# A new set of DNA methylation variants in the human genome show predominant tissue-specificity and sensitivity to reprogramming with a potential for disease susceptibility

**DOI:** 10.64898/2026.07.18.738653

**Authors:** Anuhya Anne, Lov Kumar, Monica Singh, Sumana Choudhury, Satarupa Das, Geraldine Zimmer-Bensch, Debashree Bandyopadhyay, K. Naga Mohan

## Abstract

Analyses of 3,370 normal human tissues of ectodermal, endodermal and mesodermal origins identified 12,587 regions averaging ∼585 bp with significant differences in DNA methylation levels within identical tissues. These methylation variants (MeVars) occurred in 8,037 genes enriched in neurological disorders and cancers of which, majority were tissue-specific rather than being systemic. This somatic variation was reduced by reprogramming *in vitro* into iPSCs and *in vivo* during spermatogenesis. Analysis of prefrontal cortices showed a higher incidence of MeVars in the candidate genes in controls than schizophrenia patients wherein a subset showed significantly altered transcript levels. Similar effects were observed for oral tissues and skin fibroblast cells. MeVars showed significant association with SINE1, simple and low complexity repeats, H3K27me3, H3k9me3 and H3K4me1 modifications and EZH2, SUZ12 and REST binding sites. Collectively, MeVars have postzygotic origins with an ability to ‘reset’ during reprogramming, adding a new dimension in the form of epigenetic diversity and its relevance to disease susceptibility in humans.

## Introduction

Genetic and epigenetic variation are the main contributory factors for interindividual differences in disease susceptibility. While the range of genetic variation is well-documented, information on epigenetic variation and its impact on disease susceptibility needs to be studied in similar detail. Methylation of cytosines at the fifth carbons is the first epigenetic modification identified and occurs mainly in the CpG dinucleotides that are significantly underrepresented in mammalian genomes^1,2^. This CpG suppression is not observed in ∼50% of the genes that have CpG islands (CGIs) as promoters^3,4^. CGIs are generally unmethylated, and their methylated state can influence gene expression^5,6^. For instance, in the female eutherian mammals, many genes that are silenced in the inactive X chromosome have methylated CGI promoters^7^. In several autosomal genes with CGI promoters, the strength of gene repression is directly proportional to the density of methylated CpGs^8,9^. Even for genes with non-CGI promoters, methylation has been shown to affect expression^10^. In contrast, methylation of CGIs that occur in gene bodies (between the transcription start and termination sites), has opposite effects i.e., presence of methylation is often correlated with expression^11^.

Abnormalities of DNA methylation have been reported in different disease conditions such as cancer susceptibility, imprinting disorders, etc. For instance, in some individuals with retinoblastoma susceptibility, one of the promoters of the two alleles is methylated and is associated with reduced expression of the tumor suppressor gene^12^. These individuals need somatic mutation of just one allele with the unmethylated promoter to become deficient of the tumor suppressor protein. As a second example, there are about 230-260 genes in the human and mouse, that have only one of the two alleles expressed according to their parental origin^13–14^. Expression of these imprinted genes is dependent in many cases on the methylated state of ∼50 imprinting control regions (ICRs) in human^15^. ICRs show parental allele-specific methylation patterns that are established during gametogenesis after erasure in primordial germ cells^16^. ICRs influence expression of genes located in *cis* and as a result, only one of the two parental alleles is expressed^17^. Abnormalities in methylation of ICRs cause imprinting disorders such as Prader-Willi syndrome^18^.

Results from the studies mentioned above indicate the possibility that individuals with methylation abnormality in a candidate gene for disease become susceptible despite no mutation. On this basis, it was proposed that increased methylation levels of one of the two promoters of a candidate gene decreasing its transcript levels would render the individual functionally heterozygous resulting in haploinsufficiency^19^ or an increased susceptibility for a recessive disorder through somatic mutations of the normal allele^12^. As an example, inactivation due to promoter hypermethylation of one of the two alleles of the mismatch repair gene *MLH* causes Lynch syndrome. These individuals have an increased susceptibility to develop cancers when the second allele with unmethylated promoter acquires a somatic mutation^20^. On the other hand, loss of methylation of the promoters that results in increased transcript levels is similar to triplosensitivity or ectopic expression^19^.

Genome-scale investigations on interindividual DNA methylation variation have been recently reported^21–24^. This variation is due to differences in methylation levels between individuals within the same tissues unlike majority of the studies that compared affected tissues and normal tissue counterparts or two different tissues. In one study, methylation comparisons were made using four brain regions at post-mortem and blood at pre-mortem conditions of the same individuals. In this report, sites with interindividual variation did not correlate well between brain tissues and blood whereas better correlations were obtained in different cortical regions than with cerebellum^21^. In the second study^22^ comparing fibroblasts, B-cells and T-cells from newborn and neurons and glial cells from post-mortem tissues variably methylated regions (VMRs) were identified that were mostly specific to the cell type and the others were present in all the cell types. Based on their presence in more than one cell type, the authors concluded that the VMRs are highly heritable. In the third study^23^, brain, thyroid and heart that originate from the three different germ layers were studied, resulting in identification of nearly 10,000 correlated sites of systemic interindividual variation (CoRSIVs) that are conserved across ethnic groups. Some of these variants were significantly correlated with altered transcript levels, suggesting their functional influence. Subsequently, CoRSIVs were shown to be displaying allelic bias and localization near the LINE1 and LTR transposable elements^24^. Bioinformatic analyses showed association of the genes containing CoRSIVs with conditions like cancer, mental health disease and many other disease conditions of different organ systems.

While these results constitute important foundations for epigenetic basis of interindividual differences in disease susceptibility, a detailed analysis of DNA methylation variants is needed to study whether there are regions (<u>M</u>ethylation <u>V</u>ariants or MeVars), each with successive CpG sites showing significant interindividual variation in methylation levels within the same tissue type originating from the three germ layers and if the MeVars in genes are (i) Systemic or tissue-specific, (ii) Susceptible to reprogramming events and originate during early developmental stages (iii) Conserved in different ethnic groups, and (iv) Associated with altered transcript levels of the genes of interest. These questions were addressed in this manuscript.

## Materials and Methods

### Identification of genes with MeVars

The general principle and the basis of identifying regions with hypermethylation or hypomethylation was described previously^19^. Methylation data on different samples of the same tissue type were downloaded from the NCBI GEO repository from published reports (Table 1). In the case of cortex samples from 30 controls that were from Harvard and Maryland brain banks, data using Infinium Human Methylation 450K arrays were generated at the Department of Molecular and Human Genetics, Baylor College of Medicine (USA) (GSE280053). DNA samples from six normal oral tissues were used to generate methylation data by using Infinium EPIC arrays (V1 and V2) (GSE304711). Each GEO submission was handled separately for identification of MeVars. Data on all samples for each accession number were quantile normalized by using the Minfi Bioconductor package^25^. After normalization, MeVars were identified using a custom-designed Python^®^ Script. In case of Asian population, data were pooled from different batches and pre-processed for normalization before MeVar identification. From each dataset for a given tissue, an average methylation profile was generated by calculating mean values of each CpG site among all the normal individuals. These averages were used as ‘reference’ for identifying methylation values that deviate for each CpG site in each individual (the differences referred were represented as delta-beta values). As an initial requirement for screening, only

**Table 1.**
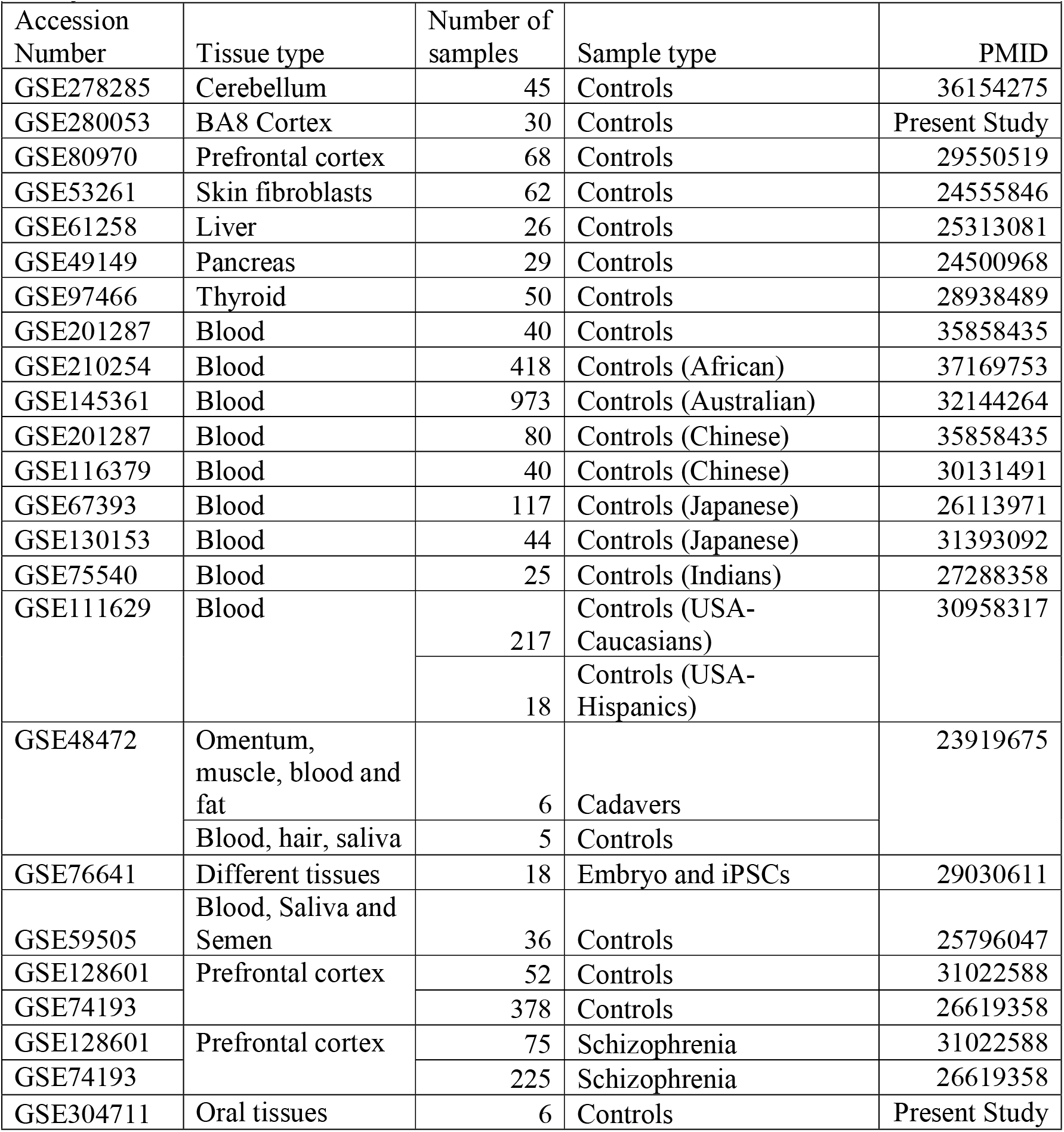
Details of DNA samples analyzed in identification and characterization of methylation variants.

CpG sites showing a minimum 20% difference in the delta-beta values were considered^19^. A Python^®^ script was used to identify regions showing the ≥ 20% difference over a stretch containing at least three probes spanning ≤ 182 bp in the 450K arrays and ≤ 75bp in the EPIC arrays. These cutoffs were used based on the distribution of probes representing the CpG sites in the promoters and gene bodies in the manifest files of Infinium HumanMethylation 450K and EPIC arrays^26^. Intergenic regions were not considered. Statistical significances in the observed methylation differences in these stretches were determined by Fisher’s exact tests to obtain initial *p* values that were then subjected to Benjamini-Hochberg corrections^27^ and regions with adjusted *p* values ≤ 0.05 were taken as Methylation Variants (MeVars). MeVars were classified using the information on the manifest files from Illumina^26^ into genic or non-genic initially and further as associated with CGIs, non-CGIs, promoters (± 1500 bp from the transcription start sites or TSS), 1^st^ Exons, 5’UTR, gene bodies and 3’UTRs. Expected proportions of these different categories were calculated from the Illumina manifest files to compare with the observed proportions among the classifications mentioned above. Bonferroni cutoff *p* value^28^ of 0.025 (0.05 ÷ 2) was used for CGI versus non-CGI comparisons, 0.003 (0.05 ÷ 15) in the case of genic Mevars spanning CGIs and 0.01(0.05 ÷ 5) for genic MeVars without CGIs.

Using the information on the coordinates of MeVars in different tissues, duplicate regions were identified and removed. Unique regions occurring within a length of 500 bp, were combined together to obtain longer coordinates resulting in a list of 12,587 coordinates in which the MeVars identified from all the samples analyzed (Supplementary Table 1).

### Graphical presentation of data

Data on methylation values of the MeVars, their frequencies, distribution among different groups (e.g., ethnic groups, tissue-specific versus systemic, controls versus patients, etc.) and correlation plots were generated using Microsoft Excel. Online tools were used for display of Venn diagrams^29^, Violin plots^30^ and protein-protein interactions (STRING^31^). Principal Component Analyses (PCA) and Hierarchical clustering data were generated using SR plots^32^ (https://www.bioinformatics.com.cn/en).

### Bioinformatic Analyses

DisGeNet and Gene Ontology terms, enrichment of transcription factor binding sites among the promoters and chromatin features (repressive and active histone modification marks) of the genes identified were studied using Enrichr^33^ and those with Benjamini-Hochberg adjusted *p* values ≤ 0.05 were taken as significant. Wherever necessary, the obtained *p* values were converted to -log_10_ *p* values and cutoff values of 1.30103 corresponding to 0.05 were taken as significant. Association of repeat sequences (LINEs, SINEs, LTR, DNA, Simple, Low-complexity, Satellite, RNA and Others) with the MeVar containing genes was studied by obtaining the information from the UCSC genome browser (https://genome.ucsc.edu/) (hg19 assembly) using a customized Python^®^ script. In this case, sequences upstream (−500 bp) and downstream (+500) to the identified MeVar coordinates were used. The proportion of MeVars containing each category of the repeat was compared with the corresponding proportion estimated for the human genome^2^ by χ^2^ two-tailed tests with Yate’s corrections. Repeats with *p* values lesser than or equal to the Bonferroni corrected threshold *p* value of 0.0056 (0.05 ÷ 9) were taken as significant. For comparing the lengths of genes with MeVars showing association with mental health disorders, cancers and others (remainder), the lengths of individual genes from the transcription start and stop sites were extracted from the UCSC genome browser using a Python^®^ Script. The gene lengths of each category were compared by student’s two-tailed *t*-test using a Bonferroni corrected cutoff *p* value of 0.0166 (0.05 ÷ 3).

### Oral tissue Methylome and Transcriptome Data

This part of the work was approved by the Institutional Human Ethics Committees of BITS Pilani Hyderabad Campus and ESIC Medical College and Hospital, Hyderabad. After obtaining informed consent from all the participants, normal oral tissues were sampled from six individuals diagnosed with oral leukoplakias. Histopathological examinations confirmed absence of any abnormal cells in these tissues. Experiments described here and the research involving the human subjects were performed following the guidelines and regulations of the Institutional Biosafety Committee, BITS Pilani Hyderabad Campus and in accordance with the guidelines of Indian Council of Medical Research (India) and the Declaration of Helsinki. Genomic DNAs and total RNAs were isolated by using AllPrep DNA/RNA minikits (Qiagen, Germany). DNAs were subjected to methylation profiling by using Illumina HumanEpic^®^ Methylation Arrays. RNA sequencing libraries were generated using NEBNext^®^ Ultra^TM^ II Directional RNA library kit (New England Biolabs, USA) and were sequenced using Illumina HiSeq 2500 (Illumina, USA) to obtain 50 million paired-end reads. After adapter trimming, the sequence reads were mapped to hg19 genome assembly using Partek^®^ Flow^®^ software v.4.0 (Illumina, USA). Log_2_-fold changes were calculated as per published protocols and differentially expressed genes were identified using FDR-adjusted *p* ≤ 0.05 (Poisson regression) and log_2_ fold-changes of ≥ 1.0 or ≤ -1.0. RNA sequencing data were deposited under the accession number in NCBI SRA (PRJNA926011).

### Methylation and expression studies

DNA and RNA samples from peripheral bloods of normal individuals were isolated using AllPrep^®^ Kit (Qiagen, Germany). Genomic DNAs were treated with bisulfite using Epitect Bisulfite conversion kit (Qiagen, Germany) and amplified with primers designed using MethPrimer^34^ (https://www.urogene.org/cgi-bin/methprimer/methprimer.cgi) following the PCR conditions as described in other reports^35^. All primer sequences used in this study were given in Supplementary Table 2. The PCR products were digested with *Taq*1α restriction enzyme (NEB, USA) at 65^0^C for 1 hour and the digested products were resolved on agarose gels to estimate the amount of digested products by using ImageJ^36^ and determine the degree of methylation in the individual samples^37^. All experiments were done in at least triplicates or six biological replicates. RNA samples were converted to cDNAs using SuperScriptIV cDNA synthesis kit (ThermoFisher, USA), amplified with primers for *AURKC* and *NAPRT1* transcripts using SYBR Green mix (BioRad, USA) and QantStudio3 real-time PCR system (ThermoFisher, USA). Primers for *β-ACTIN* were used as reference controls. Relative fold-changes in the transcript levels were calculated by 2(–ΔΔCt) method^38^. All qRT-PCR experiments were performed in triplicates for each sample. RNA sequencing data from a previously published report^39^ was used to study the relationship between methylation levels of genes containing MeVars and their transcript levels in normal oral tissues.

### Statistical analyses

For estimating the significance of the observed number of MeVars for each tissue, the total number of MeVars identified was divided by seven (the number of different tissues analyzed) to obtain an expected number. The expected and observed numbers and the corresponding remainders from the total number of MeVars were used to calculate the Fishers exact two-tailed test to obtain *p* values. A Bonferroni threshold of 0.0071 (= 0.05 **÷** 7) was then applied as a cutoff. In the case of MeVars observed in multiple tissues from cadavers, the same threshold value was used as the number of variables is seven. However, in the case of analyses of the proportion of MeVars observed in healthy individuals, a Bonferroni cutoff value of 0.01 (= 0.05 ÷ 5) was used. For determining if the ranges of methylation values of MeVars observed in multiple tissues from the embryos were significantly different from those observed in the iPSC lines derived the same tissues, median absolute deviation (MAD) values for each MeVar were determined. For this purpose, median values of the methylation levels of each MeVar were calculated. Absolute differences of each methylation value and the corresponding median value were calculated, and the median value of each difference was calculated to obtain MAD values for every MeVar. The MAD values obtained for each MeVar among the tissues and the corresponding values from the iPSCs were compared by students two-tailed *t*-tests. A Bonferroni adjusted *p* value cutoff of 4.38E-05 (= 0.05 **÷** 1141) was used. In the case of body fluids and semen, a similar approach was used except that the threshold cutoff *p* value used was 1.01E-04 (= 0.05 **÷** 495). Correlation values obtained for the respective data using Microsoft Excel scatter plots using the trendline function along with the sample sizes were used to determine the *p* values using Pearson (R) Calculator tool from Social Science Statistics^40^ and those with values ≤ 0.05 were taken as significant. To determine whether representation of MeVars is significantly different between the frontal cortex samples from controls and patients, the number of MeVars observed in each group and the remainder from the total MeVars in both groups were used to conduct a Fisher’s exact two-tailed test using the GraphPad^®^ online tool^41^ and *p* value ≤ 0.05 was taken as significant. The numbers of individuals with a specific MeVar in each group and the number of individuals without it were used to perform two-tailed χ^2^ tests of significance with Yate’s corrections using the GraphPad^®^ online tool. This yielded 112 genes with MeVars with *p* values ≤ 0.05 due to their difference in their occurrence between controls and patients. These were subjected to Bonferroni correction that gave a threshold *p* value of 4.46E-04 (= 0.05 **÷** 112). Accordingly, genic MeVars with *p* values ≤ 4.46E-04 were taken as significant. In the case of qRT-PCR experiments, Student’s two-tailed *t-*tests were performed using the ‘Data analysis plugin’ in Microsoft Excel, and *p* values ≤ 0.05 were taken as significant^37^.

## Results

Using an initial set of 45 cerebellum samples, we generated a reference methylation dataset that served to identify CpG sites showing significant differences in the methylation level (≥ 20%; *p* ≤ 0.05) in each sample. These data were then used to identify differentially methylated regions with at least three successive CpG sites with the same direction of methylation difference (either all with an increased level or all with a decreased level) and an average density of a CpG site per ≤ 182 bp^19^ (see Materials and Methods; Figure 1A). On this basis, 2,528 regions were identified as showing methylation levels significantly different from the reference of which, 571 were non-genic. Of the remaining 1,957 genic regions, 1,081 were unique and were taken as methylation variants or MeVars. Within the samples studied, there was a higher incidence of these variants at younger age (Supplementary Figure 1A). Nearly 50% occurred in just 1-2 samples each whereas 64 occurred in 5-20 individuals (Supplementary Figure 1B). When two sets of 30 and 68 frontal cortex samples were analyzed, 206 and 1,769 unique genic MeVars were identified, respectively indicating a relationship between the sample size and the number of genic MeVars identified. Some examples of MeVars identified in the two tissues were shown in Figures 1B and 1C. Only 79 MeVars were common to both cerebellum and cortex, in 203 genes (Supplementary Figure 2). In contrast to the two MeVars illustrated in Figures 1B and 1C, *TBC1D9B* contained a region that showed a tissue-specific difference or differentially methylated region (DMR) in the cerebellum and is hypomethylated (Figure 1D). This DMR did not show any individual with recognizable methylation difference in cerebellum. Comparative analysis of MeVars identified in seven different tissues of ectodermal (cerebellum, cortex and skin fibroblasts), mesodermal (blood) and endodermal (thyroid, pancreas and liver) origins suggested that the skin fibroblasts contained significantly higher proportion of MeVars whereas tissues like cerebellum, cortex, thyroid and blood had significantly lower proportions (Figure 1E). Accordingly, the number of MeVars (4,434) in skin was about three times higher than that in the frontal cortex (1,543) despite the number of samples being lesser than the former (Table 1), suggesting an association with the tissue type with the number of MeVars rather than the sample sizes (Supplementary Figure 3A). Among the seven tissues studied, only one genic MeVar present in the gene body of *HOOK2* was found among all the tissues (Figure 1F). Overall, a significantly higher proportion of MeVars occurred in CGI than non-CGI regions (Figure 1G), but there was no bias in their occurrence in the context of gene structure (Figures 1I-K).

**Figure 1.**
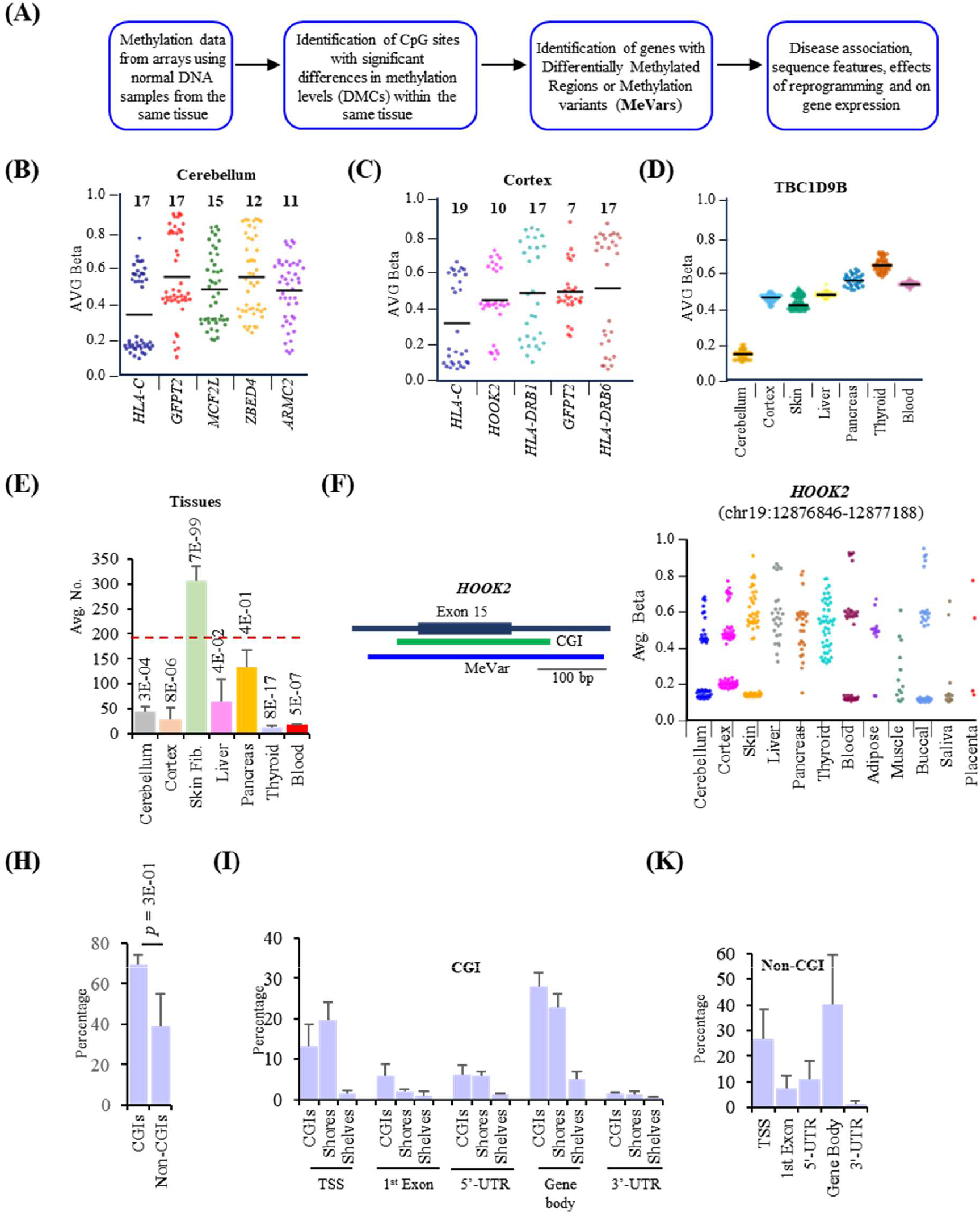
Identification of sequences showing DNA methylation variation in normal individuals. (A) Schematic outlining the method of identification of Methylation Variants (MeVars) within the same tissue in each individual. The procedure employed is given in Materials and Methods. (B-C) Examples of genic MeVars in cerebellum and cortex AVG Beta: average β-values that represent the methylation levels ranging from 0 (0%) – 1 (100%). For instance, methylation levels of the HLA-C MeVar segregates the cerebellum samples into hypomethylated (AVG Beta ∼0.2) and hypermethylated groups (AVG Beta ∼0.6). Numbers above each violin graph indicates the number of samples that showed ≥ 20% difference from the average methylation value for each sequence (shown as horizontal black line). (D) Example of a tissue specific differentially methylated region (DMR). The DMR in *TBC1D9B* is hypomethylated in cerebellum compared to the other tissues. Within these tissues, there is no individual with a methylation significantly different from the average methylation value. (E) Average number of MeVars identified in seven sets of tissues from different individuals. Bonferroni adjusted cutoff *p* value is 71.4E-03 (see Materials and Methods) is shown as dashed red line. (F) Methylation levels of MeVar identified in *HOOK2* in 12 different tissues. Left panel: Schematic showing the location of the MeVar. Right panel: Distribution of individuals with different methylation values for the MeVar in each tissue. (F) Proportion of MeVars in CpG islands (CGIs) and non-CGIs. (G-H) Distribution of MeVars in different genic regions containing CGIs (G) and non-CGIs (H).

After analysis of a total of 2,304 normal DNA samples from the seven different tissues, a set of 12,587 coordinates were derived (see Materials and Methods) that contain MeVars in 8,037 genes which when analyzed together, a statistically significant proportion were found associated with neurological/neurodegenerative and cancer disorders (Figure 2A, Supplementary Table 3). When the gene sizes of these two categories disorder were compared along with the remainder, no significant difference was observed (Supplementary Figure 4), suggesting the overall enrichment was not due to differences in disease-associated gene sizes. Pathway analysis of all the 8,037 genes showed involvement of pathways such as Neuroactive ligand-receptor interaction, Calcium signaling, cAMP signaling, Cell adhesion molecules, Pathways in cancer, Human T-cell leukemia virus 1 infection, Type I diabetes mellitus, PI3K-AKT signaling pathway, Antigen processing and presentation, Graft-versus-host disease, Serotonergic synapse, Phospholipase D signaling pathway, Amphetamine addiction, Synaptic vesicle cycle and MAPK signaling pathways (Figure 2B). STRING analysis of the proteins encoded by the total collection of genes confirmed protein-protein interactions associated with some of the significant biological processes and pathways identified through Gene Ontology analyses (Figures 2C-2D). However, it may be noted that the type of disorders and pathways associated with the MeVars varied based on the tissue type when analyzed separately (Supplementary Figure 5; Supplementary Table 4). In contrast to the 8,037 genes, analysis of the 14,291 genes without MeVars gave only one significant term each for DisGeNet (abnormal hair quantity) and the pathway terms (olfactory transduction) (Supplementary Table 5). Mapping of the genes containing MeVars and genes without MeVars to the human genome did not reveal any gross difference in their chromosomal distribution (Supplementary Figure 6).

**Figure 2.**
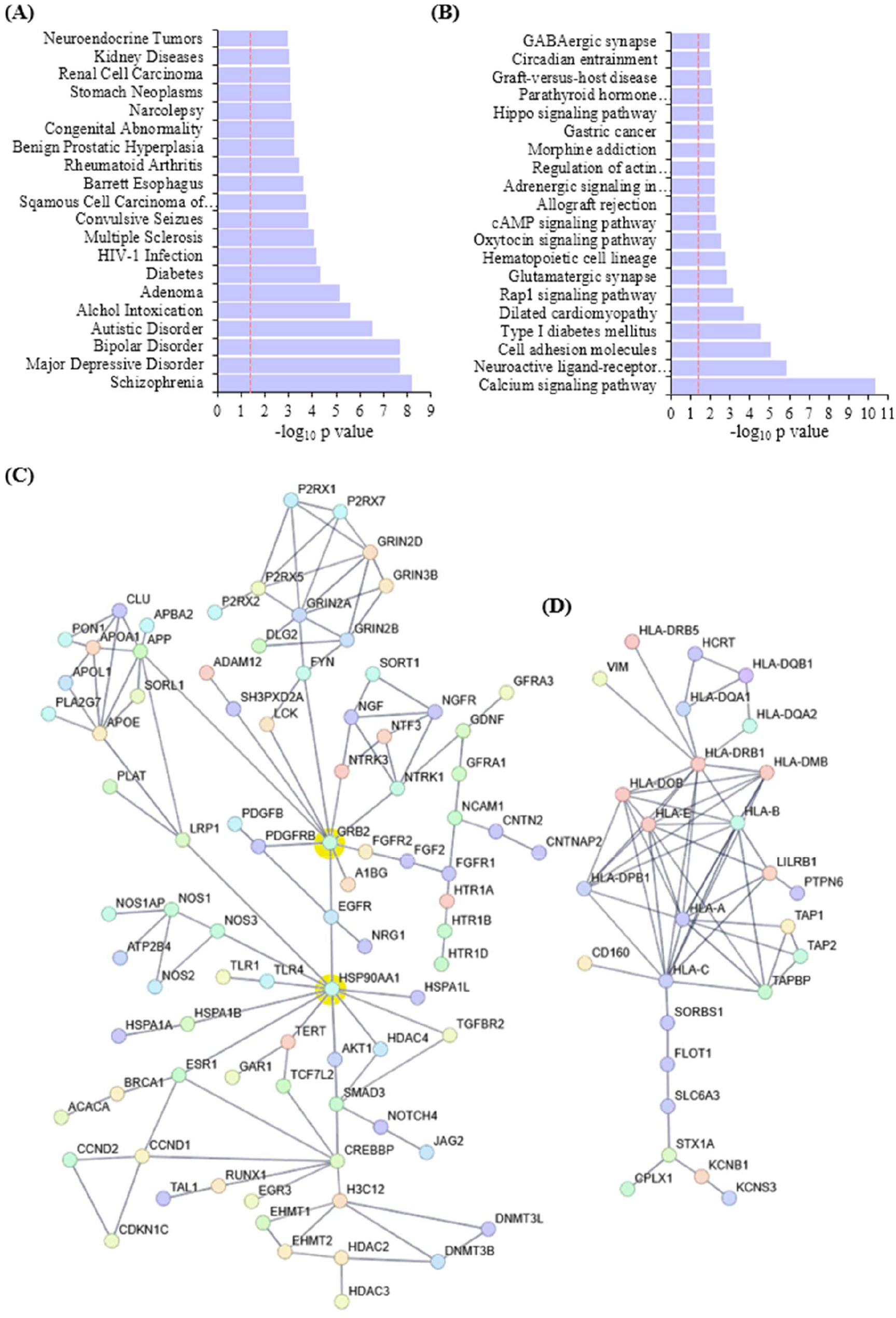
Disease ontology and KEGG pathway analysis of genes containing MeVars. (A-B) DisGeNet (A) and biological pathway analyses (B) of 8,037 genes containing MeVars. Red dotted line indicates the cutoff adjusted *p* value of 0.05 that corresponds to -log_10_ value of ∼1.3. (C-D) Protein-protein interaction analysis. In the interaction map generated using STRING in ‘C’, proteins related to neurodevelopment neurotransmitter synthesis, chromatin modification, etc. were identified. GRB2 and HSP90AA1 involved in growth factor signaling and protein folding, respectively had the highest interacting nodes. A separate cluster of proteins involved in histocompatibility was also identified (D).

## MeVars are shared in different ethnic groups

To test whether MeVars vary in different ethnic groups, blood DNA methylation data were collected on seven different populations (Table 1). We reasoned that if the initial set of samples is large from one of the three ethnic groups (Caucasians, Africans and Asians) a higher number of MeVars can be identified so that there would be better probability of identifying shared MeVars even if the sample sizes of the other two ethnic groups are smaller. Accordingly, data on 901 normal individuals from an Australian study on Caucasians were used to obtain 5,801 genic MeVars that were then compared with those identified within 418 African and 237 Asian (Chinese and Japanese) groups (Supplementary Table 1). Overall, ∼52 – 68% of the MeVars among the non-Caucasian samples were shared with the Caucasians (Supplementary Figure 7) involving a large fraction of MeVar-containing shared genes (∼82% and ∼93 % among the Africans and Asians, respectively) (Figure 3A). When a wider comparison was made using PCA by including additional 260 samples (25 Indians, 18 USA-Hispanics and 217 USA-Caucasians), in agreement with the genetic data, three distinct clusters separating Caucasian, African and Asian populations were obtained. Furthermore, Spanish and Indian data showed a closer relationship with the Caucasians (Figure 3B). This data also concurred with the hierarchical clustering of the CpG sites that showed the highest variance among the data from the seven different populations studied (Figure 3C). Bioinformatic analyses of the genes with MeVars shared between the ethnic groups revealed pathways involved in glucuronidation, morphogenesis, synaptic transmission/signaling, etc. (Supplementary Table 6). The data on methylation variation in the seven populations for *UGT2B15+17* encoding glycosyltransferase involved in detoxification^42^ indicates that some variants can be detected even in smaller sample sizes (Figure 3D).

**Figure 3.**
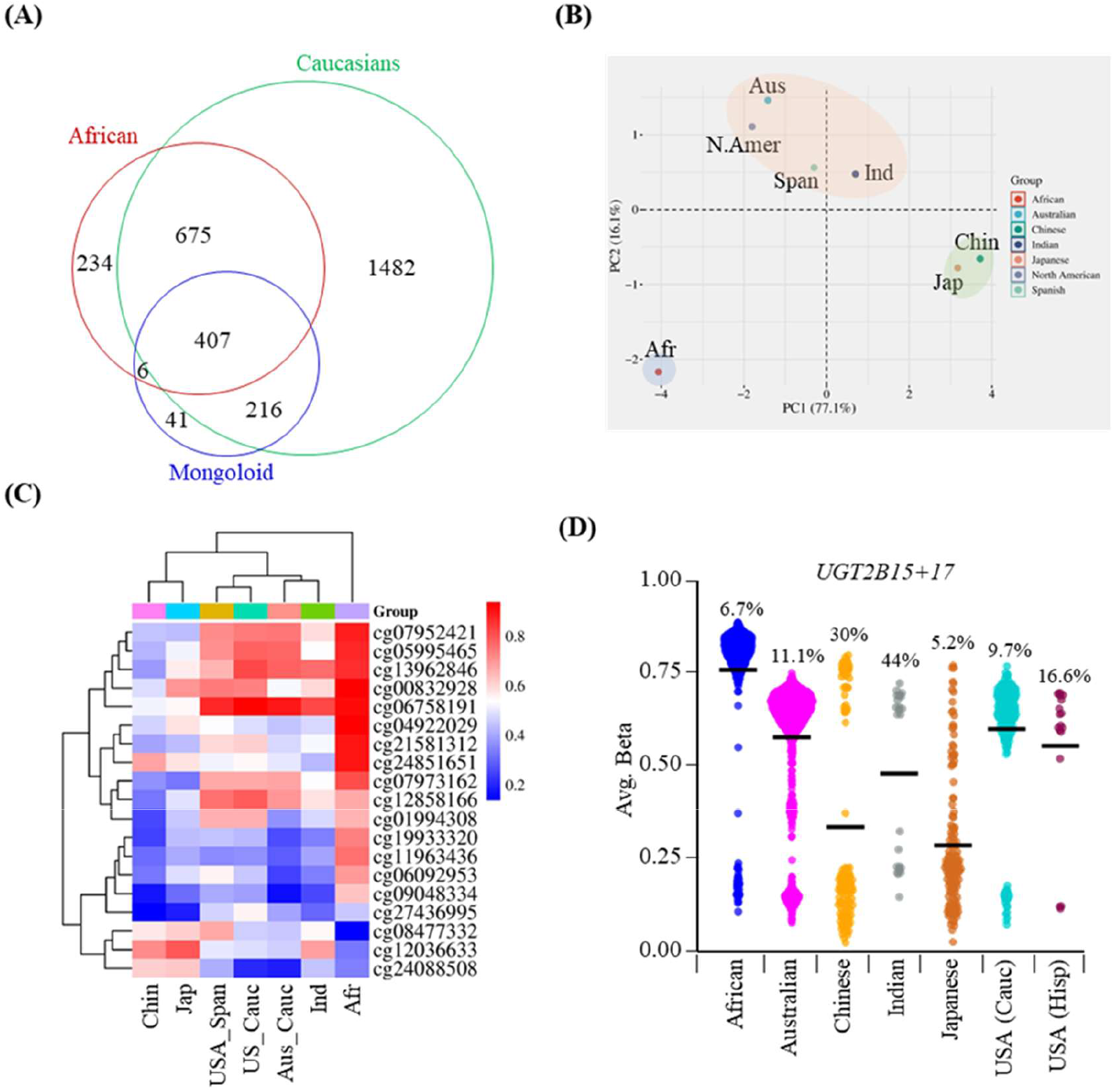
Analysis of MeVars in the three major ethnic groups (Caucasian, Asian and African). (A) Shared and ethnic group – specific MeVars identified among 901 Australians (Caucasian), 234 Chinese and Japanese (Asian) and 418 African blood DNA samples. (B) PCA analysis of the blood DNA samples in ‘A’, along with 235 Caucasians from the USA (217 North Americans and 18 Hispanics) and 25 Indians. (C) Hierarchical clustering of the CpG sites showing highest variance among the samples studied in ‘B’. Cauc: Caucasians, Span: Hispanics/Spanish, Chin: Chinese, Jap: Japanese, Ind: Indians, Afr: Africans. (D) Violin plots showing methylation levels of UGT2B15+17 in the seven population groups analyzed in ‘B’. Hisp: Hispanics. Numbers in percentage indicate the proportion of the individuals with the MeVars.

## MeVars are predominantly tissue-specific

Methylation data on multiple tissues from the same individual helps ascertain whether the MeVars are systemic or tissue-specific. When data on four different tissues from six cadaver samples were analyzed, blood and omentum showed significantly higher number of MeVars than muscle and fat tissues (Supplementary Table 7). Only 12 out of the total 722 variants (∼ 1.6%) were present in all tissues (Figure 4A). Similarly, only ∼5.0% of the MeVars were systemic among the DNAs from three different tissues from healthy individuals analyzed (Supplementary Table 8; Figure 4B). In general, the tissue-specific MeVars were significantly higher (∼98%) than systemic counterparts (∼2%; Figure 4C; *p* =4.5E-04). Examples of systemic and tissue-specific MeVars are shown in Figures 4D and 4E, respectively.

**Figure 4.**
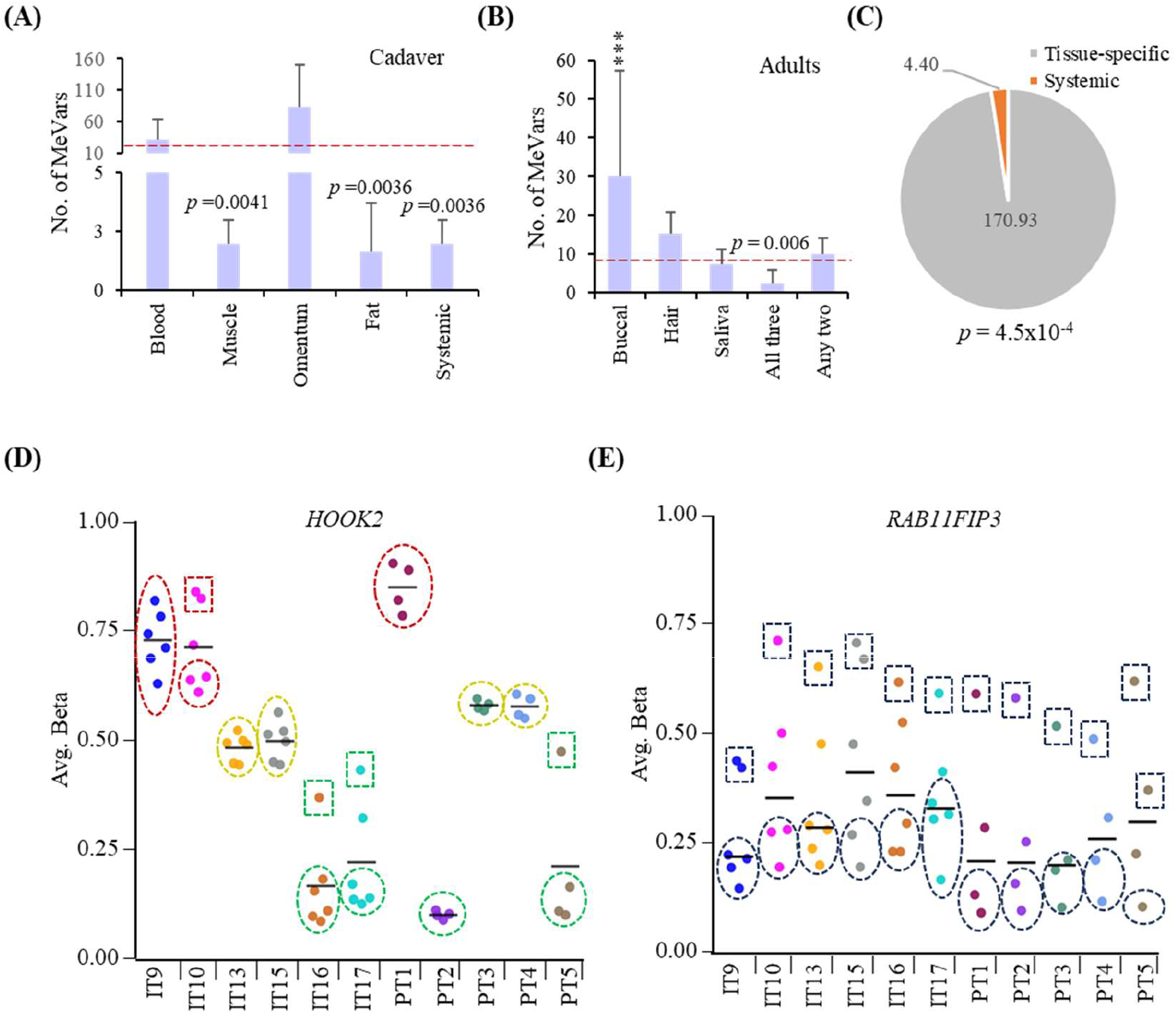
A majority of MeVars are tissue specific. (A-B) Distribution of MeVars in the DNAs from blood, muscle, omentum and fat from the same cadavers (A) and, buccal cells, hair and saliva of the same healthy individuals. The horizontal dotted line represents the average expected numbers of MeVars. (C) Pie chart showing the average number of systemic and tissue-specific MeVars identified among the samples studied in ‘A’ and ‘B’. (D-E) Methylation levels of *HOOK2*, a systemic MeVar (D) and *RAB11FIP3*, a tissue-specific MeVar (E) in the samples studied in ‘A’ and ‘B’. Black horizontal lines show the average methylation levels in each individual on the X-axes. In ‘D’, Red (hypermethylated), green (hypomethylated) and yellow (average methylation) circles or ovals indicate high, medium and low methylation levels of majority of the tissues, respectively. Squares represent the outliers. In ‘E’, average methylation levels per individual are shown as circles or ovals with the outliers as squares.

## Reprogramming reduces the tissue-specific diversity due to MeVars

In the context that a high-proportion of MeVars are tissue-specific, it is reasonable to expect that they originate during the developmental stages that are associated with tissue differentiation and/or lineage specification. If these origins were true, the differences between tissues observed should be reduced at developmental stages prior to the establishment of the tissue-specific MeVars. *In vitro* reprogramming by conversion of terminally differentiated cells into induced pluripotent stem cells (iPSCs) is one model that was used to address this question. Hierarchical clustering analysis of MeVars from six tissues from the same embryo and their iPSC derivatives (Supplementary Table 9) placed the iPSCs into one group. In addition, there was no grouping observed among tissues originating from the same germ layer. For example, kidney and muscle, despite their mesodermal origins, are more dissimilar than kidney and pancreas (endodermal origin) or muscle and skin (ectodermal origin). In addition, the distances between the iPSCs derived from the different tissues are closer than between the tissues (Figure 5A). Median deviation of about median (MAD) analyses (Figure 5B) as well as PCA (Supplementary Figure 8A) further confirmed a higher degree of similarity in the methylation profiles of the MeVars among the iPSCs than the tissues from which they were derived. The reduced heterogeneity in the methylation levels in iPSCs was accompanied by hypermethylation of a subset (e.g. *ZNF238* and *B3GNTL1*) and hypomethylation of the remaining (e.g. *TBX5* and *LTBP4*) MeVars (Figure 5C). In contrast, when tissue-specific differentially methylated regions (DMRs) that do not contain MeVars were analyzed, although an effect of the *in vitro* reprogramming was observed, the extent of methylation heterogeneity between the embryonic tissues and their iPSCs were similar (Supplementary Figure 8A).

**Figure 5.**
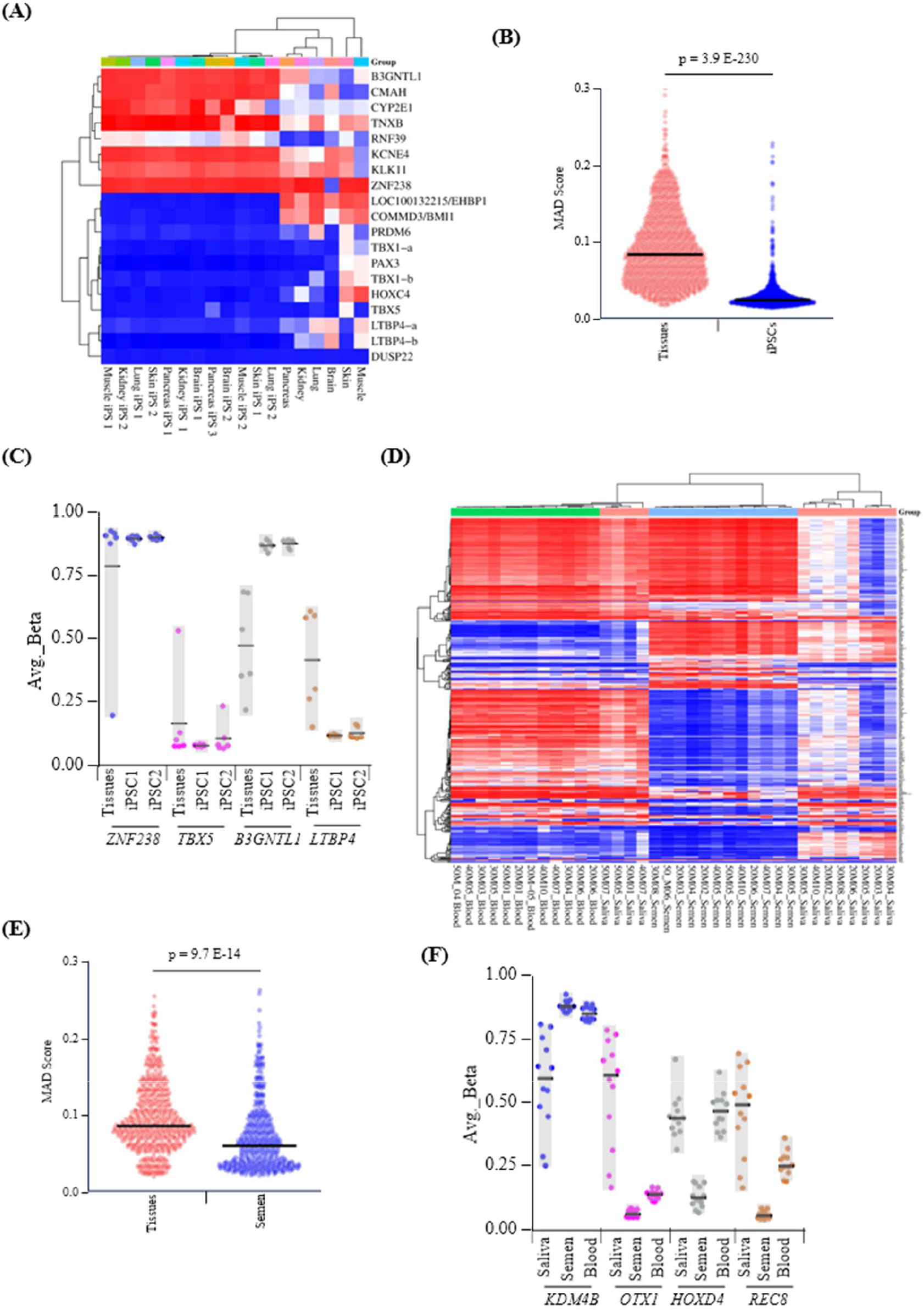
Reduction of heterogeneity in the methylation levels of MeVars in different tissues of the same individual after reprogramming. (A) Hierarchical clustering analysis of 19 MeVars showing highest variance among the different embryonic tissues and their corresponding iPSC lines. Some genes (*TBX1* and *LTBP4*) had more than one MeVar and are denoted as ‘-a’ and ‘-b’, respectively. (B) Violin plots of Median absolute deviations (MADs) of the identified MeVars in tissues and their corresponding values in the iPSCs derived from them. (C) Examples of MeVars undergoing hypomethylation or hypermethylation after reprogramming. In each case, the span of the gray box indicates the range of methylation levels observed in each cell type. (D) Hierarchical clustering of MeVars identified among the body fluids from normal healthy individuals. (E) Violin plots of MADs of MeVars detected in tissues and their corresponding values in the semen. (F) Examples of MeVars undergoing hypomethylation or hypermethylation in the sperm. Information on the gray boxes is the same as in ‘C’.

*In vivo* reprogramming occurs in gametogenesis and preimplantation development of mammals. Of the two, it is not possible to know the methylation states of the MeVars in developmental stages prior to the blastocyst as it requires syngenic pre-blastocyst stage human cells. However, comparisons of the methylation levels of tissue-specific MeVars in individuals and their gametes also provide useful information on the consequences of *in vivo* reprogramming. Of the two types of gametes, methylation data on mature oocytes of females and their somatic tissues was not available and therefore this analysis was limited to male somatic tissues and sperm. As in the case of iPSCs, analysis of data on twelve males by hierarchical clustering using data from saliva, blood and semen revealed higher degree of heterogeneity for saliva than semen (Figure 5D; Supplementary Table 10). In agreement with the previous observation (Figure 1D), blood showed the lowest range of differences in methylation levels among the MeVars. MAD analyses (Figure 5E) as well as PCA (Supplementary Figure 8B) of the methylation data on the somatic MeVars nevertheless confirmed the observed higher degree of variance among the saliva samples and its reduction in the semen samples. As in case of iPSCs, the observed reduction in the degree of differences in methylation levels of the MeVars in sperm involved hypomethylation (e.g. *OTX1*, *HOXD4* and *REC8*) as well as hypermethylation (*KDM4B*) events (Figure 5F). When tissue-specific DMRs devoid of MeVars were analyzed further, as in the case of the iPSCs, the extent of methylation differences between the tissues and the semen was similar (Supplementary Figure 8B).

## Candidate gene MeVars occur at different frequencies in both controls and schizophrenia patients

Schizophrenia (SZ), major depressive disorder, bipolar and autism spectrum disorder (ASD) were among the most significant disease ontology terms among the overall collection of genes containing MeVars identified in all the seven tissue types (Figure 2). We chose to analyze data on SZ because of high discordance rates in monozygotic twins implicating an important role of epigenetic mechanisms^27,43^. In addition, aberrant methylation patterns of candidate genes, dysregulation of DNA methylation machinery have been reported in schizophrenia^38^. Data on SZ were analyzed to test whether there is a difference in the proportion of disease – associated genes containing the genic MeVars between prefrontal cortex tissues from 547 controls and 321 patients (Supplementary Table 1). Overall, there were 7,338 MeVars in the controls and 4,088 in patients (Supplementary Table 11), indicating similar average numbers per sample (16.02 per control and 12.70 per patient; *p* = 0.0714; Figure 6A). The identified genes with MeVars yielded similar terms for DisGeNet but slightly different gene ontology (e.g. mesenchymal to epithelial cell transition in controls; neurodevelopment in patients) and pathway (e.g. Hippo signaling in controls; AMPK signaling and dopaminergic synapse in patients) terms for controls and patients (Figures 6B-C; Supplementary Table 12).

**Figure 6.**
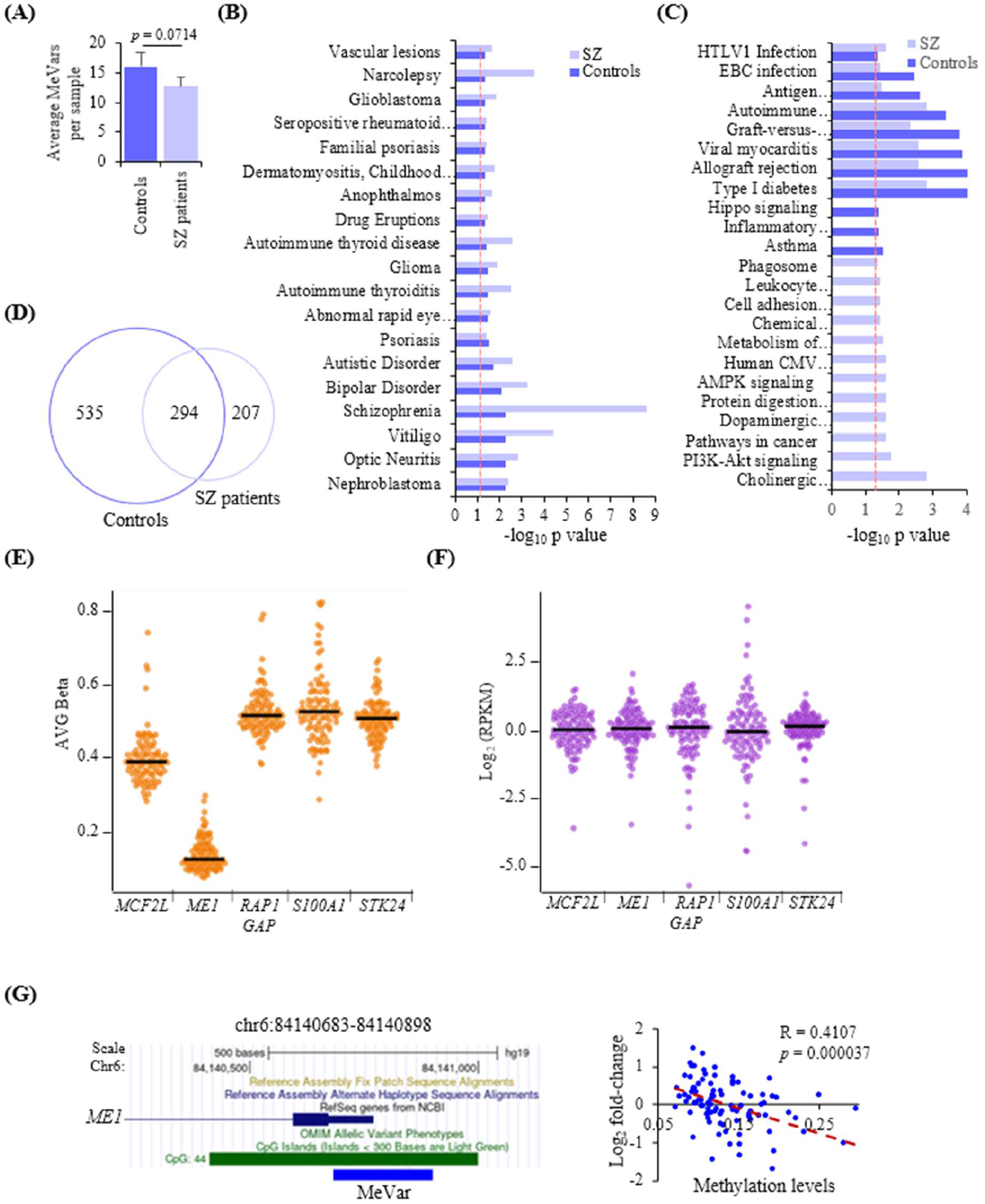
Analyses of MeVars in schizophrenia candidate genes among prefrontal cortex samples. (A) Average number of MeVars detected per each sample among prefrontal cortex tissues from controls and schizophrenia (SZ) patients. (B-C) DisGeNet (B) and Biological pathway (C) analyses of MeVars identified in controls and patients. (D) Shared and unique MeVars in controls and patients. Genes reported as dysregulated in SZ were used for generating the Venn diagram. (E-F) Methylation (E) and transcript levels reported in the BrainSeq PhaseII database for the genes shown on the X-axes. (G) Location of MeVars in the CGI promoter of *ME1* (left) coordinates of the identified MeVars is given above. (H) Correlation between the methylation levels of the indicated region and the changes from the mean value of *ME1* transcript levels.

Genes with MeVars identified in both control and patient brain samples were then compared with a list of 11,650 candidate genes listed in the SZDB2.0 database^27,44^. Of the 1,035 SZ-associated genes with MeVars identified, 534 were present in only in controls, 207 only in patients and 294 in both categories indicating significant difference (*p* = < 0.0001; Figure 6D).

A comparative analysis of the 294 candidate genes with MeVars in both controls and patients was carried out using the frequencies of individuals observed for each MeVar by two-tailed χ^2^ tests with Yate’s corrections. This analysis yielded 39 genes with significant differences between controls and patients after Bonferroni correction *p* value cutoff of 4.46 x 10^-4^ (Supplementary Table 13).

## A subset of SZ candidate genes with MeVars are associated with altered transcript levels in frontal cortex

Methylation and expression data on the same prefrontal cortex samples available for 95 controls were obtained from the BrainSeq database^45^ (https://github.com/LieberInstitute/brainseq_phase2/blob/master/BrainSeq_Phase2_phenotype_data_small_n900.csv <u>and GSE74193</u>). Transcript data on the 39 significant genes mentioned above revealed seven with a significant association between methylation and transcript level changes (Figures 6E-F; Supplementary Table 14). Four genes possessed MeVars in gene bodies wherein *FGFR2*, and *RAP1GAP* with negative correlation whereas *GLI3*, and *NUAK1* showed positive correlation. Three genes had MeVars in the promoters of which *IQSEC1* showed positive correlation whereas *ME1* and *ZBTB16* showed negative correlation (e.g., Figures 6G).

## A subset of genes with MeVars are also associated with altered transcript levels in oral tissues and skin fibroblasts

Genome-scale methylation and expression data from six normal oral tissues were analyzed to further compare the proportion of genes containing MeVars and altered transcript levels. Overall, 62 genes with MeVars were identified in three tissues (Supplementary Table 15) of which 30 showed a negative correlation (*p* < 1E-05; Figure 7A) between methylation levels and transcript level changes whereas 32 showed a positive correlation (*p* < 1E-05; Figure 7B). Among the 62, the transcript levels of 20 genes were significantly altered. When individual genes were analyzed, two showed significant negative correlation whereas four showed significant positive correlation between methylation levels and transcript levels. (e.g., Figure 7C; Supplementary Table 15; Supplementary Figure 9). This proportion (6 out of 62), when compared with that from prefrontal cortex described above (7 out of 39) showed no significant difference (*p* = 0.24).

**Figure 7.**
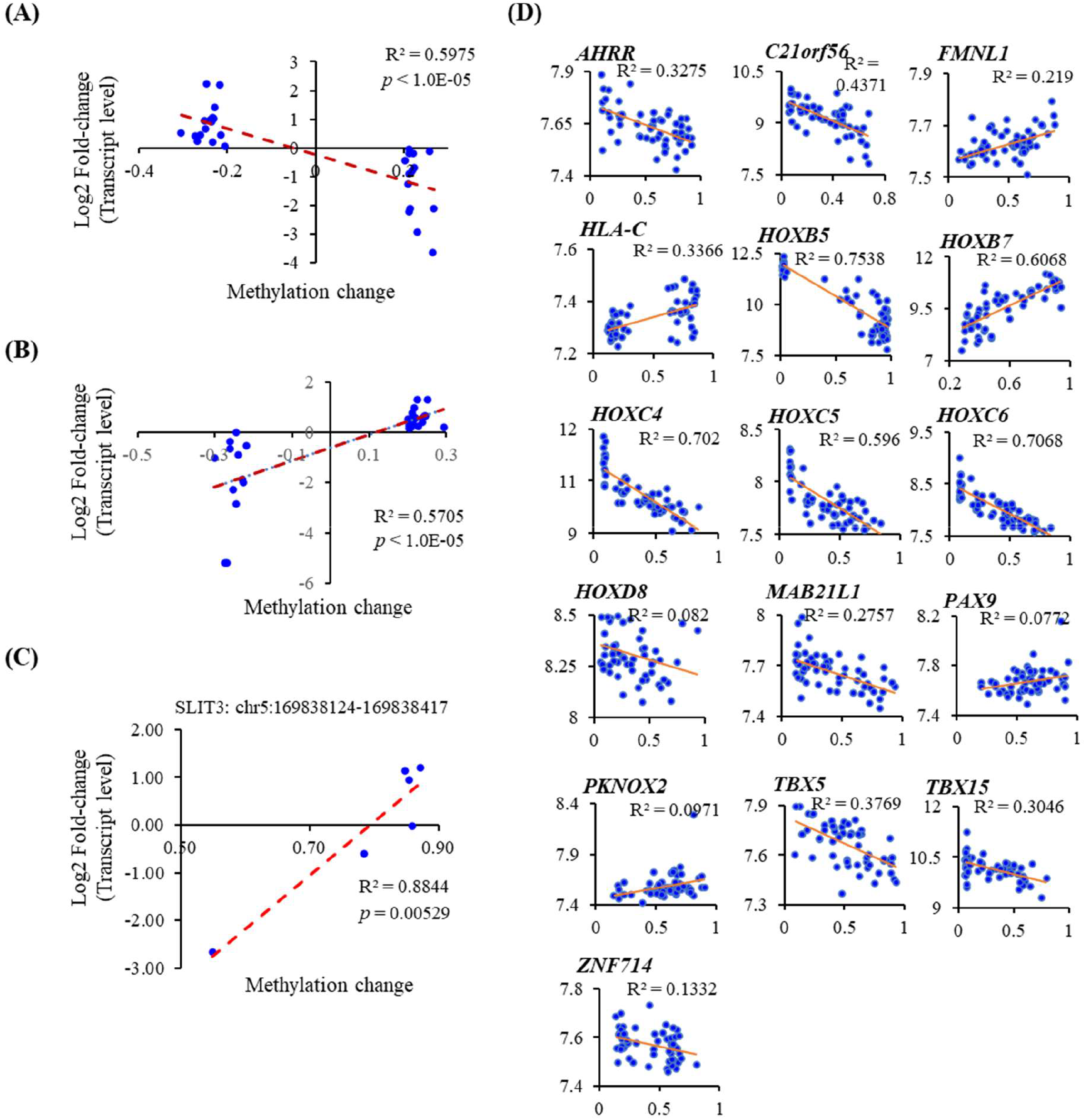
Analysis of methylation levels and transcript levels of genic MeVars. (A-B) Correlation between methylation levels of genic MeVars and transcript level changes in oral tissues. (C) Relationship between methylation levels *SLIT3* MeVar and the corresponding transcript levels. (D) Significant correlations observed between methylation transcript levels of 16 MeVar genes identified in cultured skin fibroblasts.

To further examine the relationship between MeVar methylation and expression, data on a third set of samples of 62 untransformed skin fibroblasts^46^ were analyzed. Of 7,775 unique genic MeVars identified, 59 were present in 10 or more samples (Supplementary Table 17) and were taken further for transcript level comparisons. Transcript data is available for 28 genes for which, 16 showed significant correlation between methylation and transcript levels (p = 0.035 - < 0.0001; (Supplementary Table 18; Figure 7D). Of these, 11 showed significant negative correlation.

## Replication of MeVars in *NAPRT1* and *AURKC* in Indian population

The replication potential of variable methylation of *NAPRT1* and *AURKC* was tested experimentally in a set of 30 normal Indian blood DNA samples. In both cases, the identified MeVar regions downstream to the promoter start sites are associated with the promoter CGIs.

These two genes were chosen because recessive mutations in *AURKC* cause male infertility^47^ (Mendelian disorder) whereas *NAPRT1* variant was reported to be associated with SZ in an Indian study^48^ (Complex disorder).

Methylation analyses of *NAPRT1* by COBRA assays identified 25 and five individuals with unmethylated and methylated CGI promoters, respectively. When compared with the six having low methylation, the transcript levels were significantly lower in the five samples with increased methylation (Figures 8A; *p* = 0.032).

**Figure 8.**
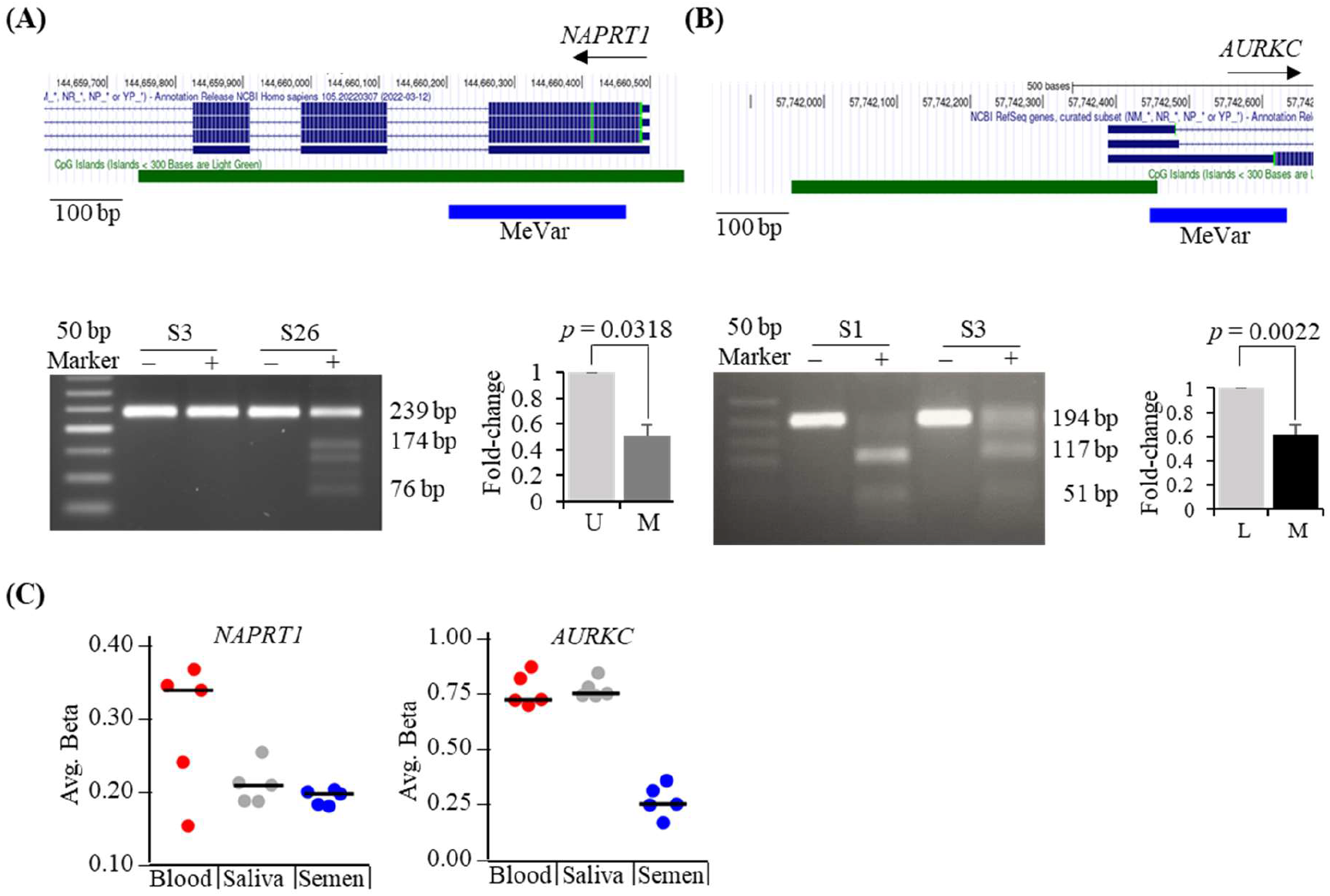
Analysis of methylation levels and transcript levels of genic MeVars in Indian samples. (A) Analysis of MeVars in *NAPRT1* promoter CpG island. Top: Location of the *NAPRT1* MeVar. Bottom Left: Identification of samples with decreased and increased methylation levels in the identified MeVar region by COBRA. ‘-’ indicates no enzyme control whereas ‘+’ indicates reactions carried out in presence of *Taq*1α restriction enzyme. Bottom Right: Transcript levels observed in samples with unmethylated (U) and methylated (M) *NAPRT1* MeVar. (B) Analysis of MeVars in *AURKC* promoter CpG island. Top: Location of the *AURKC* MeVar. Bottom Left: COBRA analysis to identify samples with moderate and high-level methlation. ‘-’ and ‘+’ signs indicate the same as in ‘E’. Bottom Right: Transcript levels measured in samples with low methylation (L) and high (H) methylation levels of the *AURKC* MeVar. (C) Methylation levels of MeVars in *NAPRT1* (Left) and *AURKC* in blood, saliva and semen of five healthy individuals.

In the case of *AURKC* CGI promoter, 12 and seven individuals showed moderate, and high methylation levels, respectively (Figure 8B). As in the case of *NAPRT1,* increased methylation levels were inversely correlated with the *AURKC* transcript levels (*p* = 0.002).

Analysis of methylation levels of the two MeVars in body fluids of the five individuals analyzed before for reprogramming effects showed that in the sperm, only *NAPRT1* showed a detectable reduction in the range of methylation values in semen compared to the two other body fluids indicating gametic reprogramming effects (Figure 8C).

## Chromatin modifications, transcription factors, repeat sequences and SNPs associated with MeVar genes

To gain insights into the type of chromatin modifications associated with the MeVar gene promoters, the gene list was mined against datasets containing histone modifications (ENCODE histone modifications, 2015 and Epigenomics Roadmap), transcription factor binding sites (ENCODE Transcription factor ChIP seq) and repeat sequences (UCSC Genome Browser, hg19 assembly). There was a highly significant association of the MeVar genes with H3K27me3 (*p* = 3.41E-145), followed by H3K4me1 (4.90E-41) and H3k9me3 (1.05E-25) among the histone modifications (Figure 9A) and four significantly associated transcription factors at the promoters mainly involved in processes related to different forms of cancer and pluripotency (Figure 9B). Consistent with the association of H3K27me3, enrichment of binding sites for SUZ12 (*p* = 7.1E-40), EZH2 (2.06E-35) and CBX2 (*p* = 0.015; not significant) were observed, REST (*p* = 2.84E-08) was the other repressor molecule that is associated with histone deacetylation.

**Figure 9.**
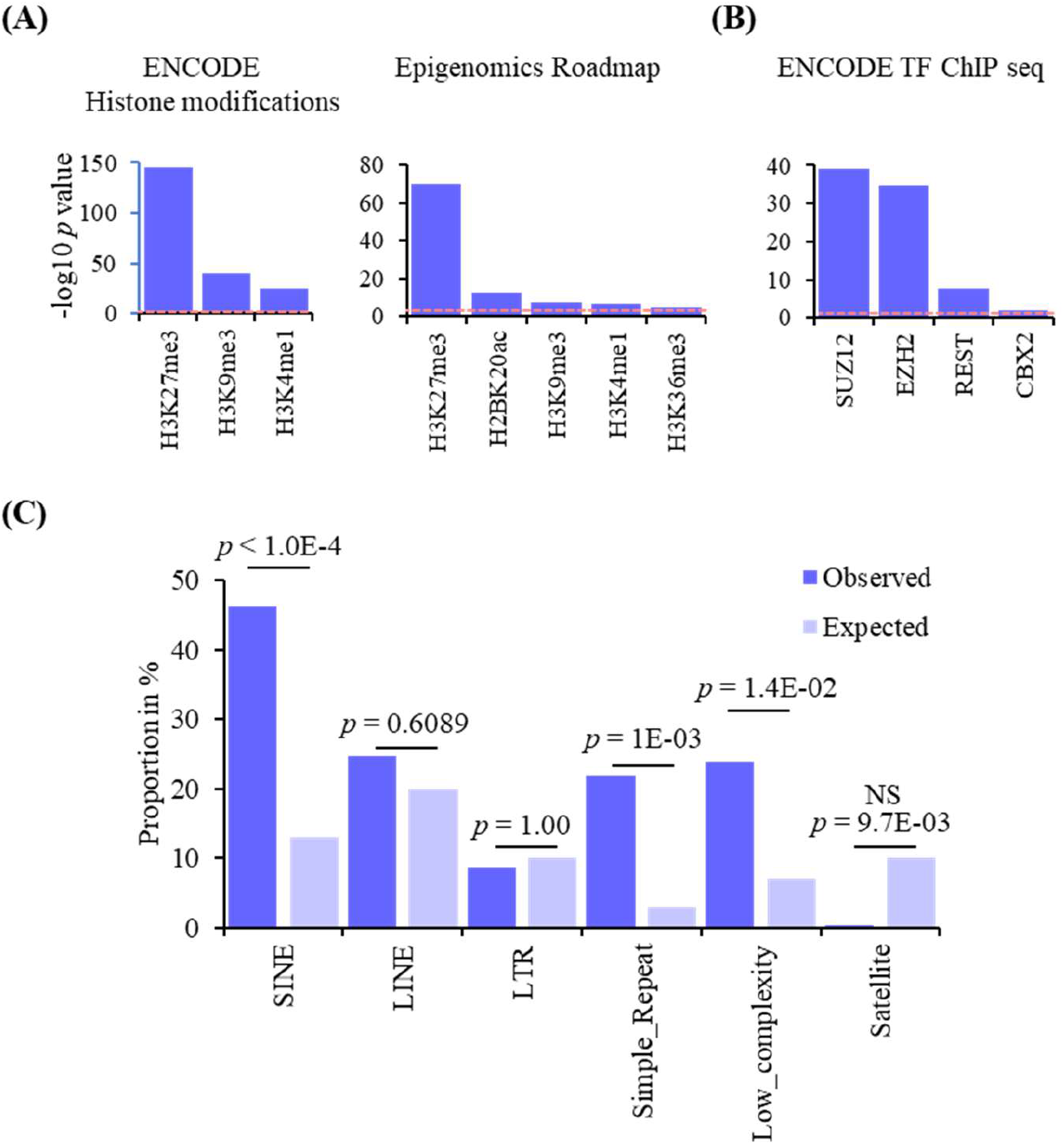
Chromatin modification and repeat analysis of MeVar genes. (A) Histone modifications predicted to be significantly enriched in the MeVar genes using the ENCODE (left) and Epigenomics Roadmap (right) databases. (B) ENCODE transcription factor ChIP seq analysis of the MeVar genes. In both ‘A’ and ‘B’, dashed red line indicates the threshold adjusted *p* value of 0.05. (C) Enrichment of different repeat sequences in the genic MeVars. For details on the *p* values obtained, refer the ‘Bioinformatic Analyses’ section under Materials and Methods.

Among the repeat sequences identified (Figure 9C; Supplementary Table 1), only three categories occurred at significantly different frequencies after Bonferroni correction: SINEs (*p* < 1E-04), Simple Repeats (*p* = 1E-03) and Satellites (*p* = 9.7E-03). When the frequencies of MeVars were searched against the number of SNPs in the dbSNP155, no significant correlation was observed. However, between tissues we observed differences in the degrees of correlation, suggesting that the tissue source is more likely reason than SNP association with the occurrence of MeVars (Supplementary Table 19).

## Discussion

Most studies comparing DNA methylation involved paired/unpaired normal and affected tissues or different normal tissues that identified differentially methylated cytosines (DMCs and differentially methylated regions (DMRs) in the literature^49,50^. Another context in which the term DMR is often used to refer to those regions that are gamete specific or imprinted^51^. The current study and a few others compared the DNA methylation patterns within the same tissue, but from only normal individuals. Prior to this report, the studies done so far focused primarily on individual CpG sites showing methylation differences^21–24^ whereas there is one that focused on regions containing three or more successive CpGs showing similar levels of methylation difference^22^. Thus, the category of the sequences identified here are distinct from those referred to as DMRs and because they occur among normal individuals in identical tissues, we refer to them as Methylation Variants (MeVars). To the best of our knowledge till date, this is the first report on comprehensive analysis of MeVars as regions showing significant differences in methylation levels among normal individuals with insights into their developmental origins, reprogramming and effects on transcript levels. The results described here provided some novel insights into these questions through the identification of MeVars in a large set of normal individuals in multiple tissues.

Analysis of multiple tissues per each of the six cadavers and five normal individuals showed that the number of MeVars varied with the type of tissue in the same person. Further, the proportion of MeVars shared among all the tissues from the sample was significantly lower. For example, the proportion of MeVars in muscle was significantly lower than the two other mesoderm-derived tissues (blood and fat). This observation indicated that a very large proportion of MeVars have likely originated much after lineage specification and maybe even at later stages that accompany terminal differentiation and doesn’t support their germline or early developmental origins. Although more than one lineage is involved in the development of the buccal cells and hair follicles than the salivary gland cells from whom the collected saliva samples were analyzed, the significantly low representation of regions showing systemic variation adds support to the later developmental origins of a majority of the identified MeVars. From the viewpoint of diagnosis based on MeVars, their widespread tissue-specific over systemic occurrence limits the utility of easily accessible biological materials (blood, saliva, urine, etc.) and necessitates specific/appropriate tissues by biopsies. This requirement is also in agreement with Hannon *et al*^21^. who reported disagreements in methylation levels between the blood and brain tissues from the same individuals. Another observation related to the occurrence of MeVars is their significant likelihood to be individual specific. This implies that DNA methylation-based testing for candidate genes is most likely to be personalized.

The presence of tissue-specific MeVars adds to the methylation differences between the tissues of the same individual. However, susceptibility to the reprogramming processes distinguishes the two categories of sequences. The extent of diversity observed in the methylation levels of the MeVars in the embryonic and adult tissues were significantly reduced when reprogrammed to iPSCs and sperm. Non-MeVar sequences showing tissue-specific differences also underwent reprogramming-associated methylation changes, but the resultant iPSCs or sperm showed similar extent of differences as in the tissues. In this regard, MeVars appear to be distinct from the non-MeVar sequences. However, viewed in the context of the possibility that MeVars might originate at latter stages of differentiation, their susceptibility to reprogramming and their origins during preimplantation or early post-implantation development cannot be ruled out. At present, it may not be feasible to address this question because of lack of possibility of having data on blastocysts or prior preimplantation stages of development of the same human embryo.

One of the main features of MeVars is their statistically significant association with genes implicated in certain neurodevelopmental disorders and cancers. Even more notable is the effect on transcript levels of an estimated ∼20-40% of the MeVar-containing genes displaying both positive and negative correlations with methylation changes. This absence of 100% correlation between the genes with MeVars and their transcript levels is consistent with an earlier report^46^. Given the limited amount of information on the genome-scale methylation and transcriptome data on the same tissues, more in-depth analyses are required to estimate whether a higher proportion of candidate genes with MeVars are associated with significant transcript level changes. For instance, when prefrontal cortex samples with MeVars in *NAPRT1* (a SZ candidate gene) were considered, transcript levels of samples with lower methylation levels were not available for correlation. However, qRT-PCR data on an independent set of blood samples showed a significant negative correlation. Similarly, a significant negative correlation was observed in case of *AURKC*, a candidate gene for autosomal recessive male infertility. In both cases, individuals with increased methylation at these loci are physiologically inequivalent to those with lesser methylation. The occurrence of a somatic or inherited heterozygous mutation affecting the unmethylated allele has the potential to reduce the overall normal protein levels, conferring an increased risk. However, as noted above, germline reprogramming confines the *NAPRT1* and *AURKC* MeVars to somatic tissues making them less likely to be heritable. However, more data is needed to rule out the possibility that gametic reprogramming reduces the variability of methylation levels of these genes.

A comparison of the frequencies of MeVars in controls and schizophrenia patients did not show any significant difference, but the difference was significant in case of the schizophrenia candidate genes. We speculate that a higher number of genes with methylation differences in controls can be viewed as a higher range of epigenetic variation with a probable protective effect similar to populations having increased genetic variation that confers a selective advantage. This possibility needs to be tested through careful analyses of cases and controls for schizophrenia as well as other neurodevelopmental disorders (bipolar, epilepsy, autism spectrum disorder, etc.) wherein the contribution of epigenetic phenomena is well-recognized^27,43^.

As noted under results, the MeVars were predicted to be enriched in sequences with chromatin features associated with members of the polycomb repression group 2 complex (SUZ12 and EZH2) that mediates the repressive H3K27me3 chromatin modification. The third protein is REST, a repressor protein associated with regulation of neurogenesis. These results of association indicate the possibility that the chromatin binding sites for these proteins might also be variable as the DNA methylation states in MeVars between different individuals. More genome-scale studies need to be conducted to establish this notion.

In summary, the identification of genes with MeVars contributes to the tissue-specific variation between individuals with an ability to impact expression levels of a subset of the genes identified in both Mendelian and complex disorders. These variants thus add to the genetic variants in the human genome (SNPs, indels, CNVs, etc.^2^) that may help explain individual differences in disease susceptibility. Future studies involving families with genome-scale methylation and transcriptome data on the same tissues would enable ascertain whether there is any heritability and/or allelic bias associated with the MeVar-containing genes.

## Funding Declaration

This work did not receive any specific funding from any agency. AA received a fellowship from the Council of Scientific and Industrial Research (Government of India). SC received a fellowship from the Indian Council of Medical Research, Government of India.

## Data Availability Statement

RNA sequencing data are deposited under the accession number in NCBI SRA (PRJNA926011).

## Acknowledgements

The authors thank the administrative and infrastructural support extended by the Centre for Human Disease Research (BITS Pilani Hyderabad Campus) and acknowledge the help of Koyena Das and Sohini Mukhopadhyay for compiling the summary data on MeVars identified in the seven tissues mentioned in this manuscript.

## Notes

### Competing Interest Statement

The authors have declared no competing interest.

