## Supplemental tables for "A new set of DNA methylation variants in the human genome show predominant tissue-specificity and sensitivity to reprogramming with a potential for disease susceptibility"

### Supplementary Figures

**Supplementary Figure 1. MeVars identified in 45 normal cerebellum samples. (A)** Distribution of MeVars according to age of the individuals. **(B)** Frequency of occurrence of unique MeVars.

**(A)**

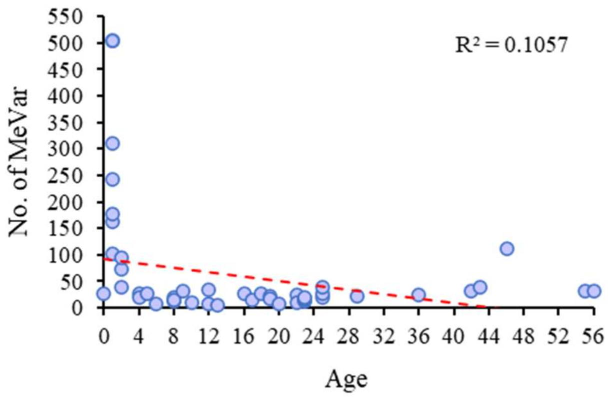

**(B)**

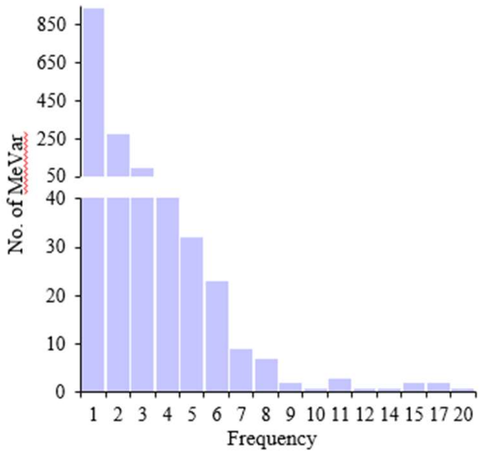

**Supplementary Figure 2. Comparison of MeVars between cerebellum and cortex.** (A) Venn diagram of coordinates containing 1,081 MeVars identified in 45 cerebellum samples and 1,769 identified in 68 cortex samples. (B) Venn diagram of genes containing MeVars in cerebellum and cortex. About 2% of the MeVar coordinates and ~ 10% of the genes with MeVars were shared between the two tissues.

**(A)**

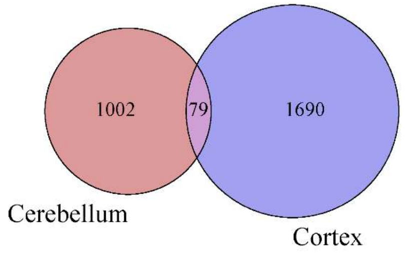

**(B)**

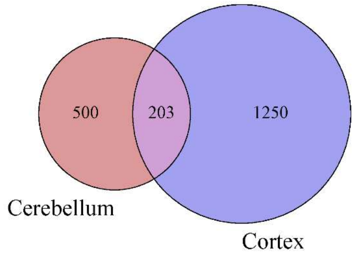

**Supplementary Figure 3. Correlation between the number of genic MeVars observed and the sample sizes.** Number of MeVars observed in different **(A)** tissues with varying sample sizes. **(B)** Blood data sets of varying sample sizes.

**(A)**

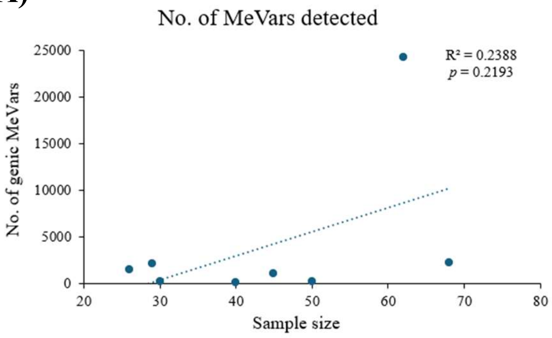

**(B)**

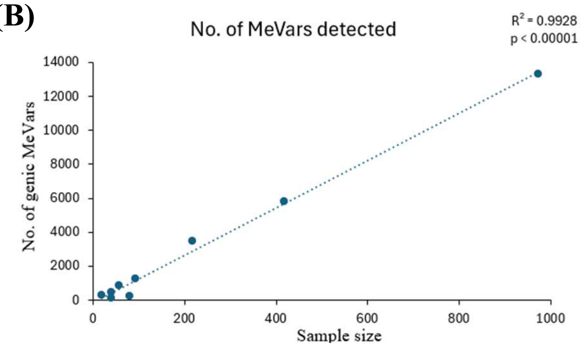

**Supplementary Figure 4. Length of genes containing MeVars and associated with cancers, mental health disorders and others (non-cancer and non-mental health disorder – associated genes.**

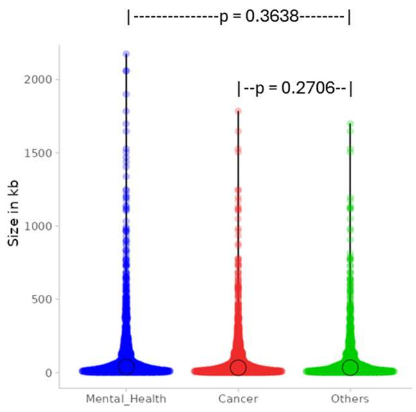

**Supplementary Figure 5. Genes with MeVars among tissues of ectodermal (cerebellum, cortex and skin), endodermal and mesodermal origins.** (A) DisGenet analyses. In each case, horizontal dashed red line indicates cutoff of significance (Benjamini-Hochberg adjusted *p* value of 0.05).

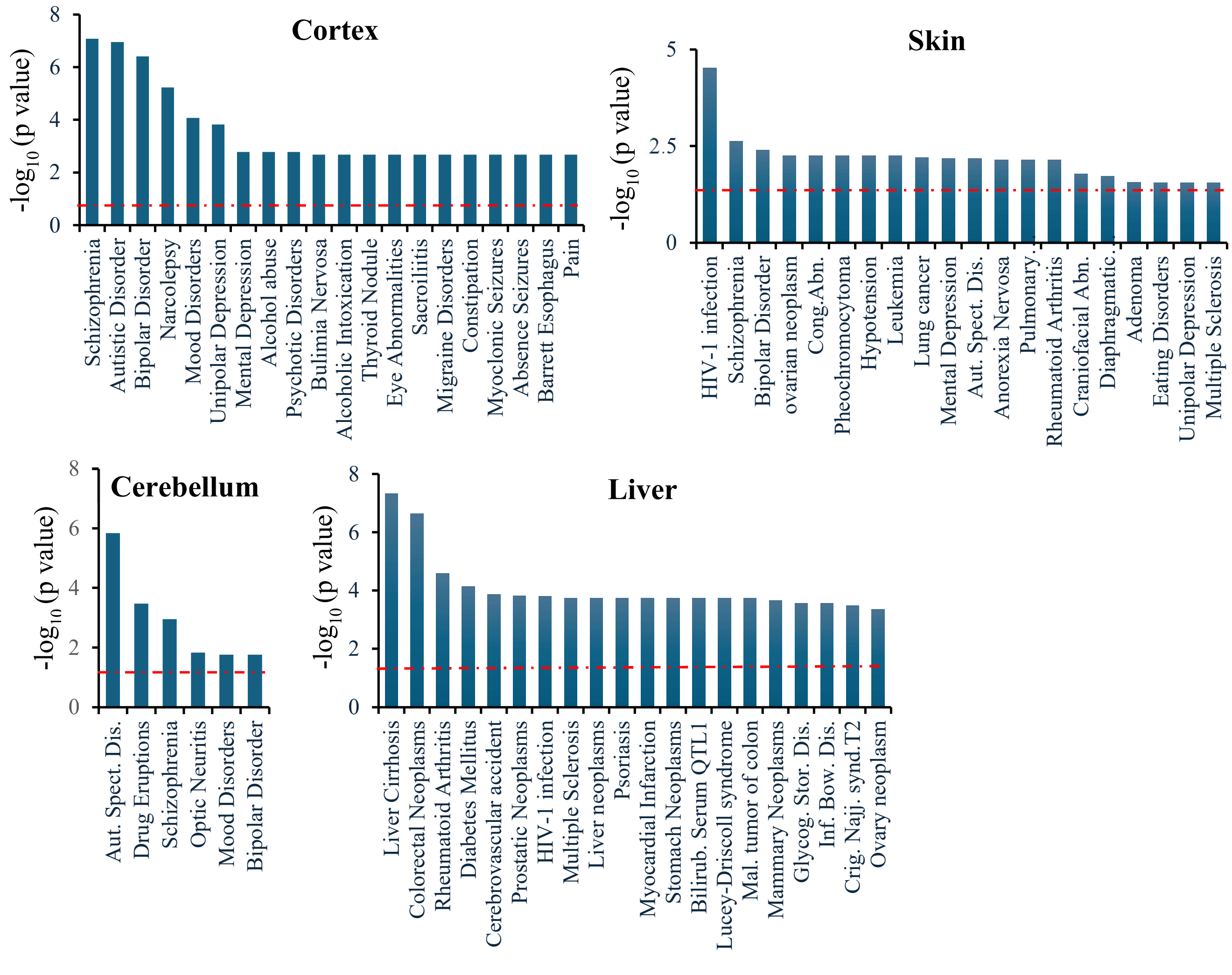

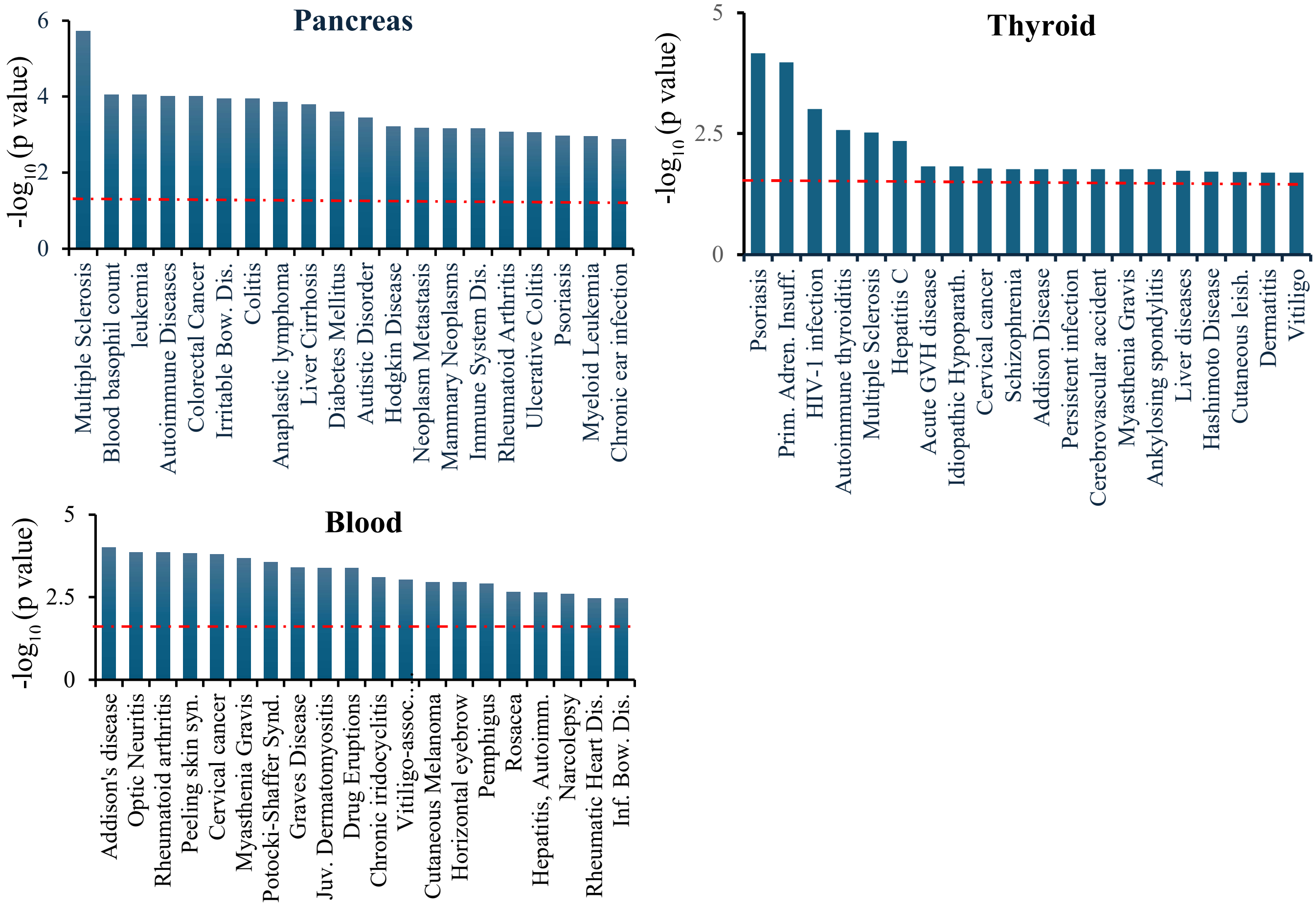

**Supplementary Figure 6. Distribution of genes with MeVars (left, blue) and without MeVars (right, red).** Ideograms showing the locations of MeVars and non-MeVars.

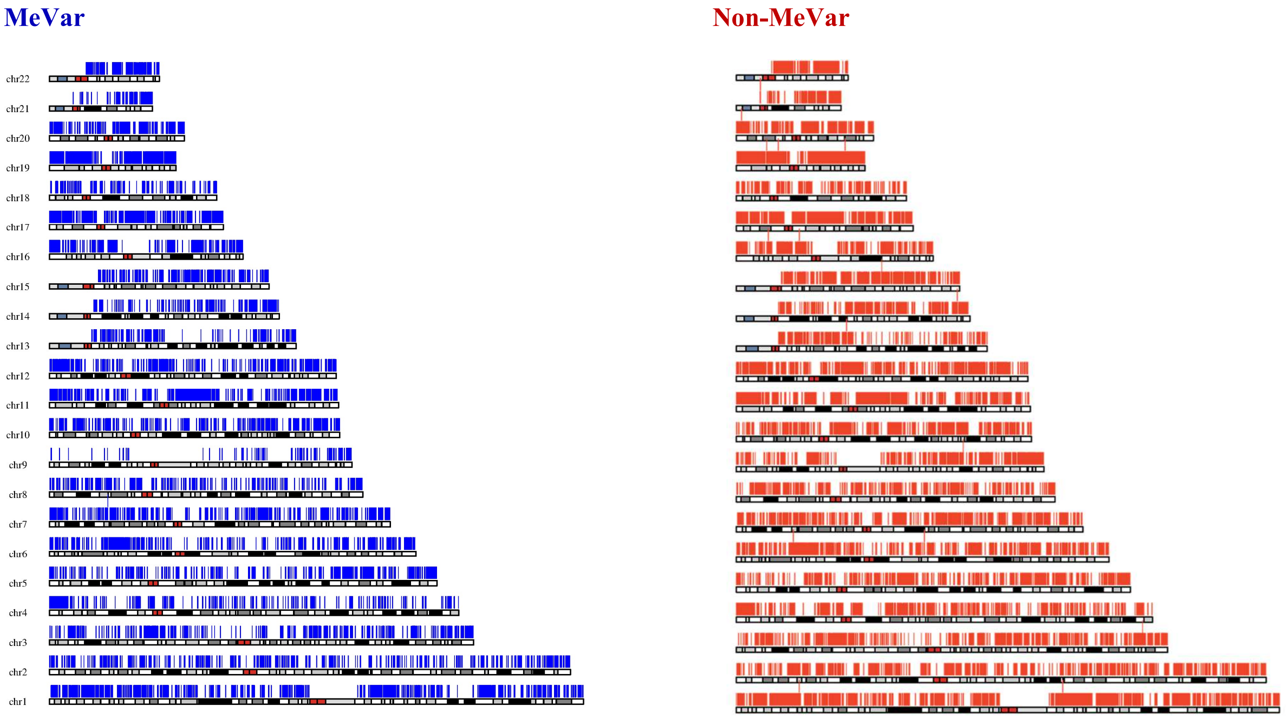

**Supplementary Figure 7. Shared and distinct MeVars among different ethnic groups.** Venn diagram showing MeVars identified in Caucasian (Australian), Asian (Chinese + Japanese) and African populations. About 50% of the MeVars in Africans are shared with Caucasians whereas ~14% with Africans. Among Asians, ~63% were shared with Caucasians and ~ 37% with Africans. For Caucasians, ~22% are shared with Africans and ~10% with Mongoloids.

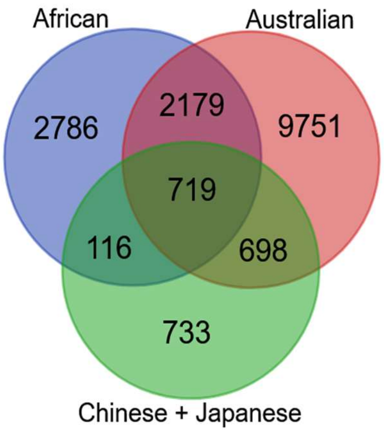

**Supplementary Figure 8. Principal Component Analysis of methylation levels of MeVars and non-MeVars.** PCA of data on MeVars (A) and non-MeVars (B) from six different tissues from the same embryo and the corresponding iPSC lines. PCA of data on MeVars (C) and non-MeVars (D) from saliva, blood and semen of five individuals (S1, S6 and S11; S2, S7 and S12; S3, S8 and S13; S4, S9 and S14; S5, S10 and S15, respectively).

**(A)** **(B)**

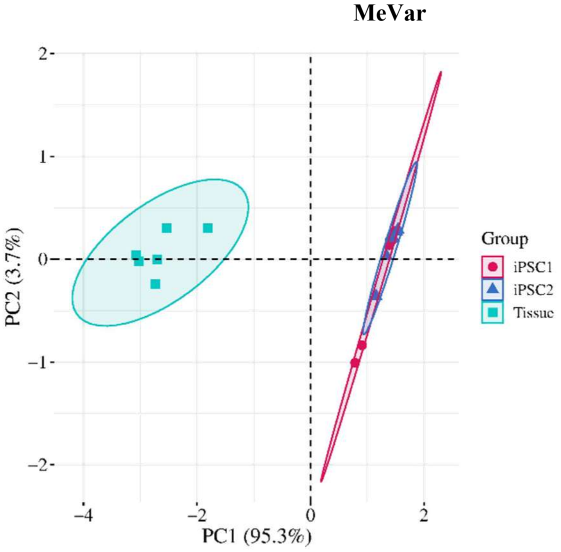

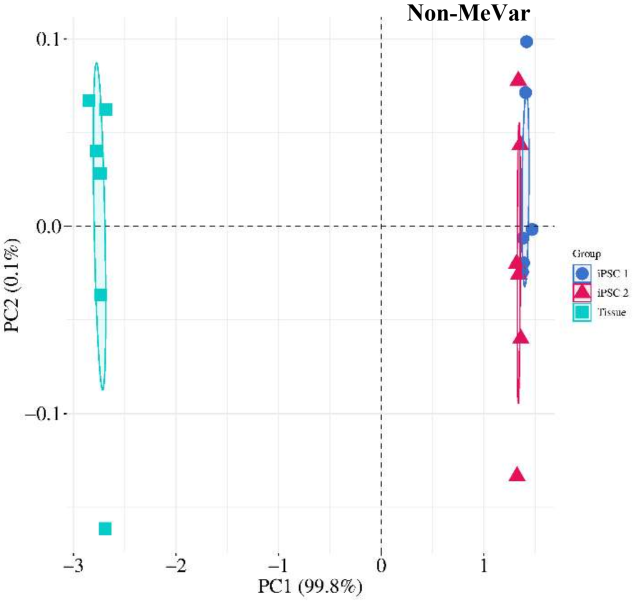

(C)

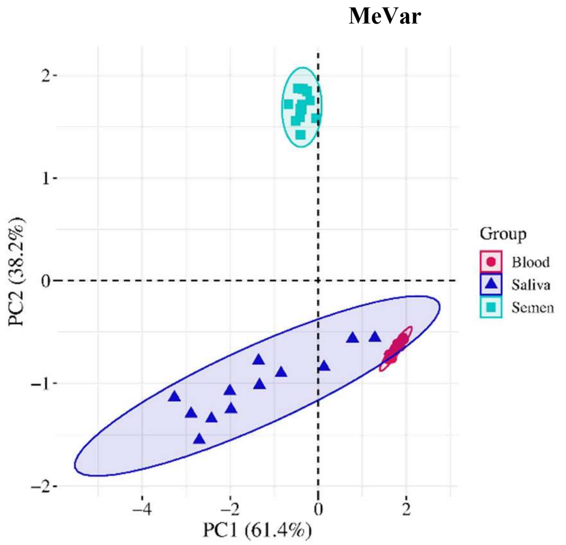

(D)

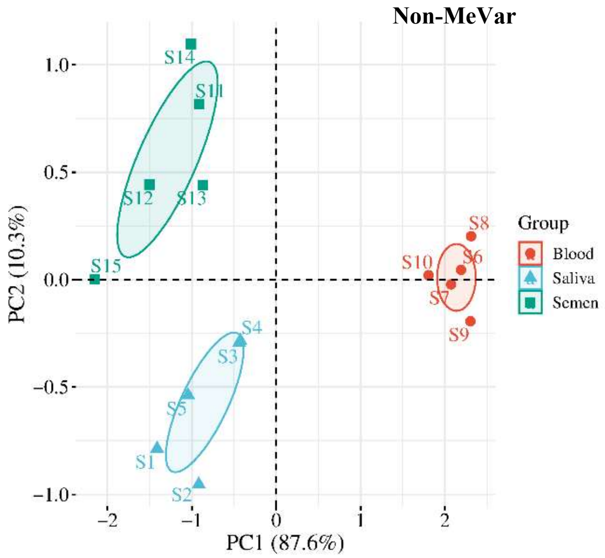

**Supplementary Figure 9. Genes with MeVars in oral mucosa showing significant correlation with changes in transcript levels.** In each case, methylation levels are shown on the X-axes and the $\log_2$ fold-changes in transcript levels on the Y-axes. $R^2$ values of 0.66 and higher correspond the $p$ values ≤ 0.05 and the genes names with significant correlations are given in blue.

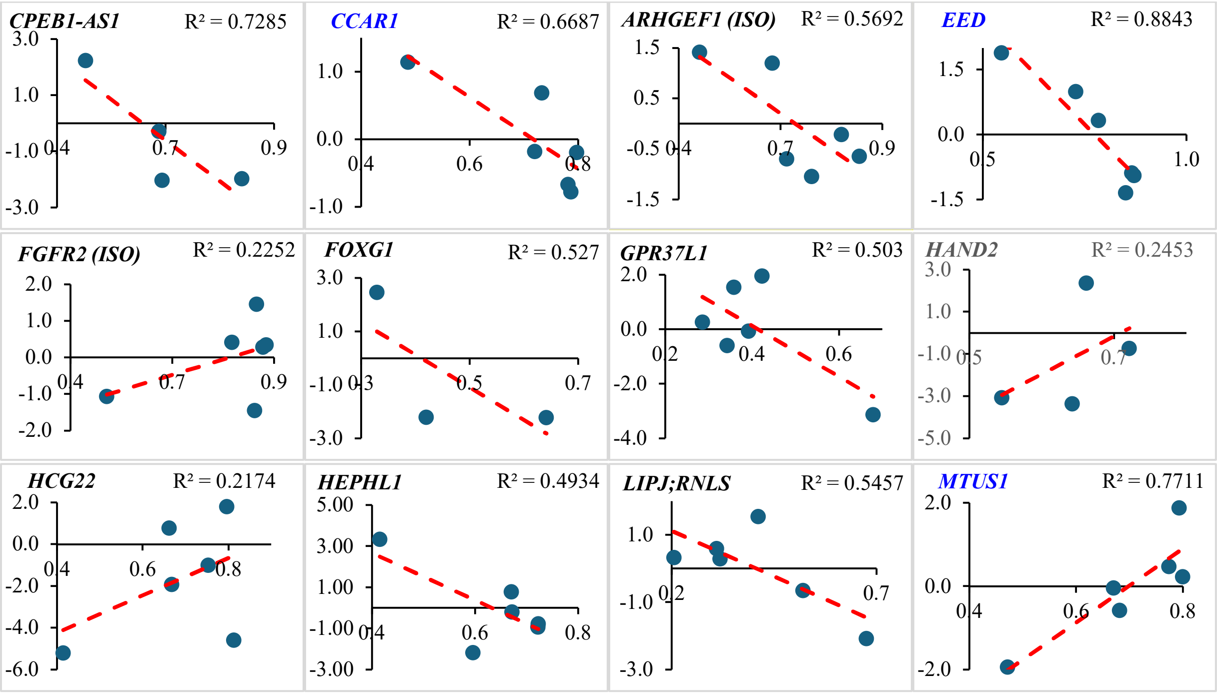

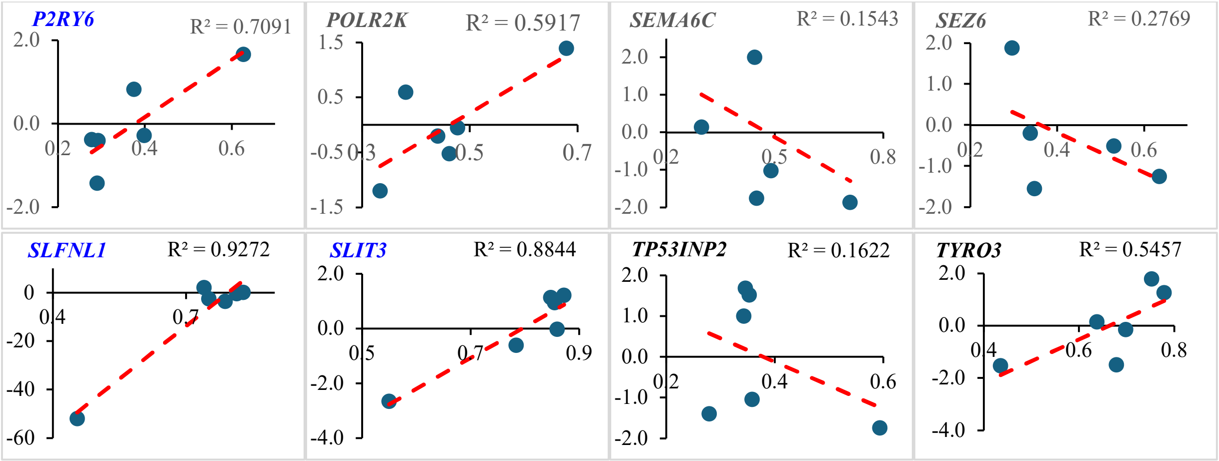
P2RY6
R² = 0.7091
POLR2K
R² = 0.5917
SEMA6C
R² = 0.1543
SEZ6
R² = 0.2769
SLFNL1
R² = 0.9272
SLIT3
R² = 0.8844
TP53INP2
R² = 0.1622
TYRO3
R² = 0.5457

**Supplementary Table 1. Genic Mevars identified in tissues of ectodermal, endodermal and mesodermal origins and their association with different categories of repeat sequences.**

| UCSC_REFGENE_NAME | ELATION_TO_UCSC_CPG_ISLAN | UCSC_REFGENE_GROUP | Coordinates | Length | n MeVar C | MeVar Coo | ast MeVar | leVar Coor | MeVar Co | MeVar Coo | leVar Coo | od MeVar ( | ood MeVa | lood MeVa | od MeVar ( | Blood Me\ | Blood MeV | SINE | LINE | LTR | DNA | mple_Repe | w_complex | Satellite | RNA | Other | Unknown |
|---|---|---|---|---|---|---|---|---|---|---|---|---|---|---|---|---|---|---|---|---|---|---|---|---|---|---|---|
| FAM41C | NA | TSS200 | chr1:812248-812679 | 431 | | | | | | | | | | | | | | | | | | yes | | | | | |
| FLJ39609 | Island;S_Shore | TSS200;TSS1500;Body | chr1:854766-855649 | 883 | | | 17 | | | | | | 2 | 1 | | | | | yes | | | | | | | | |
| SAMD11 | Island | 5'UTR | chr1:861265-861886 | 621 | | | | | | | | | | | | | | | | | | | yes | | | | |
| SAMD11 | Island | Body | chr1:870791-871980 | 1189 | | | 17 | | | | | 0 | 8 | 4 | 1 | 6 | | yes | | | | | | | | | |
| SAMD11 | S_Shore | Body | chr1:879375-879958 | 583 | | | | | | | | | 1 | | | | | | | | | yes | yes | | | | |
| KLHL17 | Island | Body | chr1:897500-897949 | 449 | | | | | | | | | 1 | | | | | | | | | yes | | | | | |
| PLEKHN1;KLHL17 | Island | TSS1500;3'UTR | chr1:901062-901725 | 663 | | | 5 | | | | | | | | | | | | | | | yes | | | | | |
| PLEKHN1 | S_Shore | Body | chr1:906965-907708 | 743 | | | | | | | | | | | | | | yes | | | | | | | | | |
| C1orf170 | N_Shore | Body | chr1:911601-912055 | 454 | | | | | | | | 0 | 1 | 1 | | 2 | | | | | yes | yes | yes | | | | |
| C1orf170 | N_Shelf | Body;TSS200 | chr1:917118-917819 | 701 | | | 11 | | | | | | | | | | | | | | | | | | | | |
| ISG15 | N_Shore;Island | TSS1500;TSS200;Body | chr1:948625-949893 | 1268 | | | 15 | | | | | 0 | 3 | | | 1 | | yes | | | | | | | | | |
| AGRN | Island | Body | chr1:968558-970294 | 1736 | | | | 2 | 1 | | | | | 1 | 1 | | | | | | | yes | yes | | | | |
| AGRN | Island | Body | chr1:975551-976227 | 676 | | | | 2 | | | | | | | | | | | | | | yes | | | | | |
| AGRN | Island | Body | chr1:983386-983611 | 225 | | | | | | | | | | | | | | yes | | | | | | | | | |
| C1orf159 | N_Shore | Body | chr1:1022642-1023319 | 677 | | | 10 | | | | | | | | | | | | | | | | | | | | |
| C1orf159 | Island | 5'UTR | chr1:1046086-1046230 | 144 | | | | | | | | | | | | | | yes | yes | yes | | | | | | | |
| C1orf159 | Island | 5'UTR | chr1:1050579-1050737 | 158 | | | | | | | | 1 | | | | | | | yes | | | | yes | | | | |
| C1orf159 | S_Shore | TSS1500 | chr1:1052600-1052949 | 349 | | | 6 | | | | | | | | | | | yes | | | | | | | | | |
| MIR200A;MIR429;MIR200B | S_Shelf;S_Shore | TSS200;TSS1500;Body | chr1:1101302-1105518 | 4216 | | | 17 | | | | | | 1 | | | | | | | | | yes | yes | | | | |
| TTLL10 | N_Shore;Island | TSS1500;TSS200;5'UTR | chr1:1108521-1110107 | 1586 | | 1 | 14 | 3 | | 2 | | 3 | 7 | | | 3 | | yes | yes | yes | | | | | | | |
| TTLL10 | N_Shore | TSS1500;5'UTR;Body;TSS200 | chr1:1114579-1115575 | 996 | | | 6 | | 1 | | | | 3 | | | | | | | | | | | | | | |
| TTLL10 | S_Shelf | Body | chr1:1118157-1118391 | 234 | | | | | 1 | | | | 1 | | | | | | | | | yes | | | | | |
| TNFRSF18 | N_Shore;Island;S_Shore | Body;1stExon;TSS200 | chr1:1141617-1142617 | 1000 | | | 7 | | | | | 2 | | | | | | | | | | | | | | | |
| TNFRSF4 | S_Shore;N_Shore;Island | TSS1500;Body | chr1:1148973-1150936 | 1963 | | | 10 | | | | | | 3 | | | | | yes | | yes | | | yes | | | | |
| FAM132A | Island | Body | chr1:1178039-1178245 | 206 | | | | | | | | | 1 | | | | | | | | | | | | | | |
| SCNN1D | NA | TSS1500;5'UTR;TSS200;1stExon | chr1:1217103-1217664 | 561 | | | 4 | | | | | | | | | | | yes | | | | | | | | | |
| SCNN1D | N_Shore;N_Shelf | Body | chr1:1226776-1227307 | 531 | | | 3 | | | | | | 2 | | | | | | | | | | | | | | |
| ACAP3 | Island | Body | chr1:1228990-1229589 | 599 | | | | | | | | 2 | 1 | | | 1 | | | | | | yes | | | | | |
| ACAP3 | N_Shelf | Body | chr1:1238436-1238702 | 266 | | | | | | | | | 2 | | | | | | | | | | | | | | |
| PUSL1 | Island | Body | chr1:1245014-1245184 | 170 | | | 4 | | | | | | | | | | | | | | | | | | | | |
| GLTPD1;CPSF3L | N_Shore | 5'UTR;TSS1500 | chr1:1261347-1261797 | 450 | | | | | | | | | 1 | | | | | | | | | | | | | | |
| TAS1R3 | N_Shore;Island | TSS200;Body;TSS1500 | chr1:1266178-1269399 | 3221 | | | 25 | | | | | | 7 | | | | | | | | | | yes | | | | |
| DVL1 | S_Shore | Body | chr1:1273375-1273707 | 332 | | | | | | | | | | | | | | | | | | yes | | | | | |
| MXRA8 | Island | Body | chr1:1289806-1290712 | 906 | | | | | | | | 4 | 7 | 1 | | 8 | 1 | | | | | | yes | | | | |
| MXRA8 | S_Shelf | 1stExon;TSS200;TSS1500 | chr1:1293868-1295077 | 1209 | | | 2 | 1 | | | | | | | | | | | | | | | | | | | |
| AURKAIP1 | Island | Body | chr1:1309331-1309641 | 310 | | | 2 | | | | | | | | | | | | | | | | | | | | |
| CCNL2 | Island | Body;3'UTR | chr1:1322783-1322923 | 140 | | | | | | | | | | | | | | | | | | | | | | | |
| LOC441869 | S_Shore | 1stExon;Body;5'UTR | chr1:1356565-1356902 | 337 | | | | | | | | | 1 | | | | | | | | | yes | | | | | |
| TMEM88B | N_Shore;Island | TSS1500;1stExon | chr1:1361216-1361729 | 513 | | | | | | | | 2 | | | | | | yes | | | | | | | | | |
| VWA1 | N_Shore | TSS1500 | chr1:1369466-1370775 | 1309 | | | | 1 | | | | | | | | | | | | | | yes | yes | | | | |
| VWA1 | Island | 3'UTR | chr1:1375448-1375967 | 519 | | | | | | | | | 4 | | | 1 | | | | | | | yes | | | | |
| ATAD3C | NA | TSS200 | chr1:1385047-1385371 | 324 | | | 1 | | | | | | | | | | | yes | | yes | | | | | | | |
| ATAD3C | N_Shore;Island | Body | chr1:1395945-1396169 | 224 | | | | | 2 | | | 1 | 1 | 1 | | | | yes | | | | | | | | | |
| ATAD3C | S_Shore | Body | chr1:1397412-1398348 | 936 | | | 3 | | | | | 1 | 2 | | | | | | | | | | | | | | |
| C1orf70 | Island;N_Shore | Body;nan;1stExon | chr1:1475143-1476269 | 1126 | | 2 | | | 1 | | | | | | 1 | | | yes | | | | yes | yes | | | | |
| SSU72 | S_Shore | Body | chr1:1478170-1478827 | 657 | | | | 1 | | | | | | | | | | | | | | | | | | | |
| SSU72 | S_Shore | Body | chr1:1490676-1490906 | 230 | | | 3 | | | | | | | | | | | yes | | | | | | | | | |
| MIB2 | N_Shore | TSS1500 | chr1:1549615-1549818 | 203 | | | 1 | | | | | | 1 | | | | | yes | yes | yes | | | | | | | |
| MIB2 | N_Shore | Body | chr1:1562771-1563096 | 325 | | | | | 1 | | | | | | | | | | | | | | yes | | | | |
| MIB2;MMP23A;MMP23B | Island | Body;3'UTR;TSS1500 | chr1:1564422-1567257 | 2835 | | | | | | | | | | | | | | | | | | | | | | | |
| LOC728661;CDK11B | N_Shore | Body;5'UTR | chr1:1606206-1606557 | 351 | | | | | | | | | 1 | | | | | yes | | | | | | | | | |
| MMP23A;CDK11B | N_Shore | TSS1500;Body;5'UTR | chr1:1630742-1631100 | 358 | | | 2 | | | | | | 1 | 1 | | | | | | | | | yes | | | | |
| CDK11B;CDK11A | Island | TSS200 | chr1:1655803-1655942 | 139 | | | 1 | | | | | | | | | | | yes | | | | | | | | | |
| SLC35E2 | N_Shore | Body | chr1:1669123-1669413 | 290 | | | | | | | | | 1 | | | | | yes | | | | | | | | | |
| NADK | N_Shore | 3'UTR | chr1:1683738-1684399 | 661 | 5 | | | | | | | | | | | | | | | | | yes | | | | | |
| CALML6 | NA | TSS1500 | chr1:1845940-1846340 | 400 | 1 | | | 1 | | | | | 2 | | | | | | | | | | | | | | |
| TMEM52 | Island;S_Shore | TSS1500 | chr1:1851372-1852238 | 866 | | | | | | | | | | | | | | | | | | | | | | | |
| KIAA1751 | Island | 3'UTR | chr1:1886501-1886865 | 364 | | | 1 | | | | | | | | | | | yes | | | | | | | | | |
| KIAA1751 | Island | Body | chr1:1897585-1897785 | 200 | | | | | | | | | | | | | | yes | | | | | yes | | | | |
| KIAA1751 | NA | Body | chr1:1920339-1920798 | 459 | | | 2 | | | | | | | | | | | yes | | yes | yes | yes | | | | | |
| KIAA1751 | Island | TSS200 | chr1:1935289-1935561 | 272 | | | | | | | | | 1 | | | | | yes | | | yes | | yes | | | | |
| GABRD | Island;N_Shore | Body | chr1:1957073-1957805 | 732 | | | 12 | | | | | 1 | 1 | | | | | yes | yes | | | | | | | | |
| GABRD | Island;N_Shore | 3'UTR;Body | chr1:1959442-1962082 | 2640 | 12 | | | | | | 1 | 6 | | 1 | | | | | | | | | | | | | |
| PRKCZ | N_Shore | TSS1500 | chr1:1981217-1982791 | 1574 | | | | | | | | | | 1 | | | | | | | yes | | yes | | | | |
| PRKCZ | Island | Body | chr1:1990780-1990986 | 206 | | | | | | | | | | | | | | yes | | | | | | | | | |
| PRKCZ | S_Shore | Body | chr1:1992389-1992687 | 298 | | | | | | | | | | | | | | | yes | | | | | | | | |
| PRKCZ | N_Shelf;N_Shore;Island;chr1:2 | Body;TSS1500;1stExon;5'UTR;Islar | chr1:2002722-2006032 | 3310 | 2 | | 28 | | 4 | | | | 3 | | 1 | | | yes | | yes | yes | | yes | | | | |
| PRKCZ | NA | Body;5'UTR | chr1:2014371-2014628 | 257 | 2 | | | | | | | | | | | | | | | | yes | | yes | | | | |
| PRKCZ | N_Shelf | Body;5'UTR | chr1:2022951-2023258 | 307 | | | | | | | | | | | | | | yes | yes | | | | yes | | | | |
| PRKCZ | Island | Body;5'UTR | chr1:2025769-2026193 | 424 | | 1 | | | | | | | | | | | | | | | | | | | | | |
| PRKCZ | N_Shelf | 1stExon;5'UTR;Body | chr1:2036283-2036643 | 360 | 2 | | | | | | | | 1 | | | | | yes | | | | | | | | | |
| PRKCZ | Island | 5'UTR;Body | chr1:2038593-2038940 | 347 | 6 | | | | | | | | | | | | | | | | | | | | | | |
| PRKCZ | N_Shelf | 5'UTR;Body | chr1:2043761-2043853 | 92 | | | | | 1 | | 1 | | 1 | | | 2 | | yes | | | | | | | | | |
| PRKCZ | Island | 5'UTR;Body | chr1:2046506-2046928 | 422 | | | 21 | | | | | | | | | | | yes | | yes | | | | | | | |
| PRKCZ | NA | 5'UTR;Body;nan | chr1:2058417-2059086 | 669 | | | 10 | | 1 | | | | 1 | | 1 | | | | | | | | | | | | |
| PRKCZ | Island | 5'UTR;Body | chr1:2063799-2064691 | 892 | | | | 1 | | | | 1 | 1 | | | 1 | | yes | yes | yes | | | yes | | | | |
| PRKCZ | chr1:2082314-2082529;Island | Island;Body | chr1:2082315-2082522 | 207 | | | | | | | | 1 | | | 1 | | | | | | | | | | | | |
| PRKCZ | S_Shore | Body | chr1:2084319-2084595 | 276 | 11 | | | | | | | | | | | | | yes | yes | | yes | | | | | | |
| PRKCZ | NA | Body | chr1:2099745-2100984 | 1239 | 6 | 2 | | | | | | | | | | | | yes | | | yes | yes | | | | | |
| PRKCZ | NA | Body | chr1:2104310-2106732 | 2422 | | | 6 | | | | | 2 | 3 | | | 1 | | | | | | yes | | | | | |
| FAAP20 | N_Shore;Island | Body;3'UTR | chr1:2120985-2121724 | 739 | | 7 | 10 | 1 | 3 | 3 | 1 | 7 | 8 | 4 | | 26 | 1 | | | | | | | | | | |
| FAAP20;C1orf86;LOC100128003 | N_Shore | Body;TSS200 | chr1:2125049-2125243 | 194 | | | | | | | | | | | | | | | | | | | | | | | |
| C1orf86 | Island | 5'UTR | chr1:2137307-2137665 | 358 | | | | | | | | 1 | 1 | | | | | | | | | | | | | | |
| C1orf86 | S_Shore | 5'UTR | chr1:2138757-2139905 | 1148 | | | 5 | | | | | | 1 | 1 | 1 | | | yes | | | | yes | | | | | |
| SKI | S_Shelf | Body | chr1:2163437-2163529 | 92 | | | | | 1 | | | | | | | | | | | | | | | | | | |
| SKI | NA | Body | chr1:2170956-2171253 | 297 | | | 2 | | 1 | | | | | 1 | | | | | | | | | | | | | |
| SKI | NA | Body | chr1:2174725-2174895 | 170 | | | 2 | | | | | | | | | | | yes | | | | | | | | | |
| SKI | NA | Body | chr1:2182126-2182612 | 486 | | 1 | | | 1 | | | | | | | | | yes | | | | | | | | | |
| SKI | NA | Body | chr1:2188767-2189027 | 260 | | | 14 | | | | | | | | | | | yes | yes | | | | | | | | |
| SKI | NA | Body | chr1:2190850-2191658 | 808 | | | 14 | | | | | | | | | | | yes | yes | | yes | | | | | | |
| SKI | NA | Body | chr1:2202931-2203648 | 717 | 1 | | | | | | | | | | | | | | | | | | | | | | |
| SKI | NA | Body | chr1:2205770-2205902 | 132 | 1 | | | | | | | | | | | | | | | | | | | | | | |
| SKI | N_Shelf | Body | chr1:2209140-2209341 | 201 | | | | | | | | | 1 | | | | | yes | | yes | | yes | | | | | |
| SKI | N_Shore;Island | Body | chr1:2221807-2222674 | 867 | | 1 | | 1 | 2 | 1 | | | | | | | | | | | | | | | | | |
| SKI | S_Shelf | Body | chr1:2224678-2225139 | 461 | | 1 | | | | | | | | | | | | | | | | | | | | | |
| SKI | NA | Body | chr1:2228160-2228319 | 159 | | | 4 | | | | | | | | | | | yes | yes | | | | | | | | |
| SKI | N_Shore | Body | chr1:2230640-2230923 | 283 | | | | | | | | | 1 | 1 | | | | | | | | | | | | | |

| | | | | | | | | | | | | | | | | | | | | | | | | | | | |
|---|---|---|---|---|---|---|---|---|---|---|---|---|---|---|---|---|---|---|---|---|---|---|---|---|---|---|---|
|SKI|N_Shore;Island|Body|chr1:2231925-2232527|602|||5|1|1||||2|||||||||||||||
|SKI|Island|3'UTR|chr1:2238720-2238932|212||1||||||||||||||||||||||
|MORN1|N_Shore|3'UTR|chr1:2252794-2252995|201|||2|||||||||||||||||||||
|MORN1|Island|Body|chr1:2254199-2254757|558||||||||||||||||||||||||
|MORN1|N_Shore;Island|Body|chr1:2257894-2259325|1431|||14|||||||||||||||yes||||||
|MORN1|N_Shelf|Body|chr1:2262232-2262923|691|||||||||2||1|||||||||||||
|MORN1|N_Shore;Island;chr1:2266007-|Body;Island|chr1:2265881-2266933|1052|||||||||2|1|1|||||||||||||
|MORN1;LOC100129534|NA|Body;TSS200;TSS1500|chr1:2284167-2285115|948|||||||||1|1||||yes||yes||||||||
|MORN1|Island|Body|chr1:2304264-2304375|111|1||||||||1|||||||||||||||
|MORN1|N_Shelf|Body|chr1:2319506-2319796|290|||||||||1|||||yes||||||||||
|MORN1;RER1|S_Shore|TSS1500;5'UTR|chr1:2324205-2324396|191|||7||||||||||||yes|||||||||
|PEX10|N_Shelf|3'UTR;Body|chr1:2337198-2337711|513|||5|||||||||||||||||||||
|PEX10|Island|Body|chr1:2339890-2340342|452||||1|||||||||||yes|||||||||
|PEX10|N_Shore|TSS1500|chr1:2344979-2345475|496|||25|1||||2|3|||||||||||||||
|PLCH2|N_Shelf|TSS1500|chr1:2406721-2406795|74||||||||||||||||||||||||
|PLCH2|S_Shore;Island|Body|chr1:2411240-2411643|403|||4|||||||||||||||||||||
|PLCH2|Island|Body|chr1:2415199-2415453|254|||||||||2|||||||||||||||
|PLCH2|NA|Body|chr1:2425860-2426035|175||1|9|||||||||||||||||||||
|PLCH2|NA|Body|chr1:2428864-2429005|141|||2||||||2|||||||||||||||
|PLCH2|Island|Body|chr1:2434058-2435827|1769|||2||||||||||||||||yes|||||
|PLCH2|S_Shore|3'UTR|chr1:2436883-2437658|775|||10||1||||||||||||||||||
|PANK4|N_Shore|Body|chr1:2444426-2444644|218|1|||||||||||||||||||||||
|MMEL1|Island|Body|chr1:2527256-2527502|246||||||||||||||||||||||||
|MMEL1|N_Shore|Body|chr1:2559822-2559930|108||||||||1||||||||||||||||
|MMEL1|S_Shelf|TSS1500|chr1:2564880-2565265|385||||||||||||||||||||||||
|ACTRT2|Island|1stExon|chr1:2938646-2939463|817||||||||||||||||||||||||
|FLJ42875|Island|Body|chr1:2979311-2980937|1626|||||||||4|1||||yes|||||yes|||||
|FLJ42875;PRDM16|N_Shore;Island|Body;TSS1500;TSS200|chr1:2982592-2987961|5369||2|15|||||||||||||||yes|yes|||||
|PRDM16|Island|Body|chr1:2990031-2990678|647||||||2||||||||yes||||yes||||||
|PRDM16|N_Shore|Body|chr1:2996522-2997409|887|||4|1||||1||||||||||||||||
|PRDM16|NA|Body|chr1:3006329-3006755|426||||||||||||||||||yes||||||
|PRDM16|NA|Body|chr1:3017674-3018024|350||||||||||||||||||||||||
|PRDM16|Island;S_Shore|Body|chr1:3028230-3029070|840|||21||||||||||||||||yes|||||
|PRDM16|S_Shelf;NA|Body|chr1:3032302-3032965|663||||||||||||||||||||||||
|PRDM16|S_Shore;N_Shore;Island|Body|chr1:3036916-3039210|2294|||9||||||1|||||||||yes||||||
|PRDM16|S_Shelf|Body|chr1:3042085-3042459|374||||||||||||||||||||||||
|PRDM16|NA|Body|chr1:3044768-3045052|284||||||||||||||||||||||||
|PRDM16|NA|Body|chr1:3047342-3047677|335|||||||||||||||||||yes|||||
|PRDM16|NA|Body|chr1:3050837-3052175|1338|||3|||||||||||||||||||||
|PRDM16|Island|Body|chr1:3056640-3056742|102|||||||||1||||||||||||||yes|
|PRDM16|N_Shore;Island;S_Shore|Body|chr1:3057810-3059820|2010|||20||||||1|||||yes||||yes||||||
|PRDM16|N_Shore|Body|chr1:3070603-3070786|183|||||||||1|||||||||||||||
|PRDM16|Island|Body|chr1:3071963-3072235|272|||||1||||||||||||||||||
|PRDM16|N_Shelf;N_Shore|Body|chr1:3077356-3081481|4125|||10|1|1|||1|1||1|1||yes|yes|yes||||||||
|PRDM16|S_Shore|Body|chr1:3082862-3083173|311||||||||||||||yes||||||||||
|PRDM16|NA|Body|chr1:3094566-3094859|293||||||||||||||yes|||yes|||||||
|PRDM16|N_Shelf|Body|chr1:3099917-3100497|580|||8|||||||||||||||||||||
|PRDM16|Island|Body|chr1:3104999-3105326|327|||2||||2|2|2|1||5||||||||||||
|PRDM16|S_Shore|Body|chr1:3106929-3107290|361||||||||||||||||||||||||
|PRDM16|S_Shelf|Body|chr1:3114820-3115309|489||||||||||||||||||||||||
|PRDM16|NA|Body|chr1:3123054-3123854|800|||3|||||||||||||||||||||
|PRDM16|NA|Body|chr1:3129005-3130550|1545|||8|||||||||||yes||||||||||
|PRDM16|NA|Body|chr1:3133763-3133857|94||||1||||||||||||||||||||
|PRDM16|NA|Body|chr1:3140299-3140906|607|||||||||||||||||||yes|||||
|PRDM16|NA;N_Shelf|Body|chr1:3142925-3144710|1785||||||||1|1|||||yes|||||yes|||||
|PRDM16|N_Shore;Island|Body|chr1:3147329-3148357|1028|||4||||||||||||||yes|||||||
|PRDM16|N_Shore;S_Shore|Body|chr1:3157321-3159645|2324|||7|||||||||||yes|yes|||||||||
|PRDM16|Island;N_Shelf;N_Shore|Body|chr1:3160739-3165191|4452||1|14|1|2|||2|3||1|||||||||||||
|PRDM16|N_Shore|Body|chr1:3182736-3183165|429||||||||||||||yes|yes|||||||||
|PRDM16|N_Shore;Island|Body|chr1:3184434-3185042|608|||||||||||||||yes|||||||||
|PRDM16|N_Shelf|Body|chr1:3191481-3192542|1061|||4|||||||||||||||||||||
|PRDM16|Island|Body|chr1:3195570-3196029|459|||||||||1|||||||||||||||
|PRDM16|S_Shelf|Body|chr1:3207502-3207588|86||||||||||||||||||||||||
|PRDM16|NA|Body|chr1:3209519-3210424|905|||||2||||||||||yes|||||||||
|PRDM16|NA|Body|chr1:3228966-3230647|1681|||10|1||||1|1||||||||||yes|||||
|PRDM16|N_Shore;Island|Body|chr1:3239712-3240227|515|||6|||||||||||||||||||||
|PRDM16|NA|Body|chr1:3269252-3269478|226|||14||||||1|||||||||||||||
|PRDM16|Island|Body|chr1:3275052-3275283|231||||||||||||||yes||||||||||
|PRDM16|S_Shore;S_Shelf|Body|chr1:3276413-3277623|1210|||26||||||1|||||||||||||||
|PRDM16|NA|Body|chr1:3282449-3282870|421|||3|||||||||||||||yes|yes|||||
|PRDM16|NA|Body|chr1:3289798-3289994|196|||||||||1|||||||||||||||
|PRDM16|NA;N_Shelf|Body|chr1:3302968-3304131|1163|||2||||||1||||||||yes|||||||
|PRDM16|N_Shore|Body|chr1:3308286-3308568|282||||||||||||||||||||||||
|PRDM16|N_Shore;Island|Body|chr1:3309864-3310413|549|||||3||||||||||||||yes|||||
|PRDM16|N_Shore;Island|Body|chr1:3320431-3322363|1932|||6||||||1||||||yes|||||||||
|PRDM16|Island|Body|chr1:3331137-3332705|1568|||15||||||1|||||||||||||||
|PRDM16|N_Shelf;N_Shore|Body|chr1:3339870-3340187|317|||7|||||2|2|||2||||||||||||
|PRDM16|N_Shore|Body|chr1:3341944-3342307|363||||||||||||||yes||||||||||
|PRDM16|N_Shelf|Body|chr1:3347577-3347994|417|||1||||||||||||||||yes|||||
|PRDM16|S_Shore|3'UTR|chr1:3352257-3352884|627|||3|||||||||||||||||||||
|ARHGEF16|NA|5'UTR|chr1:3379492-3379884|392||||||||||||||yes||||||||||
|ARHGEF16|NA|Body|chr1:3382237-3382565|328|||||||||1||||||yes|||||||||
|ARHGEF16|NA|Body|chr1:3387913-3388196|283|||2|||||||||||||||yes||||||
|ARHGEF16|N_Shore|Body|chr1:3394013-3394871|858|||||||||||||||yes||||yes|||||
|ARHGEF16|S_Shore|Body|chr1:3397061-3397125|64|||1||1||||||||||||||||||
|MEGF6|Island|Body|chr1:3427407-3427598|191|||||||||1|||||||||||||||
|MEGF6|Island|Body|chr1:3459949-3460467|518|||19||||||||1||||||||||||
|MEGF6|NA|Body|chr1:3473923-3474376|453||||||||||||||||yes||||||||
|MEGF6|NA|Body|chr1:3477168-3477442|274|||4||||||1||||||yes|||||||||
|MEGF6|N_Shelf|Body|chr1:3496027-3496139|112||||||||||||||||||||||||
|MEGF6|Island|Body|chr1:3511710-3512048|338|||3|||||||||||||||||||||
|MEGF6|S_Shelf|Body|chr1:3514921-3515194|273|2||6|||||1|1|||||||yes||yes|yes|||||
|WDR8|N_Shelf;N_Shore|Body|chr1:3554769-3555737|968|||1||||||1|||||yes|yes|||||||||
|WDR8|N_Shore|Body|chr1:3559559-3559966|407|1|||||||||||||yes||||||||||
|WDR8;TP73|Island|TSS1500|chr1:3566950-3568245|1295|3||4||||||5|9|1||3|||||yes|yes|||||
|TP73|NA|5'UTR|chr1:3574507-3575147|640|||11||||||1|1|||||||yes|yes||||||
|TP73|NA|Body|chr1:3600735-3600879|144||||||||1||||1||||yes||||||||
|TP73|N_Shore;Island|TSS1500;Body;TSS200|chr1:3605747-3607425|1678|||15|||||1|3|1||||||||||||||
|TP73|NA|Body|chr1:3613723-3614190|467|||4||||||||||||yes|||||||||

| | | | | | | | | | | | | | | | | | | | | | | | |
|---|---|---|---|---|---|---|---|---|---|---|---|---|---|---|---|---|---|---|---|---|---|---|---|
| TP73 | NA | Body | chr1:3615390-3615762 | 372 | | | | | | | | | 1 | | | | | | | | | | |
| TP73 | N_Shelf | Body | chr1:3620218-3621445 | 1227 | | | 1 | | | | | 3 | 6 | | | | | yes | yes | | yes | | |
| TP73 | Island | Body | chr1:3623500-3623859 | 359 | | | 2 | | | | | | 1 | 1 | | | | | | | | | |
| TP73 | Island | Body | chr1:3634867-3635210 | 343 | | | 3 | | | | | 2 | | | | | | | | | | yes | |
| TP73 | N_Shore;Island;S_Shore | Body;3'UTR | chr1:3649250-3649971 | 721 | | | 4 | | | | | 1 | | | | | | | | | | | |
| KIAA0495 | N_Shelf | 3'UTR | chr1:3659644-3659702 | 58 | | | | | | | | | | | | | | yes | | | | | |
| KIAA0495 | Island;S_Shore | 1stExon;5'UTR;TSS1500 | chr1:3663339-3664680 | 1341 | | 1 | | | 2 | | | 1 | | | | | | yes | | yes | | | yes |
| CCDC27;LOC388588 | N_Shore;Island | Body;TSS200 | chr1:3687804-3689201 | 1397 | | | | | | | | | | | | | | | | | | | |
| LRRC47 | Island;S_Shore | Body | chr1:3703792-3704559 | 767 | | | | | | | | | 2 | | | | | | | | | | yes |
| KIAA0562;DFFB | N_Shore;Island | TSS1500;Body | chr1:3774827-3775206 | 379 | | | 5 | | | | | 2 | 2 | | | 1 | | yes | | | | | |
| LOC100133612 | Island | Body | chr1:3824319-3824447 | 128 | | | | | | | | | | | | | | | | | | yes | |
| LOC100133612 | S_Shore;S_Shelf | Body | chr1:3826425-3827428 | 1003 | | 1 | 4 | | 1 | | | | 1 | | | | | yes | | yes | | | yes |
| LOC284661 | NA | TSS1500 | chr1:4470956-4471758 | 802 | | | 4 | | | | | | | | | | | | yes | yes | | | yes |
| AJAP1 | Island | TSS1500 | chr1:4714008-4714991 | 983 | | 1 | | | | | | | 2 | | | | | | | | | | yes |
| AJAP1 | S_Shore | Body | chr1:4718212-4718573 | 361 | | | | | | | | | 1 | | | | | | | | | yes | |
| AJAP1 | NA | Body | chr1:4736099-4736187 | 88 | | | | | | | | | | | | | | | | yes | | | |
| AJAP1 | N_Shore | Body | chr1:4770676-4770935 | 259 | | | | | | | | | | | | | | | | | | yes | |
| AJAP1 | NA | Body | chr1:4794912-4795114 | 202 | | | 3 | | | | | | | | | | | yes | | | | | |
| AJAP1 | Island | Body | chr1:4832352-4832865 | 513 | | | 9 | | | | | | | | | | | | | | | | yes |
| AJAP1 | S_Shelf | 3'UTR | chr1:4834960-4835412 | 452 | | | 8 | | | | | | | | | | | | | | | | |
| AJAP1 | NA | 3'UTR | chr1:4841913-4842898 | 985 | | | 4 | | | | | | | | | | | | | | yes | yes | |
| KCNAB2 | N_Shore;Island | TSS1500;TSS200 | chr1:6085799-6086652 | 853 | | | 19 | | | | | | | | | 1 | | yes | | | | | |
| KCNAB2 | NA | 5'UTR | chr1:6093770-6094316 | 546 | | | 6 | | | | | | | | | | | yes | yes | | | yes | |
| KCNAB2 | NA | Body | chr1:6105859-6106358 | 499 | | | | | 1 | | | | | | | | | | | | | | |
| KCNAB2 | N_Shore | Body | chr1:6111480-6111671 | 191 | | | | | | | | | | | | | | yes | | | | | |
| KCNAB2 | NA | Body | chr1:6149049-6149444 | 395 | | | | | | | | | | | | | | yes | | | | yes | |
| CHD5 | NA | Body | chr1:6169988-6170117 | 129 | | | | | | | | | | | | | | | | | | | |
| CHD5 | Island | Body | chr1:6187886-6188120 | 234 | 1 | | | | | | | | 1 | | | | | yes | | | | | |
| CHD5 | S_Shelf | Body | chr1:6191821-6192121 | 300 | | | | | | | | | | | | | | | yes | yes | | | |
| CHD5 | Island | Body | chr1:6208717-6209266 | 549 | 3 | | | | | | | | | | | | | yes | | | | yes | yes |
| CHD5 | Island | TSS1500 | chr1:6240455-6240508 | 53 | | | | | | | | | 1 | | | | | | | | | | yes |
| RPL22 | S_Shore | TSS1500 | chr1:6260181-6260616 | 435 | | | 4 | | | | | | | | | | | | yes | yes | yes | | |
| RNF207 | N_Shore;Island | TSS1500 | chr1:6265071-6266360 | 1289 | | | | | 1 | | | | 1 | | | | | yes | yes | | | yes | |
| RNF207 | N_Shore;Island | Body | chr1:6268709-6269635 | 926 | | 1 | | 1 | | | | 0 | 3 | 2 | | | | yes | yes | | | | yes |
| ICMT | N_Shore | Body | chr1:6294656-6294850 | 194 | | | 4 | | | | | | | | | | | yes | | | | | |
| ICMT | S_Shore | TSS1500 | chr1:6296335-6297128 | 793 | | | 4 | | | | | | 1 | | | | | yes | yes | | | | yes |
| HES3 | N_Shore | TSS200;TSS1500 | chr1:6303793-6304272 | 479 | 4 | | 2 | | | | | | 1 | | | | | | | | | | |
| GPR153 | Island | 3'UTR | chr1:6309300-6309724 | 424 | | | | | | | | | | | | | | yes | | | | | yes |
| ACOT7 | NA | Body | chr1:6333502-6333843 | 341 | | | | | 1 | | | | | | | | | | | | | | |
| ACOT7 | Island | Body | chr1:6341230-6341327 | 97 | | | 2 | | | | | | | | | | | | yes | | | | |
| ACOT7 | NA | Body | chr1:6363562-6363886 | 324 | | | | | | | | | 1 | | | | | | | | | yes | |
| ACOT7 | NA | Body | chr1:6390053-6390830 | 777 | | 1 | | | | | | | 1 | | | | | | | yes | | yes | |
| ACOT7 | NA | Body;TSS1500 | chr1:6419606-6419906 | 300 | | | 2 | | | | | | 1 | | | | | | | | | | |
| ACOT7 | NA | Body;TSS1500 | chr1:6421246-6421502 | 256 | | | | | 1 | | | | | | | | | yes | yes | | | | |
| ACOT7 | NA | Body;1stExon;5'UTR;TSS1500 | chr1:6445632-6446787 | 1155 | | 1 | 3 | | | | | | 1 | 1 | | | | yes | yes | | yes | | |
| ESPN | Island | Body | chr1:6507979-6508035 | 56 | | | | 1 | | | | | | | | | | | | | | | |
| ESPN | Island | Body | chr1:6514605-6515624 | 1019 | | | 1 | | 1 | | | 0 | | | | | | | | | | yes | yes |
| ESPN | N_Shore | Body | chr1:6519923-6520354 | 431 | 2 | | 2 | | | | | | | | | | | yes | | | yes | | |
| TNFRSF25;PLEKHG5 | N_Shore;Island;S_Shore | Body;1stExon;3'UTR;5'UTR;TSS20 | chr1:6524642-6526531 | 1889 | 2 | | 35 | | | | | 0 | 5 | 6 | 1 | | | yes | | | | | yes |
| PLEKHG5 | Island | Body | chr1:6531717-6531813 | 96 | | | | | | | | | 1 | | | | | yes | yes | | | | |
| PLEKHG5 | S_Shore | 5'UTR;TSS1500;Body | chr1:6546574-6546839 | 265 | | | 2 | | | | | | | | | | | yes | | | | | |
| PLEKHG5 | S_Shore | Body;TSS1500 | chr1:6552273-6552403 | 130 | | | | | | | | | | | | | | yes | yes | | | | |
| PLEKHG5 | S_Shore | Body;TSS1500 | chr1:6558085-6558497 | 412 | | | 14 | | | | | | | | | | | | yes | | | | |
| TAS1R1;NOL9 | S_Shore | TSS200;TSS1500 | chr1:6615264-6615418 | 154 | | | 12 | | | | | | | | | | | | | | | | |
| KLHL21 | Island | Body | chr1:6659205-6659504 | 299 | | | | | | | | | 3 | 1 | | | | yes | | | | | |
| DNAJC11 | S_Shore | TSS1500 | chr1:6762215-6762265 | 50 | | | | | | | | | | | | | | yes | | | | | |
| CAMTA1 | chr1:7022001-7022308;Island | Island;Body | chr1:7022041-7022284 | 243 | | | 7 | | | | | | 1 | | 1 | | | | | | | yes | yes |
| CAMTA1 | NA | Body | chr1:7111184-7111316 | 132 | | | | | | | | | | | | | | | | | | | |
| CAMTA1 | N_Shore | Body | chr1:7120893-7121245 | 352 | 1 | | 6 | | | | | | | | | | | | | yes | yes | | |
| CAMTA1 | Island | Body | chr1:7122667-7123191 | 524 | | | 4 | | | | | 0 | | | | | | yes | yes | yes | yes | | yes |
| CAMTA1 | NA | Body | chr1:7209666-7210116 | 450 | | | 2 | | | | | | | | | | | | | | yes | | |
| CAMTA1 | NA | Body | chr1:7308012-7308117 | 105 | | | | | 1 | | | 0 | 1 | | | | | | | | | | |
| CAMTA1 | NA | Body | chr1:7408793-7409003 | 210 | | | 2 | | | | | | | | | | | | | yes | | | |
| CAMTA1 | NA | Body | chr1:7692321-7692367 | 46 | | | 6 | | 1 | | | | 1 | 1 | 1 | | | | | | | | |
| CAMTA1 | NA | Body | chr1:7710719-7711079 | 360 | | | 1 | | | | | | | | | | | | | | | | |
| CAMTA1 | Island | Body | chr1:7724097-7724533 | 436 | | | | | | | | | | | | | | | | | | | |
| PER3 | Island | 5'UTR;1stExon;TSS200 | chr1:7844735-7845237 | 502 | 2 | | | 1 | 1 | | | | 1 | | | | | | | | | | yes |
| PER3 | Island | Body | chr1:7887199-7887560 | 361 | | | | | | | | 0 | 3 | 2 | | | | | | | | | |
| TNFRSF9 | N_Shore | TSS200 | chr1:8001027-8001252 | 225 | | | | | | | | | | | | | | yes | | | | | yes |
| ERRFI1 | Island;S_Shore | TSS1500 | chr1:8086721-8087590 | 869 | | | 2 | | | | | | | | | | | | | | | yes | |
| SLC45A1 | N_Shore | TSS1500;TSS200 | chr1:8383899-8384718 | 819 | | | 2 | | | | | | | | | | | yes | | | | | |
| SLC45A1 | N_Shore | Body | chr1:8403631-8404050 | 419 | | | 42 | 1 | | | | | | | | | | | | | yes | | |
| SLC2A7 | NA | TSS200 | chr1:9086460-9086648 | 188 | | | 1 | | | | | | 1 | | | | | yes | | | | | |
| SLC2A5 | Island | Body | chr1:9099281-9100506 | 1225 | | | 5 | | | | | | | | | | | yes | yes | | | | |
| SLC2A5 | N_Shelf;N_Shore;Island | Body;5'UTR;1stExon;TSS1500 | chr1:9129487-9131783 | 2296 | | | 25 | 1 | 2 | | | | | | | | | yes | | | yes | yes | |
| GPR157 | Island | TSS200 | chr1:9189312-9189507 | 195 | | | | | | | | | 2 | | | | | | | | | yes | |
| SPSB1 | NA | 5'UTR | chr1:9400692-9400775 | 83 | | | 3 | | | | | | | | | | | | | | | | |
| SPSB1 | NA | 5'UTR | chr1:9409349-9409810 | 461 | | | 7 | | | | | | | | | | | | | | | yes | |
| SLC25A33 | Island | Body | chr1:9600345-9600721 | 376 | | | | | | | | | | | | | | yes | | | | | |
| C1orf200;PIK3CD | Island;S_Shore | Body;5'UTR;TSS200 | chr1:9712928-9714845 | 1917 | | | 1 | | 2 | 1 | | | 1 | 1 | | | | yes | | | | | |
| PIK3CD | Island | Body | chr1:9775755-9776133 | 378 | | | | | 1 | | | | 1 | | | | | | | | | | |
| PIK3CD;CLSTN1 | N_Shore | 3'UTR | chr1:9788517-9789699 | 1182 | | | | | 1 | | | | 4 | | | | | yes | | | yes | | yes |
| CTNNBIP1 | Island | TSS1500 | chr1:9970639-9970659 | 20 | | | | | | | | | 1 | | | | | | | | | yes | yes |
| RBP7 | N_Shore | TSS1500 | chr1:10056837-10057059 | 222 | | | 12 | | | | | | 2 | | | | | yes | | yes | | | yes |
| APITD1 | N_Shore | TSS200;TSS1500 | chr1:10489962-10490063 | 101 | | | 17 | | | | | 3 | 1 | 1 | | | | yes | yes | | | | yes |
| PEX14 | NA | Body | chr1:10601389-10601595 | 206 | | 1 | | | | | | | | | | | | | | | yes | | |
| CASZ1 | Island | 3'UTR | chr1:10698313-10698794 | 481 | | | | | | | | 1 | 1 | | | 2 | | | | | | yes | yes |
| CASZ1 | Island | Body | chr1:10707917-10708091 | 174 | | | | | | | | | | 1 | | | | | | | | | |
| CASZ1 | S_Shore | Body | chr1:10709185-10710134 | 949 | | | | | | | | | | | | 1 | | | | | | yes | yes |
| CASZ1 | N_Shelf | Body | chr1:10711457-10711695 | 238 | | | 1 | | | | | | | | | | | | | | | yes | |
| CASZ1 | NA | Body | chr1:10732063-10733030 | 967 | | | 9 | 1 | | | | | | | | | | | | | | | yes |
| CASZ1 | NA | Body | chr1:10738569-10738769 | 200 | | | 1 | | | | | | | | | | | | | | | | |
| CASZ1 | Island | 5'UTR | chr1:10757892-10757954 | 62 | | | | | | | | | 1 | 1 | | | | | | | | | |
| CASZ1 | Island | 5'UTR | chr1:10764512-10764896 | 384 | | | | | | | | 1 | 1 | | | | | | | | | | |
| CASZ1 | NA | 5'UTR | chr1:10839450-10839574 | 124 | | | | | | | | | 1 | 1 | | | | | | | | yes | |
| CASZ1 | Island | TSS200 | chr1:10856707-10856943 | 236 | | | | | | | | | 1 | | | | | yes | yes | | | yes | |
| C1orf127 | NA | TSS200 | chr1:11042161-11042381 | 220 | | | | | | | | | 1 | | | | | yes | | | | | |
| MASP2 | NA | Body | chr1:11107045-11107452 | 407 | | | | 1 | | | | | | | | | | yes | yes | | | | |
| SRM | N_Shore | Body | chr1:11118880-11119225 | 345 | | | | | | | | | | | | | | yes | | | | | |

| | | | | | | | | | | | | | | | | | | | | | | | | | | | |
|---|---|---|---|---|---|---|---|---|---|---|---|---|---|---|---|---|---|---|---|---|---|---|---|---|---|---|---|
| MTOR;ANGPTL7 | NA | Body;TSS1500 | chr1:11249037-11249057 | 20 | | | | 1 | | | 1 | | | | | | | | | | | | | | | | |
| PTCHD2 | Island | Body | chr1:11561497-11562134 | 637 | | | 14 | | | | | 1 | 1 | | | 1 | | | | | | | | | | | |
| PTCHD2 | Island | Body | chr1:11589839-11590043 | 204 | | | | | | | | | | | | | | yes | | | | | | | | | |
| FBXO2 | Island | Body | chr1:11708949-11709271 | 322 | 1 | | | | 1 | | | | | | | | | yes | yes | | | yes | | | | | |
| FBXO44;FBXO2 | N_Shore;Island | TSS1500;Body | chr1:11713123-11714254 | 1131 | | | 8 | | 1 | | | 1 | 1 | | | | | yes | yes | | | yes | | | | | |
| FBXO44;FBXO2 | S_Shore | Body;TSS1500 | chr1:11716015-11716171 | 156 | | | | | | | | | | | | | | yes | | | | | | | | | |
| MAD2L2;C1orf187 | Island | TSS1500;5'UTR | chr1:11752342-11752772 | 430 | 1 | | | | | | | | | | | | | yes | | | | yes | | | | | |
| C1orf187 | NA | 5'UTR | chr1:11760856-11761766 | 910 | | | 10 | | | | | | 1 | | | | | yes | | | | yes | yes | | | | |
| AGTRAP | N_Shore | TSS1500 | chr1:11795897-11795985 | 88 | | | | | 1 | | | | 2 | 1 | | | | yes | | yes | | yes | | | | | |
| NPPA | NA | TSS200;TSS1500 | chr1:11907986-11908347 | 361 | 4 | | 5 | 1 | | | | | | | | | | | | | | | | | | | |
| TNFRSF8 | Island | 5'UTR;1stExon | chr1:12123554-12123934 | 380 | | 1 | | | | | | | | | | | | | | | | | | | | | |
| TNFRSF1B | Island | Body | chr1:12227700-12227927 | 227 | | | 6 | | | | | | | | | | | yes | | | | | | | | | |
| TNFRSF1B | N_Shore;Island | Body | chr1:12251718-12251934 | 216 | | | | | | | | | 2 | | | | | | yes | | | yes | | | | | |
| VPS13D | NA | Body | chr1:12538341-12538678 | 337 | | | 1 | | 1 | | | | 1 | | | | | | | | | | | | | | |
| DHRS3 | Island | Body | chr1:12656114-12656315 | 201 | | | | | | | | | | 1 | | | | yes | | | | | yes | | | | |
| AADACL4 | NA | TSS1500 | chr1:12703864-12704007 | 143 | | | | | | | | | | | | | | yes | | yes | | | | | | | |
| AADACL3 | NA | TSS1500 | chr1:12775848-12776160 | 312 | | | | | | | | | | | | | | | yes | | | | yes | | | | |
| PRAMEF1 | NA | 5'UTR;1stExon | chr1:12851605-12851682 | 77 | | | | | | | | | | | | | | yes | | | | | yes | | | | |
| HNRNPCL1;LOC649330 | NA | Body;1stExon;5'UTR;TSS1500 | chr1:12907837-12908861 | 1024 | | | 2 | | | | | | | | | | | yes | | | | | | | | | |
| LOC440563 | NA | Body | chr1:13183557-13184214 | 657 | | | 2 | | | | | | | | | | | yes | | | | | | | | | |
| PRAMEF20 | NA | TSS200 | chr1:13736832-13737267 | 435 | | | | | | | | | | | | | | yes | | | | yes | | | | | |
| PDPN | Island;N_Shore | TSS1500;TSS200 | chr1:13909522-13910555 | 1033 | | | | | | | | | 2 | | | 1 | | | | | yes | | yes | | | | |
| PRDM2 | N_Shore | TSS1500 | chr1:14026054-14026360 | 306 | | | | | | | | | | | | | | yes | yes | | | yes | | | | | |
| PRDM2 | S_Shore | 5'UTR | chr1:14027378-14027581 | 203 | | | 17 | | | | | 1 | | | | | | | | | | yes | yes | | | | |
| PRDM2 | S_Shelf | TSS1500;5'UTR | chr1:14029478-14030141 | 663 | | | | | | | | | 2 | | | | | | | | yes | | | | | | |
| KIAA1026 | Island | TSS200;1stExon | chr1:14925135-14925992 | 857 | | | | | | | | | 2 | | | | | | | | | yes | yes | | | | |
| KIAA1026 | NA | Body | chr1:15175204-15175356 | 152 | | | 6 | | | | | | | | | | | yes | | | | | yes | | | | |
| KIAA1026 | Island | Body;TSS1500 | chr1:15250374-15250388 | 14 | | | 1 | | | | | | | | | | | yes | | | | yes | yes | | | | |
| KIAA1026 | S_Shelf;NA | Body;TSS1500;TSS200 | chr1:15255438-15256481 | 1043 | | | 34 | | | | | | 2 | | | | | yes | | | | | yes | | | | |
| KIAA1026 | NA | Body;5'UTR;TSS1500;nan | chr1:15271830-15272326 | 496 | | 1 | | | | | | | 2 | | 1 | | | yes | | | | | | | | | |
| KIAA1026 | Island | Body | chr1:15375148-15375386 | 238 | | | | 1 | | | | | | | | | | | yes | | yes | yes | | | | | |
| KIAA1026 | NA | Body;3'UTR | chr1:15392793-15392907 | 114 | | | | | | | | 1 | 1 | | | | | | | | | yes | yes | | | | |
| TMEM51;C1orf126 | N_Shore | TSS1500;Body | chr1:15478206-15478401 | 195 | | | | | | | | | | | | | | yes | yes | | | | | | | | |
| TMEM51 | Island | 5'UTR | chr1:15480844-15481367 | 523 | | | | | | | | | 1 | | | | | yes | | | | | yes | | | | |
| TMEM51 | Island | 5'UTR | chr1:15541182-15541446 | 264 | | | 2 | | | | | | | | | | | | | | | | | | | | |
| FHAD1 | N_Shore | TSS1500 | chr1:15573050-15573566 | 516 | | | 3 | | | | | | | | | | | yes | | yes | | | | | | | |
| FHAD1 | NA | Body | chr1:15655764-15656086 | 322 | | | | | | | | | | | | | | yes | | | | | | | | | |
| CASP9 | N_Shore | TSS200 | chr1:15850916-15851444 | 528 | | | | | | | | | 2 | | | | | | | | | yes | yes | | | | |
| PLEKHM2 | Island | Body | chr1:16058422-16058504 | 82 | | 1 | | | | | | | | | | | | | | | | | | | | | |
| SLC25A34 | S_Shelf;NA | TSS1500;TSS200 | chr1:16061408-16063471 | 2063 | | | | 1 | 1 | | | | 2 | | | | | yes | | | | | | | | | |
| TMEM82 | N_Shore | Body | chr1:16069388-16069700 | 312 | | | | 1 | | | | | | | | | | yes | | | | | yes | | | | |
| FBLIM1 | N_Shore | TSS1500 | chr1:16084445-16084939 | 494 | | | 7 | | 1 | | | | | | | | | yes | | | | | yes | | | | |
| UQCRHL | NA | 3'UTR;1stExon | chr1:16133715-16134184 | 469 | | | | | | | | | | | | | | yes | | | | yes | | | | | |
| FLJ37453 | N_Shore;Island | Body | chr1:16163479-16164122 | 643 | | | 7 | 1 | 2 | | | 3 | 3 | 6 | 1 | 1 | | yes | | | | | | | | | |
| HSPB7 | NA | 1stExon;5'UTR;TSS200;TSS1500 | chr1:16344977-16346317 | 1340 | | | 10 | | | | | | | | | | | yes | yes | | yes | | | | | | |
| CLCNKA | NA | TSS1500 | chr1:16347984-16348397 | 413 | | | | | | | | | | | | | | yes | | | | | | | | | |
| CLCNKB | NA | TSS1500;TSS200 | chr1:16369395-16370265 | 870 | | | 9 | | | | | | 2 | | 1 | | | yes | | | | | | | | | |
| ARHGEF19 | Island | Body | chr1:16533891-16534579 | 688 | | | 8 | 1 | | | | | | | | | | yes | | | | yes | yes | | | | |
| ARHGEF19 | NA | 5'UTR;1stExon | chr1:16539013-16539119 | 106 | | | | | | | | | | | | | | yes | | | | | | | | | |
| FBXO42 | S_Shore | TSS1500 | chr1:16679388-16679780 | 392 | | | | | | | | | | | | | | yes | | yes | | yes | yes | | | | |
| CROCCL2 | N_Shore;Island | Body | chr1:16810147-16810776 | 629 | 3 | | | | | | | 2 | | | | | | yes | | | | | | | | | |
| MST1P2 | N_Shore | Body | chr1:16972460-16972917 | 457 | | | | 1 | | | | | | | | | | | | | | | | | | | |
| MST1P2 | Island | Body | chr1:16974532-16975005 | 473 | | | | | | | | | | | | | | | | | | | | | | | |
| ESPNP | Island | Body | chr1:17023008-17023283 | 275 | | | | | | | | 1 | | | | | | yes | | | | | | | | | |
| ESPNP | Island | Body | chr1:17027832-17027941 | 109 | 2 | | | 1 | | | | | | | | | | | | | | | | | | | |
| MST1P9 | Island | Body | chr1:17085860-17086558 | 698 | | | | | | | | | | | | | | | | | | | | | | | |
| MST1P9 | S_Shelf;NA | Body;TSS1500 | chr1:17090834-17092279 | 1445 | | | 9 | 1 | 1 | | | | 1 | | | | | | yes | | | yes | | | | | |
| CROCC | NA | TSS1500 | chr1:17247699-17248394 | 695 | | | | | | | | | | | | | | yes | | | | | yes | | | | |
| CROCC | NA | Body | chr1:17265239-17265567 | 328 | | | | | | | | | | | | | | | | | | yes | | | | | |
| MFAP2 | S_Shore | TSS200 | chr1:17308032-17308303 | 271 | | | | | | | | | | | 1 | | | yes | yes | yes | | | | | | | |
| SDHB | S_Shore | TSS1500 | chr1:17381800-17382130 | 330 | | | 2 | | | | | | 1 | | | | | yes | yes | | | yes | | | | | |
| PADI2 | Island | TSS200 | chr1:17446068-17446087 | 19 | | | 1 | | | | | | | | | | | | yes | | | | yes | | | | |
| PADI3 | NA | TSS1500;TSS200 | chr1:17575210-17575440 | 230 | | | 1 | | | | | | 2 | | | | | yes | | | yes | | | | | | |
| PADI4 | NA | TSS200 | chr1:17634543-17634716 | 173 | | | | | | | | | | | | | | yes | | | | | | | | | |
| PADI6 | NA | TSS200 | chr1:17698725-17699021 | 296 | | | | | | | | | | | | | | yes | | | | yes | | | | | |
| RCC2 | Island | Body | chr1:17746438-17746840 | 402 | | | 3 | 1 | | | | | | | | | | yes | | | | | | | | | |
| ARHGEF10L | NA | Body | chr1:17944356-17944660 | 304 | | | 18 | | | | | | | | | | | yes | yes | | | yes | | | | | |
| ARHGEF10L | N_Shore | Body | chr1:17949212-17949604 | 392 | | 1 | | | | | | | | | | | | yes | | yes | | | | | | | |
| ACTL8 | N_Shore | Body | chr1:18152144-18152639 | 495 | | | | | | | | | | | | | | | | | | | yes | | | | |
| IGSF21 | Island | Body | chr1:18435619-18435858 | 239 | | 1 | | | | | | | | | | | | | | | | yes | | | | | |
| IGSF21 | Island | Body | chr1:18691875-18692019 | 144 | | | | | | | | | | | | | | yes | | | | | | | | | |
| IGSF21 | N_Shelf | Body | chr1:18700603-18700899 | 296 | | | | | | | | 1 | 1 | | | 1 | | yes | | | | | | | | | |
| KLHDC7A | N_Shelf | TSS1500 | chr1:18806517-18807388 | 871 | | | | | | | | | 1 | | | | | yes | | | yes | | | | | | |
| KLHDC7A | N_Shore;Island | 1stExon | chr1:18808559-18809656 | 1097 | | | 4 | | | | | 2 | 4 | | | | | | | | | | | | | | |
| PAX7 | N_Shore;Island | TSS1500;TSS200 | chr1:18956762-18958084 | 1322 | | 1 | | | | | | | 2 | | | | | | | | | yes | | | | | |
| PAX7 | Island | Body | chr1:18963477-18963958 | 481 | | | | | | | | | 1 | | | | | | | | | | yes | | | | |
| PAX7 | NA | Body | chr1:19051735-19052235 | 500 | | | 2 | | | | | 1 | 1 | | | | | yes | | | yes | yes | yes | | | | |
| TAS1R2 | Island | Body | chr1:19181150-19181459 | 309 | | | | | | | | | 3 | | | 1 | | yes | yes | | | | | | | | |
| ALDH4A1 | NA | TSS1500;Body | chr1:19217794-19218212 | 418 | | | | | | | | | 1 | | | | | yes | | yes | | | | | | | |
| AKR7L | Island | 1stExon;TSS200;TSS1500 | chr1:19600454-19601069 | 615 | | | 32 | | 1 | 1 | | 18 | 8 | 3 | | 3 | | yes | yes | | | | | | | | |
| AKR7A3 | Island;N_Shore;chr1:19614998 | TSS200;TSS1500;Body;1stExon;Isla | chr1:19614851-19615702 | 851 | 1 | | 10 | | | | 1 | 5 | 2 | 2 | 1 | 3 | | yes | yes | | | | yes | | | | |
| CAPZB | S_Shore | 3'UTR | chr1:19665446-19665757 | 311 | | | | 1 | | | | | | | | | | | | | | | | | | | |
| CAPZB | NA | Body | chr1:19764480-19764821 | 341 | | | | 1 | | | | | | | | | | | | | | | | | | | |
| NBL1 | N_Shore | TSS1500;Body | chr1:19970204-19970337 | 133 | | | | | | | | 1 | | | | | | | yes | | | | yes | | | | |
| HTR6 | Island | TSS200;1stExon | chr1:19991676-19992771 | 1095 | | | | | | | | | 1 | | | | | | yes | | | | yes | | | | |
| RNF186 | NA | 5'UTR;1stExon;TSS1500 | chr1:20141662-20142706 | 1044 | | | 5 | 1 | | | | | 2 | | | | | yes | | yes | | | | | | | yes |
| PLA2G2E | NA | TSS200;TSS1500 | chr1:20250186-20251013 | 827 | | | 6 | | | | | | | | | | | yes | | | yes | | | | | | |
| PLA2G5 | NA | TSS200 | chr1:20396560-20396750 | 190 | | | | | | | | | | | | | | yes | yes | | | | | | | | |
| PLA2G2F | NA | TSS1500 | chr1:20465286-20465716 | 430 | | | | | | | | | | | | | | yes | | yes | | | | | | | |
| VWA5B1 | NA | TSS1500;TSS200 | chr1:20616951-20617456 | 505 | | | 15 | | | | | | 1 | | | | | yes | | | | yes | yes | | | | |
| VWA5B1 | Island | Body | chr1:20669724-20669975 | 251 | 2 | | | | | | | | | | | | | | | | | | yes | | | | |
| FAM43B | Island | 1stExon;3'UTR | chr1:20880461-20880663 | 202 | | | 1 | | | | | | | | | | | | | | | yes | yes | | | | |
| DDOST | S_Shore | TSS1500 | chr1:20988590-20988763 | 173 | | | 7 | | | | | | | | | | | | | | | | | | | | |
| KIF17 | Island | 1stExon;TSS1500 | chr1:21044031-21044945 | 914 | | | 2 | | | | | | 1 | | | | | yes | yes | | | | yes | | | | |
| ECE1 | NA | Body | chr1:21586831-21587174 | 343 | 3 | | 7 | | | | | | | | | | | yes | | | | yes | | | | | |
| ECE1 | NA | Body;5'UTR;1stExon | chr1:21606051-21606767 | 716 | | | | 1 | | | | | | | | | | yes | yes | | | | | | | | |
| ECE1 | Island | 1stExon;Body | chr1:21616619-21616825 | 206 | | | | | 1 | | | | | | | | | | | | yes | | yes | | | | |
| ALPL | Island | 5'UTR | chr1:21836133-21836326 | 193 | | 1 | | | | | | | | | | | | | | | | | yes | | | | |
| ALPL | NA | 5'UTR;TSS1500 | chr1:21877322-21877781 | 459 | | 1 | | 1 | | | | | | | | | | yes | yes | | | | | | | | |

| | | | | | | | | | | | | | | | | | | | | | | | | | | | |
|---|---|---|---|---|---|---|---|---|---|---|---|---|---|---|---|---|---|---|---|---|---|---|---|---|---|---|---|
| ALPL | NA | Body | chr1:21900987-21901260 | 273 | | | | 1 | | | | | | | | | | yes | | | | | | | | | |
| RAP1GAP | Island | Body | chr1:21930180-21930336 | 156 | | | | | | | | | 2 | | | | | | | | | | yes | | | | |
| RAP1GAP | S_Shelf | Body | chr1:21934312-21934726 | 414 | | | 1 | | | | | | | | | 1 | | yes | | | | | | | | | |
| RAP1GAP | NA | Body | chr1:21966310-21966500 | 190 | | | | | | | | | | | 1 | | | yes | yes | | | | | | | | |
| RAP1GAP | Island | 5'UTR;TSS1500 | chr1:21978698-21978863 | 165 | | | | | 1 | | | | | | | | | yes | | | | yes | yes | | | | |
| CELA3A | NA | TSS200 | chr1:22328026-22328339 | 313 | | | | | | | | | | | | | | yes | yes | | | | | | | | |
| HSPC157 | N_Shore | TSS1500;TSS200 | chr1:22351389-22352348 | 959 | | | | | 7 | | | | | | | | | yes | | | | | | | | | |
| WNT4 | Island | Body | chr1:22446661-22446854 | 193 | | | | | | | | | 1 | | | | | | | | | | | | | | |
| WNT4 | S_Shore | TSS1500 | chr1:22470290-22470343 | 53 | | | | | | | | | | | | | | yes | | | | | | | | | |
| EPHA8 | Island | TSS1500 | chr1:22889611-22890264 | 653 | | | | | | | | | 2 | | | | | yes | | | | yes | yes | | | | |
| EPHA8 | Island | Body | chr1:22919873-22920169 | 296 | | | | | | | | 1 | 3 | 1 | | 2 | | yes | | | | | | | | | |
| C1QA | NA | TSS1500 | chr1:22962506-22962986 | 480 | | | 6 | | | | | | | | | | | yes | | yes | | | | | | | |
| C1QC | Island | Body | chr1:22974149-22974243 | 94 | | | | | | | | | | | | | | | | | | | yes | | | | |
| C1QB | NA | TSS1500 | chr1:22978607-22978981 | 374 | 1 | | | | | | | | | | | | | yes | yes | | | | | | | | |
| LUZP1 | NA | TSS200 | chr1:23495492-23495577 | 85 | | | | | | | | | 2 | | | | | yes | | | | | yes | | | | |
| HTR1D | NA | TSS200 | chr1:23521272-23521815 | 543 | | | 27 | | | | | | | | | | | yes | | yes | | yes | | | | | yes |
| C1orf213;ZNF436 | S_Shore | 3'UTR;Body;1stExon;TSS1500 | chr1:23697258-23697656 | 398 | | | 4 | | | 1 | | | 1 | | | | | yes | | | yes | | | | | | |
| TCEA3 | Island | 1stExon;5'UTR | chr1:23751195-23751210 | 15 | | | | | | | | | 1 | | | | | | | | | yes | | | | | |
| ASAP3 | N_Shore;Island | Body | chr1:23763081-23763767 | 686 | | | | | 1 | | | 4 | 5 | | | 4 | | yes | | | | | | | | | |
| ASAP3 | S_Shore | TSS200 | chr1:23810871-23810942 | 71 | | | | | | | | | 1 | | | | | yes | | | | yes | yes | | | | |
| ID3 | S_Shore | TSS1500 | chr1:23887324-23887816 | 492 | | | 4 | | | | | | | | | | | yes | | | | yes | | | | | |
| MDS2 | NA | TSS1500 | chr1:23953480-23953778 | 298 | | | | | | | | | 1 | | | | | | | | | | | | | | |
| GALE | N_Shore | 5'UTR;1stExon | chr1:24126017-24126194 | 177 | | | | | | | | | | | | | | yes | | | | | | | | | |
| HMGCL | NA | TSS1500 | chr1:24152579-24152774 | 195 | | | 5 | | | | | | 2 | | | | | yes | yes | | yes | yes | | | | | |
| FUCA1 | S_Shore;Island | TSS1500;TSS200 | chr1:24194974-24195659 | 685 | | | 12 | | | | 2 | 1 | 3 | 1 | 1 | 3 | | yes | | | | yes | | | | | |
| CNR2 | Island | 5'UTR | chr1:24229232-24229682 | 450 | | | | | | | | 1 | 1 | 3 | | | | yes | | | | | yes | | | | |
| CNR2 | NA | 5'UTR;1stExon;TSS200 | chr1:24239771-24240069 | 298 | | | | | 1 | | | | 1 | 1 | | | | yes | | | yes | yes | | | | | |
| SFRS13A | S_Shore | TSS1500 | chr1:24307291-24307535 | 244 | | | 5 | | | | | | | | | | | yes | | | yes | | | | | | |
| MYOM3 | NA | 1stExon;5'UTR;TSS200;TSS1500 | chr1:24438607-24439418 | 811 | | | 13 | | | | | | | | | | | yes | | yes | | yes | | | | | |
| IL22RA1 | NA | 1stExon;TSS200 | chr1:24469564-24470205 | 641 | | | | | 1 | | | | 2 | | 1 | | | | yes | | | | | | | | |
| IL28RA | Island | TSS1500 | chr1:24514412-24514557 | 145 | | | 3 | | | | | | | | | | | yes | yes | | yes | yes | yes | | | | |
| GRHL3 | N_Shelf | TSS200;1stExon;5'UTR;Body;TSS15 | chr1:24645184-24646392 | 1208 | | | 18 | 3 | | | | | 1 | | | | | yes | yes | | | | | | | | |
| RCAN3 | N_Shore | TSS1500 | chr1:24828096-24828586 | 490 | | | | | | | | | | | | | | yes | | | | yes | yes | | | | |
| RCAN3 | N_Shore;Island | Body;3'UTR | chr1:24861604-24861919 | 315 | | | | | 1 | | | | 3 | | | | | yes | | | | | | | | | |
| RUNX3 | Island | 3'UTR | chr1:25228582-25228690 | 108 | | | | | | | | | | | | | | | | | | | | | | | |
| RUNX3 | NA | Body | chr1:25240865-25240938 | 73 | | | | | | | | | | | | | | yes | yes | | | | | | | | |
| RUNX3 | N_Shore | Body | chr1:25254088-25254149 | 61 | | | | | | | | | 1 | | | | | | | | | | | | | | |
| RUNX3 | Island;chr1:25255527-2525900 | TSS1500;Body;Island | chr1:25257505-25258679 | 1174 | | | 4 | | 3 | | | | 6 | | 1 | | | | | | | | | | | | |
| RUNX3 | NA | TSS200;TSS1500;1stExon;5'UTR | chr1:25291385-25292274 | 889 | | | | | 1 | | | | 11 | 1 | | 1 | | yes | yes | | | | yes | | | | |
| MAN1C1 | Island | 1stExon | chr1:25944514-25945042 | 528 | | | 18 | 1 | 1 | | | | 1 | | | 1 | | | | | | | yes | | | | |
| MAN1C1 | Island | Body | chr1:26097918-26098214 | 296 | | | | | | | | | | | | | | yes | | | | | | | | | |
| FAM54B | N_Shore | TSS1500 | chr1:26145328-26145437 | 109 | | | | | | | | | 1 | 1 | | | | | | yes | yes | yes | | | | | |
| PAQR7 | N_Shelf | TSS200 | chr1:26197855-26198360 | 505 | | | | | | | | | 1 | | | | | yes | yes | | | yes | | | | | |
| STMN1 | Island | 1stExon;TSS1500;5'UTR;TSS200 | chr1:26233332-26234015 | 683 | 2 | | 21 | 1 | | | | 3 | 2 | | | 4 | | | | | | | | | | | |
| EXTL1 | NA | TSS1500 | chr1:26346812-26347541 | 729 | 1 | | | | | | | | | | | | | yes | yes | | | | | | | | |
| TRIM63 | NA | 1stExon;5'UTR | chr1:26394009-26394300 | 291 | | | 7 | | | | | | | | | | | yes | yes | | | | | | | | |
| GRRP1 | N_Shore | 5'UTR | chr1:26487591-26488053 | 462 | | | | | | | | | | | | 1 | | yes | | | | | yes | | | | |
| CNKSR1 | NA | TSS1500;TSS200 | chr1:26503623-26504548 | 925 | | | 13 | 1 | | | | 1 | 6 | 1 | | 4 | | yes | yes | | yes | | | | | | |
| CATSPER4 | NA | TSS1500 | chr1:26516478-26516508 | 30 | | | | | | | | | 1 | | | | | | | | | | | | | | |
| CATSPER4 | NA | Body | chr1:26527870-26527928 | 58 | 1 | | | 1 | | | 1 | 1 | 1 | 1 | | 16 | 1 | yes | | | | | | | | | |
| SH3BGRL3 | N_Shore | TSS1500 | chr1:26605913-26606110 | 197 | | | | 1 | | | | | | | | | | | | | | | | | | | |
| UBXN11 | NA | Body | chr1:26611311-26611453 | 142 | | | | 1 | | | | | 1 | 1 | | | | | | | | | | | | | |
| UBXN11 | Island | 5'UTR | chr1:26632948-26633338 | 390 | | | | | | | | 1 | | | | | | | | | | | | | | | |
| UBXN11;CD52 | NA | TSS200;Body;5'UTR;1stExon | chr1:26644515-26645313 | 798 | | | | | 1 | | | | 1 | 1 | | | | yes | | | | | | | | | |
| LIN28 | Island | TSS200 | chr1:26737262-26737711 | 449 | | 1 | | | | | | | | | | | | | | | | yes | | | | | |
| DHDDS | N_Shore | TSS1500 | chr1:26758228-26758308 | 80 | | | | | | | | | 1 | | | | | | | | | | | | | | |
| DHDDS;HMGN2 | N_Shore | 3'UTR;TSS1500 | chr1:26797145-26797619 | 474 | | | 4 | | | | | | | | | | | | | | | | | | | | |
| MIR1976;RPS6KA1 | NA | TSS1500;Body | chr1:26880547-26880807 | 260 | | | | | | | | 1 | | | | | | yes | | | | | | | | | |
| SFN | N_Shore;Island | TSS1500;TSS200;1stExon;5'UTR | chr1:27188587-27190639 | 2052 | | 1 | 1 | | 4 | | | 1 | 4 | 1 | | | | yes | | | | yes | | | | | |
| NR0B2 | NA | 1stExon | chr1:27240319-27240694 | 375 | | | | | 3 | | | | | | | | | yes | | | | | | | | | |
| C1orf172 | Island;S_Shore | TSS200;TSS1500 | chr1:27287082-27287541 | 459 | | | 6 | | | | | 1 | 1 | | | | | yes | yes | | | | | | | | |
| FAM46B | N_Shore | Body | chr1:27338787-27339287 | 500 | | | | | | | | | | | | | | yes | | | | yes | yes | | | | |
| SLC9A1 | N_Shore | 1stExon | chr1:27480543-27480668 | 125 | | | | | | | | | | | | | | | | | | | | | | | |
| SYTL1 | NA | TSS1500;TSS200 | chr1:27668058-27668610 | 552 | | | 5 | | | | | 1 | | | | | | yes | | | | | | | | | |
| SYTL1 | Island | Body | chr1:27676205-27676652 | 447 | | | | | | | | | 1 | | | 1 | | yes | | | | | yes | | | | |
| SYTL1 | Island | Body | chr1:27677860-27678138 | 278 | | | 6 | | | | | | | | | | | yes | yes | | | | | | | | |
| MAP3K6 | N_Shore | Body | chr1:27683139-27683501 | 362 | | | 1 | | | | | | 2 | | | 1 | | | | | | | | | | | |
| MAP3K6 | N_Shore;Island | Body | chr1:27686768-27687695 | 927 | | | 1 | | | | | 2 | 1 | | | 3 | | | | | | | | | | | |
| MAP3K6 | S_Shore | TSS1500 | chr1:27693795-27693942 | 147 | | | | | | | | | | | | | | yes | | | | | | | | | |
| CD164L2 | Island;S_Shore;N_Shore | 1stExon;5'UTR;TSS200;Body | chr1:27709575-27709928 | 353 | | | 3 | | | | | 2 | 3 | | | 1 | | yes | | | | yes | | | | | |
| GPR3 | N_Shore | TSS1500 | chr1:27718221-27718277 | 56 | | | 2 | | | | | 1 | | | | | | | | | | | | | | | |
| FGR | N_Shore;Island;S_Shore | 5'UTR;1stExon;TSS200;TSS1500 | chr1:27961334-27962412 | 1078 | | | 11 | | | | | 2 | 5 | 1 | | | | yes | | | | | yes | | | | |
| C1orf38 | N_Shore | TSS1500 | chr1:28198451-28198813 | 362 | | | | | | | | | | | | | | yes | | | | yes | | | | | |
| RPA2 | Island | TSS1500 | chr1:28241495-28241577 | 82 | | | | | | | | | 1 | | | | | yes | yes | | | | | | | | |
| PHACTR4 | NA | TSS200;Body | chr1:28764523-28765031 | 508 | | | | | | | | | 1 | | | | | yes | | yes | | | yes | | | | |
| SNHG3-RCC1;RCC1 | N_Shore | Body;TSS1500;5'UTR | chr1:28843736-28844028 | 292 | | | | | | | | | 1 | | | | | yes | yes | | | | | | | | |
| SNHG12;SNORD99;SNORA61 | N_Shore | Body;TSS1500 | chr1:28906382-28906538 | 156 | | | | | | | | | 1 | | | | | yes | | | | | | | | | |
| OPRD1 | N_Shore;Island | TSS1500;1stExon | chr1:29138043-29139121 | 1078 | | | | | | | | | 1 | | | | | yes | yes | | | | yes | | | | |
| OPRD1 | Island | Body | chr1:29189486-29189789 | 303 | | | | | | | | | 1 | | | | | | yes | | | | | | | | |
| EPB41 | N_Shore | TSS1500 | chr1:29213106-29213367 | 261 | | | | | | | | 1 | | | | | | yes | yes | | | | | | | | |
| EPB41 | N_Shore | 5'UTR;TSS1500 | chr1:29240256-29240554 | 298 | | | | | | | | | | | | | | yes | | | | | yes | | | | |
| SFRS4 | NA | 3'UTR | chr1:29474864-29475256 | 392 | | | | 1 | | | | | | | | | | | | | | | yes | | | | |
| SFRS4 | S_Shore | TSS200 | chr1:29508776-29508888 | 112 | | | | | | | | | 1 | | | | | yes | | | | yes | | | | | |
| PTPRU | Island | Body | chr1:29564615-29565306 | 691 | | | 5 | | | | | | | | | | | yes | | | | yes | yes | | | | |
| PTPRU | Island;N_Shore | Body | chr1:29585853-29587357 | 1504 | 2 | | | | 3 | | | | | | | | | yes | | | yes | | yes | | | | |
| NKAIN1 | N_Shore;Island | 3'UTR | chr1:31654355-31655009 | 654 | | | 6 | | | | | 2 | 4 | | | 7 | | yes | | | | | | | | | |
| NKAIN1 | NA | TSS1500 | chr1:31661978-31662035 | 57 | | | | | | | | | | | | | | yes | | yes | | | | | | | |
| FABP3 | NA | 1stExon;5'UTR;TSS200 | chr1:31845904-31846110 | 206 | | | | 1 | 2 | | | | | | | | | yes | | | | | | | | | |
| LOC284551 | NA | Body | chr1:31988357-31988579 | 222 | | | 2 | | | | | | | | | | | yes | | | | | | | | | |
| TINAGL1 | NA | TSS1500;TSS200 | chr1:32041052-32042921 | 1869 | | | 13 | 3 | | | | | | | | | | yes | | | | yes | | | | | |
| TINAGL1 | N_Shore | Body | chr1:32052246-32052651 | 405 | | | 1 | | | | | | | | | | | yes | yes | | | | yes | | | | |
| COL16A1 | Island | 1stExon;5'UTR | chr1:32169568-32170507 | 939 | 5 | | | | | | | | 1 | | | | | yes | | | | | | | | | |
| SPOCD1 | NA | 5'UTR | chr1:32281460-32281665 | 205 | | | 4 | | | | | | | | | | | yes | yes | | | | | | | | |
| CCDC28B | N_Shore | TSS1500 | chr1:32665450-32665584 | 134 | | | 1 | | 1 | | | | 1 | | | | | yes | | | | | yes | | | | |
| MTMR9L | S_Shore | TSS1500 | chr1:32708234-32708485 | 251 | | | 4 | 1 | | | | 1 | 2 | | | | | yes | yes | | | yes | | | | | |
| FAM167B | Island | Body | chr1:32714023-32714308 | 285 | | | 12 | 1 | | | | | 1 | | | | | yes | | | yes | | | | | | |
| LCK | N_Shore | 5'UTR;TSS1500 | chr1:32739049-32739752 | 703 | | | | | | | | | 1 | | | | | yes | | | | yes | | | | | |
| LCK | Island | Body | chr1:32741152-32742037 | 885 | | | 12 | | | | | | 1 | | | | | yes | | | | | | | | | |
| MARCKSL1 | Island;S_Shore | TSS1500 | chr1:32802133-32802312 | 179 | | | 12 | | 1 | | | 1 | | | | | | | | | | | yes | | | | |

| | | | | | | | | | | | | | | | | | | | | | | | |
|---|---|---|---|---|---|---|---|---|---|---|---|---|---|---|---|---|---|---|---|---|---|---|---|
| LOC100128071;TSSK3 | Island | 3'UTR;TSS1500;1stExon;5'UTR;TSS | chr1:32826969-32828016 | 1047 | | | 1 | | | | | 2 | 2 | 1 | | | | | | | | yes | yes |
| ZBTB8B | Island | TSS200 | chr1:32930511-32930544 | 33 | | | | | | | | | 1 | | | | | | yes | | | | |
| ZBTB8B | NA | Body | chr1:32936780-32936996 | 216 | | | | | | | | 1 | 1 | | | 1 | | | | | | yes | |
| SYNC | S_Shore | TSS1500 | chr1:33169558-33169850 | 292 | 1 | | 6 | | | | | | | | | | | yes | yes | | | yes | |
| KIAA1522 | NA | Body | chr1:33231190-33231382 | 192 | | | 3 | | | | | | | | | | | yes | | | | | |
| KIAA1522 | N_Shore | Body | chr1:33235324-33235619 | 295 | | | 1 | | | | | | | | | | | yes | | | yes | | |
| TRIM62 | S_Shore | TSS1500 | chr1:33648648-33648819 | 171 | | | 3 | | | | | | 2 | | | | | | yes | yes | | yes | |
| HMGB4;CSMD2 | NA | 5'UTR;Body;TSS1500 | chr1:34329069-34329788 | 719 | | | | | | | | | 1 | | | | | yes | | | | | |
| C1orf94;CSMD2 | S_Shore | TSS200;TSS1500 | chr1:34632573-34633066 | 493 | 2 | | 2 | | | | | | 1 | | | | | | yes | | | | |
| C1orf94 | Island;N_Shore | 5'UTR;1stExon;TSS1500;TSS200 | chr1:34642000-34643106 | 1106 | | 1 | 10 | | | | | 2 | 6 | | | 2 | | yes | yes | | yes | | |
| GJB5 | NA | TSS1500;TSS200 | chr1:35220224-35220839 | 615 | | | 7 | | | | | | 2 | 2 | | | | yes | yes | | | | |
| GJB4 | Island | Body | chr1:35227029-35227158 | 129 | | | | | | | | | | | | | | yes | | | | | |
| GJB3 | N_Shore | TSS1500;TSS200 | chr1:35245692-35246759 | 1067 | | | | | | | | 1 | | | | | | yes | yes | | yes | | |
| GJA4 | N_Shore | TSS1500 | chr1:35257475-35258061 | 586 | | | | | | | | | 1 | | | | | yes | yes | | | | yes |
| GJA4 | S_Shore | 5'UTR | chr1:35259746-35259937 | 191 | | | | | | | | | 1 | | | | | yes | | | | | |
| DLGAP3 | N_Shore | Body | chr1:35333886-35334529 | 643 | | | 1 | | | | | | | | | | | | | | yes | yes | |
| DLGAP3 | Island | Body | chr1:35351340-35351627 | 287 | | | | | 1 | | | | | | | | | | | | | yes | |
| NCDN | S_Shelf | Body | chr1:36026194-36026611 | 417 | | | | | | | | | | | | | | | yes | | | | |
| TFAP2E | N_Shore;Island | TSS1500;TSS200;Body | chr1:36038468-36039885 | 1417 | 1 | | 19 | | | | | 5 | | | | 1 | | yes | | | | | yes |
| TFAP2E | Island | Body | chr1:36042433-36043633 | 1200 | 4 | | 5 | 1 | 1 | | | | 1 | | 1 | | | yes | yes | | | yes | |
| PSMB2 | S_Shore | TSS1500 | chr1:36108206-36108233 | 27 | | | 6 | | | | | 2 | 2 | | | 1 | | yes | yes | | | | |
| TRAPPC3 | S_Shore | TSS1500 | chr1:36615845-36616030 | 185 | | | 10 | | | | | | | | | | | yes | yes | | | | |
| C1orf113 | N_Shore | TSS1500 | chr1:36771714-36771844 | 130 | | | 1 | 1 | | | | | 3 | | | | | | | | | | |
| FAM176B | Island | 3'UTR;Body | chr1:36787678-36788436 | 758 | | | | 1 | 2 | | | 2 | 1 | | | | | | | | | | yes |
| FAM176B | S_Shore | 1stExon;5'UTR | chr1:36789496-36789937 | 441 | | | | | | | | | | | | | | yes | | | | yes | yes |
| OSCP1 | S_Shore | TSS1500 | chr1:36916605-36916744 | 139 | | | | | | | | | 1 | | | 1 | | yes | | | | | |
| CSF3R | NA | TSS200;5'UTR | chr1:36948313-36948981 | 668 | | | 3 | | | | | | 1 | | | | | | | | | | |
| GRIK3 | Island | Body | chr1:37499309-37499649 | 340 | | 1 | | | | | | | | | | | | | | | | | yes |
| SNIP1 | S_Shore | TSS200 | chr1:38020091-38020505 | 414 | | | | | | | | | | | | | | yes | yes | yes | | | |
| DNALI1 | Island | 1stExon | chr1:38022586-38022718 | 132 | | | 7 | | 1 | | | | 1 | | | | | yes | yes | | | | |
| C1orf109 | N_Shore | TSS1500 | chr1:38156575-38156939 | 364 | | | | | | | | | | | | | | | | | | | |
| EPHA10 | Island | Body | chr1:38200920-38201123 | 203 | | | | | | | | 1 | 1 | | | | | | yes | | yes | | |
| EPHA10 | N_Shore | Body | chr1:38218080-38218194 | 114 | | | | | | | | | | | | | | | yes | | | | yes |
| EPHA10 | Island | 1stExon;5'UTR | chr1:38230769-38230887 | 118 | | 1 | | | | | | | | | | | | | | | | yes | |
| MANEAL | N_Shore | TSS1500 | chr1:38258709-38259096 | 387 | | | | | | | | | | | | | | yes | yes | yes | | yes | |
| MANEAL | S_Shore | Body;TSS200 | chr1:38260988-38261192 | 204 | | | 2 | | | | | | 2 | | | 1 | | | | | | | |
| INPP5B | NA | 5'UTR;1stExon | chr1:38412519-38412818 | 299 | | 1 | 1 | | | | | | 3 | | | | | yes | | | | | |
| MYCBP;GJA9 | S_Shore | TSS1500;Body | chr1:39340248-39340512 | 264 | | | | | | | | | | | | | | | | | | | |
| RHBDL2 | NA | 1stExon;5'UTR | chr1:39407387-39407651 | 264 | | | 2 | | | | | | | | | | | yes | yes | | | | |
| NDUFS5 | NA | TSS1500 | chr1:39491318-39491459 | 141 | | | 1 | 1 | | | | | | | | | | yes | | | | | yes |
| MACF1 | N_Shore | TSS1500 | chr1:39546511-39546656 | 145 | | | 2 | | | | | | | | | | | yes | yes | | | yes | yes |
| MACF1 | Island | 5'UTR | chr1:39547881-39548001 | 120 | | | 1 | | | | | | | | | | | | | | | | yes |
| BMP8A | N_Shore;Island | TSS1500;1stExon;5'UTR | chr1:39956204-39957400 | 1196 | | | | | | | | | 1 | | | | | yes | yes | | | | yes |
| LOC728448 | NA | Body | chr1:40024971-40025415 | 444 | | 2 | 5 | | | | | | 2 | | | 2 | | yes | | | | | |
| SNORA55;PABPC4 | NA | TSS1500;Body | chr1:40034481-40034535 | 54 | | | | | 1 | | | | | | | | | yes | | | yes | | |
| HEYL | Island | TSS1500 | chr1:40105664-40105764 | 100 | | | 22 | | | | | | 2 | | | | | | | | | | yes |
| NT5C1A | Island | TSS1500 | chr1:40137966-40138052 | 86 | | | | | | | | | 1 | | | | | | | | | yes | yes |
| HPCAL4 | Island | Body | chr1:40149432-40149837 | 405 | | | | | | | | | 1 | | | | | | yes | | | | |
| BMP8B | S_Shore | TSS1500 | chr1:40255321-40255555 | 234 | | | 4 | | | | | | | | | | | yes | yes | | | | yes |
| TRIT1 | S_Shore | TSS1500 | chr1:40349894-40350052 | 158 | | | | | | | | | 1 | | | | | yes | yes | | | | |
| MYCL1 | Island | TSS1500 | chr1:40368189-40368366 | 177 | | | | | | | | | 1 | | | | | yes | | | | | |
| COL9A2 | Island | Body | chr1:40769669-40769744 | 75 | | | 3 | | | | | | 1 | | | | | | | | | | yes |
| COL9A2 | Island;S_Shore | TSS200;TSS1500 | chr1:40783060-40783264 | 204 | | | 2 | | | | | 1 | 1 | | | | | | | | | | yes |
| RIMS3 | Island;N_Shore | 5'UTR | chr1:41119634-41119988 | 354 | | | | | | | | | 2 | | | | | | | | yes | yes | |
| NFYC;LOC100130557 | N_Shore | TSS1500;Body | chr1:41156730-41156988 | 258 | | | 1 | | | | | 2 | 3 | | | | | yes | | | | | |
| KCNQ4 | Island | 1stExon | chr1:41250062-41250117 | 55 | | | 18 | 1 | | | | | | | | | | | | | | | yes |
| CTPS | N_Shore | TSS1500;TSS200 | chr1:41444651-41444854 | 203 | | | 6 | | | | | | | | | 1 | | yes | yes | | | yes | yes |
| SLFNL1 | S_Shelf;NA | 5'UTR;TSS1500 | chr1:41487093-41488323 | 1230 | | | 3 | | | | | | | | | | | yes | | | yes | | |
| EDN2 | NA | 1stExon | chr1:41950237-41950362 | 125 | | | | | | | | 1 | 1 | | | 1 | | yes | | | | | |
| HIVEP3 | N_Shore;Island;S_Shore | 5'UTR;1stExon;TSS1500 | chr1:42383598-42385941 | 2343 | | | 9 | | 2 | | | 2 | 6 | 2 | 1 | 12 | | | | | | yes | |
| GUCA2B | NA | TSS1500;TSS200 | chr1:42618041-42619118 | 1077 | | | | 2 | 1 | | | | 1 | | | | | yes | yes | | | | |
| GUCA2A | NA | TSS1500;TSS200 | chr1:42630436-42631835 | 1399 | | | | 1 | | | | | | | | | | yes | yes | | | yes | |
| CCDC23;ERMAP | S_Shore | TSS1500;5'UTR | chr1:43283685-43283884 | 199 | | | | 1 | | | | | | | | | | yes | yes | | | | yes |
| ZNF691 | NA | Body | chr1:43317080-43317191 | 111 | | | | | | | | | 1 | | | | | | | | | | |
| FAM183A | N_Shore;Island | TSS1500;TSS200 | chr1:43613173-43613627 | 454 | | | | | | | | 0 | 1 | | | | | yes | yes | | yes | | yes |
| TMEM125 | N_Shelf | 5'UTR | chr1:43736290-43736600 | 310 | | | | | | | | 0 | | | | | | yes | | | | yes | |
| TMEM125 | Island | Body | chr1:43738485-43738659 | 174 | | | | | | | | | | | | | | | | | | | |
| C1orf210 | NA | TSS200;TSS1500;5'UTR;1stExon | chr1:43750693-43752185 | 1492 | | 1 | 23 | 1 | 1 | | | | 4 | | | 1 | | yes | yes | | | | |
| TIE1 | NA | TSS1500 | chr1:43765617-43766752 | 1135 | | | | | 1 | | | | | | | | | yes | yes | | | | |
| TIE1 | Island | Body | chr1:43770523-43770851 | 328 | | | 4 | | | | | | 1 | | | | | | yes | | | | |
| MPL | N_Shore;Island | Body | chr1:43814169-43815465 | 1296 | | | | | 1 | | | 0 | 3 | 1 | | 2 | | | yes | | | yes | |
| HYI | S_Shore | TSS200;TSS1500 | chr1:43920056-43920264 | 208 | | | 2 | | | | | | 1 | | | | | | yes | yes | | | |
| PTPRF | NA | Body | chr1:44016014-44016049 | 35 | | | 11 | | | | | | | | | | | yes | yes | | | | |
| PTPRF | Island | Body | chr1:44031300-44032202 | 902 | | | | 1 | 1 | | | 0 | 1 | | | | | yes | | | | | |
| PTPRF | NA | 3'UTR | chr1:44087716-44087800 | 84 | | | 2 | | | | | | | | | | | | | | | | |
| ARTN | Island;N_Shore | 1stExon;5'UTR;TSS1500 | chr1:44399363-44399894 | 531 | | | 4 | | | | | | | | | | | | | | | | |
| ARTN | Island | 1stExon;Body | chr1:44401760-44402522 | 762 | | | 3 | | | | | 0 | 1 | | 1 | | | | | | | | yes |
| B4GALT2 | N_Shore | TSS1500 | chr1:44444217-44444603 | 386 | | | | | | | | | 1 | | | | | | | | | yes | yes |
| SLC6A9 | NA | Body | chr1:44482420-44482918 | 498 | 1 | | | | | | | | | | | | | | | | | yes | |
| SLC6A9 | S_Shore | TSS1500 | chr1:44497352-44497694 | 342 | | | | | | | | | 1 | | | | | yes | | | | yes | |
| KLF17 | NA | TSS200;TSS1500 | chr1:44583886-44584446 | 560 | | | 2 | | | | | | 1 | | | | | yes | yes | | | | yes |
| ERI3 | NA | Body | chr1:44703750-44704073 | 323 | | | 6 | | 1 | | | | | | 1 | | | | | | | | |
| ERI3 | NA | Body | chr1:44715958-44716296 | 338 | | | | | 1 | | | | | | | | | | | | | | |
| ERI3 | NA | Body | chr1:44726943-44727313 | 370 | | 1 | | | | | | | 2 | 1 | | | | | yes | | | | yes |
| RNF220 | Island | 5'UTR | chr1:44871429-44871634 | 205 | | | | | | | | | 1 | | | | | | | | | yes | |
| RNF220 | S_Shore | 5'UTR | chr1:44874524-44874649 | 125 | | | | | | | | | | | | | | | | | | | |
| RNF220 | Island | Body | chr1:44883990-44884150 | 160 | | | 4 | | | | | | 1 | | | | | | | | | | |
| RNF220 | N_Shore;chr1:45082739-4508 | Body;N_Shore | chr1:45082704-45083278 | 574 | | | | | 2 | | | | 2 | 1 | 1 | | | yes | | | | | yes |
| TMEM53;C1orf228 | Island | TSS1500;TSS200 | chr1:45140355-45140465 | 110 | | | | | | | | | | | | | | | | | | yes | |
| C1orf228 | N_Shelf | Body | chr1:45187551-45187946 | 395 | | | 8 | | | | | | | | | | | yes | yes | | | | |
| C1orf228 | Island | Body | chr1:45189743-45190519 | 776 | | | 2 | | | | | | | | | | | yes | | | | | |
| SNORD46;RPS8;SNORD38A | S_Shore | TSS200;Body;TSS1500 | chr1:45242073-45242191 | 118 | | | 3 | | | | | | | | | | | yes | | | | | |
| BEST4 | S_Shore | TSS1500 | chr1:45254092-45254530 | 438 | | | | | | | | | | | | | | yes | | | | | |
| TCTEX1D4 | Island | Body | chr1:45272255-45272637 | 382 | | | 11 | | | | | | 1 | | | | | | | | | | |
| BTBD19 | Island | Body | chr1:45278971-45279684 | 713 | | | 7 | 1 | | | | 0 | 1 | | | | | yes | | | | yes | |
| PTCH2 | NA | 3'UTR | chr1:45285821-45286062 | 241 | | | | | | | | 0 | | | | | | yes | yes | | | | |
| PTCH2 | N_Shore | Body | chr1:45308248-45308593 | 345 | | | 1 | | | | | | | | | | | yes | | | | | |
| LOC400752 | Island | TSS1500 | chr1:45769308-45769554 | 246 | | | | | | | | | 1 | | | | | yes | | | | | yes |
| HPDL | Island | 1stExon;5'UTR | chr1:45792660-45792733 | 73 | | | | | | | | 0 | 1 | | | | | | | | | yes | |

| | | | | | | | | | | | | | | | | | | | | | | | |
|---|---|---|---|---|---|---|---|---|---|---|---|---|---|---|---|---|---|---|---|---|---|---|---|
| TESK2 | N_Shore | 5'UTR | chr1:45955733-45957456 | 1723 | | | | | | | | 0 | 1 | | | | | yes | yes | | | | |
| CCDC163P;MMACHC | N_Shore;chr1:45965586-45966 | Body;TSS1500;N_Shore | chr1:45965020-45966115 | 1095 | | | | | | | | 0 | 1 | | 1 | | | yes | | | | | |
| CCDC17 | Island;S_Shore | Body;5'UTR;1stExon | chr1:46088336-46089804 | 1468 | | | 6 | | | | | 0 | 4 | | | 1 | | | | | | | |
| TSPAN1 | NA | TSS1500 | chr1:46639632-46640020 | 388 | 1 | | | | | | | | | | | | | | | | | yes | |
| C1orf190;POMGNT1 | Island | TSS200;Body | chr1:46668822-46669221 | 399 | | | | 1 | 2 | | | | | | | | | yes | yes | | yes | yes | yes |
| NSUN4 | S_Shore | Body | chr1:46807113-46807522 | 409 | | | | | | | | | | | | | | | | | | | yes |
| FAAH | N_Shore;Island | TSS1500;TSS200 | chr1:46859671-46860511 | 840 | | 1 | 1 | | | | | 0 | 5 | | | | | yes | | yes | | | |
| MOBKL2C | Island | Body | chr1:47078581-47078966 | 385 | | | | | 2 | | | | | | | | | yes | | | | | |
| CYP4A11 | NA | 1stExon | chr1:47406950-47407158 | 208 | | | | | | | | | 1 | | | | | | yes | | | | |
| CYP4X1 | N_Shore;Island | TSS200;1stExon;5'UTR | chr1:47489195-47489776 | 581 | | | | 1 | | | | | 1 | | | | | | yes | | yes | | yes |
| PDZK1IP1 | NA | TSS1500 | chr1:47656003-47656140 | 137 | 1 | | | | | | | | 1 | | | | | | | | | | |
| TAL1 | Island | Body | chr1:47691158-47691550 | 392 | | | | | | | | | | 1 | 1 | | | | | | | yes | yes |
| TAL1 | N_Shore | 5'UTR;1stExon;TSS1500 | chr1:47694919-47696505 | 1586 | | | 6 | | | | | | | | | | | | | | | | yes |
| FOXE3 | Island | 1stExon | chr1:47882265-47882739 | 474 | | | | | | | | | 2 | | | | | | | | | yes | yes |
| MGC12982;FOXD2 | N_Shore;Island | Body;TSS1500 | chr1:47898239-47900630 | 2391 | | 1 | | 1 | 2 | | | 0 | 3 | 2 | | | | yes | | | | | |
| FOXD2 | N_Shore | 1stExon;5'UTR | chr1:47902269-47902371 | 102 | | | 3 | | | | | | | | | | | yes | | | | | |
| SLC5A9 | NA | TSS1500 | chr1:48687359-48687989 | 630 | | | | | | | | | | | | | | yes | | | | yes | |
| AGBL4 | N_Shore;Island | Body;TSS200 | chr1:50489319-50489954 | 635 | | | | | | | | | 2 | | | | | | yes | yes | | | yes |
| ELAVL4 | Island | TSS200 | chr1:50513645-50514319 | 674 | | 1 | | | 2 | | | 0 | 5 | | | | | | | | | | |
| ELAVL4 | NA | Body;TSS1500;TSS200 | chr1:50573970-50574837 | 867 | | | 9 | | | | | 0 | | | | | | | | | | | |
| DMRTA2 | Island | 3'UTR | chr1:50884242-50884544 | 302 | | | | | | | | | 1 | 1 | | | | | | | | yes | |
| DMRTA2 | Island | Body;5'UTR;1stExon;TSS200;TSS15 | chr1:50886393-50890952 | 4559 | | | 4 | | | | | | 3 | 1 | | | | | | | | yes | yes |
| TTC39A | S_Shore | TSS200;TSS1500 | chr1:51810836-51811558 | 722 | | | 5 | | | | | | 1 | | | | | yes | | | | | yes |
| EPS15 | NA | TSS1500;Body | chr1:51888289-51888961 | 672 | | | 6 | | | | | | | | | | | | | | | | |
| OSBPL9 | Island;N_Shore | TSS1500;Body;5'UTR | chr1:52194898-52195686 | 788 | | | 9 | | | | | | | | | | | yes | | | | | |
| NRD1 | Island | TSS1500 | chr1:52344844-52345196 | 352 | | | | | | | | | | | | 1 | | yes | yes | | | | |
| TXNDC12;KTI12 | Island | Body;1stExon | chr1:52498502-52499228 | 726 | | | | | | | | | | 1 | | | | | | | | | |
| GPX7 | N_Shore | TSS1500 | chr1:53067365-53067717 | 352 | | | | | | | | | 1 | | | | | yes | yes | yes | | | |
| ZYG11A | Island | 1stExon | chr1:53308597-53309261 | 664 | | | | | | | | | 1 | | | | | yes | | | | yes | yes |
| PODN | Island | Body | chr1:53528612-53528991 | 379 | | | 2 | | | | | | | | | | | yes | | | | | |
| SLC1A7 | Island | Body | chr1:53558392-53558511 | 119 | | | | | | | | | 1 | | | | | | | | | | |
| SLC1A7 | NA | Body | chr1:53600741-53600889 | 148 | | | 7 | | | | | | | | | | | yes | yes | | yes | | |
| CPT2 | N_Shore | TSS1500 | chr1:53661346-53661861 | 515 | | | | 1 | | | | | | | | | | yes | yes | | yes | | |
| MAGOH | Island | TSS1500 | chr1:53704498-53704978 | 480 | | | 6 | | | | | 0 | 2 | 1 | | | | yes | | | yes | | |
| LRP8 | N_Shore | Body | chr1:53742252-53742380 | 128 | | | | | | | | | 1 | | | | | yes | yes | | | | |
| LRP8 | S_Shore | TSS1500 | chr1:53794150-53794760 | 610 | | | | | | | | | | | | | | | yes | | yes | yes | yes |
| FLJ40434 | N_Shore | Body | chr1:53904065-53904319 | 254 | | | 1 | | | | | | | | | | | | | | | | |
| DMRTB1 | N_Shore | TSS1500;1stExon | chr1:53924164-53925368 | 1204 | | | | | | | | | 1 | | | | | yes | | | | yes | yes |
| GLIS1 | NA | 5'UTR | chr1:54194919-54195078 | 159 | | | 3 | | | | | | | | | | | | | | yes | | |
| GLIS1 | N_Shelf | 5'UTR;1stExon;TSS1500 | chr1:54199696-54200587 | 891 | | | 25 | | | | | | | | | | | yes | | | | | |
| C1orf83;TMEM59 | S_Shore | Body;TSS1500 | chr1:54520114-54520450 | 336 | | | 1 | | | | | | 1 | | | | | yes | yes | | | yes | |
| SSBP3 | N_Shore | Body | chr1:54723213-54723426 | 213 | | | | | 1 | | | | | | | | | | | | | | |
| SSBP3 | N_Shore;Island;chr1:54821990 | Body;Island | chr1:54821853-54822503 | 650 | | | | 1 | | | | 0 | 1 | | 1 | | | | | | | | |
| SSBP3 | N_Shore | Body | chr1:54856035-54856511 | 476 | | | 27 | | | | | | | | | | | yes | | | | | |
| ACOT11 | NA | TSS1500;TSS200 | chr1:55013419-55014160 | 741 | | | 7 | | | | | | | | | | | yes | | | | | |
| ACOT11;FAM151A | NA | Body;1stExon;5'UTR | chr1:55089104-55089372 | 268 | | | 6 | | | | | | 1 | | 1 | | | yes | | | | | |
| TTC4 | N_Shore | TSS1500 | chr1:55181235-55181269 | 34 | | | 2 | | | | | | | | | | | | yes | | | | |
| TTC22 | Island | 3'UTR;Body | chr1:55246867-55247408 | 541 | | 1 | | | | | | 0 | 3 | | | 2 | | yes | | | | | |
| TTC22 | Island;S_Shore | TSS200;TSS1500 | chr1:55267046-55267293 | 247 | | | 16 | | | | | | | | | | | yes | | | | | |
| C1orf177 | N_Shore;Island | TSS1500;TSS200 | chr1:55271295-55271921 | 626 | | | 5 | | | | 1 | | 1 | 1 | | | | yes | yes | | | | |
| TMEM61 | N_Shore;Island | TSS1500;1stExon;5'UTR | chr1:55445834-55446960 | 1126 | 2 | | | | | | | | 1 | | | | | yes | yes | | | yes | |
| BSND | S_Shore | 1stExon;5'UTR | chr1:55464825-55464927 | 102 | | | | | | | | | | | | | | | | | | yes | |
| PCSK9 | N_Shore;Island | TSS1500;TSS200;1stExon | chr1:55504649-55506014 | 1365 | 2 | | 7 | | | | | | 1 | 1 | | | | yes | yes | | | yes | |
| PCSK9 | NA | Body | chr1:55522013-55522543 | 530 | | | | | | | | | 1 | | | | | | yes | yes | | | yes |
| PRKAA2 | Island | TSS1500;TSS200;1stExon | chr1:57110686-57111199 | 513 | | | | | | | | 0 | 4 | | | 2 | | yes | | | | yes | yes |
| C1orf168 | NA | 1stExon;5'UTR | chr1:57285156-57285466 | 310 | | | 4 | 1 | | | | | 1 | | | | | yes | | | | | |
| DAB1 | N_Shore | 5'UTR | chr1:58713941-58714452 | 511 | | | | | | | | 0 | | | | | | yes | | | | | yes |
| TACSTD2 | Island | 1stExon;5'UTR;TSS200 | chr1:59042065-59043370 | 1305 | | | | | 1 | | | 0 | 12 | 2 | 1 | 32 | | yes | | | | yes | |
| MYSM1 | NA | Body | chr1:59165508-59165868 | 360 | | | | | | | | 0 | 1 | | | | | yes | | | | | |
| LOC729467 | Island | Body | chr1:59612408-59612710 | 302 | | | 4 | | | | | | | | | | | yes | yes | | yes | | yes |
| HOOK1 | N_Shore;Island | TSS1500;1stExon | chr1:60280088-60281077 | 989 | | | | | | | | 0 | 2 | | | | | yes | | | | yes | yes |
| CYP2J2 | Island;N_Shore | 1stExon;Body | chr1:60392140-60392933 | 793 | | | | | | | | 0 | 2 | | | | | yes | yes | | | yes | |
| C1orf87 | Island;S_Shore | 1stExon;5'UTR;TSS200 | chr1:60539362-60539675 | 313 | | 1 | 10 | | | | | | 2 | | | 1 | | yes | | | | | |
| NFIA | N_Shore | TSS200;Body | chr1:61547859-61547969 | 110 | | | | | | | | | 1 | | | | | | | | | yes | yes |
| L1TD1 | Island | 1stExon;5'UTR | chr1:62660556-62660861 | 305 | | 1 | | | | | | | 1 | | | | | yes | yes | | | | |
| KANK4 | Island | 5'UTR | chr1:62784762-62785079 | 317 | | | | | | | | | 1 | | | | | yes | | | | | |
| ATG4C | NA | TSS1500 | chr1:63249197-63249213 | 16 | | | 29 | | 1 | | 1 | 0 | 3 | 1 | | 11 | | yes | | | | yes | |
| PGM1 | N_Shore | TSS1500 | chr1:64058180-64058672 | 492 | | | | 1 | 1 | | | | 1 | | | | | yes | | | | | |
| UBE2U | NA | TSS1500;TSS200 | chr1:64668576-64669599 | 1023 | | | | | | | | | 5 | | | | | | yes | | | | |
| CACHD1 | S_Shore | Body | chr1:64937345-64937528 | 183 | | | | 1 | | | | | | | | | | | | | | | yes |
| RAVER2 | N_Shore | TSS1500 | chr1:65209468-65209572 | 104 | | | 1 | | | | | | | | | | | yes | yes | | | yes | |
| AK3L1 | N_Shore | TSS1500 | chr1:65612850-65613042 | 192 | | | | | | | | | | | | | | yes | yes | | | yes | yes |
| DNAJC6 | N_Shore | TSS200 | chr1:65730339-65731107 | 768 | | | | | | | | | 2 | | | | | yes | yes | | | | |
| DNAJC6 | Island | Body | chr1:65775527-65775734 | 207 | | | | | | | | | 1 | | | | | | | | | | yes |
| LEPR | Island | 5'UTR | chr1:65991176-65991765 | 589 | | | | | | | | 0 | 2 | | | | | | yes | | | | yes |
| PDE4B | N_Shore;Island | TSS1500;5'UTR;TSS200 | chr1:66257732-66259084 | 1352 | | | | | 1 | | | | 3 | | 1 | 1 | | | yes | | | yes | yes |
| PDE4B | NA | TSS200;Body | chr1:66458361-66458803 | 442 | | 1 | | | | | | | | | | | | | | | | | |
| SGIP1 | N_Shore;S_Shore | TSS1500;TSS200 | chr1:66998484-66999929 | 1445 | | | 3 | | | | | | | | | | | | | | | yes | |
| TCTEX1D1 | N_Shore | TSS1500;TSS200 | chr1:67217673-67218505 | 832 | | 1 | 6 | 1 | | | | | 2 | | | | | yes | yes | | | yes | |
| IL12RB2 | N_Shore | TSS200 | chr1:67772896-67773175 | 279 | | | 14 | | | | | | 1 | | | | | yes | | | yes | | |
| GNG12 | S_Shore | TSS1500 | chr1:68299388-68299511 | 123 | | | | 1 | | | | 0 | 2 | | | | | | yes | | | yes | yes |
| DIRAS3 | N_Shore;Island | Body | chr1:68512539-68513063 | 524 | | | | | | | | | 3 | 1 | | | | | | | | | |
| DIRAS3 | N_Shore;Island | 5'UTR;1stExon;TSS200;TSS1500 | chr1:68515788-68517273 | 1485 | | | 4 | | | | | 0 | 5 | 1 | | | | yes | yes | | | | |
| DEPDC1 | N_Shore;Island | Body;TSS200 | chr1:68961838-68962972 | 1134 | | | | | | | | | 2 | | | | | yes | | | | yes | |
| NEGR1 | Island | TSS1500 | chr1:72749275-72749735 | 460 | | | | | | | | | 1 | | | | | | | | | | yes |
| LRRIQ3;TNNI3K;FPGT | N_Shore | 5'UTR;TSS1500 | chr1:74663583-74663750 | 167 | | | | | | | | 0 | 1 | | | | | yes | | | | | |
| C1orf173 | NA | Body;1stExon;5'UTR;TSS200;TSS15 | chr1:75139139-75140289 | 1150 | | | 2 | | | | | 0 | 3 | | | | | yes | yes | | | yes | yes |
| TYW3;CRYZ | NA | TSS1500;5'UTR;TSS200;1stExon;Bo | chr1:75198211-75199496 | 1285 | | | 6 | | 2 | | | 0 | 7 | | | 4 | | | yes | | yes | | yes |
| LHX8 | N_Shelf;N_Shore;S_Shore | TSS1500;TSS200;1stExon;5'UTR | chr1:75592950-75594592 | 1642 | 1 | 1 | 31 | | 1 | 1 | | 0 | | | | | | yes | yes | | | | yes |
| LHX8 | Island;S_Shore | 5'UTR;Body | chr1:75595919-75598070 | 2151 | | 2 | 4 | | | | | | 2 | | | | | | | | | yes | |
| LHX8 | N_Shore | Body | chr1:75599532-75600685 | 1153 | 1 | | 11 | | | | | | | | | | | | | | | | |
| LHX8 | Island | Body | chr1:75602451-75602925 | 474 | | 1 | | | | | | | | | | | | | | | | | yes |
| SLC44A5 | N_Shelf | TSS1500 | chr1:76077832-76078230 | 398 | | | | | | | | | | | | | | | yes | | | | |
| ACADM | N_Shore | TSS1500 | chr1:76189192-76189801 | 609 | | | 17 | | | 1 | 1 | 0 | 1 | 1 | | 12 | 1 | | | yes | | yes | |
| MSH4 | Island;N_Shore | 1stExon;5'UTR;TSS1500;TSS200 | chr1:76262302-76262984 | 682 | | | 12 | 1 | | 1 | | | 4 | 1 | | 1 | | yes | | | | | |
| ST6GALNAC3 | Island | TSS200 | chr1:76540222-76540574 | 352 | | | | | | | | | 1 | | | | | | | | | yes | yes |
| AK5 | Island | TSS1500 | chr1:77747374-77747875 | 501 | | | 3 | | | | | | 2 | | | | | | | | | | |
| DNAJB4 | NA | TSS200 | chr1:78470449-78470640 | 191 | | | | | | | | | 1 | | | | | | | | | | yes |
| GIPC2 | N_Shore | TSS1500;TSS200 | chr1:78511140-78511600 | 460 | | | 13 | | | | | 0 | 1 | | | 2 | | | yes | | | yes | |

| | | | | | | | | | | | | | | | | | | | | | | | |
|---|---|---|---|---|---|---|---|---|---|---|---|---|---|---|---|---|---|---|---|---|---|---|---|
| PTGFR | Island | 5'UTR;1stExon | chr1:78956845-78957203 | 358 | | | | | 1 | | | | | | | | | | | | | | |
| IFI44L | NA | TSS1500 | chr1:79085162-79085765 | 603 | | | | | | | | 0 | 4 | | | 1 | | | yes | yes | | | |
| ELTD1 | Island | 5'UTR;1stExon;TSS200;Body | chr1:79472282-79472643 | 361 | | 1 | 6 | | 3 | | | 0 | 6 | | 1 | 4 | | | yes | | | | |
| LPHN2 | Island | 5'UTR | chr1:82267682-82268801 | 1119 | | | | | | | | | 2 | | | | | | | | | | |
| CTBS | Island | TSS200 | chr1:85040254-85040339 | 85 | | | | | | | | | 1 | | | | | yes | yes | | | yes | |
| LPAR3 | Island | TSS1500 | chr1:85359119-85359334 | 215 | | | | | | | | | 1 | | | | | | | | | yes | |
| MCOLN2 | Island | TSS1500 | chr1:85464000-85464261 | 261 | | | | | | | | | 1 | | | | | yes | yes | | yes | | |
| MCOLN3 | N_Shore | 5'UTR | chr1:85512957-85513315 | 358 | | | | | | | | | | | | | | | | | | | |
| MCOLN3 | Island;S_Shore | TSS1500 | chr1:85514554-85514708 | 154 | | | 9 | | | | | | | | | | | | | | | | yes |
| DDAH1 | Island | TSS200;5'UTR | chr1:85931072-85931116 | 44 | | | | | | | | | 1 | | | | | | yes | | | yes | |
| DDAH1 | S_Shore | 1stExon;5'UTR;TSS200 | chr1:86043795-86044230 | 435 | | | | | | | | 0 | 2 | 1 | | | | | | | yes | | |
| CYR61 | Island;chr1:86046362-8604724 | Body;S_Shore | chr1:86047227-86048585 | 1358 | | | 3 | | | | | | 1 | | 1 | | | | | | | | |
| ZNHIT6 | Island | TSS200 | chr1:86174161-86174240 | 79 | | | | | | | | 0 | | | | | | yes | | | | | |
| COL24A1 | Island | TSS1500 | chr1:86622624-86623101 | 477 | | | 6 | | | | | | | | | | | | yes | | | yes | |
| ODF2L | N_Shore | 5'UTR | chr1:86861147-86861232 | 85 | | | | | | | | 0 | | | | | | yes | yes | | | | |
| CLCA2 | NA | TSS1500 | chr1:86888585-86888629 | 44 | | | | | | | | | 1 | 1 | | 2 | | yes | yes | | | | |
| LOC339524 | Island | Body | chr1:87598069-87598514 | 445 | | | 1 | | | | | | | | | | | yes | | | | | |
| LOC339524 | Island | Body | chr1:87617296-87617972 | 676 | | | 17 | | | | | | | | | | | | | | yes | | |
| LMO4 | N_Shore | TSS1500 | chr1:87793364-87793510 | 146 | | | | | | | | | 1 | | | | | | | | | | |
| GTF2B | S_Shore | TSS1500 | chr1:89357908-89358043 | 135 | | | 9 | 1 | 1 | | | | 1 | | | | | | | yes | | yes | |
| CCBL2;RBMXL1 | S_Shore | TSS1500 | chr1:89459504-89459658 | 154 | | | 1 | | | | | 0 | 1 | | | 1 | | | yes | yes | | | |
| GBP4 | NA | Body | chr1:89663868-89664582 | 714 | 1 | | 29 | | | | | | | | | | | | yes | yes | | | |
| GBP6 | NA | TSS200;5'UTR | chr1:89829308-89829951 | 643 | | | 3 | | | | | | | 1 | | | | | yes | | yes | yes | yes |
| LRRC8C | S_Shore | 5'UTR | chr1:90099704-90100070 | 366 | | | 1 | | | | | | 1 | | | | | | | | | | yes |
| LRRC8D | Island | 5'UTR | chr1:90309559-90309998 | 439 | | | | | | | | 0 | | | | | | | | | | | |
| BARHL2 | Island;N_Shore | 1stExon;5'UTR;TSS1500 | chr1:91182128-91185441 | 3313 | | 2 | | | | | | | 3 | | | | | | | | | yes | yes |
| TGFBR3 | S_Shore | TSS1500 | chr1:92352293-92352481 | 188 | | | 6 | | | | | 0 | | | | | | yes | yes | | | | yes |
| BRDT | N_Shore | TSS1500 | chr1:92414221-92414782 | 561 | | | 26 | 1 | 1 | | | 0 | 1 | | | 1 | | yes | | | | | |
| BTBD8 | N_Shore | TSS1500 | chr1:92545580-92545662 | 82 | | | 5 | | | | | | | | | | | yes | | | | | |
| C1orf146 | N_Shore | TSS200 | chr1:92683403-92683821 | 418 | | | | | | | | | 1 | | | | | yes | yes | | | | |
| GFI1 | Island | Body | chr1:92946132-92947035 | 903 | | | | | | | | 0 | 3 | | 1 | 1 | | | | | | | yes |
| GFI1 | Island | 5'UTR;TSS1500 | chr1:92950123-92950244 | 121 | | | 1 | | | | | | | | | | | | | | | | |
| GFI1 | Island;S_Shore | TSS1500;TSS200 | chr1:92952467-92953037 | 570 | | | 12 | | | | | | 2 | | | | | | | | | | yes |
| EVI5 | S_Shore | Body | chr1:93251184-93251218 | 34 | | | | | | | | 0 | 1 | | | 1 | | yes | | | | | |
| EVI5 | NA | TSS1500 | chr1:93258288-93258319 | 31 | | | | | | | | | 1 | | | | | yes | | | | | yes |
| ABCA4 | NA | Body | chr1:94560697-94561088 | 391 | | 1 | | 1 | | | | | | | | | | | | | | | yes |
| ABCA4 | NA | 1stExon;5'UTR | chr1:94586670-94587076 | 406 | | | | | | | | | 1 | | | | | | | | yes | yes | |
| ARHGAP29 | Island | 1stExon;5'UTR | chr1:94703176-94703378 | 202 | | | | | | | | | 1 | | | | | yes | | | | yes | yes |
| ABCD3 | N_Shore;Island | TSS1500;1stExon | chr1:94883469-94884541 | 1072 | | 2 | | | | | | | 1 | | | | | | | | yes | yes | yes |
| F3 | N_Shore | Body | chr1:95006322-95006977 | 655 | | 1 | 12 | | 1 | | | 0 | 1 | | | | | | | | | | yes |
| SLC44A3 | N_Shore | TSS1500 | chr1:95285252-95285531 | 279 | | | 5 | | | | | | | | | | | yes | | | | yes | |
| CNN3 | Island | Body;TSS200 | chr1:95392372-95393138 | 766 | | | | | 1 | | | | 2 | | | | | | | | | | yes |
| RWDD3 | N_Shore;Island | TSS1500;1stExon | chr1:95698378-95699865 | 1487 | | | 7 | | 1 | | | 0 | 5 | 2 | | 1 | 1 | yes | yes | | | yes | yes |
| MIR137 | Island | Body | chr1:98511629-98511792 | 163 | | | | | | | | | 1 | | | | | | | | | yes | |
| SNX7 | N_Shore;Island | TSS1500;TSS200 | chr1:99126704-99127250 | 546 | | | | | | | | | 3 | | | | | | | | | | |
| LPPR5 | N_Shore;Island | Body;TSS200 | chr1:99469577-99470967 | 1390 | | 1 | | | | | | | | | | | | | | | | yes | yes |
| LPPR4 | N_Shore | TSS200;TSS1500 | chr1:99729504-99730221 | 717 | | | | | | | | 0 | 5 | 1 | | 1 | | | yes | | | | |
| PALMD | NA | TSS200;1stExon;5'UTR | chr1:100111387-100111949 | 562 | | | | | 2 | | | 2 | 2 | | | 3 | | | | | | | |
| FRRS1 | NA | TSS1500 | chr1:100231977-100232077 | 100 | | | | | | | | 1 | | | | | | yes | | | | | yes |
| AGL | Island | TSS1500 | chr1:100315421-100316030 | 609 | | | | | | | | 1 | | | | | | | | | | | |
| OLFM3 | NA | Body | chr1:102312608-102312671 | 63 | | | | | | | | 1 | 1 | | | 1 | | | | | | | yes |
| OLFM3 | NA | Body;5'UTR;1stExon;TSS200 | chr1:102462043-102462877 | 834 | | | | | | | | | 2 | | | | | | | | | yes | |
| COL11A1 | NA | TSS1500 | chr1:103574469-103574958 | 489 | | | 5 | | | | | 1 | | | | | | | yes | | | | yes |
| NTNG1 | N_Shore | TSS1500 | chr1:107682283-107682552 | 269 | | | | | | | | | 1 | | | | | | yes | yes | | yes | |
| NTNG1 | N_Shore;Island | Body | chr1:108022767-108023482 | 715 | 1 | | 4 | | | | 1 | 4 | 4 | 4 | | 10 | | | | | | | |
| SLC25A24 | N_Shore;Island | Body;1stExon;5'UTR | chr1:108734992-108735549 | 557 | 1 | 7 | 5 | 1 | | | 1 | 1 | 1 | 4 | 2 | 9 | 1 | | yes | | | | |
| C1orf59 | Island | 1stExon;5'UTR;TSS200;TSS1500 | chr1:109203610-109204325 | 715 | | | | | 1 | | | 1 | 1 | | | 1 | | yes | yes | | | | |
| CLCC1 | S_Shore | TSS1500 | chr1:109506696-109506963 | 267 | | | | | | | | | | | | | | yes | yes | | yes | | |
| WDR47 | Island | TSS200 | chr1:109584988-109585031 | 43 | | | | | | | | | 1 | | | | | yes | | | | yes | |
| SARS | N_Shore | TSS1500 | chr1:109756023-109756191 | 168 | | | 1 | | | | | | | | | | | | | yes | | | |
| MYBPHL | NA | TSS1500;TSS200 | chr1:109849705-109850837 | 1132 | | | | | | | | | 1 | | | | | yes | yes | | yes | | |
| SORT1 | N_Shore | Body | chr1:109939800-109940029 | 229 | | | | | | | | | | | | | | | | | | | yes |
| SORT1 | S_Shore | TSS1500 | chr1:109941043-109941201 | 158 | | | 13 | | | 1 | | 2 | 3 | | | 1 | | | | | | | yes |
| SYPL2 | N_Shore | TSS1500 | chr1:110008870-110009509 | 639 | | | | | | | | | 1 | | | | | yes | | | | yes | |
| CYB561D1 | N_Shore | TSS1500 | chr1:110036141-110036417 | 276 | | | 4 | | | | | | | | | | | | | | | | |
| GPR61 | NA | TSS200 | chr1:110082294-110082688 | 394 | | | | | | | | | | | | | | yes | | | yes | | |
| AMPD2 | S_Shelf | Body;TSS1500 | chr1:110166563-110166664 | 101 | | | 10 | | | | | | | | | | | | | | | | |
| GSTM2 | N_Shore | TSS1500 | chr1:110210439-110210641 | 202 | | | | 1 | | | | | | | | | | yes | | | | | |
| GSTM1 | Island | TSS200 | chr1:110230252-110230633 | 381 | 5 | 1 | 32 | 1 | | | 3 | | 1 | 3 | 1 | 20 | 2 | | | | | | |
| GSTM5 | NA | TSS1500;TSS200;1stExon;5'UTR | chr1:110253977-110255096 | 1119 | 7 | 7 | 38 | 1 | 1 | | 4 | 9 | 13 | 9 | 5 | 71 | 4 | | | yes | | | |
| EPS8L3 | NA | 5'UTR;1stExon | chr1:110306507-110306642 | 135 | | | | 1 | | | | | | | | | | | | | | | |
| CSF1 | N_Shore | TSS1500 | chr1:110452372-110452616 | 244 | | | 7 | | | | | | | | | | | | yes | | | | |
| ALX3 | N_Shore;Island | Body | chr1:110609973-110612474 | 2501 | | 1 | 27 | | | | | | | | 1 | | | | | | | | |
| UBL4B | NA | TSS1500 | chr1:110654497-110655000 | 503 | | | | | | | | | 1 | | | | | yes | yes | yes | | | |
| KCNA10 | NA | TSS200 | chr1:111061829-111062127 | 298 | | | | | | | | | 1 | | 1 | | | yes | yes | | | yes | |
| KCNA3 | N_Shore;Island;chr1:11121624 | 1stExon;Island;TSS200;TSS1500 | chr1:111216000-111218287 | 2287 | | 1 | 20 | | 2 | | | 2 | 3 | | 1 | 1 | | | | | | | yes |
| DENND2D | N_Shelf | 1stExon;5'UTR | chr1:111743020-111743411 | 391 | | | | 1 | 1 | | | | 1 | | | | | | | | | | |
| CHI3L2 | NA | TSS200 | chr1:111770111-111770291 | 180 | | | | | | | | | | | | | | yes | | | | | |
| CHIA | NA | TSS1500 | chr1:111832109-111832593 | 484 | | | | | | | | | 2 | | | | | | | yes | | | |
| C1orf88 | Island | Body | chr1:111889406-111889533 | 127 | | | | | | | | | 1 | | | | | | | | | | |
| C1orf88 | S_Shore | Body | chr1:111890580-111890880 | 300 | | | | | | | | | | | | | | | | | | | yes |
| ADORA3 | NA | Body;TSS200 | chr1:112046915-112047365 | 450 | | | | | | | | | 1 | | | | | | | | | yes | |
| ADORA3 | Island | Body | chr1:112058236-112058750 | 514 | | | | 1 | | | | | 1 | | | | | yes | | | | | |
| RAP1A | N_Shore | TSS1500 | chr1:112161618-112162068 | 450 | | | | | | | | 1 | | | | | | | yes | | | yes | |
| KCND3 | NA | Body | chr1:112438321-112438797 | 476 | | | | | | | | | | | | | | yes | | | | | yes |
| KCND3 | N_Shore | 5'UTR | chr1:112531263-112531653 | 390 | | | 3 | | | | | | | | | | | | | | | | yes |
| WNT2B | Island | Body;1stExon | chr1:113051925-113052167 | 242 | | | | | | | | | | | | | | | | | | | |
| FAM19A3 | Island | TSS1500 | chr1:113261711-113261764 | 53 | | | | | | | | | | | | | | | | | yes | | |
| FAM19A3 | Island | Body | chr1:113265679-113265883 | 204 | | | 2 | | | | | | | | | | | | | | | | |
| RSBN1 | S_Shore | TSS1500 | chr1:114355571-114355693 | 122 | | | | | | | | | 1 | | | | | yes | | | | | |
| PTPN22 | NA | Body;5'UTR;1stExon | chr1:114414042-114414408 | 366 | | | | | 1 | | | | 1 | 2 | | | | yes | | | | | yes |
| BCL2L15 | NA | 1stExon;Body | chr1:114429815-114430280 | 465 | | | | | 1 | | | | 2 | 1 | | | | yes | yes | | | | |
| SYT6 | Island | 5'UTR;1stExon | chr1:114696350-114696641 | 291 | | 1 | | | | | | | 1 | | | | | yes | | | | | |
| DENND2C | S_Shore | TSS1500 | chr1:115213243-115213259 | 16 | | | 1 | | | | | | 1 | | | | | | | | | yes | |
| AMPD1 | NA | TSS200;TSS1500 | chr1:115238213-115238559 | 346 | | | 21 | | | | | | 1 | | | | | yes | | | | yes | |
| SYCP1 | N_Shore;Island | TSS200;1stExon;5'UTR | chr1:115397374-115398123 | 749 | | 1 | 9 | | | | | 2 | 7 | | | 5 | | yes | yes | | | | |
| NGF | Island | TSS200;TSS1500 | chr1:115881044-115881259 | 215 | | | 9 | | | | | | | | | | | | | | yes | | |
| VANGL1 | N_Shore | TSS1500 | chr1:116184219-116184387 | 168 | | | 1 | | | | | | | | | | | yes | | | | yes | yes |
| CASQ2 | NA | 5'UTR;1stExon | chr1:116311341-116311561 | 220 | | | | | | | | | | | | | | yes | | | | yes | |

| | | | | | | | | | | | | | | | | | | | | | | | |
|---|---|---|---|---|---|---|---|---|---|---|---|---|---|---|---|---|---|---|---|---|---|---|---|
| NHLH2 | Island | 5'UTR | chr1:116381475-116381660 | 185 |||||||||1|||||||||yes||
| C1orf161 | NA | TSS1500 | chr1:116653300-116653561 | 261 ||||||||||||||||||||
| ATP1A1 | NA | TSS1500;Body | chr1:116925411-116925753 | 342 |||||||||1|||||yes||||||
| CD58 | N_Shore | Body | chr1:117112616-117112854 | 238 ||||||||||||||yes|||||yes|
| MIR320B1 | S_Shelf | TSS1500 | chr1:117213980-117214228 | 248 |||5|||||||||||yes|||yes|yes|yes|
| PTGFRN | Island | TSS200 | chr1:117452517-117452580 | 63 |||||||||1|||||yes||||yes|yes|
| PTGFRN | Island | Body | chr1:117487269-117487539 | 270 |||||||||1|||||yes||||||
| PTGFRN | NA | Body;3'UTR | chr1:117529478-117530009 | 531 |||||||||4||1||||||||yes|
| VTCN1 | NA | Body;TSS200;nan | chr1:117753313-117753965 | 652 ||1|4|1|||||2|1|1||||yes|||yes||
| FAM46C | NA | Body | chr1:118165598-118166402 | 804 |||||2|||||||||yes||||||
| SPAG17 | N_Shore;Island | Body;5'UTR;1stExon;TSS200 | chr1:118727658-118728226 | 568 |||1|||||6|1|||||yes|yes|yes||||
| TBX15 | N_Shore;Island | 5'UTR | chr1:119522188-119522855 | 667 ||1||||||||||||||||yes||
| TBX15 | N_Shore;Island;S_Shore | 5'UTR | chr1:119526060-119530144 | 4084 ||||||||||||||||||||
| TBX15 | N_Shore;chr1:119531991-1195 | 5'UTR;N_Shore;1stExon;TSS200;T | chr1:119531625-119534088 | 2463 ||||||||||||||||||||
| ZNF697 | NA | 5'UTR | chr1:120173989-120174570 | 581 ||||||||1|1|||||yes|||||yes|
| HMGCS2 | NA | 1stExon;5'UTR | chr1:120311542-120311653 | 111 |||||1||||1|1||||yes||||||
| REG4 | NA | TSS1500 | chr1:120354493-120354772 | 279 |||||||||1|||||yes||||||
| ADAM30 | N_Shore | 1stExon | chr1:120438925-120439036 | 111 |||||||||1|||||||||||
| HIST2H2BA | N_Shore | TSS1500 | chr1:120905306-120905767 | 461 ||||||||1||||||||||||
| LOC647121 | N_Shore | TSS1500 | chr1:121260116-121260727 | 611 |||1||||||1|||||||||yes||
| FAM72D | Island | TSS1500 | chr1:143913409-143913552 | 143 ||1||||||||||||||||||
| PDE4DIP | NA | Body | chr1:144994318-144994715 | 397 ||1|||||||||||yes|||yes||||
| HFE2 | N_Shelf | TSS1500 | chr1:145412774-145413193 | 419 ||||||||||||||yes||||yes||
| HFE2 | Island | 5'UTR;Body | chr1:145415412-145415659 | 247 ||1|||||||||||||||||yes|
| CD160 | Island | 5'UTR | chr1:145714010-145714143 | 133 |||||||||1|||1||yes||||||
| CD160 | S_Shore | TSS200 | chr1:145715571-145715719 | 148 |||||1||||3|||||yes|||yes|||
| PDZK1 | NA | TSS1500 | chr1:145726979-145727762 | 783 |||||1||||1|||||||||||
| PDIA3P | Island | Body | chr1:146649635-146649851 | 216 |||2|||||1|2|||2|||yes|||||
| CHD1L | N_Shore | TSS1500 | chr1:146713493-146714019 | 526 |||13||||||1|||1||yes|yes|yes||||
| GJA5 | NA | 5'UTR | chr1:147245168-147245626 | 458 |||2||||||||||||||||yes|
| LOC645166 | NA | TSS1500;Body;nan | chr1:148929617-148930220 | 603 ||1|||||||||1|||||||||
| LOC388692 | Island | Body | chr1:149287076-149288007 | 931 |||||||||||||||yes|||||
| HIST2H2BF | N_Shore;Island | TSS1500 | chr1:149784571-149785211 | 640 |||||||||1|||1||||||||
| MTMR11 | NA | Body;TSS200;5'UTR;1stExon | chr1:149908085-149908781 | 696 ||||||||1|3|1|||||yes||||yes|
| PLEKHO1 | Island | Body | chr1:150123037-150123734 | 697 ||||1|1||||||||||||||yes|
| C1orf51 | S_Shore | TSS1500;TSS200 | chr1:150254869-150255503 | 634 |||||||||3||||||||||yes|
| ECM1 | NA | 1stExon;5'UTR | chr1:150480534-150480856 | 322 |||||||||1|||||yes||||||
| MCL1 | S_Shore | TSS1500 | chr1:150552537-150552817 | 280 ||||||||||||||yes||yes||yes||
| LASS2 | Island | TSS1500 | chr1:150947832-150948030 | 198 |||16|1|1|||1|1|||2||yes||||||
| FAM63A | N_Shelf | 5'UTR;Body | chr1:150978385-150979364 | 979 |||||||||1|||||yes||||||
| TNFAIP8L2 | NA | TSS200 | chr1:151129062-151129298 | 236 |||||1|||||||||yes||||||
| SCNM1;LYSMD1 | NA | TSS1500;Body | chr1:151137376-151137676 | 300 ||||||||1||||1||yes||||||
| SELENBP1 | NA | Body;5'UTR;1stExon;TSS200 | chr1:151344806-151345248 | 442 |||2||1||||2|1||||yes||||yes||
| TUFT1 | Island;S_Shore | Body | chr1:151513141-151513502 | 361 |||1||||||1|||||||||||
| TNRC4 | NA | TSS1500 | chr1:151689587-151690075 | 488 |||3||||||1|||1|||yes||||yes|
| C1orf230 | N_Shore | TSS1500 | chr1:151693222-151693503 | 281 |||1||||||||||||||yes||yes|
| OAZ3;MRPL9 | N_Shore | TSS1500;Body | chr1:151734928-151735194 | 266 |||1|||||||||||||||||
| LINGO4 | NA | TSS1500 | chr1:151778255-151778657 | 402 ||||||||||||||||||yes|yes|
| LOC100132111;C2CD4D | Island;N_Shore | TSS1500;Body;TSS200;5'UTR | chr1:151810485-151812710 | 2225 ||||||||||||||||||||
| THEM5 | NA | 1stExon;5'UTR | chr1:151826123-151826533 | 410 |||||||||1|||||yes||||||
| THEM4 | Island;S_Shore | TSS200;TSS1500 | chr1:151882284-151882779 | 495 ||1||||||1||||||yes|||yes|||
| S100A10 | Island | TSS200 | chr1:151966892-151967023 | 131 |||||1|||||||||yes|||||yes|
| TCHH | Island | Body | chr1:152080354-152080543 | 189 ||||||||||||||||||||
| TCHH | S_Shore | TSS200;TSS1500 | chr1:152086623-152087315 | 692 |||||||||2|||||||||||
| HRNR | NA | 5'UTR | chr1:152196106-152196674 | 568 |||7||||||1|||||yes|yes|||||
| FLG | NA | TSS1500 | chr1:152298884-152299078 | 194 |||||||||||||||yes|||||
| LCE5A | NA | TSS200 | chr1:152483199-152483675 | 476 ||||||||||||||yes||||yes||
| CRCT1 | N_Shore;Island | 5'UTR;Body;3'UTR | chr1:152487844-152488368 | 524 ||1|1||||||1|||||yes|yes||||yes|
| LCE3E | NA | 3'UTR;Body | chr1:152538380-152538888 | 508 |||||||||1|||||||||||
| LCE3A | NA | 1stExon;TSS1500 | chr1:152595322-152596413 | 1091 |||5|||||1|2|1||1||||||||
| LCE2B | NA | TSS1500 | chr1:152657950-152658407 | 457 |||||||||1|||||yes|||yes|||
| KPRP | N_Shelf | TSS1500 | chr1:152730027-152730426 | 399 |||||||||2|||||yes|yes|||||
| KPRP | Island | Body | chr1:152733025-152733395 | 370 |||||||||1|||||||||||
| LCE1F | NA | TSS1500 | chr1:152747472-152747562 | 90 ||||||||||||||yes||||yes||
| LCE1D | NA | TSS1500 | chr1:152768267-152768727 | 460 ||||||||1||||6||||yes||||
| LCE6A | NA | TSS1500 | chr1:152815068-152815459 | 391 |||||||||||||||yes|||||
| SPRR4 | NA | 5'UTR;TSS1500 | chr1:152942401-152943352 | 951 ||||||||||||||yes|||yes|||
| SPRR3 | NA | TSS1500 | chr1:152973720-152974279 | 559 ||||||||||1||||yes|||||yes|
| SPRR2A | NA | TSS1500 | chr1:153030514-153030751 | 237 |||3|||||||||||yes||||||
| SPRR2F | NA | 5'UTR | chr1:153085649-153085998 | 349 ||||||||||||||||||yes||
| LELP1 | NA | TSS1500 | chr1:153174654-153175262 | 608 ||||||||||||||||yes||||
| LOR | NA | TSS200 | chr1:153232076-153232460 | 384 ||||||||||||||||||||
| LOR | NA | Body | chr1:153234037-153234809 | 772 ||||||||1|2|||1||||||yes||
| PGLYRP3 | NA | TSS200;TSS1500 | chr1:153283317-153283994 | 677 |||||||||1|||||yes||||||
| S100A9 | NA | TSS1500;TSS200 | chr1:153329512-153330776 | 1264 |||8|||||||||||yes|yes|||||
| S100A8 | NA | 5'UTR;Body | chr1:153362927-153363580 | 653 |||||||||1|1|||||||yes|||
| S100A7L2 | NA | TSS1500 | chr1:153413641-153413940 | 299 |||||||||1|1||||yes|yes|||yes||
| S100A6 | Island;S_Shore | 5'UTR;1stExon;TSS1500 | chr1:153508511-153509635 | 1124 ||1|1|||||||||2||||||||
| S100A5 | NA | 1stExon;5'UTR | chr1:153514114-153514442 | 328 |||3|||||||||||||||||
| S100A3 | NA | 5'UTR;1stExon | chr1:153521712-153521909 | 197 |||||||||1||1|||yes||||||
| S100A2 | NA | TSS200;TSS1500 | chr1:153538406-153539404 | 998 |4|||1|||||||||||yes|yes||||
| S100A16 | NA | 5'UTR | chr1:153581494-153581793 | 299 |||32||||||1|||||||||||
| S100A13;S100A1 | NA | 5'UTR;1stExon;TSS1500;TSS200 | chr1:153599479-153600156 | 677 |||||1||2|3|7|2|2|29|2|||||||
| C1orf77;S100A13 | S_Shore | 5'UTR;TSS1500 | chr1:153607617-153608034 | 417 |||||||||1|||||yes|||yes|yes||
| ILF2 | Island | TSS200 | chr1:153643515-153643582 | 67 ||||||||||||||yes||||||
| NPR1 | Island | 1stExon;5'UTR | chr1:153651568-153651938 | 370 ||||||||||||||||||yes|yes|
| SLC27A3 | N_Shore | TSS1500 | chr1:153747551-153747609 | 58 |||||||||1|||||||||yes||
| JTB | S_Shore | TSS1500 | chr1:153950843-153951279 | 436 |||1|1||||||||||yes||||||
| NUP210L;TPM3 | NA | Body;1stExon;5'UTR;TSS200;3'UTR | chr1:154127138-154128041 | 903 ||1|15|1||||1|5|||||yes||||||
| TPM3 | Island | Body | chr1:154155295-154156180 | 885 ||||||||2||||||yes||||||
| C1orf189;C1orf43 | NA | TSS1500;3'UTR | chr1:154179856-154179996 | 140 ||||||||||||||yes||||||
| UBAP2L;C1orf43 | N_Shore | TSS1500;Body | chr1:154192205-154192370 | 165 |||||||||||||||yes|||||
| AQP10 | NA | TSS200 | chr1:154293482-154293651 | 169 ||||||||||||||yes||yes|||yes|
| ATP8B2 | Island | 5'UTR | chr1:154298335-154298543 | 208 |||||||||1|||||||||yes|yes|
| ATP8B2 | Island | Body | chr1:154301468-154301964 | 496 |||1||||||||||||||||yes|
| IL6R | N_Shore | TSS1500 | chr1:154377091-154377403 | 312 ||||||||||||||yes|||yes||yes|
| TDRD10;SHE | Island | TSS1500;1stExon;TSS200;5'UTR | chr1:154474108-154475269 | 1161 ||1||1||||1|3||||||||||yes|
| TDRD10 | NA | Body | chr1:154519846-154520187 | 341 ||||1|||||1|||||yes||||||
| CHRNB2 | N_Shore | TSS1500 | chr1:154539974-154540074 | 100 |||||||||1|||1|||||||yes|

| | | | | | | | | | | | | | | | | | | | | | | | |
|---|---|---|---|---|---|---|---|---|---|---|---|---|---|---|---|---|---|---|---|---|---|---|---|
| ADAR | S_Shore | TSS1500 | chr1:154600682-154600712 | 30 | | | 1 | | | | | | | | | | | | yes | | | | |
| KCNN3 | NA | Body | chr1:154839813-154839983 | 170 | | | | | | | 1 | 1 | 1 | 1 | 1 | 14 | 1 | | | | | | |
| PBXIP1;PYGO2 | NA | TSS1500;3'UTR | chr1:154929317-154929791 | 474 | | | 4 | | | | | | 1 | 1 | 1 | | | | | | | | |
| SHC1 | N_Shelf | Body;1stExon | chr1:154942509-154942910 | 401 | | 1 | | | | | | | 1 | | | | | | yes | | | | |
| DCST1;DCST2 | NA | Body;TSS1500 | chr1:155007014-155007323 | 309 | | | | | | | | | 1 | | | | | yes | | | | yes | |
| EFNA4 | Island | TSS1500 | chr1:155035388-155035748 | 360 | | | | | | | | 1 | | | | | | | | | | | |
| MUC1 | N_Shelf;N_Shore | Body;TSS200 | chr1:155161211-155162871 | 1660 | | | 3 | | 1 | | | | | | | | | | | | | | yes |
| SCAMP3;CLK2 | S_Shore | TSS1500;3'UTR | chr1:155232677-155233025 | 348 | | | | | | | | | | | | | | | | | | | |
| PKLR | Island | Body | chr1:155265033-155265378 | 345 | | 1 | | | | | | | | | | | | | | | | | |
| PKLR | NA | TSS200;Body | chr1:155270866-155271149 | 283 | | | | 1 | | | | | | | | | | yes | | yes | | | |
| RUSC1;FDPS;C1orf104 | N_Shore;Island | TSS1500;Body;3'UTR;TSS200;5'UT | chr1:155290236-155291224 | 988 | | | 6 | 1 | | | | | | | | | | | | | yes | | |
| KIAA0907 | S_Shore | TSS1500 | chr1:155904702-155904916 | 214 | | | 4 | | | | | | | | | | | yes | yes | | | | |
| RXFP4 | NA | TSS1500 | chr1:155910523-155911361 | 838 | | | | 1 | | | | | 1 | | | | | yes | | | | | |
| ARHGEF2 | NA | Body | chr1:155931629-155931858 | 229 | | | 1 | | | | | | | | | | | yes | | | | | |
| RAB25 | NA | TSS200 | chr1:156030809-156031182 | 373 | | 1 | | | 1 | | | | 1 | 1 | | | | yes | | | | | |
| MEX3A | Island | 3'UTR | chr1:156046344-156046832 | 488 | | | | 1 | | | | | 1 | | | 2 | | | | | | yes | |
| SEMA4A | NA | TSS200 | chr1:156123352-156123500 | 148 | | | | | | | | | 1 | 1 | | | | | | | | | yes |
| PMF1;SLC25A44 | N_Shore | TSS1500;3'UTR | chr1:156181616-156182246 | 630 | | | | | | | | | | | | | | | | | | | |
| BGLAP | N_Shelf | TSS1500;TSS200 | chr1:156211409-156211896 | 487 | | | 13 | | | | | 1 | 1 | | | | | | | | | | |
| PAQR6 | Island | Body;5'UTR | chr1:156215418-156215951 | 533 | | | 8 | | | | | | | | | | | | | | | | |
| PAQR6 | S_Shore;S_Shelf | 5'UTR;TSS200 | chr1:156217178-156218009 | 831 | | | | | | | | | 2 | | | | | yes | | | | yes | |
| TMEM79 | N_Shore;Island | Body | chr1:156260918-156261841 | 923 | | | 6 | | | | | 3 | 2 | 1 | | 2 | | yes | | | | yes | |
| CCT3;C1orf182 | N_Shelf | Body;TSS1500 | chr1:156305656-156305725 | 69 | | | | | | | | | 1 | | | | | | yes | | yes | | |
| RHBG | N_Shore;Island | TSS1500 | chr1:156338639-156338826 | 187 | | | | | | | | | | | | | | yes | | | | yes | |
| MIR9-1;C1orf61 | N_Shore | TSS200;5'UTR | chr1:156390237-156390523 | 286 | | | 1 | | | | | | | | | | | | | | | yes | |
| MEF2D | NA | 5'UTR | chr1:156460453-156460747 | 294 | | | | 1 | | | | | | | | | | yes | | | | | |
| TTC24 | NA | TSS1500 | chr1:156549200-156549431 | 231 | | | | | | | | | 1 | | | | | yes | | | | | |
| BCAN | N_Shore | TSS1500 | chr1:156610966-156611425 | 459 | | | 2 | | | | | | | | | | | yes | | | yes | | yes |
| BCAN | N_Shore;Island | Body | chr1:156615962-156617074 | 1112 | | | | | | | | 4 | 1 | | | | | yes | yes | | | yes | |
| BCAN | N_Shore | Body | chr1:156626749-156627099 | 350 | | 1 | | | | | | | | | | | | yes | | | | yes | yes |
| NES | Island | 1stExon | chr1:156646293-156646635 | 342 | | | | | | | | 1 | | | | | | | | | | | yes |
| ISG20L2;C1orf66 | N_Shore | Body;1stExon;TSS1500 | chr1:156696467-156697041 | 574 | | | 10 | | | | | | 1 | | | | | yes | | | | | |
| SH2D2A;NTRK1 | S_Shore;S_Shelf | Body;TSS200;5'UTR;1stExon;TSS1 | chr1:156784869-156786653 | 1784 | | | 13 | | | | | | | | | | | yes | | | | yes | |
| NTRK1;INSRR | Island | Body | chr1:156814901-156815529 | 628 | | 1 | | | | | | | | | | | | | | | | | |
| NTRK1;INSRR | Island | Body | chr1:156818749-156819191 | 442 | | | 2 | | | | | | | | | | | yes | yes | | | | yes |
| PEAR1 | N_Shore | TSS1500 | chr1:156863129-156863732 | 603 | | | 1 | | | | | | | | | | | yes | | | | | |
| C1orf92 | N_Shore | TSS1500 | chr1:156889826-156889964 | 138 | | | 1 | 1 | | | | | | | | | | yes | | | yes | | |
| ETV3L | NA | 1stExon;5'UTR | chr1:157069325-157069611 | 286 | | | 1 | | | | | | | | | | | yes | | | | | yes |
| FCRL5 | NA | Body | chr1:157490546-157490968 | 422 | | | 1 | | | | | | | | | | | | yes | | | | |
| FCRL3 | NA | 5'UTR;TSS200 | chr1:157670328-157670869 | 541 | | | 6 | | 1 | | 1 | | 1 | 1 | | | | | | | yes | | |
| CD5L | NA | Body;TSS200 | chr1:157811303-157811905 | 602 | | 1 | | | | | | | | | | | | yes | | | | | |
| CD1D | N_Shore | TSS200 | chr1:158149705-158149974 | 269 | | | 1 | | | | | | 1 | | | | | yes | yes | | | | |
| CD1A | NA | TSS1500 | chr1:158223513-158223934 | 421 | | | 4 | | | | | | | | | | | | yes | | | yes | |
| OR10K2 | NA | 1stExon;TSS200 | chr1:158390649-158391119 | 470 | | | 2 | | | | | | | | | | | | yes | | | | |
| OR6K2 | NA | 1stExon | chr1:158670124-158670617 | 493 | | | | | | | | | | | | | | | | | yes | | |
| OR6K3 | NA | TSS1500 | chr1:158688775-158689141 | 366 | | | | | | | | | | | | | | yes | yes | | | | yes |
| OR6K6 | NA | TSS1500 | chr1:158723939-158724362 | 423 | | | | | | | | | | | | | | yes | | | | | |
| PYHIN1 | NA | TSS1500 | chr1:158900088-158900644 | 556 | | | | | | | | | 2 | | | | | | | yes | | | |
| AIM2 | NA | 5'UTR;TSS200 | chr1:159046391-159047034 | 643 | | | 1 | | | | | | 1 | | | | | | | yes | | | |
| CADM3 | N_Shore | TSS1500 | chr1:159140372-159140702 | 330 | | | 4 | | | | | | | | | | | yes | | | | | |
| DARC | NA | TSS1500 | chr1:159174076-159174728 | 652 | | | | | | | | | | | | | | | | | yes | | yes |
| CRP | NA | TSS200;Body;1stExon;5'UTR | chr1:159683347-159684684 | 1337 | | 1 | 8 | 1 | | | | | | | | | | yes | yes | | | yes | |
| DUSP23 | Island | 5'UTR | chr1:159750836-159751352 | 516 | | | | | | | | | 1 | | | | | | yes | | yes | | |
| FCRL6 | NA | TSS1500 | chr1:159771711-159772194 | 483 | | | | | | | | | 1 | | | | | | yes | | | | |
| SLAMF8 | NA | TSS1500;TSS200;5'UTR;1stExon | chr1:159796231-159796658 | 427 | | | 18 | 1 | | | | 1 | 1 | | | 1 | | yes | yes | | | | |
| VSIG8;C1orf204 | S_Shore | Body;TSS1500 | chr1:159825552-159825761 | 209 | | | | 1 | | | | | | | | | | | yes | | | yes | |
| CCDC19 | Island | TSS200 | chr1:159869927-159870915 | 988 | | | 5 | | 1 | 1 | | | 3 | | | 1 | | yes | | | yes | | yes |
| IGSF9 | Island | Body | chr1:159899530-159899714 | 184 | | | 1 | | | | | | 1 | | | | | | | | | | |
| IGSF9 | S_Shore | TSS1500 | chr1:159915970-159916077 | 107 | | | | | | | | | | | | | | yes | | | | | |
| SLAMF9 | NA | Body | chr1:159923525-159923861 | 336 | | | | | | | | | | | | | | | | | | | |
| KCNJ9 | N_Shelf | TSS1500 | chr1:160050877-160051087 | 210 | | | | | | | | | | | | | | yes | yes | | | | yes |
| KCNJ9 | N_Shore;Island | 5'UTR;Body | chr1:160053689-160054321 | 632 | | | 10 | | | | | | | | | | | yes | | | | | yes |
| IGSF8 | S_Shore | TSS200 | chr1:160068607-160068842 | 235 | | | | | | | | | 3 | | | | | yes | | | | yes | yes |
| ATP1A2 | NA | TSS1500;TSS200 | chr1:160084398-160085581 | 1183 | | | | | | | | | 1 | | | | | yes | | | | | |
| CASQ1 | NA | TSS200 | chr1:160160106-160160312 | 206 | | | | | | | | | | | | | | | | | | | |
| NCSTN;COPA | S_Shore | Body;TSS1500 | chr1:160314452-160314746 | 294 | | | | | | | | | 1 | | | | | yes | | | | | |
| NHLH1 | NA | TSS1500 | chr1:160336590-160336994 | 404 | | | | | | | | | 1 | | | | | | | | | | |
| CD84 | NA | Body | chr1:160520346-160520625 | 279 | | | | | | | | | | | | | | yes | | | | | |
| SLAMF1 | NA | Body;5'UTR;1stExon;TSS200 | chr1:160616604-160617974 | 1370 | | | | 1 | | | | | | | | | | | yes | | | yes | |
| CD48 | NA | 1stExon | chr1:160681530-160682008 | 478 | | | | | | | | | | | | | | yes | | yes | yes | | |
| LY9 | NA | TSS1500 | chr1:160764664-160765349 | 685 | | | | | | | | | | | | | | yes | | yes | yes | | |
| ITLN1 | NA | TSS200 | chr1:160854954-160855148 | 194 | | | | | | | | 1 | | | 1 | | | yes | | | | | |
| F11R | Island | TSS200 | chr1:160991154-160991191 | 37 | | | | | | | | | | | | | | yes | | | | | |
| TSTD1 | N_Shore;Island;chr1:16100837 | Body;5'UTR;1stExon;TSS200;Island | chr1:161008127-161008977 | 850 | 9 | 1 | 29 | 1 | | | | 2 | 2 | | 1 | 1 | | | | | | yes | |
| ARHGAP30 | NA | 1stExon;5'UTR | chr1:161039681-161040044 | 363 | | | | | | | | | 2 | 1 | | | | yes | | | | | |
| PVRL4 | N_Shore;Island | Body | chr1:161049147-161050227 | 1080 | | | 15 | | | | | | 2 | | | 1 | | yes | yes | | | | |
| NECTIN4 | NA | TSS200;5'UTR;1stExon | chr1:161059267-161059654 | 387 | | | | | 2 | | | | 2 | | | | | yes | | | | yes | |
| KLHDC9 | N_Shore;Island | TSS1500;TSS200;5'UTR;1stExon | chr1:161067927-161068781 | 854 | | | 7 | | | | | | | | | | | yes | | | | yes | |
| PFDN2;NIT1 | S_Shore | TSS1500;Body | chr1:161088957-161089118 | 161 | | | | | | | | | 1 | | | | | | yes | | | | |
| USP21;UFC1 | N_Shore | TSS1500;3'UTR | chr1:161128567-161128720 | 153 | | | 3 | | | | | | | | | | | | | | | | |
| NDUFS2;ADAMTS4 | N_Shelf;chr1:161171809-1611 | TSS1500;1stExon;TSS200;N_Shelf; | chr1:161168145-161169574 | 1429 | | 1 | | | | | | | 1 | | 1 | | | | | | | yes | |
| NDUFS2 | N_Shore | 5'UTR;TSS1500 | chr1:161171211-161171632 | 421 | | | | 1 | | | | 1 | 1 | | | | | | | | | | |
| FCER1G | NA | TSS200 | chr1:161184921-161185092 | 171 | | | | | | | | | 1 | 1 | | | | yes | | | | | |
| APOA2 | NA | TSS1500 | chr1:161193933-161193991 | 58 | | | | | | | | | 1 | | | | | | yes | | | | |
| C1orf192 | NA | Body | chr1:161337603-161337703 | 100 | | | 1 | | | | | 1 | | | | | | yes | | | yes | | |
| FCGR2A | NA | TSS1500 | chr1:161474586-161474976 | 390 | | | | | | | | | | | | | | yes | yes | | | | |
| HSPA6 | N_Shore | TSS1500 | chr1:161492666-161493011 | 345 | | | 2 | | | | | | | | | | | | | | | | |
| HSPA7 | Island | Body;TSS200 | chr1:161575716-161576956 | 1240 | | | 2 | | | | | 2 | | | | | | | | | | | |
| FCRLA | NA | TSS1500 | chr1:161676266-161676581 | 315 | | | | | | | | | | | 1 | | | yes | | | | | |
| FCRLB | Island | Body | chr1:161695659-161695839 | 180 | | | | | | | | | | | | | | yes | | | | | |
| NOS1AP | Island | Body | chr1:162336877-162337375 | 498 | | | 12 | | | | | | 2 | | | | | | | | | | |
| UHMK1 | N_Shore | TSS1500 | chr1:162467180-162467363 | 183 | | | 6 | | | | | 1 | 2 | | | | | | | yes | | | |
| DDR2 | NA | TSS1500;TSS200 | chr1:162601807-162602559 | 752 | | | | | | | | 1 | 1 | | | | | | | | | | |
| HSD17B7 | N_Shore | TSS1500 | chr1:162759914-162760659 | 745 | | | | | | | | 1 | 1 | 3 | | 1 | | | | yes | yes | | |
| RGS4 | NA | 1stExon;Body;TSS200;5'UTR | chr1:163039012-163039184 | 172 | | 1 | | | | | | | 1 | | | | | | | | | | |
| RGS5 | NA | TSS200 | chr1:163172914-163173262 | 348 | | | 5 | | | | | | | | | | | yes | yes | | | | |
| PBX1 | Island | Body | chr1:164545553-164546143 | 590 | | 1 | 11 | | | | | | 3 | | 1 | 1 | | | | | | | |
| PBX1 | NA | Body | chr1:164581169-164587676 | 6507 | | | | | | | | | | | | | | yes | yes | | yes | | |
| LMX1A | N_Shore;Island | Body | chr1:165204410-165205199 | 789 | | | 6 | | | | | 2 | 3 | | | | | | | | yes | yes | yes |

| | | | | | | | | | | | | | | | | | | | | | | | |
|---|---|---|---|---|---|---|---|---|---|---|---|---|---|---|---|---|---|---|---|---|---|---|---|
| LMX1A | Island | Body | chr1:165321741-165322199 | 458 | | | 3 | | | | | | | | | | | | | | | | |
| LMX1A | N_Shore;Island | Body | chr1:165323232-165325136 | 1904 | | 1 | 4 | | | | | | 1 | | | | | | | | | yes | |
| LMX1A | Island | TSS1500 | chr1:165326269-165326345 | 76 | | | | | | | | | | | 1 | | | yes | | | | | |
| LOC400794;LRRC52 | NA | Body;TSS200 | chr1:165513318-165513361 | 43 | | | | | | | | 2 | | | | 2 | | | | | | | |
| FAM78B | NA | 3'UTR;Body | chr1:166039444-166039858 | 414 | | | 4 | | | | | | | | | | | | | | | | |
| MIR921;FAM78B | NA | TSS1500;Body | chr1:166124419-166124903 | 484 | | | | | | | | | | | | | | yes | | | | | |
| FAM78B | S_Shore | TSS1500 | chr1:166136630-166136939 | 309 | | | 1 | | | | | | | | | | | | yes | | | | |
| ILDR2 | Island | Body | chr1:166890380-166890436 | 56 | | | | | | | | | 1 | | | | | | | | | | |
| MAEL | Island | TSS1500;TSS200 | chr1:166958233-166958583 | 350 | | | 3 | | | | | | 1 | | | | | | yes | | | | |
| STYXL2 | Island | Body | chr1:167090618-167090974 | 356 | | | | | | | | 1 | 1 | | | | | yes | | yes | | yes | |
| CD247 | Island | Body | chr1:167408709-167409198 | 489 | | | | | | | | | 1 | 1 | | | | yes | yes | | | | |
| CD247 | NA | Body | chr1:167487295-167487633 | 338 | | | | | | | | | 1 | 1 | 1 | | | yes | | | | | |
| RCSD1 | N_Shore | TSS1500 | chr1:167598958-167599124 | 166 | | | | | | | | | 1 | | 1 | | | | | | | yes | yes |
| MPZL1 | N_Shore | TSS1500 | chr1:167690438-167690788 | 350 | | | | | | | | 1 | 1 | | | | | | | | | | |
| ADCY10 | Island | Body | chr1:167789455-167789646 | 191 | | | 1 | | | | | 1 | 1 | | | 1 | | yes | | | yes | | |
| XCL2 | NA | Body | chr1:168513067-168513310 | 243 | | | | | | | | | 1 | | | | | yes | | | | | |
| DPT | NA | TSS1500 | chr1:168698867-168698977 | 110 | | | | | | | | | 1 | | | | | | yes | | | | |
| C1orf114 | Island | 1stExon;5'UTR;TSS200 | chr1:169396635-169396909 | 274 | | 1 | | | | | | 1 | 1 | | | | | | | | | | |
| METTL11B | NA | TSS1500;TSS200 | chr1:170114946-170115351 | 405 | | | | | | | | 1 | 2 | | | 1 | | | yes | yes | | | |
| PRRX1 | S_Shelf | Body | chr1:170637600-170637796 | 196 | | | | | | | | | | | | | | | | | | | |
| PRRX1 | NA | Body | chr1:170638807-170639289 | 482 | | | 4 | | | | | | 1 | | | | | | | | | yes | |
| FMO6P | NA | TSS1500 | chr1:171105781-171106397 | 616 | | | | | | | | | | | | | | | yes | yes | | | |
| FMO2 | NA | 5'UTR;1stExon | chr1:171154475-171154802 | 327 | | | | | | | | | | | | | | | yes | | | | |
| FMO1 | NA | TSS1500 | chr1:171217189-171217592 | 403 | | | | | | | | | 1 | | | | | yes | yes | | | | yes |
| MYOC | NA | TSS200;1stExon | chr1:171621279-171622329 | 1050 | | | | | | | | | | | | | | | yes | | yes | yes | |
| DNM3;MIR199A2 | NA | Body;TSS200;TSS1500 | chr1:172113920-172114419 | 499 | | | | | | | | 1 | 1 | | | | | | | | | | yes |
| C1orf105;PIGC | N_Shore;S_Shore | Body;5'UTR;TSS1500 | chr1:172412837-172413612 | 775 | | | 7 | | | | | | | 1 | | 1 | | yes | yes | | | | |
| FASLG | NA | TSS200;nan | chr1:172628020-172628514 | 494 | | | | | | | | 1 | 2 | | 1 | | | | | | | yes | |
| TNFSF4 | NA | 5'UTR;1stExon | chr1:173176362-173176523 | 161 | | | 35 | | | | | | 1 | 1 | | | | yes | | | | | |
| GAS5;SNORD78;SNORD76;SNORD77;SN | N_Shore | Body;TSS1500;TSS200 | chr1:173835989-173836307 | 318 | | | 2 | | | | | | | | | | | yes | | | | | |
| RABGAP1L | NA | TSS1500;Body | chr1:174843754-174844490 | 736 | | 1 | 5 | | | | | 1 | 1 | | | 3 | | yes | | | | yes | |
| CACYBP | N_Shore | TSS1500 | chr1:174967663-174968144 | 481 | | | 6 | | | | | | 1 | | | | | | yes | yes | | | |
| KIAA0040 | S_Shore | TSS1500;TSS200 | chr1:175162397-175163045 | 648 | | | 8 | | | | | | 2 | | | | | yes | | | | yes | |
| TNR | NA | Body | chr1:175355212-175355362 | 150 | | | | | | | | | | | | | | yes | | | yes | | |
| TNR | N_Shore;Island | 5'UTR | chr1:175568216-175568710 | 494 | | 1 | | | | | | 1 | | | | 1 | 1 | yes | yes | | | | |
| ASTN1 | NA | Body | chr1:177001793-177002228 | 435 | | | | | | | | | 1 | | | | | yes | | | | yes | |
| FAM5B | Island | TSS1500 | chr1:177140322-177140680 | 358 | | | | | | | | | 1 | | | | | | | | | yes | |
| FAM5B | NA | Body | chr1:177225466-177225888 | 422 | | | 2 | 1 | | | | 1 | 1 | | | 2 | | | | | | | |
| SEC16B | NA | TSS1500 | chr1:177939251-177939546 | 295 | | | | | 1 | | | | 1 | | | | | yes | yes | | | | |
| RASAL2 | NA | Body;TSS200 | chr1:178310534-178310650 | 116 | 2 | | | 1 | | | | | 1 | | 1 | | | | | | | | yes |
| RALGPS2 | N_Shore;Island | TSS1500 | chr1:178693595-178694013 | 418 | | | 8 | | | | | | | | | | | | | | | yes | yes |
| FAM20B | N_Shore | TSS1500 | chr1:178994243-178994355 | 112 | | | | | | | | | 1 | | | | | yes | | | | | |
| TOR3A | N_Shore | TSS1500 | chr1:179050404-179050691 | 287 | | | 2 | | | | | | | | | | | | | | | | |
| ABL2 | S_Shore | TSS1500 | chr1:179199424-179199939 | 515 | | | | | | | | | 1 | | | | | | | | yes | yes | |
| NPHS2 | Island;S_Shore | Body;1stExon;5'UTR;TSS1500 | chr1:179544721-179545790 | 1069 | | 1 | | | | | | 1 | 1 | | | | | yes | | | | yes | yes |
| TDRD5 | N_Shore;Island | TSS1500;TSS200 | chr1:179560700-179561111 | 411 | | | 2 | | | | | 1 | 5 | | | | | | | | yes | | |
| FAM163A | Island | TSS200;5'UTR | chr1:179712281-179713950 | 1669 | | | 10 | | | | | 1 | 2 | | | 3 | | | | | | | yes |
| FAM163A | NA | 5'UTR | chr1:179781780-179782240 | 460 | | 1 | | | | | | | | | | | | yes | | | | | |
| KIAA1614 | N_Shore | TSS1500 | chr1:180881086-180881163 | 77 | | | | 1 | | | | | | | | | | | | yes | | | |
| KIAA1614 | Island | Body | chr1:180882591-180882681 | 90 | | 1 | | | | | | | | | | | | yes | | | | | yes |
| KIAA1614 | Island | Body | chr1:180904677-180904914 | 237 | | | | 1 | | | | | | | | | | | | | | | |
| KIAA1614 | NA | Body | chr1:180911407-180911763 | 356 | | | | | | | | | 1 | | | | | yes | yes | | | | |
| STX6 | Island | TSS200 | chr1:180992098-180992117 | 19 | | | | | | | | | 1 | | | | | yes | | | | yes | |
| MR1 | NA | TSS200 | chr1:181002976-181003203 | 227 | | | | | | | | | 1 | | | | | | | | | | yes |
| IER5 | N_Shore | TSS1500 | chr1:181057360-181057458 | 98 | | | 1 | | | | | | | | | | | | | | | | |
| CACNA1E | Island | TSS1500 | chr1:181451906-181454164 | 2258 | | 1 | | | | | | | | | | | | | yes | | | yes | yes |
| CACNA1E | Island | Body | chr1:181767609-181767774 | 165 | | | | | | | | | | | | | | | yes | | yes | | |
| ZNF648 | Island | Body | chr1:182026435-182026795 | 360 | | | | | | | | | | | | | | | | | | | |
| RGS16 | Island | TSS200 | chr1:182573731-182573759 | 28 | | | | | 1 | | | 1 | | | | | | yes | yes | | | | |
| RGS8 | NA | 5'UTR;1stExon | chr1:182642036-182642214 | 178 | | | | | | | | | 1 | | | | | yes | | | yes | | |
| C1orf14 | N_Shore;Island | Body | chr1:182921660-182922502 | 842 | | | 12 | | | | | | 3 | 1 | | 2 | | yes | | | | yes | |
| LAMC2 | NA | TSS1500;TSS200;1stExon;5'UTR | chr1:183154778-183155644 | 866 | | | 16 | | | | | | 1 | | | | | yes | yes | | | yes | |
| NMNAT2 | NA | TSS200;Body | chr1:183274124-183274661 | 537 | | | | | | | | | 1 | | | | | yes | | | | | |
| HMCN1 | N_Shore;Island | TSS1500;TSS200 | chr1:185703335-185703557 | 222 | | | | | | | | | 2 | 1 | | | | | | | | | |
| PRG4;MIR548F1 | NA | TSS200;Body | chr1:186265270-186265765 | 495 | | | | 1 | | | | | | | | | | | | | | | |
| PTGS2 | Island | TSS1500 | chr1:186649812-186649909 | 97 | | | | | | | | | 1 | | | | | | | | | | |
| FAM5C | NA | 5'UTR | chr1:190444666-190445197 | 531 | | 1 | | | | | | | | | | | | | | | | yes | yes |
| DENND1B | Island;S_Shore | TSS200;TSS1500 | chr1:197744757-197745238 | 481 | | | 13 | | | | | | | | | | | yes | | | | | yes |
| LHX9 | Island;S_Shore | Body | chr1:197887649-197888880 | 1231 | | 1 | | | | | | | 1 | | | | | | | | | | |
| LHX9 | Island | Body | chr1:197890479-197890812 | 333 | | | 1 | | | | | | | | | | | | | | | | yes |
| NR5A2 | N_Shore;Island;S_Shore | Body | chr1:200003267-200005635 | 2368 | | | 2 | 1 | | | | | 1 | | | | | | | | | yes | |
| NR5A2 | Island | Body | chr1:200010626-200011976 | 1350 | | 1 | | | | | | | 1 | | | | | | | | | | |
| DDX59 | S_Shore | TSS1500 | chr1:200640177-200640542 | 365 | | | | | | | | | 1 | | | | | | yes | | yes | yes | |
| GPR25 | Island | 1stExon | chr1:200842282-200842756 | 474 | | | 1 | | | | | | | | | | | | | | | | yes |
| C1orf106 | Island | TSS1500 | chr1:200860363-200860534 | 171 | | | | | | | | | 1 | | | | | | yes | | | | |
| KIF21B | Island | Body | chr1:200969751-200969906 | 155 | | | 1 | | | | | | | | | | | yes | yes | | | | |
| KIF21B | S_Shore | TSS200 | chr1:200992964-200993200 | 236 | | | 1 | | | | | | | | | | | yes | | | | yes | yes |
| CACNA1S | N_Shelf | 1stExon | chr1:201081382-201081686 | 304 | | | | | | | | | | | | | | | | | | yes | |
| TMEM9 | S_Shore | TSS1500 | chr1:201123894-201123992 | 98 | | | | | | | | | 1 | | | 1 | | yes | | | | | |
| IGFN1 | NA | TSS200 | chr1:201159756-201159988 | 232 | | | | | | | | | | | | | | yes | | | | | yes |
| PKP1 | Island | 1stExon;5'UTR | chr1:201252734-201253537 | 803 | | | 10 | | | | | | 1 | | | 1 | | | | | | | |
| TNNT2 | NA | 5'UTR | chr1:201346647-201346876 | 229 | | | | | | | | | 1 | | | | | yes | | | | | |
| LAD1 | Island | TSS200;TSS1500 | chr1:201368843-201369031 | 188 | | | 14 | | | | | | 1 | | | | | | | | | | yes |
| CSRP1 | S_Shore | TSS200 | chr1:201476557-201476775 | 218 | | | 4 | | | | | | | | | | | yes | | | | | |
| NAV1 | Island | 1stExon;TSS200 | chr1:201617355-201618284 | 929 | | | 1 | | | | | 1 | 2 | | | 1 | | | | | | yes | yes |
| NAV1 | Island | Body | chr1:201619295-201619900 | 605 | | | | | | | | 1 | 1 | | | | | | | | | yes | |
| NAV1 | N_Shore;Island | TSS1500;Body;TSS200;1stExon;5'U | chr1:201708500-201709675 | 1175 | | | 24 | 1 | | | | | 1 | | | 1 | | | | | | | |
| SHISA4 | NA | TSS1500 | chr1:201857591-201857787 | 196 | | | | | | | | | 1 | | | | | | | | | | yes |
| RNPEP | N_Shore | TSS1500 | chr1:201951076-201951234 | 158 | | | 1 | | | | | | | | | | | yes | yes | | yes | | yes |
| ELF3 | NA | TSS1500;nan;TSS200;5'UTR;1stExo | chr1:201979128-201979938 | 810 | | | 2 | 1 | 1 | | | | 4 | 1 | 1 | | | yes | | | | | |
| GPR37L1 | NA | TSS1500;TSS200 | chr1:202090924-202092005 | 1081 | | | 19 | 1 | | | | | 3 | | | | | yes | yes | | | | |
| PTPN7 | NA | 5'UTR;1stExon;TSS200;TSS1500;na | chr1:202128682-202131184 | 2502 | | | 5 | 1 | | | | | 4 | 1 | 1 | | | | | | | yes | |
| LGR6 | NA | TSS200;Body;nan | chr1:202172778-202172912 | 134 | | | | | | | 1 | 1 | 2 | 2 | 2 | 15 | | | | | | yes | |
| LGR6 | N_Shore;Island | TSS1500;Body;TSS200 | chr1:202182417-202183387 | 970 | 4 | | 2 | | | | | | | | | | | yes | yes | yes | | yes | yes |
| UBE2T | Island | 1stExon;5'UTR | chr1:202311052-202311228 | 176 | | | | | | | | | | | | | | yes | yes | | | | |
| KDM5B | N_Shore | TSS1500 | chr1:202778061-202778869 | 808 | | | 3 | | | | | | 2 | | | | | yes | yes | | | | yes |
| RABIF | S_Shore | TSS1500 | chr1:202858761-202858826 | 65 | | | | | | | | | 1 | | | | | | yes | | | | |
| PPFIA4 | N_Shore;Island | Body | chr1:203044677-203045229 | 552 | | | 13 | | | | | 4 | 4 | | | 1 | | yes | | | | | |
| MYOG | NA | TSS200 | chr1:203055479-203055662 | 183 | | | 1 | | | | | | | | | | | | yes | | | | |

| | | | | | | | | | | | | | | | | | | | | | | | |
|---|---|---|---|---|---|---|---|---|---|---|---|---|---|---|---|---|---|---|---|---|---|---|---|
| ADORA1 | N_Shore | TSS1500 | chr1:203095747-203096683 | 936 | | | 5 | | | | | | | | | | | yes | | | | | yes |
| CHI3L1 | NA | 1stExon;TSS200;TSS1500 | chr1:203155737-203156764 | 1027 | | | 6 | | | | | 2 | 1 | 1 | | | | | yes | yes | | yes | |
| FMOD | NA | 1stExon;5'UTR;TSS200 | chr1:203320190-203320541 | 351 | | 1 | | | | | | 4 | 8 | 2 | | 13 | | yes | yes | | | yes | |
| PRELP | NA | TSS1500 | chr1:203444518-203444945 | 427 | | | | | 1 | | | | 1 | | | | | yes | | | | yes | |
| ATP2B4 | N_Shelf | TSS1500 | chr1:203595048-203595283 | 235 | | | 5 | 1 | | | | | | | | | | yes | | | | yes | yes |
| ATP2B4 | Island | 5'UTR | chr1:203598594-203599089 | 495 | | | | 1 | | | | | | | | | | | | | | yes | |
| LAX1 | NA | TSS200;1stExon;5'UTR | chr1:203734256-203734559 | 303 | | | 1 | | | | | | 3 | 1 | | | | | | | | | |
| SNRPE | N_Shore | TSS1500 | chr1:203829481-203830200 | 719 | | | 8 | | 1 | | | | 1 | | | | | yes | | | | | |
| ETNK2 | Island | Body | chr1:204120607-204120825 | 218 | | | | | | | | | 1 | | | | | | | | | | yes |
| ETNK2 | S_Shore | TSS1500 | chr1:204121899-204121925 | 26 | | | | | | | | | | | | 1 | | | | | yes | | |
| KISS1 | N_Shore | 3'UTR | chr1:204159498-204159832 | 334 | 1 | | | | | | | | | | | | | | yes | | yes | | yes |
| KISS1 | NA | 5'UTR | chr1:204165307-204165688 | 381 | | | 4 | | | | | | | | | | | | | | | yes | yes |
| GOLT1A | NA | 5'UTR;1stExon;TSS200;TSS1500 | chr1:204183060-204183655 | 595 | | 1 | 1 | | | | | | 1 | | | | | yes | | | yes | | |
| PLEKHA6 | NA | 5'UTR;TSS200 | chr1:204328377-204329307 | 930 | | 1 | | 1 | | | | | | | | | | | | | | | yes |
| PIK3C2B | NA | 5'UTR | chr1:204459139-204459584 | 445 | | | | | | | | | | | | | | yes | yes | | | | |
| LRRN2 | S_Shore | TSS1500 | chr1:204655147-204655678 | 531 | | | 3 | | | | | | | | | | | | | | | | |
| NFASC | NA | Body | chr1:204966197-204966338 | 141 | | | 4 | | | | | | | | | | | | | | | yes | |
| CNTN2 | NA | TSS1500 | chr1:205011906-205012569 | 663 | | | | 1 | | | | | | | | | | yes | | | | yes | yes |
| CNTN2 | N_Shore;Island | Body | chr1:205038308-205039083 | 775 | | 1 | 2 | | | | | | | | | | | yes | | | | yes | |
| DSTYK | S_Shore | TSS1500 | chr1:205181237-205181581 | 344 | | | | | | | | 1 | 1 | | | | | yes | | | | | |
| KLHDC8A | N_Shore;Island | Body;5'UTR | chr1:205312199-205313833 | 1634 | | | | | | | | | | | | | | yes | | | | | yes |
| KLHDC8A | NA | TSS200 | chr1:205326059-205326437 | 378 | | | | | | | | | 2 | | 1 | | | yes | | | | | |
| CDK18 | N_Shore;Island | Body | chr1:205497713-205498551 | 838 | | | | | | | | 1 | 1 | | | | | yes | | | yes | | |
| MFSD4 | Island | TSS1500 | chr1:205537752-205537837 | 85 | | | | | 1 | | | | | | | | | | | | yes | | yes |
| MFSD4 | N_Shore | Body | chr1:205560943-205561401 | 458 | | 1 | | | | | | | | | | | | yes | yes | | | | |
| ELK4 | S_Shore | TSS1500 | chr1:205602864-205603464 | 600 | | | 4 | 1 | | | | | | | | | | yes | | | | | |
| SLC45A3 | Island | Body | chr1:205631978-205632101 | 123 | | | | | | | | | 1 | | | | | yes | | | | | |
| RAB7L1 | S_Shore | TSS1500 | chr1:205745431-205745560 | 129 | | | | | | | | | 1 | | | | | | yes | yes | | | |
| PM20D1 | N_Shore;Island;S_Shore | Body;1stExon;5'UTR;TSS200 | chr1:205818484-205819609 | 1125 | 7 | 23 | 47 | 3 | 4 | 16 | 1 | 6 | 16 | 3 | 4 | 91 | 4 | yes | | | | | |
| AVPR1B | N_Shore | TSS1500 | chr1:206223241-206223499 | 258 | | | 3 | 1 | | | | | | | | | | yes | | | | | |
| IKBKE | NA | TSS1500 | chr1:206643301-206643377 | 76 | | | 2 | | | | | | 1 | | | | | | | | | | |
| RASSF5 | Island | TSS1500 | chr1:206680250-206680628 | 378 | | | 1 | | | | | | | | | | | | | | yes | | |
| RASSF5 | Island;N_Shore | Body;5'UTR;1stExon;TSS1500 | chr1:206729034-206731166 | 2132 | | 1 | 11 | 1 | | | | | 2 | | | | | yes | | | | | yes |
| LGTN | S_Shore | TSS1500 | chr1:206786170-206786222 | 52 | | 1 | 26 | | | | | 2 | 2 | | | 2 | | yes | | | | | |
| IL10 | NA | TSS200 | chr1:206945924-206946187 | 263 | | | | | | | | 1 | 1 | | 1 | | | yes | | | | | |
| IL19 | NA | TSS1500;TSS200 | chr1:206972002-206972262 | 260 | | | | | | | | | | | | | | | | | yes | | |
| IL20 | NA | TSS1500 | chr1:207038584-207039008 | 424 | | | | | | | | | 1 | | | | | | | | yes | | |
| IL24 | NA | TSS1500 | chr1:207070539-207071109 | 570 | | | | | | | | | 2 | | | | | yes | | | | | |
| FAIM3 | N_Shore;Island | Body | chr1:207082479-207083129 | 650 | | | | | | | | 1 | 2 | | | | | | yes | | | | yes |
| FAIM3 | NA | Body;5'UTR;1stExon;TSS1500 | chr1:207095153-207096833 | 1680 | | 1 | 3 | | | | | | 1 | | | | | yes | | | | yes | |
| PIGR | NA | TSS200 | chr1:207119987-207120407 | 420 | | | 1 | | | | | | | | | | | yes | | | | | |
| YOD1 | N_Shore;Island | TSS200;TSS1500;1stExon | chr1:207224090-207225647 | 1557 | | | 7 | | | | | 1 | 4 | 1 | | 2 | | yes | yes | | | | |
| C4BPB | NA | TSS1500;TSS200 | chr1:207262190-207262706 | 516 | | | | 1 | 1 | | | | | | | | | | | yes | | | |
| C4BPA | NA | TSS1500;TSS200 | chr1:207277018-207277581 | 563 | | 1 | 2 | 1 | | | | | 2 | | | | | | | | | | |
| CD55 | N_Shore | TSS1500 | chr1:207494213-207494472 | 259 | | | | 1 | | | | | | | | | | | yes | | yes | yes | |
| CR1 | N_Shore | TSS1500 | chr1:207669116-207669334 | 218 | | | 5 | | | | | | | | | | | | yes | | | | |
| CR1L | N_Shore;Island | TSS1500;TSS200 | chr1:207817630-207818818 | 1188 | | | 2 | | | | | 3 | | | | 1 | | | | | | | |
| CR1L | Island | Body | chr1:207842833-207843084 | 251 | | 2 | 1 | | | | 1 | 3 | 3 | 3 | | 8 | 1 | | | | | | |
| LOC148696 | NA | TSS1500;TSS200;Body | chr1:207990895-207992548 | 1653 | 16 | | 21 | | 2 | | | | | | 1 | | | | | | | | |
| CD34 | N_Shore;S_Shore | Body;TSS1500 | chr1:208083913-208085455 | 1542 | | | 6 | | | | | | 1 | 1 | | | | | | | | yes | yes |
| LOC642587 | NA | TSS1500 | chr1:209600926-209601066 | 140 | | | | | | | | | 1 | | | | | yes | | | | | |
| LOC642587;MIR205 | NA | Body;TSS1500 | chr1:209604378-209605304 | 926 | | | | | | | | | 2 | | | | | | | | | yes | |
| CAMK1G | NA | 5'UTR | chr1:209759145-209759344 | 199 | | | 1 | | | | | | | | | | | yes | | | yes | | |
| LAMB3 | N_Shore;Island | Body | chr1:209798721-209799353 | 632 | 4 | | 13 | | | | | | 2 | 1 | | | | yes | | | | yes | |
| LAMB3 | NA | TSS1500;5'UTR;1stExon;TSS200 | chr1:209825672-209826863 | 1191 | | | | | | | | | 2 | 1 | | | | yes | | | | | |
| G0S2 | N_Shore;Island | TSS1500;TSS200;1stExon;5'UTR | chr1:209848402-209849445 | 1043 | | | | | | | | 6 | 3 | | | 1 | | | | | | yes | yes |
| HSD11B1 | NA | 5'UTR;TSS1500 | chr1:209877941-209878178 | 237 | | | 2 | | | | | | | | | | | | | | | | |
| TRAF3IP3 | NA | TSS200;nan | chr1:209929496-209943131 | 13635 | | | | | | | | | 1 | 1 | 1 | | | yes | yes | yes | yes | yes | |
| IRF6 | N_Shore;Island | 5'UTR;TSS200 | chr1:209979111-209979779 | 668 | | | 2 | | | | | 2 | 2 | | | 1 | | yes | | | | | |
| C1orf107 | N_Shore | TSS1500 | chr1:210000883-210001082 | 199 | | | | | | | | 1 | 1 | | | | | yes | | | | | |
| SYT14 | N_Shore;Island | TSS1500;1stExon;5'UTR;Body | chr1:210110998-210112031 | 1033 | | 1 | | | | | | | 1 | | | | | | | | | yes | yes |
| C1orf133;SERTAD4 | Island;S_Shore | TSS200;5'UTR;TSS1500 | chr1:210407473-210407973 | 500 | 1 | | 21 | | | | | | 1 | | | | | | | | | | yes |
| HHAT | N_Shore | TSS1500 | chr1:210501368-210501634 | 266 | | | 1 | | | | | | | | | | | yes | | | yes | | yes |
| KCNH1 | N_Shore | Body | chr1:210856989-210857380 | 391 | | | 2 | | | | | | | | | | | | | | yes | | |
| KCNH1 | S_Shore | TSS1500 | chr1:211307761-211308115 | 354 | | | | | | | | | | | | | | yes | | | | | |
| TRAF5 | N_Shore | TSS1500 | chr1:211498962-211499648 | 686 | | | 7 | | | | | | 1 | | | | | yes | | | yes | yes | |
| RD3 | N_Shore;Island | 3'UTR;Body | chr1:211652276-211652741 | 465 | | | 12 | | | | | 2 | 2 | | | 2 | | | | | | | |
| RD3 | NA | TSS1500;TSS200 | chr1:211666304-211667682 | 1378 | 1 | | | 1 | | | | | | | | | | | | | | | |
| TMEM206 | S_Shore | TSS1500 | chr1:212588536-212588848 | 312 | | | | | | | | 1 | | | | | | | | | | | |
| FAM71A | NA | TSS1500 | chr1:212797322-212797670 | 348 | | | | | | | | | 1 | | | | | | | | | | |
| PROX1 | Island | TSS1500 | chr1:214161306-214162039 | 733 | | | | | | | | | 1 | | | | | | | | | yes | yes |
| PROX1 | NA | Body | chr1:214169912-214170376 | 464 | | | 4 | | | | | | | | | | | | | | | | |
| SMYD2 | NA | Body | chr1:214477063-214477280 | 217 | | | 6 | | | | | | 1 | | | 1 | | yes | | | | | |
| PTPN14 | Island;S_Shore | Body | chr1:214557399-214557928 | 529 | | | | | | | | | 3 | | | | | | | | | | |
| PTPN14 | Island | TSS1500 | chr1:214724875-214725246 | 371 | | | | | | | | | 1 | | | | | | | | | | yes |
| KCNK2 | NA | TSS1500 | chr1:215178601-215178854 | 253 | | | | | | | | | 1 | | | | | | | | | yes | |
| KCNK2 | N_Shore;Island;S_Shore | TSS1500;Body | chr1:215255110-215257222 | 2112 | | 1 | 5 | | | | | | 2 | 1 | | | | | | | | yes | yes |
| ESRRG | NA | 5'UTR;TSS200 | chr1:216896852-216896863 | 11 | | | | | | | | | 1 | | | | | | | | | | |
| ESRRG | Island;S_Shore | TSS1500;5'UTR | chr1:217263213-217263847 | 634 | | | 6 | | | | | | | | | | | | | | | | yes |
| ESRRG | N_Shore | 5'UTR | chr1:217306590-217306763 | 173 | | | 5 | 1 | | | | | 1 | | | 2 | | | | | | yes | yes |
| ESRRG | N_Shore;Island;S_Shore | 5'UTR;1stExon;TSS200;TSS1500 | chr1:217309568-217312478 | 2910 | | | | | | | | | | | | | | | | | | | |
| SLC30A10 | chr1:220101132-220102147;S_ | Island;TSS1500 | chr1:220101698-220102517 | 819 | | | | 1 | | | | 1 | | | 1 | | | yes | | | | | yes |
| RAB3GAP2 | N_Shore | Body | chr1:220445206-220446043 | 837 | | | | | | | | | 1 | | | | | | yes | | yes | yes | |
| C1orf115 | N_Shore | TSS1500 | chr1:220863263-220863407 | 144 | | | 2 | | | | | 1 | 1 | | | 1 | | yes | yes | | | | |
| MOSC2 | Island | TSS200;1stExon;5'UTR | chr1:220921611-220922217 | 606 | | | | | | | | 2 | 3 | | | | | yes | | yes | | | |
| MOSC1 | N_Shore | TSS1500 | chr1:220959586-220959931 | 345 | | | | | | | | | 1 | | | | | | | yes | | | |
| HLX | N_Shore | TSS1500 | chr1:221051758-221052143 | 385 | | | 11 | | | | | | | | | | | | | | | | yes |
| HLX | N_Shore;Island | Body | chr1:221053841-221055964 | 2123 | | | 53 | | | | | | 2 | | | | | | | | | yes | |
| HLX | N_Shore;Island | Body | chr1:221057236-221058198 | 962 | | | | | 1 | 1 | | | 4 | | | | | | | | yes | yes | |
| HHIPL2 | NA | 1stExon | chr1:222721340-222721509 | 169 | | | | | | | | 1 | | | | | | | | | | | yes |
| TLR5 | Island | 5'UTR | chr1:223316219-223316272 | 53 | | | | | | | | | | | | | | | | | | | |
| SUSD4 | Island | TSS1500 | chr1:223538174-223538379 | 205 | | | | | | | | | 2 | | | | | | | | | | yes |
| C1orf65 | N_Shore;Island | TSS1500;TSS200;5'UTR;1stExon | chr1:223566127-223567173 | 1046 | 1 | | 33 | 1 | | | | 10 | 9 | 1 | | 12 | | yes | | | | | |
| CAPN8 | NA | TSS1500 | chr1:223854447-223854704 | 257 | | | | | | | | | | | | | | yes | | yes | yes | | |
| CAPN2 | NA | TSS1500;nan | chr1:223888431-223889345 | 914 | | | | 1 | | | | | 1 | 1 | 1 | | | yes | yes | | | yes | yes |
| CAPN2 | N_Shore | TSS1500;Body | chr1:223899685-223899845 | 160 | | | | | | | | 1 | | | | | | yes | | | yes | | |
| CAPN2 | Island | Body | chr1:223936343-223937043 | 700 | | | 14 | | 3 | | | | 2 | | 1 | | | yes | | | | | |
| DNAH14 | Island | 1stExon;5'UTR | chr1:225117389-225117817 | 428 | | | | | | | | | 1 | | | | | yes | | | yes | | yes |
| EPHX1 | S_Shore | 5'UTR | chr1:225998730-225999183 | 453 | | | | | | | | | | | | | | | yes | yes | yes | | |

| | | | | | | | | | | | | | | | | | | | | | | | |
|---|---|---|---|---|---|---|---|---|---|---|---|---|---|---|---|---|---|---|---|---|---|---|---|
| EPHX1 | NA | TSS1500;5'UTR | chr1:226011953-226013212 | 1259 | | | | 1 | | | | | | | | | | | yes | | | | |
| LEFTY1 | Island | Body | chr1:226075151-226075679 | 528 | | | | | | | | 1 | | | | | | | | | | | |
| LEFTY2 | S_Shore | TSS1500 | chr1:226129481-226129824 | 343 | | | | | | | | | | | | | | yes | | | | yes | |
| C1orf55 | S_Shore | TSS1500 | chr1:226187852-226188006 | 154 | | | 4 | | 1 | | | | 1 | 1 | 1 | | | | | | | | |
| MIXL1 | Island;N_Shore | TSS1500 | chr1:226410871-226411057 | 186 | | | | | | | | 1 | | | | | | | | | | | yes |
| C1orf95 | N_Shore | TSS1500 | chr1:226735384-226735660 | 276 | | | | | | | | | | | | | | yes | yes | | | yes | |
| ITPKB | NA | Body | chr1:226825435-226825606 | 171 | | | 18 | | | | | | | | | | | | | | | | |
| ITPKB | N_Shore;Island | Body;5'UTR | chr1:226924257-226925702 | 1445 | | | 2 | | 4 | | | | | | 1 | | | | | | | yes | |
| CDC42BPA | Island | TSS1500 | chr1:227506513-227506639 | 126 | | | | | | | | | 1 | | | | | | | | | | yes |
| SNAP47 | NA | Body | chr1:227962723-227963296 | 573 | | | 22 | | | | | | | | | | | | | | | | |
| WNT9A | N_Shore | Body | chr1:228112806-228113207 | 401 | | | | | | | | | 2 | | | | | yes | | | | | |
| WNT3A | Island | Body | chr1:228196198-228196376 | 178 | | | | | | | | | 1 | | | | | | | | | | |
| WNT3A | Island | Body | chr1:228225533-228225659 | 126 | | | | | | | | | 1 | | | | | | | | | | yes |
| WNT3A | Island | Body | chr1:228246632-228246937 | 305 | | | 5 | | | | | 1 | | | | | | | | | | | |
| C1orf35 | S_Shore | TSS1500 | chr1:228291486-228291705 | 219 | | | 19 | | | | | 3 | 5 | | | 7 | | yes | | yes | | | |
| MRPL55 | S_Shore | TSS1500 | chr1:228297864-228297975 | 111 | | | 4 | 1 | | | | 1 | 1 | | | 1 | | | | yes | | | |
| GUK1 | S_Shelf | Body;5'UTR;TSS1500 | chr1:228332058-228332502 | 444 | | | | | | | | | 1 | | | | | | | | | yes | |
| GJC2;GUK1 | NA | TSS200;TSS1500;3'UTR | chr1:228336474-228337776 | 1302 | | | 10 | 1 | | | | | 2 | 1 | | | | | | | | | |
| GJC2 | Island | Body | chr1:228346014-228346347 | 333 | | | 12 | 1 | 1 | | | | | | | | | | | | | yes | yes |
| C1orf69 | N_Shore | Body | chr1:228362309-228362509 | 200 | | | | | | | | | 1 | | | 1 | | | | | | yes | |
| OBSCN | NA;N_Shelf | TSS1500;TSS200 | chr1:228394509-228396009 | 1500 | | | 18 | | | | | | | | | | | | | yes | | | |
| OBSCN | Island | Body | chr1:228400217-228400628 | 411 | | | | | | | | 1 | | | | | | | | | | | yes |
| OBSCN | Island | Body | chr1:228463634-228464232 | 598 | | | | 1 | | | | 1 | | | | | | | | | | | yes |
| OBSCN | NA | Body | chr1:228494821-228495111 | 290 | | | 2 | | | | | | | | | | | | | | | | |
| OBSCN | Island | Body | chr1:228503882-228504579 | 697 | | | | | | | | | | | | | | | | | | | |
| OBSCN | S_Shore | 3'UTR;Body | chr1:228548644-228548842 | 198 | | | | | | | | | | | | | | | | | | | |
| OBSCN | Island | Body | chr1:228558987-228559321 | 334 | | | | | | | | | 1 | | | | | | | | | | |
| OBSCN | Island | Body;3'UTR | chr1:228566080-228566824 | 744 | | | | | 1 | | | | | 2 | | | | | | | | | yes |
| TRIM17 | Island | 1stExon;5'UTR | chr1:228604566-228604800 | 234 | | | | | | | | | 1 | | | | | | | yes | | | yes |
| HIST3H3 | Island | TSS200 | chr1:228613100-228613207 | 107 | | | | | | | | | | | | | | yes | | | | | |
| HIST3H2BB;HIST3H2A | N_Shore;Island | TSS1500;1stExon | chr1:228644750-228645627 | 877 | | 1 | 9 | | | | | | | | | | | | | | | | yes |
| DUSP5P | Island | Body | chr1:228785987-228786295 | 308 | | | | | | | | | | | | 1 | | | | yes | | | |
| RHOU | N_Shore | TSS1500 | chr1:228870454-228870637 | 183 | | | | | | | | | | | | | | yes | | | yes | | yes |
| RAB4A | N_Shore | TSS1500 | chr1:229406360-229406661 | 301 | | | 2 | | | | | | 1 | | | | | yes | | | | | yes |
| ACTA1 | Island | Body | chr1:229568077-229568360 | 283 | | 1 | | | | | | | | | | | | | | | | yes | yes |
| ACTA1 | Island | TSS200;5'UTR;1stExon | chr1:229569795-229569902 | 107 | 1 | | | | | | | | | | | 1 | | | | | | | |
| GALNT2 | NA | Body | chr1:230272306-230272656 | 350 | | | | | | | | | | | | | | yes | | | | | |
| GALNT2 | N_Shore;Island | Body;3'UTR | chr1:230415185-230415902 | 717 | | | | | 1 | | | 2 | 7 | | | 2 | | | | | | | |
| PGBD5 | NA | 3'UTR | chr1:230457979-230458393 | 414 | 1 | | | | | | | | | | | | | | | | | | yes |
| PGBD5 | N_Shore;Island | Body | chr1:230468464-230469330 | 866 | | | | | | | | | 2 | | | 2 | | yes | | | | | |
| PGBD5 | NA | 1stExon;TSS200 | chr1:230513277-230514361 | 1084 | | | | | | | | | | | | | | yes | | | | | |
| COG2 | N_Shore | TSS1500 | chr1:230777685-230777875 | 190 | | | | 1 | 1 | | | | | | | | | | | | | | |
| CAPN9 | NA | TSS1500;TSS200 | chr1:230882387-230883552 | 1165 | | | | | | | | | 1 | | | | | yes | yes | | yes | yes | |
| C1orf198 | NA | Body;TSS1500 | chr1:230992820-230992990 | 170 | | | 1 | | | | | | | | | | | yes | yes | | | | |
| FAM89A;MIR1182 | NA | Body;TSS200;TSS1500 | chr1:231155632-231155983 | 351 | | | 4 | | | | | | 4 | | | 1 | | | | | | | |
| FAM89A | Island | TSS1500 | chr1:231176985-231177201 | 216 | | | | | | | | | | 1 | | | | | | | | | yes |
| TRIM67 | Island | TSS1500 | chr1:231298177-231298651 | 474 | | | | | | | | | 1 | | | | | | | | | | |
| EXOC8 | N_Shore | 1stExon | chr1:231472152-231472445 | 293 | | | 1 | | 1 | | | | | | | | | | | | | | |
| TSNAX-DISC1;DISC1 | N_Shore | Body;TSS1500 | chr1:231761688-231761845 | 157 | | | 1 | | | | | | | | | | | yes | yes | | | | |
| SIPA1L2 | NA | 1stExon;5'UTR | chr1:232651092-232651427 | 335 | | | | | | | | | 1 | | | | | | yes | | | | |
| KIAA1383 | Island | 1stExon | chr1:232941468-232941775 | 307 | | | | | | | | | 1 | | | | | | | | | yes | |
| KIAA1804 | N_Shore | TSS1500 | chr1:233462575-233462803 | 228 | | | 5 | | | | | | | | | | | yes | | | yes | yes | |
| KIAA1804 | N_Shore | Body | chr1:233497542-233497916 | 374 | | | 2 | | | | | | | | | | | | | | | | |
| KCNK1 | Island | TSS200 | chr1:233749633-233749666 | 33 | | | | | | | | | 1 | | | | | | | | | yes | |
| KCNK1 | NA | Body | chr1:233765199-233765350 | 151 | | | | | | | | | 1 | | | | | | | | | | |
| SLC35F3 | N_Shore;Island | TSS1500;1stExon;5'UTR | chr1:234039761-234041306 | 1545 | | | 2 | | | | | 1 | 1 | | | 1 | | | yes | | | yes | yes |
| SLC35F3 | S_Shore | Body | chr1:234350834-234351342 | 508 | | | 5 | | | | | 1 | | | | | | yes | | | | | |
| SLC35F3 | Island | Body | chr1:234367145-234367586 | 441 | | | | 1 | 1 | 1 | 1 | 5 | 5 | 3 | | 7 | | | | | yes | | |
| GNG4 | Island | 5'UTR | chr1:235805403-235806070 | 667 | | | 4 | | 1 | | | 4 | 1 | | | | | yes | yes | | | | |
| GNG4 | N_Shore;Island | TSS1500;TSS200 | chr1:235814145-235814814 | 669 | | | 9 | | | | | 2 | 1 | | | | | yes | | | yes | | yes |
| NID1 | Island | TSS200;TSS1500 | chr1:236228625-236228755 | 130 | | | | | | | | 1 | 2 | 1 | | 1 | | | | | | | yes |
| GPR137B | Island | Body | chr1:236306640-236306767 | 127 | | | | | 1 | | | | | | | | | | | | | | |
| EDARADD | N_Shore | TSS200;TSS1500 | chr1:236557473-236557758 | 285 | | | 4 | | 1 | | | | | | | | | yes | | | | | |
| EDARADD | S_Shore | Body | chr1:236559438-236559675 | 237 | | | 3 | | | | | 1 | | | | | | | | | | | |
| LGALS8 | N_Shore | TSS200;5'UTR;TSS1500 | chr1:236686012-236687053 | 1041 | 1 | | 21 | | 2 | | | 2 | 6 | 1 | | 3 | | yes | | | | | |
| ACTN2 | Island;chr1:236849472-236850 | TSS1500;TSS200;Island;1stExon;5' | chr1:236849473-236850361 | 888 | | 1 | | | | | | | 3 | 1 | 1 | 1 | | | | yes | | yes | yes |
| ACTN2 | NA | Body | chr1:236902424-236902765 | 341 | | | 1 | | | | | | | | | | | | yes | | | | |
| ACTN2 | NA | Body | chr1:236925843-236925965 | 122 | | | 16 | 1 | | | | 1 | 2 | | | 3 | | | | | | | |
| RYR2 | NA | Body | chr1:237628962-237629007 | 45 | | | 3 | | | | | | | | | | | yes | yes | | yes | | yes |
| RYR2 | NA | Body | chr1:237729917-237730049 | 132 | | | | | | | | | | | | | | | yes | | | | |
| RYR2 | Island | Body | chr1:237947246-237947435 | 189 | | | | | | | | | | | | | | | | | | | |
| LOC100130331 | NA | TSS1500 | chr1:238023986-238024671 | 685 | | | 1 | | | | | | 2 | | | | | yes | yes | | | | |
| ZP4;LOC100130331 | NA | 1stExon;Body | chr1:238053828-238053991 | 163 | | | 2 | | | | | | | | | | | yes | | | | | |
| FMN2 | Island | 1stExon | chr1:240255627-240256019 | 392 | | | | | | | | | 1 | | | | | | | | | | |
| GREM2 | N_Shore;Island | 3'UTR;Body | chr1:240656217-240656737 | 520 | | | | | | | | 6 | 8 | | | 8 | | | | | | | |
| GREM2 | NA | TSS1500 | chr1:240776162-240776316 | 154 | | | 34 | 1 | | | | | 1 | | | | | | | | | | |
| RGS7 | Island | 5'UTR;1stExon | chr1:241520286-241520331 | 45 | | | | | | | | | | | | 1 | | | | | | yes | yes |
| OPN3 | S_Shore | TSS1500 | chr1:241804036-241804939 | 903 | | | | | | | | | 1 | | | | | yes | yes | | | yes | yes |
| WDR64 | NA | Body | chr1:241912510-241912768 | 258 | | | 21 | | | | | | 1 | 1 | | | | yes | | | | | |
| MAP1LC3C | NA | Body | chr1:242161780-242161881 | 101 | | | 2 | | | | | | | | | | | yes | | | | | |
| SDCCAG8 | Island | Body | chr1:243637838-243638141 | 303 | | | | | | | | | 1 | | | | | | | | | | |
| SDCCAG8 | N_Shore;Island | Body | chr1:243645911-243647204 | 1293 | | | 22 | 1 | 2 | | | | | | 1 | | | yes | yes | | | | |
| SDCCAG8; AKT3 | Island | Body | chr1:243651346-243651920 | 574 | | | 3 | | | | | | | | | | | | | | | | |
| ZNF238 | N_Shore;Island | TSS1500 | chr1:244213358-244214679 | 1321 | | | 5 | | | | | 2 | 5 | | | 1 | | | | | | | |
| C1orf101 | N_Shore | TSS200 | chr1:244624752-244624883 | 131 | | | | | | | | | | | 1 | | | yes | | | | | yes |
| KIF26B | Island | TSS1500 | chr1:245317950-245317980 | 30 | | | | | | | | | 1 | | | | | | | | | | yes |
| KIF26B | NA | Body | chr1:245710332-245710401 | 69 | | | | | | | | 1 | 1 | 1 | | 8 | | yes | | | | | |
| KIF26B | N_Shore;Island | Body | chr1:245753302-245753635 | 333 | | | 10 | | | | | | | | | | | yes | | | | | |
| CNST;TFB2M | N_Shore | TSS1500;Body | chr1:246728663-246728727 | 64 | | | | | | | | 1 | | | | 2 | | yes | | | | | |
| LOC149134 | chr1:246952214-246952511;Isl | Body;TSS1500 | chr1:246952118-246952508 | 390 | | 1 | | | | | | 1 | 1 | | 1 | 1 | | yes | | | | | |
| ZNF695 | S_Shore | TSS1500 | chr1:247171572-247171838 | 266 | | | | | | | | | 1 | | | | | yes | | | yes | yes | |
| ZNF496 | Island;S_Shore | TSS1500 | chr1:247495818-247496575 | 757 | | | | | | | | | 2 | 1 | | 1 | | yes | | | | | yes |
| NLRP3 | NA | TSS1500;5'UTR | chr1:247578859-247580565 | 1706 | | | 6 | | 1 | | | | 2 | | | | | yes | | | | | |
| NLRP3 | N_Shelf | Body;3'UTR | chr1:247611448-247611931 | 483 | | | 1 | | 1 | | | 1 | 4 | 1 | | 4 | | | yes | | | yes | yes |
| OR2B11 | Island;S_Shore | 1stExon | chr1:247614727-247615683 | 956 | | | | | | | | | | | | | | yes | | | | | |
| LOC148824;OR2C3 | Island | Body;3'UTR;TSS200;TSS1500 | chr1:247694041-247694557 | 516 | | | 24 | | | | 1 | 2 | 6 | 3 | | 8 | | | | yes | yes | | |
| C1orf150 | NA | TSS1500;TSS200 | chr1:247711595-247712591 | 996 | | | | | 1 | | | | 3 | 1 | 1 | | | yes | | | yes | yes | |
| OR14A16 | NA | 1stExon | chr1:247978690-247979157 | 467 | | | | | | | | | | | | | | | | | | | yes |
| OR11L1 | NA | 1stExon | chr1:248004886-248005715 | 829 | | | 3 | | | | | | | | | | | yes | yes | | | yes | |

| | | | | | | | | | | | | | | | | | | | | | | | | |
|---|---|---|---|---|---|---|---|---|---|---|---|---|---|---|---|---|---|---|---|---|---|---|---|---|
| TRIM58 | Island | TSS200;1stExon | chr1:248020350-248021091 | 741 | | | | | 4 | | | | 3 | | 1 | | | yes | yes | | | | yes | |
| OR2L13 | N_Shore;Island | TSS1500;TSS200;1stExon;5'UTR | chr1:248099880-248101009 | 1129 | 2 | 1 | 34 | 1 | 1 | | 2 | 12 | 19 | 14 | 5 | 59 | 4 | | yes | | | | | |
| OR2M5 | NA | TSS1500 | chr1:248307068-248307316 | 248 | | | | | | | | | | | | | | | yes | yes | | | yes | |
| LOC646627 | NA | 1stExon;TSS200;3'UTR | chr1:248902767-248903467 | 700 | | | | | | | | 5 | 2 | 1 | 1 | | | yes | | | | | | |
| SH3BP5L | Island | Body | chr1:249106207-249106516 | 309 | | | | 1 | | | | | | | | | | | | | | | | |
| TUBB8 | S_Shore;N_Shore;Island | TSS200;1stExon;5'UTR;Body | chr10:94136-95661 | 1525 | | | 7 | | | | | 0 | | | | | | | | yes | | yes | | |
| DIP2C | NA | Body | chr10:334537-334773 | 236 | | | | | | | | | 1 | | | | | yes | | yes | | | | |
| DIP2C | N_Shelf | Body | chr10:384286-384652 | 366 | | | 16 | 1 | | | | | 1 | 1 | | | | | | | yes | yes | | |
| DIP2C | N_Shore | Body | chr10:407721-407935 | 214 | | | | | | | | | 1 | | | | | | | | | | | |
| DIP2C | NA | Body | chr10:438227-438517 | 290 | | | 10 | | | | | | | | | | | | | | | yes | | |
| DIP2C | Island | Body | chr10:459697-459968 | 271 | | | | | | | | | 1 | | | 1 | | yes | | | | | | |
| DIP2C | N_Shelf | Body | chr10:463561-463801 | 240 | 1 | | | | | | | 0 | | | | | | | | | | | | |
| DIP2C | S_Shore | Body | chr10:466737-466878 | 141 | | | | | | | | | 1 | | | | | | | | | | | |
| DIP2C | S_Shore | Body | chr10:468347-468412 | 65 | | | | | | | | | 1 | | | | | | | | | | | |
| DIP2C | N_Shore;Island;S_Shore | Body | chr10:530635-533102 | 2467 | | | 43 | | | | | 0 | 24 | 8 | | 13 | | yes | | | | | | |
| DIP2C | NA | Body | chr10:556470-557936 | 1466 | | 1 | 5 | | | | | | 3 | 1 | | 1 | | yes | | | | | | |
| DIP2C | N_Shore | Body | chr10:574291-574361 | 70 | | | | | | | | | 1 | | | | | | | yes | | | | |
| DIP2C | NA | Body | chr10:601759-601816 | 57 | | | | | | | | | 1 | 1 | | | | | | | | | | |
| DIP2C | NA | Body | chr10:612266-612391 | 125 | | | | | 1 | | | | | | | | | | | | | | | |
| DIP2C | NA | Body | chr10:616959-617105 | 146 | | | 10 | | | | | | | | | | | | | | | | yes | |
| DIP2C | NA | Body | chr10:636140-636331 | 191 | | | | | | | | | 1 | | | | | | | | | | | |
| DIP2C | NA | Body | chr10:639935-640085 | 150 | 3 | | 9 | 1 | 1 | | | | | | | | | | yes | | | | | |
| DIP2C | NA | Body | chr10:641615-641641 | 26 | | | | | | | | | 1 | | | | | | yes | | yes | | | |
| DIP2C | NA | Body | chr10:661009-661730 | 721 | | | 38 | | | | | | | | | | | | | | | yes | | |
| DIP2C | NA | Body | chr10:664719-664994 | 275 | | | 5 | | | | | | | | | | | | | yes | | | | |
| DIP2C | S_Shore | Body | chr10:670562-671628 | 1066 | | | 16 | | | | | 0 | 3 | | | | | | | | | | | |
| DIP2C | Island | Body | chr10:679322-679514 | 192 | | 1 | | | | | | | | | | | | | | | | | | |
| DIP2C | N_Shelf | Body | chr10:682125-682871 | 746 | | | 10 | | | | | 0 | 1 | | | | | | yes | yes | | | | |
| C10orf108;DIP2C | NA | TSS1500;Body;TSS200 | chr10:695403-696356 | 953 | 1 | | | 1 | 1 | | | 0 | 8 | 1 | | 3 | | | | | | | | |
| DIP2C;C10orf108 | N_Shelf | Body | chr10:709839-710082 | 243 | 1 | 1 | | | | | | | | | | | | | yes | | | | | |
| DIP2C | N_Shore;Island | Body | chr10:711556-712442 | 886 | | | 5 | 1 | | | | | 1 | | | 2 | | | | | | | | |
| DIP2C | NA | Body | chr10:729001-729956 | 955 | 3 | 1 | 4 | | | | | 0 | 6 | 1 | 1 | | | | | | | | yes | |
| LARP4B | S_Shelf | 3'UTR | chr10:858817-858923 | 106 | | | 1 | | | | | | | | | | | | | | | | yes | |
| C10orf110;IDI2 | NA | TSS1500;Body | chr10:1067372-1067538 | 166 | | | | | | | | | 1 | | | | | yes | | | | | | |
| C10orf110;IDI2 | NA | Body | chr10:1068667-1068763 | 96 | 1 | | | | | | | | | | | | | yes | yes | | | | | |
| C10orf110 | NA | Body | chr10:1083192-1083304 | 112 | | 1 | | | | | | | | | | | | | yes | yes | | | | |
| WDR37 | NA | Body | chr10:1156113-1156594 | 481 | | | | | 2 | | | | 1 | | | | | yes | | | | | | |
| WDR37 | NA | Body | chr10:1166782-1167544 | 762 | 1 | | 1 | | | | | | | | | | | | | | | | | |
| NCRNA00200 | Island;N_Shore | TSS200;TSS1500 | chr10:1205222-1206555 | 1333 | | | 13 | | | | | 0 | 14 | 2 | | 4 | | | | yes | yes | | | |
| NCRNA00200 | Island | Body | chr10:1209379-1209463 | 84 | | | 3 | | | | | 0 | | | | | | | | | | | | |
| ADARB2 | N_Shelf;N_Shore | 3'UTR | chr10:1228316-1229080 | 764 | | | 4 | | | | | | | | | | | | | yes | | | | |
| ADARB2 | S_Shelf | Body | chr10:1234237-1234378 | 141 | | | | 1 | | | | | | | | | | | | | | | yes | |
| ADARB2 | NA | Body | chr10:1245261-1246035 | 774 | | | 3 | | | | | | 1 | | | 1 | | | | | | | yes | |
| ADARB2 | N_Shelf | Body | chr10:1248770-1248818 | 48 | | | | | | | | | 1 | | | | | yes | | | | | | |
| ADARB2 | N_Shore | Body | chr10:1251771-1252146 | 375 | | | | | | | | | | | | | | | | yes | | | | |
| ADARB2 | S_Shelf | Body | chr10:1256339-1256487 | 148 | | | 7 | | | | | | | | | | | | | | | | | |
| ADARB2 | S_Shore | Body | chr10:1285332-1285535 | 203 | | | | | | | | | | | | | | | yes | | | | | |
| ADARB2 | NA | Body | chr10:1410519-1411072 | 553 | | | 2 | | | | | | | | | | | yes | | | | | | |
| ADARB2 | Island;S_Shore | Body | chr10:1416008-1416920 | 912 | | | 17 | | | | 1 | | 2 | | | 1 | | | | | | yes | | |
| ADARB2 | NA | Body | chr10:1444557-1445134 | 577 | | | 5 | | | | | | | | | | | | | | yes | | | |
| ADARB2 | NA | Body | chr10:1451381-1451460 | 79 | | | | | | | | | | | | | | | | | | | | |
| ADARB2 | NA | Body | chr10:1453917-1454848 | 931 | | | | 1 | 2 | | | | | | | 1 | | yes | | | | | yes | |
| ADARB2 | NA | Body | chr10:1462906-1463108 | 202 | | | | | | | | | 1 | | | | | | | | | | | |
| ADARB2 | N_Shore | Body | chr10:1505595-1506175 | 580 | | | 1 | | | | | | 2 | | | 1 | | | yes | | | | | |
| ADARB2 | S_Shore | Body | chr10:1507708-1508061 | 353 | | | | | | | | | 1 | 1 | 1 | | | | yes | yes | | | | |
| ADARB2 | S_Shelf;NA | Body | chr10:1509224-1511277 | 2053 | | 7 | 11 | | | | | | | | | | | yes | | | | yes | yes | |
| ADARB2 | Island | Body | chr10:1517083-1517340 | 257 | | | 2 | | | | | | 3 | | | 1 | | | | | | | | |
| ADARB2 | NA | Body | chr10:1531034-1531529 | 495 | | | 3 | | | | | | 2 | | | | | | | | | | | |
| ADARB2 | NA | Body | chr10:1547459-1547983 | 524 | | | 2 | | | | | | | | | | | | | | yes | yes | | |
| ADARB2 | NA | Body | chr10:1551251-1551326 | 75 | | | 2 | | | | | | 1 | | | | | yes | | | | | | |
| ADARB2 | NA | Body | chr10:1558762-1559249 | 487 | | 1 | | | | | | 0 | | | | | | | | | | yes | | |
| ADARB2 | NA | Body | chr10:1560471-1560628 | 157 | | | | | | | | | | | | | | | yes | | | | | |
| ADARB2 | NA | Body | chr10:1567746-1567897 | 151 | | | | | | | | | | | | | | | | | | yes | | |
| ADARB2 | Island | Body | chr10:1585223-1585345 | 122 | | | | | | | | | 1 | | | | | | | | | | | |
| ADARB2 | Island;S_Shore | Body | chr10:1595704-1596108 | 404 | | | 13 | | | | | | | | 1 | | | | | yes | | | | |
| ADARB2 | NA | Body | chr10:1609383-1609753 | 370 | | | | | | | | | | | | | | | yes | | | | | |
| ADARB2 | NA | Body | chr10:1651238-1651391 | 153 | | | | | | | | 0 | | | | 1 | | | yes | | | yes | | |
| ADARB2 | NA | Body | chr10:1668625-1669312 | 687 | | | 2 | | | | | | | | | | | | | | yes | | | |
| ADARB2 | NA | Body | chr10:1707523-1707576 | 53 | | | | | | | | | | | | | | | | | | | | |
| ADARB2 | NA | Body | chr10:1712918-1713900 | 982 | | 1 | 14 | | | | | | 1 | | | | | | | | yes | | | |
| ADARB2 | NA | Body | chr10:1725096-1725190 | 94 | | | | | | | | | | | | | | yes | | | | | | |
| ADARB2 | N_Shelf | Body | chr10:1739346-1739665 | 319 | | | | | | | | | | | | | | | | | yes | | | |
| ADARB2 | NA | Body | chr10:1767482-1767830 | 348 | | | 19 | | | | | | | | | | | yes | | | | | | |
| ADARB2 | NA | Body | chr10:1769480-1769577 | 97 | | | | | | | | | | | | | | | yes | | | | | |
| PFKP | NA | Body | chr10:3138418-3138534 | 116 | | | 16 | | | | | 0 | 1 | 1 | | | | yes | | | yes | | | |
| PFKP | N_Shelf | Body | chr10:3143354-3143783 | 429 | | 1 | | | | | | | | | | | | yes | | | | yes | | |
| PFKP | N_Shore | Body | chr10:3146486-3147056 | 570 | | | 2 | | | | | | | | | | | | | | | | | |
| PFKP | N_Shore | Body | chr10:3148096-3148613 | 517 | | | | | | | | | 1 | | | | | | | | | | yes | |
| PFKP | S_Shore | Body | chr10:3149673-3149852 | 179 | 1 | | 23 | 1 | 1 | 1 | | | 2 | 1 | | | | yes | | | | | yes | |
| PFKP | Island | Body | chr10:3160926-3160982 | 56 | | | | | 1 | | | | | | | | | yes | | | | | | |
| PFKP | S_Shelf | Body | chr10:3164740-3165100 | 360 | | | | | | | | | | | | | | yes | | | yes | | | |
| PFKP | NA | Body | chr10:3168776-3169516 | 740 | 1 | | | | 1 | | | | 1 | 1 | | | | | | | | | | |
| KLF6 | Island;S_Shore | Body | chr10:3823790-3824687 | 897 | | | 5 | 1 | | | | 0 | 5 | | | 2 | | | | | | | | |
| LOC100216001 | NA | TSS1500 | chr10:4720653-4720912 | 259 | | | | | | | | | | | | | | yes | | yes | | | | |
| AKR1E2 | Island | TSS200 | chr10:4868328-4868892 | 564 | | | 10 | 1 | | | | 0 | 3 | 2 | | 1 | | yes | yes | | | | yes | |
| AKR1C2 | NA | TSS200 | chr10:5060291-5060566 | 275 | | | 4 | | | | | | 1 | | | | | yes | | yes | | | | |
| UCN3 | NA | TSS1500;TSS200 | chr10:5406423-5407119 | 696 | | 2 | 26 | | 1 | | | 0 | 2 | | | 3 | | yes | yes | | | yes | | |
| NET1 | N_Shore;Island | Body;TSS200 | chr10:5488346-5488628 | 282 | | 1 | | | | | | | 3 | | | | | | | | | | | |
| CALML5 | S_Shore | TSS200 | chr10:5541568-5541726 | 158 | | | | | | | | | | | | | | | | | | | | |
| CALML3 | N_Shore;Island | TSS1500;1stExon | chr10:5566543-5567827 | 1284 | | 1 | | | 1 | | | | 1 | | | | | | yes | | | | | |
| ASB13 | NA | Body | chr10:5688470-5688620 | 150 | | 1 | | | | | | 0 | 1 | 1 | | 1 | | yes | | yes | | | | |
| C10orf18 | NA | 5'UTR | chr10:5734571-5735135 | 564 | | | | | | | | | | 1 | | | | yes | | | | | yes | |
| IL15RA | S_Shore | TSS1500 | chr10:6020369-6020585 | 216 | | | | | | | | | | | | | | yes | yes | | | | yes | |
| IL2RA | NA | 1stExon;5'UTR;TSS200 | chr10:6104137-6104630 | 493 | | | 4 | | 2 | | | | | | | | | | | | | yes | | |
| RBM17 | N_Shore | TSS1500 | chr10:6130479-6130580 | 101 | | | | | | | | 0 | 1 | | | | | | yes | | | yes | | yes |
| PFKFB3 | Island | Body | chr10:6205688-6206210 | 522 | | | | | | | | 0 | | | | 1 | | | yes | yes | | | yes | |
| PFKFB3 | NA | Body | chr10:6214016-6214079 | 63 | | | 3 | | | | | | 1 | 1 | | | | yes | | | yes | | | |
| PFKFB3 | S_Shore | Body | chr10:6264625-6264776 | 151 | | | | | | | | | | | | | | | yes | | | | | |
| PRKCQ | NA | Body | chr10:6472750-6472833 | 83 | | | | | | | | | | | | | | yes | | | | yes | | |

| | | | | | | | | | | | | | | | | | | | | | | | |
|---|---|---|---|---|---|---|---|---|---|---|---|---|---|---|---|---|---|---|---|---|---|---|---|
| PRKCQ | Island | 5'UTR | chr10:6621965-6622297 | 332 | | 1 | | | | | | | | | | | | | | | | | yes |
| SFMBT2 | NA | Body | chr10:7221679-7221713 | 34 | | | | | | | | | | | | | | yes | | yes | | yes | |
| SFMBT2 | NA | Body | chr10:7290492-7290545 | 53 | | | | | | | | | | | | | | | | | | | |
| SFMBT2 | NA | Body | chr10:7376656-7376726 | 70 | | | | | | | | | | | | | | | yes | | | | |
| SFMBT2 | Island | Body | chr10:7409618-7410045 | 427 | | | | | | | | | | | | | | | | | yes | | yes |
| ITIH5 | NA | Body;TSS200 | chr10:7661651-7661796 | 145 | | | | | | | | | | | | | | yes | yes | yes | | | |
| ITIH5 | NA | Body | chr10:7670683-7671259 | 576 | | | | | | | | | 3 | | | | | | yes | | | | |
| ITIH5 | NA | Body | chr10:7708240-7708290 | 50 | | | | | | | | | 1 | | | | | yes | | | | | yes |
| ITIH5 | NA | TSS1500 | chr10:7709848-7710179 | 331 | | | | | | | | | | | | | | | yes | | | | yes |
| TAF3 | N_Shore | TSS1500 | chr10:7859623-7859768 | 145 | | | | | | | | | 1 | | | | | | yes | | yes | yes | |
| FLJ45983;GATA3 | Island | Body;TSS200;TSS1500;5'UTR;1stE› | chr10:8094093-8098005 | 3912 | | 2 | 7 | | | | | 0 | 9 | | 1 | 1 | | | | | | | yes |
| GATA3 | S_Shore;N_Shore | Body | chr10:8100114-8101566 | 1452 | | | 7 | | | | | | | | | | | | | | | | |
| GATA3 | NA | Body | chr10:8106003-8106035 | 32 | | | 2 | | | | | | | | | | | | | | | yes | |
| CUGBP2 | NA | 1stExon;5'UTR | chr10:11047271-11047354 | 83 | | | | | 1 | | | | | | | | | | | | | yes | yes |
| CUGBP2 | N_Shore;Island | Body;TSS1500;5'UTR | chr10:11206221-11208339 | 2118 | | | | | | | | | | | | | | | | | | | |
| CUGBP2 | NA | Body | chr10:11246612-11246763 | 151 | | | | | | | | | 1 | | | | | | | | | | |
| USP6NL | NA | Body;TSS200 | chr10:11574300-11574616 | 316 | | | | | | | | | | | | | | | | | | | |
| ECHDC3 | N_Shore | TSS1500 | chr10:11783823-11785161 | 1338 | | | 7 | | 1 | | | 0 | | | | | | yes | | yes | yes | yes | yes |
| C10orf47 | Island | Body | chr10:11912126-11912597 | 471 | | | 10 | | | | | | 1 | | | | | | | | | | yes |
| UPF2 | S_Shore | TSS1500 | chr10:12085555-12086036 | 481 | | | 3 | | | | | | 2 | | | 1 | | yes | | | | | yes |
| CAMK1D | NA | Body | chr10:12648032-12648526 | 494 | | | 19 | | | | | | | | | | | yes | | | | | |
| LOC283070 | NA | TSS1500 | chr10:12874207-12874395 | 188 | | | | | 1 | | | | | | | | | yes | | | | | |
| CCDC3 | NA | Body | chr10:12999550-12999618 | 68 | | | 1 | | | | | | 1 | | | | | yes | | yes | | | |
| UCMA | NA | 5'UTR;1stExon | chr10:13276279-13276892 | 613 | | | | | | | | | | | | | | yes | | | | | yes |
| SEPHS1 | NA | 5'UTR | chr10:13388442-13388567 | 125 | | 1 | | | | | | | 1 | | | | | | | | | | |
| BEND7 | NA | Body | chr10:13481846-13481944 | 98 | | | | | | | | 0 | 1 | | | 1 | | | | | | yes | |
| BEND7 | N_Shore | Body | chr10:13514864-13515334 | 470 | | | | | | | | | 1 | | | | | | | | | | |
| PRPF18 | S_Shore | Body | chr10:13629624-13629984 | 360 | | | 1 | | | | | | | | | | | yes | | yes | | | |
| FRMD4A | NA | 3'UTR | chr10:13688030-13688106 | 76 | 2 | | | | | | | | | | | | | | | | | | yes |
| FRMD4A | NA | Body | chr10:13749193-13749625 | 432 | | | 25 | | | | | | | | | | | | | | | yes | |
| FRMD4A | N_Shore | Body | chr10:13932842-13933452 | 610 | | | 30 | | | 1 | | | | | | | | | | | | yes | yes |
| FRMD4A | NA | Body | chr10:14014553-14014651 | 98 | | | | | | | | | 1 | | | | | yes | | | | | |
| FRMD4A | NA | Body | chr10:14050305-14050521 | 216 | | | | | 2 | | | 0 | 1 | | | | | | | | | yes | |
| FRMD4A | NA | Body;nan | chr10:14051636-14052028 | 392 | 3 | | 1 | 1 | 1 | | | 0 | 7 | | 1 | 5 | 1 | | | | yes | | |
| FRMD4A | NA | 5'UTR;TSS200 | chr10:14372431-14372913 | 482 | | | 5 | | | | | | 2 | | | | | | | | | yes | |
| MIR1265 | NA | TSS200 | chr10:14478428-14478750 | 322 | | | | | | | | | | | | | | | | | yes | | |
| FAM107B | NA | Body | chr10:14575319-14575436 | 117 | | | | | | | | | 1 | | | | | | yes | | | | |
| DCLRE1C | N_Shore | Body;5'UTR | chr10:14995476-14995625 | 149 | | | | 1 | | | | | | | | | | | | yes | | | |
| FAM171A1 | N_Shore | Body | chr10:15411709-15412027 | 318 | | | | | | | | | | | | | | | | | | yes | |
| ITGA8 | Island | TSS200;TSS1500 | chr10:15761854-15762312 | 458 | | 1 | 1 | | 1 | | | | 1 | | | | | yes | | | | yes | |
| FAM188A | S_Shore | TSS1500 | chr10:15902731-15902887 | 156 | | | | | | | | | | | | | | yes | | | | | |
| PTER | N_Shore | TSS1500 | chr10:16478471-16478613 | 142 | | | 1 | | | | | | | | | | | | | yes | | | |
| C1QL3 | N_Shore | Body | chr10:16561202-16561605 | 403 | 3 | | | | | | | | 3 | 1 | | | | | | | | | yes |
| VIM | N_Shore | TSS1500 | chr10:17269274-17269775 | 501 | | | | | | | 1 | | | | | | | | | | | | |
| CACNB2 | Island | TSS200;1stExon;5'UTR | chr10:18429643-18429760 | 117 | | 1 | | | | | | 0 | 3 | | | | | | | | | | |
| CACNB2 | NA | Body;TSS200;1stExon;5'UTR | chr10:18549413-18550223 | 810 | | | 15 | | | | | 0 | 3 | | | 1 | | | | | | | |
| CACNB2 | NA | Body;TSS1500;TSS200 | chr10:18628543-18629677 | 1134 | 2 | 1 | | | | | | | 2 | | | | | yes | yes | | yes | | |
| CACNB2 | NA | Body;TSS200;TSS1500 | chr10:18689035-18689503 | 468 | | | 5 | | | | | 0 | 2 | | | 3 | | yes | | | | | |
| PLXDC2 | N_Shore;Island | TSS1500;1stExon;5'UTR | chr10:20104333-20106089 | 1756 | | | | | | | | | 2 | | | | | yes | | | | | yes |
| NEBL;C10orf113 | NA | Body;TSS200 | chr10:21435534-21435895 | 361 | | | | | | | | | 1 | | | | | | yes | | | | |
| C10orf140 | S_Shore | 5'UTR | chr10:21807166-21807252 | 86 | | | 1 | | | | | | | | | | | | | | | | |
| COMMD3 | Island;S_Shore | TSS200;Body | chr10:22605173-22606072 | 899 | | | 18 | | | | | 0 | 1 | | | | | | | | | | yes |
| SPAG6 | Island | TSS200;Body;TSS1500 | chr10:22634038-22634733 | 695 | | | 4 | | | | | 0 | 6 | | | | | | | | | | |
| PIP4K2A | NA | Body | chr10:22843607-22843938 | 331 | 7 | | | | | | | | | | | | | yes | | | | | |
| ARMC3 | NA | TSS1500;TSS200 | chr10:23216637-23216847 | 210 | | | 3 | | | | | | 1 | | | | | yes | | | | | |
| PTF1A | N_Shore;Island | TSS1500 | chr10:23479993-23481421 | 1428 | | | 1 | | | | | | 2 | | | | | | | | | yes | yes |
| KIAA1217 | Island;N_Shore | TSS1500 | chr10:23982480-23983498 | 1018 | | 2 | | | | | | | | | | | | | | | | | |
| PRTFDC1 | Island | TSS1500 | chr10:25241883-25242031 | 148 | | | | | | | | | 1 | | | | | | | | | yes | yes |
| ENKUR;THNSL1 | Island | TSS1500;TSS200 | chr10:25305418-25305874 | 456 | | | | | | | | | 1 | | | | | | | | | | |
| LOC100128811;GPR158 | Island | Body;TSS1500;1stExon;5'UTR;TSS2 | chr10:25463757-25466027 | 2270 | | 2 | | | | | | | 4 | | | | | yes | | | | | yes |
| GPR158 | NA | Body | chr10:25507099-25507202 | 103 | | | | | | | | | 1 | | | | | | yes | | | | |
| MYO3A | Island | TSS200;1stExon;5'UTR | chr10:26222922-26223310 | 388 | | | | | | | | 0 | 2 | | | | | | | | | | |
| MYO3A | Island | Body | chr10:26500780-26501065 | 285 | | 1 | | | | | | | | | | | | | | | yes | | |
| GAD2 | Island | 5'UTR;1stExon | chr10:26505440-26505503 | 63 | | | | | | | | | 1 | | | | | | | | | | |
| GAD2 | Island | Body | chr10:26506933-26507383 | 450 | | | | | | | | | 1 | | | | | | | | | yes | yes |
| APBB1IP | N_Shore;Island | TSS1500;TSS200 | chr10:26726877-26727925 | 1048 | | 1 | 6 | | 1 | | | | | | | | | | yes | | | yes | |
| APBB1IP | Island | Body | chr10:26855907-26856454 | 547 | | | 6 | 1 | 2 | 1 | | | 3 | 1 | 1 | | | yes | | | | yes | yes |
| C10orf50 | NA | TSS1500 | chr10:26877515-26877803 | 288 | | | | | | | | | | | | | | | | | | | |
| C10orf50 | NA | Body | chr10:26883013-26883165 | 152 | | | 2 | 1 | | | | | 1 | | | | | yes | yes | | yes | yes | yes |
| LOC731789 | NA | TSS200 | chr10:26931969-26931991 | 22 | | | | | | | | | 1 | | | | | | | | | | |
| ABI1 | S_Shore | TSS1500 | chr10:27150407-27150529 | 122 | | | 3 | | 1 | | | | | | | | | yes | yes | | | | |
| ANKRD26 | Island | TSS200 | chr10:27389600-27389908 | 308 | | 1 | | | | | | | | | | | | yes | | | | | |
| LOC387646 | Island | TSS200 | chr10:27541298-27541913 | 615 | | | 21 | | | | | 0 | 1 | | | | | yes | | | | | yes |
| PTCHD3 | N_Shore;Island | 1stExon;5'UTR;TSS200 | chr10:27702309-27703547 | 1238 | | | 7 | | | | | 0 | 3 | | | 1 | | | yes | | | | |
| MKX | Island | Body | chr10:28030398-28031091 | 693 | | 1 | 10 | | | | | | | | | | | | | | yes | | yes |
| MKX | Island;chr10:28030182-280352 | 5'UTR;1stExon;TSS200;Island;TSS1 | chr10:28033840-28036151 | 2311 | | | | | | | | | | | | | | | | | | | |
| ARMC4 | Island;S_Shore | TSS200 | chr10:28288003-28288192 | 189 | | | 1 | | | | | | | 1 | | | | | | | | | yes |
| MPP7 | NA | Body | chr10:28525513-28525755 | 242 | | | | | | | | | | | | | | | yes | yes | | | |
| MPP7 | NA | TSS200;TSS1500 | chr10:28571186-28571848 | 662 | | | 1 | | | | | | 1 | | | | | yes | | yes | | | |
| LOC387647 | N_Shore;Island | TSS1500;TSS200 | chr10:29698152-29699043 | 891 | | | 5 | 1 | | | | 0 | 8 | | | 4 | | yes | yes | | | | |
| SVIL | NA | Body | chr10:29759227-29759454 | 227 | | | | | | | | | | | | | | | | | | | |
| SVIL | NA | Body | chr10:29811530-29811659 | 129 | | | 13 | | | | | | | | | | | | | | | | |
| SVIL;MIR604 | NA | Body;TSS200;TSS1500 | chr10:29834018-29834523 | 505 | | | 4 | | | | | 0 | | | | | | yes | | | | | |
| SVIL | NA | Body | chr10:29839837-29840083 | 246 | | | | | | | | | | | | | | yes | | | | | |
| SVIL | NA | 5'UTR;1stExon | chr10:29923736-29924258 | 522 | | | | | | | | | 1 | | | | | | | | | yes | |
| SVIL | Island | TSS1500 | chr10:30026028-30026143 | 115 | | | | | | | | | 1 | | | | | | | yes | yes | | yes |
| KIAA1462 | Island | 5'UTR | chr10:30347964-30348638 | 674 | | | | | | | | | 1 | | | | | | | | | yes | yes |
| ZNF438 | NA | 5'UTR;Body | chr10:31191938-31192123 | 185 | | | | | | | | | | | | | | | | yes | | | |
| ZNF438 | NA | 5'UTR;TSS200;Body | chr10:31288634-31288997 | 363 | | | | | | | | | 1 | | | | | | yes | | yes | | |
| ZNF438 | Island | TSS200 | chr10:31321002-31321013 | 11 | | | | | | | | | 1 | | | | | | | | | | yes |
| ZEB1;LOC220930 | N_Shore | TSS1500;Body | chr10:31607150-31607160 | 10 | | | 8 | | | | | | | | | | | yes | | | | | |
| ARHGAP12 | N_Shore | 5'UTR | chr10:32216031-32216359 | 328 | | | | | 2 | | | 0 | 1 | | | 1 | | | | | | | |
| PARD3 | NA | Body | chr10:34408530-34408654 | 124 | | | | | | | | | 1 | | | 1 | | yes | | | | | |
| PARD3 | NA | Body | chr10:35080979-35081084 | 105 | | | | | | | | 0 | | | | 1 | | yes | | | | | |
| GJD4 | N_Shelf | TSS1500;TSS200 | chr10:35893624-35894430 | 806 | | | 7 | | | | | 0 | 1 | | | | | yes | | | | | |
| FZD8 | Island | 1stExon | chr10:35929139-35929369 | 230 | | | | | | | | 0 | | | | | | | | | | yes | yes |
| ANKRD30A | N_Shore;Island | TSS1500;TSS200 | chr10:37413782-37414802 | 1020 | | | 4 | | | | | | 2 | | | | | yes | | | | | |
| ZNF37A | N_Shore | TSS1500 | chr10:38382849-38383098 | 249 | | | 1 | | | | | | | | | | | yes | yes | | | | |
| HSD17B7P2 | N_Shore;Island | TSS1500;TSS200;Body | chr10:38644991-38645740 | 749 | | | 8 | | | | 1 | 0 | 14 | 5 | | 14 | 1 | | | | yes | | |

| | | | | | | | | | | | | | | | | | | | | | | | | |
|---|---|---|---|---|---|---|---|---|---|---|---|---|---|---|---|---|---|---|---|---|---|---|---|---|
| CDC10L | N_Shore;Island;S_Shore | Body;TSS200;TSS1500 | chr10:38691306-38692317 | 1011 | | | 7 | 1 | | | 2 | 0 | 5 | 4 | | 7 | 1 | yes | | | | | | |
| LOC441666 | N_Shore;Island;S_Shore | Body;TSS200 | chr10:42862876-42863594 | 718 | 1 | 2 | 23 | 1 | 4 | 1 | 1 | 0 | 16 | 8 | 1 | 19 | 1 | yes | | yes | yes | | | |
| LOC84856 | N_Shore;Island | TSS1500;TSS200;Body | chr10:42970351-42971732 | 1381 | | | 7 | | | | | 0 | 2 | 1 | | 1 | | | yes | | yes | | | yes |
| LOC84856 | Island | Body | chr10:42972753-42973001 | 248 | | | | | | | | | | | | | | yes | yes | | | | | |
| ZNF37B | N_Shore;Island;S_Shore | Body;TSS200;TSS1500 | chr10:43047784-43048734 | 950 | | | | | | | | 0 | | 3 | | | | yes | yes | | | | | |
| ZNF33B | N_Shore | 5'UTR | chr10:43133692-43134302 | 610 | | | | | | | | 0 | | 2 | | | | yes | | | | | | |
| RASGEF1A | Island | Body | chr10:43698834-43699070 | 236 | | | | | | | | 0 | 1 | | | | | | | | | yes | yes | |
| RASGEF1A | Island | 5'UTR | chr10:43725376-43725741 | 365 | | | | | | | | | 1 | | | | | | yes | | | yes | | |
| FXYD4 | NA | TSS200 | chr10:43867064-43867104 | 40 | | | | | | | | | 1 | | | | | yes | | | | yes | | |
| ZNF239 | Island | TSS200 | chr10:44070173-44070200 | 27 | | | | | | | | | | | | 1 | | | yes | | | | | |
| ZNF485 | N_Shore | TSS1500 | chr10:44101569-44101728 | 159 | | | 10 | | | | | 0 | | | | | | | | | | | | |
| CXCL12 | Island | TSS200;TSS1500 | chr10:44880708-44881934 | 1226 | | | 3 | | | | | | 1 | 1 | | | | yes | | | | yes | | |
| TMEM72 | NA | TSS1500;TSS200 | chr10:45406295-45406847 | 552 | | | 1 | | | | | 0 | 4 | 1 | | 1 | | yes | | | yes | | | |
| C10orf10;RASSF4 | S_Shelf | 5'UTR;Body | chr10:45473712-45474372 | 660 | | | 10 | | | | | | | | | | | yes | yes | | | | | |
| RASSF4 | NA | TSS1500;Body | chr10:45477550-45477807 | 257 | | | | | | | | 0 | | | | | | | | | | | | |
| C10orf25;ZNF22 | N_Shore | 3'UTR;TSS1500 | chr10:45495435-45496087 | 652 | | | 38 | 1 | | | | | 2 | | | | | yes | yes | | | | yes | |
| OR13A1 | NA | 3'UTR | chr10:45799211-45799430 | 219 | | | | | | | | | | | | | | | | | | | | |
| ALOX5 | N_Shelf;N_Shore;Island | Body | chr10:45914276-45914949 | 673 | 1 | | | 2 | 4 | | | | | | | | | yes | | | | | | |
| ALOX5 | N_Shore | Body | chr10:45922877-45923705 | 828 | | | 2 | | | | | | | | | | | yes | | yes | | | | |
| MARCH8 | NA | 3'UTR | chr10:45958759-45958885 | 126 | | | | | | | | 0 | 1 | 1 | | | | yes | | | | | | |
| MARCH8 | NA | Body | chr10:45967484-45967498 | 14 | | | | | | | | | 1 | | | | | yes | yes | | yes | | | |
| SYT15 | Island | Body;TSS1500 | chr10:46970584-46971495 | 911 | | | 1 | | | | | 0 | 1 | | | | | yes | | | | | | |
| GPRIN2 | N_Shore;Island;S_Shore | TSS1500;TSS200 | chr10:46992448-46993924 | 1476 | | | 15 | | | | | | | | | | | yes | | | | | yes | |
| PPYR1 | N_Shore;Island | TSS1500;TSS200 | chr10:47083283-47083504 | 221 | | | 1 | | | | | | 4 | | | | | yes | yes | | | | yes | |
| ANXA8L2 | NA | TSS1500 | chr10:47746733-47747039 | 306 | | | | | | | | | | | | | | yes | yes | | | | | |
| RBP3 | S_Shore | 1stExon;5'UTR | chr10:48390936-48391548 | 612 | | | | | | | | | | | | | | yes | | | | yes | | |
| GDF2 | N_Shore | 3'UTR | chr10:48413742-48413850 | 108 | | | | | | | | | | | | | | | | | | | | |
| GDF2 | S_Shelf | 1stExon;TSS200 | chr10:48416463-48417338 | 875 | | | 6 | | 1 | | | | 1 | | | | | yes | | | yes | | yes | |
| GDF10 | Island | TSS200 | chr10:48439430-48439452 | 22 | | | | | | | | | 1 | | | | | | | | yes | | yes | |
| FRMPD2 | NA | Body | chr10:49459800-49460249 | 449 | | | | | | | | | 1 | | | | | | yes | | | | yes | |
| FRMPD2 | NA | Body | chr10:49482695-49482826 | 131 | | | | | | | | | | | | | | yes | | | | | | |
| ARHGAP22 | N_Shore;Island;chr10:4967424 | Body;Island | chr10:49674251-49674883 | 632 | | | 6 | | | | | | 2 | | 1 | | | yes | | | | | yes | |
| ARHGAP22 | Island;N_Shore | TSS200;Body | chr10:49812686-49813434 | 748 | | | | | | | | 0 | 2 | | | | | yes | | | | yes | | |
| WDFY4 | NA | TSS1500;nan | chr10:49892741-49893549 | 808 | | | 4 | | 2 | | | 0 | 1 | 3 | 1 | | | yes | yes | | | | | |
| WDFY4 | NA | TSS1500;Body | chr10:50143337-50143517 | 180 | 1 | | 6 | | 1 | | | | | | | | | | | | | | | |
| C10orf72 | Island | TSS200 | chr10:50323708-50323832 | 124 | | | 16 | | | | | | | | | | | yes | | | | | yes | |
| FAM170B | N_Shelf;N_Shore;Island | nan;3'UTR;Body | chr10:50339356-50340173 | 817 | | | 19 | | | | | | | | | | | yes | yes | | yes | | | |
| FAM170B | Island;S_Shore | Body;1stExon;5'UTR | chr10:50341989-50342145 | 156 | | | 6 | | | | | | | | | | | | | yes | yes | | | |
| C10orf128 | NA | Body;TSS200 | chr10:50396425-50396833 | 408 | | | | | | | | | | | | | | yes | | | yes | | | |
| C10orf71 | NA | TSS1500;TSS200 | chr10:50506748-50507298 | 550 | | | 35 | | 2 | | | | 3 | 1 | 1 | | | | | | | | | |
| C10orf71 | NA | 5'UTR | chr10:50530557-50530868 | 311 | | | 1 | | | | | | | | | | | | yes | yes | | | | |
| SLC18A3;CHAT | N_Shore;Island | TSS1500;1stExon;5'UTR | chr10:50816963-50818652 | 1689 | 3 | | | | | | | | 2 | | | | | yes | | | | | yes | |
| CHAT | Island | 1stExon;5'UTR | chr10:50822606-50822752 | 146 | | | | | | | | | 1 | | | | | | | | | | yes | |
| C10orf53 | N_Shore | TSS1500 | chr10:50886999-50887731 | 732 | | | 1 | | | | | 0 | 4 | | | 2 | | yes | yes | yes | yes | | | |
| AGAP7 | S_Shelf | Body | chr10:51487558-51487759 | 201 | | | | | | | | | | | | | | yes | | | | | | |
| MSMB | NA | TSS1500 | chr10:51548895-51549656 | 761 | | | 2 | | | | | | 2 | | | | | yes | | yes | yes | | | |
| NCOA4 | S_Shore;chr10:51572198-5157 | 5'UTR;Body;S_Shore | chr10:51575592-51576452 | 860 | | | | 1 | | | | 0 | 1 | 1 | 1 | 1 | | yes | | | | | | |
| SGMS1 | N_Shore | 5'UTR | chr10:52177823-52178225 | 402 | | | | | 1 | | | | | | | | | yes | | | yes | | | |
| DKK1 | Island;N_Shelf;N_Shore | 5'UTR;1stExon;nan;TSS1500;TSS20 | chr10:54073047-54074524 | 1477 | | 1 | 17 | | | | | | 2 | | | | | | | | | | yes | |
| MBL2 | NA | nan | chr10:54531345-54531653 | 308 | | | | | | | | | | | | | | | | yes | | | | |
| PCDH15 | NA | 5'UTR;1stExon;TSS200 | chr10:56561096-56561213 | 117 | | | | | | | | | 2 | | | 1 | | | | | | yes | | |
| UBE2D1 | N_Shelf | nan | chr10:60094109-60094511 | 402 | | | 1 | | | | | | | | | | | yes | yes | yes | yes | | yes | |
| SLC16A9 | N_Shelf;N_Shore | 5'UTR | chr10:61468158-61468757 | 599 | | | 2 | | | | | | | | | | | | | | | | | |
| ANK3 | NA | 1stExon;5'UTR | chr10:62149329-62149642 | 313 | | 1 | | | | | | | | | | | | | | | | | | |
| RHOBTB1 | NA | Body | chr10:62671136-62671228 | 92 | | | 6 | | | | | | | | | | | | | | | | | |
| RHOBTB1 | Island | TSS200;TSS1500 | chr10:62761500-62761819 | 319 | | | 2 | | | | | | | | | | | | | | | | yes | |
| TMEM26 | N_Shore | Body | chr10:63212206-63212265 | 59 | | | | | | | | | | | | | | | | | | yes | | |
| TMEM26 | S_Shore | 1stExon;5'UTR;TSS200 | chr10:63213339-63213510 | 171 | | 1 | 22 | | | | | | 1 | | | 1 | | | | | yes | | | |
| C10orf107 | N_Shore;Island | TSS1500;TSS200 | chr10:63422511-63423079 | 568 | | | | | 1 | | | | 2 | 1 | | 3 | | yes | yes | | | | yes | |
| ARID5B | NA | Body;nan | chr10:63809073-63809170 | 97 | | | | | | | | | 3 | | 1 | | | | | | | | | |
| ZNF365 | N_Shore | TSS1500 | chr10:64133686-64133801 | 115 | | | | | | | | | 1 | | | | | | | | | yes | | |
| ADO | Island | 1stExon;3'UTR | chr10:64565642-64565772 | 130 | | | | | | | | 0 | 1 | | | | | | | | | | | |
| EGR2 | N_Shore | 3'UTR | chr10:64572927-64573476 | 549 | | | | | | | | | 2 | | | 1 | | | | | | | | |
| EGR2 | Island | 5'UTR | chr10:64578919-64579140 | 221 | | | | | | | | | | | | | | yes | | | | | | |
| NRBF2 | N_Shore | TSS1500 | chr10:64892673-64892844 | 171 | | | | | | | | | | | | | | yes | | | | | yes | |
| REEP3 | N_Shelf | nan | chr10:65280473-65280795 | 322 | | | 2 | | | | | | | | | | | yes | | | | | yes | |
| CTNNA3;LRRTM3 | NA | Body | chr10:68687951-68688167 | 216 | | 1 | 7 | | | | | | | | | | | | | | | | | |
| ATOH7 | S_Shore | 5'UTR;1stExon | chr10:69991859-69992156 | 297 | | | | | | | | | 1 | | | | | | | | | | yes | |
| DNA2 | S_Shore | TSS200 | chr10:70232027-70232188 | 161 | | | | | | | | | | | | | | yes | yes | | | | | |
| TET1 | S_Shore | 5'UTR | chr10:70321383-70322442 | 1059 | | | | 1 | | | | 0 | 1 | | | | | | | | | | yes | |
| SRGN | NA | nan;TSS1500 | chr10:70847350-70847882 | 532 | | | 2 | 1 | 1 | | | | 1 | | | | | yes | | | | | | |
| SRGN | NA | Body | chr10:70863699-70863740 | 41 | | | | | | | | | 1 | | | | | yes | | | | | | |
| HKDC1 | NA | nan;TSS1500 | chr10:70979777-70980112 | 335 | | | | 1 | | | | | | | | | | yes | | | | | | |
| HK1 | NA | Body | chr10:71087834-71088038 | 204 | | | 14 | | | | | 0 | 2 | | | 1 | | yes | yes | | | | | |
| HK1 | NA | Body | chr10:71142347-71142526 | 179 | 1 | | | | | | | | | | | | | yes | | | | | | |
| TACR2 | NA | 1stExon;5'UTR;TSS200 | chr10:71176501-71177172 | 671 | | | 7 | | | | | | | | | | | yes | | | yes | | | |
| TSPAN15 | N_Shelf | nan | chr10:71210704-71211006 | 302 | | | | | | | | | | | | | | yes | | | | | | |
| NEUROG3 | Island | Body | chr10:71332776-71333037 | 261 | | 1 | | | | | | | | | | | | yes | | | | | | |
| C10orf35 | N_Shore;Island | TSS1500;TSS200 | chr10:71389696-71390385 | 689 | | | | | | | | | 3 | | | | | yes | yes | | | | yes | |
| H2AFY2 | N_Shore | TSS200 | chr10:71812319-71812330 | 11 | | | | | | | | | | | | | | | | | | | | |
| NODAL | Island;S_Shore | Body;1stExon;TSS200 | chr10:72201417-72201943 | 526 | | | 12 | | | | | | | | | | | | | | | | | |
| KIAA1274 | Island | TSS200 | chr10:72238622-72238844 | 222 | | | 1 | | | | | | | | | | | | | | | | yes | |
| KIAA1274 | S_Shelf | 5'UTR | chr10:72254314-72254335 | 21 | | | 11 | | | | | 0 | 1 | 1 | | | | yes | yes | | | | | |
| KIAA1274 | NA | Body | chr10:72311914-72312093 | 179 | | | 27 | | | | | | | | | | | | yes | | | | | |
| PRF1 | N_Shore | Body | chr10:72360348-72360448 | 100 | | | | | | | | | 1 | | | | | yes | | | | | | |
| PRF1 | S_Shore;S_Shelf | 1stExon;5'UTR;TSS200 | chr10:72362618-72362866 | 248 | | | | | | | | | 2 | 1 | | | | yes | | | | | | |
| ADAMTS14 | Island | TSS200;1stExon | chr10:72432540-72433218 | 678 | | | 1 | | | | | | | | | 1 | | | | | | yes | | |
| ADAMTS14 | NA | Body | chr10:72453975-72454026 | 51 | 6 | | | | | | | | | | | | | | yes | | | | | |
| UNC5B | NA | Body | chr10:73004558-73004610 | 52 | | | | 1 | | | | | | | | | | | yes | | yes | | | |
| C10orf105;CDH23 | NA | 5'UTR;Body;1stExon | chr10:73497514-73497772 | 258 | 1 | | 18 | | | | | | | | | | | yes | | | yes | | | |
| CDH23 | N_Shore | Body | chr10:73565625-73565966 | 341 | | | 1 | | | | | 0 | | | | | | yes | yes | | | | | |
| SPOCK2 | chr10:73846806-73848233;S_S | S_Shore;5'UTR;1stExon;TSS200;Bo | chr10:73848153-73849167 | 1014 | | | 18 | | 1 | | | | 3 | 1 | 1 | | | yes | | | | | yes | |
| ASCC1;C10orf104 | S_Shore | TSS1500;5'UTR | chr10:73976790-73976821 | 31 | | | 6 | 1 | | | | | 1 | | 1 | | | yes | | | | | | |
| MYOZ1 | S_Shelf | nan | chr10:75401236-75401315 | 79 | | | | | | | | | | | | | | | | | | | | |
| SYNPO2L | S_Shelf | TSS1500;Body | chr10:75415704-75415875 | 171 | | | | | | | | | 1 | | | | | yes | | | | | yes | |
| CAMK2G | NA | Body | chr10:75631588-75632066 | 478 | | | | | | | | | 1 | 1 | | | | | yes | | | | | |
| DUPD1 | N_Shore | Body | chr10:76803270-76803925 | 655 | | | | | | | | 0 | 1 | | | 1 | | | | yes | | yes | | |
| DUPD1 | NA | Body;1stExon | chr10:76818207-76818797 | 590 | | | | | | | | | | | | | | yes | | yes | | | | |
| DUSP13 | NA | Body;3'UTR;TSS200 | chr10:76859350-76859441 | 91 | | | | | | | | | | | | | | yes | | | | | | |

| | | | | | | | | | | | | | | | | | | | | | | | | | | | |
|---|---|---|---|---|---|---|---|---|---|---|---|---|---|---|---|---|---|---|---|---|---|---|---|---|---|---|---|
| COMTD1 | Island | TSS200 | chr10:76996280-76996533 | 253 | | | | | | | | 0 | | | | | | yes | | | | | | | | | |
| C10orf41 | Island | Body;TSS200 | chr10:77163368-77166318 | 2950 | | | 2 | 1 | | | | 0 | 7 | | | 1 | | | | | | | yes | | | | |
| C10orf11 | NA | TSS1500;TSS200 | chr10:77542302-77542585 | 283 | | | 4 | | | | | 0 | 11 | | | 3 | | | | | | | | | | | |
| C10orf11 | N_Shelf;N_Shore;Island | Body | chr10:77871618-77872458 | 840 | | | 32 | | | | | 0 | 5 | | | 1 | | | yes | | | | | | | | |
| KCNMA1 | NA | Body | chr10:78870177-78870221 | 44 | | | 4 | | | | | | | | | | | | | | | | | | | | |
| ZMIZ1 | NA | 5'UTR | chr10:80853937-80854035 | 98 | | | 7 | | | | | | 1 | 1 | | | | yes | | | | | | | | | |
| ZMIZ1 | Island | Body | chr10:81002480-81003020 | 540 | | | | | 1 | | | | | | | | | | | | | yes | yes | | | | |
| ZMIZ1 | NA | Body | chr10:81056358-81056385 | 27 | | | | | | | | | 1 | | | | | | yes | | | | yes | | | | |
| ZMIZ1 | NA | 3'UTR | chr10:81075012-81075492 | 480 | 2 | | 3 | | | | | | 1 | 1 | | | | | | | | | | | | | |
| ZCCHC24 | NA | Body | chr10:81154142-81154192 | 50 | | | | 1 | | | | | | | | | | | | | | | | | | | |
| MBL1P | NA | TSS1500 | chr10:81679675-81680307 | 632 | | | | | | | | | 1 | | | | | yes | yes | | | | | | | | |
| PLAC9 | N_Shore | TSS1500;TSS200 | chr10:81892109-81892161 | 52 | | | | | | | | 0 | 1 | | | | | yes | | yes | | | yes | | | | |
| PLAC9 | NA | Body | chr10:81904160-81904331 | 171 | | | | | | | | | 1 | | | | | | | | | | | | | | |
| MAT1A | NA | Body;1stExon;5'UTR | chr10:82049201-82049858 | 657 | | | 7 | | | | | | 1 | | | | | | yes | | | yes | | | | | |
| DYDC2;DYDC1 | Island | TSS1500;5'UTR;TSS200 | chr10:82116365-82116596 | 231 | | | | 1 | | | | | 1 | | | | | | | | | | | | | | |
| C10orf58 | N_Shelf;N_Shore | nan;TSS1500 | chr10:82167757-82168077 | 320 | | | 25 | | | | | 0 | 2 | 1 | | 2 | | yes | | | | | | | | | |
| C10orf58 | S_Shelf;S_Shore | 5'UTR;TSS1500 | chr10:82172262-82173570 | 1308 | | | | | | | | | | | | | | yes | yes | | | | | | | | |
| SH2D4B | S_Shore | TSS1500 | chr10:82297397-82297920 | 523 | | | | | | | | | | | | | | yes | | | | | | | | | |
| SH2D4B | S_Shore | 5'UTR;1stExon | chr10:82299606-82299737 | 131 | | | | | | | | | 1 | | | | | yes | | | | | | | | | |
| SH2D4B | NA | Body | chr10:82345699-82345955 | 256 | | | 2 | | | | | | | | | | | yes | | yes | | | | | | | |
| SH2D4B | NA | Body | chr10:82362642-82363434 | 792 | | | 2 | | | | | | 1 | | | | | yes | | | yes | | | | | | |
| PCDH21 | NA | Body | chr10:85974018-85974623 | 605 | | | 2 | | | | | 0 | 2 | 1 | | 2 | | | | | yes | | | | | | |
| LRIT2 | NA | Body;1stExon;TSS200 | chr10:85985222-85985779 | 557 | | | 3 | | | | | 0 | 3 | | | | | yes | yes | | | | | | | | |
| LRIT1 | N_Shore | Body | chr10:85997177-85997426 | 249 | | | | | | | | | | | | 1 | | yes | | | yes | | | | | | |
| LRIT1 | S_Shelf | 1stExon;TSS200 | chr10:86001241-86001560 | 319 | | | | | | | | | 1 | | | | | yes | yes | yes | | | | | | | |
| RGR | NA | TSS1500;TSS200 | chr10:86004660-86005121 | 461 | | | | | | | | | 1 | | | | | yes | | | | | | | | | |
| GRID1;MIR346 | N_Shore;Island;S_Shore | Body;TSS200;TSS1500 | chr10:88022641-88025642 | 3001 | | | | | | | | | | | | | | | | | | | | | | | |
| GRID1 | Island | TSS200 | chr10:88126291-88126306 | 15 | | | | | | | | | | 1 | | | | | | | | yes | yes | | | | |
| LDB3 | NA | 3'UTR;TSS1500;TSS200;1stExon;5' | chr10:88427400-88428295 | 895 | | | | | | | | | | | | | | yes | yes | | | yes | | | | | |
| MMRN2 | N_Shore | Body | chr10:88702316-88702795 | 479 | | | 5 | | | | | | | | | | | | | | | | | | | | |
| MMRN2;SNCG | NA | Body;TSS1500;TSS200 | chr10:88717364-88718393 | 1029 | | | | 1 | | | | | 2 | | | | | yes | | | | yes | | | | | |
| C10orf116 | N_Shore;Island | TSS1500;TSS200 | chr10:88728073-88728572 | 499 | | | | | | | | 0 | 1 | 2 | | | | yes | | | | | | | | | |
| AGAP11;C10orf116 | N_Shore | TSS1500;Body | chr10:88730324-88730599 | 275 | | | | | | | | 0 | 1 | | | | | | | | | | | | | | |
| MINPP1 | N_Shelf;N_Shore | nan;TSS1500 | chr10:89263900-89264128 | 228 | | | 14 | | | | | 0 | 2 | | | | | | | | yes | | | | | | |
| PAPSS2 | N_Shore | TSS1500 | chr10:89418898-89419292 | 394 | | | 1 | | | | | | | | | | | yes | | | | yes | | | | | |
| RNLS | Island | TSS200 | chr10:90343192-90343435 | 243 | | | 2 | | | | | | | | | | | yes | yes | | | | | | | | |
| ANKRD22 | NA | TSS200;Body;1stExon;5'UTR | chr10:90611604-90612232 | 628 | | | 4 | 1 | 2 | | | 0 | 5 | 1 | | 3 | | | | yes | | | | | | | |
| ACTA2 | NA | Body;5'UTR;TSS200 | chr10:90712478-90712948 | 470 | | | 35 | | | | | | 1 | | | | | | | yes | | | | | | | |
| SLC16A12 | N_Shore;Island | 5'UTR;TSS200;TSS1500 | chr10:91295045-91296457 | 1412 | | | 1 | | | | | | 2 | | | 1 | | yes | | | yes | | | | | | |
| RPP30 | NA | TSS1500 | chr10:92631135-92631194 | 59 | | 1 | | | | | | | | | | | | | | | yes | | | | | | |
| LOC100188947 | NA | Body | chr10:93334974-93335196 | 222 | | | | 1 | | | | | 1 | | | | | yes | | | yes | | | | | | |
| PPP1R3C | N_Shelf;N_Shore;Island | Body;1stExon;5'UTR;TSS200;TSS1 | chr10:93391963-93393509 | 1546 | | | | | | | | | | | | | | | | | | | | | | | |
| CPEB3 | S_Shelf | 3'UTR | chr10:93811958-93812043 | 85 | | | | | | | | | 1 | | | | | yes | | | | yes | | | | | |
| HHEX | N_Shore;Island | Body;1stExon | chr10:94450531-94452554 | 2023 | | | 13 | | 1 | | | 0 | 3 | 2 | 1 | | | | | | | yes | | | | | |
| EXOC6 | NA | TSS1500 | chr10:94594240-94594494 | 254 | | | 1 | | | | | | | | | | | | | | | | yes | | | | |
| CYP26C1 | N_Shore;Island | TSS1500;TSS200 | chr10:94819802-94821085 | 1283 | | | 8 | | | | | 0 | 3 | | | 1 | | yes | | yes | | | | | | | |
| CYP26C1 | N_Shore;Island | Body | chr10:94825547-94827745 | 2198 | | | | | | | | | 3 | | | | | yes | | | | yes | yes | | | | |
| CYP26A1 | Island | Body | chr10:94834557-94834763 | 206 | | 1 | | | | | | | | | | | | | | | | | | | | | |
| MYOF | NA | Body | chr10:95123718-95123774 | 56 | | | | | | | | | 1 | | | | | yes | | | | | | | | | |
| MYOF | NA | Body | chr10:95198200-95198374 | 174 | | | 9 | | | | | | 1 | | | | | | | yes | | | | | | | |
| GPR120 | Island;S_Shore | 1stExon;Body | chr10:95327107-95327884 | 777 | | 1 | 2 | | | | | 0 | 2 | | | | | yes | | | | | | | | | |
| PDE6C | S_Shelf;NA | nan;TSS1500 | chr10:95372040-95372445 | 405 | | | | | | | | | | | | | | | | yes | | | | | | | |
| C10orf4 | N_Shelf;N_Shore;Island;S_Sho | Body;5'UTR;TSS200;TSS1500 | chr10:95461526-95463124 | 1598 | | | | | | | | 0 | 5 | | | 2 | | yes | yes | | | yes | yes | | | | |
| LGI1 | NA | TSS1500;TSS200 | chr10:95516861-95517895 | 1034 | | | 1 | 1 | | | | | | | | | | | | | | yes | | | | | |
| PLCE1 | NA | Body;TSS1500;TSS200 | chr10:95848649-95848841 | 192 | | | 11 | | | | | | | | | | | | | | | | | | | | |
| CYP2C18 | NA | TSS1500;TSS200 | chr10:96443071-96443383 | 312 | | | | 1 | | | | | | | | | | yes | yes | yes | yes | yes | | | | | |
| SORBS1 | NA | Body | chr10:97175272-97175479 | 207 | | | 19 | | | | | | | | | | | | | | | | | | | | |
| SORBS1 | NA | Body | chr10:97200911-97200987 | 76 | | | | 1 | | | | | | | | | | yes | yes | | | | | | | | |
| ENTPD1 | NA | TSS1500;TSS200;Body | chr10:97515222-97515753 | 531 | | | 9 | | | | | 0 | 4 | 2 | | 1 | | yes | yes | | | | yes | | | | |
| C10orf131 | N_Shelf;Island | nan;TSS1500 | chr10:97666894-97667731 | 837 | 1 | | | | | | | | 1 | | | | | | | | | | | | | | |
| ZNF518A | N_Shore | TSS200 | chr10:97889379-97889448 | 69 | | | 3 | | | | | | | | | | | | | | | | yes | | | | |
| BLNK | NA | Body;1stExon;TSS200 | chr10:98031125-98031964 | 839 | | | 5 | 1 | | | | | 3 | | | | | | | | | | | | | | |
| OPALIN | NA | TSS200;Body | chr10:98118853-98120242 | 1389 | | | | | | | | | 1 | | | | | | yes | | yes | | | | | | |
| TLL2 | NA | 3'UTR | chr10:98129751-98130127 | 376 | | | | | 1 | | | | | | | | | yes | yes | | | | | | | | |
| TLL2 | Island | Body | chr10:98144406-98144452 | 46 | | | | | | | | | 1 | | | | | yes | | yes | yes | | yes | | | | |
| PIK3AP1 | NA | Body | chr10:98469437-98469552 | 115 | | | | | | | | | | | | | | yes | yes | | | | | | | | |
| SLIT1 | NA | Body | chr10:98908755-98909007 | 252 | | | 5 | | | | | | | | | | | yes | | | yes | | yes | | | | |
| SLIT1 | Island | TSS1500 | chr10:98946111-98946778 | 667 | | | 1 | | | | | | | | | | | | | | | yes | | | | | yes |
| FRAT1 | Island | 1stExon;3'UTR | chr10:99080594-99081016 | 422 | | | 12 | | | | | | 1 | | | | | | | | | | | | | | |
| RRP12 | N_Shelf | Body | chr10:99160096-99160200 | 104 | | | | | | | | 0 | 1 | | | | | yes | | | | | | | | | |
| PGAM1 | N_Shelf | nan | chr10:99185264-99185307 | 43 | | | | | | | | 0 | | | | | | yes | | | | yes | | | | | |
| UBTD1 | Island | Body | chr10:99330056-99330133 | 77 | | | | | | | | | 1 | | | | | yes | | yes | | | | | | | |
| ANKRD2 | S_Shore | 3'UTR | chr10:99331332-99331808 | 476 | | | | | | | | | 1 | 1 | | | | yes | | | | | | | | | |
| ANKRD2 | Island;N_Shelf;N_Shore | Body | chr10:99337800-99338240 | 440 | | | 16 | | | 1 | | 0 | 3 | | | 2 | | yes | | | yes | | | | | | |
| C10orf62;DHDPSL | NA | TSS1500;Body | chr10:99349003-99349395 | 392 | 1 | | | | | | | | | | | | | | | | yes | | | | | | |
| MORN4 | Island;S_Shore | 1stExon;5'UTR;TSS200 | chr10:99393374-99393886 | 512 | | | | | | | | 0 | 1 | | | 1 | | yes | | yes | | | | | | | |
| MARVELD1 | N_Shore | TSS1500 | chr10:99473085-99473224 | 139 | | | | 1 | 1 | | | | 1 | | | | | yes | | | | yes | yes | | | | |
| LOC100270710 | S_Shelf | Body;3'UTR | chr10:99477939-99478572 | 633 | | | | | | | | | | | | | | yes | | | | | | | | | |
| SFRP5 | Island | Body;TSS200 | chr10:99531164-99532367 | 1203 | | | 1 | | 1 | | | | | | | | | yes | yes | | | yes | yes | | | | |
| CRTAC1 | NA | Body | chr10:99734081-99735202 | 1121 | 1 | 1 | 19 | | | | 1 | 0 | 17 | 2 | | 8 | | | | | yes | | yes | | | | |
| CRTAC1 | Island | Body | chr10:99789971-99790910 | 939 | | | | | | | | | 1 | | | | | | | | | yes | yes | | | | |
| C10orf28 | N_Shelf | nan | chr10:99894013-99894087 | 74 | | | 9 | | | | | | 1 | | | | | yes | yes | | | | | | | | |
| PYROXD2;MIR1287 | NA | Body | chr10:100155182-100155583 | 401 | | | | | | | | | | | | | | yes | yes | yes | | | | | | | |
| PYROXD2 | NA | Body;1stExon;TSS200 | chr10:100174853-100175560 | 707 | | | | 1 | 1 | | | | | | | | | yes | | | | | | | | | |
| HPSE2 | N_Shelf;N_Shore;Island | Body | chr10:100991745-100994478 | 2733 | | | 19 | | 5 | | | | 2 | | | 1 | | yes | | | | yes | yes | | | | |
| HPSE2 | S_Shore;S_Shelf | TSS1500 | chr10:100996070-100997023 | 953 | | 1 | 2 | | 1 | | | | 1 | | | | | | | | | yes | | | | | |
| CNNM1 | N_Shore | TSS200 | chr10:101088811-101089592 | 781 | | | | | | | | | 1 | | | | | yes | yes | | | yes | yes | | | | |
| NKX2-3 | N_Shore;Island | TSS1500;TSS200 | chr10:101292403-101293089 | 686 | | | 3 | | | | | | | | | | | yes | yes | | | | | | | | |
| NKX2-3 | N_Shore | Body | chr10:101294300-101294643 | 343 | | | | | | | | | 1 | | | | | | | | | | yes | | | | |
| ENTPD7 | N_Shore | TSS1500 | chr10:101418483-101418880 | 397 | | | | | | | | | 1 | | | | | | yes | yes | | | yes | | | | |
| NCRNA00093;DNMBP | NA | TSS1500;Body | chr10:101686538-101686925 | 387 | | | | | | | | | | | | | | yes | yes | | | | | | | | |
| CPN1 | Island;N_Shore | Body | chr10:101825029-101825185 | 156 | | | | 1 | | | | | | | | 1 | | | yes | | yes | yes | | | | | |
| CPN1 | NA | Body | chr10:101841253-101841973 | 720 | | | | 1 | 2 | | | | | | | | | yes | yes | | | | | | | | |
| BLOC1S2 | Island | TSS1500;TSS200 | chr10:102046685-102046768 | 83 | | | | | | | | 0 | | | | | | | | | | | | | | | |
| PKD2L1 | NA;S_Shelf | TSS200;Body;5'UTR;1stExon | chr10:102089681-102090589 | 908 | | | 8 | | | | | | 1 | | | | | | | | | | yes | | | | |
| SEC31B | N_Shore;Island | 5'UTR;TSS200 | chr10:102278918-102279758 | 840 | | 1 | | | | | | 0 | 2 | | | | | yes | | | | | | | | | |
| PAX2 | Island | Body | chr10:102507979-102508529 | 550 | | | | | | | | | 1 | | | | | | | | | | | | | | |
| PAX2 | N_Shore | 3'UTR | chr10:102589467-102589840 | 373 | | 1 | | | | | | | 1 | | | | | | | | | | yes | | | | |
| SEMA4G | S_Shelf | TSS200 | chr10:102732224-102732605 | 381 | | | | | | | | | 1 | | | | | | | | yes | | | | | | |

| | | | | | | | | | | | | | | | | | | | | | | | | | | | |
|---|---|---|---|---|---|---|---|---|---|---|---|---|---|---|---|---|---|---|---|---|---|---|---|---|---|---|---|
| LZTS2 | Island | 5'UTR | chr10:102760724-102760970 | 246 | | | | 1 | | | | | | | | | | | | | | | | | | | |
| PDZD7 | Island | Body | chr10:102778743-102779095 | 352 | | | | | | | | 0 | 1 | | | 2 | | yes | | | | yes | | | | | |
| KAZALD1 | Island | 1stExon;5'UTR | chr10:102821552-102822249 | 697 | | | 3 | | | | | 0 | 1 | | | | | | | | | yes | | | | | |
| TLX1NB | N_Shore | 5'UTR | chr10:102882978-102883550 | 572 | | | | 1 | | | | | | | | | | | | | | | | | | | |
| TLX1;TLX1NB | N_Shore;Island | TSS1500;5'UTR;TSS200 | chr10:102890145-102891280 | 1135 | | | | | | | | | 1 | | | | | | yes | | | yes | yes | | | | |
| TLX1NB;TLX1 | N_Shore | TSS1500;Body | chr10:102892391-102892787 | 396 | | | 2 | | | | | | | | | | | | | | | | yes | | | | |
| TLX1 | Island | Body | chr10:102893925-102894120 | 195 | | 1 | | | | | | | | | | | | | | | | | yes | | | | |
| TLX1 | N_Shore;Island | Body | chr10:102895708-102896869 | 1161 | | | 7 | | | | | | 1 | | | | | | | | | | | | | | |
| LBX1 | Island | nan | chr10:102986808-102987257 | 449 | | | | | | | | | 1 | | | | | | | | | | yes | | | | |
| FLJ41350;LBX1 | Island | TSS200;TSS1500;1stExon | chr10:102988831-102989329 | 498 | | | | | | | | | 2 | | | | | | | | | | yes | | | | |
| FLJ41350 | S_Shore | Body | chr10:102997382-102998762 | 1380 | | 1 | 16 | | | | | | | | | | | yes | | | | yes | yes | | | | |
| FGF8 | NA | nan;3'UTR | chr10:103530053-103530255 | 202 | | | | | | | | 0 | 2 | | | | | | | | | yes | | | | | |
| FGF8 | N_Shore | Body | chr10:103534501-103534934 | 433 | | | 1 | | | | | | | | | | | yes | | | | | | | | | |
| KCNIP2 | N_Shore | Body | chr10:103588818-103589245 | 427 | | | 5 | | | | | | 1 | | | 1 | | | | | | yes | | | | | |
| KCNIP2 | S_Shelf | Body | chr10:103599520-103599705 | 185 | | | | | | | | | | | | | | | | | | | yes | | | | |
| KCNIP2 | S_Shore;Island | TSS200;1stExon;5'UTR | chr10:103603563-103604222 | 659 | | | 4 | 1 | 2 | | | | | | | | | | | | | | yes | | | | |
| LDB1 | S_Shore | TSS1500 | chr10:103880810-103881480 | 670 | | | | 1 | 1 | | | | | | | | | | | | | | yes | | | | |
| ELOVL3 | Island | TSS200 | chr10:103986245-103986736 | 491 | | | | | | | | | | | | | | yes | | | | | yes | | | | |
| PITX3 | Island | 1stExon;5'UTR | chr10:104001240-104001261 | 21 | | | | | | | | | 1 | | | | | | | | | | yes | | | | |
| PSD | Island | Body | chr10:104170217-104170513 | 296 | | 1 | | | | | | | | | | | | yes | | | | | | | | | |
| FBXL15;PSD | Island | TSS200;TSS1500 | chr10:104179054-104179484 | 430 | | | | | | | | | 2 | | | | | yes | yes | | | | yes | | | | |
| MIR146B | S_Shelf | TSS1500;TSS200 | chr10:104196206-104196541 | 335 | | | | 1 | | | | | 3 | | | | | yes | | | | | | | | | |
| C10orf95 | Island;S_Shore | Body;5'UTR;1stExon;TSS200 | chr10:104211296-104211488 | 192 | | | 9 | | | | | | | | | | | yes | | | | | | | | | |
| C10orf26 | NA | TSS1500;Body;TSS200;5'UTR;1stE | chr10:104535521-104536695 | 1174 | 1 | 1 | 1 | | | | | 0 | 4 | | | | | | | | | | | | | | |
| CYP17A1 | NA | Body;1stExon | chr10:104596890-104597435 | 545 | | | 1 | | | | | | | | | | | yes | | yes | | | | | | | |
| C10orf32 | N_Shore | TSS1500 | chr10:104613767-104613787 | 20 | | | 3 | 1 | | | | | 1 | | | 2 | | yes | | | | yes | | | | | |
| NT5C2 | NA | Body | chr10:104905387-104915551 | 10164 | | | | | | | | | | | | | | yes | yes | | yes | | yes | | | | |
| INA | Island | TSS200 | chr10:105037349-105037661 | 312 | | | | | | | | | | | | 1 | | | | | | | yes | | | | |
| CALHM2 | NA | Body | chr10:105209452-105209552 | 100 | | | | | | | | | | | | | | | | | | | | | | | |
| CALHM2 | N_Shore;Island;S_Shore | 5'UTR;Body;1stExon;TSS200;TSS1 | chr10:105212056-105212813 | 757 | | | 2 | | | | | 0 | 2 | | | 1 | | yes | yes | | | | yes | | | | |
| CALHM1 | N_Shore | 3'UTR | chr10:105215053-105215306 | 253 | | | | | | | | | | | | | | yes | | | | | | | | | |
| CALHM1 | S_Shore | 1stExon;TSS200 | chr10:105218524-105219172 | 648 | | | | | | | | | 1 | | | | | | | | | yes | | | | | |
| CALHM3 | N_Shelf;N_Shore;Island | nan;3'UTR;Body | chr10:105232723-105233423 | 700 | | | 17 | | | | | | | | | | | yes | | | yes | yes | | | | | |
| CALHM3 | NA | 1stExon;5'UTR;nan;TSS200 | chr10:105238873-105239205 | 332 | | | 8 | | | | | | 1 | | 1 | | | | yes | | | yes | | | | | |
| NEURL | N_Shelf | nan | chr10:105252720-105253152 | 432 | | | | | | | | 0 | | | | | | | yes | | | | yes | | | | |
| NEURL | N_Shore;Island | Body | chr10:105344174-105344807 | 633 | | | 7 | | | | | 0 | | | | | | yes | yes | | | | yes | | | | |
| SH3PXD2A | N_Shore | 3'UTR | chr10:105361785-105362057 | 272 | 1 | | | | | | | | | | | | | | | | | | | | | | |
| SH3PXD2A | Island;N_Shore | Body | chr10:105420747-105421480 | 733 | | | | 1 | | | | 0 | 3 | | | | | | | | | | yes | | | | |
| SH3PXD2A | N_Shelf;N_Shore | Body | chr10:105428385-105428818 | 433 | | | 4 | | | | | 0 | 3 | | | 4 | | yes | | | | | | | | | |
| SH3PXD2A | Island | Body | chr10:105452885-105453229 | 344 | | | | 1 | | | | | | | 1 | | | | | | | yes | | | | | |
| MIR609;C10orf79 | NA | Body;TSS200 | chr10:105978651-105978956 | 305 | | | 13 | | 1 | | | | 3 | 1 | 1 | | | | yes | | yes | | | | | | |
| GSTO2 | N_Shelf;N_Shore;Island | Body;TSS1500;TSS200;5'UTR;1stEx | chr10:106028206-106029358 | 1152 | | | 31 | 1 | | | | 0 | | | | | | yes | | | yes | | | | | | |
| ITPRIP | NA | 5'UTR | chr10:106088702-106089203 | 501 | | | 14 | | 1 | | | | 1 | | | | | yes | yes | | | yes | | | | | |
| ITPRIP | N_Shelf | TSS200 | chr10:106093718-106094166 | 448 | | | | | | | | | 2 | | | | | yes | | | | | | | | | yes |
| CCDC147 | NA | TSS1500 | chr10:106113175-106113270 | 95 | | | | | | | | | 1 | | | | | | yes | | | | | | | | |
| SORCS3 | Island | TSS200 | chr10:106400686-106401479 | 793 | | | | | 1 | | | | | | | 1 | | | | | | | | | | | |
| SORCS3 | S_Shore | Body | chr10:106425960-106426210 | 250 | | | 4 | | | | | | | | | | | yes | | | | | | | | | |
| SORCS3 | NA | Body | chr10:106440778-106441036 | 258 | | | | | | | | | 1 | | | | | yes | yes | | | | | | | | |
| SORCS1 | Island | 1stExon;TSS200 | chr10:108924366-108924867 | 501 | | 1 | | | | | | 0 | 3 | 1 | | | | | yes | | | yes | yes | | | | |
| XPNPEP1 | Island | 1stExon;Body;5'UTR;TSS200 | chr10:111683342-111683654 | 312 | | | 6 | | | | | 0 | 2 | | | | | | yes | | | | yes | | | | |
| ADD3 | N_Shelf | nan | chr10:111765673-111766049 | 376 | | | | | 1 | | | | | | | | | | | | | yes | yes | | | | |
| MXI1 | N_Shelf | nan | chr10:111966029-111966094 | 65 | | | 1 | | | | | | 1 | | | 1 | | yes | | | | | | | | | |
| DUSP5 | N_Shelf;N_Shore | nan;TSS1500 | chr10:112256517-112256825 | 308 | | | | 1 | | | | | 1 | | | | | | yes | | | | | | | | |
| RBM20 | S_Shore;N_Shelf;N_Shore | 1stExon;Body;nan;TSS1500 | chr10:112402748-112404997 | 2249 | | | 10 | | | | | | | | | | | | | | | yes | yes | | | | |
| RBM20 | Island | Body | chr10:112431906-112432418 | 512 | | 1 | | | | | | | | | | | | | | | | yes | | | | | |
| RBM20 | NA | Body | chr10:112562640-112582591 | 19951 | | | | | | | | | | | | | | yes | yes | yes | yes | yes | yes | | | | |
| LOC282997;PDCD4 | N_Shelf | Body | chr10:112630726-112630835 | 109 | | | | | 1 | | | | | | | | | yes | | | | yes | | | | | |
| GPAM | N_Shore | Body | chr10:113943149-113944114 | 965 | | | | 1 | | | | | | | | | | yes | yes | | | | yes | | | | |
| TECTB | NA | TSS200 | chr10:114043412-114043548 | 136 | | | | | | | | | 1 | | | | | | | | | | | | | | |
| ACSL5 | NA | TSS1500;5'UTR;TSS200 | chr10:114134878-114136073 | 1195 | | | | | 2 | | | | 1 | | | | | yes | | | | | | | | | |
| LOC143188 | NA | TSS200 | chr10:114610560-114610635 | 75 | | | | | 1 | | | | | | | | | | | | yes | | | | | | |
| TCF7L2 | N_Shelf | TSS1500 | chr10:114709739-114709957 | 218 | | | | | | | | | 1 | | | | | | | | | | yes | | | | |
| TCF7L2 | Island;S_Shore | Body | chr10:114712695-114713187 | 492 | | | 4 | | | | | 0 | 2 | | | 2 | | | | | | | | | | | |
| TCF7L2 | NA | Body | chr10:114877616-114877689 | 73 | | | 38 | 1 | | | | | | | | | | | | | | yes | yes | | | | |
| TCF7L2 | NA | Body | chr10:114886413-114886678 | 265 | | | | | | | | | | | | | | | | | | yes | yes | | | | |
| NRAP | NA | Body | chr10:115370894-115371055 | 161 | | | 4 | 1 | | | | | | | | | | | | | | | | | | | |
| CASP7 | NA | nan | chr10:115438129-115438230 | 101 | | | | | | | | | | | | | | yes | | | | yes | | | | | |
| C10orf81 | NA | TSS1500 | chr10:115510928-115511153 | 225 | | | | | | | | | | | | | | yes | | | | | | | | | |
| TDRD1 | Island;N_Shore | TSS200;TSS1500 | chr10:115938729-115939717 | 988 | | | 7 | | | | | | 4 | | | | | yes | yes | yes | | | | | | | |
| VWA2 | N_Shore | TSS1500;TSS200 | chr10:115998493-115999174 | 681 | | | | | | | | | | | | | | yes | yes | | | | | | | | |
| VWA2 | N_Shelf | Body | chr10:116045562-116046030 | 468 | | | 1 | | | | | | | | | | | yes | yes | | | | | | | | |
| AFAP1L2 | Island;S_Shore | TSS200 | chr10:116164640-116165190 | 550 | | | 6 | | | | | | | | | | | yes | | | | | yes | | | | |
| ABLIM1 | NA | Body | chr10:116307486-116307714 | 228 | 6 | | | | | | | | 1 | | | | | yes | | | yes | | | | | | |
| ABLIM1 | NA | Body;1stExon;TSS200 | chr10:116418181-116418797 | 616 | | | | | | | | | 1 | | | | | yes | yes | | | | | | | | |
| GFRA1 | Island | 5'UTR;1stExon;TSS1500;TSS200 | chr10:118031654-118034031 | 2377 | | 4 | | | | | | | 3 | | | | | | | | | yes | | | | | |
| C10orf96 | NA | 5'UTR;TSS200 | chr10:118083918-118084587 | 669 | | | 4 | | | | | 0 | 1 | | | | | | | | | | yes | | | | |
| PNLIP | NA | TSS1500 | chr10:118305388-118305444 | 56 | | | | | | | | | | | | | | yes | | | yes | | | | | | |
| PNLIPRP1 | NA | TSS1500;TSS200 | chr10:118350377-118350608 | 231 | | | 6 | | | | | | | | | | | | | | | yes | | | | | |
| PNLIPRP2 | N_Shelf;Island;N_Shore | Body | chr10:118386951-118387518 | 567 | | 1 | | | 1 | | | 0 | 1 | 1 | | 3 | | yes | | | | | yes | | | | |
| C10orf82 | N_Shore | 5'UTR | chr10:118429452-118429658 | 206 | | | 3 | | | | | | | | | | | yes | | | | | | | | | |
| HSPA12A | Island | TSS1500 | chr10:118502705-118502947 | 242 | | | | | | | | | 1 | | | | | yes | | | | | | | | | |
| KIAA1598 | Island | TSS200 | chr10:118765174-118765255 | 81 | | | | | | | | | | | | | | yes | | | | yes | | | | | |
| VAX1 | N_Shelf;N_Shore | 3'UTR | chr10:118890977-118891706 | 729 | | | 5 | | | | | | 1 | | | | | | | | | | yes | | | | |
| VAX1 | Island;N_Shore | Body;3'UTR | chr10:118892869-118893602 | 733 | | | 6 | | | | | | | | | | | | | | | yes | yes | | | | |
| VAX1 | Island | Body | chr10:118896755-118897280 | 525 | | | | | | | | 0 | 1 | | | | | | | | | | yes | | | | |
| VAX1 | Island;N_Shore | TSS1500 | chr10:118899024-118899407 | 383 | | 1 | | | | | | | | | | 1 | | | | | | | yes | | | | |
| KCNK18 | S_Shelf;NA | nan;TSS1500;TSS200 | chr10:118956011-118957259 | 1248 | | | 7 | | | | | | | | | | | yes | | yes | | | | | | | |
| SLC18A2 | N_Shelf | nan | chr10:119000083-119000236 | 153 | | | 9 | | | | | | | | | | | | | | | | yes | | | | |
| EMX2OS | NA;N_Shore;Island | Body | chr10:119291766-119296942 | 5176 | | | | | | | | | | | | | | | | | | | | | | | |
| EMX2;EMX2OS | Island | TSS200;Body | chr10:119302050-119302062 | 12 | | | | | | | | | 1 | | | | | | | | | yes | yes | | | | |
| EMX2;EMX2OS | Island;S_Shore | Body;TSS200;TSS1500 | chr10:119304708-119306659 | 1951 | | | 6 | | | | | | 1 | | | | | | | | | | | | | | |
| C10orf84 | S_Shore | TSS200 | chr10:120102302-120102359 | 57 | | | 2 | | | | | | | | | | | yes | yes | | yes | | yes | | | | |
| PRLHR | N_Shore;Island | 3'UTR;5'UTR;1stExon;TSS200 | chr10:120354081-120356072 | 1991 | | 2 | | | | | | | 3 | | | | | | | | | | | | | | |
| NANOS1 | NA | nan | chr10:120788366-120788696 | 330 | | | 1 | | | | | 0 | | | | 2 | | yes | | | | yes | yes | | | | |
| FAM45B;FAM45A | N_Shore | TSS1500 | chr10:120863603-120863723 | 120 | | | | | | | | | | | | 1 | | | yes | | | yes | yes | | | | |
| GRK5 | NA | Body | chr10:121138105-121138206 | 101 | | | | | | | | | | | | | | yes | yes | yes | | | | | | | |
| GRK5 | NA | Body | chr10:121171859-121172898 | 1039 | | | | | | | | | | | | | | yes | yes | | | | | | | | |
| RGS10 | NA | Body | chr10:121265587-121265893 | 306 | | | 9 | | | | | | | | | | | yes | | | | | | | | | |
| RGS10 | N_Shelf | Body | chr10:121300857-121301343 | 486 | | | 1 | | | | | | | | | | | yes | | | yes | | | | | | |

| | | | | | | | | | | | | | | | | | | | | | | | |
|---|---|---|---|---|---|---|---|---|---|---|---|---|---|---|---|---|---|---|---|---|---|---|---|
| TIAL1 | Island | TSS1500 | chr10:121357400-121357524 | 124 | | | | | | | | | 1 | | | | | | | yes | | | |
| PPAPDC1A | N_Shelf;Island | nan;Body | chr10:122215735-122217470 | 1735 | | 1 | 2 | | | | | 0 | | | | | | | yes | | | | yes |
| FGFR2 | NA;N_Shelf;N_Shore;chr10:12 | Body;1stExon;5'UTR;TSS200;TSS1 | chr10:123353298-123356336 | 3038 | | | 13 | | | | 1 | 0 | 8 | 5 | 2 | 12 | | | | | | yes | |
| TACC2 | NA | 5'UTR | chr10:123778998-123779362 | 364 | | | | 1 | | | | | 1 | | | | | | | | | yes | |
| TACC2 | NA | 5'UTR | chr10:123781519-123781602 | 83 | | | 2 | | | | | | | | | | | yes | | | | | |
| TACC2 | NA | Body | chr10:123909356-123909456 | 100 | | | | | | | | 0 | 1 | | | 1 | | | | | | | |
| TACC2 | N_Shelf;chr10:123922850-1239 | Body;N_Shore;TSS1500;TSS200;1s | chr10:123922620-123924085 | 1465 | | | 8 | | 3 | | | | 3 | 1 | 2 | | | | | | | | yes |
| BTBD16 | NA | TSS1500 | chr10:124030740-124030801 | 61 | | | 5 | | | | | | | | | | | yes | | | | | |
| PLEKHA1 | N_Shelf | nan | chr10:124133785-124133822 | 37 | | | | | | | | | 1 | | | | | | yes | | | | yes |
| ARMS2 | NA | TSS1500 | chr10:124213760-124214151 | 391 | | | | | | | | | 1 | | | | | yes | yes | | yes | | |
| HTRA1 | N_Shore;Island | TSS1500 | chr10:124220359-124220874 | 515 | | | 3 | | | | | 0 | 2 | 2 | | | | | | | | yes | yes |
| HTRA1 | NA | Body | chr10:124248943-124249010 | 67 | | | | | | | | | 1 | 1 | | | | | | | | | |
| DMBT1 | NA | TSS1500 | chr10:124319733-124320064 | 331 | | | 5 | | | | | | | | | | | | | | | | |
| DMBT1 | N_Shore | Body | chr10:124402673-124402857 | 184 | | | | | | | | | | | | | | yes | yes | | | | |
| C10orf120 | S_Shelf;NA | nan;Body | chr10:124459039-124459681 | 642 | | | | | | | | | | | | | | yes | yes | | | | |
| FAM24B;LOC399815 | N_Shore;Island | 5'UTR;TSS1500;TSS200;1stExon | chr10:124638756-124639221 | 465 | | | 5 | | | | 1 | 0 | 7 | 1 | | 8 | | | | | | | |
| C10orf88 | Island | TSS200 | chr10:124714098-124714384 | 286 | | | 3 | | | | | | | | | | | | | | | | |
| PSTK | N_Shore | TSS1500 | chr10:124739466-124739553 | 87 | | | 4 | | | | | 0 | | | | | | yes | | | | | |
| IKZF5;ACADSB | Island | 5'UTR;1stExon;TSS200 | chr10:124768267-124768276 | 9 | | | | | | | | 0 | | | | | | | | | | yes | |
| HMX3 | Island | Body | chr10:124896659-124897048 | 389 | | | | | | | | | 1 | | | | | | | | | | yes |
| HMX2 | N_Shore | TSS1500 | chr10:124906565-124906859 | 294 | | | | | | | | | | | | | | yes | | | | | |
| HMX2 | Island | 1stExon | chr10:124908427-124908747 | 320 | | | | | | | | | 1 | | | | | | | | | | |
| HMX2 | Island | 3'UTR | chr10:124909889-124910217 | 328 | | | 3 | | | | | 0 | 1 | | | | | | | | | | yes |
| GPR26 | N_Shelf;Island | nan;TSS1500 | chr10:125425262-125426048 | 786 | | 1 | | | | | | 0 | 3 | | | 1 | | | | | | yes | yes |
| CPXM2 | N_Shore;Island;S_Shore | Body;1stExon;5'UTR;TSS200 | chr10:125650957-125651726 | 769 | | | 13 | | | | | | 1 | | | 1 | | yes | | yes | | yes | yes |
| CHST15 | NA | nan | chr10:125767421-125768098 | 677 | | | | | | | | | | | | | | | | | | yes | |
| CHST15 | N_Shelf;N_Shore | Body | chr10:125803767-125804378 | 611 | | | 8 | 1 | | | | | | | | | | | | | | yes | |
| CHST15 | Island;S_Shore | Body;1stExon;5'UTR | chr10:125805493-125807047 | 1554 | | | 8 | | | | | | 2 | | | | | yes | | | | | |
| LHPP | NA | Body | chr10:126211704-126211892 | 188 | | | | | | | | | 1 | | | 1 | | | | | | | |
| LHPP | NA | nan | chr10:126222996-126223401 | 405 | | | | | | | | | | | 1 | | | yes | yes | | | | |
| LHPP | NA | Body | chr10:126262805-126262876 | 71 | | | 4 | | | | | | | | | | | yes | yes | | | | |
| LHPP | N_Shelf;N_Shore;Island | Body | chr10:126278272-126278502 | 230 | 1 | | 30 | | 1 | | | | 2 | 1 | | | | | | | | | |
| LHPP | S_Shelf | Body | chr10:126289835-126290192 | 357 | | | | | | | | | 1 | | | | | | yes | | yes | | |
| FAM53B | S_Shore;S_Shelf | nan;3'UTR | chr10:126308034-126308552 | 518 | | | 3 | | 1 | | | | 1 | | | | | yes | | | | | |
| FAM53B | NA | Body | chr10:126314680-126314798 | 118 | | | | | | | | | 1 | | | | | | | yes | | | |
| FAM53B | NA | Body | chr10:126330845-126331425 | 580 | | | 5 | | | | | | | | | | | yes | | | | | |
| CTBP2 | NA | Body | chr10:126703576-126704259 | 683 | | | 3 | 1 | | | | 0 | 2 | | | | 1 | | | | | | |
| CTBP2 | NA | Body | chr10:126712074-126712482 | 408 | | | 2 | | | | | | | | | | | | | | | | |
| CTBP2 | NA | 5'UTR | chr10:126751950-126752037 | 87 | | | 5 | | | | | | | | | | | yes | | | | | |
| CTBP2 | NA | 5'UTR | chr10:126782326-126782592 | 266 | | | 5 | | | | | | | | | | | | | | | | |
| LOC100169752 | NA | nan;TSS1500 | chr10:127262250-127262774 | 524 | | | 2 | | | | | | 1 | | | | | yes | | yes | yes | | |
| C10orf122 | NA | Body | chr10:127371542-127371728 | 186 | | | 1 | | | | | | | | | | | yes | yes | | | | |
| MMP21 | S_Shore | TSS200 | chr10:127464557-127464762 | 205 | | | 2 | | | | | | | | | | | | | | | | |
| DHX32 | NA | 5'UTR;1stExon | chr10:127569751-127569973 | 222 | | | | | | | | | 1 | | | 1 | | yes | yes | | | | yes |
| ADAM12 | Island | TSS200;Body;1stExon;5'UTR | chr10:128076910-128077376 | 466 | | | 8 | | | | | | 3 | | | | | | | | | | |
| C10orf90 | NA | Body | chr10:128152075-128152247 | 172 | | | 7 | | | | | 0 | 1 | | | | | yes | yes | | | | |
| C10orf90 | N_Shore | Body | chr10:128193219-128193400 | 181 | | | | | | | | | 1 | | | | | | | | | | |
| C10orf90 | NA | Body | chr10:128209980-128210076 | 96 | | | 2 | | | | | | | | | | | yes | | yes | | | |
| DOCK1 | Island | TSS1500;TSS200 | chr10:128593922-128594417 | 495 | | | | | | | | 0 | 6 | 2 | | 4 | | | yes | | | | yes |
| DOCK1 | NA | Body | chr10:128810484-128810904 | 420 | | | | | | | | 0 | 1 | | | | | | | | | yes | |
| FAM196A;DOCK1 | N_Shore;Island | 5'UTR;Body;1stExon;TSS200 | chr10:128993810-128995192 | 1382 | | | | | | | | 0 | 7 | 3 | | 5 | | | | | | yes | yes |
| DOCK1 | NA | Body | chr10:129144210-129144306 | 96 | | 1 | | | | | | | | | | | | yes | | | | | |
| DOCK1 | NA | Body | chr10:129245294-129245398 | 104 | | 1 | | | | | | | | | | | | | | | | | |
| FOXI2 | Island;N_Shore | TSS1500;TSS200;1stExon | chr10:129534182-129537308 | 3126 | | | | | | | | | | | | | | | | | | | |
| PTPRE | N_Shelf | nan | chr10:129704488-129704949 | 461 | | | | | | | | | | | | | | | yes | | | | |
| PTPRE | NA | 5'UTR | chr10:129785053-129785537 | 484 | | | 3 | | | | | | | | | | | | | | | | |
| PTPRE | NA | 5'UTR | chr10:129797760-129797840 | 80 | | | | 1 | | | | 0 | 1 | | | 6 | | | yes | | | | |
| PTPRE | NA | 5'UTR | chr10:129828720-129828798 | 78 | | | 6 | | | | | | | | | | | | | | | | yes |
| PTPRE | NA | Body;TSS1500 | chr10:129845660-129845903 | 243 | | | | 1 | | | | | | | 1 | | | | | | | | |
| PTPRE | NA | Body | chr10:129868087-129868324 | 237 | | | | | | | | | | | | | | | | | | | |
| MGMT | N_Shore;Island | TSS1500;TSS200;1stExon;5'UTR | chr10:131264786-131265796 | 1010 | | | 8 | | | | | 0 | 1 | | | | | | | yes | | | yes |
| MGMT | NA | nan;Body | chr10:131323903-131324189 | 286 | | | | | | | | | 2 | | 1 | | | | | | | | |
| MGMT | NA | Body | chr10:131355112-131355302 | 190 | | | | | | | | | 3 | | | | | | | yes | yes | | |
| MGMT | NA | Body | chr10:131367501-131367623 | 122 | | | | | | | | | 1 | | | | | yes | | | | | |
| MGMT | NA | Body | chr10:131412507-131412968 | 461 | | | 9 | | | | | | | | | | | | yes | | | | |
| MGMT | NA | Body | chr10:131505762-131506261 | 499 | | | | 1 | | | | | | | | | | yes | | | | | |
| EBF3 | S_Shore | Body | chr10:131640006-131640303 | 297 | | | | | | | | | | | | | | yes | | | | | |
| EBF3 | NA | Body | chr10:131647800-131648297 | 497 | | | | | | | | | | | | | | | | | | | |
| EBF3 | N_Shore | Body | chr10:131665635-131666017 | 382 | | | 6 | | | | | | | | | | | | | | | | |
| EBF3 | S_Shore | Body | chr10:131669406-131669630 | 224 | | | 21 | | | | 1 | 0 | 1 | 1 | 1 | 25 | 1 | | | | | | |
| EBF3 | NA | Body | chr10:131671373-131671776 | 403 | | | | | | | | | 1 | | | | | | | | | | |
| EBF3 | NA | Body | chr10:131682251-131682452 | 201 | | | 3 | | | | | | 1 | 1 | | | | | yes | | | | |
| EBF3 | S_Shelf | Body | chr10:131690763-131691217 | 454 | | | | | | | | | | | | | | | | | | | |
| EBF3 | N_Shore;Island;S_Shore | Body | chr10:131697080-131697980 | 900 | 6 | | 13 | | | | | 0 | 3 | | | 4 | | | | | | | |
| EBF3 | Island | Body | chr10:131757191-131758786 | 1595 | | | 14 | | | | | | 2 | | | | | | | | | yes | yes |
| EBF3 | Island;N_Shore | Body | chr10:131760778-131761660 | 882 | | | 18 | | | | | | | | | | | | | | | | yes |
| TCERG1L | NA;N_Shelf;N_Shore | nan;3'UTR;Body | chr10:132890839-132892510 | 1671 | | | 15 | | | | | | | | | | | | yes | | | | yes |
| TCERG1L | NA | Body | chr10:132909113-132909289 | 176 | | | | | | | | | | 1 | | 2 | | | | | | | |
| TCERG1L | NA | Body | chr10:132910868-132911152 | 284 | | | | 1 | | | | | | 1 | | 1 | | | | yes | | | |
| TCERG1L | NA | Body | chr10:132912784-132913233 | 449 | | | | | | | | | | | | | | | | yes | | | |
| TCERG1L | NA | Body | chr10:132977221-132977408 | 187 | 1 | | 3 | | | | | | | | | | | | | | | | |
| TCERG1L | NA | Body | chr10:132999647-133000471 | 824 | | | 22 | | | | | | | | | | | yes | yes | yes | yes | | yes |
| TCERG1L | NA | Body | chr10:133019987-133020227 | 240 | | | | | | | | | | | | | | | | | | | |
| TCERG1L | NA | Body | chr10:133036065-133036507 | 442 | | | 2 | | | | | | | | | | | | yes | | | | |
| TCERG1L | NA | Body | chr10:133038915-133039348 | 433 | | | 3 | | | | | | | | | | | | yes | | | yes | |
| TCERG1L | NA | Body | chr10:133051487-133051839 | 352 | | | | | | | | | | | | | | | yes | | | | |
| TCERG1L | N_Shore | Body | chr10:133058339-133058861 | 522 | | | 1 | | | | | | | | | | | | | | | yes | |
| TCERG1L | S_Shelf | Body | chr10:133087013-133087135 | 122 | | | | | | | | | | | | | | | yes | yes | | | |
| TCERG1L | N_Shore | Body | chr10:133109013-133109194 | 181 | | | | | | | | 0 | | | | | | | | | | yes | yes |
| PPP2R2D | S_Shelf | 5'UTR;Body | chr10:133758719-133758887 | 168 | 1 | | | | | | | | | | | | | yes | yes | | | | |
| BNIP3 | N_Shore;Island | Body | chr10:133793442-133793734 | 292 | | 1 | 2 | | | | | 0 | 4 | 1 | | 5 | | yes | | | | | |
| BNIP3 | Island | TSS1500 | chr10:133796164-133796494 | 330 | | | 3 | | | | | 0 | 1 | | | | | yes | | yes | | | yes |
| JAKMIP3 | NA | TSS1500 | chr10:133918102-133918267 | 165 | | | | | | | | | | | | | | | yes | | | | |
| JAKMIP3 | N_Shore | Body | chr10:133929429-133929519 | 90 | 1 | | 4 | | | | | | | | | | | | | | | | |
| JAKMIP3 | S_Shore | Body | chr10:133932254-133932445 | 191 | | | | | | | | | | | | | | | yes | | | | |
| JAKMIP3 | Island;S_Shore | Body | chr10:133938156-133938970 | 814 | | | 8 | | | | | | 1 | | | | | yes | yes | | | | |
| JAKMIP3 | N_Shelf;N_Shore | Body | chr10:133946674-133948362 | 1688 | 1 | | 9 | | | | | | | | | | | | | | yes | | |
| JAKMIP3 | Island;S_Shore;S_Shelf | Body | chr10:133955398-133957726 | 2328 | | | 4 | | | | | 0 | 2 | | | 1 | | yes | | | | | yes |
| JAKMIP3 | S_Shelf;NA | Body | chr10:133960386-133961306 | 920 | | | 4 | | | | | | 1 | | | | | | | | | | |

| | | | | | | | | | | | | | | | | | | | | | | | |
|---|---|---|---|---|---|---|---|---|---|---|---|---|---|---|---|---|---|---|---|---|---|---|---|
| JAKMIP3 | NA | 3'UTR | chr10:133981394-133982031 | 637 | | | | | | | | | | | | | | | | | | | yes |
| JAKMIP3 | NA | 3'UTR | chr10:133985911-133985960 | 49 | | 1 | 9 | | | | | | | | | | | yes | yes | | | | yes |
| JAKMIP3 | NA | 3'UTR | chr10:133995286-133995387 | 101 | | | 5 | | | | 1 | 0 | 1 | | | | | yes | | | | | |
| DPYSL4 | N_Shore;chr10:133998912-134 | 3'UTR;Island;TSS1500 | chr10:133998913-134000377 | 1464 | | 1 | | | | | | | 2 | | 1 | | | | | | | yes | yes |
| DPYSL4 | N_Shore | 3'UTR | chr10:134019250-134019775 | 525 | | | | | | | | | | | | | | | | | | | |
| STK32C | Island | Body | chr10:134041506-134041783 | 277 | | | | | | | | | 1 | | | | | yes | | yes | | yes | |
| STK32C | Island;S_Shore | Body | chr10:134043362-134043897 | 535 | 1 | | 8 | 1 | | | | 0 | 5 | 1 | | 7 | 1 | yes | | yes | | | |
| STK32C | S_Shelf;chr10:134040198-1340 | Body;S_Shelf | chr10:134045514-134046066 | 552 | | | | | 1 | | 2 | 0 | 4 | 4 | 3 | 17 | | | | yes | | | |
| STK32C | NA | Body | chr10:134053802-134055144 | 1342 | 3 | | 3 | | | | | | | | | | | yes | yes | | | | |
| STK32C | NA | Body | chr10:134061604-134062614 | 1010 | | | 13 | | | | | | 1 | | | | | | yes | | | | yes |
| STK32C | S_Shelf | Body | chr10:134079311-134079425 | 114 | | | 1 | | | | | | | | | | | | yes | | | | |
| STK32C | N_Shore;S_Shore | Body | chr10:134095321-134096880 | 1559 | | | 6 | | | | | | | | | | | | | | | | |
| STK32C | NA | Body | chr10:134109919-134110326 | 407 | | | | | | | | 0 | 1 | | | | | | yes | | yes | | |
| STK32C | NA | Body | chr10:134113312-134113511 | 199 | | | | | | | | | | | | | | | | yes | | | |
| STK32C | N_Shore | Body | chr10:134119433-134120029 | 596 | 1 | | 1 | | | | | | | | | | | | | | | | |
| LRRC27 | S_Shelf;NA | Body;TSS1500;TSS200 | chr10:134150451-134150760 | 309 | | | 5 | | 1 | | | 0 | 6 | 1 | | 14 | 1 | yes | | | | | |
| PWWP2B | S_Shore | Body | chr10:134211620-134211908 | 288 | | | 3 | | | | | | 1 | 1 | | | | | | | | | |
| PWWP2B | S_Shore;S_Shelf | 3'UTR;Body | chr10:134221503-134222507 | 1004 | | | 1 | | 1 | | | 0 | 3 | | | | | | | | | | |
| PWWP2B | NA | 3'UTR;Body | chr10:134226361-134226547 | 186 | | | 6 | | | | | | | | | | | | | | | | |
| C10orf91 | NA | TSS1500 | chr10:134258421-134258819 | 398 | | | | | | | | | | | | | | | | | yes | yes | |
| C10orf91 | NA | Body | chr10:134260981-134261480 | 499 | | | | | | | | | | | | | | | | | | | yes |
| INPP5A | S_Shore | TSS200 | chr10:134351165-134351197 | 32 | | 1 | | | | | | | | | | | | | | | | yes | yes |
| INPP5A | N_Shelf;N_Shore | Body | chr10:134358030-134358685 | 655 | | | 7 | 1 | 2 | | | | 2 | | 1 | | | | yes | | | | |
| INPP5A | Island;S_Shore | Body | chr10:134360912-134362170 | 1258 | | | | | | | | | 4 | | | 2 | | yes | yes | | | | |
| INPP5A | NA | Body | chr10:134377343-134377481 | 138 | | | 2 | | | | | | 1 | | | | | | | | | yes | |
| INPP5A | NA | Body | chr10:134399880-134400191 | 311 | | | 1 | | | | | | | | | | | | yes | | | | |
| INPP5A | NA | Body | chr10:134404873-134404945 | 72 | | | | | | | | | 1 | | | | | | yes | | | | |
| INPP5A | N_Shelf;Island | Body | chr10:134411295-134411480 | 185 | | | | | | | | | 2 | | | | | | yes | | | | |
| INPP5A | NA | Body | chr10:134420036-134420623 | 587 | 2 | | 1 | | | | | | | | | | | | yes | | | | |
| INPP5A | NA | Body | chr10:134455814-134456012 | 198 | | | 3 | | | | | | | | | | | | | | | | |
| INPP5A | NA | Body | chr10:134498830-134498960 | 130 | 1 | | | | | | | | | | | | | | | | | | yes |
| INPP5A | NA | Body | chr10:134511494-134511968 | 474 | | | | | | | | | | | | | | | | | | | |
| INPP5A | NA | Body | chr10:134527172-134528050 | 878 | | | | | | | | | 1 | | | | | | | | | yes | |
| INPP5A | Island;N_Shore | Body | chr10:134536030-134536498 | 468 | 1 | 1 | | | | | | | | | | | | | | | | | |
| INPP5A | N_Shore | Body | chr10:134548998-134549665 | 667 | | | | | | | | | | | | | | | | | | | |
| INPP5A | S_Shore | Body | chr10:134556463-134556533 | 70 | | | 1 | | | | | | | | | | | | | | | | |
| INPP5A | S_Shelf | Body | chr10:134559553-134561116 | 1563 | | | | | | | | | | | | | | yes | | | | | yes |
| INPP5A | N_Shore | Body | chr10:134566575-134566915 | 340 | | | | | | | | | | | | | | | | | | yes | |
| INPP5A | NA | Body | chr10:134578261-134578481 | 220 | | | 1 | | | | | | 1 | | | | | | | | | | |
| INPP5A | N_Shore | 3'UTR | chr10:134596873-134596989 | 116 | | | | | | | | | 1 | 1 | | | | | | | | | yes |
| NKX6-2 | Island | TSS1500;Body | chr10:134598927-134602163 | 3236 | | 5 | | | | | | 0 | 12 | 1 | | 1 | | | | | | yes | yes |
| C10orf93 | NA;N_Shelf | Body | chr10:134751598-134752472 | 874 | | | 4 | 1 | | | | | | | | | | | yes | | | | |
| C10orf93 | N_Shelf;N_Shore;Island | Body;1stExon;TSS200;TSS1500 | chr10:134754286-134756994 | 2708 | | 1 | 3 | | 1 | | 2 | 0 | 2 | | | | | yes | | | | yes | yes |
| GPR123 | Island | TSS200;5'UTR | chr10:134901279-134902278 | 999 | | 1 | | | | | | | 1 | 1 | | | | | | | | | yes |
| GPR123 | N_Shore | Body | chr10:134910285-134912411 | 2126 | | | 8 | | | | | | 1 | | | | | yes | | yes | | | |
| GPR123 | Island | Body | chr10:134913608-134913922 | 314 | | | | | | | | | 1 | | | | | | yes | | | | |
| GPR123 | S_Shore | Body | chr10:134915746-134916087 | 341 | | | | | | | | | | | | | | | | | | | |
| GPR123 | N_Shore | Body | chr10:134918671-134918974 | 303 | | | 4 | | | | | | | | | | | | | | | | |
| GPR123 | S_Shore | Body | chr10:134928138-134928457 | 319 | | | 1 | | | | | | | | | | | | | | | | |
| GPR123 | Island;N_Shore | Body | chr10:134937798-134938528 | 730 | | | 11 | | | | | | | | | | | yes | yes | yes | | | yes |
| GPR123 | Island | Body | chr10:134943246-134943309 | 63 | | | | | | | | 0 | | | | | | | | | | | |
| GPR123 | N_Shore | 3'UTR | chr10:134944512-134944737 | 225 | | | | | | | | | | | | | | | | | | yes | |
| KNDC1 | N_Shore | nan | chr10:134972886-134972969 | 83 | | | | | | | | | | | | | | | yes | | | | yes |
| KNDC1 | N_Shore;Island | Body | chr10:134977981-134978147 | 166 | | | 4 | | | | | | 2 | | | | | | yes | | | | |
| KNDC1 | S_Shore;S_Shelf | Body | chr10:134983383-134984426 | 1043 | | | | | | | | | | | | | | yes | yes | | yes | | |
| KNDC1 | N_Shore;S_Shore | Body | chr10:134992030-134994455 | 2425 | 1 | | | | | | | | | | | | | | yes | | | yes | |
| KNDC1 | Island;S_Shore;N_Shore | Body | chr10:134997074-134998521 | 1447 | 1 | | 5 | | | | | | 1 | | | | | | | | | yes | yes |
| KNDC1 | N_Shore | Body | chr10:135002718-135002976 | 258 | | | 1 | | | | | | | | | | | | | | yes | | yes |
| KNDC1 | Island | Body | chr10:135020822-135021100 | 278 | | 1 | | | | | | | 1 | 1 | | | | | | | | | |
| KNDC1 | N_Shore | Body | chr10:135029097-135029365 | 268 | | | | | | | | | | | | | | | | | | | |
| KNDC1 | S_Shore | Body | chr10:135030439-135030554 | 115 | | | | | | | | | | | | | | yes | yes | | | | |
| KNDC1 | N_Shore;S_Shore | Body | chr10:135032154-135033590 | 1436 | | | 1 | | | | | 0 | | | | | | | | | | | yes |
| KNDC1 | N_Shore | Body | chr10:135038107-135038446 | 339 | | | | | | 1 | | | | | | | | | | | | | |
| UTF1 | Island | TSS1500;1stExon | chr10:135043492-135045023 | 1531 | | 1 | | | | | | 0 | | | | | | | yes | | | yes | yes |
| VENTX | Island | nan;TSS1500 | chr10:135049991-135051270 | 1279 | | 3 | 1 | | | | | 0 | 10 | | | 1 | | | | | | | |
| MIR202 | NA | nan;Body;TSS200 | chr10:135061015-135061670 | 655 | | | 17 | | | | | | | | | | | | | | | | yes |
| ADAM8 | N_Shore | Body | chr10:135082008-135082125 | 117 | | | | | | | | | | | | | | | | | | | |
| ADAM8 | N_Shelf | Body | chr10:135087441-135087497 | 56 | | | | | 1 | | | | 1 | | | | | | | | | | yes |
| ADAM8 | Island | Body | chr10:135090255-135090565 | 310 | | | | 1 | 1 | | | 0 | | | | | | yes | | | | | yes |
| TUBGCP2 | Island | Body | chr10:135098662-135099129 | 467 | | | 1 | | | | | | | | | | | yes | | | | | |
| TUBGCP2 | NA | Body | chr10:135113191-135113384 | 193 | | | | | | | | | 1 | | | | | yes | | | | | |
| ZNF511;TUBGCP2 | Island | Body;TSS1500;5'UTR;1stExon | chr10:135122230-135123421 | 1191 | 2 | 1 | 2 | 1 | 2 | | | | 1 | 1 | | | | | | | | | yes |
| PRAP1 | NA | TSS1500;TSS200 | chr10:135160697-135161047 | 350 | | | 4 | | | | | 0 | 3 | | | 4 | | yes | | | | | |
| C10orf125 | Island | TSS200 | chr10:135171713-135172096 | 383 | | | 11 | | | | | | 1 | | | | | | | | | yes | |
| ECHS1 | NA | Body | chr10:135178626-135178849 | 223 | | | | | | | | | | 1 | | | | yes | | | | | |
| PAOX | N_Shore;Island | TSS1500 | chr10:135191333-135191798 | 465 | | | 2 | 2 | | | | 0 | 2 | | | | | | | | | | |
| PAOX | N_Shelf;N_Shore | Body;3'UTR | chr10:135202522-135203200 | 678 | | | | | 1 | 1 | | 0 | 5 | 1 | | 2 | | | yes | | yes | | |
| LOC619207 | N_Shore;Island | Body | chr10:135278717-135279147 | 430 | | | 17 | | | | | 0 | 5 | 1 | | 2 | 1 | | | | | | |
| CYP2E1 | N_Shelf;N_Shore;Island;S_Sho | nan;TSS1500;TSS200;Body;1stExo | chr10:135340445-135343280 | 2835 | 11 | 6 | 44 | | 2 | 4 | | 0 | 15 | 12 | 1 | 76 | 3 | | yes | | | | |
| SYCE1 | N_Shore;Island | Body;5'UTR;TSS200 | chr10:135379213-135379754 | 541 | | | 12 | | | | | 0 | 2 | | | | | | | | | yes | |
| FRG2B | S_Shore | nan | chr10:135440923-135441372 | 449 | | | | | | | | | | | | | | | | | | | yes |
| SCGB1C1;LOC653486 | NA | TSS200 | chr11:192996-193112 | 116 | | | | | | | | | | | | | | | | | | yes | |
| NLRP6 | N_Shore | TSS1500;TSS200 | chr11:278092-278450 | 358 | | | | | | | | | | | | | | | yes | yes | | | |
| ATHL1 | N_Shore | TSS1500 | chr11:287723-288305 | 582 | | | | | | | | | 1 | | | 5 | | | yes | yes | | | |
| ATHL1 | Island | 5'UTR;Body | chr11:289774-290447 | 673 | | | | | | | 1 | 0 | 2 | 1 | 1 | 6 | 1 | | | | | | yes |
| IFITM5 | S_Shore;Island | TSS1500;1stExon;5'UTR;TSS200 | chr11:299503-300772 | 1269 | | | 10 | | 1 | | | 0 | 2 | | | | | yes | yes | yes | | | |
| IFITM1 | S_Shore;N_Shelf;N_Shore | TSS1500;1stExon;5'UTR;Body | chr11:312841-316456 | 3615 | | | 15 | 2 | 4 | | | 0 | 6 | 1 | | | | yes | | yes | | | |
| B4GALNT4 | NA | TSS1500 | chr11:368366-368966 | 600 | | | 2 | | | | | 0 | | | | | 1 | yes | | yes | | yes | yes |
| B4GALNT4 | S_Shore | Body | chr11:381709-381875 | 166 | | | | | | | | | | | | | | | | | | | yes |
| PKP3 | N_Shore;Island | TSS1500;1stExon;5'UTR | chr11:392903-394545 | 1642 | | 1 | 9 | | | 1 | | | 3 | 1 | 1 | | | | | | | | |
| PKP3 | Island | Body | chr11:396686-397077 | 391 | | | 4 | | | | | 0 | 1 | 1 | 1 | 10 | | | | | | | |
| PKP3 | N_Shore | 3'UTR | chr11:404596-404762 | 166 | | | | | | | | | 1 | | | | | | | | | | |
| SIGIRR | N_Shore | 3'UTR;Body | chr11:405846-406901 | 1055 | | | 19 | 1 | | | | | 1 | 1 | | | | | | | | yes | yes |
| SIGIRR | N_Shore | 1stExon;5'UTR;TSS200;TSS1500 | chr11:414951-415457 | 506 | | | 9 | 1 | 2 | | | | 1 | 1 | 1 | | | | | | | yes | |
| ANO9 | N_Shore | Body | chr11:430036-430292 | 256 | | | | | | | | | 1 | | | | | | | | | yes | yes |
| ANO9 | NA | TSS200 | chr11:442189-442649 | 460 | | | | | | | | | | | | | | | | | | yes | |
| PTDSS2 | N_Shore | Body | chr11:457256-457304 | 48 | | | | | 1 | | | | 1 | | | | | yes | yes | | | | |
| PTDSS2 | Island | Body | chr11:460469-460914 | 445 | | | | | | | | | 2 | | | | | | | | | | |
| RNH1 | Island;N_Shore | 5'UTR | chr11:504301-504937 | 636 | | | | | | | | 0 | 5 | 1 | | | | | | yes | | | yes |

| | | | | | | | | | | | | | | | | | | | | | | | | | | | |
|---|---|---|---|---|---|---|---|---|---|---|---|---|---|---|---|---|---|---|---|---|---|---|---|---|---|---|---|
| LRRC56 | Island | 1stExon;5'UTR;TSS200 | chr11:537432-537737 | 305 | | | 5 | | | | | | | | | | | | | | | | | | | | |
| PHRF1;IRF7 | N_Shore;S_Shore | Body;3'UTR | chr11:611114-612837 | 1723 | | | | | | | | | | | | | | | | | | | | | | | |
| IRF7 | Island | 5'UTR;1stExon;TSS1500;TSS200 | chr11:615519-616112 | 593 | | | | | 2 | | | | | | | | | yes | | | | | yes | | | | |
| MUPCDH | S_Shore | Body | chr11:618993-619080 | 87 | | | | | | | | | 2 | | | | | | | | | | | | | | |
| MUPCDH | N_Shore;Island | Body;5'UTR;1stExon;TSS200;TSS1 | chr11:624457-626128 | 1671 | | | 12 | 1 | | | | | | | | | | yes | yes | | | | yes | | | | |
| SCT | Island;S_Shore | TSS1500 | chr11:627582-628677 | 1095 | | | 20 | | | | | 0 | 1 | | | 1 | | yes | yes | | | yes | yes | | | | |
| DRD4 | Island | Body;TSS1500;TSS200 | chr11:636659-638939 | 2280 | | 1 | 13 | | 4 | | | 0 | 7 | 1 | 1 | 6 | 1 | yes | | | | | yes | | | | |
| DRD4 | Island | Body | chr11:640328-640606 | 278 | | | | | | | | | | | | | | | | | | | yes | | | | |
| DEAF1 | NA | Body | chr11:655445-655611 | 166 | | | | 1 | | | | | | | | | | yes | | | | | | | | | |
| DEAF1 | NA | Body | chr11:674556-674687 | 131 | | 1 | | | | | | | | | | | | yes | | | | | | | | | |
| DEAF1 | NA | Body | chr11:679706-679841 | 135 | | | | 1 | 1 | | | | | | | | | yes | | | yes | | | | | | |
| DEAF1 | N_Shore | Body | chr11:692963-693032 | 69 | | | | | | | | | 1 | 1 | | | | yes | yes | | | | | | | | |
| EPS8L2 | N_Shore | TSS1500 | chr11:705692-705892 | 200 | | | | | | | | | | | | | | | | | | yes | yes | | | | |
| CEND1 | N_Shore | 3'UTR | chr11:787971-788714 | 743 | | | 2 | | | | | | | | | | | yes | | | | | | | | | |
| CEND1 | N_Shore | TSS200 | chr11:790293-790795 | 502 | | | 6 | | | | | | | | | | | | | | | yes | yes | | | | |
| SLC25A22 | N_Shelf | Body | chr11:794433-794867 | 434 | | | 1 | | 1 | | | | | | | | | | | | | | | | | | |
| RPLP2;SNORA52 | S_Shore | Body;TSS200 | chr11:811545-811791 | 246 | | | | | | | | | 1 | | | | | yes | | | | | | | | | |
| PNPLA2 | N_Shore | TSS1500;TSS200 | chr11:818466-818917 | 451 | | | 11 | | | 1 | | 0 | 3 | 1 | | | | yes | | | | | | | | | |
| EFCAB4A | Island | TSS200 | chr11:827488-827613 | 125 | | | | | | | | | | | | | | | | | | | yes | | | | |
| EFCAB4A | Island | Body | chr11:830190-830527 | 337 | | | | | 1 | | | | | | | | | | | | | | | | | | |
| POLR2L | N_Shore | 3'UTR | chr11:840367-840487 | 120 | | | 3 | | | | | | | | | | | yes | | | | | | | | | |
| TSPAN4;POLR2L | S_Shore | 5'UTR;TSS1500;1stExon;TSS200 | chr11:843897-844400 | 503 | 1 | | | | | | | | 2 | | | | | | yes | | | | | | | | |
| TSPAN4 | N_Shore | 5'UTR;Body | chr11:856942-857071 | 129 | | | 5 | | | | | | 1 | | | | | yes | | | | | | | | | |
| TSPAN4 | S_Shore | 5'UTR;Body | chr11:859532-860157 | 625 | | | 2 | | | | | | | | | | | | | | | yes | | | | | |
| TSPAN4 | NA | Body | chr11:865673-865988 | 315 | | | | | | | | | | | | | | | yes | | | | | | | | |
| CHID1 | S_Shore | Body | chr11:882636-882687 | 51 | | | 3 | | | | | | | | | | | yes | yes | | | | | | | | |
| CHID1 | N_Shore | Body | chr11:889148-889231 | 83 | 2 | | | | | | | | | | | | | | | | | | | | | | |
| CHID1 | S_Shelf | 1stExon;5'UTR;TSS200 | chr11:914849-915239 | 390 | | | 3 | | | | | | 1 | | | | | yes | | | | | | | | | |
| AP2A2 | NA | Body | chr11:968423-968956 | 533 | | | 1 | | | | | | | | | | | yes | | yes | | | yes | | | | |
| AP2A2 | NA | Body | chr11:970389-971527 | 1138 | | | | | | | | | | | | | | | yes | | | | | | | | |
| AP2A2 | S_Shore | 3'UTR | chr11:1010983-1011415 | 432 | | | | | 3 | | | | | | | | | | | | | | | | | | |
| MUC6 | Island | Body | chr11:1024747-1025969 | 1222 | | | 4 | | | | | | | | | | | | | | yes | | | | | | |
| MUC6 | S_Shore;N_Shore;Island | Body | chr11:1028291-1030347 | 2056 | | | 9 | | | | | | | | | | | | | | | yes | | | | | |
| MUC6 | Island;N_Shore | TSS200;Body;1stExon;5'UTR | chr11:1036307-1036938 | 631 | | | 20 | | | | | 0 | 3 | | | | | | | | | | | | | | |
| MUC2 | NA | TSS200 | chr11:1074682-1074873 | 191 | | | 5 | | | | | | | | | | | | | | | | | | | | |
| MUC2 | Island;S_Shore | Body | chr11:1083178-1084842 | 1664 | | 1 | | | | | | | 1 | | | | | | | | | | yes | | | | |
| MUC2 | S_Shelf | Body | chr11:1086413-1086863 | 450 | | | | | | | | | | | | | | | | | | yes | | | | | |
| MUC2 | Island;S_Shore | Body | chr11:1095156-1097783 | 2627 | | | | | | | | | 1 | | | | | | | | | | | | | | |
| MUC2 | N_Shelf | Body | chr11:1104049-1104492 | 443 | | | 1 | | | | | | | | | | | | | | | yes | | | | | |
| MUC5B | N_Shelf | TSS1500 | chr11:1243947-1244397 | 450 | | | 5 | | | | | | | | | | | | | | | | | | | | |
| MUC5B | S_Shore;Island | Body | chr11:1247948-1248971 | 1023 | | | 4 | | | | | | | | | | | | | | | | | | | | |
| MUC5B | N_Shore | Body | chr11:1274018-1274140 | 122 | | | | | | | | | | | | | | | | | | yes | | | | | |
| MUC5B | S_Shore | Body | chr11:1275741-1276661 | 920 | | | | | | | | | | | | | | | | | | | yes | | | | |
| TOLLIP | N_Shore | 3'UTR | chr11:1297250-1297558 | 308 | | | | | | | | 0 | | | | | | | | | | | | | | | |
| TOLLIP | N_Shore | Body | chr11:1312565-1313100 | 535 | | | | | | | | | 1 | | | | | | yes | | | | | | | | |
| TOLLIP | S_Shelf | Body | chr11:1319648-1321009 | 1361 | | 1 | 7 | | 1 | 1 | | 0 | 3 | 1 | | | | | yes | | | | | | | | yes |
| TOLLIP | N_Shelf;N_Shore;NA | Body | chr11:1325718-1328622 | 2904 | | | 2 | 1 | 1 | | | | 2 | | | | | yes | yes | yes | | | yes | | | | |
| LOC255512;TOLLIP | S_Shore | Body;TSS1500 | chr11:1331736-1331831 | 95 | | | | 1 | | | | | 1 | | | | | | yes | yes | | | | | | | |
| BRSK2 | S_Shore | Body | chr11:1413093-1413315 | 222 | | | | | | | 2 | 0 | 1 | 2 | 1 | 16 | | | | | | | | | | | |
| BRSK2 | NA | Body | chr11:1422319-1422804 | 485 | | | 2 | | | | | | | | | | | | | | | | | | | | |
| BRSK2 | NA | Body | chr11:1444768-1445041 | 273 | | | 1 | | | | | | | | | | | | | yes | | | | | | | |
| BRSK2 | NA | Body | chr11:1446247-1446386 | 139 | | | | | | | | | | | | | | | | yes | | | | | | | |
| BRSK2 | NA | Body | chr11:1456891-1457346 | 455 | | | | | | | 1 | 0 | 1 | 1 | | 4 | | | | | | | yes | | | | |
| BRSK2 | N_Shore;Island | Body | chr11:1463541-1464792 | 1251 | 8 | 1 | | | | | 2 | 0 | 3 | 2 | | 18 | | | | | | | yes | | | | |
| BRSK2 | Island | Body | chr11:1471410-1473279 | 1869 | | | | | | | | | | | | | | | | | | | | | | | |
| BRSK2 | S_Shelf;NA | Body | chr11:1474386-1475899 | 1513 | | | | | | | 1 | 0 | 2 | | | | | | | yes | | yes | | | | | |
| BRSK2 | Island;N_Shore | Body;3'UTR | chr11:1481779-1483973 | 2194 | | | 4 | | | | | | | | | | | | | | | yes | yes | | | | |
| HCCA2 | NA | Body | chr11:1506936-1507347 | 411 | | | | | | | | | 3 | | 1 | | | | | | | yes | | | | | |
| HCCA2 | NA | Body | chr11:1510644-1510747 | 103 | | | | | 1 | | | | | | | | | yes | yes | | | | | | | | |
| HCCA2 | NA | Body | chr11:1526031-1526270 | 239 | 1 | | | | | | | | | | | | | yes | | | | | | | | | |
| HCCA2 | N_Shore | Body | chr11:1536428-1536810 | 382 | | | | | | | | | 1 | | | | | yes | | | | | | | | | |
| HCCA2 | NA | Body | chr11:1542313-1542394 | 81 | | | 1 | | | | | | 1 | | | | | yes | yes | | | | | | | | |
| HCCA2 | S_Shore | Body | chr11:1569783-1569812 | 29 | | | | | | | | | 1 | | | | | | | | | | | | | | |
| HCCA2;KRTAP5-4 | NA | Body;TSS200 | chr11:1643498-1643842 | 344 | 2 | | | | | | | 0 | 1 | | | | | yes | | | | yes | | | | | |
| HCCA2;KRTAP5-5 | NA | Body;TSS1500 | chr11:1649947-1650595 | 648 | | | 2 | | | | | | | | | | | | | | yes | yes | | | | | |
| HCCA2 | NA | Body | chr11:1660513-1661201 | 688 | | | 2 | | | | | | 1 | | | | | yes | yes | | | | | | | | |
| HCCA2 | NA | Body | chr11:1669278-1669376 | 98 | | | | | | | | | | | | | | yes | | | | yes | | | | | |
| HCCA2 | N_Shore | Body | chr11:1673766-1673899 | 133 | | | 1 | | | | | | | | | | | | | yes | | | | | | | |
| HCCA2 | NA | Body | chr11:1681556-1682139 | 583 | | | | 1 | | | | | 1 | | | | | | yes | yes | | | yes | | | | |
| HCCA2;FAM99A | NA | Body;TSS1500 | chr11:1686117-1686642 | 525 | | | | | | | | | | | | | | | yes | | | | yes | | | | |
| HCCA2;FAM99B | NA | Body | chr11:1704507-1704707 | 200 | | | 1 | | | | | | | | | | | | yes | | | | yes | | | | |
| HCCA2;FAM99B | NA | Body;TSS200 | chr11:1707044-1707647 | 603 | | | | | | | | | | | | | | | | | | | yes | | | | |
| HCCA2 | Island | Body | chr11:1715569-1715850 | 281 | | | | | | | | | | | | | | yes | | | yes | | | | | | |
| HCCA2;KRTAP5-6 | S_Shore | Body;TSS1500 | chr11:1717919-1718086 | 167 | | | | | | | | | | | | | | | | yes | | yes | | | | | |
| HCCA2 | NA | Body | chr11:1750302-1750939 | 637 | | | 22 | | 1 | | | | | | | | | | yes | | | | | | | | |
| HCCA2;LOC402778 | Island | Body;5'UTR;1stExon;TSS200 | chr11:1769152-1769645 | 493 | | | 22 | | | | | 0 | 8 | | | 11 | | | | | | | | | | | |
| CTSD;HCCA2 | S_Shore | TSS1500;TSS200 | chr11:1785618-1785695 | 77 | | | 2 | | | | | | | | | | | yes | | yes | | yes | yes | | | | |
| SYT8 | NA | TSS1500;TSS200;1stExon;5'UTR | chr11:1854174-1856025 | 1851 | | 1 | 7 | 1 | 1 | | | | 5 | 1 | | | | yes | | | yes | yes | | | | | |
| SYT8 | NA | Body | chr11:1857442-1858135 | 693 | | | | | | | | | 2 | | | | | | | | | yes | | | | | |
| TNNI2 | NA | Body | chr11:1862020-1862477 | 457 | | | | | | | | | 1 | | | | | | | | | | yes | | | | |
| LSP1 | NA | TSS1500;TSS200;nan;5'UTR;1stExo | chr11:1873757-1874445 | 688 | | | 5 | | 1 | | | | 2 | 1 | 1 | | | | | | | | | | | | |
| LSP1 | N_Shore;Island | 5'UTR;TSS1500;Body;TSS200;1stE | chr11:1890901-1892608 | 1707 | | | 6 | | | | | | 2 | | | | | yes | | | | | yes | | | | |
| LSP1 | Island;N_Shore | Body | chr11:1902923-1903333 | 410 | | | 4 | | | | | | 1 | | | | | | | | | | | | | | |
| TNNT3 | NA | TSS200;TSS1500 | chr11:1940459-1940912 | 453 | | | 4 | | | | | | | | | | | | | | | | | | | | |
| TNNT3 | N_Shore | Body | chr11:1945564-1945918 | 354 | | 1 | | | | | | | 1 | 1 | | | | | | | | yes | | | | | |
| TNNT3 | Island | Body | chr11:1947743-1949113 | 1370 | | | 3 | | | | | 0 | 3 | | | 1 | | | | | | | | | | | |
| TNNT3 | S_Shore | Body | chr11:1952638-1952756 | 118 | | | | | | | | | 1 | 1 | | | | | yes | | | | yes | | | | |
| MRPL23 | Island | Body | chr11:1974132-1974332 | 200 | | | | 1 | | | | | | | | | | yes | | | | | | | | | |
| LOC100133545 | N_Shelf;N_Shore | TSS200;TSS1500 | chr11:2011216-2012411 | 1195 | | 1 | 6 | | | | | | 2 | | | | | | | | | yes | yes | | | | |
| H19;MIR675 | N_Shore;Island;S_Shore | Body;TSS200;TSS1500 | chr11:2016513-2020556 | 4043 | | | 29 | | | | | | 6 | | | | | | | | | | | | | | |
| INS-IGF2;IGF2 | N_Shelf | Body;3'UTR | chr11:2151698-2151725 | 27 | | | 21 | | | | | | | | | | | | | | | yes | | | | | |
| INS-IGF2;IGF2 | N_Shore;Island;S_Shore | Body;3'UTR | chr11:2153991-2154952 | 961 | | | | | | | | 0 | 1 | | | 1 | | | | | | | | | | | |
| MIR483;INS-IGF2;IGF2 | S_Shore;N_Shelf | TSS1500;Body | chr11:2156282-2156699 | 417 | | | | | | | | | | | | | | | | | | | | | | | |
| INS-IGF2;IGF2;IGF2AS | Island;S_Shore | Body;5'UTR;1stExon;TSS1500;TSS2 | chr11:2160129-2163513 | 3384 | | | | | | | | | | | | | | | | | | | | | | | |
| IGF2AS;INS-IGF2;IGF2 | Island | Body;5'UTR | chr11:2165656-2165961 | 305 | | | 1 | | | | | | | | | | | | | | | | | | | | |
| INS-IGF2;IGF2 | NA | Body;1stExon;5'UTR;TSS200;TSS1 | chr11:2170412-2172098 | 1686 | | | 14 | | | | | | 2 | | | | | | | | | | | | | | |
| INS-IGF2 | Island | Body | chr11:2176877-2177498 | 621 | | | 2 | 1 | | | | | | | | | | | | | | | | | | | |
| INS-IGF2;INS | NA | 5'UTR;Body;TSS200;TSS1500 | chr11:2182300-2183708 | 1408 | | 1 | 5 | | | | | 0 | 1 | | | | | | | | | | yes | | | | |
| TH | Island | Body | chr11:2187921-2188244 | 323 | | | 2 | | | | | | | | | | | | | | | | | | | | |

| | | | | | | | | | | | | | | | | | | | | | | | |
|---|---|---|---|---|---|---|---|---|---|---|---|---|---|---|---|---|---|---|---|---|---|---|---|
| TH | S_Shore | Body | chr11:2189713-2190111 | 398 | | | | | | | | | | | | | | | | | | yes | |
| TH | S_Shelf | Body | chr11:2191147-2191495 | 348 | | | 1 | | | | | | | | | | | | | | | | |
| ASCL2 | Island;S_Shore | 3'UTR;1stExon;5'UTR;TSS1500 | chr11:2290953-2293665 | 2712 | | | | | | | | | | | | | | | | | | | |
| C11orf21;TSPAN32 | NA | Body;TSS1500;nan;1stExon;TSS20 | chr11:2321770-2323938 | 2168 | | | 20 | | 4 | | | | 11 | 5 | 5 | | | | | | | yes | |
| CD81 | N_Shore;Island | TSS1500;1stExon;5'UTR | chr11:2397201-2398776 | 1575 | | 1 | | 1 | 2 | | | | | | | | | | yes | | yes | yes | yes |
| CD81 | N_Shore | Body | chr11:2405835-2407008 | 1173 | | | 4 | | | | | | 1 | | | | | | yes | | | yes | |
| TSSC4 | S_Shore | TSS1500;TSS200 | chr11:2422730-2423571 | 841 | | | 18 | | | | | | | | | | | yes | | | | | |
| TRPM5 | NA | Body | chr11:2431174-2431603 | 429 | | | | | | | | | | | | | | | yes | | | | |
| TRPM5 | N_Shore | Body | chr11:2441045-2441141 | 96 | | | 2 | | | | | 0 | | | | | | yes | | | | | |
| TRPM5 | S_Shelf | TSS200;TSS1500 | chr11:2444462-2445316 | 854 | | 1 | 12 | | | | | | 1 | | | | | | | yes | | | |
| KCNQ1 | N_Shore;Island | TSS1500;1stExon;Body | chr11:2464822-2466909 | 2087 | | 2 | 26 | | | | | 0 | 1 | | | | | | yes | | | yes | yes |
| KCNQ1 | S_Shelf;NA | Body | chr11:2470675-2471097 | 422 | | | 4 | | | | | | | | | | | | | | | | |
| KCNQ1 | NA | TSS1500;Body | chr11:2481545-2482531 | 986 | | | 18 | | 3 | | | | 2 | | | | | | | | | yes | |
| KCNQ1 | NA | Body | chr11:2548282-2548955 | 673 | | | | | 1 | | | | | | | | | | yes | | | | |
| KCNQ1 | N_Shore | Body | chr11:2554410-2555576 | 1166 | | 1 | 3 | | | | | | | | | | | | yes | | | yes | |
| KCNQ1 | NA | Body | chr11:2569380-2569788 | 408 | | | 1 | | | | | | | | | | | | yes | | | | |
| KCNQ1 | NA | Body | chr11:2573972-2574042 | 70 | | | | | | | | | | | | | | | yes | | | | |
| KCNQ1 | NA | Body | chr11:2584446-2584917 | 471 | | | | | | | | | | | | | | | yes | | | | |
| KCNQ1 | N_Shore | Body | chr11:2593857-2594099 | 242 | | | | | | | | | | | | | | | | | yes | | |
| KCNQ1 | N_Shore;Island | Body | chr11:2595605-2596606 | 1001 | | | | | | | | | 1 | | | | | | | yes | | | |
| KCNQ1 | N_Shore | Body | chr11:2602926-2603011 | 85 | | | | | | | | | | | | | | | | | | | |
| KCNQ1 | S_Shore | Body | chr11:2604365-2604499 | 134 | | | | | | | | | | | | | | | | | | | |
| KCNQ1 | NA | Body | chr11:2610026-2610741 | 715 | | | 2 | | | | | | | | | | | | yes | | | | |
| KCNQ1 | NA | Body | chr11:2612781-2612876 | 95 | | | | | | | | | | | | | | | | | | | |
| KCNQ1OT1;KCNQ1 | Island;S_Shore | TSS200;Body;TSS1500 | chr11:2721366-2722713 | 1347 | | | 17 | 1 | | | | 0 | 11 | | | | | yes | | | yes | | yes |
| KCNQ1 | NA | Body | chr11:2772759-2772852 | 93 | | 1 | | | | | | | | | | | | | | | | | |
| KCNQ1 | Island | Body | chr11:2812674-2813298 | 624 | | | 2 | | | | | 0 | 2 | 1 | | 1 | | | yes | | | | |
| KCNQ1 | N_Shore;Island | Body | chr11:2828214-2829241 | 1027 | | | 15 | | | | | | 5 | 1 | 1 | 1 | | yes | | | | | yes |
| KCNQ1 | NA | Body | chr11:2846681-2847258 | 577 | | 1 | 2 | 1 | | | | | 1 | | | | | | yes | | | | |
| KCNQ1DN | N_Shore;Island;S_Shore | TSS1500;TSS200;Body | chr11:2889809-2891495 | 1686 | | 2 | 62 | 1 | 3 | | | 0 | 15 | 2 | | 8 | 1 | | | | | yes | yes |
| CDKN1C | N_Shore | 3'UTR | chr11:2904591-2905372 | 781 | | | 6 | 1 | | | | | | | | | | | | | | yes | yes |
| CDKN1C | Island;S_Shore | TSS1500 | chr11:2907670-2908447 | 777 | | | 24 | 1 | | | | 0 | 1 | | | | | | | | | yes | yes |
| SLC22A18;SLC22A18AS | N_Shelf | TSS200;Body;TSS1500 | chr11:2919689-2920819 | 1130 | | | 7 | 1 | 1 | | | 0 | 5 | 3 | | | | | yes | | | | |
| SLC22A18AS;SLC22A18 | N_Shore | 5'UTR;TSS1500 | chr11:2922334-2922517 | 183 | | | 3 | | | | | | | | | | | yes | | yes | | | |
| SLC22A18AS;SLC22A18 | S_Shore | 5'UTR;Body;TSS1500 | chr11:2924523-2925969 | 1446 | | | 19 | 1 | 2 | | | | 4 | 1 | | | | yes | yes | | | | |
| SLC22A18 | Island | Body | chr11:2930322-2930995 | 673 | | | 5 | | | | | 0 | 2 | | | 4 | | | | | | | |
| PHLDA2 | Island;S_Shore | TSS1500 | chr11:2951168-2951844 | 676 | | | | | | | | 0 | 1 | | | | | yes | | | yes | yes | yes |
| NAP1L4 | S_Shore | TSS1500 | chr11:3014225-3014291 | 66 | | | | | | | | | | | | | | yes | | | | | |
| CARS | NA | Body;5'UTR | chr11:3073136-3073488 | 352 | | | 3 | | | | | 0 | 2 | | | 2 | | yes | yes | | | | yes |
| OSBPL5 | NA | Body | chr11:3122933-3122963 | 30 | | 1 | | | | | | | | | | | | yes | | | | | |
| OSBPL5 | NA | Body | chr11:3127422-3127663 | 241 | | | | | | | | | 1 | | | | | | yes | yes | | | |
| OSBPL5 | S_Shelf | Body | chr11:3144685-3144774 | 89 | | | 4 | | | | | | 1 | | | | | yes | | | yes | | |
| OSBPL5 | NA | 5'UTR | chr11:3164631-3164851 | 220 | | | 4 | | | | | | | | | | | | yes | | | yes | yes |
| OSBPL5 | NA | 5'UTR | chr11:3168788-3169027 | 239 | | | | | | | | | | | | | | | | | | | |
| OSBPL5 | NA | 5'UTR | chr11:3175007-3175636 | 629 | | | | | | | | 0 | 2 | 1 | | | | yes | yes | yes | | | |
| OSBPL5 | N_Shelf | 5'UTR | chr11:3177622-3178024 | 402 | | | 1 | | | | | | 2 | 1 | | | | yes | yes | | | | |
| OSBPL5 | Island | 5'UTR | chr11:3181822-3182119 | 297 | | | | | | | | | 1 | | | | | | | | | | |
| OSBPL5 | S_Shore | TSS1500 | chr11:3187271-3187803 | 532 | | | | | | | | | 4 | 1 | | | | | | | | | |
| MRGPRG;C11orf36 | Island | 1stExon;TSS1500 | chr11:3239218-3239509 | 291 | | | 1 | | | | | | | | | | | | | | | | |
| MRGPRE | Island | TSS1500;TSS200 | chr11:3253769-3254065 | 296 | | | 35 | | | | | | 3 | | | | | | | yes | | | |
| TRPC2 | NA | TSS1500;TSS200 | chr11:3647419-3648013 | 594 | | | | | 1 | | | | 1 | | | | | yes | yes | | yes | | yes |
| ART1 | S_Shelf | TSS200 | chr11:3666271-3666662 | 391 | | | | | | | | | | | | | | | yes | yes | | | |
| NUP98;PGAP2 | N_Shore | 5'UTR;TSS1500 | chr11:3817793-3817849 | 56 | | | | | | | | 0 | | | | | | yes | | | | | |
| OR52B4 | NA | 1stExon;TSS200;TSS1500 | chr11:4389408-4390263 | 855 | | | | | | | | | 1 | | | | | yes | | | | | |
| TRIM21 | NA | TSS1500 | chr11:4415193-4415245 | 52 | | | 5 | | | | | 0 | 1 | | | 1 | | | yes | | | | yes |
| C11orf40 | NA | TSS1500 | chr11:4599637-4600120 | 483 | | | 1 | | | | | | 2 | | | | | | yes | yes | | yes | |
| TRIM68 | N_Shore;Island;S_Shore | 5'UTR;1stExon;TSS200 | chr11:4628823-4629667 | 844 | | | 1 | | | | | 0 | 5 | | | | | | yes | | | yes | |
| OR51T1 | NA | TSS1500 | chr11:4902464-4902583 | 119 | | | | | | | | | | | | | | yes | | | | | |
| OR51A7 | NA | TSS200 | chr11:4928572-4928760 | 188 | | | | | | | | | | | | | | yes | | | | | |
| OR51G1 | NA | 1stExon | chr11:4945422-4945892 | 470 | | | | | | | | | 1 | | | | | yes | | | | | |
| MMP26 | NA | TSS1500 | chr11:5008936-5009311 | 375 | | | | | | | | | | | | | | yes | | | yes | yes | |
| HBB | NA | 1stExon | chr11:5248218-5248566 | 348 | | | | | | | | | | | | | | yes | | | | yes | |
| HBE1 | NA | 1stExon;5'UTR | chr11:5291214-5291692 | 478 | | | 4 | | | | | | | | 1 | | | | | | | | |
| OR51Q1 | NA | TSS1500 | chr11:5442434-5442511 | 77 | | 1 | | | | | | | 1 | | | 1 | | yes | yes | | | | |
| UBQLN3 | NA | TSS200 | chr11:5531235-5531524 | 289 | | | | | | | | | | | | | | | yes | | | | |
| TRIM6-TRIM34;TRIM6 | NA | TSS1500;TSS200;1stExon;5'UTR | chr11:5616902-5618408 | 1506 | 2 | 3 | 7 | | | | | 0 | 7 | | 1 | 27 | 1 | yes | | | | | yes |
| TRIM34;TRIM6-TRIM34 | NA | 5'UTR;TSS1500;Body | chr11:5653254-5653405 | 151 | | | | | 1 | | | | 1 | | | | | | yes | | | | |
| TRIM5 | NA | 5'UTR | chr11:5705995-5706435 | 440 | | | 2 | | | | | | | | | | | yes | | yes | | | |
| CCKBR | Island;N_Shore | Body | chr11:6290951-6292511 | 1560 | | | 21 | | 1 | | 1 | 0 | 9 | 2 | | 11 | | | yes | | | | |
| PRKCDBP | Island;S_Shore | 5'UTR;1stExon;TSS200;TSS1500 | chr11:6341717-6342510 | 793 | | 1 | 4 | 1 | 3 | 1 | 1 | 0 | | | | | | yes | | | | | |
| APBB1 | N_Shore;S_Shore | 5'UTR;TSS1500 | chr11:6439806-6440968 | 1162 | | | | 1 | | | | 0 | | | | | | yes | yes | | | | yes |
| DNHD1 | NA | TSS1500;TSS200;5'UTR;1stExon | chr11:6518108-6518907 | 799 | | | 11 | | | | | 0 | 1 | | | | | yes | yes | | | | |
| DNHD1 | Island | Body | chr11:6592066-6592745 | 679 | 1 | | | | | 1 | 1 | 0 | 2 | 2 | | 1 | 1 | | | | | | |
| DCHS1 | S_Shore | TSS1500 | chr11:6677919-6678143 | 224 | | | 6 | | | | | | | | | | | | yes | | yes | | yes |
| OR2AG2 | NA | TSS1500 | chr11:6790768-6790970 | 202 | | | | | | | | | | | | | | | yes | | | | yes |
| ZNF214;NLRP14 | Island | 5'UTR;TSS1500;1stExon;TSS200 | chr11:7041491-7041987 | 496 | | 1 | 2 | | | | | | | | | | | yes | | | | | |
| NLRP14 | NA | 5'UTR | chr11:7052239-7052637 | 398 | | | | | | | | | | | | | | | yes | | | | |
| RBMXL2 | N_Shore;Island | TSS200 | chr11:7110074-7111072 | 998 | | | 10 | | | | | 0 | 6 | 1 | | 2 | | | yes | | | | yes |
| SYT9 | N_Shore;Island | TSS1500;TSS200 | chr11:7272653-7273498 | 845 | | | | | | | | | 5 | | | | | yes | yes | | | yes | yes |
| PPFIBP2 | S_Shore | 5'UTR | chr11:7535887-7535983 | 96 | | | | | | | | | 1 | | | | | | | | | | |
| PPFIBP2 | NA | Body | chr11:7597349-7597983 | 634 | | | 6 | | | | | 0 | 1 | | | 1 | | | | | | | |
| CYB5R2 | Island | TSS1500 | chr11:7695245-7695528 | 283 | | | | 1 | | | | 0 | | | | | | | | yes | | | yes |
| NLRP10 | NA | TSS200 | chr11:7985166-7985764 | 598 | | | | | | | | | | | | | | yes | | | | | yes |
| TUB | NA | TSS200 | chr11:8060031-8060468 | 437 | | | | | | | | | | | | | | | | | | | |
| TUB | N_Shore | TSS1500;Body | chr11:8101793-8102510 | 717 | | | | 1 | | | | | 1 | | | | | yes | | | | yes | yes |
| STK33 | chr11:8615397-8615889;S_Sho | Island;TSS1500 | chr11:8615579-8616172 | 593 | | | 3 | | | | | | | | 2 | | | yes | yes | | | | yes |
| TRIM66 | NA | 5'UTR;1stExon;TSS200 | chr11:8679885-8680517 | 632 | | | 9 | | | | | | | | | | | | | | yes | | |
| ST5 | NA | 5'UTR;1stExon;TSS1500;TSS200 | chr11:8832111-8833095 | 984 | | | 6 | | | | | 0 | 1 | | | | 1 | yes | | | | yes | |
| C11orf16 | NA | 5'UTR;TSS200 | chr11:8954206-8954598 | 392 | | 1 | | | | | | | | | | | | yes | | | | | |
| ASCL3 | NA | 5'UTR;1stExon | chr11:8964505-8964967 | 462 | | | | | | | | | 2 | | | | | yes | | | | | yes |
| IPO7 | N_Shore | TSS1500 | chr11:9405796-9405884 | 88 | | | | | | | | 0 | | | | | | yes | yes | | | | |
| SWAP70 | N_Shore | TSS1500 | chr11:9684886-9685299 | 413 | | | 2 | | | | | | | | | | | yes | | | | | |
| ADM | N_Shore | TSS1500 | chr11:10325567-10326100 | 533 | | | 23 | | | | | | | | | | | yes | | | | | |
| AMPD3 | Island | TSS200 | chr11:10472149-10472161 | 12 | | | 2 | | | | | | | | | | | | yes | | | yes | yes |
| AMPD3 | S_Shelf;NA | TSS1500;Body;TSS200;1stExon;5'U | chr11:10476023-10477847 | 1824 | 5 | | 23 | 1 | | | | 0 | 5 | 3 | | | | | | | yes | | |
| LYVE1 | NA | 1stExon;5'UTR | chr11:10590159-10590587 | 428 | | | | | | | | | 1 | | | | | yes | | | | | |
| MRVI1 | NA | 5'UTR;1stExon | chr11:10673844-10673997 | 153 | | | 1 | | | | | | | | | | | yes | | | | | |
| MRVI1 | NA | 1stExon;5'UTR;TSS200;TSS1500 | chr11:10715175-10716164 | 989 | | | 25 | | | | | | 2 | | | | | yes | | | yes | | |
| GALNTL4 | N_Shore | Body | chr11:11642011-11642360 | 349 | | | | | | | | 0 | 1 | | | 1 | | | | | | yes | |

| | | | | | | | | | | | | | | | | | | | | | | | |
|---|---|---|---|---|---|---|---|---|---|---|---|---|---|---|---|---|---|---|---|---|---|---|---|
| USP47 | N_Shore | TSS1500 | chr11:11862470-11862879 | 409 | | | 7 | | | | | 0 | 3 | | | | | | | | yes | | |
| DKK3 | Island;S_Shore | Body;TSS200;5'UTR;TSS1500 | chr11:12029738-12031508 | 1770 | | | 29 | | | | | | 3 | | | 2 | | yes | yes | | | yes | |
| PARVA | Island | Body;TSS200 | chr11:12399109-12399973 | 864 | | | | | | | | | 2 | | | | | | yes | yes | | yes | yes |
| RASSF10 | N_Shore;Island | TSS1500;TSS200 | chr11:13030309-13030876 | 567 | | | | | 6 | | | | 1 | | | | | | | | | | |
| PTH | NA | TSS1500 | chr11:13518676-13519025 | 349 | | | | | | | | | 1 | | | | | yes | yes | | | | |
| FAR1 | Island | TSS1500 | chr11:13689686-13690194 | 508 | | | | | | | | 0 | | | | | | | yes | | | yes | yes |
| SPON1 | N_Shore | TSS1500;TSS200 | chr11:13983009-13984067 | 1058 | | | 9 | | 1 | | | 0 | 6 | 1 | | 3 | | yes | yes | | yes | yes | |
| SPON1 | Island | Body | chr11:13985911-13986348 | 437 | | | 13 | | | | | | | | | | | yes | | | | | |
| SPON1 | Island | Body | chr11:14280742-14281053 | 311 | | | | | 1 | | | | 3 | 2 | | | | | | | | | |
| CALCA | N_Shore;Island | 5'UTR;TSS200;TSS1500 | chr11:14993378-14995129 | 1751 | | | 20 | | | | | 0 | 6 | 6 | | 1 | | yes | | | | yes | yes |
| CALCB | Island | TSS200 | chr11:15095017-15095978 | 961 | | | 1 | | | | | | 1 | | | 2 | | yes | | | | | yes |
| INSC | Island | TSS200;Body;TSS1500 | chr11:15136150-15136505 | 355 | | | | | | | | | 2 | | | | | | | | | | yes |
| C11orf58 | S_Shore | Body;5'UTR | chr11:16761290-16761533 | 243 | | | 5 | | 1 | | | | | | | | | yes | | | | | |
| PLEKHA7 | NA | Body | chr11:16838596-16838675 | 79 | | 1 | | | | | | | | | | | | yes | | | | | |
| KCNJ11 | Island | TSS1500 | chr11:17411270-17411766 | 496 | | | 1 | | | | | | | | | | | | yes | | yes | yes | yes |
| ABCC8 | N_Shore | Body | chr11:17496264-17496725 | 461 | | | | | | | | | | | | | | yes | yes | | | | |
| ABCC8 | Island | TSS200 | chr11:17498517-17498541 | 24 | | | | | | | | | 1 | | | | | yes | | | | yes | |
| USH1C | NA | Body | chr11:17555142-17555319 | 177 | | | | | | | | | | | | | | | yes | | | | |
| USH1C | N_Shore;Island | Body;TSS200 | chr11:17565744-17566051 | 307 | | | | | | | | | 3 | | | | | yes | | | | | |
| MYOD1 | N_Shore;Island | TSS1500 | chr11:17740430-17741070 | 640 | | | | | | | | | 2 | | | | | yes | | | | | |
| MYOD1 | Island | 3'UTR | chr11:17743282-17743667 | 385 | | | | | | | | | 1 | | | | | | | | | | |
| KCNC1 | S_Shelf | Body | chr11:17760574-17760956 | 382 | | | 1 | | | | | | 1 | | | | | yes | | | | yes | |
| SAAL1 | S_Shore | TSS1500 | chr11:18127958-18128177 | 219 | | | 1 | | | | | | | | | | | yes | | | | | |
| MRGPRX4 | NA | TSS1500 | chr11:18193360-18193674 | 314 | | | | | | | | | | | | | | yes | yes | | | | |
| LOC494141 | Island;N_Shore | TSS200;TSS1500;Body | chr11:18230494-18231021 | 527 | | | 11 | 1 | | | | 0 | 3 | 1 | | 8 | | | yes | yes | | | |
| SAA4 | NA | 5'UTR | chr11:18258055-18258323 | 268 | | | | | | | | | | | | | | yes | | | | yes | |
| SAA2 | NA | TSS200 | chr11:18270324-18270548 | 224 | | | | | | | | 0 | | | | | | | | | | | |
| SAA1 | NA | TSS1500 | chr11:18287542-18287689 | 147 | | | | | | | | | | | | | | yes | | | | | |
| LDHC | NA | TSS1500;TSS200 | chr11:18433500-18434354 | 854 | 1 | 5 | 27 | 3 | 1 | | 1 | 0 | 16 | 6 | 1 | 16 | | yes | | | | | yes |
| LDHAL6A | N_Shore;Island | TSS200;1stExon;5'UTR;TSS1500 | chr11:18477153-18477753 | 600 | | 2 | 33 | | | | | 0 | 16 | 1 | 1 | 25 | | yes | | | | | |
| IGSF22 | Island | Body | chr11:18727321-18727774 | 453 | | | | | | | | 0 | 1 | 1 | | 4 | | | | | yes | | |
| IGSF22 | S_Shelf;NA | 5'UTR;1stExon;TSS200 | chr11:18747315-18748167 | 852 | | | 6 | | | | | | | | | | | yes | | | | | |
| CSRP3 | NA | 1stExon;5'UTR;TSS200 | chr11:19223517-19224117 | 600 | | | | | | | | | 1 | | | | | yes | | | | | |
| NAV2 | S_Shelf | TSS1500 | chr11:19371800-19372014 | 214 | | | | | | | | | 1 | | | | | | | | | yes | |
| LOC100126784;NAV2 | S_Shore | TSS200;Body;TSS1500 | chr11:19736150-19737237 | 1087 | | | | | | 1 | | 0 | 4 | | | | | | | | | yes | |
| NAV2 | NA | Body;TSS200;TSS1500 | chr11:20044428-20044639 | 211 | | | | 1 | 1 | | | | 1 | | | | | | | | | | |
| DBX1 | Island | Body | chr11:20178040-20178496 | 456 | | | | | | | | | 1 | | | | | yes | | | | | |
| DBX1 | N_Shore;Island | Body;1stExon;TSS200 | chr11:20180437-20182021 | 1584 | | 1 | | | 2 | | | | 4 | | | 1 | | | | | | | |
| HTATIP2 | N_Shore | TSS1500 | chr11:20384886-20385172 | 286 | | | 1 | | | | | | | | | | | | | | | yes | |
| SLC6A5 | N_Shore;S_Shore | Body;TSS1500;TSS200 | chr11:20620002-20622198 | 2196 | | | | | | | | | 6 | | | | | yes | | | | yes | |
| SLC6A5 | S_Shelf | Body | chr11:20627327-20627781 | 454 | | | | | | | | | | | | | | | yes | | | | |
| NELL1 | Island | TSS1500;TSS200 | chr11:20690682-20690957 | 275 | | | | | | | | | 2 | | | | | | | | | | yes |
| NELL1 | S_Shore | Body | chr11:20692709-20692867 | 158 | | 1 | 8 | | | | | | 1 | | | | | | yes | | | yes | |
| ANO5 | Island | TSS200;5'UTR;1stExon | chr11:22214554-22215306 | 752 | | | | | | | | | 2 | | | | | | | | | yes | |
| GAS2 | N_Shore | TSS1500 | chr11:22687589-22687660 | 71 | | | | | | | | | | | | | | | | | | | |
| GAS2 | NA | 5'UTR;TSS1500 | chr11:22696087-22696308 | 221 | | | | | 1 | | | | 1 | | | | | yes | | | | yes | |
| SVIP | S_Shore | TSS200 | chr11:22851416-22851502 | 86 | | | | | | | | | 1 | | | | | | | | | yes | |
| LUZP2 | Island | TSS200 | chr11:24518367-24518737 | 370 | | | | | | | | | 1 | | | | | | | | | | |
| ANO3 | N_Shore;Island | TSS200;1stExon;5'UTR | chr11:26353484-26354057 | 573 | | | 1 | | | | | 0 | 4 | | | 1 | | yes | | | | | yes |
| SLC5A12 | NA | 5'UTR;1stExon | chr11:26743403-26743888 | 485 | | | | | | | | | | | | | | yes | | | | | yes |
| FIBIN | NA | 5'UTR;1stExon;TSS200 | chr11:27015519-27017005 | 1486 | 1 | | 6 | | | | | 0 | 3 | | | 4 | | | | | | | |
| BBOX1 | NA | TSS1500 | chr11:27061533-27061899 | 366 | | | | | | | | | | | | | | | yes | | | | |
| BDNFOS;LIN7C | N_Shore;Island | TSS1500;Body;TSS200 | chr11:27527687-27528592 | 905 | | | 2 | | | | | 0 | | | | | | yes | | | | | |
| BDNF | N_Shore | Body;5'UTR;TSS200 | chr11:27721270-27721350 | 80 | | | | | | | | | 1 | | | | | | | | | | |
| BDNF | Island | Body;5'UTR | chr11:27740813-27741077 | 264 | | | | | | | | 0 | 1 | | | 1 | | | | | | | yes |
| BDNF | N_Shore | Body;TSS1500;TSS200 | chr11:27742435-27742832 | 397 | | | | | | | | | 1 | | | | | | | | | | yes |
| KCNA4 | N_Shelf | 5'UTR | chr11:30034396-30034490 | 94 | | | | | | | | | 1 | | | | | | | | | | |
| KCNA4 | Island;N_Shore | 5'UTR;1stExon;TSS200;TSS1500 | chr11:30037833-30039113 | 1280 | | 1 | | | 1 | | | 0 | 10 | 2 | 1 | 2 | | | | yes | | yes | yes |
| C11orf46 | N_Shore | TSS1500 | chr11:30344200-30344399 | 199 | | | | | | | | | 1 | | | | | | yes | | | | |
| MPPED2 | N_Shelf | TSS1500;5'UTR | chr11:30602788-30603030 | 242 | | | | | | | | | 1 | | | | | | | | | | |
| MPPED2 | N_Shore;Island | 5'UTR | chr11:30605411-30606165 | 754 | | 1 | 2 | | | | | | 1 | | | | | | | | | yes | |
| ELP4;IMMP1L | Island | 1stExon;TSS1500 | chr11:31531492-31531759 | 267 | | | | | | | 1 | | | 1 | | | | | | | | | |
| PAX6 | N_Shelf | Body | chr11:31817810-31817881 | 71 | | | | | | | | 0 | | | | | | | | | | | |
| PAX6 | Island;S_Shore;N_Shore | Body | chr11:31819148-31828715 | 9567 | | | | | | | | | | | | | | | | | | | |
| PAX6 | Island | 5'UTR;1stExon;TSS200 | chr11:31832710-31833016 | 306 | | | | | | | | | 1 | | | | | | | | | | yes |
| PAX6 | Island | 5'UTR | chr11:31835929-31837237 | 1308 | 1 | | 23 | | | | | | | | | | | | | | | yes | |
| PAX6 | Island;N_Shore | TSS200;TSS1500 | chr11:31839521-31840862 | 1341 | | 1 | 3 | | | | | | 2 | | | | | | | | | yes | yes |
| WT1 | NA | Body | chr11:32421514-32421845 | 331 | | | 6 | | 1 | | | | 3 | 1 | | | | yes | | | | | |
| WT1 | N_Shore;Island;S_Shore | Body | chr11:32447944-32450692 | 2748 | | 2 | 4 | | | | | 0 | 2 | | | | | | | | | | |
| WT1 | N_Shore;Island | Body | chr11:32452038-32452839 | 801 | | 1 | | | | | | | 1 | | | | | yes | | | | yes | |
| WT1 | N_Shore | Body | chr11:32454718-32454768 | 50 | | | 1 | | | | | | 1 | | | | | | | | | | |
| WT1;WIT1 | Island | TSS200;1stExon;TSS1500 | chr11:32456340-32457162 | 822 | | 2 | | | | | | | 1 | | | | | | | | | yes | |
| WIT1 | N_Shore;Island | Body | chr11:32460375-32461240 | 865 | | | | | | | | | 3 | | | | | yes | | | | yes | |
| C11orf41 | NA | TSS1500 | chr11:33563060-33563720 | 660 | | | | | | | | | | | | | | | yes | | | | |
| C11orf41 | Island | Body | chr11:33680273-33680427 | 154 | | | | | | | | | 1 | | | | | | yes | | | | |
| CD59 | S_Shore | TSS200;TSS1500 | chr11:33758214-33758529 | 315 | | | | | | | | 0 | 1 | | | | | yes | | yes | | yes | |
| LMO2 | NA | 5'UTR;1stExon | chr11:33913187-33914088 | 901 | | | | | | | | | 3 | | | | | yes | | | | yes | |
| ABTB2 | N_Shore | Body | chr11:34195914-34196220 | 306 | | | | | | | | | 1 | 1 | | | | yes | | | | | |
| CAT | N_Shore;Island | TSS1500;TSS200;1stExon;5'UTR | chr11:34460107-34460789 | 682 | | | 4 | | | | | 0 | 4 | 4 | 1 | 5 | | | yes | | | | |
| ELF5 | NA | 5'UTR;TSS200 | chr11:34535085-34535579 | 494 | | 1 | | | 2 | | | | 1 | | | | | yes | | | | | |
| CD44 | N_Shore | TSS1500 | chr11:35159087-35159382 | 295 | | | | | | | | | 1 | | | | | yes | yes | | | yes | |
| SLC1A2 | N_Shore;chr11:35440375-3544 | Body;Island;TSS1500 | chr11:35439994-35442225 | 2231 | | | 8 | | | | | | 1 | 1 | 1 | | | yes | | | | | yes |
| PAMR1 | Island | TSS200 | chr11:35547311-35547519 | 208 | | | | | | | | | 1 | | | | | | | | | | |
| PRR5L | NA | 5'UTR;TSS200;Body;nan | chr11:36422377-36422615 | 238 | | | | | 1 | | | | 1 | | 1 | | | yes | yes | | | | |
| RAG2;C11orf74 | NA | 5'UTR;TSS200 | chr11:36615998-36616246 | 248 | | | | | | | | | 1 | | | | | | | | | | |
| LRRC4C | NA | 5'UTR;1stExon | chr11:40314618-40315861 | 1243 | | 1 | | | 1 | | | | 3 | | | | | yes | | | | yes | yes |
| MIR670 | NA | TSS200 | chr11:43581108-43581175 | 67 | | | | | | | | | 1 | | | | | | | | yes | | |
| MIR129-2 | chr11:43602545-43603215;Isla | Island;TSS200 | chr11:43602845-43602965 | 120 | | | | | 3 | | | | | | 1 | | | | | | | | |
| ALKBH3 | N_Shore | TSS1500 | chr11:43902134-43902557 | 423 | | | | | | | | 0 | 1 | | | | | yes | | | yes | | |
| ALX4 | Island;N_Shore | Body | chr11:44324976-44327869 | 2893 | | | 18 | | | | | | | | | 1 | | | yes | | | | |
| ALX4 | S_Shore;N_Shore;Island | TSS1500 | chr11:44332195-44335843 | 3648 | | 2 | 103 | | | | | 0 | 5 | | 1 | | | yes | yes | | | | yes |
| CD82 | Island | TSS200 | chr11:44587044-44587138 | 94 | | | | | | | | | 1 | | | | | yes | | | | | |
| TSPAN18 | NA | 5'UTR;TSS1500 | chr11:44926927-44927222 | 295 | | | | | | | | | 1 | | | | | yes | yes | | | | yes |
| TP53I11 | Island;S_Shore | TSS200 | chr11:44972627-44972890 | 218 | | | 8 | 1 | | | | | | | | | | yes | | | | yes | |
| LOC221122 | NA | TSS1500 | chr11:44994308-44994687 | 379 | 1 | | | | | | | | | | | | | yes | | yes | | | |
| PRDM11 | N_Shelf | Body | chr11:45230723-45231545 | 822 | 3 | | | | | | | | | | | | | yes | yes | | yes | | yes |
| SLC35C1 | N_Shore | TSS1500;TSS200 | chr11:45825356-45825518 | 162 | | | 18 | | | | | | 2 | | | | | yes | yes | | | | |
| SLC35C1 | S_Shore | Body;1stExon | chr11:45827475-45828080 | 605 | | | 4 | | | | | | | | | | | yes | | | | | yes |
| CRY2 | N_Shore | TSS1500 | chr11:45868030-45868501 | 471 | | | | | | | | 0 | 3 | | | | | yes | yes | | | yes | |

| | | | | | | | | | | | | | | | | | | | | | | | | | | | |
|---|---|---|---|---|---|---|---|---|---|---|---|---|---|---|---|---|---|---|---|---|---|---|---|---|---|---|---|
| C11orf94 | S_Shelf;NA | Body;5'UTR;1stExon;TSS200;TSS15 | chr11:45928370-45929541 | 1171 | | | 7 | | | | | | 1 | 1 | | | | yes | yes | | yes | | | | | | |
| CREB3L1 | Island | Body | chr11:46317036-46317577 | 541 | | | | | 1 | | | | | | | | | | | | | | | | | | |
| DGKZ | Island | 1stExon | chr11:46354997-46355283 | 286 | | | 1 | | | | | | | | | | | | | | | | yes | | | | |
| DGKZ | N_Shore;Island | TSS1500;Body;TSS200 | chr11:46366643-46367100 | 457 | | | | | 1 | | | | 4 | | | | | yes | | | | | | | | | |
| DGKZ | NA | Body;TSS1500;TSS200;nan | chr11:46382544-46383924 | 1380 | | | 4 | 1 | 2 | | | | 2 | | 1 | | | yes | | | | | | | | | |
| CKAP5 | NA | 5'UTR | chr11:46848312-46848442 | 130 | | | | | | | | | | | | 1 | | | | | | | | | | | |
| LRP4 | S_Shore | TSS1500 | chr11:46940434-46940517 | 83 | 1 | | | | | | | | | | | | | | | | | | yes | | | | |
| DDB2 | N_Shore | TSS1500 | chr11:47235900-47236131 | 231 | | | | | | | | 0 | 1 | | | | | yes | | | | | | | | | |
| NR1H3 | NA | 5'UTR;TSS1500;TSS200 | chr11:47278977-47279586 | 609 | 1 | | | | | | | | 2 | | | | | yes | yes | | | | | | | | |
| MADD;NR1H3 | N_Shore | TSS1500;Body | chr11:47290188-47290613 | 425 | | | | | | | | | | | | | | | | | | | | | | | |
| SPI1 | Island | 3'UTR | chr11:47376754-47377212 | 458 | | | | | 1 | | | | | | | | | | | | | yes | yes | | | | |
| SPI1 | N_Shore;Island | Body;1stExon;5'UTR | chr11:47399213-47400146 | 933 | | | | | 1 | | | 0 | 3 | 2 | | | | | yes | | | yes | | | | | |
| RAPSN | NA | TSS200;1stExon;5'UTR | chr11:47470674-47470803 | 129 | | | 7 | | 1 | | | | 1 | | | | | yes | | | | | | | | | |
| CUGBP1 | S_Shore | TSS1500 | chr11:47574990-47575283 | 293 | | | | | | | | | | | | | | | yes | | | yes | yes | | | | |
| FAM180B | N_Shelf | TSS1500;TSS200 | chr11:47607809-47608310 | 501 | | | | | | | | | | | | | | yes | | | | | yes | | | | |
| FOLH1 | Island | Body | chr11:49229833-49230168 | 335 | | | | | | | | | 1 | | | | | yes | | | | yes | | | | | |
| LOC440040 | NA | TSS200 | chr11:49579997-49580074 | 77 | | 1 | | | | | | 0 | 1 | | | | | | | | | yes | | | | | |
| LOC441601 | Island | Body;TSS200 | chr11:50257256-50257979 | 723 | 1 | | 11 | | | | | 0 | | | | 1 | | yes | | | | | | | | | |
| OR4A15 | NA | TSS200 | chr11:55135222-55135656 | 434 | | | | | | | | | | | | | | | yes | | | | | | | | |
| OR10AG1 | NA | TSS1500 | chr11:55736968-55737072 | 104 | | | | | | | | | 1 | | | | | | yes | | | | | | | | |
| OR8H2 | NA | TSS200 | chr11:55872467-55872882 | 415 | | | | | | | | | 1 | | | | | yes | | | | | | | | | |
| OR8H3 | NA | TSS200 | chr11:55889658-55890274 | 616 | | | | | | | | | | | | | | yes | | | | | | | | | |
| OR5M1;OR8U8 | NA | 1stExon;Body | chr11:56380614-56381124 | 510 | | | | | | | | | 1 | | | | | | | | | | | | | | |
| APLNR | NA | TSS1500 | chr11:57005876-57006273 | 397 | | | | | | | | | | | | | | yes | | | yes | | | | | | |
| TNKS1BP1 | N_Shelf | 5'UTR | chr11:57089366-57089744 | 378 | | | | | 1 | | | | 1 | 1 | | | | yes | | | | | yes | | | | |
| TNKS1BP1 | S_Shore | TSS1500 | chr11:57093322-57093618 | 296 | | | 4 | | | | | 0 | 1 | | | | | | | | | | | | | | |
| P2RX3 | S_Shelf | TSS1500 | chr11:57105500-57105814 | 314 | | | | | | | | | | | | | | | yes | | | | | | | | |
| SLC43A3 | N_Shelf | Body | chr11:57192251-57194120 | 1869 | | | | | | | | | | | | | | yes | | | | | | | | | |
| RTN4RL2 | Island | Body | chr11:57232331-57232742 | 411 | | | 3 | | | | | | | | | | | yes | | | | | | | | | |
| SLC43A1 | Island | Body | chr11:57267101-57267445 | 344 | | | | | | | | 0 | | | | | | yes | yes | | | | | | | | |
| SERPING1 | NA | TSS1500;TSS200;1stExon;5'UTR | chr11:57364890-57365948 | 1058 | 1 | | | | | | | | 1 | | | | | yes | | | | yes | yes | | | | |
| YPEL4 | Island | Body | chr11:57414401-57414908 | 507 | | | | | | | | | 1 | | | | | | | | | | | | | | |
| CLP1 | N_Shore | TSS1500 | chr11:57424284-57424513 | 229 | | | | | | | | | | | | | | yes | yes | | | | | | | | |
| TMX2;C11orf31 | N_Shore | 3'UTR;Body;TSS1500 | chr11:57508416-57508545 | 129 | | | 18 | | | | | 0 | | | | 1 | | | | | | | | | | | |
| BTBD18 | N_Shore | TSS200 | chr11:57519282-57519584 | 302 | | | 4 | | 2 | | | | 3 | | | | | yes | | | | | | | | | |
| CTNND1 | NA | TSS1500;5'UTR;1stExon | chr11:57528838-57548688 | 19850 | | | | | | | | | 4 | 1 | | | | yes | yes | | yes | yes | yes | | | | yes |
| OR9Q2 | NA | TSS1500;5'UTR;1stExon | chr11:57957575-57958248 | 673 | | | 3 | | | | | | | | | | | yes | | | | | | | | | |
| OR10Q1 | NA | 1stExon | chr11:57995971-57996371 | 400 | | | | | | | | | | | | | | yes | | | | | | | | | |
| FAM111B | Island;S_Shore | TSS1500;TSS200 | chr11:58873955-58874733 | 778 | | | | | | | | 0 | | | | | | | | | | | | | | | |
| DTX4 | Island | Body | chr11:58940964-58941415 | 451 | | | | | | | | 0 | | | | | | | | | yes | | | | | | |
| MPEG1 | NA | TSS200;1stExon | chr11:58980157-58981095 | 938 | | | 9 | | | | | | 1 | | 1 | | | | | | | | | | | | |
| OR5A2 | NA | TSS1500 | chr11:59191289-59191832 | 543 | | | | | | | | | 1 | | | | | yes | yes | | | yes | | | | | |
| MRPL16 | S_Shore | TSS200 | chr11:59578490-59578540 | 50 | | | 1 | | | | | | | | | | | yes | | yes | | | | | | | |
| TCN1 | NA | TSS200 | chr11:59634191-59634276 | 85 | | | | | | | | | 1 | | | | | | yes | | | | | | | | |
| PLAC1L | NA | TSS200;TSS1500 | chr11:59806898-59808049 | 1151 | | | 6 | | | | | | 4 | | 1 | | | | | | | | | | | | |
| MS4A3 | NA | TSS200;1stExon;5'UTR | chr11:59823993-59824541 | 548 | | | 6 | | | | | 0 | 10 | 4 | 1 | 5 | | yes | | | | yes | | | | | |
| MS4A6A | NA | TSS1500 | chr11:59951557-59952153 | 596 | | | | | | | | | 2 | | | | | yes | | | yes | | yes | | | | |
| MS4A4A | NA | Body;TSS1500;TSS200 | chr11:60048004-60049475 | 1471 | | | | | | | | | | | | | | yes | | | yes | | | | | | |
| MS4A13 | N_Shore;Island | TSS1500;TSS200;5'UTR;1stExon | chr11:60282420-60282972 | 552 | | | 9 | | | | | | | | | | | | | | | | | | | | |
| C11orf64 | NA | TSS1500;TSS200 | chr11:60382707-60383361 | 654 | | | | | | | | | 1 | | | | | yes | yes | | | | | | | | |
| C11orf64 | N_Shore;Island | Body | chr11:60414615-60415111 | 496 | | 1 | 12 | | | 1 | | | 2 | | | | | yes | yes | | | | | | | | |
| MS4A15 | NA | TSS1500;TSS200 | chr11:60524105-60524575 | 470 | | | | | | | | | | | | | | yes | | | | | | | | | |
| MS4A15 | Island | 5'UTR;Body | chr11:60534911-60535066 | 155 | | | 12 | | | | | | 1 | | | 1 | | yes | | | | | | | | | |
| GPR44 | S_Shelf | 5'UTR;TSS200;TSS1500 | chr11:60623144-60624162 | 1018 | | 1 | 3 | | | | | | | | | | | yes | | | | | | | | | |
| ZP1 | NA | TSS1500;TSS200 | chr11:60634606-60635029 | 423 | 1 | | | | | | | | 2 | | | | | yes | yes | | yes | | | | | | |
| TMEM109 | N_Shore | TSS1500 | chr11:60680006-60680515 | 509 | | | 3 | | | | | | | | | | | yes | | yes | | | yes | | | | |
| SLC15A3 | Island | 1stExon;Body | chr11:60718568-60718882 | 314 | | | | | | | | | | | | 1 | | yes | | | | | yes | | | | |
| SLC15A3 | S_Shore | TSS1500 | chr11:60719947-60720229 | 282 | | | 7 | | | | | | | | | | | yes | yes | | | | | | | | |
| CD6 | NA | TSS200;5'UTR;1stExon | chr11:60738971-60739183 | 212 | | | 4 | | 1 | | | | 3 | 2 | 1 | | | | yes | | | | | | | | |
| CD6 | Island | Body | chr11:60775715-60776172 | 457 | | | | | | | | | 3 | | | 3 | | | yes | | | | | | | | |
| CD5 | NA | TSS1500;TSS200 | chr11:60869601-60869969 | 368 | | | 1 | | 2 | | | | 1 | | | | | | yes | | | yes | | | | | |
| VWCE | NA | Body | chr11:61049230-61049520 | 290 | | | | | | | | 0 | 1 | | | | | | | | | yes | | | | | |
| VWCE | chr11:61061751-61063063;Isla | Island;TSS200;TSS1500 | chr11:61062902-61063144 | 242 | | | 6 | | | | | | | | 1 | | | | yes | | | | | | | | |
| DAK;DDB1 | S_Shore | 5'UTR;TSS1500 | chr11:61102045-61102165 | 120 | | | | 1 | | | | | | | | | | yes | | | | | | | | | |
| DAK | S_Shelf | Body | chr11:61110107-61110297 | 190 | | | | | | | | | | | | | | yes | yes | | | | | | | | |
| TMEM216 | N_Shore | TSS1500;TSS200 | chr11:61159602-61159837 | 235 | | | 12 | | | | | 0 | 1 | | | | | yes | | | | | | | | | |
| C11orf66 | NA | TSS1500;TSS200 | chr11:61248217-61248818 | 601 | | | 5 | | | | | | | | | | | yes | | | | | | | | | |
| SYT7 | Island | 3'UTR | chr11:61283872-61284417 | 545 | | 1 | | | | | | | | | | | | yes | | | | | | | | | |
| SYT7 | NA | Body | chr11:61309911-61310204 | 293 | | | 1 | | | | | | | | | | | | | | | | | | | | |
| SYT7 | NA | Body | chr11:61314680-61314936 | 256 | | | | | | | | | | | | | | yes | | | | yes | yes | | | | |
| SYT7 | N_Shore;Island | Body | chr11:61334859-61335600 | 741 | 1 | | | | | | | | 2 | | | | | yes | | | | | | | | | |
| DAGLA | S_Shore | 5'UTR | chr11:61449737-61450294 | 557 | | | 6 | | 1 | | | | | | | | | yes | | | | | | | | | |
| DAGLA | Island | Body | chr11:61511275-61511973 | 698 | 1 | | | | | | | | | | | | | yes | | | | | | | | | |
| C11orf9 | Island | Body | chr11:61544854-61545348 | 494 | | | 3 | | | | | 0 | 1 | | | | | yes | | | | | yes | | | | |
| RAB3IL1 | N_Shore | Body | chr11:61684526-61684937 | 411 | | | | | | | | | 1 | | | | | | | | | | yes | | | | |
| BEST1 | NA | TSS200;TSS1500 | chr11:61716821-61717766 | 945 | | | 3 | | 1 | | | | 1 | | | | | yes | yes | | yes | | | | | | |
| SCGB1D1 | NA | TSS1500 | chr11:61956390-61956860 | 470 | | | 5 | | | | | | | | | | | yes | | | yes | yes | | | | | |
| SCGB2A2 | NA | TSS1500 | chr11:62037191-62037564 | 373 | | | | | | | | | | | | | | yes | | | | | yes | | | | |
| SCGB1D4 | NA | 5'UTR;1stExon;TSS200 | chr11:62066533-62067245 | 712 | | | 6 | | | | | | | | | | | yes | | | yes | | | | | | |
| AHNAK | N_Shore;Island | Body | chr11:62211788-62212431 | 643 | | | | 1 | 1 | | | 0 | | | | | | yes | | | | | | | | | |
| AHNAK | NA | Body | chr11:62272972-62273418 | 446 | | | 1 | | | | | | 1 | | | | | yes | | | yes | | | | | | |
| EML3;MTA2 | Island | 3'UTR;TSS1500 | chr11:62369728-62370246 | 518 | | | | | | | | 0 | 2 | 1 | | | | | | | | | yes | | | | |
| GANAB;INTS5 | S_Shore | TSS1500;Body | chr11:62414700-62415024 | 324 | | | | | | | | | | | | | | | | | | | | | | | |
| LRRN4CL | S_Shore | 5'UTR;1stExon;TSS200 | chr11:62456939-62457345 | 406 | | | | | | | | | 2 | 1 | | | | yes | | | | | yes | | | | |
| BSCL2;GNG3 | S_Shore | 5'UTR;TSS1500;1stExon | chr11:62474735-62474759 | 24 | 1 | | | | | | | | | | | | | | | | | | | | | | |
| SNORD30;SNORD22;SNORD31;SNHG1;S | N_Shelf;N_Shore | TSS200;TSS1500;Body | chr11:62620843-62622234 | 1391 | | | 11 | | 1 | | | | 2 | | | 1 | | | | | | | | | | | |
| CHRM1 | N_Shelf | 5'UTR;1stExon | chr11:62688748-62689081 | 333 | | | | | 1 | | | | 1 | | | | | | | | | | yes | | | | |
| SLC22A8 | NA | 1stExon;5'UTR;TSS200;TSS1500 | chr11:62783272-62783573 | 301 | | | 12 | | | | | 0 | | | | 1 | | | | | | | yes | | | | |
| SLC22A25 | NA | 1stExon | chr11:62996801-62997285 | 484 | | | | | | | | | | | | | | | | | yes | | | | | | |
| SLC22A9 | NA | TSS1500 | chr11:63136783-63137555 | 772 | | | | 1 | | | | | | | | | | | yes | | | | | | | | |
| HRASLS5 | S_Shore | TSS1500 | chr11:63259187-63259603 | 416 | | | | | | | | | | | | | | | yes | yes | | | yes | | | | |
| LGALS12 | NA | TSS200 | chr11:63273331-63273565 | 234 | | | | | | | | | 1 | | | | | yes | | | | | | | | | |
| RTN3 | N_Shore | TSS1500 | chr11:63448356-63448455 | 99 | | | 7 | | | | | | | | | | | yes | | | | | | | | | |
| MARK2 | NA | TSS1500;Body;TSS200 | chr11:63655748-63656090 | 342 | | | 12 | | | | | | | | | | | | | | | | | | | | |
| MARK2 | Island | Body | chr11:63676370-63676760 | 390 | | | | | | | | | | | | | | | | | | | yes | | | | |
| OTUB1 | N_Shore | TSS200;TSS1500 | chr11:63752561-63753251 | 690 | | | 15 | 1 | | | | 0 | 1 | | | | | yes | yes | | | | | | | | |
| MACROD1 | Island | Body | chr11:63768032-63768794 | 762 | | | | | 1 | | | | | | | | | yes | | | | | | | | | |
| MACROD1 | N_Shore;Island | Body | chr11:63775226-63775612 | 386 | | | 24 | | | | | 0 | 1 | 1 | | | | | yes | | | | yes | | | | |
| MACROD1 | NA | Body | chr11:63783267-63784126 | 859 | 3 | | 9 | | | | | | | | | | | yes | | | | | | | | | |

| | | | | | | | | | | | | | | | | | | | | | | | |
|---|---|---|---|---|---|---|---|---|---|---|---|---|---|---|---|---|---|---|---|---|---|---|---|
| MACROD1 | NA | Body | chr11:63827361-63828762 | 1401 | | | 4 | 1 | 6 | | | 0 | 2 | 1 | | 1 | | yes | | | yes | yes | yes |
| MACROD1 | Island | Body | chr11:63852804-63853037 | 233 | | | | 1 | 2 | | | | | | | | | yes | | | | | |
| FLRT1;MACROD1 | NA | TSS1500;Body;TSS200 | chr11:63870619-63871265 | 646 | | | 2 | | | | | | | | | | | yes | | | | | |
| MACROD1 | NA | Body | chr11:63906050-63906104 | 54 | 1 | | | | | | | | | | | | | yes | | | | | |
| FERMT3 | N_Shore;chr11:63974829-6397 | TSS1500;N_Shore;TSS200;1stExon | chr11:63973536-63975481 | 1945 | | | 13 | | 3 | 1 | | | 4 | 3 | 1 | | | yes | | | | | |
| FERMT3 | N_Shelf | Body | chr11:63990138-63990629 | 491 | 5 | | | | | | | | 1 | 1 | | | | yes | | | | | |
| PLCB3 | N_Shelf | Body | chr11:64023086-64023183 | 97 | | | | | | | | | | | | | | yes | | | | | |
| PLCB3 | Island | Body | chr11:64034861-64035061 | 200 | | | | | | | | | 1 | | | | | yes | | | | | yes |
| KCNK4 | N_Shore | TSS1500 | chr11:64058550-64058878 | 328 | | | 9 | | | | | | | | | | | yes | | | | | yes |
| KCNK4 | N_Shelf | Body | chr11:64064727-64065371 | 644 | | | 4 | | | | | | | | | | | | | | | | |
| KCNK4;C11orf20 | N_Shore;Island | Body;TSS1500;TSS200 | chr11:64066625-64068491 | 1866 | | | | | | | | | 1 | | | | | yes | | | yes | | yes |
| ESRRA | N_Shelf | Body | chr11:64081430-64081874 | 444 | | | 20 | | | | | | 2 | | | | | | yes | | | | |
| CCDC88B | N_Shelf;N_Shore | TSS200;Body | chr11:64107517-64109158 | 1641 | | | 7 | | | | | | 2 | 2 | 1 | | | yes | yes | | | | yes |
| CCDC88B | N_Shore | Body | chr11:64110657-64110785 | 128 | | 1 | 4 | | | | | | | | | | | | | | | | |
| CCDC88B | S_Shelf | Body | chr11:64115989-64116279 | 290 | | | 1 | | | | | 0 | 1 | 1 | | | | yes | | | | | |
| CCDC88B | NA | Body | chr11:64120286-64120728 | 442 | | 1 | 1 | | | | | | | | | | | | | | | | |
| SLC22A11 | NA | TSS1500 | chr11:64322890-64323456 | 566 | | | | 1 | | | | | | | | | | yes | | | | | |
| SLC22A11 | Island | Body | chr11:64335733-64335895 | 162 | | | | | | | | | | | | | | | | | | | |
| SLC22A12 | NA | Body;5'UTR | chr11:64360724-64361020 | 296 | | | | | | | | | | | | | | yes | | | | | yes |
| NRXN2 | N_Shore | Body | chr11:64397333-64397807 | 474 | | | 1 | | | | | | | | | | | | yes | | | | yes |
| NRXN2 | N_Shore;Island | Body | chr11:64405346-64406051 | 705 | | | | | | | | 0 | 4 | | | | | | | | | | |
| NRXN2 | Island | Body | chr11:64428252-64428434 | 182 | | | 11 | | | | | | | | | | | | | | | | |
| NRXN2 | N_Shore | Body | chr11:64478186-64478374 | 188 | 1 | | | | | | | | | | | | | | | | | yes | |
| RASGRP2 | N_Shore | Body | chr11:64508056-64508509 | 453 | | | 3 | | | | | | | | | | | | yes | | | | |
| PYGM | NA | 1stExon | chr11:64527240-64527414 | 174 | | | | | | | | | | | | | | | | | | | yes |
| CDC42BPG | N_Shore | Body | chr11:64594467-64594862 | 395 | | | | | | | | | | | | | | | | | | | |
| CDC42BPG | S_Shore | Body | chr11:64599019-64599084 | 65 | | | 1 | | | | | | | | | | | yes | yes | | | | |
| EHD1 | Island | 3'UTR | chr11:64621386-64622176 | 790 | | | | | | | | | | | | | | yes | | | | | |
| EHD1 | NA | Body | chr11:64635728-64635846 | 118 | | | | | 1 | | | | | | | | | yes | yes | | | | |
| EHD1 | N_Shelf | Body | chr11:64642316-64642558 | 242 | | | | | 1 | | | | | | | | | yes | | | | | |
| MIR192;MIR194-2 | N_Shelf | TSS200;TSS1500 | chr11:64658726-64659065 | 339 | | | | | 1 | | | | | | | | | yes | yes | | | | |
| GPHA2 | NA | Body;5'UTR;TSS200 | chr11:64702921-64703637 | 716 | | | | | | | | | 1 | | | 1 | | yes | | | | yes | |
| BATF2 | NA | Body | chr11:64764297-64764490 | 193 | | | | | | | | | | | | | | yes | | | | | |
| NAALADL1 | Island | Body | chr11:64815189-64815744 | 555 | | | 1 | | | | | | | | | | | yes | | | | yes | |
| CAPN1 | N_Shore | Body | chr11:64951536-64952354 | 818 | | | | | | | | | 1 | 1 | | | | yes | | yes | | yes | |
| SLC22A20 | NA | TSS1500;TSS200;nan | chr11:64980819-64981596 | 777 | | | 8 | 1 | | | | | 1 | | 1 | | | yes | | | | | |
| SLC22A20 | N_Shore | Body | chr11:64993076-64993335 | 259 | | 1 | | 1 | | | | | | | | | | yes | | | | | |
| TIGD3 | N_Shore;Island | 5'UTR;Body | chr11:65123257-65123609 | 352 | | | 4 | | | | | 0 | | | | | | | | | | | |
| NEAT1 | Island | Body | chr11:65190825-65191707 | 882 | | | | | | | | 0 | 1 | | | | | | | | | | |
| MIR612 | NA | TSS1500 | chr11:65210600-65211084 | 484 | | | | | 1 | | | 0 | | | | | | yes | | | | | |
| LTBP3 | Island | Body | chr11:65306731-65307006 | 275 | | | | | | | | | 1 | | | | | | | | | | yes |
| LTBP3 | Island | Body | chr11:65314021-65315625 | 1604 | | | | | | | | 0 | 4 | | | | | yes | | | | | |
| LTBP3 | Island | Body | chr11:65321226-65321806 | 580 | | | | | | | | 0 | 1 | | | 1 | | yes | | | | | |
| EHBP1L1 | N_Shore | Body | chr11:65352168-65352327 | 159 | | 1 | | | | | | | | | | | | | | | | | |
| KCNK7 | Island | 3'UTR | chr11:65360327-65360509 | 182 | | | | | | | | | 1 | | | | | | | | | | |
| KCNK7 | N_Shelf | Body | chr11:65362891-65363503 | 612 | | | | | | | | | 1 | | | | | | yes | | | yes | |
| MAP3K11 | N_Shore;chr11:65374699-6537 | Body;N_Shore | chr11:65374663-65375203 | 540 | | | | | | | | | 2 | | 1 | | | | | | | | |
| SIPA1 | N_Shelf | 5'UTR | chr11:65405760-65406254 | 494 | | | | | 1 | | | | | | | | | yes | | | | | yes |
| SIPA1 | N_Shore;Island | 5'UTR;Body;TSS200;1stExon | chr11:65407399-65408566 | 1167 | | | | | | | | | 4 | 5 | 1 | | | yes | | | | | |
| SIPA1 | Island | Body | chr11:65409760-65410126 | 366 | | | 5 | | | | | | | | | 1 | | yes | | | yes | | |
| DKFZp761E198 | N_Shore;Island | 1stExon | chr11:65546557-65547889 | 1332 | | | | | 1 | | | 0 | 2 | | | 1 | | | | | | | yes |
| OVOL1 | NA | Body | chr11:65560174-65562214 | 2040 | | | 5 | | 1 | | | | 1 | | | | | | | | | | |
| SNX32 | N_Shore | TSS1500 | chr11:65600888-65601417 | 529 | | 1 | | | | | | | | | | | | | yes | | | yes | yes |
| CTSW | NA | TSS1500;TSS200 | chr11:65646791-65647532 | 741 | | | 10 | 1 | | | | | 1 | | 1 | | | yes | yes | | | | |
| C11orf68;DRAP1 | Island | Body;TSS1500 | chr11:65685108-65685874 | 766 | | | 5 | | | | | | 1 | 1 | | | | yes | | | | | |
| CST6 | N_Shore | TSS1500 | chr11:65778967-65779263 | 296 | | | | | | | | | | | | | | yes | | | | | |
| GAL3ST3 | Island;S_Shore | 5'UTR;TSS200;TSS1500 | chr11:65816463-65816985 | 522 | | | | | | | | | 2 | | | | | | | | | | yes |
| KLC2 | N_Shore | TSS1500 | chr11:66024375-66024593 | 218 | | | | | | | | | | | | | | yes | yes | | | | |
| KLC2;RAB1B | Island | 3'UTR;TSS1500 | chr11:66034784-66034926 | 142 | | | | 1 | | | | | | | | | | | | | | yes | |
| YIF1A | N_Shore | Body | chr11:66055170-66055327 | 157 | | | 1 | | | | | | | | | | | yes | | | | | |
| TMEM151A | N_Shore | TSS1500 | chr11:66059064-66059164 | 100 | | | | | | | | | 1 | | | | | yes | | | | yes | yes |
| CD248 | Island;S_Shore;N_Shore | 1stExon;TSS1500 | chr11:66083541-66085299 | 1758 | 7 | | | 1 | | | | | | | | | | yes | | | | | yes |
| RIN1 | Island;S_Shore;chr11:6610202 | Body;TSS200;S_Shelf;TSS1500 | chr11:66102055-66105148 | 3093 | | | | 1 | 1 | | | | 4 | | 1 | | | | | | | | yes |
| BRMS1;B3GNT1 | N_Shore | TSS1500;Body | chr11:66113666-66114071 | 405 | | | | | 1 | | | | | | | | | yes | | | | yes | |
| SLC29A2 | N_Shelf | Body | chr11:66135554-66135627 | 73 | | | | | | | | | | | | | | yes | | | | | |
| NPAS4 | N_Shore | TSS1500 | chr11:66188019-66188433 | 414 | | 1 | | | | | | | | | | | | | | | | yes | |
| NPAS4 | NA | 3'UTR | chr11:66193931-66194068 | 137 | | | 4 | | | | | | 1 | | | | | | | | | yes | |
| MRPL11 | NA | TSS1500 | chr11:66206894-66207458 | 564 | | | 9 | | | | | | | | | | | yes | | yes | | | |
| ZDHHC24 | N_Shore | Body | chr11:66311334-66311422 | 88 | | | 10 | 1 | 1 | | | 0 | 1 | 1 | | 2 | | yes | yes | | | | |
| ACTN3 | Island | Body | chr11:66326652-66327250 | 598 | | | | | | | | 0 | 1 | | | 3 | | yes | | | | | yes |
| LRFN4;PC | Island | TSS1500;Body | chr11:66624256-66624853 | 597 | | | | 1 | | | | | | | | | | | | | | yes | yes |
| PC;LRFN4 | Island;S_Shore | Body;3'UTR | chr11:66627706-66628295 | 589 | 2 | | | | | | | | 1 | | | | | | | | | | yes |
| PC | Island | Body | chr11:66631259-66631894 | 635 | | 1 | | | | | | | | | | | | yes | | | yes | yes | |
| PC | NA | Body | chr11:66639308-66639327 | 19 | | | | | | | | | | | | | | | | | | | |
| PC | NA | 5'UTR | chr11:66643359-66643572 | 213 | | | | 1 | | | | | | | | | | | yes | | | | |
| PC | NA | 5'UTR | chr11:66650500-66650560 | 60 | | | 11 | | | | | | 1 | | | | | | yes | | | yes | |
| SYT12 | Island | TSS1500 | chr11:66790671-66791156 | 485 | | | | | | | | | 1 | | | | | yes | | | | yes | yes |
| RHOD | N_Shore;Island | TSS1500;TSS200 | chr11:66823587-66824189 | 602 | | | 3 | | | | | 0 | 2 | | | | | yes | yes | | | yes | yes |
| RHOD | Island | Body;3'UTR | chr11:66838934-66839363 | 429 | | | 10 | | | | | 0 | 1 | | | | | yes | | | | | |
| KDM2A | N_Shore | TSS1500 | chr11:66885240-66885281 | 41 | | | 19 | | 1 | | | | | | | | | yes | | | | | yes |
| ADRBK1 | S_Shore | Body | chr11:67035965-67039119 | 3154 | | | | | | | | | | | | | | yes | | | | | yes |
| ADRBK1 | Island | Body | chr11:67052656-67052992 | 336 | | | | | | | | | 1 | | | | | yes | | | | | |
| SSH3;ANKRD13D | N_Shore;Island | TSS1500;Body;3'UTR;TSS200 | chr11:67069814-67071885 | 2071 | | | | | | | | | | | | | | | | | | | |
| LOC100130987;CLCF1 | S_Shore | Body;TSS1500 | chr11:67142001-67142268 | 267 | | | | | | | | | 1 | | | | | | | | | | yes |
| PPP1CA;TBC1D10C | S_Shore;S_Shelf | TSS1500 | chr11:67170610-67171585 | 975 | | | | 1 | 1 | | | | 1 | 1 | 1 | | | yes | | | | | |
| TBC1D10C | N_Shore;Island | Body | chr11:67176914-67177296 | 382 | | | 1 | | 1 | | | | 1 | 3 | 1 | | | | | | | | |
| ATPGD1 | S_Shore | 5'UTR | chr11:67183498-67183616 | 118 | | | | | | | | | 1 | | | | | | | | | yes | yes |
| RPS6KB2 | N_Shore | 3'UTR | chr11:67202808-67203661 | 853 | | | | | | | | | 1 | | | | | | yes | | | yes | yes |
| PTPRCAP;CORO1B | chr11:67203319-67203662;S_S | S_Shore;Body;5'UTR;1stExon;TSS2 | chr11:67204799-67206434 | 1635 | | | | | 1 | | | | 3 | 1 | 1 | | | | yes | | | | |
| CORO1B | NA | Body;5'UTR | chr11:67209355-67210282 | 927 | | | | | | | | | 2 | | | | | yes | | | | | |
| GPR152 | NA | 1stExon;TSS200 | chr11:67219295-67220318 | 1023 | | 1 | | | | | | | 1 | | | | 1 | yes | | | | | |
| TMEM134 | N_Shore;Island | 3'UTR;Body | chr11:67231878-67232511 | 633 | | | 25 | | | | | | | | 1 | | | yes | yes | | | | yes |
| CABP2 | S_Shelf | 1stExon;5'UTR | chr11:67290850-67291026 | 176 | | | | | | | | | | | | | | yes | | | | | |
| GSTP1 | N_Shore;Island | TSS1500;1stExon;5'UTR | chr11:67350153-67352041 | 1888 | | | 8 | 2 | | | | 0 | | | | | | yes | yes | | | yes | |
| NDUFV1 | N_Shore | TSS1500 | chr11:67373625-67373841 | 216 | | | 4 | | | | | | | | | | | yes | | | | | |
| TBX10 | NA | 5'UTR;1stExon;TSS200 | chr11:67407029-67407224 | 195 | | | 2 | | | | | | | | | | | yes | | | | | |
| ACY3 | NA | 5'UTR;1stExon;TSS200 | chr11:67417958-67418365 | 407 | | | 12 | | | | | 0 | 5 | 3 | | 6 | | yes | | yes | | | |
| ALDH3B2 | NA | 1stExon;5'UTR;TSS200 | chr11:67442067-67442697 | 630 | | | 5 | 1 | | | | | 1 | | | | | yes | yes | | | | |
| UNC93B1 | S_Shore | TSS200 | chr11:67771726-67772273 | 547 | | | 3 | | | | | | | | | | | yes | | | | yes | yes |

| | | | | | | | | | | | | | | | | | | | | | | | | | | | |
|---|---|---|---|---|---|---|---|---|---|---|---|---|---|---|---|---|---|---|---|---|---|---|---|---|---|---|---|
| ALDH3B1 | NA | TSS200 | chr11:67775857-67776030 | 173 | | | 4 | | | | | | | | | | | yes | | | | | | | | | |
| NDUFS8 | N_Shelf | Body | chr11:67803951-67804112 | 161 | | | | | | | | | 1 | | | | | yes | | | | | | | | | |
| TCIRG1 | Island | TSS200 | chr11:67806337-67806668 | 331 | | | | | | | | | 1 | | | | | yes | | | | | | | | | |
| TCIRG1 | NA | Body;1stExon;5'UTR | chr11:67810742-67811641 | 899 | | | 20 | | | | | | 1 | | | | | | | | | | | | | | |
| SUV420H1 | S_Shore | TSS1500 | chr11:67982143-67982703 | 560 | | | | | | | | | | | | | | yes | | | | | yes | | | | |
| LRP5 | Island;S_Shore | Body | chr11:68080928-68081686 | 758 | | | | | | | | 0 | 3 | | | | | | | | | | yes | | | | |
| LRP5 | NA | Body | chr11:68095623-68096049 | 426 | | | 1 | | | | | | | | | | | yes | | | | | | | | | |
| LRP5 | NA | Body | chr11:68138269-68139350 | 1081 | | | 6 | | | | | | 2 | | | | | | | | | | | | | | |
| LRP5 | NA | Body | chr11:68142198-68142385 | 187 | | | | | | | | | | | | | | | yes | | | | | | | | |
| LRP5 | NA | Body | chr11:68147880-68148079 | 199 | | | 21 | | | | | | | | | | | yes | yes | yes | yes | | | | | | |
| LRP5 | NA | Body | chr11:68174077-68174286 | 209 | | | | 1 | | | | | | | | | | yes | | | yes | | | | | | |
| LRP5 | N_Shore | Body | chr11:68178500-68179097 | 597 | | 1 | | | | | | | | | | | | yes | | | | | | | | | |
| LRP5 | N_Shore;Island | Body | chr11:68180584-68181383 | 799 | | | 3 | 1 | | | | | 1 | | | | | yes | | | | | | | | | |
| LRP5 | N_Shore | Body | chr11:68199656-68199794 | 138 | | | 5 | | | | | | | | | | | | | | | | | | | | |
| LRP5 | N_Shore | Body | chr11:68205840-68206027 | 187 | | | | | | | | | | | | | | | | | yes | | | | | | |
| SAPS3 | NA | 5'UTR | chr11:68249304-68271518 | 22214 | | | | | | | | | | | | | | yes | yes | yes | yes | yes | yes | | | | |
| GAL | Island | TSS200 | chr11:68451849-68451858 | 9 | | | | | | | | | 1 | | | | | | | | | | | | | | |
| MTL5 | S_Shore | TSS200 | chr11:68519060-68519174 | 114 | | | | | | | | | 2 | | | | | yes | | | | yes | yes | | | | |
| CPT1A | NA | Body | chr11:68579970-68580041 | 71 | | | 7 | | 1 | | | | | | | | | | yes | | | | | | | | |
| CPT1A | N_Shore | TSS1500 | chr11:68609747-68611596 | 1849 | | | | | | | | | | | | | | | | | | | | | | | |
| MRGPRD | NA | TSS200;1stExon | chr11:68747554-68748907 | 1353 | | 1 | | | | | | | | | | | | yes | yes | | | | | | | | |
| MRGPRF | S_Shelf;Island;S_Shore | TSS1500;5'UTR;1stExon | chr11:68778570-68782211 | 3641 | 2 | | 31 | 2 | | | | 0 | 1 | | | 2 | | yes | | yes | | yes | yes | | | | |
| TPCN2 | S_Shore | Body | chr11:68851580-68851641 | 61 | 2 | | | | | | | | 1 | | | | | | | | | yes | | | | | |
| TPCN2 | S_Shelf | Body | chr11:68853686-68854196 | 510 | | | 2 | | | | | | | | | | | yes | | | | | | | | | |
| MYEOV | NA | TSS200 | chr11:69061454-69061634 | 180 | | | | | | | | | | | | | | yes | yes | | | yes | | | | | |
| CCND1 | Island | TSS1500 | chr11:69454696-69455217 | 521 | | | 3 | | | | | | | | | | | | | | | | yes | | | | |
| CCND1 | N_Shore;Island;S_Shore | 3'UTR | chr11:69468789-69469349 | 560 | | 1 | 2 | | | | | | 1 | | | | | | | | yes | yes | | | | | |
| ORAOV1 | NA | 3'UTR | chr11:69481734-69482540 | 806 | | | | | | | | | | 1 | | | | yes | | | | | yes | | | | |
| ORAOV1 | NA | Body | chr11:69484433-69484574 | 141 | | | | | | | | | | 1 | | | | | | | yes | yes | | | | | |
| FGF19 | Island | Body | chr11:69516939-69518545 | 1606 | | 1 | 2 | | | | | | | | | | | yes | | | | | yes | | | | |
| FGF19 | Island;S_Shore | TSS1500 | chr11:69519845-69520418 | 573 | 1 | | 8 | 1 | | | | | 1 | 1 | | | | yes | | | | | | | | | yes |
| FGF3 | Island | Body | chr11:69632117-69632335 | 218 | | 1 | | | | | | | | | | | | | | | | | | | | | |
| FGF3 | Island | TSS200;TSS1500 | chr11:69634296-69634648 | 352 | 6 | 2 | | | | | | 0 | 3 | | | | | | | | | yes | yes | | | | |
| ANO1 | Island;N_Shore | TSS200;TSS1500 | chr11:69924156-69924724 | 568 | | | | 2 | | | | 0 | 1 | | | | | yes | yes | | | | yes | | | | |
| ANO1 | N_Shelf | TSS200;Body | chr11:69931337-69931514 | 177 | | | 2 | | | | | | | | | | | yes | | | | | | | | | |
| ANO1 | Island | Body | chr11:69933978-69934128 | 150 | | | 5 | | | | | 0 | 2 | | | 4 | | yes | yes | | | | | | | | |
| ANO1 | NA | Body | chr11:69949353-69949774 | 421 | | | 3 | | | | | | | | | | | yes | | | | | | | | | |
| ANO1 | NA | Body | chr11:69962358-69962465 | 107 | | | | | | | | | | | | | | yes | | | | | | | | | |
| ANO1 | NA | Body | chr11:69970464-69970618 | 154 | | | | | | | | | 1 | | | | | | yes | | | | | | | | |
| ANO1 | NA | Body | chr11:69982916-69982996 | 80 | 1 | 1 | | | | | | | 1 | | 1 | 1 | 1 | yes | yes | | | yes | | | | | |
| ANO1 | NA | Body | chr11:69987299-69987633 | 334 | | | 1 | | | | | | | | | | | yes | | yes | | | | | | | |
| ANO1 | Island;S_Shore | Body | chr11:70001754-70002831 | 1077 | | | 11 | | | | | | 1 | | | | | yes | | | | | | | | | |
| ANO1 | N_Shore | Body | chr11:70023032-70023142 | 110 | | | | | | | | | 1 | | | | | yes | | yes | yes | | | | | | |
| FADD | N_Shore | TSS1500 | chr11:70048796-70048921 | 125 | | | 17 | 1 | | | | 0 | 5 | | | 4 | | yes | yes | | | | | | | | |
| PPFIA1 | NA | Body | chr11:70211301-70211531 | 230 | | | | | | | | | 1 | 1 | 1 | | | | | | | | | | | | |
| CTTN | N_Shore | Body | chr11:70257035-70257149 | 114 | | | | | | | | 0 | 1 | 1 | 1 | | | | | | | | | | | | |
| CTTN | S_Shelf | Body | chr11:70261100-70261373 | 273 | 1 | | 1 | | | | | | | | | | | yes | | | | | | | | | |
| CTTN | NA | Body | chr11:70280382-70281091 | 709 | | | | | | | | | 2 | | | | | | yes | | | yes | | | | | |
| SHANK2 | NA | Body | chr11:70368624-70368768 | 144 | | | 1 | | | | | | | | | | | | yes | | | | | | | | |
| SHANK2 | NA | Body | chr11:70385301-70385511 | 210 | 7 | | 1 | | | | | | | | | | | | | yes | yes | yes | | | | | |
| SHANK2 | NA | Body | chr11:70391215-70391706 | 491 | | | 5 | | | | | | | | | | | yes | | yes | | | yes | | | | |
| SHANK2 | NA | Body | chr11:70458719-70459053 | 334 | 5 | | 12 | | | | | | | | | | | yes | yes | | | yes | | | | | |
| SHANK2 | N_Shore;Island | 1stExon;Body;TSS200;TSS1500 | chr11:70507825-70508659 | 834 | | 1 | | 1 | 3 | | | 0 | 8 | 2 | 1 | 4 | 1 | | | | | yes | | | | | |
| SHANK2 | NA | Body | chr11:70516627-70517411 | 784 | | | | 1 | | | | | | | | | | yes | | | | | | | | | |
| SHANK2 | NA | Body | chr11:70557445-70557881 | 436 | | | 8 | | | | | | 2 | 1 | | | | | | yes | | | yes | | | | |
| SHANK2 | NA | Body | chr11:70559053-70559616 | 563 | | | 12 | | | | | | | | | | | yes | yes | | | | | | | | |
| SHANK2 | N_Shore;Island;S_Shore | Body | chr11:70563131-70565226 | 2095 | | | 16 | | | | | 0 | 1 | | | | | yes | | | | | | | | | |
| SHANK2 | NA | Body | chr11:70584165-70584425 | 260 | | | | | | | | | | | | | | | | | | | | | | | |
| SHANK2 | NA | Body | chr11:70601781-70601971 | 190 | | | | 1 | | | | | | | | | | yes | | | | | | | | | |
| SHANK2 | NA | Body | chr11:70628938-70628992 | 54 | | 1 | | | | | | | | | | | | yes | yes | yes | yes | | | | | | |
| SHANK2 | NA | Body | chr11:70643833-70644532 | 699 | | | | | | | | | 1 | | | | | | | yes | | yes | | | | | |
| SHANK2 | Island | Body | chr11:70666452-70666748 | 296 | | | | | | | | | | | | | | | | yes | | | | | | | |
| SHANK2 | N_Shore;Island | Body | chr11:70672365-70673256 | 891 | | | 1 | | 1 | | | 0 | 9 | 2 | | 5 | | | | | | | yes | | | | |
| SHANK2 | NA | Body | chr11:70691944-70692038 | 94 | | | | 1 | | | | 0 | 1 | | | | | yes | | | | | | | | | |
| SHANK2 | NA | Body | chr11:70805455-70805627 | 172 | | | | | | | | | | | | | | | | yes | | | | | | | |
| SHANK2 | NA | Body | chr11:70829968-70830058 | 90 | | | 1 | | | | | | | | | | | yes | | yes | | | | | | | |
| SHANK2 | NA | Body | chr11:70858342-70858572 | 230 | | | | | | | | | | | | | | | yes | | | | | | | | |
| SHANK2 | NA | TSS200 | chr11:70935863-70936231 | 368 | | | 1 | | | | | | | | | | | | yes | | | | | | | | |
| DHCR7 | S_Shore | TSS1500 | chr11:71159853-71160219 | 366 | | | | | | | | | | | | | | yes | yes | yes | | | yes | | | | |
| NADSYN1 | N_Shore | TSS1500 | chr11:71163428-71163743 | 315 | | | 2 | | | | | | | | | | | | yes | yes | | | yes | | | | |
| KRTAP5-8 | NA | TSS1500 | chr11:71248741-71249212 | 471 | | | | | | | | | | | | | | | | | | | yes | | | | |
| KRTAP5-9 | NA | TSS1500;TSS200 | chr11:71259251-71259581 | 330 | | | | | | | | | | | | | | yes | | | | | | | | | |
| IL18BP | NA | TSS200;TSS1500;5'UTR;1stExon | chr11:71709725-71711087 | 1362 | | | 12 | 1 | 1 | | | | 6 | 1 | 1 | | | yes | | | yes | | | | | | |
| NUMA1 | NA | Body | chr11:71715003-71715168 | 165 | | | | | | | | | | | | | | | | | | | yes | | | | |
| NUMA1 | NA | Body | chr11:71724287-71725590 | 1303 | | | 27 | | | | | | 1 | | | | | | | | | | | | | | |
| NUMA1 | NA | 5'UTR | chr11:71752919-71752997 | 78 | | | | | | | | 0 | | | | | | | yes | | | | | | | | |
| FOLR3 | NA | TSS1500 | chr11:71846494-71846788 | 294 | | | | | | | | | 1 | 1 | | | | yes | yes | | | | | | | | |
| PHOX2A | Island | Body | chr11:71952131-71952431 | 300 | | | 10 | | | | | | | | | | | | | | | | yes | | | | |
| PHOX2A | Island | TSS200;1stExon | chr11:71954982-71955599 | 617 | | 2 | | | | | | | 2 | | | | | | | | | yes | yes | | | | |
| PDE2A | Island | Body | chr11:72295168-72295716 | 548 | | | 13 | | | | | | | | | | | yes | yes | | | | yes | | | | |
| PDE2A | N_Shore;Island | Body | chr11:72301235-72301724 | 489 | | | 2 | | | | | | | | 1 | | | yes | | | yes | | yes | | | | |
| ARAP1 | Island | Body | chr11:72408186-72408788 | 602 | | | 5 | | | | | | | | | | | | | | | | | | | | |
| ARAP1 | NA | Body;TSS1500 | chr11:72433619-72433866 | 247 | | | | | | | | | | | | | | yes | | | | yes | | | | | |
| STARD10 | N_Shore | Body | chr11:72492171-72492461 | 290 | | | 2 | | | | | | | | | | | yes | | | | | yes | | | | |
| ATG16L2 | N_Shore;Island | Body | chr11:72532555-72533664 | 1109 | | | | | | | | 0 | 2 | 1 | | 1 | | yes | | | | | | | | | |
| P2RY6 | N_Shore | TSS200 | chr11:72975404-72975702 | 298 | | | | | | | | 0 | | | | 2 | | yes | | | | | | | | | |
| ARHGEF17 | Island | 1stExon | chr11:73020351-73020729 | 378 | | | 1 | | | | | | | | | | | | | | | | | | | | |
| ARHGEF17 | Island | Body | chr11:73053649-73054401 | 752 | | | | | | | | 0 | 5 | | | | | | | | | yes | yes | | | | |
| ARHGEF17 | Island | Body | chr11:73073153-73073652 | 499 | | | | | | | | | | | | | | yes | | | | | | | | | |
| RELT | N_Shore | TSS1500 | chr11:73086934-73087205 | 271 | | | | | | | | | | | | | | | | | | | | | | | |
| FAM168A | Island | TSS200 | chr11:73309256-73309264 | 8 | | | | | | | | 0 | | | | | | | | | | yes | | | | | |
| PLEKHB1 | NA | TSS200;TSS1500 | chr11:73356316-73357396 | 1080 | 1 | | 12 | 1 | | | | 0 | 3 | | | 1 | | yes | | | yes | | yes | | | | |
| PLEKHB1 | NA | TSS200;TSS1500;Body | chr11:73358572-73358657 | 85 | | | 1 | | | | | | | | | | | yes | yes | | | | | | | | |
| PLEKHB1 | NA | 5'UTR;TSS200;Body | chr11:73359662-73359987 | 325 | | | | | | | | | 1 | | | | | yes | yes | | yes | | | | | | |
| PLEKHB1 | N_Shore | Body | chr11:73371753-73371940 | 187 | | | | | | | | | | | | | | | | | yes | | | | | | |
| MRPL48 | N_Shore | TSS1500 | chr11:73498278-73498718 | 440 | | | 1 | | | | | | | | | | | yes | | | | | | | | | |
| DNAJB13 | NA | Body | chr11:73668626-73669449 | 823 | | | 3 | | | | | | 2 | | | | | | | yes | | | | | | | |
| P4HA3 | S_Shore | TSS200;TSS1500 | chr11:74022875-74023050 | 175 | | | 9 | | | | | | 1 | | | 1 | | | yes | | | yes | yes | | | | |
| KCNE3 | Island | TSS200 | chr11:74178684-74178883 | 199 | | | 23 | 1 | | | | 0 | 1 | | | | | yes | | | | | | | | | |

| | | | | | | | | | | | | | | | | | | | | | | | |
|---|---|---|---|---|---|---|---|---|---|---|---|---|---|---|---|---|---|---|---|---|---|---|---|
| CHRDL2 | Island | TSS1500 | chr11:74442740-74442900 | 160 | | | 1 | | | | | | | | | | | yes | | | | yes | yes |
| SLCO2B1 | NA | Body;TSS1500 | chr11:74870268-74870640 | 372 | | | | | 1 | | | | | | | | | yes | | | | | |
| ARRB1;MIR326 | NA | Body;TSS1500 | chr11:75046147-75047180 | 1033 | | | | | | | | | 1 | | | | | yes | | | yes | | |
| KLHL35 | Island;N_Shore | Body | chr11:75139390-75139736 | 346 | 3 | 2 | 14 | | | 4 | 1 | 0 | 2 | 2 | | 18 | | yes | yes | | | | |
| KLHL35 | Island;S_Shore | TSS200;TSS1500 | chr11:75141835-75142449 | 614 | | | 7 | | | | | | | | | | | yes | yes | yes | | | yes |
| GDPD5 | NA | Body | chr11:75162105-75162492 | 387 | | | | | | | | | 1 | | | | | yes | | | | | |
| GDPD5 | Island | TSS1500 | chr11:75237636-75237739 | 103 | | | 1 | | | | | | | | | | | yes | | | | | |
| WNT11 | NA | Body | chr11:75898132-75898203 | 71 | | | | | | | | | | | | | | yes | | | | | |
| WNT11 | N_Shore | Body | chr11:75917251-75918089 | 838 | | | 1 | 1 | | | | | | | | | | yes | | | | | yes |
| LRRC32 | N_Shore | 5'UTR;TSS200 | chr11:76380605-76381236 | 631 | | | | 1 | | | | | 2 | | | | | yes | | | | yes | |
| ACER3 | N_Shore | TSS1500 | chr11:76571458-76571534 | 76 | | | | | | | | | | | | | | yes | | | | | yes |
| B3GNT6 | Island | Body | chr11:76751369-76751777 | 408 | | | | | | | | | 1 | | | | | | | | yes | | yes |
| MYO7A | NA | TSS1500;TSS200 | chr11:76838749-76839476 | 727 | 5 | | | | | | | 0 | 2 | | | 1 | | yes | | | | | |
| MYO7A | Island | Body | chr11:76849101-76849564 | 463 | | | 1 | | | | | 0 | | | | 1 | | yes | | | | | yes |
| MYO7A | N_Shore;Island | Body | chr11:76858648-76859387 | 739 | 4 | | | 1 | | | 1 | 0 | 1 | 1 | | | | yes | yes | | yes | yes | |
| MYO7A | NA | Body | chr11:76902992-76903183 | 191 | | | | | | | | | 1 | | | 1 | | | | | | | |
| GDPD4 | NA | TSS1500 | chr11:76998821-76998857 | 36 | | | | | | | | 0 | | | 1 | | | yes | yes | yes | | | |
| PAK1 | N_Shore | 5'UTR | chr11:77122506-77123047 | 541 | | 1 | | 1 | | | | | | | | | | yes | | | | | |
| AQP11 | N_Shore | TSS1500 | chr11:77299805-77299962 | 157 | | | 15 | | | | 1 | 0 | 1 | | | 2 | | yes | yes | | | yes | |
| KCTD14 | NA | 1stExon | chr11:77734206-77734490 | 284 | | | 4 | | | | | | | | | 1 | | yes | | yes | | | yes |
| USP35 | Island | 5'UTR | chr11:77907224-77907656 | 432 | | | 5 | | | | | 0 | 1 | | | | | yes | yes | | | | yes |
| NARS2 | S_Shore | TSS1500 | chr11:78286207-78286346 | 139 | | | | | | | | | | | | 1 | | yes | yes | | yes | yes | |
| ODZ4 | Island | Body | chr11:78614239-78614522 | 283 | | | | | | | | | 1 | 1 | | 3 | | yes | yes | | yes | | |
| ODZ4 | N_Shore;Island | Body | chr11:78672632-78673290 | 658 | | | 8 | | | | | | 1 | 1 | | | | yes | | | | | |
| ODZ4 | Island | Body | chr11:78780700-78780983 | 283 | | | | | | | | | 1 | | | 1 | | yes | yes | | | | |
| ODZ4 | Island | 5'UTR | chr11:79151202-79151719 | 517 | | | | | | | | | 1 | | | | | | | | | yes | yes |
| FAM181B | N_Shore;Island | 1stExon;3'UTR | chr11:82443149-82443614 | 465 | | 1 | | | | | | 0 | | | | | | | | | | | |
| DLG2 | NA | nan;Body;TSS200 | chr11:83393062-83394074 | 1012 | | | | | | | | | 1 | | 1 | | | yes | | | | | |
| DLG2 | NA | Body | chr11:83501743-83502111 | 368 | | | | | | | | | | | | | | | | | | | |
| DLG2 | NA | 1stExon;Body;5'UTR;TSS200 | chr11:84634126-84634620 | 494 | | 1 | 4 | | 1 | | | 0 | 6 | | | 6 | | | | | | | |
| CCDC89 | NA | nan;1stExon;5'UTR | chr11:85397109-85397684 | 575 | | | 3 | | | | | | | | 1 | | | | yes | | | | |
| SYTL2 | NA | 5'UTR;Body;1stExon;TSS1500 | chr11:85430070-85430657 | 587 | | | 22 | | | | | | | | | | | | yes | | | | |
| CCDC81 | N_Shore;Island | TSS1500;TSS200 | chr11:86085366-86086101 | 735 | | 1 | | | | | | 0 | | | | | | | yes | | | | |
| ME3 | Island;S_Shore;chr11:8638269 | TSS200;TSS1500;1stExon;5'UTR;Isl | chr11:86382900-86383809 | 909 | | | 24 | 1 | | | | 0 | 1 | | 1 | 1 | | yes | yes | | yes | | |
| RAB38 | Island;S_Shore | 1stExon;5'UTR;TSS200;TSS1500 | chr11:87908558-87908938 | 380 | | | 23 | | | | | 0 | 4 | | | | | yes | | | | | yes |
| GRM5 | Island;N_Shore | Body;3'UTR | chr11:88241594-88242604 | 1010 | | | 2 | | | | | | 2 | | | | | | | | | yes | yes |
| NOX4 | Island;S_Shore | 5'UTR;TSS200;1stExon;Body | chr11:89224506-89225042 | 536 | | | | | | | | | 4 | | | | | | | | | yes | |
| NAALAD2 | NA | TSS1500;TSS200 | chr11:89867385-89867976 | 591 | | 1 | | | | | | 0 | 5 | | | 2 | | | yes | yes | | yes | |
| NAALAD2 | NA | Body | chr11:89901285-89901450 | 165 | | | | | | | | | | | | | | yes | | yes | yes | | |
| CHORDC1 | S_Shore | TSS200 | chr11:89956704-89957033 | 329 | | | | | | | | | 2 | | | | | | yes | | | | |
| FAT3 | Island | Body | chr11:92599942-92600238 | 296 | | | | | | | | | 1 | | | | | | yes | | | | |
| FAT3 | Island | Body | chr11:92615932-92616492 | 560 | | | | | | | | | 2 | | | | | yes | | | yes | | |
| MTNR1B | N_Shore;Island | TSS200;1stExon;Body;TSS1500 | chr11:92702373-92703536 | 1163 | | 1 | 5 | | | | | 0 | 4 | | | 3 | | | yes | | | | |
| CCDC67 | N_Shore;Island | TSS1500;1stExon;5'UTR | chr11:93063516-93064092 | 576 | | | | | | | | 0 | 2 | | | | | yes | yes | | | | yes |
| C11orf75 | S_Shore | TSS1500 | chr11:93277161-93277191 | 30 | | | 13 | | | | | | 1 | | | | | yes | yes | | | | |
| KIAA1731 | N_Shore | TSS1500 | chr11:93394605-93394624 | 19 | | | 2 | | | | | | | | | | | yes | | | | | |
| SNORA32;SNORA25 | NA | TSS200;TSS1500 | chr11:93464342-93464769 | 427 | | | | | | | | | 1 | | | | | | | | | | |
| SNORA1;SNORA8;SNORD5 | NA | TSS1500;Body | chr11:93466424-93466913 | 489 | | | 3 | | | | | | | | | | | yes | | | | | |
| C11orf90 | N_Shore;Island | Body;TSS200 | chr11:93583253-93583763 | 510 | | | | | | | | 0 | 3 | 1 | | | | yes | | | | yes | yes |
| HEPHL1 | NA | TSS1500;TSS200 | chr11:93753891-93754390 | 499 | | | 5 | | | | | | 1 | | | | | yes | | | | | |
| HEPHL1 | N_Shore;Island | Body | chr11:93822022-93822294 | 272 | | | | | | | | 0 | 2 | 1 | | 2 | | | | | | | |
| GPR83 | Island | 1stExon;5'UTR | chr11:94134580-94134666 | 86 | | | | | | | | | 1 | | | | | | | | | | |
| FUT4 | Island | TSS200 | chr11:94276895-94276903 | 8 | | | | | | | | | 1 | | | | | | | | | yes | |
| FUT4 | Island | 1stExon | chr11:94278324-94279068 | 744 | | | 28 | | | | | | 4 | 6 | | | | | | | | | |
| AMOTL1 | Island | Body | chr11:94502388-94502824 | 436 | | | | | | | | | 1 | | | | | | | | | | yes |
| SESN3 | Island | TSS1500 | chr11:94965364-94965610 | 246 | | | 4 | | | | | 0 | | | | 1 | | | | | | | |
| CNTN5 | N_Shore;Island | TSS1500;TSS200 | chr11:98891492-98891811 | 319 | | 1 | | | | | | | 2 | | | | | | | | | | yes |
| PGR | Island | 1stExon | chr11:100998943-100999077 | 134 | | 1 | | | | | | | | | | | | | | | | | |
| TRPC6 | S_Shore | TSS200 | chr11:101454733-101454996 | 263 | | 1 | | | | | | | | | | | | | yes | | | | |
| KIAA1377;ANGPTL5 | Island | TSS200;5'UTR | chr11:101785659-101785698 | 39 | | 1 | | | | | | | 1 | | | | | | | | | | |
| MMP7 | NA | 1stExon;5'UTR | chr11:102401438-102401616 | 178 | | | | 1 | | | | | | | | | | yes | yes | | | | yes |
| MMP20 | NA | TSS200 | chr11:102496104-102496159 | 55 | | | 4 | | | | | | | | | | | | | | yes | | |
| MMP3 | NA | 1stExon | chr11:102714244-102714358 | 114 | | | | | | | | | 1 | | | | | yes | | | | | |
| MMP12 | NA | 5'UTR;1stExon | chr11:102745682-102745934 | 252 | | | | | | | | | 1 | | | | | | | | | yes | |
| MMP13 | NA | TSS200 | chr11:102826565-102826680 | 115 | | | | | | | | | 1 | | | | | yes | | | | | |
| PDGFD | Island;S_Shore | 1stExon;TSS1500 | chr11:104034619-104035942 | 1323 | | 1 | | 1 | | | | | | | | | | yes | | | | | |
| CASP1;CARD16 | NA | Body | chr11:104915759-104916075 | 316 | | | | | 1 | | | | | | | | | yes | | | | | |
| GRIA4 | N_Shore | TSS1500;1stExon;5'UTR | chr11:105480347-105481509 | 1162 | | 1 | | | | | | 0 | 1 | | | | | | | | | yes | |
| KIAA1826 | Island | 1stExon;5'UTR | chr11:105892855-105893048 | 193 | | | | | | | | | | | | 1 | | | yes | | | | |
| GUCY1A2 | Island | 5'UTR;1stExon | chr11:106888787-106889320 | 533 | | | | | | | | | 1 | | | | | | | | | yes | yes |
| ELMOD1;LOC643923 | Island | 5'UTR;TSS200 | chr11:107462430-107462445 | 15 | | | | | | | | | | | | | | | | | | | |
| SLN | NA | 3'UTR;Body | chr11:107578208-107578793 | 585 | | | | | | | | | 2 | | | 2 | | | | yes | yes | | |
| SLN | NA | 5'UTR;1stExon;TSS200;TSS1500 | chr11:107582734-107583406 | 672 | | | 10 | | | | | | 3 | | | | | | | | yes | yes | |
| KDELC2 | S_Shore | TSS1500 | chr11:108369884-108370005 | 121 | | | 2 | | | | | | 1 | | | | | yes | | | | yes | |
| EXPH5 | NA | Body;TSS200;1stExon;5'UTR | chr11:108408907-108409365 | 458 | | | 6 | | | | | | 1 | | | | | | | | | | |
| EXPH5 | NA | nan;5'UTR;Body;TSS200 | chr11:108422663-108423325 | 662 | | | 9 | | 2 | | | | 3 | 1 | 1 | | | yes | | | | | |
| C11orf87 | N_Shore;Island | TSS200;5'UTR | chr11:109292789-109294584 | 1795 | | | 7 | | 1 | | | 0 | 1 | | | | | | | | | yes | yes |
| ZC3H12C | Island | TSS1500 | chr11:109963338-109963583 | 245 | | | | | | | | | 1 | | | | | | | | | | yes |
| ARHGAP20 | S_Shore | TSS1500 | chr11:110584063-110584327 | 264 | | | | | | | | | 1 | | | | | yes | yes | | | | |
| C11orf53 | NA | TSS200 | chr11:111126546-111126790 | 244 | | | | | | | | | | | | | | yes | yes | | | | |
| C11orf53 | NA | Body | chr11:111154826-111155021 | 195 | | | 5 | | | | | 0 | 2 | | | 1 | | | yes | | | | |
| C11orf93;C11orf92 | N_Shore | TSS1500;5'UTR | chr11:111169284-111169473 | 189 | | | | 1 | | | | | | | | | | | | | | | |
| POU2AF1 | NA | Body;1stExon;5'UTR;TSS200 | chr11:111249659-111250431 | 772 | | | 14 | | 2 | | | 0 | 3 | | | 1 | | | | | | | |
| C11orf88 | Island | TSS200 | chr11:111385338-111385496 | 158 | | 1 | | | | | | | | | | 1 | | | | | | | |
| LAYN | Island | TSS1500 | chr11:111410935-111411076 | 141 | | | 12 | | | | | | | | | | | | | | | | |
| PPP2R1B | S_Shore | TSS1500 | chr11:111637395-111637613 | 218 | | | | | | | | 0 | | | | | | | | | yes | | yes |
| HSPB2;CRYAB | NA | TSS1500;1stExon;TSS200 | chr11:111782364-111783563 | 1199 | | 1 | | | 1 | | | 0 | | | | | | | | | | yes | |
| C11orf52 | NA | TSS200 | chr11:111789452-111789693 | 241 | | | 6 | | 1 | | | | 1 | | | | | yes | | | | | |
| DIXDC1 | N_Shore;Island | Body;TSS1500;TSS200;1stExon;5'U | chr11:111846087-111848638 | 2551 | | 1 | 1 | | | | | 0 | 5 | | | | | | yes | | | | yes |
| DLAT | N_Shore | TSS1500 | chr11:111895234-111895455 | 221 | | | 4 | | | | | | | | | | | yes | yes | | | | |
| TIMM8B;SDHD | N_Shore | Body;TSS1500 | chr11:111956362-111956829 | 467 | | | | | | | | | 1 | | | | | yes | | | yes | | |
| IL18 | NA | 5'UTR;1stExon;TSS200 | chr11:112034801-112035175 | 374 | | | | | | | | | 2 | | | | | | | | yes | | yes |
| BCO2 | NA | TSS1500 | chr11:112045965-112046088 | 123 | | | | | | | | 0 | | | | | | | yes | | | | |
| NCAM1 | N_Shore | TSS1500;TSS200 | chr11:112831501-112831915 | 414 | | | 14 | | | | | 0 | 2 | | | 1 | | yes | | | | | |
| NCAM1 | NA | Body | chr11:112916105-112916250 | 145 | | | | | | | | | 1 | | | | | yes | | | | | |
| NCAM1 | NA | Body | chr11:113143664-113143807 | 143 | | 1 | | | | | | | 2 | | | | | | | | | | |
| TTC12 | N_Shore;Island | TSS1500;5'UTR | chr11:113184936-113185879 | 943 | | | 11 | | | | | | | | | | | yes | | yes | yes | | yes |
| ANKK1 | N_Shore;Island | TSS1500;TSS200 | chr11:113258210-113258908 | 698 | | | 6 | | | | | 0 | 1 | | | | | | | | | | |
| DRD2 | S_Shore;Island | TSS1500;5'UTR | chr11:113345686-113346491 | 805 | | | | | | | | | 2 | | | 1 | | | | | | yes | yes |

| | | | | | | | | | | | | | | | | | | | | | | | |
|---|---|---|---|---|---|---|---|---|---|---|---|---|---|---|---|---|---|---|---|---|---|---|---|
| TMPRSS5 | NA | Body;TSS200;TSS1500 | chr11:113576693-113577528 | 835 | 4 | | 6 | | | | | | | | | | | yes | | | | yes | |
| HTR3B | NA | TSS200 | chr11:113775400-113775900 | 500 | | | 3 | | | | | | | | | | | | | | | | |
| HTR3A | NA | TSS1500;nan;Body | chr11:113845487-113846937 | 1450 | | | | | 1 | | | 0 | 1 | 1 | 1 | 2 | | yes | yes | | | yes | yes |
| ZBTB16 | N_Shore | TSS1500 | chr11:113929164-113929287 | 123 | | | 2 | | | | | 0 | 1 | | | | | | | | | | |
| ZBTB16 | chr11:113953620-113953839;I | Island;Body | chr11:113953621-113954170 | 549 | | | | | | | | | 1 | | 1 | | | yes | yes | | | | |
| ZBTB16 | NA | Body | chr11:114112973-114113047 | 74 | | | | | | | | | | | | | | yes | | | yes | | |
| NNMT | NA | TSS200 | chr11:114166440-114166636 | 196 | | | | | | | | | 1 | | 1 | | | yes | | yes | | | yes |
| FAM55B | NA | TSS1500 | chr11:114548759-114549291 | 532 | | | | | | | | | 2 | | | | | | | | yes | yes | |
| CADM1 | NA | Body | chr11:115088649-115088907 | 258 | | | | | | | | | 1 | | | | | yes | | | | | |
| CADM1 | N_Shore;Island | Body | chr11:115372874-115375226 | 2352 | | | 8 | | | | | | 2 | | | | | | | | | yes | yes |
| APOA5 | Island | Body | chr11:116661043-116661678 | 635 | | | | | | | | | 2 | | | | | | | | | | |
| APOA4 | S_Shore | 1stExon;5'UTR;TSS200 | chr11:116694005-116694341 | 336 | | | | | | | | | 2 | | | | | yes | | | yes | | |
| APOC3 | NA | TSS1500;TSS200 | chr11:116700049-116700483 | 434 | | | | | | | | | | | | | | yes | | | | | yes |
| APOA1 | S_Shore | 5'UTR | chr11:116708299-116708517 | 218 | | | | 1 | | | | | | | | | | | | | yes | | |
| TAGLN | NA | TSS1500;TSS200 | chr11:117069780-117070046 | 266 | | | | 1 | 1 | | | 0 | 7 | 4 | 1 | 4 | | yes | | | | | yes |
| DSCAML1 | NA | Body | chr11:117314243-117314759 | 516 | 2 | | | | | | | | | | | | | | | | | | |
| DSCAML1 | Island | Body | chr11:117352692-117352790 | 98 | | | | | | | | | 1 | | | | | | | | yes | | yes |
| DSCAML1 | Island | Body | chr11:117389445-117389755 | 310 | | | 2 | | | | | | | | | | | yes | | | | | |
| DSCAML1 | S_Shelf | Body | chr11:117391914-117392002 | 88 | 2 | | | | | | | | | | | | | yes | | | | | |
| DSCAML1 | N_Shore | Body | chr11:117665395-117666889 | 1494 | 3 | | | | | | | 0 | | | | | | | | | | | |
| DSCAML1 | Island | TSS200 | chr11:117668140-117668336 | 196 | | | 1 | | | | | | | | | | | | yes | | | | yes |
| FXYD2 | NA | Body;TSS200 | chr11:117695591-117696015 | 424 | | | | | | | | 0 | | | | | | yes | | | | | |
| FXYD6 | S_Shore | 5'UTR;TSS200;TSS1500;1stExon | chr11:117747934-117748512 | 578 | 1 | | 23 | | | | | 0 | 1 | | | | | yes | yes | | | | |
| TMPRSS13 | NA | 5'UTR;1stExon;TSS200 | chr11:117800062-117800571 | 509 | | | 3 | 1 | | | | | 2 | | | | | yes | yes | | | | |
| TMPRSS4 | NA | TSS200;TSS1500 | chr11:117947214-117947997 | 783 | | 1 | 5 | | 1 | | | | 2 | 1 | | | | yes | | | | | |
| SCN4B | N_Shore;Island | Body;1stExon;5'UTR | chr11:118022607-118023765 | 1158 | | | 6 | | | | | | 1 | | | | | | | | | | |
| AMICA1 | NA | 1stExon;5'UTR;Body;TSS1500 | chr11:118083938-118085281 | 1343 | | | | | | | | | 1 | | | | | | yes | | | | |
| MPZL2 | NA | Body;TSS1500 | chr11:118134801-118135407 | 606 | | | | | 1 | | | | 2 | | | | | yes | | | | | |
| CD3E | NA | TSS200;5'UTR;1stExon | chr11:118175172-118175675 | 503 | | | | | 1 | | | | | | | | | yes | yes | | yes | | |
| CD3D | NA | 1stExon | chr11:118213272-118213414 | 142 | | | | | | | | | 1 | | | | | yes | | | | | |
| CD3G;CD3D | NA | TSS1500 | chr11:118214810-118215112 | 302 | | | | | | | | | 2 | | | | | yes | yes | | | yes | |
| UBE4A | N_Shore | TSS1500 | chr11:118230048-118230095 | 47 | | | 5 | | | | | 0 | | | | 1 | | yes | yes | | | | |
| TTC36 | N_Shelf | TSS1500 | chr11:118397867-118398318 | 451 | | | | | | | | | | | | | | | | | | | |
| TMEM25 | S_Shore | 5'UTR | chr11:118402280-118402540 | 260 | | | | | | | | | | | | | | | | | | | yes |
| PHLDB1 | N_Shore | TSS200;TSS1500 | chr11:118477034-118477147 | 113 | | | | | | | | | | | | | | | | | | | |
| TREH | NA | 1stExon;5'UTR;TSS200 | chr11:118550379-118550644 | 265 | | | 4 | | | | | | 1 | 1 | | | | yes | | yes | | | yes |
| CXCR5 | NA | TSS200 | chr11:118754507-118754593 | 86 | | | | | 2 | | | | | | | | | yes | | | yes | | |
| CXCR5 | NA | Body;TSS1500 | chr11:118763859-118764076 | 217 | | | | | | | | | 3 | | | | | yes | | | | | |
| BCL9L | NA | Body | chr11:118773657-118773983 | 326 | | | | | | | | | 1 | | | | | yes | | | | yes | |
| BCL9L | Island;S_Shore | 1stExon;5'UTR;TSS200 | chr11:118781150-118781978 | 828 | | | | | | | | 0 | 3 | | | | | yes | | | | | |
| UPK2 | NA | TSS1500 | chr11:118826709-118827082 | 373 | | | | | | | | | | | | | | | | | | | |
| FOXR1 | N_Shore;Island | TSS200;5'UTR;1stExon | chr11:118842295-118842650 | 355 | | | 16 | | | | | 0 | 6 | 1 | | 4 | | yes | yes | | | | yes |
| SLC37A4 | S_Shore | TSS1500 | chr11:118901842-118902351 | 509 | | | | | | | | | | | | 1 | | yes | | yes | | | |
| ABCG4 | N_Shore | TSS1500 | chr11:119019361-119019412 | 51 | | | 4 | | | | | | | | | | | | | | | yes | yes |
| MFRP;C1QTNF5 | NA | TSS1500 | chr11:119217884-119218238 | 354 | | | 1 | | | | | | | | | | | yes | | | | | |
| USP2 | N_Shore;Island | 3'UTR | chr11:119227020-119227876 | 856 | | | | 2 | 5 | | | | | | | | | | | | yes | | yes |
| THY1 | S_Shore;Island;chr11:1192933 | TSS200;5'UTR;Island | chr11:119293601-119294705 | 1104 | | 1 | 11 | | | | | | | | 1 | | | | | | | | |
| TRIM29 | NA | Body | chr11:119993722-119995687 | 1965 | | | | | | | | | 1 | 1 | | | | yes | | | yes | | |
| TRIM29 | Island;S_Shore | 1stExon;5'UTR;TSS200 | chr11:120008220-120009124 | 904 | | | 2 | | | | | 0 | 2 | | | 1 | | yes | | | | yes | |
| GRIK4 | NA | TSS1500;TSS200 | chr11:120530774-120531034 | 260 | | | 1 | | | | | | 1 | | | 1 | | yes | | | | | |
| GRIK4 | Island | Body | chr11:120856801-120857123 | 322 | | | | | | | | 0 | 1 | | | | | | yes | | | yes | yes |
| TECTA | NA | TSS200 | chr11:120973197-120973274 | 77 | | 1 | | | | | | | 1 | | | | | | | | | | |
| TECTA | NA | Body | chr11:121023627-121023720 | 93 | | 1 | | | | | | | | | | | | | | | | | |
| SORL1 | NA | Body | chr11:121440849-121440892 | 43 | | | | | 1 | | | | | | | | | | | yes | | | yes |
| LOC399959;MIR125B1 | NA | Body;TSS200;TSS1500 | chr11:121970285-121973246 | 2961 | | | | | | | | | | | | | | | | | | | |
| LOC399959;BLID | NA | Body;1stExon;5'UTR;TSS200 | chr11:121986736-121987203 | 467 | | | 5 | | | | | | 4 | | | | | | | | | | |
| MIR100;LOC399959 | NA | Body | chr11:122022999-122023504 | 505 | | | | | | | | | 1 | | | | | | | | | | |
| LOC399959 | NA | TSS1500 | chr11:122074011-122074433 | 422 | | | 2 | | | | | | | | | | | yes | | | | yes | |
| UBASH3B | Island | TSS200 | chr11:122526279-122526311 | 32 | | | | | | | | | | | | | | | | | | | yes |
| BSX | S_Shore | Body | chr11:122850140-122851122 | 982 | | 1 | | | | | | | | | | | | yes | | | | | yes |
| BSX | Island | TSS200 | chr11:122852412-122852698 | 286 | 2 | | | | | | | | 1 | | | 1 | | | | | | | |
| LOC341056 | NA | TSS1500 | chr11:122887526-122887993 | 467 | | | 1 | | | | | | | | | | | yes | | | | | |
| ASAM | Island;chr11:123066517-12306 | TSS200;TSS1500;Island | chr11:123066179-123067275 | 1096 | | | 11 | | | | | | 1 | | 1 | | | yes | yes | | | | yes |
| GRAMD1B | NA | TSS200;TSS1500;nan | chr11:123396039-123396593 | 554 | | 1 | 16 | | 1 | | | | 2 | 1 | 1 | | | | | | | | |
| GRAMD1B | NA | Body | chr11:123430575-123431162 | 587 | | | | | 1 | | | 0 | 3 | 2 | | 3 | | yes | | | | yes | yes |
| OR6X1 | NA | TSS1500 | chr11:123626485-123626544 | 59 | | | | | | | | | | | | | | | yes | | | yes | |
| TMEM225 | NA | 5'UTR;1stExon | chr11:123756314-123756564 | 250 | | | | | | | | | | | | | | | | | | | |
| OR6T1 | NA | TSS200;TSS1500 | chr11:123814620-123815202 | 582 | | | 7 | | | | | | 1 | | | | | yes | | | | | |
| OR10G8 | NA | TSS1500 | chr11:123899655-123900083 | 428 | | | | | | | | | | | | | | yes | | yes | | | |
| OR8B8 | NA | 1stExon;TSS1500 | chr11:124310939-124311423 | 484 | | | 5 | | | | | | | | | | | yes | | | | | |
| OR8A1 | NA | TSS200 | chr11:124439882-124440132 | 250 | | | | | | | | | 1 | | | | | | | | yes | | |
| PANX3 | NA | TSS1500 | chr11:124480990-124481218 | 228 | | | | | | | | | | | | | | yes | | | | | |
| VSIG2 | NA | Body;5'UTR;1stExon;TSS1500 | chr11:124621829-124622815 | 986 | | | | 1 | | | | 0 | 4 | 1 | | | | | | | | | |
| ESAM | Island | Body | chr11:124629580-124629925 | 345 | | | 4 | | 1 | | | | | | | | | | | | | | |
| ROBO3 | Island | 5'UTR;1stExon;TSS200 | chr11:124735105-124735994 | 889 | | | | | | | | | 3 | | | | | | | | | | |
| ROBO3 | S_Shore | Body | chr11:124747075-124747263 | 188 | | | 5 | | | | | 0 | | | | | | | | | yes | | |
| ROBO4 | NA | Body;5'UTR;1stExon;TSS200;TSS1 | chr11:124767455-124768950 | 1495 | | | 16 | 1 | 2 | | | | 3 | | | | | yes | yes | | | yes | |
| HEPACAM | Island | 3'UTR | chr11:124790815-124791389 | 574 | | | | | | | | | 1 | | | | | yes | yes | | yes | yes | |
| HEPACAM | NA | 5'UTR;1stExon | chr11:124805912-124805981 | 69 | | | | 1 | | | | | 1 | | | | | | | | | | |
| PKNOX2 | Island;S_Shore | 5'UTR | chr11:125036088-125037348 | 1260 | | | 13 | | 1 | | | | | | | | | | | | | | yes |
| PKNOX2 | NA | 5'UTR | chr11:125106056-125106135 | 79 | | | | | | | | | 1 | | | | | | yes | | | | |
| FEZ1 | Island;S_Shore | 5'UTR;TSS1500 | chr11:125365478-125366540 | 1062 | | 1 | | | | | | | 1 | | | | | | | | | yes | |
| PATE1 | NA | TSS200 | chr11:125616118-125616168 | 50 | | | | | | | | | | | | | | | yes | | | | |
| PATE4 | NA | TSS1500 | chr11:125702592-125703230 | 638 | | | | | | | | | | | | | | yes | yes | yes | | yes | |
| DDX25;PUS3 | N_Shore | TSS200;TSS1500 | chr11:125774082-125774447 | 365 | | 1 | | | 1 | | | | | | | | | | | | | yes | yes |
| TIRAP | Island | TSS200 | chr11:126152840-126152981 | 141 | | | | | | | | | | | | 1 | | yes | | | | | yes |
| DCPS | Island;N_Shore | TSS200;1stExon;5'UTR;TSS1500 | chr11:126173246-126174329 | 1083 | | | 3 | | | | | 0 | 2 | 1 | | 2 | | yes | yes | | | yes | |
| DCPS | NA | Body | chr11:126197277-126198043 | 766 | | | | 1 | | | | | | | | | | yes | yes | | | | |
| KIRREL3 | NA | Body | chr11:126310323-126310475 | 152 | | | | | 1 | | | | | | | | | | yes | | | | |
| KIRREL3 | NA | Body | chr11:126456008-126456119 | 111 | 1 | | | | | | 1 | 0 | 1 | 1 | | | | yes | | | | | |
| KIRREL3 | NA | Body | chr11:126581947-126582317 | 370 | | | 9 | | | | | 0 | | 1 | | | | yes | | | yes | yes | |
| KIRREL3 | NA | Body | chr11:126763432-126763487 | 55 | | | | | | | | | 1 | | | | | | | | | | |
| ETS1 | N_Shore | Body | chr11:128391271-128391494 | 223 | | | | | 1 | | | | 1 | | | | | | | | | yes | |
| ETS1 | N_Shore;Island | Body | chr11:128418692-128419512 | 820 | | 1 | | 1 | | | | | | | | | | yes | | | | | |
| ETS1 | NA | 1stExon;5'UTR | chr11:128457400-128457682 | 282 | | | | | 1 | | | | | | | | | | | | | | |
| FLI1 | N_Shore | TSS200 | chr11:128562245-128562408 | 163 | | | | | | | | | 2 | | | | | | | | | | |
| FLI1 | NA | Body | chr11:128646953-128647107 | 154 | | | 17 | | | | | | 1 | | | | | yes | | | | | |
| KCNJ1 | NA | 5'UTR;TSS200 | chr11:128712399-128712462 | 63 | | | | | | | | | | | | | | | | | | | |
| KCNJ1 | NA | 5'UTR | chr11:128729652-128730011 | 359 | | 1 | 18 | | | | | | 1 | 1 | | | | yes | | | | | |

| | | | | | | | | | | | | | | | | | | | | | | | | | | | |
|---|---|---|---|---|---|---|---|---|---|---|---|---|---|---|---|---|---|---|---|---|---|---|---|---|---|---|---|
| KCNJ1 | NA | TSS200 | chr11:128737300-128737415 | 115 | | | | | 1 | | | | | | | | | | | | yes | yes | | | | | |
| KCNJ5 | N_Shore;Island | TSS1500;TSS200 | chr11:128760367-128761419 | 1052 | | | | | | | | | 2 | | | | | | | | | yes | yes | | | | yes |
| KCNJ5;C11orf45 | NA | 5'UTR;Body | chr11:128774411-128774649 | 238 | | | | | | | | | 1 | | | | | yes | yes | | yes | | | | | | |
| C11orf45;KCNJ5 | NA | TSS1500;5'UTR | chr11:128776059-128776216 | 157 | | | | | | | | | | | | | | yes | | yes | | | | | | | |
| TP53AIP1 | NA | 5'UTR;TSS200;nan;TSS1500 | chr11:128812462-128813583 | 1121 | 4 | | 29 | | | | | | 3 | 1 | 1 | | | yes | yes | | yes | | | | | | |
| BARX2 | Island | TSS1500;TSS200 | chr11:129245558-129246394 | 836 | | | | | | | | | 2 | | | | | | | | | | yes | | | | |
| BARX2 | N_Shore;Island | Body | chr11:129306366-129306881 | 515 | | | 5 | | | | | | 1 | | | | | yes | | | yes | | | | | | |
| PRDM10 | NA | TSS200;Body;5'UTR;1stExon | chr11:129817418-129817717 | 299 | | | | | | | | | 2 | | | | | yes | | yes | | | | | | | |
| APLP2 | Island | TSS200 | chr11:129939695-129939713 | 18 | | | | | | | | | 1 | | | | | | | | | yes | yes | | | | |
| APLP2 | N_Shelf;N_Shore | Body | chr11:129991445-129992368 | 923 | | | 32 | | 2 | | | 0 | 1 | | | | | | yes | | | yes | yes | | | | |
| APLP2 | NA | Body;3'UTR | chr11:130013355-130013410 | 55 | 3 | | | | | | | | | | | | | | yes | | | | | | | | |
| ST14 | N_Shore | TSS1500;TSS200 | chr11:130028844-130029684 | 840 | | | 10 | | | | | | | | | | | | yes | | | | yes | | | | |
| ST14 | Island | Body | chr11:130059549-130059791 | 242 | | 1 | | | 2 | | | | 1 | | | | | yes | | | | | | | | | |
| ST14 | NA | Body | chr11:130068175-130068448 | 273 | | | | | | | | | | | | | | | | | | | | | | | |
| ST14 | N_Shelf | Body | chr11:130076148-130076290 | 142 | | | | | | | | | 1 | | | | | yes | yes | | | | | | | | |
| ZBTB44 | S_Shore | TSS1500 | chr11:130185384-130185670 | 286 | | | | | | | | 0 | 1 | 1 | | 3 | | yes | | | | | yes | | | | |
| ADAMTS8 | S_Shore | TSS1500 | chr11:130298880-130299333 | 453 | | | | | | | | | | | | | | | | | | | | | | | |
| NTM | NA | TSS1500;1stExon;5'UTR | chr11:131239289-131240660 | 1371 | | | | | 1 | | | | 4 | | | | | yes | yes | | | yes | yes | | | | |
| NTM | NA | Body | chr11:131410868-131411235 | 367 | | | | | | | | | | | | | | yes | | | | | | | | | |
| NTM | NA | Body | chr11:131506338-131506539 | 201 | | | | | | | | | 1 | | | | | yes | | | yes | | | | | | |
| NTM | NA | Body | chr11:131630464-131630648 | 184 | | | | | | | | | 1 | | | 1 | | yes | | | | | | | | | |
| NTM | N_Shore;Island | TSS1500;Body | chr11:131779255-131780946 | 1691 | 4 | | 14 | | | | | 0 | 5 | 1 | | 1 | | | | | | yes | yes | | | | |
| OPCML | NA | Body | chr11:132526947-132527438 | 491 | | | 6 | | | | | | | | | | | | | | | | | | | | |
| OPCML | NA | Body | chr11:132662455-132662963 | 508 | | | 8 | 1 | | | 1 | 0 | 3 | 1 | | 4 | 1 | | | | | | yes | | | | |
| OPCML | Island | TSS1500;Body | chr11:132813563-132814206 | 643 | | | | | | | | | 1 | | | | | | | | | | yes | | | | |
| OPCML | NA | Body | chr11:132912205-132912368 | 163 | | | 1 | | | | | | | | | | | | | | | | | | | | |
| OPCML | N_Shelf | Body | chr11:132947723-132948177 | 454 | | | 2 | | | | | | | | | | | | yes | | | | | | | | |
| OPCML | Island | Body | chr11:132951838-132951950 | 112 | | | 5 | | | | 1 | | 1 | 1 | | 3 | | | | | | | yes | | | | |
| OPCML | NA | Body | chr11:133098424-133098499 | 75 | | | | | | | | 0 | 1 | | | | | yes | | | | | | | | | |
| OPCML | N_Shore | Body | chr11:133229994-133230391 | 397 | | | | | | | | | | | | | | yes | yes | | | | yes | | | | |
| OPCML | NA | TSS200 | chr11:133402545-133402919 | 374 | | | | | | | | | 1 | | | | | | | | | yes | | | | | |
| SPATA19 | NA | Body | chr11:133715076-133715511 | 435 | | | | | | | | | 1 | | | | | | | yes | | | | | | | |
| LOC283174 | NA | Body | chr11:133770764-133771191 | 427 | | | 2 | | | | | | | | | | | | | | | yes | | | | | |
| LOC283174 | NA | TSS1500 | chr11:133772615-133773069 | 454 | | | | | | | | | 1 | | | | | | | | | yes | | | | | |
| IGSF9B | N_Shore;Island | Body | chr11:133800241-133801225 | 984 | 9 | | 11 | | | | | | 2 | | | | | yes | | | | | | | | | |
| IGSF9B | S_Shelf | Body | chr11:133804526-133805057 | 531 | | | 6 | | | | | | 1 | | | | | | | | | | | | | | |
| IGSF9B | NA | Body | chr11:133820837-133821252 | 415 | | | 14 | | | | | | | | | | | yes | | | | yes | | | | | |
| IGSF9B | N_Shore;Island | Body;TSS200 | chr11:133825319-133827520 | 2201 | | 1 | | | | | | | 1 | | | | | | yes | | | | yes | | | | |
| LOC100128239 | NA | TSS200 | chr11:133902108-133902371 | 263 | | | | | | | | 0 | | | | | | | | | | | | | | | |
| JAM3 | N_Shore;Island | Body | chr11:133994642-133995182 | 540 | | 1 | 7 | | | | | 0 | 1 | | | | | | | | | | | | | | |
| ACAD8 | S_Shelf | Body | chr11:134126284-134126504 | 220 | | | | | | | 1 | 0 | 3 | 3 | 1 | 3 | | yes | | | | | | | | | |
| GLB1L3 | S_Shore | Body | chr11:134147327-134147634 | 307 | | | | | | | | 0 | | | | | | | | | | | | | | | |
| GLB1L3 | NA | Body | chr11:134176692-134176888 | 196 | | | 5 | | | | | | | | | | | | yes | | | | | | | | |
| GLB1L2 | Island | 1stExon;5'UTR | chr11:134201952-134201993 | 41 | | 1 | | | | | | | 1 | | | | | | | | | | yes | | | | |
| GLB1L2 | Island | Body | chr11:134211952-134212324 | 372 | | | | | | | | 0 | | | | | | | | | | | yes | | | | |
| GLB1L2 | S_Shelf | Body | chr11:134216065-134216220 | 155 | | | | | | | | | 1 | | | | | yes | | | | | | | | | |
| GLB1L2 | N_Shelf | Body | chr11:134228227-134228370 | 143 | | | | | | | | | | | | | | | | | | | | | | | |
| GLB1L2 | N_Shore;S_Shore | Body | chr11:134231621-134232313 | 692 | | | 2 | | | | | | | | | | | yes | | | | yes | | | | | |
| B3GAT1 | Island;S_Shore | Body | chr11:134253609-134254877 | 1268 | | | | | 1 | | | | 1 | | | 1 | | yes | yes | | | | | | | | |
| B3GAT1 | Island | Body | chr11:134257429-134257617 | 188 | | | | | | | | | | | | | | yes | | yes | | | | | | | |
| B3GAT1 | NA | 5'UTR | chr11:134262988-134263349 | 361 | | | | | | | | | | | | | | yes | yes | | yes | | | | | | |
| IQSEC3 | Island | TSS1500 | chr12:175809-176424 | 615 | | | | | | | | | 1 | | | | | yes | | | | | yes | | | | |
| IQSEC3 | Island | Body;5'UTR | chr12:213776-214224 | 448 | | | 9 | | | | | 0 | 1 | | | 1 | | yes | yes | | | | | | | | |
| IQSEC3 | N_Shore | Body;5'UTR | chr12:247224-247459 | 235 | | | | | | | | | 1 | | | | | yes | yes | | | yes | yes | | | | |
| IQSEC3 | S_Shore | Body | chr12:248889-249286 | 397 | | | | | | | | | 1 | | | | | yes | yes | | yes | | | | | | |
| IQSEC3 | NA | Body | chr12:258355-258436 | 81 | 1 | | | | | | | | 1 | 1 | | | | | yes | | | | | | | | |
| SLC6A12 | Island | Body | chr12:312592-313104 | 512 | | | | 1 | | | 1 | 0 | 1 | 1 | | 8 | | yes | | | | | yes | | | | |
| SLC6A12 | NA | 5'UTR;TSS200;TSS1500 | chr12:322514-323242 | 728 | | 1 | 10 | 1 | | | | | 3 | | | | | | | | | | yes | | | | |
| SLC6A13 | NA | 5'UTR;TSS200 | chr12:371916-372259 | 343 | 2 | | 4 | 1 | | | | 0 | 3 | | | | | yes | yes | | | yes | | | | | |
| B4GALNT3 | N_Shore | TSS1500 | chr12:568543-568826 | 283 | | | | | | | | | | | | | | yes | | | | yes | yes | | | | |
| B4GALNT3 | NA | Body | chr12:656703-657046 | 343 | | 1 | 1 | | | | | | | | | | | yes | yes | | | | | | | | |
| B4GALNT3 | NA | Body;3'UTR | chr12:670520-671027 | 507 | | | 4 | | | | 1 | | 2 | 1 | | | | yes | | | | | | | | | |
| NINJ2 | N_Shore;Island | Body | chr12:678853-679668 | 815 | 1 | 2 | 2 | | | | | 0 | 1 | | 1 | 1 | | | | | | | | | | | |
| NINJ2 | NA | Body | chr12:739980-740338 | 358 | | | 2 | | | | | 0 | 1 | 1 | 1 | 6 | | yes | | | | | | | | | |
| NINJ2 | NA | 1stExon;5'UTR;TSS200 | chr12:772529-772982 | 453 | | | 18 | 1 | | | | | | | | | | | | | | | | | | | |
| RAD52 | NA | Body | chr12:1025529-1025755 | 226 | | | | | 1 | | | | 1 | 1 | | 1 | | yes | | | | yes | | | | | |
| LOC100292680 | NA | TSS200;TSS1500 | chr12:1609034-1609549 | 515 | | 1 | 27 | | | | | | 2 | | | | | | | | | | | | | | |
| FBXL14 | NA | 3'UTR | chr12:1675846-1676077 | 231 | | | | | | | | | 1 | | | | | yes | yes | | | yes | | | | | |
| WNT5B | N_Shore | TSS200;5'UTR;TSS1500 | chr12:1738110-1738446 | 336 | | | | | | | | | | | | | | | | | | yes | yes | | | | |
| WNT5B | Island | 5'UTR | chr12:1739687-1739917 | 230 | | | 7 | | | | | | | | | | | | | | | | | | | | |
| WNT5B | Island | Body | chr12:1748816-1749159 | 343 | | | | | | | | | 1 | | | | | yes | yes | | | yes | | | | | |
| CACNA2D4 | N_Shore | Body | chr12:1904502-1904847 | 345 | | | 4 | | | | | | | | | | | | | yes | yes | | | | | | |
| LRTM2;CACNA2D4 | NA | TSS200;Body | chr12:1929265-1929588 | 323 | | | | | | | | 0 | 1 | 2 | | 1 | | | | | | | yes | | | | |
| LRTM2;CACNA2D4 | NA | Body | chr12:1937877-1938156 | 279 | | | | | | | | | | | | | | yes | | | | | | | | | |
| LRTM2;CACNA2D4 | NA | nan;Body | chr12:1940380-1940618 | 238 | | | | | | | | 0 | | 1 | 1 | | | | | | | | | | | | |
| CACNA2D4 | NA | Body | chr12:1948435-1948648 | 213 | | | 2 | | | | | | | | | | | yes | | | | | | | | | |
| CACNA2D4 | Island | Body | chr12:1973872-1974168 | 296 | | | | | | | | 0 | 1 | | | 2 | | | | | | | | | | | |
| CACNA2D4 | N_Shore | Body | chr12:2017093-2017649 | 556 | | | 7 | | | | | 0 | 1 | | | 1 | | | | | | | | | | | |
| CACNA1C | Island | Body | chr12:2224410-2224755 | 345 | 2 | | | | | | | | | | | | | yes | yes | | yes | | | | | | |
| CACNA1C | Island | Body | chr12:2339439-2339614 | 175 | | | 9 | | 1 | | | 0 | 1 | | | 2 | | yes | yes | | | | | | | | |
| CACNA1C | NA | Body | chr12:2734150-2734591 | 441 | | | | | | | | | 1 | | | | | yes | | | | | | | | | |
| CACNA1C | N_Shore;Island;chr12:2800139 | Body;3'UTR;Island | chr12:2800055-2801584 | 1529 | 4 | | 41 | | | | | | | | 2 | | | yes | | | | | yes | | | | |
| NRIP2 | NA | TSS1500;1stExon;5'UTR;TSS200 | chr12:2943902-2944493 | 591 | | | | | | | | 0 | 10 | 1 | | 3 | | | yes | | | yes | | | | | |
| FOXM1;C12orf32 | S_Shore | TSS1500;Body | chr12:2986879-2987138 | 259 | | | 1 | | | | | | | | | | | yes | yes | | | | | | | | |
| TSPAN9 | NA | 5'UTR | chr12:3240109-3240217 | 108 | | | | 1 | | | | | | | | | | yes | | | | | | | | | |
| TSPAN9 | Island | 5'UTR | chr12:3309264-3310269 | 1005 | | | | 1 | | | | | | | | | | | | | | | yes | | | | |
| TSPAN9 | NA | Body | chr12:3371886-3371908 | 22 | | | | | 1 | | | | | | | | | yes | | | | | | | | | |
| TSPAN9 | NA | Body | chr12:3373534-3373764 | 230 | | | | | | | | | | | | | | yes | | | | | | | | | |
| PRMT8 | Island | TSS1500 | chr12:3600087-3600764 | 677 | | | | | | | | | 1 | | | | | | | | | yes | yes | | | | |
| EFCAB4B | N_Shore;Island | 5'UTR;1stExon;TSS200;TSS1500 | chr12:3861567-3862810 | 1243 | 1 | | 4 | 1 | 7 | | | 0 | 2 | 1 | | 1 | | yes | yes | | | yes | yes | | | | |
| CCND2 | Island | TSS1500 | chr12:4381435-4382188 | 753 | | 3 | 2 | | | | | | | | | | | | | | | yes | | | | | |
| FGF23 | NA | TSS200 | chr12:4489075-4489221 | 146 | | | | | | | | | 1 | | | 1 | | | | | | | | | | | |
| FGF6 | Island;N_Shore | 1stExon;Body | chr12:4554295-4554786 | 491 | | | | | | | | | 1 | | | | | | | | | | | | | | |
| DYRK4 | NA | TSS200 | chr12:4699085-4699618 | 533 | | | | | | | | 0 | 1 | 1 | | | | yes | yes | | | | | | | | |
| GALNT8 | NA | TSS200 | chr12:4829617-4829636 | 19 | | | 3 | | | | | | | | | | | | | yes | | | | | | | |
| KCNA6 | Island | 5'UTR;1stExon | chr12:4918391-4919591 | 1200 | | 1 | | | 1 | | | 0 | 1 | 1 | 1 | 7 | 1 | | | | | | | | | | |
| KCNA6 | S_Shelf | 3'UTR;1stExon | chr12:4923430-4923529 | 99 | | | | | | | | | 1 | | | | | | | | | | | | | | |
| KCNA1 | N_Shore;Island | TSS1500;TSS200 | chr12:5018229-5019933 | 1704 | | 1 | | | | | | | 1 | | | | | | | | | | | | | | |
| KCNA1 | Island | Body | chr12:5021087-5021871 | 784 | 3 | | | | | | | | | | | | | | | | | | | | | | |

| | | | | | | | | | | | | | | | | | | | | | | | |
|---|---|---|---|---|---|---|---|---|---|---|---|---|---|---|---|---|---|---|---|---|---|---|---|
| KCNA5 | Island | 5'UTR;1stExon | chr12:5153174-5154148 | 974 | | 1 | | | | | | | | | | | | | | | | | yes |
| NTF3 | N_Shore;Island | TSS1500 | chr12:5539832-5540426 | 594 | 1 | | 11 | 1 | | | | | | | | | | | | | | | |
| NTF3 | N_Shore | TSS200;Body | chr12:5603131-5603180 | 49 | | | | | 2 | | | | | | | | | | | | | | |
| ANO2 | Island | Body | chr12:5936980-5937409 | 429 | | | 12 | | | 1 | | | 3 | | | | | yes | | | | | |
| ANO2 | NA | Body | chr12:6054362-6054749 | 387 | | | | | | | | 0 | | | | | | | | | | yes | yes |
| VWF | NA | Body | chr12:6125715-6125828 | 113 | | | | | | | | | | | | | | yes | | | | | |
| VWF | N_Shore;Island | Body | chr12:6165683-6167147 | 1464 | | | 10 | 1 | | | | | | | | | | yes | yes | | | yes | |
| VWF | NA | Body | chr12:6184176-6184516 | 340 | | | 1 | | | | | | | | | | | | | | | yes | |
| CD9 | N_Shore | TSS1500 | chr12:6308758-6309025 | 267 | | | | 1 | | | | | | | | | | | yes | | | | yes |
| PLEKHG6 | N_Shore;S_Shore | TSS1500;TSS200;5'UTR | chr12:6419409-6420252 | 843 | | | 3 | | | | | | 3 | | | | | yes | | | yes | | yes |
| PLEKHG6 | S_Shore | TSS200;Body | chr12:6421667-6421806 | 139 | | | | | | | | | 1 | | | | | | | | | | |
| TNFRSF1A | S_Shelf;NA | Body | chr12:6442873-6443282 | 409 | | | 35 | | | | | | 1 | | | | | yes | | | | | |
| SCNN1A | Island | Body | chr12:6473299-6473643 | 344 | | | | | | | | | | | | | | yes | yes | | yes | | |
| SCNN1A | NA | 5'UTR;Body;1stExon;TSS200 | chr12:6484219-6484713 | 494 | | | | 1 | | | | | | | | | | | | | | | yes |
| SCNN1A | NA | TSS200 | chr12:6486598-6487080 | 482 | | | 2 | | | | | 0 | 1 | | | | | yes | yes | | | | |
| LTBR | Island | TSS1500;TSS200 | chr12:6492890-6493521 | 631 | | | | | | | | | 2 | 1 | | | | yes | | | | | |
| CD27;LOC678655 | NA | TSS1500;Body | chr12:6553145-6553585 | 440 | | | | | | | | | 1 | | | | | | | | yes | | |
| MRPL51;NCAPD2 | N_Shore | Body;TSS1500 | chr12:6602187-6602447 | 260 | | | | | | | | | 1 | | | | | yes | | | | | |
| IFFO1 | Island | Body | chr12:6649733-6649994 | 261 | | | | 1 | 2 | | | 0 | | | | | | yes | | | | | |
| IFFO1 | Island | Body;1stExon | chr12:6657744-6658164 | 420 | | | 3 | | | | | | | | | | | | | | | | yes |
| IFFO1 | Island;chr12:6664425-6665336 | 1stExon;Island;TSS200 | chr12:6664536-6665447 | 911 | | | | | 3 | | | | 1 | | 1 | | | yes | | | | | yes |
| LPAR5 | NA | 5'UTR;TSS1500 | chr12:6741649-6741882 | 233 | | | | | 1 | | | | | | | | | yes | yes | | | | |
| LPAR5 | NA | 5'UTR;1stExon | chr12:6745057-6745568 | 511 | | | 2 | | | | | | | | | | | | yes | | yes | yes | |
| ACRBP | Island | Body;1stExon;5'UTR | chr12:6756377-6756751 | 374 | | | 4 | | | | | 0 | | | | | | yes | | | | | |
| ZNF384 | N_Shore | 5'UTR;TSS200 | chr12:6797909-6798118 | 209 | | | | | | | | | 1 | | | | | | | | | | |
| LAG3 | N_Shore | TSS200 | chr12:6881590-6881997 | 407 | | | 5 | | 1 | 1 | | | 1 | 1 | 1 | | | yes | | | | | |
| CD4 | NA | TSS200 | chr12:6898500-6898785 | 285 | | | 2 | 1 | | | | | 1 | | | | | | yes | | | | yes |
| GPR162 | Island | Body | chr12:6933322-6933689 | 367 | | | 2 | | | | | | | | | | | | | | | | |
| GNB3;LEPREL2 | NA | TSS1500;3'UTR | chr12:6948846-6949472 | 626 | | | | | 1 | | | | 3 | | | | | | | | | yes | |
| CDCA3 | N_Shore | Body | chr12:6959471-6959657 | 186 | | | | | | | | | | | | | | yes | | | | yes | |
| ENO2;ATN1 | NA | 3'UTR;TSS1500 | chr12:7032793-7033146 | 353 | | | | | | | | | 1 | | | | | | | | | yes | |
| PTPN6;C12orf57 | S_Shore | TSS1500;3'UTR;TSS200 | chr12:7055153-7056008 | 855 | | 1 | | 1 | | | | | | | | | | | yes | | | | |
| PTPN6 | NA | TSS1500;Body;TSS200 | chr12:7060187-7060569 | 382 | | | 11 | | | | | | 1 | 1 | | | | yes | | | | | |
| PTPN6 | NA | Body | chr12:7066563-7066664 | 101 | | | | | | | | | 1 | | | | | yes | yes | | | | |
| PTPN6;MIR200C;MIR141 | NA | 3'UTR;TSS1500;TSS200 | chr12:7070317-7073264 | 2947 | | | | | | | | | | | | | | | | | | | |
| C1S | NA | TSS1500 | chr12:7167584-7167900 | 316 | | | | | | | | | 2 | | | | | yes | | | yes | | |
| C1RL;LOC283314 | NA | Body;TSS1500 | chr12:7260546-7260776 | 230 | | | 3 | | | | | | | | | | | yes | | | | | yes |
| RBP5 | NA | 3'UTR | chr12:7276360-7276714 | 354 | | | | | | | | | | | | | | | | | | | |
| RBP5;CLSTN3 | NA | 1stExon;TSS1500;TSS200;nan | chr12:7281343-7283120 | 1777 | | | 12 | | | | | | | | 1 | | | | | | | | |
| PEX5 | N_Shore | TSS1500;TSS200 | chr12:7341252-7341666 | 414 | | | | 1 | | | | | | | | | | yes | | | | | |
| CD163L1 | Island | Body | chr12:7592509-7593007 | 498 | | 1 | | | | | | 0 | 1 | | | | | yes | yes | | | | |
| APOBEC1 | NA | TSS200 | chr12:7818585-7818654 | 69 | | 1 | | | | | | | 1 | | | | | yes | | | | yes | |
| GDF3 | NA | Body | chr12:7847919-7848348 | 429 | | | 1 | | | | | | | | | | | yes | yes | | yes | | |
| DPPA3 | NA | TSS1500;TSS200 | chr12:7863853-7864122 | 269 | | | 4 | | | | | | | | | | | yes | | | | yes | |
| CLEC4C | NA | TSS1500 | chr12:7902721-7903170 | 449 | | | 6 | | | | | | | | | | | yes | | yes | | yes | |
| NANOG | NA | TSS1500 | chr12:7940546-7941120 | 574 | | | | | 1 | | | | 3 | | | | | yes | yes | | | | |
| SLC2A14 | Island | 5'UTR;TSS200 | chr12:8025394-8025646 | 252 | | 1 | 1 | | 1 | | | 0 | 1 | | | 2 | | yes | yes | | | | yes |
| C3AR1 | NA | 1stExon;5'UTR | chr12:8218922-8219358 | 436 | | | 1 | | | | | | | | | | | | yes | | | yes | |
| FAM66C | Island | TSS200 | chr12:8332802-8333010 | 208 | | | | | | | | | | | | | | | yes | | | yes | |
| MFAP5 | NA | 1stExon;5'UTR | chr12:8815308-8815634 | 326 | 2 | | 7 | | | | | | | | | | | yes | | | | | |
| A2ML1 | NA | 1stExon;5'UTR | chr12:8975196-8975309 | 113 | | | | | | | | | 1 | | | | | yes | yes | | | | |
| LOC144571 | S_Shore;N_Shore;Island | TSS200;Body;TSS1500 | chr12:9217079-9218039 | 960 | | | 3 | | | 1 | | 0 | 24 | 3 | 2 | 22 | | yes | | yes | | yes | |
| A2M | NA | Body | chr12:9227173-9227237 | 64 | | 1 | | 1 | | | | | | | | | | yes | | | | | |
| PZP | NA | 1stExon | chr12:9360854-9360986 | 132 | | | | 1 | | | | | | | | | | | | | | | |
| CLEC2B | NA | 1stExon;5'UTR | chr12:10022435-10022688 | 253 | | | | | | | | | | | | | | | yes | | yes | | |
| CLEC2A | NA | 5'UTR;1stExon | chr12:10084936-10085174 | 238 | | | 18 | | | | | | 1 | | | | | yes | yes | | | | |
| CLEC9A | NA | TSS200 | chr12:10183167-10183364 | 197 | | | | | | | 1 | 0 | 4 | 7 | | 1 | | | yes | yes | | | |
| CLEC1A | NA | 5'UTR;1stExon | chr12:10251536-10252119 | 583 | | | 15 | | | | | | | | 1 | | | yes | | yes | | | |
| KLRD1 | NA | TSS1500;5'UTR | chr12:10459923-10460639 | 716 | | | | | | | | | 1 | | | | | yes | yes | | | | |
| STYK1 | S_Shore | TSS200 | chr12:10827012-10827583 | 571 | | | 3 | | | | | | | | | | | yes | yes | | | yes | yes |
| PRR4 | NA | TSS200;Body;TSS1500 | chr12:11002197-11002609 | 412 | | | | | | | | 0 | 1 | | | | | | yes | | | | yes |
| PRR4;TAS2R20;PRH1 | NA | Body;TSS1500;5'UTR | chr12:11151430-11151622 | 192 | | | 5 | | | | | | | | | | | | | | | yes | yes |
| TAS2R30;PRR4;PRH1 | NA | TSS1500;5'UTR | chr12:11287271-11287815 | 544 | | | | | | | | | 1 | | | | | | | | | | yes |
| BCL2L14 | NA | TSS1500;TSS200;5'UTR;1stExon | chr12:12223661-12224457 | 796 | | 1 | | | | | | 0 | 2 | | | 1 | | yes | | | | | |
| LRP6 | N_Shore | Body | chr12:12419053-12419209 | 156 | | | | | | | | | 1 | | | | | | | | | | yes |
| GPR19 | Island;S_Shore;N_Shore | TSS200;TSS1500;5'UTR;1stExon | chr12:12848977-12850082 | 1105 | | | 1 | | | | | 0 | 4 | | 1 | 5 | | | | | | yes | |
| APOLD1 | N_Shelf;N_Shore | Body;TSS1500 | chr12:12937632-12938425 | 793 | | 1 | 4 | | | | | | | | | | | | | | | | |
| DDX47 | N_Shore | TSS1500 | chr12:12965884-12966031 | 147 | | | 3 | | | | | 0 | | | | | | yes | yes | | | | |
| GPRC5A | N_Shore | TSS1500 | chr12:13043502-13043755 | 253 | | | 4 | | | | | | | | | | | yes | | | | | |
| KIAA1467 | N_Shore | TSS1500 | chr12:13196876-13196954 | 78 | 2 | | 1 | | | | | | 1 | | | | | yes | | | | | |
| GSG1 | S_Shelf | TSS200;TSS1500 | chr12:13256639-13257892 | 1253 | | | | | | | | | 2 | | | | | | yes | | yes | | |
| GRIN2B | N_Shore | 3'UTR | chr12:13715497-13715942 | 445 | | | | | | | | | | | | | | | | | | | |
| GRIN2B | NA | 5'UTR | chr12:14109211-14109584 | 373 | | | 2 | | | | | | | | | | | | yes | | | | |
| GRIN2B | N_Shore | TSS200 | chr12:14133129-14133336 | 207 | | 1 | | | | | | | | | | | | | | | | yes | yes |
| PLBD1 | Island | TSS200 | chr12:14720938-14721288 | 350 | | | 12 | | | | | 0 | 1 | | | 1 | | yes | yes | | | yes | yes |
| GUCY2C | NA | 1stExon;5'UTR | chr12:14849268-14849579 | 311 | | | 4 | | | | | | 1 | | | | | yes | yes | | | yes | |
| H2AFJ | N_Shore | TSS1500 | chr12:14926572-14928083 | 1511 | | | | | | | | | | | | | | | | | | | |
| ART4 | NA | 5'UTR;1stExon | chr12:14996143-14996776 | 633 | | | | 1 | | | | | | | | | | | yes | | | | |
| MGP | NA | TSS200;TSS1500 | chr12:15039025-15039560 | 535 | | | 1 | | | | | | 1 | | | | | | | | | | yes |
| ARHGDIB | NA | 5'UTR;1stExon;TSS200 | chr12:15114233-15114904 | 671 | | | 42 | 1 | | | | | | | | | | yes | | | yes | | |
| MGST1 | NA | TSS1500;TSS200 | chr12:16499963-16500403 | 440 | | | | | | | | | | | | | | | yes | | | | |
| MGST1 | NA | Body | chr12:16512895-16513009 | 114 | | | 2 | | | | | | 1 | | | | | | | | | | yes |
| LMO3 | NA | 5'UTR | chr12:16757954-16758715 | 761 | | | 4 | | | | | | 3 | | | | | | | | | | |
| LMO3 | NA | 5'UTR;TSS1500;1stExon | chr12:16759782-16761118 | 1336 | | | 3 | 1 | | | | 0 | | | | 1 | | | | yes | | | yes |
| PDE3A | Island;N_Shore | TSS1500 | chr12:20521087-20523143 | 2056 | | | | | | | | | 2 | | | | | | | | | yes | yes |
| C12orf39 | Island | Body | chr12:21680409-21680508 | 99 | | | | | | | | | 1 | | | | | | | | | | |
| LDHB | N_Shore | 5'UTR | chr12:21809973-21810139 | 166 | | | | 1 | | | | | | | | | | | | | | | |
| KCNJ8 | NA | Body | chr12:21926447-21926524 | 77 | | | 5 | | | | | 0 | 1 | | | 2 | | | | | | | |
| ST8SIA1 | N_Shore | Body | chr12:22486029-22486484 | 455 | | | | | | | | | 1 | | | | | yes | | | | | |
| ETNK1 | N_Shore | TSS1500 | chr12:22777409-22777465 | 56 | | | 6 | | | | | | 1 | | | | | yes | | | | | |
| SOX5 | NA | Body | chr12:23998921-23999034 | 113 | | | | | | | | | | | | | | | | | | | |
| SOX5 | NA | 5'UTR;1stExon;TSS200 | chr12:24102581-24102832 | 251 | | | 3 | | | | | | 1 | | | | | | | | | yes | yes |
| SOX5 | NA | nan | chr12:24104007-24104357 | 350 | | | | | | | | | | | 1 | | | | | | | yes | |
| C12orf67 | NA | Body | chr12:24737079-24737251 | 172 | | | | | | | | | 1 | | | | | | | | yes | yes | |
| BCAT1 | N_Shore;Island | Body | chr12:25054873-25056243 | 1370 | | | 44 | | 2 | | | | 1 | | | | | | | | | | |
| BCAT1 | S_Shore | 1stExon;5'UTR;TSS200 | chr12:25102274-25102521 | 247 | | | | | | | | 0 | | | | | | yes | | | | | |
| C12orf77 | NA | TSS200 | chr12:25150402-25150626 | 224 | | | 8 | | | | | | | | | | | yes | | | | | |
| LRMP | NA | TSS1500 | chr12:25204113-25204672 | 559 | | | 2 | | | | | | | | | | | yes | yes | | | yes | |
| KRAS | NA | TSS1500 | chr12:25404502-25404596 | 94 | | | | | | | | | 1 | | | | | | | | | | yes |

| | | | | | | | | | | | | | | | | | | | | | | | |
|---|---|---|---|---|---|---|---|---|---|---|---|---|---|---|---|---|---|---|---|---|---|---|---|
| IFLTD1 | NA | 1stExon;TSS200 | chr12:25801455-25801621 | 166 | | | 15 | | | | | 0 | 2 | | | 1 | | | yes | | yes | | |
| SSPN | NA | Body | chr12:26379588-26379889 | 301 | | | 14 | | | | | | | | | | | | | | | | |
| ITPR2 | S_Shore | TSS1500 | chr12:26986825-26987045 | 220 | | | 1 | | | | | | | | | | | yes | | | | | yes |
| STK38L | N_Shore | TSS1500 | chr12:27396229-27396937 | 708 | | | | | | | | | 3 | | 1 | 1 | | yes | | | yes | | |
| PTHLH | S_Shore | TSS1500;5'UTR | chr12:28124011-28124665 | 654 | | | 13 | | 1 | | | | | | | | | | | | | yes | |
| FAR2 | NA | TSS200 | chr12:29376421-29376872 | 451 | | 1 | | | | | | | | | | | | yes | | | | | |
| OVCH1 | NA | 1stExon;TSS200 | chr12:29650574-29651041 | 467 | | | | | | | | | | | | | | yes | | | | | |
| TMTC1 | Island | 5'UTR | chr12:29936407-29936845 | 438 | | | | | | | | | 1 | | | | | | | | | yes | yes |
| FAM60A | S_Shore | TSS1500 | chr12:31479853-31480300 | 447 | | | | | 1 | | | | | | | | | yes | yes | | | | |
| C12orf72 | NA | TSS1500 | chr12:31798865-31799118 | 253 | | | | | | | | 0 | | | | | | yes | | | | | |
| AMN1 | Island | Body;5'UTR;1stExon | chr12:31881778-31882198 | 420 | | | | | | | | 0 | 1 | 1 | | | | yes | | | | | |
| H3F3C | N_Shore;Island | 1stExon;3'UTR | chr12:31944556-31945286 | 730 | | | | | | | | | | | | | | yes | yes | | | | |
| FGD4 | NA | TSS200 | chr12:32654929-32655192 | 263 | | | 4 | 1 | 1 | | 1 | 0 | 1 | 1 | | 1 | | | | | | | |
| PKP2 | NA | Body | chr12:33030976-33031216 | 240 | | | | | | | | | | | | | | | | | | | |
| SYT10 | Island | Body | chr12:33592078-33592642 | 564 | | | | | | | | | 1 | | | | | yes | | | | yes | |
| ALG10 | Island | 5'UTR;1stExon | chr12:34175443-34175454 | 11 | | 1 | | | | | | | | | | | | | | | | yes | |
| ALG10B | Island | 1stExon;5'UTR | chr12:38710604-38710700 | 96 | | 1 | | | | | | | | | | | | | | | | | |
| CPNE8 | Island;N_Shore | 1stExon;5'UTR;TSS200;Body | chr12:39299132-39299726 | 594 | | 1 | | | 2 | | | | 1 | | | | | | | | | | |
| ABCD2 | NA | 1stExon;5'UTR | chr12:40013596-40014292 | 696 | | | | | | | | 0 | | | | | | | | | | | |
| CNTN1 | Island;S_Shore | 5'UTR | chr12:41086680-41087584 | 904 | | 1 | | | | | | | 2 | | | | | | | | | | |
| CNTN1 | NA | 5'UTR | chr12:41221639-41221855 | 216 | | | 1 | | | | | | 1 | 1 | | 2 | | | | | | | |
| PDZRN4 | N_Shore | TSS1500 | chr12:41581414-41582067 | 653 | | | 5 | | | | | | 1 | | | | | | yes | | | | |
| PRICKLE1 | N_Shore | 5'UTR | chr12:42875816-42876128 | 312 | | | | | | | | | | | | | | | | | | | |
| ADAMTS20 | Island;S_Shore | TSS200;TSS1500 | chr12:43945892-43946880 | 988 | | | 2 | | | | | | 1 | | | | | | | | yes | | |
| PUS7L;IRAK4 | N_Shore;Island | 1stExon;5'UTR;TSS1500;TSS200 | chr12:44152117-44153217 | 1100 | | | 5 | | | | | 0 | 1 | | | | | yes | yes | | | yes | |
| NELL2 | Island | Body;TSS1500;TSS200 | chr12:45270707-45270818 | 111 | | | | | | | | | 1 | | | | | | | | | yes | |
| DBX2 | N_Shore;Island | Body;1stExon;TSS1500 | chr12:45443808-45445385 | 1577 | | 2 | 7 | | | | | | 1 | | | | | | | | | | |
| SFRS2IP | NA | Body | chr12:46319990-46320036 | 46 | | | | | | | | | | | | | | | | | | | |
| SLC38A1 | NA | Body | chr12:46599923-46599935 | 12 | | | | | | | | | 1 | | | | | | | | | | yes |
| SLC38A1 | Island;S_Shore | TSS200;TSS1500 | chr12:46663270-46664096 | 826 | | | 2 | | 1 | | | | | | | | | yes | | | | | |
| SLC38A2 | S_Shore | TSS1500 | chr12:46767665-46768016 | 351 | | 1 | 6 | | | | | | 1 | | | | | | | | | | |
| SLC38A4 | NA | TSS200;5'UTR;nan;1stExon | chr12:47219626-47220092 | 466 | | 1 | 31 | | 1 | | | 0 | 16 | 5 | 1 | 20 | 2 | yes | | | | | |
| AMIGO2 | N_Shore | Body | chr12:47472743-47472810 | 67 | | | | | | | | | | | | 1 | | | | | | yes | |
| FAM113B;LOC100233209 | NA | 5'UTR;1stExon;Body | chr12:47610220-47610734 | 514 | | | | | | | | | 1 | 1 | | | | yes | | | | yes | |
| RAPGEF3 | N_Shore | 1stExon;TSS200;5'UTR;Body | chr12:48152204-48152428 | 224 | | | 7 | | | | | | 1 | | | | | | | | | | |
| COL2A1 | Island;N_Shore | TSS1500 | chr12:48398730-48399493 | 763 | | | 9 | | | | | | | | | | | | | | | | |
| H1FNT | N_Shore | TSS1500;TSS200;5'UTR;1stExon | chr12:48722169-48723631 | 1462 | | | 6 | | | | | | | | | | | yes | yes | | | | |
| ANP32D | NA | TSS1500 | chr12:48866138-48866481 | 343 | | | | | | | | | | | | | | | yes | | | | yes |
| CCNT1 | S_Shore | TSS1500 | chr12:49111321-49111433 | 112 | | | 2 | | | | | 0 | 1 | | | 1 | | yes | yes | | | | |
| ADCY6 | N_Shore | TSS200 | chr12:49182837-49182908 | 71 | | | | | | | | | 1 | | | | | | | | | | yes |
| CCDC65 | NA | TSS200 | chr12:49297822-49298070 | 248 | | 1 | | | | | | | | | | | | yes | | | | | |
| FKBP11 | Island | Body;TSS200;TSS1500;1stExon;5'U | chr12:49318735-49319475 | 740 | | | | | | | | 0 | 1 | | | 4 | | yes | | | | yes | yes |
| WNT10B | S_Shore | TSS1500 | chr12:49366152-49366538 | 386 | | | | | | | | | | | | | | | | | | yes | |
| DHH | Island | Body | chr12:49484932-49485151 | 219 | 2 | | 4 | | | | | | | | | | | yes | | | | yes | |
| LMBR1L | S_Shore | TSS1500 | chr12:49504967-49505053 | 86 | | | 4 | | | | | 0 | 1 | | | 1 | | yes | | | | | |
| TUBA1C | Island | 5'UTR;1stExon;TSS200 | chr12:49658703-49659154 | 451 | 2 | | | | | | | | | | | | | | | | | | yes |
| PRPH | N_Shore;Island | TSS200;5'UTR;1stExon | chr12:49688844-49689463 | 619 | | | 3 | | | | | | | | | | | | | | | | |
| C1QL4 | N_Shore;Island | 3'UTR | chr12:49726500-49727110 | 610 | | | 6 | | | | | 0 | 2 | | | 2 | | yes | | | | | |
| C1QL4 | S_Shore | TSS1500 | chr12:49731393-49731885 | 492 | | | 1 | | | | | | | | | | | | | | | yes | yes |
| DNAJC22 | S_Shore | TSS200 | chr12:49740942-49740996 | 54 | | | 1 | | | | | | | | | | | yes | | | | | |
| SPATS2 | Island | 5'UTR | chr12:49782966-49783222 | 256 | | | | | | | | 0 | 4 | 1 | | 10 | | yes | yes | | | | |
| KCNH3 | Island | Body | chr12:49933539-49934029 | 490 | | | | | | | | | | | | | | | | | | | yes |
| KCNH3 | N_Shore;Island | Body | chr12:49942757-49943340 | 583 | | | | 1 | 1 | | | | | | | | | | | | | | |
| AQP2 | NA | TSS200 | chr12:50344463-50344548 | 85 | | | | | | | | | | | | | | | | | | | |
| AQP2 | Island | Body | chr12:50349173-50349522 | 349 | | | 3 | | | | | 0 | 1 | | | | | yes | | | | | |
| AQP6 | NA | TSS1500 | chr12:50365827-50366624 | 797 | | | | | | | | | 2 | | | | | | yes | yes | | | |
| ACCN2 | Island | Body | chr12:50475035-50475372 | 337 | | | 9 | | | | | | | | | | | | | | | | |
| GPD1 | NA | TSS200 | chr12:50497732-50498016 | 284 | | | | | | | | | 2 | | | | | | | | | | |
| TMPRSS12 | N_Shore;Island | TSS200 | chr12:51236582-51236768 | 186 | | | 3 | | | | | | 3 | | | | | yes | | yes | | | |
| TFCP2 | N_Shore;Island | 5'UTR;1stExon;TSS200 | chr12:51566379-51567112 | 733 | | | | | | | | 0 | 1 | 2 | | 1 | | yes | | | | | |
| BIN2 | NA | TSS200;Body;1stExon | chr12:51717674-51718251 | 577 | | | | | 2 | | | | 3 | 9 | 1 | | | yes | | | | | |
| GALNT6 | S_Shore | TSS1500 | chr12:51786038-51786547 | 509 | | | | | | | | 0 | 1 | | | 3 | | yes | | | yes | | yes |
| SCN8A | Island;N_Shore | Body | chr12:52115238-52115678 | 440 | | | | 1 | | | | 0 | 3 | 2 | | 6 | 1 | yes | | | | yes | |
| FIGNL2 | Island | 1stExon | chr12:52214320-52215216 | 896 | | | 1 | | | | | 0 | 2 | 1 | | 2 | | | | | | | yes |
| ANKRD33 | NA | TSS200;1stExon;5'UTR | chr12:52281652-52282079 | 427 | | | 12 | | | | | 0 | 3 | | | 1 | | yes | | | | | |
| ACVRL1 | N_Shelf | TSS1500;5'UTR | chr12:52305328-52305945 | 617 | | | | | | | | | 1 | | | | | yes | yes | | | | |
| GRASP | S_Shelf | Body | chr12:52404134-52404422 | 288 | | | | 1 | | | | | 2 | | | | | | | | yes | | |
| KRT80 | NA | 1stExon;5'UTR;TSS200 | chr12:52585695-52585892 | 197 | | 1 | | 2 | | | | | 2 | | | | | yes | | | | | |
| KRT7 | Island;chr12:52626793-526275 | TSS200;Island;5'UTR;1stExon;TSS1 | chr12:52626427-52627576 | 1149 | | | 13 | | | | | | | | 1 | | | | | | | | |
| KRT86 | N_Shore | TSS200;TSS1500 | chr12:52695412-52695728 | 316 | | | 6 | | | | | | | | | | | | | | | yes | |
| KRT85 | Island | 1stExon | chr12:52760975-52761153 | 178 | | | | | | | | 0 | | | | | | | | | | yes | |
| KRT75 | NA | TSS200 | chr12:52828203-52828633 | 430 | | | 3 | | | | | | | | | | | yes | yes | | | | |
| KRT6B | NA | TSS200 | chr12:52845930-52846088 | 158 | | | | | | | | | 1 | 1 | | | | | yes | | | | |
| KRT5 | NA | 1stExon | chr12:52913824-52914229 | 405 | | | 2 | | | | | | 1 | | | | | | | | | yes | |
| KRT74 | NA | 1stExon;TSS200 | chr12:52967554-52968145 | 591 | | | | | | | | | 1 | | | | | yes | yes | | | | |
| KRT72 | Island;S_Shore | 1stExon;5'UTR;TSS200 | chr12:52994933-52995446 | 513 | | | 13 | | | | | 0 | 2 | | | 1 | | yes | | yes | | | |
| KRT1 | NA | 1stExon;TSS200 | chr12:53074023-53074328 | 305 | | | | | | | | | 1 | | | | | | | | | yes | |
| KRT1 | NA | TSS1500 | chr12:53075359-53075482 | 123 | | | 3 | | | | | | 1 | | | | | yes | yes | | | | |
| KRT76 | NA | 1stExon | chr12:53170909-53171191 | 282 | | | | | | | | | | | | | | yes | | | | yes | |
| KRT3 | Island;S_Shore | 3'UTR;Body | chr12:53183773-53184086 | 313 | | | 6 | | | | | 0 | 4 | 2 | | 6 | | | | | | yes | |
| KRT4 | NA | 1stExon;5'UTR | chr12:53208325-53208752 | 427 | | | | | | | | | | | | | | yes | | yes | yes | | |
| KRT78 | NA | 5'UTR;1stExon;TSS200;TSS1500 | chr12:53242733-53243489 | 756 | | | 5 | | | | | | 2 | 1 | | | | yes | yes | | | | |
| KRT8 | S_Shore | Body;1stExon;5'UTR;TSS200 | chr12:53297987-53299310 | 1323 | | | | 1 | 1 | | | | 2 | | | | | yes | | | | yes | |
| KRT18 | S_Shore | Body | chr12:53343514-53343849 | 335 | | 1 | | 1 | | 1 | | 0 | 1 | | | | | | | | | | |
| TENC1 | S_Shore | 5'UTR;TSS1500 | chr12:53442322-53443485 | 1163 | | | | | | | | | | | | | | | | | | | |
| TENC1 | Island | Body | chr12:53448085-53448399 | 314 | | | 7 | 1 | | | | | | | | | | yes | | | | | |
| IGFBP6 | N_Shore | TSS1500 | chr12:53490352-53490632 | 280 | | | | | | | | | 1 | | | | | | | | | | |
| SOAT2 | NA | TSS200 | chr12:53497138-53497306 | 168 | | | | | 1 | | | | | | | | | yes | yes | | | | |
| ITGB7 | Island | Body | chr12:53591398-53592259 | 861 | | 1 | 12 | | | | | 0 | 2 | | | | | yes | | | | | |
| RARG | N_Shore | Body | chr12:53612641-53613154 | 513 | | | | 1 | | | | | | | | | | | | | | | yes |
| C12orf10;PFDN5 | Island;N_Shore | 1stExon;5'UTR;Body;TSS1500;TSS2 | chr12:53693054-53694123 | 1069 | | | | | | | | 0 | 1 | | | 2 | | yes | | | | | |
| SP7 | NA | TSS200;TSS1500 | chr12:53729629-53730277 | 648 | | | 10 | | | | | | | | | | | | | | | yes | yes |
| ATP5G2 | N_Shore;Island | TSS1500 | chr12:54070891-54071194 | 303 | 1 | | | 1 | 4 | | | 0 | 2 | | | 1 | | | yes | | | | yes |
| HOXC13 | Island | 1stExon | chr12:54333323-54333823 | 500 | | | | | | | | | 1 | | | | | | | | | | yes |
| HOXC12 | N_Shore | TSS200 | chr12:54348629-54348719 | 90 | | | | | | | | | | | | | | | | | | yes | |
| HOTAIR | Island;N_Shore | Body | chr12:54359761-54360836 | 1075 | | | 11 | | | | | | | | | | | | | | | | yes |
| HOXC11 | Island | TSS200;Body | chr12:54366816-54368332 | 1516 | | | | | | | | | 2 | | | 1 | | | | | | | yes |
| HOXC10;MIR196A2 | S_Shelf;N_Shelf | 3'UTR;TSS1500;TSS200 | chr12:54383639-54385625 | 1986 | | | | | | | | | | | | | | | | | | | |
| HOXC9 | N_Shore;Island | TSS1500;1stExon | chr12:54392734-54394212 | 1478 | | 1 | 10 | | | | | 0 | 4 | | | 2 | | | | | | | yes |

| | | | | | | | | | | | | | | | | | | | | | | | |
|---|---|---|---|---|---|---|---|---|---|---|---|---|---|---|---|---|---|---|---|---|---|---|---|
| HOXC4;HOXC6;HOXC5 | S_Shore;N_Shore | TSS1500;TSS200;5'UTR;Body;1stEx | chr12:54409207-54410671 | 1464 | | | 34 | | | | | 0 | 5 | | | 3 | | | | | | yes | yes |
| HOXC4;HOXC5;HOXC6 | N_Shore;Island | 5'UTR;Body | chr12:54412269-54413345 | 1076 | | | 10 | | | | | 0 | 2 | | | 2 | | | | | | | |
| HOXC4;HOXC5;HOXC6 | N_Shelf | 5'UTR;Body;TSS1500 | chr12:54420837-54421335 | 498 | | | 6 | | | | | | | | | | | | | | | | yes |
| HOXC4;HOXC6;HOXC5 | Island;N_Shore | 5'UTR;Body;3'UTR | chr12:54423428-54425418 | 1990 | | | 15 | | 1 | | | | 1 | | | | | | | | | yes | yes |
| HOXC4;HOXC5 | 5'UTR;Body;1stExon | nan | chr12:54427293-54427528 | 235 | | | | | | | | | | | | | | | | | | | |
| HOXC4 | Island | 5'UTR | chr12:54440773-54441509 | 736 | | 1 | 29 | | | | | | | | | | | | | | | | |
| HOXC4 | N_Shore;Island;S_Shore | 5'UTR;TSS1500;TSS200;Body | chr12:54444985-54449761 | 4776 | | | | | | | | | | | | | | | | | | | |
| FLJ12825 | N_Shelf | TSS200 | chr12:54451877-54452069 | 192 | | | | | | | | | 1 | | | 1 | | | | | | | |
| FLJ12825 | N_Shore;Island | Body | chr12:54454163-54454445 | 282 | | 1 | 3 | | | | | | | 1 | | 2 | | | | | | | |
| FLJ12825;LOC100240735 | N_Shore;Island | Body | chr12:54475010-54476240 | 1230 | | | | | | | | | | | | | | | | | | yes | yes |
| CBX5 | NA | 1stExon;5'UTR;TSS200 | chr12:54653065-54653427 | 362 | | | | | 2 | | | 0 | 1 | | | | | | | | | | |
| ZNF385A | Island | 3'UTR | chr12:54763081-54763433 | 352 | | | 6 | | | | | 0 | 1 | 1 | | 1 | | | | | | | |
| ZNF385A | NA | Body;TSS1500 | chr12:54778750-54779175 | 425 | | | | | 1 | | | | | | | | | | | | | | |
| ZNF385A | chr12:54784900-54785238;Isla | N_Shore;Body;5'UTR;1stExon;TSS | chr12:54784747-54785320 | 573 | | | 11 | 1 | | | | | | | 1 | | | | | | | | yes |
| GTSF1 | NA | 5'UTR;TSS200 | chr12:54867312-54867653 | 341 | | | | | | | | 0 | 4 | | | 1 | | yes | | | | yes | |
| NCKAP1L | NA | TSS1500;TSS200 | chr12:54891213-54891636 | 423 | | | | | 1 | | | | 1 | 1 | 1 | | | yes | yes | | | yes | yes |
| PDE1B | NA | TSS1500 | chr12:54942782-54943009 | 227 | | | 5 | | | | | | | | | | | yes | yes | | | yes | yes |
| PDE1B | NA | TSS1500;Body | chr12:54954337-54954941 | 604 | | | 1 | | | | | 0 | 2 | 1 | | 1 | | yes | yes | | | | |
| LACRT | NA | Body;TSS200 | chr12:55028545-55029147 | 602 | | | | | | | | | | | | | | yes | yes | | yes | | |
| DCD | NA | Body | chr12:55041948-55042383 | 435 | | | | | | | | | | | | | | | yes | | | | |
| KIAA0748 | NA | TSS200;1stExon;5'UTR | chr12:55378411-55379228 | 817 | | | 4 | | 2 | | | | 1 | 1 | | | | | | | | | |
| OR10P1 | NA | TSS1500 | chr12:56030242-56030437 | 195 | | | 6 | | | | | | | | | | | yes | yes | | | | |
| METTL7B | NA | TSS200 | chr12:56075321-56075511 | 190 | | | | | | | | 0 | | | | | | yes | yes | | | yes | |
| ITGA7 | N_Shelf | Body | chr12:56105981-56106386 | 405 | | | 1 | | | | | | | | | | | | yes | | | | yes |
| BLOC1S1 | N_Shore | TSS1500 | chr12:56109593-56109624 | 31 | | | 1 | | | | | | | | | | | yes | | | | | |
| RDH5 | S_Shelf | TSS1500 | chr12:56113808-56114269 | 461 | | | | 1 | 1 | | | | 1 | | | | | yes | | | | | |
| DGKA | N_Shelf | TSS1500 | chr12:56323639-56323816 | 177 | | | | | | | | | 1 | | | | | yes | | | | | |
| DGKA | N_Shore | TSS1500;TSS200 | chr12:56324915-56325149 | 234 | | | 5 | | | | | | | | | | | yes | | | | | |
| DGKA | N_Shore;Island | 5'UTR | chr12:56329296-56329903 | 607 | | | 15 | | | | | | 3 | | | | | yes | yes | | | | yes |
| CDK2 | N_Shore | 3'UTR | chr12:56365823-56365978 | 155 | | | 1 | | | | | | | | | | | | | | | | yes |
| SUOX | NA | TSS1500 | chr12:56390805-56390878 | 73 | | | | | | | | 0 | | | | | | yes | | | | | |
| IKZF4 | NA | TSS1500;TSS200 | chr12:56414442-56415591 | 1149 | | | 8 | 1 | | | | | 1 | | | | | | | | | yes | |
| ERBB3 | S_Shore | Body | chr12:56474569-56475064 | 495 | | | | | | | | | | | | | | | yes | | | | |
| MYL6B | N_Shore | TSS1500 | chr12:56545428-56545649 | 221 | | | | | | | | | | | | | | yes | | | | | |
| MYL6B;MYL6 | NA | 3'UTR;TSS1500 | chr12:56551637-56551679 | 42 | | | | | | | | | 1 | | | | | yes | | | | | yes |
| OBFC2B | N_Shore | TSS1500 | chr12:56617576-56617737 | 161 | | | | | | | | 0 | 1 | | | 2 | | yes | | | | | |
| CS | S_Shore | TSS1500 | chr12:56694490-56694667 | 177 | | | | | | | | 0 | | | | | | | | | | | |
| PAN2 | N_Shore | 5'UTR;1stExon;TSS200 | chr12:56727304-56728493 | 1189 | | | | | | | | 0 | | | | | | yes | | | | | |
| IL23A | NA | TSS1500;TSS200;5'UTR;1stExon | chr12:56732427-56733052 | 625 | | | 14 | | 1 | | | | | | | | | | | | | | |
| APOF | S_Shelf | TSS1500 | chr12:56757753-56757836 | 83 | | | | | | | | | | | | | | | yes | | | | |
| MIP | NA | TSS1500 | chr12:56849120-56849618 | 498 | | | | | | | | | | | | | | | yes | | | | |
| TAC3 | NA | 5'UTR;1stExon | chr12:57410233-57410604 | 371 | | | 3 | | | | | | | | | | | yes | | yes | | | |
| LRP1 | NA | Body | chr12:57559728-57559901 | 173 | | | | | | | | | 1 | | | | | yes | | | | | |
| LRP1 | N_Shore;Island | Body | chr12:57569262-57569940 | 678 | | | 6 | | | | | 0 | 1 | | | | | | yes | | | | |
| LRP1;MIR1228 | NA | Body;TSS200 | chr12:57588144-57588350 | 206 | 1 | | | | | | | | 1 | | | | | | | | | | |
| NXPH4 | S_Shelf | Body | chr12:57614100-57614182 | 82 | | 1 | | | | | | | | | | | | | | | | yes | yes |
| NXPH4 | Island | Body | chr12:57618943-57619401 | 458 | | | 5 | | | | | | 2 | | 1 | | | yes | | | | | yes |
| SHMT2 | N_Shore | TSS1500 | chr12:57622801-57623095 | 294 | | | | | | | | | | | | | | yes | | | yes | | yes |
| NDUFA4L2 | N_Shore | TSS1500 | chr12:57634677-57635479 | 802 | | | | | | | | | | | | | | | | | | | |
| INHBC | S_Shelf | TSS1500 | chr12:57828180-57828515 | 335 | | | | | | 1 | | | | | | | | yes | | | | | |
| INHBE | NA | TSS1500 | chr12:57848560-57849029 | 469 | | | | | | | | | | | | | | yes | | | | | |
| ARHGAP9 | N_Shore | Body | chr12:57869031-57869490 | 459 | | | | | | | | | 1 | | | | | | | | | | |
| MARS | S_Shore | Body | chr12:57882891-57883315 | 424 | | | | | | | | | | | | | | yes | | | | yes | |
| MBD6;DDIT3 | N_Shore | TSS1500 | chr12:57915555-57915636 | 81 | | | 13 | | | | | 0 | | | | | | yes | | | | | |
| DCTN2 | N_Shore | Body | chr12:57940444-57940828 | 384 | | | | | | | | | 1 | | | | | | | | | | |
| DTX3;GEFT | N_Shore;Island | 3'UTR;TSS1500;TSS200 | chr12:58003151-58003925 | 774 | 5 | | 7 | 1 | | | | | | | 1 | | | | | | | | yes |
| SLC26A10 | N_Shore;Island;chr12:5801317 | TSS1500;Island;TSS200 | chr12:58013109-58013915 | 806 | | | 50 | | | | | | 1 | | 1 | | | yes | | | | | |
| B4GALNT1 | Island;S_Shore | Body | chr12:58021295-58022620 | 1325 | | | 21 | | | | | 0 | | | 1 | 1 | | yes | yes | | | | yes |
| LOC100130776;AGAP2 | N_Shore;Island | TSS1500;3'UTR;TSS200 | chr12:58119543-58121004 | 1461 | | | | | | | | | | | | | | | | | | | |
| AGAP2 | Island | Body | chr12:58130154-58131011 | 857 | | | | | | | | | | | | | | | | | | yes | |
| AGAP2 | N_Shore | TSS200;Body | chr12:58132093-58132733 | 640 | | | | | | | | 0 | 2 | | | 3 | | | | | | yes | yes |
| AGAP2 | S_Shelf | 1stExon;TSS200 | chr12:58135833-58136245 | 412 | | | 4 | | | | | | | | | | | yes | | | | | |
| AVIL | NA | TSS1500 | chr12:58210661-58211193 | 532 | | | 3 | 1 | | | | 0 | 1 | | | | | yes | yes | | | | |
| LRIG3 | Island | 1stExon;TSS1500 | chr12:59313817-59314175 | 358 | | | | | | | | | 1 | | | | | | | | | | |
| FAM19A2 | N_Shore;Island | 5'UTR | chr12:62583974-62585877 | 1903 | | 1 | 14 | | | | | | | | | | | | | | | yes | |
| DPY19L2 | Island | TSS200 | chr12:64062498-64062526 | 28 | | | | | | | | | 1 | | | | | | | | | | yes |
| C12orf56 | N_Shore;Island | Body;1stExon | chr12:64782908-64784686 | 1778 | | | | | | | | | 2 | | | 1 | | yes | yes | | | | |
| WIF1 | Island | 5'UTR;1stExon | chr12:65515021-65515470 | 449 | | 1 | | | | | | | | | | | | | | | | yes | yes |
| MSRB3 | N_Shore | TSS1500 | chr12:65671664-65672174 | 510 | | | 1 | | | | | | 1 | | | | | | | | | yes | yes |
| LLPH | N_Shore | 5'UTR | chr12:66524311-66524791 | 480 | | | | | | | | | 1 | | | 1 | | yes | yes | | | | |
| IRAK3 | N_Shore | TSS1500 | chr12:66582582-66582780 | 198 | | | | | | | | | 1 | | | | | yes | | | | | yes |
| IRAK3 | NA | Body | chr12:66627900-66628232 | 332 | | | | | | | | | | | | | | | | | | yes | |
| GRIP1 | NA | 1stExon;5'UTR;TSS200 | chr12:67072769-67073760 | 991 | | 1 | 13 | | | | | | 1 | | | | | | | | | yes | |
| DYRK2 | N_Shore | TSS1500 | chr12:68041257-68041428 | 171 | | | 2 | | | | | | 1 | | | | | yes | | | | yes | |
| IFNG | NA | 1stExon;5'UTR | chr12:68553396-68553819 | 423 | | | 2 | | | | | | 1 | | | | | yes | | | yes | | |
| IL22 | NA | Body | chr12:68647015-68647289 | 274 | | | | | | | | | | | | | | | | | | | |
| SLC35E3 | Island | 1stExon;5'UTR | chr12:69140014-69140079 | 65 | | | | | | | | 0 | | | | | | | | | | | |
| CPM | Island | TSS1500;5'UTR | chr12:69327449-69327531 | 82 | | | | | | | | | 1 | | | | | | yes | | | | yes |
| YEATS4 | N_Shore | TSS1500 | chr12:69753119-69753446 | 327 | | | | | | | | | | | | | | | yes | | | | |
| LRRC10 | NA | TSS200 | chr12:70005043-70005609 | 566 | | | | | | | | | | | | | | yes | yes | | | | |
| BEST3 | NA | Body;TSS200 | chr12:70083090-70083105 | 15 | | 1 | | 1 | | | | | | | | | | | | | yes | | |
| BEST3 | NA | 5'UTR | chr12:70092840-70093350 | 510 | | | | | | | | | 1 | | | | | yes | | | | | |
| PTPRR | NA | 1stExon;5'UTR | chr12:71314144-71314752 | 608 | | 1 | | | | | | | 1 | | | | | | | | | yes | |
| TSPAN8 | NA | TSS200;TSS1500 | chr12:71551926-71552569 | 643 | | | 5 | | | | | | | | | | | | | | | | |
| TBC1D15 | N_Shore | TSS1500 | chr12:72232889-72233266 | 377 | | | 9 | | | | | | | | | | | yes | | | | | |
| TPH2 | NA | TSS1500 | chr12:72332386-72332608 | 222 | | | | | | | | | 1 | | | | | | yes | | | | |
| LOC283392;TRHDE | Island | Body;TSS1500 | chr12:72665880-72666281 | 401 | | | | | | | | | 1 | | | | | | | | | yes | yes |
| KCNC2 | Island | Body | chr12:75601157-75601928 | 771 | | | | | 1 | | | | 1 | | | | | | | | | yes | yes |
| GLIPR1L1 | S_Shelf | TSS1500 | chr12:75728104-75728752 | 648 | | 1 | | | | | | | | | | | | | | | | | |
| GLIPR1L2 | Island;N_Shore | TSS200;1stExon;Body;TSS1500 | chr12:75783428-75785295 | 1867 | | | 5 | | | | | 0 | 11 | 2 | | 5 | | yes | yes | | yes | yes | yes |
| SYT1 | NA | 5'UTR;TSS200 | chr12:79258313-79258407 | 94 | | | | | | | | | 1 | | | | | | | | | yes | |
| MYF6 | N_Shore;Island | TSS1500;Body | chr12:81101022-81102749 | 1727 | | | 13 | | | | | 0 | 2 | | | | | | | | | | |
| MYF5 | N_Shore;Island | TSS200;1stExon;5'UTR | chr12:81110706-81111012 | 306 | | | 10 | | | | | 0 | | | | 1 | | | | | | | |
| MIR618;LIN7A | Island | TSS1500;Body;1stExon | chr12:81331078-81331718 | 640 | | | | | | | | | 4 | | | | | | | | | | yes |
| ACSS3 | Island | 1stExon;5'UTR | chr12:81471867-81471938 | 71 | | | | | | | | | 2 | | | | | | | | | | |
| PPFIA2 | S_Shore | 1stExon;5'UTR | chr12:82153070-82153227 | 157 | | | | | | | | | 1 | | | | | | | | | yes | yes |
| TMTC2 | Island | TSS200 | chr12:83080830-83080838 | 8 | | | | | | | | | 1 | | | | | | | | | yes | |
| SLC6A15 | N_Shore;Island;S_Shore | 5'UTR;TSS200;TSS1500 | chr12:85306021-85307424 | 1403 | | | 17 | | | | | | 3 | | | | | | | | | yes | |
| TSPAN19;LRRIQ1 | NA | TSS200;1stExon;5'UTR | chr12:85430123-85430174 | 51 | | | | | | | | | | | | | | | | | | | yes |

| | | | | | | | | | | | | | | | | | | | | | | | |
|---|---|---|---|---|---|---|---|---|---|---|---|---|---|---|---|---|---|---|---|---|---|---|---|
| ALX1 | N_Shore;Island | TSS1500;TSS200;Body | chr12:85673200-85674531 | 1331 | | 3 | 2 | | | | | | 2 | | | | | | | | | | |
| RASSF9 | NA | Body;5'UTR;1stExon | chr12:86229804-86230557 | 753 | | | | | | | | 0 | 2 | | | 6 | | | | | | yes | |
| MGAT4C | NA | 1stExon;5'UTR | chr12:87232511-87232821 | 310 | | | | | | | | | | | | | | | | | | | |
| KITLG | Island | TSS200 | chr12:88974433-88974521 | 88 | | | | | | | | | 1 | | | | | yes | | | | | |
| DUSP6 | N_Shelf;N_Shore | 3'UTR;Body | chr12:89742886-89744877 | 1991 | | | 45 | | 1 | | | 0 | 4 | | | 1 | | | | | | | |
| DCN | NA | 5'UTR;TSS1500;1stExon;Body | chr12:91572142-91573372 | 1230 | | | 2 | | | | | | 2 | | | 2 | | yes | yes | | | yes | |
| C12orf74 | NA | TSS200 | chr12:93096780-93096844 | 64 | | | | 1 | 1 | | | | 1 | | | | | yes | | | | | |
| SOCS2 | Island | Body | chr12:93966978-93967711 | 733 | | | | | 2 | | | | | | | | | | | | | yes | |
| CRADD | N_Shore | TSS1500 | chr12:94070475-94070836 | 361 | | | | | | | | | 1 | | | 1 | | | yes | yes | | | |
| TMCC3 | NA | Body | chr12:95009858-95010113 | 255 | | | | | 1 | | | | | | | | | | yes | | | | |
| MIR492 | NA | TSS1500 | chr12:95226786-95226993 | 207 | 2 | | | | | | | | | | | | | yes | yes | | | | |
| MIR492 | NA | TSS200 | chr12:95228096-95228478 | 382 | | | | | | | | | | | | | | yes | | | | | |
| NDUFA12 | Island | TSS200 | chr12:95397608-95397732 | 124 | | | | | | | | | | | | | | yes | yes | | | | |
| MIR331 | NA | TSS200 | chr12:95702071-95702491 | 420 | | | | | | | | | 1 | | | | | | | | | | |
| USP44 | Island | TSS1500;5'UTR | chr12:95942907-95943114 | 207 | | 1 | | | | | | | | | | | | | | | | | yes |
| USP44 | S_Shelf | 1stExon;5'UTR;TSS200 | chr12:95945082-95945767 | 685 | | | 1 | | | | | 0 | 3 | | | 1 | | | | | | | |
| SNRPF | N_Shore | TSS1500 | chr12:96251729-96252105 | 376 | | | 3 | | | | | 0 | 3 | | | 4 | | yes | | yes | | | |
| AMDHD1 | Island | Body | chr12:96350519-96350795 | 276 | 1 | | | | | | | | | | | | | yes | | | yes | | |
| HAL | Island | Body | chr12:96389483-96389694 | 211 | | | | 1 | | | | | | | | 1 | | yes | yes | | | | |
| NEDD1 | N_Shore | TSS1500 | chr12:97300410-97300582 | 172 | | | 22 | | | | | 0 | 1 | | | 2 | | | yes | | | | |
| RMST;MIR1251 | NA | Body;TSS1500 | chr12:97884975-97886312 | 1337 | | | | | | | | | | | | | | | | | | yes | yes |
| SLC25A3;SNORA53 | S_Shelf | Body;TSS1500 | chr12:98992088-98992188 | 100 | | | | | | | | | 1 | | | | | yes | | | | | |
| ANKS1B | Island | 3'UTR;Body | chr12:99139387-99139867 | 480 | | | | | | | | 0 | 2 | | | | | | yes | | | yes | |
| ANKS1B | N_Shore | Body | chr12:99287129-99287793 | 664 | | | | | | | | | | | | | | | yes | | | | |
| ANKS1B | NA | 1stExon;5'UTR;Body;TSS200;TSS15 | chr12:99548329-99549023 | 694 | | 1 | 14 | | 2 | | | | 3 | | | 1 | | | yes | | | | yes |
| SLC17A8 | NA | TSS200;TSS1500 | chr12:100750473-100750899 | 426 | | | | | | | | 0 | 2 | | | 1 | | yes | yes | | | | |
| NR1H4 | NA | TSS1500;TSS200;nan | chr12:100867409-100867758 | 349 | | | | | | | | 0 | 1 | | 1 | 1 | | yes | | | | | |
| GAS2L3 | Island | TSS200;5'UTR;1stExon | chr12:100967381-100967609 | 228 | | | | | | | | | 2 | | | | | | | | | yes | |
| SLC5A8 | Island;S_Shore | 1stExon;5'UTR | chr12:101603453-101604147 | 694 | | 1 | | | | | | 0 | 1 | | | | | yes | | | | | |
| MYBPC1 | NA | TSS200 | chr12:101988604-101988803 | 199 | | | 2 | | | | | 0 | | | | | | | yes | | | | |
| MYBPC1 | Island;N_Shore | Body | chr12:102036140-102036563 | 423 | | | 4 | | | | | | 1 | | | 1 | | | | | | | |
| PAH | S_Shore | 5'UTR;1stExon | chr12:103311300-103311500 | 200 | | | | | | | | | 1 | | | | | yes | yes | | | | |
| ASCL1 | Island;S_Shore;N_Shore | 1stExon;3'UTR;TSS1500 | chr12:103351188-103353362 | 2174 | | | | | 1 | | | | 1 | | | | | yes | | | | yes | yes |
| C12orf42 | Island | Body | chr12:103696126-103696381 | 255 | | 1 | | | | | | | 1 | | | 1 | | | | | | | |
| C12orf42 | Island | 5'UTR;TSS200 | chr12:103889307-103889960 | 653 | | | | | 2 | | | 0 | 1 | | | | | | | | | yes | |
| STAB2 | NA | TSS1500 | chr12:103980767-103981113 | 346 | | | 1 | | | | | | | | | | | yes | yes | | | | |
| NT5DC3 | Island | TSS1500 | chr12:104235298-104235554 | 256 | | | 12 | | 1 | | | | | | | | | | | yes | | yes | yes |
| NFYB | Island | TSS1500 | chr12:104532339-104532524 | 185 | | | | | | | | | | | | | | | | | | yes | yes |
| TXNRD1 | Island | 1stExon;TSS200 | chr12:104609432-104610064 | 632 | | 1 | 6 | | | | | | | | | | | yes | yes | | | | |
| TXNRD1;EID3 | N_Shore;Island | 5'UTR;Body;TSS1500;TSS200;1stEx | chr12:104697193-104697983 | 790 | | 1 | | 1 | | | | 0 | 8 | | | 2 | | | | | | | yes |
| CHST11 | Island | Body | chr12:104852151-104852446 | 295 | | | | | 1 | | | | | | | | | | | | | | |
| CHST11 | NA | Body | chr12:105114162-105114399 | 237 | | | 2 | 1 | 1 | | | | 1 | 1 | 1 | | | yes | | | | | |
| ALDH1L2 | N_Shore | Body | chr12:105477526-105477694 | 168 | | | | | | | | 0 | | | | | | | | | | yes | |
| NUAK1 | NA | Body | chr12:106480546-106480771 | 225 | 1 | 1 | | 1 | | | | | | | | | | | yes | | | | |
| NUAK1 | Island;S_Shore | 5'UTR;1stExon;TSS200;TSS1500 | chr12:106533667-106534620 | 953 | | | 3 | 1 | 1 | | | | 2 | 1 | | | | yes | | yes | | | yes |
| RFX4 | Island;N_Shore | Body | chr12:106978920-106979482 | 562 | | | 4 | | | | | | 1 | | | | | | | | | | |
| RFX4 | NA | Body | chr12:106988340-106988413 | 73 | | | 9 | | | | | 0 | 1 | | | 2 | | yes | | | | | yes |
| RFX4 | NA | Body | chr12:107039017-107039145 | 128 | 1 | | | | | | | | | | | | | | yes | | | | |
| RFX4 | NA | Body;1stExon | chr12:107078609-107078901 | 292 | | | 7 | | | | | | | | | | | yes | yes | | yes | | |
| C12orf23 | N_Shore | TSS1500 | chr12:107348689-107348944 | 255 | | | 18 | | | | | | 2 | 1 | | | | yes | yes | | | | |
| BTBD11 | Island | TSS200 | chr12:107712120-107712210 | 90 | | | | | | | | | 1 | | | | | | | | | | yes |
| BTBD11 | N_Shore | TSS1500;Body;TSS200 | chr12:107974296-107975299 | 1003 | | | 2 | | | | | | 2 | | | | | yes | | | yes | | yes |
| BTBD11 | NA | Body | chr12:108039569-108039856 | 287 | | | 8 | 1 | | | | | | | 1 | | | yes | yes | | yes | | |
| ASCL4 | N_Shore | 1stExon;5'UTR;TSS1500 | chr12:108167448-108169020 | 1572 | | | | | | | | 0 | 1 | | | | | yes | yes | | | | |
| WSCD2 | Island | TSS1500 | chr12:108523043-108523395 | 352 | | 1 | | | | | | | | | | | | yes | yes | | | | yes |
| WSCD2 | S_Shore | 5'UTR | chr12:108525241-108525514 | 273 | | | | | | | | | | | | | | yes | | | | yes | |
| WSCD2 | NA | Body | chr12:108603943-108604021 | 78 | | | | | | | | | | | | | | yes | | | | | |
| WSCD2 | NA | Body | chr12:108634147-108634275 | 128 | | | | | | | | 0 | 1 | 1 | | | | yes | yes | | | yes | |
| CMKLR1 | NA | 1stExon;5'UTR;TSS200;TSS1500 | chr12:108732994-108733791 | 797 | 5 | | | | 2 | | | | 1 | | | | | yes | yes | | | | yes |
| SSH1 | NA | 3'UTR | chr12:109181748-109182032 | 284 | | | 3 | | | | | | | | | | | | | | | | |
| SSH1 | NA | 1stExon;5'UTR;Body | chr12:109221232-109221600 | 368 | | | 2 | | | | | | | | | | | | | | | | |
| SSH1 | NA | Body | chr12:109234959-109235071 | 112 | | | | | | | | | | | | | | yes | | | | | |
| SSH1 | NA | Body | chr12:109246503-109246512 | 9 | 8 | | | | | | | | | | | | | yes | | | | | |
| DAO | NA | TSS200 | chr12:109273766-109274241 | 475 | | | | 1 | | | | | | | | | | yes | | | | | |
| FOXN4 | N_Shore | Body | chr12:109728152-109728241 | 89 | | | | | | | | | | | | | | yes | | | yes | | |
| FOXN4 | N_Shore | Body | chr12:109729478-109729809 | 331 | | | | 1 | | | | | | | | | | yes | | | | | yes |
| MVK | NA | Body | chr12:110026737-110026822 | 85 | | | | 1 | | | | | | | | | | yes | yes | | | | |
| C12orf34 | N_Shore;Island | 5'UTR | chr12:110156245-110156459 | 214 | | 1 | 5 | | | | | | | | | | | yes | | | | | yes |
| MGC14436;C12orf34 | NA | Body;5'UTR | chr12:110172208-110172346 | 138 | 1 | | 8 | | | | | | 1 | | | | | yes | yes | | | yes | |
| MGC14436 | NA | TSS200 | chr12:110211321-110211701 | 380 | | | 12 | | | | | | 1 | | | | | yes | | | | | |
| TRPV4 | NA | 5'UTR | chr12:110252616-110252831 | 215 | | | | | | | | | | | | | | yes | | | | yes | yes |
| TRPV4 | Island | TSS1500 | chr12:110271439-110271581 | 142 | | | | | | | | | | | | | | yes | | | | | yes |
| TCHP | NA | Body | chr12:110352548-110352941 | 393 | | | | | | | | | | 1 | | | | | | | | yes | |
| C12orf76 | NA | TSS1500 | chr12:110506167-110506707 | 540 | | | | | | | | | 1 | | | | | yes | | yes | yes | yes | |
| HVCN1 | S_Shore | TSS1500 | chr12:111128005-111128247 | 242 | | | | | | | | | 1 | | | | | yes | | yes | | | |
| PPP1CC | S_Shore | TSS1500 | chr12:111181439-111181673 | 234 | | | | | | | | 0 | | | | 1 | | | | | | | |
| CCDC63 | NA | TSS1500 | chr12:111284228-111284585 | 357 | | | 2 | | | | | | | | | | | yes | | | yes | yes | yes |
| MYL2 | NA | TSS200;Body;1stExon;5'UTR | chr12:111358188-111358639 | 451 | | | 4 | | | | | | | | | | | yes | yes | | | | |
| CUX2 | N_Shore | Body | chr12:111473641-111474088 | 447 | | | | 2 | | | | | | | | | | | | | | yes | |
| CUX2 | NA | Body;nan | chr12:111514551-111515293 | 742 | | | | | | | | | | | 1 | | | yes | | | | | |
| CUX2 | NA | Body | chr12:111536664-111537195 | 531 | | | | | | | | | 1 | 1 | | | | yes | yes | | | | |
| CUX2 | NA | Body | chr12:111618925-111620313 | 1388 | 2 | | | 1 | 1 | | | | 1 | | | | | yes | | | | | |
| CUX2 | NA | Body | chr12:111664582-111664785 | 203 | | | 6 | | 1 | | | | | | | | | yes | yes | | | | |
| CUX2 | NA | Body | chr12:111725167-111725516 | 349 | | 1 | | | | | | | | | | | | yes | yes | | | | |
| SH2B3 | N_Shore | 5'UTR | chr12:111855848-111855911 | 63 | | | | | | | | | | | | | | yes | yes | | | | yes |
| RPH3A | NA | TSS200;5'UTR;1stExon | chr12:113229534-113230065 | 531 | | | 13 | | | | | 0 | 4 | | | | | | | yes | | | |
| OAS1 | NA | TSS1500 | chr12:113343409-113343603 | 194 | | | | | | | | | 1 | | | | | yes | yes | yes | | | |
| OAS2 | NA | TSS1500 | chr12:113415883-113417284 | 1401 | | | | | | | | | | | | | | | | | | | |
| DTX1 | Island;chr12:113494389-11349 | TSS1500;S_Shore;TSS200;1stExon | chr12:113495386-113496168 | 782 | | | | | 1 | | | | 2 | | 1 | | | yes | | | | | |
| DTX1 | Island | Body | chr12:113515332-113515969 | 637 | | | 1 | | | | | | | | | | | yes | | | | yes | yes |
| DTX1 | N_Shelf | Body | chr12:113528764-113528845 | 81 | | | | | 1 | | | | | | | | | yes | | | | yes | |
| DTX1 | N_Shore | Body | chr12:113530566-113531060 | 494 | | | | | | | | | 1 | | | | | yes | | | yes | | |
| RASAL1 | Island | Body | chr12:113541698-113541993 | 295 | | | | 1 | 1 | | | | | | | | | yes | | | | | |
| RASAL1 | N_Shore | Body | chr12:113549832-113550113 | 281 | | | | | | | | | | | | | | yes | yes | | | | |
| RASAL1 | S_Shore | TSS200 | chr12:113574084-113574276 | 192 | | | | | | | | | | | | | | yes | | | | | |
| CCDC42B | N_Shelf | TSS1500 | chr12:113587240-113587340 | 100 | | | | | | | | | | | | | | yes | | | | | |
| CCDC42B | Island | Body | chr12:113592319-113592619 | 300 | | | | | | 1 | | | | | | | | yes | | | | | |
| DDX54 | NA | Body | chr12:113599272-113599683 | 411 | | | | | | | | | 1 | | | | | yes | | | | yes | |
| RBM19 | NA | Body | chr12:114312919-114313256 | 337 | | | 1 | | | | | | | | | | | | | | | | |

| | | | | | | | | | | | | | | | | | | | | | | | |
|---|---|---|---|---|---|---|---|---|---|---|---|---|---|---|---|---|---|---|---|---|---|---|---|
| RBM19 | NA | Body | chr12:114337676-114337927 | 251 | 1 | | | | | | | | 1 | 1 | | | | yes | yes | | yes | | |
| TBX5 | N_Shore | Body | chr12:114833837-114834201 | 364 | | | 5 | | | | | | | | | | | | | | | | yes |
| TBX5 | N_Shore;Island | Body | chr12:114836487-114839007 | 2520 | | 2 | 23 | | | | | | | | | | | yes | | | | | yes |
| TBX5 | Island;N_Shore | Body;5'UTR;1stExon;TSS200 | chr12:114840854-114842031 | 1177 | | | | | | | | | 2 | | | | | | | | | yes | |
| TBX5 | N_Shore;Island | TSS1500;5'UTR;1stExon;TSS200 | chr12:114843188-114847578 | 4390 | | | | | | | | | | | | | | | | | | | |
| TBX3 | Island;N_Shore | 3'UTR | chr12:115109452-115109934 | 482 | | | 3 | | | | | | | | | | | yes | | | | | |
| TBX3 | Island | Body | chr12:115112189-115112626 | 437 | | 1 | | | 1 | | | | 1 | | | 1 | | yes | | | | | |
| TBX3 | S_Shelf | Body | chr12:115115415-115117551 | 2136 | | | | | | | | | | | | | | | | | | yes | |
| NCRNA00173 | NA | TSS1500 | chr12:116970856-116971031 | 175 | | | | | | | | | | | | | | yes | yes | | | | |
| MAP1LC3B2 | NA | TSS1500;TSS200;5'UTR;1stExon | chr12:116996773-116997535 | 762 | | | 31 | | 5 | | | | | | | | | yes | | | | | |
| C12orf49 | NA | 3'UTR | chr12:117155382-117173003 | 17621 | | | 4 | | | | | | | | | | | yes | yes | yes | yes | | yes |
| HRK | N_Shore | 3'UTR | chr12:117316230-117316665 | 435 | | | 3 | | | | | | | | | | | yes | | | | | |
| HRK | S_Shore | TSS1500 | chr12:117320134-117320316 | 182 | | | 1 | | | | | 0 | | | | | | yes | | | | | |
| FBXW8 | NA | Body | chr12:117407879-117408163 | 284 | | | 18 | | | | | | | | | | | | | | | | |
| TESC | Island | Body | chr12:117477000-117477424 | 424 | | | | | | | | | | | | | | yes | | | | | |
| TESC | N_Shore | Body | chr12:117482953-117483310 | 357 | 1 | | | | 1 | | | 0 | 1 | 1 | | | | yes | | | | | |
| TESC | N_Shore | Body | chr12:117536182-117536531 | 349 | | | 1 | | | | | | | | | | | yes | | | | | yes |
| FBXO21 | N_Shore | Body | chr12:117626777-117627478 | 701 | | 1 | | | | | | | 1 | | | 1 | | yes | | | | yes | yes |
| NOS1 | N_Shore;Island | 5'UTR | chr12:117797056-117798748 | 1692 | | | 4 | | | | | | 3 | | | | | yes | | | | yes | yes |
| KSR2 | NA | Body | chr12:118313657-118313924 | 267 | | | 6 | | | | | | 1 | | | | | yes | yes | | | | yes |
| KSR2 | N_Shore | Body | chr12:118405665-118406158 | 493 | | | | | 1 | | | 0 | 1 | 2 | | | | | | | | yes | |
| RFC5 | N_Shore | TSS1500 | chr12:118453982-118454133 | 151 | | | | | | | | 0 | | | | | | yes | | | | | |
| PEBP1 | NA | 3'UTR | chr12:118583229-118583290 | 61 | | | | | | | | | 1 | | | | | | | | | yes | |
| SRRM4 | Island | TSS1500 | chr12:119418676-119419143 | 467 | | | | | | | | | 1 | | | | | | | | | | |
| SRRM4 | N_Shore | Body | chr12:119591664-119592343 | 679 | | | 2 | | | | 1 | | 3 | 2 | | 1 | | yes | | | | yes | |
| CCDC60 | Island | TSS200 | chr12:119772354-119772421 | 67 | | 1 | | | | | | 0 | | | | | | | yes | | | yes | |
| MIR1178;CIT | NA | Body | chr12:120151527-120151706 | 179 | | | | 1 | | | | | 1 | | | | | | yes | | | | |
| CIT | N_Shore;chr12:120241921-120 | Body;Island | chr12:120241287-120242513 | 1226 | | | | | 2 | | | | 4 | 1 | 1 | 2 | | | | | | | |
| RAB35 | S_Shore | Body | chr12:120538681-120539137 | 456 | | | | | | | | | | | | | | yes | | | | | |
| PXN | S_Shore | Body | chr12:120652604-120652844 | 240 | | | | | | | | | 1 | | | | | yes | | | | | |
| SIRT4 | NA | Body | chr12:120741565-120741653 | 88 | | | | | | | | | | | | | | yes | | yes | | | |
| PLA2G1B | NA | Body | chr12:120763591-120763869 | 278 | | | | | | | | | | 1 | | | | yes | yes | | | yes | yes |
| MSI1 | N_Shore;Island | Body | chr12:120799332-120800066 | 734 | | | | 1 | 1 | | | | | | | | | yes | | | | yes | yes |
| COQ5 | Island;S_Shore | 1stExon;5'UTR;TSS1500 | chr12:120966954-120967582 | 628 | | | | | | | | | | 3 | | | | yes | | | yes | | |
| CABP1 | NA | TSS1500;Body;TSS200 | chr12:121087382-121088408 | 1026 | 2 | | 2 | | | | | 0 | 2 | | | 1 | | | | | | yes | |
| CABP1 | NA | Body | chr12:121093625-121093709 | 84 | | | | | | | | | 1 | | | | | yes | | | yes | yes | |
| CABP1 | NA | 3'UTR | chr12:121104809-121104909 | 100 | | | | | | | | | | | | | | yes | yes | | | | |
| UNC119B | N_Shore | TSS1500 | chr12:121147222-121147730 | 508 | | | | | | | | | 2 | | | | | yes | yes | | yes | | yes |
| ACADS | S_Shore | Body | chr12:121164194-121164371 | 177 | | | | 1 | | | | | | | | | | yes | | | yes | | |
| HNF1A | N_Shore;Island | TSS1500;TSS200;1stExon;5'UTR | chr12:121416315-121416796 | 481 | | | | 1 | 1 | | | | | | | | | yes | | | | | |
| OASL | NA | Body | chr12:121476549-121476792 | 243 | | | | | | | | | | | | | | yes | yes | | | | |
| P2RX7 | NA | TSS1500 | chr12:121569924-121570081 | 157 | | | | 1 | | | | | | | | | | | yes | yes | | | |
| P2RX4 | N_Shore | TSS1500 | chr12:121647177-121647217 | 40 | | | | | | | | | | | | | | yes | | | | | |
| CAMKK2 | NA | 3'UTR | chr12:121678474-121678720 | 246 | | | 6 | | | | | | | | | | | | yes | | | | |
| CAMKK2 | Island | Body | chr12:121711847-121712245 | 398 | 1 | 1 | | | | | | | | | | | | yes | | yes | | | |
| KDM2B | N_Shore | Body | chr12:121882033-121882415 | 382 | | | | | | | | | 1 | | | | | yes | | | | | |
| KDM2B | N_Shore | Body | chr12:121890163-121890503 | 340 | | | 4 | | | | | | 1 | | | | | yes | yes | | | | |
| KDM2B | NA | Body | chr12:121958763-121958832 | 69 | | | | | | | | | 1 | | | | | | | | | | |
| KDM2B | N_Shore | Body | chr12:121973871-121974305 | 434 | | | | | | | | | 1 | | | 1 | | | yes | | | yes | |
| KDM2B | S_Shore | 1stExon;TSS1500;5'UTR;TSS200 | chr12:122018770-122019760 | 990 | | | 20 | 1 | | | | 0 | | | | | | yes | | | | yes | yes |
| RHOF | Island | 1stExon | chr12:122231455-122231626 | 171 | | 1 | | | | | | | | | | | | | | | | | |
| SETD1B | N_Shore | Body | chr12:122260498-122261162 | 664 | | | | | | | | | 1 | | | | | | yes | | yes | | yes |
| HPD | NA | TSS1500;Body;TSS200 | chr12:122296680-122297408 | 728 | | | 5 | 1 | | | | | | | | | | yes | | | | | |
| WDR66 | N_Shore;Island | TSS1500;TSS200;1stExon;5'UTR | chr12:122355842-122356852 | 1010 | | 1 | 42 | | | 2 | | 0 | 5 | | 1 | 14 | 1 | yes | | | | yes | |
| WDR66 | NA | Body | chr12:122396633-122396687 | 54 | | 1 | | | | | | | | | | | | yes | yes | | | | |
| LRRC43 | NA | TSS1500 | chr12:122651623-122652224 | 601 | | | | | | | | | 1 | | | | | yes | | yes | yes | | yes |
| IL31;LRRC43 | NA | TSS200;5'UTR;TSS1500 | chr12:122658897-122659211 | 314 | | | | | | | | | 1 | | | | | yes | | | | | |
| LRRC43 | N_Shore | 5'UTR;TSS1500 | chr12:122666575-122666672 | 97 | | | | | | | | | 1 | | | | | yes | | yes | | | |
| LRRC43;B3GNT4 | Island | 3'UTR;TSS1500 | chr12:122687992-122688969 | 977 | | | | | | | | | 1 | | | | | yes | | | | | yes |
| DIABLO | S_Shore | TSS200 | chr12:122712075-122712925 | 850 | 3 | | | | 1 | | | | 3 | | | | | | | | | | |
| GPR81 | NA | 1stExon;5'UTR;TSS200;TSS1500 | chr12:123215010-123215684 | 674 | | 1 | 18 | | | | | 0 | 9 | | | 3 | | yes | | | | | |
| HIP1R | Island | TSS200 | chr12:123319894-123319981 | 87 | | | | | | | | | 1 | | | | | | | | | yes | yes |
| HIP1R | NA | Body | chr12:123335357-123335687 | 330 | | | | | | | | | | | | | | yes | | | | | |
| VPS37B | S_Shore | Body | chr12:123352600-123352774 | 174 | | | | | | | | | 1 | | | | | | | | | | |
| VPS37B | Island | TSS200 | chr12:123380870-123381003 | 133 | | 1 | 13 | | | | | | 1 | | | | | | yes | | | | yes |
| ABCB9 | NA | 3'UTR | chr12:123414345-123414638 | 293 | 1 | | | | | | | | | | | | | yes | | | | | |
| PITPNM2 | NA | Body | chr12:123477107-123477147 | 40 | | | 1 | | | | | | 1 | | | | | yes | | | | | |
| CDK2AP1 | N_Shelf | Body | chr12:123750782-123750864 | 82 | | | | | | | | | 1 | | | | | | | | | | |
| CDK2AP1 | N_Shore;Island | Body | chr12:123753186-123754328 | 1142 | | | 6 | 1 | | | | | | | | | | yes | | | | | |
| CDK2AP1 | S_Shore | TSS1500 | chr12:123757070-123758062 | 992 | 4 | | 6 | | | | | | | | | | | yes | yes | | yes | | yes |
| DDX55 | Island | TSS200 | chr12:124086654-124086660 | 6 | | | | | | | | 0 | | | | | | yes | | | | | |
| DNAH10 | Island;N_Shore | TSS200;TSS1500 | chr12:124246391-124247223 | 832 | 10 | | | | 1 | | | 0 | | | | | | yes | | | yes | | |
| DNAH10 | Island | Body | chr12:124272266-124272433 | 167 | | | | | | | | | | | | | | | yes | | | yes | |
| DNAH10 | NA | Body | chr12:124343780-124344211 | 431 | | | | | | | | | 1 | | | | | yes | yes | | | | |
| DNAH10 | NA | Body | chr12:124371660-124371801 | 141 | | 1 | | | 1 | | | | 1 | | | | | yes | | | | | |
| CCDC92 | NA | 5'UTR | chr12:124429228-124429745 | 517 | | | | | | | | | 1 | | | | | | yes | | | | |
| FAM101A | NA | TSS1500;TSS200;1stExon;5'UTR | chr12:124773236-124774078 | 842 | | | 36 | | 1 | | | | 1 | | 1 | | | | yes | yes | | yes | |
| FAM101A | N_Shore | 5'UTR | chr12:124787364-124787465 | 101 | | | | 1 | | | | | | | | | | | | | | | |
| FAM101A | Island | 5'UTR | chr12:124788627-124788814 | 187 | 1 | | | | | | | | | | | | | | | | | | |
| FAM101A | Island | Body | chr12:124798815-124799011 | 196 | 3 | | | | | | | | | | | | | | yes | | | | |
| NCOR2 | N_Shore | Body | chr12:124856376-124856587 | 211 | | | | | | | | | 1 | | | | | | yes | yes | | | |
| NCOR2 | NA | Body | chr12:124864528-124865130 | 602 | | 1 | | 1 | | | | 0 | 2 | 2 | | | | yes | yes | yes | | | |
| NCOR2 | N_Shelf | Body | chr12:124870112-124870394 | 282 | | | 13 | | | | | | | | | | | | | | | | yes |
| NCOR2 | N_Shelf | Body | chr12:124908601-124908653 | 52 | | | | | | | | | 1 | | 1 | | | yes | | | | | |
| NCOR2 | S_Shore | Body | chr12:124911924-124912182 | 258 | 2 | | 1 | | | | | | | | | | | | | | | yes | |
| NCOR2 | N_Shore | Body | chr12:124929590-124930160 | 570 | | | 1 | | | | | | | | | | | yes | yes | | | | |
| NCOR2 | N_Shelf | Body | chr12:124938116-124938573 | 457 | | | | | | | | 0 | 1 | | | | | | yes | | | | |
| NCOR2 | Island;S_Shore | Body | chr12:124941423-124942248 | 825 | | 1 | 10 | 1 | 1 | | | | 1 | 1 | | | | yes | | | | yes | |
| NCOR2 | N_Shore;Island | Body | chr12:124950669-124950990 | 321 | | | 5 | | 2 | | | | 3 | 3 | 1 | | | yes | | | | | |
| NCOR2 | NA | 5'UTR | chr12:124985272-124985889 | 617 | | | | | | | | | 1 | | | | | | yes | yes | | yes | |
| NCOR2 | Island | 5'UTR | chr12:125002007-125003407 | 1400 | | | | | 1 | 1 | | | 2 | | | | | yes | | | | yes | |
| SCARB1 | NA | Body | chr12:125333267-125333358 | 91 | | | 2 | | | | | | | | | | | yes | yes | | | | |
| DHX37 | NA | Body | chr12:125450326-125450666 | 340 | | | | | | | | | | | | | | yes | yes | | | | |
| DHX37 | Island | TSS200 | chr12:125473823-125474021 | 198 | | | 1 | | | | | | | | | | | yes | | | | | |
| TMEM132B | NA | TSS1500 | chr12:125810688-125811164 | 476 | | | | | | | | | | | | | | | yes | | | yes | |
| LOC100128554 | NA | TSS1500;TSS200 | chr12:126926510-126926925 | 415 | | | | | | | | | | | | | | yes | yes | | | yes | |
| TMEM132C | Island | TSS1500 | chr12:128751460-128752058 | 598 | | | | | | | | | 1 | | | | | | | | | yes | yes |
| TMEM132C | NA | Body | chr12:128866172-128866735 | 563 | 1 | | | | | | | 0 | 2 | | | 2 | | | | | yes | | |
| TMEM132C | NA | Body | chr12:129028586-129028733 | 147 | | | | | | | | | | | | | | | yes | | yes | | |

| | | | | | | | | | | | | | | | | | | | | | | | | |
|---|---|---|---|---|---|---|---|---|---|---|---|---|---|---|---|---|---|---|---|---|---|---|---|---|
| SLC15A4 | N_Shore;S_Shore | Body | chr12:129280133-129282057 | 1924 | | | | | | | | | 4 | | | | | yes | | | | | | |
| SLC15A4;MGC16384 | NA | Body | chr12:129299088-129299605 | 517 | | | 14 | | | | | | | | | | | yes | | | yes | | | |
| SLC15A4 | S_Shore | TSS1500 | chr12:129309117-129309376 | 259 | | | 3 | | | | | | 1 | | | | | yes | yes | | yes | | | |
| GLT1D1 | N_Shore;Island | TSS1500;Body | chr12:129337339-129338841 | 1502 | | | 2 | | | | | 0 | | | | | | yes | | | | yes | yes | |
| GLT1D1 | NA | Body | chr12:129416689-129417818 | 1129 | | | 8 | | | | | | 1 | | | | | | yes | | yes | yes | | |
| GLT1D1 | Island | Body | chr12:129429698-129430047 | 349 | | | 1 | | | | | | | | | | | | yes | yes | | yes | | |
| GLT1D1 | NA | Body | chr12:129460559-129461043 | 484 | | 1 | 2 | 1 | | | | 0 | 2 | | | | | yes | | | yes | | | |
| TMEM132D | Island | Body | chr12:130184740-130185016 | 276 | | | | | | | | | | | | | | | | | yes | | | |
| TMEM132D | NA | Body | chr12:130195357-130195777 | 420 | | | 2 | | | | | | | | | | | yes | yes | | | | | |
| TMEM132D | NA | Body | chr12:130315506-130316058 | 552 | | | 16 | | | | | | 1 | | | | | yes | | yes | | | | |
| TMEM132D | Island | 1stExon;5'UTR;TSS200 | chr12:130387861-130389138 | 1277 | | | | | | | | | 3 | | | | | | | | | yes | yes | |
| FZD10 | Island | TSS200;5'UTR;1stExon | chr12:130647004-130647580 | 576 | | | | | | | | | 1 | | | 1 | | | | | | | yes | |
| PIWIL1 | S_Shore;N_Shore;Island | 5'UTR;TSS1500;TSS200 | chr12:130821530-130824831 | 3301 | 1 | | 28 | | 2 | 1 | 1 | 0 | 16 | 2 | 1 | 1 | | yes | | | yes | yes | yes | |
| RIMBP2 | NA | Body | chr12:130918514-130919323 | 809 | | | | | | | | | | | | | | yes | | yes | | | | |
| RIMBP2 | N_Shore | Body | chr12:130933962-130934389 | 427 | | | 2 | | | | | | | | | | | | | yes | | | | |
| RIMBP2 | Island;S_Shore | Body | chr12:130935923-130936634 | 711 | | | 6 | | | | | | 1 | | | | | | | yes | | | | |
| RIMBP2 | S_Shelf | Body | chr12:130939700-130940065 | 365 | | | | | | | | | | | | | | | | yes | | | | |
| RIMBP2 | NA | Body | chr12:130958085-130959214 | 1129 | | | 6 | | | | | | | | | | | yes | yes | | | | | |
| RIMBP2 | NA | Body | chr12:130961424-130961759 | 335 | | | 5 | | | | | 0 | 2 | | | | | | yes | | | | | |
| RIMBP2 | NA | 5'UTR | chr12:130976260-130976551 | 291 | | | 5 | | | | | | 1 | | | | | yes | | | | | yes | |
| STX2 | Island;N_Shore | Body | chr12:131303016-131303993 | 977 | 2 | | 26 | | | | | 0 | 4 | | | 3 | 1 | yes | | | yes | yes | | |
| RAN | N_Shore | TSS1500 | chr12:131355340-131355834 | 494 | | | 4 | | | | | | 1 | | | | | yes | yes | | | | | |
| GPR133 | NA | TSS1500 | chr12:131438016-131438283 | 267 | | | 1 | | 1 | | | | | | | | | yes | | | | yes | | |
| GPR133 | NA | Body | chr12:131443383-131443486 | 103 | | | | | 1 | | | | | | | | | yes | yes | | | | | |
| GPR133 | NA | Body | chr12:131455513-131455619 | 106 | | | | | | | | | 1 | | | | | | | | | | yes | |
| GPR133 | N_Shore;Island | Body | chr12:131488390-131488926 | 536 | 1 | | | | | | | 0 | 3 | | | | | | | | yes | | | |
| GPR133 | NA | Body | chr12:131496914-131497012 | 98 | | | | 1 | 1 | | | | | | | | | | | | yes | | | |
| GPR133 | NA | Body | chr12:131505834-131506092 | 258 | | | | | | | | | | | | | | yes | yes | yes | yes | | | |
| GPR133 | NA | Body | chr12:131561080-131561270 | 190 | | | 1 | | | | | | | | | | | | yes | | | | | |
| GPR133 | N_Shelf;NA | Body | chr12:131568441-131569407 | 966 | | | 2 | | | | | | | | | | | yes | yes | | | | | |
| GPR133 | NA | Body | chr12:131589353-131590596 | 1243 | | | 2 | | 1 | | | | | | | | | yes | yes | | | | | |
| GPR133 | NA | Body | chr12:131617044-131617092 | 48 | | | 3 | | | | | | | | | | 1 | yes | | yes | | | | |
| GPR133 | N_Shore | Body | chr12:131622584-131623508 | 924 | | | 13 | | 1 | 1 | | 0 | 1 | | | 2 | | | | | | yes | | |
| LOC116437 | NA | Body | chr12:131687337-131687695 | 358 | | | | | | | | | | | | | | | | | | | | |
| SFRS8 | S_Shore | Body | chr12:132196975-132197229 | 254 | | | 1 | | | | | | | | | | | | | | | | yes | |
| SFRS8 | NA;N_Shelf | Body | chr12:132259591-132259908 | 317 | | | | | | | | | 2 | | | | | yes | | | | | | |
| SFRS8 | NA | Body | chr12:132269563-132270218 | 655 | | | | | | | | | | | | | | | | | | | | |
| SFRS8 | S_Shore | Body | chr12:132278430-132278510 | 80 | | | 8 | | | | | | | 1 | | | | yes | yes | | | | | |
| ULK1 | Island | Body | chr12:132380696-132380903 | 207 | | | | | | | | | 1 | | | | | | | | | | yes | |
| SNORA49;EP400 | Island | TSS1500;Body | chr12:132514403-132514508 | 105 | | 1 | | | | | | | | | | | | | | | | | | |
| EP400 | NA | Body | chr12:132536833-132537775 | 942 | | | | | | | | | 3 | | | | | | | yes | | yes | | |
| EP400 | NA | Body | chr12:132546772-132547245 | 473 | | | | | | | | | | | | | | | | | yes | yes | | |
| EP400 | NA | Body | chr12:132549292-132549404 | 112 | | | | | | | | | 1 | | | | | | | | | | | |
| EP400 | NA | Body | chr12:132551398-132551592 | 194 | | | | | | | | | 1 | | | | | | | | | | | |
| EP400 | NA | Body | chr12:132561971-132562125 | 154 | | 1 | | | | | | | | | | | | | | | | | | |
| EP400NL | N_Shore | Body | chr12:132603220-132603671 | 451 | | | | | | | | 0 | | | | | | yes | yes | | | | | |
| GALNT9 | N_Shore | 3'UTR | chr12:132681397-132681641 | 244 | | | | | | | | | | | | | | yes | | | yes | | | |
| GALNT9 | S_Shore;Island | Body;TSS200;TSS1500;5'UTR;1stEx | chr12:132689907-132692072 | 2165 | | | 13 | | | | | 0 | 8 | 2 | 1 | 5 | 1 | yes | yes | yes | | | | |
| GALNT9 | NA | Body | chr12:132703301-132704101 | 800 | | | 2 | | | | | | 1 | | | | | | | | yes | | | |
| GALNT9 | Island;N_Shore | Body | chr12:132834399-132836216 | 1817 | | | 4 | | | | | | | | | | | | yes | | | | yes | |
| GALNT9 | Island | Body | chr12:132837412-132837679 | 267 | | 1 | | | | | | | | | | | | | yes | yes | | yes | | |
| GALNT9 | S_Shore;N_Shelf | Body | chr12:132839125-132840868 | 1743 | | | 1 | | | | | | 1 | | | | | | yes | yes | | | | |
| GALNT9 | N_Shore | Body | chr12:132845336-132845530 | 194 | | | 3 | | | | | | | | | | | | yes | | | yes | | |
| GALNT9 | Island | Body | chr12:132847907-132848554 | 647 | | | | | | | | | | | | | | | | yes | | | yes | |
| GALNT9;LOC100130238 | N_Shelf | Body;TSS1500 | chr12:132851518-132851967 | 449 | | | | | | | | | | | | | | | | yes | | yes | yes | |
| GALNT9;LOC100130238 | Island;N_Shore | Body | chr12:132853954-132855381 | 1427 | | | 11 | | | | | | 2 | | | | | | | yes | | yes | | |
| GALNT9 | N_Shore;Island | Body | chr12:132859441-132859951 | 510 | | | 11 | | | | | 0 | 2 | | | | | | yes | yes | | yes | | |
| GALNT9 | N_Shore;Island | Body | chr12:132862024-132865307 | 3283 | | | 7 | | | | | 0 | 3 | | | 1 | | yes | yes | | | | yes | |
| GALNT9 | N_Shelf | Body | chr12:132869000-132869299 | 299 | 3 | | | | | | | | | | | | | | | | | | | |
| GALNT9 | N_Shore | Body | chr12:132870378-132870448 | 70 | | | 6 | | | | | | | | | | | | | | yes | | | |
| GALNT9 | N_Shore | Body | chr12:132879809-132880128 | 319 | 6 | | | | | | | | | | | | | | | | | yes | | |
| GALNT9 | S_Shore | Body | chr12:132885871-132887175 | 1304 | 1 | | 2 | | | | | | | | | 1 | | | yes | yes | yes | | | |
| GALNT9 | N_Shore;Island;S_Shore | Body | chr12:132895346-132897840 | 2494 | 1 | | 13 | | | | | | | | | | | yes | | yes | | | | |
| GALNT9 | N_Shore;Island;S_Shore | Body | chr12:132899065-132901800 | 2735 | | | | | | | | | | | | | | | | | | | | |
| GALNT9 | N_Shore;Island | Body;1stExon | chr12:132904123-132906010 | 1887 | | 1 | | | | | | 0 | 3 | | | | | | | | | yes | yes | |
| FBRSL1 | N_Shore;Island | TSS1500 | chr12:133065724-133066054 | 330 | 11 | | 12 | 1 | 8 | | | | | | | | | | yes | | | yes | | |
| FBRSL1 | N_Shore | Body | chr12:133101168-133101612 | 444 | | | | | | | | | 4 | 1 | | | | yes | | | | | | |
| FBRSL1 | NA | Body | chr12:133120548-133120760 | 212 | | | | | | | | | 1 | | | | | | | | yes | yes | | |
| FBRSL1 | N_Shore;Island | Body | chr12:133134938-133135698 | 760 | | | 5 | | 1 | | | | | | | | | yes | | | | | | |
| FBRSL1 | N_Shelf | Body | chr12:133156510-133156565 | 55 | 1 | | | | | | | | | | | | | yes | yes | | | | | |
| FBRSL1 | Island | Body | chr12:133159472-133159939 | 467 | | | 1 | | | | | | | | | | | | | | | | yes | |
| FBRSL1 | N_Shore | 3'UTR | chr12:133161320-133161839 | 519 | | | 1 | | | | | | | | | | | | | | | yes | | |
| P2RX2 | Island | TSS1500;Body | chr12:133195061-133196647 | 1586 | | 1 | | | | | | 0 | | | | 1 | | | yes | | | yes | yes | |
| P2RX2 | S_Shore | Body | chr12:133197668-133198063 | 395 | | | | | | | | | | | | | | | | | | | | |
| POLE | N_Shore | Body | chr12:133248929-133249308 | 379 | | | | 1 | | | | | | | | | | | | | | | | |
| ANKLE2 | N_Shore | Body | chr12:133306722-133306893 | 171 | 1 | | | | | | | | | | | | | yes | | | | | yes | |
| ANKLE2 | S_Shore | Body | chr12:133312126-133312361 | 235 | | | | | | | | | | | | | | yes | | | | | | |
| ANKLE2 | N_Shore;Island | Body | chr12:133335468-133335817 | 349 | 3 | | | | | | | | | | | | | yes | yes | | | yes | | |
| GOLGA3 | N_Shelf | 3'UTR | chr12:133346264-133346306 | 42 | | | | | | | | | 1 | | | | | yes | | | | | | |
| GOLGA3 | N_Shore;Island | 3'UTR;Body | chr12:133349325-133349952 | 627 | 2 | | | | | | | | 1 | | | | | yes | | | | | | |
| GOLGA3 | S_Shelf;S_Shore | Body | chr12:133380384-133381504 | 1120 | 1 | | 1 | | | | | | | | | | | | | | | | | |
| CHFR | S_Shore | Body | chr12:133419166-133419256 | 90 | | | | | | | | | | | | | | yes | | | | | | |
| CHFR | S_Shore | TSS1500 | chr12:133464892-133465592 | 700 | | | 4 | | | | | 0 | | | | | | yes | | yes | | yes | | |
| ZNF268 | Island | TSS200 | chr12:133757947-133758599 | 652 | | 1 | | | | | | | | | | | | | yes | | yes | | | |
| DKFZp686A1627 | NA | Body | chr13:19692335-19692444 | 109 | | | | | | | | | | | | | | | | | | | | yes |
| TUBA3C | NA | 5'UTR;1stExon;TSS200 | chr13:19755916-19756905 | 989 | | | 13 | | | | | | | | | | | yes | | yes | yes | | | |
| LOC100101938 | N_Shore;Island | Body;TSS200 | chr13:19918525-19919394 | 869 | | | 9 | | | | | 0 | 3 | | | | | yes | yes | | | | | |
| GJA3 | Island | Body | chr13:20716432-20717597 | 1165 | 1 | | 4 | | | | | 0 | 3 | 1 | | 1 | | | | | yes | | yes | |
| GJA3 | Island | TSS1500 | chr13:20735967-20736342 | 375 | | | 8 | | | | | | | | | | | | yes | | | | yes | |
| GJB2 | S_Shore | TSS1500 | chr13:20767968-20768560 | 592 | | | 4 | 1 | | | | | | | | | | | | | | | yes | |
| GJB6 | NA | Body | chr13:20797356-20797693 | 337 | | | 3 | | | | | | | | | | | yes | | | | | | |
| GJB6 | N_Shore;Island | 5'UTR;1stExon;TSS200;TSS1500 | chr13:20805094-20806813 | 1719 | | 1 | 32 | 2 | | | | 0 | 3 | | | | | | | | | | yes | |
| CRYL1 | Island | Body | chr13:20989142-20989818 | 676 | | | | 1 | | | | 0 | 2 | | | 1 | | | yes | | | | | |
| CRYL1 | Island | Body | chr13:21009861-21010223 | 362 | | | | | | | | | 1 | | | | | | yes | | | | | |
| IL17D | NA | Body | chr13:21287237-21287304 | 67 | | | | | 1 | | | | | | | | | yes | | | | | yes | |
| IL17D | NA | Body | chr13:21291173-21291467 | 294 | | | | | | | | | 1 | | | | | yes | | | | | | |
| IL17D | N_Shore;Island | Body | chr13:21295590-21296465 | 875 | | | | | | | | 0 | | 1 | | 1 | | | | | | | | |
| SKA3;MRP63 | N_Shore;Island | Body;TSS200;TSS1500 | chr13:21750229-21751373 | 1144 | | | | | | | | | 2 | | | | | yes | | | | | | |
| ZDHHC20 | Island | Body | chr13:21951294-21951631 | 337 | | | | | | | | 0 | | | | | | | | | | | | |

| | | | | | | | | | | | | | | | | | | | | | | | |
|---|---|---|---|---|---|---|---|---|---|---|---|---|---|---|---|---|---|---|---|---|---|---|---|
| SGCG | NA | Body | chr13:23898364-23898864 | 500 | | | 2 | | | | | 0 | 1 | | | | | yes | | | | | |
| SACS | NA | 5'UTR | chr13:24007434-24007965 | 531 | | | | | | | | | | | | | | yes | | | | | |
| TNFRSF19 | NA | TSS1500 | chr13:24144257-24144682 | 425 | | | | | | | | 0 | 2 | | | 1 | | yes | | | | | |
| TNFRSF19 | N_Shore | 5'UTR | chr13:24152135-24152779 | 644 | | | | | | | | | | | | | | | yes | | | | |
| SPATA13 | N_Shore | TSS200 | chr13:24734742-24734795 | 53 | | | | | | | | | 1 | | | | | yes | | | | | |
| SPATA13;MIR2276 | S_Shore | 5'UTR;TSS1500;TSS200 | chr13:24736095-24736569 | 474 | | | | 1 | 2 | | | | 2 | | | | | yes | | | | | |
| SPATA13 | NA | Body | chr13:24844846-24844938 | 92 | | 1 | | | | | | | 1 | | | | | | yes | | | yes | |
| PARP4 | N_Shore | 5'UTR | chr13:25085301-25085776 | 475 | | 1 | | | 2 | | | | | | | | | yes | yes | | | | |
| RNF17 | NA | TSS200 | chr13:25338113-25338621 | 508 | | | 5 | | | | | | 3 | | | | | yes | | | | yes | |
| CENPJ | Island | Body | chr13:25469147-25469406 | 259 | | | 1 | | | | | | | | | | | yes | yes | | | | yes |
| LOC646405 | Island | Body | chr13:25506131-25506383 | 252 | | | 8 | | | | | 0 | 2 | 2 | | | | yes | yes | | | | |
| PABPC3 | Island | TSS1500;TSS200;1stExon;5'UTR | chr13:25670047-25670530 | 483 | | | 44 | | | | 1 | 0 | 11 | 2 | | 24 | | yes | | | | yes | yes |
| FAM123A | Island;S_Shore | TSS1500 | chr13:25746120-25746644 | 524 | | | 5 | | | | | 0 | 2 | | | | | | | | | yes | |
| NUPL1 | Island | TSS1500 | chr13:25875020-25875436 | 416 | | 1 | | | | | | | 1 | | | | | | yes | yes | | | |
| ATP8A2 | N_Shore | Body | chr13:26042598-26043066 | 468 | | 1 | | | | | | | | | | | | | | | | yes | yes |
| ATP8A2 | N_Shore;Island | Body | chr13:26586254-26587011 | 757 | 2 | 1 | 13 | | 1 | | | 0 | 2 | 1 | | 3 | | yes | | | | | yes |
| SHISA2 | NA | 3'UTR | chr13:26620612-26620952 | 340 | | | 2 | | | | | | | | | | | yes | | | | | |
| SHISA2 | Island | TSS200 | chr13:26625307-26625325 | 18 | | | | | | | | | 1 | | | | | | | | | | |
| WASF3 | Island | TSS200 | chr13:27131676-27131724 | 48 | | | | | | | | | 1 | | | | | | | | | | yes |
| GPR12 | N_Shore | Body | chr13:27332979-27333506 | 527 | 2 | | | | | | | | | | | | | | | | | | |
| GPR12 | Island | TSS200;TSS1500;1stExon;5'UTR | chr13:27334723-27335204 | 481 | | | 4 | | | | | | 2 | | | | | | | | | | yes |
| SNORA27;SNORD102;RPL21P28;RPL21 | S_Shelf | TSS1500;TSS200;Body | chr13:27829014-27829259 | 245 | | | | | | | | | 1 | | | | | yes | | | | | |
| GTF3A | N_Shore | TSS1500 | chr13:27998438-27998564 | 126 | | | | | | | | | 1 | | | | | yes | | | yes | yes | |
| GSX1 | Island | TSS1500;TSS200 | chr13:28365638-28367741 | 2103 | | | | | | | | | 2 | | | | | | | | | yes | yes |
| PDX1 | N_Shore | TSS1500 | chr13:28492971-28494004 | 1033 | | 1 | 2 | 1 | | | | | | | | | | | | | | | yes |
| PDX1 | N_Shore;Island | Body;3'UTR | chr13:28496641-28499045 | 2404 | | | | 2 | 4 | | | | | | | | | | | | yes | | yes |
| CDX2 | Island | TSS1500 | chr13:28544592-28545566 | 974 | | 1 | | | 4 | | | | | | | | | yes | yes | | | | |
| PRHOXNB | N_Shore;Island | Body | chr13:28553501-28555387 | 1886 | 6 | 1 | | | 2 | | | | 2 | | | 1 | | yes | yes | | | yes | |
| PRHOXNB | NA | TSS200;1stExon | chr13:28562668-28563651 | 983 | | | 14 | 1 | | | | | | | | | | yes | yes | | | yes | |
| FLT1 | Island | TSS1500 | chr13:29069623-29069941 | 318 | | | | | | | | | | | | | | | | | | | |
| SLC46A3 | chr13:29292670-29293329 | Island | chr13:29293133-29293213 | 80 | | | | | | | | | | | 1 | | | yes | | yes | | | yes |
| MTUS2 | Island | Body | chr13:29913935-29914200 | 265 | | | | | | | | | | | | | | | | | | | |
| MTUS2 | NA | Body | chr13:29933441-29933565 | 124 | | 1 | | | | | | | | | | | | | | | | | |
| MTUS2 | Island | Body | chr13:30077315-30077489 | 174 | | | 15 | | | | | 0 | 1 | 1 | | 9 | | | | | | | |
| SLC7A1 | Island | 3'UTR | chr13:30088615-30088771 | 156 | | | 28 | | | | | | 1 | | | | | | | | | | |
| LOC100188949 | NA | Body;TSS200 | chr13:30947713-30948716 | 1003 | | | | | | | | | 1 | | | | | yes | yes | | | yes | yes |
| C13orf33 | N_Shore;Island | TSS1500;TSS200;1stExon;5'UTR;Bo | chr13:31479935-31481184 | 1249 | 1 | | 12 | | | | | 0 | 2 | | | | | | | | | | yes |
| C13orf26 | NA | TSS200;1stExon;5'UTR | chr13:31506685-31507139 | 454 | | 1 | 11 | | | | 1 | 0 | 5 | 1 | | 4 | | | | | | | |
| FRY | N_Shore;Island | TSS1500;TSS200;1stExon;5'UTR | chr13:32605187-32605952 | 765 | | | | | 7 | | | | | | | | | | | | | yes | |
| KL | Island | TSS1500;TSS200 | chr13:33589962-33590495 | 533 | | | 10 | | | | | | | 1 | | | | | | | | yes | yes |
| STARD13 | NA | 1stExon;5'UTR;Body | chr13:33760215-33760693 | 478 | | | 1 | | | | | | | | | | | | | | | | |
| STARD13 | NA | 1stExon;5'UTR;Body;TSS200;TSS15 | chr13:33780078-33781288 | 1210 | | 1 | 18 | 1 | 2 | | | | | | | | | | yes | yes | | yes | |
| STARD13 | NA | 1stExon | chr13:33859636-33859885 | 249 | | | | | | | | | 1 | | | | | | | | | | |
| RFC3 | N_Shore | TSS1500 | chr13:34391985-34392073 | 88 | | | | | | | | | 1 | | | | | | yes | | | yes | |
| NBEA | N_Shore;Island | Body | chr13:36044768-36045352 | 584 | | 1 | 19 | | | | | | 1 | | | | | | | | | | |
| MAB21L1;MIR548F5;NBEA | N_Shore;Island;S_Shore | 1stExon;Body;3'UTR;5'UTR;TSS200 | chr13:36048892-36053118 | 4226 | | | | | | | | | | | | | | | | | | | |
| MIR548F5;DCLK1 | NA | Body | chr13:36421844-36421949 | 105 | | | | | | | | | 1 | | | | | | | | | yes | |
| MIR548F5;DCLK1 | NA | Body | chr13:36429936-36430134 | 198 | | | 3 | | | | | 0 | | | | | | | | | | | |
| DCLK1 | Island | 5'UTR | chr13:36705041-36705475 | 434 | | | | | | | | 0 | | | | | | | | | | yes | yes |
| SOHLH2 | N_Shore;Island | Body;TSS200 | chr13:36787885-36788859 | 974 | | | | | | | | | | | | | | yes | | | | | |
| C13orf38 | N_Shore;Island;S_Shore | Body;5'UTR;1stExon;TSS200 | chr13:36871646-36872346 | 700 | | | 6 | | | | | 0 | 6 | | | 3 | | | | yes | | | |
| SPG20 | Island | TSS200;5'UTR | chr13:36920685-36920813 | 128 | | | | | | | | | 1 | | | | | | | | | | yes |
| SPG20 | NA | 1stExon;5'UTR;TSS200 | chr13:36944294-36944471 | 177 | 2 | 3 | | | | | | | | | | | | yes | yes | | | | |
| CCNA1 | N_Shore;Island | TSS1500;1stExon;5'UTR | chr13:37004536-37006127 | 1591 | | | 38 | | 1 | | | | 6 | | | 2 | | yes | | | | | yes |
| C13orf36 | N_Shore | TSS1500 | chr13:37247048-37247708 | 660 | | | 3 | | | | | | | | | | | yes | | | yes | | |
| C13orf36 | S_Shore | 5'UTR | chr13:37248953-37249481 | 528 | | | 2 | | | | | | | | | 1 | | | | | | | |
| C13orf36 | NA | 5'UTR | chr13:37269125-37269461 | 336 | | | | | | | | | | | | | | | | | | | |
| SMAD9 | NA | Body | chr13:37453429-37454023 | 594 | | | | | | | | 0 | 2 | | | 1 | | yes | | | | | |
| POSTN | NA | 1stExon | chr13:38172803-38173002 | 199 | | | 6 | | | | | | | | | | | | | | | | yes |
| TRPC4 | N_Shore | 5'UTR | chr13:38443634-38443809 | 175 | | | | | | | | | 1 | | | | | | | | | | |
| FREM2 | Island | 1stExon;5'UTR | chr13:39261425-39262298 | 873 | | | | | | | | | 2 | | | | | | | | | yes | |
| FREM2 | NA | Body | chr13:39358777-39358893 | 116 | | | 2 | 1 | 1 | | | | | | | | | yes | | | | | |
| LOC646982 | NA | TSS200 | chr13:41055185-41055588 | 403 | | | | | | | | | 1 | | | | | | yes | | yes | | |
| SUGT1L1 | Island;N_Shore | TSS1500 | chr13:41496184-41496979 | 795 | | | 15 | 1 | | | | | 1 | | | 1 | | yes | yes | | | yes | |
| ELF1 | NA | 5'UTR;1stExon | chr13:41593385-41593582 | 197 | | | 28 | | | | | | | | | | | | | | | | yes |
| C13orf15 | Island | TSS1500 | chr13:42031103-42031291 | 188 | | | | | | | | | | | | | | yes | yes | | | | yes |
| TNFSF11 | NA | TSS200 | chr13:43136804-43136933 | 129 | | | | | | | | | | | | | | | yes | yes | | | |
| TNFSF11 | N_Shore;Island | TSS200;5'UTR | chr13:43148214-43148436 | 222 | | | 3 | | | | | | 1 | | | | | yes | | | | | |
| C13orf30 | NA | TSS200;1stExon;5'UTR | chr13:43355514-43356116 | 602 | | | | | | | | | 1 | | | | | yes | | | | | |
| EPSTI1 | S_Shore | TSS1500 | chr13:43566618-43566902 | 284 | | | 9 | | | | | | | | | | | | yes | | yes | | |
| DNAJC15 | N_Shore;Island | TSS200;5'UTR;1stExon | chr13:43597297-43598038 | 741 | | | 6 | | | 1 | | 0 | 1 | 1 | | 6 | 1 | yes | yes | | yes | | |
| LOC121838 | NA | TSS1500 | chr13:44595496-44595905 | 409 | | | | | | | | | 1 | | | | | | | | yes | | yes |
| SERP2 | Island;N_Shore | TSS200;TSS1500 | chr13:44947247-44948413 | 1166 | | | | | | | | | 1 | | | | | | yes | | | yes | |
| TSC22D1 | N_Shore | 1stExon | chr13:45149661-45149706 | 45 | | | 8 | | | | | 0 | 1 | | | | | | | | | yes | |
| NUFIP1;KIAA1704 | Island | TSS200;1stExon | chr13:45563807-45564352 | 545 | | | | | | | | 0 | | | | | | yes | yes | | | | |
| LOC100190939 | NA | Body | chr13:45945197-45945362 | 165 | | 1 | 13 | | | | | | | | | | | yes | | | yes | | |
| FAM194B | NA | 5'UTR;1stExon | chr13:46189838-46189906 | 68 | | | | | | | | | 1 | | | | | | | | yes | | yes |
| SIAH3 | NA | 5'UTR;1stExon;TSS200;TSS1500 | chr13:46425712-46426263 | 551 | | 1 | 10 | | | | | 0 | 5 | | | 2 | | | | | | | |
| LCP1 | NA | 5'UTR | chr13:46742691-46742802 | 111 | | | 3 | | | | | | | | | | | yes | yes | | | | |
| LCP1 | NA | TSS1500 | chr13:46757318-46757415 | 97 | | | | | | | | | 1 | | | | | | | | | | |
| LRCH1 | N_Shore | TSS1500 | chr13:47125801-47126551 | 750 | | | 7 | | | | | | 3 | | | 2 | | yes | yes | | | | |
| ESD | Island | 5'UTR | chr13:47370918-47371523 | 605 | | | | | | | | 0 | 1 | | | | | yes | | | | | |
| HTR2A | NA | 5'UTR;Body;1stExon;TSS200;TSS15 | chr13:47470350-47472349 | 1999 | | 1 | | 1 | | | | 0 | 3 | 1 | | | | yes | | | | | |
| SUCLA2 | S_Shore | TSS1500 | chr13:48575859-48575927 | 68 | | | 18 | | | | | 0 | 1 | | | | | yes | yes | yes | | yes | |
| RB1 | N_Shore | TSS1500 | chr13:48876866-48876950 | 84 | | | 2 | | | | | | | | | | | yes | yes | | yes | | |
| RB1 | S_Shore;Island | Body | chr13:48892948-48894595 | 1647 | | | | | | | | | 1 | | | | | yes | | | | | |
| CYSLTR2 | NA | TSS1500 | chr13:49280257-49280411 | 154 | | | | | | | | | | | | | | yes | yes | | | yes | |
| FNDC3A | NA | TSS200;Body | chr13:49684383-49684404 | 21 | | | | | 1 | | | 0 | | | | | | | | | | | |
| MLNR | N_Shore | TSS1500 | chr13:49793054-49793170 | 116 | | | | | | | | | 1 | | | | | yes | yes | | | | |
| CAB39L | NA | 5'UTR;1stExon | chr13:49975383-49975643 | 260 | | | | 1 | | | | | 1 | | | | | | | | | | |
| ARL11 | NA | TSS200 | chr13:50202547-50202713 | 166 | | | 1 | 1 | 1 | | | | | | | | | yes | | | | yes | |
| TRIM13;DLEU2 | N_Shore | TSS1500;Body | chr13:50569769-50570250 | 481 | | | | | | | | 0 | 2 | | | | | yes | | | | | |
| DLEU2 | Island | TSS1500 | chr13:50700845-50701501 | 656 | | | | 1 | 1 | | | 0 | 1 | | | | | | | | | | |
| DLEU7 | Island | 1stExon;5'UTR;TSS200 | chr13:51417469-51418221 | 752 | | 1 | 8 | | | | | 0 | 2 | | | 2 | | | yes | | | | yes |
| GUCY1B2 | NA | TSS200;Body | chr13:51640142-51640442 | 300 | | | | | | | | 0 | 2 | 1 | | | | yes | | | | | |
| FLJ37307 | NA | TSS200 | chr13:52419380-52419483 | 103 | | | | | | | | 0 | 1 | | | 1 | | | yes | | yes | | |
| NEK3 | S_Shore | TSS1500;TSS200 | chr13:52734018-52734071 | 53 | | | | | | | | | | | | | | yes | | yes | | | yes |
| THSD1P | Island;N_Shore | TSS1500 | chr13:52769289-52769951 | 662 | 1 | | 2 | | | | | 0 | 1 | | | 1 | | yes | | | yes | | |
| LECT1 | S_Shore | TSS200 | chr13:53314142-53314226 | 84 | | | | | | | | | | | | | | | | | | yes | |

| | | | | | | | | | | | | | | | | | | | | | | | |
|---|---|---|---|---|---|---|---|---|---|---|---|---|---|---|---|---|---|---|---|---|---|---|---|
| PCDH8 | N_Shore | Body | chr13:53419596-53419729 | 133 | | | | | | | | | 1 | | | | | | | | | | yes |
| PCDH8 | Island;N_Shore | TSS200;TSS1500 | chr13:53422777-53424127 | 1350 | | 1 | | | | 3 | | | 2 | | | | | | | | | | |
| OLFM4 | NA | TSS200 | chr13:53602856-53603286 | 430 | | | | | | | | | | | | | | | | | yes | yes | |
| PCDH20 | NA | 5'UTR;1stExon;TSS200 | chr13:61989597-61989859 | 262 | | | 4 | | | | | | 1 | | | 2 | | | yes | | | | yes |
| KLHL1;ATXN8OS | N_Shore;Island;S_Shore | Body;TSS200;1stExon;5'UTR | chr13:70681276-70683127 | 1851 | | | 7 | | | | | 0 | 3 | | | 1 | | | | | | yes | yes |
| COMMD6 | S_Shore | TSS1500 | chr13:76112973-76113014 | 41 | | | | | | | | | 1 | | | | | yes | yes | yes | | | |
| LMO7 | NA | TSS1500;Body;TSS200 | chr13:76334583-76334866 | 283 | | | 27 | | | | | 0 | | | | | | | | | | | yes |
| KCTD12 | S_Shore | TSS1500 | chr13:77461279-77461426 | 147 | | | 1 | | | | | 0 | 1 | | | 4 | | | | | | | |
| EDNRB | N_Shore;Island;S_Shore | Body;1stExon;5'UTR;TSS200;TSS1! | chr13:78491982-78494462 | 2480 | | 1 | 18 | | 9 | | | 0 | 4 | 1 | | 2 | | yes | | | yes | | yes |
| EDNRB | NA | TSS1500 | chr13:78550069-78550384 | 315 | | | | | | | | | | | | | | | yes | yes | | | |
| POU4F1 | Island | 5'UTR;1stExon;TSS200 | chr13:79177467-79177925 | 458 | | | 1 | | | | | | 4 | | | | | | | | | yes | yes |
| RNF219 | S_Shore | TSS1500 | chr13:79234251-79234847 | 596 | | | 10 | | | | | | 1 | | | | | yes | yes | | | | |
| NDFIP2 | N_Shore | TSS1500 | chr13:80054976-80055067 | 91 | | | | | | | | 0 | | | | | | yes | yes | | | | |
| SPRY2 | N_Shore | Body | chr13:80911626-80911808 | 182 | | 1 | 4 | | | | | | 1 | | | | | | | | | | |
| SLITRK1 | S_Shelf | 1stExon;5'UTR | chr13:84456127-84456308 | 181 | | | | | 1 | | | | 1 | | | | | | | | | yes | yes |
| SLITRK5 | Island | TSS1500 | chr13:88324432-88324597 | 165 | | | | | | | | | 1 | | | | | | | | | yes | yes |
| SLITRK5 | N_Shore;S_Shore | Body | chr13:88328009-88330614 | 2605 | | | | | | | | | | | | | | | | | | | |
| MIR19A;MIR17HG;MIR17;MIR18A | S_Shore | TSS1500;Body | chr13:92001764-92002454 | 690 | 2 | | | | | | | | | | | | | | | | | | yes |
| GPC5 | N_Shore;Island | TSS1500;1stExon;Body | chr13:92050675-92051786 | 1111 | | 1 | 4 | | | | | 0 | 3 | | | | | | | | | yes | |
| GPC6 | Island | 1stExon;5'UTR | chr13:93879303-93879769 | 466 | | | | | | | | 0 | 1 | | | | | | | | | | |
| TGDS | S_Shore | TSS1500 | chr13:95248771-95248888 | 117 | | | 7 | | | | | | | | | | | yes | | | | | |
| GPR180 | N_Shore | TSS1500 | chr13:95253641-95253676 | 35 | | | | | | | | | 1 | | | | | | yes | | | | |
| SOX21 | N_Shore | 1stExon;3'UTR | chr13:95362829-95363007 | 178 | | | 1 | | | | | | | | | | | | | | | | yes |
| SOX21 | Island | TSS200 | chr13:95364514-95364633 | 119 | | | | | | | | | 2 | | | | | | | | | yes | |
| ABCC4 | NA | Body | chr13:95826930-95827029 | 99 | | | | | | | | | | | | | | | yes | yes | | | |
| CLDN10 | Island;S_Shore | TSS200;Body;5'UTR;1stExon | chr13:96204917-96206104 | 1187 | 1 | | 4 | | 2 | | 1 | 0 | 3 | 2 | | 8 | | | | | | | yes |
| DZIP1 | S_Shore | TSS1500 | chr13:96297197-96297338 | 141 | | | 3 | | | | | | 2 | | | 1 | | | | | | yes | |
| UGGT2 | N_Shore;Island | Body | chr13:96705123-96705786 | 663 | | | | | | | | 0 | 1 | | | | | yes | yes | | yes | | |
| OXGR1 | N_Shore;Island;S_Shore | 5'UTR;TSS200 | chr13:97646341-97647222 | 881 | | | | | | | | 0 | 4 | 2 | | 3 | | yes | yes | | | | |
| MBNL2 | NA | TSS1500 | chr13:97873877-97874279 | 402 | | | 8 | 1 | | | | 0 | | | | | | | | | | yes | |
| MBNL2 | NA | 5'UTR | chr13:97928122-97928270 | 148 | | | | 1 | | | | | | | | | | | | | | | |
| IPO5 | NA | TSS200 | chr13:98605770-98606196 | 426 | | | 19 | | | | | | 1 | | | | | yes | | | | | |
| FARP1 | N_Shore | TSS1500 | chr13:98794353-98794593 | 240 | | | 9 | | | | | | | | | | | | | | yes | | |
| FARP1 | NA | Body | chr13:98960562-98960620 | 58 | | | | 1 | | | | | 1 | | | | | yes | yes | | yes | yes | |
| FARP1 | NA | Body;3'UTR | chr13:99100506-99101389 | 883 | | | 5 | | | | | | 2 | | | | | yes | | | | | |
| STK24 | NA | Body | chr13:99135543-99136580 | 1037 | | | 5 | 1 | 1 | | | | 2 | 1 | 1 | | | | | | | | |
| STK24 | NA | Body | chr13:99158698-99158901 | 203 | | | | | | | | | | | | | | | | | | | |
| STK24 | NA | Body;5'UTR;1stExon;TSS200 | chr13:99174312-99174494 | 182 | | | 15 | | | | | | 1 | | | | | | yes | | | | |
| DOCK9 | NA | Body;TSS200 | chr13:99629970-99630583 | 613 | | | | | 1 | | | | 1 | | 1 | | | | yes | | | | |
| UBAC2 | NA | Body | chr13:99867950-99868015 | 65 | | | | | | | | 0 | 1 | | | | | yes | | | yes | yes | |
| UBAC2;MIR623 | NA | Body;TSS1500 | chr13:100008030-100008450 | 420 | | 1 | | | | | | | | | | | | | yes | | | | |
| CLYBL | N_Shore;Island | Body | chr13:100310162-100310841 | 679 | | 1 | 21 | 1 | 1 | | | 0 | 4 | | | | | | | | | | |
| CLYBL | NA | Body | chr13:100506703-100506823 | 120 | 1 | | | | | | | | | | | | | | | | | | |
| CLYBL | NA | Body | chr13:100517009-100517033 | 24 | | | 2 | | | | | | | | | | | | | | | yes | |
| CLYBL | Island | 3'UTR | chr13:100547192-100548780 | 1588 | 6 | | 7 | | | | | | 2 | | | | | | yes | | | | yes |
| ZIC5 | Island | Body | chr13:100622253-100622582 | 329 | | | | | | | | | 1 | | | | | | | | | yes | |
| ZIC5 | Island;S_Shore | 1stExon;5'UTR;TSS200;TSS1500 | chr13:100623942-100624965 | 1023 | | 1 | 4 | | | | | | 1 | | | | | | | | | | yes |
| ZIC2 | N_Shore;Island | Body | chr13:100636252-100637560 | 1308 | | | 9 | | | | | | 1 | | | | | | | | | yes | |
| PCCA | N_Shore | TSS1500 | chr13:100740910-100741026 | 116 | | | | | | | | | 1 | | | | | yes | | | | | |
| PCCA | NA | Body | chr13:101173617-101174420 | 803 | 1 | | | | 1 | | | 0 | 12 | | | 5 | | | | | | | |
| PCCA | NA | Body | chr13:101175894-101175948 | 54 | | | | 1 | | | | | | | | | | | | | | | |
| A2LD1 | S_Shore | 5'UTR;1stExon;TSS200 | chr13:101185965-101187390 | 1425 | | | | | | | | | | | | | | | | | | | |
| NALCN | NA | Body | chr13:101910744-101910857 | 113 | | 1 | | | | | | | | | | | | | | | | | |
| NALCN | Island | 5'UTR | chr13:102068432-102068650 | 218 | | | | | | | | | 1 | | | | | | | | | | yes |
| ITGBL1 | N_Shore | TSS1500;TSS200;1stExon;5'UTR | chr13:102104573-102105440 | 867 | | | | | | | | | 3 | | | 1 | | | yes | | | | |
| FGF14 | NA | Body | chr13:103019936-103020405 | 469 | | | 11 | | | | | | | | | | | yes | | yes | | | |
| C13orf39 | NA | TSS200;1stExon | chr13:103346811-103346962 | 151 | | 4 | | | | | | | | | | | | | | | | yes | |
| LOC121952 | NA | TSS1500 | chr13:103532159-103532304 | 145 | | | | | | | | | 1 | | | | | yes | | | | | |
| DAOA | NA | TSS1500 | chr13:106116963-106117425 | 462 | | | 2 | | | | | | | | | | | | yes | | yes | | |
| EFNB2 | N_Shore | 3'UTR | chr13:107143980-107144036 | 56 | | | | | | | | | 1 | | 1 | | | | | | | | |
| EFNB2 | S_Shore | Body | chr13:107145933-107146164 | 231 | | | | | | | | | 1 | 1 | 1 | | | | | | | yes | |
| EFNB2 | Island | Body | chr13:107186870-107187232 | 362 | | | | | | | | | 1 | | | | | | | | | | yes |
| LIG4 | N_Shore;Island | 5'UTR;1stExon;TSS200 | chr13:108866993-108868194 | 1201 | | | | | | 1 | 2 | 0 | 4 | 1 | | 5 | | yes | yes | | | yes | |
| TNFSF13B | NA | TSS1500 | chr13:108920801-108921052 | 251 | | | | | | | | | 1 | | | | | yes | yes | | | | |
| TNFSF13B | NA | 1stExon | chr13:108922361-108922745 | 384 | | | | | | | | | | | | | | | | | | | |
| MYO16 | NA | TSS1500;TSS200 | chr13:109247864-109248503 | 639 | | | 4 | | | | | | 1 | 1 | | | | | | | | yes | |
| MYO16 | NA | Body | chr13:109538794-109539022 | 228 | | | | | | | | | 1 | | | | | yes | | yes | yes | | |
| IRS2 | N_Shore | Body | chr13:110433636-110434016 | 380 | | | | 1 | | | | | 1 | | | | | | | | yes | | |
| IRS2 | Island | 1stExon;5'UTR;TSS200 | chr13:110438256-110439004 | 748 | | | | | | | | 0 | 4 | 1 | | 4 | | | | | | yes | yes |
| COL4A1 | NA | Body | chr13:110838110-110838288 | 178 | | | 2 | | | | | | | | | | | yes | yes | | | | |
| COL4A1 | NA | Body | chr13:110874267-110874326 | 59 | | | 5 | | | | | | 1 | | | | | | | | | | |
| COL4A1 | NA | nan | chr13:110885345-110885926 | 581 | | | | | | | | | | | 1 | | | yes | | | | | |
| COL4A1 | Island | Body | chr13:110918331-110918683 | 352 | | 1 | 22 | | | | | | 1 | | | | | | yes | | | | |
| COL4A1 | NA | Body | chr13:110951025-110951418 | 393 | | 1 | 1 | | | | | | 1 | | | | | yes | | | yes | | |
| COL4A2;COL4A1 | Island;S_Shore | TSS1500;1stExon;5'UTR;TSS200;B | chr13:110959356-110961606 | 2250 | | | 14 | | | | | 0 | 4 | 2 | | | | yes | | | | | yes |
| COL4A2 | NA | Body | chr13:110992996-110993558 | 562 | | | 33 | 1 | | | | | 1 | 1 | | | | yes | | | | | |
| COL4A2 | S_Shore | Body | chr13:111039873-111040579 | 706 | | 1 | | | | | | 0 | 1 | | | 2 | 1 | yes | | | | | |
| COL4A2 | NA | Body | chr13:111061780-111062287 | 507 | | | 7 | | | | | 0 | 1 | | | 4 | | | | | | yes | |
| COL4A2 | NA | Body | chr13:111128054-111128102 | 48 | | | | | | | | | 1 | | | | | | | | | | |
| COL4A2 | NA | Body | chr13:111141830-111141841 | 11 | | | | | | | | | 1 | | | | | yes | | | | | |
| COL4A2 | Island | 3'UTR | chr13:111164546-111164966 | 420 | | | | | | | | | 1 | | | | | | | | | | |
| CARKD | N_Shore | Body | chr13:111277190-111278058 | 868 | | 1 | | | | | | | | | | | | yes | | | | | |
| CARKD | S_Shelf | Body | chr13:111280738-111281129 | 391 | | | | | | | | | | | | | | yes | | | | | |
| CARKD | N_Shelf | Body | chr13:111288257-111288583 | 326 | | | | | | | | | | | | | | | | | yes | yes | |
| CARS2 | N_Shelf | Body | chr13:111297400-111297551 | 151 | | | 5 | | | | 1 | 0 | | | | | | | | | | | |
| CARS2 | Island | Body | chr13:111301317-111301487 | 170 | | | 3 | | | | | | | | | | | | | | | | |
| CARS2 | NA | Body | chr13:111305805-111306312 | 507 | | | | | | | | | 2 | | | | | | | | yes | yes | |
| CARS2 | NA | Body | chr13:111317935-111318640 | 705 | | | | | 1 | | | 0 | 4 | 1 | | | | yes | | | | | |
| C13orf29 | NA | Body;TSS200;TSS1500 | chr13:111521981-111522985 | 1004 | | 1 | 24 | 1 | 1 | | | 0 | 4 | 1 | 1 | | | yes | yes | yes | yes | | |
| ANKRD10 | Island | TSS200 | chr13:111567603-111567833 | 230 | | | | | | | | | 1 | | | | | | | | | | yes |
| ARHGEF7 | NA | Body | chr13:111773936-111774127 | 191 | | 1 | | | | | | | | | | | | yes | | | | | |
| ARHGEF7 | NA | TSS200;Body;5'UTR | chr13:111839104-111839335 | 231 | | | | | 1 | | | | | | | | | | | yes | | | |
| ARHGEF7 | S_Shore | Body | chr13:111936044-111936071 | 27 | 1 | | | | | | | | | | | | | | | | | | |
| C13orf16 | NA | TSS200;1stExon;5'UTR | chr13:111972854-111973143 | 289 | | 1 | 5 | | | | | | 1 | | | | | | yes | | | | |
| C13orf16 | N_Shore | Body | chr13:111977248-111978024 | 776 | | | 3 | | | | | | | | | | | yes | | | | | |
| SOX1 | Island | TSS1500 | chr13:112721325-112721725 | 400 | | | | | | | | | 1 | | | | | | | | | yes | yes |
| SOX1 | N_Shore | 1stExon;3'UTR | chr13:112725605-112726124 | 519 | | | 8 | | | | | | | | | | | | | | | yes | |
| TUBGCP3 | S_Shore | TSS1500 | chr13:113242878-113243141 | 263 | | 1 | 3 | | 1 | | | 0 | 1 | | | 1 | | | | yes | | | yes |
| ATP11A | NA | Body | chr13:113356724-113357112 | 388 | | | 4 | | | | | | | | | | | yes | | yes | | | yes |

| | | | | | | | | | | | | | | | | | | | | | | | |
|---|---|---|---|---|---|---|---|---|---|---|---|---|---|---|---|---|---|---|---|---|---|---|---|
| ATP11A | S_Shore | Body | chr13:113411497-113411977 | 480 | | | | | | | | | 2 | | | | | yes | | | yes | | |
| ATP11A | S_Shore;N_Shelf;N_Shore | Body | chr13:113422162-113423268 | 1106 | | | | | | | | 0 | 6 | | | | | | | | | | |
| ATP11A | Island;S_Shore | Body | chr13:113424938-113426343 | 1405 | | 2 | | 1 | 2 | 2 | | | 2 | | | | | | yes | yes | | | |
| ATP11A | N_Shore | Body | chr13:113427738-113428172 | 434 | | | | | | | | | 1 | 1 | | | | | | yes | | yes | |
| ATP11A | S_Shore | Body | chr13:113430153-113430397 | 244 | | | 1 | | | | | | | | | | | | yes | yes | | | |
| ATP11A | NA;N_Shelf | Body | chr13:113435942-113437214 | 1272 | | 1 | 32 | | | | | | | | | | | yes | | yes | | | |
| ATP11A | N_Shelf | Body | chr13:113461305-113461652 | 347 | | 1 | | | | | | | | | | | | | yes | | yes | yes | |
| ATP11A | N_Shore | Body | chr13:113491766-113492103 | 337 | | | 2 | | | | | | | | | | | | | | | | |
| ATP11A | N_Shore | Body | chr13:113496400-113496719 | 319 | | | | 1 | | | | | 1 | | | | | | | | | | |
| ATP11A | S_Shore | Body | chr13:113498128-113498450 | 322 | | | | | 1 | | | | | | | | | | yes | | | | |
| ATP11A | NA | Body | chr13:113516304-113516424 | 120 | 1 | | | | | | | | | | | | | | yes | | | | |
| ATP11A | NA | Body | chr13:113529568-113529737 | 169 | | | | | | | | | 1 | 1 | | | | | yes | | | | |
| ATP11A | N_Shore;Island | 3'UTR | chr13:113540189-113540631 | 442 | | | 26 | | | | | 0 | 6 | | | 8 | 2 | | | | | yes | |
| MCF2L | N_Shore;Island | TSS1500;TSS200 | chr13:113622102-113622750 | 648 | | | 11 | | | | | | | | | 1 | | | yes | | | yes | |
| MCF2L | S_Shelf;NA | Body | chr13:113627484-113628068 | 584 | | 1 | 2 | | | 1 | | | | | | | | | | | | yes | |
| MCF2L | N_Shore | Body;TSS1500;TSS200 | chr13:113633379-113633654 | 275 | 2 | | | 1 | | | | | 1 | | | | | yes | | | | | |
| MCF2L | S_Shore;Island | Body | chr13:113635690-113637903 | 2213 | 1 | | 2 | 1 | | | | | | | 1 | | | | | | | yes | |
| MCF2L | NA | Body | chr13:113642457-113642703 | 246 | 1 | | | 1 | | | | | | | | | | | | | | | |
| MCF2L | N_Shore | Body | chr13:113646732-113647549 | 817 | 3 | | 2 | | | | | | 1 | | | | | | yes | | yes | | yes |
| MCF2L | Island | Body | chr13:113648607-113650155 | 1548 | | 2 | | | | | | | | | | | | yes | | | | | |
| MCF2L | N_Shelf | Body | chr13:113652980-113653020 | 40 | | | | | | | | | | | | | | | | | | | |
| MCF2L | N_Shore;Island | Body | chr13:113655468-113656424 | 956 | | | 10 | 1 | | | | 0 | | | | | | | | | | | |
| MCF2L | N_Shelf | Body | chr13:113659340-113659625 | 285 | | | | | | | | | | | | | | yes | yes | | | | |
| MCF2L | N_Shelf | Body | chr13:113674587-113674654 | 67 | | | | | | | | | | | | | | | | yes | | yes | |
| MCF2L | Island;N_Shore;chr13:1136767 | Body;Island | chr13:113676726-113677777 | 1051 | | | 5 | 1 | | | | | 3 | 1 | 1 | | | | | yes | | | |
| MCF2L | Island | Body | chr13:113684606-113684951 | 345 | | | 3 | | | | | | | | | | | | yes | yes | | | yes |
| MCF2L | S_Shore | Body | chr13:113689422-113689776 | 354 | | | | | | | | 0 | 1 | 1 | | 2 | | yes | | | | | yes |
| MCF2L | N_Shelf | Body | chr13:113698408-113699526 | 1118 | | 1 | 38 | | | | | | | | | | | yes | | | | | |
| MCF2L | Island;N_Shore | Body | chr13:113700818-113702478 | 1660 | 19 | | 7 | | | | | | | | | | | yes | | yes | | | |
| MCF2L | S_Shore;S_Shelf | Body | chr13:113707471-113708529 | 1058 | 9 | | | | | | | | | | | | | | yes | | | yes | |
| MCF2L | N_Shore | Body | chr13:113713970-113714917 | 947 | | | 8 | | | | | | | | | | | yes | | yes | | yes | |
| MCF2L | N_Shelf | Body | chr13:113718119-113719440 | 1321 | 16 | | | | | | | | | | | | | | | | | | |
| MCF2L | S_Shelf | Body | chr13:113724295-113724878 | 583 | 1 | | 1 | | | | | | | | | | | yes | yes | | | | |
| MCF2L | N_Shore | Body | chr13:113743564-113744010 | 446 | | | 2 | | | | | | | | | | | | yes | | | | yes |
| MCF2L | N_Shelf | Body | chr13:113748321-113748721 | 400 | | | | | | | | | | | | | | | | | | | |
| F7 | N_Shore | TSS1500;TSS200 | chr13:113759771-113760286 | 515 | | | 4 | 1 | | | | | 2 | | | | | yes | | | | yes | yes |
| F10 | S_Shelf | TSS1500;TSS200 | chr13:113776873-113777160 | 287 | 2 | | | | | | | 0 | 1 | | | | | yes | | | | yes | |
| F10 | Island | Body | chr13:113783975-113784663 | 688 | | | 16 | | | | | 0 | 1 | | | | | | | yes | | | |
| PROZ | NA | TSS1500 | chr13:113812524-113812911 | 387 | | | | | | | | | | | | | | | | yes | yes | | yes |
| PROZ | NA | Body | chr13:113819073-113819409 | 336 | | | | | | | | | 1 | | | | | yes | yes | | | | |
| PROZ | NA | Body | chr13:113825781-113826053 | 272 | | 1 | | | | | | | | | | | | yes | | | | | |
| PCID2 | NA | 3'UTR | chr13:113832355-113832595 | 240 | | | 4 | | | | | | | | | | | | | | yes | | |
| LAMP1 | Island | Body | chr13:113975974-113976115 | 141 | | | | 1 | | | | | | | | | | yes | | | | | yes |
| GRTP1 | S_Shelf | Body | chr13:113980035-113980343 | 308 | | | | | | | | | | | | | | yes | | | | | |
| ADPRHL1 | NA | 3'UTR | chr13:114077033-114077172 | 139 | | | 7 | | | | | | | | | | | yes | | | | | |
| ADPRHL1 | NA | 1stExon;5'UTR;Body;TSS200;TSS1! | chr13:114103452-114104057 | 605 | | 2 | 13 | | | | | | 4 | 1 | | 4 | | yes | | | | | |
| ADPRHL1 | NA | TSS200;TSS1500;1stExon;5'UTR | chr13:114107554-114108713 | 1159 | 2 | | 56 | | | | | | 1 | | | | | yes | yes | | | | |
| DCUN1D2 | NA | 3'UTR | chr13:114111864-114112218 | 354 | | | | | | | 1 | | 2 | 2 | | | | yes | | | | | yes |
| TMCO3;DCUN1D2 | Island | 5'UTR;TSS1500 | chr13:114145973-114146138 | 165 | | | | | | | | | 1 | | | | | yes | yes | | | | yes |
| TMCO3 | N_Shore | Body | chr13:114164151-114164654 | 503 | | | | | | | | 0 | 1 | 1 | | | | yes | | | | | |
| TMCO3 | Island | Body | chr13:114187104-114187548 | 444 | | | | | | | | | | | | | | | | | | | |
| TMCO3 | N_Shelf | Body | chr13:114193150-114193689 | 539 | 1 | | | | | | | | | | | | | | yes | | | | |
| TMCO3 | N_Shore;Island;S_Shore | Body | chr13:114202067-114204079 | 2012 | 1 | | 3 | | | | | | | | | | | | | | | | yes |
| TFDP1 | S_Shore | Body | chr13:114261869-114261986 | 117 | | | | | | | | | 1 | | | | | | | | | | |
| TFDP1 | N_Shelf | Body | chr13:114284038-114284318 | 280 | | 1 | | | | | | | | | | | | | | | yes | yes | |
| TFDP1 | S_Shore | Body | chr13:114294418-114294515 | 97 | | | | | | | | | | | | | | | | | | yes | |
| ATP4B | NA | Body | chr13:114308619-114308749 | 130 | | | 17 | | | | | | | | | | | | | | | | |
| ATP4B | NA | Body;TSS200 | chr13:114312105-114312686 | 581 | | | 2 | 1 | | | | | | | | | | | | | yes | | |
| GRK1 | NA | TSS1500;TSS200 | chr13:114321214-114321749 | 535 | | | 13 | | | | | | | | | | | | | | | | |
| GRK1 | NA | Body | chr13:114325250-114325720 | 470 | | | | | | | | | | | | | | yes | | | | | |
| RASA3 | S_Shelf | Body | chr13:114750991-114751280 | 289 | | 1 | | | | | | | | | | | | | | | | | |
| RASA3 | N_Shelf | Body | chr13:114760667-114761210 | 543 | | | | | | | | | | | | | | yes | | | | | |
| RASA3 | Island | Body | chr13:114764394-114764621 | 227 | | | | | | | | | | | | | | | yes | | | | |
| RASA3 | N_Shelf | Body | chr13:114770495-114770710 | 215 | | | | | 1 | | | | | | | | | | yes | | | | |
| RASA3 | Island | Body | chr13:114780772-114780992 | 220 | | | | | | | | | 1 | | | | | | | | | | |
| RASA3 | S_Shore | Body | chr13:114786825-114786904 | 79 | | 1 | | | | | | | 1 | | | | | | | | yes | | |
| RASA3 | NA | Body | chr13:114793125-114793396 | 271 | | | | | | | | | 1 | | | | | | | | | | yes |
| RASA3 | N_Shelf | Body | chr13:114800309-114801587 | 1278 | | | 6 | | | | | | 1 | | | | | yes | yes | | | | |
| RASA3 | Island | Body | chr13:114803626-114803843 | 217 | | | | | | | | | | | | | | | yes | | | | |
| RASA3 | N_Shelf | Body | chr13:114810631-114810794 | 163 | | 1 | | | | | | | | | | | | | yes | | | | |
| RASA3 | N_Shore | Body | chr13:114812780-114813015 | 235 | | 1 | | | | | | | 1 | | 1 | | | yes | yes | | | | |
| RASA3 | Island;S_Shore | Body | chr13:114814024-114815139 | 1115 | | | | | 4 | 1 | | 0 | 4 | | | 2 | | yes | | | | | |
| RASA3 | NA | Body | chr13:114828067-114828455 | 388 | | | 4 | | | | | | | | | | | | yes | | | yes | |
| RASA3 | NA | Body | chr13:114831705-114833435 | 1730 | | | | | | | | | | | | | | yes | | | | | |
| RASA3 | N_Shore | Body | chr13:114837593-114837807 | 214 | | | 2 | | | | | | | | | | | | | | | | |
| RASA3 | S_Shelf | Body | chr13:114843594-114845297 | 1703 | | | | | | | | | 1 | | | | | | | | | | |
| RASA3 | S_Shelf | Body | chr13:114855198-114855572 | 374 | | | | | 1 | | | | 1 | 1 | | | | | yes | | | | yes |
| RASA3 | S_Shore | Body | chr13:114871422-114871575 | 153 | | | 3 | | | | | | | | | | | | | | | | |
| RASA3 | N_Shelf | Body | chr13:114872599-114872787 | 188 | | | | | | | | | | | | | | | | | | | |
| RASA3 | N_Shore;Island | Body | chr13:114875001-114876521 | 1520 | | | 13 | | | | | | 1 | | | | | yes | | | yes | | |
| RASA3 | S_Shore | Body | chr13:114880761-114881556 | 795 | | | 29 | | | | | | 1 | | | | | yes | | | | | |
| RASA3 | NA | Body | chr13:114886700-114887133 | 433 | | | 4 | | | | | | 1 | | | | | | | | | | |
| RASA3 | NA | Body | chr13:114890515-114890805 | 290 | | | | | | | | | 1 | | | | | | | yes | | | |
| RASA3 | NA | Body | chr13:114892556-114892618 | 62 | | | | | | | | | 1 | | | | | yes | | | | | |
| CDC16 | Island | TSS1500;TSS200 | chr13:115000149-115000234 | 85 | | | | | | | | | 2 | | | 1 | | | yes | | | | |
| OR11H4 | NA | TSS200 | chr14:20710881-20711344 | 463 | | | | | | | | | | | | | | | | yes | | | |
| KLHL33 | Island;S_Shore | 5'UTR;TSS200 | chr14:20903410-20904169 | 759 | | | | | | | | | 1 | | | | | yes | | | | yes | |
| PNP | NA | TSS1500 | chr14:20937221-20937572 | 351 | | | | | | | | | | 1 | | | | | | | | yes | yes |
| ANG;RNASE4 | N_Shore | TSS1500;TSS200;5'UTR;1stExon | chr14:21151782-21152735 | 953 | | 2 | | | | | | | | | | | | | | | | | |
| FAM12B | NA | 5'UTR | chr14:21238111-21238436 | 325 | | | | | | | | | | | | | | | yes | | | | |
| RNASE1 | NA | 5'UTR;1stExon;TSS200 | chr14:21270407-21271355 | 948 | | | | | 2 | | | | | | | | | | | | | | |
| SLC39A2 | NA | TSS200 | chr14:21467273-21467375 | 102 | | | | | 1 | | | | | | | | | | | | | | |
| NDRG2 | Island | TSS1500;TSS200 | chr14:21493964-21494161 | 197 | | | | | | | | | 1 | | | | | | | | | yes | |
| RNASE7 | NA | TSS1500 | chr14:21510056-21510479 | 423 | | | | | | | | | 1 | | | | | | | | | | |
| ZNF219;C14orf176 | S_Shore | 5'UTR;1stExon;TSS1500 | chr14:21567164-21567520 | 356 | | | | | | | | 0 | | | | | | yes | yes | | | yes | yes |
| OR10G2 | NA | TSS200;TSS1500 | chr14:22103031-22103726 | 695 | | | 8 | | | | | | | | | | | | yes | | | | |
| MRPL52 | N_Shore | TSS200 | chr14:23298926-23299469 | 543 | | | | | | | | 0 | | | | | | yes | | | | | |
| MMP14 | N_Shore | TSS1500 | chr14:23305548-23305957 | 409 | | | | | 1 | 1 | | 0 | 3 | 1 | | | | | | | | | |
| MMP14 | Island | Body | chr14:23315028-23315249 | 221 | | | | | | | | | | | | | | yes | | | | | |

| | | | | | | | | | | | | | | | | | | | | | | | |
|---|---|---|---|---|---|---|---|---|---|---|---|---|---|---|---|---|---|---|---|---|---|---|---|
| REM2 | NA;N_Shelf | TSS1500 | chr14:23351487-23352488 | 1001 | | | 2 | | | | | | 1 | | | | | yes | | | yes | | |
| JUB | N_Shelf | Body;TSS1500 | chr14:23447078-23447792 | 714 | | | | | | | | | | | | | | | yes | | | | |
| JUB | Island | Body | chr14:23450238-23450527 | 289 | | | | | | | | | | | | | | | | | | | yes |
| JUB | Island | TSS200 | chr14:23451983-23452408 | 425 | | | | | | | | | 1 | | | | | | | | | | yes |
| C14orf93 | S_Shore | TSS1500 | chr14:23479657-23479736 | 79 | | | | | | | | | | | | | | yes | | | | | yes |
| PSMB11 | NA | TSS1500 | chr14:23510934-23511306 | 372 | | | 9 | | | | | | | | | | | yes | yes | | | | |
| PPP1R3E | S_Shore | TSS1500 | chr14:23772548-23772695 | 147 | | | | | | | | | 1 | | | | | yes | yes | | | yes | |
| SLC22A17 | Island | Body | chr14:23815887-23816088 | 201 | | | | | | | | | | | | | | yes | | | | | |
| EFS | Island | TSS200;TSS1500 | chr14:23834909-23836012 | 1103 | | | | | | | | 0 | 2 | | | | | | | | yes | | yes |
| CMTM5 | NA | TSS200;1stExon;5'UTR | chr14:23845819-23846906 | 1087 | | | 5 | | | | | | | | | | | | | | yes | | |
| MYH6 | NA | TSS1500 | chr14:23878195-23878796 | 601 | | | | | | | | | 1 | | | | | | | | | | |
| MYH7;MIR208B | N_Shore | Body;TSS200;TSS1500 | chr14:23887330-23888492 | 1162 | | | | | | | | | | | | | | yes | | | | | |
| MYH7 | NA | 5'UTR | chr14:23904678-23905273 | 595 | | | 5 | | | | | | 1 | | 1 | | | yes | | | | yes | |
| NGDN | NA | TSS1500 | chr14:23938105-23938207 | 102 | | | | | | | | 0 | | | | | | yes | | | yes | | |
| JPH4 | Island | Body | chr14:24046374-24047021 | 647 | | | 1 | | | | | 0 | 2 | 1 | | 1 | | | | | | | yes |
| DHRS4;C14orf167 | Island;N_Shore | TSS200;Body;1stExon;TSS1500 | chr14:24422368-24423199 | 831 | 2 | | 2 | | | | | 0 | 4 | 3 | | 5 | | yes | | | yes | | |
| DHRS4L2 | N_Shore;Island;S_Shore | TSS1500;TSS200;1stExon | chr14:24457509-24458345 | 836 | 3 | | 28 | 1 | | | | 0 | 3 | | 1 | 11 | 1 | yes | | | yes | | |
| DHRS4L1 | N_Shore | TSS200 | chr14:24505537-24505671 | 134 | | | 4 | | | | | 0 | 1 | | | 4 | | | | | | | yes |
| CPNE6 | NA | TSS1500 | chr14:24540174-24540773 | 599 | | | 3 | | | | | | | | | | | | yes | | | yes | |
| CPNE6 | N_Shelf | Body | chr14:24546420-24546847 | 427 | | | | | | | | | | | | | | | | | | | |
| FITM1 | N_Shore | TSS200 | chr14:24600569-24601001 | 432 | | | | | | | | | | | | | | | | | | | yes |
| REC8 | N_Shore;Island | TSS1500;TSS200;1stExon;5'UTR | chr14:24641021-24642317 | 1296 | | 1 | 29 | | 2 | | | 0 | 3 | 1 | | | | yes | | | | | |
| TGM1 | NA | Body | chr14:24731242-24731542 | 300 | | | | | | | | | | | | | | | | | | | |
| TGM1 | NA | TSS1500 | chr14:24733439-24733757 | 318 | | | | | | | | | | | | | | yes | | | | | |
| LTB4R2;LTB4R;CIDEB | N_Shore;Island | 1stExon;5'UTR;TSS1500;Body;TSS2 | chr14:24779793-24780926 | 1133 | 1 | | 47 | | | | | 0 | 12 | 5 | 1 | 21 | | | | | | | yes |
| LTB4R | Island | Body | chr14:24785310-24785933 | 623 | | | 3 | | | | | | 2 | | | | | | | | | | yes |
| ADCY4 | N_Shore;Island | Body | chr14:24801073-24802150 | 1077 | 1 | | 8 | | | | | | 1 | | | | | yes | | | | | |
| RIPK3 | S_Shore;Island | 5'UTR;1stExon;Body | chr14:24808650-24809442 | 792 | | | 6 | 1 | | | | | | | | | | yes | | | | | |
| NFATC4 | N_Shore | TSS1500 | chr14:24835104-24835609 | 505 | | 1 | | | | | | | | | | | | | | | | yes | |
| NFATC4 | Island | Body | chr14:24838828-24839259 | 431 | | | | | | | | | | | | | | | | | | yes | |
| NYNRIN | N_Shore | TSS1500 | chr14:24867164-24867954 | 790 | 4 | | 3 | 1 | | | | | | 1 | | | | | | | yes | | yes |
| NOVA1 | S_Shore | TSS1500 | chr14:27067707-27068354 | 647 | | | | | | | | | 1 | | | | | yes | yes | | | yes | |
| FOXG1 | Island;N_Shore | TSS1500 | chr14:29234890-29236221 | 1331 | | | 4 | | | | | 0 | 5 | | | 1 | | | | | | yes | yes |
| C14orf23 | N_Shore | Body | chr14:29243173-29243504 | 331 | | 1 | | | 1 | | | | 1 | | | | | | | | | | |
| C14orf23 | Island | Body | chr14:29254530-29254961 | 431 | | | | | | | | | 1 | | | 1 | | | yes | | | | |
| PRKD1 | S_Shore | TSS1500 | chr14:30397464-30397763 | 299 | | | | | | | | 0 | | | | | | | | | yes | | yes |
| SCFD1 | NA | TSS1500 | chr14:31090535-31090974 | 439 | | | 1 | | | | | | | | | | | yes | | | | | |
| COCH | N_Shore | TSS1500 | chr14:31343283-31343427 | 144 | | | | | | | | | | | | | | | yes | | | | yes |
| NUBPL | NA | TSS200 | chr14:32030546-32030916 | 370 | | | 1 | | | | | | | | | | | yes | yes | | | | yes |
| ARHGAP5;C14orf128 | Island | TSS200;TSS1500 | chr14:32546448-32546466 | 18 | | | | | | | | | 1 | | | | | | | | | yes | yes |
| NPAS3 | S_Shelf | TSS1500 | chr14:33407334-33407424 | 90 | | | | | | | | | 1 | | | | | | | | | yes | |
| NPAS3 | Island | Body | chr14:34269416-34270437 | 1021 | | | 9 | | | | | 0 | 3 | | | 4 | | | | | | yes | yes |
| EGLN3 | Island | Body;1stExon;5'UTR | chr14:34419546-34420292 | 746 | | | | | | | | | 2 | | | | | | yes | | | | |
| NKX2-1 | Island | Body;1stExon | chr14:36987301-36988446 | 1145 | | | | | | | | | 2 | | | | | | | | | yes | |
| NKX2-1 | Island | TSS1500 | chr14:36990144-36990534 | 390 | | | | | | | | | 1 | | | | | | | | | | |
| NKX2-8 | Island;N_Shore | TSS1500;Body | chr14:37051417-37053169 | 1752 | | | | 1 | | | | | 2 | | | | | | | | | yes | |
| PAX9 | Island;N_Shore | TSS1500 | chr14:37125292-37126582 | 1290 | | | 27 | | | | | | | | | | | yes | | | | | |
| PAX9 | S_Shore;Island | 5'UTR | chr14:37127671-37128790 | 1119 | | | | | | | | | | | | | | | | | | | |
| PAX9 | N_Shore | 5'UTR | chr14:37130212-37130762 | 550 | | | | | | | | | 1 | | | | | | | | | | |
| PAX9 | Island | Body | chr14:37132490-37132764 | 274 | | | | | | | | | 1 | | | | | | | | | | |
| PAX9 | N_Shore;Island | Body | chr14:37135435-37136391 | 956 | | | 19 | | | | | | | | | 1 | | | | | | | |
| SLC25A21;LOC100129794 | N_Shore;Island | Body;TSS200 | chr14:37641134-37641964 | 830 | | | | | | | | 0 | 2 | | | | | | | | yes | | yes |
| MIPOL1 | Island | TSS1500 | chr14:37666893-37666931 | 38 | | | | | | | | | 1 | | | | | | | | | yes | |
| FOXA1 | Island | 5'UTR;1stExon;TSS200 | chr14:38064112-38064513 | 401 | | | | | | | | 0 | 2 | | | | | | | | | yes | yes |
| SSTR1 | Island | 5'UTR;Body | chr14:38678537-38679656 | 1119 | | | | 1 | | | | 0 | 2 | | | | | | | | | | |
| CLEC14A | Island;S_Shore | 1stExon;5'UTR;TSS200 | chr14:38724648-38726189 | 1541 | | 1 | 19 | | 1 | | | | 3 | 2 | | | | | | | | | yes |
| MIA2 | NA | TSS1500 | chr14:39702922-39703302 | 380 | | | | 1 | | | | | | | | | | yes | | yes | | | |
| CTAGE5 | N_Shore | TSS1500;Body | chr14:39734964-39735625 | 661 | | | | | 1 | | | 0 | 2 | | | 1 | | | | | | | |
| LRFN5 | S_Shore | TSS200 | chr14:42076710-42076759 | 49 | | | | | | | | | | | | 1 | | | | | | yes | |
| C14orf28 | S_Shelf | 5'UTR | chr14:45369477-45369753 | 276 | | | | | | | | | 1 | | | | | yes | | | yes | | |
| KLHL28;FAM179B | Island;S_Shore | TSS1500;1stExon | chr14:45431685-45432516 | 831 | | | 5 | | | | | 0 | 2 | 1 | | 1 | | | | | | | |
| C14orf106 | Island | TSS200 | chr14:45722704-45722987 | 283 | | | | | | | | | 1 | | | 1 | | yes | | | | | |
| RPL10L | N_Shore;Island | 1stExon;5'UTR;TSS200 | chr14:47120507-47121158 | 651 | | | 5 | | | | | | | | | | | yes | | | | | yes |
| C14orf182 | S_Shelf | 1stExon;5'UTR | chr14:50473602-50474530 | 928 | | | | | | | | | 2 | 1 | | | | yes | | yes | | | |
| NIN | N_Shore | 5'UTR | chr14:51296586-51296728 | 142 | | | 1 | | | | | | | | | | | | | | | | |
| GNG2 | NA | TSS1500 | chr14:52325747-52326169 | 422 | | | 2 | | | | | | | | | | | yes | | | | | |
| NID2 | Island | TSS200;TSS1500 | chr14:52536066-52536893 | 827 | | | | | | | | | 1 | | | | | yes | | yes | | | |
| PTGDR | Island;S_Shore | TSS200;1stExon;5'UTR;Body | chr14:52734251-52736038 | 1787 | | 1 | 4 | | | | | 0 | 1 | | | 2 | | yes | | | | | |
| GNPNAT1 | S_Shore | TSS200 | chr14:53258566-53258601 | 35 | | | | | | | | 0 | 1 | | | 1 | | | | | yes | | yes |
| BMP4 | Island | Body | chr14:54418728-54418851 | 123 | | | 33 | | | | | 0 | | | | | | | | | | | |
| BMP4 | Island | TSS200;5'UTR | chr14:54421393-54421413 | 20 | | 1 | | | | | | | | | | | | | | | | | |
| BMP4 | S_Shore | TSS1500 | chr14:54423807-54424149 | 342 | | | 20 | | 1 | | | | | | | | | | | | | yes | yes |
| LGALS3 | N_Shore | TSS1500 | chr14:55595666-55595768 | 102 | | | | 1 | | | | | | | | | | | | yes | | | yes |
| TBPL2 | N_Shore;Island | 1stExon;TSS200;Body | chr14:55907007-55907501 | 494 | | | 24 | | | | | 0 | 8 | 1 | | 5 | | yes | | | | yes | |
| KTN1;C14orf33 | N_Shore | TSS1500;Body | chr14:56045827-56046098 | 271 | | | 1 | | | | | 0 | 3 | | | 4 | | yes | | | | | |
| OTX2 | N_Shelf | 5'UTR;1stExon | chr14:57272244-57272858 | 614 | | 1 | | | | | | | | | | | | | | | | yes | |
| OTX2;OTX2OS1 | Island;N_Shore | TSS1500;5'UTR;1stExon | chr14:57274763-57279496 | 4733 | | | | | | | | | | | | | | | | | | | |
| OTX2OS1 | Island | Body | chr14:57284096-57284528 | 432 | 1 | | | | | | | | | | | | | | | | | yes | |
| C14orf105 | NA | 1stExon;5'UTR;TSS200 | chr14:57960464-57960985 | 521 | | | | 1 | | | | | 2 | | | | | yes | | | | | |
| SLC35F4 | NA | TSS1500 | chr14:58064661-58064926 | 265 | | | | | | | | | 1 | | | | | | yes | | | | |
| TOMM20L | N_Shore | TSS1500 | chr14:58862417-58862503 | 86 | | | 6 | | | | | | | | | | | yes | | | | | yes |
| GPR135 | chr14:59930634-59932251;Isla | Island;TSS200;TSS1500 | chr14:59931541-59932321 | 780 | | | 5 | | | | | | | | 1 | | | | | | yes | yes | yes |
| C14orf38 | S_Shore | TSS200;TSS1500 | chr14:60043704-60043906 | 202 | | | 4 | | | | | | | | | | | yes | yes | | | | |
| RTN1 | chr14:60193831-60194302 | S_Shore | chr14:60194823-60195307 | 484 | | | | | | | | | | | 1 | | | | yes | | | yes | |
| C14orf39 | Island | 5'UTR | chr14:60952097-60952599 | 502 | | 1 | | | | | | | | | | | | | | | | | |
| SIX6 | Island | TSS200 | chr14:60975837-60975964 | 127 | | | | | | | | | 1 | | | | | | | | | | |
| SIX6 | Island | Body | chr14:60977856-60978275 | 419 | | 1 | | | | | | | | | | | | | | | | yes | |
| SIX1 | Island | TSS200 | chr14:61116216-61116382 | 166 | | | | | | | | | 1 | | | | | | | | | | |
| TMEM30B | S_Shore | 1stExon;5'UTR | chr14:61748243-61748440 | 197 | | | 8 | 1 | 1 | | | 0 | 1 | | | | | | | | | yes | yes |
| PRKCH | Island;S_Shore | 5'UTR;1stExon;Body | chr14:61788810-61790037 | 1227 | | | 14 | | | | | | 1 | | | | | yes | | | | | yes |
| FLJ43390 | Island | Body | chr14:62584278-62584565 | 287 | | | | | | | | | 1 | | | | | | | | | | |
| KCNH5 | NA | 5'UTR;TSS200 | chr14:63568195-63568809 | 614 | | | | | | | | | 4 | | | | | yes | yes | | | | |
| GPHB5 | NA | TSS1500 | chr14:63785287-63785523 | 236 | | | | | | | | | | | | | | | | | | | |
| SYNE2 | N_Shore | TSS1500 | chr14:64319089-64319306 | 217 | | | 6 | | | | | | | | | | | yes | | | | | yes |
| SYNE2 | NA | Body | chr14:64676792-64677007 | 215 | | | 12 | | | | | | 1 | | | | | | | | | | |
| AKAP5 | S_Shore | 5'UTR | chr14:64932678-64932782 | 104 | | | | | | | | 0 | 1 | | | | | | | | | | yes |
| HSPA2 | N_Shore | TSS1500 | chr14:65005951-65006222 | 271 | | | 14 | 1 | | | | 0 | 1 | | | | | yes | | | | | |
| HSPA2 | Island | 1stExon | chr14:65009101-65009505 | 404 | | | | | | | | | | | | 1 | | | | | | yes | |
| C14orf50 | N_Shore;Island | TSS1500;TSS200 | chr14:65015998-65016802 | 804 | | | 9 | | | | | | | | | | | | | | | | yes |

| | | | | | | | | | | | | | | | | | | | | | | | |
|---|---|---|---|---|---|---|---|---|---|---|---|---|---|---|---|---|---|---|---|---|---|---|---|
| GPX2 | NA | 1stExon | chr14:65409354-65409659 | 305 | | | | | | | | | 1 | | | | | yes | yes | | | | |
| FAM71D | NA | TSS200 | chr14:67656029-67656241 | 212 | | | | | | | | 0 | 1 | 1 | | | | yes | | | | | yes |
| PLEK2 | Island | TSS1500 | chr14:67879101-67879385 | 284 | | | 4 | | | | | 0 | | | | | | yes | | | yes | | |
| RAD51L1 | NA | Body | chr14:69052728-69052887 | 159 | | | | | | | | | | | | | | | | | | | |
| ZFP36L1 | Island | Body | chr14:69256677-69256977 | 300 | | | | | | | 1 | 0 | 4 | | | | | | | | | | |
| ACTN1 | N_Shore;Island | 3'UTR;Body | chr14:69341139-69341905 | 766 | | | 4 | | | | | | 3 | | | 2 | | | | | yes | | |
| FLJ44817 | Island | TSS200 | chr14:69951305-69951322 | 17 | | | | | | | | | 1 | | | | | | | | | | |
| C14orf162 | Island | TSS1500 | chr14:70038505-70039262 | 757 | | 1 | | | | | | 0 | 1 | | | | | | | | | | |
| SLC8A3 | N_Shore | 5'UTR | chr14:70653719-70653964 | 245 | | | | | 1 | | | | | | | | | | yes | | | yes | yes |
| COX16 | S_Shore | TSS1500 | chr14:70826997-70827336 | 339 | | | | | | | | 0 | 1 | | | | | yes | yes | | | | |
| MAP3K9 | S_Shore | TSS1500 | chr14:71276874-71277192 | 318 | | | 4 | | | | | | 1 | | | | | yes | | | | yes | |
| DPF3 | NA | 3'UTR | chr14:73136973-73137242 | 269 | | | 2 | | | | | | | | | | | | | | | yes | |
| DPF3 | S_Shore | TSS1500 | chr14:73361747-73362214 | 467 | | | 7 | | | | | | | | | | | | | | | | |
| PAPLN | Island | 5'UTR | chr14:73706449-73706676 | 227 | | | | 1 | | | | | | | | | | yes | | | yes | | |
| PAPLN | N_Shore;Island | Body | chr14:73712256-73712967 | 711 | | | 25 | 1 | | | | 0 | 2 | | | 2 | | yes | | | | yes | yes |
| ACOT6 | S_Shelf | TSS200 | chr14:74083455-74083501 | 46 | | | 1 | | | | | | | | | | | yes | yes | | | | |
| C14orf43 | N_Shore | 3'UTR | chr14:74184843-74185398 | 555 | | | | | | | | | | | | | | | | | | | |
| VSX2 | N_Shore | TSS1500 | chr14:74705942-74706031 | 89 | | | 2 | | | | | | | | | | | | yes | | | | |
| VSX2 | S_Shore | Body | chr14:74727001-74727464 | 463 | | | | | | | | | | | | | | | | | | | |
| C14orf115 | NA | TSS200 | chr14:74815085-74815187 | 102 | | | | | | | | | | | | | | | | | | yes | |
| C14orf115 | Island | Body | chr14:74824136-74824568 | 432 | | | | | | | | | 1 | | | | | | | | | | |
| TMEM90A | N_Shore;Island | 5'UTR | chr14:74892331-74893169 | 838 | | 1 | | | | | | 0 | | | | | | yes | | | | yes | yes |
| LTBP2 | Island | Body | chr14:75052674-75052717 | 43 | | | 1 | | | | | | | | | | | yes | yes | | | | |
| RPS6KL1 | N_Shore | TSS1500;TSS200 | chr14:75389222-75390007 | 785 | | | 4 | | | | | | | | | | | yes | yes | | | | |
| MLH3 | Island | 5'UTR | chr14:75517797-75518217 | 420 | | | | | | | | | 1 | | | | | yes | yes | | | | yes |
| FAM164C | Island | 1stExon;5'UTR | chr14:75536360-75536561 | 201 | | | | | | | | 0 | | | | | | yes | yes | | | | |
| NEK9 | Island | TSS200 | chr14:75593943-75594058 | 115 | | | | | | | | | 1 | | | | | yes | | | | | yes |
| JDP2 | S_Shore | 5'UTR;TSS1500 | chr14:75897342-75897841 | 499 | | | | | | | | | 1 | | | | | | | | | yes | |
| JDP2 | NA | 3'UTR | chr14:75936582-75936789 | 207 | | | 6 | 1 | | | | | 1 | | | | | yes | | | | | yes |
| BATF | NA | TSS1500;TSS200;5'UTR;1stExon | chr14:75988356-75989203 | 847 | | | 2 | | 2 | | | | 2 | 1 | 1 | | | yes | | | | yes | yes |
| BATF | Island | Body | chr14:76012927-76013186 | 259 | | | | | | | | | | | | | | yes | | | yes | yes | |
| TGFB3 | N_Shore | 1stExon;5'UTR | chr14:76447327-76447431 | 104 | | | 1 | | 1 | | | | 1 | | | | | | | | | | yes |
| ESRRB | NA | TSS1500 | chr14:76837428-76837569 | 141 | | | | | | | | | | | | | | | yes | | | | yes |
| ESRRB | N_Shore | 5'UTR | chr14:76905532-76906056 | 524 | | | | | | | | | | | | | | yes | | | | | |
| TMEM63C | N_Shore | TSS1500 | chr14:77647482-77647955 | 473 | | | | | | | | | | | | | | yes | | | | | |
| MIR1260;NGB | NA | TSS200;3'UTR | chr14:77732468-77732641 | 173 | | | 2 | | | | | | | | | | | yes | | | | | |
| NGB | Island | TSS200 | chr14:77737749-77738080 | 331 | | | 7 | | | | | | | | | | | yes | | | | | yes |
| POMT2 | NA | Body | chr14:77767644-77767735 | 91 | | | | | | | | | | | | | | yes | yes | | yes | | |
| ISM2 | S_Shore | TSS1500 | chr14:77965757-77965869 | 112 | | | | | | | | | | | | | | yes | | | | | yes |
| NRXN3 | NA | TSS200 | chr14:78869963-78870232 | 269 | | | | | | | | 0 | | | | | | | | | | | |
| NRXN3 | NA | 5'UTR | chr14:79111512-79111719 | 207 | | | 1 | | | | | | | | | | | | | | | | |
| NRXN3 | N_Shore;S_Shore | Body;TSS1500;1stExon;5'UTR | chr14:79744991-79746359 | 1368 | | | | | | | | | 2 | | | | | | | | | | |
| NRXN3 | Island | Body | chr14:80328189-80328595 | 406 | | | 1 | | | | | | | | | | | | | | | | yes |
| DIO2 | NA | 1stExon;Body | chr14:80677608-80678065 | 457 | | | 8 | | | | | | 1 | | | | | | | | | yes | |
| TSHR | S_Shore | 5'UTR;1stExon | chr14:81421983-81422160 | 177 | | | 12 | | | | | | | | | 2 | | | | | | yes | |
| TSHR | NA | Body | chr14:81425912-81426577 | 665 | | | 8 | | 3 | | | 0 | | | | | | | | yes | yes | yes | yes |
| SEL1L | Island | TSS200 | chr14:82000246-82000256 | 10 | | | | | | | | | 1 | | | | | | | | | | yes |
| FLRT2 | N_Shore;Island | TSS1500;TSS200;1stExon;5'UTR | chr14:85995655-85996873 | 1218 | | 2 | | | 1 | | | | 3 | | | | | | | | | | |
| FLRT2 | Island | 5'UTR | chr14:85999933-86000477 | 544 | | | | | 1 | | | | | | | | | | | | | yes | yes |
| GALC | N_Shore | Body | chr14:88458917-88459688 | 771 | | | | | 1 | | | | | | | | | yes | | yes | | | |
| KCNK10 | N_Shore | Body | chr14:88788343-88788931 | 588 | | | 2 | | | | | 0 | 3 | | | | | | yes | | | yes | |
| KCNK10 | Island | TSS200 | chr14:88793305-88793634 | 329 | | | 1 | | | | | | | | | | | | | | | yes | |
| ZC3H14 | N_Shore | TSS1500 | chr14:89028922-89029076 | 154 | | | | | | | | | | | | | | yes | | yes | | | |
| FOXN3 | N_Shore;S_Shore | 3'UTR;Body | chr14:89628141-89629974 | 1833 | | 1 | 2 | | | | | | | | | | | yes | | | | yes | yes |
| FOXN3 | N_Shelf | Body | chr14:89878675-89878733 | 58 | | | | | | | | | 1 | | | | | yes | | | | | yes |
| PRO1768;FOXN3 | NA | TSS1500;5'UTR | chr14:90042201-90042580 | 379 | | | 2 | | | | | | | | | | | | | | | | |
| TTC7B | N_Shore | Body | chr14:91281552-91281948 | 396 | | | | | | | | | | | | | | | yes | | | | yes |
| C14orf159 | NA | TSS1500 | chr14:91579859-91580142 | 283 | | | | | | | | 0 | | | | | | yes | | | yes | | |
| GPR68 | Island | 1stExon;5'UTR | chr14:91720173-91720527 | 354 | | | | | | | | | | | | | | yes | | | | | |
| CCDC88C | NA | Body | chr14:91744409-91744445 | 36 | | | | | | | | | 1 | | | | | | yes | | yes | | |
| CCDC88C | N_Shelf | Body | chr14:91751397-91752093 | 696 | 1 | | 5 | | | | | | 1 | | | | | | | | yes | | |
| CCDC88C | Island | Body | chr14:91755331-91755616 | 285 | | | | | | | | | 1 | | | | | yes | | | | | |
| CCDC88C | N_Shelf | Body | chr14:91766170-91766373 | 203 | | 1 | | | | | | | | | | | | | | | yes | | |
| CCDC88C | N_Shore;Island | Body | chr14:91770052-91770418 | 366 | | | 11 | | | | | | | | | | | yes | | | | | yes |
| CCDC88C | NA | Body | chr14:91805692-91806143 | 451 | | | | | | | | | | | | | | | | | | | |
| CCDC88C | NA | Body | chr14:91818497-91818668 | 171 | | | | | | | | | | | | | | | | | | | |
| CCDC88C | NA | Body | chr14:91862813-91862907 | 94 | | | | | | | | | 1 | | | | | yes | | | | | |
| CCDC88C | NA | nan;Body | chr14:91866325-91866751 | 426 | | | 1 | | 1 | | | | | | 1 | | | yes | yes | | | | |
| C14orf184 | S_Shore;Island | TSS1500;3'UTR;1stExon;5'UTR | chr14:92040499-92042333 | 1834 | | | | | 2 | | | | 2 | | | | | yes | | yes | | | yes |
| TC2N | S_Shore | TSS1500;5'UTR | chr14:92303622-92303710 | 88 | | | | | | | | | | | | | | yes | | | yes | | |
| TC2N | NA | 1stExon;5'UTR | chr14:92333765-92334271 | 506 | | | | | | | | 0 | 3 | | | | | | | yes | yes | | |
| RIN3 | Island | Body | chr14:93154317-93154758 | 441 | | | 3 | | | | | | | | | | | | | | | yes | |
| LGMN | NA | 3'UTR;Body | chr14:93170534-93170970 | 436 | | | | | 1 | | | 0 | 1 | 1 | 1 | | | | yes | | yes | | |
| LGMN | NA | TSS1500 | chr14:93215447-93215837 | 390 | | | | | | | | | 1 | | | | | yes | | yes | yes | | |
| CHGA | N_Shore;Island | TSS1500;TSS200 | chr14:93389102-93389891 | 789 | | 1 | | | | | | 0 | | | | | | yes | yes | | | | yes |
| ITPK1 | NA | Body | chr14:93405937-93406082 | 145 | | | | | | | | | 1 | 1 | | | | | | | | | |
| ITPK1 | NA | Body | chr14:93412562-93412768 | 206 | | | | | | | | | 1 | | | | | yes | | | | | |
| ITPK1 | NA | Body | chr14:93419016-93419554 | 538 | 1 | | | | | | | | | | | | | | | | | | |
| KIAA1409;COX8C | Island | 5'UTR;TSS200 | chr14:93813520-93813611 | 91 | | | | | | | | | 1 | | | | | | | | | yes | |
| KIAA1409 | NA | Body | chr14:94172979-94173261 | 282 | | | | | | | | | | | | | | yes | | | | | |
| PRIMA1 | Island | 5'UTR | chr14:94254131-94254586 | 455 | | | 1 | | | | | | | | | | | | | | | | |
| C14orf86;FAM181A | NA | Body;TSS200 | chr14:94385236-94385395 | 159 | | | | | | | | | | | | | | yes | | | yes | | |
| FAM181A;C14orf86 | NA | Body;TSS1500 | chr14:94393497-94393816 | 319 | | | | | | | | | 2 | | | | | yes | | | | yes | yes |
| ASB2 | Island | Body | chr14:94405267-94405681 | 414 | | | | | | | | 0 | | | | 1 | | yes | | | | | |
| ASB2 | NA | 5'UTR;1stExon | chr14:94423704-94423950 | 246 | | | | | | | | | 1 | | | | | yes | | | | | |
| C14orf48 | NA | TSS1500 | chr14:94462309-94462607 | 298 | | | 5 | | | | | | | | | | | | yes | | | yes | |
| IFI27 | NA | TSS1500;TSS200 | chr14:94576831-94577514 | 683 | | | 5 | | 2 | | | | 1 | 1 | 1 | | | yes | yes | | | | |
| SERPINA10 | NA | TSS1500;TSS200 | chr14:94759646-94759873 | 227 | | | 6 | | | | | | 1 | | | | | | yes | | yes | | |
| SERPINA6 | NA | 5'UTR;1stExon | chr14:94789681-94790112 | 431 | | | 2 | 1 | | | | | | | | | | yes | yes | yes | | | |
| SERPINA1 | NA | 1stExon;5'UTR | chr14:94856984-94857151 | 167 | | | | | | | | | | | | | | yes | | | | | |
| SERPINA4 | NA | TSS1500;TSS200 | chr14:95027382-95028105 | 723 | | 1 | 4 | | | | | | 1 | | | | | yes | yes | | yes | | |
| SERPINA5 | NA | TSS200 | chr14:95047591-95048035 | 444 | | | | | | | | | | | | | | yes | | | yes | | |
| GSC | Island | 3'UTR;Body | chr14:95234658-95235402 | 744 | | | 30 | | | | | | 2 | | | | | | | | | yes | |
| GSC | N_Shore | TSS200;TSS1500 | chr14:95236615-95237637 | 1022 | | | | | | | | | 2 | | | | | | | | | | yes |
| CLMN | NA | Body | chr14:95675591-95675657 | 66 | | | | | | | | | | | | | | yes | | | yes | | |
| CLMN | NA | Body | chr14:95693784-95722106 | 28322 | | | 3 | | 1 | | | | | | | | | yes | yes | yes | yes | yes | yes |
| C14orf49 | NA | Body | chr14:95927445-95927930 | 485 | | | | | | | | | 1 | | | | | | | yes | | | |
| TCL1B | NA | TSS1500;TSS200 | chr14:96152109-96152766 | 657 | | | | | | | | 0 | 2 | | | | | | yes | | | yes | yes |
| TCL1A | Island | TSS200;TSS1500;5'UTR;1stExon | chr14:96180406-96180847 | 441 | | | 3 | | 1 | | | 0 | 5 | | | | | | yes | | | | |

| | | | | | | | | | | | | | | | | | | | | | | | |
|---|---|---|---|---|---|---|---|---|---|---|---|---|---|---|---|---|---|---|---|---|---|---|---|
| C14orf132 | Island | Body | chr14:96505722-96505874 | 152 | | | | | | | | | 1 | | | | | yes | | | | yes | |
| BDKRB2 | NA | TSS1500 | chr14:96670628-96670862 | 234 | | | 6 | | | | | 0 | 1 | | | 1 | | | | | | | |
| BDKRB1 | NA | Body | chr14:96730482-96730669 | 187 | 2 | | | | | | | | | | | | | | | | yes | | |
| ATG2B | NA | 3'UTR | chr14:96750481-96750614 | 133 | | | | 1 | | | | | | | | | | | | | | | |
| ATG2B | NA | 3'UTR | chr14:96752011-96752271 | 260 | | | 3 | | | | | | | | | | | | | | | | |
| AK7 | N_Shore;Island | Body | chr14:96890120-96890632 | 512 | | | | | | | | 0 | 2 | | | | | yes | | | | | |
| C14orf64 | NA | Body;TSS200 | chr14:98444151-98444533 | 382 | | | | 2 | 1 | | | 0 | 6 | | | | | | | | yes | | |
| BCL11B | Island | 3'UTR | chr14:99640471-99641151 | 680 | | | | | | | | | 1 | | | | | | | | | yes | yes |
| BCL11B | NA | Body | chr14:99655593-99655747 | 154 | | | 2 | | 1 | | | | 1 | | | | | | | | | | |
| BCL11B | NA | Body | chr14:99664135-99664279 | 144 | | | | | | | | | 1 | | | | | | yes | | | | |
| BCL11B | NA | Body | chr14:99707650-99708008 | 358 | | | 2 | | | | | | | | | | | | | | | | yes |
| BCL11B | NA | Body | chr14:99728984-99729088 | 104 | | | 13 | 1 | | | | | 1 | | | 2 | | | | | | | |
| CCDC85C | Island | 1stExon | chr14:100069535-100070043 | 508 | | | 2 | | | | | 0 | 2 | | | 2 | | | | | | yes | yes |
| HHIPL1 | N_Shore;Island | TSS1500;TSS200 | chr14:100110845-100111354 | 509 | | | | | | | | | 2 | | | | | | yes | yes | | | yes |
| HHIPL1 | N_Shore;Island | Body | chr14:100125548-100126741 | 1193 | | | 8 | | | | | | | | | | | yes | yes | | | | |
| HHIPL1 | N_Shore;Island | Body | chr14:100141423-100142298 | 875 | | | 10 | | | | | | 2 | | | | | | | | | | yes |
| CYP46A1 | Island | TSS1500 | chr14:100149967-100149990 | 23 | | | | | | | | | 1 | | | | | | | yes | | yes | |
| EML1 | N_Shore | TSS1500 | chr14:100258773-100259020 | 247 | | | | | | | | | | | | | | | | | | | yes |
| EVL | NA | TSS200 | chr14:100531591-100532036 | 445 | | | | | | | | | 1 | | | | | yes | | | | | |
| EVL | N_Shore | Body | chr14:100610071-100610570 | 499 | | 1 | | | | | | | | | | | | | | | | | |
| DEGS2 | N_Shelf | Body | chr14:100613226-100613692 | 466 | | | | | 1 | | | | | | | | | | | | | | |
| C14orf68 | NA | TSS1500 | chr14:100788291-100788777 | 486 | | | | | | | | | | | | | | yes | yes | | | | |
| C14orf68 | Island | Body | chr14:100795151-100795297 | 146 | | | | 1 | | | | | | | | | | yes | | | | | |
| BEGAIN | S_Shore | TSS1500;5'UTR;TSS200 | chr14:101035818-101036470 | 652 | | | 15 | | | | | 0 | 2 | | | 2 | | yes | yes | | | | yes |
| C14orf70 | S_Shore;S_Shelf | TSS1500 | chr14:101123263-101123682 | 419 | | | 4 | | | | | | | | | | | yes | | | | | |
| DLK1 | N_Shore;Island | TSS1500 | chr14:101192634-101193432 | 798 | | 1 | 2 | | | | | 0 | 2 | | | 1 | | | | | | yes | |
| MEG3 | N_Shore | TSS1500 | chr14:101291068-101291997 | 929 | | | 20 | | | | | | 1 | | | | | yes | | | | | |
| MIR770;MEG3 | NA | TSS1500;Body | chr14:101317620-101317819 | 199 | | | | | | | | | | | | | | yes | | | | | yes |
| MIR136;RTL1;MIR432 | S_Shore | TSS1500;1stExon;TSS200 | chr14:101350694-101350872 | 178 | | | 1 | | | | | | | | | | | | | | | | |
| MEG8 | NA | TSS1500 | chr14:101360088-101360188 | 100 | | | | | | | | | 1 | | | | | yes | | | | | |
| SNORD114-12;SNORD114-13 | NA | TSS200;TSS1500 | chr14:101435254-101435465 | 211 | | | | | | | | | 1 | | | | | | | | | | yes |
| SNORD114-14;SNORD114-15;SNORD114 | NA | Body;TSS1500 | chr14:101438461-101438663 | 202 | | | | | | | | | 1 | | | | | | | | | | |
| SNORD114-27;SNORD114-26 | NA | TSS1500 | chr14:101453070-101453431 | 361 | | | | | | | | | 1 | | | | | | | | | | |
| MIR299;MIR380 | NA | TSS200;TSS1500;Body | chr14:101490118-101490676 | 558 | | | | | | | | | 1 | | | | | | | | | | |
| MIR494 | NA | TSS1500 | chr14:101495604-101495907 | 303 | 1 | | 1 | | | | | | | | | | | | | | | | |
| MIR495 | NA | TSS200 | chr14:101499893-101500125 | 232 | | 1 | | | | | | | 1 | | | | | | | yes | | | |
| MIR654;MIR376B;MIR376A1;MIR376A2; | NA | TSS1500;Body | chr14:101506087-101506323 | 236 | | | | | | | | | | | | | | | | | | | |
| MIR300 | NA | TSS200 | chr14:101507570-101507727 | 157 | | | | | | | | | 1 | | | | | yes | | | | | |
| MIR1185-1 | NA | TSS1500 | chr14:101508789-101509144 | 355 | | | 1 | | | | | | | | | | | yes | | | | | |
| MIR487B;MIR539;MIR889 | NA | TSS200;TSS1500 | chr14:101512732-101514051 | 1319 | | | 2 | | | | | | 1 | | | | | yes | | | | yes | |
| MIR134;MIR668;MIR382;MIR485;MIR45 | NA | TSS1500;TSS200;Body | chr14:101520412-101521809 | 1397 | | | 8 | | | | | | | | | | | | yes | | | | |
| MIR496;MIR154 | NA | TSS1500;TSS200;Body | chr14:101525465-101527144 | 1679 | | 1 | | | | | | | | | | | | yes | yes | | | | |
| MIR377 | N_Shelf | TSS200 | chr14:101528234-101528671 | 437 | 1 | | 4 | | | | | | 1 | | | | | | yes | | | yes | |
| MIR409;MIR410;MIR412;MIR369;MIR65 | N_Shore;Island | TSS200;TSS1500;Body | chr14:101531617-101531755 | 138 | | | 1 | | | | | | 1 | | | | | | | | | | |
| DIO3OS | N_Shelf | TSS1500 | chr14:102022513-102022947 | 434 | | | | | | | | | | | | | | yes | | | | yes | |
| DIO3;MIR1247 | Island | TSS1500;TSS200 | chr14:102026230-102027760 | 1530 | | 2 | | | | | | 0 | 3 | | | | | | | | | yes | yes |
| DIO3 | Island | 3'UTR;1stExon | chr14:102028994-102029375 | 381 | | | | | | | | | 1 | | | | | | | | | | |
| PPP2R5C | N_Shore | Body | chr14:102247610-102248253 | 643 | | | | | | | | 0 | | | | | | yes | | | | | yes |
| PPP2R5C | N_Shore | 3'UTR | chr14:102394274-102394448 | 174 | | | | | | | | | 1 | | | | | | | | | | |
| DYNC1H1 | NA | Body | chr14:102504880-102505062 | 182 | | | | | | | | | | | | | | yes | | | yes | | |
| DYNC1H1 | NA | Body | chr14:102510281-102510478 | 197 | | | 3 | | | | | | | | | | | | | | | | |
| HSP90AA1 | S_Shore | TSS1500;Body | chr14:102554826-102554977 | 151 | 1 | | 19 | 1 | 1 | | | | | | | | | yes | | | | | |
| WDR20 | N_Shelf | Body | chr14:102676957-102677294 | 337 | | | | | | | | 0 | 1 | 1 | 1 | | | | | | | | |
| RAGE | N_Shore | 3'UTR | chr14:102695264-102695768 | 504 | | | | | | | | | 1 | | | | | yes | | | | | |
| TECPR2 | NA | Body | chr14:102918789-102918977 | 188 | 2 | | | | | | | | | | | | | | | | | | |
| RCOR1 | NA | Body | chr14:103150012-103150300 | 288 | | | | 1 | | | | | | | | | | | | | | | |
| TRAF3 | NA | 5'UTR | chr14:103294363-103294740 | 377 | 1 | | | | 1 | | | | | | | | | | | | yes | | |
| TRAF3 | NA | Body | chr14:103367306-103367858 | 552 | | | | | 2 | | 1 | 0 | 1 | | | 2 | | yes | | | | | |
| AMN | N_Shore | TSS200;TSS1500 | chr14:103388656-103389022 | 366 | | | | | | | | 0 | 2 | | | | | yes | | | | yes | |
| CDC42BPB | NA | Body | chr14:103415458-103416389 | 931 | | | 5 | | | | | | | | | | | | | | | | yes |
| CDC42BPB | S_Shore | TSS1500 | chr14:103524584-103524901 | 317 | | | | | | | | | 1 | | | | | yes | yes | | yes | yes | |
| C14orf73 | N_Shelf | TSS1500 | chr14:103565867-103566172 | 305 | | | | | | | | | | | | | | | | | | | |
| C14orf73 | Island | Body | chr14:103570781-103571370 | 589 | | | | | | | | 0 | | | | | | | | | | yes | |
| C14orf73 | NA | Body | chr14:103576399-103576818 | 419 | | | | | | | | | | | | | | | yes | | | | yes |
| TNFAIP2 | N_Shore;Island | TSS200;5'UTR;1stExon;Body | chr14:103592526-103593995 | 1469 | | | 37 | | 1 | | | | 3 | | 1 | | | yes | yes | | | yes | |
| CKB | N_Shore | Body | chr14:103986227-103986467 | 240 | | | | | | | | | 1 | | | | | | | | | | |
| TRMT61A | N_Shore | TSS1500 | chr14:103995143-103995361 | 218 | | | | | | | | 0 | | | | | | yes | | | | | yes |
| TRMT61A | N_Shore | Body | chr14:104000447-104000505 | 58 | | | 1 | | | | | | | | | | | yes | | | | | |
| XRCC3;KLC1 | Island | Body;3'UTR | chr14:104165052-104165475 | 423 | | | | 1 | | | | | 1 | | | | | | | | | | |
| XRCC3 | NA | Body | chr14:104171695-104172109 | 414 | | | | | | | | | 2 | 1 | | | | | yes | | | | |
| XRCC3 | NA | Body | chr14:104175555-104175847 | 292 | | | | | | | | | 1 | | | | | | | | | | |
| XRCC3 | N_Shelf | 5'UTR | chr14:104178670-104179160 | 490 | | | 8 | | | | | | 1 | | | | | | | yes | | | |
| ZFYVE21 | N_Shelf | Body | chr14:104190678-104190829 | 151 | | | | | 1 | | | | | | | | | | | | | | |
| ZFYVE21 | S_Shore | Body | chr14:104195181-104195666 | 485 | | | | | | 1 | | | 1 | 1 | | | | | | | | | |
| PPP1R13B | N_Shore | Body | chr14:104204601-104205071 | 470 | | | 5 | | | | | | | | | | | | yes | | | | |
| PPP1R13B | S_Shore | TSS1500 | chr14:104315145-104315223 | 78 | 1 | | 1 | | | | | | | | | | | yes | | | | | |
| TDRD9 | chr14:104394529-104395234;I | N_Shore;TSS1500 | chr14:104394430-104395485 | 1055 | | | | | | | | 0 | 2 | | 1 | | | yes | yes | | | yes | yes |
| ASPG | N_Shore;Island | TSS200 | chr14:104551867-104552397 | 530 | 3 | 1 | | | | | | 0 | 3 | 1 | 1 | 2 | | | | | | yes | yes |
| ASPG | Island | Body | chr14:104569519-104569653 | 134 | | | | | | | | 0 | 2 | | | 3 | | | | | | | |
| MIR203 | Island;N_Shore | TSS1500 | chr14:104582743-104583740 | 997 | | | | | | | | 0 | 2 | | | | | yes | | | | | |
| KIF26A | NA | Body | chr14:104611591-104611819 | 228 | | | | | | | | | | | | | | | yes | | | | |
| KIF26A | N_Shore;Island | Body | chr14:104621176-104622128 | 952 | | | 18 | | | | | | 3 | | | | | | | | yes | | |
| KIF26A | N_Shore | Body | chr14:104623477-104623651 | 174 | | | | | | | | | 1 | | | | | | | | | | |
| KIF26A | N_Shore | Body | chr14:104636633-104636933 | 300 | | | | | | | | | | | | | | | | | | | |
| KIF26A | Island | Body | chr14:104638170-104638843 | 673 | | | | | | | | | 1 | | | | | | | | | | |
| KIF26A | Island | Body | chr14:104644850-104645063 | 213 | | | | | | 1 | | | 1 | | | | | | | | | | |
| KIF26A | S_Shore;Island | 3'UTR | chr14:104646441-104647094 | 653 | | | | | | | | | | | | | | | | | | yes | |
| C14orf180 | S_Shelf;NA | TSS1500;TSS200 | chr14:105044984-105046247 | 1263 | | | 7 | | | | | | 1 | | | | | yes | | | | | |
| C14orf180 | Island | Body | chr14:105054940-105055171 | 231 | | | | | | | | | | | | | | | | | | | |
| INF2 | N_Shore | TSS1500 | chr14:105154557-105155327 | 770 | | | 15 | | | 1 | | | | | | | | yes | | | | | |
| INF2 | Island | 5'UTR | chr14:105167300-105167607 | 307 | | | | | | | | 0 | 1 | | | | | | yes | | | | |
| INF2 | NA | Body | chr14:105173643-105173980 | 337 | | | 6 | | | | | | 1 | | | | | | | | | yes | yes |
| INF2 | N_Shelf | Body;3'UTR | chr14:105184057-105184176 | 119 | | | | | | | | | 1 | | | | | | yes | | | | |
| AKT1 | N_Shelf | 3'UTR | chr14:105236538-105236654 | 116 | | | | | | | | | | | | | | | | | | | |
| AKT1 | S_Shelf | Body | chr14:105255285-105255472 | 187 | | | | 1 | | | | | | | | | | | | | yes | | |
| ZBTB42 | N_Shore | TSS1500 | chr14:105265453-105265909 | 456 | | | 2 | | | | | | | | | | | yes | yes | | | | |
| MGC23270 | S_Shelf;NA | TSS1500 | chr14:105286870-105287427 | 557 | | | 5 | | | | | 0 | 1 | | | 1 | | yes | | yes | yes | yes | |
| KIAA0284 | Island | TSS1500 | chr14:105330630-105331124 | 494 | | | | | | | | | 1 | | | | | | | | | yes | yes |
| KIAA0284 | S_Shore | Body | chr14:105352845-105352991 | 146 | | | | | | | | | | | | | | | | | | | |

| | | | | | | | | | | | | | | | | | | | | | | | |
|---|---|---|---|---|---|---|---|---|---|---|---|---|---|---|---|---|---|---|---|---|---|---|---|
| KIAA0284 | Island | Body | chr14:105359836-105360085 | 249 | | | | | | | | | | | | | | | | | | | |
| AHNAK2 | NA | Body | chr14:105436973-105437141 | 168 | | | | | | | | | | | | | | yes | | | | | |
| AHNAK2 | Island | TSS1500 | chr14:105444897-105444910 | 13 | | | | | | | | | 1 | | | | | | | | | yes | |
| GPR132 | NA | 5'UTR | chr14:105521842-105522348 | 506 | | | | | | | | | | | | | | yes | yes | | | | yes |
| GPR132 | NA | TSS200;TSS1500;nan | chr14:105531893-105532252 | 359 | | | | | | | | | 2 | | 1 | | | | | | | | |
| JAG2 | Island | Body | chr14:105616988-105617308 | 320 | | | 3 | | | | | | | | | | | | | | | | |
| JAG2 | S_Shore;N_Shelf | Body | chr14:105618482-105619701 | 1219 | | | | | | | | | 2 | | | 1 | | yes | | | | | |
| JAG2 | NA | TSS1500 | chr14:105635706-105635769 | 63 | | | 12 | | 1 | | | | | | | | | | | | | | yes |
| BRF1 | S_Shelf | Body | chr14:105688129-105688421 | 292 | 1 | | | | 2 | | | | | | | | | | yes | | | | |
| BRF1 | NA | Body | chr14:105707628-105707747 | 119 | | | | | | | | | | | | | | | | | | | |
| BTBD6;BRF1 | Island;N_Shore | Body;TSS1500;1stExon;5'UTR;TSS2 | chr14:105714153-105716052 | 1899 | | | 8 | | 1 | | | | 1 | | | | | yes | yes | | | yes | yes |
| BRF1 | NA | Body | chr14:105747951-105748448 | 497 | | | | | | | | | | | | | | | yes | | | | |
| PACS2 | N_Shore | TSS1500 | chr14:105779826-105780137 | 311 | | | | 1 | | | | 0 | 2 | | | | | | yes | | | | |
| PACS2 | Island | Body | chr14:105796713-105796964 | 251 | | | | | | | | | | | | | | yes | | | | | |
| PACS2 | N_Shore;Island | Body | chr14:105830509-105830859 | 350 | 5 | 3 | | | 1 | | 1 | 0 | 3 | 3 | 1 | 6 | 2 | yes | | | yes | | |
| PACS2 | N_Shore | Body | chr14:105851564-105851981 | 417 | | | | | | | | | 1 | | | | | | | | | | |
| MTA1 | NA | TSS200 | chr14:105886024-105886028 | 4 | | | | | | | | | 1 | | | | | | | | | yes | yes |
| CRIP2 | N_Shore | TSS1500 | chr14:105940179-105940391 | 212 | | | | 1 | | | | | | | | | | | | | | | yes |
| CRIP2 | Island | Body | chr14:105944604-105945699 | 1095 | | | 7 | 1 | | | | 0 | 6 | 1 | | 2 | | | | | | yes | yes |
| C14orf80 | N_Shore;Island | Body | chr14:105964243-105965379 | 1136 | 2 | | | | | | | | | | | | | | | | | | |
| TMEM121 | Island | Body | chr14:105995251-105995755 | 504 | | | 3 | | | | | 0 | | | | 1 | 1 | | | | | yes | yes |
| ADAM6 | NA | Body | chr14:106438219-106438519 | 300 | | | | | | | | | 1 | | | | | | | yes | | | |
| LOC100133469 | Island;N_Shore | Body;TSS1500;TSS200 | chr14:106937310-106938915 | 1605 | | | 21 | | | | | | 2 | | | | | yes | yes | yes | | | yes |
| TUBGCP5 | N_Shore;chr15:22833282-2283 | TSS1500;TSS200;N_Shore;Body | chr15:22833149-22833975 | 826 | 1 | | 4 | | | | | 0 | 7 | 6 | 1 | 4 | | yes | | | | | yes |
| CYFIP1 | NA | 5'UTR | chr15:22922243-22922392 | 149 | | | 1 | | | | | | | | | | | | yes | | yes | | |
| CYFIP1 | NA | 5'UTR | chr15:22923501-22923823 | 322 | | | | | | | | | 1 | | | | | yes | yes | | | | |
| CYFIP1 | NA | Body | chr15:22929886-22929993 | 107 | | 1 | | | | | | | | | | | | | | | | | |
| CYFIP1 | NA | Body | chr15:22968612-22968689 | 77 | | | | | | | | | 1 | | | | | | yes | | yes | | |
| CYFIP1 | N_Shore | Body | chr15:22980991-22981419 | 428 | 2 | | | | | | | | | | | | | yes | | | | | |
| CYFIP1 | NA | Body | chr15:22992842-22993172 | 330 | | | | | | | | | 1 | | | | | | yes | | yes | yes | |
| NIPA1 | Island | Body;5'UTR;TSS200 | chr15:23085583-23086888 | 1305 | | | | | | | | | | 2 | | | | yes | | | | yes | yes |
| WHAMML1 | N_Shore | Body | chr15:23207925-23208349 | 424 | | | | | | | | | 1 | | | | | | yes | | | | |
| HERC2P2 | Island | TSS200;TSS1500 | chr15:23378429-23378814 | 385 | | 2 | 2 | | | | 2 | 0 | 3 | 1 | 1 | 5 | | yes | yes | | | | yes |
| MKRN3 | NA | TSS1500;TSS200;5'UTR;1stExon | chr15:23810163-23811369 | 1206 | | | 10 | 1 | | | | 0 | 8 | | | 2 | | | yes | | | | |
| MAGEL2 | NA | 1stExon;TSS1500 | chr15:23892770-23894248 | 1478 | | | 1 | | | | | | 1 | | | | | | yes | | yes | | |
| NDN | N_Shore;Island;S_Shore | 3'UTR;1stExon;5'UTR;TSS1500 | chr15:23930952-23933620 | 2668 | | | 7 | | | | | | 2 | | | 1 | | | yes | | | | yes |
| C15orf2 | N_Shore | TSS1500;TSS200 | chr15:24919600-24921054 | 1454 | | | 5 | | | | | | | | | | | yes | | | | | yes |
| SNRPN | NA | TSS1500;TSS200 | chr15:25068564-25068850 | 286 | | | 3 | | | | | 0 | 2 | | | | | | yes | | | | |
| SNRPN | Island | 5'UTR | chr15:25093244-25093521 | 277 | | | | | | | | 0 | 2 | | | 1 | | yes | | | | yes | |
| SNRPN | N_Shore;Island | 5'UTR | chr15:25123287-25124213 | 926 | | | 2 | | | | | | 6 | | | 1 | | yes | | | | yes | yes |
| SNRPN;SNURF | N_Shore;Island | 5'UTR;TSS1500;Body | chr15:25198793-25200490 | 1697 | | | 5 | | | | | | 1 | | | | | yes | | | yes | | |
| PAR5;SNORD64 | NA | Body;TSS200 | chr15:25230200-25230707 | 507 | | | | | | | | | 2 | | | | | yes | | | | yes | |
| SNORD116-1 | NA | TSS1500;TSS200 | chr15:25296404-25296693 | 289 | | | | | | | | 0 | | | | | | | | | | yes | |
| SNORD115-1 | NA | TSS1500 | chr15:25414716-25414781 | 65 | | | 3 | | | | | | | | | | | | | | | | yes |
| SNORD115-2 | NA | TSS1500;Body | chr15:25417453-25418074 | 621 | | | | | | | | | | | | | | | | | | | |
| SNORD115-9;SNORD115-5;SNORD115-1 | NA | TSS1500 | chr15:25422945-25423404 | 459 | | | | | | | | | | | | | | | | | | | |
| SNORD115-6 | NA | TSS1500 | chr15:25425326-25425818 | 492 | | | 6 | | | | | | | | | | | | | | | | |
| SNORD115-8;SNORD115-5;SNORD115-1 | NA | TSS200;TSS1500 | chr15:25427394-25429408 | 2014 | | | | | | | | | | | | | | | | | | | |
| SNORD115-9;SNORD115-10;SNORD115- | NA | TSS200 | chr15:25430614-25430671 | 57 | 2 | | | | | | | | | | | | | | | | | | |
| SNORD115-10 | NA | TSS1500 | chr15:25432236-25432321 | 85 | | | 4 | | | | | | | | | | | | | | | | |
| SNORD115-29;SNORD115-36;SNORD115 | NA | TSS1500 | chr15:25434030-25434533 | 503 | | | 6 | | | | | | 1 | | | | | | | | | | |
| SNORD115-9;SNORD115-10;SNORD115- | NA | TSS1500 | chr15:25436179-25436246 | 67 | | | | | | | | | | | | | | | | | | | |
| SNORD115-13 | NA | TSS1500 | chr15:25437871-25438443 | 572 | | | | | | | | 0 | 1 | | | | | | | | | | |
| SNORD115-16 | NA | TSS1500 | chr15:25443938-25444498 | 560 | | | | | | | | | | | | | | | | | | | |
| SNORD115-19;SNORD115-17;SNORD115 | NA | TSS200;TSS1500 | chr15:25448194-25448352 | 158 | | | | | | | | | | | | | | | | | | | |
| SNORD115-17;SNORD115-18;SNORD115 | NA | TSS200;TSS1500 | chr15:25449371-25450131 | 760 | | | | | | | | | | | | | | | | | | | |
| SNORD115-15;SNORD115-21;HBII-52-24 | NA | Body;TSS1500 | chr15:25457439-25457690 | 251 | | | | | | | | | 1 | | | | | | | | | | |
| SNORD115-15;SNORD115-21;HBII-52-27 | NA | Body;TSS1500 | chr15:25465087-25465466 | 379 | | | | | | | | | | | | | | | | | | | |
| SNORD115-15;SNORD115-21;SNORD115 | NA | Body;TSS1500 | chr15:25472950-25473398 | 448 | | | | | | | | | | | | | | | | | | | |
| SNORD115-15;SNORD115-21;SNORD115 | NA | Body;TSS200 | chr15:25475833-25476386 | 553 | | | | | | | | | | | | | | | | | | | |
| SNORD115-29;SNORD115-11;SNORD115 | NA | TSS1500 | chr15:25480975-25481539 | 564 | | | | | | | | | | | | | | | | | | | |
| SNORD115-39 | NA | TSS1500 | chr15:25486613-25486871 | 258 | | | | | | | | | | | | | | | | | | | |
| SNORD115-10;SNORD115-42 | NA | TSS1500;TSS200 | chr15:25491315-25492931 | 1616 | | | | | | | | | | | | | | | | | | | |
| UBE3A | S_Shore | TSS1500 | chr15:25684804-25684849 | 45 | | | | | | | | 0 | 2 | | | | | yes | | | | yes | |
| ATP10A | NA | Body | chr15:26044050-26044289 | 239 | | | 2 | | 1 | | | 0 | 1 | | | | | yes | | yes | | | |
| ATP10A | NA | Body | chr15:26047382-26047653 | 271 | | | 12 | | | | | | | | | | | | | yes | | | |
| ATP10A | S_Shore;Island | TSS1500;1stExon;5'UTR;TSS200 | chr15:26108263-26109614 | 1351 | | 2 | 46 | | 7 | | | | 7 | | | | | | | | | yes | yes |
| GABRB3 | N_Shore;Island | Body | chr15:26873818-26874788 | 970 | | | | | | | | 0 | 3 | | | 4 | | | yes | | yes | yes | |
| GABRB3 | Island | Body | chr15:26915257-26915752 | 495 | | | | | | | | | 1 | | | | | yes | | | | yes | |
| GABRB3 | S_Shore | TSS1500 | chr15:27019248-27019697 | 449 | | | | | | | | | | | | 1 | | | | | | yes | |
| GABRA5 | N_Shore;Island | TSS1500;TSS200;5'UTR;1stExon | chr15:27111388-27113511 | 2123 | | 1 | 30 | | 2 | | | 0 | 6 | 1 | | 1 | | | | | | yes | |
| GABRA5 | Island | Body | chr15:27128279-27128934 | 655 | | | | | | | | | | | | | | yes | | | | | yes |
| GABRA5 | NA | Body | chr15:27184766-27185073 | 307 | | | 5 | | | | | | | | | | | | | | yes | | |
| GABRA5 | NA | Body | chr15:27189784-27189846 | 62 | | | | | | | | | | | | | | | | | | | |
| GABRG3 | N_Shore;Island | TSS1500;TSS200 | chr15:27215448-27216819 | 1371 | | 1 | 22 | | 2 | | | 0 | 10 | 1 | | 3 | | yes | | | | | yes |
| OCA2 | N_Shore;Island | Body | chr15:28014040-28016654 | 2614 | | | 3 | | | | | | 1 | | | | | | yes | yes | | yes | yes |
| OCA2 | N_Shore;Island | Body | chr15:28050147-28050623 | 476 | | | 3 | | | | | | | | | | | yes | | | | | |
| OCA2 | Island | Body | chr15:28147928-28148431 | 503 | | | | | | | | | 1 | | | | | | | | | | |
| OCA2 | Island | Body | chr15:28248727-28249165 | 438 | | | | | | | | | | | | | | | | | | yes | |
| OCA2 | NA | Body | chr15:28272345-28272656 | 311 | | | | | | | | | 1 | | | | | | | | | | |
| OCA2 | Island | 5'UTR | chr15:28339873-28340459 | 586 | | | 12 | | | | | | | | | | | yes | | | | | |
| OCA2 | S_Shore | TSS1500;TSS200 | chr15:28344513-28344730 | 217 | | | 2 | | | | | | 2 | | | | | | | | | yes | |
| APBA2 | NA | TSS1500;TSS200 | chr15:29213250-29213860 | 610 | | | 12 | | | | | | | | | | | yes | | | | | |
| APBA2 | Island | Body | chr15:29395685-29396267 | 582 | | | | | | | | 0 | | | | | 1 | | | | | | |
| APBA2 | Island | Body | chr15:29407792-29408145 | 353 | | | | | | | | 0 | 1 | | | | | yes | | | | | |
| APBA2 | S_Shore | 3'UTR | chr15:29409410-29410245 | 835 | | | 3 | | | | | | | | | | | | | | | | |
| FAM189A1 | NA | Body | chr15:29431453-29432511 | 1058 | | | 1 | | | | | | 1 | | | | | | | | | | |
| FAM189A1 | NA | Body | chr15:29499242-29499526 | 284 | | | | | | | | | | | | | | | yes | yes | | | |
| FAM189A1 | Island | Body | chr15:29862036-29862568 | 532 | | | | | | | | | 1 | | | | | | | | | | yes |
| TJP1 | Island | 5'UTR;1stExon | chr15:30114295-30114816 | 521 | | | | | | | | | 3 | | | | | | | | | yes | yes |
| MTMR15 | N_Shore | TSS1500 | chr15:31195612-31195741 | 129 | | | | | | | | 0 | | | | | | yes | | | yes | | |
| TRPM1 | NA | Body | chr15:31355304-31355430 | 126 | | | | | | | | | | | | | | yes | | | | | |
| TRPM1 | NA | TSS200 | chr15:31394021-31394285 | 264 | | | | | | | | | | | | | | yes | | | | | |
| SCG5 | NA | TSS1500;TSS200;1stExon;5'UTR | chr15:32933661-32934836 | 1175 | | | 5 | | | | | 0 | 2 | | | 2 | | yes | | | | | |
| SCG5 | NA | Body | chr15:32964155-32964500 | 345 | | | 41 | | | | | | 1 | | | | | yes | | | | | |
| GREM1 | N_Shore;Island | TSS1500;TSS200 | chr15:33009409-33010536 | 1127 | | | 5 | | 1 | | | | 1 | | | | | | | | | | yes |
| FMN1 | NA | 1stExon;TSS200 | chr15:33359788-33360901 | 1113 | | | | 2 | | | | 0 | 1 | | | | | | | | | | |
| RYR3 | Island | TSS1500 | chr15:33602817-33602824 | 7 | | | | | 1 | | | | 1 | | | | | | | | | yes | yes |
| AVEN;CHRM5 | NA | Body;TSS1500 | chr15:34260719-34261172 | 453 | | | | | | | | | 1 | | | | | | | | | yes | |

| | | | | | | | | | | | | | | | | | | | | | | | |
|---|---|---|---|---|---|---|---|---|---|---|---|---|---|---|---|---|---|---|---|---|---|---|---|
| TMEM85 | NA | TSS1500 | chr15:34516488-34516777 | 289 | | | 11 | | | | | 0 | 1 | | | 2 | | yes | | | | yes | |
| SLC12A6 | NA | Body;1stExon | chr15:34610829-34611069 | 240 | | | 4 | | | | | 0 | | | | | | yes | | | | | |
| SLC12A6 | N_Shore | 1stExon;5'UTR;Body | chr15:34628731-34629056 | 325 | | | | | 1 | | | | 2 | | | | | | | | | | |
| GOLGA8B | Island | TSS1500 | chr15:34876295-34876471 | 176 | | | 1 | | | | | | | | | | | | | | | | |
| GJD2 | Island | Body;1stExon;TSS200 | chr15:35046610-35047203 | 593 | | | | | 1 | | | | 2 | | | | | | | | | | |
| ACTC1 | NA | 5'UTR;TSS200;Body | chr15:35086890-35088261 | 1371 | 5 | | 7 | | | | | 0 | 3 | | | 1 | | | | | | | yes |
| ATPBD4 | N_Shore | Body | chr15:35837840-35838691 | 851 | | | | | | | | | 1 | | | | | | yes | | yes | | |
| C15orf41 | N_Shore;Island | TSS1500 | chr15:36871420-36872349 | 929 | | | 4 | | | | | | | | | | | yes | yes | | | | |
| CSNK1A1P | S_Shore | TSS1500 | chr15:37111929-37112149 | 220 | | | | | | | | | | | | | | yes | | | | | yes |
| LOC145845 | N_Shelf | Body | chr15:37169665-37171175 | 1510 | | | 20 | | | | | | 1 | | | | | | | | | yes | yes |
| LOC145845 | Island | Body | chr15:37172509-37173017 | 508 | | | 8 | | | | | | | | | | | | | | | | |
| LOC145845 | S_Shore | Body | chr15:37174661-37175117 | 456 | | 1 | 4 | | | | | | 1 | | | | | | | | | yes | |
| LOC145845 | Island | TSS1500 | chr15:37179603-37180325 | 722 | | | 9 | | | | | | | | | | | | | | | | yes |
| MEIS2 | chr15:37387386-37387614 | Island | chr15:37387523-37387577 | 54 | | | | | | | | | | | 1 | | | | | | | yes | |
| MEIS2 | N_Shore | Body | chr15:37389489-37390284 | 795 | | | | 1 | | | | | | | | | | | | | | | yes |
| MEIS2 | S_Shore | TSS1500 | chr15:37394840-37395171 | 331 | | | | | | | | | 1 | | | | | yes | | | | yes | yes |
| C15orf53 | NA | TSS1500 | chr15:38988533-38988860 | 327 | | | 3 | | | | | | 1 | | | | | | | | | | |
| THBS1 | N_Shore | TSS1500 | chr15:39871808-39872186 | 378 | | | | | | 1 | | 0 | 7 | 1 | 1 | 7 | | | | | | yes | |
| EIF2AK4 | Island;N_Shore | Body | chr15:40268421-40269214 | 793 | 8 | 3 | 28 | | | | 1 | | 2 | 1 | | | | yes | | | yes | yes | yes |
| BMF | N_Shelf | TSS200;5'UTR;TSS1500;1stExon | chr15:40398307-40399131 | 824 | | | | | | | | | 2 | | | | | | | | | yes | |
| BUB1B;PAK6 | NA | Body;TSS200 | chr15:40509472-40509703 | 231 | | | | | | | | | | | | | | yes | | | | | yes |
| PAK6 | S_Shore | 5'UTR;TSS200 | chr15:40531871-40532180 | 309 | | | | | | | | | | | | | | | | | | | |
| PAK6;C15orf56 | N_Shore;chr15:40544352-4054 | 5'UTR;3'UTR;Island | chr15:40544174-40545215 | 1041 | | | 14 | | | | | | | | 1 | | | | | | | | yes |
| PLCB2 | Island | Body | chr15:40583227-40583422 | 195 | | | | | | | | 0 | 1 | | 1 | 1 | | | | | | yes | |
| PLCB2 | NA | Body;1stExon;5'UTR;TSS200 | chr15:40599680-40600635 | 955 | | | | | 2 | | | | 3 | 2 | | | | yes | | | | yes | |
| C15orf52 | NA | 1stExon | chr15:40633124-40633294 | 170 | | | | | | | | | 1 | 1 | | | | | | | yes | | |
| C15orf23 | N_Shore | TSS1500;TSS200 | chr15:40674438-40674910 | 472 | | | | | | | | | 1 | | | 1 | | yes | yes | | | | |
| DNAJC17;C15orf62 | NA | Body;TSS1500;TSS200 | chr15:41061384-41062273 | 889 | | | 3 | | 1 | | | 0 | 3 | | | | | | | | | | |
| SPINT1 | N_Shore;Island | TSS1500 | chr15:41135432-41135900 | 468 | | | 12 | | | | | 0 | 1 | | | | | yes | | | | | yes |
| VPS18 | N_Shore | TSS1500 | chr15:41185800-41186038 | 238 | | | | | | | | | | | | | | | yes | | | | |
| NDUFAF1 | Island | 1stExon;5'UTR | chr15:41694502-41694790 | 288 | | 1 | | 1 | | | | | | | | | | yes | | | | | |
| ITPKA | Island | Body | chr15:41795087-41795475 | 388 | | | 3 | | | | | | | | | | | | | | | | |
| LTK | N_Shore;Island | Body | chr15:41803428-41803850 | 422 | | | 8 | | | | | 0 | 1 | | | | | yes | | | yes | yes | yes |
| LTK | S_Shore | TSS200 | chr15:41806135-41806588 | 453 | | | 5 | 1 | | | | | 1 | | | | | yes | | | | | |
| TYRO3 | N_Shore | TSS1500;TSS200 | chr15:41850280-41851231 | 951 | | | 3 | | | | | | 1 | | | | | | yes | | | yes | yes |
| MGA | N_Shore | TSS1500 | chr15:41951943-41952429 | 486 | | | | | | | | | | | | | | | | | | | |
| JMJD7-PLA2G4B;JMJD7;PLA2G4B | NA | Body;3'UTR;TSS1500 | chr15:42129464-42130735 | 1271 | | | | | 1 | | | | | | | | | yes | | | | | |
| EHD4 | Island | Body | chr15:42211437-42211581 | 144 | | | | | 2 | | | | | | | | | yes | | | | | |
| PLA2G4E | NA | Body | chr15:42289618-42289939 | 321 | | | | | | | | | 1 | | | | | yes | yes | | | | |
| PLA2G4E | NA | 1stExon | chr15:42302379-42303032 | 653 | | | | | | | | | | | | | | | yes | | | yes | |
| PLA2G4D | N_Shore;Island | Body | chr15:42371511-42371967 | 456 | | | 18 | | | | | 0 | 7 | 1 | | 1 | | yes | | | yes | | |
| PLA2G4D | NA | Body | chr15:42376692-42376765 | 73 | | | | | | | | 0 | | | | | | | | | | | |
| PLA2G4D | NA | TSS200 | chr15:42386816-42387193 | 377 | | | | | | | | | 1 | | | | | yes | | | | | yes |
| PLA2G4F | NA | Body | chr15:42447989-42449015 | 1026 | | | | | | | | | 2 | 1 | | | | | yes | | | yes | |
| VPS39 | S_Shore | TSS200 | chr15:42500697-42500843 | 146 | | | 1 | | | | | | | | | 1 | | yes | | | yes | | |
| TMEM87A;GANC | Island | TSS1500;TSS200 | chr15:42566203-42566390 | 187 | | | | | | | | 0 | 4 | | | | | yes | yes | | | | |
| TGM5 | NA | Body;nan | chr15:43531947-43532243 | 296 | | | | | | | | | 1 | 1 | 1 | | | yes | | yes | | | |
| TGM5 | NA | TSS200 | chr15:43559170-43559486 | 316 | | | 2 | | | | | | | | | | | yes | | | | | |
| TUBGCP4;ZSCAN29 | N_Shore | TSS1500 | chr15:43662581-43662934 | 353 | | | | | | | | | | 1 | | | | | | | | yes | |
| CATSPER2 | Island | 5'UTR;1stExon | chr15:43941034-43941407 | 373 | | | | | | | | | | | | | | | | | | | yes |
| SERF2;MIR1282 | S_Shore | Body;3'UTR;TSS1500 | chr15:44085947-44087089 | 1142 | | | 16 | | | | | | | | | | | | | | | | |
| SERINC4;C15orf63 | N_Shore | TSS1500;TSS200 | chr15:44092606-44092937 | 331 | | | 1 | | | | | | | | | | | | yes | | | | |
| SPG11 | S_Shore | TSS1500 | chr15:44956098-44956192 | 94 | | | | | | | | | | | | | | | yes | | | | |
| PATL2 | NA | 5'UTR | chr15:44968935-44969244 | 309 | | | | | | | | | 1 | | | | | yes | | | | yes | |
| TRIM69 | NA | TSS1500 | chr15:45028083-45028270 | 187 | | | | 1 | 1 | | | | | | | | | | | | yes | yes | yes |
| C15orf43 | NA | TSS200 | chr15:45248744-45248878 | 134 | | | | | | | | | | | | | | yes | yes | | | | |
| SORD | N_Shore;Island | TSS1500;TSS200;5'UTR;1stExon | chr15:45314789-45315642 | 853 | | 1 | 28 | | | 1 | 1 | 0 | 3 | | | 6 | | yes | yes | | yes | yes | |
| DUOXA2;DUOXA1 | Island;S_Shore | Body | chr15:45408861-45410236 | 1375 | | | | | | | | | 1 | | | | | yes | | | | | |
| DUOX1;DUOXA1 | Island | TSS200;1stExon;5'UTR;TSS1500 | chr15:45421998-45422553 | 555 | | | 1 | 1 | 3 | | | | 2 | | | | | | | | | | |
| DUOX1 | N_Shore;Island | Body | chr15:45427034-45427734 | 700 | | | | | | | | 0 | | | | | | | | | | | |
| SHF | N_Shore | Body | chr15:45479462-45479755 | 293 | | | | | 1 | | | | | | | | | | | | | yes | yes |
| LOC145663;GATM | S_Shore;Island | Body;TSS1500;TSS200 | chr15:45670068-45671708 | 1640 | | 1 | 15 | 1 | 5 | | | 0 | 5 | | | | | yes | | | | | yes |
| SLC24A5 | NA | TSS200 | chr15:48413102-48413344 | 242 | | | | | | | | | 1 | | | | | | yes | | | | |
| SLC24A5 | NA | Body | chr15:48428985-48429067 | 82 | | | | | | | | | 1 | | | | | | yes | | | | yes |
| MYEF2 | S_Shore | TSS200 | chr15:48470678-48470973 | 295 | | | | | | | | | | | | | | | | | | | |
| CTXN2 | NA | 1stExon;5'UTR | chr15:48483948-48484302 | 354 | | 1 | | | | | | | | | | | | | | | | | |
| GALK2 | NA | TSS1500;Body | chr15:49462118-49462287 | 169 | | | | | | | | | 1 | | | | | yes | | | | yes | |
| C15orf33;FGF7 | NA | Body;TSS200;5'UTR | chr15:49715342-49716645 | 1303 | | | 4 | | | | | | | | | | | yes | | | | yes | |
| ATP8B4 | NA | TSS1500 | chr15:50412376-50412855 | 479 | | | | | | | | | 1 | | | | | yes | yes | | | yes | yes |
| SLC27A2 | N_Shore | TSS1500 | chr15:50474069-50474232 | 163 | | | 5 | | 1 | | | 0 | 2 | | | 2 | | yes | | | | | |
| HDC | NA | 5'UTR;1stExon | chr15:50557873-50558051 | 178 | | | | | | | | | | | | | | | | | | | |
| TNFAIP8L3 | NA | Body | chr15:51369862-51370019 | 157 | | | 10 | | | | | | | | | 6 | | | | yes | | | |
| TNFAIP8L3 | N_Shore;Island | Body | chr15:51385476-51386254 | 778 | | | 15 | | | | | | | | | | | yes | | | | | |
| TNFAIP8L3 | Island | Body | chr15:51387651-51387921 | 270 | | | 12 | | | | | | | | | | | yes | | | | | yes |
| GLDN | N_Shore | TSS1500 | chr15:51633430-51633821 | 391 | | | 6 | | | | | | | | | | | yes | | | | | |
| SCG3 | Island;N_Shore | 5'UTR;1stExon;TSS1500 | chr15:51972699-51973920 | 1221 | | | | | | | | | 2 | | | | | yes | | | | | |
| LYSMD2 | S_Shore | TSS200;5'UTR | chr15:52030519-52030746 | 227 | | | | | | | | | 1 | | | | | | yes | | | | |
| BCL2L10 | Island;S_Shore | 1stExon;TSS200 | chr15:52404463-52405162 | 699 | | | 8 | | | | | 0 | | | | | | yes | | yes | | | yes |
| GNB5 | NA | TSS1500 | chr15:52484443-52484920 | 477 | | | | | | | | | 1 | | | | | yes | yes | | | | yes |
| MIR1266;MYO5C | NA | TSS1500;Body | chr15:52570517-52570612 | 95 | | | 5 | | | | | | | | | | | yes | | | | | |
| KIAA1370 | N_Shore | 5'UTR | chr15:52970418-52971181 | 763 | | | | | | | | | 1 | | | | | | | | | | |
| ONECUT1 | N_Shore | Body | chr15:53075421-53075533 | 112 | | | 2 | | | | | | 1 | | | | | | | | | | |
| ONECUT1 | Island | Body | chr15:53077609-53078063 | 454 | | 1 | | | | | | | | | | | | | | | | | |
| ONECUT1 | Island | TSS1500 | chr15:53083238-53083978 | 740 | | | | | 2 | | | | | | | | | | | | | | |
| PIGB | Island | 5'UTR;1stExon | chr15:55611305-55611440 | 135 | | | | | | | | 0 | | | | | | yes | | | yes | | |
| PYGO1 | Island;S_Shore | TSS200;TSS1500 | chr15:55881075-55881706 | 631 | | | 18 | | | | | | 2 | | | | | yes | yes | | | yes | yes |
| PRTG | N_Shore | Body | chr15:56033606-56034245 | 639 | | | 4 | | | | | | | | | | | | | | | | yes |
| ZNF280D | Island | TSS200;TSS1500 | chr15:57025816-57026315 | 499 | | | 4 | | | | | | | | | 1 | | | | yes | | yes | yes |
| TCF12 | NA | TSS1500;Body | chr15:57510284-57510575 | 291 | | | | | | | | | 1 | | | | | | | | | | |
| LOC283663 | NA | TSS1500 | chr15:57592040-57592222 | 182 | | | | 1 | | | | | | | 1 | | | yes | | | | | |
| GCOM1 | N_Shore | TSS200 | chr15:57883962-57884357 | 395 | | | | | | | | | 1 | | | | | yes | yes | | | yes | yes |
| ALDH1A2 | Island | 5'UTR;1stExon;TSS1500 | chr15:58357874-58358879 | 1005 | | | 8 | | | | | | 1 | | | | | yes | | | | | yes |
| AQP9 | NA | TSS200 | chr15:58430391-58430682 | 291 | | | | | | | | | | | | | | | | | | | |
| LIPC | NA | TSS1500 | chr15:58723851-58724041 | 190 | | | | | 1 | | | | | | | | | | yes | | | | |
| LIPC | NA | Body | chr15:58844359-58844583 | 224 | | | 11 | 1 | | | | | | | | | | yes | | | | | yes |
| RNF111 | N_Shore | TSS1500 | chr15:59279406-59279556 | 150 | | | | | | | | | | | | | | yes | | | | yes | |
| GCNT3 | NA | 5'UTR | chr15:59908160-59908652 | 492 | | | | | | | | | | | | | | yes | | yes | | | |
| FOXB1 | N_Shore | TSS1500 | chr15:60295158-60295312 | 154 | | | | | | | | | 1 | | | | | | | | | yes | yes |
| FOXB1 | Island | 5'UTR;1stExon | chr15:60296809-60296996 | 187 | | | | | | | | | 1 | | | | | | | | | yes | |

| | | | | | | | | | | | | | | | | | | | | | | | |
|---|---|---|---|---|---|---|---|---|---|---|---|---|---|---|---|---|---|---|---|---|---|---|---|
| ANXA2 | N_Shore | TSS1500 | chr15:60690667-60690974 | 307 |  |  |  | 1 | 1 |  |  |  |  |  |  |  |  |  |  |  |  | yes | yes |
| RORA | S_Shore | TSS1500 | chr15:61521923-61521987 | 64 |  | 1 |  |  |  |  |  |  |  |  |  |  |  |  |  |  |  | yes |  |
| C2CD4A | N_Shore | TSS1500;TSS200 | chr15:62358531-62359275 | 744 |  |  | 21 |  |  |  |  |  |  |  |  |  |  | yes | yes |  |  |  |  |
| C2CD4B | Island | 5'UTR;1stExon;TSS200 | chr15:62457209-62457883 | 674 |  |  | 5 |  |  |  |  | 0 | 1 |  |  |  |  | yes | yes |  |  |  | yes |
| MIR190;TLN2 | NA | TSS1500;Body | chr15:63114897-63116022 | 1125 |  |  |  |  |  |  |  |  | 2 |  |  |  |  | yes | yes |  |  |  |  |
| TPM1 | N_Shore;Island | TSS1500;TSS200 | chr15:63333476-63334714 | 1238 |  |  |  |  |  |  |  |  | 2 |  |  |  |  | yes | yes |  |  |  | yes |
| TPM1 | chr15:63340256-63341464 | Island | chr15:63340515-63340702 | 187 |  |  |  |  |  |  |  |  |  |  | 1 |  |  | yes |  |  |  |  |  |
| RAB8B | S_Shore | Body | chr15:63483946-63504871 | 20925 |  |  |  |  |  |  |  |  |  |  |  |  |  | yes | yes |  | yes | yes | yes |
| APH1B | Island | TSS200 | chr15:63569620-63569686 | 66 |  |  |  |  |  |  |  |  | 1 |  |  |  |  |  |  |  | yes |  | yes |
| FBXL22 | N_Shelf | TSS1500 | chr15:63889093-63889406 | 313 |  |  |  |  |  |  |  |  |  |  |  |  |  |  |  |  |  | yes |  |
| SNX1 | N_Shore | TSS1500 | chr15:64387786-64387986 | 200 |  |  |  |  |  |  |  |  |  |  |  |  |  |  | yes |  |  |  |  |
| KIAA0101 | NA | 1stExon;5'UTR;Body | chr15:64673455-64673872 | 417 |  |  | 2 |  |  |  |  |  |  |  |  |  |  | yes |  |  |  |  |  |
| RBPMS2 | S_Shore | TSS1500 | chr15:65068554-65068885 | 331 |  |  | 1 |  |  |  |  |  |  |  |  |  |  | yes | yes |  |  |  |  |
| PIF1 | Island | Body | chr15:65116194-65116557 | 363 |  |  | 1 | 1 |  |  |  |  | 1 |  |  |  |  | yes |  |  |  | yes | yes |
| ANKDD1A | Island | Body | chr15:65204370-65204428 | 58 |  |  | 1 |  |  |  |  |  |  |  |  |  |  | yes |  |  |  |  | yes |
| OSTBETA | NA | 5'UTR | chr15:65341703-65342634 | 931 |  |  | 9 | 1 |  |  |  |  |  |  |  |  |  | yes | yes |  | yes | yes |  |
| RASL12 | Island | 5'UTR;1stExon | chr15:65360370-65360486 | 116 |  |  |  | 1 |  |  |  |  |  |  |  |  |  |  | yes |  |  |  |  |
| LOC390594 | N_Shore;Island | TSS200 | chr15:65368982-65369213 | 231 |  |  |  |  |  |  |  |  | 2 |  |  |  |  |  |  | yes |  |  |  |
| CILP | NA | 5'UTR;1stExon | chr15:65503447-65503889 | 442 |  |  |  |  |  |  |  |  |  |  |  |  |  |  |  |  |  |  |  |
| IGDCC3 | S_Shore | TSS1500 | chr15:65670863-65670881 | 18 |  |  | 2 |  |  |  |  | 0 |  |  |  |  |  | yes |  |  |  |  | yes |
| IGDCC4 | chr15:65688039-65688539;Isla | Island;Body | chr15:65688530-65689325 | 795 |  |  |  |  |  |  |  |  | 1 |  | 1 |  |  | yes | yes |  |  |  |  |
| IGDCC4 | S_Shore | TSS1500 | chr15:65715664-65715951 | 287 |  |  | 1 |  |  |  |  |  |  |  |  |  |  | yes | yes |  |  | yes | yes |
| MEGF11 | Island | Body | chr15:66274614-66274821 | 207 |  |  |  |  | 3 |  |  |  |  |  |  |  |  |  |  |  | yes | yes |  |
| DIS3L | N_Shore | TSS1500 | chr15:66584218-66584493 | 275 |  |  |  |  |  |  |  |  |  |  |  |  |  | yes |  |  |  | yes |  |
| SNORD16;RPL4;SNORD18B | NA | Body;TSS1500 | chr15:66795208-66795737 | 529 |  |  |  |  |  |  |  |  | 2 |  |  |  |  |  | yes |  |  |  | yes |
| LCTL | NA | TSS1500 | chr15:66858212-66858446 | 234 |  |  | 4 |  |  |  |  |  |  |  |  |  |  |  | yes |  |  |  |  |
| SMAD6 | S_Shelf | Body | chr15:66998926-66998998 | 72 |  |  | 2 |  |  |  |  |  |  |  |  |  |  | yes |  |  |  |  |  |
| SMAD6 | NA | Body;TSS200 | chr15:67000790-67000955 | 165 |  |  |  |  |  |  |  |  | 2 |  |  |  |  | yes | yes |  |  |  |  |
| SMAD3 | NA | Body;TSS1500;TSS200 | chr15:67417607-67418428 | 821 |  |  |  | 1 |  |  |  |  | 2 | 1 |  |  |  | yes | yes |  |  |  |  |
| SMAD3 | NA | Body;TSS1500 | chr15:67457584-67458522 | 938 |  |  | 3 |  |  |  |  |  |  |  |  |  |  |  | yes |  |  |  |  |
| IQCH;AAGAB | N_Shore | TSS1500;Body | chr15:67546365-67546624 | 259 |  |  |  |  |  |  |  |  |  |  |  |  |  | yes |  |  |  |  |  |
| C15orf61 | N_Shore | TSS1500 | chr15:67813132-67813145 | 13 | 1 |  | 1 |  |  |  |  |  |  |  |  |  |  | yes | yes |  |  |  |  |
| MAP2K5 | NA | Body | chr15:68098991-68099144 | 153 |  |  |  |  |  |  |  |  |  |  |  |  |  | yes |  |  |  |  |  |
| LBXCOR1 | Island | TSS1500;TSS200 | chr15:68117551-68118127 | 576 |  | 1 |  |  |  |  |  |  | 2 |  |  |  |  | yes |  |  |  |  | yes |
| LBXCOR1 | Island | Body | chr15:68120393-68121563 | 1170 |  | 1 |  |  |  |  |  |  | 1 |  |  |  |  |  |  |  |  | yes | yes |
| LBXCOR1 | N_Shore | Body | chr15:68125197-68125599 | 402 |  |  | 5 |  | 1 |  |  |  |  |  |  |  |  | yes |  |  |  |  | yes |
| CLN6 | S_Shore | TSS1500 | chr15:68522777-68523075 | 298 |  |  |  |  |  |  |  |  | 1 |  |  |  |  | yes |  |  |  |  |  |
| ITGA11 | Island | 1stExon | chr15:68724370-68724752 | 382 |  |  |  |  |  |  |  |  | 1 |  |  |  |  |  |  |  |  |  |  |
| CORO2B | NA | Body | chr15:68924114-68924220 | 106 |  |  |  |  |  |  |  |  | 1 |  |  |  |  | yes |  | yes |  |  |  |
| C15orf28;ANP32A | NA | TSS1500;Body | chr15:69099678-69099834 | 156 |  |  |  |  |  |  |  |  | 1 | 1 |  |  |  | yes |  |  |  |  |  |
| MIR548H4;SPESP1;NOX5 | Island | Body;TSS1500;5'UTR;1stExon | chr15:69222592-69223368 | 776 |  |  | 2 |  |  |  |  | 0 | 2 |  |  |  |  | yes | yes |  | yes |  | yes |
| MIR548H4;NOX5 | Island;N_Shore | Body | chr15:69323876-69324156 | 280 |  |  | 1 |  |  |  |  | 0 | 2 |  |  | 2 |  | yes | yes |  |  | yes |  |
| MIR548H4;NOX5 | N_Shore;Island | Body | chr15:69325271-69325560 | 289 |  |  | 1 |  |  |  |  | 0 | 4 |  |  | 2 |  | yes |  |  |  |  | yes |
| MIR548H4 | N_Shore;Island | Body | chr15:69366047-69366554 | 507 |  |  | 1 |  |  |  |  |  | 1 |  |  |  |  |  | yes | yes |  |  |  |
| RPLP1 | N_Shore | TSS1500 | chr15:69744390-69744850 | 460 |  |  | 15 |  |  |  |  | 0 | 3 | 2 |  | 3 |  | yes |  |  |  |  |  |
| LOC145837 | NA | TSS1500 | chr15:69853608-69854024 | 416 |  | 1 |  |  |  |  |  |  |  |  |  |  |  | yes | yes |  |  |  |  |
| TLE3 | NA | Body | chr15:70354647-70354825 | 178 | 7 |  | 16 |  |  |  |  |  |  |  |  |  |  |  | yes |  |  |  |  |
| MIR629;TLE3 | NA | TSS200;Body | chr15:70371992-70372614 | 622 |  |  |  |  | 1 |  |  |  |  |  |  |  |  | yes |  |  | yes | yes |  |
| UACA | NA | 5'UTR;Body;1stExon;TSS200 | chr15:70994435-70995014 | 579 |  |  | 25 | 1 |  |  |  |  |  |  |  |  |  |  |  |  |  |  |  |
| UACA | Island | 1stExon;5'UTR | chr15:71055835-71056115 | 280 |  |  |  |  |  |  |  |  | 1 |  |  |  |  |  |  |  |  |  |  |
| LRRC49;THAP10 | Island | TSS1500;1stExon | chr15:71184476-71184618 | 142 |  |  |  |  |  |  |  | 0 | 1 |  |  |  |  | yes |  |  |  |  |  |
| CT62 | S_Shore | TSS1500 | chr15:71408603-71408914 | 311 |  |  | 19 |  |  |  |  |  | 1 |  |  |  |  | yes |  |  |  |  |  |
| THSD4 | N_Shore | Body | chr15:71507300-71507695 | 395 |  |  |  |  |  |  |  | 0 |  |  |  |  |  | yes |  | yes | yes |  | yes |
| GRAMD2 | S_Shore | TSS1500 | chr15:72490380-72491032 | 652 |  |  |  |  |  |  |  |  | 1 |  |  |  |  |  |  |  |  |  | yes |
| PKM2 | N_Shelf | 5'UTR | chr15:72519739-72520804 | 1065 |  |  |  |  |  |  |  |  |  |  |  |  |  | yes |  |  |  |  |  |
| PARP6 | N_Shore;Island | TSS1500 | chr15:72564179-72565415 | 1236 |  |  | 5 |  | 1 |  |  | 0 | 1 |  |  |  |  | yes |  |  | yes |  | yes |
| C15orf60 | N_Shore;Island | TSS1500;TSS200 | chr15:73735098-73735531 | 433 |  |  | 10 |  |  |  |  |  |  |  |  |  |  | yes | yes |  |  |  |  |
| TBC1D21 | NA | TSS1500 | chr15:74164753-74164906 | 153 |  |  |  |  |  |  |  |  |  |  |  |  |  | yes | yes |  |  |  |  |
| LOXL1 | Island;N_Shore | TSS200;TSS1500 | chr15:74218418-74218921 | 503 | 2 |  |  |  |  |  |  | 0 | 4 |  |  |  |  |  |  |  |  |  |  |
| PML | Island | Body | chr15:74290466-74290733 | 267 |  |  |  |  |  |  |  |  | 1 | 1 |  |  |  |  | yes |  |  |  |  |
| LOC283731;ISLR2 | Island;chr15:74419870-744230 | Body;TSS1500;TSS200;Island;5'UT | chr15:74420796-74423077 | 2281 |  |  | 9 |  |  |  |  |  | 1 |  | 1 |  |  |  |  |  |  |  | yes |
| ISLR2 | chr15:74425091-74428821;Isla | Island;Body;5'UTR | chr15:74424640-74427499 | 2859 |  |  | 6 |  |  |  |  |  | 1 |  | 1 |  |  | yes |  |  |  |  | yes |
| ISLR | N_Shore;chr15:74467613-744€ | TSS200;TSS1500;1stExon;5'UTR;N_ | chr15:74465888-74467626 | 1738 |  |  | 1 |  |  |  |  |  | 1 |  | 1 |  |  | yes |  |  |  | yes |  |
| STRA6 | NA | 5'UTR | chr15:74494781-74495354 | 573 |  |  | 17 |  |  |  |  | 0 | 2 |  |  |  |  | yes |  |  |  |  | yes |
| STRA6 | NA | 1stExon;5'UTR;TSS200 | chr15:74501317-74501771 | 454 |  |  |  |  |  |  |  | 0 |  |  |  |  |  |  |  |  |  |  |  |
| CCDC33 | NA | TSS200 | chr15:74528522-74528719 | 197 |  |  |  | 1 |  |  |  |  |  | 1 |  |  |  | yes | yes |  |  |  |  |
| CCDC33 | NA | Body | chr15:74592566-74592786 | 220 |  |  | 3 |  |  |  |  | 0 | 1 |  |  |  |  |  |  |  |  |  | yes |
| CYP11A1 | S_Shore | 1stExon;TSS1500 | chr15:74659901-74660110 | 209 |  |  |  |  |  |  |  |  | 1 |  |  |  |  | yes |  |  | yes | yes |  |
| SEMA7A | Island | TSS200;TSS1500 | chr15:74726469-74726614 | 145 |  |  |  | 1 | 2 |  |  |  |  |  |  |  |  |  |  |  |  | yes |  |
| CYP1A1 | Island | TSS1500 | chr15:75019070-75019376 | 306 |  |  | 13 |  |  |  |  | 0 | 8 | 2 |  | 17 |  | yes | yes |  |  |  |  |
| CSK | S_Shore | 5'UTR | chr15:75076251-75081572 | 5321 |  |  |  |  |  |  |  |  |  |  |  |  |  | yes | yes |  |  | yes |  |
| LMAN1L | NA | TSS1500 | chr15:75104625-75104987 | 362 |  |  |  |  |  |  |  |  |  |  |  |  |  | yes | yes |  |  |  |  |
| SCAMP2 | NA | Body | chr15:75146514-75146663 | 149 |  |  |  |  |  |  |  |  | 1 |  |  |  |  |  |  |  |  |  |  |
| C15orf39 | Island | Body | chr15:75500859-75501277 | 418 | 2 |  |  |  |  |  |  |  |  |  |  |  |  |  | yes |  |  | yes |  |
| COMMD4 | Island | TSS200 | chr15:75628299-75628367 | 68 |  |  |  |  |  |  |  |  |  | 1 |  |  |  |  | yes |  |  |  |  |
| NEIL1 | N_Shore | TSS200 | chr15:75639163-75639433 | 270 |  |  |  |  |  |  |  |  |  |  |  |  |  | yes | yes |  |  |  |  |
| NEIL1;MIR631 | N_Shelf | Body;TSS200 | chr15:75646093-75646181 | 88 |  |  |  | 1 |  |  |  |  |  |  |  |  |  | yes |  |  |  | yes |  |
| CSPG4 | NA | TSS1500 | chr15:76005861-76006143 | 282 |  |  | 10 |  |  |  |  |  |  |  |  |  |  | yes | yes |  |  |  |  |
| C15orf27 | Island | Body | chr15:76484102-76484625 | 523 |  |  |  |  |  |  |  | 0 |  |  |  | 1 |  | yes |  |  |  |  |  |
| ISL2 | N_Shore | TSS1500 | chr15:76628385-76628531 | 146 |  |  | 1 |  |  |  |  |  |  |  |  |  |  |  |  |  |  | yes | yes |
| ISL2 | N_Shore;Island | Body | chr15:76629838-76630962 | 1124 |  | 1 |  |  |  |  |  |  | 1 |  |  | 1 |  |  |  |  |  |  | yes |
| ISL2 | Island | Body | chr15:76632287-76632813 | 526 |  |  |  |  |  |  |  |  |  |  |  |  |  |  |  |  |  |  |  |
| PSTPIP1 | NA | TSS1500 | chr15:77287243-77287899 | 656 |  |  |  |  | 1 |  |  |  |  |  |  |  |  | yes | yes |  |  | yes |  |
| PSTPIP1 | Island | Body | chr15:77320384-77320787 | 403 |  |  |  |  | 1 |  |  |  | 1 |  |  |  |  | yes |  |  |  |  |  |
| SGK269 | NA | TSS1500 | chr15:77577533-77577655 | 122 |  |  |  |  |  |  |  |  | 1 |  |  |  |  | yes | yes |  |  | yes |  |
| LINGO1 | N_Shore | Body | chr15:77924281-77924697 | 416 |  |  |  |  |  |  |  |  |  |  |  |  |  |  |  |  |  |  | yes |
| LOC91450 | NA | Body | chr15:78286521-78286905 | 384 |  |  |  | 1 |  |  |  |  |  |  |  |  |  | yes |  |  |  |  |  |
| SH2D7 | NA | TSS1500 | chr15:78384712-78384868 | 156 |  |  |  |  |  |  |  |  |  |  |  |  |  |  | yes |  |  |  |  |
| ACSBG1 | NA | 1stExon | chr15:78526804-78527275 | 471 |  |  |  |  |  |  |  |  | 1 |  |  |  |  | yes |  | yes |  |  |  |
| DNAJA4 | Island | TSS1500;TSS200;5'UTR;1stExon | chr15:78556423-78556534 | 111 |  | 1 | 1 |  |  |  |  |  |  |  |  |  |  |  |  |  |  |  | yes |
| CRABP1 | Island;N_Shore | 1stExon;5'UTR;Body;TSS1500;TSS2 | chr15:78631179-78633662 | 2483 |  |  | 11 |  |  |  |  | 0 | 3 |  |  |  |  | yes | yes |  | yes | yes | yes |
| CHRNA5 | N_Shore | TSS1500 | chr15:78856949-78857577 | 628 |  |  | 3 |  |  |  |  |  |  |  |  |  |  | yes |  |  |  |  | yes |
| CHRNA3 | Island;S_Shore | Body;TSS200;TSS1500 | chr15:78912658-78914439 | 1781 |  |  |  |  |  |  |  |  | 2 |  |  |  |  | yes | yes | yes |  | yes |  |
| CHRNB4 | Island | TSS1500 | chr15:78933807-78934280 | 473 |  |  |  |  |  |  |  |  |  |  |  | 1 |  |  |  |  |  |  |  |
| ADAMTS7 | Island | TSS1500 | chr15:79104124-79104481 | 357 |  |  |  |  |  |  |  |  |  |  |  |  |  |  | yes |  |  | yes | yes |
| MORF4L1 | N_Shore | TSS1500 | chr15:79164541-79164806 | 265 |  |  |  |  |  |  |  |  | 1 |  |  |  |  | yes |  |  |  |  | yes |
| RASGRF1 | NA | Body;5'UTR | chr15:79296377-79296585 | 208 |  |  |  |  |  |  |  |  | 1 |  |  |  |  | yes |  |  |  |  |  |
| MIR184 | NA | TSS200 | chr15:79502065-79502137 | 72 | 3 |  |  |  |  |  |  |  |  |  |  |  |  |  |  |  |  |  | yes |
| ANKRD34C | Island | 5'UTR | chr15:79576060-79576216 | 156 |  |  |  |  |  |  |  |  | 1 |  |  |  |  | yes |  |  |  |  |  |

| | | | | | | | | | | | | | | | | | | | | | | | |
|---|---|---|---|---|---|---|---|---|---|---|---|---|---|---|---|---|---|---|---|---|---|---|---|
| KIAA1024 | Island | 5'UTR | chr15:79725284-79725737 | 453 | | | | | | | | | | | | | | | | | |yes| |
| MTHFS | Island | TSS200 | chr15:80189496-80189782 | 286 | | | | | | | | |1| | | | | | | |yes| |yes|
| MIR549;KIAA1199 | NA | TSS200;5'UTR | chr15:81134479-81134721 | 242 | | | | | | | | |1| | | | |yes| | | | | |
| C15orf26 | N_Shore;Island | TSS1500;TSS200 | chr15:81426347-81426669 | 322 | | | | | | | |0|7|2| |10|1| | | | |yes| |
| IL16 | NA | TSS200 | chr15:81489075-81489338 | 263 | |1| | | | | | | | | | | |yes| | | | | |
| TMC3 | NA | 1stExon;TSS200 | chr15:81666393-81666847 | 454 | | |11| | | | | | | | | | |yes|yes| | | | |
| MEX3B | Island | Body | chr15:82335638-82336202 | 564 | | | | | | | | |1| | | | | | | | | |yes|
| CPEB1 | NA | 5'UTR;1stExon;Body;TSS200 | chr15:83240550-83240791 | 241 | | | | | | | |0|1| | | | | | | | | | |
| CPEB1 | Island | Body;TSS1500 | chr15:83315615-83317632 | 2017 | |1|7| | | | | |2| | | | | | | | | |yes|
| AP3B2 | NA | Body | chr15:83331441-83332065 | 624 | | | | | | | | | | | | | | | | | | | |
| WHAMM | N_Shore | TSS1500;TSS200 | chr15:83477608-83477965 | 357 | | |7| | | | |0|3| | |1| |yes|yes| | | | |
| FAM103A1 | N_Shore | TSS1500 | chr15:83654517-83654607 | 90 | | |2| | | | | |1| | | | | |yes| | | | |
| C15orf40 | S_Shore | TSS1500 | chr15:83680832-83681092 | 260 | | |1| | | | | |1| | | | | |yes| | | | |
| TM6SF1 | Island | 5'UTR;1stExon;Body | chr15:83776420-83777082 | 662 | | |3| |1| | | | | | | | |yes| | | | |yes|
| HDGFRP3 | N_Shore | Body | chr15:83875537-83876098 | 561 | | | | | | | | |1| | | | | | | | |yes|yes|
| BNC1 | Island;S_Shore | TSS1500 | chr15:83953690-83954392 | 702 | |1|30| | | | |0|6| | |1| |yes| | | |yes|yes|
| SH3GL3 | Island;N_Shore | TSS200;TSS1500 | chr15:84115433-84116920 | 1487 | |2|5| | | | | |1| | | | |yes|yes| | |yes|yes|
| ADAMTSL3 | N_Shore;Island | TSS1500 | chr15:84322277-84323154 | 877 | |1| | | | | | |2| | | | | | | | | |yes|
| NMB | S_Shore | TSS200 | chr15:85201891-85201913 | 22 | | | | | | | | | | | | | |yes|yes| | | |yes|
| ALPK3 | S_Shore;Island | 1stExon | chr15:85360217-85360691 | 474 | | |6| | | | |0|1| | |2| |yes|yes| | | |yes|
| AKAP13 | NA | Body | chr15:86233214-86233236 | 22 | |1| |1|1| | | | | | | | | | | | | | |
| KLHL25;MIR1276 | S_Shore | 5'UTR;TSS200 | chr15:86313372-86313973 | 601 |5| | |1| | | | |2|1| | | |yes| | | |yes| |
| KLHL25;MIR1276 | S_Shelf | 5'UTR;TSS1500 | chr15:86315115-86315211 | 96 | | |3| | |1| | | | | | | | | | | | | |
| NTRK3 | NA | Body | chr15:88576296-88576320 | 24 | | |38| | | | | | | | | | |yes|yes| | |yes| |
| NTRK3 | Island | TSS1500 | chr15:88800567-88801004 | 437 | |1| | | | | | | | | | | | | | | |yes|yes|
| MIR7-2 | NA | TSS1500 | chr15:89153667-89154196 | 529 | | |7|1|1| | |0|1| | | | |yes| | | | | |
| AEN | N_Shore | TSS1500 | chr15:89163928-89164106 | 178 | | | | | | | |0| | | | | |yes| | | | | |
| ISG20 | NA | TSS1500 | chr15:89181425-89181732 | 307 | | |11| | | | | | | | | | | |yes| | | | |
| ACAN | Island | TSS1500 | chr15:89346205-89346795 | 590 | | | | | | | | |1| | | | |yes| | | | |yes|
| ACAN | NA | Body | chr15:89386741-89386809 | 68 | |1| | | | | | | | | | | |yes| | | | | |
| HAPLN3 | S_Shore | TSS1500 | chr15:89440036-89440192 | 156 | | | | | | | | | | | | | | | | | |yes| |
| RLBP1 | NA | 1stExon;5'UTR;TSS200 | chr15:89764836-89765425 | 589 | | | | | | | | |1| | | | |yes|yes| |yes| | |
| FANCI | N_Shore | TSS1500 | chr15:89786188-89786622 | 434 | | | | | | | | | | | | | | |yes| | | | |
| POLG | S_Shore | TSS1500 | chr15:89879097-89879515 | 418 | | | | | | | | | | | | | |yes|yes| |yes| | |
| LOC254559 | N_Shore;Island | TSS1500;TSS200;Body | chr15:89920348-89922989 | 2641 | | |26| | |1| |0|1| | |2| |yes|yes| | | |yes|
| LOC254559 | N_Shelf | Body | chr15:89939252-89939709 | 457 | | | | | | | | | | | | | | |yes| | | | |
| RHCG | Island | Body | chr15:90030016-90030410 | 394 | | |5| |1| | |0|1| | |2| |yes| | | |yes| |
| RHCG | Island | TSS200 | chr15:90039805-90039822 | 17 | | | | | | | | |1| | | | | |yes| | |yes| |
| LOC283761 | NA | TSS1500 | chr15:90068044-90068496 | 452 | | | | | | | | |1| | | | |yes|yes|yes|yes| | |
| KIF7 | N_Shore | 3'UTR | chr15:90171643-90171768 | 125 | | | | | | | | | | | | | | | | | | | |
| KIF7 | N_Shore | Body | chr15:90190822-90191316 | 494 | | | | | | | | | | | | | |yes| | | | |yes|
| PEX11A;WDR93 | S_Shore | TSS1500;5'UTR | chr15:90234207-90234232 | 25 | | |1| | | | | | | | | | |yes| | | | | |
| MESP2 | N_Shore | TSS1500;TSS200 | chr15:90319155-90319516 | 361 | | |3| | | | | |2| | | | |yes| | | | | |
| ANPEP | NA | Body | chr15:90349511-90349979 | 468 | | | |1|1| | | | | | | | | | | | | | |
| C15orf38 | S_Shore | TSS1500 | chr15:90456541-90456686 | 145 | | |3| | | | | | | | | | |yes|yes| |yes|yes| |
| ZNF710 | N_Shore | TSS1500 | chr15:90543392-90543610 | 218 | | | | | | | | |1| | | | | |yes| | |yes|yes|
| ZNF710 | S_Shore | 5'UTR | chr15:90547692-90548043 | 351 | | | |1| | | | | |1| | | |yes|yes| | | | |
| ZNF710 | Island | Body | chr15:90610983-90611414 | 431 | | | | | | | | |1| | | | | | | | | | |
| SEMA4B | N_Shore | TSS1500 | chr15:90727560-90728020 | 460 |2| | | | | | | |2| | | | |yes| | | |yes| |
| SEMA4B | NA | Body | chr15:90755932-90756121 | 189 | | | | | | | | |1| | | | |yes| | | |yes| |
| TTLL13 | Island | TSS200 | chr15:90792609-90793056 | 447 | | | | | | | |0| | | | | |yes| | | | | |
| NGRN | N_Shore | TSS1500 | chr15:90808551-90808716 | 165 | | | | | | | | | | | |1| | | | | | | |
| FURIN | Island | Body | chr15:91423375-91423574 | 199 |1| | | | | | | | | | | | | | | |yes| | |
| FES | N_Shore;Island | TSS1500;TSS200;5'UTR | chr15:91427184-91428056 | 872 |1| |3| |1| | |0|2| | | | | | | | | |yes|
| MAN2A2 | S_Shore | TSS1500 | chr15:91447195-91447540 | 345 | | | | | | | | |1| | | | |yes| | | | | |
| UNC45A | Island;N_Shore | TSS200;5'UTR;1stExon;TSS1500 | chr15:91473059-91473569 | 510 | | |11|1| | | |0|3|1| |9| |yes|yes| | | | |
| HDDC3;UNC45A | N_Shore | TSS1500;5'UTR | chr15:91477073-91477210 | 137 | | | | | | | | | | | | | | | | | | | |
| RCCD1 | N_Shore | TSS1500 | chr15:91497863-91498497 | 634 | |1| | | | | | | | | | | |yes|yes| | | |yes|
| RCCD1 | Island | Body | chr15:91500781-91500893 | 112 | | |6| | | | | |1| | | | | | | | | | |
| SLCO3A1 | S_Shore | Body | chr15:92399594-92407388 | 7794 | | | | | | | | | | | | | |yes| | | |yes| |
| ST8SIA2 | N_Shore | TSS1500 | chr15:92936232-92936465 | 233 | | | | | | | | |1| | | | | | | | | |yes|
| ST8SIA2 | NA | Body | chr15:92944664-92944756 | 92 | | |9| | | | | |1| | |1| | | | |yes| | |
| FAM174B | S_Shore | TSS1500 | chr15:93199458-93199536 | 78 | | | | | | | |0| | | | | |yes| |yes| | | |
| RGMA | N_Shore | Body | chr15:93614758-93615146 | 388 | | | | | | | |0|3| | |3| | | | | | | |
| RGMA | Island;S_Shore | 1stExon;Body;5'UTR;TSS1500;TSS2 | chr15:93616894-93618028 | 1134 |1| |28| | | | |0| | | | | | | | | | |yes|
| RGMA | Island | TSS1500 | chr15:93632896-93633408 | 512 | | |25| | | | | | | | | | |yes|yes| | |yes|yes|
| MCTP2 | NA | TSS1500 | chr15:94840793-94841334 | 541 | | | |1|1| | | |2|1|1| | | | | |yes| |yes|
| NR2F2 | S_Shore;S_Shelf | TSS1500;TSS200 | chr15:96868114-96869220 | 1106 |1| |33|1|1| | | |1| | | | | | | | | | |
| NR2F2;MIR1469 | Island | TSS200;Body;TSS1500;1stExon;5'U | chr15:96874031-96877855 | 3824 | | | | | | | | |2| | |1| | | | | |yes|yes|
| LOC91948 | NA | TSS200 | chr15:98417731-98418214 | 483 | | | | | | | | | | | | | |yes| | |yes| |yes|
| ARRDC4 | Island | TSS200;1stExon;5'UTR | chr15:98503857-98504537 | 680 | | | | | | | |0|4| | |2| | | | | |yes| |
| FAM169B | NA | 5'UTR;1stExon | chr15:99057419-99058098 | 679 | | | |1|1| | | |1| | | | |yes| | | | | |
| IGF1R | NA | Body | chr15:99408636-99409957 | 1321 | | |20| | | |1|0|12|2| |3| | | | | |yes| |
| LOC145814 | NA | 5'UTR;1stExon | chr15:99548859-99549047 | 188 | | | | | | | | |1| | | | |yes| | | | | |
| SYNM | N_Shore;Island | TSS1500;TSS200 | chr15:99644973-99645663 | 690 | | | | | | | | |2| | | | |yes| | | |yes|yes|
| SYNM | NA | Body | chr15:99670386-99670731 | 345 | | | | | | | | |1| | | | | | | | | | |
| TTC23 | N_Shore | 1stExon;5'UTR | chr15:99789622-99789855 | 233 | | |3| | | | |0|5| | |12|1| | | | | | |
| ADAMTS17 | NA | Body | chr15:100516580-100516630 | 50 | | | | | | | | | | | | | | | |yes| | | |
| ADAMTS17 | Island | Body | chr15:100533202-100533253 | 51 | | | | | | | | | | | | | | |yes| | | | |
| ADAMTS17 | NA;S_Shelf | Body | chr15:100537304-100538145 | 841 | | |13| | | | | | | | | | |yes| | | | |yes|
| ADAMTS17 | NA | Body | chr15:100654734-100654807 | 73 | | |2| | | | |0|1| | | | | | | | |yes| |
| ADAMTS17 | NA | Body | chr15:100664651-100664844 | 193 | | | | | | | | | | | | | |yes|yes| | | | |
| ADAMTS17 | NA | Body | chr15:100672282-100672379 | 97 | | | | | | | |0|1| | |7| | | | | | | |
| ADAMTS17 | NA | Body | chr15:100804081-100804162 | 81 | | | | | | | | | | | | | |yes| | | | | |
| ADAMTS17 | N_Shore | Body | chr15:100831929-100832329 | 400 | | | | | | | | | | | | | | |yes| | | | |
| ADAMTS17 | N_Shore | Body | chr15:100879199-100879500 | 301 | | | | | | | | | | | |1| | | | |yes|yes| |
| ADAMTS17 | Island | Body | chr15:100881458-100882231 | 773 | | | | | | | | |1| | | | |yes| | | | |yes|
| FLJ42289 | N_Shore;Island;S_Shore | Body;TSS200;TSS1500 | chr15:100889683-100892205 | 2522 | | |6| | | | |0|2|2| | | | | |yes|yes| | |
| LASS3 | Island | 5'UTR;TSS200 | chr15:101084442-101085710 | 1268 |1| |7|1|1|2|3|0|9|3| |19|1|yes| | | | |yes|
| ALDH1A3 | Island;chr15:101419261-10142 | TSS1500;Body;N_Shore | chr15:101419011-101420989 | 1978 |2| |39| | | | | |4| |1| | | | | | | |yes|
| CHSY1 | NA | Body | chr15:101728611-101729526 | 915 | | |11| | | | | |1| | | | | | | |yes| | |
| CHSY1 | NA | Body | chr15:101738274-101738616 | 342 |1| |2| | | | | | | | | | | | | |yes| |yes|
| PCSK6 | NA | Body | chr15:101869493-101869560 | 67 | | |1| | | | | | | | | | | | | | | | |
| PCSK6 | NA | Body | chr15:101926307-101927126 | 819 | | |3| | | | | | | | | | | |yes| |yes| | |
| PCSK6 | NA | Body | chr15:101991512-101991886 | 374 | | |8| | | | |0|3| | |1| | | | | | | |
| PCSK6 | NA | Body | chr15:102009795-102010195 | 400 | | |3|1| | | | | | | | | | | | | | | |
| TM2D3 | Island | TSS1500 | chr15:102193228-102193416 | 188 | | |3| | | | | | | | | | |yes| | | | | |
| POLR3K;SNRNP25 | Island | 1stExon;TSS1500;TSS200 | chr16:103568-104183 | 615 | | | | | | | |0|3| | | | |yes|yes| |yes| | |
| RHBDF1 | S_Shore | TSS1500 | chr16:123246-123676 | 430 | | | | | | | | | | | | | | | | | |yes| |

| | | | | | | | | | | | | | | | | | | | | | | | |
|---|---|---|---|---|---|---|---|---|---|---|---|---|---|---|---|---|---|---|---|---|---|---|---|
| MPG | N_Shore | 5'UTR;TSS1500;1stExon | chr16:127072-127772 | 700 | | | 1 | 1 | | | | | | | | | | | | | | | yes |
| HBZ | N_Shore | TSS1500 | chr16:202311-202974 | 663 | | | | | | | | | 1 | | | | | yes | | | | | yes |
| HBM | Island | Body | chr16:216626-216757 | 131 | | | 1 | | | | | | | | | | | | | | | | |
| HBA1 | Island | Body | chr16:227203-227517 | 314 | | | | | | | | | 1 | | | | | | | | | | |
| HBQ1 | N_Shore | TSS1500 | chr16:229407-229528 | 121 | | | | | | | | | | | | | | yes | yes | | | yes | |
| ITFG3 | NA | 5'UTR | chr16:302852-303192 | 340 | | | | 1 | 2 | | | 0 | 2 | 2 | | 6 | | yes | yes | yes | | | yes |
| RGS11 | Island | 3'UTR | chr16:318694-318900 | 206 | | | | | | | | | | | | | | | | | | yes | |
| RGS11 | Island;S_Shore | Body;TSS1500 | chr16:325184-326728 | 1544 | | | | | | | | | | | | 1 | | yes | | | | yes | yes |
| ARHGDIG;PDIA2 | N_Shore | Body;TSS1500 | chr16:331727-331892 | 165 | | | 2 | | | | | | | | | | | | | | | | |
| PDIA2 | N_Shore | TSS200 | chr16:333104-333155 | 51 | | | 5 | | | | | | | | | | | | | | | | |
| AXIN1 | Island | Body | chr16:374924-375327 | 403 | | | | | 1 | | | | | | 1 | | | | | | | | |
| MRPL28 | N_Shore;Island | Body | chr16:419800-420230 | 430 | 4 | 2 | 10 | 1 | | 1 | 1 | 0 | 5 | 3 | 1 | 14 | | | | | | | yes |
| TMEM8A | Island | Body | chr16:427684-427912 | 228 | 2 | | | | | | | | | | | | | | | | | | yes |
| TMEM8A;LOC100134368 | S_Shore | TSS1500;Body | chr16:433439-434356 | 917 | | | | 1 | | | | 0 | 1 | | | | | yes | | | | | yes |
| DECR2;NME4 | N_Shore;Island | TSS1500;3'UTR | chr16:450562-451495 | 933 | | | 9 | | | | | 0 | 1 | | | 1 | | yes | | | | | yes |
| RAB11FIP3 | N_Shore | TSS1500 | chr16:474271-474528 | 257 | | | | | | | | 0 | 3 | 2 | 1 | | | yes | | | | yes | |
| RAB11FIP3 | NA | Body | chr16:570765-571091 | 326 | | | | | | | | | 1 | | | | | | | | | | |
| C16orf11 | N_Shore | 5'UTR | chr16:613006-613526 | 520 | | 1 | | | 1 | | | | | | | | | | | | yes | yes | |
| C16orf11;NHLRC4 | N_Shore;Island;S_Shore | Body;3'UTR;TSS1500 | chr16:614645-616935 | 2290 | | | | | | | | | | | | | | | | | | | |
| PIGQ | N_Shore;Island | 3'UTR | chr16:633722-634271 | 549 | | | 6 | | | | | 0 | 1 | | | 6 | | | | | | | |
| RAB40C | S_Shore;N_Shore | Body | chr16:671905-673115 | 1210 | | | | | | | | | | | | | | yes | yes | | | | |
| RAB40C | Island | Body;3'UTR | chr16:677526-678050 | 524 | | | 2 | | | | | | | | | | | | | | | | |
| RAB40C;WFIKKN1 | S_Shore;N_Shore | 3'UTR;TSS1500;TSS200 | chr16:679189-681022 | 1833 | | | | | | | | | | | | | | | | | | | |
| C16orf13 | Island | 3'UTR | chr16:684431-684669 | 238 | | | | | | | | | | | | | | | | | | | |
| FAM195A | N_Shore | TSS1500 | chr16:691444-691570 | 126 | | | 1 | | | | | | | | | | | | | | | | yes |
| WDR90 | Island | TSS200 | chr16:699218-699355 | 137 | | | | | | | | 0 | | | | | | | | | | | |
| WDR90 | N_Shore | Body | chr16:710674-711235 | 561 | | | 9 | | | | | 0 | 1 | | | | | | | | | | yes |
| RHOT2;WDR90 | N_Shore | TSS1500;Body | chr16:717419-717707 | 288 | | | | | | | | | 2 | 1 | | | | yes | | | | | yes |
| FBXL16 | N_Shore;Island;S_Shore | 3'UTR;Body | chr16:743338-747310 | 3972 | | | 11 | | | | | 0 | 5 | 3 | 1 | 4 | | yes | | | yes | yes | yes |
| FBXL16 | Island | 5'UTR | chr16:751056-751450 | 394 | | | | | | | | 0 | | | | 1 | | | | | | | yes |
| NARFL | S_Shore | Body | chr16:787799-788184 | 385 | | | | | 1 | | | 0 | 3 | | | 11 | | yes | | | | | |
| MSLN | NA | TSS1500;TSS200 | chr16:809679-811744 | 2065 | | 1 | 7 | | | | | | 3 | | | | | yes | | yes | | | |
| MSLN | NA | Body | chr16:815553-816149 | 596 | | | 2 | | | | | | | | | | | yes | | | | | |
| MSLNL;MIR662 | NA | Body;TSS200 | chr16:820122-820212 | 90 | | | | | | | | | | | | | | | | | | yes | |
| MSLNL | Island;S_Shore | Body;TSS200 | chr16:832828-833250 | 422 | | | 17 | | | | | | 4 | | 1 | | | | | | | | |
| GNG13 | Island;S_Shore | 5'UTR;TSS1500 | chr16:850371-851298 | 927 | | | | | | | | | 2 | | | 2 | | | | | | | |
| PRR25 | N_Shelf | TSS1500 | chr16:854168-854640 | 472 | | | 2 | | | | | | 1 | | | | | yes | yes | | | | |
| PRR25 | Island | Body | chr16:857454-858499 | 1045 | 2 | 2 | 36 | | 1 | 2 | 1 | 0 | 3 | 5 | | 15 | 2 | yes | | yes | | | |
| LMF1 | S_Shore | Body | chr16:928778-929020 | 242 | | | | | | | | | | | | | | | yes | | | | yes |
| LMF1 | NA | Body | chr16:936993-937477 | 484 | | | | | | | | | | | | | | | | | | | |
| LMF1 | Island | Body | chr16:962818-962909 | 91 | | | | | 1 | | | | | | | | | | yes | | | | |
| LMF1 | N_Shore;Island;S_Shore | Body | chr16:966593-968682 | 2089 | | 1 | 4 | | | | | | | | | | | | | | | | |
| LMF1 | Island;N_Shore | Body | chr16:971186-972115 | 929 | | | 10 | | | | | | | | | | | | yes | | | | |
| LMF1 | Island | Body | chr16:979488-979898 | 410 | | | | | | | | 0 | 2 | | | 1 | | yes | | | | | |
| LMF1 | S_Shore | Body | chr16:981631-982031 | 400 | | 1 | | | | | | | | | | | | yes | | | yes | | |
| LMF1 | N_Shore | Body | chr16:1014592-1014935 | 343 | | | | | | | | | 1 | | | | | yes | | | | | |
| SOX8 | Island | TSS1500;1stExon | chr16:1030383-1032491 | 2108 | | 2 | 8 | | 1 | | | | | | | | | | | | | yes | yes |
| LOC146336 | Island | Body | chr16:1116010-1116361 | 351 | | | | | | | | | | | | | | | yes | yes | | | |
| SSTR5;LOC146336 | Island | TSS200;Body | chr16:1128689-1128865 | 176 | | | 15 | | | 1 | | | 2 | | | | | | | | | | |
| SSTR5;LOC146336 | Island | 3'UTR;TSS1500;1stExon | chr16:1129990-1130449 | 459 | | | | | | | | | | | | | | | | | | | |
| C1QTNF8 | N_Shore | 3'UTR | chr16:1138344-1139166 | 822 | | 1 | 4 | | 1 | | | | 1 | | | 1 | | yes | yes | | | | |
| C1QTNF8 | Island | Body | chr16:1143566-1143887 | 321 | | | 4 | | | | | | | | | | | yes | | | | yes | |
| CACNA1H | Island;S_Shore | Body | chr16:1204617-1205342 | 725 | | 1 | 1 | 1 | | | | | | | | | | | | | | | |
| CACNA1H | N_Shore | Body | chr16:1206750-1207075 | 325 | | | 16 | | | | | | 1 | | | | | | | | | | |
| CACNA1H | Island | Body | chr16:1209990-1210484 | 494 | | | | | | | | | | | | | | | | | | | |
| CACNA1H | Island | Body | chr16:1217171-1217857 | 686 | | | | | | | | | | | | | | | yes | | | | yes |
| CACNA1H | Island;N_Shore | Body | chr16:1219430-1220524 | 1094 | | | | | | | | | | | | | | | | | | yes | |
| CACNA1H | Island | Body | chr16:1221943-1222626 | 683 | | | 5 | | | | | | | | | | | | | | | | |
| CACNA1H | S_Shore | Body | chr16:1230997-1231464 | 467 | | | | | | | | | | | | 1 | | | | | | yes | |
| CACNA1H | S_Shore | Body | chr16:1245911-1246033 | 122 | | | | | | | | | | | | | | | | | | | |
| CACNA1H | N_Shore | Body | chr16:1248836-1249204 | 368 | | | | | | | | | 1 | | | | | | yes | | | | |
| CACNA1H | N_Shore | Body | chr16:1267474-1267876 | 402 | | | | | | | | | | | | | | | | | | | |
| CACNA1H | S_Shelf | 3'UTR | chr16:1271505-1271998 | 493 | | | | | | | | | | | | | | | | | | | |
| TPSG1 | NA | TSS200;TSS1500 | chr16:1275450-1276278 | 828 | | | 4 | | | | | | | | | | | | | | | | yes |
| TPSD1 | N_Shore | Body | chr16:1307050-1308233 | 1183 | | | | | | | | | | | | | | yes | | | | | yes |
| UBE2I | S_Shore | Body | chr16:1365877-1366304 | 427 | | | 2 | | | | | | | | | | | yes | yes | | | | |
| UBE2I | NA | Body | chr16:1373608-1373786 | 178 | | | | | | | | | | | | | | yes | | | | | |
| BAIAP3 | Island | Body | chr16:1393584-1394089 | 505 | | | 3 | | | | | 0 | | | | | | | | | | | yes |
| BAIAP3 | N_Shore | Body | chr16:1396337-1396608 | 271 | | | | | | | | | | | | | | | | | | | |
| UNKL | Island | 3'UTR | chr16:1414422-1414718 | 296 | | | | | | | | | 1 | | | | | yes | | | | | |
| UNKL | Island | TSS200 | chr16:1429863-1430103 | 240 | | | 5 | | | | | | | | | | | | | | | yes | yes |
| C16orf91 | N_Shore | Body | chr16:1476899-1477230 | 331 | | | | | | | | | | | | | | | | yes | | | |
| C16orf91 | N_Shore | TSS1500 | chr16:1480453-1480992 | 539 | | | 5 | | | | | | 2 | | | | | | | yes | | | |
| CCDC154 | Island;N_Shore | Body;TSS200 | chr16:1493540-1494876 | 1336 | | | 6 | | | | | | | | | | | | | | yes | | |
| CLCN7 | S_Shelf | Body | chr16:1518249-1518751 | 502 | | | 2 | | | | | | | | | | | yes | yes | | | | |
| CLCN7 | N_Shelf | Body | chr16:1521617-1521676 | 59 | | | | | | | | 0 | | | | 1 | | yes | | | | | |
| CLCN7 | N_Shore | Body | chr16:1524174-1524521 | 347 | | | 2 | | | | | | | | | | | yes | | | | | yes |
| C16orf38 | S_Shore | 1stExon | chr16:1538347-1538740 | 393 | | | | | | | | | | | | | | | yes | | | | yes |
| TELO2 | S_Shore | Body | chr16:1545088-1545405 | 317 | | | | | | | | | 1 | | | 1 | | | yes | | yes | | |
| TELO2 | N_Shelf | Body | chr16:1557366-1557733 | 367 | | | 1 | | | | | | | | | | | yes | | | | yes | |
| IFT140 | S_Shore | Body | chr16:1575979-1576146 | 167 | | | | | 1 | | | | | | | | | | | | | | |
| IFT140 | NA | Body | chr16:1579439-1579962 | 523 | | | | | | | | | 1 | | | | | yes | | | | | |
| IFT140 | N_Shore | Body | chr16:1581834-1582581 | 747 | | | 17 | | | | | 0 | 1 | 1 | 1 | | | | | | | | |
| IFT140;TMEM204 | Island | Body;TSS1500;TSS200 | chr16:1583810-1584516 | 706 | | 1 | 3 | | 1 | | | 0 | 6 | 5 | 1 | 11 | 1 | yes | | | | | |
| IFT140;TMEM204 | S_Shore | Body | chr16:1585644-1586043 | 399 | | | 2 | | | | | | 1 | | | | | yes | | | | | |
| IFT140;TMEM204 | Island | Body | chr16:1587684-1587842 | 158 | | | | | | | | | 1 | | | | | | | | | | |
| IFT140;TMEM204 | S_Shore | Body | chr16:1589653-1590085 | 432 | | | | | | | | | | | | | | | | | | yes | |
| IFT140;TMEM204 | S_Shore | Body | chr16:1593178-1593766 | 588 | | | 3 | | | | | | | | | | | yes | | | | | |
| IFT140 | S_Shelf | Body | chr16:1608794-1608904 | 110 | | | 1 | | | | | | | | | | | | yes | | | | |
| IFT140 | NA | Body | chr16:1630488-1630880 | 392 | | 1 | | | | | | | | | | | | yes | | yes | yes | | |
| MAPK8IP3 | NA | Body | chr16:1764111-1764432 | 321 | | | | | 1 | | | | 1 | | | | | | yes | | | | |
| MAPK8IP3 | NA | Body | chr16:1782523-1782905 | 382 | | | | | | | | | 1 | | | | | | | | | | |
| MAPK8IP3 | Island | Body | chr16:1790400-1790700 | 300 | 4 | | | | | | | | | | | | | | | | | yes | |
| MAPK8IP3 | N_Shore | Body | chr16:1813796-1814481 | 685 | | | | | | | | | 1 | | | | | | | | | yes | |
| SPSB3 | N_Shore | 3'UTR | chr16:1826832-1827257 | 425 | | | | | | | | | | | | | | | | | | | |
| IGFALS | Island | Body | chr16:1841732-1842146 | 414 | | | | 1 | | | | | | | | | | | | | | | |
| IGFALS | S_Shore;S_Shelf | Body;5'UTR;1stExon;TSS200;TSS1 | chr16:1843424-1845460 | 2036 | | | 16 | 1 | 1 | 1 | 1 | | 2 | 4 | | | | yes | | | | | |
| HAGH | N_Shore | Body | chr16:1869043-1869611 | 568 | 2 | | | | | | | | | | | | | | | | | | |
| C16orf73 | Island | TSS200;TSS1500 | chr16:1922207-1922425 | 218 | | 1 | | | | | | | 2 | | | 1 | | yes | | | | | |

| | | | | | | | | | | | | | | | | | | | | | | | |
|---|---|---|---|---|---|---|---|---|---|---|---|---|---|---|---|---|---|---|---|---|---|---|---|
| HS3ST6 | Island | Body | chr16:1961953-1962215 | 262 | | | | | | | | | 1 | | | | | | | | | | |
| HS3ST6 | Island | TSS1500 | chr16:1968576-1968598 | 22 | | | | | | | | | 1 | | | | | | | | | yes | yes |
| SEPX1 | Island | TSS200 | chr16:1993336-1993412 | 76 | | | | | | | | | 1 | | | | | yes | | yes | | | |
| RPL3L | NA | TSS200 | chr16:2004686-2005062 | 376 | | | | | | | | | 1 | | | | | yes | | | | | |
| SNORA64;SNORA10;RPS2 | N_Shore | Body;TSS1500 | chr16:2013058-2013573 | 515 | | | | | | | | | 1 | | | | | yes | | | | | |
| NOXO1 | S_Shore | Body | chr16:2030892-2031217 | 325 | | | | | | | | | | | | | | yes | | | | | |
| SYNGR3 | N_Shore | TSS1500 | chr16:2039030-2039411 | 381 | | | | | | | | | | | | | | yes | yes | | | | yes |
| ZNF598 | Island | Body | chr16:2048236-2048574 | 338 | | 1 | 3 | | | | | | | | | | | | | | | | |
| ZNF598 | S_Shore | Body | chr16:2054944-2055684 | 740 | | | 3 | | | | | | | | | | | | | | | | |
| SLC9A3R2 | N_Shelf | Body | chr16:2082689-2083168 | 479 | | | 17 | | | | | | | | | | | yes | | | | | |
| NTHL1 | N_Shelf | Body | chr16:2094602-2094700 | 98 | | | | | | | | | | | | | | yes | | | | | |
| TSC2 | Island | Body | chr16:2114026-2114243 | 217 | | | | | | | | | | | | | | | | | | yes | |
| TSC2 | Island | Body | chr16:2128872-2129354 | 482 | | | | | | | | | 1 | | | | | | | | | | |
| TSC2 | N_Shelf;N_Shore | Body | chr16:2131718-2132723 | 1005 | | | 2 | | | | | | 2 | 1 | | | | | | | | | |
| TSC2 | Island | Body | chr16:2134631-2135281 | 650 | 1 | | | | | | | | | | | | | | yes | | | | |
| PKD1 | S_Shelf | Body | chr16:2145598-2145820 | 222 | 1 | | | | | | | | | | | | | yes | yes | | yes | | |
| MLST8 | S_Shore | Body | chr16:2256384-2256716 | 332 | | | 1 | | | | | | | | | | | yes | | yes | | | |
| C16orf79 | N_Shelf | 1stExon;5'UTR;TSS200 | chr16:2260806-2261168 | 362 | | | | | | | | | | | | | | yes | yes | | | yes | |
| DNASE1L2;E4F1 | Island | TSS1500;Body | chr16:2285495-2285775 | 280 | 2 | | | | | | | | | | | | | | | | | | |
| DNASE1L2 | Island | 5'UTR | chr16:2286786-2287294 | 508 | | | | | | | | 0 | | | | | | | | | | | yes |
| ABCA3 | Island | Body | chr16:2334163-2334596 | 433 | | | | | | | | | 1 | 1 | | | | | | yes | | | |
| TBC1D24 | Island | Body | chr16:2546265-2546596 | 331 | | | | | | | | | 1 | | | | | yes | | | | | |
| CEMP1 | N_Shore | 1stExon;5'UTR | chr16:2581122-2581570 | 448 | | | | | | | | | | | | | | | yes | | | | |
| LOC652276 | Island | TSS200;Body | chr16:2653222-2653987 | 765 | 1 | | 3 | | 1 | | 1 | 0 | 3 | 1 | | 4 | 1 | | | | | | |
| KCTD5 | NA | Body | chr16:2737220-2737356 | 136 | | | | | | | | | 1 | 1 | | | | | | | yes | | |
| KCTD5 | NA | Body | chr16:2740348-2740814 | 466 | | | | | | | | | 1 | | | | | yes | | | | | |
| PRSS27 | Island | Body | chr16:2765620-2766247 | 627 | | | 1 | | | | | 0 | | | | | | | | | | | |
| LOC100128788;SRRM2 | N_Shore | Body;TSS1500 | chr16:2801092-2801952 | 860 | | | 3 | | | | | | 2 | | | | | yes | yes | | | | |
| TESSP1 | Island;N_Shore | Body;TSS1500;TSS200;1stExon | chr16:2848223-2849144 | 921 | | | 13 | | | | | 0 | 7 | | | 1 | | yes | yes | | | yes | |
| PRSS21 | N_Shore;Island | TSS1500;TSS200;Body | chr16:2866901-2868001 | 1100 | | | 18 | | | | 2 | | 7 | 3 | | 6 | | yes | yes | | | | yes |
| ZG16B | NA | TSS1500;TSS200 | chr16:2879544-2880326 | 782 | | | | | | | | | | | | | | yes | | | | | |
| TMPRSS8 | chr16:2892579-2892810;Island | Island;TSS200;Body | chr16:2892518-2892912 | 394 | | | 20 | | | | | 0 | 3 | 1 | 2 | 1 | | yes | yes | | | | |
| PRSS22 | Island | 3'UTR;Body | chr16:2902830-2903392 | 562 | | | 4 | | | | | | | | | | | | | | | | |
| PRSS22 | N_Shore;Island;S_Shore | Body;1stExon;5'UTR;TSS200;TSS1 | chr16:2907517-2908934 | 1417 | | | 36 | | | | | 0 | 11 | 2 | | 8 | | yes | | | | | yes |
| FLYWCH2 | Island | TSS200 | chr16:2932999-2933047 | 48 | | | | | | | | 0 | | | | | | | | | | | |
| KREMEN2;PAQR4 | Island | Body;TSS1500;3'UTR | chr16:3017593-3018470 | 877 | | | 4 | | | | | | 2 | | | 1 | | | | | | yes | |
| CLDN9 | N_Shore;chr16:3063679-3063 | 1stExon;5'UTR;Island;TSS200 | chr16:3062349-3063894 | 1545 | | | | 1 | | 1 | | 0 | 10 | | 1 | 4 | | | | | | | |
| CLDN6 | N_Shore;Island | 5'UTR;TSS200;TSS1500 | chr16:3067500-3068707 | 1207 | | 1 | 7 | | | | | | | | | | | yes | | | | | |
| CCDC64B | N_Shore;Island | Body | chr16:3079423-3079953 | 530 | | 1 | | 1 | 1 | | | 0 | 3 | | | 1 | | yes | yes | | | | |
| CCDC64B | Island | 1stExon | chr16:3085246-3085571 | 325 | | | | | | | | | | | | | | yes | | | | yes | |
| IL32 | NA | TSS1500;TSS200;5'UTR | chr16:3115053-3115809 | 756 | | | 4 | | | | | | 1 | | | | | yes | | yes | yes | yes | |
| ZSCAN10 | S_Shelf | 1stExon;TSS200 | chr16:3142645-3143041 | 396 | | | 8 | | | | | | | | | | | yes | | | | | |
| OR1F2P | NA | TSS1500 | chr16:3265228-3265467 | 239 | | | 1 | | | | | | | | | | | yes | | | | | |
| MEFV | S_Shore | TSS200 | chr16:3306685-3307037 | 352 | | | | | | | | | | | | | | yes | yes | | | | |
| ZNF75A | Island | 5'UTR;1stExon | chr16:3355553-3356081 | 528 | | | | | | | | 0 | 1 | | | | 1 | yes | | | | | |
| ZNF597;NAT15 | Island;chr16:3493098-3493569 | 5'UTR;TSS1500;1stExon;Island | chr16:3493336-3494155 | 819 | | | | | | | | 0 | 2 | | 1 | 1 | | yes | | yes | | | |
| NLRC3 | NA | 1stExon;5'UTR | chr16:3627243-3628416 | 1173 | | | | | | | | | 1 | | | | | yes | | | | | yes |
| TRAP1 | Island | Body | chr16:3724734-3724848 | 114 | | | 1 | | | | | | | | | | | | yes | | | | |
| CREBBP | S_Shelf | Body | chr16:3784620-3784955 | 335 | | | | | | | | | 1 | | | | | yes | | | | | |
| CREBBP | NA | Body | chr16:3843012-3843415 | 403 | | | 3 | | | | | | | | | | | | | | yes | | |
| ADCY9 | N_Shore | 3'UTR | chr16:4013337-4014161 | 824 | | | | | | | | | 2 | | | | | | | | | yes | |
| ADCY9 | NA | Body | chr16:4102293-4103533 | 1240 | | | 9 | 1 | | | | | 2 | | | | | | yes | yes | | | |
| ADCY9 | N_Shore | Body | chr16:4163819-4164087 | 268 | | | 6 | | | | | | | | | | | yes | | | | | |
| TFAP4 | N_Shelf | 3'UTR | chr16:4307906-4308192 | 286 | | | 2 | | | | | | | | | | | yes | | | | | |
| TFAP4 | N_Shore;Island | Body | chr16:4311628-4311941 | 313 | | | 9 | | | | | | | | | | | yes | yes | | | | |
| VASN;CORO7 | N_Shore | TSS1500;Body | chr16:4421603-4422228 | 625 | | 1 | | | | | | | 1 | | | | | yes | | | yes | | yes |
| FAM100A | Island | TSS1500 | chr16:4665298-4665704 | 406 | | | 4 | | | | | | | | | | | | yes | | | | yes |
| MGRN1 | NA | Body | chr16:4689950-4690020 | 70 | | | 17 | | | | | | | | | | | | | | | | |
| MGRN1 | NA | Body | chr16:4714080-4714815 | 735 | 6 | | | 1 | | | | 0 | 2 | | | | | | yes | | | | |
| MGRN1 | N_Shore | Body | chr16:4729905-4730657 | 752 | | | 4 | | | | | 0 | 3 | 2 | | 1 | | | | | yes | | |
| ANKS3 | NA | Body | chr16:4751267-4751569 | 302 | | | | | | | | | 1 | | | | | | yes | | | | |
| SEPT12;LOC440335 | NA | TSS200;Body | chr16:4838559-4838722 | 163 | | 1 | 4 | | | | | | | | | | | yes | | | yes | | |
| A2BP1 | N_Shore | TSS1500 | chr16:6068831-6070292 | 1461 | | 1 | | | | | | | 1 | | | | | yes | | | | yes | |
| A2BP1 | N_Shore;Island | 5'UTR | chr16:6532764-6533789 | 1025 | | | | | | | | | 3 | | | | | | | | | | yes |
| A2BP1 | NA | 5'UTR | chr16:6623995-6624242 | 247 | | | | | | | | | | | | | | yes | | | | | |
| A2BP1 | NA | Body;TSS1500 | chr16:7382187-7382864 | 677 | | | 2 | | | | | | 3 | | | | | yes | | | | yes | yes |
| A2BP1 | NA | Body | chr16:7615516-7615568 | 52 | | | | | | | | | | | | | | | yes | | | | |
| A2BP1 | Island | Body | chr16:7703812-7703947 | 135 | | | 1 | | | | | | 1 | | | 1 | | yes | | | | | yes |
| TMEM114 | N_Shore | 3'UTR | chr16:8619531-8619943 | 412 | | | 5 | | | | | 0 | 1 | | | | | yes | yes | | yes | | |
| C16orf68 | NA | Body | chr16:8738240-8738608 | 368 | | | | | | | | 0 | 2 | 1 | 1 | | | yes | yes | | | | |
| ABAT | NA | 5'UTR;TSS200;1stExon;TSS1500 | chr16:8806359-8807043 | 684 | | | 19 | 2 | | | | 0 | 6 | 1 | | 3 | 1 | yes | yes | | | | |
| PMM2 | NA | 3'UTR | chr16:8942340-8942906 | 566 | | | | | | | | | | | | | | | | | | | |
| USP7 | NA | Body | chr16:8995607-8995926 | 319 | | | 1 | | | | | | | | | | | | | | yes | yes | |
| GRIN2A | NA | Body;3'UTR | chr16:9857151-9857804 | 653 | | | 5 | | | | | | | | | | | | | | | | |
| GRIN2A | Island | 5'UTR | chr16:10274315-10274415 | 100 | | | | | | | | | | | | | | | | | | | |
| GRIN2A | Island | 5'UTR;TSS200;1stExon;TSS1500 | chr16:10276092-10276799 | 707 | | 2 | | | | | | | | | | | | | | | | | yes |
| EMP2 | NA | 5'UTR | chr16:10647084-10647275 | 191 | | | | 1 | 1 | | | 0 | 1 | 1 | | 1 | | | yes | yes | yes | | |
| EMP2 | Island | 5'UTR | chr16:10674326-10674457 | 131 | | | | | | | | | 1 | | | | | yes | | | | | yes |
| FAM18A | Island | Body;1stExon;5'UTR;TSS200 | chr16:10912331-10912718 | 387 | | | 7 | | | | 1 | 0 | 2 | | | | | yes | yes | | yes | yes | |
| CIITA | N_Shore | 1stExon;5'UTR | chr16:10971105-10971250 | 145 | | | | | | | | | | | | | | | | | | | |
| CIITA | Island | Body | chr16:10972898-10973382 | 484 | | | 1 | | | | | 0 | | | | | | yes | | | | | |
| CIITA | Island | Body | chr16:11001757-11001871 | 114 | 1 | | | | | | | | | | | | | yes | yes | | | | |
| CLEC16A | NA | Body | chr16:11219954-11220376 | 422 | | | | | | | | | 1 | | | | | | | | | | |
| SOCS1 | Island | 3'UTR | chr16:11348611-11348684 | 73 | | | | | | | | | 1 | | | | | | | | | | yes |
| PRM1 | NA | 3'UTR | chr16:11374723-11375333 | 610 | | | 1 | | | | | | | | | | | yes | yes | | | | yes |
| LITAF | S_Shore | TSS200;TSS1500 | chr16:11681482-11681599 | 117 | | | | | | | | | 1 | | | | | yes | | | | | |
| SNN | N_Shore | TSS1500 | chr16:11761803-11761964 | 161 | | | 1 | | | | | | | | | | | yes | yes | | | yes | yes |
| TXNDC11 | N_Shore | Body | chr16:11834960-11835562 | 602 | | | 3 | | | | | | | | | | | yes | yes | | yes | | |
| ZC3H7A | NA | Body;5'UTR;1stExon | chr16:11876111-11876524 | 413 | | | | | | | | | 2 | | | | | yes | yes | | | | |
| TNFRSF17 | NA | TSS1500 | chr16:12058616-12059019 | 403 | | | | | | | | | | | | | | yes | yes | | | | |
| SNX29 | NA | Body | chr16:12183835-12184133 | 298 | | | 14 | 1 | | | | | 1 | | | 1 | | yes | yes | | | | |
| SNX29 | NA | Body | chr16:12192263-12192430 | 167 | | 1 | | | | | | | | | | | | | yes | | | | yes |
| SNX29 | Island | Body | chr16:12210656-12211186 | 530 | | | | | | | | | 2 | 1 | | | | yes | yes | | | | |
| SHISA9 | Island | TSS200;1stExon | chr16:12995292-12996234 | 942 | | | | | 1 | | | | 2 | | | | | | | | | yes | yes |
| MIR193B | S_Shore | TSS1500 | chr16:14397576-14397756 | 180 | | | | | | | | 0 | | | | | | | | | | | |
| MIR365-1 | NA | TSS200 | chr16:14403004-14403425 | 421 | | | 14 | 1 | 1 | | | | 1 | 1 | | | | yes | | | | | |
| C16orf45 | N_Shore | TSS1500 | chr16:15527940-15528133 | 193 | | | 3 | | | | | | 1 | | | | | yes | | | | yes | yes |
| C16orf45 | NA | TSS1500;Body;TSS200;nan | chr16:15595889-15596215 | 326 | | | 20 | | | | | | | | 1 | | | yes | | | yes | | |
| ABCC1 | NA | Body | chr16:16215875-16216108 | 233 | | | | | | | | | 1 | | | | | yes | | | | | |

| | | | | | | | | | | | | | | | | | | | | | | | |
|---|---|---|---|---|---|---|---|---|---|---|---|---|---|---|---|---|---|---|---|---|---|---|---|
| XYLT1 | NA | Body | chr16:17562960-17563300 | 340 | | 1 | | | 1 | | | | | | | | | | | | | | |
| TMC7 | Island | TSS1500;1stExon | chr16:18995391-18995637 | 246 | | | | | | | | | 1 | | | | | yes | yes | | | yes | |
| ITPRIPL2 | N_Shore | TSS1500 | chr16:19124925-19124984 | 59 | | | | | 1 | | | | 1 | | | | | yes | yes | | | | yes |
| ITPRIPL2 | Island | 1stExon;Body | chr16:19127132-19127305 | 173 | | | 1 | | | | | | | | | | | | | | | | |
| SYT17 | S_Shore | Body | chr16:19180726-19181143 | 417 | | | | | | | | | | | | | | yes | yes | | yes | | yes |
| SYT17 | Island | Body | chr16:19194989-19195208 | 219 | | | | 1 | | | | | | | | | | yes | yes | | | | |
| GPRC5B | Island | TSS1500 | chr16:19896644-19897553 | 909 | | | | | | | | 0 | 1 | | | 1 | | yes | yes | | | | yes |
| GPR139 | Island | 1stExon | chr16:20084851-20085047 | 196 | | 1 | | | | | | | | | | | | | | | | yes | |
| UMOD | Island | Body | chr16:20360206-20360497 | 291 | | | | | | | | 0 | | | | 1 | | yes | | | | | |
| ACSM5 | NA | TSS1500 | chr16:20420222-20420716 | 494 | | | | | | | | | | | | | | yes | yes | | | | yes |
| ACSM3 | NA | TSS1500 | chr16:20774873-20775166 | 293 | | | 11 | | | | | | | | | | | | yes | | | | |
| DNAH3 | NA | Body | chr16:20996560-20996704 | 144 | | | 1 | | | | | | | | | | | yes | | | | | |
| TMEM159;DNAH3 | S_Shore | 5'UTR;TSS200;TSS1500 | chr16:21170927-21171239 | 312 | | | | | | | | | 3 | | | | | | | | | | yes |
| ANKS4B | NA | TSS200 | chr16:21244898-21245235 | 337 | | | | | 1 | | | | | | | | | | yes | | | | yes |
| SLC7A5P2 | S_Shore | TSS1500 | chr16:21532721-21532834 | 113 | | | | | | | | | | | | | | | | | | | |
| POLR3E | NA | Body | chr16:22326203-22326535 | 332 | | | | | | | | | 1 | 1 | | | | yes | | | | | |
| HS3ST2 | Island | TSS200;1stExon | chr16:22825749-22826243 | 494 | | 1 | | | | | | | 2 | | | | | | | | | | yes |
| SCNN1G | Island | 5'UTR | chr16:23194674-23194733 | 59 | | | 1 | | | | | | | | | | | yes | | | | | |
| SCNN1G | Island | 5'UTR | chr16:23197569-23197740 | 171 | | | | | | | | | | | | | | yes | yes | | | | |
| SCNN1B | N_Shore;Island | TSS1500 | chr16:23313148-23313480 | 332 | | 1 | | | | | | | 1 | | | 2 | | yes | | | | | yes |
| COG7 | S_Shore | TSS1500 | chr16:23464705-23464715 | 10 | | | | | | | | | | 1 | | | | yes | | | | | |
| ERN2 | Island;S_Shore | 1stExon;5'UTR;TSS200 | chr16:23724428-23725375 | 947 | | 1 | 11 | | | | | | 2 | | | | | yes | yes | | yes | | |
| CHP2 | N_Shore;Island | TSS1500;1stExon;5'UTR | chr16:23765258-23766525 | 1267 | | 1 | 3 | | | | | | | | | | | yes | | | yes | | |
| PRKCB | N_Shore | TSS1500 | chr16:23846893-23848003 | 1110 | | 1 | | | | | | | | | | | | | | | yes | yes | yes |
| CACNG3 | Island | 1stExon;5'UTR | chr16:24267399-24267559 | 160 | | 1 | | | | | | | | | | | | | | | | yes | |
| CACNG3 | NA | Body | chr16:24372426-24372874 | 448 | | | | | | | | | | | | | | | yes | | | | |
| SLC5A11 | NA | TSS200 | chr16:24857497-24857601 | 104 | | | | 1 | | | | | | | | | | yes | | | | | |
| AQP8 | NA | Body | chr16:25228569-25228709 | 140 | | | | | 1 | | | | | | | | | yes | | | | | |
| NSMCE1 | Island | 5'UTR | chr16:27279887-27280434 | 547 | | | | | | | | | 1 | | | | | yes | | | yes | | |
| IL21R | NA | TSS1500;1stExon;5'UTR | chr16:27413904-27414281 | 377 | | | | 1 | | | | | | | | | | yes | | | | | |
| IL21R | NA | 3'UTR | chr16:27461333-27461638 | 305 | | | 2 | 1 | | | | | | | | | | yes | yes | | | | |
| KIAA0556 | NA | 3'UTR | chr16:27791167-27791248 | 81 | | | 4 | | | | | | | | | | | | | | | | |
| GSG1L | Island | TSS1500 | chr16:28075269-28075295 | 26 | | | | | | | | | | | | | | | | | | | yes |
| APOB48R | S_Shore;chr16:28509237-2850 | TSS1500;N_Shelf | chr16:28505320-28505959 | 639 | | | | | 1 | | | | 1 | 1 | 1 | | | yes | yes | | | | |
| APOB48R | Island | Body | chr16:28509314-28509637 | 323 | | | | | | | | | | | | | | yes | | | | | |
| NUPR1 | NA | 1stExon | chr16:28550171-28550525 | 354 | | | | | | | | | 1 | | | | | yes | | | | yes | |
| CCDC101 | N_Shore | TSS1500 | chr16:28564487-28564997 | 510 | | | | | | | | | 1 | | | | | yes | yes | | | yes | |
| SULT1A2 | N_Shelf | TSS1500 | chr16:28608894-28609220 | 326 | | | | | | | | | | | | | | | | yes | | | |
| SULT1A1 | Island | 5'UTR;1stExon | chr16:28634766-28635053 | 287 | | | 2 | | | | | 0 | 1 | | | 1 | | yes | yes | | | yes | |
| ATP2A1 | N_Shelf;N_Shore | TSS1500;TSS200 | chr16:28888844-28890100 | 1256 | | | 3 | | | | | | | | | | | | | | | | |
| CD19 | NA | TSS1500;TSS200 | chr16:28943019-28943288 | 269 | | | | | 1 | | | | | | | | | yes | yes | | | | |
| CD19 | Island | Body | chr16:28948179-28948319 | 140 | | | | | | | | | | | | | | | yes | | | | |
| SPNS1;LAT | NA | 3'UTR;TSS1500;TSS200;nan;Body | chr16:28995843-28996533 | 690 | | | 13 | 2 | 2 | | | | 3 | 5 | 1 | | | | | | | | |
| RRN3P2 | Island | Body | chr16:29086271-29086456 | 185 | | | 10 | | | | | | 1 | | | | | yes | | | | | |
| RUNDC2C | NA | Body | chr16:29329433-29329559 | 126 | | | 2 | | | | | | | | | | | yes | | | | | yes |
| SLC7A5P1 | S_Shore;Island | TSS1500;TSS200 | chr16:29625216-29626439 | 1223 | | | 6 | 1 | 1 | | | | | | | | | yes | | | | | yes |
| SPN | N_Shore;Island | 1stExon;5'UTR;Body;TSS1500;TSS2 | chr16:29673933-29676071 | 2138 | | | | | 3 | 1 | | | 2 | 4 | 1 | | | yes | | | | | |
| QPRT | NA | TSS1500 | chr16:29690035-29690524 | 489 | | | | | | | | | | | | | | yes | | yes | | | yes |
| QPRT | N_Shelf | Body | chr16:29703361-29703480 | 119 | | | | | | | | 0 | 1 | | | 5 | | yes | yes | | | yes | |
| QPRT | Island | Body | chr16:29706275-29706381 | 106 | | | | 1 | | | | | | | | | | yes | | | | yes | |
| C16orf54 | NA | 5'UTR | chr16:29756951-29757461 | 510 | | | | | 1 | | | | 2 | 1 | 1 | | | yes | | | | yes | |
| MAZ | Island;S_Shore | Body | chr16:29818771-29820429 | 1658 | | | 1 | 1 | 4 | | | | | | | | | | | | | yes | yes |
| LOC440356;CDIPT | S_Shore | Body;TSS200;TSS1500 | chr16:29875145-29875187 | 42 | | | | | | | | 0 | | | | | | | | | | | yes |
| SEZ6L2 | N_Shore;Island;S_Shore | Body | chr16:29887661-29888733 | 1072 | | 1 | | | 1 | | | 0 | 1 | | | | | yes | yes | | | | |
| ASPHD1 | N_Shore | 1stExon | chr16:29912460-29913007 | 547 | | | 6 | | | | | 0 | | | | | | | | | yes | | yes |
| INO80E;DOC2A | NA | 3'UTR;Body | chr16:30016775-30018193 | 1418 | | | | | | | | 0 | | | | | | | | | | | |
| ALDOA | N_Shore;chr16:30076310-3007 | TSS1500;5'UTR;N_Shore | chr16:30074702-30075547 | 845 | | | | | | | | | | | 1 | | | yes | | | | | |
| YPEL3 | S_Shore | TSS1500 | chr16:30108092-30108841 | 749 | | 1 | | | | | | | | | | | | yes | | yes | | | yes |
| GDPD3;LOC100271831 | NA | TSS200;TSS1500;Body;1stExon;5'U | chr16:30124293-30125400 | 1107 | | | 8 | | | | | | 1 | | | | | | | | yes | | |
| MAPK3 | N_Shore | Body | chr16:30134068-30134350 | 282 | | | | | | | | | 1 | | | | | | | | | | yes |
| CORO1A;LOC606724 | S_Shelf | Body;TSS1500 | chr16:30198505-30198517 | 12 | | | | | | | | | | | | | | yes | | | | | |
| MYLPF | N_Shelf | TSS1500;TSS200 | chr16:30385284-30386150 | 866 | | | 4 | | | | | | | | | | | yes | yes | | | | yes |
| MYLPF | N_Shore | Body | chr16:30388914-30389231 | 317 | | | | 1 | | | | | | | | | | yes | | | | | |
| SEPHS2 | S_Shore | TSS200 | chr16:30457367-30457493 | 126 | | | | | | | | | | | | | | yes | | yes | | | yes |
| ITGAL | Island | Body | chr16:30485383-30485966 | 583 | | | 3 | 1 | | 1 | | | 1 | | | | | yes | yes | | | | |
| ZNF689 | Island | Body | chr16:30615808-30616285 | 477 | | | | 1 | | | | | | | | | | yes | | | | | |
| FBRS | NA | TSS1500;TSS200 | chr16:30675233-30675880 | 647 | | | | | | | | | 2 | | | | | yes | | | yes | yes | yes |
| ZNF629 | Island | TSS200 | chr16:30798716-30798776 | 60 | | | | | | | | | 1 | | | | | yes | | | | | yes |
| CTF1 | S_Shore | TSS1500 | chr16:30907246-30907679 | 433 | | | | | | | | | 1 | | | | | yes | | | | yes | |
| FBXL19;ORAI3 | Island | 3'UTR;TSS1500 | chr16:30959315-30959496 | 181 | | 1 | 5 | | | | | | | | | | | | | | | | |
| STX1B | NA | TSS1500 | chr16:31022351-31022521 | 170 | | | 4 | | | | | | | | | | | yes | yes | | | | yes |
| ZNF668 | Island | Body | chr16:31075547-31075842 | 295 | | | | | | | | | 1 | | | | | | | | | | |
| POL3S | NA | 1stExon | chr16:31100095-31100428 | 333 | | | | | | | | | | | | | | yes | | | | | |
| BCKDK | N_Shore | TSS1500 | chr16:31119067-31119484 | 417 | | | | | | | | | 1 | | | | | yes | yes | | | | |
| MYST1 | N_Shore | TSS1500 | chr16:31128310-31128414 | 104 | | | | | | | | | | | | | | yes | | | | | |
| PRSS8 | NA | Body;1stExon;5'UTR | chr16:31146682-31147199 | 517 | | | 18 | | 1 | | | | 1 | 1 | 1 | | | yes | yes | | | yes | |
| PRSS36 | N_Shore | Body | chr16:31159558-31159920 | 362 | | | | | | | | 0 | 2 | | | 3 | | yes | yes | | | | yes |
| PYCARD | S_Shore | TSS200;TSS1500 | chr16:31214307-31214993 | 686 | | | 5 | 1 | 4 | | | | 1 | | | | | yes | yes | | | | |
| TRIM72;PYDC1 | S_Shore | Body;TSS200 | chr16:31228444-31228555 | 111 | | | | | | | | | | | | | | yes | | | | | |
| TRIM72 | Island | Body | chr16:31235535-31235824 | 289 | | | 2 | | | | | | | | | | | yes | | | | yes | yes |
| ITGAM | Island | Body | chr16:31342453-31343056 | 603 | | | | | | | | 0 | | | | | | yes | | | | yes | yes |
| ITGAX | NA | TSS200 | chr16:31366377-31366429 | 52 | | | 1 | | | | | | | | | | | | yes | | | yes | |
| ITGAX | Island;S_Shore | Body | chr16:31382930-31383800 | 870 | | | 3 | | | | | | 1 | | | | | yes | yes | | | yes | |
| COX6A2 | Island | Body;1stExon;5'UTR | chr16:31439106-31439752 | 646 | | 2 | 10 | | | 1 | | 0 | 2 | 1 | | 3 | | yes | yes | yes | | | yes |
| ZNF843 | Island | 5'UTR | chr16:31453406-31453591 | 185 | | | 2 | | | | | | 1 | | | | | yes | | | | | |
| AHSP | NA | TSS1500 | chr16:31538556-31539234 | 678 | | | 8 | | | | | | 1 | | | | | yes | yes | | | | |
| CSDAP1 | Island;S_Shore | Body;TSS1500 | chr16:31580560-31581309 | 749 | | | 2 | | 1 | | | | 1 | | | | | yes | | | | yes | yes |
| MIR1826 | Island | TSS1500 | chr16:33964783-33965435 | 652 | | | 1 | | | | | | | | | | | | | | yes | | |
| ANKRD26P1 | NA | Body;TSS200 | chr16:46602737-46603259 | 522 | | | 9 | | | | | 0 | 1 | | | 1 | | yes | yes | | | yes | |
| MYLK3 | NA | 5'UTR;1stExon | chr16:46782176-46782542 | 366 | | | | | | | | | | | | | | | | | | | |
| GPT2 | Island;S_Shore | Body;TSS200 | chr16:46918956-46919194 | 238 | | | 6 | | | | | 0 | 1 | | | 1 | | | yes | | | | |
| CBLN1 | Island | 3'UTR | chr16:49312238-49312560 | 322 | | | 1 | | | | | | | | | | | | | | | | yes |
| CBLN1 | Island | TSS200 | chr16:49315822-49316197 | 375 | | 1 | | | | | | | | | | | | | yes | | | yes | yes |
| C16orf78 | NA | TSS1500;TSS200 | chr16:49407465-49407997 | 532 | | | | | | | | | | | | | | | yes | yes | | | |
| ZNF423 | N_Shore | 3'UTR | chr16:49525139-49525807 | 668 | | | 9 | | | | | | | | | | | | | | | yes | yes |
| ZNF423 | NA | Body | chr16:49686567-49686662 | 95 | | | | | | | | | | | | | | | | | | | |
| ZNF423 | N_Shore;Island | Body | chr16:49732224-49732623 | 399 | 2 | | | | 1 | | 1 | 0 | 3 | 3 | | 22 | 1 | | | | | | |
| ADCY7 | NA | Body | chr16:50347766-50347895 | 129 | | | 5 | | | | | | 1 | | | | | yes | | | | yes | |
| SNX20 | NA | 1stExon;5'UTR | chr16:50715135-50715529 | 394 | | | | | 2 | | | | 2 | 1 | 1 | | | yes | yes | | | | |

| | | | | | | | | | | | | | | | | | | | | | | | |
|---|---|---|---|---|---|---|---|---|---|---|---|---|---|---|---|---|---|---|---|---|---|---|---|
| NOD2 | NA | TSS1500 | chr16:50730385-50730881 | 496 | | | | | | | | | | | | | | | yes | | | | |
| NOD2 | NA | Body | chr16:50745944-50746063 | 119 | | | 6 | | | | | | | | | | | yes | | | | | |
| SALL1 | S_Shore | 3'UTR | chr16:51170800-51171260 | 460 | | | | 1 | | | | | | | | | | | | | | | yes |
| SALL1 | NA | Body | chr16:51174069-51174177 | 108 | | | | | | | | | | | | | | | | | | | |
| SALL1 | N_Shore;Island | Body;5'UTR;1stExon;TSS1500 | chr16:51183363-51185461 | 2098 | | | 20 | | 1 | | | 0 | 2 | | 1 | | | | | | | yes | yes |
| TOX3 | Island;S_Shore | 5'UTR;TSS1500;TSS200 | chr16:52581551-52582262 | 711 | | | 10 | | | | | 0 | 3 | | | 1 | | | | | | yes | yes |
| IRX3 | Island;N_Shore | Body;5'UTR;1stExon;3'UTR | chr16:54317413-54320430 | 3017 | | | | | | | | | | | | | | | | | | | |
| IRX5 | Island | TSS200 | chr16:54965084-54965089 | 5 | | | | | | | | | 1 | | | | | | | | | yes | yes |
| IRX6 | Island;S_Shore;N_Shore | Body | chr16:55360182-55364614 | 4432 | 1 | 1 | | | | | | | | | | | | yes | yes | | | yes | |
| MMP2 | N_Shore | TSS1500 | chr16:55512822-55512870 | 48 | | | | | | | | | 1 | | | | | | | yes | | yes | yes |
| LPCAT2 | N_Shore;Island | TSS200;1stExon;5'UTR | chr16:55542747-55543166 | 419 | | 1 | | | | | | | | | | | | yes | | yes | | | |
| SLC6A2 | Island | TSS200 | chr16:55690378-55690564 | 186 | | | | | 1 | | | | 2 | | | | | | | | | | yes |
| CES4 | N_Shore | TSS1500;TSS200;Body | chr16:55794031-55794910 | 879 | 4 | 3 | 13 | 1 | | | | 0 | 5 | | | 8 | | yes | yes | yes | | | |
| CES1 | NA | Body | chr16:55850985-55851398 | 413 | | | 2 | | | | | | | | | | | yes | | yes | | yes | |
| CES1 | Island;N_Shore | Body | chr16:55866522-55867072 | 550 | 2 | | 10 | | | | 2 | 0 | 7 | | | 2 | 1 | yes | yes | yes | | | |
| CES1 | S_Shore | TSS1500 | chr16:55868196-55868432 | 236 | | | 4 | | | | | | 1 | | | | | yes | yes | | | | |
| CES7 | NA | 1stExon;5'UTR | chr16:55909166-55909732 | 566 | | | | | | | | | | | | | | | yes | | yes | | |
| DKFZP434H168;GNAO1 | S_Shelf | TSS1500;Body | chr16:56228744-56229180 | 436 | | | | | | | | | | | | | | | | | | | yes |
| AMFR | Island | Body | chr16:56397917-56398105 | 188 | | | | | | | | | | | | | | yes | | | yes | | |
| BBS2 | S_Shore | TSS1500 | chr16:56554249-56554313 | 64 | | | 3 | | | | | | 1 | | | | | | yes | | | | |
| MT1L | Island | TSS1500 | chr16:56651049-56651456 | 407 | | | | | | | | 0 | | | | | | yes | | | | | |
| MT1M | N_Shore;Island | TSS1500 | chr16:56665855-56666334 | 479 | | | | | | | | | 1 | | | 1 | | yes | | | | | |
| MT1DP | Island;S_Shore | TSS200;Body | chr16:56677585-56677855 | 270 | | | 2 | | | | | | | | | | | yes | | | | | |
| MT1B | NA | TSS200 | chr16:56685637-56685902 | 265 | | | | | | | | | | | | | | yes | | | | | |
| MT1G | Island | Body | chr16:56701868-56702158 | 290 | | | 4 | | | | | | | | | | | | | | | | |
| MT1G;MT1H | N_Shore;Island | TSS1500;TSS200 | chr16:56703257-56703833 | 576 | | | 15 | | | | | | 1 | | | | | yes | | | | | |
| MT1IP | Island | Body | chr16:56709853-56710256 | 403 | | | 12 | | | | | | | | | | | yes | | | | | |
| CETP | NA | TSS1500;TSS200 | chr16:56995201-56995856 | 655 | | | | | | | | 0 | 3 | | | 1 | | | | | yes | | |
| PLLP | N_Shore | Body | chr16:57317679-57318087 | 408 | | | 2 | | | | | | | | | | | | | | yes | | |
| CX3CL1 | NA | TSS1500;TSS200 | chr16:57406044-57406425 | 381 | | | | | | | | | 2 | | | | | yes | yes | | | | |
| POLR2C | N_Shore | TSS1500 | chr16:57495877-57496328 | 451 | | | | | | | | | | | | | | | yes | | | | yes |
| CCDC102A | Island | Body | chr16:57562452-57562841 | 389 | | | | | 1 | | | 0 | 1 | | | 1 | 1 | yes | yes | | | | yes |
| GPR114 | NA | TSS1500;nan | chr16:57576285-57576862 | 577 | | | | | | | | | 1 | 1 | 1 | | | yes | yes | | | | |
| GPR56 | NA | TSS200 | chr16:57653904-57654347 | 443 | | | | 1 | | | | | 1 | | | | | yes | | | | | |
| GPR56 | NA | TSS1500;5'UTR;TSS200 | chr16:57662008-57662690 | 682 | | 1 | | 1 | 1 | | | | | | | | | | | | | yes | |
| GPR97 | NA | TSS1500 | chr16:57701455-57701726 | 271 | | | | | | | | 0 | 1 | | | 1 | | yes | | | | | |
| CCDC135 | NA | TSS200 | chr16:57728632-57729107 | 475 | | | | | | | | | | | | | | yes | | | yes | | |
| KATNB1 | S_Shore | 5'UTR | chr16:57770390-57770913 | 523 | | | | | | | | 0 | | | | 1 | | yes | | | | | |
| KIFC3 | NA;N_Shelf | Body;1stExon;5'UTR;TSS200;TSS15 | chr16:57831745-57832760 | 1015 | | | | 1 | 1 | | | 0 | 2 | | | | | | yes | | | | |
| CNGB1 | Island | 3'UTR | chr16:57917946-57918095 | 149 | | | 1 | | | | | 0 | 1 | | | | | yes | | | | | |
| CNGB1 | N_Shore | Body | chr16:57937614-57937756 | 142 | | | | | | | | | 1 | | | | | yes | | | | | |
| TEPP | Island | Body | chr16:58018972-58019984 | 1012 | | | 8 | | | | | | 1 | | | | | | yes | | yes | | |
| CCDC113 | N_Shore | TSS1500 | chr16:58282922-58283189 | 267 | | | 6 | | | | | | | | | | | yes | yes | yes | | | |
| KLKBL4 | NA | TSS200 | chr16:58328961-58329456 | 495 | | 1 | | | | | | | | | | | | yes | | | | yes | |
| NDRG4 | Island | TSS1500 | chr16:58497230-58497239 | 9 | | | 1 | | | | | | | | | | | yes | yes | | | | |
| NDRG4 | N_Shore;Island | Body | chr16:58533743-58535556 | 1813 | | | 24 | | 1 | | 1 | 0 | 4 | 3 | | 2 | | yes | | | | | |
| SETD6 | Island | TSS200 | chr16:58549212-58549251 | 39 | | | 3 | | | | | | | | | | | yes | | | | | |
| LOC644649 | N_Shore;Island | Body | chr16:59788728-59789485 | 757 | | | | | | | | | | | | | | yes | | yes | | | yes |
| CDH8 | Island | 5'UTR;TSS200 | chr16:62069317-62070234 | 917 | | | | | 3 | | | 0 | 4 | | | 1 | | | | | | | yes |
| CDH11 | NA | 5'UTR | chr16:65105516-65105714 | 198 | | | 11 | | | | | | | | | | | | | | | yes | |
| CDH5 | NA | TSS200;TSS1500 | chr16:66400320-66400599 | 279 | | | 5 | | | | | | 3 | 1 | | | | yes | | | | | |
| TK2 | N_Shore | Body | chr16:66582592-66583081 | 489 | | | | | | | | | 1 | | | | | | | | | | |
| CMTM1 | Island | TSS200 | chr16:66600275-66600700 | 425 | | | | | | | | | | 1 | | | | | yes | | | | |
| CMTM1;CMTM2 | Island;chr16:66612749-666134 | 3'UTR;TSS1500;Island;TSS200 | chr16:66612955-66613582 | 627 | | | 35 | | 2 | | | 0 | 2 | 1 | 2 | 4 | | | | | | | |
| CMTM3 | N_Shore;Island | TSS1500;5'UTR;1stExon;TSS200 | chr16:66637919-66638438 | 519 | 1 | | | | 1 | | | | | | | | | yes | yes | yes | | yes | yes |
| CCDC79 | N_Shore;Island | 5'UTR;TSS200 | chr16:66834841-66835860 | 1019 | | | | | | | | | | 1 | | | | yes | | | | | |
| CA7 | chr16:66878172-66879072 | N_Shore | chr16:66876847-66876938 | 91 | | | | | | | | | | | 1 | | | yes | | | | | |
| CDH16 | N_Shelf | 1stExon;5'UTR | chr16:66952714-66953432 | 718 | | | | | | | | | | | | | | | yes | | | | |
| RRAD | Island | Body | chr16:66955988-66956242 | 254 | | | | | | | | 0 | 3 | | | 1 | | yes | | | | | |
| RRAD | S_Shore | TSS1500 | chr16:66959729-66959833 | 104 | | | | | | | | | | | | | | | | | | | |
| CES8 | NA | TSS200 | chr16:67022472-67022719 | 247 | | | | | | | | | 1 | 1 | | | | yes | yes | | | | yes |
| CES8 | Island | Body | chr16:67034882-67035388 | 506 | | | | 1 | | | | | | | | | | yes | | | | yes | yes |
| HSF4;FBXL8 | N_Shore;Island;chr16:6719679 | TSS1500;Body;TSS200;Island;5'UT | chr16:67196699-67198367 | 1668 | | | 35 | | 2 | | | | | | 1 | | | yes | | | | yes | |
| KIAA0895L;EXOC3L | Island | TSS1500;Body | chr16:67219193-67219218 | 25 | | | | | | | | | | | | | | yes | yes | | | | |
| EXOC3L | NA | 5'UTR;1stExon;TSS200 | chr16:67223733-67224629 | 896 | | | 6 | | | | | | | | | | | yes | | yes | | | |
| ELMO3;E2F4 | N_Shore;Island | TSS1500;3'UTR;TSS200;Body | chr16:67232180-67233983 | 1803 | | | | | 1 | | | | 4 | | | | | | | | | | yes |
| PLEKHG4 | Island | 1stExon;5'UTR | chr16:67313600-67314073 | 473 | | | 3 | | | | | | | | | | | | | | | | yes |
| LRRC36;KCTD19 | NA | TSS1500;Body | chr16:67360129-67360300 | 171 | | | | | | | | | | | | | | yes | | | | | yes |
| TPPP3 | Island | TSS1500 | chr16:67428155-67428898 | 743 | | | 12 | | | | 1 | | 1 | 3 | | 1 | | | | | | | yes |
| ZDHHC1 | Island | Body | chr16:67433184-67433636 | 452 | | | 15 | | | | | | | | | | | | | | | | |
| HSD11B2 | N_Shore;Island | TSS1500 | chr16:67464331-67464610 | 279 | | | 2 | | | | | | | | | | | | | | | yes | yes |
| ATP6V0D1 | S_Shore | TSS1500 | chr16:67515916-67516546 | 630 | | | 11 | 1 | 1 | | | | | | | | | | | yes | | | |
| FAM65A | N_Shore | TSS1500 | chr16:67562128-67562517 | 389 | | | 1 | | | | | | | | | | | yes | | | | yes | yes |
| RLTPR | N_Shore;Island | Body | chr16:67686832-67687754 | 922 | | | | | 1 | 1 | | 0 | 6 | | | 1 | | | | | | | yes |
| PARD6A;ACD | Island;S_Shore | Body;TSS1500 | chr16:67695257-67696181 | 924 | | 1 | 4 | 1 | | | | | | | | | | | | | | | yes |
| NRN1L | N_Shore;Island | TSS1500;TSS200 | chr16:67918484-67918965 | 481 | | | 2 | | | | | | 2 | 1 | | | | | | | | | |
| CTRL | N_Shelf | 5'UTR;1stExon | chr16:67965756-67965867 | 111 | | | | | | | | | | | | | | | yes | | | | |
| SLC12A4;LCAT | NA | 3'UTR;1stExon;TSS1500;Body | chr16:67977865-67980139 | 2274 | | | 11 | | | | | | | | | | | | | | | | |
| SLC12A4 | NA | Body;1stExon | chr16:67997858-67998164 | 306 | | | 4 | | | | | 0 | 1 | 1 | | | | | | | | | |
| DPEP3 | Island | Body;1stExon | chr16:68013968-68014808 | 840 | | | 1 | | | | | | 3 | | | 2 | | yes | | | | yes | |
| DPEP2 | NA | 1stExon;5'UTR;TSS200 | chr16:68033301-68034002 | 701 | | | 3 | | | | | | | | | | | yes | | yes | | | |
| NFATC3 | N_Shore | TSS1500 | chr16:68118441-68118822 | 381 | | | | | 1 | | | | | | | | | | | | | yes | yes |
| ESRP2 | Island | Body;1stExon;5'UTR;TSS200;TSS15 | chr16:68269041-68270882 | 1841 | | | 29 | | | | | 0 | 3 | 1 | | | | | | | | | yes |
| PRMT7 | NA | Body | chr16:68378352-68378420 | 68 | | | 1 | | | | | | | | | | | yes | yes | | | yes | |
| SMPD3 | Island | 5'UTR | chr16:68481049-68481813 | 764 | | | 9 | | | | | 0 | 2 | | | 3 | | | | | | | yes |
| SMPD3 | S_Shore | TSS1500 | chr16:68482941-68483194 | 253 | | | | | | | | | | | | | | yes | yes | | | | |
| CDH3 | N_Shore | TSS200 | chr16:68677997-68678092 | 95 | | | 5 | | | | | | | | | | | yes | | | yes | | |
| CDH1 | N_Shore | TSS1500 | chr16:68770944-68771035 | 91 | | | | | | | | | 1 | | | | | yes | | | | | yes |
| CDH1 | Island | Body | chr16:68772225-68772469 | 244 | | | | | 1 | | 1 | 0 | 1 | 1 | | 1 | | yes | | | | | |
| HAS3 | Island | 5'UTR;TSS200 | chr16:69141250-69141478 | 228 | | | | | | | | 0 | | | | 1 | | | | | | | |
| TMED6 | NA | 1stExon | chr16:69385547-69385827 | 280 | | | | 1 | 1 | | | | | | | | | yes | | | yes | | |
| NFAT5 | N_Shore | TSS1500 | chr16:69597925-69598417 | 492 | | | 3 | | 1 | | | | | | | | | yes | | | | yes | |
| NQO1 | S_Shore | TSS200 | chr16:69760564-69760966 | 402 | | | | 1 | | | | | | | | | | yes | | | | | |
| WWP2 | NA | TSS1500;Body;TSS200 | chr16:69958189-69958964 | 775 | | | 22 | | | | | | | | | | | | | | | yes | |
| MIR140;WWP2 | NA | TSS200;Body | chr16:69966902-69967212 | 310 | | | | | | | | 0 | 4 | | | 4 | | | | | yes | | |
| WWP2 | NA | 3'UTR | chr16:69975149-69976228 | 1079 | | | 4 | 1 | | | | | | | | | | | | | | | |
| CLEC18C | NA | TSS1500 | chr16:70207297-70207722 | 425 | | | | | | | | | | | | | | yes | | | | | |
| EXOSC6 | Island | 3'UTR;1stExon | chr16:70284893-70285955 | 1062 | | | | | 1 | | | | | | | | | yes | | | | | yes |
| IL34 | NA | TSS1500;TSS200;1stExon;5'UTR | chr16:70680104-70680817 | 713 | 1 | | 18 | | | | | | | | | | | yes | | | | | |

| | | | | | | | | | | | | | | | | | | | | | | | |
|---|---|---|---|---|---|---|---|---|---|---|---|---|---|---|---|---|---|---|---|---|---|---|---|
| MTSS1L | Island | Body | chr16:70713933-70714378 | 445 | | | | | | | | | | | | | | | yes | | | | yes |
| MTSS1L | Island | TSS200 | chr16:70720079-70720149 | 70 | | | | | | | | | 1 | | | | | | | | | yes | yes |
| VAC14 | NA | Body | chr16:70759914-70760142 | 228 | | | | | | | | | 1 | | | | | yes | | | | | |
| HYDIN | N_Shore | 5'UTR | chr16:71264282-71264678 | 396 | | | | | | | | | 1 | | | | | | yes | yes | yes | | |
| ZNF19 | NA | TSS200 | chr16:71523432-71523560 | 128 | | | | | | | | | 1 | | | | | yes | | | | | |
| CHST4 | NA | TSS1500 | chr16:71559579-71560205 | 626 | | | | | | | | 0 | 1 | | | | | yes | | | | | |
| CHST4 | NA | Body | chr16:71570932-71571146 | 214 | | 1 | | | | | | | | | | | | | | | | | |
| MARVELD3 | Island | TSS200 | chr16:71660049-71660103 | 54 | | | | | | | | | 1 | | | | | | | | | yes | |
| PMFBP1 | NA | TSS200;TSS1500 | chr16:72206536-72206863 | 327 | | | | | | | | | 1 | | | | | yes | | yes | yes | | |
| ZFHX3 | N_Shore;Island | 3'UTR;Body | chr16:72820921-72821665 | 744 | | | 7 | 1 | 2 | | | | | | | | | | | | | yes | yes |
| ZFHX3 | NA | Body | chr16:72981203-72982398 | 1195 | | | 1 | 1 | 1 | | | | | | | | | yes | | | | | |
| HTA | NA | TSS1500;Body | chr16:73125573-73126909 | 1336 | | | 5 | | | | | | | | | | | yes | | | | | |
| MLKL | Island;S_Shore | 5'UTR;TSS1500 | chr16:74734230-74735780 | 1550 | | | | | 1 | | | | 1 | | | | | yes | yes | yes | | yes | yes |
| LDHD | N_Shore;S_Shore | Body;1stExon;TSS200 | chr16:75148330-75150880 | 2550 | | | 8 | | | | | 0 | 10 | 1 | | 3 | | yes | | yes | | yes | |
| CTRB2 | NA | Body;TSS200 | chr16:75240536-75241216 | 680 | 1 | | | | 1 | | | | | | | | | yes | | | | | |
| CTRB1 | NA | TSS1500 | chr16:75252211-75252619 | 408 | | | | | | | | | 1 | | | | | yes | | | | | |
| BCAR1 | S_Shelf | Body;TSS1500 | chr16:75273344-75273486 | 142 | | | | | | | | | | | | | | | | | | | |
| BCAR1 | Island;S_Shore | TSS200;Body;TSS1500 | chr16:75300023-75301362 | 1339 | | | 1 | | | | | | 1 | | | | | | yes | | | | yes |
| CFDP1 | S_Shore | TSS1500 | chr16:75468057-75468318 | 261 | | | 17 | | | | | 0 | 2 | | | 1 | | yes | | | yes | | |
| CHST5 | NA | 5'UTR;TSS200 | chr16:75568999-75569527 | 528 | | | 8 | | | | | 0 | | | | | | yes | | | | | |
| CNTNAP4 | NA | Body;TSS1500 | chr16:76342377-76342678 | 301 | | | | | | | | | | | | | | yes | | yes | | | |
| ADAMTS18 | S_Shore | TSS200;TSS1500 | chr16:77469167-77469998 | 831 | | | 6 | | | | | 0 | 3 | | | 1 | | yes | yes | | | yes | yes |
| CLEC3A | NA | TSS200 | chr16:78056379-78056736 | 357 | | | 1 | | | | | | 1 | | | | | | | | | | |
| WWOX | NA | nan | chr16:78539954-78540172 | 218 | | | | | | | | | | | 1 | | | yes | | | | | |
| WWOX | NA | Body | chr16:78601767-78601911 | 144 | 1 | | | | | | | | | | | | | yes | | | | | yes |
| WWOX | NA | Body | chr16:79027453-79027744 | 291 | | 1 | | | | | | | | | | | | | | | | | |
| CDYL2 | NA | Body | chr16:80654711-80654763 | 52 | 3 | | | | | | | | | | | | | | | yes | yes | | |
| CENPN;C16orf61 | N_Shore | TSS1500;5'UTR | chr16:81039227-81039458 | 231 | | | | | | | | 0 | | | | | | yes | | | | | yes |
| PKD1L2 | NA | Body | chr16:81190494-81190750 | 256 | | | | | | | | | | | | | | yes | | | | | yes |
| PKD1L2 | NA | 1stExon;TSS200 | chr16:81253692-81254209 | 517 | | | | | | | | | 2 | | | | | yes | yes | | | | |
| CMIP | NA | Body | chr16:81526954-81527504 | 550 | | | 16 | | | | | | 1 | 1 | | | | yes | yes | | | | |
| CMIP | NA | Body | chr16:81564161-81564553 | 392 | | | | | | | | | 2 | 1 | | | | | | yes | | | |
| CMIP | Island | Body | chr16:81711865-81712102 | 237 | | | | 1 | | | | | 2 | | | | | yes | yes | | | | yes |
| CMIP | NA | Body | chr16:81726721-81727056 | 335 | | | 4 | | | | | | 1 | | | | | | | | | | |
| PLCG2 | NA | Body | chr16:81924972-81925132 | 160 | | | | | | | | | | | | | | yes | yes | | | | |
| PLCG2 | NA | Body | chr16:81969883-81970170 | 287 | | | | | | | | | | | | | | | yes | | | | |
| SDR42E1 | N_Shore;Island | 5'UTR;1stExon;TSS200 | chr16:82044738-82045758 | 1020 | | | 7 | | | | | 0 | 6 | | | 4 | | | yes | yes | | | |
| HSD17B2 | NA | TSS200 | chr16:82068702-82068980 | 278 | | | | 1 | | | | | | | | | | | | | | yes | |
| CDH13 | N_Shore | TSS1500;TSS200 | chr16:82659960-82660505 | 545 | | | 8 | | | | | 0 | 1 | | | | | | yes | | | yes | |
| CDH13 | NA | Body | chr16:83171068-83171314 | 246 | | | 6 | 1 | 1 | | 1 | 0 | 3 | 2 | | 6 | | | | | yes | | |
| OSGIN1 | NA | TSS200 | chr16:83982645-83982696 | 51 | | | | 1 | | | | | | | | | | | | | yes | | |
| NECAB2 | Island | 1stExon | chr16:84002370-84002843 | 473 | | | | 1 | | | | | | | | | | | | | | | yes |
| NECAB2 | Island | Body | chr16:84029457-84029838 | 381 | | | 3 | 1 | | | | 0 | 1 | | | | | yes | | | yes | | |
| SLC38A8 | NA | TSS1500 | chr16:84076582-84076941 | 359 | | 1 | | | | | | | 1 | | | | | | | | yes | | yes |
| ADAD2 | N_Shore | TSS200 | chr16:84224564-84225080 | 516 | | | | | | | | | | | | | | yes | | | | | yes |
| KCNG4 | N_Shore | Body | chr16:84256036-84256194 | 158 | | | 1 | | | | | | | | | | | | | | | | |
| WFDC1 | N_Shore | TSS1500 | chr16:84327808-84328208 | 400 | | | 7 | | | | | | 1 | | | | | yes | | | | yes | |
| WFDC1 | NA | nan;Body | chr16:84336112-84336445 | 333 | | | | | | | | | | | 1 | | | | yes | | | yes | |
| WFDC1 | NA | Body | chr16:84341323-84341364 | 41 | | | 4 | | | | | | | | | | | | | | | | |
| WFDC1 | Island | Body | chr16:84346722-84346930 | 208 | | | 19 | | | | | 0 | 1 | | | 1 | | yes | | | | yes | yes |
| ATP2C2 | N_Shore | TSS1500 | chr16:84401247-84402496 | 1249 | | | | | | | | | | | | 1 | | yes | | | | | |
| ATP2C2 | NA | Body | chr16:84475885-84476154 | 269 | | | | | | | | | | | | | | yes | yes | | | | |
| COTL1 | N_Shore | Body | chr16:84650202-84650415 | 213 | | | | | | | | | | | | | | yes | yes | | | | |
| CRISPLD2 | N_Shore | TSS1500 | chr16:84852577-84852909 | 332 | | | | | | | | | 1 | | | | | yes | yes | | | | |
| CRISPLD2 | NA | 5'UTR | chr16:84860766-84860918 | 152 | | | 3 | | | | | | | | | | | yes | yes | | | | |
| KIAA0513 | NA | 5'UTR | chr16:85074359-85075249 | 890 | | | 16 | 1 | | | | 0 | 2 | | | 1 | | yes | yes | | yes | | |
| KIAA0513 | NA | Body | chr16:85116332-85116513 | 181 | | | | | | | 1 | | | | | | | | | | | | |
| FAM92B | NA | 5'UTR;1stExon | chr16:85146088-85146180 | 92 | | | | | | | | | | | | | | | | | | | |
| KIAA0182 | chr16:85644947-85649027 | S_Shore | chr16:85650374-85650522 | 148 | | | | | | | | | | | 1 | | | | yes | | | | |
| KIAA0182 | S_Shore | 5'UTR;Body | chr16:85659946-85660534 | 588 | | | 3 | | 1 | | | | | | | | | | | | | yes | |
| KIAA0182 | Island | 5'UTR;Body | chr16:85670135-85670262 | 127 | | | | 1 | | | | | | | | | | | yes | | | | |
| KIAA0182 | S_Shore | 5'UTR;Body | chr16:85671457-85672175 | 718 | | | 3 | | | | | | | | | | | yes | | | | | yes |
| KIAA0182 | Island | Body | chr16:85684353-85684853 | 500 | | | | | | | | | 1 | | | | | | | | | yes | |
| GINS2 | S_Shore | TSS1500 | chr16:85723150-85723539 | 389 | | | 3 | | | | | | | | | | | yes | | | yes | yes | yes |
| IRF8 | Island | TSS200 | chr16:85932591-85932853 | 262 | | 1 | | | | | | 0 | | | | | | | | | | | |
| IRF8 | S_Shelf | 5'UTR | chr16:85936211-85936666 | 455 | | | | | | | | | | | | | | yes | | | | | |
| LOC732275 | NA | Body | chr16:86370962-86371442 | 480 | | | 7 | | | | | | 2 | | | 1 | | yes | | | | | |
| FOXF1 | N_Shore;chr16:86549069-8655 | 3'UTR;N_Shore;Body | chr16:86546374-86548323 | 1949 | | | 36 | | | | | 0 | 1 | | 2 | 1 | | | | | | yes | yes |
| MTHFSD;FLJ30679 | Island | TSS200;Body | chr16:86588937-86589076 | 139 | | | | | | | | | | 2 | | | | | yes | | | | |
| FOXL1 | chr16:86612188-86613821;Isla | Island;1stExon | chr16:86612590-86613731 | 1141 | | | 1 | | | | | | | | 1 | | | | | | | | |
| FBXO31 | NA | Body | chr16:87376314-87376795 | 481 | | 1 | | | | | | | | | | | | yes | | | | | |
| FBXO31 | NA | Body | chr16:87391861-87392076 | 215 | 1 | 1 | | | | | | | | | | | | yes | | | | | |
| ZCCHC14 | Island | 3'UTR | chr16:87441441-87441659 | 218 | | | 17 | | | | | | 1 | | | | | | | | | | |
| ZCCHC14 | Island | Body | chr16:87447756-87448114 | 358 | | | | | | | | | 1 | | 1 | | | yes | | | | | |
| ZCCHC14 | NA | Body | chr16:87468092-87469329 | 1237 | | | 9 | | | | | | | | | | | yes | | | yes | | |
| ZCCHC14 | N_Shore | Body | chr16:87511017-87511500 | 483 | | | | | | | | | 1 | | | | | yes | yes | | | | |
| ZCCHC14 | NA | 1stExon | chr16:87525307-87525539 | 232 | | | | | | | | 0 | | | | | | | | | | yes | yes |
| JPH3 | S_Shore | Body | chr16:87637999-87638484 | 485 | | | 2 | | | | | | | | | | | | yes | | | yes | |
| JPH3 | S_Shelf | Body | chr16:87641133-87641681 | 548 | 1 | | | | | | | | 1 | | | | | | | | | yes | |
| JPH3 | S_Shelf | Body | chr16:87682536-87683014 | 478 | | | | | | | | | | | | | | | | | yes | | |
| LOC100129637 | N_Shore;Island | Body | chr16:87734816-87737091 | 2275 | | | | | 1 | | | 0 | 3 | | | | | | | | | | yes |
| LOC100129637 | S_Shore | TSS200 | chr16:87739377-87739453 | 76 | | | | | 1 | | | | | | | | | | | | | yes | |
| KLHDC4 | Island | Body | chr16:87743106-87743286 | 180 | | | | | | | | | 1 | | | | | | | | | | |
| KLHDC4 | Island | Body | chr16:87754019-87754207 | 188 | | | | | | | | | 1 | | | | | | | | | | |
| KLHDC4 | NA | Body | chr16:87769432-87769974 | 542 | | | 7 | | | | | | | | | | | yes | | | yes | | |
| KLHDC4 | NA | Body | chr16:87778876-87778976 | 100 | | | | | | | | | 1 | | | | | | yes | | | yes | |
| SLC7A5 | NA | 3'UTR | chr16:87863785-87863856 | 71 | | | | | 1 | | | | 1 | | | | | | | | | | |
| SLC7A5 | NA | Body | chr16:87866696-87866833 | 137 | | | | | | | | | 1 | | | | | | | | | | yes |
| SLC7A5 | NA | Body | chr16:87879502-87879698 | 196 | | | | 1 | | | | | | | | | | | | yes | | yes | |
| SLC7A5 | NA | Body | chr16:87887331-87887504 | 173 | | | | | | | | | 1 | | 1 | | | | | | | | |
| SLC7A5 | S_Shore | TSS1500 | chr16:87904491-87904580 | 89 | | | 5 | | | | | | | | | | | yes | | | | | |
| CA5A | NA | Body | chr16:87943038-87943182 | 144 | | | | 1 | | | | | | | | | | yes | | | | | |
| CA5A | NA | Body | chr16:87958145-87958493 | 348 | | 1 | | | | | | | 1 | | 1 | | | | | yes | | | |
| CA5A | NA | 1stExon | chr16:87969962-87970376 | 414 | | | | 1 | | | | | | | | | | yes | | | | | |
| BANP | NA | 5'UTR | chr16:87996731-87996870 | 139 | | | 3 | | | | | | | | | | | | yes | | | | |
| BANP | N_Shelf | Body | chr16:88016208-88016632 | 424 | | | | 1 | | | | | | | | | | | | | | | |
| BANP | N_Shore | Body | chr16:88038949-88039257 | 308 | | | | | 1 | | | 0 | 2 | | | | | | | | | | |
| BANP | Island | Body | chr16:88052922-88053339 | 417 | | | 18 | | | | | | | | | | | | yes | | yes | | |
| BANP | N_Shelf | Body | chr16:88057792-88058326 | 534 | | | 10 | | | | | | | | | | | yes | | | | | yes |

| | | | | | | | | | | | | | | | | | | | | | | | |
|---|---|---|---|---|---|---|---|---|---|---|---|---|---|---|---|---|---|---|---|---|---|---|---|
| BANP | NA | Body | chr16:88079739-88079797 | 58 | | | | | 1 | | | | | | | | | yes | | | | | yes |
| BANP | S_Shore | Body | chr16:88103035-88103629 | 594 | | | | | 1 | | | | 1 | | | | | | | | yes | yes | |
| ZNF469 | Island | 1stExon | chr16:88496788-88497293 | 505 | 1 | | 10 | | | | | | | | | | | | | | | | yes |
| ZFPM1 | N_Shelf | Body | chr16:88534398-88534585 | 187 | | | | | | | | | 1 | | | | | yes | | | | | yes |
| ZFPM1 | S_Shelf | Body | chr16:88543440-88543627 | 187 | | | | 1 | | | | | | | | | | yes | | | | | |
| ZFPM1 | NA | Body | chr16:88547533-88547743 | 210 | | | | | | | | | 1 | | | | | yes | | | | | |
| ZFPM1 | N_Shelf | Body | chr16:88550961-88551489 | 528 | | | | | | | | | 3 | | | | | yes | | | yes | | |
| ZFPM1 | Island | Body | chr16:88558065-88558237 | 172 | | | | 1 | | | | | | | | | | yes | | | | | yes |
| ZFPM1 | Island | Body | chr16:88564366-88565080 | 714 | | | | 1 | | | | | | | | | | | | | | | yes |
| ZFPM1 | N_Shore | Body | chr16:88569083-88569268 | 185 | | | | | | | | | | | | | | | yes | | | | yes |
| ZFPM1 | N_Shore;Island | Body | chr16:88589918-88590467 | 549 | | | 17 | | | | | | 2 | 1 | | | | yes | | | | | |
| IL17C | N_Shore;Island | Body | chr16:88705716-88706426 | 710 | | | 9 | | | | | | 2 | | | 1 | | yes | | | | | |
| CYBA | Island;S_Shore | Body;1stExon;TSS200;TSS1500 | chr16:88717134-88717989 | 855 | 4 | | | 1 | | | | | | | | | | | | | | | |
| MGC23284;SNAI3 | N_Shelf | Body;3'UTR | chr16:88744522-88744842 | 320 | | | 4 | | | | | | | | | | | | | | | | |
| MGC23284;SNAI3 | Island;N_Shore | Body | chr16:88747591-88747919 | 328 | | | | | | | | 0 | 3 | | | | | yes | | yes | | | |
| RNF166 | Island | Body | chr16:88765071-88765233 | 162 | | | | | | | | | 1 | | | | | | | | | | |
| RNF166 | S_Shore | Body | chr16:88767124-88767723 | 599 | | | | | | | | | 1 | 1 | 1 | | | | | | | | |
| RNF166 | N_Shelf | Body | chr16:88769549-88769970 | 421 | | | | | | | | | 1 | | | | | | yes | | | | |
| FAM38A | S_Shelf | Body | chr16:88807658-88807707 | 49 | | | | | | | | | 1 | | | | | | | | | yes | yes |
| FAM38A | Island | Body | chr16:88821292-88821905 | 613 | | | | | | | | | 1 | | | | | | | yes | | | yes |
| FAM38A | NA | Body | chr16:88832476-88832667 | 191 | | | | | | | | | 3 | | | | | yes | | yes | | | |
| FAM38A | Island | Body | chr16:88844969-88845204 | 235 | 3 | | | | | | | | | | | | | yes | | | | | |
| FAM38A | S_Shore | Body | chr16:88846234-88846723 | 489 | | | 5 | | | | | | | | | | | yes | | | | | |
| FAM38A | N_Shore;Island | Body | chr16:88849875-88850755 | 880 | | | | | 1 | | | 0 | | | | | | | | | yes | | |
| CDT1 | N_Shore | Body | chr16:88871329-88871987 | 658 | | | | | | | | | 1 | | | | | | | | | | |
| GALNS | S_Shore | Body | chr16:88884836-88885124 | 288 | | | | 1 | | | | | | | | | | | yes | | | | |
| PABPN1L | NA;N_Shelf | 1stExon;TSS200 | chr16:88932955-88933182 | 227 | 1 | | 4 | | | | | | | | | | | yes | | | | | yes |
| CBFA2T3 | N_Shore | 3'UTR | chr16:88943319-88943542 | 223 | | | 1 | | | | | | | | | | | | | | | | yes |
| CBFA2T3 | Island | Body | chr16:88949985-88950197 | 212 | | | | | | | | | 1 | | | | | yes | | yes | | | |
| CBFA2T3 | N_Shore | 5'UTR;Body | chr16:88974840-88975259 | 419 | | | 3 | | | | | | | | | | | yes | | | | | |
| CBFA2T3 | S_Shore | 5'UTR;Body | chr16:88977768-88978463 | 695 | | | | | | | | | 2 | | | | | yes | yes | | | | |
| CBFA2T3 | NA | 5'UTR;Body | chr16:88988174-88988340 | 166 | | | | | | | | | | | | | | yes | | | | | |
| CBFA2T3 | N_Shelf | 5'UTR;Body | chr16:88990161-88990746 | 585 | | | 4 | 1 | 1 | | | | | | | | | yes | | | | | |
| CBFA2T3 | N_Shelf | 5'UTR;Body | chr16:89003037-89003641 | 604 | | | | | 2 | | | | | | | | | yes | | | | | |
| CBFA2T3 | N_Shore | 5'UTR;Body | chr16:89004858-89006455 | 1597 | | | 3 | | | | | | 1 | | | | | | | | | | |
| CBFA2T3 | S_Shore | Body | chr16:89009557-89010147 | 590 | | | 1 | | | | | | 1 | | | | | yes | | | yes | | |
| CBFA2T3 | NA | Body | chr16:89019403-89019632 | 229 | | | 1 | | | | | | | | | | | | | | | | |
| CBFA2T3 | NA | Body | chr16:89023389-89024657 | 1268 | | | | | | | | 0 | 2 | | 1 | 2 | | yes | yes | | | | |
| CBFA2T3 | Island;N_Shore | Body | chr16:89033692-89034797 | 1105 | | | 9 | | | | | 0 | | | | 2 | | yes | | | | | |
| CBFA2T3 | NA | Body;1stExon;5'UTR | chr16:89041793-89043707 | 1914 | | | 7 | | | | | | 1 | | | | | yes | | | | | |
| ACSF3 | S_Shore;N_Shelf | Body;5'UTR | chr16:89162196-89163625 | 1429 | | | | | | | | | 1 | | | | | | yes | | | | |
| ACSF3 | N_Shelf | Body;5'UTR | chr16:89164953-89165300 | 347 | | | | | | | | 0 | | | | | | yes | | | | | |
| ACSF3 | N_Shore;S_Shore | Body;5'UTR | chr16:89167018-89168963 | 1945 | | | | | | | | | 2 | | | 1 | | yes | | | yes | | |
| ACSF3 | NA | Body | chr16:89180285-89180843 | 558 | | | | | | | | | 2 | | | | | | | | | | |
| ACSF3 | N_Shelf | Body | chr16:89183643-89183728 | 85 | | | | | | | | | 1 | | | | | yes | | | | | |
| C16orf81 | N_Shore | TSS200;Body | chr16:89225495-89226194 | 699 | | | 5 | | | | | | | | | | | | | | | | |
| C16orf81 | N_Shelf | Body | chr16:89228590-89229439 | 849 | | | 4 | | | | | | 1 | | | | | | yes | | | yes | |
| CDH15 | N_Shore | TSS1500 | chr16:89237789-89237997 | 208 | | | | | | | | | 1 | | | | | | | yes | | yes | |
| CDH15 | Island | Body | chr16:89245659-89245911 | 252 | | | 1 | | | | | | | | | | | | | | | | |
| ANKRD11 | N_Shelf | Body | chr16:89342065-89342399 | 334 | | | | | | | | | | | | | | | yes | | | | |
| ANKRD11 | NA | Body | chr16:89363830-89364035 | 205 | | | 2 | | | | | | | | | | | | | | | yes | |
| ANKRD11 | NA | Body | chr16:89381904-89382131 | 227 | | | 4 | | | | | | | | | | | yes | | | | | |
| ANKRD11 | NA | 5'UTR | chr16:89387225-89387805 | 580 | | | 9 | | | | | | 1 | 1 | | | | | | | yes | yes | |
| ANKRD11 | NA | 5'UTR | chr16:89390609-89390968 | 359 | | | | | | | | | 1 | | | | | yes | yes | yes | | | |
| ANKRD11 | Island | 5'UTR | chr16:89408248-89408403 | 155 | | | | | | | | | 2 | | | | | | | | | | |
| ANKRD11 | S_Shore | 5'UTR | chr16:89410196-89410474 | 278 | | | 4 | | | | | | | | | | | yes | | | | | |
| ANKRD11 | NA | 5'UTR | chr16:89421700-89421770 | 70 | 1 | | | | | | | | | | | | | yes | | | | | |
| ANKRD11 | S_Shore;S_Shelf | 5'UTR | chr16:89439771-89441654 | 1883 | | | 4 | | | | | | 2 | | | | | yes | | | yes | | |
| ANKRD11 | NA | 5'UTR | chr16:89445433-89445503 | 70 | | | 7 | | | | | | 1 | | | | | yes | | | | | yes |
| ANKRD11 | NA | 5'UTR | chr16:89450023-89450094 | 71 | | | | 1 | | | | | | | | | | yes | | | | | |
| ANKRD11 | NA | 5'UTR | chr16:89461734-89462661 | 927 | | | 14 | | | | | | 1 | 1 | | | | yes | | | yes | | |
| ANKRD11 | Island | 5'UTR | chr16:89500152-89500225 | 73 | | | | | | | | | 1 | | | | | | | | | | |
| SPG7 | N_Shore | TSS1500 | chr16:89573312-89573955 | 643 | | | 12 | | | | | | 1 | | 1 | | | yes | | | | | |
| CPNE7 | N_Shore | TSS1500 | chr16:89640906-89641296 | 390 | | | 1 | | | | | | | | | | | | yes | | | | |
| DPEP1 | NA | 5'UTR;TSS1500;TSS200 | chr16:89686618-89687052 | 434 | | | 5 | | | | | | | | | | | | | | | | yes |
| DPEP1 | NA | 5'UTR | chr16:89688145-89688353 | 208 | | 1 | | | | | | | | | | | | | | | | | |
| DPEP1 | NA | 5'UTR | chr16:89692360-89692491 | 131 | | | | | | | | | | | | | | yes | | yes | yes | | |
| CDK10 | Island | Body | chr16:89761017-89761424 | 407 | | | 1 | | | | | | | | | | | | | | | | yes |
| FANCA | NA | Body | chr16:89822293-89822399 | 106 | | | | | | | | | 1 | | | | | yes | yes | | yes | | |
| SPIRE2 | N_Shelf | Body | chr16:89913618-89913972 | 354 | | | | | | 1 | | | | | | | | yes | | | | | |
| SPIRE2 | NA | Body | chr16:89925609-89926151 | 542 | | | | | | | | | 1 | | | | | | | | | | yes |
| TCF25 | NA | Body | chr16:89959984-89960457 | 473 | 1 | | | | | | | | | | | | | | | | | | |
| DEF8 | N_Shore | TSS1500 | chr16:90013969-90014316 | 347 | | | 3 | | | | | | 1 | | | | | yes | | yes | | | |
| DEF8 | Island | 5'UTR;Body | chr16:90015832-90016551 | 719 | 1 | | 5 | | | | 1 | 0 | 2 | 1 | | 4 | | yes | | | | yes | |
| AFG3L1 | NA | Body | chr16:90050102-90050151 | 49 | | | | | | | | | 1 | | | | | yes | | | | | |
| LOC100130015 | NA | Body;TSS200;TSS1500 | chr16:90113893-90115048 | 1155 | | | 11 | | | | | 0 | 4 | | | | | yes | | yes | | | yes |
| PRDM7 | Island | TSS1500 | chr16:90143751-90144788 | 1037 | | | 3 | | | | 1 | 0 | 1 | 3 | 1 | 3 | 1 | | yes | yes | | | |
| DOC2B | Island | TSS200 | chr17:31528-31545 | 17 | | | | | | | | | | 1 | | | | | | | | | yes |
| RPH3AL | Island | Body | chr17:85064-85492 | 428 | | | | | | | | | | | | | | | | | | | |
| RPH3AL | N_Shelf | Body | chr17:95730-96796 | 1066 | | | 7 | | | | | | | | | | | yes | | | | yes | |
| RPH3AL | NA | Body | chr17:109444-110138 | 694 | | | | | | | | 0 | | | | 1 | | | yes | | | | |
| RPH3AL | N_Shore | Body | chr17:113262-113799 | 537 | | | | | | | | | 1 | | | | | | | yes | | | |
| RPH3AL | N_Shore | Body | chr17:122697-123058 | 361 | | | | | | | | | | | | | | | yes | | | yes | |
| RPH3AL | NA | Body | chr17:146404-147234 | 830 | | | 8 | | | | | | | | | | | | | yes | | | |
| RPH3AL | N_Shelf | Body | chr17:149484-149582 | 98 | | 1 | 2 | | | | | | | | | | | yes | | | | | |
| RPH3AL | N_Shore;Island | Body | chr17:151914-152350 | 436 | | 1 | 5 | 1 | | | | 0 | 4 | | | 2 | | yes | | | | | |
| RPH3AL | S_Shore;N_Shelf | Body | chr17:154410-155045 | 635 | 6 | 4 | 21 | | 1 | | 1 | 0 | 3 | 2 | | 9 | | yes | | | yes | | yes |
| RPH3AL | Island | Body | chr17:171006-172159 | 1153 | | | | | | | | 0 | 5 | 1 | | 4 | | yes | | | | | |
| RPH3AL | NA | 5'UTR | chr17:180856-181288 | 432 | | | | | | | | | 1 | | | | | yes | | | yes | yes | |
| RPH3AL | N_Shelf;N_Shore;Island | 5'UTR | chr17:182656-185263 | 2607 | 2 | | 10 | | 1 | | 1 | 0 | 5 | 1 | 1 | 72 | 1 | | yes | | yes | | yes |
| RPH3AL | N_Shelf | TSS200 | chr17:202588-203087 | 499 | | | | | 1 | | | 0 | 2 | | | 1 | | | | | yes | | |
| C17orf97 | Island | TSS1500;TSS200 | chr17:259755-260292 | 537 | | | | | | | | 0 | 3 | 3 | | 4 | | yes | yes | | | | yes |
| GEMIN4 | NA | Body | chr17:650796-651216 | 420 | | | | | | | | | | | | | | | yes | | | | |
| ELP2P;GEMIN4 | Island | TSS1500;TSS200 | chr17:655677-655848 | 171 | | | | | | | | | 1 | | | | | | | | | | |
| RNMTL1 | NA | Body | chr17:693137-693388 | 251 | | | 7 | | | | | | | 1 | | | | | yes | | | | |
| NXN | N_Shore | Body | chr17:707032-707141 | 109 | | | | | | | | 0 | 1 | | | 1 | | yes | | | | | |
| NXN | NA | Body | chr17:727706-727791 | 85 | | | | | | | | | | | | | | yes | | | yes | | |
| NXN | Island | Body | chr17:738869-739405 | 536 | | | 17 | | | | | | | | | | | yes | | | | | |
| NXN | S_Shelf;S_Shore | Body | chr17:746502-747203 | 701 | | | 6 | | | | | | 1 | | | | | yes | | | | yes | |

| | | | | | | | | | | | | | | | | | | | | | | | |
|---|---|---|---|---|---|---|---|---|---|---|---|---|---|---|---|---|---|---|---|---|---|---|---|
| NXN | NA | Body | chr17:750241-751287 | 1046 | 1 | 1 | 1 | 1 | | | | 0 | 2 | | | | | yes | | yes | | | |
| NXN | NA | Body | chr17:767301-767380 | 79 | | | | | | | | 0 | 1 | | | | | yes | yes | | yes | | |
| NXN | NA | Body | chr17:773023-773283 | 260 | | | 2 | | | | | | | | | | | yes | | | yes | | |
| NXN | NA | Body | chr17:790628-791324 | 696 | | | 6 | | | | | | 1 | | | | | yes | | | | | |
| NXN | NA | Body | chr17:800084-800749 | 665 | | | 11 | 1 | 1 | | | 0 | 1 | 1 | 1 | 1 | | yes | | | | | |
| NXN | NA | Body | chr17:805469-805576 | 107 | | | 2 | | | | | | | | | | | yes | | | | | |
| NXN | NA | Body | chr17:807121-807555 | 434 | | | 1 | | | | | | | | | | | yes | | | | | |
| NXN | NA | Body | chr17:821546-821621 | 75 | | | | | | | | | | | | 1 | | yes | | | | | |
| NXN | S_Shore | Body | chr17:851785-851933 | 148 | | | 1 | | | | | | 1 | | | | | yes | yes | | yes | | yes |
| NXN | S_Shelf | Body | chr17:853971-854068 | 97 | | | 6 | | | | | | | | | | | yes | | | | | |
| NXN | NA | Body | chr17:855187-855312 | 125 | | 1 | 5 | | | | | | | | | | | | | | | yes | |
| NXN | NA | Body | chr17:872762-873452 | 690 | | 1 | 25 | | | | | 0 | 1 | 1 | | 3 | | yes | | | | | |
| NXN | N_Shore | Body | chr17:881647-881743 | 96 | | | 1 | | | | | | | | | | | | | | | yes | |
| ABR | S_Shore | Body | chr17:927457-927652 | 195 | | | | | | | | | 1 | | | | | yes | | | | | |
| ABR | N_Shore;Island | Body | chr17:932700-933179 | 479 | 2 | | 18 | | | | | | | | | | | | | | | | yes |
| ABR | Island;S_Shore | Body | chr17:934534-935884 | 1350 | | | | | | | | | 1 | | | | | yes | | | | | yes |
| ABR | NA | Body | chr17:943108-943342 | 234 | | | 4 | | | | | | | | | | | | | | | | |
| ABR | Island | Body | chr17:981518-981730 | 212 | 10 | 1 | | 1 | | | | | | | | | | | | | | yes | |
| ABR | N_Shore;Island | Body;5'UTR | chr17:1028394-1029394 | 1000 | 2 | | 26 | | | | | | | | | | | | | | | | yes |
| ABR | NA | Body;5'UTR | chr17:1040748-1041006 | 258 | | | | | | | | | 1 | | | | | yes | yes | | yes | | |
| ABR | N_Shore;Island | Body;5'UTR | chr17:1062865-1064210 | 1345 | | | 1 | 1 | | | | 0 | 1 | | | | | yes | yes | | yes | yes | |
| BHLHA9 | Island | TSS200 | chr17:1173760-1174148 | 388 | | | | | | | | | 1 | | | | | yes | | | | | yes |
| TUSC5 | NA | TSS1500 | chr17:1182702-1183159 | 457 | | | 1 | | | | | | | | | | | | | | | yes | |
| TUSC5 | NA | 3'UTR | chr17:1202015-1202252 | 237 | | | 3 | | | | | | | | | | | | | | | | |
| MYO1C | N_Shelf | Body | chr17:1372403-1372751 | 348 | | | | 1 | | | | | | | | | | yes | | | yes | yes | |
| MYO1C | N_Shore | 5'UTR;Body;TSS1500 | chr17:1390182-1390671 | 489 | | | 1 | 1 | 1 | | | | | | | | | yes | | | | | yes |
| MYO1C | Island;S_Shore | Body;TSS1500;1stExon;5'UTR;TSS2 | chr17:1395371-1396395 | 1024 | | | 10 | | | | | 0 | | | | | | | | | | yes | yes |
| PITPNA | S_Shore | TSS1500 | chr17:1466709-1466715 | 6 | | | 6 | | | | | 0 | | | | | | yes | yes | | | | |
| SLC43A2 | Island;N_Shore | 3'UTR | chr17:1477843-1479213 | 1370 | | | | | | | | 0 | | | | | | yes | | | | | |
| SLC43A2 | N_Shore | Body | chr17:1508261-1508609 | 348 | | | | | | | | 0 | 1 | | | 4 | 1 | | | | | yes | |
| SLC43A2 | Island | Body | chr17:1509883-1510666 | 783 | | | 2 | 1 | | | | | 2 | | 1 | | | yes | | | yes | | |
| SLC43A2 | NA | Body | chr17:1521312-1521483 | 171 | | | 6 | | | | | 0 | 1 | | | 2 | | yes | yes | | | | |
| SCARF1 | S_Shore | Body | chr17:1548881-1549098 | 217 | | | | | | | | 0 | 1 | | | | | yes | | | | | |
| RILP | S_Shore | 5'UTR;1stExon;TSS200 | chr17:1553341-1554577 | 1236 | | | 13 | 1 | 2 | 1 | | | 2 | 1 | | | | | | | | | yes |
| TLCD2 | Island | TSS200 | chr17:1613831-1613851 | 20 | | | | | | | | | 1 | | | | | | | | | | yes |
| C17orf91;MIR22 | N_Shore | Body;TSS200 | chr17:1617363-1617465 | 102 | | | | | | | | | | | | | | | | | | | |
| WDR81 | N_Shore | TSS1500;Body;5'UTR | chr17:1626477-1626575 | 98 | | | 1 | 1 | 1 | | | | 1 | 1 | | | | yes | | | | yes | |
| WDR81 | N_Shore | TSS200;Body;TSS1500;5'UTR | chr17:1627652-1627746 | 94 | | | | | | | | | | | | | | yes | | | | | yes |
| WDR81 | N_Shelf | Body | chr17:1633714-1634335 | 621 | | | 11 | | | | | | | | | | | yes | yes | | | yes | |
| SERPINF2 | S_Shore | 3'UTR | chr17:1658265-1658686 | 421 | | | | 1 | | | | | | | | | | yes | | | | | |
| SERPINF1 | NA | TSS1500;TSS200 | chr17:1664077-1665303 | 1226 | | | 7 | 1 | | | | | | | | | | yes | yes | | yes | | |
| SMYD4 | NA | Body | chr17:1686627-1686737 | 110 | | | 8 | | | | | 0 | 1 | | | | | yes | | | | | |
| RTN4RL1 | N_Shore;Island | Body | chr17:1880631-1881333 | 702 | | | 5 | | | | | 0 | 1 | | | 1 | | yes | | | yes | | yes |
| OVCA2;DPH1 | N_Shore | TSS1500;Body | chr17:1944594-1945076 | 482 | | | | | | | | | 1 | | | | | yes | yes | | | | yes |
| HIC1 | Island | TSS1500 | chr17:1957370-1957926 | 556 | | | 4 | | | | | | 1 | | | | | | | | | | yes |
| HIC1 | Island | Body;3'UTR | chr17:1961109-1962627 | 1518 | 1 | | 2 | | | | | | 1 | | | 1 | | | yes | | | | yes |
| SMG6 | NA | Body | chr17:2057082-2057234 | 152 | | | | | | | | 0 | 1 | | | | | | yes | | | | |
| MNT | NA | Body | chr17:2291793-2292039 | 246 | | | | | | | | | 1 | | | | | | | | | | |
| METT10D | NA | Body | chr17:2324794-2325143 | 349 | | | 2 | | 2 | | | | | | | | | yes | | | | | |
| KIAA0664 | Island | Body | chr17:2607736-2607986 | 250 | | | | 1 | | | | | | | | | | yes | yes | | | | |
| RAP1GAP2 | NA | TSS1500;TSS200;nan | chr17:2699122-2700093 | 971 | 7 | | 16 | 1 | 4 | | | | 3 | 1 | 1 | | | | | | | yes | |
| RAP1GAP2 | NA | Body | chr17:2797964-2798327 | 363 | | | | | | | 1 | 0 | 1 | 1 | | 1 | | yes | | | yes | | |
| RAP1GAP2 | NA | Body | chr17:2831573-2831903 | 330 | | | 1 | | | | | | | | | | | yes | yes | | | | |
| RAP1GAP2 | NA | Body | chr17:2839005-2839187 | 182 | | | | 1 | 1 | | | | 1 | | 1 | | | yes | yes | | | | |
| RAP1GAP2 | NA | Body | chr17:2843149-2843257 | 108 | | | | | | | | | 1 | | | | | yes | | | | | |
| RAP1GAP2 | NA | Body | chr17:2886250-2886453 | 203 | | | 1 | | | | | | | | | | | | | | | | |
| RAP1GAP2 | NA | Body | chr17:2907678-2908369 | 691 | | | | | | | | | 1 | | | | | yes | | | yes | | |
| OR1G1 | NA | TSS1500 | chr17:3031354-3031874 | 520 | | | | | | | | | | | | | | yes | yes | | yes | | |
| OR1A1 | NA | TSS1500 | chr17:3117953-3118291 | 338 | | | 1 | | | | | | | | | | | yes | yes | | | yes | |
| OR3A1 | NA | 1stExon | chr17:3195808-3196049 | 241 | | | | | | | | | | | | | | | yes | | | | |
| OR1E2 | NA | 1stExon | chr17:3336736-3337334 | 598 | | | | | | | | | | | | | | | yes | | | | |
| SPATA22 | Island | TSS200 | chr17:3375223-3375480 | 257 | | | | | | | | | | | | | | yes | | | | | |
| ASPA | chr17:3375006-3375237 | S_Shelf | chr17:3378804-3379283 | 479 | | | | | | | | | | | 1 | | | yes | | | | | |
| TRPV3 | N_Shelf | Body | chr17:3433428-3433970 | 542 | | | | | | | | | 1 | | | | | yes | | | yes | | |
| TRPV3 | Island;S_Shore | Body | chr17:3438857-3439372 | 515 | | 1 | 4 | 2 | | | | | 1 | | | | | yes | yes | | | | yes |
| TRPV3 | NA | 1stExon;5'UTR;TSS200 | chr17:3461140-3461480 | 340 | | 1 | 13 | 1 | | | | | 2 | | 1 | | | yes | | | | | |
| TRPV1 | N_Shore | Body | chr17:3474461-3474736 | 275 | | | | | | | | | 2 | | | | | yes | | | | | |
| TRPV1 | NA | Body | chr17:3493352-3493666 | 314 | | | | | | | | | 2 | | | | | | | | | | |
| TRPV1 | NA | TSS1500;5'UTR;TSS200 | chr17:3500390-3500417 | 27 | | | | | | | | | | | | | | yes | | | yes | | |
| CTNS | NA | Body | chr17:3558426-3558808 | 382 | | | | | | | | | | | | | | yes | | | | | |
| CTNS | NA | Body | chr17:3563641-3563924 | 283 | | | | | | | | | 1 | | | | | | | | | | |
| P2RX5 | NA | Body | chr17:3584925-3585166 | 241 | | | | | | | | | | | | | | yes | yes | | | | |
| P2RX5 | S_Shore | TSS1500 | chr17:3600356-3600514 | 158 | | | 5 | | | | | | 3 | | | | | yes | | | yes | | |
| ITGAE | NA | Body | chr17:3697993-3698239 | 246 | | | | | | | | | | | | | | yes | | | | | |
| ITGAE | NA | 1stExon;5'UTR;TSS200;nan | chr17:3704471-3704621 | 150 | | | 7 | | | | | 0 | 8 | 5 | 2 | 6 | | | | | | | |
| ITGAE | NA | TSS1500 | chr17:3705631-3705875 | 244 | | | 10 | 1 | | | | | | | | | | | | | yes | | |
| C17orf85 | NA | 3'UTR | chr17:3716214-3716426 | 212 | | | | | | | | | | | | | | | | | | | |
| CAMKK1 | N_Shore | 5'UTR | chr17:3791902-3792074 | 172 | | | 1 | | 1 | | | | | | | | | yes | | | | yes | |
| CAMKK1 | N_Shore | 5'UTR;TSS200;TSS1500 | chr17:3794118-3794828 | 710 | | | 4 | | | | | | 2 | | | | | | yes | | | | |
| P2RX1 | NA | 1stExon | chr17:3819429-3820150 | 721 | | | | | 1 | | | | | | | | | yes | | | | | |
| ATP2A3 | Island | 3'UTR | chr17:3827877-3828268 | 391 | | | 2 | | | | | | | | | | | | | | | | |
| ANKFY1 | NA | Body | chr17:4079262-4079652 | 390 | 7 | | | | | | | | 1 | | | | | yes | yes | | | | |
| SPNS3 | NA | TSS1500;TSS200 | chr17:4336494-4337231 | 737 | | | | | 1 | | | | 2 | 1 | | | | yes | yes | | | | |
| SPNS2 | Island | Body | chr17:4439394-4439623 | 229 | | | 6 | | | | | | | | | | | | | | | | yes |
| GGT6 | NA | TSS200;TSS1500;Body | chr17:4463371-4464400 | 1029 | | | | 1 | | | | | 1 | 1 | | | | yes | yes | | | | |
| SMTNL2 | N_Shore;Island | TSS1500;TSS200;1stExon;5'UTR | chr17:4486934-4488345 | 1411 | | | 9 | | | | | 0 | 1 | | | | | | | | | | yes |
| SMTNL2 | Island | Body | chr17:4500150-4500399 | 249 | | | | | | | | | | | | | | yes | yes | | | | |
| ALOX15 | S_Shore | TSS200 | chr17:4545074-4545721 | 647 | | 1 | | | | | | | | | | | | yes | | | | yes | |
| ARRB2 | NA | Body | chr17:4621205-4621252 | 47 | | | | | | | | 0 | 1 | | | 4 | | | | | | | |
| ZMYND15 | NA | Body | chr17:4648949-4649333 | 384 | | | 1 | | | | | 0 | 2 | | | | | yes | | | | | yes |
| TM4SF5 | NA | TSS1500 | chr17:4674843-4675193 | 350 | | | 1 | | | | | | | | | | | yes | | | | yes | |
| VMO1 | S_Shore;Island | TSS200;Body;3'UTR;1stExon;5'UT | chr17:4689165-4689893 | 728 | | | | | 1 | | | | 3 | | | 1 | | yes | | | | yes | |
| CHRNE;C17orf107 | Island | Body;TSS200;5'UTR;1stExon;3'UT | chr17:4802847-4804838 | 1991 | | | 11 | | | | | 0 | 2 | | | 2 | | | | | | | yes |
| GP1BA | NA | TSS1500 | chr17:4835341-4835627 | 286 | | | 1 | | | | | 0 | 1 | | | 2 | | yes | yes | | | | |
| CAMTA2;SPAG7 | S_Shore | Body;TSS1500 | chr17:4872326-4872603 | 277 | | | 18 | | | | | | | | | | | | | | | | |
| INCA1;CAMTA2 | S_Shore | 3'UTR;TSS1500 | chr17:4891542-4891775 | 233 | | | | 1 | | | | | | | | | | | | | | | |
| ZNF232 | N_Shore;Island | 5'UTR | chr17:5018984-5019989 | 1005 | | | 8 | | | | | 0 | 5 | | | 6 | | yes | yes | | | yes | |
| ZNF232 | Island | 5'UTR;TSS200;TSS1500 | chr17:5025970-5027051 | 1081 | | | 3 | | | | | 0 | | | | 2 | | | yes | | | yes | |
| C17orf87 | NA | Body;TSS200 | chr17:5137892-5138696 | 804 | | | | | 1 | | | | 1 | | | | | yes | | | | | |

| | | | | | | | | | | | | | | | | | | | | | | | |
|---|---|---|---|---|---|---|---|---|---|---|---|---|---|---|---|---|---|---|---|---|---|---|---|
| DERL2;MIS12 | Island | TSS200;TSS1500 | chr17:5389550-5389625 | 75 | | | | | | | | | 1 | | | | | | | | | | yes |
| LOC728392 | N_Shore;Island;chr17:5403131 | 3'UTR;1stExon;5'UTR;Island | chr17:5402883-5404581 | 1698 | | 1 | 7 | 1 | 1 | | 2 | 0 | 5 | 3 | 2 | 4 | | | yes | | | | yes |
| WSCD1 | Island;S_Shore | 5'UTR | chr17:5974064-5974609 | 545 | | | | | | | | | 2 | | | | | | | | | | yes |
| WSCD1 | NA | Body | chr17:5993686-5994006 | 320 | 1 | | 1 | | 1 | | | | | | | | | yes | yes | | | | |
| WSCD1 | Island | Body | chr17:6023675-6024454 | 779 | | | | | | | | | | | | | | | | | | | |
| AIPL1 | NA | Body;1stExon;5'UTR;TSS200 | chr17:6338181-6339302 | 1121 | | | 13 | | | | | | 2 | | | | | yes | | | | yes | |
| FAM64A | Island | 5'UTR | chr17:6347992-6348200 | 208 | | | | | | | | 0 | 2 | | | | | yes | | | | yes | |
| PITPNM3 | N_Shore;Island | 3'UTR | chr17:6358363-6358928 | 565 | | | 19 | | | | | | | | | | | yes | | | | | yes |
| SLC13A5 | Island | TSS200 | chr17:6616867-6617030 | 163 | | | | | | | | | 1 | | | | | | | | | | yes |
| XAF1 | NA | 5'UTR;1stExon | chr17:6659164-6659562 | 398 | | 1 | | 1 | | | | | | | | | | | | | yes | | |
| FBXO39 | Island | TSS1500;TSS200 | chr17:6679264-6679597 | 333 | | 1 | | | | | | | 6 | | | 2 | | yes | yes | | | | |
| TEKT1 | NA | TSS200 | chr17:6735152-6735272 | 120 | | | | | | | | | 1 | | | | | yes | yes | | | | |
| ALOX12P2 | NA | TSS1500;TSS200 | chr17:6756633-6756891 | 258 | | | 2 | | | | | | 1 | | | | | yes | yes | yes | | | yes |
| ALOX12P2 | N_Shore;Island | Body | chr17:6796745-6797771 | 1026 | | 1 | 3 | | | | | 0 | 6 | | | 6 | | | | | yes | | |
| ALOX12 | N_Shore;Island | TSS1500;TSS200;Body | chr17:6898738-6899888 | 1150 | 1 | | 11 | 1 | | | | 0 | 16 | 4 | 1 | 7 | | | | | | | |
| ALOX12 | N_Shelf | Body | chr17:6912197-6912673 | 476 | | | | | | | | | 1 | | | | | yes | | | | | |
| RNASEK;C17orf49 | N_Shore | Body;TSS1500 | chr17:6916926-6917393 | 467 | | | | | | | | | 1 | | | | | yes | | | | | |
| C17orf49;MIR195;MIR497 | S_Shelf | 3'UTR;Body;TSS1500 | chr17:6920698-6921465 | 767 | | | | | | | | | | | | | | | | | | | |
| BCL6B | Island | TSS200 | chr17:6926269-6926742 | 473 | | 1 | | 1 | | | | | | | | | | | | | | | |
| SLC16A11 | Island | Body;TSS200 | chr17:6945510-6948023 | 2513 | | | 9 | | | | | 0 | 1 | | | | | yes | | | | | |
| CLEC10A | NA | TSS1500 | chr17:6983991-6984187 | 196 | | | 5 | | | | | | | | | | | yes | yes | | | | |
| ASGR2 | NA | Body | chr17:7017474-7017968 | 494 | | | | 1 | | | | | | | | | | | | | | yes | yes |
| ASGR1 | Island | Body | chr17:7077300-7077709 | 409 | | | 4 | | | | | 0 | | | | | | | | | | yes | |
| ASGR1 | NA | 5'UTR;1stExon;TSS200 | chr17:7082562-7083413 | 851 | | | 7 | | | | | | | | | | | yes | | | | yes | |
| DLG4 | N_Shore | Body | chr17:7116792-7117241 | 449 | | | | | | | | 0 | | | | | | yes | | | | | yes |
| ACADVL;DLG4 | N_Shore | TSS1500;Body | chr17:7121848-7122063 | 215 | | | | | | | | | | | | | | | yes | | | | |
| CLDN7 | Island | TSS200 | chr17:7165842-7165849 | 7 | | | | | | | | | 1 | | | | | | | | | | yes |
| NEURL4 | Island | Body | chr17:7227499-7227737 | 238 | | | 2 | | | | | 0 | 1 | | | | | | | | | | |
| ACAP1 | NA | TSS200 | chr17:7239771-7240223 | 452 | | | | | 1 | | | | | | | | | yes | | | | | |
| ACAP1 | N_Shore;Island | Body | chr17:7253189-7253720 | 531 | | | | | | | | | 1 | 1 | | | | yes | | | | | |
| KCTD11 | S_Shore | 1stExon | chr17:7256361-7256673 | 312 | | | 3 | | | | | | | | | | | | | | | | |
| TMEM95;KCTD11 | S_Shelf | TSS200;3'UTR;TSS1500;1stExon | chr17:7257885-7258501 | 616 | | | 4 | | | | | | | | | | | | | | | yes | |
| TNK1 | N_Shore;Island | TSS1500;TSS200 | chr17:7283817-7284351 | 534 | | | | | | | | 0 | 2 | | | 1 | | | yes | | | | |
| TNK1 | N_Shore;Island | Body | chr17:7287100-7288001 | 901 | | 1 | 29 | 1 | 3 | | | 0 | 1 | 3 | | | | yes | yes | | | | |
| NLGN2 | N_Shore;Island | TSS200;1stExon | chr17:7311314-7312081 | 767 | | | | | | | | | 3 | | | 1 | | | | | | | yes |
| SPEM1 | S_Shelf | TSS200 | chr17:7323605-7324093 | 488 | | | | | | | | | | | | | | | | | | | |
| C17orf74 | NA | TSS1500 | chr17:7328520-7328814 | 294 | | | | | | | | | | | | | | yes | | | | | |
| TMEM102;FGF11 | Island;N_Shore | 3'UTR;TSS1500 | chr17:7340867-7341936 | 1069 | | | | | | | | | | | | | | | | | | | |
| CHRNB1 | Island | TSS200 | chr17:7348316-7348490 | 174 | | | 21 | | | | | | 1 | | | | | | | | | | |
| CHRNB1 | S_Shore | Body | chr17:7350001-7350413 | 412 | | | 1 | 1 | | | | 0 | | | | | | yes | | | yes | yes | yes |
| ZBTB4 | N_Shore | 5'UTR | chr17:7381288-7381769 | 481 | | | | 1 | | | | | 1 | | | | | | | | | | |
| TNFSF13;TNFSF12-TNFSF13 | N_Shelf | TSS1500;Body | chr17:7461260-7461638 | 378 | | | | | | | | 0 | 2 | | | | | | | | | | |
| SENP3 | S_Shore | Body | chr17:7466447-7466821 | 374 | | | | | | | | | | | | | | | | | | yes | yes |
| EIF4A1;SNORA48 | S_Shore | Body;TSS1500 | chr17:7477617-7477972 | 355 | | | | | | | | | | | | | | | | | | | |
| EIF4A1;CD68 | NA | 3'UTR;TSS1500 | chr17:7482003-7482475 | 472 | | | | | | | | | | | | | | | | | | | |
| MPDU1 | Island | TSS1500 | chr17:7486551-7486821 | 270 | | | | | | | | | | | | | | | | | | | |
| SOX15 | S_Shore | 1stExon;5'UTR | chr17:7493150-7494151 | 1001 | | | | | | | | | 2 | 1 | | | | | | | | | |
| SHBG;FXR2 | N_Shore | TSS1500;Body;TSS200 | chr17:7516888-7517258 | 370 | | | 7 | | 1 | | | | | | | | | | | | | | |
| TP53 | NA | Body;TSS1500 | chr17:7579263-7579546 | 283 | | | | | | | | | | | | | | yes | | | | | |
| DNAH2 | Island | Body | chr17:7643798-7644241 | 443 | | | 2 | | | | | | 3 | | | 1 | | yes | | | | | yes |
| DNAH2 | Island | Body | chr17:7690327-7690529 | 202 | | | | | | 1 | | | 1 | | | | | yes | | | | | |
| DNAH2 | NA | Body | chr17:7701546-7701636 | 90 | | | 6 | | | | | | | | | | | | | | | | |
| KDM6B | Island | Body | chr17:7756609-7756882 | 273 | | | | | | | | | 1 | | | | | | | | | | yes |
| TMEM88 | Island | Body | chr17:7758692-7759140 | 448 | | | 3 | | | | | | | | | | | | | | | | |
| CHD3 | S_Shelf | Body;TSS1500;TSS200 | chr17:7791630-7792191 | 561 | | | 14 | 1 | | | | | 2 | 1 | | | | | | | | | |
| CHD3;SCARNA21 | NA | Body;TSS1500 | chr17:7808598-7808829 | 231 | | | | | | | | | | | | | | | | | | | |
| KCNAB3 | Island | 3'UTR | chr17:7826155-7826623 | 468 | | | | | | | | 0 | | | | | | | | | | | yes |
| KCNAB3 | Island | 1stExon;TSS200 | chr17:7832680-7833237 | 557 | | 1 | 33 | | | | | 0 | 6 | 1 | | 4 | | | | | yes | | yes |
| TRAPPC1;CNTROB | S_Shore | TSS1500;1stExon | chr17:7836533-7836945 | 412 | | | 3 | 1 | | | | 0 | 1 | | | 3 | | | | | | | |
| GUCY2D | Island;N_Shore | TSS200;TSS1500 | chr17:7905636-7906098 | 462 | | 1 | 4 | | | | | | | | | | | yes | yes | | | yes | yes |
| ALOX15B | NA | TSS1500;TSS200;5'UTR;1stExon | chr17:7942137-7942743 | 606 | | | 1 | | | | | 0 | | | | | | yes | yes | | | | |
| PER1 | Island | 5'UTR;1stExon | chr17:8055733-8055834 | 101 | | | | | 1 | | | | | | | | | | | | | | |
| AURKB | S_Shore | TSS1500 | chr17:8114504-8114626 | 122 | | | 4 | | | | | | 1 | | | | | yes | yes | yes | | | |
| C17orf68 | S_Shelf | 3'UTR | chr17:8129997-8130356 | 359 | | 1 | | 1 | | | | | 1 | | | 4 | 1 | | | | | | |
| SLC25A35;RANGRF | Island | 3'UTR;Body | chr17:8192194-8192371 | 177 | | | | | 3 | | | | | | | | | yes | | | | yes | |
| ARHGEF15 | NA | Body | chr17:8218709-8219095 | 386 | | | | | | | | | 1 | | | | | | | | | | |
| RNF222 | Island | 3'UTR | chr17:8295971-8296226 | 255 | | 1 | | | | | | | | | | | | | | | | yes | |
| RNF222 | NA | 1stExon;5'UTR;TSS200 | chr17:8301118-8301192 | 74 | | 1 | 7 | | | | | 0 | 2 | | | 2 | | | yes | | | yes | |
| MYH10 | S_Shelf | Body | chr17:8383621-8383800 | 179 | | | 15 | | | | | | | | | | | | yes | | | | |
| CCDC42 | N_Shore | 1stExon;5'UTR | chr17:8647944-8648562 | 618 | | 1 | | 1 | | | | | | | | | | yes | | | | | |
| MFSD6L | Island | 1stExon;5'UTR;TSS200 | chr17:8702369-8702896 | 527 | | | | | | | 1 | 0 | 11 | 1 | 1 | 10 | 2 | | yes | yes | | yes | |
| PIK3R6 | NA | 5'UTR;1stExon;TSS200 | chr17:8770908-8771331 | 423 | | | 5 | 1 | | | | | 3 | 1 | | | | yes | yes | | | | |
| PIK3R5 | Island | Body | chr17:8791763-8792092 | 329 | | | | | | | | | | | | | | yes | | | | yes | |
| PIK3R5 | NA | 5'UTR;TSS200 | chr17:8815262-8816266 | 1004 | | | | | | | | | 2 | | 1 | | | yes | yes | | | | |
| PIK3R5 | Island | 5'UTR | chr17:8868541-8868834 | 293 | | | 4 | | | | | | 1 | | | | | | | | | yes | |
| PIK3R5 | S_Shore | TSS1500 | chr17:8869864-8869961 | 97 | | | | | | | | | | | | | | yes | | | | | |
| NTN1 | Island;S_Shore | Body | chr17:8926755-8928451 | 1696 | | | | 1 | | | | 0 | 1 | | | | | | | | | | yes |
| NTN1 | N_Shore;Island | Body | chr17:9018806-9019336 | 530 | | | | 2 | | | | 0 | | | | 1 | | yes | | | | yes | |
| NTN1 | Island | Body | chr17:9066156-9066686 | 530 | | | | | | | | | 2 | | | | | | | yes | | yes | |
| NTN1 | NA | Body | chr17:9129824-9130163 | 339 | | | | | | | | | 1 | | | 1 | | yes | | | | yes | |
| USP43 | S_Shore | Body | chr17:9550137-9550545 | 408 | | | 9 | | | | | 0 | 1 | | | 1 | | yes | yes | | | yes | |
| USP43 | NA | Body | chr17:9558932-9559589 | 657 | | | 2 | | | | | | | | | | | | yes | | | | |
| DHRS7C | N_Shore | Body | chr17:9674817-9675101 | 284 | | | | | | | | | | | | | | yes | | | yes | | |
| DHRS7C | NA | 1stExon;TSS200 | chr17:9694472-9695137 | 665 | | | 8 | | | | | | | | | | | yes | | yes | yes | | |
| GLP2R | NA | TSS1500;TSS200 | chr17:9729077-9729424 | 347 | | | 12 | | | | | | | | | 1 | | yes | yes | | | | |
| RCVRN | NA | Body | chr17:9802783-9802860 | 77 | | | 6 | | | | | 0 | 1 | 1 | | | | | yes | | yes | | |
| RCVRN | N_Shelf | Body | chr17:9804355-9804605 | 250 | | | | | | | | | 1 | | | | | | yes | yes | yes | | |
| GAS7 | NA | Body;5'UTR;1stExon;TSS200 | chr17:9862752-9863081 | 329 | | 1 | 13 | | | | | 0 | | | | | | yes | | | | | |
| GAS7 | NA | Body;TSS200 | chr17:9929698-9930101 | 403 | | | 3 | | | | | | | | | | | | yes | | | yes | |
| GAS7 | NA | Body;1stExon;5'UTR;TSS200 | chr17:9940004-9940321 | 317 | | | | | | | | 0 | 1 | 1 | | | | | | | | yes | |
| GAS7 | NA | Body | chr17:9967417-9967499 | 82 | | | 7 | | | | | | 1 | | | | | | | yes | | | |
| MYH1 | NA | TSS200;5'UTR | chr17:10421635-10422189 | 554 | | | 8 | | | | | | | | | | | | | | | yes | |
| MYH2 | NA | 5'UTR;TSS200;1stExon | chr17:10452851-10453296 | 445 | | | 3 | | | | | | 1 | | | | | | | | | | |
| SHISA6 | Island | TSS1500 | chr17:11143830-11144509 | 679 | | | | | | | | 0 | 3 | 1 | | | | | | | | yes | yes |
| SHISA6 | Island | Body | chr17:11461103-11461665 | 562 | | | 4 | | | | | | 2 | | | | | | | | yes | yes | |
| DNAH9 | Island | TSS200;1stExon | chr17:11501632-11502254 | 622 | | | | | | | | 0 | 2 | | | 1 | | | | | yes | | |
| MYOCD | N_Shore;Island | TSS1500;TSS200 | chr17:12567873-12569135 | 1262 | | | 2 | | | | | | 4 | | | | | | | yes | | | |
| RICH2 | S_Shore | Body | chr17:12694449-12694843 | 394 | | | | | 1 | | | | 1 | | | | | | yes | | | | |
| RICH2 | N_Shore;Island | Body | chr17:12877227-12877772 | 545 | | | | 2 | 1 | | | | | | | | | yes | | | yes | | |

| | | | | | | | | | | | | | | | | | | | | | | | | |
|---|---|---|---|---|---|---|---|---|---|---|---|---|---|---|---|---|---|---|---|---|---|---|---|---|
| RICH2 | NA | Body | chr17:12893389-12893648 | 259 | | | | 1 | | | | | | | | | | | | | | | | |
| HS3ST3A1 | S_Shore | TSS1500 | chr17:13505751-13506284 | 533 | | | | | | | | | 3 | | | 3 | | | | | | yes | yes | |
| HS3ST3B1 | Island | TSS1500 | chr17:14203157-14203711 | 554 | | | 1 | | | | | | | | | | | | | | | yes | | |
| HS3ST3B1;MGC12916 | Island;S_Shore | Body;TSS1500;TSS200 | chr17:14206572-14208363 | 1791 | | | 44 | 1 | | | | 0 | 2 | | | 1 | | yes | | | | | | |
| HS3ST3B1 | S_Shore | Body | chr17:14213225-14214081 | 856 | | | | | | | | | 2 | | 1 | | | yes | | | | | yes | |
| PMP22 | Island | 5'UTR;1stExon | chr17:15164050-15164106 | 56 | | | | | | | | | 1 | | | | | | | | | | | |
| PMP22 | Island | 5'UTR;1stExon | chr17:15165803-15165922 | 119 | | | | | | | | | 1 | | | | | | | | | | yes | |
| TEKT3 | N_Shore;Island | 5'UTR;1stExon;TSS200 | chr17:15244441-15245237 | 796 | | | 31 | | | | | | | | | | | yes | yes | | | | | |
| MEIS3P1 | Island | TSS1500 | chr17:15689920-15690132 | 212 | | | | | | | | | | | | | | | | | | yes | yes | |
| NCOR1;PIGL | Island;S_Shore | TSS1500;TSS200 | chr17:16119344-16120890 | 1546 | | | 2 | | | | | | | | | | | yes | | | | | | |
| CENPV | Island | TSS1500 | chr17:16257111-16257223 | 112 | | | | | | | | | | | | | | yes | | | | yes | yes | |
| TRPV2 | NA | TSS1500;TSS200 | chr17:16318334-16318972 | 638 | | | 12 | 1 | 1 | | | | | | | | | yes | yes | | | | | |
| CCDC144A | Island | TSS1500;TSS200 | chr17:16593426-16593885 | 459 | | | 22 | | | | | 0 | 1 | 1 | | 3 | | yes | | | | | | |
| TNFRSF13B | NA | Body;TSS200 | chr17:16875129-16876525 | 1396 | | | | | 1 | | | | 3 | | | | | yes | yes | | | | | |
| PLD6 | Island;N_Shore | Body;1stExon;TSS200;TSS1500 | chr17:17108846-17110353 | 1507 | | 3 | 6 | 1 | 1 | | 1 | 0 | 10 | 6 | | 19 | 2 | | yes | | | yes | yes | |
| FLCN | N_Shore;Island | Body;3'UTR | chr17:17124521-17124828 | 307 | | | | | 1 | | | | | | | | | | yes | | | | | |
| PEMT | NA | Body | chr17:17457745-17457836 | 91 | 2 | | | | | | | | | | | | | yes | yes | | | | | |
| PEMT | Island | Body | chr17:17465162-17465682 | 520 | | | 5 | | | | | | 3 | | | | | yes | | | | | | |
| RAI1 | N_Shore;Island | 5'UTR | chr17:17603531-17604184 | 653 | | | 14 | | | | | 0 | 5 | | | 1 | | yes | | | | yes | yes | |
| RAI1 | Island | Body;3'UTR | chr17:17713061-17714414 | 1353 | | | | 1 | | | | | 2 | | | | | | | | | yes | | |
| MIR33B;SREBF1 | S_Shore | TSS1500;Body | chr17:17717894-17718525 | 631 | | | | | | | | | 2 | | | | | | | | | | | |
| LRRC48 | Island | Body | chr17:17910235-17910425 | 190 | | | 1 | | | | | | 1 | | | | | yes | yes | | | | | |
| MYO15A | NA | TSS200;TSS1500 | chr17:18011513-18012014 | 501 | | | 2 | | | | | | | | | | | yes | | | | | | |
| SHMT1 | N_Shore | 5'UTR | chr17:18266135-18266219 | 84 | | | 1 | | | | | | | | | | | yes | | | | | | |
| CCDC144B | Island | 1stExon | chr17:18528568-18529187 | 619 | | | 12 | | | | | | 1 | | | 1 | | yes | yes | | | yes | | |
| TBC1D28 | NA | 5'UTR;1stExon | chr17:18547711-18547986 | 275 | | | 1 | | | | | | | | | | | | | | | yes | | |
| SLC5A10 | NA | TSS1500 | chr17:18854715-18854799 | 84 | | | | | | | | | | | | | | yes | | | | | | |
| MIR1180;B9D1 | Island | TSS200;Body | chr17:19247837-19248009 | 172 | | | 1 | | | | | | 3 | | | | | yes | | | | | | |
| RNF112 | NA | TSS1500;TSS200 | chr17:19313024-19314618 | 1594 | | | 14 | | | | | 0 | 2 | | | 2 | | yes | yes | | | | yes | |
| SLC47A1 | Island;N_Shore | TSS1500 | chr17:19436770-19436900 | 130 | | | | | | | | | 1 | | | 1 | | yes | | | | | yes | |
| SLC47A1 | S_Shore | Body | chr17:19437929-19438253 | 324 | | | | | | | | | | | | | | yes | | | | | | |
| SLC47A2 | S_Shelf | 1stExon;5'UTR;TSS1500 | chr17:19619905-19620595 | 690 | | | | | | | | | | | | | | yes | | | | | | |
| ALDH3A1 | S_Shore;N_Shore;Island | 1stExon;5'UTR;TSS200;TSS1500;B | chr17:19647973-19650069 | 2096 | | | 16 | | 1 | | | | 3 | 1 | | | | yes | yes | | | yes | yes | |
| ALDH3A1 | S_Shelf | 5'UTR;1stExon;TSS200 | chr17:19651653-19652056 | 403 | | | | | | | | 0 | 3 | | | | | yes | yes | | | | | |
| AKAP10 | Island | TSS1500 | chr17:19881632-19881925 | 293 | | | 3 | | | | | | | | | | | | | yes | | | | |
| CCDC144C | Island | TSS1500 | chr17:20224230-20224414 | 184 | | | | | | | | 0 | | | | | | | | | | | | |
| CCDC144NL | Island | 1stExon;5'UTR;TSS200 | chr17:20799162-20799769 | 607 | | 1 | 19 | | | 2 | | 0 | 9 | | | 5 | | yes | | | | | | |
| USP22 | NA | Body | chr17:20920717-20920818 | 101 | 1 | | | | | | | | | | | | | | yes | | yes | | | |
| DHRS7B | N_Shore;Island | TSS1500;TSS200 | chr17:21030030-21030859 | 829 | | | | | | | | | 2 | | | 1 | | yes | yes | | | | | |
| MAP2K3 | NA | Body;TSS200 | chr17:21194718-21194806 | 88 | | | | 1 | | | | | 1 | | | | | yes | | | | yes | | |
| KCNJ12 | Island | 5'UTR | chr17:21281017-21281507 | 490 | | | | | | | | | | | | 1 | | | | | | | | |
| KCNJ12 | NA | 5'UTR | chr17:21300735-21300791 | 56 | | | 28 | | | | | | | | | | | yes | | | | | | |
| C17orf51 | N_Shore;Island | Body;1stExon;5'UTR | chr17:21454238-21455185 | 947 | | | 2 | | | | | 0 | 2 | 1 | | | | yes | | | yes | yes | | |
| FAM27L | NA | TSS200 | chr17:21825321-21825983 | 662 | | | | | | | | | | | | | | | yes | | | yes | | yes |
| FLJ36000 | NA | Body | chr17:21911362-21911491 | 129 | | | | | | | | | | | | | | | | | | yes | | yes |
| WSB1 | N_Shore | TSS1500 | chr17:25620596-25620677 | 81 | | 1 | | | | | | | | | | | | yes | | | | | | |
| KSR1 | NA | TSS200 | chr17:25798878-25799212 | 334 | | | | | | | | | 2 | | | | | yes | yes | | | | | |
| LGALS9 | NA | TSS1500;Body;1stExon;5'UTR | chr17:25957771-25958303 | 532 | | | 6 | | | | | | | | | | | yes | | | | yes | | |
| NOS2 | NA | 1stExon;5'UTR | chr17:26127438-26127542 | 104 | | | 4 | 1 | 1 | | | | | | | | | yes | | | | yes | | |
| PPY2 | N_Shelf | TSS200 | chr17:26574271-26574374 | 103 | | | | | | | | | | | | | | | | | | yes | | |
| FLJ40504 | Island | Body | chr17:26633978-26634469 | 491 | | | | | | | | | | | | | | yes | | yes | | | | |
| TMEM97 | N_Shore | TSS1500 | chr17:26645200-26645523 | 323 | | | | | | | | | | 1 | | | | yes | | | | | | |
| SEBOX | NA | Body;TSS200 | chr17:26692284-26692844 | 560 | | | | 1 | | | | | | | | | | yes | | | | | | |
| SARM1 | Island | 1stExon | chr17:26699169-26699551 | 382 | | | | 1 | | | | | | | | | | yes | | | | | yes | |
| SARM1 | N_Shore;Island | Body | chr17:26711117-26712412 | 1295 | | | 10 | 1 | | | | 0 | 1 | | | | | | yes | | | yes | | |
| SLC46A1 | N_Shore;Island | Body | chr17:26732120-26732374 | 254 | | | 15 | | 1 | | | | | | | | | | | | | | | |
| FOXN1 | NA | TSS1500;TSS200 | chr17:26850360-26851153 | 793 | | | | | | | | | | | | | | yes | | | | yes | | |
| PIGS | S_Shore | TSS1500 | chr17:26899172-26899428 | 256 | | | | | | | | | | | | | | yes | | | | | | |
| FLJ25006;LOC645851 | NA | Body;TSS1500 | chr17:26939298-26939704 | 406 | 1 | | 2 | | | | | | | | | | | | | | | | yes | |
| FLJ25006;KIAA0100 | NA | Body;TSS1500;3'UTR | chr17:26940926-26942215 | 1289 | | | 6 | | | | | | | | | | | | yes | | | yes | | |
| PROCA1 | Island;S_Shore | 5'UTR;1stExon;TSS200;TSS1500 | chr17:27038686-27039523 | 837 | | 1 | 3 | 1 | 2 | | | | 2 | | | | | yes | | | | yes | | |
| RAB34 | Island;N_Shore | Body;5'UTR;1stExon;TSS200;TSS1 | chr17:27044629-27045302 | 673 | | | | 1 | | | | 0 | 3 | 1 | | | | | | | | | yes | |
| RPL23A;SNORD42A;SNORD4B;SNORD4A | N_Shelf;S_Shore | Body;TSS1500;TSS200 | chr17:27048708-27050042 | 1334 | | | 1 | | 1 | | | | 2 | | | | | yes | | | | | | |
| TRAF4 | Island | Body | chr17:27071569-27072015 | 446 | | | | | | | | | | | | | | yes | | | | | yes | |
| C17orf63 | S_Shore | TSS200 | chr17:27170014-27170043 | 29 | | | | | | | | | 1 | | | | | yes | | | | | | |
| MIR451;MIR144 | NA | Body;TSS200 | chr17:27188437-27188758 | 321 | | | 2 | | | | | | | | | | | yes | | | | | | |
| DHRS13 | S_Shore | TSS200 | chr17:27230278-27230393 | 115 | | | | | | | | | | | | | | yes | | | | yes | | |
| PHF12 | NA | Body | chr17:27252884-27253274 | 390 | | | | 1 | | | | | 1 | | | | | yes | | | | | | |
| SEZ6 | S_Shore | TSS1500 | chr17:27333463-27333617 | 154 | | | | | | | | | | | | | | yes | | | | yes | | |
| PIPOX | NA | TSS200 | chr17:27369851-27370117 | 266 | | | 3 | | | | | | | | | | | | | | yes | | | |
| ABHD15 | chr17:27893086-27896078 | Island | chr17:27893087-27893542 | 455 | | | | | | | | | | | 1 | | | yes | | | | | yes | |
| TP53I13 | N_Shore | Body | chr17:27899465-27899966 | 501 | | | | 1 | | | | | | | | | | | | | | | | |
| ANKRD13B;CORO6 | N_Shore | 3'UTR | chr17:27941776-27942773 | 997 | | | | | | | | | | | | | | | | | | | | |
| SSH2 | Island;N_Shore | Body | chr17:28087898-28088606 | 708 | | | | | 3 | | | | | | | | | yes | | | | yes | yes | |
| CPD | N_Shore | TSS1500 | chr17:28705147-28705364 | 217 | | | 1 | | | | | | | | | | | | yes | | | | | |
| GOSR1 | N_Shore | TSS1500 | chr17:28803808-28803927 | 119 | | | 6 | | | | | | | | | | | | yes | | | | | |
| LRRC37B2 | N_Shore | TSS1500 | chr17:28902652-28903005 | 353 | | | 1 | | | | | | | | | | | | yes | | | | | yes |
| LRRC37B2 | Island;N_Shore | Body | chr17:28927070-28928406 | 1336 | | | | | | | | 0 | 3 | | | 1 | | yes | | | | | yes | |
| C17orf42 | Island | 5'UTR;1stExon | chr17:29233276-29233864 | 588 | | | | | | | | 0 | | | | | | yes | | | | | | |
| ADAP2 | N_Shore | TSS200 | chr17:29248604-29248848 | 244 | | | 7 | | | | | | | | | | | yes | yes | | | | | |
| RNF135 | N_Shore | TSS1500 | chr17:29297380-29297604 | 224 | 8 | 3 | 16 | | | | | 0 | 4 | | | 7 | | yes | | | yes | yes | yes | |
| MIR193A | Island | TSS200;TSS1500 | chr17:29886672-29887376 | 704 | | 1 | | | | | | 0 | 2 | | | 1 | | | | | | | yes | |
| RHBDL3 | N_Shore | TSS1500;TSS200 | chr17:30591837-30593009 | 1172 | | | 4 | | | | | | 3 | | | | | | | | | yes | yes | |
| PSMD11 | N_Shore | TSS1500 | chr17:30770932-30771154 | 222 | | | | | | | | 0 | | | | | | yes | yes | | | yes | yes | |
| MYO1D | N_Shore;Island | Body | chr17:30822561-30823145 | 584 | | 1 | 19 | | | | | | | | | | | | | | | | | |
| TMEM98 | N_Shore | TSS200;TSS1500 | chr17:31254544-31254878 | 334 | | | | | | | | 0 | 1 | | | | | yes | | yes | yes | | yes | |
| ACCN1 | N_Shore;Island | 1stExon;5'UTR;TSS200;TSS1500 | chr17:32483560-32484259 | 699 | | 1 | | | 2 | | | | 1 | | | | | | | | | | | |
| CCL8 | NA | TSS1500 | chr17:32645748-32646006 | 258 | | | | | | | | | 1 | | | | | | yes | | yes | | | |
| CCL1 | NA | 1stExon | chr17:32690127-32690367 | 240 | | 1 | 5 | | | | | | | | | | | yes | | | | | | |
| C17orf102 | Island | 1stExon | chr17:32905966-32906205 | 239 | | 1 | | | | | | | | | | | | | | | | | | |
| TMEM132E;C17orf102 | Island | TSS1500;TSS200 | chr17:32907388-32907705 | 317 | | 1 | | | | | | | 1 | | | | | | | | | | yes | |
| TMEM132E | Island | Body | chr17:32909042-32909375 | 333 | | | 3 | | | | | | | | | | | | | | | | yes | |
| TMEM132E | Island | Body | chr17:32953351-32953678 | 327 | | | | | | | | 0 | | | | | | | | | | | yes | |
| TMEM132E | N_Shore;Island | Body | chr17:32964265-32965387 | 1122 | | | 6 | | | | | | 1 | | | | | | yes | | | | | |
| RFFL | NA | 1stExon;5'UTR;TSS1500 | chr17:33390736-33391960 | 1224 | | | | | | | | | 1 | | | | | yes | | | | | | |
| RFFL | S_Shore | TSS1500 | chr17:33416834-33417113 | 279 | | | 6 | | 1 | | | | | | | | | yes | | | | yes | yes | |
| UNC45B | NA | TSS1500;TSS200 | chr17:33474376-33475031 | 655 | | | 3 | | | | | | | | | | | | | | | | | |
| SLFN12 | NA | 1stExon;5'UTR;TSS200;TSS1500 | chr17:33759484-33760527 | 1043 | 2 | 8 | 48 | | | | | 0 | 12 | 4 | 1 | 29 | 1 | | | yes | | | | |
| SLFN13 | N_Shore | TSS200;TSS1500 | chr17:33775860-33776683 | 823 | | | 8 | | | | | 0 | 4 | 3 | | | | yes | yes | | | yes | | |

| | | | | | | | | | | | | | | | | | | | | | | | |
|---|---|---|---|---|---|---|---|---|---|---|---|---|---|---|---|---|---|---|---|---|---|---|---|
| SLFN12L | N_Shore | Body | chr17:33814163-33815523 | 1360 | | | | | | | | | 1 | | | 1 | | yes | | | | | |
| SLFN12L | S_Shelf | Body | chr17:33817136-33817217 | 81 | | | | | | | | | 1 | | | | | | yes | yes | | | |
| SLFN12L | NA | Body | chr17:33825172-33825374 | 202 | | | 1 | | | | | | | | | | | yes | | | | | yes |
| SLFN12L | NA | Body | chr17:33842181-33842301 | 120 | | | 6 | | | | | 0 | 1 | | | 2 | | yes | | | | | |
| SLFN12L | NA | TSS1500 | chr17:33866067-33866073 | 6 | | | 1 | | | | | | | | | | | yes | | yes | yes | yes | |
| RASL10B | Island | Body | chr17:34062219-34062321 | 102 | | | | | | | | | | | | | | | yes | | | | |
| GAS2L2 | NA | Body | chr17:34077141-34077293 | 152 | | | | | | | | | | | | 1 | | yes | | | | | |
| GAS2L2 | NA | TSS200 | chr17:34079987-34080240 | 253 | | | 2 | | | | | | | | | | | yes | | | | | |
| C17orf50 | N_Shelf | TSS1500 | chr17:34087153-34087579 | 426 | | | | | | | | | | | | | | yes | | yes | | | |
| MMP28 | Island | Body | chr17:34093631-34093754 | 123 | | | | | | | | | | | | | | | yes | | | | |
| MMP28 | S_Shore | TSS1500 | chr17:34122866-34123272 | 406 | | | | | | | | | | | | | | yes | | | | | |
| CCL5 | NA | 1stExon;5'UTR | chr17:34207332-34207663 | 331 | | | 1 | | | | | | 1 | | 1 | | | yes | | | | | |
| CCL16 | NA | TSS1500 | chr17:34309129-34309577 | 448 | | | | | | | | | | | | | | yes | yes | | | | yes |
| CCL23 | NA | 1stExon;5'UTR;TSS200 | chr17:34344995-34345392 | 397 | | | 5 | 1 | | | | | 1 | 1 | | | | | yes | yes | | | |
| CCL3 | NA | Body;5'UTR;1stExon | chr17:34417074-34417575 | 501 | | | | | | | | | | | | | | | | | | | |
| LHX1 | Island | TSS1500;TSS200;1stExon;Body | chr17:35293930-35297146 | 3216 | | | | | | | | | 4 | | | | | | | | | yes | yes |
| LHX1 | Island | Body | chr17:35300010-35300868 | 858 | | | | | | | | | 1 | | | | | | | | | yes | yes |
| ACACA | NA | 5'UTR;Body;1stExon;TSS200 | chr17:35656660-35657046 | 386 | | | 9 | | 1 | | | | 1 | | | | | | | | | | |
| DDX52 | S_Shore | TSS1500 | chr17:36004446-36004794 | 348 | | | | | 1 | | | | | | | | | yes | yes | | yes | | |
| HNF1B | Island | Body | chr17:36103978-36105517 | 1539 | 1 | | | | | | | | | | | | | | | | | | |
| GPR179 | NA | TSS200 | chr17:36499751-36500237 | 486 | | | 2 | | | | | | | | | | | yes | | | | | |
| ARHGAP23 | S_Shore;S_Shelf | TSS1500 | chr17:36612298-36612909 | 611 | | | | | | | | | 1 | 1 | | | | yes | yes | | | | |
| ARHGAP23 | S_Shore | Body | chr17:36622522-36623107 | 585 | | | 4 | | | | | | | | | | | | | | | | |
| ARHGAP23 | Island | Body | chr17:36666154-36666642 | 488 | | | | | | | | 0 | 1 | | | 2 | | | | | | | yes |
| SRCIN1 | Island | Body | chr17:36715787-36715953 | 166 | | | 7 | | | | | | | | | | | yes | | | | | |
| SRCIN1 | Island;N_Shore | Body | chr17:36717733-36719937 | 2204 | | | | | | | | | 1 | | | | | yes | | | | | yes |
| SRCIN1 | N_Shore | Body | chr17:36728456-36728859 | 403 | | | 1 | 1 | | | | | | | | | | | | | | yes | yes |
| SRCIN1 | N_Shore;Island | Body | chr17:36734475-36735584 | 1109 | | | | 1 | | | | | 1 | | | | | | | | | | yes |
| C17orf98 | N_Shore;Island;S_Shore | Body;1stExon;TSS200 | chr17:36997420-36998108 | 688 | 1 | | 19 | | | | | 0 | 7 | | | 2 | | yes | | | | | |
| RPL23 | S_Shore | TSS1500 | chr17:37011491-37011626 | 135 | | | | | | | | 0 | | | | | | yes | yes | | | yes | |
| FBXO47 | NA | 1stExon;5'UTR;TSS200 | chr17:37123638-37124558 | 920 | | 1 | 35 | 1 | | | | 0 | 12 | 4 | 1 | 16 | 1 | yes | yes | | | yes | |
| PLXDC1 | S_Shore | TSS1500 | chr17:37308828-37309118 | 290 | | | | | 1 | | | | | | | | | | | | | yes | |
| ARL5C | S_Shore;N_Shore;Island | 1stExon;5'UTR;TSS1500;Body | chr17:37321201-37323734 | 2533 | | 1 | 5 | | | | | 0 | 1 | | | | | | yes | yes | | yes | |
| CACNB1 | N_Shore;Island | 3'UTR;Body | chr17:37330925-37331628 | 703 | | | | 1 | | | | | 1 | | | 1 | | | | | | | |
| NEUROD2 | Island | 5'UTR;1stExon | chr17:37764118-37764158 | 40 | | | | | | | | | 1 | | | | | | | | | | yes |
| PPP1R1B | N_Shore | TSS1500 | chr17:37782302-37782685 | 383 | | | | | | | | | 1 | | | | | | | | | | |
| PPP1R1B | S_Shore | TSS200;Body | chr17:37784694-37784895 | 201 | | | 6 | | | | | | 1 | | | | | yes | | | | | |
| TCAP;STARD3 | N_Shelf | TSS1500;3'UTR;TSS200 | chr17:37820135-37822045 | 1910 | | | 5 | | | | | | 1 | | | | | yes | | | | yes | yes |
| ERBB2;PGAP3 | N_Shore | TSS1500;Body | chr17:37843264-37843818 | 554 | | | 3 | | | | | | | | | | | yes | | | | | |
| ERBB2 | N_Shore | 5'UTR;TSS200 | chr17:37856133-37856786 | 653 | | | | | | | | | 2 | | | 1 | | yes | | | yes | yes | yes |
| ERBB2 | Island | Body | chr17:37879654-37879740 | 86 | | | | | | | | | 1 | | | | | yes | | | | | |
| GRB7 | NA | TSS1500;TSS200;5'UTR;1stExon | chr17:37893764-37896175 | 2411 | | | 24 | 1 | | | | 0 | 12 | 2 | | | | yes | | | | | |
| ZPBP2 | N_Shore | TSS1500 | chr17:38023480-38024394 | 914 | | | | | | | | | 2 | | | | | | yes | | | | |
| ORMDL3 | S_Shore | TSS1500 | chr17:38084377-38084459 | 82 | | | | | | | | 0 | 1 | | | | | yes | | | | | |
| GSDMA | NA | TSS1500 | chr17:38118574-38119207 | 633 | | | | | | | | | 1 | | | | | yes | | | | yes | |
| CSF3 | NA | TSS1500 | chr17:38170862-38171530 | 668 | | 1 | | | | | | | 1 | | | | | yes | | | | | |
| THRA | S_Shore | 5'UTR | chr17:38220298-38221639 | 1341 | | | | | | | | | | | | | | | yes | | | | yes |
| NR1D1;THRA | Island | 3'UTR;Body | chr17:38249266-38249729 | 463 | | | 19 | | | | | | | | | | | | yes | | | | |
| RAPGEFL1 | Island | TSS200 | chr17:38334161-38334439 | 278 | | | 9 | 1 | | | | | 2 | | | | | | | | | yes | yes |
| RAPGEFL1 | Island | Body | chr17:38347603-38347710 | 107 | | | | 1 | | | | | | | | | | yes | | | yes | | yes |
| RARA | NA | TSS200;TSS1500 | chr17:38464730-38465393 | 663 | | | | 1 | | | | | 2 | | | | | | | | | | |
| RARA | Island | 5'UTR;TSS1500 | chr17:38472961-38473382 | 421 | | | | | | | | 0 | 2 | | | | | yes | | | | | |
| RARA | Island | Body | chr17:38501397-38501773 | 376 | | | | | | | | 0 | 1 | | | | | | | | | | |
| RARA | NA | Body | chr17:38510570-38512762 | 2192 | | | | 1 | | | | | 1 | | | | | yes | yes | | yes | yes | yes |
| GJD3 | N_Shore | 1stExon;3'UTR | chr17:38517641-38518714 | 1073 | 3 | | 1 | | | | | 0 | | | | | | | yes | | | | |
| TNS4 | NA | 5'UTR;TSS200 | chr17:38657310-38657934 | 624 | | 1 | | | | | | | 1 | 1 | | | | yes | | | | | |
| CCR7 | NA | Body | chr17:38717206-38717275 | 69 | | | 3 | | | | | 0 | 1 | | | | | | | | | | |
| CCR7 | NA | 5'UTR;1stExon | chr17:38721675-38721942 | 267 | | | | | | | | | 1 | | | | | yes | yes | yes | | | yes |
| KRT222 | NA | Body;1stExon;5'UTR | chr17:38821155-38821487 | 332 | | | 2 | | | | | 0 | 4 | | | 3 | | yes | | | | | |
| KRT24 | NA | 1stExon;5'UTR | chr17:38859934-38860097 | 163 | | | | | | | | | 1 | 1 | | | | | | | | | |
| KRT27 | Island | Body | chr17:38935846-38936092 | 246 | | | | | | | | 0 | 2 | | | 2 | | yes | yes | | | | |
| KRT27 | S_Shelf | TSS1500 | chr17:38939673-38940118 | 445 | | | | | | | | | | | | | | yes | | | yes | | |
| KRT28 | Island | Body | chr17:38953285-38953512 | 227 | | | | | | | | | 1 | | | | | yes | | | | | |
| KRT12 | Island | Body | chr17:39019545-39019978 | 433 | | | | | | | | 0 | 4 | | | | | yes | | | yes | yes | |
| KRT20 | NA | 1stExon | chr17:39041199-39041602 | 403 | | | 1 | | | | | | | | | | | | | | | | yes |
| KRT40 | NA | 5'UTR | chr17:39143239-39143567 | 328 | | | | | | | | | 1 | | | | | | yes | yes | | | |
| KRTAP3-1 | NA | TSS200 | chr17:39165425-39165697 | 272 | | | | | | | | | 1 | | | | | | yes | | yes | | |
| KRTAP1-5 | NA | 1stExon | chr17:39183037-39183560 | 523 | | | | | | | | | 1 | | | | | | | | | | |
| KRTAP2-1 | NA | TSS1500 | chr17:39204767-39204922 | 155 | | | 2 | | | | | | | | | | | yes | | | | | |
| KRTAP2-2 | NA | TSS1500 | chr17:39212513-39212764 | 251 | | | | | | | | | 1 | | | | | yes | | | yes | | |
| KRTAP4-8 | NA | TSS200 | chr17:39254427-39254879 | 452 | | | | | | | | | 2 | | | | | yes | | | | yes | |
| KRTAP17-1 | NA | 1stExon;5'UTR;TSS200;TSS1500 | chr17:39471738-39472498 | 760 | 3 | | 21 | | | | | | | | | | | | | | | yes | yes |
| KRT33A | NA | TSS200 | chr17:39507077-39507134 | 57 | | | | | | | | | | | | | | | | | | yes | |
| KRT38 | NA | TSS1500 | chr17:39597813-39598430 | 617 | | | | | | | | | 1 | | | | | yes | yes | | | | |
| KRT32 | NA | TSS200;TSS1500 | chr17:39623639-39624088 | 449 | | | 2 | | | | | | | | | | | | | | | yes | |
| KRT36 | NA | 1stExon | chr17:39645876-39646338 | 462 | | | | | | | | | | | | | | yes | | | | | |
| KRT13 | NA | 1stExon;5'UTR;TSS1500 | chr17:39661855-39662723 | 868 | | | | | | | | | 2 | | | | | yes | yes | | yes | yes | |
| KRT15 | NA | 5'UTR;1stExon | chr17:39675154-39675414 | 260 | | 1 | | | | | | | | | | | | | | | | yes | |
| KRT19 | N_Shelf | Body | chr17:39680217-39680662 | 445 | | | | | | | | | 1 | | | | | | | | | | |
| KRT19 | S_Shore | TSS1500 | chr17:39685482-39685912 | 430 | | | | | | | | | | | | | | yes | | | | | |
| KRT9 | NA | 1stExon;5'UTR;TSS200 | chr17:39728221-39728962 | 741 | | 1 | 8 | | | | | 0 | 3 | | | | | yes | | | | yes | |
| KRT14 | NA | Body | chr17:39739598-39739912 | 314 | | | | | | | | | | 1 | | | | yes | | | | yes | |
| KRT14 | NA | 1stExon;TSS200 | chr17:39742965-39744055 | 1090 | | | 1 | | | | | | | 1 | | | | yes | | | | yes | |
| KRT16 | NA | 1stExon;5'UTR | chr17:39769012-39769213 | 201 | | | | | | | | | 1 | | | | | yes | | | | yes | |
| KRT17 | NA | 5'UTR;1stExon | chr17:39780836-39781130 | 294 | | | 2 | | | | | | | | | | | yes | | | | | |
| JUP | Island | Body | chr17:39913900-39914156 | 256 | | | | | | | | | | | | | | | yes | | yes | | |
| JUP | Island;S_Shore | 5'UTR;TSS1500 | chr17:39942812-39943545 | 733 | | | | | | | | | 1 | | | | | yes | | | | | yes |
| CNP | Island | Body | chr17:40119847-40120035 | 188 | | | 2 | | | | | | | | | | | | | | | | |
| DHX58 | Island | Body | chr17:40259547-40260053 | 506 | | | 5 | | | | | 0 | 1 | | | | | yes | | yes | | | yes |
| KAT2A;HSPB9 | Island | TSS1500;TSS200;1stExon | chr17:40274524-40275359 | 835 | | 1 | 11 | 2 | 1 | | | | 5 | 1 | | | | yes | | | | | |
| HCRT | S_Shore | TSS200 | chr17:40337534-40337892 | 358 | | | 1 | | | | | | 1 | | | | | yes | | | | | |
| GHDC | NA | TSS200;5'UTR | chr17:40346279-40346448 | 169 | | | | | | | | | | | | | | yes | | | | | |
| STAT5A | N_Shore;Island | TSS200;5'UTR;TSS1500 | chr17:40438312-40441155 | 2843 | | | 7 | | 2 | | | | 2 | | 1 | | | yes | | | | yes | yes |
| STAT3 | NA | Body | chr17:40489513-40489785 | 272 | | | 25 | 1 | 2 | | | | 2 | | | | | yes | | | | | |
| PTRF | N_Shore | Body | chr17:40573643-40574031 | 388 | | | | | 1 | | | | | | | | | | | | | yes | |
| PTRF | Island | TSS200 | chr17:40575479-40575494 | 15 | | | | | | | | | 1 | | | | | yes | | | | | |
| ATP6V0A1 | N_Shore | TSS1500 | chr17:40609380-40609826 | 446 | | | 2 | | | | | | | | | | | yes | | | | | |
| NAGLU | N_Shore | TSS1500;TSS200 | chr17:40687496-40687885 | 289 | | | 11 | | | | | | 1 | | | | | | yes | yes | | | yes |
| HSD17B1 | Island | Body | chr17:40705273-40706061 | 788 | | | 4 | | | | | | 1 | | | | | yes | | | | yes | |

| | | | | | | | | | | | | | | | | | | | | | | | |
|---|---|---|---|---|---|---|---|---|---|---|---|---|---|---|---|---|---|---|---|---|---|---|---|
| COASY | N_Shore | TSS1500 | chr17:40713676-40713781 | 105 | | | 1 | | | | | | 1 | | | | | yes | | | | | |
| COASY | Island | 1stExon;Body;5'UTR | chr17:40715222-40715281 | 59 | | 1 | 3 | 1 | | | | | 2 | | | | | | | | yes | | |
| PLEKHH3 | Island;S_Shore | Body | chr17:40822424-40824361 | 1937 | | | | | | | | 0 | 3 | | | 1 | | yes | | | yes | yes | yes |
| CNTNAP1 | Island | Body | chr17:40835849-40836125 | 276 | | | 12 | | | | | 0 | | | | 1 | | | yes | | | yes | |
| CNTNAP1 | Island | Body | chr17:40838861-40839766 | 905 | | | 2 | 1 | | | | 0 | 3 | | | 1 | | yes | | | | yes | |
| WNK4 | N_Shore;Island | TSS1500;TSS200 | chr17:40932199-40932520 | 321 | | | | | | | | 0 | | | | | | | | | | | yes |
| WNK4 | Island;N_Shore | Body | chr17:40935998-40937365 | 1367 | | | 24 | 1 | | | | 0 | 5 | 1 | | 4 | | yes | yes | | | | yes |
| AOC2 | NA | TSS200 | chr17:40996565-40996886 | 321 | | | 2 | | | | | | | | | | | yes | | | | | |
| AOC3 | NA | 1stExon;5'UTR | chr17:41003272-41003399 | 127 | | | 1 | | | | | | | | | | | | | | | | |
| G6PC | NA | TSS200 | chr17:41052658-41053228 | 570 | | | | 1 | | | | | | | | | | yes | | | | | |
| AARSD1 | N_Shore | Body | chr17:41116197-41116704 | 507 | | | | | | | | 0 | | | | | | yes | yes | | | yes | |
| RUNDC1;AARSD1 | N_Shore | TSS1500;1stExon;TSS200;Body;5'U | chr17:41131552-41132169 | 617 | | | 3 | | | | | | | | | | | | yes | | | | |
| BRCA1;NBR2 | N_Shore;Island;S_Shore | TSS1500;Body | chr17:41277974-41278655 | 681 | | | 35 | 1 | | 1 | 1 | 0 | 20 | 8 | | 7 | | | | yes | | | |
| NBR1;TMEM106A | N_Shore;Island | 3'UTR;TSS1500;TSS200 | chr17:41363597-41364121 | 524 | 16 | | 2 | 1 | | | | | | | | | | | | | | yes | |
| ARL4D | Island | Body | chr17:41477271-41477948 | 677 | | | 4 | 1 | | | | | 1 | | | | | | | | yes | | |
| MIR2117 | NA | TSS1500 | chr17:41521655-41522193 | 538 | | | | | | | | | 1 | | | | | yes | | | yes | | yes |
| MEOX1 | N_Shore | Body | chr17:41723125-41723953 | 828 | | | 1 | | | | | | | | | | | yes | | | | | yes |
| MEOX1 | NA | TSS1500;1stExon;5'UTR;TSS200;na | chr17:41738893-41740208 | 1315 | | | 13 | | | | | | 1 | | 1 | | | yes | | | | yes | |
| SOST | N_Shore;Island | 3'UTR;Body | chr17:41832466-41833324 | 858 | | | 16 | | | | | | | | 1 | | | yes | | | | yes | yes |
| SOST | S_Shelf | 1stExon;TSS200 | chr17:41835937-41836563 | 626 | | 1 | | | | | | | | | | | | | | | | | |
| MPP3 | N_Shore | 5'UTR;Body | chr17:41909709-41909716 | 7 | | | 6 | | 1 | | | | 1 | | | | | yes | | | | yes | |
| CD300LG | NA | TSS1500;TSS200 | chr17:41924123-41924523 | 400 | | | 6 | | | | | | | | | | | | | | yes | | |
| MPP2 | Island;S_Shore | 5'UTR;1stExon;TSS200 | chr17:41984994-41985720 | 726 | | | | 1 | 1 | | | | | | | | | yes | | | | | |
| NAGS;PYY | Island | TSS200;TSS1500;Body | chr17:42082028-42083306 | 1278 | | | 10 | | | | | | | | | | | | | | | | yes |
| NAGS | Island | Body | chr17:42084612-42084953 | 341 | | | | 1 | 1 | | | | | | | | | | | | | | |
| TMEM101 | S_Shore | TSS200 | chr17:42092472-42092588 | 116 | | | | | | | | | | | | | | yes | | | | | |
| G6PC3 | N_Shore | TSS1500 | chr17:42147465-42147617 | 152 | | | | | 1 | | | 0 | | | | | | yes | | | | yes | |
| HDAC5 | Island | 5'UTR;1stExon | chr17:42200988-42201019 | 31 | | | | | | | | | 1 | | | | | | | | | yes | yes |
| C17orf53 | N_Shore | TSS1500 | chr17:42218908-42218924 | 16 | | | 7 | | | | | | 1 | | | | | yes | yes | | | | |
| ASB16 | NA | TSS200 | chr17:42248062-42248358 | 296 | | | | | | | | | 1 | | | | | yes | | | | | yes |
| ATXN7L3 | N_Shore | Body | chr17:42275334-42275551 | 217 | | | 1 | | | | | | | | | | | | | | | | |
| UBTF | Island | Body | chr17:42287715-42288203 | 488 | | | 1 | | | | | | | | | | | yes | | | | | |
| UBTF | S_Shore | Body | chr17:42289769-42290419 | 650 | | | | | | | | | | | | | | yes | | | | | |
| RUNDC3A | Island | 1stExon | chr17:42386208-42386419 | 211 | | | | | | | | 0 | | | | | | | | | | yes | |
| FAM171A2 | Island | Body | chr17:42431109-42432488 | 1379 | | 1 | | | | | | 0 | 3 | 1 | 1 | 9 | | | yes | | | | yes |
| C17orf104 | Island | TSS1500;TSS200 | chr17:42733527-42733976 | 449 | | 1 | | | | | | | 1 | | | | | yes | | | | | yes |
| ADAM11 | N_Shore | Body | chr17:42854741-42855087 | 346 | | | | | | | | | 1 | | | | | | | | | | |
| ADAM11 | Island | Body | chr17:42856773-42857157 | 384 | | | 3 | | | | | | | | | | | yes | | | yes | | |
| HIGD1B | NA | TSS200 | chr17:42925127-42925536 | 409 | | | | | | | | | | | | | | yes | | | | | |
| GFAP | S_Shelf | 1stExon;5'UTR;TSS200 | chr17:42992567-42992976 | 409 | 6 | | 4 | | | | | | 1 | | | | | yes | yes | | | | |
| GFAP | NA | TSS1500 | chr17:42994008-42994269 | 261 | | | 3 | | | | | | 1 | | | | | yes | | | | | |
| KIF18B | S_Shore | TSS1500 | chr17:43025585-43025655 | 70 | | | 1 | | | | | 0 | | | | 1 | | yes | | | | | |
| C1QL1 | N_Shore | 3'UTR | chr17:43037114-43037427 | 313 | | | 1 | 1 | 1 | | | | | | | | | yes | | | | | yes |
| C1QL1 | Island;N_Shore | TSS1500 | chr17:43046275-43047548 | 1273 | 7 | | 6 | | | | | | | | | | | yes | | | | | yes |
| NMT1 | NA | 3'UTR | chr17:43183150-43183199 | 49 | | | 2 | | | | | | | | | | | | | | | yes | |
| PLCD3 | Island | Body | chr17:43198278-43198610 | 332 | | | | | 1 | | | 0 | 1 | | | 2 | | yes | | | | | |
| ACBD4 | S_Shelf | 5'UTR;1stExon | chr17:43213013-43213429 | 416 | | | | | | | | | | | | 1 | | yes | | | | | |
| ACBD4 | N_Shore | 3'UTR | chr17:43221220-43222106 | 886 | | | | | | | | 0 | 1 | | | 1 | | | | | | yes | yes |
| FMNL1 | NA | Body | chr17:43306344-43306802 | 458 | | | 1 | | | | | | | | | | | yes | yes | | | | |
| FMNL1 | N_Shore;Island | Body | chr17:43318045-43319382 | 1337 | | | 33 | 1 | 2 | 2 | | | 4 | 1 | 1 | | | | yes | | | yes | yes |
| LOC100133991 | Island | TSS1500;TSS200 | chr17:43324793-43325657 | 864 | | | | | | | | 0 | | | | | | | | | | | yes |
| C17orf46;LOC100133991 | N_Shore;Island | Body;TSS1500 | chr17:43339040-43340592 | 1552 | | | | | | | | | | | | | | | | | | | |
| MAP3K14 | N_Shelf | 5'UTR | chr17:43391599-43392332 | 733 | | | | | | | | | | | | | | | | | | | yes |
| ARHGAP27 | Island;N_Shore | 1stExon;5'UTR;Body | chr17:43482735-43483527 | 792 | | | 15 | | | | | | | | | | | yes | yes | | | | yes |
| ARHGAP27 | N_Shelf | 1stExon;5'UTR;TSS200 | chr17:43502999-43503401 | 402 | | | 2 | | | | | 0 | 1 | 1 | | | | yes | yes | | | | |
| LOC644172 | Island | Body | chr17:43679177-43679667 | 490 | | | | | | | | | | | | | | | | | | | yes |
| C17orf69;MGC57346 | NA | TSS1500;Body | chr17:43714888-43715162 | 274 | | | 1 | | | | | | | | | | | yes | | | | | |
| C17orf69 | NA | Body | chr17:43716515-43716617 | 102 | | | | | | | | | 1 | | | | | yes | yes | | | yes | |
| CRHR1 | Island | Body | chr17:43862910-43863356 | 446 | | | 4 | | | | | | | | | | | yes | | | | | |
| LOC100128977;IMP5 | NA | Body;TSS1500;TSS200;1stExon | chr17:43921485-43923255 | 1770 | | 1 | | | | | | | 1 | | | | | yes | | | | | |
| MAPT;LOC100128977;LOC100130148 | Island | 5'UTR;TSS1500;Body | chr17:43974344-43975063 | 719 | | | 25 | | | | | | 1 | | | | | | | | | | yes |
| MAPT | S_Shelf | 5'UTR | chr17:43978251-43978756 | 505 | | | | | | | | | | | | | | yes | | | yes | | |
| MAPT | NA | Body | chr17:44087629-44087784 | 155 | | | | | | | | | | | | | | | | | yes | | |
| KIAA1267 | NA | 1stExon;5'UTR | chr17:44249543-44249866 | 323 | 2 | | | | | | | | | | | | | | yes | | yes | | |
| WNT3 | Island | Body | chr17:44847220-44847427 | 207 | | | 14 | | | | | 0 | 1 | 1 | | 1 | | yes | | | | | |
| WNT3 | chr17:44895800-44897025;Isla | Island;TSS1500 | chr17:44896212-44897512 | 1300 | | | 1 | | | | | | 3 | | 1 | | | | | | | | yes |
| WNT9B | Island | 5'UTR;1stExon | chr17:44928967-44928995 | 28 | | | | | | | | 0 | 1 | | | | | | | | | | yes |
| LOC100272146;C17orf57 | Island | Body | chr17:45501030-45501697 | 667 | | | 11 | | | | | | | | | | | | yes | | | | |
| KPNB1 | Island | TSS1500 | chr17:45726922-45727082 | 160 | | | | | | | | | 1 | | | | | yes | | | | yes | yes |
| TBKBP1 | Island | 1stExon;5'UTR | chr17:45773468-45773570 | 102 | | | | | | | | 0 | 1 | | | | | | | | | | |
| TBKBP1 | N_Shore;Island | Body | chr17:45786010-45786737 | 727 | | | 4 | | | | | | 1 | | | | | yes | yes | | yes | | yes |
| OSBPL7 | NA | TSS1500 | chr17:45899469-45899862 | 393 | | | 2 | | | | | | 1 | | | | | | | | | | |
| SP6 | S_Shore | 5'UTR | chr17:45929714-45930106 | 392 | | | | | | | | 0 | 1 | | | 2 | | yes | | | | | |
| PNPO | NA | TSS1500;TSS200 | chr17:46018654-46019184 | 530 | | | 1 | | | | | 0 | 2 | | | 1 | | yes | | | | | |
| PRR15L | NA | 1stExon;5'UTR;TSS200;TSS1500 | chr17:46035108-46036419 | 1311 | | | | | | | | | | | | | | | | | | | |
| SKAP1 | NA | Body | chr17:46233714-46233945 | 231 | | | 2 | | | | | 0 | 1 | | | | | yes | yes | | | | |
| SKAP1 | NA | Body | chr17:46388390-46388465 | 75 | | | | | 1 | | | 0 | 1 | | 1 | | | | | | | | |
| SKAP1 | chr17:46507344-46507778;Isla | Island;TSS200;TSS1500 | chr17:46507705-46508097 | 392 | | | 9 | | | | | | | | 1 | | | yes | yes | | | | yes |
| HOXB1 | N_Shore;Island;S_Shore | Body;1stExon;TSS200;TSS1500 | chr17:46607103-46608874 | 1771 | | 2 | 18 | | 2 | 1 | | | 3 | 1 | 1 | | | | yes | | | | |
| HOXB2 | S_Shore | Body;1stExon;TSS200 | chr17:46621709-46622715 | 1006 | | | 26 | | | | | 0 | 6 | 1 | | | | | | | | yes | yes |
| HOXB2 | S_Shelf | TSS1500 | chr17:46623729-46624268 | 539 | | | 5 | | | | | | | | | | | | | | | yes | yes |
| HOXB3 | N_Shore;Island;chr17:4662955 | 3'UTR;Body;N_Shore | chr17:46627384-46629804 | 2420 | | | | | | | | | | | | | | | | | | | |
| HOXB3 | Island | 5'UTR | chr17:46631910-46632154 | 244 | | | 3 | | 1 | | | | | | | | | | | | | yes | yes |
| HOXB3 | Island;N_Shore | 5'UTR | chr17:46641504-46642104 | 600 | | | | | | | | 0 | 3 | 1 | | | | | | | | yes | yes |
| HOXB3 | S_Shelf | 5'UTR | chr17:46645972-46646444 | 472 | | | 15 | | | | | 0 | 2 | 1 | | | | | | | | yes | yes |
| HOXB3 | NA | 5'UTR | chr17:46648014-46648582 | 568 | | | 5 | | 1 | | | | | | | | | | | | | | |
| HOXB3;HOXB4 | N_Shelf;N_Shore | 5'UTR;1stExon;TSS200;3'UTR | chr17:46651186-46654330 | 3144 | | | | | | | | | | | | | | | | | | | |
| HOXB4;MIR10A | S_Shore;N_Shore | TSS1500;Body;TSS200 | chr17:46656543-46658170 | 1627 | | | | | | | | | | | | | | | | | | | |
| LOC404266;HOXB5 | N_Shelf;N_Shore;Island | TSS1500;TSS200;3'UTR;Body | chr17:46666926-46670029 | 3103 | | | 65 | | | | | 0 | 12 | | 1 | | | | | | | yes | yes |
| LOC404266;HOXB5 | N_Shore | Body;TSS1500 | chr17:46671635-46672255 | 620 | | | 4 | | | | | 0 | 1 | | | | | | | | | | |
| LOC404266;HOXB6 | N_Shore;Island | Body;3'UTR | chr17:46673442-46674514 | 1072 | | | 26 | | | | | | | | | | | | | | | yes | yes |
| LOC404266;HOXB6 | S_Shore | Body;5'UTR | chr17:46675892-46676375 | 483 | | | 19 | | | | | 0 | 1 | | | | | | | | | | |
| LOC404266;HOXB6;HOXB7 | N_Shelf;N_Shore;Island;NA | Body;5'UTR;1stExon;TSS200;TSS1 | chr17:46680288-46685448 | 5160 | | | | | | | | | | | | | | | | | | | |
| HOXB7 | N_Shore | TSS1500 | chr17:46689426-46689639 | 213 | | | 4 | | | | | | | | | | | | | | | yes | |
| HOXB8 | Island;S_Shore | Body;TSS200;TSS1500 | chr17:46691543-46692592 | 1049 | | | 8 | | | | | | | | | | | | | | | yes | yes |
| HOXB9 | Island | 3'UTR | chr17:46698881-46699155 | 274 | | | | | | | | | 2 | | 1 | | | | | | | | |
| HOXB9 | Island | 1stExon;5'UTR | chr17:46703646-46703870 | 224 | | | 1 | | | | | | | | | | | | | | | | yes |
| C17orf93;PRAC | N_Shore;Island | TSS1500;Body;1stExon;5'UTR;TSS2 | chr17:46799479-46801314 | 1835 | | 1 | 15 | | | | | | 2 | | | | | yes | | | | | yes |
| HOXB13 | N_Shore;Island | 3'UTR;Body | chr17:46803802-46804414 | 612 | | 1 | 1 | | | | | | | | | | | | | | | yes | |

| | | | | | | | | | | | | | | | | | | | | | | | |
|---|---|---|---|---|---|---|---|---|---|---|---|---|---|---|---|---|---|---|---|---|---|---|---|
| TTLL6 | NA | Body;TSS200 | chr17:46871867-46871905 | 38 | | | | | | | | | | | | | | yes | | | | | |
| IGF2BP1 | Island | Body | chr17:47075758-47075880 | 122 | | 1 | | | | | | | | | | | | | | | | | yes |
| IGF2BP1 | Island | Body | chr17:47091339-47092272 | 933 | | | 18 | | | | | 0 | 2 | 4 | | 13 | | yes | | yes | | | yes |
| IGF2BP1 | S_Shore | Body | chr17:47093955-47094252 | 297 | | | | | | | | 0 | | | | | | yes | yes | | yes | yes | yes |
| IGF2BP1 | N_Shore;Island | Body | chr17:47113141-47113596 | 455 | | | 11 | | | | | | | | | | | yes | yes | | yes | yes | |
| B4GALNT2 | N_Shore;Island | TSS1500;TSS200;Body;5'UTR | chr17:47209216-47210318 | 1102 | | | 10 | | | | | | 3 | | | | | yes | yes | | | yes | |
| GNGT2;ABI3 | NA | 1stExon;TSS1500;5'UTR;TSS200 | chr17:47286719-47287577 | 858 | | | 22 | 1 | | | | | 2 | | | | | | yes | | | | |
| ABI3 | Island | Body | chr17:47296970-47297512 | 542 | | | 6 | | | | | | | | | | | yes | | | | | |
| NGFR | N_Shore | TSS1500 | chr17:47571963-47572132 | 169 | | | | | | | | | | | | | | | yes | | | yes | yes |
| NGFR | Island | Body | chr17:47583654-47583854 | 200 | | | | | | | | | 1 | | | | | | | | | | |
| NGFR | Island | Body | chr17:47590179-47590326 | 147 | | 1 | | | | | | | | | | | | | | | | | |
| FAM117A | Island | Body | chr17:47840835-47841033 | 198 | | | | | | | | | 1 | | | | | | | | | yes | yes |
| DLX4 | Island | Body | chr17:48047123-48047608 | 485 | | | | | | | | | | | | | | | | | | | yes |
| DLX4 | Island | TSS200;Body;5'UTR;1stExon | chr17:48049952-48051089 | 1137 | | | 4 | | | | | 0 | 1 | 2 | | | | | | | | | |
| DLX3 | N_Shore;Island | Body | chr17:48070354-48071845 | 1491 | | 1 | 6 | | | | | | | | | | | | yes | | | | yes |
| ITGA3 | NA | Body | chr17:48153934-48154356 | 422 | | | | | | | 1 | 0 | 1 | | | | | | | | | | |
| SAMD14 | Island | Body | chr17:48194635-48195107 | 472 | | | 2 | | | | | | | | | | | yes | yes | | | | |
| SAMD14 | N_Shore | 5'UTR | chr17:48206283-48206749 | 466 | | | 1 | | | | | | | | | | | | | | | yes | yes |
| HILS1;SGCA | NA | TSS1500;Body | chr17:48250505-48250645 | 140 | | | 3 | | | | | | | | | | | | | | | | |
| COL1A1 | NA | Body | chr17:48262937-48263204 | 267 | | | 16 | | | | | | | | | | | | | | | yes | |
| COL1A1 | NA | Body | chr17:48266770-48267239 | 469 | | | | | | | | | | | | | | | | | | | |
| COL1A1 | S_Shore | TSS200 | chr17:48279072-48279265 | 193 | | | | | | | | | 1 | | | | | | | | | | |
| XYLT2 | N_Shore | TSS1500 | chr17:48422933-48423021 | 88 | | | 1 | | | | | | | | | | | | | | yes | | yes |
| LRRC59 | N_Shore | Body | chr17:48473757-48474021 | 264 | | | | | | | | | 1 | | | | | yes | | | | | |
| ACSF2;CHAD | Island | Body;1stExon;5'UTR;TSS200;TSS1 | chr17:48545805-48546620 | 815 | | 1 | | 2 | 2 | | | 0 | 2 | | 1 | 3 | | | | | | | |
| MYCBPAP | N_Shore | TSS1500 | chr17:48585216-48585470 | 254 | | | | | | | | | 1 | | | | | yes | | | | | yes |
| MYCBPAP;EPN3 | NA | Body;TSS1500 | chr17:48608677-48609332 | 655 | | | | | | | | | | | | | | yes | | | | | |
| SPATA20 | S_Shelf | Body | chr17:48628397-48628573 | 176 | | | | | | | | | | | | | | yes | | | | | |
| CACNA1G | Island | Body | chr17:48649272-48649370 | 98 | | | | | | | | | | | | | | yes | | | yes | | |
| CACNA1G | NA | Body | chr17:48672432-48672483 | 51 | 2 | | | | | | | | | | | | | yes | | | | | |
| CACNA1G | Island | Body | chr17:48680410-48680554 | 144 | | 1 | | | | | | | | | | | | | | | | | |
| ABCC3 | Island | TSS200 | chr17:48712051-48712219 | 168 | | | | | | | | | 2 | | | | | yes | | | | | |
| C17orf73 | NA | TSS1500 | chr17:48845563-48846189 | 626 | | | | | | | | | 1 | 1 | | | | yes | yes | | yes | yes | |
| WFIKKN2 | NA | TSS200;1stExon;5'UTR;TSS1500 | chr17:48911141-48913424 | 2283 | | | 14 | | | | | 0 | 5 | | | 2 | | yes | | | | yes | |
| SPAG9 | S_Shore | TSS1500 | chr17:49199280-49199381 | 101 | 2 | | | | | | | | | | | | | | yes | | | | |
| NME2;NME1-NME2 | N_Shore;Island | TSS200;TSS1500;Body;5'UTR;1stE | chr17:49242714-49244717 | 2003 | | 1 | 4 | | 1 | | | 0 | 1 | | | | | yes | | | | yes | |
| CA10 | Island;S_Shore | 1stExon;5'UTR;TSS1500;TSS200 | chr17:50235279-50237604 | 2325 | | 2 | 2 | | | | | | 3 | | | | | | | | | yes | yes |
| KIF2B | NA | TSS200;TSS1500 | chr17:51899837-51900238 | 401 | | | | | | | | | | | | | | | | | | | |
| TOM1L1 | N_Shore | TSS1500 | chr17:52977213-52977769 | 556 | | | | | | | | | 1 | | | | | | | | | | |
| HLF | N_Shore | TSS1500 | chr17:53341098-53342029 | 931 | | | 13 | | | | | | 2 | | | | | | yes | | yes | | yes |
| TMEM100 | NA | 5'UTR;1stExon;TSS200 | chr17:53800041-53800554 | 513 | | 1 | | 1 | | | | 0 | 2 | | | 2 | | yes | | | yes | | |
| TMEM100 | NA | TSS1500 | chr17:53810321-53810813 | 492 | | | 2 | | | | | | | | | | | yes | yes | | yes | yes | |
| PCTP | N_Shore | TSS200;1stExon;5'UTR | chr17:53828263-53828850 | 587 | | | | | | | | 0 | 1 | | | | | | | | yes | | |
| ANKFN1 | NA | TSS1500;TSS200 | chr17:54230419-54230996 | 577 | | | | | | | | | 1 | | | | | | yes | | | | |
| C17orf67 | NA | 1stExon;TSS200 | chr17:54893150-54893317 | 167 | | | 7 | | | | | | | | | | | yes | | | | | yes |
| DGKE | Island | 5'UTR | chr17:54911857-54912286 | 429 | | | | | | | | | | | | | | | | | | | yes |
| RNF126P1 | N_Shore;Island | TSS1500;TSS200 | chr17:55122538-55123356 | 818 | | | | | | | | 0 | 1 | | | | | yes | | | yes | | |
| MSI2 | NA | Body | chr17:55533142-55533389 | 247 | | | | | | | | | 1 | | 1 | | | yes | yes | | | | |
| MSI2 | NA | Body | chr17:55663225-55663284 | 59 | 2 | | 3 | | | | | | | | | | | yes | | | | | |
| MRPS23 | Island | TSS1500 | chr17:55927852-55927919 | 67 | | | | | | | | | 1 | 1 | | | | | yes | | | | |
| CUEDC1 | NA | Body | chr17:55952063-55952217 | 154 | | 1 | | 1 | 1 | | | | 1 | 1 | | | | yes | | | | | yes |
| CUEDC1 | N_Shore;Island | Body | chr17:55962522-55963102 | 580 | | | | | | | | 0 | 5 | | | 1 | | yes | yes | | yes | | yes |
| SFRS1 | N_Shore | 3'UTR;Body | chr17:56082780-56083289 | 509 | | | 2 | | | | | | 1 | | | | | | | | | | |
| OR4D1 | N_Shelf;N_Shore | TSS1500;TSS200 | chr17:56231747-56232955 | 1208 | | | 8 | | | | | | 1 | | | | | yes | yes | | yes | yes | |
| MSX2P1 | Island | Body | chr17:56234476-56234897 | 421 | | 1 | | | | | | | | | | | | yes | | | | | |
| OR4D2 | NA | TSS1500 | chr17:56246435-56247036 | 601 | | | | | | | | | 1 | | | | | yes | | | | | |
| MKS1 | S_Shore | TSS200;TSS1500 | chr17:56296980-56297443 | 463 | | | 10 | | | | | | | | | | | yes | | | | | |
| MPO | Island | Body | chr17:56355299-56355431 | 132 | 2 | | | | | | | | 1 | 1 | | | | | | | yes | | yes |
| BZRAP1 | Island;N_Shore | Body | chr17:56401623-56402238 | 615 | | | | | | | | | 2 | 1 | | | | yes | | | | | yes |
| MIR142 | NA | TSS200;TSS1500 | chr17:56408688-56409518 | 830 | | | 1 | | 1 | | | | 2 | 2 | 1 | | | | | | | yes | yes |
| HSF5 | Island | 1stExon;Body | chr17:56564855-56565644 | 789 | | | 12 | | 1 | | | 0 | 3 | | | 8 | | yes | | | | | yes |
| TEX14 | NA | 5'UTR | chr17:56744332-56744490 | 158 | | | 9 | | | | | 0 | 1 | 1 | | 2 | | yes | | | | | |
| TEX14;RAD51C | N_Shore | TSS1500;TSS200;1stExon;5'UTR | chr17:56769387-56769904 | 517 | | | 7 | | | | | 0 | 2 | | | | | yes | | | | | |
| DHX40 | N_Shore;Island | TSS1500;TSS200 | chr17:57642628-57642763 | 135 | | 1 | 15 | | | | | 0 | 3 | | | 2 | | yes | yes | | | | |
| MIR21 | NA | TSS1500;TSS200 | chr17:57918262-57918682 | 420 | | | 3 | | | | | | 1 | | | | | | | | | yes | |
| LOC645638 | NA | TSS200 | chr17:58165865-58166296 | 431 | | | | | | | | | 1 | | | | | yes | yes | | | | |
| CA4 | Island | TSS200 | chr17:58227267-58227745 | 478 | | | | | | | | | 1 | | | | | yes | | | | yes | |
| USP32 | Island | 1stExon | chr17:58469253-58469452 | 199 | | | | | | | | | 1 | | | | | | | | | | yes |
| C17orf64 | S_Shore | TSS200 | chr17:58499679-58499911 | 232 | 4 | | 2 | | | | | 0 | 4 | | | | | | | | | | |
| TBX2 | Island | Body | chr17:59478068-59478579 | 511 | | | 3 | | | | | | | | | | | | | | | yes | yes |
| TBX2 | Island;chr17:59473060-594832 | Body;Island | chr17:59480470-59482956 | 2486 | | | | | | | | | | | | | | | | | | | |
| C17orf82 | Island | TSS1500 | chr17:59488116-59488635 | 519 | | | 8 | | | | | | | | | | | | | | | | |
| TBX4 | Island | TSS1500 | chr17:59532306-59533081 | 775 | | | 2 | | | | | | 1 | | 1 | | | | | | | | |
| TBX4 | Island | Body | chr17:59534637-59534998 | 361 | | | | | | | | | | | | | | | | | | | |
| TBX4 | N_Shore | Body | chr17:59539251-59539434 | 183 | | | | | | | | | | | | | | | yes | | | | yes |
| TBX4 | NA | Body | chr17:59554672-59554806 | 134 | | | 8 | | | | | | 1 | | | | | yes | | | | | |
| NACA2 | NA | TSS200;TSS1500 | chr17:59668604-59669395 | 791 | | | | | | | | | 1 | | | | | | yes | | yes | | |
| MRC2 | S_Shelf | Body | chr17:60747393-60747678 | 285 | | | | | | | | | 1 | 1 | | | | yes | | | | | |
| MRC2 | Island | Body | chr17:60758021-60758455 | 434 | | | | | | | | | 1 | | | | | yes | | | | | |
| MARCH10 | NA | Body | chr17:60828157-60828406 | 249 | | | 15 | | | | | | 1 | | | | | yes | | | yes | | |
| 40238;MARCH10 | S_Shore;N_Shore;Island | TSS200;5'UTR;TSS1500 | chr17:60885250-60886079 | 829 | | | | | | | | | | | | | | | | | | | |
| CYB561 | Island | 3'UTR | chr17:61511683-61511829 | 146 | | | 5 | | | | | | | | | | | | | yes | | | |
| CYB561 | Island | Body;5'UTR | chr17:61514753-61515312 | 559 | | | 6 | | | | | | | | | | | | | | | | yes |
| CYB561 | Island | TSS1500 | chr17:61524073-61524304 | 231 | | | | 1 | 1 | | | | | | | | | | | | | | yes |
| ACE | Island | TSS1500 | chr17:61553938-61554106 | 168 | | | | | 1 | | | | | | | | | yes | | | | yes | yes |
| LIMD2 | Island | TSS1500 | chr17:61778366-61778813 | 447 | | | | 1 | 1 | | | | | | | | | yes | | | yes | | |
| GH2 | NA | TSS200 | chr17:61959416-61959527 | 111 | | | | | | | | | | | | | | | | | | | yes |
| CSHL1 | NA | 5'UTR;Body;TSS200 | chr17:61988297-61989275 | 978 | | | 3 | | | | | | | | | | | yes | yes | | | | |
| CD79B | NA | 1stExon | chr17:62009607-62009835 | 228 | | | | | 1 | | | | 1 | | 1 | | | | | | | | |
| SCN4A | NA | TSS1500 | chr17:62051709-62051773 | 64 | | 1 | 5 | | | | | | 1 | | | | | | | | | | |
| ICAM2 | NA | Body;5'UTR;1stExon;TSS200 | chr17:62083407-62084359 | 952 | | | 7 | | | | | | | | | | | yes | yes | | | | |
| ICAM2 | NA | 5'UTR;1stExon | chr17:62097918-62098230 | 312 | | | 10 | | | | | | 1 | | | | | yes | yes | | | | |
| ERN1 | Island | TSS1500 | chr17:62208049-62208434 | 385 | | | | | | | | | | | | | | | | | | | yes |
| POLG2 | S_Shore | TSS1500 | chr17:62493599-62493832 | 233 | | | 1 | | | | | | | | | | | | | | yes | | |
| LOC146880 | Island;chr17:62774366-62777 | Body;TSS200;Island | chr17:62774939-62777777 | 2838 | | | 23 | | 1 | | | | | | 1 | | | yes | | | | | yes |
| AXIN2 | N_Shore | Body | chr17:63532898-63533303 | 405 | | | | | | | | | | | | | | | | | | | |
| APOH | NA | Body | chr17:64225346-64225731 | 385 | | | | | 1 | | | | | | | | | yes | | | | | |
| PRKCA | NA | Body | chr17:64672129-64672341 | 212 | | | | | | | | | 1 | 1 | | | | | | | | | |
| PRKCA | NA | Body | chr17:64787379-64787784 | 405 | | | 2 | | | | | | | | | | | | yes | | yes | | |

| | | | | | | | | | | | | | | | | | | | | | | | |
|---|---|---|---|---|---|---|---|---|---|---|---|---|---|---|---|---|---|---|---|---|---|---|---|
| CACNG5 | NA | TSS1500;TSS200 | chr17:64872169-64873594 | 1425 | | | 2 | | | | | | | | | | | yes | yes | | yes | yes | |
| CACNG4 | Island | 1stExon | chr17:64961140-64961442 | 302 | | | | | | | | | 1 | | | | | | | | | yes | yes |
| CACNG1 | N_Shore | TSS1500;TSS200 | chr17:65039695-65040616 | 921 | | | 20 | | | | | | 1 | | 1 | | | yes | | | yes | | |
| HELZ | Island;S_Shore | TSS1500 | chr17:65242413-65242892 | 479 | | | | | | | | 0 | | | | | | | | | yes | | yes |
| PITPNC1 | NA | Body | chr17:65471303-65471507 | 204 | | 1 | 7 | | | | | 0 | 2 | | 1 | 2 | | yes | | | | | |
| BPTF | NA | Body | chr17:65850328-65850433 | 105 | | | | | | | | | 1 | | | | | | | | | | yes |
| C17orf58 | S_Shore | TSS1500 | chr17:65990431-65990817 | 386 | | | 2 | | | | | | | | | | | | | | | yes | yes |
| LOC440461 | Island | TSS1500;TSS200 | chr17:66194407-66195134 | 727 | 9 | | 2 | | | | | | | | | | | yes | | | | | |
| LOC440461 | N_Shore | Body | chr17:66196160-66196681 | 521 | 2 | | | | | | | | | | | | | yes | | | yes | | |
| ARSG | N_Shore;Island | TSS1500;TSS200 | chr17:66254550-66255444 | 894 | | | 3 | | | | | | 1 | | | | | | | | yes | | |
| ARSG;SLC16A6 | N_Shore | 5'UTR | chr17:66286298-66286394 | 96 | | | | | | | | | | | | | | | | | | | |
| FAM20A | Island | 5'UTR;1stExon | chr17:66596999-66597500 | 501 | | | | | | | | 0 | | | | | | | yes | | | | yes |
| KCNJ16 | NA | TSS200;TSS1500 | chr17:68070499-68071397 | 902 | | 1 | | 1 | | | | | | | | | | yes | | | | | |
| KCNJ16 | NA | 5'UTR;TSS200 | chr17:68100915-68100975 | 60 | | | | | | | | | 1 | | | | | | | | | | |
| KCNJ2 | NA | Body | chr17:68171276-68171750 | 474 | | | | | | | | | | | | | | | | | | | |
| SOX9 | N_Shore | TSS1500 | chr17:70115862-70116434 | 572 | | | 3 | | | | | | | | | | | | | | | yes | |
| SOX9 | Island | Body | chr17:70118912-70123165 | 4253 | | | | | | | | | | | | | | | | | | | |
| COG1 | Island | TSS200 | chr17:71189054-71189068 | 14 | | | | | | | | | | | | | | | | | | | yes |
| CPSF4L | NA | TSS1500 | chr17:71258589-71259036 | 447 | 2 | | | | | | | | | | | | | | | | | | |
| SDK2 | NA | Body | chr17:71361857-71361901 | 44 | | | | | | | | 0 | 1 | 1 | | 1 | | | | | | | |
| SDK2 | S_Shelf | Body | chr17:71434003-71434052 | 49 | | | 3 | | | | | | | | | | | yes | | | | | |
| SDK2 | NA | Body | chr17:71533286-71533396 | 110 | | | | | | | | | 1 | | | | | | | | | | |
| SDK2 | NA | Body | chr17:71548560-71548861 | 301 | 1 | | | | | | | | | | | | | yes | | | | | |
| C17orf54 | NA | Body | chr17:71772827-71772869 | 42 | | | 11 | | | | | | | | | | | | yes | | | | |
| TTYH2;MGC16275 | N_Shore | TSS1500;Body | chr17:72208213-72208433 | 220 | | | | | | | | | | | | | | | yes | | | | |
| TTYH2 | NA | Body;TSS1500 | chr17:72243979-72244939 | 960 | | | | | | | | | | | | | | yes | | | | yes | |
| DNAI2 | N_Shore | TSS1500 | chr17:72269945-72270433 | 488 | | | 5 | | | | | | 1 | | | | | yes | | | | | |
| KIF19 | N_Shore;Island | TSS1500 | chr17:72321767-72322154 | 387 | | 1 | 2 | | | | | 0 | | | | | | yes | | | | | yes |
| KIF19 | Island | Body | chr17:72347925-72348295 | 370 | | | | | | | | 0 | | | | | | yes | | | | | |
| KIF19 | Island | Body | chr17:72350354-72350710 | 356 | | | 4 | | | | | 0 | | | | 2 | | yes | | | | | |
| BTBD17 | S_Shore | TSS200 | chr17:72357992-72358370 | 378 | 1 | | | | | | | 0 | | | | | | | | | | | |
| GPR142 | NA | TSS1500;TSS200 | chr17:72362302-72363634 | 1332 | | | | | | | | | 1 | | | | | yes | | | | | |
| GPR142 | Island | Body | chr17:72368048-72368259 | 211 | | | | | | | | | | | | | | yes | | | | | |
| GPRC5C | N_Shore;Island | TSS1500;TSS200 | chr17:72426469-72427693 | 1224 | | | | | | | | | 2 | | | | | | | | | | |
| GPRC5C | NA;N_Shelf | Body | chr17:72438885-72439217 | 332 | | | 9 | | | | | | | | | | | yes | yes | | | | |
| GPRC5C | Island | Body | chr17:72442928-72443401 | 473 | | | | | | | | | | | | | | yes | | | | | |
| CD300A | NA | TSS1500;TSS200;5'UTR;1stExon | chr17:72462234-72463080 | 846 | | | 3 | 1 | 1 | | | 0 | 4 | 1 | | | | | | | | yes | |
| CD300LB | NA | 1stExon;5'UTR | chr17:72527607-72527813 | 206 | | | | | | | | | 1 | | | | | yes | | yes | | | |
| CD300C | NA | 1stExon;5'UTR | chr17:72541932-72542430 | 498 | | | | | | | | | | | | | | | | | | yes | |
| C17orf77;CD300LD | NA | TSS200;Body;TSS1500 | chr17:72580458-72581081 | 623 | 3 | | 7 | | | | | | | | | | | yes | yes | yes | | | |
| CD300E | NA | Body;1stExon;5'UTR | chr17:72619556-72620274 | 718 | | | 6 | | | | | | | | | | | yes | yes | | | | |
| RAB37 | N_Shore;Island | TSS1500;TSS200;5'UTR;1stExon | chr17:72666692-72667743 | 1051 | | 1 | 4 | | | | | 0 | 3 | | | | | yes | | | yes | | |
| RAB37;CD300LF | NA | Body;TSS1500;5'UTR;1stExon | chr17:72709026-72709731 | 705 | | | | | | | | | | | | | | yes | | | | | |
| SLC9A3R1 | Island | Body | chr17:72745576-72745994 | 418 | | | | | | | | | | | | | | | | | | | |
| NAT9;TMEM104 | S_Shore | TSS1500;5'UTR | chr17:72772726-72772745 | 19 | | | | | | | | 0 | | | | | | | yes | | | | yes |
| GRIN2C | Island | Body | chr17:72848197-72848918 | 721 | | | | | | | | | 1 | | | | | yes | yes | | yes | | |
| FDXR | Island | TSS200 | chr17:72869219-72869237 | 18 | | | | | | | | | 1 | | | | | | | | | | |
| FADS6 | N_Shore;Island | Body;TSS200 | chr17:72889323-72889973 | 650 | | | | | | | | | 2 | | | | | | yes | | | | |
| USH1G | Island | Body | chr17:72915784-72916509 | 725 | | | 2 | | | | | | 1 | | | | | | | | yes | | |
| OTOP2;USH1G | Island | TSS1500 | chr17:72919704-72920187 | 483 | | | | | | | | | 1 | | | | | | | | | | yes |
| OTOP3 | Island | TSS200 | chr17:72931767-72931792 | 25 | 3 | | | | | | | 0 | | | | | | | | | | | |
| C17orf28 | NA | Body | chr17:72948349-72948705 | 356 | | | | | | | | 0 | 1 | | | | | yes | | yes | | | yes |
| C17orf28 | NA | Body | chr17:72951218-72951370 | 152 | | | | | | | | | 1 | | | | | yes | yes | | | | |
| KCTD2 | NA | Body | chr17:73055792-73056797 | 1005 | | | 4 | | | | | 0 | 1 | | | | | | | | | | |
| SLC16A5 | N_Shore;Island | TSS1500;5'UTR | chr17:73083632-73084494 | 862 | | | | 1 | | | | | | | | | | yes | yes | | | | |
| ARMC7 | Island | Body | chr17:73124988-73125153 | 165 | | 1 | | | | | | | | | | | | | | | | | |
| GGA3 | NA | Body | chr17:73238441-73238513 | 72 | | | 3 | | | | | | | | | | | yes | | | | | |
| GRB2 | NA | 3'UTR;Body;nan | chr17:73316280-73316889 | 609 | | | 8 | | 1 | | | | | | 1 | | | yes | | | | | |
| CASKIN2 | Island | Body | chr17:73498127-73498302 | 175 | | | 3 | | | | | | | | | | | | | | | | yes |
| CASKIN2 | Island | Body | chr17:73501109-73501230 | 121 | | | | 1 | | | | | | | | | | yes | | | | | |
| CASKIN2 | NA | Body;TSS200 | chr17:73506140-73506230 | 90 | | | | | | | | | 1 | | | | | yes | | | yes | | |
| LLGL2 | Island | 5'UTR | chr17:73522123-73522598 | 475 | | | | | | | | | | | | | | yes | | | | | yes |
| LLGL2 | Island | Body | chr17:73559363-73559562 | 199 | | | 1 | | | | | | | | | | | yes | | | | | |
| MYO15B | Island | TSS1500;TSS200 | chr17:73583839-73584617 | 778 | 1 | | | 1 | 3 | | | | | | | 1 | | yes | yes | | | | yes |
| LOC100130933;RECQL5 | NA | TSS1500;Body;TSS200 | chr17:73642080-73642773 | 693 | | | 1 | | | | | | 1 | | | | | | yes | | yes | | |
| ITGB4 | S_Shore;S_Shelf | TSS1500;5'UTR;TSS200 | chr17:73719693-73720840 | 1147 | | | | | | | | | 1 | | | | | yes | | | yes | | |
| ITGB4 | S_Shore | Body | chr17:73727879-73728061 | 182 | | 1 | | | | | | | | | | | | yes | | | | | |
| UNK | N_Shore | TSS1500 | chr17:73780035-73780495 | 460 | | 1 | 9 | | | | | | | | | | | | | | | | |
| UNC13D | NA | Body | chr17:73824354-73824760 | 406 | | | | | | | | 0 | 1 | | | | | yes | | | | | |
| UNC13D | NA | 5'UTR;1stExon;TSS200 | chr17:73840659-73841288 | 629 | | | 8 | 1 | 1 | | | | | | | | | | | | | yes | |
| TRIM47 | Island | 1stExon | chr17:73874465-73874637 | 172 | | | | | | | | | 1 | | | | | | | | | | yes |
| TRIM65 | Island | Body | chr17:73887208-73887651 | 443 | | | | | 2 | | | | | | | | | yes | | | | yes | |
| TRIM65 | Island | TSS200 | chr17:73893198-73893424 | 226 | | 1 | | 1 | | | | | | | | | | | | yes | | yes | yes |
| CDK3;C17orf106 | NA | TSS1500;Body | chr17:73996245-73996656 | 411 | | | | 1 | | | | | | | | | | | | | | | |
| EVPL | Island | Body | chr17:74011063-74011254 | 191 | | | 1 | | | | | | 1 | | | | | yes | | | | | |
| EVPL | NA | 5'UTR;1stExon;TSS200 | chr17:74023465-74023633 | 168 | | | | | | | | 0 | 1 | | | 1 | | yes | | | | | |
| ZACN | S_Shore | TSS200 | chr17:74075092-74075189 | 97 | | | 2 | | | | | 0 | 1 | | | | | yes | | | | | |
| FOXJ1 | S_Shore | TSS1500 | chr17:74138356-74138738 | 382 | | | | | | | | | | | | | | | | | | | |
| QRICH2 | NA | Body | chr17:74274956-74275137 | 181 | | | | | | | | | | | | | | | | | | | |
| UBE2O | NA | Body | chr17:74387399-74387581 | 182 | | | | | | | | | | | | | | | | | | | |
| AANAT | NA | TSS200;Body | chr17:74463560-74463767 | 207 | | | 4 | | | | | | | | | | | yes | | | | | yes |
| CYGB | NA | 3'UTR | chr17:74524371-74525125 | 754 | | | | | | | | | 2 | | | | | yes | | | | yes | |
| CYGB | Island | TSS200 | chr17:74533951-74533976 | 25 | | | | | | | | | 1 | | | | | | | | | yes | yes |
| SNORD1A | S_Shelf | TSS1500 | chr17:74557494-74557703 | 209 | | | 3 | 1 | | | | | 1 | 1 | | 1 | | yes | | | | | yes |
| ST6GALNAC2 | NA | Body | chr17:74564821-74565009 | 188 | | | | | | | | | | | | | | | | | yes | | |
| ST6GALNAC1 | NA | 5'UTR;1stExon | chr17:74639731-74639928 | 197 | | 1 | | | | | | | 1 | | | | | yes | | | | | |
| MXRA7 | NA | Body | chr17:74696737-74696872 | 135 | | | | 1 | | | | | | | | | | yes | yes | | | | |
| MGAT5B | N_Shore | TSS1500 | chr17:74864041-74864205 | 164 | | | 6 | | | | | | | | | | | yes | | | | | yes |
| MGAT5B | S_Shore;S_Shelf | TSS1500;Body;TSS200 | chr17:74868245-74869001 | 756 | | | 19 | | | | | | 1 | | | | | | yes | | yes | | |
| MGAT5B | Island | Body | chr17:74878309-74878420 | 111 | | | | | | | | | | | | | | yes | yes | | | | |
| MGAT5B | NA | Body | chr17:74908599-74908800 | 201 | | | | | | | | | | | | | | | | yes | | | |
| MGAT5B | NA | Body | chr17:74911481-74912348 | 867 | | | | | | | | | | | | | | yes | yes | yes | | | |
| MGAT5B | NA | Body | chr17:74913669-74913881 | 212 | | | 1 | | | | | | | | | | | | | | yes | | |
| MGAT5B | Island | Body | chr17:74928737-74928833 | 96 | | | 2 | | | | | | | | | | | yes | | | | | |
| MGAT5B | Island | Body | chr17:74944775-74945219 | 444 | | | | | | | | | | | | | | | | | | | |
| SCARNA16;C17orf86 | Island | TSS1500;TSS200 | chr17:75084623-75085182 | 559 | | | | | | | | 0 | | | | 1 | | yes | yes | | | | |
| SEC14L1 | Island | TSS200 | chr17:75136973-75136989 | 16 | | | | | | | | | 1 | | | | | | | | | | yes |
| SEC14L1 | NA | Body | chr17:75143118-75143600 | 482 | 3 | | | | | | | | | | | | | yes | yes | | | | yes |
| SEC14L1 | NA | Body;TSS1500;TSS200 | chr17:75180760-75181836 | 1076 | | | 8 | | | | | | | | | | | | yes | | | | |

| | | | | | | | | | | | | | | | | | | | | | | | |
|---|---|---|---|---|---|---|---|---|---|---|---|---|---|---|---|---|---|---|---|---|---|---|---|
| SEC14L1 | N_Shelf;N_Shore;Island | Body | chr17:75207115-75209644 | 2529 | | | | | 1 | | | | 2 | | | | | | | | | | |
| 40057;SEPT9 | S_Shelf;NA | Body;TSS1500;TSS200 | chr17:75282040-75284193 | 2153 | | | | | | | | | | | | | | | | | | | |
| SEPT9 | NA | 5'UTR;Body;TSS1500;TSS200;1stEx | chr17:75312494-75316784 | 4290 | | | | 1 | 3 | | | 0 | 4 | 4 | | | | yes | yes | yes | yes | yes | |
| SEPT9 | NA | 5'UTR;Body | chr17:75318655-75320035 | 1380 | | | 5 | | | | | | | | | | | yes | | yes | | | |
| SEPT9 | chr17:75368688-75370506;Isla | Island;5'UTR;Body;1stExon | chr17:75369091-75369939 | 848 | | | | | 1 | | | | | | 1 | | | | | | | | yes |
| SEPT9 | S_Shore | 5'UTR;Body;TSS1500 | chr17:75371476-75371764 | 288 | | | 4 | 1 | | | | | | | | | | | | | | | |
| SEPT9 | S_Shelf | 5'UTR;Body | chr17:75373081-75373457 | 376 | | | 3 | 1 | 1 | | | | | | | | | | | | | | |
| SEPT9 | NA | 5'UTR;Body | chr17:75385086-75385432 | 346 | | | 6 | | | | | | | | | | | | yes | | | yes | |
| SEPT9 | N_Shelf;N_Shore;Island | Body;TSS1500;TSS200;5'UTR | chr17:75445261-75448089 | 2828 | | | | | 2 | | | 0 | 3 | | | | | yes | | | yes | yes | |
| SEPT9 | S_Shelf | Body;5'UTR | chr17:75451779-75452044 | 265 | 2 | | | | | | | | 1 | | | | | | yes | | | | |
| SEPT9 | NA | Body;TSS200;5'UTR | chr17:75471193-75471491 | 298 | | | 3 | 1 | 1 | | | | 1 | | | | | yes | | | | | |
| SEPT9 | NA | Body;5'UTR | chr17:75473895-75474070 | 175 | | | | | | | | | 1 | | | | | yes | yes | | yes | | |
| TNRC6C | N_Shore;Island | 5'UTR | chr17:76036514-76037562 | 1048 | | | | 1 | | | | 0 | 4 | 2 | | 1 | | | | | | | yes |
| TMC6 | NA | Body | chr17:76117687-76117767 | 80 | | | | | | | | | | | | | | | | | | | |
| TMC8;TMC6 | N_Shore | TSS1500;5'UTR;TSS200 | chr17:76126301-76127217 | 916 | | | | | | | | | 1 | | | | | yes | | | | | |
| TMC8;TMC6 | S_Shore;chr17:76127521-7612 | Body;5'UTR;1stExon;TSS200;S_Sh | chr17:76128481-76130496 | 2015 | | | | | | | | | | | | | | | | | | | |
| SYNGR2 | N_Shore | TSS1500 | chr17:76164111-76164144 | 33 | | | 1 | | | | | | 1 | | | 1 | | yes | | | | | |
| TK1 | NA | Body | chr17:76171191-76171794 | 603 | | | | 1 | | | | | | | | | | yes | | | | | |
| AFMID;TK1 | S_Shore | Body;TSS1500 | chr17:76183551-76183641 | 90 | | | | | | | | | | | | | | yes | | | | | |
| BIRC5;EPR1 | NA | 3'UTR;Body;TSS200 | chr17:76220608-76220955 | 347 | | 1 | 10 | | | | | | | | | | | | | | | yes | |
| LOC283999 | N_Shore;chr17:76228110-7622 | TSS200;5'UTR;N_Shore | chr17:76227255-76228233 | 978 | 3 | | | | | | | | | | 1 | | | yes | | | | | yes |
| SOCS3 | N_Shore;Island | Body | chr17:76354621-76355288 | 667 | | | 4 | | 1 | | | | | | | | | | | | | | yes |
| PGS1 | Island | TSS200 | chr17:76374567-76374572 | 5 | | | | | | | | | | 1 | | | | yes | | | | | |
| DNAH17 | Island | Body | chr17:76446778-76446851 | 73 | | 1 | | | | | | | | | | | | | | | | | |
| DNAH17 | NA | Body | chr17:76464687-76464784 | 97 | | | | | | | | | 1 | | | | | yes | | | yes | | yes |
| DNAH17 | S_Shelf | Body | chr17:76501381-76501582 | 201 | | | | | | | | | 1 | | | | | yes | | | | | |
| DNAH17 | NA | Body | chr17:76503526-76503643 | 117 | | | | | | | | | | | | | | | yes | | | | |
| DNAH17 | NA | Body | chr17:76506650-76506855 | 205 | | | | | | | | | 2 | | | | | yes | | yes | | | |
| DNAH17 | NA | Body | chr17:76565037-76565232 | 195 | | | | | | | | | | | | | | yes | | | | | |
| DNAH17 | NA | TSS200 | chr17:76573499-76573574 | 75 | | | | | 1 | | | | | | | | | yes | yes | | | | |
| CYTH1 | NA | Body | chr17:76719591-76719943 | 352 | | | | | | | | | 1 | 1 | | | | | | | yes | | |
| USP36 | N_Shelf | Body | chr17:76794506-76795028 | 522 | | | 3 | | | | | | | | | | | | | | | yes | |
| USP36 | chr17:76798154-76798461 | S_Shelf | chr17:76801579-76801775 | 196 | | | | | | | | | | | 1 | | | yes | | | | | |
| USP36 | N_Shore | 5'UTR | chr17:76834778-76834989 | 211 | | 1 | | 1 | 1 | | | | | | | | | yes | | | yes | | |
| TIMP2 | N_Shore | 3'UTR | chr17:76850052-76850277 | 225 | | 1 | | | 1 | | | 0 | 1 | 1 | 1 | 1 | | | | | | | |
| TIMP2 | NA | Body | chr17:76875678-76876239 | 561 | | | | 1 | | | 1 | 0 | 3 | 1 | 1 | 10 | 1 | yes | yes | | | | |
| TIMP2 | NA | Body | chr17:76886742-76886912 | 170 | | | 1 | | | | | | | | 1 | | | yes | | | | | |
| TIMP2 | S_Shore;S_Shelf | Body | chr17:76899257-76899627 | 370 | 1 | | | | | | | | 2 | | | | | yes | | | | | |
| TIMP2 | N_Shore | Body | chr17:76920397-76920464 | 67 | 4 | | | 1 | | | | | | | | | | | | | | | |
| LGALS3BP | NA | Body | chr17:76967757-76968215 | 458 | | | | | | | | | | | | | | | yes | | | | |
| LGALS3BP | NA | Body | chr17:76970857-76970933 | 76 | | | | | | | | | | | | | | | | | | | |
| LGALS3BP | NA | 1stExon;5'UTR;TSS200;TSS1500 | chr17:76975944-76976357 | 413 | 1 | | 8 | | | | | 0 | 2 | | | | | yes | | yes | | | |
| CANT1 | Island | Body | chr17:76993506-76993698 | 192 | | | | | | | | | 1 | 1 | | | | | yes | | | | |
| C1QTNF1 | NA | 5'UTR;TSS1500;TSS200 | chr17:77029308-77030546 | 1238 | | | | | | | | | 1 | 1 | | | | yes | | | yes | | |
| C1QTNF1 | NA | 5'UTR | chr17:77037916-77038216 | 300 | | | 20 | | | | | | | | 1 | | | yes | | | | | |
| C1QTNF1 | N_Shore | Body | chr17:77042810-77043544 | 734 | 3 | | 2 | | | | | | | | | | | | | | | yes | |
| ENGASE | NA | Body | chr17:77078091-77079031 | 940 | | | 4 | | | | | | 1 | | | | | | | | | yes | |
| HRNBP3 | NA | 3'UTR | chr17:77086350-77086810 | 460 | | | 9 | | | | | | 1 | | 1 | | | | | | | yes | |
| HRNBP3 | Island | Body | chr17:77093406-77093866 | 460 | | | | | | | | 0 | 2 | | | | | | | | | | |
| HRNBP3 | N_Shore;Island | Body | chr17:77099486-77100187 | 701 | | | | | | | | 0 | 2 | | | | | | | | | | yes |
| HRNBP3 | N_Shore;S_Shore | 5'UTR | chr17:77126866-77128623 | 1757 | | 1 | 5 | | | | | | 2 | | | | | | yes | | yes | yes | |
| HRNBP3 | N_Shore;Island | 5'UTR | chr17:77133777-77134303 | 526 | | | 2 | | | | | | 1 | | | | | | | | yes | yes | |
| HRNBP3 | NA | 5'UTR | chr17:77141853-77142186 | 333 | | | | | | | | | | | | | | | | | yes | | |
| HRNBP3 | NA | 5'UTR | chr17:77147257-77147396 | 139 | | | 2 | | | | | | | | | | | yes | | | | | yes |
| HRNBP3 | NA | 5'UTR | chr17:77151165-77151243 | 78 | | | 2 | | | | | | | | | | | | | | | | |
| HRNBP3 | NA;N_Shelf | 5'UTR | chr17:77174120-77175465 | 1345 | 3 | | 3 | | | | | | | | | | | yes | | | | | |
| HRNBP3 | N_Shore | 5'UTR | chr17:77194008-77194175 | 167 | | | | | | | | | | | | 1 | | yes | | | | | |
| HRNBP3 | NA | 5'UTR | chr17:77302162-77302351 | 189 | | | | | | | | 0 | 1 | | | | | yes | | | | | |
| HRNBP3 | NA | 5'UTR | chr17:77311577-77311812 | 235 | | | 1 | | | | | 0 | 2 | | | 1 | | | | | | | |
| HRNBP3 | NA | 5'UTR | chr17:77351040-77351122 | 82 | | | | | | | | | 1 | | | | | | | | | | |
| HRNBP3 | NA | 5'UTR | chr17:77375461-77375842 | 381 | 2 | | | | | | | | 1 | | | | | | | | | | |
| HRNBP3 | NA | 5'UTR | chr17:77389727-77390045 | 318 | 5 | | 3 | | | | | | | | | | | | | | | | |
| HRNBP3 | NA | 5'UTR | chr17:77419249-77419534 | 285 | | | | | | | | | | | | | | | yes | yes | | | yes |
| HRNBP3 | Island;S_Shore | 5'UTR | chr17:77460269-77461480 | 1211 | | | 13 | | | | | | | | | | | yes | | | | yes | |
| HRNBP3 | NA | TSS200 | chr17:77478569-77478927 | 358 | | | 1 | | | | | 0 | 1 | | | | | | yes | | | | |
| ENPP7 | NA | TSS200 | chr17:77704760-77705025 | 265 | | | | | | | | | | | | | | | | | | | |
| ENPP7 | N_Shore | Body | chr17:77706946-77707066 | 120 | | | | | | | | 0 | 1 | | | 3 | | | yes | yes | | | |
| CBX4 | S_Shore;N_Shore | Body | chr17:77809912-77810912 | 1000 | 3 | | 3 | | 1 | | | | 2 | | 1 | | | | | | | | |
| TBC1D16 | Island | Body | chr17:77916733-77916892 | 159 | | | | | | | | 0 | 1 | 1 | | 2 | 1 | yes | | | | yes | |
| TBC1D16 | Island;S_Shore | Body | chr17:77924180-77925938 | 1758 | | | | | | | | | | | | | | | | | | | |
| TBC1D16 | N_Shelf | Body | chr17:77948865-77949135 | 270 | | | | | 1 | | | | | | | | | | yes | | yes | | |
| TBC1D16 | N_Shore;Island;S_Shore | Body | chr17:77951408-77953437 | 2029 | 1 | 1 | 40 | | | | | | 2 | | | 1 | | yes | | | | | |
| TBC1D16 | NA | Body | chr17:77959804-77960007 | 203 | | | | | 1 | | | | | | | | | | | | | | |
| TBC1D16 | N_Shore | Body | chr17:77963988-77964112 | 124 | 3 | 1 | | | | | | | | | | | | | yes | | | | |
| TBC1D16 | N_Shore;Island | 5'UTR | chr17:77997198-77997997 | 799 | | | 10 | | | | | 0 | | | | | | yes | | | | yes | |
| CCDC40 | N_Shore;Island | Body | chr17:78038002-78039528 | 1526 | | | | | | | | | 1 | 1 | | | | yes | yes | | | | yes |
| GAA | N_Shore;Island | Body | chr17:78078605-78079795 | 1190 | | 1 | 12 | | | | | | 1 | 1 | | | | yes | | | yes | | |
| CARD14 | N_Shelf | TSS200 | chr17:78152219-78152466 | 247 | | 1 | | | 1 | | | | | | | | | yes | | yes | | | |
| CARD14 | Island | Body | chr17:78157950-78158248 | 298 | | | | | | | | | | | | | | | yes | yes | | yes | |
| CARD14 | N_Shelf | TSS1500;Body;TSS200 | chr17:78161110-78161706 | 596 | | 1 | 18 | | 1 | | | 0 | 1 | 1 | 1 | | | yes | yes | | | | |
| CARD14 | S_Shore;Island | Body | chr17:78165188-78166385 | 1197 | | 1 | 6 | 1 | | | | 0 | 2 | | | 1 | | yes | | yes | yes | | |
| CARD14 | Island | Body | chr17:78172177-78172538 | 361 | | | | | | | | | | | | | | yes | | | | | |
| SLC26A11;SGSH | S_Shore | Body;TSS1500 | chr17:78195442-78195518 | 76 | | | | | | | | | | | | | | yes | | | | | |
| RNF213 | S_Shelf;S_Shore | 5'UTR | chr17:78237117-78237543 | 426 | 1 | | 3 | | 2 | 1 | | | | | | | | yes | | yes | | | |
| LOC100294362;FLJ35220 | N_Shore;Island | Body;TSS1500 | chr17:78387145-78388593 | 1448 | | | | | 1 | | | 0 | 2 | | | | | yes | yes | yes | | | |
| NPTX1 | N_Shore | 3'UTR | chr17:78441956-78442827 | 871 | 1 | | | | | | | | | | | | | | | | | yes | |
| NPTX1 | N_Shore | 3'UTR | chr17:78444348-78444608 | 260 | | | | | | | | | | | | | | | | | | yes | |
| RPTOR | NA | Body | chr17:78549324-78549371 | 47 | | | | | | | | 0 | 1 | | | | | yes | yes | | yes | | |
| RPTOR | NA | Body | chr17:78560478-78560916 | 438 | | 1 | 4 | | 1 | | | | | | | | | | yes | | | | |
| RPTOR | NA | Body | chr17:78607095-78607368 | 273 | | 1 | | | | | | | | | | | | yes | yes | | | | |
| RPTOR | NA | Body | chr17:78682785-78683457 | 672 | 1 | | | | | | | | 1 | | | | | yes | | | | | |
| RPTOR | NA | Body | chr17:78694592-78694856 | 264 | | | | | | | | 0 | | | | | | yes | | | yes | | |
| RPTOR | Island | Body | chr17:78700906-78701257 | 351 | | | | | | | | | 1 | | | | | | yes | yes | | | |
| RPTOR | NA | Body | chr17:78724995-78725470 | 475 | | | 3 | | | | | | | | | | | yes | yes | | | | yes |
| RPTOR | NA | Body | chr17:78735268-78735596 | 328 | | 1 | 21 | | 1 | | | | 2 | 1 | | 1 | | | | | | | |
| RPTOR | NA | Body | chr17:78747934-78748494 | 560 | 4 | | | | | | | | | | | | | yes | | | | | |
| RPTOR | NA | Body | chr17:78753756-78754372 | 616 | | | 5 | | 1 | | | 0 | | | | | | yes | | | | | |
| RPTOR | NA | Body | chr17:78755379-78755442 | 63 | 1 | | | | | | | 0 | 1 | | 1 | | | yes | | | | | |
| RPTOR | N_Shore;N_Shelf | Body | chr17:78773453-78774058 | 605 | 2 | 1 | | | | | | | 1 | | | | | | yes | | | | |

| | | | | | | | | | | | | | | | | | | | | | | | |
|---|---|---|---|---|---|---|---|---|---|---|---|---|---|---|---|---|---|---|---|---|---|---|---|
| RPTOR | S_Shore | Body | chr17:78800321-78800806 | 485 | | | 4 | | 1 | | | 0 | 3 | 1 | | 1 | | | | yes | | | |
| RPTOR | NA | Body | chr17:78806493-78806891 | 398 | | | | 1 | 1 | | | | | | | | | | | | | | |
| RPTOR | NA | Body | chr17:78808060-78808570 | 510 | | | 3 | | | | | | | | | | | yes | | | | yes | |
| RPTOR | N_Shore | Body | chr17:78817642-78818045 | 403 | | 1 | | | | | | | | | | | | yes | | | | | |
| RPTOR | S_Shelf | Body | chr17:78821604-78821928 | 324 | | | | | 1 | | | | | | | | | yes | | | | | |
| RPTOR | S_Shore | Body | chr17:78832948-78833484 | 536 | | 1 | | | | | | | | | | | | | | | | yes | |
| RPTOR | S_Shore | Body | chr17:78851149-78851503 | 354 | | | | | 1 | 1 | | 0 | 3 | 1 | 1 | 1 | | | | | | | |
| RPTOR | N_Shore | Body | chr17:78856861-78857121 | 260 | 1 | | | | | | | | | | | | | | | | | | |
| RPTOR | S_Shore;N_Shelf | Body | chr17:78859745-78860605 | 860 | | 1 | | | | | | | 3 | | | | | yes | yes | | | | |
| RPTOR | Island;S_Shore | Body | chr17:78863570-78865514 | 1944 | | | 22 | | | | | 0 | 2 | | | 1 | | | yes | | | | |
| RPTOR | N_Shelf | Body | chr17:78876656-78878579 | 1923 | 3 | | | | | | | | | | | | | yes | | | yes | | |
| RPTOR | N_Shelf | Body | chr17:78894364-78894477 | 113 | | | 4 | | | | | | | | | | | | | | | | |
| RPTOR | Island | Body | chr17:78915655-78915881 | 226 | | | 7 | | | | | | | | | | | | | | | | |
| FLJ90757 | N_Shore | Body | chr17:79004850-79006166 | 1316 | | 1 | | | | | | 0 | 3 | | | 3 | | | | | | | |
| BAIAP2 | S_Shore | Body | chr17:79010458-79011542 | 1084 | 1 | | 10 | | 1 | | | | 1 | | | | | | | | | | |
| BAIAP2 | S_Shore;Island | Body | chr17:79026049-79027207 | 1158 | | | 2 | 1 | | | | | 1 | | | | | yes | | | | | |
| BAIAP2 | NA | Body | chr17:79030796-79031712 | 916 | | | 11 | 2 | | | | | 1 | | | | | | | | | yes | |
| BAIAP2 | Island | Body | chr17:79045695-79045877 | 182 | | | | | | | 1 | | | | | | | | | | | | |
| BAIAP2 | S_Shore | Body | chr17:79053905-79054073 | 168 | | | | | | | | | | | | | | yes | | | | | |
| BAIAP2 | S_Shelf | Body | chr17:79056061-79056183 | 122 | | | | | | | | | 1 | 1 | | | | | | | | yes | |
| BAIAP2 | NA | Body | chr17:79058303-79058859 | 556 | | | | 1 | | | | | 1 | | | | | | | | | | |
| BAIAP2 | NA | Body | chr17:79066515-79066577 | 62 | | | | | | | | | 1 | | | | | | | | | yes | |
| BAIAP2 | NA | Body | chr17:79068856-79069042 | 186 | | | | | | | | 0 | 1 | | | | | | | | | | |
| AATK | Island;S_Shore | Body | chr17:79095700-79097243 | 1543 | | | 8 | | | | | | | | | | | | | | | | yes |
| MIR657;AATK;MIR338 | N_Shelf | TSS1500;Body;TSS200 | chr17:79099246-79100682 | 1436 | | | 29 | | | | | | 2 | 1 | 2 | | | | | | | | yes |
| AATK;MIR1250 | N_Shore | Body;TSS1500 | chr17:79108051-79108224 | 173 | | | | | | | | | | | | | | | | | | | |
| AATK | Island | Body | chr17:79109729-79110023 | 294 | | | 20 | | | | | | 2 | 1 | | | | | | | | | |
| AATK | NA | nan | chr17:79122243-79122455 | 212 | | | | | | | | | | | 1 | | | | | | | yes | |
| AATK | N_Shelf | Body | chr17:79128885-79129078 | 193 | | | | | | | 1 | 0 | 5 | 3 | 2 | 5 | | yes | | | yes | yes | |
| AATK | Island | Body | chr17:79131893-79132083 | 190 | | | | | | | | | 1 | | | | | | | | | | |
| LOC388428;AATK | N_Shore;S_Shore | TSS1500;Body | chr17:79138873-79140589 | 1716 | | | 7 | | | | | | | | | | | yes | | | | yes | yes |
| AZI1 | Island | Body | chr17:79183386-79183914 | 528 | | | | | | | 1 | 0 | 1 | 1 | | | | yes | | | | | |
| AZI1 | S_Shore | TSS1500 | chr17:79197669-79198101 | 432 | | | | | | | | | | | | | | yes | yes | | | | |
| C17orf56 | N_Shelf | Body | chr17:79210394-79210852 | 458 | | | | | | | | | | | | | | | | | | | |
| SLC38A10 | NA | Body | chr17:79224065-79225974 | 1909 | 6 | | | | 2 | | | | 1 | | | | | | | | | | yes |
| SLC38A10 | NA | Body | chr17:79228937-79229385 | 448 | | | | 1 | | | | | 1 | 1 | | | | yes | | | | | |
| SLC38A10 | N_Shelf | Body | chr17:79251255-79251354 | 99 | | | 4 | 1 | | | | | 1 | 1 | | | | | | | | | |
| SLC38A10 | S_Shore | Body | chr17:79257021-79257217 | 196 | | | | | | | | | 1 | | | | | yes | | | | | |
| SLC38A10 | N_Shore;Island | Body | chr17:79258687-79259963 | 1276 | | | | | | | | | | | | | | | | | | | |
| SLC38A10 | N_Shelf | Body | chr17:79265454-79265759 | 305 | | | | | | | | | | | | | | yes | yes | | | yes | |
| C17orf55 | N_Shore | TSS1500 | chr17:79283300-79283959 | 659 | | | 2 | | | | | | 2 | | | 1 | | | | | | | |
| TMEM105 | NA | 5'UTR | chr17:79297435-79297618 | 183 | | | | | | | | | 1 | 1 | | | | | | | | | |
| TMEM105 | NA | 5'UTR;1stExon;TSS200 | chr17:79304420-79304683 | 263 | | | | 1 | 1 | | | | 2 | | | | | | | | | | |
| BAHCC1 | Island | TSS1500;TSS200;Body | chr17:79372242-79374741 | 2499 | | 2 | 23 | 1 | 1 | | | 0 | 2 | 1 | | 1 | | | | | | yes | yes |
| BAHCC1 | N_Shore;Island | Body | chr17:79377272-79378207 | 935 | | | 5 | 1 | | | | | 1 | 1 | | | | | | | | | yes |
| BAHCC1 | Island | Body | chr17:79380400-79380706 | 306 | | 1 | | | | | | 0 | 5 | | | 9 | | | | | | | |
| BAHCC1 | Island | Body | chr17:79388538-79388814 | 276 | 1 | | | | | | | | | | | | | | | | | | |
| BAHCC1 | S_Shore | Body | chr17:79390726-79391071 | 345 | | | 1 | | | | | | | | | | | | | | | yes | |
| BAHCC1 | Island | Body | chr17:79393463-79393718 | 255 | | | | | | 1 | | 0 | 2 | | | 1 | | | | | | yes | |
| BAHCC1 | N_Shore | Body | chr17:79419739-79420279 | 540 | | | | | 2 | | | | 2 | | | | | | | | | | |
| BAHCC1 | Island | Body | chr17:79428036-79428750 | 714 | | 1 | | | | | | | 1 | | | | | | | | | | yes |
| BAHCC1 | Island | Body;3'UTR | chr17:79430641-79431331 | 690 | | 2 | | | | | | | 1 | | | | | | | | | yes | |
| ACTG1 | Island | TSS1500 | chr17:79480921-79481108 | 187 | | | | | 1 | | | | | | | | | | | | | | |
| FSCN2 | Island | TSS1500;TSS200 | chr17:79495006-79495664 | 658 | 1 | | 8 | | | | | 0 | 2 | | | | | | | | | yes | |
| FSCN2 | Island | Body | chr17:79503642-79503877 | 235 | 1 | | 3 | | | | | 0 | 1 | | | | | | | | | | |
| C17orf70 | S_Shore | TSS1500 | chr17:79520615-79521089 | 474 | | | 1 | 1 | | | | | | | | | | yes | | | | | |
| TSPAN10 | NA | TSS1500;TSS200 | chr17:79609095-79609346 | 251 | | | 3 | | | | | | 2 | 1 | | | | yes | | | | | |
| TSPAN10 | Island;S_Shore | 3'UTR | chr17:79615552-79615861 | 309 | | | 5 | | | | | | | | | | | yes | | | | | |
| PDE6G | NA | TSS200 | chr17:79623661-79624018 | 357 | | | | | | | | | | | | | | | yes | yes | | | |
| C17orf90;CCDC137 | N_Shore | Body;TSS1500 | chr17:79632335-79632662 | 327 | | | | | | | | | | | | | | yes | yes | | | | |
| C17orf90;CCDC137 | S_Shore | TSS1500;Body | chr17:79634844-79635073 | 229 | | | | | | | | | 1 | | | | | yes | yes | | | | |
| CCDC137 | NA | Body | chr17:79639690-79640096 | 406 | | | | | | | | | 1 | | | | | yes | | yes | | | |
| ARL16;HGS | S_Shore | TSS1500;Body | chr17:79652024-79652389 | 365 | | | 7 | | | | | | 1 | 1 | 1 | | | yes | | | | | |
| HGS | Island | Body | chr17:79662881-79663039 | 158 | | | | | | | | | 1 | | | | | yes | | | | yes | |
| SLC25A10 | N_Shore | Body | chr17:79680718-79681052 | 334 | | | | | 1 | | | | | | | | | | | | | | |
| DYSFIP1 | S_Shore | TSS200 | chr17:79793082-79793208 | 126 | | | | | | | | | | | | | | yes | | | | | |
| P4HB | NA | Body | chr17:79813410-79813507 | 97 | 2 | | | | | | | | 1 | 1 | | | | yes | | | | | |
| P4HB | N_Shore | Body | chr17:79816504-79816678 | 174 | | | 6 | | | | | | 1 | 1 | | | | yes | | | | | |
| NPB | Island | TSS1500 | chr17:79859809-79859830 | 21 | | | | | | | | 0 | | | | | | | | yes | | | |
| PCYT2 | Island | TSS1500 | chr17:79869748-79869861 | 113 | | 1 | 1 | | | | | 0 | 1 | | | | | | | | | yes | yes |
| MAFG | S_Shore | 5'UTR;TSS200 | chr17:79881066-79882042 | 976 | | 1 | 24 | 1 | | 1 | | 0 | 6 | 5 | | 5 | | | | | | | yes |
| LOC92659;MAFG | Island | Body;TSS1500 | chr17:79886614-79887046 | 432 | | | 6 | | | | | | | | | | | | | | | | |
| MYADML2 | Island | Body | chr17:79899075-79899490 | 415 | | 1 | | | | | | | | | | | | | | | | | |
| MYADML2 | NA | TSS200 | chr17:79905152-79906039 | 887 | | 1 | 52 | 1 | | | 1 | | 5 | 2 | | 14 | 1 | yes | yes | yes | yes | | |
| NOTUM | Island | TSS1500 | chr17:79920530-79920672 | 142 | | | | | | | | | 1 | | | | | | | | | | |
| ASPSCR1 | S_Shore | Body | chr17:79936477-79936561 | 84 | 1 | | | 1 | 1 | | | | | | | | | yes | yes | | | | |
| ASPSCR1 | N_Shelf | Body | chr17:79950061-79950366 | 305 | | | 2 | | | | | | 1 | | | | | | | | | | |
| ASPSCR1 | N_Shore | Body | chr17:79952786-79952915 | 129 | | | | | | | | | | | | | | | | | | | |
| ASPSCR1 | S_Shore | Body | chr17:79963012-79963651 | 639 | | | | | | | | | 1 | | | | | | | | | yes | |
| GPS1 | N_Shore | Body | chr17:80011196-80011568 | 372 | | | | | | | | | | | | | | | | | | | |
| DUS1L | N_Shore | Body | chr17:80022616-80023016 | 400 | | | 3 | | | | | | | | | | | | | | | | |
| CCDC57 | NA | Body | chr17:80066134-80066314 | 180 | | | | 1 | | | | | | | | | | | | | | | |
| CCDC57 | N_Shore | Body | chr17:80084554-80084596 | 42 | | | | | | | | | 1 | | | | | yes | yes | | | | |
| CCDC57 | NA | Body | chr17:80145540-80145575 | 35 | | | | | | | | | 1 | | | | | yes | yes | | | | yes |
| CCDC57 | NA | Body | chr17:80159399-80159595 | 196 | | | | | 1 | | | | | | | | | yes | | | | | yes |
| SLC16A3 | Island;N_Shore | TSS200;TSS1500 | chr17:80186080-80186336 | 256 | | | 12 | | 2 | | | | | | | | | | | | | | yes |
| SLC16A3 | Island;S_Shore | 5'UTR;TSS1500;TSS200 | chr17:80189381-80190154 | 773 | | | 20 | | | | | | 1 | | | | | | | | | | yes |
| SLC16A3 | Island | Body | chr17:80194552-80194706 | 154 | | | | | | | | | | | | | | | | | | | |
| SLC16A3 | S_Shore | Body;3'UTR | chr17:80196719-80197412 | 693 | | | 8 | | 1 | | | | 1 | | | | | | | | | | |
| CD7 | N_Shelf | Body | chr17:80273242-80274000 | 758 | | | | 1 | 1 | | | | 2 | | | | | | yes | | | | |
| CD7 | N_Shore | 1stExon;5'UTR;TSS200 | chr17:80275401-80275880 | 479 | | | 3 | | | | | | | | | | | | | | | yes | |
| SECTM1 | Island | 3'UTR | chr17:80278914-80279562 | 648 | | | 9 | | | | | | 2 | 1 | | | | yes | yes | | | | |
| SECTM1 | N_Shore | 5'UTR | chr17:80289500-80290217 | 717 | | 1 | 18 | | 2 | | | | 1 | 1 | | | | yes | | | yes | | |
| SECTM1 | Island | TSS200 | chr17:80292079-80292252 | 173 | | | 19 | | 1 | | | | | | | | | | | | | | |
| C17orf101 | N_Shelf | Body | chr17:80358829-80358876 | 47 | | | 26 | | | | | | 1 | 1 | 1 | | | yes | | | | | |
| C17orf101 | N_Shore | Body | chr17:80372534-80372912 | 378 | | 1 | | | | | | 0 | 2 | | | | | yes | | | | | |
| HEXDC | S_Shore | Body | chr17:80394416-80394588 | 172 | | | | 1 | | | | | | | | | | | | | | | |
| C17orf62 | N_Shore | 3'UTR | chr17:80401500-80401857 | 357 | | | | | | | | 0 | 1 | | | | | yes | | | | | |
| C17orf62 | N_Shore | 5'UTR;Body | chr17:80407031-80407779 | 748 | | | 2 | | 1 | | | 0 | 1 | 2 | 1 | | | yes | | | | | |

| | | | | | | | | | | | | | | | | | | | | | | | |
|---|---|---|---|---|---|---|---|---|---|---|---|---|---|---|---|---|---|---|---|---|---|---|---|
| FOXK2 | Island | Body | chr17:80541549-80542006 | 457 | | | 1 | | | 1 | | | 2 | | | | | yes | | | | | |
| FOXK2 | N_Shore | Body | chr17:80545175-80545454 | 279 | | | | | | | | 0 | 7 | 2 | | 1 | | yes | | | | | |
| FOXK2 | N_Shore | Body | chr17:80553970-80554772 | 802 | | | | | | | | | 2 | | | | | yes | yes | | | | |
| FOXK2 | S_Shore | 3'UTR | chr17:80560980-80561461 | 481 | | | 1 | | | | | | 1 | | | | | | | | | | |
| WDR45L | NA | Body | chr17:80581701-80581925 | 224 | | | | | | | | | 1 | | | | | yes | yes | | | | |
| WDR45L | S_Shore | TSS1500 | chr17:80606880-80606947 | 67 | | 1 | | | | | | | | | | | | | | | | yes | |
| RAB40B | NA | Body | chr17:80635520-80635688 | 168 | | | | | | | | | 1 | | | | | yes | | | | | yes |
| FN3K | Island | Body | chr17:80708279-80708632 | 353 | 1 | | 7 | | | | 2 | 0 | 3 | 2 | 1 | 21 | 1 | yes | | yes | | | yes |
| TBCD | S_Shelf | Body | chr17:80744833-80745235 | 402 | | | 4 | | | | | | | | | | | | | | | | |
| TBCD | NA | Body | chr17:80747189-80747795 | 606 | | | | | | | | | 1 | | | | | | | | | | |
| TBCD | NA | Body | chr17:80753237-80753497 | 260 | | 1 | | | | | | | | | | | | | | yes | | | |
| TBCD | N_Shore | Body | chr17:80763035-80763117 | 82 | | | 8 | | | | | | | | | | | yes | | yes | | | |
| TBCD;ZNF750 | Island | Body;5'UTR | chr17:80790369-80790580 | 211 | | | | | | 5 | | | | | | | | | | | | | |
| TBCD;ZNF750 | S_Shelf | Body;5'UTR | chr17:80794081-80794274 | 193 | | | | | | | | | 1 | | | | | yes | | | | yes | |
| TBCD;ZNF750 | NA | Body;5'UTR;1stExon;TSS200 | chr17:80795569-80798023 | 2454 | | | | | | | | | 1 | 1 | | | | | | | | yes | |
| TBCD | NA | Body | chr17:80806135-80806435 | 300 | | | 19 | 1 | | | | | 1 | | | | | yes | | | | | yes |
| TBCD | Island | Body | chr17:80818599-80819208 | 609 | | | | | 1 | | | | 1 | | | | | | | | | | |
| TBCD | N_Shelf | Body | chr17:80825471-80825525 | 54 | | | | | | | | | 1 | | | | | | | | | | |
| TBCD | N_Shore;chr17:80829533-8082 | Body;N_Shore | chr17:80829028-80829718 | 690 | | | | | 1 | | | | 1 | | 1 | | | | | | | | |
| TBCD | S_Shelf | Body | chr17:80833054-80833638 | 584 | | 1 | | | | | | | | | | | | | | | | | yes |
| TBCD | NA | Body | chr17:80839243-80839375 | 132 | | | 10 | | | | | | | | | | | | | | | | yes |
| TBCD | NA | Body | chr17:80842262-80842469 | 207 | | | | | | | | | 1 | | | | | | | | | | |
| TBCD | NA | Body | chr17:80845107-80845337 | 230 | | | | | | | | | 1 | | | | | yes | yes | | | yes | |
| TBCD | NA | Body | chr17:80847137-80847662 | 525 | | | | 1 | 2 | | | | 1 | 1 | | | | | | | | | |
| TBCD | N_Shelf;S_Shore | Body | chr17:80858406-80860250 | 1844 | | | 4 | | | | | | | | | | | | yes | | yes | | |
| TBCD | S_Shelf | Body | chr17:80866063-80866232 | 169 | | | | | | | | | | | | | | yes | | | yes | | |
| TBCD | N_Shore | Body | chr17:80869706-80870577 | 871 | | | | | 1 | | | 0 | 1 | | | | | | | | | | |
| TBCD | S_Shore | Body | chr17:80873120-80873656 | 536 | | | | | 1 | | | | | | | | | yes | yes | yes | | | |
| TBCD | S_Shelf;NA | Body | chr17:80875493-80876074 | 581 | | | 23 | | | | | | 1 | | | | | yes | yes | | | | |
| B3GNTL1 | N_Shelf | Body | chr17:80916125-80916271 | 146 | | | | | | | | | | | | | | | | | | yes | |
| B3GNTL1 | Island;S_Shore | Body | chr17:80922369-80923777 | 1408 | | | | | | | | | | | | | | yes | yes | | | | |
| B3GNTL1 | Island | Body | chr17:80929076-80929274 | 198 | | | 1 | | | | | | | | | | | | | yes | yes | | |
| B3GNTL1 | NA | Body | chr17:80943851-80944193 | 342 | | | 8 | 1 | 1 | | | | 1 | | | | | | | | | | |
| B3GNTL1 | S_Shore;N_Shore;Island | Body | chr17:80969109-80971776 | 2667 | | | | | | | | | | | | | | | | | | | |
| B3GNTL1 | N_Shelf | Body | chr17:80986552-80986746 | 194 | | | | | | | | | 1 | | | | | yes | yes | | | | |
| B3GNTL1 | N_Shelf | Body | chr17:81006485-81006524 | 39 | | | 2 | 1 | | | | | | | | | | yes | | | | | |
| B3GNTL1 | Island | TSS200 | chr17:81009832-81009998 | 166 | | 1 | | | | | | | | | | | | | | | | yes | |
| METRNL | Island;S_Shore;N_Shore | Body | chr17:81039406-81040906 | 1500 | | | 7 | | | | | | 1 | | | | | | | | | | |
| METRNL | N_Shore;Island | Body | chr17:81042554-81043668 | 1114 | | 1 | 19 | | | | | | | | | | | | | | | | |
| METRNL | Island | Body | chr17:81045495-81045863 | 368 | | | 12 | | | | | | 1 | | | | | | | | | | |
| METRNL | N_Shore;Island | Body | chr17:81047183-81047941 | 758 | | 2 | 4 | 1 | | | | | 2 | | | | | | | | | | |
| COLEC12 | S_Shore | TSS1500 | chr18:500979-501234 | 255 | | | 1 | | | | | | | | | | | | | | | | yes |
| CLUL1 | NA | TSS1500 | chr18:596703-596759 | 56 | | | | | | | | | 1 | | | | | | | | yes | | yes |
| ENOSF1 | Island | TSS200;TSS1500 | chr18:712731-712737 | 6 | | | | | | | | | 1 | | | | | yes | | | | | |
| ADCYAP1 | Island | 5'UTR;TSS200;1stExon | chr18:905156-905611 | 455 | | | | | | | | | 2 | | | | | | | | | | |
| ADCYAP1 | Island | Body | chr18:908745-909154 | 409 | | | | | | | | | | | | | | | | | | | yes |
| EMILIN2 | Island | Body | chr18:2907240-2907472 | 232 | | | 8 | | | | | | | | | | | yes | | | | | |
| LPIN2 | N_Shore | TSS200 | chr18:3011965-3012017 | 52 | | | 4 | | | | | | | | | | | yes | | | | | |
| MYOM1 | NA | 5'UTR;1stExon;TSS200 | chr18:3219966-3220273 | 307 | | | | 1 | 1 | | | | | | | | | yes | | | | | |
| TGIF1 | NA | TSS1500;TSS200 | chr18:3411743-3412088 | 345 | | | 10 | | | | | 0 | | | | | | | | yes | | | |
| TGIF1 | Island | 5'UTR;Body;1stExon;TSS1500 | chr18:3452302-3452443 | 141 | | | | | | | | | 1 | | | 1 | | | | | | | |
| DLGAP1 | NA | TSS200;Body | chr18:3845348-3845703 | 355 | | 1 | | | | | | | 1 | | | | | | | | | yes | |
| DLGAP1 | Island | 1stExon;5'UTR | chr18:3880086-3880558 | 472 | | | | | | | | | | | | | | yes | | | | | |
| LOC642597 | Island | Body;1stExon;5'UTR;TSS200 | chr18:5196397-5197462 | 1065 | | 1 | | | | | | | 2 | | | | | | | | | | yes |
| EPB41L3 | Island | 5'UTR | chr18:5543548-5543878 | 330 | | | | | | | | | | 1 | | | | | | | | | yes |
| TMEM200C | Island;S_Shore | 1stExon;TSS200 | chr18:5890293-5892753 | 2460 | | 1 | | | | | | 0 | 3 | | | 2 | | | | | | yes | yes |
| L3MBTL4 | Island | Body | chr18:6284382-6284498 | 116 | | | | | | | | | | | | | | | | | | | |
| L3MBTL4 | Island;S_Shore | 5'UTR;TSS200 | chr18:6414330-6414978 | 648 | | | | | | | | 0 | 3 | 2 | | 5 | | | yes | | | | yes |
| LAMA1 | N_Shore;Island | Body | chr18:7011217-7011463 | 246 | | | 17 | | | 1 | | 0 | 2 | 1 | | 2 | | | | | yes | | |
| LAMA1 | Island | Body | chr18:7117321-7117790 | 469 | | | | | | | | | 1 | | | | | | | | | yes | yes |
| LRRC30 | NA | TSS200 | chr18:7231037-7231534 | 497 | | | | | | | | | | | | | | | | | | | |
| PTPRM | Island | 1stExon;5'UTR;Body | chr18:7567799-7568539 | 740 | | | | | | | | | 2 | | | | | | | | | | yes |
| LOC100192426;PTPRM | Island | TSS1500;Body | chr18:8367314-8367425 | 111 | | | | | | | | | 1 | | | | | | | | | | |
| KIAA0802 | NA | TSS1500 | chr18:8717011-8717113 | 102 | 1 | | | | | | | | | | | | | | yes | | | yes | |
| TWSG1 | N_Shore | TSS1500 | chr18:9334488-9334699 | 211 | | | | | | | | | 1 | | | | | | | | yes | | yes |
| TXNDC2 | NA | 5'UTR;1stExon;TSS200 | chr18:9885797-9886064 | 267 | | | 4 | | | | | | | | | | | yes | | | | yes | |
| APCDD1 | N_Shore | TSS1500 | chr18:10453624-10454085 | 461 | | | | | | | | | | | | | | yes | | | | | yes |
| APCDD1 | Island | Body | chr18:10471657-10471866 | 209 | | | | | | | | | | | | | | yes | | | | | |
| FAM38B | Island | Body | chr18:10726260-10726863 | 603 | | | 1 | | | | | 0 | 1 | | | | | | | | | | |
| FAM38B | N_Shore;Island | Body;TSS200;TSS1500 | chr18:11147785-11150043 | 2258 | | | | | | | | | | | | | | | | | | | |
| GNAL | Island | TSS200;Body | chr18:11751516-11752538 | 1022 | | 1 | | | | | | | | | | | | | | | | | |
| CHMP1B;GNAL | N_Shore | TSS1500;Body | chr18:11850139-11850153 | 14 | | | 7 | | | | | 0 | | | | 1 | | | yes | | yes | | |
| CIDEA | Island | TSS1500;TSS200;Body | chr18:12254173-12254566 | 393 | | 1 | | | 2 | | | 0 | 3 | | | 1 | | | | | | | yes |
| TUBB6 | N_Shore | TSS1500 | chr18:12307329-12307649 | 320 | | 1 | | | 1 | | | | 1 | | | | | | | | | | yes |
| SPIRE1 | N_Shore | TSS200;Body | chr18:12656824-12657056 | 232 | | | | | | | | | 1 | | | | | | | | | | |
| C18orf1 | Island | TSS1500 | chr18:13218443-13218607 | 164 | | | | | | | | | 1 | | | | | | | | | yes | yes |
| C18orf1 | Island | 5'UTR | chr18:13262344-13262750 | 406 | | | | | | | | | 1 | | | | | | | | | | |
| C18orf1 | NA | TSS1500;Body | chr18:13611370-13611824 | 454 | | 1 | | 1 | | | | 0 | 4 | | | 1 | | | | | | | |
| C18orf1 | N_Shore;Island | Body | chr18:13641529-13641894 | 365 | 2 | | 2 | 1 | 1 | | | 0 | 2 | 2 | | 3 | | yes | | | | yes | |
| ZNF519 | Island | Body | chr18:14132207-14132481 | 274 | | | | | | | | | | | | 1 | | yes | | yes | yes | | |
| LOC284233 | NA | TSS1500 | chr18:14336021-14336385 | 364 | | | | | | | | | | | | | | | | | | | |
| ANKRD30B | Island;S_Shore | TSS1500;TSS200;1stExon;5'UTR | chr18:14747661-14748439 | 778 | | 1 | 11 | | 1 | | | 0 | 5 | | | 5 | 1 | yes | | | | | yes |
| ABHD3 | Island | TSS1500 | chr18:19285008-19285336 | 328 | | | 1 | | | | | | | | | | | | | | | | |
| GATA6 | Island | 5'UTR;Body | chr18:19750865-19752414 | 1549 | | | | | | | | | 1 | | | | | | yes | | | yes | yes |
| GATA6 | N_Shore;Island | Body | chr18:19756582-19757022 | 440 | | 1 | 18 | | | | | 0 | | 1 | 1 | | | | | | | | yes |
| GATA6 | N_Shore | Body | chr18:19780587-19780728 | 141 | | | 6 | | | | | | | | | | | | | | | yes | |
| CTAGE1 | NA | 1stExon;5'UTR;TSS200 | chr18:19997803-19998008 | 205 | | | 8 | | | | | | | | | | | | | yes | | | |
| CABLES1 | N_Shore | TSS1500;TSS200 | chr18:20713759-20715040 | 1281 | | | 20 | 1 | | | | | | | | | | yes | | | | yes | yes |
| CABLES1 | NA | Body;TSS1500;TSS200 | chr18:20735005-20735693 | 688 | 2 | | 8 | | | | | | | | | | | yes | | | | | |
| C18orf45 | S_Shore | TSS1500 | chr18:21018217-21018724 | 507 | | | | | | | | | 1 | | | | | | | yes | | yes | yes |
| LAMA3 | NA | TSS1500;Body;TSS200 | chr18:21452730-21453131 | 401 | | 1 | | | | | | | 1 | | | | | | | | | | |
| TTC39C | NA | TSS200 | chr18:21572622-21572748 | 126 | 1 | | 6 | 1 | | | | | 2 | | | | | | yes | | | | |
| CABYR | Island | TSS200;TSS1500;5'UTR | chr18:21718863-21719433 | 570 | | | | | | | | 0 | 2 | | | 2 | | yes | | | yes | | |
| IMPACT | Island | TSS1500;TSS200 | chr18:22006316-22006496 | 180 | | | 3 | | | | | | | | | | | | | | | | |
| PSMA8 | N_Shore;Island | TSS1500;TSS200 | chr18:23713407-23713928 | 521 | | | 8 | | | | | 0 | 6 | | | 1 | | yes | | | | yes | |
| KCTD1 | Island | 5'UTR;TSS1500 | chr18:24130839-24131604 | 765 | 1 | | 9 | | 1 | | | | | | | | | | | | | yes | |
| AQP4;C18orf16 | Island;S_Shore | TSS1500;Body;TSS200 | chr18:24443262-24445692 | 2430 | | 1 | 11 | | | | | 0 | | | | | | | | | | | yes |
| CHST9 | Island | TSS1500;TSS200 | chr18:24765485-24765781 | 296 | | | 3 | | | | | | | | | | | yes | | | | yes | |
| CDH2 | N_Shore | Body | chr18:25755311-25755847 | 536 | | | | | | | | | 1 | | | | | | | | | | |

| | | | | | | | | | | | | | | | | | | | | | | | |
|---|---|---|---|---|---|---|---|---|---|---|---|---|---|---|---|---|---|---|---|---|---|---|---|
| DSC3 | Island | Body;TSS200;TSS1500 | chr18:28621490-28623090 | 1600 | | 3 | | | | | | 0 | 4 | 1 | | | | | | | | | yes |
| DSG1 | NA | TSS1500 | chr18:28897113-28897372 | 259 | | | | | | | | | 1 | | | | | | | | | | yes |
| DSG3 | NA | TSS200 | chr18:29027658-29027885 | 227 | | | | | | | | | 1 | | | | | | | | | | |
| DSG2 | Island | TSS1500 | chr18:29077698-29078241 | 543 | | | | | | | | | 1 | | | | | yes | | | | | yes |
| B4GALT6 | N_Shore | Body | chr18:29263741-29264252 | 511 | | | | | | | | | 1 | | | | | | | | | | yes |
| MCART2 | NA | TSS200;1stExon;5'UTR | chr18:29340788-29340919 | 131 | | | 4 | | | | | 0 | 2 | | | 2 | | yes | | | | | |
| FAM59A | S_Shore;Island | TSS1500 | chr18:30051320-30051572 | 252 | | | 21 | | | | | | 1 | | | | | yes | | | | yes | yes |
| KLHL14 | S_Shore | 5'UTR;1stExon;TSS200;TSS1500 | chr18:30352505-30353699 | 1194 | | 1 | 26 | | 1 | | | | 4 | 1 | | | | | | | | yes | |
| C18orf34 | Island | TSS200;TSS1500 | chr18:31020754-31020973 | 219 | | | | | | | | | 1 | | | | | yes | | | | | yes |
| NOL4 | Island | Body | chr18:31739202-31739521 | 319 | | 1 | | | 1 | | | | 1 | | | | | | | | | | |
| DTNA | NA | TSS200;5'UTR;TSS1500 | chr18:32172486-32173227 | 741 | | | | | | | | 0 | 4 | | | 1 | | | | | | yes | yes |
| DTNA | NA | Body;TSS1500 | chr18:32398024-32398287 | 263 | | 1 | | | | | | | 1 | | | | | | | | | | |
| ZNF397OS | N_Shore;Island | 5'UTR | chr18:32847251-32848016 | 765 | | | | | | | | 0 | 1 | | | 4 | | yes | | | | yes | |
| ZNF396 | S_Shore | TSS200 | chr18:32957463-32957579 | 116 | | | | | | | | | | | | | | yes | | | | | |
| MIR187 | NA | Body;TSS200 | chr18:33484837-33485300 | 463 | | | | | | | | | | | | | | | | | | | |
| KIAA1328;C18orf10 | S_Shore | Body;TSS1500 | chr18:34409391-34409524 | 133 | | | | | | | 1 | | | 1 | | | | | | | | | yes |
| BRUNOL4 | Island | Body | chr18:34854300-34854936 | 636 | | | | | | | | | | | | | | yes | | | | | yes |
| BRUNOL4 | Island | 1stExon;5'UTR | chr18:35145650-35146048 | 398 | | | | | | | | | 1 | | | | | | | | | yes | yes |
| LOC647946 | NA | TSS1500 | chr18:37332566-37333028 | 462 | | | 6 | | | | | | | | | | | | yes | | | | |
| RIT2 | NA | 5'UTR;1stExon | chr18:40695592-40696075 | 483 | | | 4 | | | | | | 2 | | | | | | | | | yes | |
| SLC14A1 | NA | TSS1500;TSS200 | chr18:43303777-43304452 | 675 | | | 2 | | | | | 0 | | | | | | | | | | yes | |
| SIGLEC15 | Island | Body | chr18:43418148-43419102 | 954 | | | | | | | | | 1 | | | | | yes | yes | | yes | | |
| PSTPIP2 | N_Shore | Body | chr18:43608105-43608509 | 404 | | | | | | | | 0 | | | | 1 | | yes | yes | | | | |
| PSTPIP2 | Island | TSS200 | chr18:43652366-43652454 | 88 | | | | | | | | | 2 | | | | | | | | | | |
| RNF165 | N_Shore | TSS1500 | chr18:43912991-43913346 | 355 | | | | | | | | | | | | | | | | | | | |
| LOXHD1 | NA | Body;5'UTR;TSS200 | chr18:44098826-44099221 | 395 | | | | | | | | | 1 | | | | | yes | yes | | | | |
| LOXHD1 | Island;S_Shore | TSS200;TSS1500 | chr18:44237019-44237801 | 782 | | 1 | | | | | | | 1 | | | | | | | | yes | | |
| ST8SIA5 | N_Shore | 3'UTR | chr18:44259787-44260048 | 261 | | | | | | | | | | | | | | | | | | | |
| ST8SIA5 | Island | TSS1500 | chr18:44337853-44338147 | 294 | | 1 | 10 | | | | | | 2 | | | | | yes | | | yes | | |
| KATNAL2 | N_Shore | TSS1500 | chr18:44526430-44527026 | 596 | | | | 1 | | | | | | | | | | yes | yes | | | | |
| TCEB3CL;TCEB3C;KATNAL2 | S_Shore | TSS1500;5'UTR | chr18:44557635-44557673 | 38 | | | | | | | | | | | | | | yes | | | yes | | |
| TCEB3B;KATNAL2 | N_Shore;Island;S_Shore | 1stExon;5'UTR;TSS200;TSS1500 | chr18:44560277-44562918 | 2641 | | | | | | | | | | | | | | | | | | | |
| SMAD2 | Island;S_Shore | TSS1500 | chr18:45458021-45458698 | 677 | | 1 | | | | | | | | | | | | | | | | yes | yes |
| ZBTB7C | NA | TSS1500 | chr18:45568004-45568503 | 499 | | | | | | | | | | | | | | | yes | | yes | | yes |
| KIAA0427 | Island | 5'UTR | chr18:46067528-46067903 | 375 | | | | | | | | | | | | | | yes | | | | | yes |
| KIAA0427 | Island | 3'UTR | chr18:46386267-46386453 | 186 | | | 14 | | | | | | | | | | | | | | | | |
| SMAD7 | N_Shore;Island | 3'UTR | chr18:46447634-46447961 | 327 | | | 9 | | | | | | | | | | | | | | | yes | |
| CXXC1 | S_Shore | TSS1500 | chr18:47815407-47815430 | 23 | | | | | | | | | 1 | | | | | yes | yes | yes | | yes | |
| SKA1 | N_Shore | TSS200 | chr18:47901210-47901635 | 425 | | | | | | | | 0 | | 1 | | | | | | yes | yes | | |
| MAPK4 | Island | Body | chr18:48190466-48190637 | 171 | | | | | | | | | | | | | | yes | | | | | |
| MAPK4 | N_Shore | Body | chr18:48255506-48256237 | 731 | | | | | | | | | 2 | | | | | yes | yes | | yes | | yes |
| MRO | NA | TSS1500;5'UTR | chr18:48346660-48346843 | 183 | | | 1 | | | | | | | | | | | yes | | | | | |
| DCC | N_Shelf;N_Shore;Island | TSS1500;5'UTR;1stExon;Body | chr18:49865270-49868788 | 3518 | | | | | | | | | | | | | | | | | | | |
| MBD2;SNORA37 | N_Shore | Body;TSS1500 | chr18:51750159-51750188 | 29 | | | | | | | | | 1 | | | | | | | | yes | yes | yes |
| CCDC68 | N_Shore | TSS200;5'UTR | chr18:52625365-52625444 | 79 | | | 7 | | 1 | | | | | | | | | | | | yes | yes | |
| CCDC68 | N_Shore;Island | TSS1500;5'UTR;1stExon | chr18:52626476-52626848 | 372 | | | | | | | | 0 | | 1 | | | | yes | yes | | | | yes |
| ST8SIA3 | N_Shore | TSS1500 | chr18:55019307-55019708 | 401 | | | | | | | | | | | | 1 | | | | | | yes | yes |
| ST8SIA3 | Island | Body | chr18:55021277-55021669 | 392 | | | | | | | | | 1 | | | | | | | | | yes | |
| ONECUT2 | Island | 1stExon | chr18:55103734-55104078 | 344 | | 1 | | | | | | | | | | | | | | | | yes | yes |
| ONECUT2 | Island | Body | chr18:55108537-55109119 | 582 | 1 | | | 1 | | | | | | | | | | | | | | | |
| NEDD4L | N_Shore;Island | 5'UTR;Body;TSS1500;TSS200;1stE | chr18:55862111-55863675 | 1564 | | | 16 | 1 | | | | | | | | | | | | | | yes | |
| MIR122 | NA | TSS1500 | chr18:56117016-56117327 | 311 | | | | | | | | | | | | | | | | | | | yes |
| ALPK2 | NA | TSS200;TSS1500 | chr18:56296243-56296544 | 301 | | | 9 | | | | | | | | | | | | | | | yes | |
| ZNF532 | N_Shore | TSS1500;1stExon;5'UTR | chr18:56529784-56530420 | 636 | | | 6 | | 1 | | | | | | | | | | | | | yes | yes |
| GRP | N_Shore;Island | TSS1500;TSS200 | chr18:56886915-56887310 | 395 | | | 1 | | | | | | 1 | | | | | | | | | yes | yes |
| RAX | Island | 3'UTR;Body | chr18:56934834-56936882 | 2048 | | | 29 | | | | | | | | | | | | yes | | | yes | yes |
| CCBE1 | N_Shore | Body | chr18:57363691-57364136 | 445 | | | | | | | | | 1 | | | | | | | | yes | | |
| PMAIP1 | N_Shore | TSS1500 | chr18:57566144-57566460 | 316 | | | | | | | | | | | | | | yes | | | yes | | |
| MC4R | NA | 1stExon;5'UTR | chr18:58039932-58040213 | 281 | | | | | | | | 0 | 3 | | | 5 | | | | | | | |
| CDH20 | N_Shore;Island | Body | chr18:59221320-59222050 | 730 | | | | | | | | 0 | 6 | 2 | | 6 | | | | | | | |
| TNFRSF11A | Island | TSS200 | chr18:59992350-59992387 | 37 | | | | | | | | | 1 | | | | | | | | | yes | yes |
| TNFRSF11A | Island | Body;3'UTR | chr18:60051870-60052464 | 594 | | | | | | | | | 4 | | | 3 | | yes | | | | | yes |
| BCL2 | NA | Body | chr18:60903834-60904418 | 584 | | | 12 | | | | | | 1 | | | | | yes | yes | yes | | | |
| BCL2 | Island | TSS1500 | chr18:60988017-60988465 | 448 | | | 4 | | | | | | | | | | | yes | | | | yes | yes |
| SERPINB5 | NA | TSS200;TSS1500 | chr18:61143926-61144339 | 413 | | 1 | | | 4 | | | | 2 | 1 | | | | yes | | | | yes | |
| SERPINB5 | NA | Body | chr18:61157478-61157738 | 260 | | | | | | | | | 1 | | | | | yes | yes | | | | |
| SERPINB13 | NA | TSS1500 | chr18:61253847-61254561 | 714 | | | | | | | | | 1 | | | | | | yes | | yes | yes | |
| HMSD | NA | TSS1500;TSS200 | chr18:61616369-61616862 | 493 | | | 15 | | | | | 0 | | | | | | | yes | | | | |
| C18orf20 | NA | TSS1500 | chr18:61816543-61817320 | 777 | | | 7 | | | | | | | | | | | | yes | | yes | | |
| CDH7 | N_Shore | TSS1500;TSS200 | chr18:63417385-63417605 | 220 | | | | | | | | | 1 | | | | | | | | | yes | yes |
| DOK6 | Island | TSS200 | chr18:67068268-67068730 | 462 | | | | | | | | | 1 | | | | | | | | | | yes |
| DOK6 | Island | Body | chr18:67069959-67070324 | 365 | | | 25 | | | | | | 1 | | | 1 | | | | | | | |
| CD226 | NA | 5'UTR;1stExon | chr18:67624071-67624393 | 322 | | | | | | | | 0 | 1 | | | | | | | | yes | | |
| SOCS6 | Island | TSS1500 | chr18:67955861-67955882 | 21 | | | | | | | | | | 1 | | | | | | | | yes | yes |
| CBLN2 | S_Shore | TSS200;TSS1500 | chr18:70211916-70212474 | 558 | | | 4 | | | | | | 2 | | | 1 | | | | | | yes | |
| NETO1 | Island | TSS1500;Body | chr18:70534110-70535005 | 895 | | | | | 3 | | | 0 | | | | | | | | | | yes | yes |
| CNDP2 | S_Shore | TSS1500;5'UTR | chr18:72166016-72166324 | 308 | | | | 1 | | | | | | | | | | | | | yes | | |
| CNDP1 | NA | TSS200;1stExon;5'UTR | chr18:72201590-72201785 | 195 | | 1 | 18 | | | | | | | | | | | | | yes | | | |
| ZADH2 | N_Shore;Island | Body | chr18:72916012-72917387 | 1375 | | | | | | | | 0 | 5 | 1 | | 10 | 2 | | | | | yes | yes |
| ZNF516 | Island | Body | chr18:74114570-74114728 | 158 | | | | | | | | 0 | 1 | 1 | 1 | 1 | | | | | | | |
| ZNF516 | NA | TSS200 | chr18:74175082-74175208 | 126 | | | | | | | | | 1 | | | | | | | | | | |
| LOC284276 | NA | TSS1500 | chr18:74240034-74240476 | 442 | | | | | | | | | 1 | | | | | yes | | | | yes | |
| ZNF236 | Island;S_Shore | TSS1500;TSS200 | chr18:74535233-74536226 | 993 | | | 9 | | | | | 0 | 2 | | | | | | yes | | | | yes |
| MBP | S_Shore | 3'UTR | chr18:74692116-74692235 | 119 | | | | | | | | | | | | | | | | | | | |
| MBP | NA | 1stExon;Body;5'UTR;TSS200;TSS1 | chr18:74728931-74729551 | 620 | | | 8 | | | | | | | | | | | | | | | yes | |
| MBP | Island | Body | chr18:74770402-74770815 | 413 | | | 12 | | | | | | | | | | | yes | yes | | | yes | |
| MBP | Island | Body | chr18:74799250-74799909 | 659 | | | | | | | 1 | 0 | 2 | 2 | 1 | 5 | 1 | | | yes | | yes | |
| MBP | Island | 5'UTR | chr18:74824154-74824413 | 259 | | | | | | | | 0 | 1 | | 1 | | | yes | | yes | | yes | |
| MBP | Island | 5'UTR | chr18:74832171-74832372 | 201 | | | | | | | | | 1 | | | | | | | | | yes | |
| MBP | Island | TSS1500 | chr18:74845422-74845829 | 407 | | | | | | | | | | | | | | | | | | | |
| GALR1 | N_Shore;Island | TSS1500;TSS200 | chr18:74960652-74962672 | 2020 | | | 22 | | 3 | | | 0 | 10 | | 1 | 6 | | | yes | | | yes | |
| SALL3 | Island | TSS200;TSS1500 | chr18:76738999-76740284 | 1285 | | | | | 1 | | | | 3 | | | | | | | | | yes | yes |
| NFATC1 | Island | Body;5'UTR | chr18:77157760-77158180 | 420 | | | | 1 | | | | | 1 | | | | | | | | | | yes |
| NFATC1 | Island | Body;5'UTR | chr18:77186389-77186511 | 122 | | | | | | | | | 1 | | | | | yes | | | | | |
| NFATC1 | Island | Body;5'UTR | chr18:77193464-77193581 | 117 | | | | | | | | | 1 | | | | | | | | yes | | |
| NFATC1 | Island | Body;5'UTR | chr18:77203371-77203542 | 171 | | | | | | | | | 1 | | | | | | | | | | |
| NFATC1 | chr18:77218227-77218866 | Island | chr18:77218378-77218862 | 484 | | | | | | | | | | | 1 | | | | | | | | |
| NFATC1 | Island;N_Shore | Body | chr18:77230484-77230795 | 311 | | 1 | | | | | | | 1 | 1 | | | | | | | | | |
| NFATC1 | Island | Body | chr18:77240530-77240640 | 110 | | 1 | | | | | | | | | | | | yes | | | | | |

| | | | | | | | | | | | | | | | | | | | | | | | |
|---|---|---|---|---|---|---|---|---|---|---|---|---|---|---|---|---|---|---|---|---|---|---|---|
| NFATC1 | Island | Body | chr18:77280171-77280586 | 415 | | | 7 | | | | | 0 | 2 | | | 6 | | | | | | yes | |
| NFATC1 | N_Shore | Body | chr18:77283493-77284742 | 1249 | | | | | | | | | | | | | | yes | | | | | |
| NFATC1 | Island | Body | chr18:77286052-77286260 | 208 | | | | | | | | | 1 | | | | | | | | | | |
| KCNG2 | Island | TSS200;TSS1500 | chr18:77623409-77623598 | 189 | | | 10 | | | | 1 | | 3 | | | | | | yes | | yes | | yes |
| KCNG2 | Island | Body | chr18:77631625-77631756 | 131 | | | | | | | | | | | | | | | | yes | | | |
| KCNG2 | Island | Body | chr18:77638264-77638890 | 626 | | | | | | | | | | | | | | | yes | | | | |
| KCNG2 | Island | Body | chr18:77659572-77659949 | 377 | | | 7 | | | | | | | | | | | | | | | | |
| HSBP1L1 | N_Shore;Island | TSS1500;TSS200 | chr18:77723226-77724447 | 1221 | | 1 | | | | | | | 1 | | | | | yes | yes | yes | | yes | |
| LOC100130522 | N_Shore;Island;S_Shore | TSS1500;TSS200 | chr18:77904435-77905947 | 1512 | | | 36 | 1 | | 1 | 2 | 0 | 8 | 3 | | 24 | | yes | yes | yes | | | |
| LOC100130522;PARD6G | N_Shore | Body;3'UTR | chr18:77917615-77918142 | 527 | | | | 1 | | | | 0 | 1 | 1 | 1 | 6 | | yes | | | | | |
| PARD6G | Island;S_Shore | Body;1stExon;TSS200 | chr18:78004045-78005665 | 1620 | | | | 1 | | | | | 2 | | | | | yes | yes | | | yes | yes |
| PPAP2C | NA | 3'UTR | chr19:281291-281467 | 176 | | | 5 | | | | | | 1 | | | | | | | yes | | | |
| PPAP2C | Island;S_Shore | Body;5'UTR;TSS200;TSS1500 | chr19:291265-292887 | 1622 | | | 21 | | | | | | 2 | | | 1 | | yes | | | | | yes |
| MIER2 | Island | Body | chr19:310296-310512 | 216 | 3 | | 1 | | | | | 0 | 1 | | 1 | 1 | | | | | | | |
| MIER2 | N_Shore;Island | TSS200;TSS1500 | chr19:344981-345778 | 797 | | | 2 | | | | | | 2 | | | | | yes | yes | | | | yes |
| THEG | N_Shore;S_Shore | 1stExon;TSS200 | chr19:375746-376649 | 903 | | | 3 | | | | | | | | | | | | | yes | | | yes |
| C2CD4C | Island | 3'UTR | chr19:406256-406560 | 304 | | | | | | | | | | | | | | | | | | yes | |
| SHC2 | N_Shore;Island | 3'UTR | chr19:418205-419190 | 985 | | | 5 | | | | | | | | | | | | yes | yes | | | |
| SHC2 | Island | Body | chr19:427022-427263 | 241 | | | 14 | | | | | | | | | | | | | | | | yes |
| ODF3L2 | NA | TSS200;1stExon;5'UTR | chr19:474973-475587 | 614 | | 1 | | | | | | | 2 | | | | | yes | yes | | | | |
| MADCAM1 | Island | TSS200 | chr19:496419-496428 | 9 | | 1 | | | | | | | | | | 2 | | yes | | | | | yes |
| GZMM | S_Shore | TSS200 | chr19:543909-544349 | 440 | | | 3 | | | | | | | | | | | yes | yes | | | yes | |
| HCN2 | N_Shore;Island | Body | chr19:610955-611506 | 551 | | | | 1 | | | | 0 | 4 | | | 1 | | yes | yes | yes | | | |
| HCN2 | Island | Body | chr19:613111-613818 | 707 | | | | | | | | 0 | 3 | 1 | | 3 | | yes | | | | | yes |
| FSTL3 | N_Shore | TSS1500 | chr19:675390-675678 | 288 | | | 3 | | | | | | | | | | | | yes | | | yes | |
| FSTL3 | Island | Body | chr19:681420-681815 | 395 | | | | | | | | | 1 | | | | | | | | | | |
| PRSSL1 | N_Shore | Body | chr19:690417-690763 | 346 | | 1 | | | | | | | | | | | | yes | | | | | |
| PRSSL1 | NA | 5'UTR;1stExon | chr19:695447-695642 | 195 | | | 5 | | | | | | 1 | | 1 | | | yes | yes | | | | |
| PRSSL1 | NA | TSS1500 | chr19:696832-696946 | 114 | | | | | | | | | | | | | | yes | | yes | | | |
| PALM | N_Shore | TSS1500 | chr19:707791-708670 | 879 | | | | | | | | | 1 | | | | | yes | yes | | yes | yes | yes |
| PALM | Island | Body | chr19:710823-711001 | 178 | | | 2 | | | | | | 1 | | | | | yes | | | yes | | |
| PALM | Island | Body | chr19:719323-719632 | 309 | | | | | | | | 0 | | | | | | yes | | | | | |
| PALM | Island | Body | chr19:720740-720993 | 253 | | | 2 | | | | | | | | | | | yes | | | | yes | yes |
| PALM | Island | Body | chr19:728040-728385 | 345 | | 1 | | | | | 1 | 0 | 1 | 1 | | 4 | | yes | | | | | |
| PALM | Island | Body | chr19:736164-736425 | 261 | | | | | | | | | 1 | | | | | yes | yes | | | | |
| PALM | Island | Body | chr19:740313-740608 | 295 | | | | 1 | | | | | | | | | | yes | | | | | |
| C19orf21 | S_Shelf | TSS1500;TSS200 | chr19:750790-751241 | 451 | | | 7 | | | | | | | 1 | | | | yes | | | | | yes |
| PTBP1 | Island | Body | chr19:807322-807355 | 33 | | | | | | | | | | | | | | | | | | yes | |
| AZU1 | NA | TSS200;TSS1500 | chr19:827429-828170 | 741 | | | 22 | | | | | | 1 | 2 | | | | yes | | | | yes | yes |
| PRTN3 | N_Shelf | TSS1500;TSS200 | chr19:840447-841082 | 635 | | | 8 | | | | | | | | | | | yes | | | yes | yes | |
| PRTN3 | Island | Body | chr19:846117-846354 | 237 | | | | | | | | | | | | 1 | | yes | | | | | |
| PRTN3 | Island | Body | chr19:847943-848071 | 128 | | | | | | | | | 1 | 1 | | | | yes | | | | yes | |
| ELANE | N_Shore;Island | TSS200;Body | chr19:852114-853540 | 1426 | | | 6 | | | | | 0 | 3 | 1 | 1 | 2 | | yes | | | yes | | yes |
| ELANE | Island | Body | chr19:855899-856145 | 246 | | | | | | | | | | | | | | yes | | | | yes | |
| CFD | N_Shore | TSS200;TSS1500 | chr19:858846-859983 | 1137 | | | 14 | | | | | | | | | | | yes | | yes | | | |
| CFD | Island | Body | chr19:863054-863244 | 190 | | | 6 | | | | | 0 | 2 | 1 | | 1 | 1 | yes | | | | | |
| ARID3A | Island | Body | chr19:940724-941571 | 847 | | | | | | | | 0 | 3 | 1 | 1 | 3 | | yes | | | yes | yes | |
| ARID3A | chr19:947547-947981;Island | Island;Body | chr19:947612-947936 | 324 | | | | | | | | | 1 | | 1 | | | yes | yes | | | | |
| GRIN3B | Island | Body | chr19:1008643-1009949 | 1306 | | | 4 | | | | | | 1 | | | | | | | | | yes | yes |
| HMHA1 | N_Shore | TSS1500 | chr19:1065688-1065728 | 40 | | | 2 | | | | | | | | | | | | | | | yes | |
| HMHA1 | Island | Body | chr19:1070986-1071622 | 636 | | | | | | | | | 1 | | | | | yes | | | | | yes |
| HMHA1 | Island;N_Shore | Body | chr19:1074425-1075104 | 679 | | | | | | | | | 2 | 2 | 1 | | | yes | | | | | yes |
| SBNO2 | Island | Body | chr19:1117289-1117571 | 282 | | | | | | | | | 1 | | | | | yes | | | | | |
| SBNO2 | NA | Body;1stExon;5'UTR;TSS200 | chr19:1130866-1132966 | 2100 | | | | 1 | 6 | | | | | | | | | yes | | | | | |
| SBNO2 | Island | Body | chr19:1140909-1141150 | 241 | | | | | | | | | 3 | | | | | yes | | | | | yes |
| SBNO2 | N_Shore;Island | Body | chr19:1151960-1152189 | 229 | | | 9 | | | | | | | | | | | yes | yes | | | | |
| SBNO2 | Island | 5'UTR | chr19:1155030-1155738 | 708 | | | | | | 1 | | | 3 | 2 | | | | yes | | | | | |
| SBNO2 | Island | 5'UTR | chr19:1168733-1169138 | 405 | | | 1 | 1 | | | | | | | | | | | | | | | |
| C19orf26 | Island | Body | chr19:1230988-1231292 | 304 | | | | | 1 | | | | | | | | | | | | | | |
| C19orf23;CIRBP | S_Shore | TSS1500;Body | chr19:1271019-1271371 | 352 | | | 9 | | | | | | 2 | | | | | yes | yes | | | | |
| C19orf24 | Island | TSS1500 | chr19:1275121-1275266 | 145 | | | | | | | | | 1 | | | | | | | | | yes | |
| EFNA2 | Island | Body | chr19:1287428-1287884 | 456 | | | | 1 | | | | | 1 | | | | | yes | | | | | |
| MUM1 | S_Shore | 5'UTR;TSS1500;TSS200 | chr19:1355910-1356444 | 534 | | 1 | 2 | | 1 | 2 | | 0 | 4 | | | | | yes | | | | | |
| MUM1 | Island | Body | chr19:1371027-1371281 | 254 | | 1 | | | | | | | | | | | | | | | yes | | |
| MUM1 | Island | Body;3'UTR | chr19:1377966-1378265 | 299 | | | | | | | | | | | | | | yes | | | | | |
| NDUFS7 | Island;N_Shore | Body | chr19:1387394-1387894 | 500 | 3 | | | | | | | | 2 | | | 1 | | | yes | yes | | | |
| NDUFS7 | Island | Body | chr19:1395365-1395572 | 207 | | | | | | | | | | | | | | | | | | yes | |
| GAMT | Island | TSS1500 | chr19:1401767-1401781 | 14 | | | 1 | | | | | | | | | | | yes | | yes | | yes | yes |
| DAZAP1 | Island | Body | chr19:1428919-1429261 | 342 | | | | | | | | | 1 | | | | | | | | | | |
| RPS15 | N_Shore | TSS1500 | chr19:1438017-1438084 | 67 | | | | | | | | 0 | 1 | | | 2 | | yes | yes | | | | |
| RPS15 | Island | Body | chr19:1439980-1440260 | 280 | | | | | | | | | | | | | | yes | | | | | |
| APC2 | Island | Body | chr19:1466162-1468359 | 2197 | | | 13 | 1 | | | | 0 | 1 | | 1 | 2 | | | | | | | yes |
| APC2 | Island | 3'UTR | chr19:1472936-1473179 | 243 | | | | | | | | 0 | | | | | | | | | | | |
| REEP6;PCSK4 | Island | Body;TSS1500 | chr19:1491701-1492043 | 342 | | | | | | | | | | | | | | yes | | | | | |
| REEP6 | Island | Body | chr19:1496416-1496580 | 164 | | | | | | | | | 1 | | | | | | | | | yes | |
| ADAMTSL5 | Island | Body | chr19:1510494-1510692 | 198 | | | | | | | | 0 | | | | | | yes | | | | | |
| PLK5P | Island | TSS200;Body;TSS1500 | chr19:1523856-1525705 | 1849 | | 1 | | | | | | 0 | 2 | | | | | | | | | yes | yes |
| ONECUT3 | Island | Body | chr19:1775211-1775422 | 211 | | 1 | | | | | | | | | | | | | | | | yes | yes |
| ATP8B3 | Island | Body | chr19:1799935-1800150 | 215 | | 1 | | | | | | | | | | | | yes | | | | | |
| ATP8B3 | Island | TSS1500 | chr19:1812664-1812854 | 190 | | | | | | | | | 1 | | | | | | yes | | | yes | yes |
| KLF16 | Island | Body | chr19:1854549-1854819 | 270 | | | 12 | | | | | | 1 | | | | | | | | | | |
| ADAT3;SCAMP4 | N_Shore | 5'UTR | chr19:1907677-1908388 | 711 | | | | | | | | | 2 | | | | | yes | | | | | |
| BTBD2 | N_Shore | Body | chr19:1992866-1993100 | 234 | | | | | | | | | 1 | | | | | yes | | | | | yes |
| BTBD2 | Island | Body | chr19:1998801-1999322 | 521 | | | | | | | | | 2 | | | | | yes | | | | | |
| MKNK2 | Island | Body | chr19:2041905-2042592 | 687 | | | 10 | | | 1 | | 0 | 1 | | 1 | 1 | | | | | | yes | yes |
| MKNK2 | Island | Body | chr19:2046085-2046303 | 218 | | | | | | | | | 1 | | | | | | | | | | |
| MOBKL2A | NA | 5'UTR | chr19:2085286-2085483 | 197 | | | 2 | | | | | | | | | | | yes | | | | | |
| MOBKL2A | N_Shore | 5'UTR | chr19:2095364-2095560 | 196 | | | 1 | | | | | 0 | | | | | | yes | | | | yes | |
| AMH | Island | Body;3'UTR | chr19:2250561-2252432 | 1871 | 1 | | 7 | | 1 | | 1 | 0 | 3 | | 1 | 3 | | | | | | | yes |
| JSRP1 | Island | Body | chr19:2253492-2253854 | 362 | | | 7 | | | | | | | | | | | | | | | | yes |
| JSRP1 | S_Shore | TSS200;TSS1500 | chr19:2255419-2255848 | 429 | 1 | | | | | | | | 2 | | | | | yes | | | | | |
| OAZ1 | Island | Body | chr19:2272716-2273216 | 500 | | | | | | | | | 1 | | | | | yes | | | | | yes |
| C19orf35 | Island | Body | chr19:2276322-2276471 | 149 | | | 7 | | | | | | | | | | | yes | | | | | |
| C19orf35 | Island | Body | chr19:2278618-2278773 | 155 | | | | | 1 | | | | | | | | | yes | | | | | |
| C19orf35 | S_Shelf | 5'UTR;1stExon;TSS200;TSS1500 | chr19:2281919-2282729 | 810 | 1 | | 26 | 1 | 1 | | | | | | | | | yes | | | | | |
| LINGO3 | Island;chr19:2289895-2291904 | Body;Island | chr19:2291020-2291872 | 852 | | | | | | | | | 3 | | 1 | | | yes | | | | | yes |
| LINGO3 | N_Shore;Island | 5'UTR | chr19:2302401-2303032 | 631 | 2 | | 4 | | | | | 0 | | 1 | | | | yes | yes | | | | yes |
| LINGO3 | Island | 5'UTR | chr19:2307592-2308475 | 883 | | | | | | | | 0 | 1 | | | 1 | | yes | | | yes | yes | yes |
| LSM7 | Island | Body | chr19:2323913-2324338 | 425 | | | | | 1 | | | | | | | | | yes | yes | | yes | | |

| | | | | | | | | | | | | | | | | | | | | | | | |
|---|---|---|---|---|---|---|---|---|---|---|---|---|---|---|---|---|---|---|---|---|---|---|---|
| TMPRSS9 | NA | TSS200 | chr19:2389655-2389813 | 158 | | | | | | | | | | | | | | yes | | | | | |
| LMNB2;TIMM13 | Island | 3'UTR;TSS1500 | chr19:2428350-2429209 | 859 | | | 15 | | | | | 0 | 2 | | | 4 | | | | | | | |
| GNG7 | N_Shore | 3'UTR | chr19:2512999-2513436 | 437 | | | 6 | | | | | | 2 | | | | | yes | | | | | |
| GNG7 | Island | 5'UTR | chr19:2525290-2525634 | 344 | | | 7 | | | 1 | | | | | | | | yes | yes | | | yes | |
| GNG7 | Island | 5'UTR | chr19:2540907-2541104 | 197 | | | | | | 2 | | | 1 | 1 | | | | yes | | | | | |
| GNG7 | N_Shore | 5'UTR | chr19:2543602-2544100 | 498 | | | | | | | | | 1 | | | | | yes | | | | | |
| GNG7 | N_Shore;Island | 5'UTR | chr19:2546598-2546938 | 340 | | | 6 | | | | 1 | 0 | 2 | | | | | | yes | yes | | | |
| GNG7 | N_Shore;Island | 5'UTR | chr19:2588241-2588629 | 388 | | 1 | 9 | 1 | 2 | | | | | | | | | yes | yes | | | | yes |
| GNG7 | Island | 5'UTR | chr19:2607726-2608495 | 769 | | 1 | 4 | | | | | | 1 | | | | | yes | yes | | | yes | |
| GNG7 | Island | 5'UTR | chr19:2650755-2650863 | 108 | | | 1 | | | | | | | | | | | yes | | | yes | yes | |
| GNG7 | Island | TSS1500 | chr19:2702986-2703086 | 100 | | | | | | | | | | | | | | yes | | | yes | yes | yes |
| DIRAS1 | Island | Body | chr19:2717293-2717932 | 639 | | | | | | | | | 4 | | | | | yes | | | | | yes |
| SGTA | Island | 5'UTR | chr19:2774197-2774549 | 352 | | | | | | | | | 1 | | | | | yes | | yes | yes | | |
| THOP1 | Island | Body | chr19:2790475-2790716 | 241 | | | | 1 | | | | | | | | | | yes | | | yes | | |
| THOP1 | Island | Body | chr19:2811654-2811736 | 82 | 2 | | | 1 | 1 | | | | | | | | | | | | | | |
| ZNF555 | Island | TSS1500 | chr19:2840865-2841094 | 229 | | | | | | | | 0 | | | | | | yes | | | | | |
| ZNF556 | NA | 1stExon;5'UTR;TSS1500;TSS200 | chr19:2866899-2867753 | 854 | | | | | | | | | 1 | | | | | yes | | yes | | | |
| ZNF57 | Island | TSS1500;5'UTR;1stExon | chr19:2900365-2901147 | 782 | | | | | | | 1 | 0 | 3 | | | 1 | | yes | | yes | | | |
| ZNF77 | Island | 1stExon;5'UTR;TSS200 | chr19:2944943-2945492 | 549 | | | | | | | | 0 | | | | | | yes | | yes | | | |
| TLE6 | S_Shore | 1stExon;5'UTR | chr19:2977603-2977843 | 240 | | | 1 | | | | | | | | | | | | yes | | | yes | |
| TLE2 | S_Shore | TSS200;Body;TSS1500 | chr19:3029785-3030324 | 539 | | | 1 | | | | | | | | | | | yes | yes | | | | yes |
| GNA11 | Island | Body | chr19:3097565-3097728 | 163 | | | 7 | | 1 | | | | 2 | | | | | | | | yes | | |
| GNA15 | NA | nan;TSS200 | chr19:3136019-3136097 | 78 | | | | | | | | | | | 1 | | | yes | | | | yes | |
| S1PR4 | Island;N_Shore | 1stExon;TSS1500;TSS200 | chr19:3178413-3180815 | 2402 | | | | | | | | | | | | | | | | | | | |
| NCLN | Island | Body | chr19:3196206-3196387 | 181 | | | | | | | | | | | | | | yes | | yes | | | |
| BRUNOL5 | Island | Body | chr19:3285973-3286244 | 271 | | | | | | | | | | | | | | yes | | | | | |
| NFIC | Island | Body | chr19:3369714-3370244 | 530 | | | | | | | | | 2 | 1 | | | | yes | | | | yes | yes |
| NFIC | Island | Body | chr19:3398706-3398810 | 104 | | | | | | 1 | | | 1 | 1 | | | | yes | yes | | | | |
| NFIC | Island | Body | chr19:3434930-3435350 | 420 | | | | 1 | 1 | | | | | | | | | yes | | | | | |
| C19orf77 | N_Shore | Body;5'UTR;1stExon;TSS200 | chr19:3480363-3480672 | 309 | | | 4 | | | | | 0 | 5 | | | 5 | | yes | | | | | |
| C19orf77 | Island | TSS1500 | chr19:3481788-3481985 | 197 | | | | | | | | | | | | | | yes | | | | | |
| C19orf71;C19orf28 | NA | TSS1500;3'UTR | chr19:3538705-3539158 | 453 | | | | | | | | | | | | | | | yes | yes | | yes | |
| GIPC3 | Island | Body | chr19:3586844-3587038 | 194 | | | 1 | | | | | | 1 | | | | | yes | | | | | |
| TBXA2R | Island | Body | chr19:3595702-3596441 | 739 | | | | | | | | | | | | | | yes | | | | | yes |
| TBXA2R | Island | TSS1500 | chr19:3607139-3607608 | 469 | | | | | | | | 0 | | | | | | | | | | | |
| PIP5K1C | Island | Body | chr19:3671903-3672411 | 508 | | | 2 | | | | | | | | | | | yes | | | | | |
| PIP5K1C | Island | Body | chr19:3688036-3688477 | 441 | | | | | | | | 0 | 1 | 1 | 1 | | | | | | | yes | yes |
| PIP5K1C | Island | Body | chr19:3697922-3698223 | 301 | | | 4 | | | | | | | | | | | | yes | | yes | | yes |
| RAX2 | S_Shore | TSS1500 | chr19:3772751-3773230 | 479 | | | | | | | | | | | | | | yes | yes | | | | |
| MATK | Island | Body;TSS200;TSS1500 | chr19:3786583-3786957 | 374 | | | 3 | | | | | | | | | | | yes | yes | | | yes | yes |
| ZFR2 | Island | Body | chr19:3810782-3811194 | 412 | | 1 | | | | | | | | | | | | yes | | | | | |
| ZFR2 | Island | Body | chr19:3821095-3822189 | 1094 | | | | | | | | 0 | | | | | | yes | | | | | |
| ZFR2 | Island | Body | chr19:3825330-3825630 | 300 | | 1 | | | | | | | | | | | | yes | | | | | |
| ZFR2 | N_Shore;Island | Body | chr19:3831293-3831859 | 566 | | | 2 | | | | | | | | | | | yes | | | | yes | |
| ZFR2 | N_Shore | Body | chr19:3834199-3834519 | 320 | | | | | | | | | | | | | | | | | | | |
| ZFR2 | Island | Body | chr19:3838098-3838222 | 124 | | | | | | | | | 1 | | | | | | | | | | |
| ZFR2 | Island | Body;TSS200 | chr19:3868624-3869525 | 901 | | 1 | | | | | | | 1 | | | 1 | | yes | | | | | yes |
| ATCAY | Island | Body | chr19:3907913-3908397 | 484 | | | 4 | | | | | | | | | | | | yes | | | | yes |
| MIR637;DAPK3 | S_Shore | TSS200;Body | chr19:3961645-3961702 | 57 | | | 4 | | | | | | | | | | | yes | yes | | | | |
| DAPK3 | chr19:3970536-3970746;N_Sh | N_Shore;1stExon;5'UTR | chr19:3969522-3969950 | 428 | | | | | | | | | | | 1 | | | yes | | | | | |
| CREB3L3 | NA | TSS1500;TSS200;1stExon;5'UTR | chr19:4153364-4153713 | 349 | | | 9 | | 1 | | | 0 | | | 1 | | 1 | yes | | | | | |
| FSD1;TMIGD2 | N_Shore;Island | TSS1500;Body | chr19:4303817-4305239 | 1422 | | | | | 1 | | | | 1 | | | | | yes | | | | | yes |
| FSD1 | Island | Body | chr19:4323034-4323200 | 166 | | 1 | | 1 | | | | | | | | | | | | | | | |
| STAP2 | Island | Body | chr19:4326975-4327350 | 375 | | | | | 1 | | | | | 1 | | | | | | | | | |
| STAP2 | NA | 1stExon;5'UTR | chr19:4338769-4338932 | 163 | | | 3 | | | | | | | | | 1 | | yes | | yes | | yes | |
| UBXN6 | N_Shelf;N_Shore | 5'UTR;1stExon;Body;TSS1500 | chr19:4455239-4455957 | 718 | | | 2 | | | | | | 2 | | | | | yes | yes | | | | |
| PLIN5 | Island | Body | chr19:4531638-4531790 | 152 | | | | | | | | | | | | | | yes | | | | | |
| PLIN5 | N_Shore;Island | 5'UTR | chr19:4534986-4535211 | 225 | | | | | | | | | 1 | | | 1 | | yes | | | | | yes |
| LRG1 | N_Shelf | Body | chr19:4539943-4540065 | 122 | | | | | | | | | 1 | 1 | | | | yes | yes | | | | |
| SEMA6B | Island | 3'UTR | chr19:4543487-4544095 | 608 | | | | | | | | | 1 | | | | | | | | | yes | yes |
| TNFAIP8L1 | S_Shore | TSS200;5'UTR | chr19:4639844-4640159 | 315 | | | 3 | | | | | | | | | | | yes | | | | | yes |
| C19orf30;MIR7-3 | NA | TSS200;TSS1500;Body | chr19:4769001-4770775 | 1774 | 1 | | 5 | | | | | 0 | 2 | | | | | yes | | | | | yes |
| UHRF1 | N_Shore | TSS1500 | chr19:4909175-4909263 | 88 | | | | | | | | | | | | | | | yes | | | | |
| UHRF1 | Island | Body | chr19:4916593-4916983 | 390 | | | | | 1 | | | | 1 | 1 | 1 | | | yes | | | yes | | |
| KDM4B | N_Shore;Island | Body | chr19:5036609-5037219 | 610 | | | 8 | | | | | | 1 | 1 | | | | | | | yes | | |
| KDM4B | Island | Body | chr19:5048645-5049135 | 490 | | | | | | | | | 1 | | | | | | | | | yes | |
| KDM4B | Island | Body | chr19:5060985-5061205 | 220 | | | | | | | | | | | | | | yes | | | | | |
| KDM4B | Island | Body | chr19:5074616-5074758 | 142 | | | | | | | | | 1 | | | | | | | | | | |
| KDM4B | N_Shore;Island | Body | chr19:5093370-5093659 | 289 | | | 8 | | | | | | | | | | | | | yes | | | |
| KDM4B | Island | Body | chr19:5147855-5148409 | 554 | | | 24 | | | | | | 1 | | | | | yes | | | | | |
| PTPRS | N_Shore;Island | Body | chr19:5250376-5251120 | 744 | | | | | 1 | 3 | | | 1 | | | | | | yes | | | | yes |
| PTPRS | Island | 5'UTR | chr19:5294093-5294252 | 159 | | | | | | | | | 1 | | | | | yes | | | | yes | |
| PTPRS | N_Shore;Island | 5'UTR | chr19:5334979-5335260 | 281 | 4 | | 1 | | | | | | | | | | | yes | | | | | |
| PTPRS | Island | 5'UTR | chr19:5338881-5339092 | 211 | | | 1 | | | | | | | | | | | | | | | | |
| ZNRF4 | N_Shore;Island | TSS1500;TSS200;5'UTR;1stExon | chr19:5455217-5455638 | 421 | 9 | 1 | 2 | | | | | | | | | | | | yes | | | | |
| PLAC2 | Island;S_Shore | Body;TSS200 | chr19:5567758-5568218 | 460 | | | 16 | | | | | 0 | 3 | | | | | yes | yes | | | | yes |
| SAFB2 | N_Shore | 3'UTR | chr19:5587239-5587909 | 670 | | 1 | | | | | | | | | | | | yes | yes | | | | yes |
| LONP1 | Island | Body | chr19:5699060-5699271 | 211 | | | | | | | | | 1 | | | | | yes | | yes | | | |
| NRTN | Island | TSS1500 | chr19:5822318-5822630 | 312 | | | 4 | | | | | | 1 | | | | | yes | yes | | | | yes |
| NRTN | N_Shore | TSS200 | chr19:5823786-5824065 | 279 | | | | 1 | | | | | | | | | | yes | yes | | | | |
| NRTN | Island | Body | chr19:5827811-5828301 | 490 | | | | 1 | 2 | | | | | 1 | | | | | | | | | yes |
| FUT6 | NA | 1stExon;5'UTR | chr19:5838729-5838779 | 50 | | | 4 | | | | | 0 | | | | | | | yes | | | | |
| FUT3 | NA | 1stExon;5'UTR | chr19:5851465-5851762 | 297 | | 1 | | | | | | | 1 | | | | | yes | yes | yes | | | |
| CAPS | N_Shore | TSS200;TSS1500 | chr19:5913990-5914669 | 679 | | | 11 | | | 1 | | | 1 | 1 | | | | yes | | | | | |
| RFX2 | Island | Body | chr19:6008033-6008127 | 94 | | | | | | | | | | | | | | yes | | | | | |
| RFX2 | Island | Body | chr19:6039991-6040198 | 207 | | | | | | | | | 1 | | | | | yes | | | | | |
| ACSBG2 | NA | TSS1500 | chr19:6135279-6135519 | 240 | | | 3 | | | | | | | | | | | yes | yes | | | | |
| ACER1 | NA | 1stExon;5'UTR | chr19:6333547-6333761 | 214 | | 1 | | | | | | | 1 | 1 | | | | yes | yes | | | | |
| ALKBH7 | Island | Body | chr19:6373131-6373627 | 496 | | | | | | | | | 1 | | | | | | | | | | yes |
| SLC25A41 | NA | 1stExon;5'UTR | chr19:6433768-6433858 | 90 | | | | | | | | | 1 | | | | | yes | | yes | | | yes |
| CRB3 | N_Shore;Island | TSS1500 | chr19:6463949-6464080 | 131 | | | | | | | | | | | | | | yes | | | | yes | |
| DENND1C | Island | Body | chr19:6475456-6477198 | 1742 | | | | | 1 | | | | 1 | | 1 | | | yes | | | | yes | |
| DENND1C | NA | Body;1stExon;5'UTR;TSS200;TSS1 | chr19:6481540-6482169 | 629 | | | 15 | | 1 | | | | 3 | 1 | | | | yes | | | | yes | |
| TNFSF9 | Island | Body | chr19:6534760-6535078 | 318 | | | | 1 | 1 | | | | 1 | | | | | yes | | | | yes | |
| TNFSF14 | NA | Body | chr19:6669849-6670390 | 541 | | | | | 1 | | | | | | | | | yes | yes | | | yes | |
| C3 | Island | Body | chr19:6710658-6711092 | 434 | | | | | | | | 0 | | | | | | | | yes | | | yes |
| C3 | S_Shore | Body | chr19:6712321-6712479 | 158 | | | | 1 | | | | | 1 | | | | | | | | | | |
| TRIP10;GPR108 | N_Shore | TSS1500 | chr19:6738762-6739192 | 430 | | | | | | | | | | | | | | | | | | yes | |
| TRIP10 | Island | Body | chr19:6740812-6741181 | 369 | | | | 2 | | | | | | | | 1 | | yes | | | | | yes |

| | | | | | | | | | | | | | | | | | | | | | | | |
|---|---|---|---|---|---|---|---|---|---|---|---|---|---|---|---|---|---|---|---|---|---|---|---|
| SH2D3A | Island | 3'UTR | chr19:6752577-6753024 | 447 | | | | 1 | | | | | | | | | | yes | | | | | |
| VAV1 | NA | TSS1500;TSS200;5'UTR;1stExon | chr19:6772370-6773074 | 704 | | | 4 | | | | | | 1 | | 1 | | | | yes | | | | |
| PEX11G | N_Shore | 3'UTR | chr19:7541760-7542191 | 431 | | | | | | | | | | | | | | | | | | | |
| ZNF358 | N_Shore | TSS1500 | chr19:7580209-7580275 | 66 | | | 3 | 1 | | | | | 1 | | | | | yes | | yes | yes | yes | |
| KIAA1543 | N_Shore | Body | chr19:7680053-7680105 | 52 | | | 1 | | | | | | | | | | | | yes | | | | |
| STXBP2 | Island | Body | chr19:7712095-7712539 | 444 | | | | | | | | | 1 | | | | | yes | | | | | yes |
| C19orf59 | N_Shelf | TSS1500;TSS200 | chr19:7741602-7741955 | 353 | | | 4 | 1 | 1 | | | | 2 | | | | | yes | | | | yes | |
| TRAPPC5 | Island | 5'UTR | chr19:7746413-7747220 | 807 | | 1 | | | | | | | | | | | | | | | | | |
| FCER2 | NA | 1stExon;5'UTR;TSS200;nan | chr19:7766717-7767584 | 867 | | | 8 | | | | | | 2 | | 2 | | | yes | yes | | | yes | |
| CLEC4G | S_Shore | 5'UTR;1stExon | chr19:7797031-7797100 | 69 | | | | | | | | | | | | | | yes | | | | yes | |
| CLEC4M | NA | TSS1500 | chr19:7827560-7827802 | 242 | | | | | | | | | | | | | | yes | yes | | | | |
| CLEC4GP1 | N_Shore | TSS1500;TSS200 | chr19:7852051-7852739 | 688 | 1 | | 13 | | | | | 0 | 1 | | | | | yes | | | | yes | |
| EVI5L | N_Shore | 5'UTR;TSS1500 | chr19:7910696-7911206 | 510 | | | | | | | | | | | | | | yes | | | | | |
| EVI5L | Island | Body | chr19:7926970-7927374 | 404 | | | 8 | | | | | | | | | | | yes | | | | | yes |
| LRRC8E | N_Shore;Island | TSS1500;1stExon;5'UTR | chr19:7953054-7953440 | 386 | | | 7 | | | | | | 2 | | | | | yes | | | | | yes |
| CCL25 | S_Shelf | TSS1500;TSS200 | chr19:8117671-8117924 | 253 | | | | | 1 | | | | | | | | | yes | yes | | | | |
| FBN3 | N_Shore;Island | Body | chr19:8200860-8201419 | 559 | | 1 | | | | | | | | | | | | yes | | | | | |
| FBN3 | N_Shelf | 1stExon | chr19:8212234-8212385 | 151 | | | | | | | | | | | | | | | | | | yes | |
| LASS4 | Island | 5'UTR | chr19:8274853-8275083 | 230 | | | | | | | | | | | | | | | | | | | yes |
| CD320 | Island | TSS200 | chr19:8373286-8373296 | 10 | | | | | | | | | | 1 | | | | | yes | | | | yes |
| KANK3 | Island | Body | chr19:8398300-8399119 | 819 | | | | | | | | | 1 | | | 1 | | yes | | | | | |
| ANGPTL4 | N_Shore | TSS1500 | chr19:8428787-8428856 | 69 | | | | | 1 | | | | | | | | | yes | | | | | yes |
| PRAM1 | Island | Body | chr19:8555465-8555649 | 184 | | | | | | | | | | | | | | yes | | | | | |
| PRAM1 | N_Shore | Body | chr19:8563373-8563755 | 382 | | | | | | | | | 1 | | | | | yes | | | | | |
| MYO1F | Island | Body | chr19:8591364-8591776 | 412 | | | | | | | | | 2 | | 1 | | | yes | | | | | yes |
| MYO1F | N_Shore | Body | chr19:8601136-8601336 | 200 | | | | | | | | | 1 | | | | | yes | | | | | yes |
| MYO1F | NA | 5'UTR;1stExon;TSS200 | chr19:8642301-8642370 | 69 | | | 12 | | | | | | | | | | | yes | | | | | |
| ACTL9 | Island;S_Shore | 1stExon | chr19:8808663-8809175 | 512 | | | | | | | | | | | | | | yes | | yes | | | |
| OR7D4 | NA | 1stExon | chr19:9325438-9325742 | 304 | | | | | | | | | | | | | | yes | | | | | |
| ZNF559 | N_Shore | TSS1500;TSS200 | chr19:9434309-9434858 | 549 | | | | | | | | | 2 | | | | | yes | | yes | | | |
| ZNF177 | N_Shore;Island | TSS200 | chr19:9473565-9474128 | 563 | | 1 | | | 2 | | | 0 | 2 | | | | | | yes | | | | |
| ZNF266 | S_Shore | TSS1500 | chr19:9546723-9546752 | 29 | | | 4 | | | | | 0 | 1 | | 1 | 4 | | yes | | | | | |
| ZNF560 | Island;S_Shore | 5'UTR;TSS200 | chr19:9609152-9609542 | 390 | | 1 | 1 | | 1 | | | | 1 | | | | | yes | | | | yes | |
| ZNF561 | N_Shore;Island | 5'UTR;1stExon | chr19:9731681-9732278 | 597 | | 1 | | | | | | 0 | 3 | 1 | | 1 | 1 | yes | | yes | | | |
| ZNF562 | N_Shore;Island | 5'UTR;TSS200 | chr19:9785295-9786115 | 820 | | 1 | 4 | | 1 | | | 0 | 4 | 7 | 1 | 3 | | yes | | yes | | | |
| FBXL12 | Island | TSS200 | chr19:9929906-9929919 | 13 | | | | | | | | | 1 | | | | | | | | | | yes |
| C3P1 | NA | TSS200 | chr19:10151833-10152171 | 338 | | | | | | | | | 1 | | | | | yes | | | | | |
| C3P1 | Island | Body | chr19:10172712-10172899 | 187 | | | | | | | | | 1 | | | 1 | | yes | yes | | | yes | |
| ANGPTL6 | N_Shelf | 5'UTR;1stExon | chr19:10213271-10213957 | 686 | | | | | | | | | 1 | | | | | yes | yes | | | yes | yes |
| PPAN-P2RY11;P2RY11 | S_Shore | Body;TSS200 | chr19:10222051-10222409 | 358 | | | | | | | | | | | | | | yes | | | | yes | |
| EIF3G | chr19:10230162-10230682 | S_Shore | chr19:10231566-10231711 | 145 | | | | | | | | | | | 1 | | | yes | | | yes | | |
| ICAM1 | Island | TSS200;TSS1500 | chr19:10381063-10381416 | 353 | | | 14 | | 1 | | | | | | | | | yes | | | | | |
| ICAM4 | N_Shore;Island | TSS1500;TSS200 | chr19:10397429-10397780 | 351 | | | 6 | | | | | | | | | | | yes | | | | | |
| ICAM5 | Island | Body | chr19:10403403-10403936 | 533 | | 1 | | | | | | | 1 | | | | | | | | | | |
| ICAM5 | Island | Body | chr19:10406935-10407538 | 603 | | 1 | | | | | | | 1 | | | 1 | | yes | | | | yes | yes |
| ICAM3 | S_Shelf;NA | Body;nan;1stExon;5'UTR | chr19:10449942-10450472 | 530 | | | | 1 | | | | | 2 | | 1 | | | yes | yes | | | yes | |
| TYK2 | Island | Body | chr19:10463831-10464320 | 489 | | | | 1 | | | | | | | | | | | | | | | |
| PDE4A | N_Shore | Body;TSS1500 | chr19:10541267-10541704 | 437 | 1 | | | | | | | 0 | | | | | | yes | | | | | yes |
| S1PR5 | N_Shore | 5'UTR | chr19:10627706-10628014 | 308 | | | 1 | | | | | | | | | | | yes | | | | | |
| ATG4D;MIR1238 | NA | Body;TSS1500 | chr19:10662582-10662619 | 37 | | | 1 | | | | | | | | | | | yes | yes | | | | yes |
| AP1M2 | N_Shore;Island | Body;5'UTR;1stExon | chr19:10697690-10698036 | 346 | | | | | 1 | | | | 2 | 1 | | | | yes | | | | yes | |
| SLC44A2 | N_Shore;Island | TSS1500;Body;TSS200;5'UTR;1stEx | chr19:10735482-10736448 | 966 | | | 1 | | 1 | 2 | | 0 | 10 | | | 3 | 1 | yes | yes | | | | |
| ILF3 | NA | Body | chr19:10792615-10792710 | 95 | | | | | | | | | 1 | | | | | yes | | | | | |
| QTRT1 | N_Shore;Island | Body | chr19:10822935-10823913 | 978 | | | | | 1 | | | | 1 | | | | | yes | yes | | yes | yes | |
| MIR199A1;DNM2 | NA | TSS200;Body | chr19:10928327-10928639 | 312 | | 1 | | | | | | | | | | | | yes | | | | yes | |
| C19orf38 | NA | TSS200 | chr19:10958952-10959157 | 205 | | | | 1 | 1 | | | | 1 | | 1 | | | yes | | | | | |
| KANK2 | Island | Body | chr19:11280655-11280913 | 258 | | | 4 | | | | | | 1 | | | | | yes | yes | | | | |
| KANK2 | Island;S_Shore | Body | chr19:11289211-11289376 | 165 | | | 24 | | | | | | | | | | | yes | yes | | | | |
| DOCK6 | N_Shore | Body | chr19:11319376-11319550 | 174 | | | 1 | | | | | | | | | | | yes | | | | | |
| LOC55908;DOCK6 | NA | TSS1500;Body | chr19:11348439-11348738 | 299 | | | | | | | | | | | | | | | | | | | |
| DOCK6 | Island | Body | chr19:11353961-11354401 | 440 | | 1 | 1 | | | | | 0 | | | | | | yes | yes | | | yes | |
| TSPAN16 | S_Shelf;S_Shore | TSS200;TSS1500 | chr19:11406114-11406993 | 879 | | | 7 | | | | | | | | | | | yes | yes | | | | |
| RAB3D | N_Shore | 5'UTR | chr19:11449933-11450198 | 265 | | | | 1 | | | | | | | | | | yes | | | | | |
| LPPR2 | Island | Body | chr19:11471954-11472397 | 443 | | | | 1 | 1 | | | 0 | | | | | | | | | | | |
| C19orf39 | N_Shore | TSS1500 | chr19:11484448-11484842 | 394 | | | 1 | | | | | | | | | | | yes | yes | | | yes | |
| RGL3 | Island | Body | chr19:11517079-11517436 | 357 | | 1 | 7 | 1 | 1 | | | 0 | 2 | 2 | | 5 | | yes | | | | | |
| RGL3 | N_Shore | TSS200 | chr19:11530021-11530065 | 44 | | | | | | | | | 1 | | | | | yes | | | | yes | |
| RGL3;CCDC151 | Island | TSS1500;3'UTR | chr19:11531294-11531469 | 175 | | | | | 1 | | | | | | | | | yes | | | | | |
| ELAVL3 | Island | Body | chr19:11565385-11565399 | 14 | | | 1 | | | | | | | | | | | yes | | | | yes | yes |
| ELAVL3 | Island | Body | chr19:11591185-11591491 | 306 | | | | | | | | 0 | | | | | | | | | | | yes |
| ACP5 | S_Shore;Island | TSS1500;TSS200;5'UTR;1stExon | chr19:11689614-11689997 | 383 | | | 17 | | 1 | | | | | | | | | yes | | | | | yes |
| ZNF833 | N_Shore;Island | TSS1500;TSS200;Body | chr19:11784246-11785337 | 1091 | | 1 | 16 | 1 | 6 | | | 0 | 6 | 1 | | 6 | | yes | | | | | |
| ZNF878 | S_Shelf | 1stExon;TSS200 | chr19:12167119-12167313 | 194 | | | 3 | | | | | | | | | | | yes | | yes | | | |
| ZNF844 | N_Shore | TSS1500 | chr19:12175107-12175268 | 161 | | | 2 | | | | | | | | | | | yes | | yes | | | |
| ZNF788 | Island | TSS200 | chr19:12203029-12203644 | 615 | | 1 | | | | | | | | | | | | yes | | yes | | yes | |
| ZNF20 | Island | Body | chr19:12250778-12251216 | 438 | | | | | | | | 0 | | | | | | yes | | yes | | | |
| ZNF44 | S_Shore | TSS1500 | chr19:12406154-12406247 | 93 | | | | | | | | | | | | | | yes | | yes | | | |
| ZNF563 | Island;S_Shore | Body;TSS200 | chr19:12444273-12444741 | 468 | | | | | | | | 0 | 3 | | | 1 | | | | | | | |
| ZNF442 | N_Shore | 5'UTR | chr19:12476027-12476246 | 219 | | | | | | | | | 1 | | | | | | | yes | | | |
| ZNF709 | Island;S_Shore | 5'UTR;1stExon;TSS200 | chr19:12624466-12624786 | 320 | | 1 | 3 | | | | | 0 | 1 | | | 1 | | yes | | | | | |
| ZNF490 | Island | Body | chr19:12707809-12708229 | 420 | | | 3 | 1 | | | | | | | | | | yes | yes | | | | |
| MAN2B1 | Island | Body | chr19:12758416-12759156 | 740 | | | 3 | | | | | | 1 | | | 3 | | yes | | | | | |
| MAN2B1;C19orf56;MORG1 | N_Shore | TSS1500;3'UTR;5'UTR;Body | chr19:12779057-12779373 | 316 | | | 3 | | | | | | | | | | | yes | | | | | |
| TNPO2 | N_Shore | Body | chr19:12831659-12832212 | 553 | | | 1 | | | | | 0 | 1 | | | | | yes | | | | | yes |
| TNPO2 | S_Shore | TSS200 | chr19:12835001-12835134 | 133 | | | 2 | | | | | | | | | | | yes | | | | | |
| HOOK2 | N_Shore;Island | Body | chr19:12876846-12877188 | 342 | 4 | 16 | 45 | 2 | 2 | 6 | 1 | 0 | 2 | 2 | 1 | 109 | 1 | yes | | | | | |
| HOOK2 | S_Shore | TSS1500 | chr19:12886980-12886986 | 6 | | 1 | | | | | | | 1 | | | 1 | | yes | | | | | |
| PRDX2 | N_Shore | Body | chr19:12911875-12912252 | 377 | | | | | | | | | | | | 1 | | yes | | | | | |
| RTBDN | S_Shore;N_Shore | Body;5'UTR;TSS200 | chr19:12940950-12941409 | 459 | | | | | | | | 0 | 1 | | | 1 | | yes | | | | yes | |
| MAST1 | N_Shore | Body | chr19:12951996-12952280 | 284 | | | | | | | | 0 | 1 | | | 1 | | | yes | | | | |
| MAST1 | N_Shore;Island | Body | chr19:12957979-12958803 | 824 | | | 3 | | | | | | 1 | | | | | yes | yes | | | | |
| MAST1 | Island | Body | chr19:12978383-12978805 | 422 | | 1 | 2 | | | | | 0 | 4 | | | 2 | | yes | | | | | yes |
| KLF1 | S_Shore | TSS200;5'UTR;1stExon | chr19:12997997-12998764 | 767 | | | 7 | | | | | | | | | | | yes | | | | | |
| CALR | N_Shelf | Body | chr19:13054427-13054718 | 291 | | | | | | | | 0 | 2 | 1 | | 2 | | yes | | | | | yes |
| NFIX | Island | Body | chr19:13112283-13113893 | 1610 | | 1 | 7 | 1 | | | | | 1 | | | | | | | | | | yes |
| NFIX | Island | Body | chr19:13124215-13125851 | 1636 | | | 1 | 1 | 6 | 1 | | | 1 | | | | | | | | | | yes |
| NFIX | S_Shelf | Body | chr19:13127389-13127873 | 484 | | | 1 | | | 2 | | | 1 | | | | | | yes | | | | yes |
| NFIX | Island | Body | chr19:13135527-13136168 | 641 | | | | 1 | 2 | | | | | | | | | | | | | | yes |
| NFIX;LYL1 | Island | 3'UTR | chr19:13207445-13210356 | 2911 | | | | | | | | | | | | | | | | | | | |

| | | | | | | | | | | | | | | | | | | | | | | | | |
|---|---|---|---|---|---|---|---|---|---|---|---|---|---|---|---|---|---|---|---|---|---|---|---|---|
| LYL1 | N_Shore | TSS1500 | chr19:13215151-13215387 | 236 |  |  | 2 |  |  |  |  |  |  |  |  |  |  | yes |  |  |  |  |  |  |
| CACNA1A | Island;N_Shore | Body | chr19:13318873-13319523 | 650 |  | 1 |  | 1 | 1 |  |  |  | 1 | 1 |  |  |  |  |  |  |  | yes | yes |  |
| CACNA1A | Island | Body | chr19:13409671-13410117 | 446 |  |  |  |  |  |  |  |  |  |  |  |  |  | yes | yes |  | yes |  | yes |  |
| CACNA1A | Island | 1stExon | chr19:13616871-13617584 | 713 |  | 1 |  |  |  |  |  |  | 2 |  |  |  |  |  |  |  |  | yes | yes |  |
| MRI1 | N_Shore | TSS1500 | chr19:13875014-13875137 | 123 |  |  | 5 |  |  |  | 1 |  | 1 |  |  |  |  | yes |  |  |  | yes |  |  |
| ZSWIM4 | Island | Body | chr19:13915659-13915877 | 218 |  |  | 2 |  |  |  |  |  |  |  |  | 1 |  | yes |  |  |  |  |  |  |
| MIR24-2;MIR27A;MIR23A | S_Shelf | TSS200;Body;TSS1500 | chr19:13947229-13947914 | 685 | 1 |  |  |  | 1 |  |  |  |  |  |  |  |  | yes |  |  | yes |  |  |  |
| NANOS3 | Island | 1stExon | chr19:13988235-13988555 | 320 |  |  |  |  |  |  |  |  |  |  |  |  |  | yes |  |  |  |  |  |  |
| C19orf57;CC2D1A | N_Shore | 5'UTR;TSS1500 | chr19:14016603-14016761 | 158 |  | 1 |  |  |  |  |  |  | 1 |  |  |  |  | yes |  |  |  |  |  |  |
| PODNL1 | Island | Body | chr19:14043646-14044197 | 551 |  | 1 |  |  |  |  |  |  |  |  |  |  |  | yes |  |  |  |  |  |  |
| PODNL1;DCAF15 | S_Shore | 5'UTR;1stExon;Body | chr19:14064176-14064374 | 198 |  | 1 |  |  |  |  |  |  |  |  |  |  |  |  | yes |  |  |  |  |  |
| RFX1 | N_Shore | Body | chr19:14089284-14089711 | 427 |  |  |  |  | 1 |  |  |  |  |  |  |  |  |  |  |  |  |  | yes |  |
| RLN3 | N_Shelf | TSS200;TSS1500 | chr19:14138763-14139137 | 374 |  |  |  |  |  |  |  |  |  |  |  |  |  | yes |  |  |  |  |  |  |
| IL27RA | N_Shore | TSS1500 | chr19:14142047-14142205 | 158 |  |  |  |  |  |  |  |  |  |  |  |  |  |  |  |  |  |  |  |  |
| PALM3 | S_Shore;S_Shelf | 1stExon;TSS1500 | chr19:14169966-14171331 | 1365 |  | 1 |  | 1 | 1 |  |  |  |  |  |  |  |  | yes |  |  |  |  |  |  |
| PRKACA | N_Shelf | TSS200;Body | chr19:14224992-14225029 | 37 |  |  |  |  |  |  |  |  |  |  |  |  |  |  |  |  |  |  |  |  |
| LPHN1 | N_Shore | 3'UTR | chr19:14260587-14260651 | 64 |  | 1 | 2 | 1 | 1 |  |  |  | 1 |  |  |  |  |  |  |  |  | yes | yes |  |
| PKN1 | Island | TSS1500 | chr19:14542769-14543048 | 279 |  |  | 1 |  |  |  |  |  | 1 |  |  |  |  | yes |  |  |  |  |  |  |
| PKN1 | N_Shore | TSS1500;Body | chr19:14550846-14550999 | 153 |  |  |  | 1 | 1 | 1 |  |  | 2 | 3 | 1 |  |  | yes |  |  |  |  |  |  |
| PKN1 | Island | Body | chr19:14552072-14552350 | 278 |  |  |  |  |  |  |  |  |  |  |  |  |  | yes |  |  |  |  |  |  |
| GIPC1 | Island | Body | chr19:14591033-14591301 | 268 |  |  |  |  | 1 |  |  |  |  |  |  |  |  | yes |  |  |  |  |  |  |
| GIPC1 | S_Shore | TSS1500 | chr19:14607597-14608092 | 495 |  |  |  |  | 1 |  |  |  | 1 |  |  |  |  |  | yes |  |  | yes |  |  |
| TECR | NA | Body | chr19:14671371-14671512 | 141 |  |  |  |  |  |  |  |  | 1 |  |  |  |  |  |  |  |  |  |  |  |
| TECR | Island | Body | chr19:14676213-14676461 | 248 |  |  |  |  |  |  |  |  |  |  |  |  |  |  |  |  |  |  |  |  |
| EMR3 | NA | 5'UTR;1stExon;TSS200 | chr19:14785593-14785849 | 256 |  |  |  |  |  |  |  | 0 |  |  |  |  |  | yes |  |  |  |  | yes |  |
| ZNF333 | N_Shore | TSS1500 | chr19:14800530-14800596 | 66 |  |  |  |  |  |  |  |  | 2 |  |  |  |  |  |  |  | yes | yes |  |  |
| OR7C2 | NA | TSS1500 | chr19:15051936-15052282 | 346 |  |  |  |  |  |  |  |  |  |  |  |  |  | yes |  |  | yes |  |  |  |
| SLC1A6 | NA | TSS200;1stExon | chr19:15083667-15084823 | 1156 |  | 1 | 13 |  |  |  |  |  |  |  |  |  |  | yes |  |  |  | yes | yes |  |
| CCDC105 | N_Shore;Island | TSS1500;TSS200;5'UTR;1stExon | chr19:15121204-15122224 | 1020 |  | 1 |  |  |  |  |  | 0 | 1 |  |  |  |  | yes | yes |  |  |  |  |  |
| CCDC105 | N_Shore | Body | chr19:15132007-15132411 | 404 |  |  | 1 |  |  |  |  |  |  |  |  |  |  | yes |  |  |  |  |  |  |
| CASP14 | NA | TSS1500 | chr19:15162515-15162971 | 456 |  |  |  |  |  |  |  |  | 1 |  |  |  |  | yes |  |  | yes |  |  |  |
| OR1I1 | NA | TSS1500 | chr19:15197635-15197788 | 153 |  |  |  |  |  |  |  |  |  |  |  |  |  | yes |  |  |  |  |  |  |
| SYDE1 | Island | TSS200 | chr19:15218139-15218169 | 30 |  |  |  |  |  |  |  |  | 1 |  |  |  |  |  |  | yes |  | yes |  |  |
| NOTCH3 | Island | Body | chr19:15292440-15292775 | 335 |  | 1 |  |  | 1 |  |  |  |  |  |  |  |  | yes |  |  | yes |  |  |  |
| NOTCH3 | Island | TSS200 | chr19:15311990-15312129 | 139 |  |  | 1 |  |  |  |  |  |  |  |  |  |  | yes |  |  |  | yes | yes |  |
| EPHX3 | Island;N_Shore | 1stExon;Body;5'UTR;TSS200;TSS1! | chr19:15342915-15344364 | 1449 |  |  | 14 |  |  |  |  |  | 2 |  |  |  |  | yes |  |  |  |  |  |  |
| BRD4 | NA | Body | chr19:15375283-15375450 | 167 |  |  | 7 |  |  |  |  |  |  |  |  |  |  |  |  |  |  |  |  |  |
| BRD4 | NA | 1stExon;5'UTR;TSS1500 | chr19:15391124-15391946 | 822 |  |  |  |  |  |  |  |  | 2 | 1 |  |  |  | yes |  |  |  |  |  |  |
| WIZ | N_Shore | Body | chr19:15550742-15550869 | 127 |  |  |  |  |  |  |  |  |  |  |  |  |  |  |  |  |  |  |  |  |
| RASAL3 | Island | Body | chr19:15563870-15564105 | 235 |  |  |  |  |  |  |  |  | 1 |  | 1 |  |  | yes |  |  |  |  |  |  |
| RASAL3 | Island | Body | chr19:15568360-15568935 | 575 |  |  |  |  | 1 |  |  |  | 2 |  | 1 |  |  |  |  |  |  |  |  |  |
| RASAL3 | NA | TSS1500 | chr19:15575612-15575685 | 73 |  |  |  |  | 1 |  |  |  |  |  |  |  |  | yes | yes |  |  |  | yes |  |
| PGLYRP2 | N_Shore;Island | Body | chr19:15580310-15580721 | 411 |  |  | 1 | 1 |  |  |  | 0 | 2 | 1 |  |  |  | yes |  |  |  | yes |  |  |
| PGLYRP2 | Island | Body | chr19:15582687-15582853 | 166 |  |  | 4 |  |  |  |  |  |  |  |  |  |  | yes |  |  |  |  | yes |  |
| PGLYRP2 | NA | Body;5'UTR;1stExon | chr19:15590069-15590368 | 299 |  |  |  | 1 |  |  |  |  |  |  |  |  |  | yes | yes |  |  |  |  |  |
| CYP4F22 | Island | Body | chr19:15662114-15662578 | 464 |  |  |  |  |  |  |  |  | 1 |  |  |  |  | yes |  |  |  |  |  |  |
| OR10H2 | N_Shore | TSS1500 | chr19:15838375-15838895 | 520 |  |  |  |  |  |  |  |  |  |  |  |  |  | yes |  | yes |  |  |  |  |
| OR10H3 | NA | TSS1500 | chr19:15851571-15852002 | 431 |  |  |  |  |  |  |  |  | 1 |  |  |  |  | yes | yes |  |  |  |  |  |
| CYP4F11 | NA | TSS200 | chr19:16045708-16046185 | 477 |  |  |  |  |  |  |  | 0 |  |  |  |  |  |  |  |  |  |  |  |  |
| OR10H4 | NA | TSS1500 | chr19:16058370-16058640 | 270 |  |  |  |  |  |  |  |  | 1 |  |  |  |  | yes | yes |  |  |  |  |  |
| FLJ25328 | NA | TSS1500 | chr19:16144282-16144499 | 217 |  |  |  |  |  |  |  |  | 1 |  |  |  |  |  |  |  |  |  |  | yes |
| TPM4 | N_Shore;Island | TSS1500;TSS200;5'UTR;1stExon | chr19:16178091-16178570 | 479 |  |  | 23 |  |  |  |  |  | 4 | 1 |  |  |  | yes | yes |  |  | yes |  |  |
| HSH2D | NA | 5'UTR | chr19:16254474-16254516 | 42 |  |  | 1 |  |  |  |  |  |  |  |  |  |  | yes | yes |  |  |  |  |  |
| CIB3 | NA | 1stExon | chr19:16284242-16284421 | 179 |  |  | 4 |  |  |  |  |  | 1 |  |  |  |  | yes |  |  |  | yes | yes |  |
| CHERP | Island | Body | chr19:16641544-16641837 | 293 |  |  |  | 1 |  |  |  |  |  |  |  |  |  |  |  |  |  |  |  |  |
| NWD1 | NA | TSS200 | chr19:16830613-16830859 | 246 |  |  |  |  |  |  |  | 0 | 3 |  | 1 | 2 |  | yes | yes |  | yes |  |  |  |
| SIN3B | NA | TSS1500 | chr19:16939845-16940037 | 192 |  |  |  |  |  |  |  |  |  |  |  | 1 |  | yes |  |  | yes | yes | yes |  |
| F2RL3 | N_Shore | TSS1500;TSS200 | chr19:16999576-17000585 | 1009 |  |  | 15 | 1 | 2 |  |  |  | 1 |  |  |  |  | yes |  |  |  |  |  |  |
| OCEL1 | N_Shore | TSS1500 | chr19:17336377-17336525 | 148 |  |  |  |  |  |  |  | 0 |  |  |  |  |  | yes |  | yes |  | yes |  |  |
| NR2F6 | N_Shore | Body | chr19:17346089-17346507 | 418 |  |  |  |  | 1 |  |  |  |  |  |  |  |  | yes |  |  |  | yes | yes |  |
| NR2F6 | Island;S_Shore | TSS1500 | chr19:17357472-17357641 | 169 |  |  |  |  |  |  |  |  |  |  |  |  |  |  | yes |  |  | yes | yes |  |
| ABHD8 | Island | 3'UTR | chr19:17403176-17403529 | 353 |  |  |  | 1 | 1 |  |  |  | 1 |  |  |  |  | yes | yes |  |  |  |  |  |
| ANO8 | chr19:17438054-17440318;Isla | Island;Body | chr19:17439208-17440072 | 864 |  |  | 1 |  |  |  |  | 0 |  |  | 1 |  |  |  |  |  |  | yes | yes |  |
| GTPBP3 | Island | Body | chr19:17448797-17449042 | 245 |  |  |  |  |  |  |  |  | 1 |  |  |  |  |  |  |  |  |  |  |  |
| PLVAP | Island | 1stExon | chr19:17487896-17488225 | 329 |  |  | 1 |  |  |  |  |  | 1 |  |  |  |  | yes |  |  |  |  |  |  |
| BST2 | NA | 1stExon;TSS200;TSS1500 | chr19:17516282-17517221 | 939 |  |  |  |  | 3 |  |  |  | 1 |  |  |  |  | yes |  |  |  |  |  |  |
| NXNL1 | Island;N_Shore | Body;3'UTR | chr19:17566260-17566626 | 366 |  |  | 1 |  |  |  |  |  | 1 |  |  |  |  | yes | yes |  |  | yes | yes |  |
| NXNL1 | NA | TSS200 | chr19:17571825-17571888 | 63 |  |  | 3 |  |  |  |  |  |  |  |  |  |  | yes | yes |  |  |  |  |  |
| SLC27A1 | N_Shore | TSS200 | chr19:17581121-17581191 | 70 |  |  |  |  |  |  |  |  | 1 |  |  |  |  |  |  |  |  |  |  |  |
| PGLS | Island | Body | chr19:17631836-17631991 | 155 |  |  |  |  |  |  |  |  |  |  |  |  |  | yes |  |  |  |  |  |  |
| FAM129C | S_Shore;S_Shelf | TSS1500;TSS200 | chr19:17633880-17634172 | 292 |  |  |  | 1 | 2 |  |  |  |  |  |  |  |  | yes | yes | yes |  |  |  |  |
| UNC13A | Island | Body | chr19:17791074-17791354 | 280 |  | 1 |  |  |  |  |  |  |  |  |  |  |  | yes |  |  |  | yes | yes |  |
| FCHO1 | N_Shore | TSS1500 | chr19:17858101-17858298 | 197 |  |  | 1 |  |  |  |  |  |  |  |  |  |  | yes | yes |  |  |  | yes |  |
| FCHO1 | S_Shelf | 5'UTR;TSS1500;TSS200 | chr19:17862017-17862385 | 368 |  |  |  |  | 2 | 1 |  |  | 2 | 1 | 1 |  |  | yes |  |  | yes |  |  |  |
| FCHO1 | N_Shore;Island | Body | chr19:17877419-17877846 | 427 |  |  |  |  | 1 |  |  | 0 | 4 | 1 |  |  |  | yes |  |  | yes |  |  |  |
| B3GNT3 | N_Shore;Island | TSS1500 | chr19:17905332-17905718 | 386 |  |  | 3 |  |  |  |  | 0 | 2 |  |  | 4 |  | yes |  |  |  |  |  |  |
| B3GNT3 | Island | Body | chr19:17918795-17919173 | 378 |  |  | 1 |  |  |  |  |  |  |  |  |  |  | yes | yes |  |  | yes |  |  |
| INSL3 | N_Shore;Island | Body;TSS200 | chr19:17931932-17932369 | 437 |  |  | 14 |  |  |  |  |  | 6 |  |  | 1 |  | yes |  |  | yes | yes |  |  |
| JAK3 | N_Shore;Island | Body | chr19:17942025-17942411 | 386 |  |  | 4 |  | 1 |  |  |  | 2 |  | 1 |  |  | yes |  |  |  |  | yes |  |
| JAK3 | Island;S_Shore | 5'UTR;TSS200 | chr19:17958736-17959082 | 346 |  |  | 16 |  | 1 |  |  |  | 2 |  |  |  |  | yes |  |  |  |  |  |  |
| SLC5A5 | Island | TSS200 | chr19:17982682-17983245 | 563 | 12 |  |  |  |  |  |  |  |  |  |  |  |  | yes | yes |  |  |  | yes |  |
| KCNN1 | Island | Body | chr19:18108895-18109484 | 589 |  |  |  |  |  |  |  | 0 | 1 |  |  |  |  | yes | yes |  |  |  |  |  |
| ARRDC2 | N_Shore;Island | Body;TSS1500 | chr19:18118666-18120819 | 2153 | 1 |  |  | 1 |  |  |  | 0 |  |  |  |  |  | yes |  |  |  |  |  |  |
| MAST3 | N_Shore | TSS1500 | chr19:18208303-18208422 | 119 |  |  |  |  |  |  |  |  |  |  |  |  |  | yes |  |  | yes | yes |  |  |
| MAST3 | Island | Body | chr19:18260330-18260479 | 149 |  |  |  |  |  |  |  |  | 1 |  |  |  |  | yes |  |  |  |  | yes |  |
| IFI30 | N_Shore | TSS1500 | chr19:18284120-18284397 | 277 |  |  | 7 |  |  |  |  |  |  |  |  |  |  | yes |  |  |  | yes |  |  |
| PDE4C | Island | Body | chr19:18331075-18331394 | 319 |  |  | 1 |  |  |  |  |  |  |  |  |  |  | yes | yes |  |  |  | yes |  |
| PDE4C | S_Shore | TSS1500;Body | chr19:18338153-18338283 | 130 |  |  |  |  |  |  |  |  |  |  |  |  |  | yes |  |  |  |  |  |  |
| PDE4C | Island | Body;5'UTR | chr19:18343901-18344520 | 619 |  | 1 | 1 |  |  |  |  |  |  |  |  |  |  |  |  |  |  |  |  |  |
| PDE4C | NA | TSS1500 | chr19:18359505-18359562 | 57 |  |  |  |  |  |  |  |  |  |  |  |  |  | yes |  |  |  |  | yes |  |
| KIAA1683 | NA | 5'UTR;1stExon;TSS1500 | chr19:18385244-18385930 | 686 |  |  |  | 1 | 1 |  |  |  |  |  |  |  |  | yes | yes |  |  |  |  |  |
| GDF15 | N_Shore | TSS1500;TSS200 | chr19:18496756-18497064 | 308 |  |  | 6 |  |  |  |  | 0 | 1 |  |  | 2 |  | yes | yes |  |  |  |  |  |
| GDF15 | Island | Body | chr19:18499098-18499688 | 590 |  |  |  | 1 | 1 |  |  |  |  |  |  |  |  | yes | yes |  |  |  |  |  |
| SSBP4 | N_Shore | Body | chr19:18539756-18540330 | 574 |  |  | 2 | 1 | 1 |  |  |  | 1 |  | 1 |  |  | yes |  |  |  |  |  |  |
| ISYNA1 | Island;S_Shore | TSS200;TSS1500 | chr19:18549131-18550061 | 930 |  |  | 3 |  |  |  |  |  | 1 |  |  |  |  |  |  | yes |  |  |  |  |
| C19orf60 | N_Shore | TSS1500 | chr19:18698825-18699444 | 619 |  | 1 | 1 |  |  |  |  | 0 | 3 |  |  |  |  | yes |  | yes |  |  | yes |  |
| CRLF1 | S_Shelf | Body | chr19:18709415-18709437 | 22 |  |  |  |  |  |  |  |  |  |  |  |  |  | yes |  |  |  |  |  |  |
| CRLF1 | Island | Body;TSS200 | chr19:18716230-18717781 | 1551 |  |  | 2 |  |  |  |  |  | 2 |  |  |  |  |  |  |  |  | yes | yes |  |
| KLHL26 | N_Shore;Island | Body | chr19:18761132-18761792 | 660 |  |  | 7 | 1 |  |  |  |  |  |  |  |  |  |  |  | yes |  |  |  |  |

| | | | | | | | | | | | | | | | | | | | | | | | | |
|---|---|---|---|---|---|---|---|---|---|---|---|---|---|---|---|---|---|---|---|---|---|---|---|---|
| KLHL26 | N_Shore;Island;chr19:1877107 | Body;Island | chr19:18770726-18771327 | 601 | | | 19 | | | | | | | | 1 | | | | | | | | | |
| CRTC1 | Island | Body | chr19:18873037-18873268 | 231 | | | | | | 1 | | | | | | | | | | | yes | | | |
| CRTC1 | NA | Body | chr19:18879431-18879450 | 19 | | | | | | | | | | | | | | yes | | | | | yes | |
| CRTC1 | Island | Body | chr19:18888081-18889003 | 922 | | | | | | | | | | | | | | | yes | | yes | | yes | |
| COMP | Island | Body | chr19:18896581-18896853 | 272 | | | | | | | | | 1 | | | | | | | | | yes | | |
| COMP | Island | Body | chr19:18899876-18900568 | 692 | | | | | | | | | 1 | | | | | | | | | | yes | |
| COMP | chr19:18899037-18902284;Isla | Island;1stExon;5'UTR;TSS200;TSS1 | chr19:18901843-18902727 | 884 | | | 9 | | | | | 0 | 1 | | 1 | | | yes | | | | yes | | |
| GDF1;LASS1 | Island | Body;3'UTR | chr19:18980103-18980744 | 641 | | | | | | | | | | | | | | | | | | | yes | |
| GDF1;LASS1 | Island | TSS1500 | chr19:19007505-19007647 | 142 | | | | | | | | | | | | | | | | | | | yes | |
| SLC25A42 | Island | Body | chr19:19221316-19221574 | 258 | | | 13 | | | | | | | | | | | yes | | | | | | |
| LOC729991-MEF2B;MEF2B | N_Shore;Island | Body;5'UTR;TSS200;TSS1500 | chr19:19280969-19281559 | 590 | | | | | | | | 0 | 1 | 1 | | | | | yes | | | yes | | |
| NR2C2AP | Island | TSS200 | chr19:19314298-19314328 | 30 | | | | | | | | | 1 | | | | | yes | | | | | | |
| NCAN | Island | TSS200 | chr19:19322676-19322778 | 102 | | | | | | | | | 1 | | | | | yes | | | | | | |
| HAPLN4 | Island | Body | chr19:19371900-19371974 | 74 | | | | | | | | | | | | | | | | | yes | | | |
| HAPLN4 | S_Shore | TSS200;5'UTR;1stExon | chr19:19373566-19374178 | 612 | | | 1 | | | | | | 1 | | | | | yes | | | | | yes | |
| TM6SF2 | Island | Body | chr19:19383838-19384165 | 327 | | | | | | | | | 1 | | | | | yes | | | | yes | | |
| KIAA0892;SF4 | Island | TSS200;TSS1500 | chr19:19431572-19431584 | 12 | | | | | | | | | 1 | | | | | | | | | yes | | |
| GATAD2A | N_Shore | Body | chr19:19611900-19612071 | 171 | | | | | | | | | | | | | | yes | | | | | | |
| TSSK6;NDUFA13 | Island | 1stExon;TSS1500;5'UTR;TSS200 | chr19:19626251-19626605 | 354 | | | 8 | | | | | | 1 | | | | | | | | | | | |
| YJEFN3 | Island | TSS200 | chr19:19639553-19640075 | 522 | | 1 | 26 | | | | | | 1 | | | | | yes | | | | | | |
| YJEFN3 | Island | Body | chr19:19643172-19643534 | 362 | | | | | | | | | 1 | | | | | yes | | | | | | |
| CILP2;YJEFN3 | Island | TSS200;TSS1500;3'UTR | chr19:19648335-19648991 | 656 | | | 3 | | | | | | 2 | | | | | yes | | | | | | |
| CILP2 | Island | Body | chr19:19651867-19652066 | 199 | | 1 | | | | | | | | | | | | | | | | yes | | |
| ZNF486 | NA | TSS200 | chr19:20278013-20278051 | 38 | | | | | | | | 0 | | | | | | | | yes | | | | |
| ZNF85 | NA | TSS200 | chr19:21106002-21106120 | 118 | | | | | 1 | | | | | | | | | yes | | yes | yes | | | |
| ZNF430 | NA | TSS200 | chr19:21203469-21203723 | 254 | | | | | | | | | 1 | | | 1 | | yes | | yes | | | | |
| ZNF714 | N_Shore;Island | TSS200;5'UTR;1stExon | chr19:21264896-21265421 | 525 | | | 55 | | | 2 | 2 | 0 | 2 | 2 | | 45 | 2 | yes | | yes | yes | | yes | |
| ZNF431 | NA | TSS200 | chr19:21324707-21324782 | 75 | | | | | | | | 0 | | | | | | | | | yes | | | |
| ZNF708 | NA | 1stExon;5'UTR | chr19:21512141-21512191 | 50 | | | | | | | | | 1 | | | | | yes | | yes | | | | |
| ZNF738 | NA | Body | chr19:21541756-21541772 | 16 | | | | | | | | | 1 | | | | | yes | | | yes | | | |
| ZNF429 | N_Shore | TSS200 | chr19:21688397-21688647 | 250 | | | | | | | | | | | | 1 | | yes | | yes | yes | | | |
| ZNF208 | Island;N_Shore | Body | chr19:22193293-22193911 | 618 | | 1 | 2 | | | | | 0 | 1 | | | | | yes | | yes | | | | |
| ZNF257 | NA | TSS1500;Body | chr19:22235022-22235850 | 828 | | 1 | | | | | | 0 | | | | | | yes | | yes | | | | |
| ZNF492 | N_Shore | TSS1500;TSS200 | chr19:22816508-22817496 | 988 | | 2 | | | 4 | | | | 2 | | | | | yes | | yes | | | | |
| ZNF681 | Island;S_Shore;N_Shore | Body;TSS200;TSS1500 | chr19:23941207-23942126 | 919 | 1 | 2 | 13 | 1 | 1 | | 1 | 0 | 3 | 5 | 1 | 10 | | yes | yes | yes | | | yes | |
| RPSAP58 | N_Shore;Island | TSS1500;TSS200;Body | chr19:23945472-23946205 | 733 | | | 9 | | | | | 0 | 7 | 2 | 1 | 6 | 2 | yes | | yes | | | | |
| ZNF254 | NA | 1stExon;5'UTR | chr19:24270007-24270468 | 461 | | | | 1 | | | | | 1 | | 1 | | | yes | | yes | | | | |
| LOC148145 | NA | TSS200;TSS1500 | chr19:29455596-29456219 | 623 | | | | | | | | | 1 | | | | | yes | yes | | | | | |
| VSTM2B | Island | Body | chr19:30019369-30019647 | 278 | | 1 | | | | | | | | | | | | | | | | yes | | |
| ZNF536 | N_Shelf | TSS200 | chr19:30863260-30863349 | 89 | | | 6 | | | | | | | | | | | yes | | | | yes | yes | |
| ZNF536 | N_Shore;Island | 5'UTR | chr19:30865486-30866365 | 879 | | 1 | | | | | | 0 | 2 | | | | | | | | | yes | yes | |
| ZNF536 | Island | Body | chr19:30941858-30942096 | 238 | | | | | | | | | 1 | | | | | | | | | | yes | |
| DKFZp566F0947 | NA;TSS200 | TSS1500;TSS200;nan | chr19:31640194-31640849 | 655 | | | 42 | | | | | | 1 | | | | | yes | | | | yes | | |
| TSHZ3 | N_Shore | Body | chr19:31839171-31839734 | 563 | | | | | | | | | 1 | | | | | | | | | yes | yes | |
| TSHZ3 | Island | TSS1500 | chr19:31841391-31842115 | 724 | | | | | | | | | 1 | | | | | | yes | | | | yes | |
| ZNF507 | N_Shore | TSS1500 | chr19:32835826-32836297 | 471 | | | 1 | | | | | | | | | | | yes | | | | | yes | |
| RGS9BP | Island | 3'UTR;1stExon | chr19:33167980-33168446 | 466 | | | | | | | | | | | | | | | | | | | | |
| TDRD12 | Island | TSS200 | chr19:33210498-33210851 | 353 | | | 3 | | | | | | 3 | | | | | yes | | yes | | | yes | |
| SLC7A9 | NA | 1stExon;5'UTR;TSS1500 | chr19:33360599-33361552 | 953 | | | | 1 | | | | | | | | | | yes | | | | | | |
| WDR88 | Island | TSS200 | chr19:33622953-33623230 | 277 | | | | | | | | | 3 | | | | | yes | | | | | yes | |
| CHST8 | Island;N_Shore | 5'UTR;TSS1500;TSS200;1stExon | chr19:34111400-34113449 | 2049 | | | 5 | | | | | 0 | 8 | 1 | | 1 | | yes | | | | yes | yes | |
| CHST8 | N_Shore;Island | 5'UTR;TSS1500;TSS200 | chr19:34174345-34175481 | 1136 | | 1 | 15 | | | | | | 5 | | | 1 | | | yes | yes | | | | |
| WTIP | Island | Body | chr19:34988832-34989280 | 448 | | | | | | | | | 1 | | | | | yes | yes | | | | | |
| LOC643719 | Island;S_Shore | Body;TSS1500 | chr19:35068555-35069691 | 1136 | | | | | 1 | | | 0 | 1 | | | | | | | yes | | | | |
| SCGBL | NA | 1stExon;5'UTR;TSS1500 | chr19:35085474-35086950 | 1476 | | | | | | | | | 1 | | | | | | yes | yes | | yes | | |
| SCN1B;HPN | N_Shore;Island | 3'UTR;TSS1500;TSS200;5'UTR | chr19:35530755-35532097 | 1342 | | | 14 | | 1 | | | | 1 | | | | | | | | | | | yes |
| FXYD3 | NA | TSS200;5'UTR;1stExon | chr19:35606534-35607221 | 687 | | 1 | 5 | 1 | 1 | | | | 2 | 1 | | | | | yes | | | | yes | |
| LGI4 | Island | 3'UTR | chr19:35615444-35615638 | 194 | | | | | | | | 0 | 1 | 1 | 1 | 3 | | | | | | | | |
| LGI4 | chr19:35617253-35617640 | Island | chr19:35617254-35617819 | 565 | | | | | | | | | | | 1 | | | yes | yes | | | | | |
| FXYD1 | N_Shelf;chr19:35632356-3563 | TSS1500;N_Shelf;5'UTR;TSS200 | chr19:35629022-35630474 | 1452 | 3 | | | | | | | 0 | 6 | 1 | 1 | 1 | | yes | | | | yes | | |
| FXYD1;FXYD7 | Island | Body;TSS1500;3'UTR | chr19:35632493-35633972 | 1479 | | | 23 | 2 | | | | | | | | | | yes | yes | | | yes | yes | |
| HAMP | S_Shelf | TSS1500;TSS200 | chr19:35772917-35773718 | 801 | | | 11 | | | | | 0 | | | | | | yes | yes | | | | | |
| MAG | N_Shelf;NA | TSS1500 | chr19:35782257-35782899 | 642 | | | | | | | | | | | | | | | yes | | | | | |
| MAG | Island | Body | chr19:35786580-35786961 | 381 | | | 7 | 1 | 1 | | | | | | | | | yes | | | yes | | | |
| MAG | Island | Body | chr19:35791192-35791409 | 217 | | | | | | | | | | | | | | yes | | | | yes | yes | |
| CD22 | NA | TSS1500 | chr19:35819775-35819985 | 210 | | | | | 1 | | | | | 1 | | | | yes | yes | | | | | |
| FFAR1 | N_Shore;Island | TSS1500;5'UTR;1stExon | chr19:35842171-35843186 | 1015 | | | 11 | | | | | | | | | | | yes | | | | | | |
| FFAR2 | NA | TSS1500;TSS200 | chr19:35940174-35940862 | 688 | | 1 | 4 | | | | | | 2 | 1 | | | | yes | | | | | | |
| KRTDAP | NA | Body;TSS200;TSS1500 | chr19:35981224-35982442 | 1218 | | 1 | 10 | | | | | | | | | | | yes | yes | | | | | |
| DMKN | NA | Body;1stExon | chr19:35992791-35992984 | 193 | | | | | | | | | | | | | | yes | | | yes | | | |
| DMKN | NA | TSS200 | chr19:36004699-36004994 | 295 | | | | | | | | | 1 | | | | | | | | | | | |
| GAPDHS | Island | TSS200 | chr19:36024217-36024282 | 65 | | | 6 | | | | | | | | | | | yes | | | | | | |
| ATP4A | Island | Body | chr19:36048757-36049672 | 915 | | 1 | | | | | | | | | | | | yes | | | | | yes | |
| ATP4A | NA | 1stExon;TSS200 | chr19:36054519-36055013 | 494 | | | 3 | | | | | | | | | | | yes | | | | | yes | |
| ETV2 | N_Shore | TSS1500;TSS200 | chr19:36132426-36132842 | 416 | | | | 1 | 4 | | | | | | | | | yes | yes | | | | | yes |
| UPK1A | NA | TSS200 | chr19:36157677-36157825 | 148 | | | 2 | | | | | | | | | | | yes | | | | yes | | |
| TMEM149 | S_Shore | 1stExon;5'UTR;TSS200 | chr19:36233297-36233469 | 172 | | | | | 1 | 1 | | | 2 | 2 | 1 | | | yes | | | | | | |
| HSPB6 | Island | 3'UTR | chr19:36246395-36246906 | 511 | 2 | 1 | | 1 | 4 | | | 0 | 8 | 4 | | 7 | | | | | | | yes | |
| NPHS1 | N_Shore | Body | chr19:36334925-36335320 | 395 | | 1 | 1 | | | | | | | | | | | yes | | | | yes | | |
| KIRREL2 | Island | Body | chr19:36352005-36352459 | 454 | | | | | | | | | 1 | | | | | | | | | | | |
| APLP1 | N_Shore | Body | chr19:36365307-36365557 | 250 | | | | | | | | | | | | | | | | | | | | |
| NFKBID | N_Shore | 5'UTR | chr19:36389500-36389833 | 333 | | | | 1 | | | | | | | | | | yes | | | | | | |
| HCST | S_Shore | TSS1500 | chr19:36393064-36393696 | 632 | | | | 1 | 2 | | | | | | | | | yes | yes | | | | | |
| TYROBP | NA | 1stExon;5'UTR;TSS1500 | chr19:36399185-36400437 | 1252 | | | | | | | | 0 | | | | | | yes | yes | | | | | |
| SDHAF1 | N_Shore;Island | TSS1500 | chr19:36484731-36485360 | 629 | 1 | 1 | 10 | | | | | 0 | 4 | 1 | | 14 | | yes | | | | yes | | |
| C19orf46 | N_Shore | Body;5'UTR;1stExon | chr19:36499326-36499839 | 513 | | | 3 | 1 | 1 | | | | 1 | | | | | yes | | | | | | |
| TBCB | Island | TSS1500 | chr19:36604588-36604764 | 176 | | 1 | | | | | | | | | | | | | | | | | | |
| ZNF146 | NA | 5'UTR | chr19:36726982-36726989 | 7 | | | | | | | | | 1 | | | | | yes | | | | | | |
| LOC100134317 | S_Shore;S_Shelf | TSS1500 | chr19:36800845-36801503 | 658 | 1 | | 12 | | | | | | | | | | | yes | | | | | | yes |
| ZFP82 | Island | 5'UTR | chr19:36909326-36909831 | 505 | | 1 | | | | | | | | | | | | yes | | | yes | | | |
| ZNF529;ZNF382 | Island | Body;5'UTR;TSS1500;1stExon;TSS2 | chr19:37096010-37096588 | 578 | | 1 | 1 | | 1 | | | | | | | 1 | | | | | | | | |
| ZNF790 | Island | TSS1500 | chr19:37329498-37329859 | 361 | | | | | | | | | 2 | | | | | yes | | | | yes | | |
| ZNF568;ZNF829 | Island | 5'UTR;1stExon;TSS200;TSS1500 | chr19:37406932-37407889 | 957 | | 2 | | | | | | | | | | | | yes | yes | yes | | | | |
| ZNF585B | NA | 1stExon;5'UTR | chr19:37701393-37701738 | 345 | | 1 | | | | | | 0 | 1 | | | | | yes | yes | | | | | |
| HKR1 | N_Shore;Island | TSS1500;TSS200 | chr19:37825009-37826008 | 999 | | 1 | 2 | | | | | 0 | 9 | 2 | | 16 | | | | yes | yes | | | |
| ZNF570;ZNF569 | N_Shore | TSS1500;TSS200 | chr19:37958522-37959104 | 582 | | | 7 | | | | | 0 | 1 | | | | | yes | | | | | | |
| ZNF793 | N_Shore | TSS1500 | chr19:37997294-37997675 | 381 | | | | | | | | | 1 | | | | | | | yes | | | | |
| ZNF540 | S_Shore | TSS200 | chr19:38042109-38042472 | 363 | | 1 | | | | | | | 1 | | | | | | yes | | | | | yes |
| ZNF781 | Island;N_Shore | TSS200;5'UTR | chr19:38182773-38183262 | 489 | | 1 | | | | | | 0 | 1 | | | 1 | | yes | yes | | | | | |

| | | | | | | | | | | | | | | | | | | | | | | | | | | |
|---|---|---|---|---|---|---|---|---|---|---|---|---|---|---|---|---|---|---|---|---|---|---|---|---|---|---|
| SIPA1L3 | Island | Body | chr19:38684237-38684328 | 91 | | 1 | | | | | | | | | | | | yes | | | | yes | | | | |
| DPF1 | Island | Body;5'UTR | chr19:38713242-38713373 | 131 | | | 5 | | | | | | 1 | 1 | | 2 | | yes | | | | yes | yes | | | |
| PPP1R14A | Island | TSS200 | chr19:38747355-38747378 | 23 | | 1 | 1 | | | | | 0 | 1 | | | 1 | | yes | yes | | | yes | | | | |
| SPINT2 | N_Shore | TSS1500 | chr19:38754529-38754768 | 239 | | | | | | | | | | | | | | yes | yes | | | | | | | |
| C19orf33;YIF1B | NA | TSS1500;3'UTR;TSS200;1stExon;5' | chr19:38794554-38795051 | 497 | | | 7 | | | | | 0 | 1 | | | 3 | | yes | | | | | yes | | | |
| GGN | Island | Body | chr19:38876250-38877220 | 970 | | | | | | | | 0 | 3 | | 1 | 1 | | | | | | yes | | | | |
| SPRED3 | Island;chr19:38886109-388866 | Body;Island | chr19:38886110-38886789 | 679 | | | 14 | | | | 1 | 0 | 3 | 2 | 1 | 4 | | | | | | yes | yes | | | |
| RYR1 | N_Shore | TSS200 | chr19:38924182-38924330 | 148 | | 1 | | | | | | | | | | | | yes | | | | | | | | |
| RYR1 | Island | Body | chr19:38943525-38944048 | 523 | | | | | | | | | 1 | | | | | yes | yes | | | | | | | |
| RYR1 | Island | Body | chr19:38973940-38974117 | 177 | | 1 | | | | | | | | | | | | | | | | | | | | |
| RYR1 | Island | Body | chr19:39055712-39056216 | 504 | | | 1 | | | | | 0 | 2 | 2 | | 10 | | yes | | | | yes | yes | | | |
| MAP4K1 | Island | Body | chr19:39086923-39087186 | 263 | | | | | | | | | 1 | | 1 | | | | | | | | | | | |
| MAP4K1;EIF3K | N_Shore | 1stExon;5'UTR;TSS1500;TSS200 | chr19:39108609-39109076 | 467 | | | 2 | | | | | | | | | | | | | | | | | | | |
| LGALS7B | N_Shelf | TSS1500 | chr19:39278997-39279703 | 706 | | | | | | | | | 1 | 1 | | | | yes | | | | yes | | | | |
| LGALS4 | NA | 5'UTR;1stExon | chr19:39303698-39303993 | 295 | | | | 1 | 1 | | | | | | | | | yes | | | | | | | | |
| RINL | Island | Body | chr19:39360313-39360961 | 648 | | | | | 1 | | | 0 | 1 | 1 | | | | | | | | | | | | |
| RINL | NA | 5'UTR;TSS200 | chr19:39368408-39369353 | 945 | | | 3 | 1 | 1 | | | | | | | | | | yes | | yes | yes | | | | |
| FBXO17 | Island | Body | chr19:39440734-39441205 | 471 | | | | | | | | | | | | | | yes | | yes | | | yes | | | |
| FBXO27 | Island;S_Shore | 5'UTR;1stExon;TSS200 | chr19:39523158-39523840 | 682 | | | 3 | | | | | 0 | 2 | | | | | yes | | yes | | | yes | | | |
| SYCN | Island;S_Shore | TSS200 | chr19:39694993-39695801 | 808 | | | 11 | | | | | 0 | 3 | | | | | yes | | | | | | | | |
| IL28B | N_Shore | TSS1500 | chr19:39736690-39736956 | 266 | | | | | | | | | | | | | | yes | | | | yes | | | | |
| IL29 | NA | TSS1500 | chr19:39785679-39785941 | 262 | | | | | | | | | 1 | | | | | yes | | yes | | | yes | | | |
| LRFN1 | Island | Body | chr19:39798481-39799379 | 898 | | | | | | | | | 2 | | | | | yes | | | | | | | | |
| GMFG | NA | Body;5'UTR;1stExon;TSS200 | chr19:39826150-39826927 | 777 | | | 3 | 1 | 1 | | | | 2 | 1 | | | | yes | | | yes | yes | | | | |
| DLL3 | Island | TSS200 | chr19:39989412-39989426 | 14 | | | | | | | | | 1 | | | | | | | | | | | | | |
| DLL3 | Island | Body | chr19:39993358-39993697 | 339 | | | | | | | | | 1 | | | | | yes | | | | | | | | |
| DLL3 | Island | Body | chr19:39997622-39998290 | 668 | | | 6 | | | | | 0 | 4 | 1 | | 5 | | | yes | | | | | | | |
| CLC | NA | TSS1500 | chr19:40229215-40229766 | 551 | | | 5 | | | | | | | | | | | | yes | yes | | | | | | |
| FCGBP | Island | Body | chr19:40421453-40421743 | 290 | | | 2 | | | | | | 1 | | | | | yes | yes | | | | | | | |
| CNTD2 | Island | Body | chr19:40729157-40729386 | 229 | | | | | | | | 0 | 1 | 1 | | 1 | 1 | yes | | | | | | | | |
| CNTD2 | S_Shore | TSS200 | chr19:40732712-40732902 | 190 | | | 7 | | | | | | 1 | | | | | yes | yes | | | | | | | |
| PLD3;C19orf47 | N_Shore | TSS1500;5'UTR | chr19:40854069-40854190 | 121 | | | | | | | | | 1 | | | | | yes | | | | | | | | |
| PLD3 | N_Shore | 5'UTR | chr19:40871557-40871823 | 266 | | 1 | | | | | | 0 | | | | 1 | | | | | | | | | | |
| HIPK4 | Island | Body | chr19:40889848-40890114 | 266 | | 1 | | | | | | | | | | | | yes | | | | | | | | |
| HIPK4 | S_Shore | TSS200 | chr19:40896163-40896506 | 343 | | | | | | | | | 1 | | | | | yes | yes | | | | | | | |
| PRX | Island | Body | chr19:40904691-40904974 | 283 | | | 14 | | | | | 0 | 1 | | | 5 | 1 | yes | yes | | | | | | | |
| PRX | NA | 1stExon;5'UTR | chr19:40919245-40919760 | 515 | | | 3 | | | | | 0 | 1 | | | | | yes | | | | | | | | |
| SPTBN4;BLVRB | Island | TSS1500 | chr19:40972323-40972683 | 360 | | | 2 | | | | | | 1 | | | | | | | | | yes | yes | | | |
| SPTBN4 | Island | Body | chr19:40995921-40996346 | 425 | | | 1 | | | | | | | | | | | yes | yes | | | | | | | |
| SPTBN4 | Island | Body | chr19:41018757-41018876 | 119 | | | 1 | | | | | | | | | | | | yes | | | | yes | | | |
| SPTBN4 | Island | Body | chr19:41025339-41025959 | 620 | | | 6 | | | | | | 2 | | | | | yes | | | yes | | yes | | | |
| SPTBN4 | S_Shore | TSS1500;Body | chr19:41035836-41036253 | 417 | | | | | | | | | | | | | | | | | | yes | yes | | | |
| SPTBN4 | Island | Body | chr19:41060190-41060564 | 374 | | | | | | | | 0 | | | | | | yes | yes | | | | | | | |
| SPTBN4 | Island | Body | chr19:41073603-41074427 | 824 | | 1 | | | | | | | 1 | | | | | yes | yes | | | | | | | |
| LTBP4 | N_Shore | Body | chr19:41115390-41115766 | 376 | 2 | | | | | | | | | | | | | yes | yes | | | | yes | | | |
| LTBP4 | chr19:41119031-41120394 | Island | chr19:41119829-41120329 | 500 | | | | | | | | | | | 1 | | | | | | | | yes | | | |
| LTBP4 | N_Shore;Island | Body | chr19:41129636-41130041 | 405 | | 1 | | 1 | | | | | | | | | | yes | | | | yes | | | | |
| LTBP4 | Island | Body | chr19:41133106-41133229 | 123 | | | | | | | | | 1 | | | | | | | | | | | | | |
| EGLN2 | Island | 1stExon;Body | chr19:41307090-41307228 | 138 | | | 1 | | | | | | | | | | | | | | | | | | | yes |
| CYP2B7P1 | NA | TSS1500 | chr19:41429786-41430189 | 403 | | | | | | | | | | | | | | yes | | yes | | | | | | |
| CYP2B6 | NA | Body | chr19:41510026-41510059 | 33 | | | 4 | | | | | | | | | | | | yes | | yes | | yes | | | |
| CYP2A13 | Island | Body | chr19:41595975-41596486 | 511 | | 1 | | | 1 | | | | 1 | | | | | | | | | yes | yes | | | |
| CYP2F1 | NA | TSS1500 | chr19:41619850-41620458 | 608 | | | | | | | | | | | | | | yes | | | | yes | yes | | | |
| CYP2F1 | Island | Body | chr19:41633902-41634046 | 144 | | | | | | | 1 | 0 | | 1 | | | | yes | | | | | | | | |
| CYP2S1 | Island | TSS1500;TSS200 | chr19:41698906-41698987 | 81 | | 1 | 1 | | | | | | 3 | | | | | yes | | | | yes | | | | |
| TMEM91 | N_Shore | TSS1500;5'UTR | chr19:41882232-41882253 | 21 | | | | | | | | | 1 | 1 | | | | yes | | | | yes | yes | | | |
| B3GNT8 | NA | 5'UTR | chr19:41934170-41934715 | 545 | | | | 1 | 1 | | | | | | | | | | | yes | | | yes | | | |
| CEACAM21 | NA | TSS1500 | chr19:42082102-42082350 | 248 | | | | | | | | | | | | | | | yes | | | | | | | |
| CEACAM5 | NA | TSS1500 | chr19:42211160-42211479 | 319 | | | | | | | | | | | | | | | yes | | | | | | | |
| LYPD4;DMRTC2 | N_Shore | TSS200;TSS1500 | chr19:42348563-42348686 | 123 | | | | | | | | | | | | | | | | | | | | | | |
| GRIK5 | Island | 3'UTR;Body | chr19:42502731-42503412 | 681 | | | | | | | | 0 | 1 | 1 | | 2 | | | yes | | | | yes | | | |
| GRIK5 | Island | Body | chr19:42509860-42510093 | 233 | | | | | | | | | | | | | | yes | yes | | | | | | | |
| GRIK5 | Island | Body | chr19:42546662-42546977 | 315 | 1 | | | | | | | | | | | | | | yes | | yes | | | | | |
| GRIK5 | NA | 1stExon;5'UTR | chr19:42569945-42570406 | 461 | | | | | | | | | | | | | | | | | | | yes | yes | | |
| POU2F2 | N_Shore | 5'UTR;1stExon;TSS200 | chr19:42636625-42637137 | 512 | | | 7 | | 2 | | | | | | | | | yes | | | | | | | | |
| DEDD2 | N_Shore;Island | 3'UTR;Body | chr19:42703228-42703852 | 624 | | | | | 2 | | | | | | 1 | | | | yes | | | | yes | | | |
| PRR19;PAFAH1B3 | Island | TSS200;Body | chr19:42806184-42806228 | 44 | | | 1 | 1 | | | | 0 | 1 | | | | | | | | | | | | | |
| PRR19 | Island | 5'UTR | chr19:42811017-42811489 | 472 | | | 1 | | | | | 0 | 3 | | | | | yes | | | | yes | | | | |
| TMEM145 | Island | Body | chr19:42817981-42818481 | 500 | | | | | | | | 0 | 1 | | | | | | | | | yes | yes | | | |
| CNFN | S_Shelf | 1stExon;5'UTR | chr19:42894425-42894693 | 268 | | | | | | | | 0 | | | | | | yes | | | | | | | | |
| CXCL17 | NA | 1stExon;5'UTR;TSS200 | chr19:42947133-42948167 | 1034 | | | | | | | | | | | | | | yes | yes | | | | | | | |
| CD177 | NA | TSS200;1stExon;5'UTR | chr19:43857717-43858174 | 457 | | | 11 | | | | | | | | | | | | | | | | yes | yes | | |
| TEX101 | Island | TSS200;Body | chr19:43918943-43919039 | 96 | | | | | | | | | | | | | | | | | | yes | | yes | | |
| LYPD3 | S_Shore | 1stExon;5'UTR;TSS200 | chr19:43969769-43970034 | 265 | | | 1 | | | | | 0 | 1 | | | 1 | | yes | yes | | | | | | | |
| ZNF575 | Island | Body | chr19:44039475-44039870 | 395 | | | | | | | | 0 | 3 | | | | | | yes | | | | yes | | | |
| CADM4 | N_Shore | TSS200 | chr19:44144149-44144337 | 188 | | | | | | | | | 1 | | | | | | | | | yes | yes | | | |
| IRGC | N_Shelf | TSS200;TSS1500 | chr19:44219675-44220214 | 539 | | | 4 | | | | | | | | | | | yes | | | | | | | | |
| KCNN4 | Island | Body | chr19:44278274-44278628 | 354 | | | | | | | | | 1 | | | | | | yes | | | | yes | | | |
| KCNN4 | NA | 1stExon;5'UTR;TSS200;TSS1500 | chr19:44285297-44286076 | 779 | | | | | | | | | 2 | | | | | yes | | | | | | yes | | |
| LYPD5 | S_Shelf | 5'UTR;TSS200;1stExon | chr19:44306565-44307056 | 491 | | | 7 | | | | | | 1 | | | | | | yes | | | yes | | yes | | |
| LYPD5 | Island | TSS200;5'UTR;1stExon | chr19:44324467-44325008 | 541 | | 1 | 4 | | | | | 0 | 10 | | | 1 | 1 | | yes | yes | yes | | | | | |
| ZNF283 | NA | TSS1500 | chr19:44330665-44331318 | 653 | | | 5 | | | | | | 1 | | | | | | yes | | yes | yes | | | | |
| ZNF233 | Island | TSS200 | chr19:44763979-44764603 | 624 | | | 2 | | | | | 0 | 1 | | | 1 | | yes | | | yes | | | | | |
| ZFP112 | Island | 5'UTR | chr19:44860658-44861154 | 496 | | | | | | | | | 1 | | | | | yes | | | | yes | | | | |
| ZNF285A | NA | 5'UTR | chr19:44905343-44905980 | 637 | | 1 | | | | | | | 1 | | | | | yes | | yes | | | | | | |
| ZNF229 | Island | TSS200;5'UTR;1stExon | chr19:44952417-44952808 | 391 | | 1 | 1 | | 1 | | | 0 | 3 | | | | | | | | | | | | | |
| CEACAM20 | NA | Body;TSS200 | chr19:45033454-45033943 | 489 | | | 4 | | | | | | 2 | | | | | yes | yes | | | yes | | | | |
| PVR | Island | Body | chr19:45150513-45150725 | 212 | | | | | | | | | 1 | | | | | yes | yes | | | | | | | |
| CEACAM19 | NA | TSS200 | chr19:45174671-45175197 | 526 | | | | | | | | | 1 | | | | | yes | yes | | yes | yes | | | | |
| CBLC | N_Shore | TSS1500;TSS200 | chr19:45279765-45281284 | 1519 | | | | | | | | 0 | 1 | | | | | yes | | | | yes | yes | yes | | |
| PVRL2 | N_Shore | TSS200 | chr19:45349207-45349558 | 351 | | | | | | | | | 1 | | | | | yes | | | | yes | yes | | | |
| APOE | N_Shelf | TSS1500 | chr19:45407868-45408121 | 253 | | | 1 | 1 | | | | | | | | | | yes | | | | | | | | |
| APOC1 | NA | TSS1500;TSS200 | chr19:45417668-45418020 | 352 | | | 2 | | | | | | | | | 1 | | | | yes | | | | | | |
| APOC1P1 | NA | TSS1500 | chr19:45429771-45430113 | 342 | | | | | | | | | | | | | | yes | | | | | | | | |
| APOC4 | NA | TSS200 | chr19:45445437-45445693 | 256 | | | | | | | | | 1 | | | | | yes | | | | yes | | | | |
| APOC2;APOC4 | NA | TSS1500;3'UTR;TSS200 | chr19:45448680-45449301 | 621 | | | | | | | | | 2 | | | | | yes | | | | yes | | yes | | |
| ZNF296 | Island | TSS1500 | chr19:45579986-45580221 | 235 | | | 2 | | | | | | | | | | | | yes | | | yes | | | | |
| GEMIN7 | Island | 3'UTR | chr19:45594410-45594670 | 260 | | | 2 | | | | | 0 | 1 | | | 1 | | | | | | | | | | |
| NKPD1 | NA | TSS200;TSS1500 | chr19:45663415-45664665 | 1250 | | | 2 | | | | | | 2 | | | | | yes | | | | | | | | |

| | | | | | | | | | | | | | | | | | | | | | | | | |
|---|---|---|---|---|---|---|---|---|---|---|---|---|---|---|---|---|---|---|---|---|---|---|---|---|
| EXOC3L2 | Island | Body | chr19:45720771-45721076 | 305 | | | 1 | | | | | 0 | 2 | | | 6 | | yes | | | | | yes | |
| EXOC3L2 | Island;N_Shore | TSS200;TSS1500;5'UTR | chr19:45737011-45738115 | 1104 | | | 10 | 1 | | | | 0 | 12 | 1 | | 7 | | yes | | | | | yes | |
| KLC3 | N_Shore | TSS1500 | chr19:45843694-45843871 | 177 | | | 2 | | | | | | 1 | | | | | yes | yes | | | | | |
| PPP1R13L | Island | Body | chr19:45885800-45885940 | 140 | | | | | | | | | 1 | | | | | yes | | | | | yes | |
| ERCC1 | S_Shore | TSS1500 | chr19:45927563-45927594 | 31 | | | | | | | | | 1 | | | | | | yes | | | | | |
| FOSB | Island;N_Shore | Body | chr19:45975694-45976195 | 501 | | | 6 | | | | | | | | | | | yes | | | | | yes | |
| RTN2 | N_Shore | Body;TSS200 | chr19:45996372-45996901 | 529 | | 1 | 2 | | | | | | | | | | | yes | | | | | | |
| RTN2;FLJ40125 | N_Shore | TSS1500;TSS200 | chr19:46001605-46001962 | 357 | | | 1 | | | | | | | | | | | yes | | | | | yes | |
| OPA3 | Island | Body | chr19:46032431-46032854 | 423 | | | | | | | | 0 | | 1 | | | | yes | | | | | | |
| OPA3 | N_Shore | 3'UTR;Body | chr19:46056723-46057298 | 575 | | | | | 1 | | | | 3 | 1 | | | | yes | | | yes | | | |
| MIR330;EML2 | N_Shelf | TSS1500;1stExon;5'UTR;TSS200;Bo | chr19:46142212-46142914 | 702 | 7 | | 2 | | | | | | | | | | | yes | | | | | yes | |
| GIPR | N_Shelf | TSS1500 | chr19:46170937-46171262 | 325 | | | | | | | | | | | | | | yes | yes | | | yes | | |
| GIPR | Island | Body | chr19:46181221-46181330 | 109 | | | 3 | | | | | | 1 | | | | | yes | yes | | | | | |
| SIX5 | Island | Body | chr19:46270244-46270700 | 456 | | | | | | | | | 1 | 1 | 1 | | | | | | | | | |
| DMPK | N_Shelf | TSS1500 | chr19:46286064-46286475 | 411 | | | | | | | | | 1 | | | | | yes | | | | yes | | |
| RSPH6A | Island | Body | chr19:46307734-46308199 | 465 | | 1 | | | | | | | | | | | | yes | | | | yes | yes | |
| RSPH6A;SYMPK | N_Shore;Island | 1stExon;5'UTR;TSS200;3'UTR;Body | chr19:46318439-46319398 | 959 | | | 18 | 1 | | | | | 4 | | | | | | | | | yes | | |
| IRF2BP1 | Island | 1stExon | chr19:46387760-46387992 | 232 | | | | | | | | | 1 | | | | | | | | | | | |
| NANOS2 | N_Shore;Island | 3'UTR;1stExon | chr19:46417255-46417734 | 479 | | | 7 | | | | | 0 | 1 | | | | | yes | | | | | | |
| NANOS2 | S_Shore | TSS1500 | chr19:46418849-46419054 | 205 | | | 1 | | | | | | | | | | | yes | | | | | | |
| NOVA2 | N_Shore;Island | Body | chr19:46443657-46444050 | 393 | | 1 | 5 | | | | | 0 | 2 | | | 4 | | yes | | | | yes | yes | |
| CCDC61 | N_Shore | Body | chr19:46503907-46504412 | 505 | | 1 | | | | | | 0 | | | | | | yes | | | | | | |
| PGLYRP1 | Island;S_Shore | 1stExon;5'UTR;TSS200 | chr19:46526321-46527410 | 1089 | | | 16 | | | | | | 1 | | | | | yes | yes | | | | yes | |
| IGFL4 | NA | 1stExon;5'UTR;TSS1500 | chr19:46544240-46544802 | 562 | | | 2 | | | | | | 1 | | | | | yes | | | | | | |
| IGFL3 | NA | 1stExon | chr19:46627896-46628367 | 471 | | | 2 | | | | | | | | | | | | | | | yes | | |
| DKFZp434J0226 | NA | TSS200 | chr19:46713446-46713976 | 530 | | | 1 | | | | | 0 | 1 | | 1 | | | | yes | yes | | | | |
| HIF3A | Island | TSS1500 | chr19:46800054-46800198 | 144 | | | | | | | | | 2 | | | | | | | | | | | |
| HIF3A | S_Shore | Body;TSS200 | chr19:46801557-46801672 | 115 | | | 1 | | | | | 0 | | | | | | | | | | | | |
| HIF3A | N_Shore;Island | 1stExon;5'UTR;Body | chr19:46806907-46807466 | 559 | | | 3 | | | | | | 2 | 1 | | | | yes | | | | yes | | |
| CCDC8 | Island | 1stExon | chr19:46915351-46915776 | 425 | | | | | | | | | 1 | | | 1 | | | | yes | | | | |
| CCDC8 | S_Shore | TSS200 | chr19:46917018-46917255 | 237 | | | | | | | | | | | | | | yes | | | | | | |
| PNMAL1 | Island | TSS200;TSS1500 | chr19:46974972-46975180 | 208 | | | 2 | | | | | 0 | 1 | | | | | | | yes | | | yes | |
| PNMAL2 | Island | 1stExon;3'UTR | chr19:46996347-46996868 | 521 | 1 | | | 1 | 3 | | | | | | | | | | | | | | yes | |
| PNMAL2 | Island;S_Shore | 1stExon;5'UTR;TSS200;TSS1500 | chr19:46998383-46999840 | 1457 | | | 20 | | | | | 0 | 1 | | | 1 | | yes | | yes | | | | |
| PTGIR | Island | Body | chr19:47127041-47127365 | 324 | | | 1 | | | | | | | | | | | yes | | | | | | |
| PTGIR | S_Shore | TSS1500 | chr19:47129337-47129637 | 300 | | | 7 | | | | | | | | | | | | | | | | | |
| GNG8 | Island | TSS1500 | chr19:47139338-47139496 | 158 | | | | | | | | 0 | 1 | | | | | | | | | yes | yes | |
| SLC1A5 | N_Shelf | Body;5'UTR;1stExon | chr19:47287778-47288263 | 485 | | | 21 | 1 | | | | | | | | | | yes | | | | | | |
| AP2S1 | S_Shore | TSS1500 | chr19:47354548-47354685 | 137 | | | | | | | | | | | | | | yes | | | | | | |
| GRLF1 | Island | 3'UTR | chr19:47507399-47507691 | 292 | | | 7 | | | | | 0 | 3 | 2 | 1 | 10 | | | | | | yes | | |
| BBC3 | S_Shore | 1stExon;5'UTR;TSS1500 | chr19:47735882-47736018 | 136 | | | | 1 | | | | | | | | | | yes | | | | | | |
| PRR24 | Island | TSS1500 | chr19:47777641-47778150 | 509 | | | | | | | | | 1 | | | | | | | | | | yes | |
| C5AR1 | NA | TSS1500 | chr19:47812519-47813130 | 611 | | | | | | | | | | | | | | yes | yes | | | | yes | |
| GPR77 | NA | TSS1500 | chr19:47840110-47840366 | 256 | | | | | | | | | | | | | | yes | | | | yes | | |
| MEIS3 | Island | Body | chr19:47910108-47910580 | 472 | | | | | | | | | | | | | | yes | | | | | yes | |
| SLC8A2 | Island | Body | chr19:47951302-47951565 | 263 | | | | | | | | | 1 | | | | | | yes | | yes | | | |
| SLC8A2 | Island | Body | chr19:47969156-47969574 | 418 | 2 | | 1 | | | | | | | | | | | yes | yes | | yes | | yes | |
| SLC8A2 | NA | 5'UTR;1stExon | chr19:47975078-47975487 | 409 | | | | | | | | | 1 | | | | | yes | | | | yes | yes | |
| EHD2 | Island | Body | chr19:48219924-48220334 | 410 | | 1 | | | | | | | | | | | | yes | | | | | | |
| EHD2 | Island | Body | chr19:48229109-48229504 | 395 | | | | | | | | | 1 | | | | | yes | | | | | yes | |
| ELSPBP1 | NA | TSS200 | chr19:48497737-48497828 | 91 | | | | | | | | | | | | | | yes | | | yes | | | |
| CABP5 | NA | 1stExon;TSS200 | chr19:48547158-48547989 | 831 | | | | | | | | 0 | 1 | | | 2 | | yes | | | | yes | yes | |
| LIG1 | N_Shore | TSS1500 | chr19:48674931-48675305 | 374 | | | 9 | | | | | | | | | | | yes | | | | | | |
| EMP3 | Island;chr19:48833394-488337 | Body;Island | chr19:48833062-48833729 | 667 | | | | | | | | | | | 1 | | | | yes | | | yes | | |
| TMEM143 | Island | Body | chr19:48837349-48837598 | 249 | | | | 1 | | | | | | | | | | yes | | | | yes | | |
| KDELR1 | N_Shore | Body | chr19:48893974-48895030 | 1056 | | | | | | | | | | | | 1 | | | | | | yes | | yes |
| GRIN2D;GRWD1 | Island | 3'UTR;TSS1500 | chr19:48947560-48948005 | 445 | | | 9 | | | | | 0 | 1 | | | | | | | yes | | | | |
| KCNJ14 | N_Shore;Island | 5'UTR;1stExon;Body | chr19:48964911-48965833 | 922 | | | 3 | | | | | | | | | | | yes | | | | | | |
| CYTH2 | N_Shore;Island | 3'UTR | chr19:48983508-48984318 | 810 | | 1 | 3 | | 1 | | | | 1 | | | | | yes | yes | | | | | |
| LMTK3 | Island | Body | chr19:49000743-49002477 | 1734 | | | 2 | | | | | 0 | 1 | 1 | | 2 | | yes | | | yes | yes | yes | |
| LMTK3 | Island | Body | chr19:49004523-49004834 | 311 | | | | | | | | 0 | | | | | | yes | | | | | yes | |
| LMTK3 | N_Shore | TSS200 | chr19:49016564-49016629 | 65 | | | 3 | | | | | 0 | | | | | | yes | | | | yes | | |
| SULT2B1 | NA | TSS1500;TSS200;1stExon;5'UTR | chr19:49054908-49055443 | 535 | | 1 | 13 | 1 | | | | | 2 | | | 1 | | yes | yes | | | yes | | |
| SULT2B1 | Island | Body | chr19:49061546-49061702 | 156 | | | | | | | | 0 | 1 | | | | | yes | | | | | | |
| SULT2B1 | NA | Body;TSS1500 | chr19:49077983-49078119 | 136 | | | | | 1 | | | | | | | | | yes | | | | yes | | |
| FAM83E | S_Shore | TSS200 | chr19:49116824-49116870 | 46 | | | | | | | | | 1 | | | | | | | | | | | |
| SPHK2 | Island | 3'UTR | chr19:49133421-49134105 | 684 | | 1 | 12 | 1 | 1 | | | 0 | 1 | 1 | | | | | | | | | | |
| FUT2 | N_Shore | TSS200 | chr19:49199172-49199234 | 62 | | | | | | | | | | | | | | yes | | | | | yes | |
| MAMSTR | N_Shelf | 1stExon;Body;5'UTR | chr19:49220102-49220235 | 133 | | | | | | 1 | | | | | | | | yes | | | | | | |
| MAMSTR;RASIP1 | N_Shore;Island | 5'UTR;TSS1500;3'UTR | chr19:49222487-49224454 | 1967 | | | | | | | | 0 | 7 | 1 | | 1 | | yes | | | | | | yes |
| RASIP1 | Island | Body | chr19:49232187-49232824 | 637 | | | 5 | | | | | | 1 | | | | | yes | | | | | | |
| RASIP1 | N_Shore | Body | chr19:49238364-49238653 | 289 | | | 1 | | | | | | 1 | | | | | yes | yes | | | | | |
| RASIP1;IZUMO1 | S_Shore | TSS200;Body;TSS1500 | chr19:49244004-49244674 | 670 | | | | | | | | 0 | 3 | | | | | | | | | | | yes |
| IZUMO1 | NA;N_Shelf | 5'UTR;1stExon | chr19:49249123-49250561 | 1438 | | 1 | | | 1 | | | | | | | | | yes | | | | | | |
| FGF21;FUT1 | S_Shelf | TSS1500;TSS200 | chr19:49258805-49259452 | 647 | | | 5 | 1 | | | | | | | | | | | | | | | | yes |
| PLEKHA4;HSD17B14 | Island;N_Shore | 3'UTR;TSS1500;Body;5'UTR;1stExo | chr19:49339776-49340765 | 989 | | | 18 | 1 | | | 1 | 0 | 4 | 2 | | 14 | 2 | | | | | | | yes |
| LHB | N_Shore | TSS200 | chr19:49520450-49520532 | 82 | | | | | | | | | | | | | | | | | | | | |
| CGB2;SNAR-G2 | NA | TSS200 | chr19:49535079-49535373 | 294 | | | | | | | | | | | | | | yes | | | | | | |
| CGB8 | NA | TSS200;1stExon;5'UTR | chr19:49552017-49552606 | 589 | | | | | | | | | 1 | | | | | | yes | | | | | |
| KCNA7 | Island;S_Shore | 1stExon;TSS1500 | chr19:49575475-49577206 | 1731 | | | | | | | | | 2 | | | | | yes | | | | yes | yes | |
| PPFIA3;C19orf73 | Island | 5'UTR;TSS1500;1stExon | chr19:49622715-49623089 | 374 | | | | | | | | | 1 | | | | | | | | | | yes | |
| PPFIA3 | Island | Body | chr19:49636270-49636594 | 324 | | | | | | | | 0 | | | | | | | | | | | | |
| HRC | N_Shelf | 1stExon;TSS200 | chr19:49658232-49659299 | 1067 | | | 2 | | | | | | 1 | | | | | | | | | yes | yes | |
| TRPM4 | Island | Body | chr19:49686020-49686254 | 234 | | | | | | | | | | | | | | yes | | | | yes | | |
| TRPM4 | N_Shore;Island | Body | chr19:49699496-49700021 | 525 | | | | | | | | 0 | 1 | | | | | yes | | | | yes | | |
| TRPM4 | N_Shore;Island | Body | chr19:49713323-49714044 | 721 | | | | 1 | | | | 0 | 4 | | | 7 | | yes | yes | | | | | |
| SLC6A16 | N_Shore;Island | 1stExon;5'UTR;TSS200 | chr19:49828347-49828787 | 440 | | | | | | | | | 5 | | | | | yes | | | yes | | | |
| CD37 | N_Shelf | TSS1500;TSS200 | chr19:49838101-49838777 | 676 | | | | | 1 | | | | 1 | | | | | yes | | | | yes | yes | |
| CD37 | Island | Body | chr19:49842655-49843160 | 505 | | | | | 1 | | | | | | | | | | | | | | | |
| CCDC155 | Island | TSS1500;TSS200 | chr19:49891270-49891603 | 333 | | | 18 | | | | | 0 | 3 | 1 | | 5 | | yes | yes | | | yes | | |
| PTH2 | S_Shore | TSS1500 | chr19:49927076-49927309 | 233 | | | | | | | | | | | | | | yes | | | | | yes | |
| SLC17A7 | Island | Body | chr19:49935823-49936274 | 451 | | | 8 | | | | | | | | 1 | | | yes | | | | | | |
| SLC17A7 | N_Shore;Island | Body | chr19:49939549-49940062 | 513 | | | 4 | | | | | | | | | | | yes | | | | | | yes |
| RPL13AP5;SNORD33;SNORD35A;SNORD | S_Shore;S_Shelf | Body;TSS1500;TSS200 | chr19:49993157-49994239 | 1082 | | | | | | | | | 2 | | | | | yes | | | | | | |
| RPS11;SNORD35B | S_Shore | Body;TSS1500 | chr19:50000586-50001058 | 472 | | | | | | | | | | | | | | | | | yes | | | |
| MIR150 | NA | TSS200 | chr19:50004152-50005045 | 893 | | | 8 | 1 | 2 | | | | | | | | | yes | yes | | | | | |
| FCGRT | N_Shore | TSS200;TSS1500 | chr19:50015523-50015780 | 257 | | | | 1 | 2 | | | 0 | 3 | 2 | | | | yes | yes | | | | | yes |
| RCN3 | Island | 1stExon;5'UTR | chr19:50031017-50031395 | 378 | | 1 | | 1 | 1 | | | | 1 | | | | | yes | | | | yes | | |
| RCN3 | N_Shore | Body | chr19:50037018-50037880 | 862 | | | | 1 | 1 | | | | | | | | | | | | | | | |

| | | | | | | | | | | | | | | | | | | | | | | | | | | |
|---|---|---|---|---|---|---|---|---|---|---|---|---|---|---|---|---|---|---|---|---|---|---|---|---|---|---|
| NOSIP | Island | Body | chr19:50059750-50060174 | 424 | | | | | 1 | | | | | | | | | yes | | | | | yes | | | |
| PRRG2;NOSIP | S_Shore | TSS1500 | chr19:50084283-50084626 | 343 | | | | | | | | | 1 | | | | | yes | | | | | | | | |
| PRR12 | Island | Body | chr19:50096610-50096876 | 266 | | | | | | | | | | 1 | | | | | | | | | | | | |
| C19orf76;PRMT1;CPT1C | Island;N_Shore | TSS1500;Body;1stExon;5'UTR;3'UT | chr19:50190825-50194252 | 3427 | | | 2 | | | | | 0 | 5 | | | 1 | | yes | yes | | yes | | | | | |
| TSKS | N_Shore;Island | Body | chr19:50249464-50249927 | 463 | | 1 | 19 | | | | 1 | 0 | 4 | 1 | 1 | 17 | | yes | | | | | yes | | | |
| FUZ | Island | Body | chr19:50312505-50313003 | 498 | | | | | | | | 0 | 1 | 1 | | 2 | | yes | | | | | | | | |
| TBC1D17 | N_Shore | Body | chr19:50391064-50391885 | 821 | | 1 | | | | | | | 2 | 1 | | | | yes | yes | | | | | | | |
| IL4I1 | NA | TSS1500;Body | chr19:50400714-50400885 | 171 | | | | 1 | | | | | | | | | | yes | | | | | | | | |
| SIGLEC11 | NA | TSS200;TSS1500 | chr19:50464453-50464887 | 434 | | | 3 | | | | | | | | | | | yes | | yes | | | | | | |
| FLJ26850 | N_Shore;Island | TSS1500;TSS200;Body | chr19:50553682-50554510 | 828 | | | | | | | | 0 | | | | | | yes | | yes | | | | | | |
| SNAR-B1;SNAR-B2;SNAR-D | NA | TSS1500;Body | chr19:50643548-50643731 | 183 | | | | | | | | | | | | | | yes | | | yes | | yes | | | yes |
| C19orf41 | Island | 1stExon;5'UTR | chr19:50666238-50666552 | 314 | | | 5 | | | | | 0 | 5 | 1 | | 1 | | yes | yes | | | | | | | |
| MYH14 | Island | TSS1500 | chr19:50706577-50707172 | 595 | | | | | | | | 0 | 1 | | | | | yes | | | | | yes | | | |
| MYH14 | Island | Body | chr19:50713660-50714060 | 400 | | | | | | | | | 1 | | | | | yes | | | yes | | | | | |
| MYH14 | Island | Body | chr19:50733985-50734287 | 302 | | | | | | | | 0 | 1 | | | 1 | | yes | | | | | | | | |
| MYH14 | N_Shore | Body | chr19:50762277-50762584 | 307 | | | | | | | | | | | | | | yes | yes | | | | | | | |
| MYH14 | N_Shore;Island | Body | chr19:50789296-50789973 | 677 | | | 2 | | | | | | | | | | | yes | yes | | | yes | | | | |
| KCNC3 | Island | 1stExon | chr19:50831901-50832037 | 136 | | | | | | | | | | | | | | | | | | yes | yes | | | |
| NAPSB | NA | TSS200 | chr19:50848024-50848840 | 816 | | | | | | | | | 2 | | | | | yes | yes | | | yes | | | | |
| NAPSA | Island | 3'UTR;Body | chr19:50861774-50862121 | 347 | | | 12 | | | | | 0 | 3 | 1 | | 3 | | yes | | | | | | | | |
| NAPSA | NA | 5'UTR;1stExon;TSS200 | chr19:50868879-50869332 | 453 | | | | | | | | | 1 | | | | | yes | yes | | | yes | | | | |
| POLD1;SPIB | NA | 3'UTR;TSS1500 | chr19:50921232-50921997 | 765 | | | | | | | | | 2 | | | | | | | | | | | | | |
| SPIB | N_Shore;Island | Body | chr19:50931222-50931875 | 653 | | | 17 | | | 1 | | 0 | 6 | 3 | | 7 | | yes | | | | | | | | |
| ASPDH | N_Shelf | 5'UTR;1stExon;Body;TSS200;TSS1 | chr19:51017123-51018365 | 1242 | | | 12 | | | | | 0 | 2 | | | 2 | | | | | | yes | | | | |
| LRRC4B | Island | Body | chr19:51051861-51052080 | 219 | | | | | | | | | 1 | | | | | yes | | | | yes | | | | |
| LRRC4B | S_Shore | 1stExon;5'UTR | chr19:51071294-51071328 | 34 | | | | | | | | | 1 | | | | | | | | | yes | yes | | | |
| SNAR-F | Island | TSS1500 | chr19:51107558-51107937 | 379 | | 1 | 2 | | | | | 0 | 1 | | | | | yes | | | | | | | | |
| SYT3 | Island | Body | chr19:51128576-51128770 | 194 | | | 1 | | | | | | | | | | | yes | | | | | yes | yes | | |
| SYT3 | N_Shore;Island | 5'UTR;1stExon;TSS1500 | chr19:51141227-51142613 | 1386 | | 1 | 3 | | | | | | | | | | | | yes | | | yes | | | | |
| SHANK1 | N_Shore;Island | 3'UTR;Body | chr19:51165207-51165845 | 638 | | | 6 | | | | | | 7 | | | | | yes | | | | yes | | | | |
| SHANK1 | Island | Body | chr19:51170816-51171247 | 431 | 1 | | | | | | | | | | | | | | | | | yes | yes | | | |
| SHANK1 | Island | Body | chr19:51189975-51190179 | 204 | | | | | | | | | | | | 1 | | | | | | yes | | | | |
| SHANK1 | Island | TSS200 | chr19:51220256-51220537 | 281 | | | | | | | | | | | | 1 | | | | | | | | | | |
| CLEC11A | N_Shore | TSS1500 | chr19:51225848-51226536 | 688 | | 1 | | | | | | 0 | 1 | | | | | | yes | | | | yes | | | |
| ACPT | N_Shore | TSS200 | chr19:51293620-51293677 | 57 | | | | | | | | | 1 | | | | | yes | | yes | | yes | | | | |
| KLK1 | N_Shelf | 1stExon | chr19:51326980-51327477 | 497 | | | | | | | | | 1 | | | | | | | | | | | | | |
| KLK15 | Island | Body | chr19:51330265-51330469 | 204 | | | 5 | | | | | 0 | 3 | | | 2 | | | | | | yes | | | | |
| KLK4 | Island | Body | chr19:51411737-51411940 | 203 | | | 6 | | | | | 0 | 2 | | | | | | | | yes | | yes | | | |
| KLK4 | N_Shore | TSS1500 | chr19:51415152-51415452 | 300 | | | 7 | | | | | | 3 | | | | | yes | yes | | | | yes | | | |
| KLK6 | NA | TSS1500;5'UTR;1stExon | chr19:51472725-51473823 | 1098 | | | 4 | | | | | | | | | | | yes | | | | | yes | yes | | |
| KLK7 | NA | 5'UTR;1stExon;TSS200;TSS1500 | chr19:51486901-51487968 | 1067 | | | 14 | | | | | 0 | 4 | | | 2 | | yes | yes | | | | | yes | | |
| KLK8;KLK9 | N_Shore;Island | TSS1500;3'UTR | chr19:51505507-51506362 | 855 | | | 5 | | | | | 0 | 2 | | | 3 | | | | | | | | yes | | |
| KLK9 | NA | TSS1500 | chr19:51513252-51513883 | 631 | | | | | | | | | 1 | | | | | yes | yes | | | yes | | | | |
| KLK10 | Island | 5'UTR | chr19:51522454-51522641 | 187 | | | | | 2 | | | | | | | | | yes | | | | | | | | |
| KLK10 | S_Shore | TSS1500 | chr19:51523736-51524828 | 1092 | | | 8 | | | | | 0 | 1 | | | 1 | | yes | | | yes | yes | | | | |
| KLK11 | NA | Body;5'UTR;TSS200 | chr19:51529902-51529999 | 97 | | | | | | | | | | | | | | | | | | yes | | | | |
| KLK11 | N_Shelf | TSS1500;TSS200 | chr19:51531297-51531484 | 187 | | | 3 | | | | | | | | | | | yes | | | yes | | | | | |
| KLK12 | Island | Body | chr19:51535339-51535405 | 66 | | | 2 | | | | | | | | | | | yes | yes | | | yes | | | | |
| KLK12 | S_Shelf | 1stExon;5'UTR;TSS200 | chr19:51538134-51538653 | 519 | | | | | | | | | 2 | | | | | yes | | | | | yes | | | |
| KLK13 | S_Shore | TSS1500 | chr19:51568863-51569106 | 243 | | 1 | | | | | | | | | | | | | | | | | yes | | | |
| KLK14 | NA | 5'UTR | chr19:51587102-51587409 | 307 | | | | | | | | | | | | | | | | | | yes | yes | | | |
| CTU1 | Island | 3'UTR | chr19:51601823-51602462 | 639 | | | | | | | | | 1 | | | | | | | | | | yes | yes | | |
| SIGLEC7 | NA | TSS1500 | chr19:51644168-51644625 | 457 | | | 1 | | | | | | | | | | | | | yes | | | | | | |
| SIGLECP3 | NA | TSS1500;TSS200 | chr19:51670251-51670552 | 301 | | | 7 | | | | | | 1 | | | | | | yes | | | yes | | | | |
| IGLON5 | N_Shore;Island | Body | chr19:51829984-51830960 | 976 | | | | | | | | 0 | 2 | | | | | | yes | | | | yes | | | |
| ETFB | Island | TSS200 | chr19:51869672-51869687 | 15 | | 1 | | | | | | | | | | | | yes | | | | | | | | |
| NKG7 | NA | TSS1500 | chr19:51876470-51876788 | 318 | | | | | | | | | | | | | | yes | | yes | | yes | | yes | | |
| SIGLEC10;LOC100129083 | NA | Body | chr19:51920435-51920976 | 541 | | | | | | | | | | | | | | yes | | | yes | yes | | | | |
| SIGLEC8 | NA | 1stExon;5'UTR | chr19:51961666-51961937 | 271 | | | | | | | | | | | | | | yes | yes | | | yes | | | | |
| CEACAM18 | NA | TSS1500 | chr19:51978789-51979244 | 455 | | | | | | | | | | | | | | yes | yes | yes | | | | | | |
| SIGLEC12 | NA | TSS200;TSS1500;nan;1stExon;5'UT | chr19:52004630-52005990 | 1360 | | | 11 | | 1 | | | | 2 | 2 | 1 | | | | yes | | yes | yes | | | | |
| SIGLEC6 | NA | TSS200 | chr19:52035161-52035287 | 126 | | | | | | | | | | | | | | | yes | | | yes | | | | |
| SIGLEC5 | NA | TSS1500 | chr19:52134289-52134393 | 104 | | | | | | | 1 | | 1 | 1 | 1 | | | yes | yes | | yes | | | | | |
| SIGLEC14 | NA | TSS200 | chr19:52150230-52150634 | 404 | | | | | | | | | | | | | | yes | yes | yes | | | | | | |
| NCRNA00085 | N_Shore;Island | Body | chr19:52206224-52206912 | 688 | | | 3 | 1 | | | | 0 | 2 | | | 2 | | yes | | | | | | | | |
| HAS1 | Island | Body | chr19:52222431-52223099 | 668 | | | | | 1 | | | | 1 | | | | | yes | | | | yes | yes | | | |
| HAS1 | NA | TSS200;TSS1500 | chr19:52227309-52228400 | 1091 | | | 6 | | | | | 0 | 1 | | | | | yes | | | | yes | yes | | | |
| FPR2 | NA | TSS1500;TSS200 | chr19:52263842-52264429 | 587 | | 1 | 12 | | | | | | 1 | | | | | yes | | yes | yes | | | | | |
| ZNF577 | N_Shore;Island;S_Shore | Body;5'UTR;1stExon;TSS200;TSS1 | chr19:52390810-52391789 | 979 | | 1 | | | | | | 0 | 13 | 1 | | 5 | | | yes | yes | | | | | | |
| ZNF432 | Island;S_Shore | TSS200;TSS1500 | chr19:52552120-52552461 | 341 | | | | | 1 | | | | 2 | | | | | | | yes | | | | | | |
| ZNF841 | Island | 5'UTR | chr19:52598653-52599256 | 603 | | | | | | | | | 1 | | | | | | yes | yes | | | | | | |
| ZNF836 | S_Shore | TSS1500 | chr19:52675203-52675470 | 267 | | | 2 | | | | | 0 | 1 | | | 2 | | yes | yes | yes | | | | | | |
| ZNF610 | Island | 5'UTR | chr19:52839924-52840042 | 118 | | | | | | | | | 1 | | | | | | | | | | | | | |
| ZNF880 | N_Shore | TSS1500 | chr19:52872916-52873426 | 510 | | 1 | | | | | | | | | | | | yes | | | yes | | yes | | | |
| ZNF528 | N_Shore;Island | TSS1500;TSS200;5'UTR;1stExon | chr19:52900469-52901639 | 1170 | | 2 | | | | | | 0 | 2 | | | 2 | | yes | | | | yes | yes | | | |
| ZNF578 | N_Shore;Island | TSS1500;TSS200 | chr19:52956622-52957409 | 787 | | 1 | | | | | | 0 | 3 | | | | | | | yes | | | | | | |
| ZNF578 | Island | 5'UTR | chr19:52996083-52996617 | 534 | | | 1 | | | | | 0 | 1 | | | 1 | | yes | | | | | | | | |
| ZNF808 | N_Shore | 5'UTR | chr19:53038972-53039444 | 472 | | 1 | | 1 | | | | 0 | 1 | | | | | yes | | | | | | | | |
| ZNF83 | S_Shore | Body;TSS1500 | chr19:53142106-53142304 | 198 | | | | | | | | 0 | 1 | | | | | yes | yes | | | | | | | |
| ZNF677 | N_Shore;Island;S_Shore | 5'UTR;TSS200 | chr19:53757677-53758609 | 932 | | 2 | | | | | | | 3 | | | | | yes | | | | | yes | | | |
| VN1R4 | NA | 1stExon | chr19:53770873-53771332 | 459 | | 1 | | | | | | | | | | | | yes | | | | yes | | | | |
| BIRC8 | Island;S_Shore | 5'UTR;1stExon | chr19:53794492-53794975 | 483 | | | | | | | | | 1 | | | | | yes | yes | | | yes | | | | |
| ZNF331 | Island | TSS1500;1stExon;5'UTR | chr19:54023893-54024834 | 941 | | | | | | | | | 2 | | | | | yes | | | | yes | | | | |
| ZNF331 | Island;N_Shore | TSS200;5'UTR;TSS1500 | chr19:54040774-54041856 | 1082 | | | | | | | | 0 | 1 | | | 1 | | | yes | | | | | | | |
| LOC284379 | NA | Body;TSS200 | chr19:54106593-54107634 | 1041 | | | 3 | | | | 2 | | 2 | 2 | | 2 | | yes | yes | | yes | yes | yes | | | |
| DPRX | NA | TSS1500;TSS200 | chr19:54134711-54135342 | 631 | | | 6 | | | | | | | | | | | yes | | | yes | | | | | |
| MIR498 | NA | TSS1500;TSS200 | chr19:54177144-54177678 | 534 | | | | | | | | | 1 | | | | | yes | yes | | | yes | | | | |
| MIR526B | NA | TSS1500 | chr19:54196849-54197391 | 542 | | | | | | | | | | | | | | yes | yes | | | yes | | | | |
| MIR520B | NA | TSS200 | chr19:54204403-54204773 | 370 | | | | | | | | | | | | | | yes | | | | | | | | |
| MIR519D;MIR517A | NA | TSS1500;TSS200 | chr19:54215487-54215577 | 90 | | | | | | | | | | | | | | yes | | | | | | | | |
| MIR521-2 | NA | TSS1500 | chr19:54219381-54219607 | 226 | | | 2 | | | | | | | | | | | yes | yes | | | | | | | |
| MIR517B;MIR520G | NA | Body;TSS1500 | chr19:54224374-54224556 | 182 | | | 2 | | | | | | | | | | | yes | | | | | | | | |
| MIR518A2 | NA | TSS1500 | chr19:54242124-54242644 | 520 | | | | | | | | | | | | | | yes | | | | | | | | |
| MIR517C;MIR520H | NA | TSS200;TSS1500 | chr19:54244371-54244435 | 64 | | | | | | | | | | | | | | yes | | | | | | | | |
| MIR522 | NA | TSS1500 | chr19:54253831-54254493 | 662 | | | 2 | | | | | | | | | | | yes | yes | | | yes | | | | |
| MIR372;MIR371;MIR373 | NA | TSS1500;TSS200 | chr19:54290011-54291017 | 1006 | | | 8 | | | | | | 1 | | | | | yes | | | | | | | | |
| NLRP12 | NA | Body;TSS1500 | chr19:54312855-54313004 | 149 | | | | | | | | | 1 | | | 1 | | yes | | | | | | | | |
| NLRP12 | NA | 1stExon;5'UTR;TSS200 | chr19:54327404-54327995 | 591 | | | | | 1 | | | | 2 | 2 | 1 | | | yes | | | | | | | | |
| MYADM | N_Shore;Island | TSS1500;TSS200 | chr19:54369133-54370839 | 1706 | | | | | | | | | | | | | | | | | | | | | | |

| | | | | | | | | | | | | | | | | | | | | | | | | |
|---|---|---|---|---|---|---|---|---|---|---|---|---|---|---|---|---|---|---|---|---|---|---|---|---|
| MYADM | Island | TSS200;5'UTR | chr19:54372625-54372927 | 302 | | | | | | | | | 2 | | | | | | | | | | | yes |
| PRKCG | Island | Body | chr19:54393040-54393394 | 354 | | | | | | | | | 1 | | | | | yes | yes | | | | | |
| CACNG7 | S_Shelf | TSS1500;TSS200 | chr19:54415736-54416165 | 429 | | | 11 | | | | | 0 | 1 | | | | | yes | | | | | | |
| CACNG8 | N_Shore | 1stExon;5'UTR | chr19:54466297-54466538 | 241 | | | | | | | | | 1 | | | | | | | | | yes | yes | |
| CACNG8 | Island | Body | chr19:54481467-54481859 | 392 | | | | | | | | | 1 | | | | | yes | | | | yes | yes | |
| CACNG8 | Island | Body | chr19:54483251-54483767 | 516 | | | | | | | | | 1 | | | | | | | | | yes | yes | |
| CACNG8;MIR935 | N_Shore;Island | Body;TSS1500;TSS200 | chr19:54484823-54486058 | 1235 | | 1 | 1 | | 1 | | | 0 | 3 | | | | | | | | | yes | yes | |
| CACNG6 | N_Shore;Island | TSS1500;TSS200;1stExon;5'UTR | chr19:54495183-54496611 | 1428 | | | 13 | | | | | 0 | 2 | | | 1 | | | | | | | yes | |
| CACNG6 | Island | Body | chr19:54515169-54515341 | 172 | | | 3 | | | | | 0 | 1 | 1 | | 2 | | yes | yes | | | | | |
| VSTM1 | NA | Body | chr19:54566838-54567279 | 441 | | | | | | | | 0 | 1 | 1 | 1 | 1 | | yes | | | | | | |
| TMC4 | Island | Body | chr19:54668230-54668565 | 335 | | 1 | | | | | | 0 | | | | | | yes | | | | | | |
| MBOAT7;TMC4 | NA | 3'UTR;TSS1500 | chr19:54677144-54677232 | 88 | | | | | | | | | | | | | | | | | | | | |
| LILRB3 | NA | 1stExon;5'UTR | chr19:54726870-54727203 | 333 | | | 3 | | | | | 0 | | | | | | yes | | | | | | |
| LILRB5 | NA | 5'UTR;1stExon | chr19:54761090-54761178 | 88 | | | | | | | | 0 | | | | | | | | | | | | |
| LILRB2 | NA | TSS200 | chr19:54785076-54785518 | 442 | | | | | | | | | | | | | | yes | | yes | | | | |
| LILRA4 | NA | 1stExon;5'UTR;Body | chr19:54849738-54850645 | 907 | | | 2 | | | | | 0 | | | | | | yes | | | | | | |
| LAIR1 | NA | TSS200;5'UTR;1stExon | chr19:54876446-54876795 | 349 | | | 22 | 1 | 2 | | | | 1 | 1 | | | | yes | yes | | | | | |
| LAIR1 | NA | TSS1500 | chr19:54877827-54878038 | 211 | | | | | | | | | 1 | | | | | yes | yes | yes | | | | |
| TTYH1 | N_Shore | TSS1500;Body | chr19:54926431-54927717 | 1286 | | 1 | | | | | | 0 | | | | | | | | | | yes | | yes |
| LENG9;CDC42EP5 | Island | TSS1500;3'UTR | chr19:54976211-54976506 | 295 | | | 9 | | | | | | | | 1 | | | yes | | | | | yes | |
| CDC42EP5 | Island | 5'UTR | chr19:54982500-54982731 | 231 | | | | | 1 | | | | | | | | | yes | | | | | | yes |
| CDC42EP5 | S_Shore | 5'UTR;1stExon | chr19:54984272-54984708 | 436 | | | | 1 | 1 | | | | | | | | | | | | | yes | | yes |
| LAIR2 | NA | TSS1500;TSS200 | chr19:55013549-55014357 | 808 | | | 6 | | | | | 0 | 3 | 1 | | 2 | 1 | yes | yes | | | | | |
| LILRB1 | NA | TSS200;5'UTR;TSS1500 | chr19:55141366-55142181 | 815 | | | 3 | | | | | | | | | | | yes | | yes | | | | |
| LILRB4 | NA | TSS1500;TSS200 | chr19:55172736-55174624 | 1888 | | | | | | | | | | | | | | | | | | | | |
| KIR2DL4 | NA | TSS1500;TSS200 | chr19:55314050-55315083 | 1033 | | | | | | | | | | | | | | | yes | | | | | |
| KIR3DL1 | NA | TSS1500 | chr19:55326840-55327711 | 871 | | | | | | | | | | | | | | yes | yes | | yes | | yes | |
| KIR3DL2 | NA | TSS1500 | chr19:55361685-55361904 | 219 | | | | | | | | | | | | | | yes | yes | | | | | |
| FCAR | NA | TSS200 | chr19:55385405-55385587 | 182 | | | | | | | | | 1 | | | | | yes | yes | | | | | |
| NCR1 | NA | TSS1500;TSS200 | chr19:55416928-55417647 | 719 | | | 7 | | | | 1 | 0 | 1 | 3 | | | 1 | | yes | yes | | yes | | |
| NLRP7 | NA | TSS200;TSS1500 | chr19:55458886-55459282 | 396 | | | 7 | | | | | | | | | | | yes | | | yes | | yes | |
| NLRP2 | NA | TSS1500;TSS200 | chr19:55476665-55478588 | 1923 | | 4 | | 1 | | | | 0 | 6 | 4 | | 8 | 1 | yes | | | yes | | | |
| NLRP2 | N_Shore | Body | chr19:55493672-55494075 | 403 | | | | | | | | | | | | | | yes | | | | | | |
| GP6 | NA | TSS200;1stExon | chr19:55549590-55549842 | 252 | | | 3 | | | | | 0 | 7 | | | 9 | | yes | yes | | | | | |
| EPS8L1 | N_Shore | TSS1500;Body | chr19:55591148-55591437 | 289 | | | 4 | | | | | | 1 | | | | | yes | yes | | | yes | | |
| EPS8L1 | Island | Body | chr19:55598443-55599320 | 877 | | | | 2 | | | | 0 | 2 | | | | | yes | yes | | | | | |
| TNNT1 | NA | 1stExon;5'UTR;TSS200 | chr19:55660514-55660625 | 111 | | | | | | | 1 | 0 | 3 | 2 | 1 | 7 | 1 | | | | | yes | | |
| TNNI3 | N_Shore;Island;S_Shore | Body | chr19:55667446-55669189 | 1743 | | 1 | 1 | | | | 1 | 0 | 5 | 1 | 1 | 3 | 2 | | | | | | | yes |
| SYT5 | N_Shore | 3'UTR | chr19:55684640-55684939 | 299 | | | | | | | | 0 | | | | | | yes | | | | | yes | |
| SYT5 | S_Shore | TSS1500 | chr19:55692397-55692711 | 314 | | | | | | | | | | | | | | yes | | | | | | |
| PTPRH | NA | 1stExon;5'UTR | chr19:55720817-55721290 | 473 | | 1 | | | | | | | | | | | | yes | yes | | | | | yes |
| HSPBP1 | Island | Body | chr19:55790889-55791101 | 212 | | | | | | | | 0 | | | | | | yes | | | | | | |
| COX6B2 | N_Shore;Island | 3'UTR | chr19:55863937-55864365 | 428 | | | | | | | | | | | | | | yes | | | | | | |
| FAM71E2 | S_Shore | 5'UTR;1stExon | chr19:55874586-55874851 | 265 | | | | | | | | | | | | | | yes | | | | yes | | |
| IL11 | S_Shore | TSS1500 | chr19:55882154-55882550 | 396 | | | | | | | | | | | | | | yes | | | | yes | yes | |
| TMEM190 | N_Shore;Island | TSS1500;TSS200;Body | chr19:55887704-55889611 | 1907 | | 1 | 7 | | | | 1 | 0 | 4 | 1 | | 2 | | yes | | yes | | | yes | yes |
| SHISA7 | Island | Body | chr19:55944864-55945233 | 369 | | | 5 | | | | | | | | | | | | | | yes | yes | | |
| ISOC2 | N_Shore;Island | 5'UTR;1stExon;TSS200;TSS1500 | chr19:55972646-55973778 | 1132 | | 1 | 7 | | 1 | | | 0 | 6 | 1 | 3 | 9 | 1 | yes | yes | | | | | |
| ZNF628 | N_Shore | TSS1500 | chr19:55987366-55987415 | 49 | | | 4 | | | | | | 1 | | | | | yes | | | yes | | | |
| NAT14;SSC5D | Island;S_Shore | Body;3'UTR;TSS1500 | chr19:55997780-55998931 | 1151 | | | 12 | | | | | | 1 | | | | | | | | | yes | | |
| SSC5D | Island | Body | chr19:56002240-56002446 | 206 | | | | | | | | | 1 | | | 1 | | yes | | | | | | |
| SSC5D | N_Shore | Body | chr19:56004925-56005437 | 512 | | | | | | | | | 1 | | | | | yes | | | yes | | | |
| SSC5D | Island | Body | chr19:56028687-56028883 | 196 | | | 5 | | | | | | | | | | | | | | | yes | yes | |
| SBK2 | Island | Body | chr19:56042469-56042579 | 110 | | | 1 | | | | | | | | | | | yes | | | | | | |
| SBK2 | N_Shore | Body | chr19:56047224-56047543 | 319 | | | | | | | | | | | | | | yes | yes | | | | | yes |
| ZNF784 | Island;S_Shore | 1stExon;TSS200 | chr19:56135916-56136215 | 299 | | | | | | | | | 3 | 1 | | 1 | | yes | | | | yes | | |
| CCDC106 | N_Shore | TSS1500;TSS200;5'UTR;1stExon | chr19:56158714-56159485 | 771 | | | | | | | | 0 | 3 | | | | | yes | yes | | | | yes | |
| EPN1 | S_Shore | 5'UTR;TSS1500 | chr19:56187475-56187915 | 440 | | | | | | | | | | | | | | yes | | | yes | yes | | |
| NLRP9 | NA | 1stExon;5'UTR | chr19:56249743-56250127 | 384 | | | | | | | | | | | | | | yes | yes | | | | | |
| NLRP11;NLRP4 | NA | 5'UTR;TSS200;1stExon | chr19:56347771-56348216 | 445 | | | 4 | | | | | | | | | | | yes | yes | | | | | |
| NLRP8 | NA | TSS1500;TSS200 | chr19:56458926-56459137 | 211 | | | | | | | | | 1 | | | | | | yes | | | | | |
| NLRP5 | NA | TSS1500 | chr19:56510833-56510965 | 132 | | | | | | | | | | | | | | yes | yes | | | | | |
| GALP | NA | TSS1500;TSS200 | chr19:56686707-56687388 | 681 | | | 2 | | | | | | | | | 1 | | yes | yes | | yes | | yes | |
| ZSCAN5A | Island | 5'UTR | chr19:56739573-56739749 | 176 | | | | | | | | | | | | | | yes | | | | | yes | |
| ZNF542 | N_Shore;Island | TSS1500;TSS200;Body | chr19:56879369-56879933 | 564 | | 1 | | | 1 | | | | 2 | | | 1 | | yes | | yes | | | | |
| ZNF582 | Island;chr19:56904636-569053 | TSS200;Island | chr19:56904901-56905383 | 482 | | 1 | | | | | | | | | 1 | 1 | | | | | | | | |
| ZNF583 | Island | 5'UTR;TSS200 | chr19:56915650-56915855 | 205 | | 1 | | | | | | | | | | | | | yes | | | | | |
| ZNF667 | Island;N_Shore | TSS200;5'UTR;Body;1stExon | chr19:56988205-56989740 | 1535 | | 2 | | | 1 | | | | 2 | | | | | yes | | | yes | yes | | |
| ZNF471 | Island | TSS1500;TSS200 | chr19:57018848-57019958 | 1110 | | 1 | | | | | | 0 | 4 | 1 | 1 | 1 | | | yes | yes | | yes | | |
| ZFP28 | Island | TSS1500;1stExon;5'UTR | chr19:57049695-57050834 | 1139 | | | | | | | | 0 | 5 | | | | | | | | | yes | | |
| ZNF470 | Island | TSS200 | chr19:57078727-57079146 | 419 | | 1 | | | | | | | 1 | | | | | | | | | | | |
| ZNF835 | N_Shore;Island | Body | chr19:57182844-57183268 | 424 | | 1 | | | 2 | | | | 1 | | | | | yes | | | | | | |
| ZIM2 | N_Shore;Island | Body | chr19:57306631-57307081 | 450 | | | | | | | | 0 | 2 | | | | | yes | yes | | yes | | | |
| PEG3;ZIM2 | NA | TSS1500;5'UTR | chr19:57337294-57337648 | 354 | | | 2 | | | | | | 1 | | | | | | yes | | | | | |
| ZIM2;PEG3;MIMT1 | N_Shore;Island;S_Shore | 1stExon;5'UTR;TSS1500;TSS200;Bo | chr19:57349204-57352746 | 3542 | | | 8 | | | | | | 6 | | | | | | yes | | | | yes | |
| USP29 | N_Shore;Island;S_Shore | TSS1500;TSS200 | chr19:57630034-57631405 | 1371 | | | 29 | | | | | 0 | 10 | 1 | | 1 | | yes | | yes | | | | |
| ZNF264 | N_Shore | TSS1500 | chr19:57702324-57702660 | 336 | | | 11 | | | | | | 2 | | | | | | yes | yes | | | | |
| AURKC | N_Shore;Island | TSS1500;TSS200;5'UTR;1stExon | chr19:57741325-57742444 | 1119 | 3 | 1 | 36 | 1 | | 3 | 1 | 0 | 15 | 2 | | 9 | 2 | yes | | yes | | | yes | |
| ZNF304 | N_Shore | TSS1500 | chr19:57862242-57863057 | 815 | | | | | | | | | 1 | | | | | | yes | yes | yes | yes | | |
| ZNF772 | NA | Body;1stExon;5'UTR;TSS200 | chr19:57988497-57989182 | 685 | | 2 | 1 | | | | | 0 | 2 | | 1 | 1 | | | yes | | | | | |
| ZNF549 | Island | TSS200 | chr19:58038573-58038588 | 15 | | | 4 | | | | | | 1 | | | | | | | yes | | | | |
| ZIK1 | Island | TSS200;TSS1500 | chr19:58095011-58095659 | 648 | | 2 | | | 1 | | | 0 | 1 | | | 1 | | yes | yes | | | | | |
| ZNF154 | N_Shore;Island;S_Shore | Body;5'UTR;1stExon;TSS200 | chr19:58220080-58220837 | 757 | 1 | 1 | | | 3 | | 1 | 0 | 2 | 1 | | | | yes | yes | | | | | |
| ZNF671 | N_Shore | Body | chr19:58238435-58239222 | 787 | | 1 | | | | | | | | | | | | yes | yes | | | | | |
| ZNF586 | N_Shore | TSS1500 | chr19:58280767-58280832 | 65 | | | | | | | | | | | | 1 | | yes | | | | | | |
| ZNF552 | Island;S_Shore | 5'UTR;1stExon;TSS200 | chr19:58326226-58326446 | 220 | | | 1 | | | | | | | | | 1 | | yes | | yes | | | | |
| ZNF587 | Island | TSS200 | chr19:58361147-58361533 | 386 | | | | | 1 | | | | | | | | | | yes | yes | | | | |
| ZNF814 | N_Shore;S_Shore | Body;TSS200 | chr19:58399865-58400574 | 709 | | 1 | 1 | 1 | | | | 0 | 3 | | | 2 | | yes | | yes | | | | |
| ZNF418 | Island | TSS200 | chr19:58446745-58446988 | 243 | | | 1 | | 1 | | | 0 | 1 | | | 1 | | | | yes | | | | |
| ZSCAN1 | N_Shore;Island | TSS1500;TSS200;5'UTR | chr19:58545001-58546307 | 1306 | | | | | | | | | | | | | | | | | | | | |
| ZSCAN1 | Island | Body | chr19:58549531-58549906 | 375 | | | | | | | | 0 | | | | 1 | | | | | | | | |
| ZSCAN1 | Island | Body | chr19:58554401-58554527 | 126 | | | 3 | | | | | | 1 | | | 1 | | yes | | | yes | | | |
| ZSCAN1 | Island | Body | chr19:58556185-58556609 | 424 | | | | | | | | | | | | | | | | | | | | |
| ZNF135 | Island | TSS200 | chr19:58570419-58570995 | 576 | | 1 | | | 1 | | | 0 | 6 | 2 | | 2 | | | | | | yes | | |
| ZSCAN18 | Island | Body;5'UTR;1stExon;TSS200 | chr19:58609361-58609987 | 626 | | 2 | | | | | | 0 | 1 | 1 | | | | | | | | | yes | |
| ZSCAN18 | Island | Body;1stExon | chr19:58629719-58630089 | 370 | | 1 | | | 1 | | | | | | | | | | | | | | yes | |
| ZNF274 | N_Shore | TSS1500 | chr19:58693421-58693616 | 195 | | | | | | | | | 1 | | | | | yes | yes | | | | | |
| ZNF274 | Island | 1stExon;5'UTR | chr19:58694770-58695028 | 258 | | 1 | | | | | | | | | | | | | | | | | | |
| ZNF274 | Island | Body;5'UTR | chr19:58715251-58716135 | 884 | | 1 | | | 2 | | | 0 | 2 | | | | | yes | | yes | | | | |

| | | | | | | | | | | | | | | | | | | | | | | | |
|---|---|---|---|---|---|---|---|---|---|---|---|---|---|---|---|---|---|---|---|---|---|---|---|
| ZNF544 | Island | 5'UTR;1stExon | chr19:58740253-58740365 | 112 | | | | | | | | 0 | | | | | | | | yes | | yes | |
| ZSCAN22 | N_Shore | TSS1500 | chr19:58837043-58837257 | 214 | | | | | | | | | | | | | | | yes | yes | | | |
| A1BG;NCRNA00181 | Island | Body;TSS1500 | chr19:58861502-58862398 | 896 | | | | 1 | | 2 | | 0 | 4 | 1 | | | | yes | | | | | yes |
| NCRNA00181;A1BG | N_Shelf | Body;TSS200;5'UTR;1stExon | chr19:58864846-58865367 | 521 | | | 1 | 1 | | | | | | | | | | | yes | | | | |
| ZNF837 | Island | 3'UTR | chr19:58879022-58879059 | 37 | | | 6 | | | | | | 1 | | | | | | | | | | |
| ZNF132 | N_Shore | Body | chr19:58951095-58952249 | 1154 | | 1 | | | | | | | | | | | | | yes | | | | |
| SLC27A5 | N_Shelf | 1stExon | chr19:59023222-59023537 | 315 | | | | | | | | | | | | | | | yes | | | yes | |
| ZBTB45 | Island | 3'UTR | chr19:59025345-59025573 | 228 | | | 3 | | | | | 0 | 1 | 1 | | 4 | | | | | | | |
| CHMP2A;UBE2M | S_Shore | TSS1500;3'UTR | chr19:59067160-59067429 | 269 | | | | | | | | | | | | | | | | | | | |
| MZF1;LOC100131691 | Island | Body | chr19:59073902-59074507 | 605 | | | | | | | | 0 | 1 | 3 | | 1 | | yes | | | | | |
| MGC2752 | Island | TSS200 | chr19:59086698-59087201 | 503 | | | | | | | | | | 1 | | 1 | | yes | | yes | | | |
| FAM110C | S_Shore | TSS1500 | chr2:47319-47889 | 570 | | | 3 | | | | | | 1 | | | 1 | | | yes | | | | yes |
| FAM150B | N_Shore | Body | chr2:286830-287395 | 565 | | | 9 | | | | | | 1 | | | | | | | | | yes | yes |
| C2orf90 | NA | TSS200;TSS1500 | chr2:906044-906974 | 930 | | | 6 | | | | | | | | | | | | | | | | |
| SNTG2 | Island;S_Shore | 1stExon;5'UTR;Body | chr2:946648-947634 | 986 | | | 18 | | | | | 0 | 2 | 1 | | 1 | | | | | | | yes |
| SNTG2 | NA | Body | chr2:983306-983636 | 330 | | | | | | | | | | | | | | | yes | yes | | | |
| SNTG2 | NA | Body | chr2:1204376-1204879 | 503 | | | | | | | | | | | | | | yes | | | | | |
| SNTG2 | NA | Body | chr2:1237808-1238426 | 618 | | | 2 | | | | | | | | | | | yes | | | | | |
| SNTG2 | N_Shore;Island | Body | chr2:1286216-1286641 | 425 | | | 11 | | | | | 0 | 3 | | | | | | | | yes | | |
| SNTG2 | NA | Body | chr2:1358283-1358730 | 447 | | | 5 | | | | | | | | | | | yes | yes | | | | |
| TPO | NA | TSS1500;TSS200 | chr2:1416855-1418294 | 1439 | | | 12 | | | | | | | | | | | yes | | | | | |
| TPO | NA | Body | chr2:1425035-1426845 | 1810 | | 1 | 9 | | | | | | | | | | | yes | | | | | |
| TPO | Island | Body | chr2:1480789-1481097 | 308 | | | | | | | | | 1 | | | | | | | | | | |
| TPO | NA | Body | chr2:1487895-1488638 | 743 | | | 12 | | | | | | | | | | | yes | yes | | | | |
| TPO | S_Shelf | Body | chr2:1497815-1498138 | 323 | | | 4 | | | | | | | | | | | yes | | | | | |
| TPO | NA | Body | chr2:1507694-1507832 | 138 | | | 4 | | | | | | | | | | | | | | | | |
| TPO | N_Shore;chr2:1544272-154448 | Body;N_Shore | chr2:1544076-1544352 | 276 | | | | | | | | | | | 1 | | | | | | | | |
| PXDN | NA | Body | chr2:1647185-1647260 | 75 | | | 5 | | | | | | | | | | | yes | | | | | |
| PXDN | NA | Body | chr2:1690592-1691062 | 470 | | 1 | | 1 | | | | | 1 | | | | | | | | | | |
| PXDN | NA | Body | chr2:1711759-1711966 | 207 | | | | | | | | 0 | | 1 | | | | | | | | | |
| PXDN | N_Shore;Island | Body;1stExon;TSS1500 | chr2:1746207-1748970 | 2763 | | 1 | 3 | | | | | | 3 | | | | | | | yes | | | yes |
| MYT1L | NA | Body | chr2:1796076-1796228 | 152 | | | | | | | | 0 | 1 | | | | | yes | | | | | |
| MYT1L | NA | Body | chr2:1817263-1817951 | 688 | | 3 | 2 | 2 | | | 2 | 0 | 3 | 1 | 1 | 12 | 1 | yes | | | yes | | |
| MYT1L | NA | Body | chr2:1820169-1820433 | 264 | | | 3 | | | | | | | | | | | yes | | | | | |
| MYT1L | Island | Body | chr2:1842860-1843184 | 324 | | | 1 | | | | | 0 | 3 | | | | | | | | | | |
| MYT1L | NA | Body | chr2:1859035-1859146 | 111 | | | | | | | | 0 | 1 | | | 2 | | | | | | | |
| MYT1L | NA | Body | chr2:1879019-1880131 | 1112 | | | | | | | | | 1 | | | | | | | | | | |
| MYT1L | NA | Body | chr2:1926366-1927209 | 843 | | | 9 | | | | | | 2 | | | | | | | | | | yes |
| MYT1L | NA | Body | chr2:1932996-1933212 | 216 | | | | | | | | | | | | | | yes | | | | | |
| MYT1L | N_Shore | Body | chr2:1940991-1941523 | 532 | | | 2 | | | | | | | | | | | | | | | | |
| MYT1L | S_Shore | Body | chr2:1942814-1942942 | 128 | | | | | | | | | | | | | | yes | | | | | |
| MYT1L | NA | Body | chr2:1983190-1983518 | 328 | | | | | | | | | | | | | | | | | | yes | |
| MYT1L | Island | 5'UTR | chr2:2193511-2193665 | 154 | | | 9 | | | | | | | | | | | yes | | | yes | | yes |
| MYT1L | NA | 5'UTR | chr2:2295451-2295969 | 518 | | | | | | | | | 1 | | | | | | | | | | |
| MYT1L;LOC730811 | NA | 5'UTR;Body | chr2:2328391-2328709 | 318 | | | | | | | | | 1 | | | | | | | | | | |
| TSSC1 | NA | Body | chr2:3245794-3245997 | 203 | | | | | | | | | | | | | | | | | | | |
| TSSC1 | NA | Body | chr2:3281036-3281426 | 390 | | 1 | | | | | | | 1 | | | | | yes | | | yes | | yes |
| TSSC1 | N_Shore;Island | Body | chr2:3285967-3286479 | 512 | | | 25 | | | | | | 1 | | | | | | | | | | |
| TSSC1 | N_Shore;Island | Body | chr2:3317469-3318039 | 570 | | | 7 | | | | | | | | | | | yes | | | | | |
| TSSC1 | S_Shore | Body | chr2:3322136-3323078 | 942 | 3 | | | | | | | | | | | | | yes | yes | yes | | | |
| TSSC1 | Island | TSS200 | chr2:3381671-3381855 | 184 | | | 1 | | 1 | | | 0 | | 3 | | | | yes | | | | | |
| TTC15 | N_Shore;Island | Body | chr2:3451506-3452919 | 1413 | | | 39 | | | | | | | | | | | yes | | yes | | | |
| TTC15 | NA | Body | chr2:3471009-3471071 | 62 | | 1 | | | | | | | | | | | | | | | | | |
| COLEC11 | N_Shore;Island | TSS1500;TSS200;1stExon;5'UTR | chr2:3642211-3642967 | 756 | | | 40 | | | | 1 | 0 | 4 | 2 | | 3 | | | yes | | yes | | |
| COLEC11 | NA | 5'UTR | chr2:3648528-3648554 | 26 | | | 4 | | | | | | 1 | | | | | yes | yes | | | | |
| COLEC11 | NA | Body | chr2:3654242-3654582 | 340 | | | 4 | | | | | | | | | | | yes | | | | | |
| COLEC11 | NA | Body | chr2:3658894-3659019 | 125 | | | | | | | | 0 | | | | | | | yes | yes | | | |
| COLEC11 | NA | Body | chr2:3660868-3660939 | 71 | | | | | | | | | 1 | | | | | yes | | | | | |
| COLEC11 | Island | Body | chr2:3683030-3683757 | 727 | | | 11 | | | | | | | | | | | yes | | | yes | | yes |
| COLEC11 | S_Shore | Body | chr2:3685161-3685430 | 269 | | | 1 | | | | | | | | | | | | | | | | |
| COLEC11 | N_Shore | Body | chr2:3689356-3689727 | 371 | | | | | | | | | | | | | | | | | | | |
| COLEC11 | Island | Body | chr2:3691419-3691651 | 232 | | | | | | | | | | | | | | | | | | | |
| ALLC | NA | TSS1500 | chr2:3704363-3704793 | 430 | 2 | | | | 1 | | 1 | 0 | 9 | 2 | | 6 | 1 | yes | | | yes | | |
| ALLC | NA | 5'UTR | chr2:3714278-3715083 | 805 | | 1 | 10 | 1 | | | | | | | | | | yes | yes | | yes | | |
| SOX11 | Island | 3'UTR;1stExon | chr2:5836713-5837057 | 344 | | | | | | | | 0 | 1 | | | | | | | | | | yes |
| LOC150622 | NA | TSS1500;TSS200 | chr2:6072139-6072981 | 842 | | | 8 | | 1 | | | | 1 | | | | | yes | | | | | |
| LOC400940 | NA | Body | chr2:6122798-6122814 | 16 | | 1 | | | | | | | 1 | | | | | | | | | | yes |
| RSAD2 | NA | TSS200 | chr2:7017787-7018153 | 366 | | | | | | | | 0 | | | | | | | | | | | |
| RNF144A | N_Shore | TSS1500 | chr2:7057153-7057649 | 496 | | | 4 | | | | | | 1 | | | | | yes | | | | yes | yes |
| RNF144A | N_Shore | Body | chr2:7171869-7172297 | 428 | | | | 1 | | | | | 3 | 2 | | | | | yes | yes | yes | | |
| LOC339788 | NA | TSS1500 | chr2:8118122-8118404 | 282 | | | 5 | | | | | | | | | | | | | | | yes | |
| ID2 | N_Shore | 3'UTR | chr2:8823649-8823756 | 107 | | | | | 1 | | | | | | | | | | | | | | yes |
| MBOAT2 | Island | TSS1500 | chr2:9144618-9144764 | 146 | | | 1 | 1 | | | | | | | | | | yes | | | | | yes |
| ASAP2 | Island;N_Shore | Body | chr2:9459065-9459596 | 531 | | | | 1 | | | | | 3 | 1 | | | | | yes | | | | |
| KLF11 | Island;N_Shore | TSS1500 | chr2:10182525-10183271 | 746 | | | 2 | | 2 | | | | | | | | | | yes | | | yes | yes |
| C2orf48 | NA | Body | chr2:10302391-10302462 | 71 | | | | | | | | | 1 | | | | | yes | | | | | |
| HPCAL1 | S_Shore | 5'UTR | chr2:10444757-10445016 | 259 | | | | | | | | | | | | | | | | | | | yes |
| HPCAL1 | NA | 5'UTR | chr2:10517323-10517361 | 38 | | | | | | | | | 1 | | | | | yes | | | | yes | |
| HPCAL1 | Island;S_Shore | 3'UTR | chr2:10566989-10567843 | 854 | | | 23 | | | | | | 1 | | | | | yes | | | | | |
| NOL10 | S_Shore | TSS1500 | chr2:10830656-10830764 | 108 | | 1 | 1 | | | | | | 3 | | | 2 | | yes | | | | | |
| C2orf50 | NA | TSS1500 | chr2:11272783-11273291 | 508 | | | | | | | | 0 | | | | | | yes | | | | yes | |
| GREB1 | NA | 5'UTR;TSS200;TSS1500 | chr2:11679605-11680144 | 539 | 7 | | 17 | 2 | 1 | | | | 2 | | | | | yes | | | | yes | yes |
| GREB1 | Island | Body | chr2:11732905-11733209 | 304 | | | | | | | | | 1 | | | | | yes | | | | | yes |
| GREB1 | Island | Body | chr2:11758793-11759194 | 401 | | | | | | | | | | | | | | | yes | | | | |
| NTSR2 | S_Shore;Island | TSS1500;1stExon;TSS200 | chr2:11809681-11810631 | 950 | | 1 | 5 | | | | | | 2 | | | | | yes | | | yes | | yes |
| LPIN1 | NA | Body | chr2:11917490-11917787 | 297 | | 1 | | | | | | | | | | | | | | | yes | | |
| LPIN1 | Island | Body | chr2:11924984-11925375 | 391 | | 1 | 1 | | | | | | 1 | | | | | | | | | | |
| FAM84A | Island | TSS1500;TSS200;5'UTR;Body | chr2:14772562-14775199 | 2637 | | | | | | | | 0 | 3 | | | | | | | | | yes | yes |
| MYCN | S_Shelf | Body | chr2:16085728-16086016 | 288 | 1 | | | | | | | | 1 | | | 1 | | yes | | | | | yes |
| MSGN1 | NA | TSS1500 | chr2:17996994-17997779 | 785 | | | | | | | | | 2 | 1 | 1 | | | yes | | | | | |
| KCNS3 | Island | TSS1500 | chr2:18059103-18059130 | 27 | | | 1 | | | | | | 1 | | | | | | | | | | |
| NT5C1B | N_Shore;Island | Body | chr2:18765443-18766294 | 851 | | | 3 | | | | | | 3 | | | | | | | | | | |
| OSR1 | chr2:19551735-19551961;Islan | N_Shore;3'UTR | chr2:19551398-19551860 | 462 | | | 5 | | | | | | | | 1 | | | | | | | | yes |
| OSR1 | Island | Body | chr2:19553237-19554237 | 1000 | | | 10 | | | | | 0 | | | | | | | | | | yes | |
| MATN3 | Island | TSS200 | chr2:20212501-20212517 | 16 | | | | | | | | | 1 | | | | | yes | | | | | yes |
| LAPTM4A | Island | Body | chr2:20251148-20251361 | 213 | | | | | | | | | 1 | | | | | | | | | | |
| GDF7 | Island | TSS1500 | chr2:20865651-20865847 | 196 | | 1 | 1 | | | | | | | | | | | | | | | | |
| GDF7 | Island | Body;3'UTR | chr2:20870087-20871401 | 1314 | | | 15 | | | | | 0 | 6 | 2 | 1 | 9 | | yes | | | | | yes |
| C2orf43 | S_Shore | TSS1500 | chr2:21023387-21023697 | 310 | | | | | | | | | 1 | | | | | yes | | yes | yes | | |

| | | | | | | | | | | | | | | | | | | | | | | | | | | | |
|---|---|---|---|---|---|---|---|---|---|---|---|---|---|---|---|---|---|---|---|---|---|---|---|---|---|---|---|
| APOB | NA | Body | chr2:21229220-21229936 | 716 | | | | | | | | | | | | | | | | | | | | | | | |
| APOB | NA | Body | chr2:21255279-21255320 | 41 | | | | | | | | | | | | | | | yes | | | | | | | | |
| APOB | N_Shore;Island;S_Shore | Body;TSS200 | chr2:21266500-21267334 | 834 | | | 20 | 1 | | | | 0 | 11 | | | 5 | | | | | | yes | | | | | |
| KLHL29 | N_Shelf | 5'UTR | chr2:23746926-23747197 | 271 | | | 12 | | | | | | | | | | | yes | yes | | | | | | | | |
| KLHL29 | NA | Body | chr2:23839842-23840763 | 921 | 6 | | 2 | 1 | | | | | | | | | | yes | | yes | | | | | | | |
| KLHL29 | Island | Body | chr2:23851841-23852064 | 223 | | | | | | | | 0 | 1 | | | 1 | | yes | | | | | | | | | |
| KLHL29 | Island | Body | chr2:23913466-23913912 | 446 | | | 1 | | | | | | | | | | | yes | | | | | | | | | |
| MFSD2B | S_Shore | Body | chr2:24233923-24234017 | 94 | | | 3 | | | | | 0 | 1 | | | 1 | | | | | | | | | | | |
| FKBP1B | N_Shore | TSS1500 | chr2:24271827-24272044 | 217 | | | 7 | | | | | | 1 | | | | | yes | yes | | | yes | | | | | |
| SF3B14 | Island | TSS1500 | chr2:24300061-24300242 | 181 | | | | 1 | | | | | | | | | | | | | | yes | | | | | |
| C2orf84 | Island | TSS1500;TSS200;5'UTR;Body | chr2:24397649-24398859 | 1210 | | | 22 | | 1 | | | 0 | 4 | 2 | | 3 | | yes | yes | | | | | | | | |
| ITSN2 | S_Shore | TSS1500 | chr2:24583953-24584081 | 128 | | | | | | | | | 1 | | | | | yes | | | | | | | | | |
| ADCY3 | N_Shore | TSS200;Body;1stExon;5'UTR | chr2:25141152-25142391 | 1239 | 5 | | 11 | 1 | 3 | | | | 1 | | | | | yes | | | | | yes | | | | |
| EFR3B | Island | Body | chr2:25354401-25354986 | 585 | | | 7 | | | | | | | | | | | | | | yes | yes | yes | | | | |
| POMC | N_Shore | 3'UTR | chr2:25383851-25384293 | 442 | | | 1 | | | | | | | | | | | yes | | | | yes | | | | | |
| POMC | N_Shore;Island | 5'UTR;1stExon;TSS200;TSS1500 | chr2:25390385-25391911 | 1526 | | | 21 | | | | | 0 | 1 | | | | | yes | | | | | | | | | |
| DNMT3A | N_Shore;Island | Body;TSS1500 | chr2:25473529-25475916 | 2387 | 4 | | | 1 | | | | 0 | | | | | | | | | | | yes | | | | |
| DNMT3A | N_Shore;Island | Body | chr2:25499619-25500416 | 797 | | | 13 | | | | | | 2 | 1 | 1 | | | yes | | | | yes | | | | | |
| DNMT3A | NA | Body | chr2:25527266-25527366 | 100 | | | 1 | | | | | | | | | | | yes | yes | | | | | | | | |
| ASXL2 | NA | Body | chr2:26045189-26045287 | 98 | | | 11 | | | | | 0 | 1 | 1 | | 6 | | | | | | | | | | | |
| FAM59B | N_Shore | TSS1500 | chr2:26395359-26395458 | 99 | | 1 | | | | | | | | | | | | yes | | | | | yes | | | | |
| FAM59B | Island;N_Shore | Body | chr2:26401640-26402318 | 678 | | 1 | 8 | | | | | | 3 | 2 | | 1 | | yes | | | | | yes | | | | |
| FAM59B | Island | Body | chr2:26407628-26408167 | 539 | | | 1 | | | | | | | 1 | | | | | | | yes | | yes | | | | |
| GPR113 | NA | 5'UTR;1stExon;TSS200;Body | chr2:26541960-26542188 | 228 | | | 8 | | | | | | | | | | | yes | | | | | yes | | | | |
| C2orf39 | Island | TSS200 | chr2:26624760-26624934 | 174 | | | | | | | | | 1 | | | | | yes | yes | yes | | | yes | | | | |
| C2orf70 | Island | TSS1500;TSS200;Body | chr2:26785120-26786046 | 926 | | | 8 | | | | | 0 | 1 | 2 | | | | | | | | yes | | | | | |
| EMILIN1 | N_Shelf;NA | TSS200;1stExon;5'UTR;TSS1500 | chr2:27301195-27301943 | 748 | | | | | | | | 0 | 3 | | | 1 | | yes | | | | | yes | | | | |
| CGREF1 | Island | 5'UTR;TSS200;1stExon | chr2:27341795-27341896 | 101 | | | | | | | | | 1 | | | | | | | | | yes | yes | | | | |
| ABHD1 | N_Shore;Island | TSS1500 | chr2:27346296-27346765 | 469 | | | | | | | | 0 | | | | | | | yes | | yes | | | | | | |
| TCF23 | N_Shore | TSS1500;TSS200 | chr2:27371537-27371891 | 354 | | | 10 | | | | | | | | | | | yes | yes | | | | | | | | |
| TRIM54 | NA | TSS1500;TSS200 | chr2:27504695-27505625 | 930 | | | 7 | | | | | | 1 | | 1 | | | | | | | | | | | | |
| TRIM54;UCN | Island | Body;3'UTR;5'UTR;TSS200 | chr2:27529325-27531535 | 2210 | | | | | | | | | | | | | | | | | | | | | | | |
| PPM1G | Island | TSS1500;TSS200 | chr2:27632687-27632926 | 239 | | | | | | | | | | | | 1 | | yes | | | | | | | | | |
| KRTCAP3;NRBP1 | N_Shore;Island | TSS1500;Body;3'UTR;TSS200;1stE | chr2:27664399-27665711 | 1312 | | | 27 | 1 | 1 | 1 | | 0 | 16 | 2 | | 16 | 1 | | | | | yes | yes | | | | |
| IFT172 | S_Shore | TSS1500 | chr2:27712902-27713092 | 190 | | | | | | | | 0 | 2 | | | | | yes | | | | | | | | | |
| C2orf16 | NA | TSS1500;TSS200 | chr2:27798724-27799411 | 687 | | | 6 | | | | | | | | | | | | | | | | | | | | |
| MRPL33 | N_Shore | TSS1500 | chr2:27994018-27994325 | 307 | | | | | | | | | 1 | 1 | | 1 | | yes | yes | | | | | | | | |
| BRE;RBKS;LOC100302650 | S_Shore | 5'UTR;TSS1500 | chr2:28114349-28114462 | 113 | | | | 1 | | | | | | | | | | | | | | | | | | | |
| PLB1 | NA | TSS1500 | chr2:28718569-28718959 | 390 | | | | | | | | | | | | | | yes | | | | | | | | | |
| SPDYA | Island | 5'UTR;1stExon | chr2:29033780-29034052 | 272 | | 1 | | | | | | | | | | | | yes | yes | | | | | | | | |
| FAM179A | NA | TSS1500;TSS200 | chr2:29202812-29204695 | 1883 | | | 9 | | | | | | 3 | 1 | | | | yes | yes | | | | | | | | |
| CLIP4 | N_Shore | TSS1500 | chr2:29337946-29337988 | 42 | | | | | | | | | | | | | | | yes | | | | | | | | |
| CLIP4 | chr2:29337983-29338909 | S_Shore | chr2:29339076-29339307 | 231 | | | | | | | | | | | 1 | | 1 | | yes | | | | | | | | |
| ALK | N_Shore | Body | chr2:30142063-30142677 | 614 | | | | | | | | | | | | | | | | | | | | | | | |
| LCLAT1 | N_Shore;Island | TSS1500;TSS200 | chr2:30669597-30670304 | 707 | 4 | 7 | 29 | 1 | 2 | 5 | 3 | 0 | 6 | 3 | 1 | 13 | 1 | yes | | | | | | | | | |
| CAPN13 | NA | TSS1500 | chr2:31031702-31031755 | 53 | | | | | | | | | 1 | | | | | yes | | | yes | | | | | | |
| GALNT14 | Island | TSS200 | chr2:31361687-31361701 | 14 | | | | | | | | | 1 | | | | | | | | | yes | | | | | |
| XDH | NA | Body | chr2:31637348-31637849 | 501 | | | | 1 | | | | | | | | | | yes | | | | | | | | | |
| SRD5A2 | Island | TSS1500;1stExon;TSS200 | chr2:31805915-31806898 | 983 | | 1 | 12 | | | | | 0 | 4 | | | 2 | | | | | | | yes | | | | |
| LOC285045 | NA | TSS200 | chr2:33050369-33057712 | 7343 | | | | | | | | | | | | | | yes | yes | yes | yes | yes | | | | | |
| LTBP1 | NA | Body;TSS1500 | chr2:33359198-33359688 | 490 | | | | | | | | 0 | 5 | 2 | | 1 | | yes | | | yes | | yes | | | | |
| MYADML | S_Shore | Body | chr2:33953195-33953370 | 175 | | | | | | | | | | | | | | yes | | yes | | | | | | | |
| FEZ2 | N_Shore | Body | chr2:36824259-36824338 | 79 | | | 4 | | | | | | | | | | | yes | yes | | | | | | | | |
| FEZ2 | S_Shore | TSS1500 | chr2:36825704-36825888 | 184 | | | | | | | | | 1 | | | | | yes | yes | yes | | yes | | | | | |
| EIF2AK2 | Island | TSS1500 | chr2:37384401-37384708 | 307 | | | | | | | | | | | | | | yes | | | | yes | | | | | |
| PRKD3 | NA | TSS1500 | chr2:37545307-37545668 | 361 | | | 4 | | | | | | 1 | | | | | yes | yes | | | | | | | | |
| QPCT | Island | TSS200 | chr2:37571677-37572102 | 425 | | | 8 | | | | | | | | | | | yes | | | | | | | | | |
| CDC42EP3 | Island | TSS1500 | chr2:37899776-37900205 | 429 | | | | | | | | | | | | | | | | | | | yes | | | | |
| CYP1B1 | N_Shore | Body | chr2:38300537-38300707 | 170 | | | | | | | | | 1 | | | | | | | | yes | | | | | | |
| CYP1B1 | Island | Body | chr2:38302487-38302674 | 187 | | | 1 | | | | | | | | | | | | | | | | | | | | |
| CYP1B1 | S_Shore | TSS1500 | chr2:38304720-38304804 | 84 | | | | | | | | | | | | | | yes | | | | | | | | | |
| GALM | NA | TSS1500 | chr2:38892846-38892901 | 55 | | | | | | | | 0 | 1 | | | | | yes | | | | yes | | | | | |
| GEMIN6 | NA | TSS200 | chr2:39005158-39005834 | 676 | | | 1 | | | | | | | | | | | yes | | | | | | | | | |
| LOC375196;LOC100271715 | Island | TSS200;Body;TSS1500 | chr2:39187533-39188006 | 473 | | 1 | | | | | | | 1 | | | 1 | | | | | | | yes | | | | |
| TMEM178 | N_Shore | TSS1500;TSS200 | chr2:39892450-39892612 | 162 | | 1 | | | 1 | | | | | | | | | | | | | | yes | | | | |
| SLC8A1 | NA | 5'UTR;1stExon | chr2:40739515-40739771 | 256 | | | | | | | | | | | | | | | | | | yes | | | | | |
| PKDCC | NA | TSS1500 | chr2:42274082-42274619 | 537 | | | | | | | | | 1 | | | | | | | | | yes | yes | | | | |
| KCNG3 | N_Shore | Body | chr2:42719678-42719933 | 255 | | | | | | | | | 1 | | | | | | | | | | yes | | | | |
| MTA3 | N_Shore | TSS1500 | chr2:42795193-42795486 | 293 | | | | | | | | | 1 | | | | | | | | | yes | yes | | | | |
| HAAO | Island | TSS1500 | chr2:43020264-43020682 | 418 | | | | | | | | | | | | | | | | | | | yes | | | | |
| ZFP36L2;LOC100129726 | S_Shore | TSS1500;Body | chr2:43455088-43455103 | 15 | 1 | | | | | | | | | | | | | yes | | | | | | | | | |
| PLEKHH2;LOC728819 | NA | Body;1stExon;TSS200;TSS1500 | chr2:43903227-43904011 | 784 | | | 24 | | | | | 0 | 6 | | | 7 | | | | | | | | | | yes | |
| ABCG5 | Island | Body | chr2:44058766-44059266 | 500 | | | | 1 | | | | | | | | | | yes | yes | | yes | | yes | | | | |
| ABCG5;ABCG8 | NA | Body;TSS1500 | chr2:44065056-44065680 | 624 | | | 9 | | | | | | | | | | | yes | | | | | | | | | |
| SLC3A1 | NA | Body | chr2:44513757-44513905 | 148 | | | | | | | | | 1 | | | | | yes | yes | | | yes | | | | | |
| SIX3 | N_Shore;Island;S_Shore | TSS1500;Body;3'UTR | chr2:45168819-45172265 | 3446 | 2 | 1 | | | | | | | 1 | | | | | | | | | yes | yes | | | | |
| SIX2 | N_Shore;Island | Body;3'UTR | chr2:45232417-45233773 | 1356 | | | | | 1 | | | | | | | | | | | | | | yes | | | | |
| SIX2 | N_Shore | Body | chr2:45235241-45235512 | 271 | | | | | | | | | 1 | | | | | | | | | | | | | | |
| SIX2 | Island | TSS200;TSS1500 | chr2:45236686-45237781 | 1095 | | | | | | | | | 2 | | | | | | | | | | yes | | | | |
| EPAS1 | Island | Body | chr2:46526785-46527098 | 313 | | | 1 | | | | | 0 | | | | | | | | | | | | | | | |
| SOCS5 | N_Shore | TSS1500 | chr2:46925129-46925381 | 252 | | | | | | | | | 1 | | | | | yes | yes | | | | | | | | |
| LOC100134259 | NA | TSS200 | chr2:47054833-47055138 | 305 | | | | | | | | | 1 | | | | | | | | | yes | yes | | | | |
| TTC7A | NA | Body | chr2:47182345-47197798 | 15453 | | | | | | | | | | | | | | yes | yes | yes | yes | | | | | | yes |
| C2orf61 | NA | Body;1stExon;5'UTR;TSS200 | chr2:47382287-47382739 | 452 | | | 10 | | | | 1 | 0 | 2 | 3 | | | | yes | | | yes | | yes | | | | |
| EPCAM | Island | Body | chr2:47596912-47597331 | 419 | | | 9 | | | | | | 1 | | | 1 | | yes | | yes | | | | | | | |
| GTF2A1L;STON1-GTF2A1L | N_Shore;Island | TSS1500;Body;TSS200 | chr2:48844195-48845068 | 873 | | | 8 | | | 2 | | 0 | 9 | | | 6 | | yes | | | yes | yes | | | | | |
| LHCGR | N_Shore;Island | Body;1stExon;5'UTR;TSS200 | chr2:48982509-48983178 | 669 | | 1 | 3 | | | | | | | | | 1 | | yes | yes | | | yes | | | | | |
| FSHR | NA | TSS1500 | chr2:49382159-49382401 | 242 | | | | | | | | | 1 | | | | | | | | | | | | | | |
| NRXN1 | NA | Body | chr2:50201382-50201511 | 129 | | | 19 | | | | | 0 | 5 | 2 | | 15 | 1 | | | | | | | | | | |
| NRXN1 | Island | Body | chr2:51254823-51255627 | 804 | | | | | | | | | 4 | | | 1 | | | | | | | | | | | |
| NRXN1 | NA | TSS200 | chr2:51259693-51259786 | 93 | | 1 | | | | | | | 1 | | | 1 | | | | | | yes | | | | | |
| GPR75;LOC100302652 | Island;S_Shore | 5'UTR;Body;1stExon;TSS200;TSS1 | chr2:54086854-54087517 | 663 | | 1 | 33 | 1 | 1 | | | 0 | 6 | | | 2 | | yes | | | | | | | | | |
| SPTBN1 | Island | TSS200;Body;TSS1500 | chr2:54785178-54786148 | 970 | | | 52 | | | | | | | | | | | | | | | | | | | | |
| C2orf63 | N_Shore;Island | 5'UTR | chr2:55450577-55450874 | 297 | | | 1 | | 1 | | 1 | 0 | 2 | 2 | 1 | 3 | | yes | | | | | yes | | | | |
| PRDXDD1P | Island | TSS1500 | chr2:55509253-55509334 | 81 | | | | | | | | 0 | | | 1 | | | yes | | | | | | | | | |
| PNPT1 | N_Shore;Island | Body;TSS200 | chr2:55920477-55921125 | 648 | | | | | | | | | 2 | | | | | yes | | | | yes | | | | | |
| MIR216A | NA | Body | chr2:56216086-56216234 | 148 | | | | | | | | | | | | | | | | | | | | | | | |
| CCDC85A | N_Shore | TSS1500 | chr2:56409774-56410300 | 526 | | | 17 | | | | | | | | | | | yes | | yes | | | | | | | |
| VRK2 | N_Shore | TSS1500 | chr2:58273420-58273552 | 132 | | | | | | | | | | | | | | yes | | | | | | | | | |

| | | | | | | | | | | | | | | | | | | | | | | | | | | | |
|---|---|---|---|---|---|---|---|---|---|---|---|---|---|---|---|---|---|---|---|---|---|---|---|---|---|---|---|
| BCL11A | NA | Body | chr2:60680616-60680761 | 145 | | | | | | | | | | | | | | | | | | yes | yes | | | | |
| BCL11A | NA | Body | chr2:60763331-60763808 | 477 | | | | | | | | | 1 | | | | | | | | | | | | | | |
| BCL11A | N_Shore | Body | chr2:60780027-60780427 | 400 | | | | | | | | | | | | | | | | | | yes | yes | | | | |
| C2orf74 | NA | TSS1500;TSS200 | chr2:61371880-61372316 | 436 | | 1 | 2 | | 1 | | | 0 | 2 | | | 2 | | yes | | | | | yes | | | | |
| EHBP1 | Island | TSS1500;5'UTR;TSS200 | chr2:62932961-62933102 | 141 | | | | | | | | | 1 | | | | | | | | | yes | | | | | |
| LOC100132215;EHBP1 | Island | Body | chr2:63272232-63272334 | 102 | 1 | | | | | | | | | | | | | | | | | | | | | | |
| LOC100132215;OTX1 | Island;chr2:63274475-6327943 | Body;Island;TSS200;TSS1500 | chr2:63274615-63276779 | 2164 | | | | | | | | | | | | | | | | | | | | | | | |
| OTX1 | Island;N_Shore | 3'UTR;5'UTR;Body | chr2:63279495-63284768 | 5273 | 2 | | 6 | | 4 | | | | 5 | 7 | 1 | | | | | | | yes | yes | | | | |
| SLC1A4 | Island;S_Shore | 1stExon;5'UTR;Body | chr2:65217211-65217623 | 412 | | | 3 | | | | | 0 | 1 | 1 | | | 1 | yes | | | | | | | | | |
| SPRED2 | NA | 5'UTR;1stExon;Body | chr2:65593761-65594021 | 260 | | | | 1 | 1 | | | 0 | 2 | | | | | | yes | | | yes | | | | | |
| MEIS1 | S_Shelf | Body | chr2:66664535-66665428 | 893 | | | | 1 | | 1 | | | | | | 1 | | | | | | yes | yes | | | | |
| MEIS1 | NA | Body | chr2:66666470-66668012 | 1542 | | | | | 1 | | | | 1 | | | | | | | | | yes | yes | | | | |
| MEIS1 | N_Shore;Island | Body | chr2:66671478-66673449 | 1971 | | | 3 | | | | | 0 | 1 | 2 | 1 | | | | | | | yes | yes | | | | |
| CNRIP1 | N_Shore | Body | chr2:68545960-68546038 | 78 | | 1 | | | | | | 0 | | | | | | | | | | | | | | | |
| PLEK | NA | TSS1500;1stExon;5'UTR;nan | chr2:68591915-68592737 | 822 | | | 9 | | 1 | | | | 1 | | 1 | | | yes | | | yes | | | | | | |
| ARHGAP25 | NA | TSS1500 | chr2:68961495-68962012 | 517 | | | | | 2 | | | | | 1 | 1 | | | yes | | | | | | | | | |
| ARHGAP25 | NA | nan;Body;TSS200;5'UTR;1stExon;T | chr2:69001665-69002378 | 713 | | | 11 | 1 | 2 | | | | 2 | 4 | 1 | | | | | | | | | | | | |
| ANTXR1 | Island | TSS200 | chr2:69240197-69240453 | 256 | | | | | | | | | 1 | | | | | | | | | | | | | | |
| GFPT1 | S_Shore | TSS1500 | chr2:69614945-69615301 | 356 | | | 2 | | | | | | | | | | | | yes | | | | | | | | |
| NFU1 | S_Shore | TSS1500 | chr2:69665094-69665232 | 138 | | | | | 1 | | | | 1 | | | | | yes | | | yes | | | | | | |
| PCBP1 | N_Shore | TSS1500 | chr2:70313163-70313417 | 254 | | | | | 1 | | | | | | | | | yes | | | | yes | | | | | |
| PCYOX1 | N_Shore | TSS1500 | chr2:70484527-70484614 | 87 | | | | 1 | | | | 0 | 1 | | | | | yes | | yes | | | yes | | | | |
| FAM136A | S_Shore | TSS1500 | chr2:70529494-70529513 | 19 | | | 2 | | | | | | 1 | | | | | yes | yes | | | | yes | | | | |
| FIGLA | Island | 1stExon;TSS200 | chr2:71017541-71018044 | 503 | | 1 | | | | | | | | | | | | | | | yes | yes | yes | | | | |
| VAX2 | N_Shore;Island | TSS1500;TSS200 | chr2:71126275-71127779 | 1504 | 2 | | 7 | | | | | | 1 | | | | | | yes | | | yes | yes | | | | |
| VAX2 | N_Shore;Island | Body | chr2:71133854-71134936 | 1082 | | | 27 | | 3 | | | | | | | | | yes | yes | | | | | | | | |
| ATP6V1B1 | NA | TSS1500 | chr2:71162693-71163123 | 430 | | | | | | | | | | | | | | | | | yes | yes | | | | | |
| ATP6V1B1 | N_Shore | Body | chr2:71192044-71192445 | 401 | | 1 | | | | | | | | | | | | | yes | | | yes | | | | | |
| ANKRD53 | NA | 3'UTR;Body | chr2:71211981-71212504 | 523 | | | 8 | | | | | 0 | 2 | | | 2 | | | | | | | | | | | |
| PAIP2B | Island | TSS200 | chr2:71454280-71454317 | 37 | | | | | | | | | 1 | | | | | | | | | | | | | | |
| DYSF | N_Shore;Island | TSS1500;TSS200 | chr2:71679675-71680761 | 1086 | | | | | 1 | | | | 1 | | | | | | yes | yes | | | | | | | |
| DYSF | Island | Body | chr2:71787615-71788134 | 519 | | | 6 | | | | | | 1 | | | | | | yes | | | | | | | | |
| CYP26B1 | Island | Body | chr2:72371255-72371838 | 583 | | | 15 | | | | | | 1 | | | | | | | | yes | yes | yes | | | | |
| EXOC6B | Island | TSS200 | chr2:73053217-73053339 | 122 | | | | | | | | | 1 | | | | | | yes | | yes | | | | | | |
| NOTO | Island | TSS200 | chr2:73429331-73429791 | 460 | | | | | | | | | 2 | | | 1 | | yes | | | | yes | yes | | | | |
| NAT8 | NA | TSS200;5'UTR;1stExon | chr2:73869516-73870105 | 589 | | | | 1 | | | | | | | | | | yes | | yes | yes | | | | | | |
| ALMS1P | NA | TSS1500 | chr2:73871230-73871849 | 619 | | | | | | | | | | | | | | | yes | | yes | yes | | | | | |
| NAT8B | NA | 1stExon | chr2:73928348-73928500 | 152 | | | | | | | | | | | | | | yes | | | | | | | | | |
| TPRKB | S_Shore | TSS1500 | chr2:73964768-73964834 | 66 | | | 5 | | | | | | 1 | | | | | yes | | yes | | | | | | | |
| C2orf78 | S_Shelf | TSS1500 | chr2:74010396-74010621 | 225 | | | | | | | | | | | | | | yes | | | | | | | | | |
| ACTG2 | NA | TSS1500 | chr2:74119781-74120092 | 311 | | | 2 | | | | | | | | | | | yes | | | | | | | | | |
| DGUOK | N_Shore | TSS1500 | chr2:74153086-74153317 | 231 | | | | 1 | | | | | 1 | 1 | | | | | | | yes | yes | | | | | |
| BOLA3 | S_Shore | TSS1500 | chr2:74375507-74375930 | 423 | | | | | | | | | | | | | | | yes | yes | | | | | | | |
| DCTN1 | NA | Body | chr2:74601431-74601926 | 495 | | | 2 | | | | | | | | | | | | | | | | | | | | |
| DCTN1 | NA | 5'UTR;1stExon | chr2:74607470-74607600 | 130 | | | | 1 | | | | | | | | | | | | | | | | | | | |
| LOC100189589 | NA | TSS1500;TSS200 | chr2:74612020-74612812 | 792 | | | | | | | | | | | | | | | | | | | | | | | |
| RTKN | Island | Body;5'UTR | chr2:74663416-74663598 | 182 | | | | | | | | | | | | | | | | | | | | | | | |
| RTKN | Island | Body;TSS1500;1stExon;5'UTR | chr2:74668072-74669516 | 1444 | | | 22 | 1 | | | | 0 | 6 | | | 2 | | yes | | | | | | | | | |
| CCDC142;MRPL53 | S_Shore | 3'UTR;TSS200 | chr2:74700039-74700246 | 207 | | | 1 | | | | | | | | | | | yes | | | | | | | | | |
| LBX2 | Island | 3'UTR | chr2:74725040-74725369 | 329 | | | | 1 | | | | | | | | | | yes | | | | | | | | | |
| LBX2 | Island | Body | chr2:74726478-74727286 | 808 | | 1 | | | | | | | | | | | | | | | | | | | | | |
| LOC151534;LBX2 | Island;S_Shore | TSS1500;Body;TSS200 | chr2:74729429-74731413 | 1984 | | | 21 | | | | | | | | | | | | | | | | | | | | |
| TLX2 | N_Shore | TSS1500 | chr2:74740238-74740746 | 508 | | 1 | | | | | | | | | | | | yes | | | | | yes | | | | |
| TLX2 | Island | Body | chr2:74742362-74742786 | 424 | | | | | | | | | | | | | | | | | | | | | | | |
| DQX1 | N_Shelf | 1stExon;5'UTR | chr2:74753281-74753759 | 478 | | | | | 2 | | | | 2 | 1 | | | | | | | | | | | | | |
| LOXL3 | N_Shore | 5'UTR | chr2:74780172-74780617 | 445 | | | 27 | | | | | | | | | | | | | | | yes | yes | | | | |
| DOK1 | Island | Body | chr2:74782684-74783039 | 355 | | | | | | | | | | | | | | | | | | | | | | | |
| C2orf65 | Island | TSS200;TSS1500 | chr2:74875227-74875932 | 705 | | | 15 | | | | | 0 | 2 | | | 1 | | yes | yes | | yes | | | | | | |
| FAM176A | Island | 5'UTR | chr2:75787193-75787340 | 147 | | | | | | | | | 1 | | | | | yes | | | | | | | | | |
| FAM176A | NA | 5'UTR;1stExon;TSS1500 | chr2:75796759-75798232 | 1473 | | | | | | | | | 1 | | | | | | yes | yes | | | yes | | | | |
| C2orf3 | S_Shore;Island | TSS200 | chr2:75938118-75938438 | 320 | | | 2 | | | | | 0 | 2 | | | 1 | | | | yes | yes | | | | | | |
| SNAR-H | NA | TSS200 | chr2:78182187-78182350 | 163 | | | | | | | | | | | | | | yes | yes | | | | | | | | |
| REG3G | NA | TSS200 | chr2:79252673-79253210 | 537 | | | 1 | | | | | | | | | | | | | | | | yes | | | | |
| REG1A | NA | TSS200 | chr2:79347474-79347699 | 225 | | | | | | | | | | | | | | | | | | | | | | | |
| CTNNA2 | Island;N_Shore | TSS200;1stExon;5'UTR;TSS1500 | chr2:79739278-79740322 | 1044 | | | | | | | | 0 | 2 | | | | | | yes | | | | yes | | | | yes |
| CTNNA2;LRRTM1 | Island | Body | chr2:80529990-80530431 | 441 | | 1 | | | | | | | | | | | | | | | | | | | | | |
| CTNNA2;LRRTM1 | Island | Body;1stExon;5'UTR;TSS200 | chr2:80531483-80531898 | 415 | | | | | | | | | 2 | | | | | | | | | yes | yes | | | | |
| CTNNA2 | Island | Body | chr2:80549579-80549973 | 394 | | 1 | | | | | | | | | | | | | | | yes | yes | yes | | | | |
| LOC388965 | N_Shore | TSS1500 | chr2:84517321-84517763 | 442 | | | 3 | | | | | | | | | | | yes | | | | | | | | | |
| DNAH6 | N_Shore;Island | TSS1500;TSS200 | chr2:84743263-84743743 | 480 | | | | | | | | 0 | 1 | | | | | yes | | | yes | | | | | | |
| C2orf89 | N_Shore | Body | chr2:85106954-85107488 | 534 | | | 6 | | | | | | | | | | | yes | | | yes | | | | | | |
| TCF7L1 | Island;chr2:85359956-8536259 | Body;Island | chr2:85361485-85362480 | 995 | | 1 | | | | | | | | | 1 | | | | | | | | | | | | |
| CAPG | N_Shelf | 5'UTR;1stExon;TSS200 | chr2:85637116-85638535 | 1419 | | | 9 | | | | | | 2 | | | | | yes | | | | | | | | | |
| VAMP5 | N_Shore;Island | TSS1500;TSS200 | chr2:85810844-85812023 | 1179 | | 1 | | | 3 | | 1 | 0 | 1 | 2 | | 2 | | yes | yes | | | | yes | | | | |
| USP39 | N_Shore | TSS1500 | chr2:85843042-85843078 | 36 | | 1 | | | | | | | | | | | | yes | | | yes | | | | | | |
| GNLY | NA | 5'UTR;1stExon;TSS1500 | chr2:85921036-85921963 | 927 | | | | | | | | | 2 | | | | | yes | yes | yes | yes | | | | | | |
| ATOH8 | Island | TSS1500 | chr2:85980500-85980609 | 109 | | | | 1 | 1 | | | | | | | | | yes | | | | | | | | | |
| MRPL35 | S_Shelf | TSS200 | chr2:86426460-86426669 | 209 | | | | | | | | | 1 | | | | | yes | | | | | | | | | |
| VPS24 | Island | TSS200 | chr2:86790614-86790627 | 13 | | | | | | | | 0 | | | | | | | | | | | | | | | |
| CD8A | Island;S_Shore | 5'UTR;1stExon;Body;TSS200 | chr2:87018104-87018966 | 862 | | | 5 | | | | | 0 | 1 | | | | | yes | yes | | | | yes | | | | |
| CD8A | Island | TSS1500 | chr2:87036626-87037038 | 412 | 1 | | 1 | | | | | 0 | 1 | | | 1 | | yes | yes | | | yes | yes | | | | |
| SMYD1 | NA | Body | chr2:88390167-88390376 | 209 | | | | | | | | | 1 | | | | | | | | | | | | | | |
| FABP1 | NA | TSS200;Body;1stExon;5'UTR | chr2:88427027-88428542 | 1515 | | | 12 | 1 | | | | | | | | | | yes | yes | | | yes | | | | | |
| C2orf51 | NA | TSS200 | chr2:88823999-88824472 | 473 | | | | | | | | | | | | | | yes | yes | | yes | | | | | | |
| FLJ40330 | Island | Body | chr2:89065442-89065605 | 163 | | | | | | | | 0 | | | | 1 | | | | | | | yes | | | | |
| LOC654342 | NA | TSS200 | chr2:91847976-91847996 | 20 | | | 7 | | | | | | | | | | | | | | | | | yes | | | |
| ANKRD20B | Island | TSS200 | chr2:95522876-95522902 | 26 | | 1 | | | | | | | | | | | | | | | | | | | | | |
| TEKT4 | N_Shore;Island | TSS200;1stExon;5'UTR | chr2:95537044-95537708 | 664 | | | 10 | 1 | | | | 0 | 2 | 1 | | 1 | | | | | | | | | | | |
| TEKT4 | N_Shore | Body | chr2:95539964-95540415 | 451 | | | | | | | | | | | | | | | | | | | | | | | |
| MAL | Island | Body | chr2:95692352-95692524 | 172 | | | | | | | | | 1 | | | | | | | | | | | | | | |
| PROM2 | NA | TSS1500;TSS200 | chr2:95939815-95940475 | 660 | | | | | | | | | | | | | | yes | yes | | | | | | | | |
| PROM2 | NA | Body | chr2:95941719-95941820 | 101 | | | | | 1 | | | | 1 | | | | | yes | | | | | | | | | |
| KCNIP3 | NA | Body | chr2:95971840-95971994 | 154 | | | 2 | | 1 | | | | | | | | | | yes | | | | | | | | |
| ASTL | NA | TSS200 | chr2:96804247-96804757 | 510 | | | | | | | 1 | | | | | | | | yes | | | | yes | | | | |
| CIAO1;TMEM127 | N_Shore | TSS1500;Body | chr2:96930546-96930860 | 314 | | | 1 | | | | | | | | | | | | yes | | | | | | | | |
| ITPRIPL1 | N_Shore | TSS1500 | chr2:96990549-96990861 | 312 | | | 3 | 1 | 1 | | | | | | | | | yes | | | | | | | | | |
| NEURL3 | N_Shore | Body | chr2:97166001-97166422 | 421 | | | | | | | | | 1 | | | | | | yes | | yes | | | | | | |
| NEURL3 | N_Shelf | Body | chr2:97171137-97171449 | 312 | | 1 | | 1 | | 1 | | | | | 1 | | | yes | | | yes | | | | | | |
| NEURL3 | S_Shore | TSS1500 | chr2:97174815-97175065 | 250 | | | 1 | | | | | | | | | | | | yes | | | | | | | | |

| | | | | | | | | | | | | | | | | | | | | | | | | |
|---|---|---|---|---|---|---|---|---|---|---|---|---|---|---|---|---|---|---|---|---|---|---|---|---|
| ARID5A | Island | Body | chr2:97215789-97216131 | 342 | | | | 1 | 1 | | | | | | | | | | | | yes | | yes | |
| FER1L5 | NA | Body;TSS200 | chr2:97361034-97361073 | 39 | | | | | | | | | | | | | | yes | | | | | | |
| CNNM4 | Island;S_Shore | 1stExon | chr2:97427027-97427975 | 948 | | | | | 2 | | | | | | | | | | | | | yes | yes | |
| ANKRD23 | Island | 3'UTR | chr2:97505275-97505708 | 433 | | | 2 | | 1 | | | 0 | 1 | 1 | | 1 | | | | | | | | |
| ANKRD39 | S_Shore | TSS1500 | chr2:97524862-97525009 | 147 | | | | | | | | | | | | | | yes | | | | | | |
| SEMA4C | Island | Body | chr2:97530602-97530930 | 328 | | 1 | | | | | | | | | | | | | | | | | | |
| ZAP70 | NA | TSS200 | chr2:98329830-98330087 | 257 | | | | | 1 | | | | | | | | | | | | | | | |
| ZAP70 | Island | Body;5'UTR | chr2:98340425-98340921 | 496 | | | 13 | | | | | | 1 | | | | | | yes | | | | | |
| TMEM131 | NA | Body | chr2:98377791-98377939 | 148 | | | | | | | | 0 | 1 | | | | | | | | yes | | | |
| VWA3B | NA | Body | chr2:98852704-98852895 | 191 | | | | | | | | | 1 | | | | | yes | | | | | | |
| VWA3B | NA | Body | chr2:98928364-98928898 | 534 | | | 10 | | | | | | 5 | | | 1 | | | | | | | | |
| CNGA3 | N_Shore;Island | TSS200;1stExon;5'UTR | chr2:98962444-98964436 | 1992 | | 1 | 4 | | | | | | 2 | | | | | | | | | yes | | |
| CNGA3 | NA | Body | chr2:98993614-98993730 | 116 | | | 4 | | | | | | | | | | | | | | | | | |
| CNGA3 | NA | Body | chr2:99013128-99013409 | 281 | | | | | | | | | | | | | | | | | | | | |
| C2orf55 | Island | Body | chr2:99439533-99439883 | 350 | | 1 | | | | | | | | | | | | yes | | | | | yes | |
| LYG2 | NA | TSS200 | chr2:99871738-99872119 | 381 | | | | | | | | | | | | | | yes | | | | | | |
| AFF3 | NA | Body | chr2:100170766-100171136 | 370 | | | 1 | | 1 | | | | 1 | | | | | yes | yes | | | | | |
| AFF3 | S_Shore | TSS200;5'UTR;1stExon | chr2:100722022-100722584 | 562 | | | 6 | | | | | | | | | | | | | | | | yes | |
| AFF3 | NA | 1stExon;5'UTR;TSS200 | chr2:100759014-100759623 | 609 | | 1 | | | 1 | | | | 1 | 1 | 1 | | | | yes | | | yes | | |
| LONRF2 | N_Shore | Body | chr2:100937072-100937584 | 512 | | | | | 1 | | | | 1 | | | | | | | | yes | | | |
| RPL31 | N_Shore | TSS1500 | chr2:101617804-101618261 | 457 | | | | | | | | | | | | | | | yes | | | | | |
| RNF149 | Island | 1stExon | chr2:101924912-101925327 | 415 | | | | | | | | | | | | | | yes | | | | | yes | |
| IL1R2 | NA | TSS200 | chr2:102608155-102608349 | 194 | | | | | | | | 0 | 3 | 2 | | | | | | yes | | | yes | |
| IL1RL2 | N_Shore | TSS200 | chr2:102803290-102803787 | 497 | | | | | 1 | | | | 2 | | | | | | | | yes | | | |
| IL1RL1 | NA | TSS1500 | chr2:102927278-102927898 | 620 | | | | | | | | | | | | | | yes | | | | yes | | |
| IL1RL1 | NA | 5'UTR;TSS1500 | chr2:102952285-102953421 | 1136 | | | | | | | | | | | | | | yes | | | | | | |
| TMEM182 | NA | TSS1500 | chr2:103378135-103378691 | 556 | | | 3 | | | | | | | | | | | | | | | | yes | |
| POU3F3 | Island | TSS1500 | chr2:105471252-105472193 | 941 | | | | | | | | | 2 | | | | | | | | | yes | | |
| MRPS9 | NA | Body | chr2:105697589-105697777 | 188 | | | | | 1 | | | | | | | | | | | | | | | |
| GPR45 | N_Shore | TSS1500;TSS200 | chr2:105857955-105858489 | 534 | | | | | | | | | 3 | | | | | | yes | | | | | |
| FHL2 | Island | TSS200;5'UTR | chr2:106015739-106015771 | 32 | | | 6 | | | | | 0 | 1 | | | 1 | | | | | | | yes | |
| FHL2 | NA | 1stExon;5'UTR;TSS1500 | chr2:106054920-106055507 | 587 | | | 21 | | | | | | | | | | | yes | yes | | | | | |
| NCK2 | S_Shore | 5'UTR | chr2:106362547-106362832 | 285 | | | 6 | | | | | | | | | | | | | | | | | |
| C2orf40 | N_Shore;Island | TSS1500;TSS200;1stExon;5'UTR | chr2:106681831-106682640 | 809 | | 1 | | | | | | | 1 | | | | | | | | | | yes | |
| UXS1 | NA | Body | chr2:106764528-106777038 | 12510 | | | | 1 | | | | | 1 | 1 | 1 | | | yes | yes | yes | yes | yes | yes | |
| ST6GAL2 | Island | TSS200;TSS1500 | chr2:107503662-107504169 | 507 | | | | | | | | | 3 | | | | | | | | | | yes | |
| LOC729121;RGPD4 | Island | TSS1500;TSS200 | chr2:108443292-108443305 | 13 | | | | | | | | | 1 | | | | | yes | | | | | | |
| SLC5A7 | N_Shore;Island | TSS1500;TSS200 | chr2:108602361-108603244 | 883 | | | | | | | | | 3 | | | | | | | | | yes | yes | |
| SULT1C2 | NA | TSS1500;5'UTR;1stExon | chr2:108904812-108905468 | 656 | | | | | | | | 0 | 2 | 2 | 1 | 2 | 1 | yes | | | | | | |
| SULT1C4 | NA | TSS1500;TSS200 | chr2:108993737-108994528 | 791 | | | 4 | | | | | | 2 | | | | | yes | yes | | | | | |
| EDAR | Island | 5'UTR | chr2:109559185-109559473 | 288 | | | | | | | | | | | | | | yes | | | | | | |
| EDAR | NA | 5'UTR;1stExon | chr2:109605518-109606018 | 500 | | | 2 | | | | | | | | | | | yes | | | | | | |
| LOC100287216;SH3RF3 | Island | Body;TSS200;1stExon | chr2:109745828-109747003 | 1175 | | | | | | | | 0 | 3 | | | 4 | | | | | | | yes | |
| SH3RF3 | NA | Body | chr2:109788051-109788583 | 532 | | | | 1 | | | | | | | | | | | | | | | | |
| SH3RF3 | NA | Body | chr2:109892050-109892206 | 156 | | 1 | | | | | | | | | | | | yes | | | yes | | | |
| SH3RF3 | NA | Body | chr2:109934728-109934860 | 132 | 2 | | | | | | | | 1 | | 1 | | | | | | | | | |
| SEPT10;ANKRD57 | Island | Body;TSS1500 | chr2:110371141-110371632 | 491 | | | | | | | | | 1 | | | | | | | yes | yes | | yes | |
| SEPT10;ANKRD57 | Island | TSS1500;1stExon | chr2:110372875-110373509 | 634 | | | 4 | | | | | | 1 | | | | | | | | | | | |
| MALL | N_Shore;Island;NA | Body | chr2:110863886-110873396 | 9510 | | | 4 | | | | | | 1 | | | | | yes | yes | | | yes | | |
| NCRNA00116 | N_Shore;Island | Body | chr2:110969641-110970249 | 608 | | 1 | 6 | | | | | 0 | 3 | 1 | | 13 | | yes | | | yes | | | |
| ACOXL | NA | Body | chr2:111556249-111556607 | 358 | | | | | | | | | | | | | | yes | | yes | | | | |
| ACOXL | Island | 3'UTR | chr2:111875749-111876361 | 612 | | | | | 3 | | | | | | | | | | | | | | | |
| FBLN7 | N_Shore | TSS1500 | chr2:112895559-112895793 | 234 | | | | | | | | 0 | | | | 1 | | yes | | yes | | | yes | |
| FBLN7 | Island | Body | chr2:112917127-112917424 | 297 | | 1 | | | | | | | 1 | | | | | yes | | | | | | |
| FBLN7 | N_Shore | Body | chr2:112939119-112939562 | 443 | | | 4 | | | | | 0 | | | | | | | yes | | | | | |
| RGPD8;RGPD5 | S_Shore | TSS1500 | chr2:113191815-113192561 | 746 | | | | | | | | | 1 | | | | | | yes | | | | | |
| IL1B | NA | Body;5'UTR;1stExon | chr2:113593785-113594654 | 869 | | | 5 | | 2 | | | | | | | | | | | | | | yes | |
| IL1F7 | NA | TSS1500 | chr2:113669419-113670582 | 1163 | | | 5 | | | | | | 1 | | | | | yes | | yes | yes | | yes | |
| IL1F7 | NA | Body;TSS200 | chr2:113672587-113672758 | 171 | | | | | | | | | | | | | | | | | yes | | | |
| IL1F9 | NA | TSS1500;TSS200 | chr2:113735377-113735612 | 235 | | | | | | | | | 2 | | 1 | | | | | | | | | |
| IL1F8 | S_Shore | 5'UTR | chr2:113810244-113810743 | 499 | | | | | | | | | | | | | | yes | | yes | | | | |
| IL1RN | NA | TSS1500 | chr2:113875226-113875723 | 497 | | | | | | | | | 2 | | | | | | yes | | yes | | | |
| PSD4;LOC440839 | NA | TSS1500;Body;TSS200 | chr2:113930819-113931565 | 746 | | | | | 1 | | | | 1 | | 1 | | | yes | | | yes | | | |
| PAX8;LOC440839;LOC654433 | N_Shore | Body;TSS1500 | chr2:113992762-113994035 | 1273 | | | 20 | 5 | | | 2 | 0 | 13 | 6 | 3 | 19 | 2 | yes | | | | | | |
| PAX8;LOC440839 | Island | Body | chr2:114033360-114033830 | 470 | | | 6 | 1 | | | | | | | | | | | | | | | | |
| LOC440839 | Island | Body | chr2:114048829-114049041 | 212 | | | | 1 | | | | 0 | 1 | | | | | yes | yes | | | | | |
| WASH2P | Island | Body | chr2:114341631-114342108 | 477 | | | | | | | | 0 | | | | | | | yes | | | | | |
| MGC13005 | NA | TSS1500;Body | chr2:114359736-114359873 | 137 | | | | | | | | 0 | | | | | | | yes | | yes | yes | | yes |
| RPL23AP7;RABL2A | Island | TSS200;5'UTR;1stExon | chr2:114384855-114384875 | 20 | | | | | | | | | 1 | | | | | yes | | | | | | |
| DPP10 | N_Shore;Island | Body | chr2:115419537-115420260 | 723 | 1 | 1 | 8 | | | | | 0 | 3 | 3 | 1 | 6 | | yes | | | | | | |
| DPP10 | Island | TSS1500;Body;1stExon;5'UTR | chr2:115918745-115920825 | 2080 | | 2 | | | | | | 0 | 3 | | | | | | yes | | | yes | yes | |
| EN1 | Island | 3'UTR | chr2:119599748-119600627 | 879 | | | | | | | | | | | | | | | | | | yes | yes | |
| EN1 | N_Shore;Island | Body | chr2:119602212-119603969 | 1757 | | | 15 | | | | | | | | | | | | | | | yes | yes | |
| EN1 | Island;N_Shore | TSS1500 | chr2:119605959-119609691 | 3732 | | | | | | | | | | | | | | | | | | | | |
| MARCO | NA | TSS200 | chr2:119699682-119700029 | 347 | | | | | | | | | | | | | | | | | yes | | | |
| TMEM37 | N_Shore;Island | Body | chr2:120189867-120190297 | 430 | | | 22 | | 2 | | | | | | | | | yes | yes | | | | | |
| SCTR | Island;N_Shore | TSS200;Body | chr2:120281526-120282272 | 746 | | | 2 | | | | | | 1 | | | | | yes | | | yes | | | |
| INHBB | Island;N_Shore | 1stExon;Body | chr2:121104029-121105535 | 1506 | | | 3 | | | | | | 1 | | | | | | | | | yes | yes | |
| LOC84931 | NA | TSS200;TSS1500;Body | chr2:121223534-121224679 | 1145 | | | 3 | | | | 1 | 0 | | | | | | yes | | | | | | |
| GLI2 | NA | TSS1500;TSS200 | chr2:121554307-121554926 | 619 | | | 32 | | | | | | 1 | | | | | | yes | | | | yes | |
| GLI2 | Island;S_Shore | Body | chr2:121624862-121625735 | 873 | 1 | 1 | | | | | | 0 | 2 | | | 1 | | yes | | | | | yes | |
| GLI2 | NA | Body | chr2:121670282-121670634 | 352 | | | 1 | | | | | | | | | | | | | | yes | | | |
| GLI2 | S_Shore | Body | chr2:121748215-121748257 | 42 | | | 6 | | | | | | | | | | | | | | | | | |
| CNTNAP5 | N_Shore;Island | TSS1500;TSS200;1stExon | chr2:124782117-124783698 | 1581 | | 1 | 2 | | | | | | 6 | | | | | | | | | | yes | |
| GYPC | N_Shore | TSS1500 | chr2:127413363-127413369 | 6 | | 1 | | | | | | 0 | 1 | | | | | | | | | yes | | |
| GYPC | S_Shore | Body | chr2:127414802-127415590 | 788 | | | 3 | | | | | | 2 | 1 | | | | yes | | | | | | |
| BIN1 | Island | Body | chr2:127818000-127818226 | 226 | | | | 1 | | | | | | | | | | | | | yes | | | |
| CYP27C1 | NA | 5'UTR;1stExon;TSS200 | chr2:127963067-127963653 | 586 | | | 4 | | | | | | 4 | | | 1 | | yes | | yes | | | | |
| PROC | Island | Body | chr2:128180488-128181325 | 837 | | | | 1 | | | | | | | | | | yes | yes | | yes | | yes | |
| MYO7B | NA | TSS1500 | chr2:128293021-128293239 | 218 | | 1 | | | | | | | | | | | | yes | | yes | | | | |
| MYO7B | NA | Body | chr2:128389103-128389207 | 104 | | | | | | | | | 1 | | | | | yes | | | | | | |
| LIMS2 | Island | Body | chr2:128410602-128410803 | 201 | 2 | | | | | | | | | | | | | | yes | | | | | |
| LIMS2 | Island;S_Shore | Body;5'UTR;1stExon;TSS200;TSS1 | chr2:128421828-128422717 | 889 | | | 10 | 1 | | | | | | | | | | yes | | | | | | |
| LIMS2 | NA | 5'UTR;1stExon | chr2:128438738-128439345 | 607 | | | 8 | 1 | | | | | 1 | | | | | yes | yes | | | | yes | |
| SFT2D3 | N_Shore | TSS1500 | chr2:128458240-128458556 | 316 | | 1 | 3 | | | | | | 2 | 1 | | | | | yes | yes | | | | |
| HS6ST1 | Island | TSS1500;TSS200 | chr2:129076335-129077561 | 1226 | | | 5 | | | | | 0 | 2 | | | | | | | | | yes | yes | |
| LOC389033 | Island | Body | chr2:130691738-130691926 | 188 | | | | | | | | 0 | | | | | | yes | | yes | | | yes | |
| RAB6C | Island | TSS200 | chr2:130737216-130737664 | 448 | | 1 | | | | | | | 1 | | | | | | | yes | | | yes | |
| LOC440905 | Island | Body | chr2:130795350-130795713 | 363 | | | | | | | | 0 | 1 | | | | | yes | | | | | | |

| | | | | | | | | | | | | | | | | | | | | | | | |
|---|---|---|---|---|---|---|---|---|---|---|---|---|---|---|---|---|---|---|---|---|---|---|---|
| POTEF | N_Shore | Body | chr2:130877827-130878313 | 486 | | | | | | | | | | | | | | | | | | yes | |
| CCDC74B | Island | 3'UTR | chr2:130897120-130897302 | 182 | | 1 | | | | | | | | | | | | | | | | | |
| CCDC74B | N_Shore | Body | chr2:130901223-130901674 | 451 | | | | | | | | | 1 | | | | | yes | | | | | yes |
| SMPD4;FAM128B | S_Shore | TSS1500;Body | chr2:130940668-130940706 | 38 | | | 1 | | | | | | 1 | | | | | yes | | | | | |
| TUBA3E | Island | TSS200 | chr2:130956098-130956199 | 101 | | | | | | | | | | 1 | | | | yes | | yes | | | |
| PTPN18 | N_Shore | Body | chr2:131129567-131130131 | 564 | | | | 1 | | | | | | | | | | yes | | | | | yes |
| FAM123C | N_Shore | TSS1500;TSS200;5'UTR;1stExon | chr2:131512651-131513724 | 1073 | 1 | | 4 | | | | | | 3 | | | | | | | | | | yes |
| ARHGEF4 | N_Shore;Island;S_Shore | TSS1500;TSS200 | chr2:131673406-131674494 | 1088 | | | 10 | 1 | | | | 0 | 6 | | | 1 | | | | | | | |
| ARHGEF4 | S_Shore | 5'UTR | chr2:131676068-131676293 | 225 | | | | | | | | | | | | | | yes | yes | | | | |
| ARHGEF4 | Island;S_Shore | Body | chr2:131721692-131722722 | 1030 | | | | 1 | | | | | 1 | 1 | | | | | | | | | yes |
| ARHGEF4 | Island | Body | chr2:131792528-131793064 | 536 | | | | | 1 | | | | 1 | | | | | yes | | | | | yes |
| ARHGEF4 | Island | Body | chr2:131797438-131798087 | 649 | | | 10 | | 1 | | 1 | 0 | 4 | 1 | | 3 | | | | | | | |
| POTEE | NA | TSS200 | chr2:131975750-131975771 | 21 | | | 5 | | | | | | 1 | | | | | | | | | yes | |
| LOC440910 | N_Shore | Body | chr2:132056834-132057166 | 332 | | | | | | | | 0 | | | | | | yes | | | | | |
| LOC150786 | Island | 5'UTR;1stExon | chr2:132121314-132121815 | 501 | | 1 | | | | | | | | | | | | | | yes | | | yes |
| LOC401010 | Island | Body;TSS200 | chr2:132202427-132202537 | 110 | | | 4 | | | | 1 | 0 | 1 | | | | | | | | | yes | |
| C2orf27B | N_Shore;Island | 5'UTR;1stExon | chr2:132558939-132559484 | 545 | | | | | | | | | | | | 1 | | yes | | | | | yes |
| NCRNA00164;MIR663B | Island | Body | chr2:133014640-133014698 | 58 | | | | | | | | | 1 | | | | | | | | | | |
| GPR39 | N_Shore;Island;S_Shore | Body | chr2:133342838-133343657 | 819 | | | 4 | | | | | 0 | 1 | | | | | | | | | yes | |
| GPR39;LYPD1 | N_Shore;Island | Body;3'UTR | chr2:133402475-133403009 | 534 | | | 17 | | | | | 0 | 4 | | | | | | | | | | |
| LYPD1 | N_Shore | 5'UTR;Body | chr2:133426129-133426934 | 805 | | | 9 | | | | | | 1 | | | | | | | | | | yes |
| LYPD1;NCKAP5 | Island;S_Shore | TSS200;5'UTR;1stExon;TSS1500;3' | chr2:133428422-133430560 | 2138 | | | 18 | | | | | 0 | 1 | | | 1 | | | | | | yes | yes |
| NCKAP5 | NA | TSS200 | chr2:134326172-134326366 | 194 | | | 4 | | | | | | 1 | | | | | | | | | | |
| MGAT5 | NA | TSS1500 | chr2:135011522-135011937 | 415 | | | | | | | | | 1 | | | | | | | | | | |
| TMEM163 | NA | Body | chr2:135226787-135227167 | 380 | | | | | | | | | | | | | | yes | | yes | | | |
| DARS | Island | TSS1500 | chr2:136743460-136743698 | 238 | | | 2 | | | | | | | | | | | | yes | | | | |
| THSD7B | NA | TSS1500;TSS200 | chr2:137747719-137748833 | 1114 | 1 | | | | | | | | 2 | | | | | yes | yes | | | | yes |
| NXPH2 | Island | Body;TSS200;TSS1500 | chr2:139537745-139539181 | 1436 | | | | | 2 | | 1 | 0 | 8 | 4 | | 22 | 2 | yes | | | | | |
| LRP1B | S_Shore | TSS200 | chr2:142889283-142889392 | 109 | | 1 | 19 | | | | | | 1 | | | 4 | | | | | | yes | |
| ARHGAP15 | NA | TSS1500 | chr2:143886326-143886940 | 614 | | | 2 | | | | | | 1 | | 1 | | | | | | | | yes |
| GTDC1 | NA | 5'UTR;TSS200 | chr2:144995072-144995448 | 376 | | | | | | | | | 1 | | | | | | yes | yes | | | |
| ZEB2 | N_Shore;Island | 3'UTR;Body | chr2:145146928-145147558 | 630 | | 1 | 6 | 1 | 2 | | | | 2 | | | | | | | | | | |
| ZEB2 | N_Shelf | TSS200;TSS1500 | chr2:145278113-145278808 | 695 | | | | | | 1 | | | | | | | | | | | | | yes |
| MBD5 | NA | TSS1500 | chr2:149215431-149215884 | 453 | | | | | | | | | 1 | | | | | | | | yes | | yes |
| KIF5C | N_Shore | TSS1500 | chr2:149632599-149632683 | 84 | | | 1 | | | | | | | | | | | | | | | yes | |
| KIF5C;MIR1978 | NA | Body;TSS200 | chr2:149639612-149640259 | 647 | 5 | | | | | | | | | | | | | yes | yes | | | | |
| KIF5C | N_Shore;Island | Body | chr2:149866270-149866889 | 619 | 2 | 1 | | | | | | | | | | | | yes | yes | | | | |
| LYPD6B | N_Shore | TSS1500 | chr2:149894539-149894879 | 340 | | | | | | | | | | | | | | | yes | | yes | | yes |
| LYPD6 | Island | TSS1500 | chr2:150186706-150186923 | 217 | | | | | | | | | 1 | | | | | | | | | | yes |
| MMADHC | NA | TSS1500 | chr2:150444569-150444834 | 265 | | | 2 | | | | | | 1 | | | | | | yes | | | | |
| CACNB4 | NA | Body;1stExon;TSS200 | chr2:152830189-152831214 | 1025 | | | 12 | 1 | | | | 0 | 3 | | | | | yes | | | | | yes |
| RPRM | N_Shore;Island;S_Shore | 3'UTR;1stExon;5'UTR;TSS1500 | chr2:154334294-154336341 | 2047 | | | | | | | | | 4 | | | | | | yes | | | yes | yes |
| GALNT13 | Island | TSS1500 | chr2:154727907-154728468 | 561 | | | | | | | | | 3 | | | | | yes | | yes | | | yes |
| NR4A2 | N_Shore;Island | Body | chr2:157184137-157187204 | 3067 | | | 7 | | 2 | | | 0 | 6 | 1 | | | | | | | | | yes |
| GPD2 | N_Shore | TSS1500;1stExon;5'UTR | chr2:157291262-157292127 | 865 | | | 6 | | | 1 | | 0 | 3 | | 1 | 1 | | yes | | yes | yes | | |
| GALNT5 | NA | TSS200 | chr2:158114300-158114424 | 124 | | | | | | | | 0 | 1 | | | | | | | | | yes | |
| ERMN | NA | TSS1500 | chr2:158184535-158184641 | 106 | | | 3 | | | | | | | | | | | yes | yes | | | | |
| CYTIP | NA | 1stExon | chr2:158300475-158301005 | 530 | | | | | 1 | | | | 1 | 1 | | | | | | | | | |
| ACVR1C | N_Shore;Island | Body;5'UTR;1stExon;TSS200 | chr2:158453759-158454243 | 484 | | 1 | 2 | | 2 | | | 0 | | | | | | | | | yes | | |
| DAPL1 | NA | TSS1500;TSS200 | chr2:159651559-159652019 | 460 | | | | | | | | 0 | 1 | | | 3 | | | yes | | | | |
| BAZ2B | S_Shore | 1stExon;5'UTR;TSS200 | chr2:160472940-160473528 | 588 | | | | | | | | 0 | 3 | | | | | | | | | | yes |
| CD302 | N_Shore;S_Shore | Body;TSS200;TSS1500 | chr2:160653686-160655081 | 1395 | 3 | | | | 1 | | | 0 | 3 | | | 1 | | | yes | | | yes | |
| LY75 | Island | Body;TSS200 | chr2:160761085-160761622 | 537 | | | 4 | | | | | 0 | | | | | | | | | | | yes |
| PLA2R1 | S_Shore | TSS200 | chr2:160919253-160919282 | 29 | | | | | | | | | 1 | | | | | yes | | | | yes | |
| ITGB6 | NA | 1stExon | chr2:161056520-161056639 | 119 | | | | | 1 | | | | 1 | | | | | | | | | | yes |
| RBMS1 | S_Shore | Body | chr2:161265247-161265366 | 119 | | | 8 | | | | | | | | | | | | | | | | |
| TBR1 | N_Shore | TSS1500;TSS200 | chr2:162271751-162272687 | 936 | | | 13 | | | | | | | | | | | | | | | yes | |
| TBR1 | N_Shore | Body | chr2:162279686-162280009 | 323 | | | | | | | | | | | | | | | | | | | yes |
| SLC4A10 | NA | TSS1500 | chr2:162480565-162480821 | 256 | | | | | | | | | | | | | | | | | | | |
| DPP4 | N_Shore;S_Shore | Body;1stExon;5'UTR | chr2:162930175-162931397 | 1222 | | 1 | | | | | | | | | | | | yes | yes | | | | |
| GCA | N_Shore | TSS200 | chr2:163200532-163200637 | 105 | | | | | | | | 0 | | | | | | | | | yes | | |
| KCNH7 | NA | TSS200 | chr2:163695408-163695793 | 385 | | | | | | | | | 1 | | | | | | | | | yes | yes |
| FIGN | N_Shore | TSS200 | chr2:164592561-164592854 | 293 | | | | | | | | | 3 | | | | | | | | | | yes |
| SLC38A11 | NA | 1stExon;5'UTR;TSS200 | chr2:165811816-165812297 | 481 | | 1 | 16 | | | | | 0 | 2 | | | 5 | | yes | | | yes | | |
| GALNT3 | Island | TSS200 | chr2:166650930-166650965 | 35 | | | | | | | | | 1 | | | 1 | | | | | | | |
| B3GALT1 | NA | TSS1500;TSS200 | chr2:168674769-168675357 | 588 | 2 | | 6 | | | | | | 1 | | | | | yes | yes | | yes | | |
| NOSTRIN | NA | 5'UTR;TSS1500;TSS200 | chr2:169658881-169659182 | 301 | | 1 | 2 | | 1 | | | | | | | | | | yes | yes | | | |
| G6PC2 | NA | TSS200 | chr2:169757561-169757780 | 219 | | | | | | | | | | | | | | yes | | | | | |
| LOC440925 | N_Shore | Body | chr2:171568984-171569282 | 298 | | | 6 | | | | | | | | | | | yes | | | | | |
| LOC440925;SP5 | Island | Body;TSS1500 | chr2:171570373-171571007 | 634 | | | | | | | | | 1 | | | | | | | | | yes | yes |
| LOC440925;SP5 | Island;S_Shore | TSS1500;Body;3'UTR | chr2:171572348-171574592 | 2244 | | 1 | 3 | 1 | 1 | | | | 2 | | | | | | | | | yes | yes |
| GAD1 | N_Shore;Island | Body | chr2:171676306-171676925 | 619 | 1 | | 6 | | | | | | 1 | | | | | yes | | | yes | yes | |
| GAD1 | Island | Body | chr2:171678751-171680337 | 1586 | | | | | | | | | 2 | | | | | | | | | yes | |
| GORASP2 | N_Shore | TSS1500 | chr2:171784674-171785124 | 450 | | | 2 | | | | | | 2 | | | 3 | | | yes | | | | yes |
| MAP1D | N_Shore | TSS1500 | chr2:172864014-172864337 | 323 | | | 2 | | | | | | | | | | | | yes | | | | |
| MAP1D | chr2:172944988-172945384 | Island | chr2:172945328-172945698 | 370 | | | | | | | | | | | 1 | | | | yes | | | | |
| DLX1 | Island | TSS1500 | chr2:172949284-172950116 | 832 | | | | | | | | | 1 | | | | | | | | | | yes |
| DLX1 | Island;N_Shore;S_Shore | Body;3'UTR | chr2:172951501-172953925 | 2424 | | | | | | | | | | | | | | | | | | | |
| DLX2 | Island | Body | chr2:172965650-172966180 | 530 | | | 7 | | | | | | | | | | | | | | | | |
| ITGA6 | Island | Body | chr2:173292978-173293392 | 414 | | | | | | | | | | | | | | | | | | | |
| ITGA6 | NA | Body | chr2:173330296-173330395 | 99 | | | 2 | | 1 | | | | 1 | | | 1 | | | | | yes | | |
| RAPGEF4;LOC91149 | S_Shore | Body;TSS200 | chr2:173601103-173601413 | 310 | | | 6 | | | | | | | | | | | | | | | | |
| RAPGEF4 | NA | Body | chr2:173792545-173792879 | 334 | | | 4 | | 1 | | | | | | | | | yes | | | | | |
| SP9 | N_Shore | TSS1500 | chr2:175199259-175199349 | 90 | | | 7 | | | | | | | | | | | | | | | | |
| WIPF1 | NA | 5'UTR;1stExon | chr2:175499113-175499639 | 526 | | | 10 | 1 | 1 | | | | 2 | 1 | | | | | | | yes | | yes |
| CHRNA1 | NA | TSS200 | chr2:175629219-175629917 | 698 | | | 1 | | | | | | | | | | | | | | | | |
| ATP5G3 | N_Shore;Island | Body | chr2:176045768-176046632 | 864 | | | | | | | | | 1 | | | 1 | | | | | | | |
| EVX2 | N_Shore;Island | TSS200;Body;5'UTR;1stExon | chr2:176947435-176950273 | 2838 | | 2 | 29 | | | | | 0 | 5 | | | 4 | | | | | | yes | yes |
| HOXD13 | N_Shore;Island | TSS1500;TSS200 | chr2:176956396-176957908 | 1512 | | 1 | 12 | | | | | | | | | | | | | | | yes | yes |
| HOXD12 | N_Shore;Island | TSS1500;TSS200 | chr2:176963315-176965729 | 2414 | | | | | | | | | | | | | | | | | | | |
| HOXD11 | N_Shore;Island;S_Shore | TSS1500;TSS200;1stExon;Body | chr2:176971304-176974060 | 2756 | | | 25 | | | | | | 3 | | | | | | | | | yes | yes |
| HOXD10 | Island;N_Shore | TSS1500;TSS200;1stExon | chr2:176980837-176982342 | 1505 | | | 30 | | | | | | 2 | | | | | | | | | | |
| HOXD10 | S_Shore | Body | chr2:176983519-176983949 | 430 | | | 6 | | | | | | | | | | | | | | | yes | |
| HOXD9 | N_Shore;Island | TSS1500;TSS200;5'UTR;1stExon;Bo | chr2:176985953-176989586 | 3633 | | 1 | 21 | | | | | 0 | 4 | | | | | | | | | yes | yes |
| HOXD8 | Island | TSS1500;TSS200;5'UTR;1stExon | chr2:176993017-176995088 | 2071 | | | 30 | | | | | | 5 | | | | | | | | | yes | yes |
| MIR10B;HOXD4 | N_Shore;Island | TSS1500;TSS200;Body;5'UTR;1stEx | chr2:177014009-177017630 | 3621 | | 2 | 43 | | | | | 0 | 9 | | | 2 | | | | | | yes | yes |
| HOXD3 | N_Shore;Island;TSS200;chr2:17 | TSS1500;nan;Island;5'UTR | chr2:177027440-177029850 | 2410 | | 1 | | | | | | | | | 1 | | | | | | | | yes |
| HOXD1 | Island;chr2:177052957-177054 | TSS200;Island | chr2:177053283-177053345 | 62 | | | | | | | | | 3 | | 1 | | | | | | | | yes |
| MIR548N | NA | Body | chr2:179387853-179388064 | 211 | | | 12 | | | | 1 | 0 | 1 | 1 | | 2 | | yes | | yes | | | |

| | | | | | | | | | | | | | | | | | | | | | | | | | | | |
|---|---|---|---|---|---|---|---|---|---|---|---|---|---|---|---|---|---|---|---|---|---|---|---|---|---|---|---|
| TTN;MIR548N | NA | Body | chr2:179395885-179396214 | 329 | | | | | 2 | | | 0 | 2 | | | 1 | | | | | | | | | | | |
| TTN | NA | 1stExon;5'UTR | chr2:179672121-179672691 | 570 | | | 1 | | | | | | 1 | | | | | | | | | | | | | | |
| ZNF385B;MIR1258 | Island | TSS200;TSS1500 | chr2:180726252-180726349 | 97 | | | | | | | | | 1 | | | | | | | | | yes | yes | | | | |
| ITGA4 | N_Shore | TSS1500 | chr2:182321354-182321496 | 142 | | | 1 | | 2 | | | | 1 | | | | | | | | | | | | | | |
| ITGA4 | Island | 1stExon | chr2:182322501-182322841 | 340 | | 1 | | | | | | | | | | | | | | | | | | | | | |
| CERKL | Island | TSS200 | chr2:182521918-182521926 | 8 | | | | | | | | | | | | | | | | | yes | | | | | | |
| PPP1R1C | NA | TSS1500 | chr2:182850059-182850269 | 210 | | | 2 | | 1 | | | | 1 | | | | | yes | | | | | | | | | |
| PDE1A | NA | 1stExon;TSS200;5'UTR | chr2:183387392-183387885 | 493 | | | | | 1 | | | | | | | | | yes | | | | | yes | | | | |
| NCKAP1 | Island;S_Shore | 1stExon;TSS1500 | chr2:183902764-183903761 | 997 | | | | | | | | | 2 | | | | | | | | | yes | yes | | | | |
| DUSP19 | NA | TSS200;1stExon;5'UTR;TSS1500 | chr2:183942982-183944048 | 1066 | | | 2 | | | | | 0 | 5 | | | 2 | | | | yes | | | | | | | |
| ZNF804A | N_Shore | TSS1500 | chr2:185462269-185462655 | 386 | | 1 | | | | | | | | | | | | | | | | | yes | | | | |
| GULP1 | Island | TSS1500;TSS200 | chr2:189156425-189157361 | 936 | | | | | | | | | 3 | | | | | | | | | | | | | | |
| COL5A2 | NA | 1stExon;5'UTR;TSS200 | chr2:190044504-190044983 | 479 | | | | | | | | 0 | 2 | | 1 | 1 | | yes | | | | | | | | | |
| SLC40A1 | Island | TSS1500 | chr2:190445880-190446792 | 912 | | | 1 | | | | | | | | | | | yes | | | | yes | | | | | |
| C2orf88 | N_Shore | TSS1500;5'UTR | chr2:191044481-191045026 | 545 | | | | | | | | | | | | | | yes | | | | | yes | | | | |
| TMEM194B | N_Shore | Body | chr2:191398360-191398806 | 446 | 1 | | 7 | | | | | | 1 | | | 1 | | yes | | yes | | | yes | | | | |
| STAT4 | NA | 1stExon;5'UTR | chr2:192015726-192015996 | 270 | | | 8 | | | | | | | | | | | yes | | | | | | | | | |
| MYO1B | Island | TSS200;TSS1500 | chr2:192109990-192110021 | 31 | | | | | | | | | 1 | | | | | | | | | | | | | | |
| OBFC2A | S_Shore | Body | chr2:192543718-192543908 | 190 | | | | 1 | | | | | | | | | | yes | | | | | yes | | | | |
| SDPR | NA | Body | chr2:192701019-192701514 | 495 | | | | | | | | | 1 | | | | | | | | | | | | | | |
| DNAH7 | NA | Body | chr2:196642630-196642673 | 43 | | | | | | | | | 1 | | | | | yes | yes | | | | | | | | |
| DNAH7 | S_Shore | TSS1500 | chr2:196933852-196934154 | 302 | | | 3 | | | | | | | | | | | | yes | | | | | | | | |
| HECW2 | Island | TSS200 | chr2:197457393-197457420 | 27 | | | | | | | | | 1 | | | | | | | | | | yes | | | | |
| CCDC150 | N_Shore | TSS1500 | chr2:197503878-197504139 | 261 | | | 2 | | | | | | 1 | | | | | yes | | | | | | | | | |
| ANKRD44 | Island | Body | chr2:198029141-198029332 | 191 | | 1 | 5 | | | | | | 1 | | | | | yes | | | | | | | | | |
| ANKRD44 | NA | TSS1500 | chr2:198063653-198063781 | 128 | | | 2 | 1 | | | | | | | | | | yes | | | | yes | | | | | |
| RFTN2 | NA | 1stExon;5'UTR | chr2:198540518-198540621 | 103 | | 1 | | | | | | | 1 | | | 1 | | | | | | yes | yes | | | | |
| BOLL | N_Shore;Island | 5'UTR;Body;1stExon;TSS200;TSS15 | chr2:198649727-198651576 | 1849 | | 1 | 8 | | | | | 0 | 10 | 1 | | 5 | | yes | | | | | | | | | |
| SATB2 | Island;N_Shore | 5'UTR;Body | chr2:200320455-200321348 | 893 | | | 33 | | | | | | | | | | | | | | | | | | | | |
| SATB2 | Island | TSS1500 | chr2:200326591-200327336 | 745 | | 1 | 7 | | | | | | 1 | | | | | | | | | | | | | | |
| FLJ32063 | S_Shore;N_Shore | TSS1500 | chr2:200331675-200332380 | 705 | | | 18 | | | | | | | | | | | | | | | | yes | | | | |
| FLJ32063 | N_Shore;Island | Body | chr2:200333401-200335874 | 2473 | | | | | | | | | | | | | | | | | | | | | | | |
| AOX1 | N_Shore | TSS1500 | chr2:201450506-201450743 | 237 | | | | | | | | 0 | | | | | | | yes | | yes | | | | | | |
| AOX2P | NA | TSS200 | chr2:201635337-201635643 | 306 | | | 2 | | | | | | | | | | | | | yes | | yes | | | | | yes |
| NDUFB3;FAM126B | Island | TSS1500;5'UTR | chr2:201936101-201936323 | 222 | | | | | | | | | 1 | | | | | | | | | | | | | | |
| CASP10 | NA | TSS1500 | chr2:202047246-202047473 | 227 | | | | | | | | | 1 | | | | | yes | | | | | yes | | | | |
| CASP8 | NA | 5'UTR;TSS1500;TSS200 | chr2:202122309-202122669 | 360 | | | | 2 | | | | | | | | | | yes | | | | yes | | | | | |
| ALS2CR11 | S_Shore;Island | TSS200;1stExon;5'UTR | chr2:202483704-202484320 | 616 | | | 6 | | | | | 0 | 2 | | | 1 | | | yes | | | | | | | | |
| ALS2CR4 | S_Shore | TSS1500 | chr2:202508694-202508813 | 119 | | | 4 | | | | | | 1 | | | | | | | yes | | yes | | | | | |
| CDK15 | NA | TSS200 | chr2:202671158-202671583 | 425 | | | 1 | | | | | | | | | | | | | | | | | | | | |
| FZD7 | Island;S_Shore | 1stExon;3'UTR | chr2:202901045-202901470 | 425 | | | 7 | | | | | 0 | 4 | 2 | 1 | 3 | | | | | | | | | | | |
| CYP20A1 | N_Shore | TSS1500 | chr2:204102418-204102500 | 82 | | | | | | | | | 1 | | | | | yes | | | yes | | | | | | |
| CD28 | NA | TSS1500 | chr2:204569955-204570067 | 112 | | | | | | | | | 1 | | | | | | yes | | | yes | yes | | | | |
| NRP2 | Island | Body | chr2:206551212-206551466 | 254 | | 1 | 3 | | | | | | | | | | | | | | | yes | | | | | |
| NRP2 | NA | Body | chr2:206628553-206628692 | 139 | | | 3 | | | | | 0 | 1 | | | 3 | | yes | | | | | | | | | |
| ZDBF2 | Island | TSS200 | chr2:207139431-207139445 | 14 | | | | | | | | | 1 | | | | | yes | | | | yes | yes | | | | |
| LOC200726 | N_Shore;Island | TSS1500;TSS200 | chr2:207506535-207507528 | 993 | | 1 | 1 | | | | | | 2 | | | | | | | | | | | | | | |
| FZD5 | S_Shore | TSS1500 | chr2:208635549-208635942 | 393 | | 1 | 1 | | | | | | | | | | | yes | yes | | | | | | | | |
| PLEKHM3 | Island | Body | chr2:208795620-208795859 | 239 | | | | 1 | 1 | | | | 1 | | | | | yes | yes | | | | yes | | | | |
| PLEKHM3 | S_Shore | TSS200 | chr2:208890468-208890802 | 334 | | | | | | | | | | | | | | yes | yes | | | | | | | | |
| CRYGD | N_Shore;Island;S_Shore | Body;1stExon;5'UTR;TSS200 | chr2:208988453-208989831 | 1378 | | 1 | 7 | | | | | 0 | 4 | | | 1 | | | | | | yes | yes | | | | |
| CRYGB | NA | TSS200 | chr2:209010890-209011053 | 163 | | | 4 | | | | | | | | | | | yes | | | | yes | | | | | |
| C2orf80 | NA | 5'UTR | chr2:209054670-209055151 | 481 | | | 2 | | | | | | | | | | | yes | | yes | | | | | | | |
| PTH2R | Island | TSS200 | chr2:209271383-209272057 | 674 | | | | | | | | | 1 | | | 1 | | yes | | | | | | | | | |
| MAP2 | NA | TSS1500;5'UTR | chr2:210444154-210444327 | 173 | | | | | 1 | | | | 1 | | | | | | | | | yes | | | | | |
| UNC80 | Island | TSS1500;TSS200 | chr2:210636398-210636748 | 350 | | | 1 | | | | | 0 | 3 | | | | | | | | | yes | | | | | |
| UNC80 | Island | Body | chr2:210673437-210674232 | 795 | | | | | | | | | | | | | | yes | | | | | | | | | |
| RPE | Island | TSS200 | chr2:210867278-210867333 | 55 | | | | | | | | | 1 | | | | | yes | | | | | | | | | |
| ACADL | S_Shore | TSS200 | chr2:211090247-211090281 | 34 | | | | | | | | | 1 | | | | | yes | | yes | yes | | | | | | |
| MYL1 | NA | Body;1stExon;TSS200 | chr2:211179560-211180386 | 826 | | | | | | | | | | | | | | | | | | yes | | | | | |
| CPS1 | NA | Body;TSS1500 | chr2:211457632-211458173 | 541 | | | 8 | | | | | | 1 | | | | | | | | | | yes | | | | |
| ERBB4 | N_Shore | Body | chr2:213401195-213401996 | 801 | | | | | | | | | 1 | | | | | | | | | yes | | | | | |
| ERBB4 | Island | 5'UTR;1stExon | chr2:213403321-213403859 | 538 | | | | | | | | | 1 | | | | | | | | | | yes | | | | |
| SPAG16 | Island;N_Shore | 5'UTR;1stExon;TSS1500 | chr2:214148856-214149735 | 879 | | | 2 | | | | | | | | | | | | | | | | | | | | |
| BARD1 | S_Shore | TSS1500 | chr2:215675105-215675637 | 532 | | | 8 | | | | | 0 | 1 | | | | | yes | yes | | | | | | | | |
| ATIC | Island | Body | chr2:216177085-216177303 | 218 | | | 1 | | | | | | | | | | | yes | | | | | | | | | |
| MARCH4 | Island | 5'UTR;1stExon | chr2:217236040-217236529 | 489 | | | | | | | | | 1 | | | | | | | | | yes | | | | | |
| SMARCAL1 | N_Shore | TSS1500 | chr2:217276744-217276821 | 77 | | | | | | | | | 1 | | | 1 | | yes | | | | | | | | | |
| IGFBP2 | S_Shore | Body | chr2:217500594-217501125 | 531 | | | | | | | | 0 | 1 | | | 1 | | | yes | | | yes | | | | | |
| TNP1 | NA | Body;3'UTR | chr2:217724380-217724794 | 414 | | | 2 | | | | | | | | | | | | | | yes | | | | | | |
| DIRC3 | N_Shore;Island | Body;TSS200 | chr2:218621258-218621552 | 294 | | 1 | | | | | | | | | | | | yes | yes | yes | yes | | | | | | |
| TNS1 | NA | 5'UTR | chr2:218770208-218770270 | 62 | | | | | | | | | 1 | | | | | yes | | | | | | | | | |
| TNS1 | NA | 1stExon;5'UTR | chr2:218808652-218808859 | 207 | | | | | | | | | 1 | | | | | yes | | | | yes | | | | | |
| RUFY4 | NA | Body | chr2:218937635-218938006 | 371 | | | | | | | | | 1 | | | | | | | | | yes | | | | | |
| CXCR2 | NA | TSS200;5'UTR | chr2:218990598-218990976 | 378 | | | | | | | | | | | | | | yes | yes | | yes | | | | | | |
| PNKD;TMBIM1 | N_Shore | Body;5'UTR | chr2:219155248-219155468 | 220 | | | 1 | | | | | | | | | | | | yes | | yes | | | | | | |
| PNKD;TMBIM1 | Island | Body;5'UTR | chr2:219157083-219157119 | 36 | | | | | | | | 0 | 3 | 2 | | 4 | 1 | | yes | | | | yes | | | | |
| SLC11A1 | NA | TSS1500;TSS200 | chr2:219246468-219247055 | 587 | | | | | | | | | 1 | 1 | | | | yes | | | | yes | | | | | |
| SLC11A1 | Island | Body | chr2:219252533-219252818 | 285 | | 1 | | | | | | | 1 | | | | | yes | yes | | | | | | | | |
| CTDSP1;MIR26B | S_Shore | Body;TSS200 | chr2:219267198-219267443 | 245 | | | | | | | | | 1 | | | | | yes | | | | | | | | | |
| PLCD4 | NA | TSS1500;TSS200 | chr2:219472218-219473002 | 784 | | | | | | | | 0 | 2 | | | 1 | | yes | yes | | yes | | | | | | |
| PRKAG3 | NA | 1stExon;TSS200 | chr2:219696472-219697159 | 687 | | | 16 | | | | | | | | | | | | | yes | | yes | | | | | |
| WNT6 | Island | Body | chr2:219724891-219725069 | 178 | | | | | | | | | 1 | | | | | | | | | | yes | | | | |
| WNT6 | N_Shore;Island | Body | chr2:219735876-219736549 | 673 | | | 11 | | | | | | 2 | 3 | 1 | | | | | | | | yes | | | | |
| WNT6 | Island | Body | chr2:219738529-219738732 | 203 | | | 12 | | | | | 0 | | | | 1 | | | | | | | | | | | |
| CDK5R2 | N_Shore | 3'UTR;1stExon | chr2:219826532-219827194 | 662 | | 1 | | | | | | 0 | 2 | | | | | | | | | | yes | | | | |
| CCDC108 | Island;S_Shore | 5'UTR;TSS200 | chr2:219906116-219906468 | 352 | | 2 | | | | | | | | | | 1 | | yes | | | | yes | | | | | |
| IHH | Island | Body | chr2:219920195-219920345 | 150 | | 1 | | | | | | | | | | | | | | | | | | | | | |
| GLB1L | Island | Body | chr2:220107847-220108267 | 420 | | | 5 | | | | | 0 | 1 | | | 1 | | yes | | | yes | yes | | | | | |
| TUBA4B;TUBA4A | N_Shore | TSS1500;Body | chr2:220117599-220117875 | 276 | | | 2 | 1 | | | | | | | | | | | | | | | | | | | |
| PTPRN | N_Shore;Island | Body;TSS200 | chr2:220173591-220174251 | 660 | | 1 | | | | | | | 1 | | | | | yes | | | | | | | | | |
| RESP18 | Island | Body | chr2:220196530-220196755 | 225 | | 1 | | | | | | 0 | 1 | | | | | | | | | | | | | | |
| RESP18 | S_Shelf | TSS1500 | chr2:220199185-220199310 | 125 | | | | | | | | | | | | | | | yes | | | | | | | | |
| DNPEP | S_Shore | TSS1500 | chr2:220253653-220254248 | 595 | | | 6 | | | | | 0 | 1 | | | | | yes | | | | yes | | | | | |
| DES | N_Shore | TSS200;1stExon;5'UTR;TSS1500 | chr2:220282165-220283442 | 1277 | | | | | | | | 0 | 3 | | | 1 | | yes | | | | | yes | | | | |
| SPEG | Island | TSS200 | chr2:220299584-220299900 | 316 | | | 3 | | | | | | 3 | | | | | yes | | | | | | | | | |
| SPEG | Island | Body | chr2:220312854-220313972 | 1118 | | | | | | | | 0 | 2 | 2 | 1 | | | | | | | | | | | | |
| SPEG | N_Shore;Island;chr2:22034895 | Body;Island | chr2:220348312-220349733 | 1421 | | | | | | | | 0 | 1 | | 1 | | | | | | | | yes | | | | |
| GMPPA | Island | TSS1500 | chr2:220362163-220362242 | 79 | | | | | | | | | | | | | | yes | | | | | | | | | |

| | | | | | | | | | | | | | | | | | | | | | | | |
|---|---|---|---|---|---|---|---|---|---|---|---|---|---|---|---|---|---|---|---|---|---|---|---|
| ACCN4 | Island | TSS1500 | chr2:220377759-220377885 | 126 | 3 | | | | | | | | | | | | | | | | | | |
| ACCN4 | S_Shore | 5'UTR;1stExon | chr2:220378996-220379109 | 113 | | | | | | | | | | | | | | | | | | yes | |
| CHPF | Island | Body | chr2:220406470-220406756 | 286 | | | 12 | | | | | | | | | | | yes | | | | | |
| OBSL1 | N_Shore | Body | chr2:220426848-220427342 | 494 | | 1 | | | | | | | | | | | | | yes | | yes | | |
| OBSL1;INHA | S_Shore | TSS1500;TSS200 | chr2:220436894-220436948 | 54 | | | 5 | | 1 | | | | | | | | | yes | | | | yes | |
| EPHA4 | N_Shore | Body | chr2:222434303-222434778 | 475 | 3 | | | 1 | | | | 0 | 1 | | | 3 | | | yes | | | | |
| PAX3 | N_Shore | Body | chr2:223154140-223154201 | 61 | | 1 | | | | | | | | | | | | | | | | | |
| PAX3 | N_Shore;Island | Body | chr2:223155306-223156136 | 830 | | | 4 | | | | | | | | | | | | | | | yes | |
| PAX3 | Island | Body | chr2:223160175-223160486 | 311 | | | 1 | | | | | | | | | | | | | | | | |
| PAX3;CCDC140 | Island;N_Shore;S_Shore | Body;TSS1500;TSS200;5'UTR;1stEx | chr2:223161771-223169965 | 8194 | | | | | | | | | | | | | | | | | | | |
| SGPP2 | N_Shore;Island;S_Shore | TSS1500;TSS200;Body | chr2:223288331-223290515 | 2184 | | | 10 | | | | | 0 | 2 | | | 1 | | yes | | | | yes | yes |
| MOGAT1 | Island | TSS200 | chr2:223536267-223536609 | 342 | | | | | | | | | 3 | | | | | | yes | | | | yes |
| KCNE4 | N_Shore;Island | TSS1500;TSS200;Body | chr2:223916502-223918000 | 1498 | | 1 | 9 | | 1 | | | | 3 | | | 2 | | yes | | | | | yes |
| SERPINE2 | Island | TSS1500 | chr2:224904626-224904837 | 211 | | | 12 | | | | | | | | | | | | yes | | | | |
| FAM124B | NA | 1stExon;5'UTR;TSS200 | chr2:225265963-225266880 | 917 | 2 | 2 | | 1 | 2 | | | 0 | 3 | 1 | 1 | | | yes | | | yes | | yes |
| KIAA1486 | NA | TSS1500 | chr2:226265132-226265475 | 343 | | | | | | | | | | | | | | | | | | | yes |
| RHBDD1 | N_Shore | TSS1500 | chr2:227700114-227700467 | 353 | | | | | | | | 0 | | | | | | yes | | | | | |
| COL4A3;COL4A4 | Island | TSS1500;5'UTR;1stExon;TSS200;Bo | chr2:228028741-228029856 | 1115 | | | | | | | | | 2 | | | | | yes | | | yes | | |
| SLC19A3 | Island | 5'UTR | chr2:228582486-228582688 | 202 | | 1 | | | | | | | 1 | | | | | yes | | | | | |
| WDR69 | N_Shore;Island | TSS1500;TSS200 | chr2:228736089-228736459 | 370 | | 1 | 4 | | | | | | 5 | | | 1 | | yes | yes | | yes | | |
| SPHKAP | NA | Body | chr2:228882340-228882605 | 265 | | | | | | | | | | | | | | | | | | | |
| SPHKAP | N_Shore | Body | chr2:229045417-229046785 | 1368 | | 1 | | | | | | | | | | | | | yes | | | | |
| PID1 | Island | 5'UTR;1stExon | chr2:230135879-230136146 | 267 | | | | | | | | | | | | 1 | | | yes | | | yes | yes |
| TRIP12 | N_Shore | 5'UTR | chr2:230785050-230785180 | 130 | | | | | | | | 0 | | | | | | | | | | | |
| SLC16A14 | Island | 1stExon;5'UTR | chr2:230933543-230934399 | 856 | | | | | | | | | | | | 1 | | yes | | yes | | | yes |
| SP140 | NA | TSS200;5'UTR;1stExon | chr2:231090329-231090745 | 416 | | | | | 1 | | | | 1 | | | 1 | | | | yes | | | |
| SP140L | NA | TSS1500 | chr2:231191662-231191916 | 254 | | | | | | | | | | | | | | | | | | | |
| GPR55 | NA | TSS1500 | chr2:231790529-231790813 | 284 | | | | | | | | | 1 | | | | | yes | | | | | |
| B3GNT7 | N_Shore;Island | TSS1500 | chr2:232259431-232260139 | 708 | | | 1 | | | | | 0 | 2 | | | | | yes | | | | | yes |
| NCL;SNORA75;SNORD20 | NA | Body;TSS1500;TSS200 | chr2:232321296-232321649 | 353 | | | | | | | | | 1 | | | | | | yes | | | | |
| NMUR1 | Island | TSS200 | chr2:232395269-232395300 | 31 | | | | 1 | | | | | | | | | | yes | | | | | |
| MIR1471 | NA | Body;TSS200 | chr2:232756980-232757414 | 434 | | | | | | | | | | | | | | yes | | | | | |
| NPPC | chr2:232791575-232792016;Isl | Island;TSS1500 | chr2:232791814-232792397 | 583 | | | | | | | | | | | 1 | | | yes | | | | | |
| ALPP | Island | Body | chr2:233244424-233244503 | 79 | | | | | | | | | 1 | | | | | | | | | | |
| ALPP | N_Shore;Island | Body | chr2:233245886-233246594 | 708 | | | | | | | | 0 | | | | 1 | | yes | | | | | |
| ECEL1P2 | Island | TSS200 | chr2:233251770-233251881 | 111 | | 1 | | | | 1 | | | 1 | | | | | | | | | | yes |
| ALPPL2 | Island | Body | chr2:233274144-233274723 | 579 | | | 4 | | | | 1 | | 2 | | | | | yes | | | | | |
| ALPI | N_Shelf | TSS1500;TSS200 | chr2:233320486-233321088 | 602 | | | | | 1 | | | | | | | | | | yes | | | | |
| ALPI | N_Shore;Island | Body | chr2:233322928-233324053 | 1125 | 1 | | 16 | | | | | 0 | 3 | | | | | yes | | | | | |
| ECEL1 | Island | TSS1500 | chr2:233352776-233352951 | 175 | | | | | | | | | | | | | | | | | | yes | yes |
| CHRND | S_Shore | TSS1500;TSS200;1stExon | chr2:233390712-233391274 | 562 | | | 5 | | 1 | | 1 | 0 | 2 | 1 | | 1 | | yes | | | | | |
| CHRNG | NA | TSS1500 | chr2:233403858-233404334 | 476 | | | | | | | | | 1 | | | | | yes | | | | yes | yes |
| EFHD1 | Island | Body | chr2:233499283-233499637 | 354 | | | 1 | | | | | | | | | | | yes | | | | | |
| NGEF | NA | Body | chr2:233749413-233750589 | 1176 | | | 7 | | | | | | | | | | | | | | | yes | |
| NGEF | Island | Body | chr2:233765872-233766118 | 246 | | | 1 | 1 | | | | | | | | | | | | | | yes | |
| NGEF | NA | 1stExon;5'UTR;TSS200 | chr2:233877857-233878714 | 857 | | 1 | 5 | 1 | | | | | | | | | | yes | yes | yes | yes | yes | yes |
| INPP5D | N_Shore | TSS1500;TSS200 | chr2:233924713-233925181 | 468 | | | 8 | 1 | | | | | 2 | 4 | 1 | | | | | | | yes | |
| INPP5D | NA | Body | chr2:233981788-233981999 | 211 | | | | | | | | | 1 | | | | | yes | | yes | | | |
| INPP5D | NA | Body | chr2:234072415-234072980 | 565 | | | 5 | | | | | | | | | | | | yes | | yes | | |
| DGKD | NA | Body;TSS1500;5'UTR;1stExon | chr2:234296280-234297039 | 759 | | 1 | | 1 | | | | | 1 | | | | | yes | | | | | yes |
| DGKD | Island | Body | chr2:234369962-234370070 | 108 | | | | 1 | | | | | 1 | | | | | yes | | | | | yes |
| UGT1A6;UGT1A10;UGT1A9;UGT1A7;UG | NA | TSS1500;Body | chr2:234599988-234600149 | 161 | | | 1 | | | | | | 1 | | | | | yes | | yes | | | |
| UGT1A10;UGT1A6;UGT1A8;UGT1A9;UG | NA | Body;TSS1500;TSS200 | chr2:234626084-234627450 | 1366 | | | | 1 | | | | | 1 | | | | | | | | | | |
| UGT1A10;UGT1A6;UGT1A8;UGT1A4;UG | NA | Body;TSS1500 | chr2:234637253-234637667 | 414 | | | 3 | | | | | | | | | | | | | | | | |
| UGT1A10;UGT1A6;UGT1A8;UGT1A4;UG | Island | Body | chr2:234663729-234664336 | 607 | | | | | | | | 0 | 1 | | | | | | | | yes | | |
| UGT1A10;UGT1A6;UGT1A8;UGT1A4;UG | NA | Body;TSS200 | chr2:234668834-234669166 | 332 | | | | | | | | | | | | | | | | | | | |
| MSL3L2 | Island | TSS200 | chr2:234777078-234777085 | 7 | | | | | | | | | 1 | | | | | yes | yes | | | | yes |
| TRPM8 | N_Shore;Island | Body | chr2:234847247-234847856 | 609 | | | 4 | | | | | 0 | 3 | 1 | | 10 | | | yes | yes | | | |
| TRPM8 | NA | Body | chr2:234891753-234891861 | 108 | | | 4 | | | | | | | | | | | | | | | | |
| ARL4C | Island | TSS1500 | chr2:235406291-235406540 | 249 | | | | | | | | 0 | | | | | | | | | | yes | |
| SH3BP4 | N_Shore | TSS1500 | chr2:235859315-235859792 | 477 | | 1 | 7 | | | | | | | | | | | yes | yes | | | | |
| SH3BP4 | NA | 5'UTR | chr2:235883549-235883680 | 131 | | | 2 | | | | | | | | | | | yes | | | | | |
| SH3BP4 | NA | 5'UTR | chr2:235907647-235907796 | 149 | | | | 1 | | | | | | | | | | | yes | | | | |
| SH3BP4 | NA | 5'UTR | chr2:235935606-235936596 | 990 | | | 16 | | | | | | 1 | | | | | | | | | | |
| AGAP1 | Island | 1stExon;5'UTR | chr2:236402993-236403031 | 38 | | | | | | | | | 1 | | | | | | | | | yes | yes |
| AGAP1 | NA | Body | chr2:236415673-236416064 | 391 | | 1 | | 1 | | | | | | | | | | yes | yes | | | | |
| AGAP1 | N_Shore;S_Shore | Body | chr2:236443773-236444284 | 511 | | | | | 1 | | | | 3 | 2 | | | | | | | | | |
| AGAP1 | NA | Body | chr2:236586191-236586744 | 553 | | | 4 | | | | | | | | | | | | yes | | | yes | |
| AGAP1 | NA | Body | chr2:236619304-236619825 | 521 | 1 | 1 | | 1 | | | | 0 | | | | | | yes | yes | | yes | | |
| AGAP1 | NA | Body | chr2:236643527-236643760 | 233 | | | 5 | | | | | | | | | | | yes | | | | | |
| AGAP1 | S_Shore | Body | chr2:236674174-236674489 | 315 | | | 9 | | | | | | | | | | | yes | | | | | |
| AGAP1 | NA | Body | chr2:236715747-236716285 | 538 | | | 2 | | | | | | | | | | | | | | | | |
| AGAP1 | NA | Body | chr2:236760940-236761574 | 634 | 1 | | | | | | | | 1 | | | | | | | | | | |
| AGAP1 | Island | Body | chr2:236773924-236774164 | 240 | | | | | | | | | | 1 | | | | | | | | | yes |
| AGAP1 | NA | Body | chr2:236826245-236826420 | 175 | | | 2 | | | | | | | | | | | | | | | | yes |
| AGAP1 | N_Shore;Island | Body | chr2:236867278-236867905 | 627 | | | 2 | | | | | | | | | | | | | | | | |
| AGAP1 | NA | Body | chr2:236877288-236877492 | 204 | | 1 | | | | | | | | | | | | | | | | | |
| AGAP1 | NA | Body | chr2:236923322-236923534 | 212 | | | | | 1 | | | | 1 | | | | | | | | | | yes |
| AGAP1 | NA | Body | chr2:237022740-237022804 | 64 | | | | | | | | | 1 | | | | | | | yes | | | |
| GBX2 | Island | 1stExon;TSS200;TSS1500 | chr2:237076164-237079099 | 2935 | | | | | | | | | | | | | | | | | | | |
| ASB18 | Island | Body | chr2:237120069-237120313 | 244 | | | | | | | | | | | | | | | yes | | | | |
| ASB18 | N_Shore | Body | chr2:237122857-237123218 | 361 | | | 3 | | | | | | 1 | | | | | yes | | | yes | yes | |
| ASB18 | NA | 1stExon | chr2:237172796-237173121 | 325 | | | 3 | | | | | | 1 | | | | | | yes | yes | | | |
| CXCR7 | S_Shore | 1stExon;5'UTR | chr2:237478526-237478664 | 138 | 1 | | | | | | | | | | | | | | | | | | |
| COL6A3 | NA | 5'UTR | chr2:238322274-238322680 | 406 | | 1 | | | | | | | | | | | | | | | | | yes |
| MLPH | N_Shore;chr2:238395061-2383 | TSS1500;Island | chr2:238394933-238395852 | 919 | | | 14 | | | | | | | | 1 | | | yes | | | | | |
| RAB17 | NA | 1stExon;5'UTR;TSS200 | chr2:238499464-238500325 | 861 | | | 4 | 1 | | | | | 2 | 1 | | | | yes | | yes | yes | | |
| LRRFIP1 | NA | Body | chr2:238578376-238578491 | 115 | | | | | 1 | | | | 1 | | | | | | yes | | | | |
| LRRFIP1 | NA | Body | chr2:238581924-238582364 | 440 | | | 3 | 1 | | | | | | | | | | yes | yes | | | | |
| LRRFIP1 | N_Shore | TSS1500;Body | chr2:238599734-238600061 | 327 | | 1 | 1 | | | | | 0 | 1 | | | 1 | | yes | | | | yes | |
| LRRFIP1 | S_Shelf | Body | chr2:238647283-238647913 | 630 | | | 5 | | | | | | | | | | | | | | | | |
| RBM44 | Island | TSS200 | chr2:238707372-238708018 | 646 | | | | | | | | | | | | | | yes | yes | | yes | | |
| RAMP1 | S_Shore | Body | chr2:238769060-238769419 | 359 | | | 1 | | | | | | | | | | | | | | | | |
| RAMP1 | NA | Body | chr2:238777587-238777806 | 219 | | | 5 | | 1 | | | | | | | | | | yes | | | yes | |
| SCLY | NA | Body | chr2:238990144-238990573 | 429 | | | | 1 | | | | | | | | | | yes | | | | | |
| ESPNL | N_Shore;Island | TSS1500;TSS200;1stExon | chr2:239008679-239009246 | 567 | 2 | 1 | | | | | | 0 | | | | | | | | | | | |
| ESPNL | NA | Body | chr2:239023538-239023741 | 203 | | | | | | | | | | | | | | | | | yes | | |
| ESPNL | N_Shore | Body | chr2:239037049-239037533 | 484 | | | | | | | | | 1 | | | 1 | | | yes | | | | |
| KLHL30 | N_Shore;Island | TSS1500;TSS200;5'UTR | chr2:239046684-239048579 | 1895 | | | 7 | | | | | | 3 | | | | | yes | | | | | |

| | | | | | | | | | | | | | | | | | | | | | | | |
|---|---|---|---|---|---|---|---|---|---|---|---|---|---|---|---|---|---|---|---|---|---|---|---|
| LOC151174;LOC643387 | Island;N_Shore | Body;TSS200;TSS1500 | chr2:239139911-239140733 | 822 | 4 | 2 | 5 | | 1 | 1 | 1 | 0 | 4 | 2 | 1 | 5 | | | yes | | | yes | yes |
| TWIST2 | NA | 3'UTR | chr2:239799440-239799954 | 514 | | | 13 | | | | | | | | | | | | | | | | |
| HDAC4 | NA | Body | chr2:239983744-239984105 | 361 | | | | | | | | | 2 | 1 | | 3 | | | | | | | |
| HDAC4 | NA | Body | chr2:240014758-240014781 | 23 | | | 4 | | | | | | | | | | | | | | | | yes |
| HDAC4 | NA | Body | chr2:240020028-240020174 | 146 | | | 22 | | | | | | 1 | 1 | | | | | | | | | |
| HDAC4 | N_Shore | Body | chr2:240031454-240031635 | 181 | | 1 | | 1 | | | | | | | | | | yes | yes | | | | |
| HDAC4 | NA | Body | chr2:240040089-240040290 | 201 | | | 1 | | | | | | | | | | | | yes | | | | |
| HDAC4 | NA | Body | chr2:240090401-240090667 | 266 | | | | | | | | | | | | | | | | | | | |
| HDAC4 | Island | Body | chr2:240112249-240112434 | 185 | 4 | | | | | | | | | | | | | | | | | | |
| HDAC4 | S_Shore | Body | chr2:240114298-240114406 | 108 | | | | | | | | 0 | 1 | 1 | | 6 | | | | | | | |
| HDAC4;MGC16025 | NA | Body;TSS200 | chr2:240117005-240117640 | 635 | | 2 | | | | | | | | | | | | | | | | | |
| HDAC4 | NA | Body | chr2:240142589-240143979 | 1390 | | 1 | | | | | | | 1 | | | | | | yes | | | yes | |
| HDAC4 | N_Shore;Island | Body | chr2:240152257-240153820 | 1563 | | | 2 | | | | | 0 | 1 | | 1 | | | | yes | yes | | yes | |
| HDAC4 | NA | Body | chr2:240161619-240162323 | 704 | | | 14 | | | | | | | | | | | | | yes | | | |
| HDAC4 | NA | Body | chr2:240168283-240169280 | 997 | | 1 | | | | | | | | | | | | | | | | | |
| HDAC4 | NA | Body | chr2:240172612-240172742 | 130 | | | | | | | | | 1 | | | | | yes | | | | | |
| HDAC4 | NA | Body | chr2:240174613-240174659 | 46 | | | | 1 | | | | 0 | 1 | | | | | | | | | | |
| HDAC4 | N_Shore;Island | Body | chr2:240196078-240197131 | 1053 | | 1 | 6 | 1 | 1 | | | 0 | 1 | | | | | | yes | | | | |
| HDAC4 | NA | Body | chr2:240212763-240213173 | 410 | | | 5 | | | | | | 1 | | | | | | | | yes | yes | |
| HDAC4 | NA | Body | chr2:240230452-240230892 | 440 | | 1 | 1 | 1 | | | | | 2 | | | | | | | | | | |
| HDAC4 | NA | Body | chr2:240233043-240233240 | 197 | | | 19 | | | | | | | | | | | | | | | | |
| HDAC4 | NA | Body | chr2:240270793-240271276 | 483 | 6 | | | | 2 | | | | | | | | | | | | | | |
| HDAC4 | NA | 5'UTR | chr2:240279976-240280217 | 241 | | | | 1 | | | | | | | | | | yes | yes | | | | |
| HDAC4 | NA | 5'UTR | chr2:240291068-240291657 | 589 | | | 20 | | 2 | | | | 1 | | | | | | | | | | |
| HDAC4 | NA | 5'UTR | chr2:240298235-240298595 | 360 | | | | | | | | | | | | | | | | | | | |
| HDAC4 | NA | 5'UTR | chr2:240302585-240302855 | 270 | | 1 | | 1 | | | | | 1 | | | | | | yes | | | | |
| HDAC4 | NA | 5'UTR | chr2:240308692-240308810 | 118 | | | | | | | | | 1 | | | | | | | | | yes | |
| OR6B2 | NA | TSS200 | chr2:240969975-240970263 | 288 | | | | | | | | | | | | | | | yes | | | | |
| OR6B3 | NA | TSS1500 | chr2:240986376-240986651 | 275 | | | | | | | | | | | | | | yes | yes | | | | |
| MYEOV2 | S_Shore | TSS1500 | chr2:241076131-241077006 | 875 | | | 18 | | | | 2 | 0 | 1 | 2 | | 2 | | | yes | yes | | | yes |
| OTOS | S_Shelf;NA | 5'UTR;1stExon;TSS200 | chr2:241079766-241080464 | 698 | 3 | | 5 | | | | | | 1 | | | | | | | yes | | | |
| MIR149;PP14571;GPC1 | Island;N_Shore | TSS200;Body;TSS1500 | chr2:241395181-241395678 | 497 | | | 12 | 1 | | | | | | | 1 | | | | | | | | |
| GPC1 | Island | Body | chr2:241401799-241401880 | 81 | | | | | | | | | 1 | | | | | | | | | | |
| ANKMY1 | N_Shore | Body | chr2:241458886-241460002 | 1116 | 5 | 1 | | 1 | | 1 | | 0 | 5 | 1 | | | | | | | | yes | yes |
| ANKMY1 | Island | 5'UTR;TSS200 | chr2:241496830-241497599 | 769 | | 1 | | | | | | 0 | 2 | 1 | | 1 | | yes | yes | | | | yes |
| CAPN10 | Island | Body | chr2:241535685-241536098 | 413 | | | | 1 | | | | | | | | | | yes | | | | | |
| AQP12A | NA | TSS200 | chr2:241631239-241631498 | 259 | | | 1 | | | | | | 1 | | | | | | | | | | |
| KIF1A | NA | Body | chr2:241696788-241697024 | 236 | | | 2 | | | | | | | | | | | | yes | | | yes | |
| KIF1A | Island | Body | chr2:241719496-241719655 | 159 | | | 4 | | | | | | | | | | | | | | | yes | |
| AGXT | NA | TSS1500;TSS200 | chr2:241807708-241808595 | 887 | | | | | | | | | | | | | | | | yes | yes | | |
| C2orf54 | NA | Body | chr2:241834752-241835740 | 988 | | | | 2 | | | | | | | | | | | | | | yes | |
| SNED1 | N_Shore | TSS1500 | chr2:241936844-241937034 | 190 | | | | | | 1 | | | | | | | | | | | | | |
| SNED1 | Island | TSS200 | chr2:241938191-241938436 | 245 | | | | | | | | | | 1 | | 1 | | | | | | yes | yes |
| SNED1 | N_Shore;Island | Body | chr2:241974858-241976658 | 1800 | | | | | | | 2 | 0 | 3 | 1 | | 4 | | | | | | | yes |
| MTERFD2;SNED1 | NA | Body | chr2:242027434-242027520 | 86 | | | | | | | | | 1 | | | | | | yes | | | | |
| MTERFD2 | N_Shore | Body | chr2:242040840-242041063 | 223 | | | 1 | | | | | | | | | | | yes | | | yes | | |
| PASK | NA | Body | chr2:242048127-242048359 | 232 | | | | 1 | | | | | | | | | | yes | | | | | |
| ANO7 | NA | TSS1500;TSS200 | chr2:242127690-242128250 | 560 | | | 3 | | | | | | 4 | | 1 | | | yes | | | | | yes |
| ANO7 | N_Shore | Body | chr2:242138408-242138658 | 250 | | | | | | | | | 1 | | | | | | | | | | |
| HDLBP | NA | Body | chr2:242170166-242170236 | 70 | | 1 | | | | | | | | | | | | yes | | | | | |
| HDLBP | S_Shore | 5'UTR;TSS1500 | chr2:242212884-242212901 | 17 | | | 8 | | 1 | | | | | | | | | | yes | | | | |
| STK25 | Island | TSS1500 | chr2:242448543-242448802 | 259 | | | 1 | | | | | | 1 | | | | | yes | yes | | | | |
| BOK | N_Shore;Island;S_Shore | TSS1500;TSS200;Body | chr2:242497595-242499542 | 1947 | | | | | 1 | | | | 3 | | | | | | yes | | | yes | yes |
| THAP4 | NA | Body | chr2:242529748-242530174 | 426 | 5 | | | | | | | | | | | | | | yes | | | | |
| THAP4 | Island | Body | chr2:242549431-242549943 | 512 | | | | | | | | | 1 | | | | | yes | | | | yes | |
| ATG4B | NA | 3'UTR | chr2:242612684-242612949 | 265 | | | | 1 | | | | | | | | | | | | | | | |
| ING5 | S_Shore | 3'UTR | chr2:242665670-242666066 | 396 | | | | | | | | | | | | | | yes | | | | | |
| D2HGDH | NA | Body | chr2:242702553-242702881 | 328 | | 1 | | | | | | | 1 | | | | | yes | yes | | yes | | |
| GAL3ST2 | S_Shelf | TSS1500 | chr2:242715019-242715549 | 530 | | | 5 | | | | | | 1 | | | | | | | | | yes | |
| GAL3ST2 | Island | Body | chr2:242743214-242743958 | 744 | | | | | | | | 0 | | | | | | | | | | | |
| NEU4 | N_Shelf | TSS1500;5'UTR;TSS200 | chr2:242751796-242752475 | 679 | | | 24 | | | | | | 1 | | | | | | | | | | |
| NEU4 | Island | Body | chr2:242757444-242757618 | 174 | | | | | | | | | | | | | | | | | | | |
| PDCD1 | Island | Body | chr2:242794854-242795282 | 428 | | 1 | | | | | | | | | | | | | | | | | |
| PDCD1 | N_Shore | Body | chr2:242797846-242798512 | 666 | | | | | | | | | | | | | | yes | yes | yes | | | |
| PDCD1 | N_Shelf | TSS1500 | chr2:242801896-242802192 | 296 | | | | | 1 | | | | 2 | | | | | | | | yes | | |
| C2orf85 | S_Shelf | TSS1500 | chr2:242810748-242811552 | 804 | | | 3 | | | | | | 5 | | | | | | yes | | | | |
| C2orf85 | N_Shore | Body | chr2:242813634-242815259 | 1625 | | | | | 1 | | | | | | | | | | | | | | yes |
| DEFB128 | NA | TSS200;TSS1500 | chr20:170364-170641 | 277 | | | | | | | | | 1 | | | | | yes | yes | | | | |
| DEFB129 | NA | TSS1500 | chr20:207144-207803 | 659 | | | 2 | | | | | | | | | | | yes | yes | | | | |
| SOX12 | Island | TSS200 | chr20:306044-306049 | 5 | | | | | | | | | | 1 | | | | | | | | | yes |
| NRSN2 | Island | 1stExon;5'UTR | chr20:327607-327623 | 16 | | | | | | | | | 1 | | | 1 | | | | | | | yes |
| TCF15 | Island | 1stExon;TSS200 | chr20:590799-591221 | 422 | | 1 | | | 1 | | | | 2 | | | | | yes | yes | | yes | yes | yes |
| SCRT2 | N_Shore;Island | 3'UTR | chr20:642436-642894 | 458 | 4 | | | | | | | 0 | 2 | | | | | | | | | yes | |
| C20orf54 | NA | 1stExon;5'UTR;TSS200;TSS1500 | chr20:749148-749620 | 472 | | | 4 | | | | | 0 | 3 | | | 2 | | yes | yes | | yes | | |
| FAM110A | N_Shore;Island | TSS200;5'UTR;1stExon | chr20:825155-826023 | 868 | | | 8 | 1 | 5 | | | | | | | | | yes | | | | yes | |
| RSPO4 | Island;N_Shore | Body;5'UTR;1stExon;TSS200 | chr20:981587-983104 | 1517 | | 1 | | | | | | | | 1 | | 1 | | yes | | | | | yes |
| PSMF1 | NA | TSS1500;TSS200 | chr20:1093633-1093802 | 169 | | | | | | | | | | | | | | yes | yes | | yes | yes | yes |
| C20orf46 | N_Shore | 3'UTR | chr20:1161338-1161820 | 482 | | | | | | | | | | | | | | | yes | | yes | | |
| C20orf46 | N_Shore | 1stExon;5'UTR | chr20:1165065-1165228 | 163 | | | | | | | | 0 | | | | | | | | | | yes | |
| SNPH | Island | TSS200 | chr20:1246769-1246779 | 10 | | | | 1 | | | | | | | | | | yes | | | | | |
| SDCBP2 | S_Shelf | TSS1500 | chr20:1310425-1310884 | 459 | | | | | | | | | | | | | | yes | yes | | | | |
| FKBP1A | S_Shore | TSS200 | chr20:1373995-1374350 | 355 | | | | | | | | | | | | | | | yes | yes | | | |
| NSFL1C | S_Shore | TSS200 | chr20:1448437-1448645 | 208 | | | | | | | | 0 | 2 | | 2 | | | | yes | yes | yes | | |
| SIRPB2 | NA | Body;1stExon;5'UTR | chr20:1471884-1472419 | 535 | | | 2 | | | | | | 2 | | | | | yes | | yes | yes | | |
| PDYN | NA | 5'UTR;1stExon;TSS200;TSS1500 | chr20:1974646-1975372 | 726 | 2 | | 7 | | | | | | 1 | | | | | yes | | | | yes | yes |
| TGM3 | NA | TSS200 | chr20:2276503-2277040 | 537 | | | | | | | | | | | | | | yes | | | | | |
| TGM3 | Island | Body | chr20:2308779-2308988 | 209 | | | 1 | | | | | | | | | | | | yes | | | yes | |
| TGM6 | NA | TSS1500;TSS200 | chr20:2360385-2361461 | 1076 | | | 3 | | | | | | | | | | | yes | | | | | |
| ZNF343 | S_Shore | TSS1500 | chr20:2490030-2490109 | 79 | | | 1 | | | | | | | | | | | yes | | yes | | | yes |
| TMC2 | NA | TSS200;TSS1500 | chr20:2516256-2517613 | 1357 | | | 9 | | | | | | | | | | | yes | | | yes | | yes |
| TMC2 | Island | Body | chr20:2539167-2539552 | 385 | | | | | 1 | | | | | | | | | yes | | yes | | | |
| NOP56;SNORD110;SNORA51 | S_Shore | Body;TSS200;TSS1500 | chr20:2634697-2635105 | 408 | | | 1 | | | | | | 1 | | | | | yes | | | | | |
| SNORD57;NOP56;SNORD56 | S_Shelf | TSS1500;Body;TSS200 | chr20:2637111-2637211 | 100 | | | | 1 | | | | | | | | | | | | | | | |
| EBF4 | Island | Body | chr20:2731102-2731560 | 458 | | | | | | | | | | 1 | | | | | | | | yes | |
| GNRH2 | N_Shelf | TSS1500;TSS200 | chr20:3022925-3024368 | 1443 | | | | 1 | | | | | | | | | | | | | | | |
| OXT | N_Shore;Island | TSS1500;TSS200 | chr20:3051954-3052483 | 529 | | | 27 | 2 | | | | 0 | 6 | 2 | | 8 | | yes | | | yes | | |
| AVP | S_Shore | TSS200 | chr20:3065559-3066014 | 455 | | | | | | | | | | | | | | | | | | | |
| ProSAPiP1 | Island | Body | chr20:3145178-3145745 | 567 | | | 17 | | | | | 0 | 4 | | | 1 | | | | | | | |
| ProSAPiP1 | Island;S_Shore | 1stExon;5'UTR;TSS200 | chr20:3148785-3149777 | 992 | | | 13 | | | | | 0 | 1 | | | 1 | | yes | | | yes | | |

| | | | | | | | | | | | | | | | | | | | | | | | | | | | |
|---|---|---|---|---|---|---|---|---|---|---|---|---|---|---|---|---|---|---|---|---|---|---|---|---|---|---|---|
| SLC4A11 | N_Shore | Body;1stExon;5'UTR;TSS200 | chr20:3218145-3218579 | 434 | 1 | | 16 | | | | | | 3 | | | 1 | | | | | | | | | | | |
| C20orf194 | S_Shore | TSS1500 | chr20:3389329-3389535 | 206 | | | | | | | | | | | | 1 | | yes | | | | | | | | | |
| GFRA4 | S_Shore | 1stExon;TSS200;Body | chr20:3643863-3644192 | 329 | | | 3 | | | | | 0 | 1 | | | | | yes | yes | | | | | | | | |
| ADAM33 | Island | Body;1stExon;5'UTR | chr20:3662512-3663186 | 674 | | | 2 | | | | | | | | | | | yes | | | | yes | yes | | | | |
| HSPA12B | N_Shore | TSS1500 | chr20:3712795-3713032 | 237 | | | | | | | | | | | | | | yes | | | | | yes | | | | |
| SPEF1 | N_Shelf | TSS1500 | chr20:3762422-3762653 | 231 | | | 2 | | | | | | | | | | | yes | | | | | | | | | |
| ADRA1D | Island | 1stExon | chr20:4229561-4230374 | 813 | | 1 | | | | | | | | | | | | yes | | | | yes | yes | | | | |
| PRND | NA | TSS200 | chr20:4702516-4702581 | 65 | | | | | | | | | | | | | | | | | | | yes | | | | |
| RASSF2 | NA | 5'UTR;nan;TSS200 | chr20:4795381-4795999 | 618 | | | 1 | | | | | | 2 | 1 | 1 | | | | | | | | | | | | |
| SLC23A2 | NA | 5'UTR;1stExon | chr20:4990828-4991028 | 200 | | | | | | | | | 1 | | | | | yes | | yes | | | | | | | |
| PROKR2 | Island | TSS1500 | chr20:5296267-5296442 | 175 | | | | | | | | | | | | | | | yes | | | | | | | | |
| LOC643406 | NA | TSS1500 | chr20:5450926-5451697 | 771 | | | 1 | | | | | | | | | | | | | | yes | yes | | | | | |
| LOC149837 | NA | TSS200;Body | chr20:5485144-5485511 | 367 | | 2 | 43 | | | | | 0 | 5 | 4 | 1 | 24 | 2 | yes | yes | | | | | | | | |
| CHGB | N_Shore | TSS200;TSS1500 | chr20:5891730-5891968 | 238 | | | 2 | | | | | | | | | | | | | | | | | | | | |
| CRLS1 | S_Shore | Body;TSS200 | chr20:5987794-5988223 | 429 | | | | | | | | | | | | | | | | | | | | | | | |
| LRRN4 | S_Shore | TSS200;5'UTR | chr20:6034446-6035139 | 693 | | | 10 | | | | | | 1 | | | | | | | yes | | | yes | | | | |
| FERMT1 | S_Shore | 1stExon;5'UTR;TSS200 | chr20:6104018-6104886 | 868 | | | 8 | | | | | 0 | 2 | | | | | yes | yes | | | yes | | | | | |
| BMP2 | Island | 5'UTR;Body | chr20:6750696-6750989 | 293 | | 1 | | 1 | | | | | | | | | | | | | | yes | | | | | |
| HAO1 | NA | TSS1500 | chr20:7921905-7922235 | 330 | | | | | | | | | 1 | | | | | yes | yes | | yes | | | | | | |
| PLCB4 | NA | TSS1500 | chr20:9075493-9075962 | 469 | | | | | | | | | 1 | | | | | | yes | | | | | | | | |
| C20orf103 | N_Shore | TSS1500 | chr20:9494838-9494997 | 159 | | | 1 | | | | | | | | | | | | | | | | | | | | |
| PAK7 | S_Shore;N_Shore;Island | TSS1500;5'UTR;TSS200 | chr20:9818397-9820894 | 2497 | | | | | | | | 0 | 1 | | | | | | yes | | yes | yes | | | | | |
| MKKS | N_Shore | 5'UTR;TSS200 | chr20:10412596-10412723 | 127 | | | | 1 | | | | | | | | | | yes | yes | yes | yes | yes | | | | | |
| BTBD3 | Island | TSS200 | chr20:11871375-11871743 | 368 | | | | | | | | 0 | 2 | 1 | | 2 | | | | | | | yes | | | | |
| BTBD3 | NA | 5'UTR;TSS200 | chr20:11898531-11898851 | 320 | | | | | | | | | | | | | | | | | | yes | | | | | |
| SPTLC3 | NA | TSS1500 | chr20:12988727-12989053 | 326 | | | | | | | | | | | | | | yes | yes | | | | | | | | |
| ISM1 | Island | TSS1500 | chr20:13200931-13201353 | 422 | | | | | | | | | 6 | | | | | | | | | | | | | | |
| MACROD2 | Island;N_Shore | TSS200;TSS1500 | chr20:13975439-13976782 | 1343 | | | 1 | | | | | 0 | 9 | 4 | 1 | 2 | | yes | yes | | | yes | | | | | |
| FLRT3;MACROD2 | NA | 5'UTR;Body;1stExon | chr20:14318152-14318231 | 79 | | | | | | | | | 1 | | | 1 | | | | | | | | | | | |
| KIF16B | Island | Body | chr20:16553861-16553921 | 60 | | | | | | | | | 1 | | | | | | | | | | yes | | | | |
| KIF16B | Island | TSS1500 | chr20:16555519-16555633 | 114 | | | | | | | | | 1 | | | | | yes | | | | | | | | | |
| OTOR | NA | TSS1500;TSS200 | chr20:16728794-16729319 | 525 | | | 10 | | | | | | | | | | | | | | | | | | | | |
| PCSK2 | N_Shore;Island | Body | chr20:17295815-17296597 | 782 | | 1 | 4 | | | | | 0 | 1 | | | 2 | | | yes | yes | | | | | | | |
| RRBP1 | Island | Body | chr20:17595355-17595472 | 117 | | | 6 | | | | | | 1 | 1 | | | | | | | | | | | | | |
| BANF2 | NA | TSS200 | chr20:17674195-17674250 | 55 | | | | | | | | | 1 | | | | | | yes | | | | | | | | |
| DTD1 | Island | TSS200 | chr20:18568437-18568511 | 74 | | | | | | | | | 1 | | | | | yes | | | | | | | | | |
| C20orf79 | NA | TSS1500 | chr20:18793817-18794303 | 486 | | | | | | | | | 1 | | | | | yes | | | | | | | | | |
| SLC24A3 | Island | TSS1500 | chr20:19193057-19193587 | 530 | | | | | | | | | 1 | | | | | | | | | yes | yes | | | | |
| RIN2 | NA | Body | chr20:19915716-19916087 | 371 | | | 7 | | 1 | | | | 3 | | | | | | | | | | | | | | |
| RIN2 | N_Shore | Body | chr20:19955436-19955868 | 432 | | | | | 1 | 1 | | | | | | | | | | | | | | | | | |
| C20orf26;CRNKL1 | S_Shelf | 5'UTR;1stExon;TSS1500;TSS200 | chr20:20036653-20037150 | 497 | | | | | | | 1 | 0 | 5 | 1 | | 1 | | yes | yes | | yes | | | | | | |
| INSM1 | Island | 1stExon | chr20:20350367-20350813 | 446 | | | 8 | | | | | | | | | | | | | | | | yes | | | | |
| RALGAPA2 | N_Shore | Body | chr20:20433062-20433328 | 266 | | | | 1 | 1 | | | | 1 | | | | | yes | | | | | | | | | |
| NKX2-4 | Island | 5'UTR;1stExon | chr20:21378045-21378200 | 155 | | | | | | | | | | | | 1 | | | | | | yes | yes | | | | |
| PAX1 | Island;N_Shore | TSS200;TSS1500 | chr20:21685401-21686548 | 1147 | | | 9 | | | | | 0 | 2 | | | | | | | | | yes | | | | | |
| PAX1 | Island | Body | chr20:21689759-21690036 | 277 | | | | | | | | | | | | | | | | | | | | | | | |
| PAX1 | N_Shore | Body | chr20:21694428-21694827 | 399 | | | | | | | | | | | | | | | | | | yes | | | | | |
| LOC284788 | NA | Body | chr20:22401074-22401407 | 333 | | | 8 | | | | | | | | | | | | | yes | | | | | | | |
| FOXA2 | N_Shore;Island | TSS1500;TSS200 | chr20:22566140-22567920 | 1780 | | 1 | | 1 | 1 | | | 0 | 2 | | | | | | | | | yes | | | | | |
| SSTR4 | Island | TSS200 | chr20:23015899-23016315 | 416 | | 1 | | | 1 | | | 0 | 5 | 1 | | 2 | | | | | | | yes | | | | |
| THBD | Island | 1stExon;5'UTR | chr20:23029287-23030446 | 1159 | | | | | | | | | 2 | | | | | | | | | | | | | | |
| CD93 | NA | TSS200;5'UTR;1stExon;nan | chr20:23066944-23067691 | 747 | 1 | | 15 | | 1 | | | | 3 | 3 | 1 | | | | | yes | | yes | | | | | |
| CST11 | NA | Body | chr20:23432977-23433608 | 631 | | | 1 | | | | | | | | 1 | | | | | | yes | | | | | | |
| CST8 | NA | TSS200 | chr20:23471693-23471737 | 44 | | | | | | | | | | | | | | | | yes | | | | | | | |
| CSTT | NA | TSS1500 | chr20:23498937-23499697 | 760 | | | 4 | | | | | | | | | | | | | | | | | | | | yes |
| CST9L | NA | TSS200;1stExon;5'UTR | chr20:23549106-23550446 | 1340 | | | | | | | | | | | | | | yes | | | | | | | | | |
| GGTLC1 | Island | 5'UTR;1stExon | chr20:23969237-23969832 | 595 | | | | | | | | | | | | | | yes | | | | | | | | | |
| TMEM90B | Island | 5'UTR;1stExon | chr20:24449865-24450353 | 488 | | | | | | | | | 1 | | | | | | | | | | yes | | | | |
| TMEM90B | Island | 5'UTR | chr20:24451890-24452131 | 241 | | | 8 | | | | | | | | | | | | yes | | | yes | | | | | |
| CST7 | NA | TSS1500;TSS200 | chr20:24929607-24929716 | 109 | | | 2 | | | | | | 1 | 1 | 1 | | | | yes | | | | | | | | |
| ACSS1 | S_Shore | TSS200;1stExon;5'UTR | chr20:25039314-25039812 | 498 | | | | 1 | | | | 0 | 2 | | | | | yes | | | | | | | | | |
| VSX1 | Island;N_Shore | Body;TSS200;TSS1500 | chr20:25061762-25064268 | 2506 | 7 | 3 | | | | | | | | | | | | | yes | | yes | yes | | | | | |
| LOC284798 | N_Shore;Island;S_Shore | TSS200;Body;TSS1500 | chr20:25128681-25130869 | 2188 | 3 | | 19 | | | | | 0 | 5 | 1 | | 3 | | yes | | | | | | | | | |
| NANP | Island | TSS200 | chr20:25604805-25605095 | 290 | | | | | | | | | 1 | | | | | yes | | | yes | | | | | | |
| MIR663 | Island | TSS200;TSS1500 | chr20:26188943-26190354 | 1411 | | 2 | | | | | | | 4 | | 1 | 3 | | | yes | | yes | | | | | | |
| FRG1B | Island | TSS1500 | chr20:29611652-29611924 | 272 | | | | | | 2 | | | | | | | | | yes | | | | | | | | |
| DEFB115 | NA | TSS1500 | chr20:29844516-29844575 | 59 | | | 12 | | | | | | | | | | | yes | yes | | | | | | | | |
| DEFB116 | NA | TSS200 | chr20:29896456-29896928 | 472 | | | | | | | | | | | | | | | yes | | | | | | | | |
| DEFB118 | NA | TSS1500 | chr20:29956028-29956330 | 302 | | | | | | | | | | | | | | yes | yes | | | | | | | | |
| DEFB123 | NA | TSS1500;TSS200 | chr20:30028062-30028418 | 356 | | | 5 | | | | | | 1 | | | | | yes | | yes | | | | | | | |
| DEFB124 | N_Shelf | TSS200 | chr20:30060822-30060884 | 62 | | | 2 | | | | | | 1 | | | | | yes | yes | | yes | yes | | | | | |
| REM1;NCRNA00028 | Island;N_Shore;S_Shore | TSS200;Body;TSS1500 | chr20:30063030-30074041 | 11011 | | | | | | | | | | | | | | | | | | | | | | | |
| HM13;PSIMCT-1 | Island;N_Shore | Body;TSS200;TSS1500 | chr20:30134929-30135362 | 433 | | | 1 | | | | | | 3 | 2 | | 1 | | yes | | yes | | | | | | | |
| COX4I2 | N_Shore;Island | TSS1500;TSS200 | chr20:30225325-30226046 | 721 | | | 30 | | | | | 0 | 9 | | | 5 | | yes | | | | | yes | | | | |
| MYLK2 | N_Shelf | TSS200;5'UTR;1stExon | chr20:30406997-30407388 | 391 | 4 | | 4 | | 2 | | | | | | | | | | | | | | yes | | | | |
| FOXS1 | Island;S_Shore;chr20:3043290 | 1stExon;5'UTR;S_Shore | chr20:30432938-30433673 | 735 | | | 8 | | | | | | 2 | | 1 | | | | | | | yes | | | | | |
| XKR7 | Island | Body | chr20:30582838-30583001 | 163 | | | 1 | | | | | | | | | | | yes | | | | | | | | | |
| C20orf160 | NA | TSS1500 | chr20:30597590-30597694 | 104 | | | 2 | | | | | | | | | | | yes | | yes | | | | | | | |
| C20orf160 | N_Shore;Island | Body | chr20:30605656-30605916 | 260 | | | | | | | | | | | | | | yes | | | | yes | | | | | |
| C20orf160 | Island | 3'UTR | chr20:30618874-30619137 | 263 | | | | | | | | 0 | 3 | | | 1 | | yes | | | | yes | | | | | |
| LOC284805 | NA | TSS200 | chr20:31261814-31261994 | 180 | | | | | | | | | | | | | | yes | yes | | | | | | | | |
| COMMD7 | S_Shore | TSS200 | chr20:31331814-31332353 | 539 | | | 2 | | | | | | | | | | | yes | yes | | yes | yes | | | | | |
| DNMT3B | NA | 5'UTR;TSS1500 | chr20:31366437-31366486 | 49 | | | 6 | | 1 | | | | | | | | | yes | | | | yes | | | | | |
| BPIL1 | NA | TSS1500;TSS200;5'UTR;1stExon | chr20:31595115-31595739 | 624 | | 1 | | 1 | | | | | 1 | | | | | | | | | | | | | | |
| C20orf185 | NA | TSS1500 | chr20:31641808-31642282 | 474 | | | | | | | | | | | | | | | yes | | | | | | | | |
| C20orf70 | NA | TSS200 | chr20:31755760-31756024 | 264 | | | 5 | | | | | | | | | | | yes | yes | | | | | | | | |
| BASE | NA | TSS1500;TSS200 | chr20:31781060-31781393 | 333 | | | 6 | | | | | | 1 | | | | | yes | | yes | | | | | | | |
| C20orf71 | NA | TSS1500 | chr20:31804901-31805291 | 390 | | | 3 | | | | | | | | | | | yes | yes | | | | | | | | |
| PLUNC | NA | TSS1500;TSS200 | chr20:31823545-31824068 | 523 | | | 2 | | | | | | | | | | | yes | yes | | yes | | | | | | |
| C20orf114 | NA | TSS1500;TSS200 | chr20:31870244-31871352 | 1108 | | | | | | | | | | | | | | yes | yes | | | | | | | | |
| NECAB3;C20orf144 | N_Shore | Body;TSS200 | chr20:32250035-32250111 | 76 | | | | | | | | | 1 | | | | | yes | | | | | | | | | |
| PXMP4 | Island;S_Shore | Body;TSS1500 | chr20:32307885-32308529 | 644 | | | | | | | | | 1 | | | 1 | | yes | yes | | | | | | | | |
| ASIP | Island | Body;3'UTR | chr20:32856747-32857227 | 480 | | 1 | 29 | 1 | | | | 0 | 5 | | 1 | 2 | | yes | | | | | yes | | | | |
| DYNLRB1 | N_Shore | TSS1500 | chr20:33103246-33103584 | 338 | | | 4 | | | | | 0 | 1 | | | | | yes | yes | | | | | | | | |
| MAP1LC3A | chr20:33146135-33147318 | Island | chr20:33146263-33146543 | 280 | | | | | | | | | | | 1 | | | | | | | | yes | | | | |
| NCOA6 | S_Shore | TSS1500 | chr20:33413664-33413713 | 49 | | | | | | | | | 1 | | | | | yes | yes | | | yes | yes | | | | |
| MYH7B;MIR499 | NA | Body;TSS200 | chr20:33578069-33578276 | 207 | | | 5 | | | | | | 1 | | | | | | | | | | | | | | |
| MYH7B | Island | Body | chr20:33585040-33585435 | 395 | | | 4 | | | | | | 1 | | | | | | | | | | | | | | |
| PROCR | N_Shelf | TSS200 | chr20:33759645-33760017 | 372 | | | | 1 | | | | | | | | | | yes | | | yes | | yes | | | | |

| | | | | | | | | | | | | | | | | | | | | | | | |
|---|---|---|---|---|---|---|---|---|---|---|---|---|---|---|---|---|---|---|---|---|---|---|---|
| PROCR | N_Shore;Island | Body | chr20:33762157-33762943 | 786 | 1 | | 8 | 1 | | | | | | | | | | yes | yes | | | yes | |
| FAM83C | S_Shore | TSS200;TSS1500 | chr20:33880337-33880935 | 598 | | | 8 | | | | | | 1 | | 1 | | | yes | | | yes | yes | |
| UQCC | NA | TSS1500 | chr20:34000170-34000481 | 311 | | | | | | | | 0 | | | | | | yes | | | | | |
| SPAG4 | Island | Body | chr20:34204988-34205182 | 194 | | | | | | | | | 1 | | | | | yes | | | | | |
| LOC647979 | S_Shore | TSS200 | chr20:34639042-34639058 | 16 | | | | | | | | 0 | | | | | | | yes | | | | |
| EPB41L1 | NA | TSS1500 | chr20:34700112-34700675 | 563 | | | | | | | | | 3 | 1 | | | | yes | yes | | | | |
| DLGAP4 | NA | TSS1500;TSS200 | chr20:34994743-34995517 | 774 | | | 2 | | | | | | 1 | | | | | yes | | | | | |
| DLGAP4 | Island | Body | chr20:35064637-35064957 | 320 | | | | | | | | | | | | | | yes | | | | | |
| MYL9 | Island;S_Shore | TSS1500;TSS200 | chr20:35169260-35169886 | 626 | | | 23 | 1 | | | | 0 | 1 | | | 4 | | | yes | | | | yes |
| C20orf24 | N_Shore | TSS1500 | chr20:35233380-35233582 | 202 | | | 2 | | | | | 0 | 1 | | | | | yes | | | yes | | |
| SLA2 | NA | TSS200;5'UTR;1stExon;nan | chr20:35274281-35274828 | 547 | | | 19 | 1 | 1 | | | | 1 | 2 | 1 | | | yes | | | yes | | |
| C20orf118 | NA | TSS1500;TSS200 | chr20:35503983-35504553 | 570 | | 1 | | | | | | | 2 | 1 | 1 | | | yes | | | | yes | |
| SAMHD1 | Island | TSS1500 | chr20:35580436-35580555 | 119 | | | 1 | | | | | | | | | | | yes | | | | | |
| SRC | N_Shore | TSS200 | chr20:35972924-35973318 | 394 | | | | 1 | 1 | | | | 2 | 1 | | | | yes | | | yes | | |
| SRC | N_Shore | 5'UTR | chr20:36011999-36012596 | 597 | | | 15 | | | | | | 1 | | | | | yes | yes | | | | |
| BLCAP;NNAT | Island;N_Shore | 5'UTR;TSS1500;1stExon;Body | chr20:36148133-36151338 | 3205 | | | 7 | 1 | 3 | | | 0 | 42 | | | 6 | | | | | | yes | yes |
| BLCAP | S_Shore;N_Shelf;N_Shore | 5'UTR;1stExon;TSS1500 | chr20:36152813-36154388 | 1575 | | | 13 | | | | | | 3 | 1 | 1 | | | yes | | | | | |
| BLCAP | S_Shore | TSS1500 | chr20:36156919-36157675 | 756 | | | 6 | | | | | | 2 | | | | | | yes | yes | | | |
| VSTM2L | N_Shore | TSS1500 | chr20:36530548-36531053 | 505 | | | | | | | | 0 | | | | | | yes | | | | yes | yes |
| TGM2 | Island;S_Shore | TSS200;TSS1500;5'UTR;1stExon | chr20:36793608-36794002 | 394 | | | 11 | | | | | 0 | 3 | | 1 | | | yes | | | | yes | yes |
| KIAA1755 | Island | TSS200 | chr20:36889364-36889389 | 25 | | | | 1 | | | | 0 | 1 | | | 1 | 1 | yes | | yes | | yes | yes |
| BPI | NA | TSS200 | chr20:36932385-36933189 | 804 | | | 8 | | 1 | | | | | 1 | | | | yes | | | | yes | |
| LBP | NA | TSS200;TSS1500 | chr20:36974445-36975083 | 638 | | | | 1 | | | | | | | | | | yes | yes | yes | | | |
| LOC388796;SNORA71A | NA | Body;TSS1500 | chr20:37057182-37057418 | 236 | | | | | | | | | | | | | | yes | | yes | | | |
| LOC388796;SNORA71C | N_Shelf | Body;TSS1500 | chr20:37059717-37059936 | 219 | | | | 1 | | | | | | | | | | yes | | | | | |
| LOC388796 | S_Shore | TSS1500 | chr20:37064274-37064504 | 230 | | | | | | | | | | | | | | | yes | | | | |
| ADIG | NA | TSS1500 | chr20:37208947-37209395 | 448 | | | 2 | | | | | | | | | | | yes | | | yes | yes | |
| C20orf95 | N_Shore | TSS1500;TSS200 | chr20:37230326-37230741 | 415 | | | | | | | | 0 | 2 | 1 | | 2 | | yes | | | | | |
| SLC32A1 | Island | TSS200 | chr20:37353032-37353700 | 668 | | | | | 3 | | | | 1 | 1 | | | | | | | | yes | |
| PPP1R16B | N_Shore | TSS1500;TSS200 | chr20:37433745-37434231 | 486 | | | 1 | 1 | 1 | | | | 3 | 1 | 1 | | | yes | | | | | |
| FAM83D | N_Shore | TSS1500 | chr20:37554591-37554897 | 306 | | | | | | | | | | | | | | yes | yes | | | yes | |
| MAFB | Island | 5'UTR;1stExon | chr20:39317833-39318137 | 304 | | 1 | | | | | | | | | | | | | | | | yes | yes |
| MAFB | Island | TSS1500 | chr20:39319235-39320005 | 770 | | 1 | | | | | | | | | | | | | | | | | |
| TOP1;PRO0628 | NA | Body;TSS1500 | chr20:39666232-39666765 | 533 | | | | | | | | | 1 | | | | | yes | | | | | |
| LPIN3 | S_Shore | TSS200 | chr20:39969481-39969604 | 123 | | | 2 | | | | | | | | | | | yes | yes | | | | |
| EMILIN3 | Island | TSS200 | chr20:39995649-39996039 | 390 | | | 13 | | | | | | 1 | | | | | | yes | | | | yes |
| PTPRT | Island | Body | chr20:41817903-41819125 | 1222 | | 1 | | | | | | | 1 | | | | | | | | | yes | yes |
| L3MBTL | N_Shore;Island | TSS1500;TSS200;5'UTR;1stExon | chr20:42142005-42143502 | 1497 | | | 6 | | | | | 0 | 10 | | | | | | | | | | yes |
| SGK2 | NA | TSS1500;TSS200 | chr20:42186621-42187837 | 1216 | | | | 1 | 1 | | | | | | | | | yes | yes | | | | yes |
| TOX2 | Island | TSS200;Body;5'UTR;1stExon;TSS1 | chr20:42544086-42545443 | 1357 | | | 3 | | | | | 0 | 1 | | | 1 | | | | | | | yes |
| TOX2 | NA | 5'UTR;TSS1500;Body | chr20:42573639-42574356 | 717 | | | | | | | | | 1 | | | | | yes | yes | | | | |
| GDAP1L1 | N_Shore;Island | TSS1500;TSS200 | chr20:42875135-42876091 | 956 | | | 1 | | 2 | | | | 1 | | | | | yes | yes | | | | |
| R3HDML | NA | TSS200 | chr20:42965428-42965753 | 325 | | | | | | | | | | | | | | yes | | | | | yes |
| HNF4A | NA | TSS1500;TSS200 | chr20:42983920-42984878 | 958 | | | 5 | 1 | 1 | | | | | | | | | yes | yes | | | | |
| HNF4A | NA | TSS1500;Body;TSS200 | chr20:43029468-43029997 | 529 | | | | 1 | 1 | | | | | | | | | yes | | | | | yes |
| PKIG | N_Shore | TSS200 | chr20:43160249-43160301 | 52 | | | 1 | | | | | | | | | | | yes | | | | | |
| WISP2 | NA | TSS1500 | chr20:43343365-43343760 | 395 | | | | | | | | | 1 | | | | | | yes | | | yes | |
| KCNK15 | N_Shore;Island | Body | chr20:43378461-43379501 | 1040 | | | | | | | | | 2 | | | 1 | | | yes | | yes | | yes |
| KCNS1 | S_Shore;N_Shore;Island | TSS200;TSS1500;5'UTR | chr20:43729410-43730241 | 831 | | | 26 | 1 | | | | | | | | | | yes | | | yes | yes | |
| WFDC5 | NA | 1stExon;5'UTR;TSS200 | chr20:43743736-43743967 | 231 | | | | | | | | | | | | | | yes | | | | yes | |
| WFDC12 | NA | Body | chr20:43752871-43753347 | 476 | | | | | | | | | | | | | | yes | | | | | |
| SLPI | NA | Body;TSS1500 | chr20:43882990-43883746 | 756 | | | 1 | | | | | | 1 | | | | | yes | yes | | | yes | |
| MATN4;RBPJL | Island | 5'UTR;TSS1500;TSS200 | chr20:43935222-43935551 | 329 | | | 30 | | | | | 0 | 5 | 1 | | 3 | | yes | | | | | |
| RBPJL;MATN4 | S_Shore | Body;5'UTR;1stExon;TSS200;TSS1 | chr20:43936663-43937467 | 804 | | | | | | | | 0 | 2 | | | | | yes | | | | | |
| RBPJL | Island | Body | chr20:43945254-43945735 | 481 | | | | | | | | | 2 | | | | | yes | | | | | |
| DBNDD2;SYS1-DBNDD2 | S_Shore | 5'UTR;TSS200;Body | chr20:44036899-44037436 | 537 | | 2 | | | | | | | | | | | | yes | | | | | yes |
| WFDC2 | N_Shore | TSS1500;TSS200 | chr20:44097320-44098287 | 967 | | | | | | | | | | | | 1 | | yes | yes | | | | |
| SPINT3 | NA | 1stExon;TSS1500 | chr20:44144225-44144711 | 486 | | | | | | | | | | | | | | yes | yes | | yes | | |
| SPINLW1 | NA | 1stExon;TSS1500 | chr20:44175957-44176575 | 618 | | | 6 | | | | | | | | | | | yes | | | | | |
| WFDC10A;WFDC9 | NA | TSS200;5'UTR | chr20:44258267-44258384 | 117 | | | | | | | | | 1 | | | | | | | | | yes | yes |
| WFDC10A;WFDC9 | NA | Body;5'UTR;3'UTR;TSS200 | chr20:44259589-44260207 | 618 | | | 6 | | | | | | | | | | | | | | | | |
| WFDC10B;WFDC13 | NA | TSS1500;3'UTR | chr20:44334749-44334913 | 164 | | | | | | | | | | | | | | yes | | | | | |
| WFDC3;DNTTIP1 | N_Shore | 5'UTR;TSS1500 | chr20:44420276-44420375 | 99 | | | | | | | | | | | | | | yes | | | | | |
| UBE2C | N_Shore;Island | TSS1500;TSS200;1stExon;5'UTR;Bo | chr20:44440824-44441818 | 994 | | | | | | | | 0 | | | | 2 | | yes | | | | | |
| TNNC2 | Island | Body | chr20:44452801-44453161 | 360 | | | 1 | | | | | | | | | | | yes | | | | | |
| CTSA;NEURL2 | N_Shore | TSS1500;Body | chr20:44518802-44519396 | 594 | | | | | | | | | 1 | | | | | yes | | | | | |
| SLC12A5 | N_Shore | TSS200 | chr20:44650220-44650449 | 229 | | | | | | | | | 1 | | | | | | yes | | | yes | yes |
| SLC12A5 | Island | TSS200;Body;1stExon;5'UTR | chr20:44657637-44658225 | 588 | | | | | 1 | | | | 2 | | | 1 | | | | | | yes | |
| CD40 | N_Shore | TSS1500;TSS200 | chr20:44746392-44747006 | 614 | | | 6 | | | | | 0 | 1 | | | 1 | | yes | | | | yes | |
| CDH22 | Island | Body | chr20:44829602-44829821 | 219 | | | 5 | | | | | 0 | 1 | | | | | yes | yes | | | | |
| CDH22 | N_Shore | Body | chr20:44838776-44839187 | 411 | | | 1 | | | | | 0 | 1 | | 1 | 1 | | yes | yes | | | | yes |
| CDH22 | Island | Body | chr20:44875334-44875586 | 252 | | | 5 | | 1 | | | | 1 | | | | | yes | | | yes | | yes |
| CDH22 | Island | 1stExon;5'UTR | chr20:44879700-44880819 | 1119 | | | 9 | | | | | | | | | | | | yes | | | yes | yes |
| C20orf123 | NA | 1stExon;TSS200 | chr20:45179157-45179413 | 256 | 5 | | 4 | | | | | 0 | 4 | | | | | yes | | | yes | | |
| SLC13A3 | S_Shore | TSS200;5'UTR | chr20:45280237-45280288 | 51 | | | 1 | | | | | | | | | | | yes | yes | | | | |
| TP53RK;SLC13A3 | NA | 3'UTR;TSS200 | chr20:45313109-45313390 | 281 | | | 7 | | | | | | | | | | | yes | yes | yes | | | |
| EYA2 | N_Shore | TSS200 | chr20:45523098-45523137 | 39 | | | | | | | | | | | | | | yes | yes | | | yes | yes |
| ZMYND8;LOC100131496 | NA | Body;TSS1500 | chr20:45947025-45947892 | 867 | | | | | | | | 0 | 2 | 1 | | | | yes | | | | | yes |
| SULF2 | Island | 1stExon;5'UTR;TSS200 | chr20:46414900-46414952 | 52 | | | | | | | | | 1 | | | | | | | | | yes | yes |
| LOC284749 | NA | TSS1500 | chr20:46988236-46988477 | 241 | | | | | | | | | 1 | | | | | | yes | yes | yes | yes | |
| SNORD12B;C20orf199;MIR1259;SNORD1 | S_Shore;chr20:47894197-4789 | TSS1500;Body;TSS200;S_Shore | chr20:47896448-47897451 | 1003 | 1 | 1 | 11 | | | | | 0 | 2 | | 1 | | | yes | | yes | | | |
| KCNB1 | Island | 1stExon;5'UTR;TSS200 | chr20:48099146-48099890 | 744 | 10 | 1 | | | | | | 0 | 2 | | | | | | | | | yes | yes |
| SPATA2 | N_Shore | 5'UTR;1stExon;TSS200 | chr20:48530082-48530654 | 572 | | 1 | 1 | | | 1 | | | | | | | | yes | | | | yes | |
| FAM65C | NA | 1stExon;5'UTR;TSS200 | chr20:49253364-49253710 | 346 | | | 24 | | | | | | | | | | | yes | yes | | yes | | |
| BCAS4 | N_Shore | TSS1500 | chr20:49410810-49410882 | 72 | | | | | | | | | | | | | | yes | | | yes | | yes |
| ADNP | Island | TSS1500 | chr20:49548515-49548661 | 146 | | | 2 | | | | | | | | | | | | | | | | |
| KCNG1 | N_Shore | 3'UTR | chr20:49620366-49620597 | 231 | 4 | | | | | | | | | | | | | | | | | | |
| KCNG1 | Island | Body | chr20:49626064-49626347 | 283 | | | | | | | | | 1 | | | | | yes | yes | | | | |
| NFATC2 | Island | Body | chr20:50108912-50109375 | 463 | | | | 1 | | | | | 1 | | | | | yes | | | | yes | |
| NFATC2 | Island;chr20:50158904-501595 | 1stExon;5'UTR;Body;TSS200;Island | chr20:50158996-50159653 | 657 | | | 18 | | | | | | | | 1 | | | | yes | | | | yes |
| ATP9A | NA | Body | chr20:50312386-50312632 | 246 | | | 3 | 1 | 1 | | | | | | | | | yes | | | | | |
| SALL4 | Island;S_Shore | Body;1stExon;5'UTR;TSS200 | chr20:50418554-50419274 | 720 | | | 8 | | | | | | 2 | | | 1 | | | | | | | yes |
| ZNF217 | Island | 1stExon | chr20:52198164-52198378 | 214 | | | | | | | | | 1 | | | | | yes | | | yes | | yes |
| ZNF217 | S_Shore | 5'UTR;1stExon;TSS200 | chr20:52199520-52199778 | 258 | | | 3 | | | | | | | | | | | | | | | | |
| BCAS1 | NA | 1stExon;5'UTR | chr20:52687007-52687394 | 387 | | | | | | | | | 1 | 1 | | | | yes | yes | | | | |
| CYP24A1 | Island | TSS200 | chr20:52790589-52790733 | 144 | | | | | | | | | 1 | | | | | yes | | | | yes | |
| DOK5 | Island | TSS1500;TSS200;Body | chr20:53092040-53093152 | 1112 | | 2 | | | | | | | 1 | | | | | | yes | | | | yes |
| CBLN4 | Island | TSS200 | chr20:54580175-54580677 | 502 | | | | | | | | | 1 | | | | | yes | | | | yes | |
| MC3R | N_Shore | TSS1500 | chr20:54823194-54823303 | 109 | | | | | | | | | 1 | | | | | | yes | | | | yes |

| | | | | | | | | | | | | | | | | | | | | | | | | |
|---|---|---|---|---|---|---|---|---|---|---|---|---|---|---|---|---|---|---|---|---|---|---|---|---|
| MC3R | Island | 1stExon | chr20:54824490-54824646 | 156 | | | 3 | | | | | 0 | 1 | | | 1 | | | | | | | | |
| C20orf108 | N_Shore | TSS1500 | chr20:54933689-54933797 | 108 | | | | | | | | | | | | | | | yes | | yes | | yes | |
| CASS4 | NA | TSS1500;nan;TSS200;5'UTR | chr20:54986793-54987662 | 869 | | 2 | | | 2 | | | | 2 | 1 | 1 | | | yes | | | | | | |
| GCNT7;C20orf106 | NA | TSS200;Body | chr20:55101013-55101313 | 300 | | | | | | | | | | | | | | yes | | | yes | | | |
| C20orf107 | NA | TSS1500 | chr20:55107833-55108156 | 323 | | | | | | | | | 1 | | | | | yes | | | yes | | | |
| TFAP2C | Island | TSS1500 | chr20:55203291-55203849 | 558 | | | 17 | | | | | | | | | | | | | | | | yes | |
| BMP7 | N_Shore;Island | Body | chr20:55837483-55839663 | 2180 | | | 9 | | | | | 0 | 3 | | | 1 | | yes | yes | | yes | | yes | |
| BMP7 | S_Shore;Island | TSS200;1stExon;5'UTR | chr20:55841342-55842071 | 729 | | | 3 | | | | | 0 | 3 | | | | | | yes | | | yes | | |
| SPO11 | Island | TSS200;5'UTR;1stExon | chr20:55904686-55905076 | 390 | 1 | | 3 | | | | | 0 | 1 | | | 1 | | | | | yes | | | |
| RAE1 | N_Shore | TSS1500 | chr20:55925570-55925603 | 33 | | | 16 | | | | | | | | | | | yes | | | yes | yes | | |
| RBM38 | Island | TSS1500 | chr20:55964998-55965497 | 499 | | | 2 | | 1 | | | | 1 | | 1 | | | | yes | | yes | | | |
| RBM38 | Island | Body;3'UTR | chr20:55982703-55982978 | 275 | | 1 | | | | | | | | | | | | | yes | | | | | |
| PCK1 | NA | TSS200;nan;TSS1500 | chr20:56135615-56136569 | 954 | | | 1 | | 1 | | | | 2 | | 1 | | | yes | yes | | | | | |
| PMEPA1 | N_Shore;Island | 5'UTR;Body | chr20:56239154-56239706 | 552 | | | | | | | | | 1 | | | | | | | | | | | |
| PMEPA1 | Island | 5'UTR;Body | chr20:56247302-56247611 | 309 | | | | 1 | | | | | | | | | | yes | | | yes | | | |
| PMEPA1 | S_Shore | 5'UTR;1stExon;TSS1500;TSS200 | chr20:56286391-56287532 | 1141 | | | 15 | 1 | | | | | | | | | | yes | yes | | yes | | | |
| C20orf85 | Island | TSS200 | chr20:56725871-56725876 | 5 | | | | | | | | | 1 | | | 1 | | | yes | | | | | |
| APCDD1L | Island | Body | chr20:57042304-57042703 | 399 | | | | | | | | 0 | | | | | | | | | yes | | | |
| APCDD1L | Island;S_Shore | TSS200;TSS1500 | chr20:57089953-57090749 | 796 | | | | | | | | 0 | 4 | | | | | | | | | | yes | |
| NPEPL1 | N_Shore | TSS1500 | chr20:57267176-57267680 | 504 | | | 2 | | | | | | | | | | | | | | | yes | yes | |
| MIR298;MIR296 | NA | TSS1500;TSS200 | chr20:57392898-57394097 | 1199 | | | 2 | | | | | | | | | | | | yes | | | | | |
| GNASAS;GNAS | N_Shore | Body;TSS1500 | chr20:57413698-57414274 | 576 | | | | | | | | | | | | | | yes | yes | | | | | |
| GNASAS;GNAS | N_Shore;Island | TSS200;3'UTR;TSS1500 | chr20:57425979-57427973 | 1994 | | | 24 | | | | | 0 | 18 | 2 | | 9 | | | | | | | yes | |
| GNAS | N_Shore;Island | 3'UTR;TSS1500;Body;TSS200 | chr20:57463325-57464002 | 677 | | | 3 | | | | | | 4 | | | | | | | | | | yes | |
| GNAS | Island | 3'UTR;TSS1500;Body | chr20:57465123-57465511 | 388 | | | 12 | | | | | | 1 | | | | | | | | | yes | yes | |
| GNAS | S_Shelf | Body;3'UTR | chr20:57471654-57471844 | 190 | | | 6 | | | 1 | | | | | | | | | | | | yes | | |
| GNAS | NA | Body;3'UTR | chr20:57485837-57486076 | 239 | | | | | | | | | 2 | | | | | | | | | | yes | |
| CTSZ | Island;S_Shore;chr20:5758190 | Body;TSS1500;S_Shore | chr20:57582006-57583709 | 1703 | | | | | 4 | | | 0 | 7 | | 1 | | | | | | yes | | yes | |
| ATP5E | Island | Body;1stExon;5'UTR;TSS200 | chr20:57607061-57607762 | 701 | | | | | | | | 0 | 1 | | | 1 | | | | | | | yes | |
| SLMO2 | N_Shore | Body | chr20:57617083-57617431 | 348 | | | 3 | | | | | | | | | | | yes | yes | | | | | |
| ZNF831 | N_Shore | TSS1500 | chr20:57765489-57766332 | 843 | | | 6 | | | | | | | | | | | yes | | | yes | | | |
| EDN3 | Island;N_Shore | TSS200;TSS1500 | chr20:57875163-57876125 | 962 | | 1 | 1 | | | | | | 3 | | | | | yes | | | | yes | | |
| PHACTR3 | N_Shore;Island | TSS1500;TSS200;5'UTR;1stExon | chr20:58179244-58180616 | 1372 | | 1 | 37 | | | | | | 2 | | | | | | | | | yes | yes | |
| SYCP2 | Island | TSS1500 | chr20:58508586-58508922 | 336 | | | | | | | | | 1 | | | | | | | | | | | |
| CDH26 | NA | Body;TSS1500 | chr20:58570133-58570523 | 390 | | | | | | | | | | | | | | | | yes | | | | |
| C20orf197 | NA | TSS1500;TSS200 | chr20:58630240-58631038 | 798 | | | 5 | | | | | | 2 | | 1 | | | yes | | | | yes | | |
| CDH4 | Island | Body | chr20:60119508-60119597 | 89 | | | 3 | | | | | | | | | | | | | | yes | | | |
| CDH4 | N_Shore;Island | Body | chr20:60347821-60348407 | 586 | | | 3 | | | | | | | | | | | yes | yes | | | | | |
| CDH4 | Island | Body | chr20:60456296-60456503 | 207 | | | | | | | | | | | | | | | | yes | | | | |
| CDH4 | Island | Body | chr20:60470081-60470267 | 186 | | | | | | | | | | | | | | | | | | | | |
| MIR1257 | NA | TSS200 | chr20:60528758-60529384 | 626 | | | | | | | | | | | | | | yes | yes | yes | | | | |
| TAF4 | Island | Body | chr20:60638679-60639721 | 1042 | | | | | | 1 | | | 2 | | | | | | | | | yes | yes | |
| C20orf151 | NA | TSS200 | chr20:61002595-61002866 | 271 | 1 | 1 | 2 | 2 | 2 | | | | 3 | | | | | yes | | | | | | |
| GATA5 | Island | 1stExon;5'UTR;TSS200;TSS1500 | chr20:61051021-61051585 | 564 | | 1 | | | | | | | 5 | | | 2 | | | | | | yes | yes | |
| C20orf166;C20orf200 | N_Shore | TSS1500;5'UTR | chr20:61147144-61147604 | 460 | | | | | | | | | | | | | | | | | | yes | | |
| C20orf166;MIR1-1 | N_Shore;Island | Body;TSS200;TSS1500;5'UTR | chr20:61150637-61151899 | 1262 | | | 23 | | | | | | 1 | | | | | | | | | | yes | |
| C20orf166;MIR133A2 | NA | Body;TSS200 | chr20:61162065-61162112 | 47 | | | | | | | | | | | | | | | | | | yes | | |
| SLCO4A1;LOC100127888 | NA | Body;TSS200;TSS1500 | chr20:61297564-61299190 | 1626 | | | 9 | | | | | | 2 | | | | | yes | | | | | yes | |
| NTSR1 | Island | TSS1500 | chr20:61339875-61340116 | 241 | | 1 | | | | | | | | | | | | | | | | | yes | |
| NTSR1 | NA | Body | chr20:61371425-61371808 | 383 | 1 | | | 1 | | | 1 | 0 | 1 | 1 | | 4 | 1 | | | | | | | |
| OGFR | Island | TSS1500 | chr20:61435974-61436024 | 50 | | | | | | | | | 1 | | | | | | | yes | | yes | yes | |
| COL9A3 | N_Shore;Island | TSS1500 | chr20:61446962-61447900 | 938 | | | 33 | | | | | 0 | 2 | | | 1 | | yes | | | | | yes | |
| DIDO1 | chr20:61560433-61560954;N_S | N_Shore;5'UTR | chr20:61559756-61561020 | 1264 | | 1 | | | | | | | | | 1 | | | yes | | | | | | |
| SLC17A9 | N_Shore;Island | TSS1500;TSS200;1stExon;5'UTR | chr20:61583537-61584850 | 1313 | | | 37 | | | | | 0 | 1 | 1 | 1 | 1 | | yes | | | | | | |
| SLC17A9 | N_Shore;Island | Body | chr20:61590751-61591209 | 458 | | 1 | | | | | | 0 | 2 | 2 | 1 | 37 | 1 | | | | yes | | | |
| BHLHE23 | Island | TSS200 | chr20:61638420-61638488 | 68 | | 1 | | | | | | | | | | | | | | | | yes | yes | |
| NCRNA00029 | S_Shore | TSS200 | chr20:61668397-61668469 | 72 | | | | | | | | | | | | | | yes | yes | | | | | |
| HAR1B;HAR1A | N_Shore | Body;TSS1500;TSS200 | chr20:61731979-61732608 | 629 | 3 | | | | | | | 0 | 2 | | | 4 | | | | | | yes | | |
| MIR124-3 | Island | TSS200 | chr20:61809724-61809841 | 117 | | | | | | | | | 2 | | | | | | | | | | yes | |
| BIRC7 | N_Shelf | TSS1500;TSS200 | chr20:61866902-61867349 | 447 | | | 10 | | | | | 0 | 1 | | | | | | | | | | yes | |
| NKAIN4 | Island | Body | chr20:61877875-61878032 | 157 | | 1 | | | | | | | 1 | | | | | yes | | | | | | |
| ARFGAP1 | Island;N_Shore | Body | chr20:61915437-61916279 | 842 | | | | | | | | | 3 | 1 | | | | | | | | | | |
| COL20A1 | N_Shelf | TSS200 | chr20:61924382-61924802 | 420 | | | 7 | | | | | | | | | | | | | | | | | |
| COL20A1 | N_Shore;Island | Body | chr20:61953801-61954258 | 457 | | | 7 | | | | | 0 | 3 | | | 1 | | yes | yes | | | | | |
| KCNQ2 | Island | 3'UTR | chr20:62037718-62038111 | 393 | | | | | | | | | | | | | | | | | | | yes | |
| KCNQ2 | Island | Body | chr20:62046270-62046582 | 312 | | | | | | | | | | | | | | | yes | | | | yes | |
| KCNQ2 | Island | Body | chr20:62059392-62059559 | 167 | | | | | | | | | | | | | | | | yes | | | | |
| KCNQ2 | Island | Body | chr20:62079638-62079911 | 273 | | | 1 | | | | | | | | | | | yes | | yes | | | | |
| KCNQ2 | Island;N_Shore | Body | chr20:62084549-62086195 | 1646 | | | 6 | | | | 1 | 0 | 5 | 1 | | 8 | 1 | yes | yes | | | | | |
| KCNQ2 | S_Shore | TSS1500 | chr20:62104476-62104613 | 137 | | | 3 | | | | | | | | | | | | | | | | yes | |
| PPDPF | Island | Body | chr20:62153067-62153431 | 364 | | 1 | 6 | | | | | | | | | | | | | | | | yes | |
| PTK6 | S_Shore | TSS200 | chr20:62168727-62168831 | 104 | | | 6 | | 1 | | | | | | | | | | | | | | | |
| SRMS | Island;S_Shore | 1stExon;TSS1500 | chr20:62178498-62179752 | 1254 | | | 2 | | | | | 0 | 1 | | | | | | | | | yes | | |
| C20orf195 | N_Shore | TSS1500;TSS200 | chr20:62183924-62184397 | 473 | | | 12 | | 1 | | | | | 1 | | | | yes | | | | | | |
| PRIC285 | Island;N_Shore | Body;1stExon;5'UTR;TSS200;TSS1 | chr20:62198872-62200854 | 1982 | | 4 | | | | 1 | | | 2 | 2 | 1 | | | yes | | | | | | |
| RTEL1 | Island | Body | chr20:62317768-62318588 | 820 | | | 34 | 1 | 1 | | | 0 | 1 | 1 | | | | yes | | | | | | |
| TNFRSF6B | Island;N_Shore | 5'UTR;1stExon;TSS1500;TSS200 | chr20:62327732-62328427 | 695 | | | 13 | | | | 1 | 0 | 4 | 3 | 1 | 20 | 1 | | | | | | | |
| ZGPAT;ARFRP1 | N_Shore | TSS1500;Body | chr20:62338235-62338338 | 103 | | | 1 | | | | | | | | | | | | yes | | | | | |
| LIME1 | N_Shore;Island | TSS1500;TSS200;Body | chr20:62367698-62370310 | 2612 | | | 10 | 1 | 7 | 3 | | 0 | 3 | 1 | 1 | 2 | | | | | | | yes | |
| ZBTB46 | Island | Body | chr20:62383275-62383481 | 206 | | 1 | | | | | | | 1 | | | | | | | | | | | |
| ZBTB46 | Island | Body | chr20:62401287-62401458 | 171 | | 1 | | | | | | | | | | | | | | | | | | |
| ZBTB46 | N_Shore | Body | chr20:62406428-62406721 | 293 | | 1 | | | | | | 0 | 1 | 1 | | 2 | | yes | | | | | | |
| ZBTB46 | Island | 5'UTR | chr20:62433998-62434140 | 142 | | | 1 | | | | | 0 | 1 | | | | | yes | | yes | | | | |
| ZBTB46 | N_Shelf | TSS200 | chr20:62436995-62437035 | 40 | | | | | | | | | 1 | | | | | yes | | yes | | | yes | |
| TPD52L2 | Island | Body | chr20:62509164-62509309 | 145 | | | | | | | | | 1 | | | | | yes | | yes | | | | |
| MIR941-1;MIR941-3;MIR941-2;DNAJC5 | N_Shore | TSS1500;5'UTR;TSS200 | chr20:62550415-62551362 | 947 | 4 | | | | | | | | | | | | | yes | yes | | | | | |
| UCKL1;MIR1914;MIR647 | N_Shore | Body;TSS1500;TSS200 | chr20:62574023-62574518 | 495 | | | | 1 | | | | | 1 | | | | | yes | | | | | yes | |
| UCKL1AS;UCKL1 | N_Shelf | TSS1500;Body | chr20:62583555-62583946 | 391 | 2 | | | | | | | | | | | | | yes | | | | | | |
| SOX18 | N_Shore | 3'UTR | chr20:62679255-62679713 | 458 | | | 6 | | | | | 0 | 1 | | 1 | 8 | | | | | | | yes | |
| SOX18 | Island | TSS1500 | chr20:62681243-62681428 | 185 | | | | | | | | | | | | | | | | | | yes | yes | |
| TCEA2 | N_Shore;Island | TSS1500;TSS200;1stExon;5'UTR | chr20:62687969-62688896 | 927 | | | 15 | | | | | 0 | 1 | | | | | yes | | | | | | |
| TCEA2 | N_Shore;Island | 5'UTR;TSS1500;TSS200 | chr20:62693657-62694005 | 348 | | | 12 | | | | | 0 | 3 | | | | | yes | yes | | | | yes | |
| OPRL1;C20orf201 | Island;N_Shore | 5'UTR;TSS200;TSS1500 | chr20:62715756-62716606 | 850 | 1 | | 14 | | | | | 0 | | | | | | | | | | | | |
| NPBWR2 | Island;S_Shore | 1stExon;TSS1500 | chr20:62737547-62739073 | 1526 | | | 5 | | | | | | 1 | | | | | | | | | yes | yes | |
| MYT1 | NA | 5'UTR;1stExon;TSS200 | chr20:62795807-62796417 | 610 | | | 9 | | | | | | 1 | | | | | | | | | | | |
| PCMTD2 | N_Shore | TSS1500 | chr20:62886664-62886904 | 240 | | | | | | | | | 1 | | | | | | | yes | | yes | | |
| BAGE;BAGE5;BAGE3;BAGE2;BAGE4 | NA | 1stExon;5'UTR | chr21:11098909-11099162 | 253 | | | | | | | | | | | | | | | | | | | | yes |
| C21orf99 | Island | TSS1500 | chr21:14410030-14410422 | 392 | | | | | | | | | | | | | | | | | | | | |
| POTED | NA | TSS1500 | chr21:14982225-14982363 | 138 | | | | | | | | | 1 | | | | | | | | | yes | | |

| | | | | | | | | | | | | | | | | | | | | | | | | |
|---|---|---|---|---|---|---|---|---|---|---|---|---|---|---|---|---|---|---|---|---|---|---|---|---|
| C21orf81 | Island;S_Shore | Body;TSS200 | chr21:15352213-15352983 | 770 | | 1 | 20 | | | | 2 | 0 | 6 | 2 | | 13 | 1 | | | | | | | yes |
| ABCC13 | NA | TSS1500;TSS200 | chr21:15645649-15646312 | 663 | | | 6 | | | | | 0 | 3 | | | 1 | | | | | | | | |
| C21orf34 | NA | TSS1500 | chr21:17442467-17442803 | 336 | | | | | | | | | | | | | | | | | | yes | | |
| C21orf34;MIR125B2 | NA | Body;TSS1500 | chr21:17961737-17962124 | 387 | | | 18 | | | | | | | | | | | | | | | | | |
| CHODL | N_Shore;Island | TSS1500;TSS200 | chr21:19616939-19617439 | 500 | | | | | | | | 0 | 1 | | | | | | | | | | | |
| NCAM2 | N_Shore;Island | TSS1500 | chr21:22369310-22370733 | 1423 | | | 2 | | | | | | 1 | | | | | | | | | yes | yes | |
| MIR155HG | Island | Body | chr21:26934576-26934885 | 309 | | | | 1 | | | | | | | | | | yes | | | | | | |
| APP | NA | Body | chr21:27372387-27372461 | 74 | | 1 | | | 2 | | | 0 | 2 | | | 10 | | | | | | | | |
| ADAMTS5 | N_Shore;Island | Body;1stExon;5'UTR | chr21:28337594-28339334 | 1740 | | | 2 | | | | | | 1 | | | | | | | | | | | |
| RNF160 | S_Shore | TSS1500 | chr21:30365886-30366250 | 364 | | | 4 | | | | | | | | | | | yes | | yes | yes | | | |
| RWDD2B | Island | Body;1stExon;5'UTR | chr21:30391383-30392188 | 805 | | | 2 | | | | | 0 | 1 | | | | | yes | yes | | yes | | yes | |
| NCRNA00189 | NA | TSS200 | chr21:30565636-30565706 | 70 | | | | | | | | 0 | | | | | | yes | | yes | | | | |
| GRIK1 | N_Shore | Body | chr21:31310508-31310920 | 412 | | | | | | | | | 1 | | | | | | | | | | yes | |
| GRIK1 | S_Shore | TSS200 | chr21:31312328-31312905 | 577 | | 1 | | | | | | | 1 | | | | | | yes | | | yes | | |
| CLDN17 | NA | 1stExon | chr21:31538714-31539057 | 343 | | | | | | | | | | | | | | | | | | | | |
| CLDN8 | NA | 1stExon | chr21:31588123-31588661 | 538 | | | | | | | | | | | | | | | | | | | | |
| KRTAP24-1 | NA | 1stExon;5'UTR;TSS200;TSS1500 | chr21:31655261-31655982 | 721 | | | 7 | | | | | | 2 | | | | | | | | | yes | yes | |
| KRTAP6-3 | NA | TSS200 | chr21:31964616-31964845 | 229 | | | | | | | | | | | | | | | yes | | | yes | | |
| KRTAP6-2;KRTAP22-1 | NA | TSS1500 | chr21:31972533-31972923 | 390 | | | | | | | | | | | | | | yes | yes | | | yes | | |
| KRTAP7-1 | NA | 1stExon | chr21:32202010-32202284 | 274 | | | | | | | | | | | | | | | yes | | | | | |
| KRTAP11-1 | NA | 1stExon | chr21:32253584-32254161 | 577 | | | | | | | | | | | | | | | | | | | | |
| TIAM1 | N_Shore | Body | chr21:32623901-32624274 | 373 | 2 | | | | | | | | | | | | | yes | yes | | yes | | | |
| TIAM1 | N_Shore | 5'UTR | chr21:32929871-32932267 | 2396 | | | | 1 | | | | | | | | | | yes | yes | | | | yes | |
| C21orf62 | NA | 5'UTR;1stExon | chr21:34185927-34186122 | 195 | | | | | | | | 0 | | | | | | | | yes | | | yes | |
| OLIG2 | Island | TSS200;TSS1500 | chr21:34396944-34398532 | 1588 | | 1 | 1 | | 2 | | | 0 | 2 | | | | | | | | | yes | yes | |
| OLIG1 | Island | TSS200 | chr21:34442350-34442377 | 27 | | | | | | | | | 1 | | | | | | yes | | | yes | yes | |
| IL10RB | N_Shore;Island | TSS1500;TSS200;Body | chr21:34638126-34639113 | 987 | 5 | | | | | | | | | | | | | | yes | | | yes | | |
| IFNGR2 | N_Shore | TSS200;TSS1500 | chr21:34774882-34775045 | 163 | 1 | | | 1 | 1 | | | | | | | | | yes | | | | | | |
| KCNE2 | NA | TSS1500 | chr21:35736023-35736383 | 360 | | | | | | | | | 1 | | | | | yes | | | | yes | | |
| KCNE1 | Island | 5'UTR;1stExon;TSS200;TSS1500 | chr21:35831871-35832364 | 493 | 1 | | 33 | | | | | 0 | 8 | 2 | | 19 | 1 | | yes | | | | | |
| KCNE1 | NA | TSS1500 | chr21:35884348-35884508 | 160 | | | | | | | | | 1 | | | | | | | yes | yes | | | |
| CLIC6 | Island | TSS200;TSS1500 | chr21:36041334-36042170 | 836 | | 1 | 6 | | 1 | | | 0 | 2 | | | | | | | | | | | |
| RUNX1 | N_Shore;Island | Body;1stExon | chr21:36258423-36259460 | 1037 | | | | | | | | 0 | 4 | 2 | | 6 | | | | | | | | |
| RUNX1 | Island | Body | chr21:36263347-36263808 | 461 | | | | | | | | 0 | | | | | | | | | | | yes | |
| RUNX1 | Island | Body | chr21:36399146-36399540 | 394 | | | 15 | 1 | 1 | | | | | | | | | | | | yes | | yes | |
| RUNX1 | NA | 5'UTR;1stExon;TSS1500 | chr21:36421467-36421955 | 488 | | | | | 2 | | | | 2 | 2 | | | | | | | | yes | | |
| CBR1 | N_Shore;Island | TSS200;1stExon;5'UTR | chr21:37442104-37442777 | 673 | 1 | 3 | 2 | 1 | 1 | | | 0 | 8 | 2 | | 7 | 1 | | yes | | | | | |
| CLDN14 | NA | 5'UTR;TSS200 | chr21:37852492-37852822 | 330 | | | | | 1 | | | | | | | | | | | | | | | |
| CLDN14 | NA | TSS200;5'UTR | chr21:37914924-37915354 | 430 | | | 2 | 1 | | | | | 3 | | | 4 | | yes | yes | | | | yes | |
| CLDN14 | NA | 1stExon;5'UTR | chr21:37948709-37949197 | 488 | | | | | | | | | | | | | | yes | | | | | yes | |
| SIM2 | Island | TSS1500 | chr21:38070524-38070724 | 200 | | | | | | | | | 1 | | | | | | | | | | | |
| SIM2 | N_Shore | Body | chr21:38074992-38075171 | 179 | | | 3 | | | | | | | | | | | yes | | | | | | |
| SIM2 | N_Shore | Body | chr21:38076709-38077258 | 549 | | | 1 | | | | | | | | | | | | | | | | | |
| SIM2 | Island | Body | chr21:38080526-38081193 | 667 | | | | | | | | | 1 | | | | | | | | | yes | | |
| SIM2 | S_Shore | Body | chr21:38082613-38083149 | 536 | | | | | | | | | 1 | | | | | yes | | | | | yes | |
| SIM2 | N_Shelf | Body | chr21:38117454-38118338 | 884 | | | | | | | | | 1 | | | | | yes | | | | | | |
| SIM2 | Island | Body;3'UTR | chr21:38119920-38120824 | 904 | | | 2 | | | | | | 2 | 3 | | | | | | | yes | | yes | |
| HLCS | Island | 5'UTR | chr21:38353114-38353318 | 204 | | | | | | | | | 2 | | | 1 | | yes | yes | | | | | |
| HLCS | Island | TSS200 | chr21:38362632-38362754 | 122 | | | 6 | | | | | 0 | 2 | | 1 | 1 | | | | yes | | | | |
| DSCR9 | NA | TSS1500;TSS200 | chr21:38580129-38580856 | 727 | | | 8 | 1 | | | | | 1 | | | | | | | | | | | |
| DSCR3 | N_Shore;Island | Body | chr21:38629859-38630728 | 869 | | | 2 | | | | | 0 | 2 | 1 | | 7 | 1 | yes | | yes | | | | |
| KCNJ6 | Island | Body | chr21:39047670-39047855 | 185 | | 1 | | | | | | | | | | | | | yes | | yes | yes | | |
| KCNJ15 | NA | 5'UTR;TSS1500;TSS200 | chr21:39643733-39644354 | 621 | | | 4 | | | | | | 2 | | | | | yes | | | | | | |
| ERG | NA | Body;1stExon;5'UTR | chr21:39870397-39870704 | 307 | | | | | 1 | | | | 1 | | 1 | | | | | | | | yes | |
| NCRNA00114 | NA | Body;TSS1500 | chr21:40145361-40146149 | 788 | | | 9 | | | | | | 2 | | 1 | | | | | | | | yes | |
| ETS2 | N_Shore | TSS1500 | chr21:40176950-40177165 | 215 | | | 2 | | | | | | | | | | | yes | | | | | yes | |
| PSMG1 | Island | TSS200 | chr21:40555470-40555479 | 9 | | | | | | | | 0 | | | | | | yes | | | | | yes | |
| WRB | S_Shore;Island | Body;TSS1500 | chr21:40757691-40758346 | 655 | | | | | | | | 0 | | | | | | yes | | | | | | |
| WRB | N_Shore | Body;TSS200 | chr21:40759534-40760258 | 724 | | | 9 | | | | | 0 | 2 | | | 2 | | yes | yes | | | | | |
| SH3BGR | NA | 5'UTR;TSS1500 | chr21:40823279-40823722 | 443 | | | 2 | | 1 | | | | | | | | | yes | | | | | | |
| C21orf88 | NA | TSS200 | chr21:40984760-40984777 | 17 | | | | | | | | | 1 | | | | | yes | | | | | yes | |
| DSCAM | Island | 1stExon;TSS200 | chr21:42218551-42219221 | 670 | | | 2 | | | | | | 3 | | | | | | | | | yes | yes | |
| FAM3B | N_Shore | TSS200 | chr21:42688468-42688675 | 207 | | | | | | | | | 2 | | | | | | | | | | yes | |
| MX2 | NA | TSS1500;TSS200 | chr21:42733729-42734266 | 537 | | | | 1 | 3 | | | | | | | | | yes | | | yes | | | |
| MX1 | Island | 5'UTR | chr21:42798386-42798747 | 361 | | | | | | | | 0 | | | | | | | | | yes | | yes | |
| TMPRSS2 | Island | 1stExon;TSS200;5'UTR;TSS1500 | chr21:42880077-42880714 | 637 | | | 2 | | | | | | 1 | | | | | yes | | yes | | | yes | |
| NCRNA00111 | NA | TSS200;TSS1500 | chr21:43098516-43099460 | 944 | | 1 | | 1 | | | | 0 | | 1 | | 1 | | yes | | yes | | | | |
| C21orf129;NCRNA00112 | NA | Body;TSS1500;TSS200 | chr21:43135854-43136563 | 709 | | | 2 | | | | | | 2 | 1 | | | | | | yes | | | | |
| RIPK4 | N_Shore | Body | chr21:43185385-43185884 | 499 | | | | | 1 | | | | 1 | 1 | | | | | | | yes | | | |
| PRDM15 | Island | Body | chr21:43255477-43255700 | 223 | | 1 | | | | | | | | | | | | | | | | | yes | |
| C2CD2 | N_Shore | Body | chr21:43372922-43373136 | 214 | | | 1 | | | | | 0 | | | | | | yes | | | | | | |
| C21orf121 | NA | TSS200 | chr21:43442027-43442254 | 227 | | | 3 | | | | | | | | | | | | | yes | | | | |
| C21orf128;UMODL1 | S_Shore | TSS200;Body | chr21:43528734-43529162 | 428 | | | | | | | 1 | 0 | 4 | 2 | 1 | 8 | 1 | | | | | | | |
| UMODL1 | Island | Body | chr21:43547872-43548126 | 254 | | | | | | 1 | | | | | | | | | | | | | yes | |
| ABCG1 | NA | TSS1500 | chr21:43619310-43619651 | 341 | | | 1 | | | | | | 1 | | | | | yes | yes | | | | | |
| ABCG1 | Island | Body | chr21:43655256-43655464 | 208 | | | | | | | | | | | | | | | | yes | | | | |
| TFF3 | NA | 1stExon;TSS1500 | chr21:43735412-43737014 | 1602 | | 1 | | | 1 | | | | | | 1 | | | yes | | | | yes | | |
| TFF2 | NA | TSS1500;5'UTR;1stExon;TSS200 | chr21:43771120-43772018 | 898 | | | 3 | | | | | | | | | | | yes | | | | | | |
| TFF1 | NA | 1stExon;5'UTR;TSS200 | chr21:43786627-43786997 | 370 | | 1 | | | | | | | | | | | | yes | yes | | | | | |
| TMPRSS3 | NA | Body;TSS200 | chr21:43809360-43809567 | 207 | | | | 1 | | | | | | | | | | | | | | | | |
| TMPRSS3 | NA | 1stExon;5'UTR | chr21:43816086-43816355 | 269 | | | | | | | | | | | | | | | | | | yes | | |
| UBASH3A | NA | TSS1500;TSS200;1stExon | chr21:43823604-43824262 | 658 | | | 12 | | | | | | 4 | | | | | | yes | | yes | | | |
| SLC37A1 | S_Shelf | TSS1500 | chr21:43919316-43919853 | 537 | | | 2 | | | | | | | | | | | yes | | | | | | |
| PDE9A | Island | Body;5'UTR | chr21:44088823-44089142 | 319 | | | 2 | | | | | | | | | | | | yes | yes | | | | |
| CBS | N_Shore | 3'UTR | chr21:44473496-44473962 | 466 | | | 1 | | | | | | 2 | | | | | yes | yes | | | | | |
| CBS | N_Shore | Body | chr21:44480336-44480711 | 375 | | | | | | | | | 1 | | | | | | | | | | | |
| CRYAA | N_Shore | TSS200 | chr21:44588950-44589128 | 178 | | | | | | | | | | | | | | | | | | | | |
| SIK1 | NA | Body | chr21:44845925-44845971 | 46 | | | | | | | | | 1 | | | | | | | | | | yes | |
| C21orf125 | NA | TSS1500 | chr21:44869682-44869785 | 103 | | | | | 1 | | | | | | | | | | | yes | | | | |
| C21orf84 | NA | Body | chr21:44898090-44898206 | 116 | | | | | | | | 0 | 1 | | | 1 | | | | | | | | |
| HSF2BP | N_Shore | Body | chr21:44985005-44985357 | 352 | | | | | | | | | | | | | | | yes | yes | | | | |
| PDXK | Island;N_Shore | Body | chr21:45148332-45149398 | 1066 | 2 | | 22 | | 4 | | | | | | | | | yes | | | | | | |
| LOC284837 | NA | Body;TSS200;TSS1500 | chr21:45232232-45233662 | 1430 | | | 7 | | | | | 0 | 1 | | | | | yes | | yes | | | | |
| AGPAT3 | N_Shore | 5'UTR | chr21:45379489-45379721 | 232 | | | 1 | | | | | | | | | | | | | | | yes | | |
| DNMT3L | NA | TSS1500 | chr21:45683146-45683528 | 382 | | | 8 | | | | | | | | | | | yes | yes | | | yes | yes | |
| AIRE | Island | TSS1500;TSS200 | chr21:45705429-45706043 | 614 | 3 | 1 | 6 | | 1 | 1 | | 0 | 7 | | | 2 | | yes | | | | | | |
| AIRE | N_Shelf | Body;TSS200 | chr21:45710206-45710296 | 90 | | | | | | | | | 1 | | | | | | yes | | | | | |
| C21orf2 | Island | Body | chr21:45754009-45754232 | 223 | | 1 | | | | | | | | | | | | yes | | | | | | |
| TRPM2 | NA | TSS200;Body;TSS1500 | chr21:45773189-45774294 | 1105 | | | 11 | | | | | 0 | 1 | 1 | 1 | 3 | | yes | | | | | | |

| | | | | | | | | | | | | | | | | | | | | | | | |
|---|---|---|---|---|---|---|---|---|---|---|---|---|---|---|---|---|---|---|---|---|---|---|---|
| TRPM2 | Island | Body | chr21:45789122-45789379 | 257 | 1 | | 6 | | | | | 0 | 1 | 1 | | | | | | | | yes | |
| TRPM2 | Island | Body | chr21:45811363-45811549 | 186 | | | 5 | | | | | | 1 | | | | | yes | | yes | | | yes |
| TRPM2 | Island | Body | chr21:45845112-45845495 | 383 | | | | | | | | | | | | | | yes | | | | | |
| C21orf29 | N_Shore | Body | chr21:45926106-45926719 | 613 | | | 7 | | | | | | | | | | | | yes | | | | |
| KRTAP10-1;C21orf29 | NA | TSS1500;Body | chr21:45960718-45961033 | 315 | | | 1 | | | | | | | | | | | yes | yes | | | | yes |
| KRTAP10-8;C21orf29 | NA | TSS1500;Body | chr21:46031707-46032026 | 319 | | | | | | | | | | | | | | | | yes | | | |
| KRTAP10-10;C21orf29 | NA | TSS1500;Body | chr21:46056911-46057082 | 171 | | | | | | | | | | | | | | | | | | yes | |
| KRTAP10-11;C21orf29 | NA | TSS1500;Body | chr21:46065587-46065782 | 195 | | | | | | | | | | | | 1 | | yes | | | | | |
| C21orf29;KRTAP12-3 | NA | Body;TSS1500;TSS200 | chr21:46077454-46078007 | 553 | | | 1 | 1 | | | | 0 | 7 | 1 | | 2 | | | | | yes | yes | |
| C21orf29;KRTAP12-2 | NA | Body;1stExon | chr21:46086578-46087026 | 448 | | | | | | | | | 1 | | | | | | | | | | |
| SUMO3 | Island | TSS1500 | chr21:46238781-46238957 | 176 | | | 1 | | | | | | | | | | | yes | | | | | yes |
| ITGB2 | NA | 5'UTR | chr21:46332181-46334322 | 2141 | | | | | | | | | | | | | | yes | | | | | |
| ITGB2 | NA | 5'UTR;1stExon;TSS200 | chr21:46340776-46341389 | 613 | | 2 | 5 | 1 | | | | | 3 | 2 | 1 | | | yes | | | | yes | |
| ITGB2 | N_Shelf | 5'UTR;1stExon;TSS1500 | chr21:46348306-46349095 | 789 | | | | | | | | | 2 | | | | | | | | yes | yes | |
| C21orf70 | N_Shore | Body | chr21:46367939-46368090 | 151 | | | 1 | | | | | | | | | | | | | | | | |
| C21orf70 | N_Shore | Body | chr21:46378243-46379089 | 846 | | | 1 | | | | | | | | | | | | | | | | |
| NCRNA00162 | N_Shore | Body | chr21:46421023-46421509 | 486 | | | | | | | | | | | | | | | yes | | | yes | |
| NCRNA00162 | S_Shelf | TSS200 | chr21:46424834-46424880 | 46 | | | | | | | | | | | | | | | yes | yes | | | |
| ADARB1;C21orf122 | N_Shore | TSS1500;TSS200 | chr21:46493156-46493436 | 280 | | | 3 | | 1 | | | | 3 | | | | | yes | | yes | | yes | yes |
| POFUT2;LOC642852 | Island | TSS1500;Body | chr21:46708497-46708729 | 232 | | | | | | | | | | | | | | | yes | yes | | yes | yes |
| COL18A1 | Island | Body | chr21:46825840-46826053 | 213 | | | 2 | | | | | | | | | | | | | | | | |
| COL18A1 | Island | Body | chr21:46847581-46847898 | 317 | | | | | | | | | 1 | | | | | | yes | | | | |
| COL18A1 | N_Shore;Island | Body | chr21:46850449-46851112 | 663 | | | 7 | | | | 1 | | 2 | | | | | | | | | | yes |
| COL18A1 | Island | Body;1stExon | chr21:46876658-46876904 | 246 | | | | | | | | | 1 | | | | | yes | | | | | |
| COL18A1 | Island | Body | chr21:46890849-46891093 | 244 | | | | | | | | 0 | 2 | | | | | yes | | | | yes | |
| COL18A1 | Island | Body | chr21:46898048-46898789 | 741 | | | | | | | | | 1 | | | | | | | | | yes | |
| COL18A1 | Island | Body | chr21:46929500-46929741 | 241 | | | | | | | | | 1 | | | | | | | | | | |
| PCBP3 | N_Shore;Island | Body | chr21:47294739-47295375 | 636 | | | 15 | | | | | 0 | 1 | | | | | yes | yes | | | | |
| COL6A1 | S_Shore;Island | TSS200;Body;TSS1500 | chr21:47401109-47402144 | 1035 | | | | | 1 | | 2 | 0 | | 3 | | | | | | | | yes | |
| COL6A1 | Island | Body | chr21:47404109-47404268 | 159 | | | 3 | | | | | | | | | | | yes | | | | yes | |
| C21orf56 | Island | 5'UTR;Body | chr21:47602432-47602702 | 270 | | 1 | | | | | | | | | | | | | | | | | |
| C21orf56 | S_Shelf;S_Shore | TSS1500;5'UTR;1stExon | chr21:47604052-47605174 | 1122 | 11 | 11 | 43 | 1 | | 6 | 1 | 0 | 12 | 6 | 1 | 57 | 4 | | yes | | | | |
| C21orf57 | Island | Body;3'UTR | chr21:47717406-47718080 | 674 | | | 2 | | | | | | | | | | | yes | | | | | yes |
| PCNT | Island | Body | chr21:47802916-47803548 | 632 | | | | | | | | 0 | 2 | 2 | | | | | | | | | |
| PCNT | N_Shore;Island | Body | chr21:47813025-47813491 | 466 | | | 10 | | | | | | | | | | | | yes | | | | |
| PCNT | Island | Body | chr21:47845610-47845863 | 253 | | 1 | | | | | | | | | | | | yes | | | | yes | |
| PCNT | Island | Body | chr21:47855893-47856020 | 127 | | | | | | | | | 1 | | | | | yes | | | | | yes |
| DIP2A | Island | Body | chr21:47918542-47918727 | 185 | | 1 | | | | | | | | | | | | | | | | | |
| S100B | NA | TSS1500 | chr21:48026043-48026298 | 255 | | | | | | | | | 1 | 1 | 1 | | | yes | | | | | |
| PRMT2 | Island | Body | chr21:48081242-48081686 | 444 | | | | | | | | 0 | 1 | | | 6 | 1 | | | | | | yes |
| psiTPTE22 | Island;N_Shore | TSS1500;TSS200;Body | chr22:17082581-17083727 | 1146 | | 1 | | | 2 | | | 0 | 4 | | | | | | | | | yes | yes |
| GAB4 | Island | TSS200 | chr22:17489112-17489225 | 113 | | | 4 | | | | | | | | | | | | | | | yes | |
| CECR7 | Island | Body | chr22:17518345-17518561 | 216 | | | | | | | | | | | | | | yes | yes | | | | |
| CECR1 | NA | TSS200;Body;1stExon;5'UTR | chr22:17680477-17680706 | 229 | | | | 2 | 2 | | | | 1 | | 1 | | | yes | | | | | |
| CECR2 | NA | TSS1500;TSS200 | chr22:17955976-17956641 | 665 | 3 | | 3 | | | | | 0 | 3 | 1 | | | | yes | | | | | |
| SLC25A18 | NA | TSS1500 | chr22:18042550-18043043 | 493 | | | 4 | | | | | | | | | | | yes | | | | | |
| SLC25A18 | N_Shore;Island | Body | chr22:18063857-18064224 | 367 | | | 6 | | | | | 0 | 1 | | | | | yes | | | | | |
| BID | Island | Body;5'UTR | chr22:18231406-18231825 | 419 | | | | | | | | | 1 | | | | | yes | | | | | |
| MICAL3 | Island | Body | chr22:18324579-18324769 | 190 | 1 | | | | | | | | | | | | | yes | | | | | |
| MICAL3;MIR648 | NA | 5'UTR;TSS1500 | chr22:18464717-18465000 | 283 | 1 | | | | | | | | | | | | | yes | | | yes | yes | |
| MICAL3 | S_Shore | TSS1500 | chr22:18507711-18508298 | 587 | | | | | | | | | | | | | | yes | | | | | |
| TUBA8 | Island | TSS1500 | chr22:18593294-18593362 | 68 | | | | | | | | | 1 | | | | | | | | | | |
| USP18 | N_Shore | TSS1500 | chr22:18632404-18632476 | 72 | | | 8 | | | | | 0 | 2 | | | 1 | | yes | | | yes | | |
| DGCR6 | Island | Body | chr22:18894280-18894389 | 109 | | | | 1 | | | | | | | | | | | yes | | | | yes |
| PRODH | S_Shore | TSS1500 | chr22:18924883-18925277 | 394 | | | 1 | | | | | | | | | | | yes | yes | yes | | | |
| DGCR9 | NA | TSS1500 | chr22:19003848-19004088 | 240 | | | 1 | | | | | | | | | | | yes | yes | | | | |
| DGCR14;TSSK2 | N_Shore | 3'UTR;TSS200;1stExon;5'UTR | chr22:19118165-19118918 | 753 | | | 11 | | | | | | | | | | | | | yes | | | |
| GSC2 | Island | TSS1500 | chr22:19138124-19138856 | 732 | | | | | | | | | | 1 | | | | yes | | | | | yes |
| SLC25A1 | Island | TSS1500 | chr22:19166691-19167120 | 429 | | | | | | | | | | | | | | | | | | | yes |
| LOC150185 | Island | TSS200 | chr22:19554366-19554560 | 194 | | | | | | | | | 1 | | | | | yes | yes | | | | |
| SEPT5;GP1BB | Island | Body;TSS1500;TSS200;3'UTR | chr22:19709696-19711051 | 1355 | | | | | | | | 0 | 3 | 1 | | | | | | | | | yes |
| TBX1 | Island | TSS1500;TSS200 | chr22:19743336-19744078 | 742 | | | 14 | | | | | | 1 | | | | | | | | | yes | yes |
| TBX1 | Island | 5'UTR | chr22:19746156-19747140 | 984 | 1 | | 27 | | | | | 0 | 1 | | | | | | | | | yes | yes |
| TBX1 | Island | Body | chr22:19748910-19749188 | 278 | | | | | | | | | | | | | | | | | | yes | yes |
| TBX1 | Island | Body | chr22:19754125-19754815 | 690 | | | 5 | | | | | | | 1 | | | | | | yes | | yes | yes |
| TXNRD2 | NA | Body | chr22:19879176-19880135 | 959 | | | 2 | | 3 | | | | | | | | | | | | | | |
| COMT;TXNRD2 | N_Shore;Island | TSS1500;Body;TSS200;1stExon | chr22:19928445-19929286 | 841 | | | 2 | | | | | 0 | 1 | 2 | | 2 | | | | yes | | | yes |
| COMT | NA | 5'UTR;TSS200 | chr22:19938916-19939061 | 145 | | | | | | | | | | | | | | yes | | yes | | | |
| COMT | NA | 5'UTR;TSS200;TSS1500 | chr22:19949585-19950166 | 581 | | | 7 | | | | | | 1 | | | | | | | | | yes | |
| MIR1286;RTN4R | NA | TSS200;Body | chr22:20236751-20236780 | 29 | | | | | | | | | | | | | | | | | | | |
| ZNF74 | Island | TSS200 | chr22:20748318-20748341 | 23 | | | | | | | | | 1 | | | | | | | | | | |
| SCARF2 | Island | Body | chr22:20783497-20783963 | 466 | | | | | | | | 0 | | | 1 | | | | | | | | yes |
| MED15 | N_Shore | TSS1500 | chr22:20861470-20861479 | 9 | | | 6 | | | | | | | | | | | | yes | | | | |
| AIFM3 | S_Shore | 5'UTR;TSS200 | chr22:21321253-21321585 | 332 | | | 1 | | | | | | 1 | | | | | | | | | | |
| MGC16703;P2RX6 | S_Shore | TSS1500;TSS200 | chr22:21369411-21369610 | 199 | | | | | | | | | | | | 1 | | | yes | | | | |
| SLC7A4 | Island | TSS200 | chr22:21386999-21387058 | 59 | | | | | | | | | | | | | | | | | | | |
| UBE2L3 | N_Shore | TSS1500 | chr22:21921269-21921839 | 570 | | | | | | | | | 1 | | | | | yes | yes | | | | |
| CCDC116 | S_Shelf | TSS200 | chr22:21986960-21987190 | 230 | | | 7 | | | | | | | | | | | yes | | | | | |
| MIR130B;MIR301B | S_Shore | TSS1500;TSS200 | chr22:22007095-22007528 | 433 | | | 7 | | | | | | | | | | | yes | | | | | |
| PPIL2 | N_Shore | TSS1500 | chr22:22020012-22020076 | 64 | | | 5 | | | | | | | | | | | yes | yes | | | | |
| YPEL1 | Island | Body | chr22:22058080-22058271 | 191 | | | 6 | | | | | | | | | | | yes | | | | | |
| YPEL1 | Island | TSS1500 | chr22:22090796-22090816 | 20 | | | 3 | | | | | 0 | 1 | | | | | yes | | | | yes | yes |
| VPREB1 | NA | TSS1500 | chr22:22598937-22599189 | 252 | | | 2 | | | | | | | | | | | yes | | yes | | | yes |
| ZNF280B | S_Shore | 1stExon;5'UTR | chr22:22863293-22863663 | 370 | | | | | | | | | | | | | | | yes | | yes | | |
| ZNF280A | NA | 5'UTR;1stExon;TSS200 | chr22:22874447-22875038 | 591 | | | 3 | | | | | | 1 | | 1 | | | yes | | | yes | | yes |
| PRAME;LOC648691 | Island;S_Shore | TSS200;5'UTR;1stExon;Body;TSS15 | chr22:22901569-22902198 | 629 | | | 12 | | | | | | | | | 2 | | | | | | | |
| POM121L1P | Island | TSS200 | chr22:22987059-22987126 | 67 | | | | | | | | | 2 | | | 1 | | yes | | | | | |
| RAB36 | N_Shore;Island | TSS1500;TSS200 | chr22:23486678-23487440 | 762 | | | | | 1 | | | | 2 | | | | | yes | yes | | | | |
| BCR | N_Shore | TSS1500 | chr22:23522307-23522374 | 67 | | | | | | | | | 1 | | | | | | | | | yes | |
| IGLL1 | NA | 1stExon;5'UTR | chr22:23922378-23922551 | 173 | | | | | | | | | | | | | | | | | yes | | |
| C22orf15 | NA | TSS200 | chr22:24105087-24105692 | 605 | | | | | 1 | | | 0 | 1 | | | | | yes | yes | | yes | | yes |
| DERL3 | Island;chr22:24180491-241816 | Body;5'UTR;1stExon;Island;TSS200 | chr22:24180995-24181665 | 670 | | | 9 | | | | | | | | 1 | | | | yes | | | yes | |
| SLC2A11 | N_Shore | TSS200;TSS1500 | chr22:24199008-24199125 | 117 | | | | | | | | 0 | 1 | | | | | | yes | | | | |
| MIF | Island | TSS200 | chr22:24236375-24236617 | 242 | | | | 2 | | | | | | | | | | | | | | | |
| GSTTP1 | NA | TSS1500 | chr22:24348549-24348715 | 166 | | 7 | 9 | 1 | | | 1 | | | 1 | 1 | 30 | 1 | | | | | | |
| LOC391322 | Island | TSS1500;TSS200;5'UTR;1stExon;Bo | chr22:24372913-24373618 | 705 | | 1 | 16 | | | | | | 1 | 2 | 1 | 3 | 1 | yes | | | yes | | |
| GSTT1 | N_Shore;Island;chr22:2438413 | Body;1stExon;5'UTR;Island;TSS200 | chr22:24384105-24384944 | 839 | 10 | 16 | 18 | 3 | | | 2 | | 2 | 8 | 1 | 28 | | yes | yes | | | | |
| SUSD2 | NA | TSS1500;TSS200 | chr22:24577142-24577448 | 306 | | | 14 | | | | | | 1 | 1 | 1 | | | yes | | | | yes | |
| ADORA2A | S_Shelf | TSS1500;TSS200 | chr22:24823050-24824362 | 1312 | | | | | 2 | 1 | | | 3 | 1 | 1 | | | | | | | | |

| | | | | | | | | | | | | | | | | | | | | | | | | |
|---|---|---|---|---|---|---|---|---|---|---|---|---|---|---|---|---|---|---|---|---|---|---|---|---|
| C22orf45;UPB1 | Island | Body;TSS1500;TSS200 | chr22:24890690-24891338 | 648 | 1 | | 6 | | | | | 0 | 7 | 1 | | 4 | | | | | | | | |
| GGT1 | NA | TSS1500;TSS200 | chr22:24979254-24979964 | 710 | | | 39 | | | | | | 2 | 1 | | | | yes | | | | | | |
| GGT1 | NA | 5'UTR;1stExon | chr22:24999188-24999195 | 7 | | | | | | | | | | | | | | yes | | | | | | |
| GGT1 | NA | 5'UTR;TSS1500;TSS200 | chr22:25003400-25003552 | 152 | | | | | | | | | | | | | | | | | | | | |
| TOP1P2;PIWIL3 | N_Shore;Island | TSS1500;5'UTR | chr22:25159336-25160406 | 1070 | | | 20 | | | | | | 6 | | | | | yes | | yes | | yes | yes | |
| PIWIL3 | NA | TSS200 | chr22:25170858-25170892 | 34 | | | | | | | | | | | | | | yes | | yes | | | | |
| SGSM1 | Island | TSS200 | chr22:25201954-25202552 | 598 | | | | | | | | | 2 | | | | | yes | | | | yes | yes | |
| KIAA1671 | N_Shore;Island | TSS1500;1stExon | chr22:25423700-25424512 | 812 | | | | | | | | | 1 | | | | | yes | | | | | | |
| KIAA1671 | Island | Body | chr22:25507318-25507612 | 294 | | | | | 1 | | | | | | | | | | | | | yes | yes | |
| KIAA1671 | Island | Body | chr22:25575426-25575559 | 133 | | | | | | | | | | | | | | | yes | | | | | |
| CRYBB3 | NA | TSS1500;TSS200 | chr22:25595352-25595955 | 603 | | | 10 | | | | | | 1 | | | | | yes | | | | | | |
| CRYBB2 | NA | TSS200;TSS1500 | chr22:25615317-25615538 | 221 | | | | | | | | | | | | | | yes | yes | | | | | |
| LRP5L | S_Shelf | 5'UTR;1stExon;TSS200 | chr22:25758455-25758749 | 294 | | | 6 | | | | | | | | | | | yes | | | | | | |
| HPS4 | N_Shelf | 1stExon;5'UTR;Body;TSS200;TSS1 | chr22:26875499-26876075 | 576 | | | 7 | 1 | | | | | | | | | | yes | yes | | | | | |
| CRYBA4 | NA | TSS200 | chr22:27017854-27017910 | 56 | | | | | | | | | | | | | | yes | yes | | | | | |
| ZNRF3 | Island | Body | chr22:29426511-29426832 | 321 | | | 1 | | | | | | | | | | | | | | | | yes | |
| KREMEN1 | Island | Body | chr22:29469461-29469719 | 258 | | | | | | | | | 1 | | | | | | | | | yes | yes | |
| GAS2L1 | Island | 3'UTR;Body | chr22:29706595-29708246 | 1651 | | | | | | | | 0 | 3 | | | | | yes | | | | | yes | |
| RASL10A | S_Shore | TSS1500 | chr22:29712045-29712472 | 427 | | | 3 | | | | | | | | | | | yes | yes | | | | | |
| RFPL1S;RFPL1 | NA | Body;TSS200 | chr22:29834388-29834550 | 162 | | | | | | | | | | | | | | yes | | | | | | |
| NEFH | Island;N_Shore | TSS1500;1stExon | chr22:29875578-29877285 | 1707 | | 1 | | | | | | 0 | 1 | | | 1 | | yes | | | | | | |
| CABP7 | N_Shore | TSS1500 | chr22:30115154-30115348 | 194 | | | | | | | | | | | | | | yes | | | | yes | | |
| CABP7 | Island | 1stExon;Body | chr22:30116489-30117159 | 670 | | | | | | | | | 2 | | | | | | | | | yes | yes | |
| HORMAD2 | Island | TSS1500;TSS200 | chr22:30476089-30476525 | 436 | | 1 | 17 | | | | | | 4 | | | 1 | | yes | | | | | | |
| LIF | S_Shore | TSS200;TSS1500 | chr22:30642980-30643376 | 396 | | | 4 | | | | | | | | | | | | | | | | | |
| OSM | NA | 1stExon;5'UTR;TSS200;TSS1500;B | chr22:30661949-30663881 | 1932 | | | | | 2 | | | | 3 | 4 | 1 | | | yes | yes | | | | | |
| SEC14L3 | NA | TSS1500;1stExon;TSS200 | chr22:30867939-30869361 | 1422 | | | 4 | | | | | | | | | | | yes | | | yes | | | |
| SEC14L4 | Island;S_Shore | Body;1stExon;5'UTR;TSS200 | chr22:30901249-30901886 | 637 | | | | | | | | 0 | 4 | 1 | | 2 | | yes | | | | | yes | |
| GAL3ST1 | NA | 5'UTR;1stExon | chr22:30960857-30961151 | 294 | | | | | | | | | 1 | | | | | yes | yes | | | | | |
| C22orf27 | N_Shore;Island | TSS1500;TSS200;Body | chr22:31317914-31318546 | 632 | 1 | 1 | 30 | | | | | 0 | 15 | 6 | | 11 | | yes | yes | | | | | |
| TUG1;MORC2 | N_Shore | TSS1500;TSS200 | chr22:31364274-31364349 | 75 | | | | | | | | | | | | | | | | | | yes | | |
| SMTN | Island | Body | chr22:31480934-31481318 | 384 | | | | | 1 | | | | | | | | | | | | | | yes | |
| SMTN | Island | Body | chr22:31494714-31495174 | 460 | | | 4 | | | | | | | | | | | yes | | | | yes | | |
| SELM | Island | 3'UTR | chr22:31500896-31501382 | 486 | | | | | | | | | 1 | | | | | | | | yes | | | |
| INPP5J | NA | TSS1500;TSS200 | chr22:31518707-31518963 | 256 | | | | | 1 | | | 0 | | | | 1 | | yes | | | | | | |
| PLA2G3 | NA | 5'UTR;1stExon;TSS200 | chr22:31536431-31536784 | 353 | | | | | | | | 0 | | | | 1 | | | | | | | | |
| LIMK2 | NA | TSS1500;Body;TSS200 | chr22:31643997-31644558 | 561 | | | 19 | 1 | | | | | 1 | | | | | yes | | | | | | |
| PISD | S_Shore | TSS1500 | chr22:32027580-32027605 | 25 | | | 15 | | | | | | | | | | | yes | | | | | | |
| DEPDC5 | S_Shore | 5'UTR;TSS200 | chr22:32150725-32150934 | 209 | | | | | | | | | 1 | | | | | yes | | | | | | |
| C22orf24;YWHAH | S_Shore | TSS200;Body;TSS1500 | chr22:32341402-32341780 | 378 | | | 2 | | | | | | | | | | | | | | yes | | | |
| SLC5A1 | N_Shore;Island | TSS200;5'UTR;1stExon | chr22:32438947-32439391 | 444 | | | 12 | | | | | 0 | 6 | 1 | | 5 | | | | | | | | |
| C22orf42 | NA | Body;1stExon;TSS200 | chr22:32554848-32555310 | 462 | | | 6 | | | | | | | | | | | | | | | | | |
| RFPL2 | NA | TSS200;5'UTR;TSS1500 | chr22:32599511-32599648 | 137 | | | 8 | | | | | 0 | 4 | 1 | | | | | | | | | | yes |
| RFPL2 | NA | TSS1500;TSS200 | chr22:32600722-32601274 | 552 | | | 4 | | | | | | 1 | | | | | | | | | | | yes |
| SLC5A4 | NA | Body;1stExon;TSS200;TSS1500 | chr22:32650997-32651776 | 779 | | | 7 | | | | | | 1 | | | | | yes | | | | | yes | |
| RFPL3 | NA | TSS200 | chr22:32750734-32750777 | 43 | | | | | | | | | | | | | | | | | | | | yes |
| C22orf28 | S_Shore | TSS1500 | chr22:32808489-32808786 | 297 | | | 3 | | | | | | 1 | | | | | yes | | | | | | |
| FBXO7 | N_Shore | TSS1500 | chr22:32869991-32870183 | 192 | | | | | | | | | | | | | | yes | | yes | | | | |
| FBXO7 | Island | TSS200;Body | chr22:32871209-32871233 | 24 | | | | | | | | | 1 | | | | | | | | | | | |
| SYN3;TIMP3 | N_Shore;Island | Body;TSS1500;1stExon;5'UTR | chr22:33195943-33198055 | 2112 | | 1 | 11 | 1 | | | | | 1 | | | | | yes | yes | | | yes | yes | |
| SYN3 | NA | 1stExon;Body | chr22:33402393-33402998 | 605 | | | | 1 | | | | | | | | | | | yes | yes | yes | | | |
| SYN3 | S_Shore;Island | TSS200;TSS1500;5'UTR | chr22:33453994-33454632 | 638 | | | 8 | | | | | 0 | 1 | | | 1 | | yes | yes | | | | yes | |
| LARGE | Island | Body | chr22:34046427-34046529 | 102 | | | | | | | | | 1 | | | | | yes | | | | | | |
| MCM5 | N_Shore | TSS1500 | chr22:35795274-35795394 | 120 | | | 1 | | | | | | 1 | | | | | yes | | yes | | | | |
| MB | NA | 1stExon;TSS200;5'UTR;TSS1500 | chr22:36013432-36014356 | 924 | | | 1 | | | | | | 2 | | | | | yes | yes | | | yes | | |
| APOL6 | NA | TSS1500 | chr22:36044151-36044234 | 83 | | | | | | | | | | | | | | yes | | | yes | yes | | |
| RBM9 | NA | 5'UTR;Body;1stExon;TSS200 | chr22:36236303-36236760 | 457 | | | | 1 | | | | 0 | 2 | | | | | | | | | yes | yes | |
| APOL3 | NA | TSS1500 | chr22:36563027-36563117 | 90 | | | | | | | | 0 | 1 | 1 | | 2 | | yes | | yes | | | | |
| APOL4 | NA | 5'UTR;1stExon | chr22:36600770-36600979 | 209 | | | | | | | | 0 | 1 | | | 3 | 1 | yes | | | | | | |
| APOL1 | NA | TSS200 | chr22:36648973-36649144 | 171 | | | 1 | | | | | | | | | | | yes | | | | | | |
| MYH9 | NA | Body | chr22:36681296-36681341 | 45 | | | 6 | 1 | | | | | | | | | | yes | | | | | | |
| MYH9 | NA | Body | chr22:36682843-36682870 | 27 | | | | | | | | | | | | | | | | | | | | |
| MYH9 | NA | Body | chr22:36691016-36691114 | 98 | | | | | | | | | | | | | | yes | | | | yes | | |
| TXN2 | NA | TSS200 | chr22:36877706-36878180 | 474 | | | | | | | | | 1 | | | 1 | | yes | | | | | yes | |
| CACNG2 | Island | Body | chr22:36960499-36960937 | 438 | | | 4 | | | | | | 1 | | | | | yes | yes | | | | yes | |
| CACNG2 | N_Shore | TSS200 | chr22:37099037-37099095 | 58 | | | | | | | | | | | | 1 | | | | | | yes | yes | |
| PVALB | S_Shelf | TSS200;TSS1500 | chr22:37215619-37216067 | 448 | | | 6 | | | | | | | | | | | yes | | | | | | |
| NCF4 | NA | TSS200 | chr22:37256900-37257404 | 504 | | | | | 1 | | | | | | | | | yes | yes | | | | | |
| CSF2RB | NA | TSS1500 | chr22:37308182-37308423 | 241 | | | 1 | | | | | | | | | | | yes | | | yes | yes | | |
| KCTD17 | N_Shore | TSS1500 | chr22:37447306-37447468 | 162 | | | 1 | | | | | | | | | | | yes | | | | yes | yes | |
| TMPRSS6 | Island;N_Shore | Body | chr22:37493456-37494173 | 717 | | | | | | | | | 1 | | | 1 | | yes | | yes | | | | |
| TMPRSS6 | NA | Body;1stExon;5'UTR | chr22:37499451-37499859 | 408 | | | 1 | | | | | | 1 | | | | | yes | | | | | | |
| IL2RB | NA | 5'UTR;1stExon;TSS200 | chr22:37545423-37546325 | 902 | | | 8 | 1 | 2 | | | | 2 | | | | | yes | yes | | | yes | | |
| C1QTNF6 | NA | TSS1500 | chr22:37585337-37585617 | 280 | | | | | | | | | 1 | | | | | | yes | yes | | | | |
| SSTR3 | NA | TSS1500 | chr22:37608611-37608819 | 208 | | | | | | | | | 1 | | | | | | | | yes | yes | yes | |
| RAC2 | NA | TSS200 | chr22:37640402-37640514 | 112 | | | 1 | | | | | | | | | | | | yes | | | | | |
| CYTH4 | NA | TSS200 | chr22:37678450-37678564 | 114 | | | 2 | | | | | | | | | | | yes | yes | | | yes | | |
| ELFN2 | Island | TSS1500 | chr22:37824118-37824617 | 499 | | | | | | | | | | | | | | yes | yes | | | yes | yes | |
| CARD10 | N_Shore | Body | chr22:37914685-37915089 | 404 | | | | | | | | | | | | | | | | | | | yes | |
| CDC42EP1 | Island | Body | chr22:37962569-37962818 | 249 | | | | | | | | | 1 | | | | | yes | | | | | | |
| LGALS2 | NA | 1stExon;5'UTR;TSS200 | chr22:37975941-37976192 | 251 | | | 8 | | | | | 0 | 1 | | | | | yes | | | | | | |
| SH3BP1 | N_Shore | TSS1500 | chr22:38035109-38035456 | 347 | | | 1 | | | | | | | | | | | | yes | | | | yes | |
| LGALS1 | N_Shore;N_Shelf | TSS200;TSS1500 | chr22:38071001-38071607 | 606 | | | | | 1 | | | | 2 | | | | | yes | | | | | | |
| LGALS1 | Island | Body | chr22:38073049-38073472 | 423 | 4 | | | | | | | | | | | | | yes | | | | | yes | |
| TRIOBP | NA | TSS1500;TSS200 | chr22:38092643-38093207 | 564 | | | | 1 | | | | 0 | 6 | | | 5 | | yes | | | | | | |
| TRIOBP | Island | TSS200;Body | chr22:38142152-38142162 | 10 | 1 | | | | | | | | | | | | | yes | | | | yes | yes | |
| H1F0 | N_Shore | TSS1500 | chr22:38199759-38200221 | 462 | | | | | | | | | | | | | | | | | | | | |
| H1F0 | Island | 1stExon | chr22:38201594-38201848 | 254 | | | | | | | | 0 | | | | 1 | | | | | | yes | | |
| GALR3 | N_Shore | TSS1500;TSS200 | chr22:38218945-38219400 | 455 | | | 7 | | | | | | | | | | | yes | | | | | | |
| GALR3 | Island | Body | chr22:38221009-38221187 | 178 | | | | | 1 | | | | | | | | | yes | | | | | yes | |
| SOX10 | Island | Body | chr22:38379218-38379749 | 531 | | | | | | | | 0 | 1 | | | | | | | | | yes | | |
| PICK1 | Island | Body | chr22:38470264-38470483 | 219 | | | | | | | | | | | | | | yes | | | | | | |
| SLC16A8 | Island;S_Shore | Body;TSS200;TSS1500 | chr22:38478322-38480061 | 1739 | | | | | | | | 0 | 2 | | | | | yes | | | | | yes | |
| BAIAP2L2 | NA | 1stExon;5'UTR;TSS200 | chr22:38506623-38506781 | 158 | | | | 1 | | | | | 1 | 1 | | | | | | | | yes | | |
| MAFF | N_Shore;Island | Body;5'UTR | chr22:38610234-38610795 | 561 | | | | | | | | | 1 | | | | | yes | yes | | | | yes | |
| CSNK1E | N_Shore;S_Shore | 5'UTR;TSS1500;TSS200 | chr22:38712210-38714466 | 2256 | | | 1 | 1 | | | | 0 | 3 | 1 | | | | yes | | | yes | yes | yes | |
| KCNJ4 | NA | 5'UTR;TSS200 | chr22:38840050-38840560 | 510 | | | | | | | | | 1 | | | | | yes | yes | | | yes | | |
| KDELR3 | Island | TSS200 | chr22:38863905-38864244 | 339 | | | | | | | | 0 | | | | | | yes | | | | | | |
| APOBEC3B | NA | TSS1500;TSS200 | chr22:39377790-39378382 | 592 | | | 1 | | | | | 0 | | 3 | | | | | yes | | | yes | | |

| | | | | | | | | | | | | | | | | | | | | | | | |
|---|---|---|---|---|---|---|---|---|---|---|---|---|---|---|---|---|---|---|---|---|---|---|---|
| APOBEC3F | NA | TSS1500 | chr22:39436141-39436644 | 503 | | | | | | | | | | | | | | yes | | yes | | | |
| PDGFB | S_Shore | TSS200 | chr22:39641127-39641536 | 409 | | | 2 | | | | | | | | | | | yes | | | | | yes |
| RPL3;SNORD83B | NA | Body;TSS200 | chr22:39709951-39710023 | 72 | | | 1 | | | | | | | | | | | | | | | | |
| RPL3;SNORD83A;RNU86 | N_Shelf | Body;TSS1500;TSS200 | chr22:39712730-39713062 | 332 | | | 3 | | | | | | 1 | | | | | yes | | | | | |
| SYNGR1 | NA | TSS1500;Body;TSS200 | chr22:39759848-39760158 | 310 | | | 3 | | | | | 0 | | | | | | yes | yes | | | | |
| MGAT3 | N_Shore | 5'UTR;TSS200 | chr22:39883190-39883347 | 157 | 2 | | | | | | | | | | | | | yes | | | | | |
| CACNA1I | NA | TSS200;TSS1500 | chr22:39965495-39966921 | 1426 | | | 6 | | | | | | 1 | | | | | yes | yes | | yes | | yes |
| CACNA1I | Island | Body | chr22:40057942-40058706 | 764 | | | 2 | 1 | 1 | | 1 | 0 | 2 | 3 | 1 | 2 | | yes | | | | | yes |
| CACNA1I | N_Shelf | Body | chr22:40079107-40079626 | 519 | | | | | | | | | 1 | | | | | yes | yes | | | | |
| CACNA1I | Island | Body | chr22:40081812-40082521 | 709 | 2 | | | | | | | | | | | | | | yes | | | | yes |
| ENTHD1 | NA | 5'UTR | chr22:40289538-40289840 | 302 | | | | | | | | | | | | | | yes | yes | | | | yes |
| GRAP2 | NA | TSS1500 | chr22:40296407-40297048 | 641 | | | 8 | | 2 | | | | 1 | | | | | yes | yes | | | | |
| FAM83F | Island | 1stExon | chr22:40391434-40391617 | 183 | | | | | | | | | 1 | | | | | yes | yes | | | yes | yes |
| MKL1 | N_Shore | Body | chr22:40814431-40815101 | 670 | | | | | | | | | 1 | | | | | yes | | | | yes | |
| MCHR1 | NA | TSS1500 | chr22:41074803-41075007 | 204 | | | | | | | | | | | | | | | yes | | | | |
| CHADL | S_Shelf | TSS200 | chr22:41636941-41637354 | 413 | | | 1 | | | | | | | | | | | yes | yes | | | | |
| TEF | NA | TSS200 | chr22:41763253-41763280 | 27 | | | 1 | | | | | | 1 | | | | | | | | | | |
| CSDC2 | NA | TSS1500;TSS200 | chr22:41956440-41957443 | 1003 | | | 2 | 1 | | | | | | | | | | yes | | | | yes | |
| NHP2L1 | N_Shore;Island;S_Shore | Body;TSS200 | chr22:42077939-42079293 | 1354 | | | 3 | | | | | | 3 | | | | | yes | yes | | | | |
| MEI1 | N_Shore;Island | TSS200 | chr22:42095347-42095773 | 426 | | 1 | | | | | | 0 | 2 | | | 1 | | yes | | | | | |
| WBP2NL | N_Shore;Island | TSS1500;TSS200 | chr22:42394382-42395067 | 685 | | | 7 | | | | | 0 | 3 | 2 | | 4 | | yes | | | | yes | |
| FAM109B | N_Shore | TSS1500 | chr22:42469976-42470123 | 147 | | | 1 | | | | | 0 | 1 | | | | | | yes | | | | |
| C22orf32;FAM109B | N_Shore | TSS1500;3'UTR | chr22:42474656-42475135 | 479 | | | | | | | | | | | | | | yes | | | | | |
| CYP2D6 | Island | Body | chr22:42524984-42525279 | 295 | | | 6 | | | | | 0 | 1 | | | | | | | | | | |
| CYP2D6 | S_Shore | TSS1500 | chr22:42527611-42528123 | 512 | | | | | | | | 0 | | | | | | yes | yes | | | yes | |
| NFAM1 | NA | Body;1stExon;5'UTR;TSS200 | chr22:42828125-42828946 | 821 | | | 14 | 1 | 1 | | | 0 | 3 | 2 | 1 | | | yes | | | | | |
| SERHL | N_Shore;Island | TSS1500;Body | chr22:42896105-42896985 | 880 | | 1 | | | | | | | 1 | | | 1 | | yes | yes | | | yes | |
| CYB5R3 | N_Shelf | 5'UTR;Body;1stExon | chr22:43042879-43043270 | 391 | | | | | | | | | | | | | | yes | | | | | |
| ARFGAP3 | Island | TSS200 | chr22:43253521-43253559 | 38 | | 1 | | | | | | 0 | 1 | | | | | yes | | | | | |
| BIK | N_Shore;Island | TSS1500 | chr22:43505892-43506033 | 141 | | | | | | | | 0 | 1 | | | 1 | | yes | | | | | |
| TTLL12 | Island | Body | chr22:43570199-43570418 | 219 | | | | | | | | | 1 | | | | | yes | | | | | |
| SCUBE1 | Island | TSS1500 | chr22:43739790-43740003 | 213 | | 1 | | | | | | | 1 | | | | | | | | | | yes |
| MPPED1 | N_Shore | TSS1500 | chr22:43807121-43807429 | 308 | | | | 1 | | | | | 1 | | | | | | yes | | | yes | yes |
| MPPED1 | Island | 5'UTR | chr22:43817258-43817384 | 126 | | | | | | | | 0 | 1 | | | | | yes | yes | | | yes | |
| MPPED1 | Island | 5'UTR;Body | chr22:43820888-43821227 | 339 | | | 2 | | | | | | 1 | | | | | yes | | | yes | | yes |
| MPPED1 | Island | Body | chr22:43831005-43831161 | 156 | | | | | | | | | | | | | | yes | yes | | yes | | |
| SULT4A1 | Island | Body;TSS200 | chr22:44257997-44258602 | 605 | | | | | | | | | 3 | | | 1 | | yes | yes | | | yes | yes |
| PNPLA5 | Island | TSS200 | chr22:44287930-44288021 | 91 | | | | | | | | | 1 | | | | | yes | | | | | |
| PNPLA3 | Island | 1stExon;5'UTR | chr22:44319666-44320173 | 507 | | | | | | | | | 1 | | | | | yes | | | | | yes |
| PARVG | NA | TSS1500;TSS200 | chr22:44568337-44568913 | 576 | | | 30 | | 2 | | | 0 | 1 | 1 | | | | yes | | | | | |
| PARVG | NA | 5'UTR;Body;TSS200;TSS1500;1stE | chr22:44576654-44577920 | 1266 | | | | | 1 | | | | 1 | 2 | | | | yes | yes | yes | | | |
| KIAA1644 | NA | 5'UTR;1stExon;TSS200 | chr22:44708669-44708908 | 239 | | | 14 | | | | | | | | | | | yes | yes | | | | |
| PRR5 | NA | 5'UTR;TSS1500;TSS200;1stExon | chr22:45072455-45073165 | 710 | | | 7 | | | | | | 1 | | 1 | 1 | | yes | yes | | | | yes |
| PRR5;PRR5-ARHGAP8 | Island | Body | chr22:45125218-45126040 | 822 | | | 21 | 1 | | | | 0 | 2 | | | 1 | | | yes | | | | |
| PRR5;PRR5-ARHGAP8 | Island | Body | chr22:45132782-45133246 | 464 | | | | 1 | | | | | 1 | | | | | yes | | | | | |
| ARHGAP8;PRR5-ARHGAP8 | NA | TSS1500;Body | chr22:45148218-45148229 | 11 | | | 1 | | | | | | 1 | | | | | yes | | | | yes | yes |
| C22orf9;MIR1249 | N_Shore | Body;TSS200 | chr22:45596948-45597767 | 819 | | | 4 | | | | | | | | | | | yes | yes | | | | |
| C22orf9 | NA | Body;TSS200;nan;TSS1500;1stExo | chr22:45608345-45609421 | 1076 | | | 12 | 2 | | | | | 5 | 1 | 1 | | | yes | | | | | |
| UPK3A | N_Shore;Island | TSS1500;TSS200 | chr22:45680498-45680702 | 204 | | | 3 | | | | | 0 | | | | | | yes | yes | | | | |
| FAM118A | N_Shore | TSS1500;TSS200 | chr22:45704988-45705265 | 277 | | | | | | | | 0 | 4 | | | 5 | 1 | yes | yes | | | | yes |
| SMC1B;RIBC2 | Island | Body;TSS1500;1stExon;TSS200;5'U | chr22:45809244-45810043 | 799 | | | 14 | | | | | 0 | 2 | | | 2 | | yes | | | | | yes |
| ATXN10 | N_Shore | TSS1500 | chr22:46067363-46067586 | 223 | | | | | | | | | 1 | | | | | | | | | | yes |
| WNT7B | Island | Body | chr22:46367617-46368025 | 408 | | 1 | | | | | | | 1 | | | | | | | | | yes | yes |
| C22orf26;LOC150381 | N_Shore;Island | Body;TSS200;1stExon;5'UTR;TSS1 | chr22:46448690-46450114 | 1424 | | | | | | | | 0 | 7 | 1 | | 9 | | yes | | | | | yes |
| LOC400931 | N_Shore;Island | TSS1500;TSS200;Body | chr22:46481248-46482023 | 775 | | | | 1 | 1 | | | | 2 | 1 | 1 | | | yes | | | | | |
| TTC38 | Island | Body | chr22:46685472-46685728 | 256 | | | 2 | | | | | | 2 | | | | | yes | yes | | | | |
| CN5H6.4;GTSE1 | Island | Body;TSS1500 | chr22:46692412-46692446 | 34 | | | | | | | | 0 | | | | | | yes | | | | | |
| TRMU | N_Shore | TSS1500 | chr22:46730289-46730765 | 476 | | | | | | | | | | | | | | yes | yes | yes | | | |
| CELSR1 | Island | Body | chr22:46760120-46760678 | 558 | | | | | | | | | 2 | | | | | | | | | | |
| CELSR1 | Island | Body | chr22:46762841-46763229 | 388 | | | 2 | | | | | 0 | 1 | | | 1 | | | | | | | |
| CELSR1 | N_Shore | Body | chr22:46770063-46770440 | 377 | | | 6 | 1 | 1 | | | | | | 1 | | | | | | | | yes |
| CELSR1 | Island | Body | chr22:46805673-46805918 | 245 | | 1 | 1 | | | | | | | | | | | | | | | | |
| CELSR1 | Island | Body | chr22:46859899-46860524 | 625 | | | 1 | | | | | | | | | | | yes | | | | | yes |
| GRAMD4 | N_Shore;Island | TSS1500;TSS200;Body | chr22:47021746-47023209 | 1463 | | | 2 | | | | | | 7 | | | | | | yes | | | | |
| GRAMD4 | N_Shore | Body | chr22:47069200-47069799 | 599 | | | | | | | | | 1 | | | | | | | | | | |
| CERK | Island | 3'UTR | chr22:47081751-47082260 | 509 | | | | 1 | | | | 0 | 1 | | | | | yes | | | | | |
| TBC1D22A | Island | Body | chr22:47370132-47370342 | 210 | | | | | | | | | | | | | | | | | | | |
| TBC1D22A | N_Shore | Body | chr22:47513051-47513521 | 470 | 3 | | | | 1 | | | | | | | | | yes | | | | | |
| FAM19A5 | Island | TSS1500 | chr22:48884945-48885446 | 501 | | | | | | | | | 1 | | | | | | | | | yes | yes |
| FAM19A5 | Island | Body | chr22:48972850-48973245 | 395 | | | 1 | | | | | | | | | | | | | | | | yes |
| FAM19A5 | Island | 3'UTR | chr22:49146166-49146584 | 418 | | | | | | | | | 1 | | | | | | | | | | |
| C22orf34 | Island | Body | chr22:50025915-50026188 | 273 | | | | | 1 | | 1 | | 1 | | | | | | | | | | |
| C22orf34 | NA | TSS1500 | chr22:50052294-50052407 | 113 | | | | | | | | 0 | 1 | 1 | 1 | | | | | yes | | | |
| BRD1;LOC90834 | N_Shelf | Body;TSS200 | chr22:50174028-50174170 | 142 | | | | | | | | | 1 | | | | | | | | | | |
| BRD1 | Island | Body | chr22:50177941-50178457 | 516 | | | 2 | | | | | | | | | | | | | | | | |
| ZBED4 | Island | 5'UTR | chr22:50250238-50250494 | 256 | 12 | | | | | | | | | | | | | | | | | | |
| PIM3 | Island | Body | chr22:50356763-50357265 | 502 | | | | | | | | | 1 | | | | | | | | | | |
| IL17REL | Island | 5'UTR | chr22:50448833-50448934 | 101 | | | 1 | | | | | | | | | | | | yes | | | | |
| IL17REL | S_Shore;N_Shore | 5'UTR;1stExon;TSS200 | chr22:50451054-50451975 | 921 | | | 2 | | | | | | 4 | 1 | | | | | | | | yes | |
| TTLL8 | Island | Body | chr22:50470022-50470419 | 397 | | | | | | | | | | | | | | yes | | | | | |
| TTLL8 | N_Shore | Body | chr22:50482925-50483545 | 620 | | | 1 | | | | | 0 | 2 | 1 | 1 | | | yes | yes | yes | | | |
| TTLL8 | N_Shelf | TSS200 | chr22:50493212-50493642 | 430 | | | | | | | | | | | | | | yes | | yes | | | |
| MLC1 | NA;N_Shelf | 5'UTR;1stExon;TSS200;TSS1500 | chr22:50523686-50524691 | 1005 | | | 17 | | 1 | | | | 4 | 1 | 1 | | | | | | | | |
| MOV10L1 | N_Shore;Island | TSS1500;TSS200;1stExon | chr22:50528179-50528752 | 573 | | 1 | 10 | | 1 | | | 0 | 9 | | | 5 | | yes | yes | | | yes | |
| MOV10L1 | NA | Body;TSS1500;TSS200 | chr22:50584184-50585427 | 1243 | | | 30 | | 1 | | | | 3 | | | | | yes | | | yes | | |
| PANX2 | Island | Body | chr22:50617003-50617460 | 457 | | | 3 | | | | | | | | | | | | | | | yes | |
| TRABD | Island | 5'UTR | chr22:50629038-50629224 | 186 | | | 14 | 1 | | | | | | | | | | | | | | | |
| TRABD | S_Shore | 5'UTR | chr22:50630587-50631079 | 492 | | | | | | | | | | | | | | yes | | | | | |
| TRABD | N_Shore | Body | chr22:50632560-50632819 | 259 | 1 | | | | | | | | | | | | | | | | | | |
| PLXNB2 | S_Shore | TSS1500 | chr22:50746891-50747878 | 987 | | | 5 | 1 | | | | | | | | | | | | | | | |
| SAPS2 | Island | Body | chr22:50845159-50845380 | 221 | 1 | | | | | | | | | | | | | yes | | | | | |
| SAPS2 | Island | Body | chr22:50873337-50873573 | 236 | 1 | | | | | | | | | | | | | | | | | | |
| ADM2 | Island | TSS1500 | chr22:50919482-50919512 | 30 | | | | | | | | 0 | | | | | | yes | | | | | |
| ADM2;MIOX | NA | 3'UTR;TSS1500;TSS200;5'UTR;1stE | chr22:50924745-50925337 | 592 | | 1 | 10 | | | | | | | | | | | | | yes | | | |
| LMF2;NCAPH2 | Island | TSS1500;TSS200 | chr22:50946451-50946470 | 19 | | | | | | | | | 1 | | | | | | | | | yes | yes |
| KLHDC7B | Island | TSS1500;1stExon | chr22:50985681-50987453 | 1772 | | | 3 | | | | | 0 | 4 | 1 | | 2 | | | | | | yes | yes |
| CPT1B;CHKB-CPT1B | Island;S_Shore | 5'UTR;Body;1stExon;TSS200;TSS1 | chr22:51016501-51017166 | 665 | | | 6 | 2 | | | 1 | 0 | 9 | 5 | | 17 | | | | | | | yes |
| MAPK8IP2 | N_Shore | Body | chr22:51043692-51044053 | 361 | | | | | | | | | | | | | | | | | | | |
| RPL23AP82;RABL2B | Island | Body;1stExon;TSS200;5'UTR | chr22:51222011-51222251 | 240 | | | | | | | | | 1 | | | | | yes | | yes | | | |

| | | | | | | | | | | | | | | | | | | | | | | | |
|---|---|---|---|---|---|---|---|---|---|---|---|---|---|---|---|---|---|---|---|---|---|---|---|
| CHL1 | N_Shore;Island | TSS1500;TSS200;5'UTR | chr3:238048-239733 | 1685 | | | | | | | | 0 | 7 | | | 1 | | yes | | | | | |
| CHL1 | N_Shore;Island | 5'UTR | chr3:350503-351003 | 500 | | | 12 | | 1 | | | 0 | 7 | 1 | | 2 | | yes | yes | | | yes | yes |
| CNTN6 | NA | TSS1500;nan | chr3:1133865-1134730 | 865 | | | 1 | | 1 | | | | | | 1 | | | | | | | | yes |
| CNTN4 | NA | Body;TSS1500 | chr3:3080197-3080621 | 424 | | | | | | | | | 1 | | | | | yes | | | | | |
| IL5RA | NA | Body;3'UTR | chr3:3133700-3134187 | 487 | | | | | | | | | 1 | | | | | yes | yes | | yes | | |
| IL5RA | NA | 5'UTR;1stExon | chr3:3151899-3152530 | 631 | | | 3 | | | | | | | | | | | | | | | | |
| LRRN1 | Island | TSS200 | chr3:3841061-3841457 | 396 | | | | | | | | | 1 | | | | | | | | | yes | yes |
| SUMF1 | S_Shore | TSS1500 | chr3:4509268-4509421 | 153 | | | 3 | | | | | | 1 | | | | | yes | yes | | yes | | |
| ITPR1;EGOT | NA | Body;TSS1500 | chr3:4794020-4794082 | 62 | | | 1 | 1 | | | | 0 | 1 | | | | | yes | yes | | | | yes |
| BHLHE40 | Island | Body | chr3:5024761-5024963 | 202 | | | | 1 | | | | | | | | | | | yes | | | | |
| GRM7 | N_Shore;Island | TSS200;1stExon;5'UTR | chr3:6902689-6903019 | 330 | | | | | 1 | | | 0 | 1 | | | | | | | | | | yes |
| C3orf32 | NA | 1stExon;5'UTR;TSS200 | chr3:8693706-8693859 | 153 | | | | 1 | | | | | | | | | | yes | | | | | |
| CAV3 | NA | TSS1500;TSS200 | chr3:8774972-8775601 | 629 | | | | | | | | | | | | | | yes | yes | | | | |
| OXTR | Island | Body | chr3:8809306-8810139 | 833 | | | 7 | | | | | 0 | 1 | | | 1 | | yes | | | | | yes |
| OXTR | S_Shore | TSS200;TSS1500 | chr3:8811437-8811758 | 321 | | 1 | | | | | | | | | | | | yes | | | | | |
| RAD18 | S_Shore | TSS1500 | chr3:9005400-9005479 | 79 | | | | | | | | | 1 | | | | | yes | | yes | | | |
| SRGAP3 | NA | Body | chr3:9258385-9258573 | 188 | | | 14 | | | | | | | | | | | yes | yes | | yes | | |
| SRGAP3 | NA | Body | chr3:9289747-9289893 | 146 | | | 2 | 1 | | | | 0 | 1 | 1 | | 1 | | | | | | | |
| THUMPD3 | N_Shore | TSS1500 | chr3:9404422-9404532 | 110 | | | | | | | | 0 | | | | | | yes | | yes | | | |
| CAMK1 | S_Shore | TSS200 | chr3:9811806-9811990 | 184 | | | | | | | | 0 | | | | | | yes | | | | | |
| CIDEC | NA | 5'UTR | chr3:9920535-9920698 | 163 | | | | 1 | | | | | | | | | | yes | | | yes | | |
| IL17RE | NA | TSS1500;TSS200 | chr3:9943828-9944537 | 709 | | | 6 | | | | | 0 | 1 | 1 | 1 | | | yes | yes | | | | |
| PRRT3 | Island | 5'UTR | chr3:9993818-9994197 | 379 | | | | | 2 | | | | | | | | | yes | yes | | | yes | |
| C3orf24 | Island | 1stExon;5'UTR;TSS200 | chr3:10149803-10150139 | 336 | 1 | | 23 | | | | | | 2 | | | | | yes | | | | | |
| VHL | N_Shore | TSS1500 | chr3:10183060-10183196 | 136 | | | | | | | | | | | | | | yes | yes | | | | |
| IRAK2 | N_Shore | TSS1500 | chr3:10206254-10206344 | 90 | | | 4 | | | | | | | | | | | yes | yes | | | | |
| GHRL;GHRLOS | NA | Body;5'UTR;TSS200 | chr3:10332170-10332589 | 419 | | | 5 | | | | | | | | | | | yes | | | | | |
| GHRLOS;GHRL | NA | Body;TSS200 | chr3:10334717-10335375 | 658 | | | | | | | | 0 | 2 | | | | | yes | | | yes | | |
| ATP2B2 | N_Shore;Island | 3'UTR;Body | chr3:10370264-10370704 | 440 | | | 3 | | | | | | 1 | | | | | | | | | yes | |
| ATP2B2 | NA | Body | chr3:10377957-10378071 | 114 | | | | | | | | | 1 | | | | | yes | | | | | |
| ATP2B2;MIR885 | NA | Body;TSS200 | chr3:10436278-10436406 | 128 | | | | | | | | | 1 | | | | | yes | | | yes | | |
| LOC285370 | NA | TSS200;TSS1500 | chr3:10805956-10806370 | 414 | 2 | | 16 | | | | | 0 | 7 | | | 5 | | | yes | | | | |
| SLC6A11 | Island | TSS1500;TSS200;Body | chr3:10857699-10858504 | 805 | | 1 | 1 | | 1 | | | | 5 | 1 | | 1 | | yes | | | yes | | yes |
| SLC6A1 | S_Shore | 5'UTR | chr3:11036537-11036898 | 361 | | | 3 | | | | | | | | | | | yes | | | | | |
| SLC6A1 | Island | 5'UTR;Body | chr3:11058836-11059480 | 644 | | | 6 | | | | | | | | | | | yes | | | | | |
| SLC6A1 | NA | 3'UTR | chr3:11078979-11079832 | 853 | | | | | | | | | 1 | 1 | | | | | | | | | |
| HRH1 | NA | TSS1500;nan;TSS200 | chr3:11178064-11178798 | 734 | | | | | | | | | 3 | | 1 | | | yes | yes | | yes | | |
| HRH1 | NA | 5'UTR;TSS1500 | chr3:11267020-11267627 | 607 | | 1 | 8 | | | | | | 1 | | | | | yes | yes | | | | |
| HRH1 | NA | 5'UTR;TSS1500 | chr3:11293681-11294188 | 507 | | | | | | | | | 1 | | | | | | | yes | | | |
| VGLL4 | Island;chr3:11610137-1161037 | Body;1stExon;5'UTR;Island | chr3:11610138-11610613 | 475 | | | 1 | | | | | | | | 1 | | | | | | | | |
| VGLL4 | NA | Body;1stExon;5'UTR | chr3:11623526-11624023 | 497 | | | | 1 | | | | | 1 | | | | | | | | | | |
| VGLL4 | NA | Body | chr3:11643368-11643630 | 262 | 1 | | | | | | | | 1 | | | | | | | | | | |
| VGLL4 | NA | Body | chr3:11651536-11651881 | 345 | | | | | | | | | | | | | | yes | | | | | |
| VGLL4 | N_Shore | 5'UTR | chr3:11746183-11746765 | 582 | | | 17 | | | | | 0 | 3 | | | 1 | | yes | yes | | yes | | |
| SYN2 | N_Shore | TSS1500 | chr3:12045449-12045672 | 223 | | | | | | | | | 1 | | | | | | | | | yes | yes |
| SYN2;TIMP4 | N_Shore;Island;S_Shore | Body;1stExon;TSS200 | chr3:12199836-12200856 | 1020 | | | 10 | | | | | 0 | 1 | | | | | yes | yes | | | yes | |
| TMEM40 | NA | 1stExon;5'UTR | chr3:12800802-12801112 | 310 | | | | | | | | | 1 | | | | | yes | yes | | | | |
| IQSEC1 | NA | Body | chr3:13028573-13028642 | 69 | | | | | 1 | | | | 1 | | | | | | yes | | | | |
| IQSEC1 | NA | Body | chr3:13060741-13060974 | 233 | | | 4 | | | | | | 2 | | | | | yes | | | | | |
| IQSEC1 | NA | Body | chr3:13082751-13083151 | 400 | | | | | | | | | | | | | | yes | yes | | | | |
| IQSEC1 | N_Shore;Island | Body;5'UTR;1stExon;TSS200 | chr3:13113959-13115168 | 1209 | | | 2 | 1 | 2 | | | | | | | 1 | | yes | yes | | | | yes |
| NUP210 | NA | Body | chr3:13396268-13397022 | 754 | | | 2 | | | | | | | | | | | yes | | yes | yes | | |
| NUP210 | S_Shore | TSS1500 | chr3:13462465-13462903 | 438 | | | | | 1 | | | | | | | | | | | | | yes | yes |
| HDAC11 | N_Shore;Island | TSS1500 | chr3:13520685-13521023 | 338 | | | | | | | | | 1 | 1 | | | | yes | | | | | |
| FBLN2 | Island | TSS1500;TSS200 | chr3:13590415-13590450 | 35 | 2 | | | | | | | 0 | 1 | | | | | | | | | yes | yes |
| FBLN2 | NA | Body | chr3:13663398-13663538 | 140 | | | 7 | | | | | | | | | | | yes | | | | | |
| FBLN2 | N_Shore;Island | Body | chr3:13678918-13679574 | 656 | | 1 | | | | | | | | | | | | yes | | yes | | | yes |
| LOC285375 | NA | TSS200 | chr3:13692141-13692212 | 71 | | 1 | | | | | | | | | | | | | yes | | yes | yes | |
| WNT7A | Island | Body | chr3:13860638-13860812 | 174 | | | | | | | | | | | | | | yes | yes | | | | |
| WNT7A | Island | Body | chr3:13895992-13896256 | 264 | | | | | | | | | | | | | | yes | yes | | | | yes |
| SLC6A6 | N_Shore | TSS1500 | chr3:14443428-14443694 | 266 | | | 11 | | | | | | 1 | 1 | | | | yes | | | | yes | yes |
| GRIP2 | NA | TSS1500 | chr3:14584745-14584788 | 43 | | | | | | | | | 1 | | | | | yes | | | yes | | |
| C3orf20 | NA | Body | chr3:14755007-14755338 | 331 | | | 3 | | | | | | | | | | | yes | yes | | yes | | |
| SH3BP5 | NA | Body | chr3:15311021-15311269 | 248 | | | 24 | 1 | 1 | | | | | | | | | yes | | yes | | | |
| COLQ | NA | Body;1stExon;5'UTR | chr3:15540137-15540542 | 405 | | | | | | | | | | | | | | | | | yes | yes | |
| ANKRD28 | S_Shore | TSS1500 | chr3:15902307-15902540 | 233 | | | | | | | | | 1 | | | | | yes | | | | yes | |
| MIR563 | NA | TSS1500 | chr3:15914712-15915340 | 628 | | | | | | | | | 2 | | | | | yes | yes | | | | yes |
| GALNTL2 | NA | TSS1500;TSS200 | chr3:16215032-16216215 | 1183 | | | 1 | | | | | 0 | 2 | | | | | | yes | | | | |
| PLCL2 | NA | Body;TSS1500 | chr3:16974160-16974681 | 521 | | | | | | | | 0 | 1 | | | 1 | | | yes | | | | yes |
| ZNF385D | NA | 1stExon;5'UTR;TSS200 | chr3:21792543-21792991 | 448 | | | | | | | | 0 | 1 | | | | | | | | | yes | |
| UBE2E2 | Island | TSS1500 | chr3:23244051-23244068 | 17 | | | 1 | | | | | | | | | | | | | | | | yes |
| THRB | Island | TSS1500;5'UTR;TSS200 | chr3:24536077-24537407 | 1330 | | | 30 | | | | | | 6 | | | 1 | | | | | | | yes |
| RARB | NA | TSS1500;5'UTR;1stExon | chr3:25469148-25469991 | 843 | | | 5 | | | | | | 1 | | | | | yes | | | | | |
| LRRC3B | N_Shore;Island | TSS1500;1stExon;5'UTR | chr3:26663551-26664741 | 1190 | | | 4 | | 1 | | | | 2 | | | | | | | | | yes | |
| LRRC3B | N_Shore | 5'UTR | chr3:26665815-26666151 | 336 | | | 5 | | | | | | | | | | | | | | | | |
| LRRC3B | NA | 5'UTR | chr3:26750855-26751141 | 286 | | | | | | | | | | | | | | | | | | | |
| NEK10 | Island | 5'UTR;TSS200 | chr3:27410627-27411065 | 438 | | | 6 | | | | | | | | | | | yes | yes | | | | |
| EOMES | N_Shore | Body | chr3:27761408-27761677 | 269 | | | 1 | | 1 | | | | | | | 1 | | | | | | yes | |
| EOMES | Island;N_Shore | Body;TSS1500 | chr3:27762805-27766009 | 3204 | 5 | 1 | | | | | | | 2 | | | | | | | | | yes | |
| RBMS3 | NA | TSS200 | chr3:29322819-29323042 | 223 | | 1 | | | | | | | | | | | | | | | | | yes |
| TGFBR2 | N_Shore | TSS1500 | chr3:30647378-30647672 | 294 | | | 3 | | | | | 0 | 3 | | | | | | | | yes | | |
| TGFBR2 | Island | Body | chr3:30648819-30649017 | 198 | | | | | | | | 0 | | | | | | yes | | | | yes | |
| GADL1 | Island | 1stExon;TSS200 | chr3:30936070-30936531 | 461 | | | 2 | | | | | 0 | 3 | | | 2 | | yes | | | | yes | |
| ZNF860;OSBPL10 | Island;S_Shore | 5'UTR;1stExon;TSS200;TSS1500 | chr3:32023351-32023722 | 371 | | | 9 | | | | | | 1 | | | | | yes | | | yes | | |
| CMTM7 | N_Shore | TSS1500 | chr3:32432642-32432942 | 300 | 1 | | | | | | | | | | | | | yes | yes | | | | yes |
| TRIM71 | N_Shore;Island | Body | chr3:32860909-32861407 | 498 | 4 | | 7 | 1 | | | | | 1 | | | | | | | | | | |
| ARPP-21 | NA | 5'UTR | chr3:35706038-35706161 | 123 | | | | 1 | | | | 0 | 1 | | | 1 | | | yes | | | | |
| ARPP-21 | NA | 5'UTR;TSS200;TSS1500 | chr3:35720528-35721284 | 756 | | | 7 | | | | | | 2 | | | | | | | | | | |
| MIR128-2;ARPP-21 | NA | TSS1500;Body;TSS200 | chr3:35785041-35786480 | 1439 | | | 10 | | | | | | | | | | | | | yes | | | |
| STAC | Island | TSS200;1stExon | chr3:36422059-36422406 | 347 | | | | | | | | 0 | 2 | | | | | yes | | | | | yes |
| TRANK1 | NA | Body | chr3:36902601-36903038 | 437 | | | | | | | | | 1 | | | | | yes | | | yes | | |
| TRANK1 | NA | Body | chr3:36949957-36950004 | 47 | | 1 | 21 | | | | | | | | | | | yes | | | | | |
| TRANK1 | Island | TSS1500 | chr3:36986805-36987040 | 235 | | | 1 | | | | | | | | | | | | | | | | |
| MLH1;EPM2AIP1 | N_Shore | TSS1500;1stExon | chr3:37033373-37033980 | 607 | | | 13 | | 1 | | | | 1 | | | 1 | | | | | | | |
| CTDSPL;MIR26A1 | NA | Body;TSS1500 | chr3:38010391-38010902 | 511 | | | 3 | | | | | | | | | | | yes | | | | | |
| VILL | N_Shore | TSS1500 | chr3:38033770-38034235 | 465 | | | | | 1 | | | | | | | | | | yes | | | | |
| VILL | N_Shore;Island | Body | chr3:38035463-38036060 | 597 | | | | | 4 | | | 0 | | | | | | | yes | | | | |
| DLEC1 | Island;N_Shore | 1stExon;TSS1500;TSS200 | chr3:38080203-38081186 | 983 | | | 22 | | 1 | | | | | | | | | yes | yes | | | | |
| ACAA1;MYD88 | Island | 1stExon;Body;5'UTR;TSS1500 | chr3:38178591-38178743 | 152 | | | | | | | | | 1 | | | | | | | | | | |

| | | | | | | | | | | | | | | | | | | | | | | | |
|---|---|---|---|---|---|---|---|---|---|---|---|---|---|---|---|---|---|---|---|---|---|---|---|
| EXOG | N_Shore | TSS1500 | chr3:38537366-38537683 | 317 | | | 3 | | | | | | | | | | | yes | yes | | | | |
| XIRP1 | NA | TSS1500 | chr3:39234924-39235366 | 442 | | | 3 | | | | | | | | | | | yes | | | | | |
| CX3CR1 | NA | 1stExon;TSS1500;5'UTR;Body | chr3:39322541-39323539 | 998 | | | 4 | | | | | | 2 | | | | | | yes | yes | yes | | |
| SLC25A38 | Island | TSS200;1stExon;5'UTR;Body | chr3:39424697-39425978 | 1281 | | | 1 | | 1 | | | 0 | 1 | | | | | yes | | | yes | | yes |
| MOBP | N_Shore;Island | 5'UTR;Body | chr3:39543515-39544326 | 811 | | | 26 | | | | 1 | 0 | 4 | 2 | | 6 | 1 | | yes | yes | | | yes |
| ENTPD3 | N_Shore | TSS200 | chr3:40428638-40428986 | 348 | | | | | | | | | | | | | | | | | | | yes |
| TRAK1 | NA | TSS1500;Body | chr3:42201314-42201898 | 584 | | | 7 | 1 | | | | 0 | 3 | | | | | | yes | | | | |
| CCK | Island;S_Shore | 5'UTR;TSS200;TSS1500 | chr3:42305191-42307866 | 2675 | | 2 | 6 | | | | | | 3 | | | | | | | | yes | yes | yes |
| LYZL4 | NA | TSS1500 | chr3:42452635-42453125 | 490 | | | | | | | | | 1 | | | | | yes | | yes | | | |
| SS18L2 | N_Shore | TSS1500 | chr3:42631429-42631642 | 213 | | | | | | | | 0 | | | | | | yes | | | yes | | |
| ZBTB47 | N_Shore | TSS1500 | chr3:42694803-42695307 | 504 | | | | | | | | | | | | | | yes | | | | yes | |
| ZBTB47 | N_Shore;Island;S_Shore | Body | chr3:42699933-42700678 | 745 | | | 18 | | | | 1 | 0 | 1 | | | | | | | | yes | yes | yes |
| ZBTB47 | NA | Body | chr3:42705828-42706106 | 278 | | | 5 | | | | | | 2 | | | | | | | | | | yes |
| KBTBD5 | N_Shore;Island | TSS200;5'UTR;1stExon | chr3:42726812-42727843 | 1031 | | | 12 | | | | | 0 | 2 | | | 2 | | yes | | | | | |
| HHATL | NA | Body;5'UTR | chr3:42742993-42743794 | 801 | | | 9 | | | | | | 1 | | | | | yes | | | | yes | |
| HIGD1A | S_Shore | TSS1500 | chr3:42846907-42847116 | 209 | | | | | | | | | 2 | | | | | | | yes | | | |
| CCBP2 | NA | TSS200 | chr3:42850848-42850974 | 126 | | | | | | | | | 1 | | | | | yes | | yes | | | |
| CYP8B1 | N_Shelf | TSS1500 | chr3:42918681-42919025 | 344 | | | 2 | | 1 | | | | | | | | | yes | yes | | yes | | |
| ZNF662 | N_Shore | TSS1500;TSS200 | chr3:42947263-42947565 | 302 | | | 7 | | | | | | 1 | | | | | | | yes | | | |
| MIR138-1 | NA | TSS1500 | chr3:44154910-44155137 | 227 | | | | | | | | | | | | | | | | | yes | | |
| C3orf23 | Island | 5'UTR;TSS200 | chr3:44379878-44380138 | 260 | | | | | | | | | | 1 | | | | | | | | | |
| ZNF445 | S_Shore | TSS1500 | chr3:44519652-44519983 | 331 | | | 2 | | | | | | | | | | | | | yes | | | |
| ZNF167 | N_Shore | TSS1500 | chr3:44596178-44596512 | 334 | | | 3 | | | | | | | | | | | yes | | | | | |
| ZNF167 | N_Shore;Island | Body | chr3:44622287-44622865 | 578 | | | 6 | | | | | | | | | | | | | | | yes | yes |
| ZNF35 | Island | TSS200 | chr3:44690187-44690229 | 42 | | | | | | | | | 1 | | | | | | yes | | | | |
| ZNF502 | N_Shore;Island | TSS200 | chr3:44753946-44754587 | 641 | | 1 | | | 1 | | | 0 | 2 | | | | | yes | yes | | | yes | |
| KIF15;KIAA1143 | N_Shore;Island | TSS1500;Body;1stExon;5'UTR;TSS2 | chr3:44802549-44803293 | 744 | | | 8 | 1 | | | | 0 | 1 | | | 4 | | yes | | | | | yes |
| CLEC3B | Island;N_Shore | Body | chr3:45076599-45078075 | 1476 | | | 24 | | | | | 0 | 3 | | | 2 | | yes | yes | yes | yes | | yes |
| CDCP1 | Island;S_Shore | 1stExon;TSS1500 | chr3:45187717-45188208 | 491 | | | 2 | | | | | | | | | | | | | | | | yes |
| LIMD1 | S_Shore | TSS1500;TSS200 | chr3:45636060-45636386 | 326 | 2 | | 2 | | | | | | | | | | | | | | | yes | yes |
| CCR9 | NA | TSS200 | chr3:45927914-45928137 | 223 | | | 3 | | | | | | | | | | | yes | yes | | yes | | |
| CXCR6;FYCO1 | NA | TSS200;Body | chr3:45984838-45985168 | 330 | | | | | | | | 0 | 1 | | | | | yes | yes | | yes | yes | |
| CCR1 | NA | 1stExon;5'UTR | chr3:46249795-46250024 | 229 | | | | | | | | | 1 | | | | | yes | | | | | |
| CCR5 | NA | TSS200 | chr3:46411447-46411541 | 94 | | | | | | | | 0 | | | | | | yes | | | yes | | |
| CCRL2 | NA | TSS1500;TSS200;5'UTR;1stExon | chr3:46448084-46449636 | 1552 | | 1 | 27 | 1 | 4 | | | | 1 | 1 | | | | yes | yes | | | | |
| LTF | N_Shore;Island;S_Shore | Body;TSS200 | chr3:46506104-46506864 | 760 | | | 10 | | | | | 0 | 1 | | | 1 | | yes | yes | | | | yes |
| LRRC2 | S_Shore | 5'UTR;TSS200 | chr3:46608056-46608179 | 123 | | | | | 1 | | | | | | | | | yes | | | | | |
| LRRC2;TDGF1 | Island | 5'UTR;TSS1500 | chr3:46618325-46619364 | 1039 | | | 9 | 1 | | | | 0 | 2 | | | 1 | | yes | | | | | |
| ALS2CL | NA | Body;5'UTR;TSS200 | chr3:46718369-46719086 | 717 | | | 8 | | | | | | | | | | | | yes | | | | |
| TMIE | N_Shore;Island | TSS200;1stExon;5'UTR | chr3:46742703-46743199 | 496 | | | | | | | | 0 | 1 | 1 | | | | yes | | | | | yes |
| PRSS50 | Island;S_Shore | TSS200;TSS1500;Body;1stExon;5'U | chr3:46759096-46759698 | 602 | | 1 | 25 | | | | | 0 | 14 | 2 | | 8 | | yes | | | | | |
| PRSS42 | Island | Body;TSS200 | chr3:46875226-46875768 | 542 | | | | | | | | | 5 | | | 2 | | | | yes | | | |
| MYL3 | NA | TSS1500 | chr3:46906118-46906385 | 267 | | | 4 | | | | | | | | | | | | yes | | | yes | |
| PTH1R | Island | Body | chr3:46933782-46934204 | 422 | | | | | | | | | | | | | | | yes | | | | |
| PTH1R | N_Shore;Island | Body | chr3:46939978-46940439 | 461 | | | | | | | | 0 | 4 | 2 | | 7 | 1 | | | | | yes | |
| NBEAL2 | NA | Body | chr3:47029335-47029475 | 140 | | | 3 | | 1 | | | | | | | | | | | | | | |
| NBEAL2 | NA | Body | chr3:47040214-47040357 | 143 | | | | | | | | | | | | | | | | | | | |
| NBEAL2 | Island | Body | chr3:47050803-47051341 | 538 | | | | | | | | | 1 | | | | | | | | | | yes |
| SCAP | S_Shore | TSS1500 | chr3:47517781-47517819 | 38 | | | | | | | | 0 | | | | | | yes | | | | yes | |
| DHX30 | Island | Body | chr3:47866555-47866946 | 391 | | | | | | | | | 1 | | | | | yes | | | | | yes |
| DHX30;MIR1226 | N_Shore | Body;TSS1500 | chr3:47889960-47891032 | 1072 | | | | | | | | | | | | | | | | | | yes | yes |
| ZNF589 | N_Shore | TSS1500 | chr3:48282209-48282311 | 102 | | | 1 | | | | | | | | | | | | | yes | yes | | |
| ZNF589 | NA | 3'UTR | chr3:48310544-48310897 | 353 | | | 2 | | | | | | 1 | | | | | | | | | | |
| FBXW12 | NA | TSS200 | chr3:48413550-48413769 | 219 | | | | | | | | | 1 | | | | | | | yes | | | |
| PLXNB1 | S_Shore | TSS1500;TSS200 | chr3:48471575-48471901 | 326 | | | 6 | | | | | | 2 | | | | | yes | | | | | |
| ATRIP | N_Shore | TSS1500 | chr3:48487347-48487808 | 461 | | | | | | | | | | | | | | | yes | | | | yes |
| TREX1 | NA | 5'UTR;1stExon | chr3:48507354-48507508 | 154 | | | | | | | 1 | | 1 | 1 | | | | | | | | yes | |
| PFKFB4 | chr3:48594022-48594853 | Island | chr3:48594205-48594229 | 24 | | | | | | | | | | | 1 | | | yes | | | | | |
| UCN2 | NA | 5'UTR | chr3:48601117-48601585 | 468 | | | 2 | | | | | | | | | | | | | | | | yes |
| COL7A1 | N_Shore;Island | Body;1stExon;TSS200 | chr3:48631834-48632783 | 949 | | | 2 | | | | | 0 | 4 | | | 3 | 2 | | | | | | |
| CELSR3 | Island | Body | chr3:48693597-48694673 | 1076 | | 1 | 11 | | | | | 0 | 2 | | | 2 | | | | | | | |
| CELSR3 | N_Shore | 1stExon | chr3:48697500-48697594 | 94 | | | | | | | | | | | | | | | yes | | | | |
| MIR425;DALRD3;NDUFAF3 | N_Shore | Body;5'UTR;TSS1500;TSS200 | chr3:49057259-49058043 | 784 | | | 16 | | | | | | | | | | | yes | | | yes | | |
| LAMB2 | NA | 1stExon;5'UTR;TSS200 | chr3:49170496-49171051 | 555 | | | | | | | | 0 | 5 | | | 6 | | | | | | | |
| CCDC71 | S_Shore | TSS200 | chr3:49203910-49204548 | 638 | | | 6 | | | | | | | | | | | | | | | | |
| CCDC36 | N_Shore;Island | TSS1500;5'UTR;TSS200 | chr3:49236376-49236946 | 570 | | 1 | | | | | 2 | 0 | 2 | 2 | | 2 | | yes | yes | | | | |
| C3orf62;USP4 | N_Shore;Island | 1stExon;3'UTR;TSS200 | chr3:49314155-49314920 | 765 | | | 2 | | | | | | | | | | | | | | | | |
| USP4 | S_Shore | TSS1500 | chr3:49377951-49378124 | 173 | | | | 1 | | | | | | | | | | yes | yes | | | | |
| GPX1 | Island | 1stExon;5'UTR;TSS200 | chr3:49395745-49396232 | 487 | | | | | 2 | | | | | | | | | | | | | yes | |
| AMT;NICN1 | NA | Body;5'UTR;1stExon;3'UTR | chr3:49459855-49460177 | 322 | | | 11 | 1 | | | | | 2 | | | | | | yes | | | | |
| APEH | N_Shore;Island | TSS1500 | chr3:49710768-49711095 | 327 | | | 1 | | | | 1 | 0 | 2 | | | | | yes | | | | yes | |
| MST1 | Island | Body | chr3:49723947-49724291 | 344 | | | | 1 | | | | | | | | | | | | | | | |
| AMIGO3;RNF123 | Island;S_Shore | 1stExon;Body;TSS200;TSS1500 | chr3:49756301-49757476 | 1175 | | | | 2 | | | | | 1 | | | 1 | | yes | | | | | |
| CDH29 | N_Shelf | TSS1500 | chr3:49837529-49837689 | 160 | | | 4 | | | | | | | | | | | yes | yes | | | | |
| CAMKV | S_Shore | TSS200 | chr3:49907556-49907692 | 136 | | | 1 | | | | | | | | | | | | | | | | yes |
| MST1R | Island;N_Shore | 1stExon;TSS200 | chr3:49940126-49941613 | 1487 | | | 15 | | 1 | | | 0 | 3 | | | | | yes | | | | | |
| MON1A | S_Shore | TSS200;TSS1500 | chr3:49967517-49967713 | 196 | | | | | 1 | | | 0 | 1 | | | 1 | | yes | | | yes | | |
| GNAI2 | N_Shore | TSS1500 | chr3:50273096-50273314 | 218 | | | | | | | | | 1 | | | | | yes | | | | | yes |
| GNAI2 | N_Shore | Body | chr3:50275112-50275694 | 582 | | | | | 1 | | | | | | | | | | | | | | |
| GNAI2 | NA | TSS1500;Body | chr3:50284010-50284305 | 295 | | | | 1 | | | | | | | | | | yes | | | | | |
| SEMA3B | NA | TSS1500 | chr3:50303807-50304463 | 656 | | | 4 | | | | | | 3 | | | | | | | | | | |
| SEMA3B | NA | 5'UTR | chr3:50306107-50306878 | 771 | | | | | | | | | | | | | | | | | | yes | |
| SEMA3B | Island | Body | chr3:50310766-50311213 | 447 | | | | | | | | | | | | | | yes | | | | yes | |
| SEMA3B | Island | Body | chr3:50312913-50313213 | 300 | | | | | | | | | | | | | | yes | | | | | |
| HYAL1 | NA | TSS200;5'UTR | chr3:50341009-50341293 | 284 | | | | 1 | | | | | 1 | | | | | yes | | | yes | yes | |
| HYAL2 | Island | 5'UTR;TSS1500 | chr3:50359363-50360097 | 734 | | | 11 | 1 | | | | 0 | 2 | | | 1 | | | | | | | yes |
| RASSF1 | Island;N_Shore | 1stExon;TSS200;TSS1500;Body;5'U | chr3:50375459-50378664 | 3205 | | | | | | | | | | | | | | | | | | | |
| ZMYND10 | Island | 1stExon;5'UTR;TSS200 | chr3:50382952-50383485 | 533 | | | | | 3 | | | 0 | | | | 1 | | yes | yes | | | | |
| CACNA2D2 | Island;S_Shore | Body | chr3:50402333-50403312 | 979 | 1 | | | | | | | | 1 | | | | | | | | | yes | yes |
| CACNA2D2 | NA | Body | chr3:50487955-50488230 | 275 | | | 26 | | | | | 0 | 3 | | | 2 | | yes | | | | | |
| CACNA2D2 | N_Shore | Body | chr3:50540137-50540224 | 87 | | | 6 | | | | | | | | | | | | | | | | yes |
| CISH | N_Shelf | Body | chr3:50645106-50645426 | 320 | | | | | | | | | 1 | | | | | | | | | | |
| GRM2 | Island | TSS1500;1stExon;5'UTR | chr3:51740741-51741473 | 732 | | | 1 | | | | | 0 | 3 | | | 3 | | | | | | yes | |
| IQCF5 | NA | TSS1500 | chr3:51910238-51910733 | 495 | | | | | | | | | | | | | | yes | yes | | | | |
| GPR62 | Island | 1stExon | chr3:51990575-51990929 | 354 | | | 3 | | | | | | | | | | | yes | | | | | |
| PCBP4 | NA | 5'UTR;TSS1500 | chr3:51996797-51997033 | 236 | | | 2 | | | | | | 1 | | | | | yes | yes | | | | |
| WDR51A | Island | TSS200 | chr3:52188725-52188732 | 7 | | | | | | | | | | 1 | | | | yes | | | | | |
| ALAS1 | N_Shore | TSS1500 | chr3:52231347-52231840 | 493 | | | 1 | | | | | | | | | | | yes | | | | | |
| TLR9 | NA | 5'UTR;1stExon;TSS200 | chr3:52260090-52260671 | 581 | | | | | | | | | 1 | | | | | | yes | | | | |

| | | | | | | | | | | | | | | | | | | | | | | | |
|---|---|---|---|---|---|---|---|---|---|---|---|---|---|---|---|---|---|---|---|---|---|---|---|
| DNAH1 | NA | 5'UTR | chr3:52351355-52351963 | 608 | | | 3 | 1 | | | | 0 | 1 | | | 2 | | yes | yes | | | | |
| SEMA3G | NA | 1stExon;TSS200 | chr3:52478965-52479108 | 143 | | | 6 | | | | | | | | | | | | | | | yes | |
| TNNC1;NISCH | N_Shore | Body;1stExon;5'UTR;TSS1500;TSS2 | chr3:52487733-52488229 | 496 | | | 13 | | | | | | | | | | | yes | | | | yes | |
| STAB1 | NA | TSS1500;TSS200 | chr3:52529085-52529393 | 308 | | | 6 | | | | | | 1 | | | | | | yes | | | | |
| STAB1 | N_Shore;Island | Body | chr3:52552593-52553433 | 840 | | | | | | | | | 1 | | | | | yes | | | | yes | |
| NT5DC2 | Island;N_Shore | Body;TSS1500;5'UTR;1stExon | chr3:52567010-52569169 | 2159 | | | 3 | 1 | | | | | 1 | 1 | | | | yes | yes | | | yes | yes |
| ITIH1 | NA | TSS200;Body | chr3:52812367-52812520 | 153 | | | | | | | | | | | | | | | | | | | |
| ITIH1 | NA | TSS200;Body | chr3:52813813-52813920 | 107 | | | 7 | | 1 | | | | | | | 1 | | yes | yes | | | | |
| ITIH3 | NA | TSS200 | chr3:52828628-52828818 | 190 | | | 2 | | | | | | | | | | | yes | yes | | | | |
| ITIH4 | NA | 5'UTR;1stExon;TSS1500 | chr3:52864659-52865917 | 1258 | | | 6 | 1 | | | | | | | | | | yes | | | | | |
| MUSTN1 | NA | 1stExon;5'UTR;TSS200 | chr3:52868999-52869738 | 739 | 1 | | 10 | | | | | | | | | | | | | | | yes | |
| SFMBT1 | Island | 5'UTR | chr3:53032818-53032988 | 170 | | 1 | | | | | | | | | | 1 | | yes | | | | | |
| TKT | S_Shore | TSS1500 | chr3:53290327-53290343 | 16 | | | 7 | | | | | | | | | | | yes | | | | yes | yes |
| CACNA1D | Island | Body | chr3:53529387-53530075 | 688 | | | | 1 | | | | | | | | | | | yes | | | yes | yes |
| CACNA1D | NA | Body | chr3:53700141-53700276 | 135 | | | | | 1 | | 1 | 0 | 1 | 1 | | 2 | | | | | | | |
| CHDH | Island | Body | chr3:53857939-53857989 | 50 | | | 1 | | | | | | | | | | | | yes | | | | |
| SELK | Island | TSS200 | chr3:53926056-53926101 | 45 | | | 2 | | | | | | | | | | | | yes | | | | |
| WNT5A | Island | Body | chr3:55508460-55508686 | 226 | | | | | | | | 0 | | | | 1 | | | | | | | yes |
| WNT5A | N_Shore;chr3:55517658-55517 | Body;N_Shore | chr3:55517101-55518798 | 1697 | 11 | 1 | 7 | 1 | | 1 | | 0 | 3 | 1 | 1 | | | yes | | | | yes | |
| C3orf51;ERC2 | NA | Body;3'UTR | chr3:55693182-55693741 | 559 | | | | | | | | | 2 | | | | | | yes | | | | |
| C3orf63 | S_Shore | TSS1500 | chr3:56717769-56717804 | 35 | | | | | | | | | | | | | | yes | | | | | |
| ARHGEF3 | NA | TSS200;Body | chr3:56809796-56810320 | 524 | | | | | | | | | 1 | | | | | | | | | yes | |
| ARHGEF3 | Island | TSS200 | chr3:57113447-57113702 | 255 | | | | | | | | | 1 | | | | 1 | yes | | | | yes | yes |
| DNAH12 | NA | 5'UTR;1stExon | chr3:57529952-57530606 | 654 | | | | | | | | | | 1 | | | | yes | yes | | | | yes |
| SLMAP | S_Shore | TSS200;5'UTR;1stExon | chr3:57743163-57743599 | 436 | | | 2 | | 1 | | | 0 | 2 | | | | | | | | | | |
| PXK | N_Shore | TSS1500 | chr3:58318260-58318305 | 45 | | | | | | | | 0 | 1 | | | | | | | | | | yes |
| ACOX2 | NA | 5'UTR | chr3:58522689-58523313 | 624 | | | | | | | | | | | | | | yes | yes | | | | |
| FAM107A | NA | 1stExon;5'UTR;TSS200 | chr3:58563097-58563907 | 810 | 1 | | 15 | | | | | | 1 | | | | | yes | yes | | | | |
| C3orf67 | Island | 5'UTR;1stExon | chr3:59035407-59035943 | 536 | | | 2 | | | | | | | | | | | yes | yes | yes | yes | | |
| FHIT | N_Shore;Island | 5'UTR | chr3:61235829-61236911 | 1082 | | | 3 | 1 | | | | | 1 | | | | | yes | | | | yes | |
| C3orf14 | S_Shore;Island | TSS200;TSS1500 | chr3:62304515-62305486 | 971 | | | | | | | | | 1 | | | | | yes | | | | | |
| FEZF2 | Island | 3'UTR | chr3:62355443-62355884 | 441 | 1 | | | | | | | | | | | | | | | | | | |
| FEZF2 | Island | Body;TSS1500 | chr3:62358234-62359456 | 1222 | | | | | | | | | 2 | | | | | | | | | yes | |
| CADPS | S_Shore | TSS1500 | chr3:62861544-62861925 | 381 | | | | | | | | | 1 | | | | | | | | | | |
| LOC285401 | NA | TSS1500 | chr3:63087235-63087266 | 31 | | | 9 | | | | | | 1 | | | | | | | yes | | | |
| SYNPR | Island;N_Shore | 5'UTR;1stExon;TSS200 | chr3:63263752-63264249 | 497 | | | | | | | | | 2 | | | | | | yes | | | | yes |
| SYNPR | NA | Body;TSS1500 | chr3:63427866-63428587 | 721 | | | | | | | | | 1 | | | | | yes | | | | yes | yes |
| SNTN | NA | TSS1500 | chr3:63637118-63637558 | 440 | | | | | | | | | | | | | | | yes | | | | yes |
| PSMD6 | N_Shore | Body | chr3:64008161-64008387 | 226 | | | | | | | | | | | | | | | | | | | |
| PRICKLE2 | NA | 1stExon;5'UTR;TSS200 | chr3:64210991-64211659 | 668 | | 1 | | 1 | 2 | | | 0 | 1 | | | 1 | | yes | yes | | | | |
| ADAMTS9 | NA | Body | chr3:64547108-64547373 | 265 | | | 24 | | | | | | | | | | | | | | | | |
| ADAMTS9 | N_Shore;Island | Body | chr3:64670013-64671764 | 1751 | | | 11 | | | | | 0 | 2 | | | | | | | | | | |
| MAGI1 | Island | Body;3'UTR | chr3:65342216-65342971 | 755 | | | 15 | 1 | 2 | 1 | | 0 | 3 | | | 5 | | | yes | | | | |
| MAGI1 | NA | Body | chr3:66002927-66002991 | 64 | 10 | | | | | | | | | | | | | | yes | | | | |
| MAGI1 | Island | TSS200 | chr3:66024549-66024561 | 12 | | | | | | | | | 1 | | | | | | | | | | yes |
| FAM19A1 | N_Shelf | TSS200 | chr3:68053273-68053484 | 211 | | | 2 | | | | | | 1 | | | | | | | | | | |
| FAM19A1 | S_Shore | Body | chr3:68057306-68057357 | 51 | | | | | | | | | 1 | | | | | yes | | | | | |
| FAM19A4 | Island | TSS200 | chr3:68981854-68982064 | 210 | | | | | | | | | 2 | 1 | | | | yes | | | | | |
| C3orf64 | N_Shore | 5'UTR | chr3:69061356-69062103 | 747 | | | | 1 | 1 | | | | 1 | | 1 | | | | | | | | yes |
| MITF | NA | Body;TSS1500;TSS200 | chr3:69915189-69915449 | 260 | | | | | 2 | | | | 1 | | | | | | | | | yes | |
| FOXP1 | N_Shore | Body | chr3:71113124-71113603 | 479 | | | 2 | | | | | | | | | | | | | | | | yes |
| FOXP1 | NA | Body | chr3:71180032-71180071 | 39 | 8 | | 4 | | | | | | | | | | | | | | | | yes |
| FOXP1 | S_Shore | TSS1500 | chr3:71633826-71634044 | 218 | | | 1 | | | | | | | | | | | | | | | yes | |
| GPR27;EIF4E3 | Island | 1stExon;TSS200;TSS1500 | chr3:71804005-71804142 | 137 | | | 8 | | | | | 0 | | | | | | | | | | | yes |
| PROK2 | N_Shore | Body | chr3:71833543-71833689 | 146 | | | 2 | | | | | | | | | | | | | | | | |
| RYBP | Island | TSS1500 | chr3:72496799-72496875 | 76 | | | | | | | | 0 | 1 | | | | | yes | | | | | yes |
| PPP4R2 | N_Shore | TSS1500 | chr3:73045440-73045686 | 246 | | 1 | 5 | | | | 1 | | 1 | 1 | | 8 | 1 | yes | | | | yes | |
| PDZRN3 | Island;S_Shore | Body;1stExon;5'UTR;TSS200;TSS1 | chr3:73673231-73674702 | 1471 | | | | | | | | 0 | 5 | | | 2 | | yes | | | yes | yes | yes |
| ZNF717 | Island | TSS1500 | chr3:75834525-75834658 | 133 | | | | | | | | 0 | | | | | | yes | | | | | |
| ROBO2 | N_Shore | TSS1500 | chr3:77087860-77088570 | 710 | | | | | | | | | 3 | | | 1 | | | | | | yes | |
| ROBO2 | N_Shore | TSS200;Body | chr3:77147070-77147279 | 209 | | | | | | | | | 1 | | | | | yes | yes | | | | |
| ROBO1 | Island | 5'UTR | chr3:79815639-79815848 | 209 | | | | | | | | | 1 | | | | | | | | | | yes |
| CADM2 | Island | 5'UTR;1stExon | chr3:85008587-85008828 | 241 | | | | | | | | 0 | 1 | | | | | | | | | yes | |
| VGLL3 | Island | Body | chr3:87039687-87039900 | 213 | | | | | | | | | 1 | | | | | | | | | yes | |
| EPHA3 | NA | TSS1500 | chr3:89156071-89156478 | 407 | | | | | | | | 0 | 1 | | | | | | | | | | |
| LOC255025 | Island | TSS200 | chr3:94657022-94657048 | 26 | | | 2 | | | | | | | | | | | | yes | | | | |
| EPHA6 | Island | 1stExon;TSS200 | chr3:96533290-96534005 | 715 | | 1 | | | | | | | 1 | | | | | | | | | yes | |
| MINA | Island | 5'UTR;1stExon | chr3:97690786-97691400 | 614 | | | | | | | | | 1 | | | | | yes | | | yes | | yes |
| ST3GAL6 | N_Shore | TSS1500 | chr3:98451116-98451287 | 171 | | | | | | | | | 1 | | | | | yes | | | | | |
| ST3GAL6 | S_Shore | 5'UTR | chr3:98452812-98453086 | 274 | | | 3 | | | | | | | | | | | | | | | | |
| DCBLD2 | Island | 5'UTR;1stExon | chr3:98620265-98620572 | 307 | | | | | | | | | 1 | | | | | | | | | | yes |
| MIR548G;COL8A1 | NA | Body;TSS200 | chr3:99357437-99357613 | 176 | | | 2 | | | | | | | | | | | | | | | | yes |
| C3orf26;FILIP1L;MIR548G | Island | Body;1stExon;5'UTR | chr3:99595026-99595145 | 119 | | | | | | | | | | 1 | | | | | | | | yes | yes |
| C3orf26;FILIP1L | NA | Body;5'UTR;1stExon;TSS1500 | chr3:99833293-99833735 | 442 | | | | | | | | | 1 | | | | | yes | yes | | | yes | |
| TMEM30C | NA | TSS200 | chr3:99904510-99904650 | 140 | | | 8 | | | | | | | | | | | yes | | | | | |
| TBC1D23 | N_Shore | TSS1500 | chr3:99979117-99979421 | 304 | | | | | | | | 0 | | | | | | | yes | yes | yes | | yes |
| NIT2 | N_Shore | TSS1500 | chr3:100052640-100053188 | 548 | | | 7 | | | | | 0 | | | | | | yes | yes | | yes | | |
| LNP1;TOMM70A | Island | 5'UTR;1stExon;TSS1500 | chr3:100120773-100120961 | 188 | | | 1 | | | | | | 1 | | | | | | | | | | |
| GPR128 | NA | TSS1500 | chr3:100327897-100328265 | 368 | | | 2 | | | | | | | | | | | | | yes | | | |
| ABI3BP | NA | Body;5'UTR;1stExon | chr3:100712058-100712345 | 287 | | | | | | | | 0 | 2 | | | 4 | | | | | | | |
| RG9MTD1 | chr3:101280545-101280952 | Island | chr3:101280570-101280610 | 40 | | | | | | | | | | | 1 | | | yes | yes | | | | |
| LOC100009676;ZBTB11 | Island | TSS1500;Body | chr3:101395051-101395095 | 44 | | | | | | | | | 1 | | | | | yes | | | | | |
| ZBTB11;LOC100009676 | S_Shore | TSS1500;Body | chr3:101396658-101396897 | 239 | | | | | | | | | 1 | | | | | yes | | | | | |
| RPL24 | S_Shore | TSS1500 | chr3:101406220-101406543 | 323 | | | 3 | | | | | 0 | 1 | | | | | yes | | | | | |
| FAM55C | N_Shore | TSS1500 | chr3:101497475-101497779 | 304 | | | 14 | | | | | | | | | | | | yes | | | | |
| CD47 | Island;S_Shore | TSS1500 | chr3:107810361-107810791 | 430 | | | 6 | | | | | | 2 | | | | | | | | | | yes |
| HHLA2 | NA | TSS1500 | chr3:108020727-108021214 | 487 | | | | | | | | | 1 | | | | | yes | | | | | |
| RETNLB | NA | 5'UTR;1stExon;Body | chr3:108475765-108476278 | 513 | | 1 | 3 | | | | | | 1 | | | | | yes | | | | | |
| MORC1 | N_Shore;Island | Body;TSS200 | chr3:108836693-108837484 | 791 | | | 4 | | | | | | | | | | | | yes | | | yes | |
| C3orf66 | NA | TSS200 | chr3:108896930-108897122 | 192 | | | | | | | | | | | | | | yes | yes | yes | | | |
| DPPA2 | NA | 1stExon;5'UTR;TSS200 | chr3:109035305-109035649 | 344 | | | 4 | | | | | | | | | | | yes | | | yes | | |
| DPPA4 | NA | 1stExon | chr3:109056349-109056897 | 548 | | | | | | | | 0 | 3 | | | 2 | | yes | | | | | |
| CD96 | NA | TSS200 | chr3:111260756-111261010 | 254 | | | 5 | 1 | | | | 0 | | | | | | | | | | | |
| PHLDB2;PLCXD2 | NA | TSS1500;Body | chr3:111451090-111451470 | 380 | | | | | 1 | | | | | | | | | yes | yes | yes | | | |
| PHLDB2 | N_Shore | TSS1500;Body;TSS200 | chr3:111577634-111577981 | 347 | | | 14 | | | | | | 1 | | | | | yes | yes | | | | |
| ABHD10 | N_Shore | TSS1500 | chr3:111697479-111697575 | 96 | | | | | 1 | | | 0 | 1 | | | 2 | | | yes | | | | |
| TAGLN3 | NA | 1stExon;5'UTR;TSS1500 | chr3:111717682-111718565 | 883 | | | 3 | | | | | 0 | 3 | | | | | yes | | | | | yes |
| C3orf52 | N_Shore;Island | TSS1500;TSS200 | chr3:111804343-111805005 | 662 | | | 4 | | | | | 0 | | | | | | yes | yes | yes | | yes | |
| SLC9A10 | NA | 5'UTR;1stExon;TSS200 | chr3:112012867-112013357 | 490 | | | 11 | | | | | 0 | 5 | 2 | | 3 | | | yes | | | | |

| | | | | | | | | | | | | | | | | | | | | | | | |
|---|---|---|---|---|---|---|---|---|---|---|---|---|---|---|---|---|---|---|---|---|---|---|---|
| CD200 | N_Shore;Island | TSS200 | chr3:112051808-112052405 | 597 | | | | | 3 | | | | | | | | | yes | | | | yes | yes |
| CCDC80 | NA | Body | chr3:112358351-112358468 | 117 | | | 12 | | | | | | | | | | | | | | | | |
| CCDC80 | NA | 5'UTR;1stExon;TSS1500 | chr3:112359488-112361234 | 1746 | | 1 | 1 | | 2 | | | 0 | 1 | | | | | yes | | | | yes | yes |
| WDR52 | N_Shore;Island | 5'UTR;1stExon;TSS200 | chr3:113160071-113160637 | 566 | 1 | 1 | 31 | | | | | 0 | 3 | | | 2 | | yes | yes | | | | |
| TIGIT | NA | TSS1500 | chr3:114012316-114012806 | 490 | | | | | | | | | 1 | | | | | | yes | | | | |
| ZBTB20 | NA | 5'UTR;TSS200 | chr3:114102565-114102615 | 50 | | | 1 | | | | | | | | | | | | | | | yes | |
| ZBTB20 | NA | 5'UTR;TSS1500;TSS200 | chr3:114343068-114343825 | 757 | | | | | | | | | 1 | | | | | yes | | | | | |
| ZBTB20 | NA | 1stExon;5'UTR | chr3:114790243-114790596 | 353 | | | 9 | | | | | | | | | | | yes | | | | | |
| IGSF11 | NA | Body | chr3:118663470-118663697 | 227 | | | | | | | | | | | | | | yes | | | | yes | |
| C3orf30;IGSF11 | NA | TSS200;1stExon;5'UTR | chr3:118864870-118865478 | 608 | | | 5 | | | | | | 1 | | | | | | | | | | |
| UPK1B | NA | TSS1500 | chr3:118892215-118892330 | 115 | | | 4 | | | | | 0 | 1 | | | 1 | | | yes | | | yes | yes |
| TMEM39A | NA | TSS1500 | chr3:119182753-119182858 | 105 | | | | | | | | | | | | | | yes | yes | | | | |
| ADPRH | N_Shore | TSS1500 | chr3:119298195-119298236 | 41 | | | 9 | | | | | 0 | | | | | | yes | yes | yes | yes | | yes |
| PLA1A | NA | nan | chr3:119316673-119316820 | 147 | | | | | | | | | | | 1 | | | yes | yes | | | | yes |
| POPDC2 | NA | TSS200 | chr3:119379414-119380012 | 598 | | | 5 | | | | | | | | | | | yes | | | | yes | |
| C3orf15 | N_Shore;Island | TSS1500;5'UTR;1stExon;Body | chr3:119421553-119422359 | 806 | | | | | | | | 0 | 1 | | | 1 | | | yes | | | | yes |
| NR1I2 | N_Shore;Island | Body | chr3:119528656-119529158 | 502 | | | | 1 | 1 | | | | | | | | | | | | | | yes |
| FSTL1 | Island | TSS1500 | chr3:120170383-120170646 | 263 | | | 2 | | | | | 0 | | | | | | | | | | | yes |
| FBXO40 | NA | TSS200 | chr3:121312127-121312241 | 114 | | | 1 | | | | | | 1 | | | | | yes | | | | | |
| HCLS1 | NA | 1stExon;5'UTR | chr3:121379745-121380233 | 488 | | | 4 | | 1 | | | | 1 | 1 | | | | | | yes | | | |
| SLC15A2 | NA | TSS1500 | chr3:121612956-121613703 | 747 | | | | | | | | | | | | | | yes | yes | | | | |
| ILDR1 | NA | Body | chr3:121726186-121726421 | 235 | | | | | | | | | | | | | | yes | yes | | | | |
| CD86 | NA | TSS1500;Body;TSS200 | chr3:121795768-121796767 | 999 | | | 5 | | 1 | | | | 1 | | 1 | | | yes | | | | | |
| CASR | N_Shore | TSS200;5'UTR;1stExon | chr3:121902497-121902622 | 125 | | | 2 | | | | | 0 | 1 | | | 1 | | | yes | | | | |
| CASR | N_Shore | 5'UTR | chr3:121949083-121949308 | 225 | | | 3 | | | | | | 1 | | | | | yes | | | | | |
| CASR | NA | Body | chr3:121980683-121980842 | 159 | | | | | | | | | | | | | | | | | | yes | |
| CSTA | NA | TSS200;TSS1500 | chr3:122043799-122044172 | 373 | | | 1 | | | | | 0 | 2 | 1 | | 3 | | yes | yes | | | | |
| PARP9;DTX3L | N_Shore | 5'UTR;TSS1500 | chr3:122281881-122281975 | 94 | | | 4 | | | | 1 | 0 | 1 | | | 2 | | | yes | | yes | | |
| PARP15 | N_Shore | TSS1500;TSS200;5'UTR;1stExon | chr3:122295656-122296778 | 1122 | | 1 | 5 | 1 | 3 | | | | 1 | | | 1 | | yes | yes | yes | | | |
| PARP15 | NA | Body;TSS1500 | chr3:122334227-122334717 | 490 | | | | | | | | | | | | | | yes | | | | | |
| DIRC2 | Island;S_Shore | Body | chr3:122514443-122514814 | 371 | | | | | | | | 0 | 1 | | | | | | | yes | | | yes |
| SEMA5B | Island;N_Shore | Body | chr3:122640625-122641316 | 691 | | | 5 | | | | | | 1 | | 1 | | | yes | | | | | |
| SEMA5B | NA | 5'UTR | chr3:122712138-122712271 | 133 | | | | | | | | 0 | 1 | | | | | | | | | | |
| ADCY5 | Island | 1stExon | chr3:123167276-123167522 | 246 | | | | | | | | | 1 | | | | | | | | | yes | yes |
| MYLK | NA | 1stExon;5'UTR;Body | chr3:123339334-123339918 | 584 | | | 5 | | | | | | 1 | | | | | yes | yes | | | | yes |
| MYLK | N_Shore | 5'UTR | chr3:123602485-123602795 | 310 | | | | | | | | | 1 | | | | | | | | | yes | yes |
| ROPN1 | NA | Body | chr3:123699208-123699389 | 181 | | | | | | | | 0 | | | | | | | yes | | | | |
| KALRN | NA | TSS200 | chr3:123813419-123813504 | 85 | 1 | | | | | | | | 1 | | | | | yes | | | | | yes |
| KALRN | NA | Body | chr3:124103021-124103696 | 675 | | | | | | | | | 2 | | | | | | | yes | | | |
| KALRN | N_Shore | Body;TSS1500 | chr3:124303268-124303404 | 136 | | | 3 | | | | | | | | | | | yes | | | | | |
| MUC13 | NA | TSS200 | chr3:124653632-124653718 | 86 | | | 1 | | | | | | | | | | | yes | | | | | |
| SLC12A8 | Island | Body | chr3:124860616-124860871 | 255 | | | | | | | | | 1 | | 1 | | | | | | | yes | |
| SLC12A8 | S_Shore;Island | TSS1500;TSS200 | chr3:124931746-124932002 | 256 | | | | | | | | 0 | 2 | | | 1 | | | | | | | yes |
| ZNF148 | Island | 5'UTR | chr3:125076233-125076480 | 247 | | | | | | | | 0 | 1 | | | | | yes | | | | yes | |
| ROPN1B | S_Shelf | 5'UTR | chr3:125690774-125691054 | 280 | | 1 | | | | | | | | | | | | | yes | | | | |
| SLC41A3 | NA | Body;5'UTR;1stExon;TSS1500 | chr3:125775413-125776338 | 925 | | | | | 1 | | | | | | | | | | yes | yes | | | |
| ALDH1L1 | S_Shore;Island | TSS1500;TSS200 | chr3:125899547-125900108 | 561 | | | 2 | | | | | 0 | 3 | | | | | | yes | | | | |
| KLF15 | N_Shore;Island | 3'UTR;Body | chr3:126061950-126062822 | 872 | | | | | | | | 0 | 2 | | | | | yes | | | | yes | |
| CCDC37 | N_Shore;Island | TSS1500;TSS200 | chr3:126113454-126113918 | 464 | | 1 | | 1 | | | | | 2 | | | | | yes | yes | | yes | | |
| CCDC37 | N_Shore | Body | chr3:126134977-126135211 | 234 | | | 5 | | | | | | 1 | | | | | | | | yes | | |
| UROC1 | NA | 1stExon | chr3:126236504-126237462 | 958 | | | | 1 | | | | | | | | | | yes | | yes | | | |
| CHCHD6 | N_Shore | TSS1500 | chr3:126422260-126422575 | 315 | | | | | | | | | | | | | | yes | yes | yes | | | |
| PLXNA1 | NA | Body | chr3:126721131-126721586 | 455 | | | | 1 | | | | | 1 | | | | | yes | | | | yes | |
| PLXNA1 | Island | Body | chr3:126748781-126749321 | 540 | | | | | | | | | 1 | | | | | | | | | | |
| GATA2 | N_Shelf | Body | chr3:128202176-128202797 | 621 | | | 2 | | | | | | | | | | | | | | | | |
| GATA2 | N_Shore | Body | chr3:128204413-128205496 | 1083 | | | 13 | | | | | | 1 | | | | | | | | | yes | |
| GATA2 | Island | 5'UTR;1stExon | chr3:128210888-128212476 | 1588 | | | 6 | | | | | | | | | | | | | | | | yes |
| LOC90246 | NA | TSS1500 | chr3:128225824-128226818 | 994 | | 1 | | | | | | | | | | | | | yes | yes | | | |
| C3orf27 | NA | Body;5'UTR | chr3:128292314-128292922 | 608 | | | 4 | 1 | | | | | | | | | | yes | | | yes | | |
| C3orf27 | NA | TSS200;nan | chr3:128295045-128295134 | 89 | | | | | | | | | 1 | | 1 | | | yes | | | | | |
| RPN1 | S_Shore | TSS1500 | chr3:128370232-128370463 | 231 | | | | | | | | | 1 | | | | | yes | yes | | | | |
| KIAA1257 | S_Shore | TSS200 | chr3:128713146-128713392 | 246 | | | | | | | | | 1 | | | | | yes | yes | | | | |
| CCDC48 | Island | Body | chr3:128722515-128723007 | 492 | | | 7 | | | | | 0 | | | | | | yes | yes | | | yes | |
| GP9 | NA | TSS200;TSS1500 | chr3:128778575-128779601 | 1026 | 1 | | | | 3 | | | 0 | 1 | 1 | | 2 | | yes | | | yes | | |
| RAB43 | S_Shore | TSS200 | chr3:128840692-128840803 | 111 | | | | | | | | | 1 | | | | | | | | | yes | yes |
| C3orf37 | N_Shore | TSS200 | chr3:128997662-128997925 | 263 | | | | | | | | | 1 | | | | | | | | yes | yes | yes |
| C3orf25 | Island | 5'UTR;1stExon | chr3:129147397-129147642 | 245 | | | | | | | | | 3 | | | | | yes | yes | | | | |
| TMCC1 | NA | TSS200;Body;1stExon;5'UTR | chr3:129407365-129407548 | 183 | | | 13 | 1 | | | | | 2 | | | | | | | | | | |
| TRH | Island | 5'UTR;1stExon | chr3:129693370-129694057 | 687 | | 1 | | | | | | | | | | | | yes | | | | yes | |
| LOC729375 | Island | TSS200 | chr3:129830460-129830607 | 147 | | | | | | | | | 1 | | | | | | | | | | yes |
| NEK11;ASTE1 | Island | TSS1500;1stExon;5'UTR;TSS200 | chr3:130745442-130745959 | 517 | | | 2 | | | | | 0 | | | | | | | yes | | yes | | |
| NUDT16P | Island;N_Shore | Body;TSS200;TSS1500 | chr3:131080516-131081228 | 712 | | | 2 | | | | | 0 | 1 | | | 1 | | | | | yes | | |
| TMEM108 | N_Shore;Island | TSS1500;TSS200 | chr3:132756721-132757267 | 546 | | | 2 | | 1 | | | 0 | 3 | | | | | | | | yes | | |
| BFSP2 | NA | Body | chr3:133191270-133191407 | 137 | | | 1 | | | | | | 1 | | | 2 | | yes | | | yes | yes | yes |
| TF | N_Shore;Island | TSS1500;TSS200 | chr3:133464651-133465360 | 709 | | 1 | | | | | | 0 | 2 | 1 | | 1 | | yes | | | yes | | |
| SRPRB | N_Shore | TSS1500 | chr3:133524006-133524362 | 356 | | | | | | | | | | | | | | | | yes | yes | | |
| AMOTL2 | Island | Body | chr3:134083080-134083286 | 206 | 1 | | | 1 | | | | | | | | | | | yes | | | | |
| KY | N_Shore | Body | chr3:134369339-134369661 | 322 | | | 1 | | | | | | | | | | | | | | | | |
| PPP2R3A | N_Shore | TSS1500 | chr3:135684107-135684163 | 56 | | | | | 1 | 1 | | | 1 | | | | | | yes | | | yes | yes |
| PCCB | S_Shore | Body | chr3:135969916-135969981 | 65 | | | 1 | | | | | 0 | 1 | | | | | | | | | | |
| TMEM22 | N_Shore | TSS1500 | chr3:136537169-136537752 | 583 | | | 8 | | | | | | 1 | | | | | | | yes | | | |
| TMEM22 | Island | 5'UTR | chr3:136538801-136538990 | 189 | | | | | | | | 0 | | | | | | | | | | | |
| SOX14 | Island | TSS1500;TSS200 | chr3:137483328-137484586 | 1258 | | | | | | | | | | | | | | | | | | | |
| CLDN18 | NA | TSS200 | chr3:137717580-137718043 | 463 | | 1 | 1 | 1 | | | | | | | | | | | | | yes | | |
| CLDN18 | N_Shore | TSS1500;Body;TSS200 | chr3:137728712-137728997 | 285 | | | 1 | 1 | | 1 | | | 1 | | | | | yes | | | | | |
| DBR1 | Island | 1stExon;5'UTR | chr3:137893707-137893743 | 36 | | | 4 | | | | | 0 | | | | | | yes | yes | | | yes | |
| TXNDC6 | S_Shore | TSS200 | chr3:138048926-138049011 | 85 | | | 3 | | | | | | | | | | | | yes | | yes | | |
| ESYT3 | N_Shore | TSS1500 | chr3:138152837-138153170 | 333 | | | 15 | | | | | 0 | 1 | | | | | | | | | | yes |
| FAIM | N_Shore | TSS200;TSS1500 | chr3:138327370-138327728 | 358 | | | | | | | | 0 | 2 | | | | | yes | yes | | | | yes |
| FOXL2;C3orf72 | Island | TSS200;5'UTR;1stExon | chr3:138665984-138666473 | 489 | | | | | | | | | 1 | | | | | | | yes | | yes | yes |
| C3orf72 | Island | Body | chr3:138668832-138669434 | 602 | | | 6 | | | | | | | | | | | | | | | | |
| PRR23A | Island | TSS200;TSS1500 | chr3:138725153-138725425 | 272 | | | 4 | | | | | | 1 | | | 1 | | | yes | | | | |
| PRR23B | Island | 1stExon;5'UTR;TSS200 | chr3:138738869-138740022 | 1153 | | | 11 | | | | | 0 | 4 | | | | | yes | | | | | |
| PRR23C | Island | 1stExon;5'UTR;TSS200 | chr3:138763424-138763910 | 486 | | 1 | 13 | | | | | 0 | 6 | | | 3 | | | | | yes | | |
| RBP2 | NA | TSS1500 | chr3:139196491-139196813 | 322 | | | | | | | | | | | | | | | yes | | | | |
| RBP1 | Island;S_Shore | Body;TSS1500 | chr3:139257713-139259435 | 1722 | | | 10 | 1 | | | | | 2 | | | | | yes | yes | | yes | | |
| NMNAT3 | NA | 5'UTR | chr3:139346734-139346955 | 221 | | | 9 | | | | | | 1 | | | | | yes | yes | | | | |
| NMNAT3 | S_Shore | TSS200 | chr3:139396924-139397363 | 439 | | | | | | | | | 1 | | | | | | yes | | yes | yes | yes |
| TRIM42 | NA | TSS200 | chr3:140396803-140397089 | 286 | | | | | | | | | | | | | | yes | | | | | |

| | | | | | | | | | | | | | | | | | | | | | | | |
|---|---|---|---|---|---|---|---|---|---|---|---|---|---|---|---|---|---|---|---|---|---|---|---|
| TRIM42 | Island | Body | chr3:140401765-140402167 | 402 | | | 6 | | | | | | | | | | | yes | | | yes | | |
| SPSB4 | Island;N_Shore | Body | chr3:140813769-140814133 | 364 | 3 | | | 1 | | | | | 1 | | | 3 | | | yes | yes | | | yes |
| ZBTB38 | NA | TSS1500;TSS200 | chr3:141042705-141043480 | 775 | | | | 1 | | | | | | | | | | | yes | | | | |
| ZBTB38 | NA | 5'UTR | chr3:141087187-141087363 | 176 | | | | 2 | | | | 0 | 2 | | | 3 | | | | | | | |
| GRK7 | N_Shore | TSS1500;TSS200 | chr3:141495859-141497111 | 1252 | | | 2 | | | | | | 1 | | | | | yes | | | | | |
| CHST2 | N_Shore | TSS1500 | chr3:142837472-142837564 | 92 | | | | | | | | 0 | 1 | | | | | | | | | | |
| CHST2 | Island | Body | chr3:142839903-142840240 | 337 | | 1 | | | | | | | | | | | | | | | | | yes |
| PLOD2 | Island;S_Shore | 5'UTR;1stExon;TSS200;TSS1500 | chr3:145878963-145879710 | 747 | | | | | 1 | | | 0 | 5 | 1 | | 3 | | | | | | | |
| PLSCR4 | Island;S_Shore | TSS200;5'UTR;TSS1500 | chr3:145968700-145969718 | 1018 | | | | | | 1 | | 0 | | | | | | yes | | yes | | | |
| PLSCR2 | Island;N_Shore | 5'UTR | chr3:146187037-146187740 | 703 | | | | | | | | 0 | 2 | | | 1 | | yes | | | | | |
| PLSCR1 | N_Shore | 5'UTR | chr3:146261941-146262761 | 820 | | | 2 | | | | | 0 | 1 | 1 | 1 | 3 | | yes | yes | | yes | | |
| ZIC4 | N_Shelf | 3'UTR;Body | chr3:147105899-147107109 | 1210 | | | | | | | | | | | | | | | | | | | |
| ZIC4 | Island;N_Shore | Body;TSS200;TSS1500 | chr3:147108512-147114406 | 5894 | | | | | | | | | | | | | | | | | | | |
| ZIC4;ZIC1 | N_Shelf;N_Shore;Island | 1stExon;5'UTR;TSS1500;TSS200 | chr3:147124363-147128123 | 3760 | | 3 | 25 | | | | | 0 | 15 | | | 1 | | | | | | yes | yes |
| ZIC1 | N_Shore | Body | chr3:147129778-147129952 | 174 | | 1 | 6 | | | | | | | | | | | | | | | | yes |
| ZIC1 | Island | Body | chr3:147131069-147131860 | 791 | | | 5 | 1 | | | | | | | | | | | | | | yes | |
| AGTR1 | S_Shore | 5'UTR | chr3:148415960-148416308 | 348 | | | | | | | | | | | | 1 | | | | | | | |
| HLTF | Island;S_Shore | TSS1500 | chr3:148804541-148804746 | 205 | | | 4 | | | | | | | | | | | | | | | | |
| HPS3 | Island | TSS200 | chr3:148847273-148847295 | 22 | | | 1 | | | | | 0 | 1 | | | 1 | | | | | | | |
| TM4SF18 | NA | Body | chr3:149051159-149051523 | 364 | | | 1 | | | | | | | | | | | yes | | | | | yes |
| TM4SF1 | NA | 1stExon;TSS200 | chr3:149095283-149096029 | 746 | | | | | | | | | | | | | | | | | | | yes |
| WWTR1 | N_Shore;Island | Body | chr3:149374066-149374914 | 848 | | | | | 2 | | | | 2 | 1 | 1 | | | | | | | yes | |
| RNF13 | Island;N_Shore | TSS200;TSS1500 | chr3:149530133-149530341 | 208 | | | | | | | | | 2 | | | | 1 | | | | | | yes |
| CLRN1;CLRN1OS | NA | 1stExon;TSS1500;TSS200 | chr3:150690251-150690585 | 334 | | | 8 | | | | | | | | | | | | | | | yes | |
| P2RY14;MED12L | NA | 1stExon;Body;5'UTR | chr3:150996078-150996563 | 485 | | | 2 | | 1 | | | | 1 | | | | | | | | | | |
| P2RY12;MED12L | NA | 1stExon;5'UTR;Body | chr3:151058527-151058837 | 310 | | | | | | | | | 1 | | | | | | | | | | yes |
| P2RY12;MED12L | NA | TSS200;Body | chr3:151102703-151102879 | 176 | | | | | | | 1 | 0 | 1 | | | | | yes | | | | | yes |
| P2RY1 | Island | TSS200;5'UTR;1stExon | chr3:152552615-152553711 | 1096 | | 2 | | | | | | | | | | | | | | | | | |
| RAP2B | Island | 1stExon | chr3:152880982-152881217 | 235 | | | | | | | | | | 1 | | | | | | | | yes | |
| SGEF | Island;S_Shore | Body | chr3:153840379-153840718 | 339 | | | | | | | | 0 | | | | 1 | | | | | | | |
| DHX36 | S_Shore;N_Shore;Island | TSS1500;Body;1stExon;5'UTR | chr3:154041542-154042778 | 1236 | | | 3 | | | | | | 1 | | | | | yes | | | | | yes |
| MME | N_Shore;Island | TSS1500;TSS200;5'UTR | chr3:154796861-154797867 | 1006 | | 1 | | | | | | | 1 | | | | | | | | | yes | yes |
| PLCH1 | NA | 5'UTR;TSS200 | chr3:155394109-155394428 | 319 | | | 27 | | 1 | | | | 1 | | 1 | | | | | | | yes | |
| PLCH1 | NA | 5'UTR;1stExon | chr3:155421970-155422103 | 133 | | | 1 | | | | | | | | | | | | | | | | |
| SLC33A1 | N_Shelf | Body | chr3:155570275-155570809 | 534 | | | | 1 | | | | | | | | | | yes | | | | | yes |
| KCNAB1 | NA | TSS200 | chr3:155838317-155838625 | 308 | | | 4 | | | | | 0 | | | | | | | | | | | yes |
| KCNAB1 | N_Shore | TSS200;Body | chr3:156008770-156009319 | 549 | | 1 | | | | | | | | | | | | | | | | | |
| SSR3 | S_Shore | TSS1500 | chr3:156273297-156273458 | 161 | | | 7 | | | | | 0 | 1 | 1 | | 9 | | yes | yes | | | | |
| VEPH1;PTX3 | Island | Body | chr3:157155401-157155998 | 597 | | | | | | | | | 1 | | | | | | | | | yes | |
| VEPH1 | NA | 5'UTR;1stExon;TSS200 | chr3:157217187-157217842 | 655 | | | 15 | | | | | | | | | | | yes | | | | yes | |
| VEPH1 | NA | nan | chr3:157221198-157221260 | 62 | | | | | | | | | | | 1 | | | | | | | | |
| C3orf55 | Island;N_Shore | TSS200;TSS1500 | chr3:157260627-157261202 | 575 | | 2 | | | | | | | 2 | | | | | yes | | yes | | yes | |
| SHOX2 | Island;N_Shore | Body | chr3:157821262-157822072 | 810 | | | 31 | | | | | | | | | | | | | | | | |
| SHOX2 | Island | Body | chr3:157823158-157823479 | 321 | | | | | | | | | 1 | | | | | | | | | yes | yes |
| SHOX2 | S_Shore | TSS1500 | chr3:157824714-157825196 | 482 | | | 2 | | | | | | | | | | | | | | | yes | |
| MLF1 | N_Shore;Island | TSS1500;TSS200 | chr3:158288680-158289160 | 480 | | | | | | | | | 3 | | | | | yes | | | yes | | yes |
| GFM1;LXN | NA | Body;1stExon;5'UTR;TSS200;TSS1 | chr3:158390074-158390821 | 747 | 1 | | 20 | 1 | | | | | 1 | | | 1 | | yes | | yes | | | |
| MFSD1 | N_Shore | TSS1500 | chr3:158518888-158519176 | 288 | | | 7 | | | | | | | | | 1 | | | yes | | | | |
| SCHIP1 | NA | TSS200 | chr3:158991537-158991638 | 101 | | | | | | | | | 1 | | | | | | | | | | |
| SCHIP1 | NA | Body | chr3:159557552-159558031 | 479 | | 1 | | | | | | 0 | 2 | 1 | | 3 | | | | | | yes | |
| TRIM59 | Island | TSS200;1stExon;5'UTR | chr3:160167507-160167789 | 282 | | 1 | | | | | | | 1 | | | | | yes | | | | | |
| PPM1L | S_Shore | Body | chr3:160475035-160475336 | 301 | | 1 | 16 | | 2 | | | 0 | 2 | | | 1 | | | | | | yes | |
| PPM1L | NA | Body | chr3:160580235-160580380 | 145 | | | | | | | | | 1 | | | | | | | yes | | yes | yes |
| B3GALNT1 | S_Shore | TSS200;TSS1500 | chr3:160823315-160823505 | 190 | | | | | | | | | | | | | | | | | | | |
| SLITRK3 | NA | 5'UTR;TSS200;TSS1500 | chr3:164914150-164915115 | 965 | | | 2 | | | | | 0 | 4 | | | | | | | | | yes | yes |
| ZBBX | Island | 5'UTR | chr3:167097928-167098267 | 339 | | 1 | | | | | | | | | | | | yes | | | | yes | |
| MECOM | N_Shore | 5'UTR;Body | chr3:168863860-168864123 | 263 | | | 2 | | | | | | | | | | | | | | | | |
| MECOM | N_Shore | Body;TSS1500 | chr3:168866705-168866814 | 109 | | | 1 | | | | | | | | | | | | | | | yes | |
| MECOM | N_Shore | Body | chr3:169377725-169379010 | 1285 | | 1 | 8 | | | 2 | | | 1 | | 1 | | | | | | | yes | yes |
| MECOM | S_Shore | TSS200;TSS1500 | chr3:169381574-169382106 | 532 | | | | | | | | | | | | | | | | | | yes | yes |
| TERC | S_Shore | TSS1500 | chr3:169483122-169483295 | 173 | | | 14 | | | | | | | | | | | yes | | | | | |
| LRRC34 | S_Shore | TSS1500 | chr3:169531663-169531864 | 201 | | | 3 | | | | | 0 | | | | 1 | | yes | | yes | | | |
| LOC100128164;SEC62 | N_Shore | Body;TSS1500 | chr3:169683976-169684239 | 263 | | | 9 | | | | | | | | | | | yes | | | yes | yes | |
| GPR160 | Island | TSS200 | chr3:169755595-169755693 | 98 | | | | | | | | 0 | | | | | | yes | | | yes | yes | |
| GPR160 | NA | 5'UTR | chr3:169781404-169781717 | 313 | | | 13 | | | | | | 1 | | | | | yes | | | | | yes |
| PHC3 | S_Shore | TSS1500 | chr3:169900418-169900473 | 55 | | | | | 1 | | | | | | | | | yes | | | | yes | |
| PRKCI | Island | TSS200 | chr3:169940172-169940769 | 597 | | | | | | | | | 1 | | | | | | | | | | |
| CLDN11 | N_Shore;Island | TSS1500;1stExon;Body | chr3:170136159-170138205 | 2046 | | 1 | 5 | | | | | 0 | 1 | | | | | yes | yes | | | yes | yes |
| SLC7A14 | N_Shore;S_Shore | 5'UTR;1stExon;TSS200 | chr3:170302404-170304045 | 1641 | | 1 | 4 | | | | | | 1 | | | | | yes | yes | | | | |
| EIF5A2 | Island | TSS1500 | chr3:170626725-170627029 | 304 | | | 2 | | | | | | | | | | | | | | | yes | |
| SLC2A2 | N_Shore;Island | TSS1500 | chr3:170746029-170746586 | 557 | | | 3 | | | | | 0 | 2 | | | 1 | | yes | yes | | | | |
| TNIK | N_Shelf;N_Shore | Body | chr3:171175713-171176016 | 303 | | | 12 | | | | | | | | | | | | yes | | | | |
| TMEM212 | NA | TSS1500 | chr3:171559851-171560011 | 160 | | | | | | | | | 1 | | | | | yes | yes | | | | |
| TMEM212 | NA | TSS200 | chr3:171561060-171561201 | 141 | | | 10 | | | | | | | | | | | | | | | | |
| GHSR | Island;N_Shore | 1stExon;TSS200;TSS1500 | chr3:172165517-172167810 | 2293 | | 2 | | | | | | 0 | 2 | | | 1 | | yes | | | | yes | yes |
| TNFSF10 | NA | 1stExon | chr3:172241069-172241384 | 315 | | | 1 | | | | | | | | | | | | | yes | | | |
| SPATA16 | NA | 1stExon;5'UTR | chr3:172858969-172859049 | 80 | | | | | | | | | | | | | | | yes | | | yes | |
| NLGN1 | Island;S_Shore | TSS1500;TSS200 | chr3:173115536-173116248 | 712 | | | | | | | | | 4 | | | | | | | | | | |
| NLGN1 | NA | 5'UTR | chr3:173302303-173303193 | 890 | | 1 | | | | | | | | | | | | | | | | yes | |
| NAALADL2 | NA | Body | chr3:175338899-175338932 | 33 | | | | | | | | | 1 | | | | | | | yes | yes | | |
| KCNMB2 | NA | TSS1500;5'UTR | chr3:178275369-178275698 | 329 | | | 2 | | | | | | 1 | | | | | yes | | | yes | | |
| KCNMB3 | N_Shore;Island | TSS1500;Body | chr3:178978849-178979013 | 164 | | | 16 | | | | | 0 | 3 | 1 | | 5 | | | | | yes | | yes |
| KCNMB3 | NA | 5'UTR;1stExon;nan;TSS200 | chr3:178984747-178984973 | 226 | | | 10 | | | | | 0 | 3 | 1 | 1 | 2 | | | yes | | yes | | yes |
| GNB4 | N_Shore | 5'UTR | chr3:179168677-179168798 | 121 | | | | | 1 | | | | | | | | | | | | | | |
| PEX5L | Island | TSS200 | chr3:179754533-179754615 | 82 | | | | | 2 | | | | | | | | | | | | | | yes |
| TTC14 | N_Shore | TSS1500 | chr3:180319597-180319840 | 243 | | | | | | | | | 1 | | | | | yes | | | yes | | |
| SOX2OT | NA | TSS1500 | chr3:181326706-181327125 | 419 | | | 2 | | | | | | | | | 1 | | yes | | | | | yes |
| SOX2OT | Island | Body | chr3:181413847-181414848 | 1001 | | | 2 | | | | | | | | | | | | | | | | yes |
| SOX2OT | NA | Body | chr3:181421427-181422217 | 790 | | | 3 | | | | | | 2 | 1 | | | | | | | | | |
| SOX2OT;SOX2 | N_Shelf;N_Shore;Island | Body;TSS1500;TSS200;1stExon | chr3:181428046-181431306 | 3260 | | 1 | 24 | | | | | | 7 | 5 | | | | | | | | | yes |
| SOX2OT | S_Shore;N_Shore | Body | chr3:181437145-181438328 | 1183 | 1 | | 2 | | | | | 0 | | | | | | yes | | | | | |
| SOX2OT | N_Shore;Island;S_Shore;N_She | Body | chr3:181441345-181445647 | 4302 | 1 | | 16 | | | | | | 4 | 1 | | | | | | | | yes | |
| MCCC1 | N_Shore;Island;S_Shore | Body;1stExon;5'UTR;TSS200;TSS1 | chr3:182816738-182817626 | 888 | | | 3 | | 1 | | | 0 | 8 | 4 | 1 | 4 | | yes | | | | | |
| MCF2L2;B3GNT5 | N_Shore | Body;TSS1500;TSS200 | chr3:182970675-182970850 | 175 | 6 | | 7 | | | | | | 1 | | | | | yes | | | | | yes |
| MCF2L2;B3GNT5 | Island | Body;5'UTR | chr3:182972625-182973032 | 407 | | | | 1 | | | | | 1 | | | | | | | | | | |
| KLHL6 | Island | 3'UTR;Body | chr3:183209713-183209993 | 280 | | | | | | | | | | | | | | yes | | | | | |
| KLHL6 | NA | 5'UTR;1stExon;TSS200 | chr3:183273245-183273894 | 649 | | | 6 | 1 | | | | | 1 | 1 | 1 | | | yes | | | | | |
| MAP6D1 | Island | Body | chr3:183542750-183543862 | 1112 | | | | 1 | | | | | | | | | | | yes | | | | yes |
| HTR3D | NA | Body;TSS1500;5'UTR | chr3:183750284-183750690 | 406 | | | | | | | | | | | | | | yes | | | | | |

| | | | | | | | | | | | | | | | | | | | | | | | | | | | |
|---|---|---|---|---|---|---|---|---|---|---|---|---|---|---|---|---|---|---|---|---|---|---|---|---|---|---|---|
| HTR3C | NA | TSS1500 | chr3:183770589-183770677 | 88 | | | | | | | | | | | | 1 | | yes | | | | | | | | | |
| HTR3E | NA | TSS200 | chr3:183817853-183818368 | 515 | | | 2 | | | | | | 1 | | | | | yes | | | yes | | yes | | | | |
| DVL3 | Island;S_Shore | Body;3'UTR | chr3:183888141-183889898 | 1757 | | | | | | | | | 2 | | | | | yes | | | | | | | | | |
| AP2M1 | N_Shore | TSS1500 | chr3:183891928-183892060 | 132 | | | | | | | | | | | | | | | yes | | | | | | | | |
| AP2M1 | S_Shore | 5'UTR | chr3:183894727-183894778 | 51 | | 1 | 25 | 1 | | | | | 1 | | | | | | | | | | | | | | |
| ABCF3 | N_Shore | TSS1500 | chr3:183903129-183903315 | 186 | | | 3 | | | | | | | | | | | yes | yes | | | | | | | | |
| VWA5B2;MIR1224 | N_Shore;Island | Body;TSS1500;TSS200 | chr3:183958478-183959853 | 1375 | | | 9 | | | | | 0 | 4 | | | 1 | | | yes | | yes | | | | | | |
| ECE2 | N_Shore | Body;TSS1500;TSS200 | chr3:183993080-183993726 | 646 | | | 4 | | | | | | 1 | | | | | | | | | yes | yes | | | | |
| EIF4G1 | S_Shore;S_Shelf | Body;5'UTR | chr3:184035178-184035527 | 349 | | | 3 | 1 | | | | | | | | | | | | | yes | | | | | | |
| EIF4G1 | NA | Body;TSS1500;5'UTR | chr3:184037459-184038317 | 858 | | | | | | | | | | | | | | | | | | yes | | | | | |
| FAM131A | N_Shore | 5'UTR;TSS200 | chr3:184055123-184055379 | 256 | | 1 | | | | | | | | | | | | | | | | | | | | | |
| FAM131A | Island | 5'UTR;Body | chr3:184056553-184056843 | 290 | | | | | | | | | 1 | | | | | yes | | | | | yes | | | | |
| THPO | N_Shelf | 5'UTR | chr3:184095505-184095976 | 471 | | | | | | | | | 1 | | | | | | | | | | | | | | |
| CHRD | N_Shore;Island | Body | chr3:184098790-184099432 | 642 | | | 2 | 1 | | | | | | | | | | yes | | | | | yes | | | | |
| CHRD | N_Shore;Island | Body | chr3:184104115-184104688 | 573 | | | | 1 | | | | 0 | 2 | | | 1 | | yes | | | | | | | | | |
| EPHB3 | N_Shelf | Body | chr3:184297380-184297522 | 142 | | | | | 1 | | | | | | | | | yes | yes | | | | | | | | |
| EHHADH | Island;S_Shore | 5'UTR;Body;1stExon;TSS200 | chr3:184971652-184972153 | 501 | | | 1 | | | | | 0 | 2 | | | 1 | | yes | yes | yes | | yes | | | | | |
| MAP3K13 | NA | TSS1500;TSS200;1stExon;5'UTR | chr3:185080218-185081403 | 1185 | | | 2 | 1 | 2 | | | | 1 | | | | | | | | | yes | | | | | |
| MAP3K13 | NA | 5'UTR | chr3:185136277-185136588 | 311 | | | | 1 | | | | | 1 | | | | | yes | yes | | | | | | | | |
| DGKG | Island;N_Shore | Body | chr3:185911316-185912486 | 1170 | | | 4 | | | | | 0 | 2 | | | 1 | | yes | yes | | | | yes | | | | |
| AHSG | NA | TSS1500 | chr3:186330026-186330067 | 41 | | | | | | | | | | | | | | | yes | | | | | | | | |
| HRG | NA | TSS1500 | chr3:186383386-186383874 | 488 | | | 5 | 1 | | | | | | | | | | yes | yes | yes | | | | | | | |
| SNORA4;EIF4A2 | S_Shelf | TSS200;Body | chr3:186505240-186505713 | 473 | | | | | | | | 0 | 1 | | | | | yes | | | | | | | | | |
| ADIPOQ | NA | TSS1500;TSS200 | chr3:186560083-186560617 | 534 | | | | | | | | | | | | | | | yes | | | | | | | | |
| RPL39L | Island | TSS200 | chr3:186857284-186857475 | 191 | | 1 | | | | | | | | | | | | yes | | | | | yes | | | | |
| RTP1 | N_Shelf | TSS1500 | chr3:186914980-186915320 | 340 | | | | | | | | | | | | | | yes | | | | | | | | | |
| RTP4 | NA | TSS200 | chr3:187086147-187086530 | 383 | | | 3 | | | | | | | | | | | | | yes | | | | | | | |
| SST | N_Shore;S_Shore | Body;TSS1500 | chr3:187387555-187389513 | 1958 | | 1 | | | | | | | 3 | | | | | | | | | yes | | | | | |
| BCL6 | N_Shore | 5'UTR;TSS1500;1stExon;TSS200 | chr3:187453721-187454482 | 761 | 3 | | 4 | | | | | | | | | | | yes | | | | | | | | | |
| LPP | N_Shore | TSS1500 | chr3:187870863-187871034 | 171 | | | | | | | | 0 | 1 | | | | | yes | | | | | | | | | |
| LPP | NA | TSS200;5'UTR | chr3:187930521-187930704 | 183 | | | 3 | | | | | | | | | | | yes | | | | | yes | | | | |
| TP63 | NA | TSS1500 | chr3:189348936-189349129 | 193 | | | 24 | | | | | | 1 | | | | | | | | | yes | | | | | |
| LEPREL1 | Island;S_Shore | 5'UTR;1stExon;TSS200 | chr3:189838376-189839095 | 719 | | | | | | | | | 2 | | | | | yes | yes | | | yes | yes | | | | |
| CLDN16 | NA | TSS200 | chr3:190105762-190105986 | 224 | | | | | | | | | 1 | | | | | yes | | | | | | | | | |
| LOC647309 | Island | TSS200 | chr3:190580644-190580923 | 279 | | 1 | | | | | | | | | | | | | yes | | | yes | yes | | | | |
| UTS2D;CCDC50 | chr3:191046948-191047710;S | S_Shore;TSS200;Body;1stExon;5'U | chr3:191048308-191049285 | 977 | | | 5 | | | | | | 2 | | 1 | | | | | | | yes | | | | | |
| FGF12 | Island | 5'UTR;1stExon;Body | chr3:192126023-192126960 | 937 | | 1 | | | 5 | | | | | | | | | | | | | yes | yes | | | | |
| FGF12 | Island | Body | chr3:192232863-192233231 | 368 | | | | | | | | | 1 | | | | | yes | | | | | yes | | | | |
| FGF12 | NA | Body | chr3:192289245-192289293 | 48 | 1 | | 22 | | | | 1 | 0 | 1 | 1 | | 12 | 1 | | | | | | yes | | | | |
| FGF12 | NA | TSS200 | chr3:192445500-192445594 | 94 | | | | | | | | | 1 | | | | | | | | | | yes | | | | |
| C3orf59 | N_Shore | Body | chr3:192634059-192634364 | 305 | | | 2 | | | | | | | | | | | | | | | yes | | | | | |
| ATP13A5 | NA | Body;1stExon;TSS200 | chr3:193096009-193097361 | 1352 | | | 4 | | | | | | 1 | 1 | | | | yes | | | | yes | | | | | |
| ATP13A4 | NA | Body;1stExon;5'UTR | chr3:193227429-193272869 | 45440 | | | 1 | | 1 | | | | 1 | 1 | | | | yes | yes | yes | yes | yes | yes | | | | |
| CPN2 | NA | TSS1500;5'UTR;TSS200 | chr3:194071770-194072416 | 646 | | | | 2 | | | | | | | | | | yes | yes | | yes | yes | yes | | | | |
| LRRC15 | NA | TSS200 | chr3:194090477-194091167 | 690 | | | | | | | | | 2 | | | | | | | | | yes | | | | | |
| GP5 | N_Shore;Island;S_Shore | 3'UTR;Body;5'UTR;TSS1500 | chr3:194117018-194121120 | 4102 | | 1 | | 1 | | | | | 1 | | | | | yes | yes | | | yes | yes | | | yes | |
| TMEM44 | NA | Body | chr3:194319463-194320017 | 554 | | | 13 | | | | | | | | | | | yes | | | | | | | | | yes |
| FAM43A | N_Shore | TSS1500 | chr3:194405973-194407045 | 1072 | | | | | | | | 0 | | | | | | | | | | | yes | | | | |
| FAM43A | chr3:194406440-194409045;Isl | Island;1stExon;3'UTR | chr3:194408516-194408965 | 449 | | | | | | | | | 2 | 2 | 1 | | | | | | | | yes | | | | |
| C3orf21 | NA | Body | chr3:194815259-194815434 | 175 | | | | | | | | | 1 | | | | | yes | | | yes | yes | | | | | |
| C3orf21 | NA | Body | chr3:194826411-194826585 | 174 | | 1 | | | | | | 0 | 1 | 1 | | | | | | | yes | | | | | | |
| C3orf21 | NA | Body | chr3:194833431-194834008 | 577 | | | 9 | | | | | | 1 | 1 | | | | yes | | | | yes | | | | | |
| C3orf21 | NA | Body | chr3:194868507-194868843 | 336 | | | 7 | | 1 | | | 0 | 1 | | | | | yes | | | | | | | | | |
| C3orf21 | NA | Body | chr3:194930170-194930312 | 142 | | | 4 | | | | | | | | | | | | | | | | | | | | |
| C3orf21 | NA | Body | chr3:194967958-194968024 | 66 | 2 | | | | | | | | | | | | | | | yes | | | | | | | |
| APOD | NA | 1stExon;5'UTR | chr3:195310883-195311591 | 708 | | | 1 | 1 | | | | | 2 | | | | | yes | yes | | | yes | | | | | |
| SDHAP2 | Island | TSS1500 | chr3:195384694-195384742 | 48 | | | | | | | | 0 | | 1 | | 1 | | yes | yes | | yes | yes | | | | | |
| MIR570 | NA | TSS1500 | chr3:195424838-195425337 | 499 | | | | | | | | | | | | | | yes | yes | yes | | | | | | | |
| MUC4 | Island | Body | chr3:195477778-195477968 | 190 | | 1 | | | | | | | | | | | | | | | | yes | | | | | |
| MUC4 | N_Shore;Island;S_Shore | Body | chr3:195488493-195490309 | 1816 | | | | | | | | | | | | | | | | | | | | | | | |
| MUC4 | NA | Body | chr3:195528460-195528709 | 249 | | | 6 | | | | | | | | | | | | yes | | | | | | | | |
| MUC4 | NA | Body | chr3:195530091-195530654 | 563 | | | 14 | | | | | | 1 | | | | | yes | | yes | | | | | | | |
| MUC4 | NA | Body | chr3:195536520-195537065 | 545 | | 1 | | | | | | | | | | | | | | | | | | | | | |
| MUC4 | NA | 1stExon;5'UTR | chr3:195538675-195539313 | 638 | | | | | | | | 0 | | | | | | yes | | | | | yes | | | | |
| TNK2 | N_Shore | Body | chr3:195599898-195600410 | 512 | 1 | | | | | | | | | | | | | yes | yes | | | | yes | | | | |
| TNK2 | N_Shore | Body | chr3:195613212-195613641 | 429 | | | | | | | | | | | | | | yes | | | yes | | | | | | |
| TNK2 | S_Shelf | Body;5'UTR | chr3:195616294-195616361 | 67 | | 1 | | | | | | | | | | | | | | | yes | | | | | | |
| TNK2 | S_Shore | TSS1500;5'UTR | chr3:195623338-195623792 | 454 | | | | 1 | | | | | | | | | | | | | | | yes | | | | |
| ZDHHC19 | NA | Body | chr3:195929389-195929544 | 155 | | | 5 | | | | | | | | | | | yes | yes | | | | | | | | |
| ZDHHC19 | Island | Body | chr3:195934213-195934604 | 391 | | 1 | 1 | | | | | | | | | | | yes | | | | yes | yes | | | | |
| ZDHHC19 | S_Shelf | 5'UTR;1stExon | chr3:195938201-195938668 | 467 | | | | | | | | | | | | | | yes | | | | | | | | | |
| OSTalpha | NA | TSS1500;TSS200;1stExon;5'UTR | chr3:195942717-195943713 | 996 | | | 7 | | | | | | 4 | | | | | yes | yes | | | | | | | | |
| OSTalpha | NA | Body | chr3:195946851-195947062 | 211 | | | | | | | | 0 | | | | | | yes | | | yes | | | | | | |
| TM4SF19 | NA | 5'UTR;1stExon;TSS200;TSS1500 | chr3:196065106-196065688 | 582 | | 1 | 31 | | | | | | 3 | | 1 | | | yes | yes | yes | | | | | | | |
| LRRC33 | NA | 5'UTR | chr3:196373543-196373751 | 208 | | | | | | | 1 | 0 | 1 | | | 1 | | | yes | | | | | | | | |
| LRRC33 | Island | Body | chr3:196386877-196387120 | 243 | | | | | | | | | | | | | | yes | | | | | | | | | |
| PIGZ | Island | 5'UTR | chr3:196693809-196694153 | 344 | | | | | | | | | 1 | | | | | | | | | | | | | | |
| MFI2 | NA;N_Shelf | Body | chr3:196750426-196751186 | 760 | | 1 | 4 | | | | | | 1 | | | | | | | yes | | | | | | | |
| DLG1 | Island | TSS1500 | chr3:197025853-197026013 | 160 | | | | | | | | | | | | | | yes | | | | yes | yes | | | | |
| MIR922;KIAA0226 | NA | Body;3'UTR;TSS200;TSS1500 | chr3:197401399-197402549 | 1150 | 1 | 1 | | | 4 | | | | 2 | | | | | yes | | | | | yes | | | | |
| FYTTD1 | NA | Body | chr3:197500745-197500859 | 114 | | | | | | | | | 1 | | | | | yes | | | | | | | | | |
| IQCG | NA | Body | chr3:197639792-197639960 | 168 | | | | | | | | | 1 | | | | | yes | | | | yes | | | | | |
| IQCG | NA | Body | chr3:197663860-197664106 | 246 | | | | | | | | | 1 | | | | | yes | | | yes | | | | | | |
| LOC348840 | Island | TSS1500 | chr3:197807842-197808169 | 327 | | | | | | | | | 1 | | | | | | | | | | | | | | |
| ZNF718;ZNF595 | N_Shore | TSS200 | chr4:53109-53146 | 37 | | | | | | | | | | | | | | | | | yes | yes | yes | | | | |
| ZNF718 | N_Shore | Body | chr4:124232-124622 | 390 | | | | | | | | 0 | 1 | | | 2 | 1 | yes | | yes | | yes | | | | | |
| ZNF876P | N_Shore;Island | TSS1500;TSS200;Body | chr4:206112-206877 | 765 | | 1 | | | | | | | 3 | | | | | | | yes | yes | | | | | | |
| PIGG | NA | Body | chr4:503198-503399 | 201 | | | | | | | | | | | | | | | yes | yes | | | | | | | |
| PDE6B | N_Shore | TSS1500;TSS200;1stExon;5'UTR | chr4:618114-619659 | 1545 | | | 11 | | | | | | 1 | | | | | | yes | | | yes | | | | | |
| PDE6B | NA | Body | chr4:640348-640594 | 246 | | | | | | | | 0 | 1 | 1 | | 2 | | yes | yes | | | | | | | | |
| PDE6B | N_Shelf | Body | chr4:642691-642885 | 194 | | | | | | | | | | | | | | | | | yes | | | | | | |
| PDE6B | N_Shore;Island | Body;TSS1500 | chr4:644805-646577 | 1772 | | | 5 | | | | 1 | 0 | 2 | | | 1 | 1 | | yes | | yes | yes | | | | | |
| PDE6B | N_Shore | Body | chr4:649503-650457 | 954 | | | 7 | | | | | | | | | | | yes | | | | | | | | | |
| PDE6B | Island | Body | chr4:652823-652907 | 84 | | | | | | | | | | | | | | | | | | yes | | | | | |
| PDE6B | Island | Body | chr4:657536-658058 | 522 | | | | | | | | | | | | 1 | | yes | | | | | | | | | |
| PDE6B | S_Shelf | Body | chr4:660376-660957 | 581 | | | | | | | | | | | | | | | | | | | | | | | |
| MYL5 | N_Shelf | TSS1500;TSS200 | chr4:671489-671851 | 362 | | | 8 | | | | | | 1 | 1 | | | | | | | yes | yes | | | | | |
| MFSD7 | N_Shore | Body | chr4:681280-681509 | 229 | | | 1 | | | | | | | | | | | | | | | | | | | | |
| MFSD7 | Island;S_Shore | 1stExon;TSS200 | chr4:682834-683240 | 406 | | | 13 | 2 | | | | | 3 | | | | | | | | | yes | | | | | |

| | | | | | | | | | | | | | | | | | | | | | | | | |
|---|---|---|---|---|---|---|---|---|---|---|---|---|---|---|---|---|---|---|---|---|---|---|---|---|
| PCGF3 | Island | 5'UTR | chr4:714341-714516 | 175 | | | | | | | | | 1 | | | | | | | | | | | |
| PCGF3 | N_Shore | 5'UTR | chr4:715865-716282 | 417 | | | 5 | | | | | | | 1 | | | | | | | | yes | | |
| PCGF3 | N_Shelf | Body | chr4:736328-736765 | 437 | | | | | | | | | 1 | | | | | | | | | | | |
| PCGF3 | S_Shelf | Body | chr4:757312-757456 | 144 | 2 | | | | | | | | | | | | | | yes | | | | | |
| CPLX1 | Island;S_Shore | Body;3'UTR | chr4:778827-781272 | 2445 | 2 | | | 1 | 2 | | | 0 | 2 | | | 1 | | | | yes | | | yes | |
| CPLX1 | S_Shelf | Body | chr4:802844-803014 | 170 | | | | | 1 | | | | | | | | | | yes | | | | | |
| CPLX1 | Island;N_Shore | Body | chr4:811981-812884 | 903 | | 1 | 3 | | | | | | | | | | | | | | | | | |
| GAK | N_Shore | Body | chr4:864597-865080 | 483 | 2 | | | | | | | | | | | | | | | | | | | |
| GAK | N_Shelf | Body | chr4:868365-868858 | 493 | | | | | | | | | | | | | | | yes | | | yes | | |
| GAK | N_Shore | Body | chr4:871223-871567 | 344 | | | | | | | | | 2 | | | | | yes | yes | | | | | |
| GAK | Island | Body | chr4:893986-894185 | 199 | | 1 | | | | | | | | | | | | | | | | | yes | |
| GAK | Island | Body | chr4:899775-899984 | 209 | 2 | | 4 | | | | | | | | | | | | | | | yes | | |
| TMEM175 | Island;S_Shore | 5'UTR;Body | chr4:940614-942342 | 1728 | | 1 | 16 | | 1 | | 1 | 0 | 3 | 2 | 2 | 32 | 2 | | | | | | | |
| TMEM175 | Island | Body | chr4:951898-952128 | 230 | | 1 | | | | | | | | | | | | | | | | | | |
| DGKQ | Island | TSS200 | chr4:967528-967541 | 13 | | | | | | | | 0 | | | | | | | | | | yes | yes | |
| SLC26A1 | N_Shore | Body | chr4:978723-979331 | 608 | 1 | | | 1 | | | | 0 | 2 | | | 1 | | yes | yes | | | | | |
| IDUA;SLC26A1 | Island | Body;3'UTR | chr4:981485-981630 | 145 | | | 4 | | | | | | 1 | | | | | | | | | | | |
| SLC26A1;IDUA | S_Shore | 1stExon;Body;5'UTR;TSS200;TSS15 | chr4:987108-987906 | 798 | | | | 1 | 1 | | | | | | | | | yes | | | | | | |
| FGFRL1 | Island | TSS1500 | chr4:1004774-1004783 | 9 | | | | | | | | | 1 | | | | | | | | | | yes | |
| FGFRL1 | Island | 5'UTR;TSS1500;TSS200 | chr4:1005994-1006278 | 284 | 1 | | 1 | | | | | | 1 | | | | | | | | | yes | yes | |
| RNF212 | Island | 1stExon;5'UTR;TSS200 | chr4:1107202-1107983 | 781 | | | 5 | | | | | 0 | 5 | | | | | yes | | | yes | yes | yes | |
| SPON2 | N_Shore;Island | 3'UTR;Body | chr4:1160783-1161653 | 870 | | | 4 | | 3 | | | 0 | 1 | | | | | | | | | | | |
| LOC100130872-SPON2;SPON2 | S_Shore;Island | Body;TSS1500;1stExon;5'UTR;TSS2 | chr4:1166531-1167860 | 1329 | 2 | | 17 | 1 | | | | 0 | 2 | | | | | yes | yes | | yes | | yes | |
| LOC100130872;LOC100130872-SPON2 | NA | Body | chr4:1195845-1196179 | 334 | | | 5 | 1 | | | | | | | | | | yes | yes | yes | | | | |
| LOC100130872;LOC100130872-SPON2 | NA | Body | chr4:1198278-1199037 | 759 | | | | 1 | | | | | | | | | | yes | yes | | | | | |
| LOC100130872;LOC100130872-SPON2 | N_Shore;Island | Body;TSS200;TSS1500 | chr4:1201315-1203653 | 2338 | | | 15 | | 2 | | | | 2 | | | | | | yes | | | yes | | |
| CTBP1 | Island;N_Shore | 3'UTR | chr4:1205447-1206150 | 703 | | | 8 | | | | | | 1 | | | | | | | | | yes | | |
| CTBP1 | N_Shore | Body | chr4:1208147-1208960 | 813 | | | 1 | | | | | | 1 | 1 | | | | | | | | yes | | |
| CTBP1 | NA;N_Shelf | Body | chr4:1214608-1216051 | 1443 | | | | | | | | | 2 | 1 | | | | | yes | | | | yes | |
| CTBP1 | S_Shore | Body | chr4:1225507-1225729 | 222 | | | | | | | | | 1 | 1 | | | | yes | | | | | yes | |
| CTBP1 | S_Shore | Body;5'UTR | chr4:1235292-1235781 | 489 | | | 1 | 1 | | | | | | | | | | | yes | | | | | |
| C4orf42;CTBP1 | Island | TSS1500;TSS200 | chr4:1243849-1244086 | 237 | | | 33 | | | | | 0 | 5 | | | 1 | | | | | | | | |
| MAEA | NA | Body | chr4:1294432-1295078 | 646 | | | 2 | | | | | 0 | 5 | 4 | | 2 | | yes | | | yes | | | |
| MAEA | N_Shore;Island | Body | chr4:1303317-1304048 | 731 | | | 2 | | | | | 0 | 1 | | | 1 | | | yes | | | | | |
| KIAA1530 | S_Shore | Body | chr4:1342459-1342894 | 435 | 2 | | | | | | | | | | | | | | | | yes | | | |
| KIAA1530 | NA | Body | chr4:1352787-1353499 | 712 | | | 8 | | | | | | | | | | | | | | | | | |
| KIAA1530 | NA | Body | chr4:1355321-1355424 | 103 | 1 | | | | | | | | | | | | | | | | | | yes | |
| KIAA1530 | N_Shore | Body | chr4:1363661-1363885 | 224 | | | | | | | | | 1 | | | | | | | | | yes | | |
| KIAA1530 | N_Shore | Body | chr4:1366121-1366597 | 476 | | | | | | | | | 1 | | | | | | yes | yes | | | | |
| FAM53A | N_Shelf | 3'UTR | chr4:1642054-1642257 | 203 | 5 | | 5 | | | | | | | | | | | | | | | | | |
| TMEM129 | N_Shelf | Body | chr4:1719983-1720184 | 201 | | | | 1 | | | | | 1 | | | | | | | | | | | |
| FGFR3 | Island;N_Shore | Body | chr4:1800154-1801294 | 1140 | | | | 1 | | | | | 1 | | | | | | | | yes | | | |
| FGFR3 | Island | Body | chr4:1804007-1804492 | 485 | | | 10 | | | | | | | | | | | | | | | | yes | |
| FGFR3 | N_Shore | Body | chr4:1806162-1806633 | 471 | | | | | | | | | | | | | | | | | | | | |
| LETM1 | S_Shore | Body | chr4:1844724-1844878 | 154 | | | 8 | | | | | | | | | | | yes | | | | | | |
| MIR943;WHSC2 | S_Shore | TSS200;Body | chr4:1988302-1988351 | 49 | | | | | | | | | 1 | | | | | | | | | | | |
| WHSC2 | N_Shore | Body | chr4:1994281-1995036 | 755 | | | 4 | | | | | | | | | | | yes | | yes | | | | |
| NAT8L | Island | 3'UTR | chr4:2066003-2066397 | 394 | | | | | | | | | | | | | | | | | | | yes | |
| MXD4 | NA | 3'UTR;Body | chr4:2252007-2252862 | 855 | 5 | | | | | | | | | | | | | | | | | yes | | |
| ZFYVE28 | S_Shore | Body | chr4:2278261-2278594 | 333 | 4 | | | | | | | | | | | | | yes | yes | | | | | |
| ZFYVE28 | N_Shelf;N_Shore | Body | chr4:2302953-2303860 | 907 | | | 17 | 1 | | 1 | | | | | | | | | yes | | | | | |
| ZFYVE28 | NA | Body | chr4:2321957-2322078 | 121 | | | | | 1 | | | | 1 | | | | | | | | | yes | | |
| ZFYVE28 | Island | Body | chr4:2341196-2341463 | 267 | | | | | | | | 0 | 2 | | | | | yes | | | | yes | | |
| ZFYVE28 | Island;N_Shore | Body | chr4:2366103-2367137 | 1034 | 1 | | 33 | 1 | | | | 0 | 3 | 2 | | 3 | | yes | | | | | | |
| ZFYVE28 | NA | Body | chr4:2395549-2395934 | 385 | | | | | | | | | | | | | | | yes | yes | | | | |
| ZFYVE28 | Island | Body | chr4:2401605-2401980 | 375 | | 1 | | | | | | | | | | | | | yes | yes | | | | |
| ZFYVE28 | S_Shore;S_Shelf;chr4:2401507 | Body;S_Shelf | chr4:2403559-2404284 | 725 | | | | 1 | | | 3 | 0 | 4 | 3 | 2 | 12 | 1 | | | | | | | |
| FAM193A | NA | TSS200 | chr4:2627014-2627142 | 128 | | | | | | | | | 1 | | | | | yes | | | | | | |
| FAM193A | NA | Body | chr4:2696631-2696786 | 155 | | | | | | | | | 1 | | | | | yes | | | | | | |
| SH3BP2 | NA | TSS1500;5'UTR | chr4:2813458-2814122 | 664 | | | 10 | 1 | 1 | | | | 2 | 2 | | | | | | | | | | |
| SH3BP2 | Island | TSS1500;Body;5'UTR;TSS200 | chr4:2819614-2820479 | 865 | | | | 1 | | | | 0 | 3 | | | 13 | | yes | | | | | yes | |
| ADD1 | S_Shore | Body;3'UTR | chr4:2930203-2930361 | 158 | | | | | | | | | | | | | | | | | | | | |
| MFSD10 | N_Shore;Island | Body | chr4:2933421-2934188 | 767 | | | | 1 | 1 | | | | 1 | 1 | | | | | | | | | | |
| C4orf10;NOP14 | NA | Body | chr4:2944550-2944692 | 142 | | | | | | | | | 1 | | | | | | | | | | | |
| GRK4;NOP14 | N_Shore | TSS1500;Body | chr4:2964211-2964239 | 28 | | | 1 | | | | | | | | | | | yes | yes | | | | yes | |
| HTT | NA | Body | chr4:3239305-3239461 | 156 | 1 | | | | | | | | 1 | | | | | yes | yes | | | | | |
| C4orf44 | Island | TSS200 | chr4:3250690-3251105 | 415 | | | 5 | | | | | | | | | | | | | | | | | |
| C4orf44 | Island | Body | chr4:3254969-3255174 | 205 | | | 2 | | | | | | | | | | | yes | yes | | | | | yes |
| C4orf44 | NA | Body | chr4:3265080-3265280 | 200 | | | 1 | | | | | | | | | | | | yes | yes | | | | |
| RGS12 | Island | Body | chr4:3365280-3365342 | 62 | | | | | | | | | 1 | | | | | | | | | yes | | |
| RGS12 | N_Shelf;N_Shore | TSS1500;Body;5'UTR;1stExon | chr4:3371236-3373405 | 2169 | | | 17 | 1 | | | | 0 | 1 | | | | | | | | | | yes | |
| RGS12 | Island | Body | chr4:3374767-3375144 | 377 | | | 5 | | | | | | | | | | | | | | | | | |
| RGS12 | NA | Body;nan | chr4:3387281-3387765 | 484 | | | | 1 | | | | | | | 1 | | | | | | | | | |
| RGS12 | NA | Body | chr4:3392103-3392318 | 215 | | | | | | | | | 1 | | | | | yes | yes | | | | | |
| RGS12 | NA | Body | chr4:3409828-3409927 | 99 | | | | | | | | | | 1 | | | | | | | | | | |
| RGS12 | N_Shore | Body | chr4:3413614-3413706 | 92 | | | 1 | | | | | | | | | | | | | | | | | |
| RGS12 | S_Shore | Body | chr4:3415812-3416370 | 558 | | | | | | | | | 2 | | | | | | yes | | | | | |
| HGFAC | NA | TSS200;TSS1500 | chr4:3443025-3444147 | 1122 | | | | 1 | | | | | 2 | | | | | | yes | | | | yes | |
| DOK7 | N_Shore;Island | TSS1500;TSS200 | chr4:3464653-3464887 | 234 | | | | | | | | | 2 | | | | | | | | | | yes | |
| DOK7 | N_Shelf | Body | chr4:3471816-3472246 | 430 | | | | | | | | | 1 | | | | | | | yes | yes | | | |
| DOK7 | Island;N_Shore | Body | chr4:3475041-3475372 | 331 | | | 1 | | | | | | 3 | | | | | | | | | yes | yes | |
| DOK7 | N_Shore;Island;S_Shore | Body | chr4:3485263-3487436 | 2173 | | | 14 | 1 | | | | | 2 | | | 1 | | yes | | | | | | |
| LRPAP1 | S_Shore | Body | chr4:3521481-3521533 | 52 | | | | | | | | | 1 | | | | | | | | | | | |
| ADRA2C | Island | TSS1500 | chr4:3768078-3768458 | 380 | | 1 | | | | | | | 1 | | | | | | | | | yes | yes | |
| ADRA2C | Island | 1stExon | chr4:3769468-3769810 | 342 | | | | | | | | | | | | | | | | | | | yes | |
| OTOP1 | Island | TSS200 | chr4:4228637-4228650 | 13 | | | | | | | | 0 | | | | | | | | | | | yes | |
| TMEM128 | S_Shore | TSS1500 | chr4:4250606-4250746 | 140 | | | 3 | | | | | 0 | 1 | | | | | yes | yes | | | | | |
| D4S234E | N_Shore | TSS1500 | chr4:4386726-4386906 | 180 | | | | | | | | | | | | | | | yes | yes | | yes | yes | |
| D4S234E | NA | Body | chr4:4397942-4398276 | 334 | | | 1 | | | | | | | | | | | | | | | yes | | |
| D4S234E | NA | Body | chr4:4405844-4406132 | 288 | | | 5 | | | | | | | | | | | yes | | yes | yes | | | |
| MSX1 | Island;chr4:4859632-4860191;I | TSS1500;Island | chr4:4859975-4860590 | 615 | | 1 | | | | | | 0 | 2 | | 1 | 1 | | | | | | yes | yes | |
| MSX1 | N_Shore;Island | Body | chr4:4863678-4864902 | 1224 | 1 | | | | | | | 0 | 7 | | | 9 | | yes | | | | yes | yes | |
| CYTL1 | N_Shore;Island | Body;1stExon;TSS200 | chr4:5020292-5021328 | 1036 | | | 31 | | | | | 0 | 6 | 1 | | 7 | | yes | yes | | | yes | | |
| STK32B | N_Shore | TSS1500 | chr4:5053152-5053504 | 352 | | | | | | | | | 1 | | | | | yes | | | | | yes | |
| C4orf6 | NA | TSS1500 | chr4:5526510-5526783 | 273 | | | | | | | | | | | | | | | | yes | | | | |
| EVC2 | chr4:5709985-5710495 | Island | chr4:5710319-5710411 | 92 | | | | | | | | | | | 1 | | | | yes | | | | | |
| CRMP1 | N_Shore | Body | chr4:5887725-5888193 | 468 | | | | | | | | | 1 | | | | | | | | | | | |
| CRMP1 | Island | Body;TSS200 | chr4:5890402-5890430 | 28 | | | | | | | | | 2 | | | | | | | | | yes | yes | |
| CRMP1 | Island | Body | chr4:5892072-5892350 | 278 | | | | | | | | | 1 | | | | | yes | | | | | | |

| | | | | | | | | | | | | | | | | | | | | | | | |
|---|---|---|---|---|---|---|---|---|---|---|---|---|---|---|---|---|---|---|---|---|---|---|---|
| CRMP1 | Island | 1stExon;TSS200 | chr4:5894691-5895098 | 407 | | | | | | | | | 1 | 1 | | 1 | | | yes | | | yes | yes |
| C4orf50 | NA | Body | chr4:5976063-5976157 | 94 | | | 12 | | | | | | | | | | | yes | | | | | |
| C4orf50 | NA | Body;TSS200 | chr4:5990039-5990590 | 551 | | | 3 | | | | | | | | | | | yes | | | | | |
| JAKMIP1 | NA | Body | chr4:6037313-6037615 | 302 | | | | | | | | | | | | | | yes | | yes | | | |
| JAKMIP1 | NA | Body | chr4:6052527-6052554 | 27 | | | 10 | | 1 | | | | | | | | | yes | | | | | |
| JAKMIP1 | NA | Body | chr4:6072170-6072234 | 64 | | | 2 | | | | | | | | | | | | | | | | |
| JAKMIP1 | N_Shore;Island | Body | chr4:6107021-6107791 | 770 | | | 11 | | | | | 0 | 2 | | | 3 | | | yes | | | yes | |
| JAKMIP1 | Island | Body | chr4:6114498-6114699 | 201 | | | 4 | | | | | | | | | | | yes | | yes | yes | | |
| JAKMIP1 | Island | 5'UTR;TSS200 | chr4:6201536-6202384 | 848 | | | | | | | | | 2 | | | | | yes | | | | yes | yes |
| PPP2R2C | NA | Body | chr4:6328369-6328508 | 139 | | | 2 | | | | | | | | | | | | yes | yes | | | |
| PPP2R2C | NA | Body | chr4:6344361-6344484 | 123 | | | | | | | | 0 | 1 | 1 | | | | yes | | | yes | yes | |
| PPP2R2C | NA | Body | chr4:6347168-6347453 | 285 | | | | | | | | | 1 | | | | | | yes | yes | | | |
| PPP2R2C | Island | Body | chr4:6380399-6380703 | 304 | | | | | | | | | | | | | | yes | | | | | |
| PPP2R2C | NA | Body | chr4:6423008-6423371 | 363 | 2 | | 5 | | | | | | | | | | | | yes | | | | |
| PPP2R2C | NA | Body | chr4:6449564-6449865 | 301 | 1 | | | | | | | | | | | | | yes | | | | yes | |
| PPP2R2C | S_Shore | TSS1500 | chr4:6474782-6475156 | 374 | | | | | | | | | | | | | | | yes | | | yes | |
| LOC93622 | N_Shore | TSS200 | chr4:6675660-6675680 | 20 | | | | | | | | 0 | | | | | | | | yes | | yes | yes |
| S100P | NA | TSS1500;TSS200 | chr4:6694923-6695698 | 775 | | | 11 | | | | | 0 | 5 | 1 | | | | | | yes | | | |
| TBC1D14 | NA | 3'UTR | chr4:7033722-7033895 | 173 | | | | | | | | | 1 | | | | | | | | | | |
| CCDC96;TADA2B | S_Shore | TSS1500;Body | chr4:7046066-7046407 | 341 | | 1 | | | | | | | | | | | | yes | | | | | |
| TADA2B | Island | Body | chr4:7052403-7052788 | 385 | | | | 1 | 2 | | | | | | | | | yes | | | | | |
| FLJ36777 | NA | Body | chr4:7104897-7105272 | 375 | | | | | | | | 0 | | | | | | yes | yes | | | yes | |
| SORCS2 | NA | Body | chr4:7228268-7228442 | 174 | 4 | | 1 | | | | | | 1 | | | 1 | | yes | | yes | | | |
| SORCS2 | NA | Body | chr4:7282819-7283184 | 365 | | | 11 | | | | | | | | | | | yes | yes | | | yes | |
| SORCS2 | NA | Body | chr4:7287492-7287737 | 245 | 2 | | 4 | | | | | | | | | | | yes | | | | | |
| SORCS2 | NA | Body | chr4:7289482-7289858 | 376 | | | 10 | | | | | | | | | | | | | yes | | yes | yes |
| SORCS2 | NA | Body | chr4:7300999-7301854 | 855 | 2 | | | | | | | | | | | | | yes | | | | | |
| SORCS2 | NA | Body | chr4:7373065-7373451 | 386 | | | 5 | | | | | | | | | | | | yes | | | | |
| SORCS2 | NA | Body | chr4:7394790-7395388 | 598 | | | | | | | | | | | | | | yes | | | | | |
| PSAPL1;SORCS2 | NA | 1stExon;Body;5'UTR;TSS200 | chr4:7436239-7437027 | 788 | | | 7 | 1 | | | | 0 | 1 | | | 1 | | | yes | | | | |
| SORCS2 | NA | Body | chr4:7462468-7463007 | 539 | | | 5 | | | | | | | | | | | | | | | | |
| SORCS2 | Island | Body | chr4:7512016-7512498 | 482 | | | | | | | | | 2 | | | 1 | | | yes | | | | |
| SORCS2 | NA | Body | chr4:7526521-7526671 | 150 | | | | | | | | | | | | | | | | yes | | | |
| SORCS2 | NA | Body | chr4:7528129-7528468 | 339 | | | | | | | | 0 | 1 | | | | | | yes | yes | yes | | yes |
| SORCS2 | Island | Body | chr4:7593370-7593493 | 123 | | | | | | | | | | | | | | | | | | | yes |
| SORCS2 | NA | Body | chr4:7637501-7637856 | 355 | | | | | | | | | 1 | | | | | yes | yes | | | yes | |
| SORCS2 | NA | Body | chr4:7657070-7657708 | 638 | 2 | | | 1 | | | | 0 | 1 | | | | | yes | | | yes | yes | |
| SORCS2 | NA | Body | chr4:7719899-7720084 | 185 | | | | | | | 1 | | | | | | | yes | yes | | | | |
| SORCS2 | NA | Body | chr4:7736302-7736837 | 535 | | | | | | | | | 1 | | | | | yes | yes | | | | |
| AFAP1;LOC84740 | NA | 3'UTR;Body | chr4:7762596-7762705 | 109 | 1 | | | | | | | | | | | | | | | | | | |
| AFAP1;LOC84740 | NA | Body | chr4:7768162-7768660 | 498 | | | | | | | | | 1 | | | | | | | | | | |
| AFAP1;LOC84740 | Island | Body | chr4:7774727-7774950 | 223 | | | 19 | | 1 | | | | 1 | | | | | | | | | | |
| AFAP1 | N_Shelf | Body | chr4:7799636-7799701 | 65 | | | | | | | | | 1 | | | | | | yes | | | | |
| AFAP1 | NA | Body | chr4:7813475-7814047 | 572 | | | 8 | | | | | | | | | | | | | yes | | | |
| AFAP1 | NA | Body | chr4:7844918-7844982 | 64 | | | | | | | | | | | | | | | | | | yes | yes |
| AFAP1 | NA | Body | chr4:7862646-7862659 | 13 | 1 | | | | | | | | 1 | 1 | | | | yes | | | yes | yes | |
| AFAP1 | N_Shore | 5'UTR | chr4:7940128-7940426 | 298 | | 1 | | | | | | | 1 | | | | | | | | | | yes |
| AFAP1 | Island;S_Shore | TSS200;TSS1500 | chr4:7941764-7942031 | 267 | | | 5 | | | | | | 4 | | | 1 | | yes | yes | | | | yes |
| ABLIM2 | NA | Body | chr4:8081109-8081417 | 308 | | | | | | | | | | | | | | | yes | | | yes | |
| ABLIM2 | N_Shore;Island | Body | chr4:8098178-8099033 | 855 | | | 8 | | | | | | 1 | | | | | | yes | | | | yes |
| ABLIM2 | NA | Body | chr4:8121992-8122233 | 241 | | | | | | | | | | | | | | | | yes | yes | | |
| ABLIM2 | NA | Body | chr4:8126305-8126646 | 341 | | | | | | | | | | | | | | | yes | | | | |
| SH3TC1 | NA | Body | chr4:8207119-8207325 | 206 | | | | | | | | 0 | 1 | | | | | yes | yes | | | | yes |
| HTRA3 | NA | Body | chr4:8292202-8292328 | 126 | | | | | | | | 0 | | | | | | yes | yes | | | | |
| ACOX3 | N_Shelf | Body | chr4:8372691-8372840 | 149 | 1 | | | | | | | | | | | | | | | | | | |
| ACOX3 | N_Shelf | Body | chr4:8386188-8386266 | 78 | | | 2 | | | | | | 1 | | | | | | | | | | |
| ACOX3 | NA | Body | chr4:8395941-8396410 | 469 | 2 | | 1 | | | | | | 2 | 1 | | | | | | | | | |
| ACOX3 | S_Shore | TSS1500 | chr4:8443412-8443777 | 365 | | | | | | | | | | | | | | | | yes | | | |
| C4orf23 | NA | Body | chr4:8477562-8477915 | 353 | | | | | | | | | | | | | | | | | | | |
| GPR78 | N_Shore | TSS1500 | chr4:8581424-8582300 | 876 | | | | | | | | | 1 | | | 1 | | yes | | | | | yes |
| GPR78 | Island | Body | chr4:8588802-8588870 | 68 | | | | | | | | | | | | | | yes | | | | | |
| CPZ | N_Shore | TSS1500 | chr4:8593507-8593933 | 426 | | | | | | | | | | | | | | | | | yes | | |
| CPZ | NA | 5'UTR;Body | chr4:8602556-8603012 | 456 | | | 7 | | | | | | | | | | | | | | | | |
| CPZ | NA | Body | chr4:8609401-8609706 | 305 | | | | | | | | | | | | | | | | | | | |
| CPZ | NA | Body | chr4:8621226-8621361 | 135 | | | | | | | | | | | | | | | yes | | | | |
| DRD5 | N_Shore;Island | TSS1500;TSS200 | chr4:9782891-9783623 | 732 | | 1 | | | | | | 0 | 5 | | 1 | 2 | | yes | | | | | |
| SLC2A9 | N_Shore | Body | chr4:10020360-10021025 | 665 | | | | 1 | | | | | | | | | | yes | | | | yes | |
| ZNF518B | Island;S_Shore | TSS200;TSS1500 | chr4:10459183-10460237 | 1054 | | | 12 | | | | | | 1 | | | | | yes | yes | | | | yes |
| CLNK | NA | TSS200;1stExon;5'UTR | chr4:10686300-10686610 | 310 | | | 7 | | 2 | | | | 2 | 1 | | | | yes | | | | | |
| MIR572 | N_Shore;Island | TSS1500;TSS200;Body | chr4:11370015-11370793 | 778 | | | | | | | | | 1 | | | | | yes | yes | | | | |
| HS3ST1 | Island | 5'UTR | chr4:11401662-11401872 | 210 | | | 1 | | | | | | | | | | | | | | yes | | |
| HS3ST1 | Island | TSS200 | chr4:11430693-11430903 | 210 | | | | | | | | 0 | 2 | | | 1 | | | | | | yes | yes |
| NKX3-2 | N_Shore | 3'UTR | chr4:13542719-13543119 | 400 | | | 4 | | | | | | | | | | | | | | yes | | |
| NKX3-2 | Island | Body;TSS200 | chr4:13545291-13546482 | 1191 | | | | | | | | | 3 | | | | | | | | | yes | yes |
| C1QTNF7 | NA | TSS1500;Body;TSS200 | chr4:15375625-15376215 | 590 | | | 41 | | | | | | | | | | | yes | | | | | |
| C1QTNF7 | NA | TSS200;Body;5'UTR | chr4:15429531-15429967 | 436 | | | | | | | | | | | | | | yes | yes | | | | |
| CC2D2A | NA | TSS1500;TSS200 | chr4:15471214-15471699 | 485 | | | 6 | | | | | | | | | | | | | yes | | yes | |
| BST1 | N_Shore | TSS200;1stExon;5'UTR | chr4:15704393-15704844 | 451 | | | 3 | 1 | | | | | | | | 1 | | yes | | | yes | yes | yes |
| CD38 | N_Shore | TSS1500;TSS200 | chr4:15779642-15780011 | 369 | | | 8 | | 1 | | | | 1 | | | | | yes | yes | | | | |
| FGFBP1 | NA | 1stExon;5'UTR | chr4:15939889-15940058 | 169 | | | | | | | | | | | | | | yes | | | | | |
| FGFBP2 | NA | TSS200 | chr4:15965016-15965102 | 86 | | | | | | | | | | | | | | | | | | | |
| PROM1 | S_Shore;Island | TSS1500;5'UTR;1stExon;TSS200 | chr4:16085401-16086085 | 684 | | | 3 | | | | | 0 | 1 | 1 | 1 | | | | yes | | | yes | |
| LDB2 | NA | 1stExon;5'UTR | chr4:16900199-16900254 | 55 | | | | | | | | | 1 | | | | | | | | | yes | yes |
| CLRN2 | S_Shelf | TSS1500 | chr4:17516550-17516739 | 189 | | | | | | | | | | | | | | | | | | | |
| FAM184B | Island | Body | chr4:17710911-17711202 | 291 | | | | | | | | | | | | | | yes | yes | yes | | | |
| FAM184B | Island | TSS200 | chr4:17783240-17783331 | 91 | | | | | | | | | 2 | | | | | | | | | | |
| KCNIP4 | NA | Body;1stExon;5'UTR | chr4:20884321-20884431 | 110 | | | | | | | | | | | | | | | | | | | |
| KCNIP4 | NA | Body;5'UTR | chr4:20985696-20985984 | 288 | | | | | | | | | 1 | | | | | | | | | | |
| KCNIP4 | NA | 5'UTR;TSS200;TSS1500;1stExon | chr4:21699209-21699988 | 779 | | | 20 | | | | | 0 | 2 | | | | | | | | | yes | |
| KCNIP4 | N_Shore;Island | 5'UTR;1stExon;TSS200;TSS1500 | chr4:21949853-21951046 | 1193 | | | | | | | | 0 | 5 | | | | | | | | | yes | |
| PPARGC1A | NA | 1stExon;5'UTR;TSS1500 | chr4:23891596-23892621 | 1025 | | | | | | | | 0 | 2 | | | 2 | | | | | | yes | yes |
| SOD3 | NA;N_Shelf | nan;TSS200;5'UTR;1stExon;TSS15( | chr4:24796689-24797176 | 487 | | | | | 2 | | | 0 | 2 | | 1 | | | yes | | | | | |
| CCDC149 | NA | Body | chr4:24818471-24818615 | 144 | | 1 | 9 | | | | | | | | | | | yes | | | | | |
| CCDC149 | NA | TSS200;5'UTR | chr4:24914777-24914941 | 164 | | | | | | | | | 1 | | | | | yes | yes | | yes | yes | yes |
| CCDC149 | NA | 1stExon;5'UTR | chr4:24981695-24981807 | 112 | | | | | | | | | | | | | | | | yes | | | |
| LGI2 | S_Shore | TSS200 | chr4:25032556-25032564 | 8 | | | | | | | | | 1 | | | 1 | | | | | | yes | yes |
| SLC34A2 | N_Shore;Island;S_Shore | TSS1500;TSS200;5'UTR | chr4:25656514-25658035 | 1521 | | | 4 | | | | | 0 | 3 | | | | | yes | | | | | yes |
| CCKAR | NA | 1stExon;5'UTR;TSS200;TSS1500 | chr4:26491963-26492870 | 907 | | | 12 | | | | | | | | | | | yes | yes | | | yes | |
| PCDH7 | Island | TSS200 | chr4:30721997-30722152 | 155 | | | | | | | | | 1 | | | | | | | | | | yes |

| | | | | | | | | | | | | | | | | | | | | | | | | | | | |
|---|---|---|---|---|---|---|---|---|---|---|---|---|---|---|---|---|---|---|---|---|---|---|---|---|---|---|---|
| DTHD1 | NA | TSS200 | chr4:36283184-36283588 | 404 | | | | | | | | | 1 | | | | | | yes | | | | | | | | |
| DTHD1 | NA | TSS1500;Body | chr4:36284867-36285698 | 831 | | | | | | | | | 1 | | | | | | yes | | | yes | | | | | |
| C4orf19 | Island | TSS200 | chr4:37455366-37455499 | 133 | | | | | | | | | 2 | | | | | | | | | yes | | | | | |
| C4orf19 | NA | TSS1500;5'UTR | chr4:37584954-37585361 | 407 | | | | | | | | | | | | | | yes | yes | | | | | | | | |
| RELL1 | Island | TSS200 | chr4:37688098-37688119 | 21 | | | | | | | | | 1 | | | | | | | | | yes | | | | | |
| TLR1 | NA | TSS1500 | chr4:38807259-38807382 | 123 | | | 2 | | | | | | | | | | | | | | | | | | | | |
| TLR6 | NA | TSS1500 | chr4:38859706-38859770 | 64 | | | | | | | | 0 | 1 | | | | | yes | | | | | | | | | |
| MIR574;FAM114A1 | Island | TSS1500;5'UTR;1stExon;TSS200 | chr4:38869436-38869707 | 271 | | | | | | | | 0 | | | | | | | | | | yes | yes | | | | |
| KLB | NA | TSS200 | chr4:39408416-39408665 | 249 | | | | 1 | 1 | | | | | | | | | | yes | | | | yes | | | | |
| KLB | N_Shore;Island | Body | chr4:39448160-39449130 | 970 | | | | 1 | | | | 0 | 2 | | 1 | 3 | | yes | yes | | | | | | | | |
| LOC401127 | S_Shore | Body | chr4:39482431-39483194 | 763 | | | 1 | | | | | 0 | 1 | | | | | yes | | | | | yes | | | | |
| RHOH | NA | TSS1500;nan;TSS200 | chr4:40197912-40198776 | 864 | | | 6 | | 1 | | | | 1 | | 1 | | | | yes | | yes | | | | | | yes |
| CHRNA9 | NA | TSS1500 | chr4:40336735-40337370 | 635 | | | | | | | | | | | | | | yes | | | | | | | | | |
| RBM47 | NA | 5'UTR;TSS200;TSS1500;nan;1stExo | chr4:40517938-40519094 | 1156 | | | 7 | 1 | | | | | 2 | | 1 | | | yes | | | | yes | yes | | | | |
| RBM47 | Island;S_Shore | TSS1500 | chr4:40632299-40633269 | 970 | | | | 1 | 2 | | | 0 | 3 | 1 | | | | yes | | | | | | | | | |
| APBB2 | Island;chr4:40858884-4085916 | Body;5'UTR;Island;1stExon;TSS20 | chr4:40858965-40859344 | 379 | | | 40 | | 1 | | | | | | 1 | | | yes | | | | | yes | | | | |
| UCHL1 | Island;N_Shore | TSS200;1stExon;5'UTR;TSS1500 | chr4:41258374-41259283 | 909 | | | | | | | | | 2 | | | | | | yes | | | | | | | | |
| LIMCH1 | N_Shore;Island | TSS1500;1stExon;5'UTR | chr4:41361628-41363229 | 1601 | | | 2 | | | | | 0 | 1 | | | | | | yes | yes | | yes | yes | | | | |
| LIMCH1 | Island | Body | chr4:41646293-41646567 | 274 | | | | | | | | 0 | | | | | | yes | | | | | yes | | | | |
| LIMCH1 | NA | Body | chr4:41694494-41694847 | 353 | | | 1 | | | | | | 1 | | | | | yes | yes | | | | | | | | |
| PHOX2B | N_Shore | Body;1stExon;5'UTR | chr4:41750362-41751001 | 639 | | 1 | 3 | | | | | | | | | | | | | | | | yes | | | | |
| BEND4 | NA | Body | chr4:42122106-42122161 | 55 | | | | | | | | | | | | | | | | | | | | | | | |
| BEND4 | Island;S_Shore | TSS200;TSS1500 | chr4:42154912-42155519 | 607 | | | | | | | | | | | | | | yes | | | | | yes | | | | |
| SHISA3 | Island;S_Shore | TSS200;Body | chr4:42399798-42401610 | 1812 | | 1 | 19 | | | | | | 1 | | 1 | | | yes | | | | | | | | | |
| GABRG1 | NA | TSS200;TSS1500;5'UTR;1stExon | chr4:46126066-46126448 | 382 | | | | | | | | 0 | 5 | | | 4 | | | | | | yes | | | | | |
| GABRA2 | Island | TSS1500 | chr4:46392526-46392657 | 131 | | | | | | | | | 1 | | | | | | | | | yes | | | | | |
| COX7B2 | NA | TSS200 | chr4:46911374-46911550 | 176 | | | | | | | | | | | | | | yes | | yes | yes | | | | | | |
| GABRA4 | Island | TSS200 | chr4:46995659-46995821 | 162 | | 1 | | | | | | | 2 | | | | | | | | | | | | | | |
| GABRB1 | N_Shore | TSS200 | chr4:47033126-47033423 | 297 | | | | | | | | 0 | 1 | | | | | | | | | yes | | | | | |
| GABRB1 | Island | Body | chr4:47427752-47428108 | 356 | | | | | | | | | | | | | | | yes | | | yes | | | | | |
| CORIN | S_Shore | 5'UTR;1stExon | chr4:47840011-47840095 | 84 | | | | | | | | 0 | 1 | | | | | | | | | | | | | | |
| TXK | NA | 5'UTR;1stExon | chr4:48136234-48136452 | 218 | | | | | 1 | | | | 1 | | | | | yes | | | | yes | | | | | |
| TEC | N_Shore | 5'UTR | chr4:48271072-48271121 | 49 | | | 1 | | | | | | | | | | | | yes | | | yes | | | | | |
| SLC10A4 | N_Shore | TSS1500 | chr4:48484788-48485090 | 302 | | | | | | | | 0 | | | | | | yes | | | | yes | | | | | |
| SLC10A4 | Island | Body;1stExon | chr4:48486154-48486785 | 631 | | | 2 | | | | | | | | | | | | | | | yes | | | | | |
| ZAR1 | Island | Body | chr4:48493350-48493846 | 496 | | | | | | | | | | | | | | yes | | | | | yes | | | | |
| OCIAD2 | Island | TSS200;5'UTR | chr4:48908469-48908902 | 433 | | | 4 | | | | | | 1 | | | | | yes | | yes | | | | | | | |
| CWH43 | Island | TSS200 | chr4:48988066-48988124 | 58 | | | | | | | | | 1 | | | | | | | | | | | | | | |
| LRRC66 | NA | 1stExon | chr4:52883367-52883909 | 542 | | | | | | | | | 1 | | | | | | | | | | | | | | |
| SPATA18 | Island;S_Shore | Body;TSS200;1stExon;5'UTR | chr4:52917539-52918489 | 950 | | | 18 | | | | | 0 | 3 | | | 4 | | yes | | | | | | | | | |
| SPATA18 | Island | Body | chr4:52942997-52943247 | 250 | | 1 | 6 | 1 | | | | | | | | | | | | yes | | | yes | | | | |
| USP46 | N_Shore | TSS1500;Body | chr4:53523482-53524296 | 814 | | | | | | | | | | | | | | yes | | | | | | | | | |
| RASL11B | N_Shore;Island | TSS1500;1stExon;5'UTR;Body | chr4:53727714-53729461 | 1747 | | | 14 | | | | | 0 | 3 | 1 | | 4 | | | | | | | yes | | | | |
| FIP1L1 | N_Shore | TSS1500 | chr4:54243152-54243328 | 176 | | | | | | | | | 1 | | | | | yes | yes | | | | | | | | |
| LNX1 | NA | TSS200;Body;1stExon;5'UTR | chr4:54424280-54425094 | 814 | | | 8 | | | | | | | | | | | yes | | | yes | | | | | | |
| LNX1 | NA | 5'UTR;1stExon | chr4:54457645-54457928 | 283 | | | | 1 | | | | | | | | | | | | | | | | | | | |
| GSX2 | N_Shore | TSS1500 | chr4:54965829-54965999 | 170 | | | 8 | | | | | | | | | | | | | | | | | | | | |
| GSX2 | Island | Body | chr4:54967423-54967809 | 386 | | | 1 | | | | | | | | | | | | | | | yes | | | | | |
| PDGFRA | N_Shore | TSS200 | chr4:55095209-55095261 | 52 | | 1 | | | | | | | | | | | | | | | | | | | | | |
| PDGFRA | Island | 5'UTR | chr4:55097420-55097576 | 156 | | | | | | | | | 1 | | | | | | | | | | | | | | |
| PDGFRA | NA | Body | chr4:55143269-55143557 | 288 | | | 12 | | | | | 0 | 1 | | | 3 | | yes | | | | | | | | | |
| KDR | Island | TSS200 | chr4:55991943-55992458 | 515 | | | | | | | | | 1 | | | | | | | | | | | | | | |
| PDCL2 | NA | Body;TSS200 | chr4:56458183-56458622 | 439 | | 1 | | | | | | | 1 | | | | | yes | | | yes | | | | | | |
| NMU | S_Shore | TSS1500 | chr4:56502849-56502898 | 49 | | | | | | | | | 1 | | | 1 | | | yes | | | yes | | | | | |
| KIAA1211 | NA | TSS1500 | chr4:57035948-57036403 | 455 | | | 8 | | | | | | | | | | | yes | | | | yes | | | | | |
| KIAA1211 | Island | Body | chr4:57181042-57181855 | 813 | | 1 | | | | | | | | | | | | | | | | | yes | | | | |
| HOPX | Island;S_Shore | 1stExon;5'UTR;Body;TSS200 | chr4:57522493-57523826 | 1333 | | 1 | 23 | | | | | | | | | | | yes | | | | | | | | | |
| HOPX | NA | 5'UTR;1stExon | chr4:57547347-57548093 | 746 | | | 12 | | | | | 0 | 5 | 1 | | 12 | | | | | | | | | | | |
| SPINK2 | S_Shore | TSS1500 | chr4:57688282-57688337 | 55 | | | | | | | | | | | | | | yes | yes | | | | yes | | | | |
| REST | N_Shore | TSS1500 | chr4:57773149-57773308 | 159 | | | | | | | | | | | | | | | yes | | yes | | | | | | |
| IGFBP7 | Island | 1stExon | chr4:57976171-57976561 | 390 | | | | | | | | | 1 | | | | | | | | | | yes | | | | |
| LPHN3 | N_Shore;Island | Body | chr4:62382932-62383240 | 308 | | | | 1 | | | | | 1 | | | 4 | | | | | yes | | yes | | | | |
| EPHA5 | Island | 1stExon;TSS1500 | chr4:66535351-66536772 | 1421 | | | 1 | | 1 | | | 0 | 1 | | | | | | | | | | yes | | | | |
| LOC100130017 | N_Shore | TSS1500 | chr4:69047758-69048156 | 398 | | | | | | | | | | | | | | yes | | | | | | | | | |
| YTHDC1 | NA | Body | chr4:69179905-69179965 | 60 | | | | | | | | | 1 | | | | | | | | | yes | yes | | | | |
| UGT2B15;UGT2B17 | NA | TSS1500 | chr4:69435250-69435601 | 351 | 4 | 12 | 7 | 1 | | | 1 | 0 | 1 | 1 | 1 | 21 | 1 | | yes | yes | | | | | | | |
| MUC7 | NA | TSS200;5'UTR | chr4:71337658-71337873 | 215 | | | | | | | | | 1 | | | | | yes | yes | yes | | | | | | | |
| RUFY3 | NA | Body;TSS200;1stExon;5'UTR | chr4:71587640-71587904 | 264 | | | 28 | | | | | | | | | | | | | | | yes | | | | | |
| NPFFR2 | N_Shore;Island | TSS1500;1stExon;5'UTR | chr4:72897281-72898212 | 931 | 1 | | 6 | 1 | | | | 0 | 5 | 3 | | 4 | | yes | | | | | | | | | |
| RASSF6 | Island;S_Shore | 1stExon;TSS200;TSS1500 | chr4:74486170-74486558 | 388 | | | 2 | | | | | | | | | | | | | | yes | yes | | | | | |
| CXCL6 | N_Shore | TSS200 | chr4:74702075-74702448 | 373 | | | | | | | | | 1 | | | | | | yes | | | | | | | | |
| PF4V1 | N_Shore;Island | TSS1500;TSS200;1stExon;5'UTR | chr4:74718399-74719597 | 1198 | | | 27 | | | | | 0 | 4 | | | 3 | | | | yes | | | yes | | | | |
| CXCL1 | N_Shore;Island | TSS1500;TSS200 | chr4:74734714-74735149 | 435 | | | | | | | | | 2 | | | | | | | | | | | | | | |
| PF4 | Island;N_Shore | 1stExon;5'UTR;TSS200;Body | chr4:74847100-74848016 | 916 | 2 | 1 | 29 | | 1 | 1 | 1 | 0 | 12 | 9 | 2 | 22 | 2 | | | | | yes | | | | | |
| CXCL5 | Island;S_Shore | Body;1stExon;TSS200 | chr4:74864165-74864725 | 560 | | 1 | 19 | | | | | 0 | 3 | | | | | | | | | | | | | | |
| CXCL2 | Island | TSS200 | chr4:74965068-74965278 | 210 | | | | | | | | | 1 | 1 | | | | | | | | yes | | | | | |
| PARM1 | NA | Body | chr4:75895596-75895858 | 262 | | | 7 | | | | | | | | | | | yes | | yes | | | | | | | |
| C4orf26 | NA | TSS1500 | chr4:76480681-76481299 | 618 | | | 16 | | | | | | 3 | | 1 | | | yes | yes | | | | | | | | |
| CDKL2 | Island | TSS200;TSS1500 | chr4:76555777-76556042 | 265 | | | | | | | | | | | | | | | | | | | | | | | |
| FAM47E | Island | TSS200;Body | chr4:77172836-77173197 | 361 | | | | | | | | | 2 | | | | | yes | | | | | yes | | | | |
| SHROOM3 | NA | TSS1500;TSS200 | chr4:77355817-77356507 | 690 | | | 6 | | | | | 0 | 2 | | | | | | | | yes | | | | | | |
| SHROOM3 | N_Shore;Island | Body | chr4:77506779-77507411 | 632 | | | 14 | | | | | | | | | | | | | | | | | | | | |
| ANKRD56 | Island | TSS1500 | chr4:77819534-77819688 | 154 | | | 2 | | | | | | | | | | | | | | | | | | | | |
| CXCL13 | NA | TSS1500 | chr4:78432011-78432403 | 392 | | | | | | | | | 1 | | | | | yes | yes | yes | | | | | | | |
| ANXA3 | N_Shore | TSS200 | chr4:79472605-79472807 | 202 | | | | | | | | | 1 | | | | | yes | yes | | yes | | | | | | |
| PAQR3 | S_Shore | TSS1500 | chr4:79861272-79861398 | 126 | | | | | | | | | | | | | | yes | yes | yes | | | | | | | |
| NAA11 | NA | 1stExon | chr4:80246713-80247171 | 458 | | | 4 | | | | | | | | | | | | yes | | | | | | | | |
| GK2 | NA | 1stExon;5'UTR | chr4:80329368-80329424 | 56 | | | | | | | | | | | | | | | yes | | | | yes | | | | |
| PRDM8 | S_Shore | 5'UTR | chr4:81106772-81107185 | 413 | | | 10 | | | | | 0 | 1 | | | 2 | | | | | | | yes | | | | |
| PRDM8 | S_Shore | 5'UTR | chr4:81111177-81111527 | 350 | | | | | | | | | 1 | | | 1 | | yes | | | | | | | | | |
| PRDM8 | N_Shore;Island | TSS1500;5'UTR;TSS200 | chr4:81117647-81119473 | 1826 | | 1 | 14 | | | | 1 | 0 | 15 | 4 | 1 | 9 | | | | | | yes | yes | | | | |
| PRDM8 | N_Shore | Body | chr4:81123306-81123380 | 74 | | | | | | | | | | | | | | | | | | yes | | | | | |
| PRDM8 | Island | Body | chr4:81124484-81124767 | 283 | | | | | | | | | | | | | | | | | | yes | yes | | | | |
| RASGEF1B | Island | TSS1500 | chr4:82393353-82393723 | 370 | | | | | | | | | | | | | | yes | yes | | | | yes | | | | |
| SCD5;MIR575 | NA | Body;TSS200 | chr4:83674615-83675306 | 691 | | | 2 | | | | | | 1 | | | | | yes | | | yes | | yes | | | | |
| NKX6-1 | Island;N_Shore | Body | chr4:85414486-85418816 | 4330 | | 1 | 2 | | | | | | 1 | | | | | | | | | yes | yes | | | | |
| CDS1 | Island;N_Shore | TSS200;TSS1500 | chr4:85503328-85504391 | 1063 | | | | | | | | | 3 | | | | | | | | | | yes | | | | |
| ARHGAP24 | NA | TSS1500 | chr4:86395991-86396103 | 112 | | | 1 | | | | | | | | | | | yes | | | | | | | | | |
| ARHGAP24 | NA | Body;5'UTR | chr4:86748944-86749040 | 96 | | | 2 | | | | | | | | | | | | | | | yes | | | | | |

| | | | | | | | | | | | | | | | | | | | | | | | |
|---|---|---|---|---|---|---|---|---|---|---|---|---|---|---|---|---|---|---|---|---|---|---|---|
| ARHGAP24 | NA | Body;TSS200 | chr4:86851225-86851425 | 200 | | | | | | | | | 2 | | | | | | | | | | |
| MAPK10 | NA | TSS200;5'UTR | chr4:87281412-87281511 | 99 | | | 7 | | | | | | | | | | | | | | | | |
| SLC10A6 | NA | nan;5'UTR;1stExon;TSS200 | chr4:87770089-87770782 | 693 | | | 2 | | | | | | 1 | 1 | 1 | | | | | | | | |
| SPP1 | NA | TSS200 | chr4:88896634-88896942 | 308 | | | 3 | | | | | | | | | | | yes | | | | | |
| ABCG2 | Island | TSS200 | chr4:89080179-89080199 | 20 | | 1 | | | | | | | | | | | | | | | | | |
| FAM13A | NA | 1stExon;5'UTR | chr4:89978251-89978566 | 315 | | | | | 1 | | | 0 | 1 | | | | | | yes | | | | |
| SNCA | N_Shore | 5'UTR | chr4:90757139-90757533 | 394 | | 1 | | | | | | | | | | | | | | | | | yes |
| MMRN1 | NA | TSS200 | chr4:90815958-90816310 | 352 | | | | | | | | | 1 | | | | | | | | | | |
| FAM190A | NA | 5'UTR;TSS200 | chr4:91156040-91156155 | 115 | | | | | | | | | | | | | | yes | | yes | | | |
| GRID2 | NA | Body | chr4:94124971-94125068 | 97 | | | 4 | | | | | | | | | | | | | yes | | | |
| ATOH1 | Island | TSS1500 | chr4:94749609-94749890 | 281 | | 1 | | | | | | | 1 | | | | | | | | | | |
| HPGDS | NA | 5'UTR;1stExon | chr4:95264019-95264373 | 354 | | | | | 1 | | | 0 | 2 | | 1 | | | | yes | | | | yes |
| UNC5C | NA | Body | chr4:96445830-96446177 | 347 | | | | | | | | | 1 | | | | | yes | | | yes | | |
| UNC5C | Island | TSS200;TSS1500 | chr4:96470286-96471105 | 819 | | 2 | | | | | | | 1 | | | | | | | | | yes | yes |
| PDHA2 | NA | TSS1500;1stExon;5'UTR | chr4:96761026-96761796 | 770 | | | 6 | | | | | | | | | | | | | | yes | | |
| C4orf37 | N_Shore;Island | TSS200;TSS1500;1stExon;5'UTR | chr4:99064322-99064904 | 582 | | | | | | | | 0 | 2 | | | | | yes | | | | | yes |
| TSPAN5 | S_Shore | TSS1500 | chr4:99580449-99580849 | 400 | | | 1 | | | | | | 1 | | | | | | | | | | |
| EIF4E | S_Shore | 5'UTR;1stExon;TSS1500 | chr4:99850801-99851281 | 480 | | | 9 | | | | 1 | 0 | 5 | 1 | | 9 | | yes | | | | yes | |
| ADH6 | NA | 5'UTR;1stExon | chr4:100140325-100140905 | 580 | | | | 1 | 1 | | | | | | | | | yes | yes | | | | |
| C4orf17 | NA | TSS1500 | chr4:100431894-100432266 | 372 | | | | | | | | | 1 | | | | | yes | | yes | | | |
| DAPP1 | NA | 1stExon;5'UTR | chr4:100738011-100768712 | 30701 | | | | | | | | | | | | | | yes | yes | yes | yes | yes | yes |
| EMCN | NA | TSS1500 | chr4:101439954-101440278 | 324 | | | 7 | | | | | | | | | | | yes | yes | | | | |
| BANK1 | N_Shore;Island | TSS200;5'UTR;1stExon | chr4:102711702-102712397 | 695 | | | 10 | | 1 | | | | 1 | | | 2 | | | | | | yes | yes |
| NHEDC1 | S_Shore;Island | TSS200;TSS1500;5'UTR;1stExon | chr4:103940711-103941300 | 589 | | | | | | | | 0 | 3 | 2 | | 3 | 1 | yes | | | yes | | |
| NHEDC2 | S_Shore | TSS200 | chr4:103998289-103998717 | 428 | | | 1 | | 1 | | | | 4 | 1 | | 1 | | yes | | | | | |
| TACR3 | N_Shore;Island;S_Shore | Body;1stExon;TSS200 | chr4:104640197-104641548 | 1351 | | 1 | 1 | | | | | 0 | 1 | | | | | | | | | | |
| FLJ20184 | NA | nan;TSS1500 | chr4:106472990-106473806 | 816 | | | | | | | | | | | 1 | | | | yes | | | | |
| GSTCD | S_Shore | TSS200;5'UTR | chr4:106631906-106631934 | 28 | | | | | | | | | 1 | | | | | yes | | | | yes | yes |
| NPNT | Island | TSS200 | chr4:106816548-106816770 | 222 | | | | | | | | | 1 | | | | | | | | | | |
| LEF1;LOC641518 | N_Shore | TSS1500;Body | chr4:109090828-109091059 | 231 | | | | | | | | 0 | | | | | | | | | | | |
| LOC641518 | N_Shore;Island | TSS1500;Body | chr4:109092312-109093158 | 846 | | | 8 | | | | | | | | | | | | | | | | |
| COL25A1 | Island | 1stExon;5'UTR | chr4:110223429-110223830 | 401 | | 1 | | | | | | | 2 | | | | | | | | | | |
| ENPEP | N_Shore;Island | TSS200;1stExon;5'UTR | chr4:111397134-111398146 | 1012 | | | 29 | 1 | | | | | 2 | | | 3 | | | yes | | | | |
| PITX2 | Island | Body | chr4:111539291-111540126 | 835 | | | 1 | | | | | | 1 | | | | | | | | | | |
| PITX2 | Island | 1stExon;5'UTR;Body;TSS200 | chr4:111542063-111545119 | 3056 | | 1 | 23 | | | | | | 2 | | | | | | | | | | |
| PITX2 | Island | Body | chr4:111549880-111550830 | 950 | | 1 | | | | | | | | | | | | | | | | | yes |
| PITX2 | Island | 5'UTR | chr4:111554966-111555503 | 537 | | | 2 | | | | | | 1 | | | | | | | | | | yes |
| PITX2 | N_Shore | TSS1500 | chr4:111558988-111559134 | 146 | | 1 | 2 | | | | | | | | | | | | | | | | |
| C4orf32 | N_Shore | TSS1500 | chr4:113066291-113066439 | 148 | | | 2 | | | | | | | | | | | yes | | | | yes | yes |
| TIFA | Island | 1stExon;5'UTR | chr4:113206875-113207097 | 222 | | | | | | | | | 1 | | | | | yes | | | | | |
| ALPK1 | N_Shore;Island | Body | chr4:113300446-113300821 | 375 | | | 7 | | | | | 0 | | | | | | | | | yes | | |
| ALPK1 | Island | Body | chr4:113333052-113333072 | 20 | | | | | | | | | | | | | | yes | | | | | |
| ANK2 | NA | TSS200;Body | chr4:113970617-113971324 | 707 | | | 17 | | | | | | 1 | | | | | | | | | yes | |
| ANK2 | NA | Body | chr4:114213914-114214093 | 179 | | | | 1 | | | | | | | | | | | | | | | yes |
| ARSJ | N_Shore | 1stExon;5'UTR | chr4:114900175-114900602 | 427 | | | | | | | | | 1 | | | | | | | | | | |
| UGT8 | Island | TSS200 | chr4:115519567-115519593 | 26 | | | | | | | | | 1 | | | | | | | | | | yes |
| NDST4 | NA | 1stExon;5'UTR;TSS200 | chr4:116034871-116035232 | 361 | | | | | 1 | | | | 1 | | | | | | | | | | |
| TRAM1L1 | N_Shore;Island | 1stExon;5'UTR;TSS200 | chr4:118006405-118006832 | 427 | | 1 | 3 | | 2 | | | 0 | 8 | 1 | | 6 | 1 | yes | | | | | |
| SNORA24;SNHG8 | Island | TSS1500;Body;TSS200 | chr4:119199621-119200210 | 589 | | | | | | | | 0 | | | | | | yes | yes | | | | |
| SYNPO2 | NA | 1stExon;5'UTR;TSS200 | chr4:119809845-119810092 | 247 | | | | | | | | | | | | | | | | | | yes | yes |
| SYNPO2 | NA | Body | chr4:119948360-119948542 | 182 | | | 29 | | | | | | | | | | | | | | | | |
| FABP2 | NA | 1stExon | chr4:120243252-120243555 | 303 | | | 2 | | | | | | 1 | | | | | yes | | yes | | | |
| C4orf31 | Island;N_Shore | 5'UTR | chr4:121991345-121992756 | 1411 | | | | | | | | | 1 | | | | | | | | | yes | |
| QRFPR | Island | 1stExon;TSS200 | chr4:122301573-122302421 | 848 | | 1 | 1 | | | | | | 2 | | | | | | yes | | | yes | |
| TMEM155;LOC100192379 | N_Shore | 5'UTR;TSS1500;TSS200 | chr4:122685352-122685675 | 323 | | 1 | 13 | | | | | 0 | 1 | | | | | | | | | | |
| BBS7 | S_Shore | TSS1500 | chr4:122791907-122792031 | 124 | | | | | | | | | 1 | | | | | | yes | | yes | | |
| TRPC3 | N_Shore;Island;S_Shore | 1stExon;Body;TSS200 | chr4:122853464-122854405 | 941 | | | 3 | | | | | | 10 | | | 2 | | yes | | | | | yes |
| ADAD1 | N_Shore;Island | TSS1500;TSS200;5'UTR | chr4:123299985-123300469 | 484 | | | 5 | | | | | | 1 | | | | | yes | | | yes | | |
| FGF2 | N_Shore | TSS1500 | chr4:123747447-123747750 | 303 | | 1 | | 1 | | | | 0 | 5 | | | | | | yes | | | yes | yes |
| SPRY1 | N_Shore | TSS200 | chr4:124317779-124318523 | 744 | | | 1 | | | | | | | | | | | | | | | | yes |
| SPRY1 | S_Shore | 1stExon;5'UTR | chr4:124320837-124321007 | 170 | | 1 | | | | | | | | | | | | yes | | | | | |
| FAT4 | Island | 1stExon | chr4:126238384-126238838 | 454 | | 1 | | | | | | | | | | | | | | | | | |
| INTU | NA | TSS1500 | chr4:128553669-128553930 | 261 | | | | | | | | | 1 | | | | | yes | | | | | |
| PCDH10 | Island | TSS1500;TSS200 | chr4:134069409-134070441 | 1032 | | | | | 1 | | | | 3 | | | | | | | | | | yes |
| PABPC4L | NA | 5'UTR | chr4:135122410-135123107 | 697 | | | | | | | | 0 | | 1 | | | | yes | yes | | | | yes |
| PCDH18 | NA | 1stExon;5'UTR | chr4:138453416-138453717 | 301 | | | | | | | | 0 | 1 | | | | | | | | | | yes |
| ELF2 | NA | TSS200 | chr4:140060774-140061152 | 378 | | | | | | | | | 1 | | | | | yes | yes | | | | yes |
| C4orf49 | S_Shore | TSS200 | chr4:140201537-140201652 | 115 | | | 19 | | | | | 0 | 1 | | | | | yes | | | | | |
| NDUFC1 | N_Shore | Body | chr4:140216130-140216277 | 147 | | | | | | | | 0 | 1 | | | | | yes | | | | | |
| SETD7 | NA | 3'UTR | chr4:140429869-140429995 | 126 | | | | | | | 1 | 0 | 1 | 1 | | 1 | | | | | | yes | |
| MGST2 | NA | TSS200;1stExon;5'UTR | chr4:140586881-140587536 | 655 | | | 6 | | | | | | | | | | | yes | | | | yes | |
| MAML3 | Island | Body | chr4:140656749-140657110 | 361 | | 1 | 1 | | 1 | | | | | | | | | | | | | yes | yes |
| MAML3 | NA | Body | chr4:141013584-141013724 | 140 | | | | | | | | | 1 | | | | | | yes | | | yes | |
| SCOC | NA | 5'UTR | chr4:141207794-141207852 | 58 | | | | | | | | | | | | | | yes | | | | yes | |
| SCOC | NA | TSS200;5'UTR | chr4:141264456-141264721 | 265 | | | 4 | | | | | | | | | | | yes | | | | yes | yes |
| SCOC | N_Shore | TSS1500;Body | chr4:141293985-141294016 | 31 | | | | | | | | | | | | | | | yes | | | | |
| UCP1 | Island | TSS200 | chr4:141490037-141490377 | 340 | | | | | | | | 0 | 3 | | | | | yes | | | | | |
| LOC441046 | Island | TSS200 | chr4:144480542-144480551 | 9 | | | | | | | | | | | | | | | | | | | |
| FREM3 | Island | TSS200 | chr4:144621898-144621945 | 47 | | | | | | | | | 1 | | | | | | | | | | |
| HHIP | N_Shore;Island | TSS1500 | chr4:145566200-145566663 | 463 | | | | | | | | 0 | | | | | | | | | | yes | yes |
| ZNF827 | Island | 3'UTR | chr4:146682572-146682755 | 183 | | 1 | | | | | | | | | | | | | | | | | yes |
| EDNRA | N_Shore | TSS1500;TSS200 | chr4:148401273-148402408 | 1135 | | 1 | 6 | | | | | | 1 | | | | | | | yes | | | |
| NR3C2 | N_Shore | 5'UTR | chr4:149362435-149362830 | 395 | | | | | | | | | 1 | | | | | yes | | | | | |
| DCLK2 | N_Shore | Body | chr4:151177137-151177553 | 416 | | | 5 | | | | | | | | | | | | | | | | yes |
| MAB21L2;LRBA | S_Shore;N_Shore;Island | TSS1500;Body | chr4:151500631-151501739 | 1108 | | | 11 | | | | | | | | | | | | | | | | |
| LRBA;MAB21L2 | N_Shore | Body;1stExon;5'UTR | chr4:151503227-151503546 | 319 | | | 1 | | | | | | | | | | | | | | | | yes |
| LRBA;MAB21L2 | Island | Body;1stExon | chr4:151504725-151505084 | 359 | | | 1 | | | | | | | | | | | | | | | | |
| FAM160A1 | Island | TSS200 | chr4:152330337-152330465 | 128 | | | | | | | | | | | | | | | | | | yes | |
| TMEM154 | NA | TSS200 | chr4:153601329-153602251 | 922 | | | 10 | | 1 | | | | | | | | | | | | yes | | |
| TRIM2 | N_Shore | TSS1500 | chr4:154073371-154073501 | 130 | | | 1 | | | | | | 1 | | | | | | | | | yes | |
| TRIM2 | NA | TSS1500;5'UTR;TSS200;1stExon | chr4:154125208-154125720 | 512 | | | 24 | | 1 | | | | 1 | | | | | yes | | | | | |
| TRIM2 | Island | Body;5'UTR | chr4:154143948-154144462 | 514 | | | | | 1 | | | | | | | | | | | | | | |
| RNF175 | Island | Body;5'UTR;1stExon | chr4:154680808-154681619 | 811 | | 1 | | | | | | | 1 | | | | | yes | | | yes | | yes |
| SFRP2 | N_Shore;Island | TSS1500;TSS200 | chr4:154710353-154712580 | 2227 | | 2 | 59 | 1 | | | | 0 | 7 | | | | | | yes | | | yes | |
| DCHS2 | N_Shore;Island | Body | chr4:155253864-155254602 | 738 | | | 12 | | | | | | | | | | | | | | yes | | yes |
| DCHS2 | S_Shore | TSS1500 | chr4:155413469-155413789 | 320 | | | 1 | | | | | | | | | | | | | | | yes | |
| LRAT | Island | TSS1500;TSS200 | chr4:155664242-155665832 | 1590 | | 1 | | | | | | | 2 | | | | | yes | | | | yes | |
| RBM46 | Island;N_Shore | 1stExon;5'UTR;TSS1500;TSS200 | chr4:155702172-155703429 | 1257 | 1 | | 17 | | | 1 | | 0 | 4 | 1 | | 2 | | yes | | | | | |
| MAP9 | S_Shore | TSS200;TSS1500 | chr4:156298204-156298427 | 223 | | | 11 | | | | | 0 | 2 | | | 2 | | yes | | | | | yes |

| | | | | | | | | | | | | | | | | | | | | | | | |
|---|---|---|---|---|---|---|---|---|---|---|---|---|---|---|---|---|---|---|---|---|---|---|---|
| GUCY1A3 | N_Shore | 5'UTR;TSS200;TSS1500 | chr4:156588121-156588683 | 562 | | | | | | | | | 1 | | | | | | | | | | yes |
| GUCY1B3 | N_Shore;Island | TSS200;1stExon;5'UTR | chr4:156680007-156680518 | 511 | | 1 | 2 | 1 | | | | 0 | 4 | | | 1 | | | | | | | |
| CTSO | S_Shore | TSS1500 | chr4:156875336-156875661 | 325 | | | 5 | | | | | | | | | | | yes | | | yes | yes | |
| PDGFC | Island | TSS1500 | chr4:157892748-157893306 | 558 | | | | | | | | 0 | | | | | | | yes | | | | yes |
| GLRB | N_Shore;Island | TSS1500;1stExon;TSS200;5'UTR | chr4:157996959-157997750 | 791 | | | 21 | | | | | 0 | 5 | | | | | yes | | | | | yes |
| GRIA2 | Island | TSS1500 | chr4:158141492-158141936 | 444 | | | | | | | | 0 | 2 | | | | | | yes | | | | yes |
| GRIA2 | Island | Body | chr4:158143439-158144318 | 879 | | | | | | | | 0 | 2 | | | | | | | | | yes | |
| FAM198B | NA | Body | chr4:159091837-159092553 | 716 | | | 6 | | 1 | | | 0 | | | | | | | | | | | yes |
| FAM198B | NA | TSS200;5'UTR;TSS1500;1stExon | chr4:159093841-159094517 | 676 | 1 | | | | | | | | | | | | | | | | | | |
| TMEM144 | N_Shore | TSS200 | chr4:159131324-159131785 | 461 | | | 3 | | | | | | 1 | | | | | yes | | | | | |
| RAPGEF2 | NA | TSS1500 | chr4:160188727-160189254 | 527 | | 1 | | | | | | | | | | | | | | | | | yes |
| TKTL2 | Island | 1stExon;5'UTR | chr4:164394898-164395149 | 251 | | | 3 | | | | | | | | | | | | | yes | | | |
| 36951 | Island | TSS200 | chr4:165304531-165305050 | 519 | | 1 | | | | | | | | | | | | | | | | | yes |
| C4orf39;TRIM61 | Island;N_Shore | TSS200;Body;1stExon;5'UTR;TSS1! | chr4:165877875-165878742 | 867 | 3 | 1 | 2 | | | | 1 | 0 | 6 | 2 | | 5 | | yes | yes | | | | |
| TRIM61 | NA | 5'UTR;1stExon;TSS200 | chr4:165898666-165898967 | 301 | 1 | | 26 | | | | | 0 | 9 | | | 8 | 1 | yes | | | | | |
| TRIM60 | NA | TSS1500;TSS200 | chr4:165952835-165953131 | 296 | | | 4 | | | | | | | | | | | yes | | | | | |
| KLHL2 | S_Shore | Body | chr4:166129561-166129640 | 79 | | | | | | | | | | | | | | | | | | | yes |
| KLHL2 | S_Shore | TSS200;Body | chr4:166131163-166131638 | 475 | | | | | | | | | | | | | | | yes | | | | |
| CPE | S_Shore | Body | chr4:166301076-166301256 | 180 | 1 | | 1 | | | 1 | | | | | | | | | | | | | |
| TLL1 | S_Shore | Body | chr4:166795401-166796183 | 782 | | | | | | | | | 1 | | | | | | | | | | yes |
| PALLD | NA | TSS200 | chr4:169418081-169418224 | 143 | | | | | | | | | | | | | | | yes | | | | |
| PALLD | N_Shore | 5'UTR;Body | chr4:169798931-169799624 | 693 | | | 5 | | | | | | 1 | 2 | | | | yes | | | | yes | yes |
| MFAP3L | NA | Body;5'UTR | chr4:170913525-170913676 | 151 | | | | | | | | | | | | | | yes | | | | | |
| GALNT7 | N_Shore | TSS200 | chr4:174089748-174089978 | 230 | | | | | | | | | 1 | | | | | | | | | | yes |
| HAND2 | N_Shore;Island | 3'UTR;Body | chr4:174447847-174448841 | 994 | | | 23 | | | | | 0 | 1 | | | | | | | | | yes | |
| HAND2;NBLA00301 | Island;S_Shore | 5'UTR;TSS1500;1stExon;TSS200;Bo | chr4:174450722-174453287 | 2565 | | 1 | 37 | | | | | | 3 | | | | | | | | | yes | yes |
| NBLA00301 | N_Shore | Body | chr4:174457224-174457692 | 468 | | | | | | | | 0 | 1 | | | | | | | | | | yes |
| HPGD | S_Shore | TSS200;TSS1500 | chr4:175444181-175444482 | 301 | | | | | | | | 0 | | | | | | yes | | | | | |
| ADAM29 | NA | TSS200 | chr4:175839468-175839610 | 142 | | | | | | | | | | | | | | | yes | | | | |
| MGC45800 | N_Shore;Island | Body | chr4:183061830-183063534 | 1704 | | 2 | 10 | | | | | | 2 | 1 | | | | | | | | | |
| ODZ3 | NA | Body | chr4:183247132-183247234 | 102 | | | 2 | | | | | | | | | | | yes | | | | | yes |
| ODZ3 | Island | Body | chr4:183721183-183721778 | 595 | | | | | | | 1 | | 2 | | | | | | | | | | yes |
| STOX2 | NA | Body;nan | chr4:184908351-184909018 | 667 | | | 11 | 1 | | | | 0 | 11 | 1 | 1 | 2 | | | | | | | |
| STOX2 | NA | Body | chr4:184922382-184922571 | 189 | | | 1 | | | | | | | | | | | | yes | | | | |
| STOX2 | NA | Body | chr4:184930579-184932237 | 1658 | | | 9 | | | | | | | | | | | | | | yes | | |
| STOX2 | NA | Body | chr4:184937961-184938470 | 509 | 1 | | 13 | 1 | | | | | | | | | | yes | | | | yes | |
| ENPP6 | Island | Body | chr4:185018391-185018682 | 291 | | | | 1 | | | | 0 | 1 | | | 2 | | | | yes | yes | | |
| ENPP6 | NA | 1stExon;5'UTR | chr4:185139078-185139478 | 400 | | | | | | | | | | | | | | | | | | | |
| HELT | Island | Body | chr4:185941265-185941625 | 360 | | | | | | | | | 1 | | | | | | | | | | |
| SNX25 | Island | TSS1500;TSS200 | chr4:186130781-186131280 | 499 | | | 4 | | | | | | | | | | | | | | | | yes |
| PDLIM3 | NA | 3'UTR | chr4:186423408-186423579 | 171 | | | 2 | | | | | | | | | | | | | | | yes | |
| PDLIM3 | NA | Body | chr4:186425697-186425754 | 57 | | | 1 | | | | | | | | | | | | yes | | | | |
| PDLIM3 | NA | Body | chr4:186434993-186435901 | 908 | | | 2 | 1 | | | | | | | | | | | | | yes | | yes |
| PDLIM3 | S_Shore | TSS1500 | chr4:186456935-186457012 | 77 | | | | | 1 | | | | | | | | | yes | | | | | yes |
| SORBS2 | NA | 3'UTR | chr4:186508724-186509045 | 321 | | | 7 | 1 | | | | 0 | | 1 | | | | | | | | | |
| SORBS2 | N_Shore;Island | Body | chr4:186559412-186560403 | 991 | | | 8 | | 1 | 1 | | 0 | 2 | 2 | | | | yes | | | | | |
| SORBS2 | NA | 5'UTR;1stExon;TSS200;TSS1500 | chr4:186732124-186734622 | 2498 | | | | | | | | | | | | | | | | | | | |
| SORBS2 | NA | 5'UTR | chr4:186741362-186742050 | 688 | 1 | | 1 | 1 | | | | | | | | | | yes | yes | | | | |
| SORBS2 | NA | 5'UTR | chr4:186847142-186847469 | 327 | | | | | 1 | | | | | | | | | | yes | | | | |
| SORBS2 | NA | 5'UTR;1stExon | chr4:186877497-186877916 | 419 | | | 10 | | | | | | 2 | | | | | yes | | | | yes | |
| FAM149A | Island | 5'UTR | chr4:187066279-187066505 | 226 | | | | | | | | | 1 | | | | | | | | | yes | yes |
| CYP4V2 | NA | Body | chr4:187125413-187126139 | 726 | | 1 | | | | | | | 1 | | | | | yes | | | | yes | |
| KLKB1 | NA | TSS1500 | chr4:187148009-187148688 | 679 | | | | 1 | | | | | 2 | | | | | | yes | | | | |
| F11 | NA | TSS200 | chr4:187186949-187187177 | 228 | | | | 1 | | | | | | | | | | yes | | | | yes | yes |
| MTNR1A | NA | Body | chr4:187459415-187459476 | 61 | 1 | | | | | | | | | | | | | | | yes | | | |
| MTNR1A | N_Shore;Island | Body;1stExon;TSS200 | chr4:187475841-187476953 | 1112 | | | | | | | | 0 | 9 | | 1 | 4 | | yes | yes | yes | | | |
| FAT1 | NA | Body | chr4:187580694-187581093 | 399 | | 1 | | | | | | | | | | | | | | | yes | | |
| FAT1 | NA | Body | chr4:187621221-187622164 | 943 | | | 23 | | | | | | 1 | | | | | yes | | | yes | | |
| FAT1 | NA | Body | chr4:187629328-187629387 | 59 | | | | | | | | | 2 | | | | | | | | | | |
| FAT1 | Island | 5'UTR;TSS200 | chr4:187644620-187645448 | 828 | | | | | | | | 0 | 5 | 3 | | | | | | | | yes | yes |
| ZFP42 | N_Shore;Island;S_Shore | TSS1500;TSS200;1stExon;5'UTR | chr4:188916496-188918218 | 1722 | | 1 | 9 | | 3 | | | 0 | 7 | 2 | | 4 | 1 | yes | yes | | yes | | |
| TRIML1 | NA | TSS1500;1stExon;5'UTR | chr4:189059510-189061030 | 1520 | | | | | | | | | 1 | | | | | yes | | | | | |
| TUBB4Q | Island | Body;1stExon;TSS200 | chr4:190905801-190906123 | 322 | | | 9 | | | | | | | | | | | | | yes | | | yes |
| PLEKHG4B | NA | TSS1500 | chr5:139859-140009 | 150 | | | | | | | | | | | | | | | | | | | |
| PLEKHG4B | NA | Body | chr5:155061-155199 | 138 | | | | | | | | | 1 | | | | | | yes | | | yes | |
| PLEKHG4B | N_Shelf | Body | chr5:161555-161852 | 297 | | | | | | | | | | | | | | | | | | | |
| PLEKHG4B | N_Shore | Body | chr5:169434-170081 | 647 | | | 2 | | | | | | | | | | | yes | | | | | |
| LRRC14B | N_Shore;Island | TSS200;1stExon;TSS1500 | chr5:191127-192470 | 1343 | | | 14 | | | | | 0 | 4 | 1 | | 4 | | | | yes | | | yes |
| LRRC14B | N_Shore | Body | chr5:194530-194879 | 349 | | | | | | | | | | | | | | | | | yes | | |
| CCDC127;SDHA | Island | 5'UTR;TSS1500 | chr5:218153-218452 | 299 | | | | | | | | | | 1 | | 1 | | | yes | | | yes | |
| SDHA | NA | Body | chr5:252410-252699 | 289 | | | | | | | | | | | | | | | | | | | |
| AHRR | S_Shore | Body | chr5:323320-323969 | 649 | | | 26 | | | | | | 1 | | | | | yes | | | | | yes |
| AHRR | Island | Body | chr5:343927-344218 | 291 | | | | | | | | | 1 | | | | | | | | | yes | |
| AHRR | S_Shore | Body | chr5:345850-346695 | 845 | | | 7 | | | | | | | | | | | | yes | yes | | yes | |
| AHRR | N_Shelf;N_Shore | Body | chr5:367976-369088 | 1112 | | | 14 | | 1 | | | | | | | | | | | | | | |
| AHRR;LOC100310782 | NA | Body;TSS200 | chr5:404264-404766 | 502 | | | | | | | | | 1 | | | | | yes | | | | | |
| AHRR | NA | Body | chr5:415575-415885 | 310 | | | 17 | | | | 1 | 0 | 1 | 1 | 1 | 2 | 1 | | | yes | | | |
| AHRR | Island | Body | chr5:421317-421733 | 416 | | | | | | 1 | | | 2 | | | 2 | | | | | | yes | |
| AHRR | S_Shelf | Body | chr5:423894-424371 | 477 | | | 15 | | | | | | | | | | | | yes | | | | yes |
| EXOC3;C5orf55 | S_Shore | 5'UTR;TSS1500 | chr5:443987-444228 | 241 | | | | | | | | | | | | | | | | | | | |
| EXOC3 | N_Shelf | Body | chr5:462141-462522 | 381 | 1 | | | | | | | | | | | | | yes | yes | | | | |
| LOC25845 | N_Shore | Body | chr5:471171-471385 | 214 | | | 1 | | | | | | | | | | | | | | | | |
| SLC9A3 | Island | Body | chr5:476497-476842 | 345 | | | | | | | | | 1 | | | | | | | | | | |
| SLC9A3 | Island | Body | chr5:488398-488966 | 568 | | | | | | | | | | | | | | yes | | | | | |
| SLC9A3 | N_Shore | Body | chr5:493262-493893 | 631 | | | 8 | | | | | | 2 | | | 1 | | | | | | | |
| SLC9A3 | Island;S_Shore;N_Shore | Body | chr5:495102-496476 | 1374 | | | 5 | | | | | | 1 | | | | | yes | yes | | | | yes |
| SLC9A3 | N_Shore | Body | chr5:497520-497639 | 119 | | | | | | | | | | | | | | yes | | | | yes | |
| SLC9A3 | N_Shore;Island;S_Shore | Body | chr5:501343-502571 | 1228 | | | 21 | | 1 | | | 0 | 1 | 1 | | 1 | | | | | yes | yes | |
| SLC9A3 | N_Shore;Island | Body | chr5:505985-506472 | 487 | | | | | | | | | 1 | | | | | yes | | | | | yes |
| SLC9A3 | S_Shore | Body | chr5:510035-510421 | 386 | | | | | | | | | 1 | 1 | | | | yes | | | | yes | |
| SLC9A3 | Island | 5'UTR;1stExon;TSS200 | chr5:524503-524848 | 345 | | | | | | | | | 2 | | | | | | yes | | | | yes |
| TPPP | N_Shore | 3'UTR | chr5:663895-664666 | 771 | | | | | | | | | 2 | | | | | | | | | | |
| TPPP | S_Shore;Island | Body | chr5:667968-669925 | 1957 | | | 4 | | | | | 0 | | | | | | | yes | | | | |
| TPPP | S_Shelf;NA | Body | chr5:672357-672910 | 553 | | | | | | | | 0 | 2 | 1 | | 6 | | | | | | | |
| TPPP | Island | Body | chr5:677840-678046 | 206 | | | 1 | | | | | | | | | | | | | | | | |
| ZDHHC11 | S_Shore;S_Shelf | 1stExon;5'UTR;TSS200;TSS1500 | chr5:850745-851892 | 1147 | 1 | 1 | 7 | | | | | | 1 | | | | | yes | | | | | |
| BRD9 | S_Shelf | Body | chr5:883880-884033 | 153 | 2 | | | | | | | | | | | | | | | | yes | | |
| TRIP13 | N_Shore;Island | Body | chr5:912383-913111 | 728 | | 1 | 4 | | | | | | 3 | 1 | 1 | | | yes | | | | | |
| NKD2 | NA | Body | chr5:1016279-1016768 | 489 | | | 9 | | | | | | 1 | 1 | | | | | | | | | |

| | | | | | | | | | | | | | | | | | | | | | | | |
|---|---|---|---|---|---|---|---|---|---|---|---|---|---|---|---|---|---|---|---|---|---|---|---|
| SLC12A7 | N_Shore;Island | Body | chr5:1052540-1054427 | 1887 | | | 31 | | | | | | 1 | 1 | | | | | | | | | |
| SLC12A7 | N_Shore | Body | chr5:1060334-1060773 | 439 | | | 1 | | | | | | | | | | | | | | | | |
| SLC12A7 | Island | Body | chr5:1068772-1068999 | 227 | | | 2 | | | | | | | | | | | yes | | | | | |
| SLC12A7 | N_Shore | Body | chr5:1076587-1076758 | 171 | | | 1 | | | | | | | | | | | | | | | | |
| SLC12A7 | S_Shore;N_Shore | Body | chr5:1090619-1092391 | 1772 | | | 11 | | | | | | 1 | | | | | | | | | | |
| SLC12A7 | Island | Body | chr5:1093580-1093797 | 217 | | | | 1 | | | | | | | | | | | | | | | |
| SLC12A7 | N_Shelf | Body | chr5:1097374-1097794 | 420 | | | 2 | | | | | | | | | | | | | | | yes | |
| SLC12A7 | Island | Body | chr5:1099476-1100544 | 1068 | | | 2 | 2 | | | | | | | | | | | yes | | | | |
| SLC12A7 | N_Shore;chr5:1108802-110905 | Body;N_Shore | chr5:1107098-1108315 | 1217 | | | | | 1 | | | | | | 1 | | | | | yes | | yes | |
| SLC6A19 | N_Shore;Island | TSS1500;TSS200 | chr5:1201359-1202449 | 1090 | | 1 | 11 | 1 | | | | | 3 | | | | | | | | | | |
| SLC6A19 | Island;N_Shore | Body | chr5:1217080-1218916 | 1836 | | 1 | 10 | | | | | | | | | | | | | | | | |
| SLC6A19 | N_Shelf | 3'UTR | chr5:1223167-1223262 | 95 | | | 5 | | | | | | | | | | | | | | | yes | |
| SLC6A19;SLC6A18 | N_Shore | 3'UTR;TSS1500;TSS200 | chr5:1225074-1225719 | 645 | | | 12 | | | | | | | | | | | | yes | | | | |
| SLC6A18 | S_Shore | Body | chr5:1227579-1228671 | 1092 | | 1 | 10 | 1 | | | | 0 | 2 | | | | | yes | yes | | | | |
| SLC6A18 | Island;S_Shore | Body | chr5:1232590-1233120 | 530 | | 1 | | | | | | | | | | | | yes | | | | | |
| SLC6A18 | N_Shore | Body | chr5:1241765-1241958 | 193 | | | | | | | | | | | | | | | | yes | yes | | |
| SLC6A18 | Island | Body | chr5:1243714-1243809 | 95 | | | 3 | | | | | | | | | | | | | | | | |
| SLC6A18 | N_Shore;Island | Body;3'UTR | chr5:1245669-1246795 | 1126 | | | 3 | | | | | 0 | 1 | | | 1 | | | | | | | yes |
| TERT | Island | 3'UTR | chr5:1253833-1254048 | 215 | | | | | | | | | | | | | | | | | | | |
| TERT | S_Shore | Body | chr5:1258644-1258994 | 350 | | | | | | | | | | | | | | | | | | | |
| TERT | NA | Body | chr5:1264435-1264568 | 133 | | 1 | | | | | | | | | | | | | | | | | |
| TERT | NA | Body | chr5:1266145-1266595 | 450 | | | | | | | | | | | | | | | | | | | |
| TERT | N_Shore;N_Shelf | Body | chr5:1268776-1271100 | 2324 | | | | | | | | | | | | | | | | | | | |
| TERT | N_Shore | Body | chr5:1277224-1277596 | 372 | | | | | | | | | | | | | | | yes | | | | |
| TERT | Island | Body | chr5:1282703-1282984 | 281 | | | | | | | | | | | | | | yes | | | | | |
| TERT | Island | TSS1500 | chr5:1295737-1296007 | 270 | | | | | | | | 0 | | | | | | | | | | yes | |
| CLPTM1L | NA | Body | chr5:1324183-1324645 | 462 | | | | | | | | | 1 | | | | | | | | | | |
| CLPTM1L | NA | Body | chr5:1325969-1326034 | 65 | | | | | | | | | 1 | | | | | | | | | | |
| CLPTM1L | S_Shore | Body | chr5:1335846-1335965 | 119 | | | | | | | | | 1 | | | | | | | | | | |
| CLPTM1L | N_Shelf | Body | chr5:1342124-1342200 | 76 | | | | | 1 | | | | 2 | | | | | | | | | yes | |
| SLC6A3 | NA | 3'UTR;Body | chr5:1393877-1395355 | 1478 | | | 11 | | | | | | | | | | | | | | | | |
| SLC6A3 | NA | Body | chr5:1405330-1405774 | 444 | | | | | | | | | | | | | | | | | yes | | |
| SLC6A3 | NA | Body | chr5:1407319-1407569 | 250 | | | | | | | | | | | | | | | yes | | | | |
| SLC6A3 | NA | Body | chr5:1409937-1410090 | 153 | | | | | | | | | | | | | | | | | | | |
| SLC6A3 | Island | Body | chr5:1416793-1417003 | 210 | | | | | | | | | | | | | | | | | | | |
| SLC6A3 | NA | Body | chr5:1430421-1430867 | 446 | | | 3 | | | | | | | | | | | | | | | | |
| SLC6A3 | NA | Body | chr5:1438168-1438531 | 363 | | | | | | | | | | | | | | | | | | | |
| SLC6A3 | Island | TSS200;TSS1500;5'UTR;1stExon | chr5:1445354-1446443 | 1089 | | 1 | | | 1 | | | | 4 | | | 2 | | | | yes | | | yes |
| LPCAT1 | Island | Body | chr5:1489818-1489889 | 71 | | | | | 1 | | 1 | | 1 | | | | | | | yes | | | |
| LPCAT1 | S_Shore | Body | chr5:1491489-1491600 | 111 | | | | | | | | | 1 | | | | | | | | yes | | |
| LPCAT1 | Island;chr5:1494853-1495287 | Body;Island | chr5:1494983-1495356 | 373 | | | | | 1 | | | 0 | | | 1 | 1 | | | | | | | |
| LPCAT1 | NA | Body | chr5:1503595-1503979 | 384 | | | 2 | | | | | 0 | | | | | | | yes | | yes | | |
| LPCAT1 | NA | Body | chr5:1510512-1511473 | 961 | | | | | | | | | 1 | | | | | | | | | | |
| LPCAT1 | NA | Body | chr5:1513791-1514140 | 349 | | | | | | | | | 1 | | | | | | | | | | |
| SDHAP3 | Island | Body;TSS200 | chr5:1594282-1595048 | 766 | 7 | 8 | 30 | 1 | 4 | 6 | 2 | 0 | 10 | 12 | 3 | 40 | 3 | yes | yes | | | yes | |
| LOC728613 | S_Shore | TSS1500 | chr5:1634460-1634505 | 45 | | | 2 | | | | | | 1 | | | 1 | | yes | | | | | |
| NDUFS6;MRPL36 | Island | TSS1500 | chr5:1800542-1800782 | 240 | | | 1 | | | | | | | | | | | | yes | | | | |
| IRX4 | Island;N_Shore | Body | chr5:1878205-1880310 | 2105 | | | 3 | | | | | 0 | 2 | 2 | 1 | | | yes | | | | yes | yes |
| IRX4 | Island | 5'UTR;1stExon | chr5:1882775-1882902 | 127 | | | | | | | | 0 | 1 | | | | | | | | | | yes |
| IRX4 | Island | TSS1500 | chr5:1883940-1884212 | 272 | | 1 | | | | | | | | | | | | | | | | | |
| IRX2;C5orf38 | Island | Body;TSS1500;1stExon;5'UTR;TSS2 | chr5:2749715-2755820 | 6105 | | | | | | | | | | | | | | | | | | | |
| IRX1 | Island | TSS200 | chr5:3595970-3596207 | 237 | | | | | | | | 0 | 1 | | | | | | | | | | yes |
| IRX1 | Island | Body | chr5:3597311-3597695 | 384 | | | | | | | | | 1 | | | | | | | | | | |
| LOC340094 | NA | TSS1500;TSS200 | chr5:5033311-5034397 | 1086 | | 1 | | | | | | | 1 | | | | | yes | | yes | | | |
| ADAMTS16 | Island | TSS1500 | chr5:5139458-5140646 | 1188 | | 1 | 3 | | | | | 0 | 11 | 1 | | 6 | | yes | | yes | | yes | yes |
| ADAMTS16 | NA | Body | chr5:5287980-5288296 | 316 | | | | | | | | | | | | | | | | | | | |
| ADAMTS16 | Island | Body | chr5:5303357-5303564 | 207 | | | | | | | | | | | | | | | yes | | | | yes |
| KIAA0947 | N_Shore | TSS1500 | chr5:5421452-5421908 | 456 | | | 1 | | | | | | | | | | | | | | | | |
| FLJ33360 | NA | TSS200 | chr5:6337408-6338017 | 609 | | | | | | | | | 1 | | | | | yes | | | yes | yes | |
| UBE2QL1 | NA | 3'UTR;Body | chr5:6491443-6491884 | 441 | | | 7 | | | | | | | | | | | | | | | | |
| LOC255167 | N_Shore | TSS1500 | chr5:6581774-6582193 | 419 | | | | | | | | | | | | | | yes | | yes | | | |
| POLS | Island | Body | chr5:6737778-6737918 | 140 | | | | | | | | | 1 | | | | | | | | | | |
| POLS | N_Shore;Island;S_Shore | Body;3'UTR | chr5:6755026-6755843 | 817 | | | | | | | | 0 | 3 | | | | | yes | | | | | |
| ADCY2 | N_Shore | TSS1500 | chr5:7394927-7395395 | 468 | | | 6 | | | | | | | | | | | | | | | | |
| ADCY2 | Island | 1stExon | chr5:7396491-7397021 | 530 | | | | | | | | | 1 | | | | | | | | | | yes |
| ADCY2 | NA | Body;3'UTR | chr5:7826900-7827964 | 1064 | | | 10 | | | | | | 1 | | | | | yes | yes | | | | |
| C5orf49 | Island | TSS200;1stExon;5'UTR | chr5:7851031-7851681 | 650 | | | 5 | | | | | | 1 | | | | | | yes | | yes | | yes |
| SNORD123;SEMA5A | S_Shore | TSS1500 | chr5:9547468-9548098 | 630 | | | 2 | | | | | | 1 | | | 2 | | yes | | | | | |
| FAM173B;CCT5 | Island | TSS1500;TSS200 | chr5:10250278-10251089 | 811 | | | | | | | | | | | | 1 | | | yes | | | yes | |
| 38777 | NA | Body | chr5:10376970-10377171 | 201 | | | | | | | | | | | | | | | | yes | | | |
| ANKRD33B | Island | TSS1500 | chr5:10563092-10563665 | 573 | | | 4 | | | | | | | | | | | | | | | | yes |
| ANKRD33B | Island | Body | chr5:10649381-10649867 | 486 | | | | | | | | | | | | | | yes | | | | yes | yes |
| CTNND2 | Island | Body | chr5:11385520-11385893 | 373 | | | | | 2 | | | | | | | | | | | | | yes | |
| CTNND2 | Island | TSS200 | chr5:11904110-11904811 | 701 | | 1 | | | | | | 0 | 1 | 1 | 1 | | | | | | | | yes |
| DNAH5 | Island | Body | chr5:13810143-13810279 | 136 | | | 9 | | | | | 0 | 1 | 1 | | 2 | | yes | | | | | |
| TRIO | NA | Body | chr5:14359544-14359614 | 70 | 1 | | | | | | | | | | | | | | | | | | yes |
| TRIO | NA | Body | chr5:14405747-14406732 | 985 | | | 7 | | 1 | | | | 1 | | 1 | | | yes | | | | | |
| TRIO | NA | Body | chr5:14410252-14410363 | 111 | | | 3 | | | | | | | | | | | yes | | | | | |
| TRIO | NA | Body | chr5:14440991-14441261 | 270 | | | | | | | | 0 | 1 | | | | | | | | | | |
| ANKH | N_Shore | Body | chr5:14870083-14870749 | 666 | | | | | | | 1 | 0 | 2 | 1 | | 5 | 1 | | | | | | yes |
| FBXL7 | N_Shore | TSS1500 | chr5:15499895-15500833 | 938 | | 1 | | | | | | | | | | | | | | | | | yes |
| 40603 | Island | TSS1500 | chr5:16180259-16181039 | 780 | | 1 | | | | | | | | | | | | yes | | | | | yes |
| FAM134B | NA | Body;1stExon;5'UTR;TSS200 | chr5:16508920-16509323 | 403 | | 1 | 11 | | 2 | | | | | | | | | | | | | | |
| MYO10 | NA | Body | chr5:16742020-16742179 | 159 | | | 6 | | | | | | | | | | | yes | | | | | yes |
| MYO10 | NA | Body | chr5:16784821-16785135 | 314 | | | | | 1 | | | | 2 | | | | | | | | | | |
| MYO10 | NA | Body | chr5:16794797-16794882 | 85 | | 1 | | | | | | | | | | | | | yes | | | yes | |
| MYO10 | Island | Body;1stExon;5'UTR | chr5:16935888-16936458 | 570 | | | | | | | | | 3 | | | | | | | | | yes | yes |
| CDH18 | NA | nan | chr5:19886414-19886600 | 186 | | | | | | | | | | | 1 | | | | yes | | yes | | |
| CDH18 | NA | 1stExon;5'UTR;TSS1500 | chr5:19988200-19988848 | 648 | | | | | 2 | | | 0 | 4 | | | | | | | | | | yes |
| CDH12 | NA | 5'UTR | chr5:22212932-22213263 | 331 | | 1 | | | 1 | | | | | | | | | | | | | | |
| CDH12 | NA | 5'UTR;1stExon | chr5:22853703-22853901 | 198 | | | 1 | | | | | | | | | | | | | | | | |
| PRDM9 | NA | TSS1500;TSS200 | chr5:23506738-23507752 | 1014 | | | 39 | 1 | | | | 0 | 10 | 2 | | 8 | | | yes | yes | | | |
| CDH10 | NA | Body | chr5:24488007-24488076 | 69 | | | 3 | | | | | | | | | | | | | | | | |
| CDH10 | NA | TSS200 | chr5:24645065-24645147 | 82 | | | | | | | | | 1 | | | | | | | | | | yes |
| CDH9 | NA | 1stExon;5'UTR;TSS200 | chr5:27038605-27038836 | 231 | | | | | 1 | | | 0 | 4 | | 1 | | 1 | | | | | | |
| CDH6 | S_Shore | 5'UTR | chr5:31194641-31194800 | 159 | | 1 | | | | | | | | | | | | | | | | | yes |
| CDH6 | NA | Body | chr5:31323107-31323297 | 190 | | | | | | | | | | | | | | | | | | | |
| PDZD2 | NA | TSS1500 | chr5:31798797-31799023 | 226 | | | | | | | | | | | | | | | yes | | | | |
| ZFR | S_Shore | TSS1500 | chr5:32445429-32445523 | 94 | | 1 | | | | | | | | | | | | | yes | | | | yes |

| | | | | | | | | | | | | | | | | | | | | | | | |
|---|---|---|---|---|---|---|---|---|---|---|---|---|---|---|---|---|---|---|---|---|---|---|---|
| NPR3 | Island | Body | chr5:32713723-32714525 | 802 | | 1 | | | | | | | | | | | | yes | | | | | |
| RXFP3 | Island | TSS1500;1stExon;5'UTR | chr5:33936171-33938225 | 2054 | | 1 | | | | | | | 2 | | | 2 | | | | | | | yes |
| AMACR | NA | Body;3'UTR | chr5:33997223-33997484 | 261 | | | | | | | | | | | | | | yes | yes | | | | |
| C1QTNF3 | NA | TSS200;Body;1stExon;5'UTR | chr5:34042976-34043509 | 533 | | | 13 | | 3 | | | | 1 | | | | | | | | | yes | |
| SPEF2 | N_Shore | TSS1500 | chr5:35617687-35618029 | 342 | | 1 | | | | | | 0 | 1 | | | | | yes | yes | | | | yes |
| UGT3A1 | NA | Body | chr5:35990997-35991624 | 627 | | | 2 | | | | | | | | | 1 | | | | | | | |
| UGT3A2 | NA | Body;1stExon;5'UTR | chr5:36066783-36067127 | 344 | | | | | | | | | 2 | | | | | | | | | | |
| SLC1A3 | NA | TSS1500 | chr5:36606102-36606293 | 191 | | 1 | | | | | | | | | | | | | | | | | yes |
| SLC1A3 | NA | Body | chr5:36662804-36662950 | 146 | | | | | | | | 0 | 1 | | | 1 | | yes | | | | | |
| C5orf42 | N_Shore;Island | Body | chr5:37208976-37209440 | 464 | | | 4 | | | | | | | | | | | yes | yes | | | yes | |
| GDNF | Island;N_Shore | 5'UTR;TSS200;TSS1500;1stExon;Bo | chr5:37833969-37836392 | 2423 | | | | | | | | | | | | | | | | | | | |
| GDNF | Island | TSS200;TSS1500 | chr5:37839918-37840839 | 921 | | | | | | | | | 2 | | | | | | | | | | |
| EGFLAM | N_Shore | TSS1500 | chr5:38257516-38257555 | 39 | | | 13 | | | | | | | | | | | | yes | | | | |
| EGFLAM | NA | nan | chr5:38403622-38403957 | 335 | | | | | | | | | | | 1 | | | | | yes | yes | | yes |
| EGFLAM | NA | Body;TSS1500;TSS200;5'UTR | chr5:38445220-38446192 | 972 | | | 10 | | | | | 0 | 4 | | | 1 | | yes | | | | | |
| OSMR | Island;S_Shore | 5'UTR;1stExon | chr5:38846100-38846973 | 873 | | | | | | | | | 3 | | | | | yes | | | | yes | |
| FYB | NA | Body | chr5:39203007-39203190 | 183 | | | | | | | | | 1 | 1 | 1 | | | | | | | | |
| FYB | NA | 5'UTR | chr5:39219245-39219698 | 453 | | | 2 | | | | | 0 | 1 | 1 | 1 | 1 | | yes | | | | | |
| DAB2 | Island;S_Shore | 1stExon;5'UTR;TSS200;TSS1500 | chr5:39425138-39425549 | 411 | | 2 | | | | | | | 1 | | | | | | | | | | |
| PTGER4 | Island | Body | chr5:40681893-40682333 | 440 | | | | | | | | | | | | | | yes | yes | | | | |
| RPL37 | Island | TSS1500 | chr5:40835740-40835760 | 20 | | | | | | | | | 1 | | | | | yes | | | | yes | |
| CARD6 | NA | TSS200;1stExon | chr5:40841298-40841590 | 292 | | | 20 | | 1 | | | | | | | | | | yes | | | | |
| C7 | NA | TSS1500 | chr5:40908228-40908780 | 552 | | | | | | | | | 2 | | | | | yes | | yes | | | |
| PLCXD3 | N_Shore;Island | Body | chr5:41509352-41510650 | 1298 | | 1 | | | | | | 0 | 4 | | | 6 | | | | | | | yes |
| OXCT1 | Island | Body | chr5:41869679-41870316 | 637 | | | | | | | | 0 | 1 | | | 1 | | | | | | | |
| GHR | Island | TSS1500;5'UTR | chr5:42423615-42425093 | 1478 | | 2 | | | | | | | | | | | | yes | | | | yes | yes |
| CCDC152 | NA | TSS200 | chr5:42756786-42757815 | 1029 | | | 19 | | | | | 0 | 5 | | | 5 | 1 | yes | | | | | |
| SEPP1 | NA | 1stExon;5'UTR | chr5:42811952-42812509 | 557 | | | | | 1 | | | | | | | | | | yes | | | | |
| C5orf39 | N_Shore | 1stExon;3'UTR | chr5:43039410-43039867 | 457 | | | | | | | | | 1 | | | | | yes | | | | | yes |
| LOC153684 | S_Shore | Body | chr5:43044573-43044703 | 130 | | 1 | 5 | 1 | | | | | | | | | | yes | | | | yes | |
| CCL28 | N_Shore;Island | Body | chr5:43396802-43397572 | 770 | | 1 | | | | | | | | | | | | yes | yes | yes | yes | yes | |
| FGF10 | NA | TSS200;TSS1500;1stExon | chr5:44388596-44389848 | 1252 | | 1 | | | 2 | | | | 1 | | | | | | | | | | yes |
| HCN1 | N_Shore;Island | Body;1stExon | chr5:45695241-45696592 | 1351 | | 2 | | | | | | | 2 | | | | | | | | | yes | yes |
| EMB | S_Shore | TSS1500 | chr5:49737492-49738016 | 524 | | | | | | | | | 1 | | | | | | | | | yes | |
| ISL1 | Island | TSS200;5'UTR;1stExon | chr5:50678913-50679290 | 377 | | | | | | | | 0 | 9 | | | 1 | | | | | | | yes |
| ISL1 | Island | Body | chr5:50683614-50683914 | 300 | | | 1 | | | | | | | | | | | | | | | | yes |
| ISL1 | Island | Body | chr5:50685512-50685716 | 204 | | | | | | | | | 1 | | | | | | | | | yes | |
| MOCS2 | Island | TSS200;TSS1500 | chr5:52405638-52405696 | 58 | | | | | | | | 0 | | | | | | | | yes | | yes | |
| NDUFS4 | N_Shore | TSS200;1stExon;5'UTR | chr5:52856402-52856530 | 128 | | | | | | | | 0 | 1 | | | | | | | | | | |
| HSPB3 | NA | TSS200 | chr5:53751272-53751493 | 221 | | | | | | | | | 1 | | 1 | | | yes | | | | | |
| ESM1 | NA | 1stExon;TSS200 | chr5:54281198-54281733 | 535 | | | 10 | | | | | 0 | 2 | | | 1 | | | yes | | | | |
| CCNO | Island | TSS1500 | chr5:54530114-54530436 | 322 | | | 1 | | | | | | | | | | | yes | | | | | |
| PPAP2A;RNF138P1 | N_Shore;Island | Body | chr5:54826547-54826768 | 221 | | | 7 | | | | | | 2 | 1 | | | | | | | | | |
| ANKRD55 | NA | 5'UTR;1stExon;TSS200;TSS1500 | chr5:55529124-55530254 | 1130 | | | 7 | | 1 | | | | 2 | | | | | | yes | yes | | | |
| C5orf35 | N_Shore | TSS1500 | chr5:56204884-56204925 | 41 | | | 1 | | | | | | | | | | | | yes | | | | |
| PLK2 | Island | 1stExon;TSS200;TSS1500 | chr5:57755773-57756539 | 766 | | 2 | | | | | | | 1 | | | | | | | | | | |
| RAB3C | Island | Body | chr5:57879168-57879417 | 249 | | | | | | | | | 1 | | | | | | | | | | |
| PDE4D | N_Shore;Island | Body;TSS1500 | chr5:59188920-59189849 | 929 | | | 1 | | | | | | 2 | | | | | | | | | yes | yes |
| PDE4D | NA | 5'UTR | chr5:59559062-59559218 | 156 | | | | | | | | 0 | | | | | | | | | | | |
| PART1;PDE4D | NA | TSS200;Body;5'UTR | chr5:59783668-59784203 | 535 | | | | | 1 | | | | | | | | | | | | | | |
| ERCC8;NDUFAF2 | N_Shore;Island | Body;TSS1500 | chr5:60240568-60241636 | 1068 | | | | | | | | | 2 | | | | | yes | yes | | | | yes |
| C5orf43 | Island | TSS200 | chr5:60458483-60458807 | 324 | | | | | | | | | 2 | | | | | | | | | | |
| HTR1A | Island | 1stExon;TSS200;TSS1500 | chr5:63257092-63257941 | 849 | | 1 | | | | | | | 2 | | | | | | | | | | yes |
| RNF180 | Island;N_Shore | TSS200;TSS1500 | chr5:63461216-63461930 | 714 | | | | | | | | 0 | 7 | | | 1 | | | yes | | | | yes |
| RGS7BP | Island | TSS200 | chr5:63802280-63802593 | 313 | | | | | | | | | 1 | | | | | | | | | | yes |
| FAM159B | N_Shore | TSS1500 | chr5:63985282-63985377 | 95 | | | | | | | | | | | | | | | yes | | yes | | |
| MAST4 | NA | TSS200;Body | chr5:66124492-66124801 | 309 | | | | | | | | | 1 | | | | | | | | | | |
| MAST4 | S_Shelf | Body | chr5:66462293-66462662 | 369 | | | 6 | 1 | | | | 0 | 1 | 1 | | 1 | | | | | | yes | |
| PIK3R1 | NA | TSS200 | chr5:67522298-67522442 | 144 | | | | | 1 | | | | | | | | | | yes | | | | |
| PIK3R1 | N_Shore;Island | Body;TSS1500;TSS200 | chr5:67583598-67584380 | 782 | 1 | 1 | 3 | | | | | | 1 | 1 | | 1 | | yes | | | | yes | |
| PIK3R1 | S_Shore | Body;TSS1500 | chr5:67586170-67586433 | 263 | | | | | | | | | 1 | | | | | | | | | | |
| MARVELD2 | Island | TSS200 | chr5:68710808-68711640 | 832 | 1 | | 34 | | 1 | | | 0 | 7 | 2 | 1 | 4 | | | | | | | yes |
| OCLN | N_Shore | TSS1500 | chr5:68787696-68787919 | 223 | | | 10 | | 2 | 1 | | | 1 | | | | | | | | | | |
| CARTPT | N_Shore | TSS1500;TSS200 | chr5:71014409-71015523 | 1114 | | 1 | | | | | | | 1 | | | | | yes | | | | | |
| MAP1B | NA | Body | chr5:71475111-71475356 | 245 | | 1 | | | | | | | 1 | | | | | | | | | | |
| TMEM171 | N_Shore;Island | TSS1500;TSS200 | chr5:72415429-72416275 | 846 | | 2 | 5 | | 1 | | | | 3 | | | | | | | | | | yes |
| FAM169A | Island;S_Shore | 1stExon;5'UTR;TSS200;TSS1500 | chr5:74162602-74162924 | 322 | | | 5 | | | | | | | | | | | | | | | | |
| SV2C | N_Shore | TSS1500 | chr5:75377860-75377968 | 108 | | 1 | | | | | | | | | | | | | | | | yes | yes |
| SV2C | S_Shore | 5'UTR | chr5:75380987-75381070 | 83 | | | | | | | | 0 | 1 | | | | | yes | | | | | yes |
| F2RL2;IQGAP2 | NA | 1stExon;5'UTR;Body | chr5:75919188-75919281 | 93 | | | | | | | | 0 | 1 | | | | | | | | | | |
| F2R | N_Shore | TSS1500 | chr5:76010444-76011138 | 694 | | | 20 | | 1 | | | | 2 | 1 | 1 | 1 | | yes | | | yes | | |
| F2RL1 | Island | Body | chr5:76115672-76116535 | 863 | | | 6 | | | | | | 1 | | | | | | | | | | |
| S100Z | NA | TSS200 | chr5:76145809-76146148 | 339 | | | 2 | | | | | | 1 | | | | | yes | yes | | | yes | |
| CRHBP | Island | Body | chr5:76250201-76250527 | 326 | | 1 | | | | | | | | | | | | | | | | | |
| ZBED3 | Island | Body | chr5:76373022-76373803 | 781 | | | | 1 | 1 | | | | | | | 1 | | | | | | | yes |
| OTP | N_Shore | 3'UTR | chr5:76925380-76925450 | 70 | | | 1 | | | | | | | | | | | | | | | yes | |
| OTP | S_Shore | Body | chr5:76928406-76928810 | 404 | | 1 | | | | | | | | | | | | | | | | | |
| OTP | N_Shore | Body | chr5:76932016-76932062 | 46 | | | 3 | | | | | | 1 | | | | | | | | | | |
| OTP | N_Shore | Body | chr5:76934327-76934634 | 307 | | | | | | | | | 1 | | | | | | | | | | yes |
| LHFPL2 | Island;chr5:77805753-7780631 | Body;5'UTR;N_Shore | chr5:77805555-77806538 | 983 | | | 29 | | | | | | | | 1 | 1 | | | | | | | yes |
| LHFPL2 | Island | TSS1500 | chr5:77945178-77945347 | 169 | | | | | 1 | | | | | | | | | | | | | | yes |
| ARSB | S_Shore | 1stExon;5'UTR;TSS200 | chr5:78281734-78282669 | 935 | | | 24 | | | 1 | | | 1 | | | | | yes | yes | yes | | | yes |
| BHMT2;DMGDH | N_Shore;Island | TSS1500;Body;TSS200 | chr5:78364769-78366076 | 1307 | | | 6 | 1 | | | | 0 | 3 | | | 2 | | yes | yes | | | | yes |
| BHMT | NA | TSS200;5'UTR;1stExon;TSS1500 | chr5:78407153-78408179 | 1026 | | 1 | 15 | | 1 | | | | 2 | | | 1 | | yes | yes | | | | |
| CMYA5 | NA | TSS1500;TSS200;Body | chr5:78985425-78986160 | 735 | | | 3 | 1 | | | | 0 | 10 | | | 8 | | yes | | | | | yes |
| THBS4 | N_Shore;Island | TSS1500;TSS200 | chr5:79330669-79330959 | 290 | | | 1 | | | | | | 1 | | | | | | | | | | yes |
| THBS4 | NA | Body | chr5:79378709-79378955 | 246 | | | | | | | | 0 | 1 | | | 1 | | | yes | | | | |
| LOC441089 | NA | TSS200 | chr5:79647805-79648261 | 456 | | | | | | | | | | | | | | yes | yes | yes | yes | | |
| ANKRD34B | Island | TSS200 | chr5:79866319-79866379 | 60 | | | | | | | | | 1 | | | | | yes | | | | | |
| RASGRF2 | NA | Body | chr5:80383034-80383224 | 190 | | | | | | | | | | | | | | | | | | | |
| RNU5E;RNU5D;CKMT2 | NA | Body;TSS200 | chr5:80529067-80529340 | 273 | | | 1 | | | | | 0 | 5 | 1 | | | | | | | | | |
| RNU5E;RNU5D;ACOT12 | Island | Body;TSS1500 | chr5:80690205-80690225 | 20 | | 1 | | | | | | | | | | | | | | | | | yes |
| HAPLN1 | N_Shore | 1stExon;5'UTR;TSS200 | chr5:83016630-83017553 | 923 | | | 7 | | | | | 0 | 6 | | | 1 | | | | | | yes | yes |
| CCNH | S_Shore | TSS1500 | chr5:86709126-86709603 | 477 | | | | | | | | | | | | | | | yes | | | | yes |
| TMEM161B | N_Shore;S_Shore | Body;TSS1500 | chr5:87564247-87565552 | 1305 | | | 2 | | | | | | | | | | | | | yes | | yes | yes |
| LOC645323 | Island | Body | chr5:87898610-87898894 | 284 | | | | | | | | | | | | | | | | | | | yes |
| LOC645323 | N_Shore | Body | chr5:87955366-87955859 | 493 | | 1 | 6 | | | | | | | | | | | | | | | | |
| LOC645323 | NA | Body | chr5:87963342-87963489 | 147 | | | | | | | | | | | | | | | | | | yes | |
| LOC645323 | N_Shore | Body | chr5:87968118-87969157 | 1039 | | | 3 | | | | | | 1 | | | | | | | | | yes | yes |

| | | | | | | | | | | | | | | | | | | | | | | | | |
|---|---|---|---|---|---|---|---|---|---|---|---|---|---|---|---|---|---|---|---|---|---|---|---|---|
| LOC645323 | N_Shore | Body;TSS200 | chr5:87974369-87974547 | 178 | | | | | | | | | 2 | | | | | | | | | | yes | |
| LOC645323 | N_Shore | Body | chr5:87975798-87975992 | 194 | | | 1 | | | | | | | | | | | | | | | yes | | |
| MEF2C | NA | Body | chr5:88018481-88018605 | 124 | | | | 1 | | | | | | | | | | | | | | | | |
| FLJ42709 | Island;S_Shore;chr5:92906239- | Body;TSS1500;S_Shore;nan | chr5:92906240-92909968 | 3728 | 3 | 1 | 12 | | | | | | | | 1 | | | | | | | | yes | |
| FLJ42709 | N_Shore | Body | chr5:92913741-92914828 | 1087 | 1 | | 1 | | 2 | | | | 2 | | | | | | | | | | | |
| FLJ42709 | Island | Body | chr5:92916862-92916935 | 73 | | | | | | | | | 1 | | | | | | | | | yes | yes | |
| NR2F1 | N_Shore | TSS200 | chr5:92918910-92918943 | 33 | | | 8 | | | | | | 1 | | | | | | | | | yes | yes | |
| NR2F1 | chr5:92929268-92929489;Islan | Island;Body | chr5:92929319-92929679 | 360 | | 1 | | 1 | | | | | | | 1 | | | | | | | | | |
| FAM172A;MIR2277 | Island | Body;3'UTR | chr5:92956474-92956884 | 410 | | | | | | | | 0 | | | | | | | | | | yes | | |
| FAM172A | S_Shore | TSS200 | chr5:93447487-93447700 | 213 | | | 1 | | | | | | | | | | | | yes | | | | | |
| MCTP1 | Island | TSS1500 | chr5:94620929-94621030 | 101 | | | | | | | | | 1 | | | | | | | | | | | |
| RFESD | N_Shore | TSS1500 | chr5:94982161-94982211 | 50 | | | | 1 | 1 | | | | | | | | | yes | | | | | | |
| GLRX | NA | TSS1500 | chr5:95159365-95159639 | 274 | | | | 1 | 1 | | | | 2 | 1 | | | | | | | yes | | | |
| ELL2 | N_Shore | Body | chr5:95295740-95296150 | 410 | | | | | | | | 0 | 2 | 1 | | | | yes | yes | | | yes | | |
| PCSK1 | N_Shore | Body | chr5:95767724-95767863 | 139 | | | | | | | | | 1 | | | | | | yes | | | | | |
| CAST | Island | TSS200;Body | chr5:96038348-96038614 | 266 | | | | | | | | | 3 | | | | | | yes | | | | yes | |
| CAST | NA | TSS1500;Body | chr5:96078430-96079175 | 745 | | | | | | | | 0 | 2 | | | 1 | | | | | yes | | | |
| FAM174A | Island;N_Shore | TSS200;TSS1500 | chr5:99870887-99871683 | 796 | | | | | | | | 0 | 1 | 2 | | | | | yes | yes | | | | |
| ST8SIA4 | S_Shore | TSS1500 | chr5:100239529-100240223 | 694 | | | 12 | | 1 | | | | 1 | | | | | | | | | yes | yes | |
| SLCO4C1 | Island | 1stExon;5'UTR;TSS200 | chr5:101632171-101632559 | 388 | | | 3 | | | | | | 1 | | | | | | | yes | | | | |
| SLCO6A1 | Island | 1stExon;5'UTR | chr5:101834472-101834780 | 308 | | | | | | | | | 1 | | | | | | | yes | | yes | | |
| PAM | N_Shore | TSS1500 | chr5:102201128-102201421 | 293 | | | | 1 | | | | | | | | | | | | | | | | |
| NUDT12 | NA | 5'UTR;1stExon;TSS200 | chr5:102898223-102898733 | 510 | 2 | 1 | 8 | | 1 | | | 0 | 4 | 1 | | 10 | | yes | | | | | yes | |
| FER | N_Shore | TSS1500;TSS200 | chr5:108082637-108083695 | 1058 | | | | | | | 1 | | 1 | 2 | | | | | yes | yes | | | | |
| TMEM232 | NA | 1stExon;5'UTR;TSS200 | chr5:110062343-110062837 | 494 | | 1 | 22 | | | | | 0 | 10 | 5 | 2 | 17 | | yes | | | yes | | yes | |
| TSLP | N_Shelf | TSS1500 | chr5:110406107-110406661 | 554 | | 1 | | | | | | | | | | | | | yes | | | | | |
| NCRNA00219;SNORA13 | N_Shore | TSS1500 | chr5:111495974-111496180 | 206 | | | 1 | | | | | | | | | | | yes | yes | | yes | | | |
| EPB41L4A;FLJ11235 | N_Shore | Body;TSS1500 | chr5:111753754-111754296 | 542 | | | 5 | | | | | | | | | | | | | | | | | |
| APC | N_Shore | TSS1500 | chr5:112042851-112042966 | 115 | | | 4 | | | | | | | | | | | | yes | | | | | |
| APC | NA | TSS1500;5'UTR;TSS200 | chr5:112073348-112074043 | 695 | | 1 | | 3 | 6 | | | | | | | | | yes | | | | | | |
| TSSK1B;MCC | NA | 1stExon;Body;TSS1500 | chr5:112769927-112771691 | 1764 | 1 | | | | | | | | | | | | | | | yes | | | | |
| MCC | Island;S_Shore | 5'UTR;1stExon;TSS200;TSS1500 | chr5:112824288-112824920 | 632 | | | 15 | | 1 | | | | 5 | | | 1 | | yes | | yes | | yes | | |
| KCNN2 | Island | TSS1500;TSS200;1stExon;5'UTR | chr5:113697632-113699683 | 2051 | | 2 | | | | | | | 1 | | | | | | | | | yes | | |
| TRIM36 | Island | Body | chr5:114514717-114515341 | 624 | | | 4 | | | | | | | | | | | | | | yes | | yes | |
| TRIM36 | S_Shore | TSS200 | chr5:114516368-114516422 | 54 | | | 5 | | | | | | | | | | | | | | | | yes | |
| CCDC112 | S_Shore | TSS1500;TSS200 | chr5:114632615-114632793 | 178 | | | | | | | | | 1 | | | | | yes | yes | | | yes | | |
| TMED7-TICAM2;TICAM2 | Island;S_Shore | Body;5'UTR;TSS1500 | chr5:114937784-114938459 | 675 | | | | | | | | 0 | 2 | | | | | | | | | yes | yes | |
| CDO1 | Island;S_Shore | 1stExon;5'UTR;TSS200;TSS1500 | chr5:115152386-115153223 | 837 | | 2 | 2 | | | | | | 1 | | | | | | | | | | yes | |
| LVRN | Island;S_Shore | TSS1500;TSS200;1stExon;Body | chr5:115297690-115299828 | 2138 | | 1 | 34 | | | | | | 1 | | | | | | yes | | yes | | | |
| TNFAIP8 | N_Shore | TSS1500 | chr5:118603742-118603938 | 196 | | | | | | | | 0 | 2 | 1 | | | | yes | yes | | | yes | | |
| TNFAIP8 | Island | Body;TSS200 | chr5:118691415-118692011 | 596 | | | | | 5 | | | | | | | | | | | | | | | |
| HSD17B4 | N_Shore | TSS1500 | chr5:118787455-118787985 | 530 | | | | | | | | | | | | | | | yes | yes | yes | | | |
| FAM170A | NA | TSS1500;TSS200 | chr5:118964760-118965347 | 587 | | | 4 | | | | | | | | | | | yes | | yes | | | | |
| PRR16 | N_Shore;Island | TSS1500;TSS200 | chr5:119799220-119800143 | 923 | | | 3 | | | | | | 1 | | | | | | | | | | yes | |
| FTMT | Island | TSS200 | chr5:121187589-121188016 | 427 | | | | | | | | | 1 | | | | | | | | | | | |
| SNCAIP | Island | TSS200 | chr5:121647630-121647858 | 228 | | | | | | | | 0 | 2 | | | | | | | | | | yes | |
| SNX2 | N_Shore | TSS1500 | chr5:122110491-122110559 | 68 | | | | | | | | | | | | | | | | | yes | yes | | |
| SNX24 | N_Shore | TSS1500 | chr5:122180440-122180522 | 82 | | | | | | | | | 1 | | | | | | | | | | | |
| PRDM6 | N_Shore | Body | chr5:122430434-122431121 | 687 | | 1 | | | | | | | 1 | | | | | | | | | yes | | |
| PRDM6 | Island | Body | chr5:122434178-122435202 | 1024 | | 1 | 6 | | | | | | 1 | | | | | | | | | yes | yes | |
| ZNF608 | NA | Body | chr5:124070989-124071107 | 118 | | | | 1 | | | | | | | | 1 | | | | | | | | |
| GRAMD3 | NA | Body;5'UTR;TSS200 | chr5:125800764-125800916 | 152 | | | | | | | | | 2 | | | | | yes | | | | | | |
| ALDH7A1 | N_Shore | Body | chr5:125929611-125930017 | 406 | | | | | | 1 | | | 1 | | | | | yes | yes | | | | | |
| 37681 | NA | 3'UTR | chr5:126205009-126205081 | 72 | | | | | | | | | | | | | | | | | | yes | | |
| 37681 | Island | TSS1500 | chr5:126367249-126367370 | 121 | | | | | | | | | | | | | | | | | | | | |
| FLJ44606 | N_Shore;Island;S_Shore | 5'UTR;1stExon;TSS200;TSS1500 | chr5:126408756-126409573 | 817 | 1 | 1 | 43 | 1 | | | | 0 | 6 | 6 | | 17 | 1 | | yes | | | | | |
| MEGF10 | N_Shore;Island | TSS1500;TSS200 | chr5:126625777-126627061 | 1284 | | 1 | 4 | 1 | | | | 0 | 4 | | | | | | | yes | | | | |
| PRRC1 | N_Shore | TSS1500 | chr5:126852955-126853064 | 109 | | | 2 | | | | | 0 | 1 | | | | | yes | yes | | | yes | | |
| CTXN3 | NA | TSS200 | chr5:126984564-126984994 | 430 | | | 8 | | | | | | | | | | | | | | | | | |
| CTXN3 | NA | 5'UTR;TSS1500 | chr5:126987825-126988029 | 204 | | | | | | | | | | | | | | | | | | | | |
| FBN2 | N_Shore;Island | Body;1stExon | chr5:127871927-127873397 | 1470 | | | 19 | 1 | 1 | | | 0 | 2 | | | | | | | | | | | yes |
| IL3 | NA | TSS200 | chr5:131396204-131396253 | 49 | | | | | | | | | | | | | | yes | | | | | | |
| CSF2 | NA | TSS200 | chr5:131409289-131409637 | 348 | | 1 | | | | | | | | | | | | | | | | | | |
| PDLIM4 | N_Shore;Island | TSS1500 | chr5:131592959-131593191 | 232 | | | 1 | | 1 | | | 0 | 1 | | | | | | yes | | | yes | yes | |
| PDLIM4 | Island | Body | chr5:131607097-131607372 | 275 | | | | | | | | | 1 | | | 1 | | | | | | | | |
| KIF3A | S_Shore | TSS1500 | chr5:132073546-132073916 | 370 | | | | | | | | | | | | | | yes | yes | | | | | |
| CCNI2 | Island | 1stExon;5'UTR;Body | chr5:132083184-132084068 | 884 | | | | | | | | 0 | 1 | | | | | yes | | | | | | |
| SEPT8 | S_Shore;Island | TSS200;TSS1500 | chr5:132113601-132114055 | 454 | | | | | 1 | | | | 3 | 1 | | | | yes | | | | | | |
| SHROOM1 | Island | Body | chr5:132158560-132159003 | 443 | | | 6 | | | | | 0 | 1 | | | | | yes | | | | | | |
| LEAP2 | NA | TSS1500 | chr5:132209089-132209201 | 112 | | | | | 1 | | | | | | | | | | | | | | | |
| FSTL4 | N_Shore | 5'UTR | chr5:132945292-132945815 | 523 | | | | | | | | | 1 | | | | | | | | | | | |
| TCF7 | Island;S_Shore | TSS200;TSS1500;Body;5'UTR | chr5:133451243-133451625 | 382 | 2 | | | | | | | 0 | | | | | | | | | | | yes | |
| CDKL3 | S_Shore;Island | TSS1500;5'UTR;1stExon | chr5:133702676-133703889 | 1213 | | | | | | | | | 2 | | | | | yes | | | yes | | | |
| CAMLG | N_Shore | TSS1500 | chr5:134073430-134073517 | 87 | | | | | | | | | 1 | | | | | yes | yes | | | | yes | |
| C5orf24 | N_Shore | TSS200;TSS1500 | chr5:134181226-134181266 | 40 | | | | | | | | | | 1 | | | | | | yes | | | | |
| PITX1 | Island | 3'UTR | chr5:134363516-134364387 | 871 | | | 4 | | 2 | | | 0 | 4 | 1 | | 1 | | | | | | yes | yes | |
| PITX1 | N_Shore;Island | Body | chr5:134365728-134367394 | 1666 | | 1 | 13 | | | | | | | | | | | | | | | | | |
| PITX1 | Island | TSS1500 | chr5:134370282-134370530 | 248 | | | 3 | | | | | | | | | | | | | | | yes | | |
| H2AFY | N_Shore | 5'UTR | chr5:134733519-134733905 | 386 | | | 2 | | | | | | | | | | | | | | | | | |
| H2AFY | S_Shore | TSS1500;5'UTR;1stExon;TSS200 | chr5:134735544-134735914 | 370 | | 1 | 2 | 1 | 1 | | | 0 | 4 | | | 3 | | yes | yes | | | | | |
| C5orf20 | NA | TSS200 | chr5:134783075-134783229 | 154 | | | | | | | | | | | | | | yes | yes | | | | | |
| TIFAB | NA | 1stExon;5'UTR | chr5:134787907-134788128 | 221 | | | 6 | | | | | | | | | | | yes | | | | | | |
| NEUROG1 | S_Shore;N_Shore;Island | TSS1500;1stExon;3'UTR;TSS200 | chr5:134870506-134872957 | 2451 | | | 19 | | | | | 0 | 6 | 1 | 1 | | | yes | | yes | | | yes | |
| CXCL14 | S_Shore | TSS200 | chr5:134915042-134915088 | 46 | | | | | | | | | 1 | | | | | | | | | | | |
| LOC340074 | NA | TSS1500 | chr5:134990285-134990707 | 422 | | | | | | | | | | | | | | yes | yes | yes | | | | |
| LOC153328 | N_Shore | TSS1500 | chr5:135170147-135170208 | 61 | | | | | | | | | 1 | | | | | | | | | | | |
| IL9 | NA | 1stExon;5'UTR;TSS200 | chr5:135231514-135231577 | 63 | | | | | | | | | 1 | | | | | | | | | | | |
| FBXL21 | Island | TSS200;1stExon;5'UTR | chr5:135265849-135266805 | 956 | | 1 | | | | | | 0 | 1 | | | | | yes | yes | | | yes | yes | |
| TGFBI | N_Shore;Island | TSS1500;TSS200;1stExon;Body | chr5:135364060-135365401 | 1341 | | | 10 | | | | | 0 | 5 | | | 1 | | yes | yes | | | | yes | |
| MIR886 | Island;chr5:135416204-135416 | TSS200;Island;TSS1500;Body | chr5:135416205-135416613 | 408 | | 25 | 28 | 3 | | 6 | | 0 | 6 | 6 | 5 | 5 | 2 | yes | | | | | | |
| TRPC7 | NA | 5'UTR;1stExon | chr5:135701050-135701240 | 190 | | 1 | | | | | | | 1 | | | | | | | | | yes | | |
| SPOCK1 | Island | Body;5'UTR | chr5:136834155-136834863 | 708 | | 1 | | | | | | | 1 | | | | | | yes | | | yes | | |
| MIR874;KLHL3 | NA | TSS200;Body | chr5:136983354-136984794 | 1440 | | | | | | | | | | 1 | | | | yes | | yes | | | yes | |
| MYOT | NA | TSS200 | chr5:137203358-137203640 | 282 | | | | | | | | | | | | | | | | | | | | |
| PKD2L2 | N_Shore;Island | TSS1500;Body | chr5:137224213-137225509 | 1296 | | | 2 | | | | | 0 | 2 | | | 2 | | yes | | | yes | | | |
| NME5 | NA | 1stExon;5'UTR | chr5:137475115-137475288 | 173 | | | 1 | | | | | | | | | | | | | | | | | |
| GFRA3 | Island | Body | chr5:137593418-137593583 | 165 | | | 3 | | | | | | | | | | | yes | yes | | | | | |
| GFRA3 | N_Shore;Island;S_Shore | Body;1stExon;5'UTR;TSS200 | chr5:137609835-137610448 | 613 | | | 2 | 1 | 1 | | | | | | | | | | | | | yes | yes | |
| EGR1 | Island | Body | chr5:137803102-137803362 | 260 | | | 1 | | | | | | | | | | | | | | | | yes | |

| | | | | | | | | | | | | | | | | | | | | | | | |
|---|---|---|---|---|---|---|---|---|---|---|---|---|---|---|---|---|---|---|---|---|---|---|---|
| CTNNA1;LRRTM2 | NA | Body;5'UTR;1stExon;TSS200 | chr5:138211043-138211184 | 141 | | 1 | | 1 | | | | | 2 | | | | | | | | | | |
| SLC23A1 | Island | Body | chr5:138713897-138714654 | 757 | | | 12 | | | | | 0 | 2 | 1 | | 1 | | yes | | | | | |
| SLC23A1 | NA | Body;1stExon | chr5:138718812-138719374 | 562 | | | | | | | | | | | | | | yes | | | yes | yes | yes |
| MGC29506 | N_Shelf | 1stExon | chr5:138725400-138725705 | 305 | | | | | 1 | 1 | | | 1 | | | | | yes | | | | yes | |
| LOC389333 | Island | 1stExon | chr5:138728367-138729283 | 916 | | | 4 | | | | | | | | | | | | | | | | |
| TMEM173 | NA | Body | chr5:138861241-138862441 | 1200 | | | | 1 | | | | | | | | | | yes | | yes | | yes | |
| CXXC5 | N_Shore;Island | 5'UTR | chr5:139040055-139041014 | 959 | | 1 | | | | | | | 1 | 3 | | | | | | | | | yes |
| CXXC5 | Island | 5'UTR | chr5:139056667-139057496 | 829 | | | 9 | | 1 | | | | 1 | 1 | | | | | | | | | yes |
| NRG2 | Island;N_Shore | Body;3'UTR | chr5:139227318-139228242 | 924 | | | 2 | | 2 | | | | 2 | 5 | | | | yes | | | | yes | yes |
| NRG2 | N_Shore | Body | chr5:139283051-139283262 | 211 | | | | | 1 | | | | | | | | | yes | | | | | |
| PURA | N_Shore | TSS1500 | chr5:139492230-139492727 | 497 | | | 1 | | | | | | | | | | | | | | | | |
| EIF4EBP3;ANKHD1-EIF4EBP3 | Island | TSS200;Body | chr5:139927191-139927471 | 280 | | | | | | | | | 1 | | | | | yes | | | | | |
| CD14 | Island | 1stExon;5'UTR | chr5:140012660-140012769 | 109 | | | | | | | | | 1 | | | 1 | | | yes | | | | |
| TMCO6 | N_Shore | TSS1500 | chr5:140018644-140018709 | 65 | | | 3 | | | 1 | | | 1 | | | | | yes | | yes | | yes | |
| DND1 | N_Shore | Body | chr5:140052017-140052475 | 458 | | | | | | | | | | | | | | yes | | | | | |
| ZMAT2 | NA | TSS200 | chr5:140079847-140080575 | 728 | | | | | | | | | 1 | | | | | yes | | | | | |
| VTRNA1-1 | NA | TSS1500 | chr5:140090300-140090746 | 446 | | | 1 | | | | | 0 | | | | | | yes | yes | | | | |
| VTRNA1-2 | NA | TSS1500;TSS200 | chr5:140097950-140098576 | 626 | | | 18 | | | | | 0 | 4 | | | 1 | | yes | yes | | | | |
| VTRNA1-3 | NA | TSS1500;TSS200 | chr5:140105271-140106029 | 758 | | 2 | 5 | | | | | 0 | 5 | | | 4 | | yes | yes | | | | |
| PCDHA1 | Island | 1stExon | chr5:140166059-140166589 | 530 | | | | | | | | | 2 | | | | | | | | | | |
| PCDHA2;PCDHA1 | N_Shore;Island | 5'UTR;Body;1stExon | chr5:140174449-140175116 | 667 | | 1 | | | | | | 0 | 2 | | | 2 | | | yes | | | | |
| PCDHA2;PCDHA1;PCDHA3 | N_Shore;Island | Body;TSS200;1stExon | chr5:140180637-140182956 | 2319 | | 1 | 2 | | | | | | 1 | | | 1 | | yes | | | | | |
| PCDHA2;PCDHA1;PCDHA4;PCDHA3 | N_Shore | Body;TSS200 | chr5:140186484-140187240 | 756 | | | | | 1 | | | | 2 | | | | | | | | | | |
| PCDHA2;PCDHA1;PCDHA3;PCDHA4;PCD | Island | Body;1stExon | chr5:140201539-140201746 | 207 | | | | | | | | | | | | | | | | | | | |
| PCDHA2;PCDHA1;PCDHA6;PCDHA5;PCD | N_Shore;Island | Body;TSS200;1stExon | chr5:140207460-140208201 | 741 | | | 7 | | | | | 0 | 3 | | | 1 | | | | | | | |
| PCDHA6;PCDHA2;PCDHA1;PCDHA7;PCD | N_Shore;Island | Body;TSS200;1stExon | chr5:140213783-140214372 | 589 | | | | | | | | 0 | 3 | | | 3 | | | | | | | |
| PCDHA6;PCDHA2;PCDHA1;PCDHA7;PCD | Island | Body;1stExon | chr5:140215762-140216145 | 383 | | | 1 | | | | | | | | | | | | | | | | |
| PCDHA6;PCDHA2;PCDHA1;PCDHA7;PCD | N_Shore | Body;TSS1500;TSS200 | chr5:140220686-140220943 | 257 | | | 9 | | | | | | 3 | | | | | | | | | | |
| PCDHA6;PCDHA2;PCDHA1;PCDHA7;PCD | Island | Body;1stExon | chr5:140223065-140223264 | 199 | | | | | | | | | | | | 1 | | | | | | | |
| PCDHA6;PCDHA2;PCDHA1;PCDHA7;PCD | Island | Body;1stExon | chr5:140229827-140230326 | 499 | | | | | | | | | 1 | | | | | | | | | | yes |
| PCDHA6;PCDHA2;PCDHA1;PCDHA9;PCD | N_Shore | Body;TSS1500;TSS200 | chr5:140234919-140235753 | 834 | | | | | | | | 0 | 4 | 1 | | 2 | | yes | | | | | |
| PCDHA6;PCDHA2;PCDHA1;PCDHA9;PCD | N_Shore | Body | chr5:140240593-140241088 | 495 | | | | | | | | | | | | | | yes | | | | | |
| PCDHA6;PCDHA2;PCDHA1;PCDHA9;PCD | Island | Body;1stExon | chr5:140250432-140250948 | 516 | | | | | | | | | 1 | | | | | | | | | | |
| PCDHA12;PCDHA7;PCDHA6;PCDHA10;PC | N_Shore | TSS200;Body | chr5:140254793-140255449 | 656 | | | | | | | | 0 | 3 | | | 2 | | | | | | | |
| PCDHA7;PCDHA12;PCDHA6;PCDHA10;PC | N_Shore | Body;TSS200 | chr5:140261653-140261930 | 277 | | | 6 | | | | | | 1 | | | | | | | | | | |
| PCDHA7;PCDHA12;PCDHA6;PCDHAC1;PC | Island | Body;TSS200;TSS1500 | chr5:140305947-140306458 | 511 | | | | | 4 | | | | 3 | | | | | | | | | | |
| PCDHA7;PCDHAC2;PCDHA12;PCDHA6;PC | N_Shore | Body;TSS1500 | chr5:140344290-140344620 | 330 | | | 4 | | | | | | | | | | | | | | | | |
| PCDHAC2;PCDHA7;PCDHA12;PCDHA6;PC | Island | 1stExon;TSS200;Body;5'UTR | chr5:140346199-140346394 | 195 | | | | | 3 | | | 0 | 4 | | | 2 | | | | | | yes | yes |
| PCDHB1 | NA | TSS1500 | chr5:140430569-140430713 | 144 | | | 1 | | | | | 0 | | | | | | yes | | | | | |
| PCDHB2 | N_Shore | TSS1500;TSS200;1stExon | chr5:140473857-140475611 | 1754 | | 1 | 7 | | | | | 0 | 1 | | | | | | yes | | | | |
| PCDHB3 | N_Shore | TSS1500;TSS200;1stExon | chr5:140479979-140481267 | 1288 | | 1 | 21 | | | | | 0 | 11 | 1 | | 1 | | | | | | | |
| PCDHB4 | N_Shore;Island | TSS1500;TSS200;1stExon | chr5:140501117-140503914 | 2797 | | 1 | 12 | | 2 | | | 0 | 5 | | | 1 | | | | | | | |
| PCDHB5 | N_Shore | TSS1500;TSS200;1stExon;5'UTR | chr5:140514381-140516095 | 1714 | | 1 | 9 | | | | 1 | 0 | 1 | | | 2 | | | yes | | | | |
| PCDHB6 | N_Shore;chr5:140531157-1405 | TSS1500;N_Shore;TSS200;1stExon | chr5:140529499-140531158 | 1659 | | 1 | 13 | 1 | | | | 0 | 7 | 3 | 2 | 3 | | | | | | | |
| PCDHB17 | Island;N_Shore | Body;TSS1500;TSS200 | chr5:140535215-140537884 | 2669 | | | | | | | | | | | | | | | | | | | |
| PCDHB7 | N_Shore | TSS200;1stExon;5'UTR | chr5:140552153-140553150 | 997 | | 3 | 8 | 2 | 1 | | 1 | 0 | 4 | | | 1 | | | | | | | |
| PCDHB8 | N_Shelf;N_Shore | TSS1500 | chr5:140556888-140557256 | 368 | | | | | | | | | 3 | | | 1 | | yes | yes | | | | |
| PCDHB16 | N_Shore | 5'UTR;1stExon | chr5:140561893-140563400 | 1507 | | 3 | 21 | 1 | | | 1 | 0 | 11 | 3 | 1 | 6 | | | | | | | |
| PCDHB9 | N_Shore | TSS1500 | chr5:140566303-140566710 | 407 | | | 1 | | | | | | | | | | | | yes | | | | |
| PCDHB9 | N_Shore;Island | 1stExon | chr5:140568075-140569144 | 1069 | | | | | | | | | | | | | | | | | | | |
| PCDHB10 | N_Shore | TSS200 | chr5:140571783-140572039 | 256 | | 1 | | | | | | | | | | | | | yes | | | | |
| PCDHB11 | N_Shore | TSS200 | chr5:140579160-140579644 | 484 | | | 12 | | | | | | 3 | | | 3 | | | | | | | |
| PCDHB12 | N_Shore | 5'UTR;1stExon | chr5:140588299-140588424 | 125 | | | | | | | | | | | | | | | | | | | |
| PCDHB13 | Island | 1stExon | chr5:140595009-140595291 | 282 | | | | | | | | | | | | | | | | | | | |
| PCDHB14 | N_Shore | TSS1500;TSS200 | chr5:140602519-140603270 | 751 | | 1 | | | | | | | 5 | | | 1 | | yes | yes | | | | yes |
| PCDHB18 | N_Shore | TSS200;Body | chr5:140613740-140615656 | 1916 | | 1 | 4 | | | | | | 5 | | | | | | | | | | |
| PCDHB19P | N_Shore | TSS1500 | chr5:140619446-140619586 | 140 | | | 1 | | | | | | 1 | | | | | | | | | | |
| PCDHB19P | Island | Body | chr5:140621344-140621754 | 410 | | | | | | | | | 1 | | | | | | | | | | |
| PCDHB15 | N_Shore | TSS1500;TSS200;1stExon | chr5:140624869-140625848 | 979 | | 2 | 3 | | | | | 0 | 2 | | | 1 | | | yes | | | | |
| SLC25A2 | Island | 1stExon;5'UTR;TSS200 | chr5:140683196-140684147 | 951 | | | 18 | | | | | 0 | 3 | | | | | yes | | | | | |
| TAF7 | Island;S_Shore | TSS200;TSS1500 | chr5:140700449-140700676 | 227 | | | | | | | | | 2 | 1 | | | | yes | | | | | |
| PCDHGA1 | N_Shore | TSS1500;TSS200;1stExon | chr5:140709986-140710688 | 702 | | | 2 | | | | | 0 | 1 | | | 1 | | | | | | yes | |
| PCDHGA1 | Island | 1stExon | chr5:140711852-140712459 | 607 | | | 4 | | | | | | 1 | | | | | | | | | | |
| PCDHGA1;PCDHGA2 | N_Shore | Body;TSS1500;TSS200;1stExon | chr5:140717961-140720212 | 2251 | | 2 | 5 | | | | | 0 | 2 | | | 4 | | | yes | | | | |
| PCDHGA2;PCDHGA3;PCDHGA1 | N_Shore;Island | Body;TSS1500;TSS200;1stExon | chr5:140723331-140725147 | 1816 | | 1 | 17 | | 1 | | | 0 | 3 | | | 1 | | | | | | | |
| PCDHGB1;PCDHGA2;PCDHGA1;PCDHGA | N_Shore | 1stExon;Body;TSS1500;TSS200 | chr5:140729605-140730516 | 911 | | 1 | 4 | | | | | 0 | 5 | | | | | | | | | | yes |
| PCDHGA4;PCDHGA2;PCDHGA1;PCDHGB | N_Shore;Island | TSS200;Body;1stExon | chr5:140734648-140735545 | 897 | | 1 | 1 | | | | | | 4 | | | 1 | | | yes | | | | |
| PCDHGA2;PCDHGA4;PCDHGA1;PCDHGB | N_Shore;Island | Body;1stExon | chr5:140736762-140737111 | 349 | | | | | | | | | | | | | | yes | | | | | |
| PCDHGA4;PCDHGA2;PCDHGB2;PCDHGA | N_Shore;Island | Body;TSS1500;TSS200;1stExon | chr5:140739501-140742017 | 2516 | | 1 | 4 | | | | | 0 | | | | 3 | | | yes | | | | |
| PCDHGA5;PCDHGA4;PCDHGA2;PCDHGB | N_Shore;Island | TSS1500;Body;TSS200;1stExon | chr5:140743575-140745340 | 1765 | | | 9 | | | | | 0 | 6 | | | 4 | | | | | | | |
| PCDHGA4;PCDHGB3;PCDHGA1;PCDHGA | N_Shore | Body;TSS1500;TSS200 | chr5:140749735-140750160 | 425 | | 1 | | | | | | 0 | 1 | | | 1 | | yes | | | | | |
| PCDHGA4;PCDHGB3;PCDHGA1;PCDHGA | Island;N_Shore | Body;1stExon;TSS1500;TSS200 | chr5:140751506-140754451 | 2945 | | | 12 | | | | | 0 | 6 | | | 1 | | yes | | | | | |
| PCDHGA4;PCDHGA6;PCDHGA1;PCDHGA | N_Shore;Island | Body;TSS1500;1stExon | chr5:140762229-140763254 | 1025 | | 1 | | | | | | 0 | 5 | | | 1 | | | | | | | |
| PCDHGA4;PCDHGA7;PCDHGA6;PCDHGA | Island | Body;1stExon | chr5:140764302-140764679 | 377 | | | | | | | | | | | | | | | | | | | |
| PCDHGB4;PCDHGA4;PCDHGA6;PCDHGA | Island;N_Shore | 1stExon;Body;TSS1500 | chr5:140766911-140769170 | 2259 | | 1 | 6 | | | | | | 6 | | | 2 | | | | | | | |
| PCDHGA4;PCDHGA6;PCDHGA1;PCDHGA | N_Shore;Island | Body;5'UTR;1stExon | chr5:140772182-140774440 | 2258 | | | 9 | | | | | 0 | 2 | | | 2 | | | | | | | |
| PCDHGA4;PCDHGA6;PCDHGA1;PCDHGA | N_Shore;Island | Body;TSS1500;TSS200;1stExon | chr5:140777344-140779461 | 2117 | | 1 | 6 | | 1 | | | 0 | 4 | | | 1 | | | | | | | |
| PCDHGA4;PCDHGA6;PCDHGA1;PCDHGA | S_Shelf | Body;TSS1500;TSS200 | chr5:140782297-140783173 | 876 | | | | | | | | | 1 | | | | | | | | | | yes |
| PCDHGA4;PCDHGA6;PCDHGA9;PCDHGA | Island | Body;TSS1500 | chr5:140787504-140787864 | 360 | | 1 | | | | | | | | | | | | | yes | | | | yes |
| PCDHGA4;PCDHGA9;PCDHGA1;PCDHGB | Island | Body;1stExon | chr5:140789247-140789745 | 498 | | | | | | | | | | | | | | | | | | | |
| PCDHGA4;PCDHGA6;PCDHGA9;PCDHGA | N_Shore | Body;TSS1500;TSS200;1stExon | chr5:140792511-140793307 | 796 | | 1 | | | | | | 0 | 2 | | | 1 | | | | | | | |
| PCDHGA4;PCDHGA6;PCDHGA9;PCDHGA | Island | Body;1stExon | chr5:140794359-140794993 | 634 | | | | | | | | | 1 | | | | | | | | | yes | |
| PCDHGA4;PCDHGA9;PCDHGA1;PCDHGB | Island;N_Shore | Body;TSS200;1stExon;TSS1500;5'U | chr5:140797172-140803130 | 5958 | | | 24 | | 1 | | | 0 | 7 | | | 4 | | | | | | | yes |
| PCDHGA4;PCDHGA11;PCDHGA9;PCDHG | N_Shore | Body;TSS200 | chr5:140805681-140806126 | 445 | | | 3 | | | | | 0 | 1 | 1 | | 2 | | | | | | | |
| PCDHGA4;PCDHGA11;PCDHGA9;PCDHG | Island | Body | chr5:140807666-140808068 | 402 | | | | | | | | | 1 | | | | | yes | | | | | |
| PCDHGA4;PCDHGA11;PCDHGA9;PCDHG | N_Shore;Island | Body;TSS200;5'UTR;1stExon | chr5:140810051-140811642 | 1591 | | 2 | 34 | | | | | 0 | 10 | | | 10 | | | | | | | |
| PCDHGA4;PCDHGA12;PCDHGA11;PCDH | N_Shore;Island | Body;1stExon | chr5:140856971-140858107 | 1136 | | | 9 | | | | | 0 | 4 | | | | | | | | | | |
| PCDHGA4;PCDHGC3;PCDHGA12;PCDHG | N_Shore;Island | Body;TSS1500;TSS200 | chr5:140864020-140865433 | 1413 | | | | | | | | | | | | | | | | | | | |
| PCDHGB5;PCDHGC3;PCDHGA6;PCDHGB | N_Shore | 3'UTR | chr5:140892370-140892822 | 452 | | | | | | | | | | | | 1 | | | | | | | yes |
| RELL2;HDAC3 | S_Shore | 5'UTR;1stExon;TSS1500 | chr5:141017773-141017903 | 130 | | | | 1 | | | | | | | | | | | | | | | |
| ARAP3 | NA | TSS1500 | chr5:141062313-141062571 | 258 | | | | 1 | 1 | | | | | | | | | yes | | | | yes | |
| KIAA0141 | N_Shore | TSS1500 | chr5:141302942-141303209 | 267 | | 1 | | | | | | | | | | | | yes | | | | | |
| PCDH12 | NA | TSS1500 | chr5:141339748-141339777 | 29 | | | | | | | | | | | | | | yes | | | | | |
| NDFIP1 | N_Shore | TSS1500 | chr5:141488016-141488100 | 84 | | | | | | | | | | | | | | | yes | | yes | yes | |
| FGF1 | NA | 5'UTR;Body;1stExon;TSS200;TSS1 | chr5:142065539-142066938 | 1399 | | | 15 | | | | | | 4 | 1 | | | | | | | yes | yes | |
| NR3C1 | NA | 1stExon;5'UTR | chr5:142814827-142815469 | 642 | | | | | | | | | | | | | | | yes | | | | |
| HMHB1 | NA | TSS200 | chr5:143191663-143192067 | 404 | | | | | 1 | | | | | | | 1 | | yes | yes | | | yes | yes |
| KCTD16 | Island | 5'UTR | chr5:143584811-143584957 | 146 | | | | | | | | | 1 | | | | | | | | | yes | yes |
| PRELID2 | S_Shore | TSS1500 | chr5:145215546-145215784 | 238 | | | | | | | | | 1 | | | | | yes | yes | | | | |

| | | | | | | | | | | | | | | | | | | | | | | | | | | | |
|---|---|---|---|---|---|---|---|---|---|---|---|---|---|---|---|---|---|---|---|---|---|---|---|---|---|---|---|
| POU4F3 | Island | TSS200 | chr5:145718392-145718491 | 99 | | 1 | | | | | | | 1 | | | | | | | | | yes | yes | | | | |
| PPP2R2B | NA | 5'UTR;1stExon | chr5:146435656-146436093 | 437 | | 1 | | | | | | | | | | | | | | | | yes | yes | | | | |
| PPP2R2B | NA | TSS1500 | chr5:146461785-146461855 | 70 | | | | | | | | | | | | | | yes | | | | | | | | | |
| STK32A | Island | TSS1500 | chr5:146614298-146614393 | 95 | | | | | | | | | 1 | | | | | | | | | yes | | yes | | | |
| DPYSL3 | N_Shore | Body | chr5:146832182-146832387 | 205 | | | 2 | | | | | 0 | | | | | | | | | | yes | | | | | |
| JAKMIP2 | NA | Body | chr5:147024842-147024959 | 117 | | | | 1 | | | | | | | | | | yes | | | yes | | | | | | |
| JAKMIP2 | NA | 5'UTR;1stExon | chr5:147161924-147162223 | 299 | | | | 1 | 1 | | | | | | | | | | | | | yes | | | | | |
| SCGB3A2 | NA | TSS1500 | chr5:147257219-147257456 | 237 | | | | | | | | | | | | | | | | yes | yes | | | | | | |
| SPINK5 | NA | TSS1500 | chr5:147443067-147443536 | 469 | | | | | | | | | 3 | | | | | yes | | | | | yes | | | | |
| SPINK7 | NA | TSS200 | chr5:147691898-147692173 | 275 | | | | | | | | | | | | | | yes | | yes | | | | | | | |
| HTR4 | NA | 3'UTR;Body | chr5:147862421-147862681 | 260 | | | 4 | | | | | | | | | | | | yes | | | | | | | | |
| HTR4 | Island | 5'UTR;1stExon;TSS200 | chr5:148033473-148034045 | 572 | | 1 | | | | | | 0 | 3 | | | | | yes | | | | yes | | | | | |
| ADRB2 | Island | TSS200 | chr5:148206096-148206180 | 84 | | 1 | | | | | | | | | | | | | | | | | yes | | | | |
| SH3TC2 | NA | TSS200;1stExon;5'UTR | chr5:148442694-148442890 | 196 | | | 9 | 1 | | | | | | | | | | | yes | | | | | | | | |
| ABLIM3 | N_Shore;Island | TSS1500;TSS200;5'UTR;1stExon | chr5:148520817-148521466 | 649 | | 1 | 9 | | | | | 0 | 2 | | | | | | yes | | | | | | | | |
| LOC728264 | NA | TSS1500 | chr5:148785250-148786302 | 1052 | | | 12 | | | | | | 1 | | | | | yes | yes | | | yes | | | | | |
| MIR143;LOC728264 | NA | nan | chr5:148808456-148808721 | 265 | | | | | | | | | | | 1 | | | yes | yes | | yes | yes | | | | | |
| LOC728264;MIR145 | NA | Body;TSS1500;TSS200 | chr5:148809826-148810203 | 377 | | | 14 | | | | | | 1 | 1 | | | | yes | | | | | | | | | yes |
| CSNK1A1 | N_Shore | Body | chr5:148928885-148929352 | 467 | | | | | | | | 0 | 1 | | | | | yes | | | | | | | | | |
| HMGXB3;TIGD6 | S_Shore | 5'UTR;TSS1500 | chr5:149381392-149381502 | 110 | | | | | | | | | | | | | | yes | | | | yes | | | | | |
| CSF1R | NA | TSS200 | chr5:149493008-149493258 | 250 | | | 7 | | | | | | | | | | | yes | | | | | yes | | | | |
| PDGFRB | NA | TSS1500 | chr5:149535646-149535765 | 119 | | | | 1 | | | | 0 | 1 | | | | | yes | yes | | | | | | | | |
| CDX1 | Island | TSS1500;TSS200;1stExon | chr5:149546073-149546946 | 873 | | | | | 2 | | | 0 | 5 | | | 4 | | yes | yes | | yes | yes | yes | | | | |
| SLC6A7 | Island | 5'UTR;1stExon | chr5:149569880-149570189 | 309 | | | 2 | | | | | | | | | | | yes | | | | | | | | | |
| CAMK2A | NA | TSS200 | chr5:149669488-149669813 | 325 | | | | | | | | | 1 | | | | | | | | | yes | | | | | |
| CD74 | NA | 1stExon;TSS200 | chr5:149792249-149792783 | 534 | | | 11 | | | | | | | | | | | yes | | yes | | | | | | | |
| NDST1 | NA | TSS1500 | chr5:149887422-149887497 | 75 | | | | | | | | 0 | 1 | | | | | yes | | | | yes | | | | | |
| SYNPO | NA | TSS200;1stExon;5'UTR | chr5:149980486-149980682 | 196 | 1 | | 44 | | | | | | 1 | | | | | yes | | | | | | | | | |
| SYNPO | NA | Body | chr5:149999033-149999249 | 216 | | | 1 | 1 | | | | | 1 | | | | | yes | yes | | | | | | | | |
| SYNPO | Island | Body | chr5:150004773-150004918 | 145 | | | 12 | | | | | | | | | | | yes | | | | | yes | | | | |
| DCTN4 | S_Shore | TSS1500 | chr5:150138866-150139148 | 282 | | | | | | | | | | | | | | yes | | yes | | | | | | | |
| IRGM | NA | 5'UTR | chr5:150227510-150227845 | 335 | | | 3 | | | | | | | | | | | yes | | | | | | | | | |
| ZNF300 | Island;N_Shore | TSS200;5'UTR;1stExon | chr5:150284302-150284796 | 494 | | | 1 | | | | | 0 | 2 | | | 2 | | | | | | | yes | | | | |
| LOC134466 | Island;S_Shore | Body;TSS200;TSS1500 | chr5:150325954-150326862 | 908 | | 1 | 10 | | 1 | | | 0 | 3 | 3 | | 5 | 1 | | | | yes | | | | | | |
| SLC36A3 | NA | Body | chr5:150677867-150678334 | 467 | | | | | | | 1 | 0 | 1 | 1 | | 2 | | yes | | | yes | yes | | | | | |
| SLC36A3 | NA | 5'UTR;1stExon | chr5:150683039-150683530 | 491 | | | | | | | | | | | | | | | yes | | | | | | | | |
| FAT2 | Island | Body | chr5:150911153-150911450 | 297 | | 1 | | | | | | | | | | | | | yes | | | yes | | | | | |
| SPARC | NA | 5'UTR;1stExon;TSS200 | chr5:151066268-151066730 | 462 | 3 | | | | | | | | 1 | | | | | yes | | | | | yes | | | | |
| G3BP1 | Island | TSS1500 | chr5:151150029-151150431 | 402 | | | | | | | | 0 | | | | | | yes | yes | | | | | | | | |
| GLRA1 | Island;S_Shore;N_Shore | TSS1500;Body;TSS200 | chr5:151303847-151305746 | 1899 | | 2 | | | 1 | | | 0 | 7 | | | 3 | | yes | | | | yes | yes | | | | |
| MFAP3;FAM114A2 | S_Shore | 5'UTR;TSS1500;Body | chr5:153418793-153418932 | 139 | | | | | | | | 0 | | | | | | yes | yes | | | | | | | | |
| HAND1 | Island | TSS1500;1stExon | chr5:153857468-153859973 | 2505 | | | | | | | | | | | | | | | | | | | | | | | |
| LARP1 | NA | TSS1500;TSS200 | chr5:154091976-154093260 | 1284 | | | | | | | | | | | | | | | | | | | | | | | |
| C5orf4 | NA | 5'UTR;1stExon;TSS200 | chr5:154230141-154230438 | 297 | | | | | | | | 0 | 1 | | | | | | yes | | | | | | | | |
| GEMIN5 | S_Shore | TSS1500 | chr5:154318039-154318104 | 65 | | | | | | | | | | | | | | yes | | | | | | | | | |
| SGCD | NA | 1stExon;5'UTR | chr5:155754009-155754265 | 256 | | | | 1 | | | | | 2 | | | | | | | | | | | | | | |
| LOC100286948;TIMD4 | NA | TSS200;Body | chr5:156351010-156351266 | 256 | | | 3 | | | | | | 1 | | | | | yes | | | | | | | | | |
| MED7 | N_Shore | 5'UTR | chr5:156569391-156569777 | 386 | | | | | | | | 0 | | | | | | yes | | | yes | | | | | | |
| FAM71B | NA | TSS1500 | chr5:156594400-156594725 | 325 | | | 1 | | | | | | 1 | | 1 | | | yes | yes | | | yes | | | | | |
| ITK | NA | TSS200 | chr5:156607836-156608118 | 282 | | | | | | | | | | 1 | | | | yes | yes | | | | | | | | |
| CYFIP2 | S_Shelf | TSS200;5'UTR | chr5:156696185-156696351 | 166 | | | | | | | | | 1 | | | | | yes | | | | | | | | | |
| CYFIP2 | NA | Body | chr5:156736991-156737228 | 237 | | | 1 | | | | | | | | | | | yes | | | | | | | | | |
| CYFIP2;C5orf40 | NA | Body;1stExon;5'UTR;TSS1500 | chr5:156772562-156773825 | 1263 | | | 3 | | | | | | | | | | | yes | | yes | | | | | | | |
| NIPAL4 | N_Shore;Island | TSS1500;TSS200 | chr5:156886415-156887393 | 978 | | | 26 | | | | | | 1 | | | | | yes | yes | | | | yes | | | | |
| ADAM19 | Island | Body | chr5:157001561-157001698 | 137 | | 1 | | | | | | | | | | | | yes | yes | | | | | | | | |
| SOX30 | Island | 1stExon;5'UTR;TSS200 | chr5:157078573-157079520 | 947 | | | 2 | | | | 1 | | 2 | 2 | | 1 | | | yes | | | | | | | | |
| EBF1 | N_Shore | Body | chr5:158522427-158522899 | 472 | 3 | | | | | | | | | | | | | | | | | | | | | | |
| EBF1 | Island | Body | chr5:158523907-158524649 | 742 | | | 6 | | | | | | | | | 1 | | | | | | yes | | | | | |
| UBLCP1 | N_Shore | TSS1500 | chr5:158689508-158689930 | 422 | | | 11 | | | | | | 1 | | | 2 | | | yes | | | | | | | | |
| ADRA1B | Island | TSS1500 | chr5:159343468-159343549 | 81 | | 1 | | | | | | | 1 | | | | | | | | | | yes | | | | |
| FABP6 | NA | 5'UTR;TSS1500 | chr5:159625590-159626112 | 522 | | | | 1 | | | | | | | | | | | yes | | | | | | | | |
| FABP6 | NA | TSS1500;Body;TSS200 | chr5:159655853-159656660 | 807 | | | | | | | | | | | | | | yes | yes | | | yes | | | | | |
| C1QTNF2 | S_Shore | TSS200 | chr5:159797753-159797775 | 22 | | | | | | | | | 1 | | | | | yes | | | | | | | | | |
| ATP10B | NA | TSS1500 | chr5:160280216-160280697 | 481 | | | | | | | | | | | | | | yes | yes | | | | | | | | |
| GABRB2 | S_Shore | TSS1500 | chr5:160975792-160975978 | 186 | | 1 | | | | | | | 1 | | | | | yes | | | | yes | | | | | |
| GABRA1 | NA | TSS1500;5'UTR;1stExon | chr5:161275138-161275449 | 311 | | 1 | | | | | | | | | | | | | | | | yes | yes | | | | |
| GABRG2 | NA | TSS1500;TSS200 | chr5:161494015-161494711 | 696 | | | | | | | | 0 | 3 | | | | | | | | | | | | | | |
| CCNG1 | N_Shore;Island | TSS1500;TSS200 | chr5:162864268-162865006 | 738 | | | | | | | | 0 | | | | 1 | | yes | | yes | | | | | | | |
| NUDCD2;HMMR | S_Shore | TSS1500;Body | chr5:162887936-162888518 | 582 | | | | | | | | | 1 | | | | | yes | yes | | | | | | | | |
| ODZ2 | NA | Body | chr5:167181519-167181836 | 317 | | | | | | | | | | | | | | | | | | | yes | | | | |
| ODZ2 | NA | Body | chr5:167614881-167615258 | 377 | | | | | | | | | | | | | | | yes | | yes | | | | | | |
| FBLL1 | Island | TSS200;Body | chr5:167956394-167957046 | 652 | | | | | | | | | 3 | | | | | | | | | yes | yes | | | | |
| SLIT3 | NA | Body | chr5:168271855-168271898 | 43 | | | | | | | | | 1 | | | | | | | | | yes | | | | | |
| SLIT3;MIR585 | NA | Body | chr5:168690633-168690818 | 185 | | | | | | | | | | | | | | yes | | yes | | | | | | | |
| SLIT3 | S_Shore;Island | TSS1500;5'UTR;1stExon | chr5:168727762-168729847 | 2085 | | | 20 | | | | | 0 | 1 | | | | | yes | | | yes | yes | | | | | |
| DOCK2 | N_Shore | TSS1500 | chr5:169063563-169064071 | 508 | | | 5 | | | | | | | | | | | yes | | | yes | | | | | | |
| FAM196B;DOCK2 | NA | 5'UTR;Body;1stExon | chr5:169407439-169407941 | 502 | | | | | | | | | 2 | | | | | | | | yes | yes | | | | | |
| FOXI1 | N_Shore | 1stExon;5'UTR;TSS1500;TSS200 | chr5:169532686-169533347 | 661 | | 1 | 11 | | 2 | | | | 2 | | | | | yes | | | | yes | yes | | | | |
| C5orf58 | N_Shore;Island | TSS200 | chr5:169659763-169659950 | 187 | | | | | | | | | 3 | | | | | yes | | | | | | | | | |
| LCP2 | NA | 1stExon;5'UTR | chr5:169724590-169725100 | 510 | | | | | 2 | | | | 2 | 2 | 1 | | | yes | | | yes | | | | | | |
| LOC257358 | NA | TSS1500 | chr5:169757754-169758055 | 301 | | | 1 | | | | | | | | | | | yes | yes | | | yes | | | | | |
| KCNIP1 | NA | TSS1500 | chr5:169780299-169780916 | 617 | | | 2 | 1 | | | | | | | | | | | yes | | yes | | | | | | |
| KCNMB1;KCNIP1 | NA | TSS200;Body | chr5:169816792-169817247 | 455 | | | | | | | | | | | | | | yes | | | | yes | | | | | |
| KCNIP1 | N_Shore | TSS200;Body | chr5:169930879-169931092 | 213 | | | | | | | | | 1 | | | | | yes | | | | yes | yes | | | | |
| GABRP | NA | TSS200;TSS1500 | chr5:170209760-170211009 | 1249 | | | 8 | | | | | | | | | | | yes | | | | | yes | | | | |
| RANBP17 | N_Shore | TSS1500 | chr5:170288255-170288766 | 511 | | | 11 | | 1 | | | 0 | 1 | | | 1 | | | yes | | | yes | | | | | |
| TLX3 | Island;N_Shore | TSS200;TSS1500 | chr5:170734856-170736572 | 1716 | | | | | | | | 0 | 4 | | | | | yes | | | | | | | | | |
| FGF18 | N_Shore | Body | chr5:170877540-170877932 | 392 | | | | 1 | | | | | | | | | | yes | | | | | yes | | | | |
| STK10 | NA | Body | chr5:171534678-171534792 | 114 | | | 8 | | | | | | 1 | | | | | yes | yes | | | | | | | | |
| STK10 | S_Shore | TSS1500 | chr5:171616253-171616574 | 321 | | 1 | | | | | | 0 | 1 | | | | | | yes | | | | yes | | | | |
| UBTD2 | N_Shore | Body | chr5:171708886-171709304 | 418 | 1 | | 1 | | | | | | | | | | | yes | | | | | | | | | |
| SH3PXD2B | N_Shore | Body | chr5:171766029-171766566 | 537 | | | | | | | | | 1 | | | | | | | | | | | | | | |
| SH3PXD2B | NA | Body | chr5:171808200-171808623 | 423 | | | 3 | | | | | | | | | | | | | | | | | | | | |
| NEURL1B | Island | Body | chr5:172096764-172097385 | 621 | | | | | | | | | 2 | | | | | yes | | | | yes | | | | | |
| NEURL1B | Island | Body | chr5:172110477-172110579 | 102 | | | 5 | | | | | | | | | | | yes | yes | | | | yes | | | | |
| DUSP1 | Island;S_Shore | TSS1500 | chr5:172199244-172199667 | 423 | 12 | | 4 | | 2 | | | | | | | | | yes | | | | | | | | | |
| ERGIC1 | N_Shore | TSS1500 | chr5:172260464-172260815 | 351 | | | | | | | | | 2 | | | | | yes | | | | yes | yes | | | | |
| ERGIC1 | NA | Body | chr5:172305889-172306136 | 247 | 4 | | 2 | | | | | | 1 | | 1 | | | yes | | | | | yes | | | | |
| LOC100268168;RPL26L1 | N_Shore;Island | Body;TSS1500;TSS200 | chr5:172385477-172385821 | 344 | | | 4 | | | | | | 1 | | | | | yes | | | | | | | | | |

| | | | | | | | | | | | | | | | | | | | | | | | |
|---|---|---|---|---|---|---|---|---|---|---|---|---|---|---|---|---|---|---|---|---|---|---|---|
| NKX2-5 | Island | 3'UTR | chr5:172659241-172659730 | 489 | | | | | | | | | | | | | | yes | yes | | | | yes |
| NKX2-5 | N_Shore;Island | TSS1500;Body;1stExon;TSS200 | chr5:172660796-172664439 | 3643 | | | 16 | | | | | | | | | | | | yes | | yes | yes | |
| STC2 | N_Shelf | Body | chr5:172750450-172751331 | 881 | | | | | | | | 0 | 2 | | | 3 | | | | | | | |
| STC2 | Island | Body | chr5:172754417-172754896 | 479 | | | 1 | | | | | | | | | | | | | | | | yes |
| CPEB4 | S_Shore | 1stExon | chr5:173316748-173317342 | 594 | | | | | | | | | | | | | | | | | | | |
| C5orf47 | Island | TSS200 | chr5:173415963-173415991 | 28 | | | | | | | | | 1 | | | | | | | | | | |
| MSX2 | N_Shore;Island | TSS1500;TSS200;5'UTR;1stExon | chr5:174151334-174152363 | 1029 | | 1 | | | | | | 0 | 3 | | | | | yes | | | | yes | |
| HRH2 | N_Shore | TSS1500 | chr5:175083981-175084718 | 737 | | | | | | | | | 1 | | | | | yes | yes | | | yes | yes |
| HRH2 | NA | 5'UTR;TSS1500;TSS200;1stExon | chr5:175107772-175109114 | 1342 | | | 16 | | | | | | 1 | | | | | yes | yes | | yes | | |
| CPLX2 | N_Shore;Island | TSS1500;TSS200;5'UTR | chr5:175223225-175224721 | 1496 | | | | | | | | | 2 | | | | | yes | yes | | | | |
| CPLX2 | N_Shore | TSS1500;5'UTR | chr5:175297349-175297573 | 224 | | | | | | | | | 1 | | | | | | yes | | | yes | |
| RNF44 | NA | 3'UTR | chr5:175955105-175955925 | 820 | | | 8 | | | | | | | | | | | | | | | | |
| PCDH24 | NA | TSS200;TSS1500 | chr5:175975901-175976402 | 501 | | | | 1 | | | | | | | | | | yes | | | | | |
| EIF4E1B;SNCB | S_Shore | 5'UTR;TSS1500;1stExon | chr5:176057813-176057996 | 183 | | | 11 | | | | | | | | | | | yes | yes | | | yes | yes |
| UNC5A | NA | Body | chr5:176270730-176270936 | 206 | | | 2 | | | | | | | | | | | | yes | | | | |
| UNC5A | NA | Body | chr5:176289125-176289552 | 427 | | | 1 | | | | | | | | | | | yes | | | | | |
| UNC5A | Island | Body | chr5:176304988-176305077 | 89 | | | 5 | | | | | | | | | | | | | | | | |
| UNC5A;HK3 | S_Shelf | 3'UTR | chr5:176307743-176308136 | 393 | | | | | | | | | | | | | | | | | | | |
| HK3 | NA | TSS200 | chr5:176326365-176326619 | 254 | | | | 1 | 1 | | | | 3 | | 1 | | | yes | | | | | |
| FGFR4 | N_Shore | Body | chr5:176519455-176520502 | 1047 | | 1 | 2 | | | | | | | | | | | yes | | | | | |
| FGFR4 | S_Shore | Body | chr5:176522599-176523370 | 771 | | 1 | | | | | | | | | | | | yes | | | | yes | |
| NSD1 | Island | TSS1500;TSS200 | chr5:176559334-176560265 | 931 | | | 5 | | 2 | | | 0 | 2 | | 1 | 3 | | | | | | yes | yes |
| MXD3 | Island | Body | chr5:176734633-176734858 | 225 | | | | | | | | | 1 | | | | | | | | yes | | |
| MXD3 | Island | 1stExon;5'UTR;TSS200 | chr5:176738847-176740039 | 1192 | | | | | 7 | | | | | | | | | yes | | | | | yes |
| RGS14 | NA | TSS1500;TSS200 | chr5:176784318-176785296 | 978 | | | 20 | | 4 | | | | | | | | | | yes | yes | | yes | |
| RGS14 | N_Shore | Body | chr5:176793941-176794405 | 464 | | | | | 1 | 1 | | | | | | | | | | | | yes | |
| RGS14 | S_Shelf | Body | chr5:176797920-176798049 | 129 | | | | | 1 | 2 | 1 | 0 | 1 | 1 | 1 | 47 | 1 | | | | | yes | |
| SLC34A1 | NA | TSS1500;TSS200 | chr5:176811016-176811305 | 289 | | | | | | | | | | | | | | yes | yes | | | | |
| SLC34A1 | N_Shore | Body | chr5:176823320-176823642 | 322 | | | 2 | | | | | | | | | | | yes | yes | | | | |
| PFN3 | Island | TSS200;1stExon;3'UTR | chr5:176827161-176827793 | 632 | | | | 1 | | | | 0 | 2 | | | | | yes | | | yes | | yes |
| PFN3 | N_Shore | TSS1500 | chr5:176828872-176829349 | 477 | | | 12 | | | | | | | | | | | yes | yes | | yes | | yes |
| F12 | NA | 1stExon | chr5:176836472-176836695 | 223 | | | | 1 | | | | | | | | | | yes | | yes | | | |
| GRK6 | Island | Body | chr5:176860863-176860947 | 84 | | | | | | | | | 1 | | | | | yes | | | | | |
| DOK3 | N_Shore | Body | chr5:176930263-176931111 | 848 | | | | | | | | | 1 | | | | | yes | yes | | | | |
| DOK3 | NA | TSS200;5'UTR;1stExon;TSS1500;Bo | chr5:176936563-176937446 | 883 | | | 10 | | | | | 0 | 1 | | | | | yes | | | | | |
| FAM193B | Island;S_Shore | TSS200;TSS1500 | chr5:176981629-176982179 | 550 | | | 9 | | | | | 0 | 1 | 1 | | 1 | | yes | yes | yes | | yes | yes |
| B4GALT7 | Island | Body | chr5:177035360-177035558 | 198 | | 1 | | | | | | | | | | | | | | | | | |
| PROP1 | NA | Body;1stExon;5'UTR;TSS200 | chr5:177422782-177423999 | 1217 | | 1 | | | | | | | 1 | | | | | yes | | yes | yes | | |
| FAM153C | S_Shore | TSS1500 | chr5:177434336-177434457 | 121 | | | 3 | | | | | | 1 | | | 1 | | yes | | | | yes | |
| N4BP3 | NA | Body | chr5:177547973-177549014 | 1041 | | | 9 | | | | | | | | | | | | | | | | |
| GMCL1L | Island | Body | chr5:177614205-177614382 | 177 | | | 4 | | | | | | | | | | | yes | | | | | |
| AGXT2L2 | S_Shore | TSS200;TSS1500 | chr5:177659971-177660351 | 380 | | | 2 | | | | | | | | | | | yes | yes | | | | yes |
| COL23A1 | NA | Body | chr5:177697400-177697669 | 269 | | | | | | | | | | | | | | yes | | yes | | | |
| COL23A1 | NA | Body | chr5:177707036-177707226 | 190 | | 1 | | | | | | 0 | 1 | | | | | | yes | | | | |
| COL23A1 | NA | Body | chr5:177708304-177708528 | 224 | | | 23 | | | | | | | | | | | yes | | | | | yes |
| COL23A1 | N_Shore;Island | Body | chr5:177764003-177764408 | 405 | | | | | | | | | | | | | | | yes | | | | yes |
| COL23A1 | NA | Body | chr5:177811198-177811648 | 450 | 1 | | | | | | | | 1 | | | | | | | | | | |
| COL23A1 | Island | Body | chr5:177823856-177824403 | 547 | | | | | | | | | | | | | | yes | | | | | |
| COL23A1 | NA | Body | chr5:177853481-177853642 | 161 | 7 | | | | | | | | | | | | | yes | yes | | | | |
| COL23A1 | NA | Body | chr5:177889945-177891221 | 1276 | | | 2 | | | | | | | | | | | yes | yes | | yes | | |
| COL23A1 | NA | Body | chr5:177913434-177913485 | 51 | | | | | | | | 0 | 1 | 1 | | | | | | | | | |
| COL23A1 | NA | Body | chr5:177980691-177980944 | 253 | | | 2 | | | | | | | | | | | yes | | | | | |
| COL23A1 | Island | Body | chr5:178003633-178004204 | 571 | | 1 | 3 | | 1 | | | 0 | 1 | | | | | yes | | yes | | | |
| ZFP2 | NA | 5'UTR | chr5:178339172-178339445 | 273 | | | | | | | | | | | | | | yes | | yes | yes | | |
| ZNF454 | N_Shore;Island | TSS1500;TSS200 | chr5:178367423-178368620 | 1197 | | 1 | | | 1 | | | 0 | 6 | | | | | | yes | | | | |
| GRM6 | Island | 1stExon;TSS200 | chr5:178421711-178422260 | 549 | | 1 | | | | | | 0 | 3 | | | | | | | | yes | yes | yes |
| ZNF879 | Island | TSS200 | chr5:178450758-178451176 | 418 | | 1 | | | | | | | | | | | | yes | | | | | |
| ZNF354C | Island | TSS200 | chr5:178487465-178487826 | 361 | | 1 | | | | | | | 1 | | | | | yes | yes | | | | |
| ADAMTS2 | Island | Body | chr5:178547422-178548700 | 1278 | | | | | | 1 | | 0 | 3 | 1 | | 3 | | | | | | | |
| ADAMTS2 | Island | Body | chr5:178565721-178565937 | 216 | | | | | | | | | 1 | | | | | | yes | | | | |
| ADAMTS2 | N_Shore | Body | chr5:178593785-178594097 | 312 | | | | | | | | | | | | | | yes | | | | | |
| ADAMTS2 | NA | Body | chr5:178622080-178623035 | 955 | | | 3 | | | | | | | | | | | | | | yes | | |
| ADAMTS2 | Island;N_Shore | Body | chr5:178683834-178684658 | 824 | | | 15 | | | | | | | | | | | yes | | | yes | yes | |
| ADAMTS2 | NA | Body | chr5:178692691-178692806 | 115 | | | | | | | | 0 | 1 | 1 | 1 | 5 | | | | yes | | | |
| ADAMTS2 | NA | Body | chr5:178741004-178741374 | 370 | | | 1 | | | | | | | | | | | yes | | | | yes | |
| RUFY1 | N_Shore;Island | TSS1500;Body;TSS200 | chr5:178986131-178986906 | 775 | | | 2 | | | | 1 | 0 | 13 | 8 | | 16 | | | yes | | yes | | |
| LTC4S | N_Shelf | TSS1500 | chr5:179220545-179221090 | 545 | | | | 1 | | | | | | | | | | | | | | | |
| SQSTM1 | Island | 5'UTR | chr5:179243916-179244452 | 536 | | | | | | | | | 1 | | | | | yes | | | | | |
| SQSTM1 | S_Shelf | Body | chr5:179250940-179251299 | 359 | | | | 1 | | | | | | | | | | yes | | | | | |
| RASGEF1C | N_Shore | Body | chr5:179553589-179554111 | 522 | | | | | | | | 0 | | | | 1 | | yes | yes | | | | |
| RASGEF1C | Island | Body | chr5:179563738-179564268 | 530 | | | | | | | | | 1 | | | | | yes | | yes | | | |
| RASGEF1C | S_Shore | 5'UTR | chr5:179565349-179565544 | 195 | | | 1 | | | | | | | | | | | | yes | yes | | | |
| RASGEF1C | N_Shore;Island | 5'UTR | chr5:179587950-179588838 | 888 | | | 8 | | | | 2 | 0 | 3 | | | 3 | | | | | | | |
| RASGEF1C | N_Shore;Island | 5'UTR | chr5:179634383-179635981 | 1598 | 1 | | 11 | | | | 1 | 0 | 4 | 2 | | 5 | | yes | yes | | | yes | yes |
| GFPT2 | Island | Body | chr5:179740743-179741120 | 377 | 17 | 21 | | 1 | 1 | 19 | 1 | 0 | 3 | 3 | 1 | 64 | 1 | yes | yes | | | | yes |
| GFPT2 | Island | Body | chr5:179779115-179779237 | 122 | | | 1 | | | | | 0 | 1 | | | | | | | | | yes | |
| SCGB3A1 | Island | TSS200 | chr5:180018503-180018564 | 61 | | | | | 1 | | | | | | | | | | | | | | |
| FLT4 | N_Shore | Body | chr5:180045597-180046052 | 455 | | | | | | | | 0 | 1 | | | 1 | | | | | | | yes |
| FLT4 | Island | Body | chr5:180047299-180048155 | 856 | | | | | | | | | | | | | | | | | | | |
| FLT4 | N_Shelf | Body | chr5:180071709-180071930 | 221 | | | 9 | | | | | | | | | | | yes | | | | | |
| OR2Y1 | NA | TSS1500 | chr5:180167566-180167965 | 399 | | | | | | | | | | | | | | yes | | | | yes | |
| MGAT1 | S_Shore;chr5:180229375-1802 | 1stExon;5'UTR;TSS1500;TSS200;S_ | chr5:180230767-180231194 | 427 | | | 11 | | 1 | 1 | | | 1 | 1 | 1 | | | yes | | | | | |
| BTNL8 | NA | TSS1500;TSS200 | chr5:180325579-180326186 | 607 | | | | 1 | 1 | | | | 1 | | | | | | yes | | | | |
| BTNL9 | NA | TSS1500;TSS200 | chr5:180466563-180467519 | 956 | | | | | | | | | | | | | | | | yes | | | |
| BTNL9 | Island | Body | chr5:180486604-180486787 | 183 | | | | | | | | | 1 | | | | | | yes | | | | yes |
| OR2V2 | NA | TSS1500;TSS200 | chr5:180581311-180581950 | 639 | | | | | | | | | | | | | | yes | | yes | | | |
| TRIM7 | Island | Body | chr5:180622276-180622643 | 367 | | | | | | | | | 1 | | | | | yes | yes | | | | |
| TRIM7 | S_Shore | TSS1500 | chr5:180632948-180633326 | 378 | | | 8 | | | | | | | | | | | | yes | | | yes | |
| TRIM41 | N_Shore | TSS1500 | chr5:180649174-180649807 | 633 | | | | | | | 1 | | | 1 | | | | | | | | | |
| GNB2L1;SNORD95 | Island | TSS200;TSS1500 | chr5:180670960-180670995 | 35 | | | | | | | | 0 | | | | | | yes | | | | | |
| DUSP22 | N_Shore;Island | TSS1500;TSS200;1stExon;5'UTR;Bo | chr6:291687-293331 | 1644 | 6 | 15 | 40 | 2 | 3 | 4 | 1 | 0 | 10 | 7 | 1 | 63 | 2 | | yes | | | yes | |
| DUSP22 | NA | Body | chr6:312026-312105 | 79 | | | 4 | | | | | | 1 | | | | | yes | | | | | |
| IRF4 | NA | Body | chr6:401311-401453 | 142 | | | | | | | | | | | | | | | yes | | yes | yes | |
| HUS1B;EXOC2 | Island | 1stExon;5'UTR;TSS200 | chr6:656504-657150 | 646 | | | | | | | | | 4 | | | 1 | | | yes | | | | |
| LOC285768 | NA | TSS1500 | chr6:1102332-1102438 | 106 | | | | | | | | | 1 | | | | | yes | yes | | | | |
| FOXQ1 | Island | TSS200;1stExon | chr6:1312517-1313495 | 978 | | | | | | | | | 2 | | | | | | | | | yes | yes |
| FOXF2 | N_Shore;Island | TSS1500 | chr6:1388806-1389367 | 561 | | | 14 | | | | | | | | | | | | | | | yes | yes |
| FOXF2 | S_Shore | Body | chr6:1394986-1395278 | 292 | | | | | | | | 0 | 1 | | | | | | | | | | |
| GMDS | Island | Body | chr6:1624442-1625574 | 1132 | | 1 | | | 2 | | | | | 1 | | | | | | | yes | yes | yes |

| | | | | | | | | | | | | | | | | | | | | | | | |
|---|---|---|---|---|---|---|---|---|---|---|---|---|---|---|---|---|---|---|---|---|---|---|---|
| GMDS | NA | Body | chr6:1635611-1635847 | 236 | | | 3 | 1 | | | | | | | | | | yes | yes | | | | |
| GMDS | NA | Body | chr6:1708566-1708713 | 147 | | | | | | | | | | | | | | | | | yes | | |
| MYLK4 | NA | Body | chr6:2698001-2698037 | 36 | | | 6 | | | | | | | | | | | | | | | | |
| MYLK4 | NA | 1stExon;5'UTR | chr6:2751133-2751402 | 269 | | | | | | | | | | | | | | | | | | | |
| SERPINB9 | Island | Body | chr6:2891973-2892152 | 179 | | | | | 1 | | | | 2 | | | | | yes | | | | yes | |
| SERPINB6 | NA | Body | chr6:2958750-2958852 | 102 | | | | 1 | | | | | 1 | | 1 | | | | | yes | | | |
| SERPINB6 | S_Shore | TSS200 | chr6:2972158-2972435 | 277 | | | | | | | | 0 | 1 | | | | | yes | yes | yes | | | yes |
| BPHL | N_Shore | Body | chr6:3140546-3140784 | 238 | | 1 | | | | | | | | | | | | yes | | | | | |
| TUBB2B | Island | TSS1500 | chr6:3229027-3229530 | 503 | | | | | 1 | | | | | | | | | yes | | | | yes | |
| SLC22A23 | NA | Body | chr6:3299629-3299689 | 60 | | | | | | | | | 1 | | | | | yes | yes | | yes | | yes |
| SLC22A23 | N_Shore | Body | chr6:3455459-3456062 | 603 | | | 2 | | | | | | | | | | | | | | | | |
| SLC22A23 | Island | TSS1500 | chr6:3457843-3458177 | 334 | | | 1 | | | | | | | | | | | | | | | yes | yes |
| C6orf145 | NA | Body | chr6:3743284-3743529 | 245 | | | | | | | | 0 | 1 | | | 1 | | | | | | | |
| FAM50B | N_Shore;Island | TSS1500;TSS200;1stExon;5'UTR | chr6:3848218-3850106 | 1888 | 1 | 1 | 22 | 1 | | | | 0 | 8 | | | 3 | | | | | yes | yes | |
| C6orf146;C6orf201 | N_Shore;Island | 5'UTR;TSS1500;TSS200;1stExon | chr6:4078970-4079528 | 558 | | | 4 | | | | | | | | | | | yes | | | | | yes |
| PECI | Island | TSS200;TSS1500 | chr6:4135844-4136202 | 358 | | | | | | | | 0 | 1 | | | | | | | | | yes | yes |
| CDYL | NA | TSS1500 | chr6:4706170-4706336 | 166 | | | 2 | | | | | | | | | | | yes | yes | | yes | | |
| CDYL | N_Shore | Body | chr6:4775063-4775203 | 140 | | | | | | | | | | | | | | | yes | | | | |
| CDYL | NA | 5'UTR;Body;TSS200 | chr6:4890079-4890278 | 199 | | | | 1 | 1 | | | 0 | 1 | | | | | yes | | | yes | | |
| PPP1R3G | Island;S_Shore | 3'UTR;1stExon | chr6:5087031-5087749 | 718 | | | | | | | | 0 | 5 | 1 | | 1 | | | | | yes | | |
| NRN1 | Island | 3'UTR | chr6:5999186-5999888 | 702 | | 1 | | | | | | | | | | | | | | | | | yes |
| NRN1 | Island | Body | chr6:6002543-6003303 | 760 | | | 6 | | | | | | | | | | | | | | | | yes |
| NRN1 | Island | Body | chr6:6004686-6005349 | 663 | | 1 | | | | | | | 1 | | | | | | | | | | |
| NRN1 | N_Shore | Body | chr6:6006398-6006917 | 519 | | | | | | | | | 1 | | | | | | | | | yes | |
| F13A1 | NA | 5'UTR;1stExon | chr6:6320324-6321061 | 737 | | | 5 | | | | | | | | | | | | | | yes | | yes |
| LOC285780 | Island | Body | chr6:6546375-6547273 | 898 | | | 14 | | | | | | | | | | | yes | | | | | |
| LY86;LOC285780 | NA | TSS200;Body | chr6:6588737-6589075 | 338 | | | | | 1 | | | | 1 | | | | | | | | | | |
| LOC285780;LY86 | NA | Body | chr6:6613573-6614798 | 1225 | | | 10 | | | | | | 1 | | | | | | | yes | | | |
| RREB1 | N_Shore | TSS1500 | chr6:7106952-7107129 | 177 | | | | | 1 | | | | | | | | | | | | | | yes |
| DSP | N_Shore | TSS1500 | chr6:7540834-7541519 | 685 | | | 7 | | | | | | 1 | | | | | | | | yes | | yes |
| SCARNA27;EEF1E1 | NA | TSS200;Body | chr6:8086867-8087102 | 235 | | | | | | | | | 1 | | | | | yes | | | | | |
| EEF1E1 | S_Shore | TSS200 | chr6:8103016-8103092 | 76 | | | 1 | | | | | | | | | | | | | | | yes | yes |
| SLC35B3 | Island;S_Shore | TSS200;TSS1500 | chr6:8435883-8436296 | 413 | | 1 | | | | 1 | | 0 | 1 | | | | | | | | | | yes |
| TFAP2A | N_Shore;Island | 3'UTR | chr6:10398494-10398754 | 260 | | | 4 | | | | | 0 | 1 | | | | | | | | | yes | yes |
| TFAP2A | Island;N_Shore | Body | chr6:10410042-10410916 | 874 | | | 6 | | | | | | | | | | | | | | | | |
| TFAP2A | Island | Body;5'UTR;1stExon | chr6:10412073-10412225 | 152 | | | | | | | | | 1 | | | | | | | | | yes | yes |
| TFAP2A | Island;N_Shore | Body;1stExon;5'UTR;TSS200;TSS15 | chr6:10415201-10418121 | 2920 | | 1 | 39 | | | | | 0 | 5 | | | 1 | | | | | | yes | |
| TFAP2A | Island;N_Shore | TSS200;TSS1500 | chr6:10419927-10422139 | 2212 | | 1 | 5 | | | | | | 1 | | | | | yes | | | | yes | yes |
| C6orf218 | NA | Body | chr6:10434761-10434907 | 146 | | | | | | | | | | | | | | yes | yes | yes | | | |
| GCNT2 | NA | TSS1500;TSS200 | chr6:10520874-10521612 | 738 | | | | | | | | | 1 | | | | | yes | yes | | | | |
| GCNT2 | NA | Body | chr6:10529935-10530035 | 100 | | | | 1 | | | | | | | | | | yes | | | | | |
| GCNT2 | NA | TSS200;Body;5'UTR;1stExon | chr6:10555808-10556523 | 715 | | | | | | | | 0 | 3 | 1 | | 1 | | | | | | | |
| GCNT2 | NA | Body;TSS1500;nan | chr6:10585683-10586167 | 484 | | 1 | | 1 | | | | | | | 1 | | | yes | | | | yes | |
| GCM2 | Island;S_Shore;N_Shore | TSS1500;Body;TSS200 | chr6:10881782-10884314 | 2532 | | 3 | 6 | | | | | | 1 | | | | | yes | | | | yes | yes |
| SYCP2L | N_Shore;Island | TSS200;1stExon;5'UTR;Body | chr6:10886999-10887980 | 981 | | | 4 | | | | | | 2 | | | | | yes | | | yes | | |
| ELOVL2 | Island | TSS1500 | chr6:11044877-11044974 | 97 | | 1 | 5 | | | | | | 1 | | | | | yes | | | | | yes |
| NEDD9 | NA | Body | chr6:11279656-11279727 | 71 | | | 14 | | | | | | | | | | | | | | yes | | |
| NEDD9 | NA | 1stExon;5'UTR | chr6:11382553-11382830 | 277 | | | | 1 | 2 | | | | 2 | 3 | 1 | | | yes | | | | | |
| C6orf105 | NA | Body | chr6:11771493-11771840 | 347 | | | 9 | | | | | | | | | | | | | yes | | | |
| PHACTR1 | NA | TSS1500;5'UTR | chr6:12717016-12718530 | 1514 | | 1 | | | 1 | | | | 1 | | | | | | | | | | |
| PHACTR1 | N_Shore;Island | Body | chr6:12749352-12750392 | 1040 | | 1 | | | | | | | 1 | | | | | | yes | | | | yes |
| PHACTR1 | Island | Body | chr6:13015005-13015221 | 216 | | | | | 1 | | | | 1 | | | | | | | | | | |
| PHACTR1 | NA | Body | chr6:13274151-13274354 | 203 | | | | | 1 | | | 0 | 2 | 1 | 1 | 1 | | | | | | yes | |
| GFOD1 | NA | Body | chr6:13428522-13428657 | 135 | | | 1 | 1 | 1 | | | | 1 | 1 | 1 | | | | | | | | |
| RNF182 | N_Shore | TSS1500 | chr6:13924355-13924425 | 70 | | | | | | | | 0 | 1 | | | | | | | | | yes | |
| CD83 | Island | 5'UTR;1stExon | chr6:14118033-14118415 | 382 | | | 1 | | | | | | | | | | | | | | | | yes |
| JARID2 | NA | Body | chr6:15504844-15505460 | 616 | | 1 | | | | | | 0 | 5 | 3 | | | | | | | | | |
| ATXN1 | NA | 3'UTR | chr6:16303089-16303169 | 80 | | 1 | | | | | | | | | | | | | | | | | |
| RBM24 | Island;S_Shore;N_Shore | TSS1500;TSS200;Body;5'UTR | chr6:17280551-17283113 | 2562 | | 2 | | | 1 | | | 0 | | | | | | | | | | | yes |
| FAM8A1 | N_Shore | TSS1500 | chr6:17599948-17600252 | 304 | | | | | | | | 0 | | | | | | | | yes | | | |
| NHLRC1 | Island;S_Shore | TSS200;TSS1500 | chr6:18122965-18123232 | 267 | | 1 | 5 | | | | | 0 | 2 | | | | | yes | | | | | |
| RNF144B | N_Shore | TSS1500 | chr6:18387077-18387407 | 330 | | | 13 | | | | | | | | | | | | | | | | |
| ID4 | Island | TSS1500 | chr6:19837095-19837421 | 326 | | | | | | | | | 1 | | | | | | | | | | yes |
| FLJ22536 | S_Shore | Body | chr6:21666939-21667815 | 876 | | | 10 | | | | | | | | | | | yes | | | | | yes |
| DCDC2;KAAG1 | N_Shore;Island | Body;TSS200;1stExon;5'UTR;3'UT | chr6:24356996-24358683 | 1687 | | | 11 | | | | | 0 | 5 | | | | | yes | yes | | | yes | |
| GPLD1 | NA | 1stExon;5'UTR | chr6:24489770-24489887 | 117 | | | | 1 | | | | | | | | | | | | | | | |
| ALDH5A1 | N_Shore | TSS1500 | chr6:24494605-24494793 | 188 | | 1 | | | | | | | | | | | | yes | | | | | yes |
| KIAA0319 | NA | Body | chr6:24583477-24583614 | 137 | 4 | 1 | 9 | | | | | 0 | | | | 1 | | | | | | | |
| FAM65B | NA | 5'UTR;1stExon | chr6:24877462-24877624 | 162 | | | | 1 | | | | | 1 | | | | | | | yes | | | |
| FAM65B | N_Shore;Island | 5'UTR;TSS200 | chr6:24910562-24911553 | 991 | | | | | | | | | 2 | | | 1 | | yes | | | | | yes |
| CMAH | N_Shelf | Body | chr6:25137864-25138169 | 305 | | | 3 | | | | | | 1 | | | | | | | | | yes | |
| SCGN | Island | TSS200 | chr6:25652381-25652531 | 150 | | | | | | | | | 2 | | | 1 | | | yes | | | | |
| HIST1H2BA;HIST1H2AA | Island;S_Shore | TSS1500;1stExon;TSS200 | chr6:25726597-25726912 | 315 | | | 4 | | | | | | | | | | | yes | | | | | |
| SLC17A4 | NA | TSS1500 | chr6:25753732-25754219 | 487 | | | | | | | | | | | | | | | yes | | yes | | |
| SLC17A3 | NA | 5'UTR | chr6:25874121-25874590 | 469 | | | | 1 | | | | | | | | | | yes | | | | yes | |
| SLC17A2 | NA | 1stExon;5'UTR | chr6:25930507-25931659 | 1152 | | | | 1 | | | | | | | | | | yes | | yes | | | |
| TRIM38 | NA | TSS1500 | chr6:25962567-25963109 | 542 | | 1 | | | | | | | | | | | | | | | | | |
| HIST1H1A | N_Shelf | 1stExon;TSS200 | chr6:26017619-26018602 | 983 | | 1 | 2 | | 1 | | | 0 | 2 | | | 2 | | | | | | | |
| HIST1H4B | S_Shore | TSS1500 | chr6:26028624-26028781 | 157 | | | | | | | | | 1 | | | | | | | yes | | | |
| HIST1H2BB;HIST1H3C | N_Shore;Island | 1stExon;TSS1500 | chr6:26043732-26044460 | 728 | | | | | | | | | 1 | | | | 1 | yes | | | | | |
| HIST1H3C | N_Shore | TSS200 | chr6:26045532-26045663 | 131 | | | | | | | | | 1 | | | | 1 | | | | yes | | |
| HIST1H2BE | N_Shore | TSS1500 | chr6:26183508-26183829 | 321 | | | 3 | | | | | 0 | 4 | | | 2 | 1 | | | | | | yes |
| HIST1H4D | N_Shore;S_Shore | 3'UTR;1stExon;TSS200 | chr6:26188969-26189597 | 628 | | | | | | | | | 2 | | | | | yes | | | | | |
| HIST1H4E | Island | TSS200 | chr6:26204765-26204823 | 58 | | | | | | | | | 1 | | | | | yes | | | | | |
| HIST1H2BG;HIST1H2AE | Island | TSS1500;1stExon | chr6:26217416-26217564 | 148 | | | | | | | | | 1 | | | | | | | | | | |
| HIST1H3E | N_Shore | TSS1500;TSS200 | chr6:26224013-26225539 | 1526 | | 1 | | | | | 1 | 0 | 3 | 1 | | 1 | 1 | | | | | | |
| HIST1H1D | NA | TSS200 | chr6:26235311-26235609 | 298 | | | 1 | | | | | | | | | | | | | | | | |
| HIST1H4F | N_Shore | TSS200;TSS1500 | chr6:26240307-26240939 | 632 | | 1 | | | | | | | 1 | | | 1 | | yes | | | | | |
| HIST1H4G | S_Shore | TSS200 | chr6:26247238-26247270 | 32 | | | 2 | | | | | | | | | | | yes | | | | | |
| BTN1A1 | Island | Body | chr6:26501812-26502192 | 380 | | 1 | 2 | | | | | | | | | | | | | yes | | | |
| HCG11 | Island | Body | chr6:26521980-26522578 | 598 | | | | 1 | | | | | | | | | | | yes | | | | |
| ABT1 | Island | Body | chr6:26598149-26598303 | 154 | | | | | | | | | | | | | | | | | | | |
| LOC100270746;C6orf41 | N_Shore;Island;S_Shore | Body;TSS200 | chr6:26987575-26988289 | 714 | | | | | | | | 0 | 4 | | | 1 | | | yes | | | | |
| HIST1H2BK;HIST1H4I | N_Shore;Island;S_Shore | 3'UTR;TSS200;1stExon;TSS1500 | chr6:27106564-27107756 | 1192 | | 2 | 38 | 1 | 4 | | | | | | | 1 | | | yes | yes | | | |
| PRSS16 | N_Shelf | TSS200 | chr6:27215328-27215675 | 347 | | | 3 | | | | | | 1 | | | 2 | | | yes | | | | |
| POM121L2 | Island | TSS200 | chr6:27280139-27280373 | 234 | | 1 | | | | | | | | | | | | | yes | | | | |
| HIST1H2AI | N_Shore | TSS1500 | chr6:27774865-27775042 | 177 | | 1 | | | | | | | | | | | | yes | | | | | |
| HIST1H4K | Island | TSS200 | chr6:27799308-27799385 | 77 | | | 4 | | | | | | | | | | | | | | | | |
| HIST1H2AL | N_Shore | TSS1500 | chr6:27831894-27832098 | 204 | | | 1 | | | | | | | | | | | yes | | | | | |

| | | | | | | | | | | | | | | | | | | | | | | | |
|---|---|---|---|---|---|---|---|---|---|---|---|---|---|---|---|---|---|---|---|---|---|---|---|
| HIST1H4L;HIST1H3I | N_Shore;S_Shore | 1stExon;TSS1500;3'UTR;TSS200 | chr6:27840957-27841865 | 908 | | | 2 | | | | | | | | | | | yes | | yes | yes | | |
| HIST1H3J | Island | 1stExon | chr6:27858279-27858545 | 266 | | | | | | | | 0 | | | | | | yes | | | | | |
| OR2B2 | NA | TSS1500 | chr6:27880822-27881242 | 420 | | | | | | | | | 1 | | | | | yes | | | yes | | |
| ZSCAN12L1 | NA | TSS1500;TSS200 | chr6:28058715-28059208 | 493 | | 1 | 14 | | | | | 0 | 11 | 2 | | 13 | | yes | | yes | | | |
| ZSCAN16 | NA | TSS1500;TSS200 | chr6:28092048-28093420 | 1372 | | | 3 | | | | | 0 | | | | | | yes | yes | | | yes | |
| ZNF389 | NA | TSS200 | chr6:28129359-28129616 | 257 | | | | | | | | 0 | 2 | | | | | | | | | | |
| NKAPL | Island;N_Shore | TSS200;5'UTR;1stExon;TSS1500 | chr6:28226885-28227482 | 597 | | 1 | 6 | | | | | 0 | 7 | | | 3 | | yes | | | | | yes |
| ZNF323;ZKSCAN3 | NA | 5'UTR;TSS200 | chr6:28322086-28322103 | 17 | | | | | | | | | | | | | | | | yes | | | |
| GPX6 | NA | Body;1stExon;5'UTR | chr6:28477957-28483691 | 5734 | | | | | | | | | | | | | | | | | | | |
| GPX5 | NA | TSS1500;TSS200;Body | chr6:28493113-28494863 | 1750 | | | 21 | 1 | 1 | | | | 4 | | | | | yes | yes | | | yes | |
| GPX5 | NA | Body | chr6:28497278-28497370 | 92 | | | 6 | | | | | | | | | | | | yes | | yes | | yes |
| GPX5 | NA | Body | chr6:28498759-28499825 | 1066 | | | 8 | | | | | | 2 | | | | | yes | yes | | yes | | |
| GPX5 | NA | Body;3'UTR | chr6:28501867-28501916 | 49 | | | 1 | | | | | | | | | | | | | | | | |
| SCAND3 | NA | Body | chr6:28540442-28541462 | 1020 | | | | | | | | | 1 | | | | | | | | yes | | |
| SCAND3 | NA | Body | chr6:28543237-28543514 | 277 | | | | | | | | | 1 | | | | | | | | | | |
| SCAND3 | NA | Body | chr6:28549832-28550423 | 591 | | | | | | | | | | | | | | yes | yes | yes | yes | | |
| SCAND3 | N_Shore;S_Shore | 1stExon;5'UTR;TSS1500 | chr6:28554262-28556338 | 2076 | | | 17 | | | | | 0 | 2 | | | 1 | | yes | | | yes | | |
| TRIM27 | NA | Body | chr6:28874479-28876377 | 1898 | | | 5 | | | | | 0 | 6 | 2 | 2 | 7 | | yes | yes | | | | yes |
| TRIM27 | NA | Body | chr6:28881365-28881574 | 209 | | | | | | | | | 1 | | | | | yes | yes | | | yes | |
| TRIM27 | N_Shelf | Body | chr6:28887141-28887284 | 143 | | 1 | | | | | | | | | | | | yes | yes | | yes | | |
| TRIM27 | S_Shore | TSS1500 | chr6:28892660-28892999 | 339 | | | | | | | | | 3 | | | | | yes | | yes | yes | | |
| ZNF311 | NA | Body | chr6:28963370-28963898 | 528 | | | 8 | | | | | | 1 | | | | | | | | | | |
| ZNF311 | NA | Body | chr6:28967766-28968397 | 631 | | | | | | | | | 1 | | | | | yes | | | | yes | |
| ZNF311 | NA | 5'UTR;1stExon;TSS1500 | chr6:28972065-28973524 | 1459 | | 1 | 7 | | | | | 0 | 3 | | | 1 | | yes | yes | | | | |
| OR2B3 | NA | 1stExon | chr6:29054325-29054566 | 241 | | | | | | | | | | | | | | yes | | | | | |
| OR2J3 | NA | 1stExon | chr6:29080040-29080437 | 397 | | | | | | | | | | | | | | | yes | | | | |
| OR2J2 | NA | 5'UTR;1stExon | chr6:29141346-29141678 | 332 | | | | | | | | | | | | | | | | | | | |
| OR14J1 | NA | TSS1500;1stExon | chr6:29274096-29275337 | 1241 | | | 21 | | | | | | 1 | | | | | yes | | | | | yes |
| OR5V1 | NA | 1stExon | chr6:29323191-29323978 | 787 | | | | | | | | | | | | | | | | | | | |
| OR12D3 | NA | 1stExon;3'UTR | chr6:29341892-29342706 | 814 | | 1 | 10 | | | | | | 2 | | | | | | yes | | | | |
| OR12D2 | NA | 1stExon;TSS1500;5'UTR | chr6:29364174-29365287 | 1113 | | | 9 | | | | | | | | | | | yes | | | | | |
| OR11A1 | NA | 1stExon;3'UTR;5'UTR | chr6:29394438-29396182 | 1744 | | | 16 | | | | | | | | | | | yes | yes | | | yes | yes |
| OR10C1 | NA | TSS200;1stExon | chr6:29407689-29408587 | 898 | | | 27 | | | | | | | | | | | yes | | | | | |
| OR2H1 | NA | TSS1500;TSS200 | chr6:29425885-29426320 | 435 | | | 2 | | | | | | 1 | | | | | yes | yes | | | | |
| OR2H1 | NA | Body;3'UTR | chr6:29430096-29431410 | 1314 | | | | | | | | | 2 | | 1 | | | | | yes | | | yes |
| MAS1L | NA | 1stExon | chr6:29454623-29455532 | 909 | | 1 | 25 | | | | | 0 | 2 | | | | | yes | | | | | |
| MAS1L | NA | TSS1500 | chr6:29456563-29457282 | 719 | | | | | | | | | 1 | | | | | yes | | | | yes | yes |
| UBD | S_Shore;S_Shelf | 3'UTR;Body | chr6:29523592-29524117 | 525 | | | 11 | | | | | | | | | | | yes | | yes | yes | | |
| UBD | NA | TSS200 | chr6:29527870-29528119 | 249 | | 1 | 2 | 1 | | | | | | | | | | | yes | | yes | | |
| SNORD32B | NA | TSS1500 | chr6:29548535-29550134 | 1599 | | | 6 | | | | | | 1 | | | | | | | yes | yes | | |
| OR2H2 | NA | TSS200;1stExon;TSS1500 | chr6:29555314-29556415 | 1101 | | 1 | 8 | | | | | | 2 | | | | | | yes | | | | |
| GABBR1 | NA | 3'UTR | chr6:29570507-29570739 | 232 | | | | | | | | | | | | | | | | | | | |
| GABBR1 | NA | Body | chr6:29572219-29572333 | 114 | | | | | | | | | 1 | | | | | | | yes | | | |
| GABBR1 | NA | Body | chr6:29577916-29579306 | 1390 | | | | | | | | | 3 | | | | | yes | yes | | | | yes |
| GABBR1 | NA | Body | chr6:29581054-29581228 | 174 | | | | | | | | | | | | | | yes | | | | | |
| GABBR1 | NA | Body | chr6:29585617-29585658 | 41 | | | | | | | | | 1 | | | | | yes | yes | | | yes | |
| GABBR1 | NA | Body | chr6:29588988-29590143 | 1155 | | | 13 | 2 | | | | | | | | | | | | | | yes | |
| GABBR1 | N_Shore | Body | chr6:29593382-29593490 | 108 | | | 1 | | | | | | | | | | | | | | yes | | |
| GABBR1 | N_Shore;Island | Body;1stExon | chr6:29594830-29595349 | 519 | | 1 | 29 | | | | | 0 | 2 | 1 | | 5 | | | | | | | yes |
| GABBR1 | S_Shore | TSS1500;Body | chr6:29596901-29597113 | 212 | | | | | | | | | | | | | | | | | yes | | |
| GABBR1 | N_Shore;Island;S_Shore | Body;5'UTR;TSS1500 | chr6:29599160-29602390 | 3230 | | | | 1 | | | | 0 | 9 | | | 3 | | yes | | | yes | yes | yes |
| MOG | NA | TSS1500 | chr6:29623721-29623992 | 271 | | | | | | | | | | | | | | yes | | | yes | | |
| MOG | NA | Body | chr6:29626990-29627395 | 405 | 1 | 3 | 2 | | | | | | 1 | | | | | yes | yes | | | | |
| MOG | NA | Body | chr6:29629187-29629988 | 801 | | | 4 | | | | | 0 | 3 | | | 3 | | yes | yes | | | | |
| MOG | NA | Body | chr6:29631227-29631447 | 220 | | | | | 1 | | | | | | | | | yes | yes | | | | |
| MOG | NA | 3'UTR;Body | chr6:29633622-29636669 | 3047 | | | 9 | | | | | | 4 | 1 | | 1 | | yes | yes | | yes | yes | |
| MOG | NA | 3'UTR;Body | chr6:29638112-29638575 | 463 | | | | | | | | | | | | | | yes | yes | | yes | | |
| ZFP57 | NA | TSS1500 | chr6:29645937-29649092 | 3155 | | | | | | | 1 | 0 | 2 | 1 | | 4 | | yes | | | yes | yes | yes |
| HLA-F | N_Shore;Island;S_Shore | TSS1500;TSS200;1stExon;Body | chr6:29690766-29693309 | 2543 | | 4 | 44 | 1 | 3 | 1 | | 0 | 6 | 1 | 1 | | | yes | | | | yes | yes |
| LOC285830 | S_Shelf | Body | chr6:29695164-29695590 | 426 | | | | | | | | | | | | | | | yes | | yes | | |
| LOC285830 | NA | Body | chr6:29697897-29698518 | 621 | | | | | | | | | 2 | | | | | | yes | yes | | yes | |
| LOC285830 | NA | Body | chr6:29705815-29706377 | 562 | | | 3 | | | | | | | | | | | | yes | yes | | | |
| LOC285830 | N_Shelf | Body | chr6:29713615-29714059 | 444 | | | | | | | | | | | | | | | yes | | | | |
| LOC285830 | Island;S_Shore | Body;TSS200;TSS1500 | chr6:29716536-29718093 | 1557 | | 3 | 5 | | | | | | 1 | | | | | | yes | | | yes | |
| HCG4 | N_Shore;Island;S_Shore | Body;TSS200 | chr6:29759718-29761075 | 1357 | | 3 | 22 | | | | | 0 | 3 | | | 1 | | | | yes | | | yes |
| HLA-G | N_Shore;Island;S_Shore | 5'UTR;Body | chr6:29795411-29798387 | 2976 | | 1 | 30 | | | | | 0 | 4 | | | 5 | | | | | | | yes |
| HLA-H | N_Shore;Island;S_Shore | TSS1500;TSS200;Body | chr6:29854631-29857569 | 2938 | | 4 | 12 | 1 | | | 4 | 0 | 8 | 8 | 1 | 15 | 1 | | yes | | | | yes |
| HCG2P7 | NA | Body | chr6:29868207-29868295 | 88 | | | | | | | | | 1 | | | | | | yes | yes | | | |
| HCG2P7 | NA | Body | chr6:29869838-29870062 | 224 | | | 1 | | | | 1 | | 1 | 1 | | 2 | 1 | yes | yes | | | | |
| HCG4P6 | Island;N_Shore | TSS1500;Body | chr6:29893168-29895260 | 2092 | | 3 | 48 | 1 | 1 | | 7 | 0 | 17 | 14 | 5 | 80 | 4 | | yes | | | | yes |
| HLA-A | Island;S_Shore | Body | chr6:29910778-29911836 | 1058 | 5 | 10 | 27 | 2 | | 6 | 4 | 0 | 6 | 1 | 1 | 46 | 1 | | | | | | |
| HCG9 | N_Shore;Island;S_Shore | TSS200;Body | chr6:29942706-29945948 | 3242 | 1 | 2 | 63 | 4 | | | | | 4 | | | | | yes | yes | | | | yes |
| HLA-J;NCRNA00171 | Island;S_Shore;N_Shore | Body | chr6:29974022-29975708 | 1686 | | | 5 | | | | | 0 | 4 | 2 | | 1 | 1 | | | | | | yes |
| NCRNA00171 | NA | Body | chr6:29981250-29981695 | 445 | | | | 1 | | | | | | | | | | yes | | | yes | | |
| NCRNA00171 | NA | Body | chr6:29987727-29988066 | 339 | | | | | | | | | 1 | | | | | | | yes | | | |
| NCRNA00171 | NA | Body | chr6:30001703-30002859 | 1156 | | | | | | | | | 1 | | | | | | yes | yes | | | |
| NCRNA00171 | NA | Body | chr6:30009745-30010145 | 400 | | | 1 | | | | | | | | | | | yes | yes | | yes | | |
| ZNRD1;NCRNA00171 | N_Shore | TSS1500;Body | chr6:30027963-30028257 | 294 | | | | | | | | | | | | | | | yes | | | | |
| ZNRD1;NCRNA00171 | S_Shore | Body;TSS1500 | chr6:30029929-30030069 | 140 | | 1 | | | | | | | | | | | | yes | | | | | |
| RNF39 | N_Shore;Island;S_Shore | 3'UTR;Body | chr6:30038712-30040291 | 1579 | 15 | 5 | 98 | 11 | | 7 | 7 | 0 | 37 | 36 | 11 | 72 | 10 | | | yes | | | |
| RNF39 | N_Shore;Island;S_Shore | Body;1stExon;TSS200;TSS1500 | chr6:30041727-30044259 | 2532 | | | 21 | | | | | 0 | 6 | | | 1 | | yes | | | | | yes |
| TRIM31 | Island;S_Shore | Body | chr6:30071373-30071612 | 239 | | | 11 | | | | | 0 | 4 | | | 1 | | | | | | | |
| TRIM31 | S_Shelf | Body | chr6:30074162-30074219 | 57 | | | | | | | | | | | | | | yes | | | | | yes |
| TRIM31 | NA | Body | chr6:30076659-30076980 | 321 | | | 5 | | | | | 0 | | | | | | yes | | | yes | | yes |
| TRIM31 | NA | Body | chr6:30078290-30080782 | 2492 | 1 | 2 | 48 | 1 | 1 | | | 0 | 15 | 3 | 1 | 32 | 2 | yes | | | yes | | |
| TRIM40 | NA | TSS1500;TSS200;1stExon;5'UTR | chr6:30103459-30104982 | 1523 | | 1 | 5 | | | | | | 1 | 1 | | | | yes | yes | yes | yes | | |
| TRIM40 | NA | Body;3'UTR | chr6:30112769-30116378 | 3609 | | 1 | 26 | | | | | | 1 | | | | | yes | yes | yes | | | |
| TRIM10 | NA | Body | chr6:30121774-30124792 | 3018 | | 10 | 54 | 4 | | | | 0 | 9 | 2 | 1 | 1 | | yes | yes | | yes | yes | |
| TRIM10 | NA | 1stExon;Body | chr6:30127152-30128686 | 1534 | | | | | | | | | 2 | | | | | yes | yes | | | yes | |
| TRIM10;TRIM15 | NA | TSS1500;TSS200;5'UTR;1stExon;B | chr6:30130137-30132471 | 2334 | | 2 | 20 | 3 | 2 | | | | 1 | | | | | yes | yes | | yes | yes | |
| TRIM15 | NA | Body | chr6:30134956-30135671 | 715 | | | | | | | | | | | | | | yes | yes | | | | |
| TRIM15 | N_Shore | Body | chr6:30138178-30138225 | 47 | | | | | | | | | | | | | | yes | | | | | |
| TRIM15 | N_Shore;Island;S_Shore | Body;3'UTR | chr6:30139421-30140417 | 996 | | | 15 | 1 | 5 | | | 0 | 2 | | | | | yes | | | yes | | |
| TRIM26 | NA | 3'UTR | chr6:30152322-30152802 | 480 | | | | | | | | | | | | | | | yes | | | | |
| TRIM26 | NA | Body | chr6:30159991-30160045 | 54 | | | | | | | | | | | | | | yes | yes | | | | |
| TRIM26 | NA | Body | chr6:30162942-30163370 | 428 | | | 7 | | | | | | 1 | | | | | yes | yes | | yes | | |
| TRIM26 | NA | Body;5'UTR | chr6:30166423-30167836 | 1413 | | | | | | | | | 2 | | | | | yes | | | yes | | |
| TRIM26 | NA | 5'UTR | chr6:30168896-30169120 | 224 | | | | 1 | | | | | 1 | | | | | yes | | | | | |
| TRIM26 | NA | 5'UTR | chr6:30173936-30176076 | 2140 | | | 36 | 1 | | | | | 8 | 1 | | | | yes | yes | | yes | | |

| | | | | | | | | | | | | | | | | | | | | | | | |
|---|---|---|---|---|---|---|---|---|---|---|---|---|---|---|---|---|---|---|---|---|---|---|---|
| TRIM26 | N_Shelf | 5'UTR | chr6:30177177-30177295 | 118 | | | | | | | | | 1 | | | | | yes | | yes | yes | yes | |
| TRIM26 | S_Shore | TSS1500 | chr6:30182349-30182563 | 214 | | | 1 | | | | | | | | | | | yes | | | | | |
| HLA-L | Island;N_Shore | Body;TSS1500 | chr6:30227001-30228431 | 1430 | | | 9 | 1 | | | | 0 | 2 | | | 2 | | | | | | | yes |
| HLA-L | S_Shelf | Body | chr6:30231639-30232015 | 376 | | | 8 | | | | | | | | | | | | yes | | | yes | |
| HCG18 | NA | Body | chr6:30285906-30286162 | 256 | | | | | | | | 0 | | | | | | yes | | yes | | | |
| HCG18 | N_Shelf;N_Shore | Body | chr6:30290408-30292481 | 2073 | | | 13 | | | | | | 6 | | | | | yes | yes | yes | yes | | yes |
| HCG18;TRIM39 | S_Shore;S_Shelf | TSS1500;5'UTR;1stExon;Body | chr6:30295328-30297627 | 2299 | | | 2 | | | | | 0 | 13 | 2 | 1 | 3 | | | | | yes | | |
| RPP21 | N_Shore | TSS1500 | chr6:30312580-30312818 | 238 | | | | | | | | | 1 | | | | | | | | | | |
| HLA-E | N_Shore | TSS1500 | chr6:30455846-30455964 | 118 | | | 5 | | | | | | | | | | | yes | yes | yes | yes | | |
| HLA-E | Island;S_Shore;S_Shelf | Body;3'UTR | chr6:30458078-30460600 | 2522 | | | 17 | 2 | 6 | 1 | | 0 | 6 | 1 | | | | yes | | | | | |
| GNL1 | NA | 3'UTR | chr6:30510021-30510662 | 641 | | | 7 | | 1 | | | | | | | | | | yes | | yes | | |
| GNL1 | N_Shore | Body | chr6:30522620-30523072 | 452 | | | | | | | | | | | | | | | | | | | |
| ABCF1 | S_Shore | Body | chr6:30540984-30541653 | 669 | | | | | | | | | 1 | | | | | yes | yes | | | | |
| ABCF1 | NA | Body | chr6:30558319-30558470 | 151 | | | | | | | | | | | | | | | | | | | |
| PPP1R10 | NA | 3'UTR | chr6:30568532-30568668 | 136 | | | | | | | | | 1 | | | | | yes | | | | | |
| PPP1R10 | NA | Body | chr6:30570103-30570163 | 60 | | | | | | | | | | | | | | | | | | yes | |
| PPP1R10 | NA | 5'UTR | chr6:30579560-30580581 | 1021 | | | | | | | | | | | | | | yes | | | | | |
| MRPS18B;PPP1R10 | S_Shore | Body;TSS1500 | chr6:30586293-30586468 | 175 | | | 2 | | | | | | | | | | | yes | | | | | |
| MRPS18B | S_Shore | Body | chr6:30587655-30587737 | 82 | | | | | | | | | | | | | | yes | | | | | |
| C6orf134 | N_Shelf | Body | chr6:30610781-30611056 | 275 | | | | | | | | | | | | | | yes | | | | | yes |
| C6orf136 | N_Shore | TSS1500 | chr6:30614168-30614422 | 254 | | | 6 | | | | | | | | | 1 | | | | | | yes | yes |
| C6orf136 | S_Shore | Body | chr6:30615944-30616140 | 196 | | | | 1 | 1 | | | | 1 | | | | | yes | | | | | |
| C6orf136 | S_Shelf | Body | chr6:30619047-30620732 | 1685 | | | | | 1 | | | | 4 | | | | | yes | yes | yes | yes | yes | |
| DHX16 | NA | Body | chr6:30624168-30624636 | 468 | | | | | | | | 0 | 4 | 2 | 1 | 2 | | | | | | | |
| KIAA1949 | S_Shelf | 3'UTR | chr6:30644693-30644832 | 139 | | | | | | | | | 1 | | | | | | | | | | |
| KIAA1949 | NA;N_Shelf;N_Shore | Body;1stExon | chr6:30646756-30653799 | 7043 | | | 44 | 3 | 5 | 1 | | | 1 | | | | | yes | yes | | | yes | yes |
| KIAA1949;NRM | S_Shore;S_Shelf | TSS1500;3'UTR;Body | chr6:30655896-30658014 | 2118 | | | | | | | | | | | | | | | | | | | |
| MDC1 | N_Shore | 5'UTR | chr6:30684278-30684736 | 458 | | 1 | 2 | | | | | | 3 | | | | | yes | yes | | | | |
| MDC1;TUBB | N_Shore | TSS1500 | chr6:30686511-30687221 | 710 | | | 3 | | | | | | 3 | | | | | yes | yes | | | | |
| TUBB | S_Shore;S_Shelf | Body | chr6:30691305-30692066 | 761 | | | 6 | | | | | | 1 | | | | | | yes | | | | |
| FLOT1 | NA | 3'UTR;Body | chr6:30695686-30696029 | 343 | | | 5 | | | | | | 1 | | | | | yes | | | | | |
| FLOT1 | NA | nan;Body | chr6:30698584-30699383 | 799 | | | 15 | | | | | | 3 | 1 | 1 | | | yes | | | | | |
| FLOT1 | N_Shelf | Body | chr6:30707619-30708009 | 390 | | | | 2 | | | | | | | | | | yes | | | | | |
| IER3;FLOT1 | Island | 3'UTR;TSS1500 | chr6:30711022-30711680 | 658 | | | 2 | | 1 | 3 | | | 1 | | | | | | | | | | |
| DDR1 | N_Shore | TSS1500 | chr6:30850405-30851753 | 1348 | 3 | 2 | 18 | | | 1 | | | 7 | 2 | 1 | | | | | | | yes | yes |
| DDR1 | S_Shore | 5'UTR | chr6:30852874-30854551 | 1677 | | | 14 | | 1 | | | 0 | 7 | | 1 | 4 | | | yes | | yes | | yes |
| DDR1 | S_Shelf | TSS1500;5'UTR;TSS200 | chr6:30855729-30856491 | 762 | | 1 | 20 | | | | | | | 1 | | | | yes | yes | | | | |
| DDR1 | NA | Body | chr6:30860014-30860905 | 891 | | | | | | | | | 3 | | | | | | | yes | yes | | |
| DDR1 | NA | Body | chr6:30865979-30866025 | 46 | | | | | | | | | | | | | | | | | | | |
| GTF2H4 | N_Shore | TSS1500 | chr6:30875586-30875592 | 6 | | | | | | | | | 1 | | | 1 | | yes | | | | | yes |
| GTF2H4 | S_Shore | Body | chr6:30876844-30876992 | 148 | | | | | | | | | | | | | | | | | yes | | |
| GTF2H4 | N_Shelf | Body | chr6:30878437-30878499 | 62 | | | | | | | | | | | | | | yes | | | | | |
| GTF2H4 | N_Shore | Body | chr6:30879800-30879889 | 89 | | | | | | | | | | | | | | | | | | | |
| VARS2;GTF2H4 | N_Shore;Island;S_Shore | TSS1500;Body;5'UTR | chr6:30881112-30884124 | 3012 | | | 21 | 2 | | | | 0 | 10 | 1 | | 2 | | yes | yes | | | yes | yes |
| VARS2 | S_Shelf | Body | chr6:30885205-30885567 | 362 | | | | | | | | | 1 | | | | | yes | | | | | |
| SFTA2 | NA | 3'UTR;Body;TSS200 | chr6:30899162-30900560 | 1398 | | | 4 | | | | | 0 | 1 | | | | | yes | | | | | |
| DPCR1 | NA | TSS200 | chr6:30908579-30908706 | 127 | | | | | | | | | 1 | | | 1 | | | | | yes | | |
| DPCR1 | N_Shelf;N_Shore | Body;3'UTR | chr6:30919516-30921757 | 2241 | | 1 | 7 | | | | | | 2 | | | | | | yes | | | | |
| MUC21 | NA | TSS1500 | chr6:30950666-30951394 | 728 | | 2 | | | | | | | 2 | | | | | | | yes | | | |
| MUC21 | NA | Body;3'UTR | chr6:30954150-30956796 | 2646 | | 1 | 32 | | | | | 0 | 5 | | | 1 | | yes | | | | | |
| HCG22 | NA | TSS1500;TSS200 | chr6:31021422-31021957 | 535 | | | 2 | | 1 | | | | | | | | | | | yes | | | |
| HCG22 | NA | Body | chr6:31023257-31023770 | 513 | | 1 | | | | | | | 2 | | | | | yes | | | | yes | |
| HCG22 | NA | Body | chr6:31025933-31026287 | 354 | | | 29 | | | | | | 1 | | | | | yes | | yes | yes | | |
| C6orf15 | NA | Body | chr6:31079306-31079617 | 311 | | 1 | | | | | | | | | | | | | | | | | |
| PSORS1C1;C6orf15 | NA | TSS1500;TSS200;5'UTR | chr6:31081323-31083322 | 1999 | 2 | | 11 | | | | | | 1 | | | | | | yes | yes | | yes | |
| CDSN;PSORS1C1 | NA | TSS200;5'UTR | chr6:31088343-31088434 | 91 | | | | | | | | 0 | 1 | 1 | | 2 | | yes | | | | | |
| PSORS1C1 | NA | 5'UTR | chr6:31090929-31091383 | 454 | | | | | | | | | 1 | | | | | yes | | | | | |
| PSORS1C1 | NA | 5'UTR | chr6:31093846-31094185 | 339 | | | | | | | | | 1 | | | | | | | | | | |
| PSORS1C1 | NA | 5'UTR | chr6:31095800-31097078 | 1278 | | | 8 | | | | 1 | | | | | | | yes | yes | | | yes | |
| PSORS1C1;PSORS1C2 | NA | Body;1stExon;5'UTR;TSS200;3'UTI | chr6:31105711-31107663 | 1952 | | 1 | 17 | | | | | | 2 | | | | | yes | yes | | | yes | |
| CCHCR1 | NA | 3'UTR;Body | chr6:31110228-31111270 | 1042 | | | 10 | | | | | | | | | | | yes | yes | | | | |
| CCHCR1;TCF19 | N_Shore | Body;TSS1500;5'UTR | chr6:31124978-31125445 | 467 | | | | | | | | | | | | | | | | | | | |
| CCHCR1;TCF19 | S_Shore | TSS1500;5'UTR;Body | chr6:31126864-31127863 | 999 | | | 5 | 2 | 3 | | | 0 | 2 | | | | | yes | | | | | |
| TCF19 | S_Shelf | Body | chr6:31130258-31130548 | 290 | 1 | 1 | | | | | | | | | | | | | | | | | |
| POU5F1 | NA | TSS1500 | chr6:31138925-31139796 | 871 | | | 3 | | | | | | 1 | | | | | yes | | | | | |
| PSORS1C3 | NA | Body | chr6:31143710-31144068 | 358 | | | 8 | | | | | | | | | | | yes | | | | | |
| PSORS1C3 | N_Shelf;N_Shore | Body;TSS1500 | chr6:31145541-31148748 | 3207 | | 1 | 8 | 2 | | | | | 6 | | | 1 | | yes | yes | | | yes | |
| HCG27 | N_Shore | TSS1500 | chr6:31164743-31164994 | 251 | | | 10 | | | 1 | | 0 | 8 | 2 | | 9 | 1 | yes | | yes | | | |
| HCG27 | S_Shore | Body | chr6:31166502-31166799 | 297 | | | | 1 | | | | 0 | 5 | | | 1 | | yes | | | | yes | |
| HCG27 | S_Shelf | Body | chr6:31168558-31168571 | 13 | | | | | | | | | 1 | | | | | yes | yes | yes | | yes | |
| HLA-C | N_Shore | Body | chr6:31237013-31239005 | 1992 | 19 | 47 | 110 | 2 | | | 3 | 0 | 5 | 3 | 2 | 147 | 1 | | | | | | |
| HLA-C | S_Shore | TSS1500 | chr6:31240651-31241294 | 643 | | | 2 | | | | 2 | 0 | 5 | 5 | | 18 | | | yes | | | | |
| HLA-B | N_Shore;Island;S_Shore | Body;5'UTR;1stExon;TSS1500 | chr6:31322298-31325817 | 3519 | | | 22 | | | 1 | 2 | 0 | 5 | 2 | | 4 | | | yes | | | | |
| MICA | S_Shore | Body | chr6:31373660-31373932 | 272 | | | | | | | | | | | | | | | yes | | | | |
| MICA | NA | Body | chr6:31379828-31380915 | 1087 | | | | | | | | | | | | | | | | | | | |
| HCP5 | NA | TSS1500;TSS200;Body;3'UTR | chr6:31430641-31431969 | 1328 | | | 7 | 2 | | | | 0 | 2 | | | | | | | yes | | | |
| HCG26 | NA | Body | chr6:31439037-31439497 | 460 | | | | | | | | | | | | | | | yes | yes | yes | | |
| MICB | N_Shore;Island;S_Shore | TSS1500;Body | chr6:31464826-31467192 | 2366 | | | 10 | | | | | 0 | 1 | | | | | yes | | | | | |
| MICB | S_Shelf;chr6:31465460-314661 | Body;S_Shelf | chr6:31468473-31468957 | 484 | | | 4 | | | | | | | | 1 | | | | yes | | | | |
| MICB | NA | Body | chr6:31474185-31476231 | 2046 | | | 3 | | | | | | | | | | | | | | | | |
| MICB | NA | 3'UTR | chr6:31478432-31478830 | 398 | | | | | | | | | 1 | 1 | | | | yes | | yes | | yes | yes |
| MCCD1 | NA | TSS200;1stExon;5'UTR;Body | chr6:31496644-31497756 | 1112 | | | 8 | | | | | | 1 | | | | | | | | | | yes |
| BAT1;SNORD117 | NA | Body;TSS200 | chr6:31504369-31504388 | 19 | | | | | | | | | | | | | | yes | | | | | |
| BAT1 | N_Shore;N_Shelf | Body | chr6:31506568-31508665 | 2097 | 1 | | 16 | 1 | | | | | 2 | | | | | yes | yes | | | | |
| BAT1 | S_Shore | TSS1500 | chr6:31510729-31510903 | 174 | | | 6 | | | | | | | | | | | yes | yes | | | | |
| ATP6V1G2;NFKBIL1 | S_Shelf | 3'UTR;Body;TSS1500 | chr6:31512618-31513680 | 1062 | | | 9 | | | | | | 2 | | | | | yes | | | | | |
| NFKBIL1 | NA | Body | chr6:31519182-31519410 | 228 | | | | | | | | | | | | | | yes | | | | | |
| LTA | NA | TSS1500;TSS200;1stExon;5'UTR | chr6:31538984-31541349 | 2365 | | | | | | | | | | | | | | | | | | | |
| TNF | NA | TSS200;1stExon | chr6:31543169-31543686 | 517 | | 1 | 14 | | 2 | | | | 1 | | 1 | 1 | | | yes | | | yes | yes |
| TNF;LTB | N_Shelf;Island;S_Shore | Body;3'UTR;TSS1500 | chr6:31544694-31550842 | 6148 | | | | | | | | | | | | | | | | | | | |
| LST1 | NA | Body;TSS1500;5'UTR;1stExon;TSS2 | chr6:31554199-31555016 | 817 | | | 10 | | 1 | | | | 1 | | | | | yes | | | | | |
| NCR3 | NA | 5'UTR;1stExon | chr6:31560624-31560871 | 247 | | | 4 | | | | | | | | | | | yes | | | | yes | |
| AIF1 | NA;N_Shelf | TSS1500;5'UTR;Body;1stExon | chr6:31583231-31584223 | 992 | | | | | 1 | | | 0 | 5 | 1 | 1 | 8 | | | | | | | |
| SNORA38;BAT2 | S_Shore;S_Shelf | TSS1500;Body;5'UTR | chr6:31589196-31591597 | 2401 | 1 | | 15 | | 1 | | | 0 | 3 | | | | | yes | | | | yes | |
| BAT2 | S_Shelf | Body | chr6:31592983-31593323 | 340 | | | | | | | | | | | | | | yes | | | | | |
| BAT2 | N_Shore | Body | chr6:31598379-31598424 | 45 | | | | | | | | | | | | | | yes | | | | | |
| BAT2 | Island;S_Shore | Body | chr6:31599520-31600560 | 1040 | | | 4 | | | | | | | | | | | | | | | | |
| BAT2 | S_Shelf | Body | chr6:31602130-31603011 | 881 | | | 6 | 1 | | | | | | | | | | | | | | | |
| BAT3 | NA | Body | chr6:31607340-31607674 | 334 | | | 19 | | | | | 0 | 1 | | | | | | | | | | |

| | | | | | | | | | | | | | | | | | | | | | | | |
|---|---|---|---|---|---|---|---|---|---|---|---|---|---|---|---|---|---|---|---|---|---|---|---|
| BAT3 | N_Shelf | Body | chr6:31615937-31616088 | 151 | | | | | | | | | 1 | | | | | yes | yes | | | | |
| BAT3 | N_Shelf | Body | chr6:31617472-31617530 | 58 | | | | | | | | | 1 | | | | | yes | | | | | |
| BAT3 | Island | 5'UTR;1stExon | chr6:31620040-31620059 | 19 | | | | | | | | | 1 | | | | | | | | | yes | |
| BAT3;APOM | S_Shore;S_Shelf | TSS1500 | chr6:31621210-31623302 | 2092 | | | | | | | | | | | | | | | | | | | |
| APOM | N_Shelf | Body | chr6:31625404-31625701 | 297 | | | | | | | | | | | | | | yes | | | | | |
| C6orf47 | N_Shore | 1stExon | chr6:31626986-31627714 | 728 | | | 23 | 1 | | | | 0 | 3 | | | | | | | | | | |
| C6orf47;BAT4 | S_Shore | TSS1500;3'UTR | chr6:31629900-31630051 | 151 | | | | | | | | | 1 | | | | | yes | yes | | | | |
| BAT4 | N_Shore | Body | chr6:31631638-31632248 | 610 | | 1 | 4 | | | | | 0 | 2 | | | 3 | | yes | | | | | |
| CSNK2B | S_Shore | Body | chr6:31634821-31634875 | 54 | | | | | | | | | | | | | | yes | | | | | |
| CSNK2B;LY6G5B | S_Shelf | Body;TSS1500 | chr6:31636239-31637619 | 1380 | | | | | | | | | 2 | 1 | | | | yes | | | | | |
| LY6G5C | N_Shelf | Body | chr6:31645540-31646976 | 1436 | | | | | | | | | 1 | | | | | yes | yes | | | | |
| LY6G5C | N_Shore | TSS1500 | chr6:31649432-31651291 | 1859 | | | 11 | 1 | | | | 0 | 4 | | | 1 | | yes | | | | yes | |
| LY6G6F | S_Shelf | TSS1500 | chr6:31673800-31673939 | 139 | | | 2 | | | | | | | | | | | yes | yes | | | | yes |
| LY6G6E;LY6G6D | NA | TSS1500;TSS200 | chr6:31683051-31683131 | 80 | | | | | | | | | 1 | | | | | | | | yes | | |
| LY6G6D | NA | Body;nan | chr6:31685392-31686784 | 1392 | | | 8 | | | | | | 2 | | 1 | | | yes | yes | | | | |
| LY6G6C | N_Shelf | Body | chr6:31688080-31688229 | 149 | | | | | | | | | 1 | | | | | yes | | | | | |
| LY6G6C;C6orf25 | N_Shore;Island;S_Shore | TSS200;TSS1500;1stExon;Body | chr6:31689610-31692375 | 2765 | | | 20 | | 1 | | | 0 | 14 | 5 | | 4 | | yes | | | | | |
| DDAH2 | N_Shore;Island | Body | chr6:31695027-31696729 | 1702 | | 1 | 42 | 1 | 1 | 4 | | 0 | 8 | 1 | 1 | 1 | | | yes | | | | |
| DDAH2;CLIC1 | Island;S_Shore | TSS200;3'UTR;TSS1500 | chr6:31698218-31698702 | 484 | | | 2 | 1 | 1 | | | | 1 | | | | | yes | | | | | |
| CLIC1 | S_Shelf | Body | chr6:31701110-31701288 | 178 | | | 1 | | | | | | 1 | | | | | yes | | | | | |
| CLIC1 | NA | Body | chr6:31702632-31703188 | 556 | | | | 1 | 1 | | | | | | | | | | | | | yes | |
| CLIC1 | N_Shelf | TSS1500 | chr6:31704705-31704844 | 139 | | | 7 | | | | | | | | | | | yes | yes | | | | |
| MSH5 | N_Shore | TSS1500 | chr6:31707019-31707202 | 183 | | | 2 | | | | | | 1 | | | | | yes | yes | | | yes | |
| C6orf27 | NA | Body | chr6:31733619-31734580 | 961 | | | 20 | | | | | 0 | 11 | | | 4 | | yes | yes | | | yes | |
| C6orf27 | NA | Body | chr6:31740787-31740978 | 191 | | | | | 1 | | | | | | | | | yes | | | | | |
| C6orf27 | NA | Body;nan;5'UTR | chr6:31743769-31744918 | 1149 | | 2 | 26 | | | | | 0 | | | 1 | | | | yes | | | | |
| VARS | NA | Body | chr6:31746808-31747237 | 429 | | | | 1 | | | | | 2 | 1 | | | | yes | | | | | |
| VARS | N_Shelf | Body | chr6:31760047-31760825 | 778 | | | 33 | | | | | 0 | 1 | | | | | yes | | | | | |
| VARS | N_Shore | Body | chr6:31762442-31762555 | 113 | | | 2 | | | | | | | | | | | yes | | | | | |
| VARS;LSM2 | S_Shore | TSS1500;Body | chr6:31765083-31765634 | 551 | | | | | | | | | | | | | | | | | | | |
| HSPA1L | N_Shelf | Body;5'UTR | chr6:31779598-31780562 | 964 | | | | | | | | | 2 | | | | | yes | yes | | | | |
| HSPA1A;HSPA1L | Island;N_Shore | TSS1500;TSS200;1stExon;5'UTR | chr6:31782675-31783455 | 780 | | | | | 1 | | | 0 | 1 | | | | | yes | | | | | |
| HSPA1B | N_Shore | TSS1500 | chr6:31794472-31794759 | 287 | | | | 1 | | | | | | | | | | yes | | | | | |
| C6orf48;SNORD52;SNORD48 | S_Shore | 5'UTR;TSS1500;Body | chr6:31803089-31804739 | 1650 | | | | | | | 1 | 0 | | 2 | | | | yes | | | | | |
| NEU1 | N_Shelf | Body | chr6:31827858-31827912 | 54 | | | | | | | | | | | | | | | | | | | |
| NEU1 | N_Shore | Body | chr6:31829644-31829960 | 316 | | | 24 | | | | | | 2 | | | | | | yes | | | | yes |
| SLC44A4;NEU1 | S_Shore;S_Shelf | 3'UTR;TSS1500;Body | chr6:31831340-31833883 | 2543 | | 1 | 3 | 1 | 3 | | | | 3 | | | | | yes | yes | | yes | | |
| SLC44A4 | NA | Body | chr6:31838402-31838774 | 372 | 1 | | 8 | | | | | | 2 | | | | | yes | yes | | | | |
| SLC44A4 | NA | Body | chr6:31842575-31842909 | 334 | | | | | | | | | | | | | | yes | | | | | |
| SLC44A4;EHMT2 | NA | Body;5'UTR;1stExon;TSS200;3'UTI | chr6:31845672-31848028 | 2356 | | 1 | 19 | | 1 | | | 0 | 5 | | | 1 | | yes | | | yes | | |
| EHMT2 | N_Shore | Body | chr6:31856329-31856773 | 444 | | | 1 | | | | | 0 | | | | | | | | | | | yes |
| EHMT2 | N_Shore;Island | Body | chr6:31864071-31865095 | 1024 | | | 4 | 1 | | | | | 1 | | | | | | yes | yes | | yes | yes |
| EHMT2;ZBTB12 | N_Shore;Island | TSS1500;Body | chr6:31866586-31868949 | 2363 | | | | | | | | | | | | | | | | | | | |
| ZBTB12 | S_Shore | TSS1500 | chr6:31870840-31870990 | 150 | | | | | | | | | | | | | | yes | yes | | | | |
| C2 | NA | TSS1500;1stExon;5'UTR | chr6:31894990-31895598 | 608 | | | 11 | 1 | | | | 0 | 2 | | | | | yes | | | yes | | |
| CFB;C2 | NA | Body;nan;1stExon;5'UTR | chr6:31913250-31915198 | 1948 | | | 23 | 2 | | | | 0 | 5 | | 1 | 4 | | | | | | | |
| RDBP | NA | 3'UTR | chr6:31919944-31920145 | 201 | | | 1 | | | | | | | | | | | | | | | | |
| RDBP;SKIV2L;MIR1236 | N_Shore | Body;TSS1500 | chr6:31926014-31926205 | 191 | | | 8 | | | | | | 1 | | | | | | yes | | | | |
| SKIV2L;STK19;DOM3Z | N_Shelf;N_Shore | Body;TSS1500;TSS200 | chr6:31935900-31939546 | 3646 | 1 | 1 | 51 | 2 | | | | 0 | 5 | | | 3 | | | | | | | |
| STK19;DOM3Z | S_Shore | Body;TSS1500 | chr6:31940692-31940855 | 163 | | | | | | | | 0 | 1 | | | | | | yes | | yes | | |
| C4B;C4A | NA | Body | chr6:31964193-31964391 | 198 | | | | | | | | 0 | 2 | | | | | | | | | | |
| CYP21A2 | NA | TSS200 | chr6:31973162-31973255 | 93 | | | 1 | | | | | | | | | | | yes | | | yes | yes | |
| CYP21A2;TNXB;TNXA | NA | 3'UTR;Body | chr6:31976548-31976720 | 172 | | | 2 | | | | | | | | | | | | | | | | |
| TNXB;STK19;TNXA | NA | 1stExon;TSS1500;Body;5'UTR;TSS2 | chr6:31980043-31980856 | 813 | | | 15 | | | | | | | | | | | | | | | | |
| C4A;C4B | NA | Body | chr6:31997026-31997067 | 41 | | | | | | | | 0 | | | | | | | | | | | |
| CYP21A2 | NA | TSS200;TSS1500 | chr6:32005786-32006196 | 410 | | | | 1 | | | | | | | | | | yes | | | yes | yes | |
| CYP21A2 | NA | Body | chr6:32007625-32007916 | 291 | | | | | | | | | | | | | | | | | | | |
| TNXB | NA | Body | chr6:32011506-32011575 | 69 | | | | | | | | | | | | | | | | | | | |
| TNXB | NA | Body;TSS1500;1stExon;5'UTR;TSS2 | chr6:32012900-32016602 | 3702 | 1 | 2 | 67 | 1 | | | | 0 | 11 | 1 | 2 | | | yes | | | | | |
| TNXB | NA | Body | chr6:32019556-32019840 | 284 | | | | | | | | | 1 | | | | | yes | yes | | yes | | |
| TNXB | NA | Body | chr6:32025416-32027528 | 2112 | | | 8 | | | | | 0 | 3 | | | | | yes | yes | | | | |
| TNXB | NA | Body | chr6:32029152-32032709 | 3557 | | 2 | 2 | | | | | | 1 | | | | | yes | yes | | | | |
| TNXB | NA | Body | chr6:32035439-32039176 | 3737 | | 4 | 15 | | | | | | 4 | | | | | yes | yes | | | yes | |
| TNXB | N_Shelf;NA | Body | chr6:32042797-32044869 | 2072 | 3 | | 13 | | | | | | | | | | | yes | yes | | | | |
| TNXB | S_Shelf;chr6:32046815-32047( | Body;S_Shelf;N_Shore | chr6:32049177-32057846 | 8669 | | 3 | 107 | | | 2 | | 0 | 16 | 1 | 4 | 13 | | yes | yes | | yes | yes | yes |
| TNXB | S_Shelf | Body | chr6:32059166-32059605 | 439 | | | | | | | | | 1 | | | | | yes | | | yes | | |
| TNXB | N_Shelf | Body | chr6:32060681-32061478 | 797 | | | 10 | | | 1 | | | | | | | | yes | yes | | | | |
| TNXB | N_Shore;Island;S_Shore | Body | chr6:32062528-32065890 | 3362 | | 5 | 22 | 3 | 3 | 1 | 3 | 0 | 83 | 27 | 4 | 93 | 4 | yes | yes | | yes | | |
| TNXB | NA | 5'UTR | chr6:32073123-32074397 | 1274 | | | | | | | | | | | | | | yes | yes | | | | |
| TNXB | NA | 5'UTR;1stExon;TSS1500 | chr6:32076079-32078574 | 2495 | 1 | 1 | | | | | | | 4 | | | 1 | | | | | | | yes |
| ATF6B | NA | Body | chr6:32084485-32084518 | 33 | | | | | | | | | | | | | | | | | | | |
| ATF6B | NA | Body | chr6:32086755-32086928 | 173 | | | | | | | | | 1 | | | | | yes | | | | | |
| ATF6B;FKBPL | N_Shelf;N_Shore;S_Shore | Body;1stExon;TSS1500;TSS200 | chr6:32094729-32098253 | 3524 | | | 24 | | | 1 | | | 6 | 1 | | | | yes | | | | | |
| PRRT1;PPT2 | Island;S_Shore;N_Shore;chr6:3 | Body;1stExon;5'UTR;TSS1500;N_S | chr6:32116207-32121471 | 5264 | 1 | 4 | 74 | 1 | 3 | 1 | | 0 | 34 | 3 | 3 | 4 | 3 | | | | | yes | yes |
| PPT2 | S_Shore | Body | chr6:32122864-32123651 | 787 | | | 10 | | | | | | 2 | | | | | yes | | | | yes | |
| EGFL8;PPT2 | N_Shelf | TSS1500;3'UTR | chr6:32130993-32131254 | 261 | | | | | | | | | | | | | | | | | | | |
| EGFL8 | S_Shore;Island | Body | chr6:32134573-32135728 | 1155 | | 2 | 9 | | | | | 0 | 2 | | | | | | | | | | |
| AGPAT1 | S_Shore;S_Shelf | Body | chr6:32137061-32138270 | 1209 | | | 8 | | | | | | | | | | | yes | | | | | |
| RNF5P1;RNF5;AGPAT1 | NA | TSS1500;5'UTR;1stExon;TSS200;B | chr6:32145342-32147891 | 2549 | | | | | | | | 0 | 1 | 1 | | 3 | | | | | | | |
| AGER | NA | Body | chr6:32150404-32151517 | 1113 | | | | | | | | | | | | | | | | | | | |
| PBX2 | NA | 3'UTR;Body | chr6:32153540-32157150 | 3610 | | 1 | 3 | 1 | | | | | 3 | | | | | yes | | | | yes | yes |
| GPSM3;PBX2 | N_Shelf;NA | Body;5'UTR;TSS1500;3'UTR | chr6:32158927-32161004 | 2077 | | | 6 | 1 | 3 | | | | 1 | | | | | yes | yes | | | | |
| GPSM3 | N_Shore | 5'UTR | chr6:32162202-32162695 | 493 | | | | | | | | | 1 | | | | | yes | yes | | yes | | |
| GPSM3;NOTCH4 | S_Shore;S_Shelf | TSS1500;Body | chr6:32164503-32166839 | 2336 | | | 24 | | | | | 0 | 5 | | | | | yes | | | yes | | |
| NOTCH4 | NA | Body | chr6:32168751-32170135 | 1384 | | | 5 | | | | | | | | | | | yes | yes | | | | |
| NOTCH4 | NA | Body | chr6:32171330-32172871 | 1541 | | 1 | | | | | | | 1 | | | | | yes | yes | | | | |
| NOTCH4 | NA | Body | chr6:32177664-32178017 | 353 | | | | | | | | | | | | | | yes | yes | | | | |
| NOTCH4 | NA | Body | chr6:32179971-32182326 | 2355 | | 2 | 12 | | | | | | 2 | | | | | yes | yes | | | yes | yes |
| NOTCH4 | NA | Body;1stExon;5'UTR;TSS200 | chr6:32184296-32192510 | 8214 | | 4 | 45 | | | | | | 6 | 1 | | | | yes | yes | | yes | yes | yes |
| C6orf10 | NA | Body | chr6:32289047-32289373 | 326 | | | | | | | | | | | | | | | yes | | yes | | |
| C6orf10 | NA | Body | chr6:32294260-32295237 | 977 | | | 9 | | | | | | 2 | | | | | yes | | | | yes | |
| C6orf10 | NA | Body | chr6:32297615-32297822 | 207 | | 1 | 11 | | | | | 0 | 3 | 1 | | | | | yes | | | | |
| C6orf10 | NA | Body | chr6:32299617-32299676 | 59 | | | | | | | | | 1 | | | | | | | yes | | | |
| C6orf10 | NA | Body | chr6:32300743-32301513 | 770 | | | | | | | | | | | | | | yes | | yes | | | |
| C6orf10 | NA | Body | chr6:32303626-32306373 | 2747 | | | 4 | | | | | | 1 | | | 1 | | yes | yes | | yes | yes | yes |
| C6orf10 | NA | Body | chr6:32308995-32309323 | 328 | | | | | | | | | | | | | | yes | | | yes | | |
| C6orf10 | NA | Body | chr6:32317028-32317528 | 500 | | 1 | | | | | | | | | | | | yes | | | | yes | |
| BTNL2 | NA | Body | chr6:32362626-32366347 | 3721 | | 1 | 20 | 1 | | | 2 | | 2 | 3 | | | | yes | yes | yes | | | |
| BTNL2 | NA | Body | chr6:32367672-32368318 | 646 | | | | | | | 1 | | 1 | 1 | | | | yes | | | yes | | yes |

| | | | | | | | | | | | | | | | | | | | | | | | |
|---|---|---|---|---|---|---|---|---|---|---|---|---|---|---|---|---|---|---|---|---|---|---|---|
| BTNL2 | NA | Body | chr6:32370637-32373089 | 2452 | | | 13 | | | | | 0 | 2 | | | | | yes | yes | | yes | yes | |
| BTNL2 | NA | Body;TSS200;TSS1500 | chr6:32374162-32376095 | 1933 | | 1 | 10 | | | | | | 1 | | | | | yes | | yes | | | yes |
| HLA-DRA | NA | TSS1500 | chr6:32406271-32406545 | 274 | | | | | | | | | | | | | | yes | | | | | |
| HLA-DRA | NA | Body;TSS200;5'UTR;1stExon | chr6:32407570-32411897 | 4327 | | | | | | | | | | | | | | | | | | | |
| HLA-DRB5 | N_Shore;Island;S_Shore | Body | chr6:32489203-32490444 | 1241 | 5 | 7 | 40 | 1 | | | 6 | | 14 | 6 | 1 | 25 | 1 | | yes | | | yes | yes |
| HLA-DRB5 | NA | TSS1500 | chr6:32498285-32498677 | 392 | | 1 | | | | | | | | | | | | yes | yes | | | | |
| HLA-DRB6 | NA | Body | chr6:32522400-32523136 | 736 | | 1 | 5 | 2 | | | 2 | | 1 | 1 | | 1 | | yes | | | | | |
| HLA-DRB6;HLA-DRB1 | NA;Body | Body;nan;3'UTR | chr6:32525805-32547167 | 21362 | | | | | | | | | | | | | | | | | | | |
| HLA-DRB1 | N_Shelf;N_Shore;Body;Island; | Body;nan;Island | chr6:32548321-32555411 | 7090 | | | | | | | | | | | | | | | | | | | |
| HLA-DRB1 | NA | Body;1stExon;TSS1500 | chr6:32557321-32558459 | 1138 | 1 | 7 | 5 | 2 | | | 2 | 0 | 5 | 3 | 1 | 33 | 4 | yes | | | yes | | |
| HLA-DQA1 | NA | TSS1500;TSS200;1stExon;Body;na | chr6:32604564-32607509 | 2945 | 3 | 12 | 25 | 1 | | | 4 | 0 | 4 | 8 | 1 | 25 | 4 | yes | yes | | yes | | yes |
| HLA-DQA1 | NA | Body;3'UTR | chr6:32608858-32610971 | 2113 | | 1 | 4 | | | | | | | 1 | | 7 | | | yes | yes | | | |
| HLA-DQB1 | NA | 3'UTR | chr6:32627845-32628305 | 460 | | | 2 | | | | | | | | | | | | | | | | |
| HLA-DQB1 | N_Shelf;N_Shore;Island;S_Sho | Body;1stExon | chr6:32629786-32634481 | 4695 | 12 | 21 | 58 | 1 | | 3 | 9 | 0 | 13 | 10 | 2 | 75 | 2 | yes | yes | | | | |
| HLA-DQA2 | NA | Body;1stExon | chr6:32709268-32714066 | 4798 | | 2 | 23 | | | | | | 1 | | | | | yes | yes | | yes | | |
| HLA-DQB2 | NA | Body | chr6:32725075-32725332 | 257 | | | | | | | | | | | | | | yes | | | | | |
| HLA-DQB2 | N_Shelf | Body | chr6:32726567-32726692 | 125 | | | | | | | | | | | | | | | yes | yes | | | |
| HLA-DQB2 | N_Shore;Island;S_Shore | Body | chr6:32728786-32730859 | 2073 | | 1 | 83 | | 1 | 6 | 1 | 0 | 20 | | 1 | 12 | | yes | yes | | | | |
| HLA-DQB2 | S_Shelf | TSS1500 | chr6:32732247-32732685 | 438 | | | 9 | | | | | | | | | | | yes | | | yes | | |
| HLA-DOB | NA | Body;3'UTR | chr6:32780861-32785141 | 4280 | | | 14 | | 1 | | | | 6 | | | | | | yes | yes | | | |
| TAP2 | NA | 3'UTR;Body | chr6:32795877-32796216 | 339 | | | | | | | | | 1 | | | | | yes | yes | | | yes | |
| TAP2 | NA | Body | chr6:32797234-32797268 | 34 | | | 1 | | | | | | | | | | | | | | | | |
| TAP2 | NA | Body | chr6:32798887-32800549 | 1662 | | | | | | | | | 2 | | | | | yes | yes | | | yes | |
| TAP2 | N_Shelf;N_Shore | Body | chr6:32803482-32805759 | 2277 | 2 | | 11 | 1 | | | | | 2 | | | 1 | | | | | yes | | yes |
| PSMB8 | S_Shore;N_Shelf;N_Shore | 3'UTR;Body | chr6:32808504-32810833 | 2329 | | | | 1 | 3 | 1 | | 0 | 9 | | | | | yes | | | | | |
| PSMB8;TAP1 | S_Shore;chr6:32811494-32811 | TSS1500;3'UTR;Body;S_Shore | chr6:32812933-32813715 | 782 | | | 1 | 2 | 1 | 2 | | | 1 | 2 | 2 | | | yes | | | | | |
| TAP1 | S_Shelf | Body | chr6:32814921-32815009 | 88 | | | | 1 | | | | | | | | | | yes | | | | yes | |
| TAP1 | N_Shelf;N_Shore;chr6:3282084 | Body;N_Shore | chr6:32818477-32820546 | 2069 | | | 8 | | 2 | 1 | | 0 | 14 | 1 | 1 | | | yes | | | | yes | |
| PSMB9;TAP1 | S_Shore | Body;TSS1500 | chr6:32822565-32823941 | 1376 | | | 1 | | | | 1 | 0 | 5 | 1 | | | | yes | | | | | |
| PPP1R2P1 | N_Shore;Island | Body | chr6:32846924-32847845 | 921 | 1 | | 51 | | | | | 0 | 15 | | | 16 | | yes | yes | | | | |
| HLA-DMB | NA | Body | chr6:32903598-32905320 | 1722 | | | 21 | | | | | 0 | 8 | 1 | | 1 | | | | | yes | | |
| HLA-DMB | NA | Body | chr6:32906532-32906857 | 325 | | | | | | | | | 2 | | | | | | yes | yes | yes | yes | |
| HLA-DMB | NA | 1stExon;5'UTR;TSS200;TSS1500;Bo | chr6:32908239-32909523 | 1284 | | | 13 | 1 | 1 | | | 0 | 2 | | | | | | yes | yes | | | yes |
| HLA-DMA | NA | 3'UTR;Body;nan;1stExon;5'UTR;TS | chr6:32916606-32921233 | 4627 | | | | | | | | | | | | | | | | | | | |
| BRD2 | N_Shore | TSS1500 | chr6:32935025-32935858 | 833 | | | 3 | | | | | | | | | | | | yes | | | | |
| BRD2 | S_Shore | Body | chr6:32942358-32943025 | 667 | | | | | | | | | 1 | | | 1 | | | | | | yes | yes |
| BRD2 | NA | Body | chr6:32945234-32945906 | 672 | | | 8 | 2 | | | | | | | | | | | | | yes | yes | |
| HLA-DOA | N_Shelf | 3'UTR | chr6:32972093-32972492 | 399 | | | | | | | | | | | | | | yes | | | | | |
| HLA-DOA | N_Shore;Island;S_Shore;S_She | 3'UTR;Body;TSS200;TSS1500 | chr6:32974122-32978548 | 4426 | | 2 | 55 | 3 | | 1 | | 0 | 6 | | | 2 | | yes | yes | yes | yes | yes | yes |
| HLA-DPA1 | NA | 3'UTR | chr6:33032830-33033484 | 654 | | | 6 | | | | 1 | | 1 | 2 | 1 | | | | yes | yes | yes | | |
| HLA-DPA1 | NA | Body | chr6:33036464-33037690 | 1226 | | 1 | 15 | | | | 1 | | 1 | 1 | 1 | | | yes | | | | | |
| HLA-DPA1 | NA | Body | chr6:33039396-33041343 | 1947 | 4 | | 25 | 1 | | | | | 11 | 1 | | 1 | | yes | yes | | | | |
| HLA-DPB1;HLA-DPA1 | NA | TSS1500;TSS200;1stExon;Body | chr6:33042551-33044510 | 1959 | | | 3 | 1 | | | | 0 | 5 | | | 1 | | yes | yes | | | | |
| HLA-DPB1 | N_Shore;Island;S_Shore | Body | chr6:33047944-33049360 | 1416 | 1 | 3 | 23 | | 1 | 2 | 5 | 0 | 58 | 17 | 6 | 24 | | | yes | | | | |
| HLA-DPB1 | S_Shelf;NA | Body;nan | chr6:33052331-33054001 | 1670 | | 1 | 10 | | | | | | 1 | 1 | 1 | | | yes | | | | yes | |
| HLA-DPB2 | N_Shore;Island;S_Shore | Body | chr6:33084420-33085800 | 1380 | | | 7 | | | | | 0 | 15 | 2 | | 10 | | yes | yes | yes | | | yes |
| HLA-DPB2 | NA | Body | chr6:33090166-33093109 | 2943 | 9 | 1 | 17 | | | | | 0 | 8 | 1 | | 9 | 1 | yes | | yes | yes | | |
| HLA-DPB2 | NA | Body | chr6:33094306-33094829 | 523 | | | | | | | | | | | | | | yes | yes | | | | |
| HLA-DPB2 | NA | Body | chr6:33095865-33096738 | 873 | | | 8 | | | | | | | | | | | yes | | | | | |
| COL11A2 | S_Shore;S_Shelf;NA | 3'UTR;Body | chr6:33131072-33139083 | 8011 | 3 | 6 | 53 | | | | | | 14 | 1 | 1 | | | | | | yes | yes | |
| COL11A2 | NA | Body | chr6:33140396-33144071 | 3675 | | 4 | 19 | | | | | | 5 | | | | | yes | | | | | |
| COL11A2 | NA | Body | chr6:33145396-33148096 | 2700 | | 1 | 6 | | | | | | 1 | | | | | | | | | | |
| COL11A2 | NA | Body | chr6:33152524-33153311 | 787 | | | 4 | 1 | | | | | | | | | | | | | | | yes |
| COL11A2 | NA | Body | chr6:33154830-33155135 | 305 | | | | | | | | | | | | | | yes | | | | yes | |
| COL11A2;RXRB | N_Shelf;N_Shore;Island;S_Sho | Body;TSS200;TSS1500;3'UTR | chr6:33156206-33164180 | 7974 | | 5 | 25 | 2 | | | | 0 | 6 | | | | | yes | yes | | | yes | yes |
| RXRB | N_Shelf | Body | chr6:33165404-33165707 | 303 | | | | | | | | | | | | | | yes | | | | | |
| RXRB;SLC39A7 | N_Shore | Body;TSS1500 | chr6:33166966-33167488 | 522 | | | 2 | | | | | 0 | | | | | | yes | | | | | yes |
| SLC39A7;RXRB | S_Shore;N_Shore | 1stExon;TSS1500;Body | chr6:33169253-33170838 | 1585 | | | 17 | | 1 | | | | 1 | | | | | | | | | | |
| HSD17B8;SLC39A7;MIR219-1;RING1 | N_Shore;chr6:33172289-33172 | TSS1500;3'UTR;Island;Body | chr6:33171911-33175225 | 3314 | | | | | | | | | | | | | | | | | | | |
| RING1 | N_Shore | 3'UTR | chr6:33180304-33180629 | 325 | | | 5 | | | | | | 1 | | | | | | | | | | yes |
| VPS52 | S_Shore | Body | chr6:33218697-33219654 | 957 | | | 11 | | | | | | 1 | | | | | yes | yes | | | | |
| VPS52 | NA | Body | chr6:33226913-33227015 | 102 | | | | | | | | | 1 | | | | | yes | yes | yes | yes | | |
| VPS52 | NA | Body | chr6:33228341-33228498 | 157 | | | 4 | | | | | | 1 | | | | | yes | yes | | | | |
| VPS52 | NA | Body | chr6:33231505-33231870 | 365 | | | 2 | | | | | | 1 | | | | | yes | yes | | | yes | |
| VPS52 | NA | Body | chr6:33234201-33234378 | 177 | | | | | | | | | 1 | | | | | yes | | | | | |
| VPS52;RPS18 | Island;S_Shore | TSS1500;Body | chr6:33240059-33241594 | 1535 | | 1 | 6 | | | | | | 1 | | | | | yes | | | | | |
| RPS18;B3GALT4;WDR46 | N_Shore;Island;chr6:33244677 | Body;TSS1500;TSS200;Island;1stEx | chr6:33243375-33247250 | 3875 | | | | | | | | | | | | | | | | | | | |
| PFDN6;WDR46 | N_Shore | TSS1500;Body | chr6:33256078-33256222 | 144 | | | 5 | | | | | | | | | | | | | | | | |
| PFDN6;WDR46 | S_Shore | Body;TSS1500 | chr6:33258045-33258581 | 536 | | | 8 | | | | | | | | | | | | yes | | | yes | |
| RGL2 | S_Shelf | Body | chr6:33260021-33260145 | 124 | | | | | | | | | | | | | | | | | | | |
| RGL2 | N_Shelf;N_Shore | Body | chr6:33262600-33265534 | 2934 | | | 28 | | | | | | 3 | 1 | | | | yes | | | | | yes |
| TAPBP;RGL2 | S_Shore | 3'UTR;TSS1500 | chr6:33267700-33267996 | 296 | | | | | 1 | | | | 2 | | | 1 | | yes | yes | | yes | | |
| TAPBP | NA | Body | chr6:33273014-33274335 | 1321 | | | | | | | | | 3 | | | | | yes | yes | | | | |
| TAPBP | N_Shore | Body | chr6:33279563-33280454 | 891 | 4 | | 15 | | | | | 0 | 3 | | | 1 | | yes | | | yes | | |
| ZBTB22;TAPBP | Island;S_Shore;N_Shore | Body;TSS1500 | chr6:33282867-33284965 | 2098 | | 2 | 54 | | | | | 0 | 14 | | | | | yes | | | | yes | yes |
| ZBTB22;DAXX | S_Shore;N_Shore;chr6:332887 | TSS1500;Body;N_Shore;5'UTR | chr6:33286692-33290379 | 3687 | | | 90 | 2 | | | | | 6 | | 1 | | | yes | | | | yes | yes |
| DAXX | S_Shore | TSS1500 | chr6:33291845-33292029 | 184 | | | 9 | | | | | | 1 | | | | | yes | | | yes | | |
| KIFC1 | N_Shore | TSS1500 | chr6:33358417-33359001 | 584 | | | 12 | | | | | | 2 | | | | | yes | yes | | | | yes |
| KIFC1 | S_Shore | Body | chr6:33360495-33361102 | 607 | | | 3 | | | | | | 1 | | | | | yes | yes | | | | |
| KIFC1 | S_Shelf | Body | chr6:33362576-33363179 | 603 | | | | | | | | | 1 | | | | | yes | yes | | | | |
| PHF1;CUTA | N_Shelf;N_Shore | Body;3'UTR | chr6:33383065-33385056 | 1991 | | | | | | | | | | | | | | | | | | | |
| CUTA;SYNGAP1 | S_Shore | TSS1500 | chr6:33386397-33387089 | 692 | | | 5 | | | | | | 1 | | | | | | | | | | |
| SYNGAP1 | S_Shore;Island | Body | chr6:33396160-33397710 | 1550 | | | 18 | 1 | | | | | | | | | | | | | | yes | yes |
| SYNGAP1 | S_Shelf;NA | Body | chr6:33398906-33401542 | 2636 | | 1 | 13 | 2 | | | | 0 | 3 | 1 | | 1 | | | yes | | | yes | yes |
| SYNGAP1 | N_Shelf | Body;3'UTR | chr6:33418814-33420227 | 1413 | | | 8 | | | | | | 1 | | | | | | | | yes | | yes |
| ZBTB9 | S_Shore | 5'UTR;Body | chr6:33422868-33424048 | 1180 | | | 19 | | | | | 0 | 6 | | | 3 | | | | | | | yes |
| C6orf227 | Island;S_Shore | Body;TSS200 | chr6:33560953-33561449 | 496 | | | 22 | | | | | 0 | 7 | 2 | 1 | 3 | | yes | yes | | | yes | |
| ITPR3 | NA | Body | chr6:33638313-33638550 | 237 | | | | | | | | | 1 | | | | | | | | | | yes |
| LEMD2 | Island | 3'UTR | chr6:33739406-33739653 | 247 | | | | | | | | 0 | | | | | | | | | | | |
| MLN | NA | TSS200 | chr6:33771981-33772366 | 385 | | | | | | | | | | | | | | yes | | | | | |
| MIR1275 | NA | TSS200 | chr6:33967924-33968234 | 310 | | | | | | | | | 1 | | | | | | | yes | | | |
| GRM4 | NA | 3'UTR | chr6:33989813-33990181 | 368 | | | | | | | | | | | | | | | | | | yes | |
| GRM4 | Island | Body | chr6:33995991-33996111 | 120 | 1 | | | | | | | | | | | | | yes | | | | | |
| GRM4 | Island | Body | chr6:34003640-34004360 | 720 | | 1 | | | | | | | | | | | | yes | | | yes | | yes |
| GRM4 | Island | Body | chr6:34024220-34024479 | 259 | | | 24 | | | | | 0 | 2 | | | | | | yes | | yes | | |
| GRM4 | NA | Body | chr6:34031060-34031694 | 634 | | | 2 | | | | | | | | | | | yes | yes | | | yes | |
| GRM4 | NA | 1stExon;5'UTR;TSS1500 | chr6:34100899-34102530 | 1631 | | 1 | 7 | | | | | | 1 | | | | | yes | | | | yes | |
| HMGA1 | Island | TSS1500 | chr6:34203153-34203887 | 734 | | | 6 | | 1 | | | | | | | | | | | | | | |
| HMGA1 | S_Shore;Island | 5'UTR;TSS200;TSS1500 | chr6:34206084-34206683 | 599 | | | 4 | | 1 | | | | | | | | | yes | | | | | yes |

| | | | | | | | | | | | | | | | | | | | | | | | | | | | |
|---|---|---|---|---|---|---|---|---|---|---|---|---|---|---|---|---|---|---|---|---|---|---|---|---|---|---|---|
| PACSIN1 | Island | Body | chr6:34495292-34495393 | 101 | | | 3 | | | | | | | | | | | | yes | | yes | yes | | | | | |
| PACSIN1 | N_Shore;Island | Body | chr6:34498918-34499504 | 586 | | | 13 | | 1 | | | 0 | 3 | 1 | | | | yes | | | yes | | | | | | |
| SPDEF | NA | 1stExon;5'UTR;TSS200;TSS1500 | chr6:34523925-34525026 | 1101 | 1 | | 9 | | | | | | 1 | | | | | | yes | | | | | | | | |
| SNRPC | N_Shore | TSS200 | chr6:34724739-34724756 | 17 | | | | | | | | 0 | 2 | 1 | | | | yes | yes | | | | | | | | |
| ANKS1A | NA | Body | chr6:35027147-35027324 | 177 | | | | | | | | | | | | | | | | | | | | | | | |
| TCP11 | Island;chr6:35108801-3510949 | Body;5'UTR;1stExon;N_Shore | chr6:35108605-35109353 | 748 | | | 19 | 1 | | | | 0 | 1 | 1 | 1 | | | yes | | | | | | | | | |
| DEF6 | NA | TSS1500;TSS200;1stExon;5'UTR | chr6:35265308-35265759 | 451 | 2 | | 40 | 2 | | | | | | | | | | yes | | | | | | | | | |
| DEF6 | NA | Body | chr6:35285967-35286360 | 393 | | | | | | | | | 1 | | | | | yes | | | | yes | | | | | |
| TEAD3 | Island | Body | chr6:35453938-35454434 | 496 | | | | | | | | | | | | | | yes | | | | | | | | | |
| TULP1 | N_Shore;Island | Body | chr6:35479350-35479910 | 560 | | 1 | | | | | | 0 | | | | | | yes | | | | yes | | | | | |
| C6orf127 | Island | Body | chr6:35754758-35754906 | 148 | | | 5 | | | | | | | | | 1 | | | yes | | | | | | | | |
| CLPS | NA | Body;5'UTR;1stExon;TSS200 | chr6:35764958-35766021 | 1063 | | | 4 | | | | | | 1 | | | | | yes | | | | yes | yes | | | | |
| SLC26A8 | N_Shelf | 5'UTR;TSS1500;TSS200 | chr6:35991802-35992685 | 883 | | | 3 | | | | | | | | | | | yes | yes | | | | | | | | |
| PNPLA1 | NA | TSS1500;5'UTR;TSS200 | chr6:36237982-36238410 | 428 | | | | | | | | | | | | | | | | | | | | | | | |
| PNPLA1 | NA | 5'UTR;Body | chr6:36253026-36253157 | 131 | | 1 | | | | | | | | | | | | | yes | | yes | | | | | | |
| C6orf222 | NA | TSS200 | chr6:36304683-36305155 | 472 | | | | | 1 | | | | 2 | | | | | yes | | | | | | | | | |
| ETV7 | Island | TSS1500 | chr6:36355732-36356162 | 430 | | | | | | | | | | | | | | yes | yes | | | yes | | | | | |
| PXT1;KCTD20 | Island | 5'UTR;TSS200;1stExon | chr6:36410485-36410534 | 49 | | | 1 | | | | | | | | | | | yes | | | | | yes | | | | |
| CDKN1A | N_Shore | TSS1500 | chr6:36644997-36645812 | 815 | | | 2 | | | | | | | | | | | yes | yes | | | | | | | | |
| CPNE5 | Island | TSS1500 | chr6:36807759-36808696 | 937 | | | | | | | | 0 | 1 | | | | | | | | | yes | yes | | | | |
| PI16 | Island | Body | chr6:36929838-36930062 | 224 | | | | | | | | 0 | 1 | | | 2 | | yes | | | | | | | | | |
| FGD2 | NA | TSS1500 | chr6:36972027-36972163 | 136 | | | 1 | | | | | | 1 | | | | | yes | yes | | | | | | | | |
| FTSJD2 | N_Shore | TSS1500 | chr6:37400392-37400668 | 276 | | | 1 | | | | | | | | | | | yes | yes | | | | | | | | |
| C6orf129 | NA | Body | chr6:37460940-37461350 | 410 | | | 6 | | 1 | | | | 1 | | | | | yes | | | | | | | | | |
| MDGA1 | Island | Body | chr6:37616410-37618514 | 2104 | | | 4 | | 1 | | 2 | 0 | 3 | 3 | 1 | 14 | 1 | yes | | yes | | | | | | | |
| MDGA1 | Island | Body | chr6:37626035-37626335 | 300 | | | | | | | | | 1 | | | | | yes | | yes | | | | | | | |
| BTBD9 | NA | Body | chr6:38145125-38145464 | 339 | | | | | 1 | | | | | | | | | | | | | | | | | | |
| GLP1R | Island | TSS1500 | chr6:39016326-39016470 | 144 | | | | | | | | | 1 | | | | | yes | | | | | | | | | |
| KCNK17 | S_Shore | TSS200 | chr6:39282316-39282532 | 216 | | | 6 | | 1 | 1 | | | 1 | | | | | | | | | | | | | | |
| KCNK16 | NA | TSS200 | chr6:39290415-39290877 | 462 | | 1 | 5 | | | | | | | | | | | yes | yes | | | | | | | | |
| KIF6 | Island;S_Shore | Body;TSS200 | chr6:39692744-39693366 | 622 | | | | | | | | | 2 | | | 1 | | yes | | | | | yes | | | | |
| DAAM2 | S_Shore | TSS200 | chr6:39760607-39760833 | 226 | | | | | | | | | | | | 1 | | | | | | yes | | | | | |
| DAAM2 | Island | Body | chr6:39869689-39869840 | 151 | | | | 1 | | | | | | | | | | | | | | | | | | | |
| MOCS1 | NA | nan;5'UTR;1stExon;Body | chr6:39895360-39895671 | 311 | | | | | | | | | 1 | | 1 | | | yes | | | | | | | | | |
| MOCS1 | S_Shore | TSS200 | chr6:39902322-39902356 | 34 | | | | | | | | | 1 | | | | | yes | | | yes | yes | | | | | |
| TDRG1 | NA | TSS1500 | chr6:40345376-40346044 | 668 | | | | | | | | | | | | | | | yes | yes | | | | | | | |
| LRFN2 | NA | Body | chr6:40400230-40400462 | 232 | | | | | | | | | | | | | | | | | | | | | | | |
| UNC5CL | N_Shore;Island | 3'UTR | chr6:40995388-40996213 | 825 | | | | 1 | 2 | | | | | | | | | yes | | | | yes | | | | | |
| UNC5CL | NA | TSS200 | chr6:41006974-41007485 | 511 | | | | | | | | | | | | | | yes | yes | | yes | | | | | | |
| APOBEC2 | NA | TSS1500;TSS200 | chr6:41020537-41021161 | 624 | | | | | 1 | | | | | | | | | yes | yes | | | | | | | | |
| NFYA;LOC221442 | Island | 3'UTR;TSS200;TSS1500 | chr6:41068553-41069048 | 495 | | | | | | | | 0 | 8 | 1 | 3 | 13 | 1 | | yes | | | | | | | | |
| TREML1 | NA | Body | chr6:41121714-41122081 | 367 | | | | | | | | | | | | | | | yes | | | | | | | | |
| TREM2 | NA | 1stExon;5'UTR | chr6:41130917-41131213 | 296 | | | | | | | | | | | | | | yes | | | | | | | | | |
| TREML2 | NA | Body;1stExon;5'UTR;TSS200 | chr6:41168185-41169171 | 986 | | | | | | | | 0 | 1 | | 1 | | | yes | | | | yes | yes | | | | |
| TREM1 | NA | 5'UTR;1stExon;TSS1500 | chr6:41254433-41254997 | 564 | | | 2 | | | | | | 1 | | | | | | yes | | | | | | | | |
| FOXP4 | N_Shore;Island;chr6:41528266 | 5'UTR;Island | chr6:41528198-41528785 | 587 | | 1 | 30 | 1 | | | | | 1 | 2 | 1 | | | yes | | | | | | | | | |
| FOXP4 | N_Shore | Body | chr6:41561085-41561472 | 387 | 1 | | | | | | | | | | | | | | | | | | | | | | |
| MDFI | Island | TSS1500 | chr6:41605343-41605784 | 441 | | | 4 | | | | | | | | | | | | | | | | yes | | | | |
| MDFI | Island | Body | chr6:41620951-41621322 | 371 | | 1 | | | 1 | | | | | | | | | yes | | | | | | | | | |
| TFEB | S_Shore | 5'UTR;TSS1500;1stExon | chr6:41703872-41704208 | 336 | | | 3 | | | | | | | | | | | | | | | | yes | | | | |
| PGC | NA | TSS200 | chr6:41715281-41715593 | 312 | | | | | | | | | | | | | | | yes | | yes | yes | | | | | |
| C6orf132 | S_Shore | TSS200 | chr6:42110740-42110867 | 127 | | | | | | | | | 1 | | | | | yes | | | | | yes | | | | |
| PTCRA | S_Shelf | TSS1500 | chr6:42883504-42883762 | 258 | | | | | 1 | | | | 1 | | | | | yes | | | | | | | | | |
| PTCRA | Island | Body | chr6:42892994-42893167 | 173 | | | | | | | | | | | | | | yes | | | | | | | | | |
| CNPY3 | N_Shore | TSS1500 | chr6:42896394-42896418 | 24 | | | 4 | | | | | | | | | | | yes | yes | | | | | | | | |
| GNMT | N_Shore;Island | TSS1500;TSS200 | chr6:42927940-42928773 | 833 | | | 16 | | 1 | 1 | | 0 | 9 | 4 | | 2 | | yes | | yes | | | | | | | yes |
| KLC4;MRPL2 | Island | TSS1500;5'UTR;TSS200 | chr6:43027568-43027608 | 40 | | | | | | | | | 1 | | | | | | | | | | | | | | |
| PTK7 | N_Shore;Island | TSS1500;1stExon | chr6:43043691-43045052 | 1361 | | | | 1 | | | | | 1 | | | | | | | | | yes | | | | | |
| CUL9 | N_Shore | TSS1500 | chr6:43149356-43149629 | 273 | | | 4 | | | | | | | | | | | | | | | yes | | | | | |
| C6orf108 | S_Shore | TSS1500 | chr6:43197544-43197675 | 131 | | | 3 | | | | | | | | | | | | yes | yes | | | | | | | |
| TTBK1 | Island | 5'UTR | chr6:43214396-43214590 | 194 | | | 2 | | | | | 0 | | | | | | | | | | | | | | | |
| TTBK1 | Island | Body | chr6:43243709-43244304 | 595 | | | 7 | 1 | 1 | | | | | | | | | | yes | | | yes | yes | | | | |
| TTBK1 | N_Shore;Island | Body | chr6:43251485-43253387 | 1902 | | 1 | | 1 | | | | 0 | 1 | | | | | | | | yes | | yes | | | | |
| CRIP3 | S_Shore | TSS1500 | chr6:43276744-43276781 | 37 | | | | | | | | | | | | 1 | | yes | | | | | | | | | |
| TJAP1 | NA | 5'UTR;Body;TSS1500;TSS200 | chr6:43456984-43457511 | 527 | | 1 | 7 | | | | | | | | | | | | | | | | | | | | |
| C6orf154 | Island | TSS200 | chr6:43478455-43478829 | 374 | | | 7 | | | | | 0 | 2 | | | | | | | | | | | | | | |
| RSPH9 | N_Shore | TSS1500;TSS200 | chr6:43611905-43613395 | 1490 | | | 12 | | | | | | 1 | | | | | yes | | | | yes | | | | | |
| VEGFA | N_Shore | TSS1500 | chr6:43736807-43737136 | 329 | | 1 | | | | | | | | | | | | | | | | yes | | | | | |
| CAPN11 | NA | TSS1500 | chr6:44125473-44126075 | 602 | | | 3 | | | | | | 1 | | | | | yes | | yes | | | | | | | |
| SLC29A1 | N_Shore;chr6:44187186-44187 | TSS1500;TSS200;Island;1stExon;5' | chr6:44187031-44187674 | 643 | 5 | | | | | | | | 1 | | 1 | | | yes | | | yes | yes | | | | | |
| NFKBIE | S_Shore | TSS1500 | chr6:44234126-44234511 | 385 | | | | | | | | | 1 | | | | | yes | | | | | | | | | |
| TMEM151B | Island | Body | chr6:44243304-44243750 | 446 | | | | | | | 1 | 0 | | | | 2 | | yes | | | | | yes | | | | |
| TCTE1 | Island | Body | chr6:44254074-44254149 | 75 | | | 1 | | | | | | | | | | | | yes | | | | | | | | |
| AARS2 | Island | TSS200 | chr6:44281184-44281197 | 13 | | | | | | | | | 1 | | | | | | yes | | | yes | | | | | |
| SPATS1 | NA | TSS1500;TSS200 | chr6:44309765-44310445 | 680 | | | | | 5 | | | | 2 | | | | | | yes | yes | | | | | | | |
| RUNX2 | N_Shore;Island | Body;TSS1500 | chr6:45387257-45388583 | 1326 | | | 27 | | | | | | | | | | | | | | | | | | | | |
| RUNX2 | Island;S_Shore | Body | chr6:45391645-45391973 | 328 | | | 11 | | | | | | | | | | | | | | | | | | | | |
| RCAN2 | NA | TSS200 | chr6:46293571-46293862 | 291 | | | | 1 | | | | | | | | | | | yes | | | | | | | | |
| TDRD6 | Island | TSS200;TSS1500 | chr6:46655306-46656612 | 1306 | | | | | | | | | | | | | | | | | | | | | | | |
| PLA2G7 | S_Shore | TSS1500;1stExon;5'UTR;TSS200 | chr6:46703422-46703670 | 248 | | | 14 | | | | | | 3 | | | | | yes | | | | | yes | | | | |
| MEP1A | NA | TSS200 | chr6:46761035-46761090 | 55 | | | 2 | 1 | 1 | | | | | | | | | | yes | | | | yes | | | | |
| GPR116 | NA | TSS1500 | chr6:46923414-46923611 | 197 | | | | | | | | | | | | | | | yes | | yes | | | | | | |
| GPR110 | NA | 5'UTR | chr6:47002973-47010177 | 7204 | | | | | | | | | 1 | | | | | yes | yes | | | | | | | | |
| GPR111 | NA | TSS1500 | chr6:47623886-47624276 | 390 | | | 12 | | | | | | 1 | | | | | | | yes | | | | | | | |
| OPN5 | NA | TSS200 | chr6:47749714-47750042 | 328 | | | 1 | | | | | | 1 | | | | | yes | | | | yes | | | | | |
| C6orf138 | N_Shore;Island;S_Shore | 1stExon;TSS200 | chr6:48035985-48036676 | 691 | | | | | 1 | | | 0 | 7 | 1 | | 2 | | | | | | yes | yes | | | | |
| C6orf141 | Island | 5'UTR;1stExon | chr6:49518449-49518986 | 537 | | | | | | | | | 1 | | | | | | | | | | | | | | |
| CRISP2 | NA | 5'UTR;1stExon;TSS200 | chr6:49681178-49681774 | 596 | | | 9 | | 2 | | | 0 | 10 | 3 | | 7 | | | | | | | | | | | |
| PGK2 | NA | 1stExon;TSS200 | chr6:49754848-49755190 | 342 | | 1 | 11 | | | | | | | | | | | | | | yes | | | | | | |
| DEFB133 | NA | 1stExon;Body | chr6:49916931-49917254 | 323 | | 1 | 11 | | | | | | | | | | | yes | | | | yes | | | | | |
| TFAP2D | N_Shore | Body | chr6:50690590-50691003 | 413 | | 1 | | | | | | | | | | | | | | | | | | | | | |
| TFAP2D | Island | Body | chr6:50692150-50692605 | 455 | | 1 | | | | | | | | | | | | yes | | | | | | | | | |
| TFAP2B | N_Shelf;N_Shore | TSS1500;TSS200 | chr6:50784944-50786670 | 1726 | | | 8 | | | | | | 1 | | | | | | | | | | yes | | | | |
| TFAP2B | NA | Body | chr6:50803820-50804174 | 354 | | | | | | | | | | | | | | yes | | | yes | yes | | | | | |
| TFAP2B | N_Shore;Island | Body | chr6:50810432-50810993 | 561 | | | 12 | | | | | | | | | | | | | | | | | | | | |
| TFAP2B | Island;S_Shore | 3'UTR | chr6:50813341-50814305 | 964 | | 1 | 5 | | | | | | 2 | | | | | | | | | | | | | | |
| PKHD1 | NA | TSS1500 | chr6:51953096-51953258 | 162 | | | | | | | | | 1 | | | | | | yes | | | | | | | | |
| MIR206 | NA | TSS1500;TSS200 | chr6:52008713-52009154 | 441 | | | 6 | | | | | | | | | | | yes | | | | | | | | | |
| GSTA4 | N_Shore | Body | chr6:52858837-52859107 | 270 | | | | | | | | 0 | 2 | | | 1 | | yes | yes | | | | | | | | |

| | | | | | | | | | | | | | | | | | | | | | | | |
|---|---|---|---|---|---|---|---|---|---|---|---|---|---|---|---|---|---|---|---|---|---|---|---|
| GCM1 | NA | 1stExon;5'UTR | chr6:53013589-53014189 | 600 | | | | | | | | | 1 | | | | | yes | | | | | |
| ELOVL5 | Island | TSS200 | chr6:53214008-53214076 | 68 | | | 1 | | | | | | | | | | | | | | | | |
| KLHL31 | NA | 1stExon;5'UTR;TSS200 | chr6:53530503-53530628 | 125 | | | 9 | 1 | | | | | 1 | | | | | | | | | yes | |
| FAM83B | Island | TSS200;TSS1500 | chr6:54711227-54711471 | 244 | | 1 | | | | | | | 1 | | | | | | | | | | yes |
| HCRTR2 | Island | 5'UTR;1stExon | chr6:55039232-55039382 | 150 | | 1 | | | | | | | | | | | | | | | | | yes |
| HMGCLL1 | Island;S_Shore | TSS200 | chr6:55444091-55444488 | 397 | | | | | | | | 0 | 1 | | | | | | | | | | |
| COL21A1 | N_Shore | 5'UTR | chr6:56111812-56112216 | 404 | | | | | | | | | 2 | | | | | | | | yes | | |
| BEND6;DST | S_Shore | 5'UTR;TSS1500 | chr6:56820386-56820869 | 483 | | | 17 | | | | | | | | | | | | | | | | |
| BEND6 | NA | Body | chr6:56878281-56878665 | 384 | | | | | | | | | 1 | 1 | | | | yes | yes | yes | | | |
| KHDRBS2 | Island;S_Shore | 1stExon;5'UTR;TSS200;TSS1500 | chr6:62995876-62996702 | 826 | | 2 | | | 2 | | | 0 | 6 | | | | | | | | | | |
| PTP4A1 | N_Shore | TSS1500 | chr6:64281439-64281771 | 332 | | | 10 | | | | | 0 | | | | | | | | | | | |
| PTP4A1 | NA | Body | chr6:64290014-64290049 | 35 | | | | | | | | | 1 | | | | | | yes | | | | |
| BAI3 | N_Shore | TSS1500 | chr6:69344406-69344423 | 17 | | | | | | | | | 1 | | | | | | | | | yes | yes |
| LMBRD1 | NA | Body | chr6:70500634-70501081 | 447 | | | | 1 | | | | | 1 | | | | | | | | yes | | |
| COL19A1 | N_Shore | TSS1500;TSS200 | chr6:70576168-70576541 | 373 | | | 19 | | | | | | | | | | | | | | | yes | |
| COL9A1 | N_Shore | Body | chr6:70990226-70991283 | 1057 | | | | | | | | 0 | 3 | 1 | | 2 | | | | | | yes | |
| COL9A1 | S_Shore | Body;TSS200 | chr6:70993107-70993440 | 333 | | | | | | | | | | | | | | yes | | | | | |
| COL9A1 | NA | 5'UTR;1stExon;TSS200 | chr6:71012765-71013139 | 374 | | | | | | | | | 1 | | | | | yes | | | | | yes |
| C6orf155 | Island | Body;TSS200;TSS1500 | chr6:72130407-72131020 | 613 | | 1 | | | 2 | | | 0 | 4 | | | 1 | | | | | | | yes |
| RIMS1 | NA | TSS1500;Body | chr6:72924771-72925046 | 275 | 1 | | 1 | | | | | | | | | | | yes | | | | | |
| RIMS1 | NA | TSS1500;TSS200;Body | chr6:72926121-72926485 | 364 | | | 3 | | | | | | | | | | | | | | | | |
| KCNQ5 | N_Shore | TSS1500 | chr6:73330106-73330208 | 102 | | | | | | | | | | | | | | | | | | | |
| KHDC1L | NA | TSS200 | chr6:73935270-73935676 | 406 | | | | 1 | | | | | | | | | | yes | | | | | |
| C6orf147 | N_Shore | Body | chr6:74008688-74009041 | 353 | | | | | | | | | | | | | | yes | | | | | |
| DPPA5 | Island | TSS200 | chr6:74063982-74064594 | 612 | | | 8 | | | | | 0 | 8 | 1 | | 4 | | | yes | | | yes | yes |
| C6orf221 | N_Shore | TSS200 | chr6:74072242-74072447 | 205 | | 1 | 3 | | | | | 0 | 3 | | | 1 | | yes | | | | yes | |
| OOEP | N_Shore;Island | 1stExon;TSS200 | chr6:74079335-74080260 | 925 | | | 3 | | | | | | | | | | | yes | yes | | | | |
| DDX43 | N_Shore;Island | TSS200;1stExon;5'UTR | chr6:74104137-74104868 | 731 | | 1 | 25 | | | 1 | 1 | 0 | 6 | 2 | | 5 | | yes | | | yes | | |
| C6orf150 | Island | 1stExon;5'UTR;TSS200 | chr6:74161911-74162142 | 231 | | | | 1 | | 1 | | | | | | | | yes | | | | | |
| CD109 | Island | 5'UTR;1stExon | chr6:74405811-74406097 | 286 | | | | | 1 | | | | | | | | | | | | | yes | yes |
| COL12A1 | NA | Body | chr6:75884800-75884860 | 60 | | | | 1 | | | | | | | | | | yes | | | | | |
| COL12A1 | Island;S_Shore | Body;5'UTR | chr6:75911716-75912711 | 995 | | | 27 | | | | | | 1 | | | | | | | | | | |
| COL12A1 | N_Shore | 5'UTR | chr6:75913786-75914046 | 260 | | | | | | | | | 1 | | | | | | yes | | | | |
| COL12A1 | Island | TSS200 | chr6:75915749-75915787 | 38 | | | | | | | | | 2 | | | | | | | | | | |
| COX7A2 | NA | TSS1500 | chr6:75953853-75954053 | 200 | | | | | | | | | 1 | | | 2 | | | | | | | |
| FILIP1 | NA | Body | chr6:76059576-76059773 | 197 | | | 3 | | | | | | 2 | | | | | | | | | yes | |
| FILIP1 | NA | 5'UTR;1stExon;TSS200 | chr6:76203225-76204308 | 1083 | | | 1 | | 1 | | | 0 | 11 | | | 2 | | | | | | yes | |
| HTR1B | Island | TSS1500;1stExon | chr6:78172950-78174065 | 1115 | | 1 | | | | | | | 1 | | | | | | | | | | yes |
| LCA5 | Island | 1stExon;5'UTR;TSS200 | chr6:80247037-80247368 | 331 | | | | | | | | 0 | 2 | | | | | | | | | yes | yes |
| ELOVL4 | Island | TSS200 | chr6:80657428-80657436 | 8 | | | | | | | | | 1 | | | | | | | | | | yes |
| TPBG | N_Shore | TSS1500;TSS200 | chr6:83072522-83073156 | 634 | | | | | | | | | 3 | | | | | | yes | | | | |
| UBE2CBP | Island | TSS1500 | chr6:83775763-83775899 | 136 | | | | | | | | | | | | | | yes | | | | | |
| PGM3;RWDD2A | S_Shore | TSS1500;5'UTR | chr6:83903927-83904091 | 164 | | | | | | | | 0 | 2 | | | | | | | | | | |
| ME1 | Island | Body;5'UTR;1stExon;TSS200 | chr6:84140564-84140997 | 433 | | | | | | | | 0 | 1 | 1 | | 2 | | | | | | yes | |
| SNAP91 | Island | TSS200;Body;5'UTR | chr6:84418659-84419360 | 701 | | | | | | | | 0 | 2 | | | | | | | | | | |
| RIPPLY2 | Island | TSS200 | chr6:84562888-84562928 | 40 | | | | | | | | | 1 | | | | | | | | | | |
| MRAP2 | N_Shore | TSS1500 | chr6:84742695-84743236 | 541 | | | 11 | | | | | 0 | | | | | | yes | yes | | | | yes |
| TBX18 | Island | TSS200 | chr6:85474028-85474093 | 65 | | | | | | | | | 1 | | | | | | | | | yes | yes |
| NT5E | N_Shore | TSS1500 | chr6:86158972-86159210 | 238 | | 1 | | | | | | | 2 | | | | | | yes | | | yes | |
| HTR1E | N_Shore;Island | 1stExon;5'UTR;TSS1500;TSS200 | chr6:87646738-87647644 | 906 | | 4 | | | 3 | | | | | | | | | | yes | | | | yes |
| CGA | NA | 1stExon;5'UTR | chr6:87804765-87805033 | 268 | | | | | | | | | 1 | | | | | | | | | | |
| SPACA1 | Island | TSS1500;TSS200 | chr6:88757302-88757878 | 576 | | | 20 | | | | | 0 | 8 | 1 | | 7 | | yes | | | | | |
| CNR1 | NA | 5'UTR;TSS200 | chr6:88855255-88855716 | 461 | | | | | | | | | | | | | | yes | | | | | |
| GABRR2 | NA | TSS200 | chr6:90025032-90025136 | 104 | | | | | | | | | | | | | | | | | | | |
| GJA10 | NA | TSS1500 | chr6:90603803-90604299 | 496 | | | | | | | | | | | | | | | | | | | |
| BACH2 | NA | Body | chr6:90660913-90661242 | 329 | | | | | | | | | | 1 | | | | | | | | | |
| BACH2 | N_Shore | 5'UTR | chr6:91004430-91004482 | 52 | | 1 | 12 | | 1 | | | | | | | | | | | | | | yes |
| BACH2 | S_Shore | TSS1500 | chr6:91007365-91007625 | 260 | | | | | | | | | 1 | | | | | | | | | | |
| FUT9 | Island | 5'UTR;1stExon | chr6:96463902-96464135 | 233 | | | | | 1 | | | | 1 | | | | | | | | | | yes |
| POU3F2 | Island | TSS1500 | chr6:99281500-99281818 | 318 | | | | | | | | | 1 | | | | | | | | | | |
| POU3F2 | Island | 1stExon | chr6:99282887-99283553 | 666 | | | | | | | | | 1 | | | | | | | | | yes | yes |
| C6orf168 | NA | Body | chr6:99790960-99791250 | 290 | | | | | | | | | | | | | | yes | | | | | |
| COQ3 | S_Shore | TSS200;TSS1500 | chr6:99842241-99842368 | 127 | | | 7 | | | | | | 1 | | | | | yes | | | | | |
| PRDM13 | N_Shore;Island;S_Shore | TSS1500;TSS200;Body | chr6:100054008-100057243 | 3235 | | 2 | | | | | | 0 | 2 | | | | | yes | | | | | |
| MCHR2 | Island | 5'UTR;1stExon;TSS200 | chr6:100441407-100442489 | 1082 | | 1 | | | | | | 0 | 3 | 1 | | 4 | | yes | | | | | |
| SIM1 | N_Shore;Island | Body | chr6:100894513-100897445 | 2932 | | | | | | | | | | | | | | | | | | | |
| SIM1 | N_Shore;Island | Body | chr6:100903227-100904376 | 1149 | | 1 | 6 | | | | | 0 | 1 | | | | | | | | | | |
| SIM1 | Island | Body | chr6:100906182-100906448 | 266 | | | | | | | | | | | | | | | | | | yes | |
| SIM1 | N_Shore;Island | 1stExon;5'UTR;TSS200;TSS1500 | chr6:100911240-100913250 | 2010 | | 2 | 58 | | | | | | 3 | | | | | | | | | | |
| GRIK2 | N_Shore;Island;S_Shore | TSS1500;TSS200;1stExon;5'UTR;Bo | chr6:101846409-101848236 | 1827 | | 2 | 70 | | | 6 | | 0 | 5 | 2 | | | | | | | | | |
| GRIK2 | Island | Body | chr6:101850881-101851354 | 473 | | | | | | | | | | | | | | | | | | | |
| POPDC3 | S_Shore | TSS200 | chr6:105628000-105628063 | 63 | | | | | | | | | 2 | | | | | yes | | | | yes | |
| PRDM1 | NA | Body;TSS200;TSS1500 | chr6:106545984-106546824 | 840 | | | 29 | 1 | | | | 0 | 6 | 1 | 1 | 7 | | | yes | | | | |
| ATG5 | S_Shore | TSS1500 | chr6:106773949-106774172 | 223 | | | | | | | | | 1 | | | | | yes | | | | | |
| SCML4 | NA | Body | chr6:108093229-108093680 | 451 | | | | | | | | | | | | | | | | | | | |
| SCML4 | NA | 1stExon;5'UTR;nan;TSS200 | chr6:108145374-108145930 | 556 | | | 4 | | 1 | | | | 2 | | 1 | | | | | | | yes | |
| NR2E1 | Island | TSS1500;TSS200 | chr6:108485837-108487100 | 1263 | | 1 | | | | | | | 1 | | | | | | | | | | |
| NR2E1 | Island | Body | chr6:108488335-108489292 | 957 | | | 7 | | 1 | | | | | | | | | | | | | | yes |
| NR2E1 | S_Shore | Body | chr6:108492358-108492769 | 411 | | | 3 | | | | | | | | | | | | | | | | yes |
| NR2E1 | N_Shore;Island | Body | chr6:108495385-108497734 | 2349 | | 2 | | | 1 | | | | | | | | | yes | | | yes | | |
| SNX3 | S_Shore | TSS1500 | chr6:108582894-108583071 | 177 | | | | | | | | | 1 | | | | | yes | | | | | |
| ARMC2 | NA | Body | chr6:109274429-109274496 | 67 | 11 | | 30 | | | | | | | | | | | | yes | | | | |
| CCDC162 | S_Shelf | TSS200 | chr6:109615419-109615579 | 160 | | | | | | | | | 1 | | | | | yes | | | | | |
| GPR6 | N_Shore;Island | TSS1500;TSS200;1stExon | chr6:110299151-110301479 | 2328 | | 1 | 3 | | | | | | 2 | | | | | yes | | | | yes | |
| C6orf186 | N_Shore | Body | chr6:110677913-110678179 | 266 | | | 7 | | | | | 0 | 1 | | | | | yes | | | | | |
| DDO | N_Shore;Island | Body | chr6:110720501-110721629 | 1128 | 1 | | 12 | | | | 1 | 0 | 3 | 2 | 1 | 17 | | | | | yes | | |
| DDO | NA | TSS200 | chr6:110736772-110737053 | 281 | | | 4 | | | | | 0 | 3 | | | 3 | | | | | yes | | |
| SLC22A16 | Island | Body;5'UTR;1stExon | chr6:110797476-110798022 | 546 | 1 | | | 1 | | | | | | | | | | yes | | | | | yes |
| SLC16A10 | Island | TSS200 | chr6:111408761-111408765 | 4 | | | | | | | | | | | | | | | | | | | yes |
| TRAF3IP2 | NA | Body;5'UTR;1stExon;TSS200 | chr6:111887834-111888840 | 1006 | | | 13 | 1 | 2 | | | | 1 | | | | | yes | yes | | | | |
| TRAF3IP2 | NA | 1stExon;Body;5'UTR | chr6:111927312-111927480 | 168 | 1 | | | | | | | | | | | | | yes | | | | yes | |
| FYN | NA | 5'UTR;TSS1500 | chr6:112081102-112081538 | 436 | | | | 1 | 1 | | | | 1 | | | | | yes | yes | | | | |
| FYN | NA | 5'UTR | chr6:112115117-112115209 | 92 | | | | 1 | | | | | 1 | 1 | | | | | | | | | |
| WISP3 | NA | TSS200;1stExon;5'UTR | chr6:112375186-112375333 | 147 | | | | | | | | | 3 | 1 | | | | yes | | | | | |
| LAMA4 | S_Shore | TSS200 | chr6:112575850-112576393 | 543 | | | 5 | 1 | | | | 0 | 3 | | | | | yes | | | | | yes |
| RFPL4B | NA | 5'UTR;Body | chr6:112670885-112671523 | 638 | | | | | | | | | | | | | | yes | | | | | yes |
| MARCKS | Island | Body | chr6:114181482-114181798 | 316 | | | 2 | | | | 1 | 0 | 1 | 1 | | 3 | | | | | | yes | yes |
| HDAC2 | Island | Body;TSS1500 | chr6:114291911-114292751 | 840 | | | 2 | | | | | | | | | | | yes | | | | yes | yes |
| FRK | NA | 1stExon;5'UTR;TSS200;nan | chr6:116381609-116382179 | 570 | | | 8 | | | | | 0 | 4 | | 1 | 1 | | | | | | yes | yes |

| | | | | | | | | | | | | | | | | | | | | | | | |
|---|---|---|---|---|---|---|---|---|---|---|---|---|---|---|---|---|---|---|---|---|---|---|---|
| DSE | N_Shore | TSS1500;5'UTR | chr6:116691382-116691685 | 303 | | | | | | | | 0 | 1 | | | | | | | | | | |
| FAM162B | Island;S_Shore | Body;TSS200 | chr6:117086040-117087083 | 1043 | | 1 | 1 | | | | | | 1 | | | | | yes | yes | | | | |
| RFX6 | Island | TSS200 | chr6:117198183-117199067 | 884 | | | | | | | | 0 | 2 | | | | | | | | | yes | |
| VGLL2 | Island | TSS200 | chr6:117586538-117587588 | 1050 | | 1 | | | | | | | | | | | | | | | | yes | |
| VGLL2 | N_Shore | Body | chr6:117591433-117592262 | 829 | | 1 | | | | | | | | | | | | | | | yes | yes | yes |
| DCBLD1 | N_Shore | TSS1500 | chr6:117802401-117803162 | 761 | | | 6 | 1 | | | | 0 | 2 | | | 1 | | | | | | yes | |
| DCBLD1 | N_Shore;Island | Body | chr6:117869027-117869857 | 830 | | 1 | 11 | | | | | 0 | 3 | | | 1 | | yes | | | | yes | |
| GOPC | Island | TSS200 | chr6:117923795-117923963 | 168 | | | | | | | | | | | | 1 | | yes | | | yes | | |
| SLC35F1 | Island | TSS200 | chr6:118228493-118228514 | 21 | | | 1 | | | | | | | | | | | | | | | yes | yes |
| SLC35F1 | N_Shore | Body | chr6:118241243-118241568 | 325 | | 1 | | | | | | | | | | | | | | | yes | yes | |
| C6orf204 | Island;S_Shore | TSS200;TSS1500 | chr6:118973136-118973597 | 461 | | | 3 | | | | | 0 | 1 | 1 | | 2 | | | | | | | yes |
| FAM184A;MIR548B | NA | 5'UTR;Body;TSS200 | chr6:119390418-119390712 | 294 | | | | | | | | | 1 | | | | | | | | yes | | yes |
| MAN1A1 | N_Shore | Body | chr6:119669112-119669372 | 260 | | | | | | 1 | | | | | | | | | | | | | |
| PKIB | Island | 5'UTR;TSS1500 | chr6:122931154-122931192 | 38 | | | 1 | | | | | | | | | | | | | | | | |
| FABP7 | NA | TSS1500;TSS200 | chr6:123100189-123100722 | 533 | | | 4 | | | | | 0 | 1 | | | | | | yes | | | | |
| SMPDL3A | N_Shore | TSS1500 | chr6:123109391-123109903 | 512 | | | 3 | | | | | 0 | 1 | | | | | | | yes | yes | | |
| CLVS2 | Island | TSS1500;TSS200 | chr6:123317184-123317875 | 691 | | 1 | | | | | | 0 | 3 | | | | | | | | | yes | |
| STL | Island | Body;TSS200 | chr6:125283245-125284250 | 1005 | | 1 | | | | | | | 2 | | | 1 | | | | | | yes | yes |
| TPD52L1 | S_Shore | 5'UTR;Body | chr6:125476162-125476407 | 245 | | | | | | | | | | | | | | yes | yes | | yes | | yes |
| HDDC2 | S_Shore | TSS200 | chr6:125623444-125623573 | 129 | | | | | | | | | | | | | | | yes | yes | | | yes |
| HEY2 | Island;N_Shore | Body | chr6:126080132-126080723 | 591 | | | | | | | | | 2 | 1 | | | | | | | | | |
| RSPO3 | Island | Body | chr6:127441605-127441993 | 388 | | | 11 | | | | | | | | | | | | | | | | |
| ECHDC1 | N_Shore | 5'UTR;Body | chr6:127663200-127663627 | 427 | | | | | | | | | | | | | | | | | | | |
| C6orf174 | Island | Body | chr6:127796287-127797286 | 999 | | | 20 | 1 | 1 | | | 0 | 7 | 1 | | 3 | | yes | | | | | |
| C6orf174 | N_Shore;Island | Body | chr6:127835750-127836159 | 409 | | | 2 | | | | | 0 | | | | 1 | | | | | | | |
| C6orf174 | S_Shelf | 5'UTR;1stExon;TSS200 | chr6:127839822-127840755 | 933 | | 2 | | | | | | | | | | | | | | | | | yes |
| THEMIS | NA | 5'UTR;1stExon | chr6:128222053-128222390 | 337 | | | | | | | | | 1 | | | 1 | | | | | | | |
| ARHGAP18 | NA | TSS200 | chr6:130031504-130031588 | 84 | | 1 | | | | | | | | | | | | yes | | | | | |
| EPB41L2 | S_Shore | TSS200 | chr6:131384607-131384652 | 45 | | 1 | | | | | | | | | | | | | | | | | yes |
| TAAR9 | NA | TSS1500 | chr6:132858438-132858597 | 159 | | | | | | | | | 1 | | | | | | | | | yes | yes |
| TAAR6 | NA | TSS1500 | chr6:132890978-132891505 | 527 | | | | | | | | | | | | | | | | | | | |
| TAAR3 | NA | TSS1500 | chr6:132931714-132931926 | 212 | | | 4 | | | | | | | | | | | | yes | | | | |
| TAAR2 | NA | TSS200 | chr6:132945515-132945592 | 77 | | | | | | | | | | | | | | yes | | | yes | | |
| VNN1 | NA | 1stExon | chr6:133035150-133035379 | 229 | | | 10 | | 1 | | | | 1 | | 1 | | | yes | yes | yes | | | |
| VNN1 | NA | TSS1500 | chr6:133036545-133036651 | 106 | | | 6 | | | | | | | | 1 | | | | yes | yes | | | yes |
| RPS12;SNORD100;SNORA33 | S_Shore | Body;TSS200;TSS1500 | chr6:133137167-133137893 | 726 | | | 3 | | | | | | | | | | | yes | | | | | |
| EYA4 | N_Shore;TSS1500;chr6:133562 | TSS1500;nan;N_Shore;TSS200 | chr6:133561224-133562776 | 1552 | | 2 | 14 | | 1 | | | 0 | 9 | | 1 | 5 | | yes | | | | | yes |
| EYA4 | S_Shore | 5'UTR | chr6:133564466-133564578 | 112 | | | 2 | | | | | | | | | | | | | | | yes | |
| EYA4 | NA | 3'UTR | chr6:133851427-133851542 | 115 | | | 1 | | | | | | | | | | | | | | | | |
| MGC34034 | N_Shore;Island | Body | chr6:134158982-134159412 | 430 | | | | | | | | | | | | | | yes | | | | yes | |
| TCF21 | N_Shore;Island | TSS1500;TSS200;1stExon;5'UTR | chr6:134209935-134211217 | 1282 | | 1 | 25 | | | | | | | | | 1 | | | | | | yes | |
| TCF21 | Island | 3'UTR | chr6:134213992-134214649 | 657 | | | 7 | | | | | | | | | | | yes | | | | | |
| SGK1 | NA | 3'UTR;Body | chr6:134491143-134491531 | 388 | | | | | 3 | | | | 1 | | | | | | | | | | |
| SGK1 | N_Shore;Island | Body;TSS200;TSS1500 | chr6:134497230-134497755 | 525 | | | 10 | | 1 | | | 0 | | | | | | yes | | | | | yes |
| SGK1 | S_Shore | Body;1stExon;5'UTR;TSS200;TSS1! | chr6:134498979-134499519 | 540 | 1 | 1 | | | 1 | | | | | | | | | | | | | | |
| SGK1 | S_Shore | TSS200 | chr6:134639355-134639482 | 127 | | | | | | | | | | | | 1 | | yes | | | | yes | yes |
| ALDH8A1 | NA | 1stExon;TSS200;TSS1500 | chr6:135271056-135271801 | 745 | | | 5 | 1 | | | | | 1 | | | | | yes | yes | | | | |
| MYB | NA | nan;Body | chr6:135516912-135517046 | 134 | | | | | | | | | 1 | | 1 | | | | | | | | |
| MAP7 | Island | TSS200;TSS1500 | chr6:136871907-136872552 | 645 | | | 5 | | | | | 0 | 3 | | | | | | | | | | |
| MAP3K5 | NA | Body | chr6:136915088-136915556 | 468 | | | | 1 | 1 | | | | | | | | | | yes | yes | yes | yes | |
| IL20RA | S_Shore | TSS200 | chr6:137366322-137366492 | 170 | | | | | | | | | 2 | 1 | | | | yes | | | yes | yes | |
| IL22RA2 | NA | 1stExon;5'UTR;TSS200 | chr6:137494767-137495383 | 616 | | | | | | | | | | | | | | yes | yes | | | | yes |
| OLIG3 | N_Shore;Island | 1stExon;3'UTR;TSS1500 | chr6:137813584-137816623 | 3039 | | 1 | 4 | | | | | | 2 | | | | | | | | | yes | |
| TNFAIP3 | NA | Body | chr6:138200071-138200387 | 316 | | | 10 | 1 | | | | | | | | | | | | | | | |
| PERP | S_Shore | TSS200 | chr6:138428833-138428906 | 73 | | | | | | | | | | | | | | | yes | | | yes | yes |
| HEBP2 | Island | Body | chr6:138725799-138726184 | 385 | | | | | | | | 0 | 1 | | | | | | | | | | |
| NHSL1 | NA | Body | chr6:138785358-138785472 | 114 | | | | | | | | | | | | | | yes | yes | | | | |
| NHSL1 | NA | Body;nan | chr6:138866865-138867125 | 260 | 1 | 1 | 2 | | | | | | | | 1 | | | yes | | | | | |
| NHSL1 | NA | 1stExon;5'UTR;TSS200 | chr6:138892911-138893718 | 807 | | | | | | | | 0 | 2 | | | 2 | | | | | | yes | yes |
| C6orf115 | S_Shore | 5'UTR | chr6:139351113-139351353 | 240 | | | 7 | | | | | | | | | | | | yes | | | | |
| TXLNB | NA | 5'UTR;1stExon | chr6:139613145-139613456 | 311 | | | 11 | 1 | 1 | | | | | | | | | | | | | yes | |
| NMBR | Island;S_Shore;N_Shore | TSS200;TSS1500;Body;5'UTR;1stE | chr6:142409202-142410235 | 1033 | | 2 | | | 1 | | | | 2 | | | 1 | | | yes | | | | yes |
| FUCA2 | Island | TSS200 | chr6:143832942-143833011 | 69 | | | | | | | | | | 1 | | | | yes | | | yes | yes | |
| PHACTR2 | NA | TSS200 | chr6:143929300-143929511 | 211 | | | | | | | | | 1 | | | | | | | | | | |
| PLAGL1;HYMAI | S_Shore | TSS1500;5'UTR;TSS200 | chr6:144329962-144330345 | 383 | | | | | | | | | 3 | | | | | yes | yes | | | | |
| PLAGL1 | S_Shore | TSS1500 | chr6:144386093-144386848 | 755 | | | 19 | 1 | 1 | | | 0 | 3 | | | 1 | | yes | | | yes | | |
| UTRN | NA | Body | chr6:145069578-145069599 | 21 | | | | | | | | | 1 | | | | | | yes | | | | |
| GRM1 | N_Shore;Island | 5'UTR;TSS200;1stExon | chr6:146348690-146350748 | 2058 | | 2 | | | 3 | | | 0 | 1 | | | 1 | | | | | | yes | yes |
| GRM1 | Island | 3'UTR;Body | chr6:146755494-146755900 | 406 | | 1 | | | | | | | | | | | | | | | | yes | |
| RAB32 | N_Shore;Island | TSS1500;TSS200;1stExon;5'UTR | chr6:146863543-146865263 | 1720 | | | 11 | 1 | | | | 0 | 9 | 1 | | 2 | | yes | yes | | | yes | yes |
| C6orf103 | N_Shore;Island | TSS1500;TSS200 | chr6:146919633-146920295 | 662 | | | | | | | | 0 | 1 | | | 1 | | yes | yes | | | | |
| C6orf103;LOC729176 | NA | Body;TSS200 | chr6:147124881-147125402 | 521 | 1 | | 10 | | | | | 0 | 6 | | | 2 | | | yes | yes | | | |
| SASH1 | NA | Body | chr6:148869461-148869895 | 434 | | | 16 | | | | | | | | | | | | yes | | | | |
| UST | NA | Body | chr6:149358743-149358802 | 59 | | | 7 | | | | | | | | | | | yes | | | yes | | |
| ZC3H12D | N_Shore;Island | 3'UTR;Body | chr6:149771807-149772892 | 1085 | | | 12 | | | | | | 3 | | | 1 | | | | | yes | | |
| ZC3H12D | NA | 5'UTR | chr6:149803087-149803444 | 357 | | 1 | 13 | | | | | | 1 | | | | | yes | | | | | |
| ZC3H12D | NA | 5'UTR;1stExon;TSS200;TSS1500 | chr6:149805995-149806732 | 737 | 1 | 2 | 44 | | 5 | | | 0 | 2 | 1 | | | | yes | yes | | | | yes |
| NUP43 | S_Shore | TSS200 | chr6:150067846-150068017 | 171 | | | | | | | | | | | | | | yes | | | | | |
| ULBP1 | N_Shore | TSS1500 | chr6:150284518-150284777 | 259 | | | | | | | | | | | | | | | | | | | |
| ULBP1 | Island | Body | chr6:150286157-150286646 | 489 | | | 2 | | | | | | | | | | | | | | | | |
| RAET1L | NA | Body;TSS200;TSS1500 | chr6:150346497-150347053 | 556 | | | | | | | | 0 | 6 | 2 | | | | yes | yes | | | | |
| IYD | NA | TSS200 | chr6:150689918-150690143 | 225 | | | | 1 | | | | | | | | | | yes | | yes | | yes | |
| PLEKHG1 | N_Shore | TSS1500 | chr6:150920223-150920533 | 310 | | | | | 1 | | | | | | | | | yes | | | | yes | yes |
| MTHFD1L | N_Shore | TSS1500 | chr6:151186411-151186511 | 100 | | | 3 | | | | | | | | | | | yes | | | yes | yes | yes |
| MTHFD1L | NA | Body | chr6:151258337-151258520 | 183 | | | | | | | | | 1 | | | 1 | | yes | | | | | |
| AKAP12 | Island | TSS1500;TSS200 | chr6:151560831-151561055 | 224 | | | 21 | | | | | | | | | 1 | | yes | | | | | |
| AKAP12 | N_Shore | TSS200;Body;TSS1500 | chr6:151646312-151647133 | 821 | | | | | 1 | | | 0 | 4 | | | 1 | | | | | | yes | |
| C6orf97 | N_Shore | TSS1500 | chr6:151814892-151814932 | 40 | | | | | | | | | 1 | | | | | yes | | | yes | | yes |
| ESR1 | NA | TSS1500 | chr6:152011103-152011666 | 563 | | | 14 | | | | | 0 | | | | | | | | | | | yes |
| ESR1 | NA | 5'UTR | chr6:152085565-152085641 | 76 | | 1 | 20 | 1 | | | | | | | | | | | | | yes | | |
| ESR1 | N_Shelf;N_Shore;Island;S_Sho | 5'UTR;TSS1500;TSS200;1stExon;B | chr6:152125861-152130332 | 4471 | | 1 | 102 | 2 | | | | 0 | 5 | 1 | | 3 | | yes | | | | yes | yes |
| SYNE1 | NA | Body | chr6:152702330-152702660 | 330 | | | | | 2 | | | | 1 | 1 | 1 | | | | | | | | |
| SYNE1 | S_Shore | TSS1500;TSS200 | chr6:152958548-152958624 | 76 | | | | | | | | | 1 | | | | | | | | | | yes |
| OPRM1 | Island;N_Shore | 1stExon;5'UTR;Body;TSS200;TSS1! | chr6:154360344-154360894 | 550 | | 1 | | | | | | | 2 | | | | | | | | | | |
| IPCEF1 | NA | 5'UTR | chr6:154677796-154678326 | 530 | | | | | | | | 0 | 1 | | | | | yes | | | | yes | |
| TIAM2 | N_Shelf | Body;TSS200 | chr6:155537901-155538047 | 146 | 2 | | | | | | | 0 | 2 | | | | | | | | | | |
| TIAM2 | NA | Body | chr6:155568918-155569443 | 525 | | | | | | | | | 1 | | | | | yes | | | | | |
| TIAM2 | NA | Body | chr6:155577805-155578034 | 229 | | | 1 | | | | | | | | | | | | | | | yes | |
| TFB1M;CLDN20 | NA | Body;TSS1500 | chr6:155584613-155584968 | 355 | | 2 | | | | | | | | | | | | | | | | | |

| | | | | | | | | | | | | | | | | | | | | | | | |
|---|---|---|---|---|---|---|---|---|---|---|---|---|---|---|---|---|---|---|---|---|---|---|---|
| NOX3 | NA | 1stExon;5'UTR | chr6:155776951-155777175 | 224 | | | | | | | | | 2 | | | | | yes | | | | | |
| ARID1B | NA | Body | chr6:157505432-157505631 | 199 | | | | | 1 | | | | | | | | | | | | | | |
| C6orf35 | S_Shore | TSS1500 | chr6:157746584-157746724 | 140 | | | | | | | | | 1 | | | | | yes | | | yes | yes | |
| ZDHHC14 | NA | Body | chr6:157931791-157932180 | 389 | | | | | 1 | | | 0 | 2 | | | | | yes | yes | | | | |
| ZDHHC14 | NA | Body | chr6:157982145-157982358 | 213 | | | | | 1 | | | | 1 | | | | | yes | yes | | | | |
| ZDHHC14 | N_Shore;Island | Body | chr6:158013621-158014655 | 1034 | | | | | | | | | 5 | | | | | | yes | | yes | | |
| ZDHHC14 | S_Shelf | Body | chr6:158018055-158018184 | 129 | | | 3 | | | | | | | | | | | yes | | | | yes | |
| SYNJ2 | Island;S_Shore | Body | chr6:158403321-158404121 | 800 | 3 | | 5 | | | | | | 1 | | | | | yes | | | | | yes |
| SYNJ2 | NA | Body | chr6:158447877-158448578 | 701 | 1 | | | | | | | | 2 | | | | | yes | yes | | | | |
| SYNJ2 | NA | Body | chr6:158464617-158464852 | 235 | | | 8 | | | | | | | | | | | | | | | | |
| SYNJ2 | NA | Body | chr6:158487541-158487608 | 67 | 4 | | | | | | | | | | | | | yes | | | | | |
| SYNJ2 | chr6:158507719-158508126;Isl | Island;Body | chr6:158507753-158508188 | 435 | | | | | | | | | 1 | 1 | 1 | 1 | | | | | | | |
| TULP4 | NA | Body | chr6:158834153-158834216 | 63 | | | | | | | | | 1 | | | | | | | | | | |
| TMEM181 | NA | Body | chr6:159052400-159052937 | 537 | | | | | | | | | 1 | 1 | | | | yes | | | | | |
| SYTL3 | NA | TSS1500;TSS200 | chr6:159070607-159071106 | 499 | | | 25 | 1 | 1 | | | | 1 | | | | | | | yes | yes | | |
| SYTL3 | NA | Body | chr6:159084370-159084599 | 229 | | | 5 | | | | | 0 | 1 | | | | | | | | | | |
| EZR | N_Shore | Body | chr6:159238463-159238744 | 281 | | | 4 | | | | | 0 | | | | | | yes | | | | | |
| TAGAP | NA | 5'UTR;Body | chr6:159462964-159463300 | 336 | | | | | 1 | | | | 1 | | | | | | | | | | |
| TAGAP | NA | 1stExon;5'UTR | chr6:159466054-159466427 | 373 | | | | | 1 | | | | | | | | | yes | yes | | | yes | |
| FNDC1 | Island | TSS1500 | chr6:159589874-159590231 | 357 | | 1 | 3 | | | | | | 1 | | | | | yes | | | | | yes |
| FNDC1 | NA | Body | chr6:159639257-159639549 | 292 | | | | | | | | | 1 | | | | | | | | | | |
| ACAT2 | Island;N_Shore | TSS1500;TSS200 | chr6:160182184-160183769 | 1585 | | | 13 | | | 1 | 1 | 0 | 3 | 2 | 1 | 15 | 1 | | yes | | | yes | yes |
| TCP1;MRPL18 | Island | TSS1500;TSS200;Body | chr6:160211355-160212070 | 715 | | | | | | | | | 1 | | | 1 | | | yes | | | | |
| PNLDC1 | N_Shore;Island | TSS1500;TSS200;5'UTR;1stExon | chr6:160221026-160221822 | 796 | | | | | | | | | 2 | | | | | | | | | | |
| PNLDC1 | NA | Body | chr6:160241105-160241556 | 451 | | 1 | | | | | | 0 | 3 | | | 13 | | | yes | | | | |
| IGF2R | N_Shore | TSS1500 | chr6:160389454-160389717 | 263 | | | | | | | | | | | | | | yes | | | | | yes |
| IGF2R | NA | Body | chr6:160500620-160500971 | 351 | | | | 1 | | | | | | | | | | | | | yes | | |
| IGF2R | Island | Body | chr6:160511937-160512464 | 527 | | 1 | | | | | | | | | | | | | | yes | | | |
| SLC22A2 | N_Shore;S_Shore | Body;1stExon;5'UTR;TSS200 | chr6:160678936-160680213 | 1277 | | | 12 | | | | | 0 | 1 | | | | | yes | | yes | | | |
| SLC22A3 | Island | TSS1500 | chr6:160768801-160769754 | 953 | | | | | | | | 0 | 1 | | | | | yes | | | | yes | yes |
| SLC22A3 | Island | Body | chr6:160783785-160784385 | 600 | | | 3 | | | | | 0 | 1 | | | | | | | | | | |
| LPA | NA | 5'UTR;1stExon | chr6:161086715-161087573 | 858 | | | | | | | | | 1 | | | | | yes | yes | | | yes | yes |
| AGPAT4 | NA | Body | chr6:161558075-161558105 | 30 | | | 1 | | | | | | | | | | | | yes | | | | |
| AGPAT4 | NA | Body | chr6:161560605-161561121 | 516 | 2 | 2 | 1 | 1 | | | | | 1 | | | | | | | | | | |
| C6orf59;AGPAT4 | NA | TSS1500;Body | chr6:161583481-161583530 | 49 | | | | | | | | | | | | | | yes | | yes | | | |
| PARK2 | NA | Body | chr6:161796785-161796854 | 69 | 2 | 1 | 22 | 1 | | | | | | | | | | | | | | | |
| PARK2 | NA | Body | chr6:161860203-161860635 | 432 | | 1 | 9 | | | | | 0 | 1 | | | | | yes | | | | | |
| PACRG;PARK2 | N_Shore | TSS1500;Body;TSS200 | chr6:163147733-163148022 | 289 | | | 7 | | | | | | 2 | | | | | | | | | | yes |
| PACRG;PARK2 | S_Shore | 5'UTR;1stExon;TSS1500 | chr6:163149167-163149453 | 286 | | | 2 | | | | | | | | | | | | | | | | yes |
| PACRG | NA | Body | chr6:163275952-163276038 | 86 | | 1 | | | | | | | 1 | | | | | yes | | | | | |
| PACRG | NA | nan | chr6:163289514-163289702 | 188 | | | | | | | | | | | 1 | | | yes | | | | | |
| PACRG | Island | Body | chr6:163521371-163521608 | 237 | | | | | | | | | | | | | | | | | | | |
| PACRG | Island | Body | chr6:163570371-163570725 | 354 | | | | | | | | | | | | | | yes | | | | yes | |
| PACRG | NA | Body | chr6:163602881-163602929 | 48 | | | | 1 | | | | | 1 | | | | | yes | | | | yes | |
| PACRG | NA | Body | chr6:163612896-163613008 | 112 | | | | | | | | | 1 | | | | | yes | | | | | |
| PACRG | NA | Body | chr6:163661301-163661641 | 340 | | | 2 | | | | | | | | | | | | | | | yes | |
| PACRG | NA | Body | chr6:163730796-163730853 | 57 | | | | | | | | 0 | 1 | | | 1 | | | | | | | |
| LOC285796 | NA | TSS1500 | chr6:163746113-163746175 | 62 | | | | | | | 1 | 0 | 1 | 1 | | | | yes | | | | | |
| C6orf118 | NA | Body | chr6:165714794-165715757 | 963 | | | 10 | | | | | | | | | | | | | | | | |
| PDE10A | Island | 1stExon;5'UTR | chr6:166075453-166075520 | 67 | | | | | | | | 0 | | | | | | | | | | yes | yes |
| C6orf176 | Island | Body | chr6:166354015-166354232 | 217 | | | 1 | | | | | | | | | | | yes | | | | | |
| T | N_Shore | 3'UTR | chr6:166571782-166571880 | 98 | | | | | | | | | | | | | | yes | | | | | |
| T | N_Shore | Body | chr6:166579057-166579199 | 142 | | 1 | | | | | | 0 | 1 | | | 1 | | | | | | | |
| T | Island;chr6:166579973-166583 | 1stExon;5'UTR;Island;TSS200 | chr6:166581929-166582393 | 464 | | 1 | | | | | | | 1 | | 1 | | | | | | | yes | |
| PRR18 | Island | TSS200 | chr6:166722053-166722060 | 7 | | | | | | | | | 1 | | | | | | | | | | yes |
| RPS6KA2 | NA | Body | chr6:166836643-166836791 | 148 | | 1 | | | | | | | | | | | | | yes | | | | |
| RPS6KA2 | NA | Body | chr6:166856056-166856094 | 38 | | | | | | | | 0 | 1 | 1 | | | | yes | | | yes | | |
| RPS6KA2 | NA | Body | chr6:166876585-166877038 | 453 | | | | | | | | 0 | 4 | 1 | 1 | | | | | | | | |
| RPS6KA2 | NA | Body | chr6:166892362-166892762 | 400 | | | | | | | | | | | | | | yes | | | | | |
| RPS6KA2 | NA | Body | chr6:166901788-166901854 | 66 | | | | 1 | 1 | | | | | | | | | | | | | | |
| RPS6KA2 | NA | Body | chr6:166908677-166908811 | 134 | 1 | | | | | | | | | | | | | yes | | | | | |
| RPS6KA2 | NA | Body | chr6:167062104-167062226 | 122 | | | 1 | | | | | | | | | | | | yes | | | yes | |
| RPS6KA2 | NA | Body | chr6:167104601-167104926 | 325 | | | | | | | | | 1 | | | | | yes | | | | | |
| RPS6KA2 | NA | Body | chr6:167178233-167178280 | 47 | | | | | | | | | 1 | | | | | | | | | | |
| RPS6KA2 | NA | Body | chr6:167190034-167190226 | 192 | | | 1 | | | | | | | | | | | | yes | | | | |
| RPS6KA2 | NA | Body | chr6:167250429-167250541 | 112 | | 1 | | | | | | | | | | | | | yes | | | | |
| RPS6KA2 | NA | Body | chr6:167263025-167263454 | 429 | | | 3 | | 1 | | | | 2 | | | | | | yes | | | yes | |
| CCR6 | NA | TSS1500 | chr6:167523797-167524238 | 441 | | | 1 | | | | | | | | | | | yes | yes | | | | |
| CCR6 | NA | 5'UTR | chr6:167529678-167530343 | 665 | | | 4 | | | | | | 1 | | | | | | | | yes | | yes |
| CCR6 | NA | TSS1500;5'UTR;TSS200 | chr6:167535171-167536521 | 1350 | | | 10 | | 2 | | | | 1 | | | | | yes | | yes | | | |
| GPR31 | NA | TSS200;TSS1500 | chr6:167571324-167571803 | 479 | | | | | | | | | 1 | | | | | | yes | | | | |
| UNC93A | NA | TSS200 | chr6:167704673-167705253 | 580 | | | 8 | 1 | | | | | | | | | | yes | | | yes | | |
| TTLL2 | NA | TSS1500 | chr6:167737937-167738425 | 488 | | | | | | | | | | | | | | yes | yes | | | | |
| TCP10 | NA | Body | chr6:167789195-167789539 | 344 | | | | | | | | | | | | | | | | | | | |
| C6orf123 | Island;N_Shore | Body;TSS200 | chr6:168196493-168197921 | 1428 | 7 | | 3 | | 3 | | | 0 | 7 | 1 | | 3 | | | | yes | | | |
| C6orf124 | N_Shore | Body | chr6:168225394-168225685 | 291 | | 1 | 3 | | | | | | | | | | | | | | | | |
| KIF25 | NA | TSS1500;TSS200 | chr6:168417574-168418632 | 1058 | | | 5 | | 3 | | | | 2 | | | | | | yes | | yes | yes | |
| KIF25 | NA | 5'UTR | chr6:168426201-168426413 | 212 | | | | | | | | | 1 | | | | | yes | yes | | | | |
| KIF25 | N_Shore;Island | Body | chr6:168435636-168436353 | 717 | 1 | 1 | 20 | | | 1 | 1 | 0 | 5 | 2 | 1 | 9 | | yes | | | | | yes |
| KIF25 | N_Shelf | Body | chr6:168445570-168445614 | 44 | | | | | | | | | | | | | | | | | | | |
| FRMD1 | Island | 5'UTR;1stExon | chr6:168479787-168479975 | 188 | | | | | | | | | | | | | | | | | | | yes |
| DACT2 | NA | Body | chr6:168714673-168714709 | 36 | | | | | | | | | 1 | | | | | | | | | | |
| DACT2 | Island | TSS200 | chr6:168720476-168720505 | 29 | | | | | | | | 0 | | | | | | yes | | | | | yes |
| SMOC2 | Island | 5'UTR;1stExon | chr6:168841907-168842302 | 395 | | | | | | | | 0 | | | | | | | | | | yes | yes |
| SMOC2 | NA | Body | chr6:168857165-168858319 | 1154 | | | | | | | | | 4 | 1 | | | | | | | | | |
| SMOC2 | NA | Body | chr6:168859941-168859997 | 56 | | | | | | | | | | | | | | yes | | | | yes | |
| SMOC2 | NA | Body | chr6:168897991-168898334 | 343 | | | | | | | | | | | | | | | | | | | |
| SMOC2 | N_Shelf | Body | chr6:168915440-168915507 | 67 | | | | | | | | | | | | | | | | | | | |
| SMOC2 | N_Shelf | Body | chr6:168968529-168968754 | 225 | | | | | | | | | | | | | | | yes | | | | |
| SMOC2 | N_Shore | Body | chr6:168970787-168972930 | 2143 | | | 11 | | | | | | 1 | | | | | yes | yes | | | | |
| SMOC2 | NA | Body | chr6:168983411-168983500 | 89 | | 1 | | | | | | | | | | | | | yes | | | | yes |
| SMOC2 | NA | Body | chr6:169013530-169013732 | 202 | | | | | | | | | | | | | | | | | | | |
| SMOC2 | NA | Body | chr6:169018751-169019307 | 556 | | | 3 | | | | | | | | | | | | | | yes | | yes |
| SMOC2 | NA | Body | chr6:169026027-169027218 | 1191 | | | 12 | | | | | | | | | | | | yes | | | yes | |
| SMOC2 | NA | Body | chr6:169050956-169051431 | 475 | | | 4 | | 1 | | | | | | | | | | yes | | | | |
| THBS2 | N_Shore | Body | chr6:169629860-169630013 | 153 | | | | | | | | 0 | 1 | | | | | | | | | | |
| THBS2 | N_Shore | Body | chr6:169641045-169641288 | 243 | | | | | | | | | | | | | | | | | | yes | yes |
| THBS2 | NA | 5'UTR | chr6:169653612-169653961 | 349 | | | | | | | | 0 | 4 | | | 1 | | | | | | | |
| WDR27 | Island | Body | chr6:169977394-169978387 | 993 | | | 5 | | | | | | 1 | | | | | yes | | yes | | | yes |

| | | | | | | | | | | | | | | | | | | | | | | | |
|---|---|---|---|---|---|---|---|---|---|---|---|---|---|---|---|---|---|---|---|---|---|---|---|
| WDR27 | NA | Body | chr6:170041920-170042028 | 108 | | | | | | | | | 1 | | | | | | | | | | |
| C6orf208;C6orf122 | N_Shore;Island;S_Shore | TSS1500;Body | chr6:170189761-170191082 | 1321 | | | 1 | | | | | 0 | 1 | | | 1 | | yes | | | yes | | |
| LOC154449 | Island | Body | chr6:170564889-170565151 | 262 | | | 1 | | | | | | 1 | | | | | | | | yes | | |
| LOC154449 | NA | Body;TSS200 | chr6:170571143-170572016 | 873 | 1 | | 22 | | | | | 0 | 4 | | | 5 | | yes | | | | | |
| FAM20C | N_Shore;Island | Body;1stExon | chr7:193229-195679 | 2450 | | 1 | | 1 | | | | | 2 | 1 | 1 | | | | | | | | yes |
| FAM20C | N_Shelf | Body | chr7:214577-214779 | 202 | | | 1 | 1 | | | | | | | | | | yes | yes | | | yes | |
| FAM20C | S_Shore | Body | chr7:226444-226959 | 515 | | | 14 | | | | | | | | | | | | | | | | |
| FAM20C | chr7:231272-231539 | N_Shore | chr7:229388-229695 | 307 | | | | | | | | | | | 1 | | | | | yes | | | |
| PDGFA | S_Shelf | 3'UTR | chr7:537175-537828 | 653 | | | 7 | | | | | | | | | | | | yes | | | yes | |
| PDGFA | NA | Body | chr7:544577-544847 | 270 | | | | | 1 | | | | | | | | | yes | | | | | |
| PDGFA | Island | Body | chr7:550508-550640 | 132 | | | | | | | | | | | | | | | | | | yes | |
| PDGFA | N_Shelf | Body | chr7:553412-553747 | 335 | | | 6 | | | | | | 1 | | | | | yes | | | | | |
| PRKAR1B | N_Shore | Body | chr7:592465-592840 | 375 | | | | 1 | | | | | | | | | | yes | | | | | yes |
| PRKAR1B | N_Shore | Body | chr7:597101-597596 | 495 | | | | | | | | | | | | | | yes | | | | yes | |
| PRKAR1B | N_Shore;Island | Body | chr7:598729-599291 | 562 | | | 2 | | | | | | | | | | | yes | yes | yes | yes | | |
| PRKAR1B | Island | Body | chr7:601283-601828 | 545 | | 1 | | 1 | 2 | | | | | | | | | | | | | yes | |
| PRKAR1B | N_Shore | Body | chr7:612458-612876 | 418 | | | 4 | | | | | | | | | | | | | | | | |
| PRKAR1B | NA | Body | chr7:621876-622318 | 442 | | | | | | | | | | | | | | yes | yes | | | | |
| PRKAR1B | NA | Body | chr7:624656-625136 | 480 | | | | | 1 | | | | 2 | | | | | yes | | | yes | | |
| PRKAR1B | N_Shore | Body | chr7:630473-630914 | 441 | | | 8 | | | | | | 2 | | | | | yes | | | | | |
| PRKAR1B | S_Shore | Body | chr7:632926-633202 | 276 | | | | | 1 | | | | | | | | | yes | yes | | | | |
| PRKAR1B | NA | Body | chr7:637132-637454 | 322 | | | 1 | | | | | | | | | | | | | | | | |
| PRKAR1B | N_Shelf | Body | chr7:640338-640820 | 482 | | | | | | | | | | | | | | yes | yes | | | | yes |
| PRKAR1B | Island | Body | chr7:644219-644417 | 198 | | | 6 | | | | | | 1 | 1 | | | | | yes | | | yes | |
| PRKAR1B | N_Shelf | Body | chr7:671717-672108 | 391 | 3 | | | | | | | | | | | | | yes | | | | | |
| PRKAR1B | Island;N_Shore;S_Shore | Body;5'UTR;TSS1500;TSS200;1stE | chr7:751009-752800 | 1791 | | | | | | 1 | | 0 | 4 | 1 | 1 | 1 | 1 | yes | | | | | yes |
| PRKAR1B;HEATR2 | Island | 5'UTR;TSS1500 | chr7:766100-766119 | 19 | | | 4 | | 1 | | 1 | 0 | 1 | 1 | | 6 | | | | yes | | | |
| HEATR2 | N_Shore | Body | chr7:778730-779089 | 359 | | 1 | | | | | | | 1 | | | | | | | | | | |
| HEATR2 | NA | Body | chr7:786420-786861 | 441 | | 1 | | | | | | | | | | | | | yes | | | | |
| HEATR2 | N_Shelf | Body | chr7:788797-788989 | 192 | | | | 1 | | | | | | | | | | yes | | | | | |
| HEATR2 | Island;S_Shore | Body | chr7:808592-809109 | 517 | | | 8 | | | | | | | | | | | yes | | | | | |
| HEATR2 | N_Shelf | Body | chr7:810958-811491 | 533 | | | 1 | | 2 | | | | | | | | | | | | yes | | |
| HEATR2 | Island | Body | chr7:814391-814621 | 230 | | | 1 | | | | | | 1 | | | | | yes | | | yes | | |
| HEATR2 | S_Shelf | Body | chr7:816960-817576 | 616 | | | 2 | | | | | | | | | | | | yes | | yes | | |
| UNC84A | Island | TSS1500 | chr7:855365-855632 | 267 | | | 2 | | | | | | | | | | | yes | | | | yes | |
| UNC84A | Island | 5'UTR | chr7:862776-862831 | 55 | | | | 1 | | | | | | | | | | | yes | | | | |
| UNC84A | S_Shelf | 5'UTR | chr7:866102-866406 | 304 | | | 7 | | | | | | | | | | | yes | | | | | |
| UNC84A | S_Shelf | Body | chr7:891044-891145 | 101 | 11 | | | | | | | | | | | | | | | | | | |
| UNC84A | NA | Body | chr7:896303-896603 | 300 | | | | 1 | | | | | | | | | | yes | | | | | |
| UNC84A | NA | Body | chr7:899806-900037 | 231 | | | | | | | | | 1 | | | | | | | yes | | | |
| UNC84A | S_Shore | Body | chr7:906726-908060 | 1334 | | | 18 | | 1 | | | | | | | | | | yes | yes | | | yes |
| C7orf20 | N_Shore;chr7:915753-916644 | TSS1500;Island | chr7:915073-915831 | 758 | | 1 | | | | | | | | | 1 | | | yes | yes | | | yes | |
| C7orf20 | Island | Body | chr7:917662-917805 | 143 | | | 5 | | | | | | 1 | | | | | yes | | | | | yes |
| C7orf20 | S_Shore | Body | chr7:919082-919518 | 436 | | | | | | | | | | | | | | yes | | | | | |
| C7orf20 | NA | Body | chr7:922051-923365 | 1314 | | | | 1 | 1 | | | | | | | | | | yes | | | | |
| ADAP1 | Island | Body | chr7:942748-942874 | 126 | | | | | | | | | | | | | | | yes | | | | |
| ADAP1 | S_Shore | Body | chr7:944607-944776 | 169 | | | 15 | | | | | | | | | | | yes | | | | | yes |
| ADAP1 | Island | Body | chr7:959759-960528 | 769 | | | | | 2 | | | | | | | | | yes | | yes | | | yes |
| ADAP1 | S_Shore | Body | chr7:968194-968552 | 358 | | | | | | | 1 | 0 | 1 | 1 | | 1 | | yes | yes | | | | |
| ADAP1 | N_Shore | Body | chr7:974745-975358 | 613 | | | 4 | | | | | | | | | | | yes | | | | | |
| ADAP1 | Island | Body | chr7:985614-986139 | 525 | | | 5 | | | | | | | | | | | yes | yes | | | | |
| COX19 | N_Shelf | 3'UTR | chr7:1004748-1005089 | 341 | 1 | | | | | | | | | | | | | yes | yes | | | | |
| COX19 | S_Shore | TSS200 | chr7:1015418-1015730 | 312 | | | 5 | | | | | 0 | | | | | | yes | | yes | | yes | |
| CYP2W1 | N_Shore | TSS1500;TSS200;1stExon | chr7:1022213-1023066 | 853 | 1 | 1 | 8 | | | | | | 4 | | | | | yes | yes | | | | |
| CYP2W1 | N_Shore;Island | Body | chr7:1024673-1026363 | 1690 | | | 3 | | | | | | 1 | | | | | yes | | | | | |
| C7orf50 | N_Shore;Island | Body | chr7:1038940-1040081 | 1141 | | | | | | | | | 1 | | | | | yes | yes | | | yes | |
| C7orf50 | Island;S_Shore | Body | chr7:1052392-1054449 | 2057 | | 1 | | | | | | | 2 | | | | | | | | | | |
| C7orf50 | Island | Body | chr7:1059627-1059784 | 157 | 1 | | | | | | | | | | | | | | | | | | |
| C7orf50;MIR339 | Island | Body;TSS200 | chr7:1062652-1064100 | 1448 | | | | | 2 | | | | 4 | | | | | yes | yes | | | yes | |
| C7orf50 | Island | Body | chr7:1067302-1068244 | 942 | | | | 1 | | | | 0 | | 2 | | 1 | | | | | yes | | yes |
| C7orf50 | Island;N_Shore | Body | chr7:1080473-1081709 | 1236 | 1 | 5 | 7 | 2 | | | | 0 | 10 | | 1 | 7 | 1 | yes | yes | | | | |
| C7orf50 | N_Shore | Body | chr7:1083072-1083473 | 401 | | | 1 | 1 | | | | | 1 | | | | | yes | | | yes | | |
| C7orf50 | N_Shelf | Body | chr7:1093854-1094263 | 409 | | | | | | 1 | | | | | | | | yes | | | | | |
| C7orf50 | N_Shore | Body | chr7:1095416-1095666 | 250 | | | | | 1 | | | | | | | | | | | | | | |
| C7orf50;GPR146 | N_Shore | Body;TSS200 | chr7:1097027-1097382 | 355 | 2 | | | | | | | | | | | | | | | | | | |
| C7orf50;GPR146 | S_Shore | Body;3'UTR;1stExon | chr7:1098625-1098703 | 78 | | | 13 | | | | | | 1 | | | | | | | | yes | | |
| C7orf50 | N_Shelf | Body | chr7:1102138-1103262 | 1124 | | | | | 1 | | | | 2 | 1 | | | | | yes | | yes | | |
| C7orf50 | NA | Body | chr7:1113014-1113237 | 223 | | | | | | | | 0 | 1 | | | | | yes | yes | | | | yes |
| C7orf50 | N_Shelf | Body | chr7:1117418-1117540 | 122 | | | | 1 | | | | | | | | | | yes | | | | | |
| C7orf50 | S_Shore | Body | chr7:1121738-1121929 | 191 | | | | | | | | | | | | | | | | | | | |
| GPER;C7orf50 | N_Shore;S_Shore | TSS1500;Body;TSS200;5'UTR | chr7:1125817-1127767 | 1950 | | | 7 | | 1 | | | | 3 | | | 1 | | yes | | | yes | | |
| GPER;C7orf50 | Island | Body | chr7:1132036-1132345 | 309 | 1 | | | | | | | | | | | | | | | | | | |
| C7orf50 | S_Shore | Body | chr7:1133730-1133979 | 249 | | | | | | | | | | | | | | | | | | | |
| C7orf50 | Island | Body | chr7:1135747-1136023 | 276 | | | 2 | | | | | | 1 | 1 | | | | yes | | | | | |
| C7orf50 | N_Shelf | Body | chr7:1139076-1139461 | 385 | | 1 | | | | | | | | | | | | | yes | yes | | | |
| C7orf50 | Island | Body | chr7:1142643-1142861 | 218 | | | | | | | | | 1 | | | | | | | yes | | | |
| C7orf50 | S_Shelf | Body | chr7:1160383-1160746 | 363 | | | | | | | | | | | | | | yes | yes | | | | |
| C7orf50 | NA | Body | chr7:1162860-1164063 | 1203 | | | 31 | 1 | | | | 0 | 1 | | | | | | yes | | | | |
| ZFAND2A | N_Shore | 5'UTR | chr7:1198522-1199455 | 933 | | | 12 | | | | 3 | | 2 | 1 | | 7 | 1 | | yes | | | | yes |
| UNCX | Island | TSS200 | chr7:1272512-1272559 | 47 | | | | | | | | | 2 | | | 1 | | | | | | yes | yes |
| UNCX | Island | Body | chr7:1273835-1274166 | 331 | | | 1 | | | | | | | | | | | | | | | | |
| MICALL2 | N_Shore;Island | Body | chr7:1491280-1492452 | 1172 | 4 | | 4 | | | | | | 2 | 1 | | | | yes | | yes | | | |
| INTS1 | Island | Body | chr7:1514031-1514623 | 592 | | | | | | | | | 1 | | | | | yes | | | | | |
| INTS1 | N_Shelf | Body | chr7:1530760-1531166 | 406 | | | | | | | | | | | | | | yes | yes | | | | |
| INTS1 | S_Shore | TSS1500 | chr7:1545385-1545819 | 434 | | 1 | 1 | | | | | 0 | | | | | | yes | | | | yes | yes |
| MAFK | S_Shore | 5'UTR | chr7:1576765-1577016 | 251 | | | | | | | | | 1 | | | | | | | | | | |
| TMEM184A | N_Shore;Island;S_Shore | Body | chr7:1587402-1589689 | 2287 | | | 2 | | | | | | 1 | | | | | | | | | yes | |
| TMEM184A | NA | TSS200 | chr7:1596103-1596260 | 157 | | 1 | 1 | 1 | | | | | | | | | | yes | yes | | | | |
| PSMG3 | N_Shelf;N_Shore | 3'UTR;Body | chr7:1607146-1607868 | 722 | | | 12 | | 1 | | | | | | | | | yes | | | | yes | |
| KIAA1908;PSMG3 | N_Shore | TSS200;5'UTR;1stExon | chr7:1609515-1610220 | 705 | | | | | | | | 0 | 1 | | | | | yes | | | | | yes |
| ELFN1 | NA | TSS1500 | chr7:1748081-1748639 | 558 | | | | 1 | | | | | | | | | | yes | | | | | |
| ELFN1 | N_Shore;Island | 5'UTR | chr7:1753396-1753909 | 513 | 1 | | 8 | | | | | | 2 | | | | | | yes | | yes | | |
| ELFN1 | NA | 5'UTR | chr7:1775904-1776076 | 172 | | | 6 | | | | | | 1 | | | | | yes | | | | | |
| ELFN1 | N_Shelf;N_Shore | 5'UTR | chr7:1782000-1782755 | 755 | | | | | | | | | 1 | | | 2 | | yes | | | | yes | |
| ELFN1 | S_Shore | 3'UTR | chr7:1787194-1787944 | 750 | | | 1 | | | | | | | | | | | | | | | yes | yes |
| MAD1L1 | S_Shore;N_Shore | Body | chr7:1881887-1883876 | 1989 | | | 4 | 1 | 1 | | | 0 | 4 | 2 | | 3 | | yes | | | | | |
| MAD1L1 | N_Shore;Island | Body | chr7:1895903-1896827 | 924 | | | | | | | | 0 | 4 | | | 1 | | | | yes | | | |
| MAD1L1 | Island | Body | chr7:1902804-1903385 | 581 | | | | 2 | | | | | | | | | | yes | | | yes | | |
| MAD1L1 | N_Shore | Body | chr7:1912065-1912602 | 537 | | | | 1 | | | | | | | | | | | yes | | | | |

| | | | | | | | | | | | | | | | | | | | | | | | |
|---|---|---|---|---|---|---|---|---|---|---|---|---|---|---|---|---|---|---|---|---|---|---|---|
| MAD1L1 | S_Shore | Body | chr7:1938450-1938791 | 341 | | | 8 | | | | | | | | | | | | | | | | |
| MAD1L1 | N_Shore | Body | chr7:1948755-1949247 | 492 | | | | | | | | | 1 | | | | | yes | yes | | | | yes |
| MAD1L1 | S_Shelf | Body | chr7:1952518-1952807 | 289 | 2 | | 24 | | | | | | 2 | | | 1 | | | | | | | |
| MAD1L1 | NA | Body | chr7:1956546-1957317 | 771 | 1 | | | | | | | | 1 | | | | | | yes | | | | |
| MAD1L1 | NA | Body | chr7:1961785-1963149 | 1364 | | | 2 | 1 | 1 | | | | 1 | | | | | | yes | | | | |
| MAD1L1 | N_Shore | Body | chr7:1967621-1968348 | 727 | | 1 | | | | | | | | | | | | | yes | | | | yes |
| MAD1L1 | S_Shore | Body | chr7:1980117-1981314 | 1197 | | | 19 | | 1 | 1 | | 0 | 2 | | | 1 | | | | | yes | | yes |
| MAD1L1 | NA | Body | chr7:1986245-1986533 | 288 | 5 | | | | | | | 0 | 1 | 1 | | | | yes | | | | | |
| MAD1L1 | N_Shore;Island;S_Shore | Body | chr7:2019608-2020583 | 975 | 2 | | | | 2 | | | | 2 | | | | | yes | | | | | |
| MAD1L1 | N_Shore;Island | Body | chr7:2025606-2027283 | 1677 | 2 | 1 | | | | | | | | | | | | | yes | | | | |
| MAD1L1 | S_Shelf | Body | chr7:2046821-2046963 | 142 | 2 | | | | | | | | | | | | | yes | yes | | | yes | |
| MAD1L1 | N_Shore | Body | chr7:2052961-2053275 | 314 | | | | | | | | | | | | | | yes | yes | | | | |
| MAD1L1 | N_Shore;Island | Body | chr7:2058865-2060179 | 1314 | | | 15 | | | | | 0 | 3 | | | | | yes | yes | | | yes | |
| MAD1L1 | NA | Body | chr7:2076998-2077345 | 347 | | | 4 | | | | | | | | | | | | yes | | yes | | |
| MAD1L1 | NA | Body | chr7:2078650-2079008 | 358 | | | | | | | | | 3 | | | | | yes | yes | | | | |
| MAD1L1 | NA | Body | chr7:2082529-2082961 | 432 | | | | | | | | | 2 | | | | | | | yes | | | |
| MAD1L1 | NA | Body | chr7:2094628-2094977 | 349 | 1 | | | | | | | | 1 | 1 | | | | yes | | | yes | | |
| MAD1L1 | S_Shore | Body | chr7:2109244-2109890 | 646 | 2 | | 16 | | | | | | | | | | | | yes | | | | |
| MAD1L1 | Island | Body | chr7:2116368-2116648 | 280 | | | | | | | | | 1 | | 1 | | | yes | | yes | | | |
| MAD1L1 | S_Shelf | Body | chr7:2119340-2119608 | 268 | | | | | | | | 0 | 1 | | | | | | | | yes | | |
| MAD1L1 | NA | Body | chr7:2124092-2124566 | 474 | | | | | | | | | 1 | | | | | yes | yes | | | | |
| MAD1L1 | NA | Body | chr7:2133441-2133480 | 39 | | | 12 | | | | | | | | | | | | yes | | | | |
| MAD1L1 | NA | Body | chr7:2139216-2140321 | 1105 | | 1 | 18 | | | | | | 2 | | | 1 | | | yes | | | | |
| MAD1L1 | NA | Body | chr7:2144559-2144767 | 208 | | | 6 | | | | | | 1 | | | | | | | | | | |
| MAD1L1 | NA;N_Shelf | Body | chr7:2149674-2152821 | 3147 | | | 4 | | 1 | | | | 3 | | | | | yes | yes | | | | |
| MAD1L1 | NA | Body | chr7:2167316-2167503 | 187 | | | 1 | | | | | | | | | | | | yes | | | | |
| MAD1L1 | NA | Body | chr7:2205691-2205745 | 54 | | | | | | | | | 1 | | | | | | | | | | |
| MAD1L1 | NA | Body | chr7:2208606-2208830 | 224 | | | | | | | | | 1 | | | | | yes | yes | | | | |
| MAD1L1 | NA | Body | chr7:2214759-2214790 | 31 | | | 3 | | | | | | | | | | | | yes | | | | |
| NUDT1 | S_Shelf | Body | chr7:2284600-2284960 | 360 | | | | | | | | | 1 | | | | | yes | | | yes | | |
| NUDT1 | N_Shelf;NA | Body | chr7:2288716-2289888 | 1172 | | | 3 | | | | | 0 | 1 | | 1 | 5 | 1 | yes | | | | yes | |
| SNX8 | N_Shore | Body | chr7:2316958-2317083 | 125 | | | | | | | | | 1 | | | | | yes | | | | | |
| SNX8 | NA | Body | chr7:2328746-2329077 | 331 | | | 4 | | | | | | 1 | | | | | yes | yes | | | | |
| SNX8 | S_Shore | TSS200 | chr7:2354203-2354395 | 192 | | | | | | | | 0 | 1 | | | | | | | | | yes | |
| EIF3B | N_Shelf | Body | chr7:2416396-2416675 | 279 | | | | 1 | | | | | | | | | | yes | | | | | |
| CHST12 | S_Shore | 5'UTR | chr7:2445727-2446566 | 839 | | | | | | | | | | | | | | yes | yes | | | | yes |
| LFNG | NA | TSS1500 | chr7:2551598-2551996 | 398 | | | | | | | | | | | | | | | | | yes | | |
| LFNG | N_Shore | Body;TSS1500;TSS200 | chr7:2557174-2557715 | 541 | | | 8 | | | | | | 1 | | | | | | yes | | | | |
| LFNG | S_Shore | Body | chr7:2560976-2561116 | 140 | | | 6 | | | | | | | | | | | yes | | | | | |
| LFNG | N_Shore;Island | Body | chr7:2563184-2564256 | 1072 | | | 25 | 2 | | | | | | | | | | | | | | | |
| LFNG | Island;S_Shore | Body | chr7:2565944-2566911 | 967 | | 1 | | | | | | | 2 | | | | | | | | | yes | |
| IQCE | NA | Body | chr7:2606716-2607424 | 708 | | | 20 | 1 | | | | | 2 | | | | | yes | | | yes | | |
| IQCE | NA | Body | chr7:2611890-2612101 | 211 | | | | 1 | | | | | | | | | | yes | | | yes | | |
| IQCE | S_Shore;chr7:2645524-264576 | Body;S_Shelf | chr7:2646478-2648785 | 2307 | | | 19 | | | | | 0 | 3 | | 1 | | | yes | | | | | |
| IQCE | NA | 3'UTR | chr7:2653651-2654530 | 879 | | | | | 1 | | | 0 | 3 | 1 | | | | yes | | | | yes | |
| TTYH3 | S_Shore | Body | chr7:2672929-2673408 | 479 | | | | 1 | | | | 0 | 1 | | | | | yes | | | | | |
| TTYH3 | NA | Body | chr7:2677986-2679546 | 1560 | 1 | | | | | | | | 1 | | | | | | | | | | |
| TTYH3 | N_Shore | Body | chr7:2699640-2699820 | 180 | | | | | | | | | | | | | | | | | | yes | |
| TTYH3 | Island | 3'UTR | chr7:2701419-2701641 | 222 | | | | 2 | | | | | 1 | | | | | | | | | | |
| AMZ1 | N_Shore | TSS1500 | chr7:2718172-2718413 | 241 | | | | | | | | | | | | | | | yes | yes | | | |
| AMZ1 | S_Shore | 5'UTR | chr7:2728776-2728912 | 136 | | | 7 | | | | | | 1 | | | | | yes | | | | | |
| AMZ1 | N_Shore | 5'UTR | chr7:2738728-2738961 | 233 | 3 | | 13 | 1 | | | | | | | | | | yes | | | | | |
| AMZ1 | Island | Body | chr7:2742325-2742484 | 159 | | | | | 2 | | | | | | | | | yes | | | | | |
| AMZ1 | NA | Body | chr7:2749541-2749751 | 210 | | | | | | | | 0 | | | | | | | | | | | |
| GNA12 | N_Shore;chr7:2774444-277465 | Body;N_Shore | chr7:2773010-2775893 | 2883 | 2 | | 1 | 1 | 1 | | | 0 | 4 | 1 | 1 | | | | yes | | | | |
| GNA12 | N_Shelf;N_Shore;Island;S_Sho | Body | chr7:2800436-2802976 | 2540 | 3 | | 59 | 1 | | | | 0 | 10 | 2 | 1 | 15 | 1 | yes | | | | yes | |
| GNA12 | NA | Body | chr7:2842593-2842746 | 153 | | | 3 | | | | | | | | | | | | | | | | |
| GNA12 | NA | Body | chr7:2847517-2847575 | 58 | 5 | | 28 | | | | | 0 | 1 | 1 | | 4 | | | | | | | |
| GNA12 | NA | Body | chr7:2855625-2855759 | 134 | | | | | | | | | | | | | | yes | yes | yes | | | |
| GNA12 | NA | Body | chr7:2874522-2874595 | 73 | | | | | | | | | 1 | | | | | | yes | | yes | | |
| CARD11 | NA | Body | chr7:2953836-2953930 | 94 | | | 4 | | | | | | | | | | | yes | | | | | yes |
| CARD11 | NA | Body | chr7:2962240-2962845 | 605 | | | 3 | | | | | | 1 | | | | | yes | | | | | |
| CARD11 | Island | Body | chr7:2968316-2968627 | 311 | | | | | | | | | 3 | | | | | yes | | | | | |
| CARD11 | S_Shelf | Body | chr7:2971898-2972392 | 494 | | | | | | | | | | | | | | yes | yes | | | | |
| CARD11 | NA | 5'UTR;nan | chr7:3018607-3020536 | 1929 | | | 7 | | | | | | 2 | | 1 | | | | | | | | |
| CARD11 | NA | 5'UTR | chr7:3025634-3026459 | 825 | | | 7 | | | | | | 1 | | | | | yes | yes | | yes | | |
| CARD11 | NA | 5'UTR | chr7:3033173-3033939 | 766 | | 1 | | | 1 | | | | 1 | 1 | | | | | yes | yes | | | |
| CARD11 | NA | 5'UTR | chr7:3082006-3082233 | 227 | | | 1 | | | | | | 1 | | | | | yes | yes | | | | |
| CARD11 | NA | 1stExon;5'UTR;TSS200 | chr7:3083541-3084077 | 536 | | | 6 | | | | | | | | | | | yes | yes | | | | yes |
| SDK1 | Island | Body | chr7:3341524-3341875 | 351 | | | | | | | | | 1 | | | | | | | | | | yes |
| SDK1 | NA | Body | chr7:3849340-3849602 | 262 | | | | | | | | | | | | | | | | yes | | | |
| SDK1 | NA | Body | chr7:3919928-3920534 | 606 | | 1 | | | | | | | 1 | | | | | | yes | | | yes | |
| SDK1 | NA | Body | chr7:3988772-3989167 | 395 | | | 5 | 1 | | | | | | | | | | | | | | | |
| SDK1 | NA | Body | chr7:3990479-3991559 | 1080 | | | 5 | | | | | | | | | | | yes | | | yes | | |
| SDK1 | NA | Body | chr7:3996856-3996923 | 67 | | | | | | | | | | | | | | | yes | | | | |
| SDK1 | NA | Body | chr7:4030038-4030413 | 375 | | | | | | | | | | | | | | yes | yes | | | | |
| SDK1 | NA | Body | chr7:4049531-4050875 | 1344 | | | 4 | | | | | | | | | | | yes | | | | | |
| SDK1 | NA | Body | chr7:4065931-4066127 | 196 | | | 17 | | | | | | | | | | | | | | | | |
| SDK1 | NA | Body | chr7:4118583-4118825 | 242 | | | | | | | | | | | | | | | | | | | |
| SDK1 | Island | Body | chr7:4153707-4153919 | 212 | 1 | | | | | | | | | | | | | | | yes | | | |
| SDK1 | N_Shore | Body;TSS1500 | chr7:4167607-4168039 | 432 | | | | | | | | | 1 | | | | | yes | yes | | | yes | |
| SDK1 | NA | Body | chr7:4174482-4175865 | 1383 | | | | | | | | | | | | | | yes | yes | | | | |
| SDK1 | NA | Body | chr7:4183528-4183976 | 448 | | | 3 | | | | | | | | | | | yes | | | | | |
| SDK1 | NA | Body | chr7:4187615-4187852 | 237 | | | | | | | | | | | | | | yes | yes | | | | |
| SDK1 | NA | Body | chr7:4201510-4201594 | 84 | | | | | | | | | | | | | | yes | | | | yes | |
| SDK1 | NA | Body | chr7:4210949-4212007 | 1058 | | | 8 | | | | | | | | | | | | | yes | | | |
| SDK1 | NA | Body | chr7:4218743-4219174 | 431 | | | | | | | | | | | | | | | | | | | |
| SDK1 | NA | Body | chr7:4242769-4244643 | 1874 | | 3 | 12 | 1 | | | 1 | 0 | 2 | 1 | | 7 | | | | | yes | | yes |
| SDK1 | NA | Body | chr7:4249825-4250180 | 355 | | | | | | | | | | | | | | | yes | | | | |
| SDK1 | NA | Body | chr7:4254497-4255011 | 514 | | | 6 | | | | | | | | | | | yes | | yes | | | |
| SDK1 | NA | Body | chr7:4264980-4265548 | 568 | | | | | | | | | | | | | | yes | yes | | | | |
| SDK1 | NA | Body | chr7:4266872-4267412 | 540 | 2 | | 12 | | | | | | | | | | | | | | | | |
| SDK1 | NA | Body | chr7:4275784-4276204 | 420 | | | 5 | | | | | | | | | | | yes | yes | | | | |
| SDK1 | Island | Body;3'UTR | chr7:4304397-4305535 | 1138 | | | | | | | | 0 | 3 | | | 1 | | | | | | | |
| SDK1 | S_Shelf | Body;3'UTR | chr7:4307392-4308372 | 980 | | | 3 | | | 2 | | | | | | | | | | | | | yes |
| FOXK1 | NA | Body | chr7:4743056-4743426 | 370 | | | | | | | | | | | | | | yes | yes | | | yes | |
| FOXK1 | NA | Body | chr7:4746131-4747261 | 1130 | | 1 | 4 | | | | | | 3 | | | | | yes | | | | yes | |
| FOXK1 | NA | Body | chr7:4751611-4753002 | 1391 | | 1 | | | | | | | 2 | 1 | | | | yes | | | | | |
| FOXK1 | N_Shelf | Body | chr7:4754566-4755415 | 849 | | | 4 | | | | | 0 | 8 | 4 | 1 | 4 | | yes | | | | | |

| | | | | | | | | | | | | | | | | | | | | | | | |
|---|---|---|---|---|---|---|---|---|---|---|---|---|---|---|---|---|---|---|---|---|---|---|---|
| FOXK1 | N_Shelf;N_Shore | Body | chr7:4761793-4763182 | 1389 | | | 10 | | | | | | 2 | | | | | yes | | | | | |
| FOXK1 | N_Shore | Body | chr7:4764443-4764941 | 498 | | | | | | | | 0 | 1 | | | | | yes | | yes | | | |
| FOXK1 | NA | Body | chr7:4778839-4779342 | 503 | 4 | | 7 | 1 | | | | 0 | 1 | | | | | yes | | | | yes | |
| FOXK1 | N_Shore | Body | chr7:4784090-4784485 | 395 | | | | 1 | | | | | | | | | | yes | yes | | | | |
| FOXK1 | Island | Body | chr7:4801923-4802774 | 851 | | | | | | | | | 2 | | | | | | | | | yes | |
| FOXK1 | N_Shore | 3'UTR | chr7:4806755-4807062 | 307 | | | 1 | | | | | | | | | | | | | | | | |
| RADIL | Island | Body | chr7:4848683-4848939 | 256 | | | 4 | | 1 | | 1 | 0 | 1 | 1 | 1 | | | yes | | | yes | | yes |
| RADIL | Island | Body | chr7:4856859-4857070 | 211 | | | 2 | | | | | | | | | | | | | | | yes | |
| RADIL | S_Shore;S_Shelf | Body | chr7:4859125-4859763 | 638 | | | 14 | | | | | | | | | | | yes | | | | | |
| RADIL | NA | Body | chr7:4869981-4870162 | 181 | | | | | | | | 0 | | | | 4 | | yes | yes | yes | | | |
| RADIL | S_Shore | Body | chr7:4877433-4877609 | 176 | | | 3 | | | | | | | | | | | | | | | yes | |
| RADIL | NA | Body | chr7:4885521-4885643 | 122 | | 1 | 15 | | | | | 0 | | | | | | yes | yes | | | | |
| RADIL;PAPOLB | Island | Body;1stExon;5'UTR;TSS200 | chr7:4901337-4901798 | 461 | | | 19 | | | | | 0 | 2 | | | 1 | | yes | yes | | | | |
| RADIL | NA | Body | chr7:4911022-4911121 | 99 | | | | | 1 | | | | | | | 1 | | | yes | | | | |
| RADIL | N_Shore | 5'UTR | chr7:4921827-4922039 | 212 | | | | | | | | | | | | 1 | | yes | | | yes | | |
| LOC389458 | Island;S_Shore;N_Shore | TSS200;Body;TSS1500 | chr7:5111488-5112654 | 1166 | | | 13 | | | | | 0 | 9 | 1 | | 3 | | yes | yes | | | yes | yes |
| WIPI2 | NA | Body | chr7:5257609-5258485 | 876 | | | | | 1 | | | | 2 | 1 | 1 | | | yes | yes | | | | |
| WIPI2 | Island | Body | chr7:5267749-5267994 | 245 | 2 | | | | | | | | | | | | | yes | | | | | |
| WIPI2 | chr7:5266921-5267306;S_Shelf | S_Shelf;3'UTR | chr7:5270567-5272900 | 2333 | | | | | 2 | | | 0 | 5 | | 1 | 3 | 1 | yes | | | | | |
| SLC29A4 | Island | Body | chr7:5336543-5336811 | 268 | | | 11 | | | | | | | | | | | | | | | yes | yes |
| SLC29A4 | Island | Body | chr7:5340232-5340536 | 304 | | | | | | | | | | | | | | yes | yes | | | | |
| TNRC18 | Island | Body | chr7:5353055-5353456 | 401 | | | | | | | | | | | | | | yes | | | | yes | |
| TNRC18 | N_Shore | Body | chr7:5390491-5390509 | 18 | 2 | | | | | | | | | | | | | | | | | | yes |
| FBXL18 | Island | Body | chr7:5527927-5528266 | 339 | | | | | | | | 0 | 1 | 1 | | 3 | | | | | | | |
| MIR589;FBXL18 | S_Shore | TSS200;Body | chr7:5534951-5535934 | 983 | | 1 | 31 | 2 | 1 | | | | 6 | | 1 | | | yes | yes | | | | |
| FBXL18 | N_Shore | Body | chr7:5552460-5552734 | 274 | | | 1 | | | | | | | | | | | yes | | | | | |
| FSCN1 | N_Shore | Body | chr7:5642705-5643021 | 316 | | | | | | | | | | | | | | yes | yes | | | | |
| FSCN1 | N_Shore | 3'UTR | chr7:5645728-5646121 | 393 | 1 | | | | | | | | | | | | | | | | | | |
| ZNF815 | N_Shore | TSS200;TSS1500 | chr7:5862449-5863510 | 1061 | | | 2 | | | | | | | | | | | yes | | | | | |
| C7orf28A | N_Shore | TSS1500 | chr7:5937190-5937486 | 296 | | | | | | | | | 1 | | | | | yes | | yes | | | |
| EIF2AK1 | S_Shore | TSS200 | chr7:6099044-6099250 | 206 | | | | | | | | | | | | | | yes | | | | yes | yes |
| USP42 | N_Shore | Body | chr7:6191807-6192000 | 193 | | | | | | | | 0 | | | | | | yes | | | yes | | |
| USP42 | N_Shore | Body | chr7:6197678-6198004 | 326 | | | | | | | | | 1 | | | | | yes | | | yes | | |
| USP42 | S_Shore | 3'UTR | chr7:6199980-6200140 | 160 | | 1 | | 1 | | | | | 1 | 1 | 1 | | | | | | | | |
| C7orf28B | S_Shore | TSS1500 | chr7:6866834-6867074 | 240 | | | 1 | | | | | | | | | | | yes | | yes | | | |
| COL28A1 | NA | TSS1500 | chr7:7576090-7576792 | 702 | | | | | | | | | 1 | | | | | yes | yes | | yes | | |
| GLCCI1 | S_Shelf | Body | chr7:8011594-8012104 | 510 | | | | | | | 1 | 0 | 1 | | | | | | | | | yes | yes |
| ICA1 | Island;S_Shore | TSS200;TSS1500 | chr7:8302229-8302899 | 670 | | | 12 | | | | | | | | | | | yes | | | | | yes |
| NXPH1 | S_Shore | Body | chr7:8476098-8476521 | 423 | | | 3 | | | | | 0 | 1 | | | | | | | | | | |
| NXPH1 | N_Shore;Island | Body | chr7:8480586-8483710 | 3124 | | | | | | | | | | | | | | | | | | | |
| NDUFA4 | N_Shore | Body | chr7:10979247-10979586 | 339 | | | 1 | | | | | | | | | | | | | | | | |
| THSD7A | NA | TSS200;TSS1500 | chr7:11872009-11872073 | 64 | | | | | | | | 0 | 1 | 1 | | | | | | | | yes | yes |
| VWDE | S_Shore | TSS200;TSS1500 | chr7:12443880-12444290 | 410 | | 1 | 5 | | | | | 0 | 1 | 1 | | 14 | 1 | yes | | | | yes | yes |
| SCIN | N_Shore;Island | TSS1500;TSS200;1stExon;Body | chr7:12609907-12610833 | 926 | | | 3 | | | | | | 2 | | | | | | | | | | |
| ETV1 | N_Shore | 1stExon;5'UTR | chr7:14029285-14029592 | 307 | | | | | | | | | 2 | | | | | | | | | | |
| MEOX2 | NA | Body | chr7:15725196-15725652 | 456 | | | | | | | | 0 | 1 | | | | | | | | | yes | |
| MEOX2 | NA | TSS1500 | chr7:15726821-15727598 | 777 | | 1 | | | | | | | | | | | | | | | | | |
| ISPD | NA | Body | chr7:16438702-16439165 | 463 | | | | | | | | | 1 | | | | | | | | | | |
| ISPD | Island | TSS200 | chr7:16460965-16461064 | 99 | | | | | | | | | 1 | | | | | | | | yes | | yes |
| SOSTDC1 | NA | 1stExon;5'UTR;TSS200 | chr7:16505094-16505664 | 570 | | | 20 | | | | | 0 | 5 | | | 10 | | | | | | | |
| PRPS1L1 | NA | TSS200 | chr7:18067533-18067537 | 4 | | | | | | | | | 1 | | | | | | yes | | | | |
| HDAC9 | NA | TSS1500;5'UTR | chr7:18535499-18535786 | 287 | | | | | | | | | | | | | | | | | | | |
| TWIST1 | Island;S_Shore | TSS1500 | chr7:19157813-19158954 | 1141 | | 3 | | | | | | | 4 | | 1 | | | | | | | | |
| FERD3L | Island;S_Shore | 1stExon;TSS200;TSS1500;3'UTR | chr7:19184427-19185407 | 980 | | | 25 | | | | | 0 | 3 | | | | | | | | | | yes |
| TMEM196 | Island;S_Shore | TSS200;TSS1500 | chr7:19812570-19813493 | 923 | | | | | | | | 0 | 2 | | | 1 | | | | | | | |
| SP8 | Island;S_Shore | Body;5'UTR | chr7:20823981-20826345 | 2364 | | | 3 | | | | | 0 | 4 | | | | | | | | | yes | yes |
| DNAH11 | NA | Body | chr7:21788358-21788477 | 119 | | | 1 | | | | | | | | | | | yes | | | | | |
| MGC87042 | Island | 5'UTR;1stExon | chr7:22539703-22540090 | 387 | | 1 | 1 | | | | | 0 | | | | | | yes | | | yes | | |
| IL6 | NA | Body | chr7:22767384-22767571 | 187 | | | 4 | | | | | | | | | | | yes | | | | | |
| SNORD93 | S_Shore | TSS1500 | chr7:22894795-22895283 | 488 | | | 5 | | | | | | | | | | | yes | | | | | |
| GPNMB | N_Shore | 5'UTR;1stExon | chr7:23286319-23286539 | 220 | | | | | | | | | 1 | | | | | yes | | | | | |
| IGF2BP3 | NA | Body | chr7:23387365-23387396 | 31 | | | | | | | | 0 | 1 | 1 | | | | | | | | | |
| STK31 | NA | TSS200;TSS1500 | chr7:23749667-23749740 | 73 | | | 3 | | | | | | | | | | | | yes | | | | |
| NPY | Island;N_Shore | 5'UTR;TSS1500;TSS200 | chr7:24323128-24325371 | 2243 | | | | | | | | | | | | | | | | | | | |
| DFNA5 | S_Shore | TSS1500;TSS200 | chr7:24797656-24798175 | 519 | | | | 1 | | | | 0 | 7 | | | | | | yes | yes | yes | | |
| OSBPL3 | N_Shore | 5'UTR | chr7:25018565-25018855 | 290 | | | 5 | | | | | | | | | | | | yes | | | | |
| C7orf31 | N_Shelf | Body | chr7:25216506-25216828 | 322 | | | 1 | | | | | | | | | | | yes | yes | | | | |
| MIR148A | N_Shore | TSS200 | chr7:25989618-25990026 | 408 | | 1 | | | | | | 0 | | | | | | | | | | | |
| NFE2L3 | N_Shore | TSS1500 | chr7:26191458-26191599 | 141 | | | 3 | | | | | | | | | | | yes | | | | | yes |
| NFE2L3 | Island;S_Shore | Body | chr7:26192756-26193805 | 1049 | | | 4 | | 4 | | | | | | | | | yes | | | | | |
| C7orf71 | NA | TSS1500 | chr7:26676191-26676482 | 291 | | | | | | | | | 1 | | | | | | yes | | yes | | |
| SKAP2 | NA | Body | chr7:26897182-26897714 | 532 | | 1 | 20 | | | | | 0 | 4 | | | 2 | | | | | | yes | |
| SKAP2 | Island | TSS1500 | chr7:26904682-26904713 | 31 | | | | | | | | | 1 | | | | | | | | yes | | |
| HOXA1 | Island | 3'UTR;Body | chr7:27134109-27134369 | 260 | | | 3 | | 1 | | | 0 | 3 | | | 6 | 1 | | | | | | |
| HOXA2;HOXA3 | N_Shore;Island;S_Shore;N_She | 1stExon;5'UTR;TSS200;TSS1500;3' | chr7:27140797-27146445 | 5648 | | | | | | | | | | | | | | | | | | | |
| HOXA3 | N_Shore | 3'UTR | chr7:27147513-27148204 | 691 | | 1 | | | | | | | 1 | | | | | | | | | yes | yes |
| HOXA3 | Island | 5'UTR | chr7:27150262-27150598 | 336 | | | 8 | | | | | | 1 | | | | | | | | | | yes |
| HOXA3 | N_Shore;Island;chr7:27154999 | 5'UTR;1stExon;TSS200;N_Shore;TS | chr7:27152899-27160960 | 8061 | | | | | | | | | | | | | | | | | | | |
| HOXA3 | N_Shore | 5'UTR | chr7:27163095-27163929 | 834 | | | | | | | | | | | | | | | | | | | |
| HOXA4;HOXA3 | Island;S_Shore;N_Shelf;N_Sho | 1stExon;5'UTR;TSS200;TSS1500;3' | chr7:27167315-27171528 | 4213 | | | | | | | | | | | | | | | | | | | |
| HOXA5;HOXA6 | N_Shore;Island | Body;1stExon;5'UTR;TSS200;TSS15 | chr7:27181480-27185512 | 4032 | | | | | | | | | | | | | | | | | | | |
| HOXA6 | Island;S_Shore;N_Shore | 1stExon;TSS200;TSS1500 | chr7:27186993-27188770 | 1777 | | | 51 | | 1 | | | 0 | 1 | | | | | | | | | | yes |
| HOXA7 | Island;N_Shore | Body;TSS200;TSS1500 | chr7:27194614-27199752 | 5138 | | | | | | | | | | | | | | | | | | | |
| HOXA9 | Island | Body;1stExon;TSS1500 | chr7:27204005-27206907 | 2902 | | 3 | 2 | | | | | 0 | 2 | | | | | | | | | | |
| MIR196B | Island | Body;TSS200 | chr7:27209131-27209582 | 451 | | | 3 | | | | | | 3 | | | | | | | | | yes | yes |
| HOXA10 | chr7:27212416-27214396;Islan | Island;Body;1stExon;TSS200 | chr7:27212935-27214383 | 1448 | | 1 | | | | | | | 2 | | 1 | | | | | | | yes | |
| HOXA10 | S_Shore | Body | chr7:27215464-27215610 | 146 | | | 1 | | | | | 0 | 1 | | | | | | | | | | yes |
| HOXA10 | S_Shore | TSS200 | chr7:27220034-27220398 | 364 | | | 7 | | | | | | | | | | | | | | | | yes |
| HOXA11 | N_Shore | Body | chr7:27222869-27223219 | 350 | | | 1 | | | | | | | | | | | | | | | yes | |
| HOXA11AS;HOXA11 | Island;N_Shore | Body;TSS1500;1stExon;TSS200 | chr7:27224568-27229163 | 4595 | | | | | | | | | | | | | | | | | | | |
| HOXA11AS | N_Shore | Body | chr7:27227358-27227847 | 489 | | | | | | | | | | | | | | | | | | | |
| HOXA13 | Island | TSS1500 | chr7:27240181-27240251 | 70 | | | 1 | | | | | | | | | | | | | | | | yes |
| EVX1 | N_Shore;Island | TSS1500;1stExon;5'UTR | chr7:27280914-27282431 | 1517 | | 2 | 35 | | | | | 0 | 4 | | | 1 | | | | | | | yes |
| EVX1 | N_Shore;Island | Body | chr7:27284348-27286075 | 1727 | 1 | | | | | | | | | | | | | | | | | | |
| LOC402644 | NA | TSS200;TSS1500 | chr7:28319499-28319676 | 177 | | | 5 | | | | | | | | | | | | | | | | |
| CREB5 | NA | TSS1500 | chr7:28338584-28338757 | 173 | | | | | | | | | 1 | | | | | | | | yes | | |
| CREB5 | N_Shore | 5'UTR | chr7:28448103-28448295 | 192 | | | | | | | | 0 | | | | | | yes | | | | | |
| CREB5 | S_Shore | TSS1500;5'UTR | chr7:28450687-28450797 | 110 | | | | | | | | | | | | | | | | | | | |
| CREB5 | S_Shelf | 5'UTR;TSS200 | chr7:28452066-28452289 | 223 | | | 19 | | | | | 0 | 5 | | | 6 | | | | | | yes | |

| | | | | | | | | | | | | | | | | | | | | | | | |
|---|---|---|---|---|---|---|---|---|---|---|---|---|---|---|---|---|---|---|---|---|---|---|---|
| CREB5 | NA | TSS1500;5'UTR;Body | chr7:28474589-28475121 | 532 | | | 6 | | | | | | | | | | | yes | | | | yes | |
| CREB5 | NA | Body | chr7:28838086-28838234 | 148 | | | 9 | | | | | | | | | | | | | | | | |
| TRIL | Island;S_Shore | 1stExon;TSS200;TSS1500 | chr7:28996491-28998860 | 2369 | 1 | | 15 | 1 | | | | 0 | 4 | | | | | | | | | | |
| CPVL | NA | TSS200;TSS1500 | chr7:29186267-29187019 | 752 | 3 | | 5 | | | | | | 2 | | | | | yes | | yes | yes | | |
| CHN2 | NA | TSS200;Body;5'UTR;1stExon | chr7:29519297-29519656 | 359 | | | 20 | | 1 | | | | | | 1 | | | | | | | | |
| PRR15 | N_Shore | TSS1500;TSS200 | chr7:29602763-29603407 | 644 | | 1 | 3 | | | | | | | | | | | yes | | | | | |
| PRR15 | Island;N_Shore | 5'UTR;Body | chr7:29605237-29606349 | 1112 | | 1 | 4 | | 5 | | | 0 | 6 | | | 9 | | | | | | | |
| WIPF3 | N_Shore;Island | Body | chr7:29923120-29924165 | 1045 | | | 4 | | | | | 0 | 1 | | | 2 | | yes | | yes | | | yes |
| SCRN1 | Island | TSS1500;1stExon;Body;5'UTR;TSS2 | chr7:30029717-30029776 | 59 | | 1 | 2 | | | | | | 1 | | | | | yes | yes | | | yes | yes |
| ZNRF2 | N_Shore | TSS1500 | chr7:30323267-30323735 | 468 | | | | | | | | | | | | | | yes | | | | yes | |
| NOD1 | NA | Body | chr7:30492141-30492171 | 30 | | | 1 | | | | | | | | | | | | | | | | |
| GGCT | Island | TSS200;1stExon;5'UTR | chr7:30544409-30544737 | 328 | | | | | | | | 0 | | | | | | yes | yes | | | | |
| GARS | S_Shore | Body | chr7:30635712-30636176 | 464 | | | | | | | | 0 | 3 | | | 5 | | yes | | | yes | | |
| CRHR2 | Island | TSS200 | chr7:30722320-30722361 | 41 | | 1 | | | | | | | | | | | | | | | | | yes |
| FAM188B | N_Shore | TSS200 | chr7:30810858-30810882 | 24 | | | | | | | | | 1 | | | | | yes | | | | | |
| AQP1 | NA | TSS1500;TSS200;1stExon;5'UTR | chr7:30950813-30951801 | 988 | 4 | | 24 | 2 | 2 | | | | 2 | 1 | | | | yes | | | | | yes |
| ADCYAP1R1 | NA | Body | chr7:31126479-31126531 | 52 | | | | | | | | | 1 | | | | | | | | | | |
| C7orf16 | NA | TSS200 | chr7:31726639-31726912 | 273 | 1 | | 11 | | | | | 0 | 2 | | | 1 | | yes | | | | | |
| PDE1C | N_Shore;S_Shore | Body;TSS200;TSS1500 | chr7:32109831-32111453 | 1622 | | 1 | 4 | | | | | 0 | 2 | | | 1 | | | | | | | yes |
| NT5C3 | NA | 1stExon;5'UTR;Body | chr7:33080496-33080615 | 119 | | | | 1 | | | | | | | | | | | yes | | | | |
| BMPER | Island | Body | chr7:34118464-34118758 | 294 | | | | | | | | | 3 | | | | | | | | | | |
| AAA1;NPSR1 | NA | Body;TSS200 | chr7:34873912-34874533 | 621 | | | | | | | | | 1 | | | | | yes | | yes | | yes | |
| DPY19L1 | S_Shore | TSS1500 | chr7:35078082-35078269 | 187 | | | | | | | | | | | | | | yes | | | | | |
| TBX20 | N_Shore;Island;chr7:35292985 | TSS1500;Body;1stExon;5'UTR;Islar | chr7:35292896-35295198 | 2302 | | 1 | 14 | | | | | | 1 | | 1 | | | | | | | | yes |
| EEPD1 | S_Shore | Body | chr7:36194253-36194575 | 322 | | | | 1 | | | | | | | | | | | | | | | |
| AOAH | NA | Body | chr7:36700744-36701016 | 272 | 1 | | 13 | | | | | | | | | | | | | | | | |
| AOAH | NA | 1stExon;5'UTR;TSS1500 | chr7:36764019-36765609 | 1590 | | | | 1 | | | | | | | | | | yes | | | | yes | yes |
| ELMO1 | NA | TSS1500;Body;1stExon;5'UTR;TSS2 | chr7:37025706-37025890 | 184 | | 1 | | | 1 | | | | | | | | | | | | | | |
| ELMO1 | NA | 5'UTR | chr7:37392880-37393045 | 165 | | | | | 1 | | | 0 | 1 | | | | | | | yes | | | |
| TXNDC3 | NA | TSS200 | chr7:37888078-37888670 | 592 | | | 6 | | | | | | | | | | | yes | | | | yes | yes |
| SFRP4 | S_Shore | TSS1500 | chr7:37956850-37956963 | 113 | | | 2 | | | | | | | | | | | | | | | | |
| EPDR1 | N_Shore;Island | TSS1500;1stExon | chr7:37959686-37960902 | 1216 | | 1 | 9 | | | | | | | | | | | yes | yes | | | | |
| AMPH | NA | Body | chr7:38468944-38469273 | 329 | | | | | | | | | | | | | | | | | | | |
| AMPH | Island | Body;5'UTR;1stExon | chr7:38670804-38671001 | 197 | | | | | 1 | | | | 3 | | | | | | | | | yes | yes |
| POU6F2 | NA | Body | chr7:39170497-39171113 | 616 | | 5 | 9 | 2 | 2 | | 2 | 0 | 6 | 3 | | 36 | | | yes | | | | yes |
| POU6F2 | NA | Body | chr7:39332739-39332884 | 145 | | | 2 | 1 | | | | 0 | 1 | | | | | | | | | | |
| POU6F2 | Island | Body | chr7:39453532-39453704 | 172 | | 1 | | | | | | | | | | | | | | | | | |
| LOC646999 | N_Shore | Body | chr7:39649224-39649443 | 219 | | 1 | | | | | | | 1 | | | | | | yes | | | | |
| INHBA;LOC285954 | NA | Body;TSS1500 | chr7:41732833-41733356 | 523 | | | | 1 | | | | | | | | | | | | | | | |
| LOC285954;INHBA | NA | Body | chr7:41739678-41739801 | 123 | | 1 | 10 | | | | | | | | | | | | | | | | yes |
| INHBA;LOC285954 | N_Shelf | 5'UTR;Body;1stExon | chr7:41742630-41743001 | 371 | | | 2 | | | | | | | | | | | | | | | yes | yes |
| GLI3 | NA | Body | chr7:42214276-42214361 | 85 | | | 1 | | | | | | | | | | | yes | | | | | |
| GLI3 | Island;chr7:42267546-4226782 | 5'UTR;N_Shore | chr7:42267269-42268109 | 840 | | 1 | 25 | | | | | | 4 | 4 | 2 | | | | | | | | yes |
| GLI3 | S_Shore | TSS1500 | chr7:42277950-42278096 | 146 | | | 3 | | | | | | 1 | | | | | | | | | yes | yes |
| HECW1 | N_Shore | TSS1500 | chr7:43151877-43152021 | 144 | | | | | | | | | 1 | | | | | | | | | | |
| HECW1 | Island | 5'UTR | chr7:43153201-43153595 | 394 | | 1 | | | | | | | | | | | | | | | | | |
| HECW1 | Island | Body | chr7:43288635-43288889 | 254 | | | 9 | | | | | | 1 | | 1 | 1 | | | | | | | yes |
| HECW1 | Island | Body | chr7:43484421-43484715 | 294 | | | | | | | | | 1 | | | | | | | | | yes | |
| URGCP | NA | Body | chr7:43918101-43918221 | 120 | | | | 1 | | | | | | | | | | | | | | | |
| FLJ35390;RASA4P | N_Shore | TSS200;Body | chr7:44078913-44079036 | 123 | | | | | | | | | | | | | | yes | | | | | |
| PGAM2 | N_Shore;Island;S_Shore | Body;1stExon;5'UTR;TSS200;TSS15 | chr7:44104584-44105764 | 1180 | | | 5 | | | | | 0 | 9 | | | 6 | | yes | | | | | |
| AEBP1 | Island | Body | chr7:44153188-44153333 | 145 | | | | | | | | | | | | | | | | | | yes | yes |
| GCK | NA | Body;5'UTR | chr7:44198380-44199106 | 726 | | | | 1 | | | | | | | | | | | yes | | | | |
| GCK | NA | 1stExon;5'UTR | chr7:44229001-44229319 | 318 | | | | | | | | | 1 | | | | | | | | | | |
| CAMK2B | S_Shore | Body | chr7:44261120-44261644 | 524 | | | 11 | | | | | | | | | | | yes | | | | yes | |
| CAMK2B | Island;S_Shore | Body | chr7:44279273-44279938 | 665 | | | 2 | | | | | | | | | | | | | | | | |
| CAMK2B | Island | Body | chr7:44349704-44349955 | 251 | | | 1 | | | | | | | | | | | yes | yes | | | | |
| NPC1L1 | NA | Body | chr7:44578565-44579271 | 706 | | | 3 | 1 | | | | | | | | | | | | | | yes | |
| DDX56 | N_Shore;Island;S_Shore | Body;TSS200;TSS1500 | chr7:44613073-44614489 | 1416 | | | 7 | | | | | 0 | 1 | 1 | | 1 | | yes | | yes | yes | | |
| ZMIZ2 | NA | TSS200;5'UTR | chr7:44795713-44796036 | 323 | | | 2 | | | | | | | | | | | | | | | | |
| MYO1G | Island | Body;3'UTR | chr7:45002287-45002919 | 632 | | | | | | | | 0 | 1 | | | | | | | | yes | | |
| MYO1G | NA | 5'UTR;1stExon;TSS200 | chr7:45018658-45019005 | 347 | | | | | | | | | 1 | 1 | 1 | | | | yes | | yes | | |
| CCM2 | NA | nan;Body;TSS1500 | chr7:45066738-45067319 | 581 | | | 6 | | 1 | | | | 2 | 1 | 1 | | | yes | | | yes | | |
| TBRG4;SNORA5A | NA | Body;TSS200 | chr7:45144116-45144171 | 55 | | | | | | | | | | | | | | | | | | | |
| ADCY1 | Island | TSS1500 | chr7:45613410-45613725 | 315 | | 1 | | | | | | | | | | | | yes | | | | yes | yes |
| IGFBP1 | N_Shore | TSS200 | chr7:45927824-45928133 | 309 | | | 11 | | | | | | | | | | | | | | | | yes |
| IGFBP3 | Island;S_Shore | 1stExon;5'UTR;TSS1500 | chr7:45960834-45962576 | 1742 | | | 7 | | | | | | 4 | | | 1 | | | | | | | yes |
| TNS3 | NA | 5'UTR | chr7:47539511-47539837 | 326 | | | | | | | | | 1 | | | | | yes | yes | | | yes | |
| TNS3 | Island | 5'UTR | chr7:47576580-47577090 | 510 | | | | | | | | 0 | 1 | | | 1 | | | | | | yes | |
| C7orf65 | NA | TSS1500;TSS200 | chr7:47694443-47694857 | 414 | | | 17 | | | | | | | | | | | yes | yes | yes | | | |
| HUS1 | S_Shore;Island | TSS200 | chr7:48019388-48019713 | 325 | | | 3 | | | | | | | | | | | yes | | | yes | yes | |
| SUNC1 | NA | Body | chr7:48031468-48031800 | 332 | | | | | | | | 0 | 1 | | | | | | yes | | | yes | |
| SUNC1 | NA | Body | chr7:48068321-48068758 | 437 | | | 4 | | | | | | | | | | | | yes | | | yes | |
| UPP1 | Island | 5'UTR | chr7:48129444-48129992 | 548 | | | 3 | | | | | 0 | 3 | 1 | | | | | | | | | yes |
| ABCA13 | N_Shore;Island | Body | chr7:48494156-48495072 | 916 | | | 14 | | | | | | 7 | | | | | | | | | | |
| VWC2 | Island | TSS1500;TSS200;5'UTR | chr7:49813031-49815117 | 2086 | | 1 | | | 4 | | | 0 | 2 | | | | | | | | | | yes |
| ZPBP | N_Shore;Island;S_Shore | Body;TSS200;TSS1500 | chr7:50132541-50133212 | 671 | | 1 | 5 | | | | | | | | | | | | | yes | | | yes |
| IKZF1 | S_Shelf | 5'UTR | chr7:50348174-50348870 | 696 | | | | | 1 | | | | | | | | | | | | | | yes |
| IKZF1 | NA | Body | chr7:50443901-50444052 | 151 | | | | | | | | | | | | | | | | | | | |
| IKZF1 | Island | 3'UTR | chr7:50468399-50468722 | 323 | | | | | | | | | | | | | | | | | | | |
| FIGNL1 | Island | TSS1500 | chr7:50518350-50518921 | 571 | | | 8 | | | | | 0 | 2 | 1 | | 2 | | | | yes | yes | yes | |
| DDC | NA | 5'UTR;1stExon;TSS1500 | chr7:50628718-50629987 | 1269 | | | 3 | | | | | | 1 | | | | | yes | yes | | | yes | |
| DDC | NA | 5'UTR;1stExon;TSS1500 | chr7:50632994-50634648 | 1654 | | | 12 | | | | | | 4 | | | | | yes | yes | yes | yes | | |
| GRB10 | NA | 5'UTR;Body;TSS1500 | chr7:50774060-50774217 | 157 | | | | 1 | | | | | | | | | | yes | yes | | | | |
| GRB10 | NA | 5'UTR;1stExon | chr7:50799978-50800202 | 224 | | | | | | | | | 1 | | | | | | | yes | | | |
| GRB10 | Island;S_Shore | TSS1500 | chr7:50861535-50861967 | 432 | | | 5 | | | | | 0 | 2 | | | 1 | | yes | yes | | | | yes |
| COBL | Island | 1stExon;5'UTR;TSS200 | chr7:51384371-51384676 | 305 | | | | | | | | | 1 | 3 | | 1 | | | | | yes | yes | yes |
| POM121L12 | N_Shore;Island | TSS200 | chr7:53103214-53103312 | 98 | | | 10 | | | | | | | | | | | | yes | | | | yes |
| VSTM2A | Island | TSS200;5'UTR;1stExon | chr7:54609953-54610320 | 367 | | | | | 1 | | | | 2 | | | | | | | | | yes | yes |
| VSTM2A | Island | Body | chr7:54612325-54612709 | 384 | | 1 | | | 1 | | | | 1 | | | | | | yes | | | | yes |
| VSTM2A | Island | Body | chr7:54614210-54614620 | 410 | | | | | | | | 0 | 1 | | | 1 | | | | | | yes | yes |
| SEC61G | S_Shore | TSS1500 | chr7:54827431-54827677 | 246 | | | | | | | | | | | | | | yes | yes | | yes | | |
| EGFR | NA | Body | chr7:55224863-55225183 | 320 | | | | | 2 | | | 0 | 2 | | | 1 | | yes | yes | | | | |
| LANCL2 | N_Shore | TSS1500 | chr7:55432402-55432692 | 290 | | | 3 | | | | | | | | | | | | yes | | | | |
| FKBP9L | NA | Body;TSS1500 | chr7:55756785-55757158 | 373 | | | | | | | | | | | | | | | | | | | |
| ZNF713 | NA | TSS200 | chr7:55980165-55980652 | 487 | | | | | | | | | | | | | | yes | | | yes | | |
| CCT6A;PSPH | N_Shore | TSS1500;5'UTR;1stExon | chr7:56119028-56119218 | 190 | | | | | | | | | | 2 | | | | yes | | | | | yes |
| PHKG1 | NA | 5'UTR;1stExon;TSS200 | chr7:56160409-56161020 | 611 | | | | | 1 | | | 0 | 3 | | | 1 | | yes | | | | | |
| LOC650226 | N_Shore;Island;S_Shore | Body;TSS200 | chr7:56514965-56516504 | 1539 | | | 35 | | | | | 0 | 19 | 2 | 1 | 8 | | yes | yes | yes | yes | | |

| | | | | | | | | | | | | | | | | | | | | | | | | |
|---|---|---|---|---|---|---|---|---|---|---|---|---|---|---|---|---|---|---|---|---|---|---|---|---|
| DKFZp434L192 | NA | TSS1500 | chr7:56562731-56563152 | 421 | | | 3 | | | | | | | | | | | | | | | | | |
| LOC642006 | Island;S_Shore | Body;TSS200 | chr7:57247707-57248455 | 748 | | | 8 | | | | | | | | | | | yes | | | | | yes | |
| ZNF727 | N_Shore | TSS1500;TSS200;1stExon;5'UTR | chr7:63505584-63506261 | 677 | | | 9 | | | | | 0 | 12 | 3 | | 11 | 1 | yes | | yes | | | | |
| ZNF735 | NA | TSS1500;TSS200 | chr7:63667212-63667594 | 382 | | | | | | | | | | | | | | yes | | yes | | | | |
| ZNF679 | NA | TSS1500 | chr7:63687600-63688007 | 407 | | | | | | | | | | | | | | | | | | | | yes |
| LOC649395 | NA | TSS200 | chr7:63893881-63894251 | 370 | | | | | | | | | | | | | | | yes | | | | | |
| LOC641746 | NA | Body | chr7:64043012-64043193 | 181 | 1 | | 3 | | 1 | | 1 | 0 | 1 | 1 | | 3 | 1 | | | yes | | | yes | |
| ZNF273 | NA | TSS200 | chr7:64363546-64363590 | 44 | | | | | | | | | 1 | | | | | yes | | yes | | | | |
| ZNF117 | NA | 1stExon;5'UTR | chr7:64450722-64450926 | 204 | | | | | | | | | 1 | | | | | yes | yes | yes | | | | |
| SNORA22;CCT6P1 | S_Shelf | TSS1500;Body | chr7:65219948-65220346 | 398 | | | | | | | | | 1 | | | | | yes | | | | | | |
| AUTS2 | Island | Body | chr7:70061135-70061616 | 481 | | | | | | | | 0 | | | | | | yes | | | | yes | | |
| AUTS2 | N_Shore | Body | chr7:70096029-70096458 | 429 | | | 26 | | | | | | | | | | | | | | | | | |
| AUTS2 | N_Shelf | Body | chr7:70252335-70252953 | 618 | 6 | | | | | | | | | | | | | | | | | | | |
| WBSCR17 | Island | TSS1500;TSS200 | chr7:70597058-70598282 | 1224 | | 1 | | | 2 | | | 0 | 4 | | | | | | | | | | yes | |
| NSUN5 | S_Shore | TSS1500 | chr7:72723150-72723603 | 453 | | | | | | | | | | | | | | yes | yes | | | yes | | |
| TRIM50;FKBP6 | Island | TSS200 | chr7:72742151-72742188 | 37 | | | | | | | | | | | | | | | | | | | | |
| TBL2 | Island | TSS200 | chr7:72993161-72993203 | 42 | | | | | | | | | 1 | | | | | yes | | | | | | |
| MLXIPL | S_Shore | TSS200 | chr7:73038982-73039553 | 571 | | | | | | | | | 1 | | | | | yes | | | | | | |
| STX1A | N_Shore;Island | Body | chr7:73118452-73118748 | 296 | | 1 | | | | | | 0 | | | | | | yes | | | | yes | | |
| CLDN4 | N_Shore;Island | TSS1500;TSS200;1stExon;5'UTR | chr7:73244836-73246044 | 1208 | 1 | | 4 | 1 | | | | | 3 | 1 | | | | | | | | | | |
| WBSCR27 | N_Shore;S_Shore | Body;TSS200 | chr7:73256360-73257083 | 723 | | | | | | | | | 1 | | | | | yes | | | | | yes | |
| WBSCR28 | NA | TSS200;TSS1500 | chr7:73275265-73275348 | 83 | | | | | | | | | 1 | | | | | yes | | yes | | | | |
| ELN | NA | TSS200;5'UTR;1stExon | chr7:73442282-73443113 | 831 | | | | | 1 | | | 0 | 1 | | | 1 | | yes | | | | yes | yes | |
| ELN | NA | Body | chr7:73474232-73476337 | 2105 | | | | | | | | | | | | | | yes | yes | | yes | | | |
| LIMK1 | NA | Body | chr7:73508618-73508994 | 376 | | | | | 1 | | | | | | | | | | | | | | | |
| LAT2 | NA | TSS1500 | chr7:73623637-73623876 | 239 | | | | | | | | | 1 | | | | | yes | | | | yes | yes | |
| CLIP2 | Island | 5'UTR | chr7:73720808-73721031 | 223 | | | 4 | 1 | 1 | | | | 1 | | | | | yes | | | | | | |
| GTF2IRD1 | N_Shore;Island | 5'UTR | chr7:73894573-73895278 | 705 | | | 8 | | | | | 0 | 2 | | | | 1 | yes | yes | | | | | |
| HIP1 | Island | Body | chr7:75189069-75189250 | 181 | | | | | | | | | | | | | | yes | | | | | yes | |
| CCL26 | NA | TSS200 | chr7:75419119-75419311 | 192 | | | 6 | | | | | | | | | | | yes | | yes | | yes | yes | |
| CCL24 | NA | TSS200 | chr7:75443075-75443246 | 171 | | | | | 1 | | | | 1 | | | | | yes | yes | | | | | |
| POR;SNORA14A | NA | 5'UTR;TSS1500;TSS200 | chr7:75572817-75573192 | 375 | | | | | | | | | 2 | | | | | yes | yes | | yes | | | |
| TMEM120A | S_Shore | TSS1500 | chr7:75624427-75624699 | 272 | | | | | | | | | | | | | | yes | yes | | | | | |
| SRRM3 | Island | Body | chr7:75911830-75912349 | 519 | 1 | | | | | | | | | | | | | | | | | | yes | |
| YWHAG | N_Shore | 3'UTR | chr7:75957040-75957123 | 83 | | | | | | | | 0 | 1 | | 1 | 1 | | | | | | | yes | |
| SRCRB4D;ZP3 | N_Shore;Island | Body;TSS1500;5'UTR;1stExon | chr7:76026556-76027101 | 545 | | | 4 | | | | | 0 | 1 | | | 1 | | yes | yes | | | | | |
| UPK3B | N_Shore;Island | 3'UTR | chr7:76145280-76145672 | 392 | | | 9 | | | | | | | | | 1 | 1 | yes | yes | | | | | |
| PTPN12 | Island | TSS200;TSS1500 | chr7:77166647-77166693 | 46 | | | | | | | | | 1 | | | | | | | | | yes | yes | |
| MAGI2 | Island | Body | chr7:77669697-77670016 | 319 | | | 4 | | | | | | 2 | | | | | | | | | | | |
| MAGI2 | Island | Body | chr7:79081666-79081791 | 125 | | | | | | | | | 1 | | | | | | | | | | yes | |
| MAGI2 | Island;S_Shore | TSS200;TSS1500 | chr7:79083054-79084196 | 1142 | | 1 | 1 | | | | | 0 | 6 | | | 1 | | yes | | | | yes | | |
| GNAI1 | Island | 1stExon;5'UTR | chr7:79764176-79764291 | 115 | | | | | | | | 0 | 1 | | | | | | | | | | yes | |
| SEMA3C | Island | 5'UTR;1stExon;TSS200 | chr7:80548456-80548725 | 269 | | | | | | | | | 2 | | | | | yes | | | | | yes | |
| CACNA2D1 | Island | TSS1500 | chr7:82073453-82073690 | 237 | | | | | | | | | 2 | | | | | | | | | yes | yes | |
| PCLO | Island | 5'UTR;1stExon | chr7:82792105-82792379 | 274 | | | | | | | | | 1 | | | | | | | | | | | |
| SEMA3E | NA | TSS200 | chr7:83278462-83278594 | 132 | | | | | | | | | 1 | | | 1 | | | | | | yes | | |
| C7orf23 | Island | TSS200 | chr7:86849173-86849187 | 14 | | | | | | | | | 2 | | | | | | yes | | | | | |
| ABCB1;RUNDC3B | Island | 5'UTR;TSS200 | chr7:87257538-87257786 | 248 | | 1 | | | | | | | | | | | | | | | | yes | yes | |
| SRI | N_Shore | Body | chr7:87848143-87848292 | 149 | | | | | | | | | | | | | | | | | | | | |
| SRI | S_Shore | Body;TSS1500 | chr7:87849886-87850377 | 491 | | | | | | | | | | | | | | | | | | | | |
| STEAP4 | NA | 5'UTR | chr7:87935979-87936225 | 246 | | | | | | | | | 1 | | | | | yes | | | | | yes | |
| ZNF804B;MGC26647 | NA | Body;TSS200;1stExon;5'UTR | chr7:88424985-88425148 | 163 | | | 3 | | | | | | | | | | | | | | | | | |
| DPY19L2P4 | Island | TSS1500;TSS200 | chr7:89747958-89748686 | 728 | | 2 | | | | | | 0 | 2 | 1 | | 1 | | | | | | | | |
| STEAP1 | N_Shore;Island | TSS1500;TSS200 | chr7:89783435-89783976 | 541 | | 1 | | | | | | 0 | | | | | | yes | | | yes | | | |
| C7orf63 | N_Shore;Island | TSS1500;TSS200 | chr7:89873797-89874465 | 668 | | | 12 | | | | | 0 | 2 | | | 2 | | yes | | yes | | | | |
| FZD1 | Island | 1stExon;3'UTR | chr7:90896329-90896701 | 372 | | | | | | | | | 1 | | | | | | | | | | yes | |
| MTERF | Island | TSS1500 | chr7:91510314-91510597 | 283 | | | 1 | | | | 1 | | 2 | | | | | yes | | | | | | |
| CYP51A1 | S_Shore;Island | TSS1500 | chr7:91764339-91764605 | 266 | | | 2 | | | | | 0 | 1 | | | 3 | | | yes | yes | | | | |
| FAM133B | S_Shore | TSS1500 | chr7:92220582-92220885 | 303 | | | | | | | | | 1 | | | | | yes | yes | | | | | |
| CDK6 | NA | 3'UTR | chr7:92237896-92238364 | 468 | | | 7 | 1 | | | 1 | 0 | 2 | 1 | | 1 | | | | | | yes | yes | |
| CDK6 | Island | 5'UTR | chr7:92462851-92463254 | 403 | | | | | | | | | 1 | | | | | | | | | yes | yes | |
| CALCR | N_Shore | TSS200 | chr7:93204046-93204092 | 46 | | | | | | | | | 1 | | | | | | | | | | | |
| CALCR | S_Shore | TSS1500 | chr7:93205237-93205295 | 58 | | | | | | | | 0 | 1 | | | | | | | | | | | |
| TFPI2 | S_Shore;N_Shore | TSS1500;Body | chr7:93519220-93520566 | 1346 | | | 7 | | | | | | 2 | | | 1 | | yes | yes | | | | yes | |
| GNG11 | NA | TSS200 | chr7:93551004-93551014 | 10 | | | | | | | | | 1 | | | | | | | | | | | |
| COL1A2 | NA | TSS1500 | chr7:94023308-94023557 | 249 | | | | | | | | 0 | 1 | 1 | | 2 | | | | | | yes | | |
| SGCE;PEG10 | N_Shore;Island;S_Shore | Body;TSS1500;1stExon;TSS200;5'U | chr7:94284380-94286936 | 2556 | | | 5 | | | | | 0 | 8 | | | 1 | | | | | | | yes | |
| PPP1R9A | S_Shore | TSS1500;5'UTR | chr7:94538165-94538894 | 729 | | | 4 | | | | | | | | | | | | | | | yes | | |
| PON1 | Island;S_Shore;N_Shore | 1stExon;5'UTR;TSS200;TSS1500;Bo | chr7:94953653-94954455 | 802 | | | 18 | | | | 1 | 0 | 6 | 1 | | 8 | | yes | yes | yes | | | | |
| PON3 | N_Shore;Island;S_Shore | Body;1stExon;TSS200;TSS1500 | chr7:95025407-95027158 | 1751 | | | 53 | 1 | | | 1 | 0 | 13 | 4 | | 5 | | yes | yes | | | | yes | |
| PON2 | S_Shore | TSS1500 | chr7:95064691-95064738 | 47 | | | | | | | | | | | | | | yes | | | | yes | | |
| ASB4 | NA | TSS1500;TSS200 | chr7:95114680-95115354 | 674 | | | 11 | | | | | | | | | | | | yes | | | yes | | |
| DYNC1I1 | Island | 5'UTR | chr7:95402431-95402733 | 302 | | | 3 | | | | | | | | | | | | | | | | | |
| DYNC1I1 | NA | Body | chr7:95546508-95546556 | 48 | | | | 1 | 1 | | | | | | | | | | | | yes | | | |
| MIR591;SLC25A13 | NA | TSS200;Body | chr7:95849192-95849578 | 386 | | | | | | | | | 1 | | | | | yes | | | yes | yes | | |
| DLX6AS | NA;N_Shelf | Body | chr7:96626151-96627700 | 1549 | | 1 | 1 | | | | | | | | | | | | | | | yes | | |
| DLX6AS | Island;S_Shore;N_Shore | Body | chr7:96631384-96633726 | 2342 | | | 12 | | | | | | 2 | | | | | | | | | | yes | |
| DLX6AS;DLX6 | S_Shore | Body | chr7:96636616-96637152 | 536 | | | | | | | | | 1 | | | | | | | | | | | |
| DLX6AS | NA;N_Shelf | Body;TSS200 | chr7:96641456-96643686 | 2230 | | 2 | | | | | | | 2 | | | | | | | | | yes | | |
| DLX5 | N_Shore;Island | 3'UTR;Body | chr7:96649956-96652894 | 2938 | | 1 | 50 | | | | | | | | | | | | | | | yes | | |
| DLX5 | S_Shore | TSS1500 | chr7:96654548-96656358 | 1810 | | 1 | 47 | 1 | | | | | 5 | | 1 | | | | | | | yes | | |
| TAC1 | N_Shore;Island | TSS1500;TSS200 | chr7:97360615-97361408 | 793 | | 1 | 2 | | | | | 0 | | | | | | | | | | | | |
| LMTK2 | NA | Body | chr7:97807643-97807741 | 98 | | | | | | | | | 1 | | | | | yes | | | | | | |
| LMTK2 | N_Shore | Body | chr7:97832830-97833154 | 324 | | | | | | | | | 1 | | | | | yes | yes | | | | | |
| BHLHA15 | N_Shore | TSS1500 | chr7:97841001-97841334 | 333 | | | 3 | | | | | | | | | | | | | | | | yes | |
| TECPR1 | NA | Body | chr7:97857492-97857971 | 479 | | | | | | | | | 1 | | | | | yes | | | | | | |
| TECPR1 | NA | Body | chr7:97875259-97875333 | 74 | | | | | 1 | | | | | | | | | yes | | | | | | |
| BAIAP2L1 | NA | Body | chr7:97923751-97923834 | 83 | | | | 1 | | | | | | | | | | yes | | | | | | |
| NPTX2 | Island | TSS1500 | chr7:98245830-98246006 | 176 | | | | | | | | | 1 | | | | | | | | | yes | | |
| NPTX2 | S_Shore | Body | chr7:98248993-98249103 | 110 | | | | | | | | | 1 | | | | | | | | | | | |
| NPTX2 | NA | Body | chr7:98255916-98256076 | 160 | | | | | | | | | | | | 1 | | yes | | | | | | |
| TRRAP | Island | Body | chr7:98579466-98579639 | 173 | | 1 | | | | | | | | | | | | yes | | | | | | |
| TRRAP | NA | Body | chr7:98586433-98586548 | 115 | 1 | 1 | | | | | | | | | | | | yes | yes | | | | | |
| TRRAP | NA | Body | chr7:98600622-98600799 | 177 | | | | | | | | | | | | | | yes | | | | | | |
| TRRAP | NA | Body | chr7:98602908-98603561 | 653 | | | | | | | | | 2 | | | | | | | | yes | | | |
| SMURF1 | S_Shelf | Body | chr7:98636086-98636282 | 196 | | | | | | | | | 1 | | | | | yes | | | | | | |
| SMURF1 | N_Shelf;chr7:98741205-987419 | Body;N_Shore | chr7:98738998-98739698 | 700 | | | | | | | | | 1 | | 1 | | | | | | | | | |
| ARPC1B | chr7:98971649-98973170 | Island | chr7:98971852-98972188 | 336 | | | | | | | | | | | 1 | | | yes | | | | | | |
| ARPC1B | Island | Body | chr7:98990626-98991138 | 512 | | | | | | | | | 1 | | | 3 | | | yes | | | | | |

| | | | | | | | | | | | | | | | | | | | | | | | |
|---|---|---|---|---|---|---|---|---|---|---|---|---|---|---|---|---|---|---|---|---|---|---|---|
| BUD31;PTCD1 | N_Shore | Body;3'UTR | chr7:99016932-99017073 | 141 | | | | 1 | | | | | | | | | | | | | | | |
| ZNF498 | NA | Body | chr7:99227370-99227515 | 145 | | | | | | | | | 1 | | | | | | | | | | |
| TRIM4 | Island | Body;TSS200 | chr7:99516603-99517460 | 857 | | | 3 | | | | | 0 | 5 | | | 7 | | yes | | yes | yes | | |
| ZNF3 | S_Shore | TSS1500 | chr7:99680278-99680324 | 46 | | | 20 | | | | | | | | | | | yes | | yes | | | |
| MCM7;MIR25 | S_Shelf;NA | Body;TSS200 | chr7:99690806-99691611 | 805 | | | | | | | | | 2 | | | | | | yes | | | | |
| MCM7 | N_Shore | Body;5'UTR | chr7:99697172-99697496 | 324 | | | | | | | | | | | | | | | | | | | |
| GAL3ST4 | N_Shelf | TSS1500 | chr7:99766655-99767035 | 380 | | | | | | | | | 1 | | | | | yes | | | | | |
| STAG3;GPC2 | Island | TSS200;TSS1500 | chr7:99775422-99775558 | 136 | | 1 | | | 1 | | | 0 | 1 | | | 1 | | | | | | | yes |
| GATS;PVRIG | NA | Body;3'UTR;TSS1500 | chr7:99816256-99816750 | 494 | | | | | | | | | | | | | | | | yes | | | |
| PILRA | NA | TSS1500 | chr7:99970448-99970869 | 421 | | | | | | | | | | | | | | | | | | | |
| TSC22D4 | Island;chr7:100075303-100075 | Body;Island | chr7:100075304-100075402 | 98 | | 1 | | | | | | | | | 1 | | | | | | | yes | yes |
| C7orf51 | Island | Body | chr7:100091181-100091786 | 605 | 2 | 1 | 16 | | | | | | | | 1 | | | yes | | | | | |
| LRCH4 | N_Shore | Body | chr7:100181701-100182085 | 384 | | | 4 | 1 | | | | | | | | | | | | | | | |
| PCOLCE | N_Shore | TSS200 | chr7:100199764-100199952 | 188 | | | | 1 | | | | | | | | | | yes | | | | yes | |
| PCOLCE | chr7:100201614-100203652 | Island | chr7:100203114-100203506 | 392 | | | | | | | | | | | 1 | | | yes | yes | | | | |
| MOSPD3 | N_Shore | TSS1500 | chr7:100208533-100208950 | 417 | | | 3 | | | | | | | | | | | yes | | | | | |
| TFR2 | N_Shore;Island | Body | chr7:100224015-100224557 | 542 | | | | | 2 | | | | | | | | | | | | | | |
| TFR2 | NA | Body | chr7:100238750-100239188 | 438 | | | 1 | | | | | | | | | | | | | | | | yes |
| ACTL6B | Island | Body | chr7:100253792-100254149 | 357 | | 1 | | | 1 | | | | 1 | | | | | yes | | | | | |
| GIGYF1 | S_Shore | Body | chr7:100285650-100285913 | 263 | | | | | | | | | 1 | | | | | | | | | | |
| POP7 | S_Shore | Body | chr7:100304669-100305063 | 394 | | | | | | | | | | | | | | yes | | | | | |
| EPO | Island | TSS1500 | chr7:100318190-100318239 | 49 | | | | | 2 | | | | | | | | | yes | | | | yes | |
| SLC12A9;TRIP6 | N_Shore;S_Shore | Body;TSS1500 | chr7:100463206-100465657 | 2451 | | | 15 | 1 | | | | | 5 | 2 | | | | yes | | | | | |
| SRRT | N_Shore | 3'UTR | chr7:100486248-100486423 | 175 | | | | | | | | | 1 | | | | | | | | | | |
| UFSP1;ACHE | N_Shore | TSS1500;Body;3'UTR | chr7:100488398-100489022 | 624 | | | 2 | 1 | | | | | | | | | | yes | | | | | yes |
| ACHE | N_Shore | Body;5'UTR | chr7:100490570-100492376 | 1806 | | | 5 | | | | | | | | | | | | | | | yes | |
| MUC12 | S_Shore;S_Shelf | TSS1500;TSS200 | chr7:100611492-100612891 | 1399 | | | 4 | | | | | 0 | | | | | | | | | | | yes |
| TRIM56 | N_Shore | 5'UTR | chr7:100730471-100730929 | 458 | | | | | | | | | 1 | | | | | yes | | | | yes | |
| C7orf52 | Island | Body | chr7:100817863-100818038 | 175 | | | | | | | | | 1 | | | | | yes | | | | | |
| MOGAT3 | N_Shelf | Body | chr7:100841473-100841554 | 81 | | | 3 | | | | | 0 | | | | | | yes | | | | | |
| MOGAT3 | N_Shore | 1stExon | chr7:100844060-100844445 | 385 | | | | | | | | | | | | | | yes | | | | | |
| CLDN15 | NA | 1stExon;5'UTR | chr7:100881007-100881429 | 422 | | | 3 | | | | | | | | | | | yes | | | | | |
| EMID2 | N_Shore;Island | TSS1500;TSS200 | chr7:101005832-101006308 | 476 | | 1 | 8 | | | | | 0 | 1 | | | | | yes | | | | | |
| MYL10 | NA | TSS1500 | chr7:101273741-101274037 | 296 | | 1 | 5 | | | | | | | | | | | yes | | yes | | | |
| CUX1 | NA | Body | chr7:101500014-101500514 | 500 | | | 5 | | 1 | | | | | | 1 | | | | | | | | |
| CUX1 | NA | Body | chr7:101512828-101513100 | 272 | | | | | | | | | 1 | | | | | yes | yes | | | | |
| CUX1 | N_Shelf | Body | chr7:101555711-101556183 | 472 | | | | | | | | | | | | | | | | | | | yes |
| CUX1 | N_Shore | Body | chr7:101558400-101558597 | 197 | | | 14 | | | | | | | | | | | yes | yes | | | | |
| CUX1 | NA | Body | chr7:101596140-101596404 | 264 | | | 2 | | | | | | | | | | | yes | | | yes | | |
| CUX1 | NA | Body | chr7:101688648-101688730 | 82 | | | | 1 | | | | 0 | 1 | | | | | | | | | | |
| CUX1 | NA | Body | chr7:101768610-101768874 | 264 | | | 16 | 1 | | | | | 2 | | 1 | | | yes | yes | | | | yes |
| CUX1 | NA | Body | chr7:101813842-101814109 | 267 | | | 1 | | | | | | | | | | | yes | | | | | |
| CUX1 | NA | Body | chr7:101901663-101901764 | 101 | | | | 1 | | | | | | | | | | yes | | | | | |
| SH2B2 | N_Shore | TSS1500 | chr7:101927846-101928206 | 360 | | | | | 1 | | | | | | | | | yes | | | | | |
| SH2B2 | Island;N_Shore | Body | chr7:101936230-101936527 | 297 | | | 3 | | | | | 0 | 1 | | | 2 | | yes | | | | yes | |
| SH2B2 | Island | Body | chr7:101943990-101944556 | 566 | | | 2 | | | | | | | | | | | yes | | | | yes | |
| SH2B2 | Island | Body;3'UTR | chr7:101961796-101962267 | 471 | | | 9 | | | | | 0 | 1 | 1 | | 1 | | | | | | yes | yes |
| PRKRIP1 | NA | 3'UTR | chr7:102066852-102067440 | 588 | | | | | | | | | | | | | | yes | | | | | |
| LRRC17;FBXL13 | NA | TSS200;Body;5'UTR;1stExon | chr7:102553293-102553457 | 164 | 1 | | | 1 | | | | 0 | 1 | | | 3 | | | | | | | |
| LRRC17;FBXL13 | NA | nan;Body | chr7:102574105-102574504 | 399 | | | | | | | | | 1 | | 1 | | | yes | yes | | | | yes |
| RPL19P12;NAPEPLD | NA | TSS1500;5'UTR | chr7:102783873-102784228 | 355 | | | | | | | | | 1 | | | | | yes | | yes | | | |
| NAPEPLD | Island | TSS1500 | chr7:102790112-102790406 | 294 | | | 2 | | | | | | | | | | | | | | | | yes |
| DPY19L2P2 | Island | Body;TSS200;TSS1500 | chr7:102920771-102920980 | 209 | | 1 | | | | | | | 3 | | | | | | | | | | |
| SLC26A5 | N_Shore | 5'UTR | chr7:103083817-103084288 | 471 | | | | | | | | | 1 | | | | | | | | | | |
| SLC26A5 | Island | 5'UTR | chr7:103085736-103085984 | 248 | | | | | | | | | | | | 1 | | | | | | | |
| RELN | Island | TSS200 | chr7:103630013-103630490 | 477 | | | | | | | | | 2 | | | | | | | | | yes | yes |
| ORC5L | NA | Body | chr7:103848079-103848368 | 289 | | | | | | | | | 1 | | | 1 | | | | | | | |
| EFCAB10 | N_Shore | TSS200 | chr7:105221985-105222024 | 39 | | | 1 | | | | | 0 | 1 | | | | | yes | | | | | |
| ATXN7L1 | NA | Body;5'UTR | chr7:105319437-105319819 | 382 | | | 7 | | | | | 0 | 1 | 2 | | | | | yes | | | | |
| ATXN7L1 | NA | Body | chr7:105331648-105332279 | 631 | 1 | | | | | | | | | | | | | | | | | | yes |
| FLJ23834 | NA | TSS1500 | chr7:105603432-105603743 | 311 | | 1 | | | 1 | | | | | | | | | | | | yes | | |
| FLJ36031 | Island | 3'UTR;1stExon;5'UTR | chr7:106300577-106301767 | 1190 | | | | | | | | 0 | 1 | 1 | | | | | | | | | yes |
| PIK3CG | N_Shelf;chr7:106508057-10650 | TSS1500;N_Shelf;TSS200 | chr7:106505688-106505961 | 273 | | | | 1 | | | | | 1 | 1 | 1 | | | | | | | | |
| COG5;DUS4L | S_Shore | TSS200;5'UTR | chr7:107205105-107205392 | 287 | | | 7 | | | | | 0 | | | | | | yes | yes | | yes | | |
| SLC26A4;LOC286002 | N_Shore | TSS1500;Body;TSS200 | chr7:107300287-107301048 | 761 | | | 11 | | 1 | | | | 1 | | | | | | | | yes | | |
| LAMB1 | Island | TSS200 | chr7:107643925-107643956 | 31 | | | | | | | | | 1 | | | | | yes | | | | | |
| LAMB4 | NA | 1stExon;5'UTR | chr7:107770791-107771148 | 357 | | | | | | | | | | 1 | | | | | | | | yes | |
| EIF3IP1 | NA | TSS200 | chr7:109600425-109600733 | 308 | | | | | | | | | | | | | | | | yes | yes | | |
| DOCK4 | Island | Body | chr7:111368367-111368847 | 480 | | | | | | | | 0 | 3 | | | 1 | | yes | | yes | | | |
| IFRD1 | NA | TSS1500 | chr7:112062297-112062419 | 122 | | | | | | | | | 1 | | | | | | | | | | |
| C7orf53 | NA | TSS200;TSS1500 | chr7:112120785-112121259 | 474 | | | 3 | | | | | | | | | | | yes | | | | yes | |
| TMEM168 | Island;S_Shore | 5'UTR;TSS1500 | chr7:112430152-112430779 | 627 | | | 6 | 1 | 3 | | | | | | | | | | yes | | | | |
| GPR85 | NA | 5'UTR | chr7:112725899-112726396 | 497 | | | 2 | | | | | | | | | 1 | | | | | | yes | yes |
| GPR85 | NA | TSS1500;5'UTR | chr7:112727423-112727914 | 491 | | | | | | | | | 1 | | | 3 | | | | | | yes | |
| FOXP2 | NA | TSS1500;1stExon;5'UTR | chr7:114053894-114056377 | 2483 | | | 40 | | 1 | | | 0 | 4 | | | 1 | | | yes | | | yes | |
| MDFIC | N_Shore | TSS1500 | chr7:114561804-114562063 | 259 | | | 7 | | | | | | | | | | | | | | | | |
| TES | NA | TSS1500;Body | chr7:115862648-115863026 | 378 | | | 22 | | | | | | | | | | | | | | | | |
| CAV2 | N_Shore | TSS200 | chr7:116139367-116139425 | 58 | | | | | | | | 0 | 1 | | | | | yes | | | | yes | |
| ST7OT4;ST7OT1;ST7 | N_Shore | TSS1500;Body | chr7:116592956-116592968 | 12 | | | 3 | | 1 | | | | | | | | | yes | | | | | yes |
| ST7 | NA | Body | chr7:116660172-116660240 | 68 | | | 1 | 1 | | | | | 1 | | | | | yes | | | | | yes |
| WNT2 | Island | TSS200 | chr7:116963492-116963502 | 10 | | | 1 | | | | | | 1 | 1 | | | | | | | | | yes |
| ASZ1 | N_Shore;Island;S_Shore | Body;1stExon;TSS200 | chr7:117067136-117067919 | 783 | | | | | | | | | 1 | 1 | | 2 | | | | | | | |
| CFTR | NA | TSS1500 | chr7:117119382-117120064 | 682 | | 1 | | | | | | 0 | 1 | | | | | | yes | | | | yes |
| KCND2 | N_Shore | 1stExon;5'UTR | chr7:119914282-119914778 | 496 | | | 4 | | | | | | | | | | | | | | | | |
| WNT16 | N_Shore | TSS1500;Body;TSS200 | chr7:120968877-120969174 | 297 | | 1 | 10 | | | | | 0 | | | | 2 | | | | | | | |
| PTPRZ1 | NA | Body | chr7:121651766-121652024 | 258 | | | | | | | | | | | | | | | | | | | |
| FEZF1 | Island | 1stExon | chr7:121944173-121944637 | 464 | | | | | | | | | 1 | | | | | | | | | yes | yes |
| FEZF1 | Island | TSS1500 | chr7:121946018-121946213 | 195 | | | | | | | | | 1 | | | | | | | | | | |
| CADPS2 | Island;S_Shore | 5'UTR;1stExon;TSS200;TSS1500 | chr7:122526408-122527499 | 1091 | | | | | | | | | 3 | | | | | | | | | | yes |
| LMOD2 | NA | TSS1500 | chr7:123295321-123295821 | 500 | | | | | | | | | | | | | | yes | yes | | | | |
| SPAM1 | NA | TSS1500 | chr7:123564016-123564379 | 363 | | | | | | | | | | | | | | | | yes | | | |
| GPR37 | Island | 1stExon | chr7:124404244-124404431 | 187 | | 1 | 1 | | | | | | | | | | | | | | | | |
| GPR37 | Island | TSS200 | chr7:124405732-124405760 | 28 | | | | | | | | | 1 | | | | | | | | | | yes |
| MIR592;GRM8 | NA | Body;TSS200 | chr7:126698156-126698829 | 673 | | | 2 | | | | | | 3 | | | | | | yes | | | | |
| GCC1 | Island | TSS1500 | chr7:127225891-127225949 | 58 | | | 9 | | | | | 0 | 1 | | | | | yes | yes | | | | |
| FSCN3 | NA | TSS200 | chr7:127233574-127233742 | 168 | | | | 1 | | | | | | | | | | yes | | | | yes | |
| PAX4 | NA | 5'UTR;1stExon;TSS1500 | chr7:127255591-127256920 | 1329 | | | 4 | | | | | | | | | | | | | | | | |
| LRRC4;SND1 | N_Shore;Island;chr7:12767115 | 5'UTR;1stExon;Body;TSS200;TSS1! | chr7:127670993-127673038 | 2045 | | | | | | | | | | | | | | | | | | | |
| LEP | Island | TSS1500;TSS200 | chr7:127880932-127881440 | 508 | | 1 | 3 | | | | | 0 | 5 | | | 1 | | yes | | | | yes | yes |

| | | | | | | | | | | | | | | | | | | | | | | | |
|---|---|---|---|---|---|---|---|---|---|---|---|---|---|---|---|---|---|---|---|---|---|---|---|
| C7orf68 | N_Shore | TSS1500 | chr7:128094710-128095027 | 317 | | | | | | | | | | | | | | yes | | | | | |
| FAM71F1 | NA | TSS200 | chr7:128355262-128355632 | 370 | | | | | | | | | | | | | | | | | | | |
| FLNC | Island | Body | chr7:128482287-128482679 | 392 | | | | | | | | | | | | | | | | | | yes | |
| FLNC | Island | Body | chr7:128494060-128494568 | 508 | | 1 | | | | | | | | | | | | | | | | | |
| KCP | Island | Body | chr7:128530795-128531479 | 684 | | | 8 | | | | | 0 | | | | | | | | | | yes | yes |
| KCP | Island | TSS200 | chr7:128550812-128550967 | 155 | | | 26 | | | | | | 3 | | | | | yes | | | | | |
| IRF5 | N_Shore | TSS1500 | chr7:128577593-128577819 | 226 | | | 3 | | | | | | | | | | | | yes | yes | | | |
| IRF5 | S_Shore | 5'UTR;TSS1500;TSS200 | chr7:128579876-128580792 | 916 | | | | | 1 | | | | 1 | | | | | | | | | | |
| LOC286016;TNPO3 | Island | Body;TSS1500 | chr7:128696150-128696229 | 79 | | | | | | | | 0 | 1 | | | | | yes | | | | | |
| LOC407835 | N_Shore | TSS200 | chr7:128766272-128766655 | 383 | | | 1 | | | | | | | | | | | yes | | | | | yes |
| AHCYL2 | NA | 1stExon;Body;5'UTR;TSS200 | chr7:129007902-129008407 | 505 | | | 2 | 1 | | | | | 1 | | | | | | | | | | |
| MIR182 | NA | Body;TSS1500 | chr7:129410227-129411088 | 861 | | | | | | | | | | | | | | | | | | | yes |
| MIR96;MIR183 | N_Shelf | TSS200;TSS1500;Body | chr7:129414527-129414894 | 327 | 6 | | 10 | | | | | | | | | | | | | | | | |
| TMEM209 | S_Shore | TSS1500 | chr7:129845574-129845832 | 258 | | | 6 | | | | | | | | | | | yes | | | | | |
| CPA2 | NA | Body | chr7:129912803-129913310 | 507 | | | | | 2 | | | | | | | | | | | yes | | yes | |
| CPA1 | N_Shore | TSS1500 | chr7:130019715-130020328 | 613 | | | 7 | | | | | | 1 | | | | | | | | | | |
| CPA1 | Island | Body | chr7:130021519-130021651 | 132 | | | | | 1 | | | | | | | | | | | | | | |
| MEST | N_Shore;Island | TSS1500;TSS200;1stExon;5'UTR | chr7:130124971-130126871 | 1900 | | | 29 | | | | | 0 | 9 | | 1 | 2 | 2 | yes | yes | | | | |
| MESTIT1;MEST | N_Shore;Island | Body;5'UTR;TSS1500;TSS200 | chr7:130129946-130133110 | 3164 | | | 5 | | | | | | 19 | 1 | | 5 | | | yes | | | | yes |
| KLF14 | Island;S_Shore | TSS1500 | chr7:130419136-130419932 | 796 | | | 3 | | | | | | | | | | | | yes | | | yes | yes |
| PODXL | N_Shore | Body | chr7:131240440-131240531 | 91 | | | 3 | | | | | | | | | | | | | | | yes | |
| PLXNA4 | Island | TSS1500;5'UTR | chr7:132262526-132262962 | 436 | | | | | | | | | | | | | | | | | | | yes |
| PLXNA4 | NA | 5'UTR | chr7:132299570-132300110 | 540 | 1 | | 4 | | | | | | | | | | | | | | yes | yes | yes |
| AKR1B15 | NA | TSS200 | chr7:134250141-134250240 | 99 | | | 5 | | | | | | | | | | | | | yes | yes | | |
| CALD1 | NA | TSS200 | chr7:134464123-134464411 | 288 | | | | | 1 | | | 0 | 5 | | | 5 | | | | | | | |
| CALD1 | NA | TSS1500;Body;nan;1stExon;5'UTR | chr7:134574947-134576386 | 1439 | | | 6 | 1 | | | | | 5 | | 1 | | | yes | | | | yes | |
| TMEM140 | NA | TSS200;1stExon;5'UTR;TSS1500 | chr7:134832221-134833299 | 1078 | | 1 | 41 | | | | | | 1 | | | | | yes | yes | | | | |
| C7orf49 | S_Shore | TSS1500 | chr7:134856562-134856681 | 119 | | | 5 | | | | | | 1 | | | | | | | yes | | | |
| SLC13A4 | NA | 5'UTR;1stExon;TSS200 | chr7:135412356-135413232 | 876 | | 2 | | | | | | | | | | | | | yes | | | | |
| FAM180A | NA | 1stExon;5'UTR;nan | chr7:135433353-135433790 | 437 | | | | | | | | | 1 | 1 | 1 | | | | | | | | |
| CHRM2 | N_Shore;Island | TSS1500;5'UTR;TSS200;1stExon | chr7:136553088-136553884 | 796 | | 1 | 1 | | | | | | 1 | | | | | | | | | yes | |
| PTN | NA | TSS200 | chr7:137028617-137029097 | 480 | | | | | | | | | | | | 1 | | | | | | | yes |
| DGKI | Island | TSS200;TSS1500 | chr7:137531697-137532374 | 677 | | 1 | | | | | | | 1 | | | | | | | | | yes | |
| AKR1D1 | NA | TSS1500 | chr7:137760128-137760331 | 203 | | | | | | | | | 1 | | | | | | | yes | | | |
| SVOPL | N_Shore;Island | Body;5'UTR;TSS200 | chr7:138347530-138349443 | 1913 | | | 3 | | | | | | | | | 4 | 1 | | | | | | yes |
| SVOPL | NA | TSS1500 | chr7:138364464-138364774 | 310 | | | | | | | | | | | | | | yes | | | | | |
| ATP6V0A4 | NA | Body | chr7:138413328-138413421 | 93 | | | | | | | | | | | | | | yes | | yes | | | yes |
| ATP6V0A4;TMEM213 | NA | 5'UTR;TSS200;1stExon | chr7:138482580-138483143 | 563 | | | | | | | | | 1 | | | | | yes | | | | | yes |
| KIAA1549 | NA | Body | chr7:138601707-138602695 | 988 | | | | | | | | | 2 | | | | | yes | | | | | |
| ZC3HAV1L | Island | TSS1500 | chr7:138720989-138721065 | 76 | | | | 1 | | | | | | | | | | yes | | | yes | | |
| KLRG2 | N_Shore;Island | Body;1stExon;5'UTR;TSS200 | chr7:139167427-139168804 | 1377 | | | | | 2 | | | | 1 | | | | | yes | | | | | |
| CLEC2L | N_Shore;Island | TSS1500;1stExon | chr7:139208223-139208979 | 756 | | | | | | | | | 2 | | | | | yes | yes | | | yes | yes |
| HIPK2 | NA | 3'UTR | chr7:139256620-139257086 | 466 | | | | | | | | | 1 | | | | | | | | | yes | yes |
| HIPK2 | NA | Body | chr7:139333296-139333410 | 114 | | | | 1 | | | | | | | | | | | yes | | | | |
| RAB19 | NA | TSS200 | chr7:140103671-140104086 | 415 | 1 | | 12 | | | | | | | | | 1 | | yes | yes | | yes | | yes |
| BRAF | S_Shore | TSS1500 | chr7:140625365-140625783 | 418 | | | | | | | | | | | | | | yes | yes | | | | |
| TAS2R38 | NA | TSS200 | chr7:141673582-141673903 | 321 | | | | | | | | | | | | | | | yes | | | | |
| MGAM | NA | TSS1500;TSS200 | chr7:141695293-141695759 | 466 | | | | | 2 | | | | | | | | | yes | | yes | | | |
| EPHB6 | NA | Body | chr7:142561918-142562153 | 235 | | | | | | | | | | | | | | | | | | yes | |
| TRPV6 | NA | TSS1500 | chr7:142583715-142583836 | 121 | | | | | | | | | 1 | | | | | | | | yes | | |
| C7orf34 | NA | TSS1500;TSS200 | chr7:142636123-142636518 | 395 | | | 7 | | | | | | | | | | | yes | | | | yes | |
| KEL | NA | 1stExon;5'UTR | chr7:142659349-142659425 | 76 | | | | | | | | | | | | | | | | yes | | | |
| OR6V1 | NA | TSS1500;TSS200 | chr7:142748962-142749542 | 580 | | | 5 | | | | | | 1 | | | | | yes | | | | yes | yes |
| PIP | NA | nan;TSS200 | chr7:142828903-142829304 | 401 | | | | | | | | | | | 1 | | | | | | | | |
| TMEM139 | N_Shelf | TSS1500 | chr7:142981689-142982463 | 774 | | | | 1 | 1 | | | | | | | | | | | | | | |
| CLCN1 | NA | TSS1500;TSS200 | chr7:143012960-143013614 | 654 | | | 3 | | | | | | | | | | | | | | | | |
| EPHA1 | NA | Body;TSS200 | chr7:143105595-143106298 | 703 | | | 3 | | | | | | | | | | | yes | | | | | yes |
| TAS2R60 | NA | TSS1500 | chr7:143140164-143140674 | 510 | | 1 | | | | | | | 1 | | | | | yes | | | | | yes |
| FAM115A | N_Shelf | 5'UTR | chr7:143579665-143580147 | 482 | | | | | 2 | | | | 1 | | 1 | | | yes | | | | yes | yes |
| FAM115A | Island | 5'UTR | chr7:143582146-143582630 | 484 | | | 1 | 1 | 1 | | | 0 | 1 | | | 1 | | | | | | | yes |
| OR6B1 | NA | TSS200 | chr7:143700925-143701215 | 290 | | | | | | | | | | | | | | yes | yes | | | yes | |
| OR2A5 | NA | TSS1500;TSS200 | chr7:143746979-143747596 | 617 | | | | | | | | | 2 | | 1 | | | | | | | yes | |
| CNTNAP2 | Island | TSS200;1stExon;5'UTR | chr7:145813413-145814306 | 893 | | | | | | | | 0 | 6 | | | 1 | | | | | | yes | |
| MIR548F3;CNTNAP2 | NA | TSS1500;Body | chr7:147591462-147591961 | 499 | | | | 1 | | | | | | | | | | yes | | | | | |
| CUL1 | N_Shore | TSS1500 | chr7:148394542-148394888 | 346 | | | | 1 | | | | | | | | | | | | | | | |
| CUL1 | N_Shore | Body | chr7:148480080-148480503 | 423 | | 1 | 4 | | | | | | | | | | | yes | | | yes | | |
| CUL1 | NA | 3'UTR | chr7:148497907-148497970 | 63 | | | | | | | | | | | | | | yes | | | | | |
| ZNF767 | S_Shore | TSS1500 | chr7:149322723-149323101 | 378 | | | | | | | | | | | | | | yes | | | yes | | |
| SSPO | S_Shore | TSS1500 | chr7:149471884-149472134 | 250 | | | | | | | | | 1 | | | | | | | | yes | | |
| SSPO | N_Shelf;N_Shore | Body | chr7:149484798-149485981 | 1183 | | | | | | | | 0 | 1 | | 1 | 13 | | | yes | | | | |
| ZNF862 | NA | Body | chr7:149543136-149543177 | 41 | 5 | | | | | | | | | | | | | | yes | | yes | | |
| ATP6V0E2;LOC401431 | N_Shore | TSS200;Body | chr7:149569998-149570071 | 73 | | | | | | | | 0 | 1 | | | 1 | | yes | yes | | | | |
| LRRC61;ACTR3C | N_Shore;Island;S_Shore | TSS1500;5'UTR;TSS200;1stExon | chr7:150019477-150021553 | 2076 | | | 12 | 1 | 1 | | | 0 | 17 | 10 | | 7 | | yes | | yes | | | yes |
| C7orf29;LRRC61 | NA | TSS1500;5'UTR;TSS200 | chr7:150026567-150027050 | 483 | 1 | | | | | | | | | | | | | | | | | | |
| RARRES2 | Island | 5'UTR | chr7:150038502-150038898 | 396 | | 1 | | | | | | 0 | 1 | | | | | | | | | | |
| REPIN1 | N_Shore | 5'UTR;Body;TSS1500 | chr7:150067712-150068240 | 528 | | | | | | | | | 2 | | | | | yes | | | | | |
| LOC728743 | N_Shore | Body | chr7:150104655-150105713 | 1058 | | | | | | | | | 3 | | | | | yes | yes | | | | yes |
| GIMAP8 | NA | TSS200;5'UTR;1stExon;TSS1500 | chr7:150147536-150148073 | 537 | | 1 | 21 | 1 | 1 | | | | 2 | | | | | yes | yes | | | | |
| GIMAP8 | NA | Body | chr7:150174507-150174632 | 125 | | | | | | | | | | | | | | | | | | | |
| GIMAP7 | NA | TSS1500;TSS200 | chr7:150210742-150212873 | 2131 | | | 10 | | | | | | 1 | | | | | yes | yes | | | | |
| GIMAP7 | NA | 5'UTR | chr7:150216489-150217081 | 592 | | | | | | | | | 2 | | | | | | yes | | | | |
| GIMAP4 | NA | TSS200 | chr7:150264284-150264767 | 483 | | | | | | | | | | | | | | | | | yes | | |
| GIMAP1 | N_Shelf;NA | TSS200;TSS1500 | chr7:150412715-150413801 | 1086 | | | | 1 | 1 | | | | 1 | | | | | | | yes | yes | | |
| GIMAP1 | Island | Body | chr7:150417431-150417775 | 344 | | | | | | | | | | | | | | | | | | | |
| GIMAP5 | NA | TSS1500;1stExon;5'UTR | chr7:150433828-150434640 | 812 | | | 21 | 1 | | | | | 3 | 1 | 1 | | | | | yes | | | |
| TMEM176B;TMEM176A | Island;S_Shore | TSS1500;5'UTR;1stExon;TSS200;Bo | chr7:150497065-150499737 | 2672 | | 1 | 4 | 1 | | | | 0 | 3 | 1 | | 1 | | | yes | | | | yes |
| ABP1 | NA | TSS1500 | chr7:150549175-150549472 | 297 | | 1 | | | | | | | | | | | | | | | | | |
| ABP1 | Island | Body | chr7:150554420-150554915 | 495 | | | | | | | | | | | | | | | | | | | |
| KCNH2 | Island | Body | chr7:150644459-150645092 | 633 | | | | | | | | | | | | | | | yes | | | yes | yes |
| KCNH2 | Island | TSS200;Body | chr7:150652948-150653115 | 167 | | 1 | | | 2 | | | | | | | | | | | | | | |
| NOS3 | NA | 5'UTR;TSS1500 | chr7:150689873-150689889 | 16 | | | 6 | | | | | | | | | | | | | | | | |
| NOS3 | Island | Body | chr7:150705760-150706355 | 595 | 1 | | 9 | | | | | 0 | 2 | | | 1 | | yes | | | | yes | yes |
| ACCN3;ABCB8 | N_Shore | TSS1500;3'UTR | chr7:150744274-150744599 | 325 | | | 2 | 1 | | | | | | | | | | | yes | | | | |
| FASTK;TMUB1 | Island | TSS1500;Body | chr7:150778724-150779710 | 986 | | | 6 | 1 | | | | | 3 | | | | | | | | | yes | yes |
| AGAP3 | S_Shore | Body | chr7:150786044-150786082 | 38 | | 1 | 5 | 1 | | | | 0 | 1 | | 1 | | | | yes | | | | |
| AGAP3 | Island | Body | chr7:150822655-150823386 | 731 | | | 2 | | | | | 0 | 1 | | | 4 | | | | | | yes | |
| ASB10 | NA | 1stExon;Body;5'UTR;TSS200;TSS1 | chr7:150884003-150885654 | 1651 | | | 8 | | | | | | 2 | | | | | yes | yes | | | | yes |
| WDR86 | Island | 3'UTR;Body | chr7:151078646-151079501 | 855 | | | 5 | | | | | 0 | 2 | 1 | | | | | | yes | yes | | |
| WDR86 | NA | Body | chr7:151087656-151087979 | 323 | | | | | | | | | | | | | | | yes | | | | |

| | | | | | | | | | | | | | | | | | | | | | | | |
|---|---|---|---|---|---|---|---|---|---|---|---|---|---|---|---|---|---|---|---|---|---|---|---|
| WDR86 | N_Shore | Body | chr7:151106022-151106245 | 223 | | | 6 | | | | | | | | | | | yes | | | | | |
| WDR86 | Island | TSS200 | chr7:151107261-151107400 | 139 | | | | | | | | | | 1 | | | | | | | | | |
| CRYGN | Island;S_Shore | Body;TSS1500 | chr7:151136940-151137882 | 942 | | | 6 | | | | | | | | | | | yes | yes | | | | yes |
| RHEB | Island | TSS1500 | chr7:151217575-151217961 | 386 | | | | 1 | | | | 0 | 1 | | | | | | | | | | |
| PRKAG2 | NA | Body;TSS1500 | chr7:151330008-151330230 | 222 | | | 6 | | | | | | | | | | | | yes | | | | yes |
| PRKAG2 | NA | Body | chr7:151442351-151442967 | 616 | | 2 | 7 | | | | | | 1 | | | | | yes | | | | | |
| PRKAG2 | NA | 5'UTR;Body | chr7:151493629-151493820 | 191 | | | 5 | | | | | | 1 | | | | | | | | | | |
| PRKAG2 | NA | 5'UTR;Body | chr7:151504844-151505116 | 272 | 1 | | | | | | | | | | | | | | | | | | |
| PRKAG2 | NA | Body | chr7:151526959-151527251 | 292 | | | | | | | | | | | | | | | yes | | | | |
| PRKAG2 | NA | Body | chr7:151553471-151553782 | 311 | | | | | | | | | | | | | | yes | yes | | | | |
| GALNTL5 | NA | TSS200 | chr7:151653454-151653890 | 436 | | | 6 | | | | | | | | | | | yes | | | | | |
| GALNTL5 | NA | 5'UTR;Body | chr7:151664041-151664190 | 149 | | | | | | | | | 1 | | | | | yes | yes | | | | |
| GALNT11 | N_Shore | TSS1500 | chr7:151722461-151722536 | 75 | | | | | | | | | 1 | | | | | | yes | | | | yes |
| MLL3 | NA | Body | chr7:152131930-152132218 | 288 | | | | | | | | | 1 | | | 1 | | | | | | | yes |
| FABP5L3;MLL3 | NA | TSS1500 | chr7:152133420-152133914 | 494 | | | | | 1 | | | 0 | 2 | | | 2 | | | | | | yes | yes |
| ACTR3B | NA | Body | chr7:152549172-152549357 | 185 | | 1 | | | | | | | | | | | | | | | | | |
| DPP6 | Island;S_Shore | TSS200;1stExon;5'UTR;Body | chr7:153584416-153586592 | 2176 | | | 3 | | 4 | | | 0 | 4 | | 1 | 3 | | | | | | yes | |
| DPP6 | Island | TSS1500;Body;TSS200 | chr7:153749206-153749756 | 550 | | | | | 1 | | | | 2 | | | | | | | | | yes | yes |
| DPP6 | S_Shore;Island | Body;TSS1500;1stExon | chr7:154002064-154003309 | 1245 | 3 | | 14 | | 1 | | | | | | | | | | | | | yes | yes |
| DPP6 | NA | Body | chr7:154473202-154474156 | 954 | 2 | | 3 | | | | | | | | | | | | yes | | | | |
| DPP6 | N_Shore | Body | chr7:154540294-154540369 | 75 | | | | | | | | | | | | | | | | | | | yes |
| DPP6 | Island | Body | chr7:154542060-154542313 | 253 | | | | | | | | | | | | | | yes | yes | | | | yes |
| DPP6 | Island | Body | chr7:154585834-154586375 | 541 | | | | | | | | 0 | | | | 2 | | | | | | | |
| DPP6 | N_Shore;Island;S_Shore | Body;3'UTR | chr7:154684051-154685726 | 1675 | | | 11 | | | | | | 7 | | | 4 | | yes | | | | yes | |
| HTR5A | N_Shore;Island | TSS1500;TSS200;1stExon | chr7:154862157-154863381 | 1224 | 2 | 1 | 8 | | | | | 0 | 1 | | | 1 | | | | | | | |
| EN2 | Island | Body | chr7:155255310-155255791 | 481 | | | | | | | | | 1 | | | | | yes | | | | | yes |
| CNPY1 | N_Shelf | Body | chr7:155299624-155299687 | 63 | | | 7 | | | | | | | | | | | | | | | yes | |
| CNPY1 | N_Shore | 5'UTR | chr7:155301734-155302342 | 608 | | | | | | | | | 1 | | | | | | | | | yes | |
| CNPY1 | N_Shore;Island | 5'UTR | chr7:155325552-155326740 | 1188 | | | | | 1 | | | 0 | | | | | | yes | | | | yes | |
| RBM33 | N_Shore | TSS1500 | chr7:155436380-155436512 | 132 | | | | | | | | | | | | | | yes | | | | | |
| SHH | N_Shore | Body | chr7:155599467-155599780 | 313 | | | | | | | | | | | | | | | | | | | |
| NOM1 | Island | TSS1500 | chr7:156741514-156741733 | 219 | | | 5 | | 1 | | | 0 | 2 | | | 2 | | yes | yes | yes | | yes | yes |
| MNX1 | Island | Body | chr7:156799148-156799393 | 245 | | | | | 1 | | | | | | | | | yes | | | yes | | |
| MNX1 | Island | TSS200;Body | chr7:156801419-156802502 | 1083 | | 1 | 1 | | | | | | 1 | | | | | | | | | yes | yes |
| DNAJB6 | NA | Body | chr7:157179773-157180124 | 351 | | | 4 | 1 | 1 | | | | 1 | | | | | yes | | | | | |
| DNAJB6 | NA | Body | chr7:157185279-157185402 | 123 | | | 6 | | | | | | | | | | | yes | | | | | |
| DNAJB6 | Island | Body | chr7:157192974-157193101 | 127 | | | | | | | | | 1 | | | | | yes | | | | | |
| DNAJB6 | N_Shelf | Body | chr7:157198482-157198869 | 387 | | | 11 | | | | | | | | | | | | | | | | |
| PTPRN2 | NA | Body;3'UTR | chr7:157332376-157335647 | 3271 | 3 | | 21 | | | | | 0 | | | | | | | | | | | |
| PTPRN2 | NA | Body | chr7:157340390-157340554 | 164 | | | | | | | | | | | | | | yes | yes | | | yes | |
| PTPRN2 | N_Shore | Body | chr7:157345090-157346224 | 1134 | | | 15 | | | | | | 1 | | | | | | | | | yes | |
| PTPRN2 | S_Shore | Body | chr7:157347704-157348266 | 562 | | | | | | | | | | | | | | | | | | | |
| PTPRN2 | N_Shelf | Body | chr7:157350867-157351042 | 175 | | | | | | | | | | | | | | | | | | | |
| PTPRN2 | Island | Body | chr7:157353097-157353191 | 94 | | | 6 | | | | | | | | | | | | | | | | |
| PTPRN2 | Island;N_Shore | Body | chr7:157361396-157362094 | 698 | 3 | | | | 2 | | | | 1 | | | | | | | | | | |
| PTPRN2;MIR153-2 | N_Shelf;N_Shore | Body;TSS1500 | chr7:157366933-157367778 | 845 | | | | | | | | | | | | | | | | | | yes | |
| PTPRN2 | N_Shore;Island;S_Shore | Body | chr7:157368881-157371224 | 2343 | | | 20 | | | | 1 | 0 | 3 | 1 | | 1 | | | | | | | |
| PTPRN2 | NA | Body | chr7:157381078-157381240 | 162 | | | | | 1 | | | | | | | | | yes | yes | | | | |
| PTPRN2 | N_Shore;Island | Body | chr7:157386714-157387361 | 647 | | | | | | | | | | | | | | yes | yes | | | | |
| PTPRN2 | S_Shelf | Body | chr7:157391030-157391297 | 267 | | | | | | | | | | | | | | yes | | | | | |
| PTPRN2 | NA | Body | chr7:157394633-157394970 | 337 | | | | | | | | | | | | | | | | | | | |
| PTPRN2 | N_Shelf | Body | chr7:157402263-157402673 | 410 | 1 | | | | | | | | | | | | | yes | | | | | |
| PTPRN2 | Island | Body | chr7:157405965-157406737 | 772 | | | 10 | | 1 | 1 | | 0 | 1 | 2 | | 15 | 1 | | | | | | yes |
| PTPRN2 | S_Shore;N_Shore;Island | Body | chr7:157407820-157410194 | 2374 | | | | | | | | | | | | | | | yes | yes | | | |
| PTPRN2 | N_Shore;Island;S_Shore | Body | chr7:157412195-157414580 | 2385 | | | 12 | | | | | | | | | | | yes | yes | | | | |
| PTPRN2 | N_Shelf | Body | chr7:157437773-157437844 | 71 | | | | | | | | | | | | | | yes | yes | | | | |
| PTPRN2 | S_Shore | Body | chr7:157443760-157444239 | 479 | | | 1 | | | | | | | | | | | | | | | | |
| PTPRN2 | N_Shore;Island | Body | chr7:157447022-157448944 | 1922 | | | 8 | | | | | | | | | | | | | | | yes | |
| PTPRN2 | N_Shelf | Body | chr7:157451260-157451906 | 646 | | | | | | | | | | | | | | | | | | | yes |
| PTPRN2 | Island | Body | chr7:157453775-157454207 | 432 | | | 10 | | | 1 | 1 | 0 | 4 | 2 | | 1 | | yes | | | | | |
| PTPRN2 | NA | Body | chr7:157460657-157464230 | 3573 | | | 8 | | | | | | | | | | | | | | | | |
| PTPRN2 | NA | Body | chr7:157467883-157468443 | 560 | 2 | | 6 | | | | | | | | | | | | | | | | |
| PTPRN2 | N_Shelf;N_Shore | Body | chr7:157474856-157477308 | 2452 | | 1 | | | | | | 0 | 2 | | | 1 | | | | | | yes | yes |
| PTPRN2 | Island | Body | chr7:157482508-157483006 | 498 | | | | | 1 | | | | | | | | | | | | | | |
| PTPRN2 | Island | Body | chr7:157484045-157484250 | 205 | | | | | | | | | | | | | | | | | | | yes |
| PTPRN2 | N_Shore;S_Shore | Body | chr7:157493703-157495969 | 2266 | | | 14 | | | | | | 1 | | | | | yes | | yes | | | |
| PTPRN2 | N_Shore | Body | chr7:157502370-157502620 | 250 | | | 5 | | | | | | 1 | | | | | yes | yes | | | | |
| PTPRN2 | NA | Body | chr7:157510257-157510462 | 205 | | | 1 | | | | | | | | | | | | | | | | |
| PTPRN2 | NA | Body | chr7:157513189-157514822 | 1633 | | | | | | | | | | | | | | | | | | | |
| PTPRN2 | N_Shore | Body | chr7:157523356-157523541 | 185 | | | | | | | | | | | | | | | yes | | | | |
| PTPRN2 | S_Shelf | Body | chr7:157529335-157530177 | 842 | | | | | | | | | | | | | | | | yes | | | |
| PTPRN2 | NA | Body | chr7:157536681-157537120 | 439 | | | | | | | | | 1 | | | | | | yes | | | | yes |
| PTPRN2 | NA | Body | chr7:157543597-157543942 | 345 | | | | | | | | | | | | | | | | | | | |
| PTPRN2 | Island | Body | chr7:157551024-157551275 | 251 | | | | | | | | | | | | | | | | yes | | | |
| PTPRN2 | Island;S_Shore | Body | chr7:157561010-157561925 | 915 | 3 | | 20 | | | | | | | | | | | | | | | | |
| PTPRN2 | Island | Body | chr7:157568170-157568342 | 172 | | | | | | | | | | | | | | | | | | | yes |
| PTPRN2 | NA | Body | chr7:157572612-157573039 | 427 | | | | | | | | | | | | | | yes | | yes | | | |
| PTPRN2 | N_Shore | Body | chr7:157585857-157586866 | 1009 | | | 5 | | | | | | 2 | | | | | | | | | | |
| PTPRN2 | S_Shore | Body | chr7:157588520-157588858 | 338 | | 1 | | | | | | | 1 | | | | | | | yes | | | |
| PTPRN2 | NA | Body | chr7:157602227-157602725 | 498 | | | | | | | | | | | | | | | | yes | yes | | |
| PTPRN2 | N_Shelf | Body | chr7:157608573-157608642 | 69 | | | 1 | | | | | | | | | | | | | yes | | | |
| PTPRN2 | S_Shore | Body | chr7:157620685-157620943 | 258 | | | | | | | | | | | | | | | | | | | |
| PTPRN2 | S_Shelf | Body | chr7:157622032-157622221 | 189 | | | 1 | | | | | | | | | | | | | | | | |
| PTPRN2 | NA | Body | chr7:157624788-157624958 | 170 | | | | | | | | | 1 | | | | | | | | | | |
| PTPRN2 | NA | Body | chr7:157625982-157626309 | 327 | | | 1 | | | | | | | | | | | | yes | | | | |
| PTPRN2 | NA | Body | chr7:157641737-157641916 | 179 | | | | | | | | | | | | | | | | | | | |
| PTPRN2 | N_Shelf;NA | Body | chr7:157643125-157644665 | 1540 | 2 | | 6 | | | | | 0 | 2 | | | | | | | | | | |
| PTPRN2 | N_Shore;Island;S_Shore | Body | chr7:157646605-157648749 | 2144 | | | | | | | | | | | | | | | | | | | |
| PTPRN2 | NA | Body | chr7:157653809-157654170 | 361 | | | | | | | | | | | | | | | | | | | |
| PTPRN2 | S_Shore;N_Shore | Body | chr7:157667762-157668141 | 379 | | | 4 | | | | | | 2 | | | | | | | | | yes | |
| PTPRN2 | S_Shore | Body | chr7:157670677-157670732 | 55 | | 1 | | | | 1 | | 0 | 1 | | | | | | | | | | |
| PTPRN2 | N_Shore;Island;S_Shore | Body | chr7:157690139-157692127 | 1988 | | | 22 | | | | | | | | 1 | | | yes | | | | | |
| PTPRN2 | S_Shore;Island | Body | chr7:157694404-157697240 | 2836 | | | 14 | | | | | | | | | | | | yes | yes | yes | | |
| PTPRN2 | N_Shelf;N_Shore | Body | chr7:157700935-157701262 | 327 | | | | | | | | | 1 | | | | | | yes | | yes | | |
| PTPRN2 | NA | Body | chr7:157709849-157710179 | 330 | | | | | | | | | | | | | | | | | | | |
| PTPRN2 | NA | Body | chr7:157738245-157738329 | 84 | | | | | | | | | 1 | | | | | | | | | | |
| PTPRN2 | NA | Body | chr7:157754600-157754746 | 146 | | | 2 | | | | | 0 | 1 | 1 | 1 | 6 | | yes | | | | | |
| PTPRN2 | NA | Body | chr7:157776346-157776679 | 333 | | | | | | | | | | | | | | | | | | yes | |
| PTPRN2 | NA | Body | chr7:157785628-157786631 | 1003 | | | 9 | | | | | | | | | | | | | | | yes | |

| | | | | | | | | | | | | | | | | | | | | | | | | |
|---|---|---|---|---|---|---|---|---|---|---|---|---|---|---|---|---|---|---|---|---|---|---|---|---|
| PTPRN2 | NA | Body | chr7:157791721-157792100 | 379 | 1 | | 2 | | | | | | | | | | | yes | | | | | | |
| PTPRN2 | NA | Body | chr7:157809235-157809596 | 361 | | | | | 1 | | | 0 | 6 | 1 | | 2 | 1 | | | | | | | |
| PTPRN2 | NA | Body | chr7:157834846-157835049 | 203 | | | | | | | | | | | | | | yes | | | | | | |
| PTPRN2 | NA | Body | chr7:157849639-157849744 | 105 | | | | | | | | | | | | | | | | | | | | |
| PTPRN2 | NA | Body | chr7:157866704-157867612 | 908 | | | 13 | | | | | | | | | | | | | | | | | |
| PTPRN2 | N_Shore | Body | chr7:157890171-157890659 | 488 | 3 | | | | | | | | | | | | | | | | | | | |
| PTPRN2 | N_Shelf | Body | chr7:157915200-157916687 | 1487 | | | 4 | | | | | | | | | | | yes | | yes | | | | |
| PTPRN2 | S_Shore | Body | chr7:157919649-157920174 | 525 | | | | | | | | | | | | | | | | | | | | |
| PTPRN2 | NA;N_Shelf | Body | chr7:157932168-157934005 | 1837 | | | 17 | | | | | | 1 | | | | | | yes | | | yes | | |
| PTPRN2 | N_Shelf | Body | chr7:157935557-157936027 | 470 | | | | | | | | | | | | | | | | yes | | | | |
| PTPRN2 | N_Shore;Island | Body | chr7:157951097-157951799 | 702 | | | | | | | | | 1 | | | | | | | | | | | |
| PTPRN2 | N_Shore;Island;S_Shore | Body | chr7:157956042-157958299 | 2257 | | | 5 | | | | | | | | | | | | | | | | | |
| PTPRN2 | Island;S_Shore | Body | chr7:157981275-157982589 | 1314 | | | | | | | | | 1 | | | | | yes | yes | | | | | |
| PTPRN2 | NA | Body | chr7:157985748-157985818 | 70 | | | | | | | | | | | | | | | | yes | | | | |
| PTPRN2 | NA | Body | chr7:158032120-158032496 | 376 | | | | | | | | | 1 | | | | | | | | | | | |
| PTPRN2 | Island | Body | chr7:158037763-158038007 | 244 | | | | | | | | | | | | | | | | | | yes | | |
| PTPRN2 | NA | Body | chr7:158045980-158046358 | 378 | | | 6 | | | | 2 | 0 | 7 | 3 | 1 | 15 | 1 | | | | | | | |
| PTPRN2 | NA | Body | chr7:158049946-158051019 | 1073 | | | | | | | | | | | | | | | | | yes | | | |
| PTPRN2 | Island;N_Shore | Body | chr7:158058752-158061820 | 3068 | 5 | | 10 | | | | | | 2 | 1 | | | | | | | | | | |
| PTPRN2 | NA | Body | chr7:158072687-158073079 | 392 | | | 2 | | | | | | | | | | | | | yes | | | | |
| PTPRN2 | NA | Body | chr7:158075388-158076567 | 1179 | | | | | | | | | | | | | | yes | | | | | | |
| PTPRN2 | NA | Body | chr7:158092647-158093277 | 630 | | | 7 | | | | | | | | | | | yes | | | yes | | | |
| PTPRN2 | N_Shore | Body | chr7:158103293-158103493 | 200 | | | | | | | | | | | | | | | | yes | | | | |
| PTPRN2 | N_Shelf;N_Shore | Body | chr7:158107269-158107723 | 454 | | | | | | | | | | | | | | | | yes | | | | |
| PTPRN2 | Island;N_Shore | Body | chr7:158109492-158110685 | 1193 | | | 6 | | 1 | | | 0 | 1 | | | 3 | | | | yes | | | yes | |
| PTPRN2 | S_Shelf | Body | chr7:158114137-158114659 | 522 | | | | | | | | | | | | | | | | | | | | |
| PTPRN2 | NA | Body | chr7:158122151-158122468 | 317 | | | | | | | | | | | | | | yes | | yes | | | | |
| PTPRN2 | NA | Body | chr7:158136312-158136485 | 173 | | | | | | | | | | | | | | | yes | | | | | |
| PTPRN2 | NA | Body | chr7:158155733-158156215 | 482 | | | | | | | | | | | | | | | yes | | | yes | | |
| PTPRN2 | NA | Body | chr7:158157881-158158401 | 520 | | | | | | | | | | | | | | | | | | yes | yes | |
| PTPRN2 | N_Shore | Body | chr7:158216857-158217673 | 816 | | | | | | | | | | | | | | yes | | yes | | | | |
| PTPRN2 | S_Shore | Body | chr7:158220856-158221370 | 514 | | | | | | | | | | | | | | yes | | | | | | |
| PTPRN2 | NA | Body | chr7:158235423-158236160 | 737 | | | 13 | | | | | | | | | | | | | | | | | |
| PTPRN2 | NA | Body | chr7:158243369-158243730 | 361 | | 1 | | | | | | | | | | 1 | | | yes | yes | | | | |
| PTPRN2 | Island;S_Shore | Body | chr7:158250129-158250978 | 849 | | | | | | | | | | | | | | yes | yes | | yes | | | |
| PTPRN2 | NA | Body | chr7:158263204-158263277 | 73 | | | | | | | | 0 | 1 | 1 | | 1 | | yes | | | | | | |
| PTPRN2 | NA | Body | chr7:158264663-158266198 | 1535 | 1 | | 8 | | | | | | | | | 1 | | yes | | | yes | | | |
| PTPRN2 | NA | Body | chr7:158267256-158267589 | 333 | | | | | | | | | | | | | | | | | | | | |
| PTPRN2 | NA | Body | chr7:158271757-158272547 | 790 | | | 6 | | | | | | | | | | | yes | yes | | yes | | | |
| PTPRN2 | Island;S_Shore;N_Shore | Body | chr7:158277852-158281613 | 3761 | 4 | | 13 | | | | | | | | | | | yes | yes | | | | yes | yes |
| PTPRN2 | NA | Body | chr7:158293463-158293697 | 234 | | | | | | | | | | | | | | | | | yes | | | |
| PTPRN2;MIR595 | NA | Body | chr7:158324928-158325586 | 658 | | | | | | | | 0 | 2 | | | | | | | | | | | |
| PTPRN2 | NA | Body | chr7:158336016-158336818 | 802 | | | 21 | | | | | | | | | | | yes | | | yes | | | |
| PTPRN2 | N_Shore;Island | Body | chr7:158342105-158342963 | 858 | 1 | | 4 | 1 | | | | | | | | | | | | | | | | |
| PTPRN2 | Island | Body | chr7:158349441-158350090 | 649 | | | | | | | | | | | | | | yes | | | | | | |
| PTPRN2 | Island;S_Shore | Body | chr7:158362339-158363642 | 1303 | | | 6 | | | | | 0 | 4 | | | 3 | | yes | | yes | | | yes | |
| PTPRN2 | N_Shore | Body | chr7:158379132-158379348 | 216 | | | 7 | | | | | 0 | 1 | | | | | | | | | | | |
| PTPRN2 | Island | TSS1500 | chr7:158380884-158381023 | 139 | | | | | | | | | 1 | | | | | | | | | yes | yes | |
| ESYT2 | NA | Body | chr7:158533877-158533927 | 50 | | | | | | | | | 1 | | | | | | | | | | | |
| ESYT2 | NA | Body | chr7:158575681-158575934 | 253 | | | | | | | | | 1 | | | | | | | | yes | | | |
| WDR60 | Island | Body | chr7:158656429-158656677 | 248 | | | 1 | | | | | | 3 | | | 1 | | yes | | yes | | | | |
| WDR60 | NA | Body | chr7:158664058-158664247 | 189 | | | | | | | | 0 | 1 | | | | | yes | | | | yes | | |
| WDR60 | S_Shelf | Body | chr7:158673837-158673979 | 142 | | | | 1 | | | | | | | | | | yes | | | | | | |
| LOC154822 | S_Shore | TSS1500;TSS200 | chr7:158799715-158800986 | 1271 | | | | | | | | | | | | | | yes | | | | | | |
| LOC154822 | NA | Body | chr7:158809018-158809804 | 786 | | | | | | | | | | | | | | | | yes | | | | |
| LOC154822 | Island;S_Shore | Body | chr7:158817612-158818164 | 552 | | | 4 | | | | | | 1 | | | | | | | | | | | |
| VIPR2 | S_Shelf | Body | chr7:158828235-158828603 | 368 | | | | | | | | | | | | | | | | yes | | | | |
| VIPR2 | Island | Body | chr7:158854447-158854856 | 409 | | 1 | 1 | | | | | | 1 | | | | | yes | yes | yes | | | | |
| VIPR2 | S_Shore | Body | chr7:158858057-158859224 | 1167 | | | 9 | | | | | | 1 | | | 1 | | | | yes | yes | | | |
| VIPR2 | N_Shore | Body | chr7:158885581-158886532 | 951 | | | | | | | | | 1 | | | | | yes | | | | | | |
| VIPR2 | N_Shore;Island | Body | chr7:158905015-158905536 | 521 | | | 5 | | | | 1 | 0 | 2 | 2 | 1 | 3 | | yes | yes | yes | | | | |
| VIPR2 | N_Shelf | Body | chr7:158933340-158934060 | 720 | | | | | | | | | | | | | | | | | yes | | | |
| VIPR2 | Island;chr7:158936507-158938 | Body;1stExon;5'UTR;Island | chr7:158936508-158938060 | 1552 | | 1 | | | | | | | 1 | | 1 | | | | yes | | | | yes | |
| ZNF596;RPL23AP53 | Island | 5'UTR;1stExon;TSS200 | chr8:182402-182686 | 284 | | | | | | | | 0 | | | | | | | | | | | yes | |
| C8orf42 | S_Shore | TSS1500 | chr8:496327-496739 | 412 | | | | | | | | | 2 | | | | | | | | | | | |
| ERICH1 | N_Shore | Body | chr8:618674-618751 | 77 | 3 | | | | | | | | | | | | | | | | | | | |
| ERICH1 | Island | Body | chr8:637909-638330 | 421 | | | | | | | | 0 | 1 | 1 | 1 | 7 | | | yes | | | yes | yes | |
| ERICH1 | NA | Body | chr8:644521-644878 | 357 | | 1 | | | | | | | | | | | | | | | | | | |
| ERICH1 | S_Shore | Body | chr8:652315-652403 | 88 | 2 | | | | | | | | | | | | | yes | yes | yes | | | | |
| ERICH1 | NA | Body | chr8:672007-672406 | 399 | | | | | | | | | 1 | | | | | | | | | | | |
| ERICH1 | NA | Body | chr8:674434-674899 | 465 | | | | | | | | 0 | 1 | 1 | 1 | 11 | | yes | | | | yes | | |
| DLGAP2 | NA | TSS1500;TSS200;5'UTR | chr8:1448361-1450031 | 1670 | | 1 | 14 | | | | | | 1 | | | | | | | | | | | |
| DLGAP2 | NA | 5'UTR | chr8:1458119-1458558 | 439 | | | | | | | | | | | | | | yes | | | yes | | | |
| DLGAP2 | NA | 5'UTR | chr8:1460569-1460718 | 149 | | | 8 | | | | | | | | | | | | | | | | | |
| DLGAP2 | NA | 5'UTR | chr8:1462903-1463125 | 222 | | | | | | | | | | | | | | | | | | yes | | |
| DLGAP2 | N_Shore;Island | 5'UTR | chr8:1494546-1495332 | 786 | | | 5 | | | | | | | | | | | | | | | | yes | |
| DLGAP2 | N_Shore;Island | 5'UTR | chr8:1496347-1497307 | 960 | | | 7 | | | | | | 1 | | | | | yes | | | | | | |
| DLGAP2 | chr8:1501407-1501671;N_Shor | N_Shore;Body | chr8:1500367-1501670 | 1303 | | | | | | | | | | | 1 | | | | yes | | | | | |
| DLGAP2 | NA | Body | chr8:1553395-1553879 | 484 | | | 6 | | | | | | | | | | | yes | | | | | | |
| DLGAP2 | NA | Body | chr8:1570635-1571105 | 470 | | | | | | | | | | | | | | | | | | | | |
| DLGAP2 | Island | Body | chr8:1579039-1579180 | 141 | | | | | | | | | | | | | | yes | | | | | | |
| DLGAP2 | N_Shore;Island | Body | chr8:1616209-1617868 | 1659 | | | 8 | | | | | | 1 | | | 1 | | | yes | | | | | |
| DLGAP2 | NA | Body | chr8:1626419-1626697 | 278 | | | 4 | | | | | | | | | | | | | | | | | |
| DLGAP2 | N_Shore | Body | chr8:1639573-1639909 | 336 | | | | | | | | | | | | | | | | | | yes | | |
| DLGAP2 | S_Shore | Body | chr8:1642380-1642577 | 197 | | | | | | | | | 1 | | | | | | | | | | | |
| DLGAP2 | NA | Body | chr8:1644982-1645455 | 473 | | | | | | | | | | | | | | yes | | | | yes | | |
| DLGAP2 | Island | Body | chr8:1649522-1649868 | 346 | | | 1 | | | | | | 1 | | | | | | | | | yes | | |
| DLGAP2 | S_Shore | 3'UTR | chr8:1651101-1651197 | 96 | | | 1 | | | | | | | | | | | | | | | | | |
| DLGAP2 | S_Shelf | 3'UTR | chr8:1653179-1653566 | 387 | | | | | | | | | | | | | | | | | | | | |
| MIR596 | Island;S_Shore | TSS1500;TSS200;Body | chr8:1764618-1765820 | 1202 | | | 39 | | | | | 0 | 3 | 1 | | 3 | | | yes | yes | | | | |
| ARHGEF10 | Island | TSS200 | chr8:1771973-1772264 | 291 | | 1 | | | | | | 0 | 2 | | | | | | | | | yes | | |
| ARHGEF10 | S_Shore | 5'UTR | chr8:1780242-1780401 | 159 | | | 4 | | | | | | | | | | | | | | yes | yes | | |
| ARHGEF10 | NA | 5'UTR | chr8:1785958-1786322 | 364 | | | 12 | | | | | | 1 | | | | | | | | | | | |
| ARHGEF10 | NA | 5'UTR | chr8:1790793-1790905 | 112 | | | | | | | | | 1 | 1 | | | | | | | | | | |
| ARHGEF10 | N_Shore | Body | chr8:1796429-1796676 | 247 | | | | | | | | | | | | | | yes | yes | | | | | |
| ARHGEF10 | NA | Body | chr8:1807061-1807575 | 514 | | | | | | | | 0 | | | | | | | | | | | | |
| ARHGEF10 | NA | Body | chr8:1809425-1809936 | 511 | | | 4 | | | | | | | | | | | | | | | | yes | |
| ARHGEF10 | NA | Body | chr8:1814745-1815153 | 408 | | | 6 | | | | | | | | | | | | | | | yes | | |
| ARHGEF10 | NA | Body | chr8:1847999-1848741 | 742 | 1 | | 32 | | | | | 0 | | | | | | yes | yes | | | yes | yes | |

| | | | | | | | | | | | | | | | | | | | | | | | |
|---|---|---|---|---|---|---|---|---|---|---|---|---|---|---|---|---|---|---|---|---|---|---|---|
| ARHGEF10 | NA | Body | chr8:1861037-1861136 | 99 | | | | | | | | | 1 | | | | | | | | yes | | |
| ARHGEF10 | NA | Body | chr8:1879551-1879786 | 235 | | | | 1 | | | | | | | | | | | | | yes | | |
| ARHGEF10 | S_Shore | Body | chr8:1894604-1895558 | 954 | | | | | | | | | | | | | | | | yes | | | yes |
| ARHGEF10 | NA | Body | chr8:1899943-1900992 | 1049 | | | 8 | | 1 | | | | | | | | | yes | | | | | |
| ARHGEF10 | NA | 3'UTR;Body | chr8:1905131-1906498 | 1367 | 1 | | | | 1 | | | | | 1 | | | | | | | | | |
| KBTBD11 | Island | 5'UTR | chr8:1949246-1950120 | 874 | | 1 | | | | | | 0 | 1 | | | | | | | | | | yes |
| KBTBD11 | N_Shelf;N_Shore | 3'UTR | chr8:1954777-1955323 | 546 | | | 21 | | | | | | | | | | | | | yes | | | |
| MYOM2 | NA | TSS1500;TSS200;5'UTR | chr8:1992139-1994574 | 2435 | | 1 | 12 | | 2 | | | 0 | 4 | | | 2 | | yes | | yes | yes | yes | |
| MYOM2 | S_Shore | Body | chr8:2005503-2005940 | 437 | | 1 | 8 | | | | | | | | | | | yes | yes | | yes | | |
| MYOM2 | NA | Body | chr8:2019935-2020707 | 772 | | | 4 | | | | | | | | | | | yes | | yes | | | |
| MYOM2 | Island | Body | chr8:2037717-2037954 | 237 | | | | | | | | | 1 | | | | | | yes | | | | |
| MYOM2 | Island | Body | chr8:2046805-2047010 | 205 | | | | | | | | | | | | | | yes | | | | | |
| MYOM2 | N_Shore;Island | Body | chr8:2075209-2075820 | 611 | 10 | 8 | 25 | 1 | | 4 | 2 | 0 | 2 | 2 | 1 | 38 | 2 | | yes | | | | |
| MYOM2 | Island | Body | chr8:2078112-2078387 | 275 | | | | | | | | | 1 | | | | | | | | | | |
| MYOM2 | NA | Body;3'UTR | chr8:2092607-2093220 | 613 | | | 6 | | | | | | | | | | | | | | | | |
| CSMD1 | NA | Body | chr8:3429382-3429480 | 98 | | | | | | | | | | | | | | | yes | | | | |
| CSMD1 | NA | Body | chr8:3855415-3855577 | 162 | | | | | | | | | | | | | | | | | | | |
| CSMD1 | NA | Body | chr8:4504848-4504999 | 151 | | | | | | | | | | | | | | | yes | | | | |
| CSMD1 | Island | Body | chr8:4849399-4849827 | 428 | | | | | | | | | 1 | | | | | | | | | | |
| CSMD1 | Island;S_Shore | Body;TSS1500 | chr8:4851751-4853437 | 1686 | | 1 | 4 | | | | | | | | | | | | | | | yes | yes |
| ANGPT2;MCPH1 | NA | Body | chr8:6400815-6401045 | 230 | | 1 | 2 | | | | | | | | | | | yes | | | | | |
| ANGPT2;MCPH1 | NA | Body | chr8:6407273-6407997 | 724 | | | 3 | | | | | | 2 | | | 1 | | | yes | | | | |
| ANGPT2;MCPH1 | NA;N_Shelf | Body;1stExon;5'UTR;TSS200 | chr8:6418978-6421171 | 2193 | 5 | | 16 | | 1 | | | 0 | 4 | | | 4 | 1 | yes | | | | | |
| MCPH1 | NA | Body | chr8:6456182-6456465 | 283 | | | 8 | | | | | | | | | | | | | | | | |
| MCPH1 | NA | Body | chr8:6466774-6466848 | 74 | | | 1 | | | | | | | | | | | | | | yes | | |
| MCPH1 | NA | Body | chr8:6484293-6484687 | 394 | | | 1 | | | | | | | | | | | yes | | | | yes | |
| XKR5 | S_Shore | TSS1500 | chr8:6693367-6693540 | 173 | | | 1 | | | | | | | | | | | | | yes | | yes | |
| DEFB1 | NA | Body;5'UTR;1stExon | chr8:6734962-6735559 | 597 | | 1 | 1 | | | | | | 2 | 1 | | | | yes | | | | | |
| FAM90A7 | NA | TSS1500 | chr8:7441657-7441706 | 49 | | | | | | | | | | | | | | | yes | | | yes | |
| PRAGMIN | Island | Body | chr8:8176225-8176714 | 489 | | | | | | | | | 1 | | | | | | yes | | | yes | |
| PRAGMIN | NA | Body | chr8:8182454-8182747 | 293 | | | 3 | | | | | | 1 | | 1 | | | yes | | | | | |
| MFHAS1 | Island | 1stExon | chr8:8749074-8749938 | 864 | | | 11 | | | 1 | | | | | | | | | | | | | yes |
| PPP1R3B | Island | TSS1500 | chr8:9009105-9009401 | 296 | | | | | | | | 0 | 1 | | | 1 | | | | | yes | | |
| MSRA | NA | Body;TSS1500;TSS200 | chr8:9952636-9953022 | 386 | | 3 | 23 | | | | | | | | | | | | | | | yes | |
| MSRA | NA | Body | chr8:10005423-10005590 | 167 | 2 | | | | | | | | | | | | | yes | | yes | | | |
| MSRA | NA | Body | chr8:10207852-10208257 | 405 | 1 | | 1 | | | | | | | | | | | yes | | | yes | | |
| MSRA | N_Shore | Body | chr8:10261798-10262020 | 222 | | | | | | | | | | | | | | yes | | | | | |
| MSRA | NA | Body | chr8:10282549-10283152 | 603 | | | | | | | | 0 | | | | | | | | | | yes | |
| RP1L1 | N_Shore;Island | Body | chr8:10469310-10470278 | 968 | | | 7 | | | | | | | | | | | | | | | | |
| RP1L1 | NA | 5'UTR | chr8:10489118-10489270 | 152 | | | | | | | | 0 | | | | | | yes | | | | yes | |
| RP1L1 | NA | 1stExon;5'UTR;TSS1500 | chr8:10512459-10513717 | 1258 | | | | | | | | | | | | | | | | | yes | | |
| C8orf74 | NA | TSS1500;TSS200 | chr8:10529811-10530148 | 337 | | | 4 | 1 | | | | | 1 | | | | | | | | | yes | |
| SOX7 | Island;N_Shore | Body | chr8:10586678-10587305 | 627 | | | 25 | 1 | | | | | | | | | | | | | yes | | yes |
| XKR6 | NA | Body | chr8:10755744-10756091 | 347 | | | | | | | | | | | | | | | | | | yes | |
| XKR6;MIR598 | NA | Body | chr8:10892734-10893100 | 366 | | | | | | | | | | | | | | yes | | | | yes | |
| XKR6 | NA | Body | chr8:10907718-10907788 | 70 | | | 8 | | | | | | | | | | | yes | yes | | | | yes |
| XKR6 | Island | Body | chr8:10928985-10929599 | 614 | | | 5 | | | | | | | | | | | yes | | | | | |
| XKR6 | NA | Body | chr8:10966192-10966251 | 59 | | | | | | | | | | | | | | | | | yes | | |
| MTMR9 | N_Shore | TSS1500 | chr8:11141365-11141680 | 315 | | 1 | 12 | | | | | 0 | 1 | | | 1 | | | | yes | | | |
| TDH | N_Shore | Body | chr8:11203882-11204132 | 250 | | | 3 | | | | | 0 | 3 | 1 | | 5 | | | | yes | yes | yes | |
| C8orf12;TDH | NA | TSS1500;Body;TSS200 | chr8:11225385-11226339 | 954 | | | | | | | | | | | | | | yes | yes | | | | |
| BLK | NA | Body | chr8:11413043-11413234 | 191 | | | | | | | | | 1 | | | | | yes | | | | | |
| GATA4 | Island | TSS1500 | chr8:11560451-11561531 | 1080 | | | 4 | | 2 | | | | | | | | | | | | | | yes |
| GATA4 | S_Shore | Body | chr8:11567691-11568356 | 665 | | | 5 | | | | | | 1 | | | | | yes | | | | | yes |
| FDFT1 | Island | TSS1500;TSS200 | chr8:11659832-11660209 | 377 | | | | | | | | 0 | 1 | 1 | | 1 | | yes | | | | | |
| FDFT1 | Island | Body | chr8:11665962-11666017 | 55 | | | | | | | | | 1 | | | | | | | yes | | | yes |
| DEFB136 | NA | Body;TSS1500 | chr8:11831823-11833039 | 1216 | | | 3 | | | | | | | | | | | yes | | | | yes | |
| DEFB135 | NA | TSS200 | chr8:11839764-11839844 | 80 | | 1 | 26 | | | | | | | | 1 | | | yes | | | | | |
| FAM66A | NA | Body | chr8:12237626-12237900 | 274 | | | 1 | | | | | | | | | | | | | | | yes | |
| DLC1 | NA | Body;TSS1500 | chr8:12974448-12974556 | 108 | | | | | | | | | 1 | | | | | yes | | | | | yes |
| DLC1 | Island | Body;1stExon;5'UTR | chr8:12990213-12990898 | 685 | | 1 | | | | | | | 1 | | | | | | | | | yes | |
| DLC1 | NA | 1stExon;5'UTR;TSS1500 | chr8:13372132-13373141 | 1009 | | | | | | | | 0 | 2 | | | | | | | | | yes | |
| C8orf48 | NA | TSS1500;TSS200 | chr8:13424080-13424419 | 339 | | 1 | | | | | | | 2 | | | | | | yes | | | yes | |
| SGCZ | N_Shore | Body | chr8:15093686-15094114 | 428 | | | 2 | | | | | | | | | | | | | | | | yes |
| TUSC3 | Island | TSS200;5'UTR;1stExon | chr8:15397637-15397899 | 262 | | | | | | | | | 2 | | | | | yes | | | | | |
| FGF20 | Island;S_Shore | 1stExon;5'UTR;TSS200;TSS1500 | chr8:16859451-16860121 | 670 | | | 26 | | | | | 0 | 1 | | | 1 | | | | | | | yes |
| MTMR7 | NA | Body | chr8:17206671-17206819 | 148 | | | | 1 | 1 | | | | 1 | | | | | yes | yes | | | | |
| MTMR7 | N_Shore;Island | Body;5'UTR;1stExon;TSS200 | chr8:17270347-17271340 | 993 | | | 24 | | | | | | 2 | | | | | | | | | | yes |
| SLC7A2 | NA | TSS1500;5'UTR | chr8:17394958-17395271 | 313 | | | | | | | | | | | | | | yes | | | | | |
| MTUS1 | NA | Body;1stExon;5'UTR | chr8:17554892-17555302 | 410 | | | 4 | | | | | | | | | | | | | | | | |
| NAT1 | NA | TSS1500 | chr8:18027713-18028071 | 358 | | | | 1 | | | | | | | | | | yes | yes | yes | | | yes |
| PSD3 | NA | Body | chr8:18541446-18541988 | 542 | | 2 | 6 | | | | | | | | | | | | | | | yes | |
| PSD3 | NA | Body;TSS200 | chr8:18666531-18666561 | 30 | 1 | | | | | | | | | | | | | | | | yes | | |
| CSGALNACT1 | Island;S_Shore | 1stExon;5'UTR;Body;TSS200 | chr8:19459996-19460243 | 247 | | | 1 | | 1 | | | | 2 | | | 1 | | | | | | yes | yes |
| CSGALNACT1 | NA | TSS200;Body;TSS1500;5'UTR;1stEx | chr8:19539991-19540734 | 743 | | | 14 | | | | | 0 | 4 | 1 | | 1 | | | | | yes | | |
| LPL | N_Shore | TSS1500;TSS200 | chr8:19796299-19796591 | 292 | | | | | | | | 0 | 1 | | | 1 | | | yes | | | | |
| ATP6V1B2 | N_Shore | TSS1500 | chr8:20054405-20054583 | 178 | | | | | | | | | 1 | | | | | yes | yes | | | yes | yes |
| LZTS1 | NA | 5'UTR;1stExon;TSS200 | chr8:20112683-20113188 | 505 | | | 19 | | | | | | 1 | | | | | yes | yes | | | | |
| GFRA2 | N_Shore;Island | Body | chr8:21640100-21640304 | 204 | | | | | | | | | 2 | | | 1 | | | | | | | yes |
| DOK2 | N_Shelf | Body | chr8:21767146-21767310 | 164 | | | | | | | | | 1 | | | | | yes | | | | | |
| DOK2 | S_Shore;chr8:21769694-21770 | TSS200;TSS1500;1stExon;S_Shore | chr8:21771059-21771668 | 609 | | | 3 | | 2 | | | | 3 | 2 | 1 | | | yes | | | | yes | |
| NPM2 | Island | 5'UTR | chr8:21882587-21882942 | 355 | | | | | | | | 0 | 1 | | | 3 | | yes | | | yes | | |
| FGF17 | Island | Body | chr8:21905462-21905960 | 498 | | | 4 | | 1 | | | | | | | | | yes | | | | | yes |
| EPB49 | S_Shelf | 1stExon;5'UTR | chr8:21916802-21917015 | 213 | | | 3 | 1 | 1 | | | | | | | | | yes | | | | | yes |
| FAM160B2 | NA | 3'UTR | chr8:21961713-21961788 | 75 | | | 12 | | | | | | | | | | | yes | | | | | |
| PHYHIP | N_Shore;Island | Body | chr8:22078982-22079330 | 348 | 6 | | 1 | | | | | | | | | | | | yes | | | | |
| PHYHIP | NA | Body | chr8:22084861-22085227 | 366 | | | 3 | | | | | | | | | | | yes | | | | | |
| PHYHIP | Island | 1stExon;5'UTR | chr8:22089444-22089615 | 171 | | | | | | | | 0 | | | | | | | | | | | |
| PIWIL2 | N_Shore;Island | TSS1500;TSS200;5'UTR;1stExon | chr8:22131899-22133375 | 1476 | 4 | 2 | 34 | | | | | | 7 | | | 3 | | yes | | yes | | | |
| SLC39A14 | N_Shore | TSS1500 | chr8:22223852-22224272 | 420 | | | 3 | | | | | 0 | 2 | | | 2 | | | | | | | yes |
| SORBS3 | Island | TSS1500;Body | chr8:22422657-22423994 | 1337 | | | 3 | | | | | | | | | | | | yes | | | yes | yes |
| PDLIM2 | NA | nan | chr8:22446721-22447181 | 460 | | | | | | | | | | | 1 | | | | yes | | | | yes |
| C8orf58 | Island | Body | chr8:22457446-22458182 | 736 | | | | 1 | | | | | | | | | | | | | | yes | yes |
| EGR3 | Island | Body | chr8:22548006-22548952 | 946 | | 1 | 8 | | | | | | 1 | | | | | | | | | | |
| PEBP4 | NA | Body | chr8:22642322-22642638 | 316 | | | 4 | | | | | | | | | | | | | | | | |
| PEBP4 | NA | Body | chr8:22735111-22735478 | 367 | | | | | | | | 0 | | | | 2 | | | | | | yes | |
| PEBP4 | NA | TSS200;TSS1500 | chr8:22785456-22786374 | 918 | | | 4 | | | | | | | | | | | yes | yes | | | | |
| RHOBTB2 | NA | TSS1500 | chr8:22844374-22844827 | 453 | | | | | | | | | 1 | | | | | yes | | | | | |
| RHOBTB2 | N_Shore | Body;TSS1500 | chr8:22855616-22855745 | 129 | | | | | | | | | 1 | | | | | yes | | | | | |

| | | | | | | | | | | | | | | | | | | | | | | | | |
|---|---|---|---|---|---|---|---|---|---|---|---|---|---|---|---|---|---|---|---|---|---|---|---|---|
| TNFRSF10C | Island;N_Shore | TSS200;TSS1500 | chr8:22959713-22960461 | 748 | | | 7 | 1 | 2 | | | | 1 | | | | | | | | | | | |
| TNFRSF10A | Island;S_Shore | TSS200;TSS1500 | chr8:23082721-23083578 | 857 | | | 8 | | 1 | | | | 1 | | | | | yes | yes | | | | | |
| LOXL2 | NA | Body | chr8:23162162-23162383 | 221 | | | 12 | | | | | | | | | | | | | | | | | |
| LOXL2 | S_Shore | TSS1500 | chr8:23261967-23262159 | 192 | | | | | | | | 0 | | | | | | | yes | | | | | |
| NKX2-6 | Island | Body | chr8:23560319-23560590 | 271 | | 1 | | | | | | | | | | | | yes | | | | | | |
| NKX2-6 | Island | Body;1stExon;TSS200 | chr8:23562918-23564591 | 1673 | | 1 | | | | | | 0 | 4 | | | | | | | | | | | |
| STC1 | NA | TSS1500;TSS200 | chr8:23712355-23713016 | 661 | | | | | | | | 0 | 2 | | | | | yes | | | | yes | | |
| ADAM28 | NA | TSS200 | chr8:24151472-24151874 | 402 | | | | | | | | 0 | | | | | | | | | | yes | yes | |
| NEFM | N_Shore;Island | TSS1500;1stExon;TSS200 | chr8:24770381-24773060 | 2679 | | | | | | | | | | | | | | | | | | | | |
| NEFL | N_Shore;Island;S_Shore | Body;5'UTR;1stExon;TSS1500 | chr8:24812784-24815367 | 2583 | | 2 | | | 1 | | | | | | | | | | | | | yes | | |
| EBF2 | NA | Body | chr8:25812278-25812500 | 222 | | | | | | | | 0 | 1 | | | | | | yes | | | | | |
| EBF2 | NA | Body | chr8:25868045-25868109 | 64 | | | | | | | | 0 | 1 | | | | | | | | | yes | | |
| EBF2 | N_Shore;Island | Body | chr8:25897763-25900020 | 2257 | 1 | 1 | 13 | | | | | | 1 | | | | | yes | | | | | | |
| EBF2 | Island | TSS200 | chr8:25902400-25902594 | 194 | | | 5 | | | | | | 1 | | | | | | | | | | | |
| DPYSL2 | NA | Body | chr8:26481666-26481822 | 156 | | | | | | | | | | | | | | | | | | | | |
| DPYSL2 | NA | 3'UTR | chr8:26513782-26513861 | 79 | 2 | | | | | | | | | | | | | | | | | | yes | |
| ADRA1A | Island | TSS200 | chr8:26722933-26723584 | 651 | | | | | | | | | 1 | | | | | | | | | | | |
| STMN4 | NA | 1stExon;5'UTR | chr8:27115867-27116213 | 346 | | | 3 | | | | | | 1 | | | 1 | | | yes | | | | | |
| PTK2B | N_Shore | TSS1500;5'UTR;TSS200 | chr8:27182871-27183873 | 1002 | | | | 1 | 1 | | | | | | | | | | yes | | | yes | | |
| CHRNA2 | NA | 5'UTR;1stExon;TSS1500 | chr8:27336626-27338248 | 1622 | 2 | | 3 | 1 | | | | | | | | | | yes | | yes | | yes | | |
| EPHX2 | Island | Body | chr8:27348882-27348940 | 58 | | | | | | | | | 1 | | | | | | | | | yes | yes | |
| CLU | N_Shelf | 1stExon;Body;5'UTR;TSS200;TSS15 | chr8:27468684-27469338 | 654 | | 1 | 8 | | | | | 0 | 1 | | 1 | | | yes | | | | | | |
| CLU | Island | TSS200 | chr8:27472446-27472513 | 67 | | | 1 | | | | | | | | | | | yes | | | yes | | | |
| CCDC25;ESCO2 | N_Shore;Island | Body;TSS1500 | chr8:27629592-27632322 | 2730 | | | | | | | | | | | | | | | | | | | | |
| SCARA5 | Island | Body | chr8:27737078-27737223 | 145 | | | | | | | | | 1 | | | | | yes | yes | | | | | |
| SCARA5 | NA | 5'UTR | chr8:27845182-27845277 | 95 | | 1 | | | | | | | 1 | | | | | yes | | | | | | |
| SCARA5 | NA | TSS200 | chr8:27850215-27850423 | 208 | | 1 | | | | | | | 1 | | | | | yes | | | yes | | | |
| PNOC | NA | TSS1500;5'UTR;1stExon | chr8:28173732-28175285 | 1553 | | | 10 | | | | | 0 | 4 | | | 4 | | yes | yes | | | | | |
| EXTL3 | S_Shore | 5'UTR | chr8:28560467-28560723 | 256 | | | 30 | 1 | 1 | | | | 1 | | 1 | | | | | | | | | |
| KIF13B | N_Shore;Island | Body | chr8:28928948-28929242 | 294 | | | 3 | | | | | | | | | | | | | | | | yes | |
| KIF13B | NA | Body | chr8:28961315-28961356 | 41 | | | | | | | | | 1 | | | | | | | | | | | |
| RBPMS | N_Shore | TSS1500 | chr8:30241167-30241280 | 113 | | | | | | | | | | | | | | yes | yes | | | | yes | |
| RBPMS | NA | Body | chr8:30413794-30413919 | 125 | | | | | | | | | 1 | | | | | | | | | | yes | |
| PURG;WRN | NA | TSS1500;5'UTR | chr8:30892276-30892490 | 214 | | | | | | | | | 1 | | | 1 | | | yes | yes | | | | |
| NRG1 | Island | TSS1500 | chr8:31496966-31498345 | 1379 | | | | | | | | 0 | 1 | | | | | | | | | yes | yes | |
| NRG1 | Island | 1stExon;5'UTR;Body | chr8:32406150-32406928 | 778 | | | | | | | | | 1 | | | | | | | | | yes | yes | |
| NRG1 | NA | Body;5'UTR;1stExon;TSS200 | chr8:32504110-32505269 | 1159 | | | | 1 | | | | | | | | | | | | | | yes | yes | |
| NRG1 | NA | Body;TSS1500 | chr8:32578937-32579253 | 316 | | | | | | | | 0 | | | | | | | | | | | | |
| UNC5D | Island | TSS1500;TSS200 | chr8:35092687-35094053 | 1366 | | | | | | | | | | | | | | | | | | | | |
| LOC728024;ERLIN2 | NA | Body;TSS200;TSS1500 | chr8:37605359-37606579 | 1220 | | | | | | | | | | | | | | | | | | | | |
| GPR124 | Island | Body | chr8:37698900-37699923 | 1023 | | | 2 | | | | | 0 | 2 | 1 | | 3 | | | | | | | yes | |
| BRF2 | S_Shelf | Body | chr8:37702396-37702581 | 185 | | | 4 | | | | | | | | | | | yes | | | | | | |
| RAB11FIP1 | NA | Body;TSS1500 | chr8:37731969-37732224 | 255 | | | | | 1 | | | | | | | | | | | | | | | |
| RAB11FIP1 | N_Shore | Body | chr8:37755160-37755555 | 395 | | 1 | | | | | | | | | | | | yes | yes | | | | | |
| RAB11FIP1 | Island | TSS1500 | chr8:37757250-37757401 | 151 | | | 1 | | | | | | | | | | | yes | | | | yes | | |
| GOT1L1 | NA | 1stExon;5'UTR;TSS200 | chr8:37797458-37798156 | 698 | | | 5 | | | | | | | | | | | yes | yes | | | yes | | |
| ADRB3 | S_Shore;Island | TSS1500;1stExon;TSS200 | chr8:37823409-37824765 | 1356 | | 2 | | | | | | 0 | 3 | | | | | yes | | | | | yes | |
| STAR | NA | 5'UTR;1stExon;TSS1500 | chr8:38008386-38009548 | 1162 | | 1 | | | | | | | 2 | 2 | | | | yes | | | yes | yes | yes | |
| WHSC1L1 | NA | Body | chr8:38175544-38175684 | 140 | | 1 | | | | | | | | | | | | | | | | | yes | |
| FGFR1 | S_Shore;Island | TSS1500;5'UTR;1stExon;TSS200 | chr8:38326334-38326968 | 634 | | | 2 | | | | | | 1 | | | | | yes | | | yes | | | |
| TACC1 | NA | TSS1500;1stExon;5'UTR | chr8:38584636-38586183 | 1547 | | 1 | | | 1 | | | | | | | | | yes | | | | yes | yes | |
| TACC1 | Island | 5'UTR | chr8:38614832-38615330 | 498 | | 1 | | | | | | | | | | | | | | | | yes | | |
| TACC1 | N_Shore | TSS1500;5'UTR | chr8:38643568-38643846 | 278 | | | | | | | | | 1 | | | | | yes | | | | | | |
| TACC1 | Island | Body;5'UTR | chr8:38645271-38645613 | 342 | | | | 1 | | | | | | | | | | | | | | | yes | |
| PLEKHA2 | N_Shore | TSS1500 | chr8:38757652-38757859 | 207 | | 1 | | | 2 | | | | | | | | | yes | | | | yes | | |
| HTRA4;PLEKHA2 | N_Shore;Island | TSS1500;3'UTR;TSS200;1stExon | chr8:38831148-38832335 | 1187 | | | | 1 | | | | 0 | 5 | 2 | | 2 | | yes | | | | | | |
| ADAM32 | N_Shore;Island | TSS200 | chr8:38964858-38965492 | 634 | | 1 | 1 | | | | | 0 | 3 | | | 2 | | yes | | yes | | | yes | |
| ADAM5P | NA | TSS1500;TSS200 | chr8:39171620-39172120 | 500 | | 1 | 16 | | 1 | | | 0 | 8 | | 1 | 11 | 1 | | yes | | yes | | | |
| ADAM3A | Island | Body | chr8:39380341-39380592 | 251 | | 2 | | | | | | 0 | | | | 6 | | | | | | yes | | |
| ADAM18 | NA | TSS1500;TSS200 | chr8:39441473-39442080 | 607 | | | | | | | | | | | | | | | yes | | | | | |
| ADAM2 | NA | TSS200 | chr8:39695848-39696062 | 214 | | | 5 | | | | | | | | | | | yes | yes | | | | | |
| IDO2 | NA | Body | chr8:39835697-39836064 | 367 | | | 1 | | | | | | 2 | | | 1 | | yes | yes | | | | | |
| C8orf4 | NA | TSS1500 | chr8:40010443-40010823 | 380 | | | 15 | | | | | | | | | | | | | | | | | |
| ZMAT4 | Island | 5'UTR | chr8:40755165-40755380 | 215 | | | | | | | | | 1 | | | | | | | | | | yes | |
| SFRP1 | Island;S_Shore;N_Shore | TSS200;TSS1500;Body;1stExon | chr8:41165699-41168481 | 2782 | | 1 | 9 | | 2 | | | 0 | 4 | | | 1 | | yes | | | | | yes | |
| NKX6-3 | Island;S_Shore | Body;1stExon;TSS200;TSS1500 | chr8:41504078-41505696 | 1618 | | | 10 | | 1 | | | | 1 | | | | | yes | | | | yes | yes | |
| ANK1 | NA | Body;1stExon;5'UTR;TSS200 | chr8:41522721-41523968 | 1247 | | | | | | | | | | | | | | | | | | | | |
| ANK1 | NA | Body | chr8:41528946-41529140 | 194 | | | | | | | | | | | | | | | | | | | | |
| ANK1 | NA | Body | chr8:41542092-41542172 | 80 | | | | | | | | | 1 | | | | | yes | | | | | yes | |
| ANK1 | N_Shore;Island | Body | chr8:41583018-41583523 | 505 | | | 13 | | | | | 0 | | | | 2 | | yes | yes | | | | | |
| ANK1 | Island | Body;1stExon;5'UTR;TSS200;TSS15 | chr8:41654967-41656079 | 1112 | | | 30 | | | | | 0 | 2 | | | | | | | | | yes | | |
| AP3M2 | Island | TSS1500 | chr8:42010142-42010169 | 27 | | 1 | | | | | | | | | | | | | | | | | yes | |
| PLAT | NA | Body | chr8:42037170-42037527 | 357 | | | 3 | | | | | | 1 | 1 | | | | yes | | | yes | | | |
| DKK4 | NA | Body;5'UTR;1stExon;TSS200 | chr8:42234243-42234975 | 732 | | | 12 | | | | | 0 | 3 | 1 | | 1 | | yes | | | | | | |
| SLC20A2 | NA | 5'UTR | chr8:42356637-42356871 | 234 | | | 8 | | | | | | 1 | 1 | | | | | | | | | | |
| CHRNA6 | NA | TSS200 | chr8:42623686-42623946 | 260 | 5 | | 6 | | 1 | | | 0 | | | | | | yes | | | | | | |
| SGK196 | Island | TSS200 | chr8:42948547-42948868 | 321 | | | | | | | | | 1 | | | | | yes | | | yes | yes | | |
| POTEA | NA | TSS1500 | chr8:43147211-43147554 | 343 | | | 1 | | | | | | | | | | | | | | | yes | | |
| BEYLA | NA | TSS1500 | chr8:47751494-47751953 | 459 | | | | | | | | | 1 | | | | | | | | | | | yes |
| KIAA0146 | N_Shore | TSS1500 | chr8:48172746-48172992 | 246 | | | 2 | | | | | 0 | 2 | | | | | | | | | | | |
| KIAA0146 | NA | Body | chr8:48572447-48572557 | 110 | 1 | | | | 1 | | | 0 | 1 | | | 3 | | | | | | | yes | |
| KIAA0146 | N_Shore | 3'UTR | chr8:48648112-48649086 | 974 | | | | | | | | | | | | | | | | | | yes | | |
| EFCAB1 | N_Shore;Island | Body;1stExon;5'UTR | chr8:49647579-49647907 | 328 | | 1 | 1 | | 1 | | | 0 | 3 | | | | | | | | yes | | | |
| C8orf22 | NA | TSS1500;nan | chr8:49984564-49985021 | 457 | | | | | | | | | 1 | | 1 | | | yes | | | | | | |
| PXDNL | Island | Body | chr8:52321814-52322341 | 527 | | | 5 | | | | | | 5 | | | | | | | | | | | |
| PXDNL | NA | Body;TSS200;TSS1500 | chr8:52721657-52722539 | 882 | 3 | | | | | | | | | | | | | | yes | | | yes | | |
| ST18 | NA | 5'UTR | chr8:53300830-53301664 | 834 | | | 4 | | | | | | | | | | | | yes | | yes | | | |
| ST18 | NA | 5'UTR;1stExon | chr8:53321751-53322510 | 759 | 1 | | 1 | | | | | | | | | | | yes | | | | | | |
| NPBWR1 | Island | TSS200 | chr8:53852274-53852661 | 387 | | 1 | | 1 | | | | | | | 1 | | | | | | | | | |
| OPRK1 | Island | 5'UTR;1stExon;TSS200 | chr8:54163622-54164757 | 1135 | | 1 | | | | | | 0 | 4 | | | | | | | | | yes | | |
| LYPLA1 | Island | TSS200 | chr8:55014643-55014671 | 28 | | | 1 | | | | | | | | | | | | | yes | | | yes | |
| SOX17 | Island | TSS1500;TSS200;Body | chr8:55370192-55372569 | 2377 | | | | | | | | | | | | | | | | | | | | |
| RP1 | Island | Body | chr8:55533829-55533955 | 126 | | | | | | | | | | | | 1 | | | | | | yes | | |
| XKR4;SBF1P1 | Island | Body | chr8:56362167-56362648 | 481 | | | | | | | | | | | | | | | | | | yes | | |
| XKR4;SBF1P1 | NA | Body | chr8:56367723-56368202 | 479 | | | | | | | | | | | | | | | yes | | | yes | | |
| XKR4 | NA | Body | chr8:56436393-56437720 | 1327 | | | 2 | | | | | | | | | | | | | | | | | |
| LYN | N_Shore | TSS1500 | chr8:56791576-56791798 | 222 | | | 5 | | | | | | | | | | | yes | yes | | | yes | | |
| LYN | NA | 5'UTR | chr8:56851846-56852290 | 444 | | | 1 | 1 | 1 | | | | | | | | | yes | | | yes | yes | | |
| SDR16C5 | S_Shore | TSS200;TSS1500 | chr8:57233269-57233874 | 605 | | | 3 | | | | | 0 | 4 | | | 1 | | | | | yes | | | |

| | | | | | | | | | | | | | | | | | | | | | | | |
|---|---|---|---|---|---|---|---|---|---|---|---|---|---|---|---|---|---|---|---|---|---|---|---|
| SDR16C6 | NA | TSS1500 | chr8:57309109-57309394 | 285 | | | | | | | | | 1 | | | | | yes | | | | | |
| PENK | Island | Body | chr8:57358130-57358651 | 521 | | | | | 2 | | | 0 | 1 | | | | | | | | | | |
| PENK | Island | TSS1500 | chr8:57359908-57360711 | 803 | | | 21 | | | | | 0 | 2 | 1 | | 1 | | yes | | | | yes | |
| C8orf71 | N_Shore;Island | TSS1500;TSS200;Body | chr8:58191386-58193284 | 1898 | 1 | | 6 | 1 | | 1 | | 0 | 2 | | | 1 | | | | | | | yes |
| FAM110B | Island | TSS200 | chr8:58907088-58907242 | 154 | | 1 | | | | | | | 1 | | | | | | | | | yes | yes |
| FAM110B | Island | 5'UTR | chr8:59058254-59059119 | 865 | | | 28 | 1 | | | | | 1 | | 1 | | | | | | | | yes |
| TOX | Island | TSS1500 | chr8:60032607-60032983 | 376 | | | 3 | | | | | | | | | | | | | | | yes | yes |
| CHD7 | Island;N_Shore | 5'UTR | chr8:61626185-61627281 | 1096 | | | | | | | | | 1 | 1 | | | | yes | yes | | | yes | yes |
| ASPH | Island | TSS200 | chr8:62627324-62627421 | 97 | | | | | | | | | 1 | | | | | | yes | | | yes | |
| NKAIN3 | N_Shore;Island | TSS1500;1stExon;5'UTR | chr8:63160127-63161676 | 1549 | | 2 | | | | | | 0 | 5 | 1 | | 3 | | | | | | yes | yes |
| NKAIN3 | NA | Body | chr8:63614431-63614552 | 121 | | | | | | | | | 1 | | | 1 | | | | | yes | | |
| NKAIN3 | NA | Body | chr8:63776642-63776762 | 120 | | | 5 | | | | | 0 | 1 | | | 4 | 1 | | | | | yes | |
| MIR124-2 | S_Shore | TSS1500;TSS200 | chr8:65291287-65291730 | 443 | | 1 | 1 | | | | | | 1 | | | 2 | | | | | | | |
| LOC401463;BHLHE22 | Island;N_Shore;chr8:65491083 | Body;TSS200;TSS1500;N_Shore | chr8:65488640-65492683 | 4043 | | | | | | | | | | | | | | | | | | | |
| CYP7B1 | Island | TSS200;TSS1500 | chr8:65711522-65711916 | 394 | | | 5 | 1 | | | | 0 | 2 | | | 3 | | | | | | | |
| PDE7A | NA | Body;TSS200 | chr8:66701377-66701906 | 529 | | | | | | | | | 1 | | | | | | yes | | | | |
| TRIM55 | NA | TSS1500 | chr8:67039041-67039461 | 420 | | 1 | | | | | | | | | | | | yes | | | | yes | yes |
| CRH | S_Shore | 1stExon;5'UTR;TSS200 | chr8:67090581-67090798 | 217 | | | 5 | | | 1 | | 0 | 1 | | | 2 | | | | | | yes | yes |
| ADHFE1 | Island | TSS200 | chr8:67344553-67345006 | 453 | | 1 | | | | | | | | | | | | yes | yes | | | | |
| C8orf46 | NA | TSS1500;TSS200 | chr8:67405257-67405742 | 485 | | | 8 | | | | | | 2 | | | | | | | | | yes | |
| C8orf45 | NA | 1stExon;TSS1500;5'UTR | chr8:67783002-67783229 | 227 | | | | | | | | 0 | | | | | | | yes | | | | |
| LRRC67 | N_Shore | 5'UTR | chr8:67940640-67941077 | 437 | | | 1 | | | | | | | | | | | | yes | | yes | | yes |
| CSPP1;COPS5 | N_Shore | TSS1500 | chr8:67975874-67975895 | 21 | | 1 | | | | | 1 | 0 | 1 | 1 | | 4 | | yes | yes | | | | |
| CPA6 | NA | TSS1500 | chr8:68658943-68659328 | 385 | | | | | | | | | 1 | | | | | yes | | | yes | yes | |
| PREX2 | N_Shore | TSS1500;TSS200 | chr8:68864012-68864908 | 896 | | 1 | | | 3 | | | 0 | | | | | | | | | | yes | yes |
| SULF1 | NA | TSS1500;TSS200 | chr8:70378380-70378994 | 614 | | | 8 | | | | | 0 | 1 | | | | | | | | | | |
| SLCO5A1 | Island | TSS200 | chr8:70747440-70747646 | 206 | | | 1 | | | | | | | | | | | | yes | | | yes | |
| PRDM14 | Island | TSS200;Body;1stExon;5'UTR | chr8:70982039-70984559 | 2520 | | | | | | | | | | | | | | | | | | | |
| XKR9;LACTB2 | Island;S_Shore | 5'UTR;1stExon;TSS200;TSS1500 | chr8:71581642-71581788 | 146 | | | | | | | | | 2 | | | | | yes | | | | yes | |
| EYA1 | NA | 5'UTR;1stExon;TSS200 | chr8:72274256-72274615 | 359 | | | 11 | | | | | | 1 | | | | | | | | | yes | |
| MSC | Island;N_Shore;S_Shore | 3'UTR;Body;1stExon;TSS200;TSS1 | chr8:72754272-72758701 | 4429 | | 1 | 11 | | 3 | | | | 1 | | | | | yes | | | | yes | yes |
| KCNB2 | NA | Body | chr8:73848252-73849114 | 862 | | | 6 | | | | | | | | | | | | | | | | |
| C8orf84 | Island;N_Shore | TSS200;Body | chr8:74004928-74005566 | 638 | | | | | | | | | 1 | 1 | | | | | | | | | yes |
| JPH1 | Island | Body | chr8:75227564-75227811 | 247 | | | 1 | | | | | | | | | | | | | | | | |
| CRISPLD1 | Island | TSS200 | chr8:75896645-75896836 | 191 | | | | | | | | | 1 | | | | | | | | | | |
| LOC100192378 | NA | Body | chr8:77585222-77585448 | 226 | | 1 | 9 | | | | | | | | | | | | | | | yes | |
| LOC100192378 | Island | Body | chr8:77590168-77590513 | 345 | | 1 | | | | | | | | | | | | | | | | yes | yes |
| ZFHX4;LOC100192378 | N_Shore | TSS1500;Body | chr8:77592946-77593362 | 416 | | 1 | | | | | | | | | | | | | | | | yes | yes |
| ZFHX4;LOC100192378 | S_Shore | 5'UTR;TSS1500 | chr8:77596069-77596382 | 313 | | | 10 | | | | | | 1 | | | | | | | | | yes | |
| ZFHX4 | NA | Body | chr8:77617694-77618401 | 707 | | | | | | | | | 1 | | | | | | | | | | |
| STMN2 | N_Shore | TSS200 | chr8:80523226-80523364 | 138 | | | 1 | | | | | | | | | | | | | | | | |
| HEY1 | N_Shore | Body | chr8:80678770-80679314 | 544 | | | 7 | 1 | | | | | 1 | | | | | | | | | | |
| ZNF704 | NA | Body | chr8:81599484-81599569 | 85 | | 1 | 2 | | | | | 0 | 1 | | | 1 | | | | | yes | | yes |
| ZNF704 | NA | Body | chr8:81639711-81665271 | 25560 | | | | | | | | | | | | | | yes | yes | yes | yes | yes | yes |
| FABP5 | chr8:82192378-82193685;Islan | Island;TSS1500;TSS200 | chr8:82192519-82192606 | 87 | | | 3 | | | | | | | | 1 | | | | | | | | yes |
| PMP2 | NA | 1stExon | chr8:82359567-82359759 | 192 | | | | | | | | | 1 | | | | | | | | | | |
| FABP4 | NA | TSS200 | chr8:82395569-82396069 | 500 | | | | | | | | | 1 | | | | | | | | | | |
| ZFAND1 | Island | TSS200 | chr8:82633648-82633727 | 79 | | | | | | | | 0 | | | | | | yes | | | | | |
| CHMP4C | N_Shore;Island | TSS1500;TSS200;1stExon;5'UTR | chr8:82644373-82645116 | 743 | | | 1 | | | | | 0 | 6 | | | | | | | | yes | | |
| RALYL | N_Shore | TSS200;TSS1500 | chr8:85095275-85095697 | 422 | | | | | | | | | 1 | | | | | | | | | yes | yes |
| RALYL | Island | TSS200;5'UTR;1stExon | chr8:85097031-85097429 | 398 | | | 5 | | | | | | 1 | | | | | | | | | yes | |
| CA3 | Island;N_Shore | TSS200;TSS1500 | chr8:86350278-86351195 | 917 | | 1 | 1 | | 2 | | | | | | | | | yes | | | | | |
| CA2 | Island | 5'UTR;1stExon | chr8:86376277-86376842 | 565 | | 1 | | | | | | | | | | | | yes | | | | | yes |
| PSKH2 | N_Shore;Island | Body;TSS200 | chr8:87081553-87082205 | 652 | | | | | | | | 0 | 2 | | | 1 | | | | yes | | yes | yes |
| SLC7A13 | NA | TSS1500 | chr8:87243677-87243983 | 306 | | | | | | | | | | | | | | yes | yes | | | yes | |
| WWP1 | N_Shore | TSS1500 | chr8:87354422-87354562 | 140 | | | 1 | | | | | | | | | | | yes | | | | | yes |
| MMP16 | N_Shore;Island | TSS200;TSS1500 | chr8:89339819-89340480 | 661 | | 1 | | | | | | | 2 | | | | | | | | | yes | yes |
| NECAB1 | N_Shore;Island | TSS1500 | chr8:91803412-91803692 | 280 | | | | | | | | | 1 | | | | | yes | yes | | | | |
| RUNX1T1 | NA | Body;5'UTR;1stExon | chr8:93029588-93030117 | 529 | | | | | | | | | 1 | | | | | | | | | | |
| RUNX1T1 | NA | Body;5'UTR;TSS1500 | chr8:93075534-93076177 | 643 | | | | | | | | | 1 | | | | | | | | | | |
| RUNX1T1 | NA | 5'UTR;1stExon;TSS200;TSS1500 | chr8:93107298-93108262 | 964 | | | 7 | | | | | | | | | | | | | | | | yes |
| TMEM67 | N_Shore | TSS1500 | chr8:94766196-94766468 | 272 | | | 1 | | | | | 0 | 1 | | | | | yes | yes | yes | yes | | |
| CDH17 | NA | TSS200 | chr8:95229664-95229872 | 208 | | | | | | | | | 1 | | 1 | | | yes | | | yes | | yes |
| RAD54B | S_Shore | TSS1500 | chr8:95487613-95487732 | 119 | | | | | | | | 0 | 1 | 1 | | | | yes | yes | | | | |
| ESRP1 | N_Shore | Body | chr8:95653874-95654019 | 145 | | | 1 | | | | | | | | | | | | | | | | |
| TP53INP1 | Island | TSS1500 | chr8:95962084-95962463 | 379 | | | 19 | | | | | 0 | 4 | 1 | | 3 | | yes | | | | yes | yes |
| C8orf38 | N_Shore | TSS1500 | chr8:96036568-96036905 | 337 | | | 3 | | | | | | | | | | | yes | | | yes | | |
| GDF6 | Island | Body | chr8:97157756-97158052 | 296 | | | 7 | | 2 | | | | | | | | | | yes | | | | |
| GDF6 | N_Shelf | Body | chr8:97167470-97167878 | 408 | | | 2 | | | | | | | | | | | | | | | | |
| GDF6 | N_Shore | Body | chr8:97169621-97170097 | 476 | | 1 | 6 | | | | | | | | | | | yes | | | | | |
| GDF6 | N_Shore;Island | Body;1stExon;5'UTR | chr8:97171332-97173335 | 2003 | | | | | | | | | 3 | | | | | | | | | | yes |
| PTDSS1 | NA | Body | chr8:97339939-97340270 | 331 | 6 | 1 | | | | | | | | | | | | | | | | | |
| PGCP | Island | TSS1500;TSS200;5'UTR;1stExon | chr8:97657259-97657833 | 574 | | 1 | | | | | | 0 | | | | | | | | | | yes | |
| TSPYL5 | Island | 1stExon;5'UTR;TSS200 | chr8:98290148-98290601 | 453 | | | 1 | 1 | | | | | 2 | | | 1 | | yes | | | | | |
| C8orf47 | N_Shore | TSS1500 | chr8:99076314-99076734 | 420 | | | 3 | | | | | | 1 | | | | | yes | yes | | | | yes |
| KCNS2 | Island | 5'UTR | chr8:99439948-99440331 | 383 | | | 3 | | | | | | | | | 1 | | | | | | | yes |
| OSR2 | N_Shore | TSS1500 | chr8:99955563-99956055 | 492 | | 1 | | | | | | | | | | | | | | | | | |
| OSR2 | N_Shore;Island;S_Shore | 5'UTR;Body | chr8:99959430-99962118 | 2688 | | | 13 | | | | | 0 | 2 | | 1 | | | | | | | yes | yes |
| RGS22 | Island | Body;TSS200 | chr8:101117949-101118899 | 950 | | | 7 | | | | | | | | 1 | 1 | | | | | | | |
| SPAG1 | N_Shore;Island | Body | chr8:101224915-101225902 | 987 | 2 | 1 | 2 | 1 | | | | 0 | 4 | 1 | | 4 | | | yes | | | | yes |
| RNF19A | NA | 5'UTR;1stExon;TSS200 | chr8:101315424-101315846 | 422 | | | | 1 | | | | | 1 | 1 | | 1 | | yes | yes | yes | | | |
| SNX31 | N_Shore;Island | Body;1stExon;5'UTR;TSS200 | chr8:101661245-101662053 | 808 | | | 11 | | | | | | 1 | | | | | | | | yes | | yes |
| GRHL2 | N_Shore;Island | TSS1500;TSS200;Body | chr8:102503892-102506635 | 2743 | | | 11 | | | | | 0 | 1 | | | 1 | | | | | | | yes |
| NCALD | NA | 5'UTR;TSS1500 | chr8:102804678-102804916 | 238 | | | | | | | | | 1 | | | | | yes | | | | | yes |
| NCALD | S_Shore | TSS1500;TSS200 | chr8:103137173-103137274 | 101 | | | 3 | 1 | | | | | 1 | | | | | | yes | | | | |
| ATP6V1C1 | N_Shore | TSS1500 | chr8:104032819-104032876 | 57 | | | 14 | | | | | 0 | 1 | | | 2 | 1 | yes | yes | | | | |
| C8orf56;BAALC | S_Shore | TSS200;Body;TSS1500 | chr8:104153592-104154149 | 557 | | | 27 | | 2 | | | | 4 | | | 1 | | | | | | yes | |
| FZD6 | N_Shore | TSS1500;TSS200 | chr8:104310380-104310886 | 506 | | | 4 | | | | | | 1 | | | | | yes | | | | yes | |
| CTHRC1 | Island | TSS1500;TSS200 | chr8:104383499-104383971 | 472 | | | | | | | | 0 | | | | 2 | | yes | | | | | |
| RIMS2 | Island | 1stExon | chr8:104513186-104513704 | 518 | | | | | | | | | 1 | | | | | | | | | | |
| RIMS2 | Island | Body | chr8:105235606-105235943 | 337 | | | | | 1 | | | | | | | | | | | | | | |
| DPYS | Island;S_Shore | Body;1stExon;5'UTR;TSS200;TSS1 | chr8:105478683-105479862 | 1179 | | 1 | 1 | | | | | | 3 | | | 1 | | yes | yes | | | | yes |
| LRP12 | Island | TSS1500 | chr8:105601615-105601862 | 247 | | | | | | | | | 2 | | | | | yes | | | | yes | yes |
| ZFPM2 | N_Shore;Island | TSS1500;Body | chr8:106330170-106332188 | 2018 | | | | | | | | 0 | 2 | | | | | | yes | | | yes | yes |
| ABRA | NA | TSS200 | chr8:107782513-107782734 | 221 | | | 3 | | | | | | | | | | | yes | | | | | |
| ANGPT1 | NA | TSS200 | chr8:108510286-108510343 | 57 | | | 4 | | | | | | 1 | 1 | | | | | | | | | yes |
| RSPO2 | Island;S_Shore | 5'UTR;TSS200 | chr8:109094880-109096007 | 1127 | | 1 | | | 1 | | | | 2 | | | | | | | | | | |
| TMEM74 | Island | 1stExon;5'UTR | chr8:109799739-109799909 | 170 | | | | | | | | | 1 | | | | | yes | | | | yes | |
| TRHR | NA | TSS1500;TSS200 | chr8:110098844-110100005 | 1161 | | | 14 | | | | | 0 | 1 | | | | | | | | | yes | |

| | | | | | | | | | | | | | | | | | | | | | | | |
|---|---|---|---|---|---|---|---|---|---|---|---|---|---|---|---|---|---|---|---|---|---|---|---|
| PKHD1L1 | Island | TSS200 | chr8:110374563-110374866 | 303 | | | | | | | | 0 | | | | 1 | | | | | | yes | |
| GOLSYN | NA | 3'UTR | chr8:110586775-110587264 | 489 | | | | | | | | | 1 | | | | | | | | | | |
| GOLSYN | N_Shore | Body;5'UTR;TSS1500 | chr8:110656096-110656245 | 149 | | | | | | | | | 1 | | | | | yes | | | | | |
| GOLSYN | NA | 5'UTR;1stExon | chr8:110693195-110693289 | 94 | | 1 | 8 | | | | | 0 | 1 | | | | | | | | | | |
| GOLSYN | S_Shore | TSS200;TSS1500 | chr8:110704080-110705093 | 1013 | | | | | | | | | 1 | | | | | yes | | | | yes | yes |
| KCNV1 | Island;S_Shore | TSS1500 | chr8:110987946-110988444 | 498 | | 1 | 9 | | | | | | | | | | | yes | | | | | |
| CSMD3 | NA | 5'UTR;Body;1stExon;TSS200 | chr8:114389356-114389479 | 123 | | | | | | | | | 3 | | | | | | | | | yes | |
| CSMD3 | NA | TSS1500 | chr8:114450347-114450647 | 300 | | 1 | | | | | | | | | | | | | | | | yes | |
| TRPS1 | Island;N_Shore | 5'UTR | chr8:116660311-116661063 | 752 | | | 27 | | | | | 0 | 1 | | | | | | | | | | |
| C8orf85 | N_Shore | TSS1500;TSS200 | chr8:117950244-117950504 | 260 | | | 3 | | | | | | | | | | | | yes | yes | | yes | |
| SLC30A8 | NA | TSS200 | chr8:118147163-118147534 | 371 | | | 1 | | | | | | 1 | | | | | | | | | | yes |
| MED30 | N_Shore;Island | TSS1500;1stExon;5'UTR | chr8:118532625-118533254 | 629 | | | | | | | | | 1 | | | 1 | | yes | yes | | | | |
| EXT1 | NA | Body | chr8:119086580-119086762 | 182 | | | | | | | | 0 | 1 | | | 2 | | yes | | | | yes | |
| MAL2 | N_Shore;Island | TSS1500;TSS200;1stExon;Body | chr8:120219902-120221797 | 1895 | | | 13 | | | | | 0 | 10 | 2 | | 5 | | yes | | | yes | yes | yes |
| NOV | Island | TSS200 | chr8:120428507-120428692 | 185 | | | | | | | | | 1 | | | | | | | | | | yes |
| COL14A1 | S_Shore | 5'UTR | chr8:121138133-121138317 | 184 | | | | | | | | | | | | 1 | | | | | yes | | |
| HAS2AS;HAS2 | N_Shore | TSS1500;5'UTR | chr8:122651271-122651370 | 99 | | | | | | | | | 1 | | | | | | | | | | |
| ZHX2 | NA | 5'UTR | chr8:123875036-123875223 | 187 | | | 2 | | | | | | 1 | 1 | | | | yes | | | yes | yes | |
| FAM83A | NA | TSS1500;TSS200;1stExon | chr8:124194312-124195402 | 1090 | | 1 | | | | | | 0 | 2 | | | | | | yes | yes | | yes | |
| FAM83A;LOC100131726 | NA | nan | chr8:124215033-124215152 | 119 | | | | | | | | | | | 1 | | | yes | yes | | yes | | |
| FAM83A | N_Shore | Body | chr8:124217698-124217896 | 198 | | | | | | | | 0 | 1 | | | | | yes | | | | | |
| FAM83A | N_Shore;Island | Body | chr8:124219077-124219725 | 648 | | | 5 | | | | | 0 | | | | 1 | | yes | | | | | yes |
| ZHX1 | N_Shelf | 5'UTR | chr8:124284176-124284393 | 217 | | 1 | | | | | | | | | | | | | | | | | yes |
| TRMT12 | NA | TSS200;1stExon;5'UTR | chr8:125462982-125463682 | 700 | | | 4 | | 1 | | | 0 | 9 | 4 | | 6 | 1 | | | yes | | | |
| MTSS1 | Island | TSS1500 | chr8:125741158-125741265 | 107 | | | | | | | | | 2 | | | | | | | | | | yes |
| SQLE | S_Shore | 1stExon;5'UTR | chr8:126011549-126011954 | 405 | | | 1 | | | | | | | | | | | | | | | | |
| FAM84B | N_Shore;Island | 3'UTR;Body;TSS200 | chr8:127568328-127570908 | 2580 | | | 1 | | | | | | 2 | 1 | | 2 | | | | | | | yes |
| POU5F1B | NA | TSS1500 | chr8:128427099-128427668 | 569 | | | | | | | | | 1 | | | | | yes | | yes | | | |
| LOC727677 | NA | TSS1500 | chr8:128494655-128494728 | 73 | | 1 | | | | | | | | | | | | | yes | | | | |
| MYC | Island | Body | chr8:128750586-128750924 | 338 | | | | 1 | | | | | | | | | | | | | | | |
| MIR1204;PVT1 | S_Shore | TSS200;Body | chr8:128808017-128808554 | 537 | | | | | | | | | 1 | | | | | | | | | | |
| MIR1207;PVT1 | NA | TSS1500;Body | chr8:129060815-129061382 | 567 | | | | | | | | | 2 | | | | | yes | yes | yes | | | |
| HPYR1 | NA | nan;TSS200;TSS1500 | chr8:133573739-133574073 | 334 | | 1 | | | | | | | 1 | | 1 | | | | | yes | | | |
| TMEM71 | NA | TSS200 | chr8:133773093-133773115 | 22 | | | 1 | | | | | | 1 | | | | | yes | | | | | |
| TG | NA | TSS1500;TSS200 | chr8:133878997-133880105 | 1108 | | 1 | | | | | | | | | | | | yes | yes | | | | yes |
| TG;SLA | NA | Body;1stExon;5'UTR | chr8:134114834-134115431 | 597 | | | | | | | | | 1 | | | | | yes | | | yes | | yes |
| WISP1 | NA | TSS1500 | chr8:134202337-134202370 | 33 | | 1 | | | 1 | | | | | | | | | yes | yes | | | | yes |
| NDRG1 | NA | 3'UTR | chr8:134249790-134250016 | 226 | | | | | | | | | | | | | | | | | | | |
| ST3GAL1 | NA | 5'UTR | chr8:134570528-134576933 | 6405 | | | | | | | | | | | | | | yes | yes | yes | yes | yes | |
| ST3GAL1 | N_Shore | 5'UTR | chr8:134580580-134581043 | 463 | | | 4 | | | | | | | | | | | yes | | | | | |
| ZFAT | N_Shore;Island | 3'UTR;Body | chr8:135490694-135491009 | 315 | | | 10 | | | | | | | | | | | yes | | | | | |
| ZFAT | NA | Body | chr8:135614730-135614834 | 104 | | | | | | | | | 1 | | | | | | | | | | |
| ZFAT | S_Shore | TSS1500 | chr8:135725874-135726595 | 721 | | 1 | | 1 | | | | | | | | | | yes | yes | | yes | | |
| LOC286094 | NA | TSS1500 | chr8:136245867-136246255 | 388 | | | | | | | | | | | | | | yes | | | | | yes |
| FAM135B | Island | 5'UTR | chr8:139508813-139509708 | 895 | | 1 | | | | | | | | | | | | | | | | | yes |
| COL22A1 | NA | Body | chr8:139613712-139614020 | 308 | | | 3 | | | | | | | | | | | | | | | | |
| COL22A1 | NA | Body | chr8:139784036-139784450 | 414 | | | | | | | | | | | | | | yes | | | | | |
| KCNK9 | S_Shore | Body | chr8:140632686-140632846 | 160 | | | | | | | | | | | | | | yes | yes | | | | |
| KCNK9 | NA | Body | chr8:140643009-140643434 | 425 | | 1 | | | | | | | | | | | | yes | | | | | |
| KCNK9 | N_Shelf | Body | chr8:140712424-140712499 | 75 | 1 | | | | | | | | | | | | | yes | | | | yes | |
| KCNK9 | N_Shore | Body | chr8:140714489-140714610 | 121 | | | 1 | | | | | | | | | | | | | | | | |
| KCNK9 | Island | TSS1500 | chr8:140715646-140715802 | 156 | | | | | | | | | 1 | | | | | | | | | yes | yes |
| TRAPPC9 | NA | Body | chr8:140749025-140749377 | 352 | | | | | | | | | | | | | | | yes | | yes | | |
| TRAPPC9 | NA | Body | chr8:140931190-140931443 | 253 | | 1 | | | | | | | | | | | | yes | yes | | | | |
| TRAPPC9 | NA | Body | chr8:140937286-140937410 | 124 | | | 3 | | | | | | | | | | | yes | yes | | | | |
| TRAPPC9 | NA | Body | chr8:140945782-140945843 | 61 | | | 3 | | 1 | | | 0 | 1 | | 1 | | | yes | yes | | | | |
| TRAPPC9 | NA | Body | chr8:141034325-141034355 | 30 | | | | | | | | | 1 | | | | | yes | | | | | |
| TRAPPC9 | Island | Body | chr8:141057285-141057827 | 542 | | | 12 | | 1 | | | | | | | | | yes | | | | | |
| TRAPPC9 | Island | Body | chr8:141110747-141111080 | 333 | | | | | | | | | 1 | | | | | yes | | | | | |
| TRAPPC9 | NA | Body | chr8:141220177-141220490 | 313 | | | 2 | 1 | | | | | | | | | | yes | | | | | |
| TRAPPC9 | NA | Body | chr8:141248582-141249094 | 512 | | 1 | 13 | | | | | | | | 1 | | | | yes | | | | |
| TRAPPC9 | Island | Body | chr8:141348866-141349194 | 328 | | | | | | | | | 1 | | | | | yes | | yes | | | |
| TRAPPC9 | Island | Body | chr8:141359539-141359786 | 247 | | | 8 | | | | 1 | 0 | 1 | | | | | | | | | | |
| TRAPPC9 | S_Shore | TSS1500;TSS200 | chr8:141468743-141468933 | 190 | | | 2 | | | | | | | | | | | yes | yes | | | | yes |
| EIF2C2 | Island | Body | chr8:141559367-141559710 | 343 | | | 1 | | | | | | | | | | | | | | | | |
| EIF2C2 | NA | Body | chr8:141565971-141566221 | 250 | | | | | | | | | 1 | | | | | yes | | | | | |
| EIF2C2 | NA | Body | chr8:141572044-141572291 | 247 | | | | | | | | | 1 | | | | | | | | | | |
| EIF2C2 | NA | Body | chr8:141577061-141577150 | 89 | 1 | | | | | | | | | | | | | | | | | | |
| EIF2C2 | NA | Body | chr8:141588118-141588242 | 124 | | | 3 | | | | | | 1 | 1 | | | | | yes | | | | |
| EIF2C2 | NA | Body | chr8:141599141-141599436 | 295 | | | | 1 | 1 | | | | 3 | | 1 | | | | | | yes | yes | |
| EIF2C2 | NA | Body | chr8:141608316-141609470 | 1154 | | | | 1 | 2 | | | 0 | 2 | | | | | yes | yes | | | | |
| DENND3 | chr8:142138208-142139342 | Island | chr8:142139341-142139839 | 498 | | | | | | | | | | | 1 | | | | | | | | |
| DENND3 | NA | Body | chr8:142157104-142157867 | 763 | | | 1 | | | | | | | | | | | | yes | | | | |
| DENND3 | NA | nan | chr8:142159992-142160235 | 243 | | | | | | | | | | | 1 | | | | | | yes | | |
| DENND3 | N_Shore;Island | Body | chr8:142183179-142183677 | 498 | | | | | | | | | 1 | | | | | yes | | | | | |
| DENND3 | NA | Body | chr8:142189535-142189600 | 65 | | 1 | | | 1 | | | | | | | | | yes | yes | | | | |
| DENND3 | NA | Body | chr8:142192487-142192718 | 231 | | | | | 1 | | | | | | | | | yes | yes | | | | |
| DENND3 | NA | Body | chr8:142200020-142201784 | 1764 | 3 | | | | | | | | 2 | | | | | | yes | | | | |
| SLC45A4 | S_Shore | Body | chr8:142232952-142233705 | 753 | | | 2 | | | 3 | | | 6 | | | | | yes | yes | | | | |
| SLC45A4 | N_Shore | Body | chr8:142235430-142235763 | 333 | | | | | | | | | 1 | 1 | | | | | yes | | | | |
| SLC45A4 | Island | Body | chr8:142237057-142237359 | 302 | | | | | | | | 0 | 1 | | | | | | | | | | |
| SLC45A4 | S_Shore | 5'UTR;1stExon;TSS200;TSS1500 | chr8:142238504-142239323 | 819 | | 1 | 4 | | 1 | | | 0 | 2 | 1 | | 1 | | | | | | | |
| GPR20 | S_Shore | 5'UTR | chr8:142368026-142368316 | 290 | | | | | | | | | 1 | | | | | yes | | | | | |
| PTP4A3 | S_Shelf | TSS1500 | chr8:142431437-142431765 | 328 | | | | | | | | | | | | | | | | | | | |
| PTP4A3 | S_Shore | Body | chr8:142437565-142437958 | 393 | | | | | | | | | | | | | | | yes | | | | |
| FLJ43860 | Island | Body | chr8:142452378-142452509 | 131 | | 1 | | | | | | | | | | | | | | | | | |
| FLJ43860 | NA | Body | chr8:142505085-142505578 | 493 | | | 3 | | | | | | | | | | | yes | | | | yes | |
| FLJ43860 | NA | TSS200 | chr8:142517341-142517806 | 465 | | | 4 | | | | | | 4 | | | | | yes | | | | | yes |
| NCRNA00051 | NA | TSS1500 | chr8:143279443-143279569 | 126 | | | | | | | | 0 | | | | | | | yes | | | | |
| TSNARE1 | NA | Body | chr8:143324660-143324707 | 47 | | | 9 | | | | | | | | | | | yes | yes | | | | |
| TSNARE1 | Island | Body | chr8:143333518-143333834 | 316 | | 1 | | | | | | | | | | | | yes | | | | | |
| TSNARE1 | Island | Body | chr8:143359953-143360290 | 337 | | | | | | | | | | | | | | | | | | | |
| TSNARE1 | NA | nan;Body | chr8:143386260-143387265 | 1005 | | | | | | | | | 1 | 1 | 1 | | | yes | | | | | |
| TSNARE1 | N_Shore | Body | chr8:143405663-143406192 | 529 | | 1 | | | | | | | | | | | | yes | yes | | | | |
| TSNARE1 | Island | Body | chr8:143407646-143408047 | 401 | | | | | | | | | 1 | | | | | | yes | | | | |
| TSNARE1 | NA | Body | chr8:143424755-143424827 | 72 | | | | | | | | | 1 | | | | | | | | | yes | |
| TSNARE1 | N_Shore | Body | chr8:143427631-143427750 | 119 | | | 1 | | | | | | | | | | | | | | yes | | |
| TSNARE1 | S_Shore | Body | chr8:143429591-143430202 | 611 | | | 6 | | | | | | | | | | | | yes | | | | yes |
| TSNARE1 | NA | 5'UTR | chr8:143437568-143437988 | 420 | | | 3 | | | | | | | | | | | yes | | | yes | | |

| | | | | | | | | | | | | | | | | | | | | | | | |
|---|---|---|---|---|---|---|---|---|---|---|---|---|---|---|---|---|---|---|---|---|---|---|---|
| TSNARE1 | NA | 5'UTR | chr8:143459774-143459992 | 218 | | | | | | | | | | | | | | | yes | | | | |
| TSNARE1 | NA | 5'UTR | chr8:143463227-143463667 | 440 | 6 | | | | | | | | 1 | 1 | 1 | | | | | | | | |
| BAI1 | N_Shore;Island;S_Shore | TSS200;5'UTR;1stExon;Body | chr8:143545214-143546893 | 1679 | | | 29 | | | | | | 1 | 1 | | 1 | | | yes | | | yes | yes |
| BAI1 | S_Shore | Body | chr8:143560651-143561039 | 388 | | | 1 | | | | | | | | | | | yes | | | | | |
| BAI1 | Island | Body | chr8:143580770-143581481 | 711 | | | 11 | | | | | 0 | 2 | 1 | | 1 | | | yes | | | | yes |
| BAI1 | NA;S_Shelf | Body | chr8:143596671-143596947 | 276 | | | 3 | | | | | | | | | | | | yes | | | | |
| BAI1 | N_Shore | Body | chr8:143625597-143625967 | 370 | | | | | | | | | 1 | | | | | | | | | yes | |
| PSCA | NA | TSS1500 | chr8:143761503-143761697 | 194 | | 1 | 1 | | | | | | 1 | | | | | | | yes | | | yes |
| PSCA | NA | Body | chr8:143762866-143763565 | 699 | | | 9 | | | | | | | | | | | | | | | yes | |
| LY6K | N_Shore;Island | TSS1500;TSS200;1stExon;5'UTR | chr8:143781297-143782367 | 1070 | | 2 | 8 | | 2 | | | 0 | 2 | | | 1 | | yes | | yes | | | yes |
| SLURP1 | S_Shelf | 5'UTR;1stExon;TSS200;Body | chr8:143823734-143824001 | 267 | | 1 | 3 | | | | | | 2 | 1 | 1 | | | | yes | | | | |
| LYPD2 | NA | TSS1500 | chr8:143834539-143835022 | 483 | | | | | | | | | 1 | | | | | yes | yes | | | | |
| LYNX1 | NA | TSS200;Body | chr8:143851287-143851427 | 140 | | | 1 | 1 | | | | | 1 | | | | | yes | | | | | |
| LYNX1 | Island;S_Shore | 5'UTR;TSS200;1stExon;TSS1500 | chr8:143858414-143859990 | 1576 | 3 | 1 | | 1 | | | | 0 | 8 | 2 | 1 | 3 | 2 | yes | yes | | yes | | |
| LY6D | NA | Body;1stExon;5'UTR;TSS200;TSS1 | chr8:143867799-143869473 | 1674 | 1 | | 11 | | | | | | 2 | | | | | | | | | | |
| CYP11B1 | NA | 1stExon | chr8:143961141-143961291 | 150 | | | | | | | | | | | | | | | yes | | | | |
| CYP11B2 | NA | 1stExon | chr8:143999155-143999729 | 574 | | | | | | | | | | | | | | | yes | | | | |
| LOC100133669 | NA | Body | chr8:144069512-144069888 | 376 | | | | | | | | | | | | | | | yes | yes | | | |
| LY6E;LOC100133669 | Island | 5'UTR;TSS1500 | chr8:144100300-144100535 | 235 | | 1 | | | | | | | | | | | | | | | | | yes |
| C8orf31 | NA | TSS200;TSS1500 | chr8:144120106-144120706 | 600 | | | | | | | | 0 | 5 | | | 4 | | yes | | yes | | yes | |
| LY6H | Island | Body | chr8:144240945-144242268 | 1323 | | 1 | | | | | | | | | | | | yes | yes | | | yes | yes |
| ZFP41 | NA | 3'UTR | chr8:144344397-144344793 | 396 | | | 6 | | | | | | | | | | | | | | | | yes |
| GLI4 | Island | Body | chr8:144358566-144359074 | 508 | | | | | | | | 0 | | | | | | | | | | | |
| TOP1MT | NA | Body | chr8:144410701-144410972 | 271 | | | | | | | | | 1 | | | | | yes | yes | | | | |
| TOP1MT | N_Shore | Body | chr8:144416327-144416485 | 158 | | | | | | | | | | | | | | | yes | | | | |
| RHPN1;C8orf51 | Island;S_Shore | Body;TSS1500 | chr8:144451413-144451924 | 511 | | | 5 | 1 | | | | 0 | 2 | | | 2 | | | | | | | yes |
| RHPN1 | S_Shore | Body | chr8:144457677-144458069 | 392 | | | | | | | | | | | | | | | | | | | |
| RHPN1 | N_Shore;Island | Body | chr8:144460945-144462631 | 1686 | | | | | | | | | 1 | | | | | | | | | | |
| MAFA | Island | TSS1500 | chr8:144513697-144513980 | 283 | | | 11 | | | | | | | | | | | | | | | | yes |
| ZC3H3 | NA | Body | chr8:144543486-144544041 | 555 | | | | | | | | | 4 | | | | | yes | yes | | | | |
| ZC3H3 | N_Shelf | Body | chr8:144561585-144561886 | 301 | | | 1 | | | | | | | | | | | | | | | | |
| ZC3H3 | Island | Body | chr8:144565277-144565335 | 58 | | | | | 1 | | | | | | | | | | | | | | |
| ZC3H3 | NA | Body | chr8:144576604-144577238 | 634 | 2 | | | | | | | | 2 | | | | | | yes | | | | yes |
| ZC3H3 | NA | Body | chr8:144589984-144590068 | 84 | | | | | | | | | 1 | | | | | | yes | | | | |
| ZC3H3 | Island | Body | chr8:144601734-144601883 | 149 | | | | 2 | | | | | | | | | | | | | | | |
| ZC3H3 | NA | Body | chr8:144610522-144610644 | 122 | | | | 1 | | | | | | | | | | | | | | | |
| GSDMD | N_Shore;Island | TSS1500;TSS200;5'UTR;1stExon | chr8:144635260-144636462 | 1202 | | 1 | 1 | | | | 2 | 0 | 13 | 12 | 1 | 9 | | | yes | | | yes | yes |
| GSDMD | N_Shore | TSS200;5'UTR | chr8:144640306-144640378 | 72 | | | | | | | | | | | | | | yes | yes | | | | |
| C8orf73 | S_Shore | TSS200;1stExon;5'UTR | chr8:144654887-144655447 | 560 | | | 3 | | | | | 0 | | 1 | | | | | | yes | | | |
| NAPRT1 | Island;S_Shore;N_Shore | Body;TSS200 | chr8:144659627-144661051 | 1424 | 4 | 3 | 28 | | 1 | 1 | | 0 | 6 | 5 | | 13 | 1 | | | | | | yes |
| EEF1D | NA | Body | chr8:144668199-144668378 | 179 | | | | | | | | | | | | | | | | yes | | | yes |
| EEF1D;TIGD5 | Island | TSS1500;1stExon | chr8:144680893-144681482 | 589 | | | | | | | | | | | | | | | | | | | yes |
| ZNF623 | NA | 5'UTR;TSS1500 | chr8:144730494-144730562 | 68 | | | | | | | | | 1 | | | | | | yes | | | | |
| BREA2 | S_Shelf | TSS200 | chr8:144779210-144779376 | 166 | 2 | | | | | | | | | | | | | | | yes | | | |
| LOC100130274 | Island | Body;1stExon;TSS200 | chr8:144789823-144790656 | 833 | | 1 | 7 | | 1 | | | | 3 | | | 3 | | | | | | yes | |
| MAPK15 | N_Shore;Island | TSS200;1stExon;5'UTR | chr8:144798477-144798631 | 154 | | | | | | | | 0 | | | | | | yes | yes | | | yes | |
| MAPK15 | Island | Body | chr8:144800997-144801223 | 226 | | | | | | | | | 1 | | | | | | | | | | |
| MAPK15 | S_Shelf | Body | chr8:144803601-144803661 | 60 | | | 6 | 1 | | | | | 1 | | | | | | | | | | yes |
| FAM83H | Island | Body | chr8:144809598-144810034 | 436 | | | | | | | | | | | | | | | | | | | yes |
| FAM83H | N_Shore | 5'UTR | chr8:144814911-144815275 | 364 | | | | | | | | | 1 | | | | | | | | | | yes |
| SCRIB | Island | Body | chr8:144891810-144891927 | 117 | | | | | | | | | 1 | | | 1 | 1 | | | | | | |
| SCRIB;MIR937 | Island;N_Shore | Body;TSS1500 | chr8:144894951-144896307 | 1356 | | | | | | | | 0 | 1 | | | 1 | | | | | | | |
| SCRIB | chr8:144897283-144898354 | Island | chr8:144897555-144897570 | 15 | | | | | | | | | | | 1 | | | | | | | | yes |
| PUF60 | NA | Body;5'UTR | chr8:144906376-144906507 | 131 | | | | 1 | | | | | | | | | | | | | | yes | |
| NRBP2 | S_Shore | TSS1500 | chr8:144924893-144925527 | 634 | | | | | | | | | | | | | | | yes | yes | yes | | |
| EPPK1 | Island | 1stExon | chr8:144942223-144943053 | 830 | | | | | | | | | | | | | | | | | | | |
| EPPK1 | S_Shore;Island | TSS200;TSS1500;1stExon | chr8:144946557-144948781 | 2224 | | | 6 | | | | | | 1 | | | | | | | yes | | | |
| PLEC1 | N_Shore;Island | Body | chr8:145002522-145004388 | 1866 | | | | | 1 | | | | 3 | | | | | | yes | | | | yes |
| PLEC1 | Island;N_Shore | Body | chr8:145008110-145009406 | 1296 | | | | 1 | 1 | | | 0 | 4 | 1 | 1 | 10 | 1 | | | | | yes | yes |
| PLEC1 | N_Shelf | Body;1stExon;5'UTR | chr8:145013249-145013932 | 683 | | | | 1 | 3 | 2 | | | | | | | | | | | | | |
| PLEC1 | N_Shore | Body;TSS1500 | chr8:145014989-145015122 | 133 | | | | | | | | | 1 | | | | | | | | | yes | |
| PLEC1;MIR661 | Island;S_Shore | Body;1stExon;TSS1500;TSS200 | chr8:145018816-145019839 | 1023 | | | 39 | | | | | 0 | | | | | | | | | | | |
| PLEC1 | Island | Body;1stExon;TSS200 | chr8:145024774-145026051 | 1277 | | | | 1 | | | | | 1 | 1 | | | | | | | | yes | yes |
| PLEC1 | Island | Body;TSS200;1stExon;5'UTR | chr8:145027948-145028461 | 513 | | | 8 | | | | | 0 | 1 | | | | | | | | | yes | yes |
| PLEC1 | NA | Body | chr8:145033104-145033743 | 639 | | | | 3 | | | | | 1 | | | | | yes | yes | | | | |
| PLEC1 | Island | Body;TSS1500 | chr8:145047469-145049472 | 2003 | | | 9 | 1 | | | | 0 | 2 | | 1 | | | | | | | yes | yes |
| PARP10 | NA | Body | chr8:145057101-145057174 | 73 | | | | | | | | | | | | | | yes | | | | | |
| PARP10 | N_Shelf | TSS1500 | chr8:145061234-145061628 | 394 | | | | | 1 | | | | 1 | | | | | yes | yes | | | yes | |
| SPATC1 | NA | TSS1500 | chr8:145085954-145086659 | 705 | | | 4 | | 1 | | | | 1 | 1 | 1 | | | | yes | | | | |
| OPLAH | Island | Body | chr8:145106246-145107199 | 953 | | | | | 1 | | | | | 1 | | | | | | | | yes | yes |
| KIAA1875;MAF1 | N_Shore | Body;3'UTR;TSS1500;TSS200 | chr8:145162300-145163136 | 836 | | | 20 | | | | | 0 | 4 | | | 1 | | | | | | | yes |
| BOP1 | N_Shore;Island | Body | chr8:145501817-145502297 | 480 | | | | | | | | | | | | | | | | | | yes | |
| HSF1 | NA | Body | chr8:145537504-145538097 | 593 | | | 1 | 2 | 1 | | | | | | | | | | | | | yes | |
| SCRT1 | Island | Body | chr8:145558810-145558967 | 157 | | | 2 | | | | | | | | | | | | | | | yes | |
| C8ORFK29 | Island;S_Shore | Body;TSS200 | chr8:145577701-145578687 | 986 | | | 2 | | | | | 0 | | | | | | | | | | yes | |
| CPSF1;MIR939 | N_Shore | Body;TSS1500 | chr8:145620185-145620299 | 114 | | | | | | | | | | | | | | | | | | | |
| CPSF1;MIR1234 | S_Shelf | Body;TSS1500 | chr8:145626588-145626898 | 310 | | | | | 1 | | | | | | | | | | | | | | |
| SLC39A4 | Island | Body | chr8:145638434-145639181 | 747 | | | 7 | 1 | | | | 0 | 1 | 2 | | 2 | 1 | | | | | yes | |
| SLC39A4 | Island;S_Shore | 1stExon;TSS200;5'UTR;TSS1500 | chr8:145641996-145642456 | 460 | | 1 | | 1 | | 1 | | | | | | | | yes | | | | | |
| VPS28;NFKBIL2 | Island | TSS1500;Body | chr8:145654565-145654854 | 289 | | | 13 | | | | | 0 | 3 | | | 1 | | yes | | | | | |
| KIFC2;CYHR1 | Island | TSS1500;TSS200 | chr8:145691038-145691254 | 216 | | | | | | | | | 1 | | | | | | | | | yes | yes |
| CYHR1;KIFC2 | Island | TSS1500;Body | chr8:145692504-145692591 | 87 | | | | | | | | | | | | | | | | | | | yes |
| PPP1R16A;GPT | Island;S_Shore;N_Shore | Body;3'UTR;TSS1500;5'UTR;1stExo | chr8:145725708-145729772 | 4064 | | | | | | | | | | | | | | | | | | | |
| LRRC14 | Island | 3'UTR | chr8:145747739-145748059 | 320 | | | | | | | | | | | | | | | | | | | yes |
| LRRC24;LRRC14 | N_Shore;Island | Body;3'UTR | chr8:145749776-145750363 | 587 | | | | | | | | 0 | 2 | | | 1 | | | | | yes | yes | |
| MGC70857;KIAA1688 | Island | TSS1500;3'UTR | chr8:145755264-145755974 | 710 | | | | | 1 | | | 0 | 3 | | | 1 | | | | | | | yes |
| KIAA1688 | Island | Body | chr8:145758486-145758650 | 164 | | 1 | | | | | | | | | | | | | | | | | |
| KIAA1688 | Island | Body | chr8:145806273-145806752 | 479 | | | | | | | | | 1 | | | | | | | | | yes | |
| KIAA1688 | S_Shore | Body | chr8:145808436-145808810 | 374 | | | | | 1 | | | | | | | | | yes | | | | | |
| ZNF251 | Island | Body | chr8:145955983-145956024 | 41 | | 2 | | | 1 | | | 0 | 1 | | | 12 | 1 | | | | | | |
| ZNF34 | S_Shore | TSS1500 | chr8:146013430-146013792 | 362 | | | | | | | | | | | | | | yes | | | | | |
| KANK1 | Island | 1stExon;5'UTR;Body | chr9:707166-707410 | 244 | | | | | 1 | | | | | | | | | | | | | | |
| DMRT3 | N_Shore | TSS1500 | chr9:975825-975997 | 172 | | | 2 | | | | | | | | | | | | | | | yes | |
| DMRT3 | Island | Body | chr9:977556-977818 | 262 | | | | | | | | | | | | | | | | | | yes | |
| DMRT3 | NA | Body | chr9:990618-991047 | 429 | | | 2 | | | | | | | | | | | | | | | | |
| C9orf68;PPAPDC2 | Island | Body;1stExon | chr9:4662858-4663107 | 249 | | | | | | | | | | | | | | | | | | | |
| AK3 | Island | TSS1500 | chr9:4741473-4741710 | 237 | | | | | | | | | 1 | | | | | | | | | | |
| PTPRD | Island | 5'UTR | chr9:8857486-8857502 | 16 | | | | | | | | | | 1 | | | | | | | | yes | |
| SLC24A2 | Island | TSS1500 | chr9:19788317-19788757 | 440 | | | | | | | | | 1 | | | | | | | | | | yes |

| | | | | | | | | | | | | | | | | | | | | | | | |
|---|---|---|---|---|---|---|---|---|---|---|---|---|---|---|---|---|---|---|---|---|---|---|---|
| TUSC1 | Island | 1stExon;3'UTR | chr9:25677495-25678015 | 520 | | | | | | | | | 1 | | | 1 | | | | | | yes | |
| TEK | NA | TSS200 | chr9:27109111-27109492 | 381 | | | 8 | | | | | | | | | | | | | | | yes | yes |
| TMEM215 | Island | TSS1500 | chr9:32783202-32783516 | 314 | | 1 | | | | | | | | | | | | | | | | | yes |
| KIAA1161 | Island | 3'UTR;Body | chr9:34370781-34371380 | 599 | | | 6 | | | | | | 3 | | | | | | | | | | |
| C9orf25 | N_Shore | Body | chr9:34457129-34457500 | 371 | | | 1 | | 1 | | | | 1 | | 1 | | | | | | | | |
| CNTFR | Island | TSS1500 | chr9:34590742-34591192 | 450 | | | 5 | | | | | | | | | | | | | | | | |
| ARID3C | Island | Body | chr9:34624080-34624587 | 507 | | | | | | | | | 1 | | | | | | | | yes | | yes |
| SPAG8 | N_Shore | Body | chr9:35811884-35812293 | 409 | | | | | | | | | | | | | | | | | | yes | |
| GNE | Island | 5'UTR;Body | chr9:36258171-36258600 | 429 | | 1 | | | | | | | | | | | | yes | | | | | |
| PAX5 | N_Shore | Body | chr9:36881908-36882152 | 244 | | | | 1 | | | | | | | | | | | | | | yes | |
| PAX5 | Island | Body | chr9:36985987-36986788 | 801 | | | | | | | | | 2 | | | | | yes | yes | | | | yes |
| TJP2 | N_Shore | TSS1500 | chr9:71735648-71735701 | 53 | | | | | | | | | | | | | | yes | yes | yes | | | |
| GDA | Island | 1stExon;5'UTR | chr9:74764428-74764504 | 76 | | 1 | | | | | | | | | | | | | yes | | | | |
| PRUNE2 | Island | TSS200 | chr9:79521093-79521359 | 266 | | | | | | | | | 1 | | | | | | | | yes | | |
| GNAQ | Island | TSS1500 | chr9:80647290-80647609 | 319 | | | | | | | | | 1 | | | | | | | | | yes | yes |
| KIF27 | N_Shore | 5'UTR | chr9:86535572-86535755 | 183 | | | | | | | | 0 | | | | | | yes | | | | | yes |
| NTRK2 | Island | 5'UTR;1stExon | chr9:87284722-87285186 | 464 | | | | | | | | | 1 | | | | | | | | | yes | |
| C9orf170 | Island | TSS200 | chr9:89763370-89763570 | 200 | | | | | | | | 0 | | | | | | | | | | | |
| DAPK1 | S_Shore | Body | chr9:90113998-90114156 | 158 | | | 1 | | | | | 0 | | | | | | | | | | | |
| CTSL1 | N_Shore | TSS1500 | chr9:90340652-90341458 | 806 | | 1 | | | | | | | | | | | | yes | | | | | |
| CDK20 | Island | 1stExon;5'UTR | chr9:90589530-90589806 | 276 | | 1 | | | | | | | 2 | | | | | yes | | | | | |
| GADD45G | Island | Body | chr9:92220388-92221282 | 894 | | | 5 | | | | | | | | | | | | | | | | |
| LOC100129066 | N_Shore | Body | chr9:92291210-92291523 | 313 | | 1 | 1 | 1 | | | | | 1 | 1 | | | | | | | yes | | |
| ROR2 | Island | Body | chr9:94486741-94487105 | 364 | | | | | | | | | | | | | | | | | | | |
| WNK2 | N_Shore | Body | chr9:96009796-96009985 | 189 | | | | | | | | | | | | | | | yes | | | | |
| BARX1 | Island | Body | chr9:96715843-96716665 | 822 | | | 3 | | | | | | 1 | | | | | | | | | | |
| FBP1 | S_Shore | 5'UTR;TSS1500 | chr9:97402166-97402555 | 389 | | 1 | | | | | | | | | | | | yes | yes | | | | |
| C9orf3 | N_Shore | Body | chr9:97810617-97811087 | 470 | | | | | | | | | | | | | | | | | | yes | |
| C9orf3;MIR24-1 | N_Shore | 3'UTR;TSS200 | chr9:97848107-97848457 | 350 | | | | | | | | | | 1 | | | | | | | | | |
| NCRNA00092 | S_Shore | TSS1500 | chr9:98784802-98785110 | 308 | | | 2 | | | | | | | | | | | yes | | | | | |
| ZNF510 | Island | 1stExon;5'UTR | chr9:99540324-99540452 | 128 | | | | | | | | 0 | | | | | | yes | | | | | |
| KIAA1529 | N_Shore | TSS1500 | chr9:100000026-100000415 | 389 | | | | | | | | | | | | | | yes | yes | | | yes | |
| KIAA1529 | Island | Body | chr9:100070007-100070423 | 416 | | | 1 | | | | | | | | | | | | yes | | | | |
| TMOD1 | Island | 5'UTR | chr9:100264941-100265181 | 240 | | | 3 | | | | | | | | | | | | | | | | |
| ANP32B | Island | Body | chr9:100747473-100747767 | 294 | | | 2 | | | | | | | | | | | yes | | | | | yes |
| GALNT12 | N_Shore | TSS1500 | chr9:101568975-101569334 | 359 | | | | | | | | | | | | | | yes | yes | yes | yes | | |
| C9orf152 | NA | 5'UTR;1stExon | chr9:112970404-112970590 | 186 | | | | 1 | 1 | | | | | | | | | | | | | | yes |
| SVEP1 | Island | 1stExon;5'UTR | chr9:113341925-113342326 | 401 | | | | | | | | | 1 | | | | | | yes | | | yes | yes |
| SUSD1 | Island | TSS200 | chr9:114937740-114938015 | 275 | | | | | | | | | | | | | | yes | | | | | yes |
| ZFP37 | N_Shore;Island | Body;1stExon;TSS200 | chr9:115818495-115819364 | 869 | | 1 | | | 1 | | | 0 | 2 | | | | | | yes | yes | | | |
| BSPRY | N_Shore | TSS1500 | chr9:116111377-116111676 | 299 | | | 1 | | | | | | | | | | | yes | yes | | | | yes |
| KIF12 | N_Shore;Island | 5'UTR;1stExon | chr9:116860437-116861442 | 1005 | | 1 | | | | | | | | | | | | yes | | | | yes | yes |
| COL27A1 | NA | Body | chr9:117068159-117068532 | 373 | | | | | | | | | | | | | | | yes | yes | | | |
| ATP6V1G1 | N_Shore | TSS1500 | chr9:117349316-117349621 | 305 | | | | | | | | | | | | | | yes | yes | | | | |
| TNFSF8 | NA | 1stExon;5'UTR | chr9:117692745-117693010 | 265 | | | | | | | | | 1 | 1 | 1 | | | | | | | yes | yes |
| TNC | NA | 1stExon;5'UTR | chr9:117880330-117880556 | 226 | | | | | | | | | | 1 | | | | | | | | | |
| C9orf27 | NA | TSS200 | chr9:118687465-118687943 | 478 | | 1 | | | | | | | 1 | | | | | | yes | | | | |
| PAPPA | N_Shore | TSS1500 | chr9:118915832-118915907 | 75 | 2 | | | | | | | | | | | | | | | | | yes | yes |
| ASTN2 | S_Shore | TSS1500 | chr9:120177700-120178198 | 498 | | 1 | | | | | | | | | | | | | | | | yes | yes |
| TLR4 | NA | Body;1stExon;5'UTR | chr9:120466493-120466837 | 344 | | | 8 | 1 | | | | | | | | | | yes | | yes | | | |
| DBC1 | Island | 5'UTR;1stExon;TSS1500 | chr9:122131497-122132379 | 882 | | 1 | | | | | | | | | | 1 | | yes | | | | yes | |
| PSMD5;LOC253039 | Island | TSS200 | chr9:123605229-123605666 | 437 | | | | | | | | 0 | | | | | | yes | | | | | |
| TRAF1 | N_Shelf | 5'UTR | chr9:123688715-123689193 | 478 | | | | | | | | | | | | 1 | | yes | | | | | |
| STOM | Island | Body;TSS1500 | chr9:124132159-124133094 | 935 | | | | | | | | 0 | | 1 | | | | yes | | yes | | | |
| DAB2IP | Island | Body | chr9:124461998-124462182 | 184 | | | 2 | | | | | | | | | | | | | | | | yes |
| DAB2IP | Island | Body | chr9:124535348-124535611 | 263 | | | | 1 | | | | | | | | | | | | | | | yes |
| LHX6 | N_Shore | Body | chr9:124979184-124979356 | 172 | | | | 1 | 1 | | | | 1 | | | | | yes | yes | | | | |
| LHX6 | Island | Body | chr9:124988720-124990456 | 1736 | | | | | | | | 0 | 3 | 1 | 1 | 2 | | yes | | | | | yes |
| CRB2 | N_Shore;Island | Body | chr9:126135169-126135998 | 829 | | | 6 | | | 1 | | 0 | 2 | 1 | | 1 | | yes | yes | | | | yes |
| ANGPTL2;RALGPS1 | NA | 5'UTR;Body | chr9:129884026-129884457 | 431 | | | | | | | | 0 | 1 | | | 2 | | yes | | | | | |
| SH2D3C | Island | Body;5'UTR | chr9:130516695-130517095 | 400 | | | 1 | | | | | | | | | | | yes | | | | | yes |
| PIP5KL1 | N_Shore | Body | chr9:130689253-130689627 | 374 | | | 1 | | | | | | | | | | | yes | | | | | |
| CIZ1 | N_Shore | TSS1500;5'UTR | chr9:130955135-130955436 | 301 | | 3 | 4 | | 1 | | 1 | 0 | 1 | 1 | 1 | 35 | 1 | yes | | | | | |
| CIZ1;DNM1 | S_Shore | TSS1500;Body | chr9:130967820-130968072 | 252 | | | | | | | | | | | | | | yes | | | | | |
| DNM1 | Island | Body | chr9:131012953-131013323 | 370 | | | | | 1 | | | 0 | 1 | | | 3 | | | | | | | |
| MIR219-2 | Island | Body | chr9:131154899-131155013 | 114 | | | | | | | | 0 | 1 | 1 | | | | | | | | yes | |
| ZDHHC12 | Island | TSS1500 | chr9:131486742-131486769 | 27 | | | | | | | | | | | | | | yes | | | | | |
| CRAT;PPP2R4 | N_Shore | Body;TSS1500 | chr9:131872005-131872542 | 537 | | | | | | | | | 1 | | | | | | | | | | yes |
| USP20 | N_Shore | Body | chr9:132636402-132636787 | 385 | | | | | | | | | | | | | | | yes | | | | |
| FREQ | Island | Body | chr9:132935542-132935664 | 122 | | | | 1 | | | | | | | | | | | | | | yes | yes |
| QRFP | NA | 1stExon | chr9:133768931-133769184 | 253 | | | | | | | | | 1 | 1 | | | | | | | | | |
| NTNG2 | Island | 5'UTR | chr9:135039836-135040296 | 460 | | | 5 | | | | | | | | | | | | | | | | yes |
| C9orf171 | Island | Body | chr9:135361993-135362480 | 487 | | | | | | | | 0 | 1 | | | | | yes | | | | | |
| BARHL1 | Island | Body | chr9:135464814-135465435 | 621 | 1 | | | | | | | | | | | | | yes | | | | | |
| GTF3C5 | N_Shore | TSS1500 | chr9:135904966-135905092 | 126 | | | 5 | | | | | | | | | | | yes | yes | | | | |
| ABO | N_Shore | Body | chr9:136149908-136150361 | 453 | | 1 | | | | | 1 | 0 | 2 | | | 3 | | yes | | yes | | | yes |
| ADAMTS13;C9orf7 | N_Shore | Body;TSS1500;3'UTR | chr9:136324375-136324586 | 211 | | | | | | | | | | | | | | yes | | | | | |
| FAM163B | S_Shore | Body | chr9:136445244-136445595 | 351 | | | 1 | | | | | | | | | | | yes | yes | | | | |
| DBH | NA | Body | chr9:136522662-136522927 | 265 | | | 1 | 1 | | | | | | | | | | | yes | | | yes | |
| SARDH | N_Shore | Body | chr9:136566991-136567339 | 348 | | | 3 | | | | | | 1 | | | | | | | | yes | | |
| VAV2 | N_Shore | Body | chr9:136844397-136844768 | 371 | | | 1 | | | | | 0 | 1 | | | | | | | | | | |
| COL5A1 | N_Shore | Body | chr9:137718814-137719300 | 486 | | | | | | | | | 1 | 1 | | | | | | | | yes | |
| FCN2 | NA | TSS200 | chr9:137772544-137772580 | 36 | | | | | | | | | | | | | | | | | | | |
| KCNT1 | Island | Body | chr9:138606811-138607307 | 496 | | | | | | | | | 1 | | | | | | | | | | yes |
| KCNT1 | Island | Body | chr9:138660958-138661146 | 188 | | | | | | | | | | | | | | | | | | | |
| KCNT1 | Island | Body | chr9:138670609-138670672 | 63 | | | | | | | | | | | | | | yes | | | | | |
| KCNT1 | Island | Body | chr9:138672837-138673031 | 194 | | | | | | | | | | | | | | | | | | | |
| KCNT1 | N_Shore | Body | chr9:138679528-138679950 | 422 | | | | | | | | | | | | | | | | | | | |
| NACC2 | Island | 5'UTR | chr9:138985942-138986379 | 437 | | | | | | | | | 1 | | | | | | | | | | yes |
| LHX3 | Island | Body | chr9:139091378-139092974 | 1596 | | | 10 | | | | 1 | 0 | 5 | | | | | | | | | | yes |
| LHX3 | Island | Body | chr9:139094185-139094581 | 396 | | | 1 | | | | | | | | | | | | | | | yes | |
| LHX3 | Island;N_Shore | Body;TSS1500 | chr9:139096080-139097029 | 949 | | 1 | 3 | | | | | | | | | | | | | | | | yes |
| GPSM1 | N_Shore | Body | chr9:139228737-139228912 | 175 | | | | | | | | | | | | | | | | | | | |
| GPSM1 | N_Shore;Island | Body | chr9:139236663-139237889 | 1226 | 1 | | 25 | | | | | | | | | | | yes | yes | | | | |
| GPSM1 | Island | Body | chr9:139239853-139240119 | 266 | | | | | | | | | 1 | | | | | | | | | | |
| SEC16A | N_Shore | Body | chr9:139345474-139345761 | 287 | | | | | | | | | | | | | | | yes | | | | |
| EGFL7;MIR126 | Island | Body | chr9:139565087-139565154 | 67 | | | 4 | | | | | | | | | | | | | | | | |
| LCN10 | Island | Body | chr9:139634063-139634272 | 209 | | | 2 | | | | | | | | | | | | yes | | | | |
| LCN6 | S_Shore | 1stExon | chr9:139642875-139643351 | 476 | | | | | | | | | 1 | | | | | | | | | yes | |

| | | | | | | | | | | | | | | | | | | | | | | | |
|---|---|---|---|---|---|---|---|---|---|---|---|---|---|---|---|---|---|---|---|---|---|---|---|
| LCN15 | S_Shelf | 1stExon | chr9:139658896-139659112 | 216 | | 1 | | | | | | | | | | | | yes | | | | | |
| C9orf86 | N_Shore;Island | Body | chr9:139715548-139716215 | 667 | | | 1 | | | | | 0 | 2 | | | | | yes | yes | | | | |
| CLIC3 | S_Shore | Body | chr9:139890918-139891058 | 140 | | | | | | | | | | | | | | | | | | | |
| ABCA2 | N_Shore | Body | chr9:139913237-139913752 | 515 | | 1 | | | | | | | | | | | | yes | | | | yes | yes |
| ABCA2;C9orf139 | N_Shore | Body;TSS200 | chr9:139921813-139921971 | 158 | | | | | | | | | | | | | | | | | | | yes |
| ENTPD2 | Island;S_Shore | TSS1500 | chr9:139948927-139949577 | 650 | | | | 1 | | | | | | | | | | | | | | | yes |
| GRIN1 | Island | Body | chr9:140051205-140051729 | 524 | | | | | | | | 0 | | | | | | | | | | | |
| LRRC26 | S_Shore | TSS1500 | chr9:140065568-140065972 | 404 | | | 1 | | | | | | | | | | | | | | | | |
| RNF208 | N_Shore;Island | 1stExon;TSS200;TSS1500 | chr9:140115474-140117443 | 1969 | | | 3 | | | | | | 2 | | | | | | | | | yes | yes |
| C9orf167 | N_Shore;Island | TSS200;Body;3'UTR | chr9:140172089-140175679 | 3590 | | | | | 1 | | | | 1 | | | | | | | | | | |
| PNPLA7 | Island | Body | chr9:140356475-140356830 | 355 | 1 | | 2 | | 1 | | | | 1 | 1 | | | | | | | | | |
| PNPLA7 | Island | Body | chr9:140401382-140401576 | 194 | | | 28 | | | | | | | | | | | yes | | yes | | | |
| MRPL41 | Island | Body | chr9:140446656-140446993 | 337 | | | | | | | | 0 | 1 | | | 3 | 1 | yes | | | | yes | yes |
| EHMT1 | Island | Body | chr9:140683212-140683797 | 585 | | | | | 2 | | | | | | | | | yes | | | | | |
| CACNA1B | N_Shore;Island | TSS1500 | chr9:140771180-140771990 | 810 | | | 1 | | | | | | 1 | | | 1 | | | | yes | | yes | yes |
| CACNA1B | N_Shore | Body | chr9:140916980-140917431 | 451 | | | 1 | | 1 | | | 0 | 1 | 1 | | 3 | | | | | | | yes |

**Supplementary Table 2: Details of primer sequences used in the present study**

| Target gene | Experiment | Forward primer | Reverse primer | Annealing temparatue | Product size |
|---|---|---|---|---|---|
| β- *ACTIN* | q-RT PCR | GCTCGTCGTCGACAACGGCTC | CAAACATGATCTGGGTCATCTTCTC | $60^0C$ | 353 bp |
| *AURKC* | q-RT PCR | GGCTCATCTACAACACCCCAAT | TCCTCTATTATCGTGGCTGTGC | $60^0C$ | 162 bp |
| *NAPRT1* | q-RT PCR | GGCTCTGACGGGTCTCCACT | CACAGCTGTCCCTGCTGGAG | $60^0C$ | 170 bp |

| Target gene | Experiment | Outer forward | Outer reverse | Inner forward | Inner reverse | Annealing Temperature | Product size |
|---|---|---|---|---|---|---|---|
| *AURKC* | COBRA | GTGATTTTGGAGAGTGGTTTGAAT | CTAAACCCTCTAAATAACTATTCCC | GGAATTTTGGATTGAGGTTTT | AATATCTCACTACTACACAAACTTC | $50^0C$ | 194 bp |
| *NAPRT1* | COBRA | GGGATATGGGGGGTTTTA | AATCCAACTATTTCAACCCCT | GTTATAGGTGGGGGTTTTT | CTACTCACTAACCTCTACCAAA | $50^0C$ | 239 bp |

**Supplementary Table 03: Disgenet, pathway and Gene Ontology analyses of 8,037 genes containing MeVars in normal tissues.**

| DISGENET Analysis | Adjusted P-value | Genes |
|---|---|---|
| Schizophrenia | 6.983830927462064E-9 | CYFIP1;RPL3;TRRAP;MYT1L;CLEC10A;EHMT2;UXS1;STMN2;EHMT1;C4BPA;ADARB1;C4BPB;HTR4;ABCA13;CRHBP;MIR432;SYNGAP1;CYP2D6;DPYSL2;GJA5;AKT1;PIP4K2A;VPS39;PKNOX2;CHGB;ANKS1B;DAXX;UNC5C;PRKCA;PDZK1;PDYN;DKK1;DKK3;DKK4;BCAN;DAAM2;FEZ1;RBP1;PADI2;CYP2E1;ASTN2;ANKK1;ACOT6;HPD;NGEF;ABCB1;DHH;TERC;SHMT1;SHOX2;PDGFB;AGAP1;AGAP3;A1BG;PLA2G7;MIR206;C2;ADAMTS16;C3;HSD11B2;GRIN2A;CYP2B6;PCDHA1;TERT;CHST11;BLOC1S1;PNPO;HCRT;HLA-DQA2;ST3GAL1;HLA-DQA1;JAG2;HLA-DRB6;NGFR;CREBBP;GSTM2;HLA-DRB5;CADM3;GSTM1;ACE;NFATC3;PBX2;NFATC2;EIF2AK4;GRIN2B;NR0B2;GRIN2D;MIR346;EHD1;GRIN3B;CPEB1;SLC6A9;FXYD1;PNOC;FXYD6;PLCH2;HLA-DRB1;XKR6;CDK15;CDKN1C;ACHE;GABRB2;MIR130B;ADM;SLC6A1;MED15;SLC6A2;SLC6A3;PANX2;C1QTNF3;CHL1;ADAMTSL3;NEFL;NEFM;DLGAP1;DLGAP3;JARID2;DLGAP2;SLC18A2;TIMP4;BRD1;TPSG1;PLLP;ELOVL5;EN1;TACR3;PRSS16;TET1;PAX6;DUSP6;CIT;GJD2;ZEB2;TUBB2B;MMP16;TAL1;PLSCR4;RARA;RARB;ANGPTL4;TKT;HLA-DQB2;HLA-DQB1;GRAMD1B;GSTP1;ASIP;KCNA3;ADCY1;ASCL1;SNX31;ADCY7;FSTL1;EGFR;SNX29;HLA-DMB;NTF3;SLC17A7;STX2;APOL1;CSMD2;CSMD1;SLC17A3;GABRA1;HSPA1L;KCNB1;GABRA5;GABRA4;GNMT;FABP3;FABP5;FABP7;HLA-DPB1;HSPA1B;HSPA1A;CHRM2;CSF2;CHRM1;EPO;CTNND2;ZBTB20;RORA;MSI2;CLU;FGF2;TNF;IPO5;BEST1;RIMS2;HHAT;HPGDS;ATCAY;ADORA1;GRB10;PITX2;HLA-DOB;KCNH1;KCNH2;MAGI1;PPP1R16B;SLC38A1;MAGI2;F2R;TAP2;DIO3;TAP1;DGKZ;FOXP2;FOXP1;NPAS4;CACNB2;MTNR1A;MAG;MTNR1B;MSH5;ADNP;TNIK;S100A9;PVALB;MAD1L1;DLX1;MAGEL2;SLC26A1;CNP;CCDC25;NOTCH4;GPR85;GSTT1;FHIT;NPAS3;SOCS2;TPO;SBNO2;ATXN1;MOBP;MBP;SRGAP3;NPTX1;SLC38A2;KIAA0319;DRD4;GRIA4;DRD5;TMEM132D;TRAPPC3;SRD5A2;CD160;HTR1D;HTR1A;HPS4;HTR1B;DCDC2;FOXN3;BTBD9;TMPRSS13;GFAP;BCL2;SLC26A8;LHX6;ZNF536;TSNAX-DISC1;ESAM;SHBG;CNTNAP2;ST6GALNAC1;PPP1R13B;DAGLA;WISP3;PIK3CD;ADRA1D;HTR2A;TREM1;PIK3CG;SYNE1;DNMT3L;SGCD;PSD3;DNMT3B;DGCR6;EMX2;HSP90AA1;GPX1;GRID1;HTR3E;APOA1;NRG1;HTR3A;HTR3B;KSR2;TNFRSF1B;ADRA2C;NGF;TNFRSF1A;POM121L2;ADORA2A;LCK;DNAJC5;NOS1AP;ITGA8;CRH;LANCL2;CRP;HDAC4;CX3CR1;ITIH4;HDAC2;ITIH3;NLGN1;HDAC3;LRP1;CCDC68;LRP4;FURIN;ITIH1;SELENBP1;FBXL21;CACNG8;P2RY1;FYN;UGT1A3;CACNG2;CACNG5;NTRK1;TBX1;SLC12A5;KCNIP1;ASCC1;TXNRD2;NTRK3;PCDH8;CABLES1;HTR5A;SORL1;TBX4;EFR3B;MYO16;MTOR;MYO1D;P2RX7;P2RX5;KLF6;MARCKS;NFASC;USH1G;APC;SST;P2RX2;IL2RA;LEP;P2RX1;PTPN6;TRIM31;MDGA1;APP;FAAH;OXTR;VIPR2;RFESD;DIXDC1;ICAM3;MLC1;AQP4;ICAM1;RASSF1;EDNRA;TRIM26;SYNPO;SEPHS1;SLC12A6;SOX5;HLA-J;SRRM2;HLA-B;SLC6A13;HLA-C;KRT7;SLC6A12;SLC6A11;CSNK1E;MYO7A;HLA-A;MIF;MAPK8IP3;HLA-E;UHMK1;RUNX1;KCNQ2;KCTD12;TAGLN3;CTXN3;ATP6V0D1;TRIM10;ALDH7A1;STX1A;NRN1;GIPR;SLC45A1;TRANK1;WNK4;KALRN;KLK8;CRHR2;SYNGR1;INPP5A;TSPAN8;MAP2;APOH;CMYA5;ST8SIA4;MAP7;APOD;APOE;KCNN3;PCK1;ZBED4;NTSR1;HIST1H3E;NTNG1;EGR1;NTNG2;EGR3;SPHKAP;BCL11B;ST8SIA2;IGF2;LIF;DCLK1;MSRA;TF;TH;STAR;PTK7;PPP1R1B;CCDC88C;PEMT;GRB2;IGSF9B;VSX1;DDR1;CD40;MIR17;GRIK5;GRIK3;GRIK4;GRIK1;GRIK2;BRCA1;KIF17;PTPRF;CCKAR;SH3PXD2A;SLC22A16;STAB1;PSD;SLC32A1;TUBB;MST1;INSRR;GFRA1;SYN3;GFRA3;PARP9;SYN2;CLDN11;TLR1;ADRB3;IL1B;ARHGEF2;ARHGEF7;TLR4;VGLL4;SDCCAG8;C1QB;C1QA;TNXB;TMPRSS5;SATB2;TRAK1;NOL4;CAMKK2;PURA;KIAA0513;ALS2CL;NPY;PAK6;MGAT1;CTNNA3;CTNNA2;PCNT;DISC1;GABRD;MICB;MPP2;NQO1;GABBR1;MOG;SORT1;GAD1;PLEKHA6;LILRB1;SORBS1;VEGFA;TPPP3;RIT2;PER3;KCNS3;QPCT;DLC1;CACHD1;B9D1;GHRL;TRIP13;APBA2;IL17C;NEUROG1;PNPLA2;PLAT;CELSR1;COMT;ACTG1;DAOA;RGS4;GRIP1;RGS5;CCND2;CCND1;TGM5;TGM2;NANOS1;PDGFRB;SLC30A8;KDM2B;LMO3;LIG4;TSC2;LMO7;CARTPT;SHISA9;HCG9;ERMN;DHDDS;ALDH3A1;TMX2;LMX1A;ADAM12;TSNARE1;RTN3;CHRNA2;CHRNA5;NTM;CHAT;CACNA1B;SLC1A2;SLC1A3;ARHGAP18;CACNA1D;ALOX12;SLC1A4;CSF2RB;CACNA1C;SLC1A5;SLC1A6;CPLX1;NALCN;ACACA;MCHR2;NKAPL;PSORS1C1;CACNA1I;RAI1;HRH1;HRH2;ALDH3B1;CDH23;OTX2;PRODH;TAC1;NDUFV1;WNT2;MARK2;TFAP2A;SMAD3;AUTS2;HMGA1;TBCB;INHBA;ESR1;LRP1B;PHOX2B;NR4A2;IL6;GNAL;DAO;DLG2;GDNF;BHLHE40;GNAS;GNB3;CNTN2;DEGS2;TGFBI;FGFR2;FGFR1;SNED1;MEGF10;OXT;LAMC2;ARHGEF10L;DBH;NR2E1;PPP1R9A;HAPLN4;GRM1;LMNB2;HK1;GHR;GRM2;GRM5;CYP26B1;GRM4;GRM7;ZIC2;NETO1;CA2;LRRTM1;GRM8;CASP1;PDE4B;FLOT1;PDE4A;NOS1;LRRFIP1;YWHAH;MIR193A;CAMK1D;PLA2G4D;DCC;PLA2G4B;PDE4D;NAV1;OLIG2;NAV2;OPRM1;UPK3B;DNM1;SSTR4;TAPBP;SSTR5;TGFBR2;CYP26C1;NDUFS7;IFNG;NDUFS6;ALDH1A2;CHDH;LTA;CHI3L1;MYH9;SCN4A;HAGH;MCTP2;FBP1;THOP1;SHANK2;RBKS;SHANK1;RBMS3;CAMK2B;DDC;LAMA1;NRXN1;PON1;NRXN3;NRXN2;SPG7;THBS1;DDO;CRYZ;ACAN;CYTH4;CNR2;CBS;GNG7;CLVS2;SLIT3;NCAM1;ATXN8OS;SNCB;OPCML;CHGA;TCF7L2;TTC12;NOS2;NOS3;GALNT2;B3GAT1;ZNF804A;ATP2B4;OPRK1;CCK;ATP2B2;DLK1;POMC;MIR144;PPP2R2B;CAPN10;RGS10;CD9;RGS12;VIM;SCLY;HCN1 |
| Unipolar Depression/Major deprssive Disorder | 2.0625111211996978E-8 | APP;FAAH;OXTR;TRIO;MYT1L;RFESD;STMN2;DIXDC1;ADARB1;HTR4;CRHBP;CYP2D6;ANPEP;PROK2;TRIM26;SOX9;PRIMA1;PRKCB;SLC6A15;SLC6A13;MIF;PDYN;UNC13D;ANKK1;ENPEP;ABCB1;TERC;KALRN;CRHR1;CRHR2;ADAMTS16;TERT;MAP2;CMYA5;APOE;HCRT;PROM1;NTSR1;S100A10;LHPP;NGFR;ACE;CADM1;DCLK2;GRIN2B;PARP15;STAR;TH;ARTN;PPP1R1B;FAT1;SSPN;CC2D1A;PTPRR;ARHGAP8;GRIK4;SLC6A2;SLC6A3;CCKAR;UCN3;SLC22A16;NEFL;IKBKE;CD34;SLC18A2;DUSP1;TESC;ANO8;MRPL48;TYK2;CIT;TAL1;IL1B;HCLS1;SGK1;HTT;ADCY7;EGFR;REM1;CUX1;NPY;NTF3;APBB2;PCNT;DISC1;CSMD1;CCL24;GABBR1;HSPA1L;PKLR;GAD1;VEGFA;PER1;PER3;GHRL;SIGLEC7;HSPA1B;HSPA1A;SPSB4;CHRM2;CSF3;CSF2;CTNND2;ZBTB20;DEFB1;RORA;COMT;FGF2;TNF;MPO;RGS4;DAOA;CCND1;DSCAM;F2R;FOXP2;CACNB2;PDE9A;PFKFB3;CNP;NTM;SLC1A2;SLC1A3;CSF2RB;CACNA1C;SLC1A6;FHIT;NPAS3;AANAT;PRR5-ARHGAP8;DEAF1;TPPP;MORC1;NDUFV1;DRD4;DRD5;GRIA4;KL;CBX5;TFAP2D;AUTS2;CNTN5;ZBTB16;RARRES2;HTR1A;HTR1B;BMP7;ESR1;GFAP;ISG20;NR4A2;IL6;DLG2;GDNF;BCL2;GNB3;TSNAX-DISC1;FGFR1;DPP10;CNTNAP2;UCN;SNED1;ADAR;LAMC2;OXT;HTR2A;TRH;DBH;GRM1;SYNE1;GRM5;GRM7;GRM8;BDKRB2;PDE4B;PDE4A;NOS1;TRPC7;HSP90AA1;DST;DCC;NRG1;HTR3A;HTR3B;KSR2;OPRM1;NGF;TNFRSF1B;C1QL1;SSTR5;ADORA2A;IFNG;OAS2;CAT;CRH;IRF7;LTA;MCTP2;THOP1;CRP;ITIH3;DDC;HLF;LRP1;PON1;ITIH1;INS;CNR2;NCAM1;SLIT3;MYO10;NOS3;NTRK3;ZNF804A;HTR5A;MUC5B;MTOR;P2RX7;POMC;NELL1;TEF;SST;IL2RA;IL2RB;CD9;KCNK2;MDGA1 |
| Bipolar Disorder | 2.0625111211996978E-8 | CPNE8;APP;FAAH;VIPR2;UXS1;RFESD;CLEC10A;DIXDC1;MLC1;ADARB1;HTR4;ABCA13;CRHBP;EDNRA;DPYSL2;AKT1;PIP4K2A;PROK2;TRIM26;SLC12A6;PRIMA1;DAXX;SLC6A13;HLA-C;CSNK1E;FNDC3A;PDYN;ISL1;HLA-G;MAPK8IP3;HLA-E;UHMK1;DKK4;SFRP1;ADCY9;RFX4;FEZ1;KCNQ2;KCTD12;ASTN2;ANKK1;ABCB1;NRN1;DHH;HPN;SHOX2;TRANK1;PRKCZ;MIR206;KLK8;CRHR1;CRHR2;ADAMTS16;TRPM2;GRIN2A;SYNGR1;TSPAN8;PCDHA1;MAP2;APOD;APOE;KCNN3;APOB;PROM1;ST3GAL1;S100A10;NTNG1;EGR1;NTNG2;CREBBP;GSTM1;EGR3;ACE;ST8SIA2;NR1D1;GRIN2B;DCLK1;MIR346;TH;STAR;PPP1R1B;FAT1;HLA-DRB1;GABRB2;CD40;ARHGAP8;GRIK5;GRIK3;GRIK4;GRIK1;ADM;GRIK2;SLC6A2;SLC6A3;AKAP10;CCKAR;CSRP1;SLC22A16;CHL1;CHMP1B;STAB1;NEFL;NEFM;ACADS;SLC18A2;PSD;BRD1;AVPR1B;MST1;SYN3;SYN2;SORCS2;DUSP6;CIT;NCOR2;TLR1;IL1B;MAPT;ARHGEF7;GPRC5C;TLR4;SDCCAG8;GRAMD1B;NFIX;TMPRSS5;KCNA3;ADCY3;ADCY2;ADCY1;EGFR;RASIP1;CAMKK2;EXOSC6;CUX2;NTF3;SLC17A7;STX2;CSMD2;PCNT;DISC1;CSMD1;GABRA1;GABRA5;RNLS;GAD1;GABRA4;NFYC;VEGFA;PER3;FABP5;FABP7;IL17C;CHRM2;CSF2;CTNND2;DEFB1;RORA;COMT;TNF;MIR149;RGS4;DAOA;HPGDS;TNR;NRTN;MACROD2;TGM2;MAGI1;DSCAM;LMO3;MAGI2;DIO2;TAP2;LMO7;CARTPT;HCG9;GPRIN2;CACNB2;LMX1A;DOK5;SCN8A;PVALB;MAD1L1;PDE9A;ABCG1;DLX1;CALCA;PFKFB3;CHRNA2;DLX4;NOTCH4;CHRNA6;CACNA1B;SLC1A2;ITPR1;CACNA1A;SLC1A3;GSTT1;CACNA1D;SLC1A4;ALOX12;CACNA1C;RASAL2;ADRB2;SLC1A6;CPLX1;MCHR2;NPAS3;TPO;PRR5-ARHGAP8;ALOX12P2;FGF20;OTX2;NPTX1;PRODH;TAC1;NDUFV1;DRD4;GRIA4;DRD5;HTR1E;CNTN6;RARRES2;HTR1A;HTR1B;TMPRSS13;ESR1;OPN5;GFAP;FGF17;NR4A2;IL6;PLCXD3;GNAL;DAO;DLG2;NBEA;GDNF;BHLHE40;BCL2;GNB3;SP8;TSNAX-DISC1;FGFR2;FGFR1;DPP10;CNTNAP2;SNED1;ST6GALNAC1;DOCK9;GADL1;LAMC2;ARHGEF10L;HTR2A;DBH;NR2E1;PPP1R9A;SYNE1;GRM5;GRM4;CASP8;GRM7;CA2;CA5A;BDKRB2;PDE4B;PDE4A;NOS1;YWHAH;TRPC7;ARHGEF10;EMX2;TRPC3;GRID1;NRG1;HTR3A;HTR3B;KSR2;ADRA2C;SSTR4;SSTR5;IMPACT;IFNG;EHHADH;CAT;NOS1AP;CRH;BPI;THOP1;RBMS3;MCPH1;CRP;ITIH4;ITIH3;NLGN1;DDC;NRXN1;PON1;NOD2;ITIH1;ADD3;INS;SELENBP1;CYTH4;CNR2;CBS;P2RY1;ERBB2;SLIT1;PTK2B;FYN;NCAM1;TRIM42;CACNG5;NTRK1;TBX1;TCF7L2;NOS2;ASCC1;NOS3;TXNRD1;ZNF804A;CABLES1;CCK;HTR5A;SORL1;P2RX7;POMC;KLF6;MARCKS;P2RX5;PPP2R2C;MIR144;SST;P2RX2;LEP;P2RX1;IL2RB;TLL2;MDGA1 |
| Mental Depression | 3.547376925029927E-8 | APP;FAAH;OXTR;VIPR2;DIXDC1;ABAT;AQP4;HTR4;ABCA13;SOX2;CRHBP;CYP2D6;DPYSL2;STMN1;AKT1;SLC12A7;CMKLR1;SRRM2;PRKCB;SERPINF1;SLC6A15;KRT7;PDYN;ANK1;DKK4;RNF123;SFRP1;SERPING1;CYP2E1;ANKK1;NEIL1;ALPK1;ALDH7A1;ABCB1;DHH;PDE1B;KLK8;CD1A;CRHR1;CRHR2;HSD11B1;GRIN2A;CYP2B6;GALR3;MIR330;PENK;APOE;HCRT;A2M;S100A10;NGFR;ACE;GSTM1;EGR3;FZD5;FZD6;SLC33A1;LIF;NR1D1;GRIN2B;GRIN2D;PEX5L;STAR;TH;ARTN;PNOC;PPP1R1B;FAT1;PIGR;GPR26;GRIK3;GRIK4;BRCA1;SERPINA6;SLC6A1;KRT20;SLC6A2;SLC6A3;CHL1;NEFL;SOSTDC1;NEFM;SLC18A2;AVPR1B;DUSP1;INSRR;TACR3;DUSP6;CIT;TAL1;IL1B;RARA;TFF3;TLR4;RAPGEF3;RAPGEF4;KCNA3;HTT;ADCY1;ADCY7;EGFR;REM1;EXOSC6;PRSS50;MTHFD1L;NPY;NTF3;CSMD2;PCNT;DISC1;NQO1;SORT1;GAD1;STAT3;VEGFA;PER1;PER3;FABP7;FOSB;F2RL3;PNPLA2;CHRM2;CSF2;CRTC1;RORA;COMT;CLU;FGF2;TNF;MPO;ETS2;DAOA;NRTN;STK32C;LMO3;CARTPT;ALDH3A1;HGS;PVALB;PFKFB3;CHRNA2;CNP;CHAT;CHRNA6;SLC1A2;GSTT1;ADRB2;CACNA1C;CPLX1;DPP4;RAI1;AANAT;DEAF1;FGF20;NXPH1;TAC1;DRD4;TFAP2B;HTR1A;HTR1B;TLX1NB;ESR1;OPN5;GFAP;FGF17;PYY;IL6;GDNF;ALPL;GNB3;SHBG;FGFR2;FGFR1;UCN;LAMC2;OXT;HTR2A;TRH;DBH;GRM1;SYNE1;SGCE;ADAMTS2;GRM5;GRM7;CASP1;PDE4B;PDE4A;NOS1;ARHGEF10;HSP90AA1;GPX1;DST;GRID1;PDE4D;NRG1;HTR3A;HTR3B;OPRM1;ADRA2C;NGF;TNFRSF1B;TNFRSF1A;C1QL1;TGFBR3;IFNG;OAS2;NOS1AP;CAT;CRH;CDH13;BPI;RBMS3;MCPH1;CRP;CX3CR1;HDAC4;HDAC2;NRXN1;PON1;MUC1;CNR2;CBS;SNCG;TNNI3;NCAM1;SNCA;NTRK1;LGI1;KCNJ6;EYA1;NOS2;NOS3;NTRK3;TXNRD1;OPRK1;P2RX7;POMC;NELL1;MARCKS;SUCLA2;USH1G;LEP;SCLY;KCNK2;HCN1 |
| Autistic Disorder | 2.886425627918438E-7 | CYFIP1;APP;FAAH;OXTR;SCT;DIXDC1;EHMT1;ABAT;STMN4;AQP4;HTR4;SYNGAP1;CYP2D6;DPYSL2;GJA5;RASSF5;PART1;AKT1;HOXA1;SLC12A6;SRRM2;MEF2C;PRKCB;HNF1B;HLA-A;MIF;PASK;UPB1;HLA-G;SND1;KATNAL2;HOXB1;ASTN2;HPD;STX1A;SLC22A3;ENPEP;LXN;PCDH10;HEPACAM;FOXG1;AGAP1;THY1;ABO;GRIN2A;PTDSS1;FXR2;ZNF827;MAP2;PRDM16;APOE;NTNG1;NTNG2;CREBBP;ACE;GSTM1;CADM1;ST8SIA2;POU6F2;IGF2;GRIN2B;EHD1;EXT1;BST1;FAM135B;TF;TG;GTF2IRD1;HLA-DRB1;VSX1;GSTM5;GABRB2;RYR2;ROBO3;NRP2;ROBO4;ELN;GRIK4;ADM;GRIK2;PCSK6;SLC6A3;RYR3;CHL1;CD38;DLGAP2;CTSD;PSD;ACAD8;GPR37;KCND2;TESC;PAX3;AMPD1;PAX6;SORCS1;SYN3;SYN2;SLC7A5;ALDH5A1;KCNMA1;ARHGEF2;RAPGEF4;GSTP1;NR1I2;PRICKLE2;HTT;HOXD11;NT5E;CUX1;NSD1;NTF3;CSMD3;DISC1;GABRD;XDH;GABRA1;GABBR1;MBD6;GABRA5;GABRA4;GAD1;VEGFA;SKI;PER1;GATM;WT1;F2RL1;APBA2;SEMA5A;CSF2;CTNND2;HDLBP;ZBTB20;RORA;COMT;CDH9;TNF;CDH8;RIMS1;GRIP1;MACROD2;PITX1;SNRPN;DSCAM;IGFBP3;TSC2;TBR1;FOXP2;FOXP1;CACNB2;CLIP2;SCN8A;ERG;ADNP;PVALB;DLX1;DLX2;MAGEL2;CALCA;ANKRD11;SLC1A2;CACNA1A;SLC1A3;CACNA1D;ADRB2;CACNA1C;SEZ6L2;ACACA;CACNA1H;MCHR2;CACNA1G;DPP4;FBXO40;RAI1;SNTG2;DEAF1;MBP;OTX1;PRODH;WNT2;DRD4;AUTS2;ZBTB16;HTR1D;HTR1B;MCC;BMP4;SDK1;DAB2;FER;IL6;DAO;NBEA;REEP3;BCL2;TUBGCP5;CNTNAP2;CNTNAP1;WISP3;SLC40A1;PIK3CD;LAMC2;OXT;HTR2A;DBH;PIK3CG;SYNE2;GHR;FLCN;GRM5;NETO1;GRM8;PDE4B;PDE4A;DGCR6;NOS1;GPX1;TRPC6;ITGA4;MIR23A;PDE4D;DNMT3A;NRG1;HTR3A;KCNAB2;OPRM1;TNFRSF1B;ARID1B;SSTR5;ALDH1A3;ADORA2A;IFNG;CDH10;CAT;LTA;CDH13;ITGA6;CNTNAP5;SHANK2;RBMS3;SHANK1;MCPH1;NLGN1;DDC;SRC;LAMA1;NRXN1;PON1;NRXN3;NRXN2;ATP10A;JAKMIP1;TSPAN32;ACAN;CNR2;SH3TC2;CBS;NCAM1;EIF4E;KCNJ1;TBX1;RPH3AL;NOS2;TXNRD1;LIMK1;ZNF804A;ATP2B2;HTR5A;MTOR;MYO1D;POMC;NELL1;PC;MLPH;APC;SST;LEP |
| Major Depressive Disorder | 3.9444713867433355E-7 | APP;FAAH;OXTR;VIPR2;MYT1L;ADARB1;HTR4;CRHBP;EDNRA;CYP2D6;ANPEP;STMN1;PROK2;SOX9;PRIMA1;PRKCB;SLC6A15;SLC6A13;PDYN;ANKK1;ALDH7A1;ENPEP;ABCB1;TERC;DHH;KALRN;CRHR1;CRHR2;GRIN2A;TERT;GALR3;MAP2;GALR1;CMYA5;APOE;HCRT;PROM1;NTSR1;S100A10;LHPP;NGFR;EGR1;ACE;GRIN2B;PARP15;STAR;TH;ARTN;PPP1R1B;CC2D1A;ACHE;PTPRR;GRIK3;GRIK4;SLC6A2;SLC6A3;CCKAR;UCN3;SLC22A16;NEFL;SOSTDC1;IKBKE;CD34;SLC18A2;ZHX2;DUSP1;ELOVL5;TESC;TYK2;CIT;ADRB3;TAL1;IL1B;HCLS1;SGK1;ADCY3;HTT;ADCY7;EGFR;REM1;EXOSC6;CUX1;NPY;NTF3;SLC17A7;PCNT;DISC1;XDH;CCL24;NQO1;GABBR1;HSPA1L;MOG;GAD1;VEGFA;FABP2;PER1;PER3;FOSB;SIGLEC7;HSPA1B;HSPA1A;CHRM2;CSF2;CRTC1;CTNND2;ZBTB20;COMT;FGF2;TNF;MPO;RGS4;DAOA;CCND1;F2R;FOXP2;MAG;PVALB;PDE9A;PFKFB3;CNP;NTM;SLC1A2;SLC1A3;CSF2RB;CACNA1C;SLC1A6;NPAS3;AANAT;GRK5;DEAF1;TPPP;TAC1;MORC1;NDUFV1;DRD4;GRIA4;KL;CBX5;ZBTB16;HTR1A;HTR1B;BMP7;ESR1;GFAP;ISG20;NR4A2;IL6;DLG2;GDNF;BCL2;GNB3;TSNAX-DISC1;FGFR2;FGFR1;CNTNAP2;UCN;ADAR;LAMC2;OXT;HTR2A;TRH;DBH;GRM2;GRM5;GRM7;GRM8;DNMT3B;BDKRB2;PDE4B;PDE4A;NOS1;TRPC7;KLF11;HSP90AA1;GPX1;DST;NRG1;HTR3A;HTR3B;OPRM1;ADRA2C;NGF;TNFRSF1B;SSTR4;C1QL1;SSTR5;IFNG;OAS2;CAT;CRH;IRF7;LTA;MCTP2;THOP1;CRP;DDC;HLF;LRP1;PON1;ITIH1;NCAM1;SLIT3;NOS3;NTRK3;ZNF804A;MTOR;P2RX7;GALC;POMC;NELL1;TEF;SST;IL2RA;LEP;IL2RB;CD9;KCNK2;MDGA1 |
| Depressive disorder | 1.5764967081174494E-6 | APP;FAAH;OXTR;VIPR2;DIXDC1;AQP4;HTR4;ABCA13;HS6ST1;SOX2;CRHBP;CYP2D6;DPYSL2;STMN1;AKT1;PROK2;SLC12A7;CMKLR1;SRRM2;EPHA4;UNC13A;PRKCB;SERPINF1;SLC6A15;KRT7;MIF;PDYN;ANK1;DKK4;RNF123;SFRP1;SERPING1;CYP2E1;ANKK1;NEIL1;KCTD17;ALPK1;ALDH7A1;ABCB1;NRN1;DHH;PDE1B;PDGFB;GATA3;KLK8;CD1A;CRHR1;CRHR2;HSD11B1;GRIN2A;CYP2B6;GALR3;MIR330;PENK;APOE;HCRT;A2M;S100A10;NGFR;ACE;GSTM1;EGR3;FZD5;CADM1;FZD6;SLC33A1;LIF;NR1D1;GRIN2B;GRIN2D;PEX5L;STAR;TH;ARTN;PNOC;PPP1R1B;FAT1;GTF2IRD1;PIGR;GPR26;ELN;GRIK3;GRIK4;BRCA1;SERPINA6;SLC6A1;KRT20;SLC6A2;SLC6A3;JPH3;CHL1;SOSTDC1;NEFM;NEFH;SLC18A2;AVPR1B;DUSP1;INSRR;TACR3;DUSP6;CIT;TAL1;IL1B;RARA;TFF3;MAPT;TLR4;RAPGEF3;RAPGEF4;TNXB;SLC20A2;IDUA;KCNA3;HTT;ADCY1;ADCY7;EGFR;REM1;EXOSC6;PRSS50;MTHFD1L;EPCAM;NPY;NTF3;CSMD2;PCNT;DISC1;SPDEF;SORT1;GAD1;STAT3;MLH1;VEGFA;PER1;PER3;FABP7;FOSB;GHRL;F2RL3;PNPLA2;CHRM2;CSF3;CSF2;CRTC1;RORA;COMT;CLU;FGF2;TNF;MPO;ETS2;DAOA;FGF8;GRAP2;RNF19A;NRTN;STK32C;PDGFRB;CASR;LMO3;CARTPT;CLIP2;ALDH3A1;HGS;SQSTM1;PVALB;CSF1R;NOTCH3;CHRNA2;CNP;CHAT;CHRNA6;SLC1A2;GSTT1;ADRB2;CACNA1C;CPLX1;CACNA1G;DPP4;RAI1;AANAT;DEAF1;FGF20;CDH23;TAC3;NXPH1;TAC1;DRD4;DRD5;PRPH;TFAP2B;HTR1D;HTR1A;HTR1B;TLX1NB;ESR1;OPN5;GFAP;NR4A2;FGF17;PYY;IL6;DAO;GDNF;BCL2;GNAS;ALPL;GNB3;SHBG;FGFR2;FGFR1;UCN;PTPN22;LAMC2;OXT;HTR2A;TRH;DBH;GRM1;SYNE1;SGCE;ADAMTS2;GRM5;GRM7;CASP1;PDE4B;PDE4A;NOS1;SGCG;KLF11;ARHGEF10;HSP90AA1;GPX1;DST;GRID1;PDE4D;NRG1;HTR3A;HTR3B;OPRM1;ADRA2C;NGF;TNFRSF1B;COASY;TNFRSF1A;C1QL1;TGFBR2;TGFBR3;ADORA2A;IFNG;OAS2;NOS1AP;CAT;CRH;CDH13;ANG;BPI;RBMS3;MCPH1;CRP;CX3CR1;PON3;HDAC2;DCTN1;NRXN1;PON1;INS;MUC1;CNR2;CBS;GNA11;TNNI3;NCAM1;SNCA;NTRK1;LGI1;KCNJ6;EYA1;NOS2;NOS3;NTRK3;LIMK1;OPRK1;P2RX7;KISS1;POMC;NELL1;MARCKS;SUCLA2;EIF3K;USH1G;PPP2R2B;LEP;SCLY;KCNK2;HCN1 |
| Autism Spectrum Disorders | 1.5764967081174494E-6 | CYFIP1;FAAH;OXTR;TRIO;EHMT2;EHMT1;EDNRA;SYNGAP1;CYP2D6;CYP1B1;HOXA1;DIP2A;TRIM26;SOX9;SOX5;SRRM2;MEF2C;SRRM4;PRKCB;CIRBP;HLA-B;HNF1B;PDZK1;HLA-A;MIF;SND1;DDIT3;HOXB1;ASTN2;TRIM10;NGEF;STX1A;FOXG1;TWIST1;C2;GRIN2A;HLA-DQA2;RREB1;NTSR1;HLA-DRB6;NGFR;HLA-DRB5;CADM1;FOXF1;ST8SIA2;PBX2;GRIN2B;ARTN;HLA-DRB1;DDR1;RYR2;ICA1;GRIK4;SLC6A2;SLC6A3;HEY1;CD38;SOSTDC1;JARID2;FEZF2;CC2D2A;PSD;SLC2A14;GPR37;TESC;TET1;PAX6;SYN2;MMP16;IL1B;HLA-DQB1;TNXB;GRP;GSTP1;SATB2;PRICKLE2;HLA-DMB;CUX1;MSX1;CSMD1;MOG;GABRA5;GAD1;ELP4;MIR656;FABP3;RIT2;FABP5;WT1;FABP7;APBA2;SEMA5A;CARS;CSF2;CTNND2;ZBTB20;RORA;IRS2;TNF;CDH8;CMIP;COMP;ZNF569;MACROD2;PITX2;KIRREL3;NANOS1;USP7;DIO2;DIO3;LIG4;TSC2;TBR1;FOXP2;FOXP1;CACNB2;MTNR1A;TMX2;MTNR1B;COL4A1;MSH5;ZNF436;ADNP;TSNARE1;MAD1L1;DLX1;DLX2;MAGEL2;CHRNA5;NOTCH4;ANKRD11;ITPR1;CACNA1A;CACNA1D;CACNA1C;SEZ6L2;FHIT;CACNA1H;PSORS1C1;CACNA1G;DPP4;CACNA1I;RAI1;DPP6;SNTG2;AANAT;CAMTA1;OTX1;NXPH1;ZNF423;KIAA0319;NDUFV1;WNT2;DRD4;DRD5;CNTN5;CNTN6;HTR1B;ARHGAP24;GFAP;FBXO31;NBEA;FGFR1;DPP10;CNTNAP2;PPP1R13B;PIK3CD;LAMC2;OXT;HTR2A;PIK3CG;SYNE1;UNC80;GRM5;GRM7;LRRTM3;GRM8;DNMT3B;CASP1;DGCR6;NOS1;TRPC6;DNMT3A;AMT;OPRM1;NGF;POMGNT1;ARID1B;DNM1;SSTR4;ALDH1A3;IMPACT;ADORA2A;IFNG;CDH10;TMEM216;CHDH;CDH11;LTA;CDH13;HAGH;MCTP2;CNTNAP5;SHANK2;RBMS3;SHANK1;MCPH1;CX3CR1;ITIH3;NLGN1;NRXN1;PON1;NRXN3;NRXN2;FURIN;ATP10A;JAKMIP1;INS;ACAN;EIF4E;TBX1;NOS2;TXNRD1;ZNF804A;ATP2B2;BAIAP2;MEIS2;SOD3;MTOR;MYO16;MAPK10;ACTC1;APC;P4HB;TRIM31;CBLN2;HCN1 |
| Alcoholic Intoxication, Chronic | 2.839904373404246E-6 | CHRM2;SEMA5A;CARS;SLC46A1;THSD7B;ABAT;COMT;CDH9;TNF;CDH8;RGS4;CRHBP;CDH5;EDNRB;DPYSL2;GRAP2;RNF19A;CDK20;PITX2;GGT1;PKNOX2;GHSR;MBNL2;LMO3;HLA-B;CARTPT;PDYN;GABRG3;HGS;RFX4;CYP2E1;TAGLN3;ANKK1;CFTR;CALCA;CHRNA5;PCDH10;SLC1A2;SLC1A3;GATA4;GSTT1;PIK3R1;ABO;KIAA0040;CRHR1;ADH6;IGSF22;HSD11B2;NKAIN1;ALDH3B2;GRIN2A;CYP2B6;GALR3;MOBP;GRK5;GALR1;PENK;MBP;CYTL1;APOE;KCNN3;TAC1;DRD4;NTSR1;LHPP;TFAP2B;ACE;GSTM1;HTR1E;SRD5A2;OSBPL5;CNTN6;EPHX1;HTR1A;HTR1B;GRIN2B;ST18;GFAP;EXT1;NR4A2;SLC6A6;BHMT;TF;IL6;SLC6A9;TH;NAT1;PNOC;PPP1R1B;HLA-DRA;GNB3;SHBG;FGFR4;HAMP;GABRB2;CNTNAP2;CD40;GRIK3;PIK3CD;GRIK1;OXT;HTR2A;TRH;DBH;SLC6A1;SLC6A2;SLC6A3;PIK3CG;GRM1;GLI2;PRSS21;ZCCHC14;SGCE;GRM2;GRM5;CCKAR;GRM7;SLC22A18;GRM8;PDE4B;NOS1;SLC18A2;KLF11;DST;VWF;SGIP1;TACR3;HTR3A;HTR3B;OPRM1;NGF;PARP9;PTP4A1;CCKBR;IL1B;CDH10;CAT;CDH11;CRH;TFF3;CDH12;CDH13;CDH15;SLC29A1;TKT;ARHGEF7;DSCAML1;TLR4;CDH18;KANK1;HDAC2;NRXN1;OPRL1;NRXN3;HTT;ADCY1;ADCY7;EGFR;CYP2A13;CNR2;NPY;PTK2B;FYN;NCAM1;CTNNA2;SNCA;GABRA1;KCNJ6;GABBR1;TTC12;MOG;GABRA5;ASCC1;NOS3;GAD1;OPRK1;CCK;POMC;FABP2;PER1;PER3;LEP;ALLC |
| Adenoma | 7.222872243585408E-6 | RB1;SERPINE2;NUDT1;HTR4;MIR425;LGALS4;SOX2;LGALS3;RASSF1;EDNRA;RASSF2;EDNRB;SOX17;CREB3L3;CYP2D6;MYC;TNFSF10;AKT1;CYP1B1;SOX9;TRIM21;DLEU7;EPHB3;SLC12A9;KRT7;ACSL5;HNF1B;PRKCA;MIF;HNF1A;DKK1;RUNX3;TRHR;DKK3;DKK4;SFRP4;CLDN4;SLC5A8;SFRP1;SFRP2;SPINT1;IFI27;SPATA13;MALL;CYP2E1;EPHA1;STX1A;ENPEP;ABCB1;CTBP1;SHMT1;CRABP1;GATA6;GATA5;TWIST1;KLK4;SLC5A1;MST1R;GATA2;CRHR1;KLK7;SLC5A5;CRHR2;KLK6;C3;HSD11B1;SULT1A1;HSD11B2;TERT;HLTF;ST8SIA4;ALX4;APOD;APOE;HCRT;PROM1;NTSR1;JAG2;GSTM1;ACE;ADHFE1;FZD6;SLC33A1;NR2F1;IGF2;LIF;MRPL28;ALOX15B;DCLK1;GADD45G;VANGL1;CYP26A1;H19;TF;GJB2;TG;CYP1A1;ANP32B;HAMP;TP73;PCSK2;CDKN1C;PIGR;COL18A1;CDKN1A;MIR130B;PCSK9;GPT;BRCA1;KRT20;UCN3;CHL1;TIMP3;CD34;APC2;APOBEC1;ACAD8;BRD2;ZHX2;MME;MIR21;PROX1;F3;SLC39A14;DUSP6;PAX2;SERPINB5;PAX8;CCKBR;CYP21A2;IL1B;TFF3;ARHGEF4;TFF1;SGK1;TLR4;COL11A1;GSTP1;NR1I2;KCNA3;NEDD9;ASCL1;ASCL2;EGFR;CYP17A1;PAK1;CYP2A13;BGLAP;EPCAM;WIF1;PLAGL1;HSD17B1;ALOX5;SPP1;SPDEF;CD74;NQO1;CDX2;SORT1;MGMT;GDF15;STAT3;SCTR;OSM;MIB2;LILRB1;MLH1;PDZK1IP1;KLK11;VEGFA;SKI;RIT2;FABP6;WT1;HLA-DPB1;TRIP10;ZNF217;GHRL;INHA;APBA2;NEUROG1;SCARB1;CSF2;MPG;IHH;ETS1;MPO;CLU;FGF2;FGF3;IGF1R;PYCARD;HPGDS;CDH3;CCND1;DMBT1;CDH1;MIR150;PITX2;PITX1;KCNH1;TGM2;KCNH2;PDGFRB;IGFBP1;MAGI1;GHSR;CPA1;SLC38A1;CASR;LHCGR;G6PC;IGFBP3;F2R;MIR27A;LIG4;TSC2;HIC1;SLC9A3;HGS;FSCN1;HHIPL1;PGR;NUPR1;PVALB;MAD1L1;ABCG2;FBN2;GCM2;CALCA;TCF7;ALOX15;GSTT1;CACNA1D;RASAL1;ALOX12;FHIT;RAP1GAP;FUT4;MCHR2;CACNA1G;SLC9A3R2;TPO;SCIN;FGF20;MYH14;NLRP3;WNT2;WNT3;SMAD2;KL;TFAP2E;TFPI2;EPHX1;HMGA1;SMAD9;TNFRSF10A;INHBB;MCC;INHBC;ESR1;IGF2R;ARHGAP24;BMP4;NR4A2;IL6;LRCH1;NAT1;CDK2;GNAS;BCL2;MIR184;GNB3;SLC26A4;FGFR4;SNED1;CRISP2;UCN;MIR10A;THRA;FLT4;ITGB2;FHL2;ATP2A3;CHD5;TRH;FOXM1;CXCL5;SYNE1;SPN;GHR;CASP8;DNMT3B;CASP1;PMEPA1;BDKRB1;INSL3;GRASP;SGCG;TGIF1;MMP7;GPX1;PROP1;ITGA4;DCC;USP44;DNMT3A;PDE4C;SORD;SSTR1;TNFRSF1B;SSTR4;SSTR5;C1QL1;TGFBR2;KLF17;GREM1;IFNG;ESRP1;CRH;LTA;BIRC5;CHI3L1;CDH13;BIRC7;CRP;HDAC2;HDAC3;LAMA3;NRXN3;SPG7;THBS1;PTHLH;THBS4;INS;MUC1;MUC2;CCL5;NNAT;ERBB2;SH3BP4;FYN;NCAM1;FSD1;EIF4E;KCNJ1;MUC4;ATOH1;SASH1;UGT1A6;MUC6;NTRK1;CHGA;TCF7L2;NTRK3;TIE1;TXNRD1;IRX3;USP22;EYA4;PCDHGC3;B3GAT1;CABLES1;CCK;MAPK15;HCRTR2;MTOR;TSHR;KISS1;POMC;NELL1;MIR145;APC;MIR144;SST;LEP;PDCD4;VIM;ZACN;BCAR1 |

| | | |
|---|---|---|
| Mood Disorders | 2.9432301192078943E-5 | CHRM2;FAAH;VIPR2;CSF2;CRTC1;ABAT;ADARB1;HTR4;COMT;FGF2;TNF;CNDP2;DAOA;CRHBP;HPGDS;CYP2D6;AKT1;PROK2;PITX2;PDYN;BSCL2;KCTD12;HPD;ABCG1;MAGEL2;PFKFB3;NOTCH4;SLC1A2;CACNA1D;PIK3R1;ADRB2;CACNA1C;PRKCZ;ABO;CRHR1;CRHR2;HSD11B2;TRPM2;GRIN2A;TPO;TSPAN8;MAP2;DEAF1;APOE;TPPP;HCRT;TAC1;NTSR1;DRD4;S100A10;ACE;TFAP2D;ST8SIA2;HTR1A;DCLK2;HTR1B;ESR1;GFAP;DAO;STAR;TH;PNOC;PPP1R1B;SSPN;GNB3;TSNAX-DISC1;VSX1;DPP10;ACHE;GABRB2;PPP1R13B;KCNK9;WISP3;PIK3CD;LAMC2;OXT;GRIK2;DBH;HTR2A;ADRA1B;SLC6A2;SLC6A3;PIK3CG;GLI2;GRM2;CHL1;STAB1;DNMT3B;NOS1;SLC18A2;HSP90AA1;AVPR1B;DST;PAX6;HTR3A;HTR3B;ADRA2C;NGF;DUSP6;SSTR5;C1QL1;ADORA2A;IFNG;IL1B;RARA;CRH;BPI;ARHGEF7;HDAC4;HDAC2;DCTN1;PON1;ADCY3;HTT;ADCY1;ADCY7;CRYZ;REM1;NPY;NTF3;P2RY1;APBB2;NCAM1;DISC1;GABRD;TBX1;NQO1;GABRA1;SLC12A5;GABRA5;NOS3;GAD1;NTRK3;ZNF804A;OPRK1;HTR5A;VEGFA;P2RX7;P2RX5;PER3;SST;P2RX2;P2RX1;FOSB;CD9;SPSB4;HCN1 |
| Mental disorders | 3.5882348988083786E-5 | CHRM2;FAAH;OXTR;CTNND2;ZBTB20;ADARB1;COMT;RGS4;DAOA;CRHBP;ATCAY;CYP2D6;DPYSL2;STMN1;AKT1;PIP4K2A;NCKAP5;IER3;TGM2;MAGI1;KCNK10;SLC6A15;LMO3;KRT7;PRKCA;CACNA2D4;CARTPT;PDYN;MYT1;THEG;CACNB2;MAG;FEZ1;SCN8A;ANKK1;ALDH7A1;RIN2;STX1A;PVALB;CALCA;MAGEL2;ABCB1;CHRNA5;CNP;GPR85;CSF2RB;ADRB2;CACNA1C;KLK8;NALCN;CRHR1;NPAS3;RAI1;GRIN2A;CYP2B6;ATXN1;MIR330;DEAF1;NDN;APOD;APOE;DRD4;S100A10;DRD5;ACE;EGR3;ST8SIA2;HTR1A;HTR1B;GRIN2B;ESR1;OPN5;DCLK1;GFAP;IL6;GDNF;DEGS2;CD22;FGFR2;GABRB2;UCN;CNTNAP2;CD40;GRIK4;OXT;DBH;HTR2A;NR2E1;TREM1;SERPINA4;SLC6A3;SGCE;GRM5;SLC22A16;DNMT3B;PDE4B;NOS1;YWHAH;PSD;SRGN;BRD1;TRPC6;NRG1;HTR3A;OPRM1;HTR3B;OLIG2;TAL1;IL1B;NOS1AP;CRH;ARHGEF7;MCPH1;DDC;ITIH3;NRXN1;KCNA3;HTT;ADCY7;REM1;EXOSC6;SLITRK1;NPY;NTF3;NCAM1;PCNT;DISC1;CSMD1;TBX1;ASCC1;GAD1;ZNF804A;OPRK1;CCK;HTR5A;BAIAP2;GALNS;POMC;KLF6;RIT2;PPP2R2B;SCLY;TRIM31;F2RL3 |
| Diabetes | 4.85432837157163E-5 | EHMT1;BACH2;LIPC;GJA5;MYC;AKT1;UBASH3A;PRSS8;MNX1;UNC13A;DAPK1;SERPINF1;DAPK3;PRKCA;PASK;ISL1;GPR162;CLPS;DDIT3;TBPL2;CYP2E1;CFB;DHH;GATA6;GATA4;GATA3;SLC5A1;SLC30A10;SLC5A4;ABO;CIAO1;C3;HSD11B1;SULT1A1;HSD11B2;CYP2B6;RNF213;KCNQ1OT1;HCRT;HLA-DQA2;HLA-DQA1;NGFR;CREBBP;GSTM1;ACE;ABCA4;NFATC3;NFATC2;SLC16A11;GP6;NR0B2;MIR338;PBX1;COL1A1;FAT1;HRC;CYP1A1;NOX5;HAMP;HLA-DRB1;CDKN1C;LAD1;CDKN1A;MIR130B;SETD7;PRKAG2;PRSS27;ADM;GPT;SLC6A2;JPH3;UCN3;ADAMTSL3;TIMP2;TIMP3;SLC18A2;TIMP4;TCF19;MIOX;LYPLA1;ANXA2;PAX4;PAX6;F3;MMP14;ZEB1;SLC7A9;KCNMA1;PXDN;HCLS1;SLC29A1;TKT;HLA-DQB2;HLA-DQB1;GSTP1;NR1I2;KCNA3;KCNA5;EGFR;SLC7A2;HLA-DMA;HLA-DMB;BGLAP;PLAGL1;NTF3;SPP1;APOL1;P2RY12;AOC3;AOC2;PKLR;STAT3;LETM1;FABP2;GATM;SLMAP;HLA-DPB1;SIGLEC7;HSPA1B;HSPA1A;CSF3;CSF2;EPO;DEFB1;IRS2;CLU;FGF2;TNF;BEST1;RIMS2;HPGDS;FCGRT;ADORA1;HLA-DOA;MAP3K5;IGFBP1;FBXW8;S100A1;IGFBP3;DIO2;BSCL2;SLC9A3;CLIP2;MTNR1B;RAB38;SLC27A2;KCNE1;SHC1;ITPR1;GSTT1;VTCN1;SOCS3;TPO;ZFP57;THEM4;ZNF300;FN3K;KL;GFPT1;MAFA;HOPX;GFAP;INS-IGF2;RHEB;CDK2;CPE;PLXNB1;SHBG;ALAS1;ITGB2;FHL2;PIK3CD;PTPN22;HTR2A;FOXM1;PIK3CG;CYR61;HHEX;ITGAX;ITGB7;SOST;INSL3;ADAMTS9;PGM1;HSP90AA1;TRIM63;GPX1;TRPC6;VWF;SORD;APOA1;TNFRSF1B;NGF;TNFRSF1A;ADORA2A;LCK;NOS1AP;LCP1;CRP;CX3CR1;HDAC2;HDAC3;LRP5;NOD1;NOD2;NTN1;TTN;CCL5;P2RY1;KCNJ1;ANGPT2;ANGPT1;KCNJ11;KCNJ9;TXNRD1;SORL1;MYO16;MTOR;TPCN2;P2RX7;KLF6;PC;SUCLA2;EIF3K;APC;SST;IL2RA;LEP;IL2RB;MGP;PTPN6;P4HB;ZACN;ERRFI1;APP;TNFRSF6B;SPARC;IFI30;AQP2;HS6ST1;ICAM1;LGALS3;EDNRA;LGALS1;ANPEP;TNFSF10;TNFSF11;SLC12A9;ENTPD1;CPT1A;SLC11A1;HLA-B;HLA-C;EBF2;HNF1B;TIMD4;MIF;HNF1A;HLA-A;DHX40;PPP1R3C;TCERG1L;IL23A;KCNQ1;SFRP5;EZR;STX1A;CFTR;GIPR;C2CD4A;C2CD4B;CD1D;PIK3R1;TRPM2;TSPAN8;APOH;ST8SIA4;APOE;FADD;A2M;APOB;PCK1;SPRR2A;NTSR1;STAT5A;EGR1;PTPRN2;TNFSF14;SLC33A1;IGF2;LIF;MIR200B;CIDEC;TH;ID3;PEMT;COL9A3;PCSK2;PTPRU;RYR2;CD40;PTPRN;ICA1;PCSK9;BRCA1;KRT20;ESM1;HNF4A;CD38;SOSTDC1;FFAR1;ANKRD2;CD34;ACAD8;CD52;NME2;SORCS1;MIR21;PROX1;MC3R;NR5A2;ADRB3;IL1B;VWA2;MAPT;SGK1;TLR4;RAPSN;TAGLN;REM1;DUOX1;CUX1;WIF1;NPY;HSF1;PCNT;NQO1;SORT1;GDF15;GAD1;LILRB1;SORBS1;CYP8B1;GCK;VEGFA;PER1;PER3;GHRL;IL17D;IL17C;FOXA2;PNPLA2;NEUROG3;CARS;CD86;KLB;SCARB1;GIMAP5;COMT;ETS1;MPO;CNDP1;IGF1R;NPPA;MIR150;GGT1;TGM2;GHSR;ABCG8;PDIA2;CASR;ABCG5;G6PC;ZGPAT;SLC30A8;LIG4;TSC2;CYBA;ELMO1;CARD16;ABCG1;ABCG2;FBN2;CALCA;ISYNA1;CHAT;TCF7;ALOX15;ALOX12;ADRB2;NME1-NME2;DPP4;ARHGAP22;HRH1;DEAF1;UBD;NLRP3;TAC1;MSC;FGF21;SMAD2;HMGA1;HSPA2;BMP7;ESR1;IGF2R;BMP4;ISG20;IL6;HOOK2;GDNF;FGF19;MYF6;MIR184;GNB3;TGFBI;FGFR4;SNED1;TREH;ATP2A3;HSPB2;TNFAIP3;SLC2A2;PKD1;CXCL5;GHR;GRM2;CASP8;BDKRB2;BDKRB1;NOS1;KLF11;MKKS;KLF14;POMGNT1;DCN;GREM1;FGR;MIR126;NDUFS7;IFNG;CHDH;CAT;LTA;MYH9;PON1;THBS1;PTHLH;CRYZ;INS;CBS;PTK2B;CHGA;TCF7L2;MAS1L;NOS2;NOS3;PDX1;POMC;MIR145;NPHS2;CAPN10;LPIN2 |
| HIV-1 infection | 6.779531133950383E-5 | DDR1;HCP5;UGT1A10;ITIH4;FBN3;OXTR;TNXB;ATF6B;HCG18;NOTCH4;LST1;PSORS1C3;SAMHD1;TNF;AIF1;RYR3;PSORS1C1;C2;GRM5;TPO;GRM8;UGT1A5;RNF39;UGT1A4;TRIM26;UGT1A3;CCHCR1;UGT1A9;HLA-DOB;UGT1A8;GPSM3;UGT1A7;MICB;UGT1A6;HLA-B;TAP2;HLA-C;MIR126;SH3RF3;NBEA;IL2RA;SKIV2L;HCG22;LTB;TRIM31;SDK2;MUC21;LY6E |
| Borderline Personality Disorder | 9.298446729656032E-5 | CHRM2;GABRB2;OXTR;OXT;GRIK2;HTR2A;COMT;SLC6A2;SLC6A3;AKT1;SLC18A2;CMKLR1;PRIMA1;BRD1;F2R;NRG1;HTR3A;CACNA2D4;NINJ2;SCN8A;KCNQ2;CRH;BPI;SCN4B;PVALB;HDAC2;NRN1;TERC;SLC1A2;KCNA3;NRXN3;CACNA1C;CRHR1;EGFR;CRHR2;NPAS3;REM1;TSPAN8;TAC1;DISC1;DRD4;ZBED4;CCL24;ANGPT1;EYA1;NOS3;GAD1;HTR1D;HTR1A;ZNF804A;HTR1B;VEGFA;P2RX7;TH;SST;BCL2;GNAS;APBA2;FGFR2 |
| Multiple Sclerosis | 9.728852116551235E-5 | EHMT2;BACH2;CYP2D6;ZMIZ1;DPYSL2;STMN1;C3AR1;UBASH3A;CHGB;DAXX;PRKCA;RFX1;UNC13D;MIR196A2;DDIT3;HOXB4;TYRO3;PADI2;IDO2;CFB;SKAP2;AIRE;AGAP2;GATA3;PLA2G7;C2;C3;ADAMTS14;CYP2B6;GJC2;HLA-DQA2;HLA-DQA1;HLA-DRB6;HLA-DRB5;GSTM1;ACE;TSLP;NFATC2;GRIN2B;NR0B2;MIR338;EHD1;GJB6;TNFSF9;PPT2;HLA-DRB1;LGMN;CDKN1C;SPEF2;IL18BP;TRIM5;TIMP2;NEFL;TIMP3;NEFM;JAK3;TCF19;BRD2;ANXA2;IFNGR2;FCRL3;POU5F1;SUMF1;ZEB1;KCNMA1;RARA;TKT;CRYAB;HLA-DQB2;HLA-DQB1;ANKRD55;NR1I2;KCNA3;EGFR;HLA-DMA;HLA-DMB;ALOX5;SPP1;STAT4;WWOX;PKLR;HSPA1L;IL10RB;STAT3;NR1H3;GPR3;GTF2H4;PSMB9;WT1;FABP7;HLA-DPB1;HSPA1B;HSPA1A;CSF3;CSF2;EPO;GTF2B;FAM107A;IL5RA;STK19;CLU;TNF;HPGDS;CDH3;FCGRT;GRAP2;RNF19A;HLA-DOA;TNFRSF4;MAP3K5;CD177;LAG3;S100A1;ZBTB38;TAP2;SARS;TAP1;MYT1;TUBA4A;FOXP1;MAG;MTNR1B;PGR;PTPRCAP;ERG;PVALB;LY6E;CSF1R;NOTCH3;CNP;NOTCH4;MAZ;SPATA2;GSTT1;AK5;KLC1;SOCS3;SBNO2;MBP;KL;RAB4A;TNFRSF10C;TNFRSF10A;TRAPPC9;GFAP;LRCH1;INS-IGF2;BCL2;HLA-DRA;XAF1;CNTNAP2;CIITA;ITGB2;DYSF;PTPN22;ITGAL;ADAMTS4;TAGAP;ITGAX;HSP90AA1;DST;ITGA4;PLEKHG5;APOA1;NRG1;TNFRSF1B;NGF;TNFRSF1A;TRAF3;TNFRSF25;CRP;CX3CR1;HDAC3;LRP1;UHRF1;ZBTB46;MIR922;NTN1;CCL5;IL21R;FYN;NTRK1;SEMA4A;SORL1;P2RX7;NFASC;APC;IL2RA;POU2AF1;LEP;IL2RB;PTPN6;HCG22;PDCD1;VIPR2;GFI1;AQP4;IFI30;ICAM1;LGALS3;DHX58;TNFSF10;TNFSF11;SOX8;KIF21B;LGALS8;TP63;EPHA4;ENTPD1;SLC11A1;EBF1;HLA-B;HLA-C;MIF;HLA-A;HLA-F;HLA-G;HLA-E;DHX40;IL23A;PXT1;SH2D2A;SP140;CD1D;MST1R;CD1A;CRHR1;KLK6;SEC14L3;PTDSS1;MAP2;CD19;APOD;APOE;A2M;PCK1;TRPM4;TNFSF14;IGF2;LIF;MIR200A;BST2;OCLN;TH;ANP32B;CD200;TP73;PTPRU;CD40;SLC44A4;MIR17;SLC44A2;PTPRN;CTSZ;ICA1;KRT20;CX3CL1;CCR7;FFAR2;CCR6;CD34;HTATIP2;MST1;MIR21;TYK2;SYN3;CYP7B1;ANO2;RGMA;VAV2;NCR1;CLDN11;TLR1;BANK1;IL1B;PITPNM2;FAM20C;CD47;ARHGEF2;MAPT;TLR4;TNXB;TMPRSS6;CXCR5;CST7;DUOX1;NT5E;CUX1;NPY;ADAD1;GPC5;MGAT1;CTNNA3;LAIR1;MICB;IL32;NQO1;MOG;CDX2;SORT1;GAD1;MIR26A1;OSM;LILRB1;KLK10;LILRB4;VEGFA;BATF;PER3;F2RL1;IL17D;IL17C;CD86;PLAT;PHRF1;COMT;ETS1;MPO;CCND1;MIR150;TGM2;EOMES;PDGFRA;ZGPAT;CD8A;SCN8A;ELMO1;MASP2;CARD16;CARD14;CARD11;ABCG2;ISYNA1;SLC1A2;SLC1A3;SLC1A4;ADRB2;LPP;PSORS1C1;ZFP36L1;DPP4;HRH1;NLRP3;TAC3;TAC1;MSC;WNT3;HSPA2;BTNL2;BMP7;ESR1;ARHGAP24;SMAD7;BMP4;ISG20;NR4A2;BMP2;IL6;IL17REL;NAT1;SKIV2L;PLEK;TNFAIP3;LAMC2;GRM1;SPRED2;GRM5;GRM4;CASP8;GRM8;ZIC1;CASP1;BLNK;BDKRB1;HORMAD2;NOS1;CYTIP;MMP7;DCN;MIR126;IFNG;CAT;LTA;CHI3L1;IRF8;IRF5;RGS14;PON1;THBS1;USP18;PTHLH;CRYZ;INS;CNR2;GNA11;NCAM1;CHGA;NOS2;IRGM;DLK1;GALC;POMC;CD4;MIR145;CD6;NPHS1;MIR142 |
| Convulsive Seizures | 1.496273225272235E-4 | ACHE;FAAH;NRP2;CHRM1;ABAT;GRIK1;OXT;TRH;ADRA1B;SLC6A1;CLU;FGF2;SLC6A2;NOS1;GAMT;TSC2;OPRM1;PDYN;NGF;SYN2;CNKSR1;ALDH5A1;ADORA2A;SCN8A;IL1B;KCNQ2;CAT;CRH;ERG;PVALB;ABCB1;CHRNA5;FOXG1;CHAT;INS;NPY;SLC17A7;APOE;NGFR;SLC12A5;KCNJ11;NOS2;GABRA5;HTR1A;HTR1B;OPRK1;CCK;LETM1;POMC;SLC8A3;IL6;SST;LEP;ALPL;GNB3;GHRL;PAM;HCN1 |
| Adenocarcinoma | 1.711975959932826E-4 | RB1;SCT;SAMD14;CYP2D6;SOX18;MYC;STMN1;AKT1;CYP1B1;PLCE1;HOXA1;DAPK1;SERPINF1;RFX1;PRAP1;DKK1;DKK3;CLDN6;CLDN4;SPINT1;RBP1;HOXB2;CYP2E1;HOXB7;SKAP2;ABCB1;TERC;SHMT1;GATA6;PDGFA;GATA5;GATA4;SLC5A1;A1BG;LYPD5;PRKCZ;SLC5A5;HSD11B1;LDHB;TERT;ALX4;SFN;PROM1;RREB1;HLA-DQA1;JAG2;NAPSA;ST14;GSTM1;ACE;CADM1;PLK2;VANGL1;H19;CPEB1;CYP1A1;TNFSF9;GTF2IRD1;NOX5;PLCH1;FAT4;PAM;CDKN1C;PIGR;CDKN1A;RARG;LPCAT1;TCF21;ADM;GLI3;BBC3;HEY1;UCN3;CHL1;B3GALT1;TIMP2;TIMP3;APC2;GPR39;SLC2A14;RIPK3;ANXA2;DUSP1;TNNC1;F3;DUSP6;POU5F1;PAX2;SERPINB5;DDB1;SLC7A5;MMP14;GPRC5A;PAX8;KIFC1;RARA;RARB;CHST4;CHST5;CHST2;HLA-DQB1;OCA2;GRP;GSTP1;COL11A1;NR1I2;ASCL1;EGFR;APOLD1;EPCAM;PLAGL1;ALOX5;HAS1;SPP1;PRAME;CSMD1;IL12RB2;SPDEF;AOC3;ZNF582;MGMT;STAT3;SCTR;GUCA2B;GUCA2A;FABP3;WT1;INHA;ZNF577;YPEL3;CSF2;CSF1;EPO;CTNND2;MSI2;FGF1;CLU;FGF2;TNF;FGF3;RPH3A;PYCARD;FGF7;HPGDS;CDH3;CDH1;GRAP2;RNF19A;PITX1;SLC16A3;LZTS1;KCNH2;USP7;S100A2;TPM3;S100A1;IGFBP3;UGT2B17;CHD1L;SRCIN1;DGKZ;FOXP1;RAB32;S100A6;FSCN1;PGR;FSCN3;ADAM8;ERG;S100A9;SQSTM1;DSC3;CSF1R;NOTCH3;PFKFB3;TACSTD2;SPATA2;GSTT1;RASAL1;VTCN1;FHIT;SLC9A3R2;SOCS3;SLC38A2;ESRRA;KL;TNK2;TFPI2;TNFRSF10C;TNFRSF10A;ESRRG;MCC;TMPRSS13;CDK6;CDK2;BCL2;PLXNB1;CLPTM1L;SHBG;XAF1;LAT;PSCA;ITGB4;SYCP1;FLT4;GADL1;PIK3CD;MSLN;FOXM1;CYR61;PIK3CG;SYNE1;ZNRD1;MB;DNMT3B;EMILIN2;PMEPA1;HGFAC;SGCG;DSP;EMX2;HSP90AA1;GPX1;ITGA4;DST;ITGA3;CTDSPL;NGB;APOA1;NRG1;TNFRSF1B;TNFRSF1A;PTP4A1;PCCA;PCCB;BIRC5;ITGA6;TNFRSF25;MCM5;ANG;BIRC7;CX3CR1;LRP1;SRC;SEMA3B;BARX2;MIR148A;SELENBP1;CLEC3B;DNAJB6;RAB25;CCL5;NNAT;FYN;FSD1;EIF4E;MEST;CNNM4;NTRK1;TBX1;SEMA4A;ANGPT2;ANGPT1;NTRK3;MTOR;P2RX7;P2RX5;KLF6;MARCKS;APC;SST;LEP;MGP;PDCD4;PDCD1;OCIAD2;INSM1;BCAR1;ERRFI1;IFITM1;TNFRSF6B;SPARC;HPSE2;AQP4;ICAM1;AQP1;LGALS4;SOX2;LGALS3;FRAT1;RASSF1;EDNRB;ANPEP;TNFSF10;TNFSF11;SOX7;TP63;SLC12A9;SRRM2;EPHA4;KRT8;MTUS1;HLA-C;KRT7;ACSL5;HLA-A;KRT5;MIF;HLA-G;RUNX3;RUNX2;SFRP4;SFRP1;SFRP2;SLPI;IFI27;KCNQ1;SFRP5;RIN1;EPHA1;CFTR;GRB7;SLC22A3;PTGFR;EPAS1;TWIST1;KLK4;MST1R;KLK8;CRHR1;KLK7;KLK6;NEU1;PENK;APOE;TRPM8;FADD;TRPM5;TSPAN1;NTSR1;STAT5A;EGR1;FOXF1;IQSEC1;XRCC3;IGF2;MIR200A;PTK6;MIR200B;ALOX15B;DCLK1;NET1;CYP26A1;OCLN;TG;PTK7;PPP1R1B;ID3;TP73;APOBEC3B;DDR1;COL18A1;NRP2;CD40;MIR17;CTSZ;BRCA1;CD3E;KRT20;BOLL;CCKAR;SLC22A18;HNF4A;HYAL2;CCR7;MIR192;CD34;CTSD;APOBEC1;ZHX2;MME;MST1;NME2;TFEB;AMPD1;MIR21;ANO7;CYP7B1;VAV1;TLR1;CDCP1;NR5A2;CTTN;CCKBR;IL1B;MIR17HG;CLDN18;TFF2;TFF1;ARHGEF2;TLR4;GUCY2C;TAGLN;CSTA;SLC43A2;SATB2;TMPRSS4;CXCR5;TMPRSS2;CAPG;CYP17A1;REM1;DUOX1;PAK1;CYP2A13;WIF1;HSD17B1;IL32;NQO1;CDX2;CDX1;KLHL1;KLK13;AKR1C2;KLK15;MLH1;VAX1;KLK10;VEGFA;KLK12;KRT18;KRT17;QPCT;F2RL1;GHRL;IL17D;APBA2;CD83;PLAT;PRDM2;COMT;ETS1;MPO;ETS2;IGF1R;LOXL2;HOXA11;HOXA10;CCND2;DMBT1;CCND1;NPPA;GGT1;TGM2;CASR;LMO3;WNT5A;HBE1;MIR27A;LIG4;TSC2;DHDDS;BRMS1;MYCN;ADAM12;CARD16;ABCG4;ABCG2;ISYNA1;CALCA;CHRNA5;DIABLO;ALOX15;LTBP4;SLC1A5;ADRB2;HOXB13;FBLN2;FUT4;ZFP36L1;NME1-NME2;DPP4;DEAF1;PODXL;NLRP4;NLRP2;NLRP3;FAM83B;FAM83A;TFAP2A;SMAD2;TFAP2B;UBE2I;TGFB3;HMGA1;ASPSCR1;LAYN;INHBA;EVX2;BMP7;ESR1;IGF2R;ARHGAP24;BMP4;NR4A2;BMP2;IL6;NAT1;CNTN1;GNAS;TGFBI;FGFR4;FGFR3;FGFR2;FGFR1;TNFAIP8;MS4A3;DYRK2;THRB;ATP2A3;HSPB2;OXT;LAMC2;LTB4R;CXCL5;FLCN;CASP8;CA2;SIX3;NOS1;MIR193A;EGFL7;MMP7;MIR23A;DCC;AXIN1;SSTR1;OPRM1;SSTR4;SSTR5;TGFBR2;ACTA2;GREM1;MIR126;TGFBR3;ALDH1A3;ELF3;IFNG;PALLD;CDH10;CAT;CDH12;LTA;CHI3L1;CDH13;MYH9;ZMYND10;HAGH;RBMS3;PAOX;DDC;LAMA3;NRXN3;THBS1;AURKC;THBS4;PTHLH;AURKB;CRYZ;MUC1;MUC2;WNT11;MARVELD2;MIR495;MXI1;ERBB2;PTK2B;SPOCK1;NCAM1;MUC4;ATOH1;MUC6;OPCML;SNCA;CHGA;TCF7L2;KDM4B;NOS2;NOS3;BNIP3;EYA4;PDX1;CCK;MUC5B;CHFR;KISS1;MIR145;MIR142;CD9;S100P;VIM |
| Seizures, Auditory | 1.711975959932826E-4 | ACHE;FAAH;NRP2;CHRM1;ABAT;GRIK1;OXT;TRH;ADRA1B;SLC6A1;CLU;FGF2;SLC6A2;NOS1;GAMT;TSC2;OPRM1;PDYN;NGF;SYN2;ADORA2A;IL1B;KCNQ2;CAT;CRH;PVALB;ABCB1;CHRNA5;FOXG1;CHAT;INS;NPY;SLC17A7;APOE;NGFR;SLC12A5;KCNJ11;NOS2;GABRA5;HTR1A;HTR1B;OPRK1;CCK;LETM1;POMC;SLC8A3;IL6;SST;LEP;ALPL;GNB3;GHRL;PAM;HCN1 |
| Seizures, Clonic | 1.711975959932826E-4 | ACHE;FAAH;NRP2;CHRM1;ABAT;GRIK1;OXT;TRH;ADRA1B;SLC6A1;CLU;FGF2;SLC6A2;NOS1;GAMT;TSC2;OPRM1;PDYN;NGF;SYN2;ADORA2A;IL1B;KCNQ2;CAT;CRH;PVALB;ABCB1;CHRNA5;FOXG1;CHAT;INS;NPY;SLC17A7;APOE;NGFR;SLC12A5;KCNJ11;NOS2;GABRA5;HTR1A;HTR1B;OPRK1;CCK;LETM1;POMC;SLC8A3;IL6;SST;LEP;ALPL;GNB3;GHRL;PAM;HCN1 |
| Seizures, Sensory | 1.711975959932826E-4 | ACHE;FAAH;NRP2;CHRM1;ABAT;GRIK1;OXT;TRH;ADRA1B;SLC6A1;CLU;FGF2;SLC6A2;NOS1;GAMT;TSC2;OPRM1;PDYN;NGF;SYN2;ADORA2A;IL1B;KCNQ2;CAT;CRH;PVALB;ABCB1;CHRNA5;FOXG1;CHAT;INS;NPY;SLC17A7;APOE;NGFR;SLC12A5;KCNJ11;NOS2;GABRA5;HTR1A;HTR1B;OPRK1;CCK;LETM1;POMC;SLC8A3;IL6;SST;LEP;ALPL;GNB3;GHRL;PAM;HCN1 |
| Seizures, Somatosensory | 1.711975959932826E-4 | ACHE;FAAH;NRP2;CHRM1;ABAT;GRIK1;OXT;TRH;ADRA1B;SLC6A1;CLU;FGF2;SLC6A2;NOS1;GAMT;TSC2;OPRM1;PDYN;NGF;SYN2;ADORA2A;IL1B;KCNQ2;CAT;CRH;PVALB;ABCB1;CHRNA5;FOXG1;CHAT;INS;NPY;SLC17A7;APOE;NGFR;SLC12A5;KCNJ11;NOS2;GABRA5;HTR1A;HTR1B;OPRK1;CCK;LETM1;POMC;SLC8A3;IL6;SST;LEP;ALPL;GNB3;GHRL;PAM;HCN1 |
| Atonic Absence Seizures | 1.711975959932826E-4 | ACHE;FAAH;NRP2;CHRM1;ABAT;GRIK1;OXT;TRH;ADRA1B;SLC6A1;CLU;FGF2;SLC6A2;NOS1;GAMT;TSC2;OPRM1;PDYN;NGF;SYN2;ADORA2A;IL1B;KCNQ2;CAT;CRH;PVALB;ABCB1;CHRNA5;FOXG1;CHAT;INS;NPY;SLC17A7;APOE;NGFR;SLC12A5;KCNJ11;NOS2;GABRA5;HTR1A;HTR1B;OPRK1;CCK;LETM1;POMC;SLC8A3;IL6;SST;LEP;ALPL;GNB3;GHRL;PAM;HCN1 |
| Generalized Absence Seizures | 1.711975959932826E-4 | ACHE;FAAH;NRP2;CHRM1;ABAT;GRIK1;OXT;TRH;ADRA1B;SLC6A1;CLU;FGF2;SLC6A2;NOS1;GAMT;TSC2;OPRM1;PDYN;NGF;SYN2;ADORA2A;IL1B;KCNQ2;CAT;CRH;PVALB;ABCB1;CHRNA5;FOXG1;CHAT;INS;NPY;SLC17A7;APOE;NGFR;SLC12A5;KCNJ11;NOS2;GABRA5;HTR1A;HTR1B;OPRK1;CCK;LETM1;POMC;SLC8A3;IL6;SST;LEP;ALPL;GNB3;GHRL;PAM;HCN1 |
| Gustatory seizure | 1.711975959932826E-4 | ACHE;FAAH;NRP2;CHRM1;ABAT;GRIK1;OXT;TRH;ADRA1B;SLC6A1;CLU;FGF2;SLC6A2;NOS1;GAMT;TSC2;OPRM1;PDYN;NGF;SYN2;ADORA2A;IL1B;KCNQ2;CAT;CRH;PVALB;ABCB1;CHRNA5;FOXG1;CHAT;INS;NPY;SLC17A7;APOE;NGFR;SLC12A5;KCNJ11;NOS2;GABRA5;HTR1A;HTR1B;OPRK1;CCK;LETM1;POMC;SLC8A3;IL6;SST;LEP;ALPL;GNB3;GHRL;PAM;HCN1 |
| Jacksonian Seizure | 1.711975959932826E-4 | ACHE;FAAH;NRP2;CHRM1;ABAT;GRIK1;OXT;TRH;ADRA1B;SLC6A1;CLU;FGF2;SLC6A2;NOS1;GAMT;TSC2;OPRM1;PDYN;NGF;SYN2;ADORA2A;IL1B;KCNQ2;CAT;CRH;PVALB;ABCB1;CHRNA5;FOXG1;CHAT;INS;NPY;SLC17A7;APOE;NGFR;SLC12A5;KCNJ11;NOS2;GABRA5;HTR1A;HTR1B;OPRK1;CCK;LETM1;POMC;SLC8A3;IL6;SST;LEP;ALPL;GNB3;GHRL;PAM;HCN1 |
| Non-epileptic convulsion | 1.711975959932826E-4 | ACHE;FAAH;NRP2;CHRM1;ABAT;GRIK1;OXT;TRH;ADRA1B;SLC6A1;CLU;FGF2;SLC6A2;NOS1;GAMT;TSC2;OPRM1;PDYN;NGF;SYN2;ADORA2A;IL1B;KCNQ2;CAT;CRH;PVALB;ABCB1;CHRNA5;FOXG1;CHAT;INS;NPY;SLC17A7;APOE;NGFR;SLC12A5;KCNJ11;NOS2;GABRA5;HTR1A;HTR1B;OPRK1;CCK;LETM1;POMC;SLC8A3;IL6;SST;LEP;ALPL;GNB3;GHRL;PAM;HCN1 |
| Nonepileptic Seizures | 1.711975959932826E-4 | ACHE;FAAH;NRP2;CHRM1;ABAT;GRIK1;OXT;TRH;ADRA1B;SLC6A1;CLU;FGF2;SLC6A2;NOS1;GAMT;TSC2;OPRM1;PDYN;NGF;SYN2;ADORA2A;IL1B;KCNQ2;CAT;CRH;PVALB;ABCB1;CHRNA5;FOXG1;CHAT;INS;NPY;SLC17A7;APOE;NGFR;SLC12A5;KCNJ11;NOS2;GABRA5;HTR1A;HTR1B;OPRK1;CCK;LETM1;POMC;SLC8A3;IL6;SST;LEP;ALPL;GNB3;GHRL;PAM;HCN1 |
| Olfactory seizure | 1.711975959932826E-4 | ACHE;FAAH;NRP2;CHRM1;ABAT;GRIK1;OXT;TRH;ADRA1B;SLC6A1;CLU;FGF2;SLC6A2;NOS1;GAMT;TSC2;OPRM1;PDYN;NGF;SYN2;ADORA2A;IL1B;KCNQ2;CAT;CRH;PVALB;ABCB1;CHRNA5;FOXG1;CHAT;INS;NPY;SLC17A7;APOE;NGFR;SLC12A5;KCNJ11;NOS2;GABRA5;HTR1A;HTR1B;OPRK1;CCK;LETM1;POMC;SLC8A3;IL6;SST;LEP;ALPL;GNB3;GHRL;PAM;HCN1 |
| Single Seizure | 1.711975959932826E-4 | ACHE;FAAH;NRP2;CHRM1;ABAT;GRIK1;OXT;TRH;ADRA1B;SLC6A1;CLU;FGF2;SLC6A2;NOS1;GAMT;TSC2;OPRM1;PDYN;NGF;SYN2;ADORA2A;IL1B;KCNQ2;CAT;CRH;PVALB;ABCB1;CHRNA5;FOXG1;CHAT;INS;NPY;SLC17A7;APOE;NGFR;SLC12A5;KCNJ11;NOS2;GABRA5;HTR1A;HTR1B;OPRK1;CCK;LETM1;POMC;SLC8A3;IL6;SST;LEP;ALPL;GNB3;GHRL;PAM;HCN1 |
| Vertiginous seizure | 1.711975959932826E-4 | ACHE;FAAH;NRP2;CHRM1;ABAT;GRIK1;OXT;TRH;ADRA1B;SLC6A1;CLU;FGF2;SLC6A2;NOS1;GAMT;TSC2;OPRM1;PDYN;NGF;SYN2;ADORA2A;IL1B;KCNQ2;CAT;CRH;PVALB;ABCB1;CHRNA5;FOXG1;CHAT;INS;NPY;SLC17A7;APOE;NGFR;SLC12A5;KCNJ11;NOS2;GABRA5;HTR1A;HTR1B;OPRK1;CCK;LETM1;POMC;SLC8A3;IL6;SST;LEP;ALPL;GNB3;GHRL;PAM;HCN1 |
| Squamous cell carcinoma of lung | 1.9639278229168498E-4 | CSF2;CSF1;FGF1;FRY;MPO;TNF;IGF1R;LOXL2;MYLK4;SOX2;HPGDS;HHAT;CCND1;MYC;PEG10;TNFSF10;AKT1;TRIM26;TP63;AP2M1;EPHB3;KCNH1;EPHA5;PDGFRB;CASR;USP7;LMO3;F2R;KRT8;HLA-C;HNF1B;HLA-A;HIC1;SERPING1;CYP2E1;ANKK1;SLC22A3;CSF1R;ABCB1;GATA6;GATA4;MIR206;FHIT;NPAS3;NME1-NME2;TERT;CYTL1;PROM1;VARS2;HLA-DQA1;FAM83B;SNHG1;CREBBP;GSTM1;XRCC3;EPHX1;IGF2R;HOPX;BMP4;IL6;FGF19;CYP1A1;MKS1;BCL2;GTF2IRD1;TGFBI;CLPTM1L;FGFR4;FGFR3;FGFR2;EIF4G1;FGFR1;DDR1;CDKN1A;MS4A3;MAST4;PIK3CD;SLC2A2;DBH;PIK3CG;TUBA1C;CHL1;DENND2D;TPSG1;GPX6;NME2;MIR21;DUSP6;SERPINB5;TGFBR2;ZEB1;PAX7;LCK;RARB;GAS7;TNXB;GSTP1;GRAMD4;AURKC;EGFR;PTHLH;TTN;MIR412;HLA-DMB;ACVR1C;EPCAM;ERBB2;P2RY1;MIR372;FYN;EIF4E;P2RY12;BRF2;NFYC;STAT3;DMRT3;MLH1;CHFR;VEGFA;RAD52;SST;NEK10;VIM |
| Barrett Esophagus | 2.4266837034891872E-4 | OXTR;SPARC;CRTC1;ICAM2;ICAM3;DEFB1;FGF1;FGF2;TNF;ICAM1;IGF1R;HPGDS;CCND1;ANPEP;MYC;TRIM21;PITX1;TP63;TGM2;KCNH2;TMOD1;DAPK1;IGFBP3;KRT5;RFX1;HNF1A;RUNX3;DKK1;FOXP1;SFRP1;SFRP2;IFI27;RBP1;PKP1;HOXB7;RIN2;S100A9;CFTR;HOXB6;GRB7;NOTCH3;GATA6;ADRB2;FHIT;CD1A;FUT4;DPP4;GJC2;TERT;APOB;PROM1;TAC1;WNT2;FOXF1;WNT3A;IGF2;INHBA;ESR1;DCLK1;GDF7;NET1;BMP4;ATP4A;CYP26A1;MSRA;IL6;CDK2;BCL2;NOX5;FGFR4;ATP2A3;HSPB2;OXT;ITGAE;KRT20;AKAP12;GHR;SLC22A18;MADCAM1;MIR192;CTSD;APOBEC1;GPX1;MMP7;DCC;AMPD1;MIR21;CCKBR;ALDH1A2;MIR17HG;CAT;TFF3;CDH13;TFF1;MIR155HG;DDC;GSTP1;NR1I2;BARX1;EGFR;CYP17A1;CRYZ;SELENBP1;MUC1;MUC2;EPCAM;WIF1;ERBB2;MXI1;MUC4;CD55;ATOH1;MUC6;NQO1;CDX2;CDX1;MGMT;GDF15;ASCC1;STAT3;EYA4;AKR1C2;MLH1;TBX5;VEGFA;SKI;NELL1;MIR145;APC;FABP6;SST;LEP;KRT14;PDCD4;ZNF217;VIM;CDO1 |

| | | |
|---|---|---|
| Epileptic drop attack | 2.991642081298461E-4 | ACHE;FAAH;NRP2;CHRM1;ABAT;GRIK1;OXT;TRH;ADRA1B;SLC6A1;CLU;FGF2;SLC6A2;NOS1;SCN1B;GAMT;TSC2;OPRM1;PDYN;NGF;SYN2;DNM1;STX1B;ADORA2A;IL1B;KCNQ2;CAT;CRH;PVALB;ABCB1;CHRNA5;FOXG1;CHAT;INS;NPY;SLC17A7;APOE;NGFR;GABRA1;SLC12A5;KCNJ11;NOS2;GABRA5;HTR1A;HTR1B;OPRK1;CCK;LETM1;POMC;MAPK10;SLC8A3;IL6;SST;LEP;ALPL;GNB3;GHRL;PAM;HCN1 |
| Epileptic Seizures | 3.33491499709502E-4 | ACHE;FAAH;NRP2;CHRM1;ABAT;GRIK1;OXT;TRH;ADRA1B;SLC6A1;CLU;FGF2;SLC6A2;NOS1;GAMT;TSC2;OPRM1;PDYN;NGF;SYN2;ADORA2A;IL1B;KCNQ2;CAT;CRH;PVALB;ABCB1;CHRNA5;FOXG1;CHAT;INS;NPY;SLC17A7;APOE;NGFR;SLC12A5;KCNJ11;NOS2;GABRA5;HTR1A;HTR1B;OPRK1;CCK;LETM1;POMC;SLC8A3;IL6;SST;LEP;ALPL;GNB3;GHRL;PAM;HCN1 |
| Absence Seizures | 3.33491499709502E-4 | ACHE;FAAH;NRP2;CHRM1;EHMT1;ABAT;GRIK1;OXT;TRH;ADRA1B;SLC6A1;CLU;FGF2;SLC6A2;NOS1;SCN1B;GAMT;TSC2;OPRM1;PDYN;NGF;SYN2;STX1B;ALDH5A1;ADORA2A;IL1B;KCNQ2;CAT;KCNMA1;CRH;NHLRC1;PVALB;ABCB1;CHRNA5;FOXG1;CHAT;INS;NPY;SLC17A7;APOE;NGFR;GABRA1;SLC12A5;KCNJ11;NOS2;GABRA5;HTR1A;HTR1B;OPRK1;CCK;LETM1;POMC;SLC8A3;IL6;SST;LEP;ALPL;GNB3;GHRL;FGFR3;PAM;HCN1 |
| Rheumatoid Arthritis | 3.873275794782257E-4 | RB1;MDC1;CNTFR;CLEC10A;EHMT2;BACH2;VPREB1;SYNGAP1;CYP2D6;MYC;AKT1;LRAT;UBASH3A;DLEU7;HOXA5;CMKLR1;CHID1;DAXX;PRKCH;UNC5B;SERPINF1;SERPINF2;UNC5C;ISL1;PRAP1;DKK1;SIGIRR;PADI3;PADI2;PRKCQ;HOXB2;CFB;ABCB1;AIRE;CRABP1;GATA3;FPR2;PRKCZ;ABO;CIAO1;C2;SLC5A5;C3;HSD11B1;HSD11B2;CYP2B6;TERT;PDGFD;HLA-DQA2;HLA-DQA1;ELANE;HOXC6;PRRT1;HLA-DRB6;GSTM2;HLA-DRB5;GSTM1;ACE;TSLP;ABCA4;NR2F1;PBX2;NFATC2;NFATC1;MIR338;MIR346;H19;TNFSF4;FAT1;CDK10;TNFSF9;HOXD4;PPT2;PAM;HAMP;HOXD9;HLA-DRB1;HOXD8;PIGR;CDKN1A;RARG;TINAGL1;GPT;ADM;MMP20;IL18BP;BBC3;ATIC;UCN3;CSRP1;UCN2;TIMP2;RSPO2;TIMP3;IKBKE;HLA-DPA1;TCF19;BRD2;DUSP1;IFNGR2;FCRL3;TACR2;TACR3;PAX5;GNL1;F3;DUSP6;MIR522;FCER2;MMP14;COL2A1;GPRC5A;MMP16;SLC7A9;TAL1;KCNMA1;RARA;RARB;HCLS1;ANGPTL4;SMTNL2;HLA-DQB2;HLA-DQB1;WDR27;GRP;GSTP1;SPESP1;ANKRD55;NR1I2;COL11A2;ASIP;KCNA3;HOXD13;HOXD11;FSTL1;HOXD10;EGFR;HLA-DMA;HLA-DMB;BGLAP;ALOX5;HAS1;SPP1;STAT4;JAM3;RUNX1T1;BRD4;CCL25;CCL23;HSPA1L;IL10RB;STAT3;SAAL1;PSMB8;PSMB9;FABP3;SPSB1;CENPJ;HLA-DPB1;FOSB;PKN1;CCL28;PF4;HLA-DPB2;CCL26;CSF2;CSF1;NCF4;FAM107A;IL5RA;F13A1;STK19;ZBTB22;FGF1;CLU;FGF2;TNF;RIMS2;PYCARD;FGF7;HPGDS;FCGRT;GRAP2;RNF19A;TNFRSF8;HLA-DOA;HLA-DOB;SLC16A3;MAP3K5;TLE3;S100A1;IGFBP3;F2R;TAP2;TAP1;MITF;LTB4R2;MTNR1B;MSH5;ZFP42;S100A6;FSCN1;PGR;ADAM8;ERG;S100A9;CSF1R;NOTCH3;PFKFB3;NOTCH4;CUL1;SPATA2;ITPR2;GSTT1;VTCN1;SOCS2;SOCS3;MOBP;MBP;ZNF423;STRA6;KL;LVRN;CD160;ZBTB16;TFPI2;TNFRSF10A;GFAP;CDK6;BCL6;CDK2;BCL2;HLA-DRA;SHBG;CIITA;FLT4;ITGB2;FHL2;WISP3;PIK3CD;PTPN22;HTR2A;LSP1;ITGAL;TREM1;CYR61;PIK3CG;ADAMTS4;ADAMTS5;PSTPIP1;SUMO3;TAGAP;ITGAX;UBTF;PHACTR3;ACP5;ADAMTS9;CR1;ITGA4;VWF;TNFRSF18;APOA1;TNFRSF1B;NGF;TNFRSF1A;ADORA2A;TRAF5;CRH;BIRC5;BPI;ITGA6;LCP2;TNFRSF25;ANG;CRP;CX3CR1;MIR155HG;NFAT5;HDAC3;LRP1;SRC;UHRF1;HCG18;NOD1;NOD2;NTN1;TTN;PARD6A;DNAJB6;CCL5;VPS52;SH3BP2;IL21R;TBX1;SEMA4A;CIZ1;TRIM40;ANGPT2;ANGPT1;ASCC1;TIE1;TXNRD1;HCG27;TBX5;TBX3;P2RX7;NELL1;MARCKS;TRIM39;APC;SST;IL2RA;LEP;IL2RB;MAB21L2;CD248;HCG22;PTPN6;PDCD1;P4HB;TRIM31;TNP1;BCAR1;APP;TNFRSF6B;VARS;TRAF3IP2;LST1;ICAM3;ICAM1;LGALS3;LGALS2;EDNRA;LGALS1;BAIAP2L1;EDNRB;RPS18;ANPEP;TNFSF10;TNFSF11;TRIM26;LGALS9;LGALS8;TRIM21;TP63;SLC12A9;ATP6V1G2;ACTN1;NFKBIL1;SLC11A1;HLA-B;HLA-C;KRT7;TIMD4;SLC6A11;EBF3;HLA-A;MIF;HLA-G;RUNX2;HLA-E;RUNX1;DHX40;SFRP1;CFDP1;IFI27;IL23A;SFRP5;TRIM15;EPHA1;TRIM10;CFTR;TELO2;EPAS1;CD1D;CRHR1;CRHR2;SYNGR1;ANKH;CD19;INPP5D;STC2;APOE;FADD;APOB;A2M;SLAMF1;STAT5A;EGR1;TNFSF14;TREX1;PDE2A;IGF2;LIF;PRG4;EXT1;BST1;CYP26A1;MSRA;TF;TH;ANP32B;TDGF1;SPAG16;HOTAIR;PTPRT;ROBO3;COL18A1;CD40;SLC44A4;PPP1R10;ICA1;GRIK4;MIR18A;CD3E;PCSK6;KRT20;B3GNTL1;CLYBL;CX3CL1;HYAL2;RAC2;CCR9;CD38;CCR6;CCHCR1;CD34;CTSD;CLIC1;AMPD1;IL16;MIR21;TYK2;CYP7B1;TLR1;ADRB3;BANK1;IL1B;MIR17HG;ARHGEF2;CD47;ARHGEF7;TLR4;TNXB;MCCD1;CXCR5;NEDD9;GDPD5;HSD17B8;AIF1;PAK1;WIF1;HSD17B1;GPC1;ADAD1;LAIR2;LAIR1;CD55;LY6G6F;MICB;IL32;GABBR1;MOG;GDF15;IL34;SCD5;OSM;GRHL3;LILRB1;GCK;SLURP1;VEGFA;BATF;QPCT;F2RL1;GHRL;IL17D;LY6G5C;IL17C;PNPLA2;ENGASE;CD86;CD83;DCLRE1C;MPG;IHH;NRM;PLAT;PRDM1;PHRF1;ETS1;MPO;ETS2;IGF1R;COMP;CCND1;CCRL2;TGM2;PDGFRA;WNT5A;ADAM11;MIR27A;LIG4;MYCN;EMCN;POLG2;CD8A;ELMO1;PRTN3;MASP2;WDFY4;ABCG2;ISYNA1;ALOX15;CHAT;HTRA1;ALOX12;ADRB2;FUT4;NKAPL;PSORS1C1;GRK1;DPP4;GRK5;CALD1;NLRP3;TAC1;VARS2;WNT2;MARK2;SMAD3;TGFB3;RARRES2;TRPV2;ISG15;INHBA;BTNL2;ESR1;BMP4;ISG20;NR4A2;BMP2;IL6;PROC;GDNF;CNTN2;SKIV2L;TGFBI;FGFR1;LGALS3BP;UCN;CRISP2;MIR10A;PLEK;TNFAIP2;TNFAIP3;LAMC2;AFF3;PKD1;LTB4R;HAPLN1;SPRED2;ACOXL;CASP8;CA3;CA2;CASP1;BLNK;BDKRB2;PDE4A;NOS1;GPSM3;TGIF1;OLIG3;DCC;PDE4D;SSTR4;TAPBP;GREM1;MIR126;ALDH1A3;ZAP70;ELF3;IFNG;DDAH2;ALDH1A2;CAT;CDH11;LTA;CHI3L1;IRF8;IRF5;HAGH;LTB;B4GALT7;EGFL8;PON1;LCAT;PRELP;AGPAT1;LTC4S;THBS1;PTHLH;INS;ACAN;MUC1;MUC2;CNR2;GNA12;ERBB2;PTK2B;SLIT3;NCAM1;MUC4;PDLIM4;SNCA;CHGA;URGCP;NOS2;NOS3;B3GAT1;SOD3;POMC;CRYGD;CD4;TEC;CD6;CD5;NPHS2;SARDH;SPRY2;RGS12;TEK;VIM;SCLY |
| Complex partial seizures | 3.873275794782257E-4 | ACHE;FAAH;NRP2;CHRM1;ABAT;GRIK1;OXT;TRH;ADRA1B;SLC6A1;CLU;FGF2;SLC6A2;NOS1;GAMT;TSC2;OPRM1;PDYN;NGF;SYN2;STX1B;ADORA2A;IFNG;IL1B;KCNQ2;CAT;CRH;PVALB;ABCB1;CHRNA5;FOXG1;CHAT;CACNA1D;INS;NPY;SLC17A7;APOE;LGI1;NGFR;SLC12A5;KCNJ11;NOS2;GABRA5;HTR1A;HTR1B;OPRK1;CCK;MTOR;LETM1;POMC;SLC8A3;IL6;SST;LEP;ALPL;GNB3;GHRL;PAM;HCN1 |
| Tonic Seizures | 4.5512731984357393E-4 | ACHE;FAAH;NRP2;CHRM1;ABAT;GRIK1;OXT;TRH;ADRA1B;SLC6A1;CLU;FGF2;SLC6A2;NOS1;GAMT;TSC2;OPRM1;PDYN;NGF;SYN2;ADORA2A;IL1B;KCNQ2;CAT;CRH;PVALB;ABCB1;CHRNA5;FOXG1;CHAT;INS;NPY;SLC17A7;APOE;NGFR;SLC12A5;KCNJ11;NOS2;GABRA5;HTR1A;HTR1B;OPRK1;CCK;LETM1;POMC;SLC8A3;IL6;SST;LEP;ALPL;GNB3;GHRL;PAM;HCN1 |
| Small cell carcinoma of lung | 4.760519094853721E-4 | RB1;OXTR;GFI1;NUDT1;SOX21;ICAM1;SOX2;LGALS3;RASSF1;EDNRB;SOX1;CYP2D6;MYC;ANPEP;TNFSF10;AKT1;HOXA1;DIP2C;TRIM21;TP63;PRIMA1;EPHB3;SLC12A9;SRRM2;SRRM4;PRKCB;HLA-B;KRT8;PRKCA;HLA-A;SFRP1;CYP2E1;SLC22A3;ABCB1;EPAS1;SLCO5A1;THY1;MST1R;SLC5A4;SULT1A1;TERT;FIGNL1;ST8SIA4;PROM1;ABCA2;NGFR;WWTR1;CREBBP;GSTM1;CADM1;ABCA4;MIR200B;NMBR;MIR339;NR0B2;ID2;PPP1R1B;CYP1A1;TP73;SFTA2;HOTAIR;ACHE;PIGR;CDKN1A;GRIK3;BRCA1;KRT20;SLC6A2;CHL1;HYAL1;HYAL2;TIMP3;HTATIP2;CBR1;ANXA2;AVPR1B;MST1;PAX3;PAX5;F3;MMP14;ZEB2;ZEB1;GPRC5A;PAX8;CCKBR;SLCO4A1;PLSCR4;MIR17HG;RARB;GPRC5C;GAS5;GRP;GSTP1;SATB2;COL11A2;KCNA3;CXCR5;TMPRSS2;ADCY1;ASCL1;FSTL1;EGFR;GLRA1;SPP1;PAK6;RUNX1T1;WWOX;NQO1;GABBR1;CDX2;MGMT;COL22A1;STAT3;KLHL1;LILRB1;MLH1;VEGFA;KRT18;HSPA1B;HSPA1A;CSF3;CSF2;CSF1;FGF1;FGF2;TNF;MPO;IGF1R;RIMS2;PYCARD;HPGDS;CCND2;CCND1;DMBT1;CDH1;NPPA;SLC16A3;IGFBP1;S100A1;TPM3;USP4;IGFBP3;TAP1;CDYL;TUBA4B;DHDDS;MYCN;COL4A2;SCG5;ERG;MAD1L1;ABCG2;FBN2;CSF1R;NOTCH3;BRSK2;CALCA;DIABLO;GSTT1;CACNA1D;FHIT;STK31;BCL2L10;STK33;NLRP3;NPTX1;TAC1;TMEM132D;KL;TGFB3;MCC;TMPRSS13;HOPX;ISG20;FER;IL6;GDNF;CDK2;BCL2;MIR184;CPE;SCGB3A1;CNTN2;DYNLRB1;FGFR1;CNTNAP2;THRB;ELAVL4;PIK3CD;OXT;PIK3CG;ELAVL3;SPN;CASP8;ZIC2;GRM8;ZIC1;DNMT3B;HSP90AA1;SSTR1;NGF;C1QL1;SSTR5;TAPBP;TGFBR2;MIR126;AIM2;IFNG;CAT;CRH;BIRC5;CHI3L1;ZMYND10;LTF;RBMS3;CRP;DDC;LRP1;SRC;SEMA3B;PON1;PTHLH;AURKB;TTN;CRYZ;DLL3;MUC1;MIR495;MIR494;NNAT;ERBB2;MXI1;PTK2B;NCAM1;EIF4E;NTRK1;CHGA;RCVRN;CIZ1;MARCKSL1;ANGPT2;B3GAT1;CCK;ZIC4;TBX5;MTOR;TBX3;POMC;APC;SST;IL2RA;CD9;CD248;INSM1;VIM |
| Ki-1+ Anaplastic Large Cell Lymphoma | 5.293223975026669E-4 | RB1;CARS;CSF2;CCNT1;MUM1;ADARB1;PRDM1;CLU;ETS1;TNF;ICAM1;RIMS2;LGALS1;CCND1;ANPEP;MYC;GRAP2;AKT1;TNFRSF8;ZNF569;MIR150;TP63;SLC12A9;PDGFRB;TMOD1;PDGFRA;CD96;TPM3;HNF1A;DOK2;SLPI;ZNF436;CARS2;SKAP1;CSF1R;RTEL1;ABCB1;TCF7;TWIST1;FUT1;PRH1;FUT4;DPP4;SOCS3;RNF213;HOXC5;RREB1;HOXC6;S100A10;TRAP1;NFATC1;NR0B2;HRNR;IL6;CDK6;BCL6;ID2;BCL2;TNFSF8;GRB2;FGFR4;CD40;CDKN1A;FLT4;PIK3CD;PIK3CG;SPN;ATIC;CASP8;NOS1;JAK3;CD52;HSP90AA1;SERPINB9;PAX5;NGF;TNFRSF1B;TNFRSF1A;DUSP22;ZAP70;IFNG;MIR17HG;MYH9;ARHGEF7;MUC1;MIR494;TP53INP1;PTK2B;NCAM1;NTRK1;NOS2;ASCC1;STAT3;GTF2H4;MTOR;VEGFA;PTPN6;IL17D;ALLC;HSPA1B;HSPA1A |
| Benign Prostatic Hyperplasia | 5.940393715944314E-4 | CSF2;WIPF1;CTNND2;COMT;CLU;FGF2;ETS1;IGF1R;PYCARD;SOX2;FGF7;HPGDS;EDNRA;RASSF1;EDNRB;MYC;CYP1B1;CD177;ARL11;NANOS2;EPHA6;S100A2;IGFBP3;SWAP70;F2R;AFAP1;DLEC1;HLA-A;NRBP1;DKK3;SFRP1;SFRP2;S100A6;ERG;STEAP1;S100A9;SLC22A3;CALCA;ISYNA1;ABCB1;HPN;ALOX15;GATA6;TWIST1;KLK4;GSTT1;ALOX12;GATA3;KLK7;PROM1;A2M;BCL2L14;MARK2;HOXC6;EGR1;ACE;GSTM1;SRD5A2;FOXF2;CBX4;FOXF1;TGFB3;SLC33A1;IGF2;SMAD9;ALOX15B;ESR1;FLI1;FGF17;TDRD1;SLC2A9;H19;BMP2;IL6;EFNA2;FXYD3;PTK7;CYP1A1;BCL2;HOXD3;FGFR4;PAM;FGFR3;FGFR2;FGFR1;HOTAIR;UCN;THRB;PSCA;ELN;FLT4;ADM;ADRA1D;MED15;CYR61;GLI2;AKAP12;HEY1;TRPS1;TIMP3;PMEPA1;CCR7;INSL3;SLC18A2;DUSP1;AXIN1;MIR21;SSTR1;PAX2;POU5F1;SSTR4;ANO1;GPRC5A;AIM2;IFNG;KCNMA1;TFF3;BPI;ITGA6;TFF1;SGK1;GRP;GSTP1;NR1I2;PON1;KCNA3;CXCR5;TMPRSS2;THBS1;EGFR;PTHLH;AURKB;CYP17A1;CRYZ;RASSF10;ALOX5;CCL5;HSD17B1;ERBB2;MXI1;SPP1;FYN;EIF4E;PPP1R14A;NQO1;CDX2;NOS2;TXNRD2;GDF15;IRX4;LIMK1;GUCA2B;KLK15;DMRT3;SULF1;MLH1;KLK11;GNMT;KLF6;SST;DLC1;PDCD4;F2RL1;SPRY2;INHA;KCNK2;IL17C;HSPA1B;HSPA1A |
| Congenital Abnormality | 5.940393715944314E-4 | GFI1;FGFRL1;SOX2;SOX17;GJA5;MYC;AKT1;CYP1B1;PROK2;HOXA1;SOX9;MNX1;TP63;SRRM2;EPHA4;MEF2C;NKX2-6;PRKCA;HNF1B;HNF1A;ISL1;RUNX2;RUNX1;SLPI;DDIT3;RPL24;COL8A1;CFTR;ABCB1;AIRE;FOXG1;GATA6;GATA5;TWIST1;GATA4;GATA3;GATA2;PRH1;KCNQ1OT1;NEU1;ALX3;NKX2-5;RREB1;BBS2;ACD;ACE;GSTM1;FOXF1;TGDS;IGF2;NFATC1;NR2F2;FLI1;PBX1;VANGL1;H19;BHMT;GJB2;STAR;PTK7;GJB6;GTF2IRD1;CDKN1C;RYR1;ACHE;DPYS;GLI3;SERPINA4;MMP20;HNF4A;CD38;CD34;TUBB;RIPK4;PAX3;PAX6;PAX2;PAX1;MMP14;ZEB2;TUBB2B;COL2A1;ZEB1;RAD51C;PAX8;KCNMA1;MIR17HG;ARHGEF2;GRP;GSTP1;SATB2;HTT;HOXD13;MTHFD1L;PLAGL1;TMEM67;GSC;DISC1;DYNC1I1;DYNC1H1;WWP2;VEGFA;SPSB1;KRT18;WT1;NEUROG3;DCLRE1C;EPO;CD81;PLAT;ETS1;HOXA13;FGF3;ACTG1;IGF1R;RIMS2;FGF7;FGF8;SALL1;KIF5C;SALL4;TNR;ZNF569;PITX2;LZTS2;FBXW8;MSX2;TPM3;F2R;WNT5A;LIG4;TSC2;FRMD4A;EDAR;CD8A;COL4A1;PGR;ZNF436;FBLN7;ANKRD11;CHAT;CHRD;GSTT1;SLC9A3R2;CDH23;CAMTA1;NLRP3;MBP;OTX2;SRGAP3;ZNF423;PRODH;WNT3;STRA6;EPHX1;ISG15;BMP7;ESR1;PHOX2B;PHOX2A;SMAD7;BMP4;BMP2;G6PC3;LHX3;NAT1;GDNF;BCL2;SLC26A4;FGFR3;LHX9;FGFR2;FGF10;FGFR1;THRB;PIK3CD;NR2E1;AFF3;PIK3CG;ZIC2;TRPS1;SIX3;ZIC1;DNMT3B;EMILIN2;ITGB6;SOST;INSL3;TGIF1;EMX2;HSP90AA1;MKKS;EGFL7;PROP1;VWF;UPK3A;ALDH1A3;GREM2;IFNG;HAND2;CHDH;HAND1;CAT;IRF6;MCTP2;RBMS3;HDAC4;NRXN1;PON1;NTN1;PTHLH;DLL3;PRTG;PDLIM3;ERBB2;SH3BP4;NTRK1;TBX1;ANGPT2;EYA1;NOS3;TXNRD1;ZIC4;ZIC5;TBX5;MTOR;TBX3;TSHR;KLF6;MEIS1;TEF;SPRY1;TEK |
| Diabetes Mellitus | 6.431598909111858E-4 | EHMT1;BACH2;LIPC;GJA5;MYC;AKT1;UBASH3A;PRSS8;MNX1;MBNL2;UNC13A;DAPK1;PRKCB;SERPINF1;DAPK3;PRKCA;PASK;ISL1;DKK1;GPR162;CLPS;DDIT3;MYL2;TBPL2;CYP2E1;MAML3;CFB;AIRE;DHH;GATA6;GATA4;GATA3;SLC5A1;SLC30A10;PLA2G7;MIR206;SLC5A4;ABO;C2;CIAO1;C3;HSD11B1;SULT1A1;HSD11B2;CYP2B6;RNF213;TERT;KCNQ1OT1;HCRT;PROM1;HLA-DQA2;HLA-DQA1;S100A10;NGFR;CREBBP;GSTM1;ACE;ABCA4;NFATC3;NFATC2;SLC16A11;GP6;NR0B2;MIR338;PBX1;COL1A1;FAT1;HRC;CYP1A1;NOX5;HAMP;HLA-DRB1;CDKN1C;LAD1;CDKN1A;MIR130B;SETD7;PRKAG2;PRSS27;ADM;GPT;SLC6A2;JPH3;UCN3;ADAMTSL3;TIMP2;TIMP3;SLC18A2;TIMP4;TCF19;MIOX;LYPLA1;ANXA2;ITPK1;PAX4;PAX6;F3;F7;MMP14;ZEB1;SLC7A9;KCNMA1;PXDN;HCLS1;SLC29A1;TKT;HLA-DQB2;HLA-DQB1;GSTP1;ASIP;NR1I2;KCNA3;KCNA5;EGFR;SLC7A2;HLA-DMA;MIR410;HLA-DMB;BGLAP;PLAGL1;NTF3;SPP1;APOL1;P2RY12;AOC3;AOC2;PKLR;STAT3;LETM1;FABP2;GATM;SLMAP;HLA-DPB1;SIGLEC7;HSPA1B;HSPA1A;SEMA5B;CSF3;CSF2;EPO;DEFB1;IRS2;CLU;FGF2;TNF;BEST1;RIMS2;HPGDS;FCGRT;GRAP2;RNF19A;ADORA1;HLA-DOA;MAP3K5;IGFBP1;FBXW8;S100A1;IGFBP3;DIO2;BSCL2;SLC9A3;CLIP2;MTNR1B;RAB38;S100A9;SLC27A2;KCNE1;SHC1;ITPR1;GSTT1;VTCN1;SOCS3;TPO;ZFP57;THEM4;ZNF300;FN3K;SLC14A1;KL;SRD5A2;GFPT1;MAFA;HOPX;GFAP;INS-IGF2;RHEB;CDK2;BCL2;CPE;PLXNB1;SHBG;ALAS1;ONECUT1;ITGB2;FHL2;PIK3CD;PTPN22;HTR2A;FOXM1;PIK3CG;CYR61;HHEX;ITGAX;ITGB7;PHACTR1;SOST;INSL3;ADAMTS9;PGM1;HSP90AA1;TRIM63;GPX1;TRPC6;VWF;SORD;APOA1;TNFRSF1B;NGF;TNFRSF1A;ADORA2A;LCK;NOS1AP;CRH;LCP1;CRP;CX3CR1;HDAC2;PLEKHH2;HDAC3;LRP5;SLC5A11;NOD1;NOD2;NTN1;TTN;SH3BP1;CCL5;P2RY1;KCNJ1;MEST;ANGPT2;ANGPT1;KCNJ11;KCNJ9;TXNRD1;SORL1;MYO16;MTOR;TPCN2;P2RX7;KLF6;PC;SUCLA2;EIF3K;APC;SST;IL2RA;LEP;IL2RB;MGP;PTPN6;P4HB;ZACN;ERRFI1;APP;TNFRSF6B;SPARC;TRAF3IP2;AQP4;IFI30;AQP2;HS6ST1;ICAM1;AQP1;LGALS3;EDNRA;LGALS1;ANPEP;TNFSF10;TNFSF11;SOX9;SLC12A9;ENTPD1;CPT1A;SLC11A1;HLA-B;HLA-C;EBF2;HNF1B;TIMD4;MIF;HNF1A;HLA-A;DHX40;PPP1R3C;TCERG1L;IL23A;KCNQ1;SFRP5;PAPPA;ALPK1;EZR;STX1A;CFTR;GIPR;C2CD4A;C2CD4B;CD1D;PIK3R1;TRPM2;TSPAN8;APOH;ST8SIA4;APOE;FADD;A2M;APOB;PCK1;SPRR2A;NTSR1;STAT5A;EGR1;PTPRN2;TNFSF14;SLC33A1;IGF2;LIF;MIR200B;CIDEC;TH;ARTN;ID3;PEMT;COL9A3;PCSK2;PTPRT;PTPRU;RYR2;CD40;PTPRN;ICA1;PCSK9;BRCA1;KRT20;ESM1;HNF4A;CD38;SOSTDC1;KIF1A;FFAR1;ANKRD2;CD34;CTSD;ACAD8;CD52;AVPR1B;GAA;NME2;SORCS1;MIR21;PROX1;MC3R;NR5A2;ADRB3;IL1B;VWA2;MAPT;SGK1;TLR4;RAPSN;SDCCAG8;TAGLN;CSTA;CST7;REM1;DUOX1;CUX2;CUX1;WIF1;NPY;HSF1;PCNT;NQO1;SORT1;GDF15;GAD1;LILRB1;SORBS1;CYP8B1;GCK;VEGFA;PER1;PER3;NBR1;GHRL;IL17D;IL17C;FOXA2;PNPLA2;NEUROG3;CARS;CD86;KLB;SCARB1;GIMAP5;COMT;ETS1;MPO;CNDP1;IGF1R;NPPA;MIR150;GGT1;TGM2;GHSR;ABCG8;PDIA2;CASR;ABCG5;G6PC;ZGPAT;SLC30A8;LIG4;TSC2;CYBA;CARTPT;ELMO1;CARD16;ABCG1;ABCG2;FBN2;CALCA;ISYNA1;CHAT;TCF7;ALOX15;ALOX12;ADRB2;NME1-NME2;DPP4;ARHGAP22;HRH1;DEAF1;NPC1L1;UBD;CDH23;NLRP3;TAC1;MSC;FGF21;SMAD2;ZFHX3;MGAM;TGFB3;HMGA1;HSPA2;BMP7;ESR1;IGF2R;BMP4;ISG20;SLC2A9;IL6;HOOK2;GDNF;FGF19;GNAS;MYF6;MIR184;GNB3;TGFBI;FGFR4;SNED1;UCN;TREH;ATP2A3;HSPB2;TNFAIP3;SLC2A2;PKD1;CXCL5;GHR;GRM2;CASP8;CA2;BDKRB2;BDKRB1;NOS1;KLF11;MKKS;PDE4D;KLF14;POMGNT1;DCN;GREM1;FGR;MIR126;NDUFS7;IFNG;DDAH2;CHDH;CAT;LTA;MYH9;PON1;ASAP2;THBS1;PTHLH;CRYZ;INS;CBS;PTK2B;CHGA;TCF7L2;MAS1L;NOS2;NOS3;B3GAT1;PDX1;CCK;POMC;MIR145;NPHS2;CAPN10;MIR142;TEK;LPIN2;RAN |
| Depressive Symptoms | 8.154628527691402E-4 | OXTR;GRIK4;RORA;BRCA1;HTR2A;COMT;TNF;SLC6A3;NKPD1;CRHBP;ADAMTS2;CYP2D6;GRAP2;RNF19A;AKT1;CYP1B1;NOS1;NCKAP5;CD34;SLC18A2;LMO3;KRT7;NRG1;OPRM1;KSR2;PDYN;NGF;TAL1;IL1B;CRH;CRP;KCNE1;ABCB1;NRN1;CHRNA5;PON1;HTT;ADRB2;CACNA1C;CRHR1;EGFR;INS;SULT1A1;HSD11B2;NCAM1;APOE;PROM1;DRD4;LGI1;TFAP2B;ACE;NOS3;TXNRD2;HTR1D;HTR1A;OPRK1;SYNJ2;ESR1;P2RX7;PER1;IL6;DAO;PER3;KCNS1;ARTN;LEP;CYP1A1;GNB3;SCLY;KCNK2 |
| Narcolepsy | 8.342895134121237E-4 | NPFFR2;CSF2;CTNND2;CHKB-CPT1B;COMT;FRY;TNF;EDNRA;SALL4;MAGI1;SWAP70;TAP2;MTUS2;TAP1;HLA-A;UNC5D;HLA-F;PDYN;CPT1B;GABRG3;FOXP2;TIAM1;SFRP1;ELMO1;SERPING1;FREM2;ABCG1;NOTCH4;AK5;KALRN;FHIT;SOCS2;GRIN2A;DPP6;NKAIN3;PENK;NXPH1;HCRT;TAC1;LRRC4C;HLA-DQA2;HLA-DQA1;HLA-DRB5;NFATC2;BTNL2;MCC;SDK1;TF;TNFSF4;HLA-DRA;DCHS1;PYROXD2;HLA-DRB1;FGF10;DPP10;RYR2;LAMC2;HTR2A;TRH;JPH1;GRM5;ADAMTSL3;GRM8;PDE4B;TMEM108;DLGAP1;DLGAP2;POLE;SLC18A2;GRID1;ITGA3;TNFRSF1B;SERPINB5;VAV1;VAV2;ANO3;TNFRSF1A;IFNG;PALLD;HLA-DQB2;HLA-DQB1;RBMS3;CRP;CCDC68;CATSPER4;NRXN1;DENND3;NOL4;BARX2;JAKMIP2;REM1;SNX29;EGFLAM;P2RY1;GPC5;CTNNA2;CSMD3;CACNG2;ACSS1;CA10;MOG;MX2;ATP2B2;HCRTR2;P2RX7;P2RX5;RIT2;P2RX2;LEP;P2RX1;HLA-DPB1 |
| Stomach Neoplasms | 8.654155022655956E-4 | RB1;ZNF177;IFITM1;TNFRSF6B;SPARC;PGAP3;ICAM2;BACH2;ICAM1;SOX2;LGALS3;RASSF1;RASSF2;SOX17;SOX18;MYC;AKT1;TNFSF11;HOXA2;PLCE1;SOX9;TNS4;DAPK1;PRKCB;KRT8;MRPS18B;PRKCA;CSNK1E;MIF;RNASE1;RUNX3;DKK3;RUNX1;CLDN4;SFRP1;SPINT1;SLPI;MIR196A2;DDIT3;RBP1;CYP2E1;NEIL1;ALDH7A1;TSPYL5;ABCB1;PCDH10;MIR196B;GATA6;GATA5;TWIST1;GATA4;GATA3;THY1;MST1R;SLC5A5;KLK6;TERT;HLTF;SYMPK;HIVEP3;ST3GAL6;APOE;FADD;PROM1;SPRR2A;B4GALNT2;NGFR;TRAP1;GSTM1;ACE;CADM1;FZD5;XRCC3;IGF2;MRPL28;MIR200B;GRIN2B;NR0B2;GJB2;TH;PPP1R1B;ID3;TNFSF9;FAT4;TDGF1;TP73;HRK;HOTAIR;COL18A1;CDKN1A;BRCA1;KRT20;MED15;WDR46;PTPRF;GLI3;AKAP12;TUBA1C;TIMP2;TIMP3;CCR7;CD34;HTATIP2;ACAD8;ANXA2;DUSP1;IFNGR2;TET1;CACYBP;PAX6;MIR21;F3;POU5F1;SERPINB5;DDB1;MMP14;KAT2A;GPRC5A;NR5A2;CCKBR;IL1B;TFF3;RARB;TFF2;CLDN18;TFF1;LRRC3B;HLA-DQB1;GSTP1;KCNA5;CST7;TRAK1;ASCL2;HOXD10;EGFR;CAMKK2;RXRB;NT5E;EPCAM;WIF1;PLAGL1;ALOX5;HSD17B2;SPP1;MSX1;CTNNA2;TP53I11;XDH;IL32;WWOX;CDX2;GDF15;STAT3;LILRB1;SULF1;MLH1;KLK10;VEGFA;KLK12;KRT18;DLC1;TACC1;FILIP1;GHRL;LIMS2;HSPA1B;UGT1A10;FCGBP;CD81;ZBTB20;RORA;PRDM2;IRS2;MPO;TNF;IGF1R;LOXL2;CNDP2;PYCARD;FGF7;CCND2;CCND1;CDH1;ARHGDIB;PITX1;KCNH1;KCNH2;PDGFRA;POSTN;TPM3;IGFBP3;F2R;WNT5A;SDHB;FOXP1;DHDDS;S100A6;HTRA1;SLC1A2;GSTT1;RASAL1;ADRB2;FHIT;SOCS3;PRR5-ARHGAP8;TAC1;NDUFV1;WNT2;ZFHX3;TFAP2C;SMAD3;CPSF1;TNK2;HTR1A;EPHX3;BMP7;ESR1;ARHGAP24;IGSF11;BMP2;IL6;BCL2;DYNLRB1;XAF1;FGFR3;FGFR2;LGALS3BP;PIWIL1;CRISP2;MIR10B;PSCA;FLT4;PIK3CD;FOXM1;MSLN;PKD1;PIK3CG;CYR61;CXCL5;FLCN;ZNRD1;CASP8;SCRN1;CA2;ZIC1;DNMT3B;BDKRB2;BDKRB1;ADAMTS9;POLE;EGFL7;MMP7;TRPC6;DCC;DNMT3A;APOA1;TNFRSF1A;TGFBR2;GREM1;ALDH1A3;ITGA8;LTA;BIRC5;BLVRB;TRIM59;FBP1;HDAC2;HDAC3;SRC;THBS1;AURKB;SELENBP1;MUC1;MUC2;CCL5;TP53INP1;ERBB2;FYN;FSD1;UGT1A9;MUC4;ATOH1;MUC6;OPCML;NOP56;TCF7L2;URGCP;ANGPT2;ANGPT1;NOS2;NOS3;BNIP3;TXNRD1;B3GAT1;PDX1;TBX3;CHFR;MTOR;KISS1;KLF6;ACTC1;MIR145;APC;LEP;PDCD4;CD9;TEK;VIM |

| | | |
|---|---|---|
| Hypertensive disease | 8.922329592863031E-4 | RB1;SERPINE2;RXFP4;XYLT1;ABAT;MIR425;MYLK;LIPC;CYP2D6;GJA5;HOXA3;AKT1;CYP1B1;PLCE1;PRSS8;DLEU7;CHGB;IER3;PRKCH;PRKCB;DAPK3;HPCAL1;PRKCA;TRHR;MIR202;MYL3;ENPEP;ABCB1;SHMT1;CLCNKB;PDGFB;MTMR9;AGAP1;GATA5;NEDD4L;GATA4;PLA2G7;MIR206;SLC5A4;ABO;C3;HSD11B1;ADAMTS16;HSD11B2;CYP2B6;TERT;NPBWR1;PNPO;NKX2-5;LYN;NGFR;GSTM2;GSTM1;ACE;TSLP;NFATC3;AZU1;NR2F2;EIF2AK4;COL1A1;H19;ZFAT;CYP1A1;MKS1;NOX5;HLA-DRB1;GSTM5;LAD1;CDKN1A;ELN;ADM;GPT;SERPINA6;SLC6A2;SLC6A3;JPH3;C1QTNF1;HEY1;NEFH;DUSP5;GPR39;DUSP1;TESC;TACR3;PAX5;F3;POU5F1;PAX2;APBB1IP;F7;CYP21A2;KCNMA1;IDUA;GSTP1;NR1I2;KCNA3;KCNA5;SLC7A1;EGFR;BGLAP;ALOX5;SPP1;APOL1;CSMD1;AOC3;AOC2;RNLS;F12;STAT3;F11;GUCA2B;NR1H3;FABP2;FABP3;WT1;SLCO4C1;FOSB;CLC;INHA;KLKB1;CCL28;CSF2;EPO;IRS2;FGF1;CLU;FGF2;TNF;BEST1;HPGDS;ADORA1;MACROD2;PITX2;S100A1;IGFBP3;F2R;USP2;DIO2;TAP1;SLC9A3;CACNB2;S100A6;SQSTM1;NOTCH3;GSTT1;REG3G;CYP2C18;SLC9A3R2;CD300LG;KL;HTR1B;ESRRG;HOPX;MAFB;STK24;BCL2;ZNF536;SHBG;SLC26A4;CNTNAP2;ITGB2;PIK3CD;ECE1;ADRA1D;HTR2A;TRH;ADRA1B;PIK3CG;WISP1;CYR61;RNPEP;EMILIN2;PHACTR1;HMGCS2;PLEKHG1;GPX1;TRPC6;TRPC3;VWF;APOA1;KCNAB1;TNFRSF1B;ADRA2C;NGF;TNFRSF1A;ADORA2A;ASPG;ITGA8;CRH;CRP;CX3CR1;NFAT5;RETNLB;PLEKHH2;LRP1;SRC;LRP5;SELENBP1;RFFL;KCNJ1;ANGPT2;KCNJ8;ANGPT1;KCNJ11;TBX5;SORL1;SGCZ;TBX3;MTOR;P2RX7;MFAP5;KLF6;APC;SST;LEP;APOC1;CDO1;CBLN2;APP;OXTR;SPARC;ACSM3;NUDT1;MLC1;AQP4;AQP2;ICAM1;AQP1;LGALS3;EDNRA;LGALS1;EDNRB;ANPEP;TNFSF11;SLC12A6;SEPHS1;SLC12A9;SLC6A19;ENTPD2;SLC6A18;HLA-B;MIF;HNF1A;SLPI;FNDC1;KCNQ1;ZNF831;ALDH7A1;SLC22A3;CASZ1;EPAS1;SLC22A2;WNK4;TWIST1;KLK4;PIK3R1;PRH1;CRHR1;CRHR2;APOH;CMYA5;APOE;KCNN3;FADD;APOB;ARSB;ATP6V0A1;BBS2;STAT5A;EGR1;RABGAP1L;TNFSF14;FOXF1;GREB1;SLC33A1;IGF2;ALOX15B;VNN1;TH;ARTN;ID2;SLC22A8;DDR1;RYR1;PTPRU;RYR2;COL18A1;ROBO4;MIR17;SLC35F3;PCSK9;BRCA1;PCSK6;RYR3;CX3CL1;RPTOR;HNF4A;RAC2;CCR7;CCHCR1;MME;EDN3;INSRR;AGTRAP;IL16;GP1BA;MIR21;RHOD;VAV2;MC3R;ADRB3;IL1B;MIR17HG;SGK1;TLR4;SDCCAG8;TAGLN;TNXB;CYP17A1;GNAI2;DUOX1;NT5E;NPY;XDH;NQO1;GABBR1;SORBS1;GCK;PLEKHA7;VEGFA;F2RL1;GHRL;RAMP1;HDLBP;PLAT;COMT;IGF1R;RGS5;CCND1;NPPA;GGT1;TGM2;PDGFRB;PDGFRA;ABCG8;CASR;POSTN;ABCG5;G6PC;MSX2;OSR1;ADAM11;KCTD1;CYBA;MAT1A;SDHB;SCNN1G;COL4A1;HGS;SCNN1B;SCNN1A;CALCA;ISYNA1;NPR3;ALOX15;CACNA1D;ALOX12;CACNA1C;ADRB2;FBLN2;CACNA1H;DPP4;GRK5;GRK4;CDH23;NLRP3;TAC3;KLHL29;SMAD3;TGFB3;RARRES2;SMAD9;SMAD6;ESR1;PHOX2A;BMP4;SLC2A9;SDK1;PYY;BMP2;IL6;PROC;CPS1;FES;TLX2;GNAS;GNB3;FGFR4;FGFR2;UCN;THRA;ATP2A3;SLC2A2;LAMC2;OXT;DBH;SLC2A5;MIR96;PKD1;CXCL5;GHR;BDKRB2;BDKRB1;NOS1;PSTK;MKKS;MYOCD;MMP7;MIR23A;PDE4D;ARID1B;SSTR4;DCN;TGFBR2;ACTA2;TGFBR3;IFNG;DDAH2;CHDH;CAT;CHI3L1;CDH13;MYH9;CDH18;LTF;MYH7;PON1;ADD3;THBS1;THBS4;CRYZ;INS;MUC1;SV2C;CBS;GNA12;PTK2B;CHGA;TCF7L2;NOS2;GALNT2;NOS3;B3GAT1;ATP2B2;SOD3;TSHR;POMC;ACTC1;NPHS1;CAPN10;SARDH;TEK;LPIN1 |
| Renal Cell Carcinoma | 9.42819141420779E-4 | MIR429;ABCA13;CYP2D6;MYC;AKT1;CYP1B1;PRSS8;PGLYRP1;PRKCH;UNC5B;DAPK1;UNC5C;PRKCA;PDZK1;UNC5D;DKK1;DKK3;DKK4;SPINT1;DAAM2;MYL2;CLDN8;RBP1;JTB;IDO2;MYL9;GNE;MEG3;ENPEP;ABCB1;CRABP1;HPN;PDGFA;GATA5;GATA3;PRKCZ;C3;LDHC;HSD11B2;ABR;LDHB;TERT;PDGFD;PROM1;S100A10;ZSCAN18;PROM2;GSTM2;ST14;ACD;GSTM1;FZD5;NR0B2;FAT1;CYP1A1;HLA-DRB1;ACHE;CDKN1A;HIF3A;SLC6A3;BBC3;ATIC;HEY1;TIMP2;TIMP3;TIMP4;RING1;ANXA2;DUSP1;PAX2;MMP14;ZEB2;ZEB1;MMP16;PAX8;KCNMA1;RARB;ANGPTL4;CRYAB;KANK2;KANK1;OCA2;GRP;AHNAK;GSTP1;COL11A2;NR1I2;EGFR;BGLAP;EPCAM;PRAME;CSMD3;APOL1;WWOX;EGLN2;F12;STAT3;DAB2IP;PSMB8;DEF6;PSMB9;MIR24-2;GATM;WT1;FABP7;NFIC;INHA;HSPA1B;HSPA1A;CSF2;EPO;FAM107A;DEFB1;PLOD2;IRS2;APEH;CDH9;CLU;FGF2;TNF;CDH8;CDH6;PYCARD;HPGDS;CDH3;CDH1;GRAP2;RNF19A;MIR1236;HLA-DOA;SLC16A3;S100A1;IGFBP3;AKR1E2;F2R;TAP2;TAP1;MITF;HIC1;MTNR1A;RBMXL1;RAB37;ZFP42;FSCN1;S100A9;SQSTM1;PVALB;MAD1L1;CSF1R;NOTCH3;ITPR1;CUL1;ITPR2;GSTT1;VTCN1;FHIT;ACAT2;RAP1GAP;SLC9A3R2;SOCS3;SMYD2;TPPP;AP1M2;KL;TNK2;TNFRSF10A;TMPRSS13;COPS5;RHEB;CDK2;BCL2;ZNF536;PLXNB1;XAF1;PIWIL1;BNC1;PIK3CD;ECE1;FOXM1;PIK3CG;CYR61;SYNE2;MB;RNPEP;DNMT3B;HGFAC;VWF;DNMT3A;SLC25A18;ESRP1;BIRC5;ANG;BIRC7;CRP;SRC;NTN1;SH3BP2;TP53INP1;IGF2BP1;MAL2;ACSS1;EIF4E;NTRK1;SPAG9;MARCKSL1;RCVRN;ANGPT2;ANGPT1;PCDH8;KCNIP4;MTOR;NELL1;APC;SST;IL2RA;MGP;PDCD4;PTPN6;CDO1;TNFRSF6B;SPARC;DNHD1;ICAM1;AQP1;LGALS3;RASSF1;EDNRA;EDNRB;RASSF5;TNFSF10;TNFSF11;DPF3;SLC12A9;SRRM2;SLC34A2;EGOT;MYO7B;HLA-B;KRT8;HLA-C;KRT7;HNF1B;DLEC1;HNF1A;HLA-A;OVOL1;RUNX3;HLA-G;SFRP4;TIAM1;SFRP1;SFRP2;IFI27;SLPI;SFRP5;MT1G;MT1H;EZR;GRB7;SLC22A3;MACF1;SPTBN4;EPAS1;HEPACAM;TWIST1;KLK4;CD1D;MST1R;PIK3R1;KLK7;NEU1;HIVEP3;APOE;FADD;BBS2;EGR1;INTS1;IGF2;LIF;MIR200A;MIR200B;BST2;GRB2;TP73;APOBEC3B;MIR129-2;RYR1;HOTAIR;PTPRU;ROBO3;COL18A1;CD40;NRP2;ROBO4;MIR17;BRCA1;CD3E;KRT20;AKAP13;VMO1;HNF4A;CHMP1B;CD38;SOSTDC1;ANKRD2;CD34;CTSD;MME;TUBB;MST1;INSRR;NME2;TFEB;AMPD1;CACYBP;MIR21;KRT32;TYK2;TLR1;CDCP1;IL1B;MIR17HG;FAM20C;DERL3;NEDD9;CAPG;PAK1;CUX1;WIF1;NPY;CD74;NQO1;GAD1;TMEM176B;SUSD2;KLHL1;LILRB1;DMRT3;MLH1;KLK10;KLK11;VEGFA;MAD2L2;NLRP12;RIT2;DLC1;QPCT;NBR1;F2RL1;SCARB1;NDUFA13;KLHL35;COMT;IGF1R;LOXL2;HOXA11;MIR149;RGS5;CCND1;MIR150;TGM5;PPFIA4;TGM2;PDGFRB;EOMES;GHSR;CASR;G6PC;LIG1;MIR27A;TSC2;SDHB;DHDDS;SCNN1B;CARD11;ABCG2;FBN2;DIABLO;HOXB13;DPP4;PODXL;TMEM127;ALDH3B1;NLRP2;PRODH;MARK2;TFAP2A;SMAD2;TFAP2B;WNT3A;HMGA1;ASPSCR1;INHBA;HSPA2;SSH2;ESR1;LRP1B;ARHGAP24;SYT6;BMP4;ISG20;PYY;BMP2;IL6;GNAS;MIR183;FGFR4;FGFR3;FGFR2;CRISP2;TNFAIP8;THRB;MIR10A;HSPB7;SLC2A2;MIR96;GRM1;FLCN;CASP8;CA2;SIX2;MCF2L;CASP1;NOS1;MIR193A;MMP7;MIR23A;DCC;UBE2QL1;DCN;TAPBP;TGFBR2;ACTA2;TGFBR3;MIR126;CCNA1;IFNG;CDH11;IRF8;CDH13;CDH15;HAGH;SHANK1;RBMS3;THBS1;PTHLH;AURKB;CRYZ;MUC1;WNT11;CNR2;GNG4;KDELC2;MIR495;MIR373;ERBB2;MIR372;PTK2B;MUC4;RBM38;TCF7L2;NOS2;BNIP3;B3GAT1;ZNF804A;DLK1;KISS1;POMC;PNKD;TEC;MIR145;SARDH;TEK;VIM;RAN |
| Visual seizure | 0.0010161895814339492 | ACHE;CNTNAP2;FAAH;NRP2;CHRM1;EPO;GRIK5;ABAT;GRIK1;OXT;TRH;SLC6A1;ADRA1B;SLC6A2;CLU;FGF2;TNF;CCND1;GRM8;CASP1;PDE4A;NOS1;MAP3K5;GAMT;BRD1;SLC38A1;DUSP1;TACR3;TSC2;SLC6A11;NRG1;OPRM1;PDYN;NGF;TNFRSF1B;DKK1;SYN2;DUSP6;TNFRSF1A;ADORA2A;IFNG;MAP1B;IL1B;KCNQ2;CAT;CRH;PVALB;HDAC2;ABCB1;CHRNA5;FOXG1;KCNA1;CHAT;PDGFB;AIF1;INS;ANKH;NPY;CCL5;SPP1;SLC17A7;APOE;WNT2;SLC38A2;EGR1;NGFR;NQO1;GABBR1;MARCKSL1;SLC12A5;KCNJ11;NOS2;GABRA5;HTR1A;HTR1B;OPRK1;CCK;LETM1;SLC8A3;POMC;IL6;MARCKS;SST;GJB6;LEP;ALPL;GNB3;GHRL;PAM;HCN2;FGFR1;HCN1 |
| Kidney Diseases | 0.001022787673490833 | SPARC;TP53RK;XYLT2;XYLT1;AQP2;AQP1;ICAM1;EDNRA;LIPC;EDNRB;AKT1;SYNPO;TRIM21;PRKCB;SERPINF1;HNF1B;HLA-A;MIF;HNF1A;DKK3;SFRP1;IFI27;DDIT3;KCNQ1;CYP2E1;EZR;HPD;CFTR;ABCB1;SLC22A2;CLCNKB;KLK4;GATA3;THY1;HSD11B2;CYP2B6;TERT;KCNQ1OT1;PDGFD;APOE;APOB;HLA-DQA1;NGFR;EGR1;CREBBP;ACE;GSTM1;TREX1;SLC33A1;MIR200A;GP6;FLI1;GADD45G;TNFSF4;FAT1;MKS1;NOX5;SLC22A8;HAMP;HLA-DRB1;CDKN1C;CDKN1A;NRP2;ADM;KRT20;HNF4A;STAB1;TIMP3;CD38;SOSTDC1;MIR192;CTSD;CC2D2A;HLA-DPA1;IFT140;ELOVL5;TESC;SORCS1;MIR21;F3;PAX2;TLR1;F7;MMP14;INF2;SLC7A9;IL1B;TFF3;SGK1;TLR4;TAGLN;GSTP1;SPESP1;HOXD11;EGFR;TMEM67;SPP1;STAT4;STX2;APOL1;XDH;BRD4;AOC3;NQO1;RNLS;GAD1;STAT3;PSMB8;GCK;HIPK2;VEGFA;FABP1;LETM1;FABP2;GATM;WT1;CLC;FMOD;CCL28;HLA-DPB2;CARS;CSF3;CSF1;EPO;IRS2;CLU;FGF2;TNF;MPO;CNDP1;HOXA11;CNDP2;PYCARD;SALL1;CCND1;FCGRT;NPPA;MIR150;ABCG8;CASR;G6PC;IGFBP3;TSC2;CYBA;ADAM19;SCNN1G;COL4A2;COL4A1;SCNN1B;SCNN1A;ELMO1;NOTCH3;PROZ;GSTT1;RASAL1;VTCN1;ADRB2;DPP4;SOCS3;CALD1;PODXL;UBD;NLRP3;ZNF423;PRODH;TFAP2A;KL;RAB4A;SMAD3;GFPT1;BMP7;ESR1;ARHGAP24;SMAD7;IL6;MAFB;DAO;BCL2;ALPL;GNB3;TGFBI;TREH;ITGB2;SLC2A2;DBH;ADRA1B;TREM1;PKD1;GHR;CA2;PSD4;SIX2;CASP1;BDKRB1;HORMAD2;MKKS;TRPC6;MMP7;DST;VWF;SORD;APOA1;DDOST;DCN;SSTR4;TGFBR2;ACTA2;IFNG;PALLD;DDAH2;TMEM216;CAT;CDH13;MYH9;IRF5;CRP;CX3CR1;PON1;LCAT;THBS1;INS;MUC1;CCL5;MXI1;PTK2B;NCAM1;TBX1;TCF7L2;ANGPT2;ANGPT1;NOS2;NOS3;TPRKB;MTOR;TBX3;MYO16;POMC;NPHS1;NPHS2;LEP;MGP;AQP11 |
| Neuroendocrine Tumors | 0.0010826879007088941 | SCT;AQP4;TNF;IGF1R;RASSF1;CCND1;MYC;STMN1;AKT1;CHGB;IGFBP1;GHSR;PDGFRA;DAXX;CARTPT;SDHB;CLDN4;SFRP1;MYCN;SLPI;PGR;MEG3;NOTCH3;CALCA;GIPR;CRABP1;MST1R;SLC1A5;FHIT;ABO;SLC5A5;TERT;MAP2;TMEM127;TRPM8;TAC1;NTSR1;NTSR2;NTNG1;IGF2;HMGA1;FOXN3;ESR1;TMPRSS13;FLI1;DCLK1;ATP4A;IL6;GDNF;BCL2;GNAS;CPE;PIK3CD;OXT;HTR2A;FOXM1;SLC6A2;PIK3CG;NEFM;SLC18A2;HSP90AA1;MMP7;MST1;GFRA1;PAX5;SSTR1;NGF;SSTR5;SLC7A5;PAX8;IL1B;CRH;RARB;CD302;GRP;ADCY2;ASCL1;EGFR;PTHLH;INS;PAK1;MUC1;WIF1;NPY;ERBB2;GNA12;PTK2B;NCAM1;MUC4;CHGA;NTRK1;ANGPT2;CDX2;MGMT;B3GAT1;PDX1;CCK;MLH1;POU4F1;MTOR;DLK1;VEGFA;POMC;KLF6;APC;SST;GHRL;INSM1;TEK;LGR6 |
| Secondary malignant neoplasm of lymph node | 0.001197424892364632 | RB1;CYFIP1;TRIO;SERPINE2;MIR429;ELK4;MIR675;SOX17;TMEM88;SOX18;MYC;DPYSL3;STMN1;AKT1;CYP1B1;PLCE1;SERPINF1;DAPK3;PRKCA;ISL1;DKK1;DKK3;SYTL2;CLDN4;SPINT1;BIN1;HOXB7;ABCB1;CTBP2;DHH;MIR196B;CRABP1;GATA6;PDGFA;GATA5;GATA3;THY1;GATA2;LYPD5;MIR206;SLC5A5;SULT1A1;CYB5R2;LDHB;CYP2B6;TERT;SFN;PROM1;HOXC6;ST14;GSTM1;ACE;CADM1;NR2F2;NR1D1;MIR338;FLI1;GJB2;CPEB1;FAT4;HOXD9;CDKN1C;LAD1;CDKN1A;LPCAT1;TCF21;ADM;MED15;MYOM2;TIMP2;NEFL;TIMP3;GPR39;ANXA2;RIPK4;PAX6;F3;POU5F1;PAX2;MMP14;ZEB2;ZEB1;MMP16;RARA;RARB;ANGPTL4;GAS5;GRP;COL11A1;GSTP1;CRCT1;NR1I2;MIR409;KCNA5;HOXD11;FSTL1;EGFR;MIR410;EPCAM;SPP1;CYP2F1;WWOX;MGMT;STAT3;SULF1;FABP5;FABP6;ZNF217;SEMA5A;CSF1;EPO;CTNND2;RAB3D;F13A1;MIR146B;DEFB1;MSI1;CLU;FGF2;TNF;FGF3;CDH6;FGF7;CDH1;LZTS1;KCNH2;S100A2;S100A1;IGFBP3;UGT2B15;F2R;MITF;FOXP2;FOXP1;MTNR1A;S100A6;FSCN1;PGR;ERG;S100A9;MAD1L1;NOTCH3;SEMA7A;PFKFB3;DLX4;TACSTD2;VTCN1;FHIT;SOCS3;SMYD2;ESRRA;KL;TFPI2;TRAPPC9;TMPRSS13;MFHAS1;DAB2;CDK2;BCL2;CPE;CLPTM1L;SHBG;PIWIL2;PIWIL1;PSCA;ITGB4;FLT4;ITGB2;WISP3;GADL1;PIK3CD;FOXM1;ITGAL;PIK3CG;CYR61;DNMT3B;INSL3;ADAMTS9;SGCG;HSP90AA1;ITGA4;DNMT3A;CTDSPL;MIR7-2;TNFRSF1B;NGF;TNFRSF1A;PTP4A1;BIRC5;ITGA6;ANG;CRP;HDAC3;UHRF1;SEMA3B;MIR148A;DNAJB6;FSD1;NTRK1;TBX1;ANGPT2;MYO10;ANGPT1;ANKRD30A;STYK1;MTOR;TPCN2;GZMM;KLF6;APC;LEP;MGP;PDCD4;OCIAD2;BCAR1;FAAH;OXTR;TNFRSF6B;SPARC;ICAM1;LGALS4;SOX2;TP53AIP1;LGALS3;RASSF1;FRAT1;LGALS1;TNFSF10;LGALS9;SEC62;TNS4;TP63;SLC12A9;EPHA5;SRRM2;EPHA6;MIR9-1;KRT7;DLEC1;KRT5;RUNX3;HLA-G;RUNX2;RUNX1;SFRP1;SFRP2;IFI27;SLPI;SFRP5;MT1G;EPHA1;RIN1;EZR;GRB7;SLC22A3;EPHA10;EPAS1;SLC22A2;TWIST2;TWIST1;NUAK1;SEC14L1;STC2;NEU1;HLTF;APOD;APOE;FADD;EGR1;TRAP1;IGF2;MIR200B;PTK7;PPP1R1B;GRB2;TDGF1;TP73;HOTAIR;ROBO3;COL18A1;CD40;NRP2;DOC2A;BRCA1;KRT20;CX3CL1;AKAP12;CCR7;CCR6;MIR192;CD34;CTSD;CLIC1;HTATIP2;PSD;FOXD2;ZHX2;PRMT7;MME;MST1;NCOA4;NME2;MIR21;PROX1;TYK2;ANO1;NR5A2;CTTN;CCKBR;IL1B;TFF3;B3GNT3;TFF1;CD47;TLR4;GUCY2C;TAGLN;TMPRSS4;NEDD9;TMPRSS2;CAPG;CYP17A1;PAK1;NT5E;NPY;GPC5;IL32;CD74;NQO1;CDX2;KLK13;KLK15;LILRB1;MLH1;KLK10;GRHL2;KLK11;VEGFA;KLK12;PER1;KRT18;DLC1;IL17D;SIPA1;RTKN;IHH;COMT;ETS1;IGF1R;LOXL2;MIR149;HOXA10;CCND1;PEBP4;MIR150;TGM3;PDGFRB;TMOD1;PDGFRA;MIR27A;TSC2;SDHB;BRMS1;MYCN;ABCG2;CALCA;ALOX15;HTRA1;ADRB2;HOXB13;NME1-NME2;CALD1;UBD;WNT2;SMAD3;MLKL;HMGA1;POU2F2;ESR1;ARHGAP24;NR4A2;BMP2;IL6;CASS4;MIR184;MIR183;FGFR4;FGFR3;FGFR2;FGFR1;TNFAIP8;MIR10B;MIR10A;ATP2A3;HSPB2;CHD5;MIR96;GHR;UPK1A;ZIC1;CASP1;FLOT1;MIR193A;MMP7;DCC;AXIN1;TGFBR2;KLF17;CCNA1;ELF3;DPPA2;CDH11;CDH13;MYH9;THOP1;LTF;RBMS3;RGS16;THBS1;PTHLH;AURKB;MUC1;MUC2;WNT11;MIR494;MIR373;MXI1;ERBB2;GNA12;PTK2B;SNCG;NCAM1;MUC4;MUC6;CHGA;RBM38;LGI1;KDM4B;CMTM3;USP22;LIMK1;CHFR;TSHR;KISS1;PNKD;MIR145;SARDH;CD9;VIM |
| Colitis | 0.001211279742206426 | APP;FAAH;ICAM1;ELK4;LGALS2;LGALS1;ANPEP;TNFSF10;PROK2;IER3;ENTPD1;UNC5B;KRT8;PDZK1;MIF;RUNX3;RUNX2;SFRP1;SFRP2;SLPI;BIN1;IL23A;EZR;CFTR;ABCB1;THY1;CD1D;PIK3R1;CRHR1;CRHR2;FADD;TRPM8;ELANE;NTSR1;STAT5A;ST14;TSLP;TNFSF14;LRBA;NFATC2;IL22RA1;FERMT1;VNN1;TNFSF4;TOLLIP;CYP1A1;TNFSF8;ANP32B;HAMP;HLA-DRB1;PIGR;CD40;PTPRS;SLC44A4;LRRC4;ADM;BRCA1;PTPRH;UCN2;HNF4A;RSPO2;CCR9;TIMP3;RSPO3;CCR7;FFAR2;CCR6;JAK3;APC2;ACAD8;CBR1;RIPK3;TPSG1;TACR2;PRSS16;IL16;MIR21;MIR19A;F3;CBFA2T3;TLR1;SLC7A5;COL2A1;ADRB3;CCKBR;IL1B;RARA;TFF3;CLDN18;TFF1;SGK1;TLR4;GUCY2C;COL11A2;NR1I2;FSTL1;EGFR;GNAI2;NT5E;PAK1;CUX1;ALOX5;SPP1;CD55;CCL25;NQO1;EGLN2;IL10RB;MGMT;STAT3;MLH1;GUCA2A;VEGFA;NLRP12;F2RL1;GHRL;IL17D;IL17C;CD86;CSF3;CSF2;CD83;EPO;DEFB1;RORA;PRDM1;GIMAP5;TNF;MPO;ETS2;RIMS2;FGF7;HPGDS;DMBT1;GRAP2;RNF19A;CAPN2;TNFRSF8;MIR150;TNFRSF4;MAP3K5;CASR;S100A1;F2R;DHDDS;SLC9A3;MTNR1B;PRTN3;S100A9;DSC3;CSF1R;CALCA;NPR3;FUT3;SLC9A3R2;SOCS3;NLRP3;TAC1;KL;SMAD3;RARRES2;TFPI2;HPS4;BTNL2;BMP7;SMAD7;ISG20;IL6;STK24;FGF19;BCL2;SKIV2L;UCN;CIITA;TNFAIP8;PIK3CD;PTPN22;TREM1;PIK3CG;WISP1;CXCL5;SYNE1;CASP8;ITGAX;CASP1;MADCAM1;LBP;MIR193A;HSP90AA1;GPX1;TRIM63;MMP7;ITGA4;GLP2R;TNFRSF1B;TNFRSF1A;TGFBR2;MYH2;ELF3;IFNG;MEP1A;CAT;LTA;CHI3L1;BPI;B4GALT7;CX3CR1;RETNLB;SRC;PON1;NOD1;NOD2;NTN1;MUC1;IL1RL2;MUC2;CNR2;CEACAM20;NTRK1;CHGA;NOS2;NOS3;LIMK2;P2RX7;PC;APC;IL2RA;LEP;PDCD4;TEK;VIM |
| Convulsions | 0.0013010984770893217 | ACHE;FAAH;NRP2;CHRM1;CSF2;ABAT;GRIK1;LAMC2;OXT;TRH;ADRA1B;SLC6A1;CLU;FGF2;SLC6A2;NOS1;GAMT;TSC2;OPRM1;PDYN;NGF;SYN2;CNKSR1;ADORA2A;IL1B;KCNQ2;CAT;CRH;PVALB;ABCB1;CHRNA5;FOXG1;CHAT;KCNA3;INS;NPY;SLC17A7;APOE;NGFR;SLC12A5;KCNJ11;NOS2;GABRA5;HTR1A;HTR1B;OPRK1;CCK;LETM1;POMC;SLC8A3;IL6;SST;LEP;ALPL;GNB3;GHRL;PAM;HCN1 |
| Pulmonary Edema | 0.0013147291282200534 | SGMS1;EPAS1;NR1I2;PRKAG2;TWIST1;ADRB2;MPO;TNF;AQP1;FGF7;HPGDS;NT5E;ALOX5;ITGB6;MARK2;ENTPD1;ACE;VWF;NOS2;NOS3;F2R;SARS;NRG1;PARP9;MUC5B;VEGFA;ISG20;CLDN4;IL6;IFNG;DDIT3;IL2RA;CAT;SCNN1A;BCL2;CPE;TEK;SLC29A1;SGK1;CFTR;FGF10;HSPA1A |
| Essential Hypertension | 0.001327586275418126 | ACSM3;XYLT2;C4BPA;C4BPB;MPO;TNF;MYLK;IGF1R;LGALS3;RGS5;EDNRA;HPGDS;GJA5;NPPA;ADORA1;TNFRSF4;KCNH2;SLC6A18;OSR1;HLA-B;CYBA;HLA-A;TRHR;SCNN1A;S100A6;PTGFR;CALCA;ISYNA1;NPR3;CLCNKB;WNK4;NEDD4L;GATA5;ALOX12;CACNA1D;GSTT1;ADRB2;CACNA1C;ABO;SULT1A1;HSD11B2;GRK4;ST8SIA4;NLRP3;APOE;APOB;KL;GSTM2;ACE;GSTM1;SLC33A1;TFPI2;IL6;TH;STK24;GNAS;GNB3;NOX5;HLA-DRB1;ELN;ADM;ECE1;DBH;HTR2A;ECE2;ADRA1B;SYNE1;BDKRB2;BDKRB1;NOS1;NEFH;GPX1;CAMK1D;TRPC3;TESC;TNFRSF1B;SSTR4;TGFBR2;MIR126;ADRB3;ADORA2A;CAT;LTA;SGK1;CRP;KCNA3;SLC7A1;CYP17A1;GNAI2;INS;NPY;SPP1;PTK2B;APOL1;CHGA;ANGPT2;HSPA1L;NOS2;GDF15;RNLS;NOS3;TXNRD1;STAT3;GUCA2B;ATP2B2;SOD3;GCK;VEGFA;POMC;FABP3;CAPN10;LEP;FKBP1B;HLA-DPB1;KLKB1;RAMP1;HSPA1B |
| Generalized seizures | 0.001327586275418126 | ACHE;FAAH;NRP2;CHRM1;CTNND2;ABAT;GRIK1;OXT;TRH;ADRA1B;SLC6A1;CLU;FGF2;SLC6A2;NOS1;GAMT;TSC2;OPRM1;PDYN;NGF;SYN2;ADORA2A;SCN8A;IL1B;KCNQ2;CAT;CRH;PVALB;ABCB1;CHRNA5;FOXG1;CHAT;CLCN1;INS;NPY;SLC17A7;APOE;LGI1;NGFR;SLC12A5;KCNJ11;NOS2;GABRA5;KCNB1;HTR1A;HTR1B;OPRK1;CCK;LETM1;POMC;SLC8A3;IL6;SST;LEP;ALPL;GNB3;CNTN2;GHRL;FGFR3;PAM;HCN1 |
| Abnormal behavior | 0.001327586275418126 | CHRM2;APP;FAAH;OXTR;SLC46A1;CSF2;CTNND2;DIXDC1;ZBTB20;ADARB1;COMT;DAOA;SYNGAP1;CYP2D6;DPYSL2;STMN1;AKT1;PIP4K2A;SOX5;IER3;MAGI1;CPT1A;S100A1;SLC6A15;LMO3;TSC2;PRKCA;CACNA2D4;PDYN;FEZ1;SCN8A;ANKK1;ALDH7A1;SQSTM1;STX1A;MMADHC;PVALB;SLC22A3;MAGEL2;CHRNA2;CHRNA5;GPR85;ADRB2;CACNA1C;KLK8;CRHR1;NPAS3;RAI1;GRIN2A;CYP2B6;MIR330;DEAF1;NDN;CAMTA1;APOD;APOE;HCRT;TAC1;DRD4;EGR1;ACE;EGR3;ST8SIA2;TREX1;HTR1A;HTR1B;GRIN2B;ESR1;OPN5;DCLK1;COL1A1;IL6;TH;GDNF;ARTN;BCL2;GTF2IRD1;DEGS2;SIM1;FGFR2;UCN;CNTNAP2;CD40;MIR10A;MEGF10;LAMC2;OXT;BRCA1;GRIK2;DBH;HTR2A;NR2E1;TREM1;SLC6A3;SGCE;GRM2;GRM5;ZIC2;ZIC1;PDE4B;NOS1;DLGAP3;PSD;SRGN;GPX1;NRG1;HTR3A;OPRM1;HTR3B;OLIG2;RGMA;TNFRSF1A;ALDH5A1;IL1B;PEX5;NOS1AP;CRH;MAPT;ARHGEF7;SHANK2;CX3CR1;HDAC4;DDC;ITIH3;DCTN1;NRXN1;PRICKLE2;HTT;INS;NSD1;NTF3;NCAM1;PCNT;DISC1;CSMD1;TBX1;ASCC1;GAD1;ZNF804A;OPRK1;PEX10;CCK;HTR5A;BAIAP2;PEX14;TBX2;SUCLA2;PPP2R2B;LEP;HCN1 |
| Colorectal Neoplasms | 0.0013289778545484676 | RB1;TNFRSF6B;SPARC;SERPINE2;ICAM5;ABCA13;FGFRL1;ICAM1;SOX2;LGALS3;RASSF1;RASSF2;SOX17;ZMIZ1;DPYSL2;RASSF5;MYC;STMN1;TNFSF10;AKT1;CYP1B1;SOX9;SOX7;TP63;SLC12A9;EPHA4;ARL11;UNC5A;PRKCB;SERPINF1;UNC5C;MIF;HNF1A;HLA-A;DKK1;RUNX3;HLA-G;DKK3;DKK4;SFRP4;TIAM1;CLDN4;SLC5A8;SFRP1;SFRP2;SPINT1;WDR81;TCERG1L;CLDN8;SFRP5;CD109;SPATA13;CYP2E1;EPHA1;RIN1;EZR;NEIL1;CFTR;SLC22A3;ABCB1;PCDH10;CTBP1;EIF5A2;PDGFB;GATA5;TWIST1;GATA4;LYPD3;MST1R;PIK3R1;SLC5A5;KLK6;HSD11B1;SULT1A1;HSD11B2;GRIN2A;TERT;KCNQ1OT1;SFN;APOE;PROM1;RREB1;NTSR1;ATP6V0A1;NTNG1;TRAP1;CREBBP;ST14;GSTM1;ACE;FUCA1;FZD7;ABCA4;XRCC3;IGF2;NFATC2;NR2F2;ALOX15B;DCLK1;NR2F6;STAG3;FXYD3;PTK7;GIPC1;GIPC2;FAT1;CYP1A1;SSPN;TNFSF9;HAMP;HOXD9;TDGF1;LGMN;TP73;ACHE;PTPRU;NRP2;CDKN1A;ROBO4;ARHGAP8;MIR130B;ADM;BRCA1;PTPRH;KRT20;BBC3;ABCC13;LFNG;AKAP10;BOLL;ESM1;JPH3;HNF4A;ADAMTSL3;TIMP2;TIMP3;CCR6;TK1;CLIC1;TIMP4;APC2;ZHX2;MME;NME2;MIR21;F3;SLC39A14;PAX2;SERPINB5;NCOR2;MMP14;NR5A2;ZEB1;RAD51C;CTTN;PAX8;IL1B;TFF3;ARHGEF4;RARB;TFF1;ARHGEF2;LRRC3B;TLR4;GUCY2C;COL11A1;GSTP1;NED D9;TMPRSS2;EGFR;PAK1;MIR410;EPCAM;WIF1;ALOX5;HSF1;FLNC;EXTL3;CSMD1;CD55;SPDEF;NQO1;PKLR;CDX2;MGMT;CDX1;GDF15;STAT3;MLH1;KLK10;GUCA2A;VEGFA;SKI;MAD2L2;PER1;KRT18;FABP6;DLC1;LGR6;CSF2;CSF1;MPG;MIR146B;ETS1;MPO;CLU;FGF2;TNF;IGF1R;LOXL2;PYCARD;CDH5;HPGDS;FGF7;CCND2;CDH3;CCND1;CDH1;ADORA3;SALL4;ZNF569;TNFRSF4;KCNH1;KCNH2;PDGFRB;IGFBP1;PDGFRA;CASR;POSTN;KDM2A;IGFBP3;TAP1;EPDR1;HIC1;DHDDS;SCG5;DPEP1;ZNF436;MASP2;PGR;S100A9;ZNF432;ABCG1;DSC3;ABCG2;FBN2;NOTCH3;DIABLO;TCF7;ALOX15;TACSTD2;GSTT1;RASAL1;ALOX12;HOXB13;FHIT;FBLN2;ZFP36L2;CDH22;FGF20;PRODH;WNT2;TFAP2A;SMAD2;CBX5;TFPI2;EPHX1;TNFRSF10A;MCC;BMP7;ESR1;ARHGAP24;SMAD7;BMP4;NR4A2;BMP2;IL6;NAT1;UBXN11;FES;GNAS;GNB4;BCL2;CPE;DYNLRB1;XAF1;FGFR3;FGFR2;FGFR1;PRR5;KCNK9;FLT4;ITGB2;PIK3CD;CHD5;LAMC2;FOXM1;PKD1;PIK3CG;HAPLN1;CXCL5;SYNE1;GHR;FLCN;CASP8;DNMT3B;CASP1;POLE;PGM1;HSP90AA1;MMP7;GRID1;DCC;DNMT3A;SORD;AXIN1;TNFRSF1B;ADAMTS20;TAPBP;TGFBR2;GREM1;AIM2;IFNG;RASA3;MEP1A;HAND1;BIRC5;CDH13;ANG;EVL;LCP1;BCAT1;CRP;CX3CR1;HDAC2;HDAC3;LRP1;SRC;LAMA1;NXN;NOD2;LTC4S;NTN1;THBS1;SELENBP1;MUC1;MUC2;CCL5;GNA11;ERBB2;SLIT1;SNCG;FSD1;UGT1A7;UGT1A6;PPP1R14A;TCF7L2;RPH3AL;KDM4B;LAMB3;NOS2;CMTM3;NOS3;BNIP3;TXNRD1;EYA4;SEMA4C;PDX1;CABLES1;TBX3;CHFR;MTOR;KISS1;KLF6;FMNL1;MIR145;APC;CAPN10;COL7A1;LEP;PDCD4;CD9;CDK2AP1;VIM |
| Diaphragmatic Hernia | 0.001426134855692423 | THRB;THRA;LRP1;ELN;GATA6;GATA4;FGF2;TNF;ICAM1;EDNRA;FGF7;EDNRB;HLX;NPPA;TIMP2;SLIT3;SOX7;HOXA5;STRA6;PDGFRA;ACE;NOS2;KCNB1;NOS3;IGF2;PAX3;NR2F2;BMP4;IL6;WT1;KCNMA1;FGF18;HOXB3;RARB;CFTR;FOXA2 |
| Seizures, Focal | 0.001426134855692423 | ACHE;FAAH;NRP2;CHRM1;CTNND2;ABAT;GRIK1;OXT;TRH;ADRA1B;SLC6A1;CLU;FGF2;SLC6A2;NOS1;GAMT;KCND3;TSC2;OPRM1;PDYN;NGF;SYN2;CYP26C1;ADORA2A;SCN8A;IL1B;KCNQ2;CAT;CRH;PVALB;ABCB1;CHRNA5;FOXG1;CHAT;INS;GRIN2A;NPY;SLC17A7;APOE;LGI1;NGFR;DYNC1H1;SLC12A5;KCNJ11;NOS2;GABRA5;HTR1A;HTR1B;OPRK1;CCK;LETM1;POMC;SLC8A3;IL6;SST;LEP;ALPL;GNB3;CNTN2;GHRL;PAM;HCN1 |

| | | |
|---|---|---|
| Endometriosis | 0.0015513942106665337 | RB1;ERRFI1;COL12A1;AQP2;ICAM1;LGALS4;SOX2;LGALS1;RASSF2;CREB3L1;TNFSF10;AKT1;CYP1B1;LGALS9;SERPINF1;ACTN1;HNF1B;MIF;DKK1;HLA-G;RUNX1;SFRP4;CLDN4;SFRP1;IFI27;SLPI;CRISPLD2;TBPL2;PAPPA;RBP1;HOXB2;ASTN2;EZR;CFD;PTGFR;MIR196B;GATA6;PDGFA;NEDD4L;MST1R;GATA2;CRHR1;C2;CRHR2;C3;HSD11B2;CYP2B6;TERT;ANKH;MAP2;STC2;APOE;LYN;EGR1;GSTM2;ACE;GSTM1;CADM1;TSLP;SPHKAP;SLC33A1;GREB1;MIR200A;MIR200B;NR2F2;GDF3;NR0B2;NR2F6;GRIN2D;COL1A1;CYP26A1;GJB2;STAR;CYP1A1;TDGF1;HLA-DRB1;COL18A1;MIR17;CAB39L;BRCA1;KRT20;CX3CL1;AKAP13;HEY1;UCN3;UCN2;TIMP2;CCR9;NEFM;PF4V1;BRD2;ZHX2;ANXA2;MME;DUSP1;FCRL3;MST1;TET1;IL16;MIR21;F3;PAX2;POU5F1;VAV1;SLC22A23;MMP14;PAX8;IL1B;RARB;AHRR;CD47;TLR4;HLA-DQB1;TAGLN;GSTP1;NR1I2;AIF1;EGFR;CYP17A1;PAK1;HSD17B1;NTF3;HSD17B2;SPP1;CD55;CCL25;NQO1;CDX1;STAT3;AKR1C2;LILRB1;MLH1;VEGFA;WT1;NBR1;HLA-DPB1;ZNF217;F2RL1;INHA;PLCD3;PF4;CSF2;DCLRE1C;CSF1;EPO;IHH;MSI1;IRS2;FGF1;COMT;FGF2;TNF;ETS1;HOXA13;MPO;HOXA11;IGF1R;HOXA10;RGS4;HPGDS;CDH3;CCND1;CDH1;GRAP2;RNF19A;SALL4;IGFBP1;PDGFRA;OSR2;SLC38A1;IGFBP3;F2R;DIO2;FOXP2;ZFP42;S100A6;PGR;CSF1R;SHC1;NOTCH4;ALOX15;TACSTD2;CACNA1A;GSTT1;DPP4;DPP6;CALD1;TAC1;MSC;SMAD2;SMAD3;SRD5A2;WNT3A;HMGA1;SYNJ2;INHBA;CALHM3;TMPRSS13;BMP7;ESR1;BMP4;ISG20;IL6;CDK6;UST;BCL2;HLA-DRA;SCGB3A1;TGFBI;FGFR2;UCN;MIR10B;THRA;SLC40A1;PIK3CD;CHD5;PTPN22;TRH;CYR61;PIK3CG;CXCL5;SYNE1;HK1;CA2;DNMT3B;NOS1;AMH;POLE;HS3ST3B1;KLF11;EMX2;MMP7;DNMT3A;OPRM1;TNFRSF1B;UTF1;DCN;TNFRSF1A;ACTA2;GREM1;MIR126;HSH2D;IFNG;CAT;CRH;LTA;BIRC5;ANG;CDH15;SMPDL3A;LTF;HDAC2;SRC;PON1;THBS1;MUC1;CCL5;ERBB2;SNCG;CCL1;NCAM1;SLIT3;MUC4;NTRK1;ANGPT2;ANGPT1;NOS2;NOS3;NTRK3;SGPP2;MTOR;MEIS1;MIR145;SST;LEP;IL2RB;TEK |
| Liver Cirrhosis | 0.001560478047501656 | RB1;SPARC;ADARB1;AQP2;AQP1;ICAM1;SPIB;LGALS3;EDNRA;RASSF1;EDNRB;LGALS1;CYP2D6;MYC;STMN1;TNFSF10;AKT1;SLC12A9;CPT1A;SERPINF1;SLC11A1;KRT8;HLA-C;HLA-A;DKK3;SPINT1;TFR2;RBP1;SERPING1;CYP2E1;HPD;ABCB1;TERC;PDE1B;PDGFB;CD1D;SLC30A10;PRH1;HSD11B2;DGUOK;GRIN2A;CYP2B6;TERT;TSPAN8;PDGFD;APOH;APOE;PROM1;A2M;APOB;HLA-DQA1;NGFR;EGR1;ACE;GSTM1;IGF2;GRIN2B;MIR338;DCLK1;COL1A1;CYP1A1;NOSTRIN;SLC22A8;HAMP;HLA-DRB1;TP73;DDR1;ACHE;CDKN1A;ELN;GPT;SERPINA6;GLI2;ESM1;HNF4A;TIMP2;TIMP3;DLAT;CD34;CC2D2A;DENND2D;MST1;MIR21;PROX1;CYP7B1;F3;F7;MMP14;ZEB1;ADRB3;IL1B;MIR17HG;TLR4;HLA-DQB1;C1QB;GSTP1;NR1I2;CD5L;EGFR;HLA-DMA;NT5E;WIF1;ALOX5;TMEM67;SPP1;STAT4;SLC17A2;NQO1;MGMT;GDF15;STAT3;OSM;MLH1;KLK10;GCK;VEGFA;KRT18;HLA-DPB1;GHRL;FMOD;IL17D;KLKB1;CSF3;CSF2;CSF1;PLOD2;CLU;FGF2;TNF;MPO;IGF1R;PYCARD;FGF7;HPGDS;DMBT1;CDH1;NPPA;TNFRSF8;TGM2;IGFBP1;NANOS1;PDGFRA;CASR;MSX2;IGFBP3;F2R;LIG4;CYBA;PCOLCE;MAT1A;HIC1;DHDDS;BSCL2;ADAM12;EHBP1L1;S100A9;ABCG2;NOTCH3;NOTCH4;GSTT1;ADRB2;FHIT;FUT1;DPP4;SOCS3;TPO;KL;SMAD3;RARRES2;TFPI2;DCDC2;BMP7;ESR1;IGF2R;ARHGAP24;IL6;COPS5;FGF19;BCL2;CPE;GNB3;SKIV2L;TGFBI;FGFR4;FGFR2;LGALS3BP;CNTNAP2;FHL2;ADAR;LAMC2;HTR2A;CXCL5;GHR;HK3;DNMT3B;CASP1;LBP;NOS1;CR1;GPX1;TRIM63;MMP7;VWF;H2AFY;DNMT3A;AXIN1;APOA1;NGF;TNFRSF1B;ACTA2;MIR126;CCNA1;AIM2;IFNG;CAT;CRH;IRF7;CDH13;CHI3L1;BPI;IRF5;CRP;CX3CR1;HDAC2;UNC93B1;PON1;NOD2;THBS1;AURKB;INS;MUC2;CNR2;CCL5;ERBB2;IL21R;SNCG;NCAM1;FSD1;UGT1A7;SNCA;ANGPT1;PRRX1;NOS3;KLF6;MARCKS;PC;APC;LEP;POU2AF1;BCL6B;CD9;SPRY2;CLNK;SPRY1;VIM |
| Diabetes Mellitus, Insulin-Dependent | 0.0015781436200927761 | HCP5;ERRFI1;APP;HPSE2;EHMT2;LST1;NUDT1;XYLT2;EHMT1;XYLT1;IKZF1;IKZF4;BACH2;ICAM1;LGALS3;EDNRA;LIPC;CYP2D6;ZMIZ1;ANPEP;MYC;TNFSF10;TNFSF11;UBASH3A;PRKCB;HLA-L;SLC11A1;HLA-B;HLA-C;HNF1B;UNC5D;MIF;HNF1A;HLA-A;RUNX1;IL23A;DDIT3;PRKCQ;CYP2E1;CFB;CFTR;SKAP2;MEG3;SLC22A3;ABCB1;AIRE;SLC22A2;SP140;GATA4;GATA3;SLC5A1;THY1;CD1D;SLC30A10;PLA2G7;CD1A;ABO;C2;C3;HSD11B2;SEC14L3;PCBP4;SCGB1D4;APOE;FADD;HCRT;SPRR2A;HLA-DQA2;HLA-DQA1;PRRT1;STAT5A;CREBBP;GSTM2;GSTM1;ACE;PTPRN2;IGF2;COL1A1;ATP4A;TG;TH;ID2;TNFSF4;ID3;COL9A3;PPT2;HLA-DRB1;HOXD8;LAD1;PTPRU;ROBO3;CD40;SLC44A4;PTPRN;SETD7;ICA1;ADM;GPT;CD3E;KRT20;IL18BP;C1QTNF8;HNF4A;CD38;TIMP3;FFAR2;SLC18A2;HLA-DPA1;TCF19;CD52;MIOX;CBR1;MME;DUSP1;IFNGR2;FCRL3;PAX4;PRSS16;TYK2;VAV1;CIT;NCR1;APBB1IP;TLR1;FCER2;MMP14;SLC7A9;IL1B;VWA2;TLR4;HLA-DQB2;HLA-DQB1;GRAMD1B;TNXB;ANKRD55;COL11A2;ADCY7;AIF1;EGFR;DUOX1;HLA-DMB;CUX2;BGLAP;NPY;SPP1;STAT4;LY6G6D;LAIR1;IL12RB2;MICB;IL32;AOC3;NQO1;AOC2;HSPA1L;RNLS;GAD1;STAT3;SORBS1;PSMB8;GCK;PSMB9;VEGFA;BATF;HLA-DPB1;IL17D;LY6G5C;IL17C;HSPA1B;HSPA1A;NEUROG3;CD86;CHRM2;RTL1;CSF2;DEFB1;STK19;GIMAP5;MPO;GIMAP7;FGF2;TNF;FGF3;CNDP1;CNDP2;HPGDS;FCGRT;NPPA;ADORA1;HLA-DOA;HLA-DOB;TNFRSF4;TGM2;G6PC;SLC30A8;TAP2;SARS;MIR27A;TAP1;KCTD1;CYBA;LMO7;HCG9;CLIP2;POLG2;MSH5;FSCN1;ELMO1;ERG;ABCG2;NOTCH3;KCNE1;CALCA;NOTCH4;TCF7;ALOX15;SPATA2;GSTT1;VTCN1;ADRB2;LPP;PSORS1C1;ZFP36L1;SOCS3;TPO;ZFP57;CALD1;DEAF1;NLRP3;MSC;VARS2;BCL2L15;FGF21;ESRRA;SLC14A1;KL;SMAD3;HTR1A;HSPA2;BTNL2;MAFA;ESR1;IGF2R;GFAP;SMAD7;ISG20;ADCY10;BMP2;IL6;INS-IGF2;UBXN11;BCL2;HLA-DRA;CPE;SKIV2L;TGFBI;MUC21;OR14J1;ITGB2;TNFAIP3;SLC2A2;PTPN22;FOXM1;AFF3;ADAMTS4;ADAMTS5;HHEX;ACOXL;CA2;TAGAP;ITGAX;FLOT1;BLNK;ITGB7;MADCAM1;NOS1;PGM1;GPSM3;MIR193A;HSP90AA1;CR1;GPX1;ITGA4;VWF;APOA1;OLIG2;UPK3B;DDOST;SSTR4;DCN;TNFRSF1A;FGR;MIR126;ADORA2A;IFNG;LCK;DDAH2;CHDH;CAT;IRF7;LTA;IRF8;CDH13;IRF5;BCAT1;CRP;DNAH2;HDAC3;HCG18;NRXN1;PON1;LRP5;NOD2;AGPAT1;USP18;CRYZ;INS;ACAN;MIR494;FYN;NCAM1;EIF4E;SNCA;CHGA;TCF7L2;TRIM40;ANGPT2;ANGPT1;KCNJ11;NOS2;NOS3;PDX1;EFR3B;TSHR;DLK1;POMC;CRYGD;APC;NPHS2;IL2RA;LEP;IL2RB;P4HB;PDCD1;TRIM31 |
| Liver Cirrhosis, Experimental | 0.0015895762779079418 | APP;IFITM1;SLC46A1;TES;SPARC;CERK;WIPF1;FAM89A;GMFG;SLC23A1;PLEKHB1;COL12A1;ICAM2;ABAT;LGALS3;LGALS2;LGALS1;CYP2D6;RASSF4;MYC;HOXA3;LRAT;SAMD11;ARL11;PRKCB;ACTN1;HPCAL1;KRT8;RNASE4;HLA-F;PPP1R3C;EZR;MYL9;SKAP2;CFD;ENPEP;ABCB1;SP140;PDGFA;ZBTB4;MTMR7;LDHB;TSPAN8;EPB41L2;INPP5D;NCKAP1L;A2M;PCK1;HLA-DQA1;S100A10;SDR42E1;ST14;STARD13;RABGAP1L;GSTM1;ADHFE1;OSGIN1;CIDEA;NR0B2;FLI1;COL1A1;CYP26A1;TF;HAL;ID2;FAT1;FXYD6;DUS1L;SLC22A8;HLA-DRB1;LGMN;ACHE;CDKN1A;SLC44A2;DPYS;CTSZ;ADM;GPT;SERPINA6;MEOX2;SERPINA4;IL18BP;GNGT2;RAC2;TIMP2;CD38;SH3BGRL3;CLIC1;APOBEC1;MME;ANXA2;TSC22D1;ELOVL5;SERPINB9;GFRA1;DUSP6;VAV1;NCOR2;APBB1IP;SLC22A23;PLSCR1;ZEB2;IL1B;PXDN;CRYAB;TLR4;SGK2;TMEM106A;JDP2;C1QB;KANK1;C1QA;SLC43A2;AHNAK;WRB;CD5L;CAPG;SLC7A1;FSTL1;AIF1;CYP17A1;GNAI2;HLA-DMA;NT5E;EXTL1;HLA-DMB;EPCAM;HSD17B2;SLC17A2;CD74;PKLR;MGMT;OSM;PSMB8;GCK;PSMB9;VEGFA;KTN1;FABP2;SMOC2;KRT18;GPAM;FABP7;B9D1;RAMP1;YPEL3;SIGLEC5;SPSB4;FOXA2;CD86;ALDH1L1;CSF3;CD83;PLOD2;PLAT;FRY;METRNL;FGF2;TNF;CNDP1;LOXL2;PYCARD;STK10;RGS5;CDH5;HPGDS;TUBB6;IYD;ADORA3;FAM198B;ARHGDIB;ZFP37;PDGFRB;IGFBP1;CSGALNACT1;PDGFRA;ABCG8;ABCG5;UNC5CL;G6PC;TPM4;TPM3;IGFBP3;F2R;TAP2;CYBA;SHISA3;RAB32;PROCR;COL4A2;COL4A1;S100A6;MYADM;IGFALS;NUPR1;S100A9;ABCG1;CSF1R;ISYNA1;CNP;MRO;SOCS2;UBD;KIAA1522;PRODH;SLC15A3;SMAD2;TGFB3;TFPI2;LY86;BMP7;INHBC;IGF2R;HOPX;ISG20;NAT8;DAO;CPS1;NBEA;HLA-DRA;TNFAIP8;EIF4E3;ITGB2;PLEK;FHL2;DYSF;ITGAL;ADRA1B;GHR;PSTPIP1;PTER;CA3;CA5A;CASP1;ACP5;PMEPA1;ITGB6;LRRFIP1;LBP;SRGN;VWF;NRG1;TGFBR2;ACTA2;FGR;ADORA2A;IFNG;TMBIM1;COL6A1;IRF8;LCP2;TRIM59;LCP1;HAGH;LTB;COLEC12;DDC;UNC93B1;ADD3;INS;PRTG;SELENBP1;ABLIM3;CHSY1;PALMD;CCL5;RNF39;FYN;GBP4;SLC25A24;PTPN18;RPH3AL;KCNJ8;NOS2;CMTM3;NOS3;LIMK2;TM6SF1;GALC;POMC;FAM167B;KLF6;FMNL1;MYO1C;PC;LEP;IL2RB;MGP;CD7;RGS10;VIM;CDO1;MYO1F;PFKP |
| Pancreatic carcinoma | 0.0016371928686281785 | RB1;SERPINE2;SCT;MIR429;MIR675;MYC;DPYSL3;STMN1;AKT1;CYP1B1;HOXA1;FAM19A5;IER3;PRIMA1;WSB1;PRKCB;SERPINF1;DAPK3;PRKCA;RNASE1;DKK1;EML2;DKK3;DKK4;MIR202;CLDN4;SLC5A8;SPINT1;ARMC3;CLDN8;RBP1;HOXB3;HOXB2;MAML3;ASTN2;IDO2;HPD;HOXB7;HOXB6;GNE;MEG3;ABCB1;DHH;TERC;MIR196B;GATA6;PDGFA;GATA5;PRDM14;GATA4;FPR2;LYPD5;MIR206;ABO;SLC5A5;SULT1A1;C3;CYP2B6;TERT;MIR330;CHST15;PROM1;RREB1;JAG2;NGFR;CREBBP;GSTM1;ICMT;CADM1;PLK2;NFATC2;MRPL28;NFATC1;NR2F2;PBX1;H19;FERMT1;GJB2;FXYD3;GIPC1;CYP1A1;HLA-DRB1;CDKN1C;PIGR;CDKN1A;MIR130B;MIR629;ADM;SLC6A2;TRIOBP;GLI3;BBC3;TIMP2;RSPO2;TK1;BRD2;RING1;ANXA2;DUSP1;PAX6;PAX5;F3;DUSP6;POU5F1;SERPINB5;MMP14;ZEB2;ZEB1;SLCO4A1;HCLS1;SLC29A1;GAS5;JDP2;KANK2;GRP;GSTP1;COL11A1;NR1I2;COL11A2;ADCY3;HOXD13;FSTL1;EGFR;ARFRP1;MIR410;RECQL5;EPCAM;BGLAP;ALOX5;NTF3;SPP1;BRD4;WWOX;PKLR;F10;MGMT;WNT7B;STAT3;SCTR;SULF1;SKI;WT1;FOSB;SIGLEC7;SEMA5A;CSF3;CSF2;CSF1;PLOD2;IRS2;CLU;FGF2;TNF;BEST1;RIMS2;CDH5;HHAT;FGF7;HPGDS;CDH3;CDH1;SLCO2B1;GRAP2;RNF19A;HLA-DOA;SLC16A3;SLC39A4;MAP3K5;KCNH2;IGFBP1;USP7;FBXW8;S100A2;S100A1;RPS8;IGFBP3;F2R;MITF;CHD1L;HIC1;FOXP1;LTB4R2;MTNR1B;S100A6;FSCN1;PGR;ADAM8;TNIK;S100A9;SQSTM1;NOTCH3;BRSK2;SHC1;TACSTD2;GSTT1;VTCN1;FHIT;RAP1GAP;SLC9A3R2;STK31;IGSF21;SOCS3;STK33;SMYD2;PLXNA1;KL;NOP14;OSBPL5;HTR1D;GFPT1;TFPI2;TNK1;TNFRSF10C;HTR1B;TNFRSF10A;MCC;TMPRSS13;HOPX;DAB2;COPS5;CDK6;BCL6;STK24;CDK2;BCL2;CPE;CLPTM1L;CIITA;CNTNAP1;PSCA;BNC1;FLT4;ONECUT1;ITGB2;FHL2;WISP3;PIK3CD;LSP1;MSLN;FOXM1;CYR61;PIK3CG;TRIM69;ITGB6;HGFAC;HSP90AA1;PEG3;KSR1;DNMT3A;NRG1;FOXL1;TNFRSF1B;NGF;DDOST;TNFRSF1A;ESRP1;BIRC5;ITGA6;MIR124-2;ANG;CX3CR1;HDAC2;HDAC3;SRC;UHRF1;MIR148A;ZNRF3;PARD6A;CCL5;TP53INP1;SH3BP4;MAL2;EIF4E;UGT1A7;NTRK1;MARCKSL1;ANGPT2;ANGPT1;PRRX1;KCNJ11;ASCC1;MIR935;TBX5;MTOR;TBX2;P2RX7;NELL1;P2RX5;NIN;APC;SST;IL2RA;LEP;IL2RB;APOC1;PDCD4;SPAG1;BCAR1;APP;FAAH;TNFRSF6B;SPARC;ICAM2;ICAM1;SOX2;LGALS3;FRAT1;RASSF1;LGALS1;RASSF2;ZNF608;TRIM29;PEG10;RPS6KA2;TNFSF10;LGALS9;TRIM27;SOX9;TNS4;SOX7;TP63;SLC12A9;SRRM2;EPHA4;ENTPD1;ENTPD2;HLA-B;KRT8;HLA-C;HNF1B;HLA-A;MIF;HNF1A;RUNX3;RUNX2;TIAM1;SFRP1;SLPI;IFI27;IL23A;KCNQ1;EZR;CFTR;GRB7;SLC22A3;EPAS1;EIF5A2;TWIST2;TWIST1;MST1R;ZBTB4;KLK7;DGUOK;NUAK1;TSPAN8;PENK;HIVEP3;TRPM8;FADD;MIR1178;TSPAN1;NTSR1;EGR1;WWTR1;IQSEC1;XRCC3;SLC33A1;IGF2;LIF;MICAL3;GDF2;MIR200A;PTK6;MIR200B;DCLK1;TF;OCLN;ARTN;ID2;SIM2;TDGF1;RGL2;VSX1;CC2D1A;TP73;APOBEC3B;MIR129-2;DDR1;HOTAIR;ROBO3;COL18A1;NRP2;CD40;MIR17;CTSZ;MIR18A;BRCA1;KRT20;KIF15;CCKAR;CCR9;MUC13;CCR7;CCR6;CD34;CTSD;HTATIP2;MAP4K1;CBR1;ZHX2;MME;MST1;TFEB;CACYBP;GFRA1;MIR21;PROX1;NME5;RHOF;MIR19A;VAV1;CNKSR1;CDCP1;NR5A2;CTTN;ADRB3;CCKBR;IL1B;MIR17HG;TFF3;CLDN18;TFF2;TFF1;ARHGEF2;CD47;ARHGEF7;TLR4;TAGLN;TMPRSS3;CXCR5;NEDD9;CYP17A1;NT5E;PAK1;CYP2A13;CUX1;WIF1;GPC1;NPY;HSF1;MICB;IL32;CD74;NQO1;GABBR1;CDX2;CDX1;GDF15;GAD1;OSM;KLHL1;MIR301B;SORBS2;LILRB1;MLH1;GRHL2;GCK;VEGFA;PER1;PER3;KRT18;KRT17;NBR1;TCEA2;F2RL1;IL17C;PNPLA2;F2RL3;FOXA2;NEUROG3;CD86;IHH;PLAT;PRDM1;COMT;ETS1;MPO;ETS2;IGF1R;CNDP2;MIR149;CCND2;CCND1;CAPN2;MIR150;NRTN;GGT1;PPFIA4;TGM2;PDGFRB;PDGFRA;CASR;ZGPAT;MSX2;LIG1;KDM2B;APLP2;WNT5A;MATK;MIR27A;LIG4;MAT1A;SCNN1G;MYCN;COL4A2;DPEP1;CARD16;NUPR1;ABCG2;FBN2;TCF7;ALOX15;ADRB2;FBLN2;FUT1;FUT4;MCHR2;FUT3;DPP6;PODXL;MIR296;NLRP2;TAC1;MSC;WNT2;MARK2;TFAP2A;SMAD2;FDPS;SMAD3;HMGA1;ISG15;HSPA2;BMP7;ESR1;IGF2R;ARHGAP24;SMAD7;BMP4;BMP2;IL6;EFNA2;NAT1;GDNF;MIR183;ALPI;TGFBI;FGFR4;FGFR3;FGFR2;FGF10;FGFR1;PNLIPRP2;CRISP2;MIR10B;MIR10A;HSPB2;CHD5;ADAR;CHD3;USP39;MIR96;PKD1;LTB4R;CXCL5;GHR;CASP8;CA2;MCF2L;CASP1;NOS1;HS3ST3B1;KLF11;MMP7;MIR23A;DCC;SSTR1;OPRM1;ARID1B;DCN;SSTR5;TGFBR2;MIR126;ALDH1A3;IFNG;PALLD;CAT;LTA;CHI3L1;CDH13;LTB;RBMS3;MCPH1;DNAH5;RGS16;AGPAT1;THBS1;PTHLH;AURKB;CRYZ;MUC1;MIR377;MUC2;MIR373;GNA12;ERBB2;MIR492;PTK2B;SPOCK1;SNCG;NCAM1;MUC4;MUC6;CHGA;TCF7L2;BNIP3;PDX1;CCK;MUC5B;SOD3;HS3ST1;KISS1;POMC;MIR145;MIR144;MIR142;CD9;VIM |
| Pheochromocytoma | 0.0016371928686281785 | RB1;APP;SCARB1;EPO;MPG;PKD2L1;PRDM2;HTR4;COMT;TNF;IGF1R;LGALS3;EDNRA;RASSF1;EDNRB;CCND1;DMBT1;MYC;RASSF5;STMN1;CAPN2;CHGB;HNF1B;FNDC3A;PDYN;SDHB;HIC1;MYCN;IFI27;DDIT3;SLC22A3;CALCA;EPAS1;TERC;GATA4;ACACA;TERT;TMEM127;PENK;HIVEP3;PROM1;FGF21;SMAD2;CREBBP;CBX4;IGF2;BMP7;PHOX2B;STAR;TH;GDNF;BCL2;GNAS;CPE;MIR183;SLC26A4;SIM2;PCSK2;PTPRU;PIK3CD;ADM;DBH;FOXM1;MIR96;SLC6A2;PTPRF;PIK3CG;RPTOR;CASP8;UCN3;CHL1;SLC18A2;ZHX2;HSP90AA1;DUSP1;TET1;SSTR1;NGF;SSTR5;IFNG;CAT;CRH;BIRC5;ANGPTL4;MCPH1;DDC;CBS;NPY;PLAGL1;CCL5;NTF3;ERBB2;MPZL2;CHGA;NTRK1;FARP1;RWDD3;EGLN2;MGMT;WNT7B;BNIP3;STAT3;SCTR;MLH1;POU4F1;MTOR;VEGFA;POMC;CRYGD;CRYGB;KRT17;SST;QPCT;SARDH;GHRL;ZACN;TNP1 |
| Adult Diffuse Large B-Cell Lymphoma | 0.0016371928686281785 | CD86;CD40;IKZF1;KRT20;PRDM11;CD79B;LGALS3;CCND1;MYC;TNFSF10;TP63;CREBBP;ASCC1;STAT3;HLA-B;ARAP3;MIR21;CDC42BPB;VEGFA;FOXP1;DDIT3;CD109;CDH11;CDK2;BCL2;IRF8;ARHGEF7;CARD11 |
| Childhood Diffuse Large B-Cell Lymphoma | 0.0016371928686281785 | CD86;CD40;IKZF1;KRT20;PRDM11;CD79B;LGALS3;CCND1;MYC;TNFSF10;TP63;CREBBP;ASCC1;STAT3;HLA-B;ARAP3;MIR21;CDC42BPB;VEGFA;FOXP1;DDIT3;CD109;CDH11;CDK2;BCL2;IRF8;ARHGEF7;CARD11 |
| Psychotic Disorders | 0.0016403047173265527 | PRDM8;FAAH;ADARB1;COMT;IGF1R;RGS4;DAOA;HPGDS;CYP2D6;DPYSL2;AKT1;SLC12A6;SEPHS1;NCKAP5;KCNH2;PDGFRB;SLC6A19;HLA-B;HLA-C;PRKCA;CSNK1E;CARTPT;HCG9;GABRG3;SRCIN1;HLA-E;DHDDS;MTNR1B;FEZ1;NHLRC1;RIN2;PVALB;SLC22A3;DLX1;RTN3;ABCB1;CNP;NOTCH4;SHOX2;PDGFB;SLC1A2;CACNA1A;CSF2RB;GSTT1;CACNA1C;CPLX1;NALCN;CRHR1;NPAS3;TRPM2;GRIN2A;HRH1;CYP2B6;APOE;TPPP;ST3GAL1;DRD4;DRD5;EGR1;HTR1A;HTR1B;GRIN2B;ESR1;IL6;DAO;SLC6A9;PPP1R1B;GNAS;SP8;FGFR3;FGFR2;GABRB2;CNTNAP2;CD40;ST6GALNAC1;GRIK3;GRIK4;OXT;HTR2A;CPZ;SLC6A3;GHR;GRM2;GRM5;CCKAR;ACADS;CD34;SLC18A2;TGIF1;BRD1;TET1;NRG1;OPRM1;OLIG2;LMBRD1;ALDH5A1;IFNG;RARB;MAPT;ARHGEF7;THOP1;SHANK1;CRP;DDC;HDAC2;SLC20A2;GSTP1;NR1I2;SELENBP1;CHN2;NPY;NTF3;GNA12;MGAT1;PCNT;DISC1;SNCA;NTRK1;TBX1;GABRA1;ASCC1;GAD1;B3GAT1;ZNF804A;HTR5A;TBX4;LEP;F2RL1;CD9;CPVL |
| Astrocytoma | 0.002171995713630953 | APP;SPARC;TNFRSF6B;SERPINE2;MLC1;AQP4;ADARB1;AQP1;ICAM1;LGALS4;SOX2;LGALS3;EDNRA;RASSF1;EDNRB;CYP2D6;MYC;RASSF5;STMN1;TNFSF10;AKT1;SLC12A9;ACTN1;HLA-C;SLC6A12;PRKCA;HLA-G;DKK3;RUNX1;GPR162;SLC5A8;SFRP1;IFI27;STX1A;SLC22A3;RTEL1;TERC;EPAS1;GATA6;PDGFB;PDGFA;GATA4;FPR2;PIK3R1;ADH6;C2;KLK6;C3;TERT;MAP2;PDGFD;PRDM16;NEU1;S100A13;CHST15;APOD;APOE;A2M;RREB1;NGFR;EGR1;NAPSA;STARD13;GSTM1;NFATC3;IGF2;MRPL28;NR0B2;FLI1;BST2;PCDHGA11;NANOG;HOXD9;HLA-DRB1;TP73;HRK;HOTAIR;ACHE;PTPRU;COL18A1;CDKN1A;CD40;MIR17;PPP1R10;ADM;AKAP12;HEY1;FASTK;HNF4A;TIMP2;NEFL;CTSD;TIMP4;BRD2;SLC2A14;ZHX2;ANXA2;TSC22D1;SPHK2;MST1;AMPD1;PAX6;PAX5;F3;POU5F1;VAV2;NCR1;SLC7A5;MMP14;MMP16;IL1B;MIR17HG;CD47;ARHGEF7;GSTP1;HOXD13;KCNA5;ASCL1;AIF1;EGFR;RASSF10;WIF1;ALOX5;NPY;NTF3;SPP1;CTNNA1;CTNNA3;CD55;MGMT;STAT3;OSM;MLH1;BLCAP;HIPK2;VEGFA;FABP5;FABP7;SEMA5A;CSF3;ALDH1L1;CSF2;CSF3R;CCNT1;PLAT;CLU;FGF2;TNF;ETS1;IGF1R;RIMS2;PYCARD;RGS4;RGS5;HPGDS;CCND1;DMBT1;CDH1;GRAP2;RNF19A;ZNF569;PITX1;SLC16A3;KCNH2;PDGFRA;TLE2;CASR;SNRPN;CD93;WNT5B;TPM3;IGFBP3;MATK;TSC2;DHDDS;BSCL2;MYCN;SCG5;PGR;ZNF436;PFKFB3;CNP;TUBGCP2;SLC1A2;GSTT1;NPAS3;HRH1;CAMTA1;MBP;TAC1;TFAP2A;CD160;TNFRSF10A;TMPRSS13;ESR1;ARHGAP24;GFAP;IL6;GDNF;CDK2;BCL2;MIR184;CNTN2;TUBGCP3;TGFBI;FGFR4;LHX9;FGFR2;FGFR1;MXD4;CRISP2;CIITA;GADL1;PIK3CD;ADAR;LSP1;TRH;PIK3CG;GRM5;PSD3;DNMT3B;MMP7;DNMT3A;OLIG2;NGF;UPK3B;ZAP70;IFNG;CAT;COL6A1;SNURF;IRF7;LTA;BIRC5;CHI3L1;BIRC7;ZMYND10;CX3CR1;ITIH4;SRC;SPG7;THBS1;PTHLH;DLL3;MUC1;CNR2;CCL5;P2RY1;ERBB2;MXI1;NCAM1;SNCA;NTRK1;SPAG9;LGI1;TCF7L2;MARCKSL1;ANGPT2;ANGPT1;ASCC1;NOS3;B3GAT1;CCK;MTOR;P2RX7;KLF6;APC;MIR144;SST;PTPN6;CD248;TEK;VIM |
| Anorexia Nervosa | 0.0023087814988327388 | UCN;FAAH;OXTR;ATP8A2;ABCB1;KCNA3;OXT;HTR2A;COMT;HTR4;CDH9;SLC6A2;TNF;SLC6A3;CYP17A1;DPP4;CYP2B6;STMN1;FAM19A2;KCNN3;DRD4;FGF21;IGFBP1;GHSR;LMO3;NTRK3;HTR1D;EBF1;HTR3A;INHBB;HTR3B;OLIG2;PARP9;SORCS2;ESR1;GRIN2B;C1QL1;POMC;MC3R;PYY;IL6;IFNG;LEP;NT5C1B;CRH;GHRL;SHBG;RAN |
| Urothelial Carcinoma | 0.002422515641090615 | CSF2;MSI1;FGF2;TNF;LGALS4;RASSF1;FGF7;HPGDS;CCND1;MYC;STMN1;TNFSF10;AKT1;CYP1B1;SRRM2;PDGFRB;DAPK1;S100A1;WNT5A;TSC2;KCNK17;COL4A1;DDIT3;CYP2E1;SLC22A3;ZNF671;EIF5A2;CUL1;GSTT1;GATA3;PIK3R1;FHIT;SULT1A1;TERT;NDN;KCNN3;TRPM8;CREBBP;GSTM2;GSTM1;SMAD3;IGF2;TRPV2;HSPA2;INHBA;TMPRSS13;DAB2;IL6;BCL2;TGFBI;FGFR3;FGFR2;FGFR1;HOTAIR;CDKN1A;MIR10A;DOC2A;NUMA1;PIK3CD;GRIK2;KRT20;FOXM1;PIK3CG;ELAVL3;CASP8;DNMT3B;SH3BGRL3;CD34;BOK;GSTO2;DUSP1;MST1;DNMT3A;EMP2;MIR21;NME5;UPK3A;POU5F1;DCN;KAT2A;ANGPTL4;RBMS3;HDAC2;GSTP1;COL11A2;THBS1;EGFR;HLA-DMA;MUC1;CYP2A13;TMEM67;ERBB2;MXI1;PTK2B;PRAME;CTNNA3;EIF4E;WWOX;DAB2IP;MLH1;MTOR;VEGFA;KISS1;FABP2;MIR145;PDCD1;HSPA1B;HSPA1A |
| Coronary heart disease | 0.002503151273221046 | NPFFR2;TRIO;TES;HPSE2;EHMT1;ICAM1;LGALS2;EDNRA;LIPC;CYP2D6;NDST4;GJA5;AKT1;SOX9;DLEU7;SLC12A9;MEF2C;SRRM4;SERPINF1;MIF;HNF1A;ISL1;SFRP4;GPR162;CFDP1;FNDC1;SOAT2;MYL2;KCNQ1;MAML3;ZNF831;ASTN2;CES1;SLC22A3;ABCB1;TERC;GATA6;PDGFA;AGAP1;GATA4;GATA2;PLA2G7;KALRN;MIR206;PRH1;ABO;C3;HSD11B1;CYP2B6;RNF213;ST8SIA4;APOD;APOE;APOB;NKX2-5;PROM1;ELANE;EGR1;GSTM2;HLA-DRB5;ST14;DNAH11;GSTM1;ACE;DTNA;SLC33A1;IGF2;MRPL28;NR2F2;GP6;ATP4B;BHMT;HAL;TNFSF4;CYP1A1;NOX5;ANP32B;HLA-DRB1;NODAL;PTPRT;ACHE;RYR2;ROBO3;CD40;MIR17;GRIK4;PCSK9;TCF21;KIF12;GPT;PEAR1;PCSK6;ASGR1;RYR3;CX3CL1;IL18BP;GLT1D1;JPH3;GNGT2;MECOM;HNF4A;CCR7;CCHCR1;CD34;PF4V1;ANXA2;BHMT2;RIPK4;ITPK1;AMPD1;IL16;GP1BA;MIR21;SORCS3;F3;POU5F1;F7;MMP14;ADRB3;TNNT1;IL1B;MIR17HG;GLCCI1;ANGPTL4;ARHGEF7;TLR4;RAPSN;HLA-DQB1;WRB;COL11A2;NR1I2;KCNA3;EGFR;CYP17A1;CUX2;MTHFD1L;BGLAP;ALOX5;NPY;SPP1;EXTL3;SLC17A3;SH3BGR;MPZL2;P2RY12;NQO1;SORT1;RNLS;GAD1;F12;ADAM33;DAB2IP;NR1H3;GCK;PLEKHA7;VEGFA;FABP2;CR1L;DLC1;ESYT3;GHRL;PLCD3;HSPA1A;F2RL3;CD86;SCARB1;SEMA5B;CSF3;CSF2;F13A1;PLAT;IRS2;COMT;ETS1;MPO;CLU;TNF;ETS2;CNDP1;RPH3A;HPGDS;ADORA3;MIR150;PITX2;IGFBP1;ABCG8;CASR;ABCG5;SLC30A8;CD93;DSCAM;OSR1;IGFBP3;F2R;ADAM11;CYBA;HCG9;FOXP1;PROCR;CACNB2;COL4A2;COL4A1;DPEP2;ELMO1;HHIPL1;S100A9;ABCG1;DPEP3;FBN2;ISYNA1;ALOX15;GSTT1;CACNA1C;ADRB2;FHIT;ACAT2;FUT3;GRK4;NPC1L1;NLRP3;TAC1;KIAA0319;MARK2;FGF21;KLHL29;KL;TFAP2B;ZFHX3;SMAD3;EPHX1;HMGA1;ESR1;SMAD7;BMP4;SLC2A9;IL6;PROC;CPS1;BCL2;GNB3;CPE;SKIV2L;SHBG;SLC26A4;THRA;DOCK9;TNFAIP3;ECE1;PIK3CG;CXCL5;CA2;CASP1;PHACTR1;NOS1;GPSM3;MKKS;MMP7;GPX1;TRPC3;VWF;MIR23A;APOA1;OPRM1;TNFRSF1B;WNT16;KLF14;TGFBR2;ACTA2;GREM1;MIR126;IFNG;PALLD;DDAH2;CHDH;HAND1;TRAF5;NOS1AP;COL6A1;CRH;LTA;CHI3L1;IRF8;NDUFS2;CDH13;IRF6;CRP;CX3CR1;PON3;HCG18;PON1;LCAT;LTC4S;NTN1;THBS1;THBS4;INS;SELENBP1;CBS;CCL5;TBX20;P2RY1;NCAM1;FSD1;TBX1;TCF7L2;ANGPT2;KCNJ8;KCNJ11;NOS2;GALNT2;ASCC1;NOS3;NTRK3;CMTM5;PCDH8;IRX4;B3GAT1;TBX5;SOD3;TBX3;ACTC1;MIR145;APC;CAPN10;LEP;MGP;APOC1;PDCD4;BCAR1 |
| Septic Shock | 0.0026238902108613247 | CRP;CSF3;ANGPT2;ANGPT1;NOS2;VWF;ADM;SLC5A1;TNFRSF1B;TNF;F3;AQP1;TNFRSF1A;VEGFA;PYCARD;FER;IL6;NLRP12;CD6;DDAH2;IL1B;CASP1;HLA-DRA;TLR4 |

| | | |
|---|---|---|
| Medulloblastoma | 0.0026640420282419883 | SPARC;GFI1;EHMT1;SOX2;RASSF1;MYC;STMN1;C3AR1;TNFSF10;AKT1;SOX9;HOXA5;SRRM2;DAPK1;UNC5B;DLEC1;EBF3;RFX1;DKK1;DKK3;SFRP1;SFRP2;IFI27;KCTD11;EZR;ABCB1;FOXG1;PDGFB;WNK4;CD1D;SLC5A5;C3;ABR;LDHB;TERT;MAP2;PDGFD;ALX4;PROM1;S100A10;JAG2;NGFR;CREBBP;GSTM2;GSTM1;CADM1;IGF2;LIF;FLI1;TF;L1TD1;COL9A3;GRB2;COL9A2;TP73;ACHE;COL18A1;NRP2;MIR17;GLI3;GLI2;AKAP12;HNF4A;TIMP3;DLAT;MIR192;MAP4K1;BRD2;TESC;PAX3;PAX6;MIR21;PAX5;MIR19A;F3;POU5F1;VAV1;MIR17HG;HCLS1;KCNA1;MIR520G;KCNA3;ASCL1;EGFR;NT5E;CUX1;NTF3;PRAME;BRD4;GABRA5;MGMT;STAT3;DAB2IP;MLH1;POU4F1;VEGFA;WT1;DLC1;SIGLEC7;NEUROG1;F2RL3;CSF3;CSF2;MPG;MPL;MSI1;IRS2;PRDM2;FGF2;TNF;IGF1R;COMP;HPGDS;CCND2;CCND1;DMBT1;CDH1;GRAP2;RNF19A;TNR;PDGFRB;PDGFRA;USP7;S100A2;S100A1;TPM3;IGFBP3;WNT5A;LIG4;TSC2;HIC1;DHDDS;MYCN;SCN8A;S100A6;SCG5;S100A9;MAD1L1;CSF1R;NPR3;SLC1A2;GSTT1;FHIT;FUT4;GRK5;NLRP2;SMYD4;OTX2;OTX1;DRD4;SMAD3;PTCH2;HMGA1;TNFRSF10A;DNAJC15;ESR1;ARHGAP24;GFAP;BMP2;IL6;CDK6;BCL2;GNAS;MIR183;MXD3;FGFR1;CRISP2;SNED1;MIR10B;TREH;PIK3CD;LAMC2;TRH;FOXM1;PIK3CG;CASP8;GRM4;ZIC2;ZIC1;HGFAC;HSP90AA1;AXIN1;OLIG2;NGF;TNFRSF1B;IFNG;HAND1;CRH;BIRC5;BCAT1;RBMS3;UHRF1;NTN1;THBS1;AURKB;MIR495;MIR494;NNAT;ERBB2;MKNK2;MXI1;PTK2B;SNCB;EIF4E;ATOH1;SNCA;NTRK1;CIZ1;MARCKSL1;KDM4B;EYA1;NTRK3;MIR935;MTOR;MEIS1;APC;SST;IL2RB;PDCD4;CCM2;INSM1;VIM |
| CNS disorder | 0.0026640420282419883 | APP;ACHE;CSF3;CSF2;EPO;AQP4;LAMC2;HTR2A;TNF;GRM1;SLC6A3;MYOM2;ICAM1;CRHBP;GRM5;CYP2D6;RAC2;CD34;MEF2C;DST;MME;MIR21;ADORA2A;OAS2;CRH;ADAM8;ABCB1;NRN1;NR1I2;KCNA3;MST1R;SAMHD1;PTHLH;KLK6;SOCS3;CYP2B6;CNR2;CD19;ERBB2;SPP1;NLRP3;MGAT1;NCAM1;APOE;TPPP;ARSB;MOG;GTF2H4;GFAP;POMC;IL6;SLC6A9;SST;BCL2;F2RL1;VIM;KCNK2;HLA-DRB1;LGMN |
| Nicotine Dependence | 0.0026640420282419883 | CHRM2;APP;CD40;CHRM1;DBH;HTR2A;COMT;SLC6A3;ZCCHC14;RPTOR;SLC18A2;CHRNB1;PLEKHG1;FRMD4A;HTR3A;CARTPT;OPRM1;HTR3B;DNM1;CRH;CYP2E1;LCP1;ANKK1;ARHGEF7;RAPGEF3;RAPGEF4;CHRNA1;MEG3;KANK1;DDC;CHRNA2;CHRNA5;NRXN1;CHAT;CHRNA6;PON1;NRXN3;CHRND;CYP2B6;TERT;CYP2A13;GALR1;NTF3;NCAM1;APBB1;HCRT;DRD4;DRD5;GABBR1;TTC12;ACE;EGLN2;ASCC1;GABRA4;HCRTR2;IL6;TH;CPD;PPP1R1B |
| Cerebrovascular accident | 0.0026705459067315096 | APP;ACSM3;HPSE2;AQP4;HTR4;ICAM1;CRHBP;EDNRA;LIPC;EDNRB;TNFSF10;AKT1;ENTPD1;PRKCH;ANK2;HLA-A;MIF;HLA-G;DHX40;ADCY9;KCNQ1;NINJ2;MAML3;NEIL1;ALDH7A1;CASZ1;ABCB1;GIPR;SHMT1;NEDD4L;KLK4;GATA4;GATA3;PIK3R1;PLA2G7;KALRN;ABO;SLC5A4;PRH1;KLK6;C3;GRIN2A;CYP2B6;PDGFD;INPP5D;ST8SIA4;APOD;APOE;NKX2-5;PROM1;A2M;APOB;ELANE;EGR1;TRAP1;ACE;GSTM1;FOXF2;TNFSF14;TREX1;LIF;GP6;GRIN2B;PM20D1;GP9;BHMT;OCLN;TNFSF4;TOLLIP;PPP1R1B;CYP1A1;GTF2IRD1;HLA-DRB1;RYR1;ACHE;ROBO3;CD40;TNFRSF13B;SLC35F3;SLC44A2;ELN;PRKAG2;PCSK9;ADM;ZCCHC14;JPH3;SH3PXD2A;TIMP2;TIMP3;CD34;SPHK2;ITPK1;GP1BA;F3;SERPINB6;F7;MMP14;IL1B;KCNMA1;CD47;TLR4;RAPGEF3;COL11A2;KCNA3;ASCL1;MIR410;ALOX5;NPY;SPP1;APOL1;RUNX1T1;AOC3;P2RY12;FARP1;NQO1;HSPA1L;MOG;RNLS;GDF15;F12;SORBS1;VEGFA;FABP2;GATM;HLA-DPB1;KLKB1;RAMP1;IL17C;HSPA1B;HSPA1A;SPSB4;F2RL3;SEMA5B;CSF3;CSF2;CSF3R;EPO;F13A1;DEFB1;PLAT;FGF1;CELSR1;TNF;MPO;HPGDS;NPPA;CAPN2;TEKT4;PITX2;MAP3K5;TGM2;S100A1;IGFBP3;F2R;CYBA;MAT1A;CARTPT;NPAS4;CLIP2;PROCR;MAG;LTB4R2;COL4A2;COL4A1;IGFALS;WDFY4;ABCG2;FBN2;NOTCH3;SHC1;NPR3;HTRA1;PROZ;CACNA1A;SLC1A4;ADRB2;FUT1;HRH1;CALD1;GRK4;MYH14;NLRP3;MBP;TPPP;TAC1;MSC;ZFHX3;SMAD3;ZBTB16;HTR1D;TFPI2;HTR1A;HSPA2;BMP7;ESR1;GFAP;NR4A2;IL6;CDK6;LRCH1;GDNF;BCL2;GNB3;TGFBI;UCN;ITGB2;HSPB2;PIK3CD;ECE1;PTPN22;LAMC2;HTR2A;CHD3;ITGAL;PIK3CG;TSPAN10;LTB4R;GHR;GRM2;CASP8;TRIM69;PHACTR1;NOS1;LRRFIP1;ZNF880;ARHGEF10;EGFL7;GPX1;VWF;PDE4D;NGB;APOA1;NRG1;NGF;UPK3B;TNFRSF1A;ACTA2;TGFBR3;DDAH2;CHDH;ASPG;CAT;CRH;LTA;CHI3L1;PAOX;CRP;CX3CR1;PON3;HDAC2;LRP1;LAMA1;PON1;NOD1;LTC4S;SAMHD1;THBS1;INS;CBS;CEACAM20;TNNI3;TCF7L2;ANGPT2;ANGPT1;NOS2;TXNRD2;GALNT2;NOS3;TXNRD1;LIMK1;PDE4DIP;TBX3;SLC25A38;CRYGD;PC;XPNPEP1;LEP;CCM2;P4HB;TEK;KCNK2 |
| Colorectal Cancer | 0.0026755838218706215 | TSKS;EHMT2;PREX2;MIR1-1;ISLR;DPYSL2;NDST4;FAM110B;AKT1;UNC5B;UNC5C;EML2;SLC5A8;CLDN4;SPINT1;MIR196A2;DDIT3;RUFY3;CYP2E1;ABCB1;SHMT1;MIR196B;CRABP1;GATA6;GATA5;GATA4;GATA3;LYPD5;GATA2;SLC5A5;SULT1A1;HSD11B2;ADAMTS14;GRIN2A;LARP1;CYP2B6;KCNQ1OT1;ADAMTS18;PDGFD;NPBWR2;LYN;NGFR;CREBBP;GSTM2;GSTM1;CADM1;FUCA1;ABCA4;MRPL28;EIF2AK4;NR2F2;MIR339;MIR338;NR0B2;NR2F6;VANGL1;IL22RA1;FXYD3;FAT1;NANOG;DCHS1;FAT4;HRG;HAMP;HLA-DRB1;LGMN;HRK;MIR130B;LRRC4;MMP21;BBC3;TUBA1C;JPH3;MECOM;UCN2;IKBKE;APC2;CHST8;DUSP1;KCND3;ELOVL5;SPHK2;F3;SLC39A14;POU5F1;F7;MMP14;GPRC5A;SLC29A1;LRRC3B;CRYAB;DSCAML1;MDFI;ZNF232;ZNF471;GSTP1;COL11A1;COL11A2;FSTL1;SLC7A1;EGFR;SLC7A2;PLAGL1;SPP1;CSMD3;FLNC;CSMD2;CSMD1;EXTL3;MEF2D;SPDEF;EGLN2;PKLR;KCNB2;P2RY14;STAT3;SCTR;DAB2IP;MIR539;PLXDC1;DCBLD1;SKI;FABP2;WT1;FABP6;ZNF217;LGR6;SIGLEC7;UGT1A10;ALDH1L1;NCF4;MSI1;FRY;HPGDS;ADORA3;ADORA1;PITX2;TNFRSF4;PITX1;KCNH1;KCNH2;UGT2B15;AKR1E2;UGT2B17;SRCIN1;ACAP1;FSCN1;S100A6;S100A9;SQSTM1;ZNF432;DSC3;PFKFB3;TACSTD2;SPATA2;GSTT1;FHIT;SOCS2;STK31;SOCS3;IGSF21;BCL2L10;NPTX1;KL;ZBTB16;HTR1D;TFPI2;HPS3;TNFRSF10A;MCC;TMPRSS13;LRCH1;HLA-DRA;CLPTM1L;SHBG;PIWIL1;CIITA;ITGB4;FLT4;ITGB2;FHL2;WISP3;PIK3CD;ECE1;FOXM1;PIK3CG;WISP1;SPN;PTER;ITGAX;UPP1;PGM1;GRID2;EMX2;HSP90AA1;PEG3;ITGA4;KSR1;GRID1;ITGA3;FOXL1;TNFRSF1B;TNFRSF1A;PCCA;PCCB;MYO3A;TMBIM1;BIRC5;LCP1;BIRC7;BCAT1;SNAP47;LRP1;SRC;UHRF1;LRP5;FOXK2;FURIN;FOXK1;NTN1;CCL5;BEND4;FSD1;EIF4E;NOP56;NTRK1;TBX1;PPP1R14A;MARCKSL1;SLC12A5;PRRX1;TIE1;NTRK3;STYK1;CABLES1;HTR5A;SORL1;TBX3;MYO16;NELL1;SST;IL2RB;ST7;MGP;PDCD4;CD248;PTPN6;PDCD1;BCAR1;SPON2;TNFRSF6B;NUDT1;ICAM5;IKZF1;ICAM1;LGALS4;LGALS3;TCL1A;RASSF1;LGALS1;RASSF2;RASSF5;TNFSF10;TRIM21;TP63;EPHB3;SLC12A9;ARL11;EPHA4;EPHA6;ANK2;HNF1B;CSNK1E;HNF1A;SFRP4;SFRP1;SFRP2;IFI27;PPP1R3C;CRISPLD2;KCNQ1;SFRP5;MT1G;ALPK2;EPHA1;EPAS1;PCDH10;MST1R;KALRN;PIK3R5;TSPAN8;PCBP4;NEU1;HLTF;ST8SIA4;SYMPK;APOD;HIVEP3;APOE;APOB;B4GALNT3;B4GALNT2;SNX8;TSPAN1;NTNG1;STAT5A;WWTR1;EGR1;TNFSF14;FOXF2;FOXF1;RCC2;ADHFE1;XRCC3;IGF2;NBEAL2;DCLK1;PEX5L;PTK7;GRB2;TP73;VSX2;PCSK2;DDR1;PTPRT;SPAG17;PTPRU;NRP2;CD40;PTPRS;GRIK3;ICA1;BRCA1;CD3E;AKAP12;AKAP10;BOLL;HNF4A;MUC12;CCR9;CD38;CCR7;CCR6;MCF2L2;CD34;CTSD;HTATIP2;ACAD8;APOBEC1;ZHX2;MME;MST1;CACYBP;IL16;NCR1;ANO1;NR5A2;CTTN;IL1B;CD47;PDZRN4;TAGLN;SATB2;MIR520G;TMPRSS4;NEDD9;CRIP2;CYP17A1;PAK1;FAM172A;HSD17B1;GPC1;HSD17B2;HSF1;HSF4;PAK6;MSX1;XDH;CD55;MICB;IL32;SORT1;TMEM176A;KLHL1;KLK10;VEGFA;MAD2L2;PER1;TFAP4;PER3;KIF26B;NEK10;IL17D;IL17C;NEK11;PNPLA2;CD86;MPG;MPL;IHH;PLAT;PRDM2;ETS1;MPO;ETS2;IGF1R;LOXL2;CCND2;DMBT1;CCND1;PEBP4;CAPN2;TGM2;PDGFRB;PDGFRA;CASR;MSX2;CD93;SCARA5;LMTK3;MIR27A;SDHB;DHDDS;ADAM19;SCNN1G;CD8A;SCNN1B;DPEP1;FBN2;NDRG4;CALCA;TCF7;SLC1A3;SLC1A5;HOXB13;SLC1A7;ACACA;FUT4;CACNA1G;FUT3;NME1-NME2;FUT6;AANAT;IRAK2;CALD1;DEAF1;CAMTA1;PRODH;TAC1;WNT2;WNT3;TFAP2A;UBE2I;TFAP2E;RARRES2;HMGA1;INHBB;INHBA;BMP7;IGF2R;SSH1;BMP4;BMP2;IL6;NAT1;FGF18;CNTN1;GNAS;GNB4;MKRN3;ALPI;LGALS3BP;LDB1;CNRIP1;TNFAIP3;CHD5;LAMC2;ARHGEF10L;GRM1;LMNB2;GHR;CPT2;ZIC1;GDAP1L1;LRRFIP1;MIR193A;MMP7;MIR23A;DDX56;NAV2;TAPBP;GREM1;CCNA1;DPPA2;MMRN2;IRF7;SNORD44;CHI3L1;ATG4B;IRF5;SHANK1;OTUB1;DNAH3;LAMA1;GOSR1;ASAP3;AURKC;PTHLH;AURKB;CRYZ;INS;CYTH2;CBS;MIR495;MIR494;ATOH8;MIR492;PTK2B;SLIT3;ATOH1;CHGA;LAMB3;BNIP3;EYA4;ZNF804A;CCK;MIR1914;CHFR;HS3ST2;GALNS;PNKD;MIR125B1;COL7A1;CD9;TEK;VIM;RAN;TRIO;SERPINE2;WIPF1;COL12A1;MIR429;ADARB2;ABCA13;MIR425;MIR675;SOX17;ZMIZ1;MYC;STMN1;CYP1B1;PLCE1;PROK2;DIP2C;HOXA5;ANKS1B;HOXA4;IER3;PRIMA1;NUP210;PRKCB;DAPK1;SERPINF1;RPL22;AFAP1;DKK1;DKK3;SULT2B1;DKK4;MIR202;GPR162;HOXB9;BIN1;MAP1B;HOXB3;PRR15;HOXB7;TSPYL5;MEG3;CTBP1;TERC;PRKCZ;MIR206;ABO;LDHB;TERT;GALR1;ALX3;ALX4;PROM1;HOXC9;ZSCAN16;NKX2-3;ZSCAN18;S100A10;AFAP1L2;STARD13;ST14;ACE;FZD7;FZD6;NFATC1;SLC6A6;GJB2;GIPC1;TNFSF4;CYP1A1;TNFSF9;HOXD9;HOXD8;CDKN1C;CDKN1A;RARG;ELN;ADM;FMN1;GLI3;MYOM2;ABCC13;CHL1;ADAMTSL3;TIMP2;RSPO2;TIMP3;RSPO3;TK1;BRD2;TESC;TET1;PAX6;SERPINB5;SERPINB6;GJD4;ZEB2;GJD3;COL2A1;ZEB1;PAX8;SLCO4A1;RARA;RARB;GAS5;GAS7;GRP;SGMS1;NR1I2;MIR409;MIR647;HOXD10;ASCL2;RASSF10;EPCAM;ALOX5;RUNX1T1;BRD4;AOC3;ZNF583;GABRA1;WWOX;BRF1;MGMT;GUCA2B;SULF1;GUCA2A;HIPK2;SMOC2;ABI1;LPCAT2;FOSB;HSPA1B;HSPA1A;DGKG;CSF3R;CSF2;CSF1;CLSTN3;GTF2B;CTNND2;IRS2;CLU;FGF2;TNF;RIMS2;RPH3A;PYCARD;CDH5;FGF7;TUBB6;FGF8;CDH3;LAMP1;FCGRT;SALL3;CDH1;GRAP2;RNF19A;SALL4;MAEL;ZNF569;SLC26A10;MACROD2;SLC39A7;LZTS2;LZTS1;STK32C;USP7;ZNF560;SNRPN;TPM3;USP4;IGFBP3;F2R;TAP2;TAP1;HIC1;CACNB1;CACNB2;OBSCN;PGR;RAB38;ADAM8;ERG;ADNP;TNIK;MAD1L1;NOTCH3;RASAL1;RASAL2;VTCN1;TPO;NDN;STRA6;ESRRA;ZNF540;TRAPPC9;BTBD9;MIR612;COPS5;RHEB;MAFG;CDK2;BCL2;CPE;MAP3K13;XAF1;EIF4G1;DIDO1;SFT2D3;EIF4E3;ZFP82;ADRA1B;TREM1;SYNE2;CYR61;DNALI1;SYNE1;SGCE;ADAMTS4;ADAMTS5;SCRN1;PHACTR3;DNMT3B;EFHD1;HMGCS2;GRASP;ADAMTS9;SGCG;GPX1;DNMT3A;AMT;NGF;DUSP22;TFDP1;LCK;ESRP2;ESRP1;EVL;ANG;CRP;HDAC2;RAB1B;NOD1;NOD2;JAKMIP1;MIR148A;JAKMIP2;SELENBP1;DNAJB6;PUF60;UGT1A9;UGT1A7;UGT1A6;SPAG9;SEMA4A;RPH3AL;ANGPT2;KCNJ11;ANGPT1;TXNRD2;TXNRD1;MTOR;P2RX7;RAD52;MARCKS;KLF6;APC;LEP;ZACN;CDO1;CBLN2;PANK4;APP;IFITM1;SLC46A1;SPARC;TP53RK;TTLL10;GFI1;NADSYN1;FERD3L;SOX2;FRAT1;EDNRB;PART1;SOX9;TNS4;SOX7;SOX5;SRRM2;CPT1A;HLA-B;MTUS1;KRT8;HLA-C;ACSL5;SLC6A12;DLEC1;HLA-A;RUNX3;RUNX2;HLA-E;RUNX1;TIAM1;TCERG1L;IL23A;CD109;EZR;CFTR;SLC22A3;MACF1;HEPACAM;SLC22A2;TWIST2;TWIST1;KLK4;KLK8;KLK7;CRHR2;KLK6;ZNF829;NUAK1;MAP2;STC2;FADD;TRPM8;SPTBN1;NTSR1;BANP;TRAP1;PTPRN2;HIST1H4K;BCL11B;HIST1H4L;PDE2A;SLC33A1;MIR200A;MIR200B;ALOX15B;GADD45G;SMTN;OCLN;VNN1;CYP2S1;HIST1H4I;ANP32B;TDGF1;NODAL;SLC22A9;HIST1H4F;HOTAIR;COL18A1;SLC44A4;MIR17;MIR18A;KRT20;ESM1;SLC22A18;SOSTDC1;MIR192;MAP4K1;PSD;PARP6;CBR1;AVPR1B;INSRR;NME2;TFEB;AMPD1;MIR21;PROX1;MIR19A;CBFA2T3;CDCP1;RAD51C;CCKBR;MIR17HG;MMP23A;TFF3;TFF2;VWA2;SGK1;TLR4;GUCY2C;CXCR5;NT5E;MTHFD1L;CUX1;WIF1;CTNNA1;CTNNA3;NQO1;CDX2;CDX1;GDF15;LILRB1;DMRT3;SORBS1;MLH1;GRHL2;CYP2W1;KRT17;DLC1;NEUROG1;APBA2;SIPA1;F2RL3;COMT;TMEM140;MIR149;MIR150;POSTN;KDM2A;LIG1;SPNS3;WNT5A;LIG4;TSC2;LMO7;SHISA3;MYCN;HGS;SCG5;CD300E;NUPR1;STEAP1;ABCG1;ABCG2;ISYNA1;ALOX15;LTBP4;ALOX12;ADRB2;LPP;FBLN2;ZFP36L2;DPP4;HRH1;PODXL;UBD;NLRP4;CDH22;NLRP2;NLRP3;LONRF2;MARK2;SMAD2;CBX5;SMAD3;CBX4;TGFB3;WNT3A;EPHX1;HSPA2;POU2F2;ESR1;ARHGAP24;SMAD7;NR4A2;GDNF;FES;MIR183;SKIV2L;FGFR4;FGFR3;FGFR2;FGFR1;KDM5B;CRISP2;MIR10B;MIR10A;HSPB6;PZP;HSPB2;MIR96;PKD1;CXCL5;HAPLN1;HK3;FLCN;CASP8;PDE4B;PDE4A;NOS1;POLE;DCC;USP44;AXIN1;ARAP3;WNT16;DCN;TGFBR2;MIR126;AIM2;IFNG;PALLD;CDH10;CDH12;LTA;CDH13;MYH9;CDH15;DOCK2;THOP1;RBMS3;MCPH1;NXN;RGS16;SPG7;THBS1;THBS4;MIR369;MUC1;MUC2;WNT11;ERICH1;GMDS;MIR373;MIR372;GNA12;ERBB2;SNCG;MUC4;MUC6;SNCA;TCF7L2;KDM4B;NOS2;NOS3;LIMK2;USP22;PCDHGC3;B3GAT1;PDX1;POMC;KISS1;MEIS1;MIR145;CAPN10;PPP2R2B;MIR142;CDK2AP1;LPIN1 |
| Constipation | 0.0028045305432814922 | FAAH;THRB;HPSE2;THRA;ELN;ECE1;BRCA1;TRH;AQP2;COMT;SLC6A3;UNC80;EDNRB;SALL1;IYD;SIX3;NRTN;PDGFRA;CASR;PROP1;EDN3;OPRM1;KCNAB2;ADRA2C;SDHB;DDOST;TRHR;FOXP1;TGFBR2;TNFRSF1A;SCNN1G;CLIP2;PAX8;PCCA;SCNN1B;PCCB;PGR;CFTR;GUCY2C;MAGEL2;DDC;TNXB;FOXG1;NRXN1;NPR3;PON1;ASCL1;NALCN;SLC5A5;RAI1;TPO;GJC2;EPCAM;DEAF1;PRDM16;CAMTA1;NXPH1;NKX2-5;TAC1;RREB1;GABRD;KCNJ1;NTSR1;CHGA;TBX1;CREBBP;CD160;LIMK1;B3GAT1;TBCD;MLH1;FLI1;PHOX2B;TSHR;VANGL1;SKI;NR4A2;POMC;PYY;TG;TH;GDNF;LHX3;SST;COL7A1;ALPL;GNB3;GTF2IRD1;SLC26A4;CCL28 |
| Pulmonary Hypertension | 0.002920678117765645 | CD40;CDKN1A;ALAS1;ADM;ECE1;HIF3A;CLU;FGF2;TNF;CX3CL1;HK1;SMPD3;EDNRA;EDNRB;HYAL1;MYC;HYAL2;NPPA;BDKRB2;CASR;TRPC6;TRPC3;VWF;TACR2;CYBA;HLA-A;F3;TGFBR2;GREM1;COL4A1;KCNMA1;BIRC5;CX3CR1;CALCA;EPAS1;GATA6;PDGFB;PDGFA;GATA4;KCNA5;PLA2G7;TERT;ALOX5;HAS1;CCL1;APOE;TAC1;BRD4;EGR1;ACE;ANGPT2;ANGPT1;NOS2;NOS3;GDF2;SMAD9;NFATC2;EIF2AK4;SMAD6;BMP7;MTOR;SMAD7;VEGFA;COL1A1;BMP2;IL6;BCL2;CBLN2;FGFR1 |
| Inflammatory Bowel Diseases | 0.003366633524431004 | TNFRSF6B;TRAF3IP2;WIPF1;SLC23A1;LST1;ICAM3;BACH2;MIR425;MYLK;ICAM1;ELK4;LGALS3;LGALS2;RASSF1;LGALS1;CYP2D6;ZMIZ1;TEPP;PROK2;KIF21B;TP63;ENTPD1;PRKCB;SLC11A1;KRT1;HLA-B;KRT8;HLA-C;PDZK1;MIF;HLA-A;RUNX3;HLA-G;CRNKL1;CLDN4;SFRP1;SFRP2;IL23A;PAPPA;ZNF831;EZR;CFB;CFTR;SKAP2;ABCB1;SP140;PDGFB;FPR2;CD1D;PRH1;C3;HSD11B2;TERT;CD19;SLAMF8;APOE;AMZ1;HLA-DQA1;NKX2-3;NTSR1;S100A10;STAT5A;EGR1;GSTM2;ST14;GSTM1;ACE;TNFSF14;BCL11B;LRBA;IGF2;NFATC1;MIR200B;UBAC2;MIR346;CCDC88B;FERMT1;OCLN;TG;VNN1;TNFSF4;FAT1;HAMP;HLA-DRB1;ACHE;PTPRU;DENND1B;CD40;CDKN1A;PTPRS;ICA1;TCF21;BRCA1;PTPRH;GLI3;CX3CL1;HNF4A;RAC2;TIMP2;RSPO2;MUC13;RSPO3;CCR6;JAK3;VENTX;APC2;ACAD8;SLC2A14;CBR1;RIPK3;IFNGR2;FCRL3;MST1;TACR2;IL16;MIR21;TYK2;SYN3;TMEM173;SLC39A14;CDYL2;CIT;TLR1;SLC22A23;MMP14;KAT2A;NR5A2;BANK1;IL1B;TFF3;TFF2;FAM20C;TFF1;CD47;ARHGEF2;TLR4;ERGIC1;HLA-DQB2;HLA-DQB1;GUCY2C;IDUA;ANKRD55;NR1I2;KCNA3;ADCY3;ADCY7;EGFR;GNAI2;FGD2;DUOX1;PAK1;CUX1;ALOX5;SPP1;STAT4;STX2;SPDEF;MICB;IL32;EGLN2;CDX2;IL10RB;IL34;STAT3;OSM;LILRB1;GPR3;MLH1;SNX20;LILRB4;VEGFA;PER3;KRT18;HLA-DPB1;F2RL1;LGR6;IL17D;IL17C;KLKB1;HSPA1B;HSPA1A;CCL26;CSF3;DCLRE1C;CSF2;CSF3R;NCF4;DEFB1;PLAT;PRDM1;ETS1;MPO;FGF2;TNF;CELSR3;IGF1R;HPGDS;FGF7;CCND1;DMBT1;ADORA3;GRAP2;RNF19A;MIR150;TNFRSF4;GHSR;CASR;UNC5CL;ZGPAT;S100A1;IGFBP3;F2R;MAGI2;FOXP1;SLC9A3;S100A9;ABCG2;NDRG4;CUL1;GSTT1;FHIT;LPP;ACACA;DPP4;SOCS3;SBNO2;FGF20;NLRP3;TAC1;SLC15A4;ZNF300;SMAD3;MLKL;CD160;RARRES2;HSPA2;BTNL2;BMP7;BTBD8;ESR1;SMAD7;ISG20;IL6;FER;G6PC3;PROC;IL17REL;STK24;BCL2;ALPI;FGFR2;CIITA;MIR10A;ITGB2;PTPN22;ITGAL;TREM1;CXCL5;PSTPIP1;ITGAX;DNMT3B;BDKRB2;CASP1;HORMAD2;HGFAC;HSP90AA1;TRIM63;MMP7;ITGA4;KSR1;OPRM1;TNFRSF1B;NGF;TNFRSF1A;AIM2;IFNG;MEP1A;TMBIM1;LANCL2;LTA;CHI3L1;BPI;IRF5;IRF6;LTB;CRP;PON3;RGS14;PON1;NOD1;NOD2;NTN1;MUC1;MUC2;CNR2;CBS;CCL5;GNA12;IL21R;PTK2B;NCAM1;MUC4;MUC6;NTRK1;CHGA;TCF7L2;NOS2;NOS3;B3GAT1;MUC5B;IRGM;MTOR;GALC;P2RX7;NELL1;PNKD;KLF6;P2RX5;PC;TRIM39;CD6;APC;IL2RA;LEP;IL2RB;PDCD1;TEK |
| Pancreatitis, Chronic | 0.003366633524431004 | SPARC;SERPINE2;SCT;IHH;EHMT1;FGF1;CLU;FGF2;TNF;ICAM1;LGALS3;LGALS1;CCND1;RASSF6;TNFSF10;GGT1;CPA2;ENTPD1;CASR;CPA1;ENTPD2;MSX2;HLA-B;KRT8;HLA-A;MIF;BHLHA15;HIC1;SFRP1;IFI27;CRISPLD2;CYP2E1;CFB;CFTR;NOTCH3;ABCB1;GSTT1;ABO;ST8SIA4;NLRP3;APOE;PROM1;TAC1;NTSR1;NGFR;TFPI2;TNFRSF10C;NFATC1;MIR200B;TNFRSF10A;HSPA2;COL1A1;ISG20;IL6;FXYD3;ARTN;HLA-DRB1;MIR10B;PSCA;ADM;KRT20;MSLN;CX3CL1;CCR6;TK1;MAP4K1;MMP7;DUSP1;TACR2;TACR3;MIR21;NGF;TNFRSF1B;F3;DCN;GREM1;CCKBR;IFNG;IL1B;CAT;LTA;HCLS1;CHI3L1;TFF1;ANG;CX3CR1;UHRF1;COL11A1;PON1;THBS1;EGFR;PTHLH;MIR148A;LMF1;MUC1;MUC2;CUX1;BGLAP;NTF3;ERBB2;SPP1;MUC4;UGT1A7;MUC6;NTRK1;CDX2;BNIP3;STAT3;KLHL1;PDX1;CCK;VEGFA;P2RX7;KRT18;IL2RA;LEP;APOC1;IL2RB;F2RL1;SIGLEC7;CCL28;HSPA1B;HSPA1A |
| Hypotension | 0.0034626198461428577 | RYR1;UCN;DDC;CLCNKB;PRKAG2;ADM;OXT;PLAT;ADRA1B;SERPINA6;FGF1;TNF;GLI2;INS;HRH1;MUC1;GNA11;NPPA;ADORA1;BDKRB2;CDH23;OTX2;TAC1;CASR;PROP1;ACE;EDN3;HTR1D;HTR1B;PDYN;GRIN2B;POMC;SCNN1G;SLC9A3;IL6;ADORA2A;IFNG;IL1B;SCNN1B;CAT;SCNN1A;PNOC;CRH;GHRL;LTF |
| Eating Disorders | 0.004102874300165641 | HDAC4;CNTFR;OXTR;CSF2;ATP8A2;LAMC2;HTR2A;COMT;SLC6A3;CRHR1;RXFP3;CRHR2;CCKAR;CUX1;CYP2D6;DRD4;ESRRA;NTRK1;GHSR;GABBR1;LMO3;NTRK3;HTR1D;IGF2;KRT7;CARTPT;NGF;TNFRSF1B;ESR1;TAL1;LEP;IL2RB;CRH;HCLS1;GHRL;SCLY;SIGLEC7;ABCG1 |
| Epithelioma | 0.00441479580898 6965 | CSF1;CLU;FGF2;TNF;IGF1R;RASSF1;LGALS1;CDH1;MYC;AKT1;TNFSF11;HOXA7;TP63;DAXX;POSTN;F2R;KRT8;ACSL5;TSC2;HNF1A;MAT1A;RUNX3;HLA-G;RUNX1;SFRP5;FEZ1;FSCN1;ZNF154;CSF1R;NOTCH3;RTEL1;ABCB1;TCF7;TACSTD2;KLK4;MST1R;LYPD5;FHIT;FUT1;RAP1GAP;BCL2L10;TERT;UBD;NEU1;PROM1;JAG2;EGR1;NGFR;ST14;HMGA1;PTK6;NR0B2;PHC3;ESR1;COL1A1;DAB2;IL6;ID2;BCL2;FGFR2;SNED1;CD40;PIK3CD;BRCA1;ITGAE;MMP21;PIK3CG;HHEX;CASP8;MB;HNF4A;PSD4;TIMP2;BRD2;EGFL7;TRIM62;MMP7;ANXA2;KSR1;MIR21;NGF;CNKSR1;MMP14;ZEB1;MMP23A;CAT;RARB;SGK1;LTF;HDAC3;NR1I2;TMPRSS2;EGFR;MIR148A;TTN;MUC1;BGLAP;EPCAM;ALOX5;HAS1;HSF1;ERBB2;SPP1;MUC4;NTRK1;MGMT;TBX5;PSMB9;VEGFA;TBX2;MEIS1;MIR145;TEC;KRT17;WT1;LEP;KRT14;NBR1;PTPN6;VIM;LGR6 |
| Mammary Neoplasms, Human | 0.0044240864772353145 | RB1;CSF3;CSF2;CSF1;MIR429;ICAM5;COMT;TNF;FGF3;ETS2;IGF1R;LOXL2;CDH5;EDNRB;CCND1;CYP2D6;CDH1;STMN1;TNFSF10;AKT1;CYP1B1;VPS39;TP63;KCNH1;TGM2;TLE3;CPT1A;FKBPL;RPS8;KRT8;DIO3;PDZK1;KRT5;MIF;DKK1;HIC1;CLDN4;SFRP1;HOXB9;SFRP2;DDIT3;SFRP5;CD109;PGR;ZNF432;DSC3;LLGL2;ABCG2;CSF1R;NOTCH3;GRB7;ABCB1;SHMT1;NOTCH4;PDGFA;DTX3;HOXB13;MIR206;SLC5A5;LDHB;TERT;STC2;BARD1;JAG2;ESRRA;STAT5A;TFAP2A;CADM1;TFPI2;PDE2A;XRCC3;NR2F1;MIR200B;SYNJ2;NMBR;ESR1;NR2F6;BMP4;H19;OCLN;BMP2;IL6;RALYL;ARTN;CYP1A1;BCL2;HRG;SIM1;FGFR2;TP73;APOBEC3B;FGF10;FGFR1;ACHE;CD40;MIR10B;PCDHGB6;MIR10A;FHL2;SLC2A2;ADAR;LSP1;BRCA1;FOXM1;SLC2A5;SYNE1;LIMD2;AKAP12;HHEX;UPK1B;CASP8;MECOM;HEY1;SLC22A18;DNMT3B;RSPO3;MRPL9;CLIC1;APC2;HSP90AA1;GPX1;MME;DNMT3A;MST1;NRG1;F3;SERPINB5;ACTA2;MIR126;TOX3;MMP14;ZEB2;ZEB1;RAD51C;IFNG;IL1B;CAT;RARA;RARB;BIRC5;MYH9;EVL;ANGPTL4;SRC;GSTP1;HIST1H2BK;HOXD11;THBS1;EGFR;PTHLH;CYP17A1;GNAI2;RXRB;PAK1;CNR2;ERBB2;SPP1;SNCG;RIBC2;WWOX;CD74;NQO1;NOS2;NOS3;STAT3;RPL23A;KLK10;TBX3;HEYL;MIR24-2;MEIS1;KRT18;PER3;MIR145;WT1;FABP7;LEP;COL7A1;KRT14;VIM;LGR6;HSPA1B |
| Sciatic Neuropathy | 0.004522288972752703 | CX3CR1;CALCA;ELAVL4;OPRL1;PLAT;CACNA1D;MIR96;TNF;ICAM1;CYP17A1;C3;CNR2;MAP2;CCL5;BDKRB2;BDKRB1;NEFL;NEFM;SLC17A7;APOE;NEFH;JAM3;PDGFRB;PDGFRA;NGFR;KCNJ8;KCNJ11;APOA1;GFRA1;NRG1;GFRA3;TNFRSF1B;MTOR;TNFRSF1A;RGMA;MAG;IL6;GDNF;IL1B;CAT;NOS1AP;CRH;RARA;RARB;TLR4;RAN;FGFR1 |
| Acute Myeloid Leukemia (AML-M2) | 0.004522288972752703 | CSF3;CSF2;SPARC;SPI1;TRIO;GFI1;EHMT2;CTSZ;FHL2;TRH;IFI30;MPO;BACH2;LYL1;HOXA9;CCND2;ANPEP;MYC;CAPN2;TNFSF10;PDE4B;BAALC;DLEU2;ANXA2;DAPK1;DNMT3A;TSC2;RUNX3;RUNX1;PXDN;RARA;ERG;SGK1;CSF1R;ABCB1;CAPG;GATA2;ADCY7;SYNGR1;CNR2;CTNNA1;NCAM1;S100A10;RUNX1T1;SVIL;STAT3;OPRK1;POU4F1;CDK6;WT1;ID2;ASXL2;BCL2;TCEA2;CD9;FXYD6;AGRN |
| Acute leukemia | 0.004978187185747897 | RB1;APP;CSF3;CSF2;CSF3R;SCT;EHMT2;MPL;C4BPA;IKZF1;TNF;ETS1;MPO;ETS2;HOXA11;RIMS2;CDH5;HOXA9;VPREB1;HPGDS;RASSF1;CYP2D6;MYC;ANPEP;STMN1;RASSF6;TNFSF10;AKT1;BAALC;MIR150;PITX2;SRRM2;PDGFRA;EBF1;HLA-B;HLA-C;LIG4;TSC2;MIR27A;HLA-A;RUNX3;UNC13D;RUNX1;DHDDS;THEG;TFR2;IL23A;DDIT3;SFRP5;ERG;MAD1L1;CFTR;ABCG2;CSF1R;RTEL1;CALCA;ABCB1;MIR196B;GSTT1;THY1;GATA2;FHIT;EVPL;LPP;ABO;ZFP36L2;FUT4;SULT1A1;SOCS2;DPP4;BCL2L10;CYP2B6;PCBP4;CD19;INPP5D;NLRP2;ZNF423;LYN;TMEM132D;STAT5A;GTF2A1L;NGFR;CREBBP;ANKRD26;GSTM1;SMAD3;BCL11B;ZBTB16;MRPL28;NR2F2;NR0B2;FLI1;PBX1;ARHGAP24;TF;OCLN;IL6;BCL6;TLX1;CYP1A1;BCL2;HLA-DRB1;CD22;CD200;TP73;FGFR1;RARG;CNTNAP1;LDB1;PRSS27;PIK3CD;BRCA1;CD3E;AFF3;PIK3CG;AKAP12;LYL1;MECOM;TIMP2;CD38;CCR7;JAK3;CD34;HSP90AA1;APOBEC2;ITGA4;MME;MIR23A;TESC;DNMT3A;IL16;PAX5;TNFRSF1B;TNFRSF1A;TAL1;MIR17HG;RARA;HCLS1;MYH9;BIRC7;TLR4;LTF;RBMS3;LAMA1;GSTP1;PON1;HOXD13;NOD2;AURKB;CRYZ;ARFRP1;RASSF10;CUX1;HLX;NPY;NNAT;MXI1;PTK2B;PRAME;NCAM1;EIF4E;EPS15;CD55;CYTH1;RUNX1T1;NQO1;MARCKSL1;PKLR;ANGPT1;CDX2;MGMT;STAT3;STYK1;SORL1;VEGFA;MIR24-2;KLF6;MEIS1;CD6;WT1;LEP;POU2AF1;IL2RB;CD7;HLA-DPB1;PTPN6;SIGLEC7 |
| Adenocarcinoma Of Esophagus | 0.004978187185747897 | OXTR;SPARC;CRTC1;FGF1;FGF2;TNF;ICAM1;IGF1R;SOX2;CCND1;CDH1;MYC;STMN1;TNFSF10;LGALS9;SOX9;PITX1;SLC12A9;SRRM2;PDGFRB;TMOD1;IGFBP3;F2R;KRT7;TSC2;HNF1A;RUNX3;DKK1;DKK3;FOXP1;SFRP1;ELMO1;PKP1;PGR;CFTR;KHDRBS2;GATA6;GATA5;GATA4;ADRB2;PRKCZ;FHIT;C3;MIR330;TPPP;PROM1;TAC1;FOXF1;XRCC3;MIR200A;TNFRSF10A;ESR1;DCLK1;ARHGAP24;GDF7;MSRA;IL6;BCL2;NOX5;FGFR4;TP73;EIF4G1;CD40;CDKN1A;HSPB2;OXT;FOXM1;AKAP12;CASP8;SLC22A18;TIMP3;MIR192;CTSD;APOBEC1;MMP7;AMPD1;MIR21;MMP14;ALDH1A2;MIR17HG;RARA;CDH13;TFF1;ARHGEF7;DOCK2;TLR4;RBMS3;UHRF1;GSTP1;ASIP;BARX1;EGFR;MIR148A;CRYZ;SELENBP1;MUC1;MUC2;EPCAM;ERBB2;SPP1;EIF4E;CD55;RBM38;NQO1;CDX2;MGMT;GDF15;NOS3;ASCC1;STAT3;AKR1C2;TBX5;MTOR;VEGFA;SKI;NELL1;MIR145;APC;LEP;GHRL;VIM |

| | | |
|---|---|---|
| Endometriosis of uterus | 0.005052459263652123 | NDUFA13;MIR10B;COMT;FGF1;FGF2;TNF;CYP17A1;PAK1;CYP2B6;TIMP2;TBX1;GSTM2;GSTM1;MMP7;ANXA2;DNMT3A;ESR1;F3;VEGFA;IL6;ZEB1;NBR1;BCL2;CYP1A1;PGR;TLR4 |
| Medullomyoblastoma | 0.005052459263652123 | MPL;EHMT1;IRS2;PIK3CG;RASSF1;MYC;ERBB2;OTX2;PRAME;SMYD4;ATOH1;S100A10;BRD4;PDGFRA;BRD2;KDM4B;STAT3;PTCH2;FLI1;HIC1;VAV1;CDK6;MYCN;APC;S100A6;SCG5 |
| Melanotic medulloblastoma | 0.005052459263652123 | MPL;EHMT1;IRS2;PIK3CG;RASSF1;MYC;ERBB2;OTX2;PRAME;SMYD4;ATOH1;S100A10;BRD4;PDGFRA;BRD2;KDM4B;STAT3;PTCH2;FLI1;HIC1;VAV1;CDK6;MYCN;APC;S100A6;SCG5 |
| Undifferentiated leukemia | 0.005360832834312189 | KDM5B;CSF1R;CNTNAP1;ABCB1;ALOX15;FOXM1;GATA2;EVPL;FUT1;GLI2;LYL1;HOXA9;MECOM;SALL4;DNMT3B;TNFSF10;CD38;CCR7;CD34;EIF4E;STAT5A;TBX1;CD96;SNRPN;CD93;F2R;KRT7;RUNX1;VEGFA;MIR126;ISG20;MEIS1;BCL6;TAL1;IL2RA;SCLY |
| Peripheral Vascular Diseases | 0.00544361065551013 | CX3CR1;ELN;PON1;MPL;PCSK9;ADM;FGF2;MPO;TNF;THBS1;ABO;ICAM1;SELENBP1;LIPC;CCND1;SPP1;APOE;CCHCR1;APOB;P2RY12;ABCG8;ABCG5;GPX1;TRIM63;ACE;NOS3;F12;DAB2IP;APOA1;TNFRSF1B;F3;VEGFA;DHX40;F7;IL6;CYP1A1;CD47;SHBG;TLR4;CFTR |
| Carcinoma, Transitional Cell | 0.005735864001710423 | CSF3;CSF2;MSI1;FGF2;TNF;LGALS4;LGALS3;FGF7;HPGDS;RASSF1;LGALS1;CCND2;CCND1;CDH1;MYC;STMN1;TNFSF10;AKT1;CYP1B1;LZTS1;TP63;SLC12A9;TGM2;SRRM2;PDGFRB;TMOD1;DAPK1;S100A1;IGFBP3;WNT5A;LIG4;TSC2;KCNK17;HLA-G;LTB4R2;COL4A1;DDIT3;CYP2E1;ZNF671;SLC22A3;ABCB1;TERC;EIF5A2;GATA6;CUL1;GSTT1;GATA3;PIK3R1;FHIT;ABO;SULT1A1;TERT;HRH2;NEU1;NDN;KCNN3;TRPM8;BCL2L14;CREBBP;GSTM2;GSTM1;SMAD3;IGF2;TRPV2;MRPL28;HSPA2;INHBA;TMPRSS13;ESR1;DAB2;IL6;BCL6;CDK2;BCL2;GNAS;TGFBI;FGFR3;LAT;CD200;FGFR2;FGFR1;HOTAIR;CDKN1A;CD40;MIR10A;NUMA1;PSCA;DOC2A;PIK3CD;BRCA1;GRIK2;FOXM1;MSLN;CD3E;KRT20;PIK3CG;LTB4R;GLI2;ELAVL3;UPK1B;CASP8;UPK1A;DNMT3B;SH3BGRL3;CD34;BOK;GPX1;MMP7;ANXA2;GSTO2;DUSP1;DCC;DNMT3A;MST1;APOA1;EMP2;MIR21;NME5;UPK3A;POU5F1;DCN;SLC7A5;KAT2A;MMP14;IL1B;ANGPTL4;RBMS3;HDAC2;GSTP1;COL11A2;THBS1;EGFR;HLA-DMA;MUC1;CYP2A13;TMEM67;ERBB2;MXI1;PTK2B;PRAME;CTNNA3;EIF4E;WWOX;MARCKSL1;DAB2IP;MLH1;MTOR;VEGFA;KISS1;FABP2;MIR145;CD9;PDCD1;HSPA1B;HSPA1A |
| Adenocarcinoma, Basal Cell | 0.005911530181914705 | DDR1;CDKN1C;SYCP1;MSLN;KRT20;TNF;HOXA10;PYCARD;RASSF1;CASP8;CCND1;CA2;MYC;NPPA;PLCE1;NOS1;TP63;DAPK1;TPM3;KRT8;UGT2B17;APOA1;CYP7B1;RUNX3;MMP14;KCNQ1;IL1B;BIRC5;RARB;PGR;CYP2E1;GRB7;NR1I2;EGFR;SELENBP1;MUC2;ERBB2;SPP1;APOE;STAT5A;TCF7L2;ACE;NOS2;NOS3;STAT3;TNFRSF10A;ESR1;MTOR;VEGFA;CYP26A1;APC;BCL2;ID3;GHRL;FAT4 |
| Carcinoma, Small Cell | 0.005911530181914705 | RB1;PIGR;THRB;PIK3CD;KRT20;PIK3CG;IGF1R;SOX2;PYCARD;LGALS3;RASSF1;CCND1;CYP2D6;NPPA;DNMT3B;SGCG;PDGFRA;CBR1;NRG1;PAX5;POU5F1;TGFBR2;FEZ1;BIRC5;CYP2E1;ERG;SLC22A3;CALCA;GRP;GSTP1;SPATA2;TMPRSS2;SEZ6L2;ASCL1;FHIT;EGFR;TERT;MAP2;ERBB2;NCAM1;SNCA;CHGA;CREBBP;GSTM1;CDX2;CCK;MLH1;NR0B2;MTOR;POMC;BCL2;CYP1A1;PDCD1;FGFR1;FOXA2 |
| Acute Undifferentiated Leukemia | 0.006287328841029159 | KDM5B;CSF1R;EPO;DERL3;ALOX15;FOXM1;GATA2;MPO;FUT1;GLI2;LYL1;HOXA9;MECOM;ANPEP;SALL4;DNMT3B;TNFSF10;CD38;BAALC;CCR7;CD34;EIF4E;STAT5A;TBX1;CD96;SNRPN;CD93;F2R;KRT7;RUNX1;VEGFA;MIR126;ISG20;MEIS1;BCL6;TAL1;IL2RA;ERG;SCLY |
| ovarian neoplasm | 0.006287328841029159 | SPARC;BACH2;ICAM1;AQP1;RASSF1;EDNRA;FRAT1;RASSF2;BAIAP2L1;MYC;TNFSF10;AKT1;CYP1B1;PRSS8;SOX9;HOXA7;TP63;HOXA5;SLC12A9;HOXA4;SRRM2;ARL11;SERPINF1;HLA-B;KRT7;HNF1B;PRKCA;DLEC1;MIF;UNC93A;RUNX3;HLA-G;RUNX1;CLDN4;SFRP1;SFRP2;HOXB9;IFI27;SLPI;PAPPA;RBP1;SKAP1;GPLD1;ABCB1;CTBP2;TERC;SHMT1;HPN;EIF5A2;GATA6;PDGFB;KLK4;GATA4;GATA3;THY1;MST1R;PIK3R1;KLK9;KLK8;CRHR1;KLK7;SLC5A5;KLK6;SULT1A1;HSD11B2;TERT;NEU1;HLTF;SFN;APOE;RREB1;HLA-DQA1;STAT5A;BARD1;NAPSA;ST14;GSTM1;GREB1;IGF2;LIF;MRPL28;PTK6;NFATC1;PBX1;IFFO1;GJB2;ID2;GNRH2;CYP1A1;GRB2;HOXD9;TDGF1;HLA-DRB1;LGMN;TP73;DDR1;DIRAS3;CDKN1C;ACHE;ROBO3;COL18A1;CD40;CDKN1A;MIR17;ADM;BRCA1;PCSK6;AKAP13;MECOM;HYAL1;TIMP2;MUC13;TK1;CD34;CTSD;BRD2;GSTO2;MME;DUSP1;TUBB;NME2;IL16;MIR21;F3;DUSP6;POU5F1;PAX2;SERPINB5;VAV2;TLR1;MMP14;ZEB1;RAD51C;CDCP1;CTTN;PAX8;KIFC1;IL1B;RARA;TFF1;SGK1;TLR4;GRP;GSTP1;TMPRSS3;TMPRSS4;NEDD9;HOXD11;EGFR;CAMKK2;PAK1;NT5E;HLA-DMB;CUX1;EPCAM;HAS1;PLAGL1;GPC1;ALOX5;HSF1;SPP1;CTNNA1;CTNNA3;SPDEF;RUNX1T1;WWOX;MGMT;GDF15;STAT3;KLK13;KLK15;SULF1;MLH1;KLK10;GRHL2;KLK11;PSMB9;VEGFA;WT1;DLC1;NBR1;ZNF217;YPEL3;CSF2;CSF1;EPO;LY75;MPL;PLAT;FGF1;ETS1;ACRBP;CLU;FGF2;TNF;ETS2;HPGDS;FGF7;CCND2;FGF8;CCND1;LAMP1;CDH1;PEBP4;NPPA;CHP2;ZNF569;PITX2;GGT1;TGM2;PDGFRB;PDGFRA;LHCGR;POSTN;IGFBP3;HIC1;FOXP1;ADAM19;LTB4R2;COL4A2;CHMP4C;ZNF436;PGR;SQSTM1;ABCG2;CSF1R;NOTCH3;CALCA;DLX4;HTRA1;CHRD;GSTT1;VTCN1;HOXB13;FHIT;NME1-NME2;TPO;MSC;ESRRA;SMAD2;UBE2I;EPHX1;TNFRSF10C;HMGA1;TNFRSF10A;ISG15;DNAJC15;ESR1;IGF2R;ARHGAP24;SMAD7;BMP4;IL6;DAB2;COPS5;CDK2;BCL2;CPE;PLXNB1;CLPTM1L;TGFBI;SHBG;FGFR4;FGFR2;LGALS3BP;USP36;THRB;FLT4;PIK3CD;CHD5;LAMC2;ITGAE;FOXM1;MSLN;PIK3CG;LTB4R;CYR61;PSD4;PSD3;DNMT3B;AMH;INSL3;MMP7;PEG3;TRPC3;DCC;NRG1;DCN;TGFBR2;TGFBR3;MIR485;IFNG;CDH11;ITGA8;CRH;BIRC5;CDH13;BCAT1;OTUB1;RBMS3;CRP;HDAC3;SRC;ZBTB46;PON1;SELENBP1;MUC1;ACVR1C;MUC2;RAB25;CBS;ERBB2;IGF2BP1;SNCG;EIF4E;MUC4;OPCML;NTRK1;MARCKSL1;ANGPT2;NOS2;CABLES1;TBX3;MTOR;KISS1;POMC;KLF6;APC;SST;ATP13A4;S100P;ZACN;BCAR1 |
| Myocardial Infarction | 0.0063660112431184485 | SPARC;HPSE2;MYLK;ICAM1;LGALS2;EDNRA;LIPC;EDNRB;CYP2D6;AKT1;TNFSF11;CYP1B1;MEF2C;ENTPD1;NUP210;NFKBIL1;SERPINF2;HLA-B;HLA-C;ANK2;MIF;HNF1A;GPR162;SFRP2;SLPI;BIN1;DDIT3;KCNQ1;PAPPA;GPR160;JTB;SPTBN4;ABCB1;AIRE;EPAS1;PDGFA;KLK4;GATA4;PLA2G7;PRH1;ABO;C2;SLC5A5;C3;HSD11B2;CYP2B6;TSPAN9;KCNQ1OT1;PDGFD;APOH;APOD;APOE;A2M;APOB;PROM1;ELANE;EGR1;GSTM2;GSTM1;ACE;PTPRN2;ABCA4;SLC33A1;IGF2;OSGIN1;LIF;GP6;COL1A1;MSRA;ID2;TNFSF4;CYP1A1;NOX5;ANP32B;HRG;HAMP;TP73;PCSK2;CDKN1C;ACHE;CD40;CDKN1A;MIR17;TCF25;PIGQ;ELN;PCSK9;ADM;GPT;BRCA1;B3GNTL1;GLI3;SERPINA4;RYR3;CX3CL1;JPH3;SLC22A18;TIMP2;TIMP3;CD34;CTSD;TIMP4;RIPK3;ELOVL5;ELOVL2;IFNGR2;MST1;ITPK1;AMPD1;CACYBP;GP1BA;MIR21;F3;SORCS2;COMMD7;NCOR2;F7;MMP14;CDCP1;ADRB3;SLC7A9;IL1B;KCNMA1;TFF1;ANGPTL4;MAPT;ARHGEF7;TLR4;ERGIC1;GSTP1;DERL3;NR1I2;KCNA3;CXCR5;FSTL1;CYP17A1;REM1;MTHFD1L;ALOX5;NPY;SPP1;APOL1;DISC1;RUNX1T1;MPZL2;P2RY12;F10;SORT1;GDF15;STAT3;F11;DAB2IP;OSM;LILRB1;VEGFA;FABP2;FABP3;PER3;F2RL1;GHRL;RAMP1;HLA-DPB2;SCARB1;CSF3;CSF2;EPO;MPL;F13A1;PLAT;COMT;MPO;FGF2;TNF;IGF1R;BEST1;HPGDS;CCND1;ADORA3;NPPA;ADORA1;MIR150;GGT1;TNFRSF4;MAP3K5;GHSR;ABCG8;CASR;USP7;POSTN;ABCG5;CD93;S100A1;SWAP70;F2R;DIO3;CYBA;DGKZ;HIC1;PROCR;COL4A2;COL4A1;HHIPL1;ADAM8;S100A9;PVALB;ABCG1;CUL9;NOTCH3;CHRNA5;TUBGCP2;CHAT;NPR3;ALOX15;GSTT1;LTBP2;ADRB2;DPP4;SOCS3;GRK4;UBD;NLRP3;SMAD2;SMAD3;GFPT2;HTR1D;EPHX1;INHBA;TRAPPC9;ESR1;IGF2R;GFAP;SLC2A9;IL6;DAB2;FES;BCL2;GNB3;ZNF536;SHBG;SLC26A4;FGFR3;DIDO1;CIITA;ST6GALNAC1;ZC3H3;HSPB2;PIK3CD;LAMC2;HTR2A;TREM1;PKD1;PIK3CG;TSPAN10;CASP8;CA3;MB;BDKRB2;CASP1;BDKRB1;PHACTR1;LRRFIP1;LBP;NOS1;GPX1;TRPC3;VWF;ITGA3;MIR23A;APOA1;NRG1;TNFRSF1B;NGF;DCN;TNFRSF1A;TGFBR2;ACTA2;MIR126;IFNG;PALLD;DDAH2;CHDH;CAT;LTA;BIRC5;CHI3L1;CDH13;MYH9;BPI;ITGA6;IRF5;ANG;LCP1;IRF6;LTB;MYH7;CRP;CX3CR1;ITIH3;LRP1;UHRF1;LAMA3;PON1;LCAT;LTC4S;NTN1;THBS1;THBS4;INS;TTN;WNT11;CNR2;CBS;CCL5;TBX20;TNNI3;FSD1;ACSS1;ATOH1;ANGPT2;ANGPT1;KCNJ11;NOS2;TXNRD2;ASCC1;NOS3;BNIP3;TXNRD1;PDX1;OPRK1;TBX5;SOD3;TBX3;MTOR;DLK1;PC;MGP;APOC1;RGS12;P4HB;TEK;TAF4;KCNK2 |
| Lymphoid leukemia | 0.00638298382936936 | COL18A1;UCN;CD40;CSF3;CNTNAP1;MPG;GFI1;MPL;PLAT;IKZF1;SERPINA6;ETS1;MPO;TNF;BACH2;SMPD3;LYL1;HPGDS;SIX6;CCND2;CASP8;MECOM;ANPEP;MYC;BLNK;CD38;BAALC;CD34;PDGFRA;HSP90AA1;MME;KDM2B;CTDSPL;KRT7;IL16;PAX5;TYK2;ATP11A;RUNX1;FCER2;ZAP70;MYCN;IFNG;TAL1;LCK;DDIT3;IL1B;RARA;MYADM;PRTN3;ERG;EPHA1;TLR4;CSF1R;NOTCH3;ABCB1;UHRF1;MIR196B;EVPL;ZFP36L2;ZFP36L1;TERT;CD19;NNAT;HOXC5;SPP1;RUNX1T1;LYN;TMEM132D;PKLR;STAT3;B3GAT1;MLH1;PBX1;ARHGAP24;ISG20;IL6;BCL6;WT1;FES;TLX1;IL2RA;CDK2;BCL2;GRB2;CLPTM1L;SCLY;CD22;TP73;FGFR1 |
| Bronchopulmonary Dysplasia | 0.006428947426150352 | CHRM2;GABRB2;ELN;PLOD2;GRIK2;COMT;FOXM1;MPO;TNF;SLC6A2;LOXL1;SLC6A3;LOXL2;FGF7;TIMP2;AKT1;SLC18A2;CMKLR1;PDGFRB;PDGFRA;POSTN;HSP90AA1;BRD1;NRG1;MIF;CACNA2D4;F3;EML2;F7;IFNG;MMP16;CRISPLD2;SCN8A;IL1B;KCNQ2;CRH;CLDN18;PRTN3;BPI;SCN4B;TLR4;PVALB;HDAC2;NRN1;GSTP1;GATA6;SLC1A2;KCNA3;CACNA1C;MIR206;EGFR;NPAS3;CYP2B6;TSPAN8;SPOCK2;NLRP3;TAC1;MSC;DISC1;DRD4;ELANE;ZBED4;NQO1;CCL24;ACE;ANGPT1;NOS3;GAD1;ZNF804A;VEGFA;P2RX7;IL6;PC;SST;BCL2;CYP1A1;GNAS;F2RL1;FGFR2;SIPA1;FGF10 |
| Alcohol abuse | 0.0064943285813166135 | FAAH;SPI1;CD81;HTR2A;COMT;TNF;SLC6A3;CCKAR;CCND1;CDK20;PDE4B;SOSTDC1;GGT1;CCHCR1;GHSR;TACR3;OPRM1;PDYN;CRH;MCM5;CYP2E1;ANKK1;TKT;TLR4;HDAC2;CHRNA5;OPRL1;GATA4;ABO;CRHR1;EGFR;SELENBP1;GALR3;CNR2;NPY;PENK;FYN;APOE;DRD4;SNCA;SMAD2;GABRA1;GABBR1;KCNJ6;HTR1A;CCK;POMC;TF;LEP;SHBG;HAMP |
| Malignant neoplasm of esophagus | 0.00674867688087647 | RB1;MDC1;ICAM2;ICAM3;ICAM1;SOX2;LGALS3;FRAT1;RASSF1;RASSF2;SOX17;MYC;PEG10;MAGOH;TNFSF10;AKT1;CYP1B1;PLCE1;HOXA1;SOX9;TP63;SLC12A9;SRRM2;NKX2-8;DAPK1;AFAP1;KRT7;DLEC1;CSNK1E;HLA-A;DKK1;RUNX3;RUNX1;MIR196A2;CRISPLD2;DDIT3;MT1G;CYP2E1;EZR;NEIL1;LLGL2;MEG3;GRB7;SLC22A3;ABCB1;TERC;GATA5;TWIST1;GATA4;THY1;SULT1A1;ADAMTS16;TERT;KCNQ1OT1;STC2;HLTF;SFN;EGR1;TRAP1;GSTM2;ACE;GSTM1;FZD5;FZD7;XRCC3;IGF2;ALOX15B;DCLK1;GADD45G;NET1;CYP26A1;GJB2;CYP1A1;FAT4;HLA-DRB1;TP73;CDKN1C;HOTAIR;CDKN1A;BRCA1;BBC3;AKAP12;AKAP13;SLC22A18;TIMP2;TIMP3;CD38;MIR192;CD34;GPR39;ZHX2;ANXA2;RIPK3;AMPD1;MIR21;PROX1;PAX2;POU5F1;SLC7A5;MMP14;CDCP1;CTTN;IL1B;MIR17HG;RARA;RARB;CRYAB;TLR4;GRP;GSTP1;NR1I2;CRCT1;EGFR;NKD2;RASSF10;EPCAM;WIF1;SPP1;IL32;WWOX;NQO1;CDX2;MGMT;STAT3;MLH1;VEGFA;GHRL;IL17D;LY6K;CSF3;ZBTB20;PRDM2;FGF2;TNF;ETS1;HOXA13;MPO;FGF3;IGF1R;LOXL2;RIMS2;FGF7;HPGDS;CCND1;DMBT1;CDH1;GRAP2;RNF19A;LZTS1;TGM3;TGM2;PDGFRB;USP7;TPM4;USP4;IGFBP3;UBE2E2;MIR27A;FSCN1;DPEP1;S100A9;DIABLO;ALOX15;LTBP4;GSTT1;VTCN1;FHIT;STK31;MIR296;SMYD2;SMAD2;TFPI2;EPHX1;TNFRSF10A;ISG15;MCC;ARHGAP24;DAB2;IL6;NAT1;HOOK2;CDK2;BCL2;CNTN1;GNAS;SCGB3A1;MIR183;CLPTM1L;TGFBI;XAF1;FGF12;FGFR1;MIR10B;THRB;HSPB2;PIK3CD;MSLN;MIR96;CYR61;PIK3CG;WISP1;ZNRD1;CASP8;DNMT3B;ADAMTS9;MMP7;DCC;TGFBR2;IFNG;BIRC5;CDH13;MCM5;RBMS3;CRP;LRP1;SRC;FOXK1;MIR148A;CRYZ;MUC1;MIR377;DNAJB6;PUF60;GNG7;MIR373;ERBB2;SNCG;FSD1;RGS22;EIF4E;MUC4;MARCKSL1;ANGPT1;PRRX1;NOS2;NOS3;TXNRD1;FBXL19;BNIP3;EYA4;CCK;MTOR;CHFR;MIR145;APC;SST |
| ESR raised | 0.007764335400995066 | MVK;ANKRD55;TREX1;PTPN22;MIF;TYK2;TMEM173;PSMB8;TNFRSF1A;IL6;IL2RA;IL2RB;STAT4;HLA-DPB1;NLRP3;PRTN3;HLA-DRB1 |
| Amphetamine Addiction | 0.008491144742444551 | ACHE;GSTP1;PLAT;PRDM2;GSTT1;AOAH;AK5;COMT;FHIT;SLC6A3;GRM2;CYP2D6;PSD3;PDE4B;AKT1;NOS1;CCHCR1;ZNF423;CSMD1;CD55;DRD4;SNCA;GSTM1;PDE4D;LMO3;EPHX1;HTR1B;NRG1;HTR3A;OPRM1;PDYN;SGCZ;POU5F1;SH3RF3;ADORA2A;GDNF;SLC6A9;CDH13;ASTN2 |
| Fever | 0.008761689647932206 | RYR1;CSF3;DCLRE1C;WIPF1;NCF4;ICAM2;PTPN22;IKZF1;HTR2A;AQP2;TNF;PSTPIP1;CCND1;BLNK;NOS1;DST;SLC11A1;GAA;MST1;HLA-B;LIG4;CYBA;MIF;TMEM173;UNC13D;RUNX1;FOXP1;TNFRSF1A;IFNG;CYP21A2;IL1B;KCNQ2;ASPG;CRH;IRF8;PRTN3;FBP1;TLR4;FBN2;NOTCH3;MVK;COL11A2;HTRA1;KCNA3;CACNA1A;PIK3R1;NOD2;GATA2;THBS1;CD79B;HMGCL;HSF1;SLC19A3;STAT4;NLRP3;KCNJ1;ELANE;TREX1;STAT3;HTR1A;CCK;BTNL2;UBAC2;COL1A1;POMC;IL6;NLRP12;BCL6;BCL2;HLA-DPB1;ALPL;LPIN1;LPIN2;HLA-DRB1;FGFR1 |
| Craniofacial Abnormalities | 0.008900429203669201 | THRB;UXS1;FOXI1;RAX;ECE1;ETS1;ETS2;FGF3;HAPLN1;GLI3;ACTG1;EDNRA;CYP26B1;FGF8;SALL3;SALL4;TRPS1;SH3PXD2B;HOXA3;DNMT3B;HOXA1;SOX9;PITX2;TP63;PDGFRA;UNC45B;MSX2;DNMT3A;PAX3;PAX6;RUNX2;FOXP2;TGFBR2;COL2A1;CHDH;HOXB1;IRF6;B4GALT7;FBLN7;DLX1;COLEC11;DLX2;DLX5;COL11A1;NRXN1;SATB2;COL11A2;CHRD;TWIST1;GATA2;EGFR;DLL3;RAI1;ANKH;GSC;ALX4;ALX1;FADD;MSX1;DCPS;STRA6;JAG2;SMAD2;TFAP2A;TBX1;TFAP2B;SMAD3;EYA1;EPHX1;ZIC5;FLI1;LETM1;SKI;TBX15;SPRY2;GTF2IRD1;SIM2;FGFR2;VSX1;FGFR1 |
| Ulcerative Colitis | 0.008900429203669201 | FAAH;TNFRSF6B;TRAF3IP2;SLC23A1;AQP8;ICAM2;IKZF1;BACH2;ICAM1;LGALS4;SOX2;LGALS3;RASSF1;ZMIZ1;PROK2;UBASH3A;LGALS9;KIF21B;SLC12A9;PRKCB;SERPINF1;NFKBIL1;SLC11A1;HLA-B;KRT8;KRT7;MIF;FNDC3A;HLA-A;RUNX3;HLA-G;CRNKL1;CLDN4;SLC5A8;SFRP1;SFRP2;SLPI;MIR196A2;BIN1;IL23A;PRKCQ;ZNF831;CFB;CFTR;SKAP2;ABCB1;MIR196B;SP140;TWIST1;FPR2;MST1R;PLA2G7;CRHR1;CRHR2;HSD11B2;CYP2B6;TERT;APOH;CD19;APOE;PROM1;AMZ1;NKX2-3;LYN;STAT5A;EGR1;ST14;GSTM1;ACE;GPR12;TSLP;FZD5;PLK2;XRCC3;IGF2;NFATC2;NFATC1;NR0B2;UBAC2;FERMT1;TG;VNN1;TOLLIP;PKIG;HLA-DRB1;DENND1B;CD40;CDKN1A;PTPRS;SLC44A4;MIR629;KRT20;CX3CL1;PANX2;HNF4A;MUC12;CCR9;MUC13;CCR7;CCR6;MIR192;CCHCR1;CD34;ACAD8;PSD;CBR1;DUSP1;IFNGR2;MST1;TACR2;IL16;MIR21;PAX5;TYK2;CBFA2T3;CIT;TLR1;SLC22A23;KAT2A;NR5A2;BANK1;IL1B;TFF3;TFF2;CLDN18;TFF1;CD47;TLR4;CHST2;HLA-DQB2;HLA-DQB1;GAS7;IDUA;GSTP1;ANKRD55;NR1I2;KCNA3;NEDD9;ADCY3;ADCY7;EGFR;PAK1;NT5E;NTF3;SPP1;STAT4;ADAD1;STX2;CD55;LAIR1;MICB;CCL25;CD74;CCL24;NQO1;HSPA1L;CDX2;IL10RB;MGMT;P2RY14;IL34;STAT3;MLH1;PDZK1IP1;LILRB4;SLURP1;VEGFA;FABP2;PER3;F2RL1;GHRL;IL17D;IL17C;HSPA1B;HSPA1A;CCL26;CD86;DEFB1;MSI1;PRDM1;APEH;ETS1;MPO;CLU;FGF2;TNF;CELSR3;HPGDS;FGF7;CDH3;CCND1;DMBT1;CDH1;ADORA3;GRAP2;RNF19A;MIR150;TGM2;PDGFRB;GHSR;ZGPAT;S100A1;TPM3;MAGI2;UBE4A;HIC1;SLC9A3;SCNN1G;PROCR;SCNN1B;MSH5;MASP2;PRTN3;S100A9;ABCG2;CALCA;ISYNA1;GSTT1;FHIT;ACAT2;ACACA;FUT3;SOCS3;SBNO2;IRAK2;NLRP3;TAC1;ZNF300;SMAD3;CD160;TGFB3;RARRES2;HSPA2;BTNL2;BMP7;ESR1;ISG20;IL6;IL17REL;INS-IGF2;NAT1;BCL2;HLA-DRA;ALPI;RNF186;FHL2;TNFAIP3;LAMC2;PTPN22;ITGAE;LSP1;ITGAL;TREM1;WISP1;CXCL5;CYP26B1;CA2;ITGAX;CASP1;MADCAM1;GAL3ST2;MIR193A;HSP90AA1;MMP7;KSR1;DCC;DNMT3A;HTR3A;TNFRSF1B;NGF;TNFRSF1A;TGFBR2;ACTA2;MIR126;ACTA1;ELF3;IFNG;MEP1A;TMBIM1;CAT;CDH11;CRH;LTA;CHI3L1;CDH13;BPI;IRF5;LTF;CRP;CX3CR1;RGS14;PON1;LRP5;NOD1;ASAP2;NOD2;NTN1;CRYZ;MUC2;WNT11;CCL5;MKNK2;MXI1;GNA12;MUC4;ATOH1;UGT1A7;CHGA;TCF7L2;NOS2;NOS3;EYA4;B3GAT1;IRGM;NELL1;PNKD;MIR145;CD6;APC;SST;IL2RA;LEP;P4HB;PDCD1 |
| Bone Diseases | 0.009071730028961465 | CSF3;CDKN1A;SPARC;TNFRSF6B;CSF1;GFI1;IHH;RORA;TCIRG1;TNF;MYOM2;BEST1;RPTOR;TRPS1;TNFSF10;TNFSF11;ACP5;SOST;SLC12A9;MMP7;SERPINF1;MIR21;ANK1;DKK1;RUNX2;COASY;TGFBR2;SFRP2;ELF3;IL1B;CHI3L1;ALDH7A1;SQSTM1;CFTR;LRP1;IDUA;COL11A1;LRP5;LRP4;EGFR;PTHLH;ANKH;BGLAP;CCL5;PRDM16;ERBB2;SH3BP2;SPP1;FYN;MSC;KL;TNFSF14;WNT3A;EPHX1;B3GAT1;HTR1B;BMP7;ESR1;BMP4;COL1A1;EXT1;CLCN7;P2RX7;BMP2;IL6;GNAS;RGS10;ALPL;TNFSF8;SP7;FGFR3;CD200 |
| Malignant neoplasm of kidney | 0.009122177707681754 | CSF2;EPO;MPG;CRTC1;DEFB1;COMT;FGF2;TNF;MYLK;ICAM1;IGF1R;CDH6;PYCARD;CRHBP;HPGDS;RASSF1;CCND1;CDH1;MYC;RASSF5;NPPA;TNFSF10;AKT1;CYP1B1;PITX2;SLC16A3;SRRM2;DAPK1;IGFBP3;F2R;HLA-C;TSC2;PRKCA;UNC5C;MITF;DLEC1;UNC5D;DKK1;SDHB;DKK3;ADAM2;SFRP1;SFRP2;SFRP5;SQSTM1;ZNF154;ISYNA1;ABCB1;EPAS1;ITPR1;GSTT1;GATA3;PIK3R1;PRKCZ;FHIT;RAP1GAP;SLC5A5;FIGNL1;HLTF;TRPM8;PROM1;PRODH;BBS2;EGR1;ACE;GSTM1;TNK2;OSGIN1;ARHGAP24;COL1A1;ISG20;IL6;COPS5;RHEB;GIPC1;CDK2;CYP1A1;HRC;BCL2;MIR183;HAMP;TP73;DDR1;PTPRU;CDKN1A;MIR17;THRB;LRRC4;PIK3CD;PKD1;PIK3CG;FLCN;ESM1;CHL1;MAP1LC3C;HNF4A;CASP1;TIMP3;SOSTDC1;HSP90AA1;ANXA2;MME;TESC;TUBB;TFEB;MIR21;PAX2;ZEB2;ZEB1;CDCP1;IFNG;MIR17HG;RARA;BIRC5;ANGPTL4;SGK1;RBMS3;KANK1;GRP;UHRF1;DERL3;NR1I2;SEMA3B;EGFR;MIR148A;PTHLH;CYP17A1;CRYZ;PAK1;MUC1;CUX1;WIF1;PLAGL1;MIR494;HSF1;ERBB2;MXI1;SPP1;FSD1;SPAG9;IL32;RBM38;NQO1;NOS2;BNIP3;STAT3;PDX1;MLH1;BLCAP;GRHL2;MTOR;VEGFA;MARCKS;MIR145;APC;WT1;IL2RA;LEP;PDCD4;SARDH;VIM |
| Pancreatic Neoplasm | 0.00971424479648048 | RB1;SPARC;SERPINE2;SCT;ICAM1;LGALS3;MYC;TNFSF10;AKT1;SOX9;TP63;IER3;SRRM2;DAXX;WSB1;KRT8;HNF1A;RUNX3;DKK3;SFRP4;TIAM1;CLDN4;SLC5A8;SFRP1;SFRP2;SLPI;DDIT3;SFRP5;IDO2;EZR;GATA6;PDGFB;PDGFA;GATA5;TWIST1;GATA4;GATA3;PRKCZ;ABO;KLK7;SLC5A5;CYP2B6;TERT;PDGFD;CHST15;SFN;APOE;PROM1;NGFR;WWTR1;XRCC3;NFATC1;NR2F2;GJB2;FXYD3;GIPC1;TP73;CDKN1C;DDR1;HOTAIR;COL18A1;RARG;NRP2;ADM;BRCA1;KRT20;CCKAR;TIMP2;CCR7;CCR6;CD34;CTSD;MAP4K1;ANXA2;MME;DUSP1;CACYBP;MIR21;PROX1;F3;DUSP6;SERPINB5;VAV1;MMP14;GPRC5A;NR5A2;CCKBR;IL1B;TFF2;TFF1;ANGPTL4;ARHGEF2;SLC29A1;TAGLN;GSTP1;COL11A2;NR1I2;EGFR;BGLAP;EPCAM;ALOX5;GPC1;SPP1;CTNNA1;WWOX;NQO1;GABBR1;CDX2;MGMT;GDF15;STAT3;SCTR;SORBS2;SULF1;LILRB4;VEGFA;PER1;WT1;F2RL1;CCL28;F2RL3;SEMA5A;CSF3;CSF2;CSF1;IHH;IRS2;PLAT;FGF2;TNF;ETS1;MPO;IGF1R;RIMS2;FGF7;CCND1;CDH1;GRAP2;RNF19A;MIR150;SLC39A4;MAP3K5;TGM2;POSTN;MSX2;IGFBP3;APLP2;F2R;WNT5A;TSC2;MIR27A;FOXP1;MAG;S100A6;ADAM8;LY6E;NOTCH3;PFKFB3;NOTCH4;ALOX15;TACSTD2;FHIT;RAP1GAP;FUT3;STK33;NLRP2;MARK2;SMAD2;OSBPL5;TFPI2;SMAD7;BMP4;IL6;COPS5;GDNF;CDK2;BHLHE40;BCL2;GNAS;CSPG4;TGFBI;FGFR2;FGF10;MIR10A;PSCA;SYCP1;FLT4;ONECUT1;PIK3CD;FOXM1;MSLN;CYR61;PIK3CG;LTB4R;CASP8;CA2;PSD4;HGFAC;HSP90AA1;MMP7;KSR1;DCC;NRG1;FOXL1;SSTR1;DCN;SSTR5;TAPBP;TGFBR2;TGFBR3;IFNG;PALLD;ESRP1;BIRC5;ITGA6;RBMS3;SRC;THBS1;PTHLH;MUC1;ZNRF3;MUC2;CNR2;ERBB2;NT5DC3;SPOCK1;NCAM1;MUC4;NTRK1;NOS2;NOS3;NTRK3;BNIP3;PDX1;CCK;MTOR;P2RX7;APC;SST;APOC1;S100P;BCAR1 |
| Colonic Neoplasms | 0.00971424479648048 | APP;TNFRSF6B;SPARC;ICAM1;LGALS3;RASSF1;MYC;TNFSF10;AKT1;CYP1B1;SOX9;SLC12A9;HLA-C;HNF1B;HNF1A;HLA-A;RNASE1;DKK1;RUNX3;HLA-G;SND1;RUNX2;RUNX1;DKK4;TIAM1;SLC5A8;SLPI;BIN1;DDIT3;CD109;MT1G;MT1H;EZR;ABCB1;EPAS1;MST1R;PLA2G7;PRKCZ;SLC5A5;SULT1A1;HLTF;MAP7;SYMPK;A2M;PROM1;EGR1;GSTM1;XRCC3;IGF2;NFATC2;PTK6;NR0B2;VANGL1;FERMT1;CYP26A1;MT1IP;TNFSF4;FAT1;NMT1;CYP1A1;ID3;SIM2;TDGF1;CD200;RCOR1;GSTM5;ACHE;PIGR;CD40;CDKN1A;ADM;KRT20;GLI3;SERPINA4;GLI2;AIFM3;MECOM;TIMP2;RSPO2;RSPO3;CCR7;MIR192;CLIC1;TCF19;CBR1;PARP4;NME2;TET1;AMPD3;SERPINB5;MMP14;COL2A1;ZEB1;B3GNT8;ADRB3;CCKBR;IL1B;MIR17HG;TFF3;RARB;TFF2;VWA2;CRYAB;TAGLN;GRP;NR1I2;ASCL2;EGFR;CYP17A1;EPCAM;ALOX5;HSF1;SPP1;CD55;SPDEF;BRD4;IL32;NQO1;CDX2;MGMT;CDX1;STAT3;SULF1;MLH1;PSMB8;GUCA2A;PSMB9;VEGFA;CYP2W1;PER3;FABP6;WT1;DLC1;F2RL1;GHRL;YPEL3;HSPA1A;F2RL3;CSF3;NDUFA13;CHRM1;CSF2;MPG;IRS2;COMT;CLU;FGF2;TNF;ACTG2;IGF1R;LOXL2;CCND2;CCND1;DMBT1;CDH1;ADORA3;KCNH2;EOMES;IGFBP1;CASR;POSTN;TPM3;IGFBP3;F2R;WNT5A;MATK;TAP1;TLE6;FOXP1;MTNR1A;FSCN1;S100A9;LY6E;ABCG2;CALCA;SHC1;DIABLO;ALOX15;TACSTD2;GSTT1;RASAL1;FHIT;FUT1;SLC9A3R2;NME1-NME2;DPP4;HRH1;PRR5-ARHGAP8;HRH2;PODXL;MARK2;STRA6;TFAP2A;FDPS;SMAD3;TNFRSF10A;HSPA2;BMP7;ESR1;BMP4;IL6;FGF19;FGF18;CDK2;GNAS;BCL2;XAF1;FGFR4;LGALS3BP;KDM5B;ITGB4;WISP3;PIK3CD;OXT;PIK3CG;WISP1;HAPLN1;GRM4;CASP8;DNMT3B;PMEPA1;ITGB6;HMGCS2;LBP;YWHAH;HSP90AA1;EGFL7;MMP7;PROP1;ITGA4;KSR1;ITGA3;DCC;DNMT3A;DCN;TGFBR2;ACTA2;AIM2;IFNG;MEP1A;PALLD;ESRP1;CAT;BIRC5;ITGA7;MCM5;LCP1;FBP1;LTF;CRP;ITIH4;RETNLB;HDAC2;HDAC3;SRC;UHRF1;NOD1;NTN1;PTHLH;MUC1;MUC2;RAB25;CCL5;ERBB2;IGF2BP1;UGT1A9;EIF4E;ATOH1;SASH1;TCF7L2;NOS2;TIE1;CABLES1;MEIS1;MIR145;APC;LEP;PDCD4;BCAR1 |

| | | |
|---|---|---|
| Psoriasis | 0.00971424479648048 | HCP5;APP;TNFRSF6B;TRAF3IP2;MYT1L;ICAM5;BACH2;ICAM1;AQP1;LGALS1;CYP2D6;ZMIZ1;TNFSF10;AKT1;UBASH3A;SOX9;PGLYRP1;TRIM21;TP63;SLC12A9;CMKLR1;PRKCB;NFKBIL1;HLA-B;HLA-C;MIF;FNDC3A;HLA-A;KRT5;DKK1;RUNX3;HLA-G;RUNX1;HLA-E;SFRP4;STX1B;IFI27;SLPI;IL23A;TRIM15;PRKCQ;ZNF831;SPATA18;CFB;SKAP2;ABCB1;EPAS1;SP140;MGST2;GATA3;CD1D;PRKCZ;KLK8;KLK7;C2;CRHR2;KLK6;C3;SYNGR1;APOE;APOB;HLA-DQA2;ELANE;HLA-DQA1;NTSR1;LYN;EGR1;NAPSA;GSTM1;ACE;PTPRN2;TSLP;FZD5;IGF2;NFATC1;NR0B2;ALOX15B;UBAC2;BST2;GJB2;VNN1;CYP2S1;CYP1A1;TNFSF8;ANP32B;PKIG;HLA-DRB1;DDR1;DENND1B;ROBO3;CD40;CDKN1A;TNFRSF13B;ELN;AATK;MED15;B3GNTL1;CX3CL1;RPTOR;TIMP3;CCR7;CCR6;CCHCR1;IKBKE;CD34;CHST8;ACAD8;TCF19;LYNX1;ANXA2;DUSP1;IFNGR2;MST1;NME2;MIR21;TYK2;POU5F1;GPRC5A;NR5A2;BANK1;IL1B;ARHGEF2;TLR4;HLA-DQB1;CSTA;TNXB;GSTP1;ANKRD55;ASIP;NR1I2;ADCY3;ADCY7;EGFR;RASIP1;DUOX1;LCE3A;HLA-DMB;SPP1;CSMD1;IL32;AOC3;CDX2;MGMT;STAT3;ADAM33;OSM;KLK13;NR1H3;GRHL3;PSMB8;KLK11;SLURP1;PSMB9;VEGFA;KRT17;KRT14;HLA-DPB1;FOSB;F2RL1;IL17D;IL17C;CSF3;CSF2;F13A1;MIR146B;DEFB1;PLAT;ETS1;TNF;IGF1R;HPGDS;FGF7;GRAP2;RNF19A;CCRL2;HLA-DOA;TGM5;HLA-DOB;TGM3;TGM2;LAG3;S100A1;USP2;WNT5A;TAP2;PIPOX;TAP1;HCG9;ELMO1;MASP2;S100A9;CARD14;ABCG2;ISYNA1;NOTCH4;ALOX15;PSORS1C3;SPATA2;GSTT1;ALOX12;GLIS1;PSORS1C1;PSORS1C2;NME1-NME2;SOCS3;SBNO2;NLRP3;TAC1;SMAD2;SMAD3;RARRES2;ISG15;ISG20;NR4A2;ADCY10;IL6;INS-IGF2;NAT1;BCL2;MUC21;FLT4;ITGB2;PIK3CD;TNFAIP3;PTPN22;ITGAE;ITGAL;PIK3CG;CYR61;CA2;TAGAP;ITGAX;CASP1;PDE4B;PDE4A;GAL3ST2;TRIM65;SGCG;HSP90AA1;TRPC6;KSR1;OLIG3;PDE4D;DNMT3A;APOA1;OPRM1;TNFRSF1B;NGF;TNFRSF1A;QTRT1;ACTA2;IFNG;OAS2;CAT;CRH;LTA;CHI3L1;CDH13;IRF5;ZNF750;CRP;CX3CR1;PON1;ASAP2;NOD2;IL1RL2;CLMN;CCL5;GNA12;MIR492;SLIT3;NCAM1;EIF4E;NTRK1;MARCKSL1;ANGPT2;ANGPT1;NOS2;NOS3;FBXL19;SOD3;IRGM;POMC;CD6;APC;MIR125B1;IL2RA;LEP;PTPN6;HCG22;PDCD1;TEK |
| Adenocarcinoma of pancreas | 0.00971424479648048 | APP;TNFRSF6B;CSF2;EPO;PLAT;IRS2;FGF1;ETS1;FGF2;MPO;ICAM1;IGF1R;LGALS4;CCND2;CCND1;GRAP2;RNF19A;AKT1;SLC39A4;SLC12A9;EPHA4;SERPINF1;F2R;WNT5A;MIR27A;HNF1A;HLA-G;DKK1;SLC5A8;MYCN;LTB4R2;JTB;HOXB7;SQSTM1;CFTR;ABCG2;GATA6;PDGFA;GSTT1;MIR206;FUT4;TERT;PENK;NLRP2;THEM4;TRPM8;RREB1;MARK2;KL;NAPSA;GSTM1;CADM1;XRCC3;TFPI2;HMGA1;NR2F2;DCLK1;ISG20;IL6;NAT1;ARTN;GIPC1;CYP1A1;BCL2;GNAS;FGFR4;FGFR1;PTPRT;PTPRU;PIWIL1;CD40;CDKN1A;NRP2;CIITA;PSCA;MIA2;PIK3CD;CHD5;ADM;LAMC2;BRCA1;LSP1;KRT20;MSLN;CX3CL1;PIK3CG;CA2;CCR7;POLE;MMP7;ANXA2;DCC;MIR21;F3;TGFBR2;ZEB1;CCKBR;SLCO4A1;PALLD;IL1B;CAT;SLC29A1;TLR4;HLA-DQB1;CX3CR1;HDAC2;SRC;NR1I2;SPG7;THBS1;EGFR;PTHLH;MIR148A;MUC1;CNR2;NTF3;ERBB2;MXI1;MIR372;SPP1;MUC4;UGT1A7;NQO1;CDX2;SCTR;KLHL1;PDX1;CCK;VEGFA;GNMT;KISS1;APC;IL2RA;NBR1;SPAG1;F2RL1;SPRY2;F2RL3 |
| Malignant neoplasm of pancreas | 0.00971424479648048 | RB1;SERPINE2;SCT;MIR429;MIR675;MYC;DPYSL3;STMN1;AKT1;CYP1B1;HOXA1;IER3;PRIMA1;DAXX;WSB1;PRKCB;SERPINF1;DAPK3;PRKCA;RNASE1;DKK1;EML2;DKK3;DKK4;MIR202;CLDN4;SLC5A8;SPINT1;ARMC3;CLDN8;RBP1;HOXB3;HOXB2;MAML3;IDO2;HPD;HOXB7;HOXB6;GNE;MEG3;ABCB1;DHH;TERC;MIR196B;GATA6;PDGFA;GATA5;PRDM14;GATA4;FPR2;LYPD5;PRKCZ;MIR206;ABO;SLC5A5;SULT1A1;C3;CYP2B6;TERT;MIR330;KCNQ1OT1;CHST15;PROM1;RREB1;JAG2;NGFR;CREBBP;GSTM1;ICMT;CADM1;PLK2;NFATC2;MRPL28;NFATC1;NR2F2;PBX1;H19;FERMT1;GJB2;FXYD3;GIPC1;CYP1A1;HLA-DRB1;CDKN1C;PIGR;CDKN1A;MIR130B;MIR629;ADM;SLC6A2;TRIOBP;GLI3;BBC3;TIMP2;RSPO2;TK1;BRD2;RING1;ANXA2;DUSP1;PAX6;PAX5;F3;DUSP6;POU5F1;SERPINB5;MMP14;ZEB2;ZEB1;SLCO4A1;HCLS1;SLC29A1;GAS5;JDP2;KANK2;GRP;GSTP1;COL11A1;NR1I2;COL11A2;ADCY3;HOXD13;FSTL1;EGFR;ARFRP1;MIR410;RECQL5;EPCAM;BGLAP;ALOX5;NTF3;SPP1;BRD4;WWOX;PKLR;F10;MGMT;WNT7B;STAT3;SCTR;SULF1;SKI;WT1;FOSB;SIGLEC7;SEMA5A;CSF3;CSF2;ATP8A2;PLOD2;IRS2;CLU;FGF2;TNF;BEST1;CDH5;HHAT;FGF7;HPGDS;CDH3;CDH1;SLCO2B1;GRAP2;RNF19A;HLA-DOA;SLC16A3;SLC39A4;MAP3K5;KCNH2;IGFBP1;USP7;FBXW8;S100A2;S100A1;RPS8;IGFBP3;F2R;MITF;CHD1L;HIC1;FOXP1;LTB4R2;MTNR1B;S100A6;FSCN1;PGR;ADAM8;TNIK;S100A9;SQSTM1;LY6E;NOTCH3;BRSK2;SHC1;TACSTD2;GSTT1;VTCN1;FHIT;RAP1GAP;SLC9A3R2;STK31;SOCS3;STK33;SMYD2;PLXNA1;KL;NOP14;OSBPL5;HTR1D;GFPT1;TFPI2;TNK1;TNFRSF10C;HTR1B;TNFRSF10A;MCC;TMPRSS13;HOPX;COPS5;CDK6;STK24;CDK2;BCL2;CPE;CLPTM1L;CNTNAP1;PSCA;BNC1;SYCP1;FLT4;ONECUT1;ITGB2;FHL2;WISP3;PIK3CD;LSP1;MSLN;FOXM1;CYR61;PIK3CG;TRIM69;HGFAC;HSP90AA1;PEG3;KSR1;DNMT3A;NRG1;FOXL1;TNFRSF1B;NGF;DDOST;TNFRSF1A;ESRP1;BIRC5;ITGA6;MIR124-2;ANG;CX3CR1;HDAC2;HDAC3;LRP1;SRC;UHRF1;MIR148A;ZNRF3;PARD6A;CCL5;TP53INP1;SH3BP4;MAL2;EIF4E;UGT1A7;NTRK1;ANGPT2;ANGPT1;PRRX1;KCNJ11;ASCC1;NTRK3;MIR935;MTOR;TBX2;P2RX7;NELL1;P2RX5;NIN;APC;SST;IL2RA;LEP;IL2RB;APOC1;PDCD4;SPAG1;BCAR1;APP;FAAH;TNFRSF6B;SPARC;ICAM2;ICAM1;SOX2;LGALS3;FRAT1;RASSF1;LGALS1;TRIM29;RPS6KA2;TNFSF10;TRIM27;SOX9;TNS4;TP63;SLC12A9;SRRM2;EPHA4;ENTPD1;ENTPD2;HLA-B;KRT8;HLA-C;HLA-A;MIF;HNF1A;RUNX3;RUNX2;TIAM1;SFRP1;IFI27;KCNQ1;EZR;CFTR;STX1A;GRB7;SLC22A3;EPAS1;EIF5A2;TWIST2;TWIST1;MST1R;ZBTB4;PRH1;KLK7;DGUOK;NUAK1;TSPAN8;PENK;HIVEP3;TRPM8;FADD;MIR1178;TSPAN1;NTSR1;EGR1;WWTR1;IQSEC1;XRCC3;SLC33A1;IGF2;LIF;GDF2;MIR200A;PTK6;MIR200B;DCLK1;GADD45G;TF;OCLN;ARTN;ID2;SIM2;TDGF1;RGL2;VSX1;CC2D1A;TP73;APOBEC3B;MIR129-2;DDR1;HOTAIR;DIRAS3;ROBO3;COL18A1;NRP2;CD40;MIR17;CTSZ;MIR18A;BRCA1;KRT20;KIF15;CCKAR;CCR9;MUC13;CCR7;CCR6;CD34;CTSD;HTATIP2;MAP4K1;CBR1;ZHX2;MME;MST1;TFEB;CACYBP;GFRA1;MIR21;PROX1;NME5;RHOF;MIR19A;VAV1;CNKSR1;CDCP1;NR5A2;CTTN;ADRB3;CCKBR;IL1B;MIR17HG;TFF3;CLDN18;TFF1;ARHGEF2;CD47;ARHGEF7;TLR4;GUCY2C;TAGLN;TMPRSS3;NEDD9;CYP17A1;NT5E;PAK1;CYP2A13;CUX1;WIF1;GPC1;NPY;HSF1;MICB;IL32;CD74;NQO1;GABBR1;CDX2;SORT1;CDX1;GDF15;GAD1;OSM;KLHL1;SORBS2;LILRB1;MLH1;GRHL2;GCK;VEGFA;PER1;PER3;KRT18;KRT17;NBR1;TCEA2;F2RL1;PNPLA2;F2RL3;FOXA2;CD86;CD83;IHH;PLAT;PRDM1;COMT;ETS1;MPO;ETS2;IGF1R;CNDP2;MIR149;CCND2;CCND1;CAPN2;MIR150;NRTN;GGT1;PPFIA4;TGM2;PDGFRB;PDGFRA;CASR;ZGPAT;MSX2;LIG1;KDM2B;APLP2;WNT5A;MATK;MIR27A;LIG4;TSC2;MAT1A;MYCN;COL4A2;DPEP1;CARD16;NUPR1;ABCG2;FBN2;TCF7;ALOX15;ADRB2;FBLN2;FUT1;FUT4;MCHR2;FUT3;PODXL;MIR296;TAC1;MSC;WNT2;MARK2;TFAP2A;SMAD2;FDPS;SMAD3;HMGA1;ISG15;BMP7;ESR1;IGF2R;ARHGAP24;SMAD7;BMP4;BMP2;IL6;EFNA2;NAT1;GDNF;MIR183;ALPI;TGFBI;FGFR4;FGFR3;FGFR2;FGF10;FGFR1;PNLIPRP2;CRISP2;MIR10B;MIR10A;HSPB2;CHD5;ADAR;USP39;MIR96;PKD1;LTB4R;CXCL5;GHR;CASP8;MCF2L;NOS1;HS3ST3B1;KLF11;MMP7;MIR23A;DCC;SSTR1;OPRM1;ARID1B;DCN;SSTR5;TGFBR2;MIR126;ALDH1A3;IFNG;PALLD;CAT;LTA;CHI3L1;CDH13;LTB;RBMS3;MCPH1;DNAH5;RGS16;THBS1;PTHLH;AURKB;CRYZ;MUC1;MIR377;MUC2;CNR2;MIR373;GNA12;ERBB2;MIR492;PTK2B;SPOCK1;SNCG;MUC4;MUC6;CHGA;TCF7L2;BNIP3;PDX1;CCK;MUC5B;SOD3;HS3ST1;KISS1;MIR145;MIR144;MIR142;CD9;VIM |
| Schizoaffective Disorder | 0.0103058663921663 | CNTNAP2;UXS1;ARHGEF10L;HTR2A;COMT;MED15;EDNRA;CYP2D6;NCKAP5;PDGFRB;GRID1;LMO7;PRKCA;KSR2;FOXP2;SSTR4;RARB;RIN2;THOP1;PVALB;MAD1L1;SDCCAG8;RTN3;ABCB1;DLX4;TMPRSS5;ITIH1;CPLX1;NALCN;SELENBP1;NPY;PRODH;STX2;DISC1;CSMD1;DRD4;DRD5;NTRK1;TBX1;TCF7L2;EGR3;GAD1;HTR1B;CABLES1;H19;TF;DAO;USH1G;SST;CAPN10;NEUROG1 |
| Neuroblastoma | 0.0103058663921663 | RB1;SERPINE2;SCT;STMN2;STMN4;HTR4;BACH2;MIR432;CYP2D6;DPYSL2;GJA5;MYC;STMN1;AKT1;MNX1;CHGB;WSB1;DAPK3;HPCAL1;PRKCA;UNC5D;RFX1;PDYN;ISL1;DKK1;DKK3;RBP7;BIN1;DDIT3;MAP1B;SERPING1;HOXB5;MEG3;ABCB1;CRABP1;PDGFA;GATA4;GATA3;THY1;GATA2;PRKCZ;SLC5A5;TERT;KCNQ1OT1;ZSCAN10;ALX3;PROM1;HOXC9;HOXC6;BARD1;ABCA2;NGFR;CREBBP;GSTM1;CADM1;FZD5;FZD6;MIR338;PBX1;FLI1;PNOC;TNFSF9;NANOG;PPT2;PAM;HOXD9;HOXD8;CDKN1C;ACHE;CDKN1A;RARG;ADM;SLC6A2;SLC6A3;MMP20;BBC3;GLI2;SHF;CHL1;TIMP2;NEFL;DLGAP1;NEFH;MYOG;DFFB;ANXA2;PAX3;POU5F1;NCOR2;MMP14;GPRC5A;RARA;RARB;HCLS1;ZNF236;OCA2;NFIX;GRP;GSTP1;NR1I2;COL11A2;LRRC17;KCNA3;HTT;ADCY2;ASCL1;GRAMD4;EGFR;MIR410;EXTL1;RASSF10;BGLAP;ALOX5;NSD1;CHN2;NTF3;SPP1;STAT4;WWOX;PKLR;MGMT;STAT3;POU4F1;HIPK2;FOXR1;FABP3;NFIC;FABP7;FOSB;SIGLEC7;CRLF1;HSPA1B;HSPA1A;CSF3;CSF2;CSF3R;CSF1;CCNT1;EPO;FAM107A;RORA;APEH;CLU;FGF2;TNF;RIMS2;PYCARD;CDH5;HHAT;HPGDS;FGF8;NAGLU;GRAP2;RNF19A;MAP3K5;SLC16A5;CD177;KCNH2;USP7;LAG3;SNRPN;TPM3;S100A1;IGFBP3;HIC1;FOXP1;NPAS4;S100A6;SQSTM1;PVALB;MAD1L1;DSC3;CUL9;NOTCH3;ITPR1;CUL1;ITPR2;KLC1;SOCS3;SBNO2;NDN;SRGAP3;ZNF423;TFPI2;TNFRSF10C;HTR1B;ADAP1;GFAP;CDK6;STK25;CPD;CDK2;BCL2;SCGB3A1;SHBG;XAF1;CIITA;FLT4;FHL2;PIK3CD;ECE1;HTR2A;TRH;CPZ;FOXM1;PIK3CG;MB;PSD4;RNPEP;DNMT3B;SGCG;UNC45A;HSP90AA1;GPX1;ITGA4;PEG3;PLEKHG5;SH2D3C;NGB;TNFRSF1B;NGF;COASY;C1QL1;LCK;CRH;SNURF;BIRC5;TNFRSF25;ANG;CRP;CX3CR1;ITIH4;HDAC2;HDAC3;LRP1;SRC;RAB1B;SEMA3B;MIR487B;GLRX;NTN1;MGAT5B;NNAT;P2RY1;IGF2BP1;NTRK1;MARCKSL1;ANGPT2;ASCC1;NTRK3;KCNIP3;TBX5;MTOR;P2RX7;P2RX5;KLF6;MARCKS;APC;SST;P2RX2;IL2RA;LEP;NOVA2;IL2RB;CD248;CCM2;PDCD1;P4HB;INSM1;BCAR1;NPFFR2;ERRFI1;APP;SPARC;CERK;ICAM2;ICAM1;SOX2;LGALS3;RASSF1;EDNRA;LGALS1;EDNRB;RASSF5;RASSF6;TNFSF10;TP63;SLC12A9;PTGDR;SRRM2;EBF1;HLA-C;ANK2;CSNK1E;HLA-A;MIF;HLA-G;RUNX3;RUNX2;RUNX1;TIAM2;TIAM1;THEG;SFRP1;SLPI;KCNQ4;RTEL1;CASZ1;EPAS1;SLC45A1;MGST1;CEND1;TWIST1;CD1D;PIK3R1;CRHR1;TRPM2;NEU4;MAP2;CD19;STC2;PENK;ST8SIA4;APOD;B4GALNT1;APOE;TRPM8;SPTBN1;BBS2;STAT5A;EGR1;SNHG1;BCL11B;ST8SIA2;IGF2;MIR200A;MIR200B;DCLK1;CYP26A1;MSRA;TH;ID2;ID3;GRB2;TP73;SPAG16;PTPRT;COL18A1;ACY3;CD40;MIR17;ICA1;MIR18A;BRCA1;PCSK6;PTPRF;CLYBL;CX3CL1;HNF4A;CD38;MIR192;KIF1A;CCR6;FFAR1;MCF2L2;CD34;CTSD;MME;PABPC4;INSRR;NME2;NME4;MIR21;PROX1;VAV1;AJAP1;NAALADL1;ADRB3;IL1B;MIR17HG;MPPED2;B3GNT3;ARHGEF2;CD47;MAPT;ARHGEF7;SGK1;CXCR5;NEDD9;CAPG;GDPD5;CAMKK2;RXRB;CUX1;WIF1;NPY;HSF1;GPC2;MSX1;DISC1;RAE1;CD55;NQO1;OSM;LILRB1;MLH1;PEX14;VEGFA;TPPP3;PER3;QPCT;RAMP1;IL17C;NEUROG1;F2RL3;CD86;RTL1;CD81;LOXL3;PLAT;PRDM2;ETS1;ACTG1;IGF1R;MIR149;RGS4;CCND2;DMBT1;CCND1;NPPA;TGM2;PDGFRB;PDGFRA;PDIA2;CASR;G6PC;LIG1;LMO3;APLP2;WNT5A;MATK;UBE4A;MAT1A;SDHB;DHDDS;MYCN;HGS;CD8A;SCNN1A;AIPL1;ABCG2;CALCA;DIABLO;TCF7;ALOX15;CHAT;ALOX12;SLC1A5;ADRB2;NME1-NME2;CAMTA1;CYTL1;TAC1;MSC;WNT3;PRPH;TFAP2B;ZFHX3;TFAP2E;WNT3A;PTCH2;HMGA1;ESR1;PHOX2B;ARHGAP24;PHOX2A;NR4A2;BMP2;IL6;RALYL;NAT1;GDNF;TLX2;FAM163A;GNAS;MIR184;ALPI;TGFBI;FGFR4;MXD3;FGFR2;FGFR1;LGALS3BP;KDM5B;CRISP2;MIR10B;MIR10A;MEGF10;HSPB6;ELAVL4;HSPB2;CHD5;OXT;LAMC2;DBH;NR2E1;SLC2A5;GRM1;GRM2;GRM5;CYP26B1;CASP8;CA2;CASP1;FLOT1;NOS1;GALNT9;DCC;NAV2;OPRM1;UTF1;POMGNT1;ARID1B;TGFBR3;IFNG;ALDH1A2;CAT;LTA;CDH13;ZMYND10;RBMS3;MCPH1;CAMK2B;DDC;RGS16;OPRL1;RRBP1;THBS1;PTHLH;AURKB;CRYZ;MUC1;CNR2;CLMN;MXI1;ERBB2;PTK2B;NCAM1;SNCB;SNCA;CHGA;LGI1;CDK11A;KDM4B;URGCP;NOS2;LIMK2;B3GAT1;CCK;CDK11B;MEIS2;DLK1;POMC;MEIS1;MIR145;XPNPEP1;MIR125B1;PPP2R2B;CD9;SPRY2;TEK;VIM;RAN |
| Squamous cell carcinoma | 0.010357028272521681 | RB1;HOXA9;ZMIZ1;CYP2D6;MYC;STMN1;AKT1;CYP1B1;HOXA2;PLCE1;HOXA1;HOXA7;HOXA5;IER3;ANKS1B;DAPK1;SERPINF1;SERPINF2;SOX14;PDYN;DKK3;CLDN4;SLC5A8;SPINT1;BIN1;DDIT3;FEZ1;RBP1;TYRO3;PADI2;CYP2E1;ENPEP;ABCB1;TERC;SHMT1;CRABP1;GATA5;GATA4;LYPD3;GATA3;THY1;ABO;WDR66;SLC5A5;SULT1A1;LDHB;TERT;S100A16;GALR1;KCNMB3;SFN;PROM1;HLA-DQA1;HOXC6;JAG2;NGFR;CREBBP;GSTM2;ST14;DNAH11;GSTM1;CADM1;PLK2;MRPL28;GRIN2B;NR0B2;PBX1;VANGL1;H19;FERMT1;GJB2;GJB5;FAT1;NMT1;MKS1;CYP1A1;NANOG;GTF2IRD1;PLCH1;HRG;HOXD9;HLA-DRB1;CDKN1C;PIGR;CDKN1A;RARG;TCF21;ADM;HOXC10;BBC3;GLI2;GPR132;MECOM;TIMP2;TIMP3;JAK3;IKBKE;SLC2A14;RIPK3;ANXA2;DUSP1;IFNGR2;TESC;RIPK4;TMC8;TMC6;PAX5;POU5F1;SERPINB5;PAX1;SLC7A5;MMP14;ZEB2;ZEB1;GPRC5A;PAX8;KIFC1;TAL1;ITPKA;RARA;RARB;HCLS1;CRYAB;HLA-DQB1;KANK1;OCA2;GRP;GSTP1;COL11A1;CRCT1;NR1I2;ASIP;KCNA3;ASCL1;EGFR;MIR410;EPCAM;PLAGL1;SPP1;PRAME;CSMD1;BRD4;WWOX;ZNF582;BRF2;PKLR;MGMT;STAT3;DEF8;DHRS4;SKI;FABP5;WT1;FOSB;LGR6;ZNF577;SIGLEC7;LY6K;UGT1A10;CSF3;CSF2;CSF1;EPO;MSI1;DEFB1;IRS2;CLU;FGF2;TNF;FGF3;PYCARD;FGF7;HPGDS;CDH3;CDH1;GRAP2;RNF19A;GRB10;PITX1;SLC16A3;LZTS1;AP2M1;KCNH1;MAP3K5;KCNH2;USP7;S100A2;S100A1;IGFBP3;F2R;CHD1L;HIC1;FOXP1;MTNR1A;S100A6;FSCN1;PKP1;PGR;ADAM8;ERG;S100A9;SQSTM1;DSC3;NOTCH3;DLX4;NOTCH4;TACSTD2;GSTT1;LIN7C;FHIT;RAP1GAP;GLIS1;RHOBTB1;SLC9A3R2;SOCS2;SOCS3;NDN;ZNF300;ESRRA;KL;TFPI2;TNFRSF10A;MCC;TMPRSS13;HOPX;GFAP;DAB2;COPS5;CDK6;CDK2;BCL2;SCGB3A1;CLPTM1L;LAT;EIF4G1;CIITA;ITGB4;BNC1;ONECUT1;GADL1;PIK3CD;ECE1;MSLN;FOXM1;CYR61;PIK3CG;WISP1;ITGAX;DNMT3B;ITGB6;DSP;EMX2;HSP90AA1;TRPC6;GPX1;ITGA4;DST;ITGA3;DNMT3A;CTDSPL;NGB;TNFRSF1B;NGF;TNFRSF1A;PTP4A1;TFDP1;TRAF3;CRH;BIRC5;ZNF750;CRP;HDAC2;SRC;SEMA3B;FURIN;GLRX;MIR148A;CCL5;NNAT;UGT1A5;UGT1A4;UGT1A3;CCL1;UGT1A9;EIF4E;UGT1A8;UGT1A7;UGT1A6;NTRK1;TBX1;SPAG9;SEMA4A;MARCKSL1;TXNRD1;MTOR;RAD52;MARCKS;APC;SST;IL2RA;LEP;IL2RB;PDCD4;CCDC63;PTPN6;PDCD1;ZACN;BCAR1;APP;HPSE2;ICAM2;ICAM5;ICAM1;AQP1;SOX2;LGALS3;RASSF1;LGALS1;RASSF2;EDNRB;SOX1;TRIM29;RPS6KA2;TNFSF10;LGALS9;SOX9;TRIM21;TP63;RPS12;EPHB3;SLC12A9;SRRM2;KRT1;KRT8;HLA-C;KRT7;DLEC1;EBF3;HLA-A;KRT5;MIF;HLA-G;RUNX3;KRT9;RUNX1;TIAM1;SFRP4;SFRP1;CFDP1;SFRP2;SLPI;IFI27;SFRP5;CD109;RIN1;NEIL1;EPHA1;EZR;CFTR;SLC22A3;EPAS1;TWIST2;MGST1;TWIST1;KLK4;MST1R;CD1A;INPP5A;MAP2;HLTF;ST8SIA4;APOE;TRPM8;TSPAN1;ATP6V1C1;NTSR1;STAT5A;EGR1;HIST1H4K;HIST1H4L;XRCC3;IGF2;LIF;PTK6;HAL;ID2;PPP1R1B;ID3;HIST1H4I;ANP32B;TDGF1;TP73;APOBEC3B;HIST1H4F;HOTAIR;ROBO3;COL18A1;CD40;MIR17;TNFRSF13B;MIA2;GRIK2;MIR18A;BRCA1;PCSK6;KRT20;KIF15;AKAP12;HYAL1;SLC22A18;HYAL2;CCR7;CCHCR1;CD34;CTSD;HTATIP2;MST1;NME2;MIR21;PROX1;MIR19A;VAV2;TLR1;ANO1;CDCP1;CTTN;IL1B;TFF3;TFF2;TFF1;SGK1;TLR4;CSTA;SLC43A2;TMPRSS4;CXCR5;NEDD9;PAK1;CYP2A13;CUX1;WIF1;HSD17B1;NPY;HSF1;CTNNA1;PAK6;IL32;CD74;NQO1;CDX2;GDF15;KLHL1;AKR1C2;CSHL1;MLH1;VAX1;GRHL2;KLK10;VEGFA;PPP1CA;KRT18;KRT17;QPCT;KRT14;F2RL1;IL17D;SIPA1;SCARB1;CD83;DCLRE1C;MPG;IHH;PLAT;COMT;ETS1;MPO;ETS2;IGF1R;LOXL2;HOXA11;HOXA10;CCND2;DMBT1;CCND1;PEBP4;CAPN2;PPFIA1;GGT1;TGM3;TGM2;WNT5B;OSR1;WNT5A;MIR27A;LIG4;TSC2;SCNN1G;BRMS1;ADAM12;AMPH;ABCG2;CALCA;DIABLO;ALOX15;CHAT;LTBP4;ALOX12;SLC1A5;ADRB2;BHLHE23;LPP;FUT1;NME1-NME2;PODXL;CHAD;NLRP2;OTX2;OTX1;TFAP2A;SMAD2;MGAM;SMAD3;MLKL;EPHX1;HMGA1;INHBA;EVX2;SMAD6;BMP7;ESR1;IGF2R;LRP1B;ARHGAP24;SMAD7;BMP2;IL6;NAT1;FGF19;GNAS;MIR184;MIR183;TGFBI;FGFR4;FGF12;FGFR3;MUC21;FGFR2;FGFR1;LGALS3BP;UCN;KDM5B;CRISP2;LDB1;ATP2A3;HSPB2;SLC2A2;OXT;LAMC2;NR2E1;CXCL5;HK1;CYP26B1;CASP8;ZIC2;CPT2;NOS1;MMP7;DCC;PDE4D;AXIN1;SSTR1;OPRM1;DCN;TGFBR2;GREM1;MIR126;TGFBR3;CCNA1;MYH2;AIM2;ELF3;MIR485;IFNG;CAT;LTA;IRF7;CHI3L1;CDH13;MYH9;ZMYND10;IRF6;SHANK2;RBMS3;MCPH1;DDC;THBS1;PTHLH;AURKB;MIR136;MUC1;MIR377;WNT11;GNA12;ERBB2;MIR492;PTK2B;SNCG;SLIT3;NCAM1;MUC4;CHGA;NOS2;LAMB3;CMTM3;GALNT2;PDE4DIP;MUC5B;CHFR;HS3ST2;GALC;KISS1;POMC;CD4;MIR145;MIR125B1;COL7A1;CDK2AP1;VIM;SCLY |
| Anxiety Disorders | 0.01056606005418811 | APP;FAAH;OXTR;CTNND2;DIXDC1;PKD2L1;COMT;TNF;RXFP3;CRHBP;CCND1;CYP2D6;STMN1;ADORA1;PITX2;EOMES;SLC38A1;MAGI2;KRT7;TSC2;MIF;CARTPT;MYT1;CYP2E1;ANKK1;PVALB;ABCB1;LXN;SLC26A1;CHRNA2;DLX4;CHRNA5;SLC1A2;SLC1A3;ADRB2;CACNA1C;FHIT;CRHR1;ACACA;CRHR2;HSD11B2;TRPM2;RAI1;CYP2B6;GALR3;MIR330;DEAF1;PENK;MBP;APOE;PRODH;TAC1;DRD4;S100A10;TMEM132D;EGR1;ACE;GSTM1;SLC33A1;HTR1A;IGF2;HTR1B;ESR1;SLC2A9;IL6;DAO;GDNF;PTK7;BCL2;ALPL;PAM;ACHE;LAD1;UCN;GPR26;FLT4;OXT;HTR2A;BRCA1;DBH;SLC6A1;SLC6A2;SLC6A3;GLI2;SGCE;ADAMTS2;GRM5;CCKAR;UCN3;GRM8;DNMT3B;CASP1;PDE4B;SOSTDC1;LBP;DGCR6;DLGAP3;IKBKE;SLC18A2;AVPR1B;COL25A1;TACR3;HTR3A;PAX5;OPRM1;NGF;C1QL1;ADORA2A;CCKBR;IFNG;TAL1;DNAJC5;IL1B;CAT;CRH;RAPGEF3;SHANK1;CRP;HDAC2;DDC;GRP;PON1;HTT;TTN;CYP17A1;CAMKK2;REM1;EXOSC6;NPY;NTF3;SLITRK5;FYN;NCAM1;DISC1;EYA1;NTRK3;GAD1;ZNF804A;CCK;BAIAP3;P2RX7;POMC;FABP3;FABP5;APC;KCNS1;FABP7;LEP;SCLY;ALLC;HSPA1B;HCN1 |
| Central neuroblastoma | 0.011012402924074568 | RB1;SERPINE2;SCT;STMN2;STMN4;HTR4;BACH2;MIR432;CYP2D6;DPYSL2;GJA5;MYC;STMN1;AKT1;MNX1;CHGB;WSB1;DAPK3;HPCAL1;PRKCA;UNC5D;RFX1;PDYN;ISL1;DKK1;DKK3;RBP7;BIN1;DDIT3;MAP1B;SERPING1;HOXB5;MEG3;ABCB1;CRABP1;PDGFA;GATA4;GATA3;THY1;GATA2;SLC5A5;TERT;KCNQ1OT1;ZSCAN10;ALX3;PROM1;HOXC9;HOXC6;BARD1;ABCA2;NGFR;CREBBP;GSTM1;CADM1;FZD5;FZD6;MIR338;PBX1;FLI1;PNOC;TNFSF9;PAM;HOXD9;HOXD8;CDKN1C;ACHE;CDKN1A;RARG;ADM;SLC6A2;SLC6A3;BBC3;GLI2;SHF;CHL1;TIMP2;NEFL;DLGAP1;NEFH;MYOG;DFFB;ANXA2;PAX3;POU5F1;NCOR2;MMP14;GPRC5A;RARA;RARB;HCLS1;ZNF236;OCA2;NFIX;GRP;GSTP1;NR1I2;COL11A2;LRRC17;KCNA3;HTT;ADCY2;ASCL1;GRAMD4;EGFR;MIR410;EXTL1;RASSF10;BGLAP;ALOX5;NSD1;CHN2;NTF3;SPP1;STAT4;WWOX;PKLR;STAT3;POU4F1;HIPK2;FOXR1;FABP3;NFIC;FABP7;FOSB;SIGLEC7;CRLF1;HSPA1B;HSPA1A;CSF3;CSF2;CSF3R;CSF1;CCNT1;EPO;FAM107A;RORA;APEH;CLU;FGF2;TNF;RIMS2;PYCARD;CDH5;HHAT;HPGDS;FGF8;NAGLU;GRAP2;RNF19A;MAP3K5;CD177;KCNH2;USP7;LAG3;SNRPN;TPM3;IGFBP3;HIC1;FOXP1;NPAS4;S100A6;SQSTM1;PVALB;MAD1L1;DSC3;CUL9;NOTCH3;ITPR1;CUL1;ITPR2;KLC1;SOCS3;SBNO2;NDN;SRGAP3;ZNF423;TFPI2;TNFRSF10C;HTR1B;ADAP1;GFAP;CDK6;STK25;CPD;CDK2;BCL2;SCGB3A1;SHBG;XAF1;CIITA;FLT4;FHL2;PIK3CD;ECE1;HTR2A;TRH;CPZ;FOXM1;PIK3CG;MB;PSD4;RNPEP;DNMT3B;SGCG;UNC45A;HSP90AA1;GPX1;ITGA4;PEG3;PLEKHG5;SH2D3C;NGB;TNFRSF1B;NGF;COASY;C1QL1;CRH;SNURF;BIRC5;TNFRSF25;ANG;CRP;CX3CR1;ITIH4;HDAC2;HDAC3;LRP1;SRC;RAB1B;MIR487B;GLRX;NTN1;MGAT5B;P2RY1;IGF2BP1;NTRK1;MARCKSL1;ANGPT2;ASCC1;NTRK3;KCNIP3;TBX5;MTOR;P2RX7;P2RX5;KLF6;MARCKS;APC;SST;P2RX2;IL2RA;LEP;P2RX1;NOVA2;IL2RB;CD248;CCM2;PDCD1;P4HB;INSM1;BCAR1;NPFFR2;ERRFI1;APP;SPARC;CERK;ICAM2;ICAM1;SOX2;LGALS3;RASSF1;EDNRA;LGALS1;EDNRB;RASSF5;RASSF6;TNFSF10;TP63;SLC12A9;PTGDR;SRRM2;EBF1;HLA-C;ANK2;CSNK1E;HLA-A;MIF;HLA-G;RUNX3;RUNX2;TIAM2;TIAM1;THEG;SFRP1;SLPI;KCNQ4;RTEL1;CASZ1;EPAS1;SLC45A1;MGST1;CEND1;TWIST1;CD1D;PIK3R1;CRHR1;TRPM2;NEU4;MAP2;CD19;STC2;PENK;ST8SIA4;APOD;B4GALNT1;APOE;TRPM8;SPTBN1;BBS2;STAT5A;EGR1;SNHG1;BCL11B;ST8SIA2;IGF2;MIR200A;MIR200B;DCLK1;CYP26A1;MSRA;TH;ARTN;ID2;ID3;GRB2;TP73;SPAG16;PTPRT;COL18A1;ACY3;CD40;MIR17;ICA1;MIR18A;BRCA1;PCSK6;PTPRF;CLYBL;CX3CL1;HNF4A;CD38;MIR192;KIF1A;CCR6;FFAR1;MCF2L2;CD34;CTSD;MME;PABPC4;INSRR;NME2;NME4;MIR21;PROX1;VAV1;AJAP1;NAALADL1;ADRB3;IL1B;MIR17HG;MPPED2;B3GNT3;ARHGEF2;CD47;MAPT;ARHGEF7;SGK1;NEDD9;CAPG;GDPD5;CAMKK2;RXRB;CUX1;WIF1;NPY;HSF1;GPC2;MSX1;DISC1;RAE1;NQO1;OSM;LILRB1;PEX14;VEGFA;TPPP3;PER3;QPCT;RAMP1;IL17C;NEUROG1;F2RL3;CD86;RTL1;CD83;CD81;LOXL3;PLAT;PRDM2;ETS1;ACTG1;IGF1R;MIR149;RGS4;CCND2;DMBT1;CCND1;NPPA;TGM2;PDGFRB;PDGFRA;PDIA2;CASR;G6PC;LIG1;LMO3;APLP2;WNT5A;MATK;UBE4A;MAT1A;SDHB;DHDDS;MYCN;HGS;CD8A;SCNN1A;AIPL1;ABCG2;CALCA;DIABLO;TCF7;ALOX15;CHAT;ALOX12;SLC1A5;ADRB2;NME1-NME2;CAMTA1;CYTL1;TAC1;MSC;WNT3;MARK2;PRPH;TFAP2B;ZFHX3;TFAP2E;WNT3A;PTCH2;HMGA1;ESR1;PHOX2B;ARHGAP24;PHOX2A;NR4A2;BMP2;IL6;RALYL;NAT1;GDNF;TLX2;FAM163A;GNAS;MIR184;ALPI;FGFR4;MXD3;FGFR2;FGFR1;LGALS3BP;KDM5B;CRISP2;MIR10B;MIR10A;MEGF10;HSPB6;ELAVL4;HSPB2;CHD5;OXT;LAMC2;DBH;NR2E1;GRM1;GRM2;GRM5;CYP26B1;CASP8;CA2;CASP1;FLOT1;NOS1;GALNT9;DCC;NAV2;OPRM1;POMGNT1;ARID1B;TGFBR3;IFNG;ALDH1A2;CAT;LTA;CDH13;ZMYND10;RBMS3;MCPH1;CAMK2B;DDC;RGS16;OPRL1;RRBP1;THBS1;PTHLH;AURKB;CRYZ;MUC1;CNR2;CLMN;MXI1;ERBB2;PTK2B;NCAM1;SNCB;SNCA;CHGA;LGI1;CDK11A;KDM4B;URGCP;NOS2;LIMK2;B3GAT1;CCK;CDK11B;MEIS2;DLK1;POMC;MEIS1;MIR145;XPNPEP1;MIR125B1;PPP2R2B;CD9;TEK;VIM;RAN |
| Drug-induced tardive dyskinesia | 0.01182341330538428 | TERC;DBH;HTR2A;COMT;TNF;SLC6A3;GLI2;CRYZ;CYP17A1;CYP2B6;CYP2D6;AKT1;PIP4K2A;NOS1;APOE;PITX2;SLC18A2;DRD4;NQO1;NOS3;F2R;GFRA1;GRIN2B;MTNR1A;GNAL;ADORA2A;MTNR1B;PPP1R1B |
| Tardive Dyskinesia | 0.01182341330538428 | TERC;DBH;HTR2A;COMT;TNF;SLC6A3;GLI2;CRYZ;CYP17A1;CYP2B6;CYP2D6;AKT1;PIP4K2A;NOS1;APOE;PITX2;SLC18A2;DRD4;NQO1;NOS3;F2R;GFRA1;GRIN2B;MTNR1A;GNAL;ADORA2A;MTNR1B;PPP1R1B |
| Carcinoma, Cribriform | 0.01182341330538428 | DDR1;CDKN1C;SYCP1;MSLN;KRT20;TNF;HOXA10;PYCARD;RASSF1;CASP8;CCND1;CA2;MYC;NPPA;PLCE1;TP63;DAPK1;TPM3;KRT8;UGT2B17;APOA1;CYP7B1;RUNX3;MMP14;KCNQ1;IL1B;BIRC5;RARB;PGR;CYP2E1;GRB7;NR1I2;EGFR;SELENBP1;MUC2;ERBB2;SPP1;APOE;STAT5A;TCF7L2;ACE;NOS3;STAT3;TNFRSF10A;ESR1;MTOR;VEGFA;CYP26A1;APC;BCL2;ID3;GHRL;FAT4 |
| Carcinoma, Granular Cell | 0.01182341330538428 | DDR1;CDKN1C;SYCP1;MSLN;KRT20;TNF;HOXA10;PYCARD;RASSF1;CASP8;CCND1;CA2;MYC;NPPA;PLCE1;TP63;DAPK1;TPM3;KRT8;UGT2B17;APOA1;CYP7B1;RUNX3;MMP14;KCNQ1;IL1B;BIRC5;RARB;PGR;CYP2E1;GRB7;NR1I2;EGFR;SELENBP1;MUC2;ERBB2;SPP1;APOE;STAT5A;TCF7L2;ACE;NOS3;STAT3;TNFRSF10A;ESR1;MTOR;VEGFA;CYP26A1;APC;BCL2;ID3;GHRL;FAT4 |
| Vascular lesions | 0.011885587491503086 | APP;COL18A1;NOTCH3;ELN;FLT4;ITGB2;LOXL3;TCF21;GSTT1;MPO;TNF;LOXL2;PAK1;CYP2D6;MYC;APOE;CD34;NQO1;EGR1;GSTM1;VWF;F10;PRRX1;EPHX1;HLA-B;IGF2;HLA-A;GP6;MTOR;IL6;AIM2;IL1B;CRH;IRF7;CYP1A1;PRTN3;CCM2;CYP2E1;TGFBI;LGMN |

| | | |
|---|---|---|
| Tumor Progression | 0.011885587491503086 | RB1;CYFIP2;MDC1;SPI1;MIR429;MIR425;ZC3H12D;MIR675;SOX17;CYP2D6;MYC;CREB3L1;STMN1;AKT1;PROK2;CYP1B1;DLEU2;HOXA5;PRKCB;DAPK1;SERPINF1;UNC5C;RFX1;RNASE1;PRAP1;DKK1;DKK3;SYTL2;DKK4;CLDN4;SLC5A8;HOXB9;SPINT1;BIN1;RBP1;JTB;PADI2;HOXB7;MEG3;ABCB1;LXN;CTBP2;DHH;TERC;HPN;GATA6;AGAP2;PDGFA;GATA4;GATA3;LYPD5;LY9;SLC5A5;LARP1;TERT;CHST11;S100A16;PDGFD;PROM1;ST3GAL1;ELANE;HOXC6;JAG2;BARD1;NGFR;CREBBP;ST14;ACE;TSLP;CADM1;PLK2;FZD7;NFATC2;MRPL28;NFATC1;NR2F2;AZU1;NR0B2;FLI1;H19;FERMT1;CPEB1;GJB6;GIPC1;NANOG;PAM;HLA-DRB1;LGMN;CDKN1C;HRK;PIGR;CDKN1A;MIR130B;LPCAT1;TCF21;ADM;MMP21;IL18BP;BBC3;GLI2;CHL1;TIMP3;NEFH;IKBKE;APC2;ANXA2;DUSP1;SPHK2;TESC;PAX6;PAX5;TMEM173;F3;POU5F1;SERPINB5;NCOR2;ITPKB;SLC7A5;MMP14;ZEB2;ZEB1;GPRC5A;MMP16;KIFC1;RARA;RARB;ANGPTL4;OCA2;GRP;GSTP1;COL11A1;NR1I2;KCNA3;HTT;KCNA5;FSTL1;HOXD10;EGFR;EPCAM;PLAGL1;ALOX5;NSD1;HAS1;SPP1;PRAME;MEF2D;SPDEF;BRD4;WWOX;PKLR;HSPA1L;MGMT;WNT7B;STAT3;DAB2IP;BLCAP;HIPK2;SKI;SPSB1;MIR661;WT1;ABI3;FABP7;VEPH1;ABI1;FOSB;ZNF217;LGR6;INHA;CSF3;CSF2;RASL11B;CSF1;EPO;RAB3D;DGKA;MSI1;IRS2;FGF1;CLU;FGF2;TNF;FGF3;BEST1;RIMS2;CDH6;MT1DP;HPGDS;CDH1;ADORA3;SALL4;GRAP2;RNF19A;PITX2;SLC16A3;KCNH1;KCNH2;MAP2K3;S100A2;TPM3;S100A1;IGFBP3;UGT2B15;F2R;UGT2B17;DIO3;MITF;CHD1L;HIC1;FOXP1;RAB34;S100A6;FSCN1;PGR;PKP3;ERG;TNIK;S100A9;MAD1L1;DSC3;NOTCH3;DLX2;DLX4;NOTCH4;MAZ;TACSTD2;GSTT1;VTCN1;FHIT;RAP1GAP;SLC9A3R2;SOCS2;SOCS3;SMYD2;THEM4;PLXNA4;ESRRA;KL;TNK2;TFPI2;GFAP;ADCY10;DAB2;COPS5;CDK2;BCL2;CPE;CSPG4;PLXNB1;XAF1;LAT;EIF4G1;PIWIL2;PIWIL1;PSCA;ITGB4;FLT4;ONECUT1;ITGB2;FHL2;WISP3;GADL1;PIK3CD;MSLN;FOXM1;TREM1;CYR61;PIK3CG;WISP1;ADAMTS4;CAPZB;MB;DNMT3B;SOST;HSP90AA1;GPX1;PEG3;VWF;DNMT3A;CTDSPL;APOA1;NGF;TNFRSF1A;TFDP1;BIRC5;TRIM59;MCM5;ANG;CRP;HDAC2;LRP1;SRC;UHRF1;HM13;SEMA3B;NTN1;MIR148A;SELENBP1;RAB25;CCL5;P2RY1;FYN;EIF4E;CNNM4;NTRK1;TBX1;SPAG9;ANGPT2;ANGPT1;PRRX1;ANKRD30A;PCDH8;CABLES1;MTOR;TBX2;P2RX7;MFAP5;NELL1;P2RX5;KLF6;PC;APC;SST;P2RX2;IL2RA;LEP;MGP;PDCD4;PTPN6;ZACN;P4HB;BCAR1;SPON2;IFITM1;TNFRSF6B;SPARC;GFI1;ICAM1;SOX2;LGALS3;RASSF1;EDNRA;LGALS1;EDNRB;TRIM29;ANPEP;RPS6KA2;TNFSF10;PART1;TNFSF11;SOX9;SOX7;TP63;SOX5;SRRM2;ENTPD2;EPHA6;KRT8;HLA-C;HLA-A;HNF1A;HLA-G;RUNX3;RUNX2;HLA-E;RUNX1;TIAM1;SFRP1;CCDC80;SFRP2;SLPI;IFI27;IL23A;ALPK2;EPHA1;EZR;STX1A;LLGL2;GRB7;SLC22A3;RTEL1;EPHA10;EPAS1;PCDH10;EIF5A2;TWIST1;MST1R;CRHR2;KLK6;NUAK1;TSPAN8;PCBP4;MAP2;STC2;HLTF;B4GALNT1;TRPM8;TSPAN1;NTSR1;BANP;STAT5A;NTNG1;EGR1;WWTR1;FOXF2;BCL11B;ST8SIA2;GREB1;IGF2;LIF;GDF2;MIR200A;PTK6;MIR200B;GADD45G;EXT1;BST1;OCLN;VNN1;PTK7;PPP1R1B;GRB2;ANP32B;TDGF1;CD200;TP73;DDR1;HOTAIR;DIRAS3;COL18A1;NRP2;CD40;MIR17;DOC2A;PPP1R10;BRCA1;LIMD1;PCSK6;KRT20;LIMD2;AKAP12;HYAL1;SLC22A18;HNF4A;HYAL2;PRX;STAB1;RAC2;CCR9;SOSTDC1;MIR192;CCR6;CTSD;HTATIP2;ACAD8;CBR1;ZHX2;EDN3;MME;MST1;AMPD1;IL16;MIR21;PROX1;AJAP1;ANO1;CDCP1;NR5A2;CTTN;CCKBR;IL1B;MIR17HG;MMP23A;TFF3;ARHGEF2;CD47;SGK1;TLR4;GUCY2C;TAGLN;CSTA;TNXB;TMPRSS6;SATB2;TMPRSS3;TMPRSS4;NEDD9;TMPRSS2;GNAI2;CAMKK2;NT5E;PAK1;CUX1;WIF1;GPC1;HSF1;GPC5;CTNNA2;MICB;IL32;GINS2;CD74;NQO1;CDX2;CDX1;GDF15;GRHL3;MLH1;PDZK1IP1;GRHL2;KLK10;VEGFA;PER1;KRT18;DLC1;F2RL1;BLID;TACC2;IL17D;F2RL3;PRDM2;ETS1;ETS2;IGF1R;LOXL2;MIR149;HOXA10;RGS4;RGS5;CCND2;CCND1;PEBP4;ARHGDIB;MIR150;TGM2;PDGFRB;PDGFRA;CASR;KDM2B;OSR1;WNT5A;LMTK3;MIR27A;TSC2;BRMS1;MYCN;ADAM12;PRTN3;CARD16;NUPR1;ABCG2;CALCA;DIABLO;TCF7;BHLHE23;FUT1;FUT4;DPP4;CALD1;PODXL;MIR296;UBD;OTX2;STAP2;TAC1;WNT3;MARK2;TFAP2A;SMAD2;FDPS;TFAP2C;CBX5;SMAD3;WNT3A;TGFB3;PTCH2;HMGA1;INHBA;BMP7;ESR1;ARHGAP24;SMAD7;ISG20;NR4A2;BMP2;IL6;FER;EFNA2;GDNF;FGF19;BHLHE40;GNAS;GNB4;MIR183;ALPI;TGFBI;FGFR4;FGFR3;ULBP1;FGFR2;FGFR1;LGALS3BP;KDM5B;CRISP2;TNFAIP8;MIR10B;PLEK;TNFAIP2;HSPB2;LAMC2;MIR96;GRM1;CXCL5;HK1;ELAVL3;CASP8;MLST8;FLOT1;NOS1;MIR193A;EGFL7;MMP7;H2AFY;MIR23A;DCC;OPRM1;DCN;SSTR4;TAPBP;TGFBR2;KLF17;GREM1;MIR126;FGR;TGFBR3;ELF1;AIM2;IFNG;ELF5;CAT;CDH11;IRF7;CHI3L1;CDH13;IRF8;ZMYND10;LTB;OTUB1;RBMS3;LAMA4;RGS16;THBS1;PTHLH;AURKB;MUC1;MUC2;GMDS;MIR495;MIR494;MXI1;GNA12;ATOH8;ERBB2;MIR372;PTK2B;SPOCK1;SNCG;NCAM1;MUC4;PDLIM4;CHGA;RWDD3;URGCP;NOS2;CMTM3;NOS3;BNIP3;USP22;EYA4;LIMK1;MUC5B;MIR1914;HS3ST1;KISS1;POMC;FMNL1;MIR145;MIR144;MIR142;LNX1;RGS10;CD9;VIM;LPIN1;RAN |
| Cirrhosis | 0.011885587491503086 | RB1;CSF3;CSF1;CLU;TNF;MPO;AQP1;ICAM1;IGF1R;PYCARD;FGF7;HPGDS;EDNRA;RASSF1;EDNRB;CCND2;DMBT1;CYP2D6;CDH1;AKT1;SLC12A9;IGFBP1;NANOS1;CASR;MSX2;SERPINF1;SLC11A1;KRT8;HLA-C;LIG4;HLA-A;MAT1A;DKK1;HIC1;DKK3;RBP1;CYP2E1;S100A9;SLC27A2;ABCG2;CHRNA1;FBN2;NOTCH3;ABCB1;TERC;NOTCH4;PDGFB;ADRB2;FUT1;DPP4;SOCS3;DGUOK;GRIN2A;TPO;CYP2B6;TERT;TSPAN8;PDGFD;APOH;APOE;APOB;TAC1;NGFR;ACE;GSTM1;TFPI2;IGF2;GRIN2B;BMP7;ESR1;DCLK1;ARHGAP24;COL1A1;IL6;COPS5;FGF19;NOSTRIN;BCL2;GNB3;TGFBI;HAMP;HLA-DRB1;FGFR2;TP73;DDR1;ACHE;CNTNAP2;CDKN1A;FHL2;GPT;HTR2A;PIK3CG;CXCL5;GLI2;GHR;HK3;HNF4A;DNMT3B;CASP1;DLAT;NOS1;DENND2D;GPX1;TRIM63;MMP7;VWF;H2AFY;DNMT3A;AXIN1;APOA1;MIR21;PROX1;ADRA2C;NGF;F3;DCN;F7;CCNA1;MMP14;ZEB1;AIM2;ADRB3;IFNG;IL1B;MIR17HG;CAT;CRH;CDH13;CHI3L1;BPI;TLR4;GSTP1;PON1;NOD2;THBS1;EGFR;AURKB;HLA-DMA;NT5E;WIF1;ALOX5;CCL5;ERBB2;UGT1A7;SNCA;ANGPT1;MGMT;GDF15;NOS3;STAT3;OSM;MLH1;KLK10;GCK;VEGFA;KLF6;MARCKS;APC;HLA-DPB1;SPRY2;CLNK;SPRY1;VIM;FMOD |
| Degenerative polyarthritis | 0.011885587491503086 | APP;SPON1;TNFRSF6B;VIPR2;SERPINE2;STMN2;ICAM3;XYLT1;ICAM1;LGALS3;MIR675;EDNRA;LGALS1;EDNRB;DPYSL2;MYC;STMN1;TNFSF10;AKT1;TNFSF11;CYP1B1;SOX9;DUS4L;TRIM21;SLC12A9;CMKLR1;UNC5B;COG5;ACTN1;SERPINF2;UNC5C;EBF3;HLA-A;KRT5;PDYN;ANK1;DKK1;RUNX2;DKK3;SFRP1;SLPI;IL23A;PADI2;EZR;EPAS1;TWIST1;FPR2;PIK3R1;PRKCZ;CRHR1;CRHR2;C3;ADAMTS16;ADAMTS14;ANKH;CHST11;INPP5D;APOE;A2M;B4GALNT3;S100A10;EGR1;TRAP1;ST14;ACE;TSLP;TNFSF14;TREX1;IGF2;LIF;PRG4;MIR200B;NR1D1;COL1A1;COL9A3;COL9A2;ANP32B;HOXD9;PAM;HLA-DRB1;NODAL;DDR1;CD40;CDKN1A;MIR17;ADM;LRRC32;PCSK6;SERPINA4;RYR3;BBC3;RPTOR;UCN3;UCN2;HYAL1;HNF4A;HYAL2;TRIM2;TIMP2;CD38;TIMP3;SOSTDC1;CCR6;IKBKE;CTSD;CLIC1;TIMP4;ANXA2;DUSP1;MYOF;NME2;TET1;IL16;MIR21;PAX5;NCOR2;TLR1;MMP14;COL2A1;IL1B;UFC1;RARB;FAM20C;ANGPTL4;CRYAB;GAS5;TLR4;TNXB;GRP;SLC20A2;COL11A1;COL11A2;ASIP;NR1I2;CXCR5;CAPG;HOXD13;FSTL1;BGLAP;HAS1;ALOX5;SPP1;LAIR2;CILP2;MSX1;APOL1;JAM3;IL32;GDF15;IL34;STAT3;OSM;SAAL1;AKR1C2;WWP2;SORBS2;SULF1;GCK;VEGFA;FOSB;F2RL1;CLC;LGR6;IL17D;CRLF1;CD86;CSF2;CSF1;IHH;PLOD2;RORA;FGF1;CLU;FGF2;TNF;LOXL2;PYCARD;COMP;HPGDS;GRAP2;RNF19A;NPPA;PITX1;SLC16A3;TGM2;KIRREL3;IGFBP1;PDGFRA;SNRPN;MSX2;S100A1;IGFBP3;F2R;WNT5A;DIO2;DIO3;LIG4;CYBA;FOXP1;LTB4R2;SCN8A;ADAM12;S100A9;CSF1R;NOTCH3;CALCA;DLX5;MAZ;CHAT;ALOX15;HTRA1;SOCS3;NLRP3;ZNF423;TAC1;MSC;NDUFV1;ISOC2;ESRRA;SMAD2;KL;SMAD3;RARRES2;TFPI2;INHBA;BTNL2;BMP7;ESR1;SMAD7;BMP4;MFHAS1;BMP2;IL6;LRCH1;BCL2;HLA-DRA;ALPL;PLXNB1;TGFBI;FGFR4;FGFR3;FGFR1;CRISP2;UCN;THRA;FLT4;ITGB2;FHL2;WISP3;DUSP19;SLC40A1;PIK3CD;TREM1;PIK3CG;WISP1;LTB4R;CYR61;HAPLN1;GHR;ADAMTS4;ADAMTS5;CASP8;SCRN1;SUMO3;TRPS1;MCF2L;BDKRB2;CASP1;ACP5;PDE4A;SOST;ADAMTS9;SGCG;HSP90AA1;MMP7;ITGA4;BUD31;DNMT3A;APOA1;TNFRSF1B;NGF;TNFRSF1A;TGFBR2;GREM1;RCN3;TGFBR3;ZAP70;ELF3;IFNG;ALDH1A2;CDH11;SNURF;CRH;CHI3L1;IRF5;ANG;B4GALT7;CNGB1;CRP;HDAC4;CAMK2B;HDAC2;LRP1;TUFT1;LRP5;GLRX;NOD1;TM2D3;NTN1;THBS1;PTHLH;MIR148A;PRTG;CLEC3A;ACAN;CLEC3B;CNR2;CHADL;CCL5;MIR373;ERBB2;IL21R;PTK2B;OPCML;CHGA;SEMA4A;ANGPT1;NOS2;TXNRD1;KCNIP3;OPRK1;TBX5;CLCN7;P2RX7;POMC;MIR145;LEP;MGP;SPRY2;PTPN7;VIM;RAN |
| Coronary Arteriosclerosis | 0.011885587491503086 | NPFFR2;TES;GMFG;EHMT1;AQP4;ICAM1;EDNRA;LGALS2;LIPC;EDNRB;CYP2D6;NDST4;AKT1;ENTPD1;SERPINF1;ANK2;MIF;HNF1A;RUNX3;RUNX1;GPR162;SFRP4;CFDP1;SOAT2;DDIT3;PAPPA;CES1;SLC22A3;ABCB1;AIRE;TERC;PDGFA;KLK4;SLC5A1;GATA2;PLA2G7;MIR206;KALRN;ABO;C3;HSD11B1;CYP2B6;RNF213;TERT;KCNQ1OT1;PDGFD;ST8SIA4;APOD;APOE;PROM1;APOB;ELANE;EGR1;GSTM2;ACE;GSTM1;ABCA4;SLC33A1;IGF2;MRPL28;GP6;ATP4B;BHMT;TNFSF4;CYP1A1;NOX5;ANP32B;HLA-DRB1;ACHE;PIGR;RYR2;ROBO3;CD40;MIR17;TCF21;GRIK4;PCSK9;PEAR1;ASGR1;RYR3;CX3CL1;IL18BP;JPH3;CCR9;CCR7;CCHCR1;CD34;PF4V1;PARP4;ANXA2;AMPD1;SERPINB9;GP1BA;IL16;MIR21;F3;F7;MMP14;ADRB3;TNNT1;IL1B;ANGPTL4;ARHGEF7;TLR4;RAPSN;HLA-DQB1;NR1I2;MTHFD1L;BGLAP;ALOX5;NPY;SPP1;EXTL3;MPZL2;P2RY12;NQO1;RNLS;SORT1;GDF15;F12;STAT3;ADAM33;DAB2IP;NR1H3;GCK;VEGFA;FABP2;ESYT3;GHRL;HSPA1A;F2RL3;CD86;SCARB1;CSF3;CSF2;EPO;F13A1;MIR146B;IRS2;COMT;FGF2;TNF;ETS1;MPO;CNDP1;HPGDS;SLN;ADORA3;ADORA1;ABCG8;CASR;ABCG5;MSX2;CD93;F2R;CYBA;PROCR;ELMO1;HHIPL1;ADAM8;S100A9;ABCG1;ISYNA1;ALOX15;GSTT1;ADRB2;CACNA1C;CYP2C18;ACAT2;FUT3;GRK5;GRK4;HRH2;NPC1L1;NLRP3;MARK2;FGF21;KL;TFAP2B;RARRES2;ESR1;BMP4;SLC2A9;BMP2;IL6;CPS1;BCL2;CPE;GNB3;SLC26A4;SHBG;FGFR1;TRIM72;UCN;THRA;HSPB6;FHL2;DYSF;HSPB2;PIK3CD;TNFAIP3;ECE1;HTR2A;CYR61;PIK3CG;CXCL5;CA2;CASP1;PHACTR1;GPX1;MMP7;TRPC3;VWF;MIR23A;APOA1;OPRM1;TNFRSF1B;KLF14;TGFBR2;ACTA2;GREM1;MIR126;ADORA2A;IFNG;DDAH2;CHDH;TRAF5;NOS1AP;CRH;LTA;CDH13;CHI3L1;IRF8;IRF6;CRP;CX3CR1;PON3;LRP1;PON1;LCAT;NOD1;LTC4S;NTN1;THBS1;THBS4;TTN;INS;SELENBP1;CBS;CCL5;P2RY1;NCAM1;FSD1;UGT1A6;TCF7L2;ANGPT1;KCNJ8;NOS2;ASCC1;GALNT2;NOS3;CMTM5;SOD3;KISS1;POMC;MIR145;CAPN10;LEP;APOC1;MGP;PDCD4;BCAR1 |
| Amphetamine-Related Disorders | 0.011885587491503086 | ACHE;GSTP1;PLAT;PRDM2;GSTT1;AOAH;AK5;COMT;FHIT;SLC6A3;GRM2;CYP2D6;PSD3;PDE4B;AKT1;NOS1;CCHCR1;ZNF423;CSMD1;CD55;DRD4;SNCA;GSTM1;PDE4D;EPHX1;HTR1B;NRG1;HTR3A;OPRM1;PDYN;SGCZ;POU5F1;SH3RF3;ADORA2A;GDNF;SLC6A9;CDH13;ASTN2 |
| Amphetamine Abuse | 0.011885587491503086 | ACHE;GSTP1;PLAT;PRDM2;GSTT1;AOAH;AK5;COMT;FHIT;SLC6A3;GRM2;CYP2D6;PSD3;PDE4B;AKT1;NOS1;CCHCR1;ZNF423;CSMD1;CD55;DRD4;SNCA;GSTM1;PDE4D;EPHX1;HTR1B;NRG1;HTR3A;OPRM1;PDYN;SGCZ;POU5F1;SH3RF3;ADORA2A;GDNF;SLC6A9;CDH13;ASTN2 |
| Diabetes Mellitus, Non-Insulin-Dependent | 0.0130217007965932 9 | PGAP3;DQX1;EHMT2;XYLT2;EHMT1;BACH2;MIR432;LIPC;ZMIZ1;STMN1;AKT1;CYP1B1;PRKCB;SERPINF1;UNC5D;ISL1;GABRG3;CLPS;DDIT3;SERPING1;CYP2E1;ANKK1;CFD;ABCB1;CTBP1;TERC;GATA6;GATA4;SLC5A1;ABCB9;SLC30A10;PRKCZ;SLC5A4;ABO;SULT1A1;C3;HSD11B1;CYP2B6;PRDM16;ALX4;HCRT;HLA-DQA2;RREB1;HLA-DQA1;HLA-DRB5;ACD;GSTM1;ACE;ABCA4;SLC16A11;NFATC1;NR1D1;MIR339;GP6;NR0B2;PBX1;EHD1;BHMT;FAT1;HRC;CYP1A1;PAM;HAMP;HLA-DRB1;CDKN1C;CDKN1A;ELN;SETD7;PRKAG2;GPT;ADM;SLC6A2;TRIOBP;C1QTNF3;C1QTNF1;JPH3;TIMP2;TIMP3;TCF19;GPR39;BRD2;PAX4;TACR3;PAX6;F3;DUSP6;POU5F1;NCOR2;GJD2;F7;MMP14;EXOC6;ANGPTL4;HLA-DQB2;HLA-DQB1;OCA2;GSTP1;SPESP1;ANKRD55;NR1I2;ASIP;KCNA3;MIR409;ADCY3;EGFR;HLA-DMA;HLA-DMB;ALOX5;CHN2;SPP1;APOL1;P2RY12;AOC3;AOC2;PKLR;KCNB1;RNLS;STAT3;NR1H3;TMEM154;FABP1;TMEM155;FABP2;FABP3;MIR661;FABP5;FABP6;SLMAP;SLCO4C1;HSPA1B;TTLL8;HSPA1A;EXOC3L2;DGKD;EPO;LY75;DEFB1;IRS2;FRY;CLU;FGF2;TNF;BEST3;BEST1;RIMS2;HPGDS;MAEA;SESN3;ADORA3;GRAP2;RNF19A;ADORA1;GRB10;SLC39A7;MAP3K5;IGFBP1;TLE3;IGFBP3;UGT2B15;USP2;MAGI2;DIO2;UBE2E2;FOXP1;CLIP2;MIR1228;MTNR1A;MTNR1B;S100A6;FSCN3;TOX;S100A9;SQSTM1;SHC1;GSTT1;LIN7C;CYP2C18;FHIT;RHOBTB1;SOCS3;SBNO2;ATXN1;THEM4;NKX6-1;CD300LG;STRA6;ESRRA;KL;GFPT2;ANKRD23;ESRRG;MCC;MAFA;INS-IGF2;MAFB;BCL2;CPE;HLA-DRA;SHBG;ONECUT1;ITGB2;PIK3CD;ECE1;PTPN22;HTR2A;TRH;TCIRG1;FOXM1;PIK3CG;HHEX;SGCD;ADAMTS9;SGCG;TRPC6;GPX1;DST;PEG3;VWF;SORD;APOA1;KSR2;TNFRSF1B;COASY;TNFRSF1A;TRAF3;NOS1AP;BPI;BCAT1;CRP;RBPJL;CX3CR1;LRP1;SRC;MIR487B;LRP5;ATP10A;NOD1;PRR5L;NOD2;SELENBP1;CCL5;TP53INP1;KCNJ1;UGT1A6;NTRK1;TUB;KCNJ6;PRRX1;KCNJ11;KCNJ9;TXNRD2;IRX3;PCDH8;KCNJ15;EXOG;SORL1;MTOR;TPCN2;P2RX7;MIR1249;PC;IL2RA;LEP;PDCD1;P4HB;CDO1;BCAR1;SLC35B3;SPON2;APP;FAAH;SPARC;AQP2;ICAM1;EDNRA;EDNRB;TNFSF10;SCN1B;SLC12A9;CPT1A;ENTPD1;HLA-C;KCNK16;ANK2;HNF1B;TIMD4;HLA-A;MIF;HNF1A;CPT1B;ANK1;RUNX2;RUNX1;DHX40;SFRP4;PPP1R3C;IL23A;KCNQ1;SFRP5;COL8A1;ALDH7A1;CFTR;STX1A;CES1;SLC22A3;MACF1;EPAS1;GIPR;SLC22A2;C2CD4A;C2CD4B;PIK3R1;CRHR1;TRPM2;TSPAN8;ST8SIA4;APOD;APOE;APOB;PCK1;TRPM5;TRPM4;STAT5A;PTPRN2;TNFSF14;FAM78B;SLC33A1;IGF2;LIF;MIR200A;NBEAL2;GDF3;EXT1;PEX5L;TF;STAR;GRB2;FBXL7;PCSK2;PTPRU;RYR2;PTPRS;PTPRN;PCSK9;CX3CL1;GNGT2;CCKAR;HNF4A;CD38;MIR192;FFAR1;CCHCR1;ACAD8;MME;TSC22D1;GAA;PABPC4;SORCS1;MIR21;PROX1;NCR1;MC3R;ADRB3;IL1B;MIR17HG;SGK1;TLR4;RAPSN;TMPRSS6;CYP17A1;CAMKK2;NPY;GPC5;CILP2;PCNT;DISC1;NQO1;CDX2;SORT1;GDF15;GAD1;OSM;LILRB1;SORBS1;CSHL1;GCK;VEGFA;PER1;RIT2;GHRL;IL17D;PNPLA2;FOXA2;NEUROG3;SCARB1;PLAT;CELSR1;COMT;CNDP1;CMIP;IGF1R;CNDP2;CCND2;DMBT1;NPPA;CAPN2;GGT1;TGM2;GHSR;ABCG8;PDIA2;CASR;ABCG5;G6PC;SLC30A8;MSX2;WNT5B;MIR27A;CYBA;CARTPT;MAT1A;SDHB;PROCR;LMX1A;DOK5;GPD2;ELMO1;NUPR1;ABCG1;ABCG2;FBN2;ISYNA1;CALCA;CCDC149;TCF7;ALOX15;NPR3;CACNA1D;ALOX12;CACNA1C;ADRB2;ARHGAP15;CACNA1E;LPP;SLC1A7;CPLX1;ACACA;DPP4;RAI1;AANAT;GRK5;NPC1L1;ALDH3B1;NLRP3;CAMTA1;MSC;FGF21;TFAP2A;TFAP2B;DARS;MGAM;RARRES2;EPHX1;HMGA1;HSPA2;POU2F2;BMP7;ESR1;IGF2R;LRP1B;PHOX2A;SMAD7;BMP4;ISG20;PYY;IL6;FGF14;IL17REL;HOOK2;FGF19;GNB3;TGFBI;FGFR4;FGFR1;GPSM1;USP36;MIR10B;TREH;PLEK;ATP2A3;TNFAIP3;SLC2A2;NR2E1;SLC2A5;PKD1;PITPNC1;CXCL5;HK1;CASP8;GDAP1L1;MAN1A1;NOS1;KLF11;MMP7;CAMK1D;MIR23A;GLP2R;RBM19;OPRM1;ARAP1;KLF14;POMGNT1;DCN;SSTR4;TGFBR2;GREM1;MIR126;TGFBR3;MIR485;IFNG;DDAH2;GORASP2;PALLD;CHDH;CAT;LTA;CHI3L1;MYH9;RBMS1;LTF;COLEC12;LAMA1;NRXN1;PON1;NRXN3;AGPAT1;CRYZ;INS;MIR377;CNR2;CBS;PTK2B;TAS1R3;CHGA;TCF7L2;CDK11A;NOS2;NOS3;GALNT2;PDX1;CCK;MUC5B;SOD3;POMC;CD4;MIR145;MIR144;CAPN10;SARDH;SPRY2;TEK;LPIN1;LPIN2 |
| Diabetes Mellitus, Experimental | 0.0130217007965932 9 | CHRM2;PCSK2;ATP2A3;SLC2A2;IRS2;MPO;TNF;CX3CL1;ICAM1;HK1;TIMP2;BDKRB1;MAP3K5;YWHAH;CPT1A;GPX1;CYBA;PRKCA;PAX6;CPT1B;TNFRSF1A;SLC9A3;ADRB3;IFNG;GPD2;IL1B;CAT;S100A6;HSD11B1;CYP2B6;NKX6-1;PCK1;S1PR5;S1PR4;FGF21;ESRRA;AOC3;NQO1;ACE;KCNJ11;NOS2;PKLR;NOS3;PDX1;GCK;VEGFA;IL6;PC;GPAM;LEP;BCL2;CYP1A1 |
| Alloxan Diabetes | 0.0130217007965932 9 | CHRM2;PCSK2;ATP2A3;SLC2A2;IRS2;MPO;TNF;CX3CL1;ICAM1;HK1;TIMP2;BDKRB1;MAP3K5;YWHAH;CPT1A;GPX1;CYBA;PRKCA;PAX6;CPT1B;TNFRSF1A;SLC9A3;ADRB3;IFNG;GPD2;IL1B;CAT;S100A6;HSD11B1;CYP2B6;NKX6-1;PCK1;S1PR5;S1PR4;FGF21;ESRRA;AOC3;NQO1;ACE;KCNJ11;NOS2;PKLR;NOS3;PDX1;GCK;VEGFA;IL6;PC;GPAM;LEP;BCL2;CYP1A1 |
| Streptozotocin Diabetes | 0.0130217007965932 9 | CHRM2;PCSK2;ATP2A3;SLC2A2;IRS2;MPO;TNF;CX3CL1;ICAM1;HK1;TIMP2;BDKRB1;MAP3K5;YWHAH;CPT1A;GPX1;CYBA;PRKCA;PAX6;CPT1B;TNFRSF1A;SLC9A3;ADRB3;IFNG;GPD2;IL1B;CAT;S100A6;HSD11B1;CYP2B6;NKX6-1;PCK1;S1PR5;S1PR4;FGF21;ESRRA;AOC3;NQO1;ACE;KCNJ11;NOS2;PKLR;NOS3;PDX1;GCK;VEGFA;IL6;PC;GPAM;LEP;BCL2;CYP1A1 |
| Noninfiltrating Intraductal Carcinoma | 0.0130217007965932 9 | CD86;IRS2;CLU;ICAM1;SOX2;PYCARD;LGALS3;RASSF1;HPGDS;CCND2;CCND1;CDH1;MYC;AKT1;SOX9;HOXA5;TP63;SLC12A9;SRRM2;STARD3;KDM2A;S100A1;IGFBP3;DAPK3;WNT5A;KRT5;RUNX3;CACNB1;SFRP1;BRMS1;RBP1;PGR;S100A9;GRB7;SLC22A3;TWIST1;GATA4;FHIT;LDHB;TERT;PCBP4;KCNQ1OT1;NEU1;STAT5A;TFAP2A;CREBBP;ESR1;TMPRSS13;ISG20;COPS5;BCL6;FAT1;BCL2;SCGB3A1;MIR183;SIM2;FGFR1;DIRAS3;HOTAIR;CDKN1A;LPCAT1;FHL2;HSPB2;PIK3CD;BRCA1;PIK3CG;TRPS1;NEFL;CD34;TRIM62;MME;ITGA4;MST1;PAX6;MIR21;NRG1;PAX5;DUSP6;SERPINB5;NCOR2;NR5A2;ALDH5A1;ELF3;CDH11;CDH13;IRF5;RBMS3;COL1A1;GSTP1;PON1;NOD2;EGFR;PTHLH;PAK1;MUC1;EPCAM;ERBB2;SNCG;EIF4E;NQO1;WWOX;BNIP3;TIE1;PDZK1IP1;SULF1;KLK10;VEGFA;P2RX5;PPP2R2B;KRT15;IL2RA;KRT14;PDCD4;VIM |
| Pain | 0.013089187767939242 | RB1;APP;CSF3;FAAH;OXTR;CSF2;CRTC1;ZBTB20;COMT;TNF;AQP1;COMP;CRHBP;EDNRA;TCL1A;EDNRB;CYP2D6;ANPEP;NPPA;AKT1;MIR150;NRTN;EPHA4;UNC13A;LMO3;KRT7;PDYN;KCNK18;DHDDS;MTNR1B;IL23A;SERPING1;PGR;SQSTM1;CFTR;CALCA;ABCB1;CHAT;SLC1A2;CACNA1A;ADRB2;CACNA1E;ACACA;HRH1;CYP2B6;NPBWR1;PENK;NLRP3;SFN;TRPM8;HCRT;TAC1;DRD4;NTSR1;PRPH;SMAD2;NGFR;TRAP1;ACE;CD160;HTR1A;LIF;TRPV3;GRIN2B;ESR1;COL1A1;IL6;DAO;CPS1;ARTN;PNOC;GNB3;COL9A2;SHBG;HLA-DRB1;FERMT3;KCNK9;DYSF;ADM;GPT;OXT;HTR2A;TRH;SERPINA6;PCSK6;SLC6A3;PIK3CG;GRM1;ADAMTS4;TMEM100;GRM2;CPT2;OPA3;MCF2L;GRM8;BDKRB2;BDKRB1;CCHCR1;NEFH;MCF2L2;SLC18A3;TACR2;TACR3;TET1;NRG1;IL16;HTR3B;OPRM1;ADRA2C;NGF;TNFRSF1B;SSTR4;TNFRSF1A;ANO3;COL2A1;IFNG;TAL1;IL1B;RARA;CRH;ANG;BPI;SCN4A;MAPT;LTB;CRP;CX3CR1;ITIH4;PON3;GRP;DCTN1;COL11A1;GSTP1;NR1I2;PON1;CXCR5;OPRL1;CAPG;ASAP2;AIF1;EGFR;SELENBP1;REM1;ACAN;ABLIM3;CNR2;CCL5;NPY;NTF3;P2RY1;ERBB2;SPP1;CACNG2;DISC1;SH2B2;P2RY12;NTRK1;CHGA;KCNJ6;GABBR1;CDX2;TXNRD2;NOS3;KCNIP3;GAD1;B3GAT1;OPRK1;MTOR;VEGFA;P2RX7;POMC;MIR145;KCNS1;KRT17;SST;LEP;F2RL1;SCLY;LGR6;KCNK2;RAMP1 |
| Memory impairment | 0.013359898375592132 | APP;CHRM1;CSF2;CSF1;CRTC1;EHMT1;COMT;MPO;TNF;PYCARD;GRAP2;RNF19A;CDK20;PDGFRB;S100A1;MAGI2;PRKCA;MIF;CARTPT;PRKAR1B;BIN1;RBP1;S100A9;SQSTM1;GNE;CSF1R;NOTCH3;ALOX15;CHAT;PDGFB;GPR85;GSTT1;CPLX1;TTBK1;TRPM2;PCBP4;DEAF1;CAMTA1;APOE;TPPP;MSC;NGFR;KL;CREBBP;GSTM1;IGF2;HTR1A;NR1D1;EIF2AK4;GRIN2B;GFAP;NR4A2;CPEB1;IL6;FGF14;PNOC;BCL2;SIM1;SIM2;ACHE;THRA;LAMC2;BRCA1;HTR2A;KIF17;SLC6A3;GRM5;HNF4A;PDE4A;NEFM;MME;FCRL5;SPHK2;PDE4D;DNMT3A;APOA1;NGF;TNFRSF1A;ADORA2A;IL1B;MAPT;SHANK1;CX3CR1;HDAC4;NLGN1;SLC20A2;SPG7;CLCN1;INS;CNR2;ALOX5;NPY;SLC17A7;NCAM1;EIF4E;SNCA;CD74;PKLR;GABRA5;GAD1;STAT3;SORL1;VEGFA;POMC;SUCLA2;QPCT;HLA-DPB1 |
| Congenital Hypothyroidism | 0.016920473712093435 | ACHE;CSF1R;THRA;TNF;SLC5A5;GHR;DUOX1;HHEX;TPO;IYD;BGLAP;NEFL;NEFM;OTX2;NKX2-5;NEFH;EGR1;NGFR;PROP1;KSR1;NOS2;PDE4D;INHBB;TRAPPC9;RUNX2;PAX2;TSHR;TRHR;CNKSR1;TG;PAX8;LHX3;GNAS;SLC26A4 |
| Encephalitis | 0.017001261822428507 | APP;CNTNAP2;CD40;CIITA;CSF2;AQP4;ADM;LAMC2;TCIRG1;FGF2;TNF;CX3CL1;ICAM1;PYCARD;CASP8;CPT2;CASP1;BLNK;CD38;F2R;NGB;LIG4;SARS;IL16;MIR21;FCER2;IFNG;IL1B;DNAJC5;CHI3L1;CD47;ARHGEF7;CRP;CX3CR1;ITIH4;ISYNA1;ABCB1;UNC93B1;NR1I2;PDGFB;KCNA3;MIR206;GRK1;MAP2;SPP1;NLRP3;APOD;MBP;APOE;PRODH;ATP6V0A1;LGI1;EGR1;GABBR1;NOS2;ASCC1;STAT3;GFAP;POMC;IL6;SST;MIR142;BCL2 |

| | | |
|---|---|---|
| Asthma | 0.017034825449315956 | SERPINE2;PGAP3;HTR4;BACH2;MYLK;ZMIZ1;C3AR1;STARD3;SERPINF2;CLEC11A;DKK1;SIGIRR;RBP7;DDIT3;TYRO3;SERPING1;CYP2E1;MAML3;PDGFB;PDGFA;NEDD4L;FPR2;GATA3;GATA2;PLA2G7;ABO;C3;HSD11B2;ADAMTS14;CYP2B6;ELANE;HLA-DQA1;CREBBP;GSTM2;HLA-DRB5;GSTM1;ACE;TSLP;LRBA;FUCA1;PBX2;BTBD11;NR0B2;COL1A1;H19;TNFSF4;PNOC;HLA-DRB1;XKR6;CDKN1C;ACHE;DENND1B;CDKN1A;ELN;ADM;LRRC32;MYOM2;GLT1D1;TIMP3;JAK3;HLA-DPA1;SLC18A3;BRD2;GSTO2;ANXA2;DUSP1;FCRL3;TACR2;TET1;F3;SERPINB6;F7;FCER2;RARA;GPRC5C;HLA-DQB1;GRP;AHNAK;GSTP1;SYNPO2;COL11A2;NR1I2;ADCY2;AOAH;FSTL1;EGFR;SLC7A2;BGLAP;ALOX5;NTF3;SPP1;STAT4;FLNC;CSMD1;P2RY12;CCL24;F10;HSPA1L;STAT3;ADAM33;HLA-DPB1;FOSB;INHA;KLKB1;CCL28;CCL26;CHRM2;CSF3;CHRM1;CSF2;FAM13A;F13A1;IL5RA;MIR146B;DEFB1;RORA;FGF2;TNF;RIMS2;HPGDS;CDH1;FCGRT;GRAP2;RNF19A;ADORA1;TNFRSF8;HLA-DOA;SLC39A2;TNFRSF4;MAP2K3;S100A1;IGFBP3;F2R;TAP2;OBSCN;LTB4R2;ADAM8;S100A9;SQSTM1;ZNF432;CSF1R;NOTCH4;ITPR1;MRI1;GSTT1;SOCS3;MBP;CD160;ZBTB16;MCC;PTPRE;CDK2;BCL2;HLA-DRA;SLC26A4;LAT;DPP10;ITGB4;ITGB2;COX4I2;FHL2;PIK3CD;PTPN22;ITGAL;ADRA1B;PIK3CG;WISP1;ITGAX;ITGB7;ITGB6;ADAMTS9;GRASP;CR1;ITGA4;GRID1;VPS13D;APOA1;NRG1;TNFRSF1B;NGF;ADORA2A;CRH;BIRC5;ANG;CNGB1;CRP;CX3CR1;RETNLB;HDAC2;HCG18;GLRX;NOD1;NOD2;JAKMIP1;CCL5;IL21R;FYN;CCL1;D2HGDH;UGT1A3;FSD1;EIF4E;NTRK1;SEMA4A;ANGPT1;KCNIP4;MTOR;P2RX7;MARCKS;PC;APC;IL2RA;LEP;IL2RB;PDCD4;PTPN6;PTPN7;OXTR;AQP4;IKZF4;ICAM1;LGALS3;EDNRA;LGALS1;EDNRB;TNFSF10;LGALS9;TP63;PTGDR;SOX5;EPHA5;SRRM2;ENTPD1;SLC11A1;SLC6A15;HLA-B;HLA-C;TIMD4;MIF;HNF1A;HLA-A;ANK1;RUNX3;HLA-G;RUNX1;SLPI;CRISPLD2;IL23A;ALDH7A1;ZFPM1;CFTR;SLC22A2;CD1D;CRHR1;CRHR2;CD19;INPP5D;APOE;TRPM8;A2M;STAT5A;EGR1;TNFSF14;HIST1H4K;LRRN2;HIST1H4L;SLC33A1;LIF;ALOX15B;PM20D1;BST1;VNN1;HIST1H4I;ANP32B;CD200;FBXL7;HIST1H4F;RYR2;CD40;TNFRSF13B;WFDC1;KRT20;AP2A2;CX3CL1;AKAP13;CCR9;CD38;CCR7;CCR6;TNFAIP8L2;MIR192;CCHCR1;CD34;CTSD;ACAD8;LYNX1;MST1;IL16;MIR21;PARP9;MIR19A;ANO3;GFRA4;TLR1;ANO1;CTTN;IL1B;TFF2;ARHGEF2;GLCCI1;TLR4;RAPSN;SDCCAG8;TAGLN;CSTA;CUX1;NPY;CTNNA1;CTNNA3;XDH;CD55;MICB;IL32;NQO1;OSM;SORBS1;PEX14;KLK11;VEGFA;BATF;KCNS3;NBR1;F2RL1;GHRL;IL17D;ALLC;RAMP1;IL17C;CD86;GIMAP5;ETS1;MPO;ETS2;RGS5;NPPA;GGT1;TGM2;PDGFRB;PDGFRA;POSTN;SLC30A8;CYBA;ERN2;ADAM12;CCDC151;CALCA;ISYNA1;CHRNA2;CHAT;ALOX15;ALOX12;CSF2RB;ADRB2;LPP;PSORS1C1;DPP6;HRH1;AANAT;GRK5;NLRP3;TAC3;TAC1;MARK2;SMAD2;SMAD3;EPHX1;TBCD;ESR1;SMAD7;ISG20;IL6;PROC;NAT1;ALPL;TGFBI;LGALS3BP;UCN;MS4A3;HSPB6;PLEK;HSPB2;TNFAIP3;OXT;LTB4R;GRM1;PITPNC1;CXCL5;CASP8;BDKRB2;PDE4B;BDKRB1;PDE4A;NOS1;MKKS;PTGIR;MMP7;PDE4D;UBE2QL1;IRAK4;SSTR4;DCN;TAPBP;TGFBR2;TGFBR3;ACTA1;AIM2;IFNG;OAS2;ELF5;CAT;PEX5;IRF7;LTA;CHI3L1;LTB;RBMS3;DNAH5;PON1;OPRL1;RRBP1;LTC4S;THBS1;AURKB;CRYZ;MUC1;MUC2;HLX;ERBB2;NOS2;NOS3;B3GAT1;OPRK1;MUC5B;SOD3;IRGM;MIR145;LPIN2;RAN |
| Adenocarcinoma, Tubular | 0.017034825449315956 | DDR1;CDKN1C;SYCP1;MSLN;KRT20;TNF;HOXA10;PYCARD;RASSF1;CASP8;CCND1;CA2;MYC;NPPA;PLCE1;TP63;DAPK1;TPM3;KRT8;UGT2B17;APOA1;CYP7B1;RUNX3;MMP14;KCNQ1;IL1B;BIRC5;RARB;PGR;CYP2E1;GRB7;NR1I2;EGFR;SELENBP1;MUC2;ERBB2;SPP1;APOE;MUC6;STAT5A;TCF7L2;ACE;CDX2;NOS3;STAT3;TNFRSF10A;ESR1;MTOR;VEGFA;CYP26A1;APC;APOC1;BCL2;ID3;GHRL;FAT4 |
| Familial M??ni??re disease | 0.017034825449315956 | CD86;CD40;CSF2;TNFAIP3;AQP4;DEFB1;PTPN22;AQP2;AQP1;ICAM1;CCND1;GPX1;PRKCB;GPX5;HLA-B;HLA-C;HLA-A;MIF;TNFRSF1A;IFNG;MTNR1B;IL1B;IRF7;KCNQ4;NDUFS2;TLR4;LTF;NOTCH3;KCNE1;KCNE3;KCNA1;NPR3;PON1;CACNA1A;KCNA4;KCNA5;ADD3;FCHO1;MARVELD2;CCL5;HLA-DQA1;CHGA;HLA-DRB5;DTNA;NOS2;NOS3;STAT3;FAM136A;ESR1;POU4F3;IL6;GJB6;IL2RA;SLC26A4;HLA-DRB1;HSPA1A |
| Esophageal carcinoma | 0.017034825449315956 | RB1;MDC1;ICAM2;ICAM3;MIR429;ICAM1;SOX2;LGALS3;FRAT1;RASSF1;RASSF2;SOX17;MYC;PEG10;TNFSF10;AKT1;CYP1B1;PLCE1;HOXA1;SOX9;TP63;SLC12A9;NKX2-8;DAPK1;KRT7;PRKCA;DLEC1;HLA-A;DKK1;RUNX3;RUNX1;MIR196A2;CRISPLD2;KCNQ2;CYP2E1;EZR;NEIL1;LLGL2;MEG3;GRB7;SLC22A3;ABCB1;TERC;GATA5;TWIST1;GATA4;THY1;ABO;SULT1A1;ADAMTS16;TERT;PCBP4;KCNQ1OT1;STC2;SFN;EGR1;TRAP1;CREBBP;GSTM2;ACE;GSTM1;FZD5;FOXF1;FZD7;XRCC3;IGF2;MRPL28;ALOX15B;DCLK1;GADD45G;GJB2;ARTN;CYP1A1;FAT4;HLA-DRB1;TP73;CDKN1C;HOTAIR;CDKN1A;BRCA1;MIR18A;AKAP12;SLC22A18;TIMP2;CD38;MIR192;CCHCR1;CD34;HTATIP2;GPR39;ZHX2;ANXA2;RIPK3;MIR21;PROX1;PAX2;POU5F1;SLC7A5;MMP14;CDCP1;CTTN;IL1B;MIR17HG;RARA;TFF3;RARB;TFF2;TFF1;TLR4;GRP;GSTP1;CRCT1;EGFR;NKD2;RASSF10;EPCAM;WIF1;SPP1;IL32;WWOX;NQO1;CDX2;MGMT;STAT3;MLH1;PSMB8;VEGFA;DLC1;INHA;IL17D;CHRM2;SEMA5B;LY6K;CSF2;ZBTB20;PRDM2;FGF2;TNF;ETS1;HOXA13;MPO;FGF3;IGF1R;LOXL2;RIMS2;FGF7;HPGDS;CCND1;CDH1;CCRL2;LZTS1;TGM3;PDGFRB;S100A1;USP4;IGFBP3;TAP2;UBE2E2;MIR27A;FSCN1;DPEP1;S100A9;ABCG2;DIABLO;ALOX15;LTBP4;ITPR2;GSTT1;VTCN1;FHIT;STK31;MIR296;TAC1;TFPI2;EPHX1;ISG15;ESR1;ARHGAP24;SLC2A9;DAB2;IL6;HOOK2;CDK2;BCL2;CNTN1;GNAS;SCGB3A1;MIR183;CLPTM1L;TGFBI;XAF1;FGF12;FGFR2;FGFR1;MIR10B;THRB;HSPB2;PIK3CD;MSLN;MIR96;CYR61;PIK3CG;DNALI1;WISP1;ZNRD1;CASP8;DNMT3B;ADAMTS9;MMP7;DCC;PDE4D;TGFBR2;IFNG;BIRC5;MCM5;CRP;SRC;GOSR1;FOXK1;CRYZ;MUC1;MIR377;DNAJB6;PUF60;MIR373;ERBB2;SNCG;CCL1;FSD1;RGS22;EIF4E;MUC4;MARCKSL1;ANGPT1;PRRX1;ASCC1;TXNRD1;FBXL19;BNIP3;EYA4;CCK;MTOR;CHFR;MIR145;APC;ST7 |
| Bulimia | 0.017034825449315956 | FAAH;ARHGAP8;DST;LMO3;HTR1A;IGF2;HTR2A;COMT;CRHR1;SLC6A3;C1QL1;CIT;POMC;PRR5-ARHGAP8;LEP;CRH;GHRL;APOB;DRD4 |
| HIV Encephalopathy | 0.017034825449315956 | APP;ITIH4;CSF2;DCLRE1C;NOS2;LIG4;LAMC2;ADRB2;OPRM1;CLU;TNF;IGF2R;TNFRSF1A;IL6;SST;SPP1;CD9;APOE;TPPP |
| Panic Disorder | 0.017034825449315956 | GRP;ELN;HTR2A;COMT;PRDM11;SLC6A2;CRHR1;MIR148A;CRHR2;INS;SGCE;DAOA;LDHB;CCKAR;DEAF1;NPY;NPPA;PDE4B;SOSTDC1;APOE;IKBKE;DRD4;TMEM132D;ACE;AVPR1B;GABRA5;GAD1;NTRK3;HLA-B;HTR1A;HTR3A;CCK;MIR339;SGCZ;ANO2;ADORA2A;CCKBR;GLRB;PTK7;CRH;PKP1;PGR;HLA-DRB1;SDK2 |
| Nonorganic psychosis | 0.017034825449315956 | GABRB2;FAAH;CD40;ST6GALNAC1;GRIK4;OXT;HTR2A;COMT;SLC6A3;IGF1R;RGS4;DAOA;GHR;GRM2;GRM5;CYP2D6;DPYSL2;AKT1;SOSTDC1;SEPHS1;CD34;KCNH2;PDGFRB;TGIF1;HLA-C;TET1;PRKCA;NRG1;CSNK1E;CARTPT;OPRM1;OLIG2;HCG9;GABRG3;SRCIN1;HLA-E;DHDDS;MTNR1B;RARB;MAPT;ARHGEF7;THOP1;SHANK1;SLC22A3;DLX1;RTN3;HDAC2;ITIH3;ABCB1;GSTP1;SHOX2;NR1I2;GSTT1;CACNA1C;CRHR1;NPAS3;SELENBP1;TRPM2;GRIN2A;CYP2B6;NPY;NTF3;GNA12;MGAT1;APOE;PCNT;DISC1;ST3GAL1;DRD4;SNCA;TBX1;GABRA1;ASCC1;GAD1;B3GAT1;ZNF804A;HTR5A;GRIN2B;ESR1;TBX4;IL6;DAO;SLC6A9;ARTN;LEP;PPP1R1B;F2RL1;SP8;FGFR3;FGFR2 |
| Cholangiocarcinoma | 0.017382676774641136 | UGT1A10;SCT;EPO;TNF;MPO;AQP1;LOXL2;LGALS3;HOXA9;RASSF1;EDNRB;LGALS1;SOX17;CCND1;DMBT1;CDH1;MYC;GRAP2;RNF19A;STMN1;TNFSF10;AKT1;MIR150;PDGFRB;ABCG8;DAPK1;MITF;DLEC1;SFRP4;SFRP1;IFI27;FSCN1;NEIL1;HOXB7;SQSTM1;CFTR;ABCG2;SLC22A3;SHC2;ABCB1;GATA6;PLA2G7;FHIT;SLC5A5;FUT3;SULT1A1;NME1-NME2;SOCS3;TERT;PODXL;NEU1;ALX4;NLRP3;SFN;APOE;PROM1;APOB;SPRR2A;TAC1;WNT2;MARK2;TFAP2A;TGFB3;LIF;MIR200A;PTK6;MIR200B;ESR1;IGF2R;ARHGAP24;IL6;PPFIBP2;NAT1;CYP1A1;FAT1;BCL2;GNAS;GNB3;HOXD9;LAT;FGFR2;TP73;FGFR1;RARG;CD40;RPLP1;FLT4;ONECUT1;PIK3CD;FOXM1;MSLN;KRT20;PIK3CG;WISP1;CXCL5;DNMT3B;TIMP3;MIR192;CD34;SGCG;ACAD8;CBR1;MMP7;PEG3;MME;DCC;NME2;MIR21;PROX1;NGF;MIR19A;POU5F1;TNFRSF1A;TGFBR2;MIR126;SLC7A5;ZEB1;IFNG;IL1B;MIR17HG;BIRC5;CLDN18;TFF1;CREB5;CRP;HDAC4;HDAC3;GSTP1;NR1I2;EGFR;CRYZ;MUC1;MUC2;EPCAM;ALOX5;MIR494;HSF1;ERBB2;SPP1;PTK2B;EIF4E;MUC4;MUC6;NTRK1;WWOX;NQO1;MARCKSL1;ANGPT2;CDX2;NOS2;MGMT;NOS3;STAT3;SCTR;KLHL1;PDX1;GNMT;VEGFA;MFAP5;GALNS;MARCKS;KRT18;APC;MIR144;SST;DLC1;PDCD4;TRIP10;F2RL1;CDO1 |
| Familial multiple trichoepitheliomata | 0.017451533786128185 | PIK3CD;TNF;PIK3CG;GHR;AKAP12;CCND1;CDH1;MYC;SLC22A18;AKT1;TIMP3;MIR192;IGFBP3;TAP2;AFAP1;MIR21;RUNX3;RUNX1;TNNT1;IL1B;ALDH1A2;ELMO1;PKP1;CDH13;PGR;TFF1;S100A9;RIN2;MIR155HG;GSTP1;BARX1;FHIT;FUT4;KLK7;EGFR;MIR148A;CYP17A1;SELENBP1;SEC14L3;GJC2;TERT;WIF1;ERBB2;SPP1;APOB;TAC1;EIF4E;CD55;WWOX;FOXF1;MGMT;IGF2;MRPL28;MIR200A;INHBA;MLH1;TBX5;ESR1;MTOR;GDF7;VEGFA;NELL1;CDK6;MIR145;APC;FABP6;LEP;GHRL;EIF4G1 |
| Esophageal Neoplasms | 0.018011573997658 | RB1;MDC1;IFITM1;ICAM2;ICAM3;ICAM1;SOX2;LGALS3;FRAT1;RASSF1;RASSF2;SOX17;MYC;PEG10;TNFSF10;AKT1;CYP1B1;PLCE1;SOX9;TP63;SLC12A9;NKX2-8;DAPK1;SLC11A1;KRT7;DLEC1;HLA-A;DKK1;RUNX3;RUNX1;MIR196A2;CRISPLD2;MT1G;CYP2E1;EZR;NEIL1;LLGL2;MEG3;GRB7;SLC22A3;ABCB1;TERC;GATA5;TWIST1;GATA4;THY1;SULT1A1;ADAMTS16;TERT;TSPAN8;KCNQ1OT1;STC2;SFN;EGR1;TRAP1;GSTM2;ACE;GSTM1;FZD5;FZD7;XRCC3;IGF2;MRPL28;ALOX15B;DCLK1;GADD45G;CYP26A1;GJB2;CYP1A1;FAT4;HLA-DRB1;TP73;CDKN1C;HOTAIR;CDKN1A;BRCA1;BBC3;AKAP12;AKAP13;SLC22A18;TIMP2;CD38;MIR192;CD34;GPR39;ZHX2;ANXA2;RIPK3;MIR21;PROX1;PAX2;POU5F1;SLC7A5;MMP14;CDCP1;CTTN;IL1B;MIR17HG;RARA;RARB;CRYAB;TLR4;GRP;GSTP1;NR1I2;CRCT1;EGFR;NKD2;RASSF10;EPCAM;WIF1;ALOX5;SPP1;IL32;WWOX;NQO1;CDX2;MGMT;STAT3;MLH1;VEGFA;GHRL;IL17D;LY6K;CSF3;ZBTB20;PRDM2;FGF2;TNF;ETS1;HOXA13;MPO;IGF1R;LOXL2;RIMS2;FGF7;HPGDS;CCND1;CDH1;LZTS1;TGM3;TGM2;PDGFRB;TPM4;USP4;IGFBP3;UBE2E2;MIR27A;ADAM12;FSCN1;DPEP1;S100A9;DIABLO;ALOX15;LTBP4;GSTT1;VTCN1;FHIT;STK31;MIR296;SMYD2;TFPI2;EPHX1;TNFRSF10A;ISG15;ARHGAP24;DAB2;IL6;HOOK2;CDK2;BCL2;CNTN1;GNAS;SCGB3A1;MIR183;CLPTM1L;TGFBI;XAF1;FGF12;MIR10B;THRB;HSPB2;PIK3CD;MSLN;MIR96;CYR61;PIK3CG;WISP1;ZNRD1;CASP8;DNMT3B;EMILIN3;ADAMTS9;MMP7;DCC;TGFBR2;IFNG;BIRC5;CDH13;CRP;SRC;FOXK1;AURKB;CRYZ;MUC1;MIR377;DNAJB6;PUF60;GNG7;MIR373;ERBB2;SNCG;FSD1;RGS22;EIF4E;MARCKSL1;ANGPT1;PRRX1;NOS2;NOS3;TXNRD1;FBXL19;BNIP3;EYA4;CCK;MTOR;CHFR;MIR145;APC;SST |
| Opiate Addiction | 0.0180316778926817 | ABCB1;OPRL1;NRXN3;COMT;GRIN2A;GRM6;GRM8;PENK;DISC1;DRD4;SNCA;MYOCD;OPRK1;HTR3A;OPRM1;PDYN;PARP9;GRIN2B;NR4A2;POMC;IL1B;PPP1R1B;CRH;TFF3;TSNAX-DISC1;ANKK1;CES1 |
| MYELODYSPLASTIC SYNDROME | 0.0180316778926817 | RB1;SPARC;SCT;GFI1;ICAM1;HOXA9;RASSF1;MYC;ANPEP;STMN1;TNFSF10;AKT1;CYP1B1;PIP4K2A;PLCE1;TP63;IER3;SLC12A9;CPT1A;DAPK1;DAPK3;RPL22;HLA-C;RUNX2;DKK3;RUNX1;HLA-E;UHMK1;SFRP4;SFRP1;IFI27;TFR2;IL23A;DDIT3;SFRP5;CYP2E1;ENPEP;ABCB1;CTBP1;TERC;MIR196B;TWIST1;THY1;GATA2;EVPL;ABO;CYP2B6;TERT;PCBP4;CD19;INPP5D;PRDM16;FADD;PROM1;ELANE;EGR1;CREBBP;GSTM1;INTS1;PLK2;LIF;MRPL28;NBEAL2;NR0B2;FLI1;GP9;GRB2;HAMP;HLA-DRB1;CD22;CD200;TP73;CDKN1C;ACHE;CDKN1A;CD40;GFM1;MIR17;PRSS27;GPT;KRT20;SMPD3;AKAP12;AKAP13;MECOM;TIMP2;CD38;CCR7;TK1;CD34;CD52;ANXA2;RING1;MME;NME4;MIR21;TLR1;SLC7A5;MMP14;IL1B;MIR17HG;RARA;CD47;SLC29A1;EEF1E1;TLR4;GSTP1;HOXD13;PURA;NT5E;PAK1;CUX1;ALOX5;NSD1;SPP1;CTNNA1;PRAME;CD55;RUNX1T1;IL32;NQO1;MGMT;STAT3;HIPK2;VEGFA;WT1;HLA-DPB1;CCL28;CRLF1;SIPA1;CSF3;CSF2;CSF3R;CSF1;EPO;CRTC1;MPL;IRS2;PRDM2;MSI2;TNF;HOXA13;MPO;FGF3;IGF1R;HPGDS;CCND1;CDH1;GRAP2;RNF19A;SALL4;BAALC;MIR150;CD177;PDGFRB;KDM2B;IGFBP3;HIC1;FOXP1;DHDDS;ERG;S100A9;MAD1L1;CSF1R;CALCA;DLX4;GSTT1;FHIT;FUT1;RAP1GAP;FUT4;TPO;BCL2L10;UBE2I;TNFRSF10C;ARHGAP24;LSM2;SMAD7;ISG20;NR4A2;DAB2;BMP2;IL6;G6PC3;BCL2;GNAS;ALPI;TGFBI;DIDO1;CNTNAP1;MIR10B;MIR10A;ITGB4;FHL2;PIK3CD;LAMC2;FOXM1;PIK3CG;SPN;SGCE;LYL1;CASP8;CPT2;NOS1;KLF11;DCC;DNMT3A;WIPI2;TNFRSF1B;TNFRSF1A;TGFBR2;IFNG;LTA;BIRC5;BLVRB;ANG;SCN4A;ZMYND10;ITIH4;CXXC5;AURKB;MXI1;NCAM1;FSD1;EIF4E;PDLIM4;ANGPT2;ANGPT1;NOS2;SPAG6;TIE1;OPRK1;SGCZ;DLK1;P2RX7;RAD52;MEIS1;MIR145;TEC;APC;CD6;MIR125B1;IL2RA;PTPN6;TEK |
| Cleft Palate | 0.01815394581371812 | CDKN1C;RARG;F13A1;FGF1;ETS1;FGF2;FGF3;GLI2;FGF7;FGF8;MYC;SOX9;TP63;PDGFRA;OSR2;SLC32A1;BHMT2;RUNX2;TGFBR2;F7;COL2A1;CAT;IRF6;DLX5;COL11A1;SATB2;COL11A2;NEDD4L;GSTT1;SPP1;MSX1;TFAP2A;TBX1;GSTM1;FOXF2;TGFB3;NOS3;GAD1;ABCA4;GRHL3;MEIS2;BMP4;SLC2A9;BMP2;MAFB;SPRY2;FGFR3;SIM2;FGFR2;FGF10;FGFR1 |
| Behcet Syndrome | 0.018291257523252752 | HCP5;TRAF3IP2;C4BPA;ADARB1;C4BPB;ETS1;MPO;TNF;BACH2;ICAM1;PYCARD;SLC11A1;HLA-B;TAP2;HLA-C;TAP1;HLA-A;MIF;HLA-F;HLA-G;RUNX1;HLA-E;IL23A;PRKCQ;TXK;PROZ;GSTT1;GATA3;PSORS1C1;APOH;NLRP3;APOB;ACE;GSTM1;UBAC2;IL6;CYP1A1;BCL2;HLA-DRB1;DDR1;CD40;CIITA;ITGB2;GADL1;TNFAIP3;PTPN22;ETFB;KRT20;ITGAL;TREM1;PSTPIP1;CASP8;CASP1;CCR7;NEFM;CCR6;CCHCR1;ANXA2;VWF;MIR23A;FCRL3;APOA1;MIR21;NRG1;NGF;TNFRSF1B;TNFRSF1A;TLR1;TGFBR3;AIM2;IFNG;IL1B;CAT;TRAF5;LTA;IRF8;TLR4;HLA-DQB1;CRP;MVK;NOD1;SPG7;NOD2;STAT4;IL12RB2;MICB;NTRK1;CCL23;MOG;PKLR;NOS2;NOS3;STAT3;VEGFA;GATM;QPCT;IL2RA;LEP;HLA-DPB1;PDCD4;PDCD1;IL17D;CPVL |
| Common Migraine | 0.01831129787230302 | CALCA;DDC;LRP1;KCNA3;CACNA1A;HTR2A;CACNA1C;COMT;TNF;CACNA1E;EDNRA;PRDM16;PHACTR1;KCNN3;TRPM8;HCRT;TAC1;MEF2D;DRD4;ACE;KCNB2;GFRA1;KCNK18;CACNB2;DLG2;LTA;ASTN2;RAMP1;HLA-DRB1 |
| Hydatidiform Mole | 0.018474229650889434 | CDKN1C;CSF1R;PPP1R13B;PSCA;HPSE2;TERC;ASCL2;EGFR;PAK1;LGALS1;ZFP57;TERT;NLRP7;PLAGL1;ERBB2;HOXC5;SPP1;NLRP2;TIMP3;HOXC4;SRGAP3;HLA-DOA;APOB;HOXC9;CHGB;HOXC6;IGFBP1;ZBTB16;STAT3;IGF2;HIC1;POU5F1;PSMB9;KISS1;BRMS1;LEP;BCL2;NANOG;ZACN;HLA-DRB1 |
| Osteoporosis | 0.018529080829285727 | CSF3;SPARC;CSF1;GTF2B;XYLT2;ZBTB20;PLOD2;RORA;IRS2;PRDM2;COMT;FGF2;TNF;HS6ST1;BEST1;ICAM1;ACTG1;IGF1R;HPGDS;FGF8;CCND1;MYC;CYP1B1;PROK2;TNFSF11;SOX9;TGM2;CASR;G6PC;TPM4;SLC34A1;UGT2B17;MIR27A;MITF;DKK1;RUNX3;RUNX2;SFRP4;MTNR1B;GPD2;DOK6;MT1G;ALDH7A1;GCM2;CSF1R;CALCA;ABCB1;TERC;DLX5;ALOX15;WNK4;TWIST1;GATA4;ALOX12;ADRB2;CRHR1;HSD11B1;CYP2B6;TERT;ANKH;CHAD;CDH23;TAC3;APOE;SPTBN1;ZNF266;STRA6;FGF21;ESRRA;KL;ACE;WNT3A;IGF2;MIR338;ESR1;COL1A1;EXT1;BMP4;FGF17;BMP2;IL6;BHLHE40;CYP1A1;BCL2;GNAS;ALPL;SP7;SHBG;FGFR1;CIITA;CD40;ELN;PLEK;WISP3;ADRA1D;PTPRF;GHR;ADAMTS2;ATIC;MECOM;CA2;SH3PXD2B;TIMP2;RSPO3;SOST;INSL3;SLC22A11;GPX1;ANXA2;TACR3;IL16;TNFRSF1B;WNT16;DUSP6;GREM2;MMP14;PCCA;IFNG;IL1B;PCCB;CAT;LTA;LTF;LRP1;NR1I2;LRP5;LRP4;SLC7A1;MIR148A;PTHLH;CYP17A1;CAMKK2;CNR2;BGLAP;CBS;NSD1;HSD17B1;SPP1;PTK2B;ZNF469;FYN;PDLIM4;WWOX;SVIL;KDM4B;KCNJ8;CDX2;TXNRD1;DMRT3;P2RX7;CLCN7;GALNS;KISS1;POMC;NELL1;FABP3;WT1;LEP;QPCT;COL7A1;MGP;MAB21L2;SPRY1;CLC;P4HB |
| Myoclonic Seizures | 0.018624328626861707 | PRDM8;ACHE;FAAH;NRP2;CHRM1;ABAT;GRIK1;OXT;TRH;ADRA1B;SLC6A1;CLU;FGF2;SLC6A2;NOS1;SCN1B;GAMT;TSC2;OPRM1;PDYN;NGF;SYN2;ALDH5A1;ADORA2A;IL1B;KCNQ2;CAT;CRH;NHLRC1;ALDH7A1;PVALB;ABCB1;CHRNA5;FOXG1;CHAT;CPLX1;INS;NPY;PLAGL1;SLC17A7;APOE;SLC25A22;NGFR;GABRA1;SLC12A5;KCNJ11;NOS2;GABRA5;STAT3;HTR1A;PDX1;HTR1B;OPRK1;CCK;TRAPPC9;GCK;LETM1;POMC;SLC8A3;IL6;SST;LEP;ALPL;GNB3;GHRL;PAM;HCN1 |
| Status Epilepticus | 0.018939272756703474 | CD86;CD40;EPO;ELAVL4;EHMT1;ABAT;AQP4;GRIK1;PLAT;ECE1;SLC6A1;TNF;CX3CL1;GRM1;LMNB2;AQP1;SLC8A2;GRM5;CASP8;CCND1;DPYSL2;CCR9;CASP1;CCR7;NOS1;CTSD;PDGFRB;GHSR;MEF2C;PRKCB;TSC2;PROX1;SSTR1;NGF;SSTR4;SSTR5;ALDH5A1;ADORA2A;DDIT3;IL1B;SCN8A;KCNMA1;CAT;CRH;ALDH7A1;TLR4;S100A9;CARS2;ABCG2;CRP;CX3CR1;PDXK;NLGN1;ABCB1;SRC;SLC1A2;ASCL1;AIF1;HSD11B1;SOCS3;ACAN;GRIN2A;NT5E;DEAF1;NPY;NTF3;PNPO;PTK2B;APOD;PRODH;NQO1;GABRA1;EGR3;SLC12A5;KCNJ11;NOS2;GABRA4;NTRK3;GRIN2B;MTOR;VEGFA;SLC8A3;GALC;GDNF;SST;GJB6;BCL2;FGF12;KCNK2;HCN1 |
| Crohn's disease of large bowel | 0.019161450015446054 | CRP;DENND1B;NCF4;TCF7;DEFB1;FPR2;NOD2;TNF;BACH2;SOCS3;FGF7;TAGAP;CCR9;STAT4;NLRP3;MADCAM1;NKX2-3;ACAD8;CCL25;SMAD3;SLC11A1;DNMT3A;MST1;STAT3;TYK2;TNFRSF1B;MLH1;NR0B2;IRGM;PHOX2B;TNFRSF1A;IL6;IFNG;IL2RA;LEP;FGF19;TLR4;HLA-DRB1 |
| Chronic Obstructive Airway Disease | 0.019436979008124095 | APP;SERPINE2;WIPF1;PITPNA;EHMT1;PKD2L1;HTR4;ICAM1;EDNRA;EDNRB;C3AR1;FAM19A2;PLCE1;SOX5;ENTPD1;PRKCB;SLC11A1;HLA-A;KRT5;MIF;HLA-F;ARMC2;SPINT1;SLPI;IL23A;DDIT3;MAP1B;RIN3;CYP2E1;MAML3;CFTR;SYCP2L;EPAS1;CD1A;HSD11B1;TRPM8;A2M;ELANE;EGR1;CREBBP;GSTM2;AFAP1L2;ACE;GSTM1;TSLP;BTBD11;COL1A1;H19;CYP1A1;CDKN1C;SPAG16;PIGR;CD40;ELN;SETD7;ADM;SMPD3;AKAP12;TIMP2;CCR7;CCR6;MIR192;IKBKE;DIRC3;LYNX1;TPSG1;GSTO2;TACR2;TFEB;IL16;MIR21;PARP9;SERPINB6;SUMF1;MMP14;TOX2;GPRC5A;IL1B;RARB;ARHGEF7;TLR4;RAPGEF3;CSTA;GSTP1;NR1I2;CD5L;KCNA3;ADCY2;EGFR;DUOX1;NT5E;NTF3;SPP1;STAT4;CSMD1;XDH;BRD4;IL32;CD74;NQO1;EGLN2;CDX2;GDF15;STAT3;ADAM33;SORBS1;VEGFA;FABP5;HLA-DPB1;F2RL1;IL17D;KLKB1;IL17C;FOXA2;CHRM2;SCARB1;CSF3;CSF2;KCNG4;CD83;EPO;NCF4;FAM13A;DEFB1;MSI1;CELSR1;FGF2;TNF;MPO;FGF7;HPGDS;CCND1;TNFRSF4;CD177;PDGFRB;F2R;MAGI2;HBE1;CYBA;CDYL;SDHB;ADAM19;CD8A;ADAM8;PRTN3;S100A9;CUL9;CHRNA1;CHRNA5;LTBP4;ADRB2;SLC1A5;SLC1A7;PSORS1C1;SOCS3;DPP6;SNTG2;TAC1;PLXNA4;MARK2;SMAD2;KL;SMAD3;EPHX1;BTBD9;SMAD6;SMAD7;IL6;DLG2;GDNF;BCL2;ESAM;SLC26A4;FGF10;ITGB2;PIK3CD;PTPN22;HTR2A;DBH;ITGAL;TREM1;PIK3CG;CXCL5;SGCD;PDE4A;NOS1;DSP;TRPC6;TRIM63;VWF;PDE4D;APOA1;NRG1;TNFRSF1B;TNFRSF1A;TGFBR3;ADORA2A;IFNG;CAT;CDH13;CHI3L1;LTB;DOCK1;CRP;HDAC2;HDAC3;SRC;DNAH5;LRP5;GLRX;NOD2;LTC4S;MUC1;MUC2;CBS;CCL5;P2RY1;CCL1;FSD1;S1PR5;MUC4;MUC6;ANGPT2;NOS2;ASCC1;NOS3;BNIP3;MUC5B;TBX4;SOD3;MYO1D;P2RX7;GALC;NELL1;P2RX5;P2RX2;LEP;P2RX1;SARDH;VIM |
| Chronic gastritis | 0.020107585614876508 | ABCB1;PCDH10;GSTP1;GATA4;GSTT1;NOD1;NOD2;TNF;THBS1;TP53AIP1;REM1;MUC1;TERT;CCL5;ERBB2;AKT1;TIMP3;MADCAM1;WNT2;GSTM1;MSX2;DCC;XRCC3;PDX1;MIR27A;MLH1;RUNX3;POMC;IL6;IFI27;CD8A;IL1B;MIR142;BCL2;TFF2;TFF1;CYP2E1;TRIM31;TLR4;CHST2;HLA-DQB1 |
| Psychotic symptom | 0.020139640686741237 | OXTR;GYPC;CCDC68;OXT;DBH;HTR2A;COMT;SLC6A3;MCHR2;SELENBP1;DAOA;ADAMTS2;MGAT1;APOE;NOS1;NOS2;LMO3;HTR1A;KRT7;NRG1;NR4A2;IL6;SLC6A9;TAL1;MAPT;SCLY |
| Elevated C-reactive protein | 0.020139640686741237 | CASR;CPA1;ANKRD55;PTPN22;MIF;TYK2;TNFRSF1A;IL6;NLRP12;IL2RA;IL2RB;STAT4;HLA-DPB1;NLRP3;PRTN3;CFTR;HLA-DRB1 |
| Elevated C-reactive protein level | 0.020139640686741237 | CASR;CPA1;ANKRD55;PTPN22;MIF;TYK2;TNFRSF1A;IL6;NLRP12;IL2RA;IL2RB;STAT4;HLA-DPB1;NLRP3;PRTN3;CFTR;HLA-DRB1 |
| Myasthenia Gravis | 0.02025306092168879 | CD86;CNTFR;ACHE;ROBO3;CD40;CD83;CSF1;TNFAIP3;AQP4;PTPN22;KRT20;TNF;IGF1R;LGALS1;GRAP2;RNF19A;DNMT3B;ITGAX;HLA-DOA;LGALS8;TNFRSF4;MYOG;CHRNB1;HLA-B;TAP2;HLA-C;HLA-A;KRT5;PAX5;HLA-G;CLEC4C;IFNG;IL1B;DOK7;LTA;IRF5;TLR4;CFB;HLA-DQB1;CHRNA1;CALCA;ABCB1;AIRE;KCNA3;LRP4;ADRB2;NTN1;C2;INS;TTN;CHRNE;CCL5;MBP;NCAM1;HLA-DQA1;NTRK1;TSLP;B3GAT1;LIF;OSM;ESR1;ISG20;POMC;IL6;MIR145;TEC;EIF3K;TNFSF4;IL2RA;IL2RB;HLA-DPB1;PDCD1;FGFR3;PAM;HLA-DRB1 |
| Recurrent tumor | 0.02047437134611975 | CERK;MIR429;PRDM2;CLU;FGF2;TNF;ETS2;FGF3;IGF1R;RIMS2;EDNRA;RASSF1;EDNRB;DMBT1;CYP2D6;MYC;PEG10;SALL4;STMN1;RASSF6;TNFSF10;AKT1;CYP1B1;LZTS1;TGM2;KCNH2;PDGFRA;S100A2;IGFBP3;F2R;CIRBP;HLA-A;KRT5;MIF;FSCN1;ERG;RIN1;MAD1L1;CSF1R;CALCA;MIR196B;HPN;SHOX2;EIF5A2;GATA3;HOXB13;FUT4;KLK6;SOCS2;TERT;MIR296;PROM1;TAC1;S100A10;TNFRSF10C;ESR1;IL6;TG;BCL6;CDK2;BCL2;CPE;MIR183;NANOG;XAF1;TDGF1;FGFR3;FGFR2;HOTAIR;COL18A1;HSPB6;MIR130B;LPCAT1;PIK3CD;BRCA1;KRT20;FOXM1;PIK3CG;CXCL5;HK1;SLC22A18;PSD4;TIMP3;CCR6;GPX1;MMP7;DCC;MYOF;MIR21;PROX1;POU5F1;SSTR4;NCOR2;FGR;MIR126;COL2A1;ZEB1;IL1B;B3GNT3;CDH13;TFF1;LCP1;GUCY2C;HDAC3;SATB2;THBS1;BARX2;EGFR;PTHLH;RASSF10;HAS1;HSF1;ERBB2;MXI1;MIR372;SPP1;GPC5;URGCP;MGMT;GDF15;STAT3;KLK13;CCK;MLH1;MTOR;VEGFA;KISS1;SPSB1;KLF6;PER3;MIR145;APC;LEP |

| | | |
|---|---|---|
| Precursor Cell Lymphoblastic Leukemia Lymphon | 0.02060049720371731 | RB1;CYFIP1;ZNF296;SPARC;SCT;GFI1;IKZF1;LGALS3;HOXA9;VPREB1;TCL1A;RASSF1;CYP2D6;ZMIZ1;MYC;ANPEP;STMN1;TNFSF10;AKT1;CYP1B1;PIP4K2A;TP63;ANKS1B;SLC12A9;MEF2C;EBF1;HLA-C;KRT7;RUNX1;UHMK1;DDIT3;CYP2E1;EPHA1;CES1;RTEL1;ABCB1;LXN;GYPC;PCDH10;MIR196B;PDE1A;TWIST2;PRDM14;GATA3;CD1D;EVPL;CRHR1;PRRC1;TERT;ADAMTS18;CD19;NKX2-5;HLA-DQA1;STAT5A;CREBBP;GSTM2;GSTM1;BCL11B;IGF2;NFATC2;MRPL28;NR0B2;PBX1;BHMT;TG;GIPC2;CYP1A1;TNFSF8;HOXD4;HLA-DRB1;CD22;TP73;CDKN1C;COL18A1;CDKN1A;CD40;MIR17;KRT20;GLI3;MECOM;CCR9;CD38;CCR7;JAK3;CD34;MME;SPHK2;TUBB;IL16;PAX5;TYK2;CBFA2T3;VAV1;SLC7A5;KAT2A;ZEB2;TAL1;IL1B;RARA;CD47;RAPGEF3;MIR128-2;HLA-DQB1;IDUA;COL11A1;GSTP1;RASSF10;SPP1;PRAME;MEF2D;RUNX1T1;CCL25;NQO1;PKLR;CDX2;STAT3;VEGFA;WT1;HLA-DPB1;BLID;CD86;CSF3;CSF2;PRDM9;CSF1;EPO;MPG;F13A1;MSI2;TNF;MPO;ACTG1;HOXA11;IGF1R;HPGDS;CCND1;SALL4;GRB10;TNFRSF8;BAALC;KCNH2;CPA2;PDGFRA;CASR;LIG4;ATP11A;MPEG1;CD8A;S100A6;ERG;STEAP1;ABCG2;NOTCH3;CALCA;ISYNA1;TCF7;GSTT1;ADRB2;FHIT;ZFP36L2;FUT4;ZFP36L1;DPP4;SOCS3;NLRP2;TAC1;TMEM132D;AUTS2;ZBTB16;EPHX1;ARHGAP24;SMAD7;CDK6;BCL6;TLX1;CDK2;BCL2;NFE2L3;CSPG4;FGFR1;UCN;CNTNAP1;PPP1R13B;FLT4;PIK3CD;AFF3;PIK3CG;LYL1;SIX6;CASP8;BLNK;HSP90AA1;ITGA4;TNFRSF1A;ZAP70;IFNG;LCK;ASPG;CDH11;BIRC7;DDC;HLF;UHRF1;SPG7;CXXC5;CRYZ;HLX;NCAM1;TTC12;TCF7L1;B3GAT1;MAP1LC3B2;OPRK1;MEIS1;CD6;MIR144;MIR125B1;IL2RA;LEP;MIR125B2;CD7;PTPN6;SCLY |
| Intestinal Cancer | 0.02066425142502819 | CDKN1A;NRP2;MSI1;MSI2;FOXM1;RASSF1;MUC2;MYC;ERBB2;HLTF;PART1;CCR7;SLC12A9;CSGALNACT1;TCF7L2;GPX1;CDX2;GDF15;WNT5A;HMGA1;PRKCA;CCK;MLH1;DCLK1;GREM1;NR5A2;CCKBR;APC;KCNQ1;BCL2 |
| Mammary Neoplasms | 0.02070640248723215 | MYLK;CYP2D6;AKT1;STARD3;PDZK1;SIGIRR;CLDN4;SLC5A8;SPINT1;DDIT3;RBP1;JTB;PADI2;CYP2E1;ABCB1;SHMT1;MIR196B;CRABP1;PDGFB;PDGFA;GATA4;GATA3;FPR2;LYPD5;SLC5A5;SULT1A1;HSD11B2;CYP2B6;SFN;JAG2;LYN;NGFR;CREBBP;GSTM1;CADM1;ABCA4;NR2F1;NR2F2;AZU1;PBX1;NR2F6;COL1A1;FERMT1;FXYD3;FAT4;HRG;HAMP;HLA-DRB1;ACHE;HOXC10;SMPD3;MECOM;HEY1;IKBKE;APC2;DUSP1;MYOF;F3;DUSP6;POU5F1;SLC7A5;F7;MMP14;CRYAB;GSTP1;COL11A2;KCNA3;HTT;EGFR;FSTL3;BGLAP;PLAGL1;SPP1;PRAME;IL12RB2;SPDEF;STAT3;ADAM33;DAB2IP;HEYL;MIR24-2;FABP3;SPSB1;FABP5;WT1;FABP7;ZNF217;LGR6;SIGLEC7;FRK;MSI1;APEH;ADORA3;TNFRSF8;PITX2;KCNH1;S100A2;S100A1;UGT2B15;SRCIN1;TUBA4A;FOXP1;S100A6;FSCN1;IGFALS;SLC27A2;SQSTM1;ZNF432;DSC3;DLX4;SHC1;MAZ;GSTT1;CYP2C18;FHIT;SLC9A3R2;SOCS2;SOCS3;KL;TFPI2;TNFRSF10C;TNFRSF10A;FBXO31;SCGB3A1;SHBG;ST6GALNAC1;NUMA1;KCNK9;ITGB4;FHL2;WISP3;PIK3CD;MSLN;FOXM1;PIK3CG;WISP1;SPN;HHEX;MB;ITGAX;ACP5;PMEPA1;MRPL9;TRIM62;HSP90AA1;SH2D3A;SH2D3C;NRG1;NRG2;TNFRSF1B;TNFRSF1A;BIRC5;ITGA6;BIRC7;LRP1;SRC;ITIH1;CXXC5;NTN1;CCL5;IGF2BP1;FYN;EIF4E;NTRK1;TBX1;CIZ1;MARCKSL1;TIE1;STYK1;RPL23A;TBX3;TBX2;GZMM;SST;IL2RA;IL2RB;ST7;PDCD4;PDCD1;P4HB;BCAR1;ERRFI1;CERK;ICAM5;ICAM1;LGALS4;LGALS3;RASSF1;RASSF2;TRIM29;TNFSF10;TNFSF11;LGALS9;SLC12A6;SLC12A7;TP63;SLC12A9;EPHA5;CSNK1E;MIF;ANK1;SFRP4;SFRP1;SFRP2;SLPI;IFI27;SFRP5;MT1G;MT1H;RIN1;RIN2;GRB7;EPAS1;CD1D;MST1R;PIK3R1;CD1A;PCBP1;NEU1;ST8SIA4;B4GALNT1;APOE;APOB;STAT5A;EGR1;WWTR1;FOXF1;IQSEC1;GREB1;XRCC3;IGF2;OSGIN1;PTK6;BST2;NET1;ARTN;PPP1R1B;NDUFAF3;PEMT;GRB2;CD200;TP73;APOBEC3B;DDR1;DIRAS3;CD40;BRCA1;PCSK6;AKAP12;AKAP13;HYAL1;HNF4A;HYAL2;CCR9;CCR7;CCR6;CTSD;LYNX1;MME;MST1;TOX3;ANO1;NR5A2;CTTN;IL1B;ARHGEF7;TMPRSS6;NEDD9;CAPG;HSD17B8;CYP17A1;GNAI2;RXRB;EXOSC6;PAK1;HSD17B1;GPC1;HSD17B2;HSF1;MGAT3;XDH;CD55;RIBC2;CD74;OSM;MIB2;AKR1C2;KLK15;CSHL1;KLK10;KLK11;VEGFA;PPP1CA;PER1;PER3;TACC2;CD86;MPL;PLAT;PRDM1;ETS1;ETS2;LOXL1;IGF1R;LOXL2;CCND2;DMBT1;CCND1;CCRL2;CAPN2;TGM3;TGM2;PDGFRA;CASR;MSX2;MAT1A;BRMS1;ADAM12;CARD16;DIABLO;HTRA1;HOXB13;CACNA1H;ACACA;NME1-NME2;CALD1;TAC1;WNT2;WNT3;TFAP2A;TFAP2B;TFAP2C;ZFHX3;UBE2I;HMGA1;INHBB;ISG15;BMP7;IGF2R;BMP4;ISG20;BMP2;IL6;NAT1;GNAS;GNB3;FGF10;PRR5;ADAR;OXT;LAMC2;AFF3;GRM1;GHR;MMP7;OLIG2;SSTR1;ACTA2;GREM1;CCNA1;ELF3;CHDH;CAT;CHI3L1;IRF5;PAOX;LAMA3;AURKC;PTHLH;AURKB;INS;CNR2;SLIT1;PTK2B;SLIT3;RBM38;EYA1;BNIP3;ATP2B2;MUC5B;MAPK15;CHFR;CD4;COL7A1;CD9;TEK;VIM;CPVL;RB1;SERPINE2;CLEC10A;MIR429;HOXA9;GJA3;SOX17;MYC;CREB3L1;STMN1;CYP1B1;VPS39;HOXA5;CHGB;PRIMA1;SERPINF1;AFAP1;PRKCA;DKK1;DKK3;HOXB9;BIN1;PRKCQ;HOXB1;HOXB7;ENPEP;CTBP2;CTBP1;PRKCZ;MIR206;LDHB;TERT;SLC19A3;PROM1;ST3GAL1;ELANE;S100A10;BARD1;ST14;ACE;STARD10;NFATC2;NFATC1;ST18;SQLE;H19;GJB2;GIPC1;CYP1A1;CDKN1C;CDKN1A;SETD7;ADM;SERPINA4;GLI2;LFNG;TIMP2;RSPO2;TIMP3;RSPO3;TIMP4;ANXA2;TET1;PAX2;SERPINB5;NCOR2;ZEB2;TUBB2B;ZEB1;RARA;RARB;HCLS1;ANGPTL4;PLCB2;HLA-DQB1;GRP;NR1I2;HOXD11;HOXD10;PRSS50;EPCAM;ALOX5;CHN2;GCNT2;BRD4;CCL25;WWOX;F10;MGMT;WWP1;SULF1;HIPK2;NFIC;ABI1;FOSB;CCL28;HSPA1B;CSF3;CSF2;CSF1;EPO;RAB3D;MIR146B;IRS2;FGF1;CLU;FGF2;TNF;FGF3;RIMS2;PYCARD;CDH5;FGF7;FGF8;CDH3;CDH1;SALL4;GRAP2;RNF19A;SLC39A2;MAP2K3;IGFBP1;TLE3;LHCGR;FKBPL;RPS8;IGFBP3;USP2;F2R;DIO3;HIC1;CACNB1;MTNR1A;MTNR1B;PGR;ADAM8;CSF1R;NOTCH3;SLC26A1;NOTCH4;RASAL2;SAFB2;NDN;ESRRA;FN3K;SLC37A1;TNK2;SLC4A11;SYNJ2;DAB2;COPS5;CDK6;CPD;CDK2;BCL2;SLC26A8;PLXNB1;PSCA;LSP1;ZFP82;CYR61;SYNE1;PSD4;DNMT3B;EMILIN2;AMH;GPX1;VWF;DNMT3A;NGF;DUSP22;LCK;ESRP1;CRH;EVL;HDAC3;GLRX;NOD2;BPHL;DNAJB6;MEST;SPAG9;IRX2;ANGPT2;ANGPT1;ASCC1;TXNRD2;ANKRD30A;TXNRD1;MTOR;P2RX7;RAD52;P2RX5;KLF6;MARCKS;APC;LEP;ZACN;SPARC;VIPR2;AQP1;SOX2;EDNRA;EDNRB;RPS6KA2;SOX9;CPT1A;HLA-B;KRT8;DLEC1;KRT5;HLA-G;RUNX3;RUNX2;RUNX1;TIAM1;CD109;EZR;CFTR;TELO2;LLGL2;SLC22A3;CASZ1;TWIST1;KLK4;CRHR1;CRHR2;KLK6;TRPM2;STC2;FADD;NTSR1;TRAP1;HIST1H4K;HIST1H4L;PDE2A;LIF;MIR200B;NMBR;ALOX15B;OCLN;MT1IP;TRMT12;HIST1H4I;SIM1;SIM2;TDGF1;HIST1H4F;HOTAIR;COL18A1;MIR17;MIR18A;LIMD1;KRT20;CX3CL1;LIMD2;SLC22A16;SLC22A18;FFAR1;CLIC1;CBR1;RAB6C;PRMT7;TSC22D1;TUBB;NME2;MIR21;PROX1;CBFA2T3;VAV1;CDCP1;RAD51C;B3GNT5;TFF3;TFF2;TFF1;MAPT;SGK1;TLR4;C1QA;CSTA;CXCR5;GDPD5;AIF1;NT5E;CUX1;WIF1;GABRP;GINS2;NQO1;GDF15;LILRB1;MLH1;PDZK1IP1;KRT18;KRT17;DLC1;KRT14;TRIP10;F2RL1;GHRL;SIPA1;COMT;GHSR;POSTN;WNT5A;MATK;TSC2;COL4A2;HGS;COL4A1;HIST1H2BE;NUPR1;STEAP1;ABCG2;ALOX15;LTBP4;ALOX12;DTX3;PSORS1C2;DPP4;PODXL;NLRP2;SMAD2;CBX5;SMAD3;TGFB3;ESR1;ARHGAP24;SMAD7;SLC2A9;PYY;RALYL;GDNF;MIR184;MIR183;TGFBI;FGFR4;FGFR3;FGFR2;FGFR1;ZSCAN1;KDM5B;MIR10B;THRB;MIR10A;PCDHGB6;THRA;HSPB2;SLC2A2;SLC2A5;PKD1;CXCL5;UPK1B;CASP8;H2AFJ;TRPS1;BDKRB1;NOS1;POLE;EGFL7;WNT16;DCN;TGFBR2;KLF17;MIR126;ALDH1A3;AIM2;IFNG;CDH11;LTA;CDH13;MYH9;DOCK1;LTF;CAMK2B;RGS16;HIST1H2BK;PON1;THBS1;MUC1;WNT11;GNA12;ERBB2;SNCG;NCAM1;SASH1;TCF7L2;KDM4B;NOS2;NOS3;LIMK1;KISS1;MEIS1;MIR145 |
| precancerous lesions | 0.020899395679272837 | CSF3;CDKN1A;CSF1;PSCA;DEFB1;BRCA1;FOXM1;TNF;ICAM1;CCND1;CDH1;STMN1;PLCE1;MUC13;SLC12A9;KCNH2;DENND2D;MMP7;ITGA4;DAPK1;HIC1;SND1;ANO1;MMP14;MYCN;SFRP2;LMX1A;IL1B;TFF2;TLR4;PCDH10;GSTP1;HPN;KCNA3;GSTT1;FHIT;ABO;EGFR;TERT;EPCAM;ALX4;EGR1;GSTM1;CDX1;EPHX1;TFPI2;IGF2;TNFRSF10C;PDX1;MLH1;ESR1;NR0B2;VEGFA;IL6;COPS5;BCL2;CYP1A1;VIM |
| Congenital Heart Defects | 0.02108640908974276 | RB1;TNF;LOXL2;ZC3H12D;GRIP1;EDNRA;HPGDS;SLN;GJA5;SALL4;NPPA;HOXA3;TNFSF11;HOXA1;SOX9;PITX2;SOX7;CD96;MEF2C;DSCAM;FOXP1;KCNQ1;MAML3;CCDC151;FREM2;FBLN7;ABCB1;AIRE;GATA6;TWIST1;GATA4;CACNA1C;RAI1;ADAP2;DEAF1;FGF20;APOE;NKX2-5;STRA6;SMAD2;CREBBP;ACE;NFATC1;NR2F2;VANGL1;G6PC3;WDR60;TDGF1;NODAL;FGFR2;FGF10;FGFR1;RYR2;ELN;DYSF;PRKAG2;ECE1;PKD1;SYNE1;SH3PXD2B;DNMT3B;JARID2;DSP;MKKS;TUBB;DNMT3A;PROX1;F3;ARID1B;POU5F1;TGFBR2;ACTA2;ALDH1A3;ZEB2;RAD51C;HAND2;COL6A1;PEX5;HAND1;ITGA8;RARB;IRF8;MYH7;MGRN1;EGFR;INS;TTN;PUF60;CBS;GNA11;TBX20;MSX1;FSD1;TBX1;KCNJ6;EYA1;KCNJ11;NOS3;NTRK3;STAT3;PDX1;ATP2B2;TBX5;GCK;VEGFA;FABP3;MEIS1;ACTC1;TEK |
| Endometrioid carcinoma ovary | 0.02108640908974276 | MAST4;MSX2;IGFBP3;TNK2;PLK2;KRT8;PIK3CD;HNF1B;FRY;MLH1;EGFR;PIK3CG;HOXA10;TUBA1C;FER;CCND1;TERT;ERBB2;NEK10;MAP3K6;POLE;FGFR2;KCNH1 |
| Thyroid Neoplasm | 0.02108640908974276 | CD86;ERRFI1;NDUFA13;CSF2;EPO;AQP4;PRDM2;APEH;CLU;FGF2;TNF;ETS1;IGF1R;CDH6;RIMS2;LGALS3;HOXA9;EDNRA;RASSF1;LGALS1;RASSF2;CCND2;CCND1;CDH1;GRAP2;RASSF5;TNFSF10;AKT1;MIR150;TP63;SLC12A9;POSTN;S100A1;TPM3;WNT5A;DIO2;LMTK3;HLA-C;HNF1B;HNF1A;HLA-G;RUNX2;MINPP1;CLDN4;IFI27;S100A6;FSCN1;MT1G;NUPR1;ABCG2;NOTCH3;CALCA;FBLIM1;NOTCH4;CRABP1;PDGFB;TACSTD2;PDGFA;GSTT1;RASAL1;FHIT;RAP1GAP;SLC5A5;DPP4;TPO;CYP2B6;TERT;HLTF;PROM1;ESRRA;EGR1;AFAP1L2;GSTM1;TGFB3;FUCA1;XRCC3;PTCH2;IGF2;HMGA1;NR1D1;ESR1;FLI1;IGF2R;ARHGAP24;SMAD7;TF;IL6;TG;BCL2;GNAS;ID3;GNB3;SLC26A4;FGFR4;FGFR2;TP73;CDKN1C;PCSK2;COL18A1;CDKN1A;CD40;THRB;THRA;ITGB4;FLT4;PIK3CD;MIR18A;PIK3CG;LIMD2;HHEX;ZIC1;TIMP3;CCR7;CCR6;INSL3;SLC2A14;MST1;NCOA4;NRG1;GFRA1;SSTR1;NGF;MIR19A;F3;SERPINB5;DCN;SSTR4;TGFBR2;MMP14;PAX8;CCKBR;IL1B;TFF3;BIRC5;MCM5;GAS5;LRP1;SRC;GSTP1;TMPRSS4;ASCL1;THBS1;EGFR;DUOX1;PAK1;MUC1;RASSF10;CNR2;ERBB2;SPP1;EIF4E;BRD4;NTRK1;SPAG9;TCF7L1;ANGPT2;MGMT;TXNRD2;SORT1;GDF15;TIE1;KCNIP3;STAT3;MLH1;TSHR;VEGFA;P2RX7;RAD52;APC;WT1;ABI3;SST;CENPJ;PDCD4;PDCD1;FOXA2 |
| Mixed Salivary Gland Tumor | 0.02108640908974276 | TNXB;CRTC1;SRC;XYLT2;HSPB2;PDGFA;FPR2;EGFR;RASSF1;CCND1;TERT;BGLAP;WIF1;ANPEP;ERBB2;TIMP2;NLRP3;TRIM27;FSD1;EIF4E;PDGFRA;MME;DAPK1;S100A1;MGMT;NTRK3;STAT3;B3GAT1;IGF2;TYK2;TBX5;RUNX3;BMP7;ARHGAP24;GUCA2A;GFAP;VEGFA;BMP2;IL6;IFNG;DDIT3;LEP;HMSD;CDH11;FSCN1;BCL2;FGFR2;TP73 |
| Thymoma | 0.02108640908974276 | RYR1;ERRFI1;RYR2;ROBO3;CIITA;FLT4;LY75;PIK3CD;AQP4;IRS2;PTPN22;CLEC14A;ETS1;TNF;TREM1;CYR61;PIK3CG;ICAM1;DPYSL4;MECOM;CCND1;CDH1;MB;MYC;DNMT3B;AKT1;CCR7;TNS4;TNFRSF4;TP63;MYOG;HLA-C;MIR27A;HLA-A;RUNX3;RUNX1;IFI27;IFNG;LCK;IL1B;LTA;ERG;S100A9;CHRNA1;CSF1R;CALCA;AIRE;THY1;FHIT;NTN1;CD1A;EGFR;AURKB;TTN;ATXN1;CHRNE;ERBB2;FSD1;HLA-DQA1;MICB;STAT5A;TMEM132D;ST14;TSLP;MGMT;ESR1;MTOR;VEGFA;ISG20;BMP2;IL6;CD4;TEC;STK24;EIF3K;SST;TNFSF4;IL2RA;CDK2;HLA-DPB1;BCL2;CNTN2;ULBP1;HLA-DRB1 |
| Uterine Cervical Neoplasm | 0.02123997804163478 | CDKN1A;CSF1;ETS1;TNF;GHR;SOX1;CASP8;CCND1;CDH1;MYC;SOX9;SLC12A9;MMP7;WNT5A;MST1;HLA-A;PAX1;IL1B;BIRC5;RARB;S100A9;EPAS1;PCDH10;GSTT1;SLC5A1;FHIT;SLC7A1;EGFR;WNT11;TERT;WIF1;WNT2;TSPAN1;SMAD2;SPAG9;CMTM7;ZNF582;GSTM1;MGMT;STAT3;PDZK1IP1;BMP7;TMPRSS13;POU4F1;MTOR;VEGFA;IL6;BCL2;CYP1A1;HLA-DPB1;PLXNB1;CLPTM1L;FGFR3;APOBEC3B;HLA-DPB2 |
| Acute Myeloid Leukemia, M1 | 0.02198422533717048 | CSF3;CSF2;SPARC;SPI1;TRIO;GFI1;EHMT2;CTSZ;FHL2;TRH;IFI30;MPO;BACH2;LYL1;HOXA9;CCND2;MYC;CAPN2;TNFSF10;PDE4B;BAALC;DLEU2;ANXA2;DAPK1;DNMT3A;TSC2;RUNX3;RUNX1;PXDN;ERG;SGK1;CSF1R;CAPG;GATA2;ADCY7;SYNGR1;CNR2;CTNNA1;S100A10;RUNX1T1;SVIL;STAT3;POU4F1;CDK6;WT1;ID2;ASXL2;BCL2;TCEA2;CD9;FXYD6;AGRN |
| Ileocolitis | 0.02209264302358777 | CRP;DENND1B;NCF4;TCF7;DEFB1;FPR2;NOD2;TNF;BACH2;TAGAP;NLRP3;NKX2-3;SMAD3;SLC11A1;DNMT3A;MST1;TYK2;IRGM;PHOX2B;IL6;IFNG;IL2RA;LEP;TLR4;HLA-DRB1 |
| Infiltrating Cervical Carcinoma | 0.02209264302358777 | PTPRR;CD83;PCDH10;CLU;MPO;THBS1;TNF;EGFR;PYCARD;ERBB2;DNMT3B;PEG3;CADM1;HLA-B;MIR21;HLA-G;POU5F1;VEGFA;CCNA1;BCL2;CYP1A1;F2RL1;FGFR2;HLA-DQB2;HLA-DQB1 |
| Regional enteritis | 0.02209264302358777 | CRP;DENND1B;NCF4;TCF7;FPR2;NOD2;TNF;BACH2;TAGAP;NLRP3;NKX2-3;SMAD3;HSPA1L;SLC11A1;DNMT3A;MST1;TYK2;IRGM;PHOX2B;IL6;IFNG;IL2RA;LEP;TLR4;HLA-DRB1 |
| Pain, Crushing | 0.022397671017361922 | CSF3;CSF2;ABCB1;ADM;OXT;TRH;COMT;TNF;AQP1;HRH1;NPPA;PENK;BDKRB2;AKT1;BDKRB1;HCRT;TAC1;DISC1;NTSR1;HTR1A;OPRK1;NRG1;HTR3B;PDYN;POMC;PNOC;CRH |
| Pain, Migratory | 0.022397671017361922 | CSF3;CSF2;ABCB1;ADM;OXT;TRH;COMT;TNF;AQP1;HRH1;NPPA;PENK;BDKRB2;AKT1;BDKRB1;HCRT;TAC1;DISC1;NTSR1;HTR1A;OPRK1;NRG1;HTR3B;PDYN;POMC;PNOC;CRH |
| Pain, Splitting | 0.022397671017361922 | CSF3;CSF2;ABCB1;ADM;OXT;TRH;COMT;TNF;AQP1;HRH1;NPPA;PENK;BDKRB2;AKT1;BDKRB1;HCRT;TAC1;DISC1;NTSR1;HTR1A;OPRK1;NRG1;HTR3B;PDYN;POMC;PNOC;CRH |
| Suffering, Physical | 0.022397671017361922 | CSF3;CSF2;ABCB1;ADM;OXT;TRH;COMT;TNF;AQP1;HRH1;NPPA;PENK;BDKRB2;AKT1;BDKRB1;HCRT;TAC1;DISC1;NTSR1;HTR1A;OPRK1;NRG1;HTR3B;PDYN;POMC;PNOC;CRH |
| Ache | 0.022397671017361922 | CSF3;CSF2;ABCB1;ADM;OXT;TRH;COMT;TNF;AQP1;HRH1;NPPA;PENK;BDKRB2;AKT1;BDKRB1;HCRT;TAC1;DISC1;NTSR1;HTR1A;OPRK1;NRG1;HTR3B;PDYN;POMC;PNOC;CRH |
| Radiating pain | 0.022397671017361922 | CSF3;CSF2;ABCB1;ADM;OXT;TRH;COMT;TNF;AQP1;HRH1;NPPA;PENK;BDKRB2;AKT1;BDKRB1;HCRT;TAC1;DISC1;NTSR1;HTR1A;OPRK1;NRG1;HTR3B;PDYN;POMC;PNOC;CRH |
| Acute lymphocytic leukemia | 0.022518773278946976 | RB1;CYFIP1;ZNF296;SPI1;TES;SPARC;HPSE2;SCT;GFI1;IKZF1;MIR425;ICAM1;LGALS3;VPREB1;RASSF1;HOXA9;CYP2D6;ANPEP;MYC;STMN1;TNFSF10;AKT1;CYP1B1;PIP4K2A;DLEU7;SLC12A9;MEF2C;PRKCB;RPL22;EBF1;HLA-C;KRT7;HLA-A;RUNX3;DKK3;RUNX1;UHMK1;TFR2;DDIT3;TYRO3;CYP2E1;EPHA1;RIN2;CFB;CES1;RTEL1;ABCB1;LXN;PCDH10;SHMT1;MIR196B;PDE1A;TWIST2;GATA3;THY1;CD1D;EVPL;CD1A;CRHR1;C3;CD79B;CYP2B6;PRRC1;TERT;CD19;INPP5D;PROM1;HLA-DQA1;S100A10;LYN;STAT5A;NGFR;CREBBP;GSTM2;GSTM1;TSLP;ZNF382;BCL11B;XRCC3;RANBP17;IGF2;MRPL28;NFATC2;NR0B2;FLI1;PBX1;BHMT;HAL;TG;GIPC2;FAT1;CYP1A1;TNFSF8;HOXD4;HLA-DRB1;CD22;TP73;DDR1;CDKN1C;COL18A1;CD40;CDKN1A;MIR17;ELN;GRIK2;KRT20;GLI3;ATIC;MECOM;CCR9;TIMP2;CD38;CD37;CCR7;JAK3;CD34;CD52;MME;ANXA2;SPHK2;TUBB;IL16;PAX5;TYK2;CBFA2T3;KAT2A;ZEB2;TAL1;IL1B;RARA;PIGL;CD47;MAPT;SLC29A1;TLR4;MIR128-2;HLA-DQB1;GSTP1;ASIP;EXOSC6;RASSF10;SPP1;PRAME;MEF2D;CD55;RUNX1T1;CCL25;NQO1;PKLR;CDX2;STAT3;KLHL1;MLH1;VEGFA;WT1;HLA-DPB1;FOSB;BLID;CD86;CSF3;CD83;PRDM9;CSF2;CSF3R;CSF1;MPG;EPO;PLAT;PRDM2;MSI2;MPO;FGF2;TNF;ACTG1;IGF1R;RIMS2;HOXA10;HPGDS;CCND2;CCND1;FCGRT;SALL4;TNFRSF8;BAALC;GGT1;CD177;KCNH2;PDGFRB;IGFBP1;PDGFRA;TLE3;CASR;TLE2;S100A1;WNT5A;ATP11A;HIC1;FOXP1;EMCN;S100A6;ERG;STEAP1;S100A9;ABCG2;CSF1R;NOTCH3;CALCA;ISYNA1;GSTT1;ADRB2;FHIT;FUT4;ZFP36L2;ZFP36L1;DPP4;SOCS3;SMYD2;NLRP2;TAC1;MSC;TMEM132D;AUTS2;CD160;WNT3A;ZBTB16;EPHX1;ESR1;ARHGAP24;SMAD7;BMP4;IL6;FER;CDK6;NAT1;BCL6;TLX1;CDK2;BCL2;ALPI;FGFR4;MXD3;FGFR1;UCN;CNTNAP1;PPP1R13B;ITGB2;PIK3CD;FOXM1;ITGAL;MSLN;AFF3;PIK3CG;SGCE;LYL1;SIX6;CASP8;BLNK;NOS1;HSP90AA1;CR1;ITGA4;DCC;DNMT3A;MIR126;CCNA1;ZAP70;IFNG;LCK;CAT;ASPG;CDH11;BIRC5;BLVRB;LTB;HDAC4;TRABD;HDAC2;DDC;HDAC3;HLF;LRP1;UHRF1;SPG7;CXXC5;AURKB;CRYZ;CBS;HLX;ERBB2;IGF2BP1;NCAM1;SPAG9;TCF7L2;MARCKSL1;TCF7L1;NOS2;NOS3;BNIP3;B3GAT1;OPRK1;SORL1;MTOR;TSPAN16;NELL1;MEIS1;CD6;MIR144;SST;CD5;MIR125B1;MIR125B2;LEP;CD7;PTPN6;SPRY1;SCLY |
| Camurati-Engelmann Syndrome | 0.02280987996770805 | RYR1;CSF2;LAMC2;MITF;IRAK4;TGFBR2;VEGFA;BGLAP;CA2;DPEP1;TNFSF11;APOE;MAPT;SOST;TLR4 |
| Complete hydatidiform mole | 0.023337323856681736 | CDKN1C;ENPEP;DOC2A;PEG3;IGF2;ZFP36L2;EGFR;PSMB9;TNFRSF1A;H19;DAB2;NLRP7;ERBB2;SPP1;BCL2;TNFRSF25;APOB;HLA-DOA;PGM1 |
| Agenesis | 0.023337323856681736 | CDKN1C;DIRAS3;CSF3;CSF2;GATA6;GATA4;ZNF22;AFF3;TNF;FGF3;GLI3;IGF1R;HNF4A;CD19;TMEM67;FGF20;PROK2;KCNN3;MSX1;TP63;TGIF1;EDARADD;PEG3;EYA1;WNT5A;PDX1;HNF1B;ISL1;MTOR;GCK;PAX1;EDAR;BMP4;GREM2;IFNG;PAX8;ALDH1A2;PAX9;IRF6;CFTR;CCNO;FGFR2;LHX9;FGF10 |
| Adenocarcinoma, Oxyphilic | 0.023337323856681736 | DDR1;CDKN1C;SYCP1;MSLN;KRT20;TNF;HOXA10;PYCARD;RASSF1;CASP8;CCND1;CA2;MYC;NPPA;PLCE1;TP63;DAPK1;TPM3;KRT8;UGT2B17;APOA1;CYP7B1;RUNX3;MMP14;KCNQ1;IL1B;BIRC5;RARB;PGR;CYP2E1;GRB7;NR1I2;EGFR;SELENBP1;MUC2;ERBB2;SPP1;APOE;STAT5A;TCF7L2;ACE;NOS3;STAT3;TNFRSF10A;ESR1;MTOR;TSHR;VEGFA;CYP26A1;APC;BCL2;ID3;GHRL;FAT4 |
| Gastrointestinal Neoplasms | 0.023337323856681736 | CIITA;FHL2;ADM;BRCA1;KRT20;SLC6A3;CDH4;LGALS1;RASSF2;SOX17;CDH1;AKT1;HOXA1;MIR192;POLE;APOBEC1;PDGFRA;MIR21;RUNX3;ISL1;SDHB;HIC1;TLE6;GREM1;CLDN4;SFRP2;SPINT1;CCKBR;IL1B;MIR17HG;PRR15;PKP3;SRC;FHIT;EGFR;MUC2;EPCAM;ERBB2;FADD;WNT2;CHGA;ST14;GDF15;BNIP3;SMAD9;LILRB1;MLH1;ARHGAP24;VEGFA;EFNA2;MIR145;APC;FGFR4;TP73 |
| Hematopoietic Neoplasms | 0.024139540086144957 | CSF2;PIK3CD;IKZF1;ETS1;FGF2;TNF;PIK3CG;IGF1R;BBC3;SPN;VPREB1;CCND2;CCND1;MYC;HNF4A;AKT1;MIR150;DLEU2;KCNH1;VENTX;PDGFRA;KDM2B;WNT5A;TET1;PAX5;DKK1;HIC1;RUNX1;FOXP1;MIR17HG;HOXB4;IRF8;IRF5;ABCG2;CSF1R;ABCB1;LXN;PON1;HTT;GSTT1;MRO;FHIT;CD79B;DPP4;TERT;PCBP4;CTNNA1;NCAM1;PRAME;EIF4E;ZNF300;STAT5A;WWOX;CREBBP;MARCKSL1;EGLN2;BCL11B;STAT3;HMGA1;KLHL1;ESR1;PBX1;ARHGAP24;P2RX7;IL6;FMNL1;CDK6;BCL6;CD6;FES;WT1;LEP;MIR142;BCL2;NFE2L3;HLA-DPB1;SPRY2;PTPN7;TP73 |
| Simple renal cyst | 0.024486670070732672 | PIGR;CASR;SPARC;SCT;DIABLO;FGF2;PKD1;TNF;EGFR;PAX2;DKK3;ANO1;TIMP2;FGFR3 |
| Memory Disorders | 0.024486670070732672 | ACHE;APP;KL;CHRM1;GABRA5;SPHK2;STAT3;MAGI2;HTR1A;IGF2;MIF;COMT;HTR4;SLC6A3;INS;VEGFA;NPY;IL1B;PNOC;BCL2;HLA-DPB1;SLC17A7;APOE;MAPT |
| Neuromyelitis Optica | 0.024486670070732672 | RASL12;CD40;CSF2;MOG;FCRL3;SLC1A2;AQP4;LAMC2;PTPN22;IFI30;FUT4;AQP1;GFAP;TNFRSF1A;IL6;CD19;SPP1;HLA-DPB1;IRF8;MBP;HLA-DQA1;HLA-DRB1;HLA-DQB1;ABCG2 |
| Dyschezia | 0.024486670070732672 | THRB;HPSE2;THRA;ELN;ECE1;BRCA1;TRH;AQP2;COMT;SLC6A3;UNC80;EDNRB;SALL1;IYD;SIX3;NRTN;PDGFRA;CASR;PROP1;EDN3;OPRM1;KCNAB2;SDHB;DDOST;TRHR;FOXP1;TGFBR2;TNFRSF1A;SCNN1G;CLIP2;PCCA;PAX8;SCNN1B;PCCB;MAGEL2;DDC;TNXB;FOXG1;NRXN1;PON1;ASCL1;NALCN;SLC5A5;RAI1;TPO;GJC2;DEAF1;EPCAM;PRDM16;CAMTA1;NKX2-5;RREB1;GABRD;KCNJ1;TBX1;CREBBP;LIMK1;TBCD;MLH1;FLI1;PHOX2B;TSHR;NR4A2;VANGL1;POMC;SKI;TG;GDNF;TH;LHX3;COL7A1;ALPL;GTF2IRD1;SLC26A4 |
| High-Grade Squamous Intraepithelial Lesions | 0.02504522154352109 | MPG;TERC;UHRF1;PCDH10;GSTT1;FHIT;PYCARD;RASSF1;TERT;ERBB2;AKT1;NOS1;BRD4;GSTM1;NOS2;DAPK1;RIPK4;SORBS1;NR0B2;HIC1;ARHGAP24;PAX1;SERPINB6;VEGFA;IFNG;SCGB3A1;PTPN6;ZMYND10;HLA-DQB2;HLA-DRB1;HLA-DQB1;TP73 |

| | | |
|---|---|---|
| Malignant neoplasm of stomach | 0.026517853973719894 | MDC1;PREX2;DPYSL3;AKT1;STARD3;BHLHA15;PDYN;ISL1;RNASE1;SND1;CLDN4;SLC5A8;SPINT1;MIR196A2;DDIT3;RUFY3;RBP1;CYP2E1;ABCB1;LXN;DHH;MIR196B;GATA6;GATA5;GATA4;GATA3;THY1;LYPD5;SLC5A5;SULT1A1;CYP2B6;KCNQ1OT1;SFN;LYN;NGFR;GSTM2;GSTM1;MRPL28;MIR339;MIR338;GRIN2B;NR0B2;FLI1;COL1A1;FAT4;HLA-DRB1;LGMN;HRK;ACHE;DPYS;MIR130B;MED15;BBC3;TUBA1C;DUSP5;GPR39;SPHK2;F3;POU5F1;DDB1;MMP14;GPRC5A;MMP16;KCNMA1;LRRC3B;RAPGEF3;ERGIC1;TMEM106A;KANK2;IDUA;GSTP1;COL11A2;EGFR;PLAGL1;SPP1;STAT4;FLNC;MEF2D;WNT7B;STAT3;ADAM33;BMP8B;MIR24-2;WT1;ZNF217;SIGLEC7;LY6K;ZBTB20;MSI1;DEFB1;RORA;HPGDS;MAP3K6;PITX1;MAP3K5;KCNH2;S100A2;S100A1;SRCIN1;S100A6;FSCN1;PTPRCAP;S100A9;LY6E;TACSTD2;GSTT1;FHIT;SLC9A3R2;SOCS2;SOCS3;BCL2L10;SMYD2;MBP;TFPI2;HTR1A;TNFRSF10A;MCC;TMPRSS13;FBXO31;IGSF11;SCGB3A1;SHBG;PIWIL1;ST6GALNAC1;ITGB4;FLT4;PIK3CD;ECE1;MSLN;FOXM1;PIK3CG;WISP1;ZNRD1;ITGAX;EMX2;HSP90AA1;ITGA3;NRG1;TNFRSF1B;TNFRSF1A;HAND1;ITGA8;BIRC5;TRIM59;SRC;UHRF1;FOXK1;CCL5;TP53INP1;FYN;FSD1;NOP56;MARCKSL1;TIE1;PCDH8;TBX3;NELL1;SST;IL2RB;PDCD4;CD248;PTPN6;P4HB;TRIM31;TNFRSF6B;SLC23A1;ICAM2;FOXI1;ICAM1;LGALS3;RASSF1;LGALS1;RASSF2;RASSF6;TNFSF10;LGALS9;SLC12A9;EPHA4;SLC34A2;MIF;HNF1A;SFRP1;SFRP2;SLPI;IFI27;SFRP5;RIN1;EPHA1;GRB7;EPAS1;PCDH10;MST1R;TSPAN8;PCBP1;NEU1;HLTF;PENK;SYMPK;APOE;B4GALNT2;TSPAN1;STAT5A;EGR1;WWTR1;TNFSF14;FOXF1;XRCC3;IGF2;PTK6;NET1;ATP4A;TH;PPP1R1B;GRB2;TP73;APOBEC3B;DIRAS3;PTPRT;CD40;CTSZ;GRIK2;BRCA1;PTPRF;AKAP12;HNF4A;MUC13;CCR7;CCR6;CD34;CTSD;ACAD8;ZHX2;MME;MST1;IL16;CACYBP;ZHX1;TOX3;ANO1;NR5A2;CTTN;IL1B;CD47;ARHGEF7;MIR128-2;TAGLN;SATB2;TMPRSS4;NEDD9;CAPG;CYP17A1;CAMKK2;RXRB;REM1;PAK1;HSD17B1;HSF1;MGAT3;GPC5;MSX1;XDH;CD55;IL32;CD74;KLK13;KLK10;KLK11;VEGFA;PPP1CA;KLK12;NEDD1;PDCL2;KIF26B;TACC1;IL17D;IL17C;FOXA2;DCLRE1C;CD81;IHH;PLAT;PRDM2;ETS1;MPO;ETS2;IGF1R;LOXL2;CCND2;DMBT1;CCND1;TGM2;PDGFRA;MSX2;OSR1;LMTK3;MIR27A;DHDDS;SPOCD1;DOK2;CD8A;SCNN1B;ADAM12;CARD16;DIABLO;TCF7;HTRA1;SLC1A2;FUT1;ACACA;FUT4;FUT3;FUT6;WNT6;PRR5-ARHGAP8;CALD1;MIR296;MSC;NDUFV1;WNT2;WNT3;TFAP2A;TFAP2C;ZFHX3;UBE2I;ISG15;INHBA;BMP7;IGF2R;BMP4;BMP2;IL6;NAT1;BHLHE40;CNTN1;GNAS;FGF10;LGALS3BP;TNFAIP2;ATP2A3;CHD5;LAMC2;GHR;CA2;ZIC1;MLST8;MIR193A;MMP7;MIR23A;SSTR1;TAPBP;GREM1;CCNA1;CAT;IRF7;BLVRB;IRF8;ZMYND10;FBP1;OTUB1;LAMA3;AURKB;INS;RAB40C;CBS;MIR495;MIR494;MXI1;PTK2B;SPOCK1;SLIT3;ATOH1;URGCP;LAMB3;BNIP3;CCK;MUC5B;MAPK15;CHFR;GALNS;PNKD;ACTC1;BCL6B;CD9;TEK;VIM;ZNF177;TES;SERPINE2;MYT1L;MIR429;BACH2;MIR425;MIR675;SOX17;SOX18;MYC;STMN1;CYP1B1;HOXA2;PLCE1;HOXA1;PRSS8;IER3;PRKCH;PRKCB;DAPK1;DAPK3;AFAP1;PRKCA;DKK3;SYTL2;DKK4;MIR202;HOXB8;CFB;HOXB5;TSPYL5;MEG3;CTBP2;CTBP1;MIR206;ABO;TERT;SLC19A3;ST3GAL6;PROM1;HLA-DQA2;HLA-DQA1;S100A10;GSDMD;AFAP1L2;STARD13;ST14;ACE;FZD5;FZD7;H19;CPEB1;GIPC2;CYP1A1;TNFSF9;PCDHB9;CDKN1A;SERPINA10;SETD7;TCF21;WDR46;GLI3;GLI2;TIMP2;TIMP3;DENND2D;ANXA2;IFNGR2;PAX3;TET1;PAX6;PAX5;SERPINB5;ZEB2;ZEB1;RARA;RARB;HCLS1;GAS5;HLA-DQB1;NFIX;GRP;NR1I2;MIR409;ADCY3;MIR647;ASCL2;HOXD10;MIR410;RASSF10;EPCAM;ALOX5;GSC;BRD4;WWOX;MGMT;NR1H3;WWP1;SULF1;NFIC;FOSB;FILIP1;INHA;HSPA1B;HSPA1A;SEMA5A;CSF3;EPO;MIR146B;IRS2;CLU;FGF2;TNF;FGF3;PYCARD;FGF7;CDH4;CDH1;SALL4;GRAP2;RNF19A;MACROD2;LZTS1;IGFBP1;SNRPN;TPM3;IGFBP3;F2R;HIC1;RAB32;PGR;ERG;MAD1L1;RASAL1;RASAL2;VTCN1;TNK2;HOPX;CDK6;BCL6;CDK2;BCL2;XAF1;PSCA;ZFP82;ADRA1B;CYR61;SGCE;DNMT3L;SCRN1;DNMT3B;INSL3;ADAMTS9;TRPC6;GPX1;VWF;DNMT3A;APOA1;NGF;COASY;ANG;MIR1266;HDAC4;TRABD;HDAC2;HDAC3;NOD1;NOD2;MIR148A;SELENBP1;MGAT5B;VPS52;UGT1A6;ANGPT2;ANGPT1;ASCC1;MTOR;KLF6;MARCKS;APC;LEP;IFITM1;SPARC;HPSE2;PKD2L1;TP53AIP1;SOX2;FRAT1;EDNRA;SOX9;TNS4;RPS12;SRRM2;SLC11A1;KRT8;MRPS18B;DLEC1;EBF3;HLA-A;HLA-G;RUNX3;RUNX2;HLA-E;RUNX1;TIAM1;THEG;IL23A;NEIL1;EZR;MIR1182;ALDH7A1;CFTR;LLGL2;SLC22A3;CASZ1;EIF5A2;TWIST1;KLK4;CRHR2;KLK6;NUAK1;STC2;SPRR2A;SPTBN1;NTSR1;TRAP1;SNHG1;GDF2;MIR200A;MIR200B;GADD45G;ANP32B;NODAL;MIR129-2;HOTAIR;ROBO3;COL18A1;MIR17;MIR18A;KRT20;ESM1;CCKAR;SOSTDC1;MIR192;TNFAIP8L2;CLIC1;CBR1;NME4;MIR21;GFRA3;MIR19A;RHOD;VAV1;VAV2;CLDN11;TLR1;RAD51C;CCKBR;MIR17HG;TFF3;CLDN18;TFF2;TFF1;MAPT;TLR4;VGLL4;CXCR5;VWCE;CST7;NKD2;NT5E;CUX1;WIF1;CTNNA1;CTNNA2;NQO1;CDX2;CDX1;GDF15;SORBS2;LILRB1;MLH1;CYP2W1;KRT18;DLC1;F2RL1;GHRL;APBA2;F2RL3;FCGBP;NDUFA13;COMT;HOXA13;CCDC28B;CMIP;HOXA11;CNDP2;MIR149;HOXA10;ARHGDIB;GHSR;POSTN;KDM2A;WNT5A;MPEG1;MYCN;LMX1A;ABCG2;ISYNA1;ALOX15;ALOX12;ADRB2;SEZ6L2;DPP4;SCIN;HRH2;PODXL;UBD;SMAD2;SMAD3;CPSF1;MLKL;WNT3A;EPHX1;EPHX3;HSPA2;DNAJC15;ESR1;LRP1B;ARHGAP24;SMAD7;NR4A2;EFNA2;MIR184;MIR183;TGFBI;FGFR4;FGFR2;FGFR1;UCN;MIR10B;THRB;THRA;HSPB2;MIR96;PKD1;CXCL5;HK1;ELAVL3;CASP8;UPK1A;CASP1;BDKRB2;POLE;EGFL7;DCC;USP44;AXIN1;TGFBR2;MIR126;ALDH1A3;MYH1;IFNG;LTA;CDH13;MYH9;DOCK1;RBMS3;DDC;THBS1;MIR136;MUC1;MUC2;MIR373;GNA12;ERBB2;MIR372;SNCG;NCAM1;MUC4;MUC6;OPCML;TCF7L2;KDM4B;TCF7L1;NOS2;CMTM3;NOS3;USP22;LIMK1;PDX1;SGPP2;KISS1;POMC;MIR145;MIR144;CAPN10;MIR142 |
| Chronic Kidney Diseases | 0.026601657433083657 | EPO;MRGPRF;IRS2;MPO;TNF;ICAM1;GRIP1;HPGDS;CYP2D6;SOX18;MIR150;CASR;LHCGR;POSTN;NKX6-2;MSX2;SLC34A1;SLC6A13;HNF1B;MITF;HLA-G;DKK3;DDIT3;DOK6;ELMO1;ALPK1;S100A9;ABCG2;ABCB1;SLC22A2;IDI2;KLK4;GSTT1;RASAL1;GATA3;SLC5A4;ACAT2;C3;DPP4;SOCS3;CYP2B6;GRK4;ST8SIA4;MYH14;NLRP3;APOE;APOB;EGR1;KL;ACE;GSTM1;SMAD3;FAM78B;RARRES2;SLC33A1;BMP7;ESR1;SMAD7;SLC2A9;OCLN;BMP2;DAB2;IL6;NAT8;CPS1;BCL2;FGFR4;HAMP;HLA-DRB1;FGFR1;CD40;MIR17;ELN;PRKAG2;AFF3;PKD1;GLI2;GHR;CASP1;NOS1;SLC13A3;MYOCD;WDR37;GPX1;TRPC6;TRPC3;VWF;PDE4D;SHROOM3;NGF;DDOST;C1QL1;F7;MYH2;IFNG;SLC7A9;DDAH2;IL1B;CHDH;CAT;TFF3;CRH;LTA;MYH9;SGK1;CRP;CX3CR1;SLC20A2;GSTP1;PON1;NTN1;EGFR;PTHLH;NT5E;CUX1;BGLAP;CCL5;APOL1;XDH;KCNJ1;AOC3;P2RY12;TCF7L2;ANGPT2;NOS3;TXNRD1;GUCA2B;MTOR;VEGFA;FABP1;P2RX7;POMC;MIR145;NPHS1;WT1;NPHS2;LEP;SLCO4C1;MGP;CD248;HSPA1B;HSPA1A;UMODL1 |
| Coughing | 0.02682904341133217 | RTEL1;CSF2;TERC;FAM13A;PTPN22;ADRB2;PIK3R1;ABO;EGFR;CPN1;CD79B;TERT;STMN1;BDKRB2;BLNK;TRPM8;TAC1;DSP;WWOX;PTGIR;SNRPN;ACE;MME;SLC11A1;STAT3;TACR2;TSC2;DLEC1;ATP11A;BTNL2;EIF2AK4;PARP9;MUC5B;TGFBR2;ATP4A;XPNPEP1;IL1B;SNURF;HLA-DPB1;PRTN3;ZMYND10;CCDC151;HLA-DRB1 |
| Leukemia, Myelocytic, Acute | 0.027372495407981288 | RB1;ZNF296;SPI1;TRIO;SCT;EHMT2;MRGPRF;BACH2;HOXA9;SOX17;CYP2D6;MYC;STMN1;AKT1;CYP1B1;PIP4K2A;PLCE1;DLEU2;HOXA7;IER5;MNX1;HOXA5;HOXA4;PRIMA1;DAXX;MEF2C;PRKCH;PRKCB;DAPK1;PRKCA;FNDC3A;PRAP1;DKK1;SLC5A8;HOXB9;TFR2;DDIT3;RBP1;HOXB4;TYRO3;HOXB3;HOXB2;CYP2E1;HOXB1;HOXB8;HOXB7;HOXB6;HOXB5;MEG3;ABCB1;DHH;TERC;SHMT1;MIR196B;CRABP1;PDGFB;GATA4;THY1;GATA2;ABO;SLC5A5;SULT1A1;CYP2B6;TERT;PRDM16;PROM1;ELANE;S100A10;BARD1;LYN;CREBBP;GSTM1;ACE;CADM1;PLK2;ABCA4;PBX2;MRPL28;NFATC1;NR0B2;PBX1;FLI1;NR2F6;TNFSF4;FAT1;CYP1A1;TNFSF9;FXYD6;NANOG;TNFSF8;HLA-DRB1;CDKN1C;ACHE;CDKN1A;RARG;ELN;PRSS27;ADM;BBC3;SMPD3;MECOM;CHL1;TIMP2;TIMP3;TK1;JARID2;JAK3;DFFB;RIPK3;ANXA2;DUSP1;TESC;TET1;PAX5;F3;POU5F1;NCOR2;ITPKB;MMP14;PLSCR1;KAT2A;ZEB2;GPRC5A;TAL1;PXDN;RARA;HCLS1;SLC29A1;GAS5;JDP2;HLA-DQB1;GAS7;GSTP1;SPESP1;COL11A2;KCNA3;HOXD13;HOXD11;ADCY7;EGFR;ALOX5;NSD1;NTF3;SPP1;PRAME;JAM3;RUNX1T1;BRD4;PKLR;MGMT;STAT3;GTF2H4;POU4F1;GNMT;HIPK2;SKI;WT1;ABI1;FOSB;CLC;AGRN;SIGLEC7;CCL28;FRK;SIGLEC5;PF4;CSF3;CSF2;CSF3R;CSF1;EPO;MSI2;FGF1;FGF2;TNF;FGF3;RIMS2;HPGDS;FCGRT;CDH1;SALL4;GRAP2;RNF19A;GRB10;HLA-DOA;CD177;KCNH2;F2R;HIC1;FOXP1;CLEC4C;S100A6;FSCN1;ERG;S100A9;SQSTM1;MAD1L1;CSF1R;DLX4;GSTT1;FHIT;SOCS2;SOCS3;BCL2L10;ATXN1;E4F1;TMEM132D;ZBTB16;TNFRSF10C;TNFRSF10A;G6PC3;CDK6;MAFB;CDK2;BCL2;SLC26A4;CIITA;CNTNAP1;SYCP1;FLT4;ITGB2;FHL2;PIK3CD;TRH;ITGAL;MSLN;FOXM1;CYR61;SYNE2;PIK3CG;HHEX;DNMT3L;ITGAX;DNMT3B;HSP90AA1;CR1;ITGA4;VWF;TNFRSF18;DNMT3A;CTDSPL;WIPI2;KSR2;TNFRSF1B;NGF;TNFRSF1A;LCK;BIRC5;ITGA6;ANG;BCAT1;HDAC2;HDAC3;LRP1;SRC;NOD2;CXXC5;BPHL;CKMT2;CCL5;NNAT;FYN;EIF4E;NTRK1;MARCKSL1;ANGPT2;ANGPT1;PRRX1;ASCC1;NTRK3;TIE1;SCRL1;SGCZ;MTOR;KLF1;NELL1;P2RX5;KLF6;IL2RA;LEP;IL2RB;ASXL2;PDCD4;PTPN6;PDCD1;P4HB;APP;SPARC;HPSE2;GFI1;MLC1;IKZF1;IFI30;ICAM1;LGALS3;TCL1A;RASSF1;ANPEP;TNFSF10;SOX7;SLC12A9;SRRM2;HLA-C;KRT7;EBF3;HLA-A;HLA-G;ANK1;RUNX3;RUNX2;UHMK1;RUNX1;TIAM1;SFRP1;SFRP2;IFI27;SFRP5;MT1G;CES1;SLC22A3;RTEL1;EPAS1;SLC22A2;TWIST2;TWIST1;CD1D;PIK3R1;EVPL;CD1A;DGUOK;SYNGR1;PCBP4;CD19;ZNF704;INPP5D;HLTF;SPTBN1;STAT5A;EGR1;RABGAP1L;SPHKAP;INTS1;BCL11B;XRCC3;IGF2;LIF;NBEAL2;BST1;PTK7;ARTN;ID2;CD200;TP73;APOBEC3B;DDR1;HOTAIR;PTPRT;COL18A1;PTPRU;NRP2;ROBO4;CD40;CTSZ;BRCA1;KRT20;AKAP12;AKAP13;SLC22A16;HNF4A;RAC2;CD38;CCR7;MIR192;DLAT;CD34;VENTX;CBR1;CD52;ZHX2;MME;TSC22D1;NME2;RHOH;MIR21;MIR19A;CBFA2T3;VAV1;NCR1;NCR3;NR5A2;IL1B;MIR17HG;ARHGEF2;CD47;ARHGEF7;SGK1;TLR4;CAPG;PURA;NT5E;PAK1;CUX1;WIF1;HSF1;CTNNA1;MSX1;EPS15;LAIR1;CD55;SVIL;CD74;NQO1;CDX2;CDX1;KLHL1;LILRB1;MLH1;LILRB4;VEGFA;TCEA2;F2RL1;IL17D;CD86;CD83;DCLRE1C;MPG;MPL;PRDM2;PRDM1;ETS1;HOXA13;MPO;ETS2;IGF1R;HOXA11;CNDP2;HOXA10;CCND2;CCND1;CAPN2;MIR150;BAALC;TGM6;GGT1;PDGFRB;PDGFRA;CD96;CASR;KDM2B;CD93;WNT5A;MATK;LIG4;TSC2;CYBA;CARTPT;DHDDS;MYCN;POLG2;PRTN3;CARD16;ABCG2;FBN2;CALCA;DIABLO;ALOX12;SLC1A5;LPP;FUT1;FUT4;ZFP36L2;CACNA1G;NME1-NME2;DPP6;NLRP2;EP400;MSC;MYH10;TFAP2A;UBE2I;SMAD3;EPHX1;POU2F2;ESR1;ARHGAP24;BMP4;ISG20;IL6;FER;EFNA2;FES;ALPI;FGFR4;ULBP1;FGFR1;DOCK6;UCN;MIR10A;THRA;SPO11;DDX43;LAMC2;CXCL5;LYL1;HK3;CASP8;TRPS1;CASP1;PDE4B;NOS1;TGIF1;HS3ST3B1;MIR193A;EGFL7;USP42;DCC;OLIG2;MIR126;FGR;CCNA1;ELF1;ELF3;IFNG;ELF5;LTA;IRF7;MYH9;BLVRB;IRF8;SCN4A;LTB;RBMS3;SPG7;SAMHD1;AURKB;CRYZ;INS;MUC1;MIR377;WNT11;CNR2;CBS;HLX;MIR495;MIR494;GNA12;ERBB2;NT5DC3;PTK2B;NCAM1;PDLIM4;CHGA;NOS2;OPRK1;CHFR;DLK1;HS3ST2;MEIS1;MIR145;CD6;MIR125B1;MIR142;CD7;CD9;TEK;VIM;SCLY |
| Sporadic Breast Carcinoma | 0.027372495407981288 | RB1;ACHE;COL18A1;CD40;MIR17;THRA;DIXDC1;BRCA1;COMT;PHRF1;TNF;IGF1R;HPGDS;CCND1;CDH1;MYC;TNFSF10;DNMT3B;FEZF2;SRRM2;ARL11;MME;DAPK1;WNT5A;LIG4;PROX1;TNFRSF1B;TGFBR2;TNFRSF1A;ANO1;RAD51C;PGR;RBMS3;MCPH1;UHRF1;GSTP1;COL11A2;GATA4;GSTT1;VTCN1;FHIT;ABO;WIF1;HSD17B1;ERBB2;HSD17B2;NLRP2;BARD1;GABRP;CA10;GSTM1;NOS3;XRCC3;HMGA1;MLH1;ESR1;ARHGAP24;VEGFA;KRT17;TNFSF4;LEP;CDK2;BCL2;SHBG;FGFR2;SIPA1;TP73 |
| Carcinoid Tumor | 0.027372495407981288 | APP;CDKN1A;MIR10A;SCT;PTPRN;PIK3CD;AQP4;OTP;SLC6A2;CLU;ETS1;FGF2;FGF3;PIK3CG;SOX2;RASSF1;MIR432;CASP8;CHL1;CA2;MYC;AKT1;TIMP3;PITX1;PDGFRB;MIR193A;PDGFRA;HSP90AA1;APLP2;MST1;DIO3;HLA-A;PAX5;CARTPT;SSTR1;NGF;CLDN4;GPRC5A;CCKBR;SLCO4A1;S100A9;CFTR;CALCA;ABCB1;GRP;MIR409;MIR487B;PDGFA;SLCO5A1;ASCL1;FHIT;THBS1;EGFR;PTHLH;MIR369;MIR136;MIR410;TERT;MAP2;MIR494;ERBB2;GNA12;TAC1;NTSR1;ATOH1;HOXC6;CHGA;MARCKSL1;FZD7;SCTR;MLH1;POU4F1;TMPRSS13;VEGFA;POMC;PYY;CDK6;APC;WT1;SST;BCL2;GNAS;PDCD4;FOSB;CPE;SIM2 |
| Dementia, Vascular | 0.0281351689251623 | CRP;ACHE;APP;NOTCH3;CSF2;EPO;PLEK;CHAT;PON1;HTRA1;KLK4;LAMC2;TNF;KLK6;PENK;APOE;APOB;CTSD;EIF4E;TIMP4;SNCA;PTCRA;GPX1;ACE;TNFSF14;F11;ESR1;MTOR;VEGFA;IL6;ADORA2A;SST;IL1B;SDHAF1;PHLDB2;HSPA1B;FGFR1;HSPA1A |
| Prostatic Neoplasms | 0.0281351689251623 | RB1;TES;SERPINE2;ELK4;NDST4;MYC;STMN1;AKT1;CYP1B1;PRSS8;STARD3;DAXX;SERPINF1;PRKCA;RNASE4;PDZK1;DKK1;UNC13D;DKK3;SULT2B1;HOXB9;SPINT1;CLDN9;BIN1;CYP2E1;ABCB1;CTBP2;CTBP1;TERC;HPN;AGAP2;GATA3;THY1;PRKCZ;ABO;SLC5A5;SULT1A1;LDHB;ABR;CYP2B6;TERT;NPBWR1;PDGFD;SFN;PLEK2;NGFR;CREBBP;ST14;GSTM1;ACE;CADM1;STARD10;NFATC3;NR0B2;PBX1;SQLE;GJB2;PPFIBP2;FXYD3;CYP1A1;NANOG;HOXD3;HLA-DRB1;HRK;ACHE;CDKN1A;ADM;MED15;GLI3;MYOM2;GLI2;HEY1;CSRP1;TIMP2;TIMP3;TIMP4;RING1;ANXA2;PGAM2;PAX6;F3;POU5F1;PAX2;SERPINB5;NCOR2;SLC7A5;MMP14;ZEB1;KCNMA1;RARA;RARB;CRYAB;GRP;GSTP1;NR1I2;COL11A2;ASCL1;SLC7A1;EGFR;EPCAM;BGLAP;ALOX5;NTF3;SPP1;CSMD1;GPR158;SPDEF;BRD4;WWOX;MGMT;STAT3;DAB2IP;NR1H3;POU4F1;GNMT;FABP5;NFIC;PKN1;HSPA1A;ALDH1L1;CSF2;CSF1;EPO;CTNND2;MIR146B;ZBTB20;DEFB1;CLU;FGF2;TNF;RIMS2;PYCARD;FGF7;HPGDS;FGF8;CDH3;SESN3;NAGLU;CDH1;PITX2;LZTS2;PITX1;LZTS1;CD177;KCNH2;IGFBP1;TLE3;USP7;TPM3;IGFBP3;UGT2B15;USP2;F2R;UGT2B17;PIPOX;TAP1;FOXP4;FOXP1;PKP1;PGR;PKP3;ERG;S100A9;MAD1L1;CSF1R;DLX4;SHC1;TACSTD2;ITPR1;GSTT1;CYP2C18;FBXO44;FHIT;ACAT2;SOCS2;SOCS3;MBP;SLC15A3;SLC38A2;ESRRA;SRD5A2;ZBTB16;TNK2;TNFRSF10C;DCDC2;TNFRSF10A;SYNJ2;TMPRSS13;DAB2;CDK6;RHEB;CDK2;BCL2;SP5;SCGB3A1;CLPTM1L;SHBG;SLC26A4;XAF1;PSCA;ITGB4;FLT4;FHL2;PIK3CD;FOXM1;CYR61;PIK3CG;ADAMTS5;TRIM68;DNMT3B;ACP5;PMEPA1;SOST;AMH;ADAMTS9;HSP90AA1;GPX1;PEG3;DNMT3A;NGF;ITGA7;BIRC5;TNFRSF25;ANG;CRP;CX3CR1;HDAC2;SRC;UHRF1;EHBP1;CCL5;VPS52;FYN;FSD1;NTRK1;CIZ1;PRRX1;TXNRD2;ANKRD30A;PCDH8;TBX3;REC8;MTOR;KLF6;APC;IL2RA;LEP;TRIM36;PTPN6;P4HB;BCAR1;SPON2;AHCYL2;SPARC;ACSM3;HPSE2;GFI1;NUDT1;ICAM1;SOX2;LGALS3;RASSF1;EDNRA;LGALS1;RASSF2;EDNRB;TRIM29;TNFSF10;TNFSF11;SOX9;SEC62;SOX7;TP63;SLC12A9;SOX5;PAQR4;KRT8;KRT7;DLEC1;HNF1B;HLA-A;MIF;RUNX2;RUNX1;TIAM1;SFRP4;SPATA18;EZR;CFTR;GRB7;SLC22A3;CASZ1;EPAS1;TWIST1;KLK4;MST1R;PIK3R1;CRHR2;TRPM2;TSPAN8;PENK;APOD;APOE;TRPM8;A2M;NTSR1;TRPM4;STAT5A;EGR1;EGR3;GREB1;IGF2;PTK6;ALOX15B;ID2;GNRH2;ID3;SIM2;TP73;DDR1;COL18A1;NRP2;CD40;MIR17;STAB2;WFDC1;BRCA1;CX3CL1;AKAP12;AKAP13;HYAL1;CCR9;CCR6;CCHCR1;CTSD;HTATIP2;CBR1;ZHX2;MME;MST1;NCOA4;IL16;MIR21;TYK2;ANO7;CYP7B1;TLR1;CDCP1;DNASE1L2;TLR4;MATN4;TAGLN;TMPRSS6;TMPRSS2;CYP17A1;CAMKK2;RXRB;PURA;NT5E;PAK1;WIF1;HSD17B1;NPY;PAK6;CD55;CD74;NQO1;GDF15;OSM;KLK15;LILRB1;MLH1;PDZK1IP1;KLK10;KLK11;VEGFA;DLC1;KRT14;F2RL1;GHRL;IL17D;F2RL3;FOXA2;NDUFA13;CRTC1;COMT;ETS1;MPO;ACRBP;IGF1R;CCND2;CCND1;NPPA;GGT1;TMOD1;PDGFRB;PDGFRA;GHSR;CASR;ABCG5;OSR1;WNT5A;GPD2;NUPR1;STEAP1;ABCG2;CALCA;DIABLO;ALOX15;NPR3;ALOX12;SLC1A5;ADRB2;HOXB13;CACNA1H;FUT4;ZFP36L2;CNN3;DPP4;PODXL;MYH14;TAC1;MSC;TFAP2A;SMAD2;ZFHX3;SMAD3;WNT3A;EPHX1;TRPV2;EPHX3;BMP7;ESR1;IGF2R;LRP1B;SMAD7;BMP4;ISG20;TDRD1;BMP2;IL6;FER;NAT1;GNAS;ALPL;CRYL1;TGFBI;FGFR4;FGFR3;FGFR2;FGF10;FGFR1;KDM5B;CRISP2;TNFAIP8;HSPB2;TNFAIP3;CHD3;CXCL5;GHR;CASP8;ZIC2;TRPS1;MMP7;DCC;PDE4D;IRAK4;SSTR1;DCN;SSTR4;TGFBR2;TGFBR3;ZAP70;CCNA1;ELF3;IFNG;ALDH1A2;EHHADH;CDH11;CDH12;CHI3L1;CDH13;ACOX3;HAGH;CREB5;DDC;PON1;THBS1;PTHLH;INS;MUC1;MUC2;WNT11;CNR2;MXI1;GNA12;ERBB2;MIR492;PTK2B;SPOCK1;EPS8L3;MUC4;MUC6;CHGA;TCF7L2;NOS3;BNIP3;LIMK1;B3GAT1;PDX1;MEIS2;MEIS1;MIR145;PPP2R2C;SARDH;CD9;SPRY2;S100P;TEK;VIM |
| Bulimia Nervosa | 0.0281351689251623 | GHSR;FAAH;OXTR;ATP8A2;DST;LMO3;HTR1A;IGF2;HTR1B;HTR3A;HTR2A;HTR3B;COMT;ESR1;SLC6A3;CIT;DPP4;NKAIN3;LEP;NT5C1B;SOSTDC1;GHRL;DRD4 |
| Congenital absence of kidney | 0.029753720240774866 | LRP4;HOXD13;SERPINA4;PUF60;SALL4;FGF20;SOX8;SOX9;INSL3;TP63;WNT3;TFAP2A;EYA1;PAX2;CIT;VANGL1;GDNF;ITGA8;MKS1;SPRY1;FGFR3;CFTR;FREM2;FGFR2;SDCCAG8;FGF10;FGFR1 |
| Desmoplastic Medulloblastoma | 0.029753720240774866 | MPL;EHMT1;IRS2;PIK3CG;RASSF1;MYC;ERBB2;OTX2;PRAME;SMYD4;ATOH1;S100A10;BRD4;PDGFRA;BRD2;KDM4B;STAT3;PTCH2;PAX5;HIC1;VAV1;FLI1;MYCN;CDK6;APC;S100A6;SCG5 |
| Invasive Lobular Breast Carcinoma | 0.029753720240774866 | SLC22A3;SPARC;THRA;PIK3CD;BRCA1;GATA3;FGF3;PIK3CG;CCND2;WNT11;CCND1;CDH1;ERBB2;HSD17B2;ITGAX;NOS1;EGFL7;NOS2;ADAM33;NR1D1;ESR1;FOXP1;VEGFA;SCGB3A1;MIR183;PGR;FGFR1 |
| Cleft palate, isolated | 0.030382583236786616 | CDKN1C;APP;RARG;F13A1;ADM;ETS1;FGF2;FGF3;GLI2;FGF7;FGF8;MYC;SOX9;TP63;SRGN;PDGFRA;OSR2;SLC32A1;BHMT2;TGFBR2;F7;COL2A1;IL1B;RARA;PGR;IRF6;PIF1;DLX5;COL11A1;SATB2;COL11A2;NEDD4L;GSTT1;ALX3;SPP1;MSX1;TBX1;SPAG9;GSTM1;FOXF2;TGFB3;NOS3;ABCA4;GRHL3;ESR1;BMP4;SLC2A9;BMP2;MAFB;BCL2;SPRY2;FGFR3;SIM2;FGFR2;FGF10;FGFR1 |
| Heart Diseases | 0.030661162450794303 | SCARB1;CSF3;EPO;PRDM6;NCF4;HTR4;FGF1;FGF2;TNF;MYLK;LGALS3;MIR675;HPGDS;MYC;NPPA;PITX2;KCNH1;ENTPD1;SERPINF1;DIO3;ANK2;CYBA;RUNX2;KCNQ1;PAPPA;PKP1;CYP2E1;PDE9A;NOTCH3;KCNE1;ABCB1;NPR3;GATA4;SLC5A1;CACNA1C;MIR206;ABO;PRH1;GRK5;NPC1L1;PRDM16;HOXC5;NLRP3;APOE;APOB;NKX2-5;TRPM4;STRA6;ACE;NFATC1;NR0B2;IL6;TH;CYP1A1;NOX5;SHBG;HOTAIR;RYR2;RARG;HSPB6;PIK3CD;ADM;MYOM2;CYR61;PIK3CG;SYNE1;GHR;SGCD;MB;HNF4A;RAC2;EMILIN2;PF4V1;DSP;UNC45B;PTGIR;TRPC3;VWF;GAA;AMPD1;MIR21;F3;LCK;IL1B;CHDH;CAT;HAND1;SGK1;MYH7;CRP;OCA2;HDAC3;IDUA;PON1;KCNA3;HTT;THBS1;EGFR;TTN;PDLIM3;CBS;TBX20;SPP1;TNNI3;FSD1;XDH;NQO1;ANGPT2;NOS2;NOS3;BNIP3;STAT3;TBX5;MTOR;DLK1;VEGFA;FABP3;NPHS2;LEP;APOC1;ASXL2;GHRL;PF4;PNPLA2 |

| | | |
|---|---|---|
| Neoplasm Metastasis | 0.031238300140012477 | MDC1;EHMT2;MYLK;CYP2D6;TMEM88;NDST4;DPYSL3;EPSTI1;AKT1;DAXX;UNC5B;PDZK1;RNASE1;PRAP1;SND1;CLDN6;CLDN4;SPINT1;DDIT3;RUFY3;CYP2E1;MYL9;SKAP2;LXN;ABCB1;DHH;MIR196B;HPN;GATA6;PDGFB;PDGFA;GATA5;LYPD3;GATA4;FPR2;THY1;GATA3;LYPD5;GATA2;SLC5A5;SULT1A1;HSD11B1;CYP2B6;MIR330;KCNQ1OT1;ADAMTS18;PDGFD;SFN;HCRT;JAG2;LYN;NGFR;GSTM1;CADM1;ABCA4;MRPL28;NR2F2;MIR339;MIR338;NR0B2;PBX1;FLI1;MIR346;COL1A1;VANGL1;FERMT1;FAT1;HRC;NANOG;FAT4;HRG;HLA-DRB1;LGMN;FERMT3;PIGR;MIR130B;LRRC4;MED15;MMP21;SHF;SMPD3;HEY1;UCN3;NEFL;NEFM;SH3BGRL3;JAK3;IKBKE;MYOG;RIPK3;DUSP1;SPHK2;MYOF;F3;DUSP6;POU5F1;MIR522;F7;SLC7A5;MMP14;GPRC5A;MMP16;KCNMA1;CRYAB;GPRC5C;RAPGEF3;TMEM106A;IDUA;GSTP1;SYNPO2;COL11A1;COL11A2;KCNA3;HTT;FSTL1;EGFR;BGLAP;PLAGL1;HAS1;NTF3;SPP1;MICALL2;PRAME;FLNC;CSMD1;SPDEF;JAM3;LRRC26;PKLR;WNT7B;STAT3;SCTR;DAB2IP;MIR539;PLXDC1;PSMB8;SKI;FABP5;WT1;FABP6;FABP7;ZNF217;PKN1;SIGLEC7;LY6K;LY75;MUM1;UBE2D1;MSI1;DEFB1;PLOD2;HPGDS;HHAT;SLN;ADORA3;ADORA1;MAP3K9;PITX2;SLC16A3;KCNH1;MAP3K5;KCNH2;S100A2;S100A1;UGT2B15;UGT2B17;ATP11A;SRCIN1;FOXP2;FOXP1;LY6D;FSCN1;S100A6;PKP1;ZNF436;PKP3;S100A9;SQSTM1;LY6E;DSC3;DLX2;DLX4;TUBGCP2;FBLIM1;DLX5;SHC1;TACSTD2;GSTT1;LIN7C;FHIT;RAP1GAP;SOCS2;SLC9A3R2;SOCS3;BCL2L10;ATXN1;KL;NOP14;SRD5A2;HTR1D;TFPI2;TNFRSF10C;TNFRSF10A;TMPRSS13;SCGB3A1;SP7;PIWIL2;CNTNAP1;ST6GALNAC1;PIWIL3;PPP1R13B;KCNK9;ITGB4;FLT4;ONECUT1;ITGB2;FHL2;WISP3;PIK3CD;ECE1;ITGAE;MSLN;FOXM1;PIK3CG;WISP1;MB;NCK2;ACP5;PMEPA1;ITGB6;UPP1;NKX3-2;UNC45A;DSP;TRIM62;HSP90AA1;ITGA4;KSR1;ITGA3;NRG1;FOXL1;NRG2;TNFRSF1B;TNFRSF1A;ITGA7;BIRC5;TNFRSF25;ITGA6;LCP1;BIRC7;BCAT1;ITIH3;LRP1;SRC;UHRF1;HM13;LRP5;FURIN;ITIH1;NTN1;PARD6A;SH3BP1;CCL5;NNAT;SLITRK3;TP53INP1;IGF2BP1;SH3BP4;FYN;FSD1;KCNJ1;EIF4E;NTRK1;TBX1;PPP1R14A;CIZ1;MARCKSL1;SLC12A5;MYO10;PRRX1;NTRK3;STYK1;PCDH8;TBX3;TBX2;NELL1;GZMM;PC;MYO1C;SST;IL2RA;IL2RB;MGP;PDCD4;CD248;PTPN6;PDCD1;P4HB;BCAR1;ERRFI1;TNFRSF6B;ICAM2;ICAM1;LGALS4;LGALS3;TCL1A;RASSF1;LGALS1;RASSF2;TRIM29;ANPEP;PEG10;RASSF6;TNFSF10;TNFSF11;LGALS9;TRIM27;LGALS8;TP63;EPHB3;SLC12A9;EPHA4;SLC34A2;ENTPD1;EPHA6;HNF1B;TM4SF5;MIF;SFRP4;SFRP1;CFDP1;SFRP2;SLPI;IFI27;CRISPLD2;KCNQ1;SFRP5;MT1G;MT1H;KCTD11;ALPK2;RIN1;EPHA1;ALPK1;STX1A;GRB7;EPHA10;EPAS1;PCDH10;MST1R;PIK3R1;CD1D;CD1A;PIK3R5;RAB11FIP1;TSPAN8;PCBP1;CD19;NEU1;HLTF;HIVEP3;APOE;KCNN3;A2M;B4GALNT2;TSPAN1;ATP6V1C1;BBS2;NTNG1;STAT5A;WWTR1;EGR1;EGR3;TNFSF14;FOXF2;FOXF1;IQSEC1;RCC2;XRCC3;IGF2;PTK6;NBEAL2;DCLK1;BST2;NET1;BST1;MSRA;TF;TG;TH;LPAR5;PTK7;ARTN;ID3;GRB2;RCOR1;CD200;CD22;TP73;APOBEC3B;PCSK2;DDR1;DIRAS3;PTPRT;PTPRU;NRP2;CD40;PTPRR;CTSZ;PCSK9;MIA2;BRCA1;PCSK6;PTPRF;AKAP12;CTSO;HYAL1;HYAL2;HNF4A;RAC2;MUC12;CCR9;CD38;CCR7;CCR6;CD34;CTSD;HTATIP2;ZHX2;MME;MST1;AJAP1;TOX3;IMMP1L;ANO1;NR5A2;CTTN;PITPNM3;IL1B;ARHGEF4;ARHGEF2;CD47;ARHGEF7;TAGLN;TMPRSS6;SATB2;MIR520G;TMPRSS4;NEDD9;TMPRSS2;CAPG;CYP17A1;GNAI2;RXRB;PAK1;HSD17B2;HSF1;PAK6;GPC5;MSX1;PCNT;EPS15;XDH;CD55;MICB;IL32;CD74;NUBPL;KLK13;OSM;AKR1C2;KLK10;KLK11;PPP1CA;KLK12;VEGFA;PER1;NEDD1;PER3;TTC4;KIF26B;FOXA2;PNPLA2;CD86;RTL1;CD83;DCLRE1C;CRTC1;CD81;MPL;IHH;PLAT;PRDM2;PRDM1;ETS1;MPO;ETS2;ACTG2;LOXL1;IGF1R;LOXL2;RGS5;CCND2;CCND1;PEBP4;NPPA;PPFIA1;TGM2;PDGFRB;PDGFRA;CASR;SCARA5;MIR27A;MAT1A;RUNDC3B;SDHB;DHDDS;BRMS1;SCNN1B;ADAM12;DPEP1;CARD16;FBN2;NDRG4;CALCA;TCF7;HTRA1;SLC1A5;ACACA;FUT4;FUT3;NME1-NME2;FUT6;GRK5;CALD1;FUT9;MIR296;CYTL1;TAC1;MSC;WNT2;WNT3;TFAP2A;ZFHX3;TFAP2C;UBE2I;CNTN5;CNTN6;HMGA1;ISG15;INHBA;BMP7;IGF2R;PHOX2B;FGF17;BMP4;ISG20;BMP2;IL6;CNTN1;GNB3;ALPI;FGF11;LGALS3BP;SNED1;TNFAIP8;DYRK2;TNFAIP2;ATP2A3;TNFAIP3;CHD5;LAMC2;ADAR;GRM1;PITPNC1;GHR;GRM5;GRM4;CA3;ZIC2;CA2;LRRFIP1;MIR193A;TGIF1;MYOCD;MMP7;MIR23A;BUD31;SHROOM3;OLIG2;SSTR1;SSTR4;TAPBP;SSTR5;ACTA2;GREM1;CCNA1;ACTA1;ELF3;MIR485;ELF5;CAT;IRF7;IRF8;CHI3L1;IRF5;OTUB1;LAMA4;LAMA3;PTHLH;AURKB;CNR2;CBS;MIR495;MIR494;MXI1;MIR492;PTK2B;SPOCK1;SLIT3;ATOH1;CHGA;RBM38;URGCP;BNIP3;CCK;MUC5B;CD4;TEF;BCL6B;SARDH;CD9;TEK;VIM;RAN;RB1;TES;SERPINE2;WIPF1;MIR429;MIR425;MIR675;HOXA9;SOX17;ZMIZ1;SOX18;MYC;CREB3L1;STMN1;CYP1B1;PIP4K2A;HOXA1;PRSS8;HOXA5;CHGB;HOXA4;WSB1;PRKCB;DAPK1;SERPINF1;DAPK3;AFAP1;PRKCA;DKK1;DKK3;SYTL2;DKK4;HOXB9;BIN1;MAP1B;GPR160;TYRO3;HOXB3;HOXB1;HOXB8;HOXB7;HOXB5;TSPYL5;MEG3;CTBP2;CTBP1;TERC;PRDM14;NEDD4L;MIR206;ABO;LDHB;TERT;CHST11;S100A16;S100A13;ALX1;PROM1;HOXC9;RREB1;ELANE;HOXC6;S100A10;STARD13;ST14;ICMT;ACE;FZD5;ZNF382;FZD7;FZD6;NFATC2;NFATC1;SQLE;H19;GJB2;CYP1A1;HOXD3;PCDHB9;HOXD9;CDKN1C;CDKN1A;TINAGL1;ELN;LPCAT1;TCF21;GPT;ADM;PRSS22;SLC6A2;SERPINA4;MYOM2;GLI2;CHL1;TIMP2;TIMP3;JARID2;TIMP4;DENND2D;BRD2;ANXA2;TNNC1;TESC;PAX3;TET1;SERPINB9;PAX5;PAX2;SERPINB5;NCOR2;ZEB2;ZEB1;ANGPTL7;PAX8;KIFC1;TAL1;PAX7;PAX9;RARA;RARB;HCLS1;ANGPTL4;TKT;GAS5;HLA-DQB1;GRP;AHNAK;NR1I2;MIR409;HOXD10;ASCL2;RASSF10;EPCAM;ALOX5;GCNT2;RUNX1T1;BRD4;CCL25;WWOX;BRF2;F10;MGMT;WWP1;ELP4;SULF1;DEF6;HIPK2;MIR661;ABI3;ABI1;FOSB;MSRB3;CCL28;HSPA1B;PF4;HSPA1A;SEMA5A;CSF3;CSF3R;CSF2;CSF1;EPO;RAB3D;CTNND2;MIR146B;IRS2;FGF1;CLU;FGF2;TNF;FGF3;BEST1;RIMS2;CDH6;PYCARD;CDH5;FGF7;MT1DP;FGF8;CDH3;LAMP1;CDH1;GRAP2;RNF19A;SLCO2B1;SALL4;ZNF569;LZTS1;SLC39A4;CD177;CSGALNACT1;USP7;LHCGR;TPM4;FKBPL;USP4;IGFBP3;F2R;MAGI2;DIO3;TAP1;MITF;CHD1L;DGKZ;HIC1;RAB32;MIR1228;MTNR1A;OBSCN;LTB4R2;RAB37;PGR;ADAM8;ERG;SNRPF;MAD1L1;CSF1R;NOTCH3;SEMA7A;NOTCH4;CUL1;RASAL2;VTCN1;TPO;NDN;ZNF300;ESRRA;SEC16B;TNK2;SYNJ2;TRAPPC9;HOPX;MIR612;GFAP;DAB2;COPS5;CDK6;BCL6;CDK2;BCL2;CPE;CSPG4;PLXNB1;ESAM;SLC26A4;XAF1;LAT;PSCA;SYNE2;CYR61;SYNE1;ADAMTS5;ADAMTS2;DNMT3B;SOST;HMGCS2;GAL3ST2;ADAMTS9;SGCG;HGFAC;CR1;GPX1;VWF;DTX3L;DNMT3A;APOA1;NGF;C1QL1;PTP4A1;TFDP1;ESRP2;ESRP1;CRH;EVL;ANG;CRP;HDAC4;NFAT5;HDAC2;HDAC3;RAB1B;SEMA3B;MIR148A;TTN;CLEC3B;RAB25;P2RY1;MAL2;UGT1A7;SPAG9;IRX2;ANGPT2;ANGPT1;TXNRD2;ANKRD30A;TXNRD1;SEMA4B;MTOR;P2RX7;MARCKS;NIN;P2RX5;KLF6;APC;LEP;RAB17;CDO1;APP;IFITM1;SPARC;ACSM3;HPSE2;GFI1;DIXDC1;AQP1;SOX2;EDNRA;EDNRB;PART1;SOX9;RPS12;SOX5;SRRM2;CPT1A;HLA-B;MTUS1;KRT8;HLA-C;KRT7;DLEC1;HLA-A;KRT5;HLA-G;RUNX3;RUNX2;RUNX1;TIAM1;IL23A;PAPPA;EZR;MIR1182;CFTR;TELO2;LLGL2;SLC22A3;MACF1;CASZ1;HEPACAM;EIF5A2;TWIST2;TWIST1;KLK4;KLK8;CRHR1;KLK7;KLK6;NUAK1;STC2;INPP5J;FADD;TRPM8;SPRR2A;NTSR1;BANP;SNHG1;TRAP1;TREX1;LIF;GDF2;MIR200A;MIR200B;GADD45G;OCLN;HAL;ANP32B;SIM1;TDGF1;NODAL;HOTAIR;ROBO3;COL18A1;GFM1;MIR17;MIR18A;KRT20;CX3CL1;ESM1;SH3PXD2A;STAB1;SOSTDC1;TNFAIP8L3;MIR192;TNFAIP8L2;CLIC1;PARP6;CBR1;PRMT7;NME2;TFEB;MIR21;TYK2;PROX1;MIR19A;GFRA3;CBFA2T3;VAV1;VAV2;CLDN10;CDCP1;B3GNT7;CCKBR;TNNT1;MIR17HG;MMP23A;TFF3;CLDN18;TFF2;TFF1;CD302;MAPT;TLR4;GUCY2C;C1QA;CSTA;VWCE;CST7;AIF1;NKD2;NT5E;CUX1;WIF1;NPY;CTNNA1;GINS2;NQO1;CDX2;GDF15;SUSD2;LILRB1;MLH1;GRHL2;KRT18;KRT17;DLC1;KRT14;NBR1;TRIP10;F2RL1;BLID;TRIP13;NEUROG1;SIPA1;F2RL3;NDUFA13;HOXA13;CNDP2;MIR149;HOXA10;ARHGDIB;MIR150;NANOS1;PDIA2;POSTN;KDM2A;WNT5B;KDM2B;APLP2;WNT5A;LIG4;TSC2;SHISA3;MPEG1;ALDH3A1;MYCN;LMX1A;COL4A2;ELMO1;NUPR1;STEAP1;ABCG2;ISYNA1;ALOX15;ARHGAP18;ALOX12;ADRB2;FBLN2;DPP4;SCIN;PODXL;HRH2;UBD;CDH23;NLRP2;MYH10;MARK2;SMAD2;KLHDC4;CBX5;SMAD3;CBX4;TGFB3;WNT3A;TRPV2;ASPSCR1;TBCB;HSPA2;ESR1;LRP1B;ARHGAP24;SMAD7;FER;GDNF;FES;MIR184;MIR183;TGFBI;FGFR4;FGFR3;FGFR2;FGFR1;UCN;KDM5B;CRISP2;MIR10B;MIR10A;THRA;HSPB2;MIR96;PKD1;CXCL5;ELAVL3;UPK1A;CASP8;CA5A;SIX2;SIX3;FLOT1;CASP1;BDKRB2;NOS1;POLE;SRGN;KLF11;EGFL7;DCC;AXIN1;OPRM1;UPK3A;DCN;TGFBR2;KLF17;M |
| Autoimmune thyroid disease | 0.031602552679344 | THRB;ICA1;TNFAIP3;PTPN22;TNF;BACH2;LGALS1;ZMIZ1;PDE4B;LGALS9;HLA-DOA;SLC30A8;TNFRSF18;FCRL3;TAP2;HLA-A;HLA-G;IFNG;IRF8;TNFRSF25;CFB;HLA-DQB2;HLA-DQB1;ANKRD55;SLC30A10;LPP;ADCY7;SLC5A5;INS;TPO;STAT4;IL21R;HLA-DQA1;BCL2L15;HLA-DRB6;SMAD3;IGF2;BTNL2;MAFA;ARHGAP24;TSHR;ISG20;IL6;TG;INS-IGF2;FABP5;ZFAT;IL2RA;GNAS;HLA-DPB1;SLC26A4;TRIM31;IL17C;HLA-DRB1;HSPA1B;TP73;HSPA1A |
| Gastroesophageal reflux disease | 0.031693511775038714 | SLC46A1;SPARC;FAM13A;ABAT;FGFRL1;IGF1R;SALL1;CCND1;TP63;F2R;TSC2;EBF3;ATP11A;DKK1;FOXP1;DKK4;SCNN1D;EZR;RTEL1;CALCA;ISYNA1;CTBP1;TERC;FOXG1;LTBP4;ADRB2;CPLX1;NALCN;RAI1;TERT;DEAF1;PRDM16;TAC1;NTNG1;TFAP2A;ACE;XRCC3;PHOX2B;COL1A1;ATP4A;IL6;GDNF;GNB3;NOX5;TECPR2;HLA-DRB1;ELN;CTSZ;ATP2A3;HSPB2;CTSW;HTR2A;SLC6A3;FLCN;ADAMTS2;SLC22A18;KIF1A;CCR6;DSP;MIR21;KCNAB2;NGF;PAX2;DDOST;TNFRSF1A;IL1B;IRF5;TLR4;GUCY2C;DDC;NRXN1;NR1I2;ASCL1;THBS1;EGFR;GLRA1;MUC1;GABRD;MUC4;NQO1;CDX2;NOS2;CDX1;MGMT;CCK;MUC5B;MEIS2;VEGFA;LETM1;SKI;P2RX7;GLRB;NPHS1;MIR142;F2RL1;RAMP1;CCL26 |
| Headache | 0.03276460107346538 | CX3CR1;APP;CSF3;CALCA;DDC;MVK;COL11A1;KCNA1;MPL;LRP5;CACNA1A;OXT;PTPN22;COMT;TNF;SLC6A3;EDNRA;ADORA1;CDH23;NLRP3;SOST;APOE;CASR;ACE;MSX2;VWF;NOS2;NOS3;HTR1A;HTR1B;OPRM1;VEGFA;VANGL1;IL6;NLRP12;ADORA2A;DOK7;GNB3;CCM2;LPIN2 |
| Herpes NOS | 0.03276460107346538 | COL18A1;CSF2;TNFRSF6B;MRGPRF;LAMC2;NOD2;TNF;ICAM1;MYLK;SELENBP1;EDNRA;CASP8;CYP2B6;ERBB2;GNA12;PENK;CASP1;TK1;APOE;CD34;MICB;USP7;DST;TNFSF14;CD160;SLC11A1;HBE1;ABCA4;HLA-C;HLA-A;PAX5;ESR1;SSTR4;VEGFA;IL6;APC;IL2RA;LTA;BCL2;MYH9 |
| Squamous cell carcinoma of esophagus | 0.03276460107346538 | RB1;APP;TNFRSF6B;SPARC;MIR429;FGFRL1;SOX2;RASSF1;RASSF2;EDNRB;SOX17;CYP2D6;RASSF5;MYC;STMN1;TNFSF10;AKT1;CYP1B1;PLCE1;TRIM27;SOX9;TP63;SLC12A9;PRIMA1;NKX2-8;CPT1A;HLA-C;DLEC1;HLA-F;RNASE1;RUNX3;DKK3;SYTL2;RUNX1;TIAM1;CLDN4;SFRP1;SFRP2;IFI27;MIR196A2;CRISPLD2;BIN1;CYP2E1;EZR;NEIL1;MYL9;HOXB7;ALDH7A1;GRB7;SLC22A3;ABCB1;CTBP2;SHMT1;CRABP1;EIF5A2;TWIST1;THY1;PIK3R1;EVPL;ABO;WDR66;ADAMTS16;LDHB;CYP2B6;TERT;PCBP4;STC2;SFN;FADD;SPRR2A;HLA-DQA1;S100A10;NGFR;EGR1;TRAP1;CREBBP;GSTM2;AFAP1L2;GSTM1;CADM1;HIST1H4K;HIST1H4L;FZD7;XRCC3;IGF2;MRPL28;PTK6;MIR200B;GRIN2B;GADD45G;FAM135B;GJB2;PTK7;PPP1R1B;FAT1;CYP1A1;FAT2;NANOG;GRB2;HIST1H4I;FAT4;TDGF1;HLA-DRB1;TP73;HIST1H4F;MIR129-2;PTPRT;HOTAIR;COL18A1;CDKN1A;MIR130B;BRCA1;MIR18A;MED15;MMP21;AKAP12;RPTOR;MECOM;CSRP1;HNF4A;SLC22A18;TIMP3;CCR7;MIR192;CCHCR1;NEFH;CLIC1;HTATIP2;GPR39;MME;ANXA2;MST1;MIR21;PROX1;TMEM173;MIR19A;DUSP6;POU5F1;PAX2;SERPINB5;VAV1;AJAP1;TLR1;MMP14;ZEB2;GPRC5A;ZEB1;CDCP1;CTTN;IL1B;PAX9;MMP23A;MIR17HG;HCLS1;RARB;TFF1;GPRC5C;TLR4;VGLL4;HLA-DQB1;TAGLN;CSTA;TNXB;GRP;GSTP1;COL11A2;NR1I2;CRIP2;FSTL1;EGFR;RASSF10;CUX1;EPCAM;WIF1;GPC1;SPP1;WWOX;NQO1;MGMT;NFYC;STAT3;KLHL1;AKR1C2;SORBS2;LILRB1;MLH1;PSMB8;PSMB9;VEGFA;KRT17;KRT14;SIGLEC7;CCL26;RAB3D;MSI1;PLAT;PRDM2;MSI2;ETS1;CLU;FGF2;TNF;HOXA13;ETS2;FGF3;IGF1R;LOXL2;RIMS2;HPGDS;CCND1;CDH1;GRAP2;RNF19A;MIR150;PITX2;TGM3;PDGFRB;S100A2;S100A1;IGFBP3;WNT5A;MIR27A;HIC1;DHDDS;FSCN1;DPEP1;MASP2;PHLDB2;ABCG2;NOTCH3;CHRNA5;DIABLO;ALOX15;CHRNA6;HTRA1;GSTT1;ALOX12;LTBP2;HOXB13;FHIT;ACACA;FUT4;AMDHD1;SMYD2;TAC1;SMAD2;SMAD3;EPHX1;TMPRSS13;ESR1;HOPX;MIR612;LRP1B;ARHGAP24;FBXO31;SMAD7;IL6;CDK6;HOOK2;FGF19;CDK2;BCL2;MIR183;CLPTM1L;TGFBI;SHBG;FGFR2;FGFR1;TNFAIP8;MIR10B;PSCA;TNFAIP2;HSPB2;PIK3CD;TNFAIP3;ADAR;LAMC2;MIR96;PIK3CG;WISP1;CYR61;GHR;CYP26B1;CASP8;UPK1A;CA2;MLST8;DNMT3B;HMGCS2;SOST;ADAMTS9;DSP;SRGN;EGFL7;MMP7;DCC;DNMT3A;TNFRSF1B;TGFBR2;PTP4A1;MIR126;IFNG;LTA;BIRC5;BLVRB;CHI3L1;CDH13;ITGA6;ZMYND10;SHANK2;ZNF750;RBMS3;FAM84B;CRP;MIR155HG;SEMA3B;THBS1;AURKB;CRYZ;JAKMIP2;MUC1;DNAJB6;RAB25;CCL5;MIR494;MIR373;ERBB2;SPOCK1;CCL1;FSD1;EIF4E;PDLIM4;MUC4;LGI1;LAMB3;BNIP3;EYA4;TNRC6C;NELL1;MEIS1;MIR145;APC;CAPN10;IL2RA;COL7A1;MIR142;IL2RB;PDCD4;VIM;CDO1;RAN |
| Hypersomnia | 0.03276460107346538 | CSF2;LAMC2;COMT;CPT1B;TSHR;SLC5A5;TPO;TG;PAX8;CYP2D6;IYD;NKX2-5;HCRT;SLC26A4;NCKAP5;HLA-DRB1 |
| Infection caused by Helicobacter pylori | 0.03283160443115206 | DEFB1;MPO;TNF;ICAM1;SOX2;TP53AIP1;CDH1;AKT1;TRIM21;F2R;MIR9-1;HLA-C;MIF;HPYR1;RUNX3;PROCR;SLPI;IL23A;AMPH;CYP2E1;SLC22A3;ABCB1;TACSTD2;GATA5;GATA4;GSTT1;GATA3;CYP2C18;FHIT;ABO;FUT3;FUT6;NME1-NME2;HLA-DQA1;ACE;GSTM1;XRCC3;TFPI2;PTK6;ARHGAP24;SMAD7;BMP4;ISG20;ATP4A;BMP2;IL6;CYP1A1;BCL2;HLA-DRB1;TP73;APOBEC3B;CDKN1C;UCN;CD40;PSCA;CTSZ;PIK3CD;FOXM1;TREM1;PIK3CG;GLI2;CASP1;MADCAM1;CCR6;MMP7;DCC;NME2;MIR21;SSTR1;VAV1;TNFRSF1A;TLR1;IFNG;IL1B;CAT;LTA;TFF2;CHI3L1;TFF1;IRF5;ARHGEF7;TLR4;HLA-DQB1;GRP;GSTP1;COL11A2;NR1I2;CXCR5;NOD1;NOD2;THBS1;EGFR;REM1;MUC1;CCL5;ERBB2;SPP1;ATOH1;MUC6;IL32;CDX2;NOS2;MGMT;ASCC1;STAT3;LILRB1;IRGM;VEGFA;TPPP3;NLRP12;APC;SST;IL2RA;LEP;FOSB;GHRL;PTPN6;IL17D |
| Malignant neoplasm of gallbladder | 0.0328562469928381 | HOTAIR;COL18A1;PSCA;PPP1R10;PIK3CD;TNF;FGF3;PIK3CG;BBC3;BEST1;RASSF1;CCKAR;CASP8;CDH1;MYC;TIMP2;CYP1B1;PLCE1;CCR7;CLIC1;ABCG8;DCC;MIR27A;HLA-A;MIF;FOXL1;POU5F1;DKK3;SYTL2;TGFBR2;SFRP4;CLDN4;ZEB1;ADRB3;BIRC5;PGR;TLR4;MEG3;ABCB1;TMPRSS4;LPP;EGFR;CYP17A1;NT5E;MUC2;EPCAM;WIF1;ERBB2;MIR372;ST8SIA4;GCNT2;SPOCK1;APOE;FADD;APOB;PROM1;BRD4;IL32;WWOX;GSTM2;CIZ1;ACE;GSTM1;CDX2;NOS2;MGMT;CCK;MLH1;ESR1;SOD3;ARHGAP24;VEGFA;MIR145;APC;DLC1;CDK2;CYP1A1;BCL2;VIM;SHBG;CDO1;TP73 |
| Thyroid carcinoma | 0.0328562469928381 | APP;ERRFI1;TRAF3IP2;FOXI1;AQP4;ICAM1;LGALS3;EDNRA;RASSF1;LGALS1;RASSF2;TRIM29;MYC;ANPEP;RASSF5;TNFSF10;AKT1;SEC62;TP63;PRIMA1;SLC12A9;HLA-C;PRKCA;HNF1A;DKK1;RUNX2;DKK3;SLC5A8;CCDC80;IFI27;IL23A;DDIT3;MT1G;PRKCQ;SLC22A3;ABCB1;PDGFA;TWIST1;THY1;MST1R;SLC5A5;TERT;PCBP4;HLTF;PROM1;RREB1;HLA-DRB6;CREBBP;GSTM2;AFAP1L2;GSTM1;FUCA1;XRCC3;IGF2;NR1D1;TG;CYP1A1;ID3;TP73;CDKN1A;LRRC34;CD40;NRP2;MIR17;TNFRSF13B;ELN;MIR18A;KRT20;HOXC10;EPB41L4A;CHL1;TIMP2;TIMP3;SOSTDC1;CCR7;CCR6;DIRC3;TIMP4;DUSP5;ZHX2;MST1;NCOA4;MIR21;PROX1;MIR19A;DUSP6;SERPINB5;MMP14;ZEB1;PAX8;CCKBR;IL1B;RARA;TFF3;RARB;CD47;GAS5;OCA2;GRP;GSTP1;COL11A2;TMPRSS4;CRIP2;EGFR;PAK1;RASSF10;EPCAM;SPP1;CTNNA1;BRD4;IL32;WWOX;CD74;MGMT;SORT1;GDF15;STAT3;MIB2;MLH1;PDZK1IP1;HIPK2;VEGFA;WT1;CENPJ;FOSB;GHRL;RAMP1;FCGBP;CSF2;CSF1;EPO;MPG;MIR146B;PRDM2;CLU;FGF2;TNF;ETS1;IGF1R;CDH6;PYCARD;CDH5;HPGDS;CCND2;CCND1;CDH1;IYD;SLCO2B1;MIR150;PITX2;GHSR;CASR;POSTN;S100A1;TPM3;DIO2;LMTK3;TSC2;UBE4A;MIR27A;SDHB;CLIP2;MYCN;S100A9;CSF1R;NOTCH3;CALCA;PFKFB3;FBLIM1;TACSTD2;GSTT1;RASAL1;LTBP2;FHIT;RAP1GAP;ACACA;DPP4;SOCS3;TPO;SMAD3;CBX5;PTCH2;HMGA1;SSH2;ESR1;HOPX;LRP1B;ARHGAP24;IL6;GDNF;CDK2;BCL2;GNB3;ALPI;SLC26A4;FGFR4;FGFR2;FGFR1;PIWIL2;THRB;THRA;PIK3CD;FOXM1;PIK3CG;FLCN;CASP8;ZIC1;INSL3;HSP90AA1;MMP7;KSR1;NRG1;NGF;KLF17;MIR126;CAT;BIRC5;RBMS1;MCM5;IRF5;CREB5;SRC;CXXC5;THBS1;AURKB;MUC1;ZNRF3;ERBB2;FYN;NCAM1;SLIT3;FSD1;EIF4E;NTRK1;SPAG9;TCF7L1;TXNRD2;NTRK3;LIMK1;CCK;SOD3;MTOR;REC8;TSHR;POMC;CRYGD;TBX15;MIR145;APC;SST;SPRY2;PDCD1;TEK;VIM |
| Ganglioneuroma | 0.03358433069313757 | CRISP2;NOTCH3;FLT4;PRDM2;ASCL1;THBS1;PTHLH;LGALS3;CCND1;HEY1;NCAM1;MSX1;NTRK1;EDN3;S100A1;SERPINF1;DAPK3;OLIG2;HIC1;PBX1;DLK1;PHOX2B;DKK3;GFAP;VEGFA;MYCN;GDNF;SST |
| Attention Deficit Disorder | 0.03358433069313757 | APP;ZNF132;DDC;CSF2;THRB;PON1;PIK3CD;LAMC2;DBH;CACNA1C;COMT;GRM1;PIK3CG;SLC6A3;INS;LGALS3;GRM5;GRM7;GRM8;ERBB2;NTF3;AKT1;APOE;CACNG2;DISC1;DRD4;DRD5;CREBBP;BTBD9;GRIN2B;VEGFA;POMC;PER1;MYCN;MIR183 |
| Nephritis, Interstitial | 0.03358433069313757 | CRP;ACE;ANGPT1;RNLS;APOA1;SERPINB9;HNF1B;NGF;FGF2;THBS1;DCN;MUC1;CYP2D6;CCL1;APOL1;XDH;HLA-DQA1;HLA-DRB1;HLA-DQB1 |
| Fibrosis | 0.033655647387035456 | CSF3;CDKN1A;SPARC;EPO;AQP2;FGF2;TNF;ACTG2;PIK3CG;ICAM1;LOXL2;FGF7;CCND2;CCND1;AKT1;BDKRB1;PDGFRB;PDGFRA;MMP7;PRKCB;ITGA3;PRKCA;MIR21;MIF;ADRA2C;F3;DCN;TGFBR2;GREM1;ADORA2A;IFNG;IL1B;CHI3L1;CYP2E1;SGK1;SLC27A2;TLR4;CFTR;CHRNA1;CX3CR1;GSTP1;PON1;PDGFB;GSTT1;FURIN;ADRB2;FHIT;THBS1;EGFR;PDGFD;GNA12;SPP1;NCAM1;TAC1;EGR1;ACE;GSTM1;ANGPT1;TGFB3;STAT3;PHOX2A;VEGFA;BMP4;COL1A1;IL6;WT1;CYP1A1;SSPN;F2RL1;VIM |
| Gastric Adenocarcinoma | 0.034331189957542675 | SPARC;RTKN;TRRAP;ETS1;FGF2;TNF;IGF1R;SOX2;RASSF1;FGF7;HPGDS;CCND1;NAGLU;MYC;STMN1;DHX58;TNFSF10;AKT1;PLCE1;MAP3K9;IER3;SLC12A9;PDGFRB;DAPK1;IGFBP3;TAP2;TAP1;MIR27A;HLA-A;ISL1;CLDN4;MAG;DOK2;PGR;CYP2E1;RIN1;ISYNA1;ABCB1;EPHA10;TWIST1;GATA5;GATA4;GSTT1;GATA3;ADRB2;PROM1;TSPAN1;HLA-DQA1;SMAD2;TFAP2A;CREBBP;GSTM1;MLKL;LRBA;TNK2;IGF2;MIR200A;TNFRSF10A;NR0B2;ESR1;DCLK1;ARHGAP24;NET1;ISG20;NR4A2;ATP4A;IL6;NAT1;PPP1R1B;CDK2;CYP1A1;BCL2;GNAS;HLA-DRB1;FGFR2;TP73;FGFR1;HOTAIR;COL18A1;CDKN1A;PSCA;FLT4;HSPB2;MIR18A;MSLN;SLC6A2;CYR61;ESM1;UPK1A;TIMP2;SPEG;CCR7;CLIC1;JAK3;POLE;GPR39;BRD2;MMP7;MIR23A;TEX14;F3;TGFBR2;ZEB2;CCKBR;ELF3;IL1B;LTA;TFF2;BIRC5;TFF1;ANG;HLA-DQB1;GUCY2C;GRP;EGFR;TTN;MUC1;PLAGL1;ERBB2;MIR372;MUC4;MICB;MUC6;CHGA;NQO1;WWOX;CDX2;NOS2;GAD1;NTRK3;STAT3;CCK;MLH1;MTOR;VEGFA;APC;SST;IL2RA;LEP;PTPN6;TRIM31;RAN;BRDT |
| Solid Neoplasm | 0.03511905512417143 | RB1;SLC46A1;ICAM2;ICAM1;TCL1A;RASSF1;HOXA9;SOX17;TRIM29;MYC;STMN1;TNFSF10;AKT1;DLEU2;SOX9;PGLYRP1;SLC12A9;SRRM2;ARL11;SERPINF1;HLA-A;ANK1;RUNX3;HLA-G;SND1;RUNX2;DKK3;RUNX1;CLDN4;SFRP1;SFRP2;BIN1;DDIT3;EZR;GRB7;ZNF154;SLC22A3;ABCB1;EPAS1;PCDH10;CTBP1;TWIST2;TWIST1;GATA2;MIR206;SLC5A5;CYP2B6;TERT;CD19;STC2;PROM1;TSPAN1;ELANE;STAT5A;EGR1;TSLP;IGF2;MRPL28;NFATC2;FLI1;DCLK1;BST2;H19;HAL;FAT1;CYP1A1;NANOG;ANP32B;SIM2;HLA-DRB1;CD200;LGMN;FERMT3;TP73;APOBEC3B;ROBO3;COL18A1;CD40;NRP2;CDKN1A;ROBO4;ELN;ADM;BRCA1;PCSK6;KRT20;MMP21;MECOM;PRX;CCR9;CD38;CCR7;ZHX2;MME;TSC22D1;DUSP1;PAX3;IL16;SERPINB9;MIR21;F3;POU5F1;VAV1;F7;MMP14;CDCP1;TNNT1;IL1B;MMP23A;MIR17HG;RARB;TFF1;CD47;CRYAB;GSTP1;COL11A2;TMPRSS2;EGFR;NT5E;EPCAM;WIF1;SPP1;PRAME;MEF2D;CD55;BRD4;CD74;NQO1;PKLR;MGMT;STAT3;MLH1;PDZK1IP1;PLXDC1;VEGFA;WT1;DLC1;NBR1;CSF2;CSF3R;CSF1;MPG;EPO;MPL;MSI1;PLAT;PRDM2;ETS1;FGF2;TNF;IGF1R;LOXL2;HOXA11;HPGDS;HHAT;CCND1;CDH1;SALL4;MIR150;HLA-DOA;CD177;PDGFRB;PDGFRA;CASR;USP7;POSTN;TPM3;IGFBP3;UGT2B17;MIR27A;CHD1L;HIC1;FOXP1;DHDDS;DOK2;BRMS1;MYCN;COL4A2;CD8A;FSCN1;ERG;ABCG2;CSF1R;NOTCH3;DLX2;ALOX15;ALOX12;HOXB13;FHIT;DPP4;SOCS3;TPO;UBD;NLRP2;MARK2;SMAD2;ZFHX3;TFPI2;TRPV2;MCC;ESR1;ARHGAP24;GFAP;ISG20;MFHAS1;IL6;FES;TLX1;CDK2;GNAS;BCL2;MIR183;FGFR4;FGFR3;FGFR2;FGFR1;MS4A3;KCNK9;FLT4;PIK3CD;TNFAIP3;DDX43;ECE1;CLEC14A;FOXM1;MSLN;PIK3CG;HK1;SPN;ADAMTS4;CASP8;MB;CASP1;UPP1;HSP90AA1;EGFL7;MMP7;VWF;DCC;PDE4D;WIPI2;NGF;PTP4A1;IMPACT;ADORA2A;IFNG;ASPG;CHI3L1;IRF8;BPI;ANG;BIRC7;RBMS3;HDAC2;SRC;FOXK1;SPG7;ITIH1;CXXC5;THBS1;PTHLH;AURKB;CRYZ;CLEC3B;MUC1;DNAJB6;CCL5;MIR373;ERBB2;PTK2B;SNCG;FSD1;EIF4E;MUC4;NTRK1;MARCKSL1;NOS3;NTRK3;BNIP3;CHFR;MTOR;P2RX7;CRYGD;KLF6;MEIS1;APC;SST;IL2RA;CD248;PDCD1;TEK;VIM |
| Intestinal Neoplasms | 0.03530428219849113 | CDKN1A;PIK3CD;FGF2;PIK3CG;HPGDS;CCND1;SOX17;MYC;AKT1;SOX9;PTGDR;SLC12A9;SRRM2;APOBEC1;PDGFRA;MMP7;WNT5A;DNMT3A;GLP2R;PRKCA;UNC5C;CARTPT;PROX1;DCN;TGFBR2;GREM1;TIAM1;NR5A2;ZEB1;IFNG;KCNQ1;PRR15;SGK1;TLR4;RBMS3;GUCY2C;HDAC2;SLC26A1;PRKCZ;NTN1;EGFR;MUC1;MUC2;TERT;ERBB2;SPP1;B4GALNT1;MUC4;TCF7L2;CDX2;GDF15;IGF2;CABLES1;MCC;MLH1;DCLK1;GUCA2A;SMAD7;VEGFA;P2RX5;APC;LEP;CDK2;BCL2;INSM1;IL17C |
| Choriocarcinoma | 0.035497383857133316 | CSF3;RORA;FGF2;TNF;ETS2;RIMS2;EDNRA;FGF7;EDNRB;LGALS1;CCND2;FGF8;GJA5;CYP1B1;PITX2;TP63;PDGFRB;USP7;F2R;KRT8;DIO2;HLA-A;HLA-G;DKK1;HIC1;HLA-E;BRMS1;SLPI;PGR;EZR;RIN2;SQSTM1;CFTR;ABCG2;SLC22A3;CSF1R;ENPEP;CALCA;DLX3;DLX4;TERC;TACSTD2;CUL1;PDGFA;CD1D;ABO;ZFP36L2;SLC5A5;HSD11B2;CYP2B6;TERT;HLCS;NLRP7;TFAP2A;TFPI2;IGF2;LIF;NR2F2;ESR1;H19;DAB2;IL6;NANOG;FGFR2;FGFR1;CDKN1C;CIITA;PPP1R13B;DOC2A;PSCA;HSPB2;PIK3CD;ADM;OXT;HTR2A;PIK3CG;BBC3;CASP8;TIMP2;TIMP3;CD34;CLIC1;SLC2A14;APOBEC2;PEG3;POU5F1;TNFRSF1A;C1QL1;SLC7A5;IMMP1L;PAX8;IFNG;CAT;CRH;ANG;MAPT;HM13;ASCL2;EGFR;ARFRP1;PAK1;MUC1;PDLIM3;PARD6A;EPCAM;HSD17B1;MGAT3;IGF2BP1;STAT3;VEGFA;MARCKS;KRT18;APC;LEP;CD9 |

| | | |
|---|---|---|
| Cardiomyopathies | 0.03560180306240989 | MYLK2;CSF3;FAAH;CCNT1;EPO;EHMT1;AQP2;TNF;BEST1;ICAM1;IGF1R;LGALS3;EDNRA;TCL1A;EDNRB;SLN;CYP2D6;ANPEP;NPPA;AKT1;SCN1B;KCNH1;MAP3K5;KCNH2;MEF2C;CASR;S100A1;ACTN2;DIO2;PRKCA;MIF;SDHB;BIN1;TFR2;COL4A1;MYL2;KCNQ1;PKP1;ALPK3;MASP2;CYP2E1;MYL9;LTBP4;GATA4;ADRB2;PRH1;DPP4;TRPM2;GRK5;KCNQ1OT1;PCBP1;PRDM16;NEU1;MYH14;KCNN3;NKX2-5;APOB;ARSB;TFAP2A;MYH7B;MGAM;ACE;PDE2A;IGF2;TRPV2;EHD1;ISG20;H19;IL6;ARTN;SSPN;GNAS;GNB3;HAMP;CDKN1C;RYR2;ELN;TSSK1B;DYSF;SLC40A1;PRKAG2;PIK3CD;ADM;CYR61;PIK3CG;SYNE2;SYNE1;SGCD;SGCA;CPT2;CHL1;SUMO3;TIMP2;ITGAX;BDKRB2;NOS1;ACADS;CD34;CTSD;SGCG;YWHAH;DSP;UNC45B;CBR1;TRIM63;TNNC1;GAA;MYOF;MST1;SHROOM3;CACYBP;MIR21;TYK2;NGF;DCN;ACTA2;MMP14;MYH2;ADORA2A;PCCA;IFNG;NDUFS7;TRAF3;NDUFS6;HAND2;IL1B;PCCB;HAND1;CAT;BIRC5;ITGA7;NDUFS2;RBM20;CRYAB;MYH7;HDAC2;LRP1;IDUA;HTT;EGFR;PTHLH;TTN;CACNG8;MUC2;NSD1;ERBB2;TBX20;XIRP1;TNNI3;D2HGDH;NCAM1;FLNC;FSD1;XDH;MARCKSL1;ANGPT2;NOS2;NOS3;TXNRD1;STAT3;IRX4;EYA4;TPCN2;TSHR;VEGFA;KISS1;POMC;ACTC1;EIF3K;IL2RA;FKBP1B;SARDH;TCAP;PDCD1;LIMS2 |
| Endemic Cretinism | 0.03595034164787955 | EGR1;NGFR;INHBB;RUNX2;TSHR;TRHR;GHR;TPO;TG;PAX8;BGLAP;NEFL;NEFM;APOE;NEFH |
| Weight decreased | 0.03595034164787955 | ANKLE2;EPO;MPL;ECE1;OXT;PTPN22;IKZF1;BRCA1;IGF1R;EDNRB;JPH3;CCND1;NRTN;GHSR;DAXX;EDN3;SLC11A1;KRT1;MST1;HLA-B;AMPD2;SDHB;FOXP1;TGFBR2;COL2A1;ADORA2A;PALLD;PEX5;CRH;PRTN3;MAPT;ALPK1;RIN2;HLA-DQB1;LTF;NFIX;DCTN1;GATA4;PIK3R1;GATA2;INS;DPP6;ZFP57;HLCS;TMEM127;EPCAM;NPY;PLAGL1;CDH23;SNCA;KCNJ11;STAT3;IGF2;LIF;PDX1;BTNL2;MLH1;GCK;TSHR;POMC;BCL6;GDNF;LEP;CENPJ;BCL2;HLA-DPB1;HLA-DRB1 |
| Meningioma | 0.03676892837331148 | RB1;CSF3;CSF2;SPARC;EPO;CD81;F13A1;DEFB1;PRDM2;FGF2;TNF;ETS1;IGF1R;HOXA10;LGALS3;HOXA9;HPGDS;RASSF1;EDNRB;CCND1;CYP2D6;CDH1;MYC;ANPEP;HOXA3;PART1;AKT1;CYP1B1;HOXA7;GGT1;TP63;TGM2;SRRM2;PDGFRB;TLE3;PDGFRA;DAPK1;S100A1;HLA-A;MIF;PRAP1;RUNX3;SDHB;SFRP4;SFRP1;MYCN;CD8A;S100A6;PGR;EZR;MEG3;CALCA;ABCB1;PDGFB;PDGFA;GSTT1;LTBP2;PLA2G7;CRHR1;KLK6;DPP4;TERT;MSC;S100A10;SMAD2;GSTM1;UBE2I;CADM1;TFPI2;XRCC3;IGF2;MRPL28;MIR200A;ESR1;ARHGAP24;GFAP;BMP4;H19;GJB2;IL6;BCL6;CDK2;BCL2;ALPL;GRB2;FGFR4;FGFR3;HLA-DRB1;EIF4G1;TP73;FGFR1;CDKN1A;PIK3CD;MIA2;LAMC2;BRCA1;MSLN;PIK3CG;CCKAR;CASP8;MB;TIMP2;TIMP3;SOSTDC1;CCR6;NOS1;JAK3;CD34;DSP;GPX1;ITGA3;DCC;APOA1;MIR21;TYK2;F3;CCKBR;KCNMA1;BIRC5;ANG;MCM5;SGK1;EEF1E1;CRYAB;TLR4;HLA-DQB1;RBMS3;GSTP1;PRELP;THBS1;EGFR;PTHLH;PAK1;CBS;ALOX5;ERBB2;SPP1;IGF2BP1;EIF4E;XDH;CD55;WWOX;NOS2;TMEM30B;MGMT;NOS3;STAT3;CCK;SOD3;MTOR;VEGFA;MLPH;MIR145;APC;CD6;WT1;SST;DLC1;CD9;CCM2;PDCD1 |
| Encephalomyelitis | 0.037110535663907186 | CD86;CSF3;SPARC;CSF2;EPO;AQP4;COMT;CLU;TNF;ICAM1;LGALS3;HPGDS;DPYSL2;GRAP2;RNF19A;C3AR1;TNFSF10;LGALS8;TNFRSF4;TP63;MAP3K5;EOMES;ENTPD1;S100A1;HLA-C;CLDN4;IL23A;MAP1B;PADI2;ABCG2;ISYNA1;CD1D;GATA3;MST1R;KLK8;KLK6;C3;TRPM2;HRH1;HRH2;NLRP3;MBP;APOE;FADD;MSC;TRPM4;EGR1;HLA-DRB5;ACE;TGFB3;LIF;MIR338;ESR1;GFAP;ISG20;NR4A2;OCLN;IL6;TH;BHLHE40;HLA-DRB1;CD40;MIR17;LAMC2;SLC6A1;KRT20;GRM1;DNALI1;ELAVL3;GRM5;CASP8;CCR7;CCR6;CD52;DUSP1;MST1;APOA1;TNFRSF1B;RGMA;TNFRSF1A;ZEB1;IFNG;IL1B;CAT;CD47;ARHGEF2;CRYAB;TLR4;CRP;CX3CR1;NR1I2;GOSR1;CAPG;NOD1;NOD2;AIF1;NTN1;PTHLH;TTN;ALOX5;NTF3;SPP1;STAT4;CCL1;SNCB;IL32;CHGA;NTRK1;SEMA4A;MOG;NOS2;SORT1;TMEM176B;MIR26A1;STAT3;ATP2B2;KLK10;IRGM;GCK;DLK1;VEGFA;P2RX7;APC;FABP7;IL2RA;LEP;IL2RB;HLA-DPB1;F2RL1;IL17D;KCNK2 |
| Lung diseases | 0.03734346208651116 | CSF3;CSF2;CTNND2;FAM13A;TNF;MPO;MYLK;IGF1R;HOXA9;FGF7;HPGDS;EDNRA;RASSF1;EDNRB;C3AR1;ADORA1;PDGFRA;SLC11A1;F2R;SARS;TSC2;MIF;RUNX1;SLC9A3;SCNN1G;SLPI;CRISPLD2;KCNQ1;SCNN1B;SCNN1A;PRTN3;EZR;CFTR;CCNO;ABCG1;TERC;SHOX2;GATA6;PDGFA;TWIST1;GSTT1;ADRB2;ABO;CRHR2;LDHC;DPP4;TERT;NLRP3;TPPP;A2M;SPRR2A;ELANE;MARK2;WNT3;STRA6;EGR1;ACD;ACE;GSTM1;TSLP;FOXF1;IGF2R;GADD45G;BMP4;ISG20;BST1;IL6;CYP1A1;BCL2;DYNLRB1;FGF10;DDR1;TRIM72;CDKN1A;ELN;ITGAE;ITGAL;WISP1;CX3CL1;CXCL5;UCN3;BDKRB2;PDE4A;CTSD;GPX1;TRPC6;MMP7;ITGA3;IFNGR2;IL16;MIR21;PARP9;SSTR4;TNFRSF1A;GREM1;GPRC5A;ADORA2A;ADRB3;IFNG;LCK;IL1B;CAT;LTA;TFF2;CHI3L1;BPI;TFF1;TLR4;CRP;RETNLB;HDAC3;GSTP1;NR1I2;SEMA3B;GLRX;EGFR;PTHLH;CCL5;P2RY1;SPP1;IL32;IRX2;NQO1;NOS2;NOS3;IRX3;STAT3;OSM;LILRB3;MUC5B;MTOR;VEGFA;P2RX7;POMC;P2RX5;APC;WT1;P2RX2;IL2RA;P2RX1;FOSB;F2RL1;CD9;SPRY2;VIM;FOXA2 |
| Severe Sepsis | 0.03806656207254084 | CSF3;PGAP3;ADM;TNF;TREM1;PRH1;MYLK;CASP1;APOE;LBP;ACE;ANGPT2;VWF;ANGPT1;HSPA1L;NOS2;NOS3;F2R;WNT5A;MIF;AZU1;HSPA2;TNFRSF1B;F3;IRGM;TNFRSF1A;TLR1;P2RX7;PROCR;PER1;IL6;PROC;PC;ADORA2A;IFNG;IL23A;BCL2;LTA;BPI;TGFBI;IL17D;HAMP;TLR4;CFB;HSPA1B;HSPA1A |
| Pain, Burning | 0.03843364491736768 | CSF3;CSF2;ABCB1;ADM;OXT;TRH;COMT;TNF;AQP1;HRH1;NPPA;PENK;BDKRB2;AKT1;BDKRB1;HCRT;TAC1;DISC1;NTSR1;HTR1A;OPRK1;NRG1;HTR3B;PDYN;POMC;PNOC;CRH |
| NEUROTICISM | 0.03843364491736768 | CNTNAP2;OXTR;LXN;CHRNA5;OXT;DBH;COMT;CLU;SLC6A3;ABO;NPY;PENK;CDH23;NOS1;APOE;DRD4;SPDEF;MAGI1;PDE4D;HTR1A;NRG1;OPRM1;P2RX7;LTA;CDH13;ANKK1;MCPH1 |
| Dental caries | 0.03843555717152528 | PGLYRP2;ITGB4;COX4I2;C4BPA;DEFB1;ADM;LAMC2;TCIRG1;COMT;TNF;FGF3;CXCL5;MMP20;CDH3;DMBT1;CDH1;CA2;RPS6KA2;TRPS1;TIMP2;TNFSF11;ATXN7L1;KIF1A;CCR6;TP63;ACTN2;S100A1;KRT75;RUNX2;PAX1;AJAP1;SFRP4;MMP14;MPPED2;CDH11;IRF5;CARD14;ABCG2;LTF;RTEL1;TERC;TUFT1;LAMA3;KLK4;PRH1;TERT;PROM1;RREB1;CNNM4;S100A10;TBX1;LAMB3;ADHFE1;TRPV3;INHBA;PDZK1IP1;GFAP;GALNS;FERMT1;IL6;KRT17;COL7A1;KRT14;ALPL;FGFR3;CCL28;FGFR2;HLA-DRB1;CRLF1;FGF10 |
| Tonic - clonic seizures | 0.039408760457800666 | ACHE;FAAH;NRP2;CHRM1;ABAT;GRIK1;OXT;TRH;ADRA1B;SLC6A1;CLU;FGF2;SLC6A2;CYP2D6;NOS1;SCN1B;GAMT;GPT2;TSC2;OPRM1;PDYN;NGF;SYN2;DNM1;STX1B;ALDH5A1;ADORA2A;SCN8A;IL1B;KCNQ2;CAT;KCNMA1;CRH;ALDH7A1;PVALB;ABCB1;CHRNA5;FOXG1;CHAT;CACNA1D;INS;NPY;SLC17A7;APOE;LGI1;NGFR;SLC12A5;KCNJ11;NOS2;GABRA5;STAT3;HTR1A;PDX1;HTR1B;OPRK1;CCK;GCK;LETM1;POMC;MAPK10;SLC8A3;IL6;SST;LEP;ALPL;GNB3;CNTN2;GHRL;INHA;PAM;HCN1 |
| Lymphatic Metastasis | 0.03997443786711218 | HOTAIR;NRP2;MIR17;FLT4;FGF1;MSLN;TNF;ICAM1;RASSF2;CDH1;DNMT3B;CCR7;PDGFRB;MMP7;F3;FOXP1;MMP14;IFI27;PGR;ANGPTL4;EZR;CRYAB;CFTR;EGFR;MUC1;MUC2;RAB25;PODXL;STC2;ERBB2;SFN;PROM1;MUC4;MUC6;JAM3;EGR1;ANGPT2;CADM1;ANGPT1;FOXF1;NOS3;STAT3;MLH1;ESR1;DCLK1;TSHR;SMAD7;VEGFA;KISS1;P2RX7;DAB2;IL6;GDNF;ARTN;BCL2;MIR183;PLXNB1;TEK;SHBG;BCAR1 |
| Thyroid Nodule | 0.04024440127978241 | APP;CALCA;GSTP1;QPRT;TMPRSS4;CLU;ETS1;SLC5A5;GHR;DPP4;LGALS3;TPO;TERT;HLTF;TNFRSF8;SFN;NTRK1;KCNJ6;ANGPT2;TPM3;NCOA4;GFRA1;TSHR;VEGFA;IL6;TG;MIR145;PAX8;LEP;BIRC5;GNB3;TNFSF8 |
| Malignant neoplasm of gastrointestinal tract | 0.04024440127978241 | SPARC;TNF;IGF1R;MIR149;RIMS2;HPGDS;DMBT1;CDH1;PLCE1;DAPK1;RUNX3;HIC1;SPINT1;KCNQ1;CYP2E1;PKP3;CFB;ABCB1;GSTT1;ABO;PRH1;FUT4;STK31;CYP2B6;TERT;ALX4;B4GALNT1;B4GALNT2;SPTBN1;ZSCAN18;SMAD2;ST14;GSTM1;NFATC3;IGF2R;NET1;BMP4;IL6;BCL2;CLPTM1L;XAF1;FGFR2;NRP2;MIR17;PSCA;FHL2;HSPB2;KRT20;GHR;SPRED2;ZIC1;ACAD8;CBR1;HSP90AA1;MMP7;DCC;MIR23A;AXIN1;KREMEN2;ZEB2;NR5A2;ZEB1;CCKBR;MIR17HG;TFF2;CHI3L1;DOCK1;CRP;GRP;RGS16;NOD2;EGFR;AURKB;MIR148A;MUC2;WIF1;CBS;ALOX5;GNG7;HSF1;ERBB2;UGT1A7;MUC6;NQO1;MARCKSL1;CDX2;CDX1;GDF15;BNIP3;STAT3;LILRB1;CCK;MLH1;MUC5B;KLK10;VEGFA;P2RX5;MIR145;APC;LEP;PDCD4;CDO1 |
| Attention deficit hyperactivity disorder | 0.04024440127978241 | CNTFR;APP;OXTR;TRIO;CSF2;SNORD115-1;COMT;LGALS3;EDNRA;CYP2D6;CDK20;PART1;AKT1;TRIM26;SLC12A6;MEF2C;SNRPN;LMO3;OSR1;F2R;LIG4;TSC2;MYT1;DHDDS;CACNB2;CLIP2;MTNR1A;MYCN;MAP1B;SCN8A;ADAM12;ASTN2;STX1A;GNE;CES1;MAGEL2;DLX4;TERC;CHRNA6;SLC1A3;GATA4;CACNA1C;KALRN;CPLX1;RAI1;GRIN2A;DPP6;ATXN1;MOBP;TSPAN8;DEAF1;ZBBX;NDN;APOE;TAC1;RREB1;DRD4;DRD5;PTPRN2;AUTS2;HTR1D;HTR1A;HTR1B;DCDC2;BTBD9;GRIN2B;FLI1;DCLK1;NET1;NR4A2;GDNF;PPP1R1B;MKRN3;MIR183;GTF2IRD1;SIM1;ZNF132;CNTNAP2;NRP2;THRB;ELN;PIK3CD;LAMC2;OXT;HTR2A;ITGAE;DBH;SLC6A1;ADRA1B;SLC6A2;ELFN1;SLC6A3;PIK3CG;GRM1;SYNE1;HK1;SPN;ADAMTS2;GRM5;ACOXL;GRM7;GRM8;NOS1;PDE4D;EMP2;IL16;OPRM1;ADRA2C;NGF;SYN3;SORCS2;SSTR4;TGFBR2;COL2A1;ADORA2A;CDH11;CDH13;MCTP2;ITIH3;NLGN1;DDC;COL11A2;PON1;ADCY2;SPG7;INS;PRTG;EPCAM;SLITRK1;NSD1;NPY;NTF3;ERBB2;NCAM1;CSMD2;CACNG2;DISC1;CSMD1;NTRK1;TBX1;KCNIP1;NOS2;NTRK3;TXNRD1;KCNIP4;LIMK1;ZNF804A;FBXL16;BAIAP2;MLH1;POMC;TRIM31 |
| Refractory anemias | 0.04044021117150097 | RB1;CSF3;CSF2;CSF1;EPO;F13A1;MPO;TNF;RIMS2;HPGDS;CCND1;LRAT;HOXA5;SLC12A9;TGM2;F2R;HLA-C;RUNX1;MYCN;IFI27;IL23A;ZFP42;S100A6;HOXB2;PRTN3;CSF1R;CRABP1;FUT4;HSD11B2;TERT;MOBP;STC2;APOE;FADD;WNT2;HOXC6;STRA6;TSLP;ZBTB16;IGF2;NR2F1;ISG15;ESR1;GFAP;ISG20;IL6;CDK6;TH;CDK2;BCL2;CDK10;HOXD9;TDGF1;HAMP;HLA-DRB1;HOXD8;ACHE;CDKN1A;RARG;PIK3CD;PTPN22;TREM1;CX3CL1;PIK3CG;CCR6;CD34;TGIF1;PDE4D;TNFRSF1B;MIR126;ALDH1A3;IFNG;IL1B;ALDH1A2;CAT;RARA;CRH;LTA;RARB;CRP;TMPRSS6;NOD1;GDPD5;EGFR;PAK1;ALOX5;MUC4;CD55;RUNX1T1;TBX1;URGCP;GDF15;NOS3;TBX3;CRYGD;FABP3;NPHS2;COL7A1;IL2RA;FOSB;PTPN6;PKN1;VIM |
| Embryonal Carcinoma | 0.04153931541585241 | APP;SPARC;TRRAP;EPO;FOXM1;FGF2;SOX2;DNMT3L;CCND2;FGF8;CASP8;SOX17;CCND1;DNMT3B;TNFRSF8;TP63;PDGFRA;WNT5A;NME2;TSC2;PAX6;HLA-A;POU5F1;HOXB9;NR5A2;IFNG;IL23A;PLSCR4;ZFP42;HOXB4;HOXB3;RARB;HOXB2;HOXB1;HOXB8;HOXB7;TLR4;HOXB6;HOXB5;SLC22A3;CALCA;GSTP1;CAPG;EGFR;FUT4;SLC5A5;RXRB;BGLAP;PODXL;NLRP7;SPP1;PRAME;OTX2;JAG2;TBX1;HIST1H4K;HIST1H4L;FZD7;LIF;NR2F1;HMGA1;TNFRSF10A;ARHGAP24;BMP4;BMP2;IL6;KLF6;KRT18;TH;BCL2;SPAG1;MIR183;TNFSF8;HIST1H4I;NANOG;VIM;FGFR4;TDGF1;NODAL;CDK15;TP73;HIST1H4F |
| Cardiovascular Diseases | 0.04153931541585241 | APP;GMFG;EHMT1;IFI30;ICAM1;LGALS3;LGALS2;EDNRA;LIPC;EDNRB;TNFSF10;AKT1;TNFSF11;CYP1B1;CHGB;SERPINF1;DAPK3;HLA-B;HLA-C;ANK2;PDZK1;HNF1A;RNASE1;DKK1;DHX40;KCNQ1;EPHA1;HPD;GIPR;TERC;SHMT1;AGAP1;NEDD4L;GATA4;PLA2G7;ABO;HSD11B1;HSD11B2;CYP2B6;APOH;PCBP1;NAALAD2;APOE;APOB;ELANE;ACD;GSTM1;ACE;BCL11B;SLC33A1;IGF2;PRG4;GP6;NR0B2;FLI1;DCLK1;MSRA;H19;BHMT;SLC6A7;VNN1;TH;HAMP;HLA-DRB1;VSX1;RYR1;RYR2;CD40;SLC44A2;ELN;PRKAG2;PCSK9;ADM;BRCA1;PEAR1;MED15;SLC6A2;RYR3;FFAR2;JAK3;CD34;EDN3;DUSP1;TUBB;AMPD1;MIR21;F3;RHOD;F7;MC3R;ADRB3;IL1B;KCNMA1;ANGPTL4;MAPT;GPRC5C;TLR4;RAPSN;TAGLN;NR1I2;KCNA3;HTT;ASCL1;APOL6;ALOX5;NPY;HSF1;SPP1;STAT4;FLNC;APOL1;XDH;P2RY12;CD74;SORT1;GDF15;F12;STAT3;F11;OSM;VEGFA;FABP1;FABP2;FABP3;FOSB;GHRL;PNPLA2;HSPA1A;PF4;F2RL3;CD86;SCARB1;CSF2;CSF1;EPO;PLAT;COMT;MPO;TNF;HPGDS;CCND1;NPPA;CCRL2;PITX2;MAP3K5;KCNH2;IGFBP1;ABCG8;CASR;ABCG5;S100A1;IGFBP3;ADAM11;CYBA;MAT1A;HIC1;COL4A2;HHIPL1;S100A9;ABCG1;NOTCH3;KCNE1;CHRNA5;SHC1;NPR3;ALOX15;HTRA1;GSTT1;ALOX12;ADRB2;SOCS2;DPP4;HRH1;GRK5;NLRP3;KL;RARRES2;TFPI2;TRPV2;ESR1;PHOX2B;HOPX;NR4A2;SLC2A9;IL6;PROC;GDNF;GNB3;SHBG;CRISP2;UCN;ITGB4;ITGB2;DYSF;WISP3;HSPB2;PIK3CD;SLC2A2;LAMC2;PTPN22;HTR2A;DBH;PIK3CG;CXCL5;SPRED2;FLCN;MB;MCF2L;BDKRB2;CASP1;PDE4A;NOS1;DSP;PTGIR;GPX1;VWF;APOA1;NRG1;NGF;WNT16;KLF14;TGFBR2;IFNG;DDAH2;CHDH;CAT;COL6A1;LTA;CHI3L1;CDH13;IRF5;MYH7;CRP;CX3CR1;PON3;HDAC3;PON1;LCAT;NOD1;NOD2;LTC4S;NTN1;THBS1;THBS4;INS;CNR2;CBS;CCL5;TNNI3;CHGA;TCF7L2;ANGPT2;ANGPT1;KCNJ11;NOS2;NOS3;TXNRD1;KCNIP3;IRX4;SORL1;SOD3;POMC;CRYGD;KLF6;MIR145;MIR144;CAPN10;LEP;MGP;APOC1;PDCD4;SPRY2;SPRY1;LPIN1 |
| Anxiety | 0.04205938187591577 | APP;FAAH;OXTR;CTNND2;PKD2L1;COMT;TNF;HS6ST1;RXFP3;IGF1R;CRHBP;FGF8;CCND1;CYP2D6;STMN1;ADORA1;PROK2;PITX2;SOX5;EPHA4;SLC38A1;SLC6A19;CASR;UNC13A;DSCAM;KRT7;TSC2;CARTPT;MYT1;DHDDS;CLIP2;PRKAR1B;CYP2E1;ANKK1;KCTD17;SQSTM1;PVALB;ABCB1;LXN;SLC26A1;CHRNA2;DLX4;PDGFB;SLC1A2;CACNA1A;SLC1A3;ADRB2;CACNA1C;FHIT;CRHR1;ACACA;CRHR2;HSD11B2;TRPM2;RAI1;CYP2B6;GALR3;MIR330;DEAF1;PENK;CDH23;MBP;TAC3;APOE;PRODH;TAC1;DRD4;S100A10;PRPH;TMEM132D;EGR1;ACE;GSTM1;SLC33A1;HTR1A;IGF2;HTR1B;ESR1;FGF17;IL6;DAO;GDNF;PTK7;CYP1A1;GNAS;ALPL;GTF2IRD1;FGFR1;ACHE;LAD1;GPR26;ELN;FLT4;OXT;HTR2A;BRCA1;DBH;SLC6A1;SLC6A2;SLC6A3;GLI2;SGCE;JPH3;CCKAR;UCN3;GRM8;CASP1;PDE4B;LBP;DGCR6;DLGAP3;NEFH;SLC18A2;AVPR1B;COL25A1;TACR3;HTR3A;OPRM1;NGF;DUSP6;C1QL1;TGFBR2;ALDH5A1;ADORA2A;CCKBR;IFNG;TAL1;IL1B;CAT;CRH;ANG;MAPT;RAPGEF3;SHANK1;PON3;DDC;NFIX;GRP;DCTN1;PON1;HTT;TTN;CYP17A1;CAMKK2;EXOSC6;EPCAM;NPY;GNA11;SLITRK5;FYN;EYA1;NTRK3;GAD1;LIMK1;ZNF804A;CCK;MLH1;BAIAP3;KISS1;POMC;FABP3;FABP5;APC;KCNS1;PPP2R2B;FABP7;LEP;SCLY;ALLC |
| TARSAL-CARPAL COALITION SYNDROME | 0.04205938187591577 | CD86;RB1;CDKN1C;HOTAIR;CDKN1A;CSF2;PSCA;DOC2A;KRT20;FGF2;HOXA13;ELAVL3;HPGDS;UPK1B;CCND1;CA2;STMN1;DNMT3B;PRSS8;SLC12A6;LZTS1;SLC12A9;DAPK1;DCC;IGFBP3;DNMT3A;KRT8;HLA-C;PAX5;TYK2;PRAP1;UPK3A;CTAGE1;FEZ1;PRTN3;ABCB1;GSTP1;GATA6;GSTT1;SPG7;FHIT;ABO;EGFR;TERT;KCNQ1OT1;NEU1;ERBB2;BCL2L14;SPAG9;MARCKSL1;GSTM1;MGMT;IGF2;MRPL28;MLH1;ESR1;BLCAP;VEGFA;ISG20;DAB2;IL6;IL2RA;CDK2;BCL2;GNAS;TGFBI;FGFR3;CD200 |
| Generalized glycogen storage disease of infants | 0.042781654419446966 | UGT1A10;MGAM;ACE;G6PC;GAA;FUT1;IGF2R;CPT2;EIF3K;UGT1A5;UGT1A4;UGT1A3;UGT1A9;UGT1A8;ARSB;UGT1A7;UGT1A6 |
| Neonatal hypoglycemia | 0.042781654419446966 | CDKN1C;PROP1;KCNJ11;TCF7;IGF2;PRKAG2;DBH;HNF1A;GCK;GALC;POMC;H19;HNF4A;KCNQ1OT1;NSD1;KCNQ1;ALDH7A1 |
| Kidney Failure, Chronic | 0.04285267873525805 | SCARB1;SLC46A1;CSF2;WIPF1;EPO;CLU;FGF2;TNF;MPO;CNDP1;ICAM1;MIR149;SOX2;LIPC;CYP2D6;ZMIZ1;PLCE1;GGT1;ABCG8;LHCGR;ENTPD1;CPT1A;CASR;POSTN;PRKCB;SLC34A1;HLA-B;HNF1B;CYBA;HLA-A;MIF;SLC9A3;MIR196A2;SOAT2;PAPPA;ELMO1;PRTN3;S100A9;ABCG1;ABCB1;SHC1;SLC22A2;KLK4;GSTT1;ADRB2;ACACA;C3;HSD11B1;HSD11B2;CYP2B6;TERT;CD19;SLC19A3;ST8SIA4;MYH14;APOE;NXPH1;APOB;MSC;RREB1;KL;ACE;GSTM1;TRPV2;BTBD9;BMP7;ESR1;COL1A1;BMP4;ISG20;IL6;ID2;CYP1A1;BCL2;GNB3;HAMP;HLA-DRB1;PCSK2;CD40;ADM;ADRA1D;ADRA1B;AFF3;PKD1;PTER;CPT2;HNF4A;TIMP2;BDKRB2;BDKRB1;NOS1;CBR1;GPX1;TRPC6;VWF;APOA1;MIR21;NGF;GFRA3;F3;PAX2;C1QL1;RGMA;F7;INF2;IFNG;IL1B;CHDH;CAT;CRH;MYH9;LTB;SGK1;MCTP2;TLR4;SDCCAG8;CRP;CX3CR1;DDC;GSTP1;SPESP1;PON1;LCAT;CAPG;PTHLH;MUC1;BGLAP;ALOX5;CCL5;NPY;SPP1;STAT4;SH3BP4;TNNI3;APOL1;KCNJ1;XDH;CHGA;NQO1;NOS2;RNLS;NOS3;BNIP3;LIMK2;B3GAT1;GUCA2B;MTOR;VEGFA;FABP1;POMC;MEIS1;NPHS1;WT1;NPHS2;IL2RA;LEP;MGP;GHRL;LPIN1;KLKB1 |
| Keloid | 0.043652435861064084 | DDR1;COL18A1;ELN;HSPB2;PIK3CD;FGF1;FGF2;PIK3CG;IGF1R;COMP;LGALS1;CASP8;RPS18;AKT1;CLIC1;TP63;ANXA2;IGFBP3;SERPINF1;F2R;NRG1;MIR21;MIF;DCN;SERPINB5;DKK3;SFRP1;MYH2;SFRP2;IL1B;ADAM12;S100A9;ABCG2;HLA-DQB1;TAGLN;COL11A1;CRABP1;NR1I2;PDGFA;CAPG;EGFR;ERBB2;MYH14;PTK2B;SFN;MSC;MARK2;HLA-DQA1;S100A10;SMAD2;CREBBP;SMAD3;ANGPT2;TGFB3;STAT3;ADAM33;INHBA;SMAD7;VEGFA;COL1A1;POMC;IL6;APC;BCL2;F2RL1;TGFBI;VIM;FGFR2;HLA-DRB1;RAN |
| Pancreatic Ductal Adenocarcinoma | 0.043652435861064084 | RB1;FAAH;SPARC;MIR429;PREX2;LGALS4;SOX2;LGALS3;FRAT1;LGALS1;SOX18;TRIM29;MYC;DPYSL3;DHX58;RASSF6;TNFSF10;AKT1;CYP1B1;SOX9;TP63;IER3;SLC12A9;EPHA4;SERPINF1;KRT7;DLEC1;BHLHA15;MIF;HNF1A;DKK1;RUNX3;RUNX2;DKK4;CLDN4;HOXB9;SLPI;MAML3;HOXB7;CFB;CFTR;SKAP2;SLC22A3;EPAS1;GATA6;PIK3R1;MIR206;ABO;SLC5A5;KLK6;CYP2B6;TERT;MIR330;PENK;CHST15;TRPM8;PROM1;JAG2;CREBBP;CADM1;IGF2;NFATC2;NFATC1;DCLK1;SQLE;FXYD3;RGL2;MIR129-2;DDR1;HOTAIR;ROBO3;CDKN1A;ROBO4;CTSZ;BRCA1;CX3CL1;GLI2;BBC3;AKAP12;MECOM;TIMP3;MUC13;HTATIP2;ZHX2;ANXA2;MME;DUSP1;MIR21;F3;SERPINB5;VAV1;ANO1;MMP14;ZEB1;NR5A2;MIR17HG;RARA;TFF1;SLC29A1;TLR4;CSTA;GRP;IDUA;GSTP1;COL11A2;NEDD9;EGFR;PAK1;CUX1;BGLAP;EPCAM;WIF1;GPC1;SPP1;BRD4;CDX2;RNLS;GDF15;GAD1;STAT3;MLH1;KLK10;GNMT;VEGFA;FABP5;WT1;DLC1;FOXA2;CSF2;EPO;IHH;TNF;ETS1;IGF1R;FGF7;HHAT;CDH3;CCND1;CDH1;GRAP2;RNF19A;MIR150;TGM2;KCNH2;MSX2;KDM2B;S100A1;IGFBP3;ADAM11;TAP1;MIR27A;MPEG1;ADAM19;S100A6;FSCN1;DPEP1;ADAM8;NUPR1;SQSTM1;ABCG1;ABCG2;VTCN1;ADRB2;FHIT;SOCS3;PODXL;UBD;SMYD2;NADK;SMAD2;TFAP2A;SMAD3;TFPI2;TNFRSF10A;ISG15;SMAD7;BMP2;IL6;CDK6;FGF19;BCL2;GNAS;CLPTM1L;FGFR4;FGFR3;FGFR2;FGFR1;MIR10B;ITGB4;ONECUT1;FHL2;PIK3CD;FOXM1;MSLN;PIK3CG;GHR;ZIC2;DNMT3B;ITGB6;MMP7;MIR23A;MIR7-3;MIR7-2;FOXL1;OPRM1;IRAK4;COASY;TGFBR2;IFNG;ESRP1;CX3CR1;HDAC2;SRC;CCDC68;THBS1;MIR148A;MUC1;MIR377;ABLIM2;MUC2;CCL5;MIR494;ERBB2;MKNK2;PTK2B;SPOCK1;EIF4E;MUC4;NTRK1;ANGPT2;MYO10;PRRX1;USP22;PDX1;PPP2R3A;TBX4;MTOR;KLF6;PNKD;MIR145;APC;MIR142;APOC1;PDCD4;CD9;VIM |
| Graves Disease | 0.043652435861064084 | APP;HCP5;CD83;HDLBP;AQP4;ETS1;TNF;BACH2;ICAM1;IGF1R;EDNRA;IYD;MYC;AKT1;PROK2;LGALS9;UBASH3A;HLA-DOA;TRIM21;TGM2;HLA-J;CASR;S100A1;HLA-B;DIO2;TAP2;HLA-C;TAP1;HLA-A;MIF;PDZK1;HLA-G;IL23A;CARD14;PHLDB2;NOTCH4;GATA3;ADRB2;LPP;ABO;PSORS1C1;SLC5A5;SOCS3;PTDSS1;TPO;CD19;HLA-DQA2;ST3GAL1;HLA-DQA1;CREBBP;GSTM1;HSPA2;BTNL2;ESR1;MIR346;ISG20;IL6;TG;INS-IGF2;ZFAT;CYP1A1;BCL2;GRB2;SLC26A4;HLA-DRB1;CD40;THRB;ITGB2;ICA1;PIK3CD;TNFAIP3;PTPN22;ITGAE;TRH;CYR61;PIK3CG;CASP8;RAC2;CCHCR1;INSL3;CD34;TCF19;FCRL5;FCRL3;IL16;TNFRSF1B;F3;GNL1;TLR1;FCER2;NCR3;PAX8;IFNG;IL1B;LTA;TNFRSF25;HLA-DQB1;HDAC2;CXCR5;NRXN3;INS;CCL5;SPP1;STAT4;IL21R;NCAM1;MOG;NOS3;GAD1;PSMB8;MTOR;TSHR;PSMB9;VEGFA;JSRP1;SST;QPCT;IL2RA;HLA-DPB1;GHRL;HCG22;PDCD1;IL17D;IL17C;HSPA1B;HSPA1A |
| Carcinoma breast stage IV | 0.043652435861064084 | RB1;APP;ERRFI1;CSF3;CSF2;MPG;PLAT;IFI30;TNF;MPO;IGF1R;LOXL2;HPGDS;RASSF1;CCND1;CYP2D6;CDH1;MYC;CREB3L1;GRAP2;RNF19A;RPS6KA2;CAPN2;AKT1;SLC16A3;MAP3K5;TGM2;MSX2;KRT1;RUNX2;RUNX1;DHDDS;BRMS1;FSCN1;PGR;CSF1R;SLC22A3;ABCB1;PFKFB3;TWIST1;KLK4;CD1D;KLK6;CYP2B6;NEU1;SFN;PROM1;JAG2;SMAD2;TFAP2A;SMAD3;CBX5;XRCC3;HMGA1;MIR200B;TMPRSS13;BMP7;ESR1;VANGL1;BST2;NET1;BMP2;IL6;CDK6;BCL6;BCL2;CSPG4;FGFR2;EIF4G1;FGFR1;HOTAIR;PTPRU;COL18A1;MIR10B;HSPB2;PIK3CD;ADM;LAMC2;BRCA1;FOXM1;HOXC10;CYR61;PIK3CG;PTPRF;GLI2;HK1;AKAP13;MECOM;TIMP3;CCR7;HSP90AA1;BUD31;MST1;MIR21;PROX1;TMEM173;DCN;TGFBR2;KLF17;ALDH1A3;IFNG;KCNMA1;MIR17HG;TFF3;RARB;BIRC5;ITGA6;MAPT;SLC29A1;TLR4;HDAC3;SRC;RAB1B;COL11A2;NR1I2;KCNA3;NTN1;EGFR;NT5E;MUC1;EPCAM;HSD17B1;NNAT;ERBB2;GCNT2;SPP1;EIF4E;SASH1;BRD4;SPDEF;NTRK1;ANGPT2;STAT3;LILRB1;MTOR;PSMB9;VEGFA;KISS1;NLRP12;MIR145;LEP;F2RL1;VIM;BCAR1;HSPA1A |
| Diabetes Mellitus, Ketosis-Prone | 0.043652435861064084 | CHRM2;CRP;KCNJ11;NOS3;SLC11A1;STAT3;IGF2;PAX4;PTPN22;HNF1A;TNF;BACH2;INS;HSD11B2;IFNG;DDIT3;IL2RA;CAT;PRKCQ;NOS1;HLA-DQA1;HLA-DRB1;FGF21;HLA-DQB1 |
| MICROPHTHALMIA, ISOLATED 1 | 0.04399904573949042 | SOX2;ALDH1A3;SIX6;RAX;OTX2;GDF3;VSX2 |
| Pancreatic intraepithelial neoplasia | 0.04467352360309664 | CDKN1A;CSF2;SCT;ONECUT1;PIK3CD;MSI2;KRT20;PIK3CG;CX3CL1;SOX2;CCND1;TRIM29;SLC12A9;KCNH2;ANXA2;MIR21;BHLHA15;RUNX2;DUSP6;ZEB1;FSCN1;CLDN18;HOXB2;TFF1;HOXB6;CX3CR1;EPAS1;GATA6;MST1R;EGFR;MIR148A;SOCS3;MUC1;BGLAP;ALOX5;PENK;ERBB2;MUC4;MUC6;CDX2;STAT3;PDX1;NFATC2;CCK;SQLE;IL6;FXYD3;FGF19;BCL2;FGFR4 |

| | | |
|---|---|---|
| Brain Neoplasms | 0.04539403658721485 | RB1;SPARC;TNFRSF6B;TRRAP;AQP4;ADARB1;ADARB2;ICAM1;SOX2;LGALS3;RASSF1;LGALS1;CYP2D6;MYC;TNFSF10;AKT1;SOX9;SLC12A9;DAPK1;CACNA2D2;EBF3;HLA-A;KRT5;MIF;DKK1;HLA-G;MAPK8IP3;THEG;IFI27;MYL2;CCNO;RTEL1;ABCB1;TTYH1;EPAS1;PDGFB;GATA4;PIK3R1;SULT1A1;CYP2B6;TERT;PDGFD;APOE;FADD;PROM1;TSPAN1;STAT5A;EGR1;GSTM1;ABCA4;XRCC3;IGF2;LIF;MRPL28;BST2;NET1;TF;CYP1A1;L1TD1;HRG;HOXD9;PAM;SIM2;TP73;DDR1;PTPRT;ACHE;COL18A1;CDKN1A;GFM1;LRRC4;ADM;BRCA1;KRT20;SLC6A2;TIMP2;TK1;SLC2A14;ZHX2;ANXA2;TSC22D1;GFRA1;MIR21;PAX5;MMP14;ZEB1;IL1B;MIR17HG;MAPT;CRYAB;CSTA;GSTP1;MIR520G;KCNA3;EGFR;WIF1;ALOX5;NPY;NTF3;SPP1;NQO1;MGMT;GDF15;STAT3;OSM;MLH1;VEGFA;FABP3;QPCT;TRIP10;PLVAP;CSF2;EPO;MSI1;CLU;TNF;ETS1;HPGDS;CCND2;CCND1;DMBT1;CDH1;TGM2;PDGFRB;GHSR;PDGFRA;CASR;G6PC;WNT5A;DIO2;MATK;LIG4;TSC2;DHDDS;MYCN;PGR;ABCG2;GSTT1;FHIT;FUT4;SOCS3;ATXN1;MIR296;MSC;SMAD3;GFPT1;TFPI2;EPHX1;DNAJC15;ESR1;IGF2R;ARHGAP24;GFAP;BMP4;IL6;BCL6;GDNF;CDK2;BCL2;MXD3;FGFR1;MXD4;ONECUT2;ITGB2;PIK3CD;ADAR;ITGAE;NR2E1;FOXM1;CYR61;PIK3CG;CASP8;PSD4;ZIC1;DNMT3B;PDE4A;POLE;DCC;NGB;OLIG2;NGF;DCN;TGFBR2;ARL4D;IFNG;CAT;CDH11;IRF7;CHI3L1;SRC;LAMA4;PON1;SPG7;CCL5;P2RY1;ERBB2;GNA12;MXI1;IGF2BP1;SPOCK1;NCAM1;OPCML;SNCA;NTRK1;LGI1;TCF7L2;SLC12A5;ANGPT1;NOS2;NTRK3;TXNRD1;CCK;SOD3;MTOR;DLK1;P2RX7;KLF6;P2RX5;NIN;MIR145;APC;SST;P2RX2;LEP;P2RX1;CD248;TEK;VIM |
| Osteopetrosis | 0.04550802838098503 | CD40;CSF1;SRC;MRGPRF;LRP5;TCIRG1;TNF;CLCN1;BEST1;GHR;PTDSS1;ANKH;CA2;BGLAP;MYC;ACP5;TNFSF11;CCDC154;ASCC1;MITF;HLA-A;GTF2H4;VEGFA;CLCN7;ADCY10;BCL2;CLC;ARHGEF7;FGFR4;ATP6V1B1;FERMT3 |
| Inflammation | 0.0465950373303628 | APP;CSF2;CD83;CSF1;RASL11B;EPO;AQP8;AQP4;PLAT;FGF1;CLU;FGF2;TNF;MPO;ICAM1;FGF7;DMBT1;CDH1;C3AR1;TNFSF10;AKT1;TGM2;PDGFRB;SERPINF1;F2R;TSC2;MIF;HNF1A;SULT2B1;PROCR;BIN1;IL23A;SERPING1;PGR;PRTN3;MASP2;S100A9;CFTR;CALCA;ABCB1;MAZ;PDGFB;PDGFA;FPR2;GATA3;CACNA1C;CRHR2;C3;SOCS3;HRH1;SCUBE1;NLRP3;APOD;TAC1;ELANE;EGR1;SMAD3;TSLP;TNFSF14;TREX1;TFPI2;LIF;SMAD7;BMP4;BST1;TF;IL6;PROC;NAT1;ALPL;HAMP;CD200;TP73;UCN;CDKN1A;CD40;KCNK9;TNFRSF13B;SETD7;TNFAIP3;ADM;ADAR;OXT;ITGAL;CD3E;TREM1;CX3CL1;LTB4R;UCN3;BDKRB2;CASP1;ACP5;CD38;BDKRB1;CCR7;MADCAM1;CCR6;TIMP4;YWHAH;HSP90AA1;VWF;MME;PLA2G4D;TNFRSF18;APOA1;NRG1;NGF;CYP7B1;TNFRSF1A;F7;COL2A1;ADRB3;ELF3;IFNG;IL1B;CAT;CRH;IRF7;BIRC5;CHI3L1;ANG;BPI;CD47;CRYAB;TLR4;LTF;CRP;PON1;NOD1;NOD2;NTN1;THBS1;MUC1;CNR2;CCL5;NTF3;SPP1;CCL1;NCAM1;IL32;ANGPT2;ANGPT1;NOS2;NOS3;F11;STAT3;LIMK1;LILRB1;CCK;VEGFA;GALNS;POMC;LEP;F2RL1;GHRL;PF4;CCL26 |
| Head and Neck Neoplasms | 0.046723940564495546 | CSF3;GRP;EPO;EPAS1;GSTP1;TCF21;FURIN;ICAM5;BRCA1;TNF;EGFR;IGF1R;MRPL11;BBC3;CCND1;ERBB2;CYP1B1;PRAME;UPP1;EIF4E;STAT5A;GSTM1;GPX1;S100A2;NOS2;IGFBP3;XRCC3;STAT3;MLH1;VEGFA;MINPP1;GJB2;IL6;CD4;DDIT3;GJB6;IL2RA;BCL2;CYP1A1;FAT1;GNAS;RARB;VIM;FGFR4;FGFR2;FGFR1;APOBEC3B |
| Sarcoma, Spindle Cell | 0.046723940564495546 | NTRK1;CSF2;NOS2;DHH;GSTP1;DERL3;NTRK3;PAX3;TNF;GFAP;VEGFA;SOX2;IL6;IFNG;CDH1;MOV10L1;PRAME;SOX9;MAML3 |
| MAJOR AFFECTIVE DISORDER 4 | 0.046723940564495546 | DDC;NOTCH4;MLC1;HTR2A;COMT;PRKCZ;SLC6A3;MCHR2;DAOA;CUX2;PIP4K2A;NCAM1;KCNN3;DRD4;DRD5;MAGI1;BRD1;GAD1;MAGI2;ZNF804A;HLA-C;HTR3A;HTR3B;DCLK1;SSTR5;P2RX7;IMPACT;DAO;TH;KCNQ2;CRH;BPI;GPRC5C;ABCG1 |
| MAJOR AFFECTIVE DISORDER 6 | 0.046723940564495546 | DDC;NOTCH4;MLC1;HTR2A;COMT;PRKCZ;SLC6A3;MCHR2;DAOA;CUX2;PIP4K2A;NCAM1;KCNN3;DRD4;DRD5;MAGI1;BRD1;GAD1;MAGI2;ZNF804A;HLA-C;HTR3A;HTR3B;DCLK1;SSTR5;P2RX7;IMPACT;DAO;TH;KCNQ2;CRH;BPI;GPRC5C;ABCG1 |
| Post-Traumatic Stress Disorder | 0.048880626767672776 | CRP;FAAH;OXTR;VIPR2;FLT4;ANKRD55;ALOX15;OPRL1;PCSK9;RORA;PKD2L1;ALOX12;OXT;ADRB2;BRCA1;DBH;COMT;TNF;CRHR1;SLC6A3;ICAM1;CRHR2;EXOSC6;ADAMTS2;NPY;STMN1;APOE;NOS1;SLC18A2;S100A10;EGR1;HSP90AA1;ACE;SRD5A2;HTR1A;PRKCA;OPRM1;CPT1B;ESR1;P2RX7;IL6;KLF6;IFNG;IL1B;NOS1AP;BCL2;SGK1;THOP1 |
| Endometrial Carcinoma | 0.0493497815950266 | RB1;ERRFI1;SPARC;AQP2;RXFP3;AQP1;SOX2;RASSF1;EDNRB;SOX1;MYC;STMN1;TNFSF10;AKT1;TNFSF11;CYP1B1;TRIM27;SOX9;SOX7;SLC12A9;ENTPD1;SERPINF1;SERPINF2;RPL22;CIRBP;HNF1B;PRKCA;PRAP1;RUNX3;DKK3;RUNX1;SULT2B1;CLDN4;IFI27;SLPI;DDIT3;ARMC3;RBP1;EZR;SLC22A3;ABCB1;LXN;CTBP2;PCDH10;CTBP1;TERC;TWIST1;KLK4;GATA4;PIK3R1;KLK8;SULT1A1;CYP2B6;TERT;PCBP4;PDGFD;NEU1;APOE;HCRT;PROM1;BBS2;BARD1;EGR1;CREBBP;STARD13;GSTM1;CADM1;IGF2;CIDEA;MRPL28;MIR200B;EIF2AK4;NR0B2;BST2;STAR;ARTN;GNRH2;CYP1A1;TP73;MIR129-2;DDR1;CDKN1C;HOTAIR;PIGR;CDKN1A;MIR130B;ADM;BRCA1;SERPINA6;KRT20;MED15;JPH4;MECOM;HYAL1;HNF4A;HYAL2;TIMP2;TIMP3;CCR7;CTSD;BRD2;ZHX2;ANXA2;MST1;EMP2;DUSP6;POU5F1;PAX2;TLR1;MMP14;GPRC5A;ZEB1;PAX8;IL1B;KCNMA1;TFF3;RARB;SGK1;GAS5;GRP;GSTP1;SATB2;COL11A2;NR1I2;FSTL1;ASCL2;EGFR;CYP17A1;PAK1;MIR410;NT5E;CUX1;EPCAM;HSD17B1;HSD17B2;HSF1;SPP1;IL32;WWOX;CDX2;MGMT;MIR26A1;STAT3;MLH1;VEGFA;PER1;NBR1;PKN1;LGR6;INHA;F2RL3;CSF2;CSF1;DGKA;MSI1;PLAT;PRDM2;FGF1;COMT;ETS1;CLU;FGF2;TNF;FGF3;IGF1R;HOXA11;HOXA10;HPGDS;FGF7;CCND1;CDH1;SALL4;MACROD1;GGT1;KCNH2;PDGFRB;IGFBP1;GHSR;LHCGR;IGFBP3;F2R;UGT2B17;SDHB;RAB32;S100A6;PGR;NUPR1;SQSTM1;CSF1R;NOTCH4;HTRA1;GSTT1;HOXB13;FHIT;LPP;RAP1GAP;ZFP36L1;SOCS2;SOCS3;PLXNA1;WNT2;ESRRA;SMAD2;ZFHX3;TFAP2C;SRD5A2;TGFB3;TNK2;TNFRSF10C;HTR1B;INHBB;ESR1;IGF2R;ARHGAP24;SMAD7;IL6;DAB2;EFNA2;CDK2;BCL2;MIR183;SHBG;FGFR4;FGFR2;FGF10;PIWIL2;CRISP2;UCN;ST6GALNAC1;FLT4;PLEK;PIK3CD;PIK3CG;CYR61;CASP8;ZIC2;DNMT3B;AMH;POLE;HSP90AA1;MMP7;ITGA4;DCC;AXIN1;DCN;TGFBR2;ELF1;GORASP2;HAND2;CAT;CRH;LTA;BIRC5;CDH13;MIR124-2;ANG;ZMYND10;OTUB1;LTF;CRP;LRP1;SRC;UHRF1;SEMA3B;NTN1;THBS1;AURKB;MIR148A;ERBB2;MIR372;SNCG;FSD1;EIF4E;MUC4;TCF7L2;MARCKSL1;KDM4B;ANGPT2;NOS3;CABLES1;CHFR;MTOR;KISS1;MIR145;APC;SST;LEP;CD9;PTPN6;VIM |
| Neonatal diabetes mellitus | 0.049464287531245286 | KLF11;KCNJ11;STAT3;GATA6;KCNA3;PDX1;PAX4;SLC2A2;GATA4;HNF1B;HNF1A;FOXM1;GCK;INS;ZFP57;INS-IGF2;HNF4A;PLAGL1;MNX1;ZNF300;NEUROG3 |

| REACTOME PATHWAY | Adjusted P-value | Genes |
|---|---|---|
| Neuronal System R-HSA-112316 | 1.1477607511141338E-11 | KCNG1;KCNG2;KCNG4;ABAT;COMT;RIMS1;RPS6KA2;PPFIA1;PPFIA4;AP2M1;KCNH1;KCNH2;KCNH3;SLC38A1;KCNK10;PRKCB;ACTN2;SLC6A13;CACNA2D2;SLC6A12;PRKCA;KCNK16;SLC6A11;KCNK17;KCNK18;GABRG3;RNASE1;CACNB1;CACNB2;ADCY9;CACNB4;PRKAR1B;KCNQ1;KCNQ2;KCNQ4;STX1A;CHRNA1;RTN3;CHRNA2;CHRNA5;SLC22A2;CHRNA6;CACNA1B;SLC1A2;CACNA1A;SLC1A3;LIN7C;SLC1A6;CACNA1E;CPLX1;SLC1A7;KCNV1;GRIN2A;EPB41L2;UBD;KCNMB3;KCNN2;KCNN3;SLC38A2;GRIA4;GNG12;GRIN2B;SYT7;GRIN2D;GRIN3B;PPFIBP2;LRFN4;GNAL;NBEA;GNB4;GNB3;ACHE;GABRB2;PTPRS;KCNC1;KCNK9;KCNC2;GRIK5;GRIK3;PRKAG2;GRIK4;GRIK1;SLC6A1;AP2A2;KIF17;PRKAG3;SLC6A3;PTPRF;GRM1;PANX2;GRM5;GNGT2;LRRTM3;LRRTM1;LRRTM2;FLOT1;NEFL;DLGAP1;DLGAP3;DLGAP2;SLC18A2;DLGAP4;SLC18A3;KCND2;SLC32A1;KCND3;HTR3E;SH2D3C;HTR3A;KCNAB1;NRG1;HTR3B;KCNAB2;KCNAB3;SYN3;SYN2;GJD2;GJD3;ALDH5A1;DNAJC5;KCNMA1;MAPT;ARHGEF7;PLCB2;SHANK2;SHANK1;CAMK2B;NLGN1;SRC;KCNA1;NRXN1;ADCY4;KCNA3;NRXN3;KCNA4;NRXN2;ADCY2;KCNA5;ADCY1;KCNA6;ADCY7;CAMKK1;NCALD;BEGAIN;CAMKK2;GNAI2;CHRND;GLRA1;CACNG8;GNG3;GNG2;SLITRK1;GNG4;CHRNE;GNG7;SLITRK3;SLITRK5;SLC17A7;CACNG2;CACNG3;KCNJ3;GABRA1;MPP2;KCNJ6;MPP3;KCNJ11;KCNJ8;GABRA5;KCNB1;KCNJ12;KCNJ9;GABRA4;KCNB2;NTRK3;GAD1;KCNJ14;KCNJ15;KCNJ16;GLRB;KCNS1;KCNS2;KCNS3;KCNK2;APBA2;HCN2;KCNK4;HCN1 |
| Potassium Channels R-HSA-1296071 | 1.0031574335546611E-5 | KCNG1;KCNG2;KCNG4;KCNC1;KCNK9;KCNC2;GNGT2;KCNH1;KCNH2;KCNH3;KCNK10;KCND2;KCND3;KCNK16;KCNAB1;KCNK17;KCNAB2;KCNK18;KCNAB3;KCNQ1;KCNQ2;KCNMA1;KCNQ4;KCNA1;KCNA3;KCNA4;KCNA5;KCNA6;KCNV1;GNG3;GNG2;GNG4;UBD;GNG7;KCNMB3;KCNN2;KCNN3;KCNJ1;KCNJ6;KCNJ11;KCNJ8;KCNB1;KCNJ12;KCNJ9;KCNB2;KCNJ14;KCNJ15;KCNJ16;GNG12;KCNS1;KCNS2;KCNS3;GNB4;GNB3;KCNK2;HCN2;HCN1;KCNK4 |
| Transmission Across Chemical Synapses R-HSA-1 | 1.4063479864932964E-4 | ABAT;COMT;RIMS1;RPS6KA2;PPFIA1;PPFIA4;AP2M1;SLC38A1;ACTN2;PRKCB;SLC6A13;CACNA2D2;SLC6A12;PRKCA;SLC6A11;GABRG3;RNASE1;CACNB1;CACNB2;ADCY9;CACNB4;PRKAR1B;STX1A;CHRNA1;CHRNA2;CHRNA5;SLC22A2;CHRNA6;CACNA1B;SLC1A2;CACNA1A;SLC1A3;LIN7C;SLC1A6;CACNA1E;CPLX1;SLC1A7;GRIN2A;UBD;SLC38A2;GRIA4;GNG12;GRIN2B;GRIN2D;GRIN3B;GNAL;NBEA;GNB4;GNB3;ACHE;GABRB2;GRIK5;GRIK3;GRIK4;PRKAG2;GRIK1;SLC6A1;AP2A2;KIF17;PRKAG3;SLC6A3;GNGT2;NEFL;SLC18A2;SLC18A3;SLC32A1;HTR3E;SH2D3C;HTR3A;NRG1;HTR3B;SYN3;SYN2;ALDH5A1;DNAJC5;MAPT;ARHGEF7;PLCB2;CAMK2B;SRC;ADCY4;ADCY2;ADCY1;ADCY7;CAMKK1;NCALD;CAMKK2;GNAI2;CHRND;GLRA1;CACNG8;GNG3;GNG2;GNG4;CHRNE;GNG7;SLC17A7;CACNG2;CACNG3;GABRA1;MPP2;MPP3;KCNJ6;GABRA5;KCNJ12;KCNJ9;GABRA4;GAD1;KCNJ15;KCNJ16;GLRB |
| Muscle Contraction R-HSA-397014 | 2.707602183513861E-4 | RYR1;RYR2;TRIM72;KCNK9;ATP2A3;DYSF;ATP2A1;RYR3;ACTG2;MYLK;SLC8A2;SLN;NPPA;SCN1B;KCNH2;NANOS1;TMOD1;MYBPC1;KCNK10;ANXA2;KCND2;TPM4;ACTN2;KCND3;TPM3;TNNC2;CACNA2D2;KCNK16;PRKCA;KCNK17;KCNK18;CACNB1;ACTA2;MYL5;CACNB2;TNNT1;KCNQ1;MYL2;SCN8A;MYL3;PXDN;TNNT3;SCN4A;MYL9;SCN4B;CES1;CAMK2B;KCNE1;KCNE3;KCNE4;ITPR1;ITPR2;GATA4;KCNA5;CACNA1C;CACNA1H;CACNA1G;CACNG6;CACNG7;CACNG8;CALD1;MYL10;TNNI3;NKX2-5;WWTR1;KCNIP1;KCNJ11;KCNIP2;KCNJ12;KCNIP3;KCNIP4;KCNJ14;ATP2B4;SORBS1;TBX5;SORBS3;HIPK2;SLC8A3;FGF14;FXYD3;FXYD1;FKBP1B;FXYD7;TCAP;RANGRF;PKN1;VIM;FGF12;KCNK2;FGF11;KCNK4 |
| Cardiac Conduction R-HSA-5576891 | 5.255559262728111E-4 | RYR1;RYR2;KCNK9;ATP2A3;ATP2A1;RYR3;SLC8A2;SLN;NPPA;SCN1B;KCNH2;NANOS1;KCNK10;KCND2;KCND3;CACNA2D2;KCNK16;PRKCA;KCNK17;KCNK18;CACNB1;CACNB2;KCNQ1;SCN8A;SCN4A;SCN4B;CES1;CAMK2B;KCNE1;KCNE3;KCNE4;ITPR1;ITPR2;GATA4;KCNA5;CACNA1C;CACNG6;CACNG7;CACNG8;TNNI3;NKX2-5;WWTR1;KCNIP1;KCNJ11;KCNIP2;KCNJ12;KCNIP3;KCNIP4;KCNJ14;ATP2B4;TBX5;HIPK2;SLC8A3;FGF14;FXYD3;FXYD1;FKBP1B;FXYD7;RANGRF;FGF12;KCNK2;FGF11;KCNK4 |
| GPCR Downstream Signaling R-HSA-388396 | 8.097543615676357E-4 | NPFFR2;APP;OXTR;TRIO;VIPR2;TAS2R20;SCT;RXFP4;HTR4;RXFP3;EDNRA;EDNRB;RPS6KA2;C3AR1;AKT1;PROK2;PTGDR;PRKCH;PRKCB;PRKCA;GPR37L1;PDYN;TRHR;TIAM2;TIAM1;ADCY9;PRKAR1B;PRKCQ;NGEF;PTGFR;PDE1C;GIPR;PDE1B;PDE1A;FPR2;PIK3R1;KALRN;CRHR1;PIK3R6;CRHR2;PIK3R5;C3;ABR;GALR3;GALR1;NPBWR1;NPBWR2;PENK;PCDHA4;HCRT;NTSR1;PCDHA6;NTSR2;PDE2A;NMBR;GNG12;LPAR5;PNOC;PPP1R1B;GNRH2;GRB2;GPR27;TAS2R60;GPR25;ADM;CX3CL1;AKAP13;GPR132;C1QTNF1;CCKAR;GNGT2;CCR9;CCR7;FFAR2;CCR6;FFAR1;GPR39;GPR37;AVPR1B;EDN3;TACR2;TACR3;VAV1;VAV2;MC3R;ADRB3;CCKBR;ARHGEF4;ARHGEF2;ARHGEF7;PLCB2;ADCY4;CXCR5;ADCY2;ADCY1;ADCY7;EGFR;CAMKK1;CAMKK2;GNAI2;FGD4;NPY;CCL25;P2RY12;CCL23;P2RY14;SCTR;PPP1CA;F2RL1;GHRL;PKN1;CCL28;RAMP1;PF4;F2RL3;CHRM2;DGKG;CHRM1;DGKD;DGKA;RGS4;RGS5;ADORA3;ADORA1;GHSR;LHCGR;CASR;F2R;DGKZ;MTNR1A;OBSCN;LTB4R2;MTNR1B;DGKI;CALCA;CALCB;SHC1;ITPR1;ITPR2;ADRB2;RASGRP2;MCHR2;GRK5;HRH2;UBD;NLRP3;TAC3;TAC1;DRD4;DRD5;HTR1E;HTR1D;HTR1B;OPN5;PYY;GNAL;NBEA;GNAS;GNB4;GNB3;PLXNB1;NMUR1;GPSM1;DAGLA;ADRA1D;ARHGEF10L;OXT;HTR2A;TRH;ADRA1B;PIK3CG;CXCL5;GRM1;LTB4R;GRM2;GRM5;GRM4;GRM7;GRM6;MCF2L;GRM8;BDKRB2;PDE4B;BDKRB1;PDE4A;INSL3;QRFPR;GPSM3;TRPC7;ARHGEF10;PTGIR;TRPC6;TRPC3;PLEKHG5;ARHGEF17;PDE4D;ARHGEF16;GLP2R;PDE4C;ARHGEF19;OPRM1;SSTR1;ADRA2C;SSTR4;SSTR5;ADORA2A;CRH;CX3CR1;CAMK2B;RGS14;SRC;RGS16;OPRL1;PTHLH;GNG3;GNG2;GNG4;CCL5;GNA11;GNG7;P2RY1;GNA12;CCL1;RGS22;TAS1R1;S1PR5;S1PR4;TAS1R3;CHGA;OPRK1;HTR5A;TSHR;KISS1;POMC;SST;RGS10;RGS12 |
| Signaling By GPCR R-HSA-372790 | 8.632885009241721E-4 | NPFFR2;APP;OXTR;TRIO;VIPR2;TAS2R20;SCT;RXFP4;HTR4;RXFP3;CRHBP;EDNRA;EDNRB;RPS6KA2;C3AR1;AKT1;PROK2;CMKLR1;PTGDR;PRKCH;PRKCB;PRKCA;GPR37L1;PDYN;TRHR;TIAM2;TIAM1;ADCY9;PRKAR1B;PRKCQ;NGEF;PTGFR;PDE1C;DHH;GIPR;PDE1B;PDE1A;FPR2;PIK3R1;KALRN;CRHR1;PIK3R6;CRHR2;PIK3R5;C3;ABR;GALR3;GALR1;NPBWR1;NPBWR2;PENK;PCDHA4;HCRT;NTSR1;PCDHA6;NTSR2;FZD7;FZD6;PDE2A;NMBR;GNG12;LPAR5;PNOC;PPP1R1B;GNRH2;GRB2;GPR27;TAS2R60;GPR25;ADM;CX3CL1;AKAP13;GPR132;C1QTNF1;CCKAR;GNGT2;UCN3;UCN2;CCR9;CCR7;FFAR2;CCR6;FFAR1;GPR39;GPR37;AVPR1B;EDN3;TACR2;TACR3;VAV1;VAV2;MC3R;ADRB3;CCKBR;ARHGEF4;ARHGEF2;ARHGEF7;PLCB2;ADCY4;CXCR5;ADCY2;ADCY1;ADCY7;EGFR;CAMKK1;CAMKK2;GNAI2;FGD4;NPY;CD55;CCL25;P2RY12;CCL23;P2RY14;WNT7B;SCTR;PPP1CA;F2RL1;GHRL;PKN1;CCL28;RAMP1;PF4;F2RL3;CHRM2;DGKG;CHRM1;DGKD;DGKA;IHH;RGS4;RGS5;ADORA3;CCRL2;ADORA1;GHSR;LHCGR;CASR;F2R;WNT5A;DGKZ;MTNR1A;OBSCN;LTB4R2;MTNR1B;DGKI;CALCA;CALCB;SHC1;ITPR1;ITPR2;ADRB2;RASGRP2;MCHR2;WNT6;GRK5;HRH2;UBD;NLRP3;TAC3;TAC1;WNT2;DRD4;WNT3;DRD5;HTR1E;WNT3A;HTR1D;PTCH2;HTR1A;HTR1B;OPN5;PYY;GNAL;NBEA;GNAS;GNB4;GNB3;PLXNB1;NMUR1;GPSM1;UCN;DAGLA;ADRA1D;ECE1;ARHGEF10L;OXT;HTR2A;TRH;ADRA1B;PIK3CG;CXCL5;GRM1;LTB4R;GRM2;GRM5;GRM4;GRM7;GRM6;MCF2L;GRM8;BDKRB2;PDE4B;PRLHR;BDKRB1;PDE4A;INSL3;QRFPR;GPSM3;TRPC7;ARHGEF10;PTGIR;TRPC6;TRPC3;PLEKHG5;ARHGEF17;PDE4D;ARHGEF16;GLP2R;PDE4C;ARHGEF19;OPRM1;SSTR1;ADRA2C;WNT16;SSTR4;SSTR5;ADORA2A;CRH;CX3CR1;CAMK2B;RGS14;SRC;RGS16;OPRL1;PTHLH;GNG3;GNG2;WNT11;GNG4;CCL5;GNA11;GNG7;P2RY1;GNA12;CCL1;RGS22;TAS1R1;S1PR5;S1PR4;TAS1R3;CHGA;OPRK1;HTR5A;TSHR;KISS1;POMC;SST;RGS10;RGS12 |
| Transcriptional Regulation Of Pluripotent Stem Ce | 0.0014084602150288927 | EOMES;SMAD2;CDX2;TSC22D1;EPAS1;STAT3;GATA6;PRDM14;HIF3A;DKK1;PBX1;FOXP1;SOX2;HHEX;SALL1;GSC;SALL4;ZSCAN10;NANOG;EPHA1;TDGF1 |
| Voltage Gated Potassium Channels R-HSA-12960 | 0.0014762381102235633 | KCNG1;KCNG2;KCNG4;KCNC1;KCNC2;KCNA1;KCNA3;KCNA4;KCNA5;KCNA6;KCNV1;KCNH1;KCNH2;KCNH3;KCND2;KCNB1;KCND3;KCNB2;KCNAB1;KCNAB2;KCNAB3;KCNS1;KCNS2;KCNQ1;KCNS3;KCNQ2;KCNQ4 |
| POU5F1 (OCT4), SOX2, NANOG Repress Genes R | 0.003260122326799425 | EOMES;SOX2;HHEX;CDX2;TSC22D1;GSC;GATA6;NANOG;DKK1 |
| GPCR Ligand Binding R-HSA-500792 | 0.014392676853671666 | CHRM2;NPFFR2;APP;OXTR;CHRM1;VIPR2;TAS2R20;SCT;RXFP4;IHH;HTR4;RXFP3;CRHBP;EDNRA;EDNRB;ADORA3;CCRL2;C3AR1;ADORA1;PROK2;CMKLR1;PTGDR;GHSR;LHCGR;CASR;F2R;WNT5A;GPR37L1;PDYN;TRHR;MTNR1A;LTB4R2;MTNR1B;PTGFR;CALCA;CALCB;DHH;GIPR;FPR2;ADRB2;CRHR1;MCHR2;CRHR2;C3;WNT6;GALR3;HRH2;GALR1;NPBWR1;UBD;NPBWR2;PENK;PCDHA4;NLRP3;TAC3;HCRT;TAC1;WNT2;DRD4;NTSR1;WNT3;PCDHA6;DRD5;NTSR2;HTR1E;WNT3A;FZD7;FZD6;HTR1D;PTCH2;HTR1A;HTR1B;NMBR;GNG12;OPN5;PYY;LPAR5;PNOC;GNRH2;GNAS;GNB4;GNB3;NMUR1;TAS2R60;UCN;ADM;ADRA1D;ECE1;OXT;HTR2A;TRH;ADRA1B;CX3CL1;CXCL5;GRM1;LTB4R;GRM2;GPR132;GRM5;C1QTNF1;CCKAR;GNGT2;GRM4;UCN3;GRM7;UCN2;GRM6;GRM8;CCR9;BDKRB2;PRLHR;BDKRB1;CCR7;FFAR2;CCR6;FFAR1;INSL3;QRFPR;GPR39;PTGIR;GPR37;AVPR1B;EDN3;GLP2R;TACR2;TACR3;OPRM1;SSTR1;ADRA2C;WNT16;SSTR4;SSTR5;MC3R;ADORA2A;ADRB3;CCKBR;CRH;CX3CR1;CXCR5;OPRL1;PTHLH;GNG3;GNG2;WNT11;GNG4;CCL5;NPY;GNG7;P2RY1;CCL1;TAS1R1;S1PR5;S1PR4;TAS1R3;CD55;CCL25;P2RY12;CHGA;CCL23;P2RY14;WNT7B;SCTR;OPRK1;HTR5A;TSHR;KISS1;POMC;SST;F2RL1;GHRL;CCL28;RAMP1;PF4;F2RL3 |
| Na� Dependent Neurotransmitter Transporters R- | 0.014609141957414504 | SLC6A19;SLC6A18;SLC22A2;SLC6A15;SLC6A13;SLC6A12;SLC6A11;SLC6A1;SLC6A2;SLC6A3;SLC6A6;SLC6A7;SLC6A9;SLC18A2 |
| NCAM1 Interactions R-HSA-419037 | 0.023003715872047168 | ST8SIA2;CACNA1D;GFRA1;CACNA1C;CACNA1H;CACNA1G;GFRA4;CACNB1;CACNB2;CACNB4;COL4A2;GDNF;ARTN;COL4A1;COL6A1;ST8SIA4;CNTN2;NCAM1;NRTN;COL9A3;COL9A2;AGRN |
| Netrin-1 Signaling R-HSA-373752 | 0.024112787872072928 | TRIO;SRC;PITPNA;AGAP2;NTN1;ABLIM1;ABLIM2;ABLIM3;FYN;SLIT3;TRPC7;TRPC6;DSCAM;TRPC3;UNC5A;MYO10;DCC;UNC5B;UNC5C;UNC5D;RNASE1;RGMA;PRKCQ;PKN1;EZR;DOCK1;DSCAML1 |
| Metabolism Of Amine-Derived Hormones R-HSA-2 | 0.02818444197836038 | CHGA;DIDO1;DDC;DIO2;DIO3;DBH;SLC5A5;DUOX1;TPO;AANAT;TH;IYD;PEMT |
| Platelet Activation, Signaling And Aggregation R-H | 0.02818444197836038 | DGKG;APP;SPARC;DGKD;DGKA;MPL;F13A1;CLU;ISLR;AKT1;CHID1;PRKCH;ACTN2;PRKCB;ACTN1;APLP2;SERPINF2;F2R;PRKCA;RNASE1;DGKZ;TUBA4A;CD109;SPNS1;SERPING1;PRKCQ;DGKI;SHC1;PDGFB;ITPR1;PDGFA;ITPR2;A1BG;PIK3R1;RASGRP2;PRKCZ;PIK3R6;PIK3R5;APOH;LYN;TGFB3;RARRES2;IGF2;GNG12;GP6;GP5;GP9;TF;GNB4;GNB3;GRB2;HRG;FERMT3;LGALS3BP;DAGLA;PLEK;CTSW;SERPINA4;PIK3CG;GNGT2;RAC2;TIMP3;PHACTR2;SRGN;TRPC7;TRPC6;TRPC3;VWF;APOA1;GP1BA;ADRA2C;VAV1;VAV2;APBB1IP;LCK;LCP2;RAPGEF3;RAPGEF4;ITIH4;ITIH3;SRC;THBS1;GNAI2;CLEC3B;GNG3;GNG2;GNG4;GNA11;GNG7;GNA12;P2RY1;FYN;LY6G6F;P2RY12;LHFPL2;SGPP2;VEGFA;CD9;PTPN6;BCAR1;F2RL3;PF4 |
| Neurotransmitter Release Cycle R-HSA-112310 | 0.0297693105979855 | SLC22A2;SLC1A2;SLC1A3;ABAT;SLC6A1;LIN7C;SLC1A6;CPLX1;SLC1A7;RIMS1;PPFIA1;SLC17A7;PPFIA4;SLC38A2;SLC18A2;SLC18A3;SLC32A1;GAD1;SH2D3C;SLC6A13;SLC6A12;SLC6A11;SYN3;SYN2;ALDH5A1;DNAJC5;STX1A |
| G Alpha (Q) Signaling Events R-HSA-416476 | 0.0297693105979855 | NPFFR2;DGKG;APP;OXTR;TRIO;CHRM1;DGKD;DAGLA;DGKA;ADRA1D;OXT;HTR2A;TRH;ADRA1B;GRM1;LTB4R;RGS4;RGS5;EDNRA;GPR132;GRM5;EDNRB;CCKAR;GNGT2;RPS6KA2;BDKRB2;PROK2;BDKRB1;FFAR2;FFAR1;QRFPR;TRPC7;GPR39;GHSR;CASR;PRKCH;TRPC6;AVPR1B;EDN3;TRPC3;F2R;TACR2;TACR3;PRKCA;DGKZ;TRHR;CCKBR;LTB4R2;PRKCQ;PLCB2;DGKI;PTGFR;RGS16;ITPR1;ITPR2;FPR2;PIK3R1;RASGRP2;KALRN;EGFR;MCHR2;GNG3;GNG2;GRK5;GNG4;GNA11;GNG7;P2RY1;NLRP3;TAC3;HCRT;TAC1;NTSR1;NTSR2;CCL23;NMBR;GNG12;KISS1;LPAR5;GNRH2;GNB4;F2RL1;GNB3;GRB2;GHRL;NMUR1;F2RL3 |

| KEGG Pathways | Adjusted P-value | Genes |
|---|---|---|
| Calcium signaling pathway | 4.1723655074381905E-11 | CHRM2;MYLK2;OXTR;CHRM1;CALML6;CALML3;HTR4;FGF1;FGF2;FGF3;MYLK;MYLK4;SLC8A2;EDNRA;FGF7;EDNRB;FGF8;PLCE1;PDGFRB;PDGFRA;LHCGR;PRKCB;F2R;PRKCA;TRHR;ADCY9;LTB4R2;ORAI3;PTGFR;PDE1C;PDE1B;PDE1A;PDGFB;CACNA1B;ITPR1;PDGFA;CACNA1A;ITPR2;CACNA1D;MST1R;ADRB2;CACNA1C;CACNA1E;CACNA1H;CACNA1G;CACNA1I;GRIN2A;HRH1;HRH2;PDGFD;FGF20;NTSR1;DRD5;FGF21;GRIN2D;FGF17;GNAL;GDNF;FGF19;FGF18;HRC;GNAS;FGFR4;FGFR3;FGFR2;FGF10;FGFR1;RYR1;RYR2;FLT4;ATP2A3;ADRA1D;ATP2A1;HTR2A;ADRA1B;RYR3;GRM1;GRM5;CCKAR;BDKRB2;CD38;BDKRB1;NOS1;CAMK1D;AVPR1B;SPHK2;TNNC1;MST1;TNNC2;TACR2;TACR3;NGF;ITPKB;ADRB3;CCKBR;ADORA2A;ITPKA;PLCB2;CAMK1G;CAMK2B;ADCY4;ADCY3;ADCY2;ADCY1;ADCY7;EGFR;PHKG1;GNA11;ERBB2;PTK2B;NTRK1;MCOLN3;NOS2;NOS3;NTRK3;ATP2B4;ATP2B2;HTR5A;TPCN2;VEGFA;SLC8A3;P2RX7;P2RX5;P2RX2;P2RX1;PLCD3 |
| Neuroactive ligand-receptor interaction | 1.3444250082593774E-6 | NPFFR2;CHRM2;OXTR;VIPR2;CHRM1;SCT;RXFP4;HTR4;RXFP3;EDNRA;EDNRB;ADORA3;C3AR1;ADORA1;PTGDR;CHRNB1;GHSR;LHCGR;F2R;PDYN;GABRG3;TRHR;MTNR1A;LTB4R2;MTNR1B;CHRNA1;PTGFR;CALCA;CALCB;CHRNA2;CHRNA5;GIPR;CHRNA6;FPR2;ADRB2;CRHR1;MCHR2;CRHR2;C3;GRIN2A;HRH1;GALR3;HRH2;GALR1;NPBWR1;NPBWR2;PENK;TAC3;HCRT;TAC1;NTSR1;DRD4;NTSR2;DRD5;GRIA4;HTR1E;HTR1D;HTR1A;HTR1B;NMBR;GRIN2B;GRIN2D;PYY;GRIN3B;GNRH2;NMUR1;GABRB2;UCN;THRB;THRA;GRIK5;GRIK3;GRIK4;ADM;GRIK1;ADRA1D;OXT;GRIK2;HTR2A;TRH;ADRA1B;GRM1;LTB4R;GHR;GRM2;GRM5;CCKAR;GRM4;UCN3;GRM7;UCN2;GRM6;GRM8;BDKRB2;BDKRB1;PRLHR;INSL3;GRID2;PTGIR;AVPR1B;EDN3;GRID1;GLP2R;TACR2;TACR3;OPRM1;SSTR1;ADRA2C;SSTR4;SSTR5;MC3R;ADRB3;CCKBR;ADORA2A;CRH;GRP;OPRL1;CHRND;GLRA1;CNR2;CHRNE;NPY;P2RY1;S1PR5;GABRD;S1PR4;GABRA1;GABRP;GABBR1;GABRA5;GABRA4;P2RY14;SCTR;OPRK1;CCK;HTR5A;SLURP1;HCRTR2;TSHR;KISS1;P2RX7;POMC;P2RX5;GLRB;P2RX2;SST;LEP;P2RX1;F2RL1;GHRL;F2RL3 |
| Cell adhesion molecules | 8.178066034891216E-6 | CD86;CNTNAP2;CD40;CNTNAP1;ITGB2;ICAM2;LRRC4;ICAM3;ITGAL;PTPRF;ICAM1;SPN;CDH5;CDH4;CDH3;CDH1;ITGB7;MADCAM1;HLA-DOA;HLA-DOB;CD34;HLA-DPA1;ITGA4;HLA-B;HLA-C;HLA-A;HLA-F;HLA-G;HLA-E;CLDN6;CLDN11;CLDN10;CLDN4;MAG;CLDN15;CD8A;CLDN9;CLDN14;CLDN8;ITGA8;CLDN18;ITGA6;CDH15;HLA-DQB1;NLGN1;NRXN1;NRXN3;NRXN2;VTCN1;HLA-DMA;HLA-DMB;NCAM1;NCAM2;LRRC4C;HLA-DQA2;HLA-DQA1;JAM3;NTNG1;NTNG2;CADM3;HLA-DRB5;CADM1;IGSF11;OCLN;CD4;NFASC;CD6;CNTN1;HLA-DPB1;CNTN2;HLA-DRA;ESAM;PDCD1;CD22;HLA-DRB1 |
| Type I diabetes mellitus | 3.001716084656284E-5 | CD86;PTPRN;ICA1;TNF;INS;HLA-DMA;HLA-DMB;HLA-DOA;HLA-DOB;HLA-DQA2;HLA-DQA1;HLA-DPA1;HLA-DRB5;PTPRN2;GAD1;HLA-B;HLA-C;HLA-A;HLA-F;HLA-G;HLA-E;IFNG;IL1B;HLA-DPB1;LTA;CPE;HLA-DRA;HLA-DRB1;HLA-DQB1 |
| Dilated cardiomyopathy | 2.1355418912702822E-4 | RYR2;ITGB4;ATP2A3;ATP2A1;TNF;ACTG1;SLC8A2;SGCD;SGCA;ITGB7;ITGB6;SGCG;ITGA4;TPM4;ITGA3;TPM3;TNNC1;CACNA2D2;CACNA2D4;CACNB1;CACNB2;ADCY9;CACNB4;MYL2;MYL3;ITGA8;ITGA7;ITGA6;MYH7;LAMA1;ADCY4;ADCY3;ADCY2;CACNA1D;ADCY1;CACNA1C;ADCY7;TTN;CACNG6;CACNG7;CACNG8;TNNI3;CACNG1;CACNG2;CACNG3;CACNG5;TGFB3;SLC8A3;ACTC1;GNAS |

| | | |
|---|---|---|
| Viral myocarditis | 2.1355418912702822E-4 | CD86;CD40;LAMA1;ITGB2;ITGAL;ACTG1;ICAM1;HLA-DMA;CASP8;SGCD;HLA-DMB;CCND1;SGCA;RAC2;FYN;HLA-DOA;HLA-DOB;HLA-DQA2;SGCG;CD55;HLA-DQA1;HLA-DPA1;HLA-DRB5;HLA-B;HLA-C;HLA-A;HLA-F;HLA-G;HLA-E;HLA-DPB1;HLA-DRA;HLA-DRB1;HLA-DQB1;MYH7;EIF4G1 |
| Maturity onset diabetes of the young | 2.758043751760726E-4 | PKLR;ONECUT1;PAX4;PDX1;SLC2A2;HNF1B;PAX6;BHLHA15;HNF1A;MAFA;GCK;INS;HHEX;NR5A2;HNF4A;NKX6-1;MNX1;FOXA2;NEUROG3 |
| Hypertrophic cardiomyopathy | 2.758043751760726E-4 | RYR2;ITGB4;LAMA1;ATP2A3;PRKAG2;ATP2A1;CACNA1D;CACNA1C;TNF;PRKAG3;ACTG1;SLC8A2;TTN;CACNG6;CACNG7;CACNG8;SGCD;SGCA;ITGB7;TNNI3;ITGB6;CACNG1;CACNG2;CACNG3;SGCG;CACNG5;ACE;ITGA4;TPM4;ITGA3;TPM3;TGFB3;TNNC1;CACNA2D2;CACNA2D4;CACNB1;SLC8A3;CACNB2;IL6;CACNB4;ACTC1;MYL2;MYL3;ITGA8;ITGA7;ITGA6;MYH7 |
| Rap1 signaling pathway | 7.21954599495214E-4 | CSF1;CALML6;FLT4;ITGB2;PIK3CD;CALML3;FGF1;ITGAL;SIPA1L3;FGF2;FGF3;ACTG1;IGF1R;FGF7;FGF8;CDH1;RASSF5;RAC2;AKT1;PLCE1;MAGI1;MAP2K3;PDGFRB;PDGFRA;PRKCB;F2R;MAGI2;ARAP3;PRKCA;NGF;VAV1;VAV2;APBB1IP;TIAM1;ADCY9;ADORA2A;PRKD3;RAPGEF2;EVL;LCP2;PLCB2;PFN3;SKAP1;RAPGEF3;RAPGEF4;CSF1R;RGS14;SRC;PDGFB;ADCY4;PDGFA;ADCY3;PARD6G;ADCY2;PIK3R1;ADCY1;RASGRP2;PRKCZ;ADCY7;THBS1;EGFR;RAP1GAP;GNAI2;INS;GRIN2A;PARD6A;PDGFD;P2RY1;FGF20;FGF21;NGFR;ANGPT2;ANGPT1;GRIN2B;VEGFA;FGF17;EFNA2;LPAR5;FGF19;FGF18;GNAS;TEK;FGFR4;FGFR3;SIPA1;FGFR2;LAT;BCAR1;F2RL3;FGF10;FGFR1 |
| Arrhythmogenic right ventricular cardiomyopathy | 0.0012195187477127153 | RYR2;ITGB4;LAMA1;TCF7;ATP2A3;ATP2A1;CACNA1D;CACNA1C;ACTG1;SLC8A2;CACNG6;CACNG7;CACNG8;SGCD;SGCA;CTNNA1;ITGB7;CTNNA3;ITGB6;CACNG1;CTNNA2;CACNG2;CACNG3;SGCG;CACNG5;DSP;TCF7L2;TCF7L1;ITGA4;ACTN2;ITGA3;CACNA2D2;CACNA2D4;CACNB1;SLC8A3;CACNB2;CACNB4;ITGA8;ITGA7;ITGA6 |
| Glutamatergic synapse | 0.0016308412254200046 | GRIK5;GRIK3;GRIK4;GRIK1;GRIK2;GRM1;GRM2;GRM5;GNGT2;GRM4;GRM7;GRM6;GRM8;DLGAP1;SLC38A1;PLA2G4D;PRKCB;PLA2G4B;PRKCA;ADCY9;JMJD7-PLA2G4B;PLCB2;SHANK2;SHANK1;ITPR1;SLC1A2;ADCY4;CACNA1A;ITPR2;SLC1A3;ADCY3;CACNA1D;ADCY2;CACNA1C;ADCY1;SLC1A6;SLC1A7;ADCY7;GNAI2;GRIN2A;GNG3;GNG2;GNG4;GNG7;SLC17A7;SLC38A2;GRIA4;GNG12;GRIN2B;GRIN2D;GRIN3B;GNAS;GNB4;GNB3 |
| Hematopoietic cell lineage | 0.0018292233192563092 | CSF1R;CSF3;CSF2;CSF3R;CSF1;EPO;IL5RA;CD1D;CD3E;TNF;CD1A;HLA-DMA;HLA-DMB;ANPEP;CD19;CD38;CD37;HLA-DOA;HLA-DOB;HLA-DQA2;CD34;HLA-DQA1;CD55;HLA-DPA1;HLA-DRB5;CR1;ITGA4;MME;ITGA3;GP1BA;GP5;GP9;FCER2;IL6;CD4;CD8A;CD5;CR1L;IL1B;IL2RA;CD7;HLA-DPB1;HLA-DRA;CD9;ITGA6;CD22;HLA-DRB1;HLA-DQB1 |
| Oxytocin signaling pathway | 0.002646409573513429 | RYR1;MYLK2;RYR2;OXTR;CDKN1A;CALML6;PRKAG2;CALML3;OXT;PIK3CG;PRKAG3;RYR3;ACTG1;MYLK;MYLK4;CCND1;NPPA;CD38;MEF2C;CAMK1D;PLA2G4D;PRKCB;PLA2G4B;CACNA2D2;PRKCA;CACNA2D4;CACNB1;CACNB2;ADCY9;CACNB4;JMJD7-PLA2G4B;MYL9;PLCB2;CAMK1G;CAMK2B;SRC;ITPR1;ADCY4;ITPR2;ADCY3;CACNA1D;ADCY2;CACNA1C;ADCY1;ADCY7;PIK3R6;EGFR;PIK3R5;CAMKK2;GNAI2;CACNG6;TRPM2;CACNG7;CACNG8;CACNG1;CACNG2;CACNG3;CACNG5;KCNJ6;KCNJ12;KCNJ9;NOS3;KCNJ14;NFATC3;NFATC2;NFATC1;PPP1CA;GNAS |
| cAMP signaling pathway | 0.005035271207689656 | CHRM2;RYR2;OXTR;VIPR2;CHRM1;CALML6;ATP2A3;PIK3CD;CALML3;ATP2A1;OXT;HTR4;GLI3;EDNRA;CREB3L3;CREB3L1;NPPA;RAC2;ADORA1;PDE4B;AKT1;PLCE1;PDE4A;FFAR2;CNGA3;SOX9;AMH;GHSR;LHCGR;EDN3;PDE4D;F2R;PDE4C;ARAP3;SSTR1;VAV1;VAV2;SSTR5;TIAM1;ADCY9;ADORA2A;ACOX3;MYL9;CFTR;RAPGEF3;RAPGEF4;CNGB1;CREB5;CAMK2B;GIPR;ADCY4;ADCY3;ADCY2;CACNA1D;PIK3R1;ADCY1;ADRB2;CACNA1C;ADCY7;GNAI2;GRIN2A;PAK1;NPY;TNNI3;DRD5;GRIA4;CREBBP;GABBR1;HTR1E;HTR1D;HTR1A;ATP2B4;HTR1B;NFATC1;ATP2B2;GRIN2B;TSHR;GRIN2D;PPP1CA;MAPK10;POMC;ADCY10;GRIN3B;SST;FXYD1;PPP1R1B;GNAS;GHRL;HCN2 |
| Allograft rejection | 0.005773213606513006 | CD86;CD40;HLA-DRB5;HLA-B;HLA-C;HLA-A;HLA-F;TNF;HLA-G;HLA-E;HLA-DMA;HLA-DMB;IFNG;HLA-DPB1;HLA-DRA;HLA-DOA;HLA-DOB;HLA-DQA2;HLA-DQA1;HLA-DRB1;HLA-DPA1;HLA-DQB1 |
| Adrenergic signaling in cardiomyocytes | 0.005773213606513006 | RYR2;CALML6;ATP2A3;CALML3;ADRA1D;ATP2A1;ADRA1B;PIK3CG;SLC8A2;CREB3L3;CREB3L1;AKT1;SCN1B;TPM4;TPM3;TNNC1;CACNA2D2;PRKCA;CACNA2D4;CACNB1;CACNB2;ADCY9;CACNB4;KCNQ1;MYL2;MYL3;PLCB2;SCN4B;RAPGEF3;CREB5;MYH7;RAPGEF4;CAMK2B;KCNE1;ATF6B;ADCY4;ADCY3;CACNA1D;ADCY2;ADRB2;CACNA1C;ADCY1;ADCY7;PIK3R6;PIK3R5;GNAI2;CACNG6;CACNG7;CACNG8;TNNI3;CACNG1;CACNG2;CACNG3;CACNG5;ATP2B4;ATP2B2;PPP2R3A;PPP1CA;SLC8A3;ACTC1;PPP2R2C;PPP2R2B;PPP2R2D;BCL2;GNAS |
| Regulation of actin cytoskeleton | 0.00582652556798775 | CYFIP2;CHRM2;CYFIP1;MYLK2;CHRM1;ITGB4;ITGB2;PIK3CD;ITGAE;FGF1;ITGAL;FGF2;FGF3;MYLK;ACTG1;MYLK4;FGF7;FGF8;RAC2;BDKRB2;ITGAX;PIP4K2A;BDKRB1;ITGB7;ITGB6;APC2;PDGFRB;PDGFRA;ITGA4;ITGA3;ACTN1;INSRR;F2R;VAV1;VAV2;MYL5;TIAM1;MYL2;SPATA13;ITGA8;ARHGEF4;ITGA7;MYH9;ITGA6;EZR;ARHGEF7;MYL9;DOCK1;PFN3;SRC;PDGFB;PDGFA;PIK3R1;ACTR3B;EGFR;INS;ACTR3C;PAK1;SCIN;PDGFD;GNA12;FGF20;MYL10;PAK6;MYH14;NCKAP1L;PIP5K1C;MYH10;FGF21;LIMK2;LIMK1;GNG12;SSH2;BAIAP2;SSH3;PPP1CA;SSH1;MYLPF;FGF17;LPAR5;APC;FGF19;FGF18;FGFR4;FGFR3;FGFR2;BCAR1;FGF10;FGFR1 |
| Morphine addiction | 0.00582652556798775 | GABRB2;PDE1C;PDE1B;PDE1A;ADCY4;CACNA1B;ADCY3;CACNA1A;ADCY2;ADCY1;ADCY7;GNAI2;GNG3;GNGT2;GNG2;GRK5;GNG4;GRK4;GNG7;ADORA1;PDE4B;PDE4A;GABRD;GABRA1;GABRP;GABBR1;KCNJ6;SLC32A1;PRKCB;GABRA5;KCNJ9;PDE4D;GABRA4;PDE4C;PDE2A;PRKCA;OPRM1;GNG12;GABRG3;ADCY9;GNAS;GNB4;GNB3 |
| Gastric cancer | 0.006603786943659869 | RB1;CDKN1A;PIK3CD;FGF1;FGF2;FGF3;FGF7;FRAT1;FGF8;CCND1;CDH1;MYC;AKT1;APC2;WNT5B;WNT5A;AXIN1;WNT16;TGFBR2;RARB;SHC2;ABCB1;SHC1;TERC;TCF7;LRP5;PIK3R1;EGFR;RXRB;WNT6;MUC2;WNT11;TERT;ERBB2;FGF20;CTNNA1;CTNNA3;CTNNA2;WNT2;WNT3;FGF21;SMAD2;TCF7L2;TCF7L1;SMAD3;FZD5;CDX2;TGFB3;WNT3A;FZD7;WNT7B;FZD6;MLH1;MTOR;GADD45G;FGF17;APC;FGF19;CDK2;FGF18;BCL2;GRB2;FGFR2;FGF10 |
| Type II diabetes mellitus | 0.006603786943659869 | CACNA1B;CACNA1A;PIK3CD;SLC2A2;CACNA1D;IRS2;CACNA1C;PIK3R1;CACNA1E;PRKCZ;TNF;CACNA1G;HK1;INS;SOCS2;HK3;SOCS3;KCNJ11;PKLR;PDX1;MAFA;GCK;MTOR;MAPK10;HKDC1 |
| Hippo signaling pathway | 0.006603786943659869 | ITGB2;FGF1;LIMD1;ACTG1;BBC3;GLI2;SOX2;RASSF1;CCND2;RASSF2;CCND1;CDH1;RASSF4;MYC;RASSF6;AMH;YWHAH;APC2;WNT5B;WNT5A;AXIN1;CSNK1E;WNT16;TGFBR2;BIRC5;LLGL2;CRB2;TCF7;PARD6G;PRKCZ;NKD2;WNT6;PAK1;PARD6A;WNT11;CTNNA1;CTNNA3;CTNNA2;WNT2;WNT3;SMAD2;WWTR1;TCF7L2;TCF7L1;SMAD3;FZD5;TGFB3;WNT3A;FZD7;WNT7B;FZD6;BMP8B;GDF6;BMP7;GDF7;SMAD7;PPP1CA;BMP4;BMP2;DLG2;APC;PPP2R2C;DCHS2;PPP2R2B;ID2;PPP2R2D;DCHS1;FAT4;TP73 |
| Parathyroid hormone synthesis, secretion and act | 0.00760510940091382 | GCM2;CDKN1A;ATF6B;ITPR1;ADCY4;LRP5;ITPR2;ADCY3;ADCY2;GATA3;ADCY1;ADCY7;EGFR;PTHLH;GNAI2;RXRB;AKAP13;CREB3L3;BGLAP;CREB3L1;GNA11;GNA12;PDE4B;TNFSF11;PDE4A;SOST;MEF2D;SLC34A2;EGR1;KL;CASR;MEF2C;PRKCB;SLC34A1;PDE4D;PDE4C;PRKCA;RUNX2;NR4A2;MMP14;ADCY9;MAFB;MMP16;BCL2;GNAS;PLCB2;FGFR1;CREB5 |
| Graft-versus-host disease | 0.008769182398449864 | CD86;HLA-DRB5;HLA-B;HLA-C;HLA-A;HLA-F;TNF;HLA-G;HLA-E;HLA-DMA;IL6;HLA-DMB;IFNG;IL1B;HLA-DPB1;HLA-DRA;HLA-DOA;HLA-DOB;HLA-DQA2;HLA-DQA1;HLA-DRB1;HLA-DPA1;HLA-DQB1 |
| Circadian entrainment | 0.0108566417318418 | RYR1;CAMK2B;RYR2;CALML6;ITPR1;ADCY4;ADCY3;CALML3;CACNA1D;ADCY2;CACNA1C;ADCY1;ADCY7;CACNA1H;RYR3;CACNA1G;GNAI2;CACNA1I;GRIN2A;GNG3;GNGT2;GNG2;GNG4;GNG7;NOS1;GRIA4;KCNJ6;PRKCB;KCNJ9;PRKCA;GNG12;GRIN2B;GRIN2D;ADCY10;PER1;MTNR1A;ADCY9;PER3;MTNR1B;NOS1AP;GNAS;GNB4;GNB3;PLCB2 |
| GABAergic synapse | 0.0108566417318418 | GABRB2;SRC;ADCY4;CACNA1B;ADCY3;CACNA1A;ABAT;ADCY2;CACNA1D;ADCY1;CACNA1C;SLC6A1;ADCY7;GNAI2;GNG3;GNGT2;GNG2;GNG4;GNG7;GABRD;SLC38A2;GABRA1;GABRP;SLC38A1;GABBR1;KCNJ6;SLC12A5;SLC32A1;PRKCB;GABRA5;GABRA4;GAD1;SLC6A13;SLC6A12;PRKCA;SLC6A11;GNG12;GABRG3;ADCY9;GNB4;GNB3 |
| Metabolism of xenobiotics by cytochrome P450 | 0.011261326795658333 | UGT1A10;GSTP1;MGST1;MGST2;GSTT1;ADH6;HSD11B1;ALDH3B2;HPGDS;AKR7A3;CYP2B6;CYP2A13;CYP2D6;ALDH3B1;UGT1A5;CYP1B1;UGT1A4;UGT1A3;CYP2F1;UGT1A9;UGT1A8;UGT1A7;UGT1A6;CBR1;GSTM2;GSTM1;GSTO2;UGT2B15;EPHX1;UGT2B17;AKR7L;ALDH3A1;CYP2S1;CYP1A1;CYP2E1;GSTM5 |
| Cholinergic synapse | 0.016999596658988314 | CHRM2;ACHE;CHRM1;PIK3CD;PIK3CG;GNGT2;CREB3L3;CREB3L1;AKT1;SLC18A3;PRKCB;PRKCA;ADCY9;KCNQ1;KCNQ2;KCNQ4;PLCB2;CREB5;CAMK2B;CHAT;CHRNA6;CACNA1B;ITPR1;ADCY4;CACNA1A;ITPR2;ADCY3;CACNA1D;ADCY2;CACNA1C;PIK3R1;ADCY1;ADCY7;PIK3R6;PIK3R5;GNAI2;GNG3;GNG2;GNG4;GNA11;GNG7;FYN;KCNJ6;KCNJ12;KCNJ14;GNG12;BCL2;GNB4;GNB3 |
| Serotonergic synapse | 0.016999596658988314 | APP;GABRB2;HTR2A;HTR4;GNGT2;CYP2D6;SLC18A2;KCND2;PLA2G4D;PRKCB;DUSP1;HTR3E;PLA2G4B;PRKCA;HTR3A;HTR3B;JMJD7-PLA2G4B;PLCB2;RAPGEF3;DDC;ALOX15;CACNA1B;ITPR1;CACNA1A;ITPR2;ALOX12;CACNA1D;CACNA1C;CYP2C18;GNAI2;GNG3;CYP4X1;GNG2;GNG4;ALOX5;GNG7;KCNN2;KCNJ6;HTR1E;KCNJ9;HTR1D;HTR1A;HTR1B;GNG12;HTR5A;ALOX15B;GNAS;GNB4;GNB3 |
| Antigen processing and presentation | 0.01806635927296001 | CIITA;IFI30;TNF;HLA-DMA;HLA-DMB;HLA-DOA;HLA-DOB;HLA-DQA2;HLA-DQA1;HLA-DPA1;CD74;HSP90AA1;HLA-DRB5;HSPA1L;NFYC;HSPA6;HLA-B;TAP2;HLA-C;TAP1;HLA-A;HSPA2;HLA-F;HLA-G;HLA-E;TAPBP;CD4;IFNG;CD8A;HLA-DPB1;HLA-DRA;HLA-DRB1;HSPA1B;HLA-DQB1;HSPA1A;LGMN |
| Phospholipase D signaling pathway | 0.02040524493191218 | DGKG;DGKD;DGKA;PIK3CD;GRM1;PIK3CG;GRM2;GRM5;GRM4;GRM7;GRM6;GRM8;AKT1;PDGFRB;PDGFRA;AVPR1B;PLA2G4D;SPHK2;PLA2G4B;F2R;TSC2;PRKCA;DGKZ;DNM1;ADCY9;JMJD7-PLA2G4B;PLCB2;DGKI;RAPGEF3;RAPGEF4;PTGFR;SHC2;SHC1;PDGFB;ADCY4;PDGFA;ADCY3;ADCY2;PIK3R1;ADCY1;AGPAT1;ADCY7;PIK3R6;AGPAT3;EGFR;PIK3R5;AGPAT4;INS;CYTH2;CYTH4;PDGFD;GNA12;PTK2B;FYN;PIP5K1C;CYTH1;MTOR;LPAR5;RHEB;GNAS;GRB2 |
| PI3K-Akt signaling pathway | 0.02040524493191218 | CHRM2;CSF3;CHRM1;CSF3R;CSF1;EPO;FGF1;FGF2;FGF3;IGF1R;COMP;TCL1A;FGF7;CCND2;FGF8;CREB3L3;CCND1;CREB3L1;MYC;AKT1;TNR;MAGI1;PDGFRB;PDGFRA;F2R;MAGI2;TSC2;PRKCA;COL4A2;COL4A1;EIF4E1B;CSF1R;PDGFB;PDGFA;PIK3R1;PIK3R6;PIK3R5;CD19;PDGFD;CHAD;FGF20;THEM4;PCK1;FGF21;NGFR;IGF2;GNG12;COL1A1;FGF17;IL6;CDK6;EFNA2;G6PC3;LPAR5;RHEB;FGF19;CDK2;FGF18;BCL2;GNB4;GNB3;COL9A3;GRB2;COL9A2;FGFR4;FGFR3;FGFR2;FGF10;FGFR1;CDKN1A;ITGB4;FLT4;PIK3CD;LAMC2;BRCA1;PIK3CG;RPTOR;GHR;GNGT2;MLST8;ITGB7;ITGB6;JAK3;YWHAH;HSP90AA1;ITGA4;VWF;ITGA3;NGF;COL2A1;COL6A1;ITGA8;ITGA7;ITGA6;SGK1;SGK2;TLR4;CREB5;TNXB;ATF6B;LAMA1;LAMA4;LAMA3;THBS1;EGFR;THBS4;INS;GNG3;GNG2;GNG4;GNG7;NTF3;ERBB2;SPP1;EIF4E;NTRK1;ANGPT2;ANGPT1;LAMB3;NOS3;OSM;PPP2R3A;MTOR;VEGFA;PPP2R2C;PPP2R2B;IL2RA;IL2RB;PPP2R2D;PKN1;TEK |
| Pathways in cancer | 0.0206014319505961 | RB1;SPI1;CSF3R;EPO;CALML6;IL5RA;CALML3;FGF1;FGF2;ETS1;FGF3;IGF1R;FGF7;EDNRA;FRAT1;RASSF1;FGF8;EDNRB;CCND2;CCND1;CDH1;MYC;RASSF5;AKT1;PDGFRB;PDGFRA;WNT5B;DAPK1;PRKCB;TPM3;DAPK3;F2R;WNT5A;PRKCA;MITF;RUNX1;ADCY9;COL4A2;IL23A;COL4A1;CSF1R;NOTCH3;CTBP2;CTBP1;TERC;EPAS1;NOTCH4;TCF7;PDGFB;CUL1;MGST1;PDGFA;MGST2;GSTT1;CSF2RB;PIK3R1;RASGRP2;WNT6;TERT;FGF20;FADD;WNT2;WNT3;FGF21;JAG2;STAT5A;SMAD2;CREBBP;GSTM2;GSTM1;SMAD3;FZD5;TGFB3;WNT3A;FZD7;ZBTB16;FZD6;PTCH2;IGF2;GNG12;ESR1;GADD45G;BMP4;FGF17;BMP2;IL6;CDK6;LPAR5;FGF19;CDK2;FGF18;BCL2;GNAS;GNB4;GNB3;GRB2;FGFR4;FGFR3;FGFR2;GSTM5;FGF10;FGFR1;CDKN1A;FLT4;PIK3CD;LAMC2;GLI3;BBC3;GLI2;GNGT2;CASP8;HEY1;MECOM;RAC2;BDKRB2;BDKRB1;JAK3;APC2;HSP90AA1;GSTO2;ITGA3;DCC;IFNGR2;PLEKHG5;NCOA4;AXIN1;WNT16;TGFBR2;CCNA1;PAX8;IFNG;TRAF3;TRAF5;RARA;RARB;BIRC5;ITGA6;BIRC7;PLCB2;CAMK2B;HDAC2;LAMA1;GSTP1;LAMA4;LAMA3;ADCY4;LRP5;ADCY3;ADCY2;ADCY1;ADCY7;EGFR;GNAI2;DLL3;RXRB;GNG3;GNG2;WNT11;GNG4;GNA11;GNG7;ERBB2;GNA12;STAT4;CTNNA1;CTNNA3;CTNNA2;IL12RB2;RUNX1T1;NTRK1;NQO1;TCF7L2;TCF7L1;EGLN2;LAMB3;NOS2;TXNRD2;WNT7B;TXNRD1;STAT3;MLH1;MTOR;VEGFA;MAPK10;HEYL;APC;IL2RA;IL2RB;F2RL3 |
| Toxoplasmosis | 0.021065691191752438 | CD40;CIITA;LAMA1;LAMA4;LAMA3;LAMC2;TNF;PIK3R6;PIK3CG;PIK3R5;GNAI2;HLA-DMA;CASP8;HLA-DMB;ALOX5;AKT1;HLA-DOA;HLA-DOB;HLA-DQA2;HLA-DQA1;HLA-DPA1;MAP2K3;HLA-DRB5;HSPA1L;LAMB3;NOS2;TGFB3;IL10RB;IFNGR2;HSPA6;STAT3;HSPA2;IRAK4;TYK2;IRGM;TNFRSF1A;MAPK10;IFNG;BCL2;HLA-DPB1;HLA-DRA;ITGA6;BIRC7;TLR4;HLA-DRB1;HSPA1B;HLA-DQB1;HSPA1A |
| Signaling pathways regulating pluripotency of ster | 0.021065691191752438 | ONECUT1;PIK3CD;FGF2;IGF1R;SOX2;MYC;AKT1;HOXA1;JARID2;JAK3;APC2;WNT5B;WNT5A;AXIN1;PAX6;WNT16;ISL1;POU5F1;HAND1;HOXB1;DLX5;TCF7;PIK3R1;WNT6;ACVR1C;WNT11;OTX1;WNT2;WNT3;SMAD2;ZFHX3;SMAD3;FZD5;WNT3A;FZD7;WNT7B;FZD6;PCGF3;STAT3;LIF;SMAD9;INHBB;INHBA;INHBC;TBX3;BMP4;MEIS1;APC;ID2;ID3;GRB2;NANOG;FGFR4;FGFR3;NEUROG1;FGFR2;NODAL;FGFR1;MYF5 |
| Inflammatory bowel disease | 0.035182389466074535 | RORA;GATA3;NOD2;TNF;HLA-DMA;HLA-DMB;IL21R;STAT4;HLA-DOA;HLA-DOB;HLA-DQA2;HLA-DQA1;HLA-DPA1;IL12RB2;SMAD2;HLA-DRB5;SMAD3;TGFB3;IFNGR2;STAT3;NFATC1;IL6;IFNG;IL23A;IL1B;HLA-DPB1;HLA-DRA;TLR4;HLA-DRB1;HLA-DQB1 |
| Gastric acid secretion | 0.03613419000939665 | CAMK2B;MYLK2;CALML6;ADCY4;ITPR1;ADCY3;ITPR2;CALML3;ADCY2;ADCY1;ADCY7;ACTG1;MYLK;MYLK4;GNAI2;CA2;HRH2;KCNJ1;KCNK10;PRKCB;KCNJ15;KCNJ16;PRKCA;ATP4B;ATP4A;ADCY9;CCKBR;SST;KCNQ1;GNAS;EZR;PLCB2;KCNK2;CFTR |
| Th17 cell differentiation | 0.03916634048378932 | RORA;GATA3;CD3E;RXRB;HLA-DMA;HLA-DMB;IL21R;HLA-DOA;HLA-DOB;HLA-DQA2;JAK3;HLA-DQA1;HLA-DPA1;SMAD2;STAT5A;HLA-DRB5;HSP90AA1;SMAD3;IFNGR2;STAT3;NFATC3;NFATC2;NFATC1;TYK2;MTOR;RUNX1;TGFBR2;MAPK10;ZAP70;IL6;CD4;IFNG;IL23A;LCK;IL1B;IL2RA;IL2RB;RARA;HLA-DPB1;HLA-DRA;PRKCQ;IL17D;HLA-DRB1;LAT;HLA-DQB1 |
| Breast cancer | 0.03916634048378932 | RB1;CDKN1A;FLT4;PIK3CD;BRCA1;FGF1;FGF2;FGF3;IGF1R;FGF7;FRAT1;FGF8;CCND1;HEY1;MYC;AKT1;TNFSF11;APC2;WNT5B;WNT5A;AXIN1;WNT16;PGR;NOTCH3;SHC2;SHC1;NOTCH4;TCF7;LRP5;PIK3R1;EGFR;DLL3;WNT6;WNT11;ERBB2;FGF20;WNT2;WNT3;FGF21;JAG2;TCF7L2;TCF7L1;FZD5;WNT3A;FZD7;WNT7B;FZD6;ESR1;MTOR;GADD45G;FGF17;HEYL;CDK6;APC;FGF19;FGF18;GRB2;FGF10;FGFR1 |
| beta-Alanine metabolism | 0.047795743428037434 | AOC3;AOC2;DPYS;GAD1;GADL1;ABAT;UPB1;CNDP1;CNDP2;ALDH3B2;ALDH3A1;ALDH3B1;EHHADH;ACOX3;ACADS;ALDH7A1 |
| Leukocyte transendothelial migration | 0.047795743428037434 | TXK;NCF4;ITGB2;PIK3CD;PIK3R1;THY1;ITGAL;ACTG1;ICAM1;GNAI2;CDH5;RASSF5;RAC2;CTNNA1;MYL10;PTK2B;CTNNA3;CTNNA2;JAM3;ITGA4;PRKCB;ACTN1;RHOH;CYBA;PRKCA;VAV1;VAV2;CLDN6;MYLPF;CLDN11;MYL5;CLDN10;CLDN4;OCLN;CLDN15;CLDN9;CLDN14;CLDN8;MYL2;CLDN18;ESAM;EZR;MYL9;RAPGEF3;BCAR1;SIPA1;RAPGEF4 |
| Intestinal immune network for IgA production | 0.047795743428037434 | CD86;CCL25;PIGR;CD40;HLA-DRB5;TNFRSF13B;ITGA4;HLA-DMA;IL6;HLA-DMB;CCR9;HLA-DPB1;HLA-DRA;ITGB7;MADCAM1;HLA-DOA;HLA-DOB;HLA-DQA2;CCL28;HLA-DQA1;HLA-DRB1;HLA-DPA1;HLA-DQB1 |

| GO: Biological Processes | Adjusted P-value | Genes |
|---|---|---|
| Chemical Synaptic Transmission (GO:0007268) | 0.00 | CHRM2;CHRM1;HTR4;CDH8;CRHBP;RPS6KA2;SLC12A6;SLC12A7;CHRNB1;UNC13A;PDYN;GABRG3;SHISA6;CACNB1;CACNB2;CACNB4;MTNR1B;KCNQ2;AMPH;CHRNA1;CHRNA2;CHRNA5;CNP;CHAT;CHRNA6;CACNA1B;SLC1A2;CACNA1A;SLC1A3;SLC1A4;LIN7C;SLC1A6;CACNA1E;CPLX1;GRIN2A;GALR3;HRH2;NPBWR1;PENK;APOE;NPTX1;HCRT;NTSR1;DRD4;DRD5;GRIA4;DTNA;PTPRN2;HTR1E;HTR1D;HTR1A;HTR1B;GRIN2B;GRIN2D;GRIN3B;DLG2;PCDHB2;SLC6A9;TH;GIPC1;PNOC;PCDHB6;PCDHB5;PCDHB4;PCDHB3;PCDHB9;GABRB2;DOC2A;GRIK5;GRIK3;GRIK4;GRIK1;GRIK2;DBH;HTR2A;SLC6A1;SLC6A2;HAPLN4;GRM2;GRM4;GRM7;DLGAP1;DLGAP2;SLC18A2;SLC18A3;GRID2;KCND2;SLC32A1;GRID1;HTR3E;HTR3A;HTR3B;SYN3;SYN2;SNPH;CRH;RAPSN;NLGN1;KCNA1;NRXN1;NRXN2;CHRND;GLRA1;CHRNE;SNCG;SLC17A7;SNCB;GABRD;PDLIM4;SNCA;GABRA1;GABRP;NQO1;GABBR1;SLC12A4;SLC12A5;GABRA5;KCNIP2;GABRA4;GAD1;PCDHB14;OPRK1;PCDHB11;SEPTIN9;HTR5A;PCDHB10;HCRTR2;RIT2;GLRB;SST;PCDHB16;RIMBP2;ZACN;APBA2;CBLN2;CBLN1 |
| Anterograde Trans-Synaptic Signaling (GO:00989 | 0.00 | CHRM2;CHRM1;HTR4;RPS6KA2;SLC12A6;SLC12A7;CHRNB1;PDYN;CACNB1;CACNB2;CACNB4;MTNR1B;KCNQ2;AMPH;CHRNA1;CHRNA2;CHRNA5;CNP;CHRNA6;CACNA1B;SLC1A2;CACNA1A;SLC1A3;SLC1A6;CACNA1E;CPLX1;GRIN2A;GALR3;HRH2;NPBWR1;PENK;NPTX1;HCRT;NTSR1;DRD4;DRD5;DTNA;HTR1E;HTR1D;HTR1A;HTR1B;GRIN2B;DLG2;PCDHB2;GIPC1;PNOC;PCDHB6;PCDHB5;PCDHB4;PCDHB3;PCDHB9;GABRB2;DOC2A;GRIK4;GRIK1;GRIK2;DBH;HTR2A;SLC6A1;SLC6A2;GRM2;GRM4;GRM7;DLGAP1;SLC18A2;SLC18A3;KCND2;HTR3E;HTR3A;HTR3B;SYN2;CRH;RAPSN;NLGN1;KCNA1;NRXN1;NRXN2;CHRND;GLRA1;CHRNE;SNCG;SNCB;GABRD;SNCA;GABRP;SLC12A4;SLC12A5;KCNIP2;GAD1;PCDHB14;OPRK1;PCDHB11;HTR5A;PCDHB10;HCRTR2;RIT2;GLRB;SST;PCDHB16;ZACN;APBA2;CBLN1 |
| Calcium-Dependent Cell-Cell Adhesion Via Plasm | 0.00 | NLGN1;PCDHGB4;CDH9;CDH8;CDH6;CDH5;CDH4;CDH3;CDH1;CDH22;CDH23;PCDHGC3;PCDHB14;PCDHB11;PCDHB10;PCDHB2;CDH10;CDH11;PCDHB16;CDH12;PCDHB6;CDH13;PCDHB5;DCHS1;PCDHB4;CDH15;PCDHB3;PCDHB9;CDH18 |
| Anterior/Posterior Pattern Specification (GO:0009 | 0.00 | LRP5;GATA4;HHEX;HOXA9;TIFAB;HEY1;SIX2;HOXA3;HOXC5;HOXC4;HOXA7;HES3;HOXC9;HOXA6;HOXA5;HOXA4;HOXC6;SMAD2;TBX1;CDX2;CDX1;SKI;HEYL;BMP2;CYP26C1;HOXB9;WT1;DDIT3;BHLHE40;HOXB4;HOXB3;HOXD4;HOXD3;HOXB7;HOXD9;NEUROG1;HOXB6;HOXB5 |
| Synapse Organization (GO:0050808) | 0.00 | APP;CYFIP1;PTPRS;CTNND2;LRRC4;SLC6A1;ELFN1;CX3CL1;PTPRF;SLC8A2;GRM5;CDH1;PPFIA1;PPFIA4;KIRREL3;UNC13A;DSCAM;F2R;DKK1;CACNB1;CACNB2;CACNB4;DOK7;MAPT;SHANK2;CHRNA1;C1QB;CX3CR1;C1QA;NLGN1;DCTN1;NRXN1;LRP4;NRXN2;SLITRK1;SPOCK2;SLITRK3;SNCG;APBB2;LRRC4C;SNCB;SNCA;NTNG1;FARP1;FZD5;WNT7B;PCDHB14;PCDHB11;PCDHB10;POU4F1;KY;SLC8A3;SDK1;LRFN4;PPFIBP2;PCDHB2;PTK7;PCDHB16;PCDHB6;CNTN2;PCDHB5;PCDHB4;PCDHB3;AGRN;PCDHB9;SDK2;CBLN2;CBLN1 |
| Cell-Cell Adhesion Via Plasma-Membrane Adhesi | 0.00 | PTPRT;ROBO3;ROBO4;PTPRS;PCDHGB4;ITGB2;LRRC4;ITGAL;CDH9;PKD1;CDH8;FGFRL1;PTPRF;ICAM1;CDH6;CDH5;CDH4;CDH3;CDH1;CD177;KIRREL3;GRID2;DSCAM;UNC5D;TGFBR2;CLDN6;CLDN11;CLDN10;CLDN4;MAG;CLDN15;CLDN9;CLDN14;PALLD;CLDN8;CDH10;CDH11;CDH12;CDH13;CLDN18;CDH15;DSCAML1;CDH18;CRB2;NLGN1;AMIGO3;GATA5;IGSF9;IGSF21;SLITRK1;CDH22;SLITRK3;CDH23;LRRC4C;PCDHA7;MPZL2;NTNG1;CADM3;CADM1;CNTN6;PCDHGC3;PCDHB14;PCDHB11;PCDHB10;IGSF11;SDK1;LRFN4;PCDHB2;CD6;PTK7;PCDHB16;PCDHB6;CNTN2;FAT2;PCDHB5;PLXNB2;DCHS1;ESAM;PCDHB4;FAT4;PCDHB3;PCDHB9;SDK2;CBLN1 |
| Skeletal System Morphogenesis (GO:0048705) | 0.00 | FBN2;SPEF2;ANKRD11;LRP5;CYP26B1;HOXA9;HOXA3;HOXC4;HOXC9;IFITM5;HOXA4;TFAP2A;OSR2;PDGFRA;WWOX;TBX4;SKI;SFRP4;HOXB9;HOXB4;HOXB3;HOXD4;HOXD3;DSCAML1;FGFR3;HOXD9;FGFR2;FGFR1;LTF |

| | | |
|---|---|---|
| Positive Regulation Of MAPK Cascade (GO:00434 | 0.00 | APP;CD81;FGF1;FGF2;TNF;ICAM1;IGF1R;SOX2;PYCARD;PLCE1;TNFSF11;LGALS9;PDGFRB;PDGFRA;MEF2C;SERPINF2;F2R;PRKCA;GPR37L1;CARTPT;DOK2;DOK3;DOK5;DOK6;ADAM8;NECAB2;CSF1R;NDRG4;SEMA7A;SHC1;ALOX15;PDGFB;PDGFA;GATA4;FPR2;MST1R;ADRB2;SLC30A10;PRKCZ;PIK3R6;ART4;PIK3R5;PDGFD;APOE;ELANE;DRD4;SLAMF1;FGF21;IGF2;LIF;BMP4;MFHAS1;BMP2;IL6;STK25;FGF19;FGF18;FGFR4;TDGF1;FGFR3;HLA-DRB1;NODAL;FGFR2;FGF10;FGFR1;DDR1;CD40;ARHGAP8;FLT4;DUSP19;ADRA1D;PTPN22;HTR2A;ADRA1B;CX3CL1;PIK3CG;GHR;GRM4;TRIM5;CCR7;TNFAIP8L3;AMH;MAP4K1;GPR37;AVPR1B;EDN3;KSR1;FCRL3;NRG1;OPRM1;ADRA2C;ADRB3;IL1B;HAND2;RAPGEF2;CHI3L1;TLR4;TMEM106A;NOD2;THBS1;EGFR;INS;CCL5;NTF3;ERBB2;P2RY1;GCNT2;PTK2B;CCL1;CCL25;NTRK1;TBX1;CCL24;CD74;CCL23;ANGPT1;GDF15;IL34;NTRK3;SEMA4C;DAB2IP;OSM;VEGFA;NELL1;CD4;RIT2;LEP;NEK10;F2RL1;SPRY2;GHRL;TEK;CCL26 |
| Embryonic Organ Morphogenesis (GO:0048562) | 0.00 | FBN2;RYR2;RARG;TCF21;HOXA9;SOX17;SIX2;TBX20;HOXA3;RBPMS2;HOXC4;ALX1;HOXC9;HOXA4;TBX1;OSR2;PDGFRA;FOXF1;MFAP5;ALDH1A3;HOXB9;ID2;HOXB4;HOXB3;HOXD4;HOXD3;DSCAML1;HOXD9;FGFR2 |
| Regulation Of Neurotransmitter Receptor Activity | 0.00 | APP;NLGN1;PSCA;BEGAIN;CACNG7;CRHBP;CACNG8;NETO1;PTK2B;DLGAP1;NPTX1;CACNG2;DLGAP3;CACNG3;DLGAP2;CACNG5;DLGAP4;PRRT1;LYNX1;MEF2C;DAPK1;OPRM1;SHISA9;SHISA7;SHISA6;CRH;SHANK1 |
| Regulation Of Cell Migration (GO:0030334) | 0.00 | SEMA5A;SEMA5B;IFITM1;SERPINE2;CSF1;ZMYND8;FAM107A;ADARB1;FGF1;FGF2;FGF3;MYLK;ACTG1;IGF1R;CDH5;FGF7;FGF8;CDH1;FAM110C;ADORA3;DPYSL3;AKT1;CYP1B1;TNR;PDGFRB;EPHA4;PDGFRA;WNT5B;LIMCH1;APLP2;F2R;MAGI2;PRKCA;MITF;KRT5;SRCIN1;TIAM1;CLDN4;SFRP1;SFRP2;BRMS1;DAAM2;RUFY3;SPATA13;MYADM;DPEP1;EPHA1;CSF1R;GRB7;MACF1;SEMA7A;MAZ;PDGFB;CHRD;TWIST2;PDGFA;TWIST1;THY1;FUT3;PODXL;PDGFD;FGF20;PLXNA1;SRGAP3;ZNF268;PLXNA4;ARSB;FGF21;NTNG1;NGFR;NTNG2;ACE;UBE2I;RCC2;OSGIN1;ALOX15B;SMAD7;COL1A1;BST2;FGF17;DAB2;BMP2;FER;STK24;FGF19;FGF18;CASS4;CNTN1;PLXNB2;PLXNB1;AMOTL2;TDGF1;FERMT3;PTPRT;PTPRU;ROBO4;LDB1;FLT4;PIK3CD;LAMC2;SYNE2;CX3CL1;FLCN;PIP5KL1;ITGAX;EMILIN2;PHACTR1;AMH;MMP7;TET1;RHOD;DUSP22;FGR;F7;MMP14;IL1B;CDH11;MMRN2;CDH13;ITGA6;SGK1;CAMK2B;KANK1;HDAC2;TNXB;SRC;SEMA3B;SEMA3G;NEDD9;PRR5L;AIF1;THBS1;EGFR;INS;PAK1;WNT11;CCL5;NTF3;TP53INP1;GNA12;GCNT2;MGAT3;PTK2B;FYN;CTNNA2;RFFL;SASH1;SPAG9;CCL24;SEMA4A;CD74;F10;NTRK3;SEMA4B;STAT3;SEMA4C;SULF1;TBX5;PODN;SORL1;MAPK15;SLURP1;VEGFA;MYO1C;APC;SST;KIF26A;F2RL1;SPRY2;EPPK1;LGR6;BCAR1;CCL26 |
| Monoatomic Cation Transmembrane Transport (G | 0.00 | KCNG1;KCNG2;SLC46A1;KCNG4;PKD2L1;CLU;SLC8A2;SLC39A7;SLC12A6;SLC39A2;SLC12A7;SCN1B;SLC39A4;KCNH1;KCNH2;KCNH3;SLC6A19;SLC6A18;SLC30A8;KCNK10;SCARA5;SLC11A1;SLC6A15;SLC6A13;SLC6A12;KCNK16;SLC6A11;CACNA2D4;KCNK17;KCNK18;CACNB1;SCNN1G;SLC9A3;SCNN1D;KCNQ1;SCNN1B;KCNQ2;SCNN1A;KCNQ4;ATP6V0D1;KCNE1;HPN;CACNA1A;SLC1A3;ABCB8;CACNA1D;CACNA1C;SLC30A10;NALCN;CACNA1G;TRPM2;KCNV1;GRIN2A;KCNN2;KCNN3;TRPM8;TRPM5;ATP6V1C1;ATP6V0A1;TRPM4;TRPV2;SLC4A11;TRPV3;GRIN2B;GRIN2D;ATP4A;SLC6A6;SLC6A7;SLC6A9;NOX5;KCNC1;KCNK9;KCNC2;ATP2A3;SLC40A1;ATP2A1;TCIRG1;SLC6A1;SLC6A2;PKD1;RYR3;SLC6A3;CNGA3;SLC36A3;TRPC6;KCND2;TRPC3;SLC39A14;TMEM175;SLC29A4;SCN4A;SCN4B;ATP6V1B1;CNGB1;KCNA1;KCNA3;KCNA4;KCNA5;KCNA6;CACNG1;LRRC26;MCOLN3;SLC12A4;KCNJ11;KCNJ8;KCNB1;KCNB2;ATP2B4;OTOP3;TPCN2;SLC8A3;P2RX7;KCNS1;KCNS2;KCNS3;KCNK2;HCN2;HCN1;KCNK4 |
| Inorganic Cation Transmembrane Transport (GO:( | 0.00 | KCNG1;KCNG2;SLC46A1;KCNG4;AQP8;PKD2L1;CLU;AQP1;SLC8A2;SLC39A7;SLC12A6;SLC39A2;SLC12A7;SCN1B;SLC39A4;KCNH1;KCNH2;KCNH3;SLC6A19;SLC6A18;SLC30A8;KCNK10;SCARA5;SLC11A1;SLC6A15;SLC6A13;SLC6A12;KCNK16;SLC6A11;CACNA2D4;KCNK17;KCNK18;CACNB1;SCNN1G;SLC9A3;SCNN1D;KCNQ1;SCNN1B;KCNQ2;SCNN1A;KCNQ4;ATP6V0D1;RHCG;KCNE1;HPN;CACNA1A;SLC1A3;ABCB8;CACNA1D;CACNA1C;SLC30A10;NALCN;CACNA1H;CACNA1G;TRPM2;KCNV1;KCNN2;KCNN3;TRPM8;TRPM5;ATP6V1C1;ATP6V0A1;TRPM4;TRPV2;SLC4A11;TRPV3;ATP4A;SLC6A6;SLC6A7;SLC6A9;NOX5;KCNC1;KCNK9;KCNC2;ATP2A3;SLC40A1;ATP2A1;TCIRG1;SLC6A1;SLC6A2;PKD1;RYR3;SLC6A3;SLC36A3;TRPC6;KCND2;TRPC3;HTR3E;HTR3A;HTR3B;SLC39A14;TMEM175;SCN4A;SCN4B;ATP6V1B1;KCNA1;KCNA3;KCNA4;KCNA5;KCNA6;CACNG1;LRRC26;MCOLN3;SLC12A4;KCNJ11;KCNJ8;KCNB1;KCNB2;ATP2B4;OTOP3;TPCN2;SLC8A3;P2RX7;KCNS1;KCNS2;KCNS3;KCNK2;HCN2;HCN1;KCNK4 |
| Potassium Ion Transport (GO:0006813) | 0.00 | KCNG1;KCNG2;KCNG4;KCNC1;KCNK9;KCNC2;PKD2L1;AQP1;SLC12A6;SLC12A7;KCNH1;KCNH2;KCNH3;KCNK10;KCND2;KCND3;KCNK16;KCNAB1;KCNK17;KCNK18;KCNAB3;TMEM175;SLC9A3;KCNQ1;KCNQ2;KCNMA1;KCNQ4;KCNE1;KCNA1;HPN;KCNA3;SLC1A3;KCNA4;KCNA5;KCNA6;NALCN;KCNV1;KCNMB2;KCNMB3;KCNN2;SLC17A7;KCNN3;LRRC26;KCNJ6;SLC12A4;KCNJ11;KCNJ8;KCNB1;KCNIP2;KCNJ12;KCNB2;KCNJ15;KCNJ16;KCNS1;KCNS2;KCNS3;KCNK2;HCN2;HCN1;KCNK4 |
| Neuropeptide Signaling Pathway (GO:0007218) | 0.00 | GRP;NXPH2;NXPH4;RXFP4;OPRL1;LTB4R;MCHR2;RXFP3;GLRA1;GALR3;GALR1;NPBWR1;NPY;NPPA;NPBWR2;PENK;NTSR1;NTSR2;GPR37;SORT1;OPRK1;SORCS1;OPRM1;SSTR1;PDYN;SORCS3;SORCS2;SORL1;SSTR4;HCRTR2;SSTR5;PYY;LTB4R2;GLRB;PNOC;RAPGEF2;CPE;NMUR1 |
| Skeletal System Development (GO:0001501) | 0.00 | COL18A1;SPEF2;IHH;XYLT1;HOXA13;PKD1;ETS2;HOXA11;GLI2;COMP;ADAMTS4;AKAP13;FGF8;HYAL1;HYAL2;TRPS1;SH3PXD2B;SOX9;PITX1;NKX3-2;CMKLR1;SOX5;TGM2;WNT5A;EN1;PAX1;KIAA1217;MMP14;COL2A1;MMP16;CDH11;CHI3L1;DLX5;ANKRD11;NPR3;SHOX2;COL11A2;CHRD;PRELP;COL19A1;PTHLH;DLL3;CLEC3A;EXTL1;WNT11;ANKH;DEAF1;BGLAP;ALX4;WWOX;ZBTB16;GDF2;BMP8B;GDF3;SULF1;BMP7;TBX4;TBX3;COL1A1;EXT1;BMP2;GNAS;ALPL;GHRL;COL9A2;HOXD3;INHA;FGFR3;FGFR2;FGFR1 |
| Chordate Embryonic Development (GO:0043009) | 0.00 | TRIM71;PTPRR;XYLT1;BRCA1;GATA3;PKD1;FLCN;WNT11;DEAF1;SOX8;ZNF568;NKX2-5;TBX1;NKX2-6;FOXF1;TGFB3;WNT7B;WNT5A;IGF2;LIG4;PROX1;SULF1;ISL1;BMP7;GRHL2;VEGFA;KIAA1217;COL1A1;BMP2;ALDH1A2;ZNF358;EGFL8;FGFR2;FGFR1 |
| Transmembrane Receptor Protein Tyrosine Kinase | 0.00 | CSF1;IRS2;PLAT;FGF1;FGF2;FGF3;IGF1R;FGF7;FGF8;STMN1;GRB10;AKT1;ANKS1B;EPHB3;EPHA5;PDGFRB;EPHA4;PDGFRA;EPHA6;TSC2;TIAM1;DOK2;DOK3;COL4A2;CD8A;DOK5;DOK6;TYRO3;EPHA1;GPLD1;NGEF;GRB7;CSF1R;SHC2;EPHA10;SHC1;PDGFB;PDGFA;MST1R;PIK3R1;ADRB2;KALRN;ART4;SOCS2;FLRT2;PDGFD;FLRT1;FGF20;FGF21;LYN;KL;SRMS;RARRES2;IGF2;PTK6;FGF17;FER;EFNA2;FGF19;FGF18;CASS4;GRB2;FGFR4;FGFR3;FGFR2;FGFR1;DDR1;PTPRT;TRIM71;FLT4;CD3E;GHR;BDKRB2;BLNK;INSRR;NRG1;NGF;SLC39A14;VAV1;VAV2;FGR;CNKSR1;ZAP70;LCK;RAPGEF2;LCP2;ARHGEF7;LTK;RGS14;SRC;NEDD9;EGFR;INS;PAK1;NTF3;ERBB2;PTK2B;FYN;SH2B2;NTRK1;ANGPT2;ANGPT1;GDF15;SORT1;NTRK3;TIE1;STAT3;CSHL1;SORBS1;SULF1;BAIAP2;VEGFA;NELL1;CD4;CD7;CLNK;TEK;FRK;BCAR1 |
| Enzyme-Linked Receptor Protein Signaling Pathwa | 0.01 | DDR1;PTPRT;PTPRU;TRIO;CSF1;FLT4;CD3E;PTPRF;NPPA;BDKRB2;BLNK;JAK3;PDGFRB;PDGFRA;INSRR;NGF;FGR;CNKSR1;ZAP70;DOK2;DOK3;CD8A;DOK5;LCK;DOK6;TYRO3;LCP2;GUCY2C;CSF1R;SHC2;LTK;GUCY2D;LRP1;NEDD9;MST1R;EGFR;ART4;ERBB2;NTF3;FYN;LYN;NTRK1;SRMS;NTRK3;TIE1;LIF;PTK6;GP6;CD4;CD7;CASS4;CLNK;TEK;FGFR4;FGFR3;FGFR2;BCAR1;FRK;FGFR1 |
| Positive Regulation Of Cell Population Proliferatio | 0.01 | CD86;CSF3;CSF2;CSF1;EPO;CD81;IL5RA;IRS2;FGF1;FGF2;TNF;FGF3;AQP1;IGF1R;FGF7;TCL1A;FGF8;CCND2;MYC;CHP2;AKT1;SOX8;SOX9;PDGFRB;EPHA4;OSR2;PDGFRA;MEF2C;NKX2-6;PRKCH;CLEC11A;SERPINF2;F2R;WNT5A;MATK;LIG4;FOXP1;TIAM1;SFRP1;SFRP2;S100A6;PRTN3;TOX;EPHA1;TSPYL5;CSF1R;PTGFR;SHC1;MAZ;PDGFB;CHRD;PDGFA;CSF2RB;REG3G;MST1R;CIAO1;DPP4;TERT;GRK5;PDGFD;S100A13;FGF20;NKX2-5;ZNF268;WNT2;ELANE;SLAMF1;TRPM4;FGF21;LYN;WWTR1;TFAP2B;TSLP;TGFB3;WNT3A;HTR1A;IGF2;LIF;BST2;BMP4;FGF17;BST1;BMP2;FER;IL6;CDK6;GDNF;TNFSF4;ID2;FGF19;CDK2;FGF18;BCL2;SPDYA;NANOG;FGFR4;TDGF1;FGFR3;NODAL;FGFR2;LGMN;FGF10;FGFR1;CDKN1A;MEGF10;FLT4;ADRA1D;LAMC2;TCIRG1;FCXM1;HOXC10;CX3CL1;CXCL5;ESM1;HYAL1;SIX2;RBPMS2;ITGAX;AMH;CR1;ACTL6B;EDN3;TSC22D1;GLP2R;NME2;EMP2;NRG1;PROX1;PAX2;TGFBR2;GREM1;TGFBR3;CCKBR;IFNG;IL1B;RARA;HCLS1;BIRC5;CDH13;CD47;SHANK2;LTF;HDAC4;HDAC2;TNXB;LRP5;NOD2;AIF1;THBS1;EGFR;PTHLH;GNAI2;INS;PAK1;RASSF10;RAB25;EPCAM;CCL5;NTF3;ERBB2;TBX20;GCNT2;PTK2B;BRD9;PRAME;SCAND3;IL12RB2;TBX1;CD74;TCF7L2;NTRK3;IL34;OSM;AKR1C2;MAPK15;MEIS2;TBX3;TSHR;TBX2;VEGFA;MEIS1;AQP11;PTPN6;INSM1;CRLF1 |
| Embryonic Skeletal System Morphogenesis (GO:0 | 0.01 | TBX1;TFAP2A;OSR2;PDGFRA;TWIST1;TGFBR2;HOXA9;COL2A1;HOXB9;HOXA3;HOXB4;HOXC4;HOXB3;HOXD4;HOXD3;HOXC9;DSCAML1;HOXD9;FGFR2;HOXA4 |
| Actin Filament Organization (GO:0007015) | 0.01 | WIPF1;AVIL;PPP1R9A;BAIAP2L2;SAMD14;BAIAP2L1;DPYSL3;RAC2;PRSS8;GHSR;TPM4;TPM3;ACTN1;MYO7B;SHROOM3;EMP2;MYO7A;PROX1;RHOF;ERMN;TLE6;OOEP;ACTA1;RUFY3;ELMO1;GAS2;HCLS1;ELMO3;EVL;TAGLN3;ARHGEF2;LCP1;CLRN1;NEDD9;CAPG;AIF1;ARHGAP12;CNN3;TTN;SCIN;SH3BP1;NRAP;XIRP1;VILL;WHAMM;GAS2L2;ZBED3;SVIL;MARCKSL1;BAIAP2;ARHGAP25;MAD2L2;MYO1D;MARCKS;ACTC1;MYO1C;MYO15A;MYO5C;CASS4;FAT1;NOSTRIN;SPIRE2;TCAP;GHRL;BCAR1;MYO1F |
| Cell-Cell Adhesion Mediated By Cadherin (GO:00 | 0.01 | PCDHGB4;CDH9;CDH8;PLEKHA7;CDH6;CDH5;CDH4;FER;CDH3;CDH1;CDH10;CDH22;CDH11;CDH12;CDH13;DCHS1;CDH15;CDH18 |
| Regulation Of Monoatomic Cation Transmembran | 0.01 | DPP10;KCNH2;KCNE1;KCNIP1;KCNIP2;KCNIP3;KCNIP4;NEDD4L;CACNA1D;KCNAB1;KCNAB2;PDZK1;KCNAB3;GRIN2B;GRIN2D;DPP6;GRIN2A;SLMAP;HRC;RANGRF;KCNN2;FGF12;HCN2;HCN1 |
| Cell Junction Assembly (GO:0034329) | 0.01 | NLGN1;PCDHGB4;CTNND2;NRXN1;NRXN2;CDH9;CDH8;CDH6;CDH5;FLCN;CDH4;CDH3;SH3BP1;CDH1;SLITRK1;SPOCK2;CDH22;KIRREL3;FARP1;DSCAM;FZD5;PCDHB14;PCDHB11;GRHL2;PCDHB10;POU4F1;SDK1;PCDHB2;CDH10;CDH11;PCDHB16;CDH12;FSCN1;PCDHB6;CDH13;PKP1;PCDHB5;ITGA6;DCHS1;PCDHB4;CDH15;PKP3;PCDHB3;PCDHB9;CDH18;SHANK2;SDK2 |
| Potassium Ion Transmembrane Transport (GO:00 | 0.01 | KCNG1;KCNG2;KCNG4;KCNC1;KCNK9;KCNC2;PKD2L1;SLC12A6;SLC12A7;KCNH1;SLC12A9;KCNH2;KCNH3;KCNK10;KCND2;KCND3;KCNK16;KCNK17;KCNK18;TMEM175;SLC9A3;KCNQ1;KCNQ2;KCNQ4;KCNE1;KCNE3;KCNE4;KCNA1;HPN;KCNA3;SLC1A3;KCNA4;KCNA5;KCNA6;NALCN;KCNV1;KCNN2;KCNN3;KCNJ1;LRRC26;KCNJ6;SLC12A4;SLC12A5;KCNJ11;KCNJ8;KCNB1;KCNIP2;KCNJ12;KCNJ9;KCNB2;KCNJ14;KCNJ15;KCNJ16;ATP4B;LETM1;ATP4A;KCNS1;KCNS2;KCNS3;KCNK2;HCN2;HCN1;KCNK4 |
| Nervous System Development (GO:0007399) | 0.01 | CHRM2;SEMA5A;CNTFR;APP;SH3GL3;ANKLE2;CHRM1;MYT1L;ZMYND8;MSI1;MSI2;CELSR1;FGF2;RAPGEFL1;EDNRB;DPYSL4;NAGLU;CDH1;DPF1;ADORA1;SOX8;NRTN;MNX1;KIRREL3;MEF2C;SRRM4;DSCAM;APLP2;SOX14;LIG4;FEZ1;NINJ2;SCN8A;FEZ2;KCNQ2;TYRO3;HOXB2;TAGLN3;LY6H;DLX5;SHOX2;MST1R;KALRN;KLK6;SNTG2;ATXN1;SCIN;MOBP;ST8SIA4;NPTX1;WNT2;SPTBN1;FZD5;CNTN6;ZBTB16;ST8SIA2;POU6F2;BTBD6;INHBA;DCLK1;PHOX2B;CHERP;FGF17;SDK1;FGF14;PCDHB2;GDNF;ARTN;FGF19;PCDHB6;PCDHB5;DCHS1;PCDHB4;FAT4;PCDHB3;LRCH4;PCDHB9;FGF12;FGF11;SDK2;TRIM71;DOC2A;LDB1;PCSK9;GRIK1;NR2E1;PKD1;ZIC2;ZIC1;JARID2;APC2;CHST8;EMX2;PROP1;ACTL6B;PAX6;NRG1;GFRA1;NAV1;PROX1;NAV2;GFRA3;GFRA4;ZEB2;ALDH5A1;CYP26C1;PAX8;ADORA2A;B3GNT5;ALDH1A2;NDUFS2;EVL;ZNF358;DSCAML1;PAFAH1B3;SHANK2;HDAC4;NLGN1;DCTN1;GSTP1;NRXN1;NRXN2;CAPG;SPG7;ASCL1;TRAK1;EGFR;ACVR1C;SLITRK1;SPOCK2;NTF3;ERBB2;IGF2BP1;SPOCK1;ATOH1;LGI1;FARP1;MARCKSL1;MOG;STAT3;LIMK1;WWP1;CABLES1;PCDHB14;ZIC4;ZIC5;PCDHB11;VAX1;POU4F1;GRHL2;PCDHB10;VAX2;VEGFA;NELL1;MARCKS;NFASC;P2RX5;APC;GLRB;FABP7;MAB21L2;PCDHB16;MDGA1;APBA2;NEUROG1;CBLN1;NEUROG3 |
| Macrophage Activation (GO:0042116) | 0.01 | APP;C1QA;CD93;SLC11A1;ITGB2;DYSF;AZU1;AIF1;CLU;TNF;CX3CL1;FOXP1;TLR1;FCER2;SBNO2;MYO1C;IFNG;EHHADH;MAPT;TLR4;TMEM106A;SNCA |
| Branching Morphogenesis Of An Epithelial Tube (( | 0.01 | NOTCH4;LRP5;TCF21;HOXD11;FGF2;PKD1;RASIP1;GLI2;HOXA11;PAK1;SALL1;MYC;TIE1;GDF2;PAX2;TGFBR2;VEGFA;BMP4;NELL1;SFRP2;GDNF;PAX8;COL4A1;WT1;FGF10 |
| Metal Ion Transport (GO:0030001) | 0.01 | KCNG1;RYR1;KCNG2;RYR2;KCNC1;KCNK9;ATP2A3;ATP2A1;CLU;PKD1;RYR3;AQP1;NIPAL4;SLN;KCNH1;KCNH3;TRPC3;KCND3;SLC11A1;SLC34A1;KCNK16;KCNAB1;KCNK17;KCNK18;KCNAB3;SCNN1G;SLC5A8;SCNN1D;SCN8A;SCNN1B;KCNMA1;KCNQ4;SCN4A;SGK1;SCN4B;CATSPER4;KCNA1;ITPR1;KCNA3;KCNA4;CACNA1D;KCNA5;SLC5A1;KCNA6;SLC5A4;SLC5A5;CACNG6;KCNV1;CACNG7;CACNG8;CCL5;KCNMB2;CDH23;KCNMB3;KCNN2;FYN;SLC17A7;CACNG3;CNNM4;SLC17A3;KCNJ6;KCNJ8;KCNIP2;KCNJ12;KCNB2;KCNJ15;SLC4A11;KCNJ16;ATP2B2;KCNS1;KCNS2;KCNS3;NMUR1;RAMP1;KCNK4 |
| Muscle Contraction (GO:0006936) | 0.01 | RYR1;MYLK2;RYR2;OXTR;ADRA1D;LTB4R;MYOM2;RYR3;CACNA1H;CLCN1;MYLK;TTN;CKMT2;CHRND;GLRA1;EDNRA;SGCA;CHRNE;BDKRB2;APBB2;ANKRD2;NOS1;APBB1;GAMT;TMOD1;CHRNB1;DTNA;TPM4;TPM3;KCNIP2;KCNJ12;MYOF;PGAM2;TACR2;TPCN2;MYH1;MYH2;SMTN;SLMAP;FXYD1;HRC;SSPN;SCN4A;NMUR1;CRYAB;MYH7 |
| Regulation Of Monoatomic Ion Transmembrane Tr | 0.01 | APP;NLGN1;NEDD4L;CACNA1D;CRHBP;CACNG7;CACNG8;PTK2B;CACNG2;CACNG3;SCN1B;YWHAH;CACNG5;PRRT1;MEF2C;TRPC6;DAPK1;SPHK2;ANK2;WWP2;OPRM1;PDZK1;SHISA9;SHISA7;SHISA6;FXYD3;FXYD1;CRH;FXYD7;RANGRF;FXYD6;SCN4B;FGF12;HCN2;SHANK1;HCN1 |
| Regulation Of Monocyte Differentiation (GO:0045 | 0.01 | CD74;CD4;CDK6;CSF1;FES;IL34;MYC;IRF7;HOXA7;HLA-DRB1;FOXP1;ZFP36L1 |
| Positive Regulation Of Transcription By RNA Polyn | 0.01 | MDC1;APP;SPI1;PWWP2B;ZMYND8;FOXI1;ABRA;SPIB;ELK4;SOX2;IL4I1;SOX1;CREB3L3;SOX17;SOX18;ZMIZ1;MYC;ZMIZ2;TNFSF11;HOXA1;SOX8;SOX9;HOXA7;IER5;TP63;HOXA5;NKX2-8;MEF2C;NKX2-6;SLC11A1;SERPINF2;SOX14;HNF1B;BHLHA15;HNF1A;ISL1;RUNX2;RUNX1;ISL2;SFRP2;TFR2;DDIT3;RFX4;HOXB4;HOXB3;HOXB2;HOXB1;MAML3;HOXB7;ZFPM1;SKAP1;CASZ1;CTBP2;AIRE;EPAS1;TXK;GATA6;PRDM15;GATA5;TWIST1;GATA4;GATA3;PIK3R1;GATA2;MED12L;GALR3;PCBP1;GALR1;DHX36;HLTF;HOXC4;ALX1;FAD;NKX2-5;ZNF268;RREB1;ZBED4;ZBED3;S100A10;STAT5A;EGR1;WWTR1;CREBBP;FOXF2;FZD5;BCL11B;BTBD18;FOXF1;NR2F1;IGF2;NFATC3;GDF2;LIF;NFATC1;NR2F2;NR1D1;PBX1;ST18;SOX30;NANOG;HOXD4;HOXD3;HOXD8;TP73;CD40;RARG;PTPRN;TCF21;BRCA1;MEOX2;MED15;HOXC10;GLI3;CX3CL1;GLI2;HEY1;HNF4A;HYAL2;HTATIP2;VENTX;MYOG;FOXD2;ANXA2;TCF15;NME2;TFEB;PAX3;TET1;PAX6;PROX1;POU5F1;PAX2;RGMA;TOX3;PLSCR1;TOX2;KAT2A;ZEB2;NR5A2;PAX8;TAL1;RARA;HCLS1;RARB;ARHGEF2;TLR4;NFIX;TMPRSS6;SATB2;NR1I2;HOXD13;ASCL1;ASCL2;EGFR;FSTL3;RXRB;EPCAM;PLAGL1;HSF1;STAT4;APBB2;APBB1;MEF2D;BRD4;WWOX;CDX2;CDX1;NFYC;STAT3;DAB2IP;OSM;NR1H3;GRHL3;LILRB1;POU4F1;GRHL2;VEGFA;HIPK2;POU4F3;SKI;FOXR1;HEYL;PER1;WT1;NFIC;F2RL1;AGRN;NEUROG1;FOXA2;PF4;NEUROG3;CSF3;CCNT1;CD81;CRTC1;RORA;PRDM2;FGF1;ETS1;FGF2;TNF;ETS2;HOXA10;SALL1;CHP2;PITX2;EOMES;OSR2;NKX6-3;ZBTB38;LMO3;WNT5A;MITF;LMO7;NPAS4;MYCN;LMX1A;PGR;ERG;SQSTM1;DLX1;DLX5;NOTCH4;BHLHE23;GLIS1;SBNO2;NLRP3;E4F1;CAMTA2;CYTL1;OTX2;OTX1;NKX6-1;WNT2;ZNF300;TFAP2A;ESRRA;SMAD2;TFAP2B;TFAP2C;ZFHX3;SMAD3;AUTS2;TGFB3;WNT3A;ANKRD23;HMGA1;SMAD9;FOXN4;ESRRG;INHBA;POU2F2;ESR1;PHOX2B;PHOX2A;BMP4;NR4A2;BMP2;IL6;DAB2;COPS5;LHX3;GDNF;TLX2;MYF6;SP7;FGFR2;CIITA;THRB;ONECUT2;LDB1;THRA;ONECUT1;FOXM1;PKD1;FLCN;HHEX;NCK2;NOS1;INSL3;MYOCD;PEG3;OLIG3;OLIG2;UTF1;DCN;TNFRSF1A;GREM1;ELF1;TFDP1;ELF3;HAND2;HAND1;IRF7;IRF8;CDH13;ITGA6;IRF5;IRF6;ZNF750;HDAC4;NFAT5;HDAC2;HDAC3;ATF6B;UHRF1;LRP5;FOXK2;NOD2;BARX2;MUC1;ATOH8;ATOH1;TBX1;TCF7L2;PRRX1;IRX3;PDX1;TBX5;MEIS2;TBX3;POMC;KLF6;MEIS1;POU2AF1;ASXL2;TAF4;LPIN1;LPIN2;SSBP3;LPIN3;SSBP4 |
| Embryonic Digestive Tract Development (GO:0048 | 0.01 | PDGFRA;FOXF1;RARRES2;TCF21;TNF;GLI3;GLI2;SALL1;ID2;SIX2;RBPMS2;RARB;FGFR2;STRA6 |
| Embryonic Skeletal System Development (GO:004 | 0.02 | OSR2;PDGFRA;WNT5A;XYLT1;SULF1;COL1A1;KIAA1217;HOXA9;WNT11;HOXB9;DEAF1;HOXA3;HOXB4;HOXC4;HOXB3;HOXD4;HOXD3;HOXC9;DSCAML1;HOXD9;HOXA4 |
| Heart Development (GO:0007507) | 0.02 | MESP2;CDKN1A;TCF25;ECE1;PKD1;GLI2;AKAP13;FGF8;SALL1;CASP8;SGCD;SOX17;SOX18;GJA5;SH3PXD2B;PLCE1;SOX9;SGCG;MEF2C;MKKS;MYOCD;NKX2-6;TSC2;CACYBP;FOXL1;ISL1;TGFBR2;TGFBR3;KAT2A;MYL2;HAND2;HAND1;MMP23A;ALPK3;ALPK2;ZFPM1;MYH7;POPDC2;NDRG4;PDGFB;GATA5;GATA4;CACNA1C;PDLIM3;ADAP2;SCUBE1;FLRT2;SMYD4;TNNI3;NKX2-5;MEF2D;PDLIM4;STRA6;TBX1;FOXF1;NTRK3;TBX5;BMP7;POU4F1;SGCZ;TBX3;TBX2;VEGFA;BMP4;BMP2;TH;WT1;ID2;TCAP;CCM2;TEK;DZIP1;FGF12;TDGF1 |
| Positive Regulation Of Protein Serine/Threonine Ki | 0.02 | CD40;FGF1;FGF2;TNF;PKD1;PIK3CG;SLC8A2;GHR;RPTOR;TCL1A;CCND2;CCND1;AKT1;TNFSF11;PDGFRB;MEF2C;EDN3;WNT5A;PROX1;IFNG;IL1B;RAPGEF2;ADAM8;TLR4;TELO2;LTF;CSF1R;SRC;PDGFB;PDGFA;ADRB2;MST1R;NOD2;THBS1;PIK3R6;EGFR;PIK3R5;PDGFD;ERBB2;NTF3;DRD4;ELANE;SNCA;IL34;NTRK3;ATP2B4;DAB2IP;IRGM;VEGFA;FGF18;SPDYA;NEK10;TDGF1;FGFR1 |
| Mesonephros Development (GO:0001823) | 0.02 | OSR2;FGF8;PAX8;ZBTB16;GATA3;FGF1;BMP7;PAX2;FGF10 |
| Immunoglobulin Mediated Immune Response (GO | 0.02 | CD74;HLA-DRB5;CD81;MPL;CSF2RB;HLA-DMA;HLA-DMB;CD19;IL2RB;IL21R;HLA-DRA;HLA-DOA;ZP3;HLA-DOB;HLA-DQA2;HLA-DQA1;HLA-DQB2;HLA-DRB1;HLA-DPA1 |
| Positive Regulation Of Intracellular Signal Transdu | 0.02 | APP;TRAF3IP2;SOX2;LGALS1;FAM110C;TNFSF10;PLCE1;TNFSF11;LGALS9;SOX9;TRIM21;PRKCB;UNC5B;GPR37L1;IL23A;DDIT3;ALPK1;TSPYL5;PDGFB;PDGFA;GATA3;MST1R;ART4;PIK3R5;PDGFD;DHX36;S100A13;FADD;LYN;TNFSF14;PLK2;IGF2;LIF;BST2;CORO7;TDGF1;HLA-DRB1;CC2D1A;NODAL;DDR1;CD40;SLC44A2;SECTM1;CX3CL1;BBC3;RPTOR;AKAP13;TRIM6;TRIM5;CCR7;TNFAIP8L3;IKBKE;MAP4K1;GPR37;AVPR1B;SPHK2;FCRL3;F7;ADRB3;IL1B;HCLS1;RAPGEF2;HCST;TLR4;TMEM106A;EGFR;GCNT2;EXTL3;BRD4;P2RY12;CD74;F10;GDF15;DAB2IP;SCTR;MIB2;OSM;AKR1C2;VEGFA;NLRP12;RIT2;F2RL1;GHRL;SEMA5A;CD86;CSF3;CD81;GPR62;FGF1;FGF2;TNF;IGF1R;PYCARD;TGM2;PDGFRB;PDGFRA;UNC5CL;F2R;WNT5A;CARTPT;EDAR;DOK2;DOK3;DOK5;DOK6;ADAM8;CARD16;NUPR1;S100A9;CARD11;SHC1;MAZ;ADRB2;UBD;NLRP3;KIAA0319;TGFB3;BMP7;BMP4;MFHAS1;DAB2;BMP2;IL6;CASS4;CNTN1;PLXNB1;FGFR3;FGFR2;FGF10;FGFR1;FLT4;SCHIP1;DUSP19;PIK3CD;ADRA1D;ADRA1B;PIK3CG;FLCN;CASP8;GRM4;TRIM68;CASP1;MLST8;AMH;BOK;HSP90AA1;TRIM62;KSR1;VWF;RFPL2;IRAK4;ADRA2C;WNT16;DCN;TNFRSF1A;FGR;LCK;NOS1AP;TRAF5;TRIM58;CDH13;CHI3L1;TNFRSF25;LTB;LTF;CX3CR1;NFAT5;NLGN1;HDAC3;SRC;NOD1;NOD2;CXXC5;THBS1;INS;WNT11;CCL5;ERBB2;SASH1;NTRK1;TBX1;TCF7L2;ANGPT1;NTRK3;POMC;P2RX5;CD4;MYO1C;P2RX2;LEP;PDCD4;SPRY2;PTPN6;TEK;TRIM34 |
| Positive Regulation Of Leukocyte Cell-Cell Adhesi | 0.02 | NFAT5;ITGB2;THY1;ETS1;TNF;PYCARD;HLA-DMA;HLA-DMB;FCHO1;TNFSF11;HLA-DOA;HLA-DOB;HLA-DQA2;HLA-DQA1;HLA-DPA1;HLA-DRB5;CCDC88B;IL6;LCK;HLA-DRA;PRKCQ;CD47;HLA-DQB2;HLA-DRB1;SKAP1 |
| Peptidyl-Tyrosine Modification (GO:0018212) | 0.02 | CSF1R;SRC;FLT4;PDGFB;EGFR;TTBK1;ERBB2;PTK2B;FYN;JAK3;PDGFRB;LYN;NTRK1;EPHA4;PDGFRA;SRMS;FGR;ZAP70;FER;LCK;ABI3;FES;PTPN6;TEK;EPHA1;FGFR4;FGFR3;FGFR2;FGFR1 |
| MHC Class II Protein Complex Assembly (GO:000 | 0.02 | HLA-DRB5;HLA-DMA;HLA-DMB;HLA-DRA;HLA-DOA;HLA-DOB;HLA-DQA2;HLA-DQA1;HLA-DQB2;HLA-DRB1;HLA-DPA1 |
| Peptide Antigen Assembly With MHC Class II Prot | 0.02 | HLA-DRB5;HLA-DMA;HLA-DMB;HLA-DRA;HLA-DOA;HLA-DOB;HLA-DQA2;HLA-DQA1;HLA-DQB2;HLA-DRB1;HLA-DPA1 |
| Regulation Of Macrophage Differentiation (GO:00 | 0.02 | CASP8;CSF1;IL34;ID2;HSF1;LIF;PRKCA;INHBA;FADD;INHA;PF4 |
| Regulation Of Non-Canonical Wnt Signaling Pathw | 0.02 | SFRP4;TIAM1;SFRP1;LBX2;RNF213;ZNRF3;SPEF1;WNT5B;DAAM2;RSPO3;CSNK1E |

| | | |
|---|---|---|
| Positive Regulation Of Synaptic Transmission (GO | 0.02 | APP;NLGN1;RGS14;CLSTN3;NRXN1;ADRA1D;GRIK2;FOXM1;TNF;NALCN;PRKCZ;SLC8A2;CRHR2;INS;RIMS2;RIMS1;CACNG7;CACNG8;GRIN2A;FLOT1;PTK2B;CACNG2;CACNG3;SLC18A3;CACNG5;LGI1;PLK2;EIF2AK4;GRIN2B;SHISA7;BAIAP3;GRIN2D;SLC8A3;STX1B;SQSTM1;RAPSN;SHANK2;LGMN |
| Aortic Valve Development (GO:0003176) | 0.02 | RB1;SMAD2;NOS3;ELN;TIE1;GATA5;TWIST1;GATA4;NFATC1;GATA3;SMAD6;TNFRSF1B;TGFBR2;TNFRSF1A;BMP4;HEYL;BMP2;HEY1;NPPA;TBX20;SLIT1;SOX9;NKX2-5 |
| Positive Regulation Of Myeloid Leukocyte Differen | 0.02 | CD74;CSF1;IL34;TESC;LIF;NEDD9;PRKCA;TNF;POU4F1;ZFP36L1;RUNX1;CD4;CASP8;IFNG;FES;IL23A;ID2;HSF1;TNFSF11;HCLS1;FADD;HLA-DRB1;PF4 |
| Ureteric Bud Morphogenesis (GO:0060675) | 0.02 | TCF21;GATA3;HOXD11;FGF2;PAX2;HOXA11;BMP4;SALL1;WNT11;GDNF;PAX8;WT1;MYC |
| Peptidyl-Tyrosine Phosphorylation (GO:0018108) | 0.02 | DDR1;CSF1R;LTK;SRC;FLT4;PDGFB;EGFR;TTBK1;IGF1R;ERBB2;PTK2B;FYN;JAK3;PDGFRB;LYN;NTRK1;EPHA4;PDGFRA;SRMS;PTK6;FGR;ZAP70;FER;LCK;ABI3;FES;LEP;PTPN6;TEK;EPHA1;FGFR4;FGFR3;FGFR2;FGFR1 |
| Regulation Of T Cell Activation (GO:0050863) | 0.02 | PTPN22;THY1;VTCN1;PYCARD;IL4I1;LGALS3;HLA-DMA;HLA-DMB;FCHO1;CCL5;TNFSF11;TNFAIP8L2;HLA-DOA;HLA-DOB;JAK3;HLA-DQA2;HLA-DQA1;HLA-DPA1;HLA-DRB5;TNFRSF1B;LAX1;CCDC88B;KAT2A;CD4;LCK;HLA-DRA;TNFSF9;PRKCQ;TNFSF8;CD47;HLA-DQB2;HLA-DRB1 |
| Positive Regulation Of Leukocyte Migration (GO:0 | 0.03 | PLVAP;CSF1;EDN3;TNFRSF18;THY1;AIF1;TNF;FUT4;ICAM1;VEGFA;LGALS3;F7;IL6;F2RL1;MADCAM1;NCKAP1L;ADAM8;ZP3;JAM3 |
| Modulation Of Chemical Synaptic Transmission (( | 0.03 | APP;TRIO;CLSTN3;GRIK5;DYSF;GRIK3;LRRC4;GRIK4;GRIK1;GRIK2;TNF;GRM1;SLC8A2;RIMS2;RIMS1;GRM2;GRM5;GRM4;GRM7;GRM6;GRM8;TMEM108;DLGAP1;DLGAP3;DLGAP2;DLGAP4;MEF2C;SLC38A1;GRID2;UNC13A;GRID1;MYOF;SYN3;DGKZ;NPAS4;IL1B;MCTP2;DGKI;CX3CR1;NLGN1;CACNA1B;CACNA1A;CPLX1;BEGAIN;NPTX1;HCRT;LRRC4C;GRIA4;CACNG5;LGI1;BTBD9;BAIAP3;SLC8A3;GRIN3B;SLC6A9 |
| Cell-Cell Junction Organization (GO:0045216) | 0.03 | CSF1R;PCDHGB4;CTNND2;CDH9;CDH8;CDH6;CDH5;FLCN;CDH4;CDH3;PARD6A;MARVELD2;CDH1;CDH22;CD177;DSP;SMAD3;TGFB3;F2R;OCLN;CDH10;CDH11;CDH12;FSCN1;MYADM;CDH13;PKP1;F2RL1;PRTN3;DCHS1;CDH15;PKP3;CDH18;LIMS2 |
| Circulatory System Development (GO:0072359) | 0.03 | CDKN1A;TCF25;FLT4;TCF21;ECE1;PKD1;GLI2;CDH5;AKAP13;SALL1;CASP8;SOX17;HEY1;SOX18;GJA5;SH3PXD2B;PLCE1;SOX9;MEF2C;MKKS;TSC2;CACYBP;FOXL1;ISL1;TGFBR2;KAT2A;MYL2;HAND2;HAND1;ALPK3;ZFPM1;CRB2;POPDC2;NXN;NOTCH4;PDGFB;CACNA1C;PDLIM3;ADAP2;SMYD4;TNNI3;PDLIM4;STRA6;TBX1;FOXF1;NTRK3;TBX5;POU4F1;HEYL;BMP2;TH;WT1;ID2;CCM2;TEK;DZIP1;FGF12;TDGF1 |
| Regulation Of Chemokine Production (GO:00326 | 0.03 | APP;CSF1R;CD74;EGR1;TSLP;AIRE;WNT5A;AIF1;TNF;ALOX15B;LILRB4;FOXP1;PYCARD;SIGIRR;IL6;IFNG;TNFSF4;F2RL1;FFAR2;LGALS9;LCP1;TLR4;ELANE |
| Lung Epithelium Development (GO:0060428) | 0.03 | BMP4;ERRFI1;WNT7B;RCAN3;GRHL2;PKD1;FGF10;STRA6 |
| Mesenchymal To Epithelial Transition (GO:00602 | 0.03 | BMP4;SALL1;PAX8;WT1;FZD7;SIX2;GATA3;PAX2 |
| Muscle Cell Fate Commitment (GO:0042693) | 0.03 | MYOG;TBX1;MEF2C;WT1;MYL2;MYF6;MYF5;TBX2 |
| Ventricular Cardiac Muscle Tissue Development ( | 0.03 | IRX3;TNNC1;PROX1;TBX5;ISL1;HOPX;SMAD7;TGFBR3;ID2;MYL2;HAND1;MYL3;TNNI3;NKX2-5;FGFR2;MYH7 |
| Cell-Cell Junction Assembly (GO:0007043) | 0.03 | CNTNAP1;PCDHGB4;CTNND2;CDH9;CDH8;ACTG1;CDH6;CDH5;FLCN;CDH4;CDH3;CDH1;GJA5;CDH22;CTNNA1;PIP5K1C;JAM3;TBCD;PRKCA;SMAD7;CLDN11;GJB2;GJD3;CDH10;GJB6;CDH11;CDH12;FSCN1;CDH13;PKP1;CD9;DCHS1;CDH15;PKP3;CDH18 |
| Synapse Assembly (GO:0007416) | 0.03 | GABRB2;NLGN1;NRXN1;LRRC4;NRXN2;HAPLN4;CDH1;SLITRK1;SPOCK2;SRGAP3;KIRREL3;FARP1;GABRA1;GRID2;DSCAM;FZD5;WNT3A;WNT5A;PCDHB14;PCDHB11;PCDHB10;POU4F1;NPAS4;SDK1;PCDHB2;PCDHB16;PCDHB6;PCDHB5;PLXNB2;PCDHB4;PLXNB1;PCDHB3;PCDHB9;SHANK2;SDK2 |
| SMAD Protein Signal Transduction (GO:0060395) | 0.04 | ZMIZ1;ZMIZ2;ATOH8;SMAD2;SMAD3;GDF15;TGFB3;MAGI2;GDF2;BMP8B;SMAD9;INHBB;GDF3;INHBA;GDF6;SMAD6;BMP7;INHBC;RUNX2;GDF7;HIPK2;SMAD7;BMP4;SKI;BMP2;INHA;NODAL |
| Generation Of Neurons (GO:0048699) | 0.04 | LDB1;MYT1L;PCSK9;OTP;CELSR1;RNF112;FGF8;SIX3;MEF2C;EMX2;WNT5B;DCC;WNT5A;EN1;NAV1;WNT16;RUNX3;RUNX2;RUNX1;CIT;TUBB2B;LMX1A;SNPH;MAP1B;RAPGEF2;MAPT;ASTN2;SDCCAG8;RTN3;BRSK2;TUBGCP2;LRP4;ASCL1;ASCL2;WNT6;WNT11;FUT9;ERBB2;SPOCK1;DISC1;KIAA0319;WNT2;MARK2;WNT3;IRX2;AUTS2;FZD5;WNT3A;IRX6;FZD7;IRX3;WNT7B;IRX4;DCDC2;ATP2B2;CCK;VAX1;VAX2;PHOX2B;NR4A2;NELL1;CDK6;LHX3;ARTN;NOVA2;LHX6;PLXNB2;INSM1;MDGA1;LHX9;FGFR1;LHX8 |
| Gland Development (GO:0048732) | 0.04 | GCM2;UGT1A10;NOTCH4;PCSK9;TCF21;GLI2;SOX2;SALL1;WNT11;SIX3;SOX9;FADD;NKX2-5;UGT1A9;UGT1A8;UGT1A7;EPHB3;STRA6;TBX1;NKX2-8;RIPK3;TGFB3;NRG1;PROX1;TBX3;DKK3;TGFBR3;TG;PAX8;WT1;HAND2;HOXD3;EZR;TDGF1;FGFR2;FGF10 |
| Regulation Of MAP Kinase Activity (GO:0043405) | 0.04 | CD40;DUSP19;FGF1;FGF2;TNF;PIK3CG;GHR;HYAL2;TNFSF11;PDGFRB;MEF2C;EDN3;KSR1;CD300A;DUSP1;LAX1;LIME1;HGS;IL1B;TLR4;RGS14;GSTP1;PDGFB;PDGFA;MST1R;NOD2;THBS1;PIK3R6;EGFR;PIK3R5;PDGFD;ERBB2;NTF3;APOE;DRD4;ELANE;LYN;IL34;NTRK3;DAB2IP;BMP7;SORL1;VEGFA;BMP4;BMP2;FGF18;NEK10;SPRY2;SPRY1;TDGF1;FGFR1 |
| Endocardial Cushion Morphogenesis (GO:00032 | 0.04 | SMAD2;NOS3;GATA5;ISL1;TBX2;TGFBR2;BMP4;ADAMTS5;HEYL;BMP2;APC;HEY1;TBX20;SOX9 |
| Positive Regulation Of Cytokine Production (GO:0 | 0.04 | CD86;APP;CSF2;RORA;TNF;PYCARD;DHX58;CYP1B1;LGALS9;TRIM27;SLC11A1;SERPINF2;F2R;WNT5A;CYBA;HLA-A;HLA-G;ISL1;HLA-E;RUNX1;FOXP1;IL23A;DDIT3;PRKCQ;TRIM15;ZFPM1;CARD11;CSF1R;AIRE;TXK;HMHB1;TWIST1;GATA4;GATA3;LY9;PRKCZ;C3;S100A13;NLRP3;FADD;ELANE;SPTBN1;SLAMF1;EGR1;AFAP1L2;CADM1;FZD5;TSLP;GDF2;ISG15;POU2F2;ALOX15B;CCDC88B;IL6;TNFSF4;CNTN1;NOX5;CD200;NODAL;CD40;FLT4;PIK3CD;PTPN22;BRCA1;CLEC9A;PIK3CG;C1QTNF3;TRIM6;HYAL2;CASP1;FLOT1;PDE4B;CCR7;FFAR2;AMH;IKBKE;HLA-DPA1;SPHK2;NFAM1;PDE4D;IL16;TYK2;TLR1;FGR;SLC7A5;KAT2A;IFNG;TRAF3;IL1B;EHHADH;RARA;IRF7;CHI3L1;IRF5;ARHGEF2;LCP1;LTB;ZP3;TLR4;TMEM106A;NOD1;NOD2;AGPAT1;AIF1;INS;CCL1;IL12RB2;CD74;NOS2;STAT3;OSM;LILRB1;SULF1;SORL1;BATF;CD4;CD6;LEP;POU2AF1;F2RL1;IL17D;HSPA1B;HSPA1A |
| Neurotransmitter Uptake (GO:0001504) | 0.04 | SLC22A3;SLC38A1;SLC22A2;SLC1A3;SLC29A4;SLC1A6;SLC29A1;SLC1A7;SLC18A2 |
| Regulation Of B Cell Differentiation (GO:0045577 | 0.04 | SPI1;FCRL3;INHBA;ZFP36L2;ZFP36L1;IL4I1;MMP14;SFRP1;ID2;INPP5D;SLAMF8;NCKAP1L;INHA |
| Aortic Valve Morphogenesis (GO:0003180) | 0.04 | RB1;SMAD2;NOS3;ELN;TIE1;GATA5;TWIST1;GATA4;NFATC1;GATA3;SMAD6;TGFBR2;BMP4;HEYL;HEY1;NPPA;TBX20;CDH11;SLIT1;SOX9;NKX2-5 |
| Muscle Tissue Morphogenesis (GO:0060415) | 0.04 | TBX1;BMP4;TGFBR3;MYLK2;BMP2;TBX20;TCAP;PROX1;NKX2-5;ZFPM1;MYLK;TTN |
| Positive Regulation Of Lymphocyte Activation (GC | 0.04 | NOD2;THY1;PYCARD;HLA-DMA;HLA-DMB;FCHO1;TNFSF11;NCKAP1L;HLA-DOA;HLA-DOB;HLA-DQA2;HLA-DQA1;HLA-DPA1;HLA-DRB5;CCDC88B;IL6;IL23A;LCK;TNFSF4;HLA-DRA;PRKCQ;CD47;HLA-DQB2;HLA-DRB1;FGF10 |
| Adenylate Cyclase-Inhibiting G Protein-Coupled F | 0.04 | CHRM2;CHRM1;ADCY4;OPRL1;FPR2;ADCY7;GNAI2;AKAP12;GRM4;GRM7;GRM8;INSL3;DRD4;P2RY12;GABBR1;GPR37;PSAPL1;HTR1E;HTR1D;PDE2A;HTR1A;HTR1B;OPRK1;GPR37L1;OPRM1;HTR5A;MTNR1A |
| Neurotransmitter Transport (GO:0006836) | 0.04 | SLC22A3;NRXN1;SLC22A2;CHAT;SLC1A2;NRXN2;SLC1A3;LIN7C;SLC1A6;SLC6A2;SLC1A7;SLC6A3;GRM4;SLC17A7;SLC38A2;SLC18A2;SLC18A3;SLC38A1;UNC13A;PTPRN2;SLC32A1;SLC6A15;SYN3;SYN2;SNPH;SLC29A4;SLC29A1 |
| Adherens Junction Organization (GO:0034332) | 0.04 | PCDHGB4;TBCD;CDH9;CDH8;SMAD7;CDH6;CDH5;CDH4;CDH3;CDH1;CDH10;CDH22;CDH11;CDH12;CDH13;DCHS1;PIP5K1C;CDH15;CDH18;JAM3 |
| Epithelium Development (GO:0060429) | 0.04 | TRIM71;ERRFI1;IHH;LOXL3;TCF21;KRT20;MEOX1;PKD1;LGALS3;UPK1B;FGF8;SALL1;UPK1A;DMBT1;SOX17;PTER;SOX18;SOX8;SOX9;CD34;CBR1;CPT1A;NKX2-6;KRT3;RIPK4;KRT32;PROX1;UPK3A;PAX2;KRT9;SPINT1;PAX8;ALDH1A2;HAND1;ZNF358;RHCG;CES1;KANK2;NDRG4;TAGLN;DLX3;GATA6;CNN3;NKX2-5;WNT2;WNT3;FN3K;TBX1;WNT7B;AKR1C2;PPP2R3A;TBX5;GRHL2;TBX2;VEGFA;BMP2;KRT17;WT1;TOLLIP;KRT15;RCAN3;KRT14;CCM2;FGFR2;FGF10 |
| Regulation Of Nucleotide-Binding Oligomerizatior | 0.04 | ZNRF4;PTPN22;SLC15A3;SLC15A4;HSPA1B;HSPA1A |
| Synaptic Transmission, Dopaminergic (GO:00019 | 0.04 | CRHBP;TH;SLC6A2;SLC6A3;DRD5;SNCA |
| Calcium Ion Transmembrane Import Into Cytosol | 0.04 | RYR1;RYR2;CACNA1B;ITPR1;CACNA1A;ITPR2;CACNA1D;HTR2A;CACNA1C;CLU;FGF2;JPH1;CACNA1E;CACNA1H;SLC8A2;CACNA1G;JPH4;CACNA1I;GRIN2A;JPH3;PLCE1;CCR7;F2R;ATP2B4;TRPV2;TRPV3;GP1BA;GP5;GRIN2B;CHERP;GRIN2D;SLC8A3;GP9;LCK;SCN8A;SCN4A;PLCH1;PLCH2;PLCB2 |
| Regulation Of Epithelial Cell Proliferation (GO:00 | 0.04 | CDKN1C;UHRF1;HPN;NOD2;FUT1;EGFR;ZFP36L1;SOX2;FGF7;HYAL1;DEAF1;MYC;ERBB2;SOX9;SCAND3;TBX1;OSR2;TCF7L2;GPX1;ANGPT1;TIE1;NME2;DAB2IP;NR1D1;MCC;RUNX3;SGPP2;PAX2;VEGFA;TGFBR3;BMP4;SFRP1;SFRP2;CDK6;LEP;FGF18;AQP11;EPPK1;NUPR1;FGFR2;NODAL;FGF10 |

| GO: Cellular Component | Adjusted P-value | Genes |
|---|---|---|
| Potassium Channel Complex (GO:0034705) | 1.0944419550000314E-6 | DPP10;KCNG1;CNTNAP2;KCNG2;KCNG4;KCNC1;KCNC2;GRIK5;GRIK3;GRIK4;GRIK1;GRIK2;KCNH1;KCNH2;KCND2;KCND3;KCNAB1;KCNAB2;KCNAB3;KCNQ1;KCNMA1;KCNQ2;KCNQ4;KCNE1;KCNA1;KCNA3;KCNA4;KCNA5;KCNA6;KCNV1;DPP6;KCNMB2;KCNMB3;LRRC26;KCNJ6;KCNIP1;KCNJ11;KCNJ8;KCNB1;KCNIP2;KCNB2;KCNIP3;KCNIP4;KCNJ14;KCNJ16;KCNS1;KCNS2;KCNS3;HCN2;KCNK4 |
| Collagen-Containing Extracellular Matrix (GO:006 | 9.022487414503026E-6 | SPON1;SPARC;COL16A1;SERPINE2;COL12A1;LOXL3;F13A1;PLAT;CLU;LOXL1;ICAM1;LOXL2;LGALS4;COMP;LGALS3;LGALS1;NPPA;TNR;LGALS9;TGM2;POSTN;WNT5B;SERPINF1;COL23A1;KRT1;SERPINF2;WNT5A;ADAM11;PCOLCE;ADAM19;SFRP1;CCDC80;SFRP2;SLPI;COL4A2;COL4A1;S100A6;COL8A1;SERPING1;PRTN3;MXRA7;S100A9;FREM2;FBN2;SEMA7A;PDGFB;HTRA1;LTBP4;LTBP2;FBLN2;APOH;APOE;A2M;WNT2;ELANE;S100A10;COL24A1;TGFB3;RARRES2;PRG4;SSC5D;HRNR;COL1A1;CNTN1;COL20A1;COL9A3;CSPG4;COL9A2;TGFBI;FGFR2;FGF10;FGFR1;LGALS3BP;ACHE;COL18A1;TINAGL1;ELN;CTSZ;PZP;CLEC14A;LAMC2;PCSK6;HAPLN1;ADAMTS4;TIMP2;TIMP3;EMILIN2;SOST;CTSD;ADAMTS9;EGFL7;ANXA2;DST;VWF;COL25A1;MST1;APOA1;SERPINB9;NRG1;DCN;SERPINB6;ADAMTS20;GREM1;F7;PLSCR1;COL2A1;ANGPTL7;PXDN;COL6A1;MMRN2;VWA1;CDH13;ANGPTL5;VWA2;ANG;ZP1;ANGPTL4;ZP3;MATN4;ZP4;C1QB;C1QA;ITIH4;TNXB;LAMA1;COL11A1;LAMA4;C17ORF58;COL11A2;LAMA3;SEMA3B;PRELP;ITIH1;THBS1;COL19A1;THBS4;ACAN;CLEC3B;MUC2;CHADL;GPC1;NCAM1;ANGPT2;ANGPT1;LAMB3;GDF15;F12;PODN;SULF1;KAZALD1;SOD3;MFAP5;SMOC2;COL7A1;MGP;CLC;AGRN;FMOD;PF4;CBLN1 |
| Voltage-Gated Potassium Channel Complex (GO: | 9.022487414503026E-6 | DPP10;KCNG1;CNTNAP2;KCNG2;KCNE1;KCNG4;KCNC1;KCNC2;KCNA1;KCNA3;KCNA4;KCNA5;KCNA6;KCNV1;DPP6;KCNMB2;KCNMB3;KCNH1;KCNH2;LRRC26;KCNJ6;KCND2;KCNIP1;KCNJ8;KCND3;KCNB1;KCNIP2;KCNB2;KCNIP3;KCNIP4;KCNJ14;KCNJ16;KCNAB1;KCNAB2;KCNAB3;KCNS1;KCNS2;KCNQ1;KCNS3;KCNMA1;KCNQ2;KCNQ4;HCN2;HCN1 |
| Postsynaptic Specialization Membrane (GO:0099 | 1.948933953732866E-5 | NLGN1;DAGLA;PTPRS;GRIK5;GRIK3;GRIK4;GRIK1;GRIK2;GRM1;CACNG7;CACNG8;GRIN2A;GRM5;NETO1;LRRTM1;LRRTM2;SLITRK3;CACNG2;CACNG3;CACNG5;GRIA4;GRID2;GRID1;GRIN2B;GRIN2D;GRIN3B;DLG2 |
| Neuron Projection (GO:0043005) | 6.279915940313465E-5 | CYFIP2;CYFIP1;APP;AVIL;CPNE6;STMN2;STMN4;HTR4;CRHBP;SAMD14;STMN1;EPHB3;EPHA5;EPHA4;UNC13A;EPHA6;ANK2;PDYN;GABRG3;ANK1;MAPK8IP3;UHMK1;BIN1;FEZ1;MAP1B;RUFY3;FEZ2;RIN3;STX1A;SPTBN4;EPHA10;HEPACAM;CRHR1;IGSF9;CRHR2;FXR2;INPP5A;MAP2;NPBWR1;NPBWR2;DHX36;PENK;MAP7;APOD;APOE;KCNN3;MAP9;BBS2;CADM1;PLK2;NR1D1;GRIN2B;SLC6A6;CPEB1;TH;PTK7;PNOC;IGSF9B;CD200;GABRB2;ROBO3;ROBO4;PTPRS;KCNC1;KCNC2;GRIK3;SLC6A1;SLC6A2;KIF17;SLC6A3;KIF13B;NEFL;NEFM;ACADM;KIF1A;NEFH;SLC18A2;PACRG;MME;SLC32A1;INSRR;CLDN11;RAPGEF2;MAPT;ARHGEF7;DSCAML1;GRP;KCNA1;KCNA3;HTT;KCNA4;ADCY2;TRAK1;CHRND;GLRA1;PAK1;CHRNE;MICALL2;CTNNA2;GABRD;FARP1;GABRA1;GABRP;GABRA5;KCNB1;GABRA4;KCNB2;DAB2IP;KLHL1;SORBS2;SLC8A3;RIT2;SARM1;GHRL;CDK5R2;CHRM2;CHRM1;FAM107A;KNDC1;COMT;SLC8A2;IGF1R;RPH3A;ATCAY;ADORA3;ADORA1;CAPN2;KIRREL3;CHRNB1;GHSR;DSCAM;MAGI2;SRCIN1;SCN8A;CHRNA1;RTN3;CHRNA2;KCNE3;CHRNA5;KLHL14;CHAT;CHRNA6;SLC1A3;HRH2;NPTX1;DRD4;MARK2;PRPH;HTR1E;CNTN5;CNTN6;HTR1D;HTR1A;HTR1B;SYT8;SYT7;DLG2;CNTN1;CNTN2;CNTNAP2;PNLIPRP2;MAST1;ELAVL4;HTR2A;PPP1R9A;ELFN1;UNC80;GRM2;GRM7;LRRTM1;TMEM108;PRLHR;ZNF385A;HTR3E;HTR3A;RTN4RL2;HTR3B;OPRM1;SSTR1;SSTR4;SSTR5;PALLD;CDH13;TNFRSF25;SHANK2;SHANK1;CX3CR1;CAMK2B;NLGN1;RGS14;DCTN1;OPRL1;LRP4;P2RY1;GNA12;PTK2B;FYN;SNCA;NTRK1;SLC12A5;NTRK3;BNIP3;LIMK1;OPRK1;CCK;HTR5A;MYO1D;KISS1;NELL1;NFASC;MLPH;GLRB;USH1G;RGS12;RAB17;ZACN;VIM;HCN2;HCN1 |
| MHC Protein Complex (GO:0042611) | 1.3018890611301299E-4 | CD74;HLA-DRB5;HLA-B;HLA-C;HLA-A;HLA-F;HLA-E;HLA-DMA;HLA-DMB;HLA-DRA;HLA-DOA;HLA-DOB;HLA-DQA2;HLA-DQA1;HLA-DQB2;HLA-DRB1;HLA-DPA1 |
| Postsynaptic Density Membrane (GO:0098839) | 3.496211229202391E-4 | GRID2;DAGLA;PTPRS;GRID1;GRIK5;GRIK3;GRIK4;GRIK1;GRIK2;GRIN2B;GRM1;GRIN2D;CACNG7;CACNG8;GRIN2A;GRM5;GRIN3B;DLG2;NETO1;CACNG2;CACNG3;CACNG5;GRIA4 |
| Catenin Complex (GO:0016342) | 3.50339818792014E-4 | APC2;PCDHGB4;CDH9;CDH8;CDH6;CDH5;CDH4;CDH3;CDH1;CDH10;CDH22;CDH11;CDH23;CTNNA1;CDH12;CDH13;DCHS1;CTNNA2;CDH15;CDH18 |
| MHC Class II Protein Complex (GO:0042613) | 0.004035363795672352 | CD74;HLA-DRB5;HLA-DMA;HLA-DMB;HLA-DRA;HLA-DOA;HLA-DOB;HLA-DQA2;HLA-DQA1;HLA-DQB2;HLA-DRB1;HLA-DPA1 |
| Axon (GO:0030424) | 0.005464935431501374 | APP;ROBO3;CNTNAP2;ROBO4;PTPRS;KCNC1;KCNC2;CPNE6;GRIK3;ELAVL4;SLC6A1;COMT;SLC6A3;IGF1R;SLC8A2;UNC80;GRM2;ATCAY;GRM7;LRRTM1;KIF13B;TMEM108;NEFL;NEFM;KIF1A;NEFH;SLC18A2;EPHA5;KIRREL3;EPHA4;MME;DSCAM;INSRR;RTN4RL2;OPRM1;MAPK8IP3;SRCIN1;UHMK1;CLDN11;BIN1;PALLD;FEZ1;MAP1B;RUFY3;SCN8A;FEZ2;RIN3;TNFRSF25;MAPT;DSCAML1;HEPACAM;DCTN1;KCNA3;HTT;KCNA4;IGSF9;PAK1;FXR2;DHX36;MAP7;CTNNA2;MAP9;PRPH;SNCA;NTRK1;CNTN5;KCNB1;CNTN6;NTRK3;DAB2IP;CCK;SYT8;SYT7;SLC8A3;MYO1D;NELL1;NFASC;TH;PTK7;CNTN1;CNTN2;GHRL;VIM;CD200;HCN2;HCN1 |
| Microvillus (GO:0005902) | 0.005464935431501374 | SPEF1;ICAM2;PDGFA;ESPN;SPN;PODXL;HYAL2;PROM1;JAM3;PROM2;SLC38A4;AOC3;WWOX;ANGPT1;STARD10;MYO7B;SYTL1;MYO7A;MYO1D;SLC7A5;SLC6A6;MYO1C;FSCN1;FSCN3;ANKS4B;TEK;EZR;RAPGEF3;MYO1F;ATP6V1B1;CLRN1 |
| Actin-Based Cell Projection (GO:0098858) | 0.006734758281240922 | SPEF1;ICAM2;PDGFA;ESPN;ACTG2;DNALI1;SPN;PODXL;HYAL2;PROM1;JAM3;PROM2;AOC3;WWOX;ANGPT1;MYO10;ACTN2;STARD10;MYO7B;UNC5C;MYO7A;ERMN;ACTA2;MYO1D;ACTC1;MYO1C;MYO15A;RUFY3;MYO3A;SPATA13;FSCN1;FSCN3;ANKS4B;CD302;LCP1;TEK;EZR;RAPGEF3;MYO1F;ATP6V1B1;CLRN1 |
| Ionotropic Glutamate Receptor Complex (GO:000 | 0.007602194839511686 | GRID2;GRIK5;GRIK3;GRIK4;GRIK1;GRIK2;SHISA9;GRIN2B;SHISA7;GRIN2D;SHISA6;CACNG7;CACNG8;GRIN2A;GRIN3B;VWC2;PTK2B;CACNG3;CACNG5;GRIA4 |
| Sodium Channel Complex (GO:0034706) | 0.007602194839511686 | GRIK5;GRIK3;GRIK4;GRIK2;SCNN1G;SCNN1D;SCN8A;SCNN1B;SCNN1A;SCN4A;SCN4B;SCN1B;TRPM4 |
| Cation Channel Complex (GO:0034703) | 0.02841814393805828 | RYR1;RYR2;ATP2A1;PKD2L1;KCNA5;KCNA6;PKD1;RYR3;UNC80;TRPM2;CNGA3;TRPM4;TRPC6;TRPC3;KCNIP2;HTR3E;PDE4D;HTR3A;KCNAB1;HTR3B;SCNN1G;SCNN1D;SCNN1B;SCNN1A;FKBP1B;CNGB1;KCNK4 |
| Calcium Channel Complex (GO:0034704) | 0.0335673597126832 | RYR1;RYR2;CACNA1B;CACNA1A;ATP2A1;PKD2L1;CACNA1D;CACNA1C;PKD1;CACNA1E;RYR3;CACNA1H;CACNA1G;CACNG6;CACNA1I;CACNG7;CACNG8;PDE4B;CACNG3;PDE4D;CACNA2D2;CACNA2D4;CACNB1;CACNB2;CACNB4;SCN8A;FKBP1B;CACHD1;SCN4A |
| Lumenal Side Of Endoplasmic Reticulum Membra | 0.0335673597126832 | CD74;HLA-DRB5;HM13;HLA-B;HLA-C;HLA-A;HLA-F;HLA-G;HLA-E;TAPBP;HLA-DRA;HLA-DQA2;HLA-DQA1;HLA-DQB2;HLA-DRB1;HLA-DPA1 |
| Membrane Raft (GO:0045121) | 0.0335673597126832 | APP;ITGB2;TNF;SLC6A3;HYAL2;CAPN2;FLOT1;NOS1;CTSD;CD177;GHSR;CR1;ANXA2;KCND2;MME;NFAM1;EMP2;RTN4RL2;TNFRSF1B;LAX1;TGFBR2;TNFRSF1A;TLR1;ZAP70;PLSCR1;CD8A;LCK;KCNQ1;MALL;MYADM;PRTN3;LCP2;MAPT;EZR;CARD11;SKAP1;ABCG2;KCNE3;AHNAK;SRC;SLC1A2;LRP4;ADCY2;FURIN;KCNA5;THY1;EGFR;LDHB;GPC1;MAL2;FYN;NTSR1;CD55;PROM2;S100A10;LYN;ANGPT1;ATP2B4;TNFRSF10A;ERLIN2;SORBS1;GP6;SULF1;CD4;MYO1C;RIT2;NPHS2;TEK;TDGF1 |
| Sarcoplasmic Reticulum (GO:0016529) | 0.04918342126623447 | RTN2;RYR1;RYR2;GSTM2;S100A1;ATP2A3;ITPR1;ITPR2;ATP2A1;JPH1;THBS1;RYR3;SYNE2;THBS4;CHERP;JSRP1;SLN;CMYA5;DHRS7C;FKBP1B;NOS1AP;HRC;NOS1 |

| GO: Molecular Function | Adjusted P-value | Genes |
|---|---|---|
| MHC Class II Receptor Activity (GO:0032395) | 0.0011538107091385163 | HLA-DQA1;HLA-DQB2;HLA-DRB1;HLA-DPA1 |
| MHC Class II Protein Complex Binding (GO:0023C | 0.0027439817032371408 | HLA-DRB5;HLA-DQA1;HLA-DQB2;HLA-DRB1;HLA-DPA1 |

| Supplementary Table 04: Pathway analyses of genes containing MeVars in from each tissues. | | |
|---|---|---|
| Tissue | KEGG Pathway | Adjusted p value |
| **Skin** | Type I diabetes mellitus | 5.94E-07 |
| | Cell adhesion molecules | 8.95E-06 |
| | Calcium signaling pathway | 4.42E-05 |
| | Allograft rejection | 1.25E-04 |
| | Viral myocarditis | 1.25E-04 |
| | Graft-versus-host disease | 2.24E-04 |
| | Dilated cardiomyopathy | 1.62E-03 |
| | Hypertrophic cardiomyopatl | 4.08E-03 |
| | Antigen processing and pres | 4.92E-03 |
| | Rheumatoid arthritis | 7.11E-03 |
| | Inflammatory bowel disease | 1.02E-02 |
| | Metabolism of xenobiotics b | 1.30E-02 |
| | Autoimmune thyroid disease | 2.99E-02 |
| | Maturity onset diabetes of th | 3.69E-02 |
| | Steroid hormone biosynthes | 3.69E-02 |
| | Th1 and Th2 cell differentia | 3.69E-02 |
| | Signaling pathways regulatir | 3.78E-02 |
| | Neuroactive ligand-receptor | 4.38E-02 |
| | Pathways in cancer | 4.52E-02 |
| **Liver** | Viral myocarditis | 1.21E-07 |
| | Th17 cell differentiation | 7.83E-06 |
| | Toxoplasmosis | 1.21E-05 |
| | Cell adhesion molecules | 1.21E-05 |
| | Metabolism of xenobiotics b | 1.21E-05 |
| | Type I diabetes mellitus | 1.21E-05 |
| | Antigen processing and pres | 1.27E-05 |
| | Allograft rejection | 1.27E-05 |
| | Graft-versus-host disease | 4.19E-05 |
| | Chemical carcinogenesis | 1.13E-04 |
| | Drug metabolism | 1.13E-04 |
| | Complement and coagulatio | 1.42E-04 |
| | Inflammatory bowel disease | 2.61E-04 |
| | Pathways in cancer | 2.72E-04 |
| | Autoimmune thyroid disease | 4.29E-04 |
| | Leishmaniasis | 4.48E-04 |
| | Pentose and glucuronate inte | 5.76E-04 |
| | Phagosome | 5.76E-04 |
| | Epstein-Barr virus infection | 5.76E-04 |
| | Human T-cell leukemia viru | 9.87E-04 |
| | Ascorbate and aldarate meta | 1.05E-03 |
| | Asthma | 1.31E-03 |
| | Steroid hormone biosynthes | 1.31E-03 |

| | | |
|---|---|---|
| | Kaposi sarcoma-associated l | 1.31E-03 |
| | Rap1 signaling pathway | 2.17E-03 |
| | Th1 and Th2 cell differentia | 2.57E-03 |
| | Staphylococcus aureus infec | 3.50E-03 |
| | Retinol metabolism | 3.50E-03 |
| | Cellular senescence | 4.06E-03 |
| | Central carbon metabolism i | 4.39E-03 |
| | Type II diabetes mellitus | 5.26E-03 |
| | Human cytomegalovirus info | 5.31E-03 |
| | MAPK signaling pathway | 5.50E-03 |
| | Maturity onset diabetes of th | 8.53E-03 |
| | Human immunodeficiency v | 9.32E-03 |
| | Tuberculosis | 9.87E-03 |
| | Porphyrin and chlorophyll m | 1.10E-02 |
| | Hematopoietic cell lineage | 1.20E-02 |
| | Regulation of actin cytoskel | 1.25E-02 |
| | Bile secretion | 1.25E-02 |
| | Cholinergic synapse | 1.63E-02 |
| | Neomycin, kanamycin and g | 1.70E-02 |
| | Apelin signaling pathway | 1.99E-02 |
| | Intestinal immune network f | 2.03E-02 |
| | Glutathione metabolism | 2.03E-02 |
| | Glycolysis / Gluconeogenes | 2.15E-02 |
| | GABAergic synapse | 2.70E-02 |
| | Hepatocellular carcinoma | 3.38E-02 |
| | Tight junction | 3.54E-02 |
| | Rheumatoid arthritis | 3.70E-02 |
| | Axon guidance | 3.74E-02 |
| | Insulin signaling pathway | 3.79E-02 |
| | Influenza A | 3.97E-02 |
| | Ras signaling pathway | 4.13E-02 |
| | Fluid shear stress and athero | 4.13E-02 |
| **Pancreas** | Th17 cell differentiation | 4.19E-03 |
| | Hematopoietic cell lineage | 4.19E-03 |
| | Type II diabetes mellitus | 4.19E-03 |
| | Yersinia infection | 4.19E-03 |
| | Cell adhesion molecules | 4.19E-03 |
| | Cholinergic synapse | 4.19E-03 |
| | GABAergic synapse | 4.19E-03 |
| | Platelet activation | 4.19E-03 |
| | Type I diabetes mellitus | 4.19E-03 |
| | Calcium signaling pathway | 4.19E-03 |
| | Th1 and Th2 cell differentia | 5.05E-03 |
| | Rap1 signaling pathway | 8.00E-03 |

| | | |
|---|---|---|
| | Phospholipase D signaling p | 1.37E-02 |
| | PI3K-Akt signaling pathway | 2.04E-02 |
| | Pathogenic Escherichia coli | 2.38E-02 |
| | Regulation of actin cytoskel | 2.38E-02 |
| | Maturity onset diabetes of th | 2.81E-02 |
| | Graft-versus-host disease | 3.30E-02 |
| | Serotonergic synapse | 3.30E-02 |
| | Antigen processing and pres | 4.06E-02 |
| | Insulin signaling pathway | 4.39E-02 |
| | Insulin resistance | 4.45E-02 |
| | Human cytomegalovirus infe | 4.87E-02 |
| | Morphine addiction | 4.87E-02 |
| | Allograft rejection | 4.87E-02 |
| **Cortex** | Neuroactive ligand-receptor | 1.34E-09 |
| | Type I diabetes mellitus | 3.11E-06 |
| | Allograft rejection | 1.19E-04 |
| | Calcium signaling pathway | 1.19E-04 |
| | Cell adhesion molecules | 1.39E-04 |
| | Graft-versus-host disease | 2.40E-04 |
| | Maturity onset diabetes of th | 2.86E-04 |
| | Autoimmune thyroid disease | 6.08E-04 |
| | Viral myocarditis | 2.41E-03 |
| | Intestinal immune network f | 3.45E-03 |
| | Herpes simplex virus 1 infec | 5.98E-03 |
| | Asthma | 5.98E-03 |
| | Taste transduction | 8.95E-03 |
| | Inflammatory bowel disease | 1.33E-02 |
| | Hippo signaling pathway | 1.33E-02 |
| | Leishmaniasis | 2.07E-02 |
| | Antigen processing and pres | 2.22E-02 |
| | Nicotine addiction | 2.99E-02 |
| | cAMP signaling pathway | 2.99E-02 |
| | Human T-cell leukemia viru | 3.48E-02 |
| | Rheumatoid arthritis | 3.71E-02 |
| | Glutamatergic synapse | 4.49E-02 |
| **Thyroid** | Type I diabetes mellitus | 1.32E-11 |
| | Allograft rejection | 1.10E-08 |
| | Viral myocarditis | 1.10E-08 |
| | Autoimmune thyroid disease | 1.99E-08 |
| | Graft-versus-host disease | 1.99E-08 |
| | Cell adhesion molecules | 3.64E-08 |
| | Antigen processing and pres | 1.46E-07 |
| | Epstein-Barr virus infection | 6.96E-07 |
| | Phagosome | 1.01E-05 |

| | | |
|---|---|---|
| | Human T-cell leukemia viru | 1.01E-05 |
| | Asthma | 4.57E-05 |
| | Intestinal immune network f | 8.79E-04 |
| | Th1 and Th2 cell differentia | 1.70E-03 |
| | Staphylococcus aureus infec | 1.95E-03 |
| | Human papillomavirus infec | 1.95E-03 |
| | Leishmaniasis | 2.24E-03 |
| | Hematopoietic cell lineage | 2.30E-03 |
| | Inflammatory bowel disease | 4.16E-03 |
| | Rheumatoid arthritis | 6.94E-03 |
| | Human cytomegalovirus inf | 7.29E-03 |
| | Influenza A | 7.91E-03 |
| | Th17 cell differentiation | 1.51E-02 |
| | Toxoplasmosis | 1.94E-02 |
| | PI3K-Akt signaling pathway | 2.07E-02 |
| | Fructose and mannose metal | 4.04E-02 |
| | Hepatitis C | 4.05E-02 |
| **Blood** | Type I diabetes mellitus | 5.01E-08 |
| | Allograft rejection | 1.71E-07 |
| | Graft-versus-host disease | 2.97E-07 |
| | Autoimmune thyroid disease | 1.94E-06 |
| | Antigen processing and pres | 3.98E-06 |
| | Viral myocarditis | 3.98E-06 |
| | Cell adhesion molecules | 2.68E-05 |
| | Asthma | 8.88E-05 |
| | Epstein-Barr virus infection | 9.83E-05 |
| | Phagosome | 1.62E-04 |
| | Intestinal immune network f | 8.87E-04 |
| | Hematopoietic cell lineage | 1.10E-03 |
| | Metabolism of xenobiotics b | 1.29E-03 |
| | Leishmaniasis | 1.30E-03 |
| | Inflammatory bowel disease | 3.61E-03 |
| | Herpes simplex virus 1 infec | 6.59E-03 |
| | Th17 cell differentiation | 8.23E-03 |
| | Th1 and Th2 cell differentia | 1.92E-02 |
| | Rheumatoid arthritis | 1.92E-02 |
| | Staphylococcus aureus infec | 2.04E-02 |
| | Influenza A | 2.55E-02 |
| | Human T-cell leukemia viru | 3.08E-02 |
| | Drug metabolism | 3.37E-02 |
| | Toxoplasmosis | 3.86E-02 |
| | Cellular senescence | 4.78E-02 |
| | GO: Biological Processes | 6.04E-04 |
| | MHC Class II Protein Comp | 6.04E-04 |

| | | |
|---|---|---|
| | Peptide Antigen Assembly W | 1.24E-03 |
| | Antigen Processing And Pre | 1.24E-03 |
| | Peptide Antigen Assembly W | 1.33E-03 |
| | Immunoglobulin Production | 1.89E-03 |
| | Antigen Processing And Pre | 4.97E-03 |
| | Antigen Processing And Pre | 6.35E-03 |
| | Antigen Processing And Pre | 8.01E-03 |
| | Immunoglobulin Mediated I | 1.25E-02 |
| | Aromatic Compound Catabo | 4.28E-02 |
| | Positive Regulation Of Leuk | 4.71E-05 |
| **Cerebellum** | Type I diabetes mellitus | 3.64E-04 |
| | Antigen processing and pres | 3.60E-03 |
| | Allograft rejection | 5.77E-03 |
| | Graft-versus-host disease | 2.48E-02 |
| | Autoimmune thyroid disease | 2.74E-02 |
| | Asthma | 4.12E-02 |
| | Viral myocarditis | 5.94E-07 |

| Supplementary Table 5: DISGENET, pathway and Gene Ontology analyses of 14,291 genes without MeVars | | | | | | | | |
|---|---|---|---|---|---|---|---|---|
| DisGeNet | Overlap | P-value | Adjusted P- | Old P-value | Old Adjuste | Odds Ratio | Combined | Genes |
| Abnormal hair quantity | 19/21 | 0.0368403 | 0.9999979 | 0 | 0 | 3.798802 | 12.54046 | DMPK;FOXE1;HPGD;BRAF;PTPN11;A2ML1;PEX1;NRAS;RIT1;RRAS;PEX6;RASA2;SLCO2A1;KRAS;NBN;RAF1;SOS1;LZTR1;SOS2 |
| KEGG Pathways | Overlap | P-value | Adjusted P- | Old P-value | Old Adjuste | Odds Ratio | Combined | Genes |
| Olfactory transduction | 355/440 | 4.5982022 | 0.0014714 | 0 | 0 | 1.685453 | 20.71395 | OR7G2;OR7G1;OR8I2;OR9K2;CALML5;OR11H4;OR2M7;OR2M5;OR52N1;OR2M4;OR2M3;OR1K1;OR2M2;OR4K17;CALML4;OR4Q3;OR52N5;OR51L1;OR52N2;OR5D18;OR2AE1;OR10W1;PRKACG;OR11H6;OR2A42;PRKACA;OR5B3;OR5D16;PRKACB;OR5B2;OR5D14;OR5D13;OR8H3;OR8H2;OR8H1;OR2L8;OR11G2;OR1J4;OR1J2;OR2L3;OR1J1;OR2L2;OR4P4;OR10AD1;OR52M1;OR10V1;OR4K15;OR4K14;OR4K13;OR2AT4;OR4C12;OR4C11;OR6C6;OR6C4;OR4C16;OR5A2;OR6C3;OR8G5;OR4C15;OR5A1;OR5AK2;OR6C2;OR6C1;OR4C13;OR8G1;OR9I1;OR13J1;OR1I1;OR2K2;OR10T2;OR6S1;OR52L1;OR5M11;OR5M10;OR6B3;OR7D4;OR6B2;OR6B1;OR7D2;OR2A25;OR8B12;OR51I2;OR2J3;OR4N5;OR2J2;OR4N4;OR10S1;OR4N2;OR5P3;OR5P2;OR51I1;OR52K2;OR52K1;GNG13;OR9G9;OR4D11;OR2A12;OR2Z1;OR4D10;CALM3;OR6A2;CALM1;OR7C2;OR9G4;CALM2;OR7C1;OR2A14;OR9G1;OR10J1;OR2A1;OR4E2;OR10J3;OR10J5;OR5W2;OR6Y1;OR51Q1;OR52B4;OR4D9;OR5AP2;OR5B12;OR52B2;OR4D6;OR4D5;OR5B17;OR52B6;OR4D2;OR1N1;OR10Z1;OR5V1;OR6X1;OR52R1;OR52A1;OR4C6;OR3A3;OR3A2;OR4A15;OR4C3;OR52A5;OR3A1;OR4A16;OR8K5;OR10H1;OR8K3;OR10H3;OR8K1;OR10H2;OR10H5;OR10H4;OR1M1;OR4S2;OR4S1;OR2AG1;OR2AG2;OR4C45;OR5AN1;OR4B1;OR4C46;OR6F1;OR8J3;OR8J1;OR14A16;OR10G4;OR10G3;OR1L6;OR51M1;OR1L3;OR5T3;OR5T2;OR5T1;OR10X1;OR10G8;OR4A5;OR10G7;OR10G9;OR5C1;OR7G3;OR1C1;OR5K4;OR5K3;OR5K2;OR51D1;OR5K1;OR6M1;OR2T12;SLC8A1;OR2T10;OR2T11;OR52W1;OR5H15;OR1S2;OR5H14;RGS2;OR1S1;OR13D1;OR2D3;OR1B1;OR2D2;OR14C36;OR52E4;OR5J2;OR5AS1;OR52E2;OR2T8;OR2T6;OR2T5;OR2T4;OR2T3;OR2T2;OR13C8;OR2T1;OR4X2;OR4X1;OR5AC2;OR9A4;OR51T1;OR9A2;OR52E8;OR52E6;OR13C3;OR13C4;OR1A2;OR6K6;OR2C1;OR51B2;OR6K3;OR5AR1;OR5I1;OR6K2;OR52D1;OR9Q1;OR1Q1;OR2S2;OR2AK2;OR51S1;OR51B6;OR4A47;OR51B |
| RNA polymerase | 28/31 | 0.0111235 | 0.9999969 | 0 | 0 | 3.733857 | 16.79747 | POLR2A;POLR2B;POLR2C;POLR2D;POLR2E;POLR2F;POLR2G;POLR2H;POLR2I;POLR2J;POLR2K;POLR2J3;POLR2J2;POLR3GL;POLR3A;POLR3B;POLR1A;POLR3C;POLR1B;POLR3D;POLR1C;POLR3E;POLR1D;POLR3F;POLR1E;POLR3G;POLR3H;POLR3K |
| Proteasome | 39/46 | 0.0276981 | 0.9999969 | 0 | 0 | 2.229041 | 7.994206 | PSMD12;PSMD11;PSMD14;PSMD13;POMP;PSMA7;PSMB10;PSMB6;PSMD8;PSMB7;PSMD9;PSMB4;PSMD6;PSMB5;PSMD7;PSMD4;PSMB3;PSMD2;PSMB1;PSMD3;PSMD1;PSMF1;ADRM1;PSMA5;PSMA6;PSMA3;PSMC5;PSMA4;PSMC6;PSMA1;PSMC3;PSMA2;PSMC4;PSMC1;PSME3;PSMC2;PSME4;PSME1;PSME2 |
| DNA replication | 31/36 | 0.0327333 | 0.9999969 | 0 | 0 | 2.48 | 8.480012 | PRIM2;RNASEH2C;FEN1;RNASEH2B;PCNA;RNASEH2A;MCM7;PRIM1;POLD3;POLD4;RNASEH1;POLD1;POLD2;RFC5;RFC3;RFC4;RFC1;RFC2;RPA1;RPA2;POLE4;POLA2;POLE2;RPA3;POLE3;MCM3;MCM4;DNA2;MCM6;SSBP1;MCM2 |
| GO: Biological Processes | Overlap | P-value | Adjusted P- | Old P-value | Old Adjuste | Odds Ratio | Combined | Genes |

| | | | | | | | | |
|---|---|---|---|---|---|---|---|---|
| rRNA Processing (GO:0006364) | 85/101 | 0.0021323 | 0.999998 | 0 | 0 | 2.128964 | 13.09425 | RPL5;DDX49;NVL;POP4;SPPL2B;WDR3;NAT10;TFB1M;RPL7;RRP9;RPS15;FBL;RPS14;SART1;RPS17;RPS16;RPS19;NOB1;RIOK2;RIOK1;RRP12;RRP15;PELP1;UTP15;WDR36;RPS7;DIS3;PIH1D1;RPS6;WDR74;DDX10;DDX54;EBNA1BP2;NSA2;MRTO4;XRN2;C1D;NHP2;SBDS;PIH1D2;RPL27;SRFBP1;RPL26;DDX27;PRKDC;NOP2;MAK16;RPL11;NOLC1;NOL8;NOC4L;MRM1;NOL6;RRP7A;EXOSC7;EMG1;EXOSC5;EXOSC10;EXOSC4;ERI1;EXOSC9;PES1;RPL14;UTP20;IGF2BP3;EXOSC3;LYAR;EXOSC2;DCAF13;NOP58;RIOK3;WDR18;RPL35A;WDR55;RPS25;RPS28;LSM6;RPS27;EIF6;NEDD4;MPHOSPH10;ESF1;NOL11;RPS24;MPHOSPH6 |
| ncRNA Processing (GO:0034470) | 84/100 | 0.0025351 | 0.999998 | 0 | 0 | 2.103769 | 12.57528 | RPL5;INTS12;POP7;DDX49;POP1;NVL;POP4;SPPL2B;INTS10;TSEN2;TSEN54;RPL7;RRP9;THG1L;FARS2;RPS15;FBL;RPS17;RPS16;RPS19;NOB1;RRP12;PELP1;UTP15;WDR36;RPS7;DIS3;PIH1D1;RPS6;WDR74;DDX10;DDX54;EBNA1BP2;MRTO4;XRN2;RPP21;SBDS;PIH1D2;TUT1;RPL27;RPL26;ZCCHC8;TSEN34;DDX27;RPL11;NOLC1;NOL8;NOC4L;NOL6;RRP7A;EXOSC7;EMG1;EXOSC5;EXOSC10;EXOSC4;EXOSC9;PES1;RPL14;UTP20;IGF2BP3;EXOSC3;LYAR;EXOSC2;DCAF13;NOP58;SSB;INTS3;WDR18;INTS2;ADAT1;RPL35A;WDR55;BYSL;RPS25;RPS28;RPS27;INTS5;INTS4;ESF1;INTS7;INTS6;INTS9;INTS8;RPS24 |
| Ribosome Biogenesis (GO:0042254) | 126/155 | 0.0032098 | 0.999998 | 0 | 0 | 1.742225 | 10.00306 | LTV1;RPL5;POP4;SPPL2B;FCF1;C1ORF131;NOC2L;PWP2;RPL7;RRP9;RPS15;FBL;RPS14;RPS17;RPS16;RPS19;RPS11;GLUL;RPS13;RPS9;RPS7;PIH1D1;RPS5;NGDN;RPS6;GTF3A;WDR74;DDX10;DNTTIP2;WDR75;TSC1;GTPBP4;EBNA1BP2;MRTO4;XRN2;SBDS;PIH1D2;RPL27;RPL26;RPS19BP1;NOLC1;RPF2;SPATA5;RRP7A;EMG1;SPATA5L1;C1ORF109;RRS1;SURF6;UTP20;LYAR;NOP16;PNO1;EIF2S1;MRPL22;WDR55;RCL1;RPS25;RPS28;TBL3;RPS27;ZNF658;MPHOSPH10;DHX29;RSL24D1;RPS24;RPS23;DDX49;NVL;PAK1IP1;WDR3;MRPS12;NAT10;WDR43;XPO1;NOB1;RIOK2;RIOK1;RRP12;PELP1;UTP15;WDR36;DIS3;DDX54;RPS3A;URB2;UTP18;CINP;DDX27;NIP7;HEATR1;RPL11;NOL7;NOL8;NOC4L;SDAD1;NOL6;EXOSC7;EXOSC5;EXOSC10;EXOSC4;RPS15A;BMS1;EXOSC9;PES1;RPL14;HEATR3;IGF2BP3;ZNF622;EXOSC3;EXOSC2;DCAF13;NOP58;NPM1;UTP6;KRR1;UTP3;WDR18;DDX31;RPL35A;WDR12;BYSL;CUL4A;BOP1;TNIP1;ESF1 |
| Cytoplasmic Translation (GO:0002181) | 78/93 | 0.0038993 | 0.999998 | 0 | 0 | 2.08322 | 11.55549 | EIF4A2;RPL4;EIF4A1;RPL5;RPL30;RPL32;RPL31;RPL34;MRPS12;RPLP0;RPL10A;RPL8;RPL9;RPL6;RPL7;RPS15;RPL7A;RPS14;RPS17;RPS16;RPL18A;RPS19;RPL36AL;RPL36;RPL35;RPLP2;RPL38;RPL37;RPS11;RPS10;RPS13;RPS9;RPL21;RPS7;RPS5;RPS6;RPL13A;RPS3A;RPSA;RPL37A;ETF1;RPL27;RPL26;RPL29;RPL28;UBA52;RPL12;RPL11;RPS15A;RPL14;RPS3;RPL13;RPL15;RPS2;RPL18;RPS27A;RPL17;ZC3H15;RPL19;RWDD1;EIF2B3;RPL41;RPL35A;RPS26;RPS25;EIF3M;RPS28;RPS27;RPS29;EIF3I;RPL27A;DRG2;RPL22L1;RPS20;FAU;RPS21;RPS24;RPS23 |

| | | | | | | | | |
|---|---|---|---|---|---|---|---|---|
| Sensory Perception Of Chemical Stimulus (GO:0007606) | 91/110 | 0.0043644 | 0.999998 | 0 | 0 | 1.919162 | 10.42924 | OR10J1;OR8I2;OR1C1;OR10J5;OR8U8;OR2M4;BEST2;OR8U1;OR5D18;OBP2A;OBP2B;OR5AP2;OR5B12;OR8D4;CD36;UGT2A1;OR5B3;OR5D16;OR8D2;OR5B2;OR5B17;OR8D1;OR5D14;BBS4;OR5D13;OR8H3;OR8H2;OR8H1;OR2D2;OMP;OR5AC2;OR7A5;OR52A1;OR6C3;OR8G5;OR5AK2;OR6C2;OR10A3;OR3A2;OR10A5;OR52A5;OR8G1;OR8K5;OR9I1;OR5M9;OR5M8;OR8K3;SLC24A4;OR1E2;OR8K1;OR5I1;OR1E1;OR52D1;C5AR1;OR5M3;OR9Q1;UBR3;OR5M1;ITPR3;OR5M11;OR1Q1;OR2S2;OR5M10;OR51B6;OR8B4;OR51B5;OR5AN1;OR8B3;OR51B4;OR8B2;DRD2;NXNL2;OR1D5;OR8J3;OR8B12;OR51I2;OR1D2;OR8J1;OR5B21;OR5L2;OR51M1;OR5L1;OR51I1;OR5T3;OR5T2;OR5T1;OR51V1;SYT10;OR8A1;OR6A2;OR9G4 |
| Ribonucleoprotein Complex Biogenesis (GO:0022613) | 97/118 | 0.0048470 | 0.999998 | 0 | 0 | 1.851003 | 9.864725 | LTV1;RPL5;SPPL2B;FCF1;C1ORF131;NOC2L;PWP2;RPL7;RRP9;RPS15;FBL;RPS14;RPS17;RPS16;RPS19;RPL38;RPS11;GLUL;RPS13;RPS9;RPS7;RPS5;NGDN;RPS6;GTF3A;WDR74;DNTTIP2;WDR75;GTPBP4;EBNA1BP2;SHQ1;MRTO4;RPL26;RPS19BP1;RPF2;SPATA5;RRP7A;EMG1;SPATA5L1;C1ORF109;RRS1;SURF6;UTP20;NOP16;PNO1;RCL1;RPS25;RPS28;TBL3;RPS27;ZNF658;MPHOSPH10;RSL24D1;RPS24;RPS23;NVL;PAK1IP1;WDR3;MRPS12;NAT10;WDR43;XPO1;RIOK2;RIOK1;UTP15;WDR36;RPS3A;URB2;UTP18;CINP;NIP7;HEATR1;RPL11;NOL7;SDAD1;NOL6;EXOSC10;RPS15A;BMS1;PES1;TGS1;RPL14;HEATR3;IGF2BP3;ZNF622;NOP58;NPM1;UTP6;KRR1;UTP3;DDX31;RPL35A;WDR12;BYSL;CUL4A;BOP1;TNIP1 |
| Mitochondrial Translation (GO:0032543) | 81/98 | 0.0073248 | 0.999998 | 0 | 0 | 1.908565 | 9.383428 | MRPS17;MRPS15;MRPS16;FASTKD2;GFM2;MRPS14;MRPS11;MRPS12;MRPL38;MRPL39;MRPS10;MRPL37;MRPL34;MRPL35;MRPL33;MRPL4;MRPL41;MRPL3;MRPL42;MTRF1;MRPL2;MRPL1;MRPL40;MTG1;CHCHD1;EARS2;RARS2;MRPS28;TSFM;MRPS26;MRPS27;MRPS24;MRPS25;MRPS22;MRPS23;MRPL49;MRPS18A;MRPS2;MRPS21;MRPL47;MRPL45;MRPS7;QRSL1;MRPL46;MRPS6;MRPL43;MRPS5;MRPS18C;TUFM;MRPL44;MRPL52;LARS2;MRPL50;MRPL51;MTRF1L;DAP3;MRPL18;MRPS33;MRPL19;MRPS34;MRPS31;MRPL17;MRPL14;MTIF2;MRPS30;MTIF3;MRPL12;MRPL13;MRPL10;MRPL54;MRPL20;IARS2;GADD45GIP1;PTCD3;NDUFA7;MRPL27;GATC;MRPL24;MRPL21;MRPL22;MRPL30 |
| Metaphase Plate Congression (GO:0051310) | 44/51 | 0.0104562 | 0.999998 | 0 | 0 | 2.515698 | 11.47297 | BECN1;SEH1L;DCTN2;CDCA5;CUL3;VPS4B;PINX1;VPS4A;KIF14;CDCA8;CCNB1;CDC23;CHMP1A;NUP62;RRS1;FAM83D;PIBF1;CEP55;NUDC;PDCD6IP;MAP1S;BOD1;KIF22;GEM;NDC80;RAB11A;EML3;EML4;CENPE;CENPF;KIF18A;PSRC1;INCENP;ZW10;CHMP2B;CHMP2A;CHMP4A;KIF2C;KIF2B;CHMP6;CHMP7;CENPQ;KPNB1;CHMP5 |
| Oligosaccharide-Lipid Intermediate Biosynthetic Process (GO:0006490) | 12/12 | 0.0176935 | 0.999998 | 0 | 0 | 68508 | 276399.2 | DPAGT1;DPM1;ALG8;ALG9;ALG6;PDCD6;SRD5A3;ALG14;ALG3;ALG12;ALG11;MPDU1 |
| Positive Regulation Of Transcription Elongation By RNA Polymerase II (GO:0032968) | 37/43 | 0.0203485 | 0.999998 | 0 | 0 | 2.467272 | 9.6094 | TCERG1;CCNK;ELL;MED19;MED16;SUPT6H;CDC73;MED18;MED17;MED11;MED30;MED10;MED31;LEO1;PWWP2A;MED1;NCBP1;NCBP2;MED8;MED9;MED27;MED4;ELL2;MED26;ELL3;MED29;MED6;MED28;MED7;CDK9;MED23;MED22;MED21;MED20;CDK12;CDK13;EAPP |

| | | | | | | | | |
|---|---|---|---|---|---|---|---|---|
| Mitochondrial Gene Expression (GO:0140053) | 83/103 | 0.0227409 | 0.999998 | 0 | 0 | 1.661694 | 6.287165 | MRPS17;MRPS15;MRPS16;FASTKD1;FASTKD2;GFM2;MRPS14;FASTKD3;MRPS11;MRPS12;MRPL38;MRPL39;MRPS10;MRPL37;MRPL34;MRPL35;MRPL33;MRPL4;MRPL41;MRPL3;MRPL42;MRPL2;MRPL1;MRPL40;MTG1;CHCHD1;RARS2;MRPS28;MRPS26;MRPS27;MRPS24;MRPS25;MRPS22;MRPS23;MRPL49;MRPS18A;MRPS2;MRPS21;MRPL47;TFB2M;POLRMT;MRPL45;MRPS7;QRSL1;MRPL46;MRPS6;MRPL43;MRPS5;MRPS18C;MRPL44;MRPL52;LARS2;MRPL50;MRPL51;FASTKD5;TFAM;DAP3;MRPL18;MRPS33;MRPL19;MRPS34;MRPS31;MRPL17;MRPL14;MRPS30;FOXO3;MRPL12;MRPL13;MRPL10;MRPL54;TBRG4;MRPL20;IARS2;GADD45GIP1;PTCD3;NDUFA7;MRPL27;GATC;MRPL24;MRPL21;MRPL22;MRPL30;SLC25A33 |
| NLS-bearing Protein Import Into Nucleus (GO:0006607) | 16/17 | 0.0256684 | 0.999998 | 0 | 0 | 6.397758 | 23.43175 | RANBP2;RGPD5;RGPD8;RGPD1;RGPD4;RGPD3;KPNA6;KPNA7;NUP35;KPNA4;NUP54;KPNA5;KPNA2;KPNA3;KPNB1;KPNA1 |
| Regulation Of Telomerase RNA Localization To Cajal Body (GO:1904872) | 16/17 | 0.0256684 | 0.999998 | 0 | 0 | 6.397758 | 23.43175 | CCT3;PARN;CCT6A;EXOSC10;SHQ1;TNIP1;RUVBL2;RUVBL1;TCP1;NHP2;CCT8;CCT7;CCT5;DCP2;NOP10;CCT4 |
| Positive Regulation Of Telomere Maintenance Via Telomerase (GO:0032212) | 28/32 | 0.0281719 | 0.999998 | 0 | 0 | 2.799902 | 9.994052 | TNKS;PNKP;PARN;HMBOX1;TNKS2;FBXO4;MAPK1;NEK2;CCT8;CCT7;HNRNPA1;MAP2K7;CCT5;MAP3K4;CCT4;MAPK3;CCT3;CCT2;XRCC5;NEK7;CCT6A;TNIP1;TCP1;CTNNB1;POT1;ATM;MAPKAPK5;ATR |
| Negative Regulation Of Anoikis (GO:2000811) | 15/16 | 0.0340892 | 0.999998 | 0 | 0 | 5.997478 | 20.26413 | ITGB1;NOTCH1;TLE1;CAV1;PIK3R3;PTK2;PIK3CA;CEACAM6;CEACAM5;PTRH2;PDK4;SNAI2;ZNF304;ITGA5;MCL1 |
| Regulation Of DNA Strand Elongation (GO:0060382) | 15/16 | 0.0340892 | 0.999998 | 0 | 0 | 5.997478 | 20.26413 | INO80C;INO80B;INO80D;ACTR5;MCRS1;NUCKS1;ACTR8;INO80;UCHL5;YY1;TFPT;RUVBL2;RUVBL1;POT1;NFRKB |
| Positive Regulation Of DNA-templated DNA Replication (GO:2000105) | 10/10 | 0.0346683 | 0.999998 | 0 | 0 | 57090 | 191932.4 | CDT1;DBF4;ENDOG;WIZ;CDC7;DBF4B;INO80;SSBP1;E2F7;E2F8 |
| Positive Regulation Of Epidermal Growth Factor-Activated Receptor Activity (GO:0045741) | 10/10 | 0.0346683 | 0.999998 | 0 | 0 | 57090 | 191932.4 | BTC;EPGN;NCF1;EGF;FBXW7;TGFA;AREG;ADRA2A;EREG;HBEGF |
| Small Nucleolar Ribonucleoprotein Complex Assembly (GO:0000491) | 10/10 | 0.0346683 | 0.999998 | 0 | 0 | 57090 | 191932.4 | NUFIP1;SHQ1;TNIP1;ZNHIT3;PIH1D1;RUVBL2;ZNHIT6;RUVBL1;TAF9;PIH1D2 |
| Translation (GO:0006412) | 180/234 | 0.0347151 | 0.999998 | 0 | 0 | 1.335837 | 4.489186 | RPL4;RPL5;RPL30;RPL32;RPL31;RPL34;RPL8;RPL10A;RPL9;YARS2;RPL6;RPL7;RPS15;EEF1B2;RPS14;RPS17;RPS16;RPL18A;RPS19;RPL36AL;RPL36;RPL35;RPL38;RPL37;RPS11;RPS10;CHCHD1;RPS13;RARS2;RPS9;RPL21;RPS7;RPS5;RPS6;MRPS18A;RPSA;GTPBP2;GTPBP1;MRPS18C;TUFM;EEF1A1;LARS2;EEF1A2;RPL27;RPL26;RPL29;RPL28;UBA52;DAP3;MRPL18;MRPL19;MRPL17;MRPL14;MRPL12;MRPL13;GSPT1;MRPL10;MRPL20;EIF4EBP2;ZC3H15;RPL41;MRPL27;EEF2;HARS2;MRPL24;MRPL21;RPL3L;MRPL22;MRPL30;RPS26;RPS25;RPS28;RPS27;RPS29;RPL27A;RPS20;RPS21;RPS24;FARSB;RPS23;MRPS17;HBS1L;MRPS15;MRPS16;FASTKD2;GFM2;MRPS14;MRPS11;MRPS12;RPLP0;MRPL38;MRPL39;MRPS10;MRPL37;MRPL34;MRPL35;FARS2;MRPL33;MRPL41;MRPL4;MRPL42;MRPL3;RPL7A;MRPL2;MRPL40;EEF2K;MRPL1;MTG1;ZAR1L;RPLP2;EIF5AL1;MRPS28;EIF5A;TSFM;MRPS26;MRPS27;MRPS24;MRPS25;MRPS22;MARS2;MRPS23;DHPS;MRPL49;MRPS2;MRPS21;RPL13A;MRPL47;RPS3A;MRPL45;MRPS7;QRSL1;MRPL46;MRPS6;MRPL43;MRPS5;MRPL44;ZAR1;MRPL52;EEF1G;MRPL50;MRPL51;EEF1D;RPL37A;ETF1;MRRF;MRPS33;RPL12;MRPS34;RPL11;SRBD1;MRPS31;MRPS30;MRPL54;RPS15A;RPL14;RPS3;RPL13;IARS2;IGF2BP3;RPL15;RPS2;RPL18;RPS27A;RPL17;RPL19;ABCF1;GADD45GIP1;PTCD3;RWDD1;NDUFA7;GATC;RPL35A;WARS2;TNIP1;RPS6KB2;DRG1;DRG2;RPL22L1;FAU;ABCE1 |

| | | | | | | | | |
|---|---|---|---|---|---|---|---|---|
| Sensory Perception Of Smell (GO:0007608) | 177/230 | 0.0351383 | 0.999998 | 0 | 0 | 1.33831 | 4.481281 | OR8I2;OR1C1;OR2M7;OR2M5;OR2M4;OR2M3;OR2M2;BEST2;OR2T12;OR2T10;OR2T11;OR5D18;OR2AE1;OR2A42;OR13D1;OR5B3;OR5D16;OR5B2;OR5D14;OR5D13;OR8H3;OR8H2;OR8H1;OR2D3;OR2D2;OR2L8;OR2L3;OR2L2;OR2T8;OR10AD1;OR2T6;OR2T5;OR2T4;OR2T3;OR2T2;OMP;OR13C8;OR2T1;OR5AC2;OR6C3;OR8G5;OR13C3;OR5AK2;OR6C2;OR13C4;OR8G1;OR9I1;OR6K6;OR2C1;OR6K3;OR13J1;OR5I1;OR6K2;OR52D1;OR10T2;OR2K2;OR9Q1;UBR3;OR5M11;OR5M10;OR1Q1;OR2S2;OR2AK2;OR51B6;OR51B5;OR51B4;DRD2;OR2B6;OR6B1;NXNL2;OR2A25;OR2B3;OR2B2;OR10K1;OR8B12;OR51I2;OR5B21;OR2J3;OR2J2;OR10S1;OR51I1;OR2A12;OR2Z1;OR6A2;OR9G4;OR2A5;OR2A4;OR2A14;OR2A2;OR10J1;OR2A1;OR4E2;OR10J3;OR10J5;OR8U8;OR10AG1;OR10R2;OR6Q1;OR8U1;OR6Y1;OBP2B;OR2Y1;OR5AP2;OR5B12;OR8D4;OR8D2;OR5B17;OR8D1;BBS4;OR13G1;OR6P1;OR10Z1;OR10A7;OR5V1;OR7A5;OR2B11;OR10A2;OR52A1;OR10A4;OR10A3;OR10A6;OR52A5;OR10A5;OR8K5;OR2G6;OR5M9;OR10H1;OR8K3;OR5M8;SLC24A4;OR10H3;OR8K1;OR2G3;OR10H2;OR1E1;OR2G2;OR10H5;OR10H4;OR5M3;OR5M1;OR2T34;OR2T33;OR2W3;OR2AG1;OR2W1;OR2AG2;OR8B4;OR13F1;OR8B3;OR8B2;OR1D5;OR2T27;OR8J3;OR8J1;OR1D2;OR2F2;OR2F1;OR10G4;OR2T29;OR10G3;OR1L6;OR5L2;OR51M1;OR5L1;OR6N2;OR6N1;OR5T3;OR5T2;OR5T1;OR51V1;OR2V2;SYT10;GPR148;OR10G8;OR8A1;OR10G7;OR10G9 |
| Positive Regulation Of Transcription Initiation By RNA Polymerase II (GO:0060261) | 43/52 | 0.0454209 | 0.999998 | 0 | 0 | 1.911379 | 5.909567 | GTF2A1;GTF2A2;PAXIP1;SRF;TAF9;MED19;MED16;MED18;MED17;MED11;MED13;ZNHIT1;MED30;MED10;MED31;MED1;TBP;TAF12;TAF13;TAF10;TAF11;MED8;MED9;MED27;MED4;MED26;MED29;MED6;MED28;MED7;MED23;MED22;PSMC6;MED21;MED20;TAF8;TAF4B;TAF7;TAF6;TAF5;TP53;TAF3;TAF2 |
| Modification-Dependent Protein Catabolic Process (GO:0019941) | 148/192 | 0.0467518 | 0.999998 | 0 | 0 | 1.34731 | 4.126676 | UBE2D4;UBE3C;UBE2D2;UBE2D3;CBLC;KEAP1;RNF19B;UBE2L6;UBE3B;FBXO22;CCAR1;UBE2L3;HERC5;HERC4;PSMD9;HERC3;NUB1;CDC23;RNF114;PSMD3;WDR91;LONP1;TRIM25;BTRC;FBXO8;FBXO7;HERC6;FBXW4;MYLIP;RNF43;ARRDC1;UBE2E1;RC3H1;UBE4B;FBXO10;FBXO11;TRPC4AP;KCTD6;RC3H2;RNF40;RBX1;RNF126;CDC34;RNF125;KCTD10;UBA52;PSMD11;PSMD14;PSMD13;UBA6;CUL3;UBR4;UBR3;FBXO45;ANAPC11;ATXN3;UBB;RNF215;RNF217;NAE1;PSMF1;NDFIP2;UBE2H;XBP1;HACE1;NDFIP1;SMURF2;UBE2B;SMURF1;UBE2C;SIAH2;SIAH1;KLHL22;BFAR;NTAN1;CNOT4;COPS3;NPLOC4;UBE2S;MDM2;MKRN2;FBXL4;RNF185;UBE2K;RNF11;PPP1R11;RNF13;RNF14;USP33;ARRB1;HERPUD1;TNFSF13B;ARIH2;ARIH1;RNF150;UBL7;TOPORS;RNF168;RNF167;UBE2R2;RNF166;RNF34;RNF8;NEDD8;RNF6;RNF7;CBL;DCST1;RCHY1;RNF5;C18ORF25;OS9;KCTD21;MARVELD3;FEM1C;RBBP6;RPS27A;RNF130;UBQLN4;RNF133;UBQLN3;RNF20;ANKIB1;FEM1A;RNF25;RNF26;SYVN1;KLHL3;UBE2G1;UBE2G2;FBXL14;MIB1;RNF146;SPSB2;ITCH;PINK1;RNF148;RNF149;UBQLNL;NEDD4;KLHL8;PSMC2;TTC3;PCYOX1L;STUB1;TRIP12;ASB2;TRIM32 |
| Maturation Of LSU-rRNA (GO:0000470) | 18/20 | 0.0472542 | 0.999998 | 0 | 0 | 3.598613 | 10.98373 | DDX18;MAK16;NOP2;URB1;RPF2;WDR12;FTSJ3;RPL7;NSA2;EIF6;ZNHIT3;NEDD4;PES1;RPL7L1;ZNHIT6;RPL35;RBM34;RRP15 |

| | | | | | | | | |
|---|---|---|---|---|---|---|---|---|
| Positive Regulation Of DNA-templated Transcription, Elongation (GO:0032786) | 46/56 | 0.0475081 | 0.999998 | 0 | 0 | 1.840323 | 5.607196 | TCERG1;CCNK;CCNT2;ELL;SUPT4H1;MED19;MED16;SUPT6H;CDC73;MED18;MED17;MED11;MED30;MED10;MED31;LEO1;PWWP2A;HMGN1;MED1;SUPT16H;NCBP1;PARP1;NCBP2;MED8;THOC1;MED9;MED27;MED4;THOC5;SUPT5H;MED26;ELL2;MED29;MED6;ELL3;MED28;MED7;MED23;CDK9;MED22;MED21;MED20;CDK12;ERCC6;EAPP;CDK13 |
| GO: Cellular component | Overlap | P-value | Adjusted P- | Old P-value | Old Adjuste | Odds Ratio | Combined | Genes |
| Vesicle Coat (GO:0030120) | 33/37 | 0.0090889 | 0.9999979 | 0 | 0 | 3.301042 | 15.51718 | CCL13;COPB2;COPA;SEC23A;SAR1A;PEF1;SAR1B;COPB1;PDCD6;CLTB;CLTA;KLHL12;CLINT1;TMED3;TMED7;SEC23B;SEC31A;SEC13;SEC24B;SEC24A;COPZ2;COPZ1;EPN1;EPN2;EPN3;NECAP2;NECAP1;COPG2;ENTHD1;MYCBPAP;SEC24D;SEC24C;COPE |
| Preribosome, Large Subunit Precursor (GO:0030687) | 17/18 | 0.0192765 | 0.9999979 | 0 | 0 | 6.798094 | 26.84477 | NIP7;MAK16;PPAN;WDR74;NOC2L;WDR12;FTSJ3;EBNA1BP2;NSA2;EIF6;NEDD4;PES1;MRTO4;RRS1;ZNF622;MDN1;RRP15 |
| Ribosome (GO:0005840) | 51/61 | 0.0207423 | 0.9999979 | 0 | 0 | 2.041074 | 7.910342 | RPL30;RPL32;RPL31;MRPS12;RPL10A;RPL8;RPL9;RPL6;RPL7A;RPS17;SERP1;RPL18A;RPS19;RPL36AL;RPL36;RPL38;RPS11;RPS10;RPS13;NCK1;RPS9;RPS7;RPS5;RPL13A;MRPS7;NR5A1;RPL27;NUFIP2;DHX9;RPL11;HSPA14;MRPL13;DNAJC21;RPS3;RPL15;RPL18;RPL17;RPL19;BTF3;RPL41;EIF2AK2;SRP68;RPL3L;RPS26;RPS25;RPS28;RPS27;RPS29;EIF3H;RPS21;RPS23 |
| Core Mediator Complex (GO:0070847) | 21/23 | 0.0221559 | 0.9999979 | 0 | 0 | 4.199264 | 15.99772 | MED8;MED19;MED9;MED16;MED27;MED4;MED26;MED18;MED29;MED6;MED17;MED28;MED7;MED23;MED11;MED22;MED30;MED10;MED21;MED20;MED31 |
| RNA Polymerase III Complex (GO:0005666) | 16/17 | 0.0256684 | 0.9999979 | 0 | 0 | 6.397758 | 23.43175 | POLR3GL;POLR3A;POLR3B;POLR3C;POLR3D;POLR1C;POLR3E;POLR1D;POLR2E;POLR3F;POLR2F;POLR3G;POLR3H;POLR2H;POLR3K;POLR2K |
| Clathrin Coat (GO:0030118) | 15/16 | 0.0340892 | 0.9999979 | 0 | 0 | 5.997478 | 20.26413 | CLTC;CLTB;CLTA;EPS15L1;CLINT1;EPN1;EPN2;EPN3;NECAP2;NECAP1;SYNJ1;ENTHD1;KCNQ5;MYCBPAP;PICALM |
| COPII Vesicle Coat (GO:0030127) | 14/15 | 0.0451394 | 0.9999979 | 0 | 0 | 5.597254 | 17.34029 | SEC13;SEC23A;SEC24B;SEC24A;SAR1A;PEF1;SAR1B;PDCD6;KLHL12;SEC24D;TMED7;SEC23B;SEC24C;SEC31A |
| Small-Subunit Processome (GO:0032040) | 59/73 | 0.0460395 | 0.9999979 | 0 | 0 | 1.686366 | 5.191064 | SPPL2B;WDR3;MRPS12;FCF1;C1ORF131;NAT10;FBLL1;PWP2;WDR43;RRP9;FBL;RPS14;RPS17;RPS16;RPS19;RPS11;RPS13;UTP14C;UTP15;RPS9;WDR36;RPS7;RPS5;NGDN;RPS6;DNTTIP2;WDR75;RPS3A;UTP18;RPS19BP1;PRKDC;HEATR1;UTP23;NOL7;NOC4L;NOL6;RRP7A;EMG1;EXOSC10;RPS15A;PDCD11;BMS1;IGF2BP3;UTP20;RPS27A;DCAF13;NOP58;UTP6;KRR1;PNO1;UTP3;XRCC5;RCL1;RPS28;TBL3;RPS27;MPHOSPH10;RPS24;RPS23 |
| Anaphase-Promoting Complex (GO:0005680) | 18/20 | 0.0472542 | 0.9999979 | 0 | 0 | 3.598613 | 10.98373 | ANAPC13;CUL7;ANAPC7;UBE2C;BUB1B;ANAPC10;ANAPC11;CDC20;CDC20B;FZR1;CDC23;UBE2S;CDC26;CDC16;CDC27;ANAPC4;ANAPC5;ANAPC2 |
| Term | Overlap | P-value | Adjusted P- | Old P-value | Old Adjuste | Odds Ratio | Combined | Genes |
| Mannosyltransferase Activity (GO:0000030) | 22/23 | 0.0044590 | 0.9999977 | 0 | 0 | 8.800617 | 47.63614 | ALG1L2;POMT2;ALG8;ALG9;PDCD6;POMT1;TMTC3;ALG3;PIGZ;TMTC2;ALG12;TMTC1;ALG11;PIGV;TMTC4;DPM1;PIGB;DPY19L1;DPY19L2;DPY19L3;DPY19L4;PIGM |

| | | | | | | | | |
|---|---|---|---|---|---|---|---|---|
| Olfactory Receptor Activity (GO:0004984) | 278/362 | 0.0123112 | 0.9999977 | 0 | 0 | 1.328486 | 5.841668 | OR7G2;OR7G1;OR8I2;OR9K2;OR2M7;OR2M5;OR52N1;OR2M4;OR2M3;OR2M2;OR52N5;OR51L1;OR52N2;OR5D18;OR2AE1;OR10W1;OR2A42;OR5B3;OR5D16;OR5B2;OR5D14;OR5D13;OR8H3;OR8H2;OR8H1;OR2L8;OR1J4;OR1J2;OR2L3;OR1J1;OR2L2;OR10AD1;OR52M1;OR10V1;OR2AT4;OR6C6;OR6C4;OR5A2;OR6C3;OR8G5;OR5A1;OR5AK2;OR6C2;OR6C1;OR8G1;OR9I1;OR13J1;OR10T2;OR1I1;OR2K2;OR52L1;OR6S1;OR5M11;OR5M10;OR6B3;OR7D4;OR6B2;OR6B1;OR7D2;OR2A25;OR8B12;OR51I2;OR2J3;OR2J2;OR10S1;OR5P3;OR5P2;OR51I1;OR52K2;OR52K1;OR2A12;OR2Z1;OR6A2;OR7C2;OR9G4;OR7C1;OR2A14;OR10J1;OR2A1;OR4E2;OR10J3;OR10J5;OR5W2;OR51Q1;OR6Y1;OR52B4;OR5AP2;OR5B12;OR52B2;OR5B17;OR52B6;OR1N1;OR10Z1;OR5V1;OR6X1;OR52R1;OR52A1;OR3A3;OR3A2;OR3A1;OR52A5;OR8K5;OR10H1;OR8K3;OR10H3;OR8K1;OR10H2;OR10H5;OR10H4;OR1M1;OR2AG1;OR2AG2;OR5AN1;OR6F1;OR8J3;OR8J1;OR14A16;OR10G4;OR10G3;OR1L6;OR51M1;OR1L3;OR5T3;OR5T2;OR5T1;OR10G8;OR10G7;OR10G9;OR5C1;OR7G3;OR1C1;OR5K4;OR5K3;OR5K2;OR51D1;OR5K1;OR6M1;OR2T12;OR2T10;OR2T11;OR52W1;OR5H15;OR1S2;OR5H14;OR1S1;OR13D1;OR2D3;OR1B1;OR2D2;OR14C36;OR52E4;OR5J2;OR52E2;OR5AS1;OR2T8;OR2T6;OR2T5;OR2T4;OR2T3;OR2T2;OR13C8;OR2T1;OR5AC2;OR51T1;OR52E8;OR52E6;OR13C3;OR13C4;OR1A2;OR6K6;OR2C1;OR51B2;OR6K3;OR5AR1;OR5I1;OR6K2;OR52D1;OR9Q1;OR1Q1;OR2S2;OR2AK2;OR51S1;OR51B6;OR51B5;OR51B4;OR2B6;OR2B3;OR5H6;OR2B2;OR10K1;OR5H2;OR5B21;OR5H1;OR51A2;OR13A1;OR2A5;OR2A4;OR51A7;OR2A2;OR7E24;OR7A10;OR52J3;OR8U8;OR4M1;OR10AG1;OR10R2;OR6Q1;OR8U1;OR2Y1;OR56B1;OR8D4;OR8D2;OR56B4;OR8D1;OR7A17;OR51G2;OR51G1;OR52I2;OR6C70;OR6C76;OR6C75;OR13G1;OR1F1;OR6C74;OR14I1;OR10Q1;OR6P1;OR10A7;OR56A5;OR7A5;OR2B11;OR10A2;OR10A4;OR56A4;OR10A3;OR56A3; |
| DNA-directed 5'-3' RNA Polymerase Activity (GO:0003899) | 30/34 | 0.0180803 | 0.9999977 | 0 | 0 | 3.000316 | 12.04006 | PRIM2;PRIM1;POLR2A;POLR2B;POLR2C;POLR2E;POLR2F;POLR2H;POLR2I;POLR2J;POLR2K;POLR2J3;POLR2J2;POLRMT;POLR3A;POLR3B;POLR1A;POLR3C;MED21;POLR1B;POLR3D;MED20;POLR1C;POLR3E;POLR1D;POLR3F;POLR1E;POLR3G;POLR3H;POLR3K |
| Single-Stranded DNA Helicase Activity (GO:0017116) | 20/22 | 0.0286176 | 0.9999977 | 0 | 0 | 3.999019 | 14.21145 | RFC5;POLQ;RFC3;HELQ;RFC4;MCM7;DHX9;MCM8;RFC2;DSCC1;MCM9;WRNIP1;HELB;CHTF8;RAD51;MCM3;MCM4;DNA2;MCM6;MCM2 |
| Damaged DNA Binding (GO:0003684) | 38/45 | 0.0333646 | 0.9999977 | 0 | 0 | 2.171733 | 7.384451 | DCLRE1B;DCLRE1A;FEN1;PCNA;OGG1;PNKP;HMGB2;XPA;XPC;HMGB1;SMC6;UNG;NTHL1;POLD1;TDG;RBBP8;RPS3;EP300;NBN;POLQ;PARP1;PARP2;RPA1;RPA2;XRCC1;SIRT6;APTX;FANCG;DDB2;NEIL3;ERCC3;ERCC4;ERCC1;RPA3;CRY2;ERCC2;ERCC5;RAD18 |
| RNA Exonuclease Activity, Producing 5'-Phosphomonoesters (GO:0016896) | 26/30 | 0.0433041 | 0.9999977 | 0 | 0 | 2.599544 | 8.16129 | EXD2;DIS3L;EXOSC7;EXOSC5;EXOSC10;EXOSC4;ERI1;ERI2;EXOSC9;EXOSC3;EXOSC2;DIS3L2;TOE1;PNPT1;DIS3;ANGEL2;CPSF3;POLRMT;REXO2;PAN2;CNOT7;XRN1;PDE12;XRN2;CNOT8;DCP2 |
| RNA Polymerase III Activity (GO:0001056) | 14/15 | 0.0451394 | 0.9999977 | 0 | 0 | 5.597254 | 17.34029 | POLR3A;POLR3B;POLR3C;POLR3D;POLR1C;POLR3E;POLR1D;POLR2E;POLR3F;POLR2F;POLR3H;POLR2H;POLR3K;POLR2K |
| Oxidized Purine DNA Binding (GO:0032357) | 9/9 | 0.0485265 | 0.9999977 | 0 | 0 | 51381 | 155460.6 | MSH6;RECQL4;WRN;MSH2;MSH3;OGG1;RPS3;POT1;MUTYH |
| 3'-5'-RNA Exonuclease Activity (GO:0000175) | 29/34 | 0.0488497 | 0.9999977 | 0 | 0 | 2.319675 | 7.00311 | EXD2;DIS3L;PARN;EXOSC7;EXOSC5;EXOSC10;EXOSC4;NLRP6;ERI1;ERI2;EXOSC9;EXOSC3;EXOSC2;DIS3L2;TOE1;CNOT6L;PNPT1;DIS3;ANGEL2;POLRMT;REXO2;PAN2;CNOT6;CNOT7;CNOT1;PDE12;CNOT2;XRN2;CNOT8 |

| Supplementary Table 6: Pathways involved for genes with MeVars shared in the three major ethnic groups and their bioinformatic analyses | | |
|---|---|---|
| DisGeNet | Adjusted P-value | Genes |
| HIV-1 infection | 1.2721524518738367E-5 | DDR1;UGT1A10;TNXB;CDSN;ATF6B;HCG18;NOTCH4;AIF1;PSORS1C1;C2;SH3RF3;CXCL12;NBEA;UGT1A5;RNF39;UGT1A4;UGT1A3;CCHCR1;UGT1A9;UGT1A8;GPSM3;UGT1A7;UGT1A6 |
| BILIRUBIN, SERUM LEVEL OF, QUANTITATIVE TRAIT LOCUS 1 | 2.9590904918858376E-5 | UGT1A10;UGT1A5;UGT1A4;UGT1A3;UGT1A9;UGT1A8;UGT1A7;UGT1A6 |
| Lucey-Driscoll syndrome (disorder) | 2.9590904918858376E-5 | UGT1A10;UGT1A5;UGT1A4;UGT1A3;UGT1A9;UGT1A8;UGT1A7;UGT1A6 |
| Calcification of coronary artery | 5.112950417880955E-5 | GABRB3;UGT1A10;TREH;ABAT;BACH2;MRPL33;SYNE1;PCDHAC2;ANKS1B;PCDHAC1;BCAS1;CSGALNACT1;NRG1;AMPD3;PCDHA13;PCDHA12;SORCS2;PCDHA11;PCDHA10;DNM3;COL4A2;EHHADH;PON1;SLC1A4;AMN1;NKAIN3;PCBP3;PCDHA1;PCDHA5;UGT1A5;PCDHA4;UGT1A4;PCDHA3;NCAM1;UGT1A3;PCDHA2;PCDHA9;UGT1A9;PCDHA8;KCNJ1;UGT1A8;PCDHA7;UGT1A7;PCDHA6;OPCML;UGT1A6;PTPRN2;CPEB1;GATM;CD6;COL9A1;GTF2IRD1;RADIL;SSBP3 |
| Increased bilirubin level (finding) | 5.112950417880955E-5 | UGT1A10;ANK1;HK1;DGUOK;PAX8;UGT1A5;STOM;UGT1A4;UGT1A3;UGT1A9;UGT1A8;UGT1A7;UGT1A6 |
| Crigler Najjar syndrome, type 2 | 5.112950417880955E-5 | UGT1A10;UGT1A5;UGT1A4;UGT1A3;UGT1A9;UGT1A8;UGT1A7;UGT1A6 |
| Autistic Disorder | 6.335021950528209E-4 | GABRB3;APP;NRP2;ELN;WISP3;PRF1;DIXDC1;PIK3CD;ABAT;HTR2A;GRIK2;HTR4;PCSK6;COMT;CDH9;C4B;CYP2D6;HOXA1;SLC12A6;SNRPN;TESC;DNMT3A;PAX3;TSC2;PAX6;NRG1;HTR3A;SORCS1;UPB1;SYN2;FOXP2;GABRG1;CACNB2;KATNAL2;LTA;SHANK2;MCPH1;GRIA2;CALCA;NRXN1;PON1;AGAP1;CACNA1A;SLC1A3;NRXN2;CACNA1D;CACNA1C;JAKMIP1;SEZ6L2;TSPAN32;MCHR2;CUX1;NSD1;PRDM16;MBP;NCAM1;EIF4E;KCNJ1;WNT2;TBX1;NTNG1;GABBR1;GSTM1;CADM2;GABRA5;BDNF;TXNRD1;POU6F2;IGF2;POMC;SDK1;GATM;NBEA;WT1;TUBGCP5;GTF2IRD1;APBA2;GSTM5 |
| Autism Spectrum Disorders | 0.001940585415293484 | GABRB3;DPP10;DDR1;CARS;EHMT2;PIK3CD;HTR2A;CMIP;SYNE1;UNC80;COMP;GRM7;CYP2D6;ZNF569;SOSTDC1;HOXA1;KIRREL3;TESC;DNMT3A;DIO3;LIG4;TSC2;PAX6;SYN2;FOXP2;DNM1;GABRG1;CACNB2;COL4A1;LTA;MCTP2;TRIM10;MAD1L1;SHANK2;MCPH1;TNXB;NOTCH4;NRXN1;PON1;ITPR1;CACNA1A;NRXN2;CACNA1D;CXCR6;CACNA1C;JAKMIP1;SEZ6L2;PSORS1C1;C2;INS;CACNA1I;DPP6;CUX1;EIF4E;WNT2;TBX1;MOG;GABRA5;BDNF;TXNRD1;BAIAP2;SOD3;SNAP91;NBEA;WT1;ARTN;APBA2 |
| Crigler Najjar syndrome, type 1 | 0.001978388642242012 | UGT1A10;UGT1A5;UGT1A4;UGT1A3;UGT1A9;UGT1A8;UGT1A7;UGT1A6 |

| | | |
|---|---|---|
| Attention deficit hyperactivity disorder | 0.00254744426165384 | APP;NRP2;ELN;PIK3CD;HTR2A;ITGAE;COMT;GRM1;SYNE1;HK1;SPN;C4B;MC4R;ADAMTS2;GRM7;CYP2D6;SLC12A6;SNRPN;GP1BB;LMO3;LIG4;TSC2;SORCS2;TGFBR2;CACNB2;MCTP2;CES1;DLX4;COL11A2;PON1;SLC1A3;CXCR6;CACNA1C;INS;DPP6;MOBP;NSD1;NCAM1;GPC6;TBX1;PTPRN2;BDNF;LAMB2;TXNRD1;KCNIP4;DCDC2;BAIAP2;DCLK1;POMC;GDNF;MKRN3;GTF2IRD1 |
| Generalized glycogen storage disease of infants | 0.005394893397946953 | UGT1A10;GAA;UGT1A5;UGT1A4;UGT1A3;UGT1A9;UGT1A8;UGT1A7;EPHA3;UGT1A6 |
| Bulimia Nervosa | 0.005741684279970669 | MC4R;GHSR;NKAIN3;BDNF;LMO3;IGF2;SOSTDC1;HTR3A;GHRL;HTR2A;COMT;ESR1 |
| Bulimia | 0.008547729261869653 | MC4R;POMC;PRR5-ARHGAP8;BDNF;LMO3;LEPR;IGF2;GHRL;HTR2A;COMT |
| Beckwith-Wiedemann Syndrome | 0.008547729261869653 | CDKN1C;CALCA;SNRPN;SMAD3;ELN;IGF2;PAX6;SOX2;WT1;SLC22A18;KCNQ1;PLAGL1;KCNQ1OT1;NSD1;GNAS |
| Psoriasis | 0.010181095731478166 | DDR1;APP;CD40;TNFRSF6B;FLT4;ELN;ITGB2;PRF1;PIK3CD;AATK;DEFB1;PLAT;PTPN22;ITGAE;BACH2;C4B;RPTOR;C4A;HPGDS;CYP2D6;RNF19A;CCRL2;TGM5;CCHCR1;CD34;SLC12A9;TGM2;CHST8;CAST;TCF19;LYNX1;S100A1;DNMT3A;FAM118A;NME2;TAP1;RUNX1;IFI27;LTA;ANGPTL2;CD226;CARD14;CFB;ZNF750;TNXB;NOTCH4;PON1;ASAP2;RNF5;PRKCZ;KLK8;EGFR;KLK7;RASIP1;PSORS1C1;C2;NME1-NME2;DUOX1;NLRP3;NCAM1;EIF4E;CAMK2G;GSTM1;ANGPT2;CDSN;PTPRN2;SMAD3;TNFSF13;PLCL2;IGF2;NFATC1;SOD3;PSMB8;SLURP1;PSMB9;UBAC2;POMC;ADCY10;CXCL12;INS-IGF2;CD6;MIR125B1;PTPN6;TNFSF8 |
| Bilirubin measurement | 0.010181095731478166 | UGT1A10;SLC22A18;UGT1A5;UGT1A4;UGT1A3;UGT1A9;UGT1A8;UGT1A7;UGT1A6 |
| Shared Paranoid Disorder | 0.012336941245781373 | GRM2;HPGDS;BDNF;NRG1;HTR2A;COMT;ESR1;GRIN2D |
| Xeroderma Pigmentosum, Complementation Group D | 0.012514661993275156 | UGT1A10;GSTM1;XRCC3;GTF2H4;EGFR;HPGDS;UGT1A5;UGT1A4;UGT1A3;UGT1A9;UGT1A8;UGT1A7;UGT1A6 |
| Bipolar Disorder | 0.0174940139892169 | GABRB3;DPP10;APP;CD40;DIXDC1;MLC1;DEFB1;ARRB2;HTR2A;GRIK2;HTR4;COMT;SYNE1;HPGDS;GRM7;NEFM;SLC12A6;TGM2;MAGI1;LMO3;NRG1;HTR3A;CSNK1E;SYN2;SORCS2;DUSP6;GABRG1;DKK4;NCOR2;CACNB2;EHHADH;KCNQ2;CRY2;MAD1L1;MCPH1;GRIA2;CALCA;DLX4;NOTCH4;NRXN1;PON1;ITPR1;CACNA1A;SLC1A3;CACNA1D;SLC1A4;CACNA1C;CXCR6;PRKCZ;KLK8;EGFR;MCHR2;RASIP1;NPAS3;INS;PRR5-ARHGAP8;PCDHA1;NCAM1;STX2;PROM1;GPC6;GRIA4;TBX1;NTNG1;TCF7L2;GSTM1;CDSN;CADM2;BDNF;CAV2;GABRA5;TXNRD1;NFYC;ESR1;DCLK1;POMC;KLF6;DLG2;NBEA;STAR;GDNF;PPP2R2C;SP8;FGFR2 |
| Adrenocortical cytomegaly | 0.0174940139892169 | CDKN1C;KCNQ1;KCNQ1OT1;NSD1;IGF2 |
| Overgrowth of external genitalia | 0.0174940139892169 | CDKN1C;KCNQ1;KCNQ1OT1;NSD1;IGF2 |
| Pancreatic hyperplasia | 0.0174940139892169 | CDKN1C;KCNQ1;KCNQ1OT1;NSD1;IGF2 |

| | | |
|---|---|---|
| Hamartoma | 0.021628284262685243 | TFAP2A;TICAM2;ELN;TESC;LAMA4;TMED7-TICAM2;PIK3CD;TSC2;GRM1;EGFR;IFI27;CD6;SLC25A21;CD34;SLC12A9 |
| Depressive Symptoms | 0.022610370245714977 | KCNE1;BDNF;TXNRD2;LMO3;PON1;NRG1;CACNA1C;BRCA1;HTR2A;COMT;ESR1;EGFR;INS;SULT1A1;ADAMTS2;CYP2D6;ARTN;RNF19A;NCAM1;PROM1;NCKAP5;CD34;KCNK2 |
| Craniofacial Abnormalities | 0.022610370245714977 | FBLN7;GABRB3;COLEC11;NRXN1;COL11A2;LTBP3;HAPLN1;EGFR;DLL3;FGF8;HOXA3;HOXA1;GPC6;STRA6;TFAP2A;TBX1;SMAD3;GP1BB;CSNK1A1;DNMT3A;PAX3;PAX6;FOXP2;TGFBR2;GTF2IRD1;MMACHC;FGFR2 |
| Neutrophil count (procedure) | 0.022610370245714977 | PTPN22;PCDHA13;PCDHA12;PCDHA11;PCDHA10;PNKD;CDK6;PCDHA1;LEPR;PCDHA5;PCDHA4;NLRP3;PCDHA3;PCDHA2;PCDHA9;PCDHA8;PCDHA7;PCDHA6;CREB5 |
| Amphetamine Addiction | 0.029645526132851492 | GSTM1;BDNF;LMO3;PLAT;NRG1;HTR3A;ARRB2;COMT;GABRG2;SH3RF3;GRM2;GDNF;CYP2D6;ZNF385D;CCHCR1 |
| ovarian neoplasm | 0.034386888699941644 | DDR1;DIRAS3;CDKN1C;COL18A1;CD40;FLT4;PIK3CD;PLAT;ITGAE;BRCA1;PCSK6;FGF1;CLU;LTB4R;BACH2;HPGDS;RASSF2;FGF8;ZNF569;PRSS8;CD34;HOXA5;SLC12A9;TGM2;HOXA4;TICAM2;NME2;NRG1;DUSP6;RUNX1;VAV2;TGFBR2;PYGO2;RAD51C;IFI27;CTTN;PAX8;LTB4R2;COL4A2;RARA;ANGPTL2;SGK1;SKAP1;OTUB1;CALCA;DLX4;CTBP2;TMPRSS4;PON1;NEDD9;PIK3R1;KLK9;KLK8;LTBP1;EGFR;KLK7;SULT1A1;NME1-NME2;CALCR;CUX1;PLAGL1;IGF2BP1;SFN;CD59;FILIP1L;EIF4E;MUC4;OPCML;GSTM1;ANGPT2;TMED7-TICAM2;IGF2;KLK15;NFATC1;SULF1;DNAJC15;KLK10;ESR1;IFFO1;PSMB9;POMC;KLF6;CXCL12;COPS5;WT1;DLC1;TDGF1;FGFR2;LGMN;TP73 |

| | | |
|---|---|---|
| Rheumatoid Arthritis | 0.042925219727293254 | APP;TNFRSF6B;EHMT2;PRF1;STK19;PLAT;ZBTB22;PRDM1;FGF1;CLU;BACH2;C4B;COMP;C4A;HPGDS;LGALS2;EDNRB;CYP2D6;FCGRT;RNF19A;CCRL2;LGALS8;HOXA5;SLC12A9;TGM2;CAST;TICAM2;S100A1;TAP1;LIG4;PRAP1;RUNX1;DHX40;SIGIRR;IFI27;LTB4R2;CD8A;CD226;ADAM8;TRIM10;EPHA3;CFB;NOTCH4;PRKCZ;PSORS1C1;C2;MOBP;CALD1;INPP5D;NLRP3;MBP;VARS2;WNT2;HOXC6;STRA6;PRRT1;GSTM1;CDSN;SMAD3;BDNF;CAV2;TREX1;TNFSF13;IGF2;NFATC1;ESR1;MIR338;CXCL12;CDK6;GDNF;TNFSF4;HOXD4;PPT2;TDGF1;COL18A1;CRISP2;CD40;FLT4;ITGB2;WISP3;FHL2;PIK3CD;PTPN22;HTR2A;PCSK6;AFF3;LTB4R;HAPLN1;IL18BP;SPRED2;HYAL2;DHX16;TNFSF12-TNFSF13;CCHCR1;CD34;TNPO3;GPSM3;TCF19;DUSP6;TAPBP;ELF3;DDAH1;RARA;LTA;HCLS1;TNFRSF25;SMTNL2;DDR2;TNXB;UHRF1;HCG18;SPESP1;PON1;COL11A2;NEDD9;CXCR6;AGPAT1;RNF5;AIF1;EGFR;INS;TTN;SH3BP2;LAIR2;CD59;NCAM1;MUC4;PDLIM4;LAIR1;BRD4;TBX1;CIZ1;GABBR1;CCL23;ANGPT2;MOG;HSPA1L;TXNRD1;TMED7-TICAM2;PLCL2;G0S2;SOD3;PSMB8;PML;SLURP1;PSMB9;BATF;POMC;TRIM39;CD6;MIR140;PTPN6;GHRL;RGS12;PKN1;CD247;PF4 |
| Squamous cell carcinoma of lung | 0.042925219727293254 | DDR1;TNXB;MS4A3;MAST4;PIK3CD;FGF1;AURKC;EGFR;NPAS3;TTN;SOX2;NME1-NME2;HPGDS;PEG10;PROM1;VARS2;EIF4E;EPHA5;P2RY12;GSTM1;BDNF;NFYC;LMO3;GPX6;XRCC3;NME2;DUSP6;HOPX;TGFBR2;PAX7;GTF2IRD1;EPHA3;FGFR2;DDR2;GAS7 |
| Abdominal Migraine | 0.042925219727293254 | CALCA;PRDM16;HTR2A;ESR1;TGFBR2 |
| Acute Confusional Migraine | 0.042925219727293254 | CALCA;PRDM16;HTR2A;ESR1;TGFBR2 |
| Cervical Migraine Syndrome | 0.042925219727293254 | CALCA;PRDM16;HTR2A;ESR1;TGFBR2 |
| Hemicrania migraine | 0.042925219727293254 | CALCA;PRDM16;HTR2A;ESR1;TGFBR2 |
| Sick Headaches | 0.042925219727293254 | CALCA;PRDM16;HTR2A;ESR1;TGFBR2 |
| Status Migrainosus | 0.042925219727293254 | CALCA;PRDM16;HTR2A;ESR1;TGFBR2 |
| Smoking Behaviors | 0.042925219727293254 | EHMT2;NRXN1;FAM13A;ITPR1;LTBP1;MED12L;CMIP;HOXA11;NPAS3;RPTOR;CALCR;NT5DC2;HOXC5;HOXC4;WNT2;MUC4;FAM19A4;HOXC6;TCF7L2;CADM2;BDNF;DNMT3A;TNNI3K;TRAPPC9;RUNX1;DNM3;CDK6;DOCK1 |

| KEGG Pathways | Adjusted P-value | Genes |
|---|---|---|
| Metabolism of xenobiotics by cytochrome P450 | 9.061006372463147E-4 | UGT1A10;CBR1;GSTM1;UGT2B15;UGT2B17;AKR7L;ALDH3A1;HPGDS;CYP2D6;UGT1A5;UGT1A4;UGT1A3;UGT1A9;UGT1A8;UGT1A7;UGT1A6;GSTM5 |
| Pentose and glucuronate interconversions | 9.061006372463147E-4 | UGT1A10;UGT2B15;UGT1A5;SORD;UGT1A4;UGT2B17;UGT1A3;UGT1A9;UGT1A8;UGT1A7;UGT1A6 |
| Ascorbate and aldarate metabolism | 0.001187613340464493 | UGT1A10;UGT2B15;UGT1A5;UGT1A4;UGT2B17;UGT1A3;UGT1A9;UGT1A8;UGT1A7;UGT1A6 |

| | | |
|---|---|---|
| Drug metabolism | 0.0014491447238800248 | UGT1A10;GSTM1;UGT2B15;NME2;UGT2B17;UPB1;NME1-NME2;ALDH3A1;HPGDS;CYP2D6;UGT1A5;UGT1A4;UGT1A3;UPP1;UGT1A9;UGT1A8;UGT1A7;CES1;GSTM5;UGT1A6 |
| Chemical carcinogenesis | 0.01160782649639662 | UGT1A10;ATF6B;CACNA1A;PIK3CD;CACNA1D;PIK3R1;CACNA1C;ARRB2;EGFR;GNAI1;DLL3;SULT1A1;RXRB;HPGDS;FGF8;RPS6KA2;UGT1A5;UGT1A4;UGT1A3;UGT1A9;UGT1A8;UGT1A7;UGT1A6;CBR1;GSTM1;UGT2B15;UGT2B17;ESR1;GNAS;GSTM5;CREB5 |
| Porphyrin and chlorophyll metabolism | 0.017207242398259232 | UGT1A10;UGT2B15;UGT1A5;UGT1A4;UGT2B17;UGT1A3;UGT1A9;UGT1A8;UGT1A7;UGT1A6 |
| Glutamatergic synapse | 0.017207242398259232 | GRIA2;PLA2G4F;ITPR1;CACNA1A;SLC1A3;CACNA1D;CACNA1C;GRIK2;GRM1;GNAI1;GRIN2D;GRM2;GRM7;GNG4;GNAS;SLC17A8;SHANK2;GRIA4 |
| Nicotine addiction | 0.03310235496111333 | GABRB3;GRIA2;GABRA5;CACNA1A;SLC17A8;GABRG2;GRIA4;GABRG1;GRIN2D |

| REACTOME Pathways | Adjusted P-value | Genes |
|---|---|---|
| Glucuronidation R-HSA-156588 | 0.0023157859718429784 | UGT1A10;UGT2B15;UGT1A5;UGT1A4;UGT2B17;UGT1A3;UGT1A9;UGT1A8;UGT1A7;UGT1A6 |
| NCAM1 Interactions R-HSA-419037 | 0.008666458244111165 | CACNB2;CACNB4;COL4A2;GDNF;ARTN;COL4A1;COL9A1;CACNA1D;NCAM1;CACNA1C;GFRA4 |
| Phase 2 - Plateau Phase R-HSA-5576893 | 0.008666458244111165 | CACNG6;CACNB2;KCNE1;CACNG8;KCNQ1;CACNA2D2;CACNA1C |
| Drug ADME R-HSA-9748784 | 0.040976530018653624 | UGT1A10;GGT6;GSTM1;UGT2B15;PON1;NME2;UGT2B17;VAV2;SULT1A1;UGT1A5;UGT1A4;UGT1A3;UGT1A9;UGT1A8;UGT1A7;CES1;UGT1A6 |

| GO: Biological Processes | Adjusted P-value | Genes |
|---|---|---|
| Cellular Glucuronidation (GO:0052695) | 0.001728101286851446 | UGT1A10;UGT2B15;UGT2B17;UGT1A4;UGT1A9;UGT1A8;UGT1A7;UGT1A6 |
| Glucuronate Metabolic Process (GO:0019585) | 0.001728101286851446 | UGT1A10;UGT2B15;UGT2B17;UGT1A4;UGT1A9;UGT1A8;UGT1A7;UGT1A6 |
| Embryonic Skeletal System Morphogenesis (GO:0048704) | 0.001728101286851446 | TBX1;TFAP2A;OSR2;HOXA3;HOXC4;HOXB3;SOX11;HOXD4;FGFR2;HOXA4;TGFBR2 |
| Chemical Synaptic Transmission (GO:0007268) | 0.00279378446826587 | GABRB3;GRIA2;NRXN1;CACNA1A;SLC1A3;NRXN2;SLC1A4;GRIK2;HTR2A;HTR4;GLRA1;GRM2;GRM7;RPS6KA2;CHRNE;PENK;SLC17A8;SLC12A6;PDLIM4;GRIA4;GABBR1;SLC12A4;DTNA;PTPRN2;GABRA5;HTR3A;SYN2;GABRG2;GABRG1;GRIN2D;CACNB2;CACNB4;DLG2;GLRB;KCNQ2;PNOC;PCDHB16;PCDHB5;APBA2 |
| Anterograde Trans-Synaptic Signaling (GO:0098916) | 0.027033887798235293 | GABRB3;GRIA2;NRXN1;CACNA1A;SLC1A3;NRXN2;GRIK2;HTR2A;HTR4;GLRA1;GRM2;GRM7;RPS6KA2;CHRNE;PENK;SLC12A6;SLC12A4;DTNA;HTR3A;SYN2;CACNB2;CACNB4;DLG2;GLRB;KCNQ2;PNOC;PCDHB16;PCDHB5;APBA2 |
| Positive Regulation Of T-helper 2 Cell Differentiation (GO:0045630) | 0.03971086617526126 | TNFSF4;RARA;NLRP3;PRKCZ |
| Anterior/Posterior Pattern Specification (GO:0009952) | 0.03971086617526126 | TBX1;WT1;HOXA3;HOXC5;HOXB3;HOXC4;HOXD4;NEUROG1;HOXA5;HOXB6;HOXA4;HOXB5;HOXC6 |
| Ear Morphogenesis (GO:0042471) | 0.03971086617526126 | TBX1;TFAP2A;OSR2;TMIE;PAX8;HOXA1;NEUROG1;FGFR2 |
| Inner Ear Morphogenesis (GO:0042472) | 0.04895586896537605 | TBX1;TFAP2A;TMIE;PAX8;MYO3A;HOXA1;NEUROG1;FGFR2 |

| GO: Cellular Component | Adjusted P-value | Genes |
|---|---|---|
| None Significant | | |

| GO: Molecular Function | Overlap | Genes |
|---|---|---|
| Retinoic Acid Binding (GO:0001972) | 7.085626048329898E-4 | UGT1A10;UGT2B15;RARA;UGT2B17;UGT1A4;UGT1A9;UGT1A8;UGT1A7;UGT1A6 |
| Monocarboxylic Acid Binding (GO:0033293) | 0.013288541232751434 | UGT1A10;UGT2B15;RARA;UGT1A4;UGT2B17;UGT1A9;UGT1A8;UGT1A7;UGT1A6 |
| Glucuronosyltransferase Activity (GO:0015020) | 0.013288541232751434 | CSGALNACT1;UGT1A10;UGT2B15;UGT1A4;UGT2B17;UGT1A9;UGT1A8;UGT1A7;UGT1A6 |
| Retinoid Binding (GO:0005501) | 0.013288541232751434 | UGT1A10;UGT2B15;RARA;UGT1A4;UGT2B17;UGT1A9;UGT1A8;UGT1A7;UGT1A6 |
| Voltage-Gated Monoatomic Cation Channel Activity (GO:0022843) | 0.013288541232751434 | RYR1;KCNE1;CACNA2D2;CACNA1A;CACNA1D;CACNA2D4;CACNA1C;KCNA6;CACNG6;CACNB2;CACNA1I;CACNG8;CACNB4;KCNQ1;KCNQ2 |
| Voltage-Gated Calcium Channel Activity (GO:0005245) | 0.018009850915996762 | RYR1;CACNG6;CACNA1I;CACNB2;CACNG8;CACNB4;CACNA1A;CACNA2D2;CACNA1D;CACNA1C;CACNA2D4 |
| Transmitter-Gated Monoatomic Ion Channel Activity (GO:0022824) | 0.018009850915996762 | GABRB3;GRIA2;GABRA5;CHRNE;HTR3A;GRIK2;GABRG2;GRIA4;GABRG1;GRIN2D |
| Glutamate Receptor Activity (GO:0008066) | 0.028973471578970574 | GRIA2;GRM2;GRM7;GRM1;GRIN2D;GRIA4 |

| Supplementary Table 7: Coordinates of MeVars identified in four internal tissues and matched blood tissues (duplicate values of coordinates retained) | | | | |
|---|---|---|---|---|
| Sample and Tissue | Coordinate | UCSC_REFGENE_NAME | RELATION_TO_UCSC_C | UCSC_REFGENE_GROUP |
| GSM1179497Blood_IT9 | chr1:111742515-111744309 | DENND2D | N_Shelf | Body |
| GSM1179497Blood_IT9 | chr1:153599479-153600762 | S100A13;S100A13;S100A13;S100A13;S100A1;S100A | | 5'UTR;1stExon;5'UTR;5'UTR;TSS1500;5'UTR;5'UTR |
| GSM1179497Blood_IT9 | chr1:159046773-159047163 | AIM2 | | TSS200 |
| GSM1179497Blood_IT9 | chr1:159796617-159796658 | SLAMF8;SLAMF8 | | 5'UTR;1stExon |
| GSM1179497Blood_IT9 | chr1:202172778-202172912 | LGR6;LGR6 | | TSS200;Body |
| GSM1179497Blood_IT9 | chr1:2120985-2121449 | C1orf86;LOC100128003 | N_Shore | Body;Body |
| GSM1179508Muscle_IT9 | chr1:2120985-2121521 | C1orf86;LOC100128003 | N_Shore | Body;Body |
| GSM1179524SC_Fat_IT9 | chr1:2120985-2121521 | C1orf86;LOC100128003 | N_Shore | Body;Body |
| GSM1179497Blood_IT9 | chr1:226012913-226013010 | EPHX1;EPHX1 | | TSS200;5'UTR |
| GSM1179508Muscle_IT9 | chr1:241912510-241912768 | WDR64 | | Body |
| GSM1179497Blood_IT9 | chr1:24861604-24861919 | RCAN3 | N_Shore | Body |
| GSM1179497Blood_IT9 | chr1:25290947-25292034 | RUNX3 | | Body |
| GSM1179497Blood_IT9 | chr1:25598878-25599066 | RHD;RHD | | TSS200;TSS200 |
| GSM1179497Blood_IT9 | chr1:26527870-26527928 | CATSPER4 | | Body |
| GSM1179497Blood_IT9 | chr1:45103896-45103967 | RNF220 | | Body |
| GSM1179497Blood_IT9 | chr1:92414221-92414782 | BRDT | N_Shore | TSS1500 |
| GSM1179497Blood_IT9 | chr1:9788517-9789174 | PIK3CD | N_Shore | 3'UTR |
| GSM1179497Blood_IT9 | chr10:12085555-12086036 | UPF2;UPF2 | S_Shore | TSS1500;TSS1500 |
| GSM1179497Blood_IT9 | chr10:131697080-131697345 | EBF3 | Island | Body |
| GSM1179514Omentum_IT9 | chr10:45958759-45958885 | MARCH8;MARCH8;MARCH8 | | Body;Body;Body |
| GSM1179497Blood_IT9 | chr10:6220879-6220943 | PFKFB3 | | Body |
| GSM1179497Blood_IT9 | chr10:99735010-99735202 | CRTAC1 | | Body |
| GSM1179514Omentum_IT9 | chr11:1320497-1320634 | TOLLIP | S_Shelf | Body |
| GSM1179497Blood_IT9 | chr11:1325718-1325852 | TOLLIP | | Body |
| GSM1179497Blood_IT9 | chr11:47399213-47400113 | SPI1;SPI1 | N_Shore | Body;Body |
| GSM1179497Blood_IT9 | chr11:5248218-5248566 | HBB | | 1stExon |
| GSM1179514Omentum_IT9 | chr11:59823993-59824161 | MS4A3;MS4A3;MS4A3 | | TSS200;TSS200;TSS200 |
| GSM1179497Blood_IT9 | chr11:64023044-64023183 | PLCB3 | N_Shelf | Body |
| GSM1179497Blood_IT9 | chr11:64107158-64107720 | CCDC88B | N_Shelf | TSS1500 |
| GSM1179497Blood_IT9 | chr11:67205195-67206418 | PTPRCAP | S_Shore | TSS200 |
| GSM1179497Blood_IT9 | chr11:71710885-71711021 | IL18BP;IL18BP;IL18BP;IL18BP;IL18BP;IL18BP | | 5'UTR;5'UTR;5'UTR;TSS200;5'UTR;1stExon |
| GSM1179497Blood_IT9 | chr12:14996272-14996776 | ART4;ART4 | | 5'UTR;1stExon |
| GSM1179497Blood_IT9 | chr12:27396229-27396937 | STK38L | N_Shore | TSS1500 |
| GSM1179497Blood_IT9 | chr12:53591490-53591766 | ITGB7 | Island | Body |
| GSM1179497Blood_IT9 | chr12:56414442-56414533 | IKZF4 | | TSS1500 |
| GSM1179497Blood_IT9 | chr12:57873704-57874033 | ARHGAP9;ARHGAP9 | S_Shelf | TSS1500;TSS200 |
| GSM1179497Blood_IT9 | chr12:6442526-6443045 | TNFRSF1A | S_Shelf | Body |
| GSM1179497Blood_IT9 | chr13:113426250-113426343 | ATP11A;ATP11A | S_Shore | Body;Body |
| GSM1179497Blood_IT9 | chr14:104190678-104190829 | ZFYVE21 | N_Shelf | Body |
| GSM1179508Muscle_IT9 | chr14:24458004-24458345 | DHRS4L2 | Island | TSS200 |
| GSM1179497Blood_IT9 | chr14:24458208-24458345 | DHRS4L2 | S_Shore | 1stExon |
| GSM1179497Blood_IT9 | chr14:69256677-69256977 | ZFP36L1 | Island | Body |
| GSM1179514Omentum_IT9 | chr14:69256677-69256977 | ZFP36L1 | Island | Body |
| GSM1179497Blood_IT9 | chr14:81425912-81426015 | TSHR;TSHR;TSHR | | Body;Body;Body |
| GSM1179497Blood_IT9 | chr15:43513055-43513504 | EPB42;EPB42;EPB42;EPB42 | | 1stExon;5'UTR;1stExon;5'UTR |
| GSM1179497Blood_IT9 | chr15:78555907-78556534 | DNAJA4;DNAJA4 | N_Shore | TSS1500;TSS1500 |
| GSM1179497Blood_IT9 | chr16:1404914-1405223 | GNPTG | Island | Body |
| GSM1179497Blood_IT9 | chr16:4714733-4714815 | MGRN1;MGRN1;MGRN1;MGRN1 | | Body;Body;Body;Body |
| GSM1179497Blood_IT9 | chr16:49732224-49732623 | ZNF423 | N_Shore | Body |
| GSM1179497Blood_IT9 | chr16:52582072-52582262 | TOX3;TOX3 | S_Shore | TSS1500;TSS1500 |
| GSM1179497Blood_IT9 | chr16:67686832-67687754 | RLTPR | N_Shore | Body |
| GSM1179497Blood_IT9 | chr16:787799-788184 | NARFL | S_Shore | Body |
| GSM1179497Blood_IT9 | chr16:88558065-88558237 | ZFPM1 | Island | Body |
| GSM1179497Blood_IT9 | chr16:89408248-89408567 | ANKRD11 | Island | 5'UTR |
| GSM1179497Blood_IT9 | chr17:1633714-1634018 | WDR81;WDR81;WDR81;WDR8 | N_Shelf | Body;Body;Body;Body |
| GSM1179497Blood_IT9 | chr17:19648846-19649004 | ALDH3A1;ALDH3A1;ALDH3A1 | S_Shore | 1stExon;5'UTR;TSS200;5'UTR |
| GSM1179497Blood_IT9 | chr17:27187966-27189285 | ERAL1 | | 3'UTR |
| GSM1179524SC_Fat_IT9 | chr17:30822561-30823087 | MYO1D | N_Shore | Body |
| GSM1179497Blood_IT9 | chr17:40489513-40489785 | STAT3;STAT3;STAT3 | | Body;Body;Body |
| GSM1179497Blood_IT9 | chr17:56401734-56402238 | BZRAP1;BZRAP1 | Island | Body;Body |
| GSM1179497Blood_IT9 | chr17:62083407-62084359 | ICAM2;ICAM2;ICAM2;ICAM2;ICAM2 | | Body;Body;Body;Body;Body |
| GSM1179497Blood_IT9 | chr17:76036514-76037562 | TNRC6C;TNRC6C | N_Shore | 5'UTR;5'UTR |
| GSM1179497Blood_IT9 | chr17:77916733-77916892 | TBC1D16 | Island | Body |
| GSM1179497Blood_IT9 | chr17:77952109-77952290 | TBC1D16 | Island | Body |

| | | | | |
|---|---|---|---|---|
| GSM1179514Omentum_IT9 | chr17:79128885-79129078 | AATK | N_Shelf | Body |
| GSM1179514Omentum_IT9 | chr17:79881523-79882042 | MAFG;MAFG | S_Shore | 5'UTR;TSS200 |
| GSM1179497Blood_IT9 | chr17:80084554-80085082 | CCDC57 | N_Shore | Body |
| GSM1179497Blood_IT9 | chr17:80290106-80290217 | SECTM1 | N_Shore | 5'UTR |
| GSM1179514Omentum_IT9 | chr18:74114570-74114728 | ZNF516 | Island | Body |
| GSM1179497Blood_IT9 | chr18:74799250-74799572 | MBP;MBP | Island | Body;Body |
| GSM1179497Blood_IT9 | chr19:12876846-12877188 | HOOK2;HOOK2 | N_Shore | Body;Body |
| GSM1179508Muscle_IT9 | chr19:12876846-12877188 | HOOK2;HOOK2 | N_Shore | Body;Body |
| GSM1179514Omentum_IT9 | chr19:12876846-12877188 | HOOK2;HOOK2 | N_Shore | Body;Body |
| GSM1179524SC_Fat_IT9 | chr19:12876846-12877188 | HOOK2;HOOK2 | N_Shore | Body;Body |
| GSM1179497Blood_IT9 | chr19:2607726-2607903 | GNG7 | Island | 5'UTR |
| GSM1179497Blood_IT9 | chr19:40421453-40421743 | FCGBP | Island | Body |
| GSM1179497Blood_IT9 | chr19:55013821-55013954 | LAIR2;LAIR2 | | TSS200;TSS200 |
| GSM1179497Blood_IT9 | chr19:55477755-55478262 | NLRP2 | | TSS200 |
| GSM1179497Blood_IT9 | chr2:113875226-113875512 | IL1RN;IL1RN;IL1RN | | TSS1500;TSS1500;TSS1500 |
| GSM1179497Blood_IT9 | chr2:157292018-157292127 | GPD2;GPD2;GPD2 | N_Shore | 1stExon;TSS1500;5'UTR |
| GSM1179497Blood_IT9 | chr2:231090329-231090745 | SP140;SP140 | | TSS200;TSS200 |
| GSM1179497Blood_IT9 | chr2:24233923-24234017 | MFSD2B | S_Shore | Body |
| GSM1179497Blood_IT9 | chr2:242702749-242702932 | D2HGDH | | Body |
| GSM1179497Blood_IT9 | chr2:65593761-65594021 | SPRED2;SPRED2;SPRED2 | | 5'UTR;1stExon;Body |
| GSM1179497Blood_IT9 | chr20:50108912-50109375 | NFATC2;NFATC2;NFATC2 | Island | Body;Body;Body |
| GSM1179497Blood_IT9 | chr21:15352213-15352641 | C21orf81 | Island | Body |
| GSM1179497Blood_IT9 | chr22:24823509-24823554 | ADORA2A | S_Shelf | TSS200 |
| GSM1179497Blood_IT9 | chr22:36044151-36044234 | APOL6 | | TSS1500 |
| GSM1179497Blood_IT9 | chr3:12994821-12994919 | IQSEC1;IQSEC1 | | Body;Body |
| GSM1179514Omentum_IT9 | chr4:1294566-1295078 | MAEA;MAEA | | Body;Body |
| GSM1179497Blood_IT9 | chr4:1294996-1295078 | MAEA;MAEA | | Body;Body |
| GSM1179497Blood_IT9 | chr4:186732837-186732932 | SORBS2;SORBS2;SORBS2;SORBS2;SORBS2;SORBS2; | | 5'UTR;5'UTR;5'UTR;1stExon;TSS1500;1stExon;TSS15 |
| GSM1179497Blood_IT9 | chr4:2275880-2276044 | ZFYVE28 | N_Shore | Body |
| GSM1179497Blood_IT9 | chr4:2321957-2322078 | ZFYVE28 | | Body |
| GSM1179497Blood_IT9 | chr5:176797920-176798049 | RGS14 | S_Shelf | Body |
| GSM1179497Blood_IT9 | chr5:179588440-179588838 | RASGEF1C | Island | 5'UTR |
| GSM1179514Omentum_IT9 | chr5:39219245-39219698 | FYB;FYB | | 5'UTR;5'UTR |
| GSM1179497Blood_IT9 | chr6:112115117-112115430 | FYN | | 5'UTR |
| GSM1179497Blood_IT9 | chr6:157876915-157877120 | ZDHHC14;ZDHHC14 | | Body;Body |
| GSM1179497Blood_IT9 | chr6:26183572-26183684 | HIST1H2BE | N_Shore | TSS1500 |
| GSM1179497Blood_IT9 | chr6:29911366-29911558 | HLA-A | Island | Body |
| GSM1179497Blood_IT9 | chr6:30956397-30956603 | MUC21 | | 3'UTR |
| GSM1179497Blood_IT9 | chr6:31238788-31238908 | HLA-C | N_Shore | Body |
| GSM1179497Blood_IT9 | chr6:31539735-31540121 | LTA;LTA | | TSS1500;TSS200 |
| GSM1179497Blood_IT9 | chr6:31543169-31543300 | TNF | | TSS200 |
| GSM1179497Blood_IT9 | chr6:31867698-31867919 | ZBTB12 | Island | 3'UTR |
| GSM1179497Blood_IT9 | chr6:31980836-31980856 | TNXB;TNXB;TNXA;STK19;STK19 | | 5'UTR;1stExon;TSS200;TSS1500;TSS1500 |
| GSM1179497Blood_IT9 | chr6:32135715-32136052 | EGFL8 | S_Shore | Body |
| GSM1179497Blood_IT9 | chr6:32819858-32820355 | TAP1 | N_Shore | Body |
| GSM1179497Blood_IT9 | chr6:33085565-33085590 | HLA-DPB2 | S_Shore | Body |
| GSM1179497Blood_IT9 | chr6:41010111-41010375 | TSPO2;TSPO2 | | TSS1500;TSS200 |
| GSM1179497Blood_IT9 | chr7:149543136-149543177 | ZNF862 | | Body |
| GSM1179497Blood_IT9 | chr7:1895903-1896384 | MAD1L1;MAD1L1;MAD1L1 | N_Shore | Body;Body;Body |
| GSM1179497Blood_IT9 | chr7:5271480-5271655 | WIPI2;WIPI2;WIPI2;WIPI2;WIP | S_Shelf | 3'UTR;3'UTR;3'UTR;3'UTR;3'UTR |
| GSM1179497Blood_IT9 | chr7:65419185-65419248 | VKORC1L1 | | Body |
| GSM1179497Blood_IT9 | chr8:133772742-133773115 | TMEM71;TMEM71;TMEM71;TMEM71 | | 1stExon;5'UTR;1stExon;5'UTR |
| GSM1179497Blood_IT9 | chr8:141312892-141312979 | TRAPPC9;TRAPPC9 | | Body;Body |
| GSM1179497Blood_IT9 | chr8:141599141-141599436 | EIF2C2;EIF2C2 | | Body;Body |
| GSM1179508Muscle_IT9 | chr8:144659831-144660157 | NAPRT1 | Island | Body |
| GSM1179497Blood_IT9 | chr8:145025059-145025128 | PLEC1;PLEC1;PLEC1;PLEC1 | Island | Body;Body;Body;TSS200 |
| GSM1179499Blood_IT10 | chr1:10490025-10490063 | APITD1;APITD1;APITD1 | N_Shore | TSS200;TSS1500;TSS200 |
| GSM1179516Omentum_IT10 | chr1:114414042-114414408 | PTPN22;PTPN22 | | Body;Body |
| GSM1179526SC_Fat_IT10 | chr1:119531718-119532925 | TBX15 | N_Shore | 5'UTR |
| GSM1179516Omentum_IT10 | chr1:151344806-151345248 | SELENBP1 | | Body |
| GSM1179499Blood_IT10 | chr1:153599479-153599831 | S100A13;S100A13;S100A13;S100A13;S100A1;S100A | | 5'UTR;1stExon;5'UTR;5'UTR;TSS1500;5'UTR;5'UTR |
| GSM1179499Blood_IT10 | chr1:161008127-161008977 | TSTD1;TSTD1;TSTD1 | N_Shore | Body;Body;Body |
| GSM1179499Blood_IT10 | chr1:179713788-179713950 | FAM163A | Island | 5'UTR |
| GSM1179516Omentum_IT10 | chr1:19764480-19764821 | CAPZB | | Body |
| GSM1179510Muscle_IT10 | chr1:2120985-2121724 | C1orf86;LOC100128003 | N_Shore | Body;Body |
| GSM1179526SC_Fat_IT10 | chr1:2120985-2121724 | C1orf86;LOC100128003 | N_Shore | Body;Body |

| | | | | |
|---|---|---|---|---|
| GSM1179499Blood_IT10 | chr1:245710332-245710401 | KIF26B | | Body |
| GSM1179516Omentum_IT10 | chr1:247611842-247611931 | NLRP3;NLRP3;NLRP3;NLRP3;I | N_Shelf | 3'UTR;3'UTR;3'UTR;3'UTR;3'UTR |
| GSM1179516Omentum_IT10 | chr1:247711595-247712591 | C1orf150 | | TSS1500 |
| GSM1179499Blood_IT10 | chr1:4718212-4718573 | AJAP1;AJAP1 | S_Shore | Body;Body |
| GSM1179516Omentum_IT10 | chr1:95698378-95699097 | RWDD3;RWDD3 | N_Shore | TSS1500;TSS1500 |
| GSM1179516Omentum_IT10 | chr10:111765673-111766049 | ADD3 | N_Shore | TSS200 |
| GSM1179516Omentum_IT10 | chr10:131323903-131324189 | MGMT | | Body |
| GSM1179516Omentum_IT10 | chr10:13388442-13388567 | SEPHS1 | | 5'UTR |
| GSM1179516Omentum_IT10 | chr10:1551251-1551326 | ADARB2 | | Body |
| GSM1179516Omentum_IT10 | chr10:18629497-18629677 | CACNB2;CACNB2;CACNB2;CACNB2;CACNB2;CACN | | Body;TSS200;Body;Body;Body;Body;Body;Body |
| GSM1179516Omentum_IT10 | chr10:50143375-50143483 | WDFY4 | | Body |
| GSM1179499Blood_IT10 | chr10:530635-531320 | DIP2C | N_Shore | Body |
| GSM1179516Omentum_IT10 | chr10:6104137-6104515 | IL2RA;IL2RA | | 1stExon;5'UTR |
| GSM1179516Omentum_IT10 | chr10:63422511-63422555 | C10orf107 | N_Shore | TSS1500 |
| GSM1179516Omentum_IT10 | chr11:121970694-121970768 | LOC399959;MIR125B1 | | Body;TSS200 |
| GSM1179499Blood_IT10 | chr11:132662455-132662963 | OPCML;OPCML | | Body;Body |
| GSM1179499Blood_IT10 | chr11:133098424-133098499 | OPCML | | Body |
| GSM1179499Blood_IT10 | chr11:18230620-18230629 | LOC494141;LOC494141;LOC | Island | TSS200;TSS200;TSS1500 |
| GSM1179516Omentum_IT10 | chr11:1873884-1874093 | LSP1 | | TSS1500 |
| GSM1179516Omentum_IT10 | chr11:3144685-3144774 | OSBPL5;OSBPL5;OSBPL5 | S_Shelf | Body;Body;Body |
| GSM1179516Omentum_IT10 | chr11:36422377-36422615 | PRR5L;PRR5L;PRR5L;PRR5L | | 5'UTR;5'UTR;TSS200;Body |
| GSM1179516Omentum_IT10 | chr11:460469-460914 | PTDSS2 | Island | Body |
| GSM1179499Blood_IT10 | chr11:4628823-4629667 | TRIM68 | N_Shore | 5'UTR |
| GSM1179516Omentum_IT10 | chr11:57195350-57195540 | SLC43A3;SLC43A3;SLC43A3 | S_Shore | TSS1500;TSS1500;TSS1500 |
| GSM1179516Omentum_IT10 | chr11:57364841-57365088 | SERPING1;SERPING1 | | TSS1500;TSS200 |
| GSM1179516Omentum_IT10 | chr11:60738032-60739183 | CD6 | | TSS1500 |
| GSM1179516Omentum_IT10 | chr11:70211301-70211531 | PPFIA1;PPFIA1 | | Body;Body |
| GSM1179516Omentum_IT10 | chr11:85397109-85397684 | CCDC89 | | 1stExon |
| GSM1179516Omentum_IT10 | chr12:105114162-105114399 | CHST11 | | Body |
| GSM1179516Omentum_IT10 | chr12:116996773-116997535 | MAP1LC3B2 | | TSS1500 |
| GSM1179499Blood_IT10 | chr12:130822286-130822674 | PIWIL1 | N_Shore | TSS1500 |
| GSM1179499Blood_IT10 | chr12:44152117-44153217 | PUS7L;PUS7L;IRAK4;IRAK4;PU | N_Shore | 1stExon;5'UTR;TSS1500;TSS1500;5'UTR;5'UTR;TSS1 |
| GSM1179516Omentum_IT10 | chr12:47610220-47610734 | FAM113B;FAM113B;LOC100233209 | | 5'UTR;1stExon;Body |
| GSM1179499Blood_IT10 | chr12:50475035-50475372 | ACCN2;ACCN2 | Island | Body;Body |
| GSM1179516Omentum_IT10 | chr12:54891491-54891636 | NCKAP1L | | TSS200 |
| GSM1179516Omentum_IT10 | chr13:107145933-107146164 | EFNB2 | S_Shore | Body |
| GSM1179516Omentum_IT10 | chr13:25085301-25085669 | PARP4 | N_Shore | 5'UTR |
| GSM1179516Omentum_IT10 | chr13:99629970-99630583 | DOCK9;DOCK9;DOCK9;DOCK9 | | Body;Body;Body;Body |
| GSM1179516Omentum_IT10 | chr14:20903410-20903611 | KLHL33 | Island | 5'UTR |
| GSM1179499Blood_IT10 | chr14:67656029-67656241 | FAM71D | | TSS200 |
| GSM1179499Blood_IT10 | chr14:69341430-69341905 | ACTN1;ACTN1;ACTN1 | Island | 3'UTR;3'UTR;3'UTR |
| GSM1179516Omentum_IT10 | chr15:37169665-37170554 | LOC145845 | N_Shelf | Body |
| GSM1179516Omentum_IT10 | chr16:14403004-14403425 | MIR365-1 | | TSS200 |
| GSM1179516Omentum_IT10 | chr16:2737220-2737356 | KCTD5 | | Body |
| GSM1179510Muscle_IT10 | chr16:419975-420230 | MRPL28 | Island | Body |
| GSM1179526SC_Fat_IT10 | chr16:419975-420230 | MRPL28 | Island | Body |
| GSM1179516Omentum_IT10 | chr16:420108-420230 | MRPL28 | Island | Body |
| GSM1179499Blood_IT10 | chr16:49732224-49732623 | ZNF423 | N_Shore | Body |
| GSM1179499Blood_IT10 | chr16:57562452-57562841 | CCDC102A | Island | Body |
| GSM1179516Omentum_IT10 | chr16:69597925-69598417 | NFAT5;NFAT5;NFAT5;NFAT5;N | N_Shore | TSS1500;TSS1500;TSS1500;TSS1500;TSS1500 |
| GSM1179516Omentum_IT10 | chr16:69958626-69958964 | WWP2;WWP2 | | TSS1500;Body |
| GSM1179499Blood_IT10 | chr17:17108846-17110353 | PLD6 | Island | Body |
| GSM1179499Blood_IT10 | chr17:259755-259942 | C17orf97 | Island | TSS1500 |
| GSM1179510Muscle_IT10 | chr17:33759484-33759648 | SLFN12;SLFN12 | | 1stExon;5'UTR |
| GSM1179499Blood_IT10 | chr17:33759484-33760419 | SLFN12;SLFN12 | | 1stExon;5'UTR |
| GSM1179516Omentum_IT10 | chr17:3433428-3433970 | TRPV3 | N_Shelf | Body |
| GSM1179499Blood_IT10 | chr17:74075092-74075195 | ZACN | S_Shore | TSS200 |
| GSM1179516Omentum_IT10 | chr17:75181131-75181373 | SEC14L1;SEC14L1;SEC14L1;SEC14L1;SEC14L1 | | Body;Body;TSS200;Body;Body |
| GSM1179499Blood_IT10 | chr17:80708279-80708632 | FN3K | Island | Body |
| GSM1179526SC_Fat_IT10 | chr17:80708279-80708632 | FN3K | Island | Body |
| GSM1179499Blood_IT10 | chr17:80870107-80870577 | TBCD | N_Shore | Body |
| GSM1179516Omentum_IT10 | chr18:20714217-20714399 | CABLES1 | N_Shore | TSS1500 |
| GSM1179499Blood_IT10 | chr18:49866065-49866097 | DCC | N_Shelf | TSS1500 |
| GSM1179516Omentum_IT10 | chr18:72166016-72166324 | CNDP2;CNDP2 | S_Shore | TSS1500;5'UTR |
| GSM1179516Omentum_IT10 | chr19:12876846-12877000 | HOOK2;HOOK2 | N_Shore | Body;Body |
| GSM1179510Muscle_IT10 | chr19:12876846-12877188 | HOOK2;HOOK2 | N_Shore | Body;Body |

| | | | | |
|---|---|---|---|---|
| GSM1179526SC_Fat_IT10 | chr19:12876846-12877188 | HOOK2;HOOK2 | N_Shore | Body;Body |
| GSM1179516Omentum_IT10 | chr19:14550846-14550999 | PKN1;PKN1 | N_Shore | TSS1500;Body |
| GSM1179510Muscle_IT10 | chr19:23941207-23941575 | ZNF681 | N_Shore | Body |
| GSM1179516Omentum_IT10 | chr19:23941207-23941575 | ZNF681 | N_Shore | Body |
| GSM1179526SC_Fat_IT10 | chr19:23941207-23941575 | ZNF681 | N_Shore | Body |
| GSM1179516Omentum_IT10 | chr19:2540907-2541104 | GNG7 | Island | 5'UTR |
| GSM1179499Blood_IT10 | chr19:34174967-34175052 | CHST8;CHST8;CHST8 | N_Shore | 5'UTR;5'UTR;TSS1500 |
| GSM1179516Omentum_IT10 | chr19:36233297-36233469 | TMEM149;TMEM149 | S_Shore | 1stExon;5'UTR |
| GSM1179516Omentum_IT10 | chr19:4305072-4305239 | FSD1 | Island | Body |
| GSM1179499Blood_IT10 | chr19:50249584-50249776 | TSKS | Island | Body |
| GSM1179516Omentum_IT10 | chr2:100759014-100759159 | AFF3;AFF3 | | 1stExon;5'UTR |
| GSM1179516Omentum_IT10 | chr2:143886326-143886940 | ARHGAP15 | | TSS1500 |
| GSM1179499Blood_IT10 | chr2:1817263-1817409 | MYT1L | | Body |
| GSM1179499Blood_IT10 | chr2:183943198-183944048 | DUSP19;DUSP19 | | TSS200;TSS200 |
| GSM1179499Blood_IT10 | chr2:1842964-1843136 | MYT1L | Island | Body |
| GSM1179499Blood_IT10 | chr2:217276744-217276821 | SMARCAL1;SMARCAL1 | N_Shore | TSS1500;TSS1500 |
| GSM1179516Omentum_IT10 | chr2:234296642-234297039 | DGKD;DGKD | | Body;TSS200 |
| GSM1179516Omentum_IT10 | chr2:240230452-240230892 | HDAC4 | | Body |
| GSM1179516Omentum_IT10 | chr2:240302585-240302855 | HDAC4 | | 5'UTR |
| GSM1179510Muscle_IT10 | chr2:43019614-43020457 | HAAO | N_Shore | 1stExon |
| GSM1179499Blood_IT10 | chr2:97505275-97505599 | ANKRD23 | Island | 3'UTR |
| GSM1179516Omentum_IT10 | chr20:17595355-17595472 | RRBP1;RRBP1 | Island | Body;Body |
| GSM1179516Omentum_IT10 | chr20:24929607-24929716 | CST7 | | TSS1500 |
| GSM1179516Omentum_IT10 | chr20:4795444-4795999 | RASSF2;RASSF2 | | 5'UTR;5'UTR |
| GSM1179516Omentum_IT10 | chr20:50159508-50159653 | NFATC2;NFATC2;NFATC2 | Island | TSS1500;Body;TSS1500 |
| GSM1179499Blood_IT10 | chr21:35831871-35832180 | KCNE1;KCNE1;KCNE1;KCNE1 | Island | 5'UTR;5'UTR;1stExon;1stExon;5'UTR |
| GSM1179516Omentum_IT10 | chr21:48026043-48026298 | S100B | | TSS1500 |
| GSM1179499Blood_IT10 | chr22:32438947-32439052 | SLC5A1 | N_Shore | TSS200 |
| GSM1179516Omentum_IT10 | chr3:135684107-135684163 | PPP2R3A | N_Shore | TSS1500 |
| GSM1179516Omentum_IT10 | chr3:145969579-145969718 | PLSCR4;PLSCR4;PLSCR4;PLS | S_Shore | TSS1500;TSS1500;TSS1500;TSS1500 |
| GSM1179499Blood_IT10 | chr3:159557552-159558031 | SCHIP1 | | Body |
| GSM1179499Blood_IT10 | chr3:195489708-195490309 | MUC4;MUC4;MUC4 | Island | Body;Body;Body |
| GSM1179499Blood_IT10 | chr3:48632484-48632672 | COL7A1 | Island | Body |
| GSM1179499Blood_IT10 | chr3:49203973-49204239 | CCDC71 | S_Shore | TSS200 |
| GSM1179516Omentum_IT10 | chr4:100737821-100738011 | DAPP1 | | TSS200 |
| GSM1179516Omentum_IT10 | chr4:116034871-116035232 | NDST4;NDST4 | | 1stExon;5'UTR |
| GSM1179516Omentum_IT10 | chr4:186559412-186560083 | SORBS2;SORBS2;SORBS2;SO | N_Shore | Body;Body;Body;Body;Body;Body;Body;Body |
| GSM1179499Blood_IT10 | chr4:21699426-21699745 | KCNIP4;KCNIP4 | | 5'UTR;TSS200 |
| GSM1179499Blood_IT10 | chr4:72897613-72898212 | NPFFR2;NPFFR2 | Island | 1stExon;5'UTR |
| GSM1179516Omentum_IT10 | chr4:90815958-90816310 | MMRN1 | | TSS200 |
| GSM1179499Blood_IT10 | chr4:940614-941054 | TMEM175 | Island | 5'UTR |
| GSM1179499Blood_IT10 | chr5:126408756-126409372 | FLJ44606;FLJ44606 | N_Shore | 5'UTR;5'UTR |
| GSM1179499Blood_IT10 | chr5:131607097-131607372 | PDLIM4;PDLIM4 | Island | Body;Body |
| GSM1179499Blood_IT10 | chr5:1594282-1594863 | SDHAP3 | Island | Body |
| GSM1179510Muscle_IT10 | chr5:1594282-1594863 | SDHAP3 | Island | Body |
| GSM1179516Omentum_IT10 | chr5:1594282-1594863 | SDHAP3 | Island | Body |
| GSM1179526SC_Fat_IT10 | chr5:1594282-1594863 | SDHAP3 | Island | Body |
| GSM1179499Blood_IT10 | chr5:176797920-176798049 | RGS14 | S_Shelf | Body |
| GSM1179516Omentum_IT10 | chr5:176797920-176798049 | RGS14 | S_Shelf | Body |
| GSM1179499Blood_IT10 | chr5:179740743-179741120 | GFPT2 | Island | Body |
| GSM1179516Omentum_IT10 | chr5:27038707-27038802 | CDH9 | | TSS200 |
| GSM1179499Blood_IT10 | chr5:5139508-5139512 | ADAMTS16 | Island | TSS1500 |
| GSM1179516Omentum_IT10 | chr5:9547468-9548098 | SNORD123;SEMA5A | S_Shore | TSS1500;TSS1500 |
| GSM1179516Omentum_IT10 | chr6:11382553-11382830 | NEDD9;NEDD9 | | 1stExon;5'UTR |
| GSM1179516Omentum_IT10 | chr6:13274151-13274354 | PHACTR1 | | Body |
| GSM1179516Omentum_IT10 | chr6:13428522-13428657 | GFOD1 | | Body |
| GSM1179516Omentum_IT10 | chr6:138866865-138867125 | NHSL1 | | Body |
| GSM1179516Omentum_IT10 | chr6:152702330-152702660 | SYNE1;SYNE1 | | Body;Body |
| GSM1179516Omentum_IT10 | chr6:15504844-15505460 | JARID2 | | Body |
| GSM1179499Blood_IT10 | chr6:291687-293331 | DUSP22 | N_Shore | TSS1500 |
| GSM1179510Muscle_IT10 | chr6:291687-293331 | DUSP22 | N_Shore | TSS1500 |
| GSM1179526SC_Fat_IT10 | chr6:291687-293331 | DUSP22 | N_Shore | TSS1500 |
| GSM1179510Muscle_IT10 | chr6:29910755-29910778 | HLA-A | Island | Body |
| GSM1179526SC_Fat_IT10 | chr6:29910755-29910796 | HLA-A | Island | Body |
| GSM1179516Omentum_IT10 | chr6:30122523-30122593 | TRIM10;TRIM10 | | Body;Body |
| GSM1179516Omentum_IT10 | chr6:31166502-31166799 | HCG27 | S_Shore | Body |

| | | | | |
|---|---|---|---|---|
| GSM1179499Blood_IT10 | chr6:32063835-32064148 | TNXB | Island | Body |
| GSM1179499Blood_IT10 | chr6:32064161-32064258 | TNXB | Island | Body |
| GSM1179526SC_Fat_IT10 | chr6:32305068-32305145 | C6orf10 | | Body |
| GSM1179516Omentum_IT10 | chr6:32634316-32634362 | HLA-DQB1 | S_Shore | 1stExon |
| GSM1179516Omentum_IT10 | chr6:34984836-34984953 | ANKS1A | | Body |
| GSM1179516Omentum_IT10 | chr6:76203530-76203675 | FILIP1 | | TSS200 |
| GSM1179516Omentum_IT10 | chr7:1163456-1163642 | C7orf50;C7orf50;C7orf50 | | Body;Body;Body |
| GSM1179499Blood_IT10 | chr7:157454062-157454207 | PTPRN2;PTPRN2;PTPRN2 | Island | Body;Body;Body |
| GSM1179499Blood_IT10 | chr7:1883139-1883760 | MAD1L1;MAD1L1;MAD1L1 | N_Shore | Body;Body;Body |
| GSM1179499Blood_IT10 | chr7:24797835-24797884 | DFNA5;DFNA5;DFNA5 | S_Shore | TSS1500;TSS200;TSS200 |
| GSM1179516Omentum_IT10 | chr7:27153580-27157855 | HOXA3;HOXA3;HOXA3;HOXA3 | Island | 5'UTR;1stExon;5'UTR;5'UTR |
| GSM1179516Omentum_IT10 | chr7:27168780-27169740 | HOXA4 | N_Shore | 3'UTR |
| GSM1179516Omentum_IT10 | chr7:36764019-36764205 | AOAH;AOAH | | 1stExon;5'UTR |
| GSM1179499Blood_IT10 | chr7:4304846-4305535 | SDK1;SDK1 | Island | Body;Body |
| GSM1179516Omentum_IT10 | chr7:45066738-45067319 | CCM2;CCM2;CCM2;CCM2;CCM2 | | Body;TSS1500;Body;Body;Body |
| GSM1179499Blood_IT10 | chr7:4764443-4764941 | FOXK1 | N_Shore | Body |
| GSM1179516Omentum_IT10 | chr7:4848683-4848939 | RADIL | Island | Body |
| GSM1179516Omentum_IT10 | chr8:140937286-140937410 | TRAPPC9;TRAPPC9 | | Body;Body |
| GSM1179516Omentum_IT10 | chr8:2075209-2075820 | MYOM2 | N_Shore | Body |
| GSM1179526SC_Fat_IT10 | chr8:2075209-2075820 | MYOM2 | N_Shore | Body |
| GSM1179499Blood_IT10 | chr8:2075469-2075820 | MYOM2 | Island | Body |
| GSM1179516Omentum_IT10 | chr9:140356475-140356830 | PNPLA7;PNPLA7 | Island | Body;Body |
| GSM1179499Blood_IT10 | chr9:34370781-34371380 | KIAA1161 | Island | 3'UTR |
| GSM1179515Omentum_IT13 | chr1:114414042-114414408 | PTPN22;PTPN22 | | Body;Body |
| GSM1179515Omentum_IT13 | chr1:11795746-11795985 | AGTRAP;AGTRAP;AGTRAP;AGT | N_Shore | TSS1500;TSS1500;TSS1500;TSS1500;TSS1500 |
| GSM1179515Omentum_IT13 | chr1:12538341-12538678 | VPS13D;VPS13D | | Body;Body |
| GSM1179515Omentum_IT13 | chr1:145715571-145715719 | CD160 | S_Shore | TSS200 |
| GSM1179515Omentum_IT13 | chr1:154943349-154943932 | SHC1;SHC1;SHC1;SHC1 | N_Shelf | 5'UTR;5'UTR;TSS200;TSS200 |
| GSM1179515Omentum_IT13 | chr1:158900384-158900644 | PYHIN1;PYHIN1;PYHIN1;PYHIN1 | | TSS1500;TSS1500;TSS1500;TSS1500 |
| GSM1179515Omentum_IT13 | chr1:159046391-159047163 | AIM2 | | 5'UTR |
| GSM1179515Omentum_IT13 | chr1:167690438-167690788 | MPZL1;MPZL1;MPZL1 | N_Shore | TSS1500;TSS1500;TSS1500 |
| GSM1179515Omentum_IT13 | chr1:172628020-172628263 | FASLG | | TSS200 |
| GSM1179515Omentum_IT13 | chr1:203733971-203734559 | LAX1;LAX1 | | TSS1500;TSS1500 |
| GSM1179515Omentum_IT13 | chr1:206729553-206730120 | RASSF5;RASSF5;RASSF5 | N_Shore | Body;TSS1500;Body |
| GSM1179515Omentum_IT13 | chr1:207070539-207070834 | IL24;IL24 | | TSS1500;TSS1500 |
| GSM1179498Blood_IT13 | chr1:207842917-207843084 | CR1L | Island | Body |
| GSM1179515Omentum_IT13 | chr1:25290947-25292274 | RUNX3 | | Body |
| GSM1179515Omentum_IT13 | chr1:28905573-28906538 | SNHG12;SNORD99 | N_Shelf | Body;TSS1500 |
| GSM1179515Omentum_IT13 | chr1:3703792-3704559 | LRRC47 | Island | Body |
| GSM1179515Omentum_IT13 | chr1:79085586-79085765 | IFI44L | | TSS1500 |
| GSM1179515Omentum_IT13 | chr1:95698378-95699097 | RWDD3;RWDD3 | N_Shore | TSS1500;TSS1500 |
| GSM1179515Omentum_IT13 | chr1:9788517-9789174 | PIK3CD | N_Shore | 3'UTR |
| GSM1179515Omentum_IT13 | chr10:102279690-102279758 | SEC31B | Island | TSS200 |
| GSM1179515Omentum_IT13 | chr10:102760724-102760970 | LZTS2 | S_Shore | 5'UTR |
| GSM1179515Omentum_IT13 | chr10:1156113-1156594 | WDR37 | | Body |
| GSM1179515Omentum_IT13 | chr10:121300857-121301343 | RGS10 | N_Shore | Body |
| GSM1179515Omentum_IT13 | chr10:123923881-123924085 | TACC2;TACC2;TACC2;TACC2 | S_Shore | Body;Body;Body;Body |
| GSM1179515Omentum_IT13 | chr10:124909889-124910217 | HMX2 | Island | 3'UTR |
| GSM1179515Omentum_IT13 | chr10:13388442-13388567 | SEPHS1 | | 5'UTR |
| GSM1179498Blood_IT13 | chr10:135341528-135342218 | CYP2E1 | Island | Body |
| GSM1179509Muscle_IT13 | chr10:135341528-135342218 | CYP2E1 | Island | Body |
| GSM1179515Omentum_IT13 | chr10:135341528-135342218 | CYP2E1 | Island | Body |
| GSM1179525SC_Fat_IT13 | chr10:135341528-135342218 | CYP2E1 | Island | Body |
| GSM1179498Blood_IT13 | chr10:135342560-135343280 | CYP2E1 | Island | Body |
| GSM1179525SC_Fat_IT13 | chr10:135342560-135343280 | CYP2E1 | Island | Body |
| GSM1179515Omentum_IT13 | chr10:135379213-135379242 | SYCE1;SYCE1;SYCE1 | Island | TSS200;TSS200;5'UTR |
| GSM1179515Omentum_IT13 | chr10:1551251-1551326 | ADARB2 | | Body |
| GSM1179498Blood_IT13 | chr10:34408530-34408654 | PARD3 | | Body |
| GSM1179515Omentum_IT13 | chr10:34408530-34408654 | PARD3 | | Body |
| GSM1179515Omentum_IT13 | chr10:6104137-6104515 | IL2RA;IL2RA | | 1stExon;5'UTR |
| GSM1179515Omentum_IT13 | chr10:639935-640085 | DIP2C | | Body |
| GSM1179515Omentum_IT13 | chr11:104769087-104769543 | CASP12 | | Body |
| GSM1179515Omentum_IT13 | chr11:11862730-11862879 | USP47 | N_Shore | TSS1500 |
| GSM1179515Omentum_IT13 | chr11:118754280-118754593 | CXCR5 | | TSS1500 |
| GSM1179515Omentum_IT13 | chr11:118763859-118763901 | CXCR5;CXCR5 | | Body;TSS1500 |
| GSM1179515Omentum_IT13 | chr11:121970694-121970768 | LOC399959;MIR125B1 | | Body;TSS200 |

| | | | | |
|---|---|---|---|---|
| GSM1179515Omentum_IT13 | chr11:1320497-1320634 | TOLLIP | S_Shelf | Body |
| GSM1179515Omentum_IT13 | chr11:2846681-2847642 | KCNQ1;KCNQ1 | | Body;Body |
| GSM1179515Omentum_IT13 | chr11:47278977-47279586 | NR1H3;NR1H3;NR1H3 | | 5'UTR;TSS1500;TSS1500 |
| GSM1179515Omentum_IT13 | chr11:504784-504937 | RNH1;RNH1;RNH1;RNH1;RNH | N_Shore | 5'UTR;5'UTR;5'UTR;5'UTR;5'UTR;5'UTR;5'UTR;5'UTR |
| GSM1179515Omentum_IT13 | chr11:57005876-57006273 | APLNR;APLNR | | TSS1500;TSS1500 |
| GSM1179515Omentum_IT13 | chr11:57364841-57365088 | SERPING1;SERPING1 | | TSS1500;TSS200 |
| GSM1179515Omentum_IT13 | chr11:60738032-60739183 | CD6 | | TSS1500 |
| GSM1179515Omentum_IT13 | chr11:62314820-62315320 | AHNAK;AHNAK | Island | TSS1500;TSS1500 |
| GSM1179515Omentum_IT13 | chr11:64107158-64108241 | CCDC88B | N_Shelf | TSS1500 |
| GSM1179515Omentum_IT13 | chr11:70211301-70211531 | PPFIA1;PPFIA1 | | Body;Body |
| GSM1179515Omentum_IT13 | chr11:843897-844686 | TSPAN4;TSPAN4;TSPAN4;TSPA | S_Shore | TSS1500;5'UTR;TSS200;5'UTR;5'UTR;TSS1500;5'UTR |
| GSM1179515Omentum_IT13 | chr11:86383696-86383809 | ME3;ME3;ME3 | S_Shore | TSS1500;TSS200;TSS1500 |
| GSM1179515Omentum_IT13 | chr12:123756945-123757070 | CDK2AP1 | S_Shore | TSS1500 |
| GSM1179515Omentum_IT13 | chr12:124864528-124865130 | NCOR2;NCOR2 | | Body;Body |
| GSM1179515Omentum_IT13 | chr12:125002007-125003558 | NCOR2;NCOR2 | Island | 5'UTR;5'UTR |
| GSM1179515Omentum_IT13 | chr12:131590393-131590596 | GPR133 | | Body |
| GSM1179515Omentum_IT13 | chr12:46663660-46664096 | SLC38A1;SLC38A1 | S_Shore | TSS1500;TSS1500 |
| GSM1179515Omentum_IT13 | chr12:48152204-48152428 | RAPGEF3;RAPGEF3;RAPGEF3; | N_Shore | 1stExon;TSS200;5'UTR;Body |
| GSM1179515Omentum_IT13 | chr12:52404134-52404422 | GRASP | S_Shelf | Body |
| GSM1179515Omentum_IT13 | chr12:53591398-53591766 | ITGB7 | Island | Body |
| GSM1179515Omentum_IT13 | chr12:54653065-54653427 | CBX5;CBX5;CBX5;CBX5 | | 1stExon;5'UTR;5'UTR;5'UTR |
| GSM1179515Omentum_IT13 | chr12:54891491-54891636 | NCKAP1L | | TSS200 |
| GSM1179515Omentum_IT13 | chr12:6492890-6493003 | LTBR | Island | TSS1500 |
| GSM1179515Omentum_IT13 | chr12:9217390-9217907 | LOC144571 | Island | TSS1500 |
| GSM1179515Omentum_IT13 | chr12:95226786-95226993 | MIR492 | | TSS1500 |
| GSM1179515Omentum_IT13 | chr13:107145933-107146164 | EFNB2 | S_Shore | Body |
| GSM1179515Omentum_IT13 | chr13:113540189-113540631 | ATP11A;ATP11A | N_Shore | 3'UTR;3'UTR |
| GSM1179515Omentum_IT13 | chr13:99629970-99630583 | DOCK9;DOCK9;DOCK9;DOCK9 | | Body;Body;Body;Body |
| GSM1179515Omentum_IT13 | chr14:23305548-23305957 | MMP14 | N_Shore | TSS1500 |
| GSM1179515Omentum_IT13 | chr14:39734964-39735625 | CTAGE5;CTAGE5;CTAGE5;CTA | N_Shore | TSS1500;TSS1500;TSS1500;Body |
| GSM1179515Omentum_IT13 | chr14:75988356-75989203 | BATF | | TSS1500 |
| GSM1179515Omentum_IT13 | chr14:81425912-81426234 | TSHR;TSHR;TSHR | | Body;Body;Body |
| GSM1179515Omentum_IT13 | chr14:99655593-99655747 | BCL11B;BCL11B | | Body;Body |
| GSM1179515Omentum_IT13 | chr15:34628731-34629056 | SLC12A6;SLC12A6;SLC12A6;S | N_Shore | 1stExon;5'UTR;Body;5'UTR;1stExon |
| GSM1179515Omentum_IT13 | chr15:38988533-38988860 | C15orf53 | | TSS1500 |
| GSM1179515Omentum_IT13 | chr15:40599680-40600635 | PLCB2 | | Body |
| GSM1179515Omentum_IT13 | chr15:40633124-40633294 | C15orf52 | | 1stExon |
| GSM1179515Omentum_IT13 | chr15:91427184-91427363 | FES;FES;FES;FES | N_Shore | TSS1500;TSS1500;TSS1500;TSS1500 |
| GSM1179515Omentum_IT13 | chr16:14403004-14403425 | MIR365-1 | | TSS200 |
| GSM1179515Omentum_IT13 | chr16:1575979-1576146 | IFT140 | S_Shore | Body |
| GSM1179515Omentum_IT13 | chr16:4729905-4730657 | MGRN1;MGRN1;MGRN1;MGR | N_Shore | Body;Body;Body;Body |
| GSM1179515Omentum_IT13 | chr16:474271-474528 | RAB11FIP3 | N_Shore | TSS1500 |
| GSM1179515Omentum_IT13 | chr16:50715135-50715529 | SNX20;SNX20;SNX20;SNX20;SNX20;SNX20 | | 1stExon;5'UTR;1stExon;5'UTR;5'UTR;1stExon |
| GSM1179515Omentum_IT13 | chr16:69597925-69598417 | NFAT5;NFAT5;NFAT5;NFAT5;N | N_Shore | TSS1500;TSS1500;TSS1500;TSS1500;TSS1500 |
| GSM1179515Omentum_IT13 | chr16:69958298-69958964 | WWP2;WWP2 | | TSS1500;Body |
| GSM1179515Omentum_IT13 | chr16:85936147-85936666 | IRF8 | S_Shelf | 5'UTR |
| GSM1179515Omentum_IT13 | chr16:88536950-88537260 | ZFPM1 | Island | Body |
| GSM1179515Omentum_IT13 | chr16:89180285-89180742 | ACSF3;ACSF3;ACSF3 | | Body;Body;Body |
| GSM1179515Omentum_IT13 | chr16:89390609-89390968 | ANKRD11 | | 5'UTR |
| GSM1179515Omentum_IT13 | chr17:34207332-34207663 | CCL5;CCL5 | | 1stExon;5'UTR |
| GSM1179515Omentum_IT13 | chr17:38023480-38024146 | ZPBP2;ZPBP2 | N_Shore | TSS1500;TSS1500 |
| GSM1179515Omentum_IT13 | chr17:38599056-38599366 | IGFBP4 | N_Shore | TSS1500 |
| GSM1179515Omentum_IT13 | chr17:40489513-40489785 | STAT3;STAT3;STAT3 | | Body;Body;Body |
| GSM1179515Omentum_IT13 | chr17:4079262-4079652 | ANKFY1 | | Body |
| GSM1179515Omentum_IT13 | chr17:57918500-57918682 | MIR21 | | TSS200 |
| GSM1179515Omentum_IT13 | chr17:66374702-66375195 | ARSG | | Body |
| GSM1179515Omentum_IT13 | chr17:7283817-7284049 | TNK1 | N_Shore | TSS1500 |
| GSM1179515Omentum_IT13 | chr17:76036514-76037562 | TNRC6C;TNRC6C | N_Shore | 5'UTR;5'UTR |
| GSM1179515Omentum_IT13 | chr17:77133777-77134159 | HRNBP3 | N_Shore | 5'UTR |
| GSM1179515Omentum_IT13 | chr17:78851149-78851503 | RPTOR;RPTOR | S_Shore | Body;Body |
| GSM1179515Omentum_IT13 | chr17:79419796-79420145 | BAHCC1 | N_Shore | Body |
| GSM1179515Omentum_IT13 | chr17:79881468-79882042 | MAFG;MAFG | S_Shore | 5'UTR;TSS200 |
| GSM1179515Omentum_IT13 | chr17:80084554-80085332 | CCDC57 | N_Shore | Body |
| GSM1179515Omentum_IT13 | chr17:80190129-80190154 | SLC16A3;SLC16A3;SLC16A3 | S_Shore | TSS200;5'UTR;5'UTR |
| GSM1179515Omentum_IT13 | chr18:13641529-13641894 | C18orf1;C18orf1;C18orf1;C18 | N_Shore | Body;Body;Body;Body;Body;Body |
| GSM1179515Omentum_IT13 | chr18:77196147-77196450 | NFATC1;NFATC1;NFATC1;NFA | Island | Body;5'UTR;Body;Body;Body |

| | | | | |
|---|---|---|---|---|
| GSM1179515Omentum_IT13 | chr18:77723226-77723518 | HSBP1L1 | N_Shore | TSS1500 |
| GSM1179515Omentum_IT13 | chr19:1155030-1155738 | SBNO2 | Island | 5'UTR |
| GSM1179515Omentum_IT13 | chr19:11665102-11665451 | ELOF1 | | Body |
| GSM1179515Omentum_IT13 | chr19:15391832-15391946 | BRD4;BRD4 | | TSS1500;TSS1500 |
| GSM1179515Omentum_IT13 | chr19:18197544-18197853 | IL12RB1;IL12RB1 | | Body;Body |
| GSM1179509Muscle_IT13 | chr19:21264896-21265421 | ZNF714 | N_Shore | TSS200 |
| GSM1179515Omentum_IT13 | chr19:4305072-4305239 | FSD1 | Island | Body |
| GSM1179515Omentum_IT13 | chr19:50015523-50015780 | FCGRT;FCGRT | N_Shore | TSS200;TSS1500 |
| GSM1179515Omentum_IT13 | chr19:50848114-50848461 | NAPSB | | TSS200 |
| GSM1179515Omentum_IT13 | chr2:100759014-100759159 | AFF3;AFF3 | | 1stExon;5'UTR |
| GSM1179515Omentum_IT13 | chr2:110873031-110873396 | MALL | Island | Body |
| GSM1179515Omentum_IT13 | chr2:127413363-127413369 | GYPC;GYPC | N_Shore | TSS1500;TSS1500 |
| GSM1179515Omentum_IT13 | chr2:143886326-143886940 | ARHGAP15 | | TSS1500 |
| GSM1179515Omentum_IT13 | chr2:157292018-157292127 | GPD2;GPD2;GPD2 | N_Shore | 1stExon;TSS1500;5'UTR |
| GSM1179515Omentum_IT13 | chr2:160472940-160473528 | BAZ2B;BAZ2B | S_Shore | 1stExon;5'UTR |
| GSM1179515Omentum_IT13 | chr2:160655036-160655081 | CD302 | S_Shore | TSS1500 |
| GSM1179515Omentum_IT13 | chr2:169658881-169659182 | NOSTRIN;NOSTRIN;NOSTRIN;NOSTRIN | | 5'UTR;TSS1500;TSS1500;TSS1500 |
| GSM1179515Omentum_IT13 | chr2:234296642-234297039 | DGKD;DGKD | | Body;TSS200 |
| GSM1179515Omentum_IT13 | chr2:240230452-240230892 | HDAC4 | | Body |
| GSM1179515Omentum_IT13 | chr2:240291426-240291657 | HDAC4 | | 5'UTR |
| GSM1179515Omentum_IT13 | chr2:38976250-38976409 | SFRS7 | N_Shore | Body |
| GSM1179515Omentum_IT13 | chr20:19955436-19955868 | RIN2 | N_Shore | Body |
| GSM1179515Omentum_IT13 | chr20:43343365-43343760 | WISP2 | | TSS1500 |
| GSM1179515Omentum_IT13 | chr20:45947025-45947892 | ZMYND8;LOC100131496;ZMYND8;ZMYND8 | | Body;TSS1500;Body;Body |
| GSM1179515Omentum_IT13 | chr20:47896448-47897451 | SNORD12B;C20orf199;MIR12 | S_Shore | TSS1500;Body;TSS1500;Body;TSS1500;Body |
| GSM1179515Omentum_IT13 | chr20:4795444-4796176 | RASSF2;RASSF2 | | 5'UTR;5'UTR |
| GSM1179515Omentum_IT13 | chr20:62367698-62367961 | LIME1 | N_Shore | TSS1500 |
| GSM1179515Omentum_IT13 | chr20:62549301-62549385 | MIR941-1;DNAJC5 | N_Shore | TSS1500;5'UTR |
| GSM1179515Omentum_IT13 | chr21:36263347-36263808 | RUNX1 | Island | Body |
| GSM1179515Omentum_IT13 | chr21:41554681-41554930 | DSCAM | Island | Body |
| GSM1179515Omentum_IT13 | chr22:17956453-17956641 | CECR2 | | TSS200 |
| GSM1179525SC_Fat_IT13 | chr22:32599511-32599648 | RFPL2;RFPL2;RFPL2 | | TSS200;5'UTR;TSS1500 |
| GSM1179515Omentum_IT13 | chr22:36236649-36236760 | RBM9;RBM9;RBM9;RBM9;RBM9;RBM9 | | Body;Body;TSS200;TSS200;TSS200;TSS200 |
| GSM1179515Omentum_IT13 | chr22:38714001-38714466 | CSNK1E;CSNK1E;CSNK1E | S_Shore | 1stExon;5'UTR;TSS1500 |
| GSM1179515Omentum_IT13 | chr22:40296407-40297048 | GRAP2 | | TSS1500 |
| GSM1179515Omentum_IT13 | chr3:112359488-112359935 | CCDC80;CCDC80;CCDC80 | | 5'UTR;1stExon;5'UTR |
| GSM1179515Omentum_IT13 | chr3:11623526-11624023 | VGLL4;VGLL4;VGLL4 | | Body;Body;Body |
| GSM1179515Omentum_IT13 | chr3:127473609-127473936 | MGLL;MGLL | | Body;Body |
| GSM1179515Omentum_IT13 | chr3:135684107-135684163 | PPP2R3A | N_Shore | TSS1500 |
| GSM1179515Omentum_IT13 | chr3:18480242-18480341 | SATB1;SATB1 | | 5'UTR;1stExon |
| GSM1179515Omentum_IT13 | chr3:42694803-42695307 | ZBTB47 | N_Shore | TSS1500 |
| GSM1179515Omentum_IT13 | chr3:45984743-45985168 | CXCR6;FYCO1 | | TSS1500;Body |
| GSM1179515Omentum_IT13 | chr3:46608025-46608179 | LRRC2;LRRC2 | S_Shore | 5'UTR;TSS200 |
| GSM1179515Omentum_IT13 | chr3:9958613-9959128 | IL17RC;IL17RC;IL17RC | S_Shore | TSS200;TSS200;TSS200 |
| GSM1179515Omentum_IT13 | chr4:186559412-186560083 | SORBS2;SORBS2;SORBS2;SO | N_Shore | Body;Body;Body;Body;Body;Body;Body;Body |
| GSM1179515Omentum_IT13 | chr4:187476455-187476599 | MTNR1A | Island | 1stExon |
| GSM1179515Omentum_IT13 | chr4:23892515-23892621 | PPARGC1A | | TSS1500 |
| GSM1179515Omentum_IT13 | chr4:3374219-3375144 | RGS12;RGS12;RGS12 | N_Shore | Body;Body;Body |
| GSM1179515Omentum_IT13 | chr4:48136234-48136452 | TXK;TXK | | 5'UTR;1stExon |
| GSM1179515Omentum_IT13 | chr4:6955660-6955953 | TBC1D14;TBC1D14 | | Body;Body |
| GSM1179515Omentum_IT13 | chr4:8230689-8230847 | SH3TC1 | | Body |
| GSM1179515Omentum_IT13 | chr4:89978251-89978566 | FAM13A;FAM13A | | 1stExon;5'UTR |
| GSM1179515Omentum_IT13 | chr5:118691007-118691126 | TNFAIP8;TNFAIP8 | N_Shore | Body;TSS1500 |
| GSM1179515Omentum_IT13 | chr5:150020212-150020338 | SYNPO;SYNPO;SYNPO;SYNPO;SYNPO;SYNPO | | 1stExon;5'UTR;5'UTR;Body;1stExon;Body |
| GSM1179515Omentum_IT13 | chr5:156696185-156696351 | CYFIP2;CYFIP2;CYFIP2 | S_Shelf | TSS200;5'UTR;5'UTR |
| GSM1179498Blood_IT13 | chr5:1594282-1594863 | SDHAP3 | Island | Body |
| GSM1179509Muscle_IT13 | chr5:1594282-1594863 | SDHAP3 | Island | Body |
| GSM1179515Omentum_IT13 | chr5:1594282-1594863 | SDHAP3 | Island | Body |
| GSM1179525SC_Fat_IT13 | chr5:1594282-1594863 | SDHAP3 | Island | Body |
| GSM1179515Omentum_IT13 | chr5:16784821-16785399 | MYO10 | | Body |
| GSM1179515Omentum_IT13 | chr5:169407439-169408301 | FAM196B;DOCK2;FAM196B | | 5'UTR;Body;1stExon |
| GSM1179498Blood_IT13 | chr5:179740743-179741120 | GFPT2 | Island | Body |
| GSM1179515Omentum_IT13 | chr5:179740743-179741120 | GFPT2 | Island | Body |
| GSM1179515Omentum_IT13 | chr5:75919188-75919481 | F2RL2;F2RL2;IQGAP2 | | 1stExon;5'UTR;Body |
| GSM1179515Omentum_IT13 | chr6:11382553-11382830 | NEDD9;NEDD9 | | 1stExon;5'UTR |
| GSM1179515Omentum_IT13 | chr6:128222053-128222390 | THEMIS;THEMIS;THEMIS | | 5'UTR;1stExon;1stExon |

| | | | | |
|---|---|---|---|---|
| GSM1179515Omentum_IT13 | chr6:13274151-13274354 | PHACTR1 | | Body |
| GSM1179515Omentum_IT13 | chr6:13428522-13428657 | GFOD1 | | Body |
| GSM1179515Omentum_IT13 | chr6:15504844-15505460 | JARID2 | | Body |
| GSM1179515Omentum_IT13 | chr6:159463103-159463300 | TAGAP;TAGAP;TAGAP | | 5'UTR;Body;Body |
| GSM1179515Omentum_IT13 | chr6:161796785-161796854 | PARK2;PARK2;PARK2 | | Body;Body;Body |
| GSM1179515Omentum_IT13 | chr6:29911334-29911558 | HLA-A | Island | Body |
| GSM1179515Omentum_IT13 | chr6:30124214-30124248 | TRIM10;TRIM10 | | Body;Body |
| GSM1179515Omentum_IT13 | chr6:30130698-30131001 | TRIM15 | | TSS1500 |
| GSM1179515Omentum_IT13 | chr6:30458158-30460798 | HLA-E | Island | Body |
| GSM1179515Omentum_IT13 | chr6:30647128-30648020 | KIAA1949;KIAA1949 | | Body;Body |
| GSM1179515Omentum_IT13 | chr6:30883762-30883959 | VARS2;VARS2;VARS2 | S_Shore | Body;Body;Body |
| GSM1179515Omentum_IT13 | chr6:31166502-31166799 | HCG27 | S_Shore | Body |
| GSM1179515Omentum_IT13 | chr6:31322298-31323261 | HLA-B | N_Shore | Body |
| GSM1179515Omentum_IT13 | chr6:31323397-31323856 | HLA-B | N_Shore | Body |
| GSM1179515Omentum_IT13 | chr6:31431312-31431969 | HCP5 | | 3'UTR |
| GSM1179515Omentum_IT13 | chr6:31539539-31540121 | LTA;LTA | | TSS1500;TSS1500 |
| GSM1179515Omentum_IT13 | chr6:31540169-31540750 | LTA;LTA;LTA | | 1stExon;5'UTR;5'UTR |
| GSM1179515Omentum_IT13 | chr6:31543647-31545473 | TNF | | 1stExon |
| GSM1179515Omentum_IT13 | chr6:32158927-32160455 | PBX2;GPSM3 | | TSS1500;3'UTR |
| GSM1179515Omentum_IT13 | chr6:32164723-32164816 | GPSM3;NOTCH4 | S_Shore | TSS1500;Body |
| GSM1179515Omentum_IT13 | chr6:32552016-32552042 | HLA-DRB1 | Island | Body |
| GSM1179515Omentum_IT13 | chr6:32634245-32634362 | HLA-DQB1 | S_Shore | Body |
| GSM1179509Muscle_IT13 | chr6:32634276-32634362 | HLA-DQB1 | S_Shore | 1stExon |
| GSM1179525SC_Fat_IT13 | chr6:32634276-32634362 | HLA-DQB1 | S_Shore | 1stExon |
| GSM1179515Omentum_IT13 | chr6:32808504-32808918 | PSMB8;PSMB8 | S_Shore | 3'UTR;3'UTR |
| GSM1179515Omentum_IT13 | chr6:32810258-32810833 | PSMB8;PSMB8 | N_Shore | Body;Body |
| GSM1179515Omentum_IT13 | chr6:32813337-32813715 | PSMB8;TAP1 | S_Shore | TSS1500;3'UTR |
| GSM1179515Omentum_IT13 | chr6:32819858-32820355 | TAP1 | N_Shore | Body |
| GSM1179515Omentum_IT13 | chr6:33267989-33268050 | TAPBP;TAPBP;RGL2;RGL2 | S_Shore | 3'UTR;3'UTR;TSS1500;TSS1500 |
| GSM1179515Omentum_IT13 | chr6:33383819-33384537 | PHF1;PHF1;PHF1 | N_Shore | Body;3'UTR;Body |
| GSM1179515Omentum_IT13 | chr6:52858998-52859107 | GSTA4 | N_Shore | Body |
| GSM1179515Omentum_IT13 | chr6:76203225-76203675 | FILIP1;FILIP1 | | 5'UTR;1stExon |
| GSM1179515Omentum_IT13 | chr7:101688648-101688730 | CUX1;CUX1;CUX1 | | Body;Body;Body |
| GSM1179515Omentum_IT13 | chr7:107300287-107300718 | SLC26A4;LOC286002 | N_Shore | TSS1500;Body |
| GSM1179515Omentum_IT13 | chr7:1127420-1127767 | GPER;C7orf50;C7orf50;C7orf5 | S_Shore | 5'UTR;Body;Body;Body;5'UTR;TSS1500 |
| GSM1179515Omentum_IT13 | chr7:150497065-150497157 | TMEM176B;TMEM176B;TMEM1 | Island | TSS1500;5'UTR;TSS1500;5'UTR;5'UTR;5'UTR |
| GSM1179515Omentum_IT13 | chr7:1961785-1961968 | MAD1L1;MAD1L1;MAD1L1 | | Body;Body;Body |
| GSM1179515Omentum_IT13 | chr7:6199980-6200140 | USP42 | S_Shore | 3'UTR |
| GSM1179515Omentum_IT13 | chr7:625028-625136 | PRKAR1B;PRKAR1B;PRKAR1B;PRKAR1B;PRKAR1B;PRK | | Body;Body;Body;Body;Body;Body |
| GSM1179515Omentum_IT13 | chr8:144543486-144544041 | ZC3H3 | | Body |
| GSM1179515Omentum_IT13 | chr8:145008110-145008957 | PLEC1;PLEC1;PLEC1;PLEC1;P | N_Shore | Body;Body;Body;Body;Body;Body;Body;Body |
| GSM1179515Omentum_IT13 | chr8:42064880-42065314 | PLAT;PLAT | | 5'UTR;5'UTR |
| GSM1179515Omentum_IT13 | chr8:42356637-42356871 | SLC20A2 | | 5'UTR |
| GSM1179515Omentum_IT13 | chr8:48572447-48572557 | KIAA0146 | | Body |
| GSM1179517Omentum_IT15 | chr1:110526158-110526368 | AHCYL1 | N_Shore | TSS1500 |
| GSM1179517Omentum_IT15 | chr1:151344806-151345248 | SELENBP1 | | Body |
| GSM1179517Omentum_IT15 | chr1:15392793-15392907 | KIAA1026;KIAA1026;KIAA1026;KIAA1026;KIAA1026 | | Body;3'UTR;3'UTR;3'UTR;3'UTR |
| GSM1179517Omentum_IT15 | chr1:203733971-203734559 | LAX1;LAX1 | | TSS1500;TSS1500 |
| GSM1179517Omentum_IT15 | chr1:23953480-23953778 | MDS2 | | TSS1500 |
| GSM1179517Omentum_IT15 | chr1:247611842-247611931 | NLRP3;NLRP3;NLRP3;NLRP3;I | N_Shelf | 3'UTR;3'UTR;3'UTR;3'UTR;3'UTR |
| GSM1179517Omentum_IT15 | chr1:247711595-247712591 | C1orf150 | | TSS1500 |
| GSM1179517Omentum_IT15 | chr1:25290947-25292034 | RUNX3 | | Body |
| GSM1179500Blood_IT15 | chr1:26527870-26527928 | CATSPER4 | | Body |
| GSM1179517Omentum_IT15 | chr1:42384056-42384564 | HIVEP3;HIVEP3 | S_Shore | 5'UTR;5'UTR |
| GSM1179500Blood_IT15 | chr1:76189707-76189801 | ACADM;ACADM | N_Shore | TSS1500;TSS1500 |
| GSM1179517Omentum_IT15 | chr1:9788517-9789174 | PIK3CD | N_Shore | 3'UTR |
| GSM1179517Omentum_IT15 | chr10:102760724-102760970 | LZTS2 | S_Shore | 5'UTR |
| GSM1179517Omentum_IT15 | chr10:13388442-13388567 | SEPHS1 | | 5'UTR |
| GSM1179517Omentum_IT15 | chr10:63422511-63422555 | C10orf107 | N_Shore | TSS1500 |
| GSM1179517Omentum_IT15 | chr11:118763674-118763984 | CXCR5;CXCR5 | | Body;TSS1500 |
| GSM1179500Blood_IT15 | chr11:1320497-1320634 | TOLLIP | S_Shelf | Body |
| GSM1179517Omentum_IT15 | chr11:1320497-1320634 | TOLLIP | S_Shelf | Body |
| GSM1179500Blood_IT15 | chr11:18433500-18433683 | LDHC;LDHC | | TSS1500;TSS1500 |
| GSM1179517Omentum_IT15 | chr11:320940-321324 | IFITM3 | S_Shore | TSS200 |
| GSM1179517Omentum_IT15 | chr11:457256-457304 | PTDSS2 | N_Shore | Body |
| GSM1179500Blood_IT15 | chr11:47279183-47279365 | NR1H3;NR1H3;NR1H3 | | 5'UTR;TSS1500;TSS1500 |

| | | | | |
|---|---|---|---|---|
| GSM1179517Omentum_IT15 | chr11:504784-504937 | RNH1;RNH1;RNH1;RNH1;RNH | N_Shore | 5'UTR;5'UTR;5'UTR;5'UTR;5'UTR;5'UTR;5'UTR;5'UTR |
| GSM1179517Omentum_IT15 | chr11:57364841-57365088 | SERPING1;SERPING1 | | TSS1500;TSS200 |
| GSM1179517Omentum_IT15 | chr11:62314820-62315320 | AHNAK;AHNAK | Island | TSS1500;TSS1500 |
| GSM1179517Omentum_IT15 | chr11:692963-693032 | DEAF1 | N_Shore | Body |
| GSM1179517Omentum_IT15 | chr12:124864528-124865130 | NCOR2;NCOR2 | | Body;Body |
| GSM1179517Omentum_IT15 | chr12:125002332-125003558 | NCOR2;NCOR2 | Island | 5'UTR;5'UTR |
| GSM1179511Muscle_IT15 | chr12:130822286-130822674 | PIWIL1 | N_Shore | TSS1500 |
| GSM1179517Omentum_IT15 | chr12:15039025-15039432 | MGP | | TSS200 |
| GSM1179517Omentum_IT15 | chr12:32654929-32655192 | FGD4 | | TSS200 |
| GSM1179517Omentum_IT15 | chr12:47610220-47610734 | FAM113B;FAM113B;LOC100233209 | | 5'UTR;1stExon;Body |
| GSM1179500Blood_IT15 | chr12:89743788-89744609 | DUSP6;DUSP6 | N_Shore | Body;Body |
| GSM1179517Omentum_IT15 | chr13:107145933-107146164 | EFNB2 | S_Shore | Body |
| GSM1179517Omentum_IT15 | chr13:25085301-25085669 | PARP4 | N_Shore | 5'UTR |
| GSM1179500Blood_IT15 | chr13:50701050-50701619 | DLEU2 | Island | TSS1500 |
| GSM1179517Omentum_IT15 | chr13:77461279-77461426 | KCTD12 | S_Shore | TSS1500 |
| GSM1179517Omentum_IT15 | chr14:105783627-105784090 | PACS2;PACS2 | S_Shore | Body;Body |
| GSM1179517Omentum_IT15 | chr14:39734964-39735496 | CTAGE5;CTAGE5;CTAGE5;CTA | N_Shore | TSS1500;TSS1500;TSS1500;Body |
| GSM1179500Blood_IT15 | chr14:69256677-69256977 | ZFP36L1 | Island | Body |
| GSM1179517Omentum_IT15 | chr14:81425912-81426234 | TSHR;TSHR;TSHR | | Body;Body;Body |
| GSM1179500Blood_IT15 | chr15:101084442-101084980 | LASS3 | Island | 5'UTR |
| GSM1179517Omentum_IT15 | chr15:40633124-40633294 | C15orf52 | | 1stExon |
| GSM1179517Omentum_IT15 | chr16:14403004-14403144 | MIR365-1 | | TSS200 |
| GSM1179517Omentum_IT15 | chr16:27414272-27414536 | IL21R;IL21R | | TSS200;5'UTR |
| GSM1179517Omentum_IT15 | chr16:474271-474528 | RAB11FIP3 | N_Shore | TSS1500 |
| GSM1179517Omentum_IT15 | chr16:57125674-57125755 | CPNE2 | N_Shore | TSS1500 |
| GSM1179517Omentum_IT15 | chr16:69597925-69598417 | NFAT5;NFAT5;NFAT5;NFAT5;N | N_Shore | TSS1500;TSS1500;TSS1500;TSS1500;TSS1500 |
| GSM1179517Omentum_IT15 | chr16:69958298-69958964 | WWP2;WWP2 | | TSS1500;Body |
| GSM1179517Omentum_IT15 | chr16:87447756-87448114 | ZCCHC14 | Island | Body |
| GSM1179517Omentum_IT15 | chr16:88536950-88537260 | ZFPM1 | Island | Body |
| GSM1179500Blood_IT15 | chr16:89390609-89390968 | ANKRD11 | | 5'UTR |
| GSM1179517Omentum_IT15 | chr16:89390609-89390968 | ANKRD11 | | 5'UTR |
| GSM1179511Muscle_IT15 | chr17:36997420-36997563 | C17orf98 | N_Shore | Body |
| GSM1179517Omentum_IT15 | chr17:36997420-36997740 | C17orf98 | N_Shore | Body |
| GSM1179527SC_Fat_IT15 | chr17:36997420-36997740 | C17orf98 | N_Shore | Body |
| GSM1179500Blood_IT15 | chr17:40489513-40489785 | STAT3;STAT3;STAT3 | | Body;Body;Body |
| GSM1179517Omentum_IT15 | chr17:66374702-66375195 | ARSG | | Body |
| GSM1179517Omentum_IT15 | chr17:77916733-77916892 | TBC1D16 | Island | Body |
| GSM1179517Omentum_IT15 | chr17:78851149-78851503 | RPTOR;RPTOR | S_Shore | Body;Body |
| GSM1179517Omentum_IT15 | chr17:80084554-80085332 | CCDC57 | N_Shore | Body |
| GSM1179517Omentum_IT15 | chr17:80190054-80190154 | SLC16A3;SLC16A3;SLC16A3 | S_Shore | TSS200;5'UTR;5'UTR |
| GSM1179517Omentum_IT15 | chr17:8129997-8130356 | C17orf68 | S_Shelf | 3'UTR |
| GSM1179500Blood_IT15 | chr18:13641529-13641894 | C18orf1;C18orf1;C18orf1;C18 | N_Shore | Body;Body;Body;Body;Body;Body |
| GSM1179517Omentum_IT15 | chr18:13641529-13642414 | C18orf1;C18orf1;C18orf1;C18 | N_Shore | Body;Body;Body;Body;Body;Body |
| GSM1179517Omentum_IT15 | chr19:1155030-1155738 | SBNO2 | Island | 5'UTR |
| GSM1179517Omentum_IT15 | chr19:11665102-11665451 | ELOF1 | | Body |
| GSM1179517Omentum_IT15 | chr19:15391832-15391946 | BRD4;BRD4 | | TSS1500;TSS1500 |
| GSM1179517Omentum_IT15 | chr19:18385672-18385930 | KIAA1683;KIAA1683;KIAA1683 | | TSS1500;TSS1500;TSS1500 |
| GSM1179517Omentum_IT15 | chr19:2540907-2541104 | GNG7 | Island | 5'UTR |
| GSM1179517Omentum_IT15 | chr19:35629791-35630474 | FXYD1;FXYD1;FXYD1 | N_Shelf | 5'UTR;1stExon;TSS1500 |
| GSM1179517Omentum_IT15 | chr19:4305072-4305239 | FSD1 | Island | Body |
| GSM1179517Omentum_IT15 | chr19:46801557-46801672 | HIF3A;HIF3A | S_Shore | Body;TSS200 |
| GSM1179517Omentum_IT15 | chr19:50015523-50015780 | FCGRT;FCGRT | N_Shore | TSS200;TSS1500 |
| GSM1179527SC_Fat_IT15 | chr19:613111-613818 | HCN2 | Island | Body |
| GSM1179517Omentum_IT15 | chr2:127413363-127413369 | GYPC;GYPC | N_Shore | TSS1500;TSS1500 |
| GSM1179517Omentum_IT15 | chr2:143886326-143886940 | ARHGAP15 | | TSS1500 |
| GSM1179517Omentum_IT15 | chr2:157291462-157291826 | GPD2;GPD2 | N_Shore | TSS1500;TSS1500 |
| GSM1179517Omentum_IT15 | chr2:160655036-160655081 | CD302 | S_Shore | TSS1500 |
| GSM1179517Omentum_IT15 | chr2:240230452-240230892 | HDAC4 | | Body |
| GSM1179511Muscle_IT15 | chr2:30669597-30669863 | LCLAT1;LCLAT1 | N_Shore | TSS1500;TSS1500 |
| GSM1179517Omentum_IT15 | chr2:30669597-30669863 | LCLAT1;LCLAT1 | N_Shore | TSS1500;TSS1500 |
| GSM1179517Omentum_IT15 | chr2:40739515-40739771 | SLC8A1;SLC8A1 | | 5'UTR;1stExon |
| GSM1179517Omentum_IT15 | chr20:19955436-19955868 | RIN2 | N_Shore | Body |
| GSM1179517Omentum_IT15 | chr20:36932385-36932479 | BPI | | TSS200 |
| GSM1179517Omentum_IT15 | chr20:45947025-45947892 | ZMYND8;LOC100131496;ZMYND8;ZMYND8 | | Body;TSS1500;Body;Body |
| GSM1179517Omentum_IT15 | chr20:62367698-62367961 | LIME1 | N_Shore | TSS1500 |
| GSM1179500Blood_IT15 | chr21:48081242-48081686 | PRMT2;PRMT2 | Island | Body;Body |

| | | | | |
|---|---|---|---|---|
| GSM1179517Omentum_IT15 | chr22:38714166-38714466 | CSNK1E;CSNK1E | S_Shore | TSS200;TSS1500 |
| GSM1179517Omentum_IT15 | chr3:111697479-111697575 | ABHD10 | N_Shore | TSS1500 |
| GSM1179517Omentum_IT15 | chr3:112012867-112013130 | SLC9A10 | | 5'UTR |
| GSM1179517Omentum_IT15 | chr3:127473609-127473936 | MGLL;MGLL | | Body;Body |
| GSM1179517Omentum_IT15 | chr3:135684107-135684163 | PPP2R3A | N_Shore | TSS1500 |
| GSM1179517Omentum_IT15 | chr3:141087187-141087363 | ZBTB38 | | 5'UTR |
| GSM1179517Omentum_IT15 | chr3:145969579-145969718 | PLSCR4;PLSCR4;PLSCR4;PLS | S_Shore | TSS1500;TSS1500;TSS1500;TSS1500 |
| GSM1179500Blood_IT15 | chr3:195489306-195489789 | MUC4;MUC4;MUC4 | Island | Body;Body;Body |
| GSM1179517Omentum_IT15 | chr4:24796919-24797007 | SOD3 | | TSS200 |
| GSM1179517Omentum_IT15 | chr4:48136234-48136452 | TXK;TXK | | 5'UTR;1stExon |
| GSM1179517Omentum_IT15 | chr4:6955660-6955953 | TBC1D14;TBC1D14 | | Body;Body |
| GSM1179500Blood_IT15 | chr4:89978251-89978566 | FAM13A;FAM13A | | 1stExon;5'UTR |
| GSM1179517Omentum_IT15 | chr4:89978251-89978566 | FAM13A;FAM13A | | 1stExon;5'UTR |
| GSM1179517Omentum_IT15 | chr5:1342124-1342200 | CLPTM1L | N_Shelf | Body |
| GSM1179500Blood_IT15 | chr5:1489818-1489889 | LPCAT1 | Island | Body |
| GSM1179517Omentum_IT15 | chr5:150020212-150020338 | SYNPO;SYNPO;SYNPO;SYNPO;SYNPO;SYNPO | | 1stExon;5'UTR;5'UTR;Body;1stExon;Body |
| GSM1179517Omentum_IT15 | chr5:169407439-169407941 | FAM196B;DOCK2;FAM196B | | 5'UTR;Body;1stExon |
| GSM1179517Omentum_IT15 | chr5:171534678-171534792 | STK10 | | Body |
| GSM1179500Blood_IT15 | chr5:176797920-176798049 | RGS14 | S_Shelf | Body |
| GSM1179500Blood_IT15 | chr5:179740743-179741120 | GFPT2 | Island | Body |
| GSM1179517Omentum_IT15 | chr5:179740743-179741120 | GFPT2 | Island | Body |
| GSM1179500Blood_IT15 | chr5:672845-672910 | TPPP | | Body |
| GSM1179517Omentum_IT15 | chr5:9547468-9548098 | SNORD123;SEMA5A | S_Shore | TSS1500;TSS1500 |
| GSM1179517Omentum_IT15 | chr6:11382553-11382830 | NEDD9;NEDD9 | | 1stExon;5'UTR |
| GSM1179517Omentum_IT15 | chr6:13274151-13274354 | PHACTR1 | | Body |
| GSM1179517Omentum_IT15 | chr6:152702330-152702660 | SYNE1;SYNE1 | | Body;Body |
| GSM1179517Omentum_IT15 | chr6:15504844-15505460 | JARID2 | | Body |
| GSM1179517Omentum_IT15 | chr6:160182438-160182554 | ACAT2 | Island | TSS1500 |
| GSM1179517Omentum_IT15 | chr6:161796785-161796854 | PARK2;PARK2;PARK2 | | Body;Body;Body |
| GSM1179517Omentum_IT15 | chr6:163149167-163149453 | PACRG;PACRG;PACRG;PACR( | S_Shore | 5'UTR;1stExon;5'UTR;5'UTR;TSS1500;TSS1500;TSS1 |
| GSM1179500Blood_IT15 | chr6:291687-293331 | DUSP22 | N_Shore | TSS1500 |
| GSM1179511Muscle_IT15 | chr6:291687-293331 | DUSP22 | N_Shore | TSS1500 |
| GSM1179517Omentum_IT15 | chr6:291687-293331 | DUSP22 | N_Shore | TSS1500 |
| GSM1179511Muscle_IT15 | chr6:29911087-29911104 | HLA-A | Island | Body |
| GSM1179500Blood_IT15 | chr6:30039025-30039206 | RNF39;RNF39 | Island | Body;Body |
| GSM1179517Omentum_IT15 | chr6:30039374-30039476 | RNF39;RNF39 | Island | Body;Body |
| GSM1179500Blood_IT15 | chr6:30039374-30039548 | RNF39;RNF39 | Island | Body;Body |
| GSM1179511Muscle_IT15 | chr6:30039403-30039442 | RNF39;RNF39 | Island | Body;Body |
| GSM1179527SC_Fat_IT15 | chr6:30039403-30039524 | RNF39;RNF39 | Island | Body;Body |
| GSM1179517Omentum_IT15 | chr6:30130698-30131001 | TRIM15 | | TSS1500 |
| GSM1179500Blood_IT15 | chr6:31166502-31166799 | HCG27 | S_Shore | Body |
| GSM1179517Omentum_IT15 | chr6:31166502-31166799 | HCG27 | S_Shore | Body |
| GSM1179511Muscle_IT15 | chr6:31237013-31237034 | HLA-C | N_Shore | Body |
| GSM1179500Blood_IT15 | chr6:31237405-31237722 | HLA-C | N_Shore | Body |
| GSM1179511Muscle_IT15 | chr6:31237405-31237722 | HLA-C | N_Shore | Body |
| GSM1179517Omentum_IT15 | chr6:31237405-31237722 | HLA-C | N_Shore | Body |
| GSM1179527SC_Fat_IT15 | chr6:31237405-31237722 | HLA-C | N_Shore | Body |
| GSM1179500Blood_IT15 | chr6:31539539-31540051 | LTA;LTA | | TSS1500;TSS1500 |
| GSM1179517Omentum_IT15 | chr6:31539539-31540121 | LTA;LTA | | TSS1500;TSS1500 |
| GSM1179517Omentum_IT15 | chr6:31540169-31540750 | LTA;LTA;LTA | | 1stExon;5'UTR;5'UTR |
| GSM1179517Omentum_IT15 | chr6:31544694-31545473 | TNF | N_Shelf | Body |
| GSM1179517Omentum_IT15 | chr6:33048469-33048558 | HLA-DPB1 | Island | Body |
| GSM1179517Omentum_IT15 | chr6:33048687-33048875 | HLA-DPB1 | Island | Body |
| GSM1179500Blood_IT15 | chr6:33052331-33052802 | HLA-DPB1 | S_Shelf | Body |
| GSM1179500Blood_IT15 | chr6:33141279-33141305 | COL11A2;COL11A2;COL11A2 | | Body;Body;Body |
| GSM1179517Omentum_IT15 | chr6:33141279-33141305 | COL11A2;COL11A2;COL11A2 | | Body;Body;Body |
| GSM1179517Omentum_IT15 | chr6:34984836-34984953 | ANKS1A | | Body |
| GSM1179517Omentum_IT15 | chr6:52858998-52859107 | GSTA4 | N_Shore | Body |
| GSM1179517Omentum_IT15 | chr7:1127420-1127767 | GPER;C7orf50;C7orf50;C7orf5 | S_Shore | 5'UTR;Body;Body;Body;5'UTR;TSS1500 |
| GSM1179517Omentum_IT15 | chr7:134464123-134464411 | CALD1;CALD1;CALD1 | | TSS200;TSS200;TSS200 |
| GSM1179517Omentum_IT15 | chr7:139320346-139320507 | HIPK2;HIPK2 | | Body;Body |
| GSM1179500Blood_IT15 | chr7:150786044-150786082 | AGAP3;AGAP3 | S_Shore | Body;Body |
| GSM1179517Omentum_IT15 | chr7:2773010-2773127 | GNA12 | N_Shore | Body |
| GSM1179517Omentum_IT15 | chr7:4848683-4848939 | RADIL | Island | Body |
| GSM1179517Omentum_IT15 | chr8:128808017-128808554 | MIR1204;PVT1 | S_Shore | TSS200;Body |
| GSM1179500Blood_IT15 | chr8:141599141-141599436 | EIF2C2;EIF2C2 | | Body;Body |

| | | | | |
|---|---|---|---|---|
| GSM1179517Omentum_IT15 | chr8:145003862-145004388 | PLEC1;PLEC1;PLEC1;PLEC1;P | Island | Body;Body;Body;Body;Body;Body;Body;Body |
| GSM1179517Omentum_IT15 | chr8:1900524-1900892 | ARHGEF10 | | Body |
| GSM1179517Omentum_IT15 | chr8:2074935-2075820 | MYOM2 | N_Shore | Body |
| GSM1179517Omentum_IT15 | chr8:48572447-48572557 | KIAA0146 | | Body |
| GSM1179518Omentum_IT16 | chr1:114414042-114414408 | PTPN22;PTPN22 | | Body;Body |
| GSM1179518Omentum_IT16 | chr1:12538341-12538541 | VPS13D;VPS13D | | Body;Body |
| GSM1179518Omentum_IT16 | chr1:151129062-151129844 | TNFAIP8L2 | | TSS200 |
| GSM1179518Omentum_IT16 | chr1:151344806-151345248 | SELENBP1 | | Body |
| GSM1179518Omentum_IT16 | chr1:15271830-15272326 | KIAA1026;KIAA1026;KIAA1026;KIAA1026;KIAA1026 | | Body;5'UTR;TSS1500;Body;Body |
| GSM1179518Omentum_IT16 | chr1:19764480-19764821 | CAPZB | | Body |
| GSM1179501Blood_IT16 | chr1:202172778-202172912 | LGR6;LGR6 | | TSS200;Body |
| GSM1179512Muscle_IT16 | chr1:205818956-205819492 | PM20D1 | Island | Body |
| GSM1179501Blood_IT16 | chr1:205818956-205819609 | PM20D1 | Island | Body |
| GSM1179528SC_Fat_IT16 | chr1:205818956-205819609 | PM20D1 | Island | Body |
| GSM1179518Omentum_IT16 | chr1:205818956-205819609 | PM20D1 | Island | Body |
| GSM1179518Omentum_IT16 | chr1:226187852-226188006 | C1orf55 | S_Shore | TSS1500 |
| GSM1179518Omentum_IT16 | chr1:24239771-24240069 | CNR2;CNR2 | | 5'UTR;1stExon |
| GSM1179518Omentum_IT16 | chr1:247611842-247611931 | NLRP3;NLRP3;NLRP3;NLRP3;I | N_Shelf | 3'UTR;3'UTR;3'UTR;3'UTR;3'UTR |
| GSM1179518Omentum_IT16 | chr1:247711595-247712591 | C1orf150 | | TSS1500 |
| GSM1179512Muscle_IT16 | chr10:11046888-11047586 | CUGBP2 | | TSS1500 |
| GSM1179518Omentum_IT16 | chr10:111765673-111766049 | ADD3 | N_Shore | TSS200 |
| GSM1179518Omentum_IT16 | chr10:13388442-13388567 | SEPHS1 | | 5'UTR |
| GSM1179518Omentum_IT16 | chr10:135202522-135203200 | PAOX;PAOX;PAOX | N_Shore | Body;Body;Body |
| GSM1179512Muscle_IT16 | chr10:135341528-135342218 | CYP2E1 | Island | Body |
| GSM1179528SC_Fat_IT16 | chr10:135341528-135342218 | CYP2E1 | Island | Body |
| GSM1179518Omentum_IT16 | chr10:18629497-18629677 | CACNB2;CACNB2;CACNB2;CACNB2;CACNB2;CACN | | Body;TSS200;Body;Body;Body;Body;Body;Body |
| GSM1179518Omentum_IT16 | chr10:63422511-63422555 | C10orf107 | N_Shore | TSS1500 |
| GSM1179518Omentum_IT16 | chr11:131779469-131779604 | NTM;NTM;NTM;NTM | N_Shore | TSS1500;TSS1500;TSS1500;Body |
| GSM1179518Omentum_IT16 | chr11:1873884-1874093 | LSP1 | | TSS1500 |
| GSM1179518Omentum_IT16 | chr11:36422377-36422615 | PRR5L;PRR5L;PRR5L;PRR5L | | 5'UTR;5'UTR;TSS200;Body |
| GSM1179518Omentum_IT16 | chr11:57195187-57195382 | SLC43A3;SLC43A3;SLC43A3 | S_Shore | TSS1500;TSS200;TSS1500 |
| GSM1179518Omentum_IT16 | chr11:57364841-57365088 | SERPING1;SERPING1 | | TSS1500;TSS200 |
| GSM1179518Omentum_IT16 | chr11:60738032-60739183 | CD6 | | TSS1500 |
| GSM1179518Omentum_IT16 | chr11:60869760-60869969 | CD5 | | TSS200 |
| GSM1179518Omentum_IT16 | chr11:70211301-70211531 | PPFIA1;PPFIA1 | | Body;Body |
| GSM1179501Blood_IT16 | chr11:75139390-75139736 | KLHL35 | N_Shore | Body |
| GSM1179528SC_Fat_IT16 | chr11:75139390-75139736 | KLHL35 | N_Shore | Body |
| GSM1179518Omentum_IT16 | chr11:85397109-85397684 | CCDC89 | | 1stExon |
| GSM1179518Omentum_IT16 | chr12:111537057-111537195 | CUX2 | | Body |
| GSM1179518Omentum_IT16 | chr12:116996773-116997535 | MAP1LC3B2 | | TSS1500 |
| GSM1179518Omentum_IT16 | chr12:2800055-2801584 | CACNA1C;CACNA1C;CACNA1 | N_Shore | Body;Body;Body;Body;Body;Body;Body;Body;Body;B |
| GSM1179518Omentum_IT16 | chr12:47610220-47610734 | FAM113B;FAM113B;LOC100233209 | | 5'UTR;1stExon;Body |
| GSM1179518Omentum_IT16 | chr12:54891213-54891636 | NCKAP1L | | TSS1500 |
| GSM1179518Omentum_IT16 | chr12:57869031-57869490 | ARHGAP9;ARHGAP9;ARHGAP | N_Shore | Body;Body;Body |
| GSM1179518Omentum_IT16 | chr12:58135833-58136214 | AGAP2 | S_Shelf | 1stExon |
| GSM1179518Omentum_IT16 | chr12:6881590-6881629 | LAG3 | N_Shore | TSS200 |
| GSM1179518Omentum_IT16 | chr12:95226786-95226993 | MIR492 | | TSS1500 |
| GSM1179518Omentum_IT16 | chr13:107145658-107146164 | EFNB2 | Island | Body |
| GSM1179518Omentum_IT16 | chr13:25085301-25085669 | PARP4 | N_Shore | 5'UTR |
| GSM1179518Omentum_IT16 | chr13:92001764-92002454 | MIR19A;MIR17HG;MIR17HG;M | S_Shore | TSS1500;Body;Body;TSS1500;TSS1500 |
| GSM1179518Omentum_IT16 | chr13:99629970-99630583 | DOCK9;DOCK9;DOCK9;DOCK9 | | Body;Body;Body;Body |
| GSM1179518Omentum_IT16 | chr14:20903410-20903611 | KLHL33 | Island | 5'UTR |
| GSM1179501Blood_IT16 | chr14:24779793-24780734 | LTB4R2;LTB4R2;LTB4R2;LTB4I | N_Shore | 1stExon;1stExon;5'UTR;TSS1500;5'UTR |
| GSM1179518Omentum_IT16 | chr14:94576977-94577101 | IFI27;IFI27 | | TSS200;TSS200 |
| GSM1179518Omentum_IT16 | chr15:37169665-37170554 | LOC145845 | N_Shelf | Body |
| GSM1179518Omentum_IT16 | chr15:39871808-39872186 | THBS1 | N_Shore | TSS1500 |
| GSM1179518Omentum_IT16 | chr15:40600279-40600635 | PLCB2 | | TSS200 |
| GSM1179518Omentum_IT16 | chr15:57592040-57592438 | LOC283663 | | TSS1500 |
| GSM1179518Omentum_IT16 | chr15:70994435-70995014 | UACA;UACA;UACA | | 5'UTR;Body;1stExon |
| GSM1179518Omentum_IT16 | chr16:14403004-14403425 | MIR365-1 | | TSS200 |
| GSM1179518Omentum_IT16 | chr16:2736874-2737356 | KCTD5 | S_Shelf | Body |
| GSM1179518Omentum_IT16 | chr16:29756951-29757461 | C16orf54 | | 5'UTR |
| GSM1179518Omentum_IT16 | chr16:3142911-3143041 | ZSCAN10 | S_Shelf | TSS200 |
| GSM1179518Omentum_IT16 | chr16:50715135-50715529 | SNX20;SNX20;SNX20;SNX20;SNX20;SNX20 | | 1stExon;5'UTR;1stExon;5'UTR;5'UTR;1stExon |
| GSM1179518Omentum_IT16 | chr16:69597925-69598417 | NFAT5;NFAT5;NFAT5;NFAT5;N | N_Shore | TSS1500;TSS1500;TSS1500;TSS1500;TSS1500 |
| GSM1179518Omentum_IT16 | chr16:89024511-89024657 | CBFA2T3 | | Body |

| | | | | |
|---|---|---|---|---|
| GSM1179518Omentum_IT16 | chr16:89225495-89225822 | C16orf81 | N_Shore | TSS200 |
| GSM1179518Omentum_IT16 | chr17:38472961-38473382 | RARA | Island | 5'UTR |
| GSM1179518Omentum_IT16 | chr17:48912265-48912545 | WFIKKN2 | | TSS1500 |
| GSM1179518Omentum_IT16 | chr17:800084-800431 | NXN | | Body |
| GSM1179518Omentum_IT16 | chr18:77218378-77218862 | NFATC1;NFATC1;NFATC1;NFA | Island | Body;Body;Body;Body;Body |
| GSM1179518Omentum_IT16 | chr18:77917615-77918142 | LOC100130522;LOC1001305 | N_Shore | Body;Body;3'UTR |
| GSM1179518Omentum_IT16 | chr19:10444815-10445516 | RAVER1;ICAM3 | Island | TSS1500;Body |
| GSM1179518Omentum_IT16 | chr19:10449942-10450323 | ICAM3 | S_Shelf | Body |
| GSM1179501Blood_IT16 | chr19:12876846-12877188 | HOOK2;HOOK2 | N_Shore | Body;Body |
| GSM1179512Muscle_IT16 | chr19:12876846-12877188 | HOOK2;HOOK2 | N_Shore | Body;Body |
| GSM1179528SC_Fat_IT16 | chr19:12876846-12877188 | HOOK2;HOOK2 | N_Shore | Body;Body |
| GSM1179518Omentum_IT16 | chr19:12876846-12877188 | HOOK2;HOOK2 | N_Shore | Body;Body |
| GSM1179518Omentum_IT16 | chr19:15391832-15391946 | BRD4;BRD4 | | TSS1500;TSS1500 |
| GSM1179518Omentum_IT16 | chr19:16178195-16178253 | TPM4 | N_Shore | TSS200 |
| GSM1179518Omentum_IT16 | chr19:2540907-2541104 | GNG7 | Island | 5'UTR |
| GSM1179518Omentum_IT16 | chr19:3136019-3136430 | GNA15 | | TSS200 |
| GSM1179518Omentum_IT16 | chr19:36233297-36233469 | TMEM149;TMEM149 | S_Shore | 1stExon;5'UTR |
| GSM1179518Omentum_IT16 | chr19:3969522-3969745 | DAPK3 | N_Shore | Body |
| GSM1179518Omentum_IT16 | chr19:49838478-49838777 | CD37;CD37 | N_Shelf | TSS200;TSS200 |
| GSM1179518Omentum_IT16 | chr2:100759014-100759159 | AFF3;AFF3 | | 1stExon;5'UTR |
| GSM1179518Omentum_IT16 | chr2:106776775-106777038 | UXS1 | | Body |
| GSM1179518Omentum_IT16 | chr2:143886326-143886940 | ARHGAP15 | | TSS1500 |
| GSM1179518Omentum_IT16 | chr2:234296642-234297039 | DGKD;DGKD | | Body;TSS200 |
| GSM1179512Muscle_IT16 | chr2:236867685-236867905 | AGAP1;AGAP1 | Island | Body;Body |
| GSM1179518Omentum_IT16 | chr2:240230452-240230892 | HDAC4 | | Body |
| GSM1179501Blood_IT16 | chr2:30669597-30669863 | LCLAT1;LCLAT1 | N_Shore | TSS1500;TSS1500 |
| GSM1179512Muscle_IT16 | chr2:30669597-30669863 | LCLAT1;LCLAT1 | N_Shore | TSS1500;TSS1500 |
| GSM1179528SC_Fat_IT16 | chr2:30669597-30669863 | LCLAT1;LCLAT1 | N_Shore | TSS1500;TSS1500 |
| GSM1179518Omentum_IT16 | chr2:30669597-30669863 | LCLAT1;LCLAT1 | N_Shore | TSS1500;TSS1500 |
| GSM1179518Omentum_IT16 | chr2:40739515-40739771 | SLC8A1;SLC8A1 | | 5'UTR;1stExon |
| GSM1179518Omentum_IT16 | chr2:68545960-68546038 | CNRIP1;CNRIP1 | N_Shore | Body;Body |
| GSM1179518Omentum_IT16 | chr20:36932385-36933189 | BPI | | TSS200 |
| GSM1179518Omentum_IT16 | chr20:4795444-4796176 | RASSF2;RASSF2 | | 5'UTR;5'UTR |
| GSM1179518Omentum_IT16 | chr20:62327732-62328427 | TNFRSF6B;TNFRSF6B | N_Shore | 5'UTR;TSS1500 |
| GSM1179501Blood_IT16 | chr20:62327968-62328427 | TNFRSF6B;TNFRSF6B | N_Shore | 5'UTR;TSS200 |
| GSM1179518Omentum_IT16 | chr21:36421467-36421955 | RUNX1;RUNX1 | | 5'UTR;1stExon |
| GSM1179518Omentum_IT16 | chr21:48026043-48026298 | S100B | | TSS1500 |
| GSM1179518Omentum_IT16 | chr22:17680477-17680706 | CECR1;CECR1;CECR1 | | Body;1stExon;5'UTR |
| GSM1179518Omentum_IT16 | chr22:45148218-45148229 | ARHGAP8;PRR5-ARHGAP8;ARHGAP8 | | TSS1500;Body;TSS1500 |
| GSM1179518Omentum_IT16 | chr22:45608345-45608713 | C22orf9;C22orf9;C22orf9 | | Body;1stExon;5'UTR |
| GSM1179518Omentum_IT16 | chr3:1134366-1134730 | CNTN6 | | TSS1500 |
| GSM1179518Omentum_IT16 | chr3:135684107-135684163 | PPP2R3A | N_Shore | TSS1500 |
| GSM1179518Omentum_IT16 | chr3:145969579-145969718 | PLSCR4;PLSCR4;PLSCR4;PLS | S_Shore | TSS1500;TSS1500;TSS1500;TSS1500 |
| GSM1179518Omentum_IT16 | chr3:184297380-184297522 | EPHB3 | N_Shelf | Body |
| GSM1179518Omentum_IT16 | chr3:50359849-50360041 | HYAL2;HYAL2 | Island | 5'UTR;TSS1500 |
| GSM1179518Omentum_IT16 | chr4:186508724-186509045 | SORBS2;SORBS2;SORBS2;SORBS2;SORBS2;SORBS2; | | 3'UTR;3'UTR;3'UTR;3'UTR;3'UTR;3'UTR;3'UTR;3'UTR |
| GSM1179518Omentum_IT16 | chr4:186559412-186560083 | SORBS2;SORBS2;SORBS2;SO | N_Shore | Body;Body;Body;Body;Body;Body;Body;Body |
| GSM1179518Omentum_IT16 | chr4:6955660-6955953 | TBC1D14;TBC1D14 | | Body;Body |
| GSM1179512Muscle_IT16 | chr5:134366162-134367394 | PITX1 | N_Shore | Body |
| GSM1179518Omentum_IT16 | chr5:170288742-170288766 | RANBP17 | N_Shore | TSS1500 |
| GSM1179518Omentum_IT16 | chr5:171534678-171534792 | STK10 | | Body |
| GSM1179501Blood_IT16 | chr5:179740743-179741120 | GFPT2 | Island | Body |
| GSM1179518Omentum_IT16 | chr5:59783668-59784203 | PART1;PART1;PDE4D;PART1 | | TSS200;Body;5'UTR;TSS200 |
| GSM1179518Omentum_IT16 | chr5:76145809-76146148 | S100Z | | TSS200 |
| GSM1179512Muscle_IT16 | chr6:100911240-100912072 | SIM1 | N_Shore | 1stExon |
| GSM1179518Omentum_IT16 | chr6:11382553-11382830 | NEDD9;NEDD9 | | 1stExon;5'UTR |
| GSM1179518Omentum_IT16 | chr6:13274151-13274354 | PHACTR1 | | Body |
| GSM1179518Omentum_IT16 | chr6:13428522-13428657 | GFOD1 | | Body |
| GSM1179518Omentum_IT16 | chr6:138866865-138867125 | NHSL1 | | Body |
| GSM1179518Omentum_IT16 | chr6:152702330-152702660 | SYNE1;SYNE1 | | Body;Body |
| GSM1179518Omentum_IT16 | chr6:15504844-15505460 | JARID2 | | Body |
| GSM1179501Blood_IT16 | chr6:29911542-29911558 | HLA-A | S_Shore | Body |
| GSM1179518Omentum_IT16 | chr6:30130819-30131001 | TRIM15 | | TSS200 |
| GSM1179512Muscle_IT16 | chr6:31237013-31237034 | HLA-C | N_Shore | Body |
| GSM1179528SC_Fat_IT16 | chr6:31237013-31237034 | HLA-C | N_Shore | Body |
| GSM1179518Omentum_IT16 | chr6:31237013-31237034 | HLA-C | N_Shore | Body |

| | | | | |
|---|---|---|---|---|
| GSM1179501Blood_IT16 | chr6:31237405-31237722 | HLA-C | N_Shore | Body |
| GSM1179512Muscle_IT16 | chr6:31237405-31237722 | HLA-C | N_Shore | Body |
| GSM1179528SC_Fat_IT16 | chr6:31237405-31237722 | HLA-C | N_Shore | Body |
| GSM1179518Omentum_IT16 | chr6:31237405-31237722 | HLA-C | N_Shore | Body |
| GSM1179518Omentum_IT16 | chr6:33047944-33048809 | HLA-DPB1 | N_Shore | Body |
| GSM1179518Omentum_IT16 | chr6:34984836-34984953 | ANKS1A | | Body |
| GSM1179518Omentum_IT16 | chr6:35285967-35286360 | DEF6 | | Body |
| GSM1179518Omentum_IT16 | chr7:1127420-1127767 | GPER;C7orf50;C7orf50;C7orf5 | S_Shore | 5'UTR;Body;Body;Body;5'UTR;TSS1500 |
| GSM1179518Omentum_IT16 | chr7:1163456-1163642 | C7orf50;C7orf50;C7orf50 | | Body;Body;Body |
| GSM1179518Omentum_IT16 | chr7:27168962-27169740 | HOXA4 | N_Shore | Body |
| GSM1179518Omentum_IT16 | chr7:36764019-36764205 | AOAH;AOAH | | 1stExon;5'UTR |
| GSM1179518Omentum_IT16 | chr7:45018658-45019005 | MYO1G;MYO1G | | 5'UTR;1stExon |
| GSM1179518Omentum_IT16 | chr7:45066738-45067319 | CCM2;CCM2;CCM2;CCM2;CCM2 | | Body;TSS1500;Body;Body;Body |
| GSM1179518Omentum_IT16 | chr7:4848683-4848939 | RADIL | Island | Body |
| GSM1179518Omentum_IT16 | chr7:6199980-6200140 | USP42 | S_Shore | 3'UTR |
| GSM1179518Omentum_IT16 | chr7:95546508-95546556 | DYNC1I1;DYNC1I1;DYNC1I1 | | Body;Body;Body |
| GSM1179518Omentum_IT16 | chr8:145002522-145002835 | PLEC1;PLEC1;PLEC1;PLEC1;P | N_Shore | Body;Body;Body;Body;Body;Body;Body;Body |
| GSM1179518Omentum_IT16 | chr8:22446721-22447181 | PDLIM2;PDLIM2;PDLIM2 | | Body;Body;Body |
| GSM1179518Omentum_IT16 | chr8:59058585-59058660 | FAM110B | Island | 5'UTR |
| GSM1179502Blood_IT17 | chr1:154839813-154839983 | KCNN3 | | Body |
| GSM1179513Muscle_IT17 | chr1:1910585-1911120 | KIAA1751 | Island | Body |
| GSM1179519Omentum_IT17 | chr1:1910585-1911120 | KIAA1751 | Island | Body |
| GSM1179502Blood_IT17 | chr1:2120985-2121521 | C1orf86;LOC100128003 | N_Shore | Body;Body |
| GSM1179519Omentum_IT17 | chr1:2120985-2121521 | C1orf86;LOC100128003 | N_Shore | Body;Body |
| GSM1179502Blood_IT17 | chr10:129797760-129797840 | PTPRE | | 5'UTR |
| GSM1179502Blood_IT17 | chr11:34460107-34460298 | CAT | N_Shore | TSS1500 |
| GSM1179502Blood_IT17 | chr11:34460336-34461028 | CAT | Island | TSS200 |
| GSM1179502Blood_IT17 | chr11:7110074-7110561 | RBMXL2 | N_Shore | TSS200 |
| GSM1179502Blood_IT17 | chr14:102695279-102695875 | RAGE | N_Shore | 3'UTR |
| GSM1179513Muscle_IT17 | chr16:86370962-86371248 | LOC732275 | | Body |
| GSM1179513Muscle_IT17 | chr17:259942-260058 | C17orf97 | Island | TSS200 |
| GSM1179519Omentum_IT17 | chr17:259942-260058 | C17orf97 | Island | TSS200 |
| GSM1179529SC_Fat_IT17 | chr17:259942-260058 | C17orf97 | Island | TSS200 |
| GSM1179502Blood_IT17 | chr17:259942-260292 | C17orf97 | Island | TSS200 |
| GSM1179502Blood_IT17 | chr17:44847220-44847427 | WNT3 | Island | Body |
| GSM1179502Blood_IT17 | chr17:56565286-56565644 | HSF5 | Island | 1stExon |
| GSM1179502Blood_IT17 | chr18:29340788-29340919 | MCART2;MCART2 | | 1stExon;5'UTR |
| GSM1179502Blood_IT17 | chr19:12876846-12877188 | HOOK2;HOOK2 | N_Shore | Body;Body |
| GSM1179513Muscle_IT17 | chr19:12876846-12877188 | HOOK2;HOOK2 | N_Shore | Body;Body |
| GSM1179519Omentum_IT17 | chr19:12876846-12877188 | HOOK2;HOOK2 | N_Shore | Body;Body |
| GSM1179529SC_Fat_IT17 | chr19:12876846-12877188 | HOOK2;HOOK2 | N_Shore | Body;Body |
| GSM1179502Blood_IT17 | chr19:55972646-55973228 | ISOC2;ISOC2;ISOC2 | N_Shore | 5'UTR;5'UTR;5'UTR |
| GSM1179519Omentum_IT17 | chr19:55972791-55973099 | ISOC2;ISOC2;ISOC2 | Island | 5'UTR;5'UTR;5'UTR |
| GSM1179502Blood_IT17 | chr2:113992762-113993313 | PAX8;PAX8;PAX8;PAX8;PAX8;L | N_Shore | Body;Body;Body;Body;Body;Body;TSS1500 |
| GSM1179519Omentum_IT17 | chr2:113992762-113993313 | PAX8;PAX8;PAX8;PAX8;PAX8;L | N_Shore | Body;Body;Body;Body;Body;Body;TSS1500 |
| GSM1179502Blood_IT17 | chr2:1817263-1817409 | MYT1L | | Body |
| GSM1179502Blood_IT17 | chr2:27665017-27665543 | KRTCAP3;NRBP1;KRTCAP3 | Island | TSS1500;3'UTR;TSS1500 |
| GSM1179502Blood_IT17 | chr2:3704501-3704773 | ALLC | | TSS1500 |
| GSM1179519Omentum_IT17 | chr20:62367884-62367893 | LIME1 | N_Shore | TSS200 |
| GSM1179502Blood_IT17 | chr20:62679255-62679713 | SOX18 | N_Shore | 3'UTR |
| GSM1179519Omentum_IT17 | chr21:30565636-30565706 | NCRNA00189 | | TSS200 |
| GSM1179502Blood_IT17 | chr21:47604052-47605174 | C21orf56;C21orf56 | S_Shore | 5'UTR;5'UTR |
| GSM1179513Muscle_IT17 | chr22:19929254-19929286 | TXNRD2;COMT | Island | 1stExon;TSS200 |
| GSM1179502Blood_IT17 | chr22:29707524-29707782 | GAS2L1;GAS2L1;GAS2L1 | Island | 3'UTR;Body;Body |
| GSM1179502Blood_IT17 | chr3:140813919-140814133 | SPSB4 | Island | Body |
| GSM1179502Blood_IT17 | chr3:145879568-145879710 | PLOD2;PLOD2 | S_Shore | TSS1500;TSS1500 |
| GSM1179502Blood_IT17 | chr3:169755595-169755693 | GPR160 | Island | TSS200 |
| GSM1179519Omentum_IT17 | chr3:50487955-50488230 | CACNA2D2;CACNA2D2 | | Body;Body |
| GSM1179502Blood_IT17 | chr4:156298325-156298427 | MAP9 | S_Shore | TSS1500 |
| GSM1179502Blood_IT17 | chr5:131607097-131607372 | PDLIM4;PDLIM4 | Island | Body;Body |
| GSM1179502Blood_IT17 | chr5:140552992-140553150 | PCDHB7 | N_Shore | 1stExon |
| GSM1179502Blood_IT17 | chr5:158689508-158689629 | UBLCP1 | N_Shore | TSS1500 |
| GSM1179502Blood_IT17 | chr5:179740743-179741120 | GFPT2 | Island | Body |
| GSM1179519Omentum_IT17 | chr5:179740743-179741120 | GFPT2 | Island | Body |
| GSM1179529SC_Fat_IT17 | chr5:179740743-179741120 | GFPT2 | Island | Body |
| GSM1179502Blood_IT17 | chr5:42756786-42757171 | CCDC152 | | TSS200 |

| | | | | |
|---|---|---|---|---|
| GSM1179502Blood_IT17 | chr5:54826611-54826768 | PPAP2A;PPAP2A;RNF138P1 | Island | Body;Body;Body |
| GSM1179519Omentum_IT17 | chr5:9547595-9548098 | SNORD123;SEMA5A | S_Shore | TSS1500;TSS1500 |
| GSM1179502Blood_IT17 | chr6:138892911-138893159 | NHSL1 | | 1stExon |
| GSM1179502Blood_IT17 | chr6:18123164-18123232 | NHLRC1 | S_Shore | TSS1500 |
| GSM1179502Blood_IT17 | chr6:30039025-30039206 | RNF39;RNF39 | Island | Body;Body |
| GSM1179502Blood_IT17 | chr6:30039374-30039476 | RNF39;RNF39 | Island | Body;Body |
| GSM1179502Blood_IT17 | chr6:30079203-30079307 | TRIM31 | | Body |
| GSM1179502Blood_IT17 | chr6:31164776-31164968 | HCG27 | N_Shore | TSS1500 |
| GSM1179519Omentum_IT17 | chr6:31164914-31164952 | HCG27 | N_Shore | TSS1500 |
| GSM1179529SC_Fat_IT17 | chr6:31164914-31164952 | HCG27 | N_Shore | TSS1500 |
| GSM1179519Omentum_IT17 | chr6:31237013-31237034 | HLA-C | N_Shore | Body |
| GSM1179502Blood_IT17 | chr7:89873797-89873891 | C7orf63;C7orf63 | N_Shore | TSS1500;TSS1500 |

| Supplementary Table 8: Coordinates of MeVars identified in three peripheral tissues from normal individuals (duplicate values of coordinates retained) | | | | |
|---|---|---|---|---|
| Sample and Tissue | Coordinate | UCSC_REFGENE_NAME | RELATION_TO_UCSC_CPG_ISLAND | UCSC_REFGENE_GROUP |
| Buccal_PT1 | chr1:207224740-207225128 | YOD1 | N_Shore | TSS1500 |
| Buccal_PT1 | chr1:2084319-2084595 | PRKCZ;PRKCZ;PRKCZ | S_Shore | Body;Body;Body |
| Buccal_PT1 | chr1:2266228-2266933 | MORN1 | Island | Body |
| Buccal_PT1 | chr1:243637838-243638141 | SDCCAG8 | Island | Body |
| Buccal_PT1 | chr1:3276413-3276535 | PRDM16;PRDM16 | S_Shore | Body;Body |
| Buccal_PT1 | chr1:38156462-38156652 | C1orf109 | N_Shore | TSS1500 |
| Buccal_PT1 | chr10:1507708-1508061 | ADARB2 | S_Shore | Body |
| Buccal_PT1 | chr10:3149673-3149852 | PFKP | S_Shore | Body |
| Buccal_PT1 | chr10:5541568-5541726 | CALML5 | S_Shore | TSS200 |
| Buccal_PT1 | chr10:75631588-75632066 | CAMK2G;CAMK2G;CAMK2G;CAMK2G;CAMK2G | N_Shelf | Body;Body;Body;Body;Body |
| Buccal_PT1 | chr10:99331332-99331808 | ANKRD2;ANKRD2 | S_Shore | TSS1500;TSS1500 |
| Buccal_PT1 | chr11:111847475-111847747 | DIXDC1;DIXDC1 | N_Shore | TSS1500;Body |
| Buccal_PT1 | chr11:129306730-129306881 | BARX2 | Island | Body |
| Buccal_PT1 | chr11:1471410-1471617 | BRSK2 | Island | Body |
| Buccal_PT1 | chr11:17803160-17803421 | KCNC1 | S_Shore | Body |
| Buccal_PT1 | chr11:3144685-3144774 | OSBPL5;OSBPL5;OSBPL5 | S_Shelf | Body;Body;Body |
| Buccal_PT1 | chr12:107974296-107974453 | BTBD11;BTBD11 | N_Shore | TSS1500;Body |
| Buccal_PT1 | chr12:121890163-121890503 | KDM2B;KDM2B | N_Shore | Body;Body |
| Buccal_PT1 | chr12:123750782-123750864 | CDK2AP1 | N_Shelf | Body |
| Buccal_PT1 | chr12:124876101-124876650 | NCOR2;NCOR2 | S_Shelf | Body;Body |
| Buccal_PT1 | chr12:130821607-130822674 | PIWIL1 | N_Shore | TSS1500 |
| Buccal_PT1 | chr12:64782908-64783110 | C12orf56;C12orf56 | N_Shore | Body;Body |
| Buccal_PT1 | chr13:114187973-114188051 | TMCO3 | S_Shore | Body |
| Buccal_PT1 | chr14:103593088-103593520 | TNFAIP2 | N_Shore | Body |
| Buccal_PT1 | chr15:39871808-39872186 | THBS1 | N_Shore | TSS1500 |
| Buccal_PT1 | chr16:743338-743584 | FBXL16 | N_Shore | 3'UTR |
| Buccal_PT1 | chr16:88807658-88807707 | FAM38A | S_Shelf | Body |
| Buccal_PT1 | chr17:149484-149582 | RPH3AL | N_Shelf | Body |
| Buccal_PT1 | chr17:3433428-3433970 | TRPV3 | N_Shelf | Body |
| Buccal_PT1 | chr17:40935998-40937365 | WNK4 | Island | Body |
| Buccal_PT1 | chr17:43318045-43319382 | FMNL1 | N_Shore | Body |
| Buccal_PT1 | chr17:7493222-7494151 | SOX15;SOX15 | S_Shore | 1stExon;5'UTR |
| Buccal_PT1 | chr17:7690327-7690529 | DNAH2 | Island | Body |
| Buccal_PT1 | chr17:78079668-78079795 | GAA;GAA;GAA | Island | Body;Body;Body |
| Buccal_PT1 | chr18:77230598-77230795 | NFATC1;NFATC1;NFATC1;NFATC1 | Island | Body;Body;Body;Body |
| Buccal_PT1 | chr19:10231566-10231711 | EIF3G | S_Shore | TSS1500 |
| Buccal_PT1 | chr19:1112327-1112679 | SBNO2;SBNO2 | Island | Body;Body |
| Buccal_PT1 | chr19:1117289-1117571 | SBNO2;SBNO2 | Island | Body;Body |
| Buccal_PT1 | chr19:12876846-12877188 | HOOK2;HOOK2 | N_Shore | Body;Body |
| Buccal_PT1 | chr19:1424658-1424952 | DAZAP1;DAZAP1 | Island | Body;Body |
| Buccal_PT1 | chr19:2540907-2541104 | GNG7 | Island | 5'UTR |
| Buccal_PT1 | chr19:3771880-3772414 | RAX2 | S_Shore | 5'UTR |
| Buccal_PT1 | chr19:45150513-45150725 | PVR;PVR;PVR;PVR | Island | Body;Body;Body;Body |
| Buccal_PT1 | chr19:5148023-5148409 | KDM4B | Island | Body |
| Buccal_PT1 | chr19:5250376-5251120 | PTPRS;PTPRS;PTPRS;PTPRS | N_Shore | Body;Body;Body;Body |
| Buccal_PT1 | chr2:10567628-10567843 | HPCAL1;HPCAL1 | S_Shore | 3'UTR;3'UTR |
| Buccal_PT1 | chr2:128458240-128458399 | SFT2D3 | N_Shore | TSS1500 |
| Buccal_PT1 | chr2:202508694-202508813 | ALS2CR4;ALS2CR4 | S_Shore | TSS1500;TSS1500 |
| Buccal_PT1 | chr2:26401640-26402318 | FAM59B | N_Shore | Body |
| Buccal_PT1 | chr2:74153086-74153317 | DGUOK;DGUOK | N_Shore | TSS1500;TSS1500 |
| Buccal_PT1 | chr2:74753281-74753358 | DQX1;DQX1 | N_Shelf | 1stExon;5'UTR |
| Buccal_PT1 | chr20:17595355-17595472 | RRBP1;RRBP1 | Island | Body;Body |
| Buccal_PT1 | chr20:35972924-35973098 | SRC | N_Shore | TSS200 |
| Buccal_PT1 | chr21:38117454-38117682 | SIM2;SIM2 | N_Shelf | Body;Body |
| Buccal_PT1 | chr22:43820888-43821115 | MPPED1 | Island | 5'UTR |
| Buccal_PT1 | chr22:47022418-47022567 | GRAMD4 | N_Shore | TSS1500 |
| Buccal_PT1 | chr3:195488725-195490169 | MUC4;MUC4;MUC4 | Island | Body;Body;Body |
| Buccal_PT1 | chr4:715865-715950 | PCGF3 | N_Shore | 5'UTR |
| Buccal_PT1 | chr5:510035-510421 | SLC9A3 | S_Shore | Body |
| Buccal_PT1 | chr6:157746584-157746724 | C6orf35 | S_Shore | TSS1500 |
| Buccal_PT1 | chr6:158018055-158018184 | ZDHHC14;ZDHHC14 | S_Shelf | Body;Body |
| Buccal_PT1 | chr6:29893955-29894141 | HCG4P6 | N_Shore | TSS1500 |
| Buccal_PT1 | chr6:29894162-29894197 | HCG4P6 | Island | TSS1500 |
| Buccal_PT1 | chr6:29911494-29911558 | HLA-A | S_Shore | Body |

| | | | | |
|---|---|---|---|---|
| Buccal_PT1 | chr6:30455885-30455964 | HLA-E | N_Shore | TSS1500 |
| Buccal_PT1 | chr6:30850868-30851638 | DDR1;DDR1 | N_Shore | TSS1500;TSS1500 |
| Buccal_PT1 | chr6:31237013-31237034 | HLA-C | N_Shore | Body |
| Buccal_PT1 | chr6:31237405-31237722 | HLA-C | N_Shore | Body |
| Buccal_PT1 | chr6:32060681-32061318 | TNXB | N_Shelf | Body |
| Buccal_PT1 | chr6:32165889-32165917 | NOTCH4 | S_Shore | Body |
| Buccal_PT1 | chr6:32633015-32633163 | HLA-DQB1 | Island | Body |
| Buccal_PT1 | chr7:1022213-1022806 | CYP2W1 | N_Shore | TSS1500 |
| Buccal_PT1 | chr7:1102344-1103262 | C7orf50;C7orf50;C7orf50 | N_Shelf | Body;Body;Body |
| Buccal_PT1 | chr7:157701024-157701262 | PTPRN2;PTPRN2;PTPRN2 | N_Shore | Body;Body;Body |
| Buccal_PT1 | chr7:158059704-158060003 | PTPRN2;PTPRN2;PTPRN2 | N_Shore | Body;Body;Body |
| Buccal_PT1 | chr7:23286319-23286539 | GPNMB;GPNMB;GPNMB;GPNMB | N_Shore | 5'UTR;1stExon;1stExon;5'UTR |
| Buccal_PT1 | chr7:41742630-41743001 | INHBA;LOC285954;INHBA;LOC285954 | N_Shelf | 5'UTR;Body;1stExon;Body |
| Buccal_PT1 | chr7:98738998-98739782 | SMURF1;SMURF1 | N_Shelf | Body;Body |
| Buccal_PT1 | chr8:143823809-143824001 | SLURP1;SLURP1 | S_Shelf | 5'UTR;1stExon |
| Buccal_PT1 | chr8:144894951-144895243 | SCRIB;SCRIB | Island | Body;Body |
| Buccal_PT1 | chr8:37699239-37699923 | GPR124 | Island | Body |
| Buccal_PT1 | chr8:38643568-38643846 | TACC1;TACC1;TACC1 | N_Shore | TSS1500;TSS1500;5'UTR |
| Buccal_PT2 | chr10:135341528-135342218 | CYP2E1 | Island | Body |
| Buccal_PT2 | chr10:135342560-135343280 | CYP2E1 | Island | Body |
| Buccal_PT2 | chr12:124876101-124876650 | NCOR2;NCOR2 | S_Shelf | Body;Body |
| Buccal_PT2 | chr17:149484-149582 | RPH3AL | N_Shelf | Body |
| Buccal_PT2 | chr19:12876846-12877188 | HOOK2;HOOK2 | N_Shore | Body;Body |
| Buccal_PT2 | chr2:26401640-26402318 | FAM59B | N_Shore | Body |
| Buccal_PT2 | chr6:31237013-31237034 | HLA-C | N_Shore | Body |
| Buccal_PT2 | chr6:32489750-32489963 | HLA-DRB5 | Island | Body |
| Buccal_PT2 | chr6:32551749-32551954 | HLA-DRB1 | N_Shore | Body |
| Buccal_PT2 | chr6:32632331-32632338 | HLA-DQB1 | Island | Body |
| Buccal_PT2 | chr6:32632694-32632731 | HLA-DQB1 | Island | Body |
| Buccal_PT2 | chr6:32633157-32633354 | HLA-DQB1 | S_Shore | Body |
| Buccal_PT3 | chr10:131323903-131324189 | MGMT | | Body |
| Buccal_PT3 | chr10:75631588-75632066 | CAMK2G;CAMK2G;CAMK2G;CAMK2G;CAMK2G | N_Shelf | Body;Body;Body;Body;Body |
| Buccal_PT3 | chr11:70211301-70211531 | PPFIA1;PPFIA1 | | Body;Body |
| Buccal_PT3 | chr12:258355-258436 | IQSEC3;IQSEC3 | | Body;Body |
| Buccal_PT3 | chr13:114187973-114188051 | TMCO3 | S_Shore | Body |
| Buccal_PT3 | chr17:149484-149582 | RPH3AL | N_Shelf | Body |
| Buccal_PT3 | chr17:3704471-3704621 | ITGAE;ITGAE | | 1stExon;5'UTR |
| Buccal_PT3 | chr17:927457-927652 | ABR;ABR;ABR | S_Shore | Body;Body;Body |
| Buccal_PT3 | chr18:21572622-21572748 | TTC39C | | TSS200 |
| Buccal_PT3 | chr18:61253847-61254561 | SERPINB13 | | TSS1500 |
| Buccal_PT3 | chr19:2540907-2541104 | GNG7 | Island | 5'UTR |
| Buccal_PT3 | chr2:161056520-161056639 | ITGB6 | | 1stExon |
| Buccal_PT3 | chr2:240174613-240174659 | HDAC4 | | Body |
| Buccal_PT3 | chr5:177913434-177913485 | COL23A1 | | Body |
| Buccal_PT3 | chr5:53751272-53751493 | HSPB3 | | TSS200 |
| Buccal_PT3 | chr6:158507753-158508188 | SYNJ2 | Island | Body |
| Buccal_PT3 | chr6:31023257-31023770 | HCG22 | | Body |
| Buccal_PT3 | chr6:31237013-31237034 | HLA-C | N_Shore | Body |
| Buccal_PT3 | chr6:31237405-31237722 | HLA-C | N_Shore | Body |
| Buccal_PT3 | chr6:32633114-32633354 | HLA-DQB1 | S_Shore | Body |
| Buccal_PT3 | chr6:8436218-8436296 | SLC35B3;SLC35B3;SLC35B3 | S_Shore | TSS1500;TSS1500;TSS1500 |
| Buccal_PT3 | chr7:23286319-23286539 | GPNMB;GPNMB;GPNMB;GPNMB | N_Shore | 5'UTR;1stExon;1stExon;5'UTR |
| Buccal_PT3 | chr8:145955983-145956024 | ZNF251 | Island | Body |
| Buccal_PT3 | chr9:140356475-140356830 | PNPLA7;PNPLA7 | Island | Body;Body |
| Buccal_PT4 | chr1:205818956-205819492 | PM20D1 | Island | Body |
| Buccal_PT4 | chr11:1325718-1325852 | TOLLIP | | Body |
| Buccal_PT4 | chr16:14403004-14403040 | MIR365-1 | | TSS200 |
| Buccal_PT4 | chr17:8129997-8130356 | C17orf68 | S_Shelf | 3'UTR |
| Buccal_PT4 | chr2:240174613-240174659 | HDAC4 | | Body |
| Buccal_PT4 | chr21:47604052-47605174 | C21orf56;C21orf56 | S_Shore | 5'UTR;5'UTR |
| Buccal_PT4 | chr22:47022418-47023209 | GRAMD4 | N_Shore | TSS1500 |
| Buccal_PT4 | chr3:134371285-134373303 | KY | S_Shore | TSS1500 |
| Buccal_PT4 | chr3:139397179-139397901 | NMNAT3 | S_Shore | TSS1500 |
| Buccal_PT4 | chr5:179740743-179741120 | GFPT2 | Island | Body |
| Buccal_PT4 | chr6:31237013-31237034 | HLA-C | N_Shore | Body |
| Buccal_PT5 | chr1:54822031-54822503 | SSBP3;SSBP3;SSBP3 | Island | Body;Body;Body |

| | | | | |
|---|---|---|---|---|
| Buccal_PT5 | chr10:13388442-13388567 | SEPHS1 | | 5'UTR |
| Buccal_PT5 | chr10:1454265-1454387 | ADARB2 | | Body |
| Buccal_PT5 | chr11:128457400-128457682 | ETS1;ETS1 | | 1stExon;5'UTR |
| Buccal_PT5 | chr11:70211301-70211531 | PPFIA1;PPFIA1 | | Body;Body |
| Buccal_PT5 | chr12:25801455-25801621 | IFLTD1 | | 1stExon |
| Buccal_PT5 | chr12:55378411-55378912 | KIAA0748;KIAA0748 | | 1stExon;5'UTR |
| Buccal_PT5 | chr14:104195181-104195666 | ZFYVE21 | S_Shore | Body |
| Buccal_PT5 | chr14:51955524-51956044 | FRMD6 | | TSS1500 |
| Buccal_PT5 | chr15:57592040-57592222 | LOC283663 | | TSS1500 |
| Buccal_PT5 | chr16:22326203-22326535 | POLR3E | | Body |
| Buccal_PT5 | chr16:711033-711235 | WDR90 | N_Shore | Body |
| Buccal_PT5 | chr17:149484-149582 | RPH3AL | N_Shelf | Body |
| Buccal_PT5 | chr17:7690327-7690529 | DNAH2 | Island | Body |
| Buccal_PT5 | chr19:10231566-10231711 | EIF3G | S_Shore | TSS1500 |
| Buccal_PT5 | chr19:3398706-3398810 | NFIC;NFIC | Island | Body;Body |
| Buccal_PT5 | chr2:172945058-172945698 | MAP1D | Island | Body |
| Buccal_PT5 | chr2:240020028-240020174 | HDAC4 | | Body |
| Buccal_PT5 | chr2:240174613-240174659 | HDAC4 | | Body |
| Buccal_PT5 | chr2:240302585-240302855 | HDAC4 | | 5'UTR |
| Buccal_PT5 | chr2:74753281-74753358 | DQX1;DQX1 | N_Shelf | 1stExon;5'UTR |
| Buccal_PT5 | chr20:17595355-17595472 | RRBP1;RRBP1 | Island | Body;Body |
| Buccal_PT5 | chr20:17674195-17674250 | BANF2 | | TSS200 |
| Buccal_PT5 | chr20:2276503-2276663 | TGM3 | | TSS200 |
| Buccal_PT5 | chr20:35972924-35973098 | SRC | N_Shore | TSS200 |
| Buccal_PT5 | chr21:43185385-43185884 | RIPK4 | N_Shore | Body |
| Buccal_PT5 | chr21:47604052-47604986 | C21orf56;C21orf56 | S_Shore | 5'UTR;5'UTR |
| Buccal_PT5 | chr4:87770357-87770782 | SLC10A6;SLC10A6 | | 5'UTR;1stExon |
| Buccal_PT5 | chr5:138211043-138211155 | CTNNA1;LRRTM2;LRRTM2 | | Body;5'UTR;1stExon |
| Buccal_PT5 | chr5:53751272-53751493 | HSPB3 | | TSS200 |
| Buccal_PT5 | chr6:163746113-163746175 | LOC285796 | | TSS1500 |
| Buccal_PT5 | chr6:31237013-31237722 | HLA-C | N_Shore | Body |
| Buccal_PT5 | chr6:32165889-32165917 | NOTCH4 | S_Shore | Body |
| Buccal_PT5 | chr6:33228341-33228498 | VPS52 | | Body |
| Buccal_PT5 | chr7:102574445-102574504 | FBXL13;LRRC17;FBXL13;LRRC17 | | Body;Body;Body;Body |
| Buccal_PT5 | chr7:135433540-135433790 | FAM180A;FAM180A | | 1stExon;5'UTR |
| Buccal_PT5 | chr8:1790793-1790905 | ARHGEF10 | | 5'UTR |
| Buccal_PT5 | chr8:1906312-1906498 | ARHGEF10 | | 3'UTR |
| Hair_PT1 | chr1:205818956-205819609 | PM20D1 | Island | Body |
| Hair_PT1 | chr1:38156462-38156652 | C1orf109 | N_Shore | TSS1500 |
| Hair_PT1 | chr12:130821962-130822674 | PIWIL1 | N_Shore | TSS1500 |
| Hair_PT1 | chr12:54412688-54413384 | HOXC4;HOXC5;HOXC6 | N_Shore | 5'UTR;Body;5'UTR |
| Hair_PT1 | chr14:34269416-34269637 | NPAS3;NPAS3;NPAS3;NPAS3 | Island | Body;Body;Body;Body |
| Hair_PT1 | chr19:12876846-12877188 | HOOK2;HOOK2 | N_Shore | Body;Body |
| Hair_PT1 | chr2:26401640-26402318 | FAM59B | N_Shore | Body |
| Hair_PT1 | chr21:47604052-47605174 | C21orf56;C21orf56 | S_Shore | 5'UTR;5'UTR |
| Hair_PT1 | chr3:195488493-195490309 | MUC4;MUC4;MUC4 | N_Shore | Body;Body;Body |
| Hair_PT1 | chr4:4863874-4864902 | MSX1 | N_Shore | Body |
| Hair_PT1 | chr6:29894005-29894141 | HCG4P6 | N_Shore | TSS1500 |
| Hair_PT1 | chr6:29894162-29894253 | HCG4P6 | Island | TSS1500 |
| Hair_PT1 | chr6:29911087-29911104 | HLA-A | Island | Body |
| Hair_PT1 | chr6:32632000-32632108 | HLA-DQB1 | N_Shore | Body |
| Hair_PT1 | chr6:32633102-32633163 | HLA-DQB1 | S_Shore | Body |
| Hair_PT1 | chr7:101005910-101006035 | EMID2 | Island | TSS1500 |
| Hair_PT2 | chr10:131425198-131425282 | MGMT | | Body |
| Hair_PT2 | chr10:135341528-135343280 | CYP2E1 | Island | Body |
| Hair_PT2 | chr11:69481734-69482104 | ORAOV1 | | 3'UTR |
| Hair_PT2 | chr12:94580329-94580657 | PLXNC1 | N_Shore | Body |
| Hair_PT2 | chr17:37123638-37123711 | FBXO47;FBXO47 | | 1stExon;5'UTR |
| Hair_PT2 | chr19:12876846-12877188 | HOOK2;HOOK2 | N_Shore | Body;Body |
| Hair_PT2 | chr19:17905589-17905718 | B3GNT3 | Island | TSS1500 |
| Hair_PT2 | chr2:240271171-240271276 | HDAC4 | | Body |
| Hair_PT2 | chr2:26401640-26402318 | FAM59B | N_Shore | Body |
| Hair_PT2 | chr20:55925570-55925603 | RAE1;RAE1 | N_Shore | TSS1500;TSS1500 |
| Hair_PT2 | chr4:187125413-187126139 | CYP4V2 | | Body |
| Hair_PT2 | chr6:168435914-168436353 | KIF25;KIF25 | Island | Body;Body |
| Hair_PT2 | chr6:29910796-29911011 | HLA-A | Island | Body |

| | | | | |
|---|---|---|---|---|
| Hair_PT2 | chr6:29911036-29911104 | HLA-A | Island | Body |
| Hair_PT2 | chr6:31237013-31237034 | HLA-C | N_Shore | Body |
| Hair_PT2 | chr6:31237405-31237722 | HLA-C | N_Shore | Body |
| Hair_PT2 | chr6:32489203-32489555 | HLA-DRB5 | N_Shore | Body |
| Hair_PT2 | chr6:32489750-32489963 | HLA-DRB5 | Island | Body |
| Hair_PT2 | chr6:32549139-32549361 | HLA-DRB1 | N_Shelf | Body |
| Hair_PT2 | chr6:32551749-32551954 | HLA-DRB1 | N_Shore | Body |
| Hair_PT2 | chr6:32633114-32633354 | HLA-DQB1 | S_Shore | Body |
| Hair_PT2 | chr6:32634245-32634362 | HLA-DQB1 | S_Shore | Body |
| Hair_PT2 | chr9:129433708-129433969 | LMX1B | N_Shore | Body |
| Hair_PT3 | chr1:968395-969257 | AGRN | Island | Body |
| Hair_PT3 | chr11:56947754-56948018 | LRRC55 | N_Shore | TSS1500 |
| Hair_PT3 | chr11:75139390-75139736 | KLHL35 | N_Shore | Body |
| Hair_PT3 | chr13:97646341-97647084 | OXGR1 | N_Shore | 5'UTR |
| Hair_PT3 | chr2:110969641-110970249 | NCRNA00116 | N_Shore | Body |
| Hair_PT3 | chr2:240271171-240271276 | HDAC4 | | Body |
| Hair_PT3 | chr5:102898223-102898733 | NUDT12 | | 5'UTR |
| Hair_PT3 | chr6:291687-293331 | DUSP22 | N_Shore | TSS1500 |
| Hair_PT3 | chr6:29911087-29911104 | HLA-A | Island | Body |
| Hair_PT3 | chr6:31237013-31237034 | HLA-C | N_Shore | Body |
| Hair_PT3 | chr6:31237405-31237722 | HLA-C | N_Shore | Body |
| Hair_PT3 | chr6:33245128-33245359 | B3GALT4;B3GALT4 | Island | 1stExon;5'UTR |
| Hair_PT3 | chr7:100091181-100091786 | C7orf51 | Island | Body |
| Hair_PT3 | chr7:2847517-2847575 | GNA12 | | Body |
| Hair_PT4 | chr1:205818956-205819492 | PM20D1 | Island | Body |
| Hair_PT4 | chr10:131425198-131425282 | MGMT | | Body |
| Hair_PT4 | chr11:69481734-69482104 | ORAOV1 | | 3'UTR |
| Hair_PT4 | chr12:50475167-50475372 | ACCN2;ACCN2 | Island | Body;Body |
| Hair_PT4 | chr16:67686832-67687392 | RLTPR | N_Shore | Body |
| Hair_PT4 | chr2:26401640-26402318 | FAM59B | N_Shore | Body |
| Hair_PT4 | chr2:30669597-30669863 | LCLAT1;LCLAT1 | N_Shore | TSS1500;TSS1500 |
| Hair_PT4 | chr21:47604052-47605174 | C21orf56;C21orf56 | S_Shore | 5'UTR;5'UTR |
| Hair_PT4 | chr5:179740743-179741120 | GFPT2 | Island | Body |
| Hair_PT4 | chr6:31237013-31237034 | HLA-C | N_Shore | Body |
| Hair_PT4 | chr6:31238788-31238908 | HLA-C | N_Shore | Body |
| Hair_PT4 | chr6:32632000-32632108 | HLA-DQB1 | N_Shore | Body |
| Hair_PT4 | chr6:32633015-32633354 | HLA-DQB1 | Island | Body |
| Hair_PT4 | chr6:32634245-32634362 | HLA-DQB1 | S_Shore | Body |
| Hair_PT4 | chr7:39170497-39170763 | POU6F2;POU6F2 | | Body;Body |
| Hair_PT5 | chr1:19600454-19600817 | AKR7L;AKR7L | Island | 1stExon;1stExon |
| Hair_PT5 | chr1:205819088-205819463 | PM20D1 | Island | 1stExon |
| Hair_PT5 | chr10:118387095-118387327 | PNLIPRP2 | Island | Body |
| Hair_PT5 | chr10:14051636-14052028 | FRMD4A | | Body |
| Hair_PT5 | chr11:35439994-35440525 | SLC1A2 | N_Shore | Body |
| Hair_PT5 | chr11:69481734-69482104 | ORAOV1 | | 3'UTR |
| Hair_PT5 | chr12:123201189-123201372 | GPR109B | | 1stExon |
| Hair_PT5 | chr12:312592-313104 | SLC6A12;SLC6A12;SLC6A12 | Island | Body;Body;Body |
| Hair_PT5 | chr12:53183773-53184086 | KRT3 | Island | 3'UTR |
| Hair_PT5 | chr13:43597565-43598038 | DNAJC15;DNAJC15 | Island | 5'UTR;1stExon |
| Hair_PT5 | chr16:74734230-74734885 | MLKL;MLKL | Island | 5'UTR;5'UTR |
| Hair_PT5 | chr16:88496788-88497293 | ZNF469 | Island | 1stExon |
| Hair_PT5 | chr19:12876846-12877188 | HOOK2;HOOK2 | N_Shore | Body;Body |
| Hair_PT5 | chr19:613111-613818 | HCN2 | Island | Body |
| Hair_PT5 | chr2:114033490-114033830 | PAX8;PAX8;PAX8;PAX8;PAX8;LOC440839 | Island | Body;Body;Body;Body;Body;Body |
| Hair_PT5 | chr2:26401640-26402098 | FAM59B | N_Shore | Body |
| Hair_PT5 | chr2:98340425-98340921 | ZAP70 | Island | 5'UTR |
| Hair_PT5 | chr20:32307885-32308343 | PXMP4;PXMP4 | Island | Body;Body |
| Hair_PT5 | chr21:47604052-47605174 | C21orf56;C21orf56 | S_Shore | 5'UTR;5'UTR |
| Hair_PT5 | chr3:195488493-195490309 | MUC4;MUC4;MUC4 | N_Shore | Body;Body;Body |
| Hair_PT5 | chr5:102898463-102898733 | NUDT12;NUDT12 | | 5'UTR;1stExon |
| Hair_PT5 | chr5:1594282-1594863 | SDHAP3 | Island | Body |
| Hair_PT5 | chr6:29894005-29894141 | HCG4P6 | N_Shore | TSS1500 |
| Hair_PT5 | chr6:29894162-29894253 | HCG4P6 | Island | TSS1500 |
| Hair_PT5 | chr6:29911087-29911104 | HLA-A | Island | Body |
| Hair_PT5 | chr6:31237013-31237034 | HLA-C | N_Shore | Body |

| | | | | |
|---|---|---|---|---|
| Hair_PT5 | chr6:31237405-31237722 | HLA-C | N_Shore | Body |
| Hair_PT5 | chr7:157405965-157406737 | PTPRN2;PTPRN2;PTPRN2 | Island | Body;Body;Body |
| Hair_PT5 | chr7:158032120-158032496 | PTPRN2;PTPRN2;PTPRN2 | | Body;Body;Body |
| Hair_PT5 | chr7:158854447-158854763 | VIPR2 | Island | Body |
| Hair_PT5 | chr8:41580834-41580932 | ANK1;ANK1;ANK1;ANK1;ANK1 | N_Shelf | Body;Body;Body;Body;Body |
| Hair_PT5 | chr9:140916980-140917431 | CACNA1B | N_Shore | Body |
| Saliva_PT1 | chr1:202172778-202172912 | LGR6;LGR6 | | TSS200;Body |
| Saliva_PT1 | chr12:130821962-130822674 | PIWIL1 | N_Shore | TSS1500 |
| Saliva_PT1 | chr18:74799250-74799572 | MBP;MBP | Island | Body;Body |
| Saliva_PT1 | chr19:12876846-12877188 | HOOK2;HOOK2 | N_Shore | Body;Body |
| Saliva_PT1 | chr21:36258423-36259460 | RUNX1;RUNX1;RUNX1 | N_Shore | Body;Body;Body |
| Saliva_PT1 | chr3:195488922-195490169 | MUC4;MUC4;MUC4 | Island | Body;Body;Body |
| Saliva_PT1 | chr5:176797920-176798049 | RGS14 | S_Shelf | Body |
| Saliva_PT1 | chr6:29893955-29894141 | HCG4P6 | N_Shore | TSS1500 |
| Saliva_PT1 | chr6:29894162-29894197 | HCG4P6 | Island | TSS1500 |
| Saliva_PT1 | chr6:29911542-29911558 | HLA-A | S_Shore | Body |
| Saliva_PT1 | chr6:31237405-31237722 | HLA-C | N_Shore | Body |
| Saliva_PT1 | chr7:2847517-2847575 | GNA12 | | Body |
| Saliva_PT1 | chr7:39170497-39170763 | POU6F2;POU6F2 | | Body;Body |
| Saliva_PT2 | chr1:110254662-110254709 | GSTM5 | | TSS1500 |
| Saliva_PT2 | chr10:135341528-135342218 | CYP2E1 | Island | Body |
| Saliva_PT2 | chr10:135342560-135343280 | CYP2E1 | Island | Body |
| Saliva_PT2 | chr11:5617273-5617703 | TRIM6-TRIM34;TRIM6;TRIM6 | | TSS1500;TSS1500;TSS200 |
| Saliva_PT2 | chr11:5617812-5617926 | TRIM6;TRIM6-TRIM34;TRIM6 | | TSS200;TSS200;5'UTR |
| Saliva_PT2 | chr17:2797964-2798327 | RAP1GAP2;RAP1GAP2 | | Body;Body |
| Saliva_PT2 | chr19:12876846-12877188 | HOOK2;HOOK2 | N_Shore | Body;Body |
| Saliva_PT2 | chr19:13875014-13875137 | MRI1;MRI1 | N_Shore | TSS1500;TSS1500 |
| Saliva_PT2 | chr5:176797920-176798049 | RGS14 | S_Shelf | Body |
| Saliva_PT2 | chr6:31237013-31237034 | HLA-C | N_Shore | Body |
| Saliva_PT2 | chr6:31237405-31237722 | HLA-C | N_Shore | Body |
| Saliva_PT2 | chr6:32551749-32551954 | HLA-DRB1 | N_Shore | Body |
| Saliva_PT2 | chr6:32632331-32632338 | HLA-DQB1 | Island | Body |
| Saliva_PT2 | chr6:32632694-32632731 | HLA-DQB1 | Island | Body |
| Saliva_PT3 | chr12:739280-741507 | NINJ2 | | Body |
| Saliva_PT3 | chr19:21264896-21265421 | ZNF714 | N_Shore | TSS200 |
| Saliva_PT3 | chr2:110969641-110970249 | NCRNA00116 | N_Shore | Body |
| Saliva_PT3 | chr2:20869434-20871401 | GDF7 | N_Shore | Body |
| Saliva_PT3 | chr5:102898223-102898733 | NUDT12 | | 5'UTR |
| Saliva_PT3 | chr6:291687-293331 | DUSP22 | N_Shore | TSS1500 |
| Saliva_PT3 | chr6:31237013-31237722 | HLA-C | N_Shore | Body |
| Saliva_PT3 | chr8:145955983-145956024 | ZNF251 | Island | Body |
| Saliva_PT3 | chr9:130955135-130955436 | CIZ1;CIZ1;CIZ1;CIZ1;CIZ1 | N_Shore | TSS1500;5'UTR;TSS1500;5'UTR;TSS1500 |
| Saliva_PT4 | chr1:205818956-205819609 | PM20D1 | Island | Body |
| Saliva_PT4 | chr21:47604052-47605174 | C21orf56;C21orf56 | S_Shore | 5'UTR;5'UTR |
| Saliva_PT4 | chr6:31237013-31237034 | HLA-C | N_Shore | Body |
| Saliva_PT4 | chr6:32632331-32632338 | HLA-DQB1 | Island | Body |
| Saliva_PT4 | chr7:29605808-29606349 | PRR15 | N_Shore | 5'UTR |
| Saliva_PT5 | chr1:110254662-110254709 | GSTM5 | | TSS1500 |
| Saliva_PT5 | chr1:205818956-205819609 | PM20D1 | Island | Body |
| Saliva_PT5 | chr1:76189707-76189801 | ACADM;ACADM | N_Shore | TSS1500;TSS1500 |
| Saliva_PT5 | chr13:30077315-30077489 | MTUS2;MTUS2 | Island | Body;Body |
| Saliva_PT5 | chr16:2867051-2867773 | PRSS21;PRSS21;PRSS21 | N_Shore | TSS200;TSS200;TSS200 |
| Saliva_PT5 | chr17:184833-185152 | RPH3AL | N_Shore | 5'UTR |
| Saliva_PT5 | chr17:74023465-74023633 | EVPL;EVPL | | 5'UTR;1stExon |
| Saliva_PT5 | chr19:12876846-12877188 | HOOK2;HOOK2 | N_Shore | Body;Body |
| Saliva_PT5 | chr2:139538356-139539001 | NXPH2 | Island | TSS1500 |
| Saliva_PT5 | chr21:47604052-47605174 | C21orf56;C21orf56 | S_Shore | 5'UTR;5'UTR |
| Saliva_PT5 | chr3:195488922-195490309 | MUC4;MUC4;MUC4 | Island | Body;Body;Body |
| Saliva_PT5 | chr6:29894005-29894141 | HCG4P6 | N_Shore | TSS1500 |
| Saliva_PT5 | chr6:31237013-31237034 | HLA-C | N_Shore | Body |
| Saliva_PT5 | chr6:31237405-31237722 | HLA-C | N_Shore | Body |
| Saliva_PT5 | chr7:157405965-157406737 | PTPRN2;PTPRN2;PTPRN2 | Island | Body;Body;Body |
| Saliva_PT5 | chr9:126135610-126135998 | CRB2 | Island | Body |
| Saliva_PT5 | chr9:140916980-140917431 | CACNA1B | N_Shore | Body |

**Supplementary Table 09: Selected coordinates containing MeVars identified in embryonic tissues and the corresponding iPSC lines.**

| Data Coordinate | Muscle | Pancreas | Brain | Skin | Kidney | Lung | Brain iPS #1 | Brain iPS #2 | Skin iPS #1 | Skin iPS #2 | Muscle iPS #1 | Muscle iPS #2 | Kidney iPS #1 | Kidney iPS #2 | Pancreas iPS #1 | Pancreas iPS #3 | Lung iPS #1 | Lung iPS #2 |
|---|---|---|---|---|---|---|---|---|---|---|---|---|---|---|---|---|---|---|
| chr1:110166563-110166664 | 0.447211333 | 0.554622441 | 0.304436183 | 0.335857898 | 0.408953154 | 0.539700732 | 0.819251521 | 0.778952317 | 0.808435456 | 0.770155461 | 0.818994369 | 0.813988292 | 0.791846029 | 0.813911478 | 0.757353468 | 0.810419144 | 0.793480601 | 0.805296632 |
| chr1:161008705-161008977 | 0.249763865 | 0.295868114 | 0.586530518 | 0.35775342 | 0.352636033 | 0.526990894 | 0.658514208 | 0.569774964 | 0.276541301 | 0.22199695 | 0.264543439 | 0.4021084 | 0.274179216 | 0.040799464 | 0.236024791 | 0.88014176 | 0.484506059 | 0.899925881 |
| chr1:204226566-204226892 | 0.802848577 | 0.827802779 | 0.332949741 | 0.798402101 | 0.824507003 | 0.840476902 | 0.873340671 | 0.85962392 | 0.862934811 | 0.854259348 | 0.889923436 | 0.882585036 | 0.858608802 | 0.879317971 | 0.853540603 | 0.873990971 | 0.867184122 | 0.863609297 |
| chr1:244218327-244218537 | 0.905857255 | 0.874591619 | 0.200168025 | 0.898022582 | 0.924214738 | 0.914758394 | 0.895584055 | 0.902077187 | 0.904801029 | 0.892814368 | 0.904477438 | 0.912665027 | 0.893818033 | 0.893578008 | 0.887518604 | 0.887381343 | 0.872720208 | 0.901503355 |
| chr1:3028877-3029051 | 0.736371761 | 0.601225532 | 0.768581162 | 0.590575516 | 0.806821047 | 0.78816783 | 0.912772838 | 0.883251178 | 0.903362585 | 0.883377908 | 0.922886935 | 0.92442797 | 0.890301307 | 0.924958099 | 0.879739132 | 0.88440315 | 0.897085306 | 0.913322101 |
| chr1:3100143-3100343 | 0.381978117 | 0.337033485 | 0.732383621 | 0.450071274 | 0.235685497 | 0.490794145 | 0.874729869 | 0.849217867 | 0.842855944 | 0.85862409 | 0.859206529 | 0.898832606 | 0.85640906 | 0.890056082 | 0.839813109 | 0.857973463 | 0.839423855 | 0.863328518 |
| chr1:7120893-7121446 | 0.618886755 | 0.828505003 | 0.408908544 | 0.799059582 | 0.643581672 | 0.717786253 | 0.880007852 | 0.860860765 | 0.874208968 | 0.857041189 | 0.889265376 | 0.892486317 | 0.886260311 | 0.880889876 | 0.863089746 | 0.889712892 | 0.871467359 | 0.868646341 |
| chr1:8403631-8404216 | 0.524484235 | 0.60165484 | 0.358468216 | 0.538649158 | 0.441890746 | 0.449974392 | 0.884182578 | 0.845372613 | 0.865461865 | 0.850915258 | 0.875931303 | 0.880032123 | 0.850731615 | 0.876933928 | 0.834659632 | 0.833329527 | 0.854024102 | 0.876229014 |
| chr10:133929429-133929519 | 0.44710425 | 0.615142775 | 0.370653472 | 0.717928853 | 0.459710873 | 0.691264813 | 0.922496063 | 0.917445783 | 0.923586592 | 0.916936069 | 0.916733216 | 0.927988664 | 0.883273853 | 0.932923709 | 0.879266967 | 0.91776504 | 0.891941502 | 0.928989152 |
| chr10:135343193-135343280 | 0.463937463 | 0.426749072 | 0.473971312 | 0.423343417 | 0.463515026 | 0.423206901 | 0.95092452 | 0.929797783 | 0.649481829 | 0.825929175 | 0.91969705 | 0.579451366 | 0.855449132 | 0.91421763 | 0.808587257 | 0.751836277 | 0.854321191 | 0.283342335 |
| chr10:22608674-22608951 | 0.785009327 | 0.723475638 | 0.481896468 | 0.728168202 | 0.683984348 | 0.845706201 | 0.082849983 | 0.071502451 | 0.089896685 | 0.082890086 | 0.091500366 | 0.101587112 | 0.082273262 | 0.087679936 | 0.073873448 | 0.072987681 | 0.097724083 | 0.069285343 |
| chr10:530805-531152 | 0.679540017 | 0.753245708 | 0.24762748 | 0.641179753 | 0.732157442 | 0.782340914 | 0.301029853 | 0.099065098 | 0.223313999 | 0.274615608 | 0.271988866 | 0.358477669 | 0.458885868 | 0.184849463 | 0.353308141 | 0.24722493 | 0.506915802 | 0.489546003 |
| chr10:94820923-94820988 | 0.364591257 | 0.502171004 | 0.717057062 | 0.233681084 | 0.230090251 | 0.477202397 | 0.103581657 | 0.080291222 | 0.090987597 | 0.078334601 | 0.097894319 | 0.088534997 | 0.087048537 | 0.081708421 | 0.098591494 | 0.076023948 | 0.084274015 | 0.087664075 |
| chr11:1892038-1892307 | 0.453568284 | 0.538840728 | 0.184961072 | 0.440856129 | 0.61713391 | 0.440011153 | 0.819205632 | 0.710633725 | 0.766466863 | 0.768972245 | 0.813873275 | 0.845837103 | 0.795476386 | 0.747724754 | 0.743803729 | 0.785087909 | 0.792422087 | 0.791663394 |
| chr11:2922635-2922807 | 0.780992159 | 0.864279723 | 0.492921324 | 0.776589997 | 0.807865183 | 0.875802299 | 0.856072917 | 0.829935471 | 0.83750657 | 0.808212862 | 0.835145127 | 0.854915963 | 0.831282446 | 0.849426121 | 0.807414773 | 0.834840712 | 0.827362101 | 0.851662552 |
| chr11:47607760-47608487 | 0.370284704 | 0.674724147 | 0.378356168 | 0.521507853 | 0.718803727 | 0.627402019 | 0.780323144 | 0.756320143 | 0.756068921 | 0.743394627 | 0.760461289 | 0.771170183 | 0.755258637 | 0.776990679 | 0.721154909 | 0.756033475 | 0.755702775 | 0.768324521 |
| chr11:70001754-70001938 | 0.630603166 | 0.707518302 | 0.444196714 | 0.41615403 | 0.691551814 | 0.684092636 | 0.773428478 | 0.756871253 | 0.788767743 | 0.749810769 | 0.806635341 | 0.795883351 | 0.772038283 | 0.785085815 | 0.750823702 | 0.760643799 | 0.782178963 | 0.791078543 |
| chr12:114833303-114834201 | 0.103940405 | 0.082157168 | 0.079152945 | 0.532494846 | 0.078193186 | 0.132040499 | 0.084875059 | 0.070504046 | 0.080890228 | 0.073223326 | 0.07680086 | 0.111549734 | 0.087232351 | 0.081288626 | 0.075713899 | 0.2362425 | 0.083328001 | 0.084208983 |
| chr12:120538681-120539137 | 0.392519221 | 0.620892714 | 0.388885122 | 0.642662314 | 0.602633258 | 0.546287305 | 0.823789194 | 0.804742017 | 0.816222442 | 0.793867392 | 0.821737168 | 0.829043688 | 0.791500042 | 0.828466112 | 0.765287756 | 0.819092191 | 0.798562644 | 0.827230786 |
| chr12:132865182-132865380 | 0.748853594 | 0.775809773 | 0.269138731 | 0.782992602 | 0.540340588 | 0.741129852 | 0.808804964 | 0.809184001 | 0.740498321 | 0.804541108 | 0.755827296 | 0.855953785 | 0.725883603 | 0.815306036 | 0.792613079 | 0.728320634 | 0.80440333 | 0.737423807 |
| chr12:54448729-54448913 | 0.837108072 | 0.182572865 | 0.0764119 | 0.703167531 | 0.471689404 | 0.198982825 | 0.082763438 | 0.04897324 | 0.08512554 | 0.062246687 | 0.077229577 | 0.068472011 | 0.070866849 | 0.061053556 | 0.07234821 | 0.068751543 | 0.080432914 | 0.078090433 |
| chr13:102104848-102104991 | 0.428099506 | 0.776672223 | 0.539543358 | 0.489655438 | 0.787847445 | 0.656508728 | 0.899199983 | 0.916289899 | 0.892511155 | 0.88651535 | 0.889042621 | 0.906059878 | 0.885529198 | 0.911029935 | 0.860518866 | 0.88833844 | 0.885800766 | 0.892215948 |
| chr13:111294884-111295144 | 0.741724725 | 0.860187158 | 0.596228043 | 0.629199018 | 0.874821417 | 0.860784042 | 0.898861707 | 0.891556382 | 0.89807466 | 0.887943702 | 0.899644501 | 0.92369788 | 0.909860361 | 0.914543013 | 0.893398971 | 0.916629239 | 0.899541315 | 0.89526484 |
| chr13:113424175-113424362 | 0.593038405 | 0.76967815 | 0.447129901 | 0.803691263 | 0.680441931 | 0.698276666 | 0.880403184 | 0.859609739 | 0.856898516 | 0.846304406 | 0.880749119 | 0.878800977 | 0.863801449 | 0.873866469 | 0.832802132 | 0.860360861 | 0.866178405 | 0.867955916 |
| chr13:113492269-113492708 | 0.783070542 | 0.87995818 | 0.518281746 | 0.873825598 | 0.897802912 | 0.885300822 | 0.925820967 | 0.906981316 | 0.914069748 | 0.905414374 | 0.916078203 | 0.900215792 | 0.930594401 | 0.92527223 | 0.901067296 | 0.939915951 | 0.912698971 | 0.908916877 |
| chr13:114203831-114203974 | 0.745845506 | 0.836506017 | 0.500310715 | 0.892975839 | 0.698248436 | 0.858178008 | 0.906986656 | 0.920210904 | 0.920776336 | 0.898650061 | 0.912735624 | 0.919156308 | 0.909462898 | 0.926013004 | 0.894595858 | 0.922876711 | 0.90849464 | 0.910588402 |
| chr14:104178670-104179160 | 0.492203004 | 0.665514451 | 0.396962679 | 0.583923884 | 0.734955639 | 0.551899844 | 0.865415027 | 0.847696126 | 0.861952809 | 0.848250914 | 0.868996684 | 0.887003187 | 0.859474487 | 0.875197003 | 0.841654225 | 0.859236508 | 0.853405021 | 0.8668311 |
| chr14:104636788-104637213 | 0.595706124 | 0.80134075 | 0.322528817 | 0.773493841 | 0.731461089 | 0.832791496 | 0.648719152 | 0.527777441 | 0.676070553 | 0.587947559 | 0.655021441 | 0.61828481 | 0.648979527 | 0.569896228 | 0.621804337 | 0.600021719 | 0.649137435 | 0.692520585 |
| chr14:105251188-105251610 | 0.71091231 | 0.469516494 | 0.164512365 | 0.747000735 | 0.737236803 | 0.502143423 | 0.879637412 | 0.85376782 | 0.896577471 | 0.865009793 | 0.88479147 | 0.868749168 | 0.869482642 | 0.884171809 | 0.846031097 | 0.871382433 | 0.872216017 | 0.872893456 |
| chr14:105783536-105784090 | 0.543963109 | 0.66297208 | 0.33729649 | 0.421947665 | 0.712863374 | 0.503513211 | 0.839552869 | 0.840769822 | 0.85757583 | 0.850540414 | 0.842574317 | 0.854274771 | 0.845280386 | 0.859212452 | 0.828403927 | 0.851904097 | 0.842976345 | 0.830769037 |
| chr16:11367459-11367509 | 0.705444373 | 0.727244921 | 0.431637739 | 0.678731396 | 0.557890352 | 0.738543095 | 0.832321455 | 0.795889835 | 0.825713964 | 0.799589407 | 0.832704694 | 0.835694357 | 0.791854224 | 0.822481448 | 0.774120352 | 0.809014002 | 0.802019412 | 0.838716657 |
| chr16:1480302-1480790 | 0.472897499 | 0.649841603 | 0.367165054 | 0.748421288 | 0.739525762 | 0.718415989 | 0.854452267 | 0.830732666 | 0.840263549 | 0.839358745 | 0.844120312 | 0.869297837 | 0.837954568 | 0.869407722 | 0.804637921 | 0.844096516 | 0.841898948 | 0.881435666 |
| chr16:29675846-29676071 | 0.806451885 | 0.747291738 | 0.414663018 | 0.861132891 | 0.817336158 | 0.879202388 | 0.844340948 | 0.835665376 | 0.831635208 | 0.821207335 | 0.842033891 | 0.860280242 | 0.828903828 | 0.861263285 | 0.800950203 | 0.833497869 | 0.835287767 | 0.858388327 |
| chr16:29706151-29706381 | 0.523950385 | 0.810139789 | 0.453863486 | 0.827038557 | 0.736084902 | 0.66630382 | 0.861092696 | 0.862911851 | 0.878966398 | 0.854816906 | 0.857244714 | 0.864057946 | 0.848616928 | 0.879815835 | 0.828212501 | 0.862451754 | 0.858109126 | 0.867659237 |
| chr16:474271-474528 | 0.418524618 | 0.697067029 | 0.2762287 | 0.215494671 | 0.66994433 | 0.475525919 | 0.763536483 | 0.676187901 | 0.771459621 | 0.771137713 | 0.788206365 | 0.820325566 | 0.759876456 | 0.772632582 | 0.723699392 | 0.792499559 | 0.771219948 | 0.796050464 |
| chr16:66876847-66876938 | 0.407772379 | 0.597506719 | 0.30654695 | 0.574929368 | 0.639449154 | 0.594272834 | 0.78008655 | 0.72558108 | 0.784430296 | 0.73953393 | 0.803931105 | 0.774095919 | 0.76190996 | 0.757128755 | 0.735482961 | 0.739994026 | 0.756500516 | 0.800685599 |
| chr16:84681507-84681584 | 0.628580835 | 0.585675328 | 0.217233546 | 0.532060485 | 0.636124737 | 0.579672801 | 0.539163218 | 0.545135797 | 0.544653724 | 0.54441753 | 0.55340741 | 0.578920408 | 0.521138591 | 0.576546159 | 0.510081557 | 0.562779305 | 0.541327924 | 0.513602027 |
| chr16:87738623-87738833 | 0.882953 | 0.882551667 | 0.1623295 | 0.885076141 | 0.871389049 | 0.837531865 | 0.902341057 | 0.87439575 | 0.90804166 | 0.887113839 | 0.908615353 | 0.919619912 | 0.907719976 | 0.904246673 | 0.900857781 | 0.893865172 | 0.894993122 | 0.908915696 |
| chr16:88991518-88992178 | 0.593889389 | 0.73008883 | 0.2384886 | 0.647864944 | 0.560574747 | 0.716617582 | 0.844636537 | 0.750339549 | 0.841473233 | 0.773598279 | 0.839330267 | 0.835810119 | 0.80392948 | 0.809908653 | 0.787596192 | 0.767324336 | 0.811911745 | 0.830751656 |
| chr16:89763816-89764255 | 0.546130749 | 0.797191428 | 0.560948756 | 0.768484807 | 0.795757187 | 0.823295375 | 0.862658051 | 0.863832986 | 0.866083084 | 0.85934818 | 0.869856256 | 0.884339533 | 0.847469227 | 0.878479028 | 0.827011241 | 0.877262199 | 0.851168444 | 0.87808537 |
| chr17:17124699-17124854 | 0.863641856 | 0.874349075 | 0.551512641 | 0.877844921 | 0.900601118 | 0.891098538 | 0.899487588 | 0.883066406 | 0.893163857 | 0.885566049 | 0.885978636 | 0.906769783 | 0.883454403 | 0.908130733 | 0.869086185 | 0.895892156 | 0.899548769 | 0.89029474 |
| chr17:30821755-30823087 | 0.475314801 | 0.658709846 | 0.246023151 | 0.753380189 | 0.625461135 | 0.738363508 | 0.831200384 | 0.818909125 | 0.83298367 | 0.816481461 | 0.838418018 | 0.868001909 | 0.825865836 | 0.831899244 | 0.796953316 | 0.821168687 | 0.814218661 | 0.839580419 |
| chr17:38249352-38249729 | 0.744800835 | 0.806628021 | 0.456791881 | 0.782118075 | 0.621860916 | 0.844525976 | 0.853482046 | 0.83250991 | 0.837783362 | 0.822049053 | 0.855094513 | 0.831677973 | 0.837280338 | 0.85925268 | 0.802391529 | 0.834893166 | 0.836465906 | 0.856293478 |
| chr17:44847220-44847427 | 0.107834735 | 0.418271625 | 0.760474422 | 0.403081276 | 0.044142541 | 0.490069123 | 0.93289636 | 0.921691999 | 0.935595457 | 0.921314726 | 0.942309933 | 0.938201128 | 0.935180483 | 0.940444809 | 0.929976577 | 0.927933002 | 0.941482021 | 0.933376852 |
| chr17:73559363-73559562 | 0.431324557 | 0.526619874 | 0.285918213 | 0.46311496 | 0.383041691 | 0.485924528 | 0.602562586 | 0.608428783 | 0.595025515 | 0.605877466 | 0.634149828 | 0.628895133 | 0.583836223 | 0.627891413 | 0.583541816 | 0.600163434 | 0.580615485 | 0.654164109 |
| chr17:76141038-76141068 | 0.431863241 | 0.399444822 | 0.38739467 | 0.670161519 | 0.464006451 | 0.502037987 | 0.766747091 | 0.781057595 | 0.764559362 | 0.758742612 | 0.776950031 | 0.800336152 | 0.751674791 | 0.791908229 | 0.724085318 | 0.79106859 | 0.743621155 | 0.796801631 |
| chr17:76446778-76446851 | 0.608348329 | 0.817881027 | 0.55427387 | 0.856676875 | 0.833168895 | 0.853071141 | 0.88609549 | 0.868354849 | 0.86368429 | 0.863115604 | 0.873403083 | 0.893193673 | 0.873587885 | 0.881652351 | 0.842640261 | 0.894298197 | 0.858764755 | 0.878033426 |
| chr17:77951408-77951858 | 0.718390544 | 0.84898497 | 0.274074252 | 0.882284179 | 0.676853853 | 0.931360291 | 0.909557309 | 0.916384905 | 0.905672246 | 0.891349112 | 0.901781729 | 0.930300713 | 0.882296407 | 0.910780495 | 0.879580534 | 0.90284163 | 0.890129335 | 0.890751419 |
| chr17:77952290-77952488 | 0.562069645 | 0.505745837 | 0.314823453 | 0.48060369 | 0.412849406 | 0.859896399 | 0.898962807 | 0.903242278 | 0.91372838 | 0.905477634 | 0.895449405 | 0.927863438 | 0.90980406 | 0.931365538 | 0.888330615 | 0.891970841 | 0.89418796 | 0.907726345 |
| chr17:78800767-78800806 | 0.517819985 | 0.803904176 | 0.350213911 | 0.63180157 | 0.859275905 | 0.879205477 | 0.816074828 | 0.837442657 | 0.837753473 | 0.820100277 | 0.831974386 | 0.835773673 | 0.792556048 | 0.850518624 | 0.769746221 | 0.837185897 | 0.800255414 | 0.768677798 |
| chr17:79006613-79006834 | 0.755802015 | 0.784453019 | 0.506742312 | 0.807404635 | 0.805082272 | 0.785449629 | 0.859314939 | 0.852006843 | 0.859183631 | 0.847006503 | 0.844296547 | 0.869985713 | 0.833952088 | 0.866625605 | 0.820642984 | 0.851618187 | 0.829629415 | 0.855309204 |
| chr17:79097051-79097243 | 0.61491808 | 0.820483593 | 0.55612813 | 0.697539538 | 0.80605363 | 0.792877853 | 0.825195802 | 0.813158952 | 0.810126022 | 0.813855988 | 0.80946075 | 0.844803505 | 0.817391213 | 0.825427623 | 0.775780996 | 0.814648447 | 0.805733404 | 0.827848655 |
| chr17:79099481-79099785 | 0.418829834 | 0.764550602 | 0.522946556 | 0.641194465 | 0.77519578 | 0.74037195 | 0.899539146 | 0.890751004 | 0.887018015 | 0.891311349 | 0.883516643 | 0.899939683 | 0.891614937 | 0.903040493 | 0.866625819 | 0.891213063 | 0.886821972 | 0.884707505 |
| chr17:80393504-80393598 | 0.788605693 | 0.806108273 | 0.479888392 | 0.825137293 | 0.479081314 | 0.82221897 | 0.890698245 | 0.807454017 | 0.884372994 | 0.845683637 | 0.873521572 | 0.880420614 | 0.8758045 | 0.8641806 | 0.848863006 | 0.854490159 | 0.873419344 | 0.903878065 |
| chr17:80970799-80970892 | 0.537500731 | 0.680813861 | 0.355347742 | 0.221540904 | 0.684082229 | 0.363836618 | 0.869753353 | 0.859917521 | 0.854933085 | 0.884676884 | 0.891133343 | 0.888013197 | 0.866231761 | 0.884745098 | 0.837407757 | 0.848708188 | 0.881252902 | 0.878666261 |
| chr17:915634-915895 | 0.696523613 | 0.786000605 | 0.534630692 | 0.813693435 | 0.822644985 | 0.704688886 | 0.884653254 | 0.923181513 | 0.909563935 | 0.863285746 | 0.866628472 | 0.925038159 | 0.855633652 | 0.928782503 | 0.824471879 | 0.888328389 | 0.864978308 | 0.862610446 |
| chr19:10464711-10464842 | 0.909393146 | 0.846085992 | 0.622634265 | 0.920693123 | 0.746296603 | 0.892942045 | 0.57892762 | 0.50871286 | 0.081202552 | 0.07692372 | 0.136597874 | 0.105651715 | 0.109983319 | 0.107737837 | 0.103325821 | 0.324998247 | 0.176549912 | 0.066980616 |
| chr19:11280655-11280913 | 0.581779803 | 0.758517481 | 0.54537418 | 0.752780093 | 0.662994047 | 0.73665929 | 0.863184803 | 0.854462259 | 0.871708485 | 0.852378401 | 0.862306749 | 0.868365989 | 0.843447368 | 0.859909075 | 0.828916748 | 0.856018297 | 0.842105308 | 0.848047964 |
| chr19:11285128-11285331 | 0.590359745 | 0.88045944 | 0.522076775 | 0.866784178 | 0.895667518 | 0.915222545 | 0.893669525 | 0.885672946 | 0.886932074 | 0.876944891 | 0.893396978 | 0.907345377 | 0.883600204 | 0.898876391 | 0.868822425 | 0.893995934 | 0.872802067 | 0.905279096 |
| chr19:1822386-1822532 | 0.868400474 | 0.848087973 | 0.490510651 | 0.755942419 | 0.87252077 | 0.870447545 | 0.840224813 | 0.863936638 | 0.849596916 | 0.832387706 | 0.864507276 | 0.870592215 | 0.823055963 | 0.860541391 | 0.785607487 | 0.853195605 | 0.831944298 | 0.871466555 |
| chr19:1924679-1924761 | 0.708805427 | 0.874774567 | 0.313871756 | 0.903378599 | 0.877080742 | 0.917744112 | 0.916106075 | 0.883472799 | 0.929695812 | 0.885383862 | 0.904203247 | 0.905565025 | 0.905417216 | 0.902750897 | 0.892024653 | 0.897340185 | 0.911565312 | 0.909583926 |
| chr19:3196026-3196387 | 0.833352507 | 0.863532055 | 0.524337935 | 0.862469349 | 0.87617462 | 0.879866788 | 0.906935558 | 0.89090772 | 0.896178082 | 0.873318959 | 0.880893319 | 0.887994978 | 0.87335918 | 0.891828018 | 0.8567281 | 0.899150924 | 0.888553312 | 0.891031263 |
| chr19:41115390-41115766 | 0.591656318 | 0.303467229 | 0.608919155 | 0.153760113 | 0.265032926 | 0.583195522 | 0.122130828 | 0.114555782 | 0.118218979 | 0.115836729 | 0.128044067 | 0.164738805 | 0.118007994 | 0.156701858 | 0.116811352 | 0.110751501 | 0.121696054 | 0.118412761 |
| chr19:41119829-41120329 | 0.535104818 | 0.075965945 | 0.674894522 | 0.104408996 | 0.379507281 | 0.316871741 | 0.03106726 | 0.053872721 | 0.037003087 | 0.049128381 | 0.033158695 | 0.050113258 | 0.036622714 | 0.054581515 | 0.036653514 | 0.037082018 | 0.03701496 | 0.0407848 |
| chr19:4371730-4371934 | 0.51962743 | 0.829597531 | 0.291487583 | 0.870261883 | 0.582557263 | 0.916022986 | 0.868930466 | 0.802265239 | 0.870971565 | 0.825413097 | 0.893733917 | 0.859954841 | 0.847507221 | 0.847704527 | 0.83702912 | 0.815296812 | 0.845170591 | 0.884156945 |
| chr19:4639923-4640159 | 0.277975363 | 0.333395361 | 0.262181846 | 0.323078985 | 0.451770411 | 0.436502782 | 0.781606017 | 0.744966327 | 0.756459159 | 0.781064607 | 0.752451646 | 0.800768113 | 0.762360563 | 0.796343718 | 0.726995466 | 0.768103596 | 0.735830863 | 0.799879577 |
| chr19:49116423-49116839 | 0.687596923 | 0.817645316 | 0.439276415 | 0.789838818 | 0.776447935 | 0.78901402 | 0.880315659 | 0.863742577 | 0.861176243 | 0.853257004 | 0.867846598 | 0.90829513 | 0.853321949 | 0.894587539 | 0.827063654 | 0.878501207 | 0.859825146 | 0.894097506 |
| chr19:5036609-5037219 | 0.467778796 | 0.618123765 | 0.172648042 | 0.423390091 | 0.719074823 | 0.506562623 | 0.847079295 | 0.844672191 | 0.831722979 | 0.828433604 | 0.838440734 | 0.852263552 | 0.813193443 | 0.852845038 | 0.79473988 | 0.840692951 | 0.811709558 | 0.862256802 |
| chr19:51141227-51141361 | 0.52128544 | 0.373886632 | 0.281789677 | 0.583817948 | 0.499326753 | 0.391247992 | 0.568155572 | 0.499002796 | 0.58658504 | 0.581363552 | 0.578918116 | 0.645180276 | 0.570585846 | 0.656579069 | 0.560538615 | 0.545216613 | 0.574130316 | 0.617794542 |
| chr19:51531318-51531484 | 0.306553277 | 0.690745928 | 0.780365689 | 0.684838844 | 0.630609991 | 0.748954992 | 0.83899528 | 0.799357869 | 0.813285422 | 0.777317276 | 0.820680508 | 0.823884364 | 0.792794241 | 0.831088787 | 0.786652179 | 0.804715024 | 0.812050966 | 0.809577235 |
| chr19:5455306-5455500 | 0.776261795 | 0.903126818 | 0.534803659 | 0.888831174 | 0.910526999 | 0.900203307 | 0.921311539 | 0.915660962 | 0.90634114 | 0.895826637 | 0.916500863 | 0.895900108 | 0.906584971 | 0.890523959 | 0.890326051 | 0.922247919 | 0.906839803 | 0.902905917 |
| chr2:223164854-223164869 | 0.542454854 | 0.063855984 | 0.142108514 | 0.499452544 | 0.072219067 | 0.069395924 | 0.050553651 | 0.036265869 | 0.046666772 | 0.04379495 | 0.054328947 | 0.053588688 | 0.038513352 | 0.032091773 | 0.036587132 | 0.048563894 | 0.04350439 | 0.058798357 |
| chr2:223917577-223918000 | 0.298027868 | 0.628324261 | 0.764066801 | 0.726155525 | 0.555622382 | 0.499217266 | 0.88940283 | 0.868020041 | 0.875962748 | 0.850255898 | 0.889138542 | 0.893608741 | 0.88359165 | 0.88971072 | 0.845357456 | 0.877114334 | 0.874695221 | 0.869146375 |
| chr2:63271537-63272334 | 0.822859067 | 0.839819521 | 0.736324431 | 0.843765139 | 0.690274347 | 0.856254935 | 0.08026812 | 0.068951662 | 0.077732113 | 0.069312039 | 0.089685301 | 0.079979036 | 0.070926771 | 0.082557723 | 0.085365962 | 0.069807592 | 0.07274626 | 0.074217806 |
| chr2:97530602-97530683 | 0.90103223 | 0.891467687 | 0.62457317 | 0.904637505 | 0.903471283 | 0.920330233 | 0.900671909 | 0.911320027 | 0.910547925 | 0.909601655 | 0.916325504 | 0.923486588 | 0.895191473 | 0.916940818 | 0.893220332 | 0.906152927 | 0.909073286 | 0.911109972 |
| chr20:61151446-61151472 | 0.516568422 | 0.615459851 | 0.409076059 | 0.604906666 | 0.510907196 | 0.489977194 | 0.872082479 | 0.832705204 | 0.861495292 | 0.826745955 | 0.87864157 | 0.876078258 | 0.856567242 | 0.861411178 | 0.852545182 | 0.857372937 | 0.853996194 | 0.88508073 |
| chr20:62200603-62200854 | 0.555814175 | 0.88270213 | 0.312322404 | 0.82310539 | 0.90687745 | 0.884497293 | 0.921898964 | 0.918182251 | 0.944504843 | 0.903920658 | 0.934695562 | 0.950514832 | 0.921717714 | 0.948768721 | 0.896445966 | 0.920695066 | 0.927226475 | 0.923221569 |
| chr21:38117454-38117682 | 0.639822188 | 0.707348621 | 0.371911748 | 0.73118072 | 0.690116913 | 0.692299415 | 0.828026688 | 0.799334906 | 0.838226563 | 0.800149354 | 0.845738137 | 0.825409441 | 0.830412438 | 0.836769669 | 0.819392138 | 0.824559253 | 0.82377716 | 0.835280365 |
| chr21:44480336-44480711 | 0.812452669 | 0.668381999 | 0.494222571 | 0.678806138 | 0.782967779 | 0.80478697 | 0.871207217 | 0.870841361 | 0.859856939 | 0.873242788 | 0.867896399 | 0.863449444 | 0.848346748 | 0.863571455 | 0.838325292 | 0.852543553 | 0.859315518 | 0.861993421 |
| chr22:19746268-19746984 | 0.361957613 | 0.087791481 | 0.092643388 | 0.536094238 | 0.131791947 | 0.111870372 | 0.066016519 | 0.070998466 | 0.075415425 | 0.072241457 | 0.086054493 | 0.096695027 | 0.059197944 | 0.070950632 | 0.076716756 | 0.085535584 | 0.064592279 | 0.09189395 |
| chr22:19754125-19754815 | 0.534386474 | 0.097735111 | 0.097677766 | 0.628167432 | 0.091932079 | 0.358415959 | 0.092065434 | 0.087553132 | 0.078440595 | 0.099850508 | 0.097135463 | 0.097306606 | 0.083027238 | 0.10674309 | 0.078765981 | 0.097282894 | 0.076719114 | 0.093501752 |
| chr22:22287697-22288559 | 0.850554999 | 0.869341411 | 0.492742552 | 0.864611218 | 0.878611974 | 0.893696104 | 0.852976185 | 0.842877922 | 0.858532815 | 0.841010388 | 0.852485809 | 0.840639634 | 0.837193595 | 0.857302992 | 0.804700529 | 0.843740749 | 0.842520748 | 0.859654884 |
| chr22:24890833-24891141 | 0.207983932 | 0.273228715 | 0.497055336 | 0.154691781 | 0.30958691 | 0.382206491 | 0.47740044 | 0.47462037 | 0.443510917 | 0.422830565 | 0.525898294 | 0.532945287 | 0.517754178 | 0.51576185 | 0.449462132 | 0.435716497 | 0.428323143 | 0.405446049 |
| chr22:29707524-29708028 | 0.429795236 | 0.776988006 | 0.333919386 | 0.793258787 | 0.80053249 | 0.765694753 | 0.884756698 | 0.85239553 | 0.875956093 | 0.859484684 | 0.863636063 | 0.859148688 | 0.863413278 | 0.848378335 | 0.850525841 | 0.873071402 | 0.859215057 | 0.882956764 |
| chr22:50177941-50178220 | 0.790310518 | 0.802915708 | 0.338306284 | 0.832813753 | 0.791686999 | 0.832984942 | 0.906084035 | 0.903582579 | 0.918038739 | 0.915397027 | 0.90930099 | 0.930862207 | 0.912989089 | 0.911909484 | 0.908926102 | 0.907513253 | 0.917165655 | 0.925250474 |
| chr3:148804720-148804746 | 0.810790423 | 0.858197328 | 0.380574092 | 0.827428771 | 0.883513439 | 0.877762954 | 0.817579826 | 0.817740858 | 0.825664077 | 0.823604974 | 0.820074591 | 0.84883462 | 0.796576593 | 0.856822621 | 0.76373879 | 0.833027314 | 0.79701957 | 0.794226119 |
| chr3:181441345-181441571 | 0.151301773 | 0.132948817 | 0.555987138 | 0.238079005 | 0.115860581 | 0.426780301 | 0.298769069 | 0.61594017 | 0.585241415 | 0.676532164 | 0.261531859 | 0.485519956 | 0.287065742 | 0.36561819 | 0.414617999 | 0.188372643 | 0.189028253 | 0.138541781 |
| chr4:1243980-1244042 | 0.612328003 | 0.647877355 | 0.324383703 | 0.537790462 | 0.733420235 | 0.770846459 | 0.856328229 | 0.853014737 | 0.845997045 | 0.846241351 | 0.860535128 | 0.869563327 | 0.839039837 | 0.869731224 | 0.802230927 | 0.85647742 | 0.84167743 | 0.684402609 |
| chr4:154711470-154711519 | 0.764750693 | 0.663668798 | 0.645664002 | 0.610007793 | 0.831098279 | 0.717813466 | 0.037792582 | 0.044734508 | 0.040452344 | 0.033159541 | 0.037554587 | 0.042526791 | 0.043456805 | 0.038570778 | 0.048526663 | 0.049126604 | 0.036975797 | 0.036861207 |
| chr4:4864293-4864883 | 0.35451842 | 0.433868112 | 0.50902471 | 0.594504386 | 0.292871469 | 0.506499437 | 0.14991119 | 0.142664121 | 0.160219675 | 0.136484509 | 0.173796529 | 0.161818687 | 0.17417717 | 0.139387674 | 0.16290663 | 0.147192052 | 0.168420513 | 0.138302146 |
| chr4:812499-812684 | 0.801575587 | 0.80858764 | 0.551893015 | 0.850153719 | 0.842473134 | 0.843271528 | 0.817215215 | 0.721736681 | 0.809628275 | 0.772990968 | 0.841839547 | 0.846178256 | 0.800377516 | 0.826047094 | 0.777651298 | 0.861410108 | 0.811636968 | 0.86539441 |
| chr5:1089507-1089722 | 0.362927665 | 0.477741939 | 0.444250279 | 0.324218577 | 0.451970296 | 0.494303933 | 0.863844921 | 0.838468921 | 0.859182916 | 0.841770646 | 0.866004775 | 0.862234146 | 0.842862839 | 0.8686902 | 0.816467755 | 0.846441307 | 0.842060627 | 0.857308728 |
| chr5:122423433-122423615 | 0.229670078 | 0.240572946 | 0.108485366 | 0.534384953 | 0.183993743 | 0.628962733 | 0.102867975 | 0.092803179 | 0.11522654 | 0.096245089 | 0.122927927 | 0.118745828 | 0.12690735 | 0.103086123 | 0.121647214 | 0.100778282 | 0.123401982 | 0.115121938 |
| chr5:171765285-171765653 | 0.548476133 | 0.825638821 | 0.487002854 | 0.728975929 | 0.819574966 | 0.863333368 | 0.850589384 | 0.859206744 | 0.839538016 | 0.843413364 | 0.870127048 | 0.861606398 | 0.832258529 | 0.862264462 | 0.815302155 | 0.847414247 | 0.829726774 | 0.869394185 |
| chr5:179220589-179221059 | 0.4166776 | 0.62092451 | 0.496349386 | 0.274249435 | 0.683345398 | 0.475617831 | 0.841265494 | 0.827567352 | 0.841949522 | 0.814053202 | 0.82621283 | 0.852718875 | 0.827214267 | 0.840237548 | 0.796176958 | 0.820230806 | 0.823687776 | 0.849602566 |
| chr5:510035-510421 | 0.717567806 | 0.815617279 | 0.402970469 | 0.534735332 | 0.693113668 | 0.702484654 | 0.924903591 | 0.903602699 | 0.913280674 | 0.910977196 | 0.928564294 | 0.934958943 | 0.936529919 | 0.922646628 | 0.899096708 | 0.91781621 | 0.915384746 | 0.918580462 |
| chr6:108495578-108496991 | 0.117522212 | 0.151253753 | 0.155151424 | 0.12416861 | 0.100969461 | 0.092056603 | 0.105656736 | 0.089435309 | 0.114910884 | 0.095502281 | 0.116711587 | 0.108816875 | 0.118146335 | 0.104066515 | 0.100565214 | 0.095311226 | 0.108843944 | 0.108496101 |
| chr6:108497717-108497995 | 0.133977055 | 0.188180272 | 0.168576584 | 0.172647336 | 0.120373025 | 0.134943646 | 0.14302319 | 0.130658787 | 0.145282295 | 0.144323566 | 0.159570944 | 0.161893207 | 0.144756539 | 0.153421323 | 0.143112486 | 0.128628939 | 0.151049128 | 0.130363252 |
| chr6:123109741-123109903 | 0.405992202 | 0.393058427 | 0.652798806 | 0.255599619 | 0.359802745 | 0.52623002 | 0.399423866 | 0.380305677 | 0.441846842 | 0.380392363 | 0.444481866 | 0.398045696 | 0.403018005 | 0.441042238 | 0.366906835 | 0.345750939 | 0.385025313 | 0.428858369 |
| chr6:25138632-25138951 | 0.205057137 | 0.501364366 | 0.689907635 | 0.178785596 | 0.450745943 | 0.254930389 | 0.882804639 | 0.870911157 | 0.885451992 | 0.83733802 | 0.893865747 | 0.903261226 | 0.857407795 | 0.894277112 | 0.847517264 | 0.869810194 | 0.881643009 | 0.861024605 |
| chr6:292329-292823 | 0.087713909 | 0.093768638 | 0.09281368 | 0.086675747 | 0.083107409 | 0.08153328 | 0.089936762 | 0.073975835 | 0.079234584 | 0.075394823 | 0.092832587 | 0.08567996 | 0.081753843 | 0.076627465 | 0.079163702 | 0.072325773 | 0.079121471 | 0.081471036 |
| chr6:29910994-29911011 | 0.440358827 | 0.445102541 | 0.559152791 | 0.447103417 | 0.444784192 | 0.434294834 | 0.647217958 | 0.658934317 | 0.63426415 | 0.623932857 | 0.668028571 | 0.669188842 | 0.635448692 | 0.644275977 | 0.607132703 | 0.64252877 | 0.649999416 | 0.637799867 |
| chr6:30039408-30039442 | 0.241003739 | 0.128740436 | 0.272703103 | 0.486023717 | 0.175847875 | 0.12284862 | 0.434015843 | 0.595567252 | 0.487569735 | 0.485647598 | 0.56241164 | 0.633733057 | 0.479598809 | 0.547707848 | 0.475790654 | 0.405071913 | 0.567309497 | 0.395234095 |
| chr6:32044104-32044203 | 0.328455768 | 0.776021422 | 0.229444096 | 0.584915196 | 0.697226684 | 0.683643005 | 0.873104298 | 0.86888127 | 0.85454998 | 0.840383428 | 0.878214018 | 0.877621953 | 0.86459488 | 0.875848021 | 0.837337489 | 0.880542505 | 0.871647213 | 0.864670989 |
| chr6:32063895-32064032 | 0.670666054 | 0.563511425 | 0.500917776 | 0.714130762 | 0.526393535 | 0.743697987 | 0.968781313 | 0.702706169 | 0.939458674 | 0.966854285 | 0.965733142 | 0.971140002 | 0.947871845 | 0.938652354 | 0.980712513 | 0.941568294 | 0.94575777 | 0.952330276 |
| chr6:32135715-32135776 | 0.694351317 | 0.439462379 | 0.58496905 | 0.752264 | 0.756274232 | 0.565033751 | 0.834548978 | 0.774558124 | 0.818562037 | 0.799389448 | 0.831152323 | 0.827254324 | 0.825496106 | 0.823423396 | 0.778879394 | 0.750881175 | 0.799682692 | 0.828164169 |
| chr6:43736807-43737136 | 0.357112443 | 0.660145486 | 0.395280531 | 0.461203718 | 0.713798469 | 0.494529313 | 0.812863965 | 0.80355187 | 0.80356341 | 0.790504593 | 0.812108293 | 0.821219835 | 0.787567805 | 0.836191388 | 0.765754487 | 0.807213754 | 0.782012065 | 0.815975113 |
| chr7:100463583-100463985 | 0.580067932 | 0.786156959 | 0.421657544 | 0.513444719 | 0.804293324 | 0.804831751 | 0.887065196 | 0.885892886 | 0.886849484 | 0.871289608 | 0.896237417 | 0.926260038 | 0.875070898 | 0.934714268 | 0.855589287 | 0.90062842 | 0.888039168 | 0.89868056 |
| chr7:101558400-101558698 | 0.497629606 | 0.81023345 | 0.3588597 | 0.66269537 | 0.88209493 | 0.765030152 | 0.872756724 | 0.862325565 | 0.867476506 | 0.841565806 | 0.867990202 | 0.862669946 | 0.841878749 | 0.864099235 | 0.829034008 | 0.858854804 | 0.850612399 | 0.857412149 |
| chr7:157561104-157561161 | 0.697747017 | 0.791572661 | 0.244586298 | 0.827498235 | 0.677960141 | 0.751354172 | 0.850443089 | 0.838728045 | 0.822235505 | 0.836253107 | 0.839564047 | 0.853985066 | 0.835758834 | 0.881723513 | 0.806610098 | 0.850521569 | 0.843353804 | 0.837426279 |
| chr7:2046821-2046963 | 0.874564996 | 0.88697783 | 0.439750326 | 0.873313425 | 0.8885052 | 0.916962917 | 0.911920655 | 0.899960223 | 0.907654155 | 0.881172367 | 0.906209626 | 0.911635437 | 0.897543505 | 0.914638183 | 0.873592687 | 0.907098861 | 0.909121101 | 0.907187706 |
| chr7:2059796-2060179 | 0.612194382 | 0.695389962 | 0.408627183 | 0.723990665 | 0.627009824 | 0.692931769 | 0.867529151 | 0.848309644 | 0.876719531 | 0.822378844 | 0.870812998 | 0.840957835 | 0.853796075 | 0.845744211 | 0.81144419 | 0.846567535 | 0.852703654 | 0.858116263 |
| chr7:229225-229695 | 0.268028511 | 0.646500743 | 0.271435647 | 0.490294419 | 0.503581516 | 0.578784662 | 0.888188386 | 0.864971394 | 0.867068813 | 0.850291375 | 0.877190609 | 0.883650605 | 0.876794525 | 0.887719139 | 0.849663405 | 0.872379389 | 0.867201509 | 0.871383364 |
| chr7:2648222-2648343 | 0.866472995 | 0.807989039 | 0.882518186 | 0.844631396 | 0.788418667 | 0.884883299 | 0.883298689 | 0.828976432 | 0.865921321 | 0.792854104 | 0.897966545 | 0.883377198 | 0.840996038 | 0.867486319 | 0.838918159 | 0.843560925 | 0.839997229 | 0.882213077 |
| chr7:4755010-4755415 | 0.721491361 | 0.818462252 | 0.371566578 | 0.744373727 | 0.628692268 | 0.375049756 | 0.918283798 | 0.899669099 | 0.921011101 | 0.912648529 | 0.907829501 | 0.915367913 | 0.909401655 | 0.91973091 | 0.916233799 | 0.909198922 | 0.905822373 | 0.907341589 |
| chr7:4757669-4757929 | 0.834751204 | 0.796885158 | 0.238734776 | 0.809520284 | 0.798883921 | 0.823000416 | 0.802938491 | 0.761138966 | 0.809683654 | 0.718872077 | 0.833640729 | 0.752449928 | 0.799863737 | 0.718428622 | 0.785560599 | 0.769391567 | 0.809411067 | 0.803354291 |
| chr7:5534951-5535748 | 0.544136352 | 0.854590544 | 0.437460381 | 0.843655452 | 0.795975275 | 0.887609851 | 0.89052238 | 0.875121487 | 0.886957567 | 0.870340498 | 0.885927494 | 0.898624413 | 0.871828938 | 0.893938906 | 0.86414854 | 0.882392897 | 0.881345023 | 0.887261154 |
| chr7:630581-630914 | 0.536773907 | 0.61234447 | 0.331121938 | 0.414357959 | 0.319712568 | 0.578361815 | 0.897406431 | 0.883744823 | 0.888622871 | 0.875707248 | 0.893068145 | 0.897271014 | 0.883307823 | 0.900116411 | 0.864573365 | 0.896145585 | 0.88505102 | 0.899648154 |
| chr7:70252335-70252662 | 0.889055852 | 0.879630709 | 0.531100573 | 0.841342329 | 0.900506306 | 0.897188092 | 0.886763762 | 0.903978486 | 0.897146746 | 0.882555038 | 0.899691972 | 0.903685227 | 0.884993806 | 0.899099663 | 0.874862753 | 0.887886215 | 0.878289548 | 0.896521408 |
| chr7:818287-818362 | 0.605200516 | 0.790356984 | 0.393202092 | 0.716994805 | 0.719437217 | 0.840218348 | 0.835665101 | 0.820318779 | 0.845015357 | 0.814093052 | 0.833875243 | 0.844437887 | 0.820200743 | 0.8382499 | 0.801265366 | 0.824379713 | 0.810462 | 0.844842081 |
| chr8:144561585-144561662 | 0.506824963 | 0.681096502 | 0.681776879 | 0.718769761 | 0.57968186 | 0.755495304 | 0.834920881 | 0.779088298 | 0.845833888 | 0.800031772 | 0.840952376 | 0.856275465 | 0.836885139 | 0.869514203 | 0.80284688 | 0.778300314 | 0.835823936 | 0.868110059 |

| | | | | | | | | | | | | | | | | | | |
|---|---|---|---|---|---|---|---|---|---|---|---|---|---|---|---|---|---|---|
| chr8:145024929-145025128 | 0.191764711 | 0.319613763 | 0.187835223 | 0.228014794 | 0.268085015 | 0.257845575 | 0.875199796 | 0.830111971 | 0.838213422 | 0.838213422 | 0.864121903 | 0.88255019 | 0.859818781 | 0.875965406 | 0.846759368 | 0.855559484 | 0.862661828 | 0.849909077 |
| chr9:132636402-132636787 | 0.478391705 | 0.706820769 | 0.45712736 | 0.480586033 | 0.607665115 | 0.607665115 | 0.846215062 | 0.823489248 | 0.808229128 | 0.812322091 | 0.854458139 | 0.846334944 | 0.819112756 | 0.847486904 | 0.825505201 | 0.828719525 | 0.83730109 | 0.841507772 |

| Supplementary Table 10: Coordinates of MeVars identified in four body fluids (duplicate values of coordinates retained) | | | | |
|---|---|---|---|---|
| Sample and Body fluid | Coordinate | UCSC_REFGENE_NAME | RELATION_TO_UCSC_CPG_ISLAND | UCSC_REFGENE_GROUP |
| 20M01_Blood | chr1:110230252-110230456 | GSTM1;GSTM1 | Island | TSS200;TSS200 |
| 20M01_Blood | chr3:195488922-195490033 | MUC4;MUC4;MUC4 | Island | Body;Body;Body |
| 20M01_Blood | chr12:739980-740338 | NINJ2 | | Body |
| 20M01_Blood | chr17:79905236-79905263 | MYADML2 | | TSS200 |
| 20M01_Blood | chr19:55417390-55417647 | NCR1;NCR1;NCR1;NCR1;NCR1 | | TSS200;TSS200;TSS200;TSS200;TSS200 |
| 20M01_Blood | chr20:61590751-61591066 | SLC17A9 | N_Shore | Body |
| 20M01_Blood | chr2:113992843-113992930 | PAX8;PAX8;PAX8;PAX8;PAX8;LO | N_Shore | Body;Body;Body;Body;Body;Body;TSS1500 |
| 20M01_Blood | chr8:144635430-144635478 | GSDMD | Island | TSS200 |
| 20M01_Blood | chr8:144635610-144636025 | GSDMD;GSDMD | Island | 5'UTR;1stExon |
| 20M05_Blood | chr1:7887346-7887560 | PER3 | Island | Body |
| 20M05_Blood | chr1:202172778-202172912 | LGR6;LGR6 | | TSS200;Body |
| 20M05_Blood | chr6:32297670-32297822 | C6orf10 | | Body |
| 20M05_Blood | chr11:64115989-64116279 | CCDC88B | S_Shelf | Body |
| 20M05_Blood | chr16:1844927-1845460 | IGFALS;IGFALS;IGFALS | S_Shelf | TSS200;TSS1500;TSS1500 |
| 20M05_Blood | chr17:80708367-80708632 | FN3K | Island | Body |
| 20M05_Blood | chr19:21264896-21265421 | ZNF714 | N_Shore | TSS200 |
| 20M05_Blood | chr19:55476665-55477810 | NLRP2 | | TSS1500 |
| 20M05_Blood | chr22:24348549-24348715 | GSTTP1 | | TSS1500 |
| 20M05_Blood | chr5:179740743-179741120 | GFPT2 | Island | Body |
| 20M05_Blood | chr11:1413093-1413282 | BRSK2 | S_Shore | Body |
| 20M05_Blood | chr19:23941408-23941575 | ZNF681 | Island | Body |
| 20M06_Blood | chr1:110230252-110230456 | GSTM1;GSTM1 | Island | TSS200;TSS200 |
| 20M06_Blood | chr1:110254662-110255096 | GSTM5 | | TSS1500 |
| 20M06_Blood | chr1:245710332-245710401 | KIF26B | | Body |
| 20M06_Blood | chr1:248100600-248101009 | OR2L13;OR2L13 | Island | 1stExon;5'UTR |
| 20M06_Blood | chr2:113992762-113992930 | PAX8;PAX8;PAX8;PAX8;PAX8;LO | N_Shore | Body;Body;Body;Body;Body;Body;TSS1500 |
| 20M06_Blood | chr2:113993142-113993313 | PAX8;PAX8;PAX8;PAX8;PAX8;LO | N_Shore | Body;Body;Body;Body;Body;Body;TSS1500 |
| 20M06_Blood | chr5:135416205-135416613 | MIR886 | Island | Body |
| 20M06_Blood | chr6:32063774-32063901 | TNXB | Island | Body |
| 20M06_Blood | chr6:32064212-32064258 | TNXB | Island | Body |
| 20M06_Blood | chr6:74104388-74104868 | DDX43;DDX43 | N_Shore | 1stExon;5'UTR |
| 20M06_Blood | chr10:99338074-99338240 | ANKRD2;ANKRD2 | Island | Body;Body |
| 20M06_Blood | chr12:312592-313104 | SLC6A12;SLC6A12;SLC6A12 | Island | Body;Body;Body |
| 20M06_Blood | chr12:123215301-123215553 | GPR81 | | TSS200 |
| 20M06_Blood | chr17:11461244-11461572 | SHISA6 | Island | Body |
| 20M06_Blood | chr17:16593601-16593885 | CCDC144A | Island | TSS200 |
| 20M06_Blood | chr17:79905236-79905263 | MYADML2 | | TSS200 |
| 20M06_Blood | chr22:32438947-32439052 | SLC5A1 | N_Shore | TSS200 |
| 20M06_Blood | chr1:153599671-153599831 | S100A13;S100A13;S100A13;S100A13;S100A1;S100A13;S100A13 | | 5'UTR;1stExon;5'UTR;TSS200;TSS1500;5'UTR;5'UTR |
| 20M06_Blood | chr5:415575-415885 | AHRR | | Body |
| 20M06_Blood | chr5:1594282-1594579 | SDHAP3 | Island | Body |
| 20M06_Blood | chr5:1594715-1594808 | SDHAP3 | Island | TSS200 |
| 20M06_Blood | chr10:3138418-3138534 | PFKP | | Body |
| 20M06_Blood | chr10:134043618-134043654 | STK32C | S_Shore | Body |
| 20M06_Blood | chr19:36485282-36485360 | SDHAF1 | Island | TSS1500 |
| 20M06_Blood | chr19:55972791-55973338 | ISOC2;ISOC2;ISOC2 | Island | 5'UTR;5'UTR;5'UTR |
| 30M03_Blood | chr1:110230252-110230456 | GSTM1;GSTM1 | Island | TSS200;TSS200 |
| 30M03_Blood | chr2:139538601-139539001 | NXPH2 | Island | TSS1500 |
| 30M03_Blood | chr5:135416205-135416613 | MIR886 | Island | Body |
| 30M03_Blood | chr6:32552026-32552350 | HLA-DRB1 | Island | Body |
| 30M03_Blood | chr10:123355268-123356041 | FGFR2;FGFR2;FGFR2;FGFR2;FGF | N_Shelf | 5'UTR;5'UTR;TSS1500;5'UTR;TSS1500;5'UTR;5'UTR;TSS1500;5'UTR |
| 30M03_Blood | chr17:184833-185152 | RPH3AL | N_Shore | 5'UTR |
| 30M03_Blood | chr22:24348549-24348715 | GSTTP1 | | TSS1500 |
| 30M03_Blood | chr2:26785290-26785460 | C2orf70 | Island | TSS200 |
| 30M03_Blood | chr4:69435250-69435601 | UGT2B15;UGT2B17 | | TSS1500;TSS1500 |
| 30M03_Blood | chr5:415575-415885 | AHRR | | Body |
| 30M03_Blood | chr5:1594676-1594733 | SDHAP3 | Island | TSS200 |
| 30M03_Blood | chr6:28303629-28304108 | ZNF323;ZNF323;ZNF323;ZNF323;ZNF323;ZNF323;ZNF323 | | 1stExon;5'UTR;Body;5'UTR;5'UTR;5'UTR;Body |
| 30M03_Blood | chr6:28304185-28304249 | ZNF323;ZNF323;ZNF323;ZNF323;ZNF323;ZNF323 | | TSS1500;TSS200;5'UTR;TSS200;5'UTR;TSS1500 |
| 30M03_Blood | chr6:29894141-29894162 | HCG4P6 | Island | TSS1500 |

| | | | | |
|---|---|---|---|---|
| 30M03_Blood | chr6:32526021-32526260 | HLA-DRB6 | | Body |
| 30M03_Blood | chr8:101225028-101225902 | SPAG1;SPAG1 | N_Shore | Body;Body |
| 30M03_Blood | chr8:144635861-144636462 | GSDMD;GSDMD | Island | 5'UTR;1stExon |
| 30M03_Blood | chr19:12876846-12877188 | HOOK2;HOOK2 | N_Shore | Body;Body |
| 30M04_Blood | chr1:108734992-108735549 | SLC25A24;SLC25A24 | N_Shore | Body;Body |
| 30M04_Blood | chr5:135416381-135416529 | MIR886 | Island | TSS200 |
| 30M04_Blood | chr6:32489963-32490012 | HLA-DRB5 | Island | Body |
| 30M04_Blood | chr6:32525960-32526021 | HLA-DRB6 | | Body |
| 30M04_Blood | chr10:135341528-135341933 | CYP2E1 | N_Shore | 1stExon |
| 30M04_Blood | chr10:135343193-135343280 | CYP2E1 | S_Shore | Body |
| 30M04_Blood | chr16:1584118-1584516 | IFT140;TMEM204 | Island | Body;TSS200 |
| 30M04_Blood | chr19:50249584-50249776 | TSKS | Island | Body |
| 30M04_Blood | chr22:24348549-24348715 | GSTTP1 | | TSS1500 |
| 30M04_Blood | chr1:110254662-110254835 | GSTM5 | | TSS1500 |
| 30M04_Blood | chr2:3642629-3642732 | COLEC11;COLEC11 | Island | TSS200;TSS200 |
| 30M04_Blood | chr4:69435250-69435601 | UGT2B15;UGT2B17 | | TSS1500;TSS1500 |
| 30M04_Blood | chr5:1594282-1594676 | SDHAP3 | Island | Body |
| 30M04_Blood | chr6:29856426-29856477 | HLA-H | Island | Body |
| 30M04_Blood | chr6:29894050-29894197 | HCG4P6 | N_Shore | TSS1500 |
| 30M04_Blood | chr6:31237013-31237034 | HLA-C | N_Shore | Body |
| 30M04_Blood | chr6:31237405-31237722 | HLA-C | N_Shore | Body |
| 30M04_Blood | chr10:134045514-134045609 | STK32C | S_Shelf | Body |
| 30M04_Blood | chr20:61371425-61371808 | NTSR1 | | Body |
| 30M04_Blood | chr22:24384397-24384525 | GSTT1 | Island | TSS200 |
| 30M05_Blood | chr2:48844892-48844916 | GTF2A1L;STON1-GTF2A1L;GTF2/ | Island | TSS200;Body;TSS200 |
| 30M05_Blood | chr6:32063874-32063901 | TNXB | Island | Body |
| 30M05_Blood | chr7:27183375-27183436 | HOXA5 | Island | TSS200 |
| 30M05_Blood | chr4:940644-941054 | TMEM175 | Island | 5'UTR |
| 30M05_Blood | chr4:69435250-69435601 | UGT2B15;UGT2B17 | | TSS1500;TSS1500 |
| 30M05_Blood | chr6:29856426-29856477 | HLA-H | Island | Body |
| 30M05_Blood | chr19:23941408-23941575 | ZNF681 | Island | Body |
| 40M05_Blood | chr1:3105151-3105252 | PRDM16;PRDM16 | Island | Body;Body |
| 40M05_Blood | chr4:140429869-140429995 | SETD7 | | 3'UTR |
| 40M05_Blood | chr5:415575-415885 | AHRR | | Body |
| 40M05_Blood | chr5:1594282-1594579 | SDHAP3 | Island | Body |
| 40M05_Blood | chr5:135416205-135416613 | MIR886 | Island | Body |
| 40M05_Blood | chr6:291687-293331 | DUSP22 | N_Shore | TSS1500 |
| 40M05_Blood | chr6:32551749-32552350 | HLA-DRB1 | N_Shore | Body |
| 40M05_Blood | chr6:32557321-32557422 | HLA-DRB1 | | Body |
| 40M05_Blood | chr17:79905236-79905263 | MYADML2 | | TSS200 |
| 40M05_Blood | chr4:165878055-165878219 | C4orf39;TRIM61 | Island | TSS200;Body |
| 40M05_Blood | chr6:32634316-32634362 | HLA-DQB1 | S_Shore | 1stExon |
| 40M05_Blood | chr8:144636025-144636462 | GSDMD | Island | 5'UTR |
| 40M05_Blood | chr11:1413093-1413282 | BRSK2 | S_Shore | Body |
| 40M05_Blood | chr17:33760230-33760293 | SLFN12 | | TSS1500 |
| 40M05_Blood | chr19:55972855-55973099 | ISOC2;ISOC2;ISOC2 | Island | 5'UTR;5'UTR;5'UTR |
| 40M05_Blood | chr19:55973234-55973338 | ISOC2;ISOC2;ISOC2 | Island | TSS200;TSS200;TSS200 |
| 40M05_Blood | chr20:42544728-42544794 | TOX2;TOX2;TOX2 | Island | TSS200;Body;TSS200 |
| 40M05_Blood | chr22:24348549-24348715 | GSTTP1 | | TSS1500 |
| 40M07_Blood | chr1:108734992-108735549 | SLC25A24;SLC25A24 | N_Shore | Body;Body |
| 40M07_Blood | chr2:113992843-113992930 | PAX8;PAX8;PAX8;PAX8;PAX8;LO( | N_Shore | Body;Body;Body;Body;Body;Body;TSS1500 |
| 40M07_Blood | chr7:158046025-158046222 | PTPRN2;PTPRN2;PTPRN2 | | Body;Body;Body |
| 40M07_Blood | chr12:131622584-131622739 | GPR133 | N_Shore | Body |
| 40M07_Blood | chr17:77706946-77707066 | ENPP7 | N_Shore | Body |
| 40M07_Blood | chr20:62084549-62084833 | KCNQ2;KCNQ2;KCNQ2;KCNQ2; | Island | Body;Body;Body;Body;Body |
| 40M07_Blood | chr22:24348549-24348715 | GSTTP1 | | TSS1500 |
| 40M07_Blood | chr1:248100228-248101009 | OR2L13 | N_Shore | TSS1500 |
| 40M07_Blood | chr5:135416529-135416613 | MIR886 | S_Shore | TSS1500 |
| 40M07_Blood | chr6:31803718-31804461 | C6orf48;C6orf48;SNORD52 | S_Shore | 5'UTR;5'UTR;TSS1500 |
| 40M07_Blood | chr7:91763989-91764605 | CYP51A1;CYP51A1;CYP51A1 | Island | 5'UTR;TSS200;1stExon |
| 40M07_Blood | chr8:81599484-81599569 | ZNF704 | | Body |
| 40M07_Blood | chr12:739980-740338 | NINJ2 | | Body |
| 40M07_Blood | chr15:91473475-91473569 | UNC45A;UNC45A | Island | 5'UTR;1stExon |
| 40M10_Blood | chr5:1594282-1594579 | SDHAP3 | Island | Body |
| 40M10_Blood | chr6:291687-293331 | DUSP22 | N_Shore | TSS1500 |
| 40M10_Blood | chr21:36259067-36259383 | RUNX1;RUNX1;RUNX1 | Island | Body;Body;Body |

| | | | | |
|---|---|---|---|---|
| 40M10_Blood | chr4:103940711-103940923 | NHEDC1;NHEDC1 | Island | 5'UTR;5'UTR |
| 40M10_Blood | chr4:103941189-103941300 | NHEDC1;NHEDC1 | S_Shore | TSS1500;TSS1500 |
| 40M10_Blood | chr5:178986291-178986412 | RUFY1;RUFY1;RUFY1 | N_Shore | TSS1500;Body;TSS1500 |
| 40M10_Blood | chr6:30039130-30039206 | RNF39;RNF39 | Island | Body;Body |
| 40M10_Blood | chr6:30039403-30039476 | RNF39;RNF39 | Island | Body;Body |
| 40M10_Blood | chr6:32063895-32063991 | TNXB | Island | Body |
| 40M10_Blood | chr9:140916980-140917431 | CACNA1B | N_Shore | Body |
| 40M10_Blood | chr10:135341528-135341933 | CYP2E1 | N_Shore | 1stExon |
| 40M10_Blood | chr10:135342560-135343193 | CYP2E1 | Island | Body |
| 40M10_Blood | chr19:17905589-17905626 | B3GNT3 | Island | TSS1500 |
| 40M10_Blood | chr19:38886110-38886658 | SPRED3 | Island | Body |
| 40M10_Blood | chr20:62327968-62328094 | TNFRSF6B;TNFRSF6B | N_Shore | 5'UTR;TSS200 |
| 40M10_Blood | chr22:24348549-24348715 | GSTTP1 | | TSS1500 |
| 50M04_Blood | chr6:29911542-29911558 | HLA-A | S_Shore | Body |
| 50M04_Blood | chr6:32551891-32552350 | HLA-DRB1 | Island | Body |
| 50M04_Blood | chr6:33048086-33048286 | HLA-DPB1 | N_Shore | Body |
| 50M04_Blood | chr6:33048555-33048760 | HLA-DPB1 | Island | Body |
| 50M04_Blood | chr4:69435250-69435601 | UGT2B15;UGT2B17 | | TSS1500;TSS1500 |
| 50M04_Blood | chr5:415575-415885 | AHRR | | Body |
| 50M04_Blood | chr6:31237013-31237034 | HLA-C | N_Shore | Body |
| 50M04_Blood | chr6:31237405-31237722 | HLA-C | N_Shore | Body |
| 50M04_Blood | chr6:32490012-32490350 | HLA-DRB5 | Island | Body |
| 50M04_Blood | chr8:144635430-144635547 | GSDMD | Island | TSS200 |
| 50M04_Blood | chr8:144636025-144636462 | GSDMD | Island | 5'UTR |
| 50M01_Blood | chr6:29645937-29648756 | ZFP57 | | TSS1500 |
| 50M01_Blood | chr7:157369895-157369960 | PTPRN2;PTPRN2;PTPRN2 | S_Shore | Body;Body;Body |
| 50M01_Blood | chr17:184833-185152 | RPH3AL | N_Shore | 5'UTR |
| 50M01_Blood | chr17:17109936-17110207 | PLD6 | Island | TSS1500 |
| 50M01_Blood | chr20:61371425-61371808 | NTSR1 | | Body |
| 50M01_Blood | chr22:24348549-24348715 | GSTTP1 | | TSS1500 |
| 50M01_Blood | chr6:29856278-29856294 | HLA-H | Island | Body |
| 50M01_Blood | chr6:29894619-29894679 | HCG4P6 | Island | TSS1500 |
| 50M01_Blood | chr9:126135610-126135998 | CRB2 | Island | Body |
| 50M01_Blood | chr11:1413093-1413282 | BRSK2 | S_Shore | Body |
| 50M06_Blood | chr1:108734992-108735549 | SLC25A24;SLC25A24 | N_Shore | Body;Body |
| 50M06_Blood | chr3:11746226-11746722 | VGLL4 | N_Shore | 5'UTR |
| 50M06_Blood | chr5:1594282-1594579 | SDHAP3 | Island | Body |
| 50M06_Blood | chr5:135416205-135416613 | MIR886 | Island | Body |
| 50M06_Blood | chr10:123355268-123356041 | FGFR2;FGFR2;FGFR2;FGFR2;FGF | N_Shelf | 5'UTR;5'UTR;TSS1500;5'UTR;TSS1500;5'UTR;5'UTR;TSS1500;5'UTR |
| 50M06_Blood | chr10:135341528-135342218 | CYP2E1 | N_Shore | 1stExon |
| 50M06_Blood | chr11:1413145-1413315 | BRSK2 | S_Shore | Body |
| 50M06_Blood | chr16:83171068-83171314 | CDH13 | | Body |
| 50M06_Blood | chr19:57742345-57742444 | AURKC;AURKC;AURKC | Island | TSS200;TSS200;TSS200 |
| 50M06_Blood | chr20:61371425-61371808 | NTSR1 | | Body |
| 50M06_Blood | chr2:3642629-3642732 | COLEC11;COLEC11 | Island | TSS200;TSS200 |
| 50M06_Blood | chr2:61372138-61372256 | C2orf74 | | TSS200 |
| 50M06_Blood | chr5:179740743-179741120 | GFPT2 | Island | Body |
| 50M06_Blood | chr6:30039130-30039206 | RNF39;RNF39 | Island | Body;Body |
| 50M06_Blood | chr6:30039374-30039476 | RNF39;RNF39 | Island | Body;Body |
| 50M06_Blood | chr6:160182438-160182554 | ACAT2 | Island | TSS1500 |
| 50M06_Blood | chr11:1463554-1463662 | BRSK2 | N_Shore | Body |
| 50M06_Blood | chr20:5485268-5485294 | LOC149837 | | TSS200 |
| 50M06_Blood | chr20:25129507-25129542 | LOC284798;LOC284798;LOC28 | Island | TSS1500;TSS200;TSS200 |
| 50M06_Blood | chr22:24348549-24348715 | GSTTP1 | | TSS1500 |
| 20M02_Saliva | chr1:108735228-108735549 | SLC25A24;SLC25A24 | Island | Body;1stExon |
| 20M02_Saliva | chr10:135341528-135342218 | CYP2E1 | N_Shore | 1stExon |
| 20M02_Saliva | chr5:176797920-176798049 | RGS14 | S_Shelf | Body |
| 20M02_Saliva | chr6:32552039-32552095 | HLA-DRB1 | Island | Body |
| 20M02_Saliva | chr12:131622584-131622739 | GPR133 | N_Shore | Body |
| 20M02_Saliva | chr14:104552032-104552209 | ASPG | Island | TSS200 |
| 20M02_Saliva | chr19:12876846-12877188 | HOOK2;HOOK2 | N_Shore | Body;Body |
| 20M03_Saliva | chr1:854918-854966 | FLJ39609 | Island | TSS200 |
| 20M03_Saliva | chr1:855425-856059 | FLJ39609 | S_Shore | TSS1500 |
| 20M03_Saliva | chr1:871441-871980 | SAMD11 | Island | Body |
| 20M03_Saliva | chr1:1245014-1245184 | PUSL1 | Island | Body |

| | | | | |
|---|---|---|---|---|
| 20M03_Saliva | chr1:1266178-1267117 | TAS1R3 | N_Shore | TSS1500 |
| 20M03_Saliva | chr1:1268144-1268942 | TAS1R3 | Island | Body |
| 20M03_Saliva | chr1:1562771-1563096 | MIB2;MIB2;MIB2;MIB2;MIB2;MIB | N_Shore | Body;Body;Body;Body;Body;Body |
| 20M03_Saliva | chr1:1564422-1564482 | MIB2;MIB2;MIB2;MIB2;MIB2;MIB | Island | Body;Body;Body;Body;Body;Body |
| 20M03_Saliva | chr1:1602585-1602685 | LOC728661;CDK11B;CDK11B;C | Island | Body;Body;Body;Body;Body;Body;5'UTR |
| 20M03_Saliva | chr1:1845940-1846340 | CALML6 | | TSS1500 |
| 20M03_Saliva | chr1:2138757-2139657 | C1orf86 | S_Shore | 5'UTR |
| 20M03_Saliva | chr1:2170956-2171253 | SKI | | Body |
| 20M03_Saliva | chr1:2230601-2230923 | SKI | N_Shore | Body |
| 20M03_Saliva | chr1:2262232-2262474 | MORN1 | N_Shelf | Body |
| 20M03_Saliva | chr1:2265881-2266591 | MORN1 | N_Shore | Body |
| 20M03_Saliva | chr1:2284167-2284821 | MORN1;LOC100129534 | | Body;TSS200 |
| 20M03_Saliva | chr1:2979311-2980937 | FLJ42875 | Island | Body |
| 20M03_Saliva | chr1:3028230-3028485 | PRDM16;PRDM16 | Island | Body;Body |
| 20M03_Saliva | chr1:3056542-3056679 | PRDM16;PRDM16 | Island | Body;Body |
| 20M03_Saliva | chr1:3160739-3162173 | PRDM16;PRDM16 | N_Shelf | Body;Body |
| 20M03_Saliva | chr1:3163341-3163545 | PRDM16;PRDM16 | N_Shore | Body;Body |
| 20M03_Saliva | chr1:3276413-3276535 | PRDM16;PRDM16 | S_Shore | Body;Body |
| 20M03_Saliva | chr1:3321869-3322309 | PRDM16;PRDM16 | Island | Body;Body |
| 20M03_Saliva | chr1:3332188-3332705 | PRDM16;PRDM16 | Island | Body;Body |
| 20M03_Saliva | chr1:3473665-3474202 | MEGF6 | | Body |
| 20M03_Saliva | chr1:3606892-3607425 | TP73;TP73;TP73;TP73 | N_Shore | TSS1500;TSS1500;Body;TSS1500 |
| 20M03_Saliva | chr1:6390473-6390830 | ACOT7;ACOT7;ACOT7;ACOT7 | | Body;Body;Body;Body |
| 20M03_Saliva | chr1:6445632-6446787 | ACOT7;ACOT7 | | Body;1stExon |
| 20M03_Saliva | chr1:6531717-6531813 | PLEKHG5;PLEKHG5;PLEKHG5;PL | Island | Body;Body;Body;Body;Body |
| 20M03_Saliva | chr1:6579837-6580525 | PLEKHG5;PLEKHG5 | | 5'UTR;1stExon |
| 20M03_Saliva | chr1:10707917-10708003 | CASZ1;CASZ1 | Island | Body;Body |
| 20M03_Saliva | chr1:10757892-10757954 | CASZ1;CASZ1 | Island | 5'UTR;5'UTR |
| 20M03_Saliva | chr1:10839450-10839574 | CASZ1;CASZ1 | | 5'UTR;5'UTR |
| 20M03_Saliva | chr1:11713123-11714254 | FBXO2;FBXO44 | N_Shore | Body;TSS1500 |
| 20M03_Saliva | chr1:15255847-15256481 | KIAA1026;KIAA1026;KIAA1026;KIAA1026 | | Body;TSS1500;Body;Body |
| 20M03_Saliva | chr1:15271830-15272326 | KIAA1026;KIAA1026;KIAA1026;KIAA1026;KIAA1026 | | Body;5'UTR;TSS1500;Body;Body |
| 20M03_Saliva | chr1:16061408-16061554 | SLC25A34 | S_Shelf | TSS1500 |
| 20M03_Saliva | chr1:16062720-16062757 | SLC25A34 | | TSS200 |
| 20M03_Saliva | chr1:16539013-16539119 | ARHGEF19;ARHGEF19 | | 5'UTR;1stExon |
| 20M03_Saliva | chr1:17381800-17381980 | SDHB | S_Shore | TSS1500 |
| 20M03_Saliva | chr1:17575420-17575440 | PADI3 | | TSS200 |
| 20M03_Saliva | chr1:20141662-20141781 | RNF186;RNF186 | | 5'UTR;1stExon |
| 20M03_Saliva | chr1:23697258-23698143 | C1orf213;C1orf213;C1orf213;ZN | S_Shore | 3'UTR;Body;1stExon;TSS1500 |
| 20M03_Saliva | chr1:23953480-23953778 | MDS2 | | TSS1500 |
| 20M03_Saliva | chr1:24469564-24470205 | IL22RA1 | | 1stExon |
| 20M03_Saliva | chr1:25254088-25254149 | RUNX3;RUNX3 | N_Shore | Body;Body |
| 20M03_Saliva | chr1:25257505-25257566 | RUNX3;RUNX3 | Island | TSS1500;Body |
| 20M03_Saliva | chr1:25257759-25258332 | RUNX3;RUNX3 | Island | TSS1500;Body |
| 20M03_Saliva | chr1:26503776-26504548 | CNKSR1;CNKSR1 | | TSS1500;TSS1500 |
| 20M03_Saliva | chr1:27676205-27676652 | SYTL1 | Island | Body |
| 20M03_Saliva | chr1:27683139-27683501 | MAP3K6 | N_Shore | Body |
| 20M03_Saliva | chr1:28843736-28844028 | SNHG3-RCC1;RCC1;SNHG3-RC | N_Shore | Body;TSS1500;Body;TSS1500;5'UTR;TSS1500 |
| 20M03_Saliva | chr1:35259746-35259937 | GJA4 | S_Shore | 5'UTR |
| 20M03_Saliva | chr1:36771714-36771793 | C1orf113;C1orf113 | N_Shore | TSS1500;TSS1500 |
| 20M03_Saliva | chr1:39407451-39407651 | RHBDL2;RHBDL2 | | 1stExon;5'UTR |
| 20M03_Saliva | chr1:40349894-40350052 | TRIT1 | S_Shore | TSS1500 |
| 20M03_Saliva | chr1:43751270-43751563 | C1orf210;C1orf210;C1orf210 | | 1stExon;5'UTR;TSS200 |
| 20M03_Saliva | chr1:43814169-43815035 | MPL | N_Shore | Body |
| 20M03_Saliva | chr1:44726943-44727313 | ERI3 | | Body |
| 20M03_Saliva | chr1:45082704-45083278 | RNF220 | N_Shore | Body |
| 20M03_Saliva | chr1:46089693-46090169 | CCDC17;CCDC17 | S_Shore | 5'UTR;1stExon |
| 20M03_Saliva | chr1:46645968-46646224 | TSPAN1 | | 5'UTR |
| 20M03_Saliva | chr1:46859671-46859774 | FAAH | N_Shore | TSS1500 |
| 20M03_Saliva | chr1:55089182-55089959 | ACOT11;FAM151A;FAM151A | | Body;1stExon;5'UTR |
| 20M03_Saliva | chr1:59043208-59043280 | TACSTD2 | Island | TSS200 |
| 20M03_Saliva | chr1:78277374-78277606 | FAM73A | Island | Body |
| 20M03_Saliva | chr1:86934325-86934607 | CLCA1 | | TSS200 |
| 20M03_Saliva | chr1:89357908-89357988 | GTF2B | S_Shore | TSS1500 |

| | | | | |
|---|---|---|---|---|
| 20M03_Saliva | chr1:110035670-110036205 | CYB561D1;CYB561D1;CYB561D | N_Shore | TSS1500;TSS1500;TSS1500;TSS1500;TSS1500 |
| 20M03_Saliva | chr1:117214010-117214228 | MIR320B1 | S_Shelf | TSS1500 |
| 20M03_Saliva | chr1:117753591-117753741 | VTCN1 | | TSS200 |
| 20M03_Saliva | chr1:120311542-120311653 | HMGCS2;HMGCS2;HMGCS2;HMGCS2 | | 1stExon;5'UTR;5'UTR;1stExon |
| 20M03_Saliva | chr1:149908085-149908313 | MTMR11;MTMR11;MTMR11 | | Body;5'UTR;1stExon |
| 20M03_Saliva | chr1:151345123-151345771 | SELENBP1;SELENBP1 | | 5'UTR;1stExon |
| 20M03_Saliva | chr1:151826123-151826533 | THEM5;THEM5 | | 1stExon;5'UTR |
| 20M03_Saliva | chr1:152730231-152730426 | KPRP | N_Shelf | TSS1500 |
| 20M03_Saliva | chr1:152747472-152747562 | LCE1F | | TSS1500 |
| 20M03_Saliva | chr1:152973720-152973957 | SPRR3;SPRR3 | | TSS1500;TSS1500 |
| 20M03_Saliva | chr1:153413871-153413940 | S100A7L2 | | TSS1500 |
| 20M03_Saliva | chr1:153536278-153536563 | S100A2 | | Body |
| 20M03_Saliva | chr1:154929734-154929791 | PBXIP1;PYGO2 | | TSS1500;3'UTR |
| 20M03_Saliva | chr1:155007166-155007323 | DCST1;DCST2;DCST1 | | Body;TSS1500;Body |
| 20M03_Saliva | chr1:156123352-156123500 | SEMA4A | | TSS200 |
| 20M03_Saliva | chr1:156217178-156218009 | PAQR6;PAQR6 | S_Shore | 5'UTR;5'UTR |
| 20M03_Saliva | chr1:161059267-161059654 | PVRL4;PVRL4 | | 5'UTR;1stExon |
| 20M03_Saliva | chr1:161088957-161089118 | PFDN2;NIT1 | S_Shore | TSS1500;Body |
| 20M03_Saliva | chr1:161168856-161169007 | NDUFS2;ADAMTS4 | N_Shelf | TSS1500;TSS200 |
| 20M03_Saliva | chr1:162838743-162838907 | C1orf110 | | TSS200 |
| 20M03_Saliva | chr1:175162550-175163045 | KIAA0040;KIAA0040;KIAA0040;K | S_Shore | TSS1500;TSS1500;TSS1500;TSS1500 |
| 20M03_Saliva | chr1:183154778-183155056 | LAMC2;LAMC2 | | TSS1500;TSS1500 |
| 20M03_Saliva | chr1:201979478-201979938 | ELF3;ELF3 | | TSS1500;TSS1500 |
| 20M03_Saliva | chr1:202182417-202182642 | LGR6;LGR6;LGR6 | N_Shore | TSS1500;Body;Body |
| 20M03_Saliva | chr1:203829775-203830200 | SNRPE | N_Shore | TSS1500 |
| 20M03_Saliva | chr1:206643301-206643377 | IKBKE | | TSS1500 |
| 20M03_Saliva | chr1:207224740-207225128 | YOD1 | N_Shore | TSS1500 |
| 20M03_Saliva | chr1:207992403-207992548 | LOC148696 | | Body |
| 20M03_Saliva | chr1:209825672-209825856 | LAMB3;LAMB3;LAMB3;LAMB3 | | TSS1500;5'UTR;1stExon;5'UTR |
| 20M03_Saliva | chr1:210501368-210501634 | HHAT;HHAT;HHAT;HHAT;HHAT;H | N_Shore | TSS1500;TSS1500;TSS1500;TSS1500;TSS1500;TSS1500 |
| 20M03_Saliva | chr1:215178907-215179216 | KCNK2;KCNK2 | | 1stExon;5'UTR |
| 20M03_Saliva | chr1:221057558-221057808 | HLX | Island | Body |
| 20M03_Saliva | chr1:228337181-228337379 | GJC2 | | TSS1500 |
| 20M03_Saliva | chr1:228394509-228394607 | OBSCN;OBSCN | | TSS1500;TSS1500 |
| 20M03_Saliva | chr1:234350834-234350988 | SLC35F3 | S_Shore | Body |
| 20M03_Saliva | chr1:236686564-236686618 | LGALS8;LGALS8;LGALS8;LGALS8 | N_Shore | TSS200;TSS200;5'UTR;TSS1500 |
| 20M03_Saliva | chr1:244615947-244616227 | ADSS | S_Shore | TSS1500 |
| 20M03_Saliva | chr1:247611842-247611931 | NLRP3;NLRP3;NLRP3;NLRP3;NLI | N_Shelf | 3'UTR;3'UTR;3'UTR;3'UTR;3'UTR |
| 20M03_Saliva | chr2:1358684-1358730 | SNTG2 | | Body |
| 20M03_Saliva | chr2:1417153-1417248 | TPO;TPO;TPO;TPO | | TSS1500;TSS200;TSS200;TSS1500 |
| 20M03_Saliva | chr2:1926671-1927209 | MYT1L | | Body |
| 20M03_Saliva | chr2:3452407-3452563 | TTC15 | Island | Body |
| 20M03_Saliva | chr2:3683030-3683283 | COLEC11;COLEC11 | Island | Body;Body |
| 20M03_Saliva | chr2:9459368-9459596 | ASAP2;ASAP2 | Island | Body;Body |
| 20M03_Saliva | chr2:24398427-24398859 | C2orf84 | Island | Body |
| 20M03_Saliva | chr2:27530829-27531535 | UCN | Island | 5'UTR |
| 20M03_Saliva | chr2:27665141-27665711 | KRTCAP3;KRTCAP3 | Island | TSS200;TSS200 |
| 20M03_Saliva | chr2:54086854-54087469 | GPR75;LOC100302652 | Island | 5'UTR;Body |
| 20M03_Saliva | chr2:72359383-72359612 | CYP26B1 | | Body |
| 20M03_Saliva | chr2:73215621-73215650 | SFXN5 | | Body |
| 20M03_Saliva | chr2:74153086-74153317 | DGUOK;DGUOK | N_Shore | TSS1500;TSS1500 |
| 20M03_Saliva | chr2:74668072-74668477 | RTKN;RTKN;RTKN | Island | Body;TSS1500;TSS1500 |
| 20M03_Saliva | chr2:74753281-74753358 | DQX1;DQX1 | N_Shelf | 1stExon;5'UTR |
| 20M03_Saliva | chr2:91847976-91847996 | LOC654342 | | TSS200 |
| 20M03_Saliva | chr2:95940031-95940705 | PROM2;PROM2;PROM2 | | TSS200;TSS200;TSS200 |
| 20M03_Saliva | chr2:97171137-97171243 | NEURL3 | N_Shelf | Body |
| 20M03_Saliva | chr2:102803290-102803787 | IL1RL2 | N_Shore | TSS200 |
| 20M03_Saliva | chr2:105857999-105858015 | GPR45 | N_Shore | TSS200 |
| 20M03_Saliva | chr2:106761673-106761805 | UXS1 | | Body |
| 20M03_Saliva | chr2:109934728-109934860 | SH3RF3 | | Body |
| 20M03_Saliva | chr2:113875226-113875723 | IL1RN;IL1RN;IL1RN | | TSS1500;TSS1500;TSS1500 |
| 20M03_Saliva | chr2:121554817-121554926 | GLI2 | | TSS200 |
| 20M03_Saliva | chr2:128458240-128458401 | SFT2D3 | N_Shore | TSS1500 |
| 20M03_Saliva | chr2:131975750-131975771 | POTEE | | TSS200 |

| | | | | |
|---|---|---|---|---|
| 20M03_Saliva | chr2:134326204-134326366 | NCKAP5;NCKAP5 | | TSS200;TSS200 |
| 20M03_Saliva | chr2:144994885-144995072 | GTDC1;GTDC1;GTDC1;GTDC1 | | 5'UTR;5'UTR;5'UTR;1stExon |
| 20M03_Saliva | chr2:145146928-145147350 | ZEB2;ZEB2 | N_Shore | 3'UTR;3'UTR |
| 20M03_Saliva | chr2:152830189-152830521 | CACNB4;CACNB4;CACNB4;CACNB4 | | Body;Body;1stExon;Body |
| 20M03_Saliva | chr2:161056520-161056639 | ITGB6 | | 1stExon |
| 20M03_Saliva | chr2:173330296-173330395 | ITGA6;ITGA6 | | Body;Body |
| 20M03_Saliva | chr2:201170198-201170426 | SPATS2L;SPATS2L;SPATS2L;SPAT | N_Shore | TSS1500;TSS1500;TSS1500;TSS1500 |
| 20M03_Saliva | chr2:219472218-219473002 | PLCD4 | | TSS1500 |
| 20M03_Saliva | chr2:220253653-220253668 | DNPEP | S_Shore | TSS1500 |
| 20M03_Saliva | chr2:220282165-220282281 | DES | N_Shore | TSS1500 |
| 20M03_Saliva | chr2:223916502-223916687 | KCNE4 | N_Shore | TSS1500 |
| 20M03_Saliva | chr2:231790529-231790813 | GPR55 | | TSS1500 |
| 20M03_Saliva | chr2:234526077-234526295 | UGT1A8 | | TSS1500 |
| 20M03_Saliva | chr2:236443439-236444259 | AGAP1;AGAP1 | N_Shore | Body;Body |
| 20M03_Saliva | chr2:236826207-236826311 | AGAP1;AGAP1 | | Body;Body |
| 20M03_Saliva | chr2:238499464-238499802 | RAB17;RAB17 | | 1stExon;5'UTR |
| 20M03_Saliva | chr2:240020028-240020174 | HDAC4 | | Body |
| 20M03_Saliva | chr2:240212763-240213173 | HDAC4 | | Body |
| 20M03_Saliva | chr2:240230587-240230892 | HDAC4 | | Body |
| 20M03_Saliva | chr2:241389021-241389208 | PP14571;GPC1 | N_Shelf | Body;Body |
| 20M03_Saliva | chr2:241719496-241719655 | KIF1A | Island | Body |
| 20M03_Saliva | chr2:241835670-241836501 | C2orf54 | | TSS200 |
| 20M03_Saliva | chr2:241974917-241975140 | SNED1 | Island | Body |
| 20M03_Saliva | chr2:242127690-242127948 | ANO7;ANO7 | | TSS1500;TSS1500 |
| 20M03_Saliva | chr2:242499464-242499542 | BOK | S_Shore | Body |
| 20M03_Saliva | chr2:242715019-242715041 | GAL3ST2 | S_Shelf | TSS1500 |
| 20M03_Saliva | chr2:242716559-242716801 | GAL3ST2 | | Body |
| 20M03_Saliva | chr2:242802009-242802192 | PDCD1 | N_Shelf | TSS1500 |
| 20M03_Saliva | chr2:242810748-242811149 | C2orf85 | S_Shelf | TSS1500 |
| 20M03_Saliva | chr3:1134528-1134730 | CNTN6 | | TSS200 |
| 20M03_Saliva | chr3:11267020-11267098 | HRH1;HRH1;HRH1 | | 5'UTR;TSS1500;5'UTR |
| 20M03_Saliva | chr3:11610239-11610595 | VGLL4;VGLL4;VGLL4;VGLL4;VGL | Island | Body;Body;1stExon;Body;5'UTR |
| 20M03_Saliva | chr3:11763045-11763340 | VGLL4 | S_Shore | TSS1500 |
| 20M03_Saliva | chr3:12800802-12801112 | TMEM40;TMEM40 | | 1stExon;5'UTR |
| 20M03_Saliva | chr3:13028573-13029200 | IQSEC1 | | Body |
| 20M03_Saliva | chr3:13060741-13060974 | IQSEC1 | | Body |
| 20M03_Saliva | chr3:14443428-14443473 | SLC6A6;SLC6A6;SLC6A6 | N_Shore | TSS1500;TSS1500;TSS1500 |
| 20M03_Saliva | chr3:16215315-16216215 | GALNTL2 | | TSS1500 |
| 20M03_Saliva | chr3:32023475-32023996 | ZNF860;OSBPL10 | S_Shore | 5'UTR;TSS1500 |
| 20M03_Saliva | chr3:36902601-36903038 | TRANK1 | | Body |
| 20M03_Saliva | chr3:50341009-50341293 | HYAL1;HYAL1;HYAL1;HYAL1;HYAL1;HYAL1 | | TSS200;TSS200;5'UTR;TSS200;TSS200;TSS200 |
| 20M03_Saliva | chr3:51740896-51741473 | GRM2;GRM2 | Island | TSS200;TSS200 |
| 20M03_Saliva | chr3:52552868-52553433 | STAB1 | Island | Body |
| 20M03_Saliva | chr3:52569026-52569169 | NT5DC2;NT5DC2;NT5DC2 | N_Shore | 5'UTR;1stExon;TSS1500 |
| 20M03_Saliva | chr3:58563618-58563674 | FAM107A;FAM107A | | TSS200;TSS200 |
| 20M03_Saliva | chr3:69061356-69061827 | C3orf64 | N_Shore | 5'UTR |
| 20M03_Saliva | chr3:122785328-122785650 | PDIA5;PDIA5 | N_Shore | TSS1500;TSS1500 |
| 20M03_Saliva | chr3:124102766-124103696 | KALRN;KALRN | | Body;Body |
| 20M03_Saliva | chr3:128226499-128226649 | LOC90246 | | TSS200 |
| 20M03_Saliva | chr3:146213648-146213845 | PLSCR2 | | 5'UTR |
| 20M03_Saliva | chr3:149374535-149374763 | WWTR1;WWTR1;WWTR1 | N_Shore | Body;Body;Body |
| 20M03_Saliva | chr3:150690251-150690403 | CLRN1;CLRN1OS | | 1stExon;TSS1500 |
| 20M03_Saliva | chr3:183959000-183959062 | VWA5B2;MIR1224 | N_Shore | Body;TSS200 |
| 20M03_Saliva | chr3:183959088-183959853 | VWA5B2;MIR1224 | Island | Body;TSS200 |
| 20M03_Saliva | chr3:184056553-184056843 | FAM131A;FAM131A | Island | 5'UTR;Body |
| 20M03_Saliva | chr3:191048308-191049285 | UTS2D;CCDC50;CCDC50;UTS2D | S_Shore | 1stExon;Body;Body;5'UTR |
| 20M03_Saliva | chr3:194833431-194834008 | C3orf21 | | Body |
| 20M03_Saliva | chr3:195942717-195943713 | OSTalpha | | TSS1500 |
| 20M03_Saliva | chr3:196065106-196065357 | TM4SF19 | | 5'UTR |
| 20M03_Saliva | chr3:196065506-196065688 | TM4SF19 | | TSS1500 |
| 20M03_Saliva | chr4:645256-645551 | PDE6B;PDE6B | Island | Body;Body |
| 20M03_Saliva | chr4:671543-671851 | MYL5 | N_Shelf | TSS200 |
| 20M03_Saliva | chr4:715865-715950 | PCGF3 | N_Shore | 5'UTR |
| 20M03_Saliva | chr4:981485-981630 | IDUA;SLC26A1;SLC26A1;SLC26A | Island | Body;3'UTR;Body;3'UTR |
| 20M03_Saliva | chr4:1166839-1166933 | LOC100130872-SPON2;SPON2;S | S_Shore | Body;TSS200;5'UTR |

| | | | | |
|---|---|---|---|---|
| 20M03_Saliva | chr4:1205914-1206150 | CTBP1;CTBP1 | Island | 3'UTR;3'UTR |
| 20M03_Saliva | chr4:1208301-1208960 | CTBP1;CTBP1 | N_Shore | Body;Body |
| 20M03_Saliva | chr4:1363182-1363661 | KIAA1530 | N_Shore | Body |
| 20M03_Saliva | chr4:1366463-1366662 | KIAA1530 | Island | Body |
| 20M03_Saliva | chr4:2627014-2627057 | FAM193A | | TSS200 |
| 20M03_Saliva | chr4:3372915-3373405 | RGS12;RGS12;RGS12 | N_Shore | Body;Body;Body |
| 20M03_Saliva | chr4:3387499-3387765 | RGS12;RGS12;RGS12 | | Body;Body;Body |
| 20M03_Saliva | chr4:3392103-3392318 | RGS12;RGS12;RGS12 | | Body;Body;Body |
| 20M03_Saliva | chr4:3409828-3409927 | RGS12;RGS12;RGS12 | | Body;Body;Body |
| 20M03_Saliva | chr4:3521512-3521738 | LRPAP1 | S_Shore | Body |
| 20M03_Saliva | chr4:5526510-5526783 | C4orf6 | | TSS1500 |
| 20M03_Saliva | chr4:7862646-7862659 | AFAP1;AFAP1 | | Body;Body |
| 20M03_Saliva | chr4:8395941-8396502 | ACOX3;ACOX3 | | Body;Body |
| 20M03_Saliva | chr4:40517938-40518041 | RBM47;RBM47;RBM47 | | 1stExon;5'UTR;5'UTR |
| 20M03_Saliva | chr4:41694796-41694847 | LIMCH1;LIMCH1;LIMCH1;LIMCH1;LIMCH1 | | Body;Body;Body;Body;Body |
| 20M03_Saliva | chr4:42122106-42122161 | BEND4;BEND4 | | Body;Body |
| 20M03_Saliva | chr4:57035948-57036298 | KIAA1211 | | TSS1500 |
| 20M03_Saliva | chr4:73177995-73178533 | ADAMTS3 | | Body |
| 20M03_Saliva | chr4:78432011-78432403 | CXCL13 | | TSS1500 |
| 20M03_Saliva | chr4:114214080-114214185 | ANK2;ANK2;ANK2 | | Body;Body;Body |
| 20M03_Saliva | chr4:169418081-169418224 | PALLD;PALLD | | TSS200;TSS200 |
| 20M03_Saliva | chr4:184930474-184930579 | STOX2 | | Body |
| 20M03_Saliva | chr4:186877497-186877657 | SORBS2;SORBS2 | | 5'UTR;1stExon |
| 20M03_Saliva | chr4:187148316-187148688 | KLKB1 | | TSS1500 |
| 20M03_Saliva | chr5:191127-191296 | LRRC14B | N_Shore | TSS1500 |
| 20M03_Saliva | chr5:510035-510421 | SLC9A3 | S_Shore | Body |
| 20M03_Saliva | chr5:912383-913111 | TRIP13 | N_Shore | Body |
| 20M03_Saliva | chr5:1016279-1016768 | NKD2 | | Body |
| 20M03_Saliva | chr5:1052828-1052919 | SLC12A7 | N_Shore | Body |
| 20M03_Saliva | chr5:1201479-1201608 | SLC6A19 | Island | TSS1500 |
| 20M03_Saliva | chr5:1282357-1282723 | TERT;TERT | N_Shore | Body;Body |
| 20M03_Saliva | chr5:1335846-1335965 | CLPTM1L | S_Shore | Body |
| 20M03_Saliva | chr5:14405632-14406021 | TRIO | | Body |
| 20M03_Saliva | chr5:27038605-27038836 | CDH9;CDH9 | | 1stExon;5'UTR |
| 20M03_Saliva | chr5:53751272-53751493 | HSPB3 | | TSS200 |
| 20M03_Saliva | chr5:71475111-71475356 | MAP1B | | Body |
| 20M03_Saliva | chr5:88017891-88018605 | MEF2C;MEF2C | | 3'UTR;3'UTR |
| 20M03_Saliva | chr5:95295740-95296096 | ELL2 | N_Shore | Body |
| 20M03_Saliva | chr5:118691007-118691126 | TNFAIP8;TNFAIP8 | N_Shore | Body;TSS1500 |
| 20M03_Saliva | chr5:125800764-125800916 | GRAMD3;GRAMD3;GRAMD3;GRAMD3;GRAMD3 | | Body;5'UTR;TSS200;Body;Body |
| 20M03_Saliva | chr5:125929132-125929671 | ALDH7A1 | N_Shore | Body |
| 20M03_Saliva | chr5:135529232-135529341 | LOC389332 | S_Shore | TSS1500 |
| 20M03_Saliva | chr5:137896757-137896863 | SNORD63;HSPA9 | | Body;Body |
| 20M03_Saliva | chr5:138210550-138211703 | LRRTM2;CTNNA1 | | Body;Body |
| 20M03_Saliva | chr5:140857941-140858046 | PCDHGA4;PCDHGA12;PCDHGA1 | Island | Body;Body;Body;Body;Body;Body;Body;Body;1stExon;Body;Body;Body;Body;Body;Body;Body;Body;Body;Body;Body;Body;1stExon;Body |
| 20M03_Saliva | chr5:142065539-142065676 | FGF1;FGF1;FGF1;FGF1;FGF1;FGF1;FGF1;FGF1 | | 5'UTR;Body;Body;5'UTR;1stExon;5'UTR;5'UTR;1stExon |
| 20M03_Saliva | chr5:142065761-142066938 | FGF1;FGF1;FGF1;FGF1;FGF1;FGF1;FGF1 | | 5'UTR;5'UTR;5'UTR;1stExon;Body;Body;TSS200 |
| 20M03_Saliva | chr5:148442740-148442890 | SH3TC2 | | TSS200 |
| 20M03_Saliva | chr5:149493008-149493258 | CSF1R | | TSS200 |
| 20M03_Saliva | chr5:149546087-149546366 | CDX1 | Island | TSS1500 |
| 20M03_Saliva | chr5:149980671-149980682 | SYNPO;SYNPO | | 1stExon;5'UTR |
| 20M03_Saliva | chr5:156351055-156351266 | LOC100286948;TIMD4;TIMD4 | | TSS200;Body;Body |
| 20M03_Saliva | chr5:159797813-159797845 | C1QTNF2 | S_Shore | TSS200 |
| 20M03_Saliva | chr5:175108269-175109114 | HRH2;HRH2 | | 5'UTR;TSS200 |
| 20M03_Saliva | chr5:175955105-175955524 | RNF44 | | 3'UTR |
| 20M03_Saliva | chr5:178565721-178565937 | ADAMTS2 | Island | Body |
| 20M03_Saliva | chr6:8436218-8436296 | SLC35B3;SLC35B3;SLC35B3 | S_Shore | TSS1500;TSS1500;TSS1500 |
| 20M03_Saliva | chr6:10555808-10556523 | GCNT2;GCNT2 | | TSS200;Body |
| 20M03_Saliva | chr6:11771473-11771680 | C6orf105;C6orf105 | | Body;Body |
| 20M03_Saliva | chr6:24877462-24877729 | FAM65B;FAM65B;FAM65B | | 5'UTR;1stExon;5'UTR |
| 20M03_Saliva | chr6:26225258-26225767 | HIST1H3E | N_Shore | TSS200 |

| | | | | |
|---|---|---|---|---|
| 20M03_Saliva | chr6:28876160-28876269 | TRIM27 | | Body |
| 20M03_Saliva | chr6:28967887-28968397 | ZNF311 | | Body |
| 20M03_Saliva | chr6:29911494-29911550 | HLA-A | S_Shore | Body |
| 20M03_Saliva | chr6:30043820-30043924 | RNF39;RNF39 | S_Shore | TSS200;TSS200 |
| 20M03_Saliva | chr6:30122432-30123034 | TRIM10;TRIM10 | | Body;Body |
| 20M03_Saliva | chr6:30136790-30137208 | TRIM15 | N_Shelf | Body |
| 20M03_Saliva | chr6:30174651-30174845 | TRIM26 | | 5'UTR |
| 20M03_Saliva | chr6:30175299-30175327 | TRIM26 | | 5'UTR |
| 20M03_Saliva | chr6:30177177-30177295 | TRIM26 | N_Shelf | 5'UTR |
| 20M03_Saliva | chr6:30656070-30656499 | KIAA1949;NRM;KIAA1949 | S_Shore | TSS1500;3'UTR;TSS1500 |
| 20M03_Saliva | chr6:30691764-30691914 | TUBB | S_Shelf | Body |
| 20M03_Saliva | chr6:30850405-30850581 | DDR1 | N_Shore | TSS1500 |
| 20M03_Saliva | chr6:30850829-30851638 | DDR1;DDR1 | N_Shore | TSS1500;TSS1500 |
| 20M03_Saliva | chr6:30852874-30856491 | DDR1;DDR1 | S_Shore | 5'UTR;5'UTR |
| 20M03_Saliva | chr6:30881549-30881571 | VARS2;VARS2;VARS2;GTF2H4 | Island | TSS1500;TSS1500;TSS1500;Body |
| 20M03_Saliva | chr6:30951225-30951394 | MUC21 | | TSS1500 |
| 20M03_Saliva | chr6:31023257-31023763 | HCG22 | | Body |
| 20M03_Saliva | chr6:31025933-31026287 | HCG22 | | Body |
| 20M03_Saliva | chr6:31082200-31082631 | PSORS1C1 | | TSS1500 |
| 20M03_Saliva | chr6:31096861-31097078 | PSORS1C1 | | 5'UTR |
| 20M03_Saliva | chr6:31138925-31138982 | POU5F1 | | TSS1500 |
| 20M03_Saliva | chr6:31146845-31146942 | PSORS1C3 | N_Shore | TSS1500 |
| 20M03_Saliva | chr6:31478432-31478830 | MICB | | 3'UTR |
| 20M03_Saliva | chr6:31513198-31513267 | ATP6V1G2;NFKBIL1;NFKBIL1;ATI | S_Shelf | Body;TSS1500;TSS1500;Body |
| 20M03_Saliva | chr6:31589597-31589994 | SNORA38;BAT2 | S_Shore | TSS1500;5'UTR |
| 20M03_Saliva | chr6:31590563-31590640 | SNORA38;BAT2 | S_Shore | TSS1500;5'UTR |
| 20M03_Saliva | chr6:31598451-31598504 | BAT2 | N_Shore | Body |
| 20M03_Saliva | chr6:31636239-31636913 | CSNK2B | S_Shelf | Body |
| 20M03_Saliva | chr6:31687926-31688229 | LY6G6C | N_Shelf | Body |
| 20M03_Saliva | chr6:31689610-31690170 | LY6G6C | N_Shore | TSS200 |
| 20M03_Saliva | chr6:31692234-31692375 | C6orf25;C6orf25;C6orf25;C6orf | S_Shore | Body;Body;Body;Body;Body;Body |
| 20M03_Saliva | chr6:31696223-31696259 | DDAH2 | Island | Body |
| 20M03_Saliva | chr6:31696423-31696509 | DDAH2 | Island | Body |
| 20M03_Saliva | chr6:31746808-31746820 | VARS | | Body |
| 20M03_Saliva | chr6:31868782-31868813 | ZBTB12 | Island | Body |
| 20M03_Saliva | chr6:31913250-31913323 | C2;C2;CFB | | 3'UTR;3'UTR;TSS1500 |
| 20M03_Saliva | chr6:31913814-31914068 | CFB;CFB | | 1stExon;5'UTR |
| 20M03_Saliva | chr6:32014189-32014381 | TNXB;TNXB | | Body;TSS1500 |
| 20M03_Saliva | chr6:32014484-32015083 | TNXB;TNXB | | Body;TSS1500 |
| 20M03_Saliva | chr6:32050905-32051108 | TNXB | S_Shelf | Body |
| 20M03_Saliva | chr6:32063835-32063895 | TNXB | Island | Body |
| 20M03_Saliva | chr6:32068181-32068240 | TNXB | S_Shelf | 5'UTR |
| 20M03_Saliva | chr6:32094845-32094915 | ATF6B;ATF6B | N_Shelf | Body;Body |
| 20M03_Saliva | chr6:32120324-32120826 | PPT2;PRRT1;PPT2 | N_Shore | TSS1500;TSS1500;TSS1500 |
| 20M03_Saliva | chr6:32120878-32120907 | PPT2;PRRT1;PPT2 | N_Shore | TSS1500;TSS1500;TSS1500 |
| 20M03_Saliva | chr6:32121113-32121156 | PPT2;PPT2;PRRT1 | N_Shore | TSS200;TSS1500;TSS1500 |
| 20M03_Saliva | chr6:32135195-32135285 | EGFL8 | Island | Body |
| 20M03_Saliva | chr6:32139504-32139812 | AGPAT1;AGPAT1 | | 5'UTR;5'UTR |
| 20M03_Saliva | chr6:32165889-32166206 | NOTCH4 | S_Shore | Body |
| 20M03_Saliva | chr6:32179567-32179971 | NOTCH4 | | Body |
| 20M03_Saliva | chr6:32186496-32186793 | NOTCH4 | | Body |
| 20M03_Saliva | chr6:32190325-32190528 | NOTCH4 | | Body |
| 20M03_Saliva | chr6:32191210-32191496 | NOTCH4 | | Body |
| 20M03_Saliva | chr6:32363084-32363167 | BTNL2 | | Body |
| 20M03_Saliva | chr6:32489750-32490043 | HLA-DRB5 | Island | Body |
| 20M03_Saliva | chr6:32526342-32526366 | HLA-DRB6 | | Body |
| 20M03_Saliva | chr6:32551749-32552152 | HLA-DRB1 | N_Shore | Body |
| 20M03_Saliva | chr6:32552246-32552453 | HLA-DRB1 | Island | Body |
| 20M03_Saliva | chr6:32712045-32712103 | HLA-DQA2 | | Body |
| 20M03_Saliva | chr6:32784327-32785141 | HLA-DOB | | Body |
| 20M03_Saliva | chr6:32800541-32800563 | TAP2;TAP2 | | Body;Body |
| 20M03_Saliva | chr6:32805398-32805570 | TAP2;TAP2 | N_Shore | Body;Body |
| 20M03_Saliva | chr6:32943839-32944195 | BRD2;BRD2 | S_Shelf | Body;Body |
| 20M03_Saliva | chr6:32976805-32976858 | HLA-DOA | S_Shore | Body |
| 20M03_Saliva | chr6:32978271-32978379 | HLA-DOA | S_Shelf | TSS1500 |
| 20M03_Saliva | chr6:33048540-33048558 | HLA-DPB1 | Island | Body |

| | | | | |
|---|---|---|---|---|
| 20M03_Saliva | chr6:33131222-33131560 | COL11A2;COL11A2;COL11A2 | S_Shore | 3'UTR;3'UTR;3'UTR |
| 20M03_Saliva | chr6:33173278-33173501 | HSD17B8 | S_Shore | Body |
| 20M03_Saliva | chr6:33226913-33227484 | VPS52 | | Body |
| 20M03_Saliva | chr6:33228341-33228461 | VPS52 | | Body |
| 20M03_Saliva | chr6:33264834-33265133 | RGL2;RGL2 | N_Shore | Body;Body |
| 20M03_Saliva | chr6:33282997-33283184 | ZBTB22;TAPBP;TAPBP;TAPBP;ZB | Island | Body;TSS1500;TSS1500;TSS1500;Body |
| 20M03_Saliva | chr6:33397122-33397511 | SYNGAP1 | S_Shore | Body |
| 20M03_Saliva | chr6:33418814-33419140 | SYNGAP1 | N_Shelf | Body |
| 20M03_Saliva | chr6:33638313-33638550 | ITPR3 | | Body |
| 20M03_Saliva | chr6:34024362-34024479 | GRM4 | Island | Body |
| 20M03_Saliva | chr6:35285967-35286360 | DEF6 | | Body |
| 20M03_Saliva | chr6:36238178-36238230 | PNPLA1;PNPLA1;PNPLA1 | | 5'UTR;5'UTR;TSS200 |
| 20M03_Saliva | chr6:41007450-41007522 | UNC5CL | | TSS1500 |
| 20M03_Saliva | chr6:43478455-43478829 | C6orf154 | Island | TSS200 |
| 20M03_Saliva | chr6:56878397-56878665 | BEND6 | | Body |
| 20M03_Saliva | chr6:87804765-87805033 | CGA;CGA | | 1stExon;5'UTR |
| 20M03_Saliva | chr6:112374876-112375200 | WISP3;WISP3 | | TSS1500;TSS1500 |
| 20M03_Saliva | chr6:112375302-112375333 | WISP3;WISP3;WISP3 | | TSS200;1stExon;5'UTR |
| 20M03_Saliva | chr6:116381609-116382179 | FRK;FRK | | 1stExon;5'UTR |
| 20M03_Saliva | chr6:116866658-116867006 | FAM26D;BET3L | | 5'UTR;1stExon |
| 20M03_Saliva | chr6:119390418-119390712 | FAM184A;FAM184A;MIR548B | | 5'UTR;Body;TSS200 |
| 20M03_Saliva | chr6:130182551-130182967 | C6orf191 | | TSS200 |
| 20M03_Saliva | chr6:135271222-135271518 | ALDH8A1;ALDH8A1;ALDH8A1;ALDH8A1 | | 1stExon;5'UTR;1stExon;5'UTR |
| 20M03_Saliva | chr6:137366322-137366346 | IL20RA | S_Shore | TSS200 |
| 20M03_Saliva | chr6:152126441-152126938 | ESR1;ESR1 | N_Shelf | 5'UTR;TSS1500 |
| 20M03_Saliva | chr6:152702330-152702660 | SYNE1;SYNE1 | | Body;Body |
| 20M03_Saliva | chr6:157746584-157746724 | C6orf35 | S_Shore | TSS1500 |
| 20M03_Saliva | chr6:158732256-158732600 | TULP4;TULP4 | | TSS1500;TSS1500 |
| 20M03_Saliva | chr6:159052400-159052937 | TMEM181 | | Body |
| 20M03_Saliva | chr6:161796785-161796854 | PARK2;PARK2;PARK2 | | Body;Body;Body |
| 20M03_Saliva | chr6:167017956-167017999 | RPS6KA2;RPS6KA2 | | Body;Body |
| 20M03_Saliva | chr6:167104601-167104926 | RPS6KA2 | | Body |
| 20M03_Saliva | chr6:168417574-168417743 | KIF25;KIF25 | | TSS1500;TSS1500 |
| 20M03_Saliva | chr6:168418371-168418439 | KIF25;KIF25 | | TSS200;TSS200 |
| 20M03_Saliva | chr6:168857542-168858319 | SMOC2;SMOC2 | | Body;Body |
| 20M03_Saliva | chr6:170191057-170191082 | C6orf122;C6orf208 | S_Shore | Body;Body |
| 20M03_Saliva | chr7:872208-872797 | UNC84A;UNC84A | | 5'UTR;Body |
| 20M03_Saliva | chr7:906450-906799 | UNC84A;UNC84A | S_Shore | Body;Body |
| 20M03_Saliva | chr7:1022213-1022806 | CYP2W1 | N_Shore | TSS1500 |
| 20M03_Saliva | chr7:1022904-1023156 | CYP2W1 | N_Shore | 1stExon |
| 20M03_Saliva | chr7:1024673-1024814 | CYP2W1 | Island | Body |
| 20M03_Saliva | chr7:1053726-1054092 | C7orf50;C7orf50;C7orf50 | S_Shore | Body;Body;Body |
| 20M03_Saliva | chr7:1102344-1103262 | C7orf50;C7orf50;C7orf50 | N_Shelf | Body;Body;Body |
| 20M03_Saliva | chr7:1491787-1492619 | MICALL2 | Island | Body |
| 20M03_Saliva | chr7:1514361-1514623 | INTS1 | Island | Body |
| 20M03_Saliva | chr7:2019874-2020131 | MAD1L1;MAD1L1;MAD1L1 | Island | Body;Body;Body |
| 20M03_Saliva | chr7:2094628-2094977 | MAD1L1;MAD1L1;MAD1L1 | | Body;Body;Body |
| 20M03_Saliva | chr7:2606716-2606824 | IQCE;IQCE | | Body;Body |
| 20M03_Saliva | chr7:2611890-2612101 | IQCE;IQCE | | Body;Body |
| 20M03_Saliva | chr7:2701419-2701641 | TTYH3 | Island | 3'UTR |
| 20M03_Saliva | chr7:2802942-2802976 | GNA12 | S_Shore | Body |
| 20M03_Saliva | chr7:2842593-2842746 | GNA12 | | Body |
| 20M03_Saliva | chr7:3033173-3033939 | CARD11 | | 5'UTR |
| 20M03_Saliva | chr7:4243286-4243604 | SDK1;SDK1 | | Body;Body |
| 20M03_Saliva | chr7:5534951-5535463 | FBXL18 | S_Shore | Body |
| 20M03_Saliva | chr7:23286319-23286539 | GPNMB;GPNMB;GPNMB;GPNMB | N_Shore | 5'UTR;1stExon;1stExon;5'UTR |
| 20M03_Saliva | chr7:24797835-24797884 | DFNA5;DFNA5;DFNA5 | S_Shore | TSS1500;TSS200;TSS200 |
| 20M03_Saliva | chr7:27140942-27141139 | HOXA2 | N_Shelf | Body |
| 20M03_Saliva | chr7:27150262-27150598 | HOXA3;HOXA3;HOXA3 | Island | 5'UTR;5'UTR;5'UTR |
| 20M03_Saliva | chr7:27152899-27154264 | HOXA3;HOXA3;HOXA3 | N_Shore | 5'UTR;5'UTR;5'UTR |
| 20M03_Saliva | chr7:27154537-27154845 | HOXA3;HOXA3;HOXA3 | N_Shore | 5'UTR;TSS1500;5'UTR |
| 20M03_Saliva | chr7:27168962-27169674 | HOXA4 | N_Shore | Body |
| 20M03_Saliva | chr7:27171051-27171203 | HOXA4 | S_Shore | TSS1500 |
| 20M03_Saliva | chr7:27183816-27183946 | HOXA5 | Island | TSS1500 |
| 20M03_Saliva | chr7:27183973-27184059 | HOXA5 | Island | TSS1500 |
| 20M03_Saliva | chr7:27184109-27184369 | HOXA5 | Island | TSS1500 |

| | | | | |
|---|---|---|---|---|
| 20M03_Saliva | chr7:27184438-27184521 | HOXA5 | Island | TSS1500 |
| 20M03_Saliva | chr7:29233211-29233476 | CHN2 | N_Shore | TSS1500 |
| 20M03_Saliva | chr7:45002287-45002736 | MYO1G | Island | 3'UTR |
| 20M03_Saliva | chr7:48129797-48129904 | UPP1;UPP1 | Island | 5'UTR;5'UTR |
| 20M03_Saliva | chr7:55147074-55147374 | EGFR;EGFR;EGFR;EGFR | | Body;Body;Body;Body |
| 20M03_Saliva | chr7:56160737-56161020 | PHKG1 | | TSS200 |
| 20M03_Saliva | chr7:92238207-92238364 | CDK6;CDK6 | | 3'UTR;3'UTR |
| 20M03_Saliva | chr7:99691314-99691486 | MCM7;MCM7;MIR25 | | Body;Body;TSS200 |
| 20M03_Saliva | chr7:102574105-102574504 | LRRC17;FBXL13;LRRC17;FBXL13 | | 5'UTR;Body;5'UTR;Body |
| 20M03_Saliva | chr7:116660172-116660240 | ST7;ST7 | | Body;Body |
| 20M03_Saliva | chr7:123484898-123485052 | HYAL4 | | TSS1500 |
| 20M03_Saliva | chr7:129912487-129912803 | CPA2 | | Body |
| 20M03_Saliva | chr7:130020011-130020130 | CPA1 | N_Shore | TSS1500 |
| 20M03_Saliva | chr7:134463454-134464129 | CALD1;CALD1;CALD1 | | TSS1500;TSS1500;TSS1500 |
| 20M03_Saliva | chr7:134574978-134575811 | CALD1;CALD1;CALD1;CALD1;CALD1 | | TSS1500;Body;TSS1500;Body;Body |
| 20M03_Saliva | chr7:135433540-135433790 | FAM180A;FAM180A | | 1stExon;5'UTR |
| 20M03_Saliva | chr7:142583296-142583731 | TRPV6;TRPV6 | | 1stExon;5'UTR |
| 20M03_Saliva | chr7:149471884-149472134 | SSPO | S_Shore | TSS1500 |
| 20M03_Saliva | chr7:150021488-150021553 | ACTR3C;LRRC61;LRRC61 | S_Shore | TSS1500;5'UTR;5'UTR |
| 20M03_Saliva | chr7:150778724-150779047 | FASTK;TMUB1;TMUB1;FASTK | Island | TSS1500;Body;Body;TSS1500 |
| 20M03_Saliva | chr7:150822902-150823386 | AGAP3 | Island | Body |
| 20M03_Saliva | chr7:151493629-151493820 | PRKAG2;PRKAG2 | | 5'UTR;Body |
| 20M03_Saliva | chr7:157192974-157193101 | DNAJB6 | Island | Body |
| 20M03_Saliva | chr7:157468321-157468443 | PTPRN2;PTPRN2;PTPRN2 | | Body;Body;Body |
| 20M03_Saliva | chr7:157588520-157588858 | PTPRN2;PTPRN2;PTPRN2 | S_Shore | Body;Body;Body |
| 20M03_Saliva | chr7:157701024-157701262 | PTPRN2;PTPRN2;PTPRN2 | N_Shore | Body;Body;Body |
| 20M03_Saliva | chr7:157738245-157738329 | PTPRN2;PTPRN2;PTPRN2 | | Body;Body;Body |
| 20M03_Saliva | chr7:157809235-157809385 | PTPRN2;PTPRN2;PTPRN2 | | Body;Body;Body |
| 20M03_Saliva | chr7:157866922-157867264 | PTPRN2;PTPRN2;PTPRN2 | | Body;Body;Body |
| 20M03_Saliva | chr7:158059704-158060003 | PTPRN2;PTPRN2;PTPRN2 | N_Shore | Body;Body;Body |
| 20M03_Saliva | chr7:158278869-158278954 | PTPRN2;PTPRN2;PTPRN2 | S_Shore | Body;Body;Body |
| 20M03_Saliva | chr8:1642380-1642577 | DLGAP2 | S_Shore | Body |
| 20M03_Saliva | chr8:1785958-1786207 | ARHGEF10 | | 5'UTR |
| 20M03_Saliva | chr8:1790793-1790905 | ARHGEF10 | | 5'UTR |
| 20M03_Saliva | chr8:1807061-1807575 | ARHGEF10 | | Body |
| 20M03_Saliva | chr8:1906312-1906498 | ARHGEF10 | | 3'UTR |
| 20M03_Saliva | chr8:8680903-8680971 | MFHAS1 | | Body |
| 20M03_Saliva | chr8:10529811-10530148 | C8orf74 | | TSS1500 |
| 20M03_Saliva | chr8:12869553-12869615 | C8orf79;C8orf79 | | Body;TSS1500 |
| 20M03_Saliva | chr8:19170080-19170432 | SH2D4A | N_Shore | TSS1500 |
| 20M03_Saliva | chr8:19459996-19460136 | CSGALNACT1;CSGALNACT1;CS( | Island | 1stExon;5'UTR;Body;5'UTR |
| 20M03_Saliva | chr8:22084861-22085227 | PHYHIP;PHYHIP | | Body;Body |
| 20M03_Saliva | chr8:37699239-37699923 | GPR124 | Island | Body |
| 20M03_Saliva | chr8:40010443-40010823 | C8orf4 | | TSS1500 |
| 20M03_Saliva | chr8:41707516-41707736 | ANK1 | | Body |
| 20M03_Saliva | chr8:53322321-53322619 | ST18;ST18 | | 5'UTR;1stExon |
| 20M03_Saliva | chr8:57233269-57233391 | SDR16C5 | S_Shore | TSS200 |
| 20M03_Saliva | chr8:67405405-67405742 | C8orf46 | | TSS200 |
| 20M03_Saliva | chr8:77767602-77767937 | ZFHX4 | | Body |
| 20M03_Saliva | chr8:124217698-124217896 | FAM83A;FAM83A | N_Shore | Body;Body |
| 20M03_Saliva | chr8:141588118-141588242 | EIF2C2;EIF2C2 | | Body;Body |
| 20M03_Saliva | chr8:141599185-141599356 | EIF2C2;EIF2C2 | | Body;Body |
| 20M03_Saliva | chr8:142235430-142235763 | SLC45A4 | N_Shore | Body |
| 20M03_Saliva | chr8:143424755-143424827 | TSNARE1 | | Body |
| 20M03_Saliva | chr8:143694010-143694305 | ARC;ARC | N_Shore | 1stExon;3'UTR |
| 20M03_Saliva | chr8:143761503-143761811 | PSCA | | TSS1500 |
| 20M03_Saliva | chr8:143823809-143824558 | SLURP1;SLURP1 | S_Shelf | 5'UTR;1stExon |
| 20M03_Saliva | chr8:143851287-143851427 | LYNX1;LYNX1 | | TSS200;Body |
| 20M03_Saliva | chr8:143867129-143868917 | LY6D | | Body |
| 20M03_Saliva | chr8:144120106-144120399 | C8orf31 | | TSS1500 |
| 20M03_Saliva | chr8:144576604-144577238 | ZC3H3 | | Body |
| 20M03_Saliva | chr8:144800997-144801223 | MAPK15 | Island | Body |
| 20M03_Saliva | chr8:144894951-144895243 | SCRIB;SCRIB | Island | Body;Body |
| 20M03_Saliva | chr8:145061234-145061628 | PARP10 | N_Shelf | TSS1500 |
| 20M03_Saliva | chr8:145727318-145728438 | PPP1R16A | S_Shore | 3'UTR |
| 20M03_Saliva | chr8:145728501-145728630 | GPT | N_Shore | TSS1500 |

| | | | | |
|---|---|---|---|---|
| 20M03_Saliva | chr8:145749776-145750363 | LRRC24;LRRC14 | N_Shore | Body;3'UTR |
| 20M03_Saliva | chr9:92291210-92291523 | LOC100129066 | N_Shore | Body |
| 20M03_Saliva | chr9:97848107-97848457 | C9orf3;MIR24-1 | N_Shore | 3'UTR;TSS200 |
| 20M03_Saliva | chr9:137718814-137719284 | COL5A1 | N_Shore | Body |
| 20M03_Saliva | chr9:139239853-139240119 | GPSM1 | Island | Body |
| 20M03_Saliva | chr10:530951-531152 | DIP2C | Island | Body |
| 20M03_Saliva | chr10:556470-556762 | DIP2C | | Body |
| 20M03_Saliva | chr10:641615-641641 | DIP2C | | Body |
| 20M03_Saliva | chr10:670562-670815 | DIP2C | S_Shore | Body |
| 20M03_Saliva | chr10:1248770-1248818 | ADARB2 | N_Shelf | Body |
| 20M03_Saliva | chr10:1507708-1508061 | ADARB2 | S_Shore | Body |
| 20M03_Saliva | chr10:1547717-1547983 | ADARB2 | | Body |
| 20M03_Saliva | chr10:3149673-3149852 | PFKP | S_Shore | Body |
| 20M03_Saliva | chr10:5406948-5406971 | UCN3 | | TSS200 |
| 20M03_Saliva | chr10:5541568-5541726 | CALML5 | S_Shore | TSS200 |
| 20M03_Saliva | chr10:5566543-5568004 | CALML3 | N_Shore | TSS1500 |
| 20M03_Saliva | chr10:13388442-13388567 | SEPHS1 | | 5'UTR |
| 20M03_Saliva | chr10:14575319-14575436 | FAM107B | | Body |
| 20M03_Saliva | chr10:18629006-18629677 | CACNB2;CACNB2;CACNB2;CACNB2;CACNB2;CACNB2;CACNB2;CAC | | Body;Body;Body;Body;Body;Body;TSS1500;Body |
| 20M03_Saliva | chr10:32216031-32216101 | ARHGAP12 | N_Shore | 5'UTR |
| 20M03_Saliva | chr10:45406295-45406847 | TMEM72 | | TSS1500 |
| 20M03_Saliva | chr10:45923374-45923705 | ALOX5 | Island | Body |
| 20M03_Saliva | chr10:47746733-47747039 | ANXA8L2 | | TSS1500 |
| 20M03_Saliva | chr10:49459800-49460249 | FRMPD2;FRMPD2 | | Body;Body |
| 20M03_Saliva | chr10:50506875-50507298 | C10orf71;C10orf71 | | TSS1500;TSS1500 |
| 20M03_Saliva | chr10:63422254-63422555 | C10orf107 | N_Shelf | |
| 20M03_Saliva | chr10:72432797-72433218 | ADAMTS14;ADAMTS14 | Island | 1stExon;1stExon |
| 20M03_Saliva | chr10:77165618-77166684 | C10orf41;C10orf41 | Island | Body;Body |
| 20M03_Saliva | chr10:77542423-77542483 | C10orf11 | | TSS200 |
| 20M03_Saliva | chr10:85985296-85985331 | LRIT2 | | 1stExon |
| 20M03_Saliva | chr10:88428117-88428177 | LDB3;LDB3;LDB3;LDB3;LDB3;LDB3 | | TSS1500;TSS1500;TSS200;TSS1500;TSS1500;TSS1500 |
| 20M03_Saliva | chr10:88718164-88718393 | SNCG;MMRN2 | | TSS1500;TSS1500 |
| 20M03_Saliva | chr10:88779109-88779163 | FAM25A | S_Shelf | |
| 20M03_Saliva | chr10:97200911-97200987 | SORBS1;SORBS1;SORBS1;SORBS1;SORBS1;SORBS1;SORBS1;SORBS1 | | Body;Body;Body;Body;Body;Body;Body |
| 20M03_Saliva | chr10:99330076-99330203 | UBTD1 | Island | Body |
| 20M03_Saliva | chr10:99331332-99331808 | ANKRD2;ANKRD2 | S_Shore | 3'UTR |
| 20M03_Saliva | chr10:104211388-104211488 | C10orf95 | S_Shore | TSS200 |
| 20M03_Saliva | chr10:105238873-105239205 | CALHM3;CALHM3 | | 1stExon |
| 20M03_Saliva | chr10:106093718-106094061 | ITPRIP | N_Shelf | TSS200 |
| 20M03_Saliva | chr10:121271663-121271840 | RGS10;RGS10 | | Body;Body |
| 20M03_Saliva | chr10:126278272-126278502 | LHPP;LHPP | N_Shelf | Body;Body |
| 20M03_Saliva | chr10:127588059-127588272 | FANK1 | S_Shelf | Body |
| 20M03_Saliva | chr10:129536661-129537308 | FOXI2 | Island | 1stExon |
| 20M03_Saliva | chr10:129796922-129797019 | PTPRE | | 5'UTR |
| 20M03_Saliva | chr10:131355152-131355243 | MGMT | | Body |
| 20M03_Saliva | chr10:132912784-132913153 | TCERG1L | | Body |
| 20M03_Saliva | chr10:133747849-133748048 | PPP2R2D;PPP2R2D;PPP2R2D | N_Shelf | TSS200;TSS200;TSS200 |
| 20M03_Saliva | chr10:134186822-134186967 | LRRC27;LRRC27 | | Body;Body |
| 20M03_Saliva | chr10:134404873-134405126 | INPP5A | | Body |
| 20M03_Saliva | chr10:134570057-134570283 | INPP5A | S_Shore | Body |
| 20M03_Saliva | chr10:134578261-134578481 | INPP5A | | Body |
| 20M03_Saliva | chr10:134978012-134978997 | KNDC1 | Island | Body |
| 20M03_Saliva | chr10:135032154-135032504 | KNDC1 | S_Shore | Body |
| 20M03_Saliva | chr10:135087441-135087497 | ADAM8;ADAM8;ADAM8 | N_Shelf | Body;Body;Body |
| 20M03_Saliva | chr10:135178626-135178849 | ECHS1 | | Body |
| 20M03_Saliva | chr10:135340445-135340539 | CYP2E1 | N_Shelf | |
| 20M03_Saliva | chr10:135341528-135342218 | CYP2E1 | N_Shore | 1stExon |
| 20M03_Saliva | chr11:312841-313267 | IFITM1 | S_Shore | TSS1500 |
| 20M03_Saliva | chr11:405846-405970 | SIGIRR;SIGIRR;SIGIRR | N_Shore | 3'UTR;3'UTR;3'UTR |
| 20M03_Saliva | chr11:460469-460914 | PTDSS2 | Island | Body |
| 20M03_Saliva | chr11:612680-612837 | IRF7;IRF7;IRF7 | S_Shore | Body;Body;Body |
| 20M03_Saliva | chr11:1029938-1030347 | MUC6 | S_Shore | Body |
| 20M03_Saliva | chr11:1036679-1036938 | MUC6;MUC6 | Island | 1stExon;5'UTR |
| 20M03_Saliva | chr11:1092135-1092171 | MUC2 | N_Shelf | Body |

| | | | | |
|---|---|---|---|---|
| 20M03_Saliva | chr11:1325718-1325852 | TOLLIP | | Body |
| 20M03_Saliva | chr11:1328419-1328486 | TOLLIP | N_Shore | Body |
| 20M03_Saliva | chr11:1471410-1471939 | BRSK2 | Island | Body |
| 20M03_Saliva | chr11:1474588-1474841 | BRSK2 | S_Shelf | Body |
| 20M03_Saliva | chr11:1507020-1507347 | HCCA2 | | Body |
| 20M03_Saliva | chr11:1660513-1660992 | HCCA2 | | Body |
| 20M03_Saliva | chr11:1769382-1769462 | HCCA2;LOC402778 | Island | Body;TSS200 |
| 20M03_Saliva | chr11:1855286-1856025 | SYT8 | | TSS1500 |
| 20M03_Saliva | chr11:1858467-1858605 | SYT8 | | Body |
| 20M03_Saliva | chr11:1891699-1892307 | LSP1;LSP1;LSP1;LSP1 | N_Shore | 5'UTR;TSS1500;TSS1500;Body |
| 20M03_Saliva | chr11:1945564-1945918 | TNNT3;TNNT3;TNNT3;TNNT3 | N_Shore | Body;Body;Body;Body |
| 20M03_Saliva | chr11:2011312-2011346 | LOC100133545 | N_Shelf | TSS200 |
| 20M03_Saliva | chr11:2017483-2018247 | H19 | Island | Body |
| 20M03_Saliva | chr11:2170831-2170880 | INS-IGF2;INS-IGF2;IGF2;IGF2 | | Body;Body;1stExon;5'UTR |
| 20M03_Saliva | chr11:2828364-2829241 | KCNQ1;KCNQ1 | N_Shore | Body;Body |
| 20M03_Saliva | chr11:2925570-2925869 | SLC22A18;SLC22A18;SLC22A18 | S_Shore | Body;Body;TSS1500 |
| 20M03_Saliva | chr11:3187678-3187803 | OSBPL5;OSBPL5;OSBPL5 | S_Shore | TSS1500;TSS1500;TSS1500 |
| 20M03_Saliva | chr11:4977001-4977540 | OR51A2 | | TSS200 |
| 20M03_Saliva | chr11:6518438-6518792 | DNHD1;DNHD1 | | TSS200;TSS200 |
| 20M03_Saliva | chr11:6592297-6592745 | DNHD1 | Island | Body |
| 20M03_Saliva | chr11:8832938-8833095 | ST5;ST5;ST5 | | TSS1500;TSS200;5'UTR |
| 20M03_Saliva | chr11:8964505-8964717 | ASCL3 | | 5'UTR |
| 20M03_Saliva | chr11:16761292-16761533 | C11orf58;C11orf58 | S_Shore | Body;5'UTR |
| 20M03_Saliva | chr11:17803160-17803962 | KCNC1 | S_Shore | Body |
| 20M03_Saliva | chr11:22696087-22696308 | GAS2;GAS2;GAS2 | | 5'UTR;5'UTR;TSS1500 |
| 20M03_Saliva | chr11:32421514-32421845 | WT1;WT1;WT1;WT1 | | Body;Body;Body;Body |
| 20M03_Saliva | chr11:45928848-45928988 | C11orf94 | | TSS200 |
| 20M03_Saliva | chr11:45943130-45943572 | GYLTL1B | N_Shore | TSS200 |
| 20M03_Saliva | chr11:47376519-47376985 | SPI1;SPI1 | N_Shore | 3'UTR;3'UTR |
| 20M03_Saliva | chr11:57519282-57519364 | BTBD18 | N_Shore | TSS200 |
| 20M03_Saliva | chr11:61049230-61049520 | VWCE | | Body |
| 20M03_Saliva | chr11:62456939-62457291 | LRRN4CL;LRRN4CL | S_Shore | 5'UTR;1stExon |
| 20M03_Saliva | chr11:62457334-62457677 | LRRN4CL | S_Shore | TSS200 |
| 20M03_Saliva | chr11:63990575-63990629 | FERMT3;FERMT3 | N_Shelf | Body;Body |
| 20M03_Saliva | chr11:64405452-64405993 | NRXN2;NRXN2;NRXN2 | Island | Body;Body;Body |
| 20M03_Saliva | chr11:64951916-64952034 | CAPN1 | Island | Body |
| 20M03_Saliva | chr11:65374663-65375067 | MAP3K11 | N_Shore | Body |
| 20M03_Saliva | chr11:65560174-65560763 | OVOL1 | | Body |
| 20M03_Saliva | chr11:66104468-66105148 | RIN1 | S_Shelf | TSS1500 |
| 20M03_Saliva | chr11:66454595-66454651 | SPTBN2 | | Body |
| 20M03_Saliva | chr11:66631356-66631536 | PC;PC;PC | Island | Body;Body;Body |
| 20M03_Saliva | chr11:66650500-66650560 | PC;PC;PC | | 5'UTR;5'UTR;5'UTR |
| 20M03_Saliva | chr11:67183498-67183616 | ATPGD1;ATPGD1 | S_Shore | 5'UTR;5'UTR |
| 20M03_Saliva | chr11:67350326-67350499 | GSTP1 | N_Shore | TSS1500 |
| 20M03_Saliva | chr11:67442067-67442408 | ALDH3B2;ALDH3B2;ALDH3B2 | | 1stExon;5'UTR;5'UTR |
| 20M03_Saliva | chr11:70002333-70002831 | ANO1;ANO1 | S_Shore | Body;Body |
| 20M03_Saliva | chr11:70282487-70282745 | CTTN;CTTN | | 3'UTR;3'UTR |
| 20M03_Saliva | chr11:71710614-71710982 | IL18BP;IL18BP;IL18BP;IL18BP;IL18BP;IL18BP | | TSS200;5'UTR;TSS1500;5'UTR;5'UTR;1stExon |
| 20M03_Saliva | chr11:71723428-71724677 | NUMA1 | | Body |
| 20M03_Saliva | chr11:73053649-73054401 | ARHGEF17 | Island | Body |
| 20M03_Saliva | chr11:75428910-75428970 | MOGAT2 | | TSS200 |
| 20M03_Saliva | chr11:79114133-79114457 | MIR708;ODZ4 | | TSS1500;5'UTR |
| 20M03_Saliva | chr11:84634581-84634620 | DLG2;DLG2 | | TSS200;Body |
| 20M03_Saliva | chr11:85430629-85430657 | SYTL2;SYTL2;SYTL2;SYTL2;SYTL2;SYTL2;SYTL2;SYTL2 | | Body;TSS1500;TSS1500;Body;TSS1500;Body;Body;Body |
| 20M03_Saliva | chr11:93277161-93277191 | C11orf75 | S_Shore | TSS1500 |
| 20M03_Saliva | chr11:111789452-111789693 | C11orf52 | | TSS200 |
| 20M03_Saliva | chr11:111847475-111847747 | DIXDC1;DIXDC1 | N_Shore | TSS1500;Body |
| 20M03_Saliva | chr11:113845638-113846017 | HTR3A;HTR3A | | TSS200;TSS200 |
| 20M03_Saliva | chr11:114548759-114549162 | FAM55B | | TSS1500 |
| 20M03_Saliva | chr11:117069780-117069891 | TAGLN;TAGLN | | TSS1500;TSS1500 |
| 20M03_Saliva | chr11:117800062-117800467 | TMPRSS13;TMPRSS13 | | 5'UTR;1stExon |
| 20M03_Saliva | chr11:117947459-117947997 | TMPRSS4;TMPRSS4 | | TSS1500;TSS1500 |
| 20M03_Saliva | chr11:118550379-118550644 | TREH;TREH | | 1stExon;5'UTR |
| 20M03_Saliva | chr11:118781728-118781978 | BCL9L | Island | TSS200 |

| | | | | |
|---|---|---|---|---|
| 20M03_Saliva | chr11:120008963-120009135 | TRIM29 | S_Shore | TSS200 |
| 20M03_Saliva | chr11:120973197-120973274 | TECTA | | TSS200 |
| 20M03_Saliva | chr11:121970694-121970768 | LOC399959;MIR125B1 | | Body;TSS200 |
| 20M03_Saliva | chr11:123396039-123396593 | GRAMD1B | | TSS1500 |
| 20M03_Saliva | chr11:123899655-123900083 | OR10G8 | | TSS1500 |
| 20M03_Saliva | chr11:124621829-124622096 | VSIG2 | | Body |
| 20M03_Saliva | chr11:128737415-128737467 | KCNJ1;KCNJ1;KCNJ1;KCNJ1 | | TSS200;TSS200;TSS200;TSS200 |
| 20M03_Saliva | chr11:128812462-128813583 | TP53AIP1 | | 5'UTR |
| 20M03_Saliva | chr11:129992359-129992368 | APLP2;APLP2;APLP2;APLP2;APLI | N_Shore | Body;Body;Body;Body;Body;Body |
| 20M03_Saliva | chr11:131239431-131239703 | NTM | | TSS1500 |
| 20M03_Saliva | chr11:133804743-133804959 | IGSF9B | S_Shelf | Body |
| 20M03_Saliva | chr11:134216076-134216220 | GLB1L2 | S_Shelf | Body |
| 20M03_Saliva | chr12:258355-258436 | IQSEC3;IQSEC3 | | Body;Body |
| 20M03_Saliva | chr12:372191-372259 | SLC6A13 | | TSS200 |
| 20M03_Saliva | chr12:5937031-5937409 | ANO2 | Island | Body |
| 20M03_Saliva | chr12:6421474-6422131 | PLEKHG6;PLEKHG6;PLEKHG6 | S_Shore | TSS1500;Body;Body |
| 20M03_Saliva | chr12:6442873-6442966 | TNFRSF1A | S_Shelf | Body |
| 20M03_Saliva | chr12:6484219-6484729 | SCNN1A;SCNN1A;SCNN1A;SCNN1A | | 5'UTR;Body;1stExon;5'UTR |
| 20M03_Saliva | chr12:6486701-6487080 | SCNN1A | | TSS200 |
| 20M03_Saliva | chr12:7071712-7073264 | MIR200C | | TSS1500 |
| 20M03_Saliva | chr12:7863229-7864058 | DPPA3 | | TSS1500 |
| 20M03_Saliva | chr12:8974844-8975309 | A2ML1 | | TSS1500 |
| 20M03_Saliva | chr12:9217669-9217907 | LOC144571 | Island | TSS200 |
| 20M03_Saliva | chr12:10459912-10459972 | KLRD1;KLRD1;KLRD1 | | TSS1500;5'UTR;TSS1500 |
| 20M03_Saliva | chr12:20848285-20848821 | SLCO1C1;SLCO1C1;SLCO1C1;SLCO1C1 | | TSS200;TSS200;TSS200;TSS200 |
| 20M03_Saliva | chr12:50497732-50498016 | GPD1 | | TSS200 |
| 20M03_Saliva | chr12:51236582-51236638 | TMPRSS12 | N_Shore | TSS200 |
| 20M03_Saliva | chr12:52585695-52585892 | KRT80;KRT80;KRT80;KRT80 | | 1stExon;1stExon;5'UTR;5'UTR |
| 20M03_Saliva | chr12:52845930-52846073 | KRT6B | | TSS200 |
| 20M03_Saliva | chr12:52867568-52867793 | KRT6C;KRT6C | | 1stExon;5'UTR |
| 20M03_Saliva | chr12:52887054-52887685 | KRT6A;KRT6A | | 1stExon;5'UTR |
| 20M03_Saliva | chr12:53073506-53074328 | KRT1 | | Body |
| 20M03_Saliva | chr12:53242733-53242922 | KRT78;KRT78 | | 5'UTR;1stExon |
| 20M03_Saliva | chr12:53297987-53298888 | KRT8 | S_Shore | Body |
| 20M03_Saliva | chr12:53343514-53343849 | KRT18;KRT18 | S_Shore | Body;Body |
| 20M03_Saliva | chr12:53612551-53612734 | RARG;RARG | N_Shore | Body;Body |
| 20M03_Saliva | chr12:56329615-56329731 | DGKA;DGKA;DGKA;DGKA | Island | 5'UTR;5'UTR;5'UTR;5'UTR |
| 20M03_Saliva | chr12:57559728-57559901 | LRP1 | | Body |
| 20M03_Saliva | chr12:57869031-57869490 | ARHGAP9;ARHGAP9;ARHGAP9 | N_Shore | Body;Body;Body |
| 20M03_Saliva | chr12:58119915-58121004 | AGAP2;LOC100130776;AGAP2 | Island | 3'UTR;TSS200;3'UTR |
| 20M03_Saliva | chr12:67072951-67073062 | GRIP1 | | TSS200 |
| 20M03_Saliva | chr12:68041257-68041322 | DYRK2;DYRK2 | N_Shore | TSS1500;TSS1500 |
| 20M03_Saliva | chr12:68647356-68647498 | IL22 | | TSS200 |
| 20M03_Saliva | chr12:97885575-97885729 | RMST;MIR1251 | | Body;TSS200 |
| 20M03_Saliva | chr12:117627133-117627286 | FBXO21;FBXO21 | N_Shore | Body;Body |
| 20M03_Saliva | chr12:120241287-120241760 | CIT | N_Shore | Body |
| 20M03_Saliva | chr12:120241957-120242156 | CIT | Island | Body |
| 20M03_Saliva | chr12:120242394-120242513 | CIT | S_Shore | Body |
| 20M03_Saliva | chr12:121882033-121882415 | KDM2B;KDM2B | N_Shore | Body;Body |
| 20M03_Saliva | chr12:121890163-121890503 | KDM2B;KDM2B | N_Shore | Body;Body |
| 20M03_Saliva | chr12:123352600-123352774 | VPS37B | S_Shore | Body |
| 20M03_Saliva | chr12:124773668-124774078 | FAM101A | | TSS200 |
| 20M03_Saliva | chr12:124856376-124856587 | NCOR2;NCOR2 | N_Shore | Body;Body |
| 20M03_Saliva | chr12:126926867-126926925 | LOC100128554 | | TSS200 |
| 20M03_Saliva | chr12:130316015-130316058 | TMEM132D | | Body |
| 20M03_Saliva | chr12:131303016-131303247 | STX2;STX2 | N_Shore | Body;Body |
| 20M03_Saliva | chr12:131622584-131622739 | GPR133 | N_Shore | Body |
| 20M03_Saliva | chr12:132259779-132259908 | SFRS8 | N_Shelf | Body |
| 20M03_Saliva | chr12:132690692-132690782 | GALNT9;GALNT9 | S_Shore | Body;TSS200 |
| 20M03_Saliva | chr12:132853954-132854189 | GALNT9;LOC100130238 | Island | Body;Body |
| 20M03_Saliva | chr12:133101168-133101612 | FBRSL1 | N_Shore | Body |
| 20M03_Saliva | chr13:21287237-21287304 | IL17D | | Body |
| 20M03_Saliva | chr13:21291173-21291467 | IL17D | | Body |
| 20M03_Saliva | chr13:24008109-24008532 | SACS | | TSS1500 |
| 20M03_Saliva | chr13:28562819-28562850 | PRHOXNB | | TSS200 |
| 20M03_Saliva | chr13:31480128-31480273 | C13orf33 | N_Shore | TSS200 |

| | | | | |
|---|---|---|---|---|
| 20M03_Saliva | chr13:48575859-48575927 | SUCLA2 | S_Shore | TSS1500 |
| 20M03_Saliva | chr13:51417977-51418092 | DLEU7 | Island | TSS200 |
| 20M03_Saliva | chr13:99135543-99136580 | STK24;STK24 | | Body;Body |
| 20M03_Saliva | chr13:99630462-99630583 | DOCK9;DOCK9;DOCK9;DOCK9 | | Body;Body;TSS200;TSS200 |
| 20M03_Saliva | chr13:110874267-110874326 | COL4A1 | | Body |
| 20M03_Saliva | chr13:111141830-111141841 | COL4A2 | | Body |
| 20M03_Saliva | chr13:111522651-111522985 | C13orf29 | | Body |
| 20M03_Saliva | chr13:113411497-113411561 | ATP11A;ATP11A | S_Shore | Body;Body |
| 20M03_Saliva | chr13:113491766-113492042 | ATP11A;ATP11A | N_Shore | Body;Body |
| 20M03_Saliva | chr13:113496400-113496682 | ATP11A;ATP11A | N_Shore | Body;Body |
| 20M03_Saliva | chr13:113759771-113760286 | F7;F7 | N_Shore | TSS1500;TSS1500 |
| 20M03_Saliva | chr13:114103477-114104057 | ADPRHL1;ADPRHL1 | | Body;TSS200 |
| 20M03_Saliva | chr13:114187973-114188051 | TMCO3 | S_Shore | Body |
| 20M03_Saliva | chr13:114830255-114830313 | RASA3 | | Body |
| 20M03_Saliva | chr13:114855198-114855572 | RASA3 | S_Shelf | Body |
| 20M03_Saliva | chr13:114872102-114872606 | RASA3 | N_Shelf | Body |
| 20M03_Saliva | chr13:114892556-114892618 | RASA3 | | Body |
| 20M03_Saliva | chr14:20903410-20903611 | KLHL33 | Island | 5'UTR |
| 20M03_Saliva | chr14:21510323-21510479 | RNASE7 | | TSS200 |
| 20M03_Saliva | chr14:22103031-22103310 | OR10G2 | | TSS200 |
| 20M03_Saliva | chr14:23352029-23352488 | REM2 | N_Shelf | TSS1500 |
| 20M03_Saliva | chr14:23623480-23623935 | SLC7A8;SLC7A8;SLC7A8 | | 5'UTR;Body;1stExon |
| 20M03_Saliva | chr14:23835595-23836012 | EFS;EFS | Island | TSS1500;TSS1500 |
| 20M03_Saliva | chr14:75897342-75897417 | JDP2;JDP2;JDP2;JDP2 | S_Shore | 5'UTR;5'UTR;5'UTR;TSS1500 |
| 20M03_Saliva | chr14:76447327-76447431 | TGFB3;TGFB3 | N_Shore | 1stExon;5'UTR |
| 20M03_Saliva | chr14:80328189-80328450 | NRXN3;NRXN3;NRXN3 | Island | Body;Body;Body |
| 20M03_Saliva | chr14:91087476-91087586 | TTC7B | Island | Body |
| 20M03_Saliva | chr14:93406016-93406498 | ITPK1 | | Body |
| 20M03_Saliva | chr14:93942094-93942540 | KIAA1409 | | 5'UTR |
| 20M03_Saliva | chr14:94393654-94393816 | FAM181A;C14orf86 | | Body;TSS1500 |
| 20M03_Saliva | chr14:94576977-94577101 | IFI27;IFI27 | | TSS200;TSS200 |
| 20M03_Saliva | chr14:101495758-101495847 | MIR494 | | TSS1500 |
| 20M03_Saliva | chr14:101508789-101509144 | MIR1185-1 | | TSS1500 |
| 20M03_Saliva | chr14:104165052-104165475 | XRCC3;XRCC3;XRCC3;KLC1;KL( | Island | 3'UTR;3'UTR;3'UTR;Body;Body |
| 20M03_Saliva | chr14:104175555-104175768 | XRCC3;XRCC3;XRCC3 | | Body;Body;Body |
| 20M03_Saliva | chr14:104195181-104195666 | ZFYVE21 | S_Shore | Body |
| 20M03_Saliva | chr14:104623477-104623651 | KIF26A | N_Shore | Body |
| 20M03_Saliva | chr14:105045972-105046247 | C14orf180 | | TSS200 |
| 20M03_Saliva | chr14:105154689-105154994 | INF2;INF2;INF2 | N_Shore | TSS1500;TSS1500;TSS1500 |
| 20M03_Saliva | chr14:105286870-105287427 | MGC23270 | S_Shelf | TSS1500 |
| 20M03_Saliva | chr14:105851883-105851981 | PACS2;PACS2 | Island | Body;Body |
| 20M03_Saliva | chr15:23810163-23810378 | MKRN3 | | TSS1500 |
| 20M03_Saliva | chr15:40531871-40532180 | PAK6;PAK6;PAK6 | S_Shore | 5'UTR;5'UTR;TSS200 |
| 20M03_Saliva | chr15:40583227-40583422 | PLCB2 | Island | Body |
| 20M03_Saliva | chr15:42130650-42130735 | JMJD7-PLA2G4B;JMJD7-PLA2G4B;PLA2G4B | | Body;Body;TSS1500 |
| 20M03_Saliva | chr15:42289766-42289939 | PLA2G4E | | Body |
| 20M03_Saliva | chr15:42386726-42387193 | PLA2G4D;PLA2G4D | | 1stExon;5'UTR |
| 20M03_Saliva | chr15:42448234-42448786 | PLA2G4F;PLA2G4F | | Body;Body |
| 20M03_Saliva | chr15:57883392-57883511 | GCOM1;GCOM1;GCOM1;GCOM | N_Shore | TSS1500;TSS1500;TSS1500;TSS1500 |
| 20M03_Saliva | chr15:59908622-59909035 | GCNT3 | | 5'UTR |
| 20M03_Saliva | chr15:67417607-67417899 | SMAD3;SMAD3 | | Body;TSS1500 |
| 20M03_Saliva | chr15:70994435-70994672 | UACA;UACA;UACA | | 5'UTR;Body;1stExon |
| 20M03_Saliva | chr15:72490380-72490517 | GRAMD2 | S_Shore | TSS1500 |
| 20M03_Saliva | chr15:74218418-74218529 | LOXL1 | N_Shore | TSS1500 |
| 20M03_Saliva | chr15:74290466-74290733 | PML;PML;PML;PML;PML;PML;PM | Island | Body;Body;Body;Body;Body;Body;Body;Body;Body |
| 20M03_Saliva | chr15:78631277-78631536 | CRABP1 | N_Shore | TSS1500 |
| 20M03_Saliva | chr15:83317306-83317526 | CPEB1 | Island | TSS1500 |
| 20M03_Saliva | chr15:86219783-86220255 | AKAP13;AKAP13;AKAP13 | | Body;Body;TSS1500 |
| 20M03_Saliva | chr15:86313851-86313941 | MIR1276;KLHL25 | S_Shore | TSS200;5'UTR |
| 20M03_Saliva | chr15:91500781-91500893 | RCCD1;RCCD1 | Island | Body;Body |
| 20M03_Saliva | chr15:101728611-101728653 | CHSY1 | | Body |
| 20M03_Saliva | chr16:419800-420112 | MRPL28 | N_Shore | Body |
| 20M03_Saliva | chr16:717556-717707 | RHOT2;WDR90 | N_Shore | TSS1500;Body |
| 20M03_Saliva | chr16:832683-834201 | MSLNL | N_Shore | Body |
| 20M03_Saliva | chr16:854168-854439 | PRR25 | N_Shelf | TSS1500 |

| | | | | |
|---|---|---|---|---|
| 20M03_Saliva | chr16:1127792-1128865 | SSTR5;LOC146336 | N_Shore | TSS1500;Body |
| 20M03_Saliva | chr16:1206750-1207075 | CACNA1H;CACNA1H | N_Shore | Body;Body |
| 20M03_Saliva | chr16:1210389-1210829 | CACNA1H;CACNA1H | Island | Body;Body |
| 20M03_Saliva | chr16:1813796-1814157 | MAPK8IP3;MAPK8IP3 | N_Shore | Body;Body |
| 20M03_Saliva | chr16:1843697-1843915 | IGFALS;IGFALS;IGFALS;IGFALS;I( | S_Shore | Body;5'UTR;1stExon;5'UTR;1stExon |
| 20M03_Saliva | chr16:2132315-2132723 | TSC2;TSC2;TSC2 | N_Shore | Body;Body;Body |
| 20M03_Saliva | chr16:3062349-3062426 | CLDN9 | N_Shore | TSS200 |
| 20M03_Saliva | chr16:3843012-3843415 | CREBBP;CREBBP | | Body;Body |
| 20M03_Saliva | chr16:22326203-22326535 | POLR3E | | Body |
| 20M03_Saliva | chr16:27829728-27829801 | GSG1L;GSG1L | | Body;Body |
| 20M03_Saliva | chr16:28550171-28550637 | NUPR1;NUPR1 | | 1stExon;1stExon |
| 20M03_Saliva | chr16:30124803-30125305 | GDPD3;GDPD3;LOC100271831 | | 1stExon;5'UTR;TSS1500 |
| 20M03_Saliva | chr16:55868196-55868432 | CES1;CES1;CES1 | S_Shore | TSS1500;TSS1500;TSS1500 |
| 20M03_Saliva | chr16:57406044-57406955 | CX3CL1 | | TSS1500 |
| 20M03_Saliva | chr16:57831745-57832230 | KIFC3;KIFC3;KIFC3;KIFC3 | | Body;Body;1stExon;5'UTR |
| 20M03_Saliva | chr16:57937614-57937756 | CNGB1 | N_Shore | Body |
| 20M03_Saliva | chr16:66969500-66969837 | FAM96B;CES2;FAM96B;CES2 | Island | TSS1500;1stExon;TSS1500;1stExon |
| 20M03_Saliva | chr16:67022472-67022719 | CES8 | | TSS200 |
| 20M03_Saliva | chr16:67232921-67233983 | ELMO3 | N_Shore | TSS200 |
| 20M03_Saliva | chr16:67686832-67687754 | RLTPR | N_Shore | Body |
| 20M03_Saliva | chr16:67918680-67919362 | NRN1L | Island | TSS200 |
| 20M03_Saliva | chr16:68772225-68772469 | CDH1 | Island | Body |
| 20M03_Saliva | chr16:70759914-70760142 | VAC14 | | Body |
| 20M03_Saliva | chr16:81526954-81527501 | CMIP | | Body |
| 20M03_Saliva | chr16:81564161-81564553 | CMIP;CMIP | | Body;Body |
| 20M03_Saliva | chr16:81678804-81678968 | CMIP;CMIP | | Body;Body |
| 20M03_Saliva | chr16:84076834-84076941 | SLC38A8 | | TSS1500 |
| 20M03_Saliva | chr16:84346910-84346930 | WFDC1 | Island | Body |
| 20M03_Saliva | chr16:84852577-84852909 | CRISPLD2 | N_Shore | TSS1500 |
| 20M03_Saliva | chr16:85669462-85669675 | KIAA0182;KIAA0182 | N_Shore | 5'UTR;Body |
| 20M03_Saliva | chr16:87864431-87864826 | SLC7A5 | | 3'UTR |
| 20M03_Saliva | chr16:87958145-87958493 | CA5A | | Body |
| 20M03_Saliva | chr16:88807658-88807707 | FAM38A | S_Shelf | Body |
| 20M03_Saliva | chr16:88832476-88832532 | FAM38A | | Body |
| 20M03_Saliva | chr16:88871329-88871987 | CDT1 | N_Shore | Body |
| 20M03_Saliva | chr16:89004858-89005054 | CBFA2T3;CBFA2T3 | N_Shore | 5'UTR;Body |
| 20M03_Saliva | chr16:89220516-89220688 | ACSF3;ACSF3;ACSF3 | S_Shore | Body;Body;Body |
| 20M03_Saliva | chr16:89387418-89387805 | ANKRD11 | | 5'UTR |
| 20M03_Saliva | chr16:89408322-89408567 | ANKRD11 | Island | 5'UTR |
| 20M03_Saliva | chr16:89462292-89462418 | ANKRD11 | | 5'UTR |
| 20M03_Saliva | chr16:89500152-89500225 | ANKRD11 | Island | 5'UTR |
| 20M03_Saliva | chr17:154420-154671 | RPH3AL | S_Shore | Body |
| 20M03_Saliva | chr17:693137-693388 | RNMTL1 | | Body |
| 20M03_Saliva | chr17:790628-791324 | NXN | | Body |
| 20M03_Saliva | chr17:1090291-1090326 | ABR;ABR | | 1stExon;5'UTR |
| 20M03_Saliva | chr17:1553415-1553484 | RILP | S_Shore | TSS200 |
| 20M03_Saliva | chr17:1553585-1553661 | RILP | S_Shore | TSS200 |
| 20M03_Saliva | chr17:2057082-2057234 | SMG6;SMG6 | | Body;Body |
| 20M03_Saliva | chr17:4463371-4464075 | GGT6;GGT6 | | Body;Body |
| 20M03_Saliva | chr17:6338181-6338554 | AIPL1;AIPL1;AIPL1 | | Body;Body;Body |
| 20M03_Saliva | chr17:6796745-6797119 | ALOX12P2 | N_Shore | Body |
| 20M03_Saliva | chr17:7219261-7219403 | NEURL4;NEURL4;GPS2 | S_Shore | 3'UTR;3'UTR;TSS1500 |
| 20M03_Saliva | chr17:7493150-7494151 | SOX15;SOX15 | S_Shore | 1stExon;5'UTR |
| 20M03_Saliva | chr17:7758171-7758228 | TMEM88 | N_Shore | TSS1500 |
| 20M03_Saliva | chr17:7832680-7833237 | KCNAB3 | Island | 1stExon |
| 20M03_Saliva | chr17:8702369-8702799 | MFSD6L | Island | 1stExon |
| 20M03_Saliva | chr17:8702823-8702896 | MFSD6L | Island | TSS200 |
| 20M03_Saliva | chr17:9929528-9929867 | GAS7;GAS7;GAS7;GAS7 | | 1stExon;Body;Body;5'UTR |
| 20M03_Saliva | chr17:14207530-14207890 | HS3ST3B1;MGC12916 | Island | Body;Body |
| 20M03_Saliva | chr17:14213579-14214081 | HS3ST3B1 | S_Shore | Body |
| 20M03_Saliva | chr17:17110180-17110353 | PLD6 | Island | TSS1500 |
| 20M03_Saliva | chr17:17465242-17465682 | PEMT;PEMT;PEMT | Island | Body;Body;Body |
| 20M03_Saliva | chr17:17717894-17718525 | MIR33B;SREBF1;SREBF1 | S_Shore | TSS1500;Body;Body |
| 20M03_Saliva | chr17:18150954-18151277 | FLII | Island | Body |
| 20M03_Saliva | chr17:19247837-19247918 | MIR1180;B9D1 | Island | Body;Body |
| 20M03_Saliva | chr17:19313024-19313529 | RNF112 | | TSS1500 |

| | | | | |
|---|---|---|---|---|
| 20M03_Saliva | chr17:19648718-19648853 | ALDH3A1;ALDH3A1;ALDH3A1;Al | S_Shore | 5'UTR;5'UTR;5'UTR;1stExon |
| 20M03_Saliva | chr17:19648925-19649004 | ALDH3A1;ALDH3A1;ALDH3A1 | S_Shore | 5'UTR;TSS200;TSS200 |
| 20M03_Saliva | chr17:21193332-21193716 | MAP2K3;MAP2K3 | | Body;TSS1500 |
| 20M03_Saliva | chr17:25798741-25799018 | KSR1 | | TSS1500 |
| 20M03_Saliva | chr17:32581035-32581466 | CCL2 | | TSS1500 |
| 20M03_Saliva | chr17:37782302-37782685 | PPP1R1B | N_Shore | TSS1500 |
| 20M03_Saliva | chr17:37921981-37922297 | IKZF3;IKZF3;IKZF3;IKZF3;IKZF3;IKZF3 | | 3'UTR;3'UTR;3'UTR;3'UTR;3'UTR;3'UTR |
| 20M03_Saliva | chr17:38657310-38657949 | TNS4 | | 5'UTR |
| 20M03_Saliva | chr17:38859934-38860097 | KRT24 | | 1stExon |
| 20M03_Saliva | chr17:39204767-39204922 | KRTAP2-1 | | TSS1500 |
| 20M03_Saliva | chr17:39680217-39680504 | KRT19 | N_Shelf | Body |
| 20M03_Saliva | chr17:39769012-39769213 | KRT16;KRT16 | | 1stExon;5'UTR |
| 20M03_Saliva | chr17:39943423-39943545 | JUP;JUP | S_Shore | TSS1500;TSS1500 |
| 20M03_Saliva | chr17:40274524-40274740 | KAT2A;HSPB9 | Island | TSS1500;TSS1500 |
| 20M03_Saliva | chr17:40823930-40824361 | PLEKHH3 | S_Shore | Body |
| 20M03_Saliva | chr17:41739195-41739806 | MEOX1;MEOX1;MEOX1 | | TSS1500;5'UTR;TSS1500 |
| 20M03_Saliva | chr17:43318045-43319382 | FMNL1 | N_Shore | Body |
| 20M03_Saliva | chr17:43716515-43716617 | C17orf69;C17orf69 | | Body;Body |
| 20M03_Saliva | chr17:46035108-46035343 | PRR15L;PRR15L | | 1stExon;5'UTR |
| 20M03_Saliva | chr17:46608277-46608329 | HOXB1 | Island | TSS200 |
| 20M03_Saliva | chr17:46627384-46628850 | HOXB3 | N_Shore | 3'UTR |
| 20M03_Saliva | chr17:46651722-46651867 | HOXB3;HOXB3 | N_Shelf | 5'UTR;1stExon |
| 20M03_Saliva | chr17:46652179-46652501 | HOXB3 | N_Shore | TSS1500 |
| 20M03_Saliva | chr17:46681111-46683606 | LOC404266;LOC404266;HOXB6 | | Body;Body;5'UTR |
| 20M03_Saliva | chr17:56246435-56246851 | OR4D2 | | TSS1500 |
| 20M03_Saliva | chr17:57918262-57918682 | MIR21 | | TSS1500 |
| 20M03_Saliva | chr17:59668895-59669395 | NACA2 | | TSS1500 |
| 20M03_Saliva | chr17:60757756-60758287 | MRC2 | Island | Body |
| 20M03_Saliva | chr17:62051709-62051773 | SCN4A | | TSS1500 |
| 20M03_Saliva | chr17:64225346-64225731 | APOH | | Body |
| 20M03_Saliva | chr17:65040535-65040616 | CACNG1 | N_Shore | TSS200 |
| 20M03_Saliva | chr17:65850328-65850433 | BPTF;BPTF | | Body;Body |
| 20M03_Saliva | chr17:70120318-70120541 | SOX9 | Island | Body |
| 20M03_Saliva | chr17:73506140-73506230 | CASKIN2;CASKIN2 | | Body;TSS200 |
| 20M03_Saliva | chr17:73996245-73996656 | CDK3;C17orf106 | | TSS1500;Body |
| 20M03_Saliva | chr17:74639731-74639928 | ST6GALNAC1;ST6GALNAC1 | | 5'UTR;1stExon |
| 20M03_Saliva | chr17:75143118-75143600 | SEC14L1;SEC14L1;SEC14L1;SEC14L1 | | Body;Body;Body;Body |
| 20M03_Saliva | chr17:75283431-75284193 | SEPT9;SEPT9 | | Body;TSS1500 |
| 20M03_Saliva | chr17:75445261-75446661 | SEPT9;SEPT9;SEPT9;SEPT9;SEPT | N_Shelf | Body;Body;Body;Body;TSS1500;Body |
| 20M03_Saliva | chr17:76501381-76501582 | DNAH17 | S_Shelf | Body |
| 20M03_Saliva | chr17:76719612-76719640 | CYTH1;CYTH1 | | Body;Body |
| 20M03_Saliva | chr17:76993506-76993698 | CANT1;CANT1;CANT1 | Island | Body;Body;Body |
| 20M03_Saliva | chr17:77030221-77030546 | C1QTNF1;C1QTNF1;C1QTNF1 | | 5'UTR;TSS1500;5'UTR |
| 20M03_Saliva | chr17:77086350-77086810 | HRNBP3 | | 3'UTR |
| 20M03_Saliva | chr17:77997833-77997997 | TBC1D16 | Island | 5'UTR |
| 20M03_Saliva | chr17:78039301-78039528 | CCDC40 | Island | Body |
| 20M03_Saliva | chr17:78079481-78079795 | GAA;GAA;GAA | N_Shore | Body;Body;Body |
| 20M03_Saliva | chr17:78161110-78161495 | CARD14;CARD14 | N_Shelf | TSS1500;Body |
| 20M03_Saliva | chr17:78166148-78166385 | CARD14;CARD14 | S_Shore | Body;Body |
| 20M03_Saliva | chr17:78346448-78346491 | RNF213;LOC100294362 | | Body;Body |
| 20M03_Saliva | chr17:78865087-78865373 | RPTOR;RPTOR | S_Shore | Body;Body |
| 20M03_Saliva | chr17:79056061-79056183 | BAIAP2;BAIAP2;BAIAP2;BAIAP2 | S_Shelf | Body;Body;Body;Body |
| 20M03_Saliva | chr17:79099481-79099785 | MIR657;AATK | N_Shelf | TSS1500;Body |
| 20M03_Saliva | chr17:79099882-79100218 | MIR657;AATK;MIR338 | N_Shelf | TSS1500;Body;TSS200 |
| 20M03_Saliva | chr17:79109729-79109910 | AATK | Island | Body |
| 20M03_Saliva | chr17:79251255-79251354 | SLC38A10;SLC38A10 | N_Shelf | Body;Body |
| 20M03_Saliva | chr17:79374515-79374741 | BAHCC1 | Island | Body |
| 20M03_Saliva | chr17:79495006-79495348 | FSCN2;FSCN2 | Island | TSS1500;TSS1500 |
| 20M03_Saliva | chr17:79652024-79652389 | ARL16;HGS | S_Shore | TSS1500;Body |
| 20M03_Saliva | chr17:79813410-79813507 | P4HB | | Body |
| 20M03_Saliva | chr17:79905152-79905263 | MYADML2 | | TSS200 |
| 20M03_Saliva | chr17:79961522-79961769 | ASPSCR1 | N_Shore | Body |
| 20M03_Saliva | chr17:80022616-80023016 | DUS1L | N_Shore | Body |
| 20M03_Saliva | chr17:80145540-80145575 | CCDC57 | | Body |
| 20M03_Saliva | chr17:80289500-80290217 | SECTM1 | N_Shore | 5'UTR |
| 20M03_Saliva | chr17:80401500-80401857 | C17orf62;C17orf62;C17orf62 | N_Shore | 3'UTR;3'UTR;3'UTR |

| | | | | |
|---|---|---|---|---|
| 20M03_Saliva | chr17:80796693-80798023 | TBCD;ZNF750 | | Body;5'UTR |
| 20M03_Saliva | chr17:80806135-80806435 | TBCD | | Body |
| 20M03_Saliva | chr17:80816702-80816995 | TBCD | N_Shore | Body |
| 20M03_Saliva | chr17:80832328-80832836 | TBCD | S_Shelf | Body |
| 20M03_Saliva | chr17:80847137-80847662 | TBCD | | Body |
| 20M03_Saliva | chr18:20735672-20735934 | CABLES1;CABLES1;CABLES1 | | Body;Body;TSS200 |
| 20M03_Saliva | chr18:21452730-21453131 | LAMA3;LAMA3;LAMA3;LAMA3 | | TSS1500;TSS1500;Body;Body |
| 20M03_Saliva | chr18:21572622-21572748 | TTC39C | | TSS200 |
| 20M03_Saliva | chr18:55888640-55888832 | NEDD4L;NEDD4L;NEDD4L;NEDD4L;NEDD4L;NEDD4L;NEDD4L;NEDD4L | | 5'UTR;Body;TSS200;5'UTR;5'UTR;Body;Body;Body;5'UTR |
| 20M03_Saliva | chr18:61143926-61144339 | SERPINB5 | | TSS1500 |
| 20M03_Saliva | chr18:61171358-61171722 | SERPINB5 | | 3'UTR |
| 20M03_Saliva | chr18:74240460-74240582 | LOC284276 | | TSS200 |
| 20M03_Saliva | chr18:77230598-77230795 | NFATC1;NFATC1;NFATC1;NFATC | Island | Body;Body;Body;Body |
| 20M03_Saliva | chr18:77286052-77286260 | NFATC1;NFATC1;NFATC1;NFATC | Island | Body;Body;Body;Body |
| 20M03_Saliva | chr19:736164-736425 | PALM;PALM | Island | Body;Body |
| 20M03_Saliva | chr19:1070986-1071208 | HMHA1 | Island | Body |
| 20M03_Saliva | chr19:1356269-1356304 | MUM1;MUM1 | S_Shore | TSS200;5'UTR |
| 20M03_Saliva | chr19:1395365-1395646 | NDUFS7 | Island | Body |
| 20M03_Saliva | chr19:1424658-1424952 | DAZAP1;DAZAP1 | Island | Body;Body |
| 20M03_Saliva | chr19:1496416-1496580 | REEP6 | Island | Body |
| 20M03_Saliva | chr19:1854549-1854819 | KLF16 | Island | Body |
| 20M03_Saliva | chr19:1907677-1908328 | ADAT3;SCAMP4 | N_Shore | 5'UTR;5'UTR |
| 20M03_Saliva | chr19:2428567-2428889 | LMNB2;TIMM13 | Island | 3'UTR;TSS1500 |
| 20M03_Saliva | chr19:2512999-2513356 | GNG7 | N_Shore | 3'UTR |
| 20M03_Saliva | chr19:2525231-2525634 | GNG7 | Island | 5'UTR |
| 20M03_Saliva | chr19:2540907-2541104 | GNG7 | Island | 5'UTR |
| 20M03_Saliva | chr19:4305072-4305239 | FSD1 | Island | Body |
| 20M03_Saliva | chr19:4455239-4455957 | UBXN6;UBXN6;UBXN6 | N_Shelf | 5'UTR;1stExon;Body |
| 20M03_Saliva | chr19:5036857-5037219 | KDM4B | N_Shore | Body |
| 20M03_Saliva | chr19:5048645-5049135 | KDM4B | Island | Body |
| 20M03_Saliva | chr19:5250376-5251120 | PTPRS;PTPRS;PTPRS;PTPRS | N_Shore | Body;Body;Body;Body |
| 20M03_Saliva | chr19:5587744-5587909 | SAFB2 | Island | Body |
| 20M03_Saliva | chr19:7580209-7580275 | ZNF358 | N_Shore | TSS1500 |
| 20M03_Saliva | chr19:8555061-8555169 | PRAM1 | Island | 3'UTR |
| 20M03_Saliva | chr19:10444815-10445516 | RAVER1;ICAM3 | Island | TSS1500;Body |
| 20M03_Saliva | chr19:10735548-10736448 | SLC44A2;SLC44A2 | N_Shore | TSS1500;Body |
| 20M03_Saliva | chr19:10792615-10792710 | ILF3;ILF3;ILF3;ILF3;ILF3 | | Body;Body;Body;Body;Body |
| 20M03_Saliva | chr19:11472110-11472397 | LPPR2;LPPR2 | Island | Body;Body |
| 20M03_Saliva | chr19:13127389-13127873 | NFIX | S_Shelf | Body |
| 20M03_Saliva | chr19:14170956-14171331 | PALM3 | S_Shelf | TSS1500 |
| 20M03_Saliva | chr19:14607597-14608092 | GIPC1;GIPC1;GIPC1;GIPC1;GIPC | S_Shore | TSS1500;TSS1500;TSS1500;TSS1500;TSS1500;TSS1500 |
| 20M03_Saliva | chr19:15568574-15568935 | RASAL3 | Island | Body |
| 20M03_Saliva | chr19:16058370-16058640 | OR10H4 | | TSS1500 |
| 20M03_Saliva | chr19:16178091-16179321 | TPM4 | N_Shore | TSS1500 |
| 20M03_Saliva | chr19:17942025-17942411 | JAK3 | N_Shore | Body |
| 20M03_Saliva | chr19:17972376-17972728 | RPL18AP3;SNORA68;RPL18A | S_Shore | Body;TSS1500;Body |
| 20M03_Saliva | chr19:18549689-18550061 | ISYNA1;ISYNA1;ISYNA1 | S_Shore | TSS1500;TSS1500;TSS1500 |
| 20M03_Saliva | chr19:31640249-31640849 | DKFZp566F0947 | | TSS1500 |
| 20M03_Saliva | chr19:39086733-39087186 | MAP4K1;MAP4K1 | N_Shore | Body;Body |
| 20M03_Saliva | chr19:39278997-39279703 | LGALS7B | N_Shelf | TSS1500 |
| 20M03_Saliva | chr19:43969650-43969884 | LYPD3 | S_Shore | 1stExon |
| 20M03_Saliva | chr19:45033565-45033577 | CEACAM20;CEACAM20;CEACAM20;CEACAM20 | | TSS200;TSS200;TSS200;TSS200 |
| 20M03_Saliva | chr19:45279765-45280184 | CBLC;CBLC | N_Shore | TSS1500;TSS1500 |
| 20M03_Saliva | chr19:45737623-45738115 | EXOC3L2 | Island | TSS200 |
| 20M03_Saliva | chr19:46270244-46270700 | SIX5 | Island | Body |
| 20M03_Saliva | chr19:46286064-46286683 | DMPK;DMPK;DMPK | N_Shelf | TSS1500;TSS1500;TSS1500 |
| 20M03_Saliva | chr19:46544240-46544711 | IGFL4;IGFL4 | | 1stExon;5'UTR |
| 20M03_Saliva | chr19:46651087-46651553 | IGFL2;IGFL2;IGFL2 | | 1stExon;5'UTR;TSS1500 |
| 20M03_Saliva | chr19:49220102-49220485 | MAMSTR;MAMSTR;MAMSTR | N_Shelf | 1stExon;Body;5'UTR |
| 20M03_Saliva | chr19:50192542-50193437 | C19orf76;C19orf76 | Island | 1stExon;5'UTR |
| 20M03_Saliva | chr19:51293620-51293677 | ACPT | N_Shore | TSS200 |
| 20M03_Saliva | chr19:51531297-51531393 | KLK11;KLK11;KLK11;KLK11 | N_Shelf | TSS1500;TSS200;TSS1500;TSS200 |
| 20M03_Saliva | chr19:51538134-51538416 | KLK12;KLK12;KLK12;KLK12;KLK1 | S_Shelf | 1stExon;1stExon;5'UTR;5'UTR;1stExon;5'UTR |

| | | | | |
|---|---|---|---|---|
| 20M03_Saliva | chr19:52227967-52228048 | HAS1 | | TSS1500 |
| 20M03_Saliva | chr19:56458926-56459108 | NLRP8 | | TSS1500 |
| 20M03_Saliva | chr20:1309722-1310000 | SDCBP2 | S_Shelf | 5'UTR |
| 20M03_Saliva | chr20:2276435-2276663 | TGM3 | | TSS200 |
| 20M03_Saliva | chr20:10412596-10412723 | MKKS;MKKS | N_Shore | 5'UTR;TSS200 |
| 20M03_Saliva | chr20:17595355-17595472 | RRBP1;RRBP1 | Island | Body;Body |
| 20M03_Saliva | chr20:30432938-30433512 | FOXS1 | Island | 1stExon |
| 20M03_Saliva | chr20:31781151-31781277 | BASE | | TSS1500 |
| 20M03_Saliva | chr20:33879989-33880825 | FAM83C | Island | 1stExon |
| 20M03_Saliva | chr20:34700112-34700675 | EPB41L1 | | TSS1500 |
| 20M03_Saliva | chr20:35503983-35504511 | C20orf118 | | TSS1500 |
| 20M03_Saliva | chr20:35972924-35973098 | SRC | N_Shore | TSS200 |
| 20M03_Saliva | chr20:36011999-36012016 | SRC;SRC | N_Shore | 5'UTR;5'UTR |
| 20M03_Saliva | chr20:36151003-36151338 | BLCAP;BLCAP;BLCAP;NNAT;BLC | Island | 5'UTR;5'UTR;5'UTR;Body;5'UTR;5'UTR;Body |
| 20M03_Saliva | chr20:36152813-36154388 | BLCAP;BLCAP;BLCAP;BLCAP;BL | S_Shore | 5'UTR;1stExon;5'UTR;5'UTR;5'UTR;5'UTR |
| 20M03_Saliva | chr20:36157405-36157651 | BLCAP;BLCAP;BLCAP;BLCAP | S_Shore | TSS1500;TSS1500;TSS1500;TSS1500 |
| 20M03_Saliva | chr20:37208947-37209395 | ADIG | | TSS1500 |
| 20M03_Saliva | chr20:43029732-43029895 | HNF4A;HNF4A;HNF4A;HNF4A;HNF4A;HNF4A | | TSS200;Body;TSS200;Body;Body;TSS200 |
| 20M03_Saliva | chr20:43937138-43937448 | MATN4;RBPJL;MATN4;MATN4 | S_Shore | TSS200;Body;TSS200;TSS200 |
| 20M03_Saliva | chr20:43977423-43977531 | SDC4 | S_Shore | TSS1500 |
| 20M03_Saliva | chr20:50418952-50419348 | SALL4;SALL4 | Island | 1stExon;5'UTR |
| 20M03_Saliva | chr20:52687007-52687365 | BCAS1;BCAS1 | | 1stExon;5'UTR |
| 20M03_Saliva | chr20:57485816-57486019 | GNAS;GNAS;GNAS;GNAS;GNAS;GNAS;GNAS;GNAS | | Body;Body;3'UTR;Body;Body;3'UTR;Body;Body |
| 20M03_Saliva | chr20:60639404-60639721 | TAF4 | Island | Body |
| 20M03_Saliva | chr20:61002595-61002866 | C20orf151 | | TSS200 |
| 20M03_Saliva | chr20:62367884-62367893 | LIME1 | N_Shore | TSS200 |
| 20M03_Saliva | chr20:62383275-62383481 | ZBTB46 | Island | Body |
| 20M03_Saliva | chr20:62406721-62406964 | ZBTB46 | Island | Body |
| 20M03_Saliva | chr20:62693951-62694000 | TCEA2;TCEA2 | Island | TSS200;5'UTR |
| 20M03_Saliva | chr21:15645787-15645988 | ABCC13;ABCC13 | | TSS1500;TSS1500 |
| 20M03_Saliva | chr21:35736023-35736383 | KCNE2 | | TSS1500 |
| 20M03_Saliva | chr21:35832141-35832364 | KCNE1;KCNE1;KCNE1 | Island | 5'UTR;TSS1500;TSS1500 |
| 20M03_Saliva | chr21:36259241-36259460 | RUNX1;RUNX1;RUNX1 | Island | 1stExon;1stExon;Body |
| 20M03_Saliva | chr21:38362727-38362754 | HLCS | Island | TSS200 |
| 20M03_Saliva | chr21:39643733-39644244 | KCNJ15;KCNJ15;KCNJ15 | | 5'UTR;5'UTR;TSS1500 |
| 20M03_Saliva | chr21:43098901-43099292 | NCRNA00111 | | TSS1500 |
| 20M03_Saliva | chr21:43135854-43136287 | C21orf129;NCRNA00112 | | Body;TSS1500 |
| 20M03_Saliva | chr21:43185674-43185884 | RIPK4 | N_Shore | Body |
| 20M03_Saliva | chr21:44898090-44898206 | C21orf84 | | Body |
| 20M03_Saliva | chr21:45683146-45683528 | DNMT3L;DNMT3L | | TSS1500;TSS1500 |
| 20M03_Saliva | chr21:45705740-45706043 | AIRE | Island | TSS200 |
| 20M03_Saliva | chr21:46077454-46077582 | C21orf29;KRTAP12-3 | | Body;TSS1500 |
| 20M03_Saliva | chr21:46890849-46890997 | COL18A1;COL18A1;COL18A1 | Island | Body;Body;Body |
| 20M03_Saliva | chr22:17489225-17489367 | GAB4 | S_Shore | TSS200 |
| 20M03_Saliva | chr22:19949873-19950166 | COMT;COMT;COMT;COMT | | 5'UTR;5'UTR;5'UTR;TSS200 |
| 20M03_Saliva | chr22:24577315-24577448 | SUSD2 | | TSS200 |
| 20M03_Saliva | chr22:24891158-24891338 | C22orf45;C22orf45;UPB1 | Island | TSS1500;TSS1500;TSS200 |
| 20M03_Saliva | chr22:24979554-24979642 | GGT1 | | TSS200 |
| 20M03_Saliva | chr22:31536585-31537064 | PLA2G3 | | TSS200 |
| 20M03_Saliva | chr22:33195343-33195655 | SYN3;TIMP3;SYN3;SYN3 | N_Shelf | Body;TSS1500;Body;Body |
| 20M03_Saliva | chr22:33195943-33196112 | SYN3;TIMP3;SYN3;SYN3 | N_Shore | Body;TSS1500;Body;Body |
| 20M03_Saliva | chr22:33402393-33402998 | SYN3;SYN3;SYN3 | | 1stExon;Body;1stExon |
| 20M03_Saliva | chr22:37608611-37608819 | SSTR3 | | TSS1500 |
| 20M03_Saliva | chr22:40417538-40417869 | FAM83F | Island | Body |
| 20M03_Saliva | chr22:42474656-42475135 | C22orf32;FAM109B | N_Shore | TSS1500;3'UTR |
| 20M03_Saliva | chr22:43820888-43821115 | MPPED1 | Island | 5'UTR |
| 20M03_Saliva | chr22:45608345-45608516 | C22orf9;C22orf9;C22orf9 | | Body;1stExon;5'UTR |
| 20M03_Saliva | chr22:47022899-47023209 | GRAMD4 | Island | Body |
| 20M03_Saliva | chr22:47054067-47054273 | GRAMD4 | Island | Body |
| 20M03_Saliva | chr22:47057191-47057502 | GRAMD4 | N_Shore | Body |
| 20M03_Saliva | chr22:50025915-50026188 | C22orf34 | Island | Body |
| 20M03_Saliva | chr22:50451054-50451245 | IL17REL;IL17REL | S_Shore | 5'UTR;1stExon |
| 20M03_Saliva | chr22:50585197-50585393 | MOV10L1;MOV10L1;MOV10L1;MOV10L1 | | Body;TSS1500;Body;Body |
| 20M03_Saliva | chr1:2082315-2082458 | PRKCZ;PRKCZ;PRKCZ | Island | Body;Body;Body |
| 20M03_Saliva | chr1:6526051-6526309 | TNFRSF25;TNFRSF25;TNFRSF25; | Island | Body;Body;Body;Body;Body;Body |

| | | | | |
|---|---|---|---|---|
| 20M03_Saliva | chr1:7692321-7692367 | CAMTA1 | | Body |
| 20M03_Saliva | chr1:17634543-17634716 | PADI4 | | TSS200 |
| 20M03_Saliva | chr1:26097918-26098214 | MAN1C1 | Island | Body |
| 20M03_Saliva | chr1:27687232-27687695 | MAP3K6 | Island | Body |
| 20M03_Saliva | chr1:54562040-54562213 | C1orf83 | | Body |
| 20M03_Saliva | chr1:151810887-151811364 | C2CD4D;LOC100132111 | Island | Body;TSS200 |
| 20M03_Saliva | chr1:165326269-165326345 | LMX1A | Island | TSS1500 |
| 20M03_Saliva | chr1:202130344-202131184 | PTPN7;PTPN7 | | TSS1500;TSS1500 |
| 20M03_Saliva | chr1:209929496-209929546 | TRAF3IP3 | | TSS200 |
| 20M03_Saliva | chr1:223566669-223566794 | C1orf65 | Island | TSS200 |
| 20M03_Saliva | chr1:223936508-223936812 | CAPN2;CAPN2 | Island | Body;Body |
| 20M03_Saliva | chr1:248903300-248903467 | LOC646627 | | TSS200 |
| 20M03_Saliva | chr2:25473529-25473895 | DNMT3A;DNMT3A;DNMT3A | N_Shore | Body;Body;Body |
| 20M03_Saliva | chr2:63283967-63284481 | OTX1 | Island | 3'UTR |
| 20M03_Saliva | chr2:69001774-69002378 | ARHGAP25;ARHGAP25;ARHGAP25;ARHGAP25 | | Body;TSS200;TSS200;Body |
| 20M03_Saliva | chr2:113931518-113931565 | PSD4;LOC440839 | | TSS200;Body |
| 20M03_Saliva | chr2:131797438-131798087 | ARHGEF4;ARHGEF4 | Island | Body;Body |
| 20M03_Saliva | chr2:219735876-219736549 | WNT6 | N_Shore | Body |
| 20M03_Saliva | chr2:219738226-219738732 | WNT6 | Island | Body |
| 20M03_Saliva | chr2:220313153-220313657 | SPEG | Island | Body |
| 20M03_Saliva | chr2:225266263-225266346 | FAM124B;FAM124B | | 1stExon;1stExon |
| 20M03_Saliva | chr2:233924713-233925030 | INPP5D;INPP5D | N_Shore | TSS1500;TSS1500 |
| 20M03_Saliva | chr3:9943828-9943937 | IL17RE;IL17RE;IL17RE | | TSS1500;TSS1500;TSS1500 |
| 20M03_Saliva | chr3:11078979-11079298 | SLC6A1 | | 3'UTR |
| 20M03_Saliva | chr3:42742993-42743203 | HHATL;HHATL | | Body;5'UTR |
| 20M03_Saliva | chr3:52529195-52529341 | STAB1 | | TSS200 |
| 20M03_Saliva | chr3:128779498-128779601 | GP9 | | TSS200 |
| 20M03_Saliva | chr3:181421703-181422217 | SOX2OT | | Body |
| 20M03_Saliva | chr3:181428298-181428462 | SOX2OT;SOX2 | N_Shore | Body;TSS1500 |
| 20M03_Saliva | chr3:181428547-181428697 | SOX2OT;SOX2 | N_Shore | Body;TSS1500 |
| 20M03_Saliva | chr3:183273245-183273894 | KLHL6 | | 1stExon |
| 20M03_Saliva | chr4:683148-683772 | MFSD7 | S_Shore | TSS200 |
| 20M03_Saliva | chr4:2933748-2934188 | MFSD10;MFSD10 | Island | Body;Body |
| 20M03_Saliva | chr4:154711280-154711347 | SFRP2 | N_Shore | TSS1500 |
| 20M03_Saliva | chr4:183061830-183061955 | MGC45800 | N_Shore | Body |
| 20M03_Saliva | chr5:1108315-1108651 | SLC12A7 | N_Shore | Body |
| 20M03_Saliva | chr5:1494980-1495356 | LPCAT1 | Island | Body |
| 20M03_Saliva | chr5:1878499-1878672 | IRX4 | Island | Body |
| 20M03_Saliva | chr5:37835111-37835506 | GDNF;GDNF;GDNF | Island | 5'UTR;TSS1500;5'UTR |
| 20M03_Saliva | chr5:134870506-134870517 | NEUROG1;NEUROG1 | N_Shore | 1stExon;3'UTR |
| 20M03_Saliva | chr5:135416381-135416529 | MIR886 | Island | TSS200 |
| 20M03_Saliva | chr5:139227979-139228153 | NRG2;NRG2;NRG2;NRG2 | Island | Body;Body;Body;Body |
| 20M03_Saliva | chr5:148810061-148810103 | MIR145;LOC728264 | | TSS200;Body |
| 20M03_Saliva | chr6:29943401-29943414 | HCG9 | N_Shore | Body |
| 20M03_Saliva | chr6:30039130-30039142 | RNF39;RNF39 | Island | Body;Body |
| 20M03_Saliva | chr6:30039175-30039239 | RNF39;RNF39 | Island | Body;Body |
| 20M03_Saliva | chr6:30039376-30039408 | RNF39;RNF39 | Island | Body;Body |
| 20M03_Saliva | chr6:30297257-30297338 | TRIM39;TRIM39 | S_Shelf | Body;Body |
| 20M03_Saliva | chr6:31554548-31555016 | LST1;LST1;LST1;LST1;LST1;LST1;LST1;LST1;LST1 | | Body;TSS1500;TSS1500;5'UTR;TSS1500;5'UTR;Body;5'UTR;1stExon |
| 20M03_Saliva | chr6:31691198-31691426 | C6orf25;C6orf25;C6orf25;C6orf2 | N_Shore | 1stExon;1stExon;1stExon;1stExon;1stExon;1stExon |
| 20M03_Saliva | chr6:32606845-32607174 | HLA-DQA1 | | Body |
| 20M03_Saliva | chr6:32813337-32813531 | PSMB8;TAP1 | S_Shore | TSS1500;3'UTR |
| 20M03_Saliva | chr6:33400021-33400505 | SYNGAP1 | S_Shelf | Body |
| 20M03_Saliva | chr6:41528395-41528785 | FOXP4;FOXP4;FOXP4 | Island | 5'UTR;5'UTR;5'UTR |
| 20M03_Saliva | chr6:157932074-157932180 | ZDHHC14;ZDHHC14 | | Body;Body |
| 20M03_Saliva | chr6:158507753-158508188 | SYNJ2 | Island | Body |
| 20M03_Saliva | chr7:1094737-1095720 | C7orf50;C7orf50;C7orf50 | N_Shelf | Body;Body;Body |
| 20M03_Saliva | chr7:1858916-1859172 | MAD1L1;MAD1L1;MAD1L1 | | Body;Body;Body |
| 20M03_Saliva | chr7:1980222-1980479 | MAD1L1;MAD1L1;MAD1L1 | S_Shore | Body;Body;Body |
| 20M03_Saliva | chr7:2116368-2116648 | MAD1L1;MAD1L1;MAD1L1 | Island | Body;Body;Body |
| 20M03_Saliva | chr7:2774195-2774654 | GNA12 | N_Shore | Body |
| 20M03_Saliva | chr7:4848683-4848934 | RADIL | Island | Body |
| 20M03_Saliva | chr7:30951272-30951498 | AQP1 | | TSS200 |
| 20M03_Saliva | chr7:42267547-42267747 | GLI3 | Island | 5'UTR |

| | | | | |
|---|---|---|---|---|
| 20M03_Saliva | chr7:45018658-45018849 | MYO1G;MYO1G | | 5'UTR;1stExon |
| 20M03_Saliva | chr7:45066858-45067319 | CCM2;CCM2;CCM2;CCM2;CCM2 | | Body;TSS1500;Body;Body;Body |
| 20M03_Saliva | chr7:150434452-150434541 | GIMAP5;GIMAP5 | | 1stExon;5'UTR |
| 20M03_Saliva | chr7:155556590-155556729 | RBM33 | | Body |
| 20M03_Saliva | chr7:157647109-157647241 | PTPRN2;PTPRN2;PTPRN2 | N_Shore | Body;Body;Body |
| 20M03_Saliva | chr8:21771252-21771278 | DOK2 | S_Shore | TSS200 |
| 20M03_Saliva | chr8:27469186-27469338 | CLU;CLU;CLU;CLU | N_Shelf | 1stExon;Body;5'UTR;TSS1500 |
| 20M03_Saliva | chr8:42037170-42037527 | PLAT;PLAT | | Body;Body |
| 20M03_Saliva | chr8:141609338-141609470 | EIF2C2;EIF2C2 | | Body;Body |
| 20M03_Saliva | chr8:144408230-144408588 | TOP1MT | | Body |
| 20M03_Saliva | chr8:144635478-144635547 | GSDMD | Island | TSS200 |
| 20M03_Saliva | chr9:96715687-96716527 | BARX1 | Island | Body |
| 20M03_Saliva | chr9:140356475-140356830 | PNPLA7;PNPLA7 | Island | Body;Body |
| 20M03_Saliva | chr10:6214016-6214079 | PFKFB3 | | Body |
| 20M03_Saliva | chr10:45958759-45958885 | MARCH8;MARCH8;MARCH8 | | 3'UTR;3'UTR;3'UTR |
| 20M03_Saliva | chr10:49892741-49892930 | WDFY4 | | TSS1500 |
| 20M03_Saliva | chr10:49892954-49893336 | WDFY4 | | TSS1500 |
| 20M03_Saliva | chr10:49893406-49893549 | WDFY4 | | TSS200 |
| 20M03_Saliva | chr10:73848764-73848830 | SPOCK2;SPOCK2;SPOCK2;SPOC | S_Shore | 5'UTR;5'UTR |
| 20M03_Saliva | chr10:104535854-104536035 | C10orf26;C10orf26 | | TSS200;Body |
| 20M03_Saliva | chr10:123923066-123923099 | TACC2;TACC2;TACC2;TACC2 | Island | Body;TSS200;TSS200;Body |
| 20M03_Saliva | chr10:134045913-134046237 | STK32C | S_Shelf | Body |
| 20M03_Saliva | chr11:1873884-1874049 | LSP1 | | TSS1500 |
| 20M03_Saliva | chr11:2321800-2321955 | C11orf21;TSPAN32;C11orf21 | | Body;TSS1500;Body |
| 20M03_Saliva | chr11:2322329-2322507 | C11orf21;TSPAN32;C11orf21 | | Body;TSS1500;Body |
| 20M03_Saliva | chr11:2322618-2322674 | C11orf21;TSPAN32;C11orf21 | | Body;TSS1500;Body |
| 20M03_Saliva | chr11:2322802-2323059 | C11orf21;TSPAN32;C11orf21 | | Body;TSS1500;Body |
| 20M03_Saliva | chr11:2323801-2323938 | TSPAN32;C11orf21;C11orf21 | | Body;TSS1500;TSS1500 |
| 20M03_Saliva | chr11:3647654-3648013 | TRPC2 | | TSS200 |
| 20M03_Saliva | chr11:35441777-35442225 | SLC1A2 | Island | TSS1500 |
| 20M03_Saliva | chr11:47399994-47400113 | SPI1;SPI1;SPI1;SPI1 | Island | 5'UTR;5'UTR;1stExon;1stExon |
| 20M03_Saliva | chr11:60738971-60739005 | CD6 | | TSS200 |
| 20M03_Saliva | chr11:63273543-63274030 | LGALS12;LGALS12;LGALS12;LGALS12;LGALS12;LGALS12 | | 5'UTR;1stExon;5'UTR;1stExon;1stExon;5'UTR |
| 20M03_Saliva | chr11:63973846-63974153 | FERMT3;FERMT3 | N_Shore | TSS1500;TSS1500 |
| 20M03_Saliva | chr11:64107374-64108241 | CCDC88B | N_Shelf | TSS1500 |
| 20M03_Saliva | chr11:64563983-64564012 | MAP4K2 | | Body |
| 20M03_Saliva | chr11:64981227-64981316 | SLC22A20 | | TSS200 |
| 20M03_Saliva | chr11:65407399-65408566 | SIPA1;SIPA1 | N_Shore | TSS200;5'UTR |
| 20M03_Saliva | chr11:66055170-66055327 | YIF1A | N_Shore | Body |
| 20M03_Saliva | chr11:66084841-66085299 | CD248 | S_Shore | TSS1500 |
| 20M03_Saliva | chr11:67171145-67171585 | TBC1D10C | S_Shelf | TSS1500 |
| 20M03_Saliva | chr11:67205113-67205291 | PTPRCAP;PTPRCAP | S_Shore | 5'UTR;1stExon |
| 20M03_Saliva | chr11:128729883-128730011 | KCNJ1;KCNJ1;KCNJ1;KCNJ1 | | 5'UTR;5'UTR;5'UTR;5'UTR |
| 20M03_Saliva | chr11:134254573-134254690 | B3GAT1;B3GAT1 | S_Shore | Body;Body |
| 20M03_Saliva | chr12:51718088-51718112 | BIN2 | | TSS200 |
| 20M03_Saliva | chr12:54891491-54891636 | NCKAP1L | | TSS200 |
| 20M03_Saliva | chr12:105114162-105114399 | CHST11 | | Body |
| 20M03_Saliva | chr12:117797056-117797385 | NOS1 | N_Shore | 5'UTR |
| 20M03_Saliva | chr12:123518833-123519157 | PITPNM2 | | Body |
| 20M03_Saliva | chr12:124950720-124950990 | NCOR2;NCOR2 | Island | Body;Body |
| 20M03_Saliva | chr13:51639953-51640315 | GUCY1B2 | | Body |
| 20M03_Saliva | chr13:107145933-107146164 | EFNB2 | S_Shore | Body |
| 20M03_Saliva | chr13:114261869-114261986 | TFDP1;TFDP1 | S_Shore | Body;Body |
| 20M03_Saliva | chr13:114814333-114814643 | RASA3 | Island | Body |
| 20M03_Saliva | chr14:37052470-37053169 | NKX2-8 | N_Shore | TSS1500 |
| 20M03_Saliva | chr14:75988800-75989203 | BATF;BATF | | 5'UTR;1stExon |
| 20M03_Saliva | chr14:105532012-105532252 | GPR132 | | TSS1500 |
| 20M03_Saliva | chr15:37170381-37170554 | LOC145845 | N_Shelf | Body |
| 20M03_Saliva | chr15:37387304-37387523 | MEIS2;MEIS2;MEIS2;MEIS2;MEIS | N_Shore | Body;Body;Body;Body;Body;Body;Body |
| 20M03_Saliva | chr15:86315115-86315211 | KLHL25;MIR1276 | S_Shelf | 5'UTR;TSS1500 |
| 20M03_Saliva | chr16:1583810-1584118 | IFT140;TMEM204 | Island | Body;TSS1500 |
| 20M03_Saliva | chr16:4714080-4714443 | MGRN1;MGRN1;MGRN1;MGRN1 | | Body;Body;Body;Body |
| 20M03_Saliva | chr16:8738240-8738441 | C16orf68 | | Body |
| 20M03_Saliva | chr16:28505659-28505959 | APOB48R | N_Shelf | TSS1500 |
| 20M03_Saliva | chr16:28995882-28996533 | LAT;LAT;LAT;LAT | | TSS1500;TSS1500;TSS1500;TSS1500 |

| | | | | |
|---|---|---|---|---|
| 20M03_Saliva | chr16:29674959-29676071 | SPN;SPN | N_Shore | 5'UTR;5'UTR |
| 20M03_Saliva | chr16:29756951-29757375 | C16orf54 | | 5'UTR |
| 20M03_Saliva | chr16:50715135-50715260 | SNX20;SNX20;SNX20;SNX20;SNX20;SNX20 | | 1stExon;5'UTR;1stExon;5'UTR;5'UTR;1stExon |
| 20M03_Saliva | chr16:57576285-57576394 | GPR114 | | TSS1500 |
| 20M03_Saliva | chr16:88767261-88767723 | RNF166 | S_Shore | Body |
| 20M03_Saliva | chr16:89024511-89024657 | CBFA2T3 | | Body |
| 20M03_Saliva | chr16:89033895-89034178 | CBFA2T3 | N_Shore | Body |
| 20M03_Saliva | chr16:89041793-89042185 | CBFA2T3 | | Body |
| 20M03_Saliva | chr17:998432-998711 | ABR;ABR;ABR | | Body;Body;Body |
| 20M03_Saliva | chr17:2699689-2700093 | RAP1GAP2;RAP1GAP2 | | TSS200;TSS200 |
| 20M03_Saliva | chr17:3704471-3704621 | ITGAE;ITGAE | | 1stExon;5'UTR |
| 20M03_Saliva | chr17:3819339-3819390 | P2RX1 | | Body |
| 20M03_Saliva | chr17:3819880-3820150 | P2RX1;P2RX1 | | 1stExon;5'UTR |
| 20M03_Saliva | chr17:4337082-4337231 | SPNS3 | | TSS200 |
| 20M03_Saliva | chr17:5138259-5138645 | C17orf87 | | TSS200 |
| 20M03_Saliva | chr17:33416834-33417113 | RFFL | S_Shore | TSS1500 |
| 20M03_Saliva | chr17:40936498-40936767 | WNK4 | Island | Body |
| 20M03_Saliva | chr17:43046859-43047115 | C1QL1 | N_Shore | TSS1500 |
| 20M03_Saliva | chr17:48049952-48050338 | DLX4;DLX4 | Island | TSS200;Body |
| 20M03_Saliva | chr17:55533142-55533389 | MSI2;MSI2 | | Body;Body |
| 20M03_Saliva | chr17:56408688-56408694 | MIR142 | | TSS200 |
| 20M03_Saliva | chr17:62009607-62009651 | CD79B;CD79B;CD79B | | 1stExon;1stExon;1stExon |
| 20M03_Saliva | chr17:72442928-72443179 | GPRC5C;GPRC5C | Island | Body;Body |
| 20M03_Saliva | chr17:72709026-72709572 | CD300LF;RAB37;CD300LF | | 5'UTR;Body;1stExon |
| 20M03_Saliva | chr17:75471193-75471246 | SEPT9;SEPT9;SEPT9;SEPT9;SEPT9;SEPT9;SEPT9 | | Body;TSS200;5'UTR;Body;Body;Body;Body |
| 20M03_Saliva | chr17:76126702-76127217 | TMC8;TMC6 | N_Shore | TSS200;5'UTR |
| 20M03_Saliva | chr17:77916733-77916892 | TBC1D16 | Island | Body |
| 20M03_Saliva | chr17:79005662-79006087 | FLJ90757 | N_Shore | Body |
| 20M03_Saliva | chr17:79259040-79259699 | SLC38A10;SLC38A10 | N_Shore | Body;Body |
| 20M03_Saliva | chr19:681420-681815 | FSTL3 | Island | Body |
| 20M03_Saliva | chr19:840737-841082 | PRTN3 | N_Shelf | TSS1500 |
| 20M03_Saliva | chr19:947712-947936 | ARID3A | Island | Body |
| 20M03_Saliva | chr19:3178759-3179364 | S1PR4 | Island | TSS200 |
| 20M03_Saliva | chr19:4916593-4916983 | UHRF1;UHRF1 | Island | Body;Body |
| 20M03_Saliva | chr19:5074224-5074717 | KDM4B | N_Shore | Body |
| 20M03_Saliva | chr19:11530021-11530060 | RGL3;RGL3 | N_Shore | TSS200;TSS200 |
| 20M03_Saliva | chr19:12876846-12877188 | HOOK2;HOOK2 | N_Shore | Body;Body |
| 20M03_Saliva | chr19:14550846-14550999 | PKN1;PKN1 | N_Shore | TSS1500;Body |
| 20M03_Saliva | chr19:17488042-17488281 | PLVAP | Island | 1stExon |
| 20M03_Saliva | chr19:17516712-17517221 | BST2 | | TSS1500 |
| 20M03_Saliva | chr19:36233324-36233469 | TMEM149;TMEM149 | S_Shore | 1stExon;5'UTR |
| 20M03_Saliva | chr19:39826150-39826677 | GMFG | | Body |
| 20M03_Saliva | chr19:46318633-46318918 | RSPH6A | Island | TSS200 |
| 20M03_Saliva | chr19:49340489-49340765 | PLEKHA4;PLEKHA4;HSD17B14 | Island | 3'UTR;3'UTR;TSS1500 |
| 20M03_Saliva | chr19:51628115-51628204 | SIGLEC9 | | TSS200 |
| 20M03_Saliva | chr19:51875451-51875986 | NKG7 | S_Shelf | Body |
| 20M03_Saliva | chr20:3065343-3065423 | AVP;AVP | S_Shore | 5'UTR;1stExon |
| 20M03_Saliva | chr20:4721490-4721683 | PRNT;PRNT;PRNT | | TSS200;TSS200;TSS200 |
| 20M03_Saliva | chr20:22566877-22566976 | FOXA2 | Island | TSS1500 |
| 20M03_Saliva | chr20:23066944-23067033 | CD93;CD93 | | 5'UTR;1stExon |
| 20M03_Saliva | chr20:35274570-35274715 | SLA2;SLA2 | | TSS200;TSS200 |
| 20M03_Saliva | chr20:37433745-37433803 | PPP1R16B | N_Shore | TSS1500 |
| 20M03_Saliva | chr20:57582856-57582918 | CTSZ | S_Shore | TSS1500 |
| 20M03_Saliva | chr20:57583009-57583195 | CTSZ | S_Shore | TSS1500 |
| 20M03_Saliva | chr20:62318426-62318588 | RTEL1;RTEL1 | Island | Body;Body |
| 20M03_Saliva | chr21:39870397-39870704 | ERG;ERG;ERG;ERG;ERG;ERG | | Body;1stExon;1stExon;5'UTR;5'UTR;Body |
| 20M03_Saliva | chr21:46340776-46341006 | ITGB2;ITGB2;ITGB2 | | 5'UTR;5'UTR;1stExon |
| 20M03_Saliva | chr22:19710880-19711007 | GP1BB;SEPT5 | Island | TSS200;3'UTR |
| 20M03_Saliva | chr22:19744062-19744078 | TBX1;TBX1;TBX1 | Island | TSS200;TSS200;TSS200 |
| 20M03_Saliva | chr22:19746156-19746368 | TBX1;TBX1;TBX1 | Island | 5'UTR;5'UTR;5'UTR |
| 20M03_Saliva | chr22:24823509-24823554 | ADORA2A | S_Shelf | TSS200 |
| 20M03_Saliva | chr22:31003127-31003283 | TCN2 | | TSS200 |
| 20M03_Saliva | chr22:37545924-37546124 | IL2RB;IL2RB | | 5'UTR;1stExon |
| 20M03_Saliva | chr22:37962569-37962818 | CDC42EP1 | Island | Body |
| 20M03_Saliva | chr22:42828386-42828516 | NFAM1;NFAM1 | | 1stExon;5'UTR |

| | | | | |
|---|---|---|---|---|
| 20M03_Saliva | chr22:45073148-45073165 | PRR5;PRR5;PRR5;PRR5 | | 5'UTR;5'UTR;5'UTR;5'UTR |
| 20M05_Saliva | chr1:855425-856059 | FLJ39609 | S_Shore | TSS1500 |
| 20M05_Saliva | chr1:1217103-1217511 | SCNN1D;SCNN1D | | TSS1500;TSS1500 |
| 20M05_Saliva | chr1:1245014-1245184 | PUSL1 | Island | Body |
| 20M05_Saliva | chr1:1266178-1266504 | TAS1R3 | N_Shore | TSS1500 |
| 20M05_Saliva | chr1:1266657-1267193 | TAS1R3 | N_Shore | TSS200 |
| 20M05_Saliva | chr1:1268144-1268942 | TAS1R3 | Island | Body |
| 20M05_Saliva | chr1:1602585-1602685 | LOC728661;CDK11B;CDK11B;C | Island | Body;Body;Body;Body;Body;Body;5'UTR |
| 20M05_Saliva | chr1:1845940-1846340 | CALML6 | | TSS1500 |
| 20M05_Saliva | chr1:2084391-2084595 | PRKCZ;PRKCZ;PRKCZ | S_Shore | Body;Body;Body |
| 20M05_Saliva | chr1:2138757-2139905 | C1orf86 | S_Shore | 5'UTR |
| 20M05_Saliva | chr1:2170956-2171253 | SKI | | Body |
| 20M05_Saliva | chr1:2230640-2230923 | SKI | N_Shore | Body |
| 20M05_Saliva | chr1:2262232-2262474 | MORN1 | N_Shelf | Body |
| 20M05_Saliva | chr1:2265881-2266933 | MORN1 | N_Shore | Body |
| 20M05_Saliva | chr1:2284167-2284821 | MORN1;LOC100129534 | | Body;TSS200 |
| 20M05_Saliva | chr1:2979311-2980399 | FLJ42875 | Island | Body |
| 20M05_Saliva | chr1:3276413-3277000 | PRDM16;PRDM16 | S_Shore | Body;Body |
| 20M05_Saliva | chr1:3606892-3607401 | TP73;TP73;TP73;TP73 | N_Shore | TSS1500;TSS1500;Body;TSS1500 |
| 20M05_Saliva | chr1:6390473-6390830 | ACOT7;ACOT7;ACOT7;ACOT7 | | Body;Body;Body;Body |
| 20M05_Saliva | chr1:6445632-6446787 | ACOT7;ACOT7 | | Body;1stExon |
| 20M05_Saliva | chr1:6579837-6580525 | PLEKHG5;PLEKHG5 | | 5'UTR;1stExon |
| 20M05_Saliva | chr1:9130576-9130886 | SLC2A5;SLC2A5;SLC2A5 | N_Shore | TSS1500;Body;TSS1500 |
| 20M05_Saliva | chr1:10707917-10708091 | CASZ1;CASZ1 | Island | Body;Body |
| 20M05_Saliva | chr1:10839450-10839574 | CASZ1;CASZ1 | | 5'UTR;5'UTR |
| 20M05_Saliva | chr1:15256054-15256222 | KIAA1026;KIAA1026;KIAA1026;KIAA1026 | | Body;TSS1500;Body;Body |
| 20M05_Saliva | chr1:15271830-15272326 | KIAA1026;KIAA1026;KIAA1026;KIAA1026;KIAA1026 | | Body;5'UTR;TSS1500;Body;Body |
| 20M05_Saliva | chr1:16061408-16061554 | SLC25A34 | S_Shelf | TSS1500 |
| 20M05_Saliva | chr1:16062720-16062757 | SLC25A34 | | TSS200 |
| 20M05_Saliva | chr1:17381800-17381980 | SDHB | S_Shore | TSS1500 |
| 20M05_Saliva | chr1:17575420-17575440 | PADI3 | | TSS200 |
| 20M05_Saliva | chr1:18806700-18806982 | KLHDC7A | N_Shelf | TSS1500 |
| 20M05_Saliva | chr1:20141662-20141781 | RNF186;RNF186 | | 5'UTR;1stExon |
| 20M05_Saliva | chr1:24152579-24152774 | HMGCL;HMGCL | | TSS1500;TSS1500 |
| 20M05_Saliva | chr1:24469564-24470205 | IL22RA1 | | 1stExon |
| 20M05_Saliva | chr1:25257978-25258146 | RUNX3;RUNX3 | Island | TSS1500;Body |
| 20M05_Saliva | chr1:25258225-25258332 | RUNX3;RUNX3 | Island | TSS1500;Body |
| 20M05_Saliva | chr1:26145328-26145437 | FAM54B;FAM54B;FAM54B | N_Shore | TSS1500;TSS1500;TSS1500 |
| 20M05_Saliva | chr1:26503776-26504548 | CNKSR1;CNKSR1 | | TSS1500;TSS1500 |
| 20M05_Saliva | chr1:27287468-27287776 | C1orf172 | S_Shore | TSS1500 |
| 20M05_Saliva | chr1:27676205-27676652 | SYTL1 | Island | Body |
| 20M05_Saliva | chr1:28843736-28844028 | SNHG3-RCC1;RCC1;SNHG3-RC | N_Shore | Body;TSS1500;Body;TSS1500;5'UTR;TSS1500 |
| 20M05_Saliva | chr1:35220224-35220839 | GJB5 | | TSS1500 |
| 20M05_Saliva | chr1:36771714-36771793 | C1orf113;C1orf113 | N_Shore | TSS1500;TSS1500 |
| 20M05_Saliva | chr1:39407387-39407651 | RHBDL2;RHBDL2 | | 1stExon;5'UTR |
| 20M05_Saliva | chr1:40349894-40350052 | TRIT1 | S_Shore | TSS1500 |
| 20M05_Saliva | chr1:43751270-43751563 | C1orf210;C1orf210;C1orf210 | | 1stExon;5'UTR;TSS200 |
| 20M05_Saliva | chr1:43814169-43815035 | MPL | N_Shore | Body |
| 20M05_Saliva | chr1:44726943-44727313 | ERI3 | | Body |
| 20M05_Saliva | chr1:45082704-45083278 | RNF220 | N_Shore | Body |
| 20M05_Saliva | chr1:46088486-46089045 | CCDC17 | Island | Body |
| 20M05_Saliva | chr1:46859671-46859774 | FAAH | N_Shore | TSS1500 |
| 20M05_Saliva | chr1:55089182-55089372 | ACOT11;FAM151A;FAM151A | | Body;1stExon;5'UTR |
| 20M05_Saliva | chr1:57285156-57285466 | C1orf168;C1orf168 | | 1stExon;5'UTR |
| 20M05_Saliva | chr1:59043208-59043280 | TACSTD2 | Island | TSS200 |
| 20M05_Saliva | chr1:100111387-100111727 | PALMD | | TSS200 |
| 20M05_Saliva | chr1:117753591-117753741 | VTCN1 | | TSS200 |
| 20M05_Saliva | chr1:149908085-149908524 | MTMR11;MTMR11;MTMR11 | | Body;5'UTR;1stExon |
| 20M05_Saliva | chr1:151826123-151826533 | THEM5;THEM5 | | 1stExon;5'UTR |
| 20M05_Saliva | chr1:152730027-152730426 | KPRP | N_Shelf | TSS1500 |
| 20M05_Saliva | chr1:152758453-152758755 | LCE1E | | TSS1500 |
| 20M05_Saliva | chr1:152973720-152973957 | SPRR3;SPRR3 | | TSS1500;TSS1500 |
| 20M05_Saliva | chr1:153413871-153413940 | S100A7L2 | | TSS1500 |
| 20M05_Saliva | chr1:153600762-153600865 | S100A13;S100A13;S100A13;S100A1;S100A13;S100A13 | | TSS1500;TSS200;TSS1500;TSS200;5'UTR;TSS1500 |

| | | | | |
|---|---|---|---|---|
| 20M05_Saliva | chr1:154519846-154520187 | TDRD10;TDRD10 | | Body;Body |
| 20M05_Saliva | chr1:154929734-154929791 | PBXIP1;PYGO2 | | TSS1500;3'UTR |
| 20M05_Saliva | chr1:155007166-155007323 | DCST1;DCST2;DCST1 | | Body;TSS1500;Body |
| 20M05_Saliva | chr1:156030809-156031182 | RAB25 | | TSS200 |
| 20M05_Saliva | chr1:156123352-156123500 | SEMA4A | | TSS200 |
| 20M05_Saliva | chr1:156217178-156218009 | PAQR6;PAQR6 | S_Shore | 5'UTR;5'UTR |
| 20M05_Saliva | chr1:161059267-161059654 | PVRL4;PVRL4 | | 5'UTR;1stExon |
| 20M05_Saliva | chr1:161088957-161089118 | PFDN2;NIT1 | S_Shore | TSS1500;Body |
| 20M05_Saliva | chr1:161168856-161169007 | NDUFS2;ADAMTS4 | N_Shelf | TSS1500;TSS200 |
| 20M05_Saliva | chr1:172413492-172413612 | C1orf105;PIGC;PIGC | S_Shore | Body;TSS1500;TSS1500 |
| 20M05_Saliva | chr1:178310534-178310650 | RASAL2;RASAL2 | | Body;TSS200 |
| 20M05_Saliva | chr1:183154778-183155056 | LAMC2;LAMC2 | | TSS1500;TSS1500 |
| 20M05_Saliva | chr1:201708500-201708558 | NAV1;NAV1 | N_Shore | TSS1500;Body |
| 20M05_Saliva | chr1:201979478-201979938 | ELF3;ELF3 | | TSS1500;TSS1500 |
| 20M05_Saliva | chr1:203829481-203830200 | SNRPE | N_Shore | TSS1500 |
| 20M05_Saliva | chr1:205602864-205603464 | ELK4;ELK4 | S_Shore | TSS1500;TSS1500 |
| 20M05_Saliva | chr1:206643301-206643377 | IKBKE | | TSS1500 |
| 20M05_Saliva | chr1:207038584-207039008 | IL20 | | TSS1500 |
| 20M05_Saliva | chr1:207224740-207225128 | YOD1 | N_Shore | TSS1500 |
| 20M05_Saliva | chr1:221057236-221057573 | HLX | N_Shore | Body |
| 20M05_Saliva | chr1:223888431-223889141 | CAPN2 | | TSS1500 |
| 20M05_Saliva | chr1:228394509-228394607 | OBSCN;OBSCN | | TSS1500;TSS1500 |
| 20M05_Saliva | chr1:234350834-234350988 | SLC35F3 | S_Shore | Body |
| 20M05_Saliva | chr1:236686564-236686618 | LGALS8;LGALS8;LGALS8;LGALS8 | N_Shore | TSS200;TSS200;5'UTR;TSS1500 |
| 20M05_Saliva | chr1:241912510-241912768 | WDR64 | | Body |
| 20M05_Saliva | chr1:247611842-247611931 | NLRP3;NLRP3;NLRP3;NLRP3;NLI | N_Shelf | 3'UTR;3'UTR;3'UTR;3'UTR;3'UTR |
| 20M05_Saliva | chr2:3452407-3452563 | TTC15 | Island | Body |
| 20M05_Saliva | chr2:3683030-3683186 | COLEC11;COLEC11 | Island | Body;Body |
| 20M05_Saliva | chr2:9459386-9459596 | ASAP2;ASAP2 | Island | Body;Body |
| 20M05_Saliva | chr2:10302391-10302462 | C2orf48 | | Body |
| 20M05_Saliva | chr2:17996994-17997660 | MSGN1 | | TSS1500 |
| 20M05_Saliva | chr2:24398427-24398859 | C2orf84 | Island | Body |
| 20M05_Saliva | chr2:27531236-27531320 | UCN | Island | TSS200 |
| 20M05_Saliva | chr2:27665079-27665711 | KRTCAP3;NRBP1;KRTCAP3 | Island | TSS200;3'UTR;TSS200 |
| 20M05_Saliva | chr2:29204057-29204200 | FAM179A | | TSS200 |
| 20M05_Saliva | chr2:42795193-42795486 | MTA3 | N_Shore | TSS1500 |
| 20M05_Saliva | chr2:54086854-54087343 | GPR75;LOC100302652 | Island | 5'UTR;Body |
| 20M05_Saliva | chr2:69665094-69665232 | NFU1;NFU1;NFU1;NFU1 | S_Shore | TSS1500;TSS1500;TSS1500;TSS1500 |
| 20M05_Saliva | chr2:73495839-73496203 | FBXO41 | N_Shore | Body |
| 20M05_Saliva | chr2:74153086-74153317 | DGUOK;DGUOK | N_Shore | TSS1500;TSS1500 |
| 20M05_Saliva | chr2:74668072-74668672 | RTKN;RTKN;RTKN | Island | Body;TSS1500;TSS1500 |
| 20M05_Saliva | chr2:74753281-74753358 | DQX1;DQX1 | N_Shelf | 1stExon;5'UTR |
| 20M05_Saliva | chr2:97171137-97171243 | NEURL3 | N_Shelf | Body |
| 20M05_Saliva | chr2:102803290-102803435 | IL1RL2 | N_Shore | TSS200 |
| 20M05_Saliva | chr2:109559412-109559734 | EDAR | S_Shore | 5'UTR |
| 20M05_Saliva | chr2:113735377-113735612 | IL1F9 | | TSS1500 |
| 20M05_Saliva | chr2:113875226-113875723 | IL1RN;IL1RN;IL1RN | | TSS1500;TSS1500;TSS1500 |
| 20M05_Saliva | chr2:128458240-128458399 | SFT2D3 | N_Shore | TSS1500 |
| 20M05_Saliva | chr2:134326172-134326366 | NCKAP5;NCKAP5 | | TSS200;TSS200 |
| 20M05_Saliva | chr2:145146928-145147350 | ZEB2;ZEB2 | N_Shore | 3'UTR;3'UTR |
| 20M05_Saliva | chr2:161056520-161056639 | ITGB6 | | 1stExon |
| 20M05_Saliva | chr2:219472218-219472607 | PLCD4 | | TSS1500 |
| 20M05_Saliva | chr2:223916502-223916687 | KCNE4 | N_Shore | TSS1500 |
| 20M05_Saliva | chr2:231790529-231790813 | GPR55 | | TSS1500 |
| 20M05_Saliva | chr2:232259281-232259623 | B3GNT7 | N_Shore | TSS1500 |
| 20M05_Saliva | chr2:236443439-236444259 | AGAP1;AGAP1 | N_Shore | Body;Body |
| 20M05_Saliva | chr2:238499464-238499802 | RAB17;RAB17 | | 1stExon;5'UTR |
| 20M05_Saliva | chr2:240020028-240020174 | HDAC4 | | Body |
| 20M05_Saliva | chr2:240212763-240213173 | HDAC4 | | Body |
| 20M05_Saliva | chr2:241458939-241459502 | ANKMY1;ANKMY1 | N_Shore | Body;Body |
| 20M05_Saliva | chr2:241835462-241835740 | C2orf54;C2orf54 | | 1stExon;5'UTR |
| 20M05_Saliva | chr2:241974858-241975140 | SNED1 | N_Shore | Body |
| 20M05_Saliva | chr2:242127690-242127948 | ANO7;ANO7 | | TSS1500;TSS1500 |
| 20M05_Saliva | chr2:242716183-242716388 | GAL3ST2 | | TSS200 |
| 20M05_Saliva | chr2:242802009-242802192 | PDCD1 | N_Shelf | TSS1500 |
| 20M05_Saliva | chr3:3080327-3080621 | CNTN4;CNTN4;CNTN4 | | Body;TSS1500;Body |

| | | | | |
|---|---|---|---|---|
| 20M05_Saliva | chr3:9289747-9289893 | SRGAP3;SRGAP3 | | Body;Body |
| 20M05_Saliva | chr3:13028573-13028642 | IQSEC1 | | Body |
| 20M05_Saliva | chr3:14443428-14443473 | SLC6A6;SLC6A6;SLC6A6 | N_Shore | TSS1500;TSS1500;TSS1500 |
| 20M05_Saliva | chr3:16216094-16216606 | GALNTL2 | | TSS200 |
| 20M05_Saliva | chr3:42850848-42850974 | CCBP2 | | TSS200 |
| 20M05_Saliva | chr3:50341009-50341293 | HYAL1;HYAL1;HYAL1;HYAL1;HYAL1;HYAL1 | | TSS200;TSS200;5'UTR;TSS200;TSS200;TSS200 |
| 20M05_Saliva | chr3:51741135-51741473 | GRM2;GRM2;GRM2;GRM2 | Island | 1stExon;5'UTR;1stExon;5'UTR |
| 20M05_Saliva | chr3:52552868-52553433 | STAB1 | Island | Body |
| 20M05_Saliva | chr3:57743171-57743599 | SLMAP | S_Shore | TSS200 |
| 20M05_Saliva | chr3:64211328-64211659 | PRICKLE2 | | TSS200 |
| 20M05_Saliva | chr3:69061356-69061827 | C3orf64 | N_Shore | 5'UTR |
| 20M05_Saliva | chr3:69915189-69915449 | MITF;MITF;MITF | | Body;Body;TSS1500 |
| 20M05_Saliva | chr3:123339334-123339568 | MYLK;MYLK;MYLK;MYLK;MYLK;MYLK;MYLK;MYLK | | 1stExon;5'UTR;Body;Body;Body;5'UTR;1stExon;Body |
| 20M05_Saliva | chr3:124102766-124103696 | KALRN;KALRN | | Body;Body |
| 20M05_Saliva | chr3:129407365-129407548 | TMCC1;TMCC1;TMCC1;TMCC1 | | 1stExon;Body;Body;5'UTR |
| 20M05_Saliva | chr3:141087187-141087363 | ZBTB38 | | 5'UTR |
| 20M05_Saliva | chr3:149374066-149374914 | WWTR1;WWTR1;WWTR1 | N_Shore | Body;Body;Body |
| 20M05_Saliva | chr3:183959088-183959853 | VWA5B2;MIR1224 | Island | Body;TSS200 |
| 20M05_Saliva | chr3:191048308-191048753 | UTS2D;CCDC50;CCDC50;UTS2D | S_Shore | 1stExon;Body;Body;5'UTR |
| 20M05_Saliva | chr3:193096486-193096836 | ATP13A5 | | 1stExon |
| 20M05_Saliva | chr3:194833887-194834008 | C3orf21 | | Body |
| 20M05_Saliva | chr3:195943198-195943636 | OSTalpha | | TSS200 |
| 20M05_Saliva | chr3:196065106-196065357 | TM4SF19 | | 5'UTR |
| 20M05_Saliva | chr3:196065506-196065688 | TM4SF19 | | TSS1500 |
| 20M05_Saliva | chr4:671543-671851 | MYL5 | N_Shelf | TSS200 |
| 20M05_Saliva | chr4:715865-715950 | PCGF3 | N_Shore | 5'UTR |
| 20M05_Saliva | chr4:981485-981602 | IDUA;SLC26A1;SLC26A1;SLC26A | Island | Body;3'UTR;Body;3'UTR |
| 20M05_Saliva | chr4:1166839-1166933 | LOC100130872-SPON2;SPON2;S | S_Shore | Body;TSS200;5'UTR |
| 20M05_Saliva | chr4:1205447-1206150 | CTBP1;CTBP1 | N_Shore | 3'UTR;3'UTR |
| 20M05_Saliva | chr4:1208699-1208960 | CTBP1;CTBP1 | N_Shore | Body;Body |
| 20M05_Saliva | chr4:1225507-1225729 | CTBP1;CTBP1 | S_Shore | Body;Body |
| 20M05_Saliva | chr4:1243980-1244086 | CTBP1;CTBP1;C4orf42 | Island | TSS1500;TSS1500;TSS200 |
| 20M05_Saliva | chr4:1719983-1720184 | TMEM129;TMEM129 | N_Shelf | Body;Body |
| 20M05_Saliva | chr4:3387499-3387765 | RGS12;RGS12;RGS12 | | Body;Body;Body |
| 20M05_Saliva | chr4:3392103-3392318 | RGS12;RGS12;RGS12 | | Body;Body;Body |
| 20M05_Saliva | chr4:3409828-3409927 | RGS12;RGS12;RGS12 | | Body;Body;Body |
| 20M05_Saliva | chr4:8395941-8396410 | ACOX3;ACOX3 | | Body;Body |
| 20M05_Saliva | chr4:24796919-24797007 | SOD3 | | TSS200 |
| 20M05_Saliva | chr4:36283184-36283275 | DTHD1 | | TSS200 |
| 20M05_Saliva | chr4:39448160-39448464 | KLB | N_Shore | Body |
| 20M05_Saliva | chr4:40517938-40518065 | RBM47;RBM47;RBM47 | | 1stExon;5'UTR;5'UTR |
| 20M05_Saliva | chr4:42122106-42122161 | BEND4;BEND4 | | Body;Body |
| 20M05_Saliva | chr4:57547525-57547699 | HOPX;HOPX;HOPX;HOPX;HOPX;HOPX | | 5'UTR;1stExon;1stExon;5'UTR;5'UTR;1stExon |
| 20M05_Saliva | chr4:87770357-87770782 | SLC10A6;SLC10A6 | | 5'UTR;1stExon |
| 20M05_Saliva | chr4:114214080-114214185 | ANK2;ANK2;ANK2 | | Body;Body;Body |
| 20M05_Saliva | chr4:145566200-145566595 | HHIP | N_Shore | TSS1500 |
| 20M05_Saliva | chr4:184908351-184909018 | STOX2 | | Body |
| 20M05_Saliva | chr4:186732932-186733428 | SORBS2;SORBS2;SORBS2;SORBS2;SORBS2;SORBS2;SORBS2 | | 5'UTR;5'UTR;5'UTR;1stExon;TSS1500;1stExon;TSS1500 |
| 20M05_Saliva | chr5:191127-191296 | LRRC14B | N_Shore | TSS1500 |
| 20M05_Saliva | chr5:476497-476842 | SLC9A3 | Island | Body |
| 20M05_Saliva | chr5:510035-510421 | SLC9A3 | S_Shore | Body |
| 20M05_Saliva | chr5:663787-664666 | TPPP | N_Shore | 3'UTR |
| 20M05_Saliva | chr5:912383-913111 | TRIP13 | N_Shore | Body |
| 20M05_Saliva | chr5:1016279-1016768 | NKD2 | | Body |
| 20M05_Saliva | chr5:1052828-1052919 | SLC12A7 | N_Shore | Body |
| 20M05_Saliva | chr5:1282357-1282723 | TERT;TERT | N_Shore | Body;Body |
| 20M05_Saliva | chr5:1594282-1594808 | SDHAP3 | Island | Body |
| 20M05_Saliva | chr5:27038605-27038782 | CDH9;CDH9 | | 1stExon;5'UTR |
| 20M05_Saliva | chr5:53751272-53751493 | HSPB3 | | TSS200 |
| 20M05_Saliva | chr5:55529124-55529228 | ANKRD55;ANKRD55 | | 5'UTR;1stExon |
| 20M05_Saliva | chr5:71475111-71475356 | MAP1B | | Body |
| 20M05_Saliva | chr5:78985425-78986160 | CMYA5 | | TSS1500 |

| | | | | |
|---|---|---|---|---|
| 20M05_Saliva | chr5:112824614-112824700 | MCC | S_Shore | TSS200 |
| 20M05_Saliva | chr5:118690955-118691126 | TNFAIP8;TNFAIP8 | N_Shore | Body;TSS1500 |
| 20M05_Saliva | chr5:125800366-125800916 | GRAMD3;GRAMD3;GRAMD3;GRAMD3;GRAMD3 | | Body;5'UTR;TSS1500;Body;Body |
| 20M05_Saliva | chr5:136983354-136984054 | MIR874;KLHL3 | | TSS200;Body |
| 20M05_Saliva | chr5:138211043-138211184 | CTNNA1;LRRTM2;LRRTM2 | | Body;5'UTR;1stExon |
| 20M05_Saliva | chr5:140857941-140858046 | PCDHGA4;PCDHGA12;PCDHGA1 | Island | Body;Body;Body;Body;Body;Body;Body;Body;1stExon;Body;Body;Body;Body;Body;Body;Body;Body;Body;Body;Body;Body;1stExon;Body |
| 20M05_Saliva | chr5:142065539-142066938 | FGF1;FGF1;FGF1;FGF1;FGF1;FGF1;FGF1;FGF1 | | 5'UTR;Body;Body;5'UTR;1stExon;5'UTR;5'UTR;1stExon |
| 20M05_Saliva | chr5:149546331-149546366 | CDX1 | Island | TSS200 |
| 20M05_Saliva | chr5:149980526-149980682 | SYNPO | | TSS200 |
| 20M05_Saliva | chr5:154091976-154093260 | LARP1 | | TSS1500 |
| 20M05_Saliva | chr5:176793941-176794207 | RGS14 | N_Shore | Body |
| 20M05_Saliva | chr5:178565721-178565937 | ADAMTS2 | Island | Body |
| 20M05_Saliva | chr6:6614447-6614798 | LOC285780;LY86 | | Body;Body |
| 20M05_Saliva | chr6:8436218-8436296 | SLC35B3;SLC35B3;SLC35B3 | S_Shore | TSS1500;TSS1500;TSS1500 |
| 20M05_Saliva | chr6:10555808-10556523 | GCNT2;GCNT2 | | TSS200;Body |
| 20M05_Saliva | chr6:11771473-11771680 | C6orf105;C6orf105 | | Body;Body |
| 20M05_Saliva | chr6:24877462-24877624 | FAM65B;FAM65B;FAM65B | | 5'UTR;1stExon;5'UTR |
| 20M05_Saliva | chr6:26225258-26225389 | HIST1H3E | N_Shore | TSS200 |
| 20M05_Saliva | chr6:30174651-30174845 | TRIM26 | | 5'UTR |
| 20M05_Saliva | chr6:30175299-30175327 | TRIM26 | | 5'UTR |
| 20M05_Saliva | chr6:30290443-30290603 | HCG18;HCG18 | N_Shelf | Body;Body |
| 20M05_Saliva | chr6:30850405-30850581 | DDR1 | N_Shore | TSS1500 |
| 20M05_Saliva | chr6:30850829-30851517 | DDR1;DDR1 | N_Shore | TSS1500;TSS1500 |
| 20M05_Saliva | chr6:30851624-30851638 | DDR1;DDR1 | N_Shore | TSS1500;TSS1500 |
| 20M05_Saliva | chr6:30853007-30854161 | DDR1;DDR1 | S_Shore | 5'UTR;5'UTR |
| 20M05_Saliva | chr6:30882994-30883203 | VARS2;VARS2;VARS2 | S_Shore | Body;5'UTR;Body |
| 20M05_Saliva | chr6:30900197-30900299 | SFTA2 | | TSS1500 |
| 20M05_Saliva | chr6:30951225-30951394 | MUC21 | | TSS1500 |
| 20M05_Saliva | chr6:31023257-31023763 | HCG22 | | Body |
| 20M05_Saliva | chr6:31025933-31026215 | HCG22 | | Body |
| 20M05_Saliva | chr6:31082200-31082631 | PSORS1C1 | | TSS1500 |
| 20M05_Saliva | chr6:31096861-31097078 | PSORS1C1 | | 5'UTR |
| 20M05_Saliva | chr6:31111076-31111270 | CCHCR1;CCHCR1;CCHCR1 | | Body;Body;Body |
| 20M05_Saliva | chr6:31138925-31138982 | POU5F1 | | TSS1500 |
| 20M05_Saliva | chr6:31590563-31590640 | SNORA38;BAT2 | S_Shore | TSS1500;5'UTR |
| 20M05_Saliva | chr6:31636239-31636368 | CSNK2B | S_Shelf | Body |
| 20M05_Saliva | chr6:31688080-31688229 | LY6G6C | N_Shelf | Body |
| 20M05_Saliva | chr6:31689610-31690170 | LY6G6C | N_Shore | TSS200 |
| 20M05_Saliva | chr6:31692234-31692375 | C6orf25;C6orf25;C6orf25;C6orf2 | S_Shore | Body;Body;Body;Body;Body;Body |
| 20M05_Saliva | chr6:31696223-31696259 | DDAH2 | Island | Body |
| 20M05_Saliva | chr6:31696333-31696509 | DDAH2 | Island | Body |
| 20M05_Saliva | chr6:31868724-31868813 | ZBTB12 | N_Shore | Body |
| 20M05_Saliva | chr6:31913250-31914305 | C2;C2;CFB | | 3'UTR;3'UTR;TSS1500 |
| 20M05_Saliva | chr6:32014189-32014381 | TNXB;TNXB | | Body;TSS1500 |
| 20M05_Saliva | chr6:32014510-32014663 | TNXB;TNXB | | Body;TSS1500 |
| 20M05_Saliva | chr6:32014739-32015083 | TNXB;TNXB | | Body;TSS1500 |
| 20M05_Saliva | chr6:32050905-32051108 | TNXB | S_Shelf | Body |
| 20M05_Saliva | chr6:32060681-32061084 | TNXB | N_Shelf | Body |
| 20M05_Saliva | chr6:32068181-32068240 | TNXB | S_Shelf | 5'UTR |
| 20M05_Saliva | chr6:32078398-32078453 | TNXB | | TSS1500 |
| 20M05_Saliva | chr6:32078574-32078624 | TNXB | | TSS1500 |
| 20M05_Saliva | chr6:32097184-32097481 | ATF6B;ATF6B;FKBPL | N_Shore | TSS1500;TSS1500;Body |
| 20M05_Saliva | chr6:32120625-32120826 | PPT2;PRRT1;PPT2 | N_Shore | TSS1500;TSS1500;TSS1500 |
| 20M05_Saliva | chr6:32120895-32120955 | PPT2;PRRT1;PPT2 | N_Shore | TSS1500;TSS1500;TSS1500 |
| 20M05_Saliva | chr6:32121113-32121156 | PPT2;PPT2;PRRT1 | N_Shore | TSS200;TSS1500;TSS1500 |
| 20M05_Saliva | chr6:32191210-32191496 | NOTCH4 | | Body |
| 20M05_Saliva | chr6:32297670-32297822 | C6orf10 | | Body |
| 20M05_Saliva | chr6:32526263-32526364 | HLA-DRB6 | | Body |
| 20M05_Saliva | chr6:32552042-32552106 | HLA-DRB1 | Island | Body |
| 20M05_Saliva | chr6:32784327-32785141 | HLA-DOB | | Body |
| 20M05_Saliva | chr6:32805398-32805570 | TAP2;TAP2 | N_Shore | Body;Body |
| 20M05_Saliva | chr6:32976805-32976897 | HLA-DOA | S_Shore | Body |

| | | | | |
|---|---|---|---|---|
| 20M05_Saliva | chr6:33090564-33091242 | HLA-DPB2 | | Body |
| 20M05_Saliva | chr6:33131222-33131560 | COL11A2;COL11A2;COL11A2 | S_Shore | 3'UTR;3'UTR;3'UTR |
| 20M05_Saliva | chr6:33173278-33173501 | HSD17B8 | S_Shore | Body |
| 20M05_Saliva | chr6:33226913-33227015 | VPS52 | | Body |
| 20M05_Saliva | chr6:33228341-33228498 | VPS52 | | Body |
| 20M05_Saliva | chr6:33288296-33288372 | DAXX;DAXX;DAXX;DAXX | N_Shore | Body;Body;Body;Body |
| 20M05_Saliva | chr6:33638313-33638550 | ITPR3 | | Body |
| 20M05_Saliva | chr6:34024362-34024479 | GRM4 | Island | Body |
| 20M05_Saliva | chr6:35285967-35286360 | DEF6 | | Body |
| 20M05_Saliva | chr6:43478530-43478829 | C6orf154 | Island | TSS200 |
| 20M05_Saliva | chr6:47009737-47010177 | GPR110;GPR110 | | 5'UTR;5'UTR |
| 20M05_Saliva | chr6:49681299-49681310 | CRISP2;CRISP2;CRISP2;CRISP2;CRISP2 | | TSS200;TSS200;TSS200;TSS200;TSS200 |
| 20M05_Saliva | chr6:49681391-49681774 | CRISP2;CRISP2;CRISP2;CRISP2;CRISP2 | | TSS200;TSS200;TSS200;TSS200;TSS200 |
| 20M05_Saliva | chr6:56878397-56878665 | BEND6 | | Body |
| 20M05_Saliva | chr6:106546704-106546824 | PRDM1;PRDM1 | | Body;TSS200 |
| 20M05_Saliva | chr6:112375186-112375333 | WISP3;WISP3 | | TSS200;TSS200 |
| 20M05_Saliva | chr6:116381609-116382179 | FRK;FRK | | 1stExon;5'UTR |
| 20M05_Saliva | chr6:137366322-137366492 | IL20RA | S_Shore | TSS200 |
| 20M05_Saliva | chr6:152702387-152702463 | SYNE1;SYNE1 | | Body;Body |
| 20M05_Saliva | chr6:155568918-155569443 | TIAM2;TIAM2 | | Body;Body |
| 20M05_Saliva | chr6:157746584-157746724 | C6orf35 | S_Shore | TSS1500 |
| 20M05_Saliva | chr6:159052400-159052937 | TMEM181 | | Body |
| 20M05_Saliva | chr6:168417574-168417646 | KIF25;KIF25 | | TSS1500;TSS1500 |
| 20M05_Saliva | chr6:168857542-168858078 | SMOC2;SMOC2 | | Body;Body |
| 20M05_Saliva | chr7:194719-194798 | FAM20C | N_Shore | Body |
| 20M05_Saliva | chr7:906726-906799 | UNC84A;UNC84A | S_Shore | Body;Body |
| 20M05_Saliva | chr7:1022213-1022797 | CYP2W1 | N_Shore | TSS1500 |
| 20M05_Saliva | chr7:1024673-1024814 | CYP2W1 | Island | Body |
| 20M05_Saliva | chr7:1102344-1103262 | C7orf50;C7orf50;C7orf50 | N_Shelf | Body;Body;Body |
| 20M05_Saliva | chr7:1491787-1492452 | MICALL2 | Island | Body |
| 20M05_Saliva | chr7:1514361-1514623 | INTS1 | Island | Body |
| 20M05_Saliva | chr7:1956900-1957317 | MAD1L1;MAD1L1;MAD1L1 | | Body;Body;Body |
| 20M05_Saliva | chr7:2019882-2020131 | MAD1L1;MAD1L1;MAD1L1 | Island | Body;Body;Body |
| 20M05_Saliva | chr7:2094628-2094977 | MAD1L1;MAD1L1;MAD1L1 | | Body;Body;Body |
| 20M05_Saliva | chr7:2208492-2208908 | MAD1L1;MAD1L1;MAD1L1 | | Body;Body;Body |
| 20M05_Saliva | chr7:2611890-2612101 | IQCE;IQCE | | Body;Body |
| 20M05_Saliva | chr7:2701419-2701641 | TTYH3 | Island | 3'UTR |
| 20M05_Saliva | chr7:2802560-2802976 | GNA12 | Island | Body |
| 20M05_Saliva | chr7:2842593-2842746 | GNA12 | | Body |
| 20M05_Saliva | chr7:3033173-3033939 | CARD11 | | 5'UTR |
| 20M05_Saliva | chr7:4243286-4243604 | SDK1;SDK1 | | Body;Body |
| 20M05_Saliva | chr7:27143287-27143403 | HOXA2 | Island | TSS1500 |
| 20M05_Saliva | chr7:27153580-27153944 | HOXA3;HOXA3;HOXA3;HOXA3 | Island | 5'UTR;1stExon;5'UTR;5'UTR |
| 20M05_Saliva | chr7:27154720-27155173 | HOXA3;HOXA3;HOXA3 | N_Shore | 5'UTR;TSS1500;5'UTR |
| 20M05_Saliva | chr7:27171391-27171528 | HOXA4 | S_Shore | TSS1500 |
| 20M05_Saliva | chr7:27183262-27183369 | HOXA5;HOXA5 | Island | 5'UTR;1stExon |
| 20M05_Saliva | chr7:27184264-27184316 | HOXA5 | Island | TSS1500 |
| 20M05_Saliva | chr7:28998540-28998860 | TRIL | Island | TSS1500 |
| 20M05_Saliva | chr7:43288635-43288889 | HECW1 | Island | Body |
| 20M05_Saliva | chr7:48129797-48129904 | UPP1;UPP1 | Island | 5'UTR;5'UTR |
| 20M05_Saliva | chr7:50628718-50628968 | DDC;DDC;DDC | | 5'UTR;5'UTR;1stExon |
| 20M05_Saliva | chr7:56160717-56161020 | PHKG1 | | TSS200 |
| 20M05_Saliva | chr7:73256360-73256416 | WBSCR27 | N_Shore | Body |
| 20M05_Saliva | chr7:76026556-76026648 | SRCRB4D;ZP3 | N_Shore | Body;TSS1500 |
| 20M05_Saliva | chr7:92238086-92238248 | CDK6;CDK6 | | 3'UTR;3'UTR |
| 20M05_Saliva | chr7:97807643-97807741 | LMTK2 | | Body |
| 20M05_Saliva | chr7:98738998-98739782 | SMURF1;SMURF1 | N_Shelf | Body;Body |
| 20M05_Saliva | chr7:99691314-99691486 | MCM7;MCM7;MIR25 | | Body;Body;TSS200 |
| 20M05_Saliva | chr7:101768610-101768874 | CUX1;CUX1;CUX1 | | Body;Body;Body |
| 20M05_Saliva | chr7:102574445-102574504 | FBXL13;LRRC17;FBXL13;LRRC17 | | Body;Body;Body;Body |
| 20M05_Saliva | chr7:130020011-130020130 | CPA1 | N_Shore | TSS1500 |
| 20M05_Saliva | chr7:134464123-134464288 | CALD1;CALD1;CALD1 | | TSS200;TSS200;TSS200 |
| 20M05_Saliva | chr7:134575271-134576049 | CALD1;CALD1;CALD1;CALD1;CALD1 | | TSS1500;Body;TSS1500;Body;Body |
| 20M05_Saliva | chr7:135433540-135433790 | FAM180A;FAM180A | | 1stExon;5'UTR |
| 20M05_Saliva | chr7:142583296-142583731 | TRPV6;TRPV6 | | 1stExon;5'UTR |
| 20M05_Saliva | chr7:143747422-143747596 | OR2A5 | | TSS200 |

| | | | | |
|---|---|---|---|---|
| 20M05_Saliva | chr7:150021488-150021553 | ACTR3C;LRRC61;LRRC61 | S_Shore | TSS1500;5'UTR;5'UTR |
| 20M05_Saliva | chr7:150067712-150067896 | REPIN1;REPIN1;REPIN1;REPIN1 | N_Shore | 5'UTR;Body;TSS1500;5'UTR |
| 20M05_Saliva | chr7:150778724-150778881 | FASTK;TMUB1;TMUB1;FASTK | Island | TSS1500;Body;Body;TSS1500 |
| 20M05_Saliva | chr7:151493629-151493820 | PRKAG2;PRKAG2 | | 5'UTR;Body |
| 20M05_Saliva | chr7:157608573-157608642 | PTPRN2;PTPRN2;PTPRN2 | N_Shelf | Body;Body;Body |
| 20M05_Saliva | chr7:157638062-157638259 | PTPRN2;PTPRN2;PTPRN2 | | Body;Body;Body |
| 20M05_Saliva | chr7:157701024-157701262 | PTPRN2;PTPRN2;PTPRN2 | N_Shore | Body;Body;Body |
| 20M05_Saliva | chr7:157809235-157809385 | PTPRN2;PTPRN2;PTPRN2 | | Body;Body;Body |
| 20M05_Saliva | chr7:158059704-158060003 | PTPRN2;PTPRN2;PTPRN2 | N_Shore | Body;Body;Body |
| 20M05_Saliva | chr7:158243546-158243730 | PTPRN2;PTPRN2;PTPRN2 | | Body;Body;Body |
| 20M05_Saliva | chr8:644521-644663 | ERICH1 | | Body |
| 20M05_Saliva | chr8:1642380-1642577 | DLGAP2 | S_Shore | Body |
| 20M05_Saliva | chr8:1807061-1807575 | ARHGEF10 | | Body |
| 20M05_Saliva | chr8:1906312-1906498 | ARHGEF10 | | 3'UTR |
| 20M05_Saliva | chr8:10512459-10512769 | RP1L1;RP1L1 | | 1stExon;5'UTR |
| 20M05_Saliva | chr8:10530053-10530148 | C8orf74 | | TSS200 |
| 20M05_Saliva | chr8:19170080-19170432 | SH2D4A | N_Shore | TSS1500 |
| 20M05_Saliva | chr8:22084861-22085227 | PHYHIP;PHYHIP | | Body;Body |
| 20M05_Saliva | chr8:37699239-37699923 | GPR124 | Island | Body |
| 20M05_Saliva | chr8:57233269-57233391 | SDR16C5 | S_Shore | TSS200 |
| 20M05_Saliva | chr8:123875036-123875223 | ZHX2 | | 5'UTR |
| 20M05_Saliva | chr8:128808017-128808554 | MIR1204;PVT1 | S_Shore | TSS200;Body |
| 20M05_Saliva | chr8:141565971-141566221 | EIF2C2;EIF2C2 | | Body;Body |
| 20M05_Saliva | chr8:141588118-141588242 | EIF2C2;EIF2C2 | | Body;Body |
| 20M05_Saliva | chr8:142235430-142235763 | SLC45A4 | N_Shore | Body |
| 20M05_Saliva | chr8:143386260-143387265 | TSNARE1 | | Body |
| 20M05_Saliva | chr8:143463227-143463667 | TSNARE1 | | 5'UTR |
| 20M05_Saliva | chr8:143761503-143761811 | PSCA | | TSS1500 |
| 20M05_Saliva | chr8:143823809-143824001 | SLURP1;SLURP1 | S_Shelf | 5'UTR;1stExon |
| 20M05_Saliva | chr8:143851287-143851427 | LYNX1;LYNX1 | | TSS200;Body |
| 20M05_Saliva | chr8:143868013-143868158 | LY6D | | TSS200 |
| 20M05_Saliva | chr8:143868530-143868917 | LY6D | | TSS1500 |
| 20M05_Saliva | chr8:144120335-144120484 | C8orf31 | | TSS1500 |
| 20M05_Saliva | chr8:144589984-144590068 | ZC3H3 | | Body |
| 20M05_Saliva | chr8:144659627-144659883 | NAPRT1 | N_Shore | Body |
| 20M05_Saliva | chr8:144814911-144815275 | FAM83H | N_Shore | 5'UTR |
| 20M05_Saliva | chr8:144894951-144895243 | SCRIB;SCRIB | Island | Body;Body |
| 20M05_Saliva | chr8:145727318-145728630 | PPP1R16A | S_Shore | 3'UTR |
| 20M05_Saliva | chr8:145729126-145729149 | GPT | N_Shore | TSS1500 |
| 20M05_Saliva | chr9:34457129-34457440 | C9orf25 | N_Shore | Body |
| 20M05_Saliva | chr9:92291210-92291523 | LOC100129066 | N_Shore | Body |
| 20M05_Saliva | chr9:97848107-97848457 | C9orf3;MIR24-1 | N_Shore | 3'UTR;TSS200 |
| 20M05_Saliva | chr9:124979184-124979356 | LHX6;LHX6 | N_Shore | Body;Body |
| 20M05_Saliva | chr9:129884026-129884457 | ANGPTL2;RALGPS1 | | 5'UTR;Body |
| 20M05_Saliva | chr9:137718814-137719284 | COL5A1 | N_Shore | Body |
| 20M05_Saliva | chr10:530714-531320 | DIP2C | Island | Body |
| 20M05_Saliva | chr10:531914-533102 | DIP2C | S_Shore | Body |
| 20M05_Saliva | chr10:556470-556762 | DIP2C | | Body |
| 20M05_Saliva | chr10:1507708-1508061 | ADARB2 | S_Shore | Body |
| 20M05_Saliva | chr10:1547717-1547983 | ADARB2 | | Body |
| 20M05_Saliva | chr10:3149673-3149852 | PFKP | S_Shore | Body |
| 20M05_Saliva | chr10:3169037-3169516 | PFKP | | Body |
| 20M05_Saliva | chr10:3824387-3824687 | KLF6;KLF6;KLF6;KLF6 | S_Shore | Body;Body;Body;Body |
| 20M05_Saliva | chr10:5541568-5541726 | CALML5 | S_Shore | TSS200 |
| 20M05_Saliva | chr10:5566543-5568004 | CALML3 | N_Shore | TSS1500 |
| 20M05_Saliva | chr10:11246612-11246763 | CUGBP2;CUGBP2;CUGBP2;CUGBP2 | | Body;Body;Body;Body |
| 20M05_Saliva | chr10:13388442-13388567 | SEPHS1 | | 5'UTR |
| 20M05_Saliva | chr10:26855907-26856454 | APBB1IP | Island | Body |
| 20M05_Saliva | chr10:32216031-32216101 | ARHGAP12 | N_Shore | 5'UTR |
| 20M05_Saliva | chr10:47746733-47747039 | ANXA8L2 | | TSS1500 |
| 20M05_Saliva | chr10:49459800-49460249 | FRMPD2;FRMPD2 | | Body;Body |
| 20M05_Saliva | chr10:49674251-49674883 | ARHGAP22 | N_Shore | Body |
| 20M05_Saliva | chr10:50506875-50507092 | C10orf71;C10orf71 | | TSS1500;TSS1500 |
| 20M05_Saliva | chr10:63422511-63422555 | C10orf107 | N_Shore | TSS1500 |
| 20M05_Saliva | chr10:63809073-63809170 | ARID5B | | Body |
| 20M05_Saliva | chr10:64565642-64565772 | ADO;ADO | Island | 1stExon |

| | | | | |
|---|---|---|---|---|
| 20M05_Saliva | chr10:75631588-75632066 | CAMK2G;CAMK2G;CAMK2G;CAMK2G;CAMK2G | | Body;Body;Body;Body;Body |
| 20M05_Saliva | chr10:77165618-77165905 | C10orf41;C10orf41 | Island | Body;Body |
| 20M05_Saliva | chr10:81679790-81679852 | MBL1P | | TSS1500 |
| 20M05_Saliva | chr10:88779109-88779163 | FAM25A | S_Shelf | |
| 20M05_Saliva | chr10:93811958-93812043 | CPEB3 | S_Shelf | 3'UTR |
| 20M05_Saliva | chr10:95372040-95372195 | PDE6C | S_Shelf | |
| 20M05_Saliva | chr10:97200911-97200987 | SORBS1;SORBS1;SORBS1;SORBS1;SORBS1;SORBS1;SORBS1;SORBS1 | | Body;Body;Body;Body;Body;Body;Body |
| 20M05_Saliva | chr10:98031125-98031261 | BLNK;BLNK | | Body;Body |
| 20M05_Saliva | chr10:99331332-99331808 | ANKRD2;ANKRD2 | S_Shore | 3'UTR |
| 20M05_Saliva | chr10:99894013-99894087 | C10orf28 | N_Shelf | |
| 20M05_Saliva | chr10:105238873-105239205 | CALHM3;CALHM3 | | 1stExon |
| 20M05_Saliva | chr10:106093778-106093830 | ITPRIP | N_Shelf | TSS200 |
| 20M05_Saliva | chr10:123353654-123355576 | FGFR2;FGFR2;FGFR2;FGFR2;FGI | N_Shelf | TSS200;TSS200;5'UTR;5'UTR;1stExon;5'UTR;5'UTR;5'UTR;5'UTR;5'UTR |
| 20M05_Saliva | chr10:126278272-126278502 | LHPP;LHPP | N_Shelf | Body;Body |
| 20M05_Saliva | chr10:129536661-129537308 | FOXI2 | Island | 1stExon |
| 20M05_Saliva | chr10:134222445-134222507 | PWWP2B;PWWP2B | S_Shelf | 3'UTR;Body |
| 20M05_Saliva | chr10:134404873-134404945 | INPP5A | | Body |
| 20M05_Saliva | chr10:134578261-134578481 | INPP5A | | Body |
| 20M05_Saliva | chr10:135013653-135014037 | KNDC1 | N_Shelf | Body |
| 20M05_Saliva | chr10:135032154-135032504 | KNDC1 | S_Shore | Body |
| 20M05_Saliva | chr10:135341528-135341933 | CYP2E1 | N_Shore | 1stExon |
| 20M05_Saliva | chr11:393029-394545 | PKP3 | N_Shore | TSS1500 |
| 20M05_Saliva | chr11:405846-406019 | SIGIRR;SIGIRR;SIGIRR | N_Shore | 3'UTR;3'UTR;3'UTR |
| 20M05_Saliva | chr11:460469-460914 | PTDSS2 | Island | Body |
| 20M05_Saliva | chr11:692963-693032 | DEAF1 | N_Shore | Body |
| 20M05_Saliva | chr11:1243947-1244533 | MUC5B | N_Shelf | TSS1500 |
| 20M05_Saliva | chr11:1325718-1325852 | TOLLIP | | Body |
| 20M05_Saliva | chr11:1327962-1328486 | TOLLIP | N_Shelf | Body |
| 20M05_Saliva | chr11:1471594-1471691 | BRSK2 | Island | Body |
| 20M05_Saliva | chr11:1507263-1507347 | HCCA2 | | Body |
| 20M05_Saliva | chr11:1536428-1536810 | HCCA2 | N_Shore | Body |
| 20M05_Saliva | chr11:1660513-1660992 | HCCA2 | | Body |
| 20M05_Saliva | chr11:1855286-1856025 | SYT8 | | TSS1500 |
| 20M05_Saliva | chr11:1892074-1892142 | LSP1;LSP1;LSP1;LSP1 | Island | 5'UTR;TSS200;TSS200;Body |
| 20M05_Saliva | chr11:1945564-1945918 | TNNT3;TNNT3;TNNT3;TNNT3 | N_Shore | Body;Body;Body;Body |
| 20M05_Saliva | chr11:2011216-2011346 | LOC100133545 | N_Shelf | TSS200 |
| 20M05_Saliva | chr11:2170831-2171007 | INS-IGF2;INS-IGF2;IGF2;IGF2 | | Body;Body;1stExon;5'UTR |
| 20M05_Saliva | chr11:2828364-2829241 | KCNQ1;KCNQ1 | N_Shore | Body;Body |
| 20M05_Saliva | chr11:2925439-2925869 | SLC22A18;SLC22A18;SLC22A18 | S_Shore | Body;Body;TSS1500 |
| 20M05_Saliva | chr11:3144685-3144774 | OSBPL5;OSBPL5;OSBPL5 | S_Shelf | Body;Body;Body |
| 20M05_Saliva | chr11:3187678-3187803 | OSBPL5;OSBPL5;OSBPL5 | S_Shore | TSS1500;TSS1500;TSS1500 |
| 20M05_Saliva | chr11:8832817-8833095 | ST5;ST5;ST5;ST5 | | 1stExon;5'UTR;TSS1500;5'UTR |
| 20M05_Saliva | chr11:17803160-17803421 | KCNC1 | S_Shore | Body |
| 20M05_Saliva | chr11:32421518-32421845 | WT1;WT1;WT1;WT1 | | Body;Body;Body;Body |
| 20M05_Saliva | chr11:34535377-34535416 | ELF5 | | TSS200 |
| 20M05_Saliva | chr11:45928805-45928988 | C11orf94;C11orf94 | | 5'UTR;1stExon |
| 20M05_Saliva | chr11:57089366-57089744 | TNKS1BP1 | N_Shelf | 5'UTR |
| 20M05_Saliva | chr11:57195187-57195540 | SLC43A3;SLC43A3;SLC43A3 | S_Shore | TSS1500;TSS200;TSS1500 |
| 20M05_Saliva | chr11:57519391-57519470 | BTBD18 | N_Shore | TSS200 |
| 20M05_Saliva | chr11:61049230-61049520 | VWCE | | Body |
| 20M05_Saliva | chr11:62272972-62273418 | AHNAK | | Body |
| 20M05_Saliva | chr11:62456939-62457345 | LRRN4CL;LRRN4CL | S_Shore | 5'UTR;1stExon |
| 20M05_Saliva | chr11:62621178-62621367 | SNORD22;SNORD31;SNORD30;S | N_Shelf | TSS1500;TSS1500;Body;Body |
| 20M05_Saliva | chr11:62688748-62689081 | CHRM1;CHRM1 | N_Shelf | 5'UTR;1stExon |
| 20M05_Saliva | chr11:64115989-64116279 | CCDC88B | S_Shelf | Body |
| 20M05_Saliva | chr11:64405452-64405919 | NRXN2;NRXN2;NRXN2 | Island | Body;Body;Body |
| 20M05_Saliva | chr11:64951536-64952354 | CAPN1 | N_Shore | Body |
| 20M05_Saliva | chr11:65352168-65352327 | EHBP1L1 | N_Shore | Body |
| 20M05_Saliva | chr11:65560174-65560763 | OVOL1 | | Body |
| 20M05_Saliva | chr11:66104468-66104485 | RIN1 | S_Shelf | TSS1500 |
| 20M05_Saliva | chr11:66650500-66650560 | PC;PC;PC | | 5'UTR;5'UTR;5'UTR |
| 20M05_Saliva | chr11:67350326-67350499 | GSTP1 | N_Shore | TSS1500 |
| 20M05_Saliva | chr11:67442067-67442408 | ALDH3B2;ALDH3B2;ALDH3B2 | | 1stExon;5'UTR;5'UTR |
| 20M05_Saliva | chr11:69469175-69469820 | CCND1 | S_Shore | 3'UTR |
| 20M05_Saliva | chr11:69481734-69482540 | ORAOV1 | | 3'UTR |

| | | | | |
|---|---|---|---|---|
| 20M05_Saliva | chr11:69484433-69484574 | ORAOV1 | | Body |
| 20M05_Saliva | chr11:69970464-69970618 | ANO1;ANO1 | | Body;Body |
| 20M05_Saliva | chr11:70002333-70002831 | ANO1;ANO1 | S_Shore | Body;Body |
| 20M05_Saliva | chr11:71710643-71711021 | IL18BP;IL18BP;IL18BP;IL18BP;IL18BP;IL18BP | | TSS200;5'UTR;TSS1500;5'UTR;5'UTR;1stExon |
| 20M05_Saliva | chr11:71723428-71724677 | NUMA1 | | Body |
| 20M05_Saliva | chr11:73053649-73054121 | ARHGEF17 | Island | Body |
| 20M05_Saliva | chr11:83393062-83393677 | DLG2;DLG2;DLG2;DLG2 | | 5'UTR;Body;Body;Body |
| 20M05_Saliva | chr11:93277161-93277191 | C11orf75 | S_Shore | TSS1500 |
| 20M05_Saliva | chr11:93754223-93754390 | HEPHL1 | | TSS200 |
| 20M05_Saliva | chr11:105010284-105010531 | CARD18 | | Body |
| 20M05_Saliva | chr11:111789452-111789693 | C11orf52 | | TSS200 |
| 20M05_Saliva | chr11:111847475-111847747 | DIXDC1;DIXDC1 | N_Shore | TSS1500;Body |
| 20M05_Saliva | chr11:113845638-113846017 | HTR3A;HTR3A | | TSS200;TSS200 |
| 20M05_Saliva | chr11:117069780-117070046 | TAGLN;TAGLN | | TSS1500;TSS1500 |
| 20M05_Saliva | chr11:117800062-117800467 | TMPRSS13;TMPRSS13 | | 5'UTR;1stExon |
| 20M05_Saliva | chr11:117947459-117947997 | TMPRSS4;TMPRSS4 | | TSS1500;TSS1500 |
| 20M05_Saliva | chr11:118550379-118550644 | TREH;TREH | | 1stExon;5'UTR |
| 20M05_Saliva | chr11:118781728-118781978 | BCL9L | Island | TSS200 |
| 20M05_Saliva | chr11:120008963-120009135 | TRIM29 | S_Shore | TSS200 |
| 20M05_Saliva | chr11:120973197-120973274 | TECTA | | TSS200 |
| 20M05_Saliva | chr11:121970694-121970768 | LOC399959;MIR125B1 | | Body;TSS200 |
| 20M05_Saliva | chr11:123396039-123396593 | GRAMD1B | | TSS1500 |
| 20M05_Saliva | chr11:124621829-124622096 | VSIG2 | | Body |
| 20M05_Saliva | chr11:126310323-126310475 | KIRREL3;KIRREL3 | | Body;Body |
| 20M05_Saliva | chr11:128457400-128457682 | ETS1;ETS1 | | 1stExon;5'UTR |
| 20M05_Saliva | chr11:128812462-128813451 | TP53AIP1 | | 5'UTR |
| 20M05_Saliva | chr11:129992359-129992368 | APLP2;APLP2;APLP2;APLP2;APLP | N_Shore | Body;Body;Body;Body;Body;Body |
| 20M05_Saliva | chr11:133804743-133804959 | IGSF9B | S_Shelf | Body |
| 20M05_Saliva | chr12:258355-258436 | IQSEC3;IQSEC3 | | Body;Body |
| 20M05_Saliva | chr12:6421437-6422131 | PLEKHG6;PLEKHG6;PLEKHG6 | S_Shore | TSS1500;Body;Body |
| 20M05_Saliva | chr12:6483772-6484729 | SCNN1A;SCNN1A;SCNN1A | | Body;Body;1stExon |
| 20M05_Saliva | chr12:6486701-6487080 | SCNN1A | | TSS200 |
| 20M05_Saliva | chr12:7071712-7073264 | MIR200C | | TSS1500 |
| 20M05_Saliva | chr12:8974786-8975309 | A2ML1 | | TSS1500 |
| 20M05_Saliva | chr12:9217669-9217907 | LOC144571 | Island | TSS200 |
| 20M05_Saliva | chr12:25801455-25801621 | IFLTD1 | | 1stExon |
| 20M05_Saliva | chr12:50497732-50497827 | GPD1 | | TSS200 |
| 20M05_Saliva | chr12:51236582-51236638 | TMPRSS12 | N_Shore | TSS200 |
| 20M05_Saliva | chr12:52585695-52585892 | KRT80;KRT80;KRT80;KRT80 | | 1stExon;1stExon;5'UTR;5'UTR |
| 20M05_Saliva | chr12:52845930-52846088 | KRT6B | | TSS200 |
| 20M05_Saliva | chr12:52867568-52867793 | KRT6C;KRT6C | | 1stExon;5'UTR |
| 20M05_Saliva | chr12:52887054-52887566 | KRT6A;KRT6A | | 1stExon;5'UTR |
| 20M05_Saliva | chr12:53242733-53242922 | KRT78;KRT78 | | 5'UTR;1stExon |
| 20M05_Saliva | chr12:53343514-53343849 | KRT18;KRT18 | S_Shore | Body;Body |
| 20M05_Saliva | chr12:53612551-53612734 | RARG;RARG | N_Shore | Body;Body |
| 20M05_Saliva | chr12:57559728-57559901 | LRP1 | | Body |
| 20M05_Saliva | chr12:64782908-64783110 | C12orf56;C12orf56 | N_Shore | Body;Body |
| 20M05_Saliva | chr12:67072769-67072988 | GRIP1;GRIP1 | | 1stExon;5'UTR |
| 20M05_Saliva | chr12:68041257-68041322 | DYRK2;DYRK2 | N_Shore | TSS1500;TSS1500 |
| 20M05_Saliva | chr12:93096780-93096844 | C12orf74 | | TSS200 |
| 20M05_Saliva | chr12:114337676-114337927 | RBM19;RBM19;RBM19 | | Body;Body;Body |
| 20M05_Saliva | chr12:120151527-120151890 | MIR1178;CIT | | Body;Body |
| 20M05_Saliva | chr12:120241287-120242156 | CIT | N_Shore | Body |
| 20M05_Saliva | chr12:120242394-120242513 | CIT | S_Shore | Body |
| 20M05_Saliva | chr12:121882033-121882415 | KDM2B;KDM2B | N_Shore | Body;Body |
| 20M05_Saliva | chr12:121890163-121890503 | KDM2B;KDM2B | N_Shore | Body;Body |
| 20M05_Saliva | chr12:123352600-123352774 | VPS37B | S_Shore | Body |
| 20M05_Saliva | chr12:123750782-123750864 | CDK2AP1 | N_Shelf | Body |
| 20M05_Saliva | chr12:124773668-124774078 | FAM101A | | TSS200 |
| 20M05_Saliva | chr12:128866668-128866735 | TMEM132C | | Body |
| 20M05_Saliva | chr12:132690643-132690782 | GALNT9;GALNT9 | S_Shore | Body;TSS200 |
| 20M05_Saliva | chr12:133101168-133101612 | FBRSL1 | N_Shore | Body |
| 20M05_Saliva | chr13:24736455-24736569 | SPATA13;MIR2276;SPATA13 | S_Shore | 5'UTR;TSS200;5'UTR |
| 20M05_Saliva | chr13:48575859-48575927 | SUCLA2 | S_Shore | TSS1500 |
| 20M05_Saliva | chr13:95131809-95132211 | DCT;DCT;DCT;DCT | | 1stExon;5'UTR;1stExon;5'UTR |

| | | | | |
|---|---|---|---|---|
| 20M05_Saliva | chr13:99135543-99136580 | STK24;STK24 | | Body;Body |
| 20M05_Saliva | chr13:99630462-99630583 | DOCK9;DOCK9;DOCK9;DOCK9 | | Body;Body;TSS200;TSS200 |
| 20M05_Saliva | chr13:102104976-102105049 | ITGBL1;ITGBL1 | N_Shore | 1stExon;5'UTR |
| 20M05_Saliva | chr13:110874267-110874326 | COL4A1 | | Body |
| 20M05_Saliva | chr13:111522651-111522985 | C13orf29 | | Body |
| 20M05_Saliva | chr13:113648607-113648882 | MCF2L;MCF2L | Island | Body;Body |
| 20M05_Saliva | chr13:113676726-113677777 | MCF2L;MCF2L | N_Shore | Body;Body |
| 20M05_Saliva | chr13:113759771-113760101 | F7;F7 | N_Shore | TSS1500;TSS1500 |
| 20M05_Saliva | chr13:114103452-114103646 | ADPRHL1;ADPRHL1;ADPRHL1 | | 1stExon;5'UTR;Body |
| 20M05_Saliva | chr13:114855198-114855572 | RASA3 | S_Shelf | Body |
| 20M05_Saliva | chr14:20903410-20903611 | KLHL33 | Island | 5'UTR |
| 20M05_Saliva | chr14:23352029-23352488 | REM2 | N_Shelf | TSS1500 |
| 20M05_Saliva | chr14:23623480-23623684 | SLC7A8;SLC7A8;SLC7A8 | | 5'UTR;Body;1stExon |
| 20M05_Saliva | chr14:23835595-23836012 | EFS;EFS | Island | TSS1500;TSS1500 |
| 20M05_Saliva | chr14:56045827-56046001 | KTN1;C14orf33;KTN1;KTN1;KTN1 | N_Shore | TSS1500;Body;TSS1500;TSS1500;TSS1500 |
| 20M05_Saliva | chr14:75936582-75936789 | JDP2;JDP2;JDP2;JDP2 | | 3'UTR;3'UTR;3'UTR;3'UTR |
| 20M05_Saliva | chr14:94393654-94393816 | FAM181A;C14orf86 | | Body;TSS1500 |
| 20M05_Saliva | chr14:98444151-98444533 | C14orf64 | | Body |
| 20M05_Saliva | chr14:103416122-103416389 | CDC42BPB | | Body |
| 20M05_Saliva | chr14:103593088-103593520 | TNFAIP2 | N_Shore | Body |
| 20M05_Saliva | chr14:104165052-104165475 | XRCC3;XRCC3;XRCC3;KLC1;KLC | Island | 3'UTR;3'UTR;3'UTR;Body;Body |
| 20M05_Saliva | chr14:104175555-104175768 | XRCC3;XRCC3;XRCC3 | | Body;Body;Body |
| 20M05_Saliva | chr14:105154689-105154994 | INF2;INF2;INF2 | N_Shore | TSS1500;TSS1500;TSS1500 |
| 20M05_Saliva | chr15:39871808-39872186 | THBS1 | N_Shore | TSS1500 |
| 20M05_Saliva | chr15:40531871-40532180 | PAK6;PAK6;PAK6 | S_Shore | 5'UTR;5'UTR;TSS200 |
| 20M05_Saliva | chr15:40583227-40583422 | PLCB2 | Island | Body |
| 20M05_Saliva | chr15:42289618-42289939 | PLA2G4E | | Body |
| 20M05_Saliva | chr15:42386726-42387193 | PLA2G4D;PLA2G4D | | 1stExon;5'UTR |
| 20M05_Saliva | chr15:42447989-42448786 | PLA2G4F;PLA2G4F | | Body;Body |
| 20M05_Saliva | chr15:59908622-59909035 | GCNT3 | | 5'UTR |
| 20M05_Saliva | chr15:65214075-65214588 | ANKDD1A | | Body |
| 20M05_Saliva | chr15:67417607-67417964 | SMAD3;SMAD3 | | Body;TSS1500 |
| 20M05_Saliva | chr15:68924114-68924220 | CORO2B | | Body |
| 20M05_Saliva | chr15:74290466-74290733 | PML;PML;PML;PML;PML;PML;PM | Island | Body;Body;Body;Body;Body;Body;Body;Body ;Body |
| 20M05_Saliva | chr15:74659966-74660110 | CYP11A1;CYP11A1;CYP11A1 | S_Shore | 5'UTR;1stExon;TSS1500 |
| 20M05_Saliva | chr15:86313372-86313973 | KLHL25 | S_Shore | 5'UTR |
| 20M05_Saliva | chr15:90727560-90727783 | SEMA4B | N_Shore | TSS1500 |
| 20M05_Saliva | chr16:419975-420112 | MRPL28 | Island | Body |
| 20M05_Saliva | chr16:670800-671171 | RAB40C | Island | Body |
| 20M05_Saliva | chr16:671905-672468 | RAB40C | N_Shore | Body |
| 20M05_Saliva | chr16:743338-743584 | FBXL16 | N_Shore | 3'UTR |
| 20M05_Saliva | chr16:809679-810733 | MSLN;MSLN | | TSS1500;TSS1500 |
| 20M05_Saliva | chr16:832683-834201 | MSLNL | N_Shore | Body |
| 20M05_Saliva | chr16:1128698-1128865 | SSTR5;LOC146336 | Island | TSS200;Body |
| 20M05_Saliva | chr16:1206750-1207075 | CACNA1H;CACNA1H | N_Shore | Body;Body |
| 20M05_Saliva | chr16:1582105-1582581 | IFT140 | N_Shore | Body |
| 20M05_Saliva | chr16:1813796-1814157 | MAPK8IP3;MAPK8IP3 | N_Shore | Body;Body |
| 20M05_Saliva | chr16:3062349-3062597 | CLDN9 | N_Shore | TSS200 |
| 20M05_Saliva | chr16:3843012-3843415 | CREBBP;CREBBP | | Body;Body |
| 20M05_Saliva | chr16:12210656-12211322 | SNX29 | Island | Body |
| 20M05_Saliva | chr16:57406044-57406511 | CX3CL1 | | TSS1500 |
| 20M05_Saliva | chr16:57831745-57832230 | KIFC3;KIFC3;KIFC3;KIFC3 | | Body;Body;1stExon;5'UTR |
| 20M05_Saliva | chr16:57937614-57937756 | CNGB1 | N_Shore | Body |
| 20M05_Saliva | chr16:66412919-66413098 | CDH5 | | 5'UTR |
| 20M05_Saliva | chr16:67022472-67022719 | CES8 | | TSS200 |
| 20M05_Saliva | chr16:67232921-67233983 | ELMO3 | N_Shore | TSS200 |
| 20M05_Saliva | chr16:67918680-67918965 | NRN1L | Island | TSS200 |
| 20M05_Saliva | chr16:68772225-68772469 | CDH1 | Island | Body |
| 20M05_Saliva | chr16:77469667-77469998 | ADAMTS18 | S_Shore | TSS1500 |
| 20M05_Saliva | chr16:81526954-81527504 | CMIP | | Body |
| 20M05_Saliva | chr16:81564161-81564553 | CMIP;CMIP | | Body;Body |
| 20M05_Saliva | chr16:84076834-84076941 | SLC38A8 | | TSS1500 |
| 20M05_Saliva | chr16:84852577-84852909 | CRISPLD2 | N_Shore | TSS1500 |
| 20M05_Saliva | chr16:85669462-85669675 | KIAA0182;KIAA0182 | N_Shore | 5'UTR;Body |
| 20M05_Saliva | chr16:87958145-87958493 | CA5A | | Body |

| | | | | |
|---|---|---|---|---|
| 20M05_Saliva | chr16:88807658-88807707 | FAM38A | S_Shelf | Body |
| 20M05_Saliva | chr16:88832476-88832572 | FAM38A | | Body |
| 20M05_Saliva | chr16:89220516-89220688 | ACSF3;ACSF3;ACSF3 | S_Shore | Body;Body;Body |
| 20M05_Saliva | chr16:89387014-89387805 | ANKRD11 | | 5'UTR |
| 20M05_Saliva | chr16:89408076-89408367 | ANKRD11 | N_Shore | 5'UTR |
| 20M05_Saliva | chr17:152089-152259 | RPH3AL | N_Shore | Body |
| 20M05_Saliva | chr17:790628-791324 | NXN | | Body |
| 20M05_Saliva | chr17:851785-851933 | NXN | S_Shore | Body |
| 20M05_Saliva | chr17:1553585-1553661 | RILP | S_Shore | TSS200 |
| 20M05_Saliva | chr17:1664077-1664339 | SERPINF1 | | TSS1500 |
| 20M05_Saliva | chr17:4463371-4464075 | GGT6;GGT6 | | Body;Body |
| 20M05_Saliva | chr17:6899207-6899332 | ALOX12 | Island | TSS200 |
| 20M05_Saliva | chr17:7493150-7494151 | SOX15;SOX15 | S_Shore | 1stExon;5'UTR |
| 20M05_Saliva | chr17:7690246-7690529 | DNAH2 | Island | Body |
| 20M05_Saliva | chr17:7832943-7833237 | KCNAB3 | Island | TSS200 |
| 20M05_Saliva | chr17:8218887-8219313 | ARHGEF15 | | Body |
| 20M05_Saliva | chr17:8702564-8702823 | MFSD6L;MFSD6L | Island | 5'UTR;1stExon |
| 20M05_Saliva | chr17:9929528-9929867 | GAS7;GAS7;GAS7;GAS7 | | 1stExon;Body;Body;5'UTR |
| 20M05_Saliva | chr17:14213225-14214081 | HS3ST3B1 | S_Shore | Body |
| 20M05_Saliva | chr17:17109936-17110207 | PLD6 | Island | TSS1500 |
| 20M05_Saliva | chr17:17465242-17465682 | PEMT;PEMT;PEMT | Island | Body;Body;Body |
| 20M05_Saliva | chr17:17717894-17718525 | MIR33B;SREBF1;SREBF1 | S_Shore | TSS1500;Body;Body |
| 20M05_Saliva | chr17:19247897-19248009 | MIR1180;B9D1 | Island | TSS200;Body |
| 20M05_Saliva | chr17:19313024-19313529 | RNF112 | | TSS1500 |
| 20M05_Saliva | chr17:19648718-19648853 | ALDH3A1;ALDH3A1;ALDH3A1;Al | S_Shore | 5'UTR;5'UTR;5'UTR;1stExon |
| 20M05_Saliva | chr17:38657310-38657949 | TNS4 | | 5'UTR |
| 20M05_Saliva | chr17:38717206-38717275 | CCR7 | | Body |
| 20M05_Saliva | chr17:38859934-38860097 | KRT24 | | 1stExon |
| 20M05_Saliva | chr17:39143239-39143567 | KRT40 | | 5'UTR |
| 20M05_Saliva | chr17:39165425-39165697 | KRTAP3-1 | | TSS200 |
| 20M05_Saliva | chr17:39183037-39183560 | KRTAP1-5 | | 1stExon |
| 20M05_Saliva | chr17:39623858-39624088 | KRT32 | | TSS1500 |
| 20M05_Saliva | chr17:39769012-39769213 | KRT16;KRT16 | | 1stExon;5'UTR |
| 20M05_Saliva | chr17:39943423-39943545 | JUP;JUP | S_Shore | TSS1500;TSS1500 |
| 20M05_Saliva | chr17:40274524-40274740 | KAT2A;HSPB9 | Island | TSS1500;TSS1500 |
| 20M05_Saliva | chr17:40823930-40824361 | PLEKHH3 | S_Shore | Body |
| 20M05_Saliva | chr17:41521655-41522193 | MIR2117 | | TSS1500 |
| 20M05_Saliva | chr17:46035108-46035343 | PRR15L;PRR15L | | 1stExon;5'UTR |
| 20M05_Saliva | chr17:46388390-46388465 | SKAP1;SKAP1 | | Body;Body |
| 20M05_Saliva | chr17:46627848-46628303 | HOXB3 | Island | Body |
| 20M05_Saliva | chr17:46651225-46651867 | HOXB3;HOXB3 | N_Shelf | 5'UTR;1stExon |
| 20M05_Saliva | chr17:48845563-48846189 | C17orf73 | | TSS1500 |
| 20M05_Saliva | chr17:60757756-60758287 | MRC2 | Island | Body |
| 20M05_Saliva | chr17:62051709-62051773 | SCN4A | | TSS1500 |
| 20M05_Saliva | chr17:71258589-71259036 | CPSF4L | | TSS1500 |
| 20M05_Saliva | chr17:73996245-73996656 | CDK3;C17orf106 | | TSS1500;Body |
| 20M05_Saliva | chr17:74639731-74639928 | ST6GALNAC1;ST6GALNAC1 | | 5'UTR;1stExon |
| 20M05_Saliva | chr17:75282776-75284193 | SEPT9;SEPT9 | | Body;TSS1500 |
| 20M05_Saliva | chr17:75446431-75446661 | SEPT9;SEPT9;SEPT9;SEPT9;SEPT | N_Shore | TSS200;Body;Body;Body;Body;Body |
| 20M05_Saliva | chr17:76501381-76501582 | DNAH17 | S_Shelf | Body |
| 20M05_Saliva | chr17:76976091-76976357 | LGALS3BP | | TSS200 |
| 20M05_Saliva | chr17:76993506-76993698 | CANT1;CANT1;CANT1 | Island | Body;Body;Body |
| 20M05_Saliva | chr17:77030221-77030546 | C1QTNF1;C1QTNF1;C1QTNF1 | | 5'UTR;TSS1500;5'UTR |
| 20M05_Saliva | chr17:77086350-77086532 | HRNBP3 | | 3'UTR |
| 20M05_Saliva | chr17:77962048-77962240 | TBC1D16 | N_Shelf | Body |
| 20M05_Saliva | chr17:78039301-78039528 | CCDC40 | Island | Body |
| 20M05_Saliva | chr17:78079615-78079795 | GAA;GAA;GAA | Island | Body;Body;Body |
| 20M05_Saliva | chr17:78161110-78161706 | CARD14;CARD14 | N_Shelf | TSS1500;Body |
| 20M05_Saliva | chr17:78166148-78166385 | CARD14;CARD14 | S_Shore | Body;Body |
| 20M05_Saliva | chr17:78801851-78802104 | RPTOR;RPTOR | S_Shelf | Body;Body |
| 20M05_Saliva | chr17:78865087-78865373 | RPTOR;RPTOR | S_Shore | Body;Body |
| 20M05_Saliva | chr17:79030796-79031712 | BAIAP2;BAIAP2;BAIAP2;BAIAP2 | | Body;Body;Body;Body |
| 20M05_Saliva | chr17:79056061-79056183 | BAIAP2;BAIAP2;BAIAP2;BAIAP2 | S_Shelf | Body;Body;Body;Body |
| 20M05_Saliva | chr17:79096582-79097243 | AATK | Island | Body |
| 20M05_Saliva | chr17:79099481-79099785 | MIR657;AATK | N_Shelf | TSS1500;Body |
| 20M05_Saliva | chr17:79099882-79100218 | MIR657;AATK;MIR338 | N_Shelf | TSS1500;Body;TSS200 |

| | | | | |
|---|---|---|---|---|
| 20M05_Saliva | chr17:79109729-79109910 | AATK | Island | Body |
| 20M05_Saliva | chr17:79251255-79251354 | SLC38A10;SLC38A10 | N_Shelf | Body;Body |
| 20M05_Saliva | chr17:79377548-79377850 | BAHCC1 | Island | Body |
| 20M05_Saliva | chr17:79393532-79393718 | BAHCC1 | Island | Body |
| 20M05_Saliva | chr17:79609272-79609346 | TSPAN10 | | TSS200 |
| 20M05_Saliva | chr17:79813410-79813507 | P4HB | | Body |
| 20M05_Saliva | chr17:79961522-79961769 | ASPSCR1 | N_Shore | Body |
| 20M05_Saliva | chr17:80289541-80290106 | SECTM1 | N_Shore | 5'UTR |
| 20M05_Saliva | chr17:80796693-80798023 | TBCD;ZNF750 | | Body;5'UTR |
| 20M05_Saliva | chr17:80806135-80806435 | TBCD | | Body |
| 20M05_Saliva | chr17:80816702-80816995 | TBCD | N_Shore | Body |
| 20M05_Saliva | chr17:80829028-80829309 | TBCD | N_Shore | Body |
| 20M05_Saliva | chr17:80832328-80832836 | TBCD | S_Shelf | Body |
| 20M05_Saliva | chr17:80847137-80847662 | TBCD | | Body |
| 20M05_Saliva | chr18:21452730-21453131 | LAMA3;LAMA3;LAMA3;LAMA3 | | TSS1500;TSS1500;Body;Body |
| 20M05_Saliva | chr18:21572622-21572748 | TTC39C | | TSS200 |
| 20M05_Saliva | chr18:29027658-29027885 | DSG3 | | TSS200 |
| 20M05_Saliva | chr18:55888640-55888832 | NEDD4L;NEDD4L;NEDD4L;NEDD4L;NEDD4L;NEDD4L;NEDD4L;NEDD4L | | 5'UTR;Body;TSS200;5'UTR;5'UTR;Body;Body;Body;5'UTR |
| 20M05_Saliva | chr18:61143926-61144339 | SERPINB5 | | TSS1500 |
| 20M05_Saliva | chr18:61253847-61254561 | SERPINB13 | | TSS1500 |
| 20M05_Saliva | chr18:77230598-77230795 | NFATC1;NFATC1;NFATC1;NFATC | Island | Body;Body;Body;Body |
| 20M05_Saliva | chr18:77286052-77286260 | NFATC1;NFATC1;NFATC1;NFATC | Island | Body;Body;Body;Body |
| 20M05_Saliva | chr19:736164-736425 | PALM;PALM | Island | Body;Body |
| 20M05_Saliva | chr19:751098-751241 | C19orf21 | S_Shelf | TSS200 |
| 20M05_Saliva | chr19:1356269-1356304 | MUM1;MUM1 | S_Shore | TSS200;5'UTR |
| 20M05_Saliva | chr19:1907677-1908328 | ADAT3;SCAMP4 | N_Shore | 5'UTR;5'UTR |
| 20M05_Saliva | chr19:2016402-2016532 | BTBD2 | | TSS1500 |
| 20M05_Saliva | chr19:2512999-2513356 | GNG7 | N_Shore | 3'UTR |
| 20M05_Saliva | chr19:2525231-2525634 | GNG7 | Island | 5'UTR |
| 20M05_Saliva | chr19:2540907-2541104 | GNG7 | Island | 5'UTR |
| 20M05_Saliva | chr19:4455239-4455413 | UBXN6;UBXN6;UBXN6 | N_Shelf | 5'UTR;1stExon;Body |
| 20M05_Saliva | chr19:5036609-5037219 | KDM4B | N_Shore | Body |
| 20M05_Saliva | chr19:5048645-5049135 | KDM4B | Island | Body |
| 20M05_Saliva | chr19:5250376-5251120 | PTPRS;PTPRS;PTPRS;PTPRS | N_Shore | Body;Body;Body;Body |
| 20M05_Saliva | chr19:5822318-5822630 | NRTN | Island | TSS1500 |
| 20M05_Saliva | chr19:6333587-6333761 | ACER1;ACER1 | | 5'UTR;1stExon |
| 20M05_Saliva | chr19:7580209-7580275 | ZNF358 | N_Shore | TSS1500 |
| 20M05_Saliva | chr19:8555061-8555169 | PRAM1 | Island | 3'UTR |
| 20M05_Saliva | chr19:10231566-10231711 | EIF3G | S_Shore | TSS1500 |
| 20M05_Saliva | chr19:10444815-10445516 | RAVER1;ICAM3 | Island | TSS1500;Body |
| 20M05_Saliva | chr19:10735548-10736059 | SLC44A2;SLC44A2 | N_Shore | TSS1500;Body |
| 20M05_Saliva | chr19:14607597-14608092 | GIPC1;GIPC1;GIPC1;GIPC1;GIPC | S_Shore | TSS1500;TSS1500;TSS1500;TSS1500;TSS1500;TSS1500 |
| 20M05_Saliva | chr19:15568574-15568935 | RASAL3 | Island | Body |
| 20M05_Saliva | chr19:15783239-15783725 | CYP4F12 | | TSS1500 |
| 20M05_Saliva | chr19:16178091-16178426 | TPM4 | N_Shore | TSS1500 |
| 20M05_Saliva | chr19:16830739-16830859 | NWD1 | | TSS200 |
| 20M05_Saliva | chr19:17942221-17942411 | JAK3 | Island | Body |
| 20M05_Saliva | chr19:21264896-21265421 | ZNF714 | N_Shore | TSS200 |
| 20M05_Saliva | chr19:30863260-30863349 | ZNF536 | N_Shelf | TSS200 |
| 20M05_Saliva | chr19:35085474-35086248 | SCGBL;SCGBL | | 1stExon;5'UTR |
| 20M05_Saliva | chr19:36499326-36499839 | C19orf46 | N_Shore | Body |
| 20M05_Saliva | chr19:39086733-39087186 | MAP4K1;MAP4K1 | N_Shore | Body;Body |
| 20M05_Saliva | chr19:43969650-43969810 | LYPD3 | S_Shore | 1stExon |
| 20M05_Saliva | chr19:45279765-45280184 | CBLC;CBLC | N_Shore | TSS1500;TSS1500 |
| 20M05_Saliva | chr19:45574917-45575142 | ZNF296 | Island | Body |
| 20M05_Saliva | chr19:45737623-45737873 | EXOC3L2 | Island | TSS200 |
| 20M05_Saliva | chr19:46270244-46270700 | SIX5 | Island | Body |
| 20M05_Saliva | chr19:49238364-49238653 | RASIP1 | N_Shore | Body |
| 20M05_Saliva | chr19:51531297-51531393 | KLK11;KLK11;KLK11;KLK11 | N_Shelf | TSS1500;TSS200;TSS1500;TSS200 |
| 20M05_Saliva | chr19:52227967-52228400 | HAS1 | | TSS1500 |
| 20M05_Saliva | chr19:59073369-59073430 | MZF1;MZF1;LOC100131691 | N_Shore | 3'UTR;3'UTR;Body |
| 20M05_Saliva | chr20:1309722-1310000 | SDCBP2 | S_Shelf | 5'UTR |
| 20M05_Saliva | chr20:17595355-17595472 | RRBP1;RRBP1 | Island | Body;Body |
| 20M05_Saliva | chr20:17674195-17674250 | BANF2 | | TSS200 |

| | | | | |
|---|---|---|---|---|
| 20M05_Saliva | chr20:30433233-30433512 | FOXS1 | Island | 1stExon |
| 20M05_Saliva | chr20:31781151-31781393 | BASE | | TSS1500 |
| 20M05_Saliva | chr20:33879989-33880825 | FAM83C | Island | 1stExon |
| 20M05_Saliva | chr20:34700112-34700675 | EPB41L1 | | TSS1500 |
| 20M05_Saliva | chr20:35503983-35504511 | C20orf118 | | TSS1500 |
| 20M05_Saliva | chr20:35972924-35973098 | SRC | N_Shore | TSS200 |
| 20M05_Saliva | chr20:36011999-36012016 | SRC;SRC | N_Shore | 5'UTR;5'UTR |
| 20M05_Saliva | chr20:36151003-36151338 | BLCAP;BLCAP;BLCAP;NNAT;BLC | Island | 5'UTR;5'UTR;5'UTR;Body;5'UTR;5'UTR;Body |
| 20M05_Saliva | chr20:36152813-36154388 | BLCAP;BLCAP;BLCAP;BLCAP;BL | S_Shore | 5'UTR;1stExon;5'UTR;5'UTR;5'UTR;5'UTR |
| 20M05_Saliva | chr20:43937138-43937289 | MATN4;RBPJL;MATN4;MATN4 | S_Shore | TSS200;Body;TSS200;TSS200 |
| 20M05_Saliva | chr20:43977423-43977531 | SDC4 | S_Shore | TSS1500 |
| 20M05_Saliva | chr20:45313200-45313390 | TP53RK;SLC13A3 | | 3'UTR;TSS200 |
| 20M05_Saliva | chr20:52687007-52687365 | BCAS1;BCAS1 | | 1stExon;5'UTR |
| 20M05_Saliva | chr20:61002595-61002866 | C20orf151 | | TSS200 |
| 20M05_Saliva | chr20:62367884-62367893 | LIME1 | N_Shore | TSS200 |
| 20M05_Saliva | chr20:62383275-62383481 | ZBTB46 | Island | Body |
| 20M05_Saliva | chr20:62679255-62679713 | SOX18 | N_Shore | 3'UTR |
| 20M05_Saliva | chr20:62693951-62694005 | TCEA2;TCEA2 | Island | TSS200;5'UTR |
| 20M05_Saliva | chr21:35736023-35736383 | KCNE2 | | TSS1500 |
| 20M05_Saliva | chr21:38117454-38117682 | SIM2;SIM2 | N_Shelf | Body;Body |
| 20M05_Saliva | chr21:38362725-38362754 | HLCS | Island | TSS200 |
| 20M05_Saliva | chr21:40823279-40823722 | SH3BGR;SH3BGR | | 5'UTR;TSS1500 |
| 20M05_Saliva | chr21:43135854-43136287 | C21orf129;NCRNA00112 | | Body;TSS1500 |
| 20M05_Saliva | chr21:43735412-43735844 | TFF3 | | 1stExon |
| 20M05_Saliva | chr21:44898090-44898206 | C21orf84 | | Body |
| 20M05_Saliva | chr22:17680531-17680706 | CECR1;CECR1;CECR1 | | Body;1stExon;5'UTR |
| 20M05_Saliva | chr22:19949873-19950166 | COMT;COMT;COMT;COMT | | 5'UTR;5'UTR;5'UTR;TSS200 |
| 20M05_Saliva | chr22:24348549-24348715 | GSTTP1 | | TSS1500 |
| 20M05_Saliva | chr22:24577288-24577448 | SUSD2 | | TSS200 |
| 20M05_Saliva | chr22:24891158-24891338 | C22orf45;C22orf45;UPB1 | Island | TSS1500;TSS1500;TSS200 |
| 20M05_Saliva | chr22:24979554-24979964 | GGT1 | | TSS200 |
| 20M05_Saliva | chr22:25423531-25423801 | KIAA1671 | N_Shore | TSS1500 |
| 20M05_Saliva | chr22:31536585-31537064 | PLA2G3 | | TSS200 |
| 20M05_Saliva | chr22:33402393-33402998 | SYN3;SYN3;SYN3 | | 1stExon;Body;1stExon |
| 20M05_Saliva | chr22:38506623-38506781 | BAIAP2L2;BAIAP2L2 | | 1stExon;5'UTR |
| 20M05_Saliva | chr22:43820888-43821115 | MPPED1 | Island | 5'UTR |
| 20M05_Saliva | chr22:45608345-45608713 | C22orf9;C22orf9;C22orf9 | | Body;1stExon;5'UTR |
| 20M05_Saliva | chr22:46685653-46685728 | TTC38 | Island | Body |
| 20M05_Saliva | chr22:47022899-47023209 | GRAMD4 | Island | Body |
| 20M05_Saliva | chr22:47054067-47054273 | GRAMD4 | Island | Body |
| 20M05_Saliva | chr22:47057191-47057502 | GRAMD4 | N_Shore | Body |
| 20M05_Saliva | chr22:50025915-50026188 | C22orf34 | Island | Body |
| 20M05_Saliva | chr22:50451054-50451151 | IL17REL;IL17REL | S_Shore | 5'UTR;1stExon |
| 20M05_Saliva | chr22:50585197-50585427 | MOV10L1;MOV10L1;MOV10L1;MOV10L1 | | Body;TSS1500;Body;Body |
| 20M05_Saliva | chr1:173176366-173176523 | TNFSF4;TNFSF4 | | 5'UTR;1stExon |
| 20M05_Saliva | chr1:247712239-247712383 | C1orf150 | | TSS1500 |
| 20M05_Saliva | chr2:26401640-26402318 | FAM59B | N_Shore | Body |
| 20M05_Saliva | chr2:63279495-63279693 | OTX1 | N_Shore | 5'UTR |
| 20M05_Saliva | chr2:66673316-66673449 | MEIS1 | Island | Body |
| 20M05_Saliva | chr2:106776775-106777038 | UXS1 | | Body |
| 20M05_Saliva | chr2:113992843-113992930 | PAX8;PAX8;PAX8;PAX8;PAX8;LO( | N_Shore | Body;Body;Body;Body;Body;Body;TSS1500 |
| 20M05_Saliva | chr2:225265963-225266656 | FAM124B;FAM124B | | 1stExon;1stExon |
| 20M05_Saliva | chr2:225266783-225266876 | FAM124B;FAM124B | | TSS200;TSS200 |
| 20M05_Saliva | chr3:9943828-9943937 | IL17RE;IL17RE;IL17RE | | TSS1500;TSS1500;TSS1500 |
| 20M05_Saliva | chr3:11078979-11079298 | SLC6A1 | | 3'UTR |
| 20M05_Saliva | chr3:128779586-128779601 | GP9 | | TSS200 |
| 20M05_Saliva | chr3:181428298-181428547 | SOX2OT;SOX2 | N_Shore | Body;TSS1500 |
| 20M05_Saliva | chr3:183273758-183273894 | KLHL6 | | TSS1500 |
| 20M05_Saliva | chr4:7657070-7657359 | SORCS2 | | Body |
| 20M05_Saliva | chr4:10686394-10686610 | CLNK | | TSS200 |
| 20M05_Saliva | chr4:169799087-169799624 | PALLD;PALLD;PALLD;PALLD | Island | Body;Body;Body;Body |
| 20M05_Saliva | chr4:186559707-186560083 | SORBS2;SORBS2;SORBS2;SORB | N_Shore | Body;Body;Body;Body;Body;Body;Body;Body |
| 20M05_Saliva | chr5:148810177-148810203 | MIR145;LOC728264 | | TSS200;Body |
| 20M05_Saliva | chr5:179740743-179741120 | GFPT2 | Island | Body |
| 20M05_Saliva | chr6:13428522-13428657 | GFOD1 | | Body |

| | | | | |
|---|---|---|---|---|
| 20M05_Saliva | chr6:30039466-30039524 | RNF39;RNF39 | Island | Body;Body |
| 20M05_Saliva | chr6:32904061-32904317 | HLA-DMB | | Body |
| 20M05_Saliva | chr6:41528345-41528531 | FOXP4;FOXP4;FOXP4 | Island | 5'UTR;5'UTR;5'UTR |
| 20M05_Saliva | chr6:112115117-112115209 | FYN | | 5'UTR |
| 20M05_Saliva | chr6:157932074-157932180 | ZDHHC14;ZDHHC14 | | Body;Body |
| 20M05_Saliva | chr6:158066609-158066900 | ZDHHC14;ZDHHC14 | | Body;Body |
| 20M05_Saliva | chr6:158447877-158448134 | SYNJ2 | | Body |
| 20M05_Saliva | chr7:4848683-4848939 | RADIL | Island | Body |
| 20M05_Saliva | chr7:30951328-30951425 | AQP1 | | TSS200 |
| 20M05_Saliva | chr8:21771252-21771278 | DOK2 | S_Shore | TSS200 |
| 20M05_Saliva | chr10:49893346-49893423 | WDFY4 | | TSS200 |
| 20M05_Saliva | chr10:105978702-105978956 | MIR609;C10orf79 | | TSS200;Body |
| 20M05_Saliva | chr10:123923881-123924085 | TACC2;TACC2;TACC2;TACC2 | Island | Body;Body;Body;Body |
| 20M05_Saliva | chr11:2321842-2322050 | C11orf21;TSPAN32;C11orf21 | | Body;TSS1500;Body |
| 20M05_Saliva | chr11:2322618-2322674 | C11orf21;TSPAN32;C11orf21 | | Body;TSS1500;Body |
| 20M05_Saliva | chr11:35441777-35442225 | SLC1A2 | Island | TSS1500 |
| 20M05_Saliva | chr11:60739019-60739183 | CD6 | | TSS200 |
| 20M05_Saliva | chr11:66084841-66085299 | CD248 | S_Shore | TSS1500 |
| 20M05_Saliva | chr11:69934031-69934128 | ANO1;ANO1 | Island | Body;Body |
| 20M05_Saliva | chr11:70211301-70211531 | PPFIA1;PPFIA1 | | Body;Body |
| 20M05_Saliva | chr11:78673073-78673212 | ODZ4 | Island | Body |
| 20M05_Saliva | chr11:113846918-113846937 | HTR3A;HTR3A;HTR3A | | Body;TSS1500;Body |
| 20M05_Saliva | chr12:4699085-4699618 | DYRK4 | | TSS200 |
| 20M05_Saliva | chr12:6881595-6881629 | LAG3 | N_Shore | TSS200 |
| 20M05_Saliva | chr12:55378621-55378654 | KIAA0748 | | TSS200 |
| 20M05_Saliva | chr12:117797056-117797385 | NOS1 | N_Shore | 5'UTR |
| 20M05_Saliva | chr12:123518833-123519157 | PITPNM2 | | Body |
| 20M05_Saliva | chr13:107145933-107146164 | EFNB2 | S_Shore | Body |
| 20M05_Saliva | chr13:110993474-110993558 | COL4A2 | | Body |
| 20M05_Saliva | chr15:101991560-101991828 | PCSK6;PCSK6;PCSK6;PCSK6;PCSK6;PCSK6;PCSK6;PCSK6 | | Body;Body;Body;Body;Body;Body;Body;Body |
| 20M05_Saliva | chr16:1583883-1583984 | IFT140;TMEM204 | Island | Body;TSS1500 |
| 20M05_Saliva | chr16:2892766-2892782 | TMPRSS8 | Island | TSS200 |
| 20M05_Saliva | chr16:14403004-14403040 | MIR365-1 | | TSS200 |
| 20M05_Saliva | chr16:28996021-28996053 | LAT;LAT;LAT;LAT | | TSS200;TSS1500;TSS1500;TSS1500 |
| 20M05_Saliva | chr16:28996329-28996533 | LAT;LAT;LAT;LAT | | Body;TSS200;TSS200;TSS200 |
| 20M05_Saliva | chr16:29674959-29675214 | SPN;SPN | N_Shore | 5'UTR;5'UTR |
| 20M05_Saliva | chr16:56659659-56659964 | MT1E;MT1E | Island | 1stExon;5'UTR |
| 20M05_Saliva | chr16:57576285-57576394 | GPR114 | | TSS1500 |
| 20M05_Saliva | chr17:3704494-3704602 | ITGAE;ITGAE | | 1stExon;5'UTR |
| 20M05_Saliva | chr17:5138259-5138645 | C17orf87 | | TSS200 |
| 20M05_Saliva | chr17:34344995-34345172 | CCL23;CCL23;CCL23;CCL23 | | 1stExon;5'UTR;1stExon;5'UTR |
| 20M05_Saliva | chr17:79005662-79006087 | FLJ90757 | N_Shore | Body |
| 20M05_Saliva | chr19:4326975-4327350 | STAP2;STAP2 | Island | Body;Body |
| 20M05_Saliva | chr19:11530021-11530060 | RGL3;RGL3 | N_Shore | TSS200;TSS200 |
| 20M05_Saliva | chr19:23941408-23941575 | ZNF681 | Island | Body |
| 20M05_Saliva | chr19:35819889-35820181 | CD22 | | TSS200 |
| 20M05_Saliva | chr19:51875451-51875986 | NKG7 | S_Shelf | Body |
| 20M05_Saliva | chr20:22566289-22566976 | FOXA2;FOXA2 | N_Shore | TSS1500;TSS200 |
| 20M05_Saliva | chr20:35274595-35274639 | SLA2;SLA2 | | TSS200;TSS200 |
| 20M05_Saliva | chr20:37433745-37433803 | PPP1R16B | N_Shore | TSS1500 |
| 20M05_Saliva | chr21:39870397-39870704 | ERG;ERG;ERG;ERG;ERG;ERG | | Body;1stExon;1stExon;5'UTR;5'UTR;Body |
| 20M05_Saliva | chr21:48026043-48026298 | S100B | | TSS1500 |
| 20M05_Saliva | chr22:45073148-45073165 | PRR5;PRR5;PRR5;PRR5 | | 5'UTR;5'UTR;5'UTR;5'UTR |
| 20M06_Saliva | chr1:110254709-110254854 | GSTM5 | | TSS200 |
| 20M06_Saliva | chr5:135416412-135416613 | MIR886 | Island | TSS200 |
| 20M06_Saliva | chr5:179740914-179741120 | GFPT2 | Island | Body |
| 20M06_Saliva | chr6:32063874-32063901 | TNXB | Island | Body |
| 20M06_Saliva | chr10:135341528-135341933 | CYP2E1 | N_Shore | 1stExon |
| 20M06_Saliva | chr12:312592-313104 | SLC6A12;SLC6A12;SLC6A12 | Island | Body;Body;Body |
| 20M06_Saliva | chr17:79905236-79905263 | MYADML2 | | TSS200 |
| 20M06_Saliva | chr22:24384397-24384525 | GSTT1 | Island | TSS200 |
| 20M06_Saliva | chr5:415575-415885 | AHRR | | Body |
| 20M06_Saliva | chr5:1594282-1594579 | SDHAP3 | Island | Body |
| 20M06_Saliva | chr17:74023625-74023633 | EVPL | | TSS200 |
| 20M06_Saliva | chr19:55972791-55973338 | ISOC2;ISOC2;ISOC2 | Island | 5'UTR;5'UTR;5'UTR |

| | | | | |
|---|---|---|---|---|
| 20M06_Saliva | chr22:24348549-24348715 | GSTTP1 | | TSS1500 |
| 30M04_Saliva | chr1:2266362-2266591 | MORN1 | Island | Body |
| 30M04_Saliva | chr1:2284167-2284501 | MORN1;LOC100129534 | | Body;TSS200 |
| 30M04_Saliva | chr1:3607110-3607401 | TP73;TP73;TP73;TP73 | Island | Body;TSS200;TSS200;TSS200 |
| 30M04_Saliva | chr1:3615390-3615762 | TP73;TP73;TP73;TP73 | | Body;Body;Body;Body |
| 30M04_Saliva | chr1:6390473-6390793 | ACOT7;ACOT7;ACOT7;ACOT7 | | Body;Body;Body;Body |
| 30M04_Saliva | chr1:6446095-6446234 | ACOT7;ACOT7 | | Body;TSS1500 |
| 30M04_Saliva | chr1:10707917-10707953 | CASZ1;CASZ1 | Island | Body;Body |
| 30M04_Saliva | chr1:10839450-10839574 | CASZ1;CASZ1 | | 5'UTR;5'UTR |
| 30M04_Saliva | chr1:15271952-15272238 | KIAA1026;KIAA1026;KIAA1026;KIAA1026;KIAA1026 | | Body;5'UTR;TSS1500;Body;Body |
| 30M04_Saliva | chr1:35220251-35220710 | GJB5 | | TSS1500 |
| 30M04_Saliva | chr1:43751270-43751497 | C1orf210;C1orf210;C1orf210 | | 1stExon;5'UTR;TSS200 |
| 30M04_Saliva | chr1:44726943-44727237 | ERI3 | | Body |
| 30M04_Saliva | chr1:45083079-45083207 | RNF220 | Island | Body |
| 30M04_Saliva | chr1:151345123-151345181 | SELENBP1;SELENBP1 | | 5'UTR;1stExon |
| 30M04_Saliva | chr1:178310534-178310650 | RASAL2;RASAL2 | | Body;TSS200 |
| 30M04_Saliva | chr1:205326071-205326437 | KLHDC8A | | TSS200 |
| 30M04_Saliva | chr2:74153195-74153317 | DGUOK;DGUOK | N_Shore | TSS1500;TSS1500 |
| 30M04_Saliva | chr3:11178683-11178798 | HRH1 | | TSS200 |
| 30M04_Saliva | chr3:123339334-123339568 | MYLK;MYLK;MYLK;MYLK;MYLK;MYLK;MYLK;MYLK | | 1stExon;5'UTR;Body;Body;Body;5'UTR;1stExon;Body |
| 30M04_Saliva | chr4:1214965-1215112 | CTBP1;CTBP1 | N_Shelf | Body;Body |
| 30M04_Saliva | chr4:184908351-184909018 | STOX2 | | Body |
| 30M04_Saliva | chr5:1016460-1016768 | NKD2 | | Body |
| 30M04_Saliva | chr5:59783629-59784203 | PART1;PART1;PDE4D;PART1 | | TSS200;Body;5'UTR;TSS200 |
| 30M04_Saliva | chr5:149546331-149546366 | CDX1 | Island | TSS200 |
| 30M04_Saliva | chr5:169532721-169532758 | FOXI1;FOXI1 | N_Shore | TSS200;TSS200 |
| 30M04_Saliva | chr6:8436218-8436296 | SLC35B3;SLC35B3;SLC35B3 | S_Shore | TSS1500;TSS1500;TSS1500 |
| 30M04_Saliva | chr6:30122544-30122593 | TRIM10;TRIM10 | | Body;Body |
| 30M04_Saliva | chr6:30850876-30850977 | DDR1;DDR1 | N_Shore | TSS1500;TSS1500 |
| 30M04_Saliva | chr6:32120625-32120783 | PPT2;PRRT1;PPT2 | N_Shore | TSS1500;TSS1500;TSS1500 |
| 30M04_Saliva | chr6:32552016-32552042 | HLA-DRB1 | Island | Body |
| 30M04_Saliva | chr6:33048310-33048469 | HLA-DPB1 | N_Shore | Body |
| 30M04_Saliva | chr6:33048529-33048558 | HLA-DPB1 | Island | Body |
| 30M04_Saliva | chr6:116381904-116382179 | FRK;FRK | | 1stExon;5'UTR |
| 30M04_Saliva | chr6:137366322-137366369 | IL20RA | S_Shore | TSS200 |
| 30M04_Saliva | chr7:1514455-1514623 | INTS1 | Island | Body |
| 30M04_Saliva | chr7:3033173-3033579 | CARD11 | | 5'UTR |
| 30M04_Saliva | chr7:27153655-27153847 | HOXA3;HOXA3;HOXA3 | N_Shore | 5'UTR;5'UTR;TSS200 |
| 30M04_Saliva | chr7:98738998-98739698 | SMURF1;SMURF1 | N_Shelf | Body;Body |
| 30M04_Saliva | chr7:135433540-135433715 | FAM180A;FAM180A | | 1stExon;5'UTR |
| 30M04_Saliva | chr7:158059944-158060003 | PTPRN2;PTPRN2;PTPRN2 | N_Shore | Body;Body;Body |
| 30M04_Saliva | chr8:145728507-145728543 | GPT | N_Shore | TSS1500 |
| 30M04_Saliva | chr10:1505595-1505654 | ADARB2 | N_Shore | Body |
| 30M04_Saliva | chr10:3149673-3149852 | PFKP | S_Shore | Body |
| 30M04_Saliva | chr10:26856095-26856128 | APBB1IP | Island | Body |
| 30M04_Saliva | chr10:49459949-49460249 | FRMPD2;FRMPD2 | | Body;Body |
| 30M04_Saliva | chr10:82299606-82299737 | SH2D4B;SH2D4B | S_Shore | 5'UTR;1stExon |
| 30M04_Saliva | chr10:99331332-99331808 | ANKRD2;ANKRD2 | S_Shore | 3'UTR |
| 30M04_Saliva | chr10:123353822-123354171 | FGFR2;FGFR2;FGFR2;FGFR2;FGF | N_Shelf | TSS200;5'UTR;5'UTR;TSS1500;5'UTR;TSS1500;5'UTR;5'UTR;5'UTR |
| 30M04_Saliva | chr11:67442195-67442408 | ALDH3B2;ALDH3B2 | | 5'UTR;TSS200 |
| 30M04_Saliva | chr11:69482104-69482597 | ORAOV1 | | 3'UTR |
| 30M04_Saliva | chr11:108422996-108423124 | EXPH5;EXPH5 | | TSS200;Body |
| 30M04_Saliva | chr11:117069780-117069849 | TAGLN;TAGLN | | TSS1500;TSS1500 |
| 30M04_Saliva | chr11:128812913-128813008 | TP53AIP1 | | TSS200 |
| 30M04_Saliva | chr12:258355-258436 | IQSEC3;IQSEC3 | | Body;Body |
| 30M04_Saliva | chr12:25801455-25801621 | IFLTD1 | | 1stExon |
| 30M04_Saliva | chr12:123750782-123750864 | CDK2AP1 | N_Shelf | Body |
| 30M04_Saliva | chr12:133101224-133101612 | FBRSL1 | N_Shore | Body |
| 30M04_Saliva | chr13:111522755-111522932 | C13orf29 | | TSS200 |
| 30M04_Saliva | chr14:69256690-69256977 | ZFP36L1 | Island | Body |
| 30M04_Saliva | chr14:94576977-94577101 | IFI27;IFI27 | | TSS200;TSS200 |
| 30M04_Saliva | chr14:103593503-103593520 | TNFAIP2 | Island | Body |
| 30M04_Saliva | chr14:104644850-104645063 | KIF26A | Island | Body |
| 30M04_Saliva | chr15:69099678-69099834 | C15orf28;ANP32A | | TSS1500;Body |

| | | | | |
|---|---|---|---|---|
| 30M04_Saliva | chr16:2770211-2770336 | PRSS27;PRSS27 | Island | 1stExon;5'UTR |
| 30M04_Saliva | chr16:87958281-87958493 | CA5A | | Body |
| 30M04_Saliva | chr17:693137-693388 | RNMTL1 | | Body |
| 30M04_Saliva | chr17:927457-927652 | ABR;ABR;ABR | S_Shore | Body;Body;Body |
| 30M04_Saliva | chr17:3791902-3792074 | CAMKK1;CAMKK1;CAMKK1 | N_Shore | 5'UTR;5'UTR;5'UTR |
| 30M04_Saliva | chr17:7832932-7833237 | KCNAB3 | Island | TSS200 |
| 30M04_Saliva | chr17:40274703-40274740 | HSPB9;KAT2A | Island | TSS200;TSS1500 |
| 30M04_Saliva | chr17:79031550-79031712 | BAIAP2;BAIAP2;BAIAP2;BAIAP2 | | Body;Body;Body;Body |
| 30M04_Saliva | chr17:80797881-80798013 | TBCD;ZNF750;ZNF750 | | Body;5'UTR;1stExon |
| 30M04_Saliva | chr18:21572622-21572748 | TTC39C | | TSS200 |
| 30M04_Saliva | chr18:61143926-61144250 | SERPINB5 | | TSS1500 |
| 30M04_Saliva | chr18:61253847-61254561 | SERPINB13 | | TSS1500 |
| 30M04_Saliva | chr19:751124-751241 | C19orf21 | S_Shelf | TSS200 |
| 30M04_Saliva | chr19:2525231-2525459 | GNG7 | Island | 5'UTR |
| 30M04_Saliva | chr19:4305072-4305239 | FSD1 | Island | Body |
| 30M04_Saliva | chr20:62367884-62367893 | LIME1 | N_Shore | TSS200 |
| 30M04_Saliva | chr20:62693951-62693973 | TCEA2;TCEA2 | Island | TSS200;5'UTR |
| 30M04_Saliva | chr21:43185674-43185884 | RIPK4 | N_Shore | Body |
| 30M04_Saliva | chr22:33402628-33402998 | SYN3;SYN3;SYN3 | | 1stExon;Body;1stExon |
| 30M04_Saliva | chr22:45608465-45608516 | C22orf9;C22orf9 | | Body;TSS200 |
| 30M04_Saliva | chr1:2082315-2082522 | PRKCZ;PRKCZ;PRKCZ | Island | Body;Body;Body |
| 30M04_Saliva | chr1:7692321-7692367 | CAMTA1 | | Body |
| 30M04_Saliva | chr1:9714810-9714845 | C1orf200;PIK3CD | S_Shore | TSS200;5'UTR |
| 30M04_Saliva | chr1:24239771-24240069 | CNR2;CNR2 | | 5'UTR;1stExon |
| 30M04_Saliva | chr1:54562040-54562213 | C1orf83 | | Body |
| 30M04_Saliva | chr1:114414042-114414408 | PTPN22;PTPN22 | | Body;Body |
| 30M04_Saliva | chr1:159796434-159796629 | SLAMF8 | | TSS200 |
| 30M04_Saliva | chr1:160616870-160617124 | SLAMF1;SLAMF1 | | 5'UTR;1stExon |
| 30M04_Saliva | chr1:202129865-202130344 | PTPN7;PTPN7 | | TSS1500;TSS200 |
| 30M04_Saliva | chr1:205181237-205181581 | DSTYK;DSTYK | S_Shore | TSS1500;TSS1500 |
| 30M04_Saliva | chr1:206945924-206946187 | IL10 | | TSS200 |
| 30M04_Saliva | chr1:207095153-207095332 | FAIM3;FAIM3;FAIM3 | | Body;Body;Body |
| 30M04_Saliva | chr1:207277466-207277581 | C4BPA | | TSS200 |
| 30M04_Saliva | chr1:208084942-208085455 | CD34;CD34 | S_Shore | TSS1500;TSS1500 |
| 30M04_Saliva | chr1:209929496-209930127 | TRAF3IP3 | | TSS200 |
| 30M04_Saliva | chr1:226187852-226188006 | C1orf55 | S_Shore | TSS1500 |
| 30M04_Saliva | chr1:247711595-247712591 | C1orf150 | | TSS1500 |
| 30M04_Saliva | chr2:63283939-63285950 | OTX1 | Island | 3'UTR |
| 30M04_Saliva | chr2:69001665-69001854 | ARHGAP25;ARHGAP25;ARHGAP25;ARHGAP25 | | TSS1500;Body;Body;TSS1500 |
| 30M04_Saliva | chr2:100759014-100759159 | AFF3;AFF3 | | 1stExon;5'UTR |
| 30M04_Saliva | chr2:106776775-106777038 | UXS1 | | Body |
| 30M04_Saliva | chr2:127413363-127413369 | GYPC;GYPC | N_Shore | TSS1500;TSS1500 |
| 30M04_Saliva | chr2:143886326-143886940 | ARHGAP15 | | TSS1500 |
| 30M04_Saliva | chr2:175499113-175499377 | WIPF1;WIPF1 | | 5'UTR;5'UTR |
| 30M04_Saliva | chr2:179395885-179395935 | TTN;TTN;TTN;MIR548N;TTN | | Body;Body;Body;Body;Body |
| 30M04_Saliva | chr2:225265963-225266880 | FAM124B;FAM124B | | 1stExon;1stExon |
| 30M04_Saliva | chr2:233924789-233924852 | INPP5D;INPP5D | N_Shore | TSS1500;TSS1500 |
| 30M04_Saliva | chr2:233924923-233925030 | INPP5D;INPP5D | N_Shore | TSS200;TSS200 |
| 30M04_Saliva | chr2:240174613-240174659 | HDAC4 | | Body |
| 30M04_Saliva | chr2:240302637-240302855 | HDAC4 | | 5'UTR |
| 30M04_Saliva | chr3:9943848-9943937 | IL17RE;IL17RE;IL17RE | | TSS1500;TSS1500;TSS1500 |
| 30M04_Saliva | chr3:10334717-10335375 | GHRLOS;GHRL;GHRL;GHRLOS;GHRLOS;GHRL;GHRL;GHRL;GHRL;GHR | | Body;TSS200;TSS200;Body;Body;TSS200;TSS200;TSS200;TSS200;TSS200;Body;TSS200;TSS200;TSS200 |
| 30M04_Saliva | chr3:46448579-46448868 | CCRL2;CCRL2 | | TSS1500;TSS200 |
| 30M04_Saliva | chr3:50376216-50376475 | RASSF1;RASSF1;RASSF1;RASSF1 | N_Shore | TSS1500;TSS1500;Body;Body |
| 30M04_Saliva | chr3:55517101-55517659 | WNT5A | N_Shore | Body |
| 30M04_Saliva | chr3:55517853-55518396 | WNT5A | Island | Body |
| 30M04_Saliva | chr3:121379745-121379791 | HCLS1;HCLS1 | | 1stExon;5'UTR |
| 30M04_Saliva | chr3:121796627-121796767 | CD86;CD86 | | TSS200;Body |
| 30M04_Saliva | chr3:128779498-128779601 | GP9 | | TSS200 |
| 30M04_Saliva | chr3:150996078-150996563 | P2RY14;MED12L;P2RY14 | | 1stExon;Body;5'UTR |
| 30M04_Saliva | chr3:151058527-151058837 | P2RY12;P2RY12;MED12L;P2RY12 | | 1stExon;5'UTR;Body;5'UTR |
| 30M04_Saliva | chr3:151102703-151102879 | P2RY12;MED12L | | TSS200;Body |
| 30M04_Saliva | chr3:181441364-181441978 | SOX2OT | N_Shelf | Body |
| 30M04_Saliva | chr3:183273245-183273894 | KLHL6 | | 1stExon |

| | | | | |
|---|---|---|---|---|
| 30M04_Saliva | chr4:1195845-1196179 | LOC100130872;LOC100130872-SPON2 | | Body;Body |
| 30M04_Saliva | chr4:2813542-2814122 | SH3BP2;SH3BP2 | | TSS1500;5'UTR |
| 30M04_Saliva | chr4:10686394-10686610 | CLNK | | TSS200 |
| 30M04_Saliva | chr4:15779642-15779879 | CD38 | N_Shore | TSS1500 |
| 30M04_Saliva | chr4:38807259-38807382 | TLR1 | | TSS1500 |
| 30M04_Saliva | chr4:40198392-40198595 | RHOH | | TSS200 |
| 30M04_Saliva | chr4:69435250-69435601 | UGT2B15;UGT2B17 | | TSS1500;TSS1500 |
| 30M04_Saliva | chr4:90815958-90816310 | MMRN1 | | TSS200 |
| 30M04_Saliva | chr4:95264019-95264127 | HPGDS;HPGDS | | 5'UTR;1stExon |
| 30M04_Saliva | chr4:154711280-154711347 | SFRP2 | N_Shore | TSS1500 |
| 30M04_Saliva | chr4:154711519-154711629 | SFRP2 | N_Shore | TSS1500 |
| 30M04_Saliva | chr5:37835109-37835168 | GDNF;GDNF;GDNF | Island | 5'UTR;TSS1500;5'UTR |
| 30M04_Saliva | chr5:37835684-37835784 | GDNF;GDNF;GDNF | N_Shore | TSS1500;5'UTR;TSS200 |
| 30M04_Saliva | chr5:39203007-39203190 | FYB;FYB | | Body;Body |
| 30M04_Saliva | chr5:76010444-76010658 | F2R | N_Shore | TSS1500 |
| 30M04_Saliva | chr5:95159365-95159639 | GLRX;GLRX | | TSS1500;TSS1500 |
| 30M04_Saliva | chr5:156696185-156696351 | CYFIP2;CYFIP2;CYFIP2 | S_Shelf | TSS200;5'UTR;5'UTR |
| 30M04_Saliva | chr5:169724590-169724791 | LCP2 | | 1stExon |
| 30M04_Saliva | chr5:176326365-176326528 | HK3 | | TSS200 |
| 30M04_Saliva | chr6:4890079-4890278 | CDYL;CDYL;CDYL;CDYL | | 5'UTR;Body;Body;TSS200 |
| 30M04_Saliva | chr6:11382553-11382769 | NEDD9;NEDD9 | | 1stExon;5'UTR |
| 30M04_Saliva | chr6:13274151-13274284 | PHACTR1 | | Body |
| 30M04_Saliva | chr6:13428522-13428657 | GFOD1 | | Body |
| 30M04_Saliva | chr6:28493601-28493664 | GPX5;GPX5 | | TSS200;TSS200 |
| 30M04_Saliva | chr6:28874479-28874702 | TRIM27 | | Body |
| 30M04_Saliva | chr6:28875354-28875370 | TRIM27 | | Body |
| 30M04_Saliva | chr6:29454557-29454755 | MAS1L | | 1stExon |
| 30M04_Saliva | chr6:30104440-30104626 | TRIM40 | | TSS200 |
| 30M04_Saliva | chr6:30297174-30297565 | TRIM39;TRIM39 | S_Shelf | Body;Body |
| 30M04_Saliva | chr6:31021422-31021796 | HCG22 | | TSS1500 |
| 30M04_Saliva | chr6:31237013-31237034 | HLA-C | N_Shore | Body |
| 30M04_Saliva | chr6:31237405-31237722 | HLA-C | N_Shore | Body |
| 30M04_Saliva | chr6:31240814-31241294 | HLA-C | S_Shore | TSS1500 |
| 30M04_Saliva | chr6:31554199-31555016 | LST1;LST1;LST1;LST1;LST1;LST1;LST1;LST1 | | Body;TSS1500;TSS1500;TSS1500;5'UTR;TSS1500;5'UTR;Body |
| 30M04_Saliva | chr6:31583915-31584223 | AIF1;AIF1;AIF1 | N_Shelf | Body;1stExon;Body |
| 30M04_Saliva | chr6:31695415-31695599 | DDAH2 | N_Shore | Body |
| 30M04_Saliva | chr6:32295097-32295237 | C6orf10 | | Body |
| 30M04_Saliva | chr6:32813337-32813582 | PSMB8;TAP1 | S_Shore | TSS1500;3'UTR |
| 30M04_Saliva | chr6:32904061-32904317 | HLA-DMB | | Body |
| 30M04_Saliva | chr6:32909282-32909523 | HLA-DMB | | TSS1500 |
| 30M04_Saliva | chr6:32920220-32920735 | HLA-DMA | | Body |
| 30M04_Saliva | chr6:33039481-33039606 | HLA-DPA1 | | Body |
| 30M04_Saliva | chr6:33040535-33041343 | HLA-DPA1 | | Body |
| 30M04_Saliva | chr6:33043841-33043868 | HLA-DPB1 | | 1stExon |
| 30M04_Saliva | chr6:108145539-108145930 | SCML4 | | TSS200 |
| 30M04_Saliva | chr6:133035150-133035379 | VNN1 | | 1stExon |
| 30M04_Saliva | chr6:135516912-135517046 | MYB;MYB;MYB;MYB;MYB;MYB;MYB;MYB | | Body;Body;Body;Body;Body;Body;Body;Body |
| 30M04_Saliva | chr6:166856056-166856094 | RPS6KA2;RPS6KA2 | | Body;Body |
| 30M04_Saliva | chr6:166876736-166877038 | RPS6KA2;RPS6KA2 | | Body;Body |
| 30M04_Saliva | chr7:1095595-1095720 | C7orf50;C7orf50;C7orf50 | N_Shore | Body;Body;Body |
| 30M04_Saliva | chr7:1980117-1980273 | MAD1L1;MAD1L1;MAD1L1 | S_Shore | Body;Body;Body |
| 30M04_Saliva | chr7:2648289-2648785 | IQCE;IQCE | S_Shelf | Body;Body |
| 30M04_Saliva | chr7:2773782-2774474 | GNA12 | N_Shore | Body |
| 30M04_Saliva | chr7:4752951-4753002 | FOXK1 | | Body |
| 30M04_Saliva | chr7:5271516-5271655 | WIPI2;WIPI2;WIPI2;WIPI2;WIPI2 | S_Shelf | 3'UTR;3'UTR;3'UTR;3'UTR;3'UTR |
| 30M04_Saliva | chr7:30951272-30951425 | AQP1 | | TSS200 |
| 30M04_Saliva | chr7:45066841-45066895 | CCM2;CCM2;CCM2;CCM2;CCM2 | | Body;TSS1500;Body;Body;Body |
| 30M04_Saliva | chr7:75443075-75443246 | CCL24 | | TSS200 |
| 30M04_Saliva | chr7:96655105-96655566 | DLX5 | S_Shore | TSS1500 |
| 30M04_Saliva | chr7:139333296-139333410 | HIPK2;HIPK2 | | Body;Body |
| 30M04_Saliva | chr7:151442351-151442481 | PRKAG2;PRKAG2 | | Body;Body |
| 30M04_Saliva | chr7:157647109-157647241 | PTPRN2;PTPRN2;PTPRN2 | N_Shore | Body;Body;Body |
| 30M04_Saliva | chr8:19540210-19540453 | CSGALNACT1;CSGALNACT1 | | TSS200;Body |
| 30M04_Saliva | chr8:21771059-21771258 | DOK2 | S_Shore | 1stExon |

| | | | | |
|---|---|---|---|---|
| 30M04_Saliva | chr8:27469186-27469338 | CLU;CLU;CLU;CLU | N_Shelf | 1stExon;Body;5'UTR;TSS1500 |
| 30M04_Saliva | chr8:41168072-41168205 | SFRP1 | S_Shore | TSS1500 |
| 30M04_Saliva | chr8:67975874-67975895 | CSPP1;CSPP1;COPS5 | N_Shore | TSS1500;TSS1500;TSS1500 |
| 30M04_Saliva | chr8:95229664-95229872 | CDH17 | | TSS200 |
| 30M04_Saliva | chr8:142192568-142192718 | DENND3 | | Body |
| 30M04_Saliva | chr8:145002522-145002835 | PLEC1;PLEC1;PLEC1;PLEC1;PLE | N_Shore | Body;Body;Body;Body;Body;Body;Body;Body |
| 30M04_Saliva | chr8:145086243-145086427 | SPATC1;SPATC1 | | TSS1500;TSS1500 |
| 30M04_Saliva | chr9:117692745-117693010 | TNFSF8;TNFSF8 | | 1stExon;5'UTR |
| 30M04_Saliva | chr9:123688715-123689193 | TRAF1 | N_Shelf | 5'UTR |
| 30M04_Saliva | chr10:6214016-6214079 | PFKFB3 | | Body |
| 30M04_Saliva | chr10:49892930-49893336 | WDFY4 | | TSS1500 |
| 30M04_Saliva | chr10:94451016-94451518 | HHEX | N_Shore | Body |
| 30M04_Saliva | chr10:105978651-105978956 | MIR609;C10orf79 | | Body |
| 30M04_Saliva | chr10:114134878-114135054 | ACSL5;ACSL5;ACSL5 | | TSS1500;TSS1500;5'UTR |
| 30M04_Saliva | chr10:123923881-123924085 | TACC2;TACC2;TACC2;TACC2 | Island | Body;Body;Body;Body |
| 30M04_Saliva | chr11:2321770-2322050 | C11orf21;TSPAN32;C11orf21 | | Body;TSS1500;Body |
| 30M04_Saliva | chr11:2322741-2322802 | C11orf21;TSPAN32;C11orf21 | | Body;TSS1500;Body |
| 30M04_Saliva | chr11:2322935-2323083 | C11orf21;TSPAN32;C11orf21 | | Body;TSS1500;Body |
| 30M04_Saliva | chr11:2919808-2920055 | SLC22A18AS;SLC22A18 | N_Shelf | Body;TSS1500 |
| 30M04_Saliva | chr11:3177622-3178024 | OSBPL5;OSBPL5;OSBPL5 | N_Shelf | 5'UTR;5'UTR;5'UTR |
| 30M04_Saliva | chr11:36422377-36422615 | PRR5L;PRR5L;PRR5L;PRR5L | | 5'UTR;5'UTR;TSS200;Body |
| 30M04_Saliva | chr11:46382544-46382725 | DGKZ;DGKZ;DGKZ;DGKZ | | Body;TSS1500;Body;Body |
| 30M04_Saliva | chr11:59634191-59634276 | TCN1 | | TSS200 |
| 30M04_Saliva | chr11:59823993-59824161 | MS4A3;MS4A3;MS4A3 | | TSS200;TSS200;TSS200 |
| 30M04_Saliva | chr11:60738971-60739183 | CD6 | | TSS200 |
| 30M04_Saliva | chr11:63273507-63274030 | LGALS12;LGALS12;LGALS12 | | TSS200;TSS200;TSS200 |
| 30M04_Saliva | chr11:63973846-63974123 | FERMT3;FERMT3 | N_Shore | TSS1500;TSS1500 |
| 30M04_Saliva | chr11:63974955-63975481 | FERMT3;FERMT3 | Island | Body;Body |
| 30M04_Saliva | chr11:64107517-64108241 | CCDC88B | N_Shelf | TSS200 |
| 30M04_Saliva | chr11:67170610-67171585 | PPP1CA;PPP1CA;TBC1D10C;PPI | S_Shore | TSS1500;TSS1500;TSS1500;TSS1500 |
| 30M04_Saliva | chr11:69520186-69520418 | FGF19 | S_Shore | TSS1500 |
| 30M04_Saliva | chr11:70211301-70211531 | PPFIA1;PPFIA1 | | Body;Body |
| 30M04_Saliva | chr11:70257035-70257149 | CTTN;CTTN | N_Shore | Body;Body |
| 30M04_Saliva | chr11:71709837-71709978 | IL18BP;IL18BP;IL18BP;IL18BP;IL18BP | | TSS200;TSS1500;TSS200;TSS1500;TSS1500 |
| 30M04_Saliva | chr11:78673073-78673212 | ODZ4 | Island | Body |
| 30M04_Saliva | chr11:113953462-113953812 | ZBTB16;ZBTB16 | N_Shore | Body;Body |
| 30M04_Saliva | chr11:118084920-118085281 | AMICA1;AMICA1 | | Body;TSS1500 |
| 30M04_Saliva | chr11:134126284-134126364 | ACAD8 | S_Shelf | Body |
| 30M04_Saliva | chr12:6881590-6881629 | LAG3 | N_Shore | TSS200 |
| 30M04_Saliva | chr12:7060263-7060386 | PTPN6;PTPN6;PTPN6 | | TSS200;Body;TSS200 |
| 30M04_Saliva | chr12:32654929-32655034 | FGD4 | | TSS200 |
| 30M04_Saliva | chr12:47610220-47610734 | FAM113B;FAM113B;LOC100233209 | | 5'UTR;1stExon;Body |
| 30M04_Saliva | chr12:54891213-54891636 | NCKAP1L | | TSS1500 |
| 30M04_Saliva | chr12:55378411-55378912 | KIAA0748;KIAA0748 | | 1stExon;5'UTR |
| 30M04_Saliva | chr12:58210661-58210878 | AVIL | | TSS1500 |
| 30M04_Saliva | chr12:124950784-124950990 | NCOR2;NCOR2 | Island | Body;Body |
| 30M04_Saliva | chr13:51639953-51640315 | GUCY1B2 | | Body |
| 30M04_Saliva | chr13:107145658-107146164 | EFNB2 | Island | Body |
| 30M04_Saliva | chr13:110951025-110951418 | COL4A1 | | Body |
| 30M04_Saliva | chr13:110993474-110993558 | COL4A2 | | Body |
| 30M04_Saliva | chr13:111318022-111318490 | CARS2 | | Body |
| 30M04_Saliva | chr13:114261869-114261986 | TFDP1;TFDP1 | S_Shore | Body;Body |
| 30M04_Saliva | chr14:21423684-21423837 | RNASE2;RNASE2 | | 1stExon;5'UTR |
| 30M04_Saliva | chr14:24641201-24641706 | REC8;REC8 | Island | TSS200;TSS200 |
| 30M04_Saliva | chr14:37053169-37053601 | NKX2-8 | Island | TSS1500 |
| 30M04_Saliva | chr14:38725650-38725750 | CLEC14A | S_Shore | TSS200 |
| 30M04_Saliva | chr14:75988800-75989203 | BATF;BATF | | 5'UTR;1stExon |
| 30M04_Saliva | chr14:93170710-93170970 | LGMN;LGMN | | Body;Body |
| 30M04_Saliva | chr14:102676957-102677294 | WDR20 | N_Shelf | Body |
| 30M04_Saliva | chr14:105531893-105532159 | GPR132 | | TSS200 |
| 30M04_Saliva | chr15:38988533-38988860 | C15orf53 | | TSS1500 |
| 30M04_Saliva | chr15:40599680-40600099 | PLCB2 | | Body |
| 30M04_Saliva | chr15:43531947-43532243 | TGM5;TGM5 | | Body;Body |
| 30M04_Saliva | chr15:94840793-94841334 | MCTP2;MCTP2 | | TSS1500;TSS1500 |

| | | | | |
|---|---|---|---|---|
| 30M04_Saliva | chr16:303165-303192 | ITFG3 | | 5'UTR |
| 30M04_Saliva | chr16:2737220-2737356 | KCTD5 | | Body |
| 30M04_Saliva | chr16:4714080-4714443 | MGRN1;MGRN1;MGRN1;MGRN1 | | Body;Body;Body;Body |
| 30M04_Saliva | chr16:8738240-8738441 | C16orf68 | | Body |
| 30M04_Saliva | chr16:10647084-10647275 | EMP2 | | 5'UTR |
| 30M04_Saliva | chr16:28505320-28505959 | APOB48R | S_Shore | TSS1500 |
| 30M04_Saliva | chr16:28518200-28518385 | IL27 | | TSS200 |
| 30M04_Saliva | chr16:29673933-29674184 | SPN;SPN | N_Shore | TSS1500;TSS1500 |
| 30M04_Saliva | chr16:29756951-29757323 | C16orf54 | | 5'UTR |
| 30M04_Saliva | chr16:50715135-50715260 | SNX20;SNX20;SNX20;SNX20;SNX20;SNX20 | | 1stExon;5'UTR;1stExon;5'UTR;5'UTR;1stExon |
| 30M04_Saliva | chr16:89024511-89024657 | CBFA2T3 | | Body |
| 30M04_Saliva | chr17:750241-750379 | NXN | | Body |
| 30M04_Saliva | chr17:2839005-2839187 | RAP1GAP2;RAP1GAP2 | | Body;Body |
| 30M04_Saliva | chr17:3704471-3704621 | ITGAE;ITGAE | | 1stExon;5'UTR |
| 30M04_Saliva | chr17:5138259-5138645 | C17orf87 | | TSS200 |
| 30M04_Saliva | chr17:8771003-8771141 | PIK3R6 | | TSS200 |
| 30M04_Saliva | chr17:9940004-9940321 | GAS7;GAS7;GAS7 | | Body;1stExon;5'UTR |
| 30M04_Saliva | chr17:27252884-27253274 | PHF12;PHF12 | | Body;Body |
| 30M04_Saliva | chr17:34344995-34345172 | CCL23;CCL23;CCL23;CCL23 | | 1stExon;5'UTR;1stExon;5'UTR |
| 30M04_Saliva | chr17:42466960-42467307 | ITGA2B | S_Shelf | TSS200 |
| 30M04_Saliva | chr17:56408694-56409518 | MIR142 | | TSS200 |
| 30M04_Saliva | chr17:78755379-78755442 | RPTOR;RPTOR | | Body;Body |
| 30M04_Saliva | chr17:79259040-79259699 | SLC38A10;SLC38A10 | N_Shore | Body;Body |
| 30M04_Saliva | chr18:74114570-74114728 | ZNF516 | Island | Body |
| 30M04_Saliva | chr19:7741602-7741809 | C19orf59 | N_Shelf | TSS1500 |
| 30M04_Saliva | chr19:10449942-10450323 | ICAM3 | S_Shelf | Body |
| 30M04_Saliva | chr19:10958952-10958970 | C19orf38 | | TSS200 |
| 30M04_Saliva | chr19:14550846-14550999 | PKN1;PKN1 | N_Shore | TSS1500;Body |
| 30M04_Saliva | chr19:14785593-14785749 | EMR3;EMR3 | | 5'UTR;1stExon |
| 30M04_Saliva | chr19:17862104-17862210 | FCHO1;FCHO1;FCHO1;FCHO1 | S_Shelf | 5'UTR;TSS200;TSS200;5'UTR |
| 30M04_Saliva | chr19:35819775-35819943 | CD22 | | TSS1500 |
| 30M04_Saliva | chr19:36233297-36233469 | TMEM149;TMEM149 | S_Shore | 1stExon;5'UTR |
| 30M04_Saliva | chr19:39826693-39826923 | GMFG;GMFG | | 5'UTR;1stExon |
| 30M04_Saliva | chr19:46713446-46713473 | DKFZp434J0226 | | TSS200 |
| 30M04_Saliva | chr19:49838478-49838777 | CD37;CD37 | N_Shelf | TSS200;TSS200 |
| 30M04_Saliva | chr19:51875451-51875986 | NKG7 | S_Shelf | Body |
| 30M04_Saliva | chr19:52004992-52005188 | SIGLEC12;SIGLEC12;SIGLEC12 | | 1stExon;5'UTR;TSS1500 |
| 30M04_Saliva | chr19:54566838-54567279 | VSTM1 | | Body |
| 30M04_Saliva | chr19:54876446-54877827 | LAIR1;LAIR1;LAIR1;LAIR1 | | 5'UTR;1stExon;1stExon;5'UTR |
| 30M04_Saliva | chr20:1448595-1448645 | NSFL1C;NSFL1C;NSFL1C | S_Shore | TSS200;TSS200;TSS200 |
| 30M04_Saliva | chr20:1472071-1472419 | SIRPB2;SIRPB2;SIRPB2;SIRPB2 | | 1stExon;1stExon;5'UTR;5'UTR |
| 30M04_Saliva | chr20:1600623-1600705 | SIRPB1;SIRPB1;SIRPB1;SIRPB1;SIRPB1;SIRPB1 | | 5'UTR;1stExon;1stExon;5'UTR;5'UTR;1stExon |
| 30M04_Saliva | chr20:4795886-4795935 | RASSF2;RASSF2 | | 5'UTR;TSS200 |
| 30M04_Saliva | chr20:20036779-20037100 | C20orf26;CRNKL1;C20orf26 | S_Shelf | 5'UTR;TSS200;TSS1500 |
| 30M04_Saliva | chr20:23066947-23067156 | CD93;CD93 | | 5'UTR;1stExon |
| 30M04_Saliva | chr20:24929607-24929716 | CST7 | | TSS1500 |
| 30M04_Saliva | chr20:35274570-35274715 | SLA2;SLA2 | | TSS200;TSS200 |
| 30M04_Saliva | chr20:45179226-45179354 | C20orf123 | | TSS200 |
| 30M04_Saliva | chr20:54987020-54987140 | CASS4;CASS4;CASS4;CASS4 | | TSS1500;TSS1500;TSS200;TSS1500 |
| 30M04_Saliva | chr20:57582856-57583195 | CTSZ | S_Shore | TSS1500 |
| 30M04_Saliva | chr20:62198872-62199190 | PRIC285;PRIC285 | Island | Body;1stExon |
| 30M04_Saliva | chr21:36421467-36421955 | RUNX1;RUNX1 | | 5'UTR;1stExon |
| 30M04_Saliva | chr21:45773189-45773294 | TRPM2 | | TSS1500 |
| 30M04_Saliva | chr21:46340776-46341006 | ITGB2;ITGB2;ITGB2 | | 5'UTR;5'UTR;1stExon |
| 30M04_Saliva | chr21:48026043-48026298 | S100B | | TSS1500 |
| 30M04_Saliva | chr22:23923255-23923610 | IGLL1;IGLL1 | | TSS1500;TSS1500 |
| 30M04_Saliva | chr22:24105186-24105692 | C22orf15 | | TSS200 |
| 30M04_Saliva | chr22:27013978-27014116 | CRYBB1;CRYBB1 | | 5'UTR;1stExon |
| 30M04_Saliva | chr22:37545924-37546124 | IL2RB;IL2RB | | 5'UTR;1stExon |
| 30M04_Saliva | chr22:44577211-44577265 | PARVG;PARVG;PARVG;PARVG;PARVG;PARVG | | 5'UTR;Body;TSS200;TSS200;Body;5'UTR |
| 30M04_Saliva | chr22:46770063-46770253 | CELSR1 | N_Shore | Body |
| 30M04_Saliva | chr22:50523686-50523703 | MLC1;MLC1;MLC1 | | 5'UTR;5'UTR;1stExon |
| 30M04_Saliva | chr22:50523955-50524691 | MLC1;MLC1;MLC1 | | 1stExon;5'UTR;TSS200 |
| 30M05_Saliva | chr4:69435250-69435601 | UGT2B15;UGT2B17 | | TSS1500;TSS1500 |

| | | | | |
|---|---|---|---|---|
| 30M05_Saliva | chr11:1874049-1874320 | LSP1 | | TSS200 |
| 30M05_Saliva | chr11:69970464-69970618 | ANO1;ANO1 | | Body;Body |
| 30M05_Saliva | chr19:23941207-23941575 | ZNF681 | N_Shore | Body |
| 30M05_Saliva | chr22:24348549-24348715 | GSTTP1 | | TSS1500 |
| 30M08_Saliva | chr1:108734992-108735549 | SLC25A24;SLC25A24 | N_Shore | Body;Body |
| 30M08_Saliva | chr10:134062171-134062614 | STK32C | | Body |
| 30M08_Saliva | chr11:64115989-64116279 | CCDC88B | S_Shelf | Body |
| 30M08_Saliva | chr2:26401640-26402008 | FAM59B | N_Shore | Body |
| 30M08_Saliva | chr2:113992843-113992930 | PAX8;PAX8;PAX8;PAX8;PAX8;LO( | N_Shore | Body;Body;Body;Body;Body;Body;TSS1500 |
| 30M08_Saliva | chr2:158300475-158301005 | CYTIP | | 1stExon |
| 30M08_Saliva | chr2:175499113-175499365 | WIPF1;WIPF1 | | 5'UTR;5'UTR |
| 30M08_Saliva | chr2:240174613-240174659 | HDAC4 | | Body |
| 30M08_Saliva | chr4:38807259-38807382 | TLR1 | | TSS1500 |
| 30M08_Saliva | chr4:69435250-69435601 | UGT2B15;UGT2B17 | | TSS1500;TSS1500 |
| 30M08_Saliva | chr5:169724590-169724768 | LCP2 | | 1stExon |
| 30M08_Saliva | chr6:41168911-41168960 | TREML2;TREML2 | | 1stExon;5'UTR |
| 30M08_Saliva | chr6:135516912-135517046 | MYB;MYB;MYB;MYB;MYB;MYB;MYB;MYB | | Body;Body;Body;Body;Body;Body;Body;Body |
| 30M08_Saliva | chr11:118084920-118085281 | AMICA1;AMICA1 | | Body;TSS1500 |
| 30M08_Saliva | chr17:56408694-56409028 | MIR142 | | TSS200 |
| 30M08_Saliva | chr19:54327780-54327995 | NLRP12 | | TSS200 |
| 30M08_Saliva | chr19:54876728-54876752 | LAIR1;LAIR1 | | TSS200;TSS200 |
| 30M08_Saliva | chr20:54987140-54987330 | CASS4;CASS4;CASS4;CASS4 | | TSS200;TSS200;TSS200;TSS200 |
| 40M07_Saliva | chr1:108734992-108735549 | SLC25A24;SLC25A24 | N_Shore | Body;Body |
| 40M07_Saliva | chr5:1594282-1594733 | SDHAP3 | Island | Body |
| 40M07_Saliva | chr12:10183208-10183267 | CLEC9A | | TSS200 |
| 40M07_Saliva | chr17:79905219-79905263 | MYADML2 | | TSS200 |
| 40M07_Saliva | chr20:5485144-5485268 | LOC149837 | | Body |
| 40M07_Saliva | chr1:855425-855544 | FLJ39609 | S_Shore | TSS1500 |
| 40M07_Saliva | chr1:1238436-1239259 | ACAP3 | N_Shelf | Body |
| 40M07_Saliva | chr1:1245014-1245184 | PUSL1 | Island | Body |
| 40M07_Saliva | chr1:1356565-1356902 | LOC441869;LOC441869;LOC44 | S_Shore | 1stExon;Body;5'UTR |
| 40M07_Saliva | chr1:2138757-2139175 | C1orf86 | S_Shore | 5'UTR |
| 40M07_Saliva | chr1:2265881-2266591 | MORN1 | N_Shore | Body |
| 40M07_Saliva | chr1:2304264-2304375 | MORN1 | Island | Body |
| 40M07_Saliva | chr1:3162066-3162173 | PRDM16;PRDM16 | N_Shore | Body;Body |
| 40M07_Saliva | chr1:3276413-3276535 | PRDM16;PRDM16 | S_Shore | Body;Body |
| 40M07_Saliva | chr1:3321869-3322309 | PRDM16;PRDM16 | Island | Body;Body |
| 40M07_Saliva | chr1:3607222-3607401 | TP73;TP73;TP73;TP73 | Island | Body;TSS200;TSS200;TSS200 |
| 40M07_Saliva | chr1:6390473-6390595 | ACOT7;ACOT7;ACOT7;ACOT7 | | Body;Body;Body;Body |
| 40M07_Saliva | chr1:6445632-6446787 | ACOT7;ACOT7 | | Body;1stExon |
| 40M07_Saliva | chr1:6531717-6531813 | PLEKHG5;PLEKHG5;PLEKHG5;PL | Island | Body;Body;Body;Body;Body |
| 40M07_Saliva | chr1:6579837-6580525 | PLEKHG5;PLEKHG5 | | 5'UTR;1stExon |
| 40M07_Saliva | chr1:10757892-10757954 | CASZ1;CASZ1 | Island | 5'UTR;5'UTR |
| 40M07_Saliva | chr1:15256054-15256127 | KIAA1026;KIAA1026;KIAA1026;KIAA1026 | | Body;TSS1500;Body;Body |
| 40M07_Saliva | chr1:15271952-15272326 | KIAA1026;KIAA1026;KIAA1026;KIAA1026;KIAA1026 | | Body;5'UTR;TSS1500;Body;Body |
| 40M07_Saliva | chr1:17575420-17575440 | PADI3 | | TSS200 |
| 40M07_Saliva | chr1:20141662-20141781 | RNF186;RNF186 | | 5'UTR;1stExon |
| 40M07_Saliva | chr1:24469564-24469791 | IL22RA1 | | 1stExon |
| 40M07_Saliva | chr1:26503847-26503933 | CNKSR1;CNKSR1 | | TSS200;TSS200 |
| 40M07_Saliva | chr1:27683139-27683501 | MAP3K6 | N_Shore | Body |
| 40M07_Saliva | chr1:35220224-35220839 | GJB5 | | TSS1500 |
| 40M07_Saliva | chr1:40349687-40350052 | TRIT1 | S_Shore | TSS1500 |
| 40M07_Saliva | chr1:43751497-43751563 | C1orf210;C1orf210 | | TSS1500;TSS1500 |
| 40M07_Saliva | chr1:43814764-43815035 | MPL | Island | Body |
| 40M07_Saliva | chr1:45083079-45083278 | RNF220 | Island | Body |
| 40M07_Saliva | chr1:89459504-89459658 | CCBL2;CCBL2;RBMXL1;RBMXL1 | S_Shore | TSS1500;TSS1500;TSS1500;TSS1500 |
| 40M07_Saliva | chr1:92946132-92946825 | GFI1;GFI1;GFI1 | Island | Body;Body;Body |
| 40M07_Saliva | chr1:100111387-100111727 | PALMD | | TSS200 |
| 40M07_Saliva | chr1:117753591-117753741 | VTCN1 | | TSS200 |
| 40M07_Saliva | chr1:120311542-120311653 | HMGCS2;HMGCS2;HMGCS2;HMGCS2 | | 1stExon;5'UTR;5'UTR;1stExon |
| 40M07_Saliva | chr1:152730231-152730426 | KPRP | N_Shelf | TSS1500 |
| 40M07_Saliva | chr1:152758453-152758755 | LCE1E | | TSS1500 |
| 40M07_Saliva | chr1:152973720-152973957 | SPRR3;SPRR3 | | TSS1500;TSS1500 |
| 40M07_Saliva | chr1:153413871-153413940 | S100A7L2 | | TSS1500 |
| 40M07_Saliva | chr1:154839813-154839983 | KCNN3 | | Body |

| | | | | |
|---|---|---|---|---|
| 40M07_Saliva | chr1:154929734-154929791 | PBXIP1;PYGO2 | | TSS1500;3'UTR |
| 40M07_Saliva | chr1:156030809-156031182 | RAB25 | | TSS200 |
| 40M07_Saliva | chr1:156123352-156123500 | SEMA4A | | TSS200 |
| 40M07_Saliva | chr1:156217801-156218009 | PAQR6;PAQR6;PAQR6 | S_Shore | 5'UTR;TSS200;1stExon |
| 40M07_Saliva | chr1:161059267-161059429 | PVRL4;PVRL4 | | 5'UTR;1stExon |
| 40M07_Saliva | chr1:161088957-161089118 | PFDN2;NIT1 | S_Shore | TSS1500;Body |
| 40M07_Saliva | chr1:201979478-201979938 | ELF3;ELF3 | | TSS1500;TSS1500 |
| 40M07_Saliva | chr1:203829481-203829890 | SNRPE | N_Shore | TSS1500 |
| 40M07_Saliva | chr1:206643301-206643377 | IKBKE | | TSS1500 |
| 40M07_Saliva | chr1:209605223-209605401 | LOC642587;MIR205 | | Body;TSS1500 |
| 40M07_Saliva | chr1:223888431-223889141 | CAPN2 | | TSS1500 |
| 40M07_Saliva | chr1:234350834-234350988 | SLC35F3 | S_Shore | Body |
| 40M07_Saliva | chr1:247611842-247611931 | NLRP3;NLRP3;NLRP3;NLRP3;NLI | N_Shelf | 3'UTR;3'UTR;3'UTR;3'UTR;3'UTR |
| 40M07_Saliva | chr1:248100228-248101009 | OR2L13 | N_Shore | TSS1500 |
| 40M07_Saliva | chr2:9459368-9459596 | ASAP2;ASAP2 | Island | Body;Body |
| 40M07_Saliva | chr2:10302391-10302462 | C2orf48 | | Body |
| 40M07_Saliva | chr2:17996994-17997138 | MSGN1 | | TSS1500 |
| 40M07_Saliva | chr2:24398427-24398695 | C2orf84 | Island | Body |
| 40M07_Saliva | chr2:27530884-27531236 | UCN | Island | 5'UTR |
| 40M07_Saliva | chr2:27531310-27531360 | UCN | Island | TSS200 |
| 40M07_Saliva | chr2:27665017-27665711 | KRTCAP3;NRBP1;KRTCAP3 | Island | TSS1500;3'UTR;TSS1500 |
| 40M07_Saliva | chr2:50201479-50201511 | NRXN1;NRXN1;NRXN1 | | Body;Body;Body |
| 40M07_Saliva | chr2:54087178-54087343 | LOC100302652;GPR75 | Island | TSS200;TSS200 |
| 40M07_Saliva | chr2:69665094-69665232 | NFU1;NFU1;NFU1;NFU1 | S_Shore | TSS1500;TSS1500;TSS1500;TSS1500 |
| 40M07_Saliva | chr2:74668072-74668477 | RTKN;RTKN;RTKN | Island | Body;TSS1500;TSS1500 |
| 40M07_Saliva | chr2:74753281-74753358 | DQX1;DQX1 | N_Shelf | 1stExon;5'UTR |
| 40M07_Saliva | chr2:95940031-95940705 | PROM2;PROM2;PROM2 | | TSS200;TSS200;TSS200 |
| 40M07_Saliva | chr2:97171137-97171243 | NEURL3 | N_Shelf | Body |
| 40M07_Saliva | chr2:102803290-102803787 | IL1RL2 | N_Shore | TSS200 |
| 40M07_Saliva | chr2:128458240-128458399 | SFT2D3 | N_Shore | TSS1500 |
| 40M07_Saliva | chr2:145147129-145147350 | ZEB2;ZEB2 | Island | Body;Body |
| 40M07_Saliva | chr2:159651559-159651918 | DAPL1 | | TSS1500 |
| 40M07_Saliva | chr2:177014849-177014959 | HOXD4;MIR10B | N_Shore | TSS1500;TSS200 |
| 40M07_Saliva | chr2:219472218-219472607 | PLCD4 | | TSS1500 |
| 40M07_Saliva | chr2:223916502-223916820 | KCNE4 | N_Shore | TSS1500 |
| 40M07_Saliva | chr2:232259281-232259623 | B3GNT7 | N_Shore | TSS1500 |
| 40M07_Saliva | chr2:240212763-240213173 | HDAC4 | | Body |
| 40M07_Saliva | chr2:241458886-241458974 | ANKMY1;ANKMY1 | N_Shore | Body;Body |
| 40M07_Saliva | chr2:241835462-241836501 | C2orf54;C2orf54 | | 1stExon;5'UTR |
| 40M07_Saliva | chr2:242127759-242127948 | ANO7;ANO7 | | TSS200;TSS200 |
| 40M07_Saliva | chr2:242715019-242715041 | GAL3ST2 | S_Shelf | TSS1500 |
| 40M07_Saliva | chr2:242802009-242802127 | PDCD1 | N_Shelf | TSS1500 |
| 40M07_Saliva | chr3:9289747-9289893 | SRGAP3;SRGAP3 | | Body;Body |
| 40M07_Saliva | chr3:11178365-11178798 | HRH1 | | TSS1500 |
| 40M07_Saliva | chr3:12800816-12801112 | TMEM40 | | TSS200 |
| 40M07_Saliva | chr3:52553049-52553433 | STAB1 | Island | Body |
| 40M07_Saliva | chr3:124103560-124103696 | KALRN;KALRN | | Body;Body |
| 40M07_Saliva | chr3:183959465-183959746 | VWA5B2 | Island | Body |
| 40M07_Saliva | chr3:191048361-191048753 | UTS2D;CCDC50;CCDC50 | S_Shore | TSS200;Body;Body |
| 40M07_Saliva | chr3:193096486-193096626 | ATP13A5 | | 1stExon |
| 40M07_Saliva | chr3:195943198-195943444 | OSTalpha | | TSS200 |
| 40M07_Saliva | chr3:196065506-196065688 | TM4SF19 | | TSS1500 |
| 40M07_Saliva | chr4:671543-671851 | MYL5 | N_Shelf | TSS200 |
| 40M07_Saliva | chr4:3392103-3392318 | RGS12;RGS12;RGS12 | | Body;Body;Body |
| 40M07_Saliva | chr4:40517938-40518041 | RBM47;RBM47;RBM47 | | 1stExon;5'UTR;5'UTR |
| 40M07_Saliva | chr4:40752838-40752947 | NSUN7 | Island | Body |
| 40M07_Saliva | chr4:184908351-184909018 | STOX2 | | Body |
| 40M07_Saliva | chr5:1016279-1016460 | NKD2 | | Body |
| 40M07_Saliva | chr5:1052828-1052919 | SLC12A7 | N_Shore | Body |
| 40M07_Saliva | chr5:59783629-59784203 | PART1;PART1;PDE4D;PART1 | | TSS200;Body;5'UTR;TSS200 |
| 40M07_Saliva | chr5:78985425-78986160 | CMYA5 | | TSS1500 |
| 40M07_Saliva | chr5:132113776-132114055 | SEPT8;SEPT8;SEPT8;SEPT8 | S_Shore | TSS1500;TSS1500;TSS1500;TSS1500 |
| 40M07_Saliva | chr5:136983354-136983373 | MIR874;KLHL3 | | TSS200;Body |
| 40M07_Saliva | chr5:142065539-142065676 | FGF1;FGF1;FGF1;FGF1;FGF1;FGF1;FGF1;FGF1 | | 5'UTR;Body;Body;5'UTR;1stExon;5'UTR;5'UTR;1stExon |
| 40M07_Saliva | chr5:154091976-154092619 | LARP1 | | TSS1500 |

| | | | | |
|---|---|---|---|---|
| 40M07_Saliva | chr5:178986291-178986620 | RUFY1;RUFY1;RUFY1 | N_Shore | TSS1500;Body;TSS1500 |
| 40M07_Saliva | chr6:291687-291909 | DUSP22 | N_Shore | TSS1500 |
| 40M07_Saliva | chr6:15505345-15505460 | JARID2 | | Body |
| 40M07_Saliva | chr6:26225258-26225389 | HIST1H3E | N_Shore | TSS200 |
| 40M07_Saliva | chr6:30850405-30850581 | DDR1 | N_Shore | TSS1500 |
| 40M07_Saliva | chr6:30853948-30854109 | DDR1;DDR1 | S_Shore | 5'UTR;5'UTR |
| 40M07_Saliva | chr6:30854186-30854551 | DDR1;DDR1 | S_Shore | 5'UTR;5'UTR |
| 40M07_Saliva | chr6:30882994-30883203 | VARS2;VARS2;VARS2 | S_Shore | Body;5'UTR;Body |
| 40M07_Saliva | chr6:31023257-31023709 | HCG22 | | Body |
| 40M07_Saliva | chr6:31636239-31636392 | CSNK2B | S_Shelf | Body |
| 40M07_Saliva | chr6:31688080-31688229 | LY6G6C | N_Shelf | Body |
| 40M07_Saliva | chr6:31689996-31690170 | C6orf25;C6orf25;C6orf25;C6orf2 | N_Shore | TSS1500;TSS1500;TSS1500;TSS1500;TSS1500;TSS1500;TSS1500 |
| 40M07_Saliva | chr6:31696223-31696298 | DDAH2 | Island | Body |
| 40M07_Saliva | chr6:31696423-31696482 | DDAH2 | Island | Body |
| 40M07_Saliva | chr6:31803378-31804078 | C6orf48;C6orf48;SNORD52 | S_Shore | 5'UTR;5'UTR;TSS1500 |
| 40M07_Saliva | chr6:31804159-31804306 | C6orf48;C6orf48;SNORD52 | S_Shore | 5'UTR;5'UTR;TSS1500 |
| 40M07_Saliva | chr6:31913995-31914265 | CFB;CFB | | 1stExon;5'UTR |
| 40M07_Saliva | chr6:32014739-32015083 | TNXB;TNXB | | Body;TSS1500 |
| 40M07_Saliva | chr6:32078398-32078453 | TNXB | | TSS1500 |
| 40M07_Saliva | chr6:32097184-32097439 | ATF6B;ATF6B;FKBPL | N_Shore | TSS1500;TSS1500;Body |
| 40M07_Saliva | chr6:32489984-32490012 | HLA-DRB5 | Island | Body |
| 40M07_Saliva | chr6:32552016-32552106 | HLA-DRB1 | Island | Body |
| 40M07_Saliva | chr6:33131319-33131450 | COL11A2;COL11A2;COL11A2 | S_Shore | 3'UTR;3'UTR;3'UTR |
| 40M07_Saliva | chr6:33173307-33173581 | HSD17B8 | S_Shore | Body |
| 40M07_Saliva | chr6:33282997-33283021 | ZBTB22;TAPBP;TAPBP;TAPBP;ZB1 | Island | Body;TSS1500;TSS1500;TSS1500;Body |
| 40M07_Saliva | chr6:33638313-33638550 | ITPR3 | | Body |
| 40M07_Saliva | chr6:36238178-36238230 | PNPLA1;PNPLA1;PNPLA1 | | 5'UTR;5'UTR;TSS200 |
| 40M07_Saliva | chr6:43478530-43478829 | C6orf154 | Island | TSS200 |
| 40M07_Saliva | chr6:106546704-106546754 | PRDM1;PRDM1 | | Body;TSS200 |
| 40M07_Saliva | chr6:112375200-112375333 | WISP3;WISP3 | | TSS200;TSS200 |
| 40M07_Saliva | chr6:152702330-152702463 | SYNE1;SYNE1 | | Body;Body |
| 40M07_Saliva | chr6:155568918-155569443 | TIAM2;TIAM2 | | Body;Body |
| 40M07_Saliva | chr6:157505558-157505996 | ARID1B;ARID1B;ARID1B | | Body;Body;Body |
| 40M07_Saliva | chr7:917662-917805 | C7orf20 | Island | Body |
| 40M07_Saliva | chr7:1022213-1022970 | CYP2W1 | N_Shore | TSS1500 |
| 40M07_Saliva | chr7:1024673-1024814 | CYP2W1 | Island | Body |
| 40M07_Saliva | chr7:1514361-1514568 | INTS1 | Island | Body |
| 40M07_Saliva | chr7:2094628-2094977 | MAD1L1;MAD1L1;MAD1L1 | | Body;Body;Body |
| 40M07_Saliva | chr7:2208606-2208830 | MAD1L1;MAD1L1;MAD1L1 | | Body;Body;Body |
| 40M07_Saliva | chr7:2701419-2701641 | TTYH3 | Island | 3'UTR |
| 40M07_Saliva | chr7:2802942-2802976 | GNA12 | S_Shore | Body |
| 40M07_Saliva | chr7:2842593-2842746 | GNA12 | | Body |
| 40M07_Saliva | chr7:3033173-3033579 | CARD11 | | 5'UTR |
| 40M07_Saliva | chr7:24797835-24797884 | DFNA5;DFNA5;DFNA5 | S_Shore | TSS1500;TSS200;TSS200 |
| 40M07_Saliva | chr7:27153580-27153847 | HOXA3;HOXA3;HOXA3;HOXA3 | Island | 5'UTR;1stExon;5'UTR;5'UTR |
| 40M07_Saliva | chr7:27154845-27154914 | HOXA3;HOXA3;HOXA3 | N_Shore | 5'UTR;TSS1500;5'UTR |
| 40M07_Saliva | chr7:27155036-27155173 | HOXA3;HOXA3;HOXA3 | Island | 5'UTR;TSS1500;5'UTR |
| 40M07_Saliva | chr7:27170241-27170412 | HOXA4 | Island | 1stExon |
| 40M07_Saliva | chr7:27170554-27170755 | HOXA4 | Island | TSS200 |
| 40M07_Saliva | chr7:27183133-27183369 | HOXA5 | Island | 1stExon |
| 40M07_Saliva | chr7:28338757-28338985 | CREB5 | | TSS200 |
| 40M07_Saliva | chr7:45002287-45002919 | MYO1G | Island | 3'UTR |
| 40M07_Saliva | chr7:50628841-50629067 | DDC;DDC | | TSS200;5'UTR |
| 40M07_Saliva | chr7:56160528-56161020 | PHKG1;PHKG1 | | 1stExon;5'UTR |
| 40M07_Saliva | chr7:91764177-91764605 | CYP51A1;CYP51A1 | Island | TSS200;TSS1500 |
| 40M07_Saliva | chr7:97735499-97735625 | LMTK2 | N_Shore | TSS1500 |
| 40M07_Saliva | chr7:99227370-99227515 | ZNF498 | | Body |
| 40M07_Saliva | chr7:130020011-130020130 | CPA1 | N_Shore | TSS1500 |
| 40M07_Saliva | chr7:134464123-134464242 | CALD1;CALD1;CALD1 | | TSS200;TSS200;TSS200 |
| 40M07_Saliva | chr7:134575306-134575524 | CALD1;CALD1;CALD1;CALD1;CALD1 | | TSS1500;Body;TSS1500;Body;Body |
| 40M07_Saliva | chr7:135433540-135433736 | FAM180A;FAM180A | | 1stExon;5'UTR |
| 40M07_Saliva | chr7:138482826-138483143 | ATP6V0A4;ATP6V0A4;TMEM213;ATP6V0A4;TMEM213;ATP6V0A4 | | 1stExon;5'UTR;1stExon;5'UTR;5'UTR;1stExon |
| 40M07_Saliva | chr7:142583296-142583555 | TRPV6;TRPV6 | | 1stExon;5'UTR |
| 40M07_Saliva | chr7:150021488-150021553 | ACTR3C;LRRC61;LRRC61 | S_Shore | TSS1500;5'UTR;5'UTR |

| | | | | |
|---|---|---|---|---|
| 40M07_Saliva | chr7:150778724-150778881 | FASTK;TMUB1;TMUB1;FASTK | Island | TSS1500;Body;Body;TSS1500 |
| 40M07_Saliva | chr7:151493629-151493820 | PRKAG2;PRKAG2 | | 5'UTR;Body |
| 40M07_Saliva | chr7:157809235-157809360 | PTPRN2;PTPRN2;PTPRN2 | | Body;Body;Body |
| 40M07_Saliva | chr7:158059704-158059944 | PTPRN2;PTPRN2;PTPRN2 | N_Shore | Body;Body;Body |
| 40M07_Saliva | chr8:1790793-1790905 | ARHGEF10 | | 5'UTR |
| 40M07_Saliva | chr8:22084861-22085227 | PHYHIP;PHYHIP | | Body;Body |
| 40M07_Saliva | chr8:61777711-61777859 | CHD7 | | Body |
| 40M07_Saliva | chr8:141565971-141566221 | EIF2C2;EIF2C2 | | Body;Body |
| 40M07_Saliva | chr8:141588118-141588242 | EIF2C2;EIF2C2 | | Body;Body |
| 40M07_Saliva | chr8:143386260-143386872 | TSNARE1 | | Body |
| 40M07_Saliva | chr8:143761503-143761811 | PSCA | | TSS1500 |
| 40M07_Saliva | chr8:143823809-143824001 | SLURP1;SLURP1 | S_Shelf | 5'UTR;1stExon |
| 40M07_Saliva | chr8:143851287-143851375 | LYNX1;LYNX1 | | TSS200;Body |
| 40M07_Saliva | chr8:143867129-143868158 | LY6D | | Body |
| 40M07_Saliva | chr8:143868844-143868917 | LY6D | | TSS1500 |
| 40M07_Saliva | chr8:143869231-143869473 | LY6D | | TSS1500 |
| 40M07_Saliva | chr8:144894951-144895243 | SCRIB;SCRIB | Island | Body;Body |
| 40M07_Saliva | chr9:136522662-136522927 | DBH | | Body |
| 40M07_Saliva | chr9:137718814-137719284 | COL5A1 | N_Shore | Body |
| 40M07_Saliva | chr10:1507708-1508061 | ADARB2 | S_Shore | Body |
| 40M07_Saliva | chr10:3149673-3149852 | PFKP | S_Shore | Body |
| 40M07_Saliva | chr10:3169037-3169516 | PFKP | | Body |
| 40M07_Saliva | chr10:5541568-5541726 | CALML5 | S_Shore | TSS200 |
| 40M07_Saliva | chr10:11246612-11246763 | CUGBP2;CUGBP2;CUGBP2;CUGBP2 | | Body;Body;Body;Body |
| 40M07_Saliva | chr10:45406644-45406847 | TMEM72 | | TSS1500 |
| 40M07_Saliva | chr10:49459800-49460249 | FRMPD2;FRMPD2 | | Body;Body |
| 40M07_Saliva | chr10:50506875-50507003 | C10orf71;C10orf71 | | TSS1500;TSS1500 |
| 40M07_Saliva | chr10:63809073-63809170 | ARID5B | | Body |
| 40M07_Saliva | chr10:72432797-72433218 | ADAMTS14;ADAMTS14 | Island | 1stExon;1stExon |
| 40M07_Saliva | chr10:88779109-88779163 | FAM25A | S_Shelf | |
| 40M07_Saliva | chr10:95198200-95198374 | MYOF;MYOF | | Body;Body |
| 40M07_Saliva | chr10:99331332-99331808 | ANKRD2;ANKRD2 | S_Shore | 3'UTR |
| 40M07_Saliva | chr10:105238873-105239205 | CALHM3;CALHM3 | | 1stExon |
| 40M07_Saliva | chr10:105420831-105421249 | SH3PXD2A | Island | Body |
| 40M07_Saliva | chr10:106093778-106093830 | ITPRIP | N_Shelf | TSS200 |
| 40M07_Saliva | chr10:126278272-126278502 | LHPP;LHPP | N_Shelf | Body;Body |
| 40M07_Saliva | chr10:135032219-135032504 | KNDC1 | N_Shore | Body |
| 40M07_Saliva | chr10:135341528-135341933 | CYP2E1 | N_Shore | 1stExon |
| 40M07_Saliva | chr11:393570-394262 | PKP3 | N_Shore | TSS1500 |
| 40M07_Saliva | chr11:612680-612837 | IRF7;IRF7;IRF7 | S_Shore | Body;Body;Body |
| 40M07_Saliva | chr11:1328419-1328486 | TOLLIP | N_Shore | Body |
| 40M07_Saliva | chr11:1507263-1507347 | HCCA2 | | Body |
| 40M07_Saliva | chr11:1855561-1856025 | SYT8 | | TSS200 |
| 40M07_Saliva | chr11:1945564-1945648 | TNNT3;TNNT3;TNNT3;TNNT3 | N_Shore | Body;Body;Body;Body |
| 40M07_Saliva | chr11:2011312-2011346 | LOC100133545 | N_Shelf | TSS200 |
| 40M07_Saliva | chr11:2828274-2828568 | KCNQ1;KCNQ1 | N_Shore | Body;Body |
| 40M07_Saliva | chr11:2925570-2925869 | SLC22A18;SLC22A18;SLC22A18 | S_Shore | Body;Body;TSS1500 |
| 40M07_Saliva | chr11:3187678-3187803 | OSBPL5;OSBPL5;OSBPL5 | S_Shore | TSS1500;TSS1500;TSS1500 |
| 40M07_Saliva | chr11:6518438-6518577 | DNHD1;DNHD1 | | TSS200;TSS200 |
| 40M07_Saliva | chr11:14280742-14281053 | SPON1 | Island | Body |
| 40M07_Saliva | chr11:26593556-26593881 | MUC15;MUC15;ANO3;MUC15 | | 5'UTR;5'UTR;Body;5'UTR |
| 40M07_Saliva | chr11:32421635-32421845 | WT1;WT1;WT1;WT1 | | Body;Body;Body;Body |
| 40M07_Saliva | chr11:32447944-32448293 | WT1;WT1;WT1;WT1 | N_Shore | Body;Body;Body;Body |
| 40M07_Saliva | chr11:34535281-34535416 | ELF5;ELF5 | | 1stExon;5'UTR |
| 40M07_Saliva | chr11:45928841-45928988 | C11orf94 | | TSS200 |
| 40M07_Saliva | chr11:62456939-62457334 | LRRN4CL;LRRN4CL | S_Shore | 5'UTR;1stExon |
| 40M07_Saliva | chr11:64951916-64952034 | CAPN1 | Island | Body |
| 40M07_Saliva | chr11:65560635-65560763 | OVOL1 | | Body |
| 40M07_Saliva | chr11:66104468-66104485 | RIN1 | S_Shelf | TSS1500 |
| 40M07_Saliva | chr11:66631356-66631536 | PC;PC;PC | Island | Body;Body;Body |
| 40M07_Saliva | chr11:67442067-67442408 | ALDH3B2;ALDH3B2;ALDH3B2 | | 1stExon;5'UTR;5'UTR |
| 40M07_Saliva | chr11:69469175-69469820 | CCND1 | S_Shore | 3'UTR |
| 40M07_Saliva | chr11:69481734-69482043 | ORAOV1 | | 3'UTR |
| 40M07_Saliva | chr11:69970464-69970618 | ANO1;ANO1 | | Body;Body |
| 40M07_Saliva | chr11:71710643-71711021 | IL18BP;IL18BP;IL18BP;IL18BP;IL18BP;IL18BP | | TSS200;5'UTR;TSS1500;5'UTR;5'UTR;1stExon |

| | | | | |
|---|---|---|---|---|
| 40M07_Saliva | chr11:72533295-72533664 | ATG16L2 | Island | Body |
| 40M07_Saliva | chr11:93754288-93754390 | HEPHL1 | | TSS200 |
| 40M07_Saliva | chr11:111789452-111789693 | C11orf52 | | TSS200 |
| 40M07_Saliva | chr11:117947611-117947997 | TMPRSS4;TMPRSS4 | | TSS200;TSS200 |
| 40M07_Saliva | chr11:118550379-118550644 | TREH;TREH | | 1stExon;5'UTR |
| 40M07_Saliva | chr11:118781728-118781978 | BCL9L | Island | TSS200 |
| 40M07_Saliva | chr11:119993780-119994008 | TRIM29 | | Body |
| 40M07_Saliva | chr11:121970694-121970768 | LOC399959;MIR125B1 | | Body;TSS200 |
| 40M07_Saliva | chr11:124621829-124622096 | VSIG2 | | Body |
| 40M07_Saliva | chr11:128812462-128813442 | TP53AIP1 | | 5'UTR |
| 40M07_Saliva | chr11:133804743-133804959 | IGSF9B | S_Shelf | Body |
| 40M07_Saliva | chr12:258355-258436 | IQSEC3;IQSEC3 | | Body;Body |
| 40M07_Saliva | chr12:6421513-6422131 | PLEKHG6;PLEKHG6;PLEKHG6 | S_Shore | TSS1500;Body;Body |
| 40M07_Saliva | chr12:6484521-6484729 | SCNN1A;SCNN1A;SCNN1A | | Body;TSS200;5'UTR |
| 40M07_Saliva | chr12:6486701-6487080 | SCNN1A | | TSS200 |
| 40M07_Saliva | chr12:8974844-8975309 | A2ML1 | | TSS1500 |
| 40M07_Saliva | chr12:50497778-50498016 | GPD1 | | TSS200 |
| 40M07_Saliva | chr12:51236582-51236638 | TMPRSS12 | N_Shore | TSS200 |
| 40M07_Saliva | chr12:52585695-52585849 | KRT80;KRT80;KRT80;KRT80 | | 1stExon;1stExon;5'UTR;5'UTR |
| 40M07_Saliva | chr12:52867568-52867793 | KRT6C;KRT6C | | 1stExon;5'UTR |
| 40M07_Saliva | chr12:52913824-52915036 | KRT5 | | 1stExon |
| 40M07_Saliva | chr12:53242733-53242922 | KRT78;KRT78 | | 5'UTR;1stExon |
| 40M07_Saliva | chr12:53612551-53612734 | RARG;RARG | N_Shore | Body;Body |
| 40M07_Saliva | chr12:58119915-58121004 | AGAP2;LOC100130776;AGAP2 | Island | 3'UTR;TSS200;3'UTR |
| 40M07_Saliva | chr12:58131345-58132105 | AGAP2;AGAP2 | Island | 1stExon;Body |
| 40M07_Saliva | chr12:64782908-64783110 | C12orf56;C12orf56 | N_Shore | Body;Body |
| 40M07_Saliva | chr12:68041257-68041322 | DYRK2;DYRK2 | N_Shore | TSS1500;TSS1500 |
| 40M07_Saliva | chr12:117627133-117627286 | FBXO21;FBXO21 | N_Shore | Body;Body |
| 40M07_Saliva | chr12:120151527-120151706 | MIR1178;CIT | | Body;Body |
| 40M07_Saliva | chr12:120241409-120242090 | CIT | N_Shore | Body |
| 40M07_Saliva | chr12:120242394-120242513 | CIT | S_Shore | Body |
| 40M07_Saliva | chr12:121882208-121882415 | KDM2B;KDM2B | Island | Body;Body |
| 40M07_Saliva | chr12:124773693-124774078 | FAM101A | | TSS200 |
| 40M07_Saliva | chr12:131303016-131303194 | STX2;STX2 | N_Shore | Body;Body |
| 40M07_Saliva | chr12:132853954-132854189 | GALNT9;LOC100130238 | Island | Body;Body |
| 40M07_Saliva | chr12:133101168-133101495 | FBRSL1 | N_Shore | Body |
| 40M07_Saliva | chr13:21291173-21291467 | IL17D | | Body |
| 40M07_Saliva | chr13:49793054-49793170 | MLNR | N_Shore | TSS1500 |
| 40M07_Saliva | chr13:99135543-99135711 | STK24;STK24 | | Body;Body |
| 40M07_Saliva | chr13:111522222-111522865 | C13orf29 | | Body |
| 40M07_Saliva | chr13:113676726-113677188 | MCF2L;MCF2L | N_Shore | Body;Body |
| 40M07_Saliva | chr13:114103713-114104057 | ADPRHL1;ADPRHL1 | | Body;TSS1500 |
| 40M07_Saliva | chr14:21510323-21510479 | RNASE7 | | TSS200 |
| 40M07_Saliva | chr14:23623480-23623935 | SLC7A8;SLC7A8;SLC7A8 | | 5'UTR;Body;1stExon |
| 40M07_Saliva | chr14:76447327-76447431 | TGFB3;TGFB3 | N_Shore | 1stExon;5'UTR |
| 40M07_Saliva | chr14:93412562-93412768 | ITPK1;ITPK1;ITPK1 | | Body;Body;Body |
| 40M07_Saliva | chr14:94393654-94393816 | FAM181A;C14orf86 | | Body;TSS1500 |
| 40M07_Saliva | chr14:103416122-103416389 | CDC42BPB | | Body |
| 40M07_Saliva | chr14:103593088-103593520 | TNFAIP2 | N_Shore | Body |
| 40M07_Saliva | chr14:104165052-104165401 | XRCC3;XRCC3;XRCC3;KLC1;KL( | Island | 3'UTR;3'UTR;3'UTR;Body;Body |
| 40M07_Saliva | chr14:104195301-104196038 | ZFYVE21 | S_Shore | Body |
| 40M07_Saliva | chr15:39871923-39872037 | THBS1 | N_Shore | TSS1500 |
| 40M07_Saliva | chr15:40268610-40268777 | EIF2AK4 | Island | Body |
| 40M07_Saliva | chr15:40583227-40583422 | PLCB2 | Island | Body |
| 40M07_Saliva | chr15:42130650-42130735 | JMJD7-PLA2G4B;JMJD7-PLA2G4B;PLA2G4B | | Body;Body;TSS1500 |
| 40M07_Saliva | chr15:42289618-42289939 | PLA2G4E | | Body |
| 40M07_Saliva | chr15:42448234-42448786 | PLA2G4F;PLA2G4F | | Body;Body |
| 40M07_Saliva | chr15:67417607-67417899 | SMAD3;SMAD3 | | Body;TSS1500 |
| 40M07_Saliva | chr15:86313851-86313941 | MIR1276;KLHL25 | S_Shore | TSS200;5'UTR |
| 40M07_Saliva | chr15:91473365-91473569 | UNC45A | Island | TSS200 |
| 40M07_Saliva | chr16:670800-671171 | RAB40C | Island | Body |
| 40M07_Saliva | chr16:743338-743584 | FBXL16 | N_Shore | 3'UTR |
| 40M07_Saliva | chr16:809972-810891 | MSLN;MSLN | | TSS1500;TSS1500 |
| 40M07_Saliva | chr16:832683-832841 | MSLNL | N_Shore | Body |
| 40M07_Saliva | chr16:1206750-1207075 | CACNA1H;CACNA1H | N_Shore | Body;Body |
| 40M07_Saliva | chr16:3843012-3843415 | CREBBP;CREBBP | | Body;Body |

| | | | | |
|---|---|---|---|---|
| 40M07_Saliva | chr16:12210656-12210910 | SNX29 | Island | Body |
| 40M07_Saliva | chr16:57937614-57937756 | CNGB1 | N_Shore | Body |
| 40M07_Saliva | chr16:67022472-67022719 | CES8 | | TSS200 |
| 40M07_Saliva | chr16:67233277-67233861 | ELMO3 | Island | 1stExon |
| 40M07_Saliva | chr16:67686832-67687754 | RLTPR | N_Shore | Body |
| 40M07_Saliva | chr16:67918680-67918699 | NRN1L | Island | TSS200 |
| 40M07_Saliva | chr16:81526954-81527360 | CMIP | | Body |
| 40M07_Saliva | chr16:81564161-81564314 | CMIP;CMIP | | Body;Body |
| 40M07_Saliva | chr16:83171068-83171314 | CDH13 | | Body |
| 40M07_Saliva | chr16:87637999-87638484 | JPH3 | S_Shore | Body |
| 40M07_Saliva | chr16:87958145-87958493 | CA5A | | Body |
| 40M07_Saliva | chr16:88832476-88832532 | FAM38A | | Body |
| 40M07_Saliva | chr16:89387418-89387805 | ANKRD11 | | 5'UTR |
| 40M07_Saliva | chr16:89462292-89462418 | ANKRD11 | | 5'UTR |
| 40M07_Saliva | chr17:693243-693388 | RNMTL1 | | Body |
| 40M07_Saliva | chr17:851785-851933 | NXN | S_Shore | Body |
| 40M07_Saliva | chr17:1553585-1553661 | RILP | S_Shore | TSS200 |
| 40M07_Saliva | chr17:3461140-3461398 | TRPV3;TRPV3 | | 1stExon;5'UTR |
| 40M07_Saliva | chr17:4463371-4464075 | GGT6;GGT6 | | Body;Body |
| 40M07_Saliva | chr17:6899207-6899310 | ALOX12 | Island | TSS200 |
| 40M07_Saliva | chr17:7493222-7494151 | SOX15;SOX15 | S_Shore | 1stExon;5'UTR |
| 40M07_Saliva | chr17:7690327-7690529 | DNAH2 | Island | Body |
| 40M07_Saliva | chr17:7832680-7832943 | KCNAB3 | Island | 1stExon |
| 40M07_Saliva | chr17:9929719-9929867 | GAS7;GAS7;GAS7 | | Body;Body;TSS200 |
| 40M07_Saliva | chr17:17109936-17110353 | PLD6 | Island | TSS1500 |
| 40M07_Saliva | chr17:17465242-17465682 | PEMT;PEMT;PEMT | Island | Body;Body;Body |
| 40M07_Saliva | chr17:17717894-17717967 | MIR33B;SREBF1;SREBF1 | S_Shore | TSS1500;Body;Body |
| 40M07_Saliva | chr17:19648925-19649004 | ALDH3A1;ALDH3A1;ALDH3A1 | S_Shore | 5'UTR;TSS200;TSS200 |
| 40M07_Saliva | chr17:25798415-25798942 | KSR1 | | TSS1500 |
| 40M07_Saliva | chr17:38657310-38657934 | TNS4 | | 5'UTR |
| 40M07_Saliva | chr17:38859934-38860042 | KRT24 | | 1stExon |
| 40M07_Saliva | chr17:39143239-39143416 | KRT40 | | 5'UTR |
| 40M07_Saliva | chr17:39661855-39662375 | KRT13;KRT13;KRT13;KRT13 | | 1stExon;5'UTR;5'UTR;1stExon |
| 40M07_Saliva | chr17:39943423-39943545 | JUP;JUP | S_Shore | TSS1500;TSS1500 |
| 40M07_Saliva | chr17:40274524-40274740 | KAT2A;HSPB9 | Island | TSS1500;TSS1500 |
| 40M07_Saliva | chr17:46035108-46035272 | PRR15L;PRR15L | | 1stExon;5'UTR |
| 40M07_Saliva | chr17:46608277-46608345 | HOXB1 | Island | TSS200 |
| 40M07_Saliva | chr17:46627848-46628303 | HOXB3 | Island | Body |
| 40M07_Saliva | chr17:46651722-46651867 | HOXB3;HOXB3 | N_Shelf | 5'UTR;1stExon |
| 40M07_Saliva | chr17:46681111-46682308 | LOC404266;LOC404266;HOXB6 | | Body;Body;5'UTR |
| 40M07_Saliva | chr17:48845722-48846189 | C17orf73 | | TSS1500 |
| 40M07_Saliva | chr17:60747393-60747678 | MRC2 | S_Shelf | Body |
| 40M07_Saliva | chr17:60757756-60758287 | MRC2 | Island | Body |
| 40M07_Saliva | chr17:64672129-64672341 | PRKCA | | Body |
| 40M07_Saliva | chr17:73996245-73996656 | CDK3;C17orf106 | | TSS1500;Body |
| 40M07_Saliva | chr17:74639849-74639928 | ST6GALNAC1;ST6GALNAC1 | | 5'UTR;1stExon |
| 40M07_Saliva | chr17:75446431-75446549 | SEPT9;SEPT9;SEPT9;SEPT9;SEPT | N_Shore | TSS200;Body;Body;Body;Body;Body |
| 40M07_Saliva | chr17:75473610-75473969 | SEPT9;SEPT9;SEPT9;SEPT9;SEPT9;SEPT9;SEPT9 | | Body;5'UTR;Body;Body;Body;Body;Body |
| 40M07_Saliva | chr17:76036514-76037364 | TNRC6C;TNRC6C | N_Shore | 5'UTR;5'UTR |
| 40M07_Saliva | chr17:76501381-76501582 | DNAH17 | S_Shelf | Body |
| 40M07_Saliva | chr17:76976091-76976267 | LGALS3BP | | TSS200 |
| 40M07_Saliva | chr17:76993506-76993698 | CANT1;CANT1;CANT1 | Island | Body;Body;Body |
| 40M07_Saliva | chr17:77030221-77030546 | C1QTNF1;C1QTNF1;C1QTNF1 | | 5'UTR;TSS1500;5'UTR |
| 40M07_Saliva | chr17:77133977-77134159 | HRNBP3 | Island | 5'UTR |
| 40M07_Saliva | chr17:78039301-78039528 | CCDC40 | Island | Body |
| 40M07_Saliva | chr17:78079615-78079768 | GAA;GAA;GAA | Island | Body;Body;Body |
| 40M07_Saliva | chr17:78161227-78161495 | CARD14;CARD14 | N_Shelf | TSS1500;Body |
| 40M07_Saliva | chr17:79030796-79031712 | BAIAP2;BAIAP2;BAIAP2;BAIAP2 | | Body;Body;Body;Body |
| 40M07_Saliva | chr17:79056061-79056183 | BAIAP2;BAIAP2;BAIAP2;BAIAP2 | S_Shelf | Body;Body;Body;Body |
| 40M07_Saliva | chr17:79096529-79097152 | AATK | Island | Body |
| 40M07_Saliva | chr17:79099481-79099785 | MIR657;AATK | N_Shelf | TSS1500;Body |
| 40M07_Saliva | chr17:79099882-79100218 | MIR657;AATK;MIR338 | N_Shelf | TSS1500;Body;TSS200 |
| 40M07_Saliva | chr17:79109729-79109910 | AATK | Island | Body |
| 40M07_Saliva | chr17:79609272-79609346 | TSPAN10 | | TSS200 |
| 40M07_Saliva | chr17:79961522-79961769 | ASPSCR1 | N_Shore | Body |
| 40M07_Saliva | chr17:80289541-80290106 | SECTM1 | N_Shore | 5'UTR |

| | | | | |
|---|---|---|---|---|
| 40M07_Saliva | chr17:80797948-80798023 | TBCD;ZNF750 | | Body;TSS200 |
| 40M07_Saliva | chr18:21452730-21453131 | LAMA3;LAMA3;LAMA3;LAMA3 | | TSS1500;TSS1500;Body;Body |
| 40M07_Saliva | chr18:61143993-61144339 | SERPINB5 | | TSS200 |
| 40M07_Saliva | chr18:61253847-61254561 | SERPINB13 | | TSS1500 |
| 40M07_Saliva | chr18:77230598-77230795 | NFATC1;NFATC1;NFATC1;NFATC | Island | Body;Body;Body;Body |
| 40M07_Saliva | chr18:77286052-77286260 | NFATC1;NFATC1;NFATC1;NFATC | Island | Body;Body;Body;Body |
| 40M07_Saliva | chr19:1395365-1395646 | NDUFS7 | Island | Body |
| 40M07_Saliva | chr19:2512999-2513356 | GNG7 | N_Shore | 3'UTR |
| 40M07_Saliva | chr19:2525231-2525634 | GNG7 | Island | 5'UTR |
| 40M07_Saliva | chr19:5036971-5037219 | KDM4B | Island | Body |
| 40M07_Saliva | chr19:5250376-5251120 | PTPRS;PTPRS;PTPRS;PTPRS | N_Shore | Body;Body;Body;Body |
| 40M07_Saliva | chr19:6333587-6333761 | ACER1;ACER1 | | 5'UTR;1stExon |
| 40M07_Saliva | chr19:10736006-10736059 | SLC44A2;SLC44A2 | Island | TSS200;Body |
| 40M07_Saliva | chr19:10736299-10736448 | SLC44A2;SLC44A2;SLC44A2 | Island | 5'UTR;1stExon;Body |
| 40M07_Saliva | chr19:16178091-16178426 | TPM4 | N_Shore | TSS1500 |
| 40M07_Saliva | chr19:16830613-16830749 | NWD1 | | TSS200 |
| 40M07_Saliva | chr19:18385409-18385672 | KIAA1683;KIAA1683;KIAA1683 | | TSS200;TSS200;TSS200 |
| 40M07_Saliva | chr19:39278997-39279703 | LGALS7B | N_Shelf | TSS1500 |
| 40M07_Saliva | chr19:46286064-46286683 | DMPK;DMPK;DMPK | N_Shelf | TSS1500;TSS1500;TSS1500 |
| 40M07_Saliva | chr19:49238364-49238653 | RASIP1 | N_Shore | Body |
| 40M07_Saliva | chr19:51487245-51487360 | KLK7;KLK7 | | TSS200;TSS200 |
| 40M07_Saliva | chr20:17595355-17595472 | RRBP1;RRBP1 | Island | Body;Body |
| 40M07_Saliva | chr20:17674195-17674250 | BANF2 | | TSS200 |
| 40M07_Saliva | chr20:30433405-30433512 | FOXS1;FOXS1 | S_Shore | 5'UTR;1stExon |
| 40M07_Saliva | chr20:33880337-33880825 | FAM83C | S_Shore | TSS200 |
| 40M07_Saliva | chr20:34700112-34700329 | EPB41L1 | | TSS1500 |
| 40M07_Saliva | chr20:35972957-35973098 | SRC | N_Shore | TSS200 |
| 40M07_Saliva | chr20:36011999-36012016 | SRC;SRC | N_Shore | 5'UTR;5'UTR |
| 40M07_Saliva | chr20:36151003-36151338 | BLCAP;BLCAP;BLCAP;NNAT;BLC | Island | 5'UTR;5'UTR;5'UTR;Body;5'UTR;5'UTR;Body |
| 40M07_Saliva | chr20:36152813-36153186 | BLCAP;BLCAP;BLCAP;BLCAP;BL | S_Shore | 5'UTR;1stExon;5'UTR;5'UTR;5'UTR;5'UTR |
| 40M07_Saliva | chr20:36153865-36154222 | BLCAP;BLCAP;BLCAP;BLCAP;BL | N_Shelf | 5'UTR;TSS1500;5'UTR;5'UTR;5'UTR |
| 40M07_Saliva | chr20:36157405-36157651 | BLCAP;BLCAP;BLCAP;BLCAP | S_Shore | TSS1500;TSS1500;TSS1500;TSS1500 |
| 40M07_Saliva | chr20:43937138-43937289 | MATN4;RBPJL;MATN4;MATN4 | S_Shore | TSS200;Body;TSS200;TSS200 |
| 40M07_Saliva | chr20:50418952-50419082 | SALL4;SALL4 | Island | 1stExon;5'UTR |
| 40M07_Saliva | chr20:62405935-62406964 | ZBTB46 | N_Shore | Body |
| 40M07_Saliva | chr21:38117454-38117682 | SIM2;SIM2 | N_Shelf | Body;Body |
| 40M07_Saliva | chr21:43135854-43135973 | C21orf129;NCRNA00112 | | Body;TSS1500 |
| 40M07_Saliva | chr21:45177538-45177726 | PDXK | Island | 3'UTR |
| 40M07_Saliva | chr22:17680477-17680578 | CECR1;CECR1;CECR1 | | Body;1stExon;5'UTR |
| 40M07_Saliva | chr22:19949873-19950026 | COMT;COMT;COMT;COMT | | 5'UTR;5'UTR;5'UTR;TSS200 |
| 40M07_Saliva | chr22:24577315-24577448 | SUSD2 | | TSS200 |
| 40M07_Saliva | chr22:24891158-24891220 | C22orf45;C22orf45;UPB1 | Island | TSS1500;TSS1500;TSS200 |
| 40M07_Saliva | chr22:24979554-24979642 | GGT1 | | TSS200 |
| 40M07_Saliva | chr22:38506623-38506781 | BAIAP2L2;BAIAP2L2 | | 1stExon;5'UTR |
| 40M07_Saliva | chr22:43820888-43821115 | MPPED1 | Island | 5'UTR |
| 40M07_Saliva | chr22:45608345-45608448 | C22orf9;C22orf9;C22orf9 | | Body;1stExon;5'UTR |
| 40M07_Saliva | chr22:47022945-47023209 | GRAMD4 | Island | Body |
| 40M07_Saliva | chr22:47057191-47057502 | GRAMD4 | N_Shore | Body |
| 40M07_Saliva | chr22:50451054-50451151 | IL17REL;IL17REL | S_Shore | 5'UTR;1stExon |
| 40M07_Saliva | chr22:51017151-51017166 | CPT1B;CPT1B;CPT1B;CPT1B;CH | S_Shore | TSS200;TSS200;TSS1500;TSS1500;Body;TSS200;TSS200;TSS1500 |
| 40M10_Saliva | chr5:1594282-1594808 | SDHAP3 | Island | Body |
| 40M10_Saliva | chr6:291687-292823 | DUSP22 | N_Shore | TSS1500 |
| 40M10_Saliva | chr11:117069780-117069849 | TAGLN;TAGLN | | TSS1500;TSS1500 |
| 40M10_Saliva | chr18:74799250-74799572 | MBP;MBP | Island | Body;Body |
| 40M10_Saliva | chr21:36259241-36259460 | RUNX1;RUNX1;RUNX1 | Island | 1stExon;1stExon;Body |
| 40M10_Saliva | chr6:32064212-32064258 | TNXB | Island | Body |
| 40M10_Saliva | chr10:118387095-118387327 | PNLIPRP2 | N_Shore | Body |
| 40M10_Saliva | chr10:135341528-135342218 | CYP2E1 | N_Shore | 1stExon |
| 40M10_Saliva | chr10:135342560-135343280 | CYP2E1 | Island | Body |
| 40M10_Saliva | chr11:27015808-27015991 | FIBIN;FIBIN | | 5'UTR;1stExon |
| 40M10_Saliva | chr12:4699085-4699618 | DYRK4 | | TSS200 |
| 40M10_Saliva | chr14:104552032-104552209 | ASPG | Island | TSS200 |
| 40M10_Saliva | chr19:38886110-38886658 | SPRED3 | Island | Body |
| 50M_07_Saliva | chr1:1108924-1109676 | TTLL10 | N_Shore | TSS1500 |
| 50M_07_Saliva | chr1:2082315-2082522 | PRKCZ;PRKCZ;PRKCZ | Island | Body;Body;Body |

| | | | | |
|---|---|---|---|---|
| 50M_07_Saliva | chr1:2232063-2232527 | SKI | N_Shore | Body |
| 50M_07_Saliva | chr1:6526049-6526073 | TNFRSF25;TNFRSF25;TNFRSF25; | Island | Body;Body;Body;Body;Body;Body |
| 50M_07_Saliva | chr1:6526182-6526398 | TNFRSF25;TNFRSF25;PLEKHG5;I | Island | 1stExon;1stExon;3'UTR;3'UTR;5'UTR;5'UTR;5'UTR;5'UTR;3'UTR;1stExon;1stExon;1stExon;3'UTR;5'UTR;3'UTR;1stExon;5'UTR |
| 50M_07_Saliva | chr1:7308012-7308117 | CAMTA1 | | Body |
| 50M_07_Saliva | chr1:7692321-7692367 | CAMTA1 | | Body |
| 50M_07_Saliva | chr1:9714810-9714845 | C1orf200;PIK3CD | S_Shore | TSS200;5'UTR |
| 50M_07_Saliva | chr1:16163479-16164122 | FLJ37453 | N_Shore | Body |
| 50M_07_Saliva | chr1:17634543-17634716 | PADI4 | | TSS200 |
| 50M_07_Saliva | chr1:24239771-24240069 | CNR2;CNR2 | | 5'UTR;1stExon |
| 50M_07_Saliva | chr1:26097918-26098214 | MAN1C1 | Island | Body |
| 50M_07_Saliva | chr1:27687074-27687695 | MAP3K6 | Island | Body |
| 50M_07_Saliva | chr1:27961563-27961868 | FGR | Island | 5'UTR |
| 50M_07_Saliva | chr1:36947785-36948313 | CSF3R;CSF3R;CSF3R;CSF3R | | 5'UTR;5'UTR;5'UTR;5'UTR |
| 50M_07_Saliva | chr1:38412519-38412744 | INPP5B | | 5'UTR |
| 50M_07_Saliva | chr1:54520114-54520450 | C1orf83;TMEM59 | S_Shore | Body;TSS1500 |
| 50M_07_Saliva | chr1:54562040-54562213 | C1orf83 | | Body |
| 50M_07_Saliva | chr1:54821853-54822503 | SSBP3;SSBP3;SSBP3 | N_Shore | Body;Body;Body |
| 50M_07_Saliva | chr1:67772896-67772986 | IL12RB2 | N_Shore | TSS200 |
| 50M_07_Saliva | chr1:151810485-151810904 | LOC100132111;C2CD4D | Island | TSS1500;Body |
| 50M_07_Saliva | chr1:156046344-156046832 | MEX3A | Island | 3'UTR |
| 50M_07_Saliva | chr1:159796434-159796629 | SLAMF8 | | TSS200 |
| 50M_07_Saliva | chr1:160616870-160617124 | SLAMF1;SLAMF1 | | 5'UTR;1stExon |
| 50M_07_Saliva | chr1:161039681-161039695 | ARHGAP30;ARHGAP30;ARHGAP30;ARHGAP30 | | 1stExon;5'UTR;5'UTR;1stExon |
| 50M_07_Saliva | chr1:170638807-170639289 | PRRX1;PRRX1 | | Body;Body |
| 50M_07_Saliva | chr1:202129865-202130344 | PTPN7;PTPN7 | | TSS1500;TSS200 |
| 50M_07_Saliva | chr1:202130728-202131184 | PTPN7 | | TSS1500 |
| 50M_07_Saliva | chr1:202172778-202172912 | LGR6;LGR6 | | TSS200;Body |
| 50M_07_Saliva | chr1:205181237-205181581 | DSTYK;DSTYK | S_Shore | TSS1500;TSS1500 |
| 50M_07_Saliva | chr1:205819463-205819609 | PM20D1 | S_Shore | TSS200 |
| 50M_07_Saliva | chr1:206945924-206946187 | IL10 | | TSS200 |
| 50M_07_Saliva | chr1:207095153-207095668 | FAIM3;FAIM3;FAIM3 | | Body;Body;Body |
| 50M_07_Saliva | chr1:208084942-208085455 | CD34;CD34 | S_Shore | TSS1500;TSS1500 |
| 50M_07_Saliva | chr1:223936508-223936812 | CAPN2;CAPN2 | Island | Body;Body |
| 50M_07_Saliva | chr1:226187852-226188006 | C1orf55 | S_Shore | TSS1500 |
| 50M_07_Saliva | chr1:247711595-247712591 | C1orf150 | | TSS1500 |
| 50M_07_Saliva | chr2:24397787-24397845 | C2orf84 | Island | TSS200 |
| 50M_07_Saliva | chr2:63279495-63286049 | OTX1 | N_Shore | 5'UTR |
| 50M_07_Saliva | chr2:66672337-66672688 | MEIS1 | N_Shore | Body |
| 50M_07_Saliva | chr2:66673316-66673449 | MEIS1 | Island | Body |
| 50M_07_Saliva | chr2:69001665-69002378 | ARHGAP25;ARHGAP25;ARHGAP25;ARHGAP25 | | TSS1500;Body;Body;TSS1500 |
| 50M_07_Saliva | chr2:73871849-73871942 | ALMS1P | | TSS200 |
| 50M_07_Saliva | chr2:74783005-74783088 | DOK1 | S_Shore | Body |
| 50M_07_Saliva | chr2:98377791-98377939 | TMEM131 | | Body |
| 50M_07_Saliva | chr2:100759014-100759159 | AFF3;AFF3 | | 1stExon;5'UTR |
| 50M_07_Saliva | chr2:106776775-106777038 | UXS1 | | Body |
| 50M_07_Saliva | chr2:108994116-108994528 | SULT1C4 | | TSS1500 |
| 50M_07_Saliva | chr2:127413363-127413369 | GYPC;GYPC | N_Shore | TSS1500;TSS1500 |
| 50M_07_Saliva | chr2:127414802-127415063 | GYPC;GYPC | S_Shore | Body;Body |
| 50M_07_Saliva | chr2:131674140-131674221 | ARHGEF4;ARHGEF4 | Island | TSS200;TSS200 |
| 50M_07_Saliva | chr2:143886326-143886940 | ARHGAP15 | | TSS1500 |
| 50M_07_Saliva | chr2:157184401-157184816 | NR4A2 | Island | Body |
| 50M_07_Saliva | chr2:172945058-172945383 | MAP1D | Island | Body |
| 50M_07_Saliva | chr2:175499113-175499377 | WIPF1;WIPF1 | | 5'UTR;5'UTR |
| 50M_07_Saliva | chr2:182321354-182321489 | ITGA4 | N_Shore | TSS1500 |
| 50M_07_Saliva | chr2:208989425-208989579 | CRYGD | S_Shore | TSS200 |
| 50M_07_Saliva | chr2:225265963-225266880 | FAM124B;FAM124B | | 1stExon;1stExon |
| 50M_07_Saliva | chr2:233924713-233925030 | INPP5D;INPP5D | N_Shore | TSS1500;TSS1500 |
| 50M_07_Saliva | chr3:9943828-9943937 | IL17RE;IL17RE;IL17RE | | TSS1500;TSS1500;TSS1500 |
| 50M_07_Saliva | chr3:10334717-10335288 | GHRLOS;GHRL;GHRL;GHRLOS;GHRLOS;GHRL;GHRL;GHRL;GHRL;GHR | | Body;TSS200;TSS200;Body;Body;TSS200;TSS200;TSS200;TSS200;TSS200;Body;TSS200;TSS200;TSS200 |
| 50M_07_Saliva | chr3:11078979-11079298 | SLC6A1 | | 3'UTR |
| 50M_07_Saliva | chr3:42742993-42743203 | HHATL;HHATL | | Body;5'UTR |
| 50M_07_Saliva | chr3:46448579-46448868 | CCRL2;CCRL2 | | TSS1500;TSS200 |

| | | | | |
|---|---|---|---|---|
| 50M_07_Saliva | chr3:49460057-49460177 | AMT;AMT;AMT;NICN1;AMT;AMT;AMT;AMT;AMT;AMT | | Body;5'UTR;1stExon;3'UTR;1stExon;1stExon;5'UTR;5'UTR;1stExon;5'UTR |
| 50M_07_Saliva | chr3:50376216-50376475 | RASSF1;RASSF1;RASSF1;RASSF | N_Shore | TSS1500;TSS1500;Body;Body |
| 50M_07_Saliva | chr3:52529195-52529524 | STAB1 | | TSS200 |
| 50M_07_Saliva | chr3:53700141-53700276 | CACNA1D;CACNA1D;CACNA1D | | Body;Body;Body |
| 50M_07_Saliva | chr3:55517496-55518383 | WNT5A | N_Shore | Body |
| 50M_07_Saliva | chr3:65342644-65342971 | MAGI1;MAGI1 | Island | Body;3'UTR |
| 50M_07_Saliva | chr3:121379745-121379791 | HCLS1;HCLS1 | | 1stExon;5'UTR |
| 50M_07_Saliva | chr3:121796627-121796767 | CD86;CD86 | | TSS200;Body |
| 50M_07_Saliva | chr3:128779498-128779601 | GP9 | | TSS200 |
| 50M_07_Saliva | chr3:136537169-136537607 | TMEM22;TMEM22;TMEM22 | N_Shore | TSS1500;TSS1500;TSS1500 |
| 50M_07_Saliva | chr3:150996078-150996563 | P2RY14;MED12L;P2RY14 | | 1stExon;Body;5'UTR |
| 50M_07_Saliva | chr3:151058527-151058837 | P2RY12;P2RY12;MED12L;P2RY12 | | 1stExon;5'UTR;Body;5'UTR |
| 50M_07_Saliva | chr3:151102703-151102879 | P2RY12;MED12L | | TSS200;Body |
| 50M_07_Saliva | chr3:181421427-181421951 | SOX2OT | | Body |
| 50M_07_Saliva | chr3:181428280-181428697 | SOX2OT;SOX2 | N_Shore | Body;TSS1500 |
| 50M_07_Saliva | chr3:183273245-183273894 | KLHL6 | | 1stExon |
| 50M_07_Saliva | chr3:194119885-194120150 | GP5 | S_Shore | 5'UTR |
| 50M_07_Saliva | chr4:683148-683240 | MFSD7 | S_Shore | TSS200 |
| 50M_07_Saliva | chr4:2813542-2814122 | SH3BP2;SH3BP2 | | TSS1500;5'UTR |
| 50M_07_Saliva | chr4:2933715-2934188 | MFSD10;MFSD10 | Island | Body;Body |
| 50M_07_Saliva | chr4:3374767-3374909 | RGS12;RGS12;RGS12 | Island | Body;Body;Body |
| 50M_07_Saliva | chr4:4397942-4398276 | D4S234E;D4S234E | | Body;Body |
| 50M_07_Saliva | chr4:8207119-8207325 | SH3TC1 | | Body |
| 50M_07_Saliva | chr4:10686394-10686610 | CLNK | | TSS200 |
| 50M_07_Saliva | chr4:15779642-15779879 | CD38 | N_Shore | TSS1500 |
| 50M_07_Saliva | chr4:25656514-25656865 | SLC34A2 | N_Shore | TSS1500 |
| 50M_07_Saliva | chr4:39482740-39483194 | LOC401127 | S_Shore | Body |
| 50M_07_Saliva | chr4:86851225-86851425 | ARHGAP24;ARHGAP24;ARHGAP24 | | Body;Body;TSS200 |
| 50M_07_Saliva | chr4:87281412-87281511 | MAPK10;MAPK10;MAPK10 | | TSS200;TSS200;5'UTR |
| 50M_07_Saliva | chr4:90815958-90816310 | MMRN1 | | TSS200 |
| 50M_07_Saliva | chr4:95264019-95264127 | HPGDS;HPGDS | | 5'UTR;1stExon |
| 50M_07_Saliva | chr4:154711512-154711906 | SFRP2 | N_Shore | TSS1500 |
| 50M_07_Saliva | chr4:169798931-169799624 | PALLD;PALLD;PALLD;PALLD | N_Shore | 5'UTR;Body;Body;Body |
| 50M_07_Saliva | chr4:183061830-183062004 | MGC45800 | N_Shore | Body |
| 50M_07_Saliva | chr5:368804-369088 | AHRR | N_Shore | Body |
| 50M_07_Saliva | chr5:1107837-1108315 | SLC12A7 | N_Shore | Body |
| 50M_07_Saliva | chr5:1108401-1108876 | SLC12A7 | N_Shore | Body |
| 50M_07_Saliva | chr5:1494983-1495356 | LPCAT1 | Island | Body |
| 50M_07_Saliva | chr5:1879525-1879698 | IRX4 | N_Shore | Body |
| 50M_07_Saliva | chr5:36662804-36662950 | SLC1A3;SLC1A3 | | Body;Body |
| 50M_07_Saliva | chr5:37834958-37835506 | GDNF;GDNF;GDNF | Island | 5'UTR;5'UTR;TSS200 |
| 50M_07_Saliva | chr5:37835684-37835965 | GDNF;GDNF;GDNF | N_Shore | TSS1500;5'UTR;TSS200 |
| 50M_07_Saliva | chr5:39203007-39203190 | FYB;FYB | | Body;Body |
| 50M_07_Saliva | chr5:74162771-74162924 | FAM169A | S_Shore | TSS200 |
| 50M_07_Saliva | chr5:76010444-76010658 | F2R | N_Shore | TSS1500 |
| 50M_07_Saliva | chr5:92914036-92914828 | FLJ42709;FLJ42709 | N_Shore | Body;Body |
| 50M_07_Saliva | chr5:95159365-95159639 | GLRX;GLRX | | TSS1500;TSS1500 |
| 50M_07_Saliva | chr5:100240048-100240223 | ST8SIA4;ST8SIA4 | S_Shore | TSS1500;TSS1500 |
| 50M_07_Saliva | chr5:119799220-119799531 | PRR16 | N_Shore | TSS1500 |
| 50M_07_Saliva | chr5:134870506-134870517 | NEUROG1;NEUROG1 | N_Shore | 1stExon;3'UTR |
| 50M_07_Saliva | chr5:137475115-137475532 | NME5;NME5 | | 1stExon;5'UTR |
| 50M_07_Saliva | chr5:139228062-139228153 | NRG2;NRG2;NRG2;NRG2 | Island | Body;Body;Body;Body |
| 50M_07_Saliva | chr5:141062313-141062571 | ARAP3 | | TSS1500 |
| 50M_07_Saliva | chr5:148808388-148808487 | MIR143;LOC728264 | | TSS200;Body |
| 50M_07_Saliva | chr5:148810061-148810203 | MIR145;LOC728264 | | TSS200;Body |
| 50M_07_Saliva | chr5:168728793-168729267 | SLIT3 | S_Shore | TSS1500 |
| 50M_07_Saliva | chr5:169724590-169724803 | LCP2 | | 1stExon |
| 50M_07_Saliva | chr5:169757430-169757859 | LOC257358 | | TSS1500 |
| 50M_07_Saliva | chr5:176326365-176326619 | HK3 | | TSS200 |
| 50M_07_Saliva | chr5:176937350-176937446 | DOK3;DOK3;DOK3;DOK3;DOK3 | | 1stExon;TSS1500;1stExon;5'UTR;5'UTR |
| 50M_07_Saliva | chr5:177913434-177913485 | COL23A1 | | Body |
| 50M_07_Saliva | chr6:291687-292823 | DUSP22 | N_Shore | TSS1500 |
| 50M_07_Saliva | chr6:4890079-4890278 | CDYL;CDYL;CDYL;CDYL | | 5'UTR;Body;Body;TSS200 |
| 50M_07_Saliva | chr6:11382553-11382769 | NEDD9;NEDD9 | | 1stExon;5'UTR |
| 50M_07_Saliva | chr6:13428522-13428657 | GFOD1 | | Body |

| | | | | |
|---|---|---|---|---|
| 50M_07_Saliva | chr6:25137864-25138169 | CMAH;CMAH | N_Shelf | Body;Body |
| 50M_07_Saliva | chr6:28493601-28493664 | GPX5;GPX5 | | TSS200;TSS200 |
| 50M_07_Saliva | chr6:28874479-28874702 | TRIM27 | | Body |
| 50M_07_Saliva | chr6:28875354-28875370 | TRIM27 | | Body |
| 50M_07_Saliva | chr6:30039175-30039432 | RNF39;RNF39 | Island | Body;Body |
| 50M_07_Saliva | chr6:30039466-30039600 | RNF39;RNF39 | Island | Body;Body |
| 50M_07_Saliva | chr6:30104440-30104626 | TRIM40 | | TSS200 |
| 50M_07_Saliva | chr6:30297174-30297627 | TRIM39;TRIM39 | S_Shelf | Body;Body |
| 50M_07_Saliva | chr6:30624467-30624636 | DHX16;DHX16 | | Body;Body |
| 50M_07_Saliva | chr6:31240784-31241294 | HLA-C | S_Shore | TSS1500 |
| 50M_07_Saliva | chr6:31519182-31519410 | NFKBIL1;NFKBIL1;NFKBIL1;NFKBIL1 | | Body;Body;Body;Body |
| 50M_07_Saliva | chr6:31554199-31555016 | LST1;LST1;LST1;LST1;LST1;LST1;LST1;LST1 | | Body;TSS1500;TSS1500;TSS1500;5'UTR;TSS1500;5'UTR;Body |
| 50M_07_Saliva | chr6:31583915-31584223 | AIF1;AIF1;AIF1 | N_Shelf | Body;1stExon;Body |
| 50M_07_Saliva | chr6:31607634-31607674 | BAT3;BAT3;BAT3;BAT3 | | Body;Body;Body;Body |
| 50M_07_Saliva | chr6:31690949-31691354 | C6orf25;C6orf25;C6orf25;C6orf2 | N_Shore | TSS1500;TSS1500;TSS1500;TSS1500;TSS1500;TSS1500;TSS1500 |
| 50M_07_Saliva | chr6:31765567-31765634 | LSM2 | S_Shore | Body |
| 50M_07_Saliva | chr6:32160909-32161004 | GPSM3 | N_Shelf | 5'UTR |
| 50M_07_Saliva | chr6:32294470-32295237 | C6orf10 | | Body |
| 50M_07_Saliva | chr6:32813337-32813582 | PSMB8;TAP1 | S_Shore | TSS1500;3'UTR |
| 50M_07_Saliva | chr6:32904061-32904317 | HLA-DMB | | Body |
| 50M_07_Saliva | chr6:32909440-32909523 | HLA-DMB | | TSS1500 |
| 50M_07_Saliva | chr6:33400021-33400528 | SYNGAP1 | S_Shelf | Body |
| 50M_07_Saliva | chr6:39282316-39282532 | KCNK17;KCNK17 | S_Shore | TSS200;TSS200 |
| 50M_07_Saliva | chr6:41528345-41528785 | FOXP4;FOXP4;FOXP4 | Island | 5'UTR;5'UTR;5'UTR |
| 50M_07_Saliva | chr6:73330106-73330358 | KCNQ5;KCNQ5;KCNQ5;KCNQ5; | N_Shore | TSS1500;TSS1500;TSS1500;TSS1500;TSS1500 |
| 50M_07_Saliva | chr6:76059726-76059773 | FILIP1 | | Body |
| 50M_07_Saliva | chr6:108145539-108145777 | SCML4 | | TSS200 |
| 50M_07_Saliva | chr6:112081102-112081538 | FYN;FYN | | 5'UTR;TSS1500 |
| 50M_07_Saliva | chr6:112115117-112115209 | FYN | | 5'UTR |
| 50M_07_Saliva | chr6:133035150-133035379 | VNN1 | | 1stExon |
| 50M_07_Saliva | chr6:135516912-135517046 | MYB;MYB;MYB;MYB;MYB;MYB;MYB;MYB | | Body;Body;Body;Body;Body;Body;Body;Body |
| 50M_07_Saliva | chr6:146864278-146864390 | RAB32 | N_Shore | TSS1500 |
| 50M_07_Saliva | chr6:149803206-149803444 | ZC3H12D | | 5'UTR |
| 50M_07_Saliva | chr6:151560831-151560927 | AKAP12 | Island | TSS1500 |
| 50M_07_Saliva | chr6:157931843-157932180 | ZDHHC14;ZDHHC14 | | Body;Body |
| 50M_07_Saliva | chr6:158066609-158066900 | ZDHHC14;ZDHHC14 | | Body;Body |
| 50M_07_Saliva | chr6:158447877-158448134 | SYNJ2 | | Body |
| 50M_07_Saliva | chr6:158507753-158508188 | SYNJ2 | Island | Body |
| 50M_07_Saliva | chr6:166856056-166856094 | RPS6KA2;RPS6KA2 | | Body;Body |
| 50M_07_Saliva | chr6:166876585-166877038 | RPS6KA2;RPS6KA2 | | Body;Body |
| 50M_07_Saliva | chr7:632926-633202 | PRKAR1B;PRKAR1B;PRKAR1B;PF | S_Shore | Body;Body;Body;Body;Body;Body |
| 50M_07_Saliva | chr7:1094737-1095720 | C7orf50;C7orf50;C7orf50 | N_Shelf | Body;Body;Body |
| 50M_07_Saliva | chr7:1980117-1980479 | MAD1L1;MAD1L1;MAD1L1 | S_Shore | Body;Body;Body |
| 50M_07_Saliva | chr7:2059796-2060116 | MAD1L1;MAD1L1;MAD1L1 | N_Shore | Body;Body;Body |
| 50M_07_Saliva | chr7:2116368-2116648 | MAD1L1;MAD1L1;MAD1L1 | Island | Body;Body;Body |
| 50M_07_Saliva | chr7:2124092-2124566 | MAD1L1;MAD1L1;MAD1L1 | | Body;Body;Body |
| 50M_07_Saliva | chr7:2152625-2152821 | MAD1L1;MAD1L1;MAD1L1 | N_Shelf | Body;Body;Body |
| 50M_07_Saliva | chr7:2289054-2289585 | NUDT1;NUDT1;NUDT1;NUDT1;NUDT1;NUDT1;NUDT1 | | Body;Body;Body;Body;Body;Body;Body |
| 50M_07_Saliva | chr7:2648289-2648785 | IQCE;IQCE | S_Shelf | Body;Body |
| 50M_07_Saliva | chr7:2773782-2774654 | GNA12 | N_Shore | Body |
| 50M_07_Saliva | chr7:3020155-3020536 | CARD11 | | 5'UTR |
| 50M_07_Saliva | chr7:4752951-4753002 | FOXK1 | | Body |
| 50M_07_Saliva | chr7:4848683-4848939 | RADIL | Island | Body |
| 50M_07_Saliva | chr7:5258357-5258485 | WIPI2;WIPI2;WIPI2;WIPI2;WIPI2 | | Body;Body;Body;Body;Body |
| 50M_07_Saliva | chr7:5271480-5271655 | WIPI2;WIPI2;WIPI2;WIPI2;WIPI2 | S_Shelf | 3'UTR;3'UTR;3'UTR;3'UTR;3'UTR |
| 50M_07_Saliva | chr7:5527927-5528179 | FBXL18 | Island | Body |
| 50M_07_Saliva | chr7:30951272-30951498 | AQP1 | | TSS200 |
| 50M_07_Saliva | chr7:42267269-42267547 | GLI3 | N_Shore | 5'UTR |
| 50M_07_Saliva | chr7:45018658-45019005 | MYO1G;MYO1G | | 5'UTR;1stExon |
| 50M_07_Saliva | chr7:45066841-45066895 | CCM2;CCM2;CCM2;CCM2;CCM2 | | Body;TSS1500;Body;Body;Body |
| 50M_07_Saliva | chr7:55224863-55224910 | EGFR;EGFR;EGFR | | Body;Body;Body |
| 50M_07_Saliva | chr7:75443075-75443246 | CCL24 | | TSS200 |

| | | | | |
|---|---|---|---|---|
| 50M_07_Saliva | chr7:75957040-75957123 | YWHAG | N_Shore | 3'UTR |
| 50M_07_Saliva | chr7:96655105-96655566 | DLX5 | S_Shore | TSS1500 |
| 50M_07_Saliva | chr7:100238750-100239188 | TFR2 | | Body |
| 50M_07_Saliva | chr7:128579876-128580582 | IRF5;IRF5;IRF5;IRF5;IRF5;IRF5;IR | S_Shore | 5'UTR;TSS1500;5'UTR;5'UTR;5'UTR;5'UTR;5'UTR |
| 50M_07_Saliva | chr7:139333296-139333410 | HIPK2;HIPK2 | | Body;Body |
| 50M_07_Saliva | chr7:150434452-150434541 | GIMAP5;GIMAP5 | | 1stExon;5'UTR |
| 50M_07_Saliva | chr7:151087534-151087726 | WDR86 | | Body |
| 50M_07_Saliva | chr7:157647109-157647241 | PTPRN2;PTPRN2;PTPRN2 | N_Shore | Body;Body;Body |
| 50M_07_Saliva | chr7:158110405-158110685 | PTPRN2;PTPRN2;PTPRN2 | N_Shore | Body;Body;Body |
| 50M_07_Saliva | chr8:17206671-17206819 | MTMR7 | | Body |
| 50M_07_Saliva | chr8:19540210-19540453 | CSGALNACT1;CSGALNACT1 | | TSS200;Body |
| 50M_07_Saliva | chr8:21771059-21771540 | DOK2 | S_Shore | 1stExon |
| 50M_07_Saliva | chr8:21905462-21905756 | FGF17 | Island | Body |
| 50M_07_Saliva | chr8:22223937-22223985 | SLC39A14;SLC39A14;SLC39A14 | N_Shore | TSS1500;TSS1500;TSS1500;TSS1500 |
| 50M_07_Saliva | chr8:27469186-27469338 | CLU;CLU;CLU;CLU | N_Shelf | 1stExon;Body;5'UTR;TSS1500 |
| 50M_07_Saliva | chr8:28174847-28175285 | PNOC | | 5'UTR |
| 50M_07_Saliva | chr8:38008386-38008630 | STAR;STAR;STAR;STAR | | 5'UTR;5'UTR;1stExon;1stExon |
| 50M_07_Saliva | chr8:42037170-42037527 | PLAT;PLAT | | Body;Body |
| 50M_07_Saliva | chr8:67975874-67975895 | CSPP1;CSPP1;COPS5 | N_Shore | TSS1500;TSS1500;TSS1500 |
| 50M_07_Saliva | chr8:86350278-86350390 | CA3 | N_Shore | TSS1500 |
| 50M_07_Saliva | chr8:101315498-101315560 | RNF19A;RNF19A | | 5'UTR;TSS200 |
| 50M_07_Saliva | chr8:103137173-103137274 | NCALD;NCALD;NCALD;NCALD;N | S_Shore | TSS1500;TSS1500;TSS1500;TSS1500;TSS1500;TSS1500;TSS200 |
| 50M_07_Saliva | chr8:104153753-104153829 | C8orf56;BAALC;BAALC | S_Shore | TSS200;Body;Body |
| 50M_07_Saliva | chr8:140945782-140945843 | TRAPPC9;TRAPPC9 | | Body;Body |
| 50M_07_Saliva | chr8:141609338-141609470 | EIF2C2;EIF2C2 | | Body;Body |
| 50M_07_Saliva | chr8:142159992-142160235 | DENND3 | | Body |
| 50M_07_Saliva | chr8:144408230-144408588 | TOP1MT | | Body |
| 50M_07_Saliva | chr8:145086243-145086427 | SPATC1;SPATC1 | | TSS1500;TSS1500 |
| 50M_07_Saliva | chr8:145755756-145755974 | MGC70857;KIAA1688 | Island | TSS1500;3'UTR |
| 50M_07_Saliva | chr9:34624080-34624587 | ARID3C | Island | Body |
| 50M_07_Saliva | chr9:96715687-96716527 | BARX1 | Island | Body |
| 50M_07_Saliva | chr9:98784802-98785110 | NCRNA00092 | S_Shore | TSS1500 |
| 50M_07_Saliva | chr9:117692745-117693010 | TNFSF8;TNFSF8 | | 1stExon;5'UTR |
| 50M_07_Saliva | chr9:123688715-123689193 | TRAF1 | N_Shelf | 5'UTR |
| 50M_07_Saliva | chr10:682693-682871 | DIP2C | N_Shelf | Body |
| 50M_07_Saliva | chr10:1454265-1454387 | ADARB2 | | Body |
| 50M_07_Saliva | chr10:6214016-6214079 | PFKFB3 | | Body |
| 50M_07_Saliva | chr10:49892741-49893549 | WDFY4 | | TSS1500 |
| 50M_07_Saliva | chr10:72362618-72362694 | PRF1;PRF1 | S_Shore | 1stExon;5'UTR;5'UTR;1stExon |
| 50M_07_Saliva | chr10:73848825-73848915 | SPOCK2;SPOCK2 | S_Shore | TSS200;TSS200 |
| 50M_07_Saliva | chr10:88717364-88717623 | MMRN2;SNCG;MMRN2 | | Body |
| 50M_07_Saliva | chr10:94450866-94451736 | HHEX | N_Shore | Body |
| 50M_07_Saliva | chr10:100996579-100997023 | HPSE2;HPSE2;HPSE2;HPSE2 | S_Shelf | TSS1500;TSS1500;TSS1500;TSS1500 |
| 50M_07_Saliva | chr10:102821565-102821684 | KAZALD1;KAZALD1 | Island | 1stExon;5'UTR |
| 50M_07_Saliva | chr10:104535521-104536035 | C10orf26;C10orf26 | | TSS1500;Body |
| 50M_07_Saliva | chr10:105978651-105978956 | MIR609;C10orf79 | | Body |
| 50M_07_Saliva | chr10:114134878-114135259 | ACSL5;ACSL5;ACSL5 | | TSS1500;TSS1500;5'UTR |
| 50M_07_Saliva | chr10:126222996-126223401 | LHPP;LHPP | | Body;Body |
| 50M_07_Saliva | chr10:128152075-128152124 | C10orf90 | | Body |
| 50M_07_Saliva | chr10:134109919-134110326 | STK32C | | Body |
| 50M_07_Saliva | chr10:134150489-134150567 | LRRC27;LRRC27;LRRC27;LRRC27;LRRC27 | | Body;TSS200;Body;Body;Body |
| 50M_07_Saliva | chr10:134358030-134358273 | INPP5A | N_Shelf | Body |
| 50M_07_Saliva | chr11:415430-415457 | SIGIRR;SIGIRR;SIGIRR | N_Shore | 5'UTR;5'UTR;TSS1500 |
| 50M_07_Saliva | chr11:457256-457304 | PTDSS2 | N_Shore | Body |
| 50M_07_Saliva | chr11:1874017-1874320 | LSP1 | | TSS200 |
| 50M_07_Saliva | chr11:2321770-2322050 | C11orf21;TSPAN32;C11orf21 | | Body;TSS1500;Body |
| 50M_07_Saliva | chr11:2322618-2322674 | C11orf21;TSPAN32;C11orf21 | | Body;TSS1500;Body |
| 50M_07_Saliva | chr11:2322741-2323272 | C11orf21;TSPAN32;C11orf21 | | Body;TSS1500;Body |
| 50M_07_Saliva | chr11:2323459-2323938 | TSPAN32;C11orf21;C11orf21 | | Body;TSS1500;TSS1500 |
| 50M_07_Saliva | chr11:3647419-3648013 | TRPC2 | | TSS1500 |
| 50M_07_Saliva | chr11:35441777-35442225 | SLC1A2 | Island | TSS1500 |
| 50M_07_Saliva | chr11:36422377-36422615 | PRR5L;PRR5L;PRR5L;PRR5L | | 5'UTR;5'UTR;TSS200;Body |
| 50M_07_Saliva | chr11:46382544-46382725 | DGKZ;DGKZ;DGKZ;DGKZ | | Body;TSS1500;Body;Body |
| 50M_07_Saliva | chr11:47399213-47400146 | SPI1;SPI1 | N_Shore | Body;Body |

| | | | | |
|---|---|---|---|---|
| 50M_07_Saliva | chr11:59634191-59634276 | TCN1 | | TSS200 |
| 50M_07_Saliva | chr11:59823993-59824161 | MS4A3;MS4A3;MS4A3 | | TSS200;TSS200;TSS200 |
| 50M_07_Saliva | chr11:60738971-60739183 | CD6 | | TSS200 |
| 50M_07_Saliva | chr11:63272909-63274030 | LGALS12;LGALS12;LGALS12 | | TSS1500;TSS1500;TSS1500 |
| 50M_07_Saliva | chr11:63827361-63828228 | MACROD1 | | Body |
| 50M_07_Saliva | chr11:63973846-63974153 | FERMT3;FERMT3 | N_Shore | TSS1500;TSS1500 |
| 50M_07_Saliva | chr11:64107374-64108241 | CCDC88B | N_Shelf | TSS1500 |
| 50M_07_Saliva | chr11:64981227-64981596 | SLC22A20 | | TSS200 |
| 50M_07_Saliva | chr11:65407734-65408566 | SIPA1;SIPA1;SIPA1 | N_Shore | 5'UTR;1stExon;5'UTR |
| 50M_07_Saliva | chr11:65647279-65647532 | CTSW | | TSS200 |
| 50M_07_Saliva | chr11:66055170-66055327 | YIF1A | N_Shore | Body |
| 50M_07_Saliva | chr11:66084841-66085299 | CD248 | S_Shore | TSS1500 |
| 50M_07_Saliva | chr11:66838934-66839191 | RHOD | Island | Body |
| 50M_07_Saliva | chr11:67170610-67171585 | PPP1CA;PPP1CA;TBC1D10C;PPI | S_Shore | TSS1500;TSS1500;TSS1500;TSS1500 |
| 50M_07_Saliva | chr11:67177034-67177296 | TBC1D10C | Island | Body |
| 50M_07_Saliva | chr11:67204799-67205291 | PTPRCAP | S_Shore | Body |
| 50M_07_Saliva | chr11:67803951-67804112 | NDUFS8 | N_Shelf | Body |
| 50M_07_Saliva | chr11:69520186-69520418 | FGF19 | S_Shore | TSS1500 |
| 50M_07_Saliva | chr11:69934031-69934128 | ANO1;ANO1 | Island | Body;Body |
| 50M_07_Saliva | chr11:70211301-70211531 | PPFIA1;PPFIA1 | | Body;Body |
| 50M_07_Saliva | chr11:70257035-70257149 | CTTN;CTTN | N_Shore | Body;Body |
| 50M_07_Saliva | chr11:70691944-70692038 | SHANK2 | | Body |
| 50M_07_Saliva | chr11:71709837-71709978 | IL18BP;IL18BP;IL18BP;IL18BP;IL18BP | | TSS200;TSS1500;TSS200;TSS1500;TSS1500 |
| 50M_07_Saliva | chr11:78673073-78673212 | ODZ4 | Island | Body |
| 50M_07_Saliva | chr11:89224799-89225042 | NOX4;NOX4;NOX4;NOX4 | S_Shore | 5'UTR;TSS200;TSS200;TSS200 |
| 50M_07_Saliva | chr11:124767455-124767840 | ROBO4 | | Body |
| 50M_07_Saliva | chr11:126581947-126582242 | KIRREL3;KIRREL3 | | Body;Body |
| 50M_07_Saliva | chr11:128391271-128391494 | ETS1;ETS1;ETS1 | N_Shore | Body;Body;Body |
| 50M_07_Saliva | chr11:134126284-134126504 | ACAD8 | S_Shelf | Body |
| 50M_07_Saliva | chr11:134254573-134254690 | B3GAT1;B3GAT1 | S_Shore | Body;Body |
| 50M_07_Saliva | chr12:4699085-4699618 | DYRK4 | | TSS200 |
| 50M_07_Saliva | chr12:6881590-6881629 | LAG3 | N_Shore | TSS200 |
| 50M_07_Saliva | chr12:32654929-32655034 | FGD4 | | TSS200 |
| 50M_07_Saliva | chr12:47610220-47610734 | FAM113B;FAM113B;LOC100233209 | | 5'UTR;1stExon;Body |
| 50M_07_Saliva | chr12:51717674-51718251 | BIN2 | | Body |
| 50M_07_Saliva | chr12:52404134-52404161 | GRASP | S_Shelf | Body |
| 50M_07_Saliva | chr12:53730072-53730277 | SP7 | | TSS1500 |
| 50M_07_Saliva | chr12:54891213-54891636 | NCKAP1L | | TSS1500 |
| 50M_07_Saliva | chr12:55378411-55378912 | KIAA0748;KIAA0748 | | 1stExon;5'UTR |
| 50M_07_Saliva | chr12:58210661-58210878 | AVIL | | TSS1500 |
| 50M_07_Saliva | chr12:105114162-105114399 | CHST11 | | Body |
| 50M_07_Saliva | chr12:106533840-106533867 | NUAK1 | S_Shore | TSS200 |
| 50M_07_Saliva | chr12:111537057-111537195 | CUX2 | | Body |
| 50M_07_Saliva | chr12:111618925-111618977 | CUX2 | | Body |
| 50M_07_Saliva | chr12:123518833-123519476 | PITPNM2 | | Body |
| 50M_07_Saliva | chr12:123753186-123753272 | CDK2AP1 | N_Shore | Body |
| 50M_07_Saliva | chr12:124876433-124876650 | NCOR2;NCOR2 | S_Shelf | Body;Body |
| 50M_07_Saliva | chr12:124938116-124938573 | NCOR2;NCOR2 | N_Shelf | Body;Body |
| 50M_07_Saliva | chr12:124941781-124942248 | NCOR2;NCOR2 | S_Shore | Body;Body |
| 50M_07_Saliva | chr12:124950669-124950990 | NCOR2;NCOR2 | N_Shore | Body;Body |
| 50M_07_Saliva | chr12:133135032-133135698 | FBRSL1 | N_Shore | Body |
| 50M_07_Saliva | chr13:30948103-30948716 | LOC100188949 | | TSS200 |
| 50M_07_Saliva | chr13:51639953-51641069 | GUCY1B2 | | Body |
| 50M_07_Saliva | chr13:107143980-107144036 | EFNB2 | N_Shore | 3'UTR |
| 50M_07_Saliva | chr13:107145933-107146164 | EFNB2 | S_Shore | Body |
| 50M_07_Saliva | chr13:110951025-110951418 | COL4A1 | | Body |
| 50M_07_Saliva | chr13:110993474-110993558 | COL4A2 | | Body |
| 50M_07_Saliva | chr13:111318022-111318555 | CARS2 | | Body |
| 50M_07_Saliva | chr13:113652069-113652368 | MCF2L;MCF2L | S_Shelf | Body;Body |
| 50M_07_Saliva | chr13:114261869-114261986 | TFDP1;TFDP1 | S_Shore | Body;Body |
| 50M_07_Saliva | chr13:114814333-114814680 | RASA3 | Island | Body |
| 50M_07_Saliva | chr13:114876061-114876390 | RASA3 | N_Shore | Body |
| 50M_07_Saliva | chr14:24641194-24641706 | REC8;REC8 | Island | TSS200;TSS200 |
| 50M_07_Saliva | chr14:38725536-38725750 | CLEC14A;CLEC14A | Island | 1stExon;5'UTR |
| 50M_07_Saliva | chr14:75988800-75989203 | BATF;BATF | | 5'UTR;1stExon |

| | | | | |
|---|---|---|---|---|
| 50M_07_Saliva | chr14:91866325-91866751 | CCDC88C | | Body |
| 50M_07_Saliva | chr14:93170710-93170970 | LGMN;LGMN | | Body;Body |
| 50M_07_Saliva | chr14:102676957-102677294 | WDR20 | N_Shelf | Body |
| 50M_07_Saliva | chr14:105532012-105532208 | GPR132 | | TSS1500 |
| 50M_07_Saliva | chr14:105944655-105945287 | CRIP2 | Island | Body |
| 50M_07_Saliva | chr15:26044050-26044289 | ATP10A | | Body |
| 50M_07_Saliva | chr15:37169665-37170554 | LOC145845 | N_Shelf | Body |
| 50M_07_Saliva | chr15:38988533-38988860 | C15orf53 | | TSS1500 |
| 50M_07_Saliva | chr15:43531947-43532243 | TGM5;TGM5 | | Body;Body |
| 50M_07_Saliva | chr15:45409942-45410236 | DUOXA1;DUOXA2 | S_Shore | Body;Body |
| 50M_07_Saliva | chr15:57592040-57592222 | LOC283663 | | TSS1500 |
| 50M_07_Saliva | chr15:86315115-86315211 | KLHL25;MIR1276 | S_Shelf | 5'UTR;TSS1500 |
| 50M_07_Saliva | chr15:94840793-94841334 | MCTP2;MCTP2 | | TSS1500;TSS1500 |
| 50M_07_Saliva | chr15:96877194-96877720 | NR2F2;NR2F2;NR2F2;NR2F2 | Island | Body;Body;5'UTR;5'UTR |
| 50M_07_Saliva | chr15:101418967-101419074 | ALDH1A3 | N_Shore | TSS1500 |
| 50M_07_Saliva | chr15:101991512-101991886 | PCSK6;PCSK6;PCSK6;PCSK6;PCSK6;PCSK6;PCSK6;PCSK6 | | Body;Body;Body;Body;Body;Body;Body;Body |
| 50M_07_Saliva | chr16:302852-303192 | ITFG3 | | 5'UTR |
| 50M_07_Saliva | chr16:1583899-1584050 | IFT140;TMEM204 | Island | Body;TSS1500 |
| 50M_07_Saliva | chr16:4714080-4714443 | MGRN1;MGRN1;MGRN1;MGRN1 | | Body;Body;Body;Body |
| 50M_07_Saliva | chr16:8738240-8738441 | C16orf68 | | Body |
| 50M_07_Saliva | chr16:10647084-10647275 | EMP2 | | 5'UTR |
| 50M_07_Saliva | chr16:14403004-14403040 | MIR365-1 | | TSS200 |
| 50M_07_Saliva | chr16:28505320-28505959 | APOB48R | S_Shore | TSS1500 |
| 50M_07_Saliva | chr16:28995882-28996533 | LAT;LAT;LAT;LAT | | TSS1500;TSS1500;TSS1500;TSS1500 |
| 50M_07_Saliva | chr16:29673933-29676071 | SPN;SPN | N_Shore | TSS1500;TSS1500 |
| 50M_07_Saliva | chr16:29756951-29757461 | C16orf54 | | 5'UTR |
| 50M_07_Saliva | chr16:30721139-30721715 | SNORA30;SRCAP | | TSS1500;Body |
| 50M_07_Saliva | chr16:49732224-49732623 | ZNF423 | N_Shore | Body |
| 50M_07_Saliva | chr16:50715135-50715260 | SNX20;SNX20;SNX20;SNX20;SNX20;SNX20 | | 1stExon;5'UTR;1stExon;5'UTR;5'UTR;1stExon |
| 50M_07_Saliva | chr16:56995739-56995856 | CETP | | TSS200 |
| 50M_07_Saliva | chr16:67428155-67428324 | TPPP3;TPPP3 | Island | TSS1500;TSS1500 |
| 50M_07_Saliva | chr16:88038949-88039185 | BANP;BANP | N_Shore | Body;Body |
| 50M_07_Saliva | chr16:88566483-88566748 | ZFPM1 | N_Shelf | Body |
| 50M_07_Saliva | chr16:88767124-88767454 | RNF166 | S_Shore | Body |
| 50M_07_Saliva | chr16:88844415-88844726 | FAM38A | N_Shore | Body |
| 50M_07_Saliva | chr16:89024511-89024657 | CBFA2T3 | | Body |
| 50M_07_Saliva | chr16:89033692-89034099 | CBFA2T3 | Island | Body |
| 50M_07_Saliva | chr16:89041793-89042185 | CBFA2T3 | | Body |
| 50M_07_Saliva | chr17:183931-184131 | RPH3AL | N_Shore | 5'UTR |
| 50M_07_Saliva | chr17:750241-750379 | NXN | | Body |
| 50M_07_Saliva | chr17:800717-800749 | NXN | | Body |
| 50M_07_Saliva | chr17:1548881-1549284 | SCARF1;SCARF1;SCARF1;SCARI | S_Shore | Body;Body;Body;Body;Body |
| 50M_07_Saliva | chr17:1626477-1626575 | WDR81;WDR81;WDR81 | N_Shore | TSS1500;Body;5'UTR |
| 50M_07_Saliva | chr17:2699553-2699706 | RAP1GAP2;RAP1GAP2 | | TSS200;TSS200 |
| 50M_07_Saliva | chr17:2839005-2839187 | RAP1GAP2;RAP1GAP2 | | Body;Body |
| 50M_07_Saliva | chr17:3704471-3704621 | ITGAE;ITGAE | | 1stExon;5'UTR |
| 50M_07_Saliva | chr17:3819339-3820518 | P2RX1 | | Body |
| 50M_07_Saliva | chr17:4337082-4337231 | SPNS3 | | TSS200 |
| 50M_07_Saliva | chr17:4835341-4835627 | GP1BA | | TSS1500 |
| 50M_07_Saliva | chr17:5138259-5138696 | C17orf87 | | TSS200 |
| 50M_07_Saliva | chr17:8771003-8771141 | PIK3R6 | | TSS200 |
| 50M_07_Saliva | chr17:8927521-8928028 | NTN1 | S_Shore | Body |
| 50M_07_Saliva | chr17:9940004-9940321 | GAS7;GAS7;GAS7 | | Body;1stExon;5'UTR |
| 50M_07_Saliva | chr17:13506186-13506284 | HS3ST3A1 | S_Shore | TSS1500 |
| 50M_07_Saliva | chr17:27039120-27039523 | PROCA1 | S_Shore | TSS1500 |
| 50M_07_Saliva | chr17:34344995-34345172 | CCL23;CCL23;CCL23;CCL23 | | 1stExon;5'UTR;1stExon;5'UTR |
| 50M_07_Saliva | chr17:37322028-37322431 | ARL5C;ARL5C | Island | 1stExon;5'UTR |
| 50M_07_Saliva | chr17:40439433-40439501 | STAT5A | N_Shore | TSS200 |
| 50M_07_Saliva | chr17:40440904-40441155 | STAT5A | Island | 5'UTR |
| 50M_07_Saliva | chr17:40935998-40936150 | WNK4 | Island | Body |
| 50M_07_Saliva | chr17:41857603-41857714 | DUSP3;C17orf105 | S_Shore | TSS1500;TSS200 |
| 50M_07_Saliva | chr17:44896923-44897512 | WNT3 | Island | TSS1500 |
| 50M_07_Saliva | chr17:46667683-46667737 | LOC404266;LOC404266 | N_Shore | TSS200;TSS200 |
| 50M_07_Saliva | chr17:48049952-48050512 | DLX4;DLX4 | Island | TSS200;Body |

| | | | | |
|---|---|---|---|---|
| 50M_07_Saliva | chr17:53341098-53341969 | HLF | N_Shore | TSS1500 |
| 50M_07_Saliva | chr17:55533142-55533389 | MSI2;MSI2 | | Body;Body |
| 50M_07_Saliva | chr17:55952063-55952217 | CUEDC1 | | Body |
| 50M_07_Saliva | chr17:59478953-59479306 | TBX2 | Island | Body |
| 50M_07_Saliva | chr17:62097951-62098230 | ICAM2;ICAM2;ICAM2;ICAM2;ICAM2;ICAM2;ICAM2;ICAM2 | | 5'UTR;5'UTR;5'UTR;1stExon;1stExon;5'UTR;1stExon;1stExon |
| 50M_07_Saliva | chr17:72443066-72443179 | GPRC5C;GPRC5C | Island | Body;Body |
| 50M_07_Saliva | chr17:73642547-73642628 | RECQL5;LOC100130933 | | Body;TSS200 |
| 50M_07_Saliva | chr17:74524238-74524611 | CYGB | | 3'UTR |
| 50M_07_Saliva | chr17:75471193-75471308 | SEPT9;SEPT9;SEPT9;SEPT9;SEPT9;SEPT9;SEPT9 | | Body;TSS200;5'UTR;Body;Body;Body;Body |
| 50M_07_Saliva | chr17:76126702-76127217 | TMC8;TMC6 | N_Shore | TSS200;5'UTR |
| 50M_07_Saliva | chr17:76886742-76886912 | TIMP2 | | Body |
| 50M_07_Saliva | chr17:77706946-77707066 | ENPP7 | N_Shore | Body |
| 50M_07_Saliva | chr17:77916733-77916892 | TBC1D16 | Island | Body |
| 50M_07_Saliva | chr17:78682785-78683082 | RPTOR;RPTOR | | Body;Body |
| 50M_07_Saliva | chr17:78735268-78735596 | RPTOR;RPTOR | | Body;Body |
| 50M_07_Saliva | chr17:79005548-79006166 | FLJ90757 | N_Shore | Body |
| 50M_07_Saliva | chr17:79128885-79129078 | AATK | N_Shelf | Body |
| 50M_07_Saliva | chr17:79258813-79259699 | SLC38A10;SLC38A10 | N_Shore | Body;Body |
| 50M_07_Saliva | chr17:80358829-80358876 | C17orf101;C17orf101 | N_Shelf | Body;Body |
| 50M_07_Saliva | chr17:80407031-80407779 | C17orf62;C17orf62;C17orf62 | N_Shore | Body;Body;Body |
| 50M_07_Saliva | chr17:80545175-80545434 | FOXK2 | N_Shore | Body |
| 50M_07_Saliva | chr17:80581701-80581925 | WDR45L | | Body |
| 50M_07_Saliva | chr17:80740731-80740950 | TBCD | Island | Body |
| 50M_07_Saliva | chr18:30352782-30353699 | KLHL14;KLHL14 | S_Shore | 5'UTR;1stExon |
| 50M_07_Saliva | chr18:74114570-74114728 | ZNF516 | Island | Body |
| 50M_07_Saliva | chr18:77917615-77918142 | LOC100130522;LOC100130522 | N_Shore | Body;Body;3'UTR |
| 50M_07_Saliva | chr19:681420-681815 | FSTL3 | Island | Body |
| 50M_07_Saliva | chr19:695447-695642 | PRSSL1;PRSSL1 | | 5'UTR;1stExon |
| 50M_07_Saliva | chr19:728040-728176 | PALM;PALM | Island | Body;Body |
| 50M_07_Saliva | chr19:840737-841082 | PRTN3 | N_Shelf | TSS1500 |
| 50M_07_Saliva | chr19:852114-852311 | ELANE | N_Shore | TSS200 |
| 50M_07_Saliva | chr19:947612-947936 | ARID3A | Island | Body |
| 50M_07_Saliva | chr19:3178512-3178844 | S1PR4 | N_Shore | TSS1500 |
| 50M_07_Saliva | chr19:4326975-4327350 | STAP2;STAP2 | Island | Body;Body |
| 50M_07_Saliva | chr19:4916593-4916983 | UHRF1;UHRF1 | Island | Body;Body |
| 50M_07_Saliva | chr19:6481945-6482169 | DENND1C | | TSS200 |
| 50M_07_Saliva | chr19:7733793-7734203 | RETN | N_Shore | TSS200 |
| 50M_07_Saliva | chr19:7741602-7741838 | C19orf59 | N_Shelf | TSS1500 |
| 50M_07_Saliva | chr19:10449942-10450323 | ICAM3 | S_Shelf | Body |
| 50M_07_Saliva | chr19:10958952-10958970 | C19orf38 | | TSS200 |
| 50M_07_Saliva | chr19:13207445-13207993 | NFIX | Island | 3'UTR |
| 50M_07_Saliva | chr19:13947212-13947262 | MIR24-2 | S_Shelf | TSS200 |
| 50M_07_Saliva | chr19:14550846-14550999 | PKN1;PKN1 | N_Shore | TSS1500;Body |
| 50M_07_Saliva | chr19:16999668-16999800 | F2RL3 | N_Shore | TSS200 |
| 50M_07_Saliva | chr19:17862104-17862210 | FCHO1;FCHO1;FCHO1;FCHO1 | S_Shelf | 5'UTR;TSS200;TSS200;5'UTR |
| 50M_07_Saliva | chr19:35531403-35531417 | HPN;HPN | N_Shore | TSS200;TSS200 |
| 50M_07_Saliva | chr19:35819775-35820181 | CD22 | | TSS1500 |
| 50M_07_Saliva | chr19:36233297-36233469 | TMEM149;TMEM149 | S_Shore | 1stExon;5'UTR |
| 50M_07_Saliva | chr19:39826668-39826927 | GMFG;GMFG | | 5'UTR;1stExon |
| 50M_07_Saliva | chr19:43856594-43857074 | CD177 | | TSS1500 |
| 50M_07_Saliva | chr19:46318633-46319398 | RSPH6A | Island | TSS200 |
| 50M_07_Saliva | chr19:46713446-46713473 | DKFZp434J0226 | | TSS200 |
| 50M_07_Saliva | chr19:49250252-49250561 | IZUMO1 | N_Shelf | TSS200 |
| 50M_07_Saliva | chr19:49340489-49340765 | PLEKHA4;PLEKHA4;HSD17B14 | Island | 3'UTR;3'UTR;TSS1500 |
| 50M_07_Saliva | chr19:49838478-49838777 | CD37;CD37 | N_Shelf | TSS200;TSS200 |
| 50M_07_Saliva | chr19:51628062-51628204 | SIGLEC9 | | TSS200 |
| 50M_07_Saliva | chr19:51875451-51875986 | NKG7 | S_Shelf | Body |
| 50M_07_Saliva | chr19:52004992-52005188 | SIGLEC12;SIGLEC12;SIGLEC12 | | 1stExon;5'UTR;TSS1500 |
| 50M_07_Saliva | chr19:54327404-54327780 | NLRP12 | | 1stExon |
| 50M_07_Saliva | chr19:54566838-54567279 | VSTM1 | | Body |
| 50M_07_Saliva | chr19:54604098-54604187 | OSCAR;OSCAR;OSCAR;OSCAR;( | N_Shore | 5'UTR;5'UTR;5'UTR;1stExon;1stExon;5'UTR;1stExon;1stExon |
| 50M_07_Saliva | chr19:55141723-55142181 | LILRB1;LILRB1;LILRB1;LILRB1 | | TSS200;5'UTR;TSS200;TSS1500 |
| 50M_07_Saliva | chr19:55549590-55549801 | GP6;GP6 | | 1stExon;1stExon |
| 50M_07_Saliva | chr20:1448498-1448645 | NSFL1C;NSFL1C;NSFL1C | S_Shore | TSS200;TSS200;TSS200 |

| | | | | |
|---|---|---|---|---|
| 50M_07_Saliva | chr20:3051954-3052345 | OXT | N_Shore | TSS1500 |
| 50M_07_Saliva | chr20:3065343-3065423 | AVP;AVP | S_Shore | 5'UTR;1stExon |
| 50M_07_Saliva | chr20:4795444-4795935 | RASSF2;RASSF2 | | 5'UTR;5'UTR |
| 50M_07_Saliva | chr20:20036684-20037150 | C20orf26;CRNKL1;CRNKL1;C20( | S_Shelf | 5'UTR;5'UTR;1stExon;TSS1500 |
| 50M_07_Saliva | chr20:22566877-22566976 | FOXA2 | Island | TSS1500 |
| 50M_07_Saliva | chr20:23066944-23067156 | CD93;CD93 | | 5'UTR;1stExon |
| 50M_07_Saliva | chr20:24929607-24929716 | CST7 | | TSS1500 |
| 50M_07_Saliva | chr20:35274570-35274941 | SLA2;SLA2 | | TSS200;TSS200 |
| 50M_07_Saliva | chr20:37433745-37434158 | PPP1R16B | N_Shore | TSS1500 |
| 50M_07_Saliva | chr20:45179157-45179354 | C20orf123 | | 1stExon |
| 50M_07_Saliva | chr20:54986888-54987330 | CASS4;CASS4;CASS4;CASS4 | | TSS1500;TSS1500;TSS1500;TSS1500 |
| 50M_07_Saliva | chr20:57582856-57583195 | CTSZ | S_Shore | TSS1500 |
| 50M_07_Saliva | chr20:62198872-62199190 | PRIC285;PRIC285 | Island | Body;1stExon |
| 50M_07_Saliva | chr20:62318426-62318588 | RTEL1;RTEL1 | Island | Body;Body |
| 50M_07_Saliva | chr21:34185927-34186122 | C21orf62;C21orf62;C21orf62;C21orf62;C21orf62;C21orf62 | | 5'UTR;5'UTR;1stExon;1stExon;1stExon;5'UTR |
| 50M_07_Saliva | chr21:36421467-36421955 | RUNX1;RUNX1 | | 5'UTR;1stExon |
| 50M_07_Saliva | chr21:38119946-38120518 | SIM2 | Island | Body |
| 50M_07_Saliva | chr21:45772638-45773294 | TRPM2 | | TSS1500 |
| 50M_07_Saliva | chr21:46898048-46898789 | COL18A1;COL18A1;COL18A1 | Island | Body;Body;Body |
| 50M_07_Saliva | chr21:48026043-48026298 | S100B | | TSS1500 |
| 50M_07_Saliva | chr22:17956453-17956641 | CECR2 | | TSS200 |
| 50M_07_Saliva | chr22:19710880-19711007 | GP1BB;SEPT5 | Island | TSS200;3'UTR |
| 50M_07_Saliva | chr22:24105087-24105692 | C22orf15 | | TSS200 |
| 50M_07_Saliva | chr22:24823509-24823519 | ADORA2A | S_Shelf | TSS200 |
| 50M_07_Saliva | chr22:27013978-27014116 | CRYBB1;CRYBB1 | | 5'UTR;1stExon |
| 50M_07_Saliva | chr22:37545924-37546124 | IL2RB;IL2RB | | 5'UTR;1stExon |
| 50M_07_Saliva | chr22:37962569-37962818 | CDC42EP1 | Island | Body |
| 50M_07_Saliva | chr22:37975975-37976192 | LGALS2;LGALS2 | | 1stExon;5'UTR |
| 50M_07_Saliva | chr22:42828386-42828516 | NFAM1;NFAM1 | | 1stExon;5'UTR |
| 50M_07_Saliva | chr22:45073148-45073165 | PRR5;PRR5;PRR5;PRR5 | | 5'UTR;5'UTR;5'UTR;5'UTR |
| 50M_07_Saliva | chr22:45609402-45609421 | C22orf9;C22orf9 | | Body;TSS1500 |
| 50M_07_Saliva | chr22:46770063-46770253 | CELSR1 | N_Shore | Body |
| 50M_07_Saliva | chr22:47081751-47082260 | CERK | Island | 3'UTR |
| 50M_07_Saliva | chr22:50482925-50483545 | TTLL8 | N_Shore | Body |
| 50M_07_Saliva | chr22:50523686-50524691 | MLC1;MLC1;MLC1 | | 5'UTR;5'UTR;1stExon |
| 50M_07_Saliva | chr1:1238436-1238678 | ACAP3 | N_Shelf | Body |
| 50M_07_Saliva | chr1:1846001-1846340 | CALML6 | | TSS1500 |
| 50M_07_Saliva | chr1:2084391-2084595 | PRKCZ;PRKCZ;PRKCZ | S_Shore | Body;Body;Body |
| 50M_07_Saliva | chr1:2266362-2266591 | MORN1 | Island | Body |
| 50M_07_Saliva | chr1:2284167-2284501 | MORN1;LOC100129534 | | Body;TSS200 |
| 50M_07_Saliva | chr1:2979817-2980163 | FLJ42875 | Island | Body |
| 50M_07_Saliva | chr1:3028230-3028457 | PRDM16;PRDM16 | Island | Body;Body |
| 50M_07_Saliva | chr1:3607110-3607401 | TP73;TP73;TP73;TP73 | Island | Body;TSS200;TSS200;TSS200 |
| 50M_07_Saliva | chr1:6390473-6390830 | ACOT7;ACOT7;ACOT7;ACOT7 | | Body;Body;Body;Body |
| 50M_07_Saliva | chr1:6446095-6446787 | ACOT7;ACOT7 | | Body;TSS1500 |
| 50M_07_Saliva | chr1:10707917-10708091 | CASZ1;CASZ1 | Island | Body;Body |
| 50M_07_Saliva | chr1:10839450-10839574 | CASZ1;CASZ1 | | 5'UTR;5'UTR |
| 50M_07_Saliva | chr1:15271830-15272326 | KIAA1026;KIAA1026;KIAA1026;KIAA1026;KIAA1026 | | Body;5'UTR;TSS1500;Body;Body |
| 50M_07_Saliva | chr1:17381800-17381980 | SDHB | S_Shore | TSS1500 |
| 50M_07_Saliva | chr1:20141662-20141781 | RNF186;RNF186 | | 5'UTR;1stExon |
| 50M_07_Saliva | chr1:23697258-23697506 | C1orf213;C1orf213;C1orf213;ZN | S_Shore | 3'UTR;Body;1stExon;TSS1500 |
| 50M_07_Saliva | chr1:24469768-24470205 | IL22RA1 | | TSS200 |
| 50M_07_Saliva | chr1:25257505-25257547 | RUNX3;RUNX3 | Island | TSS1500;Body |
| 50M_07_Saliva | chr1:25257978-25258146 | RUNX3;RUNX3 | Island | TSS1500;Body |
| 50M_07_Saliva | chr1:26145328-26145437 | FAM54B;FAM54B;FAM54B | N_Shore | TSS1500;TSS1500;TSS1500 |
| 50M_07_Saliva | chr1:27189648-27190330 | SFN;SFN | N_Shore | 1stExon;5'UTR |
| 50M_07_Saliva | chr1:35220224-35220814 | GJB5 | | TSS1500 |
| 50M_07_Saliva | chr1:43751270-43751563 | C1orf210;C1orf210;C1orf210 | | 1stExon;5'UTR;TSS200 |
| 50M_07_Saliva | chr1:44726943-44727237 | ERI3 | | Body |
| 50M_07_Saliva | chr1:45083079-45083278 | RNF220 | Island | Body |
| 50M_07_Saliva | chr1:55089182-55089372 | ACOT11;FAM151A;FAM151A | | Body;1stExon;5'UTR |
| 50M_07_Saliva | chr1:89829871-89829951 | GBP6 | | 5'UTR |
| 50M_07_Saliva | chr1:120311542-120311653 | HMGCS2;HMGCS2;HMGCS2;HMGCS2 | | 1stExon;5'UTR;5'UTR;1stExon |
| 50M_07_Saliva | chr1:151345123-151345248 | SELENBP1;SELENBP1 | | 5'UTR;1stExon |
| 50M_07_Saliva | chr1:152973720-152973957 | SPRR3;SPRR3 | | TSS1500;TSS1500 |

| | | | | |
|---|---|---|---|---|
| 50M_07_Saliva | chr1:154929734-154929791 | PBXIP1;PYGO2 | | TSS1500;3'UTR |
| 50M_07_Saliva | chr1:156030809-156031182 | RAB25 | | TSS200 |
| 50M_07_Saliva | chr1:161059458-161059654 | PVRL4 | | TSS200 |
| 50M_07_Saliva | chr1:178310534-178310650 | RASAL2;RASAL2 | | Body;TSS200 |
| 50M_07_Saliva | chr1:201979499-201979938 | ELF3;ELF3 | | TSS200;TSS200 |
| 50M_07_Saliva | chr1:203829481-203830200 | SNRPE | N_Shore | TSS1500 |
| 50M_07_Saliva | chr1:205326059-205326437 | KLHDC8A | | TSS200 |
| 50M_07_Saliva | chr1:205602864-205603068 | ELK4;ELK4 | S_Shore | TSS1500;TSS1500 |
| 50M_07_Saliva | chr1:207224740-207225128 | YOD1 | N_Shore | TSS1500 |
| 50M_07_Saliva | chr1:223888431-223889345 | CAPN2 | | TSS1500 |
| 50M_07_Saliva | chr1:241912510-241912768 | WDR64 | | Body |
| 50M_07_Saliva | chr1:245254391-245254667 | EFCAB2;EFCAB2;EFCAB2 | | Body;Body;Body |
| 50M_07_Saliva | chr2:17996994-17997138 | MSGN1 | | TSS1500 |
| 50M_07_Saliva | chr2:24398427-24398859 | C2orf84 | Island | Body |
| 50M_07_Saliva | chr2:31031702-31031755 | CAPN13 | | TSS1500 |
| 50M_07_Saliva | chr2:74153195-74153317 | DGUOK;DGUOK | N_Shore | TSS1500;TSS1500 |
| 50M_07_Saliva | chr2:74753281-74753358 | DQX1;DQX1 | N_Shelf | 1stExon;5'UTR |
| 50M_07_Saliva | chr2:95971840-95971994 | KCNIP3 | | Body |
| 50M_07_Saliva | chr2:113735377-113735503 | IL1F9 | | TSS1500 |
| 50M_07_Saliva | chr2:113875226-113875723 | IL1RN;IL1RN;IL1RN | | TSS1500;TSS1500;TSS1500 |
| 50M_07_Saliva | chr2:128458240-128458399 | SFT2D3 | N_Shore | TSS1500 |
| 50M_07_Saliva | chr2:161056520-161056639 | ITGB6 | | 1stExon |
| 50M_07_Saliva | chr2:202508694-202508813 | ALS2CR4;ALS2CR4 | S_Shore | TSS1500;TSS1500 |
| 50M_07_Saliva | chr2:220282165-220282281 | DES | N_Shore | TSS1500 |
| 50M_07_Saliva | chr2:231790529-231790813 | GPR55 | | TSS1500 |
| 50M_07_Saliva | chr2:236443773-236443932 | AGAP1;AGAP1 | N_Shore | Body;Body |
| 50M_07_Saliva | chr2:236444202-236444284 | AGAP1;AGAP1 | S_Shore | Body;Body |
| 50M_07_Saliva | chr2:240212763-240213173 | HDAC4 | | Body |
| 50M_07_Saliva | chr3:11178064-11178798 | HRH1 | | TSS1500 |
| 50M_07_Saliva | chr3:14443428-14443473 | SLC6A6;SLC6A6;SLC6A6 | N_Shore | TSS1500;TSS1500;TSS1500 |
| 50M_07_Saliva | chr3:48310544-48310897 | ZNF589 | | 3'UTR |
| 50M_07_Saliva | chr3:50341009-50341293 | HYAL1;HYAL1;HYAL1;HYAL1;HYAL1;HYAL1 | | TSS200;TSS200;5'UTR;TSS200;TSS200;TSS200 |
| 50M_07_Saliva | chr3:57743377-57743599 | SLMAP;SLMAP | S_Shore | 5'UTR;1stExon |
| 50M_07_Saliva | chr3:69061356-69061827 | C3orf64 | N_Shore | 5'UTR |
| 50M_07_Saliva | chr3:123339334-123339568 | MYLK;MYLK;MYLK;MYLK;MYLK;MYLK;MYLK;MYLK | | 1stExon;5'UTR;Body;Body;Body;5'UTR;1stExon;Body |
| 50M_07_Saliva | chr3:149374535-149374763 | WWTR1;WWTR1;WWTR1 | N_Shore | Body;Body;Body |
| 50M_07_Saliva | chr3:196065189-196065318 | TM4SF19;TM4SF19 | | 1stExon;5'UTR |
| 50M_07_Saliva | chr3:196065506-196065688 | TM4SF19 | | TSS1500 |
| 50M_07_Saliva | chr4:671550-671851 | MYL5 | N_Shelf | TSS200 |
| 50M_07_Saliva | chr4:1208699-1208960 | CTBP1;CTBP1 | N_Shore | Body;Body |
| 50M_07_Saliva | chr4:1214965-1215112 | CTBP1;CTBP1 | N_Shelf | Body;Body |
| 50M_07_Saliva | chr4:8395941-8396410 | ACOX3;ACOX3 | | Body;Body |
| 50M_07_Saliva | chr4:36283184-36283275 | DTHD1 | | TSS200 |
| 50M_07_Saliva | chr4:40517938-40518065 | RBM47;RBM47;RBM47 | | 1stExon;5'UTR;5'UTR |
| 50M_07_Saliva | chr4:42122106-42122161 | BEND4;BEND4 | | Body;Body |
| 50M_07_Saliva | chr4:87770357-87770782 | SLC10A6;SLC10A6 | | 5'UTR;1stExon |
| 50M_07_Saliva | chr4:114214080-114214185 | ANK2;ANK2;ANK2 | | Body;Body;Body |
| 50M_07_Saliva | chr4:184908351-184909018 | STOX2 | | Body |
| 50M_07_Saliva | chr5:912383-913111 | TRIP13 | N_Shore | Body |
| 50M_07_Saliva | chr5:1016460-1016768 | NKD2 | | Body |
| 50M_07_Saliva | chr5:53751272-53751493 | HSPB3 | | TSS200 |
| 50M_07_Saliva | chr5:78985495-78986160 | CMYA5 | | TSS200 |
| 50M_07_Saliva | chr5:125929132-125929671 | ALDH7A1 | N_Shore | Body |
| 50M_07_Saliva | chr5:136983354-136983373 | MIR874;KLHL3 | | TSS200;Body |
| 50M_07_Saliva | chr5:138211043-138211155 | CTNNA1;LRRTM2;LRRTM2 | | Body;5'UTR;1stExon |
| 50M_07_Saliva | chr5:142065819-142066245 | FGF1;FGF1;FGF1;FGF1;FGF1;FGF1;FGF1 | | 5'UTR;5'UTR;5'UTR;1stExon;Body;Body;TSS200 |
| 50M_07_Saliva | chr5:149546331-149546366 | CDX1 | Island | TSS200 |
| 50M_07_Saliva | chr5:169532686-169532758 | FOXI1;FOXI1 | N_Shore | TSS1500;TSS1500 |
| 50M_07_Saliva | chr6:6614447-6614798 | LOC285780;LY86 | | Body;Body |
| 50M_07_Saliva | chr6:8436218-8436296 | SLC35B3;SLC35B3;SLC35B3 | S_Shore | TSS1500;TSS1500;TSS1500 |
| 50M_07_Saliva | chr6:24877462-24877624 | FAM65B;FAM65B;FAM65B | | 5'UTR;1stExon;5'UTR |
| 50M_07_Saliva | chr6:30850868-30851086 | DDR1;DDR1 | N_Shore | TSS1500;TSS1500 |
| 50M_07_Saliva | chr6:30853333-30853959 | DDR1;DDR1 | S_Shore | 5'UTR;5'UTR |

| | | | | |
|---|---|---|---|---|
| 50M_07_Saliva | chr6:30951225-30951394 | MUC21 | | TSS1500 |
| 50M_07_Saliva | chr6:31023257-31023770 | HCG22 | | Body |
| 50M_07_Saliva | chr6:31025949-31026287 | HCG22 | | Body |
| 50M_07_Saliva | chr6:31096861-31097078 | PSORS1C1 | | 5'UTR |
| 50M_07_Saliva | chr6:31138925-31138982 | POU5F1 | | TSS1500 |
| 50M_07_Saliva | chr6:31636396-31636913 | CSNK2B | S_Shelf | Body |
| 50M_07_Saliva | chr6:31688080-31688229 | LY6G6C | N_Shelf | Body |
| 50M_07_Saliva | chr6:31689610-31690170 | LY6G6C | N_Shore | TSS200 |
| 50M_07_Saliva | chr6:31696223-31696240 | DDAH2 | Island | Body |
| 50M_07_Saliva | chr6:32120625-32120878 | PPT2;PRRT1;PPT2 | N_Shore | TSS1500;TSS1500;TSS1500 |
| 50M_07_Saliva | chr6:32120901-32120955 | PPT2;PRRT1;PPT2 | N_Shore | TSS1500;TSS1500;TSS1500 |
| 50M_07_Saliva | chr6:32165889-32165917 | NOTCH4 | S_Shore | Body |
| 50M_07_Saliva | chr6:32526021-32526342 | HLA-DRB6 | | Body |
| 50M_07_Saliva | chr6:32784327-32785141 | HLA-DOB | | Body |
| 50M_07_Saliva | chr6:33048529-33048558 | HLA-DPB1 | Island | Body |
| 50M_07_Saliva | chr6:33131222-33131319 | COL11A2;COL11A2;COL11A2 | S_Shore | 3'UTR;3'UTR;3'UTR |
| 50M_07_Saliva | chr6:33228341-33228498 | VPS52 | | Body |
| 50M_07_Saliva | chr6:33638313-33638550 | ITPR3 | | Body |
| 50M_07_Saliva | chr6:43478530-43478829 | C6orf154 | Island | TSS200 |
| 50M_07_Saliva | chr6:56878397-56878665 | BEND6 | | Body |
| 50M_07_Saliva | chr6:90660913-90661155 | BACH2;BACH2 | | Body;Body |
| 50M_07_Saliva | chr6:116381609-116382179 | FRK;FRK | | 1stExon;5'UTR |
| 50M_07_Saliva | chr6:137366322-137366492 | IL20RA | S_Shore | TSS200 |
| 50M_07_Saliva | chr6:152702387-152702463 | SYNE1;SYNE1 | | Body;Body |
| 50M_07_Saliva | chr6:155568918-155569443 | TIAM2;TIAM2 | | Body;Body |
| 50M_07_Saliva | chr6:157746584-157746724 | C6orf35 | S_Shore | TSS1500 |
| 50M_07_Saliva | chr6:159052400-159052937 | TMEM181 | | Body |
| 50M_07_Saliva | chr6:161796785-161796854 | PARK2;PARK2;PARK2 | | Body;Body;Body |
| 50M_07_Saliva | chr6:168857542-168858078 | SMOC2;SMOC2 | | Body;Body |
| 50M_07_Saliva | chr7:872208-872797 | UNC84A;UNC84A | | 5'UTR;Body |
| 50M_07_Saliva | chr7:985614-985719 | ADAP1 | Island | Body |
| 50M_07_Saliva | chr7:1491831-1491921 | MICALL2 | Island | Body |
| 50M_07_Saliva | chr7:2094628-2094977 | MAD1L1;MAD1L1;MAD1L1 | | Body;Body;Body |
| 50M_07_Saliva | chr7:2208492-2208830 | MAD1L1;MAD1L1;MAD1L1 | | Body;Body;Body |
| 50M_07_Saliva | chr7:2611890-2612101 | IQCE;IQCE | | Body;Body |
| 50M_07_Saliva | chr7:3033173-3033939 | CARD11 | | 5'UTR |
| 50M_07_Saliva | chr7:6199980-6200140 | USP42 | S_Shore | 3'UTR |
| 50M_07_Saliva | chr7:23286319-23286539 | GPNMB;GPNMB;GPNMB;GPNMB | N_Shore | 5'UTR;1stExon;1stExon;5'UTR |
| 50M_07_Saliva | chr7:27153605-27153847 | HOXA3;HOXA3;HOXA3;HOXA3 | Island | 5'UTR;1stExon;5'UTR;5'UTR |
| 50M_07_Saliva | chr7:27154720-27154911 | HOXA3;HOXA3;HOXA3 | N_Shore | 5'UTR;TSS1500;5'UTR |
| 50M_07_Saliva | chr7:27155002-27155039 | HOXA3;HOXA3;HOXA3 | Island | 5'UTR;TSS1500;5'UTR |
| 50M_07_Saliva | chr7:27169068-27169674 | HOXA4 | N_Shore | Body |
| 50M_07_Saliva | chr7:48129814-48129904 | UPP1;UPP1 | Island | 5'UTR;5'UTR |
| 50M_07_Saliva | chr7:90896329-90896701 | FZD1;FZD1 | Island | 1stExon;3'UTR |
| 50M_07_Saliva | chr7:92238156-92238248 | CDK6;CDK6 | | 3'UTR;3'UTR |
| 50M_07_Saliva | chr7:98738998-98739698 | SMURF1;SMURF1 | N_Shelf | Body;Body |
| 50M_07_Saliva | chr7:99612634-99612773 | ZKSCAN1 | N_Shore | TSS1500 |
| 50M_07_Saliva | chr7:102574445-102574504 | FBXL13;LRRC17;FBXL13;LRRC17 | | Body;Body;Body;Body |
| 50M_07_Saliva | chr7:134575271-134575811 | CALD1;CALD1;CALD1;CALD1;CALD1 | | TSS1500;Body;TSS1500;Body;Body |
| 50M_07_Saliva | chr7:135433540-135433790 | FAM180A;FAM180A | | 1stExon;5'UTR |
| 50M_07_Saliva | chr7:157701024-157701262 | PTPRN2;PTPRN2;PTPRN2 | N_Shore | Body;Body;Body |
| 50M_07_Saliva | chr7:158059790-158060003 | PTPRN2;PTPRN2;PTPRN2 | N_Shore | Body;Body;Body |
| 50M_07_Saliva | chr8:1790793-1790905 | ARHGEF10 | | 5'UTR |
| 50M_07_Saliva | chr8:19170080-19170432 | SH2D4A | N_Shore | TSS1500 |
| 50M_07_Saliva | chr8:123875036-123875223 | ZHX2 | | 5'UTR |
| 50M_07_Saliva | chr8:144814911-144815204 | FAM83H | N_Shore | 5'UTR |
| 50M_07_Saliva | chr8:144894951-144895243 | SCRIB;SCRIB | Island | Body;Body |
| 50M_07_Saliva | chr8:145728138-145728285 | GPT | S_Shore | TSS1500 |
| 50M_07_Saliva | chr8:145728507-145728543 | GPT | N_Shore | TSS1500 |
| 50M_07_Saliva | chr9:92291210-92291523 | LOC100129066 | N_Shore | Body |
| 50M_07_Saliva | chr10:384286-384652 | DIP2C | N_Shelf | Body |
| 50M_07_Saliva | chr10:531914-532791 | DIP2C | S_Shore | Body |
| 50M_07_Saliva | chr10:556470-556762 | DIP2C | | Body |
| 50M_07_Saliva | chr10:1507708-1508061 | ADARB2 | S_Shore | Body |
| 50M_07_Saliva | chr10:1547717-1547983 | ADARB2 | | Body |
| 50M_07_Saliva | chr10:3149673-3149852 | PFKP | S_Shore | Body |

| | | | | |
|---|---|---|---|---|
| 50M_07_Saliva | chr10:13388442-13388567 | SEPHS1 | | 5'UTR |
| 50M_07_Saliva | chr10:26855907-26856454 | APBB1IP | Island | Body |
| 50M_07_Saliva | chr10:29890938-29891284 | SVIL;SVIL | | 5'UTR;5'UTR |
| 50M_07_Saliva | chr10:47746733-47747039 | ANXA8L2 | | TSS1500 |
| 50M_07_Saliva | chr10:49459800-49460249 | FRMPD2;FRMPD2 | | Body;Body |
| 50M_07_Saliva | chr10:50506875-50506896 | C10orf71;C10orf71 | | TSS1500;TSS1500 |
| 50M_07_Saliva | chr10:75631588-75632066 | CAMK2G;CAMK2G;CAMK2G;CAMK2G;CAMK2G | | Body;Body;Body;Body;Body |
| 50M_07_Saliva | chr10:77165905-77166318 | C10orf41;C10orf41 | Island | Body;Body |
| 50M_07_Saliva | chr10:99331332-99331808 | ANKRD2;ANKRD2 | S_Shore | 3'UTR |
| 50M_07_Saliva | chr10:105238964-105239164 | CALHM3;CALHM3 | | 5'UTR;1stExon |
| 50M_07_Saliva | chr10:106093778-106093830 | ITPRIP | N_Shelf | TSS200 |
| 50M_07_Saliva | chr10:123353654-123355454 | FGFR2;FGFR2;FGFR2;FGFR2;FGI | N_Shelf | TSS200;TSS200;5'UTR;5'UTR;1stExon;5'UTR;5'UTR;5'UTR;5'UTR;5'UTR |
| 50M_07_Saliva | chr10:124248943-124249010 | HTRA1 | | Body |
| 50M_07_Saliva | chr10:134578261-134578481 | INPP5A | | Body |
| 50M_07_Saliva | chr11:692963-693032 | DEAF1 | N_Shore | Body |
| 50M_07_Saliva | chr11:1471410-1471594 | BRSK2 | Island | Body |
| 50M_07_Saliva | chr11:1855389-1855561 | SYT8 | | TSS1500 |
| 50M_07_Saliva | chr11:1855660-1855851 | SYT8 | | TSS200 |
| 50M_07_Saliva | chr11:2828525-2828652 | KCNQ1;KCNQ1 | Island | Body;Body |
| 50M_07_Saliva | chr11:2925473-2925847 | SLC22A18;SLC22A18;SLC22A18 | S_Shore | Body;Body;TSS1500 |
| 50M_07_Saliva | chr11:8832938-8833095 | ST5;ST5;ST5 | | TSS1500;TSS200;5'UTR |
| 50M_07_Saliva | chr11:17803160-17803421 | KCNC1 | S_Shore | Body |
| 50M_07_Saliva | chr11:20043971-20044639 | NAV2;NAV2;NAV2;NAV2 | | TSS1500;Body;Body;Body |
| 50M_07_Saliva | chr11:32421635-32421845 | WT1;WT1;WT1;WT1 | | Body;Body;Body;Body |
| 50M_07_Saliva | chr11:57089366-57089744 | TNKS1BP1 | N_Shelf | 5'UTR |
| 50M_07_Saliva | chr11:64951916-64952034 | CAPN1 | Island | Body |
| 50M_07_Saliva | chr11:65560174-65560763 | OVOL1 | | Body |
| 50M_07_Saliva | chr11:66104481-66104993 | RIN1 | S_Shelf | TSS1500 |
| 50M_07_Saliva | chr11:66454595-66454651 | SPTBN2 | | Body |
| 50M_07_Saliva | chr11:67442195-67442408 | ALDH3B2;ALDH3B2 | | 5'UTR;TSS200 |
| 50M_07_Saliva | chr11:76998821-76998857 | GDPD4 | | TSS1500 |
| 50M_07_Saliva | chr11:83393062-83393570 | DLG2;DLG2;DLG2;DLG2 | | 5'UTR;Body;Body;Body |
| 50M_07_Saliva | chr11:117069780-117069859 | TAGLN;TAGLN | | TSS1500;TSS1500 |
| 50M_07_Saliva | chr11:117800255-117800467 | TMPRSS13 | | TSS200 |
| 50M_07_Saliva | chr11:117947459-117947997 | TMPRSS4;TMPRSS4 | | TSS1500;TSS1500 |
| 50M_07_Saliva | chr11:118550379-118550485 | TREH;TREH | | 1stExon;5'UTR |
| 50M_07_Saliva | chr11:120008998-120009135 | TRIM29 | S_Shore | TSS200 |
| 50M_07_Saliva | chr11:120973197-120973274 | TECTA | | TSS200 |
| 50M_07_Saliva | chr11:121970694-121970768 | LOC399959;MIR125B1 | | Body;TSS200 |
| 50M_07_Saliva | chr11:123396133-123396486 | GRAMD1B | | TSS1500 |
| 50M_07_Saliva | chr11:126310323-126310475 | KIRREL3;KIRREL3 | | Body;Body |
| 50M_07_Saliva | chr11:128457400-128457682 | ETS1;ETS1 | | 1stExon;5'UTR |
| 50M_07_Saliva | chr11:128812846-128813470 | TP53AIP1 | | TSS200 |
| 50M_07_Saliva | chr12:258355-258436 | IQSEC3;IQSEC3 | | Body;Body |
| 50M_07_Saliva | chr12:8974844-8975309 | A2ML1 | | TSS1500 |
| 50M_07_Saliva | chr12:25801455-25801621 | IFLTD1 | | 1stExon |
| 50M_07_Saliva | chr12:52585695-52585892 | KRT80;KRT80;KRT80;KRT80 | | 1stExon;1stExon;5'UTR;5'UTR |
| 50M_07_Saliva | chr12:52845938-52846088 | KRT6B | | TSS200 |
| 50M_07_Saliva | chr12:52914229-52915064 | KRT5;KRT5 | | 1stExon;5'UTR |
| 50M_07_Saliva | chr12:53242733-53242917 | KRT78;KRT78 | | 5'UTR;1stExon |
| 50M_07_Saliva | chr12:53298383-53298678 | KRT8 | S_Shore | Body |
| 50M_07_Saliva | chr12:53612551-53612734 | RARG;RARG | N_Shore | Body;Body |
| 50M_07_Saliva | chr12:64782908-64783110 | C12orf56;C12orf56 | N_Shore | Body;Body |
| 50M_07_Saliva | chr12:70092840-70093350 | BEST3 | | 5'UTR |
| 50M_07_Saliva | chr12:114337676-114337927 | RBM19;RBM19;RBM19 | | Body;Body;Body |
| 50M_07_Saliva | chr12:120241957-120242156 | CIT | Island | Body |
| 50M_07_Saliva | chr12:120242394-120242513 | CIT | S_Shore | Body |
| 50M_07_Saliva | chr12:121882208-121882415 | KDM2B;KDM2B | Island | Body;Body |
| 50M_07_Saliva | chr12:124773668-124774078 | FAM101A | | TSS200 |
| 50M_07_Saliva | chr12:130976260-130976551 | RIMBP2 | | 5'UTR |
| 50M_07_Saliva | chr12:133101495-133101612 | FBRSL1 | N_Shore | Body |
| 50M_07_Saliva | chr13:99135543-99135770 | STK24;STK24 | | Body;Body |
| 50M_07_Saliva | chr13:110874267-110874326 | COL4A1 | | Body |
| 50M_07_Saliva | chr13:111522755-111522932 | C13orf29 | | TSS200 |
| 50M_07_Saliva | chr13:113676726-113676938 | MCF2L;MCF2L | N_Shore | Body;Body |

| | | | | |
|---|---|---|---|---|
| 50M_07_Saliva | chr13:114103578-114103646 | ADPRHL1;ADPRHL1 | | Body;TSS200 |
| 50M_07_Saliva | chr14:21510323-21510479 | RNASE7 | | TSS200 |
| 50M_07_Saliva | chr14:24733439-24733757 | TGM1 | | TSS1500 |
| 50M_07_Saliva | chr14:56045827-56046001 | KTN1;C14orf33;KTN1;KTN1;KTN1 | N_Shore | TSS1500;Body;TSS1500;TSS1500;TSS1500 |
| 50M_07_Saliva | chr14:69256791-69256977 | ZFP36L1 | Island | Body |
| 50M_07_Saliva | chr14:94393718-94393816 | FAM181A;C14orf86 | | Body;TSS1500 |
| 50M_07_Saliva | chr14:104195181-104195533 | ZFYVE21 | S_Shore | Body |
| 50M_07_Saliva | chr15:42289766-42289939 | PLA2G4E | | Body |
| 50M_07_Saliva | chr15:42448079-42448259 | PLA2G4F;PLA2G4F | | Body;Body |
| 50M_07_Saliva | chr15:59908652-59909035 | GCNT3 | | 5'UTR |
| 50M_07_Saliva | chr15:86313891-86313941 | MIR1276;KLHL25 | S_Shore | TSS200;5'UTR |
| 50M_07_Saliva | chr16:671905-672468 | RAB40C | N_Shore | Body |
| 50M_07_Saliva | chr16:1582105-1582320 | IFT140 | N_Shore | Body |
| 50M_07_Saliva | chr16:12210834-12211098 | SNX29 | Island | Body |
| 50M_07_Saliva | chr16:22326203-22326535 | POLR3E | | Body |
| 50M_07_Saliva | chr16:28550171-28550525 | NUPR1;NUPR1 | | 1stExon;1stExon |
| 50M_07_Saliva | chr16:31147017-31147054 | PRSS8;PRSS8 | | 1stExon;5'UTR |
| 50M_07_Saliva | chr16:66412919-66413050 | CDH5 | | 5'UTR |
| 50M_07_Saliva | chr16:67022472-67022719 | CES8 | | TSS200 |
| 50M_07_Saliva | chr16:81564161-81564553 | CMIP;CMIP | | Body;Body |
| 50M_07_Saliva | chr16:87958145-87958493 | CA5A | | Body |
| 50M_07_Saliva | chr16:89408322-89408567 | ANKRD11 | Island | 5'UTR |
| 50M_07_Saliva | chr17:693137-693388 | RNMTL1 | | Body |
| 50M_07_Saliva | chr17:790628-791217 | NXN | | Body |
| 50M_07_Saliva | chr17:4463371-4464075 | GGT6;GGT6 | | Body;Body |
| 50M_07_Saliva | chr17:7493150-7494095 | SOX15;SOX15 | S_Shore | 1stExon;5'UTR |
| 50M_07_Saliva | chr17:7690327-7690529 | DNAH2 | Island | Body |
| 50M_07_Saliva | chr17:7832932-7833237 | KCNAB3 | Island | TSS200 |
| 50M_07_Saliva | chr17:17110120-17110207 | PLD6 | Island | TSS1500 |
| 50M_07_Saliva | chr17:19313024-19313529 | RNF112 | | TSS1500 |
| 50M_07_Saliva | chr17:19648855-19650069 | ALDH3A1;ALDH3A1;ALDH3A1;Al | S_Shore | 1stExon;5'UTR;TSS200;5'UTR |
| 50M_07_Saliva | chr17:36612298-36612623 | ARHGAP23 | S_Shore | TSS1500 |
| 50M_07_Saliva | chr17:38657310-38657934 | TNS4 | | 5'UTR |
| 50M_07_Saliva | chr17:38859934-38860097 | KRT24 | | 1stExon |
| 50M_07_Saliva | chr17:39143239-39143567 | KRT40 | | 5'UTR |
| 50M_07_Saliva | chr17:39661855-39662375 | KRT13;KRT13;KRT13;KRT13 | | 1stExon;5'UTR;5'UTR;1stExon |
| 50M_07_Saliva | chr17:40274703-40274740 | HSPB9;KAT2A | Island | TSS200;TSS1500 |
| 50M_07_Saliva | chr17:43716515-43716617 | C17orf69;C17orf69 | | Body;Body |
| 50M_07_Saliva | chr17:46035108-46035343 | PRR15L;PRR15L | | 1stExon;5'UTR |
| 50M_07_Saliva | chr17:46629350-46629804 | HOXB3 | N_Shore | Body |
| 50M_07_Saliva | chr17:48845722-48846189 | C17orf73 | | TSS1500 |
| 50M_07_Saliva | chr17:62051709-62051773 | SCN4A | | TSS1500 |
| 50M_07_Saliva | chr17:75283431-75283832 | SEPT9;SEPT9 | | Body;TSS1500 |
| 50M_07_Saliva | chr17:75473895-75474070 | SEPT9;SEPT9;SEPT9;SEPT9;SEPT9;SEPT9;SEPT9 | | Body;5'UTR;Body;Body;Body;Body;Body |
| 50M_07_Saliva | chr17:76501381-76501582 | DNAH17 | S_Shelf | Body |
| 50M_07_Saliva | chr17:76993506-76993698 | CANT1;CANT1;CANT1 | Island | Body;Body;Body |
| 50M_07_Saliva | chr17:77086350-77086532 | HRNBP3 | | 3'UTR |
| 50M_07_Saliva | chr17:78079481-78079668 | GAA;GAA;GAA | N_Shore | Body;Body;Body |
| 50M_07_Saliva | chr17:78161110-78161479 | CARD14;CARD14 | N_Shelf | TSS1500;Body |
| 50M_07_Saliva | chr17:79030796-79031712 | BAIAP2;BAIAP2;BAIAP2;BAIAP2 | | Body;Body;Body;Body |
| 50M_07_Saliva | chr17:79056061-79056183 | BAIAP2;BAIAP2;BAIAP2;BAIAP2 | S_Shelf | Body;Body;Body;Body |
| 50M_07_Saliva | chr17:79099711-79099785 | MIR657;AATK;MIR338 | N_Shelf | TSS1500;Body;Body |
| 50M_07_Saliva | chr17:79099882-79100124 | MIR657;AATK;MIR338 | N_Shelf | TSS1500;Body;TSS200 |
| 50M_07_Saliva | chr17:79609272-79609346 | TSPAN10 | | TSS200 |
| 50M_07_Saliva | chr17:79652024-79652389 | ARL16;HGS | S_Shore | TSS1500;Body |
| 50M_07_Saliva | chr17:79813410-79813507 | P4HB | | Body |
| 50M_07_Saliva | chr17:80794081-80794274 | TBCD;ZNF750 | S_Shelf | Body;5'UTR |
| 50M_07_Saliva | chr17:80797881-80798023 | TBCD;ZNF750;ZNF750 | | Body;5'UTR;1stExon |
| 50M_07_Saliva | chr17:80806135-80806435 | TBCD | | Body |
| 50M_07_Saliva | chr17:80829157-80829309 | TBCD | N_Shore | Body |
| 50M_07_Saliva | chr17:80832470-80832836 | TBCD | N_Shelf | Body |
| 50M_07_Saliva | chr17:80847137-80847662 | TBCD | | Body |
| 50M_07_Saliva | chr18:21452730-21453131 | LAMA3;LAMA3;LAMA3;LAMA3 | | TSS1500;TSS1500;Body;Body |
| 50M_07_Saliva | chr18:21572622-21572748 | TTC39C | | TSS200 |
| 50M_07_Saliva | chr18:55888640-55888832 | NEDD4L;NEDD4L;NEDD4L;NEDD4L;NEDD4L;NEDD4L;NEDD4L;NEDD4L | | 5'UTR;Body;TSS200;5'UTR;5'UTR;Body;Body;Body;5'UTR |

| | | | | |
|---|---|---|---|---|
| 50M_07_Saliva | chr18:61143926-61144339 | SERPINB5 | | TSS1500 |
| 50M_07_Saliva | chr18:61253847-61254561 | SERPINB13 | | TSS1500 |
| 50M_07_Saliva | chr19:2525231-2525459 | GNG7 | Island | 5'UTR |
| 50M_07_Saliva | chr19:2540907-2541104 | GNG7 | Island | 5'UTR |
| 50M_07_Saliva | chr19:3097565-3097712 | GNA11 | Island | Body |
| 50M_07_Saliva | chr19:4305072-4305239 | FSD1 | Island | Body |
| 50M_07_Saliva | chr19:4455239-4455413 | UBXN6;UBXN6;UBXN6 | N_Shelf | 5'UTR;1stExon;Body |
| 50M_07_Saliva | chr19:5250376-5251120 | PTPRS;PTPRS;PTPRS;PTPRS | N_Shore | Body;Body;Body;Body |
| 50M_07_Saliva | chr19:10231566-10231711 | EIF3G | S_Shore | TSS1500 |
| 50M_07_Saliva | chr19:16178240-16178426 | TPM4 | N_Shore | TSS200 |
| 50M_07_Saliva | chr19:17942221-17942411 | JAK3 | Island | Body |
| 50M_07_Saliva | chr19:18385244-18385672 | KIAA1683;KIAA1683;KIAA1683;KIAA1683;KIAA1683;KIAA1683 | | 5'UTR;1stExon;5'UTR;5'UTR;1stExon;1stExon |
| 50M_07_Saliva | chr19:35085474-35085677 | SCGBL;SCGBL | | 1stExon;5'UTR |
| 50M_07_Saliva | chr19:45279765-45280184 | CBLC;CBLC | N_Shore | TSS1500;TSS1500 |
| 50M_07_Saliva | chr19:46270244-46270412 | SIX5 | Island | Body |
| 50M_07_Saliva | chr19:46286064-46286475 | DMPK;DMPK;DMPK | N_Shelf | TSS1500;TSS1500;TSS1500 |
| 50M_07_Saliva | chr19:46807119-46807466 | HIF3A;HIF3A;HIF3A | Island | 5'UTR;Body;Body |
| 50M_07_Saliva | chr20:2276526-2276663 | TGM3 | | TSS200 |
| 50M_07_Saliva | chr20:17674195-17674250 | BANF2 | | TSS200 |
| 50M_07_Saliva | chr20:33880337-33880370 | FAM83C | S_Shore | TSS200 |
| 50M_07_Saliva | chr20:34700112-34700376 | EPB41L1 | | TSS1500 |
| 50M_07_Saliva | chr20:35504371-35504511 | C20orf118 | | TSS200 |
| 50M_07_Saliva | chr20:35972924-35973098 | SRC | N_Shore | TSS200 |
| 50M_07_Saliva | chr20:36151003-36151184 | BLCAP;BLCAP;BLCAP;NNAT;BLC | Island | 5'UTR;5'UTR;5'UTR;Body;5'UTR;5'UTR;Body |
| 50M_07_Saliva | chr20:36152923-36153225 | BLCAP;BLCAP;BLCAP;BLCAP;BL | S_Shore | 5'UTR;1stExon;5'UTR;5'UTR;5'UTR;5'UTR |
| 50M_07_Saliva | chr20:36153947-36154388 | BLCAP;BLCAP;BLCAP;BLCAP;BL | N_Shelf | 5'UTR;TSS1500;5'UTR;5'UTR;5'UTR |
| 50M_07_Saliva | chr20:45313200-45313390 | TP53RK;SLC13A3 | | 3'UTR;TSS200 |
| 50M_07_Saliva | chr20:62367884-62367893 | LIME1 | N_Shore | TSS200 |
| 50M_07_Saliva | chr20:62383275-62383481 | ZBTB46 | Island | Body |
| 50M_07_Saliva | chr20:62693951-62694000 | TCEA2;TCEA2 | Island | TSS200;5'UTR |
| 50M_07_Saliva | chr21:35736023-35736383 | KCNE2 | | TSS1500 |
| 50M_07_Saliva | chr21:43135949-43136287 | C21orf129;NCRNA00112 | | TSS200;TSS1500 |
| 50M_07_Saliva | chr21:43185385-43185884 | RIPK4 | N_Shore | Body |
| 50M_07_Saliva | chr21:46890849-46890997 | COL18A1;COL18A1;COL18A1 | Island | Body;Body;Body |
| 50M_07_Saliva | chr22:17680563-17680578 | CECR1;CECR1 | | TSS200;Body |
| 50M_07_Saliva | chr22:24891158-24891220 | C22orf45;C22orf45;UPB1 | Island | TSS1500;TSS1500;TSS200 |
| 50M_07_Saliva | chr22:24979554-24979588 | GGT1 | | TSS200 |
| 50M_07_Saliva | chr22:33402628-33402998 | SYN3;SYN3;SYN3 | | 1stExon;Body;1stExon |
| 50M_07_Saliva | chr22:38506623-38506781 | BAIAP2L2;BAIAP2L2 | | 1stExon;5'UTR |
| 50M_07_Saliva | chr22:45608465-45608713 | C22orf9;C22orf9 | | Body;TSS200 |
| 50M_07_Saliva | chr22:46481727-46481785 | LOC400931 | N_Shore | TSS200 |
| 50M_07_Saliva | chr22:46481822-46482023 | LOC400931 | Island | TSS200 |
| 50M01_Saliva | chr1:1109012-1109676 | TTLL10 | N_Shore | TSS1500 |
| 50M01_Saliva | chr1:1117562-1118391 | TTLL10;TTLL10 | S_Shore | Body;Body |
| 50M01_Saliva | chr1:2058417-2058941 | PRKCZ;PRKCZ;PRKCZ | | 5'UTR;Body;5'UTR |
| 50M01_Saliva | chr1:2082315-2082522 | PRKCZ;PRKCZ;PRKCZ | Island | Body;Body;Body |
| 50M01_Saliva | chr1:2232063-2232527 | SKI | N_Shore | Body |
| 50M01_Saliva | chr1:3806817-3807709 | C1orf174 | N_Shore | Body |
| 50M01_Saliva | chr1:4718212-4718573 | AJAP1;AJAP1 | S_Shore | Body;Body |
| 50M01_Saliva | chr1:6526049-6526398 | TNFRSF25;TNFRSF25;TNFRSF25; | Island | Body;Body;Body;Body;Body;Body |
| 50M01_Saliva | chr1:7308012-7308117 | CAMTA1 | | Body |
| 50M01_Saliva | chr1:7692321-7692367 | CAMTA1 | | Body |
| 50M01_Saliva | chr1:16084445-16084830 | FBLIM1 | N_Shore | TSS1500 |
| 50M01_Saliva | chr1:16163479-16164122 | FLJ37453 | N_Shore | Body |
| 50M01_Saliva | chr1:17634543-17634716 | PADI4 | | TSS200 |
| 50M01_Saliva | chr1:24239771-24240069 | CNR2;CNR2 | | 5'UTR;1stExon |
| 50M01_Saliva | chr1:26097918-26098214 | MAN1C1 | Island | Body |
| 50M01_Saliva | chr1:27686768-27687695 | MAP3K6 | N_Shore | Body |
| 50M01_Saliva | chr1:27961563-27962037 | FGR | Island | 5'UTR |
| 50M01_Saliva | chr1:38412519-38412744 | INPP5B | | 5'UTR |
| 50M01_Saliva | chr1:40105667-40105706 | HEYL | Island | TSS1500 |
| 50M01_Saliva | chr1:50886393-50886782 | DMRTA2 | Island | Body |
| 50M01_Saliva | chr1:54520114-54520450 | C1orf83;TMEM59 | S_Shore | Body;TSS1500 |
| 50M01_Saliva | chr1:54562040-54562213 | C1orf83 | | Body |
| 50M01_Saliva | chr1:54821853-54822503 | SSBP3;SSBP3;SSBP3 | N_Shore | Body;Body;Body |

| | | | | |
|---|---|---|---|---|
| 50M01_Saliva | chr1:67772896-67772986 | IL12RB2 | N_Shore | TSS200 |
| 50M01_Saliva | chr1:92953037-92953279 | GFI1;GFI1 | S_Shore | TSS1500;TSS1500 |
| 50M01_Saliva | chr1:114414042-114414408 | PTPN22;PTPN22 | | Body;Body |
| 50M01_Saliva | chr1:151810485-151811923 | LOC100132111;C2CD4D | Island | TSS1500;Body |
| 50M01_Saliva | chr1:156046344-156046832 | MEX3A | Island | 3'UTR |
| 50M01_Saliva | chr1:156211801-156211851 | BGLAP | N_Shelf | TSS200 |
| 50M01_Saliva | chr1:159796434-159796629 | SLAMF8 | | TSS200 |
| 50M01_Saliva | chr1:160616870-160617124 | SLAMF1;SLAMF1 | | 5'UTR;1stExon |
| 50M01_Saliva | chr1:161039681-161039695 | ARHGAP30;ARHGAP30;ARHGAP30;ARHGAP30 | | 1stExon;5'UTR;5'UTR;1stExon |
| 50M01_Saliva | chr1:167598958-167599124 | RCSD1 | N_Shore | TSS1500 |
| 50M01_Saliva | chr1:170638807-170639289 | PRRX1;PRRX1 | | Body;Body |
| 50M01_Saliva | chr1:205181237-205181581 | DSTYK;DSTYK | S_Shore | TSS1500;TSS1500 |
| 50M01_Saliva | chr1:206945924-206946187 | IL10 | | TSS200 |
| 50M01_Saliva | chr1:207095153-207095332 | FAIM3;FAIM3;FAIM3 | | Body;Body;Body |
| 50M01_Saliva | chr1:207277450-207277547 | C4BPA | | TSS200 |
| 50M01_Saliva | chr1:208084942-208085455 | CD34;CD34 | S_Shore | TSS1500;TSS1500 |
| 50M01_Saliva | chr1:212588580-212589072 | TMEM206 | S_Shore | TSS1500 |
| 50M01_Saliva | chr1:223936508-223936838 | CAPN2;CAPN2 | Island | Body;Body |
| 50M01_Saliva | chr1:226187852-226188006 | C1orf55 | S_Shore | TSS1500 |
| 50M01_Saliva | chr1:228196198-228196376 | WNT3A | Island | Body |
| 50M01_Saliva | chr1:243646395-243646787 | SDCCAG8 | Island | Body |
| 50M01_Saliva | chr1:247711595-247712383 | C1orf150 | | TSS1500 |
| 50M01_Saliva | chr1:248902767-248903325 | LOC646627;LOC646627 | | 1stExon;3'UTR |
| 50M01_Saliva | chr2:11885266-11885560 | LPIN1 | N_Shore | TSS1500 |
| 50M01_Saliva | chr2:25473529-25473895 | DNMT3A;DNMT3A;DNMT3A | N_Shore | Body;Body;Body |
| 50M01_Saliva | chr2:42990957-42991422 | OXER1 | | 1stExon |
| 50M01_Saliva | chr2:45233390-45233773 | SIX2 | S_Shore | Body |
| 50M01_Saliva | chr2:45235241-45235512 | SIX2 | N_Shore | Body |
| 50M01_Saliva | chr2:56410197-56410300 | CCDC85A | N_Shore | TSS1500 |
| 50M01_Saliva | chr2:66672337-66672688 | MEIS1 | N_Shore | Body |
| 50M01_Saliva | chr2:66673316-66673449 | MEIS1 | Island | Body |
| 50M01_Saliva | chr2:68961597-68962012 | ARHGAP25;ARHGAP25 | | TSS1500;TSS1500 |
| 50M01_Saliva | chr2:69001665-69002378 | ARHGAP25;ARHGAP25;ARHGAP25;ARHGAP25 | | TSS1500;Body;Body;TSS1500 |
| 50M01_Saliva | chr2:73871849-73871942 | ALMS1P | | TSS200 |
| 50M01_Saliva | chr2:74782684-74783088 | DOK1 | Island | Body |
| 50M01_Saliva | chr2:100759014-100759159 | AFF3;AFF3 | | 1stExon;5'UTR |
| 50M01_Saliva | chr2:106776775-106777038 | UXS1 | | Body |
| 50M01_Saliva | chr2:108994116-108994528 | SULT1C4 | | TSS1500 |
| 50M01_Saliva | chr2:127413363-127413369 | GYPC;GYPC | N_Shore | TSS1500;TSS1500 |
| 50M01_Saliva | chr2:127414802-127415063 | GYPC;GYPC | S_Shore | Body;Body |
| 50M01_Saliva | chr2:131674175-131674221 | ARHGEF4;ARHGEF4 | S_Shore | TSS200;TSS200 |
| 50M01_Saliva | chr2:131721768-131722307 | ARHGEF4;ARHGEF4 | Island | Body;Body |
| 50M01_Saliva | chr2:143886326-143886940 | ARHGAP15 | | TSS1500 |
| 50M01_Saliva | chr2:157184499-157184880 | NR4A2 | Island | Body |
| 50M01_Saliva | chr2:158300475-158301005 | CYTIP | | 1stExon |
| 50M01_Saliva | chr2:172945058-172945383 | MAP1D | Island | Body |
| 50M01_Saliva | chr2:175499113-175499377 | WIPF1;WIPF1 | | 5'UTR;5'UTR |
| 50M01_Saliva | chr2:179395885-179395935 | TTN;TTN;TTN;MIR548N;TTN | | Body;Body;Body;Body;Body |
| 50M01_Saliva | chr2:182321354-182321496 | ITGA4 | N_Shore | TSS1500 |
| 50M01_Saliva | chr2:208989425-208989579 | CRYGD | S_Shore | TSS200 |
| 50M01_Saliva | chr2:225265963-225266880 | FAM124B;FAM124B | | 1stExon;1stExon |
| 50M01_Saliva | chr2:240174613-240174659 | HDAC4 | | Body |
| 50M01_Saliva | chr2:240302585-240302797 | HDAC4 | | 5'UTR |
| 50M01_Saliva | chr3:9943828-9943937 | IL17RE;IL17RE;IL17RE | | TSS1500;TSS1500;TSS1500 |
| 50M01_Saliva | chr3:10334717-10335288 | GHRLOS;GHRL;GHRL;GHRLOS;GHRLOS;GHRL;GHRL;GHRL;GHRL;GHR | | Body;TSS200;TSS200;Body;Body;TSS200;TSS200;TSS200;TSS200;TSS200;Body;TSS200;TSS200;TSS200 |
| 50M01_Saliva | chr3:11078979-11079298 | SLC6A1 | | 3'UTR |
| 50M01_Saliva | chr3:42742993-42743203 | HHATL;HHATL | | Body;5'UTR |
| 50M01_Saliva | chr3:46448579-46448868 | CCRL2;CCRL2 | | TSS1500;TSS200 |
| 50M01_Saliva | chr3:50376155-50376475 | RASSF1;RASSF1;RASSF1;RASSF1 | N_Shore | TSS1500;TSS1500;Body;Body |
| 50M01_Saliva | chr3:52529195-52529524 | STAB1 | | TSS200 |
| 50M01_Saliva | chr3:52812033-52812410 | ITIH1;ITIH1 | | TSS1500;Body |
| 50M01_Saliva | chr3:52813813-52813873 | ITIH1;ITIH1;ITIH1;ITIH1 | | TSS200;TSS200;Body;Body |
| 50M01_Saliva | chr3:55517101-55518396 | WNT5A | N_Shore | Body |
| 50M01_Saliva | chr3:65342216-65342843 | MAGI1;MAGI1 | Island | Body;3'UTR |

| | | | | |
|---|---|---|---|---|
| 50M01_Saliva | chr3:121379745-121379791 | HCLS1;HCLS1 | | 1stExon;5'UTR |
| 50M01_Saliva | chr3:121796627-121796767 | CD86;CD86 | | TSS200;Body |
| 50M01_Saliva | chr3:124860616-124860871 | SLC12A8 | Island | Body |
| 50M01_Saliva | chr3:128779498-128779601 | GP9 | | TSS200 |
| 50M01_Saliva | chr3:129800391-129800573 | ALG1L2 | | TSS1500 |
| 50M01_Saliva | chr3:139257713-139257907 | RBP1;RBP1;RBP1 | Island | Body;Body;Body |
| 50M01_Saliva | chr3:150996078-150996563 | P2RY14;MED12L;P2RY14 | | 1stExon;Body;5'UTR |
| 50M01_Saliva | chr3:151058527-151058837 | P2RY12;P2RY12;MED12L;P2RY12 | | 1stExon;5'UTR;Body;5'UTR |
| 50M01_Saliva | chr3:151102703-151102879 | P2RY12;MED12L | | TSS200;Body |
| 50M01_Saliva | chr3:155421970-155422129 | PLCH1;PLCH1 | | 5'UTR;1stExon |
| 50M01_Saliva | chr3:169377725-169377946 | MECOM | N_Shore | Body |
| 50M01_Saliva | chr3:181421427-181422217 | SOX2OT | | Body |
| 50M01_Saliva | chr3:181428242-181428697 | SOX2OT;SOX2 | N_Shore | Body;TSS1500 |
| 50M01_Saliva | chr3:183273245-183273894 | KLHL6 | | 1stExon |
| 50M01_Saliva | chr3:194408845-194408965 | FAM43A;FAM43A | Island | 3'UTR;1stExon |
| 50M01_Saliva | chr3:195946851-195947062 | OSTalpha | | Body |
| 50M01_Saliva | chr4:683148-683240 | MFSD7 | S_Shore | TSS200 |
| 50M01_Saliva | chr4:2813542-2814122 | SH3BP2;SH3BP2 | | TSS1500;5'UTR |
| 50M01_Saliva | chr4:2933715-2934188 | MFSD10;MFSD10 | Island | Body;Body |
| 50M01_Saliva | chr4:4397942-4398276 | D4S234E;D4S234E | | Body;Body |
| 50M01_Saliva | chr4:8207119-8207325 | SH3TC1 | | Body |
| 50M01_Saliva | chr4:10686335-10686610 | CLNK;CLNK | | 1stExon;5'UTR |
| 50M01_Saliva | chr4:15779642-15780011 | CD38 | N_Shore | TSS1500 |
| 50M01_Saliva | chr4:39482431-39483194 | LOC401127 | S_Shore | Body |
| 50M01_Saliva | chr4:52942997-52943247 | SPATA18 | Island | Body |
| 50M01_Saliva | chr4:62382932-62383240 | LPHN3 | N_Shore | Body |
| 50M01_Saliva | chr4:86851225-86851425 | ARHGAP24;ARHGAP24;ARHGAP24 | | Body;Body;TSS200 |
| 50M01_Saliva | chr4:87281412-87281511 | MAPK10;MAPK10;MAPK10 | | TSS200;TSS200;5'UTR |
| 50M01_Saliva | chr4:90815958-90816310 | MMRN1 | | TSS200 |
| 50M01_Saliva | chr4:95264019-95264127 | HPGDS;HPGDS | | 5'UTR;1stExon |
| 50M01_Saliva | chr4:140656749-140657110 | MAML3 | Island | Body |
| 50M01_Saliva | chr4:154711280-154711347 | SFRP2 | N_Shore | TSS1500 |
| 50M01_Saliva | chr4:154711512-154711906 | SFRP2 | N_Shore | TSS1500 |
| 50M01_Saliva | chr4:169798931-169799624 | PALLD;PALLD;PALLD;PALLD | N_Shore | 5'UTR;Body;Body;Body |
| 50M01_Saliva | chr4:186559707-186560083 | SORBS2;SORBS2;SORBS2;SORB | N_Shore | Body;Body;Body;Body;Body;Body;Body;Body |
| 50M01_Saliva | chr5:368804-369088 | AHRR | N_Shore | Body |
| 50M01_Saliva | chr5:1107837-1108315 | SLC12A7 | N_Shore | Body |
| 50M01_Saliva | chr5:1108401-1108876 | SLC12A7 | N_Shore | Body |
| 50M01_Saliva | chr5:1494980-1495356 | LPCAT1 | Island | Body |
| 50M01_Saliva | chr5:37834958-37835506 | GDNF;GDNF;GDNF | Island | 5'UTR;5'UTR;TSS200 |
| 50M01_Saliva | chr5:37835684-37835958 | GDNF;GDNF;GDNF | N_Shore | TSS1500;5'UTR;TSS200 |
| 50M01_Saliva | chr5:39203007-39203190 | FYB;FYB | | Body;Body |
| 50M01_Saliva | chr5:74162771-74162924 | FAM169A | S_Shore | TSS200 |
| 50M01_Saliva | chr5:76010444-76010658 | F2R | N_Shore | TSS1500 |
| 50M01_Saliva | chr5:92914036-92914828 | FLJ42709;FLJ42709 | N_Shore | Body;Body |
| 50M01_Saliva | chr5:95159365-95159639 | GLRX;GLRX | | TSS1500;TSS1500 |
| 50M01_Saliva | chr5:100240048-100240223 | ST8SIA4;ST8SIA4 | S_Shore | TSS1500;TSS1500 |
| 50M01_Saliva | chr5:134870506-134870517 | NEUROG1;NEUROG1 | N_Shore | 1stExon;3'UTR |
| 50M01_Saliva | chr5:137474700-137475288 | NME5 | | 5'UTR |
| 50M01_Saliva | chr5:139227318-139228153 | NRG2;NRG2;NRG2;NRG2 | N_Shore | 3'UTR;3'UTR;3'UTR;3'UTR |
| 50M01_Saliva | chr5:141062313-141062571 | ARAP3 | | TSS1500 |
| 50M01_Saliva | chr5:143191663-143192067 | HMHB1 | | TSS200 |
| 50M01_Saliva | chr5:148808388-148808487 | MIR143;LOC728264 | | TSS200;Body |
| 50M01_Saliva | chr5:148810061-148810203 | MIR145;LOC728264 | | TSS200;Body |
| 50M01_Saliva | chr5:156696185-156696351 | CYFIP2;CYFIP2;CYFIP2 | S_Shelf | TSS200;5'UTR;5'UTR |
| 50M01_Saliva | chr5:169724590-169724803 | LCP2 | | 1stExon |
| 50M01_Saliva | chr5:169780299-169780480 | KCNIP1 | | TSS1500 |
| 50M01_Saliva | chr5:176326365-176326619 | HK3 | | TSS200 |
| 50M01_Saliva | chr5:176937350-176937446 | DOK3;DOK3;DOK3;DOK3;DOK3 | | 1stExon;TSS1500;1stExon;5'UTR;5'UTR |
| 50M01_Saliva | chr5:177423311-177423598 | PROP1 | | TSS200 |
| 50M01_Saliva | chr6:4890079-4890278 | CDYL;CDYL;CDYL;CDYL | | 5'UTR;Body;Body;TSS200 |
| 50M01_Saliva | chr6:11382553-11382769 | NEDD9;NEDD9 | | 1stExon;5'UTR |
| 50M01_Saliva | chr6:13274151-13274354 | PHACTR1 | | Body |
| 50M01_Saliva | chr6:13428522-13428657 | GFOD1 | | Body |
| 50M01_Saliva | chr6:28493601-28493664 | GPX5;GPX5 | | TSS200;TSS200 |

| | | | | |
|---|---|---|---|---|
| 50M01_Saliva | chr6:28874479-28874702 | TRIM27 | | Body |
| 50M01_Saliva | chr6:28875354-28875370 | TRIM27 | | Body |
| 50M01_Saliva | chr6:29454557-29454755 | MAS1L | | 1stExon |
| 50M01_Saliva | chr6:29869838-29870060 | HCG2P7 | | Body |
| 50M01_Saliva | chr6:30039130-30039600 | RNF39;RNF39 | Island | Body;Body |
| 50M01_Saliva | chr6:30104440-30104709 | TRIM40 | | TSS200 |
| 50M01_Saliva | chr6:30297174-30297594 | TRIM39;TRIM39 | S_Shelf | Body;Body |
| 50M01_Saliva | chr6:30624395-30624636 | DHX16;DHX16 | | Body;Body |
| 50M01_Saliva | chr6:30860237-30860300 | DDR1;DDR1;DDR1 | | Body;Body;Body |
| 50M01_Saliva | chr6:31021422-31021796 | HCG22 | | TSS1500 |
| 50M01_Saliva | chr6:31240784-31241000 | HLA-C | S_Shore | TSS1500 |
| 50M01_Saliva | chr6:31519182-31519410 | NFKBIL1;NFKBIL1;NFKBIL1;NFKBIL1 | | Body;Body;Body;Body |
| 50M01_Saliva | chr6:31543647-31543686 | TNF | | 1stExon |
| 50M01_Saliva | chr6:31554199-31555016 | LST1;LST1;LST1;LST1;LST1;LST1;LST1;LST1 | | Body;TSS1500;TSS1500;TSS1500;5'UTR;TSS1500;5'UTR;Body |
| 50M01_Saliva | chr6:31690949-31691426 | C6orf25;C6orf25;C6orf25;C6orf2 | N_Shore | TSS1500;TSS1500;TSS1500;TSS1500;TSS1500;TSS1500;TSS1500 |
| 50M01_Saliva | chr6:31695415-31695599 | DDAH2 | N_Shore | Body |
| 50M01_Saliva | chr6:31765567-31765634 | LSM2 | S_Shore | Body |
| 50M01_Saliva | chr6:32182002-32182174 | NOTCH4 | | Body |
| 50M01_Saliva | chr6:32295097-32295237 | C6orf10 | | Body |
| 50M01_Saliva | chr6:32808504-32808619 | PSMB8;PSMB8 | S_Shore | 3'UTR;3'UTR |
| 50M01_Saliva | chr6:32813337-32813582 | PSMB8;TAP1 | S_Shore | TSS1500;3'UTR |
| 50M01_Saliva | chr6:32904061-32904317 | HLA-DMB | | Body |
| 50M01_Saliva | chr6:33041229-33041343 | HLA-DPA1 | | Body |
| 50M01_Saliva | chr6:33400021-33400528 | SYNGAP1 | S_Shelf | Body |
| 50M01_Saliva | chr6:39282316-39282532 | KCNK17;KCNK17 | S_Shore | TSS200;TSS200 |
| 50M01_Saliva | chr6:41528198-41528785 | FOXP4;FOXP4;FOXP4 | N_Shore | 5'UTR;5'UTR;5'UTR |
| 50M01_Saliva | chr6:50803820-50803903 | TFAP2B | | Body |
| 50M01_Saliva | chr6:75911956-75912547 | COL12A1;COL12A1 | Island | Body;Body |
| 50M01_Saliva | chr6:76059726-76059773 | FILIP1 | | Body |
| 50M01_Saliva | chr6:108145539-108145777 | SCML4 | | TSS200 |
| 50M01_Saliva | chr6:112115117-112115209 | FYN | | 5'UTR |
| 50M01_Saliva | chr6:133035150-133035379 | VNN1 | | 1stExon |
| 50M01_Saliva | chr6:146864278-146864390 | RAB32 | N_Shore | TSS1500 |
| 50M01_Saliva | chr6:149803206-149803444 | ZC3H12D | | 5'UTR |
| 50M01_Saliva | chr6:151258337-151258520 | MTHFD1L | | Body |
| 50M01_Saliva | chr6:151560831-151560981 | AKAP12 | Island | TSS1500 |
| 50M01_Saliva | chr6:157931843-157932180 | ZDHHC14;ZDHHC14 | | Body;Body |
| 50M01_Saliva | chr6:158066609-158066900 | ZDHHC14;ZDHHC14 | | Body;Body |
| 50M01_Saliva | chr6:158447877-158448455 | SYNJ2 | | Body |
| 50M01_Saliva | chr6:158507753-158508188 | SYNJ2 | Island | Body |
| 50M01_Saliva | chr6:163746113-163746175 | LOC285796 | | TSS1500 |
| 50M01_Saliva | chr6:166856056-166856094 | RPS6KA2;RPS6KA2 | | Body;Body |
| 50M01_Saliva | chr6:166876585-166877038 | RPS6KA2;RPS6KA2 | | Body;Body |
| 50M01_Saliva | chr6:170571686-170571786 | LOC154449 | | TSS200 |
| 50M01_Saliva | chr7:632926-633202 | PRKAR1B;PRKAR1B;PRKAR1B;PR | S_Shore | Body;Body;Body;Body;Body;Body |
| 50M01_Saliva | chr7:968194-968552 | ADAP1 | S_Shore | Body |
| 50M01_Saliva | chr7:1094737-1095720 | C7orf50;C7orf50;C7orf50 | N_Shelf | Body;Body;Body |
| 50M01_Saliva | chr7:1113014-1113237 | C7orf50;C7orf50;C7orf50 | | Body;Body;Body |
| 50M01_Saliva | chr7:1858916-1859172 | MAD1L1;MAD1L1;MAD1L1 | | Body;Body;Body |
| 50M01_Saliva | chr7:1980117-1980479 | MAD1L1;MAD1L1;MAD1L1 | S_Shore | Body;Body;Body |
| 50M01_Saliva | chr7:2059796-2060040 | MAD1L1;MAD1L1;MAD1L1 | N_Shore | Body;Body;Body |
| 50M01_Saliva | chr7:2116368-2116648 | MAD1L1;MAD1L1;MAD1L1 | Island | Body;Body;Body |
| 50M01_Saliva | chr7:2152625-2152821 | MAD1L1;MAD1L1;MAD1L1 | N_Shelf | Body;Body;Body |
| 50M01_Saliva | chr7:2551513-2551598 | LFNG | | TSS1500 |
| 50M01_Saliva | chr7:2648289-2648785 | IQCE;IQCE | S_Shelf | Body;Body |
| 50M01_Saliva | chr7:2773782-2774654 | GNA12 | N_Shore | Body |
| 50M01_Saliva | chr7:4752951-4753002 | FOXK1 | | Body |
| 50M01_Saliva | chr7:4848683-4848939 | RADIL | Island | Body |
| 50M01_Saliva | chr7:4911022-4911121 | RADIL | | Body |
| 50M01_Saliva | chr7:5258357-5258485 | WIPI2;WIPI2;WIPI2;WIPI2;WIPI2 | | Body;Body;Body;Body;Body |
| 50M01_Saliva | chr7:5271480-5271655 | WIPI2;WIPI2;WIPI2;WIPI2;WIPI2 | S_Shelf | 3'UTR;3'UTR;3'UTR;3'UTR;3'UTR |
| 50M01_Saliva | chr7:5527927-5528179 | FBXL18 | Island | Body |
| 50M01_Saliva | chr7:26192966-26193109 | NFE2L3 | S_Shore | Body |
| 50M01_Saliva | chr7:30951272-30951498 | AQP1 | | TSS200 |

| | | | | |
|---|---|---|---|---|
| 50M01_Saliva | chr7:42267383-42268072 | GLI3 | N_Shore | 5'UTR |
| 50M01_Saliva | chr7:45018658-45019005 | MYO1G;MYO1G | | 5'UTR;1stExon |
| 50M01_Saliva | chr7:45066841-45066895 | CCM2;CCM2;CCM2;CCM2;CCM2 | | Body;TSS1500;Body;Body;Body |
| 50M01_Saliva | chr7:55224863-55224910 | EGFR;EGFR;EGFR | | Body;Body;Body |
| 50M01_Saliva | chr7:75443075-75443246 | CCL24 | | TSS200 |
| 50M01_Saliva | chr7:75957040-75957123 | YWHAG | N_Shore | 3'UTR |
| 50M01_Saliva | chr7:96631384-96631902 | DLX6AS | Island | Body |
| 50M01_Saliva | chr7:96650096-96650192 | DLX5 | N_Shore | Body |
| 50M01_Saliva | chr7:96650407-96651111 | DLX5 | Island | Body |
| 50M01_Saliva | chr7:96654786-96654832 | DLX5 | S_Shore | TSS1500 |
| 50M01_Saliva | chr7:96655044-96655566 | DLX5 | S_Shore | TSS1500 |
| 50M01_Saliva | chr7:100238750-100239188 | TFR2 | | Body |
| 50M01_Saliva | chr7:128579876-128580582 | IRF5;IRF5;IRF5;IRF5;IRF5;IR | S_Shore | 5'UTR;TSS1500;5'UTR;5'UTR;5'UTR;5'UTR;5'UTR |
| 50M01_Saliva | chr7:139333296-139333410 | HIPK2;HIPK2 | | Body;Body |
| 50M01_Saliva | chr7:150147536-150147655 | GIMAP8 | | TSS1500 |
| 50M01_Saliva | chr7:150147922-150148022 | GIMAP8 | | TSS200 |
| 50M01_Saliva | chr7:150434452-150434640 | GIMAP5;GIMAP5 | | 1stExon;5'UTR |
| 50M01_Saliva | chr7:150549175-150549406 | ABP1 | | TSS1500 |
| 50M01_Saliva | chr7:151087534-151087726 | WDR86 | | Body |
| 50M01_Saliva | chr7:157647109-157647241 | PTPRN2;PTPRN2;PTPRN2 | N_Shore | Body;Body;Body |
| 50M01_Saliva | chr7:158263204-158263277 | PTPRN2;PTPRN2;PTPRN2 | | Body;Body;Body |
| 50M01_Saliva | chr8:6420858-6421171 | ANGPT2;ANGPT2;ANGPT2;MCPH | N_Shelf | TSS200;TSS200;TSS200;Body |
| 50M01_Saliva | chr8:17206671-17206819 | MTMR7 | | Body |
| 50M01_Saliva | chr8:19540210-19540453 | CSGALNACT1;CSGALNACT1 | | TSS200;Body |
| 50M01_Saliva | chr8:21771059-21771540 | DOK2 | S_Shore | 1stExon |
| 50M01_Saliva | chr8:22960177-22960257 | TNFRSF10C | N_Shore | TSS1500 |
| 50M01_Saliva | chr8:27469186-27469338 | CLU;CLU;CLU;CLU | N_Shelf | 1stExon;Body;5'UTR;TSS1500 |
| 50M01_Saliva | chr8:28174847-28175285 | PNOC | | 5'UTR |
| 50M01_Saliva | chr8:38008386-38008630 | STAR;STAR;STAR;STAR | | 5'UTR;5'UTR;1stExon;1stExon |
| 50M01_Saliva | chr8:38831148-38831508 | HTRA4;PLEKHA2 | N_Shore | TSS1500;3'UTR |
| 50M01_Saliva | chr8:59058585-59058660 | FAM110B | Island | 5'UTR |
| 50M01_Saliva | chr8:67975874-67975895 | CSPP1;CSPP1;COPS5 | N_Shore | TSS1500;TSS1500;TSS1500 |
| 50M01_Saliva | chr8:95229664-95229872 | CDH17 | | TSS200 |
| 50M01_Saliva | chr8:99959430-99959602 | OSR2;OSR2 | N_Shore | 5'UTR;5'UTR |
| 50M01_Saliva | chr8:101315498-101315560 | RNF19A;RNF19A | | 5'UTR;TSS200 |
| 50M01_Saliva | chr8:103137173-103137274 | NCALD;NCALD;NCALD;NCALD;N | S_Shore | TSS1500;TSS1500;TSS1500;TSS1500;TSS1500;TSS1500;TSS200 |
| 50M01_Saliva | chr8:104153753-104153829 | C8orf56;BAALC;BAALC | S_Shore | TSS200;Body;Body |
| 50M01_Saliva | chr8:140945782-140945843 | TRAPPC9;TRAPPC9 | | Body;Body |
| 50M01_Saliva | chr8:141609338-141609470 | EIF2C2;EIF2C2 | | Body;Body |
| 50M01_Saliva | chr8:142159992-142160235 | DENND3 | | Body |
| 50M01_Saliva | chr8:144408230-144408588 | TOP1MT | | Body |
| 50M01_Saliva | chr8:145086243-145086427 | SPATC1;SPATC1 | | TSS1500;TSS1500 |
| 50M01_Saliva | chr8:145106299-145106582 | OPLAH | Island | Body |
| 50M01_Saliva | chr9:34624080-34624587 | ARID3C | Island | Body |
| 50M01_Saliva | chr9:98784802-98785110 | NCRNA00092 | S_Shore | TSS1500 |
| 50M01_Saliva | chr9:123688715-123689193 | TRAF1 | N_Shelf | 5'UTR |
| 50M01_Saliva | chr9:133768931-133769184 | QRFP | | 1stExon |
| 50M01_Saliva | chr9:140356475-140356830 | PNPLA7;PNPLA7 | Island | Body;Body |
| 50M01_Saliva | chr10:601759-601816 | DIP2C | | Body |
| 50M01_Saliva | chr10:682693-682871 | DIP2C | N_Shelf | Body |
| 50M01_Saliva | chr10:1454265-1454387 | ADARB2 | | Body |
| 50M01_Saliva | chr10:6214016-6214079 | PFKFB3 | | Body |
| 50M01_Saliva | chr10:14050455-14050521 | FRMD4A | | Body |
| 50M01_Saliva | chr10:45495782-45495981 | C10orf25;ZNF22 | N_Shore | 3'UTR;TSS1500 |
| 50M01_Saliva | chr10:49892741-49893549 | WDFY4 | | TSS1500 |
| 50M01_Saliva | chr10:70847350-70847430 | SRGN | | |
| 50M01_Saliva | chr10:72362452-72362694 | PRF1;PRF1;PRF1;PRF1 | S_Shore | 5'UTR;5'UTR |
| 50M01_Saliva | chr10:73848764-73848915 | SPOCK2;SPOCK2;SPOCK2;SPOC | S_Shore | 5'UTR;5'UTR |
| 50M01_Saliva | chr10:88717364-88717623 | MMRN2;SNCG;MMRN2 | | Body |
| 50M01_Saliva | chr10:94450866-94451736 | HHEX | N_Shore | Body |
| 50M01_Saliva | chr10:100996579-100996825 | HPSE2;HPSE2;HPSE2;HPSE2 | S_Shelf | TSS1500;TSS1500;TSS1500;TSS1500 |
| 50M01_Saliva | chr10:102821565-102821684 | KAZALD1;KAZALD1 | Island | 1stExon;5'UTR |

| | | | | |
|---|---|---|---|---|
| 50M01_Saliva | chr10:103603722-103603812 | KCNIP2;KCNIP2;KCNIP2;KCNIP2 | Island | 1stExon;5'UTR;1stExon;1stExon;1stExon;5'UTR;5'UTR;5'UTR;5'UTR;1stExon;5'UTR;1stExon |
| 50M01_Saliva | chr10:104535521-104536035 | C10orf26;C10orf26 | | TSS1500;Body |
| 50M01_Saliva | chr10:105978651-105978956 | MIR609;C10orf79 | | Body |
| 50M01_Saliva | chr10:114134878-114135259 | ACSL5;ACSL5;ACSL5 | | TSS1500;TSS1500;5'UTR |
| 50M01_Saliva | chr10:126222996-126223401 | LHPP;LHPP | | Body;Body |
| 50M01_Saliva | chr10:128152075-128152124 | C10orf90 | | Body |
| 50M01_Saliva | chr10:134045913-134046237 | STK32C | S_Shelf | Body |
| 50M01_Saliva | chr10:134149366-134150760 | LRRC27;LRRC27;LRRC27;LRRC2 | S_Shelf | Body;Body;Body;Body;TSS1500 |
| 50M01_Saliva | chr10:134358030-134358273 | INPP5A | N_Shelf | Body |
| 50M01_Saliva | chr10:135202611-135203102 | PAOX;PAOX;PAOX | N_Shore | Body;Body;Body |
| 50M01_Saliva | chr11:415430-415457 | SIGIRR;SIGIRR;SIGIRR | N_Shore | 5'UTR;5'UTR;TSS1500 |
| 50M01_Saliva | chr11:457256-457304 | PTDSS2 | N_Shore | Body |
| 50M01_Saliva | chr11:1874017-1874445 | LSP1 | | TSS200 |
| 50M01_Saliva | chr11:2321770-2322050 | C11orf21;TSPAN32;C11orf21 | | Body;TSS1500;Body |
| 50M01_Saliva | chr11:2322329-2322517 | C11orf21;TSPAN32;C11orf21 | | Body;TSS1500;Body |
| 50M01_Saliva | chr11:2322618-2323272 | C11orf21;TSPAN32;C11orf21 | | Body;TSS1500;Body |
| 50M01_Saliva | chr11:3647419-3648013 | TRPC2 | | TSS1500 |
| 50M01_Saliva | chr11:35441777-35442225 | SLC1A2 | Island | TSS1500 |
| 50M01_Saliva | chr11:36422377-36422615 | PRR5L;PRR5L;PRR5L;PRR5L | | 5'UTR;5'UTR;TSS200;Body |
| 50M01_Saliva | chr11:46382544-46383924 | DGKZ;DGKZ;DGKZ;DGKZ | | Body;TSS1500;Body;Body |
| 50M01_Saliva | chr11:47399213-47400330 | SPI1;SPI1 | N_Shore | Body;Body |
| 50M01_Saliva | chr11:59634191-59634276 | TCN1 | | TSS200 |
| 50M01_Saliva | chr11:59823993-59824161 | MS4A3;MS4A3;MS4A3 | | TSS200;TSS200;TSS200 |
| 50M01_Saliva | chr11:60623415-60623639 | GPR44;GPR44 | S_Shelf | 5'UTR;1stExon |
| 50M01_Saliva | chr11:60738971-60739183 | CD6 | | TSS200 |
| 50M01_Saliva | chr11:60869760-60869969 | CD5 | | TSS200 |
| 50M01_Saliva | chr11:63272764-63274030 | LGALS12;LGALS12;LGALS12 | | TSS1500;TSS1500;TSS1500 |
| 50M01_Saliva | chr11:63827432-63828228 | MACROD1 | | Body |
| 50M01_Saliva | chr11:63973846-63975481 | FERMT3;FERMT3 | N_Shore | TSS1500;TSS1500 |
| 50M01_Saliva | chr11:64107517-64108241 | CCDC88B | N_Shelf | TSS200 |
| 50M01_Saliva | chr11:64563983-64564012 | MAP4K2 | | Body |
| 50M01_Saliva | chr11:64981227-64981596 | SLC22A20 | | TSS200 |
| 50M01_Saliva | chr11:65407399-65408566 | SIPA1;SIPA1 | N_Shore | TSS200;5'UTR |
| 50M01_Saliva | chr11:65647279-65647532 | CTSW | | TSS200 |
| 50M01_Saliva | chr11:66055170-66055327 | YIF1A | N_Shore | Body |
| 50M01_Saliva | chr11:66084841-66085299 | CD248 | S_Shore | TSS1500 |
| 50M01_Saliva | chr11:66839085-66839191 | RHOD | Island | 3'UTR |
| 50M01_Saliva | chr11:67170610-67171585 | PPP1CA;PPP1CA;TBC1D10C;PPP | S_Shore | TSS1500;TSS1500;TSS1500;TSS1500 |
| 50M01_Saliva | chr11:67176914-67177296 | TBC1D10C | N_Shore | Body |
| 50M01_Saliva | chr11:67204799-67205291 | PTPRCAP | S_Shore | Body |
| 50M01_Saliva | chr11:69519845-69520418 | FGF19 | Island | TSS1500 |
| 50M01_Saliva | chr11:69934031-69934128 | ANO1;ANO1 | Island | Body;Body |
| 50M01_Saliva | chr11:70211301-70211531 | PPFIA1;PPFIA1 | | Body;Body |
| 50M01_Saliva | chr11:70257035-70257149 | CTTN;CTTN | N_Shore | Body;Body |
| 50M01_Saliva | chr11:70691944-70692038 | SHANK2 | | Body |
| 50M01_Saliva | chr11:71709837-71709978 | IL18BP;IL18BP;IL18BP;IL18BP;IL18BP | | TSS200;TSS1500;TSS200;TSS1500;TSS1500 |
| 50M01_Saliva | chr11:71846494-71846788 | FOLR3 | | TSS1500 |
| 50M01_Saliva | chr11:78673073-78673290 | ODZ4 | Island | Body |
| 50M01_Saliva | chr11:124767455-124767840 | ROBO4 | | Body |
| 50M01_Saliva | chr11:125037226-125037348 | PKNOX2 | S_Shore | 5'UTR |
| 50M01_Saliva | chr11:126581947-126582242 | KIRREL3;KIRREL3 | | Body;Body |
| 50M01_Saliva | chr11:128391271-128391494 | ETS1;ETS1;ETS1 | N_Shore | Body;Body;Body |
| 50M01_Saliva | chr11:134126284-134126504 | ACAD8 | S_Shelf | Body |
| 50M01_Saliva | chr11:134254573-134254690 | B3GAT1;B3GAT1 | S_Shore | Body;Body |
| 50M01_Saliva | chr12:1025529-1025755 | RAD52 | | Body |
| 50M01_Saliva | chr12:4699085-4699618 | DYRK4 | | TSS200 |
| 50M01_Saliva | chr12:6664536-6665447 | IFFO1;IFFO1 | Island | 1stExon;1stExon |
| 50M01_Saliva | chr12:6881590-6881629 | LAG3 | N_Shore | TSS200 |
| 50M01_Saliva | chr12:22486029-22486484 | ST8SIA1 | N_Shore | Body |
| 50M01_Saliva | chr12:27396229-27396765 | STK38L | N_Shore | TSS1500 |
| 50M01_Saliva | chr12:32654929-32655034 | FGD4 | | TSS200 |
| 50M01_Saliva | chr12:47610220-47610734 | FAM113B;FAM113B;LOC100233209 | | 5'UTR;1stExon;Body |
| 50M01_Saliva | chr12:49731393-49731885 | C1QL4 | S_Shore | TSS1500 |

| | | | | |
|---|---|---|---|---|
| 50M01_Saliva | chr12:49782966-49783097 | SPATS2 | Island | 5'UTR |
| 50M01_Saliva | chr12:51717674-51718310 | BIN2 | | Body |
| 50M01_Saliva | chr12:52404134-52404161 | GRASP | S_Shelf | Body |
| 50M01_Saliva | chr12:53730072-53730277 | SP7 | | TSS1500 |
| 50M01_Saliva | chr12:54891213-54891636 | NCKAP1L | | TSS1500 |
| 50M01_Saliva | chr12:55378411-55378912 | KIAA0748;KIAA0748 | | 1stExon;5'UTR |
| 50M01_Saliva | chr12:56414442-56414533 | IKZF4 | | TSS1500 |
| 50M01_Saliva | chr12:57618943-57619199 | NXPH4 | Island | Body |
| 50M01_Saliva | chr12:58210661-58210878 | AVIL | | TSS1500 |
| 50M01_Saliva | chr12:81109337-81109806 | MYF5 | N_Shore | TSS1500 |
| 50M01_Saliva | chr12:105114162-105114399 | CHST11 | | Body |
| 50M01_Saliva | chr12:106533840-106533903 | NUAK1 | S_Shore | TSS200 |
| 50M01_Saliva | chr12:111537057-111537195 | CUX2 | | Body |
| 50M01_Saliva | chr12:111618925-111618977 | CUX2 | | Body |
| 50M01_Saliva | chr12:117797116-117797635 | NOS1 | N_Shore | 5'UTR |
| 50M01_Saliva | chr12:119591747-119592094 | SRRM4 | N_Shore | Body |
| 50M01_Saliva | chr12:123518833-123519476 | PITPNM2 | | Body |
| 50M01_Saliva | chr12:123753186-123753272 | CDK2AP1 | N_Shore | Body |
| 50M01_Saliva | chr12:124908601-124908653 | NCOR2;NCOR2 | N_Shelf | Body;Body |
| 50M01_Saliva | chr12:124938116-124938573 | NCOR2;NCOR2 | N_Shelf | Body;Body |
| 50M01_Saliva | chr12:124941423-124942248 | NCOR2;NCOR2 | Island | Body;Body |
| 50M01_Saliva | chr12:124950669-124950990 | NCOR2;NCOR2 | N_Shore | Body;Body |
| 50M01_Saliva | chr12:133135032-133135698 | FBRSL1 | N_Shore | Body |
| 50M01_Saliva | chr13:30947713-30948122 | LOC100188949 | | Body |
| 50M01_Saliva | chr13:51639953-51641069 | GUCY1B2 | | Body |
| 50M01_Saliva | chr13:100310162-100310405 | CLYBL | N_Shore | Body |
| 50M01_Saliva | chr13:107143980-107144036 | EFNB2 | N_Shore | 3'UTR |
| 50M01_Saliva | chr13:107145658-107146164 | EFNB2 | Island | Body |
| 50M01_Saliva | chr13:110951025-110951418 | COL4A1 | | Body |
| 50M01_Saliva | chr13:110993474-110993558 | COL4A2 | | Body |
| 50M01_Saliva | chr13:111318490-111318640 | CARS2 | | Body |
| 50M01_Saliva | chr13:113652069-113652368 | MCF2L;MCF2L | S_Shelf | Body;Body |
| 50M01_Saliva | chr13:114261869-114261986 | TFDP1;TFDP1 | S_Shore | Body;Body |
| 50M01_Saliva | chr13:114814333-114814706 | RASA3 | Island | Body |
| 50M01_Saliva | chr14:24641021-24641852 | REC8;REC8 | N_Shore | TSS1500;TSS1500 |
| 50M01_Saliva | chr14:38725536-38725750 | CLEC14A;CLEC14A | Island | 1stExon;5'UTR |
| 50M01_Saliva | chr14:75988747-75989203 | BATF | | TSS200 |
| 50M01_Saliva | chr14:91866325-91866751 | CCDC88C | | Body |
| 50M01_Saliva | chr14:93170710-93170970 | LGMN;LGMN | | Body;Body |
| 50M01_Saliva | chr14:102676957-102677294 | WDR20 | N_Shelf | Body |
| 50M01_Saliva | chr14:105532012-105532208 | GPR132 | | TSS1500 |
| 50M01_Saliva | chr14:105944604-105945699 | CRIP2 | Island | Body |
| 50M01_Saliva | chr15:26044050-26044289 | ATP10A | | Body |
| 50M01_Saliva | chr15:27111435-27111476 | GABRA5;GABRA5 | N_Shore | TSS1500;TSS1500 |
| 50M01_Saliva | chr15:27111572-27111604 | GABRA5;GABRA5 | N_Shore | TSS1500;TSS1500 |
| 50M01_Saliva | chr15:37387304-37387577 | MEIS2;MEIS2;MEIS2;MEIS2;MEIS | N_Shore | Body;Body;Body;Body;Body;Body;Body |
| 50M01_Saliva | chr15:38988533-38988860 | C15orf53 | | TSS1500 |
| 50M01_Saliva | chr15:43531947-43532243 | TGM5;TGM5 | | Body;Body |
| 50M01_Saliva | chr15:45409942-45410236 | DUOXA1;DUOXA2 | S_Shore | Body;Body |
| 50M01_Saliva | chr15:86315115-86315211 | KLHL25;MIR1276 | S_Shelf | 5'UTR;TSS1500 |
| 50M01_Saliva | chr15:94840793-94841334 | MCTP2;MCTP2 | | TSS1500;TSS1500 |
| 50M01_Saliva | chr15:96877194-96877720 | NR2F2;NR2F2;NR2F2;NR2F2 | Island | Body;Body;5'UTR;5'UTR |
| 50M01_Saliva | chr15:101418967-101419074 | ALDH1A3 | N_Shore | TSS1500 |
| 50M01_Saliva | chr15:101991512-101991886 | PCSK6;PCSK6;PCSK6;PCSK6;PCSK6;PCSK6;PCSK6;PCSK6 | | Body;Body;Body;Body;Body;Body;Body;Body |
| 50M01_Saliva | chr16:302852-303192 | ITFG3 | | 5'UTR |
| 50M01_Saliva | chr16:787799-788184 | NARFL | S_Shore | Body |
| 50M01_Saliva | chr16:1583810-1584516 | IFT140;TMEM204 | Island | Body;TSS1500 |
| 50M01_Saliva | chr16:3627243-3628416 | NLRC3;NLRC3 | | 1stExon;5'UTR |
| 50M01_Saliva | chr16:4714080-4714443 | MGRN1;MGRN1;MGRN1;MGRN1 | | Body;Body;Body;Body |
| 50M01_Saliva | chr16:8738240-8738441 | C16orf68 | | Body |
| 50M01_Saliva | chr16:10647084-10647275 | EMP2 | | 5'UTR |
| 50M01_Saliva | chr16:14403004-14403040 | MIR365-1 | | TSS200 |
| 50M01_Saliva | chr16:28505320-28505959 | APOB48R | S_Shore | TSS1500 |
| 50M01_Saliva | chr16:28995843-28996533 | SPNS1;LAT;LAT;LAT;SPNS1;SPNS1;LAT;SPNS1;SPNS1 | | 3'UTR;TSS1500;TSS1500;TSS1500;3'UTR;3'UTR;TSS1500;3'UTR;3'UTR |

| | | | | |
|---|---|---|---|---|
| 50M01_Saliva | chr16:29086271-29086456 | RRN3P2 | Island | Body |
| 50M01_Saliva | chr16:29673933-29676071 | SPN;SPN | N_Shore | TSS1500;TSS1500 |
| 50M01_Saliva | chr16:29756951-29757461 | C16orf54 | | 5'UTR |
| 50M01_Saliva | chr16:30721139-30721715 | SNORA30;SRCAP | | TSS1500;Body |
| 50M01_Saliva | chr16:49732224-49732623 | ZNF423 | N_Shore | Body |
| 50M01_Saliva | chr16:50347766-50347895 | ADCY7 | | Body |
| 50M01_Saliva | chr16:50715135-50715260 | SNX20;SNX20;SNX20;SNX20;SNX20;SNX20 | | 1stExon;5'UTR;1stExon;5'UTR;5'UTR;1stExon |
| 50M01_Saliva | chr16:56995813-56995856 | CETP | | TSS200 |
| 50M01_Saliva | chr16:57701455-57701726 | GPR97 | | TSS1500 |
| 50M01_Saliva | chr16:67428155-67428898 | TPPP3;TPPP3 | Island | TSS1500;TSS1500 |
| 50M01_Saliva | chr16:69966902-69967063 | MIR140;WWP2;WWP2 | | TSS200;Body;Body |
| 50M01_Saliva | chr16:88038949-88039185 | BANP;BANP | N_Shore | Body;Body |
| 50M01_Saliva | chr16:88566483-88566748 | ZFPM1 | N_Shelf | Body |
| 50M01_Saliva | chr16:88767124-88767723 | RNF166 | S_Shore | Body |
| 50M01_Saliva | chr16:88844415-88844726 | FAM38A | N_Shore | Body |
| 50M01_Saliva | chr16:89024511-89024657 | CBFA2T3 | | Body |
| 50M01_Saliva | chr16:89033692-89034292 | CBFA2T3 | Island | Body |
| 50M01_Saliva | chr17:750241-750379 | NXN | | Body |
| 50M01_Saliva | chr17:800717-800749 | NXN | | Body |
| 50M01_Saliva | chr17:943108-943342 | ABR;ABR;ABR | | Body;Body;Body |
| 50M01_Saliva | chr17:1548881-1549284 | SCARF1;SCARF1;SCARF1;SCARF | S_Shore | Body;Body;Body;Body;Body |
| 50M01_Saliva | chr17:1626477-1626575 | WDR81;WDR81;WDR81 | N_Shore | TSS1500;Body;5'UTR |
| 50M01_Saliva | chr17:2699505-2700093 | RAP1GAP2;RAP1GAP2 | | TSS1500;TSS1500 |
| 50M01_Saliva | chr17:2839005-2839187 | RAP1GAP2;RAP1GAP2 | | Body;Body |
| 50M01_Saliva | chr17:3704471-3704621 | ITGAE;ITGAE | | 1stExon;5'UTR |
| 50M01_Saliva | chr17:3819339-3820518 | P2RX1 | | Body |
| 50M01_Saliva | chr17:4337082-4337231 | SPNS3 | | TSS200 |
| 50M01_Saliva | chr17:5138259-5138696 | C17orf87 | | TSS200 |
| 50M01_Saliva | chr17:7287449-7287922 | TNK1 | Island | Body |
| 50M01_Saliva | chr17:9940004-9940321 | GAS7;GAS7;GAS7 | | Body;1stExon;5'UTR |
| 50M01_Saliva | chr17:19290353-19290762 | MFAP4 | | Body |
| 50M01_Saliva | chr17:27039120-27039523 | PROCA1 | S_Shore | TSS1500 |
| 50M01_Saliva | chr17:34344995-34345172 | CCL23;CCL23;CCL23;CCL23 | | 1stExon;5'UTR;1stExon;5'UTR |
| 50M01_Saliva | chr17:37322028-37322431 | ARL5C;ARL5C | Island | 1stExon;5'UTR |
| 50M01_Saliva | chr17:40439433-40439501 | STAT5A | N_Shore | TSS200 |
| 50M01_Saliva | chr17:40440904-40441155 | STAT5A | Island | 5'UTR |
| 50M01_Saliva | chr17:42466960-42467307 | ITGA2B | S_Shelf | TSS200 |
| 50M01_Saliva | chr17:44847220-44847427 | WNT3 | Island | Body |
| 50M01_Saliva | chr17:44896877-44897512 | WNT3 | Island | TSS1500 |
| 50M01_Saliva | chr17:46667587-46667812 | LOC404266;LOC404266 | N_Shore | TSS1500;TSS1500 |
| 50M01_Saliva | chr17:48049952-48050512 | DLX4;DLX4 | Island | TSS200;Body |
| 50M01_Saliva | chr17:53341098-53342029 | HLF | N_Shore | TSS1500 |
| 50M01_Saliva | chr17:55533142-55533389 | MSI2;MSI2 | | Body;Body |
| 50M01_Saliva | chr17:55952063-55952217 | CUEDC1 | | Body |
| 50M01_Saliva | chr17:56355299-56355431 | MPO | Island | Body |
| 50M01_Saliva | chr17:59478579-59479306 | TBX2 | Island | Body |
| 50M01_Saliva | chr17:59532519-59533081 | TBX4 | Island | TSS1500 |
| 50M01_Saliva | chr17:62097951-62098230 | ICAM2;ICAM2;ICAM2;ICAM2;ICAM2;ICAM2;ICAM2;ICAM2 | | 5'UTR;5'UTR;5'UTR;1stExon;1stExon;5'UTR;1stExon;1stExon |
| 50M01_Saliva | chr17:64787379-64787784 | PRKCA | | Body |
| 50M01_Saliva | chr17:65471321-65471461 | PITPNC1;PITPNC1 | | Body;Body |
| 50M01_Saliva | chr17:72442928-72443179 | GPRC5C;GPRC5C | Island | Body;Body |
| 50M01_Saliva | chr17:73642404-73642628 | LOC100130933;RECQL5 | | TSS1500;Body |
| 50M01_Saliva | chr17:75471193-75471308 | SEPT9;SEPT9;SEPT9;SEPT9;SEPT9;SEPT9;SEPT9 | | Body;TSS200;5'UTR;Body;Body;Body;Body |
| 50M01_Saliva | chr17:76126702-76127217 | TMC8;TMC6 | N_Shore | TSS200;5'UTR |
| 50M01_Saliva | chr17:76886742-76886912 | TIMP2 | | Body |
| 50M01_Saliva | chr17:77916733-77916892 | TBC1D16 | Island | Body |
| 50M01_Saliva | chr17:78735268-78735596 | RPTOR;RPTOR | | Body;Body |
| 50M01_Saliva | chr17:79005548-79006087 | FLJ90757 | N_Shore | Body |
| 50M01_Saliva | chr17:79128885-79129078 | AATK | N_Shelf | Body |
| 50M01_Saliva | chr17:79258813-79259699 | SLC38A10;SLC38A10 | N_Shore | Body;Body |
| 50M01_Saliva | chr17:80330719-80330901 | UTS2R | N_Shore | TSS1500 |
| 50M01_Saliva | chr17:80358829-80358876 | C17orf101;C17orf101 | N_Shelf | Body;Body |
| 50M01_Saliva | chr17:80407031-80407779 | C17orf62;C17orf62;C17orf62 | N_Shore | Body;Body;Body |
| 50M01_Saliva | chr17:80545175-80545454 | FOXK2 | N_Shore | Body |

| | | | | |
|---|---|---|---|---|
| 50M01_Saliva | chr17:80581701-80581925 | WDR45L | | Body |
| 50M01_Saliva | chr17:80740731-80740950 | TBCD | Island | Body |
| 50M01_Saliva | chr17:80944090-80944193 | B3GNTL1 | | Body |
| 50M01_Saliva | chr18:30352782-30353125 | KLHL14;KLHL14 | S_Shore | 5'UTR;1stExon |
| 50M01_Saliva | chr18:74114570-74114728 | ZNF516 | Island | Body |
| 50M01_Saliva | chr18:74824154-74824413 | MBP;MBP | Island | 5'UTR;5'UTR |
| 50M01_Saliva | chr19:681420-681815 | FSTL3 | Island | Body |
| 50M01_Saliva | chr19:695447-695642 | PRSSL1;PRSSL1 | | 5'UTR;1stExon |
| 50M01_Saliva | chr19:720740-720993 | PALM;PALM | Island | Body;Body |
| 50M01_Saliva | chr19:840737-841082 | PRTN3 | N_Shelf | TSS1500 |
| 50M01_Saliva | chr19:852114-852311 | ELANE | N_Shore | TSS200 |
| 50M01_Saliva | chr19:859330-859507 | CFD | N_Shore | TSS1500 |
| 50M01_Saliva | chr19:947712-947936 | ARID3A | Island | Body |
| 50M01_Saliva | chr19:1074425-1075104 | HMHA1 | N_Shore | Body |
| 50M01_Saliva | chr19:3178512-3179364 | S1PR4 | N_Shore | TSS1500 |
| 50M01_Saliva | chr19:4326975-4327350 | STAP2;STAP2 | Island | Body;Body |
| 50M01_Saliva | chr19:4916593-4916983 | UHRF1;UHRF1 | Island | Body;Body |
| 50M01_Saliva | chr19:5074616-5074758 | KDM4B | Island | Body |
| 50M01_Saliva | chr19:6481540-6481826 | DENND1C | | Body |
| 50M01_Saliva | chr19:6481945-6482169 | DENND1C | | TSS200 |
| 50M01_Saliva | chr19:6772811-6773074 | VAV1;VAV1 | | 5'UTR;1stExon |
| 50M01_Saliva | chr19:7733793-7734203 | RETN | N_Shore | TSS200 |
| 50M01_Saliva | chr19:7741602-7741955 | C19orf59 | N_Shelf | TSS1500 |
| 50M01_Saliva | chr19:10222051-10222409 | PPAN-P2RY11;P2RY11 | S_Shore | Body;TSS200 |
| 50M01_Saliva | chr19:10449942-10450472 | ICAM3 | S_Shelf | Body |
| 50M01_Saliva | chr19:10958952-10958970 | C19orf38 | | TSS200 |
| 50M01_Saliva | chr19:13319324-13319523 | CACNA1A;CACNA1A | Island | Body;Body |
| 50M01_Saliva | chr19:13947212-13947262 | MIR24-2 | S_Shelf | TSS200 |
| 50M01_Saliva | chr19:14550846-14550999 | PKN1;PKN1 | N_Shore | TSS1500;Body |
| 50M01_Saliva | chr19:14785593-14785749 | EMR3;EMR3 | | 5'UTR;1stExon |
| 50M01_Saliva | chr19:16999641-16999953 | F2RL3 | N_Shore | TSS200 |
| 50M01_Saliva | chr19:17488042-17488225 | PLVAP | Island | 1stExon |
| 50M01_Saliva | chr19:17862017-17862385 | FCHO1;FCHO1;FCHO1;FCHO1 | S_Shelf | 5'UTR;TSS1500;5'UTR;TSS1500 |
| 50M01_Saliva | chr19:35531403-35531639 | HPN;HPN | N_Shore | TSS200;TSS200 |
| 50M01_Saliva | chr19:35630106-35630305 | FXYD1;FXYD1 | N_Shelf | 5'UTR;TSS1500 |
| 50M01_Saliva | chr19:35819775-35820181 | CD22 | | TSS1500 |
| 50M01_Saliva | chr19:36233297-36233469 | TMEM149;TMEM149 | S_Shore | 1stExon;5'UTR |
| 50M01_Saliva | chr19:36399185-36399346 | TYROBP;TYROBP;TYROBP;TYROBP | | 1stExon;1stExon;5'UTR;5'UTR |
| 50M01_Saliva | chr19:39826668-39826927 | GMFG;GMFG | | 5'UTR;1stExon |
| 50M01_Saliva | chr19:43856594-43857074 | CD177 | | TSS1500 |
| 50M01_Saliva | chr19:45720771-45721076 | EXOC3L2 | Island | Body |
| 50M01_Saliva | chr19:46318439-46319398 | RSPH6A;RSPH6A | N_Shore | 1stExon;5'UTR |
| 50M01_Saliva | chr19:46713446-46713473 | DKFZp434J0226 | | TSS200 |
| 50M01_Saliva | chr19:49250252-49250561 | IZUMO1 | N_Shelf | TSS200 |
| 50M01_Saliva | chr19:49838478-49838777 | CD37;CD37 | N_Shelf | TSS200;TSS200 |
| 50M01_Saliva | chr19:51628062-51628204 | SIGLEC9 | | TSS200 |
| 50M01_Saliva | chr19:51875451-51875986 | NKG7 | S_Shelf | Body |
| 50M01_Saliva | chr19:52004992-52005188 | SIGLEC12;SIGLEC12;SIGLEC12 | | 1stExon;5'UTR;TSS1500 |
| 50M01_Saliva | chr19:54327404-54327995 | NLRP12 | | 1stExon |
| 50M01_Saliva | chr19:54566838-54567279 | VSTM1 | | Body |
| 50M01_Saliva | chr19:54604098-54604187 | OSCAR;OSCAR;OSCAR;OSCAR;( | N_Shore | 5'UTR;5'UTR;5'UTR;1stExon;1stExon;5'UTR;1stExon;1stExon |
| 50M01_Saliva | chr19:55013821-55013946 | LAIR2;LAIR2 | | TSS200;TSS200 |
| 50M01_Saliva | chr19:55141723-55142181 | LILRB1;LILRB1;LILRB1;LILRB1 | | TSS200;5'UTR;TSS200;TSS1500 |
| 50M01_Saliva | chr20:1448419-1448498 | NSFL1C;NSFL1C;NSFL1C | S_Shore | TSS200;TSS200;TSS200 |
| 50M01_Saliva | chr20:1448595-1448645 | NSFL1C;NSFL1C;NSFL1C | S_Shore | TSS200;TSS200;TSS200 |
| 50M01_Saliva | chr20:1471884-1472078 | SIRPB2;SIRPB2 | | Body;Body |
| 50M01_Saliva | chr20:1600623-1600705 | SIRPB1;SIRPB1;SIRPB1;SIRPB1;SIRPB1;SIRPB1 | | 5'UTR;1stExon;1stExon;5'UTR;5'UTR;1stExon |
| 50M01_Saliva | chr20:1974646-1974843 | PDYN;PDYN | | 5'UTR;1stExon |
| 50M01_Saliva | chr20:3065343-3065423 | AVP;AVP | S_Shore | 5'UTR;1stExon |
| 50M01_Saliva | chr20:4795444-4795935 | RASSF2;RASSF2 | | 5'UTR;5'UTR |
| 50M01_Saliva | chr20:23066944-23067159 | CD93;CD93 | | 5'UTR;1stExon |
| 50M01_Saliva | chr20:24929607-24929716 | CST7 | | TSS1500 |
| 50M01_Saliva | chr20:35274570-35274828 | SLA2;SLA2 | | TSS200;TSS200 |
| 50M01_Saliva | chr20:37433745-37434158 | PPP1R16B | N_Shore | TSS1500 |

| | | | | |
|---|---|---|---|---|
| 50M01_Saliva | chr20:44875334-44875586 | CDH22 | Island | Body |
| 50M01_Saliva | chr20:45179157-45179354 | C20orf123 | | 1stExon |
| 50M01_Saliva | chr20:54987020-54987140 | CASS4;CASS4;CASS4;CASS4 | | TSS1500;TSS1500;TSS200;TSS1500 |
| 50M01_Saliva | chr20:57582706-57583195 | CTSZ | S_Shore | TSS1500 |
| 50M01_Saliva | chr20:58630240-58630390 | C20orf197 | | TSS1500 |
| 50M01_Saliva | chr20:61371425-61371808 | NTSR1 | | Body |
| 50M01_Saliva | chr20:61590751-61591209 | SLC17A9 | N_Shore | Body |
| 50M01_Saliva | chr20:62318426-62318588 | RTEL1;RTEL1 | Island | Body;Body |
| 50M01_Saliva | chr20:62492074-62492233 | C20orf135 | N_Shore | TSS1500 |
| 50M01_Saliva | chr21:34185927-34186122 | C21orf62;C21orf62;C21orf62;C21orf62;C21orf62;C21orf62 | | 5'UTR;5'UTR;1stExon;1stExon;1stExon;5'UTR |
| 50M01_Saliva | chr21:35884348-35884508 | KCNE1 | | TSS1500 |
| 50M01_Saliva | chr21:36421467-36421955 | RUNX1;RUNX1 | | 5'UTR;1stExon |
| 50M01_Saliva | chr21:38120350-38120518 | SIM2 | Island | Body |
| 50M01_Saliva | chr21:46898048-46898789 | COL18A1;COL18A1;COL18A1 | Island | Body;Body;Body |
| 50M01_Saliva | chr22:17956453-17956641 | CECR2 | | TSS200 |
| 50M01_Saliva | chr22:19710880-19711007 | GP1BB;SEPT5 | Island | TSS200;3'UTR |
| 50M01_Saliva | chr22:24105087-24105186 | C22orf15 | | TSS200 |
| 50M01_Saliva | chr22:24348549-24348715 | GSTTP1 | | TSS1500 |
| 50M01_Saliva | chr22:24823455-24823554 | ADORA2A | S_Shelf | TSS200 |
| 50M01_Saliva | chr22:27013978-27014116 | CRYBB1;CRYBB1 | | 5'UTR;1stExon |
| 50M01_Saliva | chr22:30661949-30663881 | OSM | | Body |
| 50M01_Saliva | chr22:31003127-31003283 | TCN2 | | TSS200 |
| 50M01_Saliva | chr22:36563027-36563117 | APOL3;APOL3 | | TSS1500;TSS1500 |
| 50M01_Saliva | chr22:37545924-37546124 | IL2RB;IL2RB | | 5'UTR;1stExon |
| 50M01_Saliva | chr22:37962569-37962818 | CDC42EP1 | Island | Body |
| 50M01_Saliva | chr22:37975975-37976179 | LGALS2;LGALS2 | | 1stExon;5'UTR |
| 50M01_Saliva | chr22:42828386-42828516 | NFAM1;NFAM1 | | 1stExon;5'UTR |
| 50M01_Saliva | chr22:45073148-45073165 | PRR5;PRR5;PRR5;PRR5 | | 5'UTR;5'UTR;5'UTR;5'UTR |
| 50M01_Saliva | chr22:45609402-45609421 | C22orf9;C22orf9 | | Body;TSS1500 |
| 50M01_Saliva | chr22:46770063-46770253 | CELSR1 | N_Shore | Body |
| 50M01_Saliva | chr22:47081751-47082260 | CERK | Island | 3'UTR |
| 50M01_Saliva | chr22:50482925-50483545 | TTLL8 | N_Shore | Body |
| 50M01_Saliva | chr22:50523686-50524691 | MLC1;MLC1;MLC1 | | 5'UTR;5'UTR;1stExon |
| 50M01_Saliva | chr1:855425-855649 | FLJ39609 | S_Shore | TSS1500 |
| 50M01_Saliva | chr1:871308-871980 | SAMD11 | Island | Body |
| 50M01_Saliva | chr1:1238436-1238678 | ACAP3 | N_Shelf | Body |
| 50M01_Saliva | chr1:1846001-1846340 | CALML6 | | TSS1500 |
| 50M01_Saliva | chr1:2084391-2084595 | PRKCZ;PRKCZ;PRKCZ | S_Shore | Body;Body;Body |
| 50M01_Saliva | chr1:2138757-2139175 | C1orf86 | S_Shore | 5'UTR |
| 50M01_Saliva | chr1:2230601-2230668 | SKI | N_Shore | Body |
| 50M01_Saliva | chr1:2262232-2262474 | MORN1 | N_Shelf | Body |
| 50M01_Saliva | chr1:2266138-2266591 | MORN1 | Island | Body |
| 50M01_Saliva | chr1:2284167-2284821 | MORN1;LOC100129534 | | Body;TSS200 |
| 50M01_Saliva | chr1:2304264-2304375 | MORN1 | Island | Body |
| 50M01_Saliva | chr1:2979311-2980937 | FLJ42875 | Island | Body |
| 50M01_Saliva | chr1:3028230-3028485 | PRDM16;PRDM16 | Island | Body;Body |
| 50M01_Saliva | chr1:3161448-3162173 | PRDM16;PRDM16 | N_Shelf | Body;Body |
| 50M01_Saliva | chr1:3276413-3276535 | PRDM16;PRDM16 | S_Shore | Body;Body |
| 50M01_Saliva | chr1:3606892-3607401 | TP73;TP73;TP73;TP73 | N_Shore | TSS1500;TSS1500;Body;TSS1500 |
| 50M01_Saliva | chr1:6390473-6390830 | ACOT7;ACOT7;ACOT7;ACOT7 | | Body;Body;Body;Body |
| 50M01_Saliva | chr1:6445632-6446787 | ACOT7;ACOT7 | | Body;1stExon |
| 50M01_Saliva | chr1:9130576-9130886 | SLC2A5;SLC2A5;SLC2A5 | N_Shore | TSS1500;Body;TSS1500 |
| 50M01_Saliva | chr1:10707917-10708091 | CASZ1;CASZ1 | Island | Body;Body |
| 50M01_Saliva | chr1:10757892-10757954 | CASZ1;CASZ1 | Island | 5'UTR;5'UTR |
| 50M01_Saliva | chr1:10839450-10839574 | CASZ1;CASZ1 | | 5'UTR;5'UTR |
| 50M01_Saliva | chr1:15255847-15256127 | KIAA1026;KIAA1026;KIAA1026;KIAA1026 | | Body;TSS1500;Body;Body |
| 50M01_Saliva | chr1:15271830-15272326 | KIAA1026;KIAA1026;KIAA1026;KIAA1026;KIAA1026 | | Body;5'UTR;TSS1500;Body;Body |
| 50M01_Saliva | chr1:17381800-17381980 | SDHB | S_Shore | TSS1500 |
| 50M01_Saliva | chr1:20141662-20141781 | RNF186;RNF186 | | 5'UTR;1stExon |
| 50M01_Saliva | chr1:23953480-23953778 | MDS2 | | TSS1500 |
| 50M01_Saliva | chr1:24469564-24470205 | IL22RA1 | | 1stExon |
| 50M01_Saliva | chr1:25257931-25258146 | RUNX3;RUNX3 | Island | TSS1500;Body |
| 50M01_Saliva | chr1:26145328-26145437 | FAM54B;FAM54B;FAM54B | N_Shore | TSS1500;TSS1500;TSS1500 |
| 50M01_Saliva | chr1:27189985-27190330 | SFN | Island | 1stExon |
| 50M01_Saliva | chr1:39407387-39407851 | RHBDL2;RHBDL2 | | 1stExon;5'UTR |

| | | | | |
|---|---|---|---|---|
| 50M01_Saliva | chr1:40349894-40350052 | TRIT1 | S_Shore | TSS1500 |
| 50M01_Saliva | chr1:43751270-43751563 | C1orf210;C1orf210;C1orf210 | | 1stExon;5'UTR;TSS200 |
| 50M01_Saliva | chr1:43814169-43814358 | MPL | N_Shore | Body |
| 50M01_Saliva | chr1:44726943-44727313 | ERI3 | | Body |
| 50M01_Saliva | chr1:45083079-45083278 | RNF220 | Island | Body |
| 50M01_Saliva | chr1:55089182-55089372 | ACOT11;FAM151A;FAM151A | | Body;1stExon;5'UTR |
| 50M01_Saliva | chr1:89829871-89829951 | GBP6 | | 5'UTR |
| 50M01_Saliva | chr1:100111387-100111727 | PALMD | | TSS200 |
| 50M01_Saliva | chr1:117753591-117753741 | VTCN1 | | TSS200 |
| 50M01_Saliva | chr1:120311542-120311653 | HMGCS2;HMGCS2;HMGCS2;HMGCS2 | | 1stExon;5'UTR;5'UTR;1stExon |
| 50M01_Saliva | chr1:149908085-149908524 | MTMR11;MTMR11;MTMR11 | | Body;5'UTR;1stExon |
| 50M01_Saliva | chr1:151344806-151345771 | SELENBP1 | | Body |
| 50M01_Saliva | chr1:151826123-151826533 | THEM5;THEM5 | | 1stExon;5'UTR |
| 50M01_Saliva | chr1:152730027-152730426 | KPRP | N_Shelf | TSS1500 |
| 50M01_Saliva | chr1:152973720-152973957 | SPRR3;SPRR3 | | TSS1500;TSS1500 |
| 50M01_Saliva | chr1:153413871-153413940 | S100A7L2 | | TSS1500 |
| 50M01_Saliva | chr1:154519846-154520187 | TDRD10;TDRD10 | | Body;Body |
| 50M01_Saliva | chr1:154929734-154929791 | PBXIP1;PYGO2 | | TSS1500;3'UTR |
| 50M01_Saliva | chr1:156030809-156031182 | RAB25 | | TSS200 |
| 50M01_Saliva | chr1:156123352-156123500 | SEMA4A | | TSS200 |
| 50M01_Saliva | chr1:161059429-161059654 | PVRL4 | | TSS200 |
| 50M01_Saliva | chr1:161168856-161168957 | NDUFS2;ADAMTS4 | N_Shelf | TSS1500;TSS200 |
| 50M01_Saliva | chr1:178310534-178310650 | RASAL2;RASAL2 | | Body;TSS200 |
| 50M01_Saliva | chr1:201979478-201979938 | ELF3;ELF3 | | TSS1500;TSS1500 |
| 50M01_Saliva | chr1:203444518-203444945 | PRELP;PRELP | | TSS1500;TSS1500 |
| 50M01_Saliva | chr1:203829481-203830200 | SNRPE | N_Shore | TSS1500 |
| 50M01_Saliva | chr1:205326059-205326437 | KLHDC8A | | TSS200 |
| 50M01_Saliva | chr1:207224740-207225128 | YOD1 | N_Shore | TSS1500 |
| 50M01_Saliva | chr1:209799248-209799353 | LAMB3;LAMB3;LAMB3 | Island | Body;Body;Body |
| 50M01_Saliva | chr1:209825672-209825943 | LAMB3;LAMB3;LAMB3;LAMB3 | | TSS1500;5'UTR;1stExon;5'UTR |
| 50M01_Saliva | chr1:221057558-221057751 | HLX | Island | Body |
| 50M01_Saliva | chr1:228337181-228337379 | GJC2 | | TSS1500 |
| 50M01_Saliva | chr1:241912510-241912768 | WDR64 | | Body |
| 50M01_Saliva | chr1:247611842-247611931 | NLRP3;NLRP3;NLRP3;NLRP3;NLI | N_Shelf | 3'UTR;3'UTR;3'UTR;3'UTR;3'UTR |
| 50M01_Saliva | chr2:3452407-3452563 | TTC15 | Island | Body |
| 50M01_Saliva | chr2:9459318-9459596 | ASAP2;ASAP2 | Island | Body;Body |
| 50M01_Saliva | chr2:10302391-10302462 | C2orf48 | | Body |
| 50M01_Saliva | chr2:11679605-11679872 | GREB1;GREB1 | | TSS1500;5'UTR |
| 50M01_Saliva | chr2:17996994-17997660 | MSGN1 | | TSS1500 |
| 50M01_Saliva | chr2:24398427-24398859 | C2orf84 | Island | Body |
| 50M01_Saliva | chr2:27531244-27531535 | UCN | Island | TSS200 |
| 50M01_Saliva | chr2:29204057-29204200 | FAM179A | | TSS200 |
| 50M01_Saliva | chr2:74153086-74153317 | DGUOK;DGUOK | N_Shore | TSS1500;TSS1500 |
| 50M01_Saliva | chr2:74753281-74753358 | DQX1;DQX1 | N_Shelf | 1stExon;5'UTR |
| 50M01_Saliva | chr2:97171137-97171243 | NEURL3 | N_Shelf | Body |
| 50M01_Saliva | chr2:109934728-109934860 | SH3RF3 | | Body |
| 50M01_Saliva | chr2:113735377-113735503 | IL1F9 | | TSS1500 |
| 50M01_Saliva | chr2:113875226-113875723 | IL1RN;IL1RN;IL1RN | | TSS1500;TSS1500;TSS1500 |
| 50M01_Saliva | chr2:128458240-128458399 | SFT2D3 | N_Shore | TSS1500 |
| 50M01_Saliva | chr2:134326204-134326366 | NCKAP5;NCKAP5 | | TSS200;TSS200 |
| 50M01_Saliva | chr2:145146928-145147350 | ZEB2;ZEB2 | N_Shore | 3'UTR;3'UTR |
| 50M01_Saliva | chr2:161056520-161056639 | ITGB6 | | 1stExon |
| 50M01_Saliva | chr2:219472218-219472607 | PLCD4 | | TSS1500 |
| 50M01_Saliva | chr2:223916502-223916687 | KCNE4 | N_Shore | TSS1500 |
| 50M01_Saliva | chr2:231790529-231790813 | GPR55 | | TSS1500 |
| 50M01_Saliva | chr2:236443439-236444284 | AGAP1;AGAP1 | N_Shore | Body;Body |
| 50M01_Saliva | chr2:238499464-238499793 | RAB17;RAB17 | | 1stExon;5'UTR |
| 50M01_Saliva | chr2:238578376-238578491 | LRRFIP1 | | Body |
| 50M01_Saliva | chr2:240020028-240020174 | HDAC4 | | Body |
| 50M01_Saliva | chr2:240212763-240213173 | HDAC4 | | Body |
| 50M01_Saliva | chr2:242127690-242127948 | ANO7;ANO7 | | TSS1500;TSS1500 |
| 50M01_Saliva | chr2:242715019-242715041 | GAL3ST2 | S_Shelf | TSS1500 |
| 50M01_Saliva | chr2:242802009-242802192 | PDCD1 | N_Shelf | TSS1500 |
| 50M01_Saliva | chr2:242810748-242811149 | C2orf85 | S_Shelf | TSS1500 |
| 50M01_Saliva | chr3:14443428-14443473 | SLC6A6;SLC6A6;SLC6A6 | N_Shore | TSS1500;TSS1500;TSS1500 |
| 50M01_Saliva | chr3:32023475-32023996 | ZNF860;OSBPL10 | S_Shore | 5'UTR;TSS1500 |

| | | | | |
|---|---|---|---|---|
| 50M01_Saliva | chr3:50341009-50341293 | HYAL1;HYAL1;HYAL1;HYAL1;HYAL1;HYAL1 | | TSS200;TSS200;5'UTR;TSS200;TSS200;TSS200 |
| 50M01_Saliva | chr3:52552868-52553433 | STAB1 | Island | Body |
| 50M01_Saliva | chr3:52569026-52569169 | NT5DC2;NT5DC2;NT5DC2 | N_Shore | 5'UTR;1stExon;TSS1500 |
| 50M01_Saliva | chr3:57743377-57743599 | SLMAP;SLMAP | S_Shore | 5'UTR;1stExon |
| 50M01_Saliva | chr3:64211328-64211659 | PRICKLE2 | | TSS200 |
| 50M01_Saliva | chr3:69061356-69061827 | C3orf64 | N_Shore | 5'UTR |
| 50M01_Saliva | chr3:69915189-69915449 | MITF;MITF;MITF | | Body;Body;TSS1500 |
| 50M01_Saliva | chr3:123339334-123339568 | MYLK;MYLK;MYLK;MYLK;MYLK;MYLK;MYLK;MYLK | | 1stExon;5'UTR;Body;Body;Body;5'UTR;1stExon;Body |
| 50M01_Saliva | chr3:149374535-149374763 | WWTR1;WWTR1;WWTR1 | N_Shore | Body;Body;Body |
| 50M01_Saliva | chr3:193096486-193096836 | ATP13A5 | | 1stExon |
| 50M01_Saliva | chr3:194833887-194834008 | C3orf21 | | Body |
| 50M01_Saliva | chr3:195943345-195943579 | OSTalpha | | TSS200 |
| 50M01_Saliva | chr3:196065106-196065318 | TM4SF19 | | 5'UTR |
| 50M01_Saliva | chr3:196065506-196065688 | TM4SF19 | | TSS1500 |
| 50M01_Saliva | chr4:671543-671851 | MYL5 | N_Shelf | TSS200 |
| 50M01_Saliva | chr4:1206004-1206150 | CTBP1;CTBP1 | Island | 3'UTR;3'UTR |
| 50M01_Saliva | chr4:1208699-1208960 | CTBP1;CTBP1 | N_Shore | Body;Body |
| 50M01_Saliva | chr4:1225507-1225729 | CTBP1;CTBP1 | S_Shore | Body;Body |
| 50M01_Saliva | chr4:3387499-3387765 | RGS12;RGS12;RGS12 | | Body;Body;Body |
| 50M01_Saliva | chr4:3409828-3409927 | RGS12;RGS12;RGS12 | | Body;Body;Body |
| 50M01_Saliva | chr4:5887725-5888193 | CRMP1;CRMP1 | N_Shore | Body;Body |
| 50M01_Saliva | chr4:8395941-8396502 | ACOX3;ACOX3 | | Body;Body |
| 50M01_Saliva | chr4:24796803-24797007 | SOD3 | | TSS1500 |
| 50M01_Saliva | chr4:36283184-36283275 | DTHD1 | | TSS200 |
| 50M01_Saliva | chr4:40517938-40518065 | RBM47;RBM47;RBM47 | | 1stExon;5'UTR;5'UTR |
| 50M01_Saliva | chr4:42122106-42122161 | BEND4;BEND4 | | Body;Body |
| 50M01_Saliva | chr4:77355817-77356180 | SHROOM3 | | TSS1500 |
| 50M01_Saliva | chr4:87770357-87770782 | SLC10A6;SLC10A6 | | 5'UTR;1stExon |
| 50M01_Saliva | chr4:114214080-114214185 | ANK2;ANK2;ANK2 | | Body;Body;Body |
| 50M01_Saliva | chr4:145566200-145566595 | HHIP | N_Shore | TSS1500 |
| 50M01_Saliva | chr4:154125439-154125594 | TRIM2;TRIM2 | | TSS200;5'UTR |
| 50M01_Saliva | chr4:184908351-184909018 | STOX2 | | Body |
| 50M01_Saliva | chr4:186877497-186877657 | SORBS2;SORBS2 | | 5'UTR;1stExon |
| 50M01_Saliva | chr5:510035-510421 | SLC9A3 | S_Shore | Body |
| 50M01_Saliva | chr5:912383-913111 | TRIP13 | N_Shore | Body |
| 50M01_Saliva | chr5:1016460-1016768 | NKD2 | | Body |
| 50M01_Saliva | chr5:1335846-1335965 | CLPTM1L | S_Shore | Body |
| 50M01_Saliva | chr5:27038707-27038836 | CDH9 | | TSS200 |
| 50M01_Saliva | chr5:53751272-53751493 | HSPB3 | | TSS200 |
| 50M01_Saliva | chr5:55529124-55529228 | ANKRD55;ANKRD55 | | 5'UTR;1stExon |
| 50M01_Saliva | chr5:88018481-88018605 | MEF2C;MEF2C | | Body;Body |
| 50M01_Saliva | chr5:118690955-118691033 | TNFAIP8;TNFAIP8 | N_Shore | Body;TSS1500 |
| 50M01_Saliva | chr5:125800764-125800916 | GRAMD3;GRAMD3;GRAMD3;GRAMD3;GRAMD3 | | Body;5'UTR;TSS200;Body;Body |
| 50M01_Saliva | chr5:125929132-125929671 | ALDH7A1 | N_Shore | Body |
| 50M01_Saliva | chr5:126984564-126984994 | CTXN3 | | TSS200 |
| 50M01_Saliva | chr5:136983354-136984054 | MIR874;KLHL3 | | TSS200;Body |
| 50M01_Saliva | chr5:138211043-138211155 | CTNNA1;LRRTM2;LRRTM2 | | Body;5'UTR;1stExon |
| 50M01_Saliva | chr5:142065539-142065676 | FGF1;FGF1;FGF1;FGF1;FGF1;FGF1;FGF1;FGF1 | | 5'UTR;Body;Body;5'UTR;1stExon;5'UTR;5'UTR;1stExon |
| 50M01_Saliva | chr5:142065819-142066938 | FGF1;FGF1;FGF1;FGF1;FGF1;FGF1;FGF1 | | 5'UTR;5'UTR;5'UTR;1stExon;Body;Body;TSS200 |
| 50M01_Saliva | chr5:149546331-149546366 | CDX1 | Island | TSS200 |
| 50M01_Saliva | chr5:179634671-179634789 | RASGEF1C | Island | 5'UTR |
| 50M01_Saliva | chr5:180325579-180325969 | BTNL8;BTNL8;BTNL8;BTNL8;BTNL8 | | TSS1500;TSS1500;TSS1500;TSS1500;TSS1500 |
| 50M01_Saliva | chr6:6614447-6614798 | LOC285780;LY86 | | Body;Body |
| 50M01_Saliva | chr6:8436218-8436296 | SLC35B3;SLC35B3;SLC35B3 | S_Shore | TSS1500;TSS1500;TSS1500 |
| 50M01_Saliva | chr6:10555808-10556326 | GCNT2;GCNT2 | | TSS200;Body |
| 50M01_Saliva | chr6:12718370-12718530 | PHACTR1 | | 5'UTR |
| 50M01_Saliva | chr6:24877462-24877624 | FAM65B;FAM65B;FAM65B | | 5'UTR;1stExon;5'UTR |
| 50M01_Saliva | chr6:29629716-29629988 | MOG;MOG;MOG;MOG;MOG;MOG;MOG;MOG;MOG;MOG;MOG | | Body;Body;Body;Body;Body;Body;Body;Body;Body;Body;Body |
| 50M01_Saliva | chr6:29855636-29855914 | HLA-H | Island | Body |
| 50M01_Saliva | chr6:29894619-29894679 | HCG4P6 | Island | TSS1500 |

| | | | | |
|---|---|---|---|---|
| 50M01_Saliva | chr6:30122388-30122593 | TRIM10;TRIM10 | | Body;Body |
| 50M01_Saliva | chr6:30174651-30174845 | TRIM26 | | 5'UTR |
| 50M01_Saliva | chr6:30175299-30175327 | TRIM26 | | 5'UTR |
| 50M01_Saliva | chr6:30850405-30850581 | DDR1 | N_Shore | TSS1500 |
| 50M01_Saliva | chr6:30850829-30851417 | DDR1;DDR1 | N_Shore | TSS1500;TSS1500 |
| 50M01_Saliva | chr6:30853333-30853959 | DDR1;DDR1 | S_Shore | 5'UTR;5'UTR |
| 50M01_Saliva | chr6:30881316-30881562 | VARS2;VARS2;VARS2;GTF2H4 | N_Shore | TSS1500;TSS1500;TSS1500;Body |
| 50M01_Saliva | chr6:30951225-30951394 | MUC21 | | TSS1500 |
| 50M01_Saliva | chr6:31023257-31023770 | HCG22 | | Body |
| 50M01_Saliva | chr6:31025949-31026287 | HCG22 | | Body |
| 50M01_Saliva | chr6:31096861-31097078 | PSORS1C1 | | 5'UTR |
| 50M01_Saliva | chr6:31138925-31138982 | POU5F1 | | TSS1500 |
| 50M01_Saliva | chr6:31146845-31146942 | PSORS1C3 | N_Shore | TSS1500 |
| 50M01_Saliva | chr6:31636239-31636913 | CSNK2B | S_Shelf | Body |
| 50M01_Saliva | chr6:31688080-31688229 | LY6G6C | N_Shelf | Body |
| 50M01_Saliva | chr6:31689610-31690170 | LY6G6C | N_Shore | TSS200 |
| 50M01_Saliva | chr6:31696223-31696240 | DDAH2 | Island | Body |
| 50M01_Saliva | chr6:31913250-31913323 | C2;C2;CFB | | 3'UTR;3'UTR;TSS1500 |
| 50M01_Saliva | chr6:31913809-31914068 | CFB;CFB | | 1stExon;5'UTR |
| 50M01_Saliva | chr6:32060681-32061084 | TNXB | N_Shelf | Body |
| 50M01_Saliva | chr6:32063838-32063895 | TNXB | Island | Body |
| 50M01_Saliva | chr6:32120625-32120878 | PPT2;PRRT1;PPT2 | N_Shore | TSS1500;TSS1500;TSS1500 |
| 50M01_Saliva | chr6:32120899-32120955 | PPT2;PRRT1;PPT2 | N_Shore | TSS1500;TSS1500;TSS1500 |
| 50M01_Saliva | chr6:32121113-32121156 | PPT2;PPT2;PRRT1 | N_Shore | TSS200;TSS1500;TSS1500 |
| 50M01_Saliva | chr6:32165889-32165917 | NOTCH4 | S_Shore | Body |
| 50M01_Saliva | chr6:32191210-32191496 | NOTCH4 | | Body |
| 50M01_Saliva | chr6:32526263-32526414 | HLA-DRB6 | | Body |
| 50M01_Saliva | chr6:32552016-32552106 | HLA-DRB1 | Island | Body |
| 50M01_Saliva | chr6:32552350-32552453 | HLA-DRB1 | S_Shore | Body |
| 50M01_Saliva | chr6:32784327-32785141 | HLA-DOB | | Body |
| 50M01_Saliva | chr6:33048529-33048558 | HLA-DPB1 | Island | Body |
| 50M01_Saliva | chr6:33048732-33048760 | HLA-DPB1 | Island | Body |
| 50M01_Saliva | chr6:33131222-33131560 | COL11A2;COL11A2;COL11A2 | S_Shore | 3'UTR;3'UTR;3'UTR |
| 50M01_Saliva | chr6:33141279-33141305 | COL11A2;COL11A2;COL11A2 | | Body;Body;Body |
| 50M01_Saliva | chr6:33173482-33173501 | HSD17B8 | S_Shore | Body |
| 50M01_Saliva | chr6:33226913-33227015 | VPS52 | | Body |
| 50M01_Saliva | chr6:33228341-33228498 | VPS52 | | Body |
| 50M01_Saliva | chr6:33264534-33264844 | RGL2;RGL2 | N_Shore | Body;Body |
| 50M01_Saliva | chr6:33638313-33638550 | ITPR3 | | Body |
| 50M01_Saliva | chr6:35285967-35286360 | DEF6 | | Body |
| 50M01_Saliva | chr6:37460940-37461296 | C6orf129 | | Body |
| 50M01_Saliva | chr6:43251710-43251928 | TTBK1 | Island | Body |
| 50M01_Saliva | chr6:43478500-43478829 | C6orf154 | Island | TSS200 |
| 50M01_Saliva | chr6:56878397-56878665 | BEND6 | | Body |
| 50M01_Saliva | chr6:72926121-72926485 | RIMS1;RIMS1;RIMS1;RIMS1;RIMS1 | | TSS1500;TSS200;Body;Body;Body |
| 50M01_Saliva | chr6:106546704-106546824 | PRDM1;PRDM1 | | Body;TSS200 |
| 50M01_Saliva | chr6:112375206-112375333 | WISP3;WISP3 | | TSS200;TSS200 |
| 50M01_Saliva | chr6:116381609-116382179 | FRK;FRK | | 1stExon;5'UTR |
| 50M01_Saliva | chr6:137366322-137366492 | IL20RA | S_Shore | TSS200 |
| 50M01_Saliva | chr6:152702387-152702660 | SYNE1;SYNE1 | | Body;Body |
| 50M01_Saliva | chr6:155568918-155569443 | TIAM2;TIAM2 | | Body;Body |
| 50M01_Saliva | chr6:157746584-157746724 | C6orf35 | S_Shore | TSS1500 |
| 50M01_Saliva | chr6:159052400-159052937 | TMEM181 | | Body |
| 50M01_Saliva | chr6:168417574-168417743 | KIF25;KIF25 | | TSS1500;TSS1500 |
| 50M01_Saliva | chr6:168857542-168858319 | SMOC2;SMOC2 | | Body;Body |
| 50M01_Saliva | chr7:872208-872797 | UNC84A;UNC84A | | 5'UTR;Body |
| 50M01_Saliva | chr7:906726-906799 | UNC84A;UNC84A | S_Shore | Body;Body |
| 50M01_Saliva | chr7:1022213-1022797 | CYP2W1 | N_Shore | TSS1500 |
| 50M01_Saliva | chr7:1102344-1103262 | C7orf50;C7orf50;C7orf50 | N_Shelf | Body;Body;Body |
| 50M01_Saliva | chr7:1491831-1491921 | MICALL2 | Island | Body |
| 50M01_Saliva | chr7:2094628-2094977 | MAD1L1;MAD1L1;MAD1L1 | | Body;Body;Body |
| 50M01_Saliva | chr7:2208492-2208830 | MAD1L1;MAD1L1;MAD1L1 | | Body;Body;Body |
| 50M01_Saliva | chr7:2611890-2612101 | IQCE;IQCE | | Body;Body |
| 50M01_Saliva | chr7:3033173-3033939 | CARD11 | | 5'UTR |
| 50M01_Saliva | chr7:4230885-4231054 | SDK1;SDK1 | | Body;Body |
| 50M01_Saliva | chr7:4243286-4243526 | SDK1;SDK1 | | Body;Body |

| | | | | |
|---|---|---|---|---|
| 50M01_Saliva | chr7:4801993-4802351 | FOXK1 | Island | Body |
| 50M01_Saliva | chr7:5534951-5535302 | FBXL18 | S_Shore | Body |
| 50M01_Saliva | chr7:6199980-6200140 | USP42 | S_Shore | 3'UTR |
| 50M01_Saliva | chr7:27140942-27141088 | HOXA2 | N_Shelf | Body |
| 50M01_Saliva | chr7:27143252-27143334 | HOXA2 | Island | TSS1500 |
| 50M01_Saliva | chr7:27153605-27154264 | HOXA3;HOXA3;HOXA3;HOXA3 | Island | 5'UTR;1stExon;5'UTR;5'UTR |
| 50M01_Saliva | chr7:27154720-27155039 | HOXA3;HOXA3;HOXA3 | N_Shore | 5'UTR;TSS1500;5'UTR |
| 50M01_Saliva | chr7:27168962-27169674 | HOXA4 | N_Shore | Body |
| 50M01_Saliva | chr7:27171391-27171528 | HOXA4 | S_Shore | TSS1500 |
| 50M01_Saliva | chr7:27183133-27183643 | HOXA5 | Island | 1stExon |
| 50M01_Saliva | chr7:27183973-27184030 | HOXA5 | Island | TSS1500 |
| 50M01_Saliva | chr7:27184159-27184316 | HOXA5 | Island | TSS1500 |
| 50M01_Saliva | chr7:48129814-48129992 | UPP1;UPP1 | Island | 5'UTR;5'UTR |
| 50M01_Saliva | chr7:73256360-73256416 | WBSCR27 | N_Shore | Body |
| 50M01_Saliva | chr7:90896329-90896701 | FZD1;FZD1 | Island | 1stExon;3'UTR |
| 50M01_Saliva | chr7:98738998-98739782 | SMURF1;SMURF1 | N_Shelf | Body;Body |
| 50M01_Saliva | chr7:99612634-99612773 | ZKSCAN1 | N_Shore | TSS1500 |
| 50M01_Saliva | chr7:101936318-101936527 | SH2B2 | Island | Body |
| 50M01_Saliva | chr7:102574105-102574504 | LRRC17;FBXL13;LRRC17;FBXL13 | | 5'UTR;Body;5'UTR;Body |
| 50M01_Saliva | chr7:116660172-116660240 | ST7;ST7 | | Body;Body |
| 50M01_Saliva | chr7:123484898-123485052 | HYAL4 | | TSS1500 |
| 50M01_Saliva | chr7:130020011-130020130 | CPA1 | N_Shore | TSS1500 |
| 50M01_Saliva | chr7:134575187-134575811 | CALD1;CALD1;CALD1;CALD1;CALD1 | | TSS1500;Body;TSS1500;Body;Body |
| 50M01_Saliva | chr7:135433540-135433790 | FAM180A;FAM180A | | 1stExon;5'UTR |
| 50M01_Saliva | chr7:142583296-142583731 | TRPV6;TRPV6 | | 1stExon;5'UTR |
| 50M01_Saliva | chr7:143747422-143747596 | OR2A5 | | TSS200 |
| 50M01_Saliva | chr7:149471884-149472134 | SSPO | S_Shore | TSS1500 |
| 50M01_Saliva | chr7:150021488-150021553 | ACTR3C;LRRC61;LRRC61 | S_Shore | TSS1500;5'UTR;5'UTR |
| 50M01_Saliva | chr7:150778818-150779047 | FASTK;TMUB1;TMUB1;FASTK | Island | TSS1500;Body;Body;TSS1500 |
| 50M01_Saliva | chr7:151493629-151493820 | PRKAG2;PRKAG2 | | 5'UTR;Body |
| 50M01_Saliva | chr7:157624788-157624958 | PTPRN2;PTPRN2;PTPRN2 | | Body;Body;Body |
| 50M01_Saliva | chr7:157701024-157701262 | PTPRN2;PTPRN2;PTPRN2 | N_Shore | Body;Body;Body |
| 50M01_Saliva | chr7:158059704-158060003 | PTPRN2;PTPRN2;PTPRN2 | N_Shore | Body;Body;Body |
| 50M01_Saliva | chr8:1906312-1906498 | ARHGEF10 | | 3'UTR |
| 50M01_Saliva | chr8:19170080-19170432 | SH2D4A | N_Shore | TSS1500 |
| 50M01_Saliva | chr8:19459996-19460136 | CSGALNACT1;CSGALNACT1;CS( | Island | 1stExon;5'UTR;Body;5'UTR |
| 50M01_Saliva | chr8:22084861-22085227 | PHYHIP;PHYHIP | | Body;Body |
| 50M01_Saliva | chr8:22844374-22844827 | RHOBTB2 | | TSS1500 |
| 50M01_Saliva | chr8:57233269-57233391 | SDR16C5 | S_Shore | TSS200 |
| 50M01_Saliva | chr8:123875036-123875223 | ZHX2 | | 5'UTR |
| 50M01_Saliva | chr8:134114892-134115431 | TG;SLA;SLA;SLA;SLA | | Body;1stExon;1stExon;5'UTR;5'UTR |
| 50M01_Saliva | chr8:141588118-141588242 | EIF2C2;EIF2C2 | | Body;Body |
| 50M01_Saliva | chr8:143386260-143387265 | TSNARE1 | | Body |
| 50M01_Saliva | chr8:143463227-143463667 | TSNARE1 | | 5'UTR |
| 50M01_Saliva | chr8:143761503-143761585 | PSCA | | TSS1500 |
| 50M01_Saliva | chr8:143851287-143851427 | LYNX1;LYNX1 | | TSS200;Body |
| 50M01_Saliva | chr8:143868110-143868917 | LY6D | | TSS200 |
| 50M01_Saliva | chr8:144589984-144590068 | ZC3H3 | | Body |
| 50M01_Saliva | chr8:144800997-144801223 | MAPK15 | Island | Body |
| 50M01_Saliva | chr8:144894951-144895243 | SCRIB;SCRIB | Island | Body;Body |
| 50M01_Saliva | chr8:145728138-145728490 | GPT | S_Shore | TSS1500 |
| 50M01_Saliva | chr8:145728507-145728543 | GPT | N_Shore | TSS1500 |
| 50M01_Saliva | chr9:92291210-92291523 | LOC100129066 | N_Shore | Body |
| 50M01_Saliva | chr9:97848107-97848457 | C9orf3;MIR24-1 | N_Shore | 3'UTR;TSS200 |
| 50M01_Saliva | chr10:384286-384652 | DIP2C | N_Shelf | Body |
| 50M01_Saliva | chr10:530805-531152 | DIP2C | Island | Body |
| 50M01_Saliva | chr10:531465-533102 | DIP2C | S_Shore | Body |
| 50M01_Saliva | chr10:556470-556762 | DIP2C | | Body |
| 50M01_Saliva | chr10:1507708-1508061 | ADARB2 | S_Shore | Body |
| 50M01_Saliva | chr10:1547717-1547983 | ADARB2 | | Body |
| 50M01_Saliva | chr10:3149673-3149852 | PFKP | S_Shore | Body |
| 50M01_Saliva | chr10:3169037-3169516 | PFKP | | Body |
| 50M01_Saliva | chr10:5566543-5566849 | CALML3 | N_Shore | TSS1500 |
| 50M01_Saliva | chr10:13388442-13388567 | SEPHS1 | | 5'UTR |
| 50M01_Saliva | chr10:18629497-18629677 | CACNB2;CACNB2;CACNB2;CACNB2;CACNB2;CACNB2;CACNB2;CAC( | | Body;TSS200;Body;Body;Body;Body;Body;Bo dy |

| | | | | |
|---|---|---|---|---|
| 50M01_Saliva | chr10:26855907-26856454 | APBB1IP | Island | Body |
| 50M01_Saliva | chr10:32216031-32216311 | ARHGAP12 | N_Shore | 5'UTR |
| 50M01_Saliva | chr10:45406295-45406847 | TMEM72 | | TSS1500 |
| 50M01_Saliva | chr10:47746733-47747039 | ANXA8L2 | | TSS1500 |
| 50M01_Saliva | chr10:49459800-49460249 | FRMPD2;FRMPD2 | | Body;Body |
| 50M01_Saliva | chr10:49674348-49674883 | ARHGAP22 | Island | Body |
| 50M01_Saliva | chr10:50506875-50507092 | C10orf71;C10orf71 | | TSS1500;TSS1500 |
| 50M01_Saliva | chr10:63422511-63422555 | C10orf107 | N_Shore | TSS1500 |
| 50M01_Saliva | chr10:63809073-63809170 | ARID5B | | Body |
| 50M01_Saliva | chr10:75631588-75632066 | CAMK2G;CAMK2G;CAMK2G;CAMK2G;CAMK2G | | Body;Body;Body;Body;Body |
| 50M01_Saliva | chr10:77871958-77872170 | C10orf11 | N_Shore | Body |
| 50M01_Saliva | chr10:99331332-99331808 | ANKRD2;ANKRD2 | S_Shore | 3'UTR |
| 50M01_Saliva | chr10:99894013-99894087 | C10orf28 | N_Shelf | |
| 50M01_Saliva | chr10:105238964-105239205 | CALHM3;CALHM3 | | 5'UTR;1stExon |
| 50M01_Saliva | chr10:106093778-106093830 | ITPRIP | N_Shelf | TSS200 |
| 50M01_Saliva | chr10:123353654-123354194 | FGFR2;FGFR2;FGFR2;FGFR2;FGI | N_Shelf | TSS200;TSS200;5'UTR;5'UTR;1stExon;5'UTR;5'UTR;5'UTR;5'UTR;5'UTR |
| 50M01_Saliva | chr10:126278272-126278502 | LHPP;LHPP | N_Shelf | Body;Body |
| 50M01_Saliva | chr10:129536661-129537308 | FOXI2 | Island | 1stExon |
| 50M01_Saliva | chr10:134578261-134578481 | INPP5A | | Body |
| 50M01_Saliva | chr11:405846-406019 | SIGIRR;SIGIRR;SIGIRR | N_Shore | 3'UTR;3'UTR;3'UTR |
| 50M01_Saliva | chr11:692963-693032 | DEAF1 | N_Shore | Body |
| 50M01_Saliva | chr11:1036727-1036766 | MUC6 | Island | TSS200 |
| 50M01_Saliva | chr11:1471410-1471617 | BRSK2 | Island | Body |
| 50M01_Saliva | chr11:1660513-1660992 | HCCA2 | | Body |
| 50M01_Saliva | chr11:1769152-1769409 | HCCA2;LOC402778;LOC402778 | Island | Body;5'UTR;1stExon |
| 50M01_Saliva | chr11:1855286-1855561 | SYT8 | | TSS1500 |
| 50M01_Saliva | chr11:1855660-1855851 | SYT8 | | TSS200 |
| 50M01_Saliva | chr11:1892139-1892180 | LSP1;LSP1;LSP1;LSP1;LSP1 | Island | 5'UTR;TSS200;5'UTR;Body;1stExon |
| 50M01_Saliva | chr11:1945564-1945918 | TNNT3;TNNT3;TNNT3;TNNT3 | N_Shore | Body;Body;Body;Body |
| 50M01_Saliva | chr11:2828364-2828778 | KCNQ1;KCNQ1 | N_Shore | Body;Body |
| 50M01_Saliva | chr11:2925473-2925869 | SLC22A18;SLC22A18;SLC22A18 | S_Shore | Body;Body;TSS1500 |
| 50M01_Saliva | chr11:3187750-3187803 | OSBPL5;OSBPL5;OSBPL5 | S_Shore | TSS1500;TSS1500;TSS1500 |
| 50M01_Saliva | chr11:8832938-8833095 | ST5;ST5;ST5 | | TSS1500;TSS200;5'UTR |
| 50M01_Saliva | chr11:17803160-17803421 | KCNC1 | S_Shore | Body |
| 50M01_Saliva | chr11:20043971-20044639 | NAV2;NAV2;NAV2;NAV2 | | TSS1500;Body;Body;Body |
| 50M01_Saliva | chr11:32421514-32421845 | WT1;WT1;WT1;WT1 | | Body;Body;Body;Body |
| 50M01_Saliva | chr11:34535377-34535416 | ELF5 | | TSS200 |
| 50M01_Saliva | chr11:45928854-45928988 | C11orf94 | | TSS200 |
| 50M01_Saliva | chr11:57089366-57089744 | TNKS1BP1 | N_Shelf | 5'UTR |
| 50M01_Saliva | chr11:60775715-60776124 | CD6 | Island | Body |
| 50M01_Saliva | chr11:63990575-63990629 | FERMT3;FERMT3 | N_Shelf | Body;Body |
| 50M01_Saliva | chr11:64951536-64952034 | CAPN1 | N_Shore | Body |
| 50M01_Saliva | chr11:65560174-65560763 | OVOL1 | | Body |
| 50M01_Saliva | chr11:66104468-66104485 | RIN1 | S_Shelf | TSS1500 |
| 50M01_Saliva | chr11:66454595-66454651 | SPTBN2 | | Body |
| 50M01_Saliva | chr11:66631356-66631536 | PC;PC;PC | Island | Body;Body;Body |
| 50M01_Saliva | chr11:66650500-66650560 | PC;PC;PC | | 5'UTR;5'UTR;5'UTR |
| 50M01_Saliva | chr11:67442195-67442408 | ALDH3B2;ALDH3B2 | | 5'UTR;TSS200 |
| 50M01_Saliva | chr11:69970464-69970618 | ANO1;ANO1 | | Body;Body |
| 50M01_Saliva | chr11:71710614-71711021 | IL18BP;IL18BP;IL18BP;IL18BP;IL18BP;IL18BP | | TSS200;5'UTR;TSS1500;5'UTR;5'UTR;1stExon |
| 50M01_Saliva | chr11:73053649-73054121 | ARHGEF17 | Island | Body |
| 50M01_Saliva | chr11:76571458-76571534 | ACER3 | N_Shore | TSS1500 |
| 50M01_Saliva | chr11:76998821-76998857 | GDPD4 | | TSS1500 |
| 50M01_Saliva | chr11:83393062-83393677 | DLG2;DLG2;DLG2;DLG2 | | 5'UTR;Body;Body;Body |
| 50M01_Saliva | chr11:93754223-93754390 | HEPHL1 | | TSS200 |
| 50M01_Saliva | chr11:117069780-117070046 | TAGLN;TAGLN | | TSS1500;TSS1500 |
| 50M01_Saliva | chr11:117800255-117800467 | TMPRSS13 | | TSS200 |
| 50M01_Saliva | chr11:117947459-117947997 | TMPRSS4;TMPRSS4 | | TSS1500;TSS1500 |
| 50M01_Saliva | chr11:118550379-118550485 | TREH;TREH | | 1stExon;5'UTR |
| 50M01_Saliva | chr11:120008998-120009135 | TRIM29 | S_Shore | TSS200 |
| 50M01_Saliva | chr11:120973197-120973274 | TECTA | | TSS200 |
| 50M01_Saliva | chr11:121970694-121970768 | LOC399959;MIR125B1 | | Body;TSS200 |
| 50M01_Saliva | chr11:123396039-123396593 | GRAMD1B | | TSS1500 |
| 50M01_Saliva | chr11:126310323-126310475 | KIRREL3;KIRREL3 | | Body;Body |

| | | | | |
|---|---|---|---|---|
| 50M01_Saliva | chr11:128457400-128457682 | ETS1;ETS1 | | 1stExon;5'UTR |
| 50M01_Saliva | chr11:128812846-128813470 | TP53AIP1 | | TSS200 |
| 50M01_Saliva | chr11:133804743-133804959 | IGSF9B | S_Shelf | Body |
| 50M01_Saliva | chr11:134216076-134216220 | GLB1L2 | S_Shelf | Body |
| 50M01_Saliva | chr12:258355-258436 | IQSEC3;IQSEC3 | | Body;Body |
| 50M01_Saliva | chr12:6421437-6421768 | PLEKHG6;PLEKHG6;PLEKHG6 | S_Shore | TSS1500;Body;Body |
| 50M01_Saliva | chr12:6484219-6484607 | SCNN1A;SCNN1A;SCNN1A;SCNN1A | | 5'UTR;Body;1stExon;5'UTR |
| 50M01_Saliva | chr12:6486701-6487080 | SCNN1A | | TSS200 |
| 50M01_Saliva | chr12:7071814-7073242 | MIR141;MIR200C | | TSS1500;TSS1500 |
| 50M01_Saliva | chr12:7167751-7167900 | C1S;C1S | | TSS1500;TSS1500 |
| 50M01_Saliva | chr12:8974786-8975309 | A2ML1 | | TSS1500 |
| 50M01_Saliva | chr12:12223841-12224246 | BCL2L14;BCL2L14;BCL2L14 | | TSS200;TSS1500;TSS1500 |
| 50M01_Saliva | chr12:25801455-25801621 | IFLTD1 | | 1stExon |
| 50M01_Saliva | chr12:47219737-47219797 | SLC38A4;SLC38A4;SLC38A4;SLC38A4 | | 1stExon;5'UTR;1stExon;5'UTR |
| 50M01_Saliva | chr12:52585695-52585892 | KRT80;KRT80;KRT80;KRT80 | | 1stExon;1stExon;5'UTR;5'UTR |
| 50M01_Saliva | chr12:52845930-52846088 | KRT6B | | TSS200 |
| 50M01_Saliva | chr12:52867568-52867793 | KRT6C;KRT6C | | 1stExon;5'UTR |
| 50M01_Saliva | chr12:52887054-52887685 | KRT6A;KRT6A | | 1stExon;5'UTR |
| 50M01_Saliva | chr12:53242733-53242917 | KRT78;KRT78 | | 5'UTR;1stExon |
| 50M01_Saliva | chr12:53612551-53612734 | RARG;RARG | N_Shore | Body;Body |
| 50M01_Saliva | chr12:57559728-57559901 | LRP1 | | Body |
| 50M01_Saliva | chr12:58119915-58121004 | AGAP2;LOC100130776;AGAP2 | Island | 3'UTR;TSS200;3'UTR |
| 50M01_Saliva | chr12:64782908-64783110 | C12orf56;C12orf56 | N_Shore | Body;Body |
| 50M01_Saliva | chr12:68041257-68041322 | DYRK2;DYRK2 | N_Shore | TSS1500;TSS1500 |
| 50M01_Saliva | chr12:70092840-70093350 | BEST3 | | 5'UTR |
| 50M01_Saliva | chr12:93096780-93096844 | C12orf74 | | TSS200 |
| 50M01_Saliva | chr12:114337676-114337927 | RBM19;RBM19;RBM19 | | Body;Body;Body |
| 50M01_Saliva | chr12:120151527-120151706 | MIR1178;CIT | | Body;Body |
| 50M01_Saliva | chr12:120241957-120242156 | CIT | Island | Body |
| 50M01_Saliva | chr12:120242394-120242513 | CIT | S_Shore | Body |
| 50M01_Saliva | chr12:121882033-121882415 | KDM2B;KDM2B | N_Shore | Body;Body |
| 50M01_Saliva | chr12:121890163-121890328 | KDM2B;KDM2B | N_Shore | Body;Body |
| 50M01_Saliva | chr12:123750782-123750864 | CDK2AP1 | N_Shelf | Body |
| 50M01_Saliva | chr12:124773668-124774078 | FAM101A | | TSS200 |
| 50M01_Saliva | chr12:130976260-130976551 | RIMBP2 | | 5'UTR |
| 50M01_Saliva | chr12:132690643-132690782 | GALNT9;GALNT9 | S_Shore | Body;TSS200 |
| 50M01_Saliva | chr12:133101168-133101612 | FBRSL1 | N_Shore | Body |
| 50M01_Saliva | chr13:21291173-21291467 | IL17D | | Body |
| 50M01_Saliva | chr13:99135543-99136580 | STK24;STK24 | | Body;Body |
| 50M01_Saliva | chr13:99630462-99630505 | DOCK9;DOCK9;DOCK9;DOCK9 | | Body;Body;TSS200;TSS200 |
| 50M01_Saliva | chr13:110874267-110874326 | COL4A1 | | Body |
| 50M01_Saliva | chr13:111522755-111522985 | C13orf29 | | TSS200 |
| 50M01_Saliva | chr13:113676726-113677188 | MCF2L;MCF2L | N_Shore | Body;Body |
| 50M01_Saliva | chr13:113759771-113759909 | F7;F7 | N_Shore | TSS1500;TSS1500 |
| 50M01_Saliva | chr13:114103578-114104057 | ADPRHL1;ADPRHL1 | | Body;TSS200 |
| 50M01_Saliva | chr13:114843594-114844124 | RASA3 | S_Shelf | Body |
| 50M01_Saliva | chr14:23835595-23836012 | EFS;EFS | Island | TSS1500;TSS1500 |
| 50M01_Saliva | chr14:91744409-91744445 | CCDC88C | | Body |
| 50M01_Saliva | chr14:94393718-94393816 | FAM181A;C14orf86 | | Body;TSS1500 |
| 50M01_Saliva | chr14:94576977-94577101 | IFI27;IFI27 | | TSS200;TSS200 |
| 50M01_Saliva | chr14:98444151-98444513 | C14orf64 | | Body |
| 50M01_Saliva | chr14:101495758-101495847 | MIR494 | | TSS1500 |
| 50M01_Saliva | chr14:103593503-103593520 | TNFAIP2 | Island | Body |
| 50M01_Saliva | chr14:104165052-104165475 | XRCC3;XRCC3;XRCC3;KLC1;KL( | Island | 3'UTR;3'UTR;3'UTR;Body;Body |
| 50M01_Saliva | chr14:104175555-104175768 | XRCC3;XRCC3;XRCC3 | | Body;Body;Body |
| 50M01_Saliva | chr14:104195181-104195666 | ZFYVE21 | S_Shore | Body |
| 50M01_Saliva | chr15:39871808-39872186 | THBS1 | N_Shore | TSS1500 |
| 50M01_Saliva | chr15:40531871-40532180 | PAK6;PAK6;PAK6 | S_Shore | 5'UTR;5'UTR;TSS200 |
| 50M01_Saliva | chr15:42289766-42289939 | PLA2G4E | | Body |
| 50M01_Saliva | chr15:42386726-42387193 | PLA2G4D;PLA2G4D | | 1stExon;5'UTR |
| 50M01_Saliva | chr15:42447989-42448786 | PLA2G4F;PLA2G4F | | Body;Body |
| 50M01_Saliva | chr15:59908622-59909035 | GCNT3 | | 5'UTR |
| 50M01_Saliva | chr15:67417607-67417697 | SMAD3;SMAD3 | | Body;TSS1500 |
| 50M01_Saliva | chr15:69099678-69099834 | C15orf28;ANP32A | | TSS1500;Body |
| 50M01_Saliva | chr15:74659966-74660110 | CYP11A1;CYP11A1;CYP11A1 | S_Shore | 5'UTR;1stExon;TSS1500 |
| 50M01_Saliva | chr15:86313851-86313941 | MIR1276;KLHL25 | S_Shore | TSS200;5'UTR |

| | | | | |
|---|---|---|---|---|
| 50M01_Saliva | chr15:90727560-90727783 | SEMA4B | N_Shore | TSS1500 |
| 50M01_Saliva | chr16:670800-671171 | RAB40C | Island | Body |
| 50M01_Saliva | chr16:743338-743584 | FBXL16 | N_Shore | 3'UTR |
| 50M01_Saliva | chr16:832841-832963 | MSLNL | Island | Body |
| 50M01_Saliva | chr16:854168-854439 | PRR25 | N_Shelf | TSS1500 |
| 50M01_Saliva | chr16:1206750-1207075 | CACNA1H;CACNA1H | N_Shore | Body;Body |
| 50M01_Saliva | chr16:1582105-1582581 | IFT140 | N_Shore | Body |
| 50M01_Saliva | chr16:1742021-1742281 | HN1L | | Body |
| 50M01_Saliva | chr16:1813796-1814418 | MAPK8IP3;MAPK8IP3 | N_Shore | Body;Body |
| 50M01_Saliva | chr16:12210656-12211186 | SNX29 | Island | Body |
| 50M01_Saliva | chr16:22326203-22326535 | POLR3E | | Body |
| 50M01_Saliva | chr16:31147017-31147054 | PRSS8;PRSS8 | | 1stExon;5'UTR |
| 50M01_Saliva | chr16:67022472-67022719 | CES8 | | TSS200 |
| 50M01_Saliva | chr16:81564161-81564553 | CMIP;CMIP | | Body;Body |
| 50M01_Saliva | chr16:87958145-87958493 | CA5A | | Body |
| 50M01_Saliva | chr16:88832476-88832532 | FAM38A | | Body |
| 50M01_Saliva | chr16:89387418-89387805 | ANKRD11 | | 5'UTR |
| 50M01_Saliva | chr17:693137-693388 | RNMTL1 | | Body |
| 50M01_Saliva | chr17:790628-791324 | NXN | | Body |
| 50M01_Saliva | chr17:851785-851933 | NXN | S_Shore | Body |
| 50M01_Saliva | chr17:1664077-1664339 | SERPINF1 | | TSS1500 |
| 50M01_Saliva | chr17:3848156-3848324 | ATP2A3;ATP2A3;ATP2A3;ATP2A3 | Island | Body;Body;Body;Body;Body;Body;Body |
| 50M01_Saliva | chr17:4463371-4464075 | GGT6;GGT6 | | Body;Body |
| 50M01_Saliva | chr17:7493150-7494151 | SOX15;SOX15 | S_Shore | 1stExon;5'UTR |
| 50M01_Saliva | chr17:7690327-7690529 | DNAH2 | Island | Body |
| 50M01_Saliva | chr17:7792059-7792063 | CHD3;CHD3;CHD3 | S_Shelf | Body;TSS200;TSS200 |
| 50M01_Saliva | chr17:7832764-7833237 | KCNAB3 | Island | TSS200 |
| 50M01_Saliva | chr17:14213447-14214081 | HS3ST3B1 | S_Shore | Body |
| 50M01_Saliva | chr17:19247902-19247918 | MIR1180;B9D1 | Island | TSS200;Body |
| 50M01_Saliva | chr17:19313024-19313529 | RNF112 | | TSS1500 |
| 50M01_Saliva | chr17:25798741-25799018 | KSR1 | | TSS1500 |
| 50M01_Saliva | chr17:35656660-35656896 | ACACA;ACACA;ACACA;ACACA;ACACA;ACACA;ACACA | | 5'UTR;Body;5'UTR;Body;1stExon;1stExon;Body |
| 50M01_Saliva | chr17:37893764-37894014 | GRB7;GRB7 | | TSS1500;TSS1500 |
| 50M01_Saliva | chr17:38657310-38657934 | TNS4 | | 5'UTR |
| 50M01_Saliva | chr17:38859934-38860097 | KRT24 | | 1stExon |
| 50M01_Saliva | chr17:39143239-39143567 | KRT40 | | 5'UTR |
| 50M01_Saliva | chr17:39165425-39165697 | KRTAP3-1 | | TSS200 |
| 50M01_Saliva | chr17:39623858-39624088 | KRT32 | | TSS1500 |
| 50M01_Saliva | chr17:39680217-39680504 | KRT19 | N_Shelf | Body |
| 50M01_Saliva | chr17:39769064-39769213 | KRT16;KRT16 | | 1stExon;5'UTR |
| 50M01_Saliva | chr17:39943423-39943545 | JUP;JUP | S_Shore | TSS1500;TSS1500 |
| 50M01_Saliva | chr17:40274703-40274740 | HSPB9;KAT2A | Island | TSS200;TSS1500 |
| 50M01_Saliva | chr17:40705273-40705708 | HSD17B1 | Island | Body |
| 50M01_Saliva | chr17:40823930-40824361 | PLEKHH3 | S_Shore | Body |
| 50M01_Saliva | chr17:42994008-42994269 | GFAP;GFAP | | TSS1500;TSS1500 |
| 50M01_Saliva | chr17:43318045-43319382 | FMNL1 | N_Shore | Body |
| 50M01_Saliva | chr17:43716515-43716617 | C17orf69;C17orf69 | | Body;Body |
| 50M01_Saliva | chr17:46035108-46035343 | PRR15L;PRR15L | | 1stExon;5'UTR |
| 50M01_Saliva | chr17:46388390-46388465 | SKAP1;SKAP1 | | Body;Body |
| 50M01_Saliva | chr17:46607103-46608293 | HOXB1 | N_Shore | Body |
| 50M01_Saliva | chr17:46627384-46628850 | HOXB3 | N_Shore | 3'UTR |
| 50M01_Saliva | chr17:46681362-46682394 | LOC404266;LOC404266;HOXB6 | N_Shelf | Body;Body;5'UTR |
| 50M01_Saliva | chr17:48845563-48846189 | C17orf73 | | TSS1500 |
| 50M01_Saliva | chr17:53800041-53800554 | TMEM100;TMEM100;TMEM100 | | 5'UTR;1stExon;5'UTR |
| 50M01_Saliva | chr17:57918262-57918682 | MIR21 | | TSS1500 |
| 50M01_Saliva | chr17:62051709-62051773 | SCN4A | | TSS1500 |
| 50M01_Saliva | chr17:74639731-74640078 | ST6GALNAC1;ST6GALNAC1 | | 5'UTR;1stExon |
| 50M01_Saliva | chr17:75283431-75284193 | SEPT9;SEPT9 | | Body;TSS1500 |
| 50M01_Saliva | chr17:75473895-75474070 | SEPT9;SEPT9;SEPT9;SEPT9;SEPT9;SEPT9;SEPT9 | | Body;5'UTR;Body;Body;Body;Body;Body |
| 50M01_Saliva | chr17:76501381-76501582 | DNAH17 | S_Shelf | Body |
| 50M01_Saliva | chr17:76719612-76719640 | CYTH1;CYTH1 | | Body;Body |
| 50M01_Saliva | chr17:76993506-76993698 | CANT1;CANT1;CANT1 | Island | Body;Body;Body |
| 50M01_Saliva | chr17:77030221-77030546 | C1QTNF1;C1QTNF1;C1QTNF1 | | 5'UTR;TSS1500;5'UTR |
| 50M01_Saliva | chr17:77086350-77086532 | HRNBP3 | | 3'UTR |
| 50M01_Saliva | chr17:78039301-78039528 | CCDC40 | Island | Body |

| | | | | |
|---|---|---|---|---|
| 50M01_Saliva | chr17:78079608-78079668 | GAA;GAA;GAA | Island | Body;Body;Body |
| 50M01_Saliva | chr17:78161110-78161495 | CARD14;CARD14 | N_Shelf | TSS1500;Body |
| 50M01_Saliva | chr17:78166148-78166385 | CARD14;CARD14 | S_Shore | Body;Body |
| 50M01_Saliva | chr17:79056061-79056183 | BAIAP2;BAIAP2;BAIAP2;BAIAP2 | S_Shelf | Body;Body;Body;Body |
| 50M01_Saliva | chr17:79099481-79099785 | MIR657;AATK | N_Shelf | TSS1500;Body |
| 50M01_Saliva | chr17:79099882-79100218 | MIR657;AATK;MIR338 | N_Shelf | TSS1500;Body;TSS200 |
| 50M01_Saliva | chr17:79109777-79109910 | AATK | Island | Body |
| 50M01_Saliva | chr17:79251255-79251354 | SLC38A10;SLC38A10 | N_Shelf | Body;Body |
| 50M01_Saliva | chr17:79377548-79377850 | BAHCC1 | Island | Body |
| 50M01_Saliva | chr17:79609272-79609346 | TSPAN10 | | TSS200 |
| 50M01_Saliva | chr17:79652024-79652389 | ARL16;HGS | S_Shore | TSS1500;Body |
| 50M01_Saliva | chr17:79813410-79813507 | P4HB | | Body |
| 50M01_Saliva | chr17:80289541-80289701 | SECTM1 | N_Shore | 5'UTR |
| 50M01_Saliva | chr17:80796712-80798023 | TBCD;ZNF750 | | Body;5'UTR |
| 50M01_Saliva | chr17:80806135-80806435 | TBCD | | Body |
| 50M01_Saliva | chr17:80816702-80816995 | TBCD | N_Shore | Body |
| 50M01_Saliva | chr17:80847137-80847662 | TBCD | | Body |
| 50M01_Saliva | chr18:21452730-21453131 | LAMA3;LAMA3;LAMA3;LAMA3 | | TSS1500;TSS1500;Body;Body |
| 50M01_Saliva | chr18:21572622-21572748 | TTC39C | | TSS200 |
| 50M01_Saliva | chr18:29027658-29027885 | DSG3 | | TSS200 |
| 50M01_Saliva | chr18:55888640-55888832 | NEDD4L;NEDD4L;NEDD4L;NEDD4L;NEDD4L;NEDD4L;NEDD4L;NEDD4L | | 5'UTR;Body;TSS200;5'UTR;5'UTR;Body;Body;Body;5'UTR |
| 50M01_Saliva | chr18:61143926-61144339 | SERPINB5 | | TSS1500 |
| 50M01_Saliva | chr18:61253847-61254561 | SERPINB13 | | TSS1500 |
| 50M01_Saliva | chr18:77230598-77230795 | NFATC1;NFATC1;NFATC1;NFATC | Island | Body;Body;Body;Body |
| 50M01_Saliva | chr19:736164-736425 | PALM;PALM | Island | Body;Body |
| 50M01_Saliva | chr19:1424658-1424952 | DAZAP1;DAZAP1 | Island | Body;Body |
| 50M01_Saliva | chr19:2512999-2513356 | GNG7 | N_Shore | 3'UTR |
| 50M01_Saliva | chr19:2525231-2525634 | GNG7 | Island | 5'UTR |
| 50M01_Saliva | chr19:2540907-2541104 | GNG7 | Island | 5'UTR |
| 50M01_Saliva | chr19:3097565-3097728 | GNA11 | Island | Body |
| 50M01_Saliva | chr19:4305072-4305239 | FSD1 | Island | Body |
| 50M01_Saliva | chr19:4455239-4455413 | UBXN6;UBXN6;UBXN6 | N_Shelf | 5'UTR;1stExon;Body |
| 50M01_Saliva | chr19:5148023-5148409 | KDM4B | Island | Body |
| 50M01_Saliva | chr19:5250376-5251120 | PTPRS;PTPRS;PTPRS;PTPRS | N_Shore | Body;Body;Body;Body |
| 50M01_Saliva | chr19:5822318-5822514 | NRTN | Island | TSS1500 |
| 50M01_Saliva | chr19:6333587-6333761 | ACER1;ACER1 | | 5'UTR;1stExon |
| 50M01_Saliva | chr19:10231566-10231711 | EIF3G | S_Shore | TSS1500 |
| 50M01_Saliva | chr19:10407184-10407538 | ICAM5 | Island | Body |
| 50M01_Saliva | chr19:10444815-10445516 | RAVER1;ICAM3 | Island | TSS1500;Body |
| 50M01_Saliva | chr19:11472110-11472397 | LPPR2;LPPR2 | Island | Body;Body |
| 50M01_Saliva | chr19:15568574-15568935 | RASAL3 | Island | Body |
| 50M01_Saliva | chr19:16178240-16178426 | TPM4 | N_Shore | TSS200 |
| 50M01_Saliva | chr19:17942025-17942411 | JAK3 | N_Shore | Body |
| 50M01_Saliva | chr19:18385244-18385672 | KIAA1683;KIAA1683;KIAA1683;KIAA1683;KIAA1683;KIAA1683 | | 5'UTR;1stExon;5'UTR;5'UTR;1stExon;1stExon |
| 50M01_Saliva | chr19:35085474-35085677 | SCGBL;SCGBL | | 1stExon;5'UTR |
| 50M01_Saliva | chr19:36246402-36246408 | HSPB6 | Island | 3'UTR |
| 50M01_Saliva | chr19:39086733-39087186 | MAP4K1;MAP4K1 | N_Shore | Body;Body |
| 50M01_Saliva | chr19:39278997-39279703 | LGALS7B | N_Shelf | TSS1500 |
| 50M01_Saliva | chr19:45279765-45280184 | CBLC;CBLC | N_Shore | TSS1500;TSS1500 |
| 50M01_Saliva | chr19:45574917-45575134 | ZNF296 | Island | Body |
| 50M01_Saliva | chr19:46270244-46270700 | SIX5 | Island | Body |
| 50M01_Saliva | chr19:46286064-46286475 | DMPK;DMPK;DMPK | N_Shelf | TSS1500;TSS1500;TSS1500 |
| 50M01_Saliva | chr19:46544240-46544711 | IGFL4;IGFL4 | | 1stExon;5'UTR |
| 50M01_Saliva | chr19:46807119-46807466 | HIF3A;HIF3A;HIF3A | Island | 5'UTR;Body;Body |
| 50M01_Saliva | chr19:51513252-51513883 | KLK9 | | TSS1500 |
| 50M01_Saliva | chr19:51531297-51531393 | KLK11;KLK11;KLK11;KLK11 | N_Shelf | TSS1500;TSS200;TSS1500;TSS200 |
| 50M01_Saliva | chr20:2276503-2276663 | TGM3 | | TSS200 |
| 50M01_Saliva | chr20:17674195-17674250 | BANF2 | | TSS200 |
| 50M01_Saliva | chr20:30432938-30433512 | FOXS1 | Island | 1stExon |
| 50M01_Saliva | chr20:31781151-31781277 | BASE | | TSS1500 |
| 50M01_Saliva | chr20:33879989-33880214 | FAM83C | Island | 1stExon |
| 50M01_Saliva | chr20:33880337-33880825 | FAM83C | S_Shore | TSS200 |
| 50M01_Saliva | chr20:34700112-34700675 | EPB41L1 | | TSS1500 |
| 50M01_Saliva | chr20:35504371-35504511 | C20orf118 | | TSS200 |

| | | | | |
|---|---|---|---|---|
| 50M01_Saliva | chr20:35972924-35973098 | SRC | N_Shore | TSS200 |
| 50M01_Saliva | chr20:36151003-36151184 | BLCAP;BLCAP;BLCAP;NNAT;BLC | Island | 5'UTR;5'UTR;5'UTR;Body;5'UTR;5'UTR;Body |
| 50M01_Saliva | chr20:36152923-36153225 | BLCAP;BLCAP;BLCAP;BLCAP;BL | S_Shore | 5'UTR;1stExon;5'UTR;5'UTR;5'UTR;5'UTR |
| 50M01_Saliva | chr20:36153947-36154388 | BLCAP;BLCAP;BLCAP;BLCAP;BL | N_Shelf | 5'UTR;TSS1500;5'UTR;5'UTR;5'UTR |
| 50M01_Saliva | chr20:43937138-43937448 | MATN4;RBPJL;MATN4;MATN4 | S_Shore | TSS200;Body;TSS200;TSS200 |
| 50M01_Saliva | chr20:45313200-45313390 | TP53RK;SLC13A3 | | 3'UTR;TSS200 |
| 50M01_Saliva | chr20:50419193-50419231 | SALL4 | S_Shore | TSS200 |
| 50M01_Saliva | chr20:56136012-56136167 | PCK1 | | TSS200 |
| 50M01_Saliva | chr20:61002595-61002866 | C20orf151 | | TSS200 |
| 50M01_Saliva | chr20:62367884-62367893 | LIME1 | N_Shore | TSS200 |
| 50M01_Saliva | chr20:62383275-62383481 | ZBTB46 | Island | Body |
| 50M01_Saliva | chr20:62679255-62679713 | SOX18 | N_Shore | 3'UTR |
| 50M01_Saliva | chr20:62693951-62694000 | TCEA2;TCEA2 | Island | TSS200;5'UTR |
| 50M01_Saliva | chr21:35736023-35736383 | KCNE2 | | TSS1500 |
| 50M01_Saliva | chr21:38117454-38117682 | SIM2;SIM2 | N_Shelf | Body;Body |
| 50M01_Saliva | chr21:38362725-38362742 | HLCS | Island | TSS200 |
| 50M01_Saliva | chr21:39643733-39644244 | KCNJ15;KCNJ15;KCNJ15 | | 5'UTR;5'UTR;TSS1500 |
| 50M01_Saliva | chr21:43098901-43099292 | NCRNA00111 | | TSS1500 |
| 50M01_Saliva | chr21:43135854-43136287 | C21orf129;NCRNA00112 | | Body;TSS1500 |
| 50M01_Saliva | chr21:43185674-43185884 | RIPK4 | N_Shore | Body |
| 50M01_Saliva | chr21:45683146-45683513 | DNMT3L;DNMT3L | | TSS1500;TSS1500 |
| 50M01_Saliva | chr21:45789178-45789379 | TRPM2 | Island | Body |
| 50M01_Saliva | chr21:46890849-46890997 | COL18A1;COL18A1;COL18A1 | Island | Body;Body;Body |
| 50M01_Saliva | chr22:17680561-17680578 | CECR1;CECR1 | | TSS200;Body |
| 50M01_Saliva | chr22:19949965-19950166 | COMT;COMT;COMT;COMT | | 5'UTR;5'UTR;5'UTR;TSS200 |
| 50M01_Saliva | chr22:24891158-24891220 | C22orf45;C22orf45;UPB1 | Island | TSS1500;TSS1500;TSS200 |
| 50M01_Saliva | chr22:24979633-24979660 | GGT1 | | TSS200 |
| 50M01_Saliva | chr22:29706595-29706754 | GAS2L1;GAS2L1;GAS2L1 | Island | Body;Body;Body |
| 50M01_Saliva | chr22:33402393-33402998 | SYN3;SYN3;SYN3 | | 1stExon;Body;1stExon |
| 50M01_Saliva | chr22:38506623-38506781 | BAIAP2L2;BAIAP2L2 | | 1stExon;5'UTR |
| 50M01_Saliva | chr22:43820888-43821115 | MPPED1 | Island | 5'UTR |
| 50M01_Saliva | chr22:45608465-45608692 | C22orf9;C22orf9 | | Body;TSS200 |
| 50M01_Saliva | chr22:46481248-46481785 | LOC400931 | N_Shore | TSS1500 |
| 50M01_Saliva | chr22:50585197-50585386 | MOV10L1;MOV10L1;MOV10L1;MOV10L1 | | Body;TSS1500;Body;Body |
| 50M01_Saliva | chr22:50720288-50720552 | PLXNB2 | Island | Body |
| 50M01_Saliva | chr22:50721120-50721626 | PLXNB2 | Island | Body |
| 50M05_Saliva | chr1:1109192-1109676 | TTLL10 | N_Shore | TSS200 |
| 50M05_Saliva | chr1:7692321-7692367 | CAMTA1 | | Body |
| 50M05_Saliva | chr1:24239771-24240069 | CNR2;CNR2 | | 5'UTR;1stExon |
| 50M05_Saliva | chr1:26097918-26098214 | MAN1C1 | Island | Body |
| 50M05_Saliva | chr1:44703750-44704073 | ERI3 | | Body |
| 50M05_Saliva | chr1:202172778-202172912 | LGR6;LGR6 | | TSS200;Body |
| 50M05_Saliva | chr1:205181237-205181581 | DSTYK;DSTYK | S_Shore | TSS1500;TSS1500 |
| 50M05_Saliva | chr1:206945924-206946187 | IL10 | | TSS200 |
| 50M05_Saliva | chr1:247711595-247712377 | C1orf150 | | TSS1500 |
| 50M05_Saliva | chr2:24397787-24397845 | C2orf84 | Island | TSS200 |
| 50M05_Saliva | chr2:63279495-63280097 | OTX1 | N_Shore | 5'UTR |
| 50M05_Saliva | chr2:63283967-63284481 | OTX1 | Island | 3'UTR |
| 50M05_Saliva | chr2:106776775-106777038 | UXS1 | | Body |
| 50M05_Saliva | chr2:175499283-175499377 | WIPF1;WIPF1;WIPF1 | | 1stExon;5'UTR;5'UTR |
| 50M05_Saliva | chr2:225265963-225266880 | FAM124B;FAM124B | | 1stExon;1stExon |
| 50M05_Saliva | chr2:233924789-233924852 | INPP5D;INPP5D | N_Shore | TSS1500;TSS1500 |
| 50M05_Saliva | chr2:233924923-233925030 | INPP5D;INPP5D | N_Shore | TSS200;TSS200 |
| 50M05_Saliva | chr3:128779498-128779601 | GP9 | | TSS200 |
| 50M05_Saliva | chr3:181428280-181428307 | SOX2OT;SOX2 | N_Shore | Body;TSS1500 |
| 50M05_Saliva | chr3:181428377-181428466 | SOX2OT;SOX2 | N_Shore | Body;TSS1500 |
| 50M05_Saliva | chr3:181428579-181428669 | SOX2OT;SOX2 | N_Shore | Body;TSS1500 |
| 50M05_Saliva | chr3:183273245-183273894 | KLHL6 | | 1stExon |
| 50M05_Saliva | chr4:2813542-2814122 | SH3BP2;SH3BP2 | | TSS1500;5'UTR |
| 50M05_Saliva | chr4:4397942-4398276 | D4S234E;D4S234E | | Body;Body |
| 50M05_Saliva | chr4:10686394-10686610 | CLNK | | TSS200 |
| 50M05_Saliva | chr4:40198392-40198595 | RHOH | | TSS200 |
| 50M05_Saliva | chr4:183061850-183062004 | MGC45800 | N_Shore | Body |
| 50M05_Saliva | chr4:186559707-186560083 | SORBS2;SORBS2;SORBS2;SORB | N_Shore | Body;Body;Body;Body;Body;Body;Body;Body |
| 50M05_Saliva | chr5:37835109-37835168 | GDNF;GDNF;GDNF | Island | 5'UTR;TSS1500;5'UTR |

| | | | | |
|---|---|---|---|---|
| 50M05_Saliva | chr5:37835701-37835958 | GDNF;GDNF;GDNF | N_Shore | TSS1500;5'UTR;TSS200 |
| 50M05_Saliva | chr5:74162771-74162924 | FAM169A | S_Shore | TSS200 |
| 50M05_Saliva | chr5:76010444-76010658 | F2R | N_Shore | TSS1500 |
| 50M05_Saliva | chr5:92914036-92914202 | FLJ42709;FLJ42709 | N_Shore | Body;Body |
| 50M05_Saliva | chr5:137475115-137475288 | NME5;NME5 | | 1stExon;5'UTR |
| 50M05_Saliva | chr5:148810061-148810103 | MIR145;LOC728264 | | TSS200;Body |
| 50M05_Saliva | chr5:177913434-177913485 | COL23A1 | | Body |
| 50M05_Saliva | chr6:291687-292823 | DUSP22 | N_Shore | TSS1500 |
| 50M05_Saliva | chr6:4890079-4890278 | CDYL;CDYL;CDYL;CDYL | | 5'UTR;Body;Body;TSS200 |
| 50M05_Saliva | chr6:11382553-11382769 | NEDD9;NEDD9 | | 1stExon;5'UTR |
| 50M05_Saliva | chr6:28874479-28874702 | TRIM27 | | Body |
| 50M05_Saliva | chr6:30039239-30039380 | RNF39;RNF39 | Island | Body;Body |
| 50M05_Saliva | chr6:30104440-30104626 | TRIM40 | | TSS200 |
| 50M05_Saliva | chr6:31583915-31584223 | AIF1;AIF1;AIF1 | N_Shelf | Body;1stExon;Body |
| 50M05_Saliva | chr6:31690986-31691035 | C6orf25;C6orf25;C6orf25;C6orf2 | N_Shore | TSS200;TSS200;TSS200;TSS200;TSS1500;TSS200;TSS200 |
| 50M05_Saliva | chr6:32295097-32295237 | C6orf10 | | Body |
| 50M05_Saliva | chr6:32813337-32813582 | PSMB8;TAP1 | S_Shore | TSS1500;3'UTR |
| 50M05_Saliva | chr6:32904061-32904317 | HLA-DMB | | Body |
| 50M05_Saliva | chr6:33400021-33400505 | SYNGAP1 | S_Shelf | Body |
| 50M05_Saliva | chr6:112115117-112115209 | FYN | | 5'UTR |
| 50M05_Saliva | chr6:166876736-166877038 | RPS6KA2;RPS6KA2 | | Body;Body |
| 50M05_Saliva | chr7:1095416-1095720 | C7orf50;C7orf50;C7orf50 | N_Shore | Body;Body;Body |
| 50M05_Saliva | chr7:30951381-30951425 | AQP1 | | TSS200 |
| 50M05_Saliva | chr7:75443075-75443246 | CCL24 | | TSS200 |
| 50M05_Saliva | chr7:100238750-100239188 | TFR2 | | Body |
| 50M05_Saliva | chr7:106505772-106505961 | PIK3CG | N_Shelf | TSS200 |
| 50M05_Saliva | chr7:128579876-128580582 | IRF5;IRF5;IRF5;IRF5;IRF5;IR | S_Shore | 5'UTR;TSS1500;5'UTR;5'UTR;5'UTR;5'UTR;5'UTR |
| 50M05_Saliva | chr7:139333296-139333410 | HIPK2;HIPK2 | | Body;Body |
| 50M05_Saliva | chr7:150434452-150434480 | GIMAP5;GIMAP5 | | 1stExon;5'UTR |
| 50M05_Saliva | chr8:27469186-27469338 | CLU;CLU;CLU;CLU | N_Shelf | 1stExon;Body;5'UTR;TSS1500 |
| 50M05_Saliva | chr8:67975874-67975895 | CSPP1;CSPP1;COPS5 | N_Shore | TSS1500;TSS1500;TSS1500 |
| 50M05_Saliva | chr8:142159992-142160235 | DENND3 | | Body |
| 50M05_Saliva | chr9:123688715-123689193 | TRAF1 | N_Shelf | 5'UTR |
| 50M05_Saliva | chr10:49892930-49893336 | WDFY4 | | TSS1500 |
| 50M05_Saliva | chr10:94451016-94451518 | HHEX | N_Shore | Body |
| 50M05_Saliva | chr10:105978702-105978956 | MIR609;C10orf79 | | TSS200;Body |
| 50M05_Saliva | chr10:114134915-114135259 | ACSL5;ACSL5;ACSL5 | | TSS1500;TSS200;5'UTR |
| 50M05_Saliva | chr10:123923518-123924085 | TACC2;TACC2;TACC2;TACC2 | Island | Body;1stExon;Body;1stExon |
| 50M05_Saliva | chr10:134358030-134358273 | INPP5A | N_Shelf | Body |
| 50M05_Saliva | chr11:2321800-2322050 | C11orf21;TSPAN32;C11orf21 | | Body;TSS1500;Body |
| 50M05_Saliva | chr11:2322618-2322674 | C11orf21;TSPAN32;C11orf21 | | Body;TSS1500;Body |
| 50M05_Saliva | chr11:2322741-2322802 | C11orf21;TSPAN32;C11orf21 | | Body;TSS1500;Body |
| 50M05_Saliva | chr11:2322935-2323059 | C11orf21;TSPAN32;C11orf21 | | Body;TSS1500;Body |
| 50M05_Saliva | chr11:2323459-2323896 | TSPAN32;C11orf21;C11orf21 | | Body;TSS1500;TSS1500 |
| 50M05_Saliva | chr11:3177622-3178024 | OSBPL5;OSBPL5;OSBPL5 | N_Shelf | 5'UTR;5'UTR;5'UTR |
| 50M05_Saliva | chr11:35441881-35442225 | SLC1A2 | Island | TSS1500 |
| 50M05_Saliva | chr11:36422377-36422615 | PRR5L;PRR5L;PRR5L;PRR5L | | 5'UTR;5'UTR;TSS200;Body |
| 50M05_Saliva | chr11:47399980-47400146 | SPI1;SPI1;SPI1;SPI1 | Island | 5'UTR;5'UTR;1stExon;1stExon |
| 50M05_Saliva | chr11:59824089-59824161 | MS4A3;MS4A3;MS4A3 | | TSS200;TSS200;TSS200 |
| 50M05_Saliva | chr11:60738971-60739178 | CD6 | | TSS200 |
| 50M05_Saliva | chr11:63273507-63273565 | LGALS12;LGALS12;LGALS12 | | TSS200;TSS200;TSS200 |
| 50M05_Saliva | chr11:66084841-66085299 | CD248 | S_Shore | TSS1500 |
| 50M05_Saliva | chr11:66839085-66839191 | RHOD | Island | 3'UTR |
| 50M05_Saliva | chr11:67170610-67171585 | PPP1CA;PPP1CA;TBC1D10C;PPI | S_Shore | TSS1500;TSS1500;TSS1500;TSS1500 |
| 50M05_Saliva | chr11:70211301-70211531 | PPFIA1;PPFIA1 | | Body;Body |
| 50M05_Saliva | chr11:70257035-70257149 | CTTN;CTTN | N_Shore | Body;Body |
| 50M05_Saliva | chr11:78673073-78673212 | ODZ4 | Island | Body |
| 50M05_Saliva | chr11:113953462-113953812 | ZBTB16;ZBTB16 | N_Shore | Body;Body |
| 50M05_Saliva | chr12:4699085-4699618 | DYRK4 | | TSS200 |
| 50M05_Saliva | chr12:47610220-47610734 | FAM113B;FAM113B;LOC100233209 | | 5'UTR;1stExon;Body |
| 50M05_Saliva | chr12:51717978-51718088 | BIN2 | | TSS200 |
| 50M05_Saliva | chr12:54891491-54891636 | NCKAP1L | | TSS200 |
| 50M05_Saliva | chr12:55378411-55378654 | KIAA0748;KIAA0748 | | 1stExon;5'UTR |
| 50M05_Saliva | chr13:51640315-51641069 | GUCY1B2 | | TSS200 |

| | | | | |
|---|---|---|---|---|
| 50M05_Saliva | chr13:107145933-107146164 | EFNB2 | S_Shore | Body |
| 50M05_Saliva | chr13:110951025-110951418 | COL4A1 | | Body |
| 50M05_Saliva | chr14:21423684-21424117 | RNASE2;RNASE2 | | 1stExon;5'UTR |
| 50M05_Saliva | chr14:93170710-93170970 | LGMN;LGMN | | Body;Body |
| 50M05_Saliva | chr14:102676957-102677294 | WDR20 | N_Shelf | Body |
| 50M05_Saliva | chr15:26044050-26044289 | ATP10A | | Body |
| 50M05_Saliva | chr15:38988533-38988860 | C15orf53 | | TSS1500 |
| 50M05_Saliva | chr16:2737220-2737356 | KCTD5 | | Body |
| 50M05_Saliva | chr16:8738240-8738441 | C16orf68 | | Body |
| 50M05_Saliva | chr16:28995882-28996053 | LAT;LAT;LAT;LAT | | TSS1500;TSS1500;TSS1500;TSS1500 |
| 50M05_Saliva | chr16:28996362-28996533 | LAT;LAT;LAT;LAT | | Body;TSS200;TSS200;TSS200 |
| 50M05_Saliva | chr16:29673933-29674184 | SPN;SPN | N_Shore | TSS1500;TSS1500 |
| 50M05_Saliva | chr16:29674618-29676003 | SPN;SPN;SPN | N_Shore | 1stExon;5'UTR;5'UTR |
| 50M05_Saliva | chr16:29756951-29757323 | C16orf54 | | 5'UTR |
| 50M05_Saliva | chr16:50715135-50715260 | SNX20;SNX20;SNX20;SNX20;SNX20;SNX20 | | 1stExon;5'UTR;1stExon;5'UTR;5'UTR;1stExon |
| 50M05_Saliva | chr16:56995813-56995856 | CETP | | TSS200 |
| 50M05_Saliva | chr16:57576285-57576394 | GPR114 | | TSS1500 |
| 50M05_Saliva | chr16:67428155-67428324 | TPPP3;TPPP3 | Island | TSS1500;TSS1500 |
| 50M05_Saliva | chr16:89041793-89042185 | CBFA2T3 | | Body |
| 50M05_Saliva | chr17:263096-263615 | C17orf97 | Island | Body |
| 50M05_Saliva | chr17:3704574-3704621 | ITGAE | | TSS200 |
| 50M05_Saliva | chr17:4337082-4337111 | SPNS3 | | TSS200 |
| 50M05_Saliva | chr17:5138259-5138645 | C17orf87 | | TSS200 |
| 50M05_Saliva | chr17:34344995-34345172 | CCL23;CCL23;CCL23;CCL23 | | 1stExon;5'UTR;1stExon;5'UTR |
| 50M05_Saliva | chr17:62009607-62009835 | CD79B;CD79B;CD79B | | 1stExon;1stExon;1stExon |
| 50M05_Saliva | chr17:73642547-73642628 | RECQL5;LOC100130933 | | Body;TSS200 |
| 50M05_Saliva | chr17:77706946-77707066 | ENPP7 | N_Shore | Body |
| 50M05_Saliva | chr17:78755379-78755442 | RPTOR;RPTOR | | Body;Body |
| 50M05_Saliva | chr18:74114570-74114728 | ZNF516 | Island | Body |
| 50M05_Saliva | chr19:728040-728176 | PALM;PALM | Island | Body;Body |
| 50M05_Saliva | chr19:14550846-14550999 | PKN1;PKN1 | N_Shore | TSS1500;Body |
| 50M05_Saliva | chr19:17862104-17862210 | FCHO1;FCHO1;FCHO1;FCHO1 | S_Shelf | 5'UTR;TSS200;TSS200;5'UTR |
| 50M05_Saliva | chr19:36233297-36233469 | TMEM149;TMEM149 | S_Shore | 1stExon;5'UTR |
| 50M05_Saliva | chr19:54566838-54567279 | VSTM1 | | Body |
| 50M05_Saliva | chr19:54876664-54876752 | LAIR1;LAIR1;LAIR1;LAIR1 | | 5'UTR;1stExon;1stExon;5'UTR |
| 50M05_Saliva | chr20:1471884-1472078 | SIRPB2;SIRPB2 | | Body;Body |
| 50M05_Saliva | chr20:4795886-4795935 | RASSF2;RASSF2 | | 5'UTR;TSS200 |
| 50M05_Saliva | chr20:35274570-35274715 | SLA2;SLA2 | | TSS200;TSS200 |
| 50M05_Saliva | chr20:36932397-36932553 | BPI | | TSS200 |
| 50M05_Saliva | chr20:57582856-57582918 | CTSZ | S_Shore | TSS1500 |
| 50M05_Saliva | chr20:57583000-57583195 | CTSZ | S_Shore | TSS1500 |
| 50M05_Saliva | chr20:62198872-62199190 | PRIC285;PRIC285 | Island | Body;1stExon |
| 50M05_Saliva | chr21:48026043-48026298 | S100B | | TSS1500 |
| 50M05_Saliva | chr22:27013978-27014116 | CRYBB1;CRYBB1 | | 5'UTR;1stExon |
| 50M05_Saliva | chr22:37545924-37546115 | IL2RB;IL2RB | | 5'UTR;1stExon |
| 50M05_Saliva | chr22:50523686-50523703 | MLC1;MLC1;MLC1 | | 5'UTR;5'UTR;1stExon |
| 50M05_Saliva | chr1:6446095-6446234 | ACOT7;ACOT7 | | Body;TSS1500 |
| 50M05_Saliva | chr1:10707917-10707953 | CASZ1;CASZ1 | Island | Body;Body |
| 50M05_Saliva | chr1:15271952-15272108 | KIAA1026;KIAA1026;KIAA1026;KIAA1026;KIAA1026 | | Body;5'UTR;TSS1500;Body;Body |
| 50M05_Saliva | chr1:43751270-43751497 | C1orf210;C1orf210;C1orf210 | | 1stExon;5'UTR;TSS200 |
| 50M05_Saliva | chr1:153581494-153581793 | S100A16 | | 5'UTR |
| 50M05_Saliva | chr1:201979499-201979567 | ELF3;ELF3 | | TSS200;TSS200 |
| 50M05_Saliva | chr2:113735377-113735503 | IL1F9 | | TSS1500 |
| 50M05_Saliva | chr3:50306107-50306456 | SEMA3B;SEMA3B | | 5'UTR;5'UTR |
| 50M05_Saliva | chr3:196065506-196065688 | TM4SF19 | | TSS1500 |
| 50M05_Saliva | chr4:87770586-87770782 | SLC10A6 | | TSS200 |
| 50M05_Saliva | chr4:184908351-184908739 | STOX2 | | Body |
| 50M05_Saliva | chr5:912383-912821 | TRIP13 | N_Shore | Body |
| 50M05_Saliva | chr5:59783629-59784203 | PART1;PART1;PDE4D;PART1 | | TSS200;Body;5'UTR;TSS200 |
| 50M05_Saliva | chr5:169532721-169532758 | FOXI1;FOXI1 | N_Shore | TSS200;TSS200 |
| 50M05_Saliva | chr5:176797920-176798049 | RGS14 | S_Shelf | Body |
| 50M05_Saliva | chr6:8436218-8436296 | SLC35B3;SLC35B3;SLC35B3 | S_Shore | TSS1500;TSS1500;TSS1500 |
| 50M05_Saliva | chr6:32120625-32120783 | PPT2;PRRT1;PPT2 | N_Shore | TSS1500;TSS1500;TSS1500 |
| 50M05_Saliva | chr6:32526021-32526263 | HLA-DRB6 | | Body |
| 50M05_Saliva | chr6:32632331-32632338 | HLA-DQB1 | Island | Body |

| | | | | |
|---|---|---|---|---|
| 50M05_Saliva | chr6:33638313-33638550 | ITPR3 | | Body |
| 50M05_Saliva | chr6:116381904-116382179 | FRK;FRK | | 1stExon;5'UTR |
| 50M05_Saliva | chr6:137366336-137366492 | IL20RA | S_Shore | TSS200 |
| 50M05_Saliva | chr6:161796785-161796854 | PARK2;PARK2;PARK2 | | Body;Body;Body |
| 50M05_Saliva | chr7:23286319-23286539 | GPNMB;GPNMB;GPNMB;GPNMB | N_Shore | 5'UTR;1stExon;1stExon;5'UTR |
| 50M05_Saliva | chr7:27155234-27155548 | HOXA3;HOXA3 | Island | 5'UTR;5'UTR |
| 50M05_Saliva | chr7:73244836-73245178 | CLDN4 | N_Shore | TSS1500 |
| 50M05_Saliva | chr7:135433540-135433790 | FAM180A;FAM180A | | 1stExon;5'UTR |
| 50M05_Saliva | chr7:158059944-158060003 | PTPRN2;PTPRN2;PTPRN2 | N_Shore | Body;Body;Body |
| 50M05_Saliva | chr8:1906338-1906498 | ARHGEF10 | | 3'UTR |
| 50M05_Saliva | chr8:145728507-145728543 | GPT | N_Shore | TSS1500 |
| 50M05_Saliva | chr10:123353822-123354171 | FGFR2;FGFR2;FGFR2;FGFR2;FGF | N_Shelf | TSS200;5'UTR;5'UTR;TSS1500;5'UTR;TSS1500;5'UTR;5'UTR;5'UTR |
| 50M05_Saliva | chr11:393328-393570 | PKP3 | N_Shore | TSS1500 |
| 50M05_Saliva | chr11:692963-693032 | DEAF1 | N_Shore | Body |
| 50M05_Saliva | chr11:1855389-1855561 | SYT8 | | TSS1500 |
| 50M05_Saliva | chr11:67442195-67442408 | ALDH3B2;ALDH3B2 | | 5'UTR;TSS200 |
| 50M05_Saliva | chr11:76998821-76998857 | GDPD4 | | TSS1500 |
| 50M05_Saliva | chr11:108408962-108409365 | EXPH5;EXPH5;EXPH5 | | Body;Body;TSS200 |
| 50M05_Saliva | chr11:108422996-108423124 | EXPH5;EXPH5 | | TSS200;Body |
| 50M05_Saliva | chr11:117069780-117069849 | TAGLN;TAGLN | | TSS1500;TSS1500 |
| 50M05_Saliva | chr11:117947628-117947877 | TMPRSS4;TMPRSS4 | | TSS200;TSS200 |
| 50M05_Saliva | chr11:119993732-119993914 | TRIM29 | | Body |
| 50M05_Saliva | chr11:120008998-120009135 | TRIM29 | S_Shore | TSS200 |
| 50M05_Saliva | chr11:121970725-121970768 | LOC399959;MIR125B1 | | Body;TSS200 |
| 50M05_Saliva | chr11:123396299-123396486 | GRAMD1B | | TSS1500 |
| 50M05_Saliva | chr12:8974844-8975309 | A2ML1 | | TSS1500 |
| 50M05_Saliva | chr12:52585695-52585844 | KRT80;KRT80;KRT80;KRT80 | | 1stExon;1stExon;5'UTR;5'UTR |
| 50M05_Saliva | chr12:130976260-130976551 | RIMBP2 | | 5'UTR |
| 50M05_Saliva | chr12:133101495-133101612 | FBRSL1 | N_Shore | Body |
| 50M05_Saliva | chr13:80911626-80911808 | SPRY2 | N_Shore | Body |
| 50M05_Saliva | chr14:51955585-51956044 | FRMD6 | | TSS1500 |
| 50M05_Saliva | chr14:56045827-56046001 | KTN1;C14orf33;KTN1;KTN1;KTN1 | N_Shore | TSS1500;Body;TSS1500;TSS1500;TSS1500 |
| 50M05_Saliva | chr15:86313891-86313941 | MIR1276;KLHL25 | S_Shore | TSS200;5'UTR |
| 50M05_Saliva | chr17:7493572-7494095 | SOX15 | S_Shore | TSS200 |
| 50M05_Saliva | chr17:39143371-39143567 | KRT40;KRT40 | | 5'UTR;1stExon |
| 50M05_Saliva | chr17:79031550-79031712 | BAIAP2;BAIAP2;BAIAP2;BAIAP2 | | Body;Body;Body;Body |
| 50M05_Saliva | chr17:79652024-79652389 | ARL16;HGS | S_Shore | TSS1500;Body |
| 50M05_Saliva | chr17:80797881-80797983 | TBCD;ZNF750;ZNF750 | | Body;5'UTR;1stExon |
| 50M05_Saliva | chr17:80806135-80806388 | TBCD | | Body |
| 50M05_Saliva | chr17:80847137-80847662 | TBCD | | Body |
| 50M05_Saliva | chr18:61143926-61144339 | SERPINB5 | | TSS1500 |
| 50M05_Saliva | chr18:61253847-61254561 | SERPINB13 | | TSS1500 |
| 50M05_Saliva | chr19:35085557-35085677 | SCGBL | | TSS200 |
| 50M05_Saliva | chr20:36154222-36154388 | BLCAP;BLCAP;BLCAP;BLCAP;BL | N_Shore | 5'UTR;TSS1500;5'UTR;5'UTR;5'UTR |
| 50M05_Saliva | chr20:62367884-62367893 | LIME1 | N_Shore | TSS200 |
| 50M05_Saliva | chr22:38506712-38506781 | BAIAP2L2 | | TSS200 |
| 50M05_Saliva | chr22:45608465-45608516 | C22orf9;C22orf9 | | Body;TSS200 |
| 20M02_Semen | chr1:1959497-1959813 | GABRD | Island | Body |
| 20M02_Semen | chr6:29526338-29526376 | UBD | | Body |
| 20M02_Semen | chr6:32606884-32607509 | HLA-DQA1 | | Body |
| 20M02_Semen | chr6:32609130-32609212 | HLA-DQA1 | | Body |
| 20M02_Semen | chr6:150238016-150238164 | RAET1G | | 3'UTR |
| 20M02_Semen | chr7:138435480-138436009 | ATP6V0A4;ATP6V0A4;ATP6V0A4 | | Body;Body;Body |
| 20M02_Semen | chr7:157932168-157932971 | PTPRN2;PTPRN2;PTPRN2 | | Body;Body;Body |
| 20M02_Semen | chr7:157933750-157934005 | PTPRN2;PTPRN2;PTPRN2 | N_Shelf | Body;Body;Body |
| 20M02_Semen | chr8:27468684-27469001 | CLU;CLU;CLU;CLU | N_Shelf | 1stExon;Body;Body;5'UTR |
| 20M02_Semen | chr10:8100286-8100761 | GATA3;GATA3 | N_Shore | Body;Body |
| 20M02_Semen | chr13:113303307-113303719 | C13orf35 | Island | 5'UTR |
| 20M02_Semen | chr13:113333720-113333851 | C13orf35 | | Body |
| 20M02_Semen | chr17:714733-714935 | NXN | S_Shore | Body |
| 20M02_Semen | chr17:855187-855312 | NXN | | Body |
| 20M02_Semen | chr17:79905236-79905263 | MYADML2 | | TSS200 |
| 20M02_Semen | chr19:49061546-49061702 | SULT2B1 | Island | Body |
| 20M02_Semen | chr19:56042539-56042991 | SBK2 | Island | Body |
| 20M02_Semen | chr21:45705699-45705742 | AIRE | Island | TSS200 |

| | | | | |
|---|---|---|---|---|
| 20M02_Semen | chr4:1202653-1202700 | LOC100130872-SPON2;LOC100 | N_Shore | TSS200;Body |
| 20M02_Semen | chr5:173472623-173472732 | HMP19 | | TSS200 |
| 20M02_Semen | chr7:2313692-2313925 | SNX8 | | Body |
| 20M02_Semen | chr7:127669170-127669504 | SND1;LRRC4 | N_Shore | Body;Body |
| 20M02_Semen | chr8:145700190-145700499 | FOXH1 | N_Shore | Body |
| 20M02_Semen | chr9:990260-990692 | DMRT3 | | Body |
| 20M02_Semen | chr10:1595704-1596029 | ADARB2 | Island | Body |
| 20M02_Semen | chr11:128812846-128813470 | TP53AIP1 | | TSS200 |
| 20M02_Semen | chr11:131799642-131799701 | NTM;NTM;NTM;NTM | | Body;Body;Body;Body |
| 20M02_Semen | chr16:1204876-1205018 | CACNA1H;CACNA1H | S_Shore | Body;Body |
| 20M02_Semen | chr16:58533743-58534708 | NDRG4;NDRG4;NDRG4 | N_Shore | Body;Body;Body |
| 20M02_Semen | chr17:72948349-72948705 | C17orf28 | | Body |
| 20M02_Semen | chr17:79183705-79183966 | AZI1;AZI1 | Island | Body;Body |
| 20M02_Semen | chr19:808116-808699 | PTBP1;PTBP1;PTBP1;PTBP1 | Island | Body;Body;Body;Body |
| 20M02_Semen | chr19:1132256-1132420 | SBNO2;SBNO2;SBNO2 | | Body;1stExon;5'UTR |
| 20M02_Semen | chr19:45805741-45805935 | MARK4 | Island | 3'UTR |
| 20M02_Semen | chr22:43831005-43831161 | MPPED1 | Island | Body |
| 20M03_Semen | chr1:3128606-3129115 | PRDM16;PRDM16 | | Body;Body |
| 20M03_Semen | chr3:194120949-194121173 | GP5 | S_Shore | TSS1500 |
| 20M03_Semen | chr4:762885-763508 | PCGF3 | | 3'UTR |
| 20M03_Semen | chr4:1388830-1389270 | CRIPAK | Island | 1stExon |
| 20M03_Semen | chr5:667968-668501 | TPPP | Island | Body |
| 20M03_Semen | chr6:3083314-3083420 | RIPK1 | | Body |
| 20M03_Semen | chr6:29527870-29528119 | UBD | | TSS200 |
| 20M03_Semen | chr6:30954962-30955358 | MUC21 | | Body |
| 20M03_Semen | chr6:166959444-166959552 | RPS6KA2;RPS6KA2 | | Body;Body |
| 20M03_Semen | chr10:114133235-114133700 | ACSL5 | | |
| 20M03_Semen | chr15:89089587-89090033 | DET1;DET1;DET1 | | 5'UTR;5'UTR;Body |
| 20M03_Semen | chr17:855187-855312 | NXN | | Body |
| 20M03_Semen | chr17:46136650-46136875 | NFE2L1 | Island | Body |
| 20M03_Semen | chr17:79905236-79905263 | MYADML2 | | TSS200 |
| 20M03_Semen | chr19:10662582-10662782 | ATG4D;MIR1238 | | Body;TSS1500 |
| 20M03_Semen | chr22:47097389-47097600 | CERK | Island | Body |
| 20M03_Semen | chr1:3559559-3561076 | WDR8 | N_Shore | Body |
| 20M03_Semen | chr1:26527870-26527928 | CATSPER4 | | Body |
| 20M03_Semen | chr2:3307465-3307598 | TSSC1 | | Body |
| 20M03_Semen | chr3:9291212-9291334 | SRGAP3;SRGAP3;SRGAP3;SRGAP3 | | 1stExon;5'UTR;1stExon;5'UTR |
| 20M03_Semen | chr3:112359534-112359935 | CCDC80;CCDC80;CCDC80 | | 5'UTR;1stExon;5'UTR |
| 20M03_Semen | chr4:2278462-2278594 | ZFYVE28 | S_Shore | Body |
| 20M03_Semen | chr4:69435250-69435601 | UGT2B15;UGT2B17 | | TSS1500;TSS1500 |
| 20M03_Semen | chr6:31237013-31237034 | HLA-C | N_Shore | Body |
| 20M03_Semen | chr6:31237405-31237722 | HLA-C | N_Shore | Body |
| 20M03_Semen | chr6:32064842-32064956 | TNXB | Island | Body |
| 20M03_Semen | chr6:32606845-32607174 | HLA-DQA1 | | Body |
| 20M03_Semen | chr7:1065956-1066396 | C7orf50;C7orf50;C7orf50 | N_Shore | Body;Body;Body |
| 20M03_Semen | chr7:1105321-1105484 | C7orf50;C7orf50;C7orf50 | Island | Body;Body;Body |
| 20M03_Semen | chr7:100199952-100200009 | PCOLCE;PCOLCE | N_Shore | 1stExon;5'UTR |
| 20M03_Semen | chr8:674434-674560 | ERICH1 | | Body |
| 20M03_Semen | chr10:686080-686354 | DIP2C | Island | Body |
| 20M03_Semen | chr10:712394-712592 | DIP2C | Island | Body |
| 20M03_Semen | chr10:35484768-35484972 | CREM;CREM;CREM;CREM;CREM;CREM;CREM;CREM;CREM;CREM;CRE | | Body;Body;Body;Body;Body;Body;5'UTR;Body;Body;Body;5'UTR;TSS200;TSS200;Body;TSS200;Body;TSS200;Body;TSS200;Body |
| 20M03_Semen | chr12:7033470-7033628 | ATN1 | N_Shelf | TSS200 |
| 20M03_Semen | chr13:110918441-110918683 | COL4A1 | Island | Body |
| 20M03_Semen | chr16:2908892-2908934 | PRSS22 | S_Shore | TSS1500 |
| 20M03_Semen | chr19:1132256-1132537 | SBNO2;SBNO2;SBNO2 | | Body;1stExon;5'UTR |
| 20M03_Semen | chr19:35531639-35531859 | HPN;HPN | N_Shore | 5'UTR;5'UTR |
| 20M03_Semen | chr19:48823408-48823427 | CCDC114 | N_Shore | TSS200 |
| 20M03_Semen | chr19:51415152-51415450 | KLK4 | N_Shore | TSS1500 |
| 20M03_Semen | chr19:56092040-56092459 | ZNF579 | S_Shore | 5'UTR |
| 20M03_Semen | chr22:24384315-24384400 | GSTT1 | Island | TSS200 |
| 20M06_Semen | chr2:20870812-20871195 | GDF7 | Island | Body |
| 20M06_Semen | chr5:1594282-1594579 | SDHAP3 | Island | Body |
| 20M06_Semen | chr6:30131749-30131806 | TRIM15 | | 1stExon |
| 20M06_Semen | chr10:131686103-131686574 | EBF3 | N_Shore | Body |

| | | | | |
|---|---|---|---|---|
| 20M06_Semen | chr14:94088174-94088604 | KIAA1409 | | Body |
| 20M06_Semen | chr17:7832764-7832852 | KCNAB3 | Island | TSS200 |
| 20M06_Semen | chr17:46641708-46642104 | HOXB3 | Island | 5'UTR |
| 30M04_Semen | chr1:3078839-3079212 | PRDM16;PRDM16 | N_Shelf | Body;Body |
| 30M04_Semen | chr1:167487295-167487633 | CD247;CD247 | | Body;Body |
| 30M04_Semen | chr2:3307465-3307598 | TSSC1 | | Body |
| 30M04_Semen | chr3:11078979-11079832 | SLC6A1 | | 3'UTR |
| 30M04_Semen | chr4:1202653-1203001 | LOC100130872-SPON2;LOC100 | N_Shore | TSS200;Body |
| 30M04_Semen | chr4:186423408-186423579 | PDLIM3;PDLIM3 | | 3'UTR;3'UTR |
| 30M04_Semen | chr5:191500-191600 | LRRC14B | N_Shore | TSS200 |
| 30M04_Semen | chr5:496069-496476 | SLC9A3 | S_Shore | Body |
| 30M04_Semen | chr6:32134461-32134604 | EGFL8 | Island | Body |
| 30M04_Semen | chr6:32135129-32135195 | EGFL8 | Island | Body |
| 30M04_Semen | chr6:32371506-32372310 | BTNL2 | | Body |
| 30M04_Semen | chr7:105319437-105319819 | ATXN7L1;ATXN7L1 | | Body;5'UTR |
| 30M04_Semen | chr7:151506385-151506505 | PRKAG2;PRKAG2 | | 5'UTR;Body |
| 30M04_Semen | chr7:157594643-157595153 | PTPRN2;PTPRN2;PTPRN2 | S_Shore | Body;Body;Body |
| 30M04_Semen | chr8:27182909-27183342 | PTK2B;PTK2B;PTK2B;PTK2B | N_Shore | 5'UTR;TSS200;5'UTR;TSS200 |
| 30M04_Semen | chr10:1438165-1438686 | ADARB2 | N_Shore | Body |
| 30M04_Semen | chr10:1707523-1707576 | ADARB2 | | Body |
| 30M04_Semen | chr10:7763676-7763777 | ITIH2 | | Body |
| 30M04_Semen | chr10:131686103-131686574 | EBF3 | N_Shore | Body |
| 30M04_Semen | chr10:131697080-131697345 | EBF3 | N_Shore | Body |
| 30M04_Semen | chr11:1892139-1892307 | LSP1;LSP1;LSP1;LSP1;LSP1 | Island | 5'UTR;TSS200;5'UTR;Body;1stExon |
| 30M04_Semen | chr11:2441045-2441442 | TRPM5 | N_Shore | Body |
| 30M04_Semen | chr11:67183078-67183330 | ATPGD1;ATPGD1 | Island | TSS200;TSS200 |
| 30M04_Semen | chr11:125106056-125106135 | PKNOX2 | | 5'UTR |
| 30M04_Semen | chr11:134230036-134230439 | GLB1L2 | N_Shore | Body |
| 30M04_Semen | chr12:121087355-121087382 | CABP1;CABP1;CABP1 | | TSS1500;TSS1500;Body |
| 30M04_Semen | chr13:30079113-30079782 | MTUS2;MTUS2 | S_Shore | 3'UTR;3'UTR |
| 30M04_Semen | chr14:102904469-102904606 | TECPR2 | | Body |
| 30M04_Semen | chr15:29431453-29432511 | FAM189A1 | | Body |
| 30M04_Semen | chr16:4729905-4730657 | MGRN1;MGRN1;MGRN1;MGRN1 | N_Shore | Body;Body;Body;Body |
| 30M04_Semen | chr16:88534398-88534585 | ZFPM1 | N_Shelf | Body |
| 30M04_Semen | chr17:291473-291563 | FAM101B | N_Shore | 3'UTR |
| 30M04_Semen | chr17:767301-767380 | NXN | | Body |
| 30M04_Semen | chr17:79881483-79881543 | MAFG;MAFG | S_Shore | 5'UTR;TSS200 |
| 30M04_Semen | chr17:80540666-80540758 | FOXK2 | N_Shore | Body |
| 30M04_Semen | chr17:80545175-80545454 | FOXK2 | N_Shore | Body |
| 30M04_Semen | chr18:74692214-74692265 | MBP;MBP;MBP;MBP;MBP | S_Shore | 3'UTR;3'UTR;3'UTR;3'UTR;3'UTR |
| 30M04_Semen | chr19:13990960-13991187 | NANOS3 | N_Shore | Body |
| 30M04_Semen | chr19:35531639-35531859 | HPN;HPN | N_Shore | 5'UTR;5'UTR |
| 30M04_Semen | chr19:35940461-35940593 | FFAR2 | | TSS200 |
| 30M04_Semen | chr22:24348549-24348715 | GSTTP1 | | TSS1500 |
| 30M04_Semen | chr22:37640250-37640402 | RAC2;RAC2 | | 1stExon;5'UTR |
| 30M04_Semen | chr1:6085799-6086483 | KCNAB2;KCNAB2 | N_Shore | TSS1500;TSS1500 |
| 30M04_Semen | chr4:69435250-69435601 | UGT2B15;UGT2B17 | | TSS1500;TSS1500 |
| 30M04_Semen | chr6:31237013-31237034 | HLA-C | N_Shore | Body |
| 30M04_Semen | chr6:31237405-31237722 | HLA-C | N_Shore | Body |
| 30M04_Semen | chr6:32369227-32369487 | BTNL2 | | Body |
| 30M04_Semen | chr6:32490043-32490444 | HLA-DRB5 | Island | Body |
| 30M04_Semen | chr6:32526342-32526414 | HLA-DRB6 | | Body |
| 30M04_Semen | chr15:23810280-23810378 | MKRN3 | | TSS200 |
| 30M04_Semen | chr17:855187-855312 | NXN | | Body |
| 30M05_Semen | chr1:27332632-27332971 | FAM46B | Island | Body |
| 30M05_Semen | chr2:232262773-232263534 | B3GNT7 | N_Shore | Body |
| 30M05_Semen | chr4:2065749-2066320 | NAT8L | Island | Body |
| 30M05_Semen | chr11:1892074-1892307 | LSP1;LSP1;LSP1;LSP1 | Island | 5'UTR;TSS200;TSS200;Body |
| 30M05_Semen | chr11:2019090-2019167 | H19;MIR675 | N_Shore | TSS200;TSS1500 |
| 30M05_Semen | chr11:66083766-66084631 | CD248 | Island | 1stExon |
| 30M05_Semen | chr11:67183078-67183330 | ATPGD1;ATPGD1 | Island | TSS200;TSS200 |
| 30M05_Semen | chr16:88534398-88534585 | ZFPM1 | N_Shelf | Body |
| 30M05_Semen | chr16:88803803-88804373 | FAM38A | Island | Body |
| 30M05_Semen | chr17:79380493-79380585 | BAHCC1 | Island | Body |
| 30M05_Semen | chr17:79881483-79881543 | MAFG;MAFG | S_Shore | 5'UTR;TSS200 |
| 30M05_Semen | chr17:80545175-80545310 | FOXK2 | N_Shore | Body |

| | | | | |
|---|---|---|---|---|
| 30M05_Semen | chr19:44222974-44223436 | IRGC | Island | Body |
| 30M05_Semen | chr19:58549241-58549906 | ZSCAN1 | N_Shore | Body |
| 30M05_Semen | chr4:69435250-69435601 | UGT2B15;UGT2B17 | | TSS1500;TSS1500 |
| 30M05_Semen | chr8:1789463-1789670 | ARHGEF10 | | 5'UTR |
| 30M05_Semen | chr22:24348549-24348715 | GSTTP1 | | TSS1500 |
| 30M08_Semen | chr1:203320190-203320541 | FMOD;FMOD | | 1stExon;5'UTR |
| 30M08_Semen | chr1:236907957-236908064 | ACTN2 | | Body |
| 30M08_Semen | chr2:1203839-1204376 | SNTG2 | | Body |
| 30M08_Semen | chr2:88367018-88367362 | SMYD1 | | TSS1500 |
| 30M08_Semen | chr3:189349021-189349323 | TP63;TP63;TP63 | | TSS200;TSS200;TSS200 |
| 30M08_Semen | chr6:28499391-28499703 | GPX5;GPX5 | | Body;Body |
| 30M08_Semen | chr6:32297670-32297822 | C6orf10 | | Body |
| 30M08_Semen | chr6:32606845-32607174 | HLA-DQA1 | | Body |
| 30M08_Semen | chr7:138435480-138436009 | ATP6V0A4;ATP6V0A4;ATP6V0A4 | | Body;Body;Body |
| 30M08_Semen | chr10:3148096-3148613 | PFKP | N_Shore | Body |
| 30M08_Semen | chr11:472782-473003 | PTDSS2 | Island | Body |
| 30M08_Semen | chr11:131799642-131799701 | NTM;NTM;NTM;NTM | | Body;Body;Body;Body |
| 30M08_Semen | chr12:109678885-109678973 | ACACB | | Body |
| 30M08_Semen | chr17:80545310-80545434 | FOXK2 | N_Shore | Body |
| 30M08_Semen | chr19:56042469-56042579 | SBK2 | Island | Body |
| 30M08_Semen | chr22:24348549-24348715 | GSTTP1 | | TSS1500 |
| 30M08_Semen | chr4:69435250-69435601 | UGT2B15;UGT2B17 | | TSS1500;TSS1500 |
| 30M08_Semen | chr6:32064842-32064956 | TNXB | Island | Body |
| 30M08_Semen | chr7:75115302-75115714 | POM121C;POM121C | Island | 5'UTR;1stExon |
| 30M08_Semen | chr7:127669170-127669504 | SND1;LRRC4 | N_Shore | Body;Body |
| 30M08_Semen | chr10:686080-686354 | DIP2C | Island | Body |
| 30M08_Semen | chr16:863240-863884 | PRR25 | N_Shore | Body |
| 30M08_Semen | chr16:88051832-88052309 | BANP;BANP | N_Shore | Body;Body |
| 30M08_Semen | chr18:13262344-13262750 | C18orf1;C18orf1 | Island | 5'UTR;5'UTR |
| 30M08_Semen | chr19:56092040-56092459 | ZNF579 | S_Shore | 5'UTR |
| 40M05_Semen | chr10:132909113-132909289 | TCERG1L | | Body |
| 40M05_Semen | chr10:132910868-132911152 | TCERG1L | | Body |
| 40M05_Semen | chr12:120763487-120763716 | PLA2G1B | | Body |
| 40M05_Semen | chr12:124780809-124781038 | FAM101A | S_Shore | 5'UTR |
| 40M05_Semen | chr21:46131668-46131923 | C21orf29 | S_Shore | TSS200 |
| 40M05_Semen | chr4:10459836-10460237 | ZNF518B | S_Shore | TSS1500 |
| 40M05_Semen | chr5:10291212-10291308 | CMBL | | 5'UTR |
| 40M05_Semen | chr6:3083314-3083420 | RIPK1 | | Body |
| 40M05_Semen | chr6:32606845-32607174 | HLA-DQA1 | | Body |
| 40M05_Semen | chr6:168963358-168963731 | SMOC2;SMOC2 | | Body;Body |
| 40M05_Semen | chr10:46221225-46221684 | FAM21C;FAM21C;FAM21C | N_Shelf | |
| 40M05_Semen | chr10:131686208-131686486 | EBF3 | N_Shore | Body |
| 40M05_Semen | chr12:10183172-10183220 | CLEC9A | | TSS200 |
| 40M05_Semen | chr12:133335658-133335850 | ANKLE2 | Island | Body |
| 40M05_Semen | chr15:35529169-35529521 | LOC723972 | | TSS1500 |
| 40M05_Semen | chr16:72820681-72821665 | ZFHX3;ZFHX3 | N_Shore | 3'UTR;3'UTR |
| 40M07_Semen | chr1:3078839-3079212 | PRDM16;PRDM16 | N_Shelf | Body;Body |
| 40M07_Semen | chr2:906167-906675 | C2orf90 | | TSS200 |
| 40M07_Semen | chr4:1202653-1202700 | LOC100130872-SPON2;LOC100 | N_Shore | TSS200;Body |
| 40M07_Semen | chr4:1202861-1203001 | LOC100130872;LOC100130872 | N_Shore | TSS200;TSS1500 |
| 40M07_Semen | chr5:415575-415885 | AHRR | | Body |
| 40M07_Semen | chr9:136131118-136131484 | ABO | Island | Body |
| 40M07_Semen | chr11:45231175-45231545 | PRDM11 | N_Shelf | Body |
| 40M07_Semen | chr13:111039873-111040242 | COL4A2 | S_Shore | Body |
| 40M07_Semen | chr16:89225495-89226194 | C16orf81 | N_Shore | TSS200 |
| 40M07_Semen | chr17:855187-855312 | NXN | | Body |
| 40M07_Semen | chr17:5993686-5994006 | WSCD1 | | Body |
| 40M07_Semen | chr19:2543602-2543861 | GNG7 | N_Shore | 5'UTR |
| 40M07_Semen | chr20:33879989-33880195 | FAM83C | Island | 1stExon |
| 40M07_Semen | chr22:24348549-24348715 | GSTTP1 | | TSS1500 |
| 40M07_Semen | chr22:25424129-25424512 | KIAA1671 | Island | 1stExon |
| 40M07_Semen | chr1:108734992-108735334 | SLC25A24;SLC25A24 | N_Shore | Body;Body |
| 40M07_Semen | chr2:10340558-10340837 | C2orf48 | | Body |
| 40M07_Semen | chr7:157503488-157504143 | PTPRN2;PTPRN2;PTPRN2 | Island | Body;Body;Body |
| 40M07_Semen | chr14:94088346-94088604 | KIAA1409 | | Body |
| 40M07_Semen | chr19:51128350-51128640 | SYT3;SYT3;SYT3 | Island | Body;Body;Body |

| | | | | |
|---|---|---|---|---|
| 40M07_Semen | chr21:43098598-43099460 | NCRNA00111 | | TSS1500 |
| 40M10_Semen | chr1:3559559-3560965 | WDR8 | N_Shore | Body |
| 40M10_Semen | chr2:1358283-1358684 | SNTG2 | | Body |
| 40M10_Semen | chr2:1425035-1425560 | TPO;TPO;TPO;TPO | | Body;Body;Body;Body |
| 40M10_Semen | chr3:71247280-71247442 | FOXP1;FOXP1 | | Body;Body |
| 40M10_Semen | chr4:40428028-40428121 | RBM47;RBM47 | | Body;Body |
| 40M10_Semen | chr5:88018481-88018605 | MEF2C;MEF2C | | Body;Body |
| 40M10_Semen | chr6:30290408-30290596 | HCG18;HCG18 | N_Shelf | Body;Body |
| 40M10_Semen | chr6:31938773-31938847 | STK19;STK19;STK19;DOM3Z | N_Shore | TSS200;TSS1500;TSS1500;Body |
| 40M10_Semen | chr6:32064923-32064956 | TNXB | Island | Body |
| 40M10_Semen | chr6:33037640-33037690 | HLA-DPA1 | | Body |
| 40M10_Semen | chr7:2560976-2561116 | LFNG;LFNG;LFNG;LFNG | S_Shore | Body;Body;Body;Body |
| 40M10_Semen | chr7:127669170-127669504 | SND1;LRRC4 | N_Shore | Body;Body |
| 40M10_Semen | chr8:145700239-145700268 | FOXH1 | Island | Body |
| 40M10_Semen | chr10:1147675-1148121 | WDR37 | N_Shore | Body |
| 40M10_Semen | chr12:7033470-7033520 | ATN1 | N_Shelf | TSS200 |
| 40M10_Semen | chr12:120763487-120763869 | PLA2G1B | | Body |
| 40M10_Semen | chr14:74182387-74182560 | PNMA1;C14orf43;C14orf43 | S_Shore | TSS1500;3'UTR;3'UTR |
| 40M10_Semen | chr14:104026839-104026955 | BAG5;BAG5;BAG5 | N_Shore | Body;Body;Body |
| 40M10_Semen | chr17:714733-714935 | NXN | S_Shore | Body |
| 40M10_Semen | chr2:238499464-238500389 | RAB17;RAB17 | | 1stExon;5'UTR |
| 40M10_Semen | chr4:1201588-1202966 | LOC100130872;LOC100130872 | N_Shore | Body;Body |
| 40M10_Semen | chr5:191547-191793 | LRRC14B | N_Shore | TSS200 |
| 40M10_Semen | chr6:29526338-29526376 | UBD | | Body |
| 40M10_Semen | chr8:144640306-144640755 | GSDMD;GSDMD | N_Shore | TSS200;5'UTR |
| 40M10_Semen | chr10:1245261-1246336 | ADARB2 | | Body |
| 40M10_Semen | chr10:3160974-3161106 | PFKP | Island | Body |
| 40M10_Semen | chr10:131686103-131686208 | EBF3 | N_Shore | Body |
| 40M10_Semen | chr10:131686486-131686885 | EBF3 | Island | Body |
| 40M10_Semen | chr11:64335733-64335895 | SLC22A11 | Island | Body |
| 40M10_Semen | chr17:2601803-2602006 | KIAA0664 | Island | Body |
| 40M10_Semen | chr17:76100911-76101307 | TNRC6C;TNRC6C | Island | Body;Body |
| 40M10_Semen | chr17:77043815-77044488 | C1QTNF1;C1QTNF1;C1QTNF1 | Island | Body;Body;Body |
| 40M10_Semen | chr19:35530755-35531990 | SCN1B;HPN;HPN | N_Shore | 3'UTR;TSS1500;TSS1500 |
| 40M10_Semen | chr19:35617254-35617819 | LGI4 | Island | Body |
| 40M10_Semen | chr19:49061546-49061702 | SULT2B1 | Island | Body |
| 40M10_Semen | chr19:49106849-49107139 | FAM83E | N_Shore | Body |
| 40M10_Semen | chr22:47097389-47097600 | CERK | Island | Body |
| 50_M06_Semen | chr1:3099804-3100956 | PRDM16;PRDM16 | N_Shelf | Body;Body |
| 50_M06_Semen | chr2:1358283-1358730 | SNTG2 | | Body |
| 50_M06_Semen | chr2:1425035-1425319 | TPO;TPO;TPO;TPO | | Body;Body;Body;Body |
| 50_M06_Semen | chr4:529948-530543 | PIGG;PIGG | Island | Body;Body |
| 50_M06_Semen | chr5:179224751-179225039 | MGAT4B;MGAT4B | Island | 3'UTR;3'UTR |
| 50_M06_Semen | chr6:33037640-33037690 | HLA-DPA1 | | Body |
| 50_M06_Semen | chr6:33094069-33094306 | HLA-DPB2 | | Body |
| 50_M06_Semen | chr6:167177712-167178280 | RPS6KA2 | | Body |
| 50_M06_Semen | chr6:168971785-168972128 | SMOC2;SMOC2 | N_Shore | Body;Body |
| 50_M06_Semen | chr17:855187-855312 | NXN | | Body |
| 50_M06_Semen | chr19:45540766-45540955 | RELB | Island | Body |
| 50_M06_Semen | chr1:50574507-50574591 | ELAVL4;ELAVL4;ELAVL4;ELAVL4;ELAVL4 | | Body;Body;TSS200;Body;TSS200 |
| 50_M06_Semen | chr1:247615433-247615683 | OR2B11 | S_Shore | TSS200 |
| 50_M06_Semen | chr2:238499782-238499802 | RAB17 | | TSS200 |
| 50_M06_Semen | chr4:21699426-21699512 | KCNIP4;KCNIP4 | | 5'UTR;TSS200 |
| 50_M06_Semen | chr4:40198392-40198776 | RHOH | | TSS200 |
| 50_M06_Semen | chr4:186435443-186436021 | PDLIM3;PDLIM3 | | Body;Body |
| 50_M06_Semen | chr6:36929838-36930062 | PI16 | Island | Body |
| 50_M06_Semen | chr6:163742730-163742913 | LOC285796 | | Body |
| 50_M06_Semen | chr7:3026309-3026459 | CARD11 | | 5'UTR |
| 50_M06_Semen | chr11:1669278-1669926 | HCCA2 | | Body |
| 50_M06_Semen | chr11:2019090-2019167 | H19;MIR675 | N_Shore | TSS200;TSS1500 |
| 50_M06_Semen | chr15:91839976-91840307 | SV2B;SV2B | Island | 3'UTR;3'UTR |
| 50_M06_Semen | chr17:714733-714935 | NXN | S_Shore | Body |
| 50_M06_Semen | chr18:44140411-44140574 | LOXHD1;LOXHD1 | Island | TSS1500;Body |
| 50M04_Semen | chr1:108734992-108735334 | SLC25A24;SLC25A24 | N_Shore | Body;Body |
| 50M04_Semen | chr2:54087173-54087189 | LOC100302652;GPR75 | Island | TSS200;TSS200 |
| 50M04_Semen | chr2:238499464-238500389 | RAB17;RAB17 | | 1stExon;5'UTR |

| | | | | |
|---|---|---|---|---|
| 50M04_Semen | chr3:123813232-123813393 | KALRN;KALRN;KALRN | | TSS1500;TSS1500;TSS1500 |
| 50M04_Semen | chr3:123813504-123813596 | KALRN;KALRN;KALRN | | TSS200;TSS200;TSS200 |
| 50M04_Semen | chr4:698303-698616 | PCGF3 | Island | TSS1500 |
| 50M04_Semen | chr4:40428028-40428121 | RBM47;RBM47 | | Body;Body |
| 50M04_Semen | chr4:186435443-186435506 | PDLIM3;PDLIM3 | | Body;Body |
| 50M04_Semen | chr4:186435901-186436021 | PDLIM3;PDLIM3 | | Body;Body |
| 50M04_Semen | chr6:32489991-32490043 | HLA-DRB5 | Island | Body |
| 50M04_Semen | chr7:5372402-5372608 | TNRC18 | Island | Body |
| 50M04_Semen | chr10:8100114-8100761 | GATA3;GATA3 | N_Shore | Body;Body |
| 50M04_Semen | chr16:1204876-1205018 | CACNA1H;CACNA1H | S_Shore | Body;Body |
| 50M04_Semen | chr16:75150784-75150819 | LDHD;LDHD | S_Shore | TSS200;TSS200 |
| 50M04_Semen | chr17:17910235-17910374 | LRRC48;LRRC48;LRRC48;LRRC4 | Island | Body;Body;Body;Body |
| 50M04_Semen | chr19:21933455-21933540 | LOC641367;ZNF100 | Island | TSS200;Body |
| 50M04_Semen | chr22:50250238-50250494 | ZBED4 | Island | 5'UTR |
| 50M04_Semen | chr1:21877322-21877781 | ALPL;ALPL | | 5'UTR;TSS1500 |
| 50M04_Semen | chr1:227163700-227163884 | CABC1 | N_Shore | Body |
| 50M04_Semen | chr5:415575-415885 | AHRR | | Body |
| 50M04_Semen | chr6:31237013-31237034 | HLA-C | N_Shore | Body |
| 50M04_Semen | chr6:31237405-31237722 | HLA-C | N_Shore | Body |
| 50M04_Semen | chr7:100611492-100611764 | MUC12 | S_Shore | TSS1500 |
| 50M04_Semen | chr7:157809360-157809451 | PTPRN2;PTPRN2;PTPRN2 | | Body;Body;Body |
| 50M04_Semen | chr7:158294592-158294666 | PTPRN2;PTPRN2;PTPRN2 | | Body;Body;Body |
| 50M04_Semen | chr10:1206555-1206690 | NCRNA00200 | S_Shore | Body |
| 50M04_Semen | chr11:2019079-2019587 | H19;MIR675 | N_Shore | TSS200;TSS1500 |
| 50M04_Semen | chr11:128812978-128813183 | TP53AIP1 | | TSS200 |
| 50M04_Semen | chr11:128813223-128813470 | TP53AIP1 | | TSS1500 |
| 50M04_Semen | chr13:111039873-111040242 | COL4A2 | S_Shore | Body |
| 50M04_Semen | chr17:714733-714935 | NXN | S_Shore | Body |
| 50M04_Semen | chr17:855187-855312 | NXN | | Body |
| 50M04_Semen | chr17:77952453-77952488 | TBC1D16 | S_Shore | Body |
| 50M04_Semen | chr17:79380493-79380585 | BAHCC1 | Island | Body |
| 50M04_Semen | chr19:49061546-49061702 | SULT2B1 | Island | Body |
| 50M05_Semen | chr1:3028230-3028877 | PRDM16;PRDM16 | Island | Body;Body |
| 50M05_Semen | chr1:3184547-3184704 | PRDM16;PRDM16 | Island | Body;Body |
| 50M05_Semen | chr1:3559559-3559630 | WDR8 | N_Shore | Body |
| 50M05_Semen | chr6:32369227-32369487 | BTNL2 | | Body |
| 50M05_Semen | chr7:2801248-2801732 | GNA12 | N_Shore | Body |
| 50M05_Semen | chr10:43599476-43599566 | RET;RET | N_Shore | Body;Body |
| 50M05_Semen | chr10:131686103-131686885 | EBF3 | N_Shore | Body |
| 50M05_Semen | chr11:128812913-128813583 | TP53AIP1 | | TSS200 |
| 50M05_Semen | chr17:79380493-79380585 | BAHCC1 | Island | Body |
| 50M05_Semen | chr19:49061546-49061702 | SULT2B1 | Island | Body |
| 50M05_Semen | chr22:24348549-24348715 | GSTTP1 | | TSS1500 |
| 50M05_Semen | chr1:3128175-3128606 | PRDM16;PRDM16 | | Body;Body |
| 50M05_Semen | chr4:1201315-1202966 | LOC100130872;LOC100130872 | N_Shore | Body;Body |
| 50M05_Semen | chr6:30039403-30039432 | RNF39;RNF39 | Island | Body;Body |
| 50M05_Semen | chr6:32606845-32607174 | HLA-DQA1 | | Body |
| 50M05_Semen | chr7:1113014-1113854 | C7orf50;C7orf50;C7orf50 | | Body;Body;Body |
| 50M05_Semen | chr7:23454734-23455115 | IGF2BP3 | N_Shore | Body |
| 50M05_Semen | chr7:138435324-138436009 | ATP6V0A4;ATP6V0A4;ATP6V0A4 | | Body;Body;Body |
| 50M05_Semen | chr8:27182889-27183342 | PTK2B;PTK2B;PTK2B;PTK2B | N_Shore | 5'UTR;TSS200;5'UTR;TSS200 |
| 50M05_Semen | chr8:144635861-144636462 | GSDMD;GSDMD | Island | 5'UTR;1stExon |
| 50M05_Semen | chr11:5706226-5706435 | TRIM5;TRIM5;TRIM5;TRIM5;TRIM5 | | 5'UTR;5'UTR;1stExon;1stExon;5'UTR |
| 50M05_Semen | chr17:855187-855312 | NXN | | Body |
| 50M05_Semen | chr17:10533122-10533415 | MYH3 | | Body |

| Supplementary Table 11: Coordinates of MeVars identified in prefrontal cortex samples from 528 controls (duplicate values of coordinates retained) | | | | | |
|---|---|---|---|---|---|
| Sample | Coordinate | UCSC_REFGENE_NAME | RELATION_TO_UCSC_CPG_ISLAND | UCSC_REFGENE_GROUP | SZ Candidate gene ? |
| Control-Sample10_Beta | chr19:58861702-58862087 | A1BG | Island | Body | No |
| Control-Sample1_Beta | chr12:125625016-125625041 | AACS | | Body | No |
| Control-Sample11_Beta | chr1:12703968-12704481 | AADACL4 | | TSS1500 | No |
| Control-Sample1_Beta | chr17:74463439-74463681 | AANAT;AANAT | | TSS200;Body | Yes |
| Control-Sample13_Beta | chr17:74463560-74463681 | AANAT;AANAT | | TSS200;Body | Yes |
| Control-Sample1_Beta | chr17:79097051-79097243 | AATK | S_Shore | Body | Yes |
| Control-Sample19_Beta | chr17:79109729-79110023 | AATK | Island | Body | Yes |
| Control-Sample194_Be | chr17:79094415-79094934 | AATK | Island | Body | Yes |
| Control-Sample30_Beta | chr17:79098558-79099942 | AATK | S_Shore | Body | Yes |
| Control-Sample57_Beta | chr17:79109729-79109808 | AATK | Island | Body | Yes |
| Control-Sample108_Be | chr16:8806531-8806586 | ABAT;ABAT | | 5'UTR;TSS1500 | Yes |
| Control-Sample194_Be | chr16:8806359-8806569 | ABAT;ABAT | | 5'UTR;TSS1500 | Yes |
| Control-Sample21_Beta | chr16:8862362-8862653 | ABAT;ABAT;ABAT | | Body;Body;Body | Yes |
| Control-Sample67_Beta | chr16:8862362-8862626 | ABAT;ABAT;ABAT | | Body;Body;Body | Yes |
| Control-Sample18_Beta | chr1:94586592-94587076 | ABCA4 | | 1stExon | Yes |
| Control-Sample19_Beta | chr1:94560851-94561088 | ABCA4 | | Body | Yes |
| Control-Sample10_Beta | chr1:94586670-94587076 | ABCA4;ABCA4 | | 1stExon;5'UTR | Yes |
| Control-Sample1_Beta | chr13:95826930-95827029 | ABCC4;ABCC4 | | Body;Body | No |
| Control-Sample108_Be | chr12:40013596-40013961 | ABCD2;ABCD2 | | 1stExon;5'UTR | No |
| Control-Sample506_Be | chr2.0:27346434-27346593 | ABHD1 | Island | TSS1500 | Yes |
| Control-Sample80_Beta | chr2:27346296-27346593 | ABHD1 | N_Shore | TSS1500 | Yes |
| Control-Sample26_Beta | chr3:52008654-52008740 | ABHD14B;ABHD14B;ABHD14A;ABHD14B | N_Shore | TSS200;TSS200;TSS1500;TSS200 | Yes |
| Control-Sample26_Beta | chr3:52008802-52008944 | ABHD14B;ABHD14B;ABHD14A;ABHD14B | N_Shore | TSS200;TSS200;TSS1500;TSS200 | Yes |
| Control-Sample1_Beta | chr3:52008487-52008740 | ABHD14B;ABHD14B;ABHD14B;ABHD14A;ABHD14B;ABHD | N_Shore | 1stExon;Body;5'UTR;TSS1500;1stExon;5'UTR | Yes |
| Control-Sample10_Beta | chr3:52008538-52008740 | ABHD14B;ABHD14B;ABHD14B;ABHD14A;ABHD14B;ABHD | N_Shore | 1stExon;Body;5'UTR;TSS1500;1stExon;5'UTR | Yes |
| Control-Sample43_Beta | chr3:52008487-52008654 | ABHD14B;ABHD14B;ABHD14B;ABHD14A;ABHD14B;ABHD | N_Shore | 1stExon;Body;5'UTR;TSS1500;1stExon;5'UTR | Yes |
| Control-Sample10_Beta | chr17:27892866-27893407 | ABHD15 | N_Shore | Body | No |
| Control-Sample11_Beta | chr19:17403176-17403529 | ABHD8 | Island | 3'UTR | No |
| Control-Sample1_Beta | chr10:27150407-27150529 | ABI1;ABI1;ABI1;ABI1 | S_Shore | TSS1500;TSS1500;TSS1500;TSS1500 | No |
| Control-Sample108_Be | chr10:116444240-116444414 | ABLIM1;ABLIM1;ABLIM1;ABLIM1 | | Body;Body | Yes |
| Control-Sample108_Be | chr10:116286605-116286974 | ABLIM1;ABLIM1;ABLIM1;ABLIM1;ABLIM1 | | Body;1stExon;Body;5'UTR;Body | Yes |
| Control-Sample1_Beta | chr4:8107118-8107392 | ABLIM2;ABLIM2;ABLIM2;ABLIM2;ABLIM2;ABLIM2;ABLIM2 | | Body;Body;Body;Body;Body;Body;Body | No |
| Control-Sample16_Beta | chr17:1076874-1077099 | ABR;ABR | | Body;5'UTR | Yes |
| Control-Sample1_Beta | chr17:907643-907886 | ABR;ABR;ABR | N_Shelf | 3'UTR;3'UTR;3'UTR | Yes |
| Control-Sample1_Beta | chr17:981518-981730 | ABR;ABR;ABR | Island | Body;Body;Body | Yes |
| Control-Sample19_Beta | chr17:907729-907886 | ABR;ABR;ABR | N_Shelf | 3'UTR;3'UTR;3'UTR | Yes |
| Control-Sample20_Beta | chr17:981643-981730 | ABR;ABR;ABR | Island | Body;Body;Body | Yes |
| Control-Sample33_Beta | chr17:943108-943342 | ABR;ABR;ABR | | Body;Body;Body | Yes |
| Control-Sample5_Beta | chr17:907643-907732 | ABR;ABR;ABR | N_Shelf | 3'UTR;3'UTR;3'UTR | Yes |
| Control-Sample108_Be | chr17:35656660-35656896 | ACACA;ACACA;ACACA;ACACA;ACACA;ACACA;ACACA | | 5'UTR;Body;5'UTR;Body;1stExon;1stExon;Body | No |
| Control-Sample105_Be | chr1:76189707-76189801 | ACADM;ACADM | N_Shore | TSS1500;TSS1500 | No |
| Control-Sample464_Be | chr15:89386741-89386809 | ACAN;ACAN | | Body;Body | No |
| Control-Sample1_Beta | chr1:1238595-1239259 | ACAP3 | N_Shelf | Body | Yes |
| Control-Sample20_Beta | chr1:1238595-1238797 | ACAP3 | N_Shelf | Body | Yes |
| Control-Sample331_Be | chr1:1234354-1234844 | ACAP3 | | Body | Yes |
| Control-Sample68_Beta | chr1:1238595-1238702 | ACAP3 | N_Shelf | Body | Yes |
| Control-Sample41_Beta | chr11:107992005-107992034 | ACAT1 | Island | TSS1500 | No |
| Control-Sample15_Beta | chr6:160182184-160182624 | ACAT2 | N_Shore | TSS1500 | Yes |
| Control-Sample2_Beta | chr6:160182438-160182624 | ACAT2 | Island | TSS1500 | Yes |
| Control-Sample39_Beta | chr6:160182447-160182624 | ACAT2 | Island | TSS1500 | Yes |
| Control-Sample17_Beta | chr17:43213270-43213629 | ACBD4;ACBD4;ACBD4;ACBD4;ACBD4;ACBD4;ACBD4;ACE | S_Shelf | 5'UTR;5'UTR;5'UTR;1stExon;5'UTR;5'UTR;1stExon;1st | No |
| Control-Sample3_Beta | chr7:150744274-150744599 | ACCN3;ACCN3;ABCB8;ACCN3 | N_Shore | TSS1500;TSS1500;3'UTR;TSS1500 | No |
| Control-Sample16_Beta | chr2:220379635-220380542 | ACCN4;ACCN4 | S_Shore | 1stExon;1stExon | No |
| Control-Sample32_Beta | chr1:6390053-6390530 | ACOT7;ACOT7;ACOT7;ACOT7 | | Body;Body;Body;Body | Yes |
| Control-Sample69_Beta | chr1:6333502-6333867 | ACOT7;ACOT7;ACOT7;ACOT7 | | Body;Body;Body;Body | Yes |
| Control-Sample108_Be | chr4:8395941-8396300 | ACOX3;ACOX3 | | Body;Body | Yes |
| Control-Sample13_Beta | chr4:8372691-8372840 | ACOX3;ACOX3 | N_Shelf | Body;Body | Yes |
| Control-Sample1_Beta | chr15:78526804-78527410 | ACSBG1 | | 1stExon | Yes |
| Control-Sample2_Beta | chr15:78526836-78527410 | ACSBG1 | | 1stExon | Yes |
| Control-Sample11_Beta | chr19:6135279-6135519 | ACSBG2 | | TSS1500 | No |
| Control-Sample1_Beta | chr16:89202571-89203153 | ACSF3;ACSF3;ACSF3 | | Body;Body;Body | Yes |
| Control-Sample13_Beta | chr16:89202571-89202896 | ACSF3;ACSF3;ACSF3 | | Body;Body;Body | Yes |
| Control-Sample19_Beta | chr16:89202571-89202817 | ACSF3;ACSF3;ACSF3 | | Body;Body;Body | Yes |
| Control-Sample1_Beta | chr7:5567862-5568044 | ACTB | N_Shore | Body | No |
| Control-Sample1_Beta | chr7:150020945-150021553 | ACTR3C;LRRC61;LRRC61 | S_Shore | TSS200;5'UTR;5'UTR | Yes |
| Control-Sample11_Beta | chr7:150020925-150021553 | ACTR3C;LRRC61;LRRC61 | S_Shore | TSS200;5'UTR;5'UTR | Yes |
| Control-Sample13_Beta | chr7:150021488-150021553 | ACTR3C;LRRC61;LRRC61 | S_Shore | TSS1500;5'UTR;5'UTR | Yes |
| Control-Sample1_Beta | chr1:2937999-2938246 | ACTRT2 | N_Shore | TSS200 | No |
| Control-Sample12_Beta | chr11:67418315-67418405 | ACY3 | | TSS200 | Yes |
| Control-Sample13_Beta | chr11:67418310-67418365 | ACY3 | | TSS200 | Yes |
| Control-Sample14_Beta | chr11:67418291-67418315 | ACY3 | | TSS200 | Yes |
| Control-Sample18_Beta | chr11:67418310-67418358 | ACY3 | | TSS200 | Yes |
| Control-Sample21_Beta | chr11:67418291-67418358 | ACY3 | | TSS200 | Yes |
| Control-Sample3_Beta | chr11:67418310-67418405 | ACY3 | | TSS200 | Yes |
| Control-Sample35_Beta | chr11:67418291-67418405 | ACY3 | | TSS200 | Yes |
| Control-Sample7_Beta | chr11:67418291-67418365 | ACY3 | | TSS200 | Yes |
| Control-Sample11_Beta | chr2:54517167-54517382 | ACYP2 | | Body | No |
| Control-Sample19_Beta | chr17:42856773-42857157 | ADAM11 | Island | Body | No |
| Control-Sample194_Be | chr5:157003155-157003436 | ADAM19 | Island | TSS1500 | No |
| Control-Sample118_Be | chr8:39380341-39380592 | ADAM3A;ADAM3A;ADAM3A | Island | Body;Body;Body | Yes |
| Control-Sample524_Be | chr8.0:39380341-39380592 | ADAM3A;ADAM3A;ADAM3A | Island | Body;Body;Body | Yes |
| Control-Sample1_Beta | chr15:100804081-100804162 | ADAMTS17 | | Body | No |
| Control-Sample14_Beta | chr16:77469583-77469710 | ADAMTS18 | S_Shore | TSS1500 | No |
| Control-Sample158_Be | chr5:178548229-178548700 | ADAMTS2 | Island | Body | Yes |
| Control-Sample1_Beta | chr5:178594526-178594649 | ADAMTS2;ADAMTS2 | Island | Body;Body | Yes |
| Control-Sample23_Beta | chr4:73178007-73178545 | ADAMTS3 | | Body | No |
| Control-Sample10_Beta | chr7:951544-951962 | ADAP1 | S_Shelf | Body | No |
| Control-Sample15_Beta | chr1:154600682-154600712 | ADAR | S_Shore | TSS1500 | Yes |
| Control-Sample26_Beta | chr1:154600682-154600788 | ADAR | S_Shore | TSS1500 | Yes |

| | | | | | |
|---|---|---|---|---|---|
| Control-Sample1_Beta | chr10:1511173-1511277 | ADARB2 | | Body | Yes |
| Control-Sample1_Beta | chr10:1551251-1551326 | ADARB2 | | Body | Yes |
| Control-Sample10_Beta | chr10:1511024-1511277 | ADARB2 | | Body | Yes |
| Control-Sample10_Beta | chr10:1595543-1595704 | ADARB2 | Island | Body | Yes |
| Control-Sample16_Beta | chr10:1595543-1595633 | ADARB2 | Island | Body | Yes |
| Control-Sample19_Beta | chr10:1511024-1511228 | ADARB2 | | Body | Yes |
| Control-Sample19_Beta | chr10:1559112-1559249 | ADARB2 | | Body | Yes |
| Control-Sample36_Beta | chr10:1505595-1505654 | ADARB2 | N_Shore | Body | Yes |
| Control-Sample4_Beta | chr10:1505595-1506175 | ADARB2 | N_Shore | Body | Yes |
| Control-Sample464_Be | chr10:1692910-1693107 | ADARB2 | | Body | Yes |
| Control-Sample514_Be | chr10.0:1511024-1511277 | ADARB2 | | Body | Yes |
| Control-Sample552_Be | chr10.0:1511024-1511228 | ADARB2 | | Body | Yes |
| Control-Sample6_Beta | chr10:1531034-1531405 | ADARB2 | | Body | Yes |
| Control-Sample6_Beta | chr10:1595543-1595785 | ADARB2 | Island | Body | Yes |
| Control-Sample64_Beta | chr10:1595595-1595704 | ADARB2 | Island | Body | Yes |
| Control-Sample171_Be | chr19:1907972-1908388 | ADAT3;SCAMP4 | N_Shore | 5'UTR;5'UTR | No |
| Control-Sample4_Beta | chr16:50347766-50347895 | ADCY7 | | Body | No |
| Control-Sample1_Beta | chr11:10325567-10325809 | ADM | N_Shore | TSS1500 | No |
| Control-Sample11_Beta | chr13:114108433-114108515 | ADPRHL1 | | TSS1500 | No |
| Control-Sample1_Beta | chr20:60882762-60882801 | ADRM1;ADRM1 | Island | Body;Body | No |
| Control-Sample1_Beta | chr2:100170858-100171136 | AFF3;AFF3 | | Body;Body | Yes |
| Control-Sample33_Beta | chr2:100618399-100618466 | AFF3;AFF3 | | Body;Body | Yes |
| Control-Sample57_Beta | chr2:100210257-100210626 | AFF3;AFF3 | Island | Body;Body | Yes |
| Control-Sample1_Beta | chr2:236486327-236486625 | AGAP1;AGAP1 | | Body;Body | Yes |
| Control-Sample1_Beta | chr2:236619748-236619825 | AGAP1;AGAP1 | | Body;Body | Yes |
| Control-Sample1_Beta | chr2:236674174-236674489 | AGAP1;AGAP1 | S_Shore | Body;Body | Yes |
| Control-Sample1_Beta | chr2:236715747-236716372 | AGAP1;AGAP1 | | Body;Body | Yes |
| Control-Sample1_Beta | chr2:236761068-236761574 | AGAP1;AGAP1 | | Body;Body | Yes |
| Control-Sample10_Beta | chr2:236721769-236721826 | AGAP1;AGAP1 | | Body;Body | Yes |
| Control-Sample108_Be | chr2:236888269-236888769 | AGAP1;AGAP1 | | Body;Body | Yes |
| Control-Sample11_Beta | chr2:236867836-236868078 | AGAP1;AGAP1 | Island | Body;Body | Yes |
| Control-Sample12_Beta | chr2:236812587-236812946 | AGAP1;AGAP1 | | Body;Body | Yes |
| Control-Sample12_Beta | chr2:236826245-236826485 | AGAP1;AGAP1 | | Body;Body | Yes |
| Control-Sample124_Be | chr2:236760940-236761421 | AGAP1;AGAP1 | | Body;Body | Yes |
| Control-Sample15_Beta | chr2:236826311-236826485 | AGAP1;AGAP1 | | Body;Body | Yes |
| Control-Sample19_Beta | chr2:236443439-236443932 | AGAP1;AGAP1 | N_Shore | Body;Body | Yes |
| Control-Sample19_Beta | chr2:236643527-236643760 | AGAP1;AGAP1 | | Body;Body | Yes |
| Control-Sample19_Beta | chr2:236961065-236961189 | AGAP1;AGAP1 | | Body;Body | Yes |
| Control-Sample2_Beta | chr2:236715747-236716052 | AGAP1;AGAP1 | | Body;Body | Yes |
| Control-Sample20_Beta | chr2:236443439-236444259 | AGAP1;AGAP1 | N_Shore | Body;Body | Yes |
| Control-Sample20_Beta | chr2:236721253-236721472 | AGAP1;AGAP1 | | Body;Body | Yes |
| Control-Sample331_Be | chr2:236716219-236716372 | AGAP1;AGAP1 | | Body;Body | Yes |
| Control-Sample38_Beta | chr2:236619539-236619767 | AGAP1;AGAP1 | | Body;Body | Yes |
| Control-Sample57_Beta | chr2:236760940-236761574 | AGAP1;AGAP1 | | Body;Body | Yes |
| Control-Sample645_Be | chr2:236586347-236586710 | AGAP1;AGAP1 | | Body;Body | Yes |
| Control-Sample88_Beta | chr2:236444202-236444284 | AGAP1;AGAP1 | S_Shore | Body;Body | Yes |
| Control-Sample1_Beta | chr12:58119992-58120237 | AGAP2;LOC100130776;AGAP2 | Island | 3'UTR;TSS200;3'UTR | No |
| Control-Sample53_Beta | chr7:150822655-150822987 | AGAP3 | Island | Body | No |
| Control-Sample40_Beta | chr11:47736463-47736941 | AGBL2 | N_Shore | TSS200 | No |
| Control-Sample480_Be | chr1:100315729-100316030 | AGL;AGL;AGL;AGL;AGL;AGL | Island | TSS1500;TSS1500;5'UTR;1stExon;TSS1500;TSS1500 | No |
| Control-Sample395_Be | chr6:32146466-32146529 | AGPAT1;RNF5;RNF5P1 | | TSS1500;Body;Body | Yes |
| Control-Sample572_Be | chr6.0:32146466-32146529 | AGPAT1;RNF5;RNF5P1 | | TSS1500;Body;Body | Yes |
| Control-Sample1_Beta | chr2:241807924-241808259 | AGXT | | TSS1500 | Yes |
| Control-Sample11_Beta | chr2:241807924-241808113 | AGXT | | TSS1500 | Yes |
| Control-Sample15_Beta | chr2:241808004-241808259 | AGXT | | TSS200 | Yes |
| Control-Sample23_Beta | chr2:241807859-241808113 | AGXT | | TSS1500 | Yes |
| Control-Sample108_Be | chr7:129007618-129008130 | AHCYL2;AHCYL2;AHCYL2;AHCYL2 | | TSS1500;Body;TSS1500;Body | Yes |
| Control-Sample42_Beta | chr7:129007902-129008179 | AHCYL2;AHCYL2;AHCYL2;AHCYL2 | | Body;TSS200;Body;TSS200 | Yes |
| Control-Sample2_Beta | chr1:27931284-27931503 | AHDC1 | S_Shore | TSS1500 | No |
| Control-Sample1_Beta | chr11:62211788-62212431 | AHNAK | N_Shore | Body | Yes |
| Control-Sample11_Beta | chr11:62212114-62212431 | AHNAK | Island | Body | Yes |
| Control-Sample13_Beta | chr11:62211990-62212431 | AHNAK | Island | Body | Yes |
| Control-Sample30_Beta | chr11:62212169-62212431 | AHNAK | Island | Body | Yes |
| Control-Sample68_Beta | chr11:62212114-62212195 | AHNAK | Island | Body | Yes |
| Control-Sample10_Beta | chr5:415575-415885 | AHRR | | Body | Yes |
| Control-Sample509_Be | chr5.0:415575-415885 | AHRR | | Body | Yes |
| Control-Sample223_Be | chr3:186330026-186330067 | AHSG | | TSS1500 | No |
| Control-Sample11_Beta | chr21:45705694-45705742 | AIRE | Island | TSS200 | Yes |
| Control-Sample160_Be | chr21:45705721-45705742 | AIRE | Island | TSS200 | Yes |
| Control-Sample17_Beta | chr21:45705694-45705721 | AIRE | Island | TSS200 | Yes |
| Control-Sample1_Beta | chr1:4794912-4795114 | AJAP1;AJAP1 | | Body;Body | Yes |
| Control-Sample1_Beta | chr1:4837112-4837462 | AJAP1;AJAP1 | | 3'UTR;3'UTR | Yes |
| Control-Sample245_Be | chr1:65613383-65613392 | AK3L1;AK3L1;AK3L1 | Island | 5'UTR;TSS1500;TSS200 | No |
| Control-Sample23_Beta | chr14:96890445-96890632 | AK7 | Island | Body | No |
| Control-Sample20_Beta | chr6:151646312-151646552 | AKAP12;AKAP12 | N_Shore | TSS1500;Body | No |
| Control-Sample1_Beta | chr19:15530606-15530737 | AKAP8L | S_Shore | TSS1500 | Yes |
| Control-Sample20_Beta | chr19:15530606-15530870 | AKAP8L | S_Shore | TSS1500 | Yes |
| Control-Sample11_Beta | chr1:19600471-19600817 | AKR7L;AKR7L | Island | 1stExon;1stExon | Yes |
| Control-Sample12_Beta | chr1:19600702-19601026 | AKR7L;AKR7L | Island | TSS200;TSS200 | Yes |
| Control-Sample25_Beta | chr1:19600702-19600730 | AKR7L;AKR7L | Island | TSS200;TSS200 | Yes |
| Control-Sample26_Beta | chr1:19600817-19600912 | AKR7L;AKR7L | Island | TSS1500;TSS1500 | Yes |
| Control-Sample36_Beta | chr1:19600673-19600730 | AKR7L;AKR7L | Island | TSS200;TSS200 | Yes |
| Control-Sample511_Be | chr1.0:19600673-19600730 | AKR7L;AKR7L | Island | TSS200;TSS200 | Yes |
| Control-Sample6_Beta | chr1:19600471-19600909 | AKR7L;AKR7L | Island | 1stExon;1stExon | Yes |
| Control-Sample1_Beta | chr14:105236538-105236654 | AKT1;AKT1;AKT1 | N_Shelf | 3'UTR;3'UTR;3'UTR | No |
| Control-Sample1_Beta | chr15:101418967-101419074 | ALDH1A3 | N_Shore | TSS1500 | Yes |
| Control-Sample1_Beta | chr15:101419269-101419333 | ALDH1A3 | Island | TSS1500 | Yes |
| Control-Sample10_Beta | chr15:101419269-101419296 | ALDH1A3 | Island | TSS1500 | Yes |
| Control-Sample11_Beta | chr15:101419262-101419333 | ALDH1A3 | Island | TSS1500 | Yes |
| Control-Sample15_Beta | chr15:101418967-101419148 | ALDH1A3 | N_Shore | TSS1500 | Yes |
| Control-Sample19_Beta | chr15:101418967-101419017 | ALDH1A3 | N_Shore | TSS1500 | Yes |

| | | | | | |
|---|---|---|---|---|---|
| Control-Sample194_Be | chr15:101419275-101419296 | ALDH1A3 | Island | TSS1500 | Yes |
| Control-Sample2_Beta | chr15:101419262-101419275 | ALDH1A3 | Island | TSS1500 | Yes |
| Control-Sample270_Be | chr15:101419275-101419333 | ALDH1A3 | Island | TSS1500 | Yes |
| Control-Sample3_Beta | chr15:101419262-101419296 | ALDH1A3 | Island | TSS1500 | Yes |
| Control-Sample57_Beta | chr15:101419293-101419333 | ALDH1A3 | Island | TSS1500 | Yes |
| Control-Sample331_Be | chr11:67775972-67776123 | ALDH3B1 | | TSS200 | Yes |
| Control-Sample171_Be | chr11:67777770-67777952 | ALDH3B1;ALDH3B1;ALDH3B1 | | TSS200;TSS200;5'UTR | Yes |
| Control-Sample1_Beta | chr5:125929132-125929671 | ALDH7A1 | N_Shore | Body | No |
| Control-Sample10_Beta | chr2:3704589-3704793 | ALLC | | TSS1500 | Yes |
| Control-Sample11_Beta | chr2:3704530-3704793 | ALLC | | TSS1500 | Yes |
| Control-Sample13_Beta | chr2:3704751-3704793 | ALLC | | TSS1500 | Yes |
| Control-Sample171_Be | chr2:3704589-3704773 | ALLC | | TSS1500 | Yes |
| Control-Sample223_Be | chr2:3704530-3704773 | ALLC | | TSS1500 | Yes |
| Control-Sample3_Beta | chr2:3704501-3704793 | ALLC | | TSS1500 | Yes |
| Control-Sample475_Be | chr2:3704501-3704773 | ALLC | | TSS1500 | Yes |
| Control-Sample572_Be | chr2.0:3704589-3704793 | ALLC | | TSS1500 | Yes |
| Control-Sample63_Beta | chr2:3704530-3704751 | ALLC | | TSS1500 | Yes |
| Control-Sample1_Beta | chr17:6899207-6899888 | ALOX12 | Island | TSS200 | Yes |
| Control-Sample15_Beta | chr17:6899207-6899577 | ALOX12 | Island | TSS200 | Yes |
| Control-Sample26_Beta | chr17:6899207-6900356 | ALOX12 | Island | TSS200 | Yes |
| Control-Sample73_Beta | chr17:6899207-6899332 | ALOX12 | Island | TSS200 | Yes |
| Control-Sample73_Beta | chr17:6899364-6899577 | ALOX12 | Island | TSS200 | Yes |
| Control-Sample74_Beta | chr17:6899310-6899364 | ALOX12 | Island | TSS200 | Yes |
| Control-Sample75_Beta | chr17:6899332-6899577 | ALOX12 | Island | TSS200 | Yes |
| Control-Sample3_Beta | chr17:6756855-6756891 | ALOX12P2 | | TSS200 | No |
| Control-Sample1_Beta | chr4:113332969-113333550 | ALPK1;ALPK1 | Island | Body;Body | Yes |
| Control-Sample10_Beta | chr4:113218385-113218524 | ALPK1;ALPK1 | | TSS200;TSS200 | Yes |
| Control-Sample13_Beta | chr4:113333069-113333550 | ALPK1;ALPK1 | Island | Body;Body | Yes |
| Control-Sample21_Beta | chr4:113333052-113333550 | ALPK1;ALPK1 | Island | Body;Body | Yes |
| Control-Sample4_Beta | chr4:113332969-113333072 | ALPK1;ALPK1 | Island | Body;Body | Yes |
| Control-Sample51_Beta | chr4:113333072-113333550 | ALPK1;ALPK1 | Island | Body;Body | Yes |
| Control-Sample53_Beta | chr4:113332969-113333163 | ALPK1;ALPK1 | Island | Body;Body | Yes |
| Control-Sample1_Beta | chr18:56296354-56296607 | ALPK2 | | TSS200 | Yes |
| Control-Sample21_Beta | chr18:56296354-56296496 | ALPK2 | | TSS200 | Yes |
| Control-Sample1_Beta | chr1:110609973-110610452 | ALX3 | N_Shore | Body | Yes |
| Control-Sample22_Beta | chr1:110612044-110612474 | ALX3 | Island | Body | Yes |
| Control-Sample19_Beta | chr12:96350519-96350720 | AMDHD1 | Island | Body | No |
| Control-Sample16_Beta | chr16:2569920-2569967 | AMDHD2;ATP6V0C;AMDHD2 | Island | TSS1500;3'UTR;TSS1500 | Yes |
| Control-Sample29_Beta | chr16:2569701-2569925 | AMDHD2;ATP6V0C;AMDHD2 | Island | TSS1500;Body;TSS1500 | Yes |
| Control-Sample57_Beta | chr16:2569701-2569967 | AMDHD2;ATP6V0C;AMDHD2 | Island | TSS1500;Body;TSS1500 | Yes |
| Control-Sample25_Beta | chr19:2250561-2251061 | AMH | Island | Body | Yes |
| Control-Sample28_Beta | chr19:2250561-2250956 | AMH | Island | Body | Yes |
| Control-Sample37_Beta | chr19:2250901-2251061 | AMH | Island | Body | Yes |
| Control-Sample45_Beta | chr19:2251588-2251837 | AMH | Island | Body | Yes |
| Control-Sample15_Beta | chr3:49757438-49757476 | AMIGO3;RNF123 | S_Shore | TSS1500;Body | No |
| Control-Sample14_Beta | chr14:103388958-103389022 | AMN | N_Shore | TSS200 | No |
| Control-Sample170_Be | chr11:10476494-10476608 | AMPD3;AMPD3;AMPD3 | S_Shelf | TSS200;TSS1500;Body | Yes |
| Control-Sample475_Be | chr11:10476242-10476976 | AMPD3;AMPD3;AMPD3 | S_Shelf | TSS1500;TSS1500;Body | Yes |
| Control-Sample1_Beta | chr3:49459855-49459937 | AMT;AMT;AMT;NICN1;AMT;AMT | | Body;1stExon;1stExon;3'UTR;1stExon;1stExon | Yes |
| Control-Sample13_Beta | chr3:49459855-49460057 | AMT;AMT;AMT;NICN1;AMT;AMT | | Body;1stExon;1stExon;3'UTR;1stExon;1stExon | Yes |
| Control-Sample20_Beta | chr7:2753729-2754101 | AMZ1 | | 3'UTR | Yes |
| Control-Sample88_Beta | chr7:2742371-2742484 | AMZ1 | Island | Body | Yes |
| Control-Sample22_Beta | chr8:108510286-108510314 | ANGPT1 | | TSS200 | Yes |
| Control-Sample19_Beta | chr8:108510096-108510292 | ANGPT1;ANGPT1 | | 5'UTR;1stExon | Yes |
| Control-Sample1_Beta | chr8:41654967-41655154 | ANK1;ANK1;ANK1;ANK1;ANK1 | Island | Body;Body;Body;Body;Body | Yes |
| Control-Sample25_Beta | chr8:41654967-41655789 | ANK1;ANK1;ANK1;ANK1;ANK1 | Island | Body;Body;Body;Body;Body | Yes |
| Control-Sample28_Beta | chr8:41654967-41655673 | ANK1;ANK1;ANK1;ANK1;ANK1 | Island | Body;Body;Body;Body;Body | Yes |
| Control-Sample108_Be | chr4:113970820-113971324 | ANK2;ANK2;ANK2;ANK2;ANK2 | | 5'UTR;1stExon;5'UTR;1stExon;Body | No |
| Control-Sample1_Beta | chr17:54230419-54230878 | ANKFN1 | | TSS1500 | Yes |
| Control-Sample10_Beta | chr17:54230419-54230996 | ANKFN1 | | TSS1500 | Yes |
| Control-Sample12_Beta | chr17:54230419-54230730 | ANKFN1 | | TSS1500 | Yes |
| Control-Sample39_Beta | chr17:54230419-54230793 | ANKFN1 | | TSS1500 | Yes |
| Control-Sample53_Beta | chr17:54230793-54230996 | ANKFN1 | | TSS200 | Yes |
| Control-Sample6_Beta | chr17:54230419-54230787 | ANKFN1 | | TSS1500 | Yes |
| Control-Sample1_Beta | chr2:241458886-241459638 | ANKMY1;ANKMY1 | N_Shore | Body;Body | Yes |
| Control-Sample48_Beta | chr2:241458886-241459502 | ANKMY1;ANKMY1 | N_Shore | Body;Body | Yes |
| Control-Sample1_Beta | chr16:89357477-89357741 | ANKRD11 | | Body | Yes |
| Control-Sample1_Beta | chr16:89381904-89382131 | ANKRD11 | | Body | Yes |
| Control-Sample1_Beta | chr16:89439771-89440133 | ANKRD11 | S_Shore | 5'UTR | Yes |
| Control-Sample57_Beta | chr16:89421749-89421929 | ANKRD11 | | 5'UTR | Yes |
| Control-Sample58_Beta | chr16:89421700-89421929 | ANKRD11 | | 5'UTR | Yes |
| Control-Sample1_Beta | chr5:10562978-10563501 | ANKRD33B | Island | TSS1500 | Yes |
| Control-Sample28_Beta | chr5:10562978-10563358 | ANKRD33B | Island | TSS1500 | Yes |
| Control-Sample19_Beta | chr2:198063653-198063781 | ANKRD44 | | TSS1500 | No |
| Control-Sample15_Beta | chr2:71205564-71206291 | ANKRD53;ANKRD53 | Island | TSS200;TSS200 | Yes |
| Control-Sample3_Beta | chr2:71211981-71212157 | ANKRD53;ANKRD53 | | 3'UTR;Body | Yes |
| Control-Sample1_Beta | chr5:55530135-55530254 | ANKRD55 | | TSS1500 | No |
| Control-Sample67_Beta | chr12:99548504-99548550 | ANKS1B;ANKS1B;ANKS1B | | TSS200;TSS200;Body | Yes |
| Control-Sample291_Be | chr12:99548329-99548550 | ANKS1B;ANKS1B;ANKS1B;ANKS1B;ANKS1B | | 1stExon;5'UTR;5'UTR;Body;1stExon | Yes |
| Control-Sample321_Be | chr11:69982916-69982996 | ANO1;ANO1 | | Body;Body | Yes |
| Control-Sample530_Be | chr11.0:69982916-69982996 | ANO1;ANO1 | | Body;Body | Yes |
| Control-Sample15_Beta | chr12:5936877-5937105 | ANO2 | Island | Body | No |
| Control-Sample1_Beta | chr19:17439877-17440072 | ANO8 | Island | Body | Yes |
| Control-Sample26_Beta | chr19:17445963-17446137 | ANO8 | S_Shore | TSS1500 | Yes |
| Control-Sample1_Beta | chr4:80885279-80886264 | ANTXR2 | N_Shore | Body | Yes |
| Control-Sample11_Beta | chr4:80885279-80885735 | ANTXR2 | N_Shore | Body | Yes |
| Control-Sample20_Beta | chr4:80885735-80886143 | ANTXR2 | N_Shore | Body | Yes |
| Control-Sample29_Beta | chr4:80885981-80886264 | ANTXR2 | Island | Body | Yes |
| Control-Sample6_Beta | chr2:69969029-69969052 | ANXA4 | Island | TSS200 | No |
| Control-Sample1_Beta | chr7:36764019-36764205 | AOAH;AOAH | | 1stExon;5'UTR | No |
| Control-Sample3_Beta | chr11:924500-924646 | AP2A2 | N_Shore | TSS1500 | No |

| | | | | | |
|---|---|---|---|---|---|
| Control-Sample29_Beta | chr10:26856095-26856128 | APBB1IP | Island | Body | Yes |
| Control-Sample67_Beta | chr10:26855907-26856128 | APBB1IP | Island | Body | Yes |
| Control-Sample1_Beta | chr5:112042851-112042919 | APC | N_Shore | TSS1500 | No |
| Control-Sample10_Beta | chr19:1467008-1467032 | APC2 | Island | Body | No |
| Control-Sample1_Beta | chr1:10489962-10490063 | APITD1;APITD1;APITD1 | N_Shore | TSS200;TSS1500;TSS200 | Yes |
| Control-Sample10_Beta | chr1:10489962-10490043 | APITD1;APITD1;APITD1 | N_Shore | TSS200;TSS1500;TSS200 | Yes |
| Control-Sample17_Beta | chr1:10489962-10490058 | APITD1;APITD1;APITD1 | N_Shore | TSS200;TSS1500;TSS200 | Yes |
| Control-Sample270_Be | chr1:10509935-10510123 | APITD1;CORT | | Body;TSS200 | Yes |
| Control-Sample64_Beta | chr1:10509733-10510123 | APITD1;CORT | | Body;TSS1500 | Yes |
| Control-Sample72_Beta | chr1:10509906-10510054 | APITD1;CORT | | Body;TSS200 | Yes |
| Control-Sample23_Beta | chr2:21266727-21266953 | APOB | Island | Body | No |
| Control-Sample10_Beta | chr22:39377790-39377930 | APOBEC3B | | TSS1500 | No |
| Control-Sample1_Beta | chr3:195310883-195310936 | APOD;APOD | | 1stExon;5'UTR | No |
| Control-Sample1_Beta | chr22:36044151-36044486 | APOL6 | | TSS1500 | Yes |
| Control-Sample10_Beta | chr22:36044151-36044328 | APOL6 | | TSS1500 | Yes |
| Control-Sample17_Beta | chr22:36044151-36044234 | APOL6 | | TSS1500 | Yes |
| Control-Sample1_Beta | chr21:27372387-27372461 | APP;APP;APP;APP;APP | | Body;Body;Body;Body;Body | Yes |
| Control-Sample11_Beta | chr21:27372396-27372461 | APP;APP;APP;APP;APP | | Body;Body;Body;Body;Body | Yes |
| Control-Sample20_Beta | chr7:30951328-30951392 | AQP1 | | TSS200 | Yes |
| Control-Sample223_Be | chr7:30951425-30951492 | AQP1 | | TSS200 | Yes |
| Control-Sample8_Beta | chr7:30951432-30951498 | AQP1 | | TSS200 | Yes |
| Control-Sample1_Beta | chr18:24445134-24445318 | AQP4;C18orf16 | S_Shore | Body;TSS200 | Yes |
| Control-Sample10_Beta | chr18:24445240-24445318 | AQP4;C18orf16 | S_Shore | Body;TSS200 | Yes |
| Control-Sample1_Beta | chr11:72433619-72433866 | ARAP1;ARAP1;ARAP1 | | Body;TSS1500;TSS1500 | Yes |
| Control-Sample13_Beta | chr11:72408186-72408394 | ARAP1;ARAP1;ARAP1 | Island | Body;Body;Body | Yes |
| Control-Sample16_Beta | chr11:72433551-72433866 | ARAP1;ARAP1;ARAP1 | | Body;TSS200;TSS200 | Yes |
| Control-Sample1_Beta | chr8:143694010-143694547 | ARC;ARC | N_Shore | 1stExon;3'UTR | Yes |
| Control-Sample62_Beta | chr8:143694295-143694547 | ARC;ARC | N_Shore | 1stExon;3'UTR | Yes |
| Control-Sample19_Beta | chr10:32216031-32216101 | ARHGAP12 | N_Shore | 5'UTR | No |
| Control-Sample216_Be | chr17:36612298-36612909 | ARHGAP23 | S_Shore | TSS1500 | No |
| Control-Sample108_Be | chr4:86748944-86749040 | ARHGAP24;ARHGAP24 | | Body;5'UTR | Yes |
| Control-Sample108_Be | chr4:86851399-86851425 | ARHGAP24;ARHGAP24;ARHGAP24 | | Body;Body;TSS200 | Yes |
| Control-Sample1_Beta | chr2:69001854-69001991 | ARHGAP25;ARHGAP25;ARHGAP25;ARHGAP25 | | Body;TSS200;TSS200;Body | Yes |
| Control-Sample10_Beta | chr2:69001774-69001991 | ARHGAP25;ARHGAP25;ARHGAP25;ARHGAP25 | | Body;TSS200;TSS200;Body | Yes |
| Control-Sample16_Beta | chr2:69001774-69001815 | ARHGAP25;ARHGAP25;ARHGAP25;ARHGAP25 | | Body;TSS200;TSS200;Body | Yes |
| Control-Sample1_Beta | chr17:43502999-43503401 | ARHGAP27;ARHGAP27 | N_Shelf | 1stExon;5'UTR | Yes |
| Control-Sample10_Beta | chr17:43482735-43483527 | ARHGAP27;ARHGAP27 | N_Shore | Body;Body | Yes |
| Control-Sample12_Beta | chr17:43502999-43503394 | ARHGAP27;ARHGAP27 | N_Shelf | 1stExon;5'UTR | Yes |
| Control-Sample16_Beta | chr17:43482735-43483251 | ARHGAP27;ARHGAP27 | N_Shore | Body;Body | Yes |
| Control-Sample19_Beta | chr17:43502999-43503144 | ARHGAP27;ARHGAP27 | N_Shelf | 1stExon;5'UTR | Yes |
| Control-Sample1_Beta | chr8:1811760-1812138 | ARHGEF10 | | Body | Yes |
| Control-Sample151_Be | chr8:1786207-1786322 | ARHGEF10 | | 5'UTR | Yes |
| Control-Sample19_Beta | chr8:1848040-1848407 | ARHGEF10 | | Body | Yes |
| Control-Sample19_Beta | chr8:1906312-1906498 | ARHGEF10 | | 3'UTR | Yes |
| Control-Sample24_Beta | chr8:1796429-1796676 | ARHGEF10 | N_Shore | Body | Yes |
| Control-Sample27_Beta | chr8:1905131-1905343 | ARHGEF10 | | Body | Yes |
| Control-Sample3_Beta | chr8:1807061-1807575 | ARHGEF10 | | Body | Yes |
| Control-Sample464_Be | chr1:17949212-17949604 | ARHGEF10L;ARHGEF10L | N_Shore | Body;Body | Yes |
| Control-Sample465_Be | chr1:17949563-17949604 | ARHGEF10L;ARHGEF10L | Island | Body;Body | Yes |
| Control-Sample3_Beta | chr17:8218887-8219313 | ARHGEF15 | | Body | No |
| Control-Sample1_Beta | chr11:73020459-73020729 | ARHGEF17 | Island | 1stExon | No |
| Control-Sample1_Beta | chr1:155931629-155931858 | ARHGEF2;ARHGEF2;ARHGEF2 | | Body;Body;Body | No |
| Control-Sample108_Be | chr3:56809796-56810320 | ARHGEF3;ARHGEF3;ARHGEF3 | | TSS200;Body;Body | No |
| Control-Sample1_Beta | chr13:111773936-111774127 | ARHGEF7;ARHGEF7;ARHGEF7 | | Body;Body;Body | Yes |
| Control-Sample10_Beta | chr13:111773936-111774082 | ARHGEF7;ARHGEF7;ARHGEF7 | | Body;Body;Body | Yes |
| Control-Sample19_Beta | chr19:940724-941571 | ARID3A | Island | Body | Yes |
| Control-Sample20_Beta | chr19:941126-941571 | ARID3A | Island | Body | Yes |
| Control-Sample108_Be | chr10:63809098-63809149 | ARID5B | | Body | No |
| Control-Sample3_Beta | chr13:50202584-50202713 | ARL11 | | TSS200 | No |
| Control-Sample10_Beta | chr2:235406275-235406540 | ARL4C | Island | TSS1500 | Yes |
| Control-Sample11_Beta | chr2:235406275-235406363 | ARL4C | Island | TSS1500 | Yes |
| Control-Sample19_Beta | chr2:235406291-235406540 | ARL4C | Island | TSS1500 | Yes |
| Control-Sample465_Be | chr17:73124988-73125153 | ARMC7 | Island | Body | No |
| Control-Sample1_Beta | chr7:98972156-98972188 | ARPC1B | Island | TSS200 | Yes |
| Control-Sample6_Beta | chr7:98972156-98972169 | ARPC1B | Island | TSS200 | Yes |
| Control-Sample108_Be | chr3:35721037-35721166 | ARPP-21;ARPP-21;ARPP-21;ARPP-21 | | 5'UTR;TSS200;TSS1500;TSS200 | No |
| Control-Sample1_Beta | chr5:78281793-78281827 | ARSB;ARSB;ARSB | S_Shore | TSS200;1stExon;5'UTR | Yes |
| Control-Sample15_Beta | chr5:78281793-78281964 | ARSB;ARSB;ARSB | S_Shore | TSS200;1stExon;5'UTR | Yes |
| Control-Sample1_Beta | chr17:66255265-66255444 | ARSG | Island | TSS200 | No |
| Control-Sample1_Beta | chr11:123066956-123067275 | ASAM | Island | TSS1500 | No |
| Control-Sample19_Beta | chr2:237120069-237120313 | ASB18 | Island | Body | Yes |
| Control-Sample19_Beta | chr2:237172796-237173121 | ASB18 | | 1stExon | Yes |
| Control-Sample10_Beta | chr12:103353183-103353362 | ASCL1 | S_Shore | 3'UTR | No |
| Control-Sample13_Beta | chr11:2292890-2292902 | ASCL2 | Island | TSS1500 | No |
| Control-Sample82_Beta | chr17:3378804-3379283 | ASPA;ASPA | S_Shelf | TSS1500;5'UTR | No |
| Control-Sample31_Beta | chr14:104552032-104552209 | ASPG | Island | TSS200 | Yes |
| Control-Sample49_Beta | chr14:104551884-104552034 | ASPG | N_Shore | TSS200 | Yes |
| Control-Sample506_Be | chr14.0:104552032-104552209 | ASPG | Island | TSS200 | Yes |
| Control-Sample616_Be | chr14:104551867-104552034 | ASPG | N_Shore | TSS200 | Yes |
| Control-Sample270_Be | chr2:70188605-70189092 | ASPRV1 | Island | 1stExon | Yes |
| Control-Sample475_Be | chr2:70188605-70189111 | ASPRV1 | Island | 1stExon | Yes |
| Control-Sample673_Be | chr2:70188512-70189092 | ASPRV1 | Island | 1stExon | Yes |
| Control-Sample1_Beta | chr17:79970135-79970313 | ASPSCR1 | S_Shelf | Body | Yes |
| Control-Sample11_Beta | chr17:79958592-79958912 | ASPSCR1 | S_Shore | Body | Yes |
| Control-Sample1_Beta | chr2:242606122-242606353 | ATG4B;ATG4B | Island | Body;Body | Yes |
| Control-Sample19_Beta | chr2:242606122-242606531 | ATG4B;ATG4B | Island | Body;Body | Yes |
| Control-Sample108_Be | chr1:63249199-63249213 | ATG4C;ATG4C | | TSS1500;TSS1500 | Yes |
| Control-Sample24_Beta | chr1:63249197-63249213 | ATG4C;ATG4C | | TSS1500;TSS1500 | Yes |
| Control-Sample67_Beta | chr2:85999498-85999873 | ATOH8 | | Body | No |
| Control-Sample74_Beta | chr15:25961909-25961967 | ATP10A | S_Shelf | Body | No |
| Control-Sample1_Beta | chr13:113407479-113407769 | ATP11A;ATP11A | N_Shelf | Body;Body | Yes |

| | | | | | |
|---|---|---|---|---|---|
| Control-Sample1_Beta | chr13:113411110-113411200 | ATP11A;ATP11A | S_Shore | Body;Body | Yes |
| Control-Sample1_Beta | chr13:113425199-113425298 | ATP11A;ATP11A | S_Shore | Body;Body | Yes |
| Control-Sample1_Beta | chr13:113435838-113436400 | ATP11A;ATP11A | | Body;Body | Yes |
| Control-Sample1_Beta | chr13:113516304-113516424 | ATP11A;ATP11A | | Body;Body | Yes |
| Control-Sample124_Be | chr13:113461305-113461652 | ATP11A;ATP11A | N_Shelf | Body;Body | Yes |
| Control-Sample17_Beta | chr13:113420153-113420270 | ATP11A;ATP11A | Island | Body;Body | Yes |
| Control-Sample17_Beta | chr13:113491766-113492042 | ATP11A;ATP11A | N_Shore | Body;Body | Yes |
| Control-Sample19_Beta | chr13:113408267-113408566 | ATP11A;ATP11A | N_Shore | Body;Body | Yes |
| Control-Sample20_Beta | chr13:113346257-113346616 | ATP11A;ATP11A | S_Shore | Body;Body | Yes |
| Control-Sample20_Beta | chr13:113419790-113420270 | ATP11A;ATP11A | N_Shore | Body;Body | Yes |
| Control-Sample20_Beta | chr13:113498128-113498308 | ATP11A;ATP11A | S_Shore | Body;Body | Yes |
| Control-Sample27_Beta | chr13:113435838-113436309 | ATP11A;ATP11A | | Body;Body | Yes |
| Control-Sample33_Beta | chr13:113540400-113540557 | ATP11A;ATP11A | Island | 3'UTR;3'UTR | Yes |
| Control-Sample331_Be | chr13:113436309-113436651 | ATP11A;ATP11A | N_Shelf | Body;Body | Yes |
| Control-Sample464_Be | chr13:113492269-113492708 | ATP11A;ATP11A | N_Shore | Body;Body | Yes |
| Control-Sample5_Beta | chr13:113540400-113540631 | ATP11A;ATP11A | Island | 3'UTR;3'UTR | Yes |
| Control-Sample58_Beta | chr13:113540421-113540557 | ATP11A;ATP11A | Island | 3'UTR;3'UTR | Yes |
| Control-Sample637_Be | chr13:113424175-113424362 | ATP11A;ATP11A | N_Shore | Body;Body | Yes |
| Control-Sample673_Be | chr13:113416621-113417204 | ATP11A;ATP11A | N_Shelf | Body;Body | Yes |
| Control-Sample108_Be | chr3:193272655-193272869 | ATP13A4;ATP13A4 | | 5'UTR;1stExon | No |
| Control-Sample1_Beta | chr1:160085433-160085581 | ATP1A2 | | TSS200 | No |
| Control-Sample1_Beta | chr1:203595061-203595283 | ATP2B4;ATP2B4 | N_Shelf | TSS1500;TSS1500 | Yes |
| Control-Sample108_Be | chr1:203595283-203595871 | ATP2B4;ATP2B4 | N_Shelf | TSS1500;TSS1500 | Yes |
| Control-Sample1_Beta | chr18:43683922-43684054 | ATP5A1;HAUS1;HAUS1 | N_Shore | 5'UTR;TSS1500;TSS1500 | No |
| Control-Sample1_Beta | chr6:16306096-16306857 | ATXN1;ATXN1 | | 3'UTR;3'UTR | Yes |
| Control-Sample11_Beta | chr6:16306377-16306857 | ATXN1;ATXN1 | | 3'UTR;3'UTR | Yes |
| Control-Sample47_Beta | chr6:16306552-16306857 | ATXN1;ATXN1 | | 3'UTR;3'UTR | Yes |
| Control-Sample108_Be | chr7:105332220-105332311 | ATXN7L1 | | Body | Yes |
| Control-Sample20_Beta | chr7:105331690-105332311 | ATXN7L1 | | Body | Yes |
| Control-Sample1_Beta | chr7:105279391-105279882 | ATXN7L1;ATXN7L1 | | Body;Body | Yes |
| Control-Sample11_Beta | chr7:105279494-105279882 | ATXN7L1;ATXN7L1 | | Body;Body | Yes |
| Control-Sample69_Beta | chr7:105279549-105279882 | ATXN7L1;ATXN7L1 | | Body;Body | Yes |
| Control-Sample130_Be | chr19:57742260-57742394 | AURKC;AURKC;AURKC | Island | TSS200;TSS200;TSS200 | No |
| Control-Sample1_Beta | chr7:70061455-70061616 | AUTS2;AUTS2 | Island | Body;Body | Yes |
| Control-Sample1_Beta | chr7:70096029-70096378 | AUTS2;AUTS2 | N_Shore | Body;Body | Yes |
| Control-Sample24_Beta | chr7:70096029-70096791 | AUTS2;AUTS2 | N_Shore | Body;Body | Yes |
| Control-Sample19_Beta | chr15:34260433-34261172 | AVEN;CHRM5 | | Body;TSS1500 | No |
| Control-Sample10_Beta | chr12:58210661-58210878 | AVIL | | TSS1500 | Yes |
| Control-Sample34_Beta | chr12:58210661-58211193 | AVIL | | TSS1500 | Yes |
| Control-Sample67_Beta | chr16:374924-375327 | AXIN1;AXIN1 | Island | Body;Body | No |
| Control-Sample233_Be | chr17:63555244-63555717 | AXIN2 | N_Shore | 5'UTR | No |
| Control-Sample6_Beta | chr3:160823387-160823505 | B3GALNT1;B3GALNT1;B3GALNT1;B3GALNT1;B3GALNT1 | S_Shore | TSS1500;TSS1500;TSS1500;TSS1500;TSS1500 | No |
| Control-Sample1_Beta | chr17:80972336-80972373 | B3GNTL1 | N_Shelf | Body | Yes |
| Control-Sample12_Beta | chr17:80971776-80972373 | B3GNTL1 | S_Shore | Body | Yes |
| Control-Sample11_Beta | chr12:58021295-58021713 | B4GALNT1 | Island | Body | Yes |
| Control-Sample12_Beta | chr12:58021295-58021569 | B4GALNT1 | Island | Body | Yes |
| Control-Sample5_Beta | chr12:58021295-58021917 | B4GALNT1 | Island | Body | Yes |
| Control-Sample19_Beta | chr12:656703-657046 | B4GALNT3 | | Body | No |
| Control-Sample22_Beta | chr11:368351-368384 | B4GALNT4 | | TSS1500 | Yes |
| Control-Sample3_Beta | chr11:368351-368565 | B4GALNT4 | | TSS1500 | Yes |
| Control-Sample4_Beta | chr11:368384-368573 | B4GALNT4 | | TSS1500 | Yes |
| Control-Sample19_Beta | chr6:91004430-91004482 | BACH2;BACH2 | N_Shore | 5'UTR;5'UTR | No |
| Control-Sample1_Beta | chr17:79420145-79420436 | BAHCC1 | Island | Body | Yes |
| Control-Sample107_Be | chr17:79374327-79374515 | BAHCC1 | Island | Body | Yes |
| Control-Sample12_Beta | chr17:79432935-79433150 | BAHCC1 | S_Shore | 3'UTR | Yes |
| Control-Sample170_Be | chr17:79374327-79374691 | BAHCC1 | Island | Body | Yes |
| Control-Sample171_Be | chr17:79372242-79372528 | BAHCC1 | Island | TSS1500 | Yes |
| Control-Sample171_Be | chr17:79374327-79375254 | BAHCC1 | Island | Body | Yes |
| Control-Sample216_Be | chr17:79377272-79377665 | BAHCC1 | N_Shore | Body | Yes |
| Control-Sample331_Be | chr17:79377272-79377850 | BAHCC1 | N_Shore | Body | Yes |
| Control-Sample44_Beta | chr17:79420279-79420436 | BAHCC1 | Island | Body | Yes |
| Control-Sample464_Be | chr17:79430641-79431017 | BAHCC1 | Island | Body | Yes |
| Control-Sample465_Be | chr17:79430641-79430946 | BAHCC1 | Island | Body | Yes |
| Control-Sample65_Beta | chr17:79374327-79374741 | BAHCC1 | Island | Body | Yes |
| Control-Sample11_Beta | chr8:143545789-143545949 | BAI1 | Island | 1stExon | Yes |
| Control-Sample3_Beta | chr8:143544026-143544400 | BAI1 | N_Shore | TSS1500 | Yes |
| Control-Sample12_Beta | chr8:143545478-143545789 | BAI1;BAI1 | Island | 5'UTR;1stExon | Yes |
| Control-Sample3_Beta | chr8:143545478-143546524 | BAI1;BAI1 | Island | 5'UTR;1stExon | Yes |
| Control-Sample36_Beta | chr17:79010968-79011542 | BAIAP2;BAIAP2;BAIAP2;BAIAP2 | S_Shore | Body;Body;Body;Body | Yes |
| Control-Sample5_Beta | chr17:79047695-79047872 | BAIAP2;BAIAP2;BAIAP2;BAIAP2 | S_Shore | Body;Body;Body;Body | Yes |
| Control-Sample2_Beta | chr22:38506712-38506781 | BAIAP2L2 | | TSS200 | No |
| Control-Sample561_Be | chr16.0:1393584-1393797 | BAIAP3 | Island | Body | No |
| Control-Sample108_Be | chr6:33548127-33548425 | BAK1 | | TSS200 | No |
| Control-Sample25_Beta | chr16:88052922-88053104 | BANP;BANP | Island | Body;Body | Yes |
| Control-Sample28_Beta | chr16:88052922-88053094 | BANP;BANP | Island | Body;Body | Yes |
| Control-Sample51_Beta | chr16:88091073-88091471 | BANP;BANP | | Body;Body | Yes |
| Control-Sample52_Beta | chr16:88052956-88053104 | BANP;BANP | Island | Body;Body | Yes |
| Control-Sample1_Beta | chr6:31506604-31506990 | BAT1;BAT1 | N_Shelf | Body;Body | Yes |
| Control-Sample1_Beta | chr6:31507701-31508167 | BAT1;BAT1 | N_Shore | Body;Body | Yes |
| Control-Sample10_Beta | chr6:31507879-31508167 | BAT1;BAT1 | N_Shore | Body;Body | Yes |
| Control-Sample11_Beta | chr6:31508126-31508167 | BAT1;BAT1 | N_Shore | Body;Body | Yes |
| Control-Sample15_Beta | chr6:31508106-31508167 | BAT1;BAT1 | N_Shore | Body;Body | Yes |
| Control-Sample18_Beta | chr6:31507701-31507857 | BAT1;BAT1 | N_Shore | Body;Body | Yes |
| Control-Sample19_Beta | chr6:31507701-31507879 | BAT1;BAT1 | N_Shore | Body;Body | Yes |
| Control-Sample40_Beta | chr6:31506768-31506990 | BAT1;BAT1 | N_Shelf | Body;Body | Yes |
| Control-Sample67_Beta | chr6:31510729-31510903 | BAT1;BAT1 | S_Shore | TSS1500;TSS1500 | Yes |
| Control-Sample11_Beta | chr1:156610966-156611194 | BCAN;BCAN | N_Shore | TSS1500;TSS1500 | No |
| Control-Sample1_Beta | chr16:75300724-75300861 | BCAR1;BCAR1;BCAR1;BCAR1;BCAR1 | S_Shore | TSS1500;TSS200;Body;TSS1500;TSS1500 | Yes |
| Control-Sample19_Beta | chr16:75300724-75301362 | BCAR1;BCAR1;BCAR1;BCAR1;BCAR1 | S_Shore | TSS1500;TSS200;Body;TSS1500;TSS1500 | Yes |
| Control-Sample10_Beta | chr16:75282051-75282362 | BCAR1;BCAR1;BCAR1;BCAR1;BCAR1;BCAR1;BCAR1;BCAI | N_Shelf | Body;5'UTR;Body;Body;Body;1stExon;Body;Body;Bod | Yes |
| Control-Sample1_Beta | chr2:60680616-60680761 | BCL11A;BCL11A | | Body;Body | No |

| | | | | | |
|---|---|---|---|---|---|
| Control-Sample1_Beta | chr14:99728984-99729088 | BCL11B;BCL11B | | Body;Body | No |
| Control-Sample223_Be | chr20:30309466-30309717 | BCL2L1;BCL2L1 | N_Shore | Body;Body | No |
| Control-Sample10_Beta | chr3:187454439-187454482 | BCL6;BCL6 | N_Shore | 5'UTR;TSS200 | Yes |
| Control-Sample1_Beta | chr3:187453721-187454482 | BCL6;BCL6;BCL6 | N_Shore | 5'UTR;5'UTR;TSS1500 | Yes |
| Control-Sample13_Beta | chr3:187453973-187454482 | BCL6;BCL6;BCL6;BCL6 | N_Shore | 5'UTR;5'UTR;1stExon;TSS1500 | Yes |
| Control-Sample108_Be | chr16:81272080-81272500 | BCMO1 | | TSS1500 | No |
| Control-Sample108_Be | chr11:112046160-112046403 | BCO2;BCO2 | | TSS200;TSS1500 | No |
| Control-Sample19_Beta | chr14:101036192-101036322 | BEGAIN | S_Shore | TSS200 | Yes |
| Control-Sample21_Beta | chr14:101036192-101036470 | BEGAIN | S_Shore | TSS200 | Yes |
| Control-Sample1_Beta | chr14:101035818-101036470 | BEGAIN;BEGAIN | S_Shore | TSS1500;5'UTR | Yes |
| Control-Sample23_Beta | chr14:101035818-101036322 | BEGAIN;BEGAIN | S_Shore | TSS1500;5'UTR | Yes |
| Control-Sample1_Beta | chr4:42122106-42122161 | BEND4;BEND4 | | Body;Body | Yes |
| Control-Sample12_Beta | chr4:42154912-42154934 | BEND4;BEND4 | Island | TSS200;TSS200 | Yes |
| Control-Sample10_Beta | chr10:13481846-13481944 | BEND7 | | Body | No |
| Control-Sample108_Be | chr11:61716821-61717005 | BEST1;BEST1 | | TSS1500;TSS1500 | No |
| Control-Sample5_Beta | chr12:70092840-70093350 | BEST3 | | 5'UTR | No |
| Control-Sample10_Beta | chr1:156211434-156211609 | BGLAP | N_Shelf | TSS1500 | Yes |
| Control-Sample12_Beta | chr1:156211415-156211529 | BGLAP | N_Shelf | TSS1500 | Yes |
| Control-Sample15_Beta | chr1:156211415-156211609 | BGLAP | N_Shelf | TSS1500 | Yes |
| Control-Sample35_Beta | chr1:156211409-156211529 | BGLAP | N_Shelf | TSS1500 | Yes |
| Control-Sample6_Beta | chr1:156211409-156211609 | BGLAP | N_Shelf | TSS1500 | Yes |
| Control-Sample1_Beta | chr5:78407418-78407566 | BHMT | | TSS200 | Yes |
| Control-Sample11_Beta | chr5:78407418-78407562 | BHMT | | TSS200 | Yes |
| Control-Sample33_Beta | chr5:78407562-78407566 | BHMT | | TSS200 | Yes |
| Control-Sample41_Beta | chr5:78407509-78407562 | BHMT | | TSS200 | Yes |
| Control-Sample62_Beta | chr5:78407418-78407564 | BHMT | | TSS200 | Yes |
| Control-Sample66_Beta | chr5:78407509-78407566 | BHMT | | TSS200 | Yes |
| Control-Sample1_Beta | chr11:102187898-102187976 | BIRC3;BIRC3 | N_Shore | TSS1500;TSS1500 | Yes |
| Control-Sample4_Beta | chr11:102187898-102188025 | BIRC3;BIRC3 | N_Shore | TSS1500;TSS1500 | Yes |
| Control-Sample57_Beta | chr17:76220608-76220955 | BIRC5;BIRC5;BIRC5;EPR1 | | 3'UTR;3'UTR;3'UTR;Body | Yes |
| Control-Sample71_Beta | chr17:76220672-76220955 | BIRC5;BIRC5;BIRC5;EPR1 | | 3'UTR;3'UTR;3'UTR;Body | Yes |
| Control-Sample70_Beta | chr17:76220898-76220955 | BIRC5;EPR1;BIRC5;BIRC5 | | 3'UTR;TSS200;3'UTR;3'UTR | Yes |
| Control-Sample20_Beta | chr20:36149013-36149053 | BLCAP;BLCAP;BLCAP;BLCAP;NNAT;NNAT;BLCAP | Island | 5'UTR;5'UTR;5'UTR;5'UTR;TSS1500;TSS1500;5'UTR | Yes |
| Control-Sample24_Beta | chr20:36148375-36148604 | BLCAP;BLCAP;BLCAP;BLCAP;NNAT;NNAT;BLCAP | N_Shore | 5'UTR;5'UTR;5'UTR;5'UTR;TSS1500;TSS1500;5'UTR | Yes |
| Control-Sample30_Beta | chr20:36148375-36148615 | BLCAP;BLCAP;BLCAP;BLCAP;NNAT;NNAT;BLCAP | N_Shore | 5'UTR;5'UTR;5'UTR;5'UTR;TSS1500;TSS1500;5'UTR | Yes |
| Control-Sample32_Beta | chr20:36148457-36148615 | BLCAP;BLCAP;BLCAP;BLCAP;NNAT;NNAT;BLCAP | N_Shore | 5'UTR;5'UTR;5'UTR;5'UTR;TSS1500;TSS1500;5'UTR | Yes |
| Control-Sample2_Beta | chr15:40398565-40399101 | BMF;BMF;BMF;BMF;BMF | N_Shelf | TSS1500;5'UTR;TSS1500;5'UTR;1stExon | No |
| Control-Sample1_Beta | chr14:54423807-54424149 | BMP4;BMP4 | S_Shore | TSS1500;TSS1500 | Yes |
| Control-Sample11_Beta | chr14:54423807-54423845 | BMP4;BMP4 | S_Shore | TSS1500;TSS1500 | Yes |
| Control-Sample11_Beta | chr14:54418728-54418851 | BMP4;BMP4;BMP4 | Island | Body;Body;Body | Yes |
| Control-Sample1_Beta | chr20:55837483-55837692 | BMP7 | N_Shore | Body | No |
| Control-Sample1_Beta | chr15:83953880-83953929 | BNC1 | Island | TSS1500 | Yes |
| Control-Sample1_Beta | chr15:83954150-83954392 | BNC1 | S_Shore | TSS1500 | Yes |
| Control-Sample10_Beta | chr15:83954150-83954395 | BNC1 | S_Shore | TSS1500 | Yes |
| Control-Sample12_Beta | chr15:83954150-83954246 | BNC1 | S_Shore | TSS1500 | Yes |
| Control-Sample13_Beta | chr15:83953864-83953883 | BNC1 | Island | TSS1500 | Yes |
| Control-Sample15_Beta | chr15:83953864-83953929 | BNC1 | Island | TSS1500 | Yes |
| Control-Sample20_Beta | chr15:83954229-83954246 | BNC1 | S_Shore | TSS1500 | Yes |
| Control-Sample163_Be | chr2:198650985-198651076 | BOLL;BOLL | Island | TSS200;TSS1500 | Yes |
| Control-Sample514_Be | chr2.0:198650985-198651076 | BOLL;BOLL | Island | TSS200;TSS1500 | Yes |
| Control-Sample433_Be | chr2:198650880-198651076 | BOLL;BOLL;BOLL | Island | 1stExon;5'UTR;TSS1500 | Yes |
| Control-Sample221_Be | chr17:41278241-41278357 | BRCA1;NBR2;BRCA1;BRCA1;BRCA1;BRCA1 | Island | TSS1500;Body;TSS1500;TSS1500;TSS1500;TSS1500 | Yes |
| Control-Sample221_Be | chr17:41278581-41278655 | BRCA1;NBR2;BRCA1;BRCA1;BRCA1;BRCA1 | S_Shore | TSS1500;Body;TSS1500;TSS1500;TSS1500;TSS1500 | Yes |
| Control-Sample384_Be | chr17:41278135-41278281 | BRCA1;NBR2;BRCA1;BRCA1;BRCA1;BRCA1 | Island | TSS1500;Body;TSS1500;TSS1500;TSS1500;TSS1500 | Yes |
| Control-Sample616_Be | chr17:41278241-41278281 | BRCA1;NBR2;BRCA1;BRCA1;BRCA1;BRCA1 | Island | TSS1500;Body;TSS1500;TSS1500;TSS1500;TSS1500 | Yes |
| Control-Sample1_Beta | chr22:50174005-50174170 | BRD1;LOC90834 | N_Shelf | Body;TSS200 | Yes |
| Control-Sample10_Beta | chr22:50173998-50174170 | BRD1;LOC90834 | N_Shelf | Body;TSS200 | Yes |
| Control-Sample12_Beta | chr22:50174057-50174170 | BRD1;LOC90834 | N_Shelf | Body;TSS200 | Yes |
| Control-Sample31_Beta | chr22:50174005-50174065 | BRD1;LOC90834 | N_Shelf | Body;TSS200 | Yes |
| Control-Sample5_Beta | chr22:50174028-50174170 | BRD1;LOC90834 | N_Shelf | Body;TSS200 | Yes |
| Control-Sample64_Beta | chr22:50174028-50174065 | BRD1;LOC90834 | N_Shelf | Body;TSS200 | Yes |
| Control-Sample1_Beta | chr6:32945336-32945631 | BRD2;BRD2 | | Body;Body | Yes |
| Control-Sample12_Beta | chr6:32945423-32945631 | BRD2;BRD2 | | Body;Body | Yes |
| Control-Sample10_Beta | chr19:15375316-15375352 | BRD4;BRD4 | | Body;Body | Yes |
| Control-Sample12_Beta | chr19:15375418-15375450 | BRD4;BRD4 | | Body;Body | Yes |
| Control-Sample13_Beta | chr19:15375283-15375418 | BRD4;BRD4 | | Body;Body | Yes |
| Control-Sample19_Beta | chr19:15375283-15375450 | BRD4;BRD4 | | Body;Body | Yes |
| Control-Sample20_Beta | chr19:15375319-15375418 | BRD4;BRD4 | | Body;Body | Yes |
| Control-Sample57_Beta | chr19:15375283-15375352 | BRD4;BRD4 | | Body;Body | Yes |
| Control-Sample270_Be | chr5:883880-884033 | BRD9;BRD9;BRD9 | S_Shelf | Body;Body;Body | No |
| Control-Sample1_Beta | chr11:1474386-1474588 | BRSK2 | S_Shelf | Body | Yes |
| Control-Sample19_Beta | chr11:1483169-1483973 | BRSK2 | N_Shore | 3'UTR | Yes |
| Control-Sample7_Beta | chr11:1483169-1483390 | BRSK2 | N_Shore | 3'UTR | Yes |
| Control-Sample19_Beta | chr11:62474940-62475078 | BSCL2;BSCL2;BSCL2;GNG3;BSCL2 | N_Shore | 5'UTR;5'UTR;1stExon;TSS200;TSS1500 | Yes |
| Control-Sample1_Beta | chr11:62474725-62475078 | BSCL2;GNG3;BSCL2;BSCL2;BSCL2 | S_Shore | 5'UTR;TSS1500;5'UTR;1stExon;TSS1500 | Yes |
| Control-Sample15_Beta | chr11:62474725-62474744 | BSCL2;GNG3;BSCL2;BSCL2;BSCL2 | S_Shore | 5'UTR;TSS1500;5'UTR;1stExon;TSS1500 | Yes |
| Control-Sample15_Beta | chr11:62474910-62475078 | BSCL2;GNG3;BSCL2;BSCL2;BSCL2 | N_Shore | 5'UTR;TSS1500;5'UTR;1stExon;TSS1500 | Yes |
| Control-Sample19_Beta | chr11:62474725-62474759 | BSCL2;GNG3;BSCL2;BSCL2;BSCL2 | S_Shore | 5'UTR;TSS1500;5'UTR;1stExon;TSS1500 | Yes |
| Control-Sample57_Beta | chr11:122852429-122852461 | BSX | Island | TSS200 | No |
| Control-Sample5_Beta | chr12:108039569-108039856 | BTBD11;BTBD11 | | Body;Body | No |
| Control-Sample12_Beta | chr20:11898478-11899258 | BTBD3;BTBD3 | | 5'UTR;TSS200 | Yes |
| Control-Sample74_Beta | chr20:11898531-11898851 | BTBD3;BTBD3 | | 5'UTR;TSS200 | Yes |
| Control-Sample80_Beta | chr20:11898478-11898851 | BTBD3;BTBD3 | | 5'UTR;TSS200 | Yes |
| Control-Sample108_Be | chr6:26440101-26440610 | BTN3A3;BTN3A3 | | TSS1500;TSS1500 | No |
| Control-Sample223_Be | chr6:32363888-32363933 | BTNL2 | | Body | Yes |
| Control-Sample322_Be | chr6:32365553-32365785 | BTNL2 | | Body | Yes |
| Control-Sample475_Be | chr6:32363892-32363942 | BTNL2 | | Body | Yes |
| Control-Sample637_Be | chr6:32371506-32372310 | BTNL2 | | Body | Yes |
| Control-Sample1_Beta | chr15:40509472-40509703 | BUB1B;PAK6;PAK6 | | Body;TSS200;TSS200 | No |
| Control-Sample67_Beta | chr10:63422539-63422630 | C10orf107 | N_Shore | TSS1500 | No |
| Control-Sample1_Beta | chr10:1083192-1083304 | C10orf110;C10orf110;C10orf110;C10orf110 | | Body;Body;Body;Body | Yes |

| | | | | | |
|---|---|---|---|---|---|
| Control-Sample585_Be | chr10.0:1083192-1083304 | C10orf110;C10orf110;C10orf110;C10orf110 | | Body;Body;Body;Body | Yes |
| Control-Sample1_Beta | chr10:1068490-1068667 | C10orf110;C10orf110;C10orf110;C10orf110;IDI2 | | TSS200;TSS200;TSS200;TSS200;Body | Yes |
| Control-Sample17_Beta | chr10:1068567-1068703 | C10orf110;C10orf110;C10orf110;C10orf110;IDI2 | | TSS200;TSS200;TSS200;TSS200;Body | Yes |
| Control-Sample3_Beta | chr10:1068528-1068667 | C10orf110;C10orf110;C10orf110;C10orf110;IDI2 | | TSS200;TSS200;TSS200;TSS200;Body | Yes |
| Control-Sample39_Beta | chr10:1068490-1068703 | C10orf110;C10orf110;C10orf110;C10orf110;IDI2 | | TSS200;TSS200;TSS200;TSS200;Body | Yes |
| Control-Sample87_Beta | chr10:1068667-1068763 | C10orf110;C10orf110;C10orf110;C10orf110;IDI2 | | Body;Body;Body;Body;Body | Yes |
| Control-Sample24_Beta | chr10:88730407-88730823 | C10orf116;AGAP11 | N_Shore | TSS200;Body | No |
| Control-Sample97_Beta | chr10:97667594-97667731 | C10orf131 | Island | TSS200 | No |
| Control-Sample11_Beta | chr10:127452005-127452086 | C10orf137 | | Body | No |
| Control-Sample270_Be | chr10:21807166-21807252 | C10orf140 | S_Shore | 5'UTR | No |
| Control-Sample1_Beta | chr10:77165025-77166172 | C10orf41;C10orf41 | Island | Body;Body | Yes |
| Control-Sample18_Beta | chr10:77165025-77166318 | C10orf41;C10orf41 | Island | Body;Body | Yes |
| Control-Sample23_Beta | chr10:77165025-77166684 | C10orf41;C10orf41 | Island | Body;Body | Yes |
| Control-Sample27_Beta | chr10:77165618-77166318 | C10orf41;C10orf41 | Island | Body;Body | Yes |
| Control-Sample3_Beta | chr10:77165025-77165905 | C10orf41;C10orf41 | Island | Body;Body | Yes |
| Control-Sample40_Beta | chr10:77165905-77166318 | C10orf41;C10orf41 | Island | Body;Body | Yes |
| Control-Sample5_Beta | chr10:77165618-77165905 | C10orf41;C10orf41 | Island | Body;Body | Yes |
| Control-Sample61_Beta | chr10:77165618-77166684 | C10orf41;C10orf41 | Island | Body;Body | Yes |
| Control-Sample65_Beta | chr10:77165025-77165618 | C10orf41;C10orf41 | Island | Body;Body | Yes |
| Control-Sample66_Beta | chr10:77165025-77165293 | C10orf41;C10orf41 | Island | Body;Body | Yes |
| Control-Sample10_Beta | chr10:11868146-11868183 | C10orf47 | S_Shelf | 5'UTR | No |
| Control-Sample65_Beta | chr11:2322642-2322729 | C11orf21;TSPAN32;C11orf21 | | Body;TSS1500;Body | Yes |
| Control-Sample67_Beta | chr11:2321937-2322050 | C11orf21;TSPAN32;C11orf21 | | Body;TSS1500;Body | Yes |
| Control-Sample67_Beta | chr11:2322507-2322674 | C11orf21;TSPAN32;C11orf21 | | Body;TSS1500;Body | Yes |
| Control-Sample119_Be | chr11:3238777-3240201 | C11orf36 | N_Shore | TSS1500 | No |
| Control-Sample1_Beta | chr11:111789452-111789613 | C11orf52 | | TSS200 | Yes |
| Control-Sample19_Beta | chr11:111789452-111789693 | C11orf52 | | TSS200 | Yes |
| Control-Sample1_Beta | chr11:61248435-61248556 | C11orf66;C11orf66 | | TSS200;TSS200 | Yes |
| Control-Sample10_Beta | chr11:61248435-61248887 | C11orf66;C11orf66 | | TSS200;TSS200 | Yes |
| Control-Sample5_Beta | chr11:109294069-109294233 | C11orf87 | Island | 5'UTR | No |
| Control-Sample10_Beta | chr11:61522799-61523045 | C11orf9;C11orf9;DKFZP434K028 | S_Shelf | TSS200;Body;Body | Yes |
| Control-Sample151_Be | chr11:61525677-61525922 | C11orf9;C11orf9;DKFZP434K028 | | Body;Body;TSS1500 | Yes |
| Control-Sample194_Be | chr11:61522775-61523045 | C11orf9;C11orf9;DKFZP434K028 | S_Shelf | TSS200;Body;Body | Yes |
| Control-Sample331_Be | chr11:61522765-61522799 | C11orf9;C11orf9;DKFZP434K028 | S_Shelf | TSS200;Body;Body | Yes |
| Control-Sample475_Be | chr11:61522765-61523045 | C11orf9;C11orf9;DKFZP434K028 | S_Shelf | TSS200;Body;Body | Yes |
| Control-Sample103_Be | chr11:61522002-61522498 | C11orf9;DKFZP434K028;C11orf9 | S_Shelf | Body;Body;TSS1500 | Yes |
| Control-Sample426_Be | chr12:103889516-103889772 | C12orf42;C12orf42 | Island | 5'UTR;5'UTR | No |
| Control-Sample1_Beta | chr13:111980272-111980537 | C13orf16 | S_Shore | Body | Yes |
| Control-Sample10_Beta | chr13:111980272-111980655 | C13orf16 | S_Shore | Body | Yes |
| Control-Sample40_Beta | chr13:103426324-103426481 | C13orf27 | Island | TSS200 | No |
| Control-Sample1_Beta | chr13:111522932-111522985 | C13orf29 | | TSS1500 | No |
| Control-Sample4_Beta | chr13:31480646-31480942 | C13orf33;C13orf33 | Island | 1stExon;5'UTR | No |
| Control-Sample17_Beta | chr13:36871948-36872048 | C13orf38;C13orf38;C13orf38;C13orf38;C13orf38;C13orf3 | Island | 1stExon;1stExon;5'UTR;5'UTR;5'UTR;1stExon;1stExon | No |
| Control-Sample138_Be | chr13:103346900-103346962 | C13orf39 | | TSS200 | No |
| Control-Sample1_Beta | chr14:57960464-57960985 | C14orf105;C14orf105 | | 1stExon;5'UTR | Yes |
| Control-Sample15_Beta | chr14:57960464-57960976 | C14orf105;C14orf105 | | 1stExon;5'UTR | Yes |
| Control-Sample18_Beta | chr14:57960464-57960754 | C14orf105;C14orf105 | | 1stExon;5'UTR | Yes |
| Control-Sample108_Be | chr14:91579859-91580280 | C14orf159;C14orf159;C14orf159;C14orf159;C14orf159 | | TSS1500;TSS1500;TSS1500;TSS1500;TSS1500 | No |
| Control-Sample19_Beta | chr14:105045877-105045972 | C14orf180 | | TSS200 | No |
| Control-Sample10_Beta | chr14:92041829-92042045 | C14orf184 | S_Shore | TSS1500 | No |
| Control-Sample10_Beta | chr14:74185511-74185993 | C14orf43;C14orf43 | Island | 3'UTR;3'UTR | Yes |
| Control-Sample11_Beta | chr14:74184843-74185294 | C14orf43;C14orf43 | N_Shore | 3'UTR;3'UTR | Yes |
| Control-Sample26_Beta | chr15:67813132-67813145 | C15orf61 | N_Shore | TSS1500 | No |
| Control-Sample194_Be | chr17:71824806-71825281 | C17orf54 | | TSS200 | No |
| Control-Sample67_Beta | chr17:58499700-58499720 | C17orf64 | S_Shore | TSS200 | No |
| Control-Sample158_Be | chr17:8129997-8130356 | C17orf68 | S_Shelf | 3'UTR | No |
| Control-Sample27_Beta | chr17:72580962-72581014 | C17orf77;CD300LD | | TSS200;Body | Yes |
| Control-Sample63_Beta | chr17:72580901-72581081 | C17orf77;CD300LD | | TSS200;Body | Yes |
| Control-Sample13_Beta | chr17:259755-259930 | C17orf97 | Island | TSS1500 | Yes |
| Control-Sample2_Beta | chr17:259755-259942 | C17orf97 | Island | TSS1500 | Yes |
| Control-Sample38_Beta | chr17:259980-260058 | C17orf97 | Island | TSS200 | Yes |
| Control-Sample42_Beta | chr17:259755-260058 | C17orf97 | Island | TSS1500 | Yes |
| Control-Sample445_Be | chr17:259755-259991 | C17orf97 | Island | TSS1500 | Yes |
| Control-Sample5_Beta | chr17:259755-260292 | C17orf97 | Island | TSS1500 | Yes |
| Control-Sample638_Be | chr17:259839-259930 | C17orf97 | Island | TSS1500 | Yes |
| Control-Sample9_Beta | chr17:259991-260058 | C17orf97 | Island | TSS200 | Yes |
| Control-Sample23_Beta | chr17:36997420-36997563 | C17orf98 | N_Shore | Body | No |
| Control-Sample1_Beta | chr18:13641735-13641894 | C18orf1;C18orf1;C18orf1;C18orf1;C18orf1;C18orf1 | Island | Body;Body;Body;Body;Body;Body | No |
| Control-Sample4_Beta | chr19:1271174-1271371 | C19orf23;CIRBP;CIRBP;CIRBP | S_Shore | TSS1500;Body;Body;Body | Yes |
| Control-Sample67_Beta | chr19:1271019-1271275 | C19orf23;CIRBP;CIRBP;CIRBP | S_Shore | TSS1500;Body;Body;Body | Yes |
| Control-Sample69_Beta | chr19:1271275-1271312 | C19orf23;CIRBP;CIRBP;CIRBP | S_Shore | TSS1500;Body;Body;Body | Yes |
| Control-Sample1_Beta | chr19:38794554-38794635 | C19orf33;YIF1B;YIF1B;YIF1B;YIF1B;YIF1B;YIF1B;YIF1B | | TSS1500;3'UTR;3'UTR;3'UTR;3'UTR;3'UTR;3'UTR;3'UT | Yes |
| Control-Sample58_Beta | chr19:2095466-2095643 | C19orf36;MOBKL2A;C19orf36 | Island | TSS1500;5'UTR;TSS1500 | No |
| Control-Sample21_Beta | chr1:38156462-38156652 | C1orf109 | N_Shore | TSS1500 | Yes |
| Control-Sample47_Beta | chr1:38156462-38156575 | C1orf109 | N_Shore | TSS1500 | Yes |
| Control-Sample170_Be | chr1:1022900-1022995 | C1orf159 | N_Shore | Body | No |
| Control-Sample1_Beta | chr1:57285278-57285466 | C1orf168;C1orf168 | | 1stExon;5'UTR | No |
| Control-Sample9_Beta | chr1:75139581-75139958 | C1orf173 | | TSS200 | No |
| Control-Sample151_Be | chr1:3807168-3807387 | C1orf174 | N_Shore | Body | Yes |
| Control-Sample41_Beta | chr1:3807168-3808169 | C1orf174 | N_Shore | Body | Yes |
| Control-Sample56_Beta | chr1:3807168-3807709 | C1orf174 | N_Shore | Body | Yes |
| Control-Sample20_Beta | chr1:231005387-231005895 | C1orf198;C1orf198 | S_Shore | TSS200;TSS1500 | Yes |
| Control-Sample108_Be | chr1:230992820-230992990 | C1orf198;C1orf198;C1orf198 | | Body;Body;TSS1500 | Yes |
| Control-Sample1_Beta | chr1:43750693-43751563 | C1orf210;C1orf210 | | 5'UTR;5'UTR | Yes |
| Control-Sample10_Beta | chr1:43750693-43751497 | C1orf210;C1orf210 | | 5'UTR;5'UTR | Yes |
| Control-Sample15_Beta | chr1:43751363-43751563 | C1orf210;C1orf210 | | TSS200;TSS200 | Yes |
| Control-Sample16_Beta | chr1:43751363-43751497 | C1orf210;C1orf210 | | TSS200;TSS200 | Yes |
| Control-Sample19_Beta | chr1:43751370-43751469 | C1orf210;C1orf210 | | TSS200;TSS200 | Yes |
| Control-Sample21_Beta | chr1:43750693-43751376 | C1orf210;C1orf210 | | 5'UTR;5'UTR | Yes |
| Control-Sample21_Beta | chr1:43751497-43751563 | C1orf210;C1orf210 | | TSS1500;TSS1500 | Yes |
| Control-Sample31_Beta | chr1:43750693-43751504 | C1orf210;C1orf210 | | 5'UTR;5'UTR | Yes |

| | | | | | |
|---|---|---|---|---|---|
| Control-Sample50_Beta | chr1:43751363-43751376 | C1orf210;C1orf210 | | TSS200;TSS200 | Yes |
| Control-Sample22_Beta | chr1:43751270-43751563 | C1orf210;C1orf210;C1orf210 | | 1stExon;5'UTR;TSS200 | Yes |
| Control-Sample1_Beta | chr1:247275735-247275872 | C1orf229 | Island | TSS200 | No |
| Control-Sample1_Beta | chr1:226187852-226188006 | C1orf55 | S_Shore | TSS1500 | Yes |
| Control-Sample38_Beta | chr1:226187478-226188006 | C1orf55 | S_Shore | TSS1500 | Yes |
| Control-Sample475_Be | chr1:156399638-156399787 | C1orf61 | | TSS1500 | No |
| Control-Sample19_Beta | chr1:16330625-16330853 | C1orf64 | | TSS200 | No |
| Control-Sample19_Beta | chr1:54562040-54562543 | C1orf83 | | Body | No |
| Control-Sample1_Beta | chr1:2120985-2121724 | C1orf86;LOC100128003 | N_Shore | Body;Body | Yes |
| Control-Sample104_Be | chr1:2120985-2121521 | C1orf86;LOC100128003 | N_Shore | Body;Body | Yes |
| Control-Sample215_Be | chr1:2120985-2121349 | C1orf86;LOC100128003 | N_Shore | Body;Body | Yes |
| Control-Sample509_Be | chr1.0:2120985-2121521 | C1orf86;LOC100128003 | N_Shore | Body;Body | Yes |
| Control-Sample543_Be | chr1.0:2120985-2121724 | C1orf86;LOC100128003 | N_Shore | Body;Body | Yes |
| Control-Sample550_Be | chr1.0:2120985-2121349 | C1orf86;LOC100128003 | N_Shore | Body;Body | Yes |
| Control-Sample80_Beta | chr1:2120985-2121449 | C1orf86;LOC100128003 | N_Shore | Body;Body | Yes |
| Control-Sample121_Be | chr1:2121039-2121521 | C1orf86;LOC100128003;C1orf86 | Island | Body;Body;3'UTR | Yes |
| Control-Sample19_Beta | chr1:2121039-2121724 | C1orf86;LOC100128003;C1orf86 | Island | Body;Body;3'UTR | Yes |
| Control-Sample517_Be | chr1.0:2121039-2121521 | C1orf86;LOC100128003;C1orf86 | Island | Body;Body;3'UTR | Yes |
| Control-Sample6_Beta | chr1:22978556-22978981 | C1QB | | TSS1500 | No |
| Control-Sample1_Beta | chr12:49726500-49726893 | C1QL4 | N_Shore | 3'UTR | Yes |
| Control-Sample13_Beta | chr12:49726500-49726761 | C1QL4 | N_Shore | 3'UTR | Yes |
| Control-Sample2_Beta | chr12:49726500-49727110 | C1QL4 | N_Shore | 3'UTR | Yes |
| Control-Sample13_Beta | chr22:37584018-37584441 | C1QTNF6;C1QTNF6 | | Body;Body | No |
| Control-Sample36_Beta | chr16:1146277-1146445 | C1QTNF8 | S_Shelf | TSS200 | Yes |
| Control-Sample1_Beta | chr16:1146194-1146445 | C1QTNF8;C1QTNF8 | S_Shelf | 5'UTR;1stExon | Yes |
| Control-Sample6_Beta | chr16:1146194-1146421 | C1QTNF8;C1QTNF8 | S_Shelf | 5'UTR;1stExon | Yes |
| Control-Sample464_Be | chr20:35421865-35422703 | C20orf117;C20orf117 | N_Shore | Body;Body | Yes |
| Control-Sample19_Beta | chr20:35488024-35488454 | C20orf117;C20orf117;C20orf117 | N_Shelf | Body;1stExon;5'UTR | Yes |
| Control-Sample59_Beta | chr20:30605868-30606095 | C20orf160 | Island | Body | No |
| Control-Sample171_Be | chr20:62183924-62183966 | C20orf195 | N_Shore | TSS1500 | No |
| Control-Sample19_Beta | chr20:58630240-58630390 | C20orf197 | | TSS1500 | No |
| Control-Sample1_Beta | chr20:56725873-56725887 | C20orf85 | Island | TSS200 | Yes |
| Control-Sample12_Beta | chr20:56725873-56726093 | C20orf85 | Island | TSS200 | Yes |
| Control-Sample20_Beta | chr20:56725873-56725899 | C20orf85 | Island | TSS200 | Yes |
| Control-Sample21_Beta | chr20:56725876-56725899 | C20orf85 | Island | TSS200 | Yes |
| Control-Sample4_Beta | chr20:56725876-56726093 | C20orf85 | Island | TSS200 | Yes |
| Control-Sample63_Beta | chr20:56725887-56726093 | C20orf85 | Island | TSS200 | Yes |
| Control-Sample2_Beta | chr21:43442057-43442254 | C21orf121 | | TSS200 | Yes |
| Control-Sample3_Beta | chr21:43442027-43442254 | C21orf121 | | TSS200 | Yes |
| Control-Sample108_Be | chr21:17442467-17442803 | C21orf34 | | TSS1500 | Yes |
| Control-Sample1_Beta | chr21:17961737-17962124 | C21orf34;MIR125B2;C21orf34 | | Body;TSS1500;Body | Yes |
| Control-Sample10_Beta | chr21:17961829-17962124 | C21orf34;MIR125B2;C21orf34 | | Body;TSS1500;Body | Yes |
| Control-Sample107_Be | chr21:47604052-47604654 | C21orf56;C21orf56 | S_Shore | 5'UTR;5'UTR | Yes |
| Control-Sample117_Be | chr21:47604052-47605174 | C21orf56;C21orf56 | S_Shore | 5'UTR;5'UTR | Yes |
| Control-Sample125_Be | chr21:47604052-47604986 | C21orf56;C21orf56 | S_Shore | 5'UTR;5'UTR | Yes |
| Control-Sample230_Be | chr21:47604052-47604361 | C21orf56;C21orf56 | S_Shore | 5'UTR;5'UTR | Yes |
| Control-Sample280_Be | chr21:47604052-47605072 | C21orf56;C21orf56 | S_Shore | 5'UTR;5'UTR | Yes |
| Control-Sample380_Be | chr21:47604166-47604986 | C21orf56;C21orf56 | S_Shore | 5'UTR;5'UTR | Yes |
| Control-Sample443_Be | chr21:47604166-47604654 | C21orf56;C21orf56 | S_Shore | 5'UTR;5'UTR | Yes |
| Control-Sample504_Be | chr21.0:47604052-47604654 | C21orf56;C21orf56 | S_Shore | 5'UTR;5'UTR | Yes |
| Control-Sample522_Be | chr21.0:47604052-47605174 | C21orf56;C21orf56 | S_Shore | 5'UTR;5'UTR | Yes |
| Control-Sample538_Be | chr21.0:47604052-47604986 | C21orf56;C21orf56 | S_Shore | 5'UTR;5'UTR | Yes |
| Control-Sample10_Beta | chr21:47604291-47604986 | C21orf56;C21orf56;C21orf56;C21orf56 | S_Shore | 5'UTR;5'UTR;1stExon;1stExon | Yes |
| Control-Sample104_Be | chr21:47604291-47604654 | C21orf56;C21orf56;C21orf56;C21orf56 | S_Shore | 5'UTR;5'UTR;1stExon;1stExon | Yes |
| Control-Sample16_Beta | chr21:47604291-47605174 | C21orf56;C21orf56;C21orf56;C21orf56 | S_Shore | 5'UTR;5'UTR;1stExon;1stExon | Yes |
| Control-Sample511_Be | chr21.0:47604291-47604654 | C21orf56;C21orf56;C21orf56;C21orf56 | S_Shore | 5'UTR;5'UTR;1stExon;1stExon | Yes |
| Control-Sample525_Be | chr21.0:47604291-47604986 | C21orf56;C21orf56;C21orf56;C21orf56 | S_Shore | 5'UTR;5'UTR;1stExon;1stExon | Yes |
| Control-Sample571_Be | chr21.0:47604291-47605174 | C21orf56;C21orf56;C21orf56;C21orf56 | S_Shore | 5'UTR;5'UTR;1stExon;1stExon | Yes |
| Control-Sample13_Beta | chr21:34185927-34186122 | C21orf62;C21orf62;C21orf62;C21orf62;C21orf62;C21orf62 | | 5'UTR;5'UTR;1stExon;1stExon;1stExon;5'UTR | No |
| Control-Sample177_Be | chr21:15352213-15352641 | C21orf81 | Island | Body | Yes |
| Control-Sample517_Be | chr21.0:15352213-15352641 | C21orf81 | Island | Body | Yes |
| Control-Sample19_Beta | chr22:46449461-46449981 | C22orf26;LOC150381 | N_Shore | Body;TSS1500 | Yes |
| Control-Sample20_Beta | chr22:46449461-46450107 | C22orf26;LOC150381 | N_Shore | Body;TSS1500 | Yes |
| Control-Sample42_Beta | chr22:46449752-46449981 | C22orf26;LOC150381 | Island | 1stExon;Body | Yes |
| Control-Sample70_Beta | chr22:46449752-46450059 | C22orf26;LOC150381 | Island | 1stExon;Body | Yes |
| Control-Sample67_Beta | chr22:50025915-50026188 | C22orf34 | Island | Body | No |
| Control-Sample1_Beta | chr22:32555255-32555286 | C22orf42 | | TSS200 | No |
| Control-Sample1_Beta | chr22:24890794-24890831 | C22orf45;C22orf45;UPB1 | Island | TSS200;TSS200;TSS1500 | Yes |
| Control-Sample10_Beta | chr22:24890794-24890833 | C22orf45;C22orf45;UPB1 | Island | TSS200;TSS200;TSS1500 | Yes |
| Control-Sample13_Beta | chr22:24890814-24890833 | C22orf45;C22orf45;UPB1 | Island | TSS200;TSS200;TSS1500 | Yes |
| Control-Sample26_Beta | chr22:24890794-24890811 | C22orf45;C22orf45;UPB1 | Island | TSS200;TSS200;TSS1500 | Yes |
| Control-Sample44_Beta | chr22:24890794-24890814 | C22orf45;C22orf45;UPB1 | Island | TSS200;TSS200;TSS1500 | Yes |
| Control-Sample6_Beta | chr22:24890330-24890833 | C22orf45;C22orf45;UPB1 | N_Shore | Body;Body;TSS1500 | Yes |
| Control-Sample10_Beta | chr22:45608686-45608713 | C22orf9;C22orf9 | | Body;TSS1500 | Yes |
| Control-Sample11_Beta | chr22:45608492-45608713 | C22orf9;C22orf9 | | Body;TSS200 | Yes |
| Control-Sample27_Beta | chr22:45608516-45608713 | C22orf9;C22orf9 | | Body;TSS200 | Yes |
| Control-Sample67_Beta | chr22:45608345-45608448 | C22orf9;C22orf9;C22orf9 | | Body;1stExon;5'UTR | Yes |
| Control-Sample1_Beta | chr1:151810899-151811620 | C2CD4D;LOC100132111 | Island | Body;TSS200 | Yes |
| Control-Sample10_Beta | chr1:151810887-151811620 | C2CD4D;LOC100132111 | Island | Body;TSS200 | Yes |
| Control-Sample100_Be | chr1:151810887-151810899 | C2CD4D;LOC100132111 | Island | Body;TSS200 | Yes |
| Control-Sample101_Be | chr1:151810887-151811364 | C2CD4D;LOC100132111 | Island | Body;TSS200 | Yes |
| Control-Sample105_Be | chr1:151810899-151811364 | C2CD4D;LOC100132111 | Island | Body;TSS200 | Yes |
| Control-Sample13_Beta | chr1:151810887-151810904 | C2CD4D;LOC100132111 | Island | Body;TSS200 | Yes |
| Control-Sample87_Beta | chr1:151810893-151810904 | C2CD4D;LOC100132111 | Island | Body;TSS200 | Yes |
| Control-Sample89_Beta | chr1:151810893-151811620 | C2CD4D;LOC100132111 | Island | Body;TSS200 | Yes |
| Control-Sample1_Beta | chr2:75938223-75938438 | C2orf3 | S_Shore | TSS200 | Yes |
| Control-Sample3_Beta | chr2:75938223-75938289 | C2orf3 | S_Shore | TSS200 | Yes |
| Control-Sample54_Beta | chr2:75937535-75937889 | C2orf3 | N_Shore | Body | Yes |
| Control-Sample19_Beta | chr2:10302391-10302462 | C2orf48 | | Body | No |
| Control-Sample146_Be | chr2:11272783-11273229 | C2orf50 | | TSS1500 | Yes |
| Control-Sample147_Be | chr2:11272783-11273291 | C2orf50 | | TSS1500 | Yes |

| | | | | | |
|---|---|---|---|---|---|
| Control-Sample252_Be | chr2:11272899-11273229 | C2orf50 | | TSS1500 | Yes |
| Control-Sample40_Beta | chr2:11272783-11273075 | C2orf50 | | TSS1500 | Yes |
| Control-Sample22_Beta | chr2:55450737-55450874 | C2orf63;C2orf63 | Island | 5'UTR;5'UTR | No |
| Control-Sample14_Beta | chr2:209054740-209055251 | C2orf80;C2orf80 | | 1stExon;5'UTR | No |
| Control-Sample19_Beta | chr2:242811830-242811871 | C2orf85 | N_Shelf | TSS200 | Yes |
| Control-Sample3_Beta | chr2:242811552-242811871 | C2orf85 | N_Shelf | TSS1500 | Yes |
| Control-Sample15_Beta | chr3:194833887-194834008 | C3orf21 | | Body | No |
| Control-Sample64_Beta | chr3:55693182-55693631 | C3orf51;ERC2 | | Body;3'UTR | No |
| Control-Sample194_Be | chr3:49314539-49314655 | C3orf62 | Island | TSS200 | Yes |
| Control-Sample29_Beta | chr3:49314454-49314595 | C3orf62;C3orf62 | Island | 1stExon;5'UTR | Yes |
| Control-Sample220_Be | chr6:31997026-31997067 | C4A;C4B | | Body;Body | Yes |
| Control-Sample324_Be | chr6:31964288-31964391 | C4B;C4A | | Body;Body | No |
| Control-Sample1_Beta | chr4:121991345-121991672 | C4orf31 | N_Shore | 5'UTR | Yes |
| Control-Sample11_Beta | chr4:121991576-121991672 | C4orf31 | N_Shore | 5'UTR | Yes |
| Control-Sample568_Be | chr4.0:130017238-130017530 | C4orf33;C4orf33 | S_Shelf | 5'UTR;TSS200 | No |
| Control-Sample1_Beta | chr4:165878037-165878742 | C4orf39;TRIM61 | Island | TSS200;Body | Yes |
| Control-Sample146_Be | chr4:165878037-165878085 | C4orf39;TRIM61 | Island | TSS200;Body | Yes |
| Control-Sample1_Beta | chr4:1243849-1244086 | C4orf42;CTBP1;CTBP1 | Island | TSS1500;TSS1500;TSS1500 | Yes |
| Control-Sample2_Beta | chr4:1243849-1244042 | C4orf42;CTBP1;CTBP1 | Island | TSS1500;TSS1500;TSS1500 | Yes |
| Control-Sample6_Beta | chr4:1243849-1244024 | C4orf42;CTBP1;CTBP1 | Island | TSS1500;TSS1500;TSS1500 | Yes |
| Control-Sample19_Beta | chr4:3250745-3250921 | C4orf44;C4orf44 | Island | TSS200;TSS200 | No |
| Control-Sample331_Be | chr4:5526510-5526783 | C4orf6 | | TSS1500 | No |
| Control-Sample1_Beta | chr5:134181226-134181266 | C5orf24;C5orf24 | N_Shore | TSS200;TSS1500 | No |
| Control-Sample65_Beta | chr5:56204834-56205094 | C5orf35 | N_Shore | TSS1500 | No |
| Control-Sample108_Be | chr6:32297670-32297822 | C6orf10 | | Body | Yes |
| Control-Sample35_Beta | chr6:32305068-32305145 | C6orf10 | | Body | Yes |
| Control-Sample12_Beta | chr6:168188992-168189047 | C6orf123 | | Body | No |
| Control-Sample1_Beta | chr6:37460940-37461350 | C6orf129 | | Body | Yes |
| Control-Sample21_Beta | chr6:37461253-37461350 | C6orf129 | | Body | Yes |
| Control-Sample63_Beta | chr6:30614168-30614329 | C6orf136;C6orf136;C6orf136 | N_Shore | TSS1500;TSS1500;TSS1500 | No |
| Control-Sample1_Beta | chr6:3743284-3743529 | C6orf145 | | Body | No |
| Control-Sample16_Beta | chr6:130182478-130182512 | C6orf191 | | TSS200 | Yes |
| Control-Sample25_Beta | chr6:130182478-130182551 | C6orf191 | | TSS200 | Yes |
| Control-Sample52_Beta | chr6:130182510-130182620 | C6orf191 | | TSS200 | Yes |
| Control-Sample9_Beta | chr6:130182478-130182620 | C6orf191 | | TSS200 | Yes |
| Control-Sample10_Beta | chr6:130182413-130182620 | C6orf191;C6orf191 | | 5'UTR;1stExon | Yes |
| Control-Sample28_Beta | chr6:130182413-130182512 | C6orf191;C6orf191 | | 5'UTR;1stExon | Yes |
| Control-Sample3_Beta | chr6:130182413-130182551 | C6orf191;C6orf191 | | 5'UTR;1stExon | Yes |
| Control-Sample15_Beta | chr6:118973309-118973353 | C6orf204;C6orf204 | S_Shore | TSS1500;TSS1500 | No |
| Control-Sample24_Beta | chr6:170201673-170201785 | C6orf208 | | Body | No |
| Control-Sample134_Be | chr6:31734292-31734401 | C6orf27 | | Body | No |
| Control-Sample23_Beta | chr6:31804172-31804461 | C6orf48;C6orf48;SNORD52 | S_Shore | 5'UTR;5'UTR;TSS1500 | Yes |
| Control-Sample53_Beta | chr6:31804159-31804461 | C6orf48;C6orf48;SNORD52 | S_Shore | 5'UTR;5'UTR;TSS1500 | Yes |
| Control-Sample53_Beta | chr6:31804609-31804883 | C6orf48;C6orf48;SNORD52 | S_Shore | 5'UTR;5'UTR;TSS1500 | Yes |
| Control-Sample671_Be | chr6:31803089-31803103 | C6orf48;SNORD48;C6orf48 | S_Shore | 5'UTR;Body;5'UTR | Yes |
| Control-Sample10_Beta | chr7:31726692-31726912 | C7orf16;C7orf16 | | TSS200;TSS200 | Yes |
| Control-Sample19_Beta | chr7:31726639-31726912 | C7orf16;C7orf16 | | TSS200;TSS200 | Yes |
| Control-Sample22_Beta | chr7:31726654-31726912 | C7orf16;C7orf16 | | TSS200;TSS200 | Yes |
| Control-Sample310_Be | chr7:86849173-86849180 | C7orf23 | Island | TSS200 | No |
| Control-Sample53_Beta | chr7:6866834-6867074 | C7orf28B | S_Shore | TSS1500 | No |
| Control-Sample10_Beta | chr7:25216506-25216828 | C7orf31 | N_Shelf | Body | No |
| Control-Sample183_Be | chr7:23719851-23720838 | C7orf46;C7orf46;C7orf46 | Island | 1stExon;1stExon;1stExon | No |
| Control-Sample183_Be | chr7:134856562-134856681 | C7orf49;C7orf49;C7orf49 | S_Shore | TSS1500;TSS1500;TSS1500 | No |
| Control-Sample10_Beta | chr7:1095595-1095720 | C7orf50;C7orf50;C7orf50 | N_Shore | Body;Body;Body | Yes |
| Control-Sample11_Beta | chr7:1095595-1095701 | C7orf50;C7orf50;C7orf50 | N_Shore | Body;Body;Body | Yes |
| Control-Sample14_Beta | chr7:1113014-1113086 | C7orf50;C7orf50;C7orf50 | | Body;Body;Body | Yes |
| Control-Sample141_Be | chr7:1081061-1081250 | C7orf50;C7orf50;C7orf50 | Island | Body;Body;Body | Yes |
| Control-Sample145_Be | chr7:1067951-1068230 | C7orf50;C7orf50;C7orf50 | Island | Body;Body;Body | Yes |
| Control-Sample218_Be | chr7:1080836-1081250 | C7orf50;C7orf50;C7orf50 | Island | Body;Body;Body | Yes |
| Control-Sample331_Be | chr7:1121657-1121899 | C7orf50;C7orf50;C7orf50 | S_Shore | Body;Body;Body | Yes |
| Control-Sample419_Be | chr7:1067951-1068125 | C7orf50;C7orf50;C7orf50 | Island | Body;Body;Body | Yes |
| Control-Sample590_Be | chr7.0:1080836-1081068 | C7orf50;C7orf50;C7orf50 | Island | Body;Body;Body | Yes |
| Control-Sample1_Beta | chr7:1097090-1097131 | C7orf50;C7orf50;GPR146;C7orf50 | N_Shore | Body;Body;TSS200;Body | Yes |
| Control-Sample43_Beta | chr7:1095666-1095720 | C7orf50;C7orf50;GPR146;C7orf50 | N_Shore | Body;Body;TSS1500;Body | Yes |
| Control-Sample19_Beta | chr7:1098625-1098703 | C7orf50;GPR146;C7orf50;C7orf50;GPR146 | S_Shore | Body;3'UTR;Body;Body;1stExon | Yes |
| Control-Sample1_Beta | chr7:100817863-100818038 | C7orf52 | Island | Body | No |
| Control-Sample19_Beta | chr8:49984564-49985021 | C8orf22 | | TSS1500 | No |
| Control-Sample35_Beta | chr8:40010443-40010823 | C8orf4 | | TSS1500 | No |
| Control-Sample19_Beta | chr8:496440-496618 | C8orf42 | S_Shore | TSS1500 | Yes |
| Control-Sample20_Beta | chr8:496527-496739 | C8orf42 | S_Shore | TSS1500 | Yes |
| Control-Sample230_Be | chr8:143807962-143809371 | C8orf55 | Island | TSS1500 | No |
| Control-Sample225_Be | chr8:58192502-58192883 | C8orf71 | Island | Body | No |
| Control-Sample337_Be | chr8:117950395-117950473 | C8orf85 | N_Shore | TSS200 | No |
| Control-Sample3_Beta | chr9:34729282-34729535 | C9orf144B | | 1stExon | No |
| Control-Sample1_Beta | chr9:97848306-97848736 | C9orf3;MIR24-1 | Island | 3'UTR;Body | No |
| Control-Sample331_Be | chr1:150229871-150230233 | CA14 | | TSS1500 | No |
| Control-Sample17_Beta | chr13:49975383-49975643 | CAB39L;CAB39L;CAB39L | | 5'UTR;5'UTR;1stExon | No |
| Control-Sample194_Be | chr22:24552132-24552846 | CABIN1 | Island | Body | No |
| Control-Sample1_Beta | chr18:20713759-20714399 | CABLES1 | N_Shore | TSS1500 | Yes |
| Control-Sample171_Be | chr18:20714217-20714496 | CABLES1 | N_Shore | TSS1500 | Yes |
| Control-Sample19_Beta | chr18:20714217-20714399 | CABLES1 | N_Shore | TSS1500 | Yes |
| Control-Sample170_Be | chr18:20714428-20714496 | CABLES1;CABLES1 | N_Shore | TSS1500;TSS200 | Yes |
| Control-Sample3_Beta | chr18:20735005-20735537 | CABLES1;CABLES1;CABLES1 | | Body;Body;TSS1500 | Yes |
| Control-Sample245_Be | chr18:21718979-21719286 | CABYR;CABYR;CABYR;CABYR;CABYR;CABYR;CABYR;CABY | Island | 1stExon;1stExon;1stExon;5'UTR;5'UTR;5'UTR;TSS150 | No |
| Control-Sample245_Be | chr19:13319324-13319523 | CACNA1A;CACNA1A | Island | Body;Body | No |
| Control-Sample1_Beta | chr12:2734150-2734257 | CACNA1C;CACNA1C;CACNA1C;CACNA1C;CACNA1C;CACNA1C;CACNA1C;CACNA1C;CACNA1 | | Body;Body;Body;Body;Body;Body;Body;Body;Body;B | Yes |
| Control-Sample1_Beta | chr12:2800909-2801158 | CACNA1C;CACNA1C;CACNA1C;CACNA1C;CACNA1C;CA( | Island | 3'UTR;3'UTR;3'UTR;3'UTR;3'UTR;3'UTR;3'UTR;3'UTR;3 | Yes |
| Control-Sample31_Beta | chr12:2800909-2801061 | CACNA1C;CACNA1C;CACNA1C;CACNA1C;CACNA1C;CA( | Island | 3'UTR;3'UTR;3'UTR;3'UTR;3'UTR;3'UTR;3'UTR;3'UTR;3 | Yes |
| Control-Sample1_Beta | chr3:50488020-50488230 | CACNA2D2;CACNA2D2 | | Body;Body | Yes |
| Control-Sample10_Beta | chr3:50487955-50488230 | CACNA2D2;CACNA2D2 | | Body;Body | Yes |
| Control-Sample11_Beta | chr3:50487955-50488162 | CACNA2D2;CACNA2D2 | | Body;Body | Yes |

| | | | | | |
|---|---|---|---|---|---|
| Control-Sample108_Be | chr10:18629545-18629677 | CACNB2;CACNB2;CACNB2;CACNB2;CACNB2;CACNB2;CACNB2;CACNB2 | | Body;TSS200;Body;Body;Body;Body;Body;Body | Yes |
| Control-Sample20_Beta | chr10:18629497-18629568 | CACNB2;CACNB2;CACNB2;CACNB2;CACNB2;CACNB2;CACNB2;CACNB2 | | Body;TSS200;Body;Body;Body;Body;Body;Body | Yes |
| Control-Sample1_Beta | chr19:54445377-54445888 | CACNG7 | Island | Body | No |
| Control-Sample108_Be | chr7:134464123-134464411 | CALD1;CALD1;CALD1 | | TSS200;TSS200;TSS200 | Yes |
| Control-Sample291_Be | chr7:134464123-134464288 | CALD1;CALD1;CALD1 | | TSS200;TSS200;TSS200 | Yes |
| Control-Sample108_Be | chr7:134575145-134575306 | CALD1;CALD1;CALD1;CALD1;CALD1 | | TSS1500;Body;TSS1500;Body;Body | Yes |
| Control-Sample108_Be | chr7:134575524-134576049 | CALD1;CALD1;CALD1;CALD1;CALD1 | | TSS1500;Body;TSS1500;Body;Body | Yes |
| Control-Sample291_Be | chr7:134575187-134575306 | CALD1;CALD1;CALD1;CALD1;CALD1 | | TSS1500;Body;TSS1500;Body;Body | Yes |
| Control-Sample645_Be | chr7:134574965-134575145 | CALD1;CALD1;CALD1;CALD1;CALD1 | | TSS1500;Body;TSS1500;Body;Body | Yes |
| Control-Sample568_Be | chr7.0:134464242-134464295 | CALD1;CALD1;CALD1;CALD1;CALD1;CALD1 | | 5'UTR;1stExon;1stExon;1stExon;5'UTR;5'UTR | Yes |
| Control-Sample11_Beta | chr10:12648032-12648338 | CAMK1D;CAMK1D | | Body;Body | No |
| Control-Sample1_Beta | chr5:149669484-149669674 | CAMK2A;CAMK2A | | TSS200;TSS200 | Yes |
| Control-Sample5_Beta | chr5:149669484-149669813 | CAMK2A;CAMK2A | | TSS200;TSS200 | Yes |
| Control-Sample10_Beta | chr17:3791902-3792074 | CAMKK1;CAMKK1;CAMKK1 | N_Shore | 5'UTR;5'UTR;5'UTR | No |
| Control-Sample14_Beta | chr12:121678474-121678720 | CAMKK2;CAMKK2;CAMKK2;CAMKK2;CAMKK2 | | 3'UTR;3'UTR;3'UTR;3'UTR;3'UTR | Yes |
| Control-Sample464_Be | chr12:121711847-121712245 | CAMKK2;CAMKK2;CAMKK2;CAMKK2;CAMKK2;CAMKK2;CA | Island | Body;Body;Body;Body;Body;Body;Body | Yes |
| Control-Sample1_Beta | chr1:7022041-7022284 | CAMTA1 | Island | Body | Yes |
| Control-Sample1_Beta | chr1:7023625-7023864 | CAMTA1 | S_Shore | Body | Yes |
| Control-Sample10_Beta | chr1:7120893-7121245 | CAMTA1 | N_Shore | Body | Yes |
| Control-Sample108_Be | chr1:7308012-7308366 | CAMTA1 | | Body | Yes |
| Control-Sample16_Beta | chr1:7120893-7120926 | CAMTA1 | N_Shore | Body | Yes |
| Control-Sample19_Beta | chr1:7408793-7409003 | CAMTA1 | | Body | Yes |
| Control-Sample216_Be | chr1:7308091-7308366 | CAMTA1 | | Body | Yes |
| Control-Sample475_Be | chr1:7308012-7308117 | CAMTA1 | | Body | Yes |
| Control-Sample13_Beta | chr17:4872326-4872499 | CAMTA2;CAMTA2;CAMTA2;CAMTA2;SPAG7 | S_Shore | Body;Body;Body;Body;TSS1500 | Yes |
| Control-Sample57_Beta | chr17:4872326-4872603 | CAMTA2;CAMTA2;CAMTA2;CAMTA2;SPAG7 | S_Shore | Body;Body;Body;Body;TSS1500 | Yes |
| Control-Sample91_Beta | chr17:4872488-4872603 | CAMTA2;CAMTA2;CAMTA2;CAMTA2;SPAG7 | S_Shore | Body;Body;Body;Body;TSS1500 | Yes |
| Control-Sample11_Beta | chr11:64951536-64952034 | CAPN1 | N_Shore | Body | Yes |
| Control-Sample2_Beta | chr11:64951536-64951960 | CAPN1 | N_Shore | Body | Yes |
| Control-Sample5_Beta | chr11:64951536-64951916 | CAPN1 | N_Shore | Body | Yes |
| Control-Sample1_Beta | chr1:223936508-223936812 | CAPN2;CAPN2 | Island | Body;Body | No |
| Control-Sample1_Beta | chr7:2953836-2953930 | CARD11 | | Body | Yes |
| Control-Sample10_Beta | chr7:2971898-2972392 | CARD11 | S_Shelf | Body | Yes |
| Control-Sample15_Beta | chr5:40841315-40841579 | CARD6 | | TSS200 | No |
| Control-Sample4_Beta | chr13:111281037-111281144 | CARKD | S_Shelf | Body | No |
| Control-Sample464_Be | chr13:111330434-111331324 | CARS2 | N_Shore | Body | No |
| Control-Sample1_Beta | chr17:73506017-73506230 | CASKIN2;CASKIN2 | | Body;TSS200 | Yes |
| Control-Sample13_Beta | chr17:73506038-73506230 | CASKIN2;CASKIN2 | | Body;TSS200 | Yes |
| Control-Sample19_Beta | chr17:73506140-73506230 | CASKIN2;CASKIN2 | | Body;TSS200 | Yes |
| Control-Sample496_Be | chr17:73498127-73498302 | CASKIN2;CASKIN2 | Island | Body;Body | Yes |
| Control-Sample1_Beta | chr2:202122474-202122832 | CASP8;CASP8;CASP8;CASP8;CASP8 | | 5'UTR;5'UTR;5'UTR;TSS1500;5'UTR | Yes |
| Control-Sample10_Beta | chr2:202122309-202122474 | CASP8;CASP8;CASP8;CASP8;CASP8 | | 5'UTR;5'UTR;5'UTR;TSS1500;5'UTR | Yes |
| Control-Sample11_Beta | chr2:202122309-202122669 | CASP8;CASP8;CASP8;CASP8;CASP8 | | 5'UTR;5'UTR;5'UTR;TSS1500;5'UTR | Yes |
| Control-Sample124_Be | chr2:202122474-202122653 | CASP8;CASP8;CASP8;CASP8;CASP8 | | 5'UTR;5'UTR;5'UTR;TSS1500;5'UTR | Yes |
| Control-Sample13_Beta | chr2:202122309-202122653 | CASP8;CASP8;CASP8;CASP8;CASP8 | | 5'UTR;5'UTR;5'UTR;TSS1500;5'UTR | Yes |
| Control-Sample14_Beta | chr2:202122474-202122669 | CASP8;CASP8;CASP8;CASP8;CASP8 | | 5'UTR;5'UTR;5'UTR;TSS1500;5'UTR | Yes |
| Control-Sample16_Beta | chr2:202122474-202122753 | CASP8;CASP8;CASP8;CASP8;CASP8 | | 5'UTR;5'UTR;5'UTR;TSS1500;5'UTR | Yes |
| Control-Sample18_Beta | chr2:202122309-202122832 | CASP8;CASP8;CASP8;CASP8;CASP8 | | 5'UTR;5'UTR;5'UTR;TSS1500;5'UTR | Yes |
| Control-Sample20_Beta | chr2:202122587-202122669 | CASP8;CASP8;CASP8;CASP8;CASP8 | | 5'UTR;5'UTR;5'UTR;5'UTR;TSS200 | Yes |
| Control-Sample24_Beta | chr2:202122309-202122753 | CASP8;CASP8;CASP8;CASP8;CASP8 | | 5'UTR;5'UTR;5'UTR;TSS1500;5'UTR | Yes |
| Control-Sample1_Beta | chr1:10711457-10711695 | CASZ1;CASZ1 | N_Shelf | Body;Body | Yes |
| Control-Sample19_Beta | chr1:10710847-10711695 | CASZ1;CASZ1 | N_Shelf | Body;Body | Yes |
| Control-Sample19_Beta | chr1:10738569-10739184 | CASZ1;CASZ1 | | Body;Body | Yes |
| Control-Sample194_Be | chr1:10839450-10839574 | CASZ1;CASZ1 | | 5'UTR;5'UTR | Yes |
| Control-Sample20_Beta | chr1:10732464-10732558 | CASZ1;CASZ1 | | Body;Body | Yes |
| Control-Sample21_Beta | chr1:10732872-10733030 | CASZ1;CASZ1 | | Body;Body | Yes |
| Control-Sample27_Beta | chr1:10709702-10710134 | CASZ1;CASZ1 | S_Shore | Body;Body | Yes |
| Control-Sample57_Beta | chr1:10732464-10732900 | CASZ1;CASZ1 | | Body;Body | Yes |
| Control-Sample58_Beta | chr1:10732519-10732900 | CASZ1;CASZ1 | | Body;Body | Yes |
| Control-Sample100_Be | chr1:26527870-26527928 | CATSPER4 | | Body | No |
| Control-Sample24_Beta | chr16:88990161-88990232 | CBFA2T3;CBFA2T3 | N_Shelf | 5'UTR;Body | Yes |
| Control-Sample27_Beta | chr16:88990161-88990409 | CBFA2T3;CBFA2T3 | N_Shelf | 5'UTR;Body | Yes |
| Control-Sample32_Beta | chr16:88991238-88991518 | CBFA2T3;CBFA2T3 | N_Shelf | 5'UTR;Body | Yes |
| Control-Sample6_Beta | chr16:88990409-88990746 | CBFA2T3;CBFA2T3 | N_Shelf | 5'UTR;Body | Yes |
| Control-Sample1_Beta | chr18:70211959-70211974 | CBLN2 | S_Shore | TSS1500 | Yes |
| Control-Sample57_Beta | chr18:70211959-70212474 | CBLN2 | S_Shore | TSS1500 | Yes |
| Control-Sample12_Beta | chr14:24898992-24899054 | CBLN3;KHNYN | Island | TSS1500;TSS200 | Yes |
| Control-Sample13_Beta | chr14:24898992-24899050 | CBLN3;KHNYN | Island | TSS1500;TSS200 | Yes |
| Control-Sample1_Beta | chr21:37442289-37442476 | CBR1;CBR1 | Island | 1stExon;5'UTR | Yes |
| Control-Sample217_Be | chr21:37442289-37442759 | CBR1;CBR1 | Island | 1stExon;5'UTR | Yes |
| Control-Sample504_Be | chr21.0:37442289-37442476 | CBR1;CBR1 | Island | 1stExon;5'UTR | Yes |
| Control-Sample544_Be | chr21.0:37442289-37442759 | CBR1;CBR1 | Island | 1stExon;5'UTR | Yes |
| Control-Sample17_Beta | chr21:44473496-44473840 | CBS | N_Shore | 3'UTR | Yes |
| Control-Sample464_Be | chr21:44480336-44480699 | CBS | N_Shore | Body | Yes |
| Control-Sample1_Beta | chr17:77809912-77810742 | CBX4 | S_Shore | Body | Yes |
| Control-Sample13_Beta | chr17:77809912-77810912 | CBX4 | S_Shore | Body | Yes |
| Control-Sample17_Beta | chr17:77809912-77810509 | CBX4 | S_Shore | Body | Yes |
| Control-Sample39_Beta | chr17:77810269-77810742 | CBX4 | N_Shore | Body | Yes |
| Control-Sample108_Be | chr12:54653065-54653403 | CBX5;CBX5;CBX5;CBX5 | | 1stExon;5'UTR;5'UTR;5'UTR | No |
| Control-Sample306_Be | chr17:18528568-18528638 | CCDC144B | Island | 1stExon | No |
| Control-Sample186_Be | chr17:20799511-20799532 | CCDC144NL | Island | TSS200 | No |
| Control-Sample108_Be | chr4:24981949-24982540 | CCDC149 | | TSS200 | Yes |
| Control-Sample151_Be | chr4:24818471-24818615 | CCDC149 | | Body | Yes |
| Control-Sample89_Beta | chr1:45965595-45966925 | CCDC163P;CCDC163P;MMACHC | Island | 5'UTR;1stExon;TSS1500 | No |
| Control-Sample7_Beta | chr1:46088336-46089024 | CCDC17 | Island | Body | No |
| Control-Sample1_Beta | chr1:159869960-159870326 | CCDC19 | Island | TSS200 | Yes |
| Control-Sample10_Beta | chr1:159869927-159870326 | CCDC19 | Island | TSS200 | Yes |
| Control-Sample19_Beta | chr1:159870074-159870326 | CCDC19 | Island | TSS200 | Yes |
| Control-Sample31_Beta | chr1:159869927-159870086 | CCDC19 | Island | TSS200 | Yes |
| Control-Sample50_Beta | chr1:159869960-159870086 | CCDC19 | Island | TSS200 | Yes |
| Control-Sample61_Beta | chr1:159869927-159870074 | CCDC19 | Island | TSS200 | Yes |

| | | | | | |
|---|---|---|---|---|---|
| Control-Sample61_Beta | chr1:159870086-159870326 | CCDC19 | Island | TSS200 | Yes |
| Control-Sample1_Beta | chr15:74592566-74592786 | CCDC33 | | Body | No |
| Control-Sample10_Beta | chr3:49236860-49236946 | CCDC36;CCDC36 | Island | TSS200;5'UTR | Yes |
| Control-Sample155_Be | chr3:49236810-49236946 | CCDC36;CCDC36 | Island | TSS200;5'UTR | Yes |
| Control-Sample176_Be | chr3:49236810-49236918 | CCDC36;CCDC36 | Island | TSS200;5'UTR | Yes |
| Control-Sample19_Beta | chr17:78039096-78039528 | CCDC40 | N_Shore | Body | Yes |
| Control-Sample92_Beta | chr17:78031893-78032296 | CCDC40 | N_Shore | Body | Yes |
| Control-Sample14_Beta | chr17:80132787-80132900 | CCDC57 | | Body | No |
| Control-Sample108_Be | chr10:32734661-32734684 | CCDC7;CCDC7 | | TSS1500;TSS1500 | No |
| Control-Sample1_Beta | chr19:46916588-46917061 | CCDC8;CCDC8 | Island | 1stExon;5'UTR | Yes |
| Control-Sample10_Beta | chr19:46916741-46917061 | CCDC8;CCDC8 | Island | 1stExon;5'UTR | Yes |
| Control-Sample11_Beta | chr19:46916741-46917255 | CCDC8;CCDC8 | Island | 1stExon;5'UTR | Yes |
| Control-Sample12_Beta | chr19:46916588-46917255 | CCDC8;CCDC8 | Island | 1stExon;5'UTR | Yes |
| Control-Sample108_Be | chr3:112359488-112360446 | CCDC80;CCDC80;CCDC80 | | 5'UTR;1stExon;5'UTR | Yes |
| Control-Sample291_Be | chr3:112359557-112359935 | CCDC80;CCDC80;CCDC80 | | 5'UTR;1stExon;5'UTR | Yes |
| Control-Sample43_Beta | chr2:56409774-56410071 | CCDC85A | N_Shore | TSS1500 | No |
| Control-Sample1_Beta | chr14:100047630-100048149 | CCDC85C | | Body | Yes |
| Control-Sample1_Beta | chr14:100069535-100069666 | CCDC85C | Island | 1stExon | Yes |
| Control-Sample4_Beta | chr14:99988546-99988776 | CCDC85C | | Body | Yes |
| Control-Sample3_Beta | chr14:91770052-91770229 | CCDC88C | N_Shore | Body | No |
| Control-Sample108_Be | chr11:85397216-85397684 | CCDC89;CCDC89 | | 1stExon;5'UTR | No |
| Control-Sample46_Beta | chr6:31111076-31111270 | CCHCR1;CCHCR1;CCHCR1 | | Body;Body;Body | No |
| Control-Sample1_Beta | chr4:26491963-26492488 | CCKAR;CCKAR | | 1stExon;5'UTR | Yes |
| Control-Sample13_Beta | chr4:26491963-26492378 | CCKAR;CCKAR | | 1stExon;5'UTR | Yes |
| Control-Sample3_Beta | chr4:26491963-26492870 | CCKAR;CCKAR | | 1stExon;5'UTR | Yes |
| Control-Sample42_Beta | chr4:26491963-26492206 | CCKAR;CCKAR | | 1stExon;5'UTR | Yes |
| Control-Sample1_Beta | chr11:69460951-69461451 | CCND1 | N_Shore | Body | Yes |
| Control-Sample15_Beta | chr11:69460951-69461236 | CCND1 | N_Shore | Body | Yes |
| Control-Sample58_Beta | chr11:69460951-69461334 | CCND1 | N_Shore | Body | Yes |
| Control-Sample65_Beta | chr12:4399358-4399998 | CCND2 | | Body | No |
| Control-Sample1_Beta | chr5:132083532-132084068 | CCNI2 | Island | 1stExon | No |
| Control-Sample108_Be | chr3:46249795-46250024 | CCR1;CCR1 | | 1stExon;5'UTR | No |
| Control-Sample19_Beta | chr1:167425134-167425345 | CD247;CD247 | Island | Body;Body | No |
| Control-Sample10_Beta | chr19:49841252-49841517 | CD37;CD37 | Island | Body;Body | No |
| Control-Sample1_Beta | chr12:6898647-6898711 | CD4 | | TSS200 | No |
| Control-Sample18_Beta | chr1:207925081-207925112 | CD46;CD46;CD46;CD46;CD46;CD46;CD46;CD46;CD46;C | N_Shore | TSS1500;TSS1500;TSS1500;TSS1500;TSS1500;TSS15 | No |
| Control-Sample5_Beta | chr1:117077088-117077219 | CD58;CD58;CD58 | Island | Body;Body;Body | No |
| Control-Sample13_Beta | chr2:87018944-87018966 | CD8A;CD8A;CD8A;CD8A | S_Shore | TSS200;5'UTR;TSS200;TSS200 | Yes |
| Control-Sample1_Beta | chr2:87018585-87018966 | CD8A;CD8A;CD8A;CD8A;CD8A;CD8A | S_Shore | 5'UTR;1stExon;5'UTR;Body;5'UTR;1stExon | Yes |
| Control-Sample30_Beta | chr2:87018382-87018966 | CD8A;CD8A;CD8A;CD8A;CD8A;CD8A | S_Shore | 5'UTR;1stExon;5'UTR;Body;5'UTR;1stExon | Yes |
| Control-Sample1_Beta | chr12:6308758-6309025 | CD9 | N_Shore | TSS1500 | No |
| Control-Sample102_Be | chr10:38691787-38692145 | CDC10L | Island | TSS200 | Yes |
| Control-Sample106_Be | chr10:38691787-38691801 | CDC10L | Island | TSS200 | Yes |
| Control-Sample225_Be | chr10:38691791-38691801 | CDC10L | Island | TSS200 | Yes |
| Control-Sample567_Be | chr10.0:38691787-38692145 | CDC10L | Island | TSS200 | Yes |
| Control-Sample20_Beta | chr14:103416122-103416389 | CDC42BPB | | Body | No |
| Control-Sample475_Be | chr22:37962569-37962818 | CDC42EP1 | Island | Body | No |
| Control-Sample1_Beta | chr16:68772225-68772469 | CDH1 | Island | Body | No |
| Control-Sample108_Be | chr5:24645093-24645487 | CDH10 | | TSS200 | No |
| Control-Sample47_Beta | chr16:65105516-65105714 | CDH11 | | 5'UTR | No |
| Control-Sample216_Be | chr16:83171068-83171299 | CDH13 | | Body | Yes |
| Control-Sample5_Beta | chr16:83171068-83171314 | CDH13 | | Body | Yes |
| Control-Sample57_Beta | chr16:82660206-82660328 | CDH13 | N_Shore | TSS1500 | Yes |
| Control-Sample65_Beta | chr16:83171155-83171314 | CDH13 | | Body | Yes |
| Control-Sample171_Be | chr18:64271341-64271503 | CDH19 | | TSS200 | No |
| Control-Sample1_Beta | chr20:44875334-44875586 | CDH22 | Island | Body | No |
| Control-Sample51_Beta | chr16:66400320-66400411 | CDH5 | | TSS1500 | No |
| Control-Sample67_Beta | chr5:27038707-27038802 | CDH9 | | TSS200 | No |
| Control-Sample13_Beta | chr12:56359210-56359450 | CDK2;CDK2;SILV | N_Shore | TSS1500;TSS1500;Body | No |
| Control-Sample1_Beta | chr12:123757337-123758062 | CDK2AP1 | S_Shore | TSS1500 | Yes |
| Control-Sample10_Beta | chr12:123757337-123757536 | CDK2AP1 | S_Shore | TSS1500 | Yes |
| Control-Sample14_Beta | chr12:123757337-123757860 | CDK2AP1 | S_Shore | TSS1500 | Yes |
| Control-Sample15_Beta | chr12:123757892-123758062 | CDK2AP1 | S_Shore | TSS1500 | Yes |
| Control-Sample19_Beta | chr12:123753186-123753272 | CDK2AP1 | N_Shore | Body | Yes |
| Control-Sample2_Beta | chr12:123757536-123758062 | CDK2AP1 | S_Shore | TSS1500 | Yes |
| Control-Sample65_Beta | chr12:123757337-123758006 | CDK2AP1 | S_Shore | TSS1500 | Yes |
| Control-Sample19_Beta | chr6:36645100-36645608 | CDKN1A;CDKN1A | N_Shore | TSS1500;TSS1500 | Yes |
| Control-Sample70_Beta | chr6:36645500-36645648 | CDKN1A;CDKN1A | N_Shore | TSS1500;TSS1500 | Yes |
| Control-Sample1_Beta | chr11:2908102-2908233 | CDKN1C;CDKN1C;CDKN1C | S_Shore | TSS1500;TSS1500;TSS1500 | Yes |
| Control-Sample12_Beta | chr11:2907930-2908116 | CDKN1C;CDKN1C;CDKN1C | S_Shore | TSS1500;TSS1500;TSS1500 | Yes |
| Control-Sample24_Beta | chr11:2908036-2908116 | CDKN1C;CDKN1C;CDKN1C | S_Shore | TSS1500;TSS1500;TSS1500 | Yes |
| Control-Sample52_Beta | chr11:2907930-2908102 | CDKN1C;CDKN1C;CDKN1C | S_Shore | TSS1500;TSS1500;TSS1500 | Yes |
| Control-Sample1_Beta | chr5:149546331-149546366 | CDX1 | Island | TSS200 | Yes |
| Control-Sample12_Beta | chr5:149546335-149546366 | CDX1 | Island | TSS200 | Yes |
| Control-Sample19_Beta | chr5:149546087-149546946 | CDX1 | Island | TSS1500 | Yes |
| Control-Sample20_Beta | chr5:149546087-149547566 | CDX1 | Island | TSS1500 | Yes |
| Control-Sample1_Beta | chr6:4706170-4706427 | CDYL | | TSS1500 | Yes |
| Control-Sample5_Beta | chr6:4706170-4706336 | CDYL | | TSS1500 | Yes |
| Control-Sample19_Beta | chr16:80654711-80654763 | CDYL2 | | Body | No |
| Control-Sample136_Be | chr22:17956453-17956641 | CECR2 | | TSS200 | Yes |
| Control-Sample51_Beta | chr22:17956453-17956462 | CECR2 | | TSS200 | Yes |
| Control-Sample538_Be | chr22.0:17956462-17956641 | CECR2 | | TSS200 | Yes |
| Control-Sample57_Beta | chr22:17956455-17956641 | CECR2 | | TSS200 | Yes |
| Control-Sample622_Be | chr22:17956462-17956641 | CECR2 | | TSS200 | Yes |
| Control-Sample147_Be | chr22:46770063-46770253 | CELSR1 | N_Shore | Body | Yes |
| Control-Sample3_Beta | chr22:46770195-46770440 | CELSR1 | N_Shore | Body | Yes |
| Control-Sample220_Be | chr16:55866757-55867072 | CES1;CES1;CES1 | Island | Body;Body;Body | Yes |
| Control-Sample266_Be | chr16:55866757-55866997 | CES1;CES1;CES1 | Island | Body;Body;Body | Yes |
| Control-Sample124_Be | chr16:55794595-55794910 | CES4 | N_Shore | Body | Yes |
| Control-Sample297_Be | chr16:55794456-55794910 | CES4 | N_Shore | TSS200 | Yes |
| Control-Sample525_Be | chr16.0:55794478-55794910 | CES4 | N_Shore | TSS200 | Yes |

| | | | | | |
|---|---|---|---|---|---|
| Control-Sample6_Beta | chr16:55794456-55794731 | CES4 | N_Shore | TSS200 | Yes |
| Control-Sample670_Be | chr16:55794478-55794910 | CES4 | N_Shore | TSS200 | Yes |
| Control-Sample12_Beta | chr2:202004649-202004875 | CFLAR;CFLAR;CFLAR | | Body;Body;Body | No |
| Control-Sample331_Be | chr22:41637080-41637467 | CHADL | S_Shelf | TSS200 | No |
| Control-Sample13_Beta | chr1:203156374-203156764 | CHI3L1 | | TSS1500 | No |
| Control-Sample151_Be | chr11:888956-889231 | CHID1;CHID1;CHID1;CHID1;CHID1 | Island | Body;Body;Body;Body;Body | Yes |
| Control-Sample233_Be | chr11:889148-889231 | CHID1;CHID1;CHID1;CHID1;CHID1 | N_Shore | Body;Body;Body;Body;Body | Yes |
| Control-Sample108_Be | chr7:29519409-29519656 | CHN2;CHN2 | | TSS200;Body | No |
| Control-Sample61_Beta | chr11:89956718-89957033 | CHORDC1;CHORDC1 | S_Shore | TSS200;TSS200 | No |
| Control-Sample53_Beta | chr16:23765258-23765381 | CHP2 | N_Shore | TSS1500 | No |
| Control-Sample1_Beta | chr3:184099096-184099432 | CHRD | Island | Body | Yes |
| Control-Sample11_Beta | chr3:184104362-184104481 | CHRD | Island | Body | Yes |
| Control-Sample16_Beta | chr3:184104115-184104380 | CHRD | N_Shore | Body | Yes |
| Control-Sample17_Beta | chr3:184104362-184104518 | CHRD | Island | Body | Yes |
| Control-Sample25_Beta | chr3:184104362-184104688 | CHRD | Island | Body | Yes |
| Control-Sample53_Beta | chr3:184104115-184104481 | CHRD | N_Shore | Body | Yes |
| Control-Sample624_Be | chr15:34260956-34261172 | CHRM5;AVEN | | TSS200;Body | No |
| Control-Sample223_Be | chr2:175629219-175629267 | CHRNA1;CHRNA1 | | TSS200;TSS200 | No |
| Control-Sample19_Beta | chr20:61979272-61979563 | CHRNA4 | Island | Body | No |
| Control-Sample19_Beta | chr8:42623718-42623786 | CHRNA6 | | TSS200 | Yes |
| Control-Sample20_Beta | chr8:42623718-42623771 | CHRNA6 | | TSS200 | Yes |
| Control-Sample25_Beta | chr17:7350244-7350413 | CHRNB1 | S_Shore | Body | No |
| Control-Sample1_Beta | chr15:78933807-78934280 | CHRNB4 | Island | TSS1500 | Yes |
| Control-Sample12_Beta | chr15:78933807-78933902 | CHRNB4 | Island | TSS1500 | Yes |
| Control-Sample17_Beta | chr15:78933807-78934580 | CHRNB4 | Island | TSS1500 | Yes |
| Control-Sample1_Beta | chr17:4802692-4803053 | CHRNE;C17orf107 | Island | Body;TSS1500 | Yes |
| Control-Sample12_Beta | chr17:4802813-4803053 | CHRNE;C17orf107 | Island | Body;TSS200 | Yes |
| Control-Sample14_Beta | chr17:4802692-4802847 | CHRNE;C17orf107 | Island | Body;TSS1500 | Yes |
| Control-Sample14_Beta | chr17:4802906-4803053 | CHRNE;C17orf107 | Island | Body;TSS200 | Yes |
| Control-Sample152_Be | chr17:4802813-4802888 | CHRNE;C17orf107 | Island | Body;TSS200 | Yes |
| Control-Sample270_Be | chr17:4802813-4802906 | CHRNE;C17orf107 | Island | Body;TSS200 | Yes |
| Control-Sample65_Beta | chr12:105114162-105114399 | CHST11 | | Body | No |
| Control-Sample3_Beta | chr10:125767838-125768098 | CHST15 | | 3'UTR | No |
| Control-Sample36_Beta | chr19:34112306-34112325 | CHST8;CHST8 | Island | TSS1500;TSS1500 | Yes |
| Control-Sample6_Beta | chr19:34174731-34174777 | CHST8;CHST8;CHST8 | N_Shore | 5'UTR;5'UTR;TSS1500 | Yes |
| Control-Sample1_Beta | chr15:101728611-101729174 | CHSY1 | | Body | Yes |
| Control-Sample21_Beta | chr15:101728611-101728653 | CHSY1 | | Body | Yes |
| Control-Sample65_Beta | chr15:101728611-101729391 | CHSY1 | | Body | Yes |
| Control-Sample11_Beta | chr12:120241957-120242156 | CIT | Island | Body | No |
| Control-Sample200_Be | chr9:130955135-130955436 | CIZ1;CIZ1;CIZ1;CIZ1;CIZ1 | N_Shore | TSS1500;5'UTR;TSS1500;5'UTR;TSS1500 | No |
| Control-Sample20_Beta | chr14:103986227-103986332 | CKB | N_Shore | Body | No |
| Control-Sample1_Beta | chr11:67141628-67143062 | CLCF1;LOC100130987;CLCF1;CLCF1 | S_Shore | 1stExon;Body;TSS1500;5'UTR | Yes |
| Control-Sample15_Beta | chr11:67141628-67141747 | CLCF1;LOC100130987;CLCF1;CLCF1 | S_Shore | 1stExon;Body;TSS1500;5'UTR | Yes |
| Control-Sample63_Beta | chr11:67141628-67141792 | CLCF1;LOC100130987;CLCF1;CLCF1 | S_Shore | 1stExon;Body;TSS1500;5'UTR | Yes |
| Control-Sample65_Beta | chr13:96205679-96206104 | CLDN10;CLDN10;CLDN10 | S_Shore | Body;Body;Body | No |
| Control-Sample151_Be | chr21:37948941-37949239 | CLDN14 | | TSS200 | Yes |
| Control-Sample216_Be | chr21:37948943-37949239 | CLDN14 | | TSS200 | Yes |
| Control-Sample22_Beta | chr21:37948808-37948943 | CLDN14;CLDN14 | | 1stExon;5'UTR | Yes |
| Control-Sample223_Be | chr21:37948808-37949239 | CLDN14;CLDN14 | | 1stExon;5'UTR | Yes |
| Control-Sample51_Beta | chr1:43205623-43206037 | CLDN19;CLDN19 | S_Shore | 1stExon;1stExon | No |
| Control-Sample44_Beta | chr16:3067500-3067759 | CLDN6 | N_Shore | 5'UTR | No |
| Control-Sample1_Beta | chr16:3062653-3062975 | CLDN9;CLDN9 | N_Shore | 1stExon;5'UTR | No |
| Control-Sample1_Beta | chr3:45067564-45067788 | CLEC3B | | TSS200 | Yes |
| Control-Sample17_Beta | chr3:45067564-45067713 | CLEC3B | | TSS200 | Yes |
| Control-Sample1_Beta | chr6:31702785-31702915 | CLIC1 | | Body | Yes |
| Control-Sample10_Beta | chr6:31702785-31703139 | CLIC1 | | Body | Yes |
| Control-Sample14_Beta | chr6:31702774-31702915 | CLIC1 | | Body | Yes |
| Control-Sample2_Beta | chr6:31702774-31702862 | CLIC1 | | Body | Yes |
| Control-Sample25_Beta | chr6:31702632-31702915 | CLIC1 | | Body | Yes |
| Control-Sample29_Beta | chr6:31702774-31703139 | CLIC1 | | Body | Yes |
| Control-Sample3_Beta | chr6:31702632-31703139 | CLIC1 | | Body | Yes |
| Control-Sample10_Beta | chr14:95693784-95693981 | CLMN | | Body | Yes |
| Control-Sample18_Beta | chr14:95693784-95693881 | CLMN | | Body | Yes |
| Control-Sample108_Be | chr8:27468684-27469001 | CLU;CLU;CLU;CLU | N_Shelf | 1stExon;Body;Body;5'UTR | No |
| Control-Sample1_Beta | chr13:100497453-100497789 | CLYBL | | Body | Yes |
| Control-Sample10_Beta | chr13:100506703-100506823 | CLYBL | | Body | Yes |
| Control-Sample13_Beta | chr13:100547192-100547367 | CLYBL | Island | 3'UTR | Yes |
| Control-Sample152_Be | chr13:100547192-100547331 | CLYBL | Island | 3'UTR | Yes |
| Control-Sample21_Beta | chr13:100548513-100548910 | CLYBL | Island | 3'UTR | Yes |
| Control-Sample223_Be | chr13:100547192-100547728 | CLYBL | Island | 3'UTR | Yes |
| Control-Sample33_Beta | chr13:100547192-100547292 | CLYBL | Island | 3'UTR | Yes |
| Control-Sample40_Beta | chr13:100497610-100497789 | CLYBL | | Body | Yes |
| Control-Sample50_Beta | chr13:100547249-100547367 | CLYBL | Island | 3'UTR | Yes |
| Control-Sample585_Be | chr13.0:100547192-100547331 | CLYBL | Island | 3'UTR | Yes |
| Control-Sample7_Beta | chr13:100547728-100547897 | CLYBL | Island | 3'UTR | Yes |
| Control-Sample1_Beta | chr6:25138416-25138951 | CMAH;CMAH | N_Shore | TSS1500;Body | Yes |
| Control-Sample41_Beta | chr6:25138416-25138905 | CMAH;CMAH | N_Shore | TSS1500;Body | Yes |
| Control-Sample66_Beta | chr6:25138416-25138632 | CMAH;CMAH | N_Shore | TSS1500;Body | Yes |
| Control-Sample41_Beta | chr16:81520178-81520589 | CMIP | | Body | No |
| Control-Sample33_Beta | chr12:108733253-108733275 | CMKLR1;CMKLR1;CMKLR1 | | TSS200;TSS200;TSS200 | No |
| Control-Sample6_Beta | chr2:7005923-7005960 | CMPK2;CMPK2 | Island | 1stExon;5'UTR | No |
| Control-Sample1_Beta | chr16:66613096-66613334 | CMTM2 | Island | TSS1500 | Yes |
| Control-Sample100_Be | chr16:66613266-66613278 | CMTM2 | Island | TSS200 | Yes |
| Control-Sample2_Beta | chr16:66613220-66613334 | CMTM2 | Island | TSS200 | Yes |
| Control-Sample20_Beta | chr16:66613096-66613278 | CMTM2 | Island | TSS1500 | Yes |
| Control-Sample21_Beta | chr16:66613220-66613278 | CMTM2 | Island | TSS200 | Yes |
| Control-Sample54_Beta | chr16:66613206-66613334 | CMTM2 | Island | TSS200 | Yes |
| Control-Sample171_Be | chr14:23846456-23846906 | CMTM5;CMTM5;CMTM5;CMTM5 | | 1stExon;5'UTR;5'UTR;1stExon | No |
| Control-Sample1_Beta | chr18:72201663-72201716 | CNDP1 | | TSS200 | Yes |
| Control-Sample104_Be | chr18:72201663-72201785 | CNDP1 | | TSS200 | Yes |
| Control-Sample121_Be | chr18:72201665-72201785 | CNDP1 | | TSS200 | Yes |

| | | | | | |
|---|---|---|---|---|---|
| Control-Sample124_Be | chr18:72201504-72201785 | CNDP1 | | TSS200 | Yes |
| Control-Sample98_Beta | chr18:72201663-72201775 | CNDP1 | | TSS200 | Yes |
| Control-Sample103_Be | chr18:72201764-72201785 | CNDP1;CNDP1 | | 1stExon;5'UTR | Yes |
| Control-Sample12_Beta | chr1:26503847-26503862 | CNKSR1;CNKSR1 | | TSS200;TSS200 | Yes |
| Control-Sample17_Beta | chr1:26503847-26503979 | CNKSR1;CNKSR1 | | TSS200;TSS200 | Yes |
| Control-Sample27_Beta | chr1:26503862-26503979 | CNKSR1;CNKSR1 | | TSS200;TSS200 | Yes |
| Control-Sample73_Beta | chr17:40120250-40120496 | CNP | Island | Body | Yes |
| Control-Sample73_Beta | chr17:40125608-40125781 | CNP | | Body | Yes |
| Control-Sample76_Beta | chr17:40119957-40120496 | CNP | Island | Body | Yes |
| Control-Sample78_Beta | chr17:40125608-40126231 | CNP | | Body | Yes |
| Control-Sample89_Beta | chr17:40119847-40120496 | CNP | Island | Body | Yes |
| Control-Sample21_Beta | chr6:42896394-42896418 | CNPY3 | N_Shore | TSS1500 | No |
| Control-Sample9_Beta | chr6:88877216-88877413 | CNR1;CNR1;CNR1;CNR1 | Island | TSS1500;TSS1500;TSS1500;TSS1500 | No |
| Control-Sample464_Be | chr1:205038787-205039083 | CNTN2 | Island | Body | No |
| Control-Sample291_Be | chr3:1134528-1134730 | CNTN6 | | TSS200 | No |
| Control-Sample1_Beta | chr17:40838861-40840012 | CNTNAP1 | Island | Body | Yes |
| Control-Sample11_Beta | chr17:40835849-40836087 | CNTNAP1 | Island | Body | Yes |
| Control-Sample41_Beta | chr17:40839469-40840012 | CNTNAP1 | N_Shore | Body | Yes |
| Control-Sample51_Beta | chr17:40838861-40839766 | CNTNAP1 | Island | Body | Yes |
| Control-Sample59_Beta | chr17:40839469-40839766 | CNTNAP1 | N_Shore | Body | Yes |
| Control-Sample67_Beta | chr14:31343283-31343427 | COCH;COCH | N_Shore | TSS1500;TSS1500 | No |
| Control-Sample5_Beta | chr17:71189058-71189068 | COG1 | Island | TSS200 | No |
| Control-Sample394_Be | chr17:55038508-55038529 | COIL | Island | TSS200 | No |
| Control-Sample1_Beta | chr6:33131222-33131336 | COL11A2;COL11A2;COL11A2 | S_Shore | 3'UTR;3'UTR;3'UTR | Yes |
| Control-Sample11_Beta | chr6:33131274-33131336 | COL11A2;COL11A2;COL11A2 | S_Shore | 3'UTR;3'UTR;3'UTR | Yes |
| Control-Sample105_Be | chr6:33158503-33158538 | COL11A2;COL11A2;COL11A2;COL11A2 | N_Shore | Body;Body;Body;Body | Yes |
| Control-Sample19_Beta | chr6:33158503-33158597 | COL11A2;COL11A2;COL11A2;COL11A2 | N_Shore | Body;Body;Body;Body | Yes |
| Control-Sample223_Be | chr6:33158526-33158597 | COL11A2;COL11A2;COL11A2;COL11A2 | N_Shore | Body;Body;Body;Body | Yes |
| Control-Sample645_Be | chr6:33158020-33158538 | COL11A2;COL11A2;COL11A2;COL11A2 | N_Shore | Body;Body;Body;Body | Yes |
| Control-Sample71_Beta | chr6:33161290-33161328 | COL11A2;COL11A2;COL11A2;COL11A2 | Island | TSS1500;TSS1500;TSS1500;TSS1500 | Yes |
| Control-Sample1_Beta | chr6:75912410-75912633 | COL12A1;COL12A1 | S_Shore | Body;Body | No |
| Control-Sample14_Beta | chr1:32169701-32169868 | COL16A1;COL16A1 | Island | 1stExon;5'UTR | Yes |
| Control-Sample331_Be | chr1:32169568-32169706 | COL16A1;COL16A1 | Island | 1stExon;5'UTR | Yes |
| Control-Sample1_Beta | chr21:46850449-46850691 | COL18A1 | N_Shore | Body | Yes |
| Control-Sample12_Beta | chr21:46850594-46850691 | COL18A1 | Island | Body | Yes |
| Control-Sample2_Beta | chr21:46850594-46850840 | COL18A1 | Island | Body | Yes |
| Control-Sample57_Beta | chr21:46850449-46850840 | COL18A1 | N_Shore | Body | Yes |
| Control-Sample58_Beta | chr21:46858801-46859183 | COL18A1 | N_Shore | Body | Yes |
| Control-Sample1_Beta | chr21:46890849-46890997 | COL18A1;COL18A1;COL18A1 | Island | Body;Body;Body | Yes |
| Control-Sample14_Beta | chr8:139613712-139614020 | COL22A1 | | Body | Yes |
| Control-Sample23_Beta | chr8:139613764-139614020 | COL22A1 | | Body | Yes |
| Control-Sample30_Beta | chr8:139613712-139613769 | COL22A1 | | Body | Yes |
| Control-Sample105_Be | chr5:177876830-177876930 | COL23A1 | | Body | Yes |
| Control-Sample17_Beta | chr5:177707036-177707226 | COL23A1 | | Body | Yes |
| Control-Sample1_Beta | chr13:110993474-110993558 | COL4A2 | | Body | Yes |
| Control-Sample1_Beta | chr13:111128054-111128102 | COL4A2 | | Body | Yes |
| Control-Sample24_Beta | chr13:111077142-111077257 | COL4A2 | | Body | Yes |
| Control-Sample31_Beta | chr3:48632484-48632582 | COL7A1 | Island | Body | Yes |
| Control-Sample54_Beta | chr3:48632582-48632672 | COL7A1 | Island | 1stExon | Yes |
| Control-Sample22_Beta | chr1:40769669-40769744 | COL9A2 | Island | Body | No |
| Control-Sample1_Beta | chr20:61446962-61447038 | COL9A3 | N_Shore | TSS1500 | Yes |
| Control-Sample1_Beta | chr20:61447394-61447614 | COL9A3 | N_Shore | TSS1500 | Yes |
| Control-Sample13_Beta | chr20:61446962-61447053 | COL9A3 | N_Shore | TSS1500 | Yes |
| Control-Sample13_Beta | chr20:61447394-61447661 | COL9A3 | N_Shore | TSS1500 | Yes |
| Control-Sample14_Beta | chr20:61447394-61447623 | COL9A3 | N_Shore | TSS1500 | Yes |
| Control-Sample17_Beta | chr20:61447442-61447530 | COL9A3 | N_Shore | TSS1500 | Yes |
| Control-Sample19_Beta | chr20:61447174-61447272 | COL9A3 | N_Shore | TSS1500 | Yes |
| Control-Sample19_Beta | chr20:61447330-61447790 | COL9A3 | N_Shore | TSS1500 | Yes |
| Control-Sample20_Beta | chr20:61447174-61447268 | COL9A3 | N_Shore | TSS1500 | Yes |
| Control-Sample20_Beta | chr20:61447276-61447661 | COL9A3 | N_Shore | TSS1500 | Yes |
| Control-Sample21_Beta | chr20:61447036-61447053 | COL9A3 | N_Shore | TSS1500 | Yes |
| Control-Sample22_Beta | chr20:61447394-61447581 | COL9A3 | N_Shore | TSS1500 | Yes |
| Control-Sample25_Beta | chr20:61447530-61447614 | COL9A3 | N_Shore | TSS1500 | Yes |
| Control-Sample37_Beta | chr20:61447490-61447581 | COL9A3 | N_Shore | TSS1500 | Yes |
| Control-Sample38_Beta | chr20:61447442-61447581 | COL9A3 | N_Shore | TSS1500 | Yes |
| Control-Sample49_Beta | chr20:61447490-61447614 | COL9A3 | N_Shore | TSS1500 | Yes |
| Control-Sample57_Beta | chr20:61447394-61447530 | COL9A3 | N_Shore | TSS1500 | Yes |
| Control-Sample6_Beta | chr20:61447394-61447490 | COL9A3 | N_Shore | TSS1500 | Yes |
| Control-Sample20_Beta | chr2:3642211-3642447 | COLEC11;COLEC11 | N_Shore | TSS1500;TSS1500 | No |
| Control-Sample237_Be | chr10:22605173-22605182 | COMMD3 | Island | TSS200 | No |
| Control-Sample13_Beta | chr19:18902081-18902117 | COMP;COMP | Island | 1stExon;5'UTR | No |
| Control-Sample18_Beta | chr22:19928667-19929066 | COMT;TXNRD2 | N_Shore | TSS1500;Body | Yes |
| Control-Sample27_Beta | chr22:19928667-19929467 | COMT;TXNRD2 | N_Shore | TSS1500;Body | Yes |
| Control-Sample28_Beta | chr22:19928740-19929467 | COMT;TXNRD2 | N_Shore | TSS1500;Body | Yes |
| Control-Sample42_Beta | chr22:19928740-19929286 | COMT;TXNRD2 | N_Shore | TSS1500;Body | Yes |
| Control-Sample54_Beta | chr22:19928667-19929097 | COMT;TXNRD2 | N_Shore | TSS1500;Body | Yes |
| Control-Sample19_Beta | chr16:84600051-84600468 | COTL1 | | 3'UTR | No |
| Control-Sample15_Beta | chr7:1004748-1005089 | COX19 | N_Shelf | 3'UTR | No |
| Control-Sample28_Beta | chr16:31439106-31439393 | COX6A2 | Island | Body | No |
| Control-Sample19_Beta | chr4:166301076-166301256 | CPE | S_Shore | Body | No |
| Control-Sample11_Beta | chr4:778827-779480 | CPLX1 | Island | 3'UTR | Yes |
| Control-Sample15_Beta | chr4:778924-779480 | CPLX1 | Island | 3'UTR | Yes |
| Control-Sample171_Be | chr4:802844-803014 | CPLX1 | S_Shelf | Body | Yes |
| Control-Sample26_Beta | chr4:779173-779480 | CPLX1 | Island | 3'UTR | Yes |
| Control-Sample7_Beta | chr4:778827-779173 | CPLX1 | Island | 3'UTR | Yes |
| Control-Sample10_Beta | chr16:89641054-89641360 | CPNE7;CPNE7 | Island | TSS1500;TSS1500 | No |
| Control-Sample19_Beta | chr2:211457632-211458004 | CPS1;CPS1;CPS1 | | Body;Body;TSS1500 | Yes |
| Control-Sample23_Beta | chr2:211457684-211458173 | CPS1;CPS1;CPS1 | | Body;Body;TSS1500 | Yes |
| Control-Sample1_Beta | chr17:71258045-71258215 | CPSF4L | | TSS200 | Yes |
| Control-Sample21_Beta | chr17:71258101-71258589 | CPSF4L | | TSS200 | Yes |
| Control-Sample24_Beta | chr17:71258045-71258589 | CPSF4L | | TSS200 | Yes |

| | | | | | |
|---|---|---|---|---|---|
| Control-Sample56_Beta | chr17:71258045-71258936 | CPSF4L | | TSS200 | Yes |
| Control-Sample45_Beta | chr22:51016501-51016646 | CPT1B;CPT1B;CHKB-CPT1B;CPT1B;CPT1B;CPT1B;CPT1B; | Island | 5'UTR;5'UTR;Body;5'UTR;5'UTR;5'UTR;1stExon;5'UTR | Yes |
| Control-Sample100_Be | chr22:51016638-51016646 | CPT1B;CPT1B;CPT1B;CHKB-CPT1B;CPT1B;CPT1B;CPT1B; | Island | 5'UTR;5'UTR;TSS200;Body;5'UTR;5'UTR;5'UTR;5'UTR | Yes |
| Control-Sample1_Beta | chr10:125651549-125652323 | CPXM2 | S_Shore | TSS200 | Yes |
| Control-Sample22_Beta | chr10:125651549-125651621 | CPXM2 | S_Shore | TSS200 | Yes |
| Control-Sample3_Beta | chr10:125651549-125651726 | CPXM2 | S_Shore | TSS200 | Yes |
| Control-Sample65_Beta | chr10:125651549-125651697 | CPXM2 | S_Shore | TSS200 | Yes |
| Control-Sample42_Beta | chr4:8594413-8594549 | CPZ;CPZ;CPZ | Island | TSS200;TSS200;TSS200 | No |
| Control-Sample170_Be | chr1:207842833-207842927 | CR1L | Island | Body | Yes |
| Control-Sample181_Be | chr1:207842833-207843084 | CR1L | Island | Body | Yes |
| Control-Sample182_Be | chr1:207842917-207843084 | CR1L | Island | Body | Yes |
| Control-Sample10_Beta | chr9:126135169-126135408 | CRB2 | N_Shore | Body | No |
| Control-Sample104_Be | chr19:4153603-4153677 | CREB3L3 | | TSS200 | Yes |
| Control-Sample105_Be | chr19:4153445-4153713 | CREB3L3 | | TSS200 | Yes |
| Control-Sample372_Be | chr19:4153419-4153677 | CREB3L3 | | TSS1500 | Yes |
| Control-Sample379_Be | chr19:4153419-4153713 | CREB3L3 | | TSS1500 | Yes |
| Control-Sample558_Be | chr19.0:4153419-4153677 | CREB3L3 | | TSS1500 | Yes |
| Control-Sample108_Be | chr7:28338625-28338790 | CREB5 | | TSS1500 | Yes |
| Control-Sample108_Be | chr7:28452066-28452131 | CREB5;CREB5 | S_Shelf | 5'UTR;TSS200 | Yes |
| Control-Sample3_Beta | chr7:28452066-28452098 | CREB5;CREB5 | S_Shelf | 5'UTR;TSS200 | Yes |
| Control-Sample11_Beta | chr7:28474589-28474889 | CREB5;CREB5;CREB5 | | TSS1500;5'UTR;Body | Yes |
| Control-Sample19_Beta | chr7:28725358-28725676 | CREB5;CREB5;CREB5;CREB5 | | Body;Body;Body;TSS1500 | Yes |
| Control-Sample5_Beta | chr14:105944604-105944942 | CRIP2 | Island | Body | No |
| Control-Sample19_Beta | chr16:84860766-84860918 | CRISPLD2 | | 5'UTR | No |
| Control-Sample217_Be | chr1:16810147-16810677 | CROCCL2 | N_Shore | Body | Yes |
| Control-Sample389_Be | chr1:16810670-16810776 | CROCCL2 | Island | Body | Yes |
| Control-Sample561_Be | chr1.0:16810147-16810776 | CROCCL2 | N_Shore | Body | Yes |
| Control-Sample35_Beta | chr10:99734805-99735133 | CRTAC1 | | Body | No |
| Control-Sample10_Beta | chr19:18873037-18873268 | CRTC1;CRTC1 | Island | Body;Body | Yes |
| Control-Sample17_Beta | chr19:18879431-18879450 | CRTC1;CRTC1 | | Body;Body | Yes |
| Control-Sample492_Be | chr11:111783267-111783563 | CRYAB;HSPB2 | | TSS1500;TSS200 | No |
| Control-Sample1_Beta | chr22:27017854-27017910 | CRYBA4 | | TSS200 | Yes |
| Control-Sample62_Beta | chr22:27017201-27017910 | CRYBA4 | | TSS1500 | Yes |
| Control-Sample58_Beta | chr22:27014006-27014415 | CRYBB1 | | TSS200 | No |
| Control-Sample4_Beta | chr22:25615317-25615419 | CRYBB2 | | TSS1500 | Yes |
| Control-Sample71_Beta | chr22:25615317-25615355 | CRYBB2 | | TSS1500 | Yes |
| Control-Sample10_Beta | chr2:209029594-209029782 | CRYGA | | TSS1500 | No |
| Control-Sample19_Beta | chr2:209010890-209011053 | CRYGB | | TSS200 | Yes |
| Control-Sample20_Beta | chr2:209010986-209011053 | CRYGB | | TSS200 | Yes |
| Control-Sample1_Beta | chr2:208994529-208994720 | CRYGC;CRYGC | | 1stExon;5'UTR | Yes |
| Control-Sample58_Beta | chr2:208994529-208995254 | CRYGC;CRYGC | | 1stExon;5'UTR | Yes |
| Control-Sample1_Beta | chr13:21009861-21010223 | CRYL1 | Island | Body | Yes |
| Control-Sample23_Beta | chr13:20989142-20989349 | CRYL1 | Island | Body | Yes |
| Control-Sample379_Be | chr1:75199100-75199496 | CRYZ;TYW3;CRYZ;CRYZ;CRYZ;TYW3;TYW3 | | TSS200;1stExon;TSS200;TSS200;TSS200;1stExon;Bo | No |
| Control-Sample4_Beta | chr8:19539991-19540272 | CSGALNACT1;CSGALNACT1;CSGALNACT1 | | Body;5'UTR;1stExon | No |
| Control-Sample19_Beta | chr1:34175169-34175343 | CSMD2 | | Body | No |
| Control-Sample19_Beta | chr15:37111929-37112149 | CSNK1A1P | S_Shore | TSS1500 | No |
| Control-Sample1_Beta | chr15:71408603-71408914 | CT62 | S_Shore | TSS1500 | No |
| Control-Sample1_Beta | chr14:39734964-39735625 | CTAGE5;CTAGE5;CTAGE5;CTAGE5 | N_Shore | TSS1500;TSS1500;TSS1500;Body | No |
| Control-Sample1_Beta | chr4:1205914-1206150 | CTBP1;CTBP1 | Island | 3'UTR;3'UTR | Yes |
| Control-Sample10_Beta | chr4:1206004-1206150 | CTBP1;CTBP1 | Island | 3'UTR;3'UTR | Yes |
| Control-Sample19_Beta | chr4:1205914-1206045 | CTBP1;CTBP1 | Island | 3'UTR;3'UTR | Yes |
| Control-Sample17_Beta | chr3:38009911-38010391 | CTDSPL;CTDSPL;MIR26A1 | | Body;Body;TSS1500 | Yes |
| Control-Sample57_Beta | chr3:38010391-38010902 | CTDSPL;CTDSPL;MIR26A1 | | Body;Body;TSS1500 | Yes |
| Control-Sample7_Beta | chr16:30907246-30907679 | CTF1;CTF1 | S_Shore | TSS1500;TSS1500 | No |
| Control-Sample506_Be | chr8.0:104383715-104383798 | CTHRC1 | Island | TSS200 | No |
| Control-Sample291_Be | chr5:138211084-138211184 | CTNNA1;LRRTM2 | | Body;TSS200 | Yes |
| Control-Sample67_Beta | chr5:138211149-138211184 | CTNNA1;LRRTM2 | | Body;TSS200 | Yes |
| Control-Sample45_Beta | chr2:79738582-79738639 | CTNNA2;CTNNA2 | N_Shore | TSS1500;TSS1500 | No |
| Control-Sample36_Beta | chr5:11384581-11384753 | CTNND2 | N_Shore | Body | No |
| Control-Sample171_Be | chr16:75252619-75252744 | CTRB1 | | TSS1500 | No |
| Control-Sample20_Beta | chr1:15763737-15763967 | CTRC | | TSS1500 | No |
| Control-Sample24_Beta | chr11:66335649-66336125 | CTSF | Island | Body | No |
| Control-Sample1_Beta | chr11:70256556-70256681 | CTTN;CTTN | N_Shore | Body;Body | No |
| Control-Sample3_Beta | chr5:126984564-126984994 | CTXN3 | | TSS200 | No |
| Control-Sample108_Be | chr10:11047271-11047354 | CUGBP2;CUGBP2 | | 1stExon;5'UTR | Yes |
| Control-Sample13_Beta | chr10:11047284-11047368 | CUGBP2;CUGBP2 | | 1stExon;5'UTR | Yes |
| Control-Sample14_Beta | chr10:11047271-11047368 | CUGBP2;CUGBP2 | | 1stExon;5'UTR | Yes |
| Control-Sample58_Beta | chr10:11047271-11047586 | CUGBP2;CUGBP2 | | 1stExon;5'UTR | Yes |
| Control-Sample19_Beta | chr10:11317787-11317872 | CUGBP2;CUGBP2;CUGBP2;CUGBP2 | | Body;Body;Body;Body | Yes |
| Control-Sample13_Beta | chr6:43022044-43022182 | CUL7;CUL7;MRPL2 | S_Shore | TSS1500;TSS1500;Body | Yes |
| Control-Sample1_Beta | chr7:101723207-101723836 | CUX1;CUX1;CUX1 | | Body;Body;Body | Yes |
| Control-Sample12_Beta | chr7:101813842-101814050 | CUX1;CUX1;CUX1 | | Body;Body;Body | Yes |
| Control-Sample14_Beta | chr7:101603298-101603359 | CUX1;CUX1;CUX1 | | Body;Body;Body | Yes |
| Control-Sample17_Beta | chr7:101813842-101814109 | CUX1;CUX1;CUX1 | | Body;Body;Body | Yes |
| Control-Sample171_Be | chr7:101586761-101586820 | CUX1;CUX1;CUX1 | | Body;Body;Body | Yes |
| Control-Sample21_Beta | chr7:101468882-101469146 | CUX1;CUX1;CUX1 | | Body;Body;Body | Yes |
| Control-Sample30_Beta | chr7:101846993-101847657 | CUX1;CUX1;CUX1 | | Body;Body;Body | Yes |
| Control-Sample4_Beta | chr7:101723207-101723568 | CUX1;CUX1;CUX1 | | Body;Body;Body | Yes |
| Control-Sample44_Beta | chr7:101846993-101847575 | CUX1;CUX1;CUX1 | | Body;Body;Body | Yes |
| Control-Sample58_Beta | chr7:101558400-101558597 | CUX1;CUX1;CUX1 | N_Shore | Body;Body;Body | Yes |
| Control-Sample1_Beta | chr12:111618925-111619455 | CUX2 | | Body | Yes |
| Control-Sample1_Beta | chr12:111725167-111725516 | CUX2 | | Body | Yes |
| Control-Sample10_Beta | chr12:111589389-111589453 | CUX2 | | Body | Yes |
| Control-Sample10_Beta | chr12:111618925-111618977 | CUX2 | | Body | Yes |
| Control-Sample10_Beta | chr12:111619190-111619455 | CUX2 | | Body | Yes |
| Control-Sample11_Beta | chr12:111618925-111619190 | CUX2 | | Body | Yes |
| Control-Sample12_Beta | chr12:111619077-111619455 | CUX2 | | Body | Yes |
| Control-Sample20_Beta | chr12:111619406-111619455 | CUX2 | | Body | Yes |
| Control-Sample1_Beta | chr10:44881551-44882135 | CXCL12;CXCL12;CXCL12 | Island | TSS1500;TSS1500;TSS1500 | Yes |
| Control-Sample11_Beta | chr10:44881551-44881934 | CXCL12;CXCL12;CXCL12 | Island | TSS1500;TSS1500;TSS1500 | Yes |

| | | | | | |
|---|---|---|---|---|---|
| Control-Sample32_Beta | chr4:74864165-74864313 | CXCL5 | Island | Body | No |
| Control-Sample464_Be | chr5:139060556-139060922 | CXXC5 | Island | Body | No |
| Control-Sample571_Be | chr17.0:61514920-61515312 | CYB561;CYB561;CYB561 | Island | 5'UTR;5'UTR;5'UTR | Yes |
| Control-Sample92_Beta | chr17:61514920-61515312 | CYB561;CYB561;CYB561 | Island | 5'UTR;5'UTR;5'UTR | Yes |
| Control-Sample1_Beta | chr16:88717587-88717646 | CYBA | Island | TSS200 | Yes |
| Control-Sample10_Beta | chr16:88717134-88717456 | CYBA | Island | Body | Yes |
| Control-Sample10_Beta | chr16:88717537-88717646 | CYBA | Island | TSS200 | Yes |
| Control-Sample12_Beta | chr16:88717134-88717464 | CYBA | Island | Body | Yes |
| Control-Sample13_Beta | chr16:88717374-88717464 | CYBA | Island | 1stExon | Yes |
| Control-Sample270_Be | chr16:88717134-88717482 | CYBA | Island | Body | Yes |
| Control-Sample59_Beta | chr16:88717374-88717472 | CYBA | Island | 1stExon | Yes |
| Control-Sample216_Be | chr16:88717456-88717472 | CYBA;CYBA | Island | 1stExon;5'UTR | Yes |
| Control-Sample1_Beta | chr5:156772562-156772761 | CYFIP2;C5orf40;CYFIP2;CYFIP2;C5orf40 | | Body;1stExon;Body;Body;5'UTR | Yes |
| Control-Sample3_Beta | chr5:156695161-156695551 | CYFIP2;CYFIP2;CYFIP2 | S_Shore | 5'UTR;TSS1500;5'UTR | Yes |
| Control-Sample49_Beta | chr15:74657911-74658244 | CYP11A1;CYP11A1 | N_Shore | Body;5'UTR | No |
| Control-Sample1_Beta | chr2:72371255-72371552 | CYP26B1 | Island | Body | Yes |
| Control-Sample464_Be | chr2:72359383-72359612 | CYP26B1 | | Body | Yes |
| Control-Sample13_Beta | chr10:94819559-94820927 | CYP26C1 | N_Shore | 3'UTR;3'UTR | Yes |
| Control-Sample14_Beta | chr10:94819989-94820927 | CYP26C1 | N_Shore | TSS1500 | Yes |
| Control-Sample20_Beta | chr10:94820892-94820927 | CYP26C1 | Island | TSS1500 | Yes |
| Control-Sample4_Beta | chr10:94819802-94820927 | CYP26C1 | N_Shore | TSS1500 | Yes |
| Control-Sample5_Beta | chr10:94819989-94821773 | CYP26C1 | N_Shore | TSS1500 | Yes |
| Control-Sample58_Beta | chr10:94819802-94820945 | CYP26C1 | N_Shore | TSS1500 | Yes |
| Control-Sample65_Beta | chr10:94819989-94820945 | CYP26C1 | N_Shore | TSS1500 | Yes |
| Control-Sample1_Beta | chr10:135342218-135342560 | CYP2E1 | Island | Body | Yes |
| Control-Sample120_Be | chr10:135341870-135342560 | CYP2E1 | Island | Body | Yes |
| Control-Sample141_Be | chr10:135341870-135343280 | CYP2E1 | Island | Body | Yes |
| Control-Sample286_Be | chr10:135343193-135343280 | CYP2E1 | S_Shore | Body | Yes |
| Control-Sample408_Be | chr10:135342413-135342620 | CYP2E1 | Island | Body | Yes |
| Control-Sample425_Be | chr10:135341870-135342413 | CYP2E1 | Island | Body | Yes |
| Control-Sample487_Be | chr10:135341870-135342620 | CYP2E1 | Island | Body | Yes |
| Control-Sample561_Be | chr10.0:135341870-135342413 | CYP2E1 | Island | Body | Yes |
| Control-Sample194_Be | chr19:41633902-41634046 | CYP2F1 | Island | Body | No |
| Control-Sample11_Beta | chr7:1022471-1022675 | CYP2W1 | N_Shore | TSS1500 | Yes |
| Control-Sample23_Beta | chr7:1022213-1022675 | CYP2W1 | N_Shore | TSS1500 | Yes |
| Control-Sample19_Beta | chr8:65711644-65711916 | CYP7B1 | Island | TSS1500 | No |
| Control-Sample10_Beta | chr3:42918546-42919025 | CYP8B1 | N_Shelf | TSS1500 | Yes |
| Control-Sample40_Beta | chr3:42918546-42918806 | CYP8B1 | N_Shelf | TSS1500 | Yes |
| Control-Sample13_Beta | chr1:86047660-86047911 | CYR61 | S_Shore | Body | No |
| Control-Sample67_Beta | chr17:76719635-76719943 | CYTH1;CYTH1 | | Body;Body | No |
| Control-Sample201_Be | chr19:48983942-48984318 | CYTH2;CYTH2 | Island | 3'UTR;3'UTR | No |
| Control-Sample100_Be | chr4:5021113-5021236 | CYTL1 | Island | 1stExon | No |
| Control-Sample22_Beta | chr4:4386726-4386906 | D4S234E;D4S234E | N_Shore | TSS1500;TSS1500 | No |
| Control-Sample223_Be | chr6:39760776-39761304 | DAAM2 | S_Shore | TSS200 | No |
| Control-Sample1_Beta | chr9:124535348-124535611 | DAB2IP;DAB2IP | Island | Body;Body | No |
| Control-Sample3_Beta | chr11:61449737-61449898 | DAGLA | S_Shore | 5'UTR | No |
| Control-Sample108_Be | chr2:159651559-159651918 | DAPL1 | | TSS1500 | No |
| Control-Sample103_Be | chr20:44036899-44037436 | DBNDD2;DBNDD2;DBNDD2;SYS1-DBNDD2;DBNDD2;DBN | S_Shore | 5'UTR;5'UTR;TSS200;Body;5'UTR;5'UTR;TSS200 | Yes |
| Control-Sample68_Beta | chr20:44036723-44037519 | DBNDD2;DBNDD2;DBNDD2;SYS1-DBNDD2;DBNDD2;DBN | S_Shore | 5'UTR;5'UTR;TSS200;Body;5'UTR;5'UTR;TSS200 | Yes |
| Control-Sample74_Beta | chr20:44036723-44037436 | DBNDD2;DBNDD2;DBNDD2;SYS1-DBNDD2;DBNDD2;DBN | S_Shore | 5'UTR;5'UTR;TSS200;Body;5'UTR;5'UTR;TSS200 | Yes |
| Control-Sample201_Be | chr3:137893707-137893743 | DBR1;DBR1 | Island | 1stExon;5'UTR | Yes |
| Control-Sample524_Be | chr3.0:137893707-137893743 | DBR1;DBR1 | Island | 1stExon;5'UTR | Yes |
| Control-Sample1_Beta | chr6:117802588-117802725 | DCBLD1 | N_Shore | TSS1500 | Yes |
| Control-Sample2_Beta | chr6:117802640-117802725 | DCBLD1 | N_Shore | TSS1500 | Yes |
| Control-Sample66_Beta | chr6:117869098-117869453 | DCBLD1 | Island | Body | Yes |
| Control-Sample23_Beta | chr11:6677919-6678143 | DCHS1 | S_Shore | TSS1500 | No |
| Control-Sample108_Be | chr13:95131809-95132211 | DCT;DCT;DCT;DCT | | 1stExon;5'UTR;1stExon;5'UTR | No |
| Control-Sample67_Beta | chr13:114138019-114138260 | DCUN1D2 | | Body | No |
| Control-Sample30_Beta | chr1:86044103-86044230 | DDAH1 | S_Shore | TSS200 | No |
| Control-Sample1_Beta | chr6:110736422-110736865 | DDO;DDO | | Body;Body | Yes |
| Control-Sample108_Be | chr6:110736772-110737053 | DDO;DDO | | TSS200;TSS200 | Yes |
| Control-Sample19_Beta | chr6:110736422-110736941 | DDO;DDO | | Body;Body | Yes |
| Control-Sample270_Be | chr6:110736941-110737053 | DDO;DDO | | TSS200;TSS200 | Yes |
| Control-Sample1_Beta | chr6:30850868-30851517 | DDR1;DDR1 | N_Shore | TSS1500;TSS1500 | Yes |
| Control-Sample1_Beta | chr6:30851624-30851753 | DDR1;DDR1 | N_Shore | TSS1500;TSS1500 | Yes |
| Control-Sample10_Beta | chr6:30850977-30851517 | DDR1;DDR1 | N_Shore | TSS1500;TSS1500 | Yes |
| Control-Sample10_Beta | chr6:30851624-30851638 | DDR1;DDR1 | N_Shore | TSS1500;TSS1500 | Yes |
| Control-Sample11_Beta | chr6:30850829-30850876 | DDR1;DDR1 | N_Shore | TSS1500;TSS1500 | Yes |
| Control-Sample15_Beta | chr6:30850829-30850913 | DDR1;DDR1 | N_Shore | TSS1500;TSS1500 | Yes |
| Control-Sample2_Beta | chr6:30850829-30851517 | DDR1;DDR1 | N_Shore | TSS1500;TSS1500 | Yes |
| Control-Sample20_Beta | chr6:30850868-30850913 | DDR1;DDR1 | N_Shore | TSS1500;TSS1500 | Yes |
| Control-Sample20_Beta | chr6:30851051-30851517 | DDR1;DDR1 | N_Shore | TSS1500;TSS1500 | Yes |
| Control-Sample21_Beta | chr6:30851051-30851086 | DDR1;DDR1 | N_Shore | TSS1500;TSS1500 | Yes |
| Control-Sample33_Beta | chr6:30850829-30851069 | DDR1;DDR1 | N_Shore | TSS1500;TSS1500 | Yes |
| Control-Sample34_Beta | chr6:30850977-30851069 | DDR1;DDR1 | N_Shore | TSS1500;TSS1500 | Yes |
| Control-Sample38_Beta | chr6:30850829-30851086 | DDR1;DDR1 | N_Shore | TSS1500;TSS1500 | Yes |
| Control-Sample52_Beta | chr6:30850977-30851086 | DDR1;DDR1 | N_Shore | TSS1500;TSS1500 | Yes |
| Control-Sample61_Beta | chr6:30850868-30851069 | DDR1;DDR1 | N_Shore | TSS1500;TSS1500 | Yes |
| Control-Sample1_Beta | chr6:30855729-30856439 | DDR1;DDR1;DDR1 | S_Shelf | TSS1500;5'UTR;5'UTR | Yes |
| Control-Sample18_Beta | chr6:30856121-30856439 | DDR1;DDR1;DDR1 | S_Shelf | TSS1500;5'UTR;5'UTR | Yes |
| Control-Sample20_Beta | chr1:162602217-162602559 | DDR2;DDR2 | | TSS200;TSS200 | No |
| Control-Sample1_Beta | chr12:113599272-113599683 | DDX54;DDX54 | | Body;Body | No |
| Control-Sample276_Be | chr12:124086656-124086667 | DDX55 | Island | TSS200 | No |
| Control-Sample464_Be | chr11:674556-674687 | DEAF1 | | Body | No |
| Control-Sample10_Beta | chr6:35265534-35265555 | DEF6 | | TSS200 | No |
| Control-Sample15_Beta | chr16:90015832-90016551 | DEF8;DEF8 | Island | 5'UTR;5'UTR | Yes |
| Control-Sample270_Be | chr16:90016004-90016061 | DEF8;DEF8 | Island | 5'UTR;Body | Yes |
| Control-Sample108_Be | chr19:6482011-6482169 | DENND1C | | TSS1500 | Yes |
| Control-Sample11_Beta | chr19:6476756-6477033 | DENND1C | Island | Body | Yes |
| Control-Sample1_Beta | chr8:142179603-142179697 | DENND3 | N_Shelf | Body | Yes |
| Control-Sample585_Be | chr8.0:142179603-142179697 | DENND3 | N_Shelf | Body | Yes |

| | | | | | |
|---|---|---|---|---|---|
| Control-Sample10_Beta | chr22:32150725-32150934 | DEPDC5;DEPDC5;DEPDC5 | S_Shore | 5'UTR;TSS200;5'UTR | No |
| Control-Sample90_Beta | chr17:5389592-5389625 | DERL2;MIS12 | Island | TSS200;TSS1500 | No |
| Control-Sample15_Beta | chr2:220282914-220283156 | DES | N_Shore | TSS200 | Yes |
| Control-Sample25_Beta | chr2:220282914-220282977 | DES | N_Shore | TSS200 | Yes |
| Control-Sample19_Beta | chr2:234296280-234296642 | DGKD;DGKD | | Body;TSS1500 | No |
| Control-Sample1_Beta | chr3:185911316-185912486 | DGKG;DGKG;DGKG | N_Shore | Body;Body;Body | Yes |
| Control-Sample13_Beta | chr3:185911316-185912253 | DGKG;DGKG;DGKG | N_Shore | Body;Body;Body | Yes |
| Control-Sample24_Beta | chr3:185911702-185912486 | DGKG;DGKG;DGKG | Island | Body;Body;Body | Yes |
| Control-Sample69_Beta | chr3:185911316-185911702 | DGKG;DGKG;DGKG | N_Shore | Body;Body;Body | Yes |
| Control-Sample236_Be | chr4:967528-967541 | DGKQ | Island | TSS200 | No |
| Control-Sample13_Beta | chr11:46382544-46382725 | DGKZ;DGKZ;DGKZ;DGKZ | | Body;TSS1500;Body;Body | No |
| Control-Sample1_Beta | chr12:49484932-49485151 | DHH | Island | Body | No |
| Control-Sample464_Be | chr14:24422916-24422956 | DHRS4;C14orf167;C14orf167;C14orf167;C14orf167 | Island | TSS200;Body;Body;Body;Body | Yes |
| Control-Sample637_Be | chr14:24422749-24422928 | DHRS4;C14orf167;C14orf167;C14orf167;C14orf167 | N_Shore | TSS1500;Body;Body;Body;Body | Yes |
| Control-Sample1_Beta | chr14:24457893-24458000 | DHRS4L2 | Island | TSS200 | Yes |
| Control-Sample19_Beta | chr14:24457893-24457993 | DHRS4L2 | Island | TSS200 | Yes |
| Control-Sample54_Beta | chr14:24457893-24458014 | DHRS4L2 | Island | TSS200 | Yes |
| Control-Sample23_Beta | chr17:9694619-9694800 | DHRS7C | | TSS200 | Yes |
| Control-Sample32_Beta | chr17:9694739-9694800 | DHRS7C | | TSS200 | Yes |
| Control-Sample1_Beta | chr10:127569751-127570400 | DHX32;DHX32 | | 5'UTR;1stExon | Yes |
| Control-Sample15_Beta | chr10:127569751-127569973 | DHX32;DHX32 | | 5'UTR;1stExon | Yes |
| Control-Sample1_Beta | chr17:40259547-40260053 | DHX58 | Island | Body | Yes |
| Control-Sample6_Beta | chr17:40259724-40260053 | DHX58 | Island | Body | Yes |
| Control-Sample1_Beta | chr14:80677608-80677688 | DIO2;DIO2;DIO2 | | 1stExon;1stExon;Body | No |
| Control-Sample1_Beta | chr10:405182-405362 | DIP2C | | Body | Yes |
| Control-Sample1_Beta | chr10:531465-531569 | DIP2C | S_Shore | Body | Yes |
| Control-Sample1_Beta | chr10:676529-676761 | DIP2C | N_Shore | Body | Yes |
| Control-Sample1_Beta | chr10:679171-679514 | DIP2C | Island | Body | Yes |
| Control-Sample1_Beta | chr10:729711-729956 | DIP2C | | Body | Yes |
| Control-Sample10_Beta | chr10:555957-556044 | DIP2C | | Body | Yes |
| Control-Sample11_Beta | chr10:679171-679399 | DIP2C | Island | Body | Yes |
| Control-Sample14_Beta | chr10:669071-669285 | DIP2C | Island | Body | Yes |
| Control-Sample19_Beta | chr10:415359-415489 | DIP2C | S_Shore | Body | Yes |
| Control-Sample19_Beta | chr10:429943-429999 | DIP2C | | Body | Yes |
| Control-Sample19_Beta | chr10:729479-729956 | DIP2C | | Body | Yes |
| Control-Sample20_Beta | chr10:729259-729956 | DIP2C | | Body | Yes |
| Control-Sample21_Beta | chr10:407721-407935 | DIP2C | N_Shore | Body | Yes |
| Control-Sample21_Beta | chr10:679221-679399 | DIP2C | Island | Body | Yes |
| Control-Sample21_Beta | chr10:729846-729956 | DIP2C | | Body | Yes |
| Control-Sample25_Beta | chr10:468412-468755 | DIP2C | S_Shelf | Body | Yes |
| Control-Sample26_Beta | chr10:529214-529521 | DIP2C | N_Shore | Body | Yes |
| Control-Sample33_Beta | chr10:729001-729570 | DIP2C | | Body | Yes |
| Control-Sample331_Be | chr10:679221-679514 | DIP2C | Island | Body | Yes |
| Control-Sample39_Beta | chr10:729204-729259 | DIP2C | | Body | Yes |
| Control-Sample5_Beta | chr10:468051-468347 | DIP2C | S_Shore | Body | Yes |
| Control-Sample53_Beta | chr10:467839-468347 | DIP2C | S_Shore | Body | Yes |
| Control-Sample64_Beta | chr10:467839-468412 | DIP2C | S_Shore | Body | Yes |
| Control-Sample30_Beta | chr10:709564-709610 | DIP2C;C10orf108 | N_Shelf | Body;Body | Yes |
| Control-Sample1_Beta | chr10:695844-695877 | DIP2C;C10orf108;C10orf108 | | Body;TSS200;TSS200 | Yes |
| Control-Sample19_Beta | chr10:695853-695900 | DIP2C;C10orf108;C10orf108 | | Body;TSS200;TSS200 | Yes |
| Control-Sample20_Beta | chr10:695844-695900 | DIP2C;C10orf108;C10orf108 | | Body;TSS200;TSS200 | Yes |
| Control-Sample4_Beta | chr15:66584218-66584493 | DIS3L | N_Shore | TSS1500 | No |
| Control-Sample171_Be | chr11:61525251-61525922 | DKFZP434K028;C11orf9;C11orf9 | | TSS200;Body;Body | No |
| Control-Sample13_Beta | chr19:31640642-31640711 | DKFZp566F0947 | | TSS200 | Yes |
| Control-Sample20_Beta | chr19:31640706-31640849 | DKFZp566F0947 | | TSS200 | Yes |
| Control-Sample26_Beta | chr19:31640401-31640711 | DKFZp566F0947 | | TSS1500 | Yes |
| Control-Sample18_Beta | chr8:42234798-42234975 | DKK4 | | TSS200 | Yes |
| Control-Sample1_Beta | chr8:42234614-42234975 | DKK4;DKK4 | | 5'UTR;1stExon | Yes |
| Control-Sample18_Beta | chr8:42234614-42234724 | DKK4;DKK4 | | 5'UTR;1stExon | Yes |
| Control-Sample40_Beta | chr8:42234614-42234803 | DKK4;DKK4 | | 5'UTR;1stExon | Yes |
| Control-Sample13_Beta | chr8:12958083-12958533 | DLC1;DLC1;DLC1 | | Body;Body;Body | No |
| Control-Sample1_Beta | chr3:38080635-38080704 | DLEC1;DLEC1 | Island | TSS200;TSS200 | No |
| Control-Sample55_Beta | chr11:84634615-84634735 | DLG2;DLG2 | | TSS200;Body | Yes |
| Control-Sample19_Beta | chr11:83393344-83393677 | DLG2;DLG2;DLG2;DLG2;DLG2 | | 1stExon;5'UTR;Body;Body;Body | Yes |
| Control-Sample11_Beta | chr8:1624741-1624822 | DLGAP2 | | Body | Yes |
| Control-Sample19_Beta | chr8:1449313-1449497 | DLGAP2 | | TSS1500 | Yes |
| Control-Sample634_Be | chr8:1626419-1626697 | DLGAP2 | | Body | Yes |
| Control-Sample65_Beta | chr8:1653179-1653566 | DLGAP2 | S_Shelf | 3'UTR | Yes |
| Control-Sample15_Beta | chr1:35351340-35351627 | DLGAP3 | Island | Body | No |
| Control-Sample3_Beta | chr20:34995350-34995517 | DLGAP4 | | TSS200 | No |
| Control-Sample15_Beta | chr14:101198397-101198681 | DLK1 | N_Shelf | Body | Yes |
| Control-Sample58_Beta | chr14:101198475-101198783 | DLK1 | N_Shelf | Body | Yes |
| Control-Sample19_Beta | chr2:172965837-172966180 | DLX2 | Island | Body | No |
| Control-Sample11_Beta | chr7:96655044-96655444 | DLX5 | S_Shore | TSS1500 | No |
| Control-Sample1_Beta | chr19:46282571-46283080 | DMPK;DMPK;DMPK;DMPK | Island | Body;Body;Body;Body | No |
| Control-Sample10_Beta | chr1:50887633-50888024 | DMRTA2 | Island | 5'UTR | Yes |
| Control-Sample23_Beta | chr1:50884242-50884544 | DMRTA2 | Island | 3'UTR | Yes |
| Control-Sample1_Beta | chr3:52351355-52351680 | DNAH1 | | 5'UTR | No |
| Control-Sample1_Beta | chr12:124246391-124246446 | DNAH10 | N_Shore | TSS1500 | Yes |
| Control-Sample1_Beta | chr12:124246919-124247223 | DNAH10 | Island | TSS200 | Yes |
| Control-Sample464_Be | chr17:76446778-76446851 | DNAH17 | Island | Body | No |
| Control-Sample10_Beta | chr17:72306008-72306141 | DNAI2 | | Body | Yes |
| Control-Sample6_Beta | chr17:72270155-72270444 | DNAI2 | N_Shore | TSS1500 | Yes |
| Control-Sample194_Be | chr15:78556935-78556945 | DNAJA4;DNAJA4 | Island | Body;TSS200 | No |
| Control-Sample170_Be | chr7:157179789-157180005 | DNAJB6 | | Body | No |
| Control-Sample39_Beta | chr13:43597565-43598038 | DNAJC15;DNAJC15 | Island | 5'UTR;1stExon | No |
| Control-Sample10_Beta | chr15:41061410-41062017 | DNAJC17;C15orf62 | | Body;TSS1500 | Yes |
| Control-Sample16_Beta | chr15:41061384-41062017 | DNAJC17;C15orf62 | | Body;TSS1500 | Yes |
| Control-Sample25_Beta | chr15:41061384-41062224 | DNAJC17;C15orf62 | | Body;TSS1500 | Yes |
| Control-Sample3_Beta | chr15:41061384-41062113 | DNAJC17;C15orf62 | | Body;TSS1500 | Yes |
| Control-Sample55_Beta | chr15:41061384-41061527 | DNAJC17;C15orf62 | | Body;TSS1500 | Yes |

| | | | | | |
|---|---|---|---|---|---|
| Control-Sample3_Beta | chr5:140052017-140052475 | DND1 | N_Shore | Body | No |
| Control-Sample100_Be | chr11:6592297-6592745 | DNHD1 | Island | Body | Yes |
| Control-Sample19_Beta | chr11:6592066-6592745 | DNHD1 | Island | Body | Yes |
| Control-Sample54_Beta | chr11:6592066-6592585 | DNHD1 | Island | Body | Yes |
| Control-Sample1_Beta | chr11:6518287-6518331 | DNHD1;DNHD1 | | TSS1500;TSS1500 | Yes |
| Control-Sample1_Beta | chr11:6518559-6518907 | DNHD1;DNHD1;DNHD1;DNHD1 | | 5'UTR;1stExon;5'UTR;1stExon | Yes |
| Control-Sample1_Beta | chr2:25475583-25475916 | DNMT3A;DNMT3A;DNMT3A | Island | Body;TSS1500;Body | Yes |
| Control-Sample10_Beta | chr2:25526639-25526752 | DNMT3A;DNMT3A;DNMT3A | | Body;Body;Body | Yes |
| Control-Sample18_Beta | chr2:25475726-25475916 | DNMT3A;DNMT3A;DNMT3A | Island | Body;TSS1500;Body | Yes |
| Control-Sample56_Beta | chr2:25473781-25474327 | DNMT3A;DNMT3A;DNMT3A | N_Shore | Body;Body;Body | Yes |
| Control-Sample17_Beta | chr2:25475140-25475916 | DNMT3A;DNMT3A;DNMT3A;DNMT3A | Island | Body;5'UTR;Body;1stExon | Yes |
| Control-Sample19_Beta | chr2:25475140-25475583 | DNMT3A;DNMT3A;DNMT3A;DNMT3A | Island | Body;5'UTR;Body;1stExon | Yes |
| Control-Sample5_Beta | chr10:98063999-98064362 | DNTT;DNTT | | TSS200;TSS200 | No |
| Control-Sample1_Beta | chr7:111368512-111368847 | DOCK4 | Island | Body | Yes |
| Control-Sample13_Beta | chr7:111368367-111368847 | DOCK4 | Island | Body | Yes |
| Control-Sample291_Be | chr13:99630462-99630583 | DOCK9;DOCK9;DOCK9;DOCK9 | | Body;Body;TSS200;TSS200 | Yes |
| Control-Sample108_Be | chr13:99630210-99630583 | DOCK9;DOCK9;DOCK9;DOCK9;DOCK9;DOCK9 | | Body;5'UTR;1stExon;Body;5'UTR;1stExon | Yes |
| Control-Sample15_Beta | chr4:3464870-3464887 | DOK7;DOK7 | Island | TSS200;TSS200 | No |
| Control-Sample408_Be | chr19:38715064-38715198 | DPF1;DPF1;DPF1 | Island | TSS200;TSS200;5'UTR | No |
| Control-Sample11_Beta | chr14:73361825-73362214 | DPF3 | S_Shore | TSS1500 | No |
| Control-Sample16_Beta | chr7:153586022-153586207 | DPP6 | S_Shore | Body | No |
| Control-Sample2_Beta | chr19:4676492-4676623 | DPP9 | Island | 3'UTR | Yes |
| Control-Sample3_Beta | chr19:4676202-4676623 | DPP9 | N_Shore | 3'UTR | Yes |
| Control-Sample259_Be | chr7:89748674-89748686 | DPY19L2P4 | Island | TSS200 | No |
| Control-Sample100_Be | chr8:26513782-26513861 | DPYSL2 | | 3'UTR | Yes |
| Control-Sample11_Beta | chr8:26513239-26513861 | DPYSL2 | | 3'UTR | Yes |
| Control-Sample585_Be | chr8.0:26513782-26513861 | DPYSL2 | | 3'UTR | Yes |
| Control-Sample11_Beta | chr11:638404-638939 | DRD4 | Island | Body | Yes |
| Control-Sample13_Beta | chr11:636167-636659 | DRD4 | N_Shore | TSS1500 | Yes |
| Control-Sample21_Beta | chr11:638424-638939 | DRD4 | Island | Body | Yes |
| Control-Sample67_Beta | chr21:42212414-42212925 | DSCAM | N_Shore | Body | No |
| Control-Sample19_Beta | chr11:117665480-117666000 | DSCAML1 | N_Shore | Body | No |
| Control-Sample42_Beta | chr21:38630234-38630728 | DSCR3 | Island | Body | No |
| Control-Sample1_Beta | chr21:38580486-38580856 | DSCR9 | | TSS1500 | Yes |
| Control-Sample10_Beta | chr21:38580802-38580856 | DSCR9 | | TSS200 | Yes |
| Control-Sample2_Beta | chr21:38580484-38580856 | DSCR9 | | TSS1500 | Yes |
| Control-Sample20_Beta | chr21:38580484-38580812 | DSCR9 | | TSS1500 | Yes |
| Control-Sample21_Beta | chr21:38580129-38580856 | DSCR9 | | TSS1500 | Yes |
| Control-Sample171_Be | chr18:32398181-32398287 | DTNA;DTNA;DTNA;DTNA;DTNA;DTNA;DTNA;DTNA;DTNA | | Body;Body;Body;TSS200;Body;Body;TSS200;Body;Bo | Yes |
| Control-Sample20_Beta | chr18:32398024-32398246 | DTNA;DTNA;DTNA;DTNA;DTNA;DTNA;DTNA;DTNA;DTNA | | Body;TSS1500;Body;Body;Body;Body;Body;TSS1500; | Yes |
| Control-Sample1_Beta | chr12:113530566-113531060 | DTX1 | N_Shore | Body | No |
| Control-Sample41_Beta | chr15:45409698-45410047 | DUOXA1;DUOXA2 | S_Shore | 3'UTR;Body | No |
| Control-Sample10_Beta | chr5:172199642-172199667 | DUSP1 | S_Shore | TSS1500 | No |
| Control-Sample10_Beta | chr17:35872723-35873008 | DUSP14 | Island | Body | Yes |
| Control-Sample15_Beta | chr17:35872723-35872880 | DUSP14 | Island | Body | Yes |
| Control-Sample18_Beta | chr17:35872723-35873369 | DUSP14 | Island | Body | Yes |
| Control-Sample564_Be | chr2.0:96810690-96811060 | DUSP2 | Island | Body | Yes |
| Control-Sample87_Beta | chr2:96810690-96811060 | DUSP2 | Island | Body | Yes |
| Control-Sample100_Be | chr6:291687-292522 | DUSP22 | N_Shore | TSS1500 | Yes |
| Control-Sample101_Be | chr6:291687-292823 | DUSP22 | N_Shore | TSS1500 | Yes |
| Control-Sample2_Beta | chr6:312026-312105 | DUSP22 | | Body | Yes |
| Control-Sample207_Be | chr6:291687-292385 | DUSP22 | N_Shore | TSS1500 | Yes |
| Control-Sample218_Be | chr6:291882-292522 | DUSP22 | N_Shore | TSS1500 | Yes |
| Control-Sample394_Be | chr6:291882-292823 | DUSP22 | N_Shore | TSS1500 | Yes |
| Control-Sample505_Be | chr6.0:291687-292522 | DUSP22 | N_Shore | TSS1500 | Yes |
| Control-Sample539_Be | chr6.0:291687-292823 | DUSP22 | N_Shore | TSS1500 | Yes |
| Control-Sample57_Beta | chr1:228785987-228786295 | DUSP5P | Island | Body | No |
| Control-Sample331_Be | chr2:71680559-71680721 | DYSF;DYSF;DYSF;DYSF;DYSF;DYSF;DYSF | Island | TSS200;TSS200;TSS200;TSS200;TSS200;TSS200;TSS | No |
| Control-Sample15_Beta | chr1:23854847-23855149 | E2F2 | N_Shelf | Body | No |
| Control-Sample11_Beta | chr10:131713829-131713890 | EBF3 | | Body | Yes |
| Control-Sample24_Beta | chr10:131665838-131665896 | EBF3 | N_Shore | Body | Yes |
| Control-Sample10_Beta | chr1:21617442-21618050 | ECE1;ECE1;ECE1 | S_Shore | TSS1500;TSS1500;Body | No |
| Control-Sample30_Beta | chr3:183993080-183993368 | ECE2;ECE2;ECE2;ECE2 | N_Shore | Body;TSS1500;TSS1500;TSS1500 | No |
| Control-Sample30_Beta | chr2:233251770-233252170 | ECEL1P2 | Island | TSS200 | Yes |
| Control-Sample45_Beta | chr2:233251770-233251881 | ECEL1P2 | Island | TSS200 | Yes |
| Control-Sample1_Beta | chr16:67918001-67918484 | EDC4;NRN1L | N_Shore | Body;TSS1500 | No |
| Control-Sample108_Be | chr6:12289897-12290322 | EDN1;EDN1 | | TSS1500;TSS1500 | No |
| Control-Sample52_Beta | chr1:41950237-41950362 | EDN2 | | 1stExon | No |
| Control-Sample19_Beta | chr11:62342512-62342730 | EEF1G | S_Shore | TSS1500 | No |
| Control-Sample194_Be | chr7:36193854-36194575 | EEPD1 | S_Shore | 5'UTR | Yes |
| Control-Sample77_Beta | chr7:36194253-36194575 | EEPD1 | S_Shore | Body | Yes |
| Control-Sample13_Beta | chr12:3862417-3862428 | EFCAB4B;EFCAB4B;EFCAB4B | Island | TSS200;TSS200;TSS200 | Yes |
| Control-Sample17_Beta | chr12:3862423-3862457 | EFCAB4B;EFCAB4B;EFCAB4B | Island | TSS200;TSS200;TSS200 | Yes |
| Control-Sample29_Beta | chr12:3862417-3862444 | EFCAB4B;EFCAB4B;EFCAB4B | Island | TSS200;TSS200;TSS200 | Yes |
| Control-Sample39_Beta | chr12:3862423-3862444 | EFCAB4B;EFCAB4B;EFCAB4B | Island | TSS200;TSS200;TSS200 | Yes |
| Control-Sample19_Beta | chr19:1287428-1287884 | EFNA2 | Island | Body | No |
| Control-Sample1_Beta | chr1:155035375-155035748 | EFNA4;EFNA4;EFNA4 | Island | TSS1500;TSS1500;TSS1500 | Yes |
| Control-Sample13_Beta | chr1:155035388-155035748 | EFNA4;EFNA4;EFNA4 | Island | TSS1500;TSS1500;TSS1500 | Yes |
| Control-Sample13_Beta | chr13:107143980-107144036 | EFNB2 | N_Shore | 3'UTR | No |
| Control-Sample1_Beta | chr6:32131571-32131722 | EGFL8 | N_Shelf | TSS1500 | Yes |
| Control-Sample10_Beta | chr6:32135728-32135803 | EGFL8 | S_Shore | Body | Yes |
| Control-Sample14_Beta | chr6:32135718-32135803 | EGFL8 | S_Shore | Body | Yes |
| Control-Sample19_Beta | chr6:32135715-32135803 | EGFL8 | S_Shore | Body | Yes |
| Control-Sample12_Beta | chr7:55224863-55224910 | EGFR;EGFR;EGFR | | Body;Body;Body | Yes |
| Control-Sample19_Beta | chr7:55225082-55225183 | EGFR;EGFR;EGFR | | Body;Body;Body | Yes |
| Control-Sample35_Beta | chr7:55224863-55225133 | EGFR;EGFR;EGFR | | Body;Body;Body | Yes |
| Control-Sample11_Beta | chr11:65359521-65360509 | EHBP1L1 | Island | Body | Yes |
| Control-Sample13_Beta | chr11:65352168-65352327 | EHBP1L1 | N_Shore | Body | Yes |
| Control-Sample425_Be | chr15:40268610-40268777 | EIF2AK4 | Island | Body | No |
| Control-Sample1_Beta | chr8:141559683-141559946 | EIF2C2;EIF2C2 | S_Shore | Body;Body | Yes |
| Control-Sample1_Beta | chr8:141577061-141577413 | EIF2C2;EIF2C2 | | Body;Body | Yes |

| | | | | | |
|---|---|---|---|---|---|
| Control-Sample1_Beta | chr8:141608401-141608646 | EIF2C2;EIF2C2 | | Body;Body | Yes |
| Control-Sample100_Be | chr8:141559683-141559741 | EIF2C2;EIF2C2 | S_Shore | Body;Body | Yes |
| Control-Sample12_Beta | chr8:141577061-141577150 | EIF2C2;EIF2C2 | | Body;Body | Yes |
| Control-Sample16_Beta | chr8:141588118-141588242 | EIF2C2;EIF2C2 | | Body;Body | Yes |
| Control-Sample28_Beta | chr17:7477935-7478131 | EIF4A1;SNORA48 | S_Shore | Body;TSS200 | No |
| Control-Sample1_Beta | chr4:99851008-99851281 | EIF4E;EIF4E;EIF4E;EIF4E;EIF4E | S_Shore | 5'UTR;1stExon;5'UTR;TSS1500;1stExon | Yes |
| Control-Sample11_Beta | chr4:99851060-99851211 | EIF4E;EIF4E;EIF4E;EIF4E;EIF4E | S_Shore | 5'UTR;1stExon;5'UTR;TSS1500;1stExon | Yes |
| Control-Sample13_Beta | chr4:99851060-99851156 | EIF4E;EIF4E;EIF4E;EIF4E;EIF4E | S_Shore | 5'UTR;1stExon;5'UTR;TSS1500;1stExon | Yes |
| Control-Sample14_Beta | chr4:99851060-99851281 | EIF4E;EIF4E;EIF4E;EIF4E;EIF4E | S_Shore | 5'UTR;1stExon;5'UTR;TSS1500;1stExon | Yes |
| Control-Sample16_Beta | chr4:99851003-99851281 | EIF4E;EIF4E;EIF4E;EIF4E;EIF4E | S_Shore | 5'UTR;1stExon;5'UTR;TSS1500;1stExon | Yes |
| Control-Sample2_Beta | chr4:99851008-99851211 | EIF4E;EIF4E;EIF4E;EIF4E;EIF4E | S_Shore | 5'UTR;1stExon;5'UTR;TSS1500;1stExon | Yes |
| Control-Sample338_Be | chr4:99850801-99851156 | EIF4E;EIF4E;EIF4E;EIF4E;EIF4E | S_Shore | 5'UTR;1stExon;5'UTR;TSS1500;1stExon | Yes |
| Control-Sample339_Be | chr4:99851003-99851156 | EIF4E;EIF4E;EIF4E;EIF4E;EIF4E | S_Shore | 5'UTR;1stExon;5'UTR;TSS1500;1stExon | Yes |
| Control-Sample419_Be | chr4:99851115-99851211 | EIF4E;EIF4E;EIF4E;EIF4E;EIF4E | S_Shore | 5'UTR;1stExon;5'UTR;TSS1500;1stExon | Yes |
| Control-Sample1_Beta | chr1:50571816-50571943 | ELAVL4;ELAVL4;ELAVL4 | | Body;Body;TSS200 | Yes |
| Control-Sample1_Beta | chr1:50574383-50574591 | ELAVL4;ELAVL4;ELAVL4;ELAVL4;ELAVL4 | | Body;TSS1500;Body;Body;TSS1500 | Yes |
| Control-Sample19_Beta | chr1:50573970-50574837 | ELAVL4;ELAVL4;ELAVL4;ELAVL4;ELAVL4 | | Body;TSS1500;Body;Body;TSS1500 | Yes |
| Control-Sample20_Beta | chr1:50573970-50574517 | ELAVL4;ELAVL4;ELAVL4;ELAVL4;ELAVL4 | | Body;TSS1500;Body;Body;TSS1500 | Yes |
| Control-Sample22_Beta | chr1:50574383-50574837 | ELAVL4;ELAVL4;ELAVL4;ELAVL4;ELAVL4 | | Body;TSS1500;Body;Body;TSS1500 | Yes |
| Control-Sample27_Beta | chr1:50574383-50574517 | ELAVL4;ELAVL4;ELAVL4;ELAVL4;ELAVL4 | | Body;TSS1500;Body;Body;TSS1500 | Yes |
| Control-Sample68_Beta | chr1:50574507-50574591 | ELAVL4;ELAVL4;ELAVL4;ELAVL4;ELAVL4 | | Body;Body;TSS200;Body;TSS200 | Yes |
| Control-Sample19_Beta | chr1:201979478-201979567 | ELF3;ELF3 | | TSS1500;TSS1500 | Yes |
| Control-Sample2_Beta | chr1:201979533-201979764 | ELF3;ELF3 | | TSS200;TSS200 | Yes |
| Control-Sample19_Beta | chr11:34535357-34535416 | ELF5 | | TSS200 | No |
| Control-Sample1_Beta | chr7:1753447-1753659 | ELFN1 | Island | 5'UTR | Yes |
| Control-Sample13_Beta | chr7:1753447-1753637 | ELFN1 | Island | 5'UTR | Yes |
| Control-Sample15_Beta | chr7:1753452-1753659 | ELFN1 | Island | 5'UTR | Yes |
| Control-Sample16_Beta | chr7:1753573-1753659 | ELFN1 | Island | 5'UTR | Yes |
| Control-Sample3_Beta | chr7:1753396-1753659 | ELFN1 | N_Shore | 5'UTR | Yes |
| Control-Sample1_Beta | chr7:37025706-37025810 | ELMO1;ELMO1;ELMO1;ELMO1 | | TSS1500;Body;1stExon;5'UTR | No |
| Control-Sample1_Beta | chr16:67232180-67232460 | ELMO3;E2F4 | N_Shore | TSS1500;3'UTR | No |
| Control-Sample1_Beta | chr6:11044877-11044894 | ELOVL2 | Island | TSS1500 | No |
| Control-Sample4_Beta | chr5:60138761-60138883 | ELOVL7;ELOVL7 | N_Shore | 5'UTR;5'UTR | No |
| Control-Sample18_Beta | chr19:48497737-48497828 | ELSPBP1 | | TSS200 | Yes |
| Control-Sample475_Be | chr19:48497737-48498031 | ELSPBP1 | | TSS200 | Yes |
| Control-Sample585_Be | chr19.0:48497737-48497828 | ELSPBP1 | | TSS200 | Yes |
| Control-Sample12_Beta | chr19:46142212-46142707 | EML2 | N_Shelf | Body | Yes |
| Control-Sample24_Beta | chr19:46142212-46142673 | EML2 | N_Shelf | Body | Yes |
| Control-Sample1_Beta | chr19:46142291-46142673 | EML2;MIR330 | N_Shelf | Body;Body | Yes |
| Control-Sample15_Beta | chr19:46142291-46142707 | EML2;MIR330 | N_Shelf | Body;Body | Yes |
| Control-Sample108_Be | chr12:13349273-13349724 | EMP1 | | TSS1500 | No |
| Control-Sample14_Beta | chr10:119307692-119307931 | EMX2;EMX2 | S_Shore | Body;Body | No |
| Control-Sample1_Beta | chr10:119292587-119293800 | EMX2OS | N_Shore | Body | Yes |
| Control-Sample12_Beta | chr10:119292587-119293550 | EMX2OS | N_Shore | Body | Yes |
| Control-Sample16_Beta | chr10:119292300-119292547 | EMX2OS | N_Shore | Body | Yes |
| Control-Sample18_Beta | chr10:119292587-119293360 | EMX2OS | N_Shore | Body | Yes |
| Control-Sample19_Beta | chr10:119295827-119296612 | EMX2OS | Island | Body | Yes |
| Control-Sample20_Beta | chr10:119295827-119296182 | EMX2OS | Island | Body | Yes |
| Control-Sample22_Beta | chr10:119293360-119293800 | EMX2OS | N_Shore | Body | Yes |
| Control-Sample23_Beta | chr10:119292062-119292547 | EMX2OS | N_Shelf | Body | Yes |
| Control-Sample3_Beta | chr10:119292587-119294055 | EMX2OS | N_Shore | Body | Yes |
| Control-Sample37_Beta | chr10:119292130-119292547 | EMX2OS | N_Shore | Body | Yes |
| Control-Sample4_Beta | chr10:119292062-119294055 | EMX2OS | N_Shelf | Body | Yes |
| Control-Sample58_Beta | chr10:119293083-119293800 | EMX2OS | N_Shore | Body | Yes |
| Control-Sample7_Beta | chr10:119292549-119293550 | EMX2OS | N_Shore | Body | Yes |
| Control-Sample11_Beta | chr17:77084229-77084618 | ENGASE | | 3'UTR | No |
| Control-Sample270_Be | chr3:40428638-40428713 | ENTPD3 | N_Shore | TSS200 | Yes |
| Control-Sample331_Be | chr3:40428638-40428986 | ENTPD3 | N_Shore | TSS200 | Yes |
| Control-Sample194_Be | chr3:40428713-40429014 | ENTPD3;ENTPD3 | Island | 1stExon;5'UTR | Yes |
| Control-Sample10_Beta | chr3:27761408-27761677 | EOMES | N_Shore | Body | Yes |
| Control-Sample18_Beta | chr3:27761408-27762095 | EOMES | N_Shore | Body | Yes |
| Control-Sample1_Beta | chr22:41573046-41573176 | EP300 | | Body | Yes |
| Control-Sample22_Beta | chr22:41573046-41573155 | EP300 | | Body | Yes |
| Control-Sample1_Beta | chr12:132474636-132474827 | EP400 | | Body | Yes |
| Control-Sample19_Beta | chr12:132546659-132546836 | EP400 | | Body | Yes |
| Control-Sample3_Beta | chr12:132603531-132603671 | EP400NL | Island | Body | No |
| Control-Sample1_Beta | chr2:46593223-46593271 | EPAS1 | | Body | No |
| Control-Sample16_Beta | chr1:29213106-29213233 | EPB41;EPB41;EPB41;EPB41;EPB41 | N_Shore | TSS1500;TSS1500;TSS1500;TSS1500;TSS1500 | No |
| Control-Sample1_Beta | chr1:38200920-38201123 | EPHA10 | Island | Body | No |
| Control-Sample1_Beta | chr2:222434303-222434778 | EPHA4 | N_Shore | Body | No |
| Control-Sample1_Beta | chr19:56187475-56187915 | EPN1;EPN1;EPN1 | S_Shore | 5'UTR;5'UTR;TSS1500 | No |
| Control-Sample1_Beta | chr19:55598782-55599320 | EPS8L1;EPS8L1 | Island | Body;Body | Yes |
| Control-Sample10_Beta | chr19:55598874-55599320 | EPS8L1;EPS8L1 | Island | Body;Body | Yes |
| Control-Sample1_Beta | chr5:172305889-172305986 | ERGIC1 | | Body | Yes |
| Control-Sample22_Beta | chr5:172305889-172306136 | ERGIC1 | | Body | Yes |
| Control-Sample1_Beta | chr1:44726943-44727313 | ERI3 | | Body | Yes |
| Control-Sample19_Beta | chr1:44703750-44704073 | ERI3 | | Body | Yes |
| Control-Sample21_Beta | chr1:44715942-44716058 | ERI3 | | Body | Yes |
| Control-Sample47_Beta | chr1:44727010-44727313 | ERI3 | | Body | Yes |
| Control-Sample19_Beta | chr8:652315-652403 | ERICH1 | S_Shore | Body | Yes |
| Control-Sample64_Beta | chr8:618674-618751 | ERICH1 | N_Shore | Body | Yes |
| Control-Sample1_Beta | chr8:37595307-37595363 | ERLIN2;ERLIN2;ERLIN2 | S_Shore | 5'UTR;5'UTR;5'UTR | No |
| Control-Sample18_Beta | chr2:158184192-158184543 | ERMN | | TSS200 | No |
| Control-Sample1_Beta | chr16:23706242-23706666 | ERN2 | | Body | No |
| Control-Sample1_Beta | chr11:124629580-124629925 | ESAM | Island | Body | No |
| Control-Sample6_Beta | chr5:54281478-54281733 | ESM1;ESM1 | | TSS200;TSS200 | No |
| Control-Sample1_Beta | chr1:6519923-6520354 | ESPN | N_Shore | Body | No |
| Control-Sample1_Beta | chr16:68269041-68269694 | ESRP2 | Island | Body | Yes |
| Control-Sample15_Beta | chr16:68270252-68270282 | ESRP2 | Island | TSS200 | Yes |
| Control-Sample1_Beta | chr1:217306590-217306763 | ESRRG | N_Shore | 5'UTR | Yes |
| Control-Sample1_Beta | chr1:217309970-217310572 | ESRRG | N_Shore | 5'UTR | Yes |

| | | | | | |
|---|---|---|---|---|---|
| Control-Sample1_Beta | chr1:217311608-217312478 | ESRRG | Island | TSS1500 | Yes |
| Control-Sample12_Beta | chr1:217309568-217310572 | ESRRG | N_Shore | 5'UTR | Yes |
| Control-Sample12_Beta | chr1:217311982-217312478 | ESRRG | S_Shore | TSS1500 | Yes |
| Control-Sample359_Be | chr1:217309568-217309970 | ESRRG | N_Shore | 5'UTR | Yes |
| Control-Sample41_Beta | chr1:217309970-217310383 | ESRRG | N_Shore | 5'UTR | Yes |
| Control-Sample57_Beta | chr1:217311608-217312060 | ESRRG | Island | TSS1500 | Yes |
| Control-Sample63_Beta | chr1:217309794-217310572 | ESRRG | N_Shore | 5'UTR | Yes |
| Control-Sample67_Beta | chr1:217311982-217312060 | ESRRG | S_Shore | TSS1500 | Yes |
| Control-Sample57_Beta | chr7:158533877-158533927 | ESYT2 | | Body | No |
| Control-Sample66_Beta | chr3:138152837-138153127 | ESYT3 | N_Shore | TSS1500 | No |
| Control-Sample1_Beta | chr19:7926970-7927374 | EVI5L;EVI5L | Island | Body;Body | Yes |
| Control-Sample10_Beta | chr19:7911195-7911245 | EVI5L;EVI5L | N_Shore | 5'UTR;TSS200 | Yes |
| Control-Sample19_Beta | chr19:7911120-7911206 | EVI5L;EVI5L | N_Shore | 5'UTR;TSS1500 | Yes |
| Control-Sample223_Be | chr19:7910811-7911195 | EVI5L;EVI5L | N_Shore | 5'UTR;TSS1500 | Yes |
| Control-Sample3_Beta | chr19:7910696-7911206 | EVI5L;EVI5L | N_Shore | 5'UTR;TSS1500 | Yes |
| Control-Sample85_Beta | chr19:7913785-7913922 | EVI5L;EVI5L | Island | Body;Body | Yes |
| Control-Sample464_Be | chr14:100610071-100610570 | EVL | N_Shore | Body | No |
| Control-Sample151_Be | chr7:27281561-27281581 | EVX1 | N_Shore | TSS1500 | No |
| Control-Sample1_Beta | chr19:45737603-45737623 | EXOC3L2 | Island | TSS200 | Yes |
| Control-Sample10_Beta | chr19:45737610-45737880 | EXOC3L2 | Island | TSS200 | Yes |
| Control-Sample11_Beta | chr19:45737483-45737880 | EXOC3L2 | Island | TSS200 | Yes |
| Control-Sample2_Beta | chr19:45720771-45721000 | EXOC3L2 | Island | Body | Yes |
| Control-Sample25_Beta | chr19:45737603-45737880 | EXOC3L2 | Island | TSS200 | Yes |
| Control-Sample3_Beta | chr19:45737610-45737873 | EXOC3L2 | Island | TSS200 | Yes |
| Control-Sample32_Beta | chr19:45737485-45737880 | EXOC3L2 | Island | TSS200 | Yes |
| Control-Sample34_Beta | chr19:45737603-45737873 | EXOC3L2 | Island | TSS200 | Yes |
| Control-Sample49_Beta | chr19:45737483-45737623 | EXOC3L2 | Island | TSS200 | Yes |
| Control-Sample10_Beta | chr11:108422936-108423031 | EXPH5;EXPH5 | | TSS200;Body | No |
| Control-Sample1_Beta | chr6:133561614-133561814 | EYA4;EYA4;EYA4 | N_Shore | TSS1500;TSS1500;TSS1500 | Yes |
| Control-Sample11_Beta | chr6:133561614-133561647 | EYA4;EYA4;EYA4 | N_Shore | TSS1500;TSS1500;TSS1500 | Yes |
| Control-Sample11_Beta | chr6:133561660-133561814 | EYA4;EYA4;EYA4 | N_Shore | TSS1500;TSS1500;TSS1500 | Yes |
| Control-Sample13_Beta | chr6:133561647-133561814 | EYA4;EYA4;EYA4 | N_Shore | TSS1500;TSS1500;TSS1500 | Yes |
| Control-Sample2_Beta | chr6:133561649-133561814 | EYA4;EYA4;EYA4 | N_Shore | TSS1500;TSS1500;TSS1500 | Yes |
| Control-Sample20_Beta | chr6:133561614-133561649 | EYA4;EYA4;EYA4 | N_Shore | TSS1500;TSS1500;TSS1500 | Yes |
| Control-Sample20_Beta | chr6:133561677-133561814 | EYA4;EYA4;EYA4 | N_Shore | TSS1500;TSS1500;TSS1500 | Yes |
| Control-Sample65_Beta | chr6:133561677-133561719 | EYA4;EYA4;EYA4 | N_Shore | TSS1500;TSS1500;TSS1500 | Yes |
| Control-Sample66_Beta | chr6:133561756-133561814 | EYA4;EYA4;EYA4 | N_Shore | TSS1500;TSS1500;TSS1500 | Yes |
| Control-Sample1_Beta | chr6:159238463-159238744 | EZR;EZR | N_Shore | Body;Body | No |
| Control-Sample1_Beta | chr13:113776917-113777045 | F10 | S_Shelf | TSS200 | Yes |
| Control-Sample11_Beta | chr13:113776964-113777045 | F10 | S_Shelf | TSS200 | Yes |
| Control-Sample14_Beta | chr13:113776915-113777045 | F10 | S_Shelf | TSS200 | Yes |
| Control-Sample19_Beta | chr13:113777013-113777045 | F10 | | TSS200 | Yes |
| Control-Sample58_Beta | chr13:113776915-113777160 | F10 | S_Shelf | TSS200 | Yes |
| Control-Sample25_Beta | chr19:16999614-16999668 | F2RL3 | N_Shore | TSS1500 | No |
| Control-Sample1_Beta | chr16:74782152-74782640 | FA2H | | Body | No |
| Control-Sample3_Beta | chr1:46859671-46859774 | FAAH | N_Shore | TSS1500 | No |
| Control-Sample34_Beta | chr8:82192519-82192601 | FABP5 | Island | TSS1500 | No |
| Control-Sample108_Be | chr6:123100471-123100722 | FABP7 | | TSS200 | Yes |
| Control-Sample291_Be | chr6:123100189-123100722 | FABP7 | | TSS1500 | Yes |
| Control-Sample188_Be | chr3:138327370-138327519 | FAIM;FAIM;FAIM;FAIM | N_Shore | TSS200;TSS200;TSS200;TSS1500 | No |
| Control-Sample13_Beta | chr3:58563609-58563693 | FAM107A;FAM107A | | TSS200;TSS200 | Yes |
| Control-Sample14_Beta | chr3:58563666-58563693 | FAM107A;FAM107A | | TSS200;TSS200 | Yes |
| Control-Sample28_Beta | chr3:58563609-58563666 | FAM107A;FAM107A | | TSS200;TSS200 | Yes |
| Control-Sample78_Beta | chr3:58563609-58563674 | FAM107A;FAM107A | | TSS200;TSS200 | Yes |
| Control-Sample644_Be | chr10:14575319-14575436 | FAM107B | | Body | No |
| Control-Sample13_Beta | chr2:225266346-225266880 | FAM124B;FAM124B | | 1stExon;1stExon | Yes |
| Control-Sample14_Beta | chr2:225266346-225266656 | FAM124B;FAM124B | | 1stExon;1stExon | Yes |
| Control-Sample14_Beta | chr2:225266783-225266876 | FAM124B;FAM124B | | TSS200;TSS200 | Yes |
| Control-Sample15_Beta | chr2:225266783-225266880 | FAM124B;FAM124B | | TSS200;TSS200 | Yes |
| Control-Sample16_Beta | chr2:225266801-225266880 | FAM124B;FAM124B | | TSS200;TSS200 | Yes |
| Control-Sample4_Beta | chr2:225266346-225266876 | FAM124B;FAM124B | | 1stExon;1stExon | Yes |
| Control-Sample62_Beta | chr2:225266843-225266880 | FAM124B;FAM124B | | TSS200;TSS200 | Yes |
| Control-Sample91_Beta | chr2:225266801-225266876 | FAM124B;FAM124B | | TSS200;TSS200 | Yes |
| Control-Sample108_Be | chr2:225266580-225266880 | FAM124B;FAM124B;FAM124B;FAM124B | | 5'UTR;1stExon;1stExon;5'UTR | Yes |
| Control-Sample194_Be | chr2:225266580-225266733 | FAM124B;FAM124B;FAM124B;FAM124B | | 5'UTR;1stExon;1stExon;5'UTR | Yes |
| Control-Sample74_Beta | chr2:225266656-225266783 | FAM124B;FAM124B;FAM124B;FAM124B | | 5'UTR;1stExon;1stExon;5'UTR | Yes |
| Control-Sample80_Beta | chr2:225266580-225266783 | FAM124B;FAM124B;FAM124B;FAM124B | | 5'UTR;1stExon;1stExon;5'UTR | Yes |
| Control-Sample19_Beta | chr3:184055123-184055379 | FAM131A;FAM131A | N_Shore | 5'UTR;TSS200 | No |
| Control-Sample10_Beta | chr1:1181757-1181956 | FAM132A | Island | Body | No |
| Control-Sample108_Be | chr4:89978251-89978566 | FAM13A;FAM13A | | 1stExon;5'UTR | Yes |
| Control-Sample108_Be | chr4:89744363-89744743 | FAM13A;FAM13A;FAM13A | | 1stExon;Body;5'UTR | Yes |
| Control-Sample1_Beta | chr8:21961713-21961788 | FAM160B2 | | 3'UTR | No |
| Control-Sample1_Beta | chr1:179713788-179713950 | FAM163A | Island | 5'UTR | No |
| Control-Sample13_Beta | chr1:32714146-32714308 | FAM167B | Island | Body | No |
| Control-Sample24_Beta | chr15:99057794-99058098 | FAM169B | | TSS200 | Yes |
| Control-Sample29_Beta | chr15:99057732-99058098 | FAM169B | | TSS200 | Yes |
| Control-Sample1_Beta | chr1:36787932-36788416 | FAM176B | Island | Body | No |
| Control-Sample23_Beta | chr7:135433715-135433760 | FAM180A | | TSS200 | Yes |
| Control-Sample42_Beta | chr7:135433639-135433760 | FAM180A | | TSS200 | Yes |
| Control-Sample1_Beta | chr7:135433540-135433790 | FAM180A;FAM180A | | 1stExon;5'UTR | Yes |
| Control-Sample13_Beta | chr7:135433540-135433760 | FAM180A;FAM180A | | 1stExon;5'UTR | Yes |
| Control-Sample31_Beta | chr7:135433540-135433715 | FAM180A;FAM180A | | 1stExon;5'UTR | Yes |
| Control-Sample63_Beta | chr7:135433540-135433736 | FAM180A;FAM180A | | 1stExon;5'UTR | Yes |
| Control-Sample1_Beta | chr14:94392718-94393816 | FAM181A;C14orf86 | | Body;TSS200 | Yes |
| Control-Sample10_Beta | chr14:94392862-94393816 | FAM181A;C14orf86 | | Body;TSS200 | Yes |
| Control-Sample21_Beta | chr14:94393497-94393816 | FAM181A;C14orf86 | | Body;TSS1500 | Yes |
| Control-Sample3_Beta | chr14:94392408-94393816 | FAM181A;C14orf86 | | Body;Body | Yes |
| Control-Sample1_Beta | chr16:10912478-10912696 | FAM18A;FAM18A | Island | 1stExon;5'UTR | No |
| Control-Sample23_Beta | chr4:91048416-91048823 | FAM190A | Island | TSS1500 | Yes |
| Control-Sample24_Beta | chr4:91048416-91048584 | FAM190A | Island | TSS1500 | Yes |
| Control-Sample19_Beta | chr4:91156040-91156290 | FAM190A;FAM190A | | 5'UTR;TSS200 | Yes |

| | | | | | |
|---|---|---|---|---|---|
| Control-Sample1_Beta | chr4:2627057-2627246 | FAM193A | | TSS200 | Yes |
| Control-Sample10_Beta | chr4:2627142-2627246 | FAM193A | | TSS200 | Yes |
| Control-Sample12_Beta | chr4:2627118-2627246 | FAM193A | | TSS200 | Yes |
| Control-Sample3_Beta | chr5:176981910-176982116 | FAM193B;FAM193B | S_Shore | TSS1500;TSS1500 | Yes |
| Control-Sample6_Beta | chr5:176981888-176982116 | FAM193B;FAM193B | S_Shore | TSS1500;TSS1500 | Yes |
| Control-Sample23_Beta | chr4:159094213-159094517 | FAM198B;FAM198B;FAM198B | | TSS200;TSS1500;TSS200 | Yes |
| Control-Sample36_Beta | chr4:159093915-159094517 | FAM198B;FAM198B;FAM198B | | TSS200;5'UTR;5'UTR | Yes |
| Control-Sample63_Beta | chr4:159094034-159094517 | FAM198B;FAM198B;FAM198B;FAM198B;FAM198B | | TSS1500;5'UTR;1stExon;5'UTR;1stExon | Yes |
| Control-Sample5_Beta | chr1:113265602-113265679 | FAM19A3;FAM19A3 | Island | Body;Body | No |
| Control-Sample59_Beta | chr6:116832611-116832940 | FAM26E;BET3L | | TSS200;Body | No |
| Control-Sample1_Beta | chr16:88844125-88844726 | FAM38A | N_Shore | Body | Yes |
| Control-Sample13_Beta | chr16:88844125-88844499 | FAM38A | N_Shore | Body | Yes |
| Control-Sample1_Beta | chr6:3848218-3848280 | FAM50B | N_Shore | TSS1500 | No |
| Control-Sample1_Beta | chr4:1642054-1642257 | FAM53A | N_Shelf | 3'UTR | Yes |
| Control-Sample54_Beta | chr4:1642054-1642635 | FAM53A | N_Shelf | 3'UTR | Yes |
| Control-Sample59_Beta | chr4:1684739-1684825 | FAM53A | N_Shore | 5'UTR | Yes |
| Control-Sample69_Beta | chr4:1642061-1642257 | FAM53A | N_Shelf | 3'UTR | Yes |
| Control-Sample73_Beta | chr4:1642054-1642164 | FAM53A | N_Shelf | 3'UTR | Yes |
| Control-Sample1_Beta | chr10:126330876-126331180 | FAM53B | | Body | Yes |
| Control-Sample11_Beta | chr10:126330876-126331212 | FAM53B | | Body | Yes |
| Control-Sample108_Be | chr3:101546486-101546709 | FAM55C;FAM55C;NFKBIZ | | 3'UTR;3'UTR;TSS1500 | Yes |
| Control-Sample19_Beta | chr2:26407567-26407743 | FAM59B | Island | Body | Yes |
| Control-Sample448_Be | chr2:26401820-26402008 | FAM59B | Island | Body | Yes |
| Control-Sample1_Beta | chr1:177225850-177225888 | FAM5B | | Body | No |
| Control-Sample480_Be | chr17:6347533-6347637 | FAM64A | Island | TSS1500 | No |
| Control-Sample1_Beta | chr19:49116423-49116839 | FAM83E | Island | 1stExon | Yes |
| Control-Sample19_Beta | chr19:49116423-49116870 | FAM83E | Island | 1stExon | Yes |
| Control-Sample21_Beta | chr19:49116824-49116870 | FAM83E | S_Shore | TSS200 | Yes |
| Control-Sample1_Beta | chr16:85146088-85146180 | FAM92B;FAM92B | | 5'UTR;1stExon | No |
| Control-Sample26_Beta | chr16:89883212-89883253 | FANCA;FANCA | Island | TSS200;TSS200 | Yes |
| Control-Sample27_Beta | chr16:89883212-89883248 | FANCA;FANCA | Island | TSS200;TSS200 | Yes |
| Control-Sample334_Be | chr16:89883240-89883253 | FANCA;FANCA | Island | TSS200;TSS200 | Yes |
| Control-Sample564_Be | chr16.0:89883240-89883253 | FANCA;FANCA | Island | TSS200;TSS200 | Yes |
| Control-Sample12_Beta | chr13:99100506-99100586 | FARP1 | | Body | No |
| Control-Sample108_Be | chr4:187572844-187573046 | FAT1 | | Body | No |
| Control-Sample1_Beta | chr22:45899186-45899736 | FBLN1;FBLN1;FBLN1;FBLN1 | Island | Body;Body;Body;Body | Yes |
| Control-Sample31_Beta | chr22:45899292-45899736 | FBLN1;FBLN1;FBLN1;FBLN1 | Island | Body;Body;Body;Body | Yes |
| Control-Sample1_Beta | chr3:13590432-13590450 | FBLN2;FBLN2 | Island | TSS200;TSS200 | Yes |
| Control-Sample12_Beta | chr3:13590432-13590444 | FBLN2;FBLN2 | Island | TSS200;TSS200 | Yes |
| Control-Sample19_Beta | chr3:13590439-13590450 | FBLN2;FBLN2 | Island | TSS200;TSS200 | Yes |
| Control-Sample19_Beta | chr3:13664715-13665022 | FBLN2;FBLN2;FBLN2 | | Body;Body;Body | Yes |
| Control-Sample1_Beta | chr5:127871927-127872329 | FBN2 | N_Shore | Body | Yes |
| Control-Sample10_Beta | chr5:127871606-127872329 | FBN2 | N_Shore | Body | Yes |
| Control-Sample3_Beta | chr5:127871606-127872200 | FBN2 | N_Shore | Body | Yes |
| Control-Sample33_Beta | chr5:127871927-127872074 | FBN2 | N_Shore | Body | Yes |
| Control-Sample1_Beta | chr9:97402166-97402555 | FBP1;FBP1 | S_Shore | 5'UTR;TSS1500 | No |
| Control-Sample1_Beta | chr12:133102136-133102331 | FBRSL1 | N_Shore | Body | Yes |
| Control-Sample10_Beta | chr12:133118627-133118770 | FBRSL1 | | Body | Yes |
| Control-Sample57_Beta | chr12:133101612-133102331 | FBRSL1 | N_Shore | Body | Yes |
| Control-Sample6_Beta | chr12:133066505-133066520 | FBRSL1 | N_Shore | TSS1500 | Yes |
| Control-Sample1_Beta | chr12:1675846-1676077 | FBXL14 | | 3'UTR | No |
| Control-Sample270_Be | chr16:745663-745687 | FBXL16 | Island | Body | No |
| Control-Sample1_Beta | chr7:5534721-5535006 | FBXL18 | Island | Body | Yes |
| Control-Sample19_Beta | chr7:5534951-5535006 | FBXL18 | S_Shore | Body | Yes |
| Control-Sample10_Beta | chr16:30959315-30959496 | FBXL19;ORAI3 | Island | 3'UTR;TSS1500 | No |
| Control-Sample11_Beta | chr15:63889406-63889711 | FBXL22 | N_Shelf | TSS200 | Yes |
| Control-Sample12_Beta | chr15:63889527-63889711 | FBXL22 | N_Shelf | TSS200 | Yes |
| Control-Sample30_Beta | chr1:11708949-11709271 | FBXO2 | Island | Body | No |
| Control-Sample464_Be | chr16:87391861-87392029 | FBXO31;FBXO31 | | Body;Body | No |
| Control-Sample1_Beta | chr12:117407879-117408186 | FBXW8;FBXW8 | | Body;Body | Yes |
| Control-Sample108_Be | chr12:117407879-117408163 | FBXW8;FBXW8 | | Body;Body | Yes |
| Control-Sample124_Be | chr12:117408122-117408186 | FBXW8;FBXW8 | | Body;Body | Yes |
| Control-Sample19_Beta | chr12:117448155-117448211 | FBXW8;FBXW8 | | Body;Body | Yes |
| Control-Sample10_Beta | chr5:141031122-141031147 | FCHSD1 | Island | TSS200 | Yes |
| Control-Sample12_Beta | chr5:141031125-141031147 | FCHSD1 | Island | TSS200 | Yes |
| Control-Sample10_Beta | chr1:157670710-157670869 | FCRL3 | | TSS200 | No |
| Control-Sample564_Be | chr8.0:11659970-11660061 | FDFT1 | Island | TSS1500 | No |
| Control-Sample15_Beta | chr5:108083455-108083631 | FER | N_Shore | TSS200 | No |
| Control-Sample1_Beta | chr7:19184555-19184961 | FERD3L | Island | 1stExon | No |
| Control-Sample10_Beta | chr11:63974955-63975481 | FERMT3;FERMT3 | Island | Body;Body | No |
| Control-Sample108_Be | chr12:32654929-32655192 | FGD4 | | TSS200 | No |
| Control-Sample108_Be | chr5:142065671-142065737 | FGF1;FGF1;FGF1;FGF1;FGF1;FGF1;FGF1 | | 5'UTR;5'UTR;5'UTR;1stExon;Body;Body;TSS200 | No |
| Control-Sample11_Beta | chr11:69519974-69520418 | FGF19 | S_Shore | TSS1500 | Yes |
| Control-Sample46_Beta | chr11:69520186-69520418 | FGF19 | S_Shore | TSS1500 | Yes |
| Control-Sample464_Be | chr11:69634363-69634456 | FGF3 | Island | TSS200 | No |
| Control-Sample3_Beta | chr4:15939889-15940058 | FGFBP1;FGFBP1 | | 1stExon;5'UTR | No |
| Control-Sample520_Be | chr4.0:15965016-15965102 | FGFBP2 | | TSS200 | No |
| Control-Sample1_Beta | chr10:123355454-123355748 | FGFR2;FGFR2;FGFR2;FGFR2;FGFR2;FGFR2 | N_Shore | 5'UTR;5'UTR;5'UTR;5'UTR;TSS1500;5'UTR;5'UTR | Yes |
| Control-Sample11_Beta | chr10:123355454-123356205 | FGFR2;FGFR2;FGFR2;FGFR2;FGFR2;FGFR2 | N_Shore | 5'UTR;5'UTR;5'UTR;5'UTR;TSS1500;5'UTR;5'UTR | Yes |
| Control-Sample9_Beta | chr10:123355576-123356205 | FGFR2;FGFR2;FGFR2;FGFR2;FGFR2;FGFR2 | N_Shore | 5'UTR;5'UTR;5'UTR;5'UTR;5'UTR;5'UTR | Yes |
| Control-Sample22_Beta | chr10:123355268-123355576 | FGFR2;FGFR2;FGFR2;FGFR2;FGFR2;FGFR2;FGFR2 | N_Shelf | 5'UTR;5'UTR;TSS1500;5'UTR;TSS1500;5'UTR;5'UTR;T | Yes |
| Control-Sample10_Beta | chr10:123353800-123353889 | FGFR2;FGFR2;FGFR2;FGFR2;FGFR2;FGFR2;FGFR2;FGFR2 | N_Shelf | TSS200;5'UTR;5'UTR;TSS1500;5'UTR;TSS1500;5'UTR; | Yes |
| Control-Sample108_Be | chr10:123353889-123354171 | FGFR2;FGFR2;FGFR2;FGFR2;FGFR2;FGFR2;FGFR2;FGFR2 | N_Shelf | TSS200;5'UTR;5'UTR;TSS1500;5'UTR;TSS1500;5'UTR; | Yes |
| Control-Sample223_Be | chr10:123353298-123353418 | FGFR2;FGFR2;FGFR2;FGFR2;FGFR2;FGFR2;FGFR2;FGFR2;FGFR2 | | Body;Body;Body;Body;Body;Body;Body;Body;Body | Yes |
| Control-Sample223_Be | chr10:123353822-123353956 | FGFR2;FGFR2;FGFR2;FGFR2;FGFR2;FGFR2;FGFR2;FGFR2 | N_Shelf | TSS200;5'UTR;5'UTR;TSS1500;5'UTR;TSS1500;5'UTR; | Yes |
| Control-Sample6_Beta | chr10:123353465-123353654 | FGFR2;FGFR2;FGFR2;FGFR2;FGFR2;FGFR2;FGFR2;FGFR2 | N_Shelf | 1stExon;5'UTR;5'UTR;5'UTR;5'UTR;1stExon;1stExon;5 | Yes |
| Control-Sample65_Beta | chr4:1800154-1801294 | FGFR3;FGFR3;FGFR3 | Island | Body;Body;Body | No |
| Control-Sample1_Beta | chr4:1006237-1006278 | FGFRL1;FGFRL1;FGFRL1 | Island | 5'UTR;TSS200;5'UTR | Yes |
| Control-Sample15_Beta | chr4:1006240-1006278 | FGFRL1;FGFRL1;FGFRL1 | Island | 5'UTR;TSS200;5'UTR | Yes |
| Control-Sample171_Be | chr4:1008425-1008614 | FGFRL1;FGFRL1;FGFRL1 | S_Shore | Body;Body;Body | Yes |
| Control-Sample23_Beta | chr4:1006120-1006278 | FGFRL1;FGFRL1;FGFRL1 | Island | 5'UTR;TSS200;5'UTR | Yes |

| | | | | | |
|---|---|---|---|---|---|
| Control-Sample8_Beta | chr4:1008550-1008738 | FGFRL1;FGFRL1;FGFRL1 | S_Shore | Body;Body;Body | Yes |
| Control-Sample14_Beta | chr1:27961563-27961796 | FGR | Island | 5'UTR | No |
| Control-Sample19_Beta | chr1:15655764-15656086 | FHAD1 | | Body | No |
| Control-Sample13_Beta | chr3:61237712-61237884 | FHIT;FHIT | S_Shore | TSS1500;TSS1500 | No |
| Control-Sample5_Beta | chr2:106055450-106055507 | FHL2 | | TSS1500 | No |
| Control-Sample23_Beta | chr11:27015808-27016671 | FIBIN;FIBIN | | 5'UTR;1stExon | No |
| Control-Sample108_Be | chr6:76059496-76059756 | FILIP1 | | Body | Yes |
| Control-Sample19_Beta | chr6:76059726-76059773 | FILIP1 | | Body | Yes |
| Control-Sample1_Beta | chr7:100886961-100887364 | FIS1 | N_Shore | Body | No |
| Control-Sample441_Be | chr17:17124752-17124854 | FLCN;FLCN | Island | Body;Body | Yes |
| Control-Sample465_Be | chr17:17124699-17124854 | FLCN;FLCN | Island | Body;Body | Yes |
| Control-Sample1_Beta | chr4:106472990-106473233 | FLJ20184 | | TSS1500 | No |
| Control-Sample1_Beta | chr17:26939298-26939704 | FLJ25006;LOC645851 | | Body;TSS1500 | No |
| Control-Sample1_Beta | chr19:16144282-16144499 | FLJ25328;FLJ25328 | | TSS1500;TSS1500 | No |
| Control-Sample113_Be | chr19:50553682-50553708 | FLJ26850 | N_Shore | TSS1500 | Yes |
| Control-Sample113_Be | chr19:50553907-50554067 | FLJ26850 | N_Shore | TSS200 | Yes |
| Control-Sample15_Beta | chr19:50553682-50554013 | FLJ26850 | N_Shore | TSS1500 | Yes |
| Control-Sample16_Beta | chr19:50553682-50554067 | FLJ26850 | N_Shore | TSS1500 | Yes |
| Control-Sample655_Be | chr19:50553682-50553907 | FLJ26850 | N_Shore | TSS1500 | Yes |
| Control-Sample57_Beta | chr2:200331675-200332380 | FLJ32063 | S_Shore | TSS1500 | No |
| Control-Sample1_Beta | chr15:100890963-100890996 | FLJ42289;FLJ42289 | S_Shore | TSS1500;TSS200 | Yes |
| Control-Sample13_Beta | chr15:100890907-100890996 | FLJ42289;FLJ42289 | S_Shore | TSS1500;Body | Yes |
| Control-Sample19_Beta | chr5:92913741-92914319 | FLJ42709;FLJ42709 | N_Shore | Body;Body | Yes |
| Control-Sample20_Beta | chr5:92914085-92914319 | FLJ42709;FLJ42709 | N_Shore | Body;Body | Yes |
| Control-Sample24_Beta | chr5:92909429-92909968 | FLJ42709;FLJ42709 | S_Shore | Body;Body | Yes |
| Control-Sample3_Beta | chr5:92914202-92914319 | FLJ42709;FLJ42709 | N_Shore | Body;Body | Yes |
| Control-Sample65_Beta | chr5:92914036-92914319 | FLJ42709;FLJ42709 | N_Shore | Body;Body | Yes |
| Control-Sample19_Beta | chr1:2979311-2979817 | FLJ42875 | Island | Body | Yes |
| Control-Sample19_Beta | chr1:2980163-2980399 | FLJ42875 | Island | Body | Yes |
| Control-Sample12_Beta | chr1:2984218-2984631 | FLJ42875;FLJ42875 | Island | Body;Body | Yes |
| Control-Sample13_Beta | chr1:2983926-2984631 | FLJ42875;FLJ42875 | Island | Body;Body | Yes |
| Control-Sample40_Beta | chr1:2983926-2984245 | FLJ42875;FLJ42875 | Island | Body;Body | Yes |
| Control-Sample1_Beta | chr1:2984275-2984631 | FLJ42875;PRDM16;FLJ42875;PRDM16 | Island | Body;TSS1500;Body;TSS1500 | Yes |
| Control-Sample10_Beta | chr1:2984275-2984525 | FLJ42875;PRDM16;FLJ42875;PRDM16 | Island | Body;TSS1500;Body;TSS1500 | Yes |
| Control-Sample37_Beta | chr1:2984245-2984525 | FLJ42875;PRDM16;FLJ42875;PRDM16 | Island | Body;TSS1500;Body;TSS1500 | Yes |
| Control-Sample42_Beta | chr1:2984275-2984470 | FLJ42875;PRDM16;FLJ42875;PRDM16 | Island | Body;TSS1500;Body;TSS1500 | Yes |
| Control-Sample475_Be | chr14:62583388-62583795 | FLJ43390 | N_Shore | TSS1500 | No |
| Control-Sample24_Beta | chr5:126408756-126409007 | FLJ44606;FLJ44606 | N_Shore | 5'UTR;5'UTR | Yes |
| Control-Sample248_Be | chr5:126409007-126409227 | FLJ44606;FLJ44606 | Island | 5'UTR;5'UTR | Yes |
| Control-Sample280_Be | chr5:126409061-126409198 | FLJ44606;FLJ44606 | Island | 5'UTR;5'UTR | Yes |
| Control-Sample530_Be | chr5.0:126409061-126409227 | FLJ44606;FLJ44606 | Island | 5'UTR;5'UTR | Yes |
| Control-Sample63_Beta | chr5:126409061-126409227 | FLJ44606;FLJ44606 | Island | 5'UTR;5'UTR | Yes |
| Control-Sample164_Be | chr5:126409153-126409227 | FLJ44606;FLJ44606;FLJ44606;FLJ44606 | Island | 5'UTR;1stExon;5'UTR;1stExon | Yes |
| Control-Sample186_Be | chr5:126409153-126409211 | FLJ44606;FLJ44606;FLJ44606;FLJ44606 | Island | 5'UTR;1stExon;5'UTR;1stExon | Yes |
| Control-Sample552_Be | chr5.0:126409153-126409227 | FLJ44606;FLJ44606;FLJ44606;FLJ44606 | Island | 5'UTR;1stExon;5'UTR;1stExon | Yes |
| Control-Sample18_Beta | chr17:79005047-79005548 | FLJ90757 | N_Shore | Body | No |
| Control-Sample12_Beta | chr6:30711802-30711832 | FLOT1;IER3 | Island | TSS1500;Body | Yes |
| Control-Sample52_Beta | chr6:30711802-30711835 | FLOT1;IER3 | Island | TSS1500;Body | Yes |
| Control-Sample23_Beta | chr1:203320396-203320541 | FMOD | | TSS200 | No |
| Control-Sample103_Be | chr17:80708279-80708632 | FN3K | Island | Body | No |
| Control-Sample13_Beta | chr2:114256406-114256454 | FOXD4L1 | N_Shore | TSS1500 | Yes |
| Control-Sample25_Beta | chr2:114256406-114256673 | FOXD4L1 | N_Shore | TSS1500 | Yes |
| Control-Sample3_Beta | chr2:114256392-114256673 | FOXD4L1 | N_Shore | TSS1500 | Yes |
| Control-Sample1_Beta | chr6:1394758-1395164 | FOXF2 | S_Shore | Body | Yes |
| Control-Sample19_Beta | chr6:1394986-1395164 | FOXF2 | S_Shore | Body | Yes |
| Control-Sample27_Beta | chr6:1394758-1395061 | FOXF2 | S_Shore | Body | Yes |
| Control-Sample1_Beta | chr7:4755010-4755415 | FOXK1 | N_Shelf | Body | Yes |
| Control-Sample1_Beta | chr7:4762742-4763182 | FOXK1 | N_Shore | Body | Yes |
| Control-Sample1_Beta | chr7:4779225-4779342 | FOXK1 | | Body | Yes |
| Control-Sample1_Beta | chr7:4781704-4781872 | FOXK1 | N_Shelf | Body | Yes |
| Control-Sample10_Beta | chr7:4746131-4746462 | FOXK1 | | Body | Yes |
| Control-Sample11_Beta | chr7:4747166-4747745 | FOXK1 | | Body | Yes |
| Control-Sample12_Beta | chr7:4778908-4779342 | FOXK1 | | Body | Yes |
| Control-Sample14_Beta | chr7:4747166-4747261 | FOXK1 | | Body | Yes |
| Control-Sample15_Beta | chr7:4747216-4747745 | FOXK1 | | Body | Yes |
| Control-Sample20_Beta | chr7:4784485-4784963 | FOXK1 | N_Shore | Body | Yes |
| Control-Sample25_Beta | chr7:4757846-4757991 | FOXK1 | Island | Body | Yes |
| Control-Sample33_Beta | chr7:4747026-4747216 | FOXK1 | | Body | Yes |
| Control-Sample33_Beta | chr7:4778839-4779342 | FOXK1 | | Body | Yes |
| Control-Sample63_Beta | chr7:4751906-4752063 | FOXK1 | | Body | Yes |
| Control-Sample69_Beta | chr7:4784325-4784485 | FOXK1 | N_Shore | Body | Yes |
| Control-Sample80_Beta | chr7:4778839-4779225 | FOXK1 | | Body | Yes |
| Control-Sample94_Beta | chr7:4778839-4779312 | FOXK1 | | Body | Yes |
| Control-Sample1_Beta | chr3:138666473-138666732 | FOXL2;C3orf72 | S_Shore | TSS1500;Body | Yes |
| Control-Sample20_Beta | chr3:138666473-138666885 | FOXL2;C3orf72 | S_Shore | TSS1500;Body | Yes |
| Control-Sample19_Beta | chr14:89629585-89629974 | FOXN3;FOXN3 | S_Shore | Body;Body | No |
| Control-Sample81_Beta | chr3:71180032-71180064 | FOXP1 | | Body | No |
| Control-Sample568_Be | chr7.0:114055123-114055210 | FOXP2;FOXP2;FOXP2;FOXP2;FOXP2;FOXP2 | | 1stExon;5'UTR;5'UTR;1stExon;5'UTR;1stExon | Yes |
| Control-Sample67_Beta | chr7:114055123-114055137 | FOXP2;FOXP2;FOXP2;FOXP2;FOXP2;FOXP2 | | 1stExon;5'UTR;5'UTR;1stExon;5'UTR;1stExon | Yes |
| Control-Sample1_Beta | chr6:41528395-41528785 | FOXP4;FOXP4;FOXP4 | Island | 5'UTR;5'UTR;5'UTR | Yes |
| Control-Sample1_Beta | chr6:41561085-41561472 | FOXP4;FOXP4;FOXP4 | N_Shore | Body;Body;Body | Yes |
| Control-Sample11_Beta | chr6:41528198-41528785 | FOXP4;FOXP4;FOXP4 | N_Shore | 5'UTR;5'UTR;5'UTR | Yes |
| Control-Sample52_Beta | chr6:41528531-41528785 | FOXP4;FOXP4;FOXP4 | Island | 5'UTR;5'UTR;5'UTR | Yes |
| Control-Sample3_Beta | chr20:30433405-30433512 | FOXS1;FOXS1 | S_Shore | 5'UTR;1stExon | No |
| Control-Sample475_Be | chr6:116381957-116381976 | FRK | | TSS200 | No |
| Control-Sample108_Be | chr10:14051679-14052028 | FRMD4A | | Body | Yes |
| Control-Sample170_Be | chr10:13749010-13749382 | FRMD4A | | Body | Yes |
| Control-Sample171_Be | chr10:13749193-13749285 | FRMD4A | | Body | Yes |
| Control-Sample19_Beta | chr10:14050305-14050521 | FRMD4A | | Body | Yes |
| Control-Sample19_Beta | chr10:14372879-14372913 | FRMD4A | | TSS200 | Yes |
| Control-Sample194_Be | chr10:13749285-13749602 | FRMD4A | | Body | Yes |

| | | | | | |
|---|---|---|---|---|---|
| Control-Sample216_Be | chr10:13749193-13749382 | FRMD4A | | Body | Yes |
| Control-Sample23_Beta | chr10:13749010-13749245 | FRMD4A | | Body | Yes |
| Control-Sample4_Beta | chr10:13749245-13749382 | FRMD4A | | Body | Yes |
| Control-Sample496_Be | chr10:13749405-13749625 | FRMD4A | | Body | Yes |
| Control-Sample58_Beta | chr10:14051636-14051679 | FRMD4A | | Body | Yes |
| Control-Sample64_Beta | chr10:14372431-14372913 | FRMD4A | | 5'UTR | Yes |
| Control-Sample1_Beta | chr10:49482723-49483356 | FRMPD2;FRMPD2;FRMPD2 | | 5'UTR;1stExon;1stExon;5'UTR | Yes |
| Control-Sample11_Beta | chr10:49482767-49483356 | FRMPD2;FRMPD2;FRMPD2 | | 5'UTR;TSS200;1stExon | Yes |
| Control-Sample17_Beta | chr10:49482723-49482826 | FRMPD2;FRMPD2;FRMPD2 | | 5'UTR;1stExon;1stExon;5'UTR | Yes |
| Control-Sample1_Beta | chr17:79495268-79495324 | FSCN2;FSCN2 | Island | TSS200;TSS200 | Yes |
| Control-Sample23_Beta | chr17:79495268-79495348 | FSCN2;FSCN2 | Island | TSS200;TSS200 | Yes |
| Control-Sample53_Beta | chr17:79495268-79495519 | FSCN2;FSCN2 | Island | TSS200;TSS200 | Yes |
| Control-Sample32_Beta | chr19:4305072-4305239 | FSD1 | Island | Body | No |
| Control-Sample4_Beta | chr3:120170383-120170474 | FSTL1 | Island | TSS1500 | No |
| Control-Sample3_Beta | chr1:24195347-24195659 | FUCA1 | S_Shore | TSS1500 | Yes |
| Control-Sample4_Beta | chr1:24195227-24195659 | FUCA1 | S_Shore | TSS1500 | Yes |
| Control-Sample3_Beta | chr19:49199172-49199234 | FUT2;FUT2 | N_Shore | TSS200;TSS200 | No |
| Control-Sample19_Beta | chr11:94278413-94278603 | FUT4 | Island | 1stExon | Yes |
| Control-Sample2_Beta | chr11:94278457-94278603 | FUT4 | Island | 1stExon | Yes |
| Control-Sample331_Be | chr19:35630106-35630305 | FXYD1;FXYD1 | N_Shelf | 5'UTR;TSS1500 | No |
| Control-Sample13_Beta | chr19:35645598-35645620 | FXYD5;FXYD5;FXYD5;FXYD5 | Island | TSS200;TSS1500;TSS200;TSS200 | Yes |
| Control-Sample18_Beta | chr19:35645610-35645689 | FXYD5;FXYD5;FXYD5;FXYD5 | Island | TSS200;TSS1500;TSS200;TSS200 | Yes |
| Control-Sample62_Beta | chr19:35645620-35645689 | FXYD5;FXYD5;FXYD5;FXYD5 | Island | TSS200;TSS1500;TSS200;TSS200 | Yes |
| Control-Sample1_Beta | chr11:117747940-117748187 | FXYD6;FXYD6;FXYD6;FXYD6;FXYD6 | S_Shore | 5'UTR;TSS200;TSS200;TSS200;TSS200 | Yes |
| Control-Sample1_Beta | chr11:117748236-117748512 | FXYD6;FXYD6;FXYD6;FXYD6;FXYD6 | S_Shore | TSS1500;TSS1500;TSS1500;TSS1500;TSS200 | Yes |
| Control-Sample10_Beta | chr11:117747940-117748512 | FXYD6;FXYD6;FXYD6;FXYD6;FXYD6 | S_Shore | 5'UTR;TSS200;TSS200;TSS200;TSS200 | Yes |
| Control-Sample216_Be | chr11:117747934-117748187 | FXYD6;FXYD6;FXYD6;FXYD6;FXYD6 | S_Shore | 5'UTR;TSS200;TSS200;TSS200;TSS200 | Yes |
| Control-Sample223_Be | chr11:117747940-117748236 | FXYD6;FXYD6;FXYD6;FXYD6;FXYD6 | S_Shore | 5'UTR;TSS200;TSS200;TSS200;TSS200 | Yes |
| Control-Sample4_Beta | chr11:117747940-117748169 | FXYD6;FXYD6;FXYD6;FXYD6;FXYD6 | S_Shore | 5'UTR;TSS200;TSS200;TSS200;TSS200 | Yes |
| Control-Sample45_Beta | chr11:117747940-117748486 | FXYD6;FXYD6;FXYD6;FXYD6;FXYD6 | S_Shore | 5'UTR;TSS200;TSS200;TSS200;TSS200 | Yes |
| Control-Sample151_Be | chr11:117748132-117748187 | FXYD6;FXYD6;FXYD6;FXYD6;FXYD6;FXYD6 | S_Shore | TSS1500;1stExon;5'UTR;TSS1500;TSS1500;TSS1500 | Yes |
| Control-Sample170_Be | chr11:117748132-117748189 | FXYD6;FXYD6;FXYD6;FXYD6;FXYD6;FXYD6 | S_Shore | TSS1500;1stExon;5'UTR;TSS1500;TSS1500;TSS1500 | Yes |
| Control-Sample171_Be | chr11:117748132-117748236 | FXYD6;FXYD6;FXYD6;FXYD6;FXYD6;FXYD6 | S_Shore | TSS1500;1stExon;5'UTR;TSS1500;TSS1500;TSS1500 | Yes |
| Control-Sample32_Beta | chr6:112021635-112021676 | FYN;FYN;FYN | | Body;Body;Body | No |
| Control-Sample12_Beta | chr7:90896329-90896701 | FZD1;FZD1 | Island | 1stExon;3'UTR | No |
| Control-Sample58_Beta | chr2:208635549-208635637 | FZD5 | S_Shore | TSS1500 | No |
| Control-Sample26_Beta | chr2:202901045-202901470 | FZD7 | Island | 1stExon | Yes |
| Control-Sample270_Be | chr2:202901045-202901428 | FZD7 | Island | 1stExon | Yes |
| Control-Sample223_Be | chr2:202901288-202901428 | FZD7;FZD7 | S_Shore | 1stExon;3'UTR | Yes |
| Control-Sample24_Beta | chr1:209848845-209848901 | G0S2;G0S2 | Island | 1stExon;5'UTR | No |
| Control-Sample10_Beta | chr2:169757561-169757780 | G6PC2;G6PC2 | | TSS200;TSS200 | No |
| Control-Sample1_Beta | chr6:29579492-29580367 | GABBR1;GABBR1;GABBR1 | | Body;Body;Body | Yes |
| Control-Sample1_Beta | chr6:29593479-29593913 | GABBR1;GABBR1;GABBR1 | N_Shore | Body;Body;Body | Yes |
| Control-Sample10_Beta | chr6:29580340-29580367 | GABBR1;GABBR1;GABBR1 | | Body;Body;Body | Yes |
| Control-Sample10_Beta | chr6:29581117-29581228 | GABBR1;GABBR1;GABBR1 | | Body;Body;Body | Yes |
| Control-Sample12_Beta | chr6:29589415-29589573 | GABBR1;GABBR1;GABBR1 | | Body;Body;Body | Yes |
| Control-Sample3_Beta | chr6:29576329-29576987 | GABBR1;GABBR1;GABBR1 | | Body;Body;Body | Yes |
| Control-Sample35_Beta | chr6:29593382-29593913 | GABBR1;GABBR1;GABBR1 | N_Shore | Body;Body;Body | Yes |
| Control-Sample4_Beta | chr6:29581054-29581117 | GABBR1;GABBR1;GABBR1 | | Body;Body;Body | Yes |
| Control-Sample1_Beta | chr1:1957073-1957385 | GABRD | Island | Body | No |
| Control-Sample1_Beta | chr1:68149989-68150400 | GADD45A | N_Shore | TSS1500 | No |
| Control-Sample3_Beta | chr4:899882-899984 | GAK | Island | Body | No |
| Control-Sample108_Be | chr2:38892846-38892901 | GALM | | TSS1500 | Yes |
| Control-Sample568_Be | chr2.0:38892846-38892901 | GALM | | TSS1500 | Yes |
| Control-Sample14_Beta | chr1:230272306-230272656 | GALNT2 | | Body | Yes |
| Control-Sample67_Beta | chr1:230415225-230415377 | GALNT2 | N_Shore | 3'UTR | Yes |
| Control-Sample108_Be | chr2:158114268-158114424 | GALNT5 | | TSS200 | No |
| Control-Sample108_Be | chr3:16216094-16216215 | GALNTL2 | | TSS200 | No |
| Control-Sample10_Beta | chr22:38218945-38219283 | GALR3 | N_Shore | TSS1500 | No |
| Control-Sample1_Beta | chr22:29707524-29708028 | GAS2L1;GAS2L1;GAS2L1 | Island | 3'UTR;Body;Body | Yes |
| Control-Sample57_Beta | chr22:29707524-29708246 | GAS2L1;GAS2L1;GAS2L1 | Island | 3'UTR;Body;Body | Yes |
| Control-Sample13_Beta | chr10:8100563-8101566 | GATA3;GATA3 | Island | Body;Body | Yes |
| Control-Sample34_Beta | chr10:8100761-8101566 | GATA3;GATA3 | Island | Body;Body | Yes |
| Control-Sample64_Beta | chr10:8101307-8101566 | GATA3;GATA3 | S_Shore | Body;Body | Yes |
| Control-Sample65_Beta | chr10:8101307-8101513 | GATA3;GATA3 | S_Shore | Body;Body | Yes |
| Control-Sample23_Beta | chr1:89664260-89664582 | GBP4 | | Body | Yes |
| Control-Sample24_Beta | chr1:89663896-89664582 | GBP4 | | Body | Yes |
| Control-Sample30_Beta | chr1:89664338-89664545 | GBP4 | | Body | Yes |
| Control-Sample47_Beta | chr1:89664338-89664407 | GBP4 | | Body | Yes |
| Control-Sample622_Be | chr1:89664340-89664545 | GBP4 | | Body | Yes |
| Control-Sample19_Beta | chr7:44229001-44229319 | GCK;GCK | | 1stExon;5'UTR | No |
| Control-Sample17_Beta | chr6:10881891-10881949 | GCM2 | Island | Body | No |
| Control-Sample1_Beta | chr6:10521452-10521715 | GCNT2 | | TSS200 | No |
| Control-Sample10_Beta | chr5:37835784-37836246 | GDNF;GDNF;GDNF | N_Shore | TSS1500;5'UTR;TSS200 | Yes |
| Control-Sample12_Beta | chr5:37835784-37836392 | GDNF;GDNF;GDNF | N_Shore | TSS1500;5'UTR;TSS200 | Yes |
| Control-Sample19_Beta | chr5:37835701-37836246 | GDNF;GDNF;GDNF | N_Shore | TSS1500;5'UTR;TSS200 | Yes |
| Control-Sample22_Beta | chr5:37833969-37834171 | GDNF;GDNF;GDNF | N_Shore | Body;Body;Body | Yes |
| Control-Sample67_Beta | chr5:37835784-37835958 | GDNF;GDNF;GDNF | N_Shore | TSS1500;5'UTR;TSS200 | Yes |
| Control-Sample1_Beta | chr5:37835506-37836246 | GDNF;GDNF;GDNF;GDNF | N_Shore | 5'UTR;TSS1500;5'UTR;1stExon | Yes |
| Control-Sample10_Beta | chr5:37835506-37835684 | GDNF;GDNF;GDNF;GDNF | N_Shore | 5'UTR;TSS1500;5'UTR;1stExon | Yes |
| Control-Sample13_Beta | chr5:37835506-37836392 | GDNF;GDNF;GDNF;GDNF | N_Shore | 5'UTR;TSS1500;5'UTR;1stExon | Yes |
| Control-Sample62_Beta | chr5:37835548-37836392 | GDNF;GDNF;GDNF;GDNF | N_Shore | 5'UTR;TSS1500;5'UTR;1stExon | Yes |
| Control-Sample63_Beta | chr5:37835548-37836246 | GDNF;GDNF;GDNF;GDNF | N_Shore | 5'UTR;TSS1500;5'UTR;1stExon | Yes |
| Control-Sample45_Beta | chr12:58003898-58003925 | GEFT;GEFT | Island | TSS1500;TSS200 | No |
| Control-Sample11_Beta | chr17:42992917-42993050 | GFAP;GFAP | S_Shelf | TSS200;TSS200 | Yes |
| Control-Sample6_Beta | chr17:42992872-42993050 | GFAP;GFAP;GFAP;GFAP | S_Shelf | 1stExon;5'UTR;1stExon;5'UTR | Yes |
| Control-Sample1_Beta | chr1:92952581-92953037 | GFI1;GFI1 | Island | TSS1500;TSS200 | Yes |
| Control-Sample19_Beta | chr1:92952581-92952702 | GFI1;GFI1 | Island | TSS1500;TSS200 | Yes |
| Control-Sample19_Beta | chr6:13428522-13428657 | GFOD1 | | Body | No |
| Control-Sample100_Be | chr5:179740914-179741120 | GFPT2 | Island | Body | Yes |
| Control-Sample150_Be | chr5:179740743-179741120 | GFPT2 | Island | Body | Yes |

| | | | | | |
|---|---|---|---|---|---|
| Control-Sample540_Be | chr5.0:179740914-179741120 | GFPT2 | Island | Body | Yes |
| Control-Sample1_Beta | chr20:3644097-3644192 | GFRA4;GFRA4 | S_Shore | TSS200;TSS200 | Yes |
| Control-Sample27_Beta | chr20:3644023-3644192 | GFRA4;GFRA4 | S_Shore | 1stExon;1stExon | Yes |
| Control-Sample1_Beta | chr7:30544915-30545074 | GGCT | Island | TSS1500 | Yes |
| Control-Sample11_Beta | chr7:30544915-30544943 | GGCT | Island | TSS1500 | Yes |
| Control-Sample11_Beta | chr19:38876666-38877220 | GGN | Island | Body | Yes |
| Control-Sample12_Beta | chr19:38876762-38877220 | GGN | Island | Body | Yes |
| Control-Sample48_Beta | chr22:24979660-24979964 | GGT1 | | TSS200 | Yes |
| Control-Sample20_Beta | chr22:24998974-24999188 | GGT1;GGT1;GGT1 | | TSS1500;TSS1500;5'UTR | Yes |
| Control-Sample17_Beta | chr20:33451272-33451427 | GGT7 | Island | Body | Yes |
| Control-Sample63_Beta | chr20:33451149-33451367 | GGT7 | Island | Body | Yes |
| Control-Sample1_Beta | chr7:150147927-150148025 | GIMAP8 | | TSS200 | Yes |
| Control-Sample13_Beta | chr7:150147927-150148073 | GIMAP8 | | TSS200 | Yes |
| Control-Sample19_Beta | chr7:150147553-150148073 | GIMAP8 | | TSS1500 | Yes |
| Control-Sample20_Beta | chr7:150147783-150148025 | GIMAP8 | | TSS200 | Yes |
| Control-Sample21_Beta | chr7:150147655-150148073 | GIMAP8 | | TSS1500 | Yes |
| Control-Sample22_Beta | chr7:150147553-150147922 | GIMAP8 | | TSS1500 | Yes |
| Control-Sample4_Beta | chr7:150147922-150148073 | GIMAP8 | | TSS200 | Yes |
| Control-Sample57_Beta | chr7:150147655-150148025 | GIMAP8 | | TSS1500 | Yes |
| Control-Sample65_Beta | chr7:150147553-150147783 | GIMAP8 | | TSS1500 | Yes |
| Control-Sample10_Beta | chr7:150147988-150148025 | GIMAP8;GIMAP8 | | 5'UTR;1stExon | Yes |
| Control-Sample11_Beta | chr7:150147988-150148073 | GIMAP8;GIMAP8 | | 5'UTR;1stExon | Yes |
| Control-Sample252_Be | chr19:14606894-14607597 | GIPC1;GIPC1;GIPC1;GIPC1;GIPC1;GIPC1;GIPC1;GIPC1;G | Island | 5'UTR;5'UTR;1stExon;1stExon;5'UTR;5'UTR;1stExon;5 | No |
| Control-Sample1_Beta | chr13:20736009-20736342 | GJA3 | Island | TSS1500 | Yes |
| Control-Sample10_Beta | chr13:20736075-20736342 | GJA3 | Island | TSS1500 | Yes |
| Control-Sample622_Be | chr13:20716432-20716728 | GJA3 | Island | Body | Yes |
| Control-Sample12_Beta | chr13:20768523-20768560 | GJB2 | S_Shore | TSS1500 | Yes |
| Control-Sample36_Beta | chr13:20768389-20768560 | GJB2 | S_Shore | TSS1500 | Yes |
| Control-Sample1_Beta | chr13:20805487-20805895 | GJB6;GJB6;GJB6;GJB6 | N_Shore | 5'UTR;5'UTR;5'UTR;TSS200 | Yes |
| Control-Sample11_Beta | chr13:20805385-20805443 | GJB6;GJB6;GJB6;GJB6 | N_Shore | 5'UTR;5'UTR;5'UTR;TSS200 | Yes |
| Control-Sample17_Beta | chr13:20805656-20805895 | GJB6;GJB6;GJB6;GJB6 | N_Shore | 5'UTR;5'UTR;5'UTR;TSS1500 | Yes |
| Control-Sample2_Beta | chr13:20805487-20805679 | GJB6;GJB6;GJB6;GJB6 | N_Shore | 5'UTR;5'UTR;5'UTR;TSS200 | Yes |
| Control-Sample1_Beta | chr13:20805196-20805443 | GJB6;GJB6;GJB6;GJB6;GJB6 | N_Shore | 5'UTR;5'UTR;5'UTR;5'UTR;1stExon | Yes |
| Control-Sample14_Beta | chr13:20805196-20805385 | GJB6;GJB6;GJB6;GJB6;GJB6 | N_Shore | 5'UTR;5'UTR;5'UTR;5'UTR;1stExon | Yes |
| Control-Sample17_Beta | chr13:20805196-20805391 | GJB6;GJB6;GJB6;GJB6;GJB6 | N_Shore | 5'UTR;5'UTR;5'UTR;5'UTR;1stExon | Yes |
| Control-Sample1_Beta | chr11:134147327-134147837 | GLB1L3 | S_Shore | Body | Yes |
| Control-Sample21_Beta | chr11:134147327-134147634 | GLB1L3 | S_Shore | Body | Yes |
| Control-Sample1_Beta | chr2:121554696-121554926 | GLI2 | | TSS200 | Yes |
| Control-Sample15_Beta | chr2:121554672-121554926 | GLI2 | | TSS200 | Yes |
| Control-Sample20_Beta | chr2:121684467-121684676 | GLI2 | | Body | Yes |
| Control-Sample52_Beta | chr2:121554810-121554926 | GLI2 | | TSS200 | Yes |
| Control-Sample1_Beta | chr7:42278067-42278096 | GLI3 | S_Shore | TSS1500 | Yes |
| Control-Sample14_Beta | chr7:42278067-42278089 | GLI3 | S_Shore | TSS1500 | Yes |
| Control-Sample19_Beta | chr7:42214276-42214361 | GLI3 | | Body | Yes |
| Control-Sample19_Beta | chr7:42267269-42267747 | GLI3 | N_Shore | 5'UTR | Yes |
| Control-Sample20_Beta | chr7:42267269-42267383 | GLI3 | N_Shore | 5'UTR | Yes |
| Control-Sample33_Beta | chr7:42267383-42267547 | GLI3 | N_Shore | 5'UTR | Yes |
| Control-Sample46_Beta | chr7:42267269-42267430 | GLI3 | N_Shore | 5'UTR | Yes |
| Control-Sample64_Beta | chr7:42267269-42267547 | GLI3 | N_Shore | 5'UTR | Yes |
| Control-Sample19_Beta | chr1:54194919-54195078 | GLIS1 | | 5'UTR | No |
| Control-Sample223_Be | chr16:4382027-4382081 | GLIS2 | S_Shelf | TSS200 | No |
| Control-Sample108_Be | chr12:104444006-104444136 | GLT8D2 | | TSS200 | No |
| Control-Sample57_Beta | chr1:182361509-182361557 | GLUL;GLUL;GLUL | Island | TSS200;TSS1500;TSS200 | No |
| Control-Sample13_Beta | chr6:1708566-1708713 | GMDS | | Body | No |
| Control-Sample464_Be | chr19:3097565-3097728 | GNA11 | Island | Body | No |
| Control-Sample1_Beta | chr7:2774474-2774654 | GNA12 | Island | Body | Yes |
| Control-Sample1_Beta | chr7:2775587-2775893 | GNA12 | S_Shore | Body | Yes |
| Control-Sample10_Beta | chr7:2775587-2775923 | GNA12 | S_Shore | Body | Yes |
| Control-Sample13_Beta | chr7:2775674-2775923 | GNA12 | S_Shore | Body | Yes |
| Control-Sample14_Beta | chr7:2774414-2774654 | GNA12 | N_Shore | Body | Yes |
| Control-Sample17_Beta | chr7:2842593-2842746 | GNA12 | | Body | Yes |
| Control-Sample19_Beta | chr7:2802697-2802976 | GNA12 | Island | Body | Yes |
| Control-Sample21_Beta | chr7:2802560-2802976 | GNA12 | Island | Body | Yes |
| Control-Sample22_Beta | chr7:2802560-2802942 | GNA12 | Island | Body | Yes |
| Control-Sample25_Beta | chr7:2802374-2802697 | GNA12 | N_Shore | Body | Yes |
| Control-Sample57_Beta | chr7:2802374-2802976 | GNA12 | N_Shore | Body | Yes |
| Control-Sample20_Beta | chr18:11752538-11752904 | GNAL;GNAL;GNAL | Island | 1stExon;Body;Body | No |
| Control-Sample1_Beta | chr12:6949114-6949472 | GNB3 | | TSS1500 | Yes |
| Control-Sample12_Beta | chr12:6949114-6949260 | GNB3 | | TSS1500 | Yes |
| Control-Sample58_Beta | chr12:6949119-6949472 | GNB3 | | TSS1500 | Yes |
| Control-Sample19_Beta | chr1:235814326-235814365 | GNG4;GNG4;GNG4 | Island | TSS1500;TSS1500;TSS1500 | No |
| Control-Sample10_Beta | chr19:2614086-2614177 | GNG7 | Island | 5'UTR | Yes |
| Control-Sample11_Beta | chr19:2650727-2650826 | GNG7 | Island | 5'UTR | Yes |
| Control-Sample12_Beta | chr19:2540907-2541104 | GNG7 | Island | 5'UTR | Yes |
| Control-Sample1_Beta | chr19:47139338-47139761 | GNG8 | Island | TSS1500 | Yes |
| Control-Sample10_Beta | chr19:47139391-47139761 | GNG8 | Island | TSS1500 | Yes |
| Control-Sample568_Be | chr6.0:30509585-30509614 | GNL1 | | 3'UTR | No |
| Control-Sample113_Be | chr6:42928495-42928500 | GNMT | Island | TSS200 | Yes |
| Control-Sample130_Be | chr6:42928475-42928497 | GNMT | Island | TSS200 | Yes |
| Control-Sample154_Be | chr6:42928475-42928495 | GNMT | Island | TSS200 | Yes |
| Control-Sample226_Be | chr6:42928475-42928500 | GNMT | Island | TSS200 | Yes |
| Control-Sample524_Be | chr6.0:42928495-42928500 | GNMT | Island | TSS200 | Yes |
| Control-Sample71_Beta | chr6:42928485-42928500 | GNMT | Island | TSS200 | Yes |
| Control-Sample1_Beta | chr12:133351397-133351746 | GOLGA3 | S_Shore | Body | Yes |
| Control-Sample11_Beta | chr12:133351324-133351746 | GOLGA3 | S_Shore | Body | Yes |
| Control-Sample19_Beta | chr12:133351630-133351746 | GOLGA3 | S_Shore | Body | Yes |
| Control-Sample22_Beta | chr8:110693195-110693289 | GOLSYN;GOLSYN;GOLSYN;GOLSYN;GOLSYN;GOLSYN | | 5'UTR;5'UTR;1stExon;1stExon;5'UTR;5'UTR | Yes |
| Control-Sample270_Be | chr8:110656096-110656245 | GOLSYN;GOLSYN;GOLSYN;GOLSYN;GOLSYN;GOLSYN;GC | N_Shore | Body;Body;Body;Body;5'UTR;5'UTR;Body;Body;Body; | Yes |
| Control-Sample15_Beta | chr17:28803808-28803927 | GOSR1;GOSR1;GOSR1 | N_Shore | TSS1500;TSS1500;TSS1500 | No |
| Control-Sample1_Beta | chr8:37797789-37797828 | GOT1L1 | | TSS200 | No |
| Control-Sample1_Beta | chr3:194117018-194117679 | GP5 | N_Shore | 3'UTR | Yes |

| | | | | | |
|---|---|---|---|---|---|
| Control-Sample10_Beta | chr3:194117018-194117403 | GP5 | N_Shore | 3'UTR | Yes |
| Control-Sample21_Beta | chr3:194117018-194117261 | GP5 | N_Shore | 3'UTR | Yes |
| Control-Sample22_Beta | chr10:113911312-113911441 | GPAM | | | No |
| Control-Sample1_Beta | chr11:64703192-64703510 | GPHA2 | | 5'UTR | Yes |
| Control-Sample24_Beta | chr11:64703192-64703415 | GPHA2 | | 5'UTR | Yes |
| Control-Sample13_Beta | chr11:64703312-64703510 | GPHA2;GPHA2 | | 1stExon;5'UTR | Yes |
| Control-Sample57_Beta | chr2:207083231-207083603 | GPR1;GPR1 | | TSS1500;TSS1500 | No |
| Control-Sample23_Beta | chr2:26541960-26542163 | GPR113;GPR113;GPR113;GPR113 | | 5'UTR;1stExon;TSS200;Body | No |
| Control-Sample12_Beta | chr6:46889624-46889760 | GPR116;GPR116;GPR116 | | 5'UTR;1stExon;5'UTR | No |
| Control-Sample1_Beta | chr12:131443383-131443486 | GPR133 | | Body | Yes |
| Control-Sample1_Beta | chr12:131452238-131452297 | GPR133 | | Body | Yes |
| Control-Sample1_Beta | chr12:131488390-131488726 | GPR133 | N_Shore | Body | Yes |
| Control-Sample10_Beta | chr12:131443231-131443486 | GPR133 | | Body | Yes |
| Control-Sample10_Beta | chr12:131488390-131488474 | GPR133 | N_Shore | Body | Yes |
| Control-Sample19_Beta | chr12:131505834-131506092 | GPR133 | | Body | Yes |
| Control-Sample39_Beta | chr12:131443245-131443486 | GPR133 | | Body | Yes |
| Control-Sample180_Be | chr9:127215714-127216047 | GPR144 | Island | Body | No |
| Control-Sample108_Be | chr5:145895523-145895884 | GPR151 | | 1stExon | No |
| Control-Sample1_Beta | chr12:6933322-6933548 | GPR162;GPR162 | Island | Body;Body | Yes |
| Control-Sample11_Beta | chr12:6933322-6933689 | GPR162;GPR162 | Island | Body;Body | Yes |
| Control-Sample1_Beta | chr1:200842890-200843333 | GPR25 | Island | 1stExon | No |
| Control-Sample171_Be | chr1:202092005-202092296 | GPR37L1 | | TSS200 | Yes |
| Control-Sample216_Be | chr1:202091995-202092285 | GPR37L1 | | TSS200 | Yes |
| Control-Sample270_Be | chr1:202091944-202092296 | GPR37L1 | | TSS200 | Yes |
| Control-Sample331_Be | chr1:202091944-202092285 | GPR37L1 | | TSS200 | Yes |
| Control-Sample5_Beta | chr1:202091865-202091880 | GPR37L1 | | TSS200 | Yes |
| Control-Sample12_Beta | chr14:91720372-91720696 | GPR68 | S_Shore | TSS200 | No |
| Control-Sample10_Beta | chr2:54086854-54087028 | GPR75;LOC100302652 | Island | 5'UTR;Body | Yes |
| Control-Sample11_Beta | chr2:54086854-54087209 | GPR75;LOC100302652 | Island | 5'UTR;Body | Yes |
| Control-Sample17_Beta | chr2:54086854-54087066 | GPR75;LOC100302652 | Island | 5'UTR;Body | Yes |
| Control-Sample8_Beta | chr2:54087028-54087209 | GPR75;LOC100302652;GPR75 | Island | 5'UTR;1stExon;1stExon | Yes |
| Control-Sample16_Beta | chr2:54087066-54087209 | GPR75;LOC100302652;LOC100302652;GPR75 | Island | 5'UTR;1stExon;5'UTR;1stExon | Yes |
| Control-Sample19_Beta | chr12:123215248-123215301 | GPR81 | | TSS200 | Yes |
| Control-Sample26_Beta | chr12:123215248-123215308 | GPR81 | | TSS200 | Yes |
| Control-Sample1_Beta | chr12:123215010-123215308 | GPR81;GPR81 | | 1stExon;5'UTR | Yes |
| Control-Sample2_Beta | chr12:123215010-123215301 | GPR81;GPR81 | | 1stExon;5'UTR | Yes |
| Control-Sample1_Beta | chr10:46993515-46993924 | GPRIN2 | S_Shore | TSS200 | Yes |
| Control-Sample67_Beta | chr10:46993515-46993586 | GPRIN2 | S_Shore | TSS200 | Yes |
| Control-Sample11_Beta | chr9:139236663-139236777 | GPSM1 | N_Shore | Body | Yes |
| Control-Sample464_Be | chr9:139228737-139228912 | GPSM1;GPSM1 | N_Shore | Body;Body | Yes |
| Control-Sample67_Beta | chr11:123396299-123396505 | GRAMD1B | | TSS1500 | No |
| Control-Sample1_Beta | chr7:50861592-50861927 | GRB10 | Island | TSS1500 | Yes |
| Control-Sample10_Beta | chr7:50861602-50861750 | GRB10 | Island | TSS1500 | Yes |
| Control-Sample11_Beta | chr7:50861592-50861834 | GRB10 | Island | TSS1500 | Yes |
| Control-Sample12_Beta | chr7:50861607-50861834 | GRB10 | Island | TSS1500 | Yes |
| Control-Sample14_Beta | chr7:50861592-50861607 | GRB10 | Island | TSS1500 | Yes |
| Control-Sample17_Beta | chr7:50861592-50861750 | GRB10 | Island | TSS1500 | Yes |
| Control-Sample18_Beta | chr7:50861607-50861750 | GRB10 | Island | TSS1500 | Yes |
| Control-Sample19_Beta | chr7:50861654-50861927 | GRB10 | S_Shore | TSS1500 | Yes |
| Control-Sample30_Beta | chr7:50861602-50861654 | GRB10 | Island | TSS1500 | Yes |
| Control-Sample43_Beta | chr7:50861602-50861834 | GRB10 | Island | TSS1500 | Yes |
| Control-Sample55_Beta | chr7:50861739-50861834 | GRB10 | S_Shore | TSS1500 | Yes |
| Control-Sample1_Beta | chr7:50772477-50772879 | GRB10;GRB10;GRB10;GRB10 | | 5'UTR;Body;Body;5'UTR | Yes |
| Control-Sample1_Beta | chr17:37893803-37894027 | GRB7;GRB7 | | TSS1500;TSS1500 | No |
| Control-Sample1_Beta | chr1:240776162-240776316 | GREM2 | | TSS1500 | Yes |
| Control-Sample26_Beta | chr1:240776162-240776211 | GREM2 | | TSS1500 | Yes |
| Control-Sample1_Beta | chr8:102506389-102506635 | GRHL2 | Island | Body | No |
| Control-Sample108_Be | chr1:24645397-24645805 | GRHL3 | N_Shelf | TSS1500 | No |
| Control-Sample194_Be | chr19:42503207-42503412 | GRIK5 | Island | Body | No |
| Control-Sample1_Beta | chr12:14109211-14109584 | GRIN2B | | 5'UTR | Yes |
| Control-Sample108_Be | chr12:14109514-14109584 | GRIN2B | | 5'UTR | Yes |
| Control-Sample15_Beta | chr17:72848535-72848918 | GRIN2C | Island | Body | No |
| Control-Sample1_Beta | chr19:48917816-48918205 | GRIN2D | N_Shore | Body | No |
| Control-Sample108_Be | chr12:67072769-67073022 | GRIP1;GRIP1 | | 1stExon;5'UTR | No |
| Control-Sample19_Beta | chr13:114321390-114321473 | GRK1 | | TSS1500 | No |
| Control-Sample194_Be | chr19:47507409-47507691 | GRLF1 | Island | 3'UTR | No |
| Control-Sample464_Be | chr6:34003640-34004360 | GRM4 | Island | Body | Yes |
| Control-Sample464_Be | chr6:34059640-34059787 | GRM4 | | Body | Yes |
| Control-Sample1_Beta | chr11:88241594-88241946 | GRM5;GRM5 | N_Shore | 3'UTR;3'UTR | Yes |
| Control-Sample15_Beta | chr11:88242253-88242604 | GRM5;GRM5 | Island | Body;Body | Yes |
| Control-Sample4_Beta | chr11:88241594-88242488 | GRM5;GRM5 | N_Shore | 3'UTR;3'UTR | Yes |
| Control-Sample5_Beta | chr11:88241594-88242253 | GRM5;GRM5 | N_Shore | 3'UTR;3'UTR | Yes |
| Control-Sample12_Beta | chr14:95237330-95237637 | GSC | N_Shore | TSS1500 | No |
| Control-Sample143_Be | chr8:144635430-144636025 | GSDMD | Island | TSS200 | Yes |
| Control-Sample318_Be | chr8:144635260-144635496 | GSDMD | N_Shore | TSS1500 | Yes |
| Control-Sample358_Be | chr8:144635361-144635478 | GSDMD | Island | TSS200 | Yes |
| Control-Sample519_Be | chr8.0:144635316-144635444 | GSDMD | Island | TSS1500 | Yes |
| Control-Sample65_Beta | chr4:106631906-106631934 | GSTCD;GSTCD | S_Shore | TSS200;5'UTR | No |
| Control-Sample134_Be | chr1:110230401-110230473 | GSTM1;GSTM1 | Island | TSS200;TSS200 | Yes |
| Control-Sample160_Be | chr1:110230252-110230456 | GSTM1;GSTM1 | Island | TSS200;TSS200 | Yes |
| Control-Sample276_Be | chr1:110254662-110254692 | GSTM5 | | TSS1500 | Yes |
| Control-Sample46_Beta | chr1:110254828-110255096 | GSTM5 | | TSS200 | Yes |
| Control-Sample474_Be | chr1:110254662-110254854 | GSTM5 | | TSS1500 | Yes |
| Control-Sample530_Be | chr1.0:110254662-110254709 | GSTM5 | | TSS1500 | Yes |
| Control-Sample1_Beta | chr1:110254896-110255096 | GSTM5;GSTM5 | | 1stExon;5'UTR | Yes |
| Control-Sample121_Be | chr22:24384397-24384525 | GSTT1 | Island | TSS200 | Yes |
| Control-Sample127_Be | chr22:24384400-24384944 | GSTT1 | Island | TSS200 | Yes |
| Control-Sample131_Be | chr22:24384105-24384944 | GSTT1 | N_Shore | Body | Yes |
| Control-Sample134_Be | chr22:24384393-24384944 | GSTT1 | Island | TSS200 | Yes |
| Control-Sample154_Be | chr22:24384159-24384294 | GSTT1 | Island | 1stExon | Yes |
| Control-Sample154_Be | chr22:24384340-24384944 | GSTT1 | Island | TSS200 | Yes |

| | | | | | |
|---|---|---|---|---|---|
| Control-Sample155_Be | chr22:24384294-24384944 | GSTT1 | Island | TSS200 | Yes |
| Control-Sample168_Be | chr22:24384135-24384340 | GSTT1 | Island | 1stExon | Yes |
| Control-Sample168_Be | chr22:24384397-24384944 | GSTT1 | Island | TSS200 | Yes |
| Control-Sample176_Be | chr22:24384105-24384294 | GSTT1 | N_Shore | Body | Yes |
| Control-Sample176_Be | chr22:24384397-24384573 | GSTT1 | Island | TSS200 | Yes |
| Control-Sample220_Be | chr22:24384159-24384944 | GSTT1 | Island | 1stExon | Yes |
| Control-Sample252_Be | chr22:24384525-24384944 | GSTT1 | S_Shore | TSS1500 | Yes |
| Control-Sample312_Be | chr22:24384315-24384944 | GSTT1 | Island | TSS200 | Yes |
| Control-Sample327_Be | chr22:24384294-24384340 | GSTT1 | Island | TSS200 | Yes |
| Control-Sample403_Be | chr22:24384393-24384573 | GSTT1 | Island | TSS200 | Yes |
| Control-Sample404_Be | chr22:24384105-24384340 | GSTT1 | N_Shore | Body | Yes |
| Control-Sample412_Be | chr22:24384400-24384573 | GSTT1 | Island | TSS200 | Yes |
| Control-Sample464_Be | chr22:24384105-24384525 | GSTT1 | N_Shore | Body | Yes |
| Control-Sample504_Be | chr22.0:24384340-24384525 | GSTT1 | Island | TSS200 | Yes |
| Control-Sample529_Be | chr22.0:24384105-24384944 | GSTT1 | N_Shore | Body | Yes |
| Control-Sample551_Be | chr22.0:24384105-24384525 | GSTT1 | N_Shore | Body | Yes |
| Control-Sample552_Be | chr22.0:24384294-24384944 | GSTT1 | Island | TSS200 | Yes |
| Control-Sample558_Be | chr22.0:24384397-24384525 | GSTT1 | Island | TSS200 | Yes |
| Control-Sample559_Be | chr22.0:24384159-24384294 | GSTT1 | Island | 1stExon | Yes |
| Control-Sample559_Be | chr22.0:24384340-24384944 | GSTT1 | Island | TSS200 | Yes |
| Control-Sample560_Be | chr22.0:24384397-24384944 | GSTT1 | Island | TSS200 | Yes |
| Control-Sample576_Be | chr22.0:24384315-24384944 | GSTT1 | Island | TSS200 | Yes |
| Control-Sample585_Be | chr22.0:24384525-24384944 | GSTT1 | S_Shore | TSS1500 | Yes |
| Control-Sample630_Be | chr22:24384105-24384159 | GSTT1 | N_Shore | Body | Yes |
| Control-Sample637_Be | chr22:24384135-24384294 | GSTT1 | Island | 1stExon | Yes |
| Control-Sample638_Be | chr22:24384105-24384573 | GSTT1 | N_Shore | Body | Yes |
| Control-Sample649_Be | chr22:24384105-24384315 | GSTT1 | N_Shore | Body | Yes |
| Control-Sample71_Beta | chr22:24384159-24384573 | GSTT1 | Island | 1stExon | Yes |
| Control-Sample106_Be | chr22:24384245-24384944 | GSTT1;GSTT1 | Island | 5'UTR;1stExon | Yes |
| Control-Sample127_Be | chr22:24384245-24384315 | GSTT1;GSTT1 | Island | 5'UTR;1stExon | Yes |
| Control-Sample335_Be | chr22:24384245-24384340 | GSTT1;GSTT1 | Island | 5'UTR;1stExon | Yes |
| Control-Sample465_Be | chr22:24384245-24384573 | GSTT1;GSTT1 | Island | 5'UTR;1stExon | Yes |
| Control-Sample505_Be | chr22.0:24384245-24384944 | GSTT1;GSTT1 | Island | 5'UTR;1stExon | Yes |
| Control-Sample121_Be | chr22:24348549-24348715 | GSTTP1 | | TSS1500 | Yes |
| Control-Sample505_Be | chr22.0:24348549-24348715 | GSTTP1 | | TSS1500 | Yes |
| Control-Sample11_Beta | chr17:7905636-7905928 | GUCY2D | N_Shore | TSS1500 | Yes |
| Control-Sample18_Beta | chr17:7905636-7905840 | GUCY2D | N_Shore | TSS1500 | Yes |
| Control-Sample1_Beta | chr2:127414802-127415063 | GYPC;GYPC | S_Shore | Body;Body | Yes |
| Control-Sample20_Beta | chr2:127413363-127413369 | GYPC;GYPC | N_Shore | TSS1500;TSS1500 | Yes |
| Control-Sample26_Beta | chr2:127414802-127414883 | GYPC;GYPC | S_Shore | Body;Body | Yes |
| Control-Sample1_Beta | chr22:38199759-38200221 | H1F0 | N_Shore | TSS1500 | No |
| Control-Sample1_Beta | chr2:43019854-43020013 | HAAO | Island | TSS200 | Yes |
| Control-Sample1_Beta | chr2:43020169-43020457 | HAAO | Island | TSS1500 | Yes |
| Control-Sample1_Beta | chr5:83016717-83017184 | HAPLN1;HAPLN1 | N_Shore | 1stExon;5'UTR | Yes |
| Control-Sample10_Beta | chr5:83016779-83017184 | HAPLN1;HAPLN1 | N_Shore | 1stExon;5'UTR | Yes |
| Control-Sample16_Beta | chr5:83016779-83017078 | HAPLN1;HAPLN1 | N_Shore | 1stExon;5'UTR | Yes |
| Control-Sample38_Beta | chr5:83016717-83017078 | HAPLN1;HAPLN1 | N_Shore | 1stExon;5'UTR | Yes |
| Control-Sample39_Beta | chr5:83016630-83017184 | HAPLN1;HAPLN1 | N_Shore | 1stExon;5'UTR | Yes |
| Control-Sample57_Beta | chr5:83016779-83017009 | HAPLN1;HAPLN1 | N_Shore | 1stExon;5'UTR | Yes |
| Control-Sample100_Be | chr11:1510644-1510747 | HCCA2 | | Body | Yes |
| Control-Sample19_Beta | chr11:1669926-1670305 | HCCA2 | N_Shelf | Body | Yes |
| Control-Sample34_Beta | chr11:1506936-1507119 | HCCA2 | | Body | Yes |
| Control-Sample1_Beta | chr6:31021134-31021796 | HCG22 | | TSS1500 | Yes |
| Control-Sample18_Beta | chr6:31021365-31021796 | HCG22 | | TSS1500 | Yes |
| Control-Sample19_Beta | chr6:31021275-31021422 | HCG22 | | TSS1500 | Yes |
| Control-Sample19_Beta | chr6:31025933-31025967 | HCG22 | | Body | Yes |
| Control-Sample57_Beta | chr6:31021275-31021796 | HCG22 | | TSS1500 | Yes |
| Control-Sample6_Beta | chr6:31021134-31021957 | HCG22 | | TSS1500 | Yes |
| Control-Sample65_Beta | chr6:31021134-31021422 | HCG22 | | TSS1500 | Yes |
| Control-Sample102_Be | chr6:31164916-31164924 | HCG27 | N_Shore | TSS1500 | Yes |
| Control-Sample29_Beta | chr6:31164914-31164947 | HCG27 | N_Shore | TSS1500 | Yes |
| Control-Sample30_Beta | chr6:31164914-31164924 | HCG27 | N_Shore | TSS1500 | Yes |
| Control-Sample6_Beta | chr6:29869838-29870060 | HCG2P7 | | Body | No |
| Control-Sample13_Beta | chr6:29894050-29894195 | HCG4P6 | N_Shore | TSS1500 | Yes |
| Control-Sample174_Be | chr6:29893168-29893605 | HCG4P6 | N_Shore | Body | Yes |
| Control-Sample19_Beta | chr6:29894152-29894197 | HCG4P6 | Island | TSS1500 | Yes |
| Control-Sample20_Beta | chr6:29894152-29894195 | HCG4P6 | Island | TSS1500 | Yes |
| Control-Sample217_Be | chr6:29894228-29894253 | HCG4P6 | Island | TSS1500 | Yes |
| Control-Sample551_Be | chr6.0:29894228-29894253 | HCG4P6 | Island | TSS1500 | Yes |
| Control-Sample84_Beta | chr6:29894813-29894831 | HCG4P6 | Island | TSS1500 | Yes |
| Control-Sample10_Beta | chr6:29942951-29943188 | HCG9 | N_Shore | Body | Yes |
| Control-Sample13_Beta | chr6:29943401-29943425 | HCG9 | N_Shore | Body | Yes |
| Control-Sample183_Be | chr6:29943401-29943480 | HCG9 | N_Shore | Body | Yes |
| Control-Sample214_Be | chr6:29943401-29943414 | HCG9 | N_Shore | Body | Yes |
| Control-Sample214_Be | chr6:29943455-29943677 | HCG9 | N_Shore | Body | Yes |
| Control-Sample226_Be | chr6:29943455-29943480 | HCG9 | N_Shore | Body | Yes |
| Control-Sample445_Be | chr6:29943401-29943455 | HCG9 | N_Shore | Body | Yes |
| Control-Sample52_Beta | chr6:29945409-29945726 | HCG9 | S_Shore | Body | Yes |
| Control-Sample1_Beta | chr19:595705-595933 | HCN2 | Island | Body | Yes |
| Control-Sample31_Beta | chr19:613111-613505 | HCN2 | Island | Body | Yes |
| Control-Sample32_Beta | chr19:613111-613818 | HCN2 | Island | Body | Yes |
| Control-Sample20_Beta | chr17:40337590-40337892 | HCRT | S_Shore | TSS200 | Yes |
| Control-Sample1_Beta | chr17:40337411-40337892 | HCRT;HCRT | S_Shore | 1stExon;5'UTR | Yes |
| Control-Sample3_Beta | chr1:32082963-32083031 | HCRTR1 | N_Shore | TSS1500 | No |
| Control-Sample21_Beta | chr3:13520256-13520771 | HDAC11 | N_Shore | TSS1500 | Yes |
| Control-Sample88_Beta | chr3:13520256-13520763 | HDAC11 | N_Shore | TSS1500 | Yes |
| Control-Sample1_Beta | chr3:13520685-13520771 | HDAC11;HDAC11 | N_Shore | TSS1500;TSS1500 | Yes |
| Control-Sample1_Beta | chr2:240010264-240010446 | HDAC4 | S_Shore | Body | Yes |
| Control-Sample10_Beta | chr2:240174613-240174659 | HDAC4 | | Body | Yes |
| Control-Sample10_Beta | chr2:240230587-240230892 | HDAC4 | | Body | Yes |
| Control-Sample10_Beta | chr2:240302585-240302855 | HDAC4 | | 5'UTR | Yes |

| | | | | | |
|---|---|---|---|---|---|
| Control-Sample108_Be | chr2:240143798-240143979 | HDAC4 | | Body | Yes |
| Control-Sample18_Beta | chr2:240302585-240302797 | HDAC4 | | 5'UTR | Yes |
| Control-Sample194_Be | chr2:240313541-240313668 | HDAC4 | | 5'UTR | Yes |
| Control-Sample32_Beta | chr2:240196769-240196996 | HDAC4 | Island | Body | Yes |
| Control-Sample5_Beta | chr2:240161619-240162323 | HDAC4 | | Body | Yes |
| Control-Sample54_Beta | chr2:240230663-240230892 | HDAC4 | | Body | Yes |
| Control-Sample57_Beta | chr2:240112249-240112403 | HDAC4 | Island | Body | Yes |
| Control-Sample572_Be | chr2.0:240313541-240313668 | HDAC4 | | 5'UTR | Yes |
| Control-Sample16_Beta | chr2:240117005-240117232 | HDAC4;MGC16025 | | Body;Body | Yes |
| Control-Sample108_Be | chr7:18535072-18535407 | HDAC9;HDAC9;HDAC9;HDAC9;HDAC9 | | TSS1500;TSS1500;TSS1500;TSS1500;TSS1500 | No |
| Control-Sample108_Be | chr15:50557873-50558051 | HDC;HDC | | 5'UTR;1stExon | No |
| Control-Sample57_Beta | chr7:788797-788989 | HEATR2 | N_Shelf | Body | No |
| Control-Sample16_Beta | chr7:43288635-43289504 | HECW1 | Island | Body | Yes |
| Control-Sample22_Beta | chr7:43288635-43288889 | HECW1 | Island | Body | Yes |
| Control-Sample45_Beta | chr7:43288635-43288771 | HECW1 | Island | Body | Yes |
| Control-Sample6_Beta | chr17:65242740-65243119 | HELZ | S_Shore | TSS1500 | No |
| Control-Sample1_Beta | chr11:124790815-124791197 | HEPACAM | Island | 3'UTR | Yes |
| Control-Sample3_Beta | chr11:124805538-124805963 | HEPACAM | | Body | Yes |
| Control-Sample6_Beta | chr11:124790815-124791389 | HEPACAM | Island | 3'UTR | Yes |
| Control-Sample512_Be | chr15.0:28567004-28567528 | HERC2 | N_Shore | 5'UTR | No |
| Control-Sample358_Be | chr15:23378429-23378571 | HERC2P2 | Island | TSS200 | No |
| Control-Sample10_Beta | chr1:40105664-40105687 | HEYL | Island | TSS1500 | Yes |
| Control-Sample2_Beta | chr1:40105304-40105687 | HEYL;HEYL | Island | 1stExon;5'UTR | Yes |
| Control-Sample51_Beta | chr1:40105304-40105667 | HEYL;HEYL | Island | 1stExon;5'UTR | Yes |
| Control-Sample496_Be | chr7:81399587-81399747 | HGF;HGF;HGF;HGF;HGF | | TSS200;TSS200;TSS200;TSS200;TSS200 | Yes |
| Control-Sample19_Beta | chr7:81399308-81399458 | HGF;HGF;HGF;HGF;HGF;HGF;HGF;HGF;HGF;HGF | | 5'UTR;1stExon;1stExon;1stExon;5'UTR;1stExon;5'UTR | Yes |
| Control-Sample100_Be | chr3:42742993-42743203 | HHATL;HHATL | | Body;5'UTR | Yes |
| Control-Sample19_Beta | chr3:42743077-42743794 | HHATL;HHATL | | 5'UTR;TSS200 | Yes |
| Control-Sample1_Beta | chr10:94451351-94452311 | HHEX | N_Shore | Body | Yes |
| Control-Sample11_Beta | chr10:94451351-94451625 | HHEX | N_Shore | Body | Yes |
| Control-Sample12_Beta | chr10:94451518-94451736 | HHEX | Island | Body | Yes |
| Control-Sample14_Beta | chr10:94451518-94452311 | HHEX | Island | Body | Yes |
| Control-Sample2_Beta | chr10:94450866-94451351 | HHEX | N_Shore | Body | Yes |
| Control-Sample2_Beta | chr10:94451625-94452311 | HHEX | Island | Body | Yes |
| Control-Sample25_Beta | chr10:94451351-94451736 | HHEX | N_Shore | Body | Yes |
| Control-Sample6_Beta | chr10:94450994-94451736 | HHEX | N_Shore | Body | Yes |
| Control-Sample1_Beta | chr17:1959066-1959124 | HIC1;HIC1 | Island | TSS1500;5'UTR | Yes |
| Control-Sample1_Beta | chr17:1961440-1962236 | HIC1;HIC1 | Island | Body;Body | Yes |
| Control-Sample11_Beta | chr17:1961778-1962236 | HIC1;HIC1 | Island | Body;Body | Yes |
| Control-Sample2_Beta | chr17:1961286-1961510 | HIC1;HIC1 | Island | Body;Body | Yes |
| Control-Sample6_Beta | chr17:1961286-1962236 | HIC1;HIC1 | Island | Body;Body | Yes |
| Control-Sample34_Beta | chr19:46807272-46807660 | HIF3A;HIF3A;HIF3A | Island | 5'UTR;Body;Body | No |
| Control-Sample68_Beta | chr12:123333035-123333379 | HIP1R | | Body | No |
| Control-Sample338_Be | chr6:26044274-26044460 | HIST1H2BB;HIST1H3C | Island | TSS1500;TSS1500 | No |
| Control-Sample1_Beta | chr6:27107670-27107756 | HIST1H2BK | S_Shore | 3'UTR | Yes |
| Control-Sample29_Beta | chr6:27107588-27107756 | HIST1H2BK | S_Shore | 3'UTR | Yes |
| Control-Sample1_Beta | chr6:26225767-26226256 | HIST1H3E | Island | 1stExon | Yes |
| Control-Sample10_Beta | chr6:26225539-26226256 | HIST1H3E | Island | 1stExon | Yes |
| Control-Sample30_Beta | chr1:42385581-42385941 | HIVEP3;HIVEP3 | S_Shore | TSS1500;TSS1500 | Yes |
| Control-Sample67_Beta | chr1:42384310-42384474 | HIVEP3;HIVEP3;HIVEP3;HIVEP3 | S_Shore | 1stExon;1stExon;5'UTR;5'UTR | Yes |
| Control-Sample1_Beta | chr10:70979811-70980067 | HKDC1 | | TSS1500 | Yes |
| Control-Sample25_Beta | chr10:70979811-70980112 | HKDC1 | | TSS1500 | Yes |
| Control-Sample54_Beta | chr10:70979827-70980112 | HKDC1 | | TSS1500 | Yes |
| Control-Sample58_Beta | chr10:70979777-70980067 | HKDC1 | | | Yes |
| Control-Sample1_Beta | chr6:29911004-29911095 | HLA-A | Island | Body | Yes |
| Control-Sample11_Beta | chr6:29911004-29911104 | HLA-A | Island | Body | Yes |
| Control-Sample12_Beta | chr6:29910796-29911104 | HLA-A | Island | Body | Yes |
| Control-Sample13_Beta | chr6:29910912-29911104 | HLA-A | Island | Body | Yes |
| Control-Sample149_Be | chr6:29911104-29911265 | HLA-A | Island | Body | Yes |
| Control-Sample15_Beta | chr6:29911036-29911104 | HLA-A | Island | Body | Yes |
| Control-Sample159_Be | chr6:29911036-29911095 | HLA-A | Island | Body | Yes |
| Control-Sample164_Be | chr6:29911091-29911104 | HLA-A | Island | Body | Yes |
| Control-Sample176_Be | chr6:29911087-29911095 | HLA-A | Island | Body | Yes |
| Control-Sample217_Be | chr6:29911095-29911265 | HLA-A | Island | Body | Yes |
| Control-Sample27_Beta | chr6:29911004-29911036 | HLA-A | Island | Body | Yes |
| Control-Sample30_Beta | chr6:29910912-29911095 | HLA-A | Island | Body | Yes |
| Control-Sample314_Be | chr6:29911011-29911036 | HLA-A | Island | Body | Yes |
| Control-Sample318_Be | chr6:29910776-29910796 | HLA-A | Island | Body | Yes |
| Control-Sample321_Be | chr6:29911011-29911104 | HLA-A | Island | Body | Yes |
| Control-Sample33_Beta | chr6:29910755-29911104 | HLA-A | Island | Body | Yes |
| Control-Sample4_Beta | chr6:29910912-29911036 | HLA-A | Island | Body | Yes |
| Control-Sample5_Beta | chr6:29910755-29911011 | HLA-A | Island | Body | Yes |
| Control-Sample50_Beta | chr6:29910796-29911095 | HLA-A | Island | Body | Yes |
| Control-Sample525_Be | chr6.0:29911091-29911104 | HLA-A | Island | Body | Yes |
| Control-Sample564_Be | chr6.0:29911011-29911036 | HLA-A | Island | Body | Yes |
| Control-Sample565_Be | chr6.0:29911036-29911095 | HLA-A | Island | Body | Yes |
| Control-Sample585_Be | chr6.0:29911087-29911095 | HLA-A | Island | Body | Yes |
| Control-Sample588_Be | chr6.0:29911036-29911104 | HLA-A | Island | Body | Yes |
| Control-Sample6_Beta | chr6:29911494-29911550 | HLA-A | S_Shore | Body | Yes |
| Control-Sample67_Beta | chr6:29911028-29911095 | HLA-A | Island | Body | Yes |
| Control-Sample73_Beta | chr6:29910912-29911011 | HLA-A | Island | Body | Yes |
| Control-Sample83_Beta | chr6:29910994-29911036 | HLA-A | Island | Body | Yes |
| Control-Sample9_Beta | chr6:29910994-29911104 | HLA-A | Island | Body | Yes |
| Control-Sample15_Beta | chr6:31324227-31324259 | HLA-B | Island | Body | Yes |
| Control-Sample155_Be | chr6:31323985-31324024 | HLA-B | Island | Body | Yes |
| Control-Sample27_Beta | chr6:31324565-31324627 | HLA-B | Island | Body | Yes |
| Control-Sample12_Beta | chr6:31324972-31325000 | HLA-B;HLA-B | Island | 5'UTR;1stExon | Yes |
| Control-Sample233_Be | chr6:31324941-31324972 | HLA-B;HLA-B | Island | 5'UTR;1stExon | Yes |
| Control-Sample1_Beta | chr6:31237013-31237034 | HLA-C | N_Shore | Body | Yes |
| Control-Sample1_Beta | chr6:31237405-31237722 | HLA-C | N_Shore | Body | Yes |
| Control-Sample119_Be | chr6:31239295-31239324 | HLA-C | Island | Body | Yes |

| | | | | | |
|---|---|---|---|---|---|
| Control-Sample139_Be | chr6:31237034-31237664 | HLA-C | N_Shore | Body | Yes |
| Control-Sample356_Be | chr6:31238788-31238908 | HLA-C | N_Shore | Body | Yes |
| Control-Sample509_Be | chr6.0:31237013-31237034 | HLA-C | N_Shore | Body | Yes |
| Control-Sample512_Be | chr6.0:31237405-31237722 | HLA-C | N_Shore | Body | Yes |
| Control-Sample638_Be | chr6:31239243-31239259 | HLA-C | Island | Body | Yes |
| Control-Sample96_Beta | chr6:31239063-31239093 | HLA-C | Island | Body | Yes |
| Control-Sample20_Beta | chr6:32908901-32909003 | HLA-DMB | | TSS200 | Yes |
| Control-Sample108_Be | chr6:32908794-32909003 | HLA-DMB;HLA-DMB | | 1stExon;5'UTR | Yes |
| Control-Sample29_Beta | chr6:32908605-32908718 | HLA-DMB;HLA-DMB | | 1stExon;5'UTR | Yes |
| Control-Sample1_Beta | chr6:32978271-32978379 | HLA-DOA | S_Shelf | TSS1500 | Yes |
| Control-Sample10_Beta | chr6:32977674-32978379 | HLA-DOA | S_Shore | TSS1500 | Yes |
| Control-Sample11_Beta | chr6:32978140-32978379 | HLA-DOA | S_Shelf | TSS1500 | Yes |
| Control-Sample24_Beta | chr6:32978196-32978379 | HLA-DOA | S_Shelf | TSS1500 | Yes |
| Control-Sample60_Beta | chr6:32978233-32978379 | HLA-DOA | S_Shelf | TSS1500 | Yes |
| Control-Sample65_Beta | chr6:32977983-32978548 | HLA-DOA | S_Shelf | TSS1500 | Yes |
| Control-Sample86_Beta | chr6:32978053-32978140 | HLA-DOA | S_Shelf | TSS1500 | Yes |
| Control-Sample334_Be | chr6:33037662-33038032 | HLA-DPA1 | | Body | Yes |
| Control-Sample334_Be | chr6:33039412-33039481 | HLA-DPA1 | | Body | Yes |
| Control-Sample380_Be | chr6:33037640-33037690 | HLA-DPA1 | | Body | Yes |
| Control-Sample44_Beta | chr6:33032830-33033312 | HLA-DPA1 | | 3'UTR | Yes |
| Control-Sample50_Beta | chr6:33037640-33038032 | HLA-DPA1 | | Body | Yes |
| Control-Sample42_Beta | chr6:33048732-33048760 | HLA-DPB1 | Island | Body | Yes |
| Control-Sample464_Be | chr6:33048540-33048558 | HLA-DPB1 | Island | Body | Yes |
| Control-Sample67_Beta | chr6:33048086-33048687 | HLA-DPB1 | N_Shore | Body | Yes |
| Control-Sample108_Be | chr6:32607174-32607509 | HLA-DQA1 | | Body | Yes |
| Control-Sample116_Be | chr6:32606845-32607174 | HLA-DQA1 | | Body | Yes |
| Control-Sample117_Be | chr6:32606845-32607224 | HLA-DQA1 | | Body | Yes |
| Control-Sample119_Be | chr6:32610735-32610971 | HLA-DQA1 | | 3'UTR | Yes |
| Control-Sample18_Beta | chr6:32606445-32606845 | HLA-DQA1 | | Body | Yes |
| Control-Sample193_Be | chr6:32605385-32605704 | HLA-DQA1 | | Body | Yes |
| Control-Sample318_Be | chr6:32606884-32607509 | HLA-DQA1 | | Body | Yes |
| Control-Sample324_Be | chr6:32606884-32607224 | HLA-DQA1 | | Body | Yes |
| Control-Sample388_Be | chr6:32608858-32609094 | HLA-DQA1 | | Body | Yes |
| Control-Sample466_Be | chr6:32606445-32607224 | HLA-DQA1 | | Body | Yes |
| Control-Sample514_Be | chr6.0:32606845-32607224 | HLA-DQA1 | | Body | Yes |
| Control-Sample525_Be | chr6.0:32604564-32605043 | HLA-DQA1 | | TSS1500 | Yes |
| Control-Sample564_Be | chr6.0:32606445-32607224 | HLA-DQA1 | | Body | Yes |
| Control-Sample10_Beta | chr6:32634276-32634344 | HLA-DQB1 | S_Shore | 1stExon | Yes |
| Control-Sample100_Be | chr6:32634245-32634362 | HLA-DQB1 | S_Shore | Body | Yes |
| Control-Sample104_Be | chr6:32632694-32632731 | HLA-DQB1 | Island | Body | Yes |
| Control-Sample109_Be | chr6:32632565-32632639 | HLA-DQB1 | Island | Body | Yes |
| Control-Sample116_Be | chr6:32634245-32634344 | HLA-DQB1 | S_Shore | Body | Yes |
| Control-Sample12_Beta | chr6:32632338-32632568 | HLA-DQB1 | Island | Body | Yes |
| Control-Sample139_Be | chr6:32632331-32632338 | HLA-DQB1 | Island | Body | Yes |
| Control-Sample151_Be | chr6:32632036-32632108 | HLA-DQB1 | N_Shore | Body | Yes |
| Control-Sample18_Beta | chr6:32632937-32632961 | HLA-DQB1 | Island | Body | Yes |
| Control-Sample193_Be | chr6:32634316-32634362 | HLA-DQB1 | S_Shore | 1stExon | Yes |
| Control-Sample257_Be | chr6:32632788-32632800 | HLA-DQB1 | Island | Body | Yes |
| Control-Sample257_Be | chr6:32633157-32633354 | HLA-DQB1 | S_Shore | Body | Yes |
| Control-Sample26_Beta | chr6:32632715-32632859 | HLA-DQB1 | Island | Body | Yes |
| Control-Sample318_Be | chr6:32632643-32632731 | HLA-DQB1 | Island | Body | Yes |
| Control-Sample33_Beta | chr6:32634344-32634481 | HLA-DQB1 | S_Shore | 1stExon | Yes |
| Control-Sample338_Be | chr6:32634245-32634316 | HLA-DQB1 | S_Shore | Body | Yes |
| Control-Sample34_Beta | chr6:32632565-32632643 | HLA-DQB1 | Island | Body | Yes |
| Control-Sample368_Be | chr6:32632000-32632108 | HLA-DQB1 | N_Shore | Body | Yes |
| Control-Sample44_Beta | chr6:32632788-32632848 | HLA-DQB1 | Island | Body | Yes |
| Control-Sample448_Be | chr6:32632731-32632790 | HLA-DQB1 | Island | Body | Yes |
| Control-Sample457_Be | chr6:32634245-32634481 | HLA-DQB1 | S_Shore | Body | Yes |
| Control-Sample571_Be | chr6.0:32634245-32634362 | HLA-DQB1 | S_Shore | Body | Yes |
| Control-Sample576_Be | chr6.0:32634245-32634344 | HLA-DQB1 | S_Shore | Body | Yes |
| Control-Sample89_Beta | chr6:32632568-32632643 | HLA-DQB1 | Island | Body | Yes |
| Control-Sample9_Beta | chr6:32634276-32634362 | HLA-DQB1 | S_Shore | 1stExon | Yes |
| Control-Sample1_Beta | chr6:32552350-32552515 | HLA-DRB1 | S_Shore | Body | Yes |
| Control-Sample100_Be | chr6:32552152-32552246 | HLA-DRB1 | Island | Body | Yes |
| Control-Sample109_Be | chr6:32551888-32551954 | HLA-DRB1 | Island | Body | Yes |
| Control-Sample11_Beta | chr6:32552350-32552453 | HLA-DRB1 | S_Shore | Body | Yes |
| Control-Sample116_Be | chr6:32552106-32552453 | HLA-DRB1 | Island | Body | Yes |
| Control-Sample116_Be | chr6:32557970-32558459 | HLA-DRB1 | | TSS1500 | Yes |
| Control-Sample145_Be | chr6:32552205-32552453 | HLA-DRB1 | Island | Body | Yes |
| Control-Sample148_Be | chr6:32552246-32552443 | HLA-DRB1 | Island | Body | Yes |
| Control-Sample150_Be | chr6:32552246-32552453 | HLA-DRB1 | Island | Body | Yes |
| Control-Sample152_Be | chr6:32549849-32550067 | HLA-DRB1 | N_Shelf | Body | Yes |
| Control-Sample157_Be | chr6:32552042-32552106 | HLA-DRB1 | Island | Body | Yes |
| Control-Sample157_Be | chr6:32552205-32552515 | HLA-DRB1 | Island | Body | Yes |
| Control-Sample16_Beta | chr6:32551888-32551949 | HLA-DRB1 | Island | Body | Yes |
| Control-Sample178_Be | chr6:32552205-32552350 | HLA-DRB1 | Island | Body | Yes |
| Control-Sample187_Be | chr6:32551891-32551954 | HLA-DRB1 | Island | Body | Yes |
| Control-Sample192_Be | chr6:32552042-32552152 | HLA-DRB1 | Island | Body | Yes |
| Control-Sample193_Be | chr6:32552095-32552152 | HLA-DRB1 | Island | Body | Yes |
| Control-Sample206_Be | chr6:32549361-32549849 | HLA-DRB1 | N_Shelf | Body | Yes |
| Control-Sample220_Be | chr6:32551949-32552026 | HLA-DRB1 | Island | Body | Yes |
| Control-Sample235_Be | chr6:32551949-32552016 | HLA-DRB1 | Island | Body | Yes |
| Control-Sample259_Be | chr6:32552016-32552095 | HLA-DRB1 | Island | Body | Yes |
| Control-Sample29_Beta | chr6:32552443-32552515 | HLA-DRB1 | S_Shore | Body | Yes |
| Control-Sample314_Be | chr6:32552026-32552042 | HLA-DRB1 | Island | Body | Yes |
| Control-Sample322_Be | chr6:32552106-32552246 | HLA-DRB1 | Island | Body | Yes |
| Control-Sample386_Be | chr6:32546626-32547167 | HLA-DRB1 | | 3'UTR | Yes |
| Control-Sample468_Be | chr6:32552039-32552095 | HLA-DRB1 | Island | Body | Yes |
| Control-Sample474_Be | chr6:32552016-32552026 | HLA-DRB1 | Island | Body | Yes |
| Control-Sample519_Be | chr6.0:32552152-32552246 | HLA-DRB1 | Island | Body | Yes |
| Control-Sample537_Be | chr6.0:32551888-32551949 | HLA-DRB1 | Island | Body | Yes |

| | | | | | |
|---|---|---|---|---|---|
| Control-Sample537_Be | chr6.0:32552443-32552515 | HLA-DRB1 | S_Shore | Body | Yes |
| Control-Sample572_Be | chr6.0:32552246-32552515 | HLA-DRB1 | Island | Body | Yes |
| Control-Sample573_Be | chr6.0:32552246-32552453 | HLA-DRB1 | Island | Body | Yes |
| Control-Sample599_Be | chr6.0:32549631-32549935 | HLA-DRB1 | N_Shelf | Body | Yes |
| Control-Sample609_Be | chr6:32551749-32551949 | HLA-DRB1 | N_Shore | Body | Yes |
| Control-Sample635_Be | chr6:32548627-32549849 | HLA-DRB1 | N_Shelf | Body | Yes |
| Control-Sample635_Be | chr6:32552246-32552515 | HLA-DRB1 | Island | Body | Yes |
| Control-Sample88_Beta | chr6:32549139-32549361 | HLA-DRB1 | N_Shelf | Body | Yes |
| Control-Sample88_Beta | chr6:32549631-32549935 | HLA-DRB1 | N_Shelf | Body | Yes |
| Control-Sample109_Be | chr6:32489750-32489963 | HLA-DRB5 | Island | Body | Yes |
| Control-Sample120_Be | chr6:32489963-32489991 | HLA-DRB5 | Island | Body | Yes |
| Control-Sample125_Be | chr6:32489991-32490043 | HLA-DRB5 | Island | Body | Yes |
| Control-Sample128_Be | chr6:32489963-32490012 | HLA-DRB5 | Island | Body | Yes |
| Control-Sample148_Be | chr6:32489801-32489991 | HLA-DRB5 | Island | Body | Yes |
| Control-Sample150_Be | chr6:32489801-32489984 | HLA-DRB5 | Island | Body | Yes |
| Control-Sample270_Be | chr6:32489963-32490043 | HLA-DRB5 | Island | Body | Yes |
| Control-Sample285_Be | chr6:32489750-32490012 | HLA-DRB5 | Island | Body | Yes |
| Control-Sample322_Be | chr6:32489203-32489555 | HLA-DRB5 | N_Shore | Body | Yes |
| Control-Sample36_Beta | chr6:32489750-32489991 | HLA-DRB5 | Island | Body | Yes |
| Control-Sample401_Be | chr6:32489555-32489801 | HLA-DRB5 | N_Shore | Body | Yes |
| Control-Sample519_Be | chr6.0:32489991-32490043 | HLA-DRB5 | Island | Body | Yes |
| Control-Sample520_Be | chr6.0:32489963-32489991 | HLA-DRB5 | Island | Body | Yes |
| Control-Sample599_Be | chr6.0:32489801-32489991 | HLA-DRB5 | Island | Body | Yes |
| Control-Sample1_Beta | chr6:32526260-32526364 | HLA-DRB6 | | Body | Yes |
| Control-Sample10_Beta | chr6:32525979-32526364 | HLA-DRB6 | | Body | Yes |
| Control-Sample103_Be | chr6:32526027-32526263 | HLA-DRB6 | | Body | Yes |
| Control-Sample11_Beta | chr6:32526263-32526364 | HLA-DRB6 | | Body | Yes |
| Control-Sample117_Be | chr6:32526021-32526260 | HLA-DRB6 | | Body | Yes |
| Control-Sample135_Be | chr6:32526027-32526364 | HLA-DRB6 | | Body | Yes |
| Control-Sample143_Be | chr6:32525960-32526021 | HLA-DRB6 | | Body | Yes |
| Control-Sample143_Be | chr6:32526260-32526342 | HLA-DRB6 | | Body | Yes |
| Control-Sample158_Be | chr6:32525979-32526027 | HLA-DRB6 | | Body | Yes |
| Control-Sample166_Be | chr6:32526027-32526342 | HLA-DRB6 | | Body | Yes |
| Control-Sample183_Be | chr6:32526021-32526263 | HLA-DRB6 | | Body | Yes |
| Control-Sample190_Be | chr6:32526260-32526366 | HLA-DRB6 | | Body | Yes |
| Control-Sample217_Be | chr6:32525960-32526260 | HLA-DRB6 | | Body | Yes |
| Control-Sample296_Be | chr6:32525979-32526260 | HLA-DRB6 | | Body | Yes |
| Control-Sample318_Be | chr6:32525805-32526021 | HLA-DRB6 | | Body | Yes |
| Control-Sample327_Be | chr6:32525960-32526364 | HLA-DRB6 | | Body | Yes |
| Control-Sample372_Be | chr6:32526021-32526364 | HLA-DRB6 | | Body | Yes |
| Control-Sample42_Beta | chr6:32526027-32526366 | HLA-DRB6 | | Body | Yes |
| Control-Sample487_Be | chr6:32526021-32526342 | HLA-DRB6 | | Body | Yes |
| Control-Sample504_Be | chr6.0:32526263-32526364 | HLA-DRB6 | | Body | Yes |
| Control-Sample511_Be | chr6.0:32525960-32526021 | HLA-DRB6 | | Body | Yes |
| Control-Sample511_Be | chr6.0:32526260-32526364 | HLA-DRB6 | | Body | Yes |
| Control-Sample543_Be | chr6.0:32526021-32526260 | HLA-DRB6 | | Body | Yes |
| Control-Sample551_Be | chr6.0:32526027-32526263 | HLA-DRB6 | | Body | Yes |
| Control-Sample554_Be | chr6.0:32526021-32526263 | HLA-DRB6 | | Body | Yes |
| Control-Sample615_Be | chr6:32525960-32526027 | HLA-DRB6 | | Body | Yes |
| Control-Sample9_Beta | chr6:32525805-32525979 | HLA-DRB6 | | Body | Yes |
| Control-Sample146_Be | chr6:29690766-29690893 | HLA-F;HLA-F;HLA-F | N_Shore | TSS1500;TSS1500;TSS1500 | Yes |
| Control-Sample520_Be | chr6.0:29690766-29690893 | HLA-F;HLA-F;HLA-F | N_Shore | TSS1500;TSS1500;TSS1500 | Yes |
| Control-Sample612_Be | chr6:29692000-29692011 | HLA-F;HLA-F;HLA-F | Island | Body;Body;Body | Yes |
| Control-Sample13_Beta | chr6:29795411-29795501 | HLA-G | N_Shore | 5'UTR | Yes |
| Control-Sample14_Beta | chr6:29795501-29795583 | HLA-G | N_Shore | 5'UTR | Yes |
| Control-Sample11_Beta | chr6:29856193-29856210 | HLA-H | Island | Body | Yes |
| Control-Sample12_Beta | chr6:29856141-29856210 | HLA-H | Island | Body | Yes |
| Control-Sample14_Beta | chr6:29855325-29855347 | HLA-H | Island | TSS200 | Yes |
| Control-Sample20_Beta | chr6:29856182-29856210 | HLA-H | Island | Body | Yes |
| Control-Sample217_Be | chr6:29854740-29855110 | HLA-H | N_Shore | TSS1500 | Yes |
| Control-Sample217_Be | chr6:29855636-29855914 | HLA-H | Island | Body | Yes |
| Control-Sample512_Be | chr6.0:29856193-29856210 | HLA-H | Island | Body | Yes |
| Control-Sample8_Beta | chr6:29855302-29855347 | HLA-H | Island | TSS200 | Yes |
| Control-Sample151_Be | chr6:29974858-29974868 | HLA-J;NCRNA00171 | Island | Body;Body | Yes |
| Control-Sample266_Be | chr6:29974844-29974868 | HLA-J;NCRNA00171 | Island | Body;Body | Yes |
| Control-Sample585_Be | chr6.0:29974844-29974886 | HLA-J;NCRNA00171 | Island | Body;Body | Yes |
| Control-Sample1_Beta | chr1:221053841-221055964 | HLX | N_Shore | Body | Yes |
| Control-Sample10_Beta | chr1:221057236-221058198 | HLX | N_Shore | Body | Yes |
| Control-Sample13_Beta | chr1:221057236-221057573 | HLX | N_Shore | Body | Yes |
| Control-Sample29_Beta | chr1:221053841-221054365 | HLX | N_Shore | Body | Yes |
| Control-Sample29_Beta | chr1:221054707-221055964 | HLX | Island | Body | Yes |
| Control-Sample33_Beta | chr1:221053841-221055790 | HLX | N_Shore | Body | Yes |
| Control-Sample53_Beta | chr1:221053904-221055790 | HLX | N_Shore | Body | Yes |
| Control-Sample6_Beta | chr1:221053904-221055964 | HLX | N_Shore | Body | Yes |
| Control-Sample66_Beta | chr1:221054201-221054365 | HLX | N_Shore | Body | Yes |
| Control-Sample10_Beta | chr6:34206491-34206683 | HMGA1;HMGA1;HMGA1;HMGA1;HMGA1;HMGA1 | S_Shore | 5'UTR;5'UTR;5'UTR;5'UTR;5'UTR;TSS200 | No |
| Control-Sample1_Beta | chr1:24152579-24152604 | HMGCL;HMGCL | | TSS1500;TSS1500 | No |
| Control-Sample20_Beta | chr19:1074727-1075104 | HMHA1 | Island | Body | Yes |
| Control-Sample70_Beta | chr19:1074926-1075104 | HMHA1 | Island | Body | Yes |
| Control-Sample1_Beta | chr16:1730561-1730594 | HN1L | S_Shore | Body | Yes |
| Control-Sample7_Beta | chr16:1742021-1742281 | HN1L | | Body | Yes |
| Control-Sample23_Beta | chr20:43029652-43029777 | HNF4A;HNF4A;HNF4A;HNF4A;HNF4A;HNF4A | | TSS1500;Body;TSS1500;Body;Body;TSS1500 | Yes |
| Control-Sample3_Beta | chr20:43029652-43029895 | HNF4A;HNF4A;HNF4A;HNF4A;HNF4A;HNF4A | | TSS1500;Body;TSS1500;Body;Body;TSS1500 | Yes |
| Control-Sample129_Be | chr19:12876846-12877188 | HOOK2;HOOK2 | N_Shore | Body;Body | Yes |
| Control-Sample139_Be | chr19:12876846-12877000 | HOOK2;HOOK2 | N_Shore | Body;Body | Yes |
| Control-Sample529_Be | chr19.0:12876846-12877188 | HOOK2;HOOK2 | N_Shore | Body;Body | Yes |
| Control-Sample108_Be | chr4:57547525-57548093 | HOPX;HOPX;HOPX;HOPX;HOPX;HOPX | | 5'UTR;1stExon;1stExon;5'UTR;5'UTR;1stExon | Yes |
| Control-Sample291_Be | chr4:57547525-57547872 | HOPX;HOPX;HOPX;HOPX;HOPX;HOPX | | 5'UTR;1stExon;1stExon;5'UTR;5'UTR;1stExon | Yes |
| Control-Sample140_Be | chr22:30476089-30476281 | HORMAD2 | Island | TSS1500 | Yes |
| Control-Sample140_Be | chr22:30476345-30476525 | HORMAD2 | Island | TSS200 | Yes |
| Control-Sample15_Beta | chr7:27159883-27160586 | HOXA3;HOXA3 | N_Shelf | 5'UTR;TSS1500 | Yes |

| | | | | | |
|---|---|---|---|---|---|
| Control-Sample46_Beta | chr7:27160520-27160960 | HOXA3;HOXA3 | N_Shore | 5'UTR;TSS1500 | Yes |
| Control-Sample64_Beta | chr7:27160276-27160960 | HOXA3;HOXA3 | N_Shore | 5'UTR;TSS1500 | Yes |
| Control-Sample67_Beta | chr7:27146237-27146445 | HOXA3;HOXA3;HOXA3 | Island | 3'UTR;3'UTR;3'UTR | Yes |
| Control-Sample67_Beta | chr7:27154262-27154387 | HOXA3;HOXA3;HOXA3 | N_Shore | 5'UTR;TSS1500;5'UTR | Yes |
| Control-Sample67_Beta | chr7:27154911-27155234 | HOXA3;HOXA3;HOXA3 | N_Shore | 5'UTR;TSS1500;5'UTR | Yes |
| Control-Sample67_Beta | chr7:27153580-27153847 | HOXA3;HOXA3;HOXA3;HOXA3 | Island | 5'UTR;1stExon;5'UTR;5'UTR | Yes |
| Control-Sample67_Beta | chr7:27170394-27170554 | HOXA4;HOXA4 | Island | 5'UTR;1stExon | No |
| Control-Sample67_Beta | chr17:46607828-46608293 | HOXB1 | Island | 1stExon | No |
| Control-Sample10_Beta | chr17:46648525-46648582 | HOXB3 | | 5'UTR | Yes |
| Control-Sample67_Beta | chr17:46628663-46628850 | HOXB3 | N_Shore | Body | Yes |
| Control-Sample108_Be | chr17:46651360-46651745 | HOXB3;HOXB3 | N_Shelf | 5'UTR;1stExon | Yes |
| Control-Sample13_Beta | chr17:46669455-46669489 | HOXB5;LOC404266;LOC404266;LOC404266;LOC404266 | Island | 3'UTR;Body;TSS1500;TSS200;Body | Yes |
| Control-Sample20_Beta | chr17:46669455-46669492 | HOXB5;LOC404266;LOC404266;LOC404266;LOC404266 | Island | 3'UTR;Body;TSS1500;TSS200;Body | Yes |
| Control-Sample1_Beta | chr2:10566989-10567843 | HPCAL1;HPCAL1 | Island | 3'UTR;3'UTR | Yes |
| Control-Sample19_Beta | chr2:10566553-10567843 | HPCAL1;HPCAL1 | N_Shore | Body;Body | Yes |
| Control-Sample21_Beta | chr2:10566989-10567750 | HPCAL1;HPCAL1 | Island | 3'UTR;3'UTR | Yes |
| Control-Sample48_Beta | chr2:10567273-10567843 | HPCAL1;HPCAL1 | S_Shore | 3'UTR;3'UTR | Yes |
| Control-Sample58_Beta | chr2:10567273-10567750 | HPCAL1;HPCAL1 | S_Shore | 3'UTR;3'UTR | Yes |
| Control-Sample238_Be | chr1:45792660-45792714 | HPDL;HPDL | Island | 1stExon;5'UTR | No |
| Control-Sample13_Beta | chr19:35531403-35531417 | HPN;HPN | N_Shore | TSS200;TSS200 | No |
| Control-Sample108_Be | chr22:26875552-26875652 | HPS4;HPS4 | N_Shelf | TSS200;Body | Yes |
| Control-Sample12_Beta | chr22:26875652-26875823 | HPS4;HPS4 | N_Shelf | TSS200;Body | Yes |
| Control-Sample13_Beta | chr22:26875652-26876075 | HPS4;HPS4 | N_Shelf | TSS200;Body | Yes |
| Control-Sample46_Beta | chr22:26875806-26876075 | HPS4;HPS4 | N_Shelf | TSS1500;Body | Yes |
| Control-Sample20_Beta | chr22:26875499-26876075 | HPS4;HPS4;HPS4 | N_Shelf | 1stExon;5'UTR;Body | Yes |
| Control-Sample1_Beta | chr19:49658797-49659299 | HRC | N_Shelf | TSS200 | No |
| Control-Sample10_Beta | chr5:175108429-175109114 | HRH2;HRH2 | | 5'UTR;TSS200 | Yes |
| Control-Sample108_Be | chr5:175108269-175108315 | HRH2;HRH2 | | 5'UTR;TSS200 | Yes |
| Control-Sample11_Beta | chr17:14213447-14213642 | HS3ST3B1 | S_Shore | Body | Yes |
| Control-Sample3_Beta | chr17:14213447-14213798 | HS3ST3B1 | S_Shore | Body | Yes |
| Control-Sample35_Beta | chr17:14213579-14213798 | HS3ST3B1 | S_Shore | Body | Yes |
| Control-Sample1_Beta | chr17:14207927-14208363 | HS3ST3B1;MGC12916 | S_Shore | Body;Body | Yes |
| Control-Sample11_Beta | chr17:14207701-14208363 | HS3ST3B1;MGC12916 | Island | Body;Body | Yes |
| Control-Sample15_Beta | chr17:14207701-14207968 | HS3ST3B1;MGC12916 | Island | Body;Body | Yes |
| Control-Sample2_Beta | chr17:14207701-14207927 | HS3ST3B1;MGC12916 | Island | Body;Body | Yes |
| Control-Sample35_Beta | chr17:14207890-14208363 | HS3ST3B1;MGC12916 | S_Shore | Body;Body | Yes |
| Control-Sample40_Beta | chr17:14207890-14207968 | HS3ST3B1;MGC12916 | S_Shore | Body;Body | Yes |
| Control-Sample101_Be | chr10:38645346-38645378 | HSD17B7P2 | Island | Body | Yes |
| Control-Sample106_Be | chr10:38645346-38645740 | HSD17B7P2 | Island | Body | Yes |
| Control-Sample125_Be | chr10:38645113-38645346 | HSD17B7P2 | Island | TSS200 | Yes |
| Control-Sample125_Be | chr10:38645378-38645740 | HSD17B7P2 | Island | Body | Yes |
| Control-Sample126_Be | chr10:38644991-38645740 | HSD17B7P2 | N_Shore | TSS1500 | Yes |
| Control-Sample36_Beta | chr10:38645113-38645740 | HSD17B7P2 | Island | TSS200 | Yes |
| Control-Sample550_Be | chr10.0:38645346-38645378 | HSD17B7P2 | Island | Body | Yes |
| Control-Sample568_Be | chr10.0:38644991-38645378 | HSD17B7P2 | N_Shore | TSS1500 | Yes |
| Control-Sample19_Beta | chr6:33173278-33173333 | HSD17B8 | S_Shore | Body | No |
| Control-Sample225_Be | chr14:102554826-102554977 | HSP90AA1;HSP90AA1 | S_Shore | TSS1500;Body | No |
| Control-Sample38_Beta | chr6:44217559-44217925 | HSP90AB1 | S_Shore | Body | No |
| Control-Sample324_Be | chr1:161575766-161576629 | HSPA7 | Island | TSS200 | Yes |
| Control-Sample8_Beta | chr1:161575992-161576629 | HSPA7 | Island | Body | Yes |
| Control-Sample19_Beta | chr5:53751272-53751919 | HSPB3 | | TSS200 | Yes |
| Control-Sample20_Beta | chr5:53751272-53751493 | HSPB3 | | TSS200 | Yes |
| Control-Sample1_Beta | chr19:36246402-36246408 | HSPB6 | Island | 3'UTR | Yes |
| Control-Sample12_Beta | chr19:36246395-36246408 | HSPB6 | Island | 3'UTR | Yes |
| Control-Sample5_Beta | chr19:36246402-36246906 | HSPB6 | Island | 3'UTR | Yes |
| Control-Sample10_Beta | chr17:40274703-40274811 | HSPB9;KAT2A | Island | TSS200;TSS1500 | Yes |
| Control-Sample11_Beta | chr17:40274670-40274811 | HSPB9;KAT2A | Island | TSS200;TSS1500 | Yes |
| Control-Sample19_Beta | chr17:40274722-40274811 | HSPB9;KAT2A | Island | TSS200;TSS1500 | Yes |
| Control-Sample68_Beta | chr17:40274722-40274740 | HSPB9;KAT2A | Island | TSS200;TSS1500 | Yes |
| Control-Sample12_Beta | chr16:73126120-73126795 | HTA | | TSS200 | Yes |
| Control-Sample44_Beta | chr16:73125573-73126120 | HTA | | TSS1500 | Yes |
| Control-Sample20_Beta | chr11:20385301-20385389 | HTATIP2;HTATIP2;HTATIP2;HTATIP2;HTATIP2;HTATIP2;HTA | Island | 1stExon;5'UTR;TSS1500;1stExon;5'UTR;5'UTR;TSS20 | No |
| Control-Sample19_Beta | chr1:23521272-23521417 | HTR1D | | TSS200 | No |
| Control-Sample1_Beta | chr11:113845487-113846017 | HTR3A;HTR3A | | TSS1500;TSS1500 | No |
| Control-Sample1_Beta | chr1:19992167-19992564 | HTR6;HTR6 | Island | 5'UTR;1stExon | No |
| Control-Sample19_Beta | chr10:124220367-124220504 | HTRA1 | Island | TSS1500 | Yes |
| Control-Sample92_Beta | chr10:124266269-124266386 | HTRA1 | | Body | Yes |
| Control-Sample1_Beta | chr8:38831508-38832728 | HTRA4 | N_Shore | TSS200 | Yes |
| Control-Sample67_Beta | chr8:38832267-38832728 | HTRA4 | Island | Body | Yes |
| Control-Sample11_Beta | chr4:3239305-3239461 | HTT | | Body | No |
| Control-Sample11_Beta | chr3:50341009-50341183 | HYAL1;HYAL1;HYAL1;HYAL1;HYAL1;HYAL1 | | TSS200;TSS200;5'UTR;TSS200;TSS200;TSS200 | No |
| Control-Sample249_Be | chr6:144328421-144329829 | HYMAI;PLAGL1;PLAGL1;PLAGL1;PLAGL1;PLAGL1;PLAGL1; | N_Shore | Body;5'UTR;5'UTR;5'UTR;5'UTR;5'UTR;5'UTR;5'UTR;5' | Yes |
| Control-Sample53_Beta | chr19:10397612-10397780 | ICAM4;ICAM4;ICAM4 | Island | TSS200;TSS200;TSS200 | No |
| Control-Sample1_Beta | chr1:23884847-23885086 | ID3 | Island | 3'UTR | No |
| Control-Sample3_Beta | chr4:996052-996280 | IDUA | Island | Body | No |
| Control-Sample19_Beta | chr6:30711090-30711164 | IER3;FLOT1 | Island | 3'UTR;TSS1500 | Yes |
| Control-Sample19_Beta | chr6:30711580-30711636 | IER3;FLOT1 | Island | 3'UTR;TSS1500 | Yes |
| Control-Sample20_Beta | chr6:30711561-30711586 | IER3;FLOT1 | Island | 3'UTR;TSS1500 | Yes |
| Control-Sample270_Be | chr6:30711141-30711399 | IER3;FLOT1 | Island | 3'UTR;TSS1500 | Yes |
| Control-Sample1_Beta | chr12:6658378-6658625 | IFFO1;IFFO1;IFFO1 | S_Shore | Body;TSS200;Body | Yes |
| Control-Sample4_Beta | chr12:6649733-6649994 | IFFO1;IFFO1;IFFO1 | Island | Body;Body;Body | Yes |
| Control-Sample12_Beta | chr12:6658164-6658625 | IFFO1;IFFO1;IFFO1;IFFO1 | S_Shore | Body;1stExon;Body;5'UTR | Yes |
| Control-Sample50_Beta | chr12:6658164-6658557 | IFFO1;IFFO1;IFFO1;IFFO1 | S_Shore | Body;1stExon;Body;5'UTR | Yes |
| Control-Sample19_Beta | chr1:158979538-158979841 | IFI16 | | TSS200 | No |
| Control-Sample7_Beta | chr10:91152113-91152447 | IFIT1 | | TSS1500 | No |
| Control-Sample10_Beta | chr10:91061294-91061728 | IFIT2 | | TSS1500 | Yes |
| Control-Sample108_Be | chr10:91061115-91061728 | IFIT2 | | | Yes |
| Control-Sample1_Beta | chr11:313624-314074 | IFITM1 | N_Shelf | TSS1500 | No |
| Control-Sample2_Beta | chr11:300286-300490 | IFITM5 | S_Shore | TSS1500 | No |
| Control-Sample108_Be | chr16:1583883-1583984 | IFT140;TMEM204 | Island | Body;TSS1500 | No |
| Control-Sample41_Beta | chr15:65654017-65654217 | IGDCC3 | N_Shore | Body | No |

| | | | | | |
|---|---|---|---|---|---|
| Control-Sample1_Beta | chr15:65688530-65689325 | IGDCC4 | Island | Body | Yes |
| Control-Sample11_Beta | chr15:65689263-65689325 | IGDCC4 | Island | Body | Yes |
| Control-Sample14_Beta | chr15:65688530-65689433 | IGDCC4 | Island | Body | Yes |
| Control-Sample2_Beta | chr15:65689058-65689325 | IGDCC4 | N_Shore | Body | Yes |
| Control-Sample23_Beta | chr15:99409194-99409506 | IGF1R | | Body | Yes |
| Control-Sample6_Beta | chr15:99409942-99409957 | IGF1R | | Body | Yes |
| Control-Sample56_Beta | chr11:2162438-2162475 | IGF2AS;INS-IGF2;IGF2;IGF2AS;IGF2 | Island | Body;Body;5'UTR;Body;TSS200 | No |
| Control-Sample64_Beta | chr7:23387383-23387862 | IGF2BP3 | | Body | No |
| Control-Sample10_Beta | chr2:217500594-217501125 | IGFBP2 | S_Shore | Body | No |
| Control-Sample1_Beta | chr12:53490352-53490632 | IGFBP6 | N_Shore | TSS1500 | No |
| Control-Sample10_Beta | chr19:46544240-46544711 | IGFL4;IGFL4 | | 1stExon;5'UTR | No |
| Control-Sample10_Beta | chr19:51830477-51830960 | IGLON5 | Island | Body | No |
| Control-Sample23_Beta | chr1:18617630-18617791 | IGSF21 | | Body | No |
| Control-Sample1_Beta | chr12:56414442-56414508 | IKZF4 | | TSS1500 | No |
| Control-Sample3_Beta | chr1:67772972-67773020 | IL12RB2 | N_Shore | TSS200 | No |
| Control-Sample464_Be | chr15:81489075-81489338 | IL16 | | TSS200 | No |
| Control-Sample1_Beta | chr13:21291173-21291467 | IL17D | | Body | Yes |
| Control-Sample19_Beta | chr13:21295590-21295941 | IL17D | N_Shore | Body | Yes |
| Control-Sample26_Beta | chr13:21295590-21296301 | IL17D | N_Shore | Body | Yes |
| Control-Sample64_Beta | chr13:21295941-21296465 | IL17D | Island | Body | Yes |
| Control-Sample312_Be | chr22:17564750-17565149 | IL17RA | Island | TSS1500 | No |
| Control-Sample35_Beta | chr3:9944183-9944537 | IL17RE;IL17RE;IL17RE | | TSS200;TSS1500;TSS1500 | No |
| Control-Sample1_Beta | chr11:71710614-71710982 | IL18BP;IL18BP;IL18BP;IL18BP;IL18BP;IL18BP | | TSS200;5'UTR;TSS1500;5'UTR;5'UTR;1stExon | Yes |
| Control-Sample61_Beta | chr11:71710643-71710982 | IL18BP;IL18BP;IL18BP;IL18BP;IL18BP;IL18BP | | TSS200;5'UTR;TSS1500;5'UTR;5'UTR;1stExon | Yes |
| Control-Sample10_Beta | chr2:113875226-113875292 | IL1RN;IL1RN;IL1RN | | TSS1500;TSS1500;TSS1500 | No |
| Control-Sample1_Beta | chr12:56732615-56732723 | IL23A | | TSS200 | No |
| Control-Sample20_Beta | chr7:22766657-22766992 | IL6 | | TSS200 | No |
| Control-Sample3_Beta | chr1:166890293-166890429 | ILDR2 | Island | Body | Yes |
| Control-Sample65_Beta | chr1:166890293-166890436 | ILDR2 | Island | Body | Yes |
| Control-Sample19_Beta | chr14:105184057-105184176 | INF2;INF2 | N_Shelf | Body;3'UTR | Yes |
| Control-Sample18_Beta | chr14:105167574-105167820 | INF2;INF2;INF2 | N_Shore | 5'UTR;5'UTR;5'UTR | Yes |
| Control-Sample39_Beta | chr14:105166740-105168128 | INF2;INF2;INF2 | N_Shore | 5'UTR;5'UTR;5'UTR | Yes |
| Control-Sample40_Beta | chr14:105167300-105168128 | INF2;INF2;INF2 | Island | 5'UTR;5'UTR;5'UTR | Yes |
| Control-Sample59_Beta | chr14:105167300-105167820 | INF2;INF2;INF2 | Island | 5'UTR;5'UTR;5'UTR | Yes |
| Control-Sample61_Beta | chr14:105167457-105167820 | INF2;INF2;INF2 | Island | 5'UTR;5'UTR;5'UTR | Yes |
| Control-Sample9_Beta | chr14:105167574-105167727 | INF2;INF2;INF2 | N_Shore | 5'UTR;5'UTR;5'UTR | Yes |
| Control-Sample57_Beta | chr2:121107846-121108133 | INHBB | S_Shore | 3'UTR | No |
| Control-Sample10_Beta | chr2:191208116-191208232 | INPP1;INPP1 | N_Shore | TSS200;TSS200 | No |
| Control-Sample1_Beta | chr10:134420183-134420300 | INPP5A | | Body | Yes |
| Control-Sample1_Beta | chr10:134556463-134556992 | INPP5A | S_Shore | Body | Yes |
| Control-Sample10_Beta | chr10:134491287-134491509 | INPP5A | | Body | Yes |
| Control-Sample11_Beta | chr10:134556463-134556979 | INPP5A | S_Shore | Body | Yes |
| Control-Sample121_Be | chr10:134491331-134491593 | INPP5A | | Body | Yes |
| Control-Sample13_Beta | chr10:134536157-134536498 | INPP5A | Island | Body | Yes |
| Control-Sample19_Beta | chr10:134420183-134420623 | INPP5A | | Body | Yes |
| Control-Sample19_Beta | chr10:134556463-134556533 | INPP5A | S_Shore | Body | Yes |
| Control-Sample223_Be | chr10:134596943-134597358 | INPP5A | N_Shore | 3'UTR | Yes |
| Control-Sample40_Beta | chr10:134491287-134491593 | INPP5A | | Body | Yes |
| Control-Sample69_Beta | chr10:134498925-134499266 | INPP5A | | Body | Yes |
| Control-Sample35_Beta | chr10:121577971-121578561 | INPP5F;INPP5F | Island | Body;TSS1500 | No |
| Control-Sample568_Be | chr4.0:128554125-128554168 | INTU;INTU | | 5'UTR;1stExon | No |
| Control-Sample223_Be | chr2:237415899-237416370 | IQCA1 | N_Shore | 1stExon | No |
| Control-Sample19_Beta | chr7:2647927-2648785 | IQCE;IQCE | S_Shelf | Body;Body | Yes |
| Control-Sample20_Beta | chr7:2648222-2648343 | IQCE;IQCE | S_Shelf | Body;Body | Yes |
| Control-Sample331_Be | chr7:2646478-2646955 | IQCE;IQCE | S_Shore | Body;Body | Yes |
| Control-Sample65_Beta | chr7:2606716-2606824 | IQCE;IQCE | | Body;Body | Yes |
| Control-Sample10_Beta | chr3:197663828-197664106 | IQCG;IQCG | | Body;Body | Yes |
| Control-Sample13_Beta | chr3:197663828-197663873 | IQCG;IQCG | | Body;Body | Yes |
| Control-Sample2_Beta | chr3:197663860-197664106 | IQCG;IQCG | | Body;Body | Yes |
| Control-Sample1_Beta | chr3:13060915-13060974 | IQSEC1 | | Body | Yes |
| Control-Sample1_Beta | chr3:13082751-13083974 | IQSEC1 | | Body | Yes |
| Control-Sample10_Beta | chr3:13060741-13060974 | IQSEC1 | | Body | Yes |
| Control-Sample17_Beta | chr3:13082751-13083892 | IQSEC1 | | Body | Yes |
| Control-Sample19_Beta | chr3:13082751-13083151 | IQSEC1 | | Body | Yes |
| Control-Sample19_Beta | chr3:13083477-13083892 | IQSEC1 | | Body | Yes |
| Control-Sample21_Beta | chr3:13114797-13115104 | IQSEC1 | Island | TSS200 | Yes |
| Control-Sample25_Beta | chr3:13114681-13114803 | IQSEC1 | Island | TSS200 | Yes |
| Control-Sample26_Beta | chr3:13060741-13060971 | IQSEC1 | | Body | Yes |
| Control-Sample151_Be | chr3:12994821-12994919 | IQSEC1;IQSEC1 | | Body;Body | Yes |
| Control-Sample19_Beta | chr3:12940286-12940432 | IQSEC1;IQSEC1 | N_Shore | 3'UTR;3'UTR | Yes |
| Control-Sample420_Be | chr12:175809-176111 | IQSEC3 | Island | TSS1500 | Yes |
| Control-Sample38_Beta | chr12:187092-187480 | IQSEC3;IQSEC3 | Island | Body;5'UTR | Yes |
| Control-Sample3_Beta | chr11:616009-616089 | IRF7;IRF7;IRF7 | Island | TSS1500;TSS200;TSS200 | Yes |
| Control-Sample58_Beta | chr11:613478-613792 | IRF7;IRF7;IRF7 | N_Shore | Body;Body;Body | Yes |
| Control-Sample19_Beta | chr16:54321848-54322308 | IRX3 | Island | TSS1500 | Yes |
| Control-Sample20_Beta | chr16:54321848-54322494 | IRX3 | Island | TSS1500 | Yes |
| Control-Sample259_Be | chr1:949392-949893 | ISG15 | Island | Body | No |
| Control-Sample194_Be | chr15:76634516-76634887 | ISL2 | Island | 3'UTR | No |
| Control-Sample3_Beta | chr15:74465558-74465888 | ISLR;ISLR | N_Shelf | TSS1500;TSS1500 | No |
| Control-Sample19_Beta | chr15:74427234-74427499 | ISLR2;ISLR2;ISLR2;ISLR2 | Island | Body;Body;Body;Body | No |
| Control-Sample13_Beta | chr19:55972791-55973338 | ISOC2;ISOC2;ISOC2 | Island | 5'UTR;5'UTR;5'UTR | Yes |
| Control-Sample14_Beta | chr19:55972855-55973228 | ISOC2;ISOC2;ISOC2 | Island | 5'UTR;5'UTR;5'UTR | Yes |
| Control-Sample141_Be | chr19:55972855-55973338 | ISOC2;ISOC2;ISOC2 | Island | 5'UTR;5'UTR;5'UTR | Yes |
| Control-Sample146_Be | chr19:55973234-55973338 | ISOC2;ISOC2;ISOC2 | Island | TSS200;TSS200;TSS200 | Yes |
| Control-Sample147_Be | chr19:55972855-55973099 | ISOC2;ISOC2;ISOC2 | Island | 5'UTR;5'UTR;5'UTR | Yes |
| Control-Sample35_Beta | chr19:55972791-55973099 | ISOC2;ISOC2;ISOC2 | Island | 5'UTR;5'UTR;5'UTR | Yes |
| Control-Sample12_Beta | chr17:48153934-48154356 | ITGA3;ITGA3 | | Body;Body | No |
| Control-Sample46_Beta | chr12:54811762-54812085 | ITGA5 | N_Shore | Body | No |
| Control-Sample58_Beta | chr12:56105981-56106386 | ITGA7 | N_Shelf | Body | No |
| Control-Sample661_Be | chr21:46341006-46341067 | ITGB2;ITGB2 | | TSS200;5'UTR | Yes |
| Control-Sample662_Be | chr21:46341006-46341304 | ITGB2;ITGB2 | | TSS200;5'UTR | Yes |

| | | | | | |
|---|---|---|---|---|---|
| Control-Sample138_Be | chr21:46340833-46341304 | ITGB2;ITGB2;ITGB2 | | 5'UTR;5'UTR;1stExon | Yes |
| Control-Sample1_Beta | chr14:93419165-93419447 | ITPK1;ITPK1;ITPK1 | | Body;Body;Body | Yes |
| Control-Sample170_Be | chr14:93419016-93419209 | ITPK1;ITPK1;ITPK1 | | Body;Body;Body | Yes |
| Control-Sample2_Beta | chr14:93419165-93419496 | ITPK1;ITPK1;ITPK1 | | Body;Body;Body | Yes |
| Control-Sample1_Beta | chr15:41793472-41793881 | ITPKA | N_Shore | Body | Yes |
| Control-Sample19_Beta | chr15:41795087-41795475 | ITPKA | Island | Body | Yes |
| Control-Sample171_Be | chr3:4805396-4805550 | ITPR1;ITPR1;ITPR1 | | Body;Body;Body | No |
| Control-Sample108_Be | chr10:106088711-106089203 | ITPRIP | | 5'UTR | No |
| Control-Sample108_Be | chr19:49250180-49250452 | IZUMO1 | N_Shelf | TSS200 | Yes |
| Control-Sample8_Beta | chr19:49244571-49244674 | IZUMO1;RASIP1 | S_Shore | Body;TSS1500 | Yes |
| Control-Sample6_Beta | chr19:17942025-17942272 | JAK3 | N_Shore | Body | Yes |
| Control-Sample50_Beta | chr4:6052527-6052554 | JAKMIP1 | | Body | No |
| Control-Sample108_Be | chr10:65029020-65029168 | JMJD1C;JMJD1C | | Body;1stExon;5'UTR | No |
| Control-Sample11_Beta | chr19:2255419-2255462 | JSRP1 | S_Shore | TSS200 | Yes |
| Control-Sample44_Beta | chr19:2255419-2255493 | JSRP1 | S_Shore | TSS200 | Yes |
| Control-Sample63_Beta | chr1:153950482-153950867 | JTB | Island | TSS200 | No |
| Control-Sample88_Beta | chr6:24358236-24358566 | KAAG1;DCDC2;KAAG1;DCDC2 | Island | 1stExon;1stExon;3'UTR;5'UTR | Yes |
| Control-Sample87_Beta | chr6:24358308-24358566 | KAAG1;KAAG1;DCDC2 | Island | 1stExon;3'UTR;TSS200 | Yes |
| Control-Sample58_Beta | chr3:124103560-124103652 | KALRN;KALRN | | Body;Body | Yes |
| Control-Sample1_Beta | chr3:123813383-123813596 | KALRN;KALRN;KALRN | | TSS200;TSS200;TSS200 | Yes |
| Control-Sample19_Beta | chr3:123813419-123813465 | KALRN;KALRN;KALRN | | TSS200;TSS200;TSS200 | Yes |
| Control-Sample20_Beta | chr3:123813393-123813465 | KALRN;KALRN;KALRN | | TSS200;TSS200;TSS200 | Yes |
| Control-Sample67_Beta | chr3:123813465-123813596 | KALRN;KALRN;KALRN | | TSS200;TSS200;TSS200 | Yes |
| Control-Sample171_Be | chr9:707166-707410 | KANK1;KANK1;KANK1 | Island | 1stExon;5'UTR;Body | No |
| Control-Sample40_Beta | chr1:62729257-62729423 | KANK4 | | Body | No |
| Control-Sample1_Beta | chr17:40274524-40274811 | KAT2A;HSPB9 | Island | TSS1500;TSS1500 | Yes |
| Control-Sample1_Beta | chr10:102821670-102822201 | KAZALD1;KAZALD1 | Island | 1stExon;5'UTR | Yes |
| Control-Sample12_Beta | chr10:102821670-102822423 | KAZALD1;KAZALD1 | Island | 1stExon;5'UTR | Yes |
| Control-Sample15_Beta | chr10:102821670-102822249 | KAZALD1;KAZALD1 | Island | 1stExon;5'UTR | Yes |
| Control-Sample19_Beta | chr10:102821684-102822249 | KAZALD1;KAZALD1 | Island | 1stExon;5'UTR | Yes |
| Control-Sample22_Beta | chr10:102821684-102822201 | KAZALD1;KAZALD1 | Island | 1stExon;5'UTR | Yes |
| Control-Sample58_Beta | chr10:102821684-102822002 | KAZALD1;KAZALD1 | Island | 1stExon;5'UTR | Yes |
| Control-Sample19_Beta | chr8:1949246-1949342 | KBTBD11 | Island | 5'UTR | No |
| Control-Sample19_Beta | chr1:111218276-111218287 | KCNA3 | S_Shore | TSS1500 | No |
| Control-Sample11_Beta | chr1:6085995-6086148 | KCNAB2;KCNAB2 | N_Shore | TSS1500;TSS1500 | No |
| Control-Sample1_Beta | chr17:7832680-7832943 | KCNAB3 | Island | 1stExon | Yes |
| Control-Sample10_Beta | chr17:7832680-7832852 | KCNAB3 | Island | 1stExon | Yes |
| Control-Sample19_Beta | chr17:7832769-7832943 | KCNAB3 | Island | TSS200 | Yes |
| Control-Sample52_Beta | chr17:7832764-7832943 | KCNAB3 | Island | TSS200 | Yes |
| Control-Sample65_Beta | chr17:7832479-7832943 | KCNAB3 | N_Shore | Body | Yes |
| Control-Sample7_Beta | chr17:7832680-7832769 | KCNAB3 | Island | 1stExon | Yes |
| Control-Sample7_Beta | chr17:7832909-7832943 | KCNAB3 | Island | TSS200 | Yes |
| Control-Sample464_Be | chr20:48099323-48099359 | KCNB1 | Island | TSS200 | Yes |
| Control-Sample465_Be | chr20:48099248-48099359 | KCNB1 | Island | TSS200 | Yes |
| Control-Sample162_Be | chr21:35831871-35831958 | KCNE1;KCNE1;KCNE1;KCNE1;KCNE1 | Island | 5'UTR;5'UTR;1stExon;1stExon;5'UTR | No |
| Control-Sample1_Beta | chr2:42719503-42719933 | KCNG3;KCNG3 | N_Shore | Body;Body | Yes |
| Control-Sample20_Beta | chr2:42719678-42719933 | KCNG3;KCNG3 | N_Shore | Body;Body | Yes |
| Control-Sample1_Beta | chr7:150644631-150645092 | KCNH2;KCNH2 | Island | Body;Body | Yes |
| Control-Sample14_Beta | chr7:150644554-150645092 | KCNH2;KCNH2 | Island | Body;Body | Yes |
| Control-Sample34_Beta | chr7:150644715-150645092 | KCNH2;KCNH2 | Island | Body;Body | Yes |
| Control-Sample49_Beta | chr7:150655109-150655387 | KCNH2;KCNH2 | Island | Body;Body | Yes |
| Control-Sample19_Beta | chr5:169780299-169780916 | KCNIP1 | | TSS1500 | Yes |
| Control-Sample57_Beta | chr5:169780299-169780480 | KCNIP1 | | TSS1500 | Yes |
| Control-Sample11_Beta | chr5:170108187-170108303 | KCNIP1;KCNIP1;KCNIP1 | | Body;Body;Body | Yes |
| Control-Sample1_Beta | chr2:95978173-95978534 | KCNIP3 | | Body | No |
| Control-Sample19_Beta | chr4:21699373-21699988 | KCNIP4;KCNIP4 | | 5'UTR;TSS200 | Yes |
| Control-Sample20_Beta | chr4:21699426-21699534 | KCNIP4;KCNIP4 | | 5'UTR;TSS200 | Yes |
| Control-Sample108_Be | chr4:21305490-21305805 | KCNIP4;KCNIP4;KCNIP4;KCNIP4 | | Body;1stExon;5'UTR;5'UTR | Yes |
| Control-Sample19_Beta | chr17:21300735-21300791 | KCNJ12 | | 5'UTR | No |
| Control-Sample1_Beta | chr17:68100915-68100975 | KCNJ16;KCNJ16;KCNJ16 | | 5'UTR;TSS200;5'UTR | No |
| Control-Sample20_Beta | chr11:128775691-128775743 | KCNJ5;C11orf45 | | 5'UTR;TSS200 | No |
| Control-Sample331_Be | chr1:233765199-233765350 | KCNK1 | | Body | No |
| Control-Sample12_Beta | chr11:64064727-64065031 | KCNK4 | N_Shelf | Body | Yes |
| Control-Sample13_Beta | chr11:64064727-64065371 | KCNK4 | N_Shelf | Body | Yes |
| Control-Sample1_Beta | chr11:2612781-2612876 | KCNQ1;KCNQ1 | | Body;Body | Yes |
| Control-Sample1_Beta | chr11:2847258-2847642 | KCNQ1;KCNQ1 | | Body;Body | Yes |
| Control-Sample11_Beta | chr11:2828214-2828364 | KCNQ1;KCNQ1 | N_Shore | Body;Body | Yes |
| Control-Sample11_Beta | chr11:2870208-2871992 | KCNQ1;KCNQ1 | N_Shore | 3'UTR;3'UTR | Yes |
| Control-Sample13_Beta | chr11:2772759-2772852 | KCNQ1;KCNQ1 | | Body;Body | Yes |
| Control-Sample29_Beta | chr11:2828525-2828652 | KCNQ1;KCNQ1 | Island | Body;Body | Yes |
| Control-Sample63_Beta | chr11:2828214-2828652 | KCNQ1;KCNQ1 | N_Shore | Body;Body | Yes |
| Control-Sample65_Beta | chr11:2571399-2571525 | KCNQ1;KCNQ1 | | Body;Body | Yes |
| Control-Sample270_Be | chr11:2891360-2891495 | KCNQ1DN | S_Shore | Body | No |
| Control-Sample58_Beta | chr20:62084549-62084833 | KCNQ2;KCNQ2;KCNQ2;KCNQ2;KCNQ2 | Island | Body;Body;Body;Body;Body | No |
| Control-Sample1_Beta | chr1:41283773-41284135 | KCNQ4;KCNQ4 | N_Shore | Body;Body | Yes |
| Control-Sample1_Beta | chr1:41284639-41285096 | KCNQ4;KCNQ4 | N_Shore | Body;Body | Yes |
| Control-Sample10_Beta | chr1:41250062-41250117 | KCNQ4;KCNQ4 | Island | 1stExon;1stExon | Yes |
| Control-Sample3_Beta | chr20:43729874-43729910 | KCNS1 | S_Shore | TSS200 | No |
| Control-Sample55_Beta | chr2:18058900-18059109 | KCNS3 | N_Shore | TSS1500 | No |
| Control-Sample151_Be | chr8:36641571-36641818 | KCNU1 | | TSS1500 | Yes |
| Control-Sample572_Be | chr8.0:36641571-36641818 | KCNU1 | | TSS1500 | Yes |
| Control-Sample1_Beta | chr7:128550812-128550946 | KCP;KCP | Island | TSS200;TSS200 | Yes |
| Control-Sample12_Beta | chr7:128550862-128550946 | KCP;KCP | Island | TSS200;TSS200 | Yes |
| Control-Sample14_Beta | chr7:128550812-128550932 | KCP;KCP | Island | TSS200;TSS200 | Yes |
| Control-Sample18_Beta | chr7:128550860-128550946 | KCP;KCP | Island | TSS200;TSS200 | Yes |
| Control-Sample21_Beta | chr7:128550860-128550902 | KCP;KCP | Island | TSS200;TSS200 | Yes |
| Control-Sample27_Beta | chr7:128550860-128550932 | KCP;KCP | Island | TSS200;TSS200 | Yes |
| Control-Sample34_Beta | chr7:128550898-128550932 | KCP;KCP | Island | TSS200;TSS200 | Yes |
| Control-Sample37_Beta | chr7:128550898-128550946 | KCP;KCP | Island | TSS200;TSS200 | Yes |
| Control-Sample44_Beta | chr7:128550812-128550902 | KCP;KCP | Island | TSS200;TSS200 | Yes |
| Control-Sample47_Beta | chr7:128550860-128550898 | KCP;KCP | Island | TSS200;TSS200 | Yes |

| | | | | | |
|---|---|---|---|---|---|
| Control-Sample48_Beta | chr7:128550902-128550946 | KCP;KCP | Island | TSS200;TSS200 | Yes |
| Control-Sample50_Beta | chr7:128550862-128550902 | KCP;KCP | Island | TSS200;TSS200 | Yes |
| Control-Sample7_Beta | chr7:128550812-128550898 | KCP;KCP | Island | TSS200;TSS200 | Yes |
| Control-Sample1_Beta | chr18:24131115-24131604 | KCTD1 | Island | 5'UTR | No |
| Control-Sample1_Beta | chr13:77461335-77461919 | KCTD12 | S_Shore | TSS1500 | Yes |
| Control-Sample58_Beta | chr13:77461335-77461426 | KCTD12 | S_Shore | TSS1500 | Yes |
| Control-Sample1_Beta | chr17:73055792-73056187 | KCTD2 | | Body | Yes |
| Control-Sample108_Be | chr17:73056104-73056187 | KCTD2 | | Body | Yes |
| Control-Sample13_Beta | chr17:73055875-73056681 | KCTD2 | | Body | Yes |
| Control-Sample19_Beta | chr17:73055792-73056145 | KCTD2 | | Body | Yes |
| Control-Sample20_Beta | chr17:73056002-73056145 | KCTD2 | | Body | Yes |
| Control-Sample27_Beta | chr17:73055875-73056187 | KCTD2 | | Body | Yes |
| Control-Sample33_Beta | chr17:73055792-73056681 | KCTD2 | | Body | Yes |
| Control-Sample64_Beta | chr17:73055792-73056002 | KCTD2 | | Body | Yes |
| Control-Sample66_Beta | chr17:73056002-73056187 | KCTD2 | | Body | Yes |
| Control-Sample1_Beta | chr19:5074224-5074758 | KDM4B | N_Shore | Body | Yes |
| Control-Sample20_Beta | chr19:5060985-5061205 | KDM4B | Island | Body | Yes |
| Control-Sample23_Beta | chr19:5074224-5074717 | KDM4B | N_Shore | Body | Yes |
| Control-Sample1_Beta | chr17:7757933-7757969 | KDM6B;TMEM88 | N_Shore | 3'UTR;TSS1500 | Yes |
| Control-Sample1_Beta | chr17:7758102-7758204 | KDM6B;TMEM88 | N_Shore | 3'UTR;TSS1500 | Yes |
| Control-Sample2_Beta | chr17:7758102-7758291 | KDM6B;TMEM88 | N_Shore | 3'UTR;TSS1500 | Yes |
| Control-Sample108_Be | chr8:48572447-48572557 | KIAA0146 | | Body | No |
| Control-Sample20_Beta | chr16:85684108-85684468 | KIAA0182;KIAA0182 | Island | Body;Body | Yes |
| Control-Sample58_Beta | chr16:85684353-85684821 | KIAA0182;KIAA0182 | Island | Body;Body | Yes |
| Control-Sample1_Beta | chr18:46386267-46386453 | KIAA0427;KIAA0427 | Island | 3'UTR;3'UTR | No |
| Control-Sample1_Beta | chr16:85116332-85116513 | KIAA0513 | | Body | Yes |
| Control-Sample13_Beta | chr16:85116332-85116424 | KIAA0513 | | Body | Yes |
| Control-Sample64_Beta | chr16:85075076-85075249 | KIAA0513 | | 5'UTR | Yes |
| Control-Sample19_Beta | chr16:27781069-27781794 | KIAA0556 | | Body | Yes |
| Control-Sample20_Beta | chr16:27781069-27781295 | KIAA0556 | | Body | Yes |
| Control-Sample19_Beta | chr18:8717035-8717323 | KIAA0802 | | TSS1500 | No |
| Control-Sample1_Beta | chr15:79725284-79725737 | KIAA1024 | Island | 5'UTR | No |
| Control-Sample15_Beta | chr1:15175204-15175356 | KIAA1026;KIAA1026 | | Body;Body | Yes |
| Control-Sample6_Beta | chr1:15256123-15256136 | KIAA1026;KIAA1026;KIAA1026;KIAA1026 | | Body;TSS200;Body;Body | Yes |
| Control-Sample1_Beta | chr12:13196876-13196954 | KIAA1467 | N_Shore | TSS1500 | No |
| Control-Sample331_Be | chr1:33231306-33231382 | KIAA1522 | | Body | Yes |
| Control-Sample38_Beta | chr1:33231272-33231313 | KIAA1522 | | Body | Yes |
| Control-Sample57_Beta | chr1:33231272-33231382 | KIAA1522 | | Body | Yes |
| Control-Sample1_Beta | chr4:1376165-1376376 | KIAA1530 | | Body | Yes |
| Control-Sample64_Beta | chr4:1355321-1355424 | KIAA1530 | | Body | Yes |
| Control-Sample10_Beta | chr19:18385502-18385930 | KIAA1683;KIAA1683;KIAA1683 | | TSS200;TSS200;TSS200 | Yes |
| Control-Sample14_Beta | chr19:18385672-18385930 | KIAA1683;KIAA1683;KIAA1683 | | TSS1500;TSS1500;TSS1500 | Yes |
| Control-Sample40_Beta | chr19:18385502-18385724 | KIAA1683;KIAA1683;KIAA1683 | | TSS200;TSS200;TSS200 | Yes |
| Control-Sample464_Be | chr19:18368483-18368873 | KIAA1683;KIAA1683;KIAA1683 | Island | Body;Body;Body | Yes |
| Control-Sample40_Beta | chr2:61293253-61293272 | KIAA1841;KIAA1841 | Island | 5'UTR;TSS200 | No |
| Control-Sample108_Be | chr8:145163034-145163136 | KIAA1875 | N_Shore | Body | No |
| Control-Sample202_Be | chr7:1609793-1610220 | KIAA1908;KIAA1908;PSMG3;KIAA1908;PSMG3 | Island | Body;Body;TSS200;Body;TSS200 | Yes |
| Control-Sample1_Beta | chr6:30653512-30653549 | KIAA1949;KIAA1949 | N_Shore | Body;1stExon | Yes |
| Control-Sample10_Beta | chr6:30652396-30652771 | KIAA1949;KIAA1949 | N_Shore | Body;1stExon | Yes |
| Control-Sample10_Beta | chr6:30653407-30653549 | KIAA1949;KIAA1949 | N_Shore | Body;1stExon | Yes |
| Control-Sample12_Beta | chr6:30652376-30652399 | KIAA1949;KIAA1949 | N_Shelf | Body;1stExon | Yes |
| Control-Sample13_Beta | chr6:30652688-30652829 | KIAA1949;KIAA1949 | N_Shore | Body;1stExon | Yes |
| Control-Sample19_Beta | chr6:30651563-30651825 | KIAA1949;KIAA1949 | N_Shelf | Body;Body | Yes |
| Control-Sample28_Beta | chr6:30653407-30653736 | KIAA1949;KIAA1949 | N_Shore | Body;1stExon | Yes |
| Control-Sample3_Beta | chr6:30652396-30652829 | KIAA1949;KIAA1949 | N_Shore | Body;1stExon | Yes |
| Control-Sample32_Beta | chr6:30652376-30652771 | KIAA1949;KIAA1949 | N_Shelf | Body;1stExon | Yes |
| Control-Sample34_Beta | chr6:30652376-30652829 | KIAA1949;KIAA1949 | N_Shelf | Body;1stExon | Yes |
| Control-Sample51_Beta | chr6:30653407-30653530 | KIAA1949;KIAA1949 | N_Shore | Body;1stExon | Yes |
| Control-Sample2_Beta | chr6:17988748-17988937 | KIF13A;KIF13A;KIF13A;KIF13A | S_Shore | TSS1500;TSS1500;TSS1500;TSS1500 | Yes |
| Control-Sample21_Beta | chr6:17988861-17988960 | KIF13A;KIF13A;KIF13A;KIF13A | S_Shore | TSS1500;TSS1500;TSS1500;TSS1500 | Yes |
| Control-Sample475_Be | chr17:72347925-72348295 | KIF19 | Island | Body | No |
| Control-Sample108_Be | chr6:168418442-168418527 | KIF25;KIF25 | | TSS200;TSS200 | Yes |
| Control-Sample19_Beta | chr6:168418398-168418632 | KIF25;KIF25 | | TSS200;TSS200 | Yes |
| Control-Sample216_Be | chr6:168417634-168417743 | KIF25;KIF25 | | TSS1500;TSS1500 | Yes |
| Control-Sample322_Be | chr6:168436019-168436353 | KIF25;KIF25 | Island | Body;Body | Yes |
| Control-Sample620_Be | chr6:168435636-168435923 | KIF25;KIF25 | N_Shore | Body;Body | Yes |
| Control-Sample1_Beta | chr14:104623477-104623651 | KIF26A | N_Shore | Body | Yes |
| Control-Sample22_Beta | chr14:104625249-104625468 | KIF26A | Island | Body | Yes |
| Control-Sample3_Beta | chr1:245755268-245755687 | KIF26B | S_Shore | Body | No |
| Control-Sample10_Beta | chr8:145698116-145698660 | KIFC2 | Island | Body | No |
| Control-Sample10_Beta | chr16:57832005-57832309 | KIFC3;KIFC3;KIFC3 | N_Shelf | TSS200;Body;Body | Yes |
| Control-Sample19_Beta | chr16:57832005-57832230 | KIFC3;KIFC3;KIFC3 | N_Shelf | TSS200;Body;Body | Yes |
| Control-Sample1_Beta | chr16:57831745-57832309 | KIFC3;KIFC3;KIFC3;KIFC3 | | Body;Body;1stExon;5'UTR | Yes |
| Control-Sample1_Beta | chr11:126455596-126456119 | KIRREL3;KIRREL3 | | Body;Body | Yes |
| Control-Sample10_Beta | chr11:126456008-126456119 | KIRREL3;KIRREL3 | | Body;Body | Yes |
| Control-Sample171_Be | chr11:126455712-126456008 | KIRREL3;KIRREL3 | | Body;Body | Yes |
| Control-Sample216_Be | chr11:126455712-126456119 | KIRREL3;KIRREL3 | | Body;Body | Yes |
| Control-Sample1_Beta | chr10:3824516-3824687 | KLF6;KLF6;KLF6;KLF6 | S_Shore | Body;Body;Body;Body | Yes |
| Control-Sample12_Beta | chr10:3823758-3823907 | KLF6;KLF6;KLF6;KLF6 | N_Shore | Body;Body;Body;Body | Yes |
| Control-Sample13_Beta | chr10:3823758-3824016 | KLF6;KLF6;KLF6;KLF6 | N_Shore | Body;Body;Body;Body | Yes |
| Control-Sample19_Beta | chr10:3824387-3824687 | KLF6;KLF6;KLF6;KLF6 | S_Shore | Body;Body;Body;Body | Yes |
| Control-Sample69_Beta | chr16:87769432-87769944 | KLHDC4 | | Body | No |
| Control-Sample27_Beta | chr22:50986511-50986962 | KLHDC7B;KLHDC7B | Island | 5'UTR;1stExon | No |
| Control-Sample3_Beta | chr1:205326437-205326641 | KLHDC8A | | TSS1500 | Yes |
| Control-Sample4_Beta | chr1:205326437-205327054 | KLHDC8A | | TSS1500 | Yes |
| Control-Sample1_Beta | chr18:30353240-30353699 | KLHL14 | S_Shore | TSS1500 | Yes |
| Control-Sample2_Beta | chr18:30353125-30353699 | KLHL14 | S_Shore | TSS200 | Yes |
| Control-Sample34_Beta | chr18:30353000-30353699 | KLHL14 | S_Shore | TSS200 | Yes |
| Control-Sample80_Beta | chr4:166131119-166131313 | KLHL2;KLHL2;KLHL2 | S_Shore | TSS200;TSS200;Body | No |
| Control-Sample19_Beta | chr2:23646176-23646462 | KLHL29 | | 5'UTR | No |
| Control-Sample180_Be | chr11:75139390-75139736 | KLHL35 | N_Shore | Body | Yes |

| | | | | | |
|---|---|---|---|---|---|
| Control-Sample475_Be | chr11:75141793-75141835 | KLHL35 | Island | TSS200 | Yes |
| Control-Sample105_Be | chr16:58328961-58329456 | KLKBL4 | | TSS200 | No |
| Control-Sample1_Beta | chr10:134980513-134980924 | KNDC1 | Island | Body | Yes |
| Control-Sample51_Beta | chr10:134994215-134994945 | KNDC1 | S_Shore | Body | Yes |
| Control-Sample20_Beta | chr17:39019908-39020053 | KRT12 | Island | Body | No |
| Control-Sample15_Beta | chr17:39093658-39093939 | KRT23;KRT23 | | 1stExon;5'UTR | No |
| Control-Sample10_Beta | chr12:52828252-52828461 | KRT75 | | TSS200 | Yes |
| Control-Sample3_Beta | chr12:52828203-52828461 | KRT75 | | TSS200 | Yes |
| Control-Sample1_Beta | chr12:53297819-53298424 | KRT8 | Island | Body | Yes |
| Control-Sample12_Beta | chr12:53297987-53298424 | KRT8 | S_Shore | Body | Yes |
| Control-Sample466_Be | chr17:39212513-39212764 | KRTAP2-2 | | TSS1500 | No |
| Control-Sample13_Beta | chr17:39165425-39165697 | KRTAP3-1 | | TSS200 | No |
| Control-Sample18_Beta | chr2:27665141-27665306 | KRTCAP3;KRTCAP3 | Island | TSS200;TSS200 | Yes |
| Control-Sample2_Beta | chr2:27665128-27665150 | KRTCAP3;KRTCAP3 | Island | TSS200;TSS200 | Yes |
| Control-Sample29_Beta | chr2:27665139-27665150 | KRTCAP3;KRTCAP3 | Island | TSS200;TSS200 | Yes |
| Control-Sample33_Beta | chr2:27665141-27665507 | KRTCAP3;KRTCAP3 | Island | TSS200;TSS200 | Yes |
| Control-Sample48_Beta | chr2:27665150-27665507 | KRTCAP3;KRTCAP3 | Island | TSS200;TSS200 | Yes |
| Control-Sample57_Beta | chr2:27665128-27665306 | KRTCAP3;KRTCAP3 | Island | TSS200;TSS200 | Yes |
| Control-Sample15_Beta | chr2:27665017-27665507 | KRTCAP3;NRBP1;KRTCAP3 | Island | TSS1500;3'UTR;TSS1500 | Yes |
| Control-Sample16_Beta | chr2:27665017-27665139 | KRTCAP3;NRBP1;KRTCAP3 | Island | TSS1500;3'UTR;TSS1500 | Yes |
| Control-Sample18_Beta | chr2:27665017-27665128 | KRTCAP3;NRBP1;KRTCAP3 | Island | TSS1500;3'UTR;TSS1500 | Yes |
| Control-Sample4_Beta | chr2:27665017-27665150 | KRTCAP3;NRBP1;KRTCAP3 | Island | TSS1500;3'UTR;TSS1500 | Yes |
| Control-Sample10_Beta | chr17:25798942-25799018 | KSR1 | | TSS200 | No |
| Control-Sample1_Beta | chr14:56045827-56046098 | KTN1;C14orf33;KTN1;KTN1;KTN1 | N_Shore | TSS1500;Body;TSS1500;TSS1500;TSS1500 | Yes |
| Control-Sample34_Beta | chr14:56045827-56046001 | KTN1;C14orf33;KTN1;KTN1;KTN1 | N_Shore | TSS1500;Body;TSS1500;TSS1500;TSS1500 | Yes |
| Control-Sample58_Beta | chr1:62660191-62660421 | L1TD1;L1TD1 | N_Shore | TSS1500;TSS1500 | No |
| Control-Sample1_Beta | chr1:201368914-201369031 | LAD1 | Island | TSS1500 | Yes |
| Control-Sample13_Beta | chr1:201368914-201368948 | LAD1 | Island | TSS1500 | Yes |
| Control-Sample2_Beta | chr1:201368843-201368948 | LAD1 | Island | TSS200 | Yes |
| Control-Sample67_Beta | chr6:112576009-112576393 | LAMA4;LAMA4;LAMA4;LAMA4;LAMA4 | S_Shore | TSS200;TSS200;TSS200;TSS200;TSS200 | No |
| Control-Sample1_Beta | chr5:154092290-154092426 | LARP1 | | TSS200 | Yes |
| Control-Sample19_Beta | chr5:154091976-154092426 | LARP1 | | TSS1500 | Yes |
| Control-Sample26_Beta | chr5:154092321-154092426 | LARP1 | | TSS200 | Yes |
| Control-Sample15_Beta | chr6:150039666-150039728 | LATS1 | S_Shore | TSS1500 | No |
| Control-Sample11_Beta | chr15:68118521-68118985 | LBXCOR1 | Island | Body | Yes |
| Control-Sample67_Beta | chr15:68125197-68125599 | LBXCOR1 | N_Shore | Body | Yes |
| Control-Sample3_Beta | chr1:32739493-32740034 | LCK;LCK | N_Shore | 5'UTR;TSS1500 | No |
| Control-Sample10_Beta | chr2:30669597-30669863 | LCLAT1;LCLAT1 | N_Shore | TSS1500;TSS1500 | Yes |
| Control-Sample40_Beta | chr2:30670015-30670159 | LCLAT1;LCLAT1 | Island | TSS200;TSS200 | Yes |
| Control-Sample1_Beta | chr15:66858008-66858446 | LCTL | | TSS200 | Yes |
| Control-Sample14_Beta | chr15:66858008-66858375 | LCTL | | TSS200 | Yes |
| Control-Sample3_Beta | chr15:66858212-66858446 | LCTL | | TSS1500 | Yes |
| Control-Sample180_Be | chr11:18433554-18433683 | LDHC;LDHC | | TSS1500;TSS1500 | Yes |
| Control-Sample183_Be | chr11:18433500-18433745 | LDHC;LDHC | | TSS1500;TSS1500 | Yes |
| Control-Sample41_Beta | chr11:18433683-18433745 | LDHC;LDHC | | TSS200;TSS200 | Yes |
| Control-Sample419_Be | chr11:18433554-18433745 | LDHC;LDHC | | TSS1500;TSS1500 | Yes |
| Control-Sample504_Be | chr11.0:18433500-18433745 | LDHC;LDHC | | TSS1500;TSS1500 | Yes |
| Control-Sample1_Beta | chr16:75150784-75150819 | LDHD;LDHD | S_Shore | TSS200;TSS200 | Yes |
| Control-Sample23_Beta | chr16:75150456-75150833 | LDHD;LDHD | S_Shore | Body;Body | Yes |
| Control-Sample3_Beta | chr16:75150744-75150833 | LDHD;LDHD | S_Shore | TSS200;TSS200 | Yes |
| Control-Sample30_Beta | chr16:75150784-75150833 | LDHD;LDHD | S_Shore | TSS200;TSS200 | Yes |
| Control-Sample41_Beta | chr16:75150744-75150819 | LDHD;LDHD | S_Shore | TSS200;TSS200 | Yes |
| Control-Sample62_Beta | chr16:75149501-75150880 | LDHD;LDHD | S_Shore | Body;Body | Yes |
| Control-Sample1_Beta | chr13:53314142-53314226 | LECT1;LECT1 | S_Shore | TSS200;TSS200 | No |
| Control-Sample1_Beta | chr12:65562903-65562996 | LEMD3;LEMD3 | N_Shore | TSS1500;TSS1500 | Yes |
| Control-Sample25_Beta | chr12:65562903-65563034 | LEMD3;LEMD3 | N_Shore | TSS1500;TSS1500 | Yes |
| Control-Sample1_Beta | chr22:38071455-38071602 | LGALS1 | N_Shore | TSS200 | Yes |
| Control-Sample10_Beta | chr22:38071455-38071677 | LGALS1 | N_Shore | TSS200 | Yes |
| Control-Sample14_Beta | chr22:38071455-38071607 | LGALS1 | N_Shore | TSS200 | Yes |
| Control-Sample19_Beta | chr22:38071455-38071591 | LGALS1 | N_Shore | TSS200 | Yes |
| Control-Sample53_Beta | chr22:38071188-38071607 | LGALS1 | N_Shore | TSS1500 | Yes |
| Control-Sample1_Beta | chr14:55595666-55595768 | LGALS3 | N_Shore | TSS1500 | No |
| Control-Sample331_Be | chr1:236686564-236686618 | LGALS8;LGALS8;LGALS8;LGALS8 | N_Shore | TSS200;TSS200;5'UTR;TSS1500 | No |
| Control-Sample496_Be | chr10:95516908-95517498 | LGI1 | | TSS1500 | No |
| Control-Sample13_Beta | chr1:202182603-202183279 | LGR6;LGR6;LGR6 | N_Shore | TSS1500;Body;Body | Yes |
| Control-Sample23_Beta | chr1:202182417-202182642 | LGR6;LGR6;LGR6 | N_Shore | TSS1500;Body;Body | Yes |
| Control-Sample27_Beta | chr1:202182417-202183279 | LGR6;LGR6;LGR6 | N_Shore | TSS1500;Body;Body | Yes |
| Control-Sample15_Beta | chr1:206786170-206786180 | LGTN | S_Shore | TSS1500 | No |
| Control-Sample15_Beta | chr2:48982683-48982832 | LHCGR | Island | 1stExon | No |
| Control-Sample171_Be | chr9:126776767-126777160 | LHX2 | Island | Body | No |
| Control-Sample10_Beta | chr9:124979184-124980074 | LHX6;LHX6 | N_Shore | Body;Body | Yes |
| Control-Sample4_Beta | chr9:124979184-124979356 | LHX6;LHX6 | N_Shore | Body;Body | Yes |
| Control-Sample20_Beta | chr4:41361919-41362115 | LIMCH1;LIMCH1;LIMCH1 | N_Shore | TSS1500;TSS1500;TSS1500 | Yes |
| Control-Sample5_Beta | chr4:41361996-41362115 | LIMCH1;LIMCH1;LIMCH1 | N_Shore | TSS1500;TSS1500;TSS1500 | Yes |
| Control-Sample14_Beta | chr3:45636060-45636154 | LIMD1 | S_Shore | TSS1500 | Yes |
| Control-Sample57_Beta | chr3:45636190-45636386 | LIMD1 | S_Shore | TSS200 | Yes |
| Control-Sample10_Beta | chr20:62368956-62369504 | LIME1 | Island | Body | Yes |
| Control-Sample12_Beta | chr20:62368956-62369462 | LIME1 | Island | Body | Yes |
| Control-Sample16_Beta | chr20:62368956-62369605 | LIME1 | Island | Body | Yes |
| Control-Sample1_Beta | chr22:31644160-31644282 | LIMK2;LIMK2;LIMK2 | | TSS200;TSS200;Body | Yes |
| Control-Sample19_Beta | chr22:31644160-31644558 | LIMK2;LIMK2;LIMK2 | | TSS200;TSS200;Body | Yes |
| Control-Sample19_Beta | chr2:128403084-128403286 | LIMS2;GPR17;GPR17;LIMS2;LIMS2;GPR17;GPR17;LIMS2;LIMS2 | | Body;TSS1500;TSS1500;Body;Body;TSS1500;TSS150 | Yes |
| Control-Sample331_Be | chr2:128403084-128403710 | LIMS2;GPR17;GPR17;LIMS2;LIMS2;GPR17;GPR17;LIMS2;LIMS2 | | Body;TSS1500;TSS1500;Body;Body;TSS1500;TSS150 | Yes |
| Control-Sample1_Beta | chr19:2291613-2291872 | LINGO3 | Island | Body | Yes |
| Control-Sample145_Be | chr19:2307632-2308475 | LINGO3 | Island | 5'UTR | Yes |
| Control-Sample23_Beta | chr19:2289982-2290107 | LINGO3 | Island | 3'UTR | Yes |
| Control-Sample27_Beta | chr19:2302401-2302783 | LINGO3 | N_Shore | 5'UTR | Yes |
| Control-Sample28_Beta | chr19:2289982-2290024 | LINGO3 | Island | 3'UTR | Yes |
| Control-Sample380_Be | chr19:2307901-2308321 | LINGO3 | Island | 5'UTR | Yes |
| Control-Sample586_Be | chr19.0:2307901-2308321 | LINGO3 | Island | 5'UTR | Yes |
| Control-Sample61_Beta | chr19:2307901-2308423 | LINGO3 | Island | 5'UTR | Yes |

| | | | | | |
|---|---|---|---|---|---|
| Control-Sample74_Beta | chr19:2307901-2308355 | LINGO3 | Island | 5'UTR | Yes |
| Control-Sample1_Beta | chr15:58723851-58724225 | LIPC | | TSS1500 | Yes |
| Control-Sample2_Beta | chr15:58723851-58724041 | LIPC | | TSS1500 | Yes |
| Control-Sample31_Beta | chr15:58723879-58724225 | LIPC | | TSS1500 | Yes |
| Control-Sample313_Be | chr12:66524311-66524943 | LLPH | N_Shore | 5'UTR | No |
| Control-Sample19_Beta | chr16:979488-979662 | LMF1 | Island | Body | Yes |
| Control-Sample50_Beta | chr16:981766-982111 | LMF1 | S_Shore | Body | Yes |
| Control-Sample1_Beta | chr11:33913187-33913586 | LMO2 | | 5'UTR | No |
| Control-Sample1_Beta | chr12:16761002-16761406 | LMO3;LMO3 | | 1stExon;5'UTR | Yes |
| Control-Sample216_Be | chr12:16761002-16761118 | LMO3;LMO3 | | 1stExon;5'UTR | Yes |
| Control-Sample1_Beta | chr7:97807643-97807741 | LMTK2 | | Body | No |
| Control-Sample11_Beta | chr1:165204410-165204512 | LMX1A | N_Shore | Body | No |
| Control-Sample1_Beta | chr4:54457645-54457746 | LNX1;LNX1 | | 5'UTR;1stExon | Yes |
| Control-Sample10_Beta | chr4:54424476-54424787 | LNX1;LNX1 | | TSS200;Body | Yes |
| Control-Sample19_Beta | chr4:54457645-54457928 | LNX1;LNX1 | | 5'UTR;1stExon | Yes |
| Control-Sample29_Beta | chr4:54424596-54424787 | LNX1;LNX1 | | TSS200;Body | Yes |
| Control-Sample53_Beta | chr4:54424476-54424764 | LNX1;LNX1 | | TSS200;Body | Yes |
| Control-Sample19_Beta | chr4:54424280-54424596 | LNX1;LNX1;LNX1 | | 1stExon;Body;5'UTR | Yes |
| Control-Sample30_Beta | chr4:54424280-54424787 | LNX1;LNX1;LNX1 | | 1stExon;Body;5'UTR | Yes |
| Control-Sample14_Beta | chr11:19736912-19737237 | LOC100126784;NAV2;NAV2;NAV2 | S_Shore | TSS1500;Body;Body;Body | Yes |
| Control-Sample194_Be | chr11:19736248-19737237 | LOC100126784;NAV2;NAV2;NAV2 | S_Shore | TSS200;Body;Body;Body | Yes |
| Control-Sample331_Be | chr11:19736648-19737237 | LOC100126784;NAV2;NAV2;NAV2 | S_Shore | TSS1500;Body;Body;Body | Yes |
| Control-Sample14_Beta | chr20:61299106-61299190 | LOC100127888;SLCO4A1 | | TSS1500;Body | No |
| Control-Sample1_Beta | chr19:35597062-35597415 | LOC100128675;LOC100128675 | | Body;Body | Yes |
| Control-Sample171_Be | chr19:35597226-35597447 | LOC100128675;LOC100128675 | | TSS200;TSS200 | Yes |
| Control-Sample331_Be | chr19:35597062-35597447 | LOC100128675;LOC100128675 | | Body;Body | Yes |
| Control-Sample121_Be | chr9:92291210-92291523 | LOC100129066 | N_Shore | Body | Yes |
| Control-Sample573_Be | chr9.0:92291210-92291523 | LOC100129066 | N_Shore | Body | Yes |
| Control-Sample23_Beta | chr16:87739377-87739453 | LOC100129637 | S_Shore | TSS200 | No |
| Control-Sample113_Be | chr5:90679528-90679543 | LOC100129716;ARRDC3 | S_Shelf | Body;TSS1500 | No |
| Control-Sample238_Be | chr16:90113893-90114200 | LOC100130015;LOC100130015 | | Body;Body | No |
| Control-Sample125_Be | chr18:77905663-77905751 | LOC100130522;LOC100130522 | S_Shore | TSS200;TSS200 | Yes |
| Control-Sample554_Be | chr18.0:77905699-77905751 | LOC100130522;LOC100130522 | S_Shore | TSS200;TSS200 | Yes |
| Control-Sample82_Beta | chr18:77917615-77918142 | LOC100130522;LOC100130522;PARD6G | N_Shore | Body;Body;3'UTR | Yes |
| Control-Sample3_Beta | chr4:1198478-1198537 | LOC100130872;LOC100130872-SPON2 | | Body;Body | Yes |
| Control-Sample60_Beta | chr4:1202509-1202653 | LOC100130872;LOC100130872-SPON2 | N_Shore | Body;Body | Yes |
| Control-Sample1_Beta | chr4:1202694-1202780 | LOC100130872-SPON2;LOC100130872 | N_Shore | TSS200;Body | Yes |
| Control-Sample10_Beta | chr4:1202694-1202861 | LOC100130872-SPON2;LOC100130872 | N_Shore | TSS200;Body | Yes |
| Control-Sample20_Beta | chr4:1202657-1202780 | LOC100130872-SPON2;LOC100130872 | N_Shore | TSS200;Body | Yes |
| Control-Sample24_Beta | chr4:1202694-1202700 | LOC100130872-SPON2;LOC100130872 | N_Shore | TSS200;Body | Yes |
| Control-Sample25_Beta | chr4:1202657-1202861 | LOC100130872-SPON2;LOC100130872 | N_Shore | TSS200;Body | Yes |
| Control-Sample1_Beta | chr4:1166767-1166990 | LOC100130872-SPON2;SPON2;SPON2 | S_Shore | Body;TSS200;5'UTR | Yes |
| Control-Sample10_Beta | chr4:1166839-1166990 | LOC100130872-SPON2;SPON2;SPON2 | S_Shore | Body;TSS200;5'UTR | Yes |
| Control-Sample17_Beta | chr4:1166767-1167230 | LOC100130872-SPON2;SPON2;SPON2 | S_Shore | Body;TSS200;5'UTR | Yes |
| Control-Sample31_Beta | chr4:1166839-1167230 | LOC100130872-SPON2;SPON2;SPON2 | S_Shore | Body;TSS200;5'UTR | Yes |
| Control-Sample19_Beta | chr4:1166873-1166990 | LOC100130872-SPON2;SPON2;SPON2;SPON2 | S_Shore | Body;TSS1500;1stExon;5'UTR | Yes |
| Control-Sample2_Beta | chr4:1166531-1166990 | LOC100130872-SPON2;SPON2;SPON2;SPON2 | Island | Body;5'UTR;1stExon;5'UTR | Yes |
| Control-Sample1_Beta | chr17:73642080-73642503 | LOC100130933;RECQL5 | | TSS1500;Body | Yes |
| Control-Sample13_Beta | chr17:73642080-73642486 | LOC100130933;RECQL5 | | TSS1500;Body | Yes |
| Control-Sample171_Be | chr17:73642080-73642547 | LOC100130933;RECQL5 | | TSS1500;Body | Yes |
| Control-Sample223_Be | chr17:73642080-73642628 | LOC100130933;RECQL5 | | TSS1500;Body | Yes |
| Control-Sample10_Beta | chr11:67141747-67143062 | LOC100130987;CLCF1;CLCF1 | S_Shore | Body;TSS1500;TSS200 | Yes |
| Control-Sample11_Beta | chr11:67141686-67141747 | LOC100130987;CLCF1;CLCF1 | S_Shore | Body;TSS1500;TSS200 | Yes |
| Control-Sample22_Beta | chr11:67142001-67142268 | LOC100130987;CLCF1;CLCF1 | S_Shore | Body;TSS1500;TSS1500 | Yes |
| Control-Sample51_Beta | chr11:67141686-67141792 | LOC100130987;CLCF1;CLCF1 | S_Shore | Body;TSS1500;TSS200 | Yes |
| Control-Sample24_Beta | chr1:151810485-151811620 | LOC100132111;C2CD4D | Island | TSS1500;Body | No |
| Control-Sample10_Beta | chr2:63274476-63275039 | LOC100132215 | Island | Body | Yes |
| Control-Sample12_Beta | chr2:63275509-63275868 | LOC100132215 | Island | TSS1500 | Yes |
| Control-Sample15_Beta | chr2:63274709-63275039 | LOC100132215 | Island | Body | Yes |
| Control-Sample171_Be | chr2:63274615-63274857 | LOC100132215 | Island | Body | Yes |
| Control-Sample22_Beta | chr2:63274476-63274857 | LOC100132215 | Island | Body | Yes |
| Control-Sample23_Beta | chr2:63274615-63275039 | LOC100132215 | Island | Body | Yes |
| Control-Sample24_Beta | chr2:63274615-63275034 | LOC100132215 | Island | Body | Yes |
| Control-Sample25_Beta | chr2:63273684-63275039 | LOC100132215 | N_Shore | Body | Yes |
| Control-Sample29_Beta | chr2:63274709-63275034 | LOC100132215 | Island | Body | Yes |
| Control-Sample38_Beta | chr2:63274476-63275034 | LOC100132215 | Island | Body | Yes |
| Control-Sample6_Beta | chr2:63274857-63275039 | LOC100132215 | Island | TSS200 | Yes |
| Control-Sample1_Beta | chr2:63272232-63272334 | LOC100132215;EHBP1;EHBP1;EHBP1;EHBP1 | Island | Body;Body;Body;Body;Body | Yes |
| Control-Sample13_Beta | chr2:63271537-63272334 | LOC100132215;EHBP1;EHBP1;EHBP1;EHBP1 | N_Shore | Body;Body;Body;Body;Body | Yes |
| Control-Sample40_Beta | chr2:63271537-63272303 | LOC100132215;EHBP1;EHBP1;EHBP1;EHBP1 | N_Shore | Body;Body;Body;Body;Body | Yes |
| Control-Sample108_Be | chr7:154719422-154719718 | LOC100132707;LOC100132707 | | TSS1500;TSS1500 | No |
| Control-Sample11_Beta | chr11:2011238-2011312 | LOC100133545 | N_Shelf | TSS200 | Yes |
| Control-Sample15_Beta | chr11:2011312-2011346 | LOC100133545 | N_Shelf | TSS200 | Yes |
| Control-Sample7_Beta | chr11:2011238-2011346 | LOC100133545 | N_Shelf | TSS200 | Yes |
| Control-Sample108_Be | chr10:93335091-93335196 | LOC100188947 | | Body | No |
| Control-Sample11_Beta | chr2:74612020-74612222 | LOC100189589 | | TSS1500 | No |
| Control-Sample108_Be | chr10:99478368-99478719 | LOC100270710 | S_Shelf | TSS200 | No |
| Control-Sample151_Be | chr2:33151557-33152037 | LOC100271832;LOC285045;LOC285045;LOC285045 | | TSS1500;Body;Body;Body | No |
| Control-Sample19_Beta | chr12:1609463-1609549 | LOC100292680 | | TSS200 | Yes |
| Control-Sample20_Beta | chr12:1609470-1609549 | LOC100292680 | | TSS200 | Yes |
| Control-Sample10_Beta | chr2:54087189-54087209 | LOC100302652;GPR75 | Island | TSS200;TSS200 | Yes |
| Control-Sample17_Beta | chr2:54087178-54087209 | LOC100302652;GPR75 | Island | TSS200;TSS200 | Yes |
| Control-Sample18_Beta | chr2:54087184-54087200 | LOC100302652;GPR75 | Island | TSS200;TSS200 | Yes |
| Control-Sample1_Beta | chr1:2483719-2483933 | LOC115110 | | Body | Yes |
| Control-Sample22_Beta | chr1:2483719-2483786 | LOC115110 | | Body | Yes |
| Control-Sample40_Beta | chr1:2483719-2483734 | LOC115110 | | Body | Yes |
| Control-Sample207_Be | chr5:150326069-150326190 | LOC134466 | Island | Body | Yes |
| Control-Sample23_Beta | chr5:150326069-150326862 | LOC134466 | Island | Body | Yes |
| Control-Sample236_Be | chr5:150326069-150326193 | LOC134466 | Island | Body | Yes |
| Control-Sample237_Be | chr5:150326069-150326497 | LOC134466 | Island | Body | Yes |
| Control-Sample11_Beta | chr10:114610508-114610635 | LOC143188 | | TSS1500 | Yes |

| | | | | | |
|---|---|---|---|---|---|
| Control-Sample3_Beta | chr10:114610508-114610803 | LOC143188 | | TSS1500 | Yes |
| Control-Sample65_Beta | chr10:114610560-114610635 | LOC143188 | | TSS200 | Yes |
| Control-Sample22_Beta | chr12:9217583-9217688 | LOC144571 | Island | TSS200 | Yes |
| Control-Sample239_Be | chr12:9217583-9217769 | LOC144571 | Island | TSS200 | Yes |
| Control-Sample11_Beta | chr15:45671148-45671195 | LOC145663;GATM | S_Shore | Body;TSS200 | Yes |
| Control-Sample2_Beta | chr15:45671195-45671347 | LOC145663;GATM | S_Shore | Body;TSS1500 | Yes |
| Control-Sample3_Beta | chr15:45671148-45671246 | LOC145663;GATM | S_Shore | Body;TSS200 | Yes |
| Control-Sample5_Beta | chr15:45671155-45671246 | LOC145663;GATM | S_Shore | Body;TSS200 | Yes |
| Control-Sample13_Beta | chr15:69853722-69854024 | LOC145837 | | TSS1500 | No |
| Control-Sample20_Beta | chr15:37169665-37170554 | LOC145845 | N_Shelf | Body | No |
| Control-Sample1_Beta | chr17:62777693-62777777 | LOC146880;LOC146880 | Island | Body;TSS200 | Yes |
| Control-Sample64_Beta | chr17:62777744-62777777 | LOC146880;LOC146880 | Island | Body;TSS200 | Yes |
| Control-Sample128_Be | chr20:5485270-5485475 | LOC149837 | | TSS200 | Yes |
| Control-Sample433_Be | chr20:5485270-5485294 | LOC149837 | | TSS200 | Yes |
| Control-Sample112_Be | chr2:239139911-239140032 | LOC151174;LOC643387;LOC151174 | N_Shore | Body;TSS1500;Body | Yes |
| Control-Sample112_Be | chr2:239140182-239140296 | LOC151174;LOC643387;LOC151174 | Island | Body;TSS200;Body | Yes |
| Control-Sample257_Be | chr2:239139911-239140190 | LOC151174;LOC643387;LOC151174 | N_Shore | Body;TSS1500;Body | Yes |
| Control-Sample342_Be | chr2:239139911-239140296 | LOC151174;LOC643387;LOC151174 | N_Shore | Body;TSS1500;Body | Yes |
| Control-Sample568_Be | chr2.0:239140024-239140190 | LOC151174;LOC643387;LOC151174 | N_Shore | Body;TSS1500;Body | Yes |
| Control-Sample1_Beta | chr2:74730630-74731413 | LOC151534;LBX2 | Island | Body;TSS200 | Yes |
| Control-Sample11_Beta | chr2:74730586-74731413 | LOC151534;LBX2 | Island | Body;TSS200 | Yes |
| Control-Sample19_Beta | chr2:74730948-74731413 | LOC151534;LBX2 | Island | Body;TSS1500 | Yes |
| Control-Sample20_Beta | chr2:74731354-74731413 | LOC151534;LBX2 | S_Shore | Body;TSS1500 | Yes |
| Control-Sample23_Beta | chr2:74730580-74731413 | LOC151534;LBX2 | Island | Body;TSS200 | Yes |
| Control-Sample46_Beta | chr2:74728623-74729502 | LOC151534;LBX2 | N_Shore | TSS1500;Body | Yes |
| Control-Sample1_Beta | chr5:43044573-43045555 | LOC153684 | S_Shore | Body | No |
| Control-Sample1_Beta | chr6:170571763-170572016 | LOC154449 | | TSS200 | Yes |
| Control-Sample10_Beta | chr6:170571697-170572016 | LOC154449 | | TSS200 | Yes |
| Control-Sample151_Be | chr6:170571810-170572016 | LOC154449 | | TSS200 | Yes |
| Control-Sample171_Be | chr6:170571686-170571786 | LOC154449 | | TSS200 | Yes |
| Control-Sample8_Beta | chr6:170571786-170572016 | LOC154449 | | TSS200 | Yes |
| Control-Sample10_Beta | chr15:89921156-89921182 | LOC254559 | Island | TSS200 | Yes |
| Control-Sample100_Be | chr15:89920348-89920887 | LOC254559 | N_Shore | TSS1500 | Yes |
| Control-Sample3_Beta | chr15:89921083-89921182 | LOC254559 | Island | TSS200 | Yes |
| Control-Sample3_Beta | chr15:89939252-89939709 | LOC254559 | N_Shelf | Body | Yes |
| Control-Sample64_Beta | chr5:6581849-6582193 | LOC255167 | N_Shore | TSS1500 | No |
| Control-Sample171_Be | chr17:76226735-76227386 | LOC283999 | N_Shore | TSS1500 | Yes |
| Control-Sample331_Be | chr17:76226735-76227508 | LOC283999 | N_Shore | TSS1500 | Yes |
| Control-Sample3_Beta | chr18:74240460-74240582 | LOC284276 | | TSS200 | No |
| Control-Sample25_Beta | chr1:170253944-170254397 | LOC284688 | | TSS1500 | Yes |
| Control-Sample57_Beta | chr1:170253405-170253944 | LOC284688 | | TSS200 | Yes |
| Control-Sample19_Beta | chr20:25130549-25130869 | LOC284798;LOC284798 | S_Shore | TSS1500;TSS1500 | No |
| Control-Sample11_Beta | chr3:13697042-13697288 | LOC285375 | Island | Body | No |
| Control-Sample12_Beta | chr6:143891455-143891975 | LOC285740 | | TSS1500 | No |
| Control-Sample13_Beta | chr6:29717917-29718119 | LOC285830;LOC285830 | S_Shore | TSS1500;TSS1500 | Yes |
| Control-Sample23_Beta | chr6:29718049-29718119 | LOC285830;LOC285830 | S_Shore | TSS1500;TSS1500 | Yes |
| Control-Sample57_Beta | chr6:29717917-29718093 | LOC285830;LOC285830 | S_Shore | TSS1500;TSS1500 | Yes |
| Control-Sample1_Beta | chr1:87617710-87617972 | LOC339524 | Island | Body | Yes |
| Control-Sample10_Beta | chr1:87617710-87617916 | LOC339524 | Island | Body | Yes |
| Control-Sample11_Beta | chr1:87617811-87617972 | LOC339524 | Island | Body | Yes |
| Control-Sample21_Beta | chr10:29698460-29698470 | LOC387647 | Island | TSS200 | Yes |
| Control-Sample23_Beta | chr10:29698368-29698449 | LOC387647 | Island | TSS200 | Yes |
| Control-Sample24_Beta | chr10:29698449-29698470 | LOC387647 | Island | TSS200 | Yes |
| Control-Sample339_Be | chr10:29698447-29698462 | LOC387647 | Island | TSS200 | Yes |
| Control-Sample230_Be | chr2:130691738-130691926 | LOC389033 | Island | Body | No |
| Control-Sample653_Be | chr7:5111641-5111916 | LOC389458 | Island | TSS200 | No |
| Control-Sample1_Beta | chr15:65394414-65394873 | LOC390595 | Island | Body | No |
| Control-Sample131_Be | chr22:24372913-24372926 | LOC391322 | Island | TSS1500 | Yes |
| Control-Sample131_Be | chr22:24373054-24373322 | LOC391322 | Island | TSS200 | Yes |
| Control-Sample38_Beta | chr22:24373054-24373618 | LOC391322 | Island | TSS200 | Yes |
| Control-Sample559_Be | chr22.0:24373054-24373322 | LOC391322 | Island | TSS200 | Yes |
| Control-Sample170_Be | chr10:124638991-124639110 | LOC399815;FAM24B | Island | TSS200;5'UTR | Yes |
| Control-Sample44_Beta | chr10:124638976-124639110 | LOC399815;FAM24B | Island | 5'UTR;TSS1500 | Yes |
| Control-Sample58_Beta | chr11:121986961-121987054 | LOC399959;BLID | | Body;TSS200 | No |
| Control-Sample1_Beta | chr22:46480891-46482023 | LOC400931 | N_Shore | TSS1500 | Yes |
| Control-Sample3_Beta | chr22:46481603-46482023 | LOC400931 | N_Shore | TSS1500 | Yes |
| Control-Sample10_Beta | chr17:46668715-46669489 | LOC404266;HOXB5;LOC404266;LOC404266;LOC404266 | N_Shore | TSS1500;3'UTR;Body;TSS1500;Body | Yes |
| Control-Sample1_Beta | chr17:46667587-46669489 | LOC404266;LOC404266 | N_Shore | TSS1500;TSS1500 | Yes |
| Control-Sample10_Beta | chr17:46667587-46667789 | LOC404266;LOC404266 | N_Shore | TSS1500;TSS1500 | Yes |
| Control-Sample12_Beta | chr17:46667725-46669489 | LOC404266;LOC404266 | N_Shore | TSS200;TSS200 | Yes |
| Control-Sample13_Beta | chr17:46667725-46668733 | LOC404266;LOC404266 | N_Shore | TSS200;TSS200 | Yes |
| Control-Sample19_Beta | chr17:46667737-46667812 | LOC404266;LOC404266 | N_Shore | TSS200;TSS200 | Yes |
| Control-Sample2_Beta | chr17:46667587-46668733 | LOC404266;LOC404266 | N_Shore | TSS1500;TSS1500 | Yes |
| Control-Sample20_Beta | chr17:46667725-46667812 | LOC404266;LOC404266 | N_Shore | TSS200;TSS200 | Yes |
| Control-Sample23_Beta | chr17:46667683-46668733 | LOC404266;LOC404266 | N_Shore | TSS200;TSS200 | Yes |
| Control-Sample40_Beta | chr17:46667725-46669492 | LOC404266;LOC404266 | N_Shore | TSS200;TSS200 | Yes |
| Control-Sample45_Beta | chr17:46667683-46667789 | LOC404266;LOC404266 | N_Shore | TSS200;TSS200 | Yes |
| Control-Sample46_Beta | chr17:46667725-46667789 | LOC404266;LOC404266 | N_Shore | TSS200;TSS200 | Yes |
| Control-Sample1_Beta | chr2:114048868-114049041 | LOC440839 | Island | Body | No |
| Control-Sample11_Beta | chr2:171569144-171570003 | LOC440925 | N_Shore | Body | Yes |
| Control-Sample12_Beta | chr2:171569132-171570003 | LOC440925 | N_Shore | Body | Yes |
| Control-Sample18_Beta | chr2:171569132-171569187 | LOC440925 | N_Shore | Body | Yes |
| Control-Sample36_Beta | chr2:171569144-171569282 | LOC440925 | N_Shore | Body | Yes |
| Control-Sample4_Beta | chr2:171568984-171570003 | LOC440925 | N_Shore | Body | Yes |
| Control-Sample202_Be | chr11:50257496-50257633 | LOC441601 | Island | Body | No |
| Control-Sample23_Beta | chr10:42863173-42863550 | LOC441666 | Island | Body | Yes |
| Control-Sample24_Beta | chr10:42863508-42863550 | LOC441666 | S_Shore | Body | Yes |
| Control-Sample39_Beta | chr10:42862978-42863550 | LOC441666 | N_Shore | Body | Yes |
| Control-Sample602_Be | chr10:42863173-42863594 | LOC441666 | Island | Body | Yes |
| Control-Sample245_Be | chr7:64043012-64043193 | LOC641746 | | Body | No |
| Control-Sample170_Be | chr19:35068555-35068710 | LOC643719 | Island | Body | Yes |

| | | | | | |
|---|---|---|---|---|---|
| Control-Sample509_Be | chr19.0:35068555-35068788 | LOC643719 | Island | Body | Yes |
| Control-Sample57_Beta | chr13:41053543-41053747 | LOC646982;LOC646982;LOC646982 | | Body;Body;Body | No |
| Control-Sample23_Beta | chr7:39649359-39649706 | LOC646999 | Island | Body | No |
| Control-Sample98_Beta | chr7:56515510-56515846 | LOC650226 | N_Shore | Body | Yes |
| Control-Sample98_Beta | chr7:56516129-56516248 | LOC650226 | S_Shore | TSS200 | Yes |
| Control-Sample161_Be | chr16:2653238-2653839 | LOC652276 | Island | TSS200 | Yes |
| Control-Sample162_Be | chr16:2653222-2653839 | LOC652276 | Island | TSS200 | Yes |
| Control-Sample227_Be | chr16:2653222-2653987 | LOC652276 | Island | TSS200 | Yes |
| Control-Sample233_Be | chr16:2653222-2653306 | LOC652276 | Island | TSS200 | Yes |
| Control-Sample234_Be | chr16:2653250-2653306 | LOC652276 | Island | TSS200 | Yes |
| Control-Sample419_Be | chr16:2653250-2653280 | LOC652276 | Island | TSS200 | Yes |
| Control-Sample564_Be | chr16.0:2653250-2653280 | LOC652276 | Island | TSS200 | Yes |
| Control-Sample517_Be | chr1.0:40024971-40025415 | LOC728448 | | Body | No |
| Control-Sample1_Beta | chr2:121223534-121224327 | LOC84931 | | Body | Yes |
| Control-Sample11_Beta | chr2:121223534-121224009 | LOC84931 | | Body | Yes |
| Control-Sample19_Beta | chr2:121223534-121223964 | LOC84931 | | Body | Yes |
| Control-Sample57_Beta | chr2:121223534-121223847 | LOC84931 | | Body | Yes |
| Control-Sample57_Beta | chr2:121223947-121224009 | LOC84931 | | TSS200 | Yes |
| Control-Sample10_Beta | chr15:78286521-78287690 | LOC91450 | | Body | No |
| Control-Sample227_Be | chr4:6675660-6675680 | LOC93622 | N_Shore | TSS200 | Yes |
| Control-Sample276_Be | chr4:6675674-6675756 | LOC93622 | N_Shore | TSS200 | Yes |
| Control-Sample15_Beta | chr2:100937072-100937584 | LONRF2 | N_Shore | Body | No |
| Control-Sample14_Beta | chr15:74218715-74218767 | LOXL1 | Island | TSS200 | Yes |
| Control-Sample331_Be | chr15:74218733-74218767 | LOXL1 | Island | TSS200 | Yes |
| Control-Sample1_Beta | chr2:74780268-74780617 | LOXL3 | N_Shore | 5'UTR | Yes |
| Control-Sample12_Beta | chr2:74780272-74780617 | LOXL3 | N_Shore | 5'UTR | Yes |
| Control-Sample45_Beta | chr2:74780229-74780617 | LOXL3 | N_Shore | 5'UTR | Yes |
| Control-Sample1_Beta | chr20:39969481-39969604 | LPIN3 | S_Shore | TSS200 | Yes |
| Control-Sample12_Beta | chr20:39969481-39969542 | LPIN3 | S_Shore | TSS200 | Yes |
| Control-Sample13_Beta | chr20:39969481-39969536 | LPIN3 | S_Shore | TSS200 | Yes |
| Control-Sample1_Beta | chr19:11472110-11472731 | LPPR2;LPPR2 | Island | Body;Body | Yes |
| Control-Sample10_Beta | chr19:11472110-11472397 | LPPR2;LPPR2 | Island | Body;Body | Yes |
| Control-Sample52_Beta | chr19:11472261-11472731 | LPPR2;LPPR2 | Island | Body;Body | Yes |
| Control-Sample20_Beta | chr4:151500931-151501627 | LRBA | Island | Body | Yes |
| Control-Sample77_Beta | chr4:151503227-151503588 | LRBA;MAB21L2;MAB21L2 | N_Shore | Body;1stExon;5'UTR | Yes |
| Control-Sample194_Be | chr3:197517984-197518007 | LRCH3 | Island | TSS200 | No |
| Control-Sample19_Beta | chr10:86001254-86001509 | LRIT1 | S_Shelf | TSS200 | No |
| Control-Sample190_Be | chr2:170219228-170219405 | LRP2 | N_Shore | TSS200 | No |
| Control-Sample1_Beta | chr11:68081207-68081686 | LRP5 | S_Shore | Body | Yes |
| Control-Sample1_Beta | chr11:68199656-68199794 | LRP5 | N_Shore | Body | Yes |
| Control-Sample10_Beta | chr11:68081207-68081257 | LRP5 | S_Shore | Body | Yes |
| Control-Sample12_Beta | chr11:68081226-68081686 | LRP5 | S_Shore | Body | Yes |
| Control-Sample13_Beta | chr11:68081226-68081607 | LRP5 | S_Shore | Body | Yes |
| Control-Sample18_Beta | chr11:68081207-68081607 | LRP5 | S_Shore | Body | Yes |
| Control-Sample23_Beta | chr11:68134211-68134398 | LRP5 | | Body | Yes |
| Control-Sample42_Beta | chr11:68081257-68081686 | LRP5 | S_Shore | Body | Yes |
| Control-Sample19_Beta | chr1:53794150-53794313 | LRP8;LRP8;LRP8;LRP8 | S_Shore | TSS1500;TSS1500;TSS1500;TSS1500 | No |
| Control-Sample38_Beta | chr4:3526685-3526865 | LRPAP1 | Island | Body | No |
| Control-Sample194_Be | chr8:145747659-145747746 | LRRC14 | Island | 3'UTR | No |
| Control-Sample70_Beta | chr5:191500-192103 | LRRC14B | N_Shore | TSS200 | Yes |
| Control-Sample71_Beta | chr5:191500-191806 | LRRC14B | N_Shore | TSS200 | Yes |
| Control-Sample57_Beta | chr7:102553045-102553457 | LRRC17;FBXL13;FBXL13;LRRC17 | | TSS1500;Body;Body;TSS1500 | Yes |
| Control-Sample1_Beta | chr7:102553293-102553457 | LRRC17;FBXL13;LRRC17;FBXL13 | | TSS200;Body;TSS200;Body | Yes |
| Control-Sample27_Beta | chr7:102553293-102553437 | LRRC17;FBXL13;LRRC17;FBXL13 | | TSS200;Body;TSS200;Body | Yes |
| Control-Sample33_Beta | chr7:102553293-102553432 | LRRC17;FBXL13;LRRC17;FBXL13 | | TSS200;Body;TSS200;Body | Yes |
| Control-Sample108_Be | chr7:102553432-102553457 | LRRC17;FBXL13;LRRC17;LRRC17;LRRC17;FBXL13 | | 5'UTR;Body;5'UTR;1stExon;1stExon;Body | Yes |
| Control-Sample66_Beta | chr3:46618668-46619158 | LRRC2;TDGF1 | Island | 5'UTR;TSS1500 | No |
| Control-Sample4_Beta | chr10:134150489-134150567 | LRRC27;LRRC27;LRRC27;LRRC27;LRRC27 | | Body;TSS200;Body;Body;Body | No |
| Control-Sample230_Be | chr17:28902711-28904298 | LRRC37B2 | N_Shore | TSS1500 | No |
| Control-Sample1_Beta | chr17:17910235-17910425 | LRRC48;LRRC48;LRRC48;LRRC48 | Island | Body;Body;Body;Body | Yes |
| Control-Sample20_Beta | chr17:17910351-17910425 | LRRC48;LRRC48;LRRC48;LRRC48 | Island | Body;Body;Body;Body | Yes |
| Control-Sample20_Beta | chr1:90309410-90309998 | LRRC8D;LRRC8D | Island | 5'UTR;5'UTR | No |
| Control-Sample1_Beta | chr2:238599734-238600061 | LRRFIP1;LRRFIP1;LRRFIP1;LRRFIP1;LRRFIP1 | N_Shore | TSS1500;TSS1500;TSS1500;TSS1500;Body | Yes |
| Control-Sample13_Beta | chr2:238599761-238600061 | LRRFIP1;LRRFIP1;LRRFIP1;LRRFIP1;LRRFIP1 | N_Shore | TSS1500;TSS1500;TSS1500;TSS1500;Body | Yes |
| Control-Sample1_Beta | chr15:101587658-101587971 | LRRK1 | | Body | Yes |
| Control-Sample19_Beta | chr15:101587506-101587971 | LRRK1 | | Body | Yes |
| Control-Sample1_Beta | chr3:3843342-3843887 | LRRN1 | S_Shore | 5'UTR | No |
| Control-Sample1_Beta | chr11:62457291-62457345 | LRRN4CL | S_Shore | TSS200 | Yes |
| Control-Sample2_Beta | chr11:62457325-62457345 | LRRN4CL | S_Shore | TSS200 | Yes |
| Control-Sample108_Be | chr5:138210550-138211184 | LRRTM2;CTNNA1 | | Body;Body | Yes |
| Control-Sample19_Beta | chr3:54961956-54962380 | LRTM1;CACNA2D3;LRTM1 | | 5'UTR;Body;1stExon | Yes |
| Control-Sample19_Beta | chr11:1891872-1891947 | LSP1;LSP1;LSP1;LSP1 | N_Shore | 5'UTR;TSS1500;TSS1500;Body | No |
| Control-Sample108_Be | chr2:33359281-33359688 | LTBP1;LTBP1;LTBP1;LTBP1;LTBP1 | | Body;TSS1500;TSS1500;TSS1500;TSS1500 | No |
| Control-Sample1_Beta | chr11:65306731-65307006 | LTBP3;LTBP3;LTBP3 | Island | Body;Body;Body | Yes |
| Control-Sample1_Beta | chr11:65315205-65315466 | LTBP3;LTBP3;LTBP3 | Island | Body;Body;Body | Yes |
| Control-Sample10_Beta | chr11:65314021-65314556 | LTBP3;LTBP3;LTBP3 | Island | Body;Body;Body | Yes |
| Control-Sample13_Beta | chr11:65314161-65314556 | LTBP3;LTBP3;LTBP3 | Island | Body;Body;Body | Yes |
| Control-Sample20_Beta | chr11:65314161-65315030 | LTBP3;LTBP3;LTBP3 | Island | Body;Body;Body | Yes |
| Control-Sample1_Beta | chr5:179220795-179221059 | LTC4S | N_Shore | TSS200 | Yes |
| Control-Sample22_Beta | chr5:179220545-179220795 | LTC4S | N_Shelf | TSS1500 | Yes |
| Control-Sample58_Beta | chr5:179220589-179221059 | LTC4S | N_Shelf | TSS1500 | Yes |
| Control-Sample2_Beta | chr3:46506206-46506374 | LTF | Island | Body | Yes |
| Control-Sample6_Beta | chr3:46506104-46506292 | LTF | N_Shore | Body | Yes |
| Control-Sample11_Beta | chr15:41803428-41803772 | LTK;LTK;LTK | N_Shore | Body;Body;Body | No |
| Control-Sample1_Beta | chr5:115299071-115299646 | LVRN | Island | Body | Yes |
| Control-Sample12_Beta | chr5:115299071-115299475 | LVRN | Island | Body | Yes |
| Control-Sample1_Beta | chr3:158390329-158390671 | LXN;LXN;GFM1 | | 1stExon;5'UTR;Body | Yes |
| Control-Sample18_Beta | chr3:158390329-158390510 | LXN;LXN;GFM1 | | 1stExon;5'UTR;Body | Yes |
| Control-Sample24_Beta | chr3:158390329-158390525 | LXN;LXN;GFM1 | | 1stExon;5'UTR;Body | Yes |
| Control-Sample33_Beta | chr3:158390329-158390632 | LXN;LXN;GFM1 | | 1stExon;5'UTR;Body | Yes |
| Control-Sample1_Beta | chr8:143868023-143868062 | LY6D | | TSS200 | Yes |

| | | | | | |
|---|---|---|---|---|---|
| Control-Sample18_Beta | chr8:143868013-143868136 | LY6D | | TSS200 | Yes |
| Control-Sample24_Beta | chr8:143868062-143868158 | LY6D | | TSS200 | Yes |
| Control-Sample26_Beta | chr8:143868013-143868158 | LY6D | | TSS200 | Yes |
| Control-Sample38_Beta | chr8:143868013-143868110 | LY6D | | TSS200 | Yes |
| Control-Sample41_Beta | chr8:143868013-143868062 | LY6D | | TSS200 | Yes |
| Control-Sample10_Beta | chr8:143867942-143868110 | LY6D;LY6D | | 1stExon;5'UTR | Yes |
| Control-Sample11_Beta | chr8:143867942-143868136 | LY6D;LY6D | | 1stExon;5'UTR | Yes |
| Control-Sample12_Beta | chr8:143867942-143868158 | LY6D;LY6D | | 1stExon;5'UTR | Yes |
| Control-Sample34_Beta | chr8:143867942-143868062 | LY6D;LY6D | | 1stExon;5'UTR | Yes |
| Control-Sample3_Beta | chr6:31649619-31649638 | LY6G5C | N_Shore | TSS1500 | No |
| Control-Sample26_Beta | chr6:31683051-31683352 | LY6G6E;LY6G6E;LY6G6D | | TSS1500;TSS1500;TSS200 | Yes |
| Control-Sample40_Beta | chr6:31683051-31683109 | LY6G6E;LY6G6E;LY6G6D | | TSS1500;TSS1500;TSS200 | Yes |
| Control-Sample10_Beta | chr6:6588846-6589075 | LY86;LOC285780 | | TSS200;Body | No |
| Control-Sample5_Beta | chr19:13209901-13210356 | LYL1 | Island | 3'UTR | No |
| Control-Sample1_Beta | chr8:56852148-56852290 | LYN;LYN | | 5'UTR;5'UTR | Yes |
| Control-Sample19_Beta | chr8:56851846-56852290 | LYN;LYN | | 5'UTR;5'UTR | Yes |
| Control-Sample19_Beta | chr8:143859734-143859786 | LYNX1;LYNX1;LYNX1;LYNX1 | S_Shore | TSS200;TSS1500;TSS200;TSS1500 | Yes |
| Control-Sample230_Be | chr8:143858414-143859410 | LYNX1;LYNX1;LYNX1;LYNX1;LYNX1 | Island | 5'UTR;5'UTR;1stExon;5'UTR;5'UTR | Yes |
| Control-Sample108_Be | chr2:133430101-133430560 | LYPD1;NCKAP5;NCKAP5 | S_Shore | TSS1500;3'UTR;3'UTR | No |
| Control-Sample11_Beta | chr15:52030519-52030746 | LYSMD2;LYSMD2 | S_Shore | TSS200;5'UTR | No |
| Control-Sample18_Beta | chr8:20112813-20113038 | LZTS1 | | TSS200 | No |
| Control-Sample1_Beta | chr10:102765350-102765404 | LZTS2 | Island | Body | No |
| Control-Sample19_Beta | chr4:151502935-151504187 | MAB21L2;LRBA | N_Shore | TSS200;Body | Yes |
| Control-Sample68_Beta | chr4:151502935-151504138 | MAB21L2;LRBA | N_Shore | TSS200;Body | Yes |
| Control-Sample1_Beta | chr11:63784063-63784126 | MACROD1 | | Body | Yes |
| Control-Sample1_Beta | chr11:63887426-63887540 | MACROD1 | Island | Body | Yes |
| Control-Sample1_Beta | chr11:63906050-63906104 | MACROD1 | | Body | Yes |
| Control-Sample11_Beta | chr11:63827361-63828039 | MACROD1 | | Body | Yes |
| Control-Sample12_Beta | chr11:63827361-63828228 | MACROD1 | | Body | Yes |
| Control-Sample13_Beta | chr11:63827361-63827695 | MACROD1 | | Body | Yes |
| Control-Sample2_Beta | chr11:63912698-63912769 | MACROD1 | | Body | Yes |
| Control-Sample4_Beta | chr11:63827361-63827879 | MACROD1 | | Body | Yes |
| Control-Sample65_Beta | chr11:63894400-63894504 | MACROD1 | | Body | Yes |
| Control-Sample1_Beta | chr7:1893710-1894209 | MAD1L1;MAD1L1;MAD1L1 | N_Shelf | Body;Body;Body | Yes |
| Control-Sample1_Beta | chr7:1952518-1952807 | MAD1L1;MAD1L1;MAD1L1 | S_Shelf | Body;Body;Body | Yes |
| Control-Sample1_Beta | chr7:2082961-2083181 | MAD1L1;MAD1L1;MAD1L1 | | Body;Body;Body | Yes |
| Control-Sample1_Beta | chr7:2140143-2140321 | MAD1L1;MAD1L1;MAD1L1 | | Body;Body;Body | Yes |
| Control-Sample10_Beta | chr7:1939814-1940280 | MAD1L1;MAD1L1;MAD1L1 | S_Shore | Body;Body;Body | Yes |
| Control-Sample10_Beta | chr7:2151681-2151788 | MAD1L1;MAD1L1;MAD1L1 | N_Shelf | Body;Body;Body | Yes |
| Control-Sample11_Beta | chr7:1956900-1957317 | MAD1L1;MAD1L1;MAD1L1 | | Body;Body;Body | Yes |
| Control-Sample11_Beta | chr7:2082961-2083197 | MAD1L1;MAD1L1;MAD1L1 | | Body;Body;Body | Yes |
| Control-Sample16_Beta | chr7:1952561-1952807 | MAD1L1;MAD1L1;MAD1L1 | S_Shelf | Body;Body;Body | Yes |
| Control-Sample17_Beta | chr7:2004329-2004389 | MAD1L1;MAD1L1;MAD1L1 | N_Shelf | Body;Body;Body | Yes |
| Control-Sample19_Beta | chr7:2004020-2004329 | MAD1L1;MAD1L1;MAD1L1 | N_Shelf | Body;Body;Body | Yes |
| Control-Sample19_Beta | chr7:2025606-2025903 | MAD1L1;MAD1L1;MAD1L1 | N_Shore | Body;Body;Body | Yes |
| Control-Sample20_Beta | chr7:2004020-2004365 | MAD1L1;MAD1L1;MAD1L1 | N_Shelf | Body;Body;Body | Yes |
| Control-Sample20_Beta | chr7:2082961-2083583 | MAD1L1;MAD1L1;MAD1L1 | | Body;Body;Body | Yes |
| Control-Sample20_Beta | chr7:2139216-2139259 | MAD1L1;MAD1L1;MAD1L1 | | Body;Body;Body | Yes |
| Control-Sample20_Beta | chr7:2150534-2151007 | MAD1L1;MAD1L1;MAD1L1 | | Body;Body;Body | Yes |
| Control-Sample27_Beta | chr7:1952518-1952600 | MAD1L1;MAD1L1;MAD1L1 | S_Shelf | Body;Body;Body | Yes |
| Control-Sample368_Be | chr7:2106029-2106404 | MAD1L1;MAD1L1;MAD1L1 | N_Shore | Body;Body;Body | Yes |
| Control-Sample38_Beta | chr7:2059150-2059412 | MAD1L1;MAD1L1;MAD1L1 | N_Shore | Body;Body;Body | Yes |
| Control-Sample4_Beta | chr7:2105368-2106029 | MAD1L1;MAD1L1;MAD1L1 | N_Shore | Body;Body;Body | Yes |
| Control-Sample465_Be | chr7:2070226-2070409 | MAD1L1;MAD1L1;MAD1L1 | | Body;Body;Body | Yes |
| Control-Sample57_Beta | chr7:2078981-2079008 | MAD1L1;MAD1L1;MAD1L1 | | Body;Body;Body | Yes |
| Control-Sample57_Beta | chr7:2261337-2261506 | MAD1L1;MAD1L1;MAD1L1 | | Body;Body;Body | Yes |
| Control-Sample64_Beta | chr7:1881887-1882135 | MAD1L1;MAD1L1;MAD1L1 | S_Shore | Body;Body;Body | Yes |
| Control-Sample1_Beta | chr4:1294783-1295047 | MAEA;MAEA | | Body;Body | Yes |
| Control-Sample27_Beta | chr4:1294783-1295078 | MAEA;MAEA | | Body;Body | Yes |
| Control-Sample19_Beta | chr17:79881523-79881543 | MAFG;MAFG | S_Shore | 5'UTR;TSS200 | Yes |
| Control-Sample20_Beta | chr17:79881483-79881543 | MAFG;MAFG | S_Shore | 5'UTR;TSS200 | Yes |
| Control-Sample9_Beta | chr17:79881483-79881529 | MAFG;MAFG | S_Shore | 5'UTR;TSS200 | Yes |
| Control-Sample1_Beta | chr7:1576976-1577016 | MAFK | S_Shore | 5'UTR | No |
| Control-Sample108_Be | chr15:23894198-23894273 | MAGEL2 | | TSS1500 | No |
| Control-Sample1_Beta | chr7:77648567-77648624 | MAGI2 | Island | 3'UTR | No |
| Control-Sample19_Beta | chr2:110872620-110872729 | MALL | N_Shore | Body | No |
| Control-Sample34_Beta | chr19:49223845-49224165 | MAMSTR;RASIP1 | Island | TSS1500;3'UTR | Yes |
| Control-Sample23_Beta | chr2:172945274-172945698 | MAP1D | Island | 3'UTR | Yes |
| Control-Sample3_Beta | chr2:172945058-172945698 | MAP1D | Island | Body | Yes |
| Control-Sample82_Beta | chr2:172945058-172945274 | MAP1D | Island | Body | Yes |
| Control-Sample1_Beta | chr12:116996871-116997069 | MAP1LC3B2 | | TSS1500 | Yes |
| Control-Sample10_Beta | chr12:116996773-116997022 | MAP1LC3B2 | | TSS1500 | Yes |
| Control-Sample12_Beta | chr12:116996871-116997072 | MAP1LC3B2 | | TSS1500 | Yes |
| Control-Sample14_Beta | chr12:116996871-116997022 | MAP1LC3B2 | | TSS1500 | Yes |
| Control-Sample20_Beta | chr12:116996931-116997069 | MAP1LC3B2 | | TSS1500 | Yes |
| Control-Sample23_Beta | chr12:116996773-116996931 | MAP1LC3B2 | | TSS1500 | Yes |
| Control-Sample3_Beta | chr12:116996773-116997069 | MAP1LC3B2 | | TSS1500 | Yes |
| Control-Sample63_Beta | chr12:116996773-116997072 | MAP1LC3B2 | | TSS1500 | Yes |
| Control-Sample10_Beta | chr11:65374833-65375067 | MAP3K11 | Island | Body | No |
| Control-Sample1_Beta | chr3:185080218-185080423 | MAP3K13 | | TSS1500 | No |
| Control-Sample1_Beta | chr1:27687232-27687695 | MAP3K6 | Island | Body | Yes |
| Control-Sample1_Beta | chr1:27693782-27693840 | MAP3K6 | S_Shore | TSS1500 | Yes |
| Control-Sample14_Beta | chr1:27687232-27687685 | MAP3K6 | Island | Body | Yes |
| Control-Sample24_Beta | chr1:27687454-27687695 | MAP3K6 | Island | Body | Yes |
| Control-Sample40_Beta | chr1:27687074-27687695 | MAP3K6 | Island | Body | Yes |
| Control-Sample568_Be | chr4.0:87281412-87281511 | MAPK10;MAPK10;MAPK10 | | TSS200;TSS200;5'UTR | No |
| Control-Sample1_Beta | chr11:45924191-45924507 | MAPK8IP1 | Island | Body | No |
| Control-Sample29_Beta | chr17:60828157-60828406 | MARCH10;MARCH10 | | Body;Body | No |
| Control-Sample1_Beta | chr10:99473206-99473557 | MARVELD1;MARVELD1 | Island | TSS1500;TSS1500 | Yes |
| Control-Sample10_Beta | chr10:99473032-99473557 | MARVELD1;MARVELD1 | N_Shore | TSS1500;TSS1500 | Yes |
| Control-Sample19_Beta | chr10:99473085-99473557 | MARVELD1;MARVELD1 | N_Shore | TSS1500;TSS1500 | Yes |

| | | | | | |
|---|---|---|---|---|---|
| Control-Sample10_Beta | chr5:68710831-68710912 | MARVELD2;MARVELD2 | Island | TSS200;TSS200 | Yes |
| Control-Sample12_Beta | chr5:68710818-68710912 | MARVELD2;MARVELD2 | Island | TSS200;TSS200 | Yes |
| Control-Sample171_Be | chr5:68710818-68710907 | MARVELD2;MARVELD2 | Island | TSS200;TSS200 | Yes |
| Control-Sample1_Beta | chr19:12958803-12958913 | MAST1 | Island | Body | Yes |
| Control-Sample15_Beta | chr19:12958399-12958913 | MAST1 | Island | Body | Yes |
| Control-Sample1_Beta | chr20:43936981-43937155 | MATN4;RBPJL;MATN4;MATN4 | S_Shore | TSS200;Body;TSS200;TSS200 | Yes |
| Control-Sample11_Beta | chr20:43936853-43937155 | MATN4;RBPJL;MATN4;MATN4;MATN4;MATN4 | S_Shore | 1stExon;Body;5'UTR;5'UTR;1stExon;1stExon;5'UTR | Yes |
| Control-Sample17_Beta | chr20:43936853-43937138 | MATN4;RBPJL;MATN4;MATN4;MATN4;MATN4;MATN4 | S_Shore | 1stExon;Body;5'UTR;5'UTR;1stExon;1stExon;5'UTR | Yes |
| Control-Sample171_Be | chr16:29819451-29819931 | MAZ;MAZ | Island | Body;Body | Yes |
| Control-Sample223_Be | chr16:29819361-29819461 | MAZ;MAZ | Island | Body;Body | Yes |
| Control-Sample270_Be | chr16:29819361-29819931 | MAZ;MAZ | Island | Body;Body | Yes |
| Control-Sample6_Beta | chr16:29819451-29820119 | MAZ;MAZ | Island | Body;Body | Yes |
| Control-Sample43_Beta | chr22:36013560-36013678 | MB;MB;MB | | TSS200;TSS200;5'UTR | No |
| Control-Sample108_Be | chr3:152016872-152017240 | MBNL1;MBNL1;MBNL1;MBNL1;MBNL1;MBNL1 | | TSS1500;5'UTR;TSS1500;TSS1500;TSS1500;TSS1500 | No |
| Control-Sample25_Beta | chr13:97991057-97991122 | MBNL2;MBNL2 | | Body;Body | No |
| Control-Sample1_Beta | chr18:74770431-74770815 | MBP;MBP | Island | Body;Body | Yes |
| Control-Sample1_Beta | chr18:74691097-74692253 | MBP;MBP;MBP;MBP;MBP | N_Shore | 3'UTR;3'UTR;3'UTR;3'UTR;3'UTR | Yes |
| Control-Sample100_Be | chr18:74691269-74692145 | MBP;MBP;MBP;MBP;MBP | Island | 3'UTR;3'UTR;3'UTR;3'UTR;3'UTR | Yes |
| Control-Sample12_Beta | chr18:74691367-74692253 | MBP;MBP;MBP;MBP;MBP | Island | 3'UTR;3'UTR;3'UTR;3'UTR;3'UTR | Yes |
| Control-Sample171_Be | chr18:74691097-74691423 | MBP;MBP;MBP;MBP;MBP | N_Shore | 3'UTR;3'UTR;3'UTR;3'UTR;3'UTR | Yes |
| Control-Sample19_Beta | chr18:74691269-74692253 | MBP;MBP;MBP;MBP;MBP | Island | 3'UTR;3'UTR;3'UTR;3'UTR;3'UTR | Yes |
| Control-Sample223_Be | chr18:74691097-74691367 | MBP;MBP;MBP;MBP;MBP | N_Shore | 3'UTR;3'UTR;3'UTR;3'UTR;3'UTR | Yes |
| Control-Sample475_Be | chr18:74692214-74692253 | MBP;MBP;MBP;MBP;MBP | S_Shore | 3'UTR;3'UTR;3'UTR;3'UTR;3'UTR | Yes |
| Control-Sample67_Beta | chr18:74691423-74692253 | MBP;MBP;MBP;MBP;MBP | Island | 3'UTR;3'UTR;3'UTR;3'UTR;3'UTR | Yes |
| Control-Sample73_Beta | chr18:74691097-74692145 | MBP;MBP;MBP;MBP;MBP | N_Shore | 3'UTR;3'UTR;3'UTR;3'UTR;3'UTR | Yes |
| Control-Sample1_Beta | chr18:74728834-74729175 | MBP;MBP;MBP;MBP;MBP;MBP | | 1stExon;1stExon;Body;1stExon;Body;1stExon | Yes |
| Control-Sample1_Beta | chr18:74729481-74729551 | MBP;MBP;MBP;MBP;MBP;MBP | | TSS1500;TSS1500;Body;Body;TSS1500;TSS1500 | Yes |
| Control-Sample10_Beta | chr18:74728834-74729551 | MBP;MBP;MBP;MBP;MBP;MBP | | 1stExon;1stExon;Body;1stExon;Body;1stExon | Yes |
| Control-Sample103_Be | chr18:74728931-74728979 | MBP;MBP;MBP;MBP;MBP;MBP | | 1stExon;1stExon;Body;1stExon;Body;1stExon | Yes |
| Control-Sample105_Be | chr18:74728834-74729021 | MBP;MBP;MBP;MBP;MBP;MBP | | 1stExon;1stExon;Body;1stExon;Body;1stExon | Yes |
| Control-Sample12_Beta | chr18:74729074-74729551 | MBP;MBP;MBP;MBP;MBP;MBP | | TSS200;Body;TSS200;TSS200;TSS200;Body | Yes |
| Control-Sample124_Be | chr18:74728834-74728979 | MBP;MBP;MBP;MBP;MBP;MBP | | 1stExon;1stExon;Body;1stExon;Body;1stExon | Yes |
| Control-Sample13_Beta | chr18:74729074-74729175 | MBP;MBP;MBP;MBP;MBP;MBP | | TSS200;Body;TSS200;TSS200;TSS200;Body | Yes |
| Control-Sample151_Be | chr18:74728931-74729074 | MBP;MBP;MBP;MBP;MBP;MBP | | 1stExon;1stExon;Body;1stExon;Body;1stExon | Yes |
| Control-Sample170_Be | chr18:74728834-74729074 | MBP;MBP;MBP;MBP;MBP;MBP | | 1stExon;1stExon;Body;1stExon;Body;1stExon | Yes |
| Control-Sample2_Beta | chr18:74728834-74729064 | MBP;MBP;MBP;MBP;MBP;MBP | | 1stExon;1stExon;Body;1stExon;Body;1stExon | Yes |
| Control-Sample2_Beta | chr18:74729145-74729175 | MBP;MBP;MBP;MBP;MBP;MBP | | TSS200;Body;TSS200;TSS200;TSS200;Body | Yes |
| Control-Sample233_Be | chr18:74728834-74728931 | MBP;MBP;MBP;MBP;MBP;MBP | | 1stExon;1stExon;Body;1stExon;Body;1stExon | Yes |
| Control-Sample26_Beta | chr18:74728931-74729021 | MBP;MBP;MBP;MBP;MBP;MBP | | 1stExon;1stExon;Body;1stExon;Body;1stExon | Yes |
| Control-Sample26_Beta | chr18:74729074-74729151 | MBP;MBP;MBP;MBP;MBP;MBP | | TSS200;Body;TSS200;TSS200;TSS200;Body | Yes |
| Control-Sample26_Beta | chr18:74729251-74729551 | MBP;MBP;MBP;MBP;MBP;MBP | | TSS200;Body;TSS200;TSS200;TSS200;Body | Yes |
| Control-Sample463_Be | chr18:74728838-74729151 | MBP;MBP;MBP;MBP;MBP;MBP | | 1stExon;1stExon;Body;1stExon;Body;1stExon | Yes |
| Control-Sample54_Beta | chr18:74729151-74729551 | MBP;MBP;MBP;MBP;MBP;MBP | | TSS200;Body;TSS200;TSS200;TSS200;Body | Yes |
| Control-Sample58_Beta | chr18:74728834-74729151 | MBP;MBP;MBP;MBP;MBP;MBP | | 1stExon;1stExon;Body;1stExon;Body;1stExon | Yes |
| Control-Sample585_Be | chr18.0:74728931-74728979 | MBP;MBP;MBP;MBP;MBP;MBP | | 1stExon;1stExon;Body;1stExon;Body;1stExon | Yes |
| Control-Sample673_Be | chr18:74728838-74729021 | MBP;MBP;MBP;MBP;MBP;MBP | | 1stExon;1stExon;Body;1stExon;Body;1stExon | Yes |
| Control-Sample68_Beta | chr18:74728834-74729145 | MBP;MBP;MBP;MBP;MBP;MBP | | 1stExon;1stExon;Body;1stExon;Body;1stExon | Yes |
| Control-Sample10_Beta | chr18:13916113-13916355 | MC2R | | TSS1500 | No |
| Control-Sample1_Beta | chr5:112824614-112825132 | MCC | S_Shore | TSS200 | Yes |
| Control-Sample2_Beta | chr5:112824614-112824700 | MCC | S_Shore | TSS200 | Yes |
| Control-Sample2_Beta | chr5:112824765-112825132 | MCC | S_Shore | TSS1500 | Yes |
| Control-Sample5_Beta | chr3:182817190-182817381 | MCCC1 | Island | 1stExon | No |
| Control-Sample464_Be | chr13:113627866-113628068 | MCF2L | | Body | Yes |
| Control-Sample1_Beta | chr13:113642457-113642515 | MCF2L;MCF2L | | Body;Body | Yes |
| Control-Sample1_Beta | chr13:113698931-113700534 | MCF2L;MCF2L | N_Shelf | Body;Body | Yes |
| Control-Sample11_Beta | chr13:113698946-113700027 | MCF2L;MCF2L | N_Shelf | Body;Body | Yes |
| Control-Sample12_Beta | chr13:113698408-113698621 | MCF2L;MCF2L | N_Shelf | Body;Body | Yes |
| Control-Sample12_Beta | chr13:113698946-113700534 | MCF2L;MCF2L | N_Shelf | Body;Body | Yes |
| Control-Sample151_Be | chr13:113708289-113708662 | MCF2L;MCF2L | S_Shelf | Body;Body | Yes |
| Control-Sample18_Beta | chr13:113698994-113699014 | MCF2L;MCF2L | N_Shelf | Body;Body | Yes |
| Control-Sample19_Beta | chr13:113752435-113753244 | MCF2L;MCF2L | S_Shore | 3'UTR;3'UTR | Yes |
| Control-Sample23_Beta | chr13:113698994-113700027 | MCF2L;MCF2L | N_Shelf | Body;Body | Yes |
| Control-Sample36_Beta | chr13:113698408-113698688 | MCF2L;MCF2L | N_Shelf | Body;Body | Yes |
| Control-Sample4_Beta | chr13:113676829-113677034 | MCF2L;MCF2L | Island | Body;Body | Yes |
| Control-Sample4_Beta | chr13:113698994-113700534 | MCF2L;MCF2L | N_Shelf | Body;Body | Yes |
| Control-Sample60_Beta | chr13:113698606-113698688 | MCF2L;MCF2L | N_Shelf | Body;Body | Yes |
| Control-Sample7_Beta | chr13:113698946-113700470 | MCF2L;MCF2L | N_Shelf | Body;Body | Yes |
| Control-Sample1_Beta | chr3:182970719-182970850 | MCF2L2;B3GNT5 | N_Shore | Body;TSS1500 | Yes |
| Control-Sample1_Beta | chr3:182972625-182973032 | MCF2L2;B3GNT5 | Island | Body;5'UTR | Yes |
| Control-Sample10_Beta | chr3:182970833-182970850 | MCF2L2;B3GNT5 | N_Shore | Body;TSS200 | Yes |
| Control-Sample16_Beta | chr3:182970728-182970850 | MCF2L2;B3GNT5 | N_Shore | Body;TSS1500 | Yes |
| Control-Sample1_Beta | chr7:99690806-99691200 | MCM7;MCM7 | S_Shelf | Body;Body | Yes |
| Control-Sample19_Beta | chr7:99691314-99691447 | MCM7;MCM7;MIR25 | | Body;Body;TSS200 | Yes |
| Control-Sample57_Beta | chr7:99691200-99691447 | MCM7;MIR25;MCM7 | | Body;Body;Body | Yes |
| Control-Sample32_Beta | chr1:85512957-85513191 | MCOLN3 | N_Shore | 5'UTR | No |
| Control-Sample23_Beta | chr8:6456182-6456854 | MCPH1 | | Body | Yes |
| Control-Sample24_Beta | chr8:6456182-6456465 | MCPH1 | | Body | Yes |
| Control-Sample28_Beta | chr6:30685626-30685744 | MDC1 | S_Shore | TSS200 | No |
| Control-Sample1_Beta | chr6:37617864-37618009 | MDGA1 | Island | Body | Yes |
| Control-Sample21_Beta | chr6:37667101-37667555 | MDGA1 | Island | TSS1500 | Yes |
| Control-Sample5_Beta | chr6:84140683-84140898 | ME1;ME1 | Island | 5'UTR;1stExon | No |
| Control-Sample83_Beta | chr1:156466729-156467188 | MEF2D | N_Shelf | 5'UTR | No |
| Control-Sample58_Beta | chr14:101298824-101298920 | MEG3;MEG3;MEG3 | | Body;Body;Body | No |
| Control-Sample11_Beta | chr1:3477168-3477302 | MEGF6 | | Body | No |
| Control-Sample162_Be | chr22:42095430-42095443 | MEI1 | Island | TSS200 | No |
| Control-Sample1_Beta | chr2:66667852-66668012 | MEIS1 | | Body | Yes |
| Control-Sample21_Beta | chr2:66673316-66673985 | MEIS1 | Island | Body | Yes |
| Control-Sample23_Beta | chr2:66672841-66673449 | MEIS1 | Island | Body | Yes |
| Control-Sample24_Beta | chr2:66673316-66673449 | MEIS1 | Island | Body | Yes |
| Control-Sample57_Beta | chr2:66664667-66664871 | MEIS1 | S_Shelf | Body | Yes |
| Control-Sample65_Beta | chr2:66664798-66665428 | MEIS1 | S_Shelf | Body | Yes |

| | | | | | |
|---|---|---|---|---|---|
| Control-Sample15_Beta | chr17:41739130-41739806 | MEOX1;MEOX1;MEOX1 | | TSS200;5'UTR;TSS200 | Yes |
| Control-Sample19_Beta | chr17:41739130-41739926 | MEOX1;MEOX1;MEOX1 | | TSS200;5'UTR;TSS200 | Yes |
| Control-Sample20_Beta | chr17:41739195-41739926 | MEOX1;MEOX1;MEOX1 | | TSS1500;5'UTR;TSS1500 | Yes |
| Control-Sample5_Beta | chr17:41739195-41739806 | MEOX1;MEOX1;MEOX1 | | TSS1500;5'UTR;TSS1500 | Yes |
| Control-Sample56_Beta | chr7:100026424-100026446 | MEPCE;ZCWPW1 | Island | TSS1500;TSS200 | No |
| Control-Sample113_Be | chr7:130132360-130132419 | MEST;MEST;MESTIT1;MEST | Island | 5'UTR;5'UTR;TSS1500;Body | No |
| Control-Sample11_Beta | chr1:170114946-170115351 | METTL11B | | TSS1500 | Yes |
| Control-Sample48_Beta | chr1:170114481-170115351 | METTL11B | | TSS1500 | Yes |
| Control-Sample108_Be | chr12:51318430-51318640 | METTL7A | | TSS200 | No |
| Control-Sample108_Be | chr12:56075089-56075367 | METTL7B | | TSS1500 | No |
| Control-Sample22_Beta | chr1:156046832-156047356 | MEX3A | Island | Body | No |
| Control-Sample12_Beta | chr15:82335638-82336109 | MEX3B | Island | Body | No |
| Control-Sample520_Be | chr5.0:153418793-153418932 | MFAP3;FAM114A2;MFAP3;MFAP3 | S_Shore | 5'UTR;TSS1500;5'UTR;Body | No |
| Control-Sample3_Beta | chr17:19290708-19291120 | MFAP4 | | TSS1500 | Yes |
| Control-Sample49_Beta | chr17:19290353-19290520 | MFAP4 | | Body | Yes |
| Control-Sample1_Beta | chr17:8702564-8702799 | MFSD6L;MFSD6L | Island | 5'UTR;1stExon | No |
| Control-Sample1_Beta | chr4:683060-683206 | MFSD7 | Island | TSS200 | No |
| Control-Sample12_Beta | chr5:180237085-180237222 | MGAT1;MGAT1;MGAT1 | Island | 1stExon;TSS1500;5'UTR | No |
| Control-Sample13_Beta | chr17:74868604-74868725 | MGAT5B;MGAT5B | S_Shore | Body;TSS200 | No |
| Control-Sample10_Beta | chr12:110172208-110172346 | MGC14436;MGC14436;C12orf34 | | Body;Body;5'UTR | No |
| Control-Sample10_Beta | chr4:183063194-183063459 | MGC45800 | N_Shore | Body | No |
| Control-Sample50_Beta | chr7:22539703-22539820 | MGC87042;MGC87042 | Island | 5'UTR;5'UTR | No |
| Control-Sample11_Beta | chr10:131323903-131324189 | MGMT | | Body | Yes |
| Control-Sample13_Beta | chr10:131323903-131324090 | MGMT | | Body | Yes |
| Control-Sample23_Beta | chr10:131323986-131324189 | MGMT | | Body | Yes |
| Control-Sample29_Beta | chr10:131348484-131348599 | MGMT | | Body | Yes |
| Control-Sample1_Beta | chr16:4714647-4714815 | MGRN1;MGRN1;MGRN1;MGRN1 | | Body;Body;Body;Body | Yes |
| Control-Sample19_Beta | chr16:4714647-4714794 | MGRN1;MGRN1;MGRN1;MGRN1 | | Body;Body;Body;Body | Yes |
| Control-Sample19_Beta | chr16:4729905-4730410 | MGRN1;MGRN1;MGRN1;MGRN1 | N_Shore | Body;Body;Body;Body | Yes |
| Control-Sample1_Beta | chr1:1564920-1565027 | MIB2;MIB2;MIB2;MIB2;MIB2;MIB2 | Island | Body;Body;Body;Body;Body;Body | Yes |
| Control-Sample11_Beta | chr1:1565541-1565931 | MIB2;MIB2;MIB2;MIB2;MIB2;MIB2 | Island | Body;Body;Body;Body;Body;Body | Yes |
| Control-Sample14_Beta | chr1:1564920-1565931 | MIB2;MIB2;MIB2;MIB2;MIB2;MIB2 | Island | Body;Body;Body;Body;Body;Body | Yes |
| Control-Sample1_Beta | chr11:64135650-64136975 | MIR1237;RPS6KA4;RPS6KA4 | N_Shore | TSS1500;Body;Body | Yes |
| Control-Sample11_Beta | chr11:64135650-64137747 | MIR1237;RPS6KA4;RPS6KA4 | N_Shore | TSS1500;Body;Body | Yes |
| Control-Sample17_Beta | chr11:64135650-64135902 | MIR1237;RPS6KA4;RPS6KA4 | N_Shore | TSS1500;Body;Body | Yes |
| Control-Sample23_Beta | chr11:64135650-64138176 | MIR1237;RPS6KA4;RPS6KA4 | N_Shore | TSS1500;Body;Body | Yes |
| Control-Sample3_Beta | chr11:64135650-64137524 | MIR1237;RPS6KA4;RPS6KA4 | N_Shore | TSS1500;Body;Body | Yes |
| Control-Sample37_Beta | chr11:64135674-64135902 | MIR1237;RPS6KA4;RPS6KA4 | N_Shore | TSS1500;Body;Body | Yes |
| Control-Sample41_Beta | chr11:64135691-64135902 | MIR1237;RPS6KA4;RPS6KA4 | N_Shore | TSS1500;Body;Body | Yes |
| Control-Sample63_Beta | chr11:64135674-64137524 | MIR1237;RPS6KA4;RPS6KA4 | N_Shore | TSS1500;Body;Body | Yes |
| Control-Sample170_Be | chr8:65291287-65291562 | MIR124-2 | S_Shore | TSS1500 | Yes |
| Control-Sample171_Be | chr8:65290848-65291341 | MIR124-2 | Island | TSS1500 | Yes |
| Control-Sample19_Beta | chr8:65291513-65292321 | MIR124-2 | S_Shore | TSS200 | Yes |
| Control-Sample194_Be | chr8:65291287-65291542 | MIR124-2 | S_Shore | TSS1500 | Yes |
| Control-Sample20_Beta | chr8:65291703-65291990 | MIR124-2 | S_Shore | TSS200 | Yes |
| Control-Sample223_Be | chr8:65291523-65291562 | MIR124-2 | S_Shore | TSS200 | Yes |
| Control-Sample108_Be | chr3:35785890-35786030 | MIR128-2;ARPP-21 | | TSS200;Body | Yes |
| Control-Sample291_Be | chr3:35785958-35786480 | MIR128-2;ARPP-21 | | TSS200;Body | Yes |
| Control-Sample1_Beta | chr22:22007154-22007633 | MIR130B;MIR301B | S_Shore | TSS1500;TSS200 | Yes |
| Control-Sample13_Beta | chr22:22007154-22007515 | MIR130B;MIR301B | S_Shore | TSS1500;TSS200 | Yes |
| Control-Sample92_Beta | chr22:22007154-22007528 | MIR130B;MIR301B | S_Shore | TSS1500;TSS200 | Yes |
| Control-Sample67_Beta | chr5:148808388-148808487 | MIR143;LOC728264 | | TSS200;Body | Yes |
| Control-Sample64_Beta | chr5:148810177-148810203 | MIR145;LOC728264 | | TSS200;Body | Yes |
| Control-Sample1_Beta | chr2:241394528-241395351 | MIR149;PP14571;GPC1 | N_Shore | TSS1500;Body;Body | Yes |
| Control-Sample11_Beta | chr2:241395181-241395351 | MIR149;PP14571;GPC1 | N_Shore | TSS1500;Body;Body | Yes |
| Control-Sample15_Beta | chr2:241395288-241395351 | MIR149;PP14571;GPC1 | Island | TSS200;Body;Body | Yes |
| Control-Sample19_Beta | chr2:241395181-241395678 | MIR149;PP14571;GPC1 | N_Shore | TSS1500;Body;Body | Yes |
| Control-Sample24_Beta | chr2:241395288-241395678 | MIR149;PP14571;GPC1 | Island | TSS200;Body;Body | Yes |
| Control-Sample11_Beta | chr21:26934576-26934885 | MIR155HG | Island | Body | No |
| Control-Sample1_Beta | chr16:14397207-14397892 | MIR193B | S_Shore | TSS1500 | Yes |
| Control-Sample19_Beta | chr16:14397207-14397688 | MIR193B | S_Shore | TSS1500 | Yes |
| Control-Sample42_Beta | chr16:14397576-14397892 | MIR193B | S_Shore | TSS1500 | Yes |
| Control-Sample26_Beta | chr1:566570-567206 | MIR1977 | | TSS1500 | No |
| Control-Sample1_Beta | chr13:92001764-92002454 | MIR19A;MIR17HG;MIR17HG;MIR17;MIR18A | S_Shore | TSS1500;Body;Body;TSS1500;TSS1500 | No |
| Control-Sample10_Beta | chr9:131154899-131155013 | MIR219-2 | Island | Body | No |
| Control-Sample1_Beta | chr19:46142757-46142914 | MIR330;EML2 | N_Shelf | TSS1500;TSS200 | Yes |
| Control-Sample67_Beta | chr19:46142574-46142673 | MIR330;EML2 | N_Shelf | TSS1500;Body | Yes |
| Control-Sample71_Beta | chr19:46142647-46142707 | MIR330;EML2;EML2 | N_Shelf | TSS1500;1stExon;5'UTR | Yes |
| Control-Sample11_Beta | chr17:17717208-17717252 | MIR33B;SREBF1;SREBF1 | S_Shore | Body;Body;Body | Yes |
| Control-Sample4_Beta | chr17:17717183-17717252 | MIR33B;SREBF1;SREBF1 | S_Shore | Body;Body;Body | Yes |
| Control-Sample10_Beta | chr10:88024529-88025053 | MIR346;GRID1 | Island | Body | Yes |
| Control-Sample11_Beta | chr10:88024529-88024572 | MIR346;GRID1 | Island | Body | Yes |
| Control-Sample14_Beta | chr10:88024529-88024569 | MIR346;GRID1 | Island | Body | Yes |
| Control-Sample18_Beta | chr10:88024569-88025053 | MIR346;GRID1 | S_Shore | TSS200;Body | Yes |
| Control-Sample20_Beta | chr10:88024569-88024673 | MIR346;GRID1 | S_Shore | TSS200;Body | Yes |
| Control-Sample27_Beta | chr10:88024553-88024572 | MIR346;GRID1 | S_Shore | Body;Body | Yes |
| Control-Sample33_Beta | chr10:88024529-88024557 | MIR346;GRID1 | Island | Body | Yes |
| Control-Sample42_Beta | chr10:88024553-88025053 | MIR346;GRID1 | S_Shore | Body;Body | Yes |
| Control-Sample73_Beta | chr10:88024598-88025053 | MIR346;GRID1 | S_Shore | TSS200;Body | Yes |
| Control-Sample21_Beta | chr12:95226786-95226993 | MIR492 | | TSS1500 | No |
| Control-Sample3_Beta | chr14:101495758-101495847 | MIR494 | | TSS1500 | No |
| Control-Sample464_Be | chr14:101526802-101527046 | MIR496 | | TSS200 | Yes |
| Control-Sample465_Be | chr14:101526893-101527072 | MIR496;MIR377 | | TSS200;TSS1500 | Yes |
| Control-Sample291_Be | chr13:36421844-36421949 | MIR548F5;DCLK1 | | Body;Body | Yes |
| Control-Sample66_Beta | chr13:36052985-36053408 | MIR548F5;NBEA | Island | Body;Body | Yes |
| Control-Sample67_Beta | chr4:38869641-38869707 | MIR574;FAM114A1 | Island | TSS200;5'UTR | No |
| Control-Sample1_Beta | chr7:5535582-5535748 | MIR589;FBXL18 | S_Shore | TSS200;Body | Yes |
| Control-Sample10_Beta | chr7:5535641-5535748 | MIR589;FBXL18 | S_Shore | TSS200;Body | Yes |
| Control-Sample19_Beta | chr7:5535463-5535748 | MIR589;FBXL18 | S_Shore | Body;Body | Yes |
| Control-Sample216_Be | chr7:5535731-5535748 | MIR589;FBXL18 | S_Shore | TSS200;Body | Yes |
| Control-Sample270_Be | chr7:5535692-5535748 | MIR589;FBXL18 | S_Shore | TSS200;Body | Yes |

| | | | | | |
|---|---|---|---|---|---|
| Control-Sample1_Beta | chr8:1765066-1765477 | MIR596 | Island | TSS1500 | Yes |
| Control-Sample10_Beta | chr8:1765217-1765336 | MIR596 | S_Shore | TSS200 | Yes |
| Control-Sample10_Beta | chr8:1765387-1765477 | MIR596 | S_Shore | TSS200 | Yes |
| Control-Sample11_Beta | chr8:1764878-1765477 | MIR596 | Island | TSS1500 | Yes |
| Control-Sample14_Beta | chr8:1764878-1765336 | MIR596 | Island | TSS1500 | Yes |
| Control-Sample18_Beta | chr8:1765217-1765477 | MIR596 | S_Shore | TSS200 | Yes |
| Control-Sample3_Beta | chr8:1765066-1765820 | MIR596 | Island | TSS1500 | Yes |
| Control-Sample45_Beta | chr8:1764329-1765477 | MIR596 | Island | TSS1500 | Yes |
| Control-Sample68_Beta | chr8:1765309-1765336 | MIR596 | S_Shore | TSS200 | Yes |
| Control-Sample84_Beta | chr8:1764878-1765074 | MIR596 | Island | TSS1500 | Yes |
| Control-Sample26_Beta | chr19:40789857-40790165 | MIR641;AKT2 | N_Shore | TSS1500;5'UTR | No |
| Control-Sample1_Beta | chr17:79099481-79099942 | MIR657;AATK | N_Shelf | TSS1500;Body | Yes |
| Control-Sample64_Beta | chr17:79099481-79099819 | MIR657;AATK | N_Shelf | TSS1500;Body | Yes |
| Control-Sample67_Beta | chr17:79099481-79099785 | MIR657;AATK | N_Shelf | TSS1500;Body | Yes |
| Control-Sample74_Beta | chr17:79099481-79100218 | MIR657;AATK | N_Shelf | TSS1500;Body | Yes |
| Control-Sample171_Be | chr17:79099785-79100124 | MIR657;AATK;MIR338 | N_Shelf | TSS1500;Body;TSS200 | Yes |
| Control-Sample585_Be | chr17.0:79099777-79099882 | MIR657;AATK;MIR338 | N_Shelf | TSS1500;Body;TSS200 | Yes |
| Control-Sample67_Beta | chr17:79099882-79100214 | MIR657;AATK;MIR338 | N_Shelf | TSS1500;Body;TSS200 | Yes |
| Control-Sample69_Beta | chr17:79099777-79099942 | MIR657;AATK;MIR338 | N_Shelf | TSS1500;Body;TSS200 | Yes |
| Control-Sample72_Beta | chr17:79099882-79100287 | MIR657;AATK;MIR338 | N_Shelf | TSS1500;Body;TSS200 | Yes |
| Control-Sample78_Beta | chr17:79100214-79100287 | MIR657;AATK;MIR338 | N_Shelf | TSS1500;Body;TSS1500 | Yes |
| Control-Sample83_Beta | chr17:79099777-79100124 | MIR657;AATK;MIR338 | N_Shelf | TSS1500;Body;TSS200 | Yes |
| Control-Sample93_Beta | chr17:79099785-79099942 | MIR657;AATK;MIR338 | N_Shelf | TSS1500;Body;TSS200 | Yes |
| Control-Sample99_Beta | chr17:79099882-79100124 | MIR657;AATK;MIR338 | N_Shelf | TSS1500;Body;TSS200 | Yes |
| Control-Sample157_Be | chr20:26190328-26190354 | MIR663 | Island | TSS1500 | No |
| Control-Sample10_Beta | chr15:89153667-89154196 | MIR7-2 | | TSS1500 | Yes |
| Control-Sample67_Beta | chr15:89154058-89154196 | MIR7-2 | | TSS1500 | Yes |
| Control-Sample1_Beta | chr21:37092865-37093012 | MIR802 | | TSS200 | No |
| Control-Sample110_Be | chr5:135416412-135416613 | MIR886 | Island | TSS200 | Yes |
| Control-Sample113_Be | chr5:135416205-135416398 | MIR886 | Island | Body | Yes |
| Control-Sample130_Be | chr5:135416529-135416613 | MIR886 | S_Shore | TSS1500 | Yes |
| Control-Sample425_Be | chr5:135416205-135416381 | MIR886 | Island | Body | Yes |
| Control-Sample504_Be | chr5.0:135416412-135416613 | MIR886 | Island | TSS200 | Yes |
| Control-Sample506_Be | chr5.0:135416529-135416613 | MIR886 | S_Shore | TSS1500 | Yes |
| Control-Sample529_Be | chr5.0:135416205-135416398 | MIR886 | Island | Body | Yes |
| Control-Sample15_Beta | chr20:62550415-62550734 | MIR941-1;MIR941-3;MIR941-2;MIR941-2;MIR941-1;MIR94 | N_Shore | TSS1500;TSS1500;TSS1500;TSS1500;TSS1500;TSS15 | Yes |
| Control-Sample23_Beta | chr20:62550415-62550653 | MIR941-1;MIR941-3;MIR941-2;MIR941-2;MIR941-1;MIR94 | N_Shore | TSS1500;TSS1500;TSS1500;TSS1500;TSS1500;TSS15 | Yes |
| Control-Sample9_Beta | chr20:62550415-62550714 | MIR941-1;MIR941-3;MIR941-2;MIR941-2;MIR941-1;MIR94 | N_Shore | TSS1500;TSS1500;TSS1500;TSS1500;TSS1500;TSS15 | Yes |
| Control-Sample1_Beta | chr20:62550653-62550734 | MIR941-3;MIR941-2;MIR941-1;MIR941-3;MIR941-3;MIR94 | N_Shore | TSS1500;TSS1500;TSS1500;TSS200;TSS1500;TSS200 | Yes |
| Control-Sample11_Beta | chr3:69812798-69812885 | MITF;MITF | | Body;TSS200 | Yes |
| Control-Sample3_Beta | chr3:69812125-69812973 | MITF;MITF | | TSS1500;Body | Yes |
| Control-Sample32_Beta | chr3:69812798-69812973 | MITF;MITF | | Body;TSS200 | Yes |
| Control-Sample11_Beta | chr19:2041960-2042391 | MKNK2;MKNK2 | Island | Body;Body | Yes |
| Control-Sample25_Beta | chr19:2041985-2042391 | MKNK2;MKNK2 | Island | Body;Body | Yes |
| Control-Sample5_Beta | chr19:2041955-2041985 | MKNK2;MKNK2 | Island | Body;Body | Yes |
| Control-Sample1_Beta | chr17:56296980-56297515 | MKS1;MKS1 | S_Shore | TSS200;TSS1500 | Yes |
| Control-Sample234_Be | chr17:56297360-56297515 | MKS1;MKS1 | S_Shore | TSS1500;TSS1500 | Yes |
| Control-Sample51_Beta | chr17:56297193-56297515 | MKS1;MKS1 | S_Shore | TSS1500;TSS1500 | Yes |
| Control-Sample1_Beta | chr22:50524410-50524676 | MLC1;MLC1 | N_Shelf | TSS1500;TSS200 | Yes |
| Control-Sample10_Beta | chr22:50524541-50524691 | MLC1;MLC1 | N_Shelf | TSS1500;TSS200 | Yes |
| Control-Sample11_Beta | chr22:50524410-50524691 | MLC1;MLC1 | N_Shelf | TSS1500;TSS200 | Yes |
| Control-Sample3_Beta | chr22:50524374-50524691 | MLC1;MLC1 | N_Shelf | TSS1500;TSS200 | Yes |
| Control-Sample5_Beta | chr16:74734230-74734571 | MLKL;MLKL | Island | 5'UTR;5'UTR | Yes |
| Control-Sample54_Beta | chr16:74734230-74734321 | MLKL;MLKL | Island | 5'UTR;5'UTR | Yes |
| Control-Sample564_Be | chr16.0:74734230-74734321 | MLKL;MLKL | Island | 5'UTR;5'UTR | Yes |
| Control-Sample12_Beta | chr7:152063907-152064079 | MLL3 | Island | Body | No |
| Control-Sample4_Beta | chr17:36876238-36876514 | MLLT6 | | Body | No |
| Control-Sample19_Beta | chr6:33771981-33772366 | MLN;MLN | | TSS200;TSS200 | No |
| Control-Sample32_Beta | chr13:49796387-49796489 | MLNR | S_Shore | Body | No |
| Control-Sample11_Beta | chr1:1566860-1567257 | MMP23A;MMP23B | Island | TSS1500;TSS1500 | No |
| Control-Sample1_Beta | chr19:2095364-2095643 | MOBKL2A | N_Shore | 5'UTR | No |
| Control-Sample164_Be | chr3:39543776-39544192 | MOBP;MOBP | Island | Body;Body | Yes |
| Control-Sample3_Beta | chr3:39508863-39509195 | MOBP;MOBP | | TSS1500;TSS1500 | Yes |
| Control-Sample27_Beta | chr5:52405666-52405696 | MOCS2;MOCS2 | Island | TSS200;TSS1500 | No |
| Control-Sample1_Beta | chr6:29627290-29627845 | MOG;MOG;MOG;MOG;MOG;MOG;MOG;MOG;MOG;MOG;MOG | | Body;Body;Body;Body;Body;Body;Body;Body;Body;Bo | Yes |
| Control-Sample10_Beta | chr6:29635158-29635371 | MOG;MOG;MOG;MOG;MOG;MOG;MOG;MOG;MOG;MOG;MOG | | 3'UTR;Body;3'UTR;Body;Body;Body;Body;Body;Body; | Yes |
| Control-Sample100_Be | chr6:29627290-29627395 | MOG;MOG;MOG;MOG;MOG;MOG;MOG;MOG;MOG;MOG;MOG | | Body;Body;Body;Body;Body;Body;Body;Body;Body;Bo | Yes |
| Control-Sample104_Be | chr6:29626990-29627142 | MOG;MOG;MOG;MOG;MOG;MOG;MOG;MOG;MOG;MOG;MOG | | Body;Body;Body;Body;Body;Body;Body;Body;Body;Bo | Yes |
| Control-Sample104_Be | chr6:29627215-29627395 | MOG;MOG;MOG;MOG;MOG;MOG;MOG;MOG;MOG;MOG;MOG | | Body;Body;Body;Body;Body;Body;Body;Body;Body;Bo | Yes |
| Control-Sample105_Be | chr6:29627131-29627167 | MOG;MOG;MOG;MOG;MOG;MOG;MOG;MOG;MOG;MOG;MOG | | Body;Body;Body;Body;Body;Body;Body;Body;Body;Bo | Yes |
| Control-Sample108_Be | chr6:29624414-29624846 | MOG;MOG;MOG;MOG;MOG;MOG;MOG;MOG;MOG;MOG;MOG | | TSS1500;TSS1500;TSS1500;TSS1500;TSS1500;TSS15 | Yes |
| Control-Sample12_Beta | chr6:29623721-29624414 | MOG;MOG;MOG;MOG;MOG;MOG;MOG;MOG;MOG;MOG;MOG | | TSS1500;TSS1500;TSS1500;TSS1500;TSS1500;TSS15 | Yes |
| Control-Sample124_Be | chr6:29627167-29627395 | MOG;MOG;MOG;MOG;MOG;MOG;MOG;MOG;MOG;MOG;MOG | | Body;Body;Body;Body;Body;Body;Body;Body;Body;Bo | Yes |
| Control-Sample151_Be | chr6:29623646-29623821 | MOG;MOG;MOG;MOG;MOG;MOG;MOG;MOG;MOG;MOG;MOG | | TSS1500;TSS1500;TSS1500;TSS1500;TSS1500;TSS15 | Yes |
| Control-Sample170_Be | chr6:29626990-29627845 | MOG;MOG;MOG;MOG;MOG;MOG;MOG;MOG;MOG;MOG;MOG | | Body;Body;Body;Body;Body;Body;Body;Body;Body;Bo | Yes |
| Control-Sample2_Beta | chr6:29625991-29627395 | MOG;MOG;MOG;MOG;MOG;MOG;MOG;MOG;MOG;MOG;MOG | | Body;Body;Body;Body;Body;Body;Body;Body;Body;Bo | Yes |
| Control-Sample20_Beta | chr6:29626990-29627395 | MOG;MOG;MOG;MOG;MOG;MOG;MOG;MOG;MOG;MOG;MOG | | Body;Body;Body;Body;Body;Body;Body;Body;Body;Bo | Yes |
| Control-Sample37_Beta | chr6:29626990-29627167 | MOG;MOG;MOG;MOG;MOG;MOG;MOG;MOG;MOG;MOG;MOG | | Body;Body;Body;Body;Body;Body;Body;Body;Body;Bo | Yes |
| Control-Sample585_Be | chr6.0:29627290-29627845 | MOG;MOG;MOG;MOG;MOG;MOG;MOG;MOG;MOG;MOG;MOG | | Body;Body;Body;Body;Body;Body;Body;Body;Body;Bo | Yes |
| Control-Sample644_Be | chr6:29627167-29627290 | MOG;MOG;MOG;MOG;MOG;MOG;MOG;MOG;MOG;MOG;MOG | | Body;Body;Body;Body;Body;Body;Body;Body;Body;Bo | Yes |
| Control-Sample65_Beta | chr6:29623646-29624414 | MOG;MOG;MOG;MOG;MOG;MOG;MOG;MOG;MOG;MOG;MOG | | TSS1500;TSS1500;TSS1500;TSS1500;TSS1500;TSS15 | Yes |
| Control-Sample37_Beta | chr6:29624846-29625226 | MOG;MOG;MOG;MOG;MOG;MOG;MOG;MOG;MOG;MOG;MOG;MOG;MOG;MOG;MOG;MOG;MO | | 1stExon;1stExon;5'UTR;1stExon;1stExon;1stExon;5'U | Yes |
| Control-Sample1_Beta | chr3:49947934-49948429 | MON1A;MON1A | Island | Body;Body | Yes |
| Control-Sample24_Beta | chr3:49967669-49967713 | MON1A;MON1A | S_Shore | TSS1500;TSS1500 | Yes |
| Control-Sample11_Beta | chr1:2262232-2262474 | MORN1 | N_Shelf | Body | Yes |
| Control-Sample14_Beta | chr1:2273591-2273895 | MORN1 | N_Shelf | Body | Yes |
| Control-Sample19_Beta | chr1:2288167-2288405 | MORN1 | | Body | Yes |
| Control-Sample160_Be | chr22:50528282-50528298 | MOV10L1;MOV10L1;MOV10L1 | Island | TSS200;TSS200;TSS1500 | Yes |
| Control-Sample24_Beta | chr22:50585386-50585400 | MOV10L1;MOV10L1;MOV10L1;MOV10L1 | | TSS200;Body;Body;Body | Yes |
| Control-Sample3_Beta | chr22:50585216-50585393 | MOV10L1;MOV10L1;MOV10L1;MOV10L1 | | Body;TSS1500;Body;Body | Yes |
| Control-Sample36_Beta | chr22:50585216-50585386 | MOV10L1;MOV10L1;MOV10L1;MOV10L1 | | Body;TSS1500;Body;Body | Yes |

| | | | | | |
|---|---|---|---|---|---|
| Control-Sample4_Beta | chr22:50585216-50585400 | MOV10L1;MOV10L1;MOV10L1;MOV10L1 | | Body;TSS1500;Body;Body | Yes |
| Control-Sample1_Beta | chr16:126397-127072 | MPG | N_Shore | TSS1500 | Yes |
| Control-Sample12_Beta | chr16:126397-126892 | MPG | N_Shore | TSS1500 | Yes |
| Control-Sample13_Beta | chr16:126451-127072 | MPG | N_Shore | TSS1500 | Yes |
| Control-Sample15_Beta | chr16:126451-126892 | MPG | N_Shore | TSS1500 | Yes |
| Control-Sample3_Beta | chr16:125896-127072 | MPG | N_Shore | TSS1500 | Yes |
| Control-Sample50_Beta | chr16:126848-127072 | MPG;MPG;MPG | N_Shore | TSS200;TSS1500;TSS1500 | Yes |
| Control-Sample67_Beta | chr1:43814306-43814983 | MPL | Island | Body | No |
| Control-Sample11_Beta | chr11:30602788-30603030 | MPPED2;MPPED2 | N_Shelf | TSS1500;5'UTR | No |
| Control-Sample19_Beta | chr1:181002976-181003203 | MR1 | | TSS200 | No |
| Control-Sample15_Beta | chr17:60747596-60747678 | MRC2 | S_Shelf | Body | No |
| Control-Sample57_Beta | chr11:68782049-68782211 | MRGPRF;MRGPRF | S_Shelf | TSS1500;TSS1500 | No |
| Control-Sample1_Beta | chr19:13875014-13875137 | MRI1;MRI1 | N_Shore | TSS1500;TSS1500 | Yes |
| Control-Sample520_Be | chr19.0:13875014-13875137 | MRI1;MRI1 | N_Shore | TSS1500;TSS1500 | Yes |
| Control-Sample1_Beta | chr11:1975236-1975380 | MRPL23 | S_Shore | Body | Yes |
| Control-Sample10_Beta | chr11:1975236-1975791 | MRPL23 | S_Shore | Body | Yes |
| Control-Sample11_Beta | chr16:420108-420230 | MRPL28 | Island | Body | Yes |
| Control-Sample122_Be | chr16:419975-420112 | MRPL28 | Island | Body | Yes |
| Control-Sample17_Beta | chr16:419975-420230 | MRPL28 | Island | Body | Yes |
| Control-Sample243_Be | chr16:419800-420112 | MRPL28 | N_Shore | Body | Yes |
| Control-Sample560_Be | chr16.0:419800-420112 | MRPL28 | N_Shore | Body | Yes |
| Control-Sample568_Be | chr16.0:420108-420230 | MRPL28 | Island | Body | Yes |
| Control-Sample661_Be | chr16:419800-420230 | MRPL28 | N_Shore | Body | Yes |
| Control-Sample40_Beta | chr7:42972339-42972593 | MRPL32;PSMA2 | S_Shore | Body;TSS1500 | No |
| Control-Sample244_Be | chr1:228297864-228297975 | MRPL55;MRPL55;MRPL55;MRPL55;MRPL55;MRPL55;MRPI | S_Shore | TSS1500;TSS1500;TSS1500;TSS1500;TSS1500;TSS1! | No |
| Control-Sample1_Beta | chr2:105697589-105697777 | MRPS9 | | Body | No |
| Control-Sample1_Beta | chr11:10715217-10715595 | MRVI1;MRVI1;MRVI1;MRVI1;MRVI1 | | 1stExon;5'UTR;1stExon;5'UTR;TSS200 | Yes |
| Control-Sample11_Beta | chr11:10715217-10715445 | MRVI1;MRVI1;MRVI1;MRVI1;MRVI1 | | 1stExon;5'UTR;1stExon;5'UTR;TSS200 | Yes |
| Control-Sample17_Beta | chr11:10715217-10715571 | MRVI1;MRVI1;MRVI1;MRVI1;MRVI1 | | 1stExon;5'UTR;1stExon;5'UTR;TSS200 | Yes |
| Control-Sample21_Beta | chr11:10715217-10715360 | MRVI1;MRVI1;MRVI1;MRVI1;MRVI1 | | 1stExon;5'UTR;1stExon;5'UTR;TSS200 | Yes |
| Control-Sample25_Beta | chr11:10715213-10715445 | MRVI1;MRVI1;MRVI1;MRVI1;MRVI1 | | 1stExon;5'UTR;1stExon;5'UTR;TSS200 | Yes |
| Control-Sample36_Beta | chr11:10715213-10715571 | MRVI1;MRVI1;MRVI1;MRVI1;MRVI1 | | 1stExon;5'UTR;1stExon;5'UTR;TSS200 | Yes |
| Control-Sample39_Beta | chr11:10715213-10715360 | MRVI1;MRVI1;MRVI1;MRVI1;MRVI1 | | 1stExon;5'UTR;1stExon;5'UTR;TSS200 | Yes |
| Control-Sample63_Beta | chr11:10715213-10715595 | MRVI1;MRVI1;MRVI1;MRVI1;MRVI1 | | 1stExon;5'UTR;1stExon;5'UTR;TSS200 | Yes |
| Control-Sample175_Be | chr1:76262537-76262857 | MSH4 | Island | TSS200 | Yes |
| Control-Sample368_Be | chr1:76262594-76262711 | MSH4 | Island | TSS200 | Yes |
| Control-Sample382_Be | chr1:76262537-76262711 | MSH4 | Island | TSS200 | Yes |
| Control-Sample76_Beta | chr1:76262537-76262984 | MSH4 | Island | TSS200 | Yes |
| Control-Sample20_Beta | chr6:31711701-31711837 | MSH5;MSH5;MSH5;MSH5 | S_Shelf | Body;Body;Body;Body | No |
| Control-Sample8_Beta | chr16:810950-811031 | MSLN;MSLN | | TSS200;TSS200 | No |
| Control-Sample65_Beta | chr8:9952636-9952965 | MSRA;MSRA;MSRA | | Body;TSS1500;Body | No |
| Control-Sample1_Beta | chr4:4860061-4860439 | MSX1 | Island | TSS1500 | Yes |
| Control-Sample1_Beta | chr4:4860547-4860590 | MSX1 | N_Shore | TSS1500 | Yes |
| Control-Sample1_Beta | chr4:4863678-4864833 | MSX1 | N_Shore | Body | Yes |
| Control-Sample10_Beta | chr4:4860061-4860431 | MSX1 | Island | TSS1500 | Yes |
| Control-Sample10_Beta | chr4:4863356-4864833 | MSX1 | N_Shore | Body | Yes |
| Control-Sample13_Beta | chr4:4863874-4864532 | MSX1 | N_Shore | Body | Yes |
| Control-Sample15_Beta | chr4:4863356-4864532 | MSX1 | N_Shore | Body | Yes |
| Control-Sample19_Beta | chr4:4860061-4860157 | MSX1 | Island | TSS1500 | Yes |
| Control-Sample20_Beta | chr4:4863356-4863874 | MSX1 | N_Shore | Body | Yes |
| Control-Sample21_Beta | chr4:4863356-4864625 | MSX1 | N_Shore | Body | Yes |
| Control-Sample28_Beta | chr4:4863356-4866130 | MSX1 | N_Shore | Body | Yes |
| Control-Sample51_Beta | chr4:4863678-4864532 | MSX1 | N_Shore | Body | Yes |
| Control-Sample57_Beta | chr4:4863678-4864625 | MSX1 | N_Shore | Body | Yes |
| Control-Sample66_Beta | chr4:4864293-4864532 | MSX1 | N_Shore | Body | Yes |
| Control-Sample30_Beta | chr16:56676694-56677223 | MT1DP;MT1DP | N_Shore | TSS1500;TSS1500 | No |
| Control-Sample12_Beta | chr16:56701868-56701994 | MT1G | Island | Body | Yes |
| Control-Sample15_Beta | chr16:56701868-56702033 | MT1G | Island | Body | Yes |
| Control-Sample47_Beta | chr16:56702144-56702653 | MT1G | Island | TSS200 | Yes |
| Control-Sample1_Beta | chr16:56701918-56702033 | MT1G;MT1G | Island | 1stExon;5'UTR | Yes |
| Control-Sample1_Beta | chr16:56702653-56703551 | MT1G;MT1H | N_Shore | TSS1500;TSS1500 | Yes |
| Control-Sample23_Beta | chr16:56703257-56703551 | MT1G;MT1H | N_Shore | TSS1500;TSS1500 | Yes |
| Control-Sample21_Beta | chr16:56650508-56651049 | MT1L | N_Shore | TSS1500 | No |
| Control-Sample6_Beta | chr7:91510447-91510597 | MTERF | S_Shore | TSS1500 | No |
| Control-Sample20_Beta | chr2:242040840-242041063 | MTERFD2;MTERFD2;MTERFD2;MTERFD2 | N_Shore | Body;Body;Body;Body | No |
| Control-Sample1_Beta | chr8:11182872-11183014 | MTMR9 | | 3'UTR | No |
| Control-Sample42_Beta | chr6:74170937-74171350 | MTO1;MTO1;MTO1 | N_Shore | TSS1500;TSS1500;TSS1500 | Yes |
| Control-Sample44_Beta | chr6:74170937-74171354 | MTO1;MTO1;MTO1 | N_Shore | TSS1500;TSS1500;TSS1500 | Yes |
| Control-Sample12_Beta | chr16:70714357-70714606 | MTSS1L | Island | Body | No |
| Control-Sample1_Beta | chr3:195490033-195490169 | MUC4;MUC4;MUC4 | S_Shore | Body;Body;Body | Yes |
| Control-Sample28_Beta | chr3:195489782-195490169 | MUC4;MUC4;MUC4 | Island | Body;Body;Body | Yes |
| Control-Sample43_Beta | chr3:195489708-195489789 | MUC4;MUC4;MUC4 | Island | Body;Body;Body | Yes |
| Control-Sample51_Beta | chr3:195489708-195490169 | MUC4;MUC4;MUC4 | Island | Body;Body;Body | Yes |
| Control-Sample19_Beta | chr19:1360039-1360278 | MUM1;MUM1 | Island | Body;Body | No |
| Control-Sample1_Beta | chr3:52869263-52869521 | MUSTN1 | | TSS200 | Yes |
| Control-Sample21_Beta | chr3:52869288-52869521 | MUSTN1 | | TSS200 | Yes |
| Control-Sample36_Beta | chr3:52869266-52869521 | MUSTN1 | | TSS200 | Yes |
| Control-Sample13_Beta | chr3:52868999-52869521 | MUSTN1;MUSTN1 | | 1stExon;5'UTR | Yes |
| Control-Sample276_Be | chr19:54372709-54372808 | MYADM;MYADM;MYADM;MYADM;MYADM;MYADM | Island | 1stExon;TSS200;5'UTR;5'UTR;5'UTR;5'UTR | Yes |
| Control-Sample590_Be | chr19.0:54372709-54372808 | MYADM;MYADM;MYADM;MYADM;MYADM;MYADM | Island | 1stExon;TSS200;5'UTR;5'UTR;5'UTR;5'UTR | Yes |
| Control-Sample13_Beta | chr2:241076313-241076355 | MYEOV2;MYEOV2 | S_Shore | TSS1500;TSS1500 | Yes |
| Control-Sample54_Beta | chr2:241076282-241076355 | MYEOV2;MYEOV2 | S_Shore | TSS1500;TSS1500 | Yes |
| Control-Sample11_Beta | chr12:81109337-81109806 | MYF5 | N_Shore | TSS1500 | No |
| Control-Sample40_Beta | chr17:8383621-8383800 | MYH10 | S_Shelf | Body | No |
| Control-Sample48_Beta | chr19:50733985-50734287 | MYH14;MYH14;MYH14 | Island | Body;Body;Body | No |
| Control-Sample1_Beta | chr17:10452851-10452975 | MYH2;MYH2 | | 5'UTR;5'UTR | No |
| Control-Sample644_Be | chr3:123603306-123603311 | MYLK;MYLK;MYLK;MYLK | Island | TSS200;TSS200;TSS200;TSS200 | Yes |
| Control-Sample270_Be | chr3:123339334-123339568 | MYLK;MYLK;MYLK;MYLK;MYLK;MYLK;MYLK;MYLK | | 1stExon;5'UTR;Body;Body;Body;5'UTR;1stExon;Body | Yes |
| Control-Sample10_Beta | chr20:30406997-30407145 | MYLK2 | N_Shelf | TSS200 | Yes |
| Control-Sample39_Beta | chr20:30406997-30407202 | MYLK2 | N_Shelf | TSS200 | Yes |
| Control-Sample40_Beta | chr20:30407098-30407202 | MYLK2 | N_Shelf | TSS200 | Yes |

| | | | | | |
|---|---|---|---|---|---|
| Control-Sample12_Beta | chr6:2698001-2698037 | MYLK4 | | Body | No |
| Control-Sample1_Beta | chr5:16737972-16738125 | MYO10 | | Body | Yes |
| Control-Sample171_Be | chr5:16794797-16794882 | MYO10 | | Body | Yes |
| Control-Sample655_Be | chr5:16742020-16742179 | MYO10 | | Body | Yes |
| Control-Sample11_Beta | chr17:18011879-18012014 | MYO15A | | TSS200 | Yes |
| Control-Sample12_Beta | chr17:18011879-18012007 | MYO15A | | TSS200 | Yes |
| Control-Sample1_Beta | chr13:109779684-109779861 | MYO16 | Island | Body | Yes |
| Control-Sample10_Beta | chr13:109779684-109779925 | MYO16 | Island | Body | Yes |
| Control-Sample18_Beta | chr13:109779749-109779925 | MYO16 | Island | Body | Yes |
| Control-Sample19_Beta | chr13:109741030-109741176 | MYO16 | | Body | Yes |
| Control-Sample13_Beta | chr17:1375253-1375968 | MYO1C;MYO1C;MYO1C | Island | Body;Body;Body | Yes |
| Control-Sample34_Beta | chr17:1375253-1375459 | MYO1C;MYO1C;MYO1C | Island | Body;Body;Body | Yes |
| Control-Sample284_Be | chr10:26223100-26223310 | MYO3A;MYO3A | Island | 1stExon;5'UTR | No |
| Control-Sample1_Beta | chr11:76858729-76859387 | MYO7A;MYO7A;MYO7A | Island | Body;Body;Body | Yes |
| Control-Sample171_Be | chr11:76839191-76839217 | MYO7A;MYO7A;MYO7A | | TSS200;TSS200;TSS200 | Yes |
| Control-Sample19_Beta | chr11:76849324-76849564 | MYO7A;MYO7A;MYO7A | Island | Body;Body;Body | Yes |
| Control-Sample62_Beta | chr11:76858947-76859387 | MYO7A;MYO7A;MYO7A | Island | Body;Body;Body | Yes |
| Control-Sample66_Beta | chr11:76858729-76859056 | MYO7A;MYO7A;MYO7A | Island | Body;Body;Body | Yes |
| Control-Sample10_Beta | chr1:203055422-203055507 | MYOG | | TSS200 | Yes |
| Control-Sample1_Beta | chr1:203055296-203055507 | MYOG;MYOG | | 1stExon;5'UTR | Yes |
| Control-Sample20_Beta | chr1:203055296-203055662 | MYOG;MYOG | | 1stExon;5'UTR | Yes |
| Control-Sample108_Be | chr8:2075209-2075820 | MYOM2 | N_Shore | Body | Yes |
| Control-Sample123_Be | chr8:2075469-2075820 | MYOM2 | Island | Body | Yes |
| Control-Sample44_Beta | chr8:2074935-2075820 | MYOM2 | N_Shore | Body | Yes |
| Control-Sample553_Be | chr8.0:2075469-2075820 | MYOM2 | Island | Body | Yes |
| Control-Sample108_Be | chr20:62795464-62795830 | MYT1 | | TSS1500 | No |
| Control-Sample134_Be | chr2:1817263-1817409 | MYT1L | | Body | Yes |
| Control-Sample19_Beta | chr2:1886086-1886127 | MYT1L | | Body | Yes |
| Control-Sample19_Beta | chr2:1983190-1983518 | MYT1L | | Body | Yes |
| Control-Sample20_Beta | chr2:1942814-1942942 | MYT1L | S_Shore | Body | Yes |
| Control-Sample509_Be | chr2.0:1817263-1817409 | MYT1L | | Body | Yes |
| Control-Sample12_Beta | chr11:71163466-71163555 | NADSYN1 | N_Shore | TSS1500 | Yes |
| Control-Sample2_Beta | chr11:71163428-71163528 | NADSYN1 | N_Shore | TSS1500 | Yes |
| Control-Sample114_Be | chr8:144660590-144660631 | NAPRT1 | Island | TSS200 | Yes |
| Control-Sample13_Beta | chr8:144660659-144661051 | NAPRT1 | S_Shore | TSS200 | Yes |
| Control-Sample167_Be | chr8:144659831-144660157 | NAPRT1 | Island | Body | Yes |
| Control-Sample506_Be | chr8.0:144660590-144660631 | NAPRT1 | Island | TSS200 | Yes |
| Control-Sample543_Be | chr8.0:144659831-144660157 | NAPRT1 | Island | Body | Yes |
| Control-Sample313_Be | chr17:72772736-72772745 | NAT9;TMEM104 | S_Shore | TSS1500;5'UTR | Yes |
| Control-Sample314_Be | chr17:72772726-72772745 | NAT9;TMEM104 | S_Shore | TSS1500;5'UTR | Yes |
| Control-Sample1_Beta | chr11:20043971-20044639 | NAV2;NAV2;NAV2;NAV2 | | TSS1500;Body;Body;Body | No |
| Control-Sample3_Beta | chr8:102804678-102804916 | NCALD;NCALD;NCALD;NCALD;NCALD;NCALD;NCALD;NCALD | | 5'UTR;5'UTR;5'UTR;5'UTR;5'UTR;5'UTR;TSS1500;5'UT | No |
| Control-Sample1_Beta | chr11:113143664-113143736 | NCAM1;NCAM1 | | Body;Body | No |
| Control-Sample1_Beta | chr10:51575702-51576232 | NCOA4;NCOA4;NCOA4;NCOA4;NCOA4 | S_Shelf | 5'UTR;5'UTR;Body;TSS1500;Body | Yes |
| Control-Sample572_Be | chr10.0:51575702-51576232 | NCOA4;NCOA4;NCOA4;NCOA4;NCOA4 | S_Shelf | 5'UTR;5'UTR;Body;TSS1500;Body | Yes |
| Control-Sample54_Beta | chr17:16119344-16119441 | NCOR1;PIGL | Island | TSS1500;TSS1500 | No |
| Control-Sample1_Beta | chr12:124950720-124950990 | NCOR2;NCOR2 | Island | Body;Body | Yes |
| Control-Sample10_Beta | chr12:124907105-124907367 | NCOR2;NCOR2 | | Body;Body | Yes |
| Control-Sample14_Beta | chr12:124950784-124950990 | NCOR2;NCOR2 | Island | Body;Body | Yes |
| Control-Sample194_Be | chr12:124929704-124929991 | NCOR2;NCOR2 | N_Shore | Body;Body | Yes |
| Control-Sample29_Beta | chr12:124893511-124893649 | NCOR2;NCOR2 | | Body;Body | Yes |
| Control-Sample5_Beta | chr12:124990942-124991139 | NCOR2;NCOR2 | | 5'UTR;5'UTR | Yes |
| Control-Sample29_Beta | chr20:30073537-30073576 | NCRNA00028 | S_Shore | TSS200 | No |
| Control-Sample1_Beta | chr21:40141569-40141809 | NCRNA00114;NCRNA00114 | | Body;TSS1500 | No |
| Control-Sample106_Be | chr2:110969641-110970249 | NCRNA00116 | N_Shore | Body | Yes |
| Control-Sample134_Be | chr2:110969641-110970204 | NCRNA00116 | N_Shore | Body | Yes |
| Control-Sample10_Beta | chr21:33452218-33452381 | NCRNA00159 | | TSS1500 | No |
| Control-Sample32_Beta | chr16:58545346-58545502 | NDRG4;NDRG4;NDRG4 | N_Shelf | Body;Body;Body | No |
| Control-Sample1_Beta | chr12:57630107-57630422 | NDUFA4L2 | Island | Body | No |
| Control-Sample23_Beta | chr4:140216130-140216277 | NDUFC1 | N_Shore | Body | No |
| Control-Sample24_Beta | chr11:77791290-77791376 | NDUFC2 | S_Shore | TSS200 | No |
| Control-Sample84_Beta | chr1:161171211-161171632 | NDUFS2;NDUFS2 | N_Shore | 5'UTR;TSS1500 | No |
| Control-Sample1_Beta | chr12:97300410-97300582 | NEDD1;NEDD1;NEDD1;NEDD1 | N_Shore | TSS1500;TSS1500;TSS1500;TSS1500 | Yes |
| Control-Sample27_Beta | chr12:97300417-97300582 | NEDD1;NEDD1;NEDD1;NEDD1 | N_Shore | TSS1500;TSS1500;TSS1500;TSS1500 | Yes |
| Control-Sample275_Be | chr12:97300410-97300429 | NEDD1;NEDD1;NEDD1;NEDD1 | N_Shore | TSS1500;TSS1500;TSS1500;TSS1500 | Yes |
| Control-Sample19_Beta | chr18:55862627-55862678 | NEDD4L;NEDD4L;NEDD4L;NEDD4L;NEDD4L;NEDD4L;NEI | N_Shore | 5'UTR;TSS200;Body;Body;Body;5'UTR;5'UTR;Body | No |
| Control-Sample425_Be | chr8:24771114-24771466 | NEFM;NEFM | Island | TSS200;TSS1500 | Yes |
| Control-Sample425_Be | chr8:24772137-24772513 | NEFM;NEFM | Island | TSS1500;1stExon | Yes |
| Control-Sample637_Be | chr3:130745618-130745685 | NEK11;NEK11;ASTE1;ASTE1;NEK11 | Island | TSS200;TSS200;1stExon;5'UTR;TSS200 | Yes |
| Control-Sample333_Be | chr3:130745651-130745685 | NEK11;NEK11;NEK11;ASTE1 | Island | TSS200;TSS200;TSS200;TSS200 | Yes |
| Control-Sample1_Beta | chr1:156646065-156646635 | NES | N_Shore | Body | Yes |
| Control-Sample2_Beta | chr1:156646065-156646653 | NES | N_Shore | Body | Yes |
| Control-Sample5_Beta | chr10:5488436-5488628 | NET1;NET1 | N_Shore | Body;TSS200 | No |
| Control-Sample53_Beta | chr2:242749925-242749937 | NEU4;NEU4 | | TSS1500;TSS1500 | No |
| Control-Sample1_Beta | chr5:172110211-172110569 | NEURL1B | N_Shore | Body | No |
| Control-Sample1_Beta | chr5:134872649-134872957 | NEUROG1 | S_Shore | TSS1500 | Yes |
| Control-Sample24_Beta | chr5:134872639-134872659 | NEUROG1 | S_Shore | TSS1500 | Yes |
| Control-Sample57_Beta | chr5:134872639-134872957 | NEUROG1 | S_Shore | TSS1500 | Yes |
| Control-Sample1_Beta | chr22:42828125-42828447 | NFAM1 | | Body | Yes |
| Control-Sample14_Beta | chr22:42828411-42828447 | NFAM1 | | TSS200 | Yes |
| Control-Sample23_Beta | chr22:42828125-42828946 | NFAM1 | | Body | Yes |
| Control-Sample25_Beta | chr22:42828125-42828516 | NFAM1 | | Body | Yes |
| Control-Sample3_Beta | chr1:204839819-204839875 | NFASC;NFASC;NFASC;NFASC;NFASC | | 5'UTR;5'UTR;5'UTR;5'UTR;5'UTR | No |
| Control-Sample16_Beta | chr18:77157760-77158180 | NFATC1;NFATC1;NFATC1 | Island | Body;5'UTR;Body | No |
| Control-Sample1_Beta | chr14:24835104-24835609 | NFATC4 | N_Shore | TSS1500 | Yes |
| Control-Sample20_Beta | chr14:24835609-24836090 | NFATC4 | N_Shore | TSS1500 | Yes |
| Control-Sample22_Beta | chr14:24835006-24836090 | NFATC4 | N_Shore | TSS1500 | Yes |
| Control-Sample8_Beta | chr14:24836194-24836539 | NFATC4;NFATC4;NFATC4 | Island | 1stExon;5'UTR;TSS1500 | Yes |
| Control-Sample10_Beta | chr7:26192966-26193109 | NFE2L3 | S_Shore | Body | Yes |
| Control-Sample28_Beta | chr7:26192966-26193607 | NFE2L3 | S_Shore | Body | Yes |
| Control-Sample58_Beta | chr7:26192966-26193805 | NFE2L3 | S_Shore | Body | Yes |

| | | | | | |
|---|---|---|---|---|---|
| Control-Sample1_Beta | chr19:3369714-3369820 | NFIC;NFIC | Island | Body;Body | Yes |
| Control-Sample34_Beta | chr19:3369714-3370244 | NFIC;NFIC | Island | Body;Body | Yes |
| Control-Sample1_Beta | chr2:69664869-69665132 | NFU1;NFU1;NFU1;NFU1 | S_Shore | TSS200;TSS200;TSS200;TSS200 | Yes |
| Control-Sample19_Beta | chr2:69665094-69665232 | NFU1;NFU1;NFU1;NFU1 | S_Shore | TSS1500;TSS1500;TSS1500;TSS1500 | Yes |
| Control-Sample20_Beta | chr2:69664869-69665232 | NFU1;NFU1;NFU1;NFU1 | S_Shore | TSS200;TSS200;TSS200;TSS200 | Yes |
| Control-Sample1_Beta | chr6:41068646-41068722 | NFYA;NFYA;LOC221442 | Island | 3'UTR;3'UTR;TSS200 | Yes |
| Control-Sample19_Beta | chr6:41068646-41068741 | NFYA;NFYA;LOC221442 | Island | 3'UTR;3'UTR;TSS200 | Yes |
| Control-Sample20_Beta | chr3:25831802-25832283 | NGLY1;OXSM;OXSM;OXSM | | TSS1500;Body;5'UTR;5'UTR | No |
| Control-Sample568_Be | chr4.0:103940878-103940936 | NHEDC1;NHEDC1 | Island | TSS200;TSS200 | Yes |
| Control-Sample29_Beta | chr4:103940854-103940878 | NHEDC1;NHEDC1;NHEDC1;NHEDC1 | Island | 1stExon;1stExon;5'UTR;5'UTR | Yes |
| Control-Sample10_Beta | chr1:160336475-160336699 | NHLH1 | | TSS1500 | Yes |
| Control-Sample34_Beta | chr1:160336486-160336699 | NHLH1 | | TSS1500 | Yes |
| Control-Sample64_Beta | chr1:160336475-160336712 | NHLH1 | | TSS1500 | Yes |
| Control-Sample44_Beta | chr13:39612797-39613327 | NHLRC3;C13orf23;C13orf23;NHLRC3 | Island | 1stExon;TSS1500;TSS1500;1stExon | No |
| Control-Sample108_Be | chr6:138866882-138867046 | NHSL1 | | Body | Yes |
| Control-Sample20_Beta | chr6:138866882-138866898 | NHSL1 | | Body | Yes |
| Control-Sample291_Be | chr6:138893687-138893718 | NHSL1 | | TSS200 | Yes |
| Control-Sample67_Beta | chr6:138892911-138893159 | NHSL1 | | 1stExon | Yes |
| Control-Sample11_Beta | chr14:51290136-51290751 | NIN;NIN;NIN;NIN;NIN | | 5'UTR;5'UTR;5'UTR;1stExon;5'UTR | Yes |
| Control-Sample29_Beta | chr14:51290136-51290662 | NIN;NIN;NIN;NIN;NIN | | 5'UTR;5'UTR;5'UTR;1stExon;5'UTR | Yes |
| Control-Sample10_Beta | chr12:772861-772982 | NINJ2 | | TSS200 | Yes |
| Control-Sample310_Be | chr12:678960-679287 | NINJ2 | Island | Body | Yes |
| Control-Sample529_Be | chr12.0:678960-679287 | NINJ2 | Island | Body | Yes |
| Control-Sample572_Be | chr12.0:772861-772982 | NINJ2 | | TSS200 | Yes |
| Control-Sample104_Be | chr12:772688-772982 | NINJ2;NINJ2 | | 1stExon;5'UTR | Yes |
| Control-Sample29_Beta | chr6:28227068-28227085 | NKAPL | Island | TSS200 | No |
| Control-Sample65_Beta | chr5:1016356-1016768 | NKD2 | | Body | No |
| Control-Sample160_Be | chr19:55476665-55476717 | NLRP2 | | TSS1500 | Yes |
| Control-Sample170_Be | chr19:55477653-55477810 | NLRP2 | | TSS200 | Yes |
| Control-Sample178_Be | chr19:55477755-55477810 | NLRP2 | | TSS200 | Yes |
| Control-Sample258_Be | chr19:55476665-55477810 | NLRP2 | | TSS1500 | Yes |
| Control-Sample553_Be | chr19.0:55477755-55477810 | NLRP2 | | TSS200 | Yes |
| Control-Sample445_Be | chr17:49244154-49244616 | NME2;NME2;NME1-NME2;NME2;NME2;NME2 | Island | 5'UTR;1stExon;Body;5'UTR;5'UTR;5'UTR | Yes |
| Control-Sample64_Beta | chr17:49244154-49244169 | NME2;NME2;NME1-NME2;NME2;NME2;NME2 | Island | 5'UTR;1stExon;Body;5'UTR;5'UTR;5'UTR | Yes |
| Control-Sample11_Beta | chr17:49242821-49243027 | NME2;NME2;NME2;NME1-NME2;NME2 | N_Shore | TSS200;TSS1500;TSS1500;Body;TSS1500 | Yes |
| Control-Sample18_Beta | chr17:49242821-49243257 | NME2;NME2;NME2;NME1-NME2;NME2 | N_Shore | TSS200;TSS1500;TSS1500;Body;TSS1500 | Yes |
| Control-Sample19_Beta | chr17:49242732-49243257 | NME2;NME2;NME2;NME1-NME2;NME2 | N_Shore | TSS200;TSS1500;TSS1500;Body;TSS1500 | Yes |
| Control-Sample22_Beta | chr17:49242714-49242847 | NME2;NME2;NME2;NME1-NME2;NME2 | N_Shore | TSS200;TSS1500;TSS1500;Body;TSS1500 | Yes |
| Control-Sample39_Beta | chr17:49242732-49242847 | NME2;NME2;NME2;NME1-NME2;NME2 | N_Shore | TSS200;TSS1500;TSS1500;Body;TSS1500 | Yes |
| Control-Sample11_Beta | chr3:139346734-139346818 | NMNAT3 | | 5'UTR | No |
| Control-Sample223_Be | chr16:4511773-4511987 | NMRAL1 | | 3'UTR | No |
| Control-Sample145_Be | chr2:232395269-232395300 | NMUR1 | Island | TSS200 | Yes |
| Control-Sample564_Be | chr2.0:232395269-232395300 | NMUR1 | Island | TSS200 | Yes |
| Control-Sample1_Beta | chr10:72201484-72202715 | NODAL | S_Shore | 1stExon | Yes |
| Control-Sample10_Beta | chr10:72201484-72203124 | NODAL | S_Shore | 1stExon | Yes |
| Control-Sample19_Beta | chr10:72201484-72201622 | NODAL | S_Shore | 1stExon | Yes |
| Control-Sample20_Beta | chr10:72201600-72201622 | NODAL | S_Shore | TSS200 | Yes |
| Control-Sample66_Beta | chr10:72201741-72202215 | NODAL | S_Shore | TSS1500 | Yes |
| Control-Sample29_Beta | chr2:10830698-10830764 | NOL10 | S_Shore | TSS1500 | No |
| Control-Sample1_Beta | chr12:117797056-117797635 | NOS1 | N_Shore | 5'UTR | Yes |
| Control-Sample10_Beta | chr12:117797209-117797635 | NOS1 | N_Shore | 5'UTR | Yes |
| Control-Sample12_Beta | chr12:117797056-117797385 | NOS1 | N_Shore | 5'UTR | Yes |
| Control-Sample1_Beta | chr7:150690699-150690881 | NOS3;NOS3;NOS3;NOS3 | | 5'UTR;TSS200;TSS200;TSS200 | No |
| Control-Sample3_Beta | chr19:50059750-50059946 | NOSIP | Island | Body | No |
| Control-Sample12_Beta | chr19:15311990-15312129 | NOTCH3 | Island | TSS200 | No |
| Control-Sample331_Be | chr6:32165176-32165200 | NOTCH4 | S_Shore | Body | No |
| Control-Sample1_Beta | chr17:79920530-79920672 | NOTUM | Island | TSS1500 | No |
| Control-Sample345_Be | chr4:72897782-72898212 | NPFFR2 | Island | 1stExon | Yes |
| Control-Sample6_Beta | chr4:72897698-72897868 | NPFFR2 | Island | 1stExon | Yes |
| Control-Sample1_Beta | chr1:5946328-5946381 | NPHP4 | | Body | Yes |
| Control-Sample1_Beta | chr1:5956985-5957659 | NPHP4 | | Body | Yes |
| Control-Sample1_Beta | chr1:11908199-11908347 | NPPA | | TSS1500 | Yes |
| Control-Sample10_Beta | chr1:11907986-11908347 | NPPA | | TSS200 | Yes |
| Control-Sample22_Beta | chr1:11908058-11908199 | NPPA | | TSS1500 | Yes |
| Control-Sample23_Beta | chr1:11907986-11908164 | NPPA | | TSS200 | Yes |
| Control-Sample26_Beta | chr1:11907986-11908199 | NPPA | | TSS200 | Yes |
| Control-Sample12_Beta | chr1:11919215-11919464 | NPPB | Island | TSS1500 | No |
| Control-Sample157_Be | chr7:24323799-24323939 | NPY | Island | TSS200 | Yes |
| Control-Sample5_Beta | chr7:24323559-24323767 | NPY | Island | TSS1500 | Yes |
| Control-Sample588_Be | chr7.0:24323799-24323939 | NPY | Island | TSS200 | Yes |
| Control-Sample1_Beta | chr17:38249266-38249592 | NR1D1;THRA | Island | 3'UTR;Body | Yes |
| Control-Sample11_Beta | chr17:38249266-38249430 | NR1D1;THRA | Island | 3'UTR;Body | Yes |
| Control-Sample3_Beta | chr17:38249266-38249486 | NR1D1;THRA | Island | 3'UTR;Body | Yes |
| Control-Sample1_Beta | chr6:108488801-108489292 | NR2E1 | Island | Body | Yes |
| Control-Sample1_Beta | chr6:108489977-108490371 | NR2E1 | Island | Body | Yes |
| Control-Sample1_Beta | chr6:108491372-108492769 | NR2E1 | Island | Body | Yes |
| Control-Sample10_Beta | chr6:108492653-108492769 | NR2E1 | S_Shore | Body | Yes |
| Control-Sample12_Beta | chr6:108488335-108489292 | NR2E1 | Island | Body | Yes |
| Control-Sample13_Beta | chr6:108488335-108488751 | NR2E1 | Island | Body | Yes |
| Control-Sample13_Beta | chr6:108488915-108489292 | NR2E1 | Island | Body | Yes |
| Control-Sample17_Beta | chr6:108488801-108489064 | NR2E1 | Island | Body | Yes |
| Control-Sample65_Beta | chr6:108488335-108489064 | NR2E1 | Island | Body | Yes |
| Control-Sample14_Beta | chr5:92929372-92929573 | NR2F1 | Island | Body | Yes |
| Control-Sample23_Beta | chr5:92929473-92929679 | NR2F1 | Island | Body | Yes |
| Control-Sample24_Beta | chr5:92929372-92929679 | NR2F1 | Island | Body | Yes |
| Control-Sample108_Be | chr15:96868857-96869220 | NR2F2 | S_Shelf | TSS1500 | No |
| Control-Sample77_Beta | chr5:142781498-142781736 | NR3C1;NR3C1;NR3C1;NR3C1;NR3C1;NR3C1;NR3C1 | N_Shore | 5'UTR;5'UTR;5'UTR;5'UTR;5'UTR;5'UTR;5'UTR | No |
| Control-Sample20_Beta | chr2:157184614-157184880 | NR4A2 | Island | Body | Yes |
| Control-Sample21_Beta | chr2:157186312-157187204 | NR4A2 | Island | Body | Yes |
| Control-Sample1_Beta | chr10:115370894-115371055 | NRAP;NRAP | | Body;Body | No |
| Control-Sample194_Be | chr5:139228062-139228153 | NRG2;NRG2;NRG2;NRG2 | Island | Body;Body;Body;Body | No |

| | | | | | |
|---|---|---|---|---|---|
| Control-Sample1_Beta | chr2:206551466-206551987 | NRP2;NRP2;NRP2;NRP2;NRP2;NRP2 | S_Shore | Body;Body;Body;Body;Body;Body | No |
| Control-Sample1_Beta | chr2:51254021-51254170 | NRXN1;NRXN1 | N_Shore | Body;Body | No |
| Control-Sample22_Beta | chr11:64405452-64405919 | NRXN2;NRXN2;NRXN2 | Island | Body;Body;Body | No |
| Control-Sample10_Beta | chr5:176558658-176559563 | NSD1 | N_Shore | TSS1500 | Yes |
| Control-Sample17_Beta | chr5:176559334-176559563 | NSD1;NSD1 | Island | TSS1500;TSS1500 | Yes |
| Control-Sample10_Beta | chr3:52569026-52569097 | NT5DC2;NT5DC2;NT5DC2 | N_Shore | 5'UTR;1stExon;TSS1500 | Yes |
| Control-Sample19_Beta | chr3:52569026-52569169 | NT5DC2;NT5DC2;NT5DC2 | N_Shore | 5'UTR;1stExon;TSS1500 | Yes |
| Control-Sample33_Beta | chr3:52569026-52569147 | NT5DC2;NT5DC2;NT5DC2 | N_Shore | 5'UTR;1stExon;TSS1500 | Yes |
| Control-Sample1_Beta | chr11:131410868-131410981 | NTM | | Body | Yes |
| Control-Sample108_Be | chr11:131630464-131630648 | NTM | | Body | Yes |
| Control-Sample223_Be | chr11:131779469-131779604 | NTM;NTM;NTM;NTM | N_Shore | TSS1500;TSS1500;TSS1500;Body | Yes |
| Control-Sample1_Beta | chr17:8957664-8958090 | NTN1 | | Body | Yes |
| Control-Sample6_Beta | chr17:8928012-8928451 | NTN1 | S_Shore | Body | Yes |
| Control-Sample45_Beta | chr1:108023428-108023482 | NTNG1;NTNG1;NTNG1 | Island | Body;Body;Body | No |
| Control-Sample25_Beta | chr1:156828202-156828693 | NTRK1;INSRR | N_Shore | Body;Body | No |
| Control-Sample108_Be | chr12:86267839-86268154 | NTS | | TSS1500 | No |
| Control-Sample124_Be | chr20:61371425-61371808 | NTSR1 | | Body | No |
| Control-Sample1_Beta | chr12:106480546-106480771 | NUAK1 | | Body | Yes |
| Control-Sample124_Be | chr12:106480579-106480771 | NUAK1 | | Body | Yes |
| Control-Sample13_Beta | chr12:106480546-106480756 | NUAK1 | | Body | Yes |
| Control-Sample115_Be | chr5:102898463-102898733 | NUDT12;NUDT12 | | 5'UTR;1stExon | Yes |
| Control-Sample420_Be | chr5:102898478-102898729 | NUDT12;NUDT12 | | 5'UTR;1stExon | Yes |
| Control-Sample57_Beta | chr3:131100598-131100678 | NUDT16 | Island | TSS200 | No |
| Control-Sample276_Be | chr19:33182716-33182730 | NUDT19 | Island | TSS200 | No |
| Control-Sample1_Beta | chr11:71724287-71725590 | NUMA1 | | Body | Yes |
| Control-Sample32_Beta | chr11:71724287-71725340 | NUMA1 | | Body | Yes |
| Control-Sample40_Beta | chr11:71724338-71725590 | NUMA1 | | Body | Yes |
| Control-Sample162_Be | chr1:154127138-154127537 | NUP210L;NUP210L | | Body;Body | No |
| Control-Sample108_Be | chr19:16830613-16830746 | NWD1 | | TSS200 | Yes |
| Control-Sample19_Beta | chr19:16830746-16830859 | NWD1 | | TSS200 | Yes |
| Control-Sample20_Beta | chr19:16830739-16830859 | NWD1 | | TSS200 | Yes |
| Control-Sample65_Beta | chr19:16830739-16830749 | NWD1 | | TSS200 | Yes |
| Control-Sample1_Beta | chr17:873255-873452 | NXN | | Body | Yes |
| Control-Sample10_Beta | chr17:773023-773283 | NXN | | Body | Yes |
| Control-Sample15_Beta | chr17:750241-750766 | NXN | | Body | Yes |
| Control-Sample16_Beta | chr17:750241-750379 | NXN | | Body | Yes |
| Control-Sample17_Beta | chr17:746502-746641 | NXN | S_Shore | Body | Yes |
| Control-Sample19_Beta | chr17:840950-841205 | NXN | | Body | Yes |
| Control-Sample19_Beta | chr17:872762-873452 | NXN | | Body | Yes |
| Control-Sample21_Beta | chr17:800512-800745 | NXN | | Body | Yes |
| Control-Sample29_Beta | chr17:851785-851933 | NXN | S_Shore | Body | Yes |
| Control-Sample5_Beta | chr17:780872-781007 | NXN | | Body | Yes |
| Control-Sample57_Beta | chr17:707032-707141 | NXN | N_Shore | Body | Yes |
| Control-Sample57_Beta | chr17:800512-800749 | NXN | | Body | Yes |
| Control-Sample19_Beta | chr19:17571656-17571888 | NXNL1 | | 1stExon | No |
| Control-Sample1_Beta | chr14:24867164-24867491 | NYNRIN | N_Shore | TSS1500 | No |
| Control-Sample1_Beta | chr19:2272716-2273216 | OAZ1 | Island | Body | No |
| Control-Sample1_Beta | chr12:56617576-56617737 | OBFC2B | N_Shore | TSS1500 | No |
| Control-Sample1_Beta | chr1:228395660-228395760 | OBSCN;OBSCN | N_Shelf | TSS200;TSS200 | Yes |
| Control-Sample21_Beta | chr1:228395660-228395858 | OBSCN;OBSCN | N_Shelf | TSS200;TSS200 | Yes |
| Control-Sample65_Beta | chr1:228394509-228394607 | OBSCN;OBSCN | | TSS1500;TSS1500 | Yes |
| Control-Sample108_Be | chr1:102462821-102462877 | OLFM3 | | TSS200 | No |
| Control-Sample1_Beta | chr15:53075421-53075533 | ONECUT1 | N_Shore | Body | No |
| Control-Sample10_Beta | chr19:46056907-46057298 | OPA3;OPA3 | Island | Body;Body | Yes |
| Control-Sample18_Beta | chr19:46056784-46057298 | OPA3;OPA3 | Island | Body;Body | Yes |
| Control-Sample19_Beta | chr11:132582488-132582640 | OPCML;OPCML | | Body;Body | Yes |
| Control-Sample20_Beta | chr11:132582488-132582808 | OPCML;OPCML | | Body;Body | Yes |
| Control-Sample59_Beta | chr11:132662772-132662963 | OPCML;OPCML | | Body;Body | Yes |
| Control-Sample1_Beta | chr8:145106299-145107012 | OPLAH | Island | Body | Yes |
| Control-Sample10_Beta | chr8:145106299-145106582 | OPLAH | Island | Body | Yes |
| Control-Sample1_Beta | chr20:62715942-62716485 | OPRL1;C20orf201;OPRL1 | N_Shore | 5'UTR;TSS1500;5'UTR | Yes |
| Control-Sample30_Beta | chr20:62715942-62716443 | OPRL1;C20orf201;OPRL1 | N_Shore | 5'UTR;TSS1500;5'UTR | Yes |
| Control-Sample107_Be | chr1:248100407-248100585 | OR2L13 | Island | TSS200 | Yes |
| Control-Sample138_Be | chr1:248100228-248101009 | OR2L13 | N_Shore | TSS1500 | Yes |
| Control-Sample156_Be | chr1:248100276-248101009 | OR2L13 | N_Shore | TSS1500 | Yes |
| Control-Sample158_Be | chr1:248100345-248100585 | OR2L13 | Island | TSS200 | Yes |
| Control-Sample181_Be | chr1:248100427-248101009 | OR2L13 | Island | TSS200 | Yes |
| Control-Sample182_Be | chr1:248100407-248101009 | OR2L13 | Island | TSS200 | Yes |
| Control-Sample220_Be | chr1:248100345-248100614 | OR2L13 | Island | TSS200 | Yes |
| Control-Sample386_Be | chr1:248100407-248100614 | OR2L13 | Island | TSS200 | Yes |
| Control-Sample49_Beta | chr1:248100276-248100585 | OR2L13 | N_Shore | TSS1500 | Yes |
| Control-Sample567_Be | chr1.0:248100345-248101009 | OR2L13 | Island | TSS200 | Yes |
| Control-Sample568_Be | chr1.0:248100407-248101009 | OR2L13 | Island | TSS200 | Yes |
| Control-Sample616_Be | chr1:248100345-248100427 | OR2L13 | Island | TSS200 | Yes |
| Control-Sample107_Be | chr1:248100600-248101009 | OR2L13;OR2L13 | Island | 1stExon;5'UTR | Yes |
| Control-Sample221_Be | chr1:248100591-248100614 | OR2L13;OR2L13 | Island | 1stExon;5'UTR | Yes |
| Control-Sample146_Be | chr1:248790459-248790981 | OR2T11 | | TSS200 | No |
| Control-Sample41_Beta | chr11:7819812-7819960 | OR5P2 | | TSS1500 | No |
| Control-Sample331_Be | chr7:25018595-25019044 | OSBPL3;OSBPL3;OSBPL3;OSBPL3 | N_Shore | 5'UTR;5'UTR;5'UTR;5'UTR | No |
| Control-Sample1_Beta | chr11:3187921-3188016 | OSBPL5;OSBPL5;OSBPL5 | S_Shore | TSS1500;TSS1500;TSS1500 | Yes |
| Control-Sample16_Beta | chr11:3187631-3187704 | OSBPL5;OSBPL5;OSBPL5 | S_Shore | TSS1500;TSS1500;TSS1500 | Yes |
| Control-Sample3_Beta | chr11:3187923-3188016 | OSBPL5;OSBPL5;OSBPL5 | S_Shore | TSS1500;TSS1500;TSS1500 | Yes |
| Control-Sample10_Beta | chr2:179184854-179184866 | OSBPL6;OSBPL6 | | TSS200;Body | No |
| Control-Sample19_Beta | chr16:83986755-83987465 | OSGIN1;OSGIN1;OSGIN1 | | TSS200;5'UTR;Body | No |
| Control-Sample1_Beta | chr17:72931767-72931792 | OTOP3 | Island | TSS200 | Yes |
| Control-Sample57_Beta | chr17:72931569-72931664 | OTOP3 | N_Shore | TSS1500 | Yes |
| Control-Sample65_Beta | chr20:16728895-16729319 | OTOR | | TSS200 | No |
| Control-Sample1_Beta | chr2:241080057-241080193 | OTOS;OTOS | | 5'UTR;1stExon | Yes |
| Control-Sample10_Beta | chr2:241080057-241080394 | OTOS;OTOS | | 5'UTR;1stExon | Yes |
| Control-Sample1_Beta | chr11:63753203-63753251 | OTUB1;OTUB1 | N_Shore | TSS200;TSS200 | Yes |
| Control-Sample15_Beta | chr11:63753169-63753251 | OTUB1;OTUB1 | N_Shore | TSS200;TSS200 | Yes |

| | | | | | |
|---|---|---|---|---|---|
| Control-Sample1_Beta | chr2:63279495-63279693 | OTX1 | N_Shore | 5'UTR | Yes |
| Control-Sample10_Beta | chr2:63279495-63280097 | OTX1 | N_Shore | 5'UTR | Yes |
| Control-Sample10_Beta | chr2:63281844-63282211 | OTX1 | N_Shore | Body | Yes |
| Control-Sample10_Beta | chr2:63284132-63284557 | OTX1 | Island | 3'UTR | Yes |
| Control-Sample11_Beta | chr2:63279495-63280164 | OTX1 | N_Shore | 5'UTR | Yes |
| Control-Sample11_Beta | chr2:63283806-63285097 | OTX1 | N_Shore | 3'UTR | Yes |
| Control-Sample15_Beta | chr2:63283806-63284132 | OTX1 | N_Shore | 3'UTR | Yes |
| Control-Sample18_Beta | chr2:63283939-63284132 | OTX1 | Island | 3'UTR | Yes |
| Control-Sample25_Beta | chr2:63283806-63284066 | OTX1 | N_Shore | 3'UTR | Yes |
| Control-Sample28_Beta | chr2:63284132-63285097 | OTX1 | Island | 3'UTR | Yes |
| Control-Sample3_Beta | chr2:63283013-63285097 | OTX1 | Island | Body | Yes |
| Control-Sample32_Beta | chr2:63283939-63284066 | OTX1 | Island | 3'UTR | Yes |
| Control-Sample35_Beta | chr2:63283806-63284557 | OTX1 | N_Shore | 3'UTR | Yes |
| Control-Sample4_Beta | chr2:63283013-63285365 | OTX1 | Island | Body | Yes |
| Control-Sample49_Beta | chr2:63283806-63283967 | OTX1 | N_Shore | 3'UTR | Yes |
| Control-Sample61_Beta | chr2:63283939-63284557 | OTX1 | Island | 3'UTR | Yes |
| Control-Sample62_Beta | chr2:63283967-63284132 | OTX1 | Island | 3'UTR | Yes |
| Control-Sample7_Beta | chr2:63283806-63285365 | OTX1 | N_Shore | 3'UTR | Yes |
| Control-Sample8_Beta | chr2:63280625-63281139 | OTX1 | N_Shore | Body | Yes |
| Control-Sample464_Be | chr14:57278351-57278729 | OTX2 | N_Shore | TSS1500 | Yes |
| Control-Sample58_Beta | chr14:57278084-57278351 | OTX2 | Island | TSS1500 | Yes |
| Control-Sample70_Beta | chr14:57278188-57278729 | OTX2 | Island | TSS1500 | Yes |
| Control-Sample14_Beta | chr14:57277178-57277366 | OTX2;OTX2 | N_Shore | 1stExon;5'UTR | Yes |
| Control-Sample3_Beta | chr14:57277178-57278188 | OTX2;OTX2 | N_Shore | 1stExon;5'UTR | Yes |
| Control-Sample65_Beta | chr14:57277178-57277761 | OTX2;OTX2 | N_Shore | 1stExon;5'UTR | Yes |
| Control-Sample21_Beta | chr14:57279115-57279669 | OTX2OS1 | Island | TSS1500 | Yes |
| Control-Sample24_Beta | chr14:57279275-57279669 | OTX2OS1 | S_Shore | TSS1500 | Yes |
| Control-Sample17_Beta | chr13:97646639-97646765 | OXGR1 | Island | TSS200 | No |
| Control-Sample19_Beta | chr3:8811543-8811758 | OXTR | S_Shore | TSS1500 | No |
| Control-Sample10_Beta | chr12:121569924-121570081 | P2RX7 | | TSS1500 | No |
| Control-Sample1_Beta | chr11:72929167-72929173 | P2RY2;P2RY2;P2RY2 | Island | TSS1500;TSS200;TSS200 | Yes |
| Control-Sample13_Beta | chr11:72929135-72929173 | P2RY2;P2RY2;P2RY2 | Island | TSS1500;TSS1500;TSS1500 | Yes |
| Control-Sample16_Beta | chr11:72929054-72929173 | P2RY2;P2RY2;P2RY2 | Island | TSS1500;TSS1500;TSS1500 | Yes |
| Control-Sample522_Be | chr13.0:25670170-25670187 | PABPC3 | Island | TSS200 | No |
| Control-Sample1_Beta | chr6:163602881-163603588 | PACRG;PACRG;PACRG | | Body;Body;Body | Yes |
| Control-Sample12_Beta | chr6:163602881-163602929 | PACRG;PACRG;PACRG | | Body;Body;Body | Yes |
| Control-Sample585_Be | chr6.0:163602881-163602929 | PACRG;PACRG;PACRG | | Body;Body;Body | Yes |
| Control-Sample64_Beta | chr6:163275952-163276038 | PACRG;PACRG;PACRG | | Body;Body;Body | Yes |
| Control-Sample1_Beta | chr6:163147999-163148022 | PACRG;PACRG;PARK2;PACRG;PARK2;PARK2 | N_Shore | TSS200;TSS1500;Body;TSS200;Body;Body | Yes |
| Control-Sample10_Beta | chr14:105830248-105830509 | PACS2;PACS2 | N_Shore | Body;Body | Yes |
| Control-Sample127_Be | chr14:105830606-105830859 | PACS2;PACS2 | Island | Body;Body | Yes |
| Control-Sample13_Beta | chr14:105802387-105802918 | PACS2;PACS2 | | Body;Body | Yes |
| Control-Sample170_Be | chr14:105779794-105779952 | PACS2;PACS2 | N_Shore | TSS1500;TSS1500 | Yes |
| Control-Sample206_Be | chr14:105830631-105830859 | PACS2;PACS2 | Island | Body;Body | Yes |
| Control-Sample219_Be | chr14:105830509-105830859 | PACS2;PACS2 | N_Shore | Body;Body | Yes |
| Control-Sample23_Beta | chr14:105793554-105794085 | PACS2;PACS2 | N_Shelf | Body;Body | Yes |
| Control-Sample504_Be | chr14.0:105830606-105830859 | PACS2;PACS2 | Island | Body;Body | Yes |
| Control-Sample511_Be | chr14.0:105830631-105830859 | PACS2;PACS2 | Island | Body;Body | Yes |
| Control-Sample1_Beta | chr15:40545050-40545175 | PAK6;C15orf56;PAK6;PAK6 | Island | 5'UTR;1stExon;5'UTR;5'UTR | Yes |
| Control-Sample16_Beta | chr15:40544427-40545175 | PAK6;C15orf56;PAK6;PAK6 | Island | 5'UTR;3'UTR;5'UTR;5'UTR | Yes |
| Control-Sample10_Beta | chr15:40544855-40545175 | PAK6;PAK6;C15orf56;PAK6 | Island | 5'UTR;5'UTR;Body;5'UTR | Yes |
| Control-Sample52_Beta | chr15:40545143-40545175 | PAK6;PAK6;C15orf56;PAK6 | Island | 5'UTR;5'UTR;TSS200;5'UTR | Yes |
| Control-Sample10_Beta | chr19:710823-711001 | PALM;PALM | Island | Body;Body | No |
| Control-Sample108_Be | chr1:100111387-100111849 | PALMD | | TSS200 | No |
| Control-Sample15_Beta | chr20:3897997-3898136 | PANK2;MIR103-2;PANK2;PANK2 | | Body;TSS200;Body;Body | No |
| Control-Sample464_Be | chr16:67696047-67696181 | PARD6A;ACD;ACD;PARD6A;ACD | S_Shore | Body;TSS1500;TSS1500;Body;TSS1500 | No |
| Control-Sample108_Be | chr6:161860203-161860635 | PARK2;PARK2;PARK2 | | Body;Body;Body | Yes |
| Control-Sample19_Beta | chr6:161796785-161796854 | PARK2;PARK2;PARK2 | | Body;Body;Body | Yes |
| Control-Sample20_Beta | chr6:161860359-161860635 | PARK2;PARK2;PARK2 | | Body;Body;Body | Yes |
| Control-Sample13_Beta | chr1:8044991-8045238 | PARK7;PARK7 | | Body;Body | No |
| Control-Sample1_Beta | chr15:72564179-72564263 | PARP6 | N_Shore | TSS1500 | No |
| Control-Sample30_Beta | chr5:59783629-59783906 | PART1;PART1;PDE4D;PART1 | | TSS200;Body;5'UTR;TSS200 | No |
| Control-Sample18_Beta | chr22:44568770-44568812 | PARVG;PARVG | | TSS200;TSS200 | Yes |
| Control-Sample3_Beta | chr22:44568725-44568812 | PARVG;PARVG | | TSS200;TSS200 | Yes |
| Control-Sample4_Beta | chr22:44568753-44568812 | PARVG;PARVG | | TSS200;TSS200 | Yes |
| Control-Sample464_Be | chr2:223164854-223164869 | PAX3;PAX3;PAX3;PAX3;PAX3;CCDC140;PAX3;PAX3;PAX3 | S_Shore | TSS1500;TSS1500;TSS1500;TSS1500;TSS1500;5'UTR | No |
| Control-Sample20_Beta | chr9:36881908-36882152 | PAX5 | N_Shore | Body | No |
| Control-Sample1_Beta | chr11:31837104-31837144 | PAX6 | Island | 5'UTR | Yes |
| Control-Sample10_Beta | chr11:31836254-31836673 | PAX6 | Island | 5'UTR | Yes |
| Control-Sample11_Beta | chr11:31836254-31837144 | PAX6 | Island | 5'UTR | Yes |
| Control-Sample16_Beta | chr11:31835534-31837144 | PAX6 | Island | 5'UTR | Yes |
| Control-Sample35_Beta | chr11:31836673-31837144 | PAX6 | Island | 5'UTR | Yes |
| Control-Sample10_Beta | chr11:31819444-31820658 | PAX6;PAX6;PAX6 | N_Shore | Body;Body;Body | Yes |
| Control-Sample10_Beta | chr11:31821764-31822341 | PAX6;PAX6;PAX6 | S_Shore | Body;Body;Body | Yes |
| Control-Sample11_Beta | chr11:31819380-31820572 | PAX6;PAX6;PAX6 | N_Shore | Body;Body;Body | Yes |
| Control-Sample11_Beta | chr11:31823141-31823238 | PAX6;PAX6;PAX6 | S_Shore | Body;Body;Body | Yes |
| Control-Sample12_Beta | chr11:31819444-31819678 | PAX6;PAX6;PAX6 | N_Shore | Body;Body;Body | Yes |
| Control-Sample13_Beta | chr11:31825605-31826421 | PAX6;PAX6;PAX6 | N_Shore | Body;Body;Body | Yes |
| Control-Sample14_Beta | chr11:31825605-31825833 | PAX6;PAX6;PAX6 | N_Shore | Body;Body;Body | Yes |
| Control-Sample15_Beta | chr11:31824262-31825226 | PAX6;PAX6;PAX6 | N_Shore | Body;Body;Body | Yes |
| Control-Sample17_Beta | chr11:31819380-31819678 | PAX6;PAX6;PAX6 | N_Shore | Body;Body;Body | Yes |
| Control-Sample24_Beta | chr11:31818474-31819162 | PAX6;PAX6;PAX6 | N_Shore | Body;Body;Body | Yes |
| Control-Sample25_Beta | chr11:31817810-31820658 | PAX6;PAX6;PAX6 | N_Shelf | Body;Body;Body | Yes |
| Control-Sample27_Beta | chr11:31817810-31819219 | PAX6;PAX6;PAX6 | N_Shelf | Body;Body;Body | Yes |
| Control-Sample30_Beta | chr11:31819380-31820658 | PAX6;PAX6;PAX6 | N_Shore | Body;Body;Body | Yes |
| Control-Sample34_Beta | chr11:31817810-31817881 | PAX6;PAX6;PAX6 | N_Shelf | Body;Body;Body | Yes |
| Control-Sample36_Beta | chr11:31824262-31825969 | PAX6;PAX6;PAX6 | N_Shore | Body;Body;Body | Yes |
| Control-Sample37_Beta | chr11:31825214-31825969 | PAX6;PAX6;PAX6 | N_Shore | Body;Body;Body | Yes |
| Control-Sample4_Beta | chr11:31820441-31820658 | PAX6;PAX6;PAX6 | Island | Body;Body;Body | Yes |
| Control-Sample46_Beta | chr11:31817810-31818474 | PAX6;PAX6;PAX6 | N_Shelf | Body;Body;Body | Yes |
| Control-Sample55_Beta | chr11:31819380-31820209 | PAX6;PAX6;PAX6 | N_Shore | Body;Body;Body | Yes |

| | | | | | |
|---|---|---|---|---|---|
| Control-Sample9_Beta | chr11:31825756-31825969 | PAX6;PAX6;PAX6 | Island | Body;Body;Body | Yes |
| Control-Sample1_Beta | chr1:19051735-19052235 | PAX7;PAX7;PAX7 | | Body;Body;Body | Yes |
| Control-Sample21_Beta | chr1:19051735-19052204 | PAX7;PAX7;PAX7 | | Body;Body;Body | Yes |
| Control-Sample57_Beta | chr1:19052179-19052235 | PAX7;PAX7;PAX7 | | Body;Body;Body | Yes |
| Control-Sample53_Beta | chr2:114036571-114036953 | PAX8;PAX8;PAX8;LOC440839;PAX8;PAX8 | S_Shore | TSS200;TSS200;TSS200;Body;TSS200;TSS200 | Yes |
| Control-Sample31_Beta | chr2:114035983-114036685 | PAX8;PAX8;PAX8;PAX8;LOC440839;PAX8 | Island | 5'UTR;5'UTR;5'UTR;5'UTR;Body;5'UTR | Yes |
| Control-Sample1_Beta | chr2:114033360-114033830 | PAX8;PAX8;PAX8;PAX8;PAX8;LOC440839 | Island | Body;Body;Body;Body;Body;Body | Yes |
| Control-Sample10_Beta | chr2:114033490-114033830 | PAX8;PAX8;PAX8;PAX8;PAX8;LOC440839 | Island | Body;Body;Body;Body;Body;Body | Yes |
| Control-Sample161_Be | chr2:113992762-113992930 | PAX8;PAX8;PAX8;PAX8;PAX8;LOC440839;LOC654433 | N_Shore | Body;Body;Body;Body;Body;Body;TSS1500 | Yes |
| Control-Sample162_Be | chr2:113992762-113993142 | PAX8;PAX8;PAX8;PAX8;PAX8;LOC440839;LOC654433 | N_Shore | Body;Body;Body;Body;Body;Body;TSS1500 | Yes |
| Control-Sample346_Be | chr19:19729535-19729568 | PBX4 | Island | TSS200 | No |
| Control-Sample13_Beta | chr11:66643399-66643572 | PC;PC;PC | | 5'UTR;5'UTR;5'UTR | No |
| Control-Sample12_Beta | chr21:47294981-47295375 | PCBP3;PCBP3 | Island | Body;Body | No |
| Control-Sample1_Beta | chr13:101173617-101173774 | PCCA;PCCA | | Body;Body | Yes |
| Control-Sample1_Beta | chr13:101174096-101174420 | PCCA;PCCA | | Body;Body | Yes |
| Control-Sample10_Beta | chr13:101173617-101174420 | PCCA;PCCA | | Body;Body | Yes |
| Control-Sample11_Beta | chr13:101174060-101174420 | PCCA;PCCA | | Body;Body | Yes |
| Control-Sample19_Beta | chr13:101173617-101174096 | PCCA;PCCA | | Body;Body | Yes |
| Control-Sample57_Beta | chr13:101174130-101174420 | PCCA;PCCA | | Body;Body | Yes |
| Control-Sample68_Beta | chr13:101173669-101174420 | PCCA;PCCA | | Body;Body | Yes |
| Control-Sample108_Be | chr10:56561124-56561543 | PCDH15;PCDH15;PCDH15;PCDH15;PCDH15;PCDH15;PCDH15;PCDH15;PCDH15;PCDH15;PC | | TSS200;TSS200;TSS200;TSS200;TSS200;TSS200;TSS | Yes |
| Control-Sample568_Be | chr10.0:56561124-56561543 | PCDH15;PCDH15;PCDH15;PCDH15;PCDH15;PCDH15;PCDH15;PCDH15;PCDH15;PCDH15;PC | | TSS200;TSS200;TSS200;TSS200;TSS200;TSS200;TSS | Yes |
| Control-Sample12_Beta | chr5:140207609-140207760 | PCDHA2;PCDHA1;PCDHA1;PCDHA6;PCDHA5;PCDHA6;PC | N_Shore | Body;Body;Body;TSS200;Body;TSS200;Body;Body;TS | Yes |
| Control-Sample14_Beta | chr5:140180926-140181074 | PCDHA2;PCDHA1;PCDHA3;PCDHA3;PCDHA1 | Island | Body;Body;1stExon;1stExon;Body | Yes |
| Control-Sample537_Be | chr5.0:140552347-140552505 | PCDHB7;PCDHB7 | N_Shore | 1stExon;5'UTR | Yes |
| Control-Sample77_Beta | chr5:140552347-140552505 | PCDHB7;PCDHB7 | N_Shore | 1stExon;5'UTR | Yes |
| Control-Sample19_Beta | chr4:719908-720218 | PCGF3 | S_Shelf | 5'UTR | Yes |
| Control-Sample2_Beta | chr4:715865-715950 | PCGF3 | N_Shore | 5'UTR | Yes |
| Control-Sample17_Beta | chr20:56136012-56136167 | PCK1 | | TSS200 | No |
| Control-Sample12_Beta | chr21:47803078-47803548 | PCNT | Island | Body | No |
| Control-Sample1_Beta | chr15:102010125-102010195 | PCSK6;PCSK6;PCSK6;PCSK6;PCSK6;PCSK6;PCSK6;PCSK6 | | Body;Body;Body;Body;Body;Body;Body;Body | Yes |
| Control-Sample216_Be | chr15:101991580-101991886 | PCSK6;PCSK6;PCSK6;PCSK6;PCSK6;PCSK6;PCSK6;PCSK6 | | Body;Body;Body;Body;Body;Body;Body;Body | Yes |
| Control-Sample3_Beta | chr11:72379867-72380291 | PDE2A;PDE2A | | 5'UTR;Body | Yes |
| Control-Sample1_Beta | chr11:72295425-72295716 | PDE2A;PDE2A;PDE2A;PDE2A | Island | Body;Body;Body;Body | Yes |
| Control-Sample1_Beta | chr11:72300891-72301924 | PDE2A;PDE2A;PDE2A;PDE2A | N_Shore | Body;Body;Body;Body | Yes |
| Control-Sample13_Beta | chr11:72295168-72295535 | PDE2A;PDE2A;PDE2A;PDE2A | Island | Body;Body;Body;Body | Yes |
| Control-Sample21_Beta | chr11:72301306-72301924 | PDE2A;PDE2A;PDE2A;PDE2A | Island | Body;Body;Body;Body | Yes |
| Control-Sample24_Beta | chr11:72300891-72301558 | PDE2A;PDE2A;PDE2A;PDE2A | N_Shore | Body;Body;Body;Body | Yes |
| Control-Sample33_Beta | chr11:72295168-72295716 | PDE2A;PDE2A;PDE2A;PDE2A | Island | Body;Body;Body;Body | Yes |
| Control-Sample64_Beta | chr11:72301558-72301924 | PDE2A;PDE2A;PDE2A;PDE2A | Island | Body;Body;Body;Body | Yes |
| Control-Sample108_Be | chr1:66458116-66458803 | PDE4B;PDE4B;PDE4B | | Body;TSS1500;Body | Yes |
| Control-Sample22_Beta | chr1:66458361-66458803 | PDE4B;PDE4B;PDE4B | | TSS200;Body;Body | Yes |
| Control-Sample3_Beta | chr1:66797599-66798063 | PDE4B;PDE4B;PDE4B;PDE4B | | Body;TSS200;Body;Body | Yes |
| Control-Sample37_Beta | chr5:59559062-59559218 | PDE4D | | 5'UTR | No |
| Control-Sample1_Beta | chr4:619162-619272 | PDE6B;PDE6B | N_Shore | TSS200;TSS200 | Yes |
| Control-Sample561_Be | chr4.0:645440-645781 | PDE6B;PDE6B | Island | Body;Body | Yes |
| Control-Sample194_Be | chr4:649845-649901 | PDE6B;PDE6B;PDE6B | N_Shore | Body;Body;Body | Yes |
| Control-Sample108_Be | chr6:136172755-136172818 | PDE7B | | TSS200 | No |
| Control-Sample151_Be | chr7:537740-537828 | PDGFA;PDGFA | S_Shelf | 3'UTR;3'UTR | No |
| Control-Sample1_Beta | chr5:131592959-131593191 | PDLIM4;PDLIM4 | N_Shore | TSS1500;TSS1500 | Yes |
| Control-Sample13_Beta | chr5:131592959-131593106 | PDLIM4;PDLIM4 | N_Shore | TSS1500;TSS1500 | Yes |
| Control-Sample15_Beta | chr5:131593104-131593191 | PDLIM4;PDLIM4 | Island | TSS1500;TSS1500 | Yes |
| Control-Sample10_Beta | chr5:131593413-131593812 | PDLIM4;PDLIM4;PDLIM4;PDLIM4 | Island | 5'UTR;1stExon;1stExon;5'UTR | Yes |
| Control-Sample64_Beta | chr20:1974924-1975372 | PDYN | | TSS1500 | No |
| Control-Sample1_Beta | chr10:102778683-102778909 | PDZD7 | N_Shore | Body | No |
| Control-Sample8_Beta | chr1:145726979-145727247 | PDZK1 | | TSS1500 | No |
| Control-Sample12_Beta | chr8:22785567-22786374 | PEBP4 | | TSS200 | Yes |
| Control-Sample13_Beta | chr8:22785712-22786374 | PEBP4 | | TSS1500 | Yes |
| Control-Sample2_Beta | chr8:22786190-22786374 | PEBP4 | | TSS1500 | Yes |
| Control-Sample223_Be | chr8:22595307-22595422 | PEBP4 | | Body | Yes |
| Control-Sample543_Be | chr6.0:4135844-4135889 | PECI;PECI;PECI;PECI | Island | TSS200;TSS200;TSS200;TSS200 | No |
| Control-Sample18_Beta | chr17:17457745-17457836 | PEMT;PEMT;PEMT | | Body;Body;Body | No |
| Control-Sample3_Beta | chr2:239169537-239169696 | PER2 | | Body | No |
| Control-Sample26_Beta | chr1:7844446-7844735 | PER3 | Island | TSS1500 | No |
| Control-Sample464_Be | chr1:10601389-10601833 | PEX14 | | Body | Yes |
| Control-Sample465_Be | chr1:10601389-10601595 | PEX14 | | Body | Yes |
| Control-Sample23_Beta | chr4:74847761-74847829 | PF4 | Island | TSS200 | Yes |
| Control-Sample25_Beta | chr4:74847646-74847829 | PF4 | Island | 1stExon | Yes |
| Control-Sample284_Be | chr4:74847758-74847829 | PF4 | Island | TSS200 | Yes |
| Control-Sample524_Be | chr4.0:74847766-74847829 | PF4 | Island | TSS200 | Yes |
| Control-Sample65_Beta | chr4:74847766-74847829 | PF4 | Island | TSS200 | Yes |
| Control-Sample77_Beta | chr10:6214016-6214079 | PFKFB3 | | Body | Yes |
| Control-Sample1_Beta | chr10:6274873-6275077 | PFKFB3;PFKFB3 | | Body;Body | Yes |
| Control-Sample1_Beta | chr10:3149673-3149852 | PFKP | S_Shore | Body | Yes |
| Control-Sample171_Be | chr10:3149760-3149852 | PFKP | S_Shore | Body | Yes |
| Control-Sample19_Beta | chr10:3169037-3169516 | PFKP | | Body | Yes |
| Control-Sample464_Be | chr10:3143354-3143783 | PFKP | N_Shelf | Body | Yes |
| Control-Sample465_Be | chr10:3143596-3143783 | PFKP | N_Shelf | Body | Yes |
| Control-Sample55_Beta | chr10:3149673-3149796 | PFKP | S_Shore | Body | Yes |
| Control-Sample23_Beta | chr7:44104860-44105050 | PGAM2 | Island | 1stExon | No |
| Control-Sample10_Beta | chr1:230457906-230457979 | PGBD5 | | 3'UTR | Yes |
| Control-Sample14_Beta | chr1:230513522-230513809 | PGBD5 | | TSS200 | Yes |
| Control-Sample163_Be | chr1:230468464-230468773 | PGBD5 | N_Shore | Body | Yes |
| Control-Sample19_Beta | chr1:230468168-230468773 | PGBD5 | N_Shore | Body | Yes |
| Control-Sample53_Beta | chr1:230468464-230468699 | PGBD5 | N_Shore | Body | Yes |
| Control-Sample54_Beta | chr1:230468168-230468699 | PGBD5 | N_Shore | Body | Yes |
| Control-Sample224_Be | chr8:97657432-97657803 | PGCP | Island | TSS200 | No |
| Control-Sample34_Beta | chr6:12717016-12717192 | PHACTR1 | | TSS1500 | Yes |
| Control-Sample57_Beta | chr6:12716671-12717138 | PHACTR1 | | TSS1500 | Yes |
| Control-Sample58_Beta | chr6:12716655-12717192 | PHACTR1 | | TSS1500 | Yes |
| Control-Sample10_Beta | chr6:143929300-143929511 | PHACTR2;PHACTR2 | | TSS200;TSS200 | No |

| | | | | | |
|---|---|---|---|---|---|
| Control-Sample1_Beta | chr7:56160766-56161020 | PHKG1 | | TSS200 | Yes |
| Control-Sample5_Beta | chr7:56160893-56161020 | PHKG1 | | TSS1500 | Yes |
| Control-Sample19_Beta | chr7:56160687-56160770 | PHKG1;PHKG1 | | 1stExon;5'UTR | Yes |
| Control-Sample331_Be | chr11:118476352-118477055 | PHLDB1 | N_Shore | TSS1500 | No |
| Control-Sample1_Beta | chr19:44005893-44006432 | PHLDB3 | Island | Body | Yes |
| Control-Sample5_Beta | chr19:44005808-44006432 | PHLDB3 | N_Shore | Body | Yes |
| Control-Sample2_Beta | chr22:20399601-20400015 | PI4KAP1 | | TSS1500 | No |
| Control-Sample1_Beta | chr1:151298699-151298954 | PI4KB;PI4KB | N_Shore | 1stExon;5'UTR | No |
| Control-Sample10_Beta | chr1:9788715-9788769 | PIK3CD | N_Shore | 3'UTR | Yes |
| Control-Sample8_Beta | chr1:9788517-9788769 | PIK3CD | N_Shore | 3'UTR | Yes |
| Control-Sample103_Be | chr7:106505370-106505786 | PIK3CG | N_Shelf | TSS1500 | No |
| Control-Sample1_Beta | chr5:67583598-67583627 | PIK3R1;PIK3R1 | N_Shore | Body;TSS1500 | No |
| Control-Sample1_Beta | chr10:22843607-22843938 | PIP4K2A | | Body | Yes |
| Control-Sample17_Beta | chr10:22911537-22911629 | PIP4K2A | | Body | Yes |
| Control-Sample1_Beta | chr9:130689253-130689627 | PIP5KL1;PIP5KL1 | N_Shore | Body;Body | No |
| Control-Sample19_Beta | chr17:1466709-1466715 | PITPNA | S_Shore | TSS1500 | No |
| Control-Sample108_Be | chr17:65471321-65471461 | PITPNC1;PITPNC1 | | Body;Body | Yes |
| Control-Sample13_Beta | chr17:65471303-65471461 | PITPNC1;PITPNC1 | | Body;Body | Yes |
| Control-Sample62_Beta | chr17:65471303-65471347 | PITPNC1;PITPNC1 | | Body;Body | Yes |
| Control-Sample10_Beta | chr12:123469284-123469669 | PITPNM2 | N_Shore | 3'UTR | No |
| Control-Sample1_Beta | chr17:6358599-6358928 | PITPNM3;PITPNM3 | Island | 3'UTR;3'UTR | No |
| Control-Sample15_Beta | chr12:130821453-130821530 | PIWIL1 | N_Shore | TSS1500 | Yes |
| Control-Sample496_Be | chr12:130821962-130822605 | PIWIL1 | N_Shore | TSS1500 | Yes |
| Control-Sample181_Be | chr8:22132959-22133004 | PIWIL2;PIWIL2 | Island | 5'UTR;TSS200 | Yes |
| Control-Sample182_Be | chr8:22132926-22133076 | PIWIL2;PIWIL2 | Island | 5'UTR;TSS200 | Yes |
| Control-Sample467_Be | chr8:22132641-22132678 | PIWIL2;PIWIL2 | N_Shore | TSS200;TSS1500 | Yes |
| Control-Sample573_Be | chr8.0:22132926-22132959 | PIWIL2;PIWIL2 | Island | 5'UTR;TSS200 | Yes |
| Control-Sample89_Beta | chr8:22132678-22132874 | PIWIL2;PIWIL2 | N_Shore | TSS200;TSS1500 | Yes |
| Control-Sample89_Beta | chr8:22132932-22133076 | PIWIL2;PIWIL2 | Island | 5'UTR;TSS200 | Yes |
| Control-Sample1_Beta | chr16:81254051-81254209 | PKD1L2;PKD1L2 | | TSS200;TSS200 | Yes |
| Control-Sample13_Beta | chr16:81254010-81254209 | PKD1L2;PKD1L2 | | TSS200;TSS200 | Yes |
| Control-Sample10_Beta | chr6:51814354-51814455 | PKHD1;PKHD1 | | Body;Body | No |
| Control-Sample1_Beta | chr22:31535981-31536585 | PLA2G3 | | 1stExon | Yes |
| Control-Sample3_Beta | chr22:31535981-31536784 | PLA2G3 | | 1stExon | Yes |
| Control-Sample33_Beta | chr22:31536549-31536784 | PLA2G3 | | TSS200 | Yes |
| Control-Sample58_Beta | chr22:31535981-31536540 | PLA2G3 | | 1stExon | Yes |
| Control-Sample66_Beta | chr22:31536476-31536784 | PLA2G3 | | TSS200 | Yes |
| Control-Sample10_Beta | chr22:31536431-31536540 | PLA2G3;PLA2G3 | | 5'UTR;1stExon | Yes |
| Control-Sample15_Beta | chr22:31536431-31536784 | PLA2G3;PLA2G3 | | 5'UTR;1stExon | Yes |
| Control-Sample29_Beta | chr22:31536431-31536585 | PLA2G3;PLA2G3 | | 5'UTR;1stExon | Yes |
| Control-Sample113_Be | chr15:42371635-42371808 | PLA2G4D | Island | Body | Yes |
| Control-Sample114_Be | chr15:42371511-42371808 | PLA2G4D | N_Shore | Body | Yes |
| Control-Sample327_Be | chr15:42376692-42376765 | PLA2G4D | | Body | Yes |
| Control-Sample1_Beta | chr6:144386422-144386474 | PLAGL1;PLAGL1 | S_Shore | TSS1500;TSS1500 | Yes |
| Control-Sample11_Beta | chr6:144386275-144386416 | PLAGL1;PLAGL1 | S_Shore | TSS1500;TSS1500 | Yes |
| Control-Sample12_Beta | chr6:144386422-144386515 | PLAGL1;PLAGL1 | S_Shore | TSS1500;TSS1500 | Yes |
| Control-Sample24_Beta | chr6:144386422-144386528 | PLAGL1;PLAGL1 | S_Shore | TSS1500;TSS1500 | Yes |
| Control-Sample31_Beta | chr6:144386422-144386848 | PLAGL1;PLAGL1 | S_Shore | TSS1500;TSS1500 | Yes |
| Control-Sample30_Beta | chr12:14721020-14721288 | PLBD1 | Island | TSS1500 | No |
| Control-Sample108_Be | chr2:219472218-219472724 | PLCD4 | | TSS1500 | No |
| Control-Sample1_Beta | chr16:81929358-81929465 | PLCG2 | | Body | Yes |
| Control-Sample572_Be | chr16.0:81929358-81929465 | PLCG2 | | Body | Yes |
| Control-Sample10_Beta | chr3:155422754-155423168 | PLCH1 | | TSS1500 | Yes |
| Control-Sample108_Be | chr3:155421970-155422103 | PLCH1;PLCH1 | | 5'UTR;1stExon | Yes |
| Control-Sample108_Be | chr3:155394292-155394428 | PLCH1;PLCH1;PLCH1 | | 5'UTR;TSS200;TSS200 | Yes |
| Control-Sample496_Be | chr3:155394109-155394292 | PLCH1;PLCH1;PLCH1 | | 5'UTR;TSS200;TSS200 | Yes |
| Control-Sample568_Be | chr3.0:155394118-155394428 | PLCH1;PLCH1;PLCH1 | | 5'UTR;TSS200;TSS200 | Yes |
| Control-Sample24_Beta | chr5:41510076-41510325 | PLCXD3 | Island | Body | No |
| Control-Sample13_Beta | chr17:17109570-17109800 | PLD6 | Island | 1stExon | Yes |
| Control-Sample225_Be | chr17:17109570-17109651 | PLD6 | Island | 1stExon | Yes |
| Control-Sample23_Beta | chr17:17109651-17109691 | PLD6 | Island | TSS200 | Yes |
| Control-Sample463_Be | chr17:17109570-17109691 | PLD6 | Island | 1stExon | Yes |
| Control-Sample7_Beta | chr17:17109570-17109936 | PLD6 | Island | 1stExon | Yes |
| Control-Sample10_Beta | chr17:17109640-17109800 | PLD6;PLD6 | Island | 1stExon;5'UTR | Yes |
| Control-Sample106_Be | chr17:17109640-17109691 | PLD6;PLD6 | Island | 1stExon;5'UTR | Yes |
| Control-Sample29_Beta | chr17:17109640-17109740 | PLD6;PLD6 | Island | 1stExon;5'UTR | Yes |
| Control-Sample5_Beta | chr17:17109640-17109678 | PLD6;PLD6 | Island | 1stExon;5'UTR | Yes |
| Control-Sample1_Beta | chr8:145048278-145048767 | PLEC1;PLEC1 | Island | Body;TSS1500 | Yes |
| Control-Sample11_Beta | chr8:145048235-145048363 | PLEC1;PLEC1 | Island | Body;TSS1500 | Yes |
| Control-Sample224_Be | chr8:145047713-145048235 | PLEC1;PLEC1 | Island | Body;TSS200 | Yes |
| Control-Sample65_Beta | chr8:145028222-145028257 | PLEC1;PLEC1;PLEC1 | Island | Body;Body;TSS200 | Yes |
| Control-Sample1_Beta | chr8:145019032-145019074 | PLEC1;PLEC1;PLEC1;PLEC1;PLEC1;PLEC1 | Island | Body;Body;Body;TSS1500;Body;TSS200 | Yes |
| Control-Sample13_Beta | chr8:145019032-145019067 | PLEC1;PLEC1;PLEC1;PLEC1;PLEC1;PLEC1 | Island | Body;Body;Body;TSS1500;Body;TSS200 | Yes |
| Control-Sample23_Beta | chr8:145019032-145019116 | PLEC1;PLEC1;PLEC1;PLEC1;PLEC1;PLEC1 | Island | Body;Body;Body;TSS1500;Body;TSS200 | Yes |
| Control-Sample331_Be | chr8:145019041-145019116 | PLEC1;PLEC1;PLEC1;PLEC1;PLEC1;PLEC1 | Island | Body;Body;Body;TSS1500;Body;TSS200 | Yes |
| Control-Sample70_Beta | chr8:145019041-145019074 | PLEC1;PLEC1;PLEC1;PLEC1;PLEC1;PLEC1 | Island | Body;Body;Body;TSS1500;Body;TSS200 | Yes |
| Control-Sample19_Beta | chr8:145018075-145018300 | PLEC1;PLEC1;PLEC1;PLEC1;PLEC1;PLEC1;PLEC1;PLEC1 | Island | Body;5'UTR;Body;Body;Body;TSS1500;1stExon;Body | Yes |
| Control-Sample4_Beta | chr8:145013728-145013932 | PLEC1;PLEC1;PLEC1;PLEC1;PLEC1;PLEC1;PLEC1;PLEC1; | N_Shelf | Body;1stExon;5'UTR;Body;Body;Body;Body;Body;Bod | Yes |
| Control-Sample1_Beta | chr8:38757532-38757859 | PLEKHA2 | N_Shore | TSS1500 | Yes |
| Control-Sample66_Beta | chr8:38757532-38757772 | PLEKHA2 | N_Shore | TSS1500 | Yes |
| Control-Sample480_Be | chr19:49340574-49340765 | PLEKHA4;HSD17B14;PLEKHA4 | Island | 3'UTR;TSS1500;Body | No |
| Control-Sample1_Beta | chr11:73357019-73357164 | PLEKHB1;PLEKHB1 | | TSS1500;TSS1500 | Yes |
| Control-Sample17_Beta | chr11:73357019-73357276 | PLEKHB1;PLEKHB1 | | TSS1500;TSS1500 | Yes |
| Control-Sample18_Beta | chr11:73357019-73357194 | PLEKHB1;PLEKHB1 | | TSS1500;TSS1500 | Yes |
| Control-Sample19_Beta | chr11:73357019-73357396 | PLEKHB1;PLEKHB1 | | TSS1500;TSS1500 | Yes |
| Control-Sample28_Beta | chr11:73357019-73357159 | PLEKHB1;PLEKHB1 | | TSS1500;TSS1500 | Yes |
| Control-Sample13_Beta | chr11:73357095-73357164 | PLEKHB1;PLEKHB1;PLEKHB1;PLEKHB1 | | TSS200;TSS200;TSS1500;TSS1500 | Yes |
| Control-Sample20_Beta | chr11:73357159-73357396 | PLEKHB1;PLEKHB1;PLEKHB1;PLEKHB1 | | TSS200;TSS200;TSS1500;TSS1500 | Yes |
| Control-Sample21_Beta | chr11:73357159-73357276 | PLEKHB1;PLEKHB1;PLEKHB1;PLEKHB1 | | TSS200;TSS200;TSS1500;TSS1500 | Yes |
| Control-Sample30_Beta | chr11:73357095-73357276 | PLEKHB1;PLEKHB1;PLEKHB1;PLEKHB1 | | TSS200;TSS200;TSS1500;TSS1500 | Yes |
| Control-Sample70_Beta | chr11:73357159-73357194 | PLEKHB1;PLEKHB1;PLEKHB1;PLEKHB1 | | TSS200;TSS200;TSS1500;TSS1500 | Yes |

| | | | | | |
|---|---|---|---|---|---|
| Control-Sample71_Beta | chr11:73357159-73357183 | PLEKHB1;PLEKHB1;PLEKHB1;PLEKHB1 | | TSS200;TSS200;TSS1500;TSS1500 | Yes |
| Control-Sample1_Beta | chr1:6558085-6558113 | PLEKHG5;PLEKHG5 | S_Shore | Body;TSS1500 | No |
| Control-Sample11_Beta | chr12:6419409-6419509 | PLEKHG6;PLEKHG6 | N_Shore | TSS1500;TSS200 | Yes |
| Control-Sample66_Beta | chr12:6420034-6420041 | PLEKHG6;PLEKHG6 | S_Shore | 5'UTR;TSS200 | Yes |
| Control-Sample34_Beta | chr12:6421768-6422131 | PLEKHG6;PLEKHG6;PLEKHG6 | S_Shore | TSS200;Body;Body | Yes |
| Control-Sample108_Be | chr2:43903470-43903526 | PLEKHH2;LOC728819 | | Body;TSS200 | Yes |
| Control-Sample54_Beta | chr2:43903842-43904011 | PLEKHH2;LOC728819 | | Body;TSS1500 | Yes |
| Control-Sample23_Beta | chr3:146262174-146262421 | PLSCR1 | Island | 5'UTR | No |
| Control-Sample19_Beta | chr3:145969579-145969718 | PLSCR4;PLSCR4;PLSCR4;PLSCR4 | S_Shore | TSS1500;TSS1500;TSS1500;TSS1500 | No |
| Control-Sample331_Be | chr3:126721131-126721586 | PLXNA1 | | Body | No |
| Control-Sample1_Beta | chr3:48471575-48471771 | PLXNB1;PLXNB1 | S_Shore | TSS1500;TSS200 | Yes |
| Control-Sample11_Beta | chr3:48465017-48465320 | PLXNB1;PLXNB1 | | Body;Body | Yes |
| Control-Sample19_Beta | chr3:48471575-48471695 | PLXNB1;PLXNB1 | S_Shore | TSS1500;TSS200 | Yes |
| Control-Sample23_Beta | chr3:48471575-48471901 | PLXNB1;PLXNB1 | S_Shore | TSS1500;TSS200 | Yes |
| Control-Sample106_Be | chr1:205819088-205819492 | PM20D1 | Island | 1stExon | Yes |
| Control-Sample107_Be | chr1:205819088-205819463 | PM20D1 | Island | 1stExon | Yes |
| Control-Sample219_Be | chr1:205819088-205819251 | PM20D1 | Island | 1stExon | Yes |
| Control-Sample3_Beta | chr1:205819088-205819406 | PM20D1 | Island | 1stExon | Yes |
| Control-Sample460_Be | chr1:205819088-205819609 | PM20D1 | Island | 1stExon | Yes |
| Control-Sample525_Be | chr1.0:205819088-205819463 | PM20D1 | Island | 1stExon | Yes |
| Control-Sample539_Be | chr1.0:205819088-205819492 | PM20D1 | Island | 1stExon | Yes |
| Control-Sample31_Beta | chr20:56247302-56247611 | PMEPA1;PMEPA1;PMEPA1;PMEPA1 | Island | 5'UTR;5'UTR;Body;Body | No |
| Control-Sample10_Beta | chr8:82359719-82359799 | PMP2 | | TSS200 | Yes |
| Control-Sample171_Be | chr8:82359567-82359799 | PMP2 | | 1stExon | Yes |
| Control-Sample572_Be | chr8.0:82359567-82359799 | PMP2 | | 1stExon | Yes |
| Control-Sample223_Be | chr19:46996347-46996713 | PNMAL2;PNMAL2 | Island | 1stExon;3'UTR | Yes |
| Control-Sample6_Beta | chr19:46996516-46996868 | PNMAL2;PNMAL2 | Island | 1stExon;3'UTR | Yes |
| Control-Sample12_Beta | chr17:37824461-37824526 | PNMT | Island | TSS200 | No |
| Control-Sample57_Beta | chr8:28174847-28175003 | PNOC | | 5'UTR | Yes |
| Control-Sample58_Beta | chr8:28174847-28175021 | PNOC | | 5'UTR | Yes |
| Control-Sample12_Beta | chr9:140356475-140356830 | PNPLA7;PNPLA7 | Island | Body;Body | No |
| Control-Sample1_Beta | chr1:53527412-53527435 | PODN | N_Shore | TSS1500 | No |
| Control-Sample11_Beta | chr7:131240440-131240531 | PODXL;PODXL | N_Shore | Body;Body | No |
| Control-Sample1_Beta | chr2:25391666-25391711 | POMC;POMC | Island | TSS200;TSS200 | Yes |
| Control-Sample1_Beta | chr2:25391830-25391911 | POMC;POMC | Island | TSS1500;TSS1500 | Yes |
| Control-Sample10_Beta | chr2:25391656-25391711 | POMC;POMC | Island | TSS200;TSS200 | Yes |
| Control-Sample1_Beta | chr7:75596021-75596055 | POR | | Body | No |
| Control-Sample11_Beta | chr6:31139359-31139796 | POU5F1 | | TSS1500 | No |
| Control-Sample162_Be | chr7:39170497-39170763 | POU6F2;POU6F2 | | Body;Body | Yes |
| Control-Sample554_Be | chr7.0:39170497-39170763 | POU6F2;POU6F2 | | Body;Body | Yes |
| Control-Sample10_Beta | chr2:241389021-241389258 | PP14571;GPC1 | N_Shelf | Body;Body | Yes |
| Control-Sample42_Beta | chr2:241393833-241394528 | PP14571;GPC1 | N_Shore | Body;Body | Yes |
| Control-Sample496_Be | chr4:23892515-23892621 | PPARGC1A | | TSS1500 | No |
| Control-Sample1_Beta | chr19:49646157-49646246 | PPFIA3 | Island | Body | No |
| Control-Sample1_Beta | chr1:203044677-203045229 | PPFIA4 | N_Shore | Body | Yes |
| Control-Sample10_Beta | chr1:203044677-203045044 | PPFIA4 | N_Shore | Body | Yes |
| Control-Sample17_Beta | chr1:203044677-203044930 | PPFIA4 | N_Shore | Body | Yes |
| Control-Sample63_Beta | chr1:203044930-203045229 | PPFIA4 | Island | Body | Yes |
| Control-Sample3_Beta | chr11:7597349-7597983 | PPFIBP2 | | Body | No |
| Control-Sample44_Beta | chr6:30582241-30582352 | PPP1R10 | N_Shelf | 5'UTR | No |
| Control-Sample1_Beta | chr19:38747355-38747378 | PPP1R14A | Island | TSS200 | Yes |
| Control-Sample21_Beta | chr19:38747355-38747796 | PPP1R14A | Island | TSS200 | Yes |
| Control-Sample1_Beta | chr17:37784694-37784895 | PPP1R1B;PPP1R1B | S_Shore | TSS200;Body | Yes |
| Control-Sample10_Beta | chr17:37784694-37784711 | PPP1R1B;PPP1R1B | S_Shore | TSS200;Body | Yes |
| Control-Sample270_Be | chr2:182850269-182850683 | PPP1R1C | | TSS1500 | No |
| Control-Sample291_Be | chr5:146435656-146436093 | PPP2R2B;PPP2R2B;PPP2R2B;PPP2R2B;PPP2R2B;PPP2R2B | | 5'UTR;5'UTR;5'UTR;1stExon;1stExon;5'UTR | No |
| Control-Sample171_Be | chr4:6449564-6449865 | PPP2R2C | | Body | No |
| Control-Sample310_Be | chr14:102227932-102227943 | PPP2R5C;PPP2R5C | Island | TSS1500;TSS1500 | Yes |
| Control-Sample108_Be | chr14:102276033-102276087 | PPP2R5C;PPP2R5C;PPP2R5C;PPP2R5C;PPP2R5C | | Body;TSS200;TSS200;Body;TSS200 | Yes |
| Control-Sample67_Beta | chr6:32120773-32120901 | PPT2;PRRT1;PPT2 | N_Shore | TSS1500;TSS1500;TSS1500 | No |
| Control-Sample10_Beta | chr17:42019840-42019987 | PPY | S_Shelf | TSS200 | Yes |
| Control-Sample1_Beta | chr17:42019790-42019987 | PPY;PPY | S_Shelf | 1stExon;5'UTR | Yes |
| Control-Sample464_Be | chr10:47083392-47083504 | PPYR1 | Island | TSS200 | No |
| Control-Sample19_Beta | chr17:74535962-74536068 | PRCD | S_Shore | TSS200 | No |
| Control-Sample10_Beta | chr6:106545984-106546074 | PRDM1;PRDM1 | | Body;TSS1500 | No |
| Control-Sample1_Beta | chr1:3017674-3018024 | PRDM16;PRDM16 | | Body;Body | Yes |
| Control-Sample1_Beta | chr1:3028230-3028485 | PRDM16;PRDM16 | Island | Body;Body | Yes |
| Control-Sample1_Beta | chr1:3028813-3029070 | PRDM16;PRDM16 | S_Shore | Body;Body | Yes |
| Control-Sample1_Beta | chr1:3038564-3039059 | PRDM16;PRDM16 | S_Shore | Body;Body | Yes |
| Control-Sample1_Beta | chr1:3042085-3042459 | PRDM16;PRDM16 | S_Shelf | Body;Body | Yes |
| Control-Sample1_Beta | chr1:3070337-3070786 | PRDM16;PRDM16 | N_Shore | Body;Body | Yes |
| Control-Sample1_Beta | chr1:3115309-3115545 | PRDM16;PRDM16 | S_Shelf | Body;Body | Yes |
| Control-Sample1_Beta | chr1:3129579-3129969 | PRDM16;PRDM16 | | Body;Body | Yes |
| Control-Sample1_Beta | chr1:3142925-3144536 | PRDM16;PRDM16 | | Body;Body | Yes |
| Control-Sample1_Beta | chr1:3191660-3191815 | PRDM16;PRDM16 | N_Shelf | Body;Body | Yes |
| Control-Sample1_Beta | chr1:3269252-3269478 | PRDM16;PRDM16 | | Body;Body | Yes |
| Control-Sample1_Beta | chr1:3277525-3277623 | PRDM16;PRDM16 | S_Shelf | Body;Body | Yes |
| Control-Sample10_Beta | chr1:3028813-3028910 | PRDM16;PRDM16 | S_Shore | Body;Body | Yes |
| Control-Sample10_Beta | chr1:3089287-3089891 | PRDM16;PRDM16 | | Body;Body | Yes |
| Control-Sample10_Beta | chr1:3128175-3129969 | PRDM16;PRDM16 | | Body;Body | Yes |
| Control-Sample10_Beta | chr1:3133763-3134162 | PRDM16;PRDM16 | | Body;Body | Yes |
| Control-Sample10_Beta | chr1:3142925-3144113 | PRDM16;PRDM16 | | Body;Body | Yes |
| Control-Sample10_Beta | chr1:3144536-3144710 | PRDM16;PRDM16 | N_Shelf | Body;Body | Yes |
| Control-Sample10_Beta | chr1:3276413-3276687 | PRDM16;PRDM16 | S_Shore | Body;Body | Yes |
| Control-Sample11_Beta | chr1:3089597-3089891 | PRDM16;PRDM16 | | Body;Body | Yes |
| Control-Sample11_Beta | chr1:3128606-3129969 | PRDM16;PRDM16 | | Body;Body | Yes |
| Control-Sample11_Beta | chr1:3142925-3143965 | PRDM16;PRDM16 | | Body;Body | Yes |
| Control-Sample12_Beta | chr1:3133763-3133857 | PRDM16;PRDM16 | | Body;Body | Yes |
| Control-Sample13_Beta | chr1:3128606-3129115 | PRDM16;PRDM16 | | Body;Body | Yes |
| Control-Sample13_Beta | chr1:3191219-3191876 | PRDM16;PRDM16 | | Body;Body | Yes |
| Control-Sample14_Beta | chr1:3191660-3191876 | PRDM16;PRDM16 | N_Shelf | Body;Body | Yes |

| | | | | | |
|---|---|---|---|---|---|
| Control-Sample15_Beta | chr1:3276507-3276687 | PRDM16;PRDM16 | S_Shore | Body;Body | Yes |
| Control-Sample17_Beta | chr1:3276413-3276753 | PRDM16;PRDM16 | S_Shore | Body;Body | Yes |
| Control-Sample19_Beta | chr1:3115085-3115545 | PRDM16;PRDM16 | S_Shelf | Body;Body | Yes |
| Control-Sample19_Beta | chr1:3143808-3144113 | PRDM16;PRDM16 | | Body;Body | Yes |
| Control-Sample19_Beta | chr1:3331987-3332125 | PRDM16;PRDM16 | Island | Body;Body | Yes |
| Control-Sample21_Beta | chr1:3078839-3079212 | PRDM16;PRDM16 | N_Shelf | Body;Body | Yes |
| Control-Sample21_Beta | chr1:3143334-3144113 | PRDM16;PRDM16 | | Body;Body | Yes |
| Control-Sample22_Beta | chr1:2996949-2997409 | PRDM16;PRDM16 | N_Shore | Body;Body | Yes |
| Control-Sample22_Beta | chr1:3078524-3079212 | PRDM16;PRDM16 | N_Shelf | Body;Body | Yes |
| Control-Sample22_Beta | chr1:3143334-3143965 | PRDM16;PRDM16 | | Body;Body | Yes |
| Control-Sample24_Beta | chr1:2995301-2995604 | PRDM16;PRDM16 | N_Shore | Body;Body | Yes |
| Control-Sample26_Beta | chr1:3191219-3191815 | PRDM16;PRDM16 | | Body;Body | Yes |
| Control-Sample29_Beta | chr1:3089287-3089762 | PRDM16;PRDM16 | | Body;Body | Yes |
| Control-Sample3_Beta | chr1:3142925-3144710 | PRDM16;PRDM16 | | Body;Body | Yes |
| Control-Sample4_Beta | chr1:3129633-3129969 | PRDM16;PRDM16 | | Body;Body | Yes |
| Control-Sample40_Beta | chr1:3142925-3144209 | PRDM16;PRDM16 | | Body;Body | Yes |
| Control-Sample54_Beta | chr1:2995301-2996020 | PRDM16;PRDM16 | N_Shore | Body;Body | Yes |
| Control-Sample55_Beta | chr1:3191660-3191883 | PRDM16;PRDM16 | N_Shelf | Body;Body | Yes |
| Control-Sample57_Beta | chr1:3058521-3059051 | PRDM16;PRDM16 | N_Shore | Body;Body | Yes |
| Control-Sample65_Beta | chr1:3191219-3191883 | PRDM16;PRDM16 | | Body;Body | Yes |
| Control-Sample1_Beta | chr1:14113172-14113230 | PRDM2;PRDM2;PRDM2;PRDM2 | | Body;3'UTR;Body;3'UTR | No |
| Control-Sample525_Be | chr16.0:90143815-90144006 | PRDM7 | Island | TSS1500 | No |
| Control-Sample137_Be | chr5:23507573-23507656 | PRDM9 | | TSS200 | No |
| Control-Sample52_Beta | chr11:64085030-64085103 | PRDX5;PRDX5;TRMT112;TRMT112;PRDX5 | Island | TSS1500;TSS1500;5'UTR;1stExon;TSS1500 | No |
| Control-Sample108_Be | chr1:203444518-203444945 | PRELP;PRELP | | TSS1500;TSS1500 | No |
| Control-Sample105_Be | chr20:62200603-62200854 | PRIC285;PRIC285 | N_Shore | Body;TSS1500 | No |
| Control-Sample108_Be | chr3:64211328-64211659 | PRICKLE2 | | TSS200 | No |
| Control-Sample1_Beta | chr7:640409-640820 | PRKAR1B;PRKAR1B;PRKAR1B;PRKAR1B;PRKAR1B;PRKAR1 | N_Shelf | Body;Body;Body;Body;Body;Body | Yes |
| Control-Sample11_Beta | chr7:632926-633202 | PRKAR1B;PRKAR1B;PRKAR1B;PRKAR1B;PRKAR1B;PRKAR1 | S_Shore | Body;Body;Body;Body;Body;Body | Yes |
| Control-Sample11_Beta | chr7:640820-641337 | PRKAR1B;PRKAR1B;PRKAR1B;PRKAR1B;PRKAR1B;PRKAR1 | N_Shelf | Body;Body;Body;Body;Body;Body | Yes |
| Control-Sample19_Beta | chr7:612458-613222 | PRKAR1B;PRKAR1B;PRKAR1B;PRKAR1B;PRKAR1B;PRKAR1 | N_Shore | Body;Body;Body;Body;Body;Body | Yes |
| Control-Sample236_Be | chr7:676060-676291 | PRKAR1B;PRKAR1B;PRKAR1B;PRKAR1B;PRKAR1B;PRKAR1 | Island | Body;Body;Body;Body;Body;Body | Yes |
| Control-Sample237_Be | chr7:675570-676291 | PRKAR1B;PRKAR1B;PRKAR1B;PRKAR1B;PRKAR1B;PRKAR1 | Island | Body;Body;Body;Body;Body;Body | Yes |
| Control-Sample3_Beta | chr7:640338-641337 | PRKAR1B;PRKAR1B;PRKAR1B;PRKAR1B;PRKAR1B;PRKAR1 | N_Shelf | Body;Body;Body;Body;Body;Body | Yes |
| Control-Sample4_Beta | chr7:630581-630914 | PRKAR1B;PRKAR1B;PRKAR1B;PRKAR1B;PRKAR1B;PRKAR1 | N_Shore | Body;Body;Body;Body;Body;Body | Yes |
| Control-Sample6_Beta | chr7:640381-641028 | PRKAR1B;PRKAR1B;PRKAR1B;PRKAR1B;PRKAR1B;PRKAR1 | N_Shelf | Body;Body;Body;Body;Body;Body | Yes |
| Control-Sample170_Be | chr17:64672129-64672341 | PRKCA | | Body | No |
| Control-Sample7_Beta | chr11:6341850-6341888 | PRKCDBP | Island | TSS200 | No |
| Control-Sample1_Beta | chr1:1982791-1983033 | PRKCZ | Island | Body | Yes |
| Control-Sample13_Beta | chr1:1982688-1983003 | PRKCZ | Island | Body | Yes |
| Control-Sample10_Beta | chr1:2058417-2058941 | PRKCZ;PRKCZ;PRKCZ | | 5'UTR;Body;5'UTR | Yes |
| Control-Sample11_Beta | chr1:2058417-2058790 | PRKCZ;PRKCZ;PRKCZ | | 5'UTR;Body;5'UTR | Yes |
| Control-Sample19_Beta | chr1:2058790-2059086 | PRKCZ;PRKCZ;PRKCZ | | 5'UTR;Body;5'UTR | Yes |
| Control-Sample20_Beta | chr1:2106222-2106400 | PRKCZ;PRKCZ;PRKCZ | | Body;Body;Body | Yes |
| Control-Sample29_Beta | chr1:2063799-2064046 | PRKCZ;PRKCZ;PRKCZ | Island | 5'UTR;Body;5'UTR | Yes |
| Control-Sample444_Be | chr1:2063799-2063966 | PRKCZ;PRKCZ;PRKCZ | Island | 5'UTR;Body;5'UTR | Yes |
| Control-Sample464_Be | chr1:2099745-2099891 | PRKCZ;PRKCZ;PRKCZ | | Body;Body;Body | Yes |
| Control-Sample464_Be | chr1:2100365-2100984 | PRKCZ;PRKCZ;PRKCZ | | Body;Body;Body | Yes |
| Control-Sample58_Beta | chr1:2066006-2066430 | PRKCZ;PRKCZ;PRKCZ | N_Shore | 5'UTR;Body;5'UTR | Yes |
| Control-Sample3_Beta | chr16:11367459-11367509 | PRM3 | N_Shelf | TSS200 | No |
| Control-Sample20_Beta | chr17:27039120-27039523 | PROCA1 | S_Shore | TSS1500 | Yes |
| Control-Sample1_Beta | chr17:27038861-27039032 | PROCA1;PROCA1 | Island | 5'UTR;1stExon | Yes |
| Control-Sample18_Beta | chr17:27038861-27039523 | PROCA1;PROCA1 | Island | 5'UTR;1stExon | Yes |
| Control-Sample29_Beta | chr17:27038861-27039132 | PROCA1;PROCA1 | Island | 5'UTR;1stExon | Yes |
| Control-Sample30_Beta | chr17:27038686-27039032 | PROCA1;PROCA1 | Island | 5'UTR;1stExon | Yes |
| Control-Sample52_Beta | chr17:27038861-27039120 | PROCA1;PROCA1 | Island | 5'UTR;1stExon | Yes |
| Control-Sample1_Beta | chr4:16085702-16085814 | PROM1;PROM1 | Island | TSS200;TSS200 | Yes |
| Control-Sample10_Beta | chr4:16085702-16086085 | PROM1;PROM1 | Island | TSS200;TSS200 | Yes |
| Control-Sample11_Beta | chr4:16085702-16086023 | PROM1;PROM1 | Island | TSS200;TSS200 | Yes |
| Control-Sample19_Beta | chr4:16085702-16086051 | PROM1;PROM1 | Island | TSS200;TSS200 | Yes |
| Control-Sample41_Beta | chr4:16085716-16086023 | PROM1;PROM1 | Island | TSS200;TSS200 | Yes |
| Control-Sample5_Beta | chr4:16085716-16086085 | PROM1;PROM1 | Island | TSS200;TSS200 | Yes |
| Control-Sample51_Beta | chr4:16085814-16086085 | PROM1;PROM1 | S_Shore | TSS200;TSS1500 | Yes |
| Control-Sample1_Beta | chr1:214170114-214170241 | PROX1 | | Body | No |
| Control-Sample463_Be | chr7:29602763-29603327 | PRR15 | N_Shore | TSS1500 | No |
| Control-Sample1_Beta | chr17:46035272-46035343 | PRR15L | | TSS200 | Yes |
| Control-Sample57_Beta | chr17:46035108-46035343 | PRR15L;PRR15L | | 1stExon;5'UTR | Yes |
| Control-Sample1_Beta | chr22:45125218-45125480 | PRR5;PRR5;PRR5;PRR5-ARHGAP8;PRR5;PRR5 | Island | Body;Body;Body;Body;Body;Body | No |
| Control-Sample19_Beta | chr11:36476232-36476319 | PRR5L;PRR5L;PRR5L;PRR5L | | Body;Body;Body;Body | No |
| Control-Sample1_Beta | chr5:126852955-126853024 | PRRC1 | N_Shore | TSS1500 | No |
| Control-Sample1_Beta | chr6:32116317-32116548 | PRRT1 | N_Shore | 3'UTR | Yes |
| Control-Sample1_Beta | chr6:32116653-32117379 | PRRT1 | Island | 3'UTR | Yes |
| Control-Sample11_Beta | chr6:32116653-32117281 | PRRT1 | Island | 3'UTR | Yes |
| Control-Sample12_Beta | chr6:32118312-32118399 | PRRT1 | Island | Body | Yes |
| Control-Sample13_Beta | chr6:32118298-32118375 | PRRT1 | Island | Body | Yes |
| Control-Sample14_Beta | chr6:32116317-32117379 | PRRT1 | N_Shore | 3'UTR | Yes |
| Control-Sample16_Beta | chr6:32116317-32117281 | PRRT1 | N_Shore | 3'UTR | Yes |
| Control-Sample19_Beta | chr6:32116875-32117379 | PRRT1 | Island | 3'UTR | Yes |
| Control-Sample2_Beta | chr6:32118298-32118399 | PRRT1 | Island | Body | Yes |
| Control-Sample21_Beta | chr6:32118312-32118375 | PRRT1 | Island | Body | Yes |
| Control-Sample24_Beta | chr6:32116207-32117379 | PRRT1 | N_Shore | 3'UTR | Yes |
| Control-Sample33_Beta | chr6:32118315-32118399 | PRRT1 | Island | Body | Yes |
| Control-Sample38_Beta | chr6:32117049-32117379 | PRRT1 | Island | Body | Yes |
| Control-Sample66_Beta | chr6:32116853-32117281 | PRRT1 | Island | 3'UTR | Yes |
| Control-Sample67_Beta | chr6:32116207-32116317 | PRRT1 | N_Shore | 3'UTR | Yes |
| Control-Sample67_Beta | chr6:32116893-32116933 | PRRT1 | Island | 3'UTR | Yes |
| Control-Sample68_Beta | chr6:32116893-32116918 | PRRT1 | Island | 3'UTR | Yes |
| Control-Sample68_Beta | chr6:32117079-32117281 | PRRT1 | Island | Body | Yes |
| Control-Sample14_Beta | chr1:170635960-170636363 | PRRX1;PRRX1 | S_Shore | Body;Body | Yes |
| Control-Sample14_Beta | chr1:170638807-170639289 | PRRX1;PRRX1 | | Body;Body | Yes |
| Control-Sample107_Be | chr3:46759438-46759698 | PRSS50 | Island | TSS200 | No |

| | | | | | |
|---|---|---|---|---|---|
| Control-Sample1_Beta | chr16:31146682-31147054 | PRSS8 | | Body | Yes |
| Control-Sample15_Beta | chr16:31147017-31147054 | PRSS8;PRSS8 | | 1stExon;5'UTR | Yes |
| Control-Sample2_Beta | chr16:31146999-31147054 | PRSS8;PRSS8 | | 1stExon;5'UTR | Yes |
| Control-Sample57_Beta | chr19:40904691-40904974 | PRX;PRX | Island | Body;Body | Yes |
| Control-Sample1_Beta | chr19:40919245-40919485 | PRX;PRX;PRX;PRX | | 1stExon;5'UTR;1stExon;5'UTR | Yes |
| Control-Sample4_Beta | chr19:40919245-40919760 | PRX;PRX;PRX;PRX | | 1stExon;5'UTR;1stExon;5'UTR | Yes |
| Control-Sample108_Be | chr8:18541556-18541988 | PSD3;PSD3 | | Body;Body | Yes |
| Control-Sample108_Be | chr8:18666531-18666561 | PSD3;PSD3 | | Body;TSS200 | Yes |
| Control-Sample171_Be | chr8:18666313-18666561 | PSD3;PSD3;PSD3 | | Body;1stExon;5'UTR | Yes |
| Control-Sample194_Be | chr8:18666313-18666537 | PSD3;PSD3;PSD3 | | Body;1stExon;5'UTR | Yes |
| Control-Sample3_Beta | chr8:18666313-18666598 | PSD3;PSD3;PSD3 | | Body;1stExon;5'UTR | Yes |
| Control-Sample1_Beta | chr8:87082023-87082065 | PSKH2 | Island | TSS200 | Yes |
| Control-Sample17_Beta | chr8:87082008-87082065 | PSKH2 | Island | TSS200 | Yes |
| Control-Sample23_Beta | chr8:87082008-87082045 | PSKH2 | Island | TSS200 | Yes |
| Control-Sample6_Beta | chr8:87082014-87082065 | PSKH2 | Island | TSS200 | Yes |
| Control-Sample3_Beta | chr6:32822352-32822857 | PSMB9;PSMB9;TAP1 | Island | Body;Body;TSS1500 | No |
| Control-Sample551_Be | chr21.0:40555470-40555479 | PSMG1;PSMG1 | Island | TSS200;TSS200 | Yes |
| Control-Sample620_Be | chr21:40555470-40555479 | PSMG1;PSMG1 | Island | TSS200;TSS200 | Yes |
| Control-Sample466_Be | chr10:27703329-27703362 | PTCHD3 | Island | TSS200 | No |
| Control-Sample1_Beta | chr8:97339939-97340270 | PTDSS1 | | Body | Yes |
| Control-Sample331_Be | chr8:97340188-97340270 | PTDSS1 | | Body | Yes |
| Control-Sample42_Beta | chr8:97339939-97340225 | PTDSS1 | | Body | Yes |
| Control-Sample10_Beta | chr3:46940153-46940347 | PTH1R | Island | Body | Yes |
| Control-Sample17_Beta | chr3:46940153-46940439 | PTH1R | Island | Body | Yes |
| Control-Sample216_Be | chr8:27182889-27183285 | PTK2B;PTK2B;PTK2B;PTK2B | N_Shore | 5'UTR;TSS200;5'UTR;TSS200 | Yes |
| Control-Sample223_Be | chr8:27182965-27183285 | PTK2B;PTK2B;PTK2B;PTK2B | N_Shore | 5'UTR;TSS200;5'UTR;TSS200 | Yes |
| Control-Sample270_Be | chr8:27182909-27183285 | PTK2B;PTK2B;PTK2B;PTK2B | N_Shore | 5'UTR;TSS200;5'UTR;TSS200 | Yes |
| Control-Sample644_Be | chr8:27182930-27183097 | PTK2B;PTK2B;PTK2B;PTK2B | N_Shore | 5'UTR;TSS200;5'UTR;TSS200 | Yes |
| Control-Sample1_Beta | chr19:50361793-50361910 | PTOV1 | Island | Body | No |
| Control-Sample1_Beta | chr1:44016014-44016049 | PTPRF;PTPRF | | Body;Body | No |
| Control-Sample1_Beta | chr7:157367142-157367159 | PTPRN2;PTPRN2;MIR153-2;PTPRN2 | N_Shore | Body;Body;TSS200;Body | Yes |
| Control-Sample19_Beta | chr7:157367142-157369121 | PTPRN2;PTPRN2;MIR153-2;PTPRN2 | N_Shore | Body;Body;TSS200;Body | Yes |
| Control-Sample20_Beta | chr7:157367142-157367537 | PTPRN2;PTPRN2;MIR153-2;PTPRN2 | N_Shore | Body;Body;TSS200;Body | Yes |
| Control-Sample55_Beta | chr7:157367142-157367462 | PTPRN2;PTPRN2;MIR153-2;PTPRN2 | N_Shore | Body;Body;TSS200;Body | Yes |
| Control-Sample92_Beta | chr7:157367150-157367462 | PTPRN2;PTPRN2;MIR153-2;PTPRN2 | N_Shore | Body;Body;TSS200;Body | Yes |
| Control-Sample1_Beta | chr7:157333405-157333584 | PTPRN2;PTPRN2;PTPRN2 | | 3'UTR;3'UTR;3'UTR | Yes |
| Control-Sample1_Beta | chr7:157389643-157389729 | PTPRN2;PTPRN2;PTPRN2 | S_Shelf | Body;Body;Body | Yes |
| Control-Sample1_Beta | chr7:157394337-157394970 | PTPRN2;PTPRN2;PTPRN2 | | Body;Body;Body | Yes |
| Control-Sample1_Beta | chr7:157475692-157476575 | PTPRN2;PTPRN2;PTPRN2 | N_Shore | Body;Body;Body | Yes |
| Control-Sample1_Beta | chr7:157561786-157561925 | PTPRN2;PTPRN2;PTPRN2 | S_Shore | Body;Body;Body | Yes |
| Control-Sample1_Beta | chr7:157633771-157633985 | PTPRN2;PTPRN2;PTPRN2 | S_Shelf | Body;Body;Body | Yes |
| Control-Sample1_Beta | chr7:157643722-157643770 | PTPRN2;PTPRN2;PTPRN2 | | Body;Body;Body | Yes |
| Control-Sample1_Beta | chr7:157809325-157809596 | PTPRN2;PTPRN2;PTPRN2 | | Body;Body;Body | Yes |
| Control-Sample1_Beta | chr7:157890277-157890659 | PTPRN2;PTPRN2;PTPRN2 | N_Shore | Body;Body;Body | Yes |
| Control-Sample1_Beta | chr7:157982364-157982589 | PTPRN2;PTPRN2;PTPRN2 | S_Shore | Body;Body;Body | Yes |
| Control-Sample1_Beta | chr7:158111178-158111329 | PTPRN2;PTPRN2;PTPRN2 | S_Shore | Body;Body;Body | Yes |
| Control-Sample1_Beta | chr7:158280480-158280939 | PTPRN2;PTPRN2;PTPRN2 | N_Shore | Body;Body;Body | Yes |
| Control-Sample10_Beta | chr7:157366979-157367159 | PTPRN2;PTPRN2;PTPRN2 | N_Shelf | Body;Body;Body | Yes |
| Control-Sample10_Beta | chr7:157624788-157624958 | PTPRN2;PTPRN2;PTPRN2 | | Body;Body;Body | Yes |
| Control-Sample10_Beta | chr7:158280480-158281138 | PTPRN2;PTPRN2;PTPRN2 | N_Shore | Body;Body;Body | Yes |
| Control-Sample10_Beta | chr7:158336016-158336049 | PTPRN2;PTPRN2;PTPRN2 | | Body;Body;Body | Yes |
| Control-Sample11_Beta | chr7:158280410-158281138 | PTPRN2;PTPRN2;PTPRN2 | N_Shore | Body;Body;Body | Yes |
| Control-Sample12_Beta | chr7:157366979-157367462 | PTPRN2;PTPRN2;PTPRN2 | N_Shelf | Body;Body;Body | Yes |
| Control-Sample12_Beta | chr7:157809325-157809385 | PTPRN2;PTPRN2;PTPRN2 | | Body;Body;Body | Yes |
| Control-Sample13_Beta | chr7:157394582-157394970 | PTPRN2;PTPRN2;PTPRN2 | | Body;Body;Body | Yes |
| Control-Sample14_Beta | chr7:157561593-157561925 | PTPRN2;PTPRN2;PTPRN2 | S_Shore | Body;Body;Body | Yes |
| Control-Sample151_Be | chr7:157394582-157394661 | PTPRN2;PTPRN2;PTPRN2 | | Body;Body;Body | Yes |
| Control-Sample170_Be | chr7:157408294-157408632 | PTPRN2;PTPRN2;PTPRN2 | N_Shore | Body;Body;Body | Yes |
| Control-Sample18_Beta | chr7:158136312-158136485 | PTPRN2;PTPRN2;PTPRN2 | | Body;Body;Body | Yes |
| Control-Sample19_Beta | chr7:157333405-157333949 | PTPRN2;PTPRN2;PTPRN2 | | 3'UTR;3'UTR;3'UTR | Yes |
| Control-Sample19_Beta | chr7:157394633-157394970 | PTPRN2;PTPRN2;PTPRN2 | | Body;Body;Body | Yes |
| Control-Sample19_Beta | chr7:157461081-157461368 | PTPRN2;PTPRN2;PTPRN2 | | Body;Body;Body | Yes |
| Control-Sample19_Beta | chr7:157463300-157463692 | PTPRN2;PTPRN2;PTPRN2 | | Body;Body;Body | Yes |
| Control-Sample19_Beta | chr7:157622032-157622221 | PTPRN2;PTPRN2;PTPRN2 | S_Shelf | Body;Body;Body | Yes |
| Control-Sample19_Beta | chr7:157691177-157691376 | PTPRN2;PTPRN2;PTPRN2 | S_Shore | Body;Body;Body | Yes |
| Control-Sample19_Beta | chr7:157700935-157701201 | PTPRN2;PTPRN2;PTPRN2 | N_Shelf | Body;Body;Body | Yes |
| Control-Sample19_Beta | chr7:157791862-157792100 | PTPRN2;PTPRN2;PTPRN2 | | Body;Body;Body | Yes |
| Control-Sample19_Beta | chr7:158342105-158342600 | PTPRN2;PTPRN2;PTPRN2 | N_Shore | Body;Body;Body | Yes |
| Control-Sample2_Beta | chr7:158280771-158281138 | PTPRN2;PTPRN2;PTPRN2 | N_Shore | Body;Body;Body | Yes |
| Control-Sample20_Beta | chr7:157333405-157333752 | PTPRN2;PTPRN2;PTPRN2 | | 3'UTR;3'UTR;3'UTR | Yes |
| Control-Sample20_Beta | chr7:157368881-157368976 | PTPRN2;PTPRN2;PTPRN2 | N_Shore | Body;Body;Body | Yes |
| Control-Sample20_Beta | chr7:157461081-157462004 | PTPRN2;PTPRN2;PTPRN2 | | Body;Body;Body | Yes |
| Control-Sample20_Beta | chr7:157463300-157464230 | PTPRN2;PTPRN2;PTPRN2 | | Body;Body;Body | Yes |
| Control-Sample20_Beta | chr7:157561010-157561134 | PTPRN2;PTPRN2;PTPRN2 | Island | Body;Body;Body | Yes |
| Control-Sample20_Beta | chr7:157700872-157701024 | PTPRN2;PTPRN2;PTPRN2 | N_Shelf | Body;Body;Body | Yes |
| Control-Sample20_Beta | chr7:157818696-157818833 | PTPRN2;PTPRN2;PTPRN2 | | Body;Body;Body | Yes |
| Control-Sample20_Beta | chr7:158342283-158342600 | PTPRN2;PTPRN2;PTPRN2 | N_Shore | Body;Body;Body | Yes |
| Control-Sample21_Beta | chr7:157394337-157394633 | PTPRN2;PTPRN2;PTPRN2 | | Body;Body;Body | Yes |
| Control-Sample21_Beta | chr7:157468321-157468443 | PTPRN2;PTPRN2;PTPRN2 | | Body;Body;Body | Yes |
| Control-Sample21_Beta | chr7:158059704-158059944 | PTPRN2;PTPRN2;PTPRN2 | N_Shore | Body;Body;Body | Yes |
| Control-Sample230_Be | chr7:158032120-158032496 | PTPRN2;PTPRN2;PTPRN2 | | Body;Body;Body | Yes |
| Control-Sample230_Be | chr7:158110068-158110685 | PTPRN2;PTPRN2;PTPRN2 | N_Shore | Body;Body;Body | Yes |
| Control-Sample24_Beta | chr7:157809385-157809596 | PTPRN2;PTPRN2;PTPRN2 | | Body;Body;Body | Yes |
| Control-Sample29_Beta | chr7:158114558-158114659 | PTPRN2;PTPRN2;PTPRN2 | S_Shelf | Body;Body;Body | Yes |
| Control-Sample29_Beta | chr7:158157967-158158218 | PTPRN2;PTPRN2;PTPRN2 | | Body;Body;Body | Yes |
| Control-Sample3_Beta | chr7:157462914-157463355 | PTPRN2;PTPRN2;PTPRN2 | | Body;Body;Body | Yes |
| Control-Sample3_Beta | chr7:157811843-157811918 | PTPRN2;PTPRN2;PTPRN2 | | Body;Body;Body | Yes |
| Control-Sample30_Beta | chr7:157561684-157561848 | PTPRN2;PTPRN2;PTPRN2 | S_Shore | Body;Body;Body | Yes |
| Control-Sample32_Beta | chr7:157561684-157561925 | PTPRN2;PTPRN2;PTPRN2 | S_Shore | Body;Body;Body | Yes |
| Control-Sample4_Beta | chr7:157366933-157367462 | PTPRN2;PTPRN2;PTPRN2 | N_Shelf | Body;Body;Body | Yes |
| Control-Sample41_Beta | chr7:158280410-158280939 | PTPRN2;PTPRN2;PTPRN2 | N_Shore | Body;Body;Body | Yes |

| | | | | | |
|---|---|---|---|---|---|
| Control-Sample45_Beta | chr7:157561593-157561848 | PTPRN2;PTPRN2;PTPRN2 | S_Shore | Body;Body;Body | Yes |
| Control-Sample53_Beta | chr7:157643722-157644160 | PTPRN2;PTPRN2;PTPRN2 | | Body;Body;Body | Yes |
| Control-Sample58_Beta | chr7:157476329-157476575 | PTPRN2;PTPRN2;PTPRN2 | N_Shore | Body;Body;Body | Yes |
| Control-Sample585_Be | chr7.0:157394582-157394661 | PTPRN2;PTPRN2;PTPRN2 | | Body;Body;Body | Yes |
| Control-Sample65_Beta | chr7:157405965-157406737 | PTPRN2;PTPRN2;PTPRN2 | Island | Body;Body;Body | Yes |
| Control-Sample19_Beta | chr19:5250376-5250655 | PTPRS;PTPRS;PTPRS;PTPRS | N_Shore | Body;Body;Body;Body | Yes |
| Control-Sample58_Beta | chr19:5250495-5251120 | PTPRS;PTPRS;PTPRS;PTPRS | Island | Body;Body;Body;Body | Yes |
| Control-Sample1_Beta | chr1:29565099-29565306 | PTPRU;PTPRU;PTPRU | Island | Body;Body;Body | Yes |
| Control-Sample8_Beta | chr1:29587117-29587357 | PTPRU;PTPRU;PTPRU | Island | Body;Body;Body | Yes |
| Control-Sample65_Beta | chr8:129020343-129020489 | PVT1;MIR1206 | | Body;TSS1500 | No |
| Control-Sample18_Beta | chr12:108079556-108079615 | PWP1 | Island | TSS200 | Yes |
| Control-Sample49_Beta | chr12:108079503-108079615 | PWP1 | Island | TSS200 | Yes |
| Control-Sample1_Beta | chr2:1690592-1690740 | PXDN | | Body | No |
| Control-Sample1_Beta | chr8:52722023-52722185 | PXDNL | | TSS200 | Yes |
| Control-Sample12_Beta | chr8:52722023-52722146 | PXDNL | | TSS200 | Yes |
| Control-Sample22_Beta | chr8:52722140-52722185 | PXDNL | | TSS200 | Yes |
| Control-Sample1_Beta | chr11:64527298-64527417 | PYGM;PYGM | | 1stExon;1stExon | No |
| Control-Sample1_Beta | chr15:55881599-55881706 | PYGO1 | S_Shore | TSS1500 | No |
| Control-Sample2_Beta | chr10:100175236-100175560 | PYROXD2 | | TSS200 | No |
| Control-Sample1_Beta | chr17:42083195-42083306 | PYY;NAGS | Island | TSS1500;Body | No |
| Control-Sample1_Beta | chr16:29706151-29706381 | QPRT | N_Shore | Body | Yes |
| Control-Sample24_Beta | chr16:29706275-29706381 | QPRT | Island | Body | Yes |
| Control-Sample52_Beta | chr16:474430-474528 | RAB11FIP3 | N_Shore | TSS1500 | No |
| Control-Sample14_Beta | chr7:140103756-140103783 | RAB19 | | TSS200 | Yes |
| Control-Sample22_Beta | chr7:140103737-140103783 | RAB19 | | TSS200 | Yes |
| Control-Sample3_Beta | chr7:140103671-140103737 | RAB19 | | TSS200 | Yes |
| Control-Sample5_Beta | chr7:140103737-140103764 | RAB19 | | TSS200 | Yes |
| Control-Sample1_Beta | chr1:156030809-156031182 | RAB25 | | TSS200 | No |
| Control-Sample25_Beta | chr17:27045164-27045302 | RAB34;RAB34;RAB34;RAB34;RAB34;RAB34;RAB34 | N_Shore | Body;TSS1500;1stExon;TSS1500;1stExon;Body;1stEx | Yes |
| Control-Sample27_Beta | chr17:27045113-27045176 | RAB34;RAB34;RAB34;RAB34;RAB34;RAB34;RAB34 | N_Shore | Body;Body;TSS1500;Body;TSS1500;Body;Body | Yes |
| Control-Sample1_Beta | chr17:72732432-72732823 | RAB37;RAB37;RAB37;RAB37 | N_Shore | TSS1500;TSS1500;Body;TSS1500 | No |
| Control-Sample1_Beta | chr11:87908785-87908938 | RAB38 | S_Shore | TSS200 | Yes |
| Control-Sample10_Beta | chr11:87908783-87908817 | RAB38 | S_Shore | TSS200 | Yes |
| Control-Sample12_Beta | chr11:87908785-87908817 | RAB38 | S_Shore | TSS200 | Yes |
| Control-Sample21_Beta | chr11:87908783-87908805 | RAB38 | S_Shore | TSS200 | Yes |
| Control-Sample63_Beta | chr11:87908783-87908938 | RAB38 | S_Shore | TSS200 | Yes |
| Control-Sample6_Beta | chr16:671905-672468 | RAB40C | N_Shore | Body | No |
| Control-Sample1_Beta | chr1:174843754-174844490 | RABGAP1L;RABGAP1L | | TSS1500;Body | Yes |
| Control-Sample11_Beta | chr1:174843909-174844490 | RABGAP1L;RABGAP1L | | TSS1500;Body | Yes |
| Control-Sample19_Beta | chr1:174843971-174844490 | RABGAP1L;RABGAP1L | | TSS1500;Body | Yes |
| Control-Sample220_Be | chr20:1206693-1206703 | RAD21L1 | Island | TSS200 | No |
| Control-Sample1_Beta | chr14:68830704-68831098 | RAD51L1;RAD51L1;RAD51L1 | | Body;Body;Body | No |
| Control-Sample1_Beta | chr17:17713061-17713364 | RAI1 | Island | Body | Yes |
| Control-Sample10_Beta | chr17:17603749-17604184 | RAI1 | Island | 5'UTR | Yes |
| Control-Sample22_Beta | chr17:17603837-17604184 | RAI1 | Island | 5'UTR | Yes |
| Control-Sample171_Be | chr2:121036440-121036760 | RALB | | Body | No |
| Control-Sample69_Beta | chr5:170288742-170288766 | RANBP17 | N_Shore | TSS1500 | No |
| Control-Sample108_Be | chr5:36302094-36302704 | RANBP3L;RANBP3L | | TSS200;TSS200 | No |
| Control-Sample1_Beta | chr1:21934312-21934726 | RAP1GAP;RAP1GAP;RAP1GAP | S_Shelf | Body;Body;Body | Yes |
| Control-Sample194_Be | chr1:21966310-21966500 | RAP1GAP;RAP1GAP;RAP1GAP | | Body;5'UTR;5'UTR | Yes |
| Control-Sample1_Beta | chr17:2898745-2898878 | RAP1GAP2;RAP1GAP2 | | Body;Body | Yes |
| Control-Sample10_Beta | chr17:2699371-2699542 | RAP1GAP2;RAP1GAP2 | | TSS1500;TSS1500 | Yes |
| Control-Sample11_Beta | chr17:2886250-2886453 | RAP1GAP2;RAP1GAP2 | | Body;Body | Yes |
| Control-Sample19_Beta | chr17:2907678-2908068 | RAP1GAP2;RAP1GAP2 | | Body;Body | Yes |
| Control-Sample20_Beta | chr17:2907678-2908023 | RAP1GAP2;RAP1GAP2 | | Body;Body | Yes |
| Control-Sample194_Be | chr17:38334166-38334186 | RAPGEFL1 | Island | TSS200 | Yes |
| Control-Sample233_Be | chr17:38334170-38334186 | RAPGEFL1 | Island | TSS200 | Yes |
| Control-Sample1_Beta | chr17:38473119-38473222 | RARA;RARA;RARA | Island | TSS1500;5'UTR;TSS1500 | No |
| Control-Sample65_Beta | chr12:53612551-53612734 | RARG;RARG | N_Shore | Body;Body | No |
| Control-Sample1_Beta | chr11:63304098-63304614 | RARRES3 | | TSS200 | No |
| Control-Sample10_Beta | chr13:114810631-114810794 | RASA3 | N_Shelf | Body | Yes |
| Control-Sample17_Beta | chr13:114829170-114829833 | RASA3 | | Body | Yes |
| Control-Sample194_Be | chr13:114786825-114786887 | RASA3 | S_Shore | Body | Yes |
| Control-Sample194_Be | chr13:114802548-114802867 | RASA3 | N_Shore | Body | Yes |
| Control-Sample223_Be | chr13:114802548-114803271 | RASA3 | N_Shore | Body | Yes |
| Control-Sample496_Be | chr13:114786825-114786904 | RASA3 | S_Shore | Body | Yes |
| Control-Sample572_Be | chr13.0:114810631-114810794 | RASA3 | N_Shelf | Body | Yes |
| Control-Sample645_Be | chr13:114800549-114800841 | RASA3 | N_Shelf | Body | Yes |
| Control-Sample1_Beta | chr12:113574204-113574368 | RASAL1 | S_Shore | TSS200 | No |
| Control-Sample12_Beta | chr1:178310534-178310650 | RASAL2;RASAL2 | | Body;TSS200 | No |
| Control-Sample10_Beta | chr19:15568360-15568574 | RASAL3 | Island | Body | No |
| Control-Sample1_Beta | chr5:179634671-179634789 | RASGEF1C | Island | 5'UTR | No |
| Control-Sample1_Beta | chr11:64509084-64509592 | RASGRP2;RASGRP2;RASGRP2 | N_Shore | Body;Body;Body | No |
| Control-Sample1_Beta | chr3:50378611-50379032 | RASSF1;RASSF1;ZMYND10 | S_Shore | TSS1500;TSS1500;3'UTR | No |
| Control-Sample12_Beta | chr20:4795381-4795621 | RASSF2;RASSF2 | | 5'UTR;5'UTR | No |
| Control-Sample46_Beta | chr4:74486446-74486558 | RASSF6;RASSF6 | S_Shore | TSS200;TSS1500 | No |
| Control-Sample108_Be | chr12:86230093-86230367 | RASSF9;RASSF9 | | 5'UTR;1stExon | No |
| Control-Sample51_Beta | chr13:48894214-48894595 | RB1 | S_Shore | Body | No |
| Control-Sample19_Beta | chr12:114273539-114273782 | RBM19;RBM19;RBM19 | | Body;Body;Body | No |
| Control-Sample108_Be | chr22:36236649-36236760 | RBM9;RBM9;RBM9;RBM9;RBM9;RBM9 | | Body;Body;TSS200;TSS200;TSS200;TSS200 | No |
| Control-Sample1_Beta | chr2:161265247-161265366 | RBMS1;RBMS1 | S_Shore | Body;Body | Yes |
| Control-Sample11_Beta | chr2:161265247-161265289 | RBMS1;RBMS1 | S_Shore | Body;Body | Yes |
| Control-Sample108_Be | chr3:29322474-29323182 | RBMS3;RBMS3;RBMS3 | | TSS1500;TSS1500;TSS1500 | No |
| Control-Sample2_Beta | chr1:10056837-10056853 | RBP7 | N_Shore | TSS1500 | No |
| Control-Sample1_Beta | chr8:30241119-30241197 | RBPMS;RBPMS;RBPMS;RBPMS | N_Shore | TSS1500;TSS1500;TSS1500;TSS1500 | No |
| Control-Sample35_Beta | chr1:17751673-17751920 | RCC2;RCC2 | | Body;Body | No |
| Control-Sample201_Be | chr19:50031017-50031273 | RCN3;RCN3 | Island | 1stExon;5'UTR | No |
| Control-Sample63_Beta | chr1:167598958-167599124 | RCSD1 | N_Shore | TSS1500 | No |
| Control-Sample13_Beta | chr14:24641852-24642317 | REC8;REC8 | Island | Body;Body | No |
| Control-Sample10_Beta | chr7:150067712-150067896 | REPIN1;REPIN1;REPIN1;REPIN1 | N_Shore | 5'UTR;Body;TSS1500;5'UTR | Yes |
| Control-Sample18_Beta | chr7:150067807-150068240 | REPIN1;REPIN1;REPIN1;REPIN1 | N_Shore | 5'UTR;Body;TSS1500;5'UTR | Yes |

| | | | | | |
|---|---|---|---|---|---|
| Control-Sample3_Beta | chr7:150067712-150068240 | REPIN1;REPIN1;REPIN1;REPIN1 | N_Shore | 5'UTR;Body;TSS1500;5'UTR | Yes |
| Control-Sample1_Beta | chr4:57773300-57773436 | REST | N_Shore | TSS1500 | Yes |
| Control-Sample14_Beta | chr4:57773308-57773436 | REST | N_Shore | TSS1500 | Yes |
| Control-Sample19_Beta | chr4:57773149-57773436 | REST | N_Shore | TSS1500 | Yes |
| Control-Sample1_Beta | chr17:33391738-33391960 | RFFL;RFFL | | 5'UTR;TSS1500 | No |
| Control-Sample23_Beta | chr22:32600722-32601274 | RFPL2;RFPL2 | | TSS1500;TSS200 | Yes |
| Control-Sample161_Be | chr22:32599596-32599648 | RFPL2;RFPL2;RFPL2 | | TSS200;5'UTR;TSS1500 | Yes |
| Control-Sample203_Be | chr22:32599511-32599648 | RFPL2;RFPL2;RFPL2 | | TSS200;5'UTR;TSS1500 | Yes |
| Control-Sample108_Be | chr2:198540518-198540621 | RFTN2;RFTN2 | | 1stExon;5'UTR | Yes |
| Control-Sample568_Be | chr2.0:198540518-198540621 | RFTN2;RFTN2 | | 1stExon;5'UTR | Yes |
| Control-Sample3_Beta | chr12:106978920-106979078 | RFX4 | N_Shore | Body | Yes |
| Control-Sample5_Beta | chr12:106978920-106979229 | RFX4 | N_Shore | Body | Yes |
| Control-Sample65_Beta | chr12:106979062-106979229 | RFX4 | N_Shore | Body | Yes |
| Control-Sample2_Beta | chr6:33265386-33265424 | RGL2;RGL2 | N_Shore | Body;Body | Yes |
| Control-Sample5_Beta | chr6:33265243-33265313 | RGL2;RGL2 | N_Shore | Body;Body | Yes |
| Control-Sample10_Beta | chr15:93616959-93617141 | RGMA;RGMA;RGMA;RGMA;RGMA;RGMA | Island | 1stExon;Body;TSS200;TSS200;TSS1500;5'UTR | Yes |
| Control-Sample18_Beta | chr15:93616959-93617146 | RGMA;RGMA;RGMA;RGMA;RGMA;RGMA | Island | 1stExon;Body;TSS200;TSS200;TSS1500;5'UTR | Yes |
| Control-Sample25_Beta | chr15:93616959-93617168 | RGMA;RGMA;RGMA;RGMA;RGMA;RGMA | Island | 1stExon;Body;TSS200;TSS200;TSS1500;5'UTR | Yes |
| Control-Sample30_Beta | chr15:93616959-93617030 | RGMA;RGMA;RGMA;RGMA;RGMA;RGMA | Island | 1stExon;Body;TSS200;TSS200;TSS1500;5'UTR | Yes |
| Control-Sample1_Beta | chr15:93617030-93617134 | RGMA;RGMA;RGMA;RGMA;RGMA;RGMA;RGMA | Island | 1stExon;Body;5'UTR;TSS200;TSS200;TSS1500;5'UTR | Yes |
| Control-Sample11_Beta | chr15:93617030-93617168 | RGMA;RGMA;RGMA;RGMA;RGMA;RGMA;RGMA | Island | 1stExon;Body;5'UTR;TSS200;TSS200;TSS1500;5'UTR | Yes |
| Control-Sample12_Beta | chr15:93617030-93617146 | RGMA;RGMA;RGMA;RGMA;RGMA;RGMA;RGMA | Island | 1stExon;Body;5'UTR;TSS200;TSS200;TSS1500;5'UTR | Yes |
| Control-Sample15_Beta | chr15:93617030-93617141 | RGMA;RGMA;RGMA;RGMA;RGMA;RGMA;RGMA | Island | 1stExon;Body;5'UTR;TSS200;TSS200;TSS1500;5'UTR | Yes |
| Control-Sample27_Beta | chr15:93617080-93617141 | RGMA;RGMA;RGMA;RGMA;RGMA;RGMA;RGMA | Island | 1stExon;Body;5'UTR;TSS200;TSS200;TSS1500;5'UTR | Yes |
| Control-Sample35_Beta | chr15:93617080-93617168 | RGMA;RGMA;RGMA;RGMA;RGMA;RGMA;RGMA | Island | 1stExon;Body;5'UTR;TSS200;TSS200;TSS1500;5'UTR | Yes |
| Control-Sample3_Beta | chr10:86004761-86005121 | RGR;RGR;RGR | | TSS200;TSS200;TSS200 | No |
| Control-Sample1_Beta | chr4:3387281-3387341 | RGS12;RGS12;RGS12 | | Body;Body;Body | Yes |
| Control-Sample15_Beta | chr4:3372915-3373555 | RGS12;RGS12;RGS12 | N_Shore | Body;Body;Body | Yes |
| Control-Sample170_Be | chr4:3372915-3373405 | RGS12;RGS12;RGS12 | N_Shore | Body;Body;Body | Yes |
| Control-Sample21_Beta | chr4:3416370-3416881 | RGS12;RGS12;RGS12 | S_Shore | Body;Body;Body | Yes |
| Control-Sample1_Beta | chr5:176793941-176794207 | RGS14 | N_Shore | Body | Yes |
| Control-Sample10_Beta | chr5:176793941-176794405 | RGS14 | N_Shore | Body | Yes |
| Control-Sample14_Beta | chr5:176793941-176794799 | RGS14 | N_Shore | Body | Yes |
| Control-Sample15_Beta | chr5:176784719-176784965 | RGS14 | | TSS200 | Yes |
| Control-Sample6_Beta | chr5:176784688-176784719 | RGS14 | | TSS200 | Yes |
| Control-Sample11_Beta | chr1:163172914-163173262 | RGS5 | | TSS200 | No |
| Control-Sample43_Beta | chr16:111362-111871 | RHBDF1 | Island | Body | No |
| Control-Sample1_Beta | chr1:25747376-25747645 | RHCE;RHCE;RHCE;RHCE | | TSS200;TSS200;TSS200;TSS200 | No |
| Control-Sample21_Beta | chr1:25598878-25599066 | RHD;RHD | | TSS200;TSS200 | No |
| Control-Sample1_Beta | chr4:40198392-40198776 | RHOH | | TSS200 | Yes |
| Control-Sample41_Beta | chr4:40198392-40198595 | RHOH | | TSS200 | Yes |
| Control-Sample19_Beta | chr12:130976260-130976551 | RIMBP2 | | 5'UTR | No |
| Control-Sample80_Beta | chr6:72926121-72926485 | RIMS1;RIMS1;RIMS1;RIMS1;RIMS1 | | TSS1500;TSS200;Body;Body;Body | No |
| Control-Sample10_Beta | chr11:66104468-66104485 | RIN1 | S_Shelf | TSS1500 | Yes |
| Control-Sample331_Be | chr11:66104481-66104993 | RIN1 | S_Shelf | TSS1500 | Yes |
| Control-Sample1_Beta | chr20:19915794-19916087 | RIN2 | | Body | Yes |
| Control-Sample10_Beta | chr20:19915769-19916087 | RIN2 | | Body | Yes |
| Control-Sample108_Be | chr20:19915762-19915874 | RIN2 | | Body | Yes |
| Control-Sample21_Beta | chr20:19915762-19916087 | RIN2 | | Body | Yes |
| Control-Sample3_Beta | chr20:19915857-19916087 | RIN2 | | Body | Yes |
| Control-Sample475_Be | chr20:19915794-19915874 | RIN2 | | Body | Yes |
| Control-Sample57_Beta | chr20:19915769-19915874 | RIN2 | | Body | Yes |
| Control-Sample67_Beta | chr20:19915762-19915794 | RIN2 | | Body | Yes |
| Control-Sample72_Beta | chr20:19915762-19915857 | RIN2 | | Body | Yes |
| Control-Sample33_Beta | chr14:93154317-93154758 | RIN3 | Island | Body | No |
| Control-Sample1_Beta | chr14:24809068-24809759 | RIPK3;RIPK3 | S_Shore | 5'UTR;1stExon | Yes |
| Control-Sample12_Beta | chr14:24809226-24809759 | RIPK3;RIPK3 | S_Shore | 5'UTR;1stExon | Yes |
| Control-Sample19_Beta | chr14:24809226-24809442 | RIPK3;RIPK3 | S_Shore | 5'UTR;1stExon | Yes |
| Control-Sample1_Beta | chr21:43185385-43185803 | RIPK4 | N_Shore | Body | No |
| Control-Sample17_Beta | chr15:89765045-89765204 | RLBP1 | | TSS200 | Yes |
| Control-Sample11_Beta | chr15:89764895-89765204 | RLBP1;RLBP1 | | 1stExon;5'UTR | Yes |
| Control-Sample15_Beta | chr15:89764895-89765061 | RLBP1;RLBP1 | | 1stExon;5'UTR | Yes |
| Control-Sample25_Beta | chr15:89764836-89765204 | RLBP1;RLBP1 | | 1stExon;5'UTR | Yes |
| Control-Sample67_Beta | chr16:67686832-67687754 | RLTPR | N_Shore | Body | No |
| Control-Sample1_Beta | chr14:21269744-21271355 | RNASE1;RNASE1;RNASE1;RNASE1 | | 3'UTR;3'UTR;3'UTR;3'UTR | Yes |
| Control-Sample108_Be | chr14:21270133-21270575 | RNASE1;RNASE1;RNASE1;RNASE1 | | Body;Body;Body;Body | Yes |
| Control-Sample151_Be | chr14:21270133-21271018 | RNASE1;RNASE1;RNASE1;RNASE1 | | Body;Body;Body;Body | Yes |
| Control-Sample152_Be | chr14:21270850-21271018 | RNASE1;RNASE1;RNASE1;RNASE1 | | 5'UTR;TSS1500;5'UTR;5'UTR | Yes |
| Control-Sample170_Be | chr14:21270575-21271018 | RNASE1;RNASE1;RNASE1;RNASE1 | | 5'UTR;TSS200;5'UTR;5'UTR | Yes |
| Control-Sample194_Be | chr14:21270575-21270924 | RNASE1;RNASE1;RNASE1;RNASE1 | | 5'UTR;TSS200;5'UTR;5'UTR | Yes |
| Control-Sample234_Be | chr14:21270575-21271355 | RNASE1;RNASE1;RNASE1;RNASE1 | | 5'UTR;TSS200;5'UTR;5'UTR | Yes |
| Control-Sample270_Be | chr14:21270133-21271313 | RNASE1;RNASE1;RNASE1;RNASE1 | | Body;Body;Body;Body | Yes |
| Control-Sample475_Be | chr14:21269744-21271018 | RNASE1;RNASE1;RNASE1;RNASE1 | | 3'UTR;3'UTR;3'UTR;3'UTR | Yes |
| Control-Sample497_Be | chr14:21270575-21271148 | RNASE1;RNASE1;RNASE1;RNASE1 | | 5'UTR;TSS200;5'UTR;5'UTR | Yes |
| Control-Sample572_Be | chr14.0:21270575-21271313 | RNASE1;RNASE1;RNASE1;RNASE1 | | 5'UTR;TSS200;5'UTR;5'UTR | Yes |
| Control-Sample585_Be | chr14.0:21270575-21271355 | RNASE1;RNASE1;RNASE1;RNASE1 | | 5'UTR;TSS200;5'UTR;5'UTR | Yes |
| Control-Sample67_Beta | chr14:21270133-21271355 | RNASE1;RNASE1;RNASE1;RNASE1 | | Body;Body;Body;Body | Yes |
| Control-Sample71_Beta | chr14:21270850-21271355 | RNASE1;RNASE1;RNASE1;RNASE1 | | 5'UTR;TSS1500;5'UTR;5'UTR | Yes |
| Control-Sample223_Be | chr14:21270407-21271355 | RNASE1;RNASE1;RNASE1;RNASE1;RNASE1 | | 5'UTR;5'UTR;1stExon;5'UTR;5'UTR | Yes |
| Control-Sample78_Beta | chr14:21270407-21271018 | RNASE1;RNASE1;RNASE1;RNASE1;RNASE1 | | 5'UTR;5'UTR;1stExon;5'UTR;5'UTR | Yes |
| Control-Sample88_Beta | chr14:21270407-21270924 | RNASE1;RNASE1;RNASE1;RNASE1;RNASE1 | | 5'UTR;5'UTR;1stExon;5'UTR;5'UTR | Yes |
| Control-Sample15_Beta | chr15:59279406-59279556 | RNF111 | N_Shore | TSS1500 | No |
| Control-Sample15_Beta | chr3:149530294-149530341 | RNF13;RNF13 | Island | TSS200;TSS200 | No |
| Control-Sample1_Beta | chr17:29297380-29297414 | RNF135;RNF135 | N_Shore | TSS1500;TSS1500 | No |
| Control-Sample15_Beta | chr5:141348202-141348342 | RNF14;RNF14;RNF14;RNF14;RNF14 | N_Shore | TSS1500;TSS1500;TSS1500;TSS1500;5'UTR | No |
| Control-Sample10_Beta | chr6:13924355-13924425 | RNF182;RNF182;RNF182;RNF182 | N_Shore | TSS1500;TSS1500;TSS1500;TSS1500 | No |
| Control-Sample464_Be | chr1:20141662-20141781 | RNF186;RNF186 | | 5'UTR;1stExon | No |
| Control-Sample151_Be | chr8:101315498-101315560 | RNF19A;RNF19A | | 5'UTR;TSS200 | Yes |
| Control-Sample12_Beta | chr8:101315424-101315505 | RNF19A;RNF19A;RNF19A | | 5'UTR;5'UTR;1stExon | Yes |
| Control-Sample15_Beta | chr8:101315424-101315560 | RNF19A;RNF19A;RNF19A | | 5'UTR;5'UTR;1stExon | Yes |
| Control-Sample1_Beta | chr1:185013658-185014264 | RNF2 | N_Shore | TSS1500 | Yes |

| | | | | | |
|---|---|---|---|---|---|
| Control-Sample10_Beta | chr1:185013789-185014241 | RNF2 | N_Shore | TSS1500 | Yes |
| Control-Sample15_Beta | chr1:185013789-185014264 | RNF2 | N_Shore | TSS1500 | Yes |
| Control-Sample58_Beta | chr1:185013658-185014241 | RNF2 | N_Shore | TSS1500 | Yes |
| Control-Sample1_Beta | chr1:45103896-45103967 | RNF220 | | Body | Yes |
| Control-Sample11_Beta | chr1:45082704-45083278 | RNF220 | N_Shore | Body | Yes |
| Control-Sample16_Beta | chr1:45083079-45083278 | RNF220 | Island | Body | Yes |
| Control-Sample1_Beta | chr6:30038897-30038929 | RNF39;RNF39 | Island | Body;Body | Yes |
| Control-Sample1_Beta | chr6:30039006-30039027 | RNF39;RNF39 | Island | Body;Body | Yes |
| Control-Sample1_Beta | chr6:30039175-30039206 | RNF39;RNF39 | Island | Body;Body | Yes |
| Control-Sample1_Beta | chr6:30039376-30039442 | RNF39;RNF39 | Island | Body;Body | Yes |
| Control-Sample1_Beta | chr6:30039476-30039600 | RNF39;RNF39 | Island | Body;Body | Yes |
| Control-Sample1_Beta | chr6:30041727-30042188 | RNF39;RNF39 | N_Shore | Body;Body | Yes |
| Control-Sample1_Beta | chr6:30043820-30043858 | RNF39;RNF39 | S_Shore | TSS200;TSS200 | Yes |
| Control-Sample10_Beta | chr6:30038897-30039027 | RNF39;RNF39 | Island | Body;Body | Yes |
| Control-Sample10_Beta | chr6:30039374-30039442 | RNF39;RNF39 | Island | Body;Body | Yes |
| Control-Sample17_Beta | chr6:30039010-30039027 | RNF39;RNF39 | Island | Body;Body | Yes |
| Control-Sample17_Beta | chr6:30039408-30039442 | RNF39;RNF39 | Island | Body;Body | Yes |
| Control-Sample18_Beta | chr6:30038897-30038915 | RNF39;RNF39 | Island | Body;Body | Yes |
| Control-Sample18_Beta | chr6:30039010-30039025 | RNF39;RNF39 | Island | Body;Body | Yes |
| Control-Sample194_Be | chr6:30039432-30039442 | RNF39;RNF39 | Island | Body;Body | Yes |
| Control-Sample2_Beta | chr6:30038929-30038975 | RNF39;RNF39 | Island | Body;Body | Yes |
| Control-Sample248_Be | chr6:30039376-30039403 | RNF39;RNF39 | Island | Body;Body | Yes |
| Control-Sample255_Be | chr6:30039376-30039408 | RNF39;RNF39 | Island | Body;Body | Yes |
| Control-Sample27_Beta | chr6:30038897-30038922 | RNF39;RNF39 | Island | Body;Body | Yes |
| Control-Sample27_Beta | chr6:30038955-30039027 | RNF39;RNF39 | Island | Body;Body | Yes |
| Control-Sample28_Beta | chr6:30038897-30038975 | RNF39;RNF39 | Island | Body;Body | Yes |
| Control-Sample28_Beta | chr6:30039476-30039548 | RNF39;RNF39 | Island | Body;Body | Yes |
| Control-Sample43_Beta | chr6:30039374-30039380 | RNF39;RNF39 | Island | Body;Body | Yes |
| Control-Sample441_Be | chr6:30039374-30039408 | RNF39;RNF39 | Island | Body;Body | Yes |
| Control-Sample441_Be | chr6:30039466-30039548 | RNF39;RNF39 | Island | Body;Body | Yes |
| Control-Sample461_Be | chr6:30039374-30039435 | RNF39;RNF39 | Island | Body;Body | Yes |
| Control-Sample47_Beta | chr6:30039130-30039142 | RNF39;RNF39 | Island | Body;Body | Yes |
| Control-Sample563_Be | chr6.0:30039374-30039380 | RNF39;RNF39 | Island | Body;Body | Yes |
| Control-Sample57_Beta | chr6:30038955-30039025 | RNF39;RNF39 | Island | Body;Body | Yes |
| Control-Sample576_Be | chr6.0:30039175-30039206 | RNF39;RNF39 | Island | Body;Body | Yes |
| Control-Sample576_Be | chr6.0:30039374-30039435 | RNF39;RNF39 | Island | Body;Body | Yes |
| Control-Sample576_Be | chr6.0:30039466-30039548 | RNF39;RNF39 | Island | Body;Body | Yes |
| Control-Sample588_Be | chr6.0:30039374-30039408 | RNF39;RNF39 | Island | Body;Body | Yes |
| Control-Sample613_Be | chr6:30039466-30039524 | RNF39;RNF39 | Island | Body;Body | Yes |
| Control-Sample63_Beta | chr6:30043779-30043858 | RNF39;RNF39 | S_Shore | TSS200;TSS200 | Yes |
| Control-Sample64_Beta | chr6:30039013-30039027 | RNF39;RNF39 | Island | Body;Body | Yes |
| Control-Sample71_Beta | chr6:30043095-30043106 | RNF39;RNF39 | Island | 1stExon;1stExon | Yes |
| Control-Sample9_Beta | chr6:30038929-30039027 | RNF39;RNF39 | Island | Body;Body | Yes |
| Control-Sample394_Be | chr6:32146232-32146529 | RNF5;AGPAT1;RNF5P1;RNF5 | | 1stExon;TSS1500;Body;5'UTR | Yes |
| Control-Sample1_Beta | chr11:494944-495069 | RNH1;RNH1;RNH1;RNH1;RNH1;RNH1;RNH1;RNH1 | Island | Body;Body;Body;Body;Body;Body;Body;Body | Yes |
| Control-Sample24_Beta | chr11:494944-495198 | RNH1;RNH1;RNH1;RNH1;RNH1;RNH1;RNH1;RNH1 | Island | Body;Body;Body;Body;Body;Body;Body;Body | Yes |
| Control-Sample17_Beta | chr5:80529100-80529187 | RNU5E;RNU5D;CKMT2;CKMT2;CKMT2 | | Body;Body;TSS200;TSS200;TSS200 | Yes |
| Control-Sample18_Beta | chr5:80529100-80529135 | RNU5E;RNU5D;CKMT2;CKMT2;CKMT2 | | Body;Body;TSS200;TSS200;TSS200 | Yes |
| Control-Sample2_Beta | chr5:80529135-80529340 | RNU5E;RNU5D;CKMT2;CKMT2;CKMT2 | | Body;Body;TSS200;TSS200;TSS200 | Yes |
| Control-Sample1_Beta | chr8:10512459-10512769 | RP1L1;RP1L1 | | 1stExon;5'UTR | No |
| Control-Sample89_Beta | chr6:111303128-111303252 | RPF2 | Island | TSS200 | No |
| Control-Sample1_Beta | chr17:109444-109926 | RPH3AL | | Body | Yes |
| Control-Sample1_Beta | chr17:171006-171228 | RPH3AL | Island | Body | Yes |
| Control-Sample10_Beta | chr17:185756-186214 | RPH3AL | S_Shore | 5'UTR | Yes |
| Control-Sample11_Beta | chr17:154410-154671 | RPH3AL | S_Shore | Body | Yes |
| Control-Sample12_Beta | chr17:109444-110138 | RPH3AL | | Body | Yes |
| Control-Sample12_Beta | chr17:154410-154499 | RPH3AL | S_Shore | Body | Yes |
| Control-Sample133_Be | chr17:154410-155045 | RPH3AL | S_Shore | Body | Yes |
| Control-Sample174_Be | chr17:154420-154671 | RPH3AL | S_Shore | Body | Yes |
| Control-Sample183_Be | chr17:154420-155045 | RPH3AL | S_Shore | Body | Yes |
| Control-Sample26_Beta | chr17:109496-110138 | RPH3AL | | Body | Yes |
| Control-Sample32_Beta | chr17:171006-171257 | RPH3AL | Island | Body | Yes |
| Control-Sample40_Beta | chr17:109444-109545 | RPH3AL | | Body | Yes |
| Control-Sample464_Be | chr17:149484-149582 | RPH3AL | N_Shelf | Body | Yes |
| Control-Sample497_Be | chr17:154499-155045 | RPH3AL | N_Shelf | Body | Yes |
| Control-Sample506_Be | chr17.0:154410-155045 | RPH3AL | S_Shore | Body | Yes |
| Control-Sample509_Be | chr17.0:154420-154671 | RPH3AL | S_Shore | Body | Yes |
| Control-Sample543_Be | chr17.0:154410-154671 | RPH3AL | S_Shore | Body | Yes |
| Control-Sample19_Beta | chr3:101406220-101406543 | RPL24 | S_Shore | TSS1500 | No |
| Control-Sample1_Beta | chr22:39712694-39712956 | RPL3;SNORD83A;RPL3 | N_Shelf | Body;TSS1500;Body | Yes |
| Control-Sample40_Beta | chr22:39712633-39712730 | RPL3;SNORD83A;RPL3 | N_Shelf | Body;TSS1500;Body | Yes |
| Control-Sample1_Beta | chr22:39709853-39710068 | RPL3;SNORD83B;RPL3 | | Body;Body;Body | Yes |
| Control-Sample12_Beta | chr22:39709954-39710068 | RPL3;SNORD83B;RPL3 | | Body;TSS200;Body | Yes |
| Control-Sample53_Beta | chr22:39709951-39710068 | RPL3;SNORD83B;RPL3 | | Body;TSS200;Body | Yes |
| Control-Sample10_Beta | chr2:154336148-154336341 | RPRM | S_Shore | TSS1500 | No |
| Control-Sample10_Beta | chr6:167263193-167263401 | RPS6KA2 | | Body | Yes |
| Control-Sample17_Beta | chr6:167263193-167263454 | RPS6KA2 | | Body | Yes |
| Control-Sample19_Beta | chr6:167051714-167051781 | RPS6KA2 | | Body | Yes |
| Control-Sample1_Beta | chr6:166891902-166892216 | RPS6KA2;RPS6KA2 | | Body;Body | Yes |
| Control-Sample151_Be | chr6:166836643-166836791 | RPS6KA2;RPS6KA2 | | Body;Body | Yes |
| Control-Sample2_Beta | chr6:166891902-166892362 | RPS6KA2;RPS6KA2 | | Body;Body | Yes |
| Control-Sample10_Beta | chr11:64136015-64136975 | RPS6KA4;MIR1237;RPS6KA4 | Island | Body;TSS200;Body | Yes |
| Control-Sample19_Beta | chr11:64136015-64136089 | RPS6KA4;MIR1237;RPS6KA4 | Island | Body;TSS200;Body | Yes |
| Control-Sample2_Beta | chr11:64135875-64136975 | RPS6KA4;MIR1237;RPS6KA4 | Island | Body;TSS200;Body | Yes |
| Control-Sample58_Beta | chr11:64135875-64137747 | RPS6KA4;MIR1237;RPS6KA4 | Island | Body;TSS200;Body | Yes |
| Control-Sample115_Be | chr19:23945722-23945761 | RPSAP58 | Island | TSS200 | Yes |
| Control-Sample42_Beta | chr19:23945659-23945761 | RPSAP58 | N_Shore | TSS200 | Yes |
| Control-Sample1_Beta | chr17:78724995-78725470 | RPTOR;RPTOR | | Body;Body | Yes |
| Control-Sample10_Beta | chr17:78724995-78725375 | RPTOR;RPTOR | | Body;Body | Yes |
| Control-Sample10_Beta | chr17:78747934-78748494 | RPTOR;RPTOR | | Body;Body | Yes |
| Control-Sample10_Beta | chr17:78831230-78831518 | RPTOR;RPTOR | N_Shore | Body;Body | Yes |

| | | | | | |
|---|---|---|---|---|---|
| Control-Sample11_Beta | chr17:78725049-78725470 | RPTOR;RPTOR | | Body;Body | Yes |
| Control-Sample11_Beta | chr17:78748019-78748291 | RPTOR;RPTOR | | Body;Body | Yes |
| Control-Sample12_Beta | chr17:78725049-78725375 | RPTOR;RPTOR | | Body;Body | Yes |
| Control-Sample12_Beta | chr17:78747934-78748077 | RPTOR;RPTOR | | Body;Body | Yes |
| Control-Sample13_Beta | chr17:78747934-78748291 | RPTOR;RPTOR | | Body;Body | Yes |
| Control-Sample17_Beta | chr17:78724995-78725201 | RPTOR;RPTOR | | Body;Body | Yes |
| Control-Sample19_Beta | chr17:78607095-78607368 | RPTOR;RPTOR | | Body;Body | Yes |
| Control-Sample20_Beta | chr17:78817938-78818441 | RPTOR;RPTOR | N_Shore | Body;Body | Yes |
| Control-Sample216_Be | chr17:78725201-78725470 | RPTOR;RPTOR | | Body;Body | Yes |
| Control-Sample29_Beta | chr17:78872563-78872875 | RPTOR;RPTOR | S_Shelf | Body;Body | Yes |
| Control-Sample29_Beta | chr17:78915655-78915881 | RPTOR;RPTOR | Island | Body;Body | Yes |
| Control-Sample3_Beta | chr17:78748019-78748494 | RPTOR;RPTOR | | Body;Body | Yes |
| Control-Sample31_Beta | chr17:78747934-78748471 | RPTOR;RPTOR | | Body;Body | Yes |
| Control-Sample40_Beta | chr17:78878278-78878579 | RPTOR;RPTOR | N_Shore | Body;Body | Yes |
| Control-Sample464_Be | chr17:78682934-78683272 | RPTOR;RPTOR | | Body;Body | Yes |
| Control-Sample465_Be | chr17:78860483-78860605 | RPTOR;RPTOR | N_Shelf | Body;Body | Yes |
| Control-Sample57_Beta | chr17:78800767-78800806 | RPTOR;RPTOR | S_Shore | Body;Body | Yes |
| Control-Sample57_Beta | chr17:78940029-78940110 | RPTOR;RPTOR | Island | 3'UTR;3'UTR | Yes |
| Control-Sample64_Beta | chr17:78755379-78755442 | RPTOR;RPTOR | | Body;Body | Yes |
| Control-Sample65_Beta | chr17:78851149-78851503 | RPTOR;RPTOR | S_Shore | Body;Body | Yes |
| Control-Sample12_Beta | chr6:90120165-90120705 | RRAGD | N_Shore | Body | Yes |
| Control-Sample7_Beta | chr6:90119710-90120254 | RRAGD | N_Shore | Body | Yes |
| Control-Sample1_Beta | chr2:7017571-7018153 | RSAD2 | | TSS1500 | Yes |
| Control-Sample108_Be | chr2:7017571-7018020 | RSAD2 | | TSS1500 | Yes |
| Control-Sample63_Beta | chr2:7017787-7018153 | RSAD2 | | TSS200 | Yes |
| Control-Sample67_Beta | chr2:7017571-7017723 | RSAD2 | | TSS1500 | Yes |
| Control-Sample68_Beta | chr2:7017698-7018153 | RSAD2 | | TSS200 | Yes |
| Control-Sample75_Beta | chr2:7017571-7017787 | RSAD2 | | TSS1500 | Yes |
| Control-Sample80_Beta | chr2:7017698-7018020 | RSAD2 | | TSS200 | Yes |
| Control-Sample1_Beta | chr19:46318439-46318633 | RSPH6A;RSPH6A | N_Shore | 1stExon;5'UTR | Yes |
| Control-Sample12_Beta | chr19:46318762-46319153 | RSPH6A;SYMPK | Island | TSS200;3'UTR | Yes |
| Control-Sample6_Beta | chr19:46318762-46319145 | RSPH6A;SYMPK | Island | TSS200;3'UTR | Yes |
| Control-Sample1_Beta | chr10:16741464-16741619 | RSU1;RSU1 | | Body;Body | No |
| Control-Sample82_Beta | chr19:45996372-45996460 | RTN2;RTN2;RTN2 | N_Shore | Body;Body;Body | No |
| Control-Sample2_Beta | chr22:20237197-20237341 | RTN4R;MIR1286 | | Body;TSS1500 | No |
| Control-Sample1_Beta | chr3:187086147-187086530 | RTP4 | | TSS200 | No |
| Control-Sample67_Beta | chr4:71587640-71587904 | RUFY3;RUFY3;RUFY3 | | Body;TSS200;TSS200 | No |
| Control-Sample19_Beta | chr16:29329433-29329559 | RUNDC2C | | Body | Yes |
| Control-Sample25_Beta | chr16:29324559-29324813 | RUNDC2C | | Body | Yes |
| Control-Sample1_Beta | chr17:42392436-42393317 | RUNDC3A;RUNDC3A;RUNDC3A | Island | Body;Body;Body | Yes |
| Control-Sample1_Beta | chr17:42393931-42394004 | RUNDC3A;RUNDC3A;RUNDC3A | S_Shore | Body;Body;Body | Yes |
| Control-Sample10_Beta | chr17:42392436-42392697 | RUNDC3A;RUNDC3A;RUNDC3A | Island | Body;Body;Body | Yes |
| Control-Sample11_Beta | chr17:42392436-42392983 | RUNDC3A;RUNDC3A;RUNDC3A | Island | Body;Body;Body | Yes |
| Control-Sample15_Beta | chr17:42393858-42394004 | RUNDC3A;RUNDC3A;RUNDC3A | S_Shore | Body;Body;Body | Yes |
| Control-Sample16_Beta | chr17:42392436-42394004 | RUNDC3A;RUNDC3A;RUNDC3A | Island | Body;Body;Body | Yes |
| Control-Sample223_Be | chr17:42392668-42392983 | RUNDC3A;RUNDC3A;RUNDC3A | Island | Body;Body;Body | Yes |
| Control-Sample26_Beta | chr17:42393858-42393982 | RUNDC3A;RUNDC3A;RUNDC3A | S_Shore | Body;Body;Body | Yes |
| Control-Sample1_Beta | chr6:45391852-45391973 | RUNX2;RUNX2;RUNX2 | S_Shore | Body;Body;Body | Yes |
| Control-Sample10_Beta | chr6:45391645-45391973 | RUNX2;RUNX2;RUNX2 | Island | Body;Body;Body | Yes |
| Control-Sample27_Beta | chr6:45391645-45391916 | RUNX2;RUNX2;RUNX2 | Island | Body;Body;Body | Yes |
| Control-Sample10_Beta | chr1:25240865-25240938 | RUNX3;RUNX3 | | Body;Body | Yes |
| Control-Sample11_Beta | chr1:25257759-25257813 | RUNX3;RUNX3 | Island | TSS1500;Body | Yes |
| Control-Sample74_Beta | chr21:30391383-30391795 | RWDD2B | Island | Body | Yes |
| Control-Sample10_Beta | chr21:30391630-30391795 | RWDD2B;RWDD2B | Island | 1stExon;5'UTR | Yes |
| Control-Sample26_Beta | chr1:95698827-95698954 | RWDD3;RWDD3 | N_Shore | TSS1500;TSS1500 | No |
| Control-Sample10_Beta | chr6:33165658-33165899 | RXRB | N_Shelf | Body | Yes |
| Control-Sample2_Beta | chr6:33165658-33165707 | RXRB | N_Shelf | Body | Yes |
| Control-Sample18_Beta | chr1:151967021-151967449 | S100A10 | S_Shore | TSS1500 | No |
| Control-Sample1_Beta | chr1:153600762-153600972 | S100A13;S100A13;S100A13;S100A1;S100A13;S100A13 | | TSS1500;TSS200;TSS1500;TSS200;5'UTR;TSS1500 | Yes |
| Control-Sample171_Be | chr1:153600762-153600865 | S100A13;S100A13;S100A13;S100A1;S100A13;S100A13 | | TSS1500;TSS200;TSS1500;TSS200;5'UTR;TSS1500 | Yes |
| Control-Sample585_Be | chr1.0:153600762-153600865 | S100A13;S100A13;S100A13;S100A1;S100A13;S100A13 | | TSS1500;TSS200;TSS1500;TSS200;5'UTR;TSS1500 | Yes |
| Control-Sample67_Beta | chr1:153600790-153600972 | S100A13;S100A13;S100A13;S100A1;S100A13;S100A13 | | TSS1500;TSS200;TSS1500;TSS200;5'UTR;TSS1500 | Yes |
| Control-Sample33_Beta | chr1:153581494-153581655 | S100A16 | | 5'UTR | No |
| Control-Sample33_Beta | chr1:153508511-153508542 | S100A6;S100A6 | Island | 5'UTR;1stExon | No |
| Control-Sample1_Beta | chr21:48025691-48026298 | S100B | | TSS1500 | Yes |
| Control-Sample19_Beta | chr21:48018608-48018790 | S100B | Island | 3'UTR | Yes |
| Control-Sample223_Be | chr21:48026043-48026298 | S100B | | TSS1500 | Yes |
| Control-Sample331_Be | chr21:48025691-48026181 | S100B | | TSS1500 | Yes |
| Control-Sample10_Beta | chr20:50419193-50419348 | SALL4 | S_Shore | TSS200 | Yes |
| Control-Sample12_Beta | chr20:50419082-50419348 | SALL4 | S_Shore | TSS200 | Yes |
| Control-Sample19_Beta | chr20:50419214-50419274 | SALL4 | S_Shore | TSS200 | Yes |
| Control-Sample20_Beta | chr20:50419231-50419274 | SALL4 | S_Shore | TSS200 | Yes |
| Control-Sample21_Beta | chr20:50419193-50419274 | SALL4 | S_Shore | TSS200 | Yes |
| Control-Sample41_Beta | chr20:50418554-50418959 | SALL4 | Island | Body | Yes |
| Control-Sample42_Beta | chr20:50419082-50419274 | SALL4 | S_Shore | TSS200 | Yes |
| Control-Sample1_Beta | chr20:50418952-50419348 | SALL4;SALL4 | Island | 1stExon;5'UTR | Yes |
| Control-Sample24_Beta | chr20:50418952-50419274 | SALL4;SALL4 | Island | 1stExon;5'UTR | Yes |
| Control-Sample1_Beta | chr17:48194835-48195107 | SAMD14 | Island | Body | Yes |
| Control-Sample11_Beta | chr17:48194635-48195107 | SAMD14 | Island | Body | Yes |
| Control-Sample13_Beta | chr7:92747308-92747455 | SAMD9;SAMD9 | | 1stExon;5'UTR | No |
| Control-Sample1_Beta | chr6:148869461-148869895 | SASH1 | | Body | Yes |
| Control-Sample11_Beta | chr6:148869461-148869535 | SASH1 | | Body | Yes |
| Control-Sample15_Beta | chr6:148869489-148869895 | SASH1 | | Body | Yes |
| Control-Sample1_Beta | chr3:18391029-18391232 | SATB1;SATB1 | | Body;Body | No |
| Control-Sample1_Beta | chr2:200326591-200326725 | SATB2 | Island | TSS1500 | Yes |
| Control-Sample170_Be | chr2:200320455-200320871 | SATB2 | N_Shore | Body | Yes |
| Control-Sample464_Be | chr22:50897911-50898441 | SBF1 | Island | Body | No |
| Control-Sample19_Beta | chr19:1155030-1155184 | SBNO2 | Island | 5'UTR | Yes |
| Control-Sample10_Beta | chr19:1132313-1132420 | SBNO2;SBNO2 | | Body;TSS200 | Yes |
| Control-Sample19_Beta | chr19:1132313-1132389 | SBNO2;SBNO2 | | Body;TSS200 | Yes |
| Control-Sample21_Beta | chr19:1132313-1132372 | SBNO2;SBNO2 | | Body;TSS200 | Yes |

| | | | | | |
|---|---|---|---|---|---|
| Control-Sample216_Be | chr19:1132372-1132420 | SBNO2;SBNO2 | | Body;TSS200 | Yes |
| Control-Sample1_Beta | chr19:1132256-1132389 | SBNO2;SBNO2;SBNO2 | | Body;1stExon;5'UTR | Yes |
| Control-Sample11_Beta | chr19:1132256-1132420 | SBNO2;SBNO2;SBNO2 | | Body;1stExon;5'UTR | Yes |
| Control-Sample5_Beta | chr19:1132256-1132372 | SBNO2;SBNO2;SBNO2 | | Body;1stExon;5'UTR | Yes |
| Control-Sample1_Beta | chr3:47517781-47517819 | SCAP | S_Shore | TSS1500 | No |
| Control-Sample108_Be | chr12:125333267-125333358 | SCARB1;SCARB1 | | Body;Body | No |
| Control-Sample12_Beta | chr22:20783497-20783963 | SCARF2;SCARF2 | Island | Body;Body | No |
| Control-Sample12_Beta | chr15:51972699-51973224 | SCG3;SCG3 | N_Shore | TSS1500;TSS1500 | No |
| Control-Sample1_Beta | chr15:32964155-32964500 | SCG5;SCG5 | | Body;Body | Yes |
| Control-Sample20_Beta | chr15:32964208-32964500 | SCG5;SCG5 | | Body;Body | Yes |
| Control-Sample108_Be | chr19:35085557-35085677 | SCGBL | | TSS200 | No |
| Control-Sample194_Be | chr2:238990408-238990731 | SCLY | | Body | No |
| Control-Sample58_Beta | chr6:108145539-108145658 | SCML4 | | TSS200 | Yes |
| Control-Sample19_Beta | chr6:108145374-108146329 | SCML4;SCML4 | | 1stExon;5'UTR | Yes |
| Control-Sample63_Beta | chr19:35530755-35531417 | SCN1B;HPN;HPN | N_Shore | 3'UTR;TSS1500;TSS1500 | Yes |
| Control-Sample12_Beta | chr11:118022607-118023038 | SCN4B;SCN4B | N_Shore | Body;Body | No |
| Control-Sample1_Beta | chr12:6485881-6486123 | SCNN1A;SCNN1A;SCNN1A | | Body;TSS1500;TSS1500 | Yes |
| Control-Sample29_Beta | chr12:6483708-6483772 | SCNN1A;SCNN1A;SCNN1A | | Body;Body;1stExon | Yes |
| Control-Sample55_Beta | chr1:1217159-1217664 | SCNN1D;SCNN1D | | TSS1500;TSS1500 | No |
| Control-Sample57_Beta | chr4:141207794-141207852 | SCOC | | 5'UTR | No |
| Control-Sample21_Beta | chr7:30029776-30029938 | SCRN1;SCRN1;SCRN1;SCRN1 | Island | TSS1500;TSS200;Body;TSS1500 | No |
| Control-Sample57_Beta | chr11:628603-628677 | SCT | S_Shore | TSS1500 | No |
| Control-Sample1_Beta | chr1:243637838-243638141 | SDCCAG8 | Island | Body | Yes |
| Control-Sample331_Be | chr1:243646235-243646402 | SDCCAG8 | N_Shore | Body | Yes |
| Control-Sample51_Beta | chr1:1168432-1168550 | SDF4;SDF4;B3GALT6 | Island | TSS1500;TSS1500;1stExon | No |
| Control-Sample151_Be | chr5:231809-231967 | SDHA | | Body | No |
| Control-Sample1_Beta | chr5:1594676-1594715 | SDHAP3 | Island | TSS200 | Yes |
| Control-Sample106_Be | chr5:1594678-1594733 | SDHAP3 | Island | TSS200 | Yes |
| Control-Sample11_Beta | chr5:1594579-1594715 | SDHAP3 | Island | Body | Yes |
| Control-Sample111_Be | chr5:1594676-1594733 | SDHAP3 | Island | TSS200 | Yes |
| Control-Sample112_Be | chr5:1594282-1594579 | SDHAP3 | Island | Body | Yes |
| Control-Sample122_Be | chr5:1594282-1594715 | SDHAP3 | Island | Body | Yes |
| Control-Sample123_Be | chr5:1594282-1594808 | SDHAP3 | Island | Body | Yes |
| Control-Sample125_Be | chr5:1594579-1594678 | SDHAP3 | Island | Body | Yes |
| Control-Sample129_Be | chr5:1594579-1594733 | SDHAP3 | Island | Body | Yes |
| Control-Sample141_Be | chr5:1594282-1594733 | SDHAP3 | Island | Body | Yes |
| Control-Sample147_Be | chr5:1594808-1594863 | SDHAP3 | Island | TSS200 | Yes |
| Control-Sample15_Beta | chr5:1594330-1594733 | SDHAP3 | Island | Body | Yes |
| Control-Sample18_Beta | chr5:1594330-1594715 | SDHAP3 | Island | Body | Yes |
| Control-Sample194_Be | chr5:1594330-1594863 | SDHAP3 | Island | Body | Yes |
| Control-Sample210_Be | chr5:1594330-1594678 | SDHAP3 | Island | Body | Yes |
| Control-Sample226_Be | chr5:1594715-1594808 | SDHAP3 | Island | TSS200 | Yes |
| Control-Sample238_Be | chr5:1594330-1594808 | SDHAP3 | Island | Body | Yes |
| Control-Sample291_Be | chr5:1594282-1594863 | SDHAP3 | Island | Body | Yes |
| Control-Sample31_Beta | chr5:1594715-1594863 | SDHAP3 | Island | TSS200 | Yes |
| Control-Sample509_Be | chr5.0:1594676-1594715 | SDHAP3 | Island | TSS200 | Yes |
| Control-Sample511_Be | chr5.0:1594282-1594733 | SDHAP3 | Island | Body | Yes |
| Control-Sample512_Be | chr5.0:1594282-1594808 | SDHAP3 | Island | Body | Yes |
| Control-Sample519_Be | chr5.0:1594579-1594715 | SDHAP3 | Island | Body | Yes |
| Control-Sample524_Be | chr5.0:1594579-1594678 | SDHAP3 | Island | Body | Yes |
| Control-Sample530_Be | chr5.0:1594579-1594733 | SDHAP3 | Island | Body | Yes |
| Control-Sample539_Be | chr5.0:1594676-1594733 | SDHAP3 | Island | TSS200 | Yes |
| Control-Sample1_Beta | chr1:161283845-161283918 | SDHC;SDHC;LOC642502;SDHC;SDHC | Island | TSS1500;TSS1500;TSS1500;TSS1500;TSS1500 | No |
| Control-Sample10_Beta | chr7:3919928-3920082 | SDK1 | | Body | Yes |
| Control-Sample10_Beta | chr7:4049819-4050578 | SDK1 | | Body | Yes |
| Control-Sample15_Beta | chr7:3919928-3920439 | SDK1 | | Body | Yes |
| Control-Sample17_Beta | chr7:4167607-4167882 | SDK1 | N_Shore | Body | Yes |
| Control-Sample24_Beta | chr7:4050043-4050578 | SDK1 | | Body | Yes |
| Control-Sample43_Beta | chr7:4049967-4050578 | SDK1 | | Body | Yes |
| Control-Sample58_Beta | chr7:4049531-4049718 | SDK1 | | Body | Yes |
| Control-Sample1_Beta | chr7:4266872-4267364 | SDK1;SDK1 | | Body;Body | Yes |
| Control-Sample10_Beta | chr7:4174655-4175200 | SDK1;SDK1 | | Body;Body | Yes |
| Control-Sample100_Be | chr7:4244250-4244643 | SDK1;SDK1 | | Body;Body | Yes |
| Control-Sample12_Beta | chr7:4266872-4267372 | SDK1;SDK1 | | Body;Body | Yes |
| Control-Sample12_Beta | chr7:4304846-4305535 | SDK1;SDK1 | Island | Body;Body | Yes |
| Control-Sample17_Beta | chr7:4174939-4175200 | SDK1;SDK1 | | Body;Body | Yes |
| Control-Sample19_Beta | chr7:4175403-4175865 | SDK1;SDK1 | | Body;Body | Yes |
| Control-Sample19_Beta | chr7:4218743-4218990 | SDK1;SDK1 | | Body;Body | Yes |
| Control-Sample19_Beta | chr7:4265272-4265529 | SDK1;SDK1 | | Body;Body | Yes |
| Control-Sample20_Beta | chr7:4175655-4175865 | SDK1;SDK1 | | Body;Body | Yes |
| Control-Sample3_Beta | chr7:4266872-4267412 | SDK1;SDK1 | | Body;Body | Yes |
| Control-Sample64_Beta | chr7:4174655-4175403 | SDK1;SDK1 | | Body;Body | Yes |
| Control-Sample65_Beta | chr7:4243526-4243734 | SDK1;SDK1 | | Body;Body | Yes |
| Control-Sample68_Beta | chr7:4210949-4211174 | SDK1;SDK1 | | Body;Body | Yes |
| Control-Sample1_Beta | chr17:71361857-71361901 | SDK2 | | Body | No |
| Control-Sample23_Beta | chr22:30868156-30868226 | SEC14L3 | | TSS200 | No |
| Control-Sample474_Be | chr4:119757509-119757529 | SEC24D | Island | TSS200 | No |
| Control-Sample2_Beta | chr1:151345123-151345181 | SELENBP1;SELENBP1 | | 5'UTR;1stExon | No |
| Control-Sample20_Beta | chr3:53926056-53926101 | SELK | Island | TSS200 | No |
| Control-Sample108_Be | chr7:83824373-83824481 | SEMA3A | | TSS200 | No |
| Control-Sample19_Beta | chr3:50306107-50306456 | SEMA3B;SEMA3B | | 5'UTR;5'UTR | Yes |
| Control-Sample43_Beta | chr3:50312913-50313213 | SEMA3B;SEMA3B | Island | Body;Body | Yes |
| Control-Sample1_Beta | chr3:52479069-52479105 | SEMA3G | | TSS200 | No |
| Control-Sample1_Beta | chr15:90727560-90727783 | SEMA4B | N_Shore | TSS1500 | No |
| Control-Sample648_Be | chr15:74702956-74703101 | SEMA7A;SEMA7A;SEMA7A | N_Shore | 3'UTR;3'UTR;3'UTR | No |
| Control-Sample67_Beta | chr10:13388442-13388567 | SEPHS1 | | 5'UTR | No |
| Control-Sample64_Beta | chr2:110373002-110373509 | SEPT10;SEPT10;ANKRD57 | Island | TSS1500;TSS1500;1stExon | No |
| Control-Sample1_Beta | chr17:75315081-75315244 | SEPT9;SEPT9;SEPT9 | | 5'UTR;Body;TSS1500 | Yes |
| Control-Sample10_Beta | chr17:75315081-75315486 | SEPT9;SEPT9;SEPT9 | | 5'UTR;Body;TSS1500 | Yes |
| Control-Sample10_Beta | chr17:75318655-75318882 | SEPT9;SEPT9;SEPT9 | | 5'UTR;Body;Body | Yes |
| Control-Sample12_Beta | chr17:75315567-75315837 | SEPT9;SEPT9;SEPT9 | | 5'UTR;TSS200;Body | Yes |

| | | | | | |
|---|---|---|---|---|---|
| Control-Sample1_Beta | chr17:75373327-75373457 | SEPT9;SEPT9;SEPT9;SEPT9;SEPT9 | S_Shelf | 5'UTR;5'UTR;Body;Body;Body | Yes |
| Control-Sample216_Be | chr17:75445905-75446450 | SEPT9;SEPT9;SEPT9;SEPT9;SEPT9;SEPT9 | N_Shore | Body;Body;Body;Body;TSS1500;Body | Yes |
| Control-Sample30_Beta | chr17:75473577-75473667 | SEPT9;SEPT9;SEPT9;SEPT9;SEPT9;SEPT9;SEPT9 | | Body;5'UTR;Body;Body;Body;Body;Body | Yes |
| Control-Sample100_Be | chr6:158589294-158589307 | SERAC1;SERAC1;GTF2H5 | Island | 1stExon;5'UTR;TSS200 | Yes |
| Control-Sample150_Be | chr6:158589294-158589318 | SERAC1;SERAC1;GTF2H5 | Island | 1stExon;5'UTR;TSS200 | Yes |
| Control-Sample186_Be | chr6:158589194-158589307 | SERAC1;SERAC1;GTF2H5 | Island | 1stExon;5'UTR;TSS200 | Yes |
| Control-Sample26_Beta | chr6:2842239-2842255 | SERPINB1 | Island | TSS200 | No |
| Control-Sample3_Beta | chr6:2970624-2971087 | SERPINB6 | N_Shore | 5'UTR | No |
| Control-Sample108_Be | chr7:100769903-100770192 | SERPINE1;SERPINE1 | | TSS1500;TSS1500 | No |
| Control-Sample3_Beta | chr11:57364841-57364890 | SERPING1;SERPING1 | | TSS1500;TSS200 | No |
| Control-Sample65_Beta | chr16:29887661-29888096 | SEZ6L2;SEZ6L2;SEZ6L2;SEZ6L2 | N_Shore | Body;Body;Body;Body | No |
| Control-Sample19_Beta | chr8:41167802-41169484 | SFRP1 | S_Shore | TSS1500 | No |
| Control-Sample1_Beta | chr4:154711099-154711906 | SFRP2 | N_Shore | TSS1500 | Yes |
| Control-Sample10_Beta | chr4:154711099-154711183 | SFRP2 | N_Shore | TSS1500 | Yes |
| Control-Sample10_Beta | chr4:154711512-154711906 | SFRP2 | N_Shore | TSS1500 | Yes |
| Control-Sample11_Beta | chr4:154711099-154711347 | SFRP2 | N_Shore | TSS1500 | Yes |
| Control-Sample12_Beta | chr4:154711597-154711906 | SFRP2 | N_Shore | TSS1500 | Yes |
| Control-Sample15_Beta | chr4:154711597-154711675 | SFRP2 | N_Shore | TSS1500 | Yes |
| Control-Sample16_Beta | chr4:154711597-154711629 | SFRP2 | N_Shore | TSS1500 | Yes |
| Control-Sample3_Beta | chr4:154711099-154711457 | SFRP2 | N_Shore | TSS1500 | Yes |
| Control-Sample32_Beta | chr4:154711563-154711906 | SFRP2 | N_Shore | TSS1500 | Yes |
| Control-Sample39_Beta | chr4:154711629-154711906 | SFRP2 | N_Shore | TSS1500 | Yes |
| Control-Sample41_Beta | chr4:154711171-154711347 | SFRP2 | N_Shore | TSS1500 | Yes |
| Control-Sample62_Beta | chr4:154711512-154711675 | SFRP2 | N_Shore | TSS1500 | Yes |
| Control-Sample108_Be | chr12:132278430-132278510 | SFRS8 | S_Shore | Body | Yes |
| Control-Sample19_Beta | chr12:132269563-132269737 | SFRS8 | | Body | Yes |
| Control-Sample20_Beta | chr12:132269563-132270218 | SFRS8 | | Body | Yes |
| Control-Sample33_Beta | chr12:132221123-132221229 | SFRS8 | S_Shore | Body | Yes |
| Control-Sample57_Beta | chr12:132269563-132269687 | SFRS8 | | Body | Yes |
| Control-Sample19_Beta | chr2:128458240-128458281 | SFT2D3 | N_Shore | TSS1500 | No |
| Control-Sample12_Beta | chr7:94285329-94285352 | SGCE;PEG10;SGCE;PEG10;SGCE | Island | 1stExon;TSS1500;1stExon;TSS1500;1stExon | Yes |
| Control-Sample13_Beta | chr7:94285327-94285352 | SGCE;PEG10;SGCE;PEG10;SGCE | Island | 1stExon;TSS1500;1stExon;TSS1500;1stExon | Yes |
| Control-Sample15_Beta | chr7:94285327-94285343 | SGCE;PEG10;SGCE;PEG10;SGCE | Island | 1stExon;TSS1500;1stExon;TSS1500;1stExon | Yes |
| Control-Sample65_Beta | chr7:94285329-94285520 | SGCE;PEG10;SGCE;PEG10;SGCE | Island | 1stExon;TSS1500;1stExon;TSS1500;1stExon | Yes |
| Control-Sample1_Beta | chr6:134498979-134499189 | SGK1;SGK1;SGK1 | S_Shore | Body;1stExon;5'UTR | Yes |
| Control-Sample20_Beta | chr6:134498979-134499419 | SGK1;SGK1;SGK1 | S_Shore | Body;1stExon;5'UTR | Yes |
| Control-Sample1_Beta | chr6:134491143-134491531 | SGK1;SGK1;SGK1;SGK1 | | 3'UTR;3'UTR;3'UTR;3'UTR | Yes |
| Control-Sample13_Beta | chr6:134491163-134491531 | SGK1;SGK1;SGK1;SGK1 | | 3'UTR;3'UTR;3'UTR;3'UTR | Yes |
| Control-Sample1_Beta | chr20:42193947-42194649 | SGK2;SGK2 | | TSS1500;5'UTR | Yes |
| Control-Sample14_Beta | chr20:42194581-42194649 | SGK2;SGK2 | | 5'UTR;TSS200 | Yes |
| Control-Sample171_Be | chr20:42193808-42194649 | SGK2;SGK2 | | TSS1500;5'UTR | Yes |
| Control-Sample223_Be | chr20:42193746-42194649 | SGK2;SGK2 | | TSS1500;5'UTR | Yes |
| Control-Sample65_Beta | chr4:108744891-108745325 | SGMS2 | N_Shore | TSS1500 | No |
| Control-Sample53_Beta | chr10:82297397-82297793 | SH2D4B | S_Shore | TSS1500 | No |
| Control-Sample1_Beta | chr2:235859315-235859792 | SH3BP4 | N_Shore | TSS1500 | Yes |
| Control-Sample13_Beta | chr2:235859315-235859641 | SH3BP4 | N_Shore | TSS1500 | Yes |
| Control-Sample13_Beta | chr2:235907647-235907796 | SH3BP4 | | 5'UTR | Yes |
| Control-Sample19_Beta | chr3:15311021-15311269 | SH3BP5;SH3BP5 | | Body;Body | Yes |
| Control-Sample57_Beta | chr3:15311021-15311183 | SH3BP5;SH3BP5 | | Body;Body | Yes |
| Control-Sample14_Beta | chr17:43510478-43510841 | SH3D20 | S_Shore | TSS200 | No |
| Control-Sample19_Beta | chr5:171808200-171808623 | SH3PXD2B | | Body | No |
| Control-Sample19_Beta | chr2:109892050-109892206 | SH3RF3 | | Body | No |
| Control-Sample85_Beta | chr5:148442857-148443366 | SH3TC2 | | TSS200 | No |
| Control-Sample20_Beta | chr11:70713608-70713937 | SHANK2 | | Body | Yes |
| Control-Sample27_Beta | chr11:70644110-70644185 | SHANK2 | | Body | Yes |
| Control-Sample34_Beta | chr11:70563878-70564293 | SHANK2 | Island | Body | Yes |
| Control-Sample57_Beta | chr11:70563580-70563792 | SHANK2 | N_Shore | Body | Yes |
| Control-Sample58_Beta | chr11:70557445-70557519 | SHANK2 | | Body | Yes |
| Control-Sample58_Beta | chr11:70563264-70563792 | SHANK2 | N_Shore | Body | Yes |
| Control-Sample1_Beta | chr11:70368372-70368768 | SHANK2;SHANK2 | | Body;Body | Yes |
| Control-Sample1_Beta | chr11:70455140-70455304 | SHANK2;SHANK2 | | Body;Body | Yes |
| Control-Sample10_Beta | chr11:70368624-70368768 | SHANK2;SHANK2 | | Body;Body | Yes |
| Control-Sample10_Beta | chr11:70416238-70416309 | SHANK2;SHANK2 | | Body;Body | Yes |
| Control-Sample10_Beta | chr11:70458964-70459053 | SHANK2;SHANK2 | | Body;Body | Yes |
| Control-Sample19_Beta | chr11:70368264-70368372 | SHANK2;SHANK2 | | Body;Body | Yes |
| Control-Sample19_Beta | chr11:70398671-70398862 | SHANK2;SHANK2 | | Body;Body | Yes |
| Control-Sample475_Be | chr15:45479462-45479755 | SHF | N_Shore | Body | No |
| Control-Sample12_Beta | chr7:96339361-96339495 | SHFM1 | S_Shore | TSS200 | No |
| Control-Sample27_Beta | chr1:201857296-201857621 | SHISA4;SHISA4 | | TSS1500;TSS1500 | No |
| Control-Sample1_Beta | chr5:132158625-132159003 | SHROOM1 | Island | Body | Yes |
| Control-Sample12_Beta | chr5:132158625-132158887 | SHROOM1 | Island | Body | Yes |
| Control-Sample1_Beta | chr4:77355817-77356416 | SHROOM3 | | TSS1500 | Yes |
| Control-Sample26_Beta | chr4:77356175-77356416 | SHROOM3 | | TSS200 | Yes |
| Control-Sample1_Beta | chr13:46426026-46426263 | SIAH3 | | TSS200 | Yes |
| Control-Sample12_Beta | chr13:46426202-46426263 | SIAH3 | | TSS1500 | Yes |
| Control-Sample496_Be | chr13:46425929-46426202 | SIAH3 | | TSS200 | Yes |
| Control-Sample623_Be | chr19:52134289-52134393 | SIGLEC5 | | TSS1500 | No |
| Control-Sample103_Be | chr6:100894629-100895211 | SIM1 | N_Shore | Body | Yes |
| Control-Sample103_Be | chr6:100896055-100897445 | SIM1 | Island | Body | Yes |
| Control-Sample1_Beta | chr21:38074992-38075599 | SIM2;SIM2 | N_Shore | Body;Body | No |
| Control-Sample10_Beta | chr2:45233528-45233773 | SIX2 | N_Shore | Body | Yes |
| Control-Sample19_Beta | chr2:45233390-45234140 | SIX2 | S_Shore | Body | Yes |
| Control-Sample1_Beta | chr2:45170322-45170979 | SIX3 | Island | Body | Yes |
| Control-Sample57_Beta | chr2:45170322-45171097 | SIX3 | Island | Body | Yes |
| Control-Sample1_Beta | chr17:46233714-46233945 | SKAP1;SKAP1 | | Body;Body | No |
| Control-Sample1_Beta | chr1:2202931-2203045 | SKI | | Body | Yes |
| Control-Sample1_Beta | chr1:2230640-2230923 | SKI | N_Shore | Body | Yes |
| Control-Sample10_Beta | chr1:2232469-2232527 | SKI | Island | Body | Yes |
| Control-Sample19_Beta | chr1:2191299-2191402 | SKI | | Body | Yes |
| Control-Sample19_Beta | chr1:2222440-2222674 | SKI | Island | Body | Yes |
| Control-Sample36_Beta | chr1:2174725-2174895 | SKI | | Body | Yes |

| | | | | | |
|---|---|---|---|---|---|
| Control-Sample464_Be | chr1:2200962-2201246 | SKI | | Body | Yes |
| Control-Sample465_Be | chr1:2205770-2205902 | SKI | | Body | Yes |
| Control-Sample1_Beta | chr4:48484788-48485090 | SLC10A4 | N_Shore | TSS1500 | No |
| Control-Sample1_Beta | chr16:67997858-67998030 | SLC12A4;SLC12A4;SLC12A4;SLC12A4;SLC12A4 | | Body;Body;1stExon;Body;Body | Yes |
| Control-Sample17_Beta | chr16:67997858-67998164 | SLC12A4;SLC12A4;SLC12A4;SLC12A4;SLC12A4 | | Body;Body;1stExon;Body;Body | Yes |
| Control-Sample33_Beta | chr16:67998030-67998246 | SLC12A4;SLC12A4;SLC12A4;SLC12A4;SLC12A4 | N_Shelf | Body;Body;TSS200;Body;Body | Yes |
| Control-Sample65_Beta | chr16:67997858-67998246 | SLC12A4;SLC12A4;SLC12A4;SLC12A4;SLC12A4 | | Body;Body;1stExon;Body;Body | Yes |
| Control-Sample1_Beta | chr15:34610829-34611069 | SLC12A6;SLC12A6;SLC12A6;SLC12A6;SLC12A6;SLC12A6 | | Body;Body;Body;1stExon;Body;Body | No |
| Control-Sample58_Beta | chr7:100463416-100463583 | SLC12A9 | N_Shore | Body | Yes |
| Control-Sample19_Beta | chr7:100463583-100464145 | SLC12A9;TRIP6 | N_Shore | Body;TSS1500 | Yes |
| Control-Sample1_Beta | chr7:135413101-135413232 | SLC13A4 | | TSS200 | No |
| Control-Sample14_Beta | chr17:80186266-80186336 | SLC16A3;SLC16A3 | Island | TSS200;TSS1500 | Yes |
| Control-Sample1_Beta | chr17:80190129-80190154 | SLC16A3;SLC16A3;SLC16A3 | S_Shore | TSS200;5'UTR;5'UTR | Yes |
| Control-Sample11_Beta | chr17:73083876-73083887 | SLC16A5 | Island | TSS200 | No |
| Control-Sample11_Beta | chr22:38478322-38478787 | SLC16A8 | Island | Body | No |
| Control-Sample108_Be | chr12:100750473-100750702 | SLC17A8;SLC17A8 | | TSS1500;TSS1500 | No |
| Control-Sample108_Be | chr5:36606288-36606446 | SLC1A3;SLC1A3 | | TSS200;TSS200 | Yes |
| Control-Sample568_Be | chr5.0:36606293-36606446 | SLC1A3;SLC1A3 | | TSS200;TSS200 | Yes |
| Control-Sample1_Beta | chr2:65217528-65217623 | SLC1A4;SLC1A4 | S_Shore | Body;5'UTR | No |
| Control-Sample1_Beta | chr19:47288039-47288109 | SLC1A5;SLC1A5;SLC1A5;SLC1A5 | N_Shelf | 1stExon;5'UTR;Body;5'UTR | No |
| Control-Sample19_Beta | chr1:53608163-53608367 | SLC1A7;SLC1A7 | | 1stExon;5'UTR | No |
| Control-Sample44_Beta | chr11:2930485-2930995 | SLC22A18;SLC22A18 | Island | Body;Body | No |
| Control-Sample1_Beta | chr11:2920052-2920209 | SLC22A18AS;SLC22A18 | N_Shelf | Body;TSS1500 | Yes |
| Control-Sample20_Beta | chr11:2920052-2920146 | SLC22A18AS;SLC22A18 | N_Shelf | Body;TSS1500 | Yes |
| Control-Sample48_Beta | chr11:2920052-2920189 | SLC22A18AS;SLC22A18 | N_Shelf | Body;TSS1500 | Yes |
| Control-Sample97_Beta | chr11:2920146-2920209 | SLC22A18AS;SLC22A18 | N_Shelf | Body;TSS1500 | Yes |
| Control-Sample1_Beta | chr11:62783123-62783361 | SLC22A8 | | 5'UTR | No |
| Control-Sample1_Beta | chr5:138714136-138714337 | SLC23A1;SLC23A1 | Island | Body;Body | Yes |
| Control-Sample3_Beta | chr5:138713954-138714322 | SLC23A1;SLC23A1 | Island | Body;Body | Yes |
| Control-Sample5_Beta | chr5:138713954-138714337 | SLC23A1;SLC23A1 | Island | Body;Body | Yes |
| Control-Sample108_Be | chr22:18042646-18043043 | SLC25A18 | | TSS1500 | Yes |
| Control-Sample6_Beta | chr22:18042550-18042722 | SLC25A18 | | TSS1500 | Yes |
| Control-Sample1_Beta | chr11:796421-796607 | SLC25A22 | N_Shore | TSS1500 | Yes |
| Control-Sample10_Beta | chr11:794433-794764 | SLC25A22 | N_Shelf | Body | Yes |
| Control-Sample12_Beta | chr11:794433-794834 | SLC25A22 | N_Shelf | Body | Yes |
| Control-Sample20_Beta | chr11:796434-796607 | SLC25A22 | N_Shore | TSS1500 | Yes |
| Control-Sample122_Be | chr1:108734992-108735549 | SLC25A24;SLC25A24 | N_Shore | Body;Body | Yes |
| Control-Sample134_Be | chr1:108735312-108735549 | SLC25A24;SLC25A24;SLC25A24 | Island | 5'UTR;Body;1stExon | Yes |
| Control-Sample504_Be | chr1.0:108735312-108735549 | SLC25A24;SLC25A24;SLC25A24 | Island | 5'UTR;Body;1stExon | Yes |
| Control-Sample1_Beta | chr4:979022-979183 | SLC26A1 | N_Shore | Body | No |
| Control-Sample1_Beta | chr12:58013109-58013915 | SLC26A10 | N_Shore | TSS1500 | Yes |
| Control-Sample13_Beta | chr12:58013109-58013651 | SLC26A10 | N_Shore | TSS1500 | Yes |
| Control-Sample20_Beta | chr12:58013174-58013267 | SLC26A10 | Island | TSS1500 | Yes |
| Control-Sample21_Beta | chr12:58013109-58013458 | SLC26A10 | N_Shore | TSS1500 | Yes |
| Control-Sample21_Beta | chr12:58013487-58013915 | SLC26A10 | Island | TSS1500 | Yes |
| Control-Sample42_Beta | chr12:58013109-58013539 | SLC26A10 | N_Shore | TSS1500 | Yes |
| Control-Sample42_Beta | chr12:58013636-58013915 | SLC26A10 | Island | TSS200 | Yes |
| Control-Sample65_Beta | chr12:58013109-58013267 | SLC26A10 | N_Shore | TSS1500 | Yes |
| Control-Sample65_Beta | chr12:58013324-58013915 | SLC26A10 | Island | TSS1500 | Yes |
| Control-Sample1_Beta | chr7:107300939-107301048 | SLC26A4;LOC286002 | N_Shore | TSS200;Body | Yes |
| Control-Sample19_Beta | chr7:107300998-107301048 | SLC26A4;LOC286002 | N_Shore | TSS200;Body | Yes |
| Control-Sample1_Beta | chr7:103085736-103085984 | SLC26A5;SLC26A5;SLC26A5;SLC26A5;SLC26A5 | Island | 5'UTR;5'UTR;5'UTR;5'UTR;5'UTR | Yes |
| Control-Sample6_Beta | chr7:103085736-103086074 | SLC26A5;SLC26A5;SLC26A5;SLC26A5;SLC26A5 | Island | 5'UTR;5'UTR;5'UTR;5'UTR;5'UTR | Yes |
| Control-Sample171_Be | chr1:205897545-205897890 | SLC26A9;SLC26A9;SLC26A9 | | Body;Body;TSS200 | No |
| Control-Sample1_Beta | chr19:17608110-17608394 | SLC27A1 | Island | Body | No |
| Control-Sample291_Be | chr12:8088931-8089207 | SLC2A3 | | TSS200 | No |
| Control-Sample10_Beta | chr1:9129646-9129791 | SLC2A5;SLC2A5;SLC2A5;SLC2A5;SLC2A5 | N_Shore | 5'UTR;5'UTR;1stExon;1stExon;Body | No |
| Control-Sample23_Beta | chr4:25656830-25657160 | SLC34A2 | N_Shore | TSS1500 | No |
| Control-Sample3_Beta | chr1:100435035-100435258 | SLC35A3 | N_Shore | TSS1500 | No |
| Control-Sample467_Be | chr6:8436218-8436466 | SLC35B3;SLC35B3;SLC35B3 | S_Shore | TSS1500;TSS1500;TSS1500 | Yes |
| Control-Sample5_Beta | chr6:8435883-8435919 | SLC35B3;SLC35B3;SLC35B3 | Island | TSS200;TSS200;TSS200 | Yes |
| Control-Sample1_Beta | chr11:45825448-45825485 | SLC35C1;SLC35C1;SLC35C1 | N_Shore | TSS200;TSS1500;TSS200 | Yes |
| Control-Sample11_Beta | chr11:45825448-45825483 | SLC35C1;SLC35C1;SLC35C1 | N_Shore | TSS200;TSS1500;TSS200 | Yes |
| Control-Sample18_Beta | chr11:45825469-45825485 | SLC35C1;SLC35C1;SLC35C1 | N_Shore | TSS200;TSS1500;TSS200 | Yes |
| Control-Sample19_Beta | chr21:43919671-43919853 | SLC37A1 | S_Shelf | TSS200 | No |
| Control-Sample19_Beta | chr12:46663660-46664096 | SLC38A1;SLC38A1 | S_Shore | TSS1500;TSS1500 | No |
| Control-Sample10_Beta | chr17:79225309-79225857 | SLC38A10;SLC38A10 | | Body;Body | Yes |
| Control-Sample10_Beta | chr17:79253785-79253990 | SLC38A10;SLC38A10 | N_Shore | Body;Body | Yes |
| Control-Sample11_Beta | chr17:79253479-79253630 | SLC38A10;SLC38A10 | N_Shore | Body;Body | Yes |
| Control-Sample17_Beta | chr17:79253479-79254229 | SLC38A10;SLC38A10 | N_Shore | Body;Body | Yes |
| Control-Sample23_Beta | chr17:79253479-79253990 | SLC38A10;SLC38A10 | N_Shore | Body;Body | Yes |
| Control-Sample34_Beta | chr17:79225309-79225974 | SLC38A10;SLC38A10 | | Body;Body | Yes |
| Control-Sample4_Beta | chr17:79253479-79253888 | SLC38A10;SLC38A10 | N_Shore | Body;Body | Yes |
| Control-Sample65_Beta | chr17:79225573-79225857 | SLC38A10;SLC38A10 | | Body;Body | Yes |
| Control-Sample1_Beta | chr12:46767928-46768016 | SLC38A2 | S_Shore | TSS1500 | Yes |
| Control-Sample10_Beta | chr12:46767665-46768016 | SLC38A2 | S_Shore | TSS1500 | Yes |
| Control-Sample21_Beta | chr12:46767665-46767943 | SLC38A2 | S_Shore | TSS1500 | Yes |
| Control-Sample19_Beta | chr8:22223937-22223985 | SLC39A14;SLC39A14;SLC39A14;SLC39A14 | N_Shore | TSS1500;TSS1500;TSS1500;TSS1500 | Yes |
| Control-Sample20_Beta | chr8:22223852-22223985 | SLC39A14;SLC39A14;SLC39A14;SLC39A14 | N_Shore | TSS1500;TSS1500;TSS1500;TSS1500 | Yes |
| Control-Sample1_Beta | chr14:21467273-21467375 | SLC39A2 | | TSS200 | No |
| Control-Sample648_Be | chr8:145638881-145639181 | SLC39A4;SLC39A4 | Island | Body;Body | No |
| Control-Sample1_Beta | chr3:125775613-125775696 | SLC41A3;SLC41A3;SLC41A3;SLC41A3;SLC41A3;SLC41A3 | | Body;Body;5'UTR;Body;1stExon;5'UTR | Yes |
| Control-Sample11_Beta | chr3:125775413-125775696 | SLC41A3;SLC41A3;SLC41A3;SLC41A3;SLC41A3;SLC41A3 | | Body;Body;5'UTR;Body;1stExon;5'UTR | Yes |
| Control-Sample72_Beta | chr3:125775413-125775693 | SLC41A3;SLC41A3;SLC41A3;SLC41A3;SLC41A3;SLC41A3 | | Body;Body;5'UTR;Body;1stExon;5'UTR | Yes |
| Control-Sample5_Beta | chr17:1509953-1510041 | SLC43A2 | Island | Body | No |
| Control-Sample11_Beta | chr19:10736117-10736448 | SLC44A2;SLC44A2 | Island | TSS200;Body | Yes |
| Control-Sample3_Beta | chr19:10736117-10736355 | SLC44A2;SLC44A2 | Island | TSS200;Body | Yes |
| Control-Sample17_Beta | chr19:10736299-10736448 | SLC44A2;SLC44A2;SLC44A2 | Island | 5'UTR;1stExon;Body | Yes |
| Control-Sample20_Beta | chr6:31833153-31833199 | SLC44A4 | S_Shelf | Body | No |
| Control-Sample11_Beta | chr8:142238691-142238770 | SLC45A4 | S_Shore | TSS200 | Yes |
| Control-Sample1_Beta | chr8:142238504-142238770 | SLC45A4;SLC45A4 | S_Shore | 5'UTR;1stExon | Yes |

| | | | | | |
|---|---|---|---|---|---|
| Control-Sample6_Beta | chr17:26732249-26732374 | SLC46A1 | Island | Body | No |
| Control-Sample1_Beta | chr20:3218476-3218794 | SLC4A11 | N_Shore | TSS200 | Yes |
| Control-Sample12_Beta | chr20:3218393-3218579 | SLC4A11 | N_Shore | TSS200 | Yes |
| Control-Sample13_Beta | chr20:3218476-3218579 | SLC4A11 | N_Shore | TSS200 | Yes |
| Control-Sample19_Beta | chr20:3218476-3218500 | SLC4A11 | N_Shore | TSS200 | Yes |
| Control-Sample2_Beta | chr20:3218145-3218794 | SLC4A11 | N_Shore | Body | Yes |
| Control-Sample24_Beta | chr20:3218393-3218794 | SLC4A11 | N_Shore | TSS200 | Yes |
| Control-Sample56_Beta | chr20:3218145-3218579 | SLC4A11 | N_Shore | Body | Yes |
| Control-Sample20_Beta | chr20:3218331-3218500 | SLC4A11;SLC4A11 | N_Shore | 1stExon;5'UTR | Yes |
| Control-Sample3_Beta | chr22:32438947-32439391 | SLC5A1 | N_Shore | TSS200 | Yes |
| Control-Sample57_Beta | chr22:32438947-32439052 | SLC5A1 | N_Shore | TSS200 | Yes |
| Control-Sample58_Beta | chr22:32438947-32439254 | SLC5A1 | N_Shore | TSS200 | Yes |
| Control-Sample1_Beta | chr3:11078979-11079349 | SLC6A1 | | 3'UTR | Yes |
| Control-Sample11_Beta | chr3:11078979-11079298 | SLC6A1 | | 3'UTR | Yes |
| Control-Sample19_Beta | chr3:11079128-11079349 | SLC6A1 | | 3'UTR | Yes |
| Control-Sample22_Beta | chr3:11078979-11079491 | SLC6A1 | | 3'UTR | Yes |
| Control-Sample41_Beta | chr12:312592-313104 | SLC6A12;SLC6A12;SLC6A12 | Island | Body;Body;Body | No |
| Control-Sample1_Beta | chr12:372049-372191 | SLC6A13 | | TSS200 | No |
| Control-Sample20_Beta | chr12:85307131-85307424 | SLC6A15;SLC6A15;SLC6A15 | S_Shore | TSS1500;TSS1500;TSS1500 | No |
| Control-Sample1_Beta | chr1:44497537-44497694 | SLC6A9 | S_Shore | TSS1500 | Yes |
| Control-Sample10_Beta | chr1:44482420-44482918 | SLC6A9;SLC6A9;SLC6A9 | | Body;Body;Body | Yes |
| Control-Sample108_Be | chr4:139163341-139163535 | SLC7A11;SLC7A11 | | 1stExon;5'UTR | No |
| Control-Sample53_Beta | chr2:40739661-40740063 | SLC8A1 | | TSS200 | Yes |
| Control-Sample54_Beta | chr2:40739661-40739938 | SLC8A1 | | TSS200 | Yes |
| Control-Sample10_Beta | chr2:40739515-40740063 | SLC8A1;SLC8A1 | | 5'UTR;1stExon | Yes |
| Control-Sample171_Be | chr2:40739515-40739771 | SLC8A1;SLC8A1 | | 5'UTR;1stExon | Yes |
| Control-Sample52_Beta | chr2:40739515-40739938 | SLC8A1;SLC8A1 | | 5'UTR;1stExon | Yes |
| Control-Sample108_Be | chr3:112012867-112013130 | SLC9A10 | | 5'UTR | No |
| Control-Sample1_Beta | chr16:2082689-2083168 | SLC9A3R2;SLC9A3R2 | N_Shelf | Body;Body | Yes |
| Control-Sample66_Beta | chr16:2082777-2083168 | SLC9A3R2;SLC9A3R2 | N_Shelf | Body;Body | Yes |
| Control-Sample19_Beta | chr15:92570771-92570834 | SLCO3A1;SLCO3A1 | | Body;Body | No |
| Control-Sample114_Be | chr17:33759957-33759986 | SLFN12 | | TSS1500 | Yes |
| Control-Sample114_Be | chr17:33759512-33759648 | SLFN12;SLFN12 | | 1stExon;5'UTR | Yes |
| Control-Sample47_Beta | chr17:33759484-33759573 | SLFN12;SLFN12 | | 1stExon;5'UTR | Yes |
| Control-Sample68_Beta | chr17:33759484-33759648 | SLFN12;SLFN12 | | 1stExon;5'UTR | Yes |
| Control-Sample13_Beta | chr5:168271731-168271859 | SLIT3 | | Body | Yes |
| Control-Sample65_Beta | chr5:168271613-168271859 | SLIT3 | | Body | Yes |
| Control-Sample10_Beta | chr11:107583339-107583406 | SLN | | TSS1500 | Yes |
| Control-Sample43_Beta | chr11:107582884-107583406 | SLN | | TSS200 | Yes |
| Control-Sample1_Beta | chr15:67417860-67418316 | SMAD3;SMAD3 | | TSS200;Body | Yes |
| Control-Sample21_Beta | chr15:67417860-67418428 | SMAD3;SMAD3 | | TSS200;Body | Yes |
| Control-Sample23_Beta | chr15:67457967-67458398 | SMAD3;SMAD3;SMAD3;SMAD3 | | Body;Body;TSS1500;Body | Yes |
| Control-Sample34_Beta | chr15:67458079-67458398 | SMAD3;SMAD3;SMAD3;SMAD3 | | Body;Body;TSS1500;Body | Yes |
| Control-Sample1_Beta | chr18:46447737-46447840 | SMAD7 | Island | 3'UTR | No |
| Control-Sample1_Beta | chr9:2047160-2047595 | SMARCA2;SMARCA2 | N_Shore | Body;Body | No |
| Control-Sample499_Be | chr6:168963358-168963731 | SMOC2;SMOC2 | | Body;Body | Yes |
| Control-Sample96_Beta | chr6:169001953-169002120 | SMOC2;SMOC2 | | Body;Body | Yes |
| Control-Sample35_Beta | chr7:98739496-98739782 | SMURF1;SMURF1 | N_Shore | Body;Body | Yes |
| Control-Sample36_Beta | chr7:98739627-98739782 | SMURF1;SMURF1 | N_Shore | Body;Body | Yes |
| Control-Sample56_Beta | chr7:98739496-98739698 | SMURF1;SMURF1 | N_Shore | Body;Body | Yes |
| Control-Sample64_Beta | chr1:214477063-214477280 | SMYD2 | | Body | No |
| Control-Sample80_Beta | chr1:246363574-246363697 | SMYD3;SMYD3 | | Body;Body | No |
| Control-Sample124_Be | chr19:51107558-51107781 | SNAR-F | Island | TSS1500 | Yes |
| Control-Sample543_Be | chr19.0:51107558-51107781 | SNAR-F | Island | TSS1500 | Yes |
| Control-Sample216_Be | chr7:127670570-127670993 | SND1;LRRC4 | N_Shore | Body;Body | No |
| Control-Sample12_Beta | chr2:241975971-241976244 | SNED1 | Island | Body | Yes |
| Control-Sample57_Beta | chr2:241975971-241976658 | SNED1 | Island | Body | Yes |
| Control-Sample1_Beta | chr6:31590513-31590586 | SNORA38;BAT2 | S_Shore | TSS1500;5'UTR | Yes |
| Control-Sample10_Beta | chr6:31590513-31590622 | SNORA38;BAT2 | S_Shore | TSS1500;5'UTR | Yes |
| Control-Sample11_Beta | chr6:31590563-31590586 | SNORA38;BAT2 | S_Shore | TSS1500;5'UTR | Yes |
| Control-Sample17_Beta | chr6:31590513-31590580 | SNORA38;BAT2 | S_Shore | TSS1500;5'UTR | Yes |
| Control-Sample23_Beta | chr6:31590563-31590580 | SNORA38;BAT2 | S_Shore | TSS1500;5'UTR | Yes |
| Control-Sample47_Beta | chr6:31590576-31590586 | SNORA38;BAT2 | S_Shore | TSS1500;Body | Yes |
| Control-Sample62_Beta | chr6:31590513-31590576 | SNORA38;BAT2 | S_Shore | TSS1500;5'UTR | Yes |
| Control-Sample45_Beta | chr3:186505240-186505713 | SNORA4;EIF4A2 | S_Shelf | TSS200;Body | No |
| Control-Sample171_Be | chr5:9547468-9547825 | SNORD123;SEMA5A | S_Shore | TSS1500;TSS1500 | Yes |
| Control-Sample234_Be | chr5:9547595-9548098 | SNORD123;SEMA5A | S_Shore | TSS1500;TSS1500 | Yes |
| Control-Sample65_Beta | chr11:62620843-62621406 | SNORD22;SNORD31;SNHG1 | N_Shelf | TSS1500;Body;Body | Yes |
| Control-Sample5_Beta | chr11:62621178-62621406 | SNORD22;SNORD31;SNORD30;SNHG1 | N_Shelf | TSS1500;TSS1500;Body;Body | Yes |
| Control-Sample1_Beta | chr7:22894795-22894849 | SNORD93 | S_Shore | TSS1500 | No |
| Control-Sample475_Be | chr20:1246715-1246772 | SNPH | Island | TSS1500 | No |
| Control-Sample1_Beta | chr12:96251912-96252105 | SNRPF | N_Shore | TSS1500 | Yes |
| Control-Sample22_Beta | chr12:96251729-96252105 | SNRPF | N_Shore | TSS1500 | Yes |
| Control-Sample26_Beta | chr12:96252066-96252176 | SNRPF | Island | TSS1500 | Yes |
| Control-Sample259_Be | chr15:25068738-25068850 | SNRPN;SNRPN;SNRPN | | TSS200;TSS200;TSS200 | Yes |
| Control-Sample260_Be | chr15:25068738-25068769 | SNRPN;SNRPN;SNRPN | | TSS200;TSS200;TSS200 | Yes |
| Control-Sample576_Be | chr15.0:25068738-25068769 | SNRPN;SNRPN;SNRPN | | TSS200;TSS200;TSS200 | Yes |
| Control-Sample649_Be | chr15:25068564-25068850 | SNRPN;SNRPN;SNRPN | | TSS1500;TSS1500;TSS1500 | Yes |
| Control-Sample1_Beta | chr2:947515-947634 | SNTG2 | S_Shore | Body | No |
| Control-Sample1_Beta | chr16:12529907-12530246 | SNX29 | | Body | Yes |
| Control-Sample270_Be | chr16:12183850-12184067 | SNX29 | | Body | Yes |
| Control-Sample15_Beta | chr6:108582894-108583071 | SNX3;SNX3 | S_Shore | TSS1500;TSS1500 | No |
| Control-Sample1_Beta | chr8:101661245-101661533 | SNX31 | N_Shore | Body | No |
| Control-Sample19_Beta | chr7:2316906-2317059 | SNX8 | N_Shore | Body | Yes |
| Control-Sample19_Beta | chr7:2328746-2329077 | SNX8 | | Body | Yes |
| Control-Sample194_Be | chr12:93966978-93967711 | SOCS2 | Island | Body | No |
| Control-Sample3_Beta | chr10:97175272-97175479 | SORBS1;SORBS1;SORBS1;SORBS1;SORBS1;SORBS1;SORBS1 | | Body;Body;Body;Body;Body;Body;Body | No |
| Control-Sample1_Beta | chr4:186741362-186741887 | SORBS2 | | 5'UTR | Yes |
| Control-Sample13_Beta | chr4:186741447-186741887 | SORBS2 | | 5'UTR | Yes |
| Control-Sample51_Beta | chr4:186741362-186741788 | SORBS2 | | 5'UTR | Yes |
| Control-Sample19_Beta | chr4:186877497-186877916 | SORBS2;SORBS2 | | 5'UTR;1stExon | Yes |

| | | | | | |
|---|---|---|---|---|---|
| Control-Sample2_Beta | chr4:186877497-186877657 | SORBS2;SORBS2 | | 5'UTR;1stExon | Yes |
| Control-Sample24_Beta | chr4:186877497-186877601 | SORBS2;SORBS2 | | 5'UTR;1stExon | Yes |
| Control-Sample108_Be | chr4:186732124-186733060 | SORBS2;SORBS2;SORBS2;SORBS2;SORBS2;SORBS2 | | 5'UTR;5'UTR;5'UTR;5'UTR;1stExon;TSS200 | Yes |
| Control-Sample291_Be | chr4:186732942-186733331 | SORBS2;SORBS2;SORBS2;SORBS2;SORBS2;SORBS2;SORBS2 | | 5'UTR;5'UTR;5'UTR;1stExon;TSS1500;1stExon;TSS15 | Yes |
| Control-Sample11_Beta | chr4:186508724-186509045 | SORBS2;SORBS2;SORBS2;SORBS2;SORBS2;SORBS2;SORBS2;SORBS2 | | 3'UTR;3'UTR;3'UTR;3'UTR;3'UTR;3'UTR;3'UTR;3'UTR | Yes |
| Control-Sample1_Beta | chr8:22423014-22423055 | SORBS3;SORBS3 | Island | TSS200;Body | Yes |
| Control-Sample11_Beta | chr8:22423014-22423026 | SORBS3;SORBS3 | Island | TSS200;Body | Yes |
| Control-Sample194_Be | chr8:22423020-22423055 | SORBS3;SORBS3 | Island | TSS200;Body | Yes |
| Control-Sample26_Beta | chr8:22422816-22423055 | SORBS3;SORBS3 | Island | TSS1500;Body | Yes |
| Control-Sample46_Beta | chr8:22422816-22423026 | SORBS3;SORBS3 | Island | TSS1500;Body | Yes |
| Control-Sample585_Be | chr8.0:22423014-22423026 | SORBS3;SORBS3 | Island | TSS200;Body | Yes |
| Control-Sample1_Beta | chr4:7228268-7228442 | SORCS2 | | Body | Yes |
| Control-Sample1_Beta | chr4:7373022-7373088 | SORCS2 | | Body | Yes |
| Control-Sample1_Beta | chr4:7657070-7657481 | SORCS2 | | Body | Yes |
| Control-Sample17_Beta | chr4:7657070-7657359 | SORCS2 | | Body | Yes |
| Control-Sample19_Beta | chr4:7282819-7283184 | SORCS2 | | Body | Yes |
| Control-Sample19_Beta | chr4:7373022-7373451 | SORCS2 | | Body | Yes |
| Control-Sample20_Beta | chr4:7395027-7395538 | SORCS2 | | Body | Yes |
| Control-Sample21_Beta | chr4:7593437-7593585 | SORCS2 | Island | Body | Yes |
| Control-Sample24_Beta | chr4:7301491-7301594 | SORCS2 | | Body | Yes |
| Control-Sample28_Beta | chr4:7657303-7657481 | SORCS2 | | Body | Yes |
| Control-Sample29_Beta | chr4:7300999-7301594 | SORCS2 | | Body | Yes |
| Control-Sample3_Beta | chr4:7373022-7373375 | SORCS2 | | Body | Yes |
| Control-Sample30_Beta | chr4:7300999-7301529 | SORCS2 | | Body | Yes |
| Control-Sample331_Be | chr4:7374582-7374669 | SORCS2 | | Body | Yes |
| Control-Sample5_Beta | chr4:7719899-7720084 | SORCS2 | | Body | Yes |
| Control-Sample58_Beta | chr4:7287492-7287737 | SORCS2 | | Body | Yes |
| Control-Sample58_Beta | chr4:7300999-7301854 | SORCS2 | | Body | Yes |
| Control-Sample63_Beta | chr4:7657303-7657708 | SORCS2 | | Body | Yes |
| Control-Sample13_Beta | chr15:45314789-45314933 | SORD | N_Shore | TSS1500 | Yes |
| Control-Sample314_Be | chr15:45315313-45315430 | SORD | Island | TSS200 | Yes |
| Control-Sample557_Be | chr15.0:45315313-45315430 | SORD | Island | TSS200 | Yes |
| Control-Sample20_Beta | chr15:45315419-45315438 | SORD;SORD | Island | 5'UTR;1stExon | Yes |
| Control-Sample95_Beta | chr15:45315362-45315430 | SORD;SORD | Island | 5'UTR;1stExon | Yes |
| Control-Sample96_Beta | chr15:45315362-45315438 | SORD;SORD | Island | 5'UTR;1stExon | Yes |
| Control-Sample11_Beta | chr17:41836261-41836563 | SOST | S_Shelf | TSS200 | No |
| Control-Sample1_Beta | chr7:16505094-16505602 | SOSTDC1 | | 1stExon | Yes |
| Control-Sample10_Beta | chr7:16505589-16505602 | SOSTDC1 | | TSS200 | Yes |
| Control-Sample2_Beta | chr7:16505094-16505589 | SOSTDC1 | | 1stExon | Yes |
| Control-Sample38_Beta | chr7:16505409-16505602 | SOSTDC1;SOSTDC1 | | 1stExon;5'UTR | Yes |
| Control-Sample496_Be | chr7:16505409-16505664 | SOSTDC1;SOSTDC1 | | 1stExon;5'UTR | Yes |
| Control-Sample1_Beta | chr13:112725605-112726058 | SOX1;SOX1 | N_Shore | 1stExon;3'UTR | Yes |
| Control-Sample11_Beta | chr13:112725605-112726124 | SOX1;SOX1 | N_Shore | 1stExon;3'UTR | Yes |
| Control-Sample16_Beta | chr13:112725605-112726482 | SOX1;SOX1 | N_Shore | 1stExon;3'UTR | Yes |
| Control-Sample331_Be | chr22:38380439-38380983 | SOX10;SOX10 | S_Shore | 1stExon;5'UTR | No |
| Control-Sample10_Beta | chr3:181421427-181421703 | SOX2OT | | Body | Yes |
| Control-Sample18_Beta | chr3:181414139-181414848 | SOX2OT | S_Shore | Body | Yes |
| Control-Sample19_Beta | chr3:181428462-181428547 | SOX2OT;SOX2 | N_Shore | Body;TSS1500 | Yes |
| Control-Sample20_Beta | chr3:181428242-181428307 | SOX2OT;SOX2 | N_Shore | Body;TSS1500 | Yes |
| Control-Sample20_Beta | chr3:181428466-181428563 | SOX2OT;SOX2 | N_Shore | Body;TSS1500 | Yes |
| Control-Sample464_Be | chr3:181428242-181428377 | SOX2OT;SOX2 | N_Shore | Body;TSS1500 | Yes |
| Control-Sample1_Beta | chr11:16424673-16424935 | SOX6;SOX6;SOX6;SOX6 | | TSS1500;5'UTR;5'UTR;TSS1500 | No |
| Control-Sample1_Beta | chr8:10586678-10586959 | SOX7 | Island | Body | Yes |
| Control-Sample10_Beta | chr8:10586678-10587055 | SOX7 | Island | Body | Yes |
| Control-Sample13_Beta | chr8:10586796-10586959 | SOX7 | Island | Body | Yes |
| Control-Sample14_Beta | chr8:10586678-10587305 | SOX7 | Island | Body | Yes |
| Control-Sample80_Beta | chr8:10586678-10586882 | SOX7 | Island | Body | Yes |
| Control-Sample17_Beta | chr17:70120318-70120541 | SOX9 | Island | Body | No |
| Control-Sample108_Be | chr2:231084523-231084965 | SP110;SP110;SP110 | | 5'UTR;5'UTR;5'UTR | No |
| Control-Sample1_Beta | chr2:171574141-171574592 | SP5 | S_Shore | 3'UTR | No |
| Control-Sample1_Beta | chr17:45927893-45928250 | SP6 | N_Shore | 5'UTR | No |
| Control-Sample13_Beta | chr12:53729629-53730163 | SP7 | | TSS200 | Yes |
| Control-Sample14_Beta | chr12:53729629-53730277 | SP7 | | TSS200 | Yes |
| Control-Sample16_Beta | chr12:53729659-53730277 | SP7 | | TSS200 | Yes |
| Control-Sample25_Beta | chr12:53730072-53730277 | SP7 | | TSS1500 | Yes |
| Control-Sample65_Beta | chr12:53729629-53730072 | SP7 | | TSS200 | Yes |
| Control-Sample125_Be | chr8:101225078-101225344 | SPAG1;SPAG1 | Island | Body;Body | Yes |
| Control-Sample126_Be | chr8:101225078-101225902 | SPAG1;SPAG1 | Island | Body;Body | Yes |
| Control-Sample552_Be | chr8.0:101225078-101225344 | SPAG1;SPAG1 | Island | Body;Body | Yes |
| Control-Sample3_Beta | chr1:118727840-118727908 | SPAG17;SPAG17 | Island | 5'UTR;1stExon | No |
| Control-Sample38_Beta | chr17:26926738-26926868 | SPAG5 | S_Shore | TSS1500 | No |
| Control-Sample171_Be | chr4:88450824-88451496 | SPARCL1;SPARCL1 | | TSS200;TSS200 | Yes |
| Control-Sample19_Beta | chr4:88451223-88451496 | SPARCL1;SPARCL1 | | TSS1500;TSS1500 | Yes |
| Control-Sample11_Beta | chr13:24736095-24736569 | SPATA13;MIR2276;SPATA13 | S_Shore | 5'UTR;TSS1500;5'UTR | Yes |
| Control-Sample13_Beta | chr13:24736095-24736518 | SPATA13;MIR2276;SPATA13 | S_Shore | 5'UTR;TSS1500;5'UTR | Yes |
| Control-Sample108_Be | chr13:24844846-24845891 | SPATA13;SPATA13 | | Body;Body | Yes |
| Control-Sample296_Be | chr4:52918457-52918489 | SPATA18 | S_Shore | Body | No |
| Control-Sample19_Beta | chr20:48530082-48530654 | SPATA2;SPATA2;SPATA2 | N_Shore | 5'UTR;1stExon;5'UTR | Yes |
| Control-Sample80_Beta | chr20:48530082-48530454 | SPATA2;SPATA2;SPATA2 | N_Shore | 5'UTR;1stExon;5'UTR | Yes |
| Control-Sample15_Beta | chr2:220348312-220348803 | SPEG | N_Shore | Body | Yes |
| Control-Sample2_Beta | chr2:220299604-220299653 | SPEG | Island | TSS200 | Yes |
| Control-Sample5_Beta | chr2:220348312-220348592 | SPEG | N_Shore | Body | Yes |
| Control-Sample52_Beta | chr2:220299584-220299653 | SPEG | Island | TSS200 | Yes |
| Control-Sample104_Be | chr13:36944433-36944451 | SPG20 | | TSS200 | Yes |
| Control-Sample108_Be | chr13:36944433-36944471 | SPG20 | | TSS200 | Yes |
| Control-Sample124_Be | chr13:36944451-36944471 | SPG20 | | TSS200 | Yes |
| Control-Sample14_Beta | chr13:36944469-36944649 | SPG20 | | TSS200 | Yes |
| Control-Sample170_Be | chr13:36944469-36944640 | SPG20 | | TSS200 | Yes |
| Control-Sample463_Be | chr13:36944451-36944640 | SPG20 | | TSS200 | Yes |
| Control-Sample57_Beta | chr13:36944443-36944649 | SPG20 | | TSS200 | Yes |
| Control-Sample585_Be | chr13.0:36944451-36944640 | SPG20 | | TSS200 | Yes |

| | | | | | |
|---|---|---|---|---|---|
| Control-Sample151_Be | chr13:36944294-36944471 | SPG20;SPG20 | | 1stExon;5'UTR | Yes |
| Control-Sample171_Be | chr13:36944294-36944640 | SPG20;SPG20 | | 1stExon;5'UTR | Yes |
| Control-Sample475_Be | chr13:36944294-36944649 | SPG20;SPG20 | | 1stExon;5'UTR | Yes |
| Control-Sample88_Beta | chr13:36944294-36944451 | SPG20;SPG20 | | 1stExon;5'UTR | Yes |
| Control-Sample1_Beta | chr1:32281460-32281703 | SPOCD1 | | 5'UTR | Yes |
| Control-Sample12_Beta | chr1:32281460-32281665 | SPOCD1 | | 5'UTR | Yes |
| Control-Sample461_Be | chr4:1161157-1161319 | SPON2;SPON2 | Island | 3'UTR;3'UTR | Yes |
| Control-Sample467_Be | chr4:1161247-1161597 | SPON2;SPON2 | Island | 3'UTR;3'UTR | Yes |
| Control-Sample5_Beta | chr4:1161157-1161597 | SPON2;SPON2 | Island | 3'UTR;3'UTR | Yes |
| Control-Sample585_Be | chr4.0:1161319-1161653 | SPON2;SPON2 | Island | Body;Body | Yes |
| Control-Sample10_Beta | chr2:65593908-65594021 | SPRED2;SPRED2;SPRED2 | | 5'UTR;1stExon;Body | No |
| Control-Sample14_Beta | chr13:80911626-80911808 | SPRY2 | N_Shore | Body | No |
| Control-Sample62_Beta | chr1:9380566-9380800 | SPSB1 | | 5'UTR | No |
| Control-Sample10_Beta | chr3:140813769-140814055 | SPSB4 | N_Shore | Body | No |
| Control-Sample1_Beta | chr19:41018444-41018880 | SPTBN4 | N_Shore | Body | Yes |
| Control-Sample1_Beta | chr19:41025339-41025863 | SPTBN4 | Island | Body | Yes |
| Control-Sample15_Beta | chr19:41025574-41025863 | SPTBN4 | Island | Body | Yes |
| Control-Sample3_Beta | chr19:41025339-41025959 | SPTBN4 | Island | Body | Yes |
| Control-Sample61_Beta | chr19:41018759-41018880 | SPTBN4 | Island | Body | Yes |
| Control-Sample14_Beta | chr19:40972323-40972683 | SPTBN4;BLVRB | Island | TSS1500;TSS1500 | Yes |
| Control-Sample1_Beta | chr19:41054942-41055297 | SPTBN4;SPTBN4 | N_Shore | Body;Body | Yes |
| Control-Sample1_Beta | chr19:41060190-41060475 | SPTBN4;SPTBN4 | Island | Body;Body | Yes |
| Control-Sample29_Beta | chr19:41054942-41055208 | SPTBN4;SPTBN4 | N_Shore | Body;Body | Yes |
| Control-Sample1_Beta | chr15:42186830-42187036 | SPTBN5 | | TSS1500 | Yes |
| Control-Sample10_Beta | chr15:42186945-42187179 | SPTBN5 | | TSS1500 | Yes |
| Control-Sample22_Beta | chr15:42186830-42187179 | SPTBN5 | | TSS1500 | Yes |
| Control-Sample5_Beta | chr17:36715787-36715953 | SRCIN1 | Island | Body | No |
| Control-Sample2_Beta | chr7:76038785-76039223 | SRCRB4D;SRCRB4D;ZP3 | | 1stExon;5'UTR;5'UTR | No |
| Control-Sample1_Beta | chr7:75912071-75912462 | SRRM3 | Island | Body | Yes |
| Control-Sample11_Beta | chr7:75912102-75912462 | SRRM3 | Island | Body | Yes |
| Control-Sample12_Beta | chr7:75912071-75912349 | SRRM3 | Island | Body | Yes |
| Control-Sample270_Be | chr7:75911830-75912349 | SRRM3 | Island | Body | Yes |
| Control-Sample19_Beta | chr1:54821853-54822031 | SSBP3;SSBP3;SSBP3 | N_Shore | Body;Body;Body | No |
| Control-Sample53_Beta | chr19:18539841-18540208 | SSBP4;SSBP4 | Island | Body;Body | No |
| Control-Sample15_Beta | chr7:149471884-149472134 | SSPO | S_Shore | TSS1500 | Yes |
| Control-Sample216_Be | chr7:149484798-149484985 | SSPO | N_Shelf | Body | Yes |
| Control-Sample37_Beta | chr7:149487677-149488213 | SSPO | Island | Body | Yes |
| Control-Sample59_Beta | chr3:156273297-156273458 | SSR3 | S_Shore | TSS1500 | No |
| Control-Sample259_Be | chr20:23015908-23015936 | SSTR4 | Island | TSS200 | No |
| Control-Sample1_Beta | chr11:130029515-130029684 | ST14 | N_Shore | TSS200 | No |
| Control-Sample1_Beta | chr8:53321682-53322321 | ST18 | | 5'UTR | Yes |
| Control-Sample1_Beta | chr8:53322501-53322619 | ST18 | | TSS200 | Yes |
| Control-Sample10_Beta | chr8:53321682-53321751 | ST18 | | 5'UTR | Yes |
| Control-Sample105_Be | chr8:53322451-53322619 | ST18 | | TSS200 | Yes |
| Control-Sample19_Beta | chr8:53321682-53322619 | ST18 | | 5'UTR | Yes |
| Control-Sample572_Be | chr8.0:53321682-53322321 | ST18 | | 5'UTR | Yes |
| Control-Sample104_Be | chr8:53322321-53322619 | ST18;ST18 | | 5'UTR;1stExon | Yes |
| Control-Sample108_Be | chr7:116660172-116660240 | ST7;ST7 | | Body;Body | Yes |
| Control-Sample568_Be | chr7.0:116660172-116660240 | ST7;ST7 | | Body;Body | Yes |
| Control-Sample223_Be | chr15:92944664-92944756 | ST8SIA2 | | Body | No |
| Control-Sample205_Be | chr3:36422059-36422093 | STAC | Island | TSS200 | No |
| Control-Sample10_Beta | chr11:72465788-72466113 | STARD10 | N_Shore | 3'UTR | No |
| Control-Sample108_Be | chr13:33780283-33780362 | STARD13;STARD13 | | TSS200;Body | Yes |
| Control-Sample568_Be | chr13.0:33859885-33860074 | STARD13;STARD13 | | 1stExon;5'UTR | Yes |
| Control-Sample171_Be | chr17:40439433-40439501 | STAT5A | N_Shore | TSS200 | No |
| Control-Sample15_Beta | chr7:89783435-89783464 | STEAP1 | N_Shore | TSS1500 | No |
| Control-Sample1_Beta | chr5:171616388-171616574 | STK10 | S_Shore | TSS1500 | No |
| Control-Sample6_Beta | chr6:31937906-31937973 | STK19;DOM3Z | N_Shore | TSS1500;Body | Yes |
| Control-Sample10_Beta | chr6:31937462-31937582 | STK19;SKIV2L | N_Shelf | TSS1500;Body | Yes |
| Control-Sample1_Beta | chr13:99135543-99135711 | STK24;STK24 | | Body;Body | Yes |
| Control-Sample11_Beta | chr13:99135583-99135711 | STK24;STK24 | | Body;Body | Yes |
| Control-Sample19_Beta | chr13:99158698-99158901 | STK24;STK24 | | Body;Body | Yes |
| Control-Sample331_Be | chr13:99135543-99135625 | STK24;STK24 | | Body;Body | Yes |
| Control-Sample5_Beta | chr13:99174312-99174373 | STK24;STK24;STK24 | | Body;5'UTR;1stExon | Yes |
| Control-Sample3_Beta | chr10:134106591-134107111 | STK32C | | Body | Yes |
| Control-Sample65_Beta | chr10:134053894-134054189 | STK32C | | Body | Yes |
| Control-Sample26_Beta | chr1:26233435-26233506 | STMN1;STMN1;STMN1;STMN1 | Island | TSS1500;TSS1500;TSS1500;TSS200 | No |
| Control-Sample108_Be | chr8:27115867-27116213 | STMN4;STMN4 | | 1stExon;5'UTR | No |
| Control-Sample1_Beta | chr4:184938291-184938470 | STOX2 | | Body | Yes |
| Control-Sample10_Beta | chr4:184938316-184938470 | STOX2 | | Body | Yes |
| Control-Sample170_Be | chr4:184938291-184938359 | STOX2 | | Body | Yes |
| Control-Sample19_Beta | chr4:184922382-184922433 | STOX2 | | Body | Yes |
| Control-Sample572_Be | chr4.0:184938316-184938470 | STOX2 | | Body | Yes |
| Control-Sample585_Be | chr4.0:184922382-184922433 | STOX2 | | Body | Yes |
| Control-Sample13_Beta | chr15:74501549-74501771 | STRA6;STRA6;STRA6 | | TSS200;TSS200;TSS200 | Yes |
| Control-Sample1_Beta | chr15:74494781-74495354 | STRA6;STRA6;STRA6;STRA6;STRA6 | | 5'UTR;5'UTR;5'UTR;5'UTR;5'UTR | Yes |
| Control-Sample10_Beta | chr15:74494861-74495354 | STRA6;STRA6;STRA6;STRA6;STRA6 | | 5'UTR;5'UTR;5'UTR;5'UTR;5'UTR | Yes |
| Control-Sample11_Beta | chr15:74494900-74495354 | STRA6;STRA6;STRA6;STRA6;STRA6 | | 5'UTR;5'UTR;5'UTR;5'UTR;5'UTR | Yes |
| Control-Sample16_Beta | chr15:74494909-74495354 | STRA6;STRA6;STRA6;STRA6;STRA6 | | 5'UTR;5'UTR;5'UTR;5'UTR;5'UTR | Yes |
| Control-Sample2_Beta | chr15:74494854-74495354 | STRA6;STRA6;STRA6;STRA6;STRA6 | | 5'UTR;5'UTR;5'UTR;5'UTR;5'UTR | Yes |
| Control-Sample20_Beta | chr15:74494861-74495109 | STRA6;STRA6;STRA6;STRA6;STRA6 | | 5'UTR;5'UTR;5'UTR;5'UTR;5'UTR | Yes |
| Control-Sample3_Beta | chr15:74494781-74495384 | STRA6;STRA6;STRA6;STRA6;STRA6 | | 5'UTR;5'UTR;5'UTR;5'UTR;5'UTR | Yes |
| Control-Sample5_Beta | chr15:74494781-74494861 | STRA6;STRA6;STRA6;STRA6;STRA6 | | 5'UTR;5'UTR;5'UTR;5'UTR;5'UTR | Yes |
| Control-Sample5_Beta | chr15:74494909-74495384 | STRA6;STRA6;STRA6;STRA6;STRA6 | | 5'UTR;5'UTR;5'UTR;5'UTR;5'UTR | Yes |
| Control-Sample1_Beta | chr15:74501317-74501771 | STRA6;STRA6;STRA6;STRA6;STRA6;STRA6 | | 1stExon;5'UTR;5'UTR;1stExon;1stExon;5'UTR | Yes |
| Control-Sample10_Beta | chr16:31008574-31009232 | STX1B | N_Shore | Body | Yes |
| Control-Sample30_Beta | chr16:31008574-31009035 | STX1B | N_Shore | Body | Yes |
| Control-Sample194_Be | chr12:131303016-131303247 | STX2;STX2 | N_Shore | Body;Body | No |
| Control-Sample24_Beta | chr3:67704890-67705048 | SUCLG2 | Island | Body | No |
| Control-Sample655_Be | chr8:70378645-70378740 | SULF1;SULF1 | | TSS1500;TSS1500 | No |
| Control-Sample1_Beta | chr2:108905173-108905468 | SULT1C2;SULT1C2;SULT1C2;SULT1C2 | | 5'UTR;1stExon;1stExon;5'UTR | No |

| | | | | | |
|---|---|---|---|---|---|
| Control-Sample108_Be | chr2:108994325-108994528 | SULT1C4 | | TSS200 | No |
| Control-Sample497_Be | chr7:48031468-48031800 | SUNC1;SUNC1 | | Body;Body | No |
| Control-Sample1_Beta | chr22:24577288-24577448 | SUSD2 | | TSS200 | Yes |
| Control-Sample12_Beta | chr22:24577269-24577443 | SUSD2 | | TSS200 | Yes |
| Control-Sample21_Beta | chr22:24577288-24577438 | SUSD2 | | TSS200 | Yes |
| Control-Sample24_Beta | chr22:24577288-24577443 | SUSD2 | | TSS200 | Yes |
| Control-Sample37_Beta | chr22:24577269-24577448 | SUSD2 | | TSS200 | Yes |
| Control-Sample70_Beta | chr10:29811530-29811659 | SVIL;SVIL | | Body;Body | No |
| Control-Sample1_Beta | chr6:10887587-10887980 | SYCP2L | Island | Body | No |
| Control-Sample1_Beta | chr19:46318918-46319153 | SYMPK;RSPH6A | Island | Body;TSS1500 | Yes |
| Control-Sample108_Be | chr6:152702330-152702660 | SYNE1;SYNE1 | | Body;Body | No |
| Control-Sample13_Beta | chr14:64319089-64319275 | SYNE2;SYNE2 | N_Shore | TSS1500;TSS1500 | No |
| Control-Sample1_Beta | chr6:33421180-33421476 | SYNGAP1;ZBTB9 | N_Shore | 3'UTR;TSS1500 | Yes |
| Control-Sample12_Beta | chr6:33421180-33421704 | SYNGAP1;ZBTB9 | N_Shore | 3'UTR;TSS1500 | Yes |
| Control-Sample28_Beta | chr6:33421180-33421509 | SYNGAP1;ZBTB9 | N_Shore | 3'UTR;TSS1500 | Yes |
| Control-Sample1_Beta | chr6:158447877-158448578 | SYNJ2 | | Body | Yes |
| Control-Sample13_Beta | chr6:158447877-158448455 | SYNJ2 | | Body | Yes |
| Control-Sample170_Be | chr6:158404061-158404121 | SYNJ2 | S_Shore | Body | Yes |
| Control-Sample18_Beta | chr6:158447877-158448508 | SYNJ2 | | Body | Yes |
| Control-Sample19_Beta | chr6:158448455-158448578 | SYNJ2 | | Body | Yes |
| Control-Sample20_Beta | chr6:158447883-158448578 | SYNJ2 | | Body | Yes |
| Control-Sample29_Beta | chr6:158487541-158487608 | SYNJ2 | | Body | Yes |
| Control-Sample40_Beta | chr6:158447877-158448134 | SYNJ2 | | Body | Yes |
| Control-Sample171_Be | chr5:149980486-149980671 | SYNPO | | TSS200 | Yes |
| Control-Sample4_Beta | chr5:149980671-149980682 | SYNPO;SYNPO | | 1stExon;5'UTR | Yes |
| Control-Sample8_Beta | chr1:110009974-110010384 | SYPL2 | Island | Body | No |
| Control-Sample1_Beta | chr1:202613601-202614080 | SYT2;SYT2 | S_Shore | TSS1500;5'UTR | No |
| Control-Sample1_Beta | chr1:27668325-27668445 | SYTL1 | | TSS200 | Yes |
| Control-Sample12_Beta | chr1:27677860-27678138 | SYTL1 | Island | Body | Yes |
| Control-Sample2_Beta | chr1:27677872-27678138 | SYTL1 | Island | Body | Yes |
| Control-Sample3_Beta | chr1:27668325-27668610 | SYTL1 | | TSS200 | Yes |
| Control-Sample39_Beta | chr1:27668325-27668433 | SYTL1 | | TSS200 | Yes |
| Control-Sample67_Beta | chr1:27676205-27676652 | SYTL1 | Island | Body | Yes |
| Control-Sample108_Be | chr11:85430070-85430657 | SYTL2;SYTL2;SYTL2;SYTL2;SYTL2;SYTL2;SYTL2;SYTL2;SYTL2;SYTL2;SYTL2 | | 5'UTR;Body;5'UTR;1stExon;Body;1stExon;1stExon;Bo | No |
| Control-Sample10_Beta | chr6:132891457-132891738 | TAAR6 | | TSS200 | No |
| Control-Sample18_Beta | chr4:104640662-104640833 | TACR3 | Island | 1stExon | Yes |
| Control-Sample55_Beta | chr4:104640662-104641250 | TACR3 | Island | 1stExon | Yes |
| Control-Sample56_Beta | chr4:104640662-104641319 | TACR3 | Island | 1stExon | Yes |
| Control-Sample2_Beta | chr1:59043070-59043208 | TACSTD2;TACSTD2 | Island | 5'UTR;1stExon | No |
| Control-Sample1_Beta | chr11:117069780-117069859 | TAGLN;TAGLN | | TSS1500;TSS1500 | Yes |
| Control-Sample19_Beta | chr11:117069788-117069859 | TAGLN;TAGLN | | TSS1500;TSS1500 | Yes |
| Control-Sample62_Beta | chr11:117069780-117069849 | TAGLN;TAGLN | | TSS1500;TSS1500 | Yes |
| Control-Sample3_Beta | chr1:159893182-159893696 | TAGLN2 | N_Shore | 5'UTR | No |
| Control-Sample108_Be | chr3:111718135-111718565 | TAGLN3;TAGLN3;TAGLN3;TAGLN3 | | 1stExon;5'UTR;5'UTR;5'UTR | No |
| Control-Sample6_Beta | chr6:32820421-32820553 | TAP1 | N_Shore | Body | No |
| Control-Sample3_Beta | chr6:32800486-32800541 | TAP2;TAP2 | | Body;Body | Yes |
| Control-Sample6_Beta | chr6:32805554-32805692 | TAP2;TAP2 | N_Shore | Body;Body | Yes |
| Control-Sample10_Beta | chr12:6561007-6561352 | TAPBPL;LOC678655 | | TSS200;TSS1500 | No |
| Control-Sample1_Beta | chr1:6615418-6615604 | TAS1R1;TAS1R1;TAS1R1;NOL9;TAS1R1 | S_Shore | TSS200;TSS200;TSS200;TSS1500;TSS200 | No |
| Control-Sample134_Be | chr7:143140164-143140674 | TAS2R60 | | TSS1500 | No |
| Control-Sample1_Beta | chr11:67171145-67171200 | TBC1D10C | S_Shelf | TSS1500 | No |
| Control-Sample29_Beta | chr12:72233240-72233266 | TBC1D15;TBC1D15;TBC1D15;TBC1D15 | N_Shore | TSS1500;TSS1500;TSS1500;TSS1500 | No |
| Control-Sample1_Beta | chr17:77922775-77924180 | TBC1D16 | Island | Body | Yes |
| Control-Sample10_Beta | chr17:77922972-77924180 | TBC1D16 | Island | Body | Yes |
| Control-Sample10_Beta | chr17:77954963-77955530 | TBC1D16 | N_Shore | Body | Yes |
| Control-Sample108_Be | chr17:77916733-77916892 | TBC1D16 | Island | Body | Yes |
| Control-Sample11_Beta | chr17:77924275-77924582 | TBC1D16 | Island | Body | Yes |
| Control-Sample12_Beta | chr17:77922775-77923971 | TBC1D16 | Island | Body | Yes |
| Control-Sample13_Beta | chr17:77922865-77923603 | TBC1D16 | Island | Body | Yes |
| Control-Sample14_Beta | chr17:77922865-77923971 | TBC1D16 | Island | Body | Yes |
| Control-Sample15_Beta | chr17:77922865-77924180 | TBC1D16 | Island | Body | Yes |
| Control-Sample17_Beta | chr17:77922972-77923971 | TBC1D16 | Island | Body | Yes |
| Control-Sample18_Beta | chr17:77955167-77955530 | TBC1D16 | N_Shore | Body | Yes |
| Control-Sample19_Beta | chr17:77923275-77923971 | TBC1D16 | N_Shore | Body | Yes |
| Control-Sample19_Beta | chr17:77997833-77997997 | TBC1D16 | Island | 5'UTR | Yes |
| Control-Sample22_Beta | chr17:77959804-77960007 | TBC1D16 | | Body | Yes |
| Control-Sample29_Beta | chr17:77923275-77923603 | TBC1D16 | N_Shore | Body | Yes |
| Control-Sample3_Beta | chr17:77952453-77952488 | TBC1D16 | S_Shore | Body | Yes |
| Control-Sample30_Beta | chr17:77922972-77923603 | TBC1D16 | Island | Body | Yes |
| Control-Sample31_Beta | chr17:77922775-77923603 | TBC1D16 | Island | Body | Yes |
| Control-Sample4_Beta | chr17:77924371-77924733 | TBC1D16 | Island | Body | Yes |
| Control-Sample4_Beta | chr17:77952109-77952488 | TBC1D16 | Island | Body | Yes |
| Control-Sample404_Be | chr17:77952290-77952463 | TBC1D16 | Island | Body | Yes |
| Control-Sample475_Be | chr17:77922972-77923314 | TBC1D16 | Island | Body | Yes |
| Control-Sample56_Beta | chr17:77954963-77955492 | TBC1D16 | N_Shore | Body | Yes |
| Control-Sample1_Beta | chr15:78292470-78292550 | TBC1D2B;TBC1D2B | | Body;Body | No |
| Control-Sample1_Beta | chr17:80823240-80823397 | TBCD | S_Shelf | Body | Yes |
| Control-Sample1_Beta | chr17:80833393-80833638 | TBCD | S_Shelf | Body | Yes |
| Control-Sample11_Beta | chr17:80806135-80806375 | TBCD | | Body | Yes |
| Control-Sample11_Beta | chr17:80870163-80870577 | TBCD | N_Shore | Body | Yes |
| Control-Sample16_Beta | chr17:80806191-80806435 | TBCD | | Body | Yes |
| Control-Sample17_Beta | chr17:80806135-80806435 | TBCD | | Body | Yes |
| Control-Sample5_Beta | chr17:80858308-80858539 | TBCD | S_Shore | Body | Yes |
| Control-Sample53_Beta | chr17:80806135-80806388 | TBCD | | Body | Yes |
| Control-Sample21_Beta | chr17:80794081-80794274 | TBCD;ZNF750 | S_Shelf | Body;5'UTR | Yes |
| Control-Sample1_Beta | chr17:45786010-45786439 | TBKBP1 | N_Shore | Body | No |
| Control-Sample1_Beta | chr2:162271751-162272107 | TBR1 | N_Shore | TSS1500 | Yes |
| Control-Sample10_Beta | chr2:162271751-162271859 | TBR1 | N_Shore | TSS1500 | Yes |
| Control-Sample15_Beta | chr2:162279686-162280009 | TBR1 | N_Shore | Body | Yes |
| Control-Sample67_Beta | chr2:162271801-162272107 | TBR1 | N_Shore | TSS1500 | Yes |
| Control-Sample11_Beta | chr22:19746984-19747026 | TBX1;TBX1;TBX1 | Island | 5'UTR;5'UTR;5'UTR | No |

| | | | | | |
|---|---|---|---|---|---|
| Control-Sample1_Beta | chr1:119527638-119528018 | TBX15 | S_Shore | 5'UTR | Yes |
| Control-Sample1_Beta | chr1:119532195-119532655 | TBX15 | Island | TSS200 | Yes |
| Control-Sample1_Beta | chr1:119532773-119532925 | TBX15 | S_Shore | TSS1500 | Yes |
| Control-Sample2_Beta | chr1:119532195-119532352 | TBX15 | Island | TSS200 | Yes |
| Control-Sample58_Beta | chr1:119532195-119532736 | TBX15 | Island | TSS200 | Yes |
| Control-Sample1_Beta | chr1:119532044-119532116 | TBX15;TBX15 | Island | 1stExon;5'UTR | Yes |
| Control-Sample11_Beta | chr1:119532056-119532116 | TBX15;TBX15 | Island | 1stExon;5'UTR | Yes |
| Control-Sample15_Beta | chr1:119532044-119532195 | TBX15;TBX15 | Island | 1stExon;5'UTR | Yes |
| Control-Sample44_Beta | chr1:119532116-119532195 | TBX15;TBX15 | Island | 1stExon;5'UTR | Yes |
| Control-Sample18_Beta | chr17:59478579-59478958 | TBX2 | Island | Body | Yes |
| Control-Sample54_Beta | chr17:59478579-59479070 | TBX2 | Island | Body | Yes |
| Control-Sample1_Beta | chr17:59532564-59533081 | TBX4 | Island | TSS1500 | Yes |
| Control-Sample14_Beta | chr17:59532519-59533081 | TBX4 | Island | TSS1500 | Yes |
| Control-Sample233_Be | chr10:132909153-132909470 | TCERG1L | | Body | No |
| Control-Sample15_Beta | chr5:133451294-133451474 | TCF7;TCF7;TCF7;TCF7;TCF7;TCF7;TCF7 | S_Shore | TSS200;TSS1500;TSS1500;TSS200;Body;TSS200;TSS | Yes |
| Control-Sample31_Beta | chr5:133451383-133451625 | TCF7;TCF7;TCF7;TCF7;TCF7;TCF7;TCF7 | S_Shore | 5'UTR;TSS1500;TSS1500;Body;TSS200;5'UTR;5'UTR | Yes |
| Control-Sample9_Beta | chr5:133451294-133451625 | TCF7;TCF7;TCF7;TCF7;TCF7;TCF7;TCF7 | S_Shore | TSS200;TSS1500;TSS1500;TSS200;Body;TSS200;TSS | Yes |
| Control-Sample10_Beta | chr2:85361959-85362480 | TCF7L1 | Island | Body | No |
| Control-Sample3_Beta | chr6:35109121-35109298 | TCP11;TCP11;TCP11;TCP11 | Island | 5'UTR;1stExon;5'UTR;1stExon | No |
| Control-Sample53_Beta | chr1:67218135-67218505 | TCTEX1D1 | Island | TSS200 | No |
| Control-Sample223_Be | chr10:114043305-114043435 | TECTB | | TSS1500 | No |
| Control-Sample50_Beta | chr9:27109111-27109492 | TEK | | TSS200 | No |
| Control-Sample23_Beta | chr2:95537314-95537475 | TEKT4;TEKT4 | Island | 1stExon;5'UTR | No |
| Control-Sample1_Beta | chr16:58019866-58019984 | TEPP;TEPP | Island | Body;Body | No |
| Control-Sample3_Beta | chr7:115862648-115863026 | TES;TES | | TSS1500;Body | No |
| Control-Sample2_Beta | chr6:10416373-10417087 | TFAP2A;TFAP2A | N_Shore | Body;TSS1500 | No |
| Control-Sample1_Beta | chr1:36038468-36038701 | TFAP2E | N_Shore | TSS1500 | No |
| Control-Sample1_Beta | chr6:155584709-155584968 | TFB1M;CLDN20 | | Body;TSS1500 | Yes |
| Control-Sample151_Be | chr6:155584613-155584968 | TFB1M;CLDN20 | | Body;TSS1500 | Yes |
| Control-Sample585_Be | chr6.0:155584613-155584968 | TFB1M;CLDN20 | | Body;TSS1500 | Yes |
| Control-Sample20_Beta | chr13:114281582-114281677 | TFDP1;TFDP1 | S_Shore | Body;Body | No |
| Control-Sample63_Beta | chr19:41836950-41837123 | TGFB1 | N_Shore | 3'UTR | No |
| Control-Sample16_Beta | chr16:31487813-31488238 | TGFB1I1;TGFB1I1;TGFB1I1 | Island | Body;Body;Body | No |
| Control-Sample1_Beta | chr14:76445988-76446161 | TGFB3 | N_Shelf | Body | Yes |
| Control-Sample25_Beta | chr14:76445988-76446681 | TGFB3 | N_Shelf | Body | Yes |
| Control-Sample26_Beta | chr14:76445988-76446433 | TGFB3 | N_Shelf | Body | Yes |
| Control-Sample1_Beta | chr5:135364328-135364451 | TGFBI | N_Shore | TSS1500 | Yes |
| Control-Sample19_Beta | chr5:135364060-135364349 | TGFBI | N_Shore | TSS1500 | Yes |
| Control-Sample1_Beta | chr20:35201208-35201324 | TGIF2 | N_Shore | TSS1500 | No |
| Control-Sample29_Beta | chr2:85553759-85553966 | TGOLN2 | N_Shore | Body | No |
| Control-Sample223_Be | chr2:242529748-242530174 | THAP4;THAP4 | | Body;Body | No |
| Control-Sample67_Beta | chr15:39871876-39872032 | THBS1 | N_Shore | TSS1500 | No |
| Control-Sample171_Be | chr6:169629875-169630013 | THBS2 | N_Shore | Body | Yes |
| Control-Sample194_Be | chr6:169653935-169653961 | THBS2 | | 5'UTR | Yes |
| Control-Sample649_Be | chr6:169641941-169642301 | THBS2 | Island | Body | Yes |
| Control-Sample1_Beta | chr3:24537407-24537792 | THRB;THRB;THRB | Island | TSS1500;TSS1500;TSS1500 | No |
| Control-Sample24_Beta | chr13:52769289-52769768 | THSD1P | N_Shore | TSS1500 | No |
| Control-Sample655_Be | chr7:11872050-11872073 | THSD7A | | TSS1500 | No |
| Control-Sample331_Be | chr11:119294283-119294414 | THY1 | S_Shore | TSS200 | No |
| Control-Sample464_Be | chr21:32624145-32624274 | TIAM1 | Island | Body | No |
| Control-Sample63_Beta | chr6:155537595-155537930 | TIAM2;TIAM2 | N_Shelf | Body;TSS1500 | No |
| Control-Sample14_Beta | chr6:43456501-43457177 | TJAP1;TJAP1;TJAP1;TJAP1;TJAP1;TJAP1;TJAP1 | | 5'UTR;5'UTR;5'UTR;5'UTR;Body;TSS1500;5'UTR | Yes |
| Control-Sample16_Beta | chr6:43456501-43457274 | TJAP1;TJAP1;TJAP1;TJAP1;TJAP1;TJAP1;TJAP1 | | 5'UTR;5'UTR;5'UTR;5'UTR;Body;TSS1500;5'UTR | Yes |
| Control-Sample37_Beta | chr6:43456984-43457177 | TJAP1;TJAP1;TJAP1;TJAP1;TJAP1;TJAP1;TJAP1 | | 5'UTR;5'UTR;5'UTR;5'UTR;Body;TSS1500;5'UTR | Yes |
| Control-Sample65_Beta | chr6:43456984-43457274 | TJAP1;TJAP1;TJAP1;TJAP1;TJAP1;TJAP1;TJAP1 | | 5'UTR;5'UTR;5'UTR;5'UTR;Body;TSS1500;5'UTR | Yes |
| Control-Sample15_Beta | chr9:71735648-71735701 | TJP2 | N_Shore | TSS1500 | No |
| Control-Sample1_Beta | chr16:66583078-66583237 | TK2 | N_Shore | Body | No |
| Control-Sample1_Beta | chr4:186990006-186990315 | TLR3 | | TSS1500 | Yes |
| Control-Sample108_Be | chr4:186990006-186990425 | TLR3 | | TSS1500 | Yes |
| Control-Sample20_Beta | chr4:186990006-186990324 | TLR3 | | TSS1500 | Yes |
| Control-Sample10_Beta | chr1:223317364-223317548 | TLR5 | S_Shore | TSS1500 | Yes |
| Control-Sample14_Beta | chr1:223317456-223317548 | TLR5 | S_Shore | TSS1500 | Yes |
| Control-Sample1_Beta | chr4:38859339-38859770 | TLR6 | | TSS1500 | Yes |
| Control-Sample171_Be | chr4:38859706-38859770 | TLR6 | | TSS1500 | Yes |
| Control-Sample74_Beta | chr4:38859339-38859728 | TLR6 | | TSS1500 | Yes |
| Control-Sample12_Beta | chr10:102890038-102890886 | TLX1;TLX1NB | N_Shore | 5'UTR | Yes |
| Control-Sample47_Beta | chr10:102890145-102890514 | TLX1;TLX1NB | N_Shore | TSS1500;5'UTR | Yes |
| Control-Sample19_Beta | chr15:83777021-83777082 | TM6SF1;TM6SF1 | S_Shore | Body;Body | Yes |
| Control-Sample5_Beta | chr15:83776915-83777082 | TM6SF1;TM6SF1 | Island | Body;Body | Yes |
| Control-Sample3_Beta | chr17:76128634-76128683 | TMC8;TMC6 | S_Shore | Body;TSS200 | Yes |
| Control-Sample1_Beta | chr17:76128481-76128531 | TMC8;TMC6;TMC6 | S_Shore | Body;5'UTR;1stExon | Yes |
| Control-Sample27_Beta | chr17:76128481-76128556 | TMC8;TMC6;TMC6 | S_Shore | Body;5'UTR;1stExon | Yes |
| Control-Sample67_Beta | chr3:129407542-129407548 | TMCC1;TMCC1;TMCC1 | | TSS200;Body;Body | Yes |
| Control-Sample108_Be | chr3:129407365-129407542 | TMCC1;TMCC1;TMCC1;TMCC1 | | 1stExon;Body;Body;5'UTR | Yes |
| Control-Sample10_Beta | chr12:95010029-95010113 | TMCC3 | | Body | Yes |
| Control-Sample108_Be | chr12:95009858-95010113 | TMCC3 | | Body | Yes |
| Control-Sample223_Be | chr12:95010017-95010113 | TMCC3 | | Body | Yes |
| Control-Sample585_Be | chr12.0:95009858-95010113 | TMCC3 | | Body | Yes |
| Control-Sample1_Beta | chr13:114150007-114150139 | TMCO3 | S_Shelf | Body | Yes |
| Control-Sample1_Beta | chr13:114201796-114202861 | TMCO3 | N_Shore | Body | Yes |
| Control-Sample10_Beta | chr13:114202067-114202861 | TMCO3 | N_Shore | Body | Yes |
| Control-Sample35_Beta | chr13:114202067-114202741 | TMCO3 | N_Shore | Body | Yes |
| Control-Sample1_Beta | chr5:140018644-140018709 | TMCO6 | N_Shore | TSS1500 | No |
| Control-Sample64_Beta | chr17:53810321-53810813 | TMEM100 | | TSS1500 | Yes |
| Control-Sample291_Be | chr17:53800225-53800484 | TMEM100;TMEM100 | | TSS200;5'UTR | Yes |
| Control-Sample108_Be | chr17:53800041-53800372 | TMEM100;TMEM100;TMEM100 | | 5'UTR;1stExon;5'UTR | Yes |
| Control-Sample1_Beta | chr17:41363737-41364121 | TMEM106A | Island | TSS200 | Yes |
| Control-Sample10_Beta | chr17:41363737-41363789 | TMEM106A | Island | TSS200 | Yes |
| Control-Sample17_Beta | chr17:41363741-41363789 | TMEM106A | Island | TSS200 | Yes |
| Control-Sample27_Beta | chr17:41363741-41364121 | TMEM106A | Island | TSS200 | Yes |
| Control-Sample10_Beta | chr17:41363899-41364121 | TMEM106A;TMEM106A | Island | 5'UTR;1stExon | Yes |

| | | | | | |
|---|---|---|---|---|---|
| Control-Sample19_Beta | chr12:125810688-125811065 | TMEM132B | | TSS1500 | No |
| Control-Sample108_Be | chr12:128866402-128866735 | TMEM132C | | Body | Yes |
| Control-Sample291_Be | chr12:128866402-128866698 | TMEM132C | | Body | Yes |
| Control-Sample1_Beta | chr12:130316015-130316058 | TMEM132D | | Body | No |
| Control-Sample42_Beta | chr11:67232269-67232403 | TMEM134;TMEM134;TMEM134 | N_Shore | Body;Body;Body | No |
| Control-Sample1_Beta | chr7:134832221-134833299 | TMEM140 | | TSS1500 | Yes |
| Control-Sample15_Beta | chr7:134832671-134833299 | TMEM140 | | TSS200 | Yes |
| Control-Sample151_Be | chr7:134832671-134832850 | TMEM140 | | TSS200 | Yes |
| Control-Sample171_Be | chr7:134832671-134832939 | TMEM140 | | TSS200 | Yes |
| Control-Sample223_Be | chr7:134832671-134833031 | TMEM140 | | TSS200 | Yes |
| Control-Sample331_Be | chr7:134832221-134832939 | TMEM140 | | TSS1500 | Yes |
| Control-Sample585_Be | chr7.0:134832671-134832850 | TMEM140 | | TSS200 | Yes |
| Control-Sample68_Beta | chr7:134833031-134833299 | TMEM140 | | 5'UTR | Yes |
| Control-Sample69_Beta | chr7:134832221-134832850 | TMEM140 | | TSS1500 | Yes |
| Control-Sample92_Beta | chr7:134832584-134832770 | TMEM140 | | TSS200 | Yes |
| Control-Sample108_Be | chr7:134832844-134833299 | TMEM140;TMEM140 | | 1stExon;5'UTR | Yes |
| Control-Sample194_Be | chr7:134832850-134833031 | TMEM140;TMEM140 | | 1stExon;5'UTR | Yes |
| Control-Sample572_Be | chr7.0:134832850-134833031 | TMEM140;TMEM140 | | 1stExon;5'UTR | Yes |
| Control-Sample75_Beta | chr7:134832770-134832850 | TMEM140;TMEM140 | | 1stExon;5'UTR | Yes |
| Control-Sample80_Beta | chr7:134832770-134832939 | TMEM140;TMEM140 | | 1stExon;5'UTR | Yes |
| Control-Sample1_Beta | chr16:21170927-21171239 | TMEM159;DNAH3 | S_Shore | 5'UTR;TSS200 | Yes |
| Control-Sample24_Beta | chr16:21171067-21171239 | TMEM159;DNAH3 | S_Shore | 5'UTR;TSS1500 | Yes |
| Control-Sample40_Beta | chr16:21170927-21171084 | TMEM159;DNAH3 | S_Shore | 5'UTR;TSS200 | Yes |
| Control-Sample464_Be | chr4:951898-952128 | TMEM175 | Island | Body | No |
| Control-Sample13_Beta | chr7:150498493-150498843 | TMEM176B;TMEM176B;TMEM176B;TMEM176B;TMEM176A | S_Shore | TSS200;TSS1500;TSS1500;TSS1500;5'UTR | No |
| Control-Sample29_Beta | chr7:1596103-1596136 | TMEM184A | | TSS200 | Yes |
| Control-Sample39_Beta | chr7:1596118-1596217 | TMEM184A | | TSS200 | Yes |
| Control-Sample3_Beta | chr7:1596050-1596136 | TMEM184A;TMEM184A | | 1stExon;5'UTR | Yes |
| Control-Sample23_Beta | chr19:55889216-55889611 | TMEM190 | Island | Body | Yes |
| Control-Sample3_Beta | chr19:55888082-55888156 | TMEM190 | N_Shore | TSS200 | Yes |
| Control-Sample61_Beta | chr19:55888082-55888150 | TMEM190 | N_Shore | TSS200 | Yes |
| Control-Sample10_Beta | chr2:191398756-191398806 | TMEM194B | N_Shore | Body | No |
| Control-Sample1_Beta | chr18:5891948-5892245 | TMEM200C | S_Shore | 1stExon | Yes |
| Control-Sample17_Beta | chr18:5892119-5892245 | TMEM200C | S_Shore | TSS200 | Yes |
| Control-Sample10_Beta | chr1:9664205-9664395 | TMEM201;TMEM201 | | Body;3'UTR | Yes |
| Control-Sample38_Beta | chr1:9664334-9664395 | TMEM201;TMEM201 | | Body;3'UTR | Yes |
| Control-Sample1_Beta | chr1:212588536-212589072 | TMEM206 | S_Shore | TSS1500 | Yes |
| Control-Sample22_Beta | chr1:212588580-212589072 | TMEM206 | S_Shore | TSS1500 | Yes |
| Control-Sample108_Be | chr5:110062398-110062729 | TMEM232 | | TSS200 | Yes |
| Control-Sample443_Be | chr5:110062398-110062473 | TMEM232 | | TSS200 | Yes |
| Control-Sample568_Be | chr5.0:110062473-110062837 | TMEM232 | | TSS200 | Yes |
| Control-Sample616_Be | chr5:110062343-110062398 | TMEM232;TMEM232 | | 1stExon;5'UTR | Yes |
| Control-Sample15_Beta | chr2:120190031-120190297 | TMEM37 | Island | Body | Yes |
| Control-Sample16_Beta | chr2:120189867-120190297 | TMEM37 | N_Shore | Body | Yes |
| Control-Sample1_Beta | chr1:15541182-15541349 | TMEM51;TMEM51;TMEM51;TMEM51 | Island | 5'UTR;5'UTR;5'UTR;5'UTR | Yes |
| Control-Sample12_Beta | chr1:15541182-15541280 | TMEM51;TMEM51;TMEM51;TMEM51 | Island | 5'UTR;5'UTR;5'UTR;5'UTR | Yes |
| Control-Sample15_Beta | chr8:94766196-94766468 | TMEM67;TMEM67;TMEM67 | N_Shore | TSS1500;TSS1500;TSS1500 | No |
| Control-Sample11_Beta | chr8:133772657-133772889 | TMEM71;TMEM71 | | 5'UTR;5'UTR | Yes |
| Control-Sample13_Beta | chr8:133773093-133773115 | TMEM71;TMEM71 | | TSS200;TSS200 | Yes |
| Control-Sample3_Beta | chr8:133772657-133773099 | TMEM71;TMEM71 | | 5'UTR;5'UTR | Yes |
| Control-Sample1_Beta | chr10:45406644-45406847 | TMEM72 | | TSS1500 | Yes |
| Control-Sample11_Beta | chr10:45406295-45406681 | TMEM72 | | TSS1500 | Yes |
| Control-Sample13_Beta | chr10:45406187-45406681 | TMEM72 | | | Yes |
| Control-Sample14_Beta | chr10:45406187-45406847 | TMEM72 | | | Yes |
| Control-Sample23_Beta | chr10:45406295-45406847 | TMEM72 | | TSS1500 | Yes |
| Control-Sample18_Beta | chr1:156251280-156251456 | TMEM79;SMG5 | | TSS1500;Body | No |
| Control-Sample3_Beta | chr15:42566300-42566343 | TMEM87A;TMEM87A;GANC | Island | TSS1500;TSS1500;TSS200 | No |
| Control-Sample1_Beta | chr17:7758421-7758662 | TMEM88 | N_Shore | 1stExon | Yes |
| Control-Sample13_Beta | chr17:7758228-7758291 | TMEM88 | N_Shore | TSS200 | Yes |
| Control-Sample2_Beta | chr17:7758581-7758662 | TMEM88 | N_Shore | 1stExon | Yes |
| Control-Sample22_Beta | chr17:7758662-7758854 | TMEM88 | Island | Body | Yes |
| Control-Sample26_Beta | chr17:7758228-7758397 | TMEM88 | N_Shore | TSS200 | Yes |
| Control-Sample331_Be | chr1:1360970-1361216 | TMEM88B | N_Shore | TSS1500 | No |
| Control-Sample3_Beta | chr17:7258481-7258501 | TMEM95 | S_Shelf | TSS200 | No |
| Control-Sample28_Beta | chr21:42880557-42880714 | TMPRSS2;TMPRSS2 | Island | TSS1500;TSS1500 | No |
| Control-Sample1_Beta | chr21:43809360-43809567 | TMPRSS3;TMPRSS3;TMPRSS3 | | Body;TSS200;Body | No |
| Control-Sample1_Beta | chr11:113576693-113577528 | TMPRSS5 | | Body | Yes |
| Control-Sample108_Be | chr11:113577071-113577434 | TMPRSS5 | | TSS200 | Yes |
| Control-Sample124_Be | chr11:113577386-113577528 | TMPRSS5 | | TSS1500 | Yes |
| Control-Sample22_Beta | chr11:113576693-113577118 | TMPRSS5 | | Body | Yes |
| Control-Sample270_Be | chr11:113577118-113577386 | TMPRSS5 | | TSS200 | Yes |
| Control-Sample331_Be | chr11:113577071-113577528 | TMPRSS5 | | TSS200 | Yes |
| Control-Sample40_Beta | chr11:113577118-113577528 | TMPRSS5 | | TSS200 | Yes |
| Control-Sample42_Beta | chr11:113576693-113577434 | TMPRSS5 | | Body | Yes |
| Control-Sample35_Beta | chr16:2892518-2892782 | TMPRSS8 | N_Shore | Body | Yes |
| Control-Sample433_Be | chr16:2892766-2892782 | TMPRSS8 | Island | TSS200 | Yes |
| Control-Sample540_Be | chr16.0:2892766-2892782 | TMPRSS8 | Island | TSS200 | Yes |
| Control-Sample1_Beta | chr14:103593503-103593520 | TNFAIP2 | Island | Body | No |
| Control-Sample1_Beta | chr6:138200309-138200387 | TNFAIP3 | | Body | No |
| Control-Sample13_Beta | chr16:12058616-12059019 | TNFRSF17 | | TSS1500 | No |
| Control-Sample25_Beta | chr12:6451035-6451298 | TNFRSF1A;TNFRSF1A | | 5'UTR;1stExon | No |
| Control-Sample13_Beta | chr1:6526182-6526319 | TNFRSF25;TNFRSF25;PLEKHG5;PLEKHG5;TNFRSF25;TNFI | Island | 1stExon;1stExon;3'UTR;3'UTR;5'UTR;5'UTR;5'UTR;5'U | Yes |
| Control-Sample4_Beta | chr1:6526182-6526469 | TNFRSF25;TNFRSF25;PLEKHG5;PLEKHG5;TNFRSF25;TNFI | Island | 1stExon;1stExon;3'UTR;3'UTR;5'UTR;5'UTR;5'UTR;5'U | Yes |
| Control-Sample11_Beta | chr1:6526129-6526319 | TNFRSF25;TNFRSF25;TNFRSF25;TNFRSF25;TNFRSF25;TN | Island | 1stExon;1stExon;1stExon;1stExon;1stExon;1stExon | Yes |
| Control-Sample5_Beta | chr1:6526129-6526469 | TNFRSF25;TNFRSF25;TNFRSF25;TNFRSF25;TNFRSF25;TN | Island | 1stExon;1stExon;1stExon;1stExon;1stExon;1stExon | Yes |
| Control-Sample108_Be | chr3:172241069-172241384 | TNFSF10 | | 1stExon | No |
| Control-Sample1_Beta | chr13:108922032-108922274 | TNFSF13B;TNFSF13B;TNFSF13B;TNFSF13B | | 5'UTR;1stExon;5'UTR;1stExon | No |
| Control-Sample108_Be | chr1:173176362-173176501 | TNFSF4;TNFSF4 | | 5'UTR;1stExon | Yes |
| Control-Sample223_Be | chr1:173176362-173176523 | TNFSF4;TNFSF4 | | 5'UTR;1stExon | Yes |
| Control-Sample5_Beta | chr19:6534760-6534916 | TNFSF9 | Island | Body | No |
| Control-Sample1_Beta | chr3:195599898-195600608 | TNK2;TNK2 | N_Shore | Body;Body | Yes |

| | | | | | |
|---|---|---|---|---|---|
| Control-Sample1_Beta | chr3:195614967-195615577 | TNK2;TNK2 | S_Shore | Body;Body | Yes |
| Control-Sample10_Beta | chr3:195614967-195615726 | TNK2;TNK2 | S_Shore | Body;Body | Yes |
| Control-Sample12_Beta | chr3:195600311-195600608 | TNK2;TNK2 | N_Shore | Body;Body | Yes |
| Control-Sample13_Beta | chr3:195614967-195615626 | TNK2;TNK2 | S_Shore | Body;Body | Yes |
| Control-Sample13_Beta | chr3:195623338-195623792 | TNK2;TNK2 | S_Shore | TSS1500;5'UTR | Yes |
| Control-Sample1_Beta | chr3:52488037-52488097 | TNNC1;TNNC1;NISCH | N_Shore | 1stExon;5'UTR;TSS1500 | No |
| Control-Sample20_Beta | chr1:201390794-201391061 | TNNI1 | | 5'UTR | No |
| Control-Sample433_Be | chr19:55668066-55668612 | TNNI3 | Island | Body | Yes |
| Control-Sample550_Be | chr19.0:55668066-55668612 | TNNI3 | Island | Body | Yes |
| Control-Sample13_Beta | chr16:11363854-11364014 | TNP2 | | TSS1500 | No |
| Control-Sample67_Beta | chr17:76037239-76037364 | TNRC6C;TNRC6C | Island | 5'UTR;5'UTR | No |
| Control-Sample331_Be | chr2:218770208-218770270 | TNS1 | | 5'UTR | Yes |
| Control-Sample464_Be | chr2:218712737-218712871 | TNS1 | | Body | Yes |
| Control-Sample1_Beta | chr6:32031395-32031624 | TNXB | | Body | Yes |
| Control-Sample1_Beta | chr6:32060681-32061084 | TNXB | N_Shelf | Body | Yes |
| Control-Sample1_Beta | chr6:32078574-32078624 | TNXB | | TSS1500 | Yes |
| Control-Sample108_Be | chr6:32026656-32026797 | TNXB | | Body | Yes |
| Control-Sample108_Be | chr6:32044254-32044404 | TNXB | N_Shelf | Body | Yes |
| Control-Sample13_Beta | chr6:32060791-32060816 | TNXB | N_Shelf | Body | Yes |
| Control-Sample15_Beta | chr6:32064246-32064491 | TNXB | Island | Body | Yes |
| Control-Sample16_Beta | chr6:32060794-32061084 | TNXB | N_Shelf | Body | Yes |
| Control-Sample16_Beta | chr6:32064582-32064613 | TNXB | Island | Body | Yes |
| Control-Sample171_Be | chr6:32026605-32026656 | TNXB | | Body | Yes |
| Control-Sample20_Beta | chr6:32060681-32060816 | TNXB | N_Shelf | Body | Yes |
| Control-Sample23_Beta | chr6:32044179-32044254 | TNXB | N_Shelf | Body | Yes |
| Control-Sample280_Be | chr6:32064161-32064212 | TNXB | Island | Body | Yes |
| Control-Sample290_Be | chr6:32044104-32044254 | TNXB | N_Shelf | Body | Yes |
| Control-Sample35_Beta | chr6:32044104-32044179 | TNXB | N_Shelf | Body | Yes |
| Control-Sample35_Beta | chr6:32064239-32064491 | TNXB | Island | Body | Yes |
| Control-Sample36_Beta | chr6:32064212-32064258 | TNXB | Island | Body | Yes |
| Control-Sample49_Beta | chr6:32076079-32076144 | TNXB | | 5'UTR | Yes |
| Control-Sample1_Beta | chr6:32014605-32014739 | TNXB;TNXB | | Body;TSS1500 | Yes |
| Control-Sample10_Beta | chr6:32014300-32014739 | TNXB;TNXB | | Body;TSS1500 | Yes |
| Control-Sample11_Beta | chr6:32014484-32014739 | TNXB;TNXB | | Body;TSS1500 | Yes |
| Control-Sample12_Beta | chr6:32014605-32014674 | TNXB;TNXB | | Body;TSS1500 | Yes |
| Control-Sample18_Beta | chr6:32014300-32014484 | TNXB;TNXB | | Body;TSS1500 | Yes |
| Control-Sample4_Beta | chr6:32014484-32014674 | TNXB;TNXB | | Body;TSS1500 | Yes |
| Control-Sample58_Beta | chr6:32014510-32014739 | TNXB;TNXB | | Body;TSS1500 | Yes |
| Control-Sample64_Beta | chr6:32014300-32014674 | TNXB;TNXB | | Body;TSS1500 | Yes |
| Control-Sample65_Beta | chr6:32014300-32014476 | TNXB;TNXB | | Body;TSS1500 | Yes |
| Control-Sample66_Beta | chr6:32014381-32014484 | TNXB;TNXB | | Body;TSS1500 | Yes |
| Control-Sample1_Beta | chr11:1297250-1297558 | TOLLIP | N_Shore | 3'UTR | Yes |
| Control-Sample1_Beta | chr11:1319689-1320329 | TOLLIP | S_Shelf | Body | Yes |
| Control-Sample12_Beta | chr11:1319713-1320329 | TOLLIP | S_Shelf | Body | Yes |
| Control-Sample124_Be | chr11:1320497-1320634 | TOLLIP | S_Shelf | Body | Yes |
| Control-Sample170_Be | chr11:1320497-1321009 | TOLLIP | S_Shelf | Body | Yes |
| Control-Sample171_Be | chr11:1319713-1320114 | TOLLIP | S_Shelf | Body | Yes |
| Control-Sample171_Be | chr11:1320497-1321530 | TOLLIP | S_Shelf | Body | Yes |
| Control-Sample19_Beta | chr11:1325718-1325852 | TOLLIP | | Body | Yes |
| Control-Sample216_Be | chr11:1321009-1321530 | TOLLIP | S_Shelf | Body | Yes |
| Control-Sample270_Be | chr11:1320546-1321213 | TOLLIP | S_Shelf | Body | Yes |
| Control-Sample331_Be | chr11:1316436-1316833 | TOLLIP | Island | Body | Yes |
| Control-Sample57_Beta | chr11:1319713-1320497 | TOLLIP | S_Shelf | Body | Yes |
| Control-Sample58_Beta | chr11:1319689-1320497 | TOLLIP | S_Shelf | Body | Yes |
| Control-Sample50_Beta | chr22:22337369-22337468 | TOP3B | S_Shore | TSS1500 | No |
| Control-Sample41_Beta | chr16:52581719-52582072 | TOX3;TOX3 | S_Shore | TSS200;TSS1500 | No |
| Control-Sample10_Beta | chr11:128812846-128813008 | TP53AIP1 | | TSS200 | Yes |
| Control-Sample12_Beta | chr11:128812978-128813008 | TP53AIP1 | | TSS200 | Yes |
| Control-Sample19_Beta | chr11:128813442-128813583 | TP53AIP1 | | TSS1500 | Yes |
| Control-Sample12_Beta | chr20:45313109-45313215 | TP53RK;SLC13A3 | | 3'UTR;TSS200 | Yes |
| Control-Sample13_Beta | chr20:45313109-45313200 | TP53RK;SLC13A3 | | 3'UTR;TSS200 | Yes |
| Control-Sample18_Beta | chr20:45313109-45313390 | TP53RK;SLC13A3 | | 3'UTR;TSS200 | Yes |
| Control-Sample6_Beta | chr20:45313200-45313390 | TP53RK;SLC13A3 | | 3'UTR;TSS200 | Yes |
| Control-Sample1_Beta | chr3:189348936-189349021 | TP63;TP63;TP63 | | TSS1500;TSS1500;TSS1500 | No |
| Control-Sample29_Beta | chr1:3600735-3600879 | TP73 | | Body | Yes |
| Control-Sample1_Beta | chr1:3634811-3635100 | TP73;TP73;TP73;TP73 | Island | Body;Body;Body;Body | Yes |
| Control-Sample1_Beta | chr1:3649344-3649574 | TP73;TP73;TP73;TP73 | Island | Body;Body;3'UTR;3'UTR | Yes |
| Control-Sample17_Beta | chr1:3634044-3635100 | TP73;TP73;TP73;TP73 | N_Shore | Body;Body;Body;Body | Yes |
| Control-Sample66_Beta | chr1:3634903-3635100 | TP73;TP73;TP73;TP73 | Island | Body;Body;Body;Body | Yes |
| Control-Sample9_Beta | chr1:3634811-3635210 | TP73;TP73;TP73;TP73 | Island | Body;Body;Body;Body | Yes |
| Control-Sample1_Beta | chr1:154127952-154128041 | TPM3;TPM3;NUP210L;TPM3;TPM3;NUP210L | | 3'UTR;3'UTR;TSS1500;3'UTR;3'UTR;TSS1500 | No |
| Control-Sample1_Beta | chr19:16178099-16178253 | TPM4 | N_Shore | TSS1500 | Yes |
| Control-Sample16_Beta | chr19:16178097-16178253 | TPM4 | N_Shore | TSS1500 | Yes |
| Control-Sample19_Beta | chr19:16178091-16178253 | TPM4 | N_Shore | TSS1500 | Yes |
| Control-Sample20_Beta | chr19:16178099-16178240 | TPM4 | N_Shore | TSS1500 | Yes |
| Control-Sample53_Beta | chr19:16178091-16178099 | TPM4 | N_Shore | TSS1500 | Yes |
| Control-Sample464_Be | chr19:16178426-16178570 | TPM4;TPM4 | Island | 5'UTR;1stExon | Yes |
| Control-Sample64_Beta | chr2:1543996-1544120 | TPO;TPO;TPO;TPO | N_Shore | Body;Body;Body;Body | No |
| Control-Sample10_Beta | chr5:669733-669925 | TPPP | S_Shore | Body | Yes |
| Control-Sample44_Beta | chr5:669397-669837 | TPPP | S_Shore | Body | Yes |
| Control-Sample14_Beta | chr16:67428155-67428497 | TPPP3;TPPP3 | Island | TSS1500;TSS1500 | Yes |
| Control-Sample34_Beta | chr16:67428315-67428497 | TPPP3;TPPP3 | Island | TSS1500;TSS1500 | Yes |
| Control-Sample65_Beta | chr16:67428155-67428898 | TPPP3;TPPP3 | Island | TSS1500;TSS1500 | Yes |
| Control-Sample10_Beta | chr13:45911642-45912168 | TPT1;SNORA31 | N_Shelf | Body;Body | No |
| Control-Sample17_Beta | chr22:50629038-50629224 | TRABD | Island | 5'UTR | No |
| Control-Sample19_Beta | chr14:103294574-103294740 | TRAF3;TRAF3;TRAF3 | | 5'UTR;5'UTR;5'UTR | Yes |
| Control-Sample356_Be | chr14:103367489-103367858 | TRAF3;TRAF3;TRAF3 | | Body;Body;Body | Yes |
| Control-Sample607_Be | chr14:103367591-103367858 | TRAF3;TRAF3;TRAF3 | | Body;Body;Body | Yes |
| Control-Sample108_Be | chr6:111888620-111888683 | TRAF3IP2;TRAF3IP2;TRAF3IP2;TRAF3IP2;TRAF3IP2 | | Body;Body;Body;TSS200;Body | Yes |
| Control-Sample108_Be | chr6:111927123-111927323 | TRAF3IP2;TRAF3IP2;TRAF3IP2;TRAF3IP2;TRAF3IP2 | | 1stExon;Body;5'UTR;1stExon;5'UTR | Yes |
| Control-Sample61_Beta | chr4:118006619-118006750 | TRAM1L1;TRAM1L1 | Island | 1stExon;5'UTR | No |

| | | | | | |
|---|---|---|---|---|---|
| Control-Sample1_Beta | chr8:141248632-141249094 | TRAPPC9;TRAPPC9 | | Body;Body | Yes |
| Control-Sample10_Beta | chr8:141248582-141249094 | TRAPPC9;TRAPPC9 | | Body;Body | Yes |
| Control-Sample20_Beta | chr8:141248667-141249094 | TRAPPC9;TRAPPC9 | | Body;Body | Yes |
| Control-Sample223_Be | chr8:141057285-141057427 | TRAPPC9;TRAPPC9 | Island | Body;Body | Yes |
| Control-Sample55_Beta | chr8:141057285-141057460 | TRAPPC9;TRAPPC9 | Island | Body;Body | Yes |
| Control-Sample57_Beta | chr8:141248921-141249094 | TRAPPC9;TRAPPC9 | | Body;Body | Yes |
| Control-Sample64_Beta | chr8:141468762-141468933 | TRAPPC9;TRAPPC9 | S_Shore | TSS1500;TSS200 | Yes |
| Control-Sample1_Beta | chr3:129692887-129693370 | TRH | N_Shore | TSS1500 | No |
| Control-Sample44_Beta | chr2:12856702-12856844 | TRIB2;TRIB2 | Island | TSS1500;TSS1500 | No |
| Control-Sample101_Be | chr6:30122523-30122593 | TRIM10;TRIM10 | | Body;Body | Yes |
| Control-Sample109_Be | chr6:30122544-30122593 | TRIM10;TRIM10 | | Body;Body | Yes |
| Control-Sample19_Beta | chr6:30122947-30123226 | TRIM10;TRIM10 | | Body;Body | Yes |
| Control-Sample20_Beta | chr6:30123191-30123226 | TRIM10;TRIM10 | | Body;Body | Yes |
| Control-Sample233_Be | chr6:30122523-30122565 | TRIM10;TRIM10 | | Body;Body | Yes |
| Control-Sample544_Be | chr6.0:30122544-30122593 | TRIM10;TRIM10 | | Body;Body | Yes |
| Control-Sample558_Be | chr6.0:30122523-30122593 | TRIM10;TRIM10 | | Body;Body | Yes |
| Control-Sample567_Be | chr6.0:30122523-30122565 | TRIM10;TRIM10 | | Body;Body | Yes |
| Control-Sample12_Beta | chr6:30140145-30140325 | TRIM15 | Island | 3'UTR | Yes |
| Control-Sample34_Beta | chr6:30140305-30140366 | TRIM15 | S_Shore | 3'UTR | Yes |
| Control-Sample108_Be | chr4:154125374-154125720 | TRIM2;TRIM2 | | TSS1500;5'UTR | No |
| Control-Sample19_Beta | chr11:4415193-4415245 | TRIM21 | | TSS1500 | No |
| Control-Sample19_Beta | chr6:30174845-30175327 | TRIM26 | | 5'UTR | Yes |
| Control-Sample20_Beta | chr6:30175092-30175327 | TRIM26 | | 5'UTR | Yes |
| Control-Sample15_Beta | chr11:120008944-120008998 | TRIM29 | S_Shore | TSS200 | Yes |
| Control-Sample1_Beta | chr11:120008836-120008998 | TRIM29;TRIM29 | S_Shore | 1stExon;5'UTR | Yes |
| Control-Sample10_Beta | chr11:120008836-120008963 | TRIM29;TRIM29 | S_Shore | 1stExon;5'UTR | Yes |
| Control-Sample17_Beta | chr7:99517151-99517196 | TRIM4;TRIM4;TRIM4;TRIM4 | Island | 1stExon;1stExon;5'UTR;5'UTR | No |
| Control-Sample67_Beta | chr6:30104593-30104709 | TRIM40;TRIM40 | | 1stExon;5'UTR | No |
| Control-Sample108_Be | chr11:5705995-5706253 | TRIM5;TRIM5;TRIM5 | | 5'UTR;5'UTR;5'UTR | No |
| Control-Sample1_Beta | chr2:27505196-27505586 | TRIM54;TRIM54 | | TSS1500;TSS1500 | Yes |
| Control-Sample23_Beta | chr2:27505196-27505446 | TRIM54;TRIM54 | | TSS1500;TSS1500 | Yes |
| Control-Sample30_Beta | chr2:27505196-27505530 | TRIM54;TRIM54 | | TSS1500;TSS1500 | Yes |
| Control-Sample35_Beta | chr2:27505396-27505586 | TRIM54;TRIM54 | | TSS1500;TSS1500 | Yes |
| Control-Sample36_Beta | chr2:27505396-27505530 | TRIM54;TRIM54 | | TSS1500;TSS1500 | Yes |
| Control-Sample326_Be | chr11:5617812-5617926 | TRIM6;TRIM6-TRIM34;TRIM6 | | TSS200;TSS200;5'UTR | Yes |
| Control-Sample257_Be | chr11:5617367-5617919 | TRIM6;TRIM6-TRIM34;TRIM6;TRIM6 | | 1stExon;TSS1500;TSS1500;5'UTR | Yes |
| Control-Sample426_Be | chr11:5617367-5617926 | TRIM6;TRIM6-TRIM34;TRIM6;TRIM6 | | 1stExon;TSS1500;TSS1500;5'UTR | Yes |
| Control-Sample3_Beta | chr4:165898912-165898967 | TRIM61 | | TSS200 | Yes |
| Control-Sample612_Be | chr4:165898666-165898967 | TRIM61 | | 5'UTR | Yes |
| Control-Sample613_Be | chr4:165898666-165898848 | TRIM61 | | 5'UTR | Yes |
| Control-Sample70_Beta | chr4:165898835-165898967 | TRIM61 | | TSS200 | Yes |
| Control-Sample15_Beta | chr4:165878219-165878742 | TRIM61;C4orf39 | Island | Body;1stExon | Yes |
| Control-Sample146_Be | chr4:165878136-165878317 | TRIM61;C4orf39;C4orf39 | Island | Body;1stExon;5'UTR | Yes |
| Control-Sample2_Beta | chr4:165878136-165878742 | TRIM61;C4orf39;C4orf39 | Island | Body;1stExon;5'UTR | Yes |
| Control-Sample19_Beta | chr17:73887371-73887651 | TRIM65 | Island | Body | No |
| Control-Sample19_Beta | chr11:8680436-8680729 | TRIM66 | | TSS200 | No |
| Control-Sample279_Be | chr11:5617273-5617926 | TRIM6-TRIM34;TRIM6;TRIM6 | | TSS1500;TSS1500;TSS200 | Yes |
| Control-Sample485_Be | chr11:5617273-5617703 | TRIM6-TRIM34;TRIM6;TRIM6 | | TSS1500;TSS1500;TSS200 | Yes |
| Control-Sample551_Be | chr11.0:5617273-5617919 | TRIM6-TRIM34;TRIM6;TRIM6 | | TSS1500;TSS1500;TSS200 | Yes |
| Control-Sample28_Beta | chr5:180622276-180622643 | TRIM7;TRIM7;TRIM7;TRIM7;TRIM7 | Island | Body;Body;Body;Body;Body | No |
| Control-Sample1_Beta | chr3:32860467-32861407 | TRIM71 | Island | Body | Yes |
| Control-Sample11_Beta | chr3:32860909-32861407 | TRIM71 | N_Shore | Body | Yes |
| Control-Sample12_Beta | chr3:32860467-32861318 | TRIM71 | Island | Body | Yes |
| Control-Sample14_Beta | chr3:32861150-32861407 | TRIM71 | Island | Body | Yes |
| Control-Sample2_Beta | chr3:32860553-32861407 | TRIM71 | N_Shore | Body | Yes |
| Control-Sample1_Beta | chr5:14359544-14359614 | TRIO | | Body | No |
| Control-Sample25_Beta | chr5:912712-913111 | TRIP13 | Island | Body | No |
| Control-Sample111_Be | chr8:125463024-125463310 | TRMT12 | | TSS200 | Yes |
| Control-Sample194_Be | chr8:125463024-125463066 | TRMT12 | | TSS200 | Yes |
| Control-Sample505_Be | chr8.0:125463024-125463310 | TRMT12 | | TSS200 | Yes |
| Control-Sample517_Be | chr8.0:125463024-125463066 | TRMT12 | | TSS200 | Yes |
| Control-Sample10_Beta | chr14:104000460-104000955 | TRMT61A | N_Shore | Body | No |
| Control-Sample1_Beta | chr15:31394021-31394285 | TRPM1 | | TSS200 | No |
| Control-Sample19_Beta | chr11:65712826-65713122 | TSGA10IP | | TSS1500 | No |
| Control-Sample19_Beta | chr14:81425912-81426234 | TSHR;TSHR;TSHR | | Body;Body;Body | Yes |
| Control-Sample270_Be | chr14:81426012-81426234 | TSHR;TSHR;TSHR | | Body;Body;Body | Yes |
| Control-Sample29_Beta | chr14:81425912-81426577 | TSHR;TSHR;TSHR | | Body;Body;Body | Yes |
| Control-Sample19_Beta | chr8:143427631-143427921 | TSNARE1 | N_Shore | Body | Yes |
| Control-Sample19_Beta | chr8:143463227-143463667 | TSNARE1 | | 5'UTR | Yes |
| Control-Sample1_Beta | chr11:844390-844686 | TSPAN4;TSPAN4;TSPAN4;TSPAN4;TSPAN4;TSPAN4;TSPAN | S_Shore | TSS200;5'UTR;5'UTR;5'UTR;5'UTR;5'UTR;5'UTR | Yes |
| Control-Sample41_Beta | chr11:844284-844686 | TSPAN4;TSPAN4;TSPAN4;TSPAN4;TSPAN4;TSPAN4;TSPAN | S_Shore | TSS200;5'UTR;5'UTR;5'UTR;5'UTR;5'UTR;5'UTR | Yes |
| Control-Sample572_Be | chr12.0:71551926-71552336 | TSPAN8 | | TSS200 | No |
| Control-Sample19_Beta | chr12:3281520-3281658 | TSPAN9;TSPAN9 | | 5'UTR;5'UTR | Yes |
| Control-Sample464_Be | chr12:3365600-3365744 | TSPAN9;TSPAN9 | | Body;Body | Yes |
| Control-Sample1_Beta | chr2:3292757-3293115 | TSSC1 | S_Shore | Body | Yes |
| Control-Sample10_Beta | chr2:3317860-3318039 | TSSC1 | Island | Body | Yes |
| Control-Sample11_Beta | chr2:3286338-3286479 | TSSC1 | Island | Body | Yes |
| Control-Sample17_Beta | chr2:3285967-3286412 | TSSC1 | N_Shore | Body | Yes |
| Control-Sample19_Beta | chr2:3199889-3200316 | TSSC1 | | Body | Yes |
| Control-Sample19_Beta | chr2:3260677-3261367 | TSSC1 | | Body | Yes |
| Control-Sample475_Be | chr2:3260980-3261367 | TSSC1 | | Body | Yes |
| Control-Sample637_Be | chr2:3381687-3381734 | TSSC1 | Island | TSS200 | Yes |
| Control-Sample1_Beta | chr1:161007634-161008977 | TSTD1;TSTD1;TSTD1 | N_Shore | Body;3'UTR;Body | Yes |
| Control-Sample10_Beta | chr1:161008127-161008977 | TSTD1;TSTD1;TSTD1 | N_Shore | Body;Body;Body | Yes |
| Control-Sample12_Beta | chr1:161008127-161008542 | TSTD1;TSTD1;TSTD1 | N_Shore | Body;Body;Body | Yes |
| Control-Sample13_Beta | chr1:161007634-161008542 | TSTD1;TSTD1;TSTD1 | N_Shore | Body;3'UTR;Body | Yes |
| Control-Sample19_Beta | chr1:161008811-161008826 | TSTD1;TSTD1;TSTD1 | Island | TSS200;TSS200;TSS200 | Yes |
| Control-Sample23_Beta | chr1:161008297-161008542 | TSTD1;TSTD1;TSTD1 | N_Shore | Body;Body;Body | Yes |
| Control-Sample30_Beta | chr1:161008127-161008977 | TSTD1;TSTD1;TSTD1 | N_Shore | Body;Body;Body | Yes |
| Control-Sample12_Beta | chr1:161008705-161008977 | TSTD1;TSTD1;TSTD1;TSTD1;TSTD1;TSTD1 | Island | 5'UTR;1stExon;5'UTR;1stExon;1stExon;5'UTR | Yes |
| Control-Sample13_Beta | chr1:161008705-161008826 | TSTD1;TSTD1;TSTD1;TSTD1;TSTD1;TSTD1 | Island | 5'UTR;1stExon;5'UTR;1stExon;1stExon;5'UTR | Yes |

| | | | | | |
|---|---|---|---|---|---|
| Control-Sample21_Beta | chr1:161008705-161008824 | TSTD1;TSTD1;TSTD1;TSTD1;TSTD1;TSTD1 | Island | 5'UTR;1stExon;5'UTR;1stExon;1stExon;5'UTR | Yes |
| Control-Sample22_Beta | chr1:161008750-161008826 | TSTD1;TSTD1;TSTD1;TSTD1;TSTD1;TSTD1 | Island | 5'UTR;1stExon;5'UTR;1stExon;1stExon;5'UTR | Yes |
| Control-Sample23_Beta | chr1:161008750-161008977 | TSTD1;TSTD1;TSTD1;TSTD1;TSTD1;TSTD1 | Island | 5'UTR;1stExon;5'UTR;1stExon;1stExon;5'UTR | Yes |
| Control-Sample27_Beta | chr1:161008750-161008824 | TSTD1;TSTD1;TSTD1;TSTD1;TSTD1;TSTD1 | Island | 5'UTR;1stExon;5'UTR;1stExon;1stExon;5'UTR | Yes |
| Control-Sample3_Beta | chr6:43237330-43237642 | TTBK1 | Island | Body | No |
| Control-Sample10_Beta | chr2:3452919-3453196 | TTC15 | S_Shore | Body | No |
| Control-Sample496_Be | chr15:99789622-99789661 | TTC23;TTC23;TTC23;TTC23;TTC23;TTC23;TTC23;TT | N_Shore | 1stExon;5'UTR;5'UTR;1stExon;1stExon;1stExon;5'UTF | No |
| Control-Sample1_Beta | chr18:21572622-21572748 | TTC39C | | TSS200 | No |
| Control-Sample146_Be | chr1:1109315-1109676 | TTLL10 | Island | 5'UTR | Yes |
| Control-Sample251_Be | chr1:1109486-1109832 | TTLL10 | Island | 5'UTR | Yes |
| Control-Sample485_Be | chr1:1109315-1109832 | TTLL10 | Island | 5'UTR | Yes |
| Control-Sample561_Be | chr1.0:1109486-1109832 | TTLL10 | Island | 5'UTR | Yes |
| Control-Sample170_Be | chr7:2672989-2673408 | TTYH3 | S_Shore | Body | Yes |
| Control-Sample5_Beta | chr7:2686678-2686947 | TTYH3 | N_Shore | Body | Yes |
| Control-Sample8_Beta | chr7:2677986-2678248 | TTYH3 | | Body | Yes |
| Control-Sample459_Be | chr10:95363-95661 | TUBB8;TUBB8;TUBB8 | S_Shore | TSS200;1stExon;5'UTR | No |
| Control-Sample44_Beta | chr22:31364274-31364304 | TUG1;MORC2 | N_Shore | TSS1500;TSS200 | No |
| Control-Sample171_Be | chr6:158734097-158734300 | TULP4;TULP4;TULP4;TULP4 | | 1stExon;5'UTR;5'UTR;1stExon | Yes |
| Control-Sample223_Be | chr6:158734097-158734420 | TULP4;TULP4;TULP4;TULP4 | | 1stExon;5'UTR;5'UTR;1stExon | Yes |
| Control-Sample2_Beta | chr7:19158692-19158747 | TWIST1 | S_Shore | TSS1500 | No |
| Control-Sample1_Beta | chr2:239755455-239755585 | TWIST2 | Island | TSS1500 | No |
| Control-Sample52_Beta | chr12:104697419-104697526 | TXNRD1;TXNRD1;TXNRD1;EID3;TXNRD1;TXNRD1 | Island | 5'UTR;5'UTR;Body;TSS200;Body;5'UTR | No |
| Control-Sample108_Be | chr22:19929254-19929286 | TXNRD2;COMT | Island | 1stExon;TSS200 | Yes |
| Control-Sample168_Be | chr22:19929205-19929467 | TXNRD2;COMT | Island | Body;TSS200 | Yes |
| Control-Sample31_Beta | chr22:19929184-19929467 | TXNRD2;COMT | Island | Body;TSS200 | Yes |
| Control-Sample32_Beta | chr22:19929184-19929286 | TXNRD2;COMT | Island | Body;TSS200 | Yes |
| Control-Sample356_Be | chr22:19929205-19929286 | TXNRD2;COMT | Island | Body;TSS200 | Yes |
| Control-Sample538_Be | chr22.0:19929205-19929274 | TXNRD2;COMT | Island | Body;TSS200 | Yes |
| Control-Sample54_Beta | chr22:19929254-19929467 | TXNRD2;COMT | Island | 1stExon;TSS200 | Yes |
| Control-Sample21_Beta | chr3:126327437-126327472 | TXNRD3IT1 | | TSS200 | Yes |
| Control-Sample22_Beta | chr3:126327437-126328226 | TXNRD3IT1 | | TSS200 | Yes |
| Control-Sample1_Beta | chr19:10463831-10464842 | TYK2 | Island | Body | Yes |
| Control-Sample58_Beta | chr19:10464711-10464842 | TYK2 | Island | Body | Yes |
| Control-Sample379_Be | chr1:75198818-75198963 | TYW3;CRYZ;TYW3;CRYZ;CRYZ;CRYZ;TYW3;CRYZ;CRYZ;CRYZ;CRYZ | | TSS200;5'UTR;TSS200;1stExon;1stExon;5'UTR;TSS20 | No |
| Control-Sample30_Beta | chr15:70994632-70995014 | UACA;UACA | | Body;TSS200 | No |
| Control-Sample19_Beta | chr3:49851353-49851713 | UBA7;UBA7 | | 5'UTR;1stExon | No |
| Control-Sample20_Beta | chr16:1373608-1373786 | UBE2I;UBE2I;UBE2I;UBE2I | | Body;Body;Body;Body | No |
| Control-Sample65_Beta | chr22:21921269-21921743 | UBE2L3;UBE2L3;UBE2L3 | N_Shore | TSS1500;TSS1500;TSS1500 | No |
| Control-Sample511_Be | chr1.0:64669336-64669415 | UBE2U | | TSS200 | No |
| Control-Sample1_Beta | chr5:171708886-171709304 | UBTD2 | N_Shore | Body | No |
| Control-Sample22_Beta | chr2:27531310-27531535 | UCN | Island | TSS200 | Yes |
| Control-Sample36_Beta | chr2:27531310-27531360 | UCN | Island | TSS200 | Yes |
| Control-Sample11_Beta | chr4:69435250-69435601 | UGT2B15;UGT2B17 | | TSS1500;TSS1500 | Yes |
| Control-Sample522_Be | chr4.0:69435250-69435601 | UGT2B15;UGT2B17 | | TSS1500;TSS1500 | Yes |
| Control-Sample67_Beta | chr19:4916593-4916983 | UHRF1;UHRF1 | Island | Body;Body | No |
| Control-Sample35_Beta | chr6:150286508-150286838 | ULBP1 | Island | Body | No |
| Control-Sample1_Beta | chr21:43491213-43491319 | UMODL1;UMODL1 | | TSS1500;TSS1500 | Yes |
| Control-Sample19_Beta | chr21:43490967-43491319 | UMODL1;UMODL1 | | TSS1500;TSS1500 | Yes |
| Control-Sample20_Beta | chr21:43490967-43491238 | UMODL1;UMODL1 | | TSS1500;TSS1500 | Yes |
| Control-Sample13_Beta | chr17:73840822-73840922 | UNC13D | | TSS200 | No |
| Control-Sample13_Beta | chr7:907797-907934 | UNC84A;UNC84A | S_Shore | Body;Body | Yes |
| Control-Sample3_Beta | chr7:899806-900037 | UNC84A;UNC84A | | Body;Body | Yes |
| Control-Sample13_Beta | chr3:118892215-118892330 | UPK1B | | TSS1500 | No |
| Control-Sample1_Beta | chr7:48129814-48129904 | UPP1;UPP1 | Island | 5'UTR;5'UTR | Yes |
| Control-Sample10_Beta | chr7:48129797-48129904 | UPP1;UPP1 | Island | 5'UTR;5'UTR | Yes |
| Control-Sample616_Be | chr3:49314595-49314655 | USP4;USP4;C3orf62 | Island | 3'UTR;3'UTR;TSS200 | No |
| Control-Sample17_Beta | chr2:234384319-234384677 | USP40 | N_Shore | 3'UTR | No |
| Control-Sample171_Be | chr7:6198004-6198846 | USP42 | N_Shore | Body | Yes |
| Control-Sample2_Beta | chr7:6198496-6198703 | USP42 | Island | Body | Yes |
| Control-Sample216_Be | chr7:6198496-6198846 | USP42 | Island | Body | Yes |
| Control-Sample108_Be | chr12:95945277-95945384 | USP44 | S_Shelf | TSS200 | Yes |
| Control-Sample1_Beta | chr12:95945082-95945384 | USP44;USP44 | S_Shelf | 1stExon;5'UTR | Yes |
| Control-Sample10_Beta | chr12:95945120-95945384 | USP44;USP44 | S_Shelf | 1stExon;5'UTR | Yes |
| Control-Sample151_Be | chr10:11574300-11574616 | USP6NL;USP6NL | | Body;TSS200 | No |
| Control-Sample26_Beta | chr3:191048361-191048753 | UTS2D;CCDC50;CCDC50 | S_Shore | TSS200;Body;Body | Yes |
| Control-Sample62_Beta | chr3:191048411-191048753 | UTS2D;CCDC50;CCDC50 | S_Shore | TSS200;Body;Body | Yes |
| Control-Sample1_Beta | chr1:160369363-160369869 | VANGL2 | N_Shore | TSS1500 | No |
| Control-Sample2_Beta | chr6:30883762-30884124 | VARS2;VARS2;VARS2 | S_Shore | Body;Body;Body | Yes |
| Control-Sample62_Beta | chr6:30883074-30883203 | VARS2;VARS2;VARS2 | S_Shore | Body;5'UTR;Body | Yes |
| Control-Sample3_Beta | chr10:118891090-118891355 | VAX1 | N_Shore | 3'UTR | Yes |
| Control-Sample331_Be | chr10:118892423-118892869 | VAX1 | Island | Body | Yes |
| Control-Sample5_Beta | chr10:118891090-118892423 | VAX1 | N_Shore | 3'UTR | Yes |
| Control-Sample331_Be | chr10:118893174-118893430 | VAX1;VAX1 | N_Shore | 3'UTR;Body | Yes |
| Control-Sample57_Beta | chr10:118896012-118896516 | VAX1;VAX1 | N_Shore | Body;Body | Yes |
| Control-Sample1_Beta | chr2:71126446-71126524 | VAX2 | N_Shore | TSS1500 | Yes |
| Control-Sample10_Beta | chr2:71134694-71134936 | VAX2 | Island | Body | Yes |
| Control-Sample11_Beta | chr2:71126275-71126524 | VAX2 | N_Shore | TSS1500 | Yes |
| Control-Sample21_Beta | chr2:71134586-71134936 | VAX2 | Island | Body | Yes |
| Control-Sample3_Beta | chr2:71126275-71126505 | VAX2 | N_Shore | TSS1500 | Yes |
| Control-Sample33_Beta | chr2:71134524-71134936 | VAX2 | N_Shore | Body | Yes |
| Control-Sample41_Beta | chr2:71131748-71132250 | VAX2 | Island | Body | Yes |
| Control-Sample50_Beta | chr2:71131629-71132250 | VAX2 | Island | Body | Yes |
| Control-Sample6_Beta | chr2:71126505-71126691 | VAX2 | N_Shore | TSS1500 | Yes |
| Control-Sample9_Beta | chr2:71134630-71134936 | VAX2 | Island | Body | Yes |
| Control-Sample1_Beta | chr6:43736807-43737136 | VEGFA;VEGFA;VEGFA;VEGFA;VEGFA;VEGFA;VEGFA;VEGFA | N_Shore | TSS1500;TSS1500;TSS1500;TSS1500;TSS1500;TSS1500;TSS15 | No |
| Control-Sample15_Beta | chr3:157221198-157221329 | VEPH1;VEPH1;VEPH1;VEPH1 | | TSS200;5'UTR;TSS200;5'UTR | Yes |
| Control-Sample66_Beta | chr3:157221198-157221260 | VEPH1;VEPH1;VEPH1;VEPH1 | | TSS200;5'UTR;TSS200;5'UTR | Yes |
| Control-Sample1_Beta | chr3:11610138-11610239 | VGLL4;VGLL4;VGLL4;VGLL4;VGLL4 | Island | Body;Body;1stExon;Body;5'UTR | Yes |
| Control-Sample100_Be | chr3:11610142-11610239 | VGLL4;VGLL4;VGLL4;VGLL4;VGLL4 | Island | Body;Body;1stExon;Body;5'UTR | Yes |
| Control-Sample12_Beta | chr3:11610138-11610613 | VGLL4;VGLL4;VGLL4;VGLL4;VGLL4 | Island | Body;Body;1stExon;Body;5'UTR | Yes |
| Control-Sample57_Beta | chr3:11610138-11610595 | VGLL4;VGLL4;VGLL4;VGLL4;VGLL4 | Island | Body;Body;1stExon;Body;5'UTR | Yes |

| | | | | | |
|---|---|---|---|---|---|
| Control-Sample13_Beta | chr10:17269274-17269775 | VIM | N_Shore | TSS1500 | Yes |
| Control-Sample36_Beta | chr10:17269643-17270087 | VIM | N_Shore | TSS1500 | Yes |
| Control-Sample464_Be | chr10:17269775-17270091 | VIM | N_Shore | TSS1500 | Yes |
| Control-Sample108_Be | chr6:153071856-153072091 | VIP;VIP | | TSS200;TSS200 | No |
| Control-Sample497_Be | chr7:65419185-65419194 | VKORC1L1 | | Body | No |
| Control-Sample1_Beta | chr1:12538341-12538678 | VPS13D;VPS13D | | Body;Body | Yes |
| Control-Sample17_Beta | chr1:12538455-12538678 | VPS13D;VPS13D | | Body;Body | Yes |
| Control-Sample30_Beta | chr8:145654605-145654780 | VPS28;NFKBIL2;VPS28 | Island | TSS1500;Body;TSS1500 | Yes |
| Control-Sample43_Beta | chr8:145654565-145654854 | VPS28;NFKBIL2;VPS28 | Island | TSS1500;Body;TSS1500 | Yes |
| Control-Sample1_Beta | chr12:123352600-123352774 | VPS37B | S_Shore | Body | Yes |
| Control-Sample19_Beta | chr12:123349952-123350361 | VPS37B | N_Shore | 3'UTR | Yes |
| Control-Sample23_Beta | chr12:123380878-123380891 | VPS37B | Island | TSS200 | Yes |
| Control-Sample46_Beta | chr12:123380870-123380891 | VPS37B | Island | TSS200 | Yes |
| Control-Sample47_Beta | chr12:123380870-123380883 | VPS37B | Island | TSS200 | Yes |
| Control-Sample15_Beta | chr6:33240059-33240090 | VPS52;RPS18 | Island | TSS1500;Body | Yes |
| Control-Sample645_Be | chr6:33240024-33240066 | VPS52;RPS18 | Island | TSS1500;Body | Yes |
| Control-Sample66_Beta | chr11:124622373-124622815 | VSIG2 | | TSS1500 | Yes |
| Control-Sample15_Beta | chr11:124622096-124622815 | VSIG2;VSIG2 | | 5'UTR;1stExon | Yes |
| Control-Sample1_Beta | chr1:159825552-159825761 | VSIG8;C1orf204 | S_Shore | Body;TSS1500 | No |
| Control-Sample6_Beta | chr5:140098398-140098576 | VTRNA1-2 | | TSS200 | No |
| Control-Sample108_Be | chr5:140105669-140105729 | VTRNA1-3 | | TSS200 | No |
| Control-Sample108_Be | chr11:123986057-123986094 | VWA5A;VWA5A;VWA5A | | TSS200;TSS200;TSS200 | Yes |
| Control-Sample108_Be | chr11:123986121-123986166 | VWA5A;VWA5A;VWA5A;VWA5A;VWA5A;VWA5A | | 1stExon;5'UTR;1stExon;5'UTR;5'UTR;1stExon | Yes |
| Control-Sample1_Beta | chr1:20616951-20617456 | VWA5B1 | | TSS1500 | Yes |
| Control-Sample10_Beta | chr1:20669724-20669905 | VWA5B1 | Island | Body | Yes |
| Control-Sample18_Beta | chr1:20616975-20617456 | VWA5B1 | | TSS1500 | Yes |
| Control-Sample47_Beta | chr1:20617283-20617456 | VWA5B1 | | TSS200 | Yes |
| Control-Sample41_Beta | chr11:61049230-61049520 | VWCE | | Body | No |
| Control-Sample1_Beta | chr12:6166874-6167147 | VWF | Island | Body | Yes |
| Control-Sample23_Beta | chr12:6233657-6234313 | VWF;VWF | | 1stExon;5'UTR | Yes |
| Control-Sample38_Beta | chr2:114341631-114342108 | WASH2P | Island | Body | No |
| Control-Sample38_Beta | chr15:102501640-102501654 | WASH3P | Island | Body | No |
| Control-Sample108_Be | chr10:50143397-50143517 | WDFY4 | | Body | Yes |
| Control-Sample12_Beta | chr10:50143375-50143483 | WDFY4 | | Body | Yes |
| Control-Sample25_Beta | chr10:50143375-50143517 | WDFY4 | | Body | Yes |
| Control-Sample1_Beta | chr6:169901471-169901503 | WDR27 | | Body | No |
| Control-Sample171_Be | chr6:33255172-33255190 | WDR46;WDR46 | N_Shore | Body;Body | No |
| Control-Sample249_Be | chr7:158656429-158656620 | WDR60 | Island | Body | Yes |
| Control-Sample34_Beta | chr7:158656429-158656677 | WDR60 | Island | Body | Yes |
| Control-Sample512_Be | chr7.0:158656429-158656620 | WDR60 | Island | Body | Yes |
| Control-Sample12_Beta | chr12:122396633-122396687 | WDR66 | | Body | Yes |
| Control-Sample394_Be | chr12:122356316-122356400 | WDR66 | Island | TSS200 | Yes |
| Control-Sample24_Beta | chr2:228736253-228736357 | WDR69 | Island | TSS200 | No |
| Control-Sample151_Be | chr17:1626477-1626575 | WDR81;WDR81;WDR81 | N_Shore | TSS1500;Body;5'UTR | No |
| Control-Sample35_Beta | chr19:33622866-33622911 | WDR88 | N_Shore | TSS200 | No |
| Control-Sample1_Beta | chr16:84336112-84336445 | WFDC1 | | Body | Yes |
| Control-Sample12_Beta | chr16:84336130-84336445 | WFDC1 | | Body | Yes |
| Control-Sample19_Beta | chr16:84341323-84341364 | WFDC1 | | Body | Yes |
| Control-Sample108_Be | chr20:44258197-44258437 | WFDC10A;WFDC9 | | TSS200;5'UTR | No |
| Control-Sample13_Beta | chr20:44420225-44420291 | WFDC3;DNTTIP1 | N_Shore | 5'UTR;TSS1500 | No |
| Control-Sample22_Beta | chr20:43743835-43743921 | WFDC5 | | TSS200 | No |
| Control-Sample170_Be | chr17:48911141-48911181 | WFIKKN2 | | TSS1500 | Yes |
| Control-Sample19_Beta | chr17:48911141-48911410 | WFIKKN2 | | TSS1500 | Yes |
| Control-Sample4_Beta | chr17:48912483-48912545 | WFIKKN2 | | TSS200 | Yes |
| Control-Sample63_Beta | chr4:1994816-1994871 | WHSC2 | Island | Body | No |
| Control-Sample1_Beta | chr7:29923339-29923538 | WIPF3 | N_Shore | Body | No |
| Control-Sample3_Beta | chr7:5253804-5254072 | WIPI2;WIPI2;WIPI2;WIPI2;WIPI2 | | Body;Body;TSS200;Body;Body | No |
| Control-Sample1_Beta | chr2:219744597-219744634 | WNT10A | N_Shore | TSS1500 | Yes |
| Control-Sample13_Beta | chr2:219748503-219749205 | WNT10A | Island | Body | Yes |
| Control-Sample17_Beta | chr2:219748503-219748780 | WNT10A | Island | Body | Yes |
| Control-Sample19_Beta | chr17:44897024-44897512 | WNT3 | Island | TSS1500 | Yes |
| Control-Sample28_Beta | chr17:44896877-44897512 | WNT3 | Island | TSS1500 | Yes |
| Control-Sample61_Beta | chr17:44896923-44897512 | WNT3 | Island | TSS1500 | Yes |
| Control-Sample1_Beta | chr3:55516711-55518798 | WNT5A | N_Shore | Body | Yes |
| Control-Sample11_Beta | chr3:55516711-55518441 | WNT5A | N_Shore | Body | Yes |
| Control-Sample19_Beta | chr3:55516711-55517224 | WNT5A | N_Shore | Body | Yes |
| Control-Sample19_Beta | chr3:55517659-55518131 | WNT5A | Island | Body | Yes |
| Control-Sample19_Beta | chr3:55518383-55518441 | WNT5A | N_Shore | Body | Yes |
| Control-Sample20_Beta | chr3:55517101-55518441 | WNT5A | N_Shore | Body | Yes |
| Control-Sample41_Beta | chr3:55516366-55518131 | WNT5A | Island | Body | Yes |
| Control-Sample48_Beta | chr3:55516711-55518131 | WNT5A | N_Shore | Body | Yes |
| Control-Sample6_Beta | chr3:55516184-55518441 | WNT5A | Island | Body | Yes |
| Control-Sample67_Beta | chr3:55517101-55518131 | WNT5A | N_Shore | Body | Yes |
| Control-Sample403_Be | chr21:40759534-40759686 | WRB;WRB | N_Shore | Body;TSS200 | Yes |
| Control-Sample404_Be | chr21:40759534-40759659 | WRB;WRB | N_Shore | Body;TSS200 | Yes |
| Control-Sample1_Beta | chr11:32451461-32452187 | WT1;WT1;WT1;WT1 | N_Shore | Body;Body;Body;Body | Yes |
| Control-Sample10_Beta | chr11:32451777-32452187 | WT1;WT1;WT1;WT1 | N_Shore | Body;Body;Body;Body | Yes |
| Control-Sample21_Beta | chr11:32452038-32452370 | WT1;WT1;WT1;WT1 | N_Shore | Body;Body;Body;Body | Yes |
| Control-Sample26_Beta | chr11:32451777-32452370 | WT1;WT1;WT1;WT1 | N_Shore | Body;Body;Body;Body | Yes |
| Control-Sample1_Beta | chr16:78539954-78540172 | WWOX | | Body | Yes |
| Control-Sample10_Beta | chr16:78601767-78601961 | WWOX | | Body | Yes |
| Control-Sample17_Beta | chr16:79030857-79030890 | WWOX | | Body | Yes |
| Control-Sample21_Beta | chr16:78601767-78601911 | WWOX | | Body | Yes |
| Control-Sample3_Beta | chr3:149374066-149374535 | WWTR1;WWTR1;WWTR1 | N_Shore | Body;Body;Body | No |
| Control-Sample259_Be | chr17:6659164-6659326 | XAF1;XAF1;XAF1;XAF1 | | 5'UTR;5'UTR;1stExon;1stExon | Yes |
| Control-Sample260_Be | chr17:6659164-6659562 | XAF1;XAF1;XAF1;XAF1 | | 5'UTR;5'UTR;1stExon;1stExon | Yes |
| Control-Sample151_Be | chr8:10907718-10907788 | XKR6 | | Body | No |
| Control-Sample11_Beta | chr20:30582838-30583001 | XKR7 | Island | Body | No |
| Control-Sample1_Beta | chr14:104175555-104175847 | XRCC3;XRCC3;XRCC3 | | Body;Body;Body | Yes |
| Control-Sample1_Beta | chr14:104179095-104179160 | XRCC3;XRCC3;XRCC3 | N_Shelf | 5'UTR;5'UTR;5'UTR | Yes |
| Control-Sample1_Beta | chr16:17254551-17255059 | XYLT1 | | Body | No |

| | | | | | |
|---|---|---|---|---|---|
| Control-Sample551_Be | chr12.0:69753446-69753455 | YEATS4 | Island | TSS200 | No |
| Control-Sample21_Beta | chr19:38794814-38795051 | YIF1B;YIF1B;C19orf33;YIF1B;YIF1B;YIF1B;YIF1B;YIF1B;C19orf33 | | 3'UTR;3'UTR;1stExon;3'UTR;3'UTR;3'UTR;3'UTR;3'UTR | Yes |
| Control-Sample474_Be | chr22:22090796-22090816 | YPEL1 | Island | TSS1500 | No |
| Control-Sample171_Be | chr11:57414401-57414557 | YPEL4 | Island | Body | No |
| Control-Sample42_Beta | chr18:72917163-72917450 | ZADH2 | Island | Body | No |
| Control-Sample19_Beta | chr2:98329811-98330020 | ZAP70 | | TSS1500 | Yes |
| Control-Sample20_Beta | chr2:98329811-98330087 | ZAP70 | | TSS1500 | Yes |
| Control-Sample22_Beta | chr11:113929164-113929287 | ZBTB16 | N_Shore | TSS1500 | Yes |
| Control-Sample11_Beta | chr11:113953462-113953838 | ZBTB16;ZBTB16 | N_Shore | Body;Body | Yes |
| Control-Sample171_Be | chr11:113947148-113947359 | ZBTB16;ZBTB16 | | Body;Body | Yes |
| Control-Sample331_Be | chr11:113947036-113947359 | ZBTB16;ZBTB16 | | Body;Body | Yes |
| Control-Sample54_Beta | chr11:113953462-113953812 | ZBTB16;ZBTB16 | N_Shore | Body;Body | Yes |
| Control-Sample65_Beta | chr11:113953462-113954170 | ZBTB16;ZBTB16 | N_Shore | Body;Body | Yes |
| Control-Sample108_Be | chr3:114790243-114790596 | ZBTB20;ZBTB20;ZBTB20 | | 1stExon;5'UTR;5'UTR | Yes |
| Control-Sample64_Beta | chr3:114102565-114102615 | ZBTB20;ZBTB20;ZBTB20;ZBTB20;ZBTB20;ZBTB20;ZBTB20 | | 5'UTR;5'UTR;5'UTR;5'UTR;5'UTR;5'UTR;TSS200 | Yes |
| Control-Sample20_Beta | chr17:7381622-7381769 | ZBTB4;ZBTB4 | N_Shore | 5'UTR;5'UTR | No |
| Control-Sample223_Be | chr20:62436995-62437035 | ZBTB46 | N_Shelf | TSS200 | Yes |
| Control-Sample23_Beta | chr20:62436768-62437035 | ZBTB46;ZBTB46 | N_Shelf | 1stExon;5'UTR | Yes |
| Control-Sample1_Beta | chr3:42699933-42700585 | ZBTB47 | N_Shore | Body | Yes |
| Control-Sample15_Beta | chr3:42700108-42700585 | ZBTB47 | Island | Body | Yes |
| Control-Sample27_Beta | chr3:42700108-42700354 | ZBTB47 | Island | Body | Yes |
| Control-Sample40_Beta | chr3:42700108-42700285 | ZBTB47 | Island | Body | Yes |
| Control-Sample72_Beta | chr3:42700007-42700354 | ZBTB47 | N_Shore | Body | Yes |
| Control-Sample73_Beta | chr3:42699933-42700285 | ZBTB47 | N_Shore | Body | Yes |
| Control-Sample81_Beta | chr3:42699933-42700354 | ZBTB47 | N_Shore | Body | Yes |
| Control-Sample1_Beta | chr6:149803087-149803444 | ZC3H12D | | 5'UTR | Yes |
| Control-Sample11_Beta | chr6:149803206-149803444 | ZC3H12D | | 5'UTR | Yes |
| Control-Sample42_Beta | chr6:149803111-149803444 | ZC3H12D | | 5'UTR | Yes |
| Control-Sample1_Beta | chr8:144543568-144544041 | ZC3H3 | | Body | Yes |
| Control-Sample1_Beta | chr8:144561626-144561886 | ZC3H3 | N_Shelf | Body | Yes |
| Control-Sample17_Beta | chr8:144561585-144561886 | ZC3H3 | N_Shelf | Body | Yes |
| Control-Sample17_Beta | chr8:144576604-144577238 | ZC3H3 | | Body | Yes |
| Control-Sample19_Beta | chr8:144599027-144599347 | ZC3H3 | N_Shelf | Body | Yes |
| Control-Sample21_Beta | chr8:144543568-144543951 | ZC3H3 | | Body | Yes |
| Control-Sample1_Beta | chr16:11876111-11876391 | ZC3H7A | | Body | No |
| Control-Sample1_Beta | chr16:87469002-87469329 | ZCCHC14 | | Body | No |
| Control-Sample108_Be | chr6:157876915-157877120 | ZDHHC14;ZDHHC14 | | Body;Body | No |
| Control-Sample1_Beta | chr10:31607150-31607160 | ZEB1;ZEB1;ZEB1;LOC220930;ZEB1 | N_Shore | TSS1500;TSS1500;TSS1500;Body;TSS1500 | No |
| Control-Sample10_Beta | chr7:1198662-1198977 | ZFAND2A | N_Shore | 5'UTR | Yes |
| Control-Sample150_Be | chr7:1198841-1198977 | ZFAND2A | N_Shore | 5'UTR | Yes |
| Control-Sample1_Beta | chr16:72981543-72982048 | ZFHX3;ZFHX3 | | Body;Body | Yes |
| Control-Sample19_Beta | chr16:72981203-72982116 | ZFHX3;ZFHX3 | | Body;Body | Yes |
| Control-Sample4_Beta | chr16:72981599-72982048 | ZFHX3;ZFHX3 | | Body;Body | Yes |
| Control-Sample67_Beta | chr14:69256677-69256977 | ZFP36L1 | Island | Body | No |
| Control-Sample506_Be | chr4.0:188916581-188916726 | ZFP42 | N_Shore | TSS1500 | No |
| Control-Sample141_Be | chr6:29645937-29649092 | ZFP57 | | TSS1500 | No |
| Control-Sample19_Beta | chr5:32445429-32445523 | ZFR | S_Shore | TSS1500 | No |
| Control-Sample1_Beta | chr14:104195181-104195375 | ZFYVE21 | S_Shore | Body | Yes |
| Control-Sample11_Beta | chr14:104195181-104195533 | ZFYVE21 | S_Shore | Body | Yes |
| Control-Sample19_Beta | chr8:123875036-123875223 | ZHX2 | | 5'UTR | No |
| Control-Sample1_Beta | chr3:147129778-147131178 | ZIC1 | N_Shore | Body | Yes |
| Control-Sample11_Beta | chr3:147125942-147126119 | ZIC1 | N_Shore | TSS1500 | Yes |
| Control-Sample11_Beta | chr3:147129778-147130536 | ZIC1 | N_Shore | Body | Yes |
| Control-Sample12_Beta | chr3:147125955-147126119 | ZIC1 | N_Shore | TSS1500 | Yes |
| Control-Sample14_Beta | chr3:147125962-147126121 | ZIC1 | N_Shore | TSS1500 | Yes |
| Control-Sample15_Beta | chr3:147125942-147126121 | ZIC1 | N_Shore | TSS1500 | Yes |
| Control-Sample18_Beta | chr3:147129778-147130477 | ZIC1 | N_Shore | Body | Yes |
| Control-Sample194_Be | chr3:147125926-147126119 | ZIC1 | N_Shore | TSS1500 | Yes |
| Control-Sample194_Be | chr3:147128679-147129213 | ZIC1 | Island | 1stExon | Yes |
| Control-Sample21_Beta | chr3:147129778-147129952 | ZIC1 | N_Shore | Body | Yes |
| Control-Sample216_Be | chr3:147125926-147126121 | ZIC1 | N_Shore | TSS1500 | Yes |
| Control-Sample223_Be | chr3:147128679-147129184 | ZIC1 | Island | 1stExon | Yes |
| Control-Sample7_Beta | chr3:147125955-147126121 | ZIC1 | N_Shore | TSS1500 | Yes |
| Control-Sample1_Beta | chr3:147125712-147125758 | ZIC4;ZIC1 | N_Shore | TSS1500;TSS1500 | Yes |
| Control-Sample16_Beta | chr3:147125712-147125782 | ZIC4;ZIC1 | N_Shore | TSS1500;TSS1500 | Yes |
| Control-Sample270_Be | chr3:147125712-147126119 | ZIC4;ZIC1 | N_Shore | TSS1500;TSS1500 | Yes |
| Control-Sample331_Be | chr3:147125712-147126121 | ZIC4;ZIC1 | N_Shore | TSS1500;TSS1500 | Yes |
| Control-Sample148_Be | chr3:147124523-147124827 | ZIC4;ZIC4 | N_Shelf | TSS1500;TSS200 | Yes |
| Control-Sample3_Beta | chr3:147124523-147125114 | ZIC4;ZIC4 | N_Shelf | TSS1500;TSS200 | Yes |
| Control-Sample1_Beta | chr3:147114986-147115506 | ZIC4;ZIC4;ZIC4 | N_Shore | Body;Body;Body | Yes |
| Control-Sample150_Be | chr3:147124363-147125114 | ZIC4;ZIC4;ZIC4 | N_Shelf | 1stExon;5'UTR;TSS1500 | Yes |
| Control-Sample162_Be | chr3:147124363-147124827 | ZIC4;ZIC4;ZIC4 | N_Shelf | 1stExon;5'UTR;TSS1500 | Yes |
| Control-Sample23_Beta | chr3:147123199-147123457 | ZIC4;ZIC4;ZIC4 | N_Shelf | 5'UTR;TSS1500;1stExon | Yes |
| Control-Sample32_Beta | chr3:147113700-147113918 | ZIC4;ZIC4;ZIC4 | Island | Body;Body;Body | Yes |
| Control-Sample10_Beta | chr3:147105899-147106208 | ZIC4;ZIC4;ZIC4;ZIC4;ZIC4 | N_Shelf | 3'UTR;Body;3'UTR;Body;3'UTR | Yes |
| Control-Sample148_Be | chr3:147109426-147109784 | ZIC4;ZIC4;ZIC4;ZIC4;ZIC4 | Island | Body;Body;Body;Body;Body | Yes |
| Control-Sample153_Be | chr3:147109426-147109629 | ZIC4;ZIC4;ZIC4;ZIC4;ZIC4 | Island | Body;Body;Body;Body;Body | Yes |
| Control-Sample155_Be | chr3:147109490-147109784 | ZIC4;ZIC4;ZIC4;ZIC4;ZIC4 | Island | Body;Body;Body;Body;Body | Yes |
| Control-Sample564_Be | chr5.0:140079847-140080094 | ZMAT2 | | TSS200 | No |
| Control-Sample1_Beta | chr7:44795713-44796036 | ZMIZ2;ZMIZ2 | | TSS200;5'UTR | No |
| Control-Sample190_Be | chr19:58951599-58951672 | ZNF132 | Island | TSS200 | Yes |
| Control-Sample83_Beta | chr19:58951602-58951684 | ZNF132 | Island | TSS200 | Yes |
| Control-Sample205_Be | chr19:58220295-58220494 | ZNF154 | Island | Body | No |
| Control-Sample270_Be | chr1:244218327-244218537 | ZNF238;ZNF238 | Island | Body;1stExon | No |
| Control-Sample107_Be | chr8:145955983-145956024 | ZNF251 | Island | Body | No |
| Control-Sample2_Beta | chr6:43337754-43337791 | ZNF318 | Island | TSS1500 | No |
| Control-Sample47_Beta | chr6:28322086-28322103 | ZNF323;ZKSCAN3;ZNF323 | | 5'UTR;5'UTR;TSS200 | No |
| Control-Sample65_Beta | chr8:145998600-145998731 | ZNF34 | | 3'UTR | No |
| Control-Sample314_Be | chr20:2489550-2489564 | ZNF343;ZNF343 | Island | 5'UTR;1stExon | No |
| Control-Sample1_Beta | chr16:49525139-49525304 | ZNF423 | N_Shore | 3'UTR | Yes |
| Control-Sample270_Be | chr16:49525225-49525727 | ZNF423 | N_Shore | Body | Yes |

| | | | | | |
|---|---|---|---|---|---|
| Control-Sample11_Beta | chr16:88496788-88497293 | ZNF469 | Island | 1stExon | Yes |
| Control-Sample26_Beta | chr16:88497007-88497293 | ZNF469 | Island | 1stExon | Yes |
| Control-Sample1_Beta | chr19:12707809-12708229 | ZNF490 | Island | Body | Yes |
| Control-Sample23_Beta | chr19:12707977-12708229 | ZNF490 | Island | Body | Yes |
| Control-Sample24_Beta | chr19:12707809-12708001 | ZNF490 | Island | Body | Yes |
| Control-Sample124_Be | chr10:135122232-135122309 | ZNF511;TUBGCP2 | Island | TSS200;5'UTR | Yes |
| Control-Sample177_Be | chr10:135122757-135122788 | ZNF511;TUBGCP2 | N_Shore | Body;TSS200 | Yes |
| Control-Sample178_Be | chr10:135122752-135122788 | ZNF511;TUBGCP2 | N_Shore | Body;TSS200 | Yes |
| Control-Sample561_Be | chr10.0:135122232-135122309 | ZNF511;TUBGCP2 | Island | TSS200;5'UTR | Yes |
| Control-Sample644_Be | chr19:58038573-58038621 | ZNF549 | Island | TSS200 | Yes |
| Control-Sample89_Beta | chr19:58038573-58038588 | ZNF549 | Island | TSS200 | Yes |
| Control-Sample119_Be | chr19:9731901-9731957 | ZNF561;ZNF561 | Island | 5'UTR;1stExon | Yes |
| Control-Sample223_Be | chr19:9731901-9732278 | ZNF561;ZNF561 | Island | 5'UTR;1stExon | Yes |
| Control-Sample445_Be | chr19:9785295-9786077 | ZNF562;ZNF562;ZNF562 | N_Shore | 5'UTR;5'UTR;5'UTR | No |
| Control-Sample137_Be | chr19:2900708-2900762 | ZNF57 | Island | TSS200 | Yes |
| Control-Sample433_Be | chr19:2900365-2900762 | ZNF57 | Island | TSS1500 | Yes |
| Control-Sample217_Be | chr19:52390810-52391304 | ZNF577;ZNF577;ZNF577 | N_Shore | Body;5'UTR;5'UTR | Yes |
| Control-Sample223_Be | chr19:52390810-52391166 | ZNF577;ZNF577;ZNF577 | N_Shore | Body;5'UTR;5'UTR | Yes |
| Control-Sample223_Be | chr19:52391250-52391304 | ZNF577;ZNF577;ZNF577 | Island | TSS200;TSS200;TSS200 | Yes |
| Control-Sample565_Be | chr19.0:52390810-52391304 | ZNF577;ZNF577;ZNF577 | N_Shore | Body;5'UTR;5'UTR | Yes |
| Control-Sample34_Beta | chr19:52996083-52996617 | ZNF578 | Island | 5'UTR | No |
| Control-Sample19_Beta | chr16:2055033-2055684 | ZNF598 | S_Shore | Body | No |
| Control-Sample1_Beta | chr19:55987366-55987415 | ZNF628 | N_Shore | TSS1500 | No |
| Control-Sample10_Beta | chr1:182026044-182026510 | ZNF648 | Island | Body | No |
| Control-Sample123_Be | chr16:31085579-31085596 | ZNF668;ZNF646 | Island | TSS200;TSS200 | No |
| Control-Sample131_Be | chr19:23941408-23941575 | ZNF681 | Island | Body | Yes |
| Control-Sample588_Be | chr19.0:23941408-23941575 | ZNF681 | Island | Body | Yes |
| Control-Sample117_Be | chr19:21264896-21265421 | ZNF714 | N_Shore | TSS200 | Yes |
| Control-Sample118_Be | chr19:21264896-21265264 | ZNF714 | N_Shore | TSS200 | Yes |
| Control-Sample31_Beta | chr19:21264896-21265364 | ZNF714 | N_Shore | TSS200 | Yes |
| Control-Sample530_Be | chr19.0:21264896-21265264 | ZNF714 | N_Shore | TSS200 | Yes |
| Control-Sample590_Be | chr19.0:21264982-21265421 | ZNF714;ZNF714 | N_Shore | 5'UTR;1stExon | Yes |
| Control-Sample550_Be | chr7.0:63505714-63506261 | ZNF727 | N_Shore | TSS200 | No |
| Control-Sample671_Be | chr19:57988497-57989117 | ZNF772;ZNF772 | | Body;Body | Yes |
| Control-Sample672_Be | chr19:57988958-57989117 | ZNF772;ZNF772 | | TSS200;TSS200 | Yes |
| Control-Sample10_Beta | chr20:57765315-57765943 | ZNF831 | N_Shore | TSS1500 | Yes |
| Control-Sample15_Beta | chr20:57765815-57765943 | ZNF831 | N_Shore | TSS1500 | Yes |
| Control-Sample37_Beta | chr20:57765489-57765943 | ZNF831 | N_Shore | TSS1500 | Yes |
| Control-Sample15_Beta | chr19:52599226-52599329 | ZNF841 | Island | TSS1500 | No |
| Control-Sample11_Beta | chr3:32023402-32023473 | ZNF860;OSBPL10 | S_Shore | 5'UTR;TSS200 | Yes |
| Control-Sample12_Beta | chr3:32023402-32023475 | ZNF860;OSBPL10 | S_Shore | 5'UTR;TSS200 | Yes |
| Control-Sample19_Beta | chr3:32023257-32023475 | ZNF860;OSBPL10 | Island | TSS200;TSS200 | Yes |
| Control-Sample22_Beta | chr3:32023257-32023473 | ZNF860;OSBPL10 | Island | TSS200;TSS200 | Yes |
| Control-Sample1_Beta | chr3:32023353-32023475 | ZNF860;ZNF860;OSBPL10 | Island | 5'UTR;1stExon;TSS200 | Yes |
| Control-Sample10_Beta | chr3:32023353-32023473 | ZNF860;ZNF860;OSBPL10 | Island | 5'UTR;1stExon;TSS200 | Yes |
| Control-Sample28_Beta | chr3:32023353-32023452 | ZNF860;ZNF860;OSBPL10 | Island | 5'UTR;1stExon;TSS200 | Yes |
| Control-Sample199_Be | chr19:58549241-58549711 | ZSCAN1 | N_Shore | Body | No |
| Control-Sample108_Be | chr15:66797150-66797229 | ZWILCH;SNORD18A;RPL4;RPL4;ZWILCH | | TSS1500;TSS1500;1stExon;5'UTR;TSS1500 | Yes |
| Control-1070_M | chr4:24818471-24818615 | CCDC149 | | Body | No |
| Control-1070_M | chr5:135416205-135416412 | MIR886 | Island | Body | Yes |
| Control-1070_M | chr6:3848218-3848280 | FAM50B | N_Shore | TSS1500 | Yes |
| Control-1070_M | chr6:32526027-32526364 | HLA-DRB6 | | Body | Yes |
| Control-1070_M | chr6:32552246-32552515 | HLA-DRB1 | Island | Body | Yes |
| Control-1070_M | chr6:32604564-32605043 | HLA-DQA1 | | TSS1500 | Yes |
| Control-1070_M | chr6:33158020-33158538 | COL11A2;COL11A2;COL11A2;COL11A2 | N_Shore | Body;Body;Body;Body | Yes |
| Control-1070_M | chr6:155584709-155584968 | TFB1M;CLDN20 | | Body;TSS1500 | Yes |
| Control-1070_M | chr6:166836643-166836791 | RPS6KA2;RPS6KA2 | | Body;Body | Yes |
| Control-1070_M | chr9:92291210-92291523 | LOC100129066 | N_Shore | Body | No |
| Control-1070_M | chr13:36944294-36944471 | SPG20;SPG20 | | 1stExon;5'UTR | Yes |
| Control-1070_M | chr13:113461305-113461652 | ATP11A;ATP11A | N_Shelf | Body;Body | Yes |
| Control-1070_M | chr13:114786825-114786887 | RASA3 | S_Shore | Body | Yes |
| Control-1070_M | chr20:44036899-44037436 | DBNDD2;DBNDD2;DBNDD2;SYS1-DBNDD2;DBNDD2;DBN | S_Shore | 5'UTR;5'UTR;TSS200;Body;5'UTR;5'UTR;TSS200 | Yes |
| Control-1070_M | chr6:31237013-31237034 | HLA-C | N_Shore | Body | Yes |
| Control-1070_M | chr6:31237405-31237722 | HLA-C | N_Shore | Body | Yes |
| Control-1070_M | chr8:2075209-2075820 | MYOM2 | N_Shore | Body | Yes |
| Control-1070_M | chr8:65291287-65291523 | MIR124-2 | S_Shore | TSS1500 | No |
| Control-1070_M | chr10:1511024-1511277 | ADARB2 | | Body | Yes |
| Control-1070_M | chr11:1320497-1320634 | TOLLIP | S_Shelf | Body | Yes |
| Control-1070_M | chr19:57988497-57989182 | ZNF772;ZNF772 | | Body;Body | No |
| Control-1070_M | chr20:54987140-54987270 | CASS4;CASS4;CASS4;CASS4 | | TSS200;TSS200;TSS200;TSS200 | Yes |
| Control-1071_F | chr6:29627142-29627296 | MOG;MOG;MOG;MOG;MOG;MOG;MOG;MOG;MOG;MOG;MOG | | Body;Body;Body;Body;Body;Body;Body;Body;Body;B | Yes |
| Control-1071_F | chr6:30122523-30122593 | TRIM10;TRIM10 | | Body;Body | Yes |
| Control-1071_F | chr6:32489963-32489991 | HLA-DRB5 | Island | Body | Yes |
| Control-1071_F | chr6:32552152-32552246 | HLA-DRB1 | Island | Body | Yes |
| Control-1071_F | chr11:113143664-113143736 | NCAM1;NCAM1 | | Body;Body | Yes |
| Control-1071_F | chr19:12876846-12877188 | HOOK2;HOOK2 | N_Shore | Body;Body | Yes |
| Control-1071_F | chr3:138763852-138763899 | PRR23C | Island | TSS200 | No |
| Control-1071_F | chr4:74847646-74847829 | PF4 | Island | 1stExon | No |
| Control-1071_F | chr6:31237013-31237034 | HLA-C | N_Shore | Body | Yes |
| Control-1071_F | chr6:32526021-32526260 | HLA-DRB6 | | Body | Yes |
| Control-1071_F | chr6:32549631-32549935 | HLA-DRB1 | N_Shelf | Body | Yes |
| Control-1071_F | chr6:32557970-32558459 | HLA-DRB1 | | TSS1500 | Yes |
| Control-1071_F | chr7:39170497-39170763 | POU6F2;POU6F2 | | Body;Body | Yes |
| Control-1071_F | chr11:75139390-75139736 | KLHL35 | N_Shore | Body | Yes |
| Control-1071_F | chr16:31439106-31439393 | COX6A2 | Island | Body | Yes |
| Control-1074_M | chr4:69435250-69435601 | UGT2B15;UGT2B17 | | TSS1500;TSS1500 | Yes |
| Control-1074_M | chr6:32526021-32526263 | HLA-DRB6 | | Body | Yes |
| Control-1074_M | chr9:130955135-130955436 | CIZ1;CIZ1;CIZ1;CIZ1;CIZ1 | N_Shore | TSS1500;5'UTR;TSS1500;5'UTR;TSS1500 | Yes |
| Control-1074_M | chr11:5442434-5442511 | OR51Q1 | | TSS1500 | No |
| Control-1074_M | chr5:140552347-140552505 | PCDHB7;PCDHB7 | N_Shore | 1stExon;5'UTR | Yes |
| Control-1074_M | chr5:140552992-140553150 | PCDHB7 | N_Shore | 1stExon | Yes |

| | | | | | |
|---|---|---|---|---|---|
| Control-1074_M | chr5:140563213-140563400 | PCDHB16 | N_Shore | 1stExon | Yes |
| Control-1074_M | chr6:29627215-29627395 | MOG;MOG;MOG;MOG;MOG;MOG;MOG;MOG;MOG;MOG | | Body;Body;Body;Body;Body;Body;Body;Body;Body;Bo | Yes |
| Control-1074_M | chr6:31237013-31237034 | HLA-C | N_Shore | Body | Yes |
| Control-1074_M | chr6:32489963-32489991 | HLA-DRB5 | Island | Body | Yes |
| Control-1074_M | chr6:33158503-33158538 | COL11A2;COL11A2;COL11A2;COL11A2 | N_Shore | Body;Body;Body;Body | Yes |
| Control-1074_M | chr19:12876846-12877188 | HOOK2;HOOK2 | N_Shore | Body;Body | Yes |
| Control-1074_M | chr20:44036899-44037436 | DBNDD2;DBNDD2;DBNDD2;SYS1-DBNDD2;DBNDD2;DBN | S_Shore | 5'UTR;5'UTR;TSS200;Body;5'UTR;5'UTR;TSS200 | Yes |
| Control-1075_M | chr1:205818956-205819463 | PM20D1 | Island | Body | Yes |
| Control-1075_M | chr2:1817263-1817409 | MYT1L | | Body | Yes |
| Control-1075_M | chr4:187125413-187126139 | CYP4V2 | | Body | No |
| Control-1075_M | chr6:31237013-31237034 | HLA-C | N_Shore | Body | Yes |
| Control-1075_M | chr6:32526021-32526260 | HLA-DRB6 | | Body | Yes |
| Control-1075_M | chr6:32549631-32549935 | HLA-DRB1 | N_Shelf | Body | Yes |
| Control-1075_M | chr6:32606845-32607174 | HLA-DQA1 | | Body | Yes |
| Control-1075_M | chr13:103346900-103346962 | C13orf39 | | TSS200 | Yes |
| Control-1078_F | chr5:135416205-135416398 | MIR886 | Island | Body | Yes |
| Control-1078_F | chr6:31237013-31237034 | HLA-C | N_Shore | Body | Yes |
| Control-1078_F | chr6:32526021-32526263 | HLA-DRB6 | | Body | Yes |
| Control-1078_F | chr19:55476665-55477810 | NLRP2 | | TSS1500 | Yes |
| Control-1079_F | chr1:108734992-108735549 | SLC25A24;SLC25A24 | N_Shore | Body;Body | Yes |
| Control-1079_F | chr17:17110120-17110207 | PLD6 | Island | TSS1500 | No |
| Control-1079_F | chr1:2120985-2121521 | C1orf86;LOC100128003 | N_Shore | Body;Body | Yes |
| Control-1079_F | chr1:110230252-110230456 | GSTM1;GSTM1 | Island | TSS200;TSS200 | No |
| Control-1079_F | chr6:29911087-29911104 | HLA-A | Island | Body | Yes |
| Control-1079_F | chr6:32526021-32526260 | HLA-DRB6 | | Body | Yes |
| Control-1079_F | chr17:154420-155045 | RPH3AL | S_Shore | Body | Yes |
| Control-1086_M | chr6:291687-292823 | DUSP22 | N_Shore | TSS1500 | Yes |
| Control-1086_M | chr6:32549631-32549935 | HLA-DRB1 | N_Shelf | Body | Yes |
| Control-1087_F | chr6:291687-292823 | DUSP22 | N_Shore | TSS1500 | Yes |
| Control-1087_F | chr13:103346900-103346962 | C13orf39 | | TSS200 | Yes |
| Control-1087_F | chr13:114103713-114104057 | ADPRHL1;ADPRHL1 | | Body;TSS1500 | Yes |
| Control-1087_F | chr19:12876846-12877188 | HOOK2;HOOK2 | N_Shore | Body;Body | Yes |
| Control-1087_F | chr6:31237013-31237034 | HLA-C | N_Shore | Body | Yes |
| Control-1087_F | chr6:31237405-31237722 | HLA-C | N_Shore | Body | Yes |
| Control-1087_F | chr6:32526260-32526342 | HLA-DRB6 | | Body | Yes |
| Control-1088_M | chr6:31237013-31237034 | HLA-C | N_Shore | Body | Yes |
| Control-1088_M | chr6:32526021-32526260 | HLA-DRB6 | | Body | Yes |
| Control-1088_M | chr6:32549631-32549935 | HLA-DRB1 | N_Shelf | Body | Yes |
| Control-1088_M | chr8:2075469-2075820 | MYOM2 | Island | Body | Yes |
| Control-1091_M | chr6:31237013-31237034 | HLA-C | N_Shore | Body | Yes |
| Control-1091_M | chr6:31237405-31237722 | HLA-C | N_Shore | Body | Yes |
| Control-1091_M | chr6:32526021-32526260 | HLA-DRB6 | | Body | Yes |
| Control-1091_M | chr6:32549631-32549935 | HLA-DRB1 | N_Shelf | Body | Yes |
| Control-1091_M | chr8:2075209-2075820 | MYOM2 | N_Shore | Body | Yes |
| Control-1093_M | chr1:108734992-108735549 | SLC25A24;SLC25A24 | N_Shore | Body;Body | Yes |
| Control-1093_M | chr20:5485144-5485268 | LOC149837 | | Body | Yes |
| Control-1093_M | chr21:47604052-47605174 | C21orf56;C21orf56 | S_Shore | 5'UTR;5'UTR | Yes |
| Control-1099_M | chr5:135416205-135416398 | MIR886 | Island | Body | Yes |
| Control-1099_M | chr6:291687-292823 | DUSP22 | N_Shore | TSS1500 | Yes |
| Control-1099_M | chr6:31237013-31237034 | HLA-C | N_Shore | Body | Yes |
| Control-1099_M | chr7:1080558-1081250 | C7orf50;C7orf50;C7orf50 | N_Shore | Body;Body;Body | Yes |
| Control-1099_M | chr16:857454-857863 | PRR25 | Island | Body | Yes |
| Control-1099_M | chr19:12876846-12877188 | HOOK2;HOOK2 | N_Shore | Body;Body | Yes |
| Control-1099_M | chr1:205818956-205819609 | PM20D1 | Island | Body | Yes |
| Control-1099_M | chr5:179740743-179741120 | GFPT2 | Island | Body | Yes |
| Control-1099_M | chr6:32489963-32489991 | HLA-DRB5 | Island | Body | Yes |
| Control-1099_M | chr12:678960-679287 | NINJ2 | Island | Body | No |
| Control-1099_M | chr15:40268610-40269214 | EIF2AK4 | Island | Body | Yes |
| Control-1107_F | chr1:205819088-205819406 | PM20D1 | Island | 1stExon | Yes |
| Control-1107_F | chr6:291687-292823 | DUSP22 | N_Shore | TSS1500 | Yes |
| Control-1107_F | chr6:31237013-31237034 | HLA-C | N_Shore | Body | Yes |
| Control-1107_F | chr6:32526021-32526263 | HLA-DRB6 | | Body | Yes |
| Control-1107_F | chr6:32606884-32607224 | HLA-DQA1 | | Body | Yes |
| Control-1107_F | chr1:94883469-94884039 | ABCD3;ABCD3 | N_Shore | TSS1500;TSS1500 | Yes |
| Control-1107_F | chr1:94884090-94884541 | ABCD3;ABCD3 | Island | 1stExon;1stExon | Yes |
| Control-1107_F | chr2:110969641-110970249 | NCRNA00116 | N_Shore | Body | No |
| Control-1107_F | chr8:144659831-144660157 | NAPRT1 | Island | Body | No |
| Control-1107_F | chr10:42862876-42863594 | LOC441666 | N_Shore | Body | No |
| Control-1107_F | chr21:15352213-15352641 | C21orf81 | Island | Body | No |
| Control-1108_M | chr6:32522872-32523136 | HLA-DRB6 | | Body | Yes |
| Control-1108_M | chr6:32549631-32549935 | HLA-DRB1 | N_Shelf | Body | Yes |
| Control-1108_M | chr6:32606845-32607174 | HLA-DQA1 | | Body | Yes |
| Control-1108_M | chr19:55477653-55477810 | NLRP2 | | TSS200 | Yes |
| Control-1108_M | chr1:2120985-2121521 | C1orf86;LOC100128003 | N_Shore | Body;Body | Yes |
| Control-1108_M | chr3:46759449-46759475 | PRSS50 | Island | TSS200 | No |
| Control-1108_M | chr5:102898463-102898729 | NUDT12;NUDT12 | | 5'UTR;1stExon | No |
| Control-1108_M | chr6:29911036-29911104 | HLA-A | Island | Body | Yes |
| Control-1108_M | chr6:32489963-32489991 | HLA-DRB5 | Island | Body | Yes |
| Control-1108_M | chr6:32552443-32552515 | HLA-DRB1 | S_Shore | Body | Yes |
| Control-1108_M | chr6:32634245-32634362 | HLA-DQB1 | S_Shore | Body | Yes |
| Control-1108_M | chr11:18477280-18477534 | LDHAL6A;LDHAL6A | N_Shore | TSS200;TSS200 | No |
| Control-1108_M | chr17:77952290-77952488 | TBC1D16 | Island | Body | Yes |
| Control-1108_M | chr19:12876846-12877000 | HOOK2;HOOK2 | N_Shore | Body;Body | Yes |
| Control-1108_M | chr20:5485144-5485475 | LOC149837 | | Body | Yes |
| Control-1110_M | chr5:135416205-135416613 | MIR886 | Island | Body | Yes |
| Control-1110_M | chr6:291687-292823 | DUSP22 | N_Shore | TSS1500 | Yes |
| Control-1110_M | chr6:32549631-32549935 | HLA-DRB1 | N_Shelf | Body | Yes |
| Control-1110_M | chr10:1511024-1511277 | ADARB2 | | Body | Yes |
| Control-1111_F | chr6:29911036-29911104 | HLA-A | Island | Body | Yes |
| Control-1111_F | chr6:31237013-31237034 | HLA-C | N_Shore | Body | Yes |
| Control-1111_F | chr6:32498285-32498677 | HLA-DRB5 | | TSS1500 | Yes |

| | | | | | |
|---|---|---|---|---|---|
| Control-1111_F | chr6:32526021-32526342 | HLA-DRB6 | | Body | Yes |
| Control-1111_F | chr6:32557970-32558459 | HLA-DRB1 | | TSS1500 | Yes |
| Control-1111_F | chr6:32606845-32607509 | HLA-DQA1 | | Body | Yes |
| Control-1111_F | chr6:32608858-32609094 | HLA-DQA1 | | Body | Yes |
| Control-1117_M | chr1:205819088-205819463 | PM20D1 | Island | 1stExon | Yes |
| Control-1117_M | chr4:69435250-69435601 | UGT2B15;UGT2B17 | | TSS1500;TSS1500 | Yes |
| Control-1117_M | chr6:161561031-161561121 | AGPAT4 | | Body | Yes |
| Control-1117_M | chr17:33759484-33759573 | SLFN12;SLFN12 | | 1stExon;5'UTR | Yes |
| Control-1117_M | chr17:33759957-33759986 | SLFN12 | | TSS1500 | Yes |
| Control-1117_M | chr17:33760293-33760527 | SLFN12 | | TSS1500 | Yes |
| Control-1117_M | chr6:31237013-31237034 | HLA-C | N_Shore | Body | Yes |
| Control-1117_M | chr6:31237405-31237722 | HLA-C | N_Shore | Body | Yes |
| Control-1117_M | chr6:32526027-32526263 | HLA-DRB6 | | Body | Yes |
| Control-1118_M | chr1:154127138-154127537 | NUP210L;NUP210L | | Body;Body | No |
| Control-1118_M | chr10:1511024-1511277 | ADARB2 | | Body | Yes |
| Control-1118_M | chr13:114810631-114810794 | RASA3 | N_Shelf | Body | Yes |
| Control-1118_M | chr16:857454-857863 | PRR25 | Island | Body | Yes |
| Control-1118_M | chr22:24348549-24348715 | GSTTP1 | | TSS1500 | No |
| Control-1118_M | chr6:32526021-32526364 | HLA-DRB6 | | Body | Yes |
| Control-1118_M | chr6:32552246-32552515 | HLA-DRB1 | Island | Body | Yes |
| Control-1118_M | chr6:32634245-32634316 | HLA-DQB1 | S_Shore | Body | Yes |
| Control-1118_M | chr22:24384400-24384573 | GSTT1 | Island | TSS200 | Yes |
| Control-1123_F | chr5:135416205-135416398 | MIR886 | Island | Body | Yes |
| Control-1123_F | chr5:179740743-179741120 | GFPT2 | Island | Body | Yes |
| Control-1123_F | chr6:30079203-30079280 | TRIM31 | | Body | Yes |
| Control-1123_F | chr1:205818956-205819609 | PM20D1 | Island | Body | Yes |
| Control-1123_F | chr6:32526021-32526342 | HLA-DRB6 | | Body | Yes |
| Control-1124_M | chr1:110254709-110254835 | GSTM5 | | TSS200 | No |
| Control-1124_M | chr5:135416205-135416412 | MIR886 | Island | Body | Yes |
| Control-1124_M | chr6:291687-292823 | DUSP22 | N_Shore | TSS1500 | Yes |
| Control-1124_M | chr6:32525960-32526364 | HLA-DRB6 | | Body | Yes |
| Control-1124_M | chr6:32552443-32552515 | HLA-DRB1 | S_Shore | Body | Yes |
| Control-1124_M | chr8:145955983-145956024 | ZNF251 | Island | Body | Yes |
| Control-1125_M | chr3:73045440-73045686 | PPP4R2 | N_Shore | TSS1500 | No |
| Control-1125_M | chr6:32526021-32526263 | HLA-DRB6 | | Body | Yes |
| Control-1125_M | chr6:32557970-32558459 | HLA-DRB1 | | TSS1500 | Yes |
| Control-1125_M | chr6:32606845-32607174 | HLA-DQA1 | | Body | Yes |
| Control-1125_M | chr9:130955135-130955436 | CIZ1;CIZ1;CIZ1;CIZ1;CIZ1 | N_Shore | TSS1500;5'UTR;TSS1500;5'UTR;TSS1500 | Yes |
| Control-1125_M | chr6:32551888-32551954 | HLA-DRB1 | Island | Body | Yes |
| Control-1125_M | chr6:32632694-32632731 | HLA-DQB1 | Island | Body | Yes |
| Control-1125_M | chr21:47604052-47605174 | C21orf56;C21orf56 | S_Shore | 5'UTR;5'UTR | Yes |
| Control-1127_F | chr1:205818956-205819492 | PM20D1 | Island | Body | Yes |
| Control-1127_F | chr2:30669597-30669863 | LCLAT1;LCLAT1 | N_Shore | TSS1500;TSS1500 | Yes |
| Control-1127_F | chr7:39170497-39170763 | POU6F2;POU6F2 | | Body;Body | Yes |
| Control-1128_M | chr1:3164390-3164642 | PRDM16;PRDM16 | Island | Body;Body | Yes |
| Control-1128_M | chr1:108734992-108735549 | SLC25A24;SLC25A24 | N_Shore | Body;Body | Yes |
| Control-1128_M | chr2:223917577-223918000 | KCNE4 | Island | Body | Yes |
| Control-1128_M | chr2:235859442-235859792 | SH3BP4 | N_Shore | TSS1500 | No |
| Control-1128_M | chr2:238322274-238322680 | COL6A3;COL6A3;COL6A3;COL6A3;COL6A3 | | 5'UTR;5'UTR;5'UTR;5'UTR;5'UTR | Yes |
| Control-1128_M | chr2:241458886-241458974 | ANKMY1;ANKMY1 | N_Shore | Body;Body | Yes |
| Control-1128_M | chr3:36949957-36950004 | TRANK1 | | Body | Yes |
| Control-1128_M | chr3:46448753-46449100 | CCRL2;CCRL2;CCRL2 | | TSS1500;5'UTR;1stExon | Yes |
| Control-1128_M | chr3:55518383-55518441 | WNT5A | N_Shore | Body | Yes |
| Control-1128_M | chr3:147124363-147124827 | ZIC4;ZIC4;ZIC4 | N_Shelf | 1stExon;5'UTR;TSS1500 | Yes |
| Control-1128_M | chr4:148401828-148401958 | EDNRA;EDNRA;EDNRA | N_Shore | TSS1500;TSS1500;TSS1500 | No |
| Control-1128_M | chr5:92929319-92929679 | NR2F1 | Island | Body | No |
| Control-1128_M | chr6:25962567-25963109 | TRIM38 | | TSS1500 | No |
| Control-1128_M | chr6:30079203-30079280 | TRIM31 | | Body | Yes |
| Control-1128_M | chr6:32116893-32116918 | PRRT1 | Island | 3'UTR | Yes |
| Control-1128_M | chr6:108496460-108496811 | NR2E1 | N_Shore | Body | Yes |
| Control-1128_M | chr6:108497141-108497734 | NR2E1 | Island | Body | Yes |
| Control-1128_M | chr6:149806081-149806635 | ZC3H12D;ZC3H12D | | 5'UTR;1stExon | Yes |
| Control-1128_M | chr7:134832584-134832844 | TMEM140 | | TSS200 | Yes |
| Control-1128_M | chr7:135412356-135412849 | SLC13A4;SLC13A4 | | 5'UTR;1stExon | Yes |
| Control-1128_M | chr7:135413101-135413232 | SLC13A4 | | TSS200 | Yes |
| Control-1128_M | chr7:151442351-151442481 | PRKAG2;PRKAG2 | | Body;Body | Yes |
| Control-1128_M | chr7:157670677-157670732 | PTPRN2;PTPRN2;PTPRN2 | S_Shore | Body;Body;Body | Yes |
| Control-1128_M | chr8:6400815-6401045 | ANGPT2;ANGPT2;ANGPT2;MCPH1 | | Body;Body;Body;Body | Yes |
| Control-1128_M | chr8:41168072-41168264 | SFRP1 | S_Shore | TSS1500 | Yes |
| Control-1128_M | chr8:143781340-143781406 | LY6K;LY6K;LY6K | N_Shore | TSS200;TSS200;TSS200 | Yes |
| Control-1128_M | chr10:134536030-134536157 | INPP5A | Island | Body | Yes |
| Control-1128_M | chr11:2846681-2847258 | KCNQ1;KCNQ1 | | Body;Body | Yes |
| Control-1128_M | chr11:6342214-6342510 | PRKCDBP | S_Shore | TSS1500 | No |
| Control-1128_M | chr12:12938082-12938287 | APOLD1;APOLD1 | N_Shore | Body;TSS1500 | Yes |
| Control-1128_M | chr12:46767665-46767747 | SLC38A2 | S_Shore | TSS1500 | No |
| Control-1128_M | chr12:70083090-70083105 | BEST3;BEST3 | | Body;TSS200 | No |
| Control-1128_M | chr12:113416061-113416268 | OAS2;OAS2;OAS2 | | TSS1500;TSS1500;TSS1500 | Yes |
| Control-1128_M | chr12:123215471-123215684 | GPR81 | | TSS1500 | Yes |
| Control-1128_M | chr13:110951025-110951418 | COL4A1 | | Body | Yes |
| Control-1128_M | chr14:24835104-24835609 | NFATC4 | N_Shore | TSS1500 | No |
| Control-1128_M | chr14:70038884-70038949 | C14orf162 | Island | TSS1500 | No |
| Control-1128_M | chr16:1583899-1584050 | IFT140;TMEM204 | Island | Body;TSS1500 | Yes |
| Control-1128_M | chr16:55794031-55794910 | CES4 | N_Shore | TSS1500 | No |
| Control-1128_M | chr17:149484-149582 | RPH3AL | N_Shelf | Body | Yes |
| Control-1128_M | chr17:855187-855312 | NXN | | Body | Yes |
| Control-1128_M | chr17:6659164-6659562 | XAF1;XAF1;XAF1;XAF1 | | 5'UTR;5'UTR;1stExon;1stExon | No |
| Control-1128_M | chr17:78560478-78560916 | RPTOR;RPTOR | | Body;Body | Yes |
| Control-1128_M | chr17:79259383-79259699 | SLC38A10;SLC38A10 | Island | Body;Body | Yes |
| Control-1128_M | chr17:81047627-81047941 | METRNL | Island | Body | No |
| Control-1128_M | chr19:10928327-10928639 | MIR199A1;DNM2;DNM2;DNM2;DNM2 | | TSS200;Body;Body;Body;Body | Yes |
| Control-1128_M | chr19:13319324-13319523 | CACNA1A;CACNA1A | Island | Body;Body | Yes |

| | | | | | |
|---|---|---|---|---|---|
| Control-1128_M | chr20:54987020-54987270 | CASS4;CASS4;CASS4;CASS4 | | TSS1500;TSS1500;TSS200;TSS1500 | Yes |
| Control-1128_M | chr21:43735412-43735844 | TFF3 | | 1stExon | No |
| Control-1128_M | chr1:6390473-6390595 | ACOT7;ACOT7;ACOT7;ACOT7 | | Body;Body;Body;Body | Yes |
| Control-1128_M | chr1:66458361-66458803 | PDE4B;PDE4B;PDE4B | | TSS200;Body;Body | Yes |
| Control-1128_M | chr2:6122798-6122814 | LOC400940 | | Body | No |
| Control-1128_M | chr3:8811543-8811758 | OXTR | S_Shore | TSS1500 | No |
| Control-1128_M | chr4:186732837-186732932 | SORBS2;SORBS2;SORBS2;SORBS2;SORBS2;SORBS2;SORBS2 | | 5'UTR;5'UTR;5'UTR;1stExon;TSS1500;1stExon;TSS15 | Yes |
| Control-1128_M | chr5:1594282-1594733 | SDHAP3 | Island | Body | Yes |
| Control-1128_M | chr5:110062473-110062837 | TMEM232 | | TSS200 | No |
| Control-1128_M | chr5:138211149-138211184 | CTNNA1;LRRTM2 | | Body;TSS200 | Yes |
| Control-1128_M | chr6:12717016-12717790 | PHACTR1 | | TSS1500 | Yes |
| Control-1128_M | chr6:32557970-32558459 | HLA-DRB1 | | TSS1500 | Yes |
| Control-1128_M | chr6:101846797-101846905 | GRIK2;GRIK2;GRIK2 | Island | TSS200;TSS200;TSS200 | No |
| Control-1128_M | chr6:161860359-161860635 | PARK2;PARK2;PARK2 | | Body;Body;Body | Yes |
| Control-1128_M | chr7:4746837-4747261 | FOXK1 | | Body | Yes |
| Control-1128_M | chr7:37025796-37025890 | ELMO1;ELMO1;ELMO1 | | TSS200;TSS1500;Body | Yes |
| Control-1128_M | chr8:9952926-9953022 | MSRA;MSRA;MSRA | | Body;TSS200;Body | Yes |
| Control-1128_M | chr8:18541556-18541988 | PSD3;PSD3 | | Body;Body | Yes |
| Control-1128_M | chr10:729259-729570 | DIP2C | | Body | Yes |
| Control-1128_M | chr10:62149329-62149642 | ANK3;ANK3 | | 1stExon;5'UTR | Yes |
| Control-1128_M | chr12:67072951-67073062 | GRIP1 | | TSS200 | Yes |
| Control-1128_M | chr12:122356316-122356400 | WDR66 | Island | TSS200 | Yes |
| Control-1128_M | chr17:65471303-65471347 | PITPNC1;PITPNC1 | | Body;Body | Yes |
| Control-1128_M | chr18:24445240-24445318 | AQP4;C18orf16 | S_Shore | Body;TSS200 | No |
| Control-1128_M | chr19:23941408-23941575 | ZNF681 | Island | Body | No |
| Control-1129_M | chr6:31237013-31237034 | HLA-C | N_Shore | Body | Yes |
| Control-1129_M | chr6:31237405-31237722 | HLA-C | N_Shore | Body | Yes |
| Control-1129_M | chr6:32634245-32634362 | HLA-DQB1 | S_Shore | Body | Yes |
| Control-1132_F | chr6:31237013-31237034 | HLA-C | N_Shore | Body | Yes |
| Control-1132_F | chr6:32606845-32607224 | HLA-DQA1 | | Body | Yes |
| Control-1132_F | chr6:32634245-32634362 | HLA-DQB1 | S_Shore | Body | Yes |
| Control-1132_F | chr19:55972791-55973338 | ISOC2;ISOC2;ISOC2 | Island | 5'UTR;5'UTR;5'UTR | Yes |
| Control-1132_F | chr21:47604052-47605174 | C21orf56;C21orf56 | S_Shore | 5'UTR;5'UTR | Yes |
| Control-1132_F | chr22:24384393-24384525 | GSTT1 | Island | TSS200 | Yes |
| Control-1135_F | chr3:53032818-53032988 | SFMBT1;SFMBT1;SFMBT1 | Island | 5'UTR;5'UTR;5'UTR | Yes |
| Control-1135_F | chr6:31237013-31237034 | HLA-C | N_Shore | Body | Yes |
| Control-1135_F | chr6:32526021-32526260 | HLA-DRB6 | | Body | Yes |
| Control-1135_F | chr6:32548627-32549496 | HLA-DRB1 | N_Shelf | Body | Yes |
| Control-1135_F | chr6:32557970-32558459 | HLA-DRB1 | | TSS1500 | Yes |
| Control-1135_F | chr6:32606845-32607174 | HLA-DQA1 | | Body | Yes |
| Control-1135_F | chr6:32551888-32551954 | HLA-DRB1 | Island | Body | Yes |
| Control-1137_M | chr5:135416205-135416398 | MIR886 | Island | Body | Yes |
| Control-1137_M | chr5:135416412-135416613 | MIR886 | Island | TSS200 | Yes |
| Control-1137_M | chr19:12876846-12877188 | HOOK2;HOOK2 | N_Shore | Body;Body | Yes |
| Control-1137_M | chr1:205818956-205819609 | PM20D1 | Island | Body | Yes |
| Control-1137_M | chr5:1594282-1594808 | SDHAP3 | Island | Body | Yes |
| Control-1137_M | chr6:31237013-31237034 | HLA-C | N_Shore | Body | Yes |
| Control-1137_M | chr6:32489963-32489991 | HLA-DRB5 | Island | Body | Yes |
| Control-1137_M | chr22:24384397-24384525 | GSTT1 | Island | TSS200 | Yes |
| Control-1138_F | chr6:31237013-31237034 | HLA-C | N_Shore | Body | Yes |
| Control-1138_F | chr9:130955135-130955436 | CIZ1;CIZ1;CIZ1;CIZ1;CIZ1 | N_Shore | TSS1500;5'UTR;TSS1500;5'UTR;TSS1500 | Yes |
| Control-1138_F | chr11:18433554-18434354 | LDHC;LDHC | | TSS1500;TSS1500 | No |
| Control-1138_F | chr5:141302942-141303209 | KIAA0141;KIAA0141 | N_Shore | TSS1500;TSS1500 | No |
| Control-1138_F | chr6:32526021-32526260 | HLA-DRB6 | | Body | Yes |
| Control-1139_M | chr6:31237013-31237034 | HLA-C | N_Shore | Body | Yes |
| Control-1139_M | chr6:32634245-32634362 | HLA-DQB1 | S_Shore | Body | Yes |
| Control-1139_M | chr8:2075469-2075820 | MYOM2 | Island | Body | Yes |
| Control-1139_M | chr19:12876846-12877188 | HOOK2;HOOK2 | N_Shore | Body;Body | Yes |
| Control-1139_M | chr21:47604654-47605174 | C21orf56;C21orf56 | S_Shore | TSS1500;TSS1500 | Yes |
| Control-1139_M | chr22:24384105-24384400 | GSTT1 | N_Shore | Body | Yes |
| Control-1141_F | chr6:31237013-31237034 | HLA-C | N_Shore | Body | Yes |
| Control-1141_F | chr2:236619539-236619825 | AGAP1;AGAP1 | | Body;Body | Yes |
| Control-1141_F | chr2:240117005-240117640 | HDAC4;MGC16025 | | Body;Body | Yes |
| Control-1141_F | chr6:29627167-29627395 | MOG;MOG;MOG;MOG;MOG;MOG;MOG;MOG;MOG;MOG | | Body;Body;Body;Body;Body;Body;Body;Body;B | Yes |
| Control-1141_F | chr6:32063835-32064032 | TNXB | Island | Body | Yes |
| Control-1141_F | chr6:33158503-33158538 | COL11A2;COL11A2;COL11A2;COL11A2 | N_Shore | Body;Body;Body;Body | Yes |
| Control-1141_F | chr6:155584613-155584850 | TFB1M;CLDN20 | | Body;TSS1500 | Yes |
| Control-1141_F | chr8:9952926-9953022 | MSRA;MSRA;MSRA | | Body;TSS200;Body | Yes |
| Control-1141_F | chr12:106480579-106480771 | NUAK1 | | Body | Yes |
| Control-1141_F | chr13:36944294-36944451 | SPG20;SPG20 | | 1stExon;5'UTR | Yes |
| Control-1141_F | chr15:40268610-40269214 | EIF2AK4 | Island | Body | Yes |
| Control-1141_F | chr18:72201665-72201775 | CNDP1 | | TSS200 | Yes |
| Control-1141_F | chr19:12876846-12877188 | HOOK2;HOOK2 | N_Shore | Body;Body | Yes |
| Control-1141_F | chr22:24384525-24384944 | GSTT1 | S_Shore | TSS1500 | Yes |
| Control-GSM3680799_ | chr1:108734992-108735549 | SLC25A24;SLC25A24 | N_Shore | Body;Body | Yes |
| Control-GSM3680799_ | chr6:31237405-31237722 | HLA-C | N_Shore | Body | Yes |
| Control-GSM3680799_ | chr6:32526027-32526263 | HLA-DRB6 | | Body | Yes |
| Control-GSM3680799_ | chr6:32549849-32550067 | HLA-DRB1 | N_Shelf | Body | Yes |
| Control-GSM3680799_ | chr6:32557970-32558459 | HLA-DRB1 | | TSS1500 | Yes |
| Control-GSM3680799_ | chr6:32605385-32605704 | HLA-DQA1 | | Body | Yes |
| Control-GSM3680799_ | chr14:24457509-24458345 | DHRS4L2 | N_Shore | TSS1500 | No |
| Control-GSM3680799_ | chr14:24505547-24505653 | DHRS4L1 | N_Shore | TSS200 | No |
| Control-GSM3680799_ | chr1:205819088-205819463 | PM20D1 | Island | 1stExon | Yes |
| Control-GSM3680799_ | chr6:32551888-32551954 | HLA-DRB1 | Island | Body | Yes |
| Control-GSM3680799_ | chr7:39170497-39170763 | POU6F2;POU6F2 | | Body;Body | Yes |
| Control-GSM3680799_ | chr8:144660590-144660631 | NAPRT1 | Island | TSS200 | Yes |
| Control-GSM3680799_ | chr15:22833213-22833681 | TUBGCP5;TUBGCP5 | N_Shore | TSS200;TSS200 | Yes |
| Control-GSM3680799_ | chr20:30134929-30135291 | HM13;HM13;HM13;PSIMCT-1 | N_Shore | Body;Body;Body;TSS1500 | No |
| Control-GSM3680799_ | chr21:37442104-37442777 | CBR1 | N_Shore | TSS200 | Yes |
| Control-GSM3680801_ | chr1:205818956-205819463 | PM20D1 | Island | Body | Yes |
| Control-GSM3680801_ | chr6:291687-292823 | DUSP22 | N_Shore | TSS1500 | Yes |

| | | | | | |
|---|---|---|---|---|---|
| Control-GSM3680801_ | chr6:32551888-32551954 | HLA-DRB1 | Island | Body | Yes |
| Control-GSM3680801_ | chr6:32552152-32552246 | HLA-DRB1 | Island | Body | Yes |
| Control-GSM3680801_ | chr21:47604166-47604654 | C21orf56;C21orf56 | S_Shore | 5'UTR;5'UTR | Yes |
| Control-GSM3680801_ | chr1:248100407-248100585 | OR2L13 | Island | TSS200 | Yes |
| Control-GSM3680801_ | chr5:179740743-179741120 | GFPT2 | Island | Body | Yes |
| Control-GSM3680801_ | chr6:31237013-31237034 | HLA-C | N_Shore | Body | Yes |
| Control-GSM3680801_ | chr6:31237405-31237722 | HLA-C | N_Shore | Body | Yes |
| Control-GSM3680801_ | chr6:32526021-32526342 | HLA-DRB6 | | Body | Yes |
| Control-GSM3680801_ | chr6:32606845-32607224 | HLA-DQA1 | | Body | Yes |
| Control-GSM3680801_ | chr8:144659831-144660157 | NAPRT1 | Island | Body | Yes |
| Control-GSM3680801_ | chr8:144660590-144660631 | NAPRT1 | Island | TSS200 | Yes |
| Control-GSM3680801_ | chr10:135341528-135342218 | CYP2E1 | N_Shore | 1stExon | Yes |
| Control-GSM3680801_ | chr15:22833172-22833681 | TUBGCP5;TUBGCP5 | N_Shore | TSS1500;TSS1500 | Yes |
| Control-GSM3680801_ | chr16:2892766-2892782 | TMPRSS8 | Island | TSS200 | No |
| Control-GSM3680801_ | chr22:24384397-24384525 | GSTT1 | Island | TSS200 | Yes |
| Control-GSM3680803_ | chr6:30122523-30122593 | TRIM10;TRIM10 | | Body;Body | Yes |
| Control-GSM3680803_ | chr6:32526260-32526364 | HLA-DRB6 | | Body | Yes |
| Control-GSM3680803_ | chr2:110969641-110970249 | NCRNA00116 | N_Shore | Body | No |
| Control-GSM3680803_ | chr5:179740743-179741120 | GFPT2 | Island | Body | Yes |
| Control-GSM3680803_ | chr6:29910994-29911011 | HLA-A | Island | Body | Yes |
| Control-GSM3680803_ | chr6:29911036-29911104 | HLA-A | Island | Body | Yes |
| Control-GSM3680803_ | chr6:30039013-30039027 | RNF39;RNF39 | Island | Body;Body | Yes |
| Control-GSM3680803_ | chr8:144660590-144660631 | NAPRT1 | Island | TSS200 | Yes |
| Control-GSM3680803_ | chr11:5617273-5617926 | TRIM6-TRIM34;TRIM6;TRIM6 | | TSS1500;TSS1500;TSS200 | Yes |
| Control-GSM3680803_ | chr19:9785511-9785805 | ZNF562;ZNF562;ZNF562 | Island | 5'UTR;5'UTR;5'UTR | Yes |
| Control-GSM3680809_ | chr6:31237013-31237034 | HLA-C | N_Shore | Body | Yes |
| Control-GSM3680809_ | chr17:33759957-33759986 | SLFN12 | | TSS1500 | No |
| Control-GSM3680809_ | chr2:30669597-30669863 | LCLAT1;LCLAT1 | N_Shore | TSS1500;TSS1500 | Yes |
| Control-GSM3680809_ | chr6:29911087-29911095 | HLA-A | Island | Body | Yes |
| Control-GSM3680809_ | chr6:32551888-32551954 | HLA-DRB1 | Island | Body | Yes |
| Control-GSM3680809_ | chr12:678960-679287 | NINJ2 | Island | Body | Yes |
| Control-GSM3680811_ | chr5:179740743-179741120 | GFPT2 | Island | Body | Yes |
| Control-GSM3680811_ | chr6:32064212-32064239 | TNXB | Island | Body | Yes |
| Control-GSM3680811_ | chr6:32606845-32607224 | HLA-DQA1 | | Body | Yes |
| Control-GSM3680811_ | chr5:1594282-1594733 | SDHAP3 | Island | Body | Yes |
| Control-GSM3680811_ | chr5:102898223-102898733 | NUDT12 | | 5'UTR | No |
| Control-GSM3680811_ | chr6:31237013-31237034 | HLA-C | N_Shore | Body | Yes |
| Control-GSM3680811_ | chr6:32526021-32526364 | HLA-DRB6 | | Body | Yes |
| Control-GSM3680811_ | chr6:32552246-32552453 | HLA-DRB1 | Island | Body | Yes |
| Control-GSM3680811_ | chr7:99516603-99517378 | TRIM4;TRIM4 | Island | Body;Body | No |
| Control-GSM3680811_ | chr8:2075209-2075820 | MYOM2 | N_Shore | Body | Yes |
| Control-GSM3680811_ | chr19:13875014-13875289 | MRI1;MRI1 | N_Shore | TSS1500;TSS1500 | No |
| Control-GSM3680811_ | chr22:17956462-17956641 | CECR2 | | TSS200 | Yes |
| Control-GSM3680811_ | chr22:24384397-24384573 | GSTT1 | Island | TSS200 | Yes |
| Control-GSM3680814_ | chr3:39543776-39544192 | MOBP;MOBP | Island | Body;Body | Yes |
| Control-GSM3680814_ | chr6:291687-292385 | DUSP22 | N_Shore | TSS1500 | Yes |
| Control-GSM3680814_ | chr6:32551891-32551954 | HLA-DRB1 | Island | Body | Yes |
| Control-GSM3680814_ | chr6:32552152-32552246 | HLA-DRB1 | Island | Body | Yes |
| Control-GSM3680814_ | chr9:130955135-130955436 | CIZ1;CIZ1;CIZ1;CIZ1;CIZ1 | N_Shore | TSS1500;5'UTR;TSS1500;5'UTR;TSS1500 | Yes |
| Control-GSM3680814_ | chr1:205819088-205819463 | PM20D1 | Island | 1stExon | Yes |
| Control-GSM3680814_ | chr1:248100345-248100614 | OR2L13 | Island | TSS200 | Yes |
| Control-GSM3680814_ | chr6:32526021-32526260 | HLA-DRB6 | | Body | Yes |
| Control-GSM3680814_ | chr19:12876846-12877000 | HOOK2;HOOK2 | N_Shore | Body;Body | Yes |
| Control-GSM3680814_ | chr21:47604052-47605072 | C21orf56;C21orf56 | S_Shore | 5'UTR;5'UTR | Yes |
| Control-GSM3680815_ | chr4:69435250-69435601 | UGT2B15;UGT2B17 | | TSS1500;TSS1500 | Yes |
| Control-GSM3680815_ | chr6:31237013-31237034 | HLA-C | N_Shore | Body | Yes |
| Control-GSM3680815_ | chr6:32526027-32526364 | HLA-DRB6 | | Body | Yes |
| Control-GSM3680815_ | chr22:24348549-24348715 | GSTTP1 | | TSS1500 | Yes |
| Control-GSM3680815_ | chr1:205819088-205819463 | PM20D1 | Island | 1stExon | Yes |
| Control-GSM3680815_ | chr8:144635430-144635547 | GSDMD | Island | TSS200 | Yes |
| Control-GSM3680815_ | chr11:5617812-5617926 | TRIM6;TRIM6-TRIM34;TRIM6 | | TSS200;TSS200;5'UTR | Yes |
| Control-GSM3680815_ | chr21:47604052-47604986 | C21orf56;C21orf56 | S_Shore | 5'UTR;5'UTR | Yes |
| Control-GSM3680815_ | chr22:24384397-24384573 | GSTT1 | Island | TSS200 | Yes |
| Control-GSM3680817_ | chr1:223566643-223567173 | C1orf65 | Island | TSS200 | No |
| Control-GSM3680817_ | chr3:73045440-73045686 | PPP4R2 | N_Shore | TSS1500 | Yes |
| Control-GSM3680817_ | chr5:135416205-135416613 | MIR886 | Island | Body | Yes |
| Control-GSM3680817_ | chr5:179740743-179741120 | GFPT2 | Island | Body | Yes |
| Control-GSM3680817_ | chr6:32063835-32063895 | TNXB | Island | Body | Yes |
| Control-GSM3680817_ | chr6:32525960-32526364 | HLA-DRB6 | | Body | Yes |
| Control-GSM3680817_ | chr16:857454-857863 | PRR25 | Island | Body | Yes |
| Control-GSM3680817_ | chr22:24348549-24348715 | GSTTP1 | | TSS1500 | Yes |
| Control-GSM3680817_ | chr6:31237405-31237722 | HLA-C | N_Shore | Body | Yes |
| Control-GSM3680817_ | chr11:5617367-5617926 | TRIM6;TRIM6-TRIM34;TRIM6;TRIM6 | | 1stExon;TSS1500;TSS1500;5'UTR | Yes |
| Control-GSM3680817_ | chr18:6414974-6414978 | L3MBTL4 | S_Shore | TSS200 | No |
| Control-GSM3680817_ | chr22:24384294-24384525 | GSTT1 | Island | TSS200 | Yes |
| Control-GSM3680818_ | chr1:205818956-205819406 | PM20D1 | Island | Body | Yes |
| Control-GSM3680818_ | chr4:69435250-69435601 | UGT2B15;UGT2B17 | | TSS1500;TSS1500 | Yes |
| Control-GSM3680818_ | chr6:31237013-31237034 | HLA-C | N_Shore | Body | Yes |
| Control-GSM3680818_ | chr6:32552152-32552246 | HLA-DRB1 | Island | Body | Yes |
| Control-GSM3680818_ | chr16:55866757-55867072 | CES1;CES1;CES1 | Island | Body;Body;Body | Yes |
| Control-GSM3680818_ | chr1:2120985-2121521 | C1orf86;LOC100128003 | N_Shore | Body;Body | Yes |
| Control-GSM3680818_ | chr2:30669597-30669863 | LCLAT1;LCLAT1 | N_Shore | TSS1500;TSS1500 | Yes |
| Control-GSM3680818_ | chr5:1594282-1594863 | SDHAP3 | Island | Body | Yes |
| Control-GSM3680818_ | chr6:29911036-29911104 | HLA-A | Island | Body | Yes |
| Control-GSM3680818_ | chr6:29943401-29943414 | HCG9 | N_Shore | Body | Yes |
| Control-GSM3680818_ | chr6:32526021-32526263 | HLA-DRB6 | | Body | Yes |
| Control-GSM3680818_ | chr8:2075209-2075820 | MYOM2 | N_Shore | Body | Yes |
| Control-GSM3680818_ | chr10:135341528-135341933 | CYP2E1 | N_Shore | 1stExon | Yes |
| Control-GSM3680818_ | chr19:12876846-12877000 | HOOK2;HOOK2 | N_Shore | Body;Body | Yes |
| Control-GSM3680818_ | chr19:21265264-21265421 | ZNF714 | Island | 5'UTR | Yes |
| Control-GSM3680818_ | chr20:30073537-30073576 | NCRNA00028 | S_Shore | TSS200 | No |

| | | | | | |
|---|---|---|---|---|---|
| Control-GSM3680823_ | chr1:205818956-205819463 | PM20D1 | Island | Body | Yes |
| Control-GSM3680823_ | chr6:291687-291903 | DUSP22 | N_Shore | TSS1500 | Yes |
| Control-GSM3680823_ | chr6:292329-292823 | DUSP22;DUSP22 | Island | 1stExon;5'UTR | Yes |
| Control-GSM3680823_ | chr6:32551888-32551954 | HLA-DRB1 | Island | Body | Yes |
| Control-GSM3680823_ | chr6:32552152-32552246 | HLA-DRB1 | Island | Body | Yes |
| Control-GSM3680823_ | chr5:1594282-1594733 | SDHAP3 | Island | Body | Yes |
| Control-GSM3680823_ | chr6:32526021-32526263 | HLA-DRB6 | | Body | Yes |
| Control-GSM3680823_ | chr19:52391078-52391304 | ZNF577;ZNF577;ZNF577 | Island | Body;5'UTR;5'UTR | No |
| Control-GSM3680825_ | chr6:31237013-31237034 | HLA-C | N_Shore | Body | Yes |
| Control-GSM3680825_ | chr6:31237405-31237722 | HLA-C | N_Shore | Body | Yes |
| Control-GSM3680825_ | chr6:32526021-32526263 | HLA-DRB6 | | Body | Yes |
| Control-GSM3680825_ | chr6:32557970-32558459 | HLA-DRB1 | | TSS1500 | Yes |
| Control-GSM3680825_ | chr2:239140169-239140190 | LOC151174;LOC643387;LOC151174 | Island | Body;TSS200;Body | Yes |
| Control-GSM3680825_ | chr6:26421231-26421917 | BTN2A3 | | TSS1500 | No |
| Control-GSM3680825_ | chr6:26457984-26458093 | BTN2A1;BTN2A1 | N_Shore | TSS1500;TSS1500 | No |
| Control-GSM3680825_ | chr6:32551888-32551954 | HLA-DRB1 | Island | Body | Yes |
| Control-GSM3680826_ | chr6:30122523-30122593 | TRIM10;TRIM10 | | Body;Body | Yes |
| Control-GSM3680826_ | chr6:32526021-32526260 | HLA-DRB6 | | Body | Yes |
| Control-GSM3680826_ | chr6:32606845-32607224 | HLA-DQA1 | | Body | Yes |
| Control-GSM3680826_ | chr10:94351376-94351686 | KIF11 | N_Shore | | No |
| Control-GSM3680834_ | chr6:291687-292823 | DUSP22 | N_Shore | TSS1500 | Yes |
| Control-GSM3680834_ | chr6:32525960-32526021 | HLA-DRB6 | | Body | Yes |
| Control-GSM3680834_ | chr6:32526263-32526364 | HLA-DRB6 | | Body | Yes |
| Control-GSM3680834_ | chr7:1080558-1081344 | C7orf50;C7orf50;C7orf50 | N_Shore | Body;Body;Body | Yes |
| Control-GSM3680834_ | chr9:130955135-130955436 | CIZ1;CIZ1;CIZ1;CIZ1;CIZ1 | N_Shore | TSS1500;5'UTR;TSS1500;5'UTR;TSS1500 | Yes |
| Control-GSM3680834_ | chr1:2120985-2121521 | C1orf86;LOC100128003 | N_Shore | Body;Body | Yes |
| Control-GSM3680834_ | chr3:182817190-182817479 | MCCC1 | Island | 1stExon | No |
| Control-GSM3680834_ | chr5:1594282-1594863 | SDHAP3 | Island | Body | Yes |
| Control-GSM3680834_ | chr6:84140683-84140928 | ME1;ME1 | Island | 5'UTR;1stExon | No |
| Control-GSM3680834_ | chr8:2075209-2075820 | MYOM2 | N_Shore | Body | Yes |
| Control-GSM3680834_ | chr10:135341528-135342218 | CYP2E1 | N_Shore | 1stExon | Yes |
| Control-GSM3680834_ | chr12:81471882-81471938 | ACSS3;ACSS3 | Island | 1stExon;5'UTR | No |
| Control-GSM3680834_ | chr19:21265264-21265421 | ZNF714 | Island | 5'UTR | Yes |
| Control-GSM3680834_ | chr19:23941207-23941575 | ZNF681 | N_Shore | Body | Yes |
| Control-GSM3680834_ | chr21:37442289-37442476 | CBR1;CBR1 | Island | 1stExon;5'UTR | Yes |
| Control-GSM3680834_ | chr22:24384397-24384525 | GSTT1 | Island | TSS200 | Yes |
| Control-GSM3680835_ | chr5:135416205-135416613 | MIR886 | Island | Body | Yes |
| Control-GSM3680835_ | chr5:179740743-179741120 | GFPT2 | Island | Body | Yes |
| Control-GSM3680835_ | chr1:110254662-110254854 | GSTM5 | | TSS1500 | Yes |
| Control-GSM3680835_ | chr1:205819088-205819463 | PM20D1 | Island | 1stExon | Yes |
| Control-GSM3680835_ | chr6:29911036-29911104 | HLA-A | Island | Body | Yes |
| Control-GSM3680835_ | chr6:30039013-30039142 | RNF39;RNF39 | Island | Body;Body | Yes |
| Control-GSM3680835_ | chr6:30039175-30039206 | RNF39;RNF39 | Island | Body;Body | Yes |
| Control-GSM3680835_ | chr6:32526021-32526260 | HLA-DRB6 | | Body | Yes |
| Control-GSM3680835_ | chr11:5617273-5617926 | TRIM6-TRIM34;TRIM6;TRIM6 | | TSS1500;TSS1500;TSS200 | Yes |
| Control-GSM3680835_ | chr21:37442289-37442777 | CBR1;CBR1 | Island | 1stExon;5'UTR | Yes |
| Control-GSM3680839_ | chr2:240313541-240313668 | HDAC4 | | 5'UTR | Yes |
| Control-GSM3680839_ | chr6:291687-292823 | DUSP22 | N_Shore | TSS1500 | Yes |
| Control-GSM3680839_ | chr6:32522872-32523136 | HLA-DRB6 | | Body | Yes |
| Control-GSM3680839_ | chr6:32549631-32549935 | HLA-DRB1 | N_Shelf | Body | Yes |
| Control-GSM3680839_ | chr7:1080558-1081250 | C7orf50;C7orf50;C7orf50 | N_Shore | Body;Body;Body | Yes |
| Control-GSM3680839_ | chr11:5442434-5442511 | OR51Q1 | | TSS1500 | No |
| Control-GSM3680839_ | chr19:12876846-12877000 | HOOK2;HOOK2 | N_Shore | Body;Body | Yes |
| Control-GSM3680839_ | chr19:55477653-55477810 | NLRP2 | | TSS200 | Yes |
| Control-GSM3680839_ | chr6:32489963-32489991 | HLA-DRB5 | Island | Body | Yes |
| Control-GSM3680839_ | chr6:32551888-32551954 | HLA-DRB1 | Island | Body | Yes |
| Control-GSM3680839_ | chr6:32634245-32634362 | HLA-DQB1 | S_Shore | Body | Yes |
| Control-GSM3680839_ | chr6:160182447-160182624 | ACAT2 | Island | TSS1500 | No |
| Control-GSM3680839_ | chr8:2075209-2075820 | MYOM2 | N_Shore | Body | Yes |
| Control-GSM3680839_ | chr10:42862978-42863539 | LOC441666 | N_Shore | Body | Yes |
| Control-GSM3680844_ | chr11:75139544-75139736 | KLHL35 | Island | Body | Yes |
| Control-GSM3680844_ | chr1:2120985-2121724 | C1orf86;LOC100128003 | N_Shore | Body;Body | Yes |
| Control-GSM3680844_ | chr1:43814306-43814764 | MPL | Island | Body | No |
| Control-GSM3680844_ | chr6:168435914-168436099 | KIF25;KIF25 | Island | Body;Body | Yes |
| Control-GSM3680844_ | chr10:42862978-42863173 | LOC441666 | N_Shore | Body | Yes |
| Control-GSM3680844_ | chr10:42971209-42971732 | LOC84856 | N_Shore | Body | No |
| Control-GSM3680844_ | chr12:312592-313104 | SLC6A12;SLC6A12;SLC6A12 | Island | Body;Body;Body | No |
| Control-GSM3680844_ | chr15:101084509-101084980 | LASS3 | Island | 5'UTR | No |
| Control-GSM3680844_ | chr16:857454-857863 | PRR25 | Island | Body | Yes |
| Control-GSM3680845_ | chr1:50574517-50574837 | ELAVL4;ELAVL4;ELAVL4;ELAVL4;ELAVL4 | | Body;Body;TSS200;Body;TSS200 | Yes |
| Control-GSM3680845_ | chr1:205818956-205819463 | PM20D1 | Island | Body | Yes |
| Control-GSM3680845_ | chr2:10281159-10281258 | C2orf48 | | TSS1500 | No |
| Control-GSM3680845_ | chr6:32063874-32063901 | TNXB | Island | Body | Yes |
| Control-GSM3680845_ | chr6:32551891-32551954 | HLA-DRB1 | Island | Body | Yes |
| Control-GSM3680845_ | chr6:32552152-32552246 | HLA-DRB1 | Island | Body | Yes |
| Control-GSM3680845_ | chr7:157702935-157703425 | PTPRN2;PTPRN2;PTPRN2 | N_Shore | Body;Body;Body | Yes |
| Control-GSM3680845_ | chr8:65291730-65292217 | MIR124-2 | S_Shore | Body | Yes |
| Control-GSM3680845_ | chr10:574291-574361 | DIP2C | N_Shore | Body | Yes |
| Control-GSM3680845_ | chr10:18549378-18549439 | CACNB2;CACNB2;CACNB2;CACNB2;CACNB2;CACNB2;CACNB2 | | Body;TSS1500;Body;Body;Body;Body;Body | No |
| Control-GSM3680845_ | chr11:19736912-19737237 | LOC100126784;NAV2;NAV2;NAV2 | S_Shore | TSS1500;Body;Body;Body | Yes |
| Control-GSM3680845_ | chr11:70440347-70440395 | SHANK2;SHANK2 | | Body;Body | No |
| Control-GSM3680845_ | chr12:111619406-111619455 | CUX2 | | Body | No |
| Control-GSM3680845_ | chr13:99101678-99101761 | FARP1 | | 3'UTR | No |
| Control-GSM3680845_ | chr19:18313289-18313429 | RAB3A | N_Shore | Body | No |
| Control-GSM3680845_ | chr20:62367884-62367893 | LIME1 | N_Shore | TSS200 | No |
| Control-GSM3680845_ | chr22:19879320-19879787 | TXNRD2 | | Body | No |
| Control-GSM3680845_ | chr1:6304199-6304272 | HES3 | N_Shore | TSS200 | No |
| Control-GSM3680845_ | chr1:32708234-32708276 | MTMR9L | S_Shore | TSS1500 | No |
| Control-GSM3680845_ | chr1:151344806-151345181 | SELENBP1 | | Body | Yes |
| Control-GSM3680845_ | chr2:7017698-7017787 | RSAD2 | | TSS200 | No |
| Control-GSM3680845_ | chr2:26401640-26402098 | FAM59B | N_Shore | Body | No |

| | | | | | |
|---|---|---|---|---|---|
| Control-GSM3680845_ | chr2:108994116-108994325 | SULT1C4 | | TSS1500 | No |
| Control-GSM3680845_ | chr2:219848333-219848528 | FEV | N_Shore | Body | No |
| Control-GSM3680845_ | chr3:101546486-101547296 | FAM55C;FAM55C;NFKBIZ | | 3'UTR;3'UTR;TSS1500 | Yes |
| Control-GSM3680845_ | chr4:24796919-24797007 | SOD3 | | TSS200 | No |
| Control-GSM3680845_ | chr4:154125208-154125453 | TRIM2;TRIM2 | | TSS1500;5'UTR | Yes |
| Control-GSM3680845_ | chr4:154125594-154125681 | TRIM2;TRIM2 | | TSS200;5'UTR | Yes |
| Control-GSM3680845_ | chr5:16742020-16742179 | MYO10 | | Body | No |
| Control-GSM3680845_ | chr5:140529627-140529775 | PCDHB6 | N_Shore | TSS1500 | Yes |
| Control-GSM3680845_ | chr5:140530028-140530410 | PCDHB6 | N_Shore | 1stExon | Yes |
| Control-GSM3680845_ | chr5:140552347-140552505 | PCDHB7;PCDHB7 | N_Shore | 1stExon;5'UTR | No |
| Control-GSM3680845_ | chr5:140563213-140563400 | PCDHB16 | N_Shore | 1stExon | No |
| Control-GSM3680845_ | chr5:142065539-142065737 | FGF1;FGF1;FGF1;FGF1;FGF1;FGF1;FGF1;FGF1 | | 5'UTR;Body;Body;5'UTR;1stExon;5'UTR;5'UTR;1stExo | Yes |
| Control-GSM3680845_ | chr5:142065819-142066128 | FGF1;FGF1;FGF1;FGF1;FGF1;FGF1;FGF1 | | 5'UTR;5'UTR;5'UTR;1stExon;Body;Body;TSS200 | Yes |
| Control-GSM3680845_ | chr6:29627167-29627296 | MOG;MOG;MOG;MOG;MOG;MOG;MOG;MOG;MOG;MOG | | Body;Body;Body;Body;Body;Body;Body;Body;Body;B | Yes |
| Control-GSM3680845_ | chr6:30122523-30122593 | TRIM10;TRIM10 | | Body;Body | Yes |
| Control-GSM3680845_ | chr6:31237405-31237722 | HLA-C | N_Shore | Body | Yes |
| Control-GSM3680845_ | chr6:32526021-32526260 | HLA-DRB6 | | Body | Yes |
| Control-GSM3680845_ | chr6:72926121-72926485 | RIMS1;RIMS1;RIMS1;RIMS1;RIMS1 | | TSS1500;TSS200;Body;Body;Body | No |
| Control-GSM3680845_ | chr6:138866865-138867046 | NHSL1 | | Body | No |
| Control-GSM3680845_ | chr11:63828569-63828713 | MACROD1 | | Body | No |
| Control-GSM3680845_ | chr13:113411497-113411561 | ATP11A;ATP11A | S_Shore | Body;Body | Yes |
| Control-GSM3680845_ | chr14:21270133-21270924 | RNASE1;RNASE1;RNASE1;RNASE1 | | Body;Body;Body;Body | Yes |
| Control-GSM3680845_ | chr17:33390736-33390854 | RFFL;RFFL;RFFL | | 1stExon;5'UTR;5'UTR | No |
| Control-GSM3680845_ | chr17:41363737-41363789 | TMEM106A | Island | TSS200 | No |
| Control-GSM3680845_ | chr17:78166148-78166385 | CARD14;CARD14 | S_Shore | Body;Body | Yes |
| Control-GSM3680849_ | chr1:161008750-161008824 | TSTD1;TSTD1;TSTD1;TSTD1;TSTD1;TSTD1 | Island | 5'UTR;1stExon;5'UTR;1stExon;1stExon;5'UTR | Yes |
| Control-GSM3680849_ | chr6:291687-292823 | DUSP22 | N_Shore | TSS1500 | Yes |
| Control-GSM3680849_ | chr6:30122523-30122593 | TRIM10;TRIM10 | | Body;Body | Yes |
| Control-GSM3680849_ | chr6:31237013-31237034 | HLA-C | N_Shore | Body | Yes |
| Control-GSM3680849_ | chr6:32526260-32526364 | HLA-DRB6 | | Body | Yes |
| Control-GSM3680849_ | chr16:857454-857981 | PRR25 | Island | Body | Yes |
| Control-GSM3680849_ | chr1:110254692-110254854 | GSTM5 | | TSS200 | Yes |
| Control-GSM3680849_ | chr2:239140032-239140190 | LOC151174;LOC643387;LOC151174 | N_Shore | Body;TSS1500;Body | Yes |
| Control-GSM3680849_ | chr6:32606845-32607224 | HLA-DQA1 | | Body | Yes |
| Control-GSM3680851_ | chr5:126409061-126409198 | FLJ44606;FLJ44606 | Island | 5'UTR;5'UTR | No |
| Control-GSM3680851_ | chr6:30122523-30122593 | TRIM10;TRIM10 | | Body;Body | Yes |
| Control-GSM3680851_ | chr6:31237013-31237034 | HLA-C | N_Shore | Body | Yes |
| Control-GSM3680851_ | chr6:31237405-31237722 | HLA-C | N_Shore | Body | Yes |
| Control-GSM3680851_ | chr6:32549631-32549935 | HLA-DRB1 | N_Shelf | Body | Yes |
| Control-GSM3680851_ | chr13:109741030-109741176 | MYO16 | | Body | No |
| Control-GSM3680851_ | chr16:857454-857981 | PRR25 | Island | Body | Yes |
| Control-GSM3680851_ | chr18:77905663-77905751 | LOC100130522;LOC100130522 | S_Shore | TSS200;TSS200 | No |
| Control-GSM3680851_ | chr21:45705694-45705742 | AIRE | Island | TSS200 | No |
| Control-GSM3680851_ | chr1:2120985-2121724 | C1orf86;LOC100128003 | N_Shore | Body;Body | Yes |
| Control-GSM3680851_ | chr6:32551888-32551954 | HLA-DRB1 | Island | Body | Yes |
| Control-GSM3680851_ | chr6:32552443-32552515 | HLA-DRB1 | S_Shore | Body | Yes |
| Control-GSM3680851_ | chr8:2074935-2075820 | MYOM2 | N_Shore | Body | Yes |
| Control-GSM3680851_ | chr16:46654691-46655373 | SHCBP1 | N_Shore | Body | No |
| Control-GSM3680852_ | chr4:69435250-69435601 | UGT2B15;UGT2B17 | | TSS1500;TSS1500 | Yes |
| Control-GSM3680852_ | chr6:31237013-31237034 | HLA-C | N_Shore | Body | Yes |
| Control-GSM3680852_ | chr6:32526027-32526263 | HLA-DRB6 | | Body | Yes |
| Control-GSM3680852_ | chr16:857454-857981 | PRR25 | Island | Body | Yes |
| Control-GSM3680852_ | chr1:76189707-76189801 | ACADM;ACADM | N_Shore | TSS1500;TSS1500 | Yes |
| Control-GSM3680852_ | chr5:1594282-1594579 | SDHAP3 | Island | Body | Yes |
| Control-GSM3680854_ | chr4:69435250-69435601 | UGT2B15;UGT2B17 | | TSS1500;TSS1500 | Yes |
| Control-GSM3680854_ | chr6:31237013-31237034 | HLA-C | N_Shore | Body | Yes |
| Control-GSM3680854_ | chr16:857454-857981 | PRR25 | Island | Body | Yes |
| Control-GSM3680854_ | chr6:32526021-32526342 | HLA-DRB6 | | Body | Yes |
| Control-GSM3680854_ | chr6:32552246-32552515 | HLA-DRB1 | Island | Body | Yes |
| Control-GSM3680855_ | chr1:1369793-1369948 | VWA1;VWA1 | N_Shore | TSS1500;TSS1500 | No |
| Control-GSM3680855_ | chr1:3807168-3807387 | C1orf174 | N_Shore | Body | No |
| Control-GSM3680855_ | chr1:7308012-7308366 | CAMTA1 | | Body | No |
| Control-GSM3680855_ | chr1:11919215-11919464 | NPPB | Island | TSS1500 | No |
| Control-GSM3680855_ | chr1:19992167-19992564 | HTR6;HTR6 | Island | 5'UTR;1stExon | No |
| Control-GSM3680855_ | chr1:32169568-32170361 | COL16A1;COL16A1 | Island | 1stExon;5'UTR | No |
| Control-GSM3680855_ | chr1:33231306-33231382 | KIAA1522 | | Body | No |
| Control-GSM3680855_ | chr1:35351340-35351627 | DLGAP3 | Island | Body | No |
| Control-GSM3680855_ | chr1:40149381-40149650 | HPCAL4 | N_Shore | Body | No |
| Control-GSM3680855_ | chr1:113216774-113216829 | MOV10;MOV10 | N_Shore | TSS1500;TSS1500 | No |
| Control-GSM3680855_ | chr1:150229871-150230345 | CA14 | | TSS1500 | No |
| Control-GSM3680855_ | chr1:151345123-151345181 | SELENBP1;SELENBP1 | | 5'UTR;1stExon | Yes |
| Control-GSM3680855_ | chr1:153581494-153581793 | S100A16 | | 5'UTR | No |
| Control-GSM3680855_ | chr1:158688775-158688926 | OR6K3 | | TSS1500 | No |
| Control-GSM3680855_ | chr1:170253405-170253944 | LOC284688 | | TSS200 | No |
| Control-GSM3680855_ | chr1:173176362-173176523 | TNFSF4;TNFSF4 | | 5'UTR;1stExon | No |
| Control-GSM3680855_ | chr1:202091865-202091880 | GPR37L1 | | TSS200 | Yes |
| Control-GSM3680855_ | chr1:202091944-202092285 | GPR37L1 | | TSS200 | Yes |
| Control-GSM3680855_ | chr1:205818956-205819463 | PM20D1 | Island | Body | Yes |
| Control-GSM3680855_ | chr1:205897545-205897890 | SLC26A9;SLC26A9;SLC26A9 | | Body;Body;TSS200 | No |
| Control-GSM3680855_ | chr1:230849519-230850050 | AGT | | 5'UTR | No |
| Control-GSM3680855_ | chr1:230992820-230992990 | C1orf198;C1orf198;C1orf198 | | Body;Body;TSS1500 | No |
| Control-GSM3680855_ | chr2:3704501-3704589 | ALLC | | TSS1500 | No |
| Control-GSM3680855_ | chr2:27301195-27301490 | EMILIN1 | | TSS1500 | No |
| Control-GSM3680855_ | chr2:33151557-33152037 | LOC100271832;LOC285045;LOC285045;LOC285045 | | TSS1500;Body;Body;Body | No |
| Control-GSM3680855_ | chr2:63274615-63275602 | LOC100132215 | Island | Body | No |
| Control-GSM3680855_ | chr2:70188512-70189092 | ASPRV1 | Island | 1stExon | No |
| Control-GSM3680855_ | chr2:74729279-74729502 | LOC151534;LBX2 | N_Shore | TSS1500;Body | No |
| Control-GSM3680855_ | chr2:79738582-79738639 | CTNNA2;CTNNA2 | N_Shore | TSS1500;TSS1500 | No |
| Control-GSM3680855_ | chr2:114033490-114033830 | PAX8;PAX8;PAX8;PAX8;PAX8;LOC440839 | Island | Body;Body;Body;Body;Body;Body | No |
| Control-GSM3680855_ | chr2:121036440-121036760 | RALB | | Body | No |
| Control-GSM3680855_ | chr2:175629219-175629267 | CHRNA1;CHRNA1 | | TSS200;TSS200 | No |

| | | | | | |
|---|---|---|---|---|---|
| Control-GSM3680855_ | chr2:182850269-182850683 | PPP1R1C | | TSS1500 | No |
| Control-GSM3680855_ | chr2:218770208-218770270 | TNS1 | | 5'UTR | No |
| Control-GSM3680855_ | chr2:220381734-220381812 | ACCN4;ACCN4 | S_Shelf | Body;Body | No |
| Control-GSM3680855_ | chr2:225266580-225266880 | FAM124B;FAM124B;FAM124B;FAM124B | | 5'UTR;1stExon;1stExon;5'UTR | Yes |
| Control-GSM3680855_ | chr2:235907647-235907796 | SH3BP4 | | 5'UTR | No |
| Control-GSM3680855_ | chr2:238990408-238990731 | SCLY | | Body | No |
| Control-GSM3680855_ | chr2:239755455-239755585 | TWIST2 | Island | TSS1500 | No |
| Control-GSM3680855_ | chr2:240313541-240313668 | HDAC4 | | 5'UTR | Yes |
| Control-GSM3680855_ | chr3:10805960-10806133 | LOC285370 | | TSS200 | No |
| Control-GSM3680855_ | chr3:101546697-101546859 | FAM55C;FAM55C;NFKBIZ | | 3'UTR;3'UTR;TSS200 | Yes |
| Control-GSM3680855_ | chr3:112359488-112359557 | CCDC80;CCDC80;CCDC80 | | 5'UTR;1stExon;5'UTR | No |
| Control-GSM3680855_ | chr3:114790243-114790596 | ZBTB20;ZBTB20;ZBTB20 | | 1stExon;5'UTR;5'UTR | No |
| Control-GSM3680855_ | chr3:123339334-123339568 | MYLK;MYLK;MYLK;MYLK;MYLK;MYLK;MYLK;MYLK | | 1stExon;5'UTR;Body;Body;Body;5'UTR;1stExon;Body | Yes |
| Control-GSM3680855_ | chr3:141087187-141087313 | ZBTB38 | | 5'UTR | No |
| Control-GSM3680855_ | chr3:196693809-196694153 | PIGZ | Island | 5'UTR | No |
| Control-GSM3680855_ | chr3:197663828-197664106 | IQCG;IQCG | | Body;Body | No |
| Control-GSM3680855_ | chr4:645256-645781 | PDE6B;PDE6B | Island | Body;Body | No |
| Control-GSM3680855_ | chr4:5526510-5526783 | C4orf6 | | TSS1500 | No |
| Control-GSM3680855_ | chr4:38859339-38859770 | TLR6 | | TSS1500 | No |
| Control-GSM3680855_ | chr4:154125208-154125439 | TRIM2;TRIM2 | | TSS1500;5'UTR | Yes |
| Control-GSM3680855_ | chr4:154125594-154125681 | TRIM2;TRIM2 | | TSS200;5'UTR | Yes |
| Control-GSM3680855_ | chr4:184938291-184938470 | STOX2 | | Body | No |
| Control-GSM3680855_ | chr5:231809-231967 | SDHA | | Body | No |
| Control-GSM3680855_ | chr5:415575-415885 | AHRR | | Body | No |
| Control-GSM3680855_ | chr5:9547468-9547825 | SNORD123;SEMA5A | S_Shore | TSS1500;TSS1500 | Yes |
| Control-GSM3680855_ | chr5:42811952-42812844 | SEPP1;SEPP1;SEPP1;SEPP1;SEPP1;SEPP1 | | 1stExon;5'UTR;5'UTR;5'UTR;1stExon;1stExon | Yes |
| Control-GSM3680855_ | chr5:68710813-68710831 | MARVELD2;MARVELD2 | Island | TSS200;TSS200 | No |
| Control-GSM3680855_ | chr5:135416205-135416613 | MIR886 | Island | Body | Yes |
| Control-GSM3680855_ | chr5:141061863-141061943 | ARAP3 | | TSS200 | No |
| Control-GSM3680855_ | chr5:154230173-154230438 | C5orf4;C5orf4 | | 5'UTR;1stExon | No |
| Control-GSM3680855_ | chr5:177876830-177876930 | COL23A1 | | Body | No |
| Control-GSM3680855_ | chr6:291687-292823 | DUSP22 | N_Shore | TSS1500 | Yes |
| Control-GSM3680855_ | chr6:3848218-3848280 | FAM50B | N_Shore | TSS1500 | No |
| Control-GSM3680855_ | chr6:29631295-29631447 | MOG;MOG;MOG;MOG;MOG;MOG;MOG;MOG;MOG;MOG;MOG | | Body;Body;Body;Body;Body;Body;Body;Body;Body;Bo | Yes |
| Control-GSM3680855_ | chr6:29633622-29633971 | MOG;MOG;MOG;MOG;MOG;MOG;MOG;MOG;MOG;MOG;MOG | | 3'UTR;Body;3'UTR;Body;Body;Body;Body;Body;Body; | Yes |
| Control-GSM3680855_ | chr6:29795434-29795501 | HLA-G | N_Shore | 5'UTR | Yes |
| Control-GSM3680855_ | chr6:30039175-30039206 | RNF39;RNF39 | Island | Body;Body | Yes |
| Control-GSM3680855_ | chr6:30039376-30039442 | RNF39;RNF39 | Island | Body;Body | Yes |
| Control-GSM3680855_ | chr6:30175305-30175967 | TRIM26 | | 5'UTR | Yes |
| Control-GSM3680855_ | chr6:30850868-30850913 | DDR1;DDR1 | N_Shore | TSS1500;TSS1500 | Yes |
| Control-GSM3680855_ | chr6:30855729-30856439 | DDR1;DDR1;DDR1 | S_Shelf | TSS1500;5'UTR;5'UTR | Yes |
| Control-GSM3680855_ | chr6:32026605-32026656 | TNXB | | Body | Yes |
| Control-GSM3680855_ | chr6:32131571-32131722 | EGFL8 | N_Shelf | TSS1500 | No |
| Control-GSM3680855_ | chr6:32552152-32552246 | HLA-DRB1 | Island | Body | Yes |
| Control-GSM3680855_ | chr6:33157746-33158538 | COL11A2;COL11A2;COL11A2;COL11A2 | N_Shelf | Body;Body;Body;Body | No |
| Control-GSM3680855_ | chr6:33255172-33255190 | WDR46;WDR46 | N_Shore | Body;Body | No |
| Control-GSM3680855_ | chr6:51814354-51814455 | PKHD1;PKHD1 | | Body;Body | No |
| Control-GSM3680855_ | chr6:76059496-76059773 | FILIP1 | | Body | No |
| Control-GSM3680855_ | chr6:110736422-110737053 | DDO;DDO | | Body;Body | No |
| Control-GSM3680855_ | chr6:148869461-148869895 | SASH1 | | Body | No |
| Control-GSM3680855_ | chr6:155584613-155584968 | TFB1M;CLDN20 | | Body;TSS1500 | No |
| Control-GSM3680855_ | chr6:158404061-158404121 | SYNJ2 | S_Shore | Body | No |
| Control-GSM3680855_ | chr6:158734097-158734420 | TULP4;TULP4;TULP4;TULP4 | | 1stExon;5'UTR;5'UTR;1stExon | No |
| Control-GSM3680855_ | chr6:161796785-161796854 | PARK2;PARK2;PARK2 | | Body;Body;Body | No |
| Control-GSM3680855_ | chr6:166836643-166836791 | RPS6KA2;RPS6KA2 | | Body;Body | No |
| Control-GSM3680855_ | chr6:169629875-169630013 | THBS2 | N_Shore | Body | Yes |
| Control-GSM3680855_ | chr6:169901471-169901503 | WDR27 | | Body | No |
| Control-GSM3680855_ | chr6:170571686-170572016 | LOC154449 | | TSS200 | No |
| Control-GSM3680855_ | chr7:537740-537828 | PDGFA;PDGFA | S_Shelf | 3'UTR;3'UTR | No |
| Control-GSM3680855_ | chr7:4781704-4781872 | FOXK1 | N_Shelf | Body | Yes |
| Control-GSM3680855_ | chr7:4784090-4784419 | FOXK1 | N_Shore | Body | Yes |
| Control-GSM3680855_ | chr7:55980165-55980652 | ZNF713 | | TSS200 | No |
| Control-GSM3680855_ | chr7:75912071-75912462 | SRRM3 | Island | Body | No |
| Control-GSM3680855_ | chr7:134832221-134833299 | TMEM140 | | TSS1500 | Yes |
| Control-GSM3680855_ | chr7:134856562-134856681 | C7orf49;C7orf49;C7orf49 | S_Shore | TSS1500;TSS1500;TSS1500 | No |
| Control-GSM3680855_ | chr7:150652864-150653001 | KCNH2;KCNH2;KCNH2;KCNH2 | Island | 5'UTR;Body;Body;1stExon | No |
| Control-GSM3680855_ | chr7:157394582-157394661 | PTPRN2;PTPRN2;PTPRN2 | | Body;Body;Body | Yes |
| Control-GSM3680855_ | chr8:1786207-1786322 | ARHGEF10 | | 5'UTR | Yes |
| Control-GSM3680855_ | chr8:6419438-6419570 | ANGPT2;ANGPT2;ANGPT2;MCPH1 | | Body;Body;Body;Body | No |
| Control-GSM3680855_ | chr8:10907718-10907788 | XKR6 | | Body | No |
| Control-GSM3680855_ | chr8:18666313-18666561 | PSD3;PSD3;PSD3 | | Body;1stExon;5'UTR | No |
| Control-GSM3680855_ | chr8:53322501-53322619 | ST18 | | TSS200 | No |
| Control-GSM3680855_ | chr9:92291210-92291523 | LOC100129066 | N_Shore | Body | No |
| Control-GSM3680855_ | chr10:537033-537178 | DIP2C | | Body | Yes |
| Control-GSM3680855_ | chr10:88427400-88428177 | LDB3;LDB3;LDB3;LDB3;LDB3;LDB3 | | 3'UTR;3'UTR | Yes |
| Control-GSM3680855_ | chr10:88428228-88428257 | LDB3;LDB3;LDB3;LDB3;LDB3;LDB3;LDB3 | | TSS1500;TSS200;1stExon;TSS1500;5'UTR;TSS1500;T | Yes |
| Control-GSM3680855_ | chr10:93335091-93335196 | LOC100188947 | | Body | No |
| Control-GSM3680855_ | chr10:99477939-99478572 | LOC100270710 | S_Shelf | Body;Body;3'UTR | No |
| Control-GSM3680855_ | chr10:100174935-100175082 | PYROXD2;PYROXD2 | | 1stExon | No |
| Control-GSM3680855_ | chr10:102766322-102766610 | LZTS2 | Island | Body | No |
| Control-GSM3680855_ | chr10:121265587-121265893 | RGS10;RGS10 | | Body;Body | No |
| Control-GSM3680855_ | chr10:123353800-123353889 | FGFR2;FGFR2;FGFR2;FGFR2;FGFR2;FGFR2;FGFR2;FGFR2 | N_Shelf | TSS200;5'UTR;5'UTR;TSS1500;5'UTR;TSS1500;5'UTR; | No |
| Control-GSM3680855_ | chr10:131731416-131731569 | EBF3 | | Body | No |
| Control-GSM3680855_ | chr10:134462770-134462849 | INPP5A | S_Shore | Body | Yes |
| Control-GSM3680855_ | chr10:134499266-134499472 | INPP5A | | Body | Yes |
| Control-GSM3680855_ | chr10:134596873-134596989 | INPP5A | N_Shore | 3'UTR | Yes |
| Control-GSM3680855_ | chr11:2891065-2891098 | KCNQ1DN | Island | TSS200 | No |
| Control-GSM3680855_ | chr11:6341844-6341888 | PRKCDBP | Island | TSS200 | No |
| Control-GSM3680855_ | chr11:10476494-10476608 | AMPD3;AMPD3;AMPD3 | S_Shelf | TSS200;TSS1500;Body | No |
| Control-GSM3680855_ | chr11:16424673-16424935 | SOX6;SOX6;SOX6;SOX6 | | TSS1500;5'UTR;5'UTR;TSS1500 | Yes |
| Control-GSM3680855_ | chr11:61522002-61522498 | C11orf9;DKFZP434K028;C11orf9 | S_Shelf | Body;Body;TSS1500 | No |

| | | | | | |
|---|---|---|---|---|---|
| Control-GSM3680855_ | chr11:64067795-64068146 | C11orf20 | Island | TSS200 | No |
| Control-GSM3680855_ | chr11:65190825-65191707 | NEAT1 | Island | Body | No |
| Control-GSM3680855_ | chr11:67141628-67141961 | CLCF1;LOC100130987;CLCF1;CLCF1 | S_Shore | 1stExon;Body;TSS1500;5'UTR | Yes |
| Control-GSM3680855_ | chr11:67777618-67777952 | ALDH3B1;ALDH3B1;ALDH3B1 | | TSS200;TSS200;5'UTR | No |
| Control-GSM3680855_ | chr11:68096049-68096165 | LRP5 | | Body | No |
| Control-GSM3680855_ | chr11:69519974-69520418 | FGF19 | S_Shore | TSS1500 | No |
| Control-GSM3680855_ | chr11:76839191-76839476 | MYO7A;MYO7A;MYO7A | | TSS200;TSS200;TSS200 | No |
| Control-GSM3680855_ | chr11:113577071-113577528 | TMPRSS5 | | TSS200 | No |
| Control-GSM3680855_ | chr11:118476517-118477055 | PHLDB1 | N_Shore | TSS1500 | No |
| Control-GSM3680855_ | chr11:122852454-122852698 | BSX | Island | TSS200 | No |
| Control-GSM3680855_ | chr11:123986026-123986094 | VWA5A;VWA5A;VWA5A | | TSS200;TSS200;TSS200 | No |
| Control-GSM3680855_ | chr12:772861-772982 | NINJ2 | | TSS200 | Yes |
| Control-GSM3680855_ | chr12:2800278-2801061 | CACNA1C;CACNA1C;CACNA1C;CACNA1C;CACNA1C;CAC | Island | Body;Body;Body;Body;Body;Body;Body;Body;Body;Bo | No |
| Control-GSM3680855_ | chr12:51318430-51318640 | METTL7A | | TSS200 | No |
| Control-GSM3680855_ | chr12:95009858-95010113 | TMCC3 | | Body | No |
| Control-GSM3680855_ | chr12:95945277-95945384 | USP44 | S_Shelf | TSS200 | No |
| Control-GSM3680855_ | chr12:104443953-104444011 | GLT8D2 | | TSS200 | No |
| Control-GSM3680855_ | chr12:108732994-108733308 | CMKLR1;CMKLR1;CMKLR1;CMKLR1;CMKLR1;CMKLR1 | | 1stExon;1stExon;1stExon;5'UTR;5'UTR;5'UTR | No |
| Control-GSM3680855_ | chr12:132690643-132690752 | GALNT9;GALNT9 | S_Shore | Body;TSS200 | No |
| Control-GSM3680855_ | chr13:31506687-31506752 | C13orf26 | | TSS200 | No |
| Control-GSM3680855_ | chr13:36944294-36944649 | SPG20;SPG20 | | 1stExon;5'UTR | No |
| Control-GSM3680855_ | chr13:99629970-99630462 | DOCK9;DOCK9;DOCK9;DOCK9 | | Body;Body;Body;Body | No |
| Control-GSM3680855_ | chr13:113436400-113436568 | ATP11A;ATP11A | N_Shelf | Body;Body | Yes |
| Control-GSM3680855_ | chr13:113461305-113461652 | ATP11A;ATP11A | N_Shelf | Body;Body | Yes |
| Control-GSM3680855_ | chr13:113464041-113464532 | ATP11A;ATP11A | N_Shore | Body;Body | Yes |
| Control-GSM3680855_ | chr13:114786825-114786887 | RASA3 | S_Shore | Body | Yes |
| Control-GSM3680855_ | chr13:114800549-114800841 | RASA3 | N_Shelf | Body | Yes |
| Control-GSM3680855_ | chr13:114810631-114810794 | RASA3 | N_Shelf | Body | Yes |
| Control-GSM3680855_ | chr13:114830255-114830713 | RASA3 | | Body | Yes |
| Control-GSM3680855_ | chr14:21270407-21271018 | RNASE1;RNASE1;RNASE1;RNASE1;RNASE1 | | 5'UTR;5'UTR;1stExon;5'UTR;5'UTR | Yes |
| Control-GSM3680855_ | chr14:81426012-81426234 | TSHR;TSHR;TSHR | | Body;Body;Body | No |
| Control-GSM3680855_ | chr14:101198397-101198759 | DLK1 | N_Shelf | Body | No |
| Control-GSM3680855_ | chr14:105183652-105184057 | INF2;INF2 | N_Shelf | Body;3'UTR | No |
| Control-GSM3680855_ | chr14:105779794-105779952 | PACS2;PACS2 | N_Shore | TSS1500;TSS1500 | No |
| Control-GSM3680855_ | chr15:40544855-40545215 | PAK6;PAK6;C15orf56;PAK6 | Island | 5'UTR;5'UTR;Body;5'UTR | No |
| Control-GSM3680855_ | chr15:57510284-57510575 | TCF12;TCF12;TCF12;TCF12;TCF12 | | TSS1500;Body;Body;Body;Body | No |
| Control-GSM3680855_ | chr15:74218697-74218733 | LOXL1 | Island | TSS200 | No |
| Control-GSM3680855_ | chr15:83317306-83317526 | CPEB1 | Island | TSS1500 | No |
| Control-GSM3680855_ | chr15:89764895-89765045 | RLBP1;RLBP1 | | 1stExon;5'UTR | No |
| Control-GSM3680855_ | chr16:711033-711235 | WDR90 | N_Shore | Body | No |
| Control-GSM3680855_ | chr16:3070245-3070269 | TNFRSF12A | Island | TSS200 | No |
| Control-GSM3680855_ | chr16:4012936-4013337 | ADCY9 | N_Shelf | 3'UTR | No |
| Control-GSM3680855_ | chr16:58328961-58329456 | KLKBL4 | | TSS200 | No |
| Control-GSM3680855_ | chr16:67978158-67978450 | SLC12A4;SLC12A4;SLC12A4;LCAT;SLC12A4;SLC12A4 | | 3'UTR;3'UTR;3'UTR;TSS200;3'UTR;3'UTR | No |
| Control-GSM3680855_ | chr16:75252619-75252744 | CTRB1 | | TSS1500 | No |
| Control-GSM3680855_ | chr16:83171068-83171314 | CDH13 | | Body | No |
| Control-GSM3680855_ | chr17:6358599-6358928 | PITPNM3;PITPNM3 | Island | 3'UTR;3'UTR | No |
| Control-GSM3680855_ | chr17:36612298-36612623 | ARHGAP23 | S_Shore | TSS1500 | No |
| Control-GSM3680855_ | chr17:37896063-37896175 | GRB7;GRB7 | | 5'UTR;5'UTR | No |
| Control-GSM3680855_ | chr17:40439433-40439501 | STAT5A | N_Shore | TSS200 | No |
| Control-GSM3680855_ | chr17:41832753-41832975 | SOST | Island | Body | No |
| Control-GSM3680855_ | chr17:71258589-71259036 | CPSF4L | | TSS1500 | No |
| Control-GSM3680855_ | chr17:72208433-72208788 | TTYH2;MGC16275 | N_Shore | TSS1500;Body | No |
| Control-GSM3680855_ | chr17:72931767-72931792 | OTOP3 | Island | TSS200 | No |
| Control-GSM3680855_ | chr17:73631460-73631785 | RECQL5;LOC643008;LOC643008 | S_Shelf | Body;TSS1500;5'UTR | Yes |
| Control-GSM3680855_ | chr17:73642503-73642628 | RECQL5;LOC100130933 | | Body;TSS200 | Yes |
| Control-GSM3680855_ | chr17:76226735-76227255 | LOC283999 | N_Shore | TSS1500 | No |
| Control-GSM3680855_ | chr17:78165917-78166385 | CARD14;CARD14 | S_Shore | Body;Body | Yes |
| Control-GSM3680855_ | chr17:78724995-78725470 | RPTOR;RPTOR | | Body;Body | Yes |
| Control-GSM3680855_ | chr18:64271341-64271503 | CDH19 | | TSS200 | No |
| Control-GSM3680855_ | chr18:74692214-74692253 | MBP;MBP;MBP;MBP;MBP | S_Shore | 3'UTR;3'UTR;3'UTR;3'UTR;3'UTR | Yes |
| Control-GSM3680855_ | chr18:74728838-74729151 | MBP;MBP;MBP;MBP;MBP;MBP | | 1stExon;1stExon;Body;1stExon;Body;1stExon | Yes |
| Control-GSM3680855_ | chr19:2761388-2761892 | SGTA | N_Shore | Body | No |
| Control-GSM3680855_ | chr19:6482011-6482169 | DENND1C | | TSS1500 | No |
| Control-GSM3680855_ | chr19:7852207-7852238 | CLEC4GP1 | N_Shore | TSS200 | No |
| Control-GSM3680855_ | chr19:7910811-7911206 | EVI5L;EVI5L | N_Shore | 5'UTR;TSS1500 | No |
| Control-GSM3680855_ | chr19:10407184-10407538 | ICAM5 | Island | Body | No |
| Control-GSM3680855_ | chr19:10541267-10541704 | PDE4A;PDE4A | N_Shore | Body;TSS1500 | No |
| Control-GSM3680855_ | chr19:14671371-14671512 | TECR | | Body | No |
| Control-GSM3680855_ | chr19:16830739-16830859 | NWD1 | | TSS200 | No |
| Control-GSM3680855_ | chr19:18120692-18120819 | ARRDC2;ARRDC2 | Island | Body;Body | No |
| Control-GSM3680855_ | chr19:35085474-35085677 | SCGBL;SCGBL | | 1stExon;5'UTR | No |
| Control-GSM3680855_ | chr19:35531817-35532097 | HPN;HPN | Island | 5'UTR;5'UTR | No |
| Control-GSM3680855_ | chr19:35597062-35597447 | LOC100128675;LOC100128675 | | Body;Body | No |
| Control-GSM3680855_ | chr19:35786821-35787193 | MAG;MAG | Island | Body;Body | No |
| Control-GSM3680855_ | chr19:48497737-48498031 | ELSPBP1 | | TSS200 | No |
| Control-GSM3680855_ | chr20:4795381-4795621 | RASSF2;RASSF2 | | 5'UTR;5'UTR | No |
| Control-GSM3680855_ | chr20:30309466-30309717 | BCL2L1;BCL2L1 | N_Shore | Body;Body | No |
| Control-GSM3680855_ | chr20:30582838-30583001 | XKR7 | Island | Body | No |
| Control-GSM3680855_ | chr20:42193746-42194649 | SGK2;SGK2 | | TSS1500;5'UTR | No |
| Control-GSM3680855_ | chr20:56136012-56136167 | PCK1 | | TSS200 | No |
| Control-GSM3680855_ | chr20:62183924-62183966 | C20orf195 | N_Shore | TSS1500 | No |
| Control-GSM3680855_ | chr21:34638188-34638535 | IL10RB | N_Shore | TSS1500 | Yes |
| Control-GSM3680855_ | chr21:34638662-34639040 | IL10RB | Island | TSS200 | Yes |
| Control-GSM3680855_ | chr22:37955936-37956293 | CDC42EP1 | N_Shore | TSS1500 | No |
| Control-GSM3680855_ | chr22:38479236-38479288 | SLC16A8 | S_Shore | TSS200 | No |
| Control-GSM3680855_ | chr22:50523686-50523703 | MLC1;MLC1;MLC1 | | 5'UTR;5'UTR;1stExon | No |
| Control-GSM3680855_ | chr1:9788715-9788769 | PIK3CD | N_Shore | 3'UTR | No |
| Control-GSM3680855_ | chr1:50886393-50886969 | DMRTA2 | Island | Body | No |
| Control-GSM3680855_ | chr1:156594108-156595178 | HAPLN2 | Island | Body | No |
| Control-GSM3680855_ | chr1:230272306-230272656 | GALNT2 | | Body | No |

| | | | | | |
|---|---|---|---|---|---|
| Control-GSM3680855_ | chr1:243646235-243646787 | SDCCAG8 | N_Shore | Body | No |
| Control-GSM3680855_ | chr2:30669597-30669863 | LCLAT1;LCLAT1 | N_Shore | TSS1500;TSS1500 | Yes |
| Control-GSM3680855_ | chr2:40739515-40739771 | SLC8A1;SLC8A1 | | 5'UTR;1stExon | No |
| Control-GSM3680855_ | chr2:46593223-46593271 | EPAS1 | | Body | No |
| Control-GSM3680855_ | chr2:65593908-65594021 | SPRED2;SPRED2;SPRED2 | | 5'UTR;1stExon;Body | No |
| Control-GSM3680855_ | chr2:71680559-71680721 | DYSF;DYSF;DYSF;DYSF;DYSF;DYSF;DYSF | Island | TSS200;TSS200;TSS200;TSS200;TSS200;TSS200;TSS | No |
| Control-GSM3680855_ | chr2:200320750-200321016 | SATB2 | Island | Body | No |
| Control-GSM3680855_ | chr3:40428508-40429014 | ENTPD3 | N_Shore | TSS200 | No |
| Control-GSM3680855_ | chr3:147128679-147129184 | ZIC1 | Island | 1stExon | Yes |
| Control-GSM3680855_ | chr3:147129778-147129952 | ZIC1 | N_Shore | Body | Yes |
| Control-GSM3680855_ | chr4:802844-803014 | CPLX1 | S_Shelf | Body | No |
| Control-GSM3680855_ | chr4:1202657-1202696 | LOC100130872-SPON2;LOC100130872 | N_Shore | TSS200;Body | No |
| Control-GSM3680855_ | chr4:3372915-3373405 | RGS12;RGS12;RGS12 | N_Shore | Body;Body;Body | No |
| Control-GSM3680855_ | chr4:88450824-88451496 | SPARCL1;SPARCL1 | | TSS200;TSS200 | No |
| Control-GSM3680855_ | chr5:149669484-149669674 | CAMK2A;CAMK2A | | TSS200;TSS200 | No |
| Control-GSM3680855_ | chr5:157003083-157003436 | ADAM19 | Island | TSS1500 | No |
| Control-GSM3680855_ | chr6:32120584-32120625 | PPT2;PRRT1;PPT2 | N_Shore | TSS1500;TSS1500;TSS1500 | No |
| Control-GSM3680855_ | chr6:32526021-32526260 | HLA-DRB6 | | Body | Yes |
| Control-GSM3680855_ | chr6:32634245-32634344 | HLA-DQB1 | S_Shore | Body | Yes |
| Control-GSM3680855_ | chr6:91004430-91004482 | BACH2;BACH2 | N_Shore | 5'UTR;5'UTR | Yes |
| Control-GSM3680855_ | chr7:1004748-1005089 | COX19 | N_Shelf | 3'UTR | No |
| Control-GSM3680855_ | chr7:1080836-1081068 | C7orf50;C7orf50;C7orf50 | Island | Body;Body;Body | Yes |
| Control-GSM3680855_ | chr7:16505409-16505602 | SOSTDC1;SOSTDC1 | | 1stExon;5'UTR | No |
| Control-GSM3680855_ | chr7:29603121-29603407 | PRR15 | Island | TSS1500 | Yes |
| Control-GSM3680855_ | chr7:29606082-29606349 | PRR15 | Island | Body | Yes |
| Control-GSM3680855_ | chr7:117119382-117119424 | CFTR | | TSS1500 | No |
| Control-GSM3680855_ | chr8:1895007-1895431 | ARHGEF10 | S_Shore | Body | Yes |
| Control-GSM3680855_ | chr8:2075209-2075820 | MYOM2 | N_Shore | Body | Yes |
| Control-GSM3680855_ | chr8:22423014-22423091 | SORBS3;SORBS3 | Island | TSS200;Body | No |
| Control-GSM3680855_ | chr8:27182889-27183285 | PTK2B;PTK2B;PTK2B;PTK2B | N_Shore | 5'UTR;TSS200;5'UTR;TSS200 | No |
| Control-GSM3680855_ | chr8:65291287-65291562 | MIR124-2 | S_Shore | TSS1500 | Yes |
| Control-GSM3680855_ | chr8:145747659-145748059 | LRRC14 | Island | 3'UTR | No |
| Control-GSM3680855_ | chr9:707166-707410 | KANK1;KANK1;KANK1 | Island | 1stExon;5'UTR;Body | Yes |
| Control-GSM3680855_ | chr10:13749193-13749285 | FRMD4A | | Body | Yes |
| Control-GSM3680855_ | chr10:13749405-13749625 | FRMD4A | | Body | Yes |
| Control-GSM3680855_ | chr10:88471748-88471978 | LDB3;LDB3;LDB3 | Island | Body;Body;Body | Yes |
| Control-GSM3680855_ | chr10:118891601-118893224 | VAX1 | N_Shore | 3'UTR | No |
| Control-GSM3680855_ | chr10:124220359-124220396 | HTRA1 | N_Shore | TSS1500 | Yes |
| Control-GSM3680855_ | chr10:133948362-133948517 | JAKMIP3 | N_Shore | Body | No |
| Control-GSM3680855_ | chr11:1320329-1321530 | TOLLIP | S_Shelf | Body | No |
| Control-GSM3680855_ | chr11:31819444-31819678 | PAX6;PAX6;PAX6 | N_Shore | Body;Body;Body | No |
| Control-GSM3680855_ | chr11:57414401-57414557 | YPEL4 | Island | Body | No |
| Control-GSM3680855_ | chr11:117747940-117748189 | FXYD6;FXYD6;FXYD6;FXYD6;FXYD6 | S_Shore | 5'UTR;TSS200;TSS200;TSS200;TSS200 | No |
| Control-GSM3680855_ | chr11:119294283-119294408 | THY1 | S_Shore | TSS200 | No |
| Control-GSM3680855_ | chr11:126455712-126456119 | KIRREL3;KIRREL3 | | Body;Body | No |
| Control-GSM3680855_ | chr11:131779469-131779604 | NTM;NTM;NTM;NTM | N_Shore | TSS1500;TSS1500;TSS1500;Body | No |
| Control-GSM3680855_ | chr12:118500439-118500931 | WSB2 | S_Shore | TSS1500 | No |
| Control-GSM3680855_ | chr13:100547192-100547367 | CLYBL | Island | 3'UTR | No |
| Control-GSM3680855_ | chr14:79746212-79746627 | NRXN3;NRXN3;NRXN3;NRXN3;NRXN3 | S_Shore | Body;1stExon;1stExon;5'UTR;5'UTR | No |
| Control-GSM3680855_ | chr15:90208739-90208915 | PLIN1;PLIN1 | N_Shore | 3'UTR;3'UTR | No |
| Control-GSM3680855_ | chr17:8380278-8380351 | MYH10 | Island | Body | No |
| Control-GSM3680855_ | chr17:75315081-75315486 | SEPT9;SEPT9;SEPT9 | | 5'UTR;Body;TSS1500 | No |
| Control-GSM3680855_ | chr17:78894364-78894477 | RPTOR;RPTOR | N_Shelf | Body;Body | Yes |
| Control-GSM3680855_ | chr17:79372242-79372528 | BAHCC1 | Island | TSS1500 | Yes |
| Control-GSM3680855_ | chr17:79374327-79375254 | BAHCC1 | Island | Body | Yes |
| Control-GSM3680855_ | chr18:20714217-20714399 | CABLES1 | N_Shore | TSS1500 | No |
| Control-GSM3680855_ | chr19:1207124-1207644 | STK11 | Island | 1stExon | No |
| Control-GSM3680855_ | chr19:55972855-55973099 | ISOC2;ISOC2;ISOC2 | Island | 5'UTR;5'UTR;5'UTR | Yes |
| Control-GSM3680855_ | chr21:46890849-46890997 | COL18A1;COL18A1;COL18A1 | Island | Body;Body;Body | No |
| Control-GSM3680855_ | chr21:47604052-47605174 | C21orf56;C21orf56 | S_Shore | 5'UTR;5'UTR | Yes |
| Control-GSM3680856_ | chr3:73045440-73045686 | PPP4R2 | N_Shore | TSS1500 | Yes |
| Control-GSM3680856_ | chr4:69435250-69435601 | UGT2B15;UGT2B17 | | TSS1500;TSS1500 | Yes |
| Control-GSM3680856_ | chr6:31237013-31237034 | HLA-C | N_Shore | Body | Yes |
| Control-GSM3680856_ | chr16:857454-858205 | PRR25 | Island | Body | Yes |
| Control-GSM3680856_ | chr1:76189707-76189801 | ACADM;ACADM | N_Shore | TSS1500;TSS1500 | Yes |
| Control-GSM3680856_ | chr17:17109570-17109740 | PLD6 | Island | 1stExon | No |
| Control-GSM3680861_ | chr5:179740743-179741120 | GFPT2 | Island | Body | Yes |
| Control-GSM3680861_ | chr20:5485270-5485475 | LOC149837 | | TSS200 | Yes |
| Control-GSM3680861_ | chr1:110254662-110254854 | GSTM5 | | TSS1500 | Yes |
| Control-GSM3680861_ | chr1:205819088-205819463 | PM20D1 | Island | 1stExon | Yes |
| Control-GSM3680861_ | chr6:29910796-29911011 | HLA-A | Island | Body | Yes |
| Control-GSM3680861_ | chr6:29911036-29911104 | HLA-A | Island | Body | Yes |
| Control-GSM3680861_ | chr6:30039010-30039142 | RNF39;RNF39 | Island | Body;Body | Yes |
| Control-GSM3680861_ | chr6:30039175-30039206 | RNF39;RNF39 | Island | Body;Body | Yes |
| Control-GSM3680861_ | chr6:32526021-32526260 | HLA-DRB6 | | Body | Yes |
| Control-GSM3680861_ | chr11:5617273-5617926 | TRIM6-TRIM34;TRIM6;TRIM6 | | TSS1500;TSS1500;TSS200 | Yes |
| Control-GSM3680861_ | chr21:37442289-37442777 | CBR1;CBR1 | Island | 1stExon;5'UTR | Yes |
| Control-GSM3680863_ | chr21:47604052-47604654 | C21orf56;C21orf56 | S_Shore | 5'UTR;5'UTR | Yes |
| Control-GSM3680863_ | chr6:31237013-31237034 | HLA-C | N_Shore | Body | Yes |
| Control-GSM3680863_ | chr6:32526021-32526364 | HLA-DRB6 | | Body | Yes |
| Control-GSM3680863_ | chr6:32552205-32552453 | HLA-DRB1 | Island | Body | Yes |
| Control-GSM3680863_ | chr6:32634245-32634362 | HLA-DQB1 | S_Shore | Body | Yes |
| Control-GSM3680863_ | chr8:104310713-104311490 | FZD6;FZD6;FZD6;FZD6 | N_Shore | TSS1500;TSS1500;5'UTR;1stExon | No |
| Control-GSM3680864_ | chr6:31237013-31237034 | HLA-C | N_Shore | Body | Yes |
| Control-GSM3680864_ | chr6:32526021-32526364 | HLA-DRB6 | | Body | Yes |
| Control-GSM3680864_ | chr10:135341528-135341933 | CYP2E1 | N_Shore | 1stExon | Yes |
| Control-GSM3680864_ | chr19:12876846-12877188 | HOOK2;HOOK2 | N_Shore | Body;Body | Yes |
| Control-GSM3680864_ | chr21:47604052-47605174 | C21orf56;C21orf56 | S_Shore | 5'UTR;5'UTR | Yes |
| Control-GSM3680864_ | chr22:24384397-24384525 | GSTT1 | Island | TSS200 | Yes |
| Control-GSM3680865_ | chr3:39543776-39544192 | MOBP;MOBP | Island | Body;Body | Yes |
| Control-GSM3680865_ | chr6:291687-292823 | DUSP22 | N_Shore | TSS1500 | Yes |

| | | | | | |
|---|---|---|---|---|---|
| Control-GSM3680865_ | chr6:31237013-31237034 | HLA-C | N_Shore | Body | Yes |
| Control-GSM3680865_ | chr6:32526027-32526364 | HLA-DRB6 | | Body | Yes |
| Control-GSM3680865_ | chr8:104032819-104032876 | ATP6V1C1 | N_Shore | TSS1500 | No |
| Control-GSM3680865_ | chr17:41277974-41278241 | BRCA1;NBR2;BRCA1;BRCA1;BRCA1;BRCA1 | N_Shore | TSS1500;Body;TSS1500;TSS1500;TSS1500;TSS1500 | No |
| Control-GSM3680865_ | chr4:74847766-74847829 | PF4 | Island | TSS200 | No |
| Control-GSM3680865_ | chr19:9785511-9785805 | ZNF562;ZNF562;ZNF562 | Island | 5'UTR;5'UTR;5'UTR | Yes |
| Control-GSM3680869_ | chr5:100240048-100240223 | ST8SIA4;ST8SIA4 | S_Shore | TSS1500;TSS1500 | No |
| Control-GSM3680869_ | chr6:30122523-30122593 | TRIM10;TRIM10 | | Body;Body | Yes |
| Control-GSM3680869_ | chr6:31237013-31237034 | HLA-C | N_Shore | Body | Yes |
| Control-GSM3680869_ | chr6:32526021-32526263 | HLA-DRB6 | | Body | Yes |
| Control-GSM3680869_ | chr6:32557970-32558459 | HLA-DRB1 | | TSS1500 | Yes |
| Control-GSM3680869_ | chr1:110254709-110254835 | GSTM5 | | TSS200 | Yes |
| Control-GSM3680869_ | chr1:207842917-207843084 | CR1L | Island | Body | No |
| Control-GSM3680869_ | chr5:1594282-1594863 | SDHAP3 | Island | Body | Yes |
| Control-GSM3680869_ | chr20:5485270-5485475 | LOC149837 | | TSS200 | Yes |
| Control-GSM3680869_ | chr21:47604052-47605174 | C21orf56;C21orf56 | S_Shore | 5'UTR;5'UTR | Yes |
| Control-GSM3680870_ | chr1:161008705-161008826 | TSTD1;TSTD1;TSTD1;TSTD1;TSTD1;TSTD1 | Island | 5'UTR;1stExon;5'UTR;1stExon;1stExon;5'UTR | Yes |
| Control-GSM3680870_ | chr2:225266580-225266733 | FAM124B;FAM124B;FAM124B;FAM124B | | 5'UTR;1stExon;1stExon;5'UTR | Yes |
| Control-GSM3680870_ | chr6:291687-292823 | DUSP22 | N_Shore | TSS1500 | Yes |
| Control-GSM3680870_ | chr6:31237013-31237034 | HLA-C | N_Shore | Body | Yes |
| Control-GSM3680870_ | chr6:32526021-32526260 | HLA-DRB6 | | Body | Yes |
| Control-GSM3680870_ | chr7:134832939-134833299 | TMEM140;TMEM140 | | 1stExon;5'UTR | Yes |
| Control-GSM3680870_ | chr21:46341006-46341067 | ITGB2;ITGB2 | | TSS200;5'UTR | No |
| Control-GSM3680870_ | chr1:248100228-248101009 | OR2L13 | N_Shore | TSS1500 | Yes |
| Control-GSM3680870_ | chr6:32489963-32489991 | HLA-DRB5 | Island | Body | Yes |
| Control-GSM3680870_ | chr19:49340574-49340765 | PLEKHA4;HSD17B14;PLEKHA4 | Island | 3'UTR;TSS1500;Body | Yes |
| Control-GSM3680872_ | chr5:179740743-179741120 | GFPT2 | Island | Body | Yes |
| Control-GSM3680872_ | chr6:31237013-31237034 | HLA-C | N_Shore | Body | Yes |
| Control-GSM3680872_ | chr6:32551888-32551954 | HLA-DRB1 | Island | Body | Yes |
| Control-GSM3680872_ | chr6:32552152-32552246 | HLA-DRB1 | Island | Body | Yes |
| Control-GSM3680872_ | chr21:47604291-47604654 | C21orf56;C21orf56;C21orf56;C21orf56 | S_Shore | 5'UTR;5'UTR;1stExon;1stExon | Yes |
| Control-GSM3680872_ | chr1:205819088-205819463 | PM20D1 | Island | 1stExon | Yes |
| Control-GSM3680872_ | chr5:1594282-1594676 | SDHAP3 | Island | Body | Yes |
| Control-GSM3680872_ | chr6:32526021-32526260 | HLA-DRB6 | | Body | Yes |
| Control-GSM3680876_ | chr1:205819088-205819463 | PM20D1 | Island | 1stExon | Yes |
| Control-GSM3680876_ | chr5:179740743-179741120 | GFPT2 | Island | Body | Yes |
| Control-GSM3680876_ | chr6:31237013-31237034 | HLA-C | N_Shore | Body | Yes |
| Control-GSM3680876_ | chr6:32526021-32526260 | HLA-DRB6 | | Body | Yes |
| Control-GSM3680876_ | chr16:55866890-55867072 | CES1;CES1;CES1 | Island | Body;Body;Body | Yes |
| Control-GSM3680876_ | chr22:24348549-24348715 | GSTTP1 | | TSS1500 | Yes |
| Control-GSM3680876_ | chr6:32489963-32489991 | HLA-DRB5 | Island | Body | Yes |
| Control-GSM3680876_ | chr8:2075209-2075820 | MYOM2 | N_Shore | Body | Yes |
| Control-GSM3680876_ | chr10:134756503-134756707 | C10orf93 | Island | TSS1500 | No |
| Control-GSM3680876_ | chr11:5617812-5617926 | TRIM6;TRIM6-TRIM34;TRIM6 | | TSS200;TSS200;5'UTR | Yes |
| Control-GSM3680876_ | chr22:24384340-24384573 | GSTT1 | Island | TSS200 | Yes |
| Control-GSM3680880_ | chr4:69435250-69435601 | UGT2B15;UGT2B17 | | TSS1500;TSS1500 | Yes |
| Control-GSM3680880_ | chr6:31237013-31237034 | HLA-C | N_Shore | Body | Yes |
| Control-GSM3680880_ | chr16:857454-857981 | PRR25 | Island | Body | Yes |
| Control-GSM3680880_ | chr6:29943401-29943414 | HCG9 | N_Shore | Body | Yes |
| Control-GSM3680880_ | chr6:32526021-32526364 | HLA-DRB6 | | Body | Yes |
| Control-GSM3680880_ | chr6:32552205-32552515 | HLA-DRB1 | Island | Body | Yes |
| Control-GSM3680880_ | chr19:49340489-49340765 | PLEKHA4;PLEKHA4;HSD17B14 | Island | 3'UTR;3'UTR;TSS1500 | Yes |
| Control-GSM3680880_ | chr21:35831871-35832028 | KCNE1;KCNE1;KCNE1;KCNE1;KCNE1 | Island | 5'UTR;5'UTR;1stExon;1stExon;5'UTR | Yes |
| Control-GSM3680887_ | chr16:857454-857981 | PRR25 | Island | Body | Yes |
| Control-GSM3680887_ | chr19:55476665-55477810 | NLRP2 | | TSS1500 | Yes |
| Control-GSM3680887_ | chr6:32526260-32526342 | HLA-DRB6 | | Body | Yes |
| Control-GSM3680887_ | chr7:39170497-39170763 | POU6F2;POU6F2 | | Body;Body | Yes |
| Control-GSM3680891_ | chr5:135416205-135416613 | MIR886 | Island | Body | Yes |
| Control-GSM3680891_ | chr6:31237013-31237034 | HLA-C | N_Shore | Body | Yes |
| Control-GSM3680891_ | chr8:39380341-39380592 | ADAM3A;ADAM3A;ADAM3A | Island | Body;Body;Body | No |
| Control-GSM3680891_ | chr6:32526021-32526263 | HLA-DRB6 | | Body | Yes |
| Control-GSM3680891_ | chr6:32606845-32607224 | HLA-DQA1 | | Body | Yes |
| Control-GSM3680891_ | chr6:32634245-32634362 | HLA-DQB1 | S_Shore | Body | Yes |
| Control-GSM3680893_ | chr1:205818956-205819463 | PM20D1 | Island | Body | Yes |
| Control-GSM3680893_ | chr4:69435250-69435601 | UGT2B15;UGT2B17 | | TSS1500;TSS1500 | Yes |
| Control-GSM3680893_ | chr5:179740743-179741120 | GFPT2 | Island | Body | Yes |
| Control-GSM3680893_ | chr6:291687-292823 | DUSP22 | N_Shore | TSS1500 | Yes |
| Control-GSM3680893_ | chr6:30177166-30177194 | TRIM26 | N_Shelf | 5'UTR | Yes |
| Control-GSM3680893_ | chr6:30713244-30713295 | IER3 | S_Shore | TSS1500 | No |
| Control-GSM3680893_ | chr6:32522683-32523136 | HLA-DRB6 | | Body | Yes |
| Control-GSM3680893_ | chr6:32549631-32549935 | HLA-DRB1 | N_Shelf | Body | Yes |
| Control-GSM3680893_ | chr19:12876846-12877000 | HOOK2;HOOK2 | N_Shore | Body;Body | Yes |
| Control-GSM3680893_ | chr19:57742421-57742444 | AURKC;AURKC;AURKC;AURKC;AURKC | Island | 5'UTR;TSS200;1stExon;5'UTR;1stExon | Yes |
| Control-GSM3680893_ | chr21:39493489-39493543 | DSCR8;DSCR8;DSCR8;DSCR4;DSCR8;DSCR8 | | TSS200;TSS200;TSS200;TSS200;TSS200;TSS200 | Yes |
| Control-GSM3680893_ | chr6:31237405-31237722 | HLA-C | N_Shore | Body | Yes |
| Control-GSM3680893_ | chr6:32489963-32489991 | HLA-DRB5 | Island | Body | Yes |
| Control-GSM3680893_ | chr6:32552246-32552443 | HLA-DRB1 | Island | Body | Yes |
| Control-GSM3680893_ | chr6:32634316-32634362 | HLA-DQB1 | S_Shore | 1stExon | Yes |
| Control-GSM3680894_ | chr1:205818956-205819463 | PM20D1 | Island | Body | Yes |
| Control-GSM3680894_ | chr6:291687-292823 | DUSP22 | N_Shore | TSS1500 | Yes |
| Control-GSM3680894_ | chr6:30177166-30177194 | TRIM26 | N_Shelf | 5'UTR | Yes |
| Control-GSM3680894_ | chr7:64043012-64043193 | LOC641746 | | Body | No |
| Control-GSM3680894_ | chr10:4720653-4720912 | LOC100216001 | | TSS1500 | No |
| Control-GSM3680894_ | chr11:75139390-75139736 | KLHL35 | N_Shore | Body | Yes |
| Control-GSM3680894_ | chr19:12876846-12877188 | HOOK2;HOOK2 | N_Shore | Body;Body | Yes |
| Control-GSM3680894_ | chr1:2120985-2121521 | C1orf86;LOC100128003 | N_Shore | Body;Body | Yes |
| Control-GSM3680894_ | chr2:30669597-30669863 | LCLAT1;LCLAT1 | N_Shore | TSS1500;TSS1500 | Yes |
| Control-GSM3680894_ | chr6:32489963-32489991 | HLA-DRB5 | Island | Body | Yes |
| Control-GSM3680894_ | chr6:32552152-32552246 | HLA-DRB1 | Island | Body | Yes |
| Control-GSM3680894_ | chr8:2075209-2075820 | MYOM2 | N_Shore | Body | Yes |
| Control-GSM3680894_ | chr11:15095024-15095178 | CALCB | Island | TSS200 | No |

| | | | | | |
|---|---|---|---|---|---|
| Control-GSM3680894_ | chr21:47604052-47605174 | C21orf56;C21orf56 | S_Shore | 5'UTR;5'UTR | Yes |
| Control-GSM3680895_ | chr1:205819088-205819463 | PM20D1 | Island | 1stExon | Yes |
| Control-GSM3680895_ | chr6:32549631-32549935 | HLA-DRB1 | N_Shelf | Body | Yes |
| Control-GSM3680895_ | chr2:30669597-30669863 | LCLAT1;LCLAT1 | N_Shore | TSS1500;TSS1500 | Yes |
| Control-GSM3680895_ | chr5:176797920-176798049 | RGS14 | S_Shelf | Body | No |
| Control-GSM3680895_ | chr6:29796032-29796105 | HLA-G | Island | Body | Yes |
| Control-GSM3680895_ | chr6:29796351-29796379 | HLA-G | Island | Body | Yes |
| Control-GSM3680895_ | chr6:29943401-29943414 | HCG9 | N_Shore | Body | Yes |
| Control-GSM3680895_ | chr6:29943464-29943677 | HCG9 | N_Shore | Body | Yes |
| Control-GSM3680895_ | chr6:32489963-32489991 | HLA-DRB5 | Island | Body | Yes |
| Control-GSM3680895_ | chr6:32551888-32551949 | HLA-DRB1 | Island | Body | Yes |
| Control-GSM3680895_ | chr6:32552246-32552443 | HLA-DRB1 | Island | Body | Yes |
| Control-GSM3680895_ | chr8:2074935-2075820 | MYOM2 | N_Shore | Body | Yes |
| Control-GSM3680895_ | chr16:420108-420230 | MRPL28 | Island | Body | Yes |
| Control-GSM3680897_ | chr1:110254692-110254828 | GSTM5 | | TSS200 | Yes |
| Control-GSM3680897_ | chr5:179740743-179741120 | GFPT2 | Island | Body | Yes |
| Control-GSM3680897_ | chr6:291882-292823 | DUSP22 | N_Shore | TSS1500 | Yes |
| Control-GSM3680897_ | chr6:32526021-32526260 | HLA-DRB6 | | Body | Yes |
| Control-GSM3680897_ | chr6:32557970-32558459 | HLA-DRB1 | | TSS1500 | Yes |
| Control-GSM3680897_ | chr8:145955983-145956024 | ZNF251 | Island | Body | Yes |
| Control-GSM3680897_ | chr6:168435914-168436099 | KIF25;KIF25 | Island | Body;Body | Yes |
| Control-GSM3680897_ | chr7:5534548-5534721 | FBXL18 | Island | Body | No |
| Control-GSM3680897_ | chr8:144635430-144635478 | GSDMD | Island | TSS200 | Yes |
| Control-GSM3680900_ | chr5:179740743-179741120 | GFPT2 | Island | Body | Yes |
| Control-GSM3680900_ | chr17:73056681-73056797 | KCTD2 | | Body | No |
| Control-GSM3680900_ | chr22:24348549-24348715 | GSTTP1 | | TSS1500 | Yes |
| Control-GSM3680900_ | chr1:205819088-205819406 | PM20D1 | Island | 1stExon | Yes |
| Control-GSM3680900_ | chr6:32552246-32552515 | HLA-DRB1 | Island | Body | Yes |
| Control-GSM3680900_ | chr6:32634245-32634481 | HLA-DQB1 | S_Shore | Body | Yes |
| Control-GSM3680900_ | chr17:154420-154671 | RPH3AL | S_Shore | Body | No |
| Control-GSM3680901_ | chr6:32525960-32526364 | HLA-DRB6 | | Body | Yes |
| Control-GSM3680901_ | chr15:23378429-23378571 | HERC2P2 | Island | TSS200 | No |
| Control-GSM3680901_ | chr1:75199075-75199496 | CRYZ;CRYZ;CRYZ;TYW3;TYW3;CRYZ;CRYZ;CRYZ;TYW3;CRYZ;CRYZ | | 5'UTR;1stExon;1stExon;1stExon;Body;5'UTR;5'UTR;1s | No |
| Control-GSM3680901_ | chr1:205819088-205819406 | PM20D1 | Island | 1stExon | Yes |
| Control-GSM3680901_ | chr7:1198841-1198977 | ZFAND2A | N_Shore | 5'UTR | No |
| Control-GSM3680901_ | chr19:9731681-9732235 | ZNF561 | N_Shore | 5'UTR | No |
| Control-GSM3680901_ | chr19:23941207-23941575 | ZNF681 | N_Shore | Body | Yes |
| Control-GSM3680902_ | chr5:179740743-179741120 | GFPT2 | Island | Body | Yes |
| Control-GSM3680902_ | chr6:32552152-32552246 | HLA-DRB1 | Island | Body | Yes |
| Control-GSM3680902_ | chr22:24348549-24348715 | GSTTP1 | | TSS1500 | Yes |
| Control-GSM3680902_ | chr1:248100407-248100614 | OR2L13 | Island | TSS200 | Yes |
| Control-GSM3680902_ | chr5:1594282-1594676 | SDHAP3 | Island | Body | Yes |
| Control-GSM3680902_ | chr6:31734217-31734292 | C6orf27 | | Body | No |
| Control-GSM3680902_ | chr6:32526021-32526342 | HLA-DRB6 | | Body | Yes |
| Control-GSM3680902_ | chr6:32606845-32607224 | HLA-DQA1 | | Body | Yes |
| Control-GSM3680902_ | chr11:5617812-5617926 | TRIM6;TRIM6-TRIM34;TRIM6 | | TSS200;TSS200;5'UTR | Yes |
| Control-GSM3680902_ | chr22:17956462-17956641 | CECR2 | | TSS200 | Yes |
| Control-GSM3680905_ | chr5:135416205-135416613 | MIR886 | Island | Body | Yes |
| Control-GSM3680905_ | chr5:179740743-179741120 | GFPT2 | Island | Body | Yes |
| Control-GSM3680905_ | chr6:291687-292823 | DUSP22 | N_Shore | TSS1500 | Yes |
| Control-GSM3680905_ | chr6:32526021-32526260 | HLA-DRB6 | | Body | Yes |
| Control-GSM3680905_ | chr6:32549631-32549935 | HLA-DRB1 | N_Shelf | Body | Yes |
| Control-GSM3680905_ | chr6:32557970-32558459 | HLA-DRB1 | | TSS1500 | Yes |
| Control-GSM3680905_ | chr16:857576-858205 | PRR25 | Island | Body | Yes |
| Control-GSM3680905_ | chr1:110254692-110254854 | GSTM5 | | TSS200 | Yes |
| Control-GSM3680905_ | chr1:205819088-205819406 | PM20D1 | Island | 1stExon | Yes |
| Control-GSM3680905_ | chr2:239139911-239140190 | LOC151174;LOC643387;LOC151174 | N_Shore | Body;TSS1500;Body | Yes |
| Control-GSM3680905_ | chr4:1161247-1161597 | SPON2;SPON2 | Island | 3'UTR;3'UTR | No |
| Control-GSM3680905_ | chr6:29911087-29911104 | HLA-A | Island | Body | Yes |
| Control-GSM3680905_ | chr6:29943401-29943414 | HCG9 | N_Shore | Body | Yes |
| Control-GSM3680905_ | chr6:32551888-32551949 | HLA-DRB1 | Island | Body | Yes |
| Control-GSM3680905_ | chr6:32552246-32552443 | HLA-DRB1 | Island | Body | Yes |
| Control-GSM3680905_ | chr8:144635361-144635478 | GSDMD | Island | TSS200 | Yes |
| Control-GSM3680905_ | chr22:24384315-24384393 | GSTT1 | Island | TSS200 | Yes |
| Control-GSM3680906_ | chr1:205818956-205819463 | PM20D1 | Island | Body | Yes |
| Control-GSM3680906_ | chr6:291687-292823 | DUSP22 | N_Shore | TSS1500 | Yes |
| Control-GSM3680906_ | chr6:32525960-32526364 | HLA-DRB6 | | Body | Yes |
| Control-GSM3680906_ | chr11:18477299-18477379 | LDHAL6A;LDHAL6A | Island | TSS200;TSS200 | No |
| Control-GSM3680906_ | chr16:857576-857981 | PRR25 | Island | Body | Yes |
| Control-GSM3680906_ | chr17:79905176-79905319 | MYADML2 | | TSS200 | Yes |
| Control-GSM3680906_ | chr1:66257553-66257822 | PDE4B;PDE4B | N_Shore | TSS1500;TSS1500 | No |
| Control-GSM3680906_ | chr2:70995444-70995527 | ADD2;ADD2;ADD2;ADD2;ADD2 | S_Shore | TSS200;TSS200;TSS200;TSS200;TSS200 | Yes |
| Control-GSM3680906_ | chr6:29943401-29943414 | HCG9 | N_Shore | Body | Yes |
| Control-GSM3680906_ | chr11:5617812-5617926 | TRIM6;TRIM6-TRIM34;TRIM6 | | TSS200;TSS200;5'UTR | Yes |
| Control-GSM3680906_ | chr15:40268421-40269214 | EIF2AK4 | N_Shore | Body | Yes |
| Control-GSM3680907_ | chr5:135416205-135416398 | MIR886 | Island | Body | Yes |
| Control-GSM3680907_ | chr5:135416412-135416613 | MIR886 | Island | TSS200 | Yes |
| Control-GSM3680907_ | chr6:32552152-32552246 | HLA-DRB1 | Island | Body | Yes |
| Control-GSM3680907_ | chr21:39493489-39493543 | DSCR8;DSCR8;DSCR8;DSCR4;DSCR8;DSCR8 | | TSS200;TSS200;TSS200;TSS200;TSS200;TSS200 | Yes |
| Control-GSM3680907_ | chr4:186732837-186732932 | SORBS2;SORBS2;SORBS2;SORBS2;SORBS2;SORBS2;SORBS2 | | 5'UTR;5'UTR;5'UTR;1stExon;TSS1500;1stExon;TSS150 | No |
| Control-GSM3680907_ | chr5:179740743-179741120 | GFPT2 | Island | Body | Yes |
| Control-GSM3680907_ | chr6:31237405-31237722 | HLA-C | N_Shore | Body | Yes |
| Control-GSM3680907_ | chr10:135342560-135342936 | CYP2E1 | Island | Body | Yes |
| Control-GSM3680907_ | chr19:12876846-12877188 | HOOK2;HOOK2 | N_Shore | Body;Body | Yes |
| Control-GSM3680910_ | chr5:135416205-135416613 | MIR886 | Island | Body | Yes |
| Control-GSM3680910_ | chr6:30177166-30177194 | TRIM26 | N_Shelf | 5'UTR | Yes |
| Control-GSM3680910_ | chr6:32552152-32552246 | HLA-DRB1 | Island | Body | Yes |
| Control-GSM3680910_ | chr16:857454-858205 | PRR25 | Island | Body | Yes |
| Control-GSM3680910_ | chr17:79905152-79905263 | MYADML2 | | TSS200 | Yes |
| Control-GSM3680910_ | chr8:144659831-144660157 | NAPRT1 | Island | Body | Yes |
| Control-GSM3680910_ | chr15:40268610-40269214 | EIF2AK4 | Island | Body | Yes |

| | | | | | |
|---|---|---|---|---|---|
| Control-GSM3680910_ | chr15:100879199-100879500 | ADAMTS17 | N_Shore | Body | No |
| Control-GSM3680910_ | chr19:12876846-12877188 | HOOK2;HOOK2 | N_Shore | Body;Body | Yes |
| Control-GSM3680911_ | chr6:291687-292823 | DUSP22 | N_Shore | TSS1500 | Yes |
| Control-GSM3680911_ | chr6:32063835-32063991 | TNXB | Island | Body | Yes |
| Control-GSM3680911_ | chr6:32064212-32064239 | TNXB | Island | Body | Yes |
| Control-GSM3680911_ | chr6:32552152-32552246 | HLA-DRB1 | Island | Body | Yes |
| Control-GSM3680911_ | chr13:25670170-25670187 | PABPC3 | Island | TSS200 | No |
| Control-GSM3680911_ | chr19:55476665-55477810 | NLRP2 | | TSS1500 | Yes |
| Control-GSM3680911_ | chr22:24348549-24348715 | GSTTP1 | | TSS1500 | Yes |
| Control-GSM3680911_ | chr2:30669597-30669863 | LCLAT1;LCLAT1 | N_Shore | TSS1500;TSS1500 | Yes |
| Control-GSM3680911_ | chr5:179740743-179741120 | GFPT2 | Island | Body | Yes |
| Control-GSM3680911_ | chr6:32526021-32526260 | HLA-DRB6 | | Body | Yes |
| Control-GSM3680911_ | chr6:32549631-32549935 | HLA-DRB1 | N_Shelf | Body | Yes |
| Control-GSM3680911_ | chr6:49681299-49681310 | CRISP2;CRISP2;CRISP2;CRISP2;CRISP2 | | TSS200;TSS200;TSS200;TSS200;TSS200 | Yes |
| Control-GSM3680911_ | chr8:144660590-144660631 | NAPRT1 | Island | TSS200 | Yes |
| Control-GSM3680915_ | chr1:205819251-205819463 | PM20D1;PM20D1 | S_Shore | 5'UTR;1stExon | Yes |
| Control-GSM3680915_ | chr6:32525960-32526364 | HLA-DRB6 | | Body | Yes |
| Control-GSM3680915_ | chr7:1080558-1081344 | C7orf50;C7orf50;C7orf50 | N_Shore | Body;Body;Body | Yes |
| Control-GSM3680915_ | chr17:79905219-79905255 | MYADML2 | | TSS200 | Yes |
| Control-GSM3680915_ | chr1:76189707-76189801 | ACADM;ACADM | N_Shore | TSS1500;TSS1500 | Yes |
| Control-GSM3680915_ | chr6:31237013-31237034 | HLA-C | N_Shore | Body | Yes |
| Control-GSM3680915_ | chr6:31237405-31237722 | HLA-C | N_Shore | Body | Yes |
| Control-GSM3680915_ | chr6:32632565-32632639 | HLA-DQB1 | Island | Body | Yes |
| Control-GSM3680915_ | chr10:135342560-135342936 | CYP2E1 | Island | Body | Yes |
| Control-GSM3680915_ | chr11:5617812-5617926 | TRIM6;TRIM6-TRIM34;TRIM6 | | TSS200;TSS200;5'UTR | Yes |
| Control-GSM3680915_ | chr11:75139390-75139736 | KLHL35 | N_Shore | Body | Yes |
| Control-GSM3680919_ | chr3:195489306-195489782 | MUC4;MUC4;MUC4 | Island | Body;Body;Body | No |
| Control-GSM3680919_ | chr6:29627167-29627395 | MOG;MOG;MOG;MOG;MOG;MOG;MOG;MOG;MOG;MOG | | Body;Body;Body;Body;Body;Body;Body;Body;B | Yes |
| Control-GSM3680919_ | chr6:32525960-32526364 | HLA-DRB6 | | Body | Yes |
| Control-GSM3680919_ | chr6:169629875-169630013 | THBS2 | N_Shore | Body | Yes |
| Control-GSM3680919_ | chr8:145955983-145956024 | ZNF251 | Island | Body | Yes |
| Control-GSM3680919_ | chr10:1511024-1511277 | ADARB2 | | Body | No |
| Control-GSM3680919_ | chr12:122356390-122356598 | WDR66 | Island | TSS200 | No |
| Control-GSM3680919_ | chr18:74728931-74728979 | MBP;MBP;MBP;MBP;MBP;MBP | | 1stExon;1stExon;Body;1stExon;Body;1stExon | Yes |
| Control-GSM3680919_ | chr2:239139911-239140190 | LOC151174;LOC643387;LOC151174 | N_Shore | Body;TSS1500;Body | Yes |
| Control-GSM3680919_ | chr3:147125955-147126119 | ZIC1 | N_Shore | TSS1500 | Yes |
| Control-GSM3680919_ | chr5:179740743-179741120 | GFPT2 | Island | Body | Yes |
| Control-GSM3680919_ | chr6:91004430-91004482 | BACH2;BACH2 | N_Shore | 5'UTR;5'UTR | Yes |
| Control-GSM3680919_ | chr7:29603121-29603407 | PRR15 | Island | TSS1500 | Yes |
| Control-GSM3680919_ | chr7:155255310-155255791 | EN2 | Island | Body | No |
| Control-GSM3680919_ | chr9:707166-707410 | KANK1;KANK1;KANK1 | Island | 1stExon;5'UTR;Body | Yes |
| Control-GSM3680919_ | chr10:42862978-42863594 | LOC441666 | N_Shore | Body | Yes |
| Control-GSM3680919_ | chr10:124220359-124220396 | HTRA1 | N_Shore | TSS1500 | Yes |
| Control-GSM3680919_ | chr12:678853-679287 | NINJ2 | N_Shore | Body | Yes |
| Control-GSM3680919_ | chr16:419975-420230 | MRPL28 | Island | Body | Yes |
| Control-GSM3680919_ | chr19:9785511-9785805 | ZNF562;ZNF562;ZNF562 | Island | 5'UTR;5'UTR;5'UTR | Yes |
| Control-GSM3680919_ | chr22:17956453-17956641 | CECR2 | | TSS200 | Yes |
| Control-GSM3680920_ | chr6:32525960-32526364 | HLA-DRB6 | | Body | Yes |
| Control-GSM3680920_ | chr5:179740743-179741120 | GFPT2 | Island | Body | Yes |
| Control-GSM3680920_ | chr6:30122544-30122593 | TRIM10;TRIM10 | | Body;Body | Yes |
| Control-GSM3680920_ | chr8:2075209-2075820 | MYOM2 | N_Shore | Body | Yes |
| Control-GSM3680920_ | chr19:55972791-55973099 | ISOC2;ISOC2;ISOC2 | Island | 5'UTR;5'UTR;5'UTR | Yes |
| Control-GSM3680920_ | chr19:55973234-55973338 | ISOC2;ISOC2;ISOC2 | Island | TSS200;TSS200;TSS200 | Yes |
| Control-GSM3680922_ | chr1:108734992-108735549 | SLC25A24;SLC25A24 | N_Shore | Body;Body | Yes |
| Control-GSM3680922_ | chr1:205819088-205819463 | PM20D1 | Island | 1stExon | Yes |
| Control-GSM3680922_ | chr6:32526021-32526260 | HLA-DRB6 | | Body | Yes |
| Control-GSM3680922_ | chr6:32557970-32558459 | HLA-DRB1 | | TSS1500 | Yes |
| Control-GSM3680922_ | chr19:12876846-12877000 | HOOK2;HOOK2 | N_Shore | Body;Body | Yes |
| Control-GSM3680922_ | chr6:29911036-29911104 | HLA-A | Island | Body | Yes |
| Control-GSM3680922_ | chr12:13196876-13196954 | KIAA1467 | N_Shore | TSS1500 | No |

**Supplementary Table 12: Disgenet, pathway and Gene Ontology analyses of 1,343 genes containing MeVars in Frontal Cortex of 547 controls**

| DISGENET Analysis | Adjusted P-value | Genes |
|---|---|---|
| Nephroblastoma | 0.0054 | CDKN1C;MPG;PIK3CD;ADAR;BRCA1;IGF1R;CASP8;CCND1;SLC22A18;SALL4;SIX2;SIX3;SOSTDC1;SYNPO;AMH;HGF;WNT5A;DNMT3A;PAX6;HLA-A;SFRP1;PAX8;KCNQ1;COL6A3;RIN1;EGFR;FUT4;NPY;NSD1;NNAT;CTNNA1;NCAM1;MSX1;PROM1;SLC38A2;WWOX;FZD7;POU6F2;BLCAP;ARHGAP24;DLK1;REST;DLG2;DES;GDNF;WT1;VIM;NES;HLA-DRB1;TP73 |
| Optic Neuritis | 0.0054 | HLA-DRB5;GSTM1;MOG;AQP4;GSTT1;HLA-A;GFAP;MBP;CAPN1;HCRT;HLA-DQA1;HLA-DRB1;HLA-DQB1 |
| Vitiligo | 0.0054 | DDR1;TNXB;AIRE;GSTT1;COMT;AGPAT1;RNF5;BACH2;C4B;NPY;RNF19A;PLCG2;TRIM26;HLA-DOA;VARS2;HLA-DQA1;GSTM1;S100A1;MIR25;HGF;HLA-B;FANCA;HLA-C;MITF;HLA-A;BTNL2;DEF8;S100B;CBFA2T3;POMC;MSRA;SMOC2;TRIM15;HCG22;TRIM31;EGFL8;HLA-DRB1 |
| Schizophrenia | 0.0054 | APP;RNH1;RPL3;MYT1L;ZBTB20;MLC1;AQP4;CELSR1;COMT;GRIP1;SYNGAP1;CCND1;DPYSL2;GRB10;PIP4K2A;TRIM26;SYNPO;SOX6;ANKS1B;KCNH2;HLA-J;SFMBT1;LMO3;TAP2;HLA-B;HLA-C;MYO7A;ANK3;HLA-A;HCG9;FOXP2;DKK4;CACNB2;ARC;CYP2E1;KCTD12;TSNARE1;TRIM10;MAD1L1;CNP;NTM;PCDH15;TRANK1;SLC1A3;AGAP1;GSTT1;ALOX12;CACNA1C;KALRN;CPLX1;RAI1;SBNO2;ATXN1;INPP5A;MOBP;PCDHA1;HRH2;ALDH3B1;MBP;OTX2;HCRT;GAS2L1;SLC38A2;HLA-DQA1;DRD4;HIST1H3E;HLA-DRB6;HLA-DRB5;GSTM1;SMAD3;AUTS2;HPS4;DCDC2;EIF2AK4;GRIN2B;DCLK1;GFAP;NR4A2;MIR346;MSRA;CXCL12;DLG2;SLC6A9;GDNF;PPP1R1B;PNOC;GNB3;LHX6;FXYD6;PSMG1;TGFBI;NES;HLA-DRB1;FGFR2;DDR1;CDKN1C;SNED1;DOCK4;MIR130B;PIK3CD;ARHGEF10L;NR2E1;BRCA1;SLC6A1;RLBP1;CYP26B1;GRM5;GRM4;CCKAR;PSD3;FLOT1;PDE4B;NEFM;LRRFIP1;NOS1;DLGAP2;SREBF1;DUSP2;BRD1;PLA2G4D;GRID1;TACR3;PAX6;NAV2;DNM2;CYP26C1;DNAJC5;PMP2;ARHGEF7;SHANK2;VGLL4;TLR3;HLA-DQB1;SDCCAG8;HDAC4;TNXB;TMPRSS5;SATB2;CAMK2A;EGFR;DDO;CAMKK2;SELENBP1;KIAA0513;SNX29;HLA-DMB;CBS;GNG7;P2RY2;NPY;PAK6;SLIT3;NCAM1;OPCML;GABBR1;MOG;KCNIP1;KCNB1;GALNT2;TXNRD2;ATP2B4;CABLES1;S100B;TBX4;MYO16;DLK1;GNMT;POMC;REST;KLF6;TPPP3;DES;FABP7;HLA-DPB1;RGS12;VIM;TRIM31;MDGA1;NEUROG1 |
| Bipolar Disorder | 0.0085 | SNED1;APP;DOCK9;MLC1;ARHGEF10L;NR2E1;COMT;MIR149;GRM5;GRM4;CCKAR;CASP8;DPYSL2;PDE4B;PIP4K2A;NEFM;TRIM26;NOS1;ARHGEF10;DUSP2;BRD1;GRID1;LMO3;TAP2;HLA-C;ANK3;HCG9;SORCS2;HLA-G;DKK4;NCOR2;GPRIN2;CACNB2;SFRP1;RFX4;KCTD12;TLR6;ARHGEF7;TLR5;MAD1L1;SDCCAG8;MCPH1;PFKFB3;TMPRSS5;HPN;CHRNA6;CAMK2A;TRANK1;CACNA1A;SLC1A3;GSTT1;ALOX12;FBLN1;CACNA1C;PRKCZ;CPLX1;EGFR;RASIP1;CAMKK2;SELENBP1;CUX2;PCDHA1;CBS;P2RY2;PTK2B;OTX2;NCAM1;PROM1;DRD4;GSTM1;CABLES1;NR1D1;S100B;GRIN2B;DCLK1;GFAP;MIR346;NR4A2;POMC;KLF6;DLG2;NBEA;GDNF;FABP7;PPP1R1B;GNB3;PSMG1;MDGA1;HLA-DRB1;FGFR2 |
| Alcoholic Intoxication, Chronic | 0.0110 | SEMA5A;PIK3CD;ABAT;SLC6A1;COMT;GLI2;SGCE;GRM5;CCKAR;DPYSL2;RNF19A;SLC22A18;PDE4B;EP300;NOS1;GGT1;CHRNB4;VWF;LMO3;HLA-B;TACR3;ARC;RFX4;CDH13;CYP2E1;ARHGEF7;KANK1;CAMK2A;OPRL1;SLC1A3;GSTT1;EGFR;MOBP;NPY;PTK2B;MBP;NCAM1;DRD4;GABBR1;GSTM1;OSBPL5;MOG;GRIN2B;ST18;GFAP;NR4A2;POMC;BHMT;SLC6A9;PNOC;PPP1R1B;GNB3;ALLC |
| Mental Depression | 0.0133 | APP;UCN;CRTC1;ABAT;AQP4;BRCA1;SLC6A1;COMT;SGCE;SOX2;ADAMTS2;GRM5;DPYSL2;PDE4B;SOSTDC1;NEFM;NOS1;STK32C;ARHGEF10;CHRNB4;GRID1;LMO3;TACR3;ANK1;DKK4;ARL4C;SFRP1;OAS2;CDH13;CYP2E1;ALPK1;TLR3;MCPH1;HDAC4;PFKFB3;CNP;CHRNA6;CAMK2A;GSTT1;CACNA1C;LTBP3;CPLX1;EGFR;RAI1;AANAT;MIR330;CBS;NPY;TNNI3;NCAM1;HCRT;DRD4;GSTM1;NR1D1;TLX1NB;S100B;GRIN2B;GFAP;POMC;GDNF;FABP7;PNOC;PPP1R1B;SSPO;GNB3;FGFR2 |
| Rhabdoid Tumor | 0.0133 | CDKN1C;CDKN1A;TAGLN;ANGPT1;KRT8;PIK3CD;CRYBB2;EGFR;DES;CCND1;PGBD5;KCNQ1;VIM;FXYD5;PROM1;SLC12A9 |
| Autistic Disorder | 0.0183 | SEMA5A;APP;CNTNAP1;DOCK4;AKAP8L;ZBTB20;PIK3CD;ABAT;AQP4;PCSK6;COMT;C4B;GRIP1;FLCN;GRM5;SYNGAP1;DPYSL2;PDE4B;EP300;NOS1;DLGAP2;SNRPN;DNMT3A;PAX6;TBR1;ANK3;HLA-A;UPB1;HLA-G;FOXP2;CACNB2;ALDH1A3;CDH13;SHANK2;MCPH1;CUL7;LXN;HEPACAM;ANKRD11;PCDH15;AGAP1;CACNA1A;SLC1A3;CACNA1C;TSPAN32;RAI1;PTDSS1;CUX1;CBS;NSD1;PRDM16;MBP;NCAM1;OTX1;EIF4E;DRD4;RPH3AL;GABBR1;GSTM1;AUTS2;ZBTB16;POU6F2;MCC;GRIN2B;SKI;BMP4;POMC;SDK1;GATM;NBEA;WT1;TUBGCP5;HLA-DRB1;GSTM5 |
| Familial M??ni??re disease | 0.0184 | KCNE1;HLA-DRB5;DTNA;HLA-B;HLA-C;CACNA1A;AQP4;HLA-A;TFB1M;SLC8A1;AQP1;ADD2;CCND1;MARVELD2;GJB6;IRF7;KCNQ4;SLC26A4;HLA-DQA1;HLA-DRB1;TLR3;LTF |
| Keloid | 0.0184 | DDR1;COL18A1;TAGLN;PIK3CD;HSPB3;FGF1;EGFR;IGF1R;LGALS1;CASP8;RPS18;EP300;PTK2B;CLIC1;HLA-DQA1;SMAD3;ANGPT2;TGFB3;FOXL2;POMC;SFRP1;SFRP2;TGFBI;VIM;FGFR2;HLA-DRB1;HLA-DQB1 |
| Medulloblastoma | 0.0185 | COL18A1;CRISP2;SNED1;MCM7;MPG;GFI1;PIK3CD;GLI3;GLI2;IGF1R;SOX2;CASP8;GRM4;CCND1;RNF19A;HNF4A;ZIC1;EP300;BRD2;S100A1;HGF;WNT5A;PAX6;EBF3;HIC1;DNM2;SFRP1;SFRP2;SCG5;BIRC5;MAD1L1;GSTT1;NTN1;EGFR;FUT4;ABR;CUX1;NHLH1;NNAT;MKNK2;NLRP2;PTK2B;OTX2;OTX1;PROM1;EIF4E;DRD4;BRD4;CIZ1;KDM4B;GSTM1;MIR33B;SMAD3;MGMT;S100B;ARHGAP24;GFAP;MEIS1;REST;CXCL12;WT1;COL9A3;VIM;NES;NEUROG1;TP73 |
| oligodendroglioma | 0.0201 | CRISP2;CNTNAP1;CNP;MYT1L;COL11A2;EGFR;SOX2;LGALS1;CCND1;NPY;MBP;DUSP2;GSTM1;MGMT;MRPL28;ARHGAP24;GFAP;AJAP1;ZAP70;KLF6;REST;DES;GDNF;PARVG;BIRC5;PSMG1;NES;TP73 |
| Prelingual Deafness | 0.0281 | GJB2;CLDN14;GJB6;PCDH15;MITF;SERAC1;SLC26A4;SLC26A5 |
| Myasthenia Gravis, Ocular | 0.0281 | POMC;HLA-B;HLA-DPB1;HLA-A;HLA-DQA1;HLA-DRB1 |
| Psoriasis | 0.0281 | DDR1;APP;CDKN1A;TRAF3IP2;MYT1L;ITGB2;PIK3CD;AATK;ANTXR2;B3GNTL1;BACH2;IGF1R;AQP1;C4B;RPTOR;C4A;LGALS1;RNF19A;CCRL2;NFKBIZ;PDE4B;HLA-DOA;SLC12A9;CHST8;MIR193B;LYNX1;S100A1;DNMT3A;WNT5A;TAP2;HLA-B;NME2;HLA-C;HLA-A;TYK2;HCG9;RUNX3;HLA-G;STX1B;OAS2;ELMO1;CDH13;TRIM15;ZNF831;TLR6;CARD14;HLA-DQB1;ZNF750;TNXB;SPATA2;GSTT1;ALOX12;GATA3;RNF5;PRKCZ;EGFR;RASIP1;NME1-NME2;SBNO2;HLA-DMB;CLMN;GNA12;SLIT3;NCAM1;EIF4E;HLA-DQA1;LYN;GSTM1;ANGPT2;PTPRN2;SMAD3;ANGPT1;MGMT;S100B;NR4A2;POMC;GJB2;CXCL12;MIR143;HLA-DPB1;HCG22;IL17D;NAT9;HLA-DRB1 |
| Laryngeal Diseases | 0.0281 | CDKN1A;MLKL;FANCA;GSTT1;BRCA1;EGFR;SOX2;MIR145;CCND1;WT1;NLRP2;EIF4E;SLC12A9 |
| Squamous cell carcinoma of lung | 0.0300 | DDR1;CDKN1A;TNXB;PIK3CD;FGF1;AURKC;EGFR;IGF1R;SOX2;NME1-NME2;HLA-DMB;CCND1;PEG10;EP300;TRIM26;PROM1;VARS2;EIF4E;HLA-DQA1;SNHG1;GSTM1;LMO3;XRCC3;KRT8;NME2;HLA-C;HLA-A;HIC1;HOPX;BMP4;PAX7;FGF19;MKS1;CYP2E1;TGFBI;VIM;FGFR2 |
| Congenital sensorineural hearing loss | 0.0300 | KCNE1;GJB2;MYO15A;KCNQ1;GJB6;PCDH15;MITF;MYO7A;SLC26A4 |
| Microphthalmos | 0.0304 | LRP5;CRYBA4;GLI3;GLI2;SOX2;GRIP1;GJA3;SALL4;SIX3;ALX3;OTX2;WNT3;STRA6;S100A1;FANCA;CRYBB2;PAX6;MITF;TBR1;FOXL2;VAX1;S100B;BMP4;POMC;ALDH1A3;TBX15;MEIS1;DES;COL4A1;MAB21L2;MKS1 |
| Neurilemmoma | 0.0304 | ANGPT2;ANGPT1;S100A1;TNK2;PIK3CD;S100B;ARHGAP24;EGFR;IGF1R;GFAP;BMP4;ANO1;DES;CCND1;CUX1;WT1;MSH4;MBP;CYP2E1;ARHGEF7;NES;TP73 |
| Diastolic blood pressure | 0.0304 | LYN;CASZ1;ZFHX3;TNXB;NXN;ITPK1;TAP2;CACNA2D2;SLC8A1;DCBLD1;CACNB2;LIPC;COL4A2;EHBP1L1;GRB10;HOXA3;ZNF831;SOX6;SDCCAG8;MCPH1 |
| Congenital aneurysm of ascending aorta | 0.0304 | FBN2;TNXB;SMAD3;TGFB3;COL11A2;MYLK;SKI;FLCN;ADAMTS2;COL4A1;CBS;ZNF469;COL9A3 |
| Uterine Corpus Cancer | 0.0304 | PIWIL2;DDR1;CRISP2;UCN;CDKN1A;MIR130B;PIK3CD;BRCA1;HSPB3;FGF1;COMT;AQP1;IGF1R;SOX2;CASP8;CCND1;SALL4;HNF4A;MACROD1;EP300;AMH;SOX7;KCNH2;BRD2;SREBF1;CIRBP;UGT2B17;PAX8;BIRC5;CDH13;MIR124-2;SGK1;BIRC3;OTUB1;LXN;SATB2;HAAO;COL11A2;HTRA1;GSTT1;THBS2;NTN1;EGFR;RAP1GAP;P2RY2;HCRT;PROM1;EIF4E;MUC4;WWOX;STARD13;ZFHX3;KDM4B;GSTM1;MGMT;TNK2;ARHGAP24;GGN;CXCL12;MIR145;VIM;FGFR2;TP73 |
| Hypoplasia of the optic nerve | 0.0304 | SOX2;POMC;ALDH1A3;WT1;ANKRD11;SALL4;OTX2;PAX6;GLI2 |
| Systolic Pressure | 0.0304 | FBN2;LYN;CASZ1;ZFHX3;TNXB;ITPK1;WFDC1;TRAPPC9;ADARB2;FOXP2;ARHGAP24;CACNB2;LIPC;RPS6KA2;MCF2L;HOXA3;CDH13;ZNF831;SOX6;OPCML |
| Neuromyelitis Optica | 0.0313 | MOG;HLA-DPB1;AQP4;MBP;S100B;FUT4;HLA-DQA1;HLA-DRB1;AQP1;GFAP;HLA-DQB1 |
| Well Differentiated Oligodendroglioma | 0.0316 | CRISP2;CNTNAP1;CNP;MYT1L;COL11A2;EGFR;SOX2;LGALS1;CCND1;NPY;MBP;DUSP2;MGMT;MRPL28;ARHGAP24;GFAP;AJAP1;ZAP70;KLF6;REST;DES;GDNF;BIRC5;PSMG1;NES;TP73 |
| Abnormal rapid eye movement sleep | 0.0337 | MOG;TNFSF4;HCRT;HLA-DRB1;HLA-DQB1 |
| Autoimmune thyroiditis | 0.0337 | AIRE;ITGB2;NCOA4;HLA-B;PIK3CD;HLA-C;HLA-A;TSHR;GJB2;TNFRSF25;TLR3;HLA-DRB1;HLA-DQB1 |
| Differentiated Thyroid Gland Carcinoma | 0.0337 | WWOX;TRAF3IP2;MCM7;XRCC3;HLA-C;PIK3CD;GSTT1;BRCA1;HSPB3;RAP1GAP;ARHGAP24;EGFR;TSHR;IGF1R;PAX8;SPATA13;NCAM1;GGT1;HLA-DOA;PROM1;SLC26A4;TP73 |
| Malignant neoplasm of endometrium | 0.0360 | PIWIL2;DDR1;CRISP2;UCN;CDKN1A;MIR130B;PIK3CD;BRCA1;HSPB3;FGF1;COMT;AQP1;IGF1R;SOX2;CASP8;CCND1;SALL4;HNF4A;MACROD1;EP300;AMH;SOX7;KCNH2;BRD2;SREBF1;CIRBP;UGT2B17;PAX8;BIRC5;CDH13;MIR124-2;SGK1;BIRC3;OTUB1;SATB2;HAAO;COL11A2;HTRA1;GSTT1;THBS2;NTN1;EGFR;RAP1GAP;CUX1;P2RY2;HCRT;PROM1;EIF4E;MUC4;WWOX;STARD13;ZFHX3;KDM4B;GSTM1;MGMT;TNK2;ARHGAP24;CXCL12;MIR145;VIM;FGFR2;TP73 |
| Disorder of eye | 0.0360 | DGKG;WNT10A;GSTM1;PRRX1;ABCA4;COL11A2;CRYBB2;GSTT1;PAX6;HLA-A;LTBP3;PEX14;TSHR;SOX2;BMP4;POMC;HLA-DPB1;HLA-DQA1;HLA-DRB1;STRA6;HLA-DQB1 |

| | | |
|---|---|---|
| Glioma | 0.0360 | SEMA5A;APP;MPG;MYT1L;FAM107A;AQP4;FGF1;MYLK;IGF1R;AQP1;MIR149;SOX2;LGALS1;CCND1;PEBP4;RNF19A;SALL4;HOXA3;EP300;PRSS8;SOX6;SLC16A3;SLC12A9;IER3;S100A1;LMO3;HGF;WNT5A;CACNA2D3;HLA-B;CACNA2D2;HLA-C;MITF;HLA-A;RUNX3;HLA-G;HIC1;RUNX2;SFRP1;SFRP2;CD8A;COL4A1;MSH4;RAB34;ELMO1;SCG5;SPARCL1;MT1G;MT1H;CYP2E1;RIN2;MAD1L1;CASZ1;GSTT1;FUT4;SLC9A3R2;CALD1;S100A13;NLRP2;CAMTA1;MBP;PROM1;HIST1H3E;LYN;HLA-DRB6;SNHG1;GGT7;GSTM1;SMAD3;ZBTB16;FZD7;TNK2;ABCA4;XRCC3;MRPL28;ESRRG;GRIN2B;MIR338;DCLK1;HOPX;ARHGAP24;NFATC4;GFAP;BMP4;CXCL12;HOOK2;BCL6;GDNF;TNFSF4;TRIP6;MIR199A1;MKS1;PSMG1;COL9A3;TGFBI;XAF1;NES;HLA-DRB1;FGFR2;TP73;PIWIL2;DDR1;CDKN1C;PIWIL1;PTPRU;SNED1;COL18A1;CRISP2;UCN;CNTNAP1;CDKN1A;GFM1;MCM7;MIR130B;PIK3CD;ADAR;NR2E1;HSPB3;PCSK6;GLI2;RPTOR;GRM5;CCKAR;CASP8;HNF4A;SLC22A18;PSD3;SOSTDC1;NOS1;CLIC1;KDM6B;MIR193B;BRD2;DUSP2;PAX6;AJAP1;DNM2;ZAP70;ALDH1A3;KAT2A;IRF7;BIRC5;CDH13;ARHGEF7;SGK1;MCPH1;LTF;HDAC4;ZBTB46;COL11A2;SEMA3B;LRP5;THBS2;EGFR;LMF1;CUX1;NSD1;P2RY2;NPY;PTK2B;SLIT3;NCAM1;EIF4E;OPCML;WWOX;ANGPT2;PRRX1;F10;ANGPT1;MGMT;CDX1;S100B;DLK1;SKI;POMC;REST;KLF6;TPPP3;MIR145;WT1;MIR143;FABP7;NFIC;LNX1;ADAM3A;VIM;IL17D;CYB561;BCAR1;PF4 |
| Autoimmune thyroid disease | 0.0388 | HLA-DRB6;SMAD3;TAP2;HLA-A;BTNL2;HLA-G;BACH2;ARHGAP24;TSHR;LGALS1;CXCL12;PDE4B;HLA-DPB1;TNFRSF25;HLA-DOA;SLC26A4;TRIM31;HLA-DQA1;HLA-DRB1;TP73;HLA-DQB1 |
| Drug Eruptions | 0.0424 | C4B;GSTM1;HLA-B;HLA-DPB1;GSTT1;HLA-A;CYP2E1;HLA-DQB1 |
| Rectal Carcinoma | 0.0425 | CDKN1C;CDKN1A;PIK3CD;ALOX12;COMT;EGFR;SOX2;CCND1;RPS6KA2;CCRL2;TNNI3;PROM1;EIF4E;WNT3;SLC12A9;GSTM1;SMAD3;MGMT;HGF;XRCC3;UGT2B15;ARID3A;MCC;RUNX3;DCLK1;ARHGAP24;DNM2;BMP4;CXCL12;MIR145;CYP2E1;VIM;TLR3;FGFR2;TP73 |
| Anophthalmos | 0.0430 | SOX2;BMP4;SKI;GRIP1;ALDH1A3;COL4A1;PAX6;OTX2;VAX1;VAX2;GLI2;STRA6 |
| Dermatomyositis, Childhood Type | 0.0430 | CRISP2;COL18A1;WT1;HLA-B;NCAM1;MYOM2;HLA-DQA1;HLA-DRB1;TLR3;PF4 |
| Congenital adrenal hyperplasia due to 21 hydroxylase deficiency | 0.0430 | C4B;POMC;C4A;BRD2;TNXB;HLA-B;HLA-C;HLA-DOA;GGN |
| Congenital ectopic pupil | 0.0430 | CTBP1;MAB21L2;PAX6;CPLX1;FGFRL1 |
| Familial psoriasis | 0.0430 | HLA-B;HLA-C;CARD14;HLA-DRB1;ZNF750 |
| Seropositive rheumatoid arthritis | 0.0430 | HLA-C;HLA-A;VIM;HLA-DRB1;HLA-DQB1 |
| Glioblastoma | 0.0435 | APP;COL16A1;MPG;PLEKHB1;FAM107A;AQP4;ADARB2;CNDP1;IGF1R;AQP1;MIR149;SOX2;LGALS1;CCND1;RNF19A;SALL4;EP300;SLC16A3;SLC12A9;TNS1;KCNH2;ACOT7;S100A1;HGF;WNT5A;HLA-B;HLA-C;EBF3;HLA-A;RUNX3;HLA-G;RUNX2;SFRP1;SFRP2;COL4A1;MSH4;SCNN1A;SCG5;CYP2E1;CARD14;MAD1L1;CASZ1;PFKFB3;CNP;TUBGCP2;HEPACAM;MAZ;GSTT1;GATA3;PRKCZ;FUT4;MIR330;NLRP2;CAMTA1;MBP;PROM1;LYN;GGT7;GSTM1;SMAD3;TGFB3;ZBTB16;XRCC3;MRPL28;MIR338;HOPX;ARHGAP24;GFAP;BMP4;CXCL12;BCL6;GDNF;TNFSF4;GJB6;TRIP6;PNOC;MIR199A1;MKS1;GNB3;TGFBI;XAF1;NES;HLA-DRB1;FGFR2;TP73;DDR1;CDKN1C;PIWIL1;SNED1;COL18A1;CRISP2;UCN;CDKN1A;GFM1;DOCK9;PIK3CD;NR2E1;BRCA1;HSPB3;PCSK6;GLI3;GLI2;CASP8;SLC22A18;PSD3;SIX3;LRRFIP1;NOS1;AMH;KDM6B;SREBF1;MIR193B;BRD2;MIR25;DNMT3A;MIR7-2;PAX6;AJAP1;DNM2;NCOR2;PAX7;B3GNT5;IRF7;BIRC5;CDH13;AGXT;ARHGEF7;SGK1;TLR3;BIRC3;HDAC4;COL11A2;SEMA3B;NTN1;EGFR;MIR377;CHSY1;CBS;GPC1;NPY;CTNNA1;PTK2B;NCAM1;MSX1;EIF4E;BRD4;OPCML;WWOX;ANGPT2;ANGPT1;MGMT;S100B;DLK1;SKI;REST;KLF6;DES;MIR145;WT1;MIR143;FABP7;MIR125B2;HLA-DPB1;LNX1;VIM;CMTM2;CYB561;BCAR1 |
| Renal Cell Carcinoma | 0.0435 | FAM107A;DNHD1;KLHL35;COMT;IGF1R;AQP1;MIR149;CCND1;RNF19A;EP300;CAPN1;PRSS8;HLA-DOA;SLC16A3;PPFIA4;SLC12A9;EOMES;S100A1;SFMBT1;HGF;TAP2;KRT8;HLA-B;HLA-C;MITF;HLA-A;RUNX3;HLA-G;HIC1;DKK4;SFRP1;ARC;SFRP2;MT1G;MT1H;CARD11;MAD1L1;FBN2;SPTBN4;CUL7;HEPACAM;HPN;GSTT1;FBLN1;GATA3;PRKCZ;ACAT2;RAP1GAP;SLC9A3R2;LDHC;ABR;ALDH3B1;NLRP2;HIVEP3;TPPP;PROM1;WNT10A;GSTM1;TNK2;ASPSCR1;ARHGAP24;BMP4;CXCL12;PLXNB1;XAF1;HLA-DRB1;FGFR2;TP73;PIWIL1;PTPRU;COL18A1;CRISP2;CDKN1A;BNC1;PIK3CD;PTH1R;BRCA1;FLCN;CASP8;HNF4A;MCF2L;SIX2;SOSTDC1;NOS1;KDM6B;VWF;DNMT3A;NME2;TYK2;SLC25A18;PAX8;BIRC5;CDH13;TLR3;BIRC3;KANK1;AHNAK;COL11A2;NTN1;EGFR;CUX1;BGLAP;NPY;PTK2B;MUC4;EIF4E;WWOX;ANGPT2;ANGPT1;SUSD2;KCNIP4;S100B;DLK1;POMC;GATM;DES;MIR145;WT1;MIR143;CCDC8;FABP7;NFIC;VIM |
| Narcolepsy | 0.0435 | NPFFR2;FRMPD2;DENND3;ANTXR2;CHKB-CPT1B;COMT;KALRN;SNX29;GRM5;SALL4;P2RY2;PDE4B;SERAC1;HCRT;DLGAP2;HLA-DQA1;HLA-DRB5;MOG;GRID1;TAP2;HLA-A;BTNL2;HLA-F;MCC;CPT1B;FOXP2;SDK1;SFRP1;TNFSF4;HLA-DPB1;ELMO1;HLA-DRB1;HLA-DQB1 |
| Vascular lesions | 0.0439 | APP;COL18A1;GSTM1;VWF;F10;PRRX1;ITGB2;LOXL3;HLA-B;GSTT1;HLA-A;S100B;IRF7;CYP2E1;TGFBI |

| Supplementary Table 13: SZ candidate genes with MeVars showing significant differences in the observed frequencies between controls and patients | | | | | | | | |
|---|---|---|---|---|---|---|---|---|
| SZ candidate gene with MeVar | Controls_ Hyper | Controls_ Hypo | Controls_ Remaining | Patients_ Hyper | Patients_ Hypo | Patients_ Remaining | Chisquare value F=2 (5.991) | p-value |
| RPTOR | 58 | 119 | 273 | 0 | 3 | 221 | 8.09E+01 | 2.648E-18 |
| MCF2L | 95 | 71 | 284 | 0 | 1 | 223 | 8.02E+01 | 3.852E-18 |
| SPON2 | 78 | 67 | 305 | 2 | 0 | 222 | 6.74E+01 | 2.305E-15 |
| FBXL18 | 131 | 4 | 315 | 0 | 5 | 219 | 6.72E+01 | 2.519E-15 |
| NR2E1 | 0 | 126 | 324 | 1 | 0 | 223 | 6.47E+01 | 8.796E-15 |
| ST18 | 16 | 144 | 290 | 0 | 8 | 216 | 6.16E+01 | 4.29E-14 |
| IQSEC1 | 4 | 131 | 315 | 0 | 3 | 221 | 6.00E+01 | 9.314E-14 |
| ADARB2 | 33 | 117 | 300 | 3 | 7 | 214 | 5.26E+01 | 3.792E-12 |
| WWOX | 71 | 60 | 319 | 6 | 0 | 218 | 5.23E+01 | 4.334E-12 |
| NUAK1 | 98 | 5 | 347 | 0 | 3 | 221 | 4.88E+01 | 2.471E-11 |
| PACRG | 95 | 10 | 345 | 0 | 4 | 220 | 4.74E+01 | 5.03E-11 |
| CTTN | 84 | 3 | 363 | 0 | 4 | 220 | 4.36E+01 | 3.379E-10 |
| FXYD6 | 11 | 79 | 360 | 2 | 0 | 222 | 4.12E+01 | 1.136E-09 |
| SORBS3 | 4 | 83 | 363 | 0 | 1 | 223 | 4.09E+01 | 1.339E-09 |
| ME1 | 4 | 80 | 366 | 1 | 0 | 223 | 4.02E+01 | 1.849E-09 |
| SASH1 | 70 | 0 | 380 | 0 | 2 | 222 | 3.89E+01 | 3.64E-09 |
| TMEM140 | 76 | 13 | 361 | 0 | 6 | 218 | 3.79E+01 | 6.025E-09 |
| S100A1 | 65 | 12 | 373 | 0 | 1 | 223 | 3.62E+01 | 1.393E-08 |
| NCOA4 | 10 | 67 | 373 | 0 | 1 | 223 | 3.59E+01 | 1.603E-08 |
| SYNJ2 | 81 | 8 | 361 | 2 | 4 | 218 | 3.55E+01 | 1.915E-08 |
| LRRC4 | 1 | 69 | 380 | 1 | 0 | 223 | 3.46E+01 | 3.067E-08 |
| DPYSL2 | 69 | 4 | 377 | 0 | 2 | 222 | 3.43E+01 | 3.481E-08 |
| TOLLIP | 85 | 1 | 364 | 4 | 0 | 220 | 3.36E+01 | 4.986E-08 |
| SLC41A3 | 66 | 4 | 380 | 0 | 3 | 221 | 3.31E+01 | 6.347E-08 |
| GLI3 | 0 | 62 | 388 | 1 | 0 | 223 | 3.29E+01 | 7.28E-08 |
| FGFR2 | 47 | 31 | 372 | 0 | 3 | 221 | 3.25E+01 | 8.65E-08 |
| GNG7 | 31 | 39 | 380 | 0 | 1 | 223 | 3.25E+01 | 8.944E-08 |
| LRRFIP1 | 0 | 61 | 389 | 1 | 0 | 223 | 3.24E+01 | 9.337E-08 |
| EIF4E | 64 | 1 | 385 | 0 | 1 | 223 | 3.21E+01 | 1.065E-07 |
| FGFRL1 | 0 | 69 | 381 | 0 | 1 | 223 | 3.19E+01 | 1.175E-07 |
| KIRREL3 | 4 | 58 | 388 | 3 | 0 | 221 | 2.92E+01 | 4.648E-07 |
| SGK2 | 58 | 8 | 384 | 0 | 4 | 220 | 2.89E+01 | 5.379E-07 |
| PFKP | 57 | 5 | 388 | 0 | 2 | 222 | 2.84E+01 | 6.667E-07 |
| AHNAK | 61 | 0 | 389 | 1 | 0 | 223 | 2.79E+01 | 8.571E-07 |
| NFIC | 0 | 55 | 395 | 0 | 1 | 223 | 2.50E+01 | 3.80E-06 |
| ATXN7L1 | 4 | 48 | 398 | 1 | 0 | 223 | 2.43E+01 | 5.32E-06 |
| RAP1GAP | 16 | 36 | 398 | 1 | 0 | 223 | 2.37E+01 | 7.31E-06 |
| CABLES1 | 7 | 43 | 400 | 2 | 0 | 222 | 2.19E+01 | 1.76E-05 |
| MARVELD2 | 0 | 45 | 405 | 0 | 1 | 223 | 2.00E+01 | 4.54E-05 |

| Supplementary Table 14: Methylation levels and expression levels for seven genes showing significant correlation in prefrontal cortex samples | | | | | | | |
|---|---|---|---|---|---|---|---|
| Gene | Coordinate | Position | CGI | $R^2$ | R | p value | Correlation type |
| FGFR2 | chr10:123353889-123354171 | Gene body | No | 0.0463 | 0.2152 | 0.0361 | Negative |
| GLI3 | chr7:42267269-42267547 | Gene body | No | 0.0470 | 0.2168 | 0.0349 | Positive |
| IQSEC1 | chr3:13114797-13115104 | Promoter | Yes | 0.0578 | 0.2404 | 0.0189 | Positive |
| ME1 | chr6:84140683-84140898 | Promoter | Yes | 0.1687 | 0.4107 | 0.0000 | Negative |
| NUAK1 | chr12:106480546-106480771 | Gene body | No | 0.0434 | 0.2083 | 0.0428 | Positive |
| RAP1GAP | chr1:21966310-21966500 | Gene body | No | 0.0641 | 0.2532 | 0.0133 | Negative |
| ZBTB16 | chr11:113929164-113929287 | Promoter | No | 0.0449 | 0.2119 | 0.0394 | Negative |

| Supplementary Table 15: Coordinates of MeVars identified in three oral tissues (hg 38 coordinates) | | | | |
|---|---|---|---|---|
| Sample | Coordinate | UCSC_REFGENE_NAME | RELATION_TO_UCSC_CPG_ISLAND | UCSC_REFGENE_GROUP |
| P4 | chr1:150269253-150269467 | APH1A | Island | TSS1500;TSS1500;TSS1500;TSS1500;TSS1500;TSS1500;TSS1500 |
| P4 | chr19:41884217-41884249 | ARHGEF1 | Island | exon_1;TSS1500 |
| P4 | chr16:8861903-8862383 | CARHSP1 | | TSS200;TSS200;TSS200 |
| P4 | chr10:68721077-68721162 | CCAR1 | Island | TSS200;TSS200;TSS200;TSS200 |
| P5 | chr5:69234948-69234970 | CDK7 | Island | 5UTR;exon_1;5UTR;exon_1;5UTR;exon_1;5UTR;exon_1;5UTR;exon_1;5UTR;exon_1;5UTR;exon_1;exon_1 |
| P4 | chr2:240136781-240136865 | COPS9 | S_Shore | TSS1500;TSS1500 |
| P5 | chr15:68578640-68579066 | CORO2B | Island | TSS1500 |
| P3 | chr15:82646864-82647171 | CPEB1-AS1 | Island | TSS1500 |
| P5 | chr20:49046291-49046606 | CSE1L | Island | TSS200;TSS200;TSS200;TSS200;TSS1500;TSS1500 |
| P4 | chr10:43138309-43138399 | CSGALNACT2 | Island | TSS200;TSS200 |
| P5 | chr4:107931371-107931411 | CYP2U1 | Island | TSS200 |
| P3 | chr5:31532004-31532072 | DROSHA | Island | 5UTR;exon_1;5UTR;exon_1;5UTR;exon_1;TSS1500;TSS1500 |
| P4 | chr11:86244630-86244677 | EED | N_Shore;S_Shore | TSS200;TSS200;TSS200 |
| P4 | chr15:48877568-48877768 | EID1 | N_Shore | TSS1500 |
| P4 | chr6:53349278-53349413 | ELOVL5 | S_Shore | TSS1500;TSS1500;TSS1500;TSS1500;TSS1500 |
| P5 | chr20:25195578-25195743 | ENTPD6 | Island;N_Shore | TSS1500;TSS1500;TSS1500;TSS1500;TSS200;TSS200;TSS200;TSS200;TSS200;TSS200;TSS200;TSS200;TSS200;TSS200;TSS200;TSS200;TSS200;TSS200;TSS200;TSS200;TSS200;TSS200;TSS200;TSS200 |
| P4 | chr19:7863660-7863887 | EVI5L | Island | exon_20;exon_19 |
| P5 | chr11:22625998-22626096 | FANCF | S_Shore | TSS200 |
| P4 | chr10:121594308-121594442 | FGFR2 | N_Shelf | TSS1500;TSS200;TSS1500;TSS1500 |
| P5 | chr14:28766811-28767270 | FOXG1 | Island | 5UTR;exon_1;TSS1500 |
| P4 | chr17:65057209-65057236 | GNA13 | Island | TSS1500;TSS1500 |
| P4 | chr1:202122737-202122752 | GPR37L1 | | TSS200 |
| P4 | chr4:105708808-105708884 | GSTCD | Island | 5UTR;exon_1;5UTR;exon_1;TSS200;TSS200 |
| P4 | chr4:173527190-173527398 | HAND2 | N_Shore;N_Shelf | 3UTR;exon_2;TSS1500 |
| P4 | chr6:31055932-31056414 | HCG22 | | exon_3 |
| P4 | chr11:94088948-94089128 | HEPHL1 | Island | exon_12 |
| P4 | chr3:57227544-57227946 | HESX1 | Island | 5UTR;exon_1;5UTR;exon_1;5UTR;exon_1;5UTR;exon_1;exon_1;TSS200 |

| | | | | |
|---|---|---|---|---|
| P4 | chr7:27103784-27103966 | HOXA2 | Island;N_Shelf | TSS1500 |
| P5 | chr4:11428823-11429069 | HS3ST1 | S_Shore;Island | 5UTR;exon_1 |
| P4 | chr15:90102319-90102675 | IDH2 | Island | 5UTR;exon_1;exon_1;exon_1 |
| P5 | chr4:57109868-57110265 | IGFBP7-AS1 | Island | exon_1 |
| P4 | chr10:88583409-88583447 | LIPJ | Island | exon_1;TSS200;TSS200 |
| P4 | chr20:5950659-5950908 | MCM8 | Island | 5UTR;exon_1;5UTR;exon_1;5UTR;exon_1;5UTR;exon_1;5UTR;exon_1;TSS200;TSS200 |
| P3 | chr8:143573124-143573314 | MROH6 | S_Shelf;S_Shore | TSS1500 |
| P4 | chr8:17722607-17723040 | MTUS1 | | TSS1500;TSS200 |
| P4 | chr16:88663883-88664032 | MVD | Island | TSS1500 |
| P4 | chr8:143578459-143578489 | NAPRT | Island | TSS200;TSS200;TSS200;TSS200 |
| P5 | chr14:79279930-79280177 | NRXN3 | S_Shore | 5UTR;exon_1;5UTR;exon_1;5UTR;exon_1;exon_1 |
| P4 | chr11:73264657-73265078 | P2RY6 | Island | TSS1500 |
| P4 | chr15:65133955-65134333 | PDCD7 | S_Shore | TSS200 |
| P4 | chr8:100150030-100150420 | POLR2K | | TSS1500 |
| P5 | chr1:56645197-56645435 | PRKAA2 | Island | TSS200 |
| P4 | chr1:17438558-17438574 | RCC2 | Island | TSS200 |
| P4 | chr8:100336228-100336260 | RNF19A | | TSS200 |
| P4 | chr15:40568769-40569052 | RPUSD2 | N_Shore | TSS1500;TSS1500 |
| P5 | chr9:134325217-134325627 | RXRA | Island | TSS1500 |
| P4 | chr22:43954928-43955003 | SAMM50 | N_Shore | TSS1500 |
| P4 | chr1:151132573-151132993 | SEMA6C | Island | exon_20;exon_19;exon_19 |
| P5 | chr17:29005988-29006099 | SEZ6 | Island | exon_1;5UTR;exon_1;5UTR |
| P4 | chr1:41021645-41021842 | SLFNL1 | S_Shelf | 5UTR;exon_1;5UTR;exon_1;exon_1;5UTR;exon_1;5UTR;5UTR;exon_1 |
| P4 | chr5:169302550-169302843 | SLIT3 | S_Shore | TSS1500;TSS1500 |
| P5 | chr19:43754712-43754968 | SMG9 | Island | 5UTR;exon_1 |
| P4 | chr16:88663201-88663231 | SNAI3-AS1 | S_Shelf;Island | TSS200;TSS200;TSS200 |
| P4 | chr1:25232462-25232577 | SYF2 | Island | exon_1;exon_1 |
| P5 | chr6:33313728-33313737 | TAPBP | Island;N_Shore;N_Shelf | exon_2;exon_2;exon_2 |
| P4 | chr20:34704271-34704334 | TP53INP2 | Island | TSS1500;TSS200;TSS200;TSS200 |
| P4 | chr11:4393686-4393973 | TRIM21 | | 5UTR;exon_1 |
| P4 | chr15:41557765-41558305 | TYRO3 | N_Shore;N_Shore | TSS1500 |
| P4 | chr3:196432718-196432871 | UBXN7-AS1 | Island | exon_2;TSS1500 |

| | | | | |
|---|---|---|---|---|
| P5 | chr11:75814836-75815030 | UVRAG | Island;N_Shore | TSS1500;TSS1500;TSS1500;TSS1500;TSS1500;TSS1500;TSS1500;TSS200 |
| P4 | chr8:143541689-143542055 | ZC3H3 | Island | TSS1500 |
| P5 | chr5:80407574-80407727 | ZFYVE16 | N_Shore | TSS1500;TSS1500;TSS1500;TSS1500;TSS1500;TSS1500;TSS1500;TSS1500 |

| Supplementary Table 16: Methylation levels and expression levels for seven genes showing sign | | | | | | |
|---|---|---|---|---|---|---|
| Gene | Coordinate | Position | CGI | $R^2$ | R | p value |
| CCAR1 | chr10:68721077-68721162 | Promoter | Yes | 0.6687 | 0.8177 | 0.0468 |
| EED | chr11:86244630-86244677 | Promoter | No | 0.8843 | 0.9404 | 0.0052 |
| MTUS1 | chr8:17722607-17723040 | Gene body | No | 0.7711 | 0.8781 | 0.0214 |
| P2RY6 | chr11:73264657-73265078 | Promoter | Yes | 0.7019 | 0.8378 | 0.0373 |
| SLFNL1 | chr1:41021645-41021842 | Promoter | No | 0.9272 | 0.9629 | 0.0020 |
| SLIT3 | chr5:169302550-169302843 | Upstream | No | 0.8844 | 0.9404 | 0.0052 |

| ificant correlation in prefrontal cortex samples |
|---|
| Correlation type |
| Negative |
| Negative |
| Positive |
| Positive |
| Positive |
| Positive |

| Supplementary Table 17. MeVars identified in 62 untransformed skin fibroblasts (duplicates not removed) | | | | |
|---|---|---|---|---|
| Sample | Coordinate | UCSC_REFGENE_NAME | RELATION_ | UCSC_REFGENE_GROUP |
| GM00016 | chr6:31237405-31237722 | HLA-C | N_Shore | Body |
| GM00022 | chr6:31237013-31237034 | HLA-C | N_Shore | Body |
| GM00042 | chr6:31237405-31237722 | HLA-C | N_Shore | Body |
| GM00031 | chr6:31237013-31237034 | HLA-C | N_Shore | Body |
| GM00940 | chr6:31237405-31237722 | HLA-C | N_Shore | Body |
| GM00042 | chr6:31237013-31237034 | HLA-C | N_Shore | Body |
| GM00941 | chr6:31237405-31237722 | HLA-C | N_Shore | Body |
| GM00016 | chr10:114877616-114877689 | TCF7L2;TCF7L2;TCF7L2;TCF7L2;TCF7L2 | | Body;Body;Body;Body;Body;Body |
| GM00289 | chr6:31237013-31237034 | HLA-C | N_Shore | Body |
| GM01095 | chr6:31237405-31237722 | HLA-C | N_Shore | Body |
| GM00031 | chr10:114877616-114877689 | TCF7L2;TCF7L2;TCF7L2;TCF7L2;TCF7L2 | | Body;Body;Body;Body;Body;Body |
| GM00439 | chr6:31237013-31237034 | HLA-C | N_Shore | Body |
| GM01375 | chr6:31237405-31237722 | HLA-C | N_Shore | Body |
| GM00042 | chr10:114877616-114877689 | TCF7L2;TCF7L2;TCF7L2;TCF7L2;TCF7L2 | | Body;Body;Body;Body;Body;Body |
| GM00520 | chr19:12876846-12877000 | HOOK2;HOOK2 | N_Shore | Body;Body |
| GM00520 | chr6:31237013-31237034 | HLA-C | N_Shore | Body |
| GM01376 | chr6:31237405-31237722 | HLA-C | N_Shore | Body |
| GM00289 | chr10:114877616-114877689 | TCF7L2;TCF7L2;TCF7L2;TCF7L2;TCF7L2 | | Body;Body;Body;Body;Body;Body |
| GM00935 | chr19:12876846-12877000 | HOOK2;HOOK2 | N_Shore | Body;Body |
| GM00940 | chr6:31237013-31237034 | HLA-C | N_Shore | Body |
| GM01392 | chr6:31237405-31237722 | HLA-C | N_Shore | Body |
| GM00438 | chr10:114877616-114877689 | TCF7L2;TCF7L2;TCF7L2;TCF7L2;TCF7L2 | | Body;Body;Body;Body;Body;Body |
| GM01095 | chr19:12876846-12877000 | HOOK2;HOOK2 | N_Shore | Body;Body |
| GM00941 | chr6:31237013-31237034 | HLA-C | N_Shore | Body |
| GM01393 | chr6:31237405-31237722 | HLA-C | N_Shore | Body |
| GM00042 | chr1:119526060-119526437 | TBX15 | N_Shore | 5'UTR |
| GM00439 | chr10:114877616-114877689 | TCF7L2;TCF7L2;TCF7L2;TCF7L2;TCF7L2 | | Body;Body;Body;Body;Body;Body |
| GM00042 | chr14:54418728-54418851 | BMP4;BMP4;BMP4 | Island | Body;Body;Body |
| GM00016 | chr15:88576296-88576320 | NTRK3;NTRK3;NTRK3 | | Body;Body;Body |
| GM01376 | chr19:12876846-12877000 | HOOK2;HOOK2 | N_Shore | Body;Body |
| GM01095 | chr6:31237013-31237034 | HLA-C | N_Shore | Body |
| GM02173 | chr6:31237405-31237722 | HLA-C | N_Shore | Body |
| GM00022 | chr7:2847517-2847575 | GNA12 | | Body |
| GM00439 | chr1:119526060-119526437 | TBX15 | N_Shore | 5'UTR |
| GM00520 | chr10:114877616-114877689 | TCF7L2;TCF7L2;TCF7L2;TCF7L2;TCF7L2 | | Body;Body;Body;Body;Body;Body |
| GM00289 | chr14:54418728-54418851 | BMP4;BMP4;BMP4 | Island | Body;Body;Body |
| GM00438 | chr15:88576296-88576320 | NTRK3;NTRK3;NTRK3 | | Body;Body;Body |
| GM01650 | chr19:12876846-12877000 | HOOK2;HOOK2 | N_Shore | Body;Body |
| GM01375 | chr6:31237013-31237034 | HLA-C | N_Shore | Body |
| GM02317 | chr6:31237405-31237722 | HLA-C | N_Shore | Body |
| GM00042 | chr7:2847517-2847575 | GNA12 | | Body |
| GM00935 | chr1:119526060-119526437 | TBX15 | N_Shore | 5'UTR |
| GM00935 | chr10:114877616-114877689 | TCF7L2;TCF7L2;TCF7L2;TCF7L2;TCF7L2 | | Body;Body;Body;Body;Body;Body |
| GM00290 | chr14:54418728-54418851 | BMP4;BMP4;BMP4 | Island | Body;Body;Body |
| GM00439 | chr15:88576296-88576320 | NTRK3;NTRK3;NTRK3 | | Body;Body;Body |

| | | | | |
|---|---|---|---|---|
| GM01653 | chr19:12876846-12877000 | HOOK2;HOOK2 | N_Shore | Body;Body |
| GM01650 | chr6:31237013-31237034 | HLA-C | N_Shore | Body |
| GM02456 | chr6:31237405-31237722 | HLA-C | N_Shore | Body |
| GM00290 | chr7:2847517-2847575 | GNA12 | | Body |
| GM00940 | chr1:119526060-119526437 | TBX15 | N_Shore | 5'UTR |
| GM01095 | chr10:114877616-114877689 | TCF7L2;TCF7L2;TCF7L2;TCF7L2;TCF7L2 | | Body;Body;Body;Body;Body;Body |
| GM00438 | chr14:54418728-54418851 | BMP4;BMP4;BMP4 | Island | Body;Body;Body |
| GM00520 | chr15:88576296-88576320 | NTRK3;NTRK3;NTRK3 | | Body;Body;Body |
| GM00031 | chr17:80358829-80358876 | C17orf101;C17orf101 | N_Shelf | Body;Body |
| GM02173 | chr19:12876846-12877000 | HOOK2;HOOK2 | N_Shore | Body;Body |
| GM00031 | chr5:323320-323969 | AHRR | S_Shore | Body |
| GM02456 | chr6:31237013-31237034 | HLA-C | N_Shore | Body |
| GM02555 | chr6:31237405-31237722 | HLA-C | N_Shore | Body |
| GM00438 | chr7:2847517-2847575 | GNA12 | | Body |
| GM00941 | chr1:119526060-119526437 | TBX15 | N_Shore | 5'UTR |
| GM01096 | chr10:114877616-114877689 | TCF7L2;TCF7L2;TCF7L2;TCF7L2;TCF7L2 | | Body;Body;Body;Body;Body;Body |
| GM00439 | chr14:54418728-54418851 | BMP4;BMP4;BMP4 | Island | Body;Body;Body |
| GM00935 | chr15:88576296-88576320 | NTRK3;NTRK3;NTRK3 | | Body;Body;Body |
| GM00042 | chr17:80358829-80358876 | C17orf101;C17orf101 | N_Shelf | Body;Body |
| GM02175 | chr19:12876846-12877000 | HOOK2;HOOK2 | N_Shore | Body;Body |
| GM00289 | chr5:323320-323969 | AHRR | S_Shore | Body |
| GM02555 | chr6:31237013-31237034 | HLA-C | N_Shore | Body |
| GM02706 | chr6:31237405-31237722 | HLA-C | N_Shore | Body |
| GM00520 | chr7:2847517-2847575 | GNA12 | | Body |
| GM01095 | chr1:119526060-119526437 | TBX15 | N_Shore | 5'UTR |
| GM01392 | chr10:114877616-114877689 | TCF7L2;TCF7L2;TCF7L2;TCF7L2;TCF7L2 | | Body;Body;Body;Body;Body;Body |
| GM00520 | chr14:54418728-54418851 | BMP4;BMP4;BMP4 | Island | Body;Body;Body |
| GM00940 | chr15:88576296-88576320 | NTRK3;NTRK3;NTRK3 | | Body;Body;Body |
| GM00439 | chr17:80358829-80358876 | C17orf101;C17orf101 | N_Shelf | Body;Body |
| GM02317 | chr19:12876846-12877000 | HOOK2;HOOK2 | N_Shore | Body;Body |
| GM00290 | chr5:323320-323969 | AHRR | S_Shore | Body |
| GM02641 | chr6:31237013-31237034 | HLA-C | N_Shore | Body |
| GM03073 | chr6:31237405-31237722 | HLA-C | N_Shore | Body |
| GM02147 | chr7:2847517-2847575 | GNA12 | | Body |
| GM01096 | chr1:119526060-119526437 | TBX15 | N_Shore | 5'UTR |
| GM01653 | chr10:114877616-114877689 | TCF7L2;TCF7L2;TCF7L2;TCF7L2;TCF7L2 | | Body;Body;Body;Body;Body;Body |
| GM00935 | chr14:54418728-54418851 | BMP4;BMP4;BMP4 | Island | Body;Body;Body |
| GM01375 | chr15:88576296-88576320 | NTRK3;NTRK3;NTRK3 | | Body;Body;Body |
| GM00520 | chr17:80358829-80358876 | C17orf101;C17orf101 | N_Shelf | Body;Body |
| GM02456 | chr19:12876846-12877000 | HOOK2;HOOK2 | N_Shore | Body;Body |
| GM00438 | chr5:323320-323969 | AHRR | S_Shore | Body |
| GM02706 | chr6:31237013-31237034 | HLA-C | N_Shore | Body |
| GM03075 | chr6:31237405-31237722 | HLA-C | N_Shore | Body |
| GM02149 | chr7:2847517-2847575 | GNA12 | | Body |
| GM01393 | chr1:119526060-119526437 | TBX15 | N_Shore | 5'UTR |
| GM02555 | chr10:114877616-114877689 | TCF7L2;TCF7L2;TCF7L2;TCF7L2;TCF7L2 | | Body;Body;Body;Body;Body;Body |
| GM01095 | chr14:54418728-54418851 | BMP4;BMP4;BMP4 | Island | Body;Body;Body |

| | | | | |
|---|---|---|---|---|
| GM01376 | chr15:88576296-88576320 | NTRK3;NTRK3;NTRK3 | | Body;Body;Body |
| GM00935 | chr17:80358829-80358876 | C17orf101;C17orf101 | N_Shelf | Body;Body |
| GM02555 | chr19:12876846-12877000 | HOOK2;HOOK2 | N_Shore | Body;Body |
| GM00289 | chr19:21264896-21265421 | ZNF714 | N_Shore | TSS200 |
| GM00439 | chr5:323320-323969 | AHRR | S_Shore | Body |
| GM03075 | chr6:31237013-31237034 | HLA-C | N_Shore | Body |
| GM09255 | chr6:31237405-31237722 | HLA-C | N_Shore | Body |
| GM02456 | chr7:2847517-2847575 | GNA12 | | Body |
| GM02175 | chr1:119526060-119526437 | TBX15 | N_Shore | 5'UTR |
| GM00935 | chr1:119532320-119532542 | TBX15 | S_Shore | TSS200 |
| GM02640 | chr10:114877616-114877689 | TCF7L2;TCF7L2;TCF7L2;TCF7L2;TCF7L2 | | Body;Body;Body;Body;Body;Body |
| GM01096 | chr14:54418728-54418851 | BMP4;BMP4;BMP4 | Island | Body;Body;Body |
| GM02147 | chr15:88576296-88576320 | NTRK3;NTRK3;NTRK3 | | Body;Body;Body |
| GM00940 | chr17:80358829-80358876 | C17orf101;C17orf101 | N_Shelf | Body;Body |
| GM02640 | chr19:12876846-12877000 | HOOK2;HOOK2 | N_Shore | Body;Body |
| GM00935 | chr19:21264896-21265421 | ZNF714 | N_Shore | TSS200 |
| GM00016 | chr2:176964512-176964720 | HOXD12 | Island | TSS200 |
| GM00520 | chr5:323320-323969 | AHRR | S_Shore | Body |
| GM09255 | chr6:31237013-31237034 | HLA-C | N_Shore | Body |
| GM09256 | chr6:31237405-31237722 | HLA-C | N_Shore | Body |
| GM02555 | chr7:2847517-2847575 | GNA12 | | Body |
| GM02640 | chr1:119526060-119526437 | TBX15 | N_Shore | 5'UTR |
| GM00940 | chr1:119532320-119532542 | TBX15 | S_Shore | TSS200 |
| GM02641 | chr10:114877616-114877689 | TCF7L2;TCF7L2;TCF7L2;TCF7L2;TCF7L2 | | Body;Body;Body;Body;Body;Body |
| GM01375 | chr14:54418728-54418851 | BMP4;BMP4;BMP4 | Island | Body;Body;Body |
| GM02175 | chr15:88576296-88576320 | NTRK3;NTRK3;NTRK3 | | Body;Body;Body |
| GM00290 | chr17:21300735-21300791 | KCNJ12 | | 5'UTR |
| GM00941 | chr17:80358829-80358876 | C17orf101;C17orf101 | N_Shelf | Body;Body |
| GM02704 | chr19:12876846-12877000 | HOOK2;HOOK2 | N_Shore | Body;Body |
| GM00940 | chr19:21264896-21265421 | ZNF714 | N_Shore | TSS200 |
| GM00042 | chr2:176964512-176964720 | HOXD12 | Island | TSS200 |
| GM00935 | chr5:323320-323969 | AHRR | S_Shore | Body |
| WG1282 | chr6:31237013-31237034 | HLA-C | N_Shore | Body |
| WG1282 | chr6:31237405-31237722 | HLA-C | N_Shore | Body |
| GM02640 | chr7:2847517-2847575 | GNA12 | | Body |
| GM02641 | chr1:119526060-119526437 | TBX15 | N_Shore | 5'UTR |
| GM01096 | chr1:119532320-119532542 | TBX15 | S_Shore | TSS200 |
| GM00031 | chr1:240776162-240776211 | GREM2 | | TSS1500 |
| GM02704 | chr10:114877616-114877689 | TCF7L2;TCF7L2;TCF7L2;TCF7L2;TCF7L2 | | Body;Body;Body;Body;Body;Body |
| GM00022 | chr10:3138418-3138534 | PFKP | | Body |
| GM00031 | chr10:72254314-72254335 | KIAA1274 | S_Shelf | 5'UTR |
| GM02456 | chr14:54418728-54418851 | BMP4;BMP4;BMP4 | Island | Body;Body;Body |
| GM02456 | chr15:88576296-88576320 | NTRK3;NTRK3;NTRK3 | | Body;Body;Body |
| GM01096 | chr17:21300735-21300791 | KCNJ12 | | 5'UTR |
| GM01376 | chr17:80358829-80358876 | C17orf101;C17orf101 | N_Shelf | Body;Body |
| GM03075 | chr19:12876846-12877000 | HOOK2;HOOK2 | N_Shore | Body;Body |
| GM00941 | chr19:21264896-21265421 | ZNF714 | N_Shore | TSS200 |

| | | | | |
|---|---|---|---|---|
| GM00289 | chr2:176964512-176964720 | HOXD12 | Island | TSS200 |
| GM00290 | chr3:192289245-192289293 | FGF12 | | Body |
| GM00016 | chr5:149980671-149980682 | SYNPO;SYNPO | | 1stExon;5'UTR |
| GM01096 | chr5:323320-323969 | AHRR | S_Shore | Body |
| GM00290 | chr6:161796785-161796854 | PARK2;PARK2;PARK2 | | Body;Body;Body |
| WG1309 | chr6:31237013-31237034 | HLA-C | N_Shore | Body |
| WG1513 | chr6:31237405-31237722 | HLA-C | N_Shore | Body |
| GM02641 | chr7:2847517-2847575 | GNA12 | | Body |
| GM02704 | chr1:119526060-119526437 | TBX15 | N_Shore | 5'UTR |
| GM01393 | chr1:119532320-119532542 | TBX15 | S_Shore | TSS200 |
| GM00042 | chr1:240776162-240776211 | GREM2 | | TSS1500 |
| GM02706 | chr10:114877616-114877689 | TCF7L2;TCF7L2;TCF7L2;TCF7L2;TCF7L2 | | Body;Body;Body;Body;Body;Body |
| GM00289 | chr10:3138418-3138534 | PFKP | | Body |
| GM00042 | chr10:72254314-72254335 | KIAA1274 | S_Shelf | 5'UTR |
| GM02555 | chr14:54418728-54418851 | BMP4;BMP4;BMP4 | Island | Body;Body;Body |
| GM02706 | chr15:88576296-88576320 | NTRK3;NTRK3;NTRK3 | | Body;Body;Body |
| GM01650 | chr17:21300735-21300791 | KCNJ12 | | 5'UTR |
| GM00042 | chr17:43319137-43319382 | FMNL1 | Island | Body |
| GM00016 | chr17:79905236-79905263 | MYADML2 | | TSS200 |
| GM02175 | chr17:80358829-80358876 | C17orf101;C17orf101 | N_Shelf | Body;Body |
| WG1282 | chr19:12876846-12877000 | HOOK2;HOOK2 | N_Shore | Body;Body |
| GM01375 | chr19:21264896-21265421 | ZNF714 | N_Shore | TSS200 |
| GM00290 | chr2:176964512-176964720 | HOXD12 | Island | TSS200 |
| GM00289 | chr2:240020028-240020174 | HDAC4 | | Body |
| GM00438 | chr3:192289245-192289293 | FGF12 | | Body |
| GM00042 | chr3:50488020-50488230 | CACNA2D2;CACNA2D2 | | Body;Body |
| GM00022 | chr5:149980671-149980682 | SYNPO;SYNPO | | 1stExon;5'UTR |
| GM00031 | chr5:177708304-177708528 | COL23A1 | | Body |
| GM01375 | chr5:323320-323969 | AHRR | S_Shore | Body |
| GM00438 | chr6:161796785-161796854 | PARK2;PARK2;PARK2 | | Body;Body;Body |
| WG1310 | chr6:31237013-31237034 | HLA-C | N_Shore | Body |
| WG1514 | chr6:31237405-31237722 | HLA-C | N_Shore | Body |
| GM00042 | chr7:114055123-114055419 | FOXP2;FOXP2;FOXP2;FOXP2;FOXP2;FOX | | 1stExon;5'UTR;5'UTR;1stExon;5'UTR;1 |
| GM03073 | chr7:2847517-2847575 | GNA12 | | Body |
| GM02706 | chr1:119526060-119526437 | TBX15 | N_Shore | 5'UTR |
| GM02555 | chr1:119532320-119532542 | TBX15 | S_Shore | TSS200 |
| GM00520 | chr1:240776162-240776211 | GREM2 | | TSS1500 |
| GM09256 | chr10:114877616-114877689 | TCF7L2;TCF7L2;TCF7L2;TCF7L2;TCF7L2 | | Body;Body;Body;Body;Body;Body |
| GM00439 | chr10:3138418-3138534 | PFKP | | Body |
| GM00289 | chr10:72254314-72254335 | KIAA1274 | S_Shelf | 5'UTR |
| GM00290 | chr11:68147880-68148079 | LRP5 | | Body |
| GM00940 | chr12:54423428-54425418 | HOXC4;HOXC6;HOXC5;HOX | Island | 5'UTR;Body;Body;Body |
| GM02706 | chr14:54418728-54418851 | BMP4;BMP4;BMP4 | Island | Body;Body;Body |
| GM03073 | chr15:88576296-88576320 | NTRK3;NTRK3;NTRK3 | | Body;Body;Body |
| GM01653 | chr17:21300735-21300791 | KCNJ12 | | 5'UTR |
| GM00290 | chr17:43319137-43319382 | FMNL1 | Island | Body |
| GM00290 | chr17:79905236-79905263 | MYADML2 | | TSS200 |

| | | | | |
|---|---|---|---|---|
| GM02456 | chr17:80358829-80358876 | C17orf101;C17orf101 | N_Shelf | Body;Body |
| WG1283 | chr19:12876846-12877000 | HOOK2;HOOK2 | N_Shore | Body;Body |
| GM01650 | chr19:21264896-21265421 | ZNF714 | N_Shore | TSS200 |
| GM00438 | chr2:176964512-176964720 | HOXD12 | Island | TSS200 |
| GM00022 | chr2:176993017-176993115 | HOXD8 | Island | TSS1500 |
| GM00439 | chr2:240020028-240020174 | HDAC4 | | Body |
| GM00289 | chr2:3452347-3452624 | TTC15 | Island | Body |
| GM00042 | chr3:183894727-183894778 | AP2M1;AP2M1 | S_Shore | 5'UTR;5'UTR |
| GM00439 | chr3:192289245-192289293 | FGF12 | | Body |
| GM01095 | chr3:50488020-50488230 | CACNA2D2;CACNA2D2 | | Body;Body |
| GM00289 | chr4:71587722-71587904 | RUFY3;RUFY3;RUFY3;RUFY3;RUFY3 | | 1stExon;Body;5'UTR;1stExon;5'UTR |
| GM00289 | chr5:149980671-149980682 | SYNPO;SYNPO | | 1stExon;5'UTR |
| GM00439 | chr5:177708304-177708528 | COL23A1 | | Body |
| GM02456 | chr5:323320-323969 | AHRR | S_Shore | Body |
| GM00520 | chr6:161796785-161796854 | PARK2;PARK2;PARK2 | | Body;Body;Body |
| GM00439 | chr6:29894050-29894197 | HCG4P6 | N_Shore | TSS1500 |
| WG1513 | chr6:31237013-31237034 | HLA-C | N_Shore | Body |
| WG1657 | chr6:31237405-31237722 | HLA-C | N_Shore | Body |
| GM00289 | chr7:114055123-114055419 | FOXP2;FOXP2;FOXP2;FOXP2;FOXP2;FO | | 1stExon;5'UTR;5'UTR;1stExon;5'UTR;1 |
| GM09255 | chr7:2847517-2847575 | GNA12 | | Body |
| GM00031 | chr7:70096029-70096378 | AUTS2;AUTS2 | N_Shore | Body;Body |
| GM03073 | chr1:119526060-119526437 | TBX15 | N_Shore | 5'UTR |
| GM02640 | chr1:119532320-119532542 | TBX15 | S_Shore | TSS200 |
| GM01095 | chr1:240776162-240776211 | GREM2 | | TSS1500 |
| WG1513 | chr10:114877616-114877689 | TCF7L2;TCF7L2;TCF7L2;TCF7L2;TCF7L2 | | Body;Body;Body;Body;Body;Body |
| GM00935 | chr10:3138418-3138534 | PFKP | | Body |
| GM00439 | chr10:72254314-72254335 | KIAA1274 | S_Shelf | 5'UTR |
| GM00042 | chr10:72311914-72312093 | KIAA1274 | | Body |
| GM00439 | chr11:125037226-125037348 | PKNOX2 | S_Shore | 5'UTR |
| GM00289 | chr11:1750302-1750381 | HCCA2 | | Body |
| GM00439 | chr11:68147880-68148079 | LRP5 | | Body |
| GM00439 | chr12:114833837-114834713 | TBX5;TBX5;TBX5;TBX5 | N_Shore | Body;Body;Body;Body |
| GM01095 | chr12:54423428-54425418 | HOXC4;HOXC6;HOXC5;HOX | Island | 5'UTR;Body;Body;Body |
| GM00031 | chr13:110993474-110993558 | COL4A2 | | Body |
| GM00022 | chr14:37136192-37136391 | PAX9 | Island | Body |
| GM03075 | chr14:54418728-54418851 | BMP4;BMP4;BMP4 | Island | Body;Body;Body |
| GM03075 | chr15:88576296-88576320 | NTRK3;NTRK3;NTRK3 | | Body;Body;Body |
| GM02173 | chr17:21300735-21300791 | KCNJ12 | | 5'UTR |
| GM00439 | chr17:43319137-43319382 | FMNL1 | Island | Body |
| GM01096 | chr17:46685327-46685448 | HOXB7 | Island | Body |
| GM01392 | chr17:79905236-79905263 | MYADML2 | | TSS200 |
| GM02555 | chr17:80358829-80358876 | C17orf101;C17orf101 | N_Shelf | Body;Body |
| WG1657 | chr19:12876846-12877000 | HOOK2;HOOK2 | N_Shore | Body;Body |
| GM01653 | chr19:21264896-21265421 | ZNF714 | N_Shore | TSS200 |
| GM00520 | chr2:176964512-176964720 | HOXD12 | Island | TSS200 |
| GM00042 | chr2:176993017-176993115 | HOXD8 | Island | TSS1500 |
| GM01096 | chr2:240020028-240020174 | HDAC4 | | Body |

| | | | | |
|---|---|---|---|---|
| GM01376 | chr2:3452347-3452624 | TTC15 | Island | Body |
| GM00042 | chr21:47604052-47604654 | C21orf56;C21orf56 | S_Shore | 5'UTR;5'UTR |
| GM00289 | chr3:183894727-183894778 | AP2M1;AP2M1 | S_Shore | 5'UTR;5'UTR |
| GM00935 | chr3:192289245-192289293 | FGF12 | | Body |
| GM01096 | chr3:50488020-50488230 | CACNA2D2;CACNA2D2 | | Body;Body |
| GM00438 | chr4:71587722-71587904 | RUFY3;RUFY3;RUFY3;RUFY3;RUFY3 | | 1stExon;Body;5'UTR;1stExon;5'UTR |
| GM00290 | chr5:149980671-149980682 | SYNPO;SYNPO | | 1stExon;5'UTR |
| GM00520 | chr5:177708304-177708528 | COL23A1 | | Body |
| GM02555 | chr5:323320-323969 | AHRR | S_Shore | Body |
| GM00935 | chr6:161796785-161796854 | PARK2;PARK2;PARK2 | | Body;Body;Body |
| GM00031 | chr6:291882-292823 | DUSP22 | N_Shore | TSS1500 |
| GM00520 | chr6:29894050-29894197 | HCG4P6 | N_Shore | TSS1500 |
| WG1657 | chr6:31237013-31237034 | HLA-C | N_Shore | Body |
| WG1658 | chr6:31237405-31237722 | HLA-C | N_Shore | Body |
| GM00439 | chr7:114055123-114055419 | FOXP2;FOXP2;FOXP2;FOXP2;FOXP2;FO) | | 1stExon;5'UTR;5'UTR;1stExon;5'UTR;1 |
| GM09256 | chr7:2847517-2847575 | GNA12 | | Body |
| GM00289 | chr7:70096029-70096378 | AUTS2;AUTS2 | N_Shore | Body;Body |
| GM03075 | chr1:119526060-119526437 | TBX15 | N_Shore | 5'UTR |
| GM02704 | chr1:119532320-119532542 | TBX15 | S_Shore | TSS200 |
| GM00439 | chr1:15256054-15256222 | KIAA1026;KIAA1026;KIAA1026;KIAA1026 | | Body;TSS1500;Body;Body |
| GM00042 | chr1:205818956-205819492 | PM20D1 | Island | Body |
| GM01376 | chr1:240776162-240776211 | GREM2 | | TSS1500 |
| GM00016 | chr1:54856177-54856511 | SSBP3;SSBP3;SSBP3 | N_Shore | Body;Body;Body |
| WG1657 | chr10:114877616-114877689 | TCF7L2;TCF7L2;TCF7L2;TCF7L2;TCF7L2 | | Body;Body;Body;Body;Body;Body |
| GM01392 | chr10:3138418-3138534 | PFKP | | Body |
| GM00520 | chr10:72254314-72254335 | KIAA1274 | S_Shelf | 5'UTR |
| GM00290 | chr10:72311914-72312093 | KIAA1274 | | Body |
| GM00520 | chr11:125037226-125037348 | PKNOX2 | S_Shore | 5'UTR |
| GM00042 | chr11:128812978-128813008 | TP53AIP1 | | TSS200 |
| GM00438 | chr11:1750302-1750381 | HCCA2 | | Body |
| GM00520 | chr11:68147880-68148079 | LRP5 | | Body |
| GM01095 | chr12:114833837-114834713 | TBX5;TBX5;TBX5;TBX5 | N_Shore | Body;Body;Body;Body |
| GM01096 | chr12:54423428-54425418 | HOXC4;HOXC6;HOXC5;HOX | Island | 5'UTR;Body;Body;Body |
| GM00016 | chr12:62583974-62584454 | FAM19A2 | N_Shore | 5'UTR |
| GM00042 | chr13:110993474-110993558 | COL4A2 | | Body |
| GM00439 | chr13:36050901-36051073 | MIR548F5;NBEA;MAB21L1 | N_Shore | Body;Body;TSS200 |
| GM00042 | chr14:37136192-37136391 | PAX9 | Island | Body |
| GM09255 | chr14:54418728-54418851 | BMP4;BMP4;BMP4 | Island | Body;Body;Body |
| GM09256 | chr15:88576296-88576320 | NTRK3;NTRK3;NTRK3 | | Body;Body;Body |
| GM02555 | chr17:21300735-21300791 | KCNJ12 | | 5'UTR |
| GM00520 | chr17:43319137-43319382 | FMNL1 | Island | Body |
| GM00031 | chr17:46669485-46669492 | LOC404266;HOXB5;LOC404 | Island | TSS200;3'UTR;Body;TSS200;Body |
| GM00289 | chr17:46669485-46669644 | LOC404266;HOXB5;LOC404 | Island | TSS200;3'UTR;Body;TSS200;Body |
| GM02147 | chr17:46685327-46685448 | HOXB7 | Island | Body |
| GM00042 | chr17:62777693-62777777 | LOC146880;LOC146880 | Island | Body;TSS200 |
| GM02456 | chr17:79905236-79905263 | MYADML2 | | TSS200 |
| GM03075 | chr17:80358829-80358876 | C17orf101;C17orf101 | N_Shelf | Body;Body |

| | | | | |
|---|---|---|---|---|
| GM00031 | chr19:11289349-11289376 | KANK2;KANK2 | S_Shore | Body;Body |
| WG1658 | chr19:12876846-12877000 | HOOK2;HOOK2 | N_Shore | Body;Body |
| GM02173 | chr19:21264896-21265421 | ZNF714 | N_Shore | TSS200 |
| GM00022 | chr2:121554810-121554926 | GLI2 | | TSS200 |
| GM00935 | chr2:176964512-176964720 | HOXD12 | Island | TSS200 |
| GM00290 | chr2:176993017-176993115 | HOXD8 | Island | TSS1500 |
| GM01393 | chr2:240020028-240020174 | HDAC4 | | Body |
| GM00289 | chr2:240291426-240291657 | HDAC4 | | 5'UTR |
| GM00042 | chr2:3286338-3286479 | TSSC1 | Island | Body |
| GM02149 | chr2:3452347-3452624 | TTC15 | Island | Body |
| GM00520 | chr21:47604052-47604654 | C21orf56;C21orf56 | S_Shore | 5'UTR;5'UTR |
| GM00042 | chr3:155394109-155394303 | PLCH1;PLCH1;PLCH1 | | 5'UTR;TSS200;TSS200 |
| GM00438 | chr3:183894727-183894778 | AP2M1;AP2M1 | S_Shore | 5'UTR;5'UTR |
| GM01095 | chr3:192289245-192289293 | FGF12 | | Body |
| GM02149 | chr3:50488020-50488230 | CACNA2D2;CACNA2D2 | | Body;Body |
| GM00439 | chr4:71587722-71587904 | RUFY3;RUFY3;RUFY3;RUFY3;RUFY3 | | 1stExon;Body;5'UTR;1stExon;5'UTR |
| GM00439 | chr5:149980486-149980682 | SYNPO | | TSS200 |
| GM01095 | chr5:149980671-149980682 | SYNPO;SYNPO | | 1stExon;5'UTR |
| GM00935 | chr5:177708304-177708528 | COL23A1 | | Body |
| GM02706 | chr5:323320-323969 | AHRR | S_Shore | Body |
| GM00940 | chr6:161796785-161796854 | PARK2;PARK2;PARK2 | | Body;Body;Body |
| GM00042 | chr6:291882-292823 | DUSP22 | N_Shore | TSS1500 |
| GM01375 | chr6:29894050-29894197 | HCG4P6 | N_Shore | TSS1500 |
| WG1721 | chr6:31237013-31237034 | HLA-C | N_Shore | Body |
| WG1721 | chr6:31237405-31237722 | HLA-C | N_Shore | Body |
| GM00520 | chr7:114055123-114055419 | FOXP2;FOXP2;FOXP2;FOXP2;FOXP2;FOX | | 1stExon;5'UTR;5'UTR;1stExon;5'UTR;1 |
| WG1282 | chr7:2847517-2847575 | GNA12 | | Body |
| GM00290 | chr7:70096029-70096378 | AUTS2;AUTS2 | N_Shore | Body;Body |
| GM00022 | chr9:140401382-140401576 | PNPLA7;PNPLA7 | Island | Body;Body |
| WG1513 | chr1:119526060-119526437 | TBX15 | N_Shore | 5'UTR |
| GM02706 | chr1:119532320-119532542 | TBX15 | S_Shore | TSS200 |
| GM00935 | chr1:15256054-15256222 | KIAA1026;KIAA1026;KIAA1026;KIAA1026 | | Body;TSS1500;Body;Body |
| GM00438 | chr1:170638807-170639289 | PRRX1;PRRX1 | | Body;Body |
| GM00289 | chr1:205818956-205819492 | PM20D1 | Island | Body |
| GM01650 | chr1:240776162-240776211 | GREM2 | | TSS1500 |
| GM00439 | chr1:3459949-3460268 | MEGF6 | Island | Body |
| GM00289 | chr1:44016014-44016049 | PTPRF;PTPRF | | Body;Body |
| GM00289 | chr1:54856177-54856511 | SSBP3;SSBP3;SSBP3 | N_Shore | Body;Body;Body |
| WG1658 | chr10:114877616-114877689 | TCF7L2;TCF7L2;TCF7L2;TCF7L2;TCF7L2 | | Body;Body;Body;Body;Body;Body |
| GM02175 | chr10:3138418-3138534 | PFKP | | Body |
| GM00935 | chr10:72254314-72254335 | KIAA1274 | S_Shelf | 5'UTR |
| GM00439 | chr10:72311914-72312093 | KIAA1274 | | Body |
| GM00289 | chr10:90712684-90712948 | ACTA2;ACTA2 | | 5'UTR;5'UTR |
| GM00290 | chr11:12031093-12031099 | DKK3;DKK3;DKK3 | S_Shore | TSS1500;TSS200;TSS1500 |
| GM00935 | chr11:125037226-125037348 | PKNOX2 | S_Shore | 5'UTR |
| GM00289 | chr11:128812978-128813008 | TP53AIP1 | | TSS200 |
| GM00439 | chr11:1750302-1750381 | HCCA2 | | Body |

| | | | | |
|---|---|---|---|---|
| GM00042 | chr11:628603-628677 | SCT | S_Shore | TSS1500 |
| GM00935 | chr11:68147880-68148079 | LRP5 | | Body |
| GM00022 | chr11:77299805-77299962 | AQP11 | N_Shore | TSS1500 |
| GM01650 | chr12:114833837-114834713 | TBX5;TBX5;TBX5;TBX5 | N_Shore | Body;Body;Body;Body |
| GM02640 | chr12:54423428-54425418 | HOXC4;HOXC6;HOXC5;HOX | Island | 5'UTR;Body;Body;Body |
| GM00042 | chr12:56329615-56329903 | DGKA;DGKA;DGKA;DGKA | Island | 5'UTR;5'UTR;5'UTR;5'UTR |
| GM00022 | chr12:62583974-62584454 | FAM19A2 | N_Shore | 5'UTR |
| GM00438 | chr12:97300410-97300429 | NEDD1;NEDD1;NEDD1;NEDI | N_Shore | TSS1500;TSS1500;TSS1500;TSS1500 |
| GM01096 | chr13:110993474-110993558 | COL4A2 | | Body |
| GM00290 | chr13:114308619-114308749 | ATP4B | | Body |
| GM00935 | chr13:36050901-36051073 | MIR548F5;NBEA;MAB21L1 | N_Shore | Body;Body;TSS200 |
| GM00290 | chr14:37136192-37136391 | PAX9 | Island | Body |
| WG1309 | chr14:54418728-54418851 | BMP4;BMP4;BMP4 | Island | Body;Body;Body |
| WG1309 | chr15:88576296-88576320 | NTRK3;NTRK3;NTRK3 | | Body;Body;Body |
| GM02704 | chr17:21300735-21300791 | KCNJ12 | | 5'UTR |
| GM00935 | chr17:43319137-43319382 | FMNL1 | Island | Body |
| GM00031 | chr17:46648014-46648533 | HOXB3 | | 5'UTR |
| GM00290 | chr17:46669485-46669492 | LOC404266;HOXB5;LOC404 | Island | TSS200;3'UTR;Body;TSS200;Body |
| GM00438 | chr17:46669485-46669644 | LOC404266;HOXB5;LOC404 | Island | TSS200;3'UTR;Body;TSS200;Body |
| GM02640 | chr17:46685327-46685448 | HOXB7 | Island | Body |
| GM00520 | chr17:62777693-62777777 | LOC146880;LOC146880 | Island | Body;TSS200 |
| GM02640 | chr17:79905236-79905263 | MYADML2 | | TSS200 |
| GM09255 | chr17:80358829-80358876 | C17orf101;C17orf101 | N_Shelf | Body;Body |
| GM00520 | chr19:11289349-11289376 | KANK2;KANK2 | S_Shore | Body;Body |
| WG1721 | chr19:12876846-12877000 | HOOK2;HOOK2 | N_Shore | Body;Body |
| GM02316 | chr19:21264896-21265421 | ZNF714 | N_Shore | TSS200 |
| GM00935 | chr2:121554810-121554926 | GLI2 | | TSS200 |
| GM01392 | chr2:176964512-176964720 | HOXD12 | Island | TSS200 |
| GM01096 | chr2:176993017-176993115 | HOXD8 | Island | TSS1500 |
| GM01653 | chr2:240020028-240020174 | HDAC4 | | Body |
| GM00290 | chr2:240291426-240291657 | HDAC4 | | 5'UTR |
| GM00042 | chr2:30669597-30669863 | LCLAT1;LCLAT1 | N_Shore | TSS1500;TSS1500 |
| GM00935 | chr2:3286338-3286479 | TSSC1 | Island | Body |
| GM02640 | chr2:3452347-3452624 | TTC15 | Island | Body |
| GM00935 | chr21:47604052-47604654 | C21orf56;C21orf56 | S_Shore | 5'UTR;5'UTR |
| GM00289 | chr3:155394109-155394303 | PLCH1;PLCH1;PLCH1 | | 5'UTR;TSS200;TSS200 |
| GM00439 | chr3:183894727-183894778 | AP2M1;AP2M1 | S_Shore | 5'UTR;5'UTR |
| GM02149 | chr3:192289245-192289293 | FGF12 | | Body |
| GM00042 | chr3:36949957-36950004 | TRANK1 | | Body |
| GM02175 | chr3:50488020-50488230 | CACNA2D2;CACNA2D2 | | Body;Body |
| GM00520 | chr4:71587722-71587904 | RUFY3;RUFY3;RUFY3;RUFY3;RUFY3 | | 1stExon;Body;5'UTR;1stExon;5'UTR |
| GM00520 | chr5:149980486-149980682 | SYNPO | | TSS200 |
| GM01375 | chr5:149980671-149980682 | SYNPO;SYNPO | | 1stExon;5'UTR |
| GM02173 | chr5:177708304-177708528 | COL23A1 | | Body |
| WG1310 | chr5:323320-323969 | AHRR | S_Shore | Body |
| GM00438 | chr6:100911240-100912072 | SIM1 | N_Shore | 1stExon |
| GM02706 | chr6:161796785-161796854 | PARK2;PARK2;PARK2 | | Body;Body;Body |

| | | | | |
|---|---|---|---|---|
| GM00940 | chr6:291882-292823 | DUSP22 | N_Shore | TSS1500 |
| GM01653 | chr6:29894050-29894197 | HCG4P6 | N_Shore | TSS1500 |
| GM00031 | chr6:31025933-31025967 | HCG22 | | Body |
| WG1722 | chr6:31237013-31237034 | HLA-C | N_Shore | Body |
| WG1978 | chr6:31237405-31237722 | HLA-C | N_Shore | Body |
| GM00031 | chr6:32552152-32552246 | HLA-DRB1 | Island | Body |
| GM00022 | chr6:6002865-6003303 | NRN1 | Island | Body |
| GM00935 | chr7:114055123-114055419 | FOXP2;FOXP2;FOXP2;FOXP2;FOXP2;FOX | | 1stExon;5'UTR;5'UTR;1stExon;5'UTR;1 |
| GM00016 | chr7:158336016-158336049 | PTPRN2;PTPRN2;PTPRN2 | | Body;Body;Body |
| GM00520 | chr7:1952518-1952807 | MAD1L1;MAD1L1;MAD1L1 | S_Shelf | Body;Body;Body |
| WG1513 | chr7:2847517-2847575 | GNA12 | | Body |
| GM00438 | chr7:70096029-70096378 | AUTS2;AUTS2 | N_Shore | Body;Body |
| GM00289 | chr8:59058585-59058660 | FAM110B | Island | 5'UTR |
| GM00042 | chr9:140401382-140401576 | PNPLA7;PNPLA7 | Island | Body;Body |
| WG1977 | chr1:119526060-119526437 | TBX15 | N_Shore | 5'UTR |
| GM03075 | chr1:119532320-119532542 | TBX15 | S_Shore | TSS200 |
| GM01096 | chr1:15256054-15256222 | KIAA1026;KIAA1026;KIAA1026;KIAA1026 | | Body;TSS1500;Body;Body |
| GM01376 | chr1:170638807-170639289 | PRRX1;PRRX1 | | Body;Body |
| GM00289 | chr1:173176362-173176523 | TNFSF4;TNFSF4 | | 5'UTR;1stExon |
| GM00022 | chr1:205818956-205819463 | PM20D1 | Island | Body |
| GM00438 | chr1:205818956-205819492 | PM20D1 | Island | Body |
| GM02706 | chr1:240776162-240776211 | GREM2 | | TSS1500 |
| GM00520 | chr1:3459949-3460268 | MEGF6 | Island | Body |
| GM00290 | chr1:44016014-44016049 | PTPRF;PTPRF | | Body;Body |
| GM00520 | chr1:54856177-54856511 | SSBP3;SSBP3;SSBP3 | N_Shore | Body;Body;Body |
| WG1977 | chr10:114877616-114877689 | TCF7L2;TCF7L2;TCF7L2;TCF7L2;TCF7L2 | | Body;Body;Body;Body;Body;Body |
| GM02456 | chr10:3138418-3138534 | PFKP | | Body |
| GM01376 | chr10:72254314-72254335 | KIAA1274 | S_Shelf | 5'UTR |
| GM00935 | chr10:72311914-72312093 | KIAA1274 | | Body |
| GM00439 | chr10:90712684-90712948 | ACTA2;ACTA2 | | 5'UTR;5'UTR |
| GM00439 | chr11:12031093-12031099 | DKK3;DKK3;DKK3 | S_Shore | TSS1500;TSS200;TSS1500 |
| GM01096 | chr11:125037226-125037348 | PKNOX2 | S_Shore | 5'UTR |
| GM00935 | chr11:128812978-128813008 | TP53AIP1 | | TSS200 |
| GM00520 | chr11:1750302-1750381 | HCCA2 | | Body |
| GM00438 | chr11:628603-628677 | SCT | S_Shore | TSS1500 |
| GM01375 | chr11:68147880-68148079 | LRP5 | | Body |
| GM00042 | chr11:77299805-77299962 | AQP11 | N_Shore | TSS1500 |
| GM02640 | chr12:114833837-114834713 | TBX5;TBX5;TBX5;TBX5 | N_Shore | Body;Body;Body;Body |
| GM00290 | chr12:114837572-114837887 | TBX5;TBX5;TBX5;TBX5 | N_Shore | Body;Body;Body;Body |
| GM02641 | chr12:54423428-54425418 | HOXC4;HOXC6;HOXC5;HOX | Island | 5'UTR;Body;Body;Body |
| GM00439 | chr12:56329615-56329903 | DGKA;DGKA;DGKA;DGKA | Island | 5'UTR;5'UTR;5'UTR;5'UTR |
| GM00438 | chr12:62583974-62584454 | FAM19A2 | N_Shore | 5'UTR |
| GM00439 | chr12:97300410-97300429 | NEDD1;NEDD1;NEDD1;NEDD | N_Shore | TSS1500;TSS1500;TSS1500;TSS1500 |
| GM01393 | chr13:110993474-110993558 | COL4A2 | | Body |
| GM00439 | chr13:114308619-114308749 | ATP4B | | Body |
| GM00940 | chr13:36050901-36051073 | MIR548F5;NBEA;MAB21L1 | N_Shore | Body;Body;TSS200 |
| GM00439 | chr14:37136192-37136391 | PAX9 | Island | Body |

| | | | | |
|---|---|---|---|---|
| WG1513 | chr14:54418728-54418851 | BMP4;BMP4;BMP4 | Island | Body;Body;Body |
| GM00016 | chr15:70994435-70994672 | UACA;UACA;UACA | | 5'UTR;Body;1stExon |
| WG1514 | chr15:88576296-88576320 | NTRK3;NTRK3;NTRK3 | | Body;Body;Body |
| GM02706 | chr17:21300735-21300791 | KCNJ12 | | 5'UTR |
| GM00042 | chr17:30822561-30823006 | MYO1D | N_Shore | Body |
| GM01376 | chr17:43319137-43319382 | FMNL1 | Island | Body |
| GM00042 | chr17:46645972-46646359 | HOXB3 | S_Shelf | 5'UTR |
| GM00042 | chr17:46648014-46648533 | HOXB3 | | 5'UTR |
| GM00520 | chr17:46651945-46652179 | HOXB3 | N_Shelf | TSS200 |
| GM00439 | chr17:46669485-46669492 | LOC404266;HOXB5;LOC404 | Island | TSS200;3'UTR;Body;TSS200;Body |
| GM00520 | chr17:46669485-46669644 | LOC404266;HOXB5;LOC404 | Island | TSS200;3'UTR;Body;TSS200;Body |
| GM02641 | chr17:46685327-46685448 | HOXB7 | Island | Body |
| GM00935 | chr17:62777693-62777777 | LOC146880;LOC146880 | Island | Body;TSS200 |
| GM00289 | chr17:77037916-77037965 | C1QTNF1;C1QTNF1;C1QTNF1 | | 5'UTR;5'UTR;5'UTR |
| GM02704 | chr17:79905236-79905263 | MYADML2 | | TSS200 |
| GM09256 | chr17:80358829-80358876 | C17orf101;C17orf101 | N_Shelf | Body;Body |
| GM00935 | chr19:11289349-11289376 | KANK2;KANK2 | S_Shore | Body;Body |
| WG1722 | chr19:12876846-12877000 | HOOK2;HOOK2 | N_Shore | Body;Body |
| GM00042 | chr19:1854549-1854724 | KLF16 | Island | Body |
| GM02555 | chr19:21264896-21265421 | ZNF714 | N_Shore | TSS200 |
| GM00289 | chr19:54876728-54876752 | LAIR1;LAIR1 | | TSS200;TSS200 |
| GM00439 | chr2:10567628-10567843 | HPCAL1;HPCAL1 | S_Shore | 3'UTR;3'UTR |
| GM01096 | chr2:121554810-121554926 | GLI2 | | TSS200 |
| GM01393 | chr2:176964512-176964720 | HOXD12 | Island | TSS200 |
| GM00438 | chr2:176982108-176982342 | HOXD10 | Island | 1stExon |
| GM01376 | chr2:176993017-176993115 | HOXD8 | Island | TSS1500 |
| GM00042 | chr2:220406470-220406756 | CHPF | Island | Body |
| GM00022 | chr2:223161771-223162049 | PAX3;PAX3;PAX3;CCDC140;I | Island | Body;Body;Body;TSS1500;Body;Body; |
| GM02317 | chr2:240020028-240020174 | HDAC4 | | Body |
| GM00520 | chr2:240291426-240291657 | HDAC4 | | 5'UTR |
| WG1310 | chr2:30669597-30669863 | LCLAT1;LCLAT1 | N_Shore | TSS1500;TSS1500 |
| GM01096 | chr2:3286338-3286479 | TSSC1 | Island | Body |
| GM02641 | chr2:3452347-3452624 | TTC15 | Island | Body |
| GM00022 | chr2:66671478-66671727 | MEIS1 | N_Shore | Body |
| GM00439 | chr20:5485144-5485284 | LOC149837 | | Body |
| GM02149 | chr21:47604052-47604654 | C21orf56;C21orf56 | S_Shore | 5'UTR;5'UTR |
| GM00022 | chr3:15311021-15311269 | SH3BP5;SH3BP5 | | Body;Body |
| GM00438 | chr3:155394109-155394303 | PLCH1;PLCH1;PLCH1 | | 5'UTR;TSS200;TSS200 |
| GM00520 | chr3:183894727-183894778 | AP2M1;AP2M1 | S_Shore | 5'UTR;5'UTR |
| GM03073 | chr3:192289245-192289293 | FGF12 | | Body |
| GM00935 | chr3:36949957-36950004 | TRANK1 | | Body |
| GM09256 | chr3:50488020-50488230 | CACNA2D2;CACNA2D2 | | Body;Body |
| GM00022 | chr3:64547310-64547373 | ADAMTS9 | | Body |
| GM00042 | chr4:52918457-52918489 | SPATA18 | S_Shore | Body |
| GM02147 | chr4:55143269-55143557 | PDGFRA | | Body |
| GM01096 | chr4:71587722-71587904 | RUFY3;RUFY3;RUFY3;RUFY3;RUFY3 | | 1stExon;Body;5'UTR;1stExon;5'UTR |
| GM00935 | chr5:149980486-149980682 | SYNPO | | TSS200 |

| | | | | |
|---|---|---|---|---|
| GM02704 | chr5:149980671-149980682 | SYNPO;SYNPO | | 1stExon;5'UTR |
| GM02456 | chr5:177708304-177708528 | COL23A1 | | Body |
| WG1514 | chr5:323320-323969 | AHRR | S_Shore | Body |
| GM00520 | chr6:100911240-100912072 | SIM1 | N_Shore | 1stExon |
| GM00439 | chr6:109274429-109274496 | ARMC2 | | Body |
| GM00022 | chr6:152127812-152128024 | ESR1;ESR1;ESR1;ESR1 | N_Shore | TSS1500;5'UTR;TSS1500;5'UTR |
| GM09255 | chr6:161796785-161796854 | PARK2;PARK2;PARK2 | | Body;Body;Body |
| GM01650 | chr6:291882-292823 | DUSP22 | N_Shore | TSS1500 |
| WG1309 | chr6:29894050-29894197 | HCG4P6 | N_Shore | TSS1500 |
| GM00042 | chr6:31025933-31025967 | HCG22 | | Body |
| WG1977 | chr6:31237013-31237034 | HLA-C | N_Shore | Body |
| WG1983 | chr6:31237405-31237722 | HLA-C | N_Shore | Body |
| GM00016 | chr6:32489750-32489963 | HLA-DRB5 | Island | Body |
| GM00439 | chr6:32552152-32552246 | HLA-DRB1 | Island | Body |
| GM00439 | chr6:6002865-6003303 | NRN1 | Island | Body |
| GM01096 | chr7:114055123-114055419 | FOXP2;FOXP2;FOXP2;FOXP2;FOXP2;FOX | | 1stExon;5'UTR;5'UTR;1stExon;5'UTR;1 |
| GM00289 | chr7:158336016-158336049 | PTPRN2;PTPRN2;PTPRN2 | | Body;Body;Body |
| GM00935 | chr7:1952518-1952807 | MAD1L1;MAD1L1;MAD1L1 | S_Shelf | Body;Body;Body |
| WG1658 | chr7:2847517-2847575 | GNA12 | | Body |
| GM00935 | chr7:70096029-70096378 | AUTS2;AUTS2 | N_Shore | Body;Body |
| GM01096 | chr8:59058585-59058660 | FAM110B | Island | 5'UTR |
| GM00439 | chr9:140401382-140401576 | PNPLA7;PNPLA7 | Island | Body;Body |
| WG1978 | chr1:119526060-119526437 | TBX15 | N_Shore | 5'UTR |
| GM00940 | chr1:119531718-119532044 | TBX15 | N_Shore | 5'UTR |
| WG1310 | chr1:119532320-119532542 | TBX15 | S_Shore | TSS200 |
| GM02706 | chr1:15256054-15256222 | KIAA1026;KIAA1026;KIAA1026;KIAA1026 | | Body;TSS1500;Body;Body |
| GM02317 | chr1:170638807-170639289 | PRRX1;PRRX1 | | Body;Body |
| GM00290 | chr1:173176362-173176523 | TNFSF4;TNFSF4 | | 5'UTR;1stExon |
| GM00520 | chr1:205818956-205819463 | PM20D1 | Island | Body |
| GM01392 | chr1:205818956-205819492 | PM20D1 | Island | Body |
| GM01096 | chr1:2120985-2121724 | C1orf86;LOC100128003 | N_Shore | Body;Body |
| GM00031 | chr1:221053904-221055097 | HLX | N_Shore | Body |
| GM03073 | chr1:240776162-240776211 | GREM2 | | TSS1500 |
| GM02456 | chr1:3459949-3460268 | MEGF6 | Island | Body |
| GM01095 | chr1:36039655-36039885 | TFAP2E | Island | Body |
| GM00289 | chr1:40105667-40105706 | HEYL | Island | TSS1500 |
| GM00438 | chr1:44016014-44016049 | PTPRF;PTPRF | | Body;Body |
| GM00935 | chr1:54856177-54856511 | SSBP3;SSBP3;SSBP3 | N_Shore | Body;Body;Body |
| GM00042 | chr1:89664260-89664582 | GBP4 | | Body |
| WG1978 | chr10:114877616-114877689 | TCF7L2;TCF7L2;TCF7L2;TCF7L2;TCF7L2 | | Body;Body;Body;Body;Body;Body |
| GM02555 | chr10:3138418-3138534 | PFKP | | Body |
| GM01392 | chr10:72254314-72254335 | KIAA1274 | S_Shelf | 5'UTR |
| GM01375 | chr10:72311914-72312093 | KIAA1274 | | Body |
| GM00520 | chr10:90712684-90712948 | ACTA2;ACTA2 | | 5'UTR;5'UTR |
| GM00438 | chr10:97175272-97175479 | SORBS1;SORBS1;SORBS1;SORBS1;SORI | | Body;Body;Body;Body;Body;Body;Bod |
| GM00935 | chr11:12031093-12031099 | DKK3;DKK3;DKK3 | S_Shore | TSS1500;TSS200;TSS1500 |
| GM01392 | chr11:125037226-125037348 | PKNOX2 | S_Shore | 5'UTR |

| | | | | |
|---|---|---|---|---|
| GM01376 | chr11:128812978-128813008 | TP53AIP1 | | TSS200 |
| GM01096 | chr11:1750302-1750381 | HCCA2 | | Body |
| GM00940 | chr11:18433500-18433564 | LDHC;LDHC | | TSS1500;TSS1500 |
| GM00520 | chr11:628603-628677 | SCT | S_Shore | TSS1500 |
| GM01392 | chr11:68147880-68148079 | LRP5 | | Body |
| GM00289 | chr11:77299805-77299962 | AQP11 | N_Shore | TSS1500 |
| GM00290 | chr12:10084936-10085174 | CLEC2A;CLEC2A | | 5'UTR;1stExon |
| GM02641 | chr12:114833837-114834713 | TBX5;TBX5;TBX5;TBX5 | N_Shore | Body;Body;Body;Body |
| GM00439 | chr12:114837572-114837887 | TBX5;TBX5;TBX5;TBX5 | N_Shore | Body;Body;Body;Body |
| GM00022 | chr12:122356316-122356598 | WDR66 | Island | TSS200 |
| GM00289 | chr12:15114233-15114393 | ARHGDIB | | 5'UTR |
| GM01095 | chr12:54392734-54392884 | HOXC9 | N_Shore | TSS1500 |
| GM02704 | chr12:54423428-54425418 | HOXC4;HOXC6;HOXC5;HOX | Island | 5'UTR;Body;Body;Body |
| GM00022 | chr12:54448729-54448913 | HOXC4;HOXC4 | S_Shore | Body;Body |
| GM01392 | chr12:56329615-56329903 | DGKA;DGKA;DGKA;DGKA | Island | 5'UTR;5'UTR;5'UTR;5'UTR |
| GM00439 | chr12:62583974-62584454 | FAM19A2 | N_Shore | 5'UTR |
| GM02147 | chr12:6442873-6443045 | TNFRSF1A | S_Shelf | Body |
| GM00289 | chr12:772861-772982 | NINJ2 | | TSS200 |
| GM00520 | chr12:97300410-97300429 | NEDD1;NEDD1;NEDD1;NEDI | N_Shore | TSS1500;TSS1500;TSS1500;TSS1500 |
| GM02175 | chr13:110993474-110993558 | COL4A2 | | Body |
| GM02706 | chr13:114108433-114108652 | ADPRHL1 | | TSS1500 |
| GM00520 | chr13:114308619-114308749 | ATP4B | | Body |
| GM01375 | chr13:36050901-36051073 | MIR548F5;NBEA;MAB21L1 | N_Shore | Body;Body;TSS200 |
| GM00520 | chr14:37136192-37136391 | PAX9 | Island | Body |
| WG1721 | chr14:54418728-54418851 | BMP4;BMP4;BMP4 | Island | Body;Body;Body |
| GM00290 | chr14:99728984-99729088 | BCL11B;BCL11B | | Body;Body |
| GM00439 | chr15:55881599-55881706 | PYGO1 | S_Shore | TSS1500 |
| GM00031 | chr15:70994435-70994672 | UACA;UACA;UACA | | 5'UTR;Body;1stExon |
| WG1658 | chr15:88576296-88576320 | NTRK3;NTRK3;NTRK3 | | Body;Body;Body |
| GM00289 | chr16:69958626-69958964 | WWP2;WWP2 | | TSS1500;Body |
| WG1282 | chr17:21300735-21300791 | KCNJ12 | | 5'UTR |
| GM00290 | chr17:30822561-30823006 | MYO1D | N_Shore | Body |
| GM01392 | chr17:43319137-43319382 | FMNL1 | Island | Body |
| GM00290 | chr17:44847220-44847427 | WNT3 | Island | Body |
| GM00439 | chr17:46645972-46646359 | HOXB3 | S_Shelf | 5'UTR |
| GM00289 | chr17:46648014-46648533 | HOXB3 | | 5'UTR |
| GM01392 | chr17:46651945-46652179 | HOXB3 | N_Shelf | TSS200 |
| GM02149 | chr17:46669485-46669492 | LOC404266;HOXB5;LOC404 | Island | TSS200;3'UTR;Body;TSS200;Body |
| GM00935 | chr17:46669485-46669644 | LOC404266;HOXB5;LOC404 | Island | TSS200;3'UTR;Body;TSS200;Body |
| GM02706 | chr17:46685327-46685448 | HOXB7 | Island | Body |
| GM01375 | chr17:62777693-62777777 | LOC146880;LOC146880 | Island | Body;TSS200 |
| GM00935 | chr17:75283832-75283979 | SEPT9;SEPT9 | | TSS200;Body |
| GM01096 | chr17:77037916-77037965 | C1QTNF1;C1QTNF1;C1QTNF1 | | 5'UTR;5'UTR;5'UTR |
| WG1310 | chr17:79905236-79905263 | MYADML2 | | TSS200 |
| WG1514 | chr17:80358829-80358876 | C17orf101;C17orf101 | N_Shelf | Body;Body |
| GM01392 | chr19:11289349-11289376 | KANK2;KANK2 | S_Shore | Body;Body |
| WG1977 | chr19:12876846-12877000 | HOOK2;HOOK2 | N_Shore | Body;Body |

| | | | | |
|---|---|---|---|---|
| GM00520 | chr19:1854549-1854724 | KLF16 | Island | Body |
| GM02640 | chr19:21264896-21265421 | ZNF714 | N_Shore | TSS200 |
| GM00290 | chr19:21265264-21265421 | ZNF714 | Island | 5'UTR |
| GM00042 | chr19:427022-427263 | SHC2 | Island | Body |
| GM00290 | chr19:54876728-54876752 | LAIR1;LAIR1 | | TSS200;TSS200 |
| GM00520 | chr2:10567628-10567843 | HPCAL1;HPCAL1 | S_Shore | 3'UTR;3'UTR |
| GM02555 | chr2:121554810-121554926 | GLI2 | | TSS200 |
| GM00016 | chr2:121554817-121554926 | GLI2 | | TSS200 |
| GM00042 | chr2:142889283-142889392 | LRP1B | S_Shore | TSS200 |
| GM02147 | chr2:176964512-176964720 | HOXD12 | Island | TSS200 |
| GM00520 | chr2:176982108-176982342 | HOXD10 | Island | 1stExon |
| GM01393 | chr2:176993017-176993115 | HOXD8 | Island | TSS1500 |
| GM00439 | chr2:220406470-220406756 | CHPF | Island | Body |
| GM00042 | chr2:223161771-223162049 | PAX3;PAX3;PAX3;CCDC140;P | Island | Body;Body;Body;TSS1500;Body;Body; |
| GM03075 | chr2:240020028-240020174 | HDAC4 | | Body |
| GM01096 | chr2:240291426-240291657 | HDAC4 | | 5'UTR |
| WG1721 | chr2:30669597-30669863 | LCLAT1;LCLAT1 | N_Shore | TSS1500;TSS1500 |
| GM02316 | chr2:3286338-3286479 | TSSC1 | Island | Body |
| GM02704 | chr2:3452347-3452624 | TTC15 | Island | Body |
| GM00016 | chr2:54087173-54087189 | LOC100302652;GPR75 | Island | TSS200;TSS200 |
| GM00439 | chr2:66671478-66671727 | MEIS1 | N_Shore | Body |
| GM00439 | chr2:74780268-74780617 | LOXL3 | N_Shore | 5'UTR |
| GM00520 | chr20:5485144-5485284 | LOC149837 | | Body |
| GM02317 | chr21:47604052-47604654 | C21orf56;C21orf56 | S_Shore | 5'UTR;5'UTR |
| GM00042 | chr22:45609402-45609421 | C22orf9;C22orf9 | | Body;TSS1500 |
| GM00290 | chr3:15311021-15311269 | SH3BP5;SH3BP5 | | Body;Body |
| GM02173 | chr3:155394109-155394303 | PLCH1;PLCH1;PLCH1 | | 5'UTR;TSS200;TSS200 |
| GM00935 | chr3:183894727-183894778 | AP2M1;AP2M1 | S_Shore | 5'UTR;5'UTR |
| WG1977 | chr3:192289245-192289293 | FGF12 | | Body |
| GM01376 | chr3:196065357-196065506 | TM4SF19 | | TSS200 |
| GM01376 | chr3:36949957-36950004 | TRANK1 | | Body |
| WG1283 | chr3:50488020-50488230 | CACNA2D2;CACNA2D2 | | Body;Body |
| GM00520 | chr3:64547310-64547373 | ADAMTS9 | | Body |
| GM00289 | chr4:111398042-111398146 | ENPEP | Island | 1stExon |
| GM00016 | chr4:40858984-40859293 | APBB2;APBB2;APBB2;APBB2 | Island | Body;5'UTR;5'UTR;5'UTR;Body;5'UTR;B |
| GM00289 | chr4:52918457-52918489 | SPATA18 | S_Shore | Body |
| GM02555 | chr4:55143269-55143557 | PDGFRA | | Body |
| GM01650 | chr4:71587722-71587904 | RUFY3;RUFY3;RUFY3;RUFY3;RUFY3 | | 1stExon;Body;5'UTR;1stExon;5'UTR |
| GM01096 | chr5:149980486-149980682 | SYNPO | | TSS200 |
| WG1283 | chr5:149980671-149980682 | SYNPO;SYNPO | | 1stExon;5'UTR |
| GM02555 | chr5:177708304-177708528 | COL23A1 | | Body |
| WG1658 | chr5:323320-323969 | AHRR | S_Shore | Body |
| GM02175 | chr5:76010444-76010658 | F2R | N_Shore | TSS1500 |
| GM00935 | chr6:100911240-100912072 | SIM1 | N_Shore | 1stExon |
| GM01392 | chr6:101846791-101846872 | GRIK2;GRIK2;GRIK2 | Island | TSS200;TSS200;TSS200 |
| GM00935 | chr6:109274429-109274496 | ARMC2 | | Body |
| GM00022 | chr6:149803087-149803444 | ZC3H12D | | 5'UTR |

| | | | | |
|---|---|---|---|---|
| GM00031 | chr6:152127812-152128024 | ESR1;ESR1;ESR1;ESR1 | N_Shore | TSS1500;5'UTR;TSS1500;5'UTR |
| WG1282 | chr6:161796785-161796854 | PARK2;PARK2;PARK2 | | Body;Body;Body |
| GM02149 | chr6:291882-292823 | DUSP22 | N_Shore | TSS1500 |
| GM00438 | chr6:29690766-29690893 | HLA-F;HLA-F;HLA-F | N_Shore | TSS1500;TSS1500;TSS1500 |
| WG1657 | chr6:29894050-29894197 | HCG4P6 | N_Shore | TSS1500 |
| GM00016 | chr6:30039432-30039442 | RNF39;RNF39 | Island | Body;Body |
| GM00520 | chr6:31025933-31025967 | HCG22 | | Body |
| WG2120 | chr6:31237013-31237034 | HLA-C | N_Shore | Body |
| WG1984 | chr6:31237405-31237722 | HLA-C | N_Shore | Body |
| GM00940 | chr6:32489750-32489963 | HLA-DRB5 | Island | Body |
| GM00031 | chr6:32489991-32490043 | HLA-DRB5 | Island | Body |
| GM01375 | chr6:32552152-32552246 | HLA-DRB1 | Island | Body |
| GM00520 | chr6:6002865-6003303 | NRN1 | Island | Body |
| GM00022 | chr7:101500014-101500514 | CUX1;CUX1;CUX1 | | Body;Body;Body |
| GM01376 | chr7:114055123-114055419 | FOXP2;FOXP2;FOXP2;FOXP2;FOXP2;FOX | | 1stExon;5'UTR;5'UTR;1stExon;5'UTR;1 |
| GM00290 | chr7:158336016-158336049 | PTPRN2;PTPRN2;PTPRN2 | | Body;Body;Body |
| GM00940 | chr7:1952518-1952807 | MAD1L1;MAD1L1;MAD1L1 | S_Shelf | Body;Body;Body |
| WG1983 | chr7:2847517-2847575 | GNA12 | | Body |
| GM02175 | chr7:70096029-70096378 | AUTS2;AUTS2 | N_Shore | Body;Body |
| GM00289 | chr8:10586959-10587055 | SOX7 | N_Shore | Body |
| GM01095 | chr8:2075469-2075820 | MYOM2 | Island | Body |
| GM02149 | chr8:59058585-59058660 | FAM110B | Island | 5'UTR |
| GM00940 | chr9:140401382-140401576 | PNPLA7;PNPLA7 | Island | Body;Body |
| GM00438 | chr9:92220617-92221282 | GADD45G | Island | Body |
| WG2120 | chr1:119526060-119526437 | TBX15 | N_Shore | 5'UTR |
| GM01650 | chr1:119531718-119532044 | TBX15 | N_Shore | 5'UTR |
| WG1513 | chr1:119532320-119532542 | TBX15 | S_Shore | TSS200 |
| GM09256 | chr1:15256054-15256222 | KIAA1026;KIAA1026;KIAA1026;KIAA1026 | | Body;TSS1500;Body;Body |
| GM02706 | chr1:170638807-170639289 | PRRX1;PRRX1 | | Body;Body |
| GM00438 | chr1:173176362-173176523 | TNFSF4;TNFSF4 | | 5'UTR;1stExon |
| GM00289 | chr1:2005260-2005564 | PRKCZ;PRKCZ;PRKCZ | Island | 1stExon;Body;5'UTR |
| GM00935 | chr1:205818956-205819463 | PM20D1 | Island | Body |
| GM09256 | chr1:205818956-205819492 | PM20D1 | Island | Body |
| GM01393 | chr1:2120985-2121724 | C1orf86;LOC100128003 | N_Shore | Body;Body |
| GM00289 | chr1:221053904-221055097 | HLX | N_Shore | Body |
| GM02706 | chr1:221054615-221055097 | HLX | Island | Body |
| GM00289 | chr1:228291668-228291705 | C1orf35 | S_Shore | TSS1500 |
| WG1513 | chr1:240776162-240776211 | GREM2 | | TSS1500 |
| GM00289 | chr1:247694271-247694406 | LOC148824;OR2C3 | Island | TSS200;3'UTR |
| GM00940 | chr1:25944712-25945042 | MAN1C1 | Island | 1stExon |
| GM03075 | chr1:3459949-3460268 | MEGF6 | Island | Body |
| GM02555 | chr1:36039655-36039885 | TFAP2E | Island | Body |
| GM02456 | chr1:40105667-40105706 | HEYL | Island | TSS1500 |
| GM01392 | chr1:44016014-44016049 | PTPRF;PTPRF | | Body;Body |
| GM00941 | chr1:54199825-54200420 | GLIS1;GLIS1 | N_Shelf | 5'UTR;1stExon |
| GM01095 | chr1:54856177-54856511 | SSBP3;SSBP3;SSBP3 | N_Shore | Body;Body;Body |
| GM00439 | chr1:89664260-89664582 | GBP4 | | Body |

| | | | | |
|---|---|---|---|---|
| WG2120 | chr10:114877616-114877689 | TCF7L2;TCF7L2;TCF7L2;TCF7L2;TCF7L2 | | Body;Body;Body;Body;Body;Body |
| GM02173 | chr10:135341528-135342218 | CYP2E1 | N_Shore | 1stExon |
| GM01650 | chr10:13932842-13933247 | FRMD4A | N_Shore | Body |
| GM02706 | chr10:3138418-3138534 | PFKP | | Body |
| GM02456 | chr10:72254314-72254335 | KIAA1274 | S_Shelf | 5'UTR |
| GM02456 | chr10:72311914-72312093 | KIAA1274 | | Body |
| GM00935 | chr10:90712684-90712948 | ACTA2;ACTA2 | | 5'UTR;5'UTR |
| GM01096 | chr10:97175272-97175479 | SORBS1;SORBS1;SORBS1;SORBS1;SORI | | Body;Body;Body;Body;Body;Body;Bod |
| GM01375 | chr11:12031093-12031099 | DKK3;DKK3;DKK3 | S_Shore | TSS1500;TSS200;TSS1500 |
| GM02456 | chr11:125037226-125037348 | PKNOX2 | S_Shore | 5'UTR |
| GM01653 | chr11:128812978-128813008 | TP53AIP1 | | TSS200 |
| GM01392 | chr11:1750302-1750381 | HCCA2 | | Body |
| GM02175 | chr11:18433500-18433564 | LDHC;LDHC | | TSS1500;TSS1500 |
| GM00022 | chr11:31819444-31819678 | PAX6;PAX6;PAX6 | N_Shore | Body;Body;Body |
| GM00016 | chr11:31825756-31825833 | PAX6;PAX6;PAX6 | Island | Body;Body;Body |
| GM00940 | chr11:628603-628677 | SCT | S_Shore | TSS1500 |
| GM02173 | chr11:68147880-68148079 | LRP5 | | Body |
| GM00022 | chr11:68782049-68782202 | MRGPRF;MRGPRF | S_Shelf | TSS1500;TSS1500 |
| GM00520 | chr11:77299805-77299962 | AQP11 | N_Shore | TSS1500 |
| GM00042 | chr11:86383696-86383723 | ME3;ME3;ME3 | S_Shore | TSS1500;TSS200;TSS1500 |
| GM00520 | chr12:10084936-10085174 | CLEC2A;CLEC2A | | 5'UTR;1stExon |
| GM02706 | chr12:114833837-114834713 | TBX5;TBX5;TBX5;TBX5 | N_Shore | Body;Body;Body;Body |
| GM00520 | chr12:114837572-114837887 | TBX5;TBX5;TBX5;TBX5 | N_Shore | Body;Body;Body;Body |
| GM00042 | chr12:122356316-122356598 | WDR66 | Island | TSS200 |
| GM00031 | chr12:131303194-131303518 | STX2;STX2 | Island | Body;Body |
| GM00439 | chr12:15114233-15114393 | ARHGDIB | | 5'UTR |
| GM01096 | chr12:1609245-1609369 | LOC100292680 | | TSS1500 |
| GM00438 | chr12:312592-313104 | SLC6A12;SLC6A12;SLC6A12 | Island | Body;Body;Body |
| GM02175 | chr12:54392734-54392884 | HOXC9 | N_Shore | TSS1500 |
| GM02706 | chr12:54423428-54425418 | HOXC4;HOXC6;HOXC5;HOX | Island | 5'UTR;Body;Body;Body |
| GM00439 | chr12:54448729-54448913 | HOXC4;HOXC4 | S_Shore | Body;Body |
| GM01393 | chr12:56329615-56329903 | DGKA;DGKA;DGKA;DGKA | Island | 5'UTR;5'UTR;5'UTR;5'UTR |
| GM00940 | chr12:62583974-62584454 | FAM19A2 | N_Shore | 5'UTR |
| WG1309 | chr12:6442873-6443045 | TNFRSF1A | S_Shelf | Body |
| GM00439 | chr12:772861-772982 | NINJ2 | | TSS200 |
| GM01653 | chr12:97300410-97300429 | NEDD1;NEDD1;NEDD1;NEDI | N_Shore | TSS1500;TSS1500;TSS1500;TSS1500 |
| WG2120 | chr13:110993474-110993558 | COL4A2 | | Body |
| GM00438 | chr13:113436309-113436651 | ATP11A;ATP11A | N_Shelf | Body;Body |
| GM09256 | chr13:114108433-114108652 | ADPRHL1 | | TSS1500 |
| GM00935 | chr13:114308619-114308749 | ATP4B | | Body |
| GM02640 | chr13:36048892-36051073 | MAB21L1;MIR548F5;NBEA;M | N_Shore | 1stExon;Body;Body;3'UTR |
| GM02173 | chr13:36050901-36051073 | MIR548F5;NBEA;MAB21L1 | N_Shore | Body;Body;TSS200 |
| GM00042 | chr14:103593503-103593520 | TNFAIP2 | Island | Body |
| GM00935 | chr14:37136192-37136391 | PAX9 | Island | Body |
| WG1722 | chr14:54418728-54418851 | BMP4;BMP4;BMP4 | Island | Body;Body;Body |
| GM00290 | chr14:80677608-80677688 | DIO2;DIO2;DIO2 | | 1stExon;1stExon;Body |
| GM01376 | chr14:99728984-99729088 | BCL11B;BCL11B | | Body;Body |

| | | | | |
|---|---|---|---|---|
| GM01095 | chr15:32964155-32964500 | SCG5;SCG5 | | Body;Body |
| GM00520 | chr15:40544174-40544427 | PAK6;C15orf56;PAK6;PAK6 | N_Shore | 5'UTR;3'UTR;5'UTR;5'UTR |
| GM00042 | chr15:45314789-45314933 | SORD | N_Shore | TSS1500 |
| GM00520 | chr15:55881599-55881706 | PYGO1 | S_Shore | TSS1500 |
| GM00439 | chr15:70354647-70354825 | TLE3;TLE3;TLE3 | | Body;Body;Body |
| GM00439 | chr15:70994435-70994672 | UACA;UACA;UACA | | 5'UTR;Body;1stExon |
| WG1977 | chr15:88576296-88576320 | NTRK3;NTRK3;NTRK3 | | Body;Body;Body |
| GM01392 | chr16:2908554-2908934 | PRSS22 | S_Shore | TSS1500 |
| GM00940 | chr16:69958626-69958964 | WWP2;WWP2 | | TSS1500;Body |
| GM00439 | chr17:1028673-1028931 | ABR;ABR | Island | Body;5'UTR |
| WG1513 | chr17:21300735-21300791 | KCNJ12 | | 5'UTR |
| GM01096 | chr17:30822561-30823006 | MYO1D | N_Shore | Body |
| GM02317 | chr17:43319137-43319382 | FMNL1 | Island | Body |
| GM00940 | chr17:44847220-44847427 | WNT3 | Island | Body |
| GM02173 | chr17:46645972-46646359 | HOXB3 | S_Shelf | 5'UTR |
| GM00438 | chr17:46648014-46648533 | HOXB3 | | 5'UTR |
| GM02149 | chr17:46651945-46652179 | HOXB3 | N_Shelf | TSS200 |
| GM02173 | chr17:46669485-46669492 | LOC404266;HOXB5;LOC404 | Island | TSS200;3'UTR;Body;TSS200;Body |
| GM01392 | chr17:46669485-46669644 | LOC404266;HOXB5;LOC404 | Island | TSS200;3'UTR;Body;TSS200;Body |
| WG1282 | chr17:46685327-46685448 | HOXB7 | Island | Body |
| GM01392 | chr17:62777693-62777777 | LOC146880;LOC146880 | Island | Body;TSS200 |
| WG1722 | chr17:75283832-75283979 | SEPT9;SEPT9 | | TSS200;Body |
| GM02175 | chr17:77037916-77037965 | C1QTNF1;C1QTNF1;C1QTNF1 | | 5'UTR;5'UTR;5'UTR |
| GM00290 | chr17:77952290-77952488 | TBC1D16 | Island | Body |
| GM00042 | chr17:79109729-79109808 | AATK | Island | Body |
| WG1513 | chr17:79905236-79905263 | MYADML2 | | TSS200 |
| WG1977 | chr17:80358829-80358876 | C17orf101;C17orf101 | N_Shelf | Body;Body |
| GM01393 | chr19:11289349-11289376 | KANK2;KANK2 | S_Shore | Body;Body |
| WG1978 | chr19:12876846-12877000 | HOOK2;HOOK2 | N_Shore | Body;Body |
| GM00016 | chr19:12876846-12877188 | HOOK2;HOOK2 | N_Shore | Body;Body |
| GM00935 | chr19:1854549-1854724 | KLF16 | Island | Body |
| GM00031 | chr19:19639568-19639596 | YJEFN3 | Island | TSS200 |
| GM02706 | chr19:21264896-21265421 | ZNF714 | N_Shore | TSS200 |
| GM01095 | chr19:21265264-21265421 | ZNF714 | Island | 5'UTR |
| GM00438 | chr19:2282439-2282729 | C19orf35 | S_Shelf | TSS1500 |
| GM00289 | chr19:427022-427263 | SHC2 | Island | Body |
| GM02706 | chr19:48947560-48947578 | GRIN2D;GRWD1 | Island | 3'UTR;TSS1500 |
| GM00290 | chr19:49891270-49891574 | CCDC155 | Island | TSS1500 |
| GM00520 | chr19:54876728-54876752 | LAIR1;LAIR1 | | TSS200;TSS200 |
| GM01096 | chr2:10567628-10567843 | HPCAL1;HPCAL1 | S_Shore | 3'UTR;3'UTR |
| GM01393 | chr2:113992843-113993052 | PAX8;PAX8;PAX8;PAX8;PAX8; | N_Shore | Body;Body;Body;Body;Body;Body;TSS |
| GM02706 | chr2:121554810-121554926 | GLI2 | | TSS200 |
| GM00042 | chr2:121554817-121554926 | GLI2 | | TSS200 |
| GM01650 | chr2:142889283-142889392 | LRP1B | S_Shore | TSS200 |
| GM03073 | chr2:176964512-176964720 | HOXD12 | Island | TSS200 |
| GM00042 | chr2:176973275-176974060 | HOXD11 | N_Shore | Body |
| GM00940 | chr2:176982108-176982342 | HOXD10 | Island | 1stExon |

| | | | | |
|---|---|---|---|---|
| GM01653 | chr2:176993017-176993115 | HOXD8 | Island | TSS1500 |
| GM00438 | chr2:177016167-177017449 | HOXD4;HOXD4 | N_Shore | 5'UTR;1stExon |
| GM00289 | chr2:200320455-200321348 | SATB2 | N_Shore | Body |
| GM00520 | chr2:220406470-220406756 | CHPF | Island | Body |
| GM00439 | chr2:223161771-223162049 | PAX3;PAX3;PAX3;CCDC140;I | Island | Body;Body;Body;TSS1500;Body;Body; |
| GM00438 | chr2:223162666-223162800 | PAX3;PAX3;PAX3;PAX3;PAX3; | N_Shore | Body;Body;Body;Body;Body;Body;Bod |
| WG1513 | chr2:240020028-240020174 | HDAC4 | | Body |
| GM00520 | chr2:240233043-240233240 | HDAC4 | | Body |
| GM01393 | chr2:240291426-240291657 | HDAC4 | | 5'UTR |
| WG1978 | chr2:30669597-30669863 | LCLAT1;LCLAT1 | N_Shore | TSS1500;TSS1500 |
| WG1513 | chr2:3286338-3286479 | TSSC1 | Island | Body |
| GM03075 | chr2:3452347-3452624 | TTC15 | Island | Body |
| GM02149 | chr2:3714936-3715083 | ALLC | | 5'UTR |
| GM02175 | chr2:54087173-54087189 | LOC100302652;GPR75 | Island | TSS200;TSS200 |
| GM00940 | chr2:66671478-66671727 | MEIS1 | N_Shore | Body |
| GM00042 | chr2:71134524-71134630 | VAX2 | N_Shore | Body |
| GM00520 | chr2:74780268-74780617 | LOXL3 | N_Shore | 5'UTR |
| GM01096 | chr20:25128770-25128805 | LOC284798;LOC284798;LO( | Island | TSS200;Body;Body |
| GM00935 | chr20:5485144-5485284 | LOC149837 | | Body |
| GM00520 | chr20:55925570-55925603 | RAE1;RAE1 | N_Shore | TSS1500;TSS1500 |
| GM09256 | chr21:47604052-47604654 | C21orf56;C21orf56 | S_Shore | 5'UTR;5'UTR |
| GM00042 | chr22:22058080-22058271 | YPEL1 | Island | Body |
| GM00289 | chr22:32027580-32027605 | PISD | S_Shore | TSS1500 |
| GM00438 | chr22:45609402-45609421 | C22orf9;C22orf9 | | Body;TSS1500 |
| GM00438 | chr3:15311021-15311269 | SH3BP5;SH3BP5 | | Body;Body |
| GM02175 | chr3:155394109-155394303 | PLCH1;PLCH1;PLCH1 | | 5'UTR;TSS200;TSS200 |
| GM00940 | chr3:183894727-183894778 | AP2M1;AP2M1 | S_Shore | 5'UTR;5'UTR |
| WG2275 | chr3:192289245-192289293 | FGF12 | | Body |
| GM02175 | chr3:196065357-196065506 | TM4SF19 | | TSS200 |
| GM02706 | chr3:36949957-36950004 | TRANK1 | | Body |
| WG1513 | chr3:50488020-50488230 | CACNA2D2;CACNA2D2 | | Body;Body |
| GM00935 | chr3:64547310-64547373 | ADAMTS9 | | Body |
| GM00935 | chr4:111398042-111398146 | ENPEP | Island | 1stExon |
| GM00439 | chr4:119948360-119948542 | SYNPO2;SYNPO2;SYNPO2 | | Body;Body;Body |
| GM00042 | chr4:174447847-174448341 | HAND2 | N_Shore | 3'UTR |
| GM00042 | chr4:40858984-40859293 | APBB2;APBB2;APBB2;APBB2 | Island | Body;5'UTR;5'UTR;5'UTR;Body;5'UTR;I |
| GM00438 | chr4:52918457-52918489 | SPATA18 | S_Shore | Body |
| GM03075 | chr4:55143269-55143557 | PDGFRA | | Body |
| GM02175 | chr4:71587722-71587904 | RUFY3;RUFY3;RUFY3;RUFY3;RUFY3 | | 1stExon;Body;5'UTR;1stExon;5'UTR |
| WG1309 | chr5:14406585-14406732 | TRIO | | Body |
| GM01392 | chr5:149980486-149980682 | SYNPO | | TSS200 |
| WG1309 | chr5:149980671-149980682 | SYNPO;SYNPO | | 1stExon;5'UTR |
| WG1310 | chr5:177708304-177708528 | COL23A1 | | Body |
| WG1721 | chr5:323320-323969 | AHRR | S_Shore | Body |
| GM00031 | chr5:423894-424371 | AHRR | S_Shelf | Body |
| GM09256 | chr5:76010444-76010658 | F2R | N_Shore | TSS1500 |
| GM02175 | chr6:100911240-100912072 | SIM1 | N_Shore | 1stExon |

| | | | | |
|---|---|---|---|---|
| GM02641 | chr6:101846791-101846872 | GRIK2;GRIK2;GRIK2 | Island | TSS200;TSS200;TSS200 |
| GM01095 | chr6:109274429-109274496 | ARMC2 | | Body |
| GM00289 | chr6:11279656-11279727 | NEDD9 | | Body |
| GM00289 | chr6:149803087-149803444 | ZC3H12D | | 5'UTR |
| GM00290 | chr6:152127812-152128024 | ESR1;ESR1;ESR1;ESR1 | N_Shore | TSS1500;5'UTR;TSS1500;5'UTR |
| WG1721 | chr6:161796785-161796854 | PARK2;PARK2;PARK2 | | Body;Body;Body |
| GM00022 | chr6:161860203-161860359 | PARK2;PARK2;PARK2 | | Body;Body;Body |
| GM02456 | chr6:291882-292823 | DUSP22 | N_Shore | TSS1500 |
| GM00940 | chr6:29690766-29690893 | HLA-F;HLA-F;HLA-F | N_Shore | TSS1500;TSS1500;TSS1500 |
| WG1721 | chr6:29894050-29894197 | HCG4P6 | N_Shore | TSS1500 |
| GM01393 | chr6:30039432-30039442 | RNF39;RNF39 | Island | Body;Body |
| GM00935 | chr6:31025933-31025967 | HCG22 | | Body |
| WG2121 | chr6:31237013-31237034 | HLA-C | N_Shore | Body |
| WG2120 | chr6:31237405-31237722 | HLA-C | N_Shore | Body |
| GM02317 | chr6:32489750-32489963 | HLA-DRB5 | Island | Body |
| GM00042 | chr6:32489991-32490043 | HLA-DRB5 | Island | Body |
| GM00031 | chr6:32551891-32551954 | HLA-DRB1 | Island | Body |
| GM01653 | chr6:32552152-32552246 | HLA-DRB1 | Island | Body |
| GM01653 | chr6:6002865-6003303 | NRN1 | Island | Body |
| GM01096 | chr7:101500014-101500514 | CUX1;CUX1;CUX1 | | Body;Body;Body |
| GM01392 | chr7:114055123-114055419 | FOXP2;FOXP2;FOXP2;FOXP2;FOXP2;FOX | | 1stExon;5'UTR;5'UTR;1stExon;5'UTR;1 |
| GM00439 | chr7:158336016-158336049 | PTPRN2;PTPRN2;PTPRN2 | | Body;Body;Body |
| GM01375 | chr7:1952518-1952807 | MAD1L1;MAD1L1;MAD1L1 | S_Shelf | Body;Body;Body |
| GM00439 | chr7:2133441-2133480 | MAD1L1;MAD1L1;MAD1L1 | | Body;Body;Body |
| GM00935 | chr7:2563611-2563909 | LFNG;LFNG;LFNG;LFNG | Island | Body;Body;Body;Body |
| GM00022 | chr7:27182493-27183196 | HOXA5 | N_Shore | Body |
| WG2120 | chr7:2847517-2847575 | GNA12 | | Body |
| GM00289 | chr7:4065931-4066127 | SDK1 | | Body |
| GM00940 | chr7:4885521-4885643 | RADIL | | Body |
| GM00290 | chr7:5336543-5336811 | SLC29A4;SLC29A4 | Island | Body;Body |
| WG1309 | chr7:70096029-70096378 | AUTS2;AUTS2 | N_Shore | Body;Body |
| GM01096 | chr7:944607-944776 | ADAP1 | S_Shore | Body |
| GM00042 | chr8:104032819-104032876 | ATP6V1C1 | N_Shore | TSS1500 |
| GM00940 | chr8:10586959-10587055 | SOX7 | N_Shore | Body |
| GM00940 | chr8:116660311-116661063 | TRPS1 | N_Shore | 5'UTR |
| GM01653 | chr8:2075469-2075820 | MYOM2 | Island | Body |
| GM00439 | chr8:28560467-28560723 | EXTL3 | S_Shore | 5'UTR |
| GM02555 | chr8:59058585-59058660 | FAM110B | Island | 5'UTR |
| GM01392 | chr9:140401382-140401576 | PNPLA7;PNPLA7 | Island | Body;Body |
| GM00935 | chr9:92220617-92221282 | GADD45G | Island | Body |
| WG2121 | chr1:119526060-119526437 | TBX15 | N_Shore | 5'UTR |
| GM02175 | chr1:119531718-119532044 | TBX15 | N_Shore | 5'UTR |
| WG1978 | chr1:119532320-119532542 | TBX15 | S_Shore | TSS200 |
| WG1513 | chr1:15256054-15256222 | KIAA1026;KIAA1026;KIAA1026;KIAA1026 | | Body;TSS1500;Body;Body |
| GM00290 | chr1:156786601-156786653 | SH2D2A;NTRK1;SH2D2A;SH2 | S_Shelf | 5'UTR;Body;1stExon;TSS1500;5'UTR;5 |
| GM00520 | chr1:159796617-159796658 | SLAMF8;SLAMF8 | | 5'UTR;1stExon |
| GM09256 | chr1:170638807-170639289 | PRRX1;PRRX1 | | Body;Body |

| | | | | |
|---|---|---|---|---|
| GM00439 | chr1:173176362-173176523 | TNFSF4;TNFSF4 | | 5'UTR;1stExon |
| GM00290 | chr1:2005260-2005564 | PRKCZ;PRKCZ;PRKCZ | Island | 1stExon;Body;5'UTR |
| GM02555 | chr1:205818956-205819463 | PM20D1 | Island | Body |
| WG1309 | chr1:205818956-205819492 | PM20D1 | Island | Body |
| GM02149 | chr1:2120985-2121724 | C1orf86;LOC100128003 | N_Shore | Body;Body |
| GM00935 | chr1:21586831-21587174 | ECE1;ECE1;ECE1;ECE1 | | Body;Body;Body;Body |
| GM02641 | chr1:221053904-221054273 | HLX | N_Shore | Body |
| GM00438 | chr1:221053904-221055097 | HLX | N_Shore | Body |
| GM03075 | chr1:221054615-221055097 | HLX | Island | Body |
| GM00042 | chr1:226825435-226825606 | ITPKB | | Body |
| GM00520 | chr1:228291668-228291705 | C1orf35 | S_Shore | TSS1500 |
| WG1514 | chr1:240776162-240776211 | GREM2 | | TSS1500 |
| GM00438 | chr1:247694271-247694406 | LOC148824;OR2C3 | Island | TSS200;3'UTR |
| GM01096 | chr1:25944712-25945042 | MAN1C1 | Island | 1stExon |
| GM09256 | chr1:3459949-3460268 | MEGF6 | Island | Body |
| GM02704 | chr1:36039655-36039885 | TFAP2E | Island | Body |
| GM02706 | chr1:40105667-40105706 | HEYL | Island | TSS1500 |
| GM00022 | chr1:41250062-41250117 | KCNQ4;KCNQ4 | Island | 1stExon;1stExon |
| GM02175 | chr1:44016014-44016049 | PTPRF;PTPRF | | Body;Body |
| GM00031 | chr1:54199825-54200115 | GLIS1;GLIS1 | N_Shelf | 5'UTR;1stExon |
| GM02456 | chr1:54199825-54200420 | GLIS1;GLIS1 | N_Shelf | 5'UTR;1stExon |
| GM01375 | chr1:54856177-54856511 | SSBP3;SSBP3;SSBP3 | N_Shore | Body;Body;Body |
| GM00935 | chr1:6085995-6086148 | KCNAB2;KCNAB2 | N_Shore | TSS1500;TSS1500 |
| GM00520 | chr1:63249199-63249213 | ATG4C;ATG4C | | TSS1500;TSS1500 |
| GM00289 | chr1:6558085-6558113 | PLEKHG5;PLEKHG5 | S_Shore | Body;TSS1500 |
| GM01393 | chr1:89664260-89664582 | GBP4 | | Body |
| WG2121 | chr10:114877616-114877689 | TCF7L2;TCF7L2;TCF7L2;TCF7L2;TCF7L2 | | Body;Body;Body;Body;Body;Body |
| GM02704 | chr10:135341528-135342218 | CYP2E1 | N_Shore | 1stExon |
| GM02641 | chr10:13932842-13933247 | FRMD4A | N_Shore | Body |
| GM00042 | chr10:22605982-22606072 | COMMD3 | S_Shore | Body |
| WG1657 | chr10:3138418-3138534 | PFKP | | Body |
| GM00940 | chr10:45495739-45495981 | C10orf25;ZNF22 | N_Shore | 3'UTR;TSS1500 |
| GM00042 | chr10:50506748-50506896 | C10orf71;C10orf71 | | TSS1500;TSS1500 |
| WG1513 | chr10:72254314-72254335 | KIAA1274 | S_Shelf | 5'UTR |
| GM02555 | chr10:72311914-72312093 | KIAA1274 | | Body |
| GM00290 | chr10:82167757-82167774 | C10orf58 | N_Shelf | |
| GM01096 | chr10:90712684-90712948 | ACTA2;ACTA2 | | 5'UTR;5'UTR |
| GM02555 | chr10:97175272-97175479 | SORBS1;SORBS1;SORBS1;SORBS1;SORI | | Body;Body;Body;Body;Body;Body;Bod |
| GM00290 | chr10:99338056-99338240 | ANKRD2;ANKRD2 | N_Shore | Body;Body |
| GM01393 | chr11:12031093-12031099 | DKK3;DKK3;DKK3 | S_Shore | TSS1500;TSS200;TSS1500 |
| GM02555 | chr11:125037226-125037348 | PKNOX2 | S_Shore | 5'UTR |
| GM00016 | chr11:128729883-128730011 | KCNJ1;KCNJ1;KCNJ1;KCNJ1 | | 5'UTR;5'UTR;5'UTR;5'UTR |
| GM02555 | chr11:128812978-128813008 | TP53AIP1 | | TSS200 |
| GM00438 | chr11:129991445-129992368 | APLP2;APLP2;APLP2;APLP2;A | N_Shelf | Body;Body;Body;Body;Body;Body |
| GM02175 | chr11:1750302-1750381 | HCCA2 | | Body |
| GM02555 | chr11:18433500-18433564 | LDHC;LDHC | | TSS1500;TSS1500 |
| GM00439 | chr11:31819444-31819678 | PAX6;PAX6;PAX6 | N_Shore | Body;Body;Body |

| | | | | |
|---|---|---|---|---|
| GM02149 | chr11:31821590-31822302 | PAX6;PAX6;PAX6 | S_Shore | Body;Body;Body |
| GM00289 | chr11:31825756-31825833 | PAX6;PAX6;PAX6 | Island | Body;Body;Body |
| GM01096 | chr11:628603-628677 | SCT | S_Shore | TSS1500 |
| GM00941 | chr11:66885240-66885281 | KDM2A;KDM2A | N_Shore | TSS1500;TSS1500 |
| GM02456 | chr11:68147880-68148079 | LRP5 | | Body |
| GM00935 | chr11:68610706-68611016 | CPT1A;CPT1A | N_Shore | TSS1500;TSS1500 |
| GM01650 | chr11:68782049-68782202 | MRGPRF;MRGPRF | S_Shelf | TSS1500;TSS1500 |
| GM00940 | chr11:77299805-77299962 | AQP11 | N_Shore | TSS1500 |
| GM00520 | chr11:85430629-85430657 | SYTL2;SYTL2;SYTL2;SYTL2;SYTL2;SYTL2; | | Body;TSS1500;TSS1500;Body;TSS150 |
| GM02456 | chr11:86383696-86383723 | ME3;ME3;ME3 | S_Shore | TSS1500;TSS200;TSS1500 |
| GM00290 | chr11:87908785-87908938 | RAB38 | S_Shore | TSS200 |
| GM00935 | chr12:10084936-10085174 | CLEC2A;CLEC2A | | 5'UTR;1stExon |
| GM01375 | chr12:106988340-106988413 | RFX4 | | Body |
| GM01096 | chr12:114833837-114834201 | TBX5;TBX5;TBX5;TBX5 | N_Shore | Body;Body;Body;Body |
| WG1310 | chr12:114833837-114834713 | TBX5;TBX5;TBX5;TBX5 | N_Shore | Body;Body;Body;Body |
| GM00935 | chr12:114837572-114837887 | TBX5;TBX5;TBX5;TBX5 | N_Shore | Body;Body;Body;Body |
| GM00941 | chr12:122356056-122356598 | WDR66 | N_Shore | TSS1500 |
| GM02555 | chr12:122356316-122356598 | WDR66 | Island | TSS200 |
| GM00940 | chr12:131303194-131303518 | STX2;STX2 | Island | Body;Body |
| GM02175 | chr12:15114233-15114393 | ARHGDIB | | 5'UTR |
| GM09256 | chr12:15114703-15114732 | ARHGDIB | | TSS200 |
| GM03075 | chr12:1609245-1609369 | LOC100292680 | | TSS1500 |
| GM01095 | chr12:25150530-25150626 | C12orf77 | | TSS200 |
| GM00520 | chr12:312592-313104 | SLC6A12;SLC6A12;SLC6A12 | Island | Body;Body;Body |
| GM02706 | chr12:54392734-54392884 | HOXC9 | N_Shore | TSS1500 |
| GM03073 | chr12:54423428-54425418 | HOXC4;HOXC6;HOXC5;HOX | Island | 5'UTR;Body;Body;Body |
| GM02641 | chr12:54440773-54441509 | HOXC4 | Island | 5'UTR |
| GM00941 | chr12:54440806-54441213 | HOXC4 | Island | 5'UTR |
| GM02456 | chr12:54448729-54448913 | HOXC4;HOXC4 | S_Shore | Body;Body |
| GM02706 | chr12:56329615-56329903 | DGKA;DGKA;DGKA;DGKA | Island | 5'UTR;5'UTR;5'UTR;5'UTR |
| GM02706 | chr12:62583974-62584454 | FAM19A2 | N_Shore | 5'UTR |
| WG1310 | chr12:6442873-6443045 | TNFRSF1A | S_Shelf | Body |
| GM00940 | chr12:7281566-7281783 | CLSTN3;RBP5 | | TSS1500;TSS200 |
| GM00520 | chr12:772861-772982 | NINJ2 | | TSS200 |
| GM02706 | chr12:97300410-97300429 | NEDD1;NEDD1;NEDD1;NEDI | N_Shore | TSS1500;TSS1500;TSS1500;TSS1500 |
| WG2193 | chr13:110993474-110993558 | COL4A2 | | Body |
| GM00520 | chr13:113436309-113436651 | ATP11A;ATP11A | N_Shelf | Body;Body |
| WG1283 | chr13:114108433-114108652 | ADPRHL1 | | TSS1500 |
| GM02456 | chr13:114308619-114308749 | ATP4B | | Body |
| GM00042 | chr13:36045201-36045352 | NBEA | Island | Body |
| GM02704 | chr13:36048892-36051073 | MAB21L1;MIR548F5;NBEA;M | N_Shore | 1stExon;Body;Body;3'UTR |
| GM02555 | chr13:36050901-36051073 | MIR548F5;NBEA;MAB21L1 | N_Shore | Body;Body;TSS200 |
| GM00940 | chr14:103593088-103593520 | TNFAIP2 | N_Shore | Body |
| GM01376 | chr14:103593503-103593520 | TNFAIP2 | Island | Body |
| GM02706 | chr14:37136192-37136391 | PAX9 | Island | Body |
| WG1977 | chr14:54418728-54418851 | BMP4;BMP4;BMP4 | Island | Body;Body;Body |
| GM01096 | chr14:80677608-80677688 | DIO2;DIO2;DIO2 | | 1stExon;1stExon;Body |

| | | | | |
|---|---|---|---|---|
| GM02173 | chr14:99728984-99729088 | BCL11B;BCL11B | | Body;Body |
| GM01376 | chr15:101419275-101419296 | ALDH1A3 | Island | TSS1500 |
| GM00520 | chr15:101728611-101728653 | CHSY1 | | Body |
| GM02706 | chr15:32964155-32964500 | SCG5;SCG5 | | Body;Body |
| GM00016 | chr15:40268610-40269214 | EIF2AK4 | Island | Body |
| GM00935 | chr15:40544174-40544427 | PAK6;C15orf56;PAK6;PAK6 | N_Shore | 5'UTR;3'UTR;5'UTR;5'UTR |
| GM00031 | chr15:44086617-44087089 | SERF2;MIR1282 | S_Shore | 3'UTR;TSS1500 |
| GM00520 | chr15:45314789-45314933 | SORD | N_Shore | TSS1500 |
| GM00042 | chr15:45671155-45671246 | LOC145663;GATM | S_Shore | Body;TSS200 |
| GM00439 | chr15:49716247-49716645 | C15orf33;FGF7 | | Body;5'UTR |
| GM02555 | chr15:51369862-51370019 | TNFAIP8L3 | | Body |
| GM00935 | chr15:55881599-55881706 | PYGO1 | S_Shore | TSS1500 |
| GM00520 | chr15:70354647-70354825 | TLE3;TLE3;TLE3 | | Body;Body;Body |
| GM00940 | chr15:70994435-70994672 | UACA;UACA;UACA | | 5'UTR;Body;1stExon |
| WG1978 | chr15:88576296-88576320 | NTRK3;NTRK3;NTRK3 | | Body;Body;Body |
| GM01393 | chr16:2908554-2908934 | PRSS22 | S_Shore | TSS1500 |
| GM00016 | chr16:4103161-4103225 | ADCY9 | | Body |
| GM01096 | chr16:50347766-50347895 | ADCY7 | | Body |
| GM01376 | chr16:69958626-69958964 | WWP2;WWP2 | | TSS1500;Body |
| GM00935 | chr16:88706241-88706426 | IL17C | Island | Body |
| GM02175 | chr17:1028673-1028931 | ABR;ABR | Island | Body;5'UTR |
| WG1978 | chr17:21300735-21300791 | KCNJ12 | | 5'UTR |
| GM02149 | chr17:30822561-30823006 | MYO1D | N_Shore | Body |
| GM00520 | chr17:37123767-37123843 | FBXO47 | | TSS200 |
| GM00290 | chr17:41832753-41833161 | SOST | Island | Body |
| GM02555 | chr17:43319137-43319382 | FMNL1 | Island | Body |
| GM02704 | chr17:44847220-44847427 | WNT3 | Island | Body |
| GM09256 | chr17:46645972-46646359 | HOXB3 | S_Shelf | 5'UTR |
| GM01392 | chr17:46648014-46648533 | HOXB3 | | 5'UTR |
| WG1513 | chr17:46651945-46652179 | HOXB3 | N_Shelf | TSS200 |
| WG1721 | chr17:46669485-46669492 | LOC404266;HOXB5;LOC404 | Island | TSS200;3'UTR;Body;TSS200;Body |
| GM02175 | chr17:46669485-46669644 | LOC404266;HOXB5;LOC404 | Island | TSS200;3'UTR;Body;TSS200;Body |
| GM01095 | chr17:46673625-46674514 | LOC404266;LOC404266;LO( | Island | Body;Body;Body;Body;3'UTR;Body |
| GM01376 | chr17:46674041-46674438 | LOC404266;LOC404266;LO( | Island | Body;Body;Body;Body;Body;Body |
| WG1977 | chr17:46685327-46685448 | HOXB7 | Island | Body |
| GM02456 | chr17:62777693-62777777 | LOC146880;LOC146880 | Island | Body;TSS200 |
| WG1977 | chr17:75283832-75283979 | SEPT9;SEPT9 | | TSS200;Body |
| GM02456 | chr17:77037916-77037965 | C1QTNF1;C1QTNF1;C1QTNF1 | | 5'UTR;5'UTR;5'UTR |
| GM00935 | chr17:77952290-77952488 | TBC1D16 | Island | Body |
| GM01650 | chr17:79109729-79109808 | AATK | Island | Body |
| WG1514 | chr17:79905236-79905263 | MYADML2 | | TSS200 |
| WG2121 | chr17:80358829-80358876 | C17orf101;C17orf101 | N_Shelf | Body;Body |
| GM00042 | chr17:933047-933155 | ABR;ABR;ABR | Island | Body;Body;Body |
| GM00520 | chr18:19756582-19757022 | GATA6 | N_Shore | Body |
| GM00016 | chr18:30051538-30051572 | FAM59A | S_Shore | TSS1500 |
| GM01096 | chr18:56936434-56936882 | RAX | Island | Body |
| GM01392 | chr18:67069959-67070120 | DOK6 | Island | Body |

| | | | | |
|---|---|---|---|---|
| GM00438 | chr18:67070045-67070170 | DOK6 | Island | Body |
| GM00042 | chr18:74770402-74770815 | MBP;MBP | Island | Body;Body |
| GM00520 | chr18:77917615-77918142 | LOC100130522;LOC100130 | N_Shore | Body;Body;3'UTR |
| GM09255 | chr19:11289349-11289376 | KANK2;KANK2 | S_Shore | Body;Body |
| WG1984 | chr19:12876846-12877000 | HOOK2;HOOK2 | N_Shore | Body;Body |
| WG1309 | chr19:12876846-12877188 | HOOK2;HOOK2 | N_Shore | Body;Body |
| GM00438 | chr19:16178091-16178099 | TPM4 | N_Shore | TSS1500 |
| GM01376 | chr19:1854549-1854724 | KLF16 | Island | Body |
| GM00042 | chr19:19639553-19639596 | YJEFN3 | Island | TSS200 |
| GM00289 | chr19:19639568-19639596 | YJEFN3 | Island | TSS200 |
| GM09255 | chr19:21264896-21265421 | ZNF714 | N_Shore | TSS200 |
| GM02149 | chr19:21265264-21265421 | ZNF714 | Island | 5'UTR |
| GM00439 | chr19:2282439-2282729 | C19orf35 | S_Shelf | TSS1500 |
| GM00438 | chr19:40904691-40904974 | PRX;PRX | Island | Body;Body |
| GM00439 | chr19:427022-427263 | SHC2 | Island | Body |
| GM03075 | chr19:48947560-48947578 | GRIN2D;GRWD1 | Island | 3'UTR;TSS1500 |
| GM00439 | chr19:49891270-49891574 | CCDC155 | Island | TSS1500 |
| GM02175 | chr19:52264411-52264429 | FPR2 | | TSS200 |
| GM01376 | chr19:54876728-54876752 | LAIR1;LAIR1 | | TSS200;TSS200 |
| GM03075 | chr2:10567628-10567843 | HPCAL1;HPCAL1 | S_Shore | 3'UTR;3'UTR |
| GM01653 | chr2:113992843-113993052 | PAX8;PAX8;PAX8;PAX8;PAX8 | N_Shore | Body;Body;Body;Body;Body;Body;TSS |
| GM01653 | chr2:114033490-114033830 | PAX8;PAX8;PAX8;PAX8;PAX8 | Island | Body;Body;Body;Body;Body;Body |
| WG1722 | chr2:121554810-121554926 | GLI2 | | TSS200 |
| GM01095 | chr2:121554817-121554926 | GLI2 | | TSS200 |
| GM02175 | chr2:142889283-142889392 | LRP1B | S_Shore | TSS200 |
| GM00289 | chr2:171676553-171676925 | GAD1;GAD1 | Island | Body;Body |
| WG1978 | chr2:176964512-176964720 | HOXD12 | Island | TSS200 |
| GM00520 | chr2:176973275-176974060 | HOXD11 | N_Shore | Body |
| GM00940 | chr2:176973688-176973835 | HOXD11 | Island | Body |
| GM01393 | chr2:176982108-176982342 | HOXD10 | Island | 1stExon |
| GM02173 | chr2:176993017-176993115 | HOXD8 | Island | TSS1500 |
| GM00940 | chr2:177016167-177017449 | HOXD4;HOXD4 | N_Shore | 5'UTR;1stExon |
| GM00439 | chr2:200320455-200321348 | SATB2 | N_Shore | Body |
| GM00935 | chr2:220406470-220406756 | CHPF | Island | Body |
| GM00290 | chr2:223155925-223156136 | PAX3;PAX3;PAX3;PAX3;PAX3 | Island | Body;Body;Body;Body;Body;Body |
| GM00935 | chr2:223161771-223162049 | PAX3;PAX3;PAX3;CCDC140; | Island | Body;Body;Body;TSS1500;Body;Body; |
| GM01393 | chr2:223161771-223162128 | PAX3;PAX3;PAX3;CCDC140; | Island | Body;Body;Body;TSS1500;Body;Body; |
| GM02640 | chr2:223162666-223162800 | PAX3;PAX3;PAX3;PAX3;PAX3 | N_Shore | Body;Body;Body;Body;Body;Body;Bod |
| GM00439 | chr2:238394933-238395125 | MLPH;MLPH | N_Shore | TSS1500;TSS1500 |
| WG1722 | chr2:240020028-240020174 | HDAC4 | | Body |
| GM00935 | chr2:240233043-240233240 | HDAC4 | | Body |
| GM01650 | chr2:240291426-240291657 | HDAC4 | | 5'UTR |
| WG1983 | chr2:30669597-30669863 | LCLAT1;LCLAT1 | N_Shore | TSS1500;TSS1500 |
| WG1977 | chr2:3286338-3286479 | TSSC1 | Island | Body |
| WG1722 | chr2:3452347-3452624 | TTC15 | Island | Body |
| GM02555 | chr2:3714936-3715083 | ALLC | | 5'UTR |
| GM02555 | chr2:54087173-54087189 | LOC100302652;GPR75 | Island | TSS200;TSS200 |

| | | | | |
|---|---|---|---|---|
| GM00941 | chr2:66671478-66671727 | MEIS1 | N_Shore | Body |
| GM00439 | chr2:71134524-71134630 | VAX2 | N_Shore | Body |
| GM00935 | chr2:74780268-74780617 | LOXL3 | N_Shore | 5'UTR |
| GM01392 | chr20:25128770-25128805 | LOC284798;LOC284798;LO( | Island | TSS200;Body;Body |
| GM00439 | chr20:32856747-32856898 | ASIP | Island | Body |
| GM00042 | chr20:47896448-47896888 | SNORD12B;C20orf199;MIR1: | S_Shore | TSS1500;Body;TSS1500;Body;TSS150 |
| GM00289 | chr20:49253364-49253710 | FAM65C;FAM65C | | 1stExon;5'UTR |
| GM01653 | chr20:5485144-5485284 | LOC149837 | | Body |
| GM00290 | chr20:5485245-5485284 | LOC149837 | | TSS200 |
| GM00940 | chr20:55925570-55925603 | RAE1;RAE1 | N_Shore | TSS1500;TSS1500 |
| GM00042 | chr20:61447330-61447394 | COL9A3 | N_Shore | TSS1500 |
| WG1513 | chr21:47604052-47604654 | C21orf56;C21orf56 | S_Shore | 5'UTR;5'UTR |
| GM09256 | chr22:19754125-19754448 | TBX1;TBX1;TBX1 | Island | Body;Body;Body |
| WG1722 | chr22:22058080-22058271 | YPEL1 | Island | Body |
| GM02149 | chr22:24384105-24384294 | GSTT1 | N_Shore | Body |
| GM00940 | chr22:32027580-32027605 | PISD | S_Shore | TSS1500 |
| GM02706 | chr22:45609402-45609421 | C22orf9;C22orf9 | | Body;TSS1500 |
| GM00289 | chr3:11746183-11746765 | VGLL4 | N_Shore | 5'UTR |
| GM00290 | chr3:14443428-14443473 | SLC6A6;SLC6A6;SLC6A6 | N_Shore | TSS1500;TSS1500;TSS1500 |
| GM09256 | chr3:15311021-15311269 | SH3BP5;SH3BP5 | | Body;Body |
| GM02706 | chr3:155394109-155394303 | PLCH1;PLCH1;PLCH1 | | 5'UTR;TSS200;TSS200 |
| GM00520 | chr3:160475035-160475128 | PPM1L | S_Shore | Body |
| GM02149 | chr3:183894727-183894778 | AP2M1;AP2M1 | S_Shore | 5'UTR;5'UTR |
| WG2276 | chr3:192289245-192289293 | FGF12 | | Body |
| GM02706 | chr3:196065357-196065506 | TM4SF19 | | TSS200 |
| WG1722 | chr3:36949957-36950004 | TRANK1 | | Body |
| GM00042 | chr3:42742993-42743203 | HHATL;HHATL | | Body;5'UTR |
| WG1722 | chr3:50488020-50488230 | CACNA2D2;CACNA2D2 | | Body;Body |
| GM00290 | chr3:56809796-56810320 | ARHGEF3;ARHGEF3;ARHGEF3 | | TSS200;Body;Body |
| GM01096 | chr3:64547310-64547373 | ADAMTS9 | | Body |
| GM02175 | chr4:111398042-111398146 | ENPEP | Island | 1stExon |
| GM00520 | chr4:119948360-119948542 | SYNPO2;SYNPO2;SYNPO2 | | Body;Body;Body |
| GM00438 | chr4:174447847-174448341 | HAND2 | N_Shore | 3'UTR |
| GM00438 | chr4:40858984-40859293 | APBB2;APBB2;APBB2;APBB2 | Island | Body;5'UTR;5'UTR;5'UTR;Body;5'UTR;E |
| GM02175 | chr4:52918457-52918489 | SPATA18 | S_Shore | Body |
| GM01392 | chr4:54965829-54965904 | GSX2 | N_Shore | TSS1500 |
| WG1513 | chr4:55143269-55143557 | PDGFRA | | Body |
| GM01392 | chr4:6052527-6052554 | JAKMIP1 | | Body |
| WG1282 | chr4:71587722-71587904 | RUFY3;RUFY3;RUFY3;RUFY3;RUFY3 | | 1stExon;Body;5'UTR;1stExon;5'UTR |
| GM01096 | chr5:1053545-1053774 | SLC12A7 | Island | Body |
| WG1310 | chr5:14406585-14406732 | TRIO | | Body |
| GM01650 | chr5:149980486-149980682 | SYNPO | | TSS200 |
| WG1513 | chr5:149980671-149980682 | SYNPO;SYNPO | | 1stExon;5'UTR |
| GM00042 | chr5:176784688-176784719 | RGS14 | | TSS200 |
| GM00520 | chr5:177548398-177548913 | N4BP3 | | Body |
| WG1977 | chr5:177708304-177708528 | COL23A1 | | Body |
| WG1722 | chr5:323320-323969 | AHRR | S_Shore | Body |

| | | | | |
|---|---|---|---|---|
| GM02173 | chr5:40841493-40841590 | CARD6 | | 1stExon |
| GM00289 | chr5:423894-424371 | AHRR | S_Shelf | Body |
| GM00016 | chr5:68710808-68711277 | MARVELD2;MARVELD2 | Island | TSS200;TSS200 |
| WG1977 | chr5:76010444-76010658 | F2R | N_Shore | TSS1500 |
| GM00022 | chr5:78281793-78281964 | ARSB;ARSB;ARSB | S_Shore | TSS200;1stExon;5'UTR |
| GM02641 | chr6:100911240-100912072 | SIM1 | N_Shore | 1stExon |
| GM02704 | chr6:101846791-101846872 | GRIK2;GRIK2;GRIK2 | Island | TSS200;TSS200;TSS200 |
| GM02456 | chr6:109274429-109274496 | ARMC2 | | Body |
| GM00439 | chr6:11279656-11279727 | NEDD9 | | Body |
| GM09255 | chr6:139613145-139613456 | TXLNB;TXLNB | | 5'UTR;1stExon |
| GM01392 | chr6:149803087-149803444 | ZC3H12D | | 5'UTR |
| GM00289 | chr6:152085565-152085641 | ESR1 | | 5'UTR |
| GM00031 | chr6:152125965-152126092 | ESR1;ESR1 | N_Shelf | 5'UTR;TSS1500 |
| GM00941 | chr6:152127812-152127887 | ESR1;ESR1;ESR1;ESR1 | N_Shore | TSS1500;5'UTR;TSS1500;5'UTR |
| GM00935 | chr6:152127812-152128024 | ESR1;ESR1;ESR1;ESR1 | N_Shore | TSS1500;5'UTR;TSS1500;5'UTR |
| WG1722 | chr6:161796785-161796854 | PARK2;PARK2;PARK2 | | Body;Body;Body |
| WG1282 | chr6:161860203-161860359 | PARK2;PARK2;PARK2 | | Body;Body;Body |
| GM00022 | chr6:168435914-168436353 | KIF25;KIF25 | Island | Body;Body |
| GM02704 | chr6:291882-292823 | DUSP22 | N_Shore | TSS1500 |
| GM02147 | chr6:29690766-29690893 | HLA-F;HLA-F;HLA-F | N_Shore | TSS1500;TSS1500;TSS1500 |
| GM00042 | chr6:29690822-29690893 | HLA-F;HLA-F;HLA-F | N_Shore | TSS1500;TSS1500;TSS1500 |
| WG1977 | chr6:29894050-29894197 | HCG4P6 | N_Shore | TSS1500 |
| GM00520 | chr6:29943401-29943425 | HCG9 | N_Shore | Body |
| GM01653 | chr6:30039432-30039442 | RNF39;RNF39 | Island | Body;Body |
| GM00941 | chr6:30079256-30079280 | TRIM31 | | Body |
| GM01375 | chr6:31025933-31025967 | HCG22 | | Body |
| WG2194 | chr6:31237013-31237034 | HLA-C | N_Shore | Body |
| WG2121 | chr6:31237405-31237722 | HLA-C | N_Shore | Body |
| GM02555 | chr6:32489750-32489963 | HLA-DRB5 | Island | Body |
| GM00439 | chr6:32489991-32490043 | HLA-DRB5 | Island | Body |
| GM00016 | chr6:32526260-32526364 | HLA-DRB6 | | Body |
| GM00042 | chr6:32551891-32551954 | HLA-DRB1 | Island | Body |
| GM02147 | chr6:32552152-32552246 | HLA-DRB1 | Island | Body |
| GM00016 | chr6:33288366-33288569 | DAXX;DAXX;DAXX;DAXX | N_Shore | Body;Body;Body;Body |
| GM00042 | chr6:33288599-33288656 | DAXX;DAXX;DAXX;DAXX | N_Shore | Body;Body;Body;Body |
| GM02175 | chr6:6002865-6003303 | NRN1 | Island | Body |
| GM00042 | chr6:70576168-70576321 | COL19A1 | N_Shore | TSS1500 |
| GM01650 | chr7:101500014-101500514 | CUX1;CUX1;CUX1 | | Body;Body;Body |
| GM00290 | chr7:101768610-101768874 | CUX1;CUX1;CUX1 | | Body;Body;Body |
| GM00290 | chr7:114055123-114055137 | FOXP2;FOXP2;FOXP2;FOXP2;FOX | | 1stExon;5'UTR;5'UTR;1stExon;5'UTR;1 |
| WG1513 | chr7:114055123-114055419 | FOXP2;FOXP2;FOXP2;FOXP2;FOX | | 1stExon;5'UTR;5'UTR;1stExon;5'UTR;1 |
| GM02555 | chr7:158336016-158336049 | PTPRN2;PTPRN2;PTPRN2 | | Body;Body;Body |
| GM02175 | chr7:1952518-1952807 | MAD1L1;MAD1L1;MAD1L1 | S_Shelf | Body;Body;Body |
| GM00935 | chr7:2133441-2133480 | MAD1L1;MAD1L1;MAD1L1 | | Body;Body;Body |
| GM00016 | chr7:2139216-2139259 | MAD1L1;MAD1L1;MAD1L1 | | Body;Body;Body |
| GM01375 | chr7:2563611-2563909 | LFNG;LFNG;LFNG;LFNG | Island | Body;Body;Body;Body |
| GM00042 | chr7:27182493-27183196 | HOXA5 | N_Shore | Body |

| | | | | |
|---|---|---|---|---|
| GM00022 | chr7:27183274-27183436 | HOXA5;HOXA5 | Island | 5'UTR;1stExon |
| GM00042 | chr7:27187943-27188770 | HOXA6 | S_Shore | TSS1500 |
| GM00042 | chr7:27225299-27225342 | HOXA11AS;HOXA11 | Island | Body;TSS1500 |
| GM01392 | chr7:27225523-27225543 | HOXA11AS;HOXA11 | Island | Body;TSS1500 |
| WG2275 | chr7:2847517-2847575 | GNA12 | | Body |
| GM02456 | chr7:28998577-28998860 | TRIL | S_Shore | TSS1500 |
| GM00520 | chr7:4065931-4066127 | SDK1 | | Body |
| GM01393 | chr7:4885521-4885643 | RADIL | | Body |
| GM01392 | chr7:5336543-5336811 | SLC29A4;SLC29A4 | Island | Body;Body |
| GM00520 | chr7:553412-553693 | PDGFA;PDGFA | N_Shelf | Body;Body |
| WG1513 | chr7:70096029-70096378 | AUTS2;AUTS2 | N_Shore | Body;Body |
| GM09255 | chr7:944607-944776 | ADAP1 | S_Shore | Body |
| GM00438 | chr8:104032819-104032876 | ATP6V1C1 | N_Shore | TSS1500 |
| GM02706 | chr8:10586959-10587055 | SOX7 | N_Shore | Body |
| GM00289 | chr8:11141365-11141424 | MTMR9 | N_Shore | TSS1500 |
| GM00941 | chr8:116660311-116661063 | TRPS1 | N_Shore | 5'UTR |
| GM00016 | chr8:143545390-143545574 | BAI1;BAI1 | N_Shore | 5'UTR;1stExon |
| GM00016 | chr8:1847999-1848143 | ARHGEF10 | | Body |
| GM00022 | chr8:1847999-1848407 | ARHGEF10 | | Body |
| GM00289 | chr8:1848040-1848407 | ARHGEF10 | | Body |
| GM02704 | chr8:2075469-2075820 | MYOM2 | Island | Body |
| GM01392 | chr8:28560467-28560723 | EXTL3 | S_Shore | 5'UTR |
| WG1513 | chr8:59058585-59058660 | FAM110B | Island | 5'UTR |
| GM00289 | chr8:6456182-6456465 | MCPH1 | | Body |
| GM01393 | chr9:140401382-140401576 | PNPLA7;PNPLA7 | Island | Body;Body |
| GM01096 | chr9:92220617-92221282 | GADD45G | Island | Body |
| GM01095 | chr1:108734992-108735549 | SLC25A24;SLC25A24 | N_Shore | Body;Body |
| GM00290 | chr1:115881044-115881132 | NGF | Island | TSS200 |
| WG2193 | chr1:119526060-119526437 | TBX15 | N_Shore | 5'UTR |
| GM00438 | chr1:119526060-119526783 | TBX15 | N_Shore | 5'UTR |
| WG1282 | chr1:119531718-119532044 | TBX15 | N_Shore | 5'UTR |
| WG2121 | chr1:119532320-119532542 | TBX15 | S_Shore | TSS200 |
| WG1977 | chr1:15256054-15256222 | KIAA1026;KIAA1026;KIAA1026;KIAA1026 | | Body;TSS1500;Body;Body |
| GM00520 | chr1:153581494-153581793 | S100A16 | | 5'UTR |
| GM01393 | chr1:154127952-154128041 | TPM3;TPM3;NUP210L;TPM3;TPM3;NUP2 | | 3'UTR;3'UTR;TSS1500;3'UTR;3'UTR;TS |
| GM00438 | chr1:156786601-156786653 | SH2D2A;NTRK1;SH2D2A;SH2 | S_Shelf | 5'UTR;Body;1stExon;TSS1500;5'UTR;5 |
| GM00940 | chr1:159796617-159796658 | SLAMF8;SLAMF8 | | 5'UTR;1stExon |
| GM00042 | chr1:161049443-161049579 | PVRL4 | Island | Body |
| WG1283 | chr1:170638807-170639289 | PRRX1;PRRX1 | | Body;Body |
| GM00520 | chr1:173176362-173176523 | TNFSF4;TNFSF4 | | 5'UTR;1stExon |
| GM00022 | chr1:173176366-173176523 | TNFSF4;TNFSF4 | | 5'UTR;1stExon |
| GM01392 | chr1:2005260-2005564 | PRKCZ;PRKCZ;PRKCZ | Island | 1stExon;Body;5'UTR |
| GM00439 | chr1:202091865-202091880 | GPR37L1 | | TSS200 |
| GM02641 | chr1:205818956-205819463 | PM20D1 | Island | Body |
| WG1514 | chr1:205818956-205819492 | PM20D1 | Island | Body |
| WG1282 | chr1:207992403-207992548 | LOC148696 | | Body |
| GM02173 | chr1:2120985-2121724 | C1orf86;LOC100128003 | N_Shore | Body;Body |

| | | | | |
|---|---|---|---|---|
| GM00022 | chr1:214477063-214477280 | SMYD2 | | Body |
| GM02456 | chr1:21586831-21587174 | ECE1;ECE1;ECE1;ECE1 | | Body;Body;Body;Body |
| GM00520 | chr1:221053841-221055097 | HLX | N_Shore | Body |
| GM02704 | chr1:221053904-221054273 | HLX | N_Shore | Body |
| GM01096 | chr1:221053904-221055097 | HLX | N_Shore | Body |
| WG1978 | chr1:221054615-221055097 | HLX | Island | Body |
| GM01392 | chr1:223566643-223566669 | C1orf65 | Island | TSS200 |
| GM00935 | chr1:223936508-223936812 | CAPN2;CAPN2 | Island | Body;Body |
| GM09256 | chr1:226825435-226825606 | ITPKB | | Body |
| GM01376 | chr1:228291668-228291705 | C1orf35 | S_Shore | TSS1500 |
| WG1658 | chr1:23521272-23521417 | HTR1D | | TSS200 |
| WG1722 | chr1:240776162-240776211 | GREM2 | | TSS1500 |
| GM02173 | chr1:247694271-247694406 | LOC148824;OR2C3 | Island | TSS200;3'UTR |
| GM02147 | chr1:25944712-25945042 | MAN1C1 | Island | 1stExon |
| GM00439 | chr1:32714146-32714308 | FAM167B | Island | Body |
| GM01096 | chr1:3277525-3277623 | PRDM16;PRDM16 | S_Shelf | Body;Body |
| WG1513 | chr1:3459949-3460268 | MEGF6 | Island | Body |
| GM02706 | chr1:36039655-36039885 | TFAP2E | Island | Body |
| WG1722 | chr1:40105667-40105706 | HEYL | Island | TSS1500 |
| GM00042 | chr1:41250062-41250117 | KCNQ4;KCNQ4 | Island | 1stExon;1stExon |
| GM02706 | chr1:44016014-44016049 | PTPRF;PTPRF | | Body;Body |
| GM00042 | chr1:53600741-53600889 | SLC1A7 | | Body |
| GM02149 | chr1:54199825-54200115 | GLIS1;GLIS1 | N_Shelf | 5'UTR;1stExon |
| WG1513 | chr1:54199825-54200420 | GLIS1;GLIS1 | N_Shelf | 5'UTR;1stExon |
| GM01392 | chr1:54856177-54856511 | SSBP3;SSBP3;SSBP3 | N_Shore | Body;Body;Body |
| GM02175 | chr1:6085995-6086148 | KCNAB2;KCNAB2 | N_Shore | TSS1500;TSS1500 |
| GM01376 | chr1:63249199-63249213 | ATG4C;ATG4C | | TSS1500;TSS1500 |
| GM00031 | chr1:6526182-6526319 | TNFRSF25;TNFRSF25;PLEKH | Island | 1stExon;1stExon;3'UTR;3'UTR;5'UTR;5 |
| GM01096 | chr1:6558085-6558113 | PLEKHG5;PLEKHG5 | S_Shore | Body;TSS1500 |
| GM09255 | chr1:89664260-89664582 | GBP4 | | Body |
| GM00935 | chr1:9131622-9131783 | SLC2A5 | Island | Body |
| WG2276 | chr10:114877616-114877689 | TCF7L2;TCF7L2;TCF7L2;TCF7L2;TCF7L2 | | Body;Body;Body;Body;Body;Body |
| GM00935 | chr10:121265587-121265893 | RGS10;RGS10 | | Body;Body |
| WG1657 | chr10:126278272-126278502 | LHPP;LHPP | N_Shelf | Body;Body |
| GM00016 | chr10:131669406-131669630 | EBF3 | S_Shore | Body |
| GM00016 | chr10:131697080-131697345 | EBF3 | N_Shore | Body |
| GM00042 | chr10:131761206-131761386 | EBF3 | Island | Body |
| WG1513 | chr10:135341528-135342218 | CYP2E1 | N_Shore | 1stExon |
| GM00289 | chr10:13749193-13749625 | FRMD4A | | Body |
| GM02706 | chr10:13932842-13933247 | FRMD4A | N_Shore | Body |
| GM00290 | chr10:22605982-22606072 | COMMD3 | S_Shore | Body |
| GM00016 | chr10:26855907-26856105 | APBB1IP | Island | Body |
| GM00290 | chr10:29811530-29811659 | SVIL;SVIL | | Body;Body |
| WG1722 | chr10:3138418-3138534 | PFKP | | Body |
| GM00290 | chr10:384506-384652 | DIP2C | N_Shelf | Body |
| GM01375 | chr10:45495739-45495981 | C10orf25;ZNF22 | N_Shore | 3'UTR;TSS1500 |
| GM02456 | chr10:46993502-46993542 | GPRIN2 | Island | TSS1500 |

| | | | | |
|---|---|---|---|---|
| GM01376 | chr10:50506748-50506896 | C10orf71;C10orf71 | | TSS1500;TSS1500 |
| GM00016 | chr10:661009-661159 | DIP2C | | Body |
| WG1514 | chr10:72254314-72254335 | KIAA1274 | S_Shelf | 5'UTR |
| WG1977 | chr10:72311914-72312093 | KIAA1274 | | Body |
| GM00520 | chr10:73497514-73497629 | C10orf105;C10orf105;CDH23 | | 5'UTR;Body |
| GM00940 | chr10:73976790-73976821 | ASCC1;C10orf104 | S_Shore | TSS1500;5'UTR |
| GM01650 | chr10:82167757-82167774 | C10orf58 | N_Shelf | |
| GM01375 | chr10:90712684-90712948 | ACTA2;ACTA2 | | 5'UTR;5'UTR |
| GM03075 | chr10:97175272-97175479 | SORBS1;SORBS1;SORBS1;SORBS1;SORI | | Body;Body;Body;Body;Body;Body;Bod |
| GM01096 | chr10:99338056-99338240 | ANKRD2;ANKRD2 | N_Shore | Body;Body |
| GM00042 | chr10:99735010-99735202 | CRTAC1 | | Body |
| WG1514 | chr11:10325567-10325809 | ADM | N_Shore | TSS1500 |
| GM02456 | chr11:12031093-12031099 | DKK3;DKK3;DKK3 | S_Shore | TSS1500;TSS200;TSS1500 |
| GM03075 | chr11:125037226-125037348 | PKNOX2 | S_Shore | 5'UTR |
| GM00042 | chr11:128646953-128647107 | FLI1;FLI1 | | Body;Body |
| GM00290 | chr11:128729883-128730011 | KCNJ1;KCNJ1;KCNJ1;KCNJ1 | | 5'UTR;5'UTR;5'UTR;5'UTR |
| GM02706 | chr11:128812978-128813008 | TP53AIP1 | | TSS200 |
| GM00520 | chr11:129991445-129992368 | APLP2;APLP2;APLP2;APLP2;A | N_Shelf | Body;Body;Body;Body;Body;Body |
| GM00940 | chr11:129992297-129992368 | APLP2;APLP2;APLP2;APLP2;A | N_Shore | Body;Body;Body;Body;Body;Body |
| GM00289 | chr11:130319691-130320219 | ADAMTS15 | Island | 1stExon |
| GM02456 | chr11:1750302-1750381 | HCCA2 | | Body |
| WG2414 | chr11:18433500-18433564 | LDHC;LDHC | | TSS1500;TSS1500 |
| GM00935 | chr11:31819444-31819678 | PAX6;PAX6;PAX6 | N_Shore | Body;Body;Body |
| GM02175 | chr11:31821590-31822302 | PAX6;PAX6;PAX6 | S_Shore | Body;Body;Body |
| GM00940 | chr11:31824262-31825226 | PAX6;PAX6;PAX6 | N_Shore | Body;Body;Body |
| GM00940 | chr11:31825756-31825833 | PAX6;PAX6;PAX6 | Island | Body;Body;Body |
| GM00438 | chr11:31827084-31827757 | PAX6;PAX6;PAX6 | N_Shore | Body;Body;Body |
| GM02175 | chr11:31827084-31828040 | PAX6;PAX6;PAX6 | N_Shore | Body;Body;Body |
| GM00520 | chr11:31837104-31837144 | PAX6 | Island | 5'UTR |
| WG1513 | chr11:3253838-3254065 | MRGPRE | Island | TSS1500 |
| GM02704 | chr11:628603-628677 | SCT | S_Shore | TSS1500 |
| GM00289 | chr11:66650500-66650560 | PC;PC;PC | | 5'UTR;5'UTR;5'UTR |
| GM01096 | chr11:66885240-66885281 | KDM2A;KDM2A | N_Shore | TSS1500;TSS1500 |
| GM00439 | chr11:68138505-68138908 | LRP5 | | Body |
| GM02555 | chr11:68147880-68148079 | LRP5 | | Body |
| GM01650 | chr11:68610706-68611016 | CPT1A;CPT1A | N_Shore | TSS1500;TSS1500 |
| GM02317 | chr11:68782049-68782202 | MRGPRF;MRGPRF | S_Shelf | TSS1500;TSS1500 |
| GM09256 | chr11:77299805-77299962 | AQP11 | N_Shore | TSS1500 |
| GM01393 | chr11:85430629-85430657 | SYTL2;SYTL2;SYTL2;SYTL2;SYTL2;SYTL2; | | Body;TSS1500;TSS1500;Body;TSS150 |
| WG1977 | chr11:86383696-86383723 | ME3;ME3;ME3 | S_Shore | TSS1500;TSS200;TSS1500 |
| GM00289 | chr11:86383696-86383761 | ME3;ME3;ME3 | S_Shore | TSS1500;TSS200;TSS1500 |
| GM00935 | chr11:87908785-87908938 | RAB38 | S_Shore | TSS200 |
| GM02456 | chr12:10084936-10085174 | CLEC2A;CLEC2A | | 5'UTR;1stExon |
| GM00438 | chr12:10251536-10251638 | CLEC1A;CLEC1A | | 5'UTR;1stExon |
| GM09255 | chr12:106988340-106988413 | RFX4 | | Body |
| GM01376 | chr12:114833837-114834201 | TBX5;TBX5;TBX5;TBX5 | N_Shore | Body;Body;Body;Body |
| WG1721 | chr12:114833837-114834713 | TBX5;TBX5;TBX5;TBX5 | N_Shore | Body;Body;Body;Body |

| | | | | |
|---|---|---|---|---|
| WG1309 | chr12:114837572-114837887 | TBX5;TBX5;TBX5;TBX5 | N_Shore | Body;Body;Body;Body |
| GM00935 | chr12:114841536-114841708 | TBX5;TBX5;TBX5;TBX5 | N_Shore | Body;Body;5'UTR;Body |
| GM00439 | chr12:117155382-117155636 | C12orf49 | | 3'UTR |
| GM02175 | chr12:122356056-122356598 | WDR66 | N_Shore | TSS1500 |
| GM00289 | chr12:122356056-122356852 | WDR66 | N_Shore | TSS1500 |
| GM02640 | chr12:122356316-122356598 | WDR66 | Island | TSS200 |
| GM02175 | chr12:131303194-131303518 | STX2;STX2 | Island | Body;Body |
| GM09255 | chr12:15114233-15114393 | ARHGDIB | | 5'UTR |
| WG1283 | chr12:15114703-15114732 | ARHGDIB | | TSS200 |
| GM09255 | chr12:1609245-1609369 | LOC100292680 | | TSS1500 |
| GM00438 | chr12:25055262-25055421 | BCAT1 | N_Shore | Body |
| GM01096 | chr12:25150530-25150626 | C12orf77 | | TSS200 |
| GM09255 | chr12:312592-313104 | SLC6A12;SLC6A12;SLC6A12 | Island | Body;Body;Body |
| GM00042 | chr12:51786038-51786547 | GALNT6 | S_Shore | TSS1500 |
| GM03075 | chr12:54392734-54392884 | HOXC9 | N_Shore | TSS1500 |
| WG1977 | chr12:54423428-54425418 | HOXC4;HOXC6;HOXC5;HOX | Island | 5'UTR;Body;Body;Body |
| GM02704 | chr12:54440773-54441509 | HOXC4 | Island | 5'UTR |
| GM09255 | chr12:54440806-54441213 | HOXC4 | Island | 5'UTR |
| GM00016 | chr12:54446019-54446100 | HOXC4 | Island | 5'UTR |
| GM01393 | chr12:54446019-54446253 | HOXC4 | Island | 5'UTR |
| GM09255 | chr12:54448729-54448913 | HOXC4;HOXC4 | S_Shore | Body;Body |
| GM09255 | chr12:56329615-56329903 | DGKA;DGKA;DGKA;DGKA | Island | 5'UTR;5'UTR;5'UTR;5'UTR |
| WG1283 | chr12:62583974-62584454 | FAM19A2 | N_Shore | 5'UTR |
| WG1658 | chr12:6442873-6443045 | TNFRSF1A | S_Shelf | Body |
| GM02175 | chr12:7281566-7281783 | CLSTN3;RBP5 | | TSS1500;TSS200 |
| GM00935 | chr12:772861-772982 | NINJ2 | | TSS200 |
| WG1513 | chr12:97300410-97300429 | NEDD1;NEDD1;NEDD1;NEDD | N_Shore | TSS1500;TSS1500;TSS1500;TSS1500 |
| GM01095 | chr13:100310162-100310405 | CLYBL | N_Shore | Body |
| GM00290 | chr13:110874267-110874326 | COL4A1 | | Body |
| GM00935 | chr13:110918331-110918683 | COL4A1 | Island | Body |
| WG2194 | chr13:110993474-110993558 | COL4A2 | | Body |
| GM00520 | chr13:111521981-111522314 | C13orf29 | | Body |
| GM00935 | chr13:113436309-113436651 | ATP11A;ATP11A | N_Shelf | Body;Body |
| GM00031 | chr13:114108433-114108515 | ADPRHL1 | | TSS1500 |
| WG1722 | chr13:114108433-114108652 | ADPRHL1 | | TSS1500 |
| GM02555 | chr13:114308619-114308749 | ATP4B | | Body |
| GM02147 | chr13:114880761-114880889 | RASA3 | S_Shore | Body |
| GM00439 | chr13:20805196-20805385 | GJB6;GJB6;GJB6;GJB6;GJB6 | N_Shore | 5'UTR;5'UTR;5'UTR;5'UTR;1stExon |
| GM00289 | chr13:30088615-30088771 | SLC7A1 | Island | 3'UTR |
| GM00935 | chr13:36045201-36045352 | NBEA | Island | Body |
| GM02706 | chr13:36048892-36051073 | MAB21L1;MIR548F5;NBEA;M | N_Shore | 1stExon;Body;Body;3'UTR |
| GM00042 | chr13:36050901-36050993 | MIR548F5;NBEA;MAB21L1 | N_Shore | Body;Body;TSS200 |
| WG1283 | chr13:36050901-36051073 | MIR548F5;NBEA;MAB21L1 | N_Shore | Body;Body;TSS200 |
| GM00016 | chr13:79234435-79234847 | RNF219 | S_Shore | TSS1500 |
| GM00438 | chr13:98605892-98606196 | IPO5 | | TSS200 |
| GM00439 | chr13:98794353-98794593 | FARP1;FARP1 | N_Shore | TSS1500;TSS1500 |
| GM02173 | chr14:103593088-103593520 | TNFAIP2 | N_Shore | Body |

| | | | | |
|---|---|---|---|---|
| GM00520 | chr14:103593235-103593520 | TNFAIP2 | Island | Body |
| GM02316 | chr14:103593503-103593520 | TNFAIP2 | Island | Body |
| WG1309 | chr14:37136192-37136391 | PAX9 | Island | Body |
| WG2414 | chr14:54418728-54418851 | BMP4;BMP4;BMP4 | Island | Body;Body;Body |
| GM00042 | chr14:57277178-57277366 | OTX2;OTX2 | N_Shore | 1stExon;5'UTR |
| GM02175 | chr14:80677608-80677688 | DIO2;DIO2;DIO2 | | 1stExon;1stExon;Body |
| GM02456 | chr14:99728984-99729088 | BCL11B;BCL11B | | Body;Body |
| GM02456 | chr15:101419275-101419296 | ALDH1A3 | Island | TSS1500 |
| GM00935 | chr15:101728611-101728653 | CHSY1 | | Body |
| WG1283 | chr15:32964155-32964500 | SCG5;SCG5 | | Body;Body |
| GM00439 | chr15:37170381-37170554 | LOC145845 | N_Shelf | Body |
| GM00022 | chr15:40268610-40269214 | EIF2AK4 | Island | Body |
| GM02456 | chr15:40544174-40544427 | PAK6;C15orf56;PAK6;PAK6 | N_Shore | 5'UTR;3'UTR;5'UTR;5'UTR |
| GM00520 | chr15:44086617-44087089 | SERF2;MIR1282 | S_Shore | 3'UTR;TSS1500 |
| GM00935 | chr15:45314789-45314933 | SORD | N_Shore | TSS1500 |
| GM02149 | chr15:45671155-45671246 | LOC145663;GATM | S_Shore | Body;TSS200 |
| GM00935 | chr15:49716247-49716645 | C15orf33;FGF7 | | Body;5'UTR |
| GM02706 | chr15:51369862-51370019 | TNFAIP8L3 | | Body |
| GM02706 | chr15:55881599-55881706 | PYGO1 | S_Shore | TSS1500 |
| GM00935 | chr15:70354647-70354825 | TLE3;TLE3;TLE3 | | Body;Body;Body |
| GM02456 | chr15:70994435-70994672 | UACA;UACA;UACA | | 5'UTR;Body;1stExon |
| GM00935 | chr15:70994435-70995014 | UACA;UACA;UACA | | 5'UTR;Body;1stExon |
| WG1722 | chr15:71408603-71408847 | CT62 | S_Shore | TSS1500 |
| WG2120 | chr15:88576296-88576320 | NTRK3;NTRK3;NTRK3 | | Body;Body;Body |
| GM01095 | chr15:92944664-92944756 | ST8SIA2 | | Body |
| GM00290 | chr15:93632896-93633172 | RGMA;RGMA | Island | TSS1500;TSS1500 |
| GM01393 | chr16:2892766-2892782 | TMPRSS8 | Island | TSS200 |
| WG1977 | chr16:2908554-2908934 | PRSS22 | S_Shore | TSS1500 |
| WG1983 | chr16:4103161-4103225 | ADCY9 | | Body |
| GM01376 | chr16:50347766-50347895 | ADCY7 | | Body |
| GM02316 | chr16:55794595-55794910 | CES4 | N_Shore | Body |
| GM02175 | chr16:58019866-58019984 | TEPP;TEPP | Island | Body;Body |
| WG1282 | chr16:65105516-65105714 | CDH11 | | 5'UTR |
| GM00438 | chr16:67197084-67197186 | HSF4;FBXL8;HSF4 | Island | TSS1500;Body;TSS1500 |
| GM00439 | chr16:68270252-68270282 | ESRP2 | Island | TSS200 |
| GM09255 | chr16:69958626-69958964 | WWP2;WWP2 | | TSS1500;Body |
| GM09256 | chr16:81527427-81527504 | CMIP | | Body |
| GM00042 | chr16:857454-857981 | PRR25 | Island | Body |
| GM00016 | chr16:87441441-87441659 | ZCCHC14 | Island | 3'UTR |
| GM00439 | chr16:88052956-88053104 | BANP;BANP | Island | Body;Body |
| GM09255 | chr16:88706241-88706426 | IL17C | Island | Body |
| GM00022 | chr17:1028394-1028676 | ABR;ABR | N_Shore | Body;Body |
| WG1513 | chr17:1028673-1028931 | ABR;ABR | Island | Body;5'UTR |
| GM01096 | chr17:154410-154671 | RPH3AL | S_Shore | Body |
| WG1983 | chr17:21300735-21300791 | KCNJ12 | | 5'UTR |
| GM02149 | chr17:26732191-26732272 | SLC46A1 | Island | Body |
| GM02175 | chr17:30822561-30823006 | MYO1D | N_Shore | Body |

| | | | | |
|---|---|---|---|---|
| GM00289 | chr17:33759957-33759986 | SLFN12 | | TSS1500 |
| GM02175 | chr17:36715787-36715953 | SRCIN1 | Island | Body |
| WG1658 | chr17:3705631-3705875 | ITGAE | | TSS1500 |
| GM00935 | chr17:37123767-37123843 | FBXO47 | | TSS200 |
| GM00042 | chr17:40835849-40836087 | CNTNAP1 | Island | Body |
| GM00438 | chr17:41832753-41833161 | SOST | Island | Body |
| GM00439 | chr17:41909709-41909716 | MPP3;MPP3 | N_Shore | 5'UTR;Body |
| GM03075 | chr17:43319137-43319382 | FMNL1 | Island | Body |
| GM00520 | chr17:43974869-43975063 | MAPT;MAPT;MAPT;MAPT;LOC | Island | 5'UTR;5'UTR;5'UTR;5'UTR;Body;5'UTR; |
| GM02706 | chr17:43974919-43975063 | MAPT;MAPT;MAPT;MAPT;LOC | Island | 5'UTR;5'UTR;5'UTR;5'UTR;Body;5'UTR; |
| GM02706 | chr17:44847220-44847427 | WNT3 | Island | Body |
| WG1978 | chr17:46645972-46646359 | HOXB3 | S_Shelf | 5'UTR |
| GM00520 | chr17:46645972-46646444 | HOXB3 | S_Shelf | 5'UTR |
| GM02317 | chr17:46648014-46648533 | HOXB3 | | 5'UTR |
| WG1514 | chr17:46651945-46652179 | HOXB3 | N_Shelf | TSS200 |
| GM00941 | chr17:46656543-46656690 | HOXB4 | S_Shore | TSS1500 |
| WG1983 | chr17:46669485-46669492 | LOC404266;HOXB5;LOC404 | Island | TSS200;3'UTR;Body;TSS200;Body |
| GM02555 | chr17:46669485-46669644 | LOC404266;HOXB5;LOC404 | Island | TSS200;3'UTR;Body;TSS200;Body |
| GM02706 | chr17:46673625-46674514 | LOC404266;LOC404266;LOC | Island | Body;Body;Body;Body;3'UTR;Body |
| GM02456 | chr17:46674041-46674438 | LOC404266;LOC404266;LOC | Island | Body;Body;Body;Body;Body;Body |
| GM00042 | chr17:46682308-46683047 | LOC404266;LOC404266;HO | N_Shelf | Body;Body;1stExon;5'UTR |
| WG1978 | chr17:46685327-46685448 | HOXB7 | Island | Body |
| GM02706 | chr17:47209660-47209749 | B4GALNT2;B4GALNT2;B4GAL | N_Shore | TSS1500;TSS200;TSS1500 |
| GM00042 | chr17:47287492-47287526 | ABI3;GNGT2;ABI3 | | TSS200;TSS1500;TSS200 |
| GM00290 | chr17:59554672-59554806 | TBX4 | | Body |
| GM02555 | chr17:62777693-62777777 | LOC146880;LOC146880 | Island | Body;TSS200 |
| WG1978 | chr17:75283832-75283979 | SEPT9;SEPT9 | | TSS200;Body |
| GM03075 | chr17:77037916-77037965 | C1QTNF1;C1QTNF1;C1QTNF1 | | 5'UTR;5'UTR;5'UTR |
| GM00941 | chr17:77952290-77952488 | TBC1D16 | Island | Body |
| GM00290 | chr17:79010458-79010831 | BAIAP2;BAIAP2;BAIAP2;BAIAP | S_Shore | Body;Body;Body;Body |
| GM02175 | chr17:79099481-79099819 | MIR657;AATK | N_Shelf | TSS1500;Body |
| GM09256 | chr17:79109729-79109808 | AATK | Island | Body |
| WG1657 | chr17:79905236-79905263 | MYADML2 | | TSS200 |
| GM00022 | chr17:80292079-80292252 | SECTM1 | Island | TSS200 |
| WG2414 | chr17:80358829-80358876 | C17orf101;C17orf101 | N_Shelf | Body;Body |
| GM00290 | chr17:80806135-80806435 | TBCD | | Body |
| GM00940 | chr17:80875998-80876074 | TBCD | | Body |
| GM00439 | chr17:80943851-80944193 | B3GNTL1 | | Body |
| GM00439 | chr17:81045495-81045781 | METRNL | Island | Body |
| GM03075 | chr17:872762-873452 | NXN | | Body |
| GM02147 | chr17:933047-933155 | ABR;ABR;ABR | Island | Body;Body;Body |
| GM00935 | chr18:19756582-19757022 | GATA6 | N_Shore | Body |
| WG1513 | chr18:30051538-30051572 | FAM59A | S_Shore | TSS1500 |
| GM02706 | chr18:56936434-56936882 | RAX | Island | Body |
| WG1721 | chr18:67069959-67070120 | DOK6 | Island | Body |
| GM00520 | chr18:67070045-67070170 | DOK6 | Island | Body |
| GM02706 | chr18:74770402-74770815 | MBP;MBP | Island | Body;Body |

| | | | | |
|---|---|---|---|---|
| GM00438 | chr18:77905663-77905751 | LOC100130522;LOC100130 | S_Shore | TSS200;TSS200 |
| GM02175 | chr18:77917615-77918142 | LOC100130522;LOC100130 | N_Shore | Body;Body;3'UTR |
| WG1977 | chr19:11289349-11289376 | KANK2;KANK2 | S_Shore | Body;Body |
| GM02641 | chr19:11289356-11289376 | KANK2;KANK2 | S_Shore | Body;Body |
| WG2194 | chr19:12876846-12877000 | HOOK2;HOOK2 | N_Shore | Body;Body |
| WG1983 | chr19:12876846-12877188 | HOOK2;HOOK2 | N_Shore | Body;Body |
| GM00935 | chr19:1466647-1467032 | APC2 | Island | Body |
| GM00520 | chr19:16178091-16178099 | TPM4 | N_Shore | TSS1500 |
| WG1513 | chr19:1854549-1854724 | KLF16 | Island | Body |
| GM00042 | chr19:19626576-19626605 | NDUFA13;TSSK6 | Island | TSS1500;TSS200 |
| GM00438 | chr19:19639553-19639596 | YJEFN3 | Island | TSS200 |
| GM00520 | chr19:19639568-19639596 | YJEFN3 | Island | TSS200 |
| GM09256 | chr19:21264896-21265421 | ZNF714 | N_Shore | TSS200 |
| GM02456 | chr19:21265264-21265421 | ZNF714 | Island | 5'UTR |
| GM00520 | chr19:2282439-2282729 | C19orf35 | S_Shelf | TSS1500 |
| GM09256 | chr19:31640642-31640711 | DKFZp566F0947 | | TSS200 |
| GM01393 | chr19:39695011-39695065 | SYCN | S_Shore | TSS200 |
| GM00935 | chr19:40904691-40904974 | PRX;PRX | Island | Body;Body |
| GM00935 | chr19:427022-427263 | SHC2 | Island | Body |
| GM00289 | chr19:45260501-45261004 | BCL3 | Island | Body |
| WG2120 | chr19:48947560-48947578 | GRIN2D;GRWD1 | Island | 3'UTR;TSS1500 |
| GM02175 | chr19:49891270-49891574 | CCDC155 | Island | TSS1500 |
| GM02555 | chr19:52264411-52264429 | FPR2 | | TSS200 |
| GM02149 | chr19:54876728-54876752 | LAIR1;LAIR1 | | TSS200;TSS200 |
| GM00439 | chr19:827776-827843 | AZU1 | | TSS200 |
| WG1977 | chr2:10567628-10567843 | HPCAL1;HPCAL1 | S_Shore | 3'UTR;3'UTR |
| GM01096 | chr2:113992843-113992930 | PAX8;PAX8;PAX8;PAX8;PAX8 | N_Shore | Body;Body;Body;Body;Body;Body;TSS |
| GM02173 | chr2:113992843-113993052 | PAX8;PAX8;PAX8;PAX8;PAX8 | N_Shore | Body;Body;Body;Body;Body;Body;TSS |
| GM02555 | chr2:114033490-114033830 | PAX8;PAX8;PAX8;PAX8;PAX8 | Island | Body;Body;Body;Body;Body;Body |
| WG1977 | chr2:121554810-121554926 | GLI2 | | TSS200 |
| GM01375 | chr2:121554817-121554926 | GLI2 | | TSS200 |
| GM09256 | chr2:142889283-142889392 | LRP1B | S_Shore | TSS200 |
| GM00022 | chr2:142889367-142889392 | LRP1B | S_Shore | TSS200 |
| GM02175 | chr2:171676553-171676925 | GAD1;GAD1 | Island | Body;Body |
| WG2121 | chr2:176964512-176964720 | HOXD12 | Island | TSS200 |
| GM00935 | chr2:176973275-176974060 | HOXD11 | N_Shore | Body |
| WG1514 | chr2:176973688-176973835 | HOXD11 | Island | Body |
| GM02173 | chr2:176982108-176982342 | HOXD10 | Island | 1stExon |
| GM09256 | chr2:176993017-176993115 | HOXD8 | Island | TSS1500 |
| GM02175 | chr2:177016167-177017449 | HOXD4;HOXD4 | N_Shore | 5'UTR;1stExon |
| GM00520 | chr2:200320455-200321348 | SATB2 | N_Shore | Body |
| GM02175 | chr2:220406470-220406756 | CHPF | Island | Body |
| GM00439 | chr2:223155925-223156136 | PAX3;PAX3;PAX3;PAX3;PAX3 | Island | Body;Body;Body;Body;Body;Body |
| GM01650 | chr2:223161771-223162049 | PAX3;PAX3;PAX3;CCDC140;I | Island | Body;Body;Body;TSS1500;Body;Body; |
| GM02175 | chr2:223161771-223162128 | PAX3;PAX3;PAX3;CCDC140;I | Island | Body;Body;Body;TSS1500;Body;Body; |
| GM02641 | chr2:223162666-223162800 | PAX3;PAX3;PAX3;PAX3;PAX3 | N_Shore | Body;Body;Body;Body;Body;Body;Bod |
| WG1283 | chr2:23746926-23747197 | KLHL29 | N_Shelf | 5'UTR |

| | | | | |
|---|---|---|---|---|
| GM00520 | chr2:238394933-238395125 | MLPH;MLPH | N_Shore | TSS1500;TSS1500 |
| GM00289 | chr2:238647755-238647913 | LRRFIP1;LRRFIP1;LRRFIP1;LF | S_Shelf | Body;Body;Body;Body;Body |
| WG1977 | chr2:240020028-240020174 | HDAC4 | | Body |
| GM02173 | chr2:240233043-240233240 | HDAC4 | | Body |
| GM02555 | chr2:240291426-240291657 | HDAC4 | | 5'UTR |
| GM00289 | chr2:25391830-25391911 | POMC;POMC | Island | TSS1500;TSS1500 |
| WG1984 | chr2:30669597-30669863 | LCLAT1;LCLAT1 | N_Shore | TSS1500;TSS1500 |
| WG2120 | chr2:3286338-3286479 | TSSC1 | Island | Body |
| WG1978 | chr2:3452347-3452624 | TTC15 | Island | Body |
| GM01376 | chr2:3642400-3642634 | COLEC11;COLEC11 | N_Shore | TSS1500;TSS1500 |
| GM09255 | chr2:3714936-3715083 | ALLC | | 5'UTR |
| GM03075 | chr2:54087173-54087189 | LOC100302652;GPR75 | Island | TSS200;TSS200 |
| GM01375 | chr2:54087178-54087189 | LOC100302652;GPR75 | Island | TSS200;TSS200 |
| GM02175 | chr2:66671478-66671727 | MEIS1 | N_Shore | Body |
| GM01096 | chr2:71134524-71134630 | VAX2 | N_Shore | Body |
| GM01375 | chr2:74780268-74780617 | LOXL3 | N_Shore | 5'UTR |
| GM00289 | chr2:947167-947589 | SNTG2 | Island | Body |
| GM00520 | chr2:947515-947634 | SNTG2 | S_Shore | Body |
| GM02456 | chr20:25128770-25128805 | LOC284798;LOC284798;LO | Island | TSS200;Body;Body |
| GM00438 | chr20:25128770-25128807 | LOC284798;LOC284798;LO | Island | TSS200;Body;Body |
| GM01376 | chr20:32856747-32856898 | ASIP | Island | Body |
| GM00022 | chr20:35169539-35169609 | MYL9;MYL9 | Island | TSS1500;TSS1500 |
| GM00439 | chr20:36932397-36932553 | BPI | | TSS200 |
| GM00289 | chr20:43935460-43935471 | MATN4;MATN4;RBPJL;MATN4 | Island | 5'UTR;5'UTR;TSS200;5'UTR |
| GM01096 | chr20:47896448-47896888 | SNORD12B;C20orf199;MIR1 | S_Shore | TSS1500;Body;TSS1500;Body;TSS150 |
| GM00439 | chr20:49253364-49253710 | FAM65C;FAM65C | | 1stExon;5'UTR |
| GM02316 | chr20:5485144-5485284 | LOC149837 | | Body |
| GM01095 | chr20:5485245-5485284 | LOC149837 | | TSS200 |
| GM01096 | chr20:55925570-55925603 | RAE1;RAE1 | N_Shore | TSS1500;TSS1500 |
| GM00520 | chr20:61447330-61447394 | COL9A3 | N_Shore | TSS1500 |
| GM01653 | chr21:35832141-35832180 | KCNE1;KCNE1;KCNE1 | Island | 5'UTR;TSS1500;TSS1500 |
| GM00439 | chr21:36421467-36421503 | RUNX1;RUNX1 | | 5'UTR;1stExon |
| GM00520 | chr21:40145923-40146046 | NCRNA00114 | | TSS1500 |
| GM00520 | chr21:45148879-45149141 | PDXK | Island | Body |
| WG1722 | chr21:47604052-47604654 | C21orf56;C21orf56 | S_Shore | 5'UTR;5'UTR |
| GM00520 | chr22:19746984-19747140 | TBX1;TBX1;TBX1 | Island | 5'UTR;5'UTR;5'UTR |
| WG2275 | chr22:19754125-19754448 | TBX1;TBX1;TBX1 | Island | Body;Body;Body |
| WG1977 | chr22:22058080-22058271 | YPEL1 | Island | Body |
| GM00439 | chr22:24348549-24348715 | GSTTP1 | | TSS1500 |
| GM02456 | chr22:24384105-24384294 | GSTT1 | N_Shore | Body |
| GM02175 | chr22:24979554-24979588 | GGT1 | | TSS200 |
| GM02706 | chr22:32027580-32027605 | PISD | S_Shore | TSS1500 |
| GM00016 | chr22:42828411-42828418 | NFAM1 | | TSS200 |
| GM09256 | chr22:45609402-45609421 | C22orf9;C22orf9 | | Body;TSS1500 |
| GM00439 | chr22:50629038-50629224 | TRABD | Island | 5'UTR |
| GM01096 | chr3:11746183-11746765 | VGLL4 | N_Shore | 5'UTR |
| WG1513 | chr3:14443428-14443473 | SLC6A6;SLC6A6;SLC6A6 | N_Shore | TSS1500;TSS1500;TSS1500 |

| | | | | |
|---|---|---|---|---|
| WG1309 | chr3:15311021-15311269 | SH3BP5;SH3BP5 | | Body;Body |
| GM09256 | chr3:155394109-155394303 | PLCH1;PLCH1;PLCH1 | | 5'UTR;TSS200;TSS200 |
| GM00022 | chr3:157821262-157822072 | SHOX2;SHOX2;SHOX2 | Island | Body;Body;Body |
| GM01376 | chr3:160475035-160475128 | PPM1L | S_Shore | Body |
| GM02175 | chr3:183894727-183894778 | AP2M1;AP2M1 | S_Shore | 5'UTR;5'UTR |
| WG2414 | chr3:192289245-192289293 | FGF12 | | Body |
| GM00042 | chr3:194833887-194834008 | C3orf21 | | Body |
| WG1309 | chr3:196065357-196065506 | TM4SF19 | | TSS200 |
| WG1977 | chr3:36949957-36950004 | TRANK1 | | Body |
| GM00439 | chr3:42742993-42743203 | HHATL;HHATL | | Body;5'UTR |
| GM01096 | chr3:42947263-42947385 | ZNF662;ZNF662 | N_Shore | TSS1500;TSS200 |
| WG1977 | chr3:50488020-50488230 | CACNA2D2;CACNA2D2 | | Body;Body |
| GM00520 | chr3:56809796-56810320 | ARHGEF3;ARHGEF3;ARHGEF3 | | TSS200;Body;Body |
| GM00289 | chr3:64547108-64547373 | ADAMTS9 | | Body |
| GM02173 | chr3:64547310-64547373 | ADAMTS9 | | Body |
| GM02555 | chr4:111398042-111398146 | ENPEP | Island | 1stExon |
| GM00935 | chr4:119948360-119948542 | SYNPO2;SYNPO2;SYNPO2 | | Body;Body;Body |
| GM00031 | chr4:154711512-154711563 | SFRP2 | N_Shore | TSS1500 |
| WG1513 | chr4:174447847-174448341 | HAND2 | N_Shore | 3'UTR |
| GM00520 | chr4:40858984-40859293 | APBB2;APBB2;APBB2;APBB2 | Island | Body;5'UTR;5'UTR;5'UTR;Body;5'UTR; |
| GM00935 | chr4:5021268-5021328 | CYTL1 | Island | TSS200 |
| WG1722 | chr4:52918457-52918489 | SPATA18 | S_Shore | Body |
| GM09256 | chr4:54965829-54965904 | GSX2 | N_Shore | TSS1500 |
| WG1657 | chr4:55143269-55143557 | PDGFRA | | Body |
| GM00042 | chr4:57522762-57522881 | HOPX;HOPX;HOPX;HOPX;HC | S_Shore | 5'UTR;Body;Body;TSS200;TSS200 |
| GM09255 | chr4:6052527-6052554 | JAKMIP1 | | Body |
| WG1310 | chr4:71587722-71587904 | RUFY3;RUFY3;RUFY3;RUFY3;RUFY3 | | 1stExon;Body;5'UTR;1stExon;5'UTR |
| GM00290 | chr4:7282819-7283184 | SORCS2 | | Body |
| GM00940 | chr4:94124971-94125068 | GRID2 | | Body |
| GM00016 | chr5:102898463-102898733 | NUDT12;NUDT12 | | 5'UTR;1stExon |
| GM02175 | chr5:1053545-1053774 | SLC12A7 | Island | Body |
| GM01375 | chr5:1053643-1054262 | SLC12A7 | Island | Body |
| WG1514 | chr5:14406585-14406732 | TRIO | | Body |
| GM02456 | chr5:149980486-149980682 | SYNPO | | TSS200 |
| WG1514 | chr5:149980671-149980682 | SYNPO;SYNPO | | 1stExon;5'UTR |
| GM00935 | chr5:150004773-150004918 | SYNPO;SYNPO | Island | Body;Body |
| GM01095 | chr5:153858816-153858936 | HAND1 | Island | TSS1500 |
| GM00438 | chr5:156886990-156887005 | NIPAL4 | Island | TSS200 |
| GM00438 | chr5:1594282-1594863 | SDHAP3 | Island | Body |
| GM00042 | chr5:176057924-176057996 | EIF4E1B;SNCB;SNCB;EIF4E1 | S_Shore | 5'UTR;TSS1500;TSS1500;1stExon |
| GM00290 | chr5:176784688-176784719 | RGS14 | | TSS200 |
| GM01095 | chr5:177548398-177548913 | N4BP3 | | Body |
| WG2276 | chr5:177708304-177708528 | COL23A1 | | Body |
| GM02706 | chr5:191127-191296 | LRRC14B | N_Shore | TSS1500 |
| GM00439 | chr5:23507573-23507617 | PRDM9 | | TSS200 |
| WG1977 | chr5:323320-323969 | AHRR | S_Shore | Body |
| GM02149 | chr5:368804-369088 | AHRR | N_Shore | Body |

| | | | | |
|---|---|---|---|---|
| GM00439 | chr5:38257516-38257555 | EGFLAM | N_Shore | TSS1500 |
| WG1722 | chr5:40841493-40841590 | CARD6 | | 1stExon |
| GM00439 | chr5:423894-424371 | AHRR | S_Shelf | Body |
| GM00042 | chr5:42756786-42756860 | CCDC152 | | TSS200 |
| GM02706 | chr5:68710808-68711277 | MARVELD2;MARVELD2 | Island | TSS200;TSS200 |
| GM00520 | chr5:68710818-68711277 | MARVELD2;MARVELD2 | Island | TSS200;TSS200 |
| WG1978 | chr5:76010444-76010658 | F2R | N_Shore | TSS1500 |
| GM00520 | chr5:78281793-78281964 | ARSB;ARSB;ARSB | S_Shore | TSS200;1stExon;5'UTR |
| GM00290 | chr5:92908988-92909429 | FLJ42709;FLJ42709 | S_Shore | Body;Body |
| GM00520 | chr5:92908988-92909434 | FLJ42709;FLJ42709 | S_Shore | Body;Body |
| GM01096 | chr6:100911240-100911964 | SIM1 | N_Shore | 1stExon |
| GM02704 | chr6:100911240-100912072 | SIM1 | N_Shore | 1stExon |
| GM03075 | chr6:101846791-101846872 | GRIK2;GRIK2;GRIK2 | Island | TSS200;TSS200;TSS200 |
| WG1721 | chr6:109274429-109274496 | ARMC2 | | Body |
| GM00520 | chr6:11279656-11279727 | NEDD9 | | Body |
| GM09256 | chr6:139613145-139613456 | TXLNB;TXLNB | | 5'UTR;1stExon |
| GM02175 | chr6:149803087-149803444 | ZC3H12D | | 5'UTR |
| GM00935 | chr6:149805995-149806131 | ZC3H12D;ZC3H12D | | 5'UTR;1stExon |
| GM00290 | chr6:149806077-149806131 | ZC3H12D;ZC3H12D | | 5'UTR;1stExon |
| GM01096 | chr6:152085565-152085641 | ESR1 | | 5'UTR |
| GM00940 | chr6:152125861-152126080 | ESR1;ESR1 | N_Shelf | 5'UTR;TSS1500 |
| GM00438 | chr6:152125965-152126092 | ESR1;ESR1 | N_Shelf | 5'UTR;TSS1500 |
| WG1309 | chr6:152127812-152127887 | ESR1;ESR1;ESR1;ESR1 | N_Shore | TSS1500;5'UTR;TSS1500;5'UTR |
| GM01095 | chr6:152127812-152128024 | ESR1;ESR1;ESR1;ESR1 | N_Shore | TSS1500;5'UTR;TSS1500;5'UTR |
| GM00439 | chr6:158018055-158018184 | ZDHHC14;ZDHHC14 | S_Shelf | Body;Body |
| WG1977 | chr6:161796785-161796854 | PARK2;PARK2;PARK2 | | Body;Body;Body |
| WG1283 | chr6:161860203-161860359 | PARK2;PARK2;PARK2 | | Body;Body;Body |
| GM00042 | chr6:168435914-168436353 | KIF25;KIF25 | Island | Body;Body |
| GM00438 | chr6:21667166-21667407 | FLJ22536 | S_Shore | Body |
| GM00042 | chr6:24583477-24583614 | KIAA0319;KIAA0319;KIAA0319;KIAA0319 | | Body;Body;Body;Body;Body |
| GM02706 | chr6:291882-292823 | DUSP22 | N_Shore | TSS1500 |
| GM00289 | chr6:29595002-29595016 | GABBR1;GABBR1;GABBR1 | N_Shore | Body;Body;Body |
| GM02456 | chr6:29690766-29690893 | HLA-F;HLA-F;HLA-F | N_Shore | TSS1500;TSS1500;TSS1500 |
| GM02175 | chr6:29690822-29690893 | HLA-F;HLA-F;HLA-F | N_Shore | TSS1500;TSS1500;TSS1500 |
| WG1978 | chr6:29894050-29894197 | HCG4P6 | N_Shore | TSS1500 |
| GM02456 | chr6:29943401-29943414 | HCG9 | N_Shore | Body |
| WG1722 | chr6:29943401-29943425 | HCG9 | N_Shore | Body |
| GM00439 | chr6:29943401-29943455 | HCG9 | N_Shore | Body |
| GM00042 | chr6:29943455-29943480 | HCG9 | N_Shore | Body |
| GM00042 | chr6:29944093-29944146 | HCG9 | N_Shore | Body |
| GM03075 | chr6:30038712-30038754 | RNF39;RNF39 | N_Shore | 3'UTR;3'UTR |
| WG1722 | chr6:30039432-30039442 | RNF39;RNF39 | Island | Body;Body |
| GM01393 | chr6:30079256-30079280 | TRIM31 | | Body |
| GM00289 | chr6:30113978-30114011 | TRIM40 | | Body |
| GM00520 | chr6:30458158-30458237 | HLA-E | Island | Body |
| GM02456 | chr6:31025933-31025967 | HCG22 | | Body |
| WG2275 | chr6:31237013-31237034 | HLA-C | N_Shore | Body |

| | | | | |
|---|---|---|---|---|
| WG2276 | chr6:31237405-31237722 | HLA-C | N_Shore | Body |
| GM00438 | chr6:31760593-31760616 | VARS | N_Shelf | Body |
| GM01392 | chr6:31846769-31846813 | SLC44A4;SLC44A4 | | 5'UTR;1stExon |
| GM02706 | chr6:32055447-32055525 | TNXB | Island | Body |
| GM00042 | chr6:32097439-32097507 | ATF6B;ATF6B;FKBPL | N_Shore | TSS1500;TSS1500;Body |
| WG1283 | chr6:32489750-32489963 | HLA-DRB5 | Island | Body |
| GM00935 | chr6:32489991-32490043 | HLA-DRB5 | Island | Body |
| GM00290 | chr6:32526260-32526342 | HLA-DRB6 | | Body |
| GM02555 | chr6:32526260-32526364 | HLA-DRB6 | | Body |
| GM00439 | chr6:32551891-32551954 | HLA-DRB1 | Island | Body |
| WG2193 | chr6:32552152-32552246 | HLA-DRB1 | Island | Body |
| GM00438 | chr6:32552246-32552453 | HLA-DRB1 | Island | Body |
| GM00290 | chr6:32847762-32847788 | PPP1R2P1 | Island | Body |
| GM00940 | chr6:32908567-32908642 | HLA-DMB | | 1stExon |
| GM00022 | chr6:33288366-33288569 | DAXX;DAXX;DAXX;DAXX | N_Shore | Body;Body;Body;Body |
| GM01096 | chr6:33288599-33288656 | DAXX;DAXX;DAXX;DAXX | N_Shore | Body;Body;Body;Body |
| GM00290 | chr6:34024362-34024479 | GRM4 | Island | Body |
| GM00290 | chr6:35265616-35265643 | DEF6;DEF6 | | 1stExon;5'UTR |
| GM03073 | chr6:6002865-6003303 | NRN1 | Island | Body |
| GM00439 | chr6:6614447-6614798 | LOC285780;LY86 | | Body;Body |
| GM00520 | chr6:70576168-70576321 | COL19A1 | N_Shore | TSS1500 |
| GM02175 | chr7:101500014-101500514 | CUX1;CUX1;CUX1 | | Body;Body;Body |
| GM00439 | chr7:101768610-101768874 | CUX1;CUX1;CUX1 | | Body;Body;Body |
| GM01375 | chr7:114055123-114055137 | FOXP2;FOXP2;FOXP2;FOXP2;FOXP2;FOX | | 1stExon;5'UTR;5'UTR;1stExon;5'UTR;1 |
| WG2121 | chr7:114055123-114055419 | FOXP2;FOXP2;FOXP2;FOXP2;FOXP2;FOX | | 1stExon;5'UTR;5'UTR;1stExon;5'UTR;1 |
| GM02175 | chr7:115862648-115863026 | TES;TES | | TSS1500;Body |
| GM00439 | chr7:128550860-128550902 | KCP;KCP | Island | TSS200;TSS200 |
| GM00439 | chr7:157198482-157198869 | DNAJB6 | N_Shelf | Body |
| GM09256 | chr7:158336016-158336049 | PTPRN2;PTPRN2;PTPRN2 | | Body;Body;Body |
| GM02555 | chr7:1952518-1952807 | MAD1L1;MAD1L1;MAD1L1 | S_Shelf | Body;Body;Body |
| GM02317 | chr7:2109244-2109890 | MAD1L1;MAD1L1;MAD1L1 | S_Shore | Body;Body;Body |
| GM01375 | chr7:2133441-2133480 | MAD1L1;MAD1L1;MAD1L1 | | Body;Body;Body |
| GM02173 | chr7:2139216-2139259 | MAD1L1;MAD1L1;MAD1L1 | | Body;Body;Body |
| GM02456 | chr7:2563611-2563909 | LFNG;LFNG;LFNG;LFNG | Island | Body;Body;Body;Body |
| GM00520 | chr7:27182493-27183196 | HOXA5 | N_Shore | Body |
| GM00042 | chr7:27183274-27183436 | HOXA5;HOXA5 | Island | 5'UTR;1stExon |
| GM00438 | chr7:27187943-27188770 | HOXA6 | S_Shore | TSS1500 |
| GM00438 | chr7:27225070-27225222 | HOXA11AS;HOXA11 | Island | Body;TSS1500 |
| GM00940 | chr7:27225299-27225342 | HOXA11AS;HOXA11 | Island | Body;TSS1500 |
| GM02456 | chr7:27225523-27225543 | HOXA11AS;HOXA11 | Island | Body;TSS1500 |
| GM00022 | chr7:27281315-27281346 | EVX1 | N_Shore | TSS1500 |
| GM00289 | chr7:2800701-2801248 | GNA12 | N_Shore | Body |
| WG2276 | chr7:2847517-2847575 | GNA12 | | Body |
| GM02555 | chr7:28998577-28998860 | TRIL | S_Shore | TSS1500 |
| GM09256 | chr7:4065931-4066127 | SDK1 | | Body |
| GM00520 | chr7:42267547-42268109 | GLI3 | Island | 5'UTR |
| GM00022 | chr7:43288635-43289504 | HECW1 | Island | Body |

| | | | | |
|---|---|---|---|---|
| GM02175 | chr7:4885521-4885643 | RADIL | | Body |
| GM02704 | chr7:5336543-5336811 | SLC29A4;SLC29A4 | Island | Body;Body |
| GM00022 | chr7:5518392-5518918 | FBXL18 | N_Shelf | 3'UTR |
| GM00940 | chr7:553412-553693 | PDGFA;PDGFA | N_Shelf | Body;Body |
| WG1658 | chr7:70096029-70096378 | AUTS2;AUTS2 | N_Shore | Body;Body |
| WG1722 | chr7:944607-944776 | ADAP1 | S_Shore | Body |
| GM00935 | chr8:104032819-104032876 | ATP6V1C1 | N_Shore | TSS1500 |
| WG1513 | chr8:10586959-10587055 | SOX7 | N_Shore | Body |
| GM02173 | chr8:11141365-11141424 | MTMR9 | N_Shore | TSS1500 |
| GM01392 | chr8:116660311-116661063 | TRPS1 | N_Shore | 5'UTR |
| GM01096 | chr8:143545390-143545574 | BAI1;BAI1 | N_Shore | 5'UTR;1stExon |
| GM00520 | chr8:1847999-1848143 | ARHGEF10 | | Body |
| GM00290 | chr8:1847999-1848407 | ARHGEF10 | | Body |
| GM02704 | chr8:1848040-1848407 | ARHGEF10 | | Body |
| GM00439 | chr8:19539991-19540272 | CSGALNACT1;CSGALNACT1;CSGALNAC | | Body;5'UTR;1stExon |
| GM09255 | chr8:2075469-2075820 | MYOM2 | Island | Body |
| GM02173 | chr8:22132874-22132959 | PIWIL2;PIWIL2;PIWIL2 | Island | 5'UTR;1stExon;TSS1500 |
| GM02706 | chr8:28560467-28560723 | EXTL3 | S_Shore | 5'UTR |
| GM03075 | chr8:40010443-40010823 | C8orf4 | | TSS1500 |
| GM00520 | chr8:42356637-42356871 | SLC20A2 | | 5'UTR |
| WG1722 | chr8:59058585-59058660 | FAM110B | Island | 5'UTR |
| GM00439 | chr8:6456182-6456465 | MCPH1 | | Body |
| GM01095 | chr8:72754469-72755162 | MSC | Island | 3'UTR |
| GM00042 | chr8:95962371-95962463 | TP53INP1;TP53INP1 | Island | TSS1500;TSS1500 |
| GM02640 | chr8:9952926-9953022 | MSRA;MSRA;MSRA | | Body;TSS200;Body |
| GM00439 | chr9:139237461-139237889 | GPSM1 | Island | Body |
| GM00438 | chr9:140401382-140401455 | PNPLA7;PNPLA7 | Island | Body;Body |
| GM02317 | chr9:140401382-140401576 | PNPLA7;PNPLA7 | Island | Body;Body |
| GM01650 | chr9:92220617-92221282 | GADD45G | Island | Body |
| WG1514 | chr1:108734992-108735549 | SLC25A24;SLC25A24 | N_Shore | Body;Body |
| GM00031 | chr1:110254662-110254835 | GSTM5 | | TSS1500 |
| GM01392 | chr1:111218079-111218282 | KCNA3 | S_Shore | TSS1500 |
| GM02173 | chr1:115238457-115238559 | AMPD1 | | TSS1500 |
| GM01096 | chr1:115881044-115881132 | NGF | Island | TSS200 |
| WG2194 | chr1:119526060-119526437 | TBX15 | N_Shore | 5'UTR |
| GM09255 | chr1:119526060-119526783 | TBX15 | N_Shore | 5'UTR |
| GM00022 | chr1:119531625-119532925 | TBX15 | N_Shore | 5'UTR |
| WG1513 | chr1:119531718-119532044 | TBX15 | N_Shore | 5'UTR |
| WG2194 | chr1:119532320-119532542 | TBX15 | S_Shore | TSS200 |
| GM01650 | chr1:119532736-119532925 | TBX15 | S_Shore | TSS1500 |
| GM01653 | chr1:14027521-14027581 | PRDM2 | S_Shore | 5'UTR |
| GM00439 | chr1:151811923-151812435 | LOC100132111;C2CD4D | N_Shore | Body;5'UTR |
| WG2121 | chr1:15256054-15256222 | KIAA1026;KIAA1026;KIAA1026;KIAA1026 | | Body;TSS1500;Body;Body |
| GM00290 | chr1:15256123-15256136 | KIAA1026;KIAA1026;KIAA1026;KIAA1026 | | Body;TSS200;Body;Body |
| GM02704 | chr1:153581494-153581793 | S100A16 | | 5'UTR |
| GM02706 | chr1:154127952-154128041 | TPM3;TPM3;NUP210L;TPM3;TPM3;NUP2 | | 3'UTR;3'UTR;TSS1500;3'UTR;3'UTR;TS |
| GM00935 | chr1:156786601-156786653 | SH2D2A;NTRK1;SH2D2A;SH2 | S_Shelf | 5'UTR;Body;1stExon;TSS1500;5'UTR;5 |

| | | | | |
|---|---|---|---|---|
| GM01650 | chr1:159796617-159796658 | SLAMF8;SLAMF8 | | 5'UTR;1stExon |
| WG1984 | chr1:161008705-161008826 | TSTD1;TSTD1;TSTD1;TSTD1;T | Island | 5'UTR;1stExon;5'UTR;1stExon;1stExon |
| GM00439 | chr1:161049443-161049579 | PVRL4 | Island | Body |
| GM00016 | chr1:165324100-165324810 | LMX1A | N_Shore | Body |
| WG1514 | chr1:170638807-170639289 | PRRX1;PRRX1 | | Body;Body |
| GM09256 | chr1:173176362-173176523 | TNFSF4;TNFSF4 | | 5'UTR;1stExon |
| GM01376 | chr1:173176366-173176523 | TNFSF4;TNFSF4 | | 5'UTR;1stExon |
| GM01376 | chr1:17944356-17944660 | ARHGEF10L;ARHGEF10L | | Body;Body |
| GM02555 | chr1:182921851-182922263 | C1orf14 | Island | Body |
| GM02704 | chr1:19600673-19600730 | AKR7L;AKR7L | Island | TSS200;TSS200 |
| GM02175 | chr1:2005260-2005564 | PRKCZ;PRKCZ;PRKCZ | Island | 1stExon;Body;5'UTR |
| GM00290 | chr1:201709336-201709675 | NAV1;NAV1 | Island | Body;Body |
| GM02706 | chr1:202091865-202091880 | GPR37L1 | | TSS200 |
| GM00042 | chr1:2046702-2046928 | PRKCZ;PRKCZ;PRKCZ | Island | 5'UTR;Body;5'UTR |
| GM00940 | chr1:2046783-2046928 | PRKCZ;PRKCZ;PRKCZ | Island | 5'UTR;Body;5'UTR |
| WG1658 | chr1:205818956-205819463 | PM20D1 | Island | Body |
| WG1657 | chr1:205818956-205819492 | PM20D1 | Island | Body |
| GM00016 | chr1:205818956-205819609 | PM20D1 | Island | Body |
| GM01096 | chr1:20617283-20617456 | VWA5B1 | | TSS200 |
| WG1658 | chr1:207992403-207992548 | LOC148696 | | Body |
| GM09256 | chr1:20988590-20988763 | DDOST | S_Shore | TSS1500 |
| WG1283 | chr1:2120985-2121724 | C1orf86;LOC100128003 | N_Shore | Body;Body |
| GM00042 | chr1:214477063-214477280 | SMYD2 | | Body |
| GM02555 | chr1:21586831-21587174 | ECE1;ECE1;ECE1;ECE1 | | Body;Body;Body;Body |
| GM00520 | chr1:217309568-217309970 | ESRRG | N_Shore | 5'UTR |
| GM00520 | chr1:221051758-221052023 | HLX | N_Shore | TSS1500 |
| WG1513 | chr1:221053841-221055097 | HLX | N_Shore | Body |
| GM09255 | chr1:221053904-221054273 | HLX | N_Shore | Body |
| GM02147 | chr1:221053904-221055097 | HLX | N_Shore | Body |
| WG2120 | chr1:221054615-221055097 | HLX | Island | Body |
| GM02706 | chr1:223566643-223566669 | C1orf65 | Island | TSS200 |
| WG1513 | chr1:223566643-223566709 | C1orf65 | Island | TSS200 |
| GM02149 | chr1:223936508-223936812 | CAPN2;CAPN2 | Island | Body;Body |
| WG1513 | chr1:226825435-226825606 | ITPKB | | Body |
| GM01650 | chr1:228291668-228291705 | C1orf35 | S_Shore | TSS1500 |
| WG1978 | chr1:23521272-23521417 | HTR1D | | TSS200 |
| GM00042 | chr1:236925877-236925965 | ACTN2 | | Body |
| WG2414 | chr1:240776162-240776211 | GREM2 | | TSS1500 |
| GM00290 | chr1:241912510-241912768 | WDR64 | | Body |
| GM02706 | chr1:2425860-2426035 | PLCH2 | | Body |
| GM00935 | chr1:243646402-243646787 | SDCCAG8 | Island | Body |
| GM02555 | chr1:247694271-247694406 | LOC148824;OR2C3 | Island | TSS200;3'UTR |
| GM09255 | chr1:25944712-25945042 | MAN1C1 | Island | 1stExon |
| GM00439 | chr1:3028230-3028485 | PRDM16;PRDM16 | Island | Body;Body |
| GM00520 | chr1:32714146-32714308 | FAM167B | Island | Body |
| GM02149 | chr1:3277525-3277623 | PRDM16;PRDM16 | S_Shelf | Body;Body |
| WG1722 | chr1:3459949-3460268 | MEGF6 | Island | Body |

| | | | | |
|---|---|---|---|---|
| GM09255 | chr1:36039655-36039885 | TFAP2E | Island | Body |
| GM00940 | chr1:40024971-40025415 | LOC728448 | | Body |
| WG2415 | chr1:40105667-40105706 | HEYL | Island | TSS1500 |
| GM02175 | chr1:41250062-41250117 | KCNQ4;KCNQ4 | Island | 1stExon;1stExon |
| WG1977 | chr1:43751363-43751376 | C1orf210;C1orf210 | | TSS200;TSS200 |
| GM09256 | chr1:44016014-44016049 | PTPRF;PTPRF | | Body;Body |
| GM00290 | chr1:44703750-44704073 | ERI3 | | Body |
| GM02149 | chr1:4794912-4795114 | AJAP1;AJAP1 | | Body;Body |
| GM00438 | chr1:53600741-53600889 | SLC1A7 | | Body |
| WG1283 | chr1:54199825-54200115 | GLIS1;GLIS1 | N_Shelf | 5'UTR;1stExon |
| WG1722 | chr1:54199825-54200420 | GLIS1;GLIS1 | N_Shelf | 5'UTR;1stExon |
| GM02175 | chr1:54200078-54200420 | GLIS1 | N_Shelf | TSS1500 |
| GM02175 | chr1:54856177-54856511 | SSBP3;SSBP3;SSBP3 | N_Shore | Body;Body;Body |
| GM02706 | chr1:6085995-6086148 | KCNAB2;KCNAB2 | N_Shore | TSS1500;TSS1500 |
| GM00940 | chr1:63249197-63249213 | ATG4C;ATG4C | | TSS1500;TSS1500 |
| GM01650 | chr1:63249199-63249213 | ATG4C;ATG4C | | TSS1500;TSS1500 |
| GM02175 | chr1:6526182-6526319 | TNFRSF25;TNFRSF25;PLEKH | Island | 1stExon;1stExon;3'UTR;3'UTR;5'UTR;5 |
| GM02706 | chr1:6558085-6558113 | PLEKHG5;PLEKHG5 | S_Shore | Body;TSS1500 |
| GM00289 | chr1:8403631-8403925 | SLC45A1 | N_Shore | Body |
| WG1722 | chr1:89664260-89664582 | GBP4 | | Body |
| GM00940 | chr1:9131622-9131783 | SLC2A5 | Island | Body |
| GM00940 | chr10:106088702-106088755 | ITPRIP | | 5'UTR |
| WG2415 | chr10:114877616-114877689 | TCF7L2;TCF7L2;TCF7L2;TCF7L2;TCF7L2 | | Body;Body;Body;Body;Body;Body |
| GM01653 | chr10:119295400-119296182 | EMX2OS | Island | Body |
| GM02456 | chr10:121265587-121265893 | RGS10;RGS10 | | Body;Body |
| GM00289 | chr10:1248770-1248818 | ADARB2 | N_Shelf | Body |
| WG1722 | chr10:126278272-126278502 | LHPP;LHPP | N_Shelf | Body;Body |
| GM00438 | chr10:126278372-126278502 | LHPP;LHPP | Island | Body;Body |
| GM01650 | chr10:12648032-12648338 | CAMK1D;CAMK1D | | Body;Body |
| GM00022 | chr10:12648176-12648526 | CAMK1D;CAMK1D | | Body;Body |
| WG1283 | chr10:131669406-131669630 | EBF3 | S_Shore | Body |
| WG1283 | chr10:131697080-131697345 | EBF3 | N_Shore | Body |
| GM00520 | chr10:131761206-131761386 | EBF3 | Island | Body |
| GM02149 | chr10:132891319-132891371 | TCERG1L | N_Shelf | 3'UTR |
| GM00935 | chr10:133995286-133995387 | JAKMIP3 | | 3'UTR |
| WG2193 | chr10:135341528-135342218 | CYP2E1 | N_Shore | 1stExon |
| GM00439 | chr10:135379213-135379242 | SYCE1;SYCE1;SYCE1 | N_Shore | Body;5'UTR;Body |
| GM00439 | chr10:13749193-13749625 | FRMD4A | | Body |
| WG1977 | chr10:13932842-13933247 | FRMD4A | N_Shore | Body |
| GM02706 | chr10:1767482-1767511 | ADARB2 | | Body |
| GM02149 | chr10:1769480-1769577 | ADARB2 | | Body |
| GM00940 | chr10:22605982-22606072 | COMMD3 | S_Shore | Body |
| GM02173 | chr10:26855907-26856105 | APBB1IP | Island | Body |
| GM00520 | chr10:28035731-28036151 | MKX | S_Shore | TSS1500 |
| GM00520 | chr10:29811530-29811659 | SVIL;SVIL | | Body;Body |
| WG1978 | chr10:3138418-3138534 | PFKP | | Body |
| GM00935 | chr10:384506-384652 | DIP2C | N_Shelf | Body |

| | | | | |
|---|---|---|---|---|
| GM09256 | chr10:45495739-45495981 | C10orf25;ZNF22 | N_Shore | 3'UTR;TSS1500 |
| GM00289 | chr10:45495782-45495981 | C10orf25;ZNF22 | N_Shore | 3'UTR;TSS1500 |
| WG1310 | chr10:46993502-46993542 | GPRIN2 | Island | TSS1500 |
| GM01375 | chr10:4868328-4868333 | AKR1E2 | Island | TSS200 |
| GM02175 | chr10:50506748-50506896 | C10orf71;C10orf71 | | TSS1500;TSS1500 |
| GM00290 | chr10:50506875-50506896 | C10orf71;C10orf71 | | TSS1500;TSS1500 |
| GM00520 | chr10:556470-556762 | DIP2C | | Body |
| GM00520 | chr10:62671136-62671228 | RHOBTB1;RHOBTB1;RHOBTB1;RHOBTB1 | | Body;Body;Body;Body |
| GM00022 | chr10:661009-661159 | DIP2C | | Body |
| GM00520 | chr10:661009-661730 | DIP2C | | Body |
| GM01096 | chr10:682125-682152 | DIP2C | N_Shelf | Body |
| WG1658 | chr10:72254314-72254335 | KIAA1274 | S_Shelf | 5'UTR |
| WG2276 | chr10:72311914-72312093 | KIAA1274 | | Body |
| GM00935 | chr10:73497514-73497629 | C10orf105;C10orf105;CDH23 | | 5'UTR;Body |
| WG1513 | chr10:73848764-73848823 | SPOCK2;SPOCK2;SPOCK2;S | S_Shore | 5'UTR;5'UTR |
| GM01392 | chr10:73976790-73976821 | ASCC1;C10orf104 | S_Shore | TSS1500;5'UTR |
| GM00439 | chr10:77871958-77872170 | C10orf11 | N_Shore | Body |
| WG1977 | chr10:82167757-82167774 | C10orf58 | N_Shelf | |
| GM02456 | chr10:90712684-90712948 | ACTA2;ACTA2 | | 5'UTR;5'UTR |
| GM00438 | chr10:95848649-95848841 | PLCE1;PLCE1 | | Body;TSS1500 |
| WG1513 | chr10:97175272-97175479 | SORBS1;SORBS1;SORBS1;SORBS1;SORI | | Body;Body;Body;Body;Body;Body;Bod |
| GM02555 | chr10:99338056-99338240 | ANKRD2;ANKRD2 | N_Shore | Body;Body |
| GM00940 | chr10:99735010-99735202 | CRTAC1 | | Body |
| GM00520 | chr10:99894013-99894087 | C10orf28 | N_Shelf | |
| WG1658 | chr11:10325567-10325809 | ADM | N_Shore | TSS1500 |
| GM00022 | chr11:109292789-109292894 | C11orf87 | N_Shore | TSS200 |
| GM00042 | chr11:111250129-111250201 | POU2AF1;POU2AF1 | | 1stExon;5'UTR |
| GM02704 | chr11:12031093-12031099 | DKK3;DKK3;DKK3 | S_Shore | TSS1500;TSS200;TSS1500 |
| GM00031 | chr11:124767455-124767840 | ROBO4 | | Body |
| GM09256 | chr11:125037226-125037348 | PKNOX2 | S_Shore | 5'UTR |
| GM00935 | chr11:128646953-128647107 | FLI1;FLI1 | | Body;Body |
| GM00520 | chr11:128729883-128730011 | KCNJ1;KCNJ1;KCNJ1;KCNJ1 | | 5'UTR;5'UTR;5'UTR;5'UTR |
| GM09256 | chr11:128812978-128813008 | TP53AIP1 | | TSS200 |
| GM00935 | chr11:129991445-129992368 | APLP2;APLP2;APLP2;APLP2;A | N_Shelf | Body;Body;Body;Body;Body;Body |
| WG1310 | chr11:129992297-129992368 | APLP2;APLP2;APLP2;APLP2;A | N_Shore | Body;Body;Body;Body;Body;Body |
| GM00290 | chr11:130319691-130320219 | ADAMTS15 | Island | 1stExon |
| WG1310 | chr11:133994930-133995182 | JAM3 | Island | Body |
| GM02555 | chr11:1750302-1750381 | HCCA2 | | Body |
| WG2415 | chr11:18433500-18433564 | LDHC;LDHC | | TSS1500;TSS1500 |
| GM00439 | chr11:18477280-18477534 | LDHAL6A;LDHAL6A | N_Shore | TSS200;TSS200 |
| GM00520 | chr11:18477299-18477534 | LDHAL6A;LDHAL6A | Island | TSS200;TSS200 |
| GM02706 | chr11:2464822-2464851 | KCNQ1 | N_Shore | TSS1500 |
| GM02317 | chr11:31819444-31819678 | PAX6;PAX6;PAX6 | N_Shore | Body;Body;Body |
| GM02456 | chr11:31821590-31822302 | PAX6;PAX6;PAX6 | S_Shore | Body;Body;Body |
| GM00520 | chr11:31821764-31822302 | PAX6;PAX6;PAX6 | S_Shore | Body;Body;Body |
| GM02555 | chr11:31824262-31825226 | PAX6;PAX6;PAX6 | N_Shore | Body;Body;Body |
| GM00289 | chr11:31825166-31825226 | PAX6;PAX6;PAX6 | N_Shore | Body;Body;Body |

| | | | | |
|---|---|---|---|---|
| GM01095 | chr11:31825756-31825833 | PAX6;PAX6;PAX6 | Island | Body;Body;Body |
| GM01096 | chr11:31827084-31827757 | PAX6;PAX6;PAX6 | N_Shore | Body;Body;Body |
| GM00016 | chr11:31827084-31827920 | PAX6;PAX6;PAX6 | N_Shore | Body;Body;Body |
| GM02317 | chr11:31827084-31828040 | PAX6;PAX6;PAX6 | N_Shore | Body;Body;Body |
| GM00042 | chr11:31827899-31828040 | PAX6;PAX6;PAX6 | Island | Body;Body;Body |
| GM00940 | chr11:31837104-31837144 | PAX6 | Island | 5'UTR |
| WG1721 | chr11:3253838-3254065 | MRGPRE | Island | TSS1500 |
| GM00520 | chr11:44327267-44327572 | ALX4 | Island | Body |
| GM00438 | chr11:44332624-44332685 | ALX4 | Island | TSS1500 |
| GM00935 | chr11:44332636-44332664 | ALX4 | Island | TSS1500 |
| GM00438 | chr11:60282420-60282742 | MS4A13;MS4A13 | N_Shore | TSS1500;TSS1500 |
| GM01096 | chr11:60719947-60720229 | SLC15A3;SLC15A3 | S_Shore | TSS1500;TSS1500 |
| GM02706 | chr11:628603-628677 | SCT | S_Shore | TSS1500 |
| GM01095 | chr11:63775226-63775436 | MACROD1 | N_Shore | Body |
| GM00520 | chr11:66650500-66650560 | PC;PC;PC | | 5'UTR;5'UTR;5'UTR |
| GM02706 | chr11:66885240-66885281 | KDM2A;KDM2A | N_Shore | TSS1500;TSS1500 |
| GM00935 | chr11:68138505-68138908 | LRP5 | | Body |
| GM09256 | chr11:68147880-68148079 | LRP5 | | Body |
| GM02175 | chr11:68610706-68611016 | CPT1A;CPT1A | N_Shore | TSS1500;TSS1500 |
| GM00042 | chr11:68610706-68611260 | CPT1A;CPT1A | N_Shore | TSS1500;TSS1500 |
| GM02555 | chr11:68782049-68782202 | MRGPRF;MRGPRF | S_Shelf | TSS1500;TSS1500 |
| GM00438 | chr11:68782049-68782211 | MRGPRF;MRGPRF | S_Shelf | TSS1500;TSS1500 |
| GM00290 | chr11:70048796-70048852 | FADD | N_Shore | TSS1500 |
| GM02641 | chr11:74178761-74178800 | KCNE3 | Island | TSS200 |
| GM00520 | chr11:74178795-74178883 | KCNE3 | Island | TSS200 |
| GM00289 | chr11:75898132-75898203 | WNT11 | | Body |
| WG1721 | chr11:77299805-77299962 | AQP11 | N_Shore | TSS1500 |
| GM02706 | chr11:85430629-85430657 | SYTL2;SYTL2;SYTL2;SYTL2;SYTL2;SYTL2; | | Body;TSS1500;TSS1500;Body;TSS150 |
| WG1978 | chr11:86383696-86383723 | ME3;ME3;ME3 | S_Shore | TSS1500;TSS200;TSS1500 |
| GM00520 | chr11:86383696-86383761 | ME3;ME3;ME3 | S_Shore | TSS1500;TSS200;TSS1500 |
| GM02456 | chr11:87908785-87908938 | RAB38 | S_Shore | TSS200 |
| GM00520 | chr11:93277161-93277191 | C11orf75 | S_Shore | TSS1500 |
| GM02555 | chr12:10084936-10085174 | CLEC2A;CLEC2A | | 5'UTR;1stExon |
| GM02456 | chr12:10251536-10251638 | CLEC1A;CLEC1A | | 5'UTR;1stExon |
| GM00520 | chr12:10251536-10251879 | CLEC1A;CLEC1A | | 5'UTR;1stExon |
| GM09256 | chr12:106988340-106988413 | RFX4 | | Body |
| GM01096 | chr12:107348824-107348944 | C12orf23 | N_Shore | TSS1500 |
| GM00289 | chr12:110172208-110172346 | MGC14436;MGC14436;C12orf34 | | Body;Body;5'UTR |
| GM00022 | chr12:11151478-11151622 | PRR4;TAS2R20;PRH1 | | Body;TSS1500;5'UTR |
| GM03073 | chr12:114833837-114834201 | TBX5;TBX5;TBX5;TBX5 | N_Shore | Body;Body;Body;Body |
| WG1722 | chr12:114833837-114834713 | TBX5;TBX5;TBX5;TBX5 | N_Shore | Body;Body;Body;Body |
| WG1310 | chr12:114837572-114837887 | TBX5;TBX5;TBX5;TBX5 | N_Shore | Body;Body;Body;Body |
| GM02456 | chr12:114841536-114841708 | TBX5;TBX5;TBX5;TBX5 | N_Shore | Body;Body;5'UTR;Body |
| GM00022 | chr12:116997069-116997079 | MAP1LC3B2 | | TSS200 |
| GM00935 | chr12:117155382-117155636 | C12orf49 | | 3'UTR |
| WG1282 | chr12:122356056-122356598 | WDR66 | N_Shore | TSS1500 |
| GM02641 | chr12:122356056-122356852 | WDR66 | N_Shore | TSS1500 |

| | | | | |
|---|---|---|---|---|
| GM02704 | chr12:122356316-122356598 | WDR66 | Island | TSS200 |
| GM00289 | chr12:123380870-123380891 | VPS37B | Island | TSS200 |
| GM00031 | chr12:124773668-124773870 | FAM101A | | TSS200 |
| GM02706 | chr12:124773668-124774078 | FAM101A | | TSS200 |
| GM00022 | chr12:124773870-124774078 | FAM101A;FAM101A | | 1stExon;5'UTR |
| GM01096 | chr12:130316015-130316058 | TMEM132D | | Body |
| GM09255 | chr12:131303194-131303518 | STX2;STX2 | Island | Body;Body |
| WG1513 | chr12:15114233-15114393 | ARHGDIB | | 5'UTR |
| WG1657 | chr12:15114703-15114732 | ARHGDIB | | TSS200 |
| GM00439 | chr12:15114703-15114904 | ARHGDIB | | TSS200 |
| GM09256 | chr12:1609245-1609369 | LOC100292680 | | TSS1500 |
| GM00520 | chr12:1609245-1609470 | LOC100292680 | | TSS1500 |
| WG1657 | chr12:2339439-2339614 | CACNA1C;CACNA1C;CACNA | Island | Body;Body;Body;Body;Body;Body;Bod |
| GM00290 | chr12:25055262-25055381 | BCAT1 | N_Shore | Body |
| GM01095 | chr12:25055262-25055421 | BCAT1 | N_Shore | Body |
| WG1722 | chr12:25150530-25150626 | C12orf77 | | TSS200 |
| GM01376 | chr12:28124011-28124233 | PTHLH;PTHLH;PTHLH;PTHLH | S_Shore | TSS1500;5'UTR;5'UTR;TSS1500 |
| WG1282 | chr12:312592-313104 | SLC6A12;SLC6A12;SLC6A12 | Island | Body;Body;Body |
| GM00289 | chr12:46767665-46767747 | SLC38A2 | S_Shore | TSS1500 |
| GM00439 | chr12:51786038-51786547 | GALNT6 | S_Shore | TSS1500 |
| GM01393 | chr12:52281810-52282079 | ANKRD33;ANKRD33;ANKRD33;ANKRD33 | | 1stExon;5'UTR;1stExon;5'UTR |
| WG2120 | chr12:54392734-54392884 | HOXC9 | N_Shore | TSS1500 |
| GM00940 | chr12:54392734-54393100 | HOXC9 | N_Shore | TSS1500 |
| GM00016 | chr12:54412269-54412565 | HOXC4;HOXC5;HOXC6 | N_Shore | 5'UTR;Body;5'UTR |
| GM09255 | chr12:54423428-54424679 | HOXC4;HOXC6;HOXC5;HOX | Island | 5'UTR;Body;Body;Body |
| WG1978 | chr12:54423428-54425418 | HOXC4;HOXC6;HOXC5;HOX | Island | 5'UTR;Body;Body;Body |
| GM00439 | chr12:54424964-54425418 | HOXC4;HOXC5 | Island | 5'UTR;Body |
| WG1978 | chr12:54440773-54441509 | HOXC4 | Island | 5'UTR |
| WG1513 | chr12:54440806-54441213 | HOXC4 | Island | 5'UTR |
| GM00031 | chr12:54446019-54446100 | HOXC4 | Island | 5'UTR |
| GM01653 | chr12:54446019-54446253 | HOXC4 | Island | 5'UTR |
| WG1283 | chr12:54448729-54448913 | HOXC4;HOXC4 | S_Shore | Body;Body |
| GM09256 | chr12:56329615-56329903 | DGKA;DGKA;DGKA;DGKA | Island | 5'UTR;5'UTR;5'UTR;5'UTR |
| GM00042 | chr12:56732615-56732723 | IL23A | | TSS200 |
| GM01096 | chr12:58021295-58021917 | B4GALNT1 | Island | Body |
| GM00289 | chr12:5937031-5937409 | ANO2 | Island | Body |
| WG1513 | chr12:62583974-62584454 | FAM19A2 | N_Shore | 5'UTR |
| WG1977 | chr12:6442873-6443045 | TNFRSF1A | S_Shelf | Body |
| WG1722 | chr12:7281566-7281783 | CLSTN3;RBP5 | | TSS1500;TSS200 |
| GM01096 | chr12:772861-772982 | NINJ2 | | TSS200 |
| WG1984 | chr12:97300410-97300429 | NEDD1;NEDD1;NEDD1;NEDD | N_Shore | TSS1500;TSS1500;TSS1500;TSS1500 |
| GM01650 | chr13:100310162-100310405 | CLYBL | N_Shore | Body |
| GM09256 | chr13:103019936-103020405 | FGF14 | | Body |
| GM00439 | chr13:106116963-106117425 | DAOA;DAOA | | TSS1500;TSS1500 |
| GM02456 | chr13:110874267-110874326 | COL4A1 | | Body |
| GM01392 | chr13:110918331-110918683 | COL4A1 | Island | Body |
| WG2276 | chr13:110993474-110993558 | COL4A2 | | Body |

| | | | | |
|---|---|---|---|---|
| GM00439 | chr13:111062247-111062287 | COL4A2 | | Body |
| GM01392 | chr13:111521981-111522314 | C13orf29 | | Body |
| GM00940 | chr13:113436309-113436651 | ATP11A;ATP11A | N_Shelf | Body;Body |
| GM00520 | chr13:113698931-113699016 | MCF2L;MCF2L | N_Shelf | Body;Body |
| GM00289 | chr13:114108433-114108515 | ADPRHL1 | | TSS1500 |
| WG1977 | chr13:114108433-114108652 | ADPRHL1 | | TSS1500 |
| GM00438 | chr13:114108433-114108713 | ADPRHL1 | | TSS1500 |
| GM02706 | chr13:114308619-114308749 | ATP4B | | Body |
| GM01392 | chr13:114828067-114828455 | RASA3 | | Body |
| GM02704 | chr13:114880761-114880889 | RASA3 | S_Shore | Body |
| GM00935 | chr13:20805196-20805385 | GJB6;GJB6;GJB6;GJB6;GJB6 | N_Shore | 5'UTR;5'UTR;5'UTR;5'UTR;1stExon |
| GM00438 | chr13:30088615-30088771 | SLC7A1 | Island | 3'UTR |
| GM09256 | chr13:36045201-36045352 | NBEA | Island | Body |
| WG1978 | chr13:36048892-36051073 | MAB21L1;MIR548F5;NBEA;M | N_Shore | 1stExon;Body;Body;3'UTR |
| GM01376 | chr13:36050889-36051073 | MIR548F5;NBEA;MAB21L1 | N_Shore | Body;Body;TSS200 |
| GM02149 | chr13:36050901-36050993 | MIR548F5;NBEA;MAB21L1 | N_Shore | Body;Body;TSS200 |
| WG1513 | chr13:36050901-36051073 | MIR548F5;NBEA;MAB21L1 | N_Shore | Body;Body;TSS200 |
| GM00940 | chr13:43566633-43566902 | EPSTI1;EPSTI1 | S_Shore | TSS1500;TSS1500 |
| GM00941 | chr13:43597565-43598038 | DNAJC15;DNAJC15 | Island | 5'UTR;1stExon |
| GM00439 | chr13:48575859-48575927 | SUCLA2 | S_Shore | TSS1500 |
| GM00438 | chr13:79234435-79234847 | RNF219 | S_Shore | TSS1500 |
| GM00439 | chr13:98605892-98606196 | IPO5 | | TSS200 |
| GM00935 | chr13:98794353-98794593 | FARP1;FARP1 | N_Shore | TSS1500;TSS1500 |
| WG1513 | chr14:103593088-103593520 | TNFAIP2 | N_Shore | Body |
| GM01392 | chr14:103593235-103593520 | TNFAIP2 | Island | Body |
| WG1658 | chr14:103593503-103593520 | TNFAIP2 | Island | Body |
| GM00935 | chr14:104621867-104622128 | KIF26A | Island | Body |
| GM00042 | chr14:105714379-105714404 | BRF1;BTBD6;BRF1 | N_Shore | TSS200;TSS1500;Body |
| GM00941 | chr14:24780167-24780557 | LTB4R2;LTB4R2;LTB4R2;LTB4 | Island | 1stExon;1stExon;Body;TSS1500;1stEx |
| GM00290 | chr14:34269637-34270113 | NPAS3;NPAS3;NPAS3;NPAS3 | Island | Body;Body;Body;Body |
| GM02149 | chr14:37125914-37126283 | PAX9 | N_Shore | TSS1500 |
| WG2120 | chr14:37135898-37136391 | PAX9 | Island | Body |
| WG2275 | chr14:37136192-37136391 | PAX9 | Island | Body |
| GM00439 | chr14:45432309-45432516 | KLHL28;FAM179B | S_Shore | TSS1500;1stExon |
| WG2415 | chr14:54418728-54418851 | BMP4;BMP4;BMP4 | Island | Body;Body;Body |
| GM00438 | chr14:57277178-57277366 | OTX2;OTX2 | N_Shore | 1stExon;5'UTR |
| GM02555 | chr14:80677608-80677688 | DIO2;DIO2;DIO2 | | 1stExon;1stExon;Body |
| GM00520 | chr14:81422098-81422160 | TSHR;TSHR;TSHR | S_Shore | 1stExon;1stExon;1stExon |
| GM02555 | chr14:99728984-99729088 | BCL11B;BCL11B | | Body;Body |
| WG1310 | chr15:101419275-101419296 | ALDH1A3 | Island | TSS1500 |
| GM00022 | chr15:101419293-101419333 | ALDH1A3 | Island | TSS1500 |
| WG1310 | chr15:101728611-101728653 | CHSY1 | | Body |
| GM02175 | chr15:26109252-26109394 | ATP10A | S_Shore | TSS1500 |
| GM00438 | chr15:27215952-27215971 | GABRG3 | Island | TSS1500 |
| GM00016 | chr15:32964155-32964277 | SCG5;SCG5 | | Body;Body |
| WG1657 | chr15:32964155-32964500 | SCG5;SCG5 | | Body;Body |
| GM00520 | chr15:33009409-33009531 | GREM1 | N_Shore | TSS1500 |

| | | | | |
|---|---|---|---|---|
| GM01376 | chr15:37170381-37170554 | LOC145845 | N_Shelf | Body |
| GM01393 | chr15:40268610-40268777 | EIF2AK4 | Island | Body |
| GM03073 | chr15:40268610-40269214 | EIF2AK4 | Island | Body |
| WG1722 | chr15:40544174-40544427 | PAK6;C15orf56;PAK6;PAK6 | N_Shore | 5'UTR;3'UTR;5'UTR;5'UTR |
| GM00940 | chr15:44086617-44087089 | SERF2;MIR1282 | S_Shore | 3'UTR;TSS1500 |
| WG1513 | chr15:45314789-45314933 | SORD | N_Shore | TSS1500 |
| GM09256 | chr15:45671155-45671246 | LOC145663;GATM | S_Shore | Body;TSS200 |
| GM01392 | chr15:49716247-49716645 | C15orf33;FGF7 | | Body;5'UTR |
| WG2415 | chr15:51369862-51370019 | TNFAIP8L3 | | Body |
| GM03075 | chr15:55881599-55881706 | PYGO1 | S_Shore | TSS1500 |
| GM02555 | chr15:70354647-70354825 | TLE3;TLE3;TLE3 | | Body;Body;Body |
| GM02555 | chr15:70994435-70994672 | UACA;UACA;UACA | | 5'UTR;Body;1stExon |
| GM03075 | chr15:70994435-70995014 | UACA;UACA;UACA | | 5'UTR;Body;1stExon |
| WG1977 | chr15:71408603-71408847 | CT62 | S_Shore | TSS1500 |
| GM00042 | chr15:74494861-74494909 | STRA6;STRA6;STRA6;STRA6;STRA6 | | 5'UTR;5'UTR;5'UTR;5'UTR;5'UTR |
| GM00935 | chr15:76005861-76006143 | CSPG4 | | TSS1500 |
| GM00290 | chr15:83953690-83953744 | BNC1 | Island | TSS1500 |
| WG2275 | chr15:88576296-88576320 | NTRK3;NTRK3;NTRK3 | | Body;Body;Body |
| GM01096 | chr15:92944664-92944756 | ST8SIA2 | | Body |
| GM01392 | chr15:93632896-93633172 | RGMA;RGMA | Island | TSS1500;TSS1500 |
| GM00042 | chr15:96868935-96869220 | NR2F2 | S_Shelf | TSS1500 |
| GM00439 | chr15:96869027-96869220 | NR2F2 | S_Shelf | TSS200 |
| GM00439 | chr16:15595987-15596215 | C16orf45;C16orf45 | | TSS200;Body |
| GM00520 | chr16:1581834-1582219 | IFT140 | N_Shore | Body |
| GM00520 | chr16:2082777-2083168 | SLC9A3R2;SLC9A3R2 | N_Shelf | Body;Body |
| WG1978 | chr16:2892766-2892782 | TMPRSS8 | Island | TSS200 |
| WG2121 | chr16:2908554-2908934 | PRSS22 | S_Shore | TSS1500 |
| WG2276 | chr16:4103161-4103225 | ADCY9 | | Body |
| GM00438 | chr16:4163819-4164087 | ADCY9 | N_Shore | Body |
| GM00042 | chr16:46603015-46603059 | ANKRD26P1 | | TSS200 |
| GM09255 | chr16:50347766-50347895 | ADCY7 | | Body |
| GM00941 | chr16:51184355-51184392 | SALL1;SALL1;SALL1 | Island | Body;5'UTR;1stExon |
| GM02641 | chr16:55794595-55794910 | CES4 | N_Shore | Body |
| WG1513 | chr16:58019866-58019984 | TEPP;TEPP | Island | Body;Body |
| WG1657 | chr16:65105516-65105714 | CDH11 | | 5'UTR |
| GM00941 | chr16:66613266-66613278 | CMTM2 | Island | TSS200 |
| GM00439 | chr16:67197084-67197186 | HSF4;FBXL8;HSF4 | Island | TSS1500;Body;TSS1500 |
| GM00290 | chr16:67197129-67197640 | HSF4;HSF4;FBXL8 | Island | TSS200;TSS200;Body |
| WG1309 | chr16:68270252-68270282 | ESRP2 | Island | TSS200 |
| WG1722 | chr16:69958626-69958964 | WWP2;WWP2 | | TSS1500;Body |
| GM00290 | chr16:69958792-69958964 | WWP2;WWP2 | | TSS200;Body |
| WG1309 | chr16:81527427-81527504 | CMIP | | Body |
| GM01096 | chr16:857454-857981 | PRR25 | Island | Body |
| GM00042 | chr16:87441441-87441659 | ZCCHC14 | Island | 3'UTR |
| GM00042 | chr16:87468206-87468247 | ZCCHC14 | | Body |
| GM00520 | chr16:87904491-87904580 | SLC7A5 | S_Shore | TSS1500 |
| GM00935 | chr16:88052956-88053104 | BANP;BANP | Island | Body;Body |

| | | | | |
|---|---|---|---|---|
| GM00290 | chr16:88590325-88590404 | ZFPM1 | Island | Body |
| GM09256 | chr16:88706241-88706426 | IL17C | Island | Body |
| GM00439 | chr16:89445433-89445503 | ANKRD11 | | 5'UTR |
| GM00439 | chr16:90016004-90016061 | DEF8;DEF8 | Island | 5'UTR;Body |
| GM02706 | chr17:1028394-1028676 | ABR;ABR | N_Shore | Body;Body |
| WG1983 | chr17:1028673-1028931 | ABR;ABR | Island | Body;5'UTR |
| GM00042 | chr17:15244441-15244519 | TEKT3 | N_Shore | 5'UTR |
| GM02555 | chr17:154410-154671 | RPH3AL | S_Shore | Body |
| WG1310 | chr17:154499-155045 | RPH3AL | N_Shelf | Body |
| GM00290 | chr17:183931-184106 | RPH3AL | N_Shore | 5'UTR |
| GM00439 | chr17:20799470-20799511 | CCDC144NL | Island | TSS200 |
| WG2276 | chr17:21300735-21300791 | KCNJ12 | | 5'UTR |
| WG1977 | chr17:26732191-26732272 | SLC46A1 | Island | Body |
| GM00941 | chr17:29297380-29297414 | RNF135;RNF135 | N_Shore | TSS1500;TSS1500 |
| GM02316 | chr17:30822561-30823006 | MYO1D | N_Shore | Body |
| GM02175 | chr17:33759512-33759648 | SLFN12;SLFN12 | | 1stExon;5'UTR |
| GM09255 | chr17:33759929-33759965 | SLFN12 | | TSS1500 |
| GM02175 | chr17:33759957-33759986 | SLFN12 | | TSS1500 |
| GM00289 | chr17:3433428-3433970 | TRPV3 | N_Shelf | Body |
| GM09255 | chr17:36715787-36715953 | SRCIN1 | Island | Body |
| GM00290 | chr17:37028105-37028650 | LASP1 | S_Shore | Body |
| WG1977 | chr17:3705631-3705875 | ITGAE | | TSS1500 |
| GM00940 | chr17:37123767-37123843 | FBXO47 | | TSS200 |
| GM00439 | chr17:40259724-40260053 | DHX58 | Island | Body |
| GM00022 | chr17:40489584-40489785 | STAT3;STAT3;STAT3 | | Body;Body;Body |
| GM00289 | chr17:40687596-40687830 | NAGLU | N_Shore | TSS1500 |
| GM00940 | chr17:40835849-40836087 | CNTNAP1 | Island | Body |
| WG1657 | chr17:40935998-40936498 | WNK4 | Island | Body |
| GM00935 | chr17:41832753-41833161 | SOST | Island | Body |
| GM02175 | chr17:41909709-41909716 | MPP3;MPP3 | N_Shore | 5'UTR;Body |
| GM09255 | chr17:43319137-43319382 | FMNL1 | Island | Body |
| GM02555 | chr17:43483116-43483453 | ARHGAP27;ARHGAP27;ARHG | Island | 1stExon;5'UTR;5'UTR |
| GM00940 | chr17:43974869-43975063 | MAPT;MAPT;MAPT;MAPT;LOC | Island | 5'UTR;5'UTR;5'UTR;5'UTR;Body;5'UTR; |
| GM09256 | chr17:43974919-43975063 | MAPT;MAPT;MAPT;MAPT;LOC | Island | 5'UTR;5'UTR;5'UTR;5'UTR;Body;5'UTR; |
| GM09255 | chr17:44847220-44847427 | WNT3 | Island | Body |
| WG2414 | chr17:46645972-46646359 | HOXB3 | S_Shelf | 5'UTR |
| GM02149 | chr17:46645972-46646444 | HOXB3 | S_Shelf | 5'UTR |
| GM09256 | chr17:46648014-46648533 | HOXB3 | | 5'UTR |
| GM00438 | chr17:46651722-46651920 | HOXB3;HOXB3 | N_Shelf | 5'UTR;1stExon |
| WG1722 | chr17:46651945-46652179 | HOXB3 | N_Shelf | TSS200 |
| GM01376 | chr17:46656543-46656690 | HOXB4 | S_Shore | TSS1500 |
| WG2275 | chr17:46669485-46669492 | LOC404266;HOXB5;LOC404 | Island | TSS200;3'UTR;Body;TSS200;Body |
| GM00042 | chr17:46669485-46669566 | LOC404266;HOXB5;LOC404 | Island | TSS200;3'UTR;Body;TSS200;Body |
| GM09255 | chr17:46669485-46669644 | LOC404266;HOXB5;LOC404 | Island | TSS200;3'UTR;Body;TSS200;Body |
| WG1513 | chr17:46669485-46670029 | LOC404266;HOXB5;LOC404 | Island | TSS200;3'UTR;Body;TSS200;Body |
| GM01096 | chr17:46671861-46672255 | LOC404266;LOC404266;LO( | N_Shore | Body;Body;Body;TSS1500;TSS1500;Bo |
| GM02640 | chr17:46673442-46674514 | LOC404266;LOC404266;LO( | N_Shore | Body;Body;Body;Body;3'UTR;Body |

| | | | | |
|---|---|---|---|---|
| GM01375 | chr17:46673625-46674041 | LOC404266;LOC404266;LO( | Island | Body;Body;Body;Body;3'UTR;Body |
| GM09255 | chr17:46673625-46674514 | LOC404266;LOC404266;LO( | Island | Body;Body;Body;Body;3'UTR;Body |
| GM02641 | chr17:46674041-46674438 | LOC404266;LOC404266;LO( | Island | Body;Body;Body;Body;Body;Body |
| GM02704 | chr17:46674041-46674514 | LOC404266;LOC404266;LO( | Island | Body;Body;Body;Body;Body;Body |
| GM00438 | chr17:46674395-46674514 | LOC404266;LOC404266;LO( | N_Shore | Body;Body;Body;Body;Body;Body |
| GM00289 | chr17:46676099-46676375 | LOC404266;LOC404266;LO( | S_Shore | Body;Body;Body;Body;5'UTR;Body |
| GM02175 | chr17:46681111-46681154 | LOC404266;LOC404266;HOXB6 | | Body;Body;5'UTR |
| GM02706 | chr17:46681111-46683606 | LOC404266;LOC404266;HOXB6 | | Body;Body;5'UTR |
| GM00289 | chr17:46682308-46683047 | LOC404266;LOC404266;HO | N_Shelf | Body;Body;1stExon;5'UTR |
| GM02316 | chr17:46682319-46682413 | LOC404266;LOC404266;HO | N_Shelf | Body;Body;1stExon;5'UTR |
| GM02147 | chr17:46682390-46682398 | LOC404266;LOC404266;HO | N_Shelf | Body;Body;TSS200 |
| WG2120 | chr17:46685327-46685448 | HOXB7 | Island | Body |
| GM01095 | chr17:46691543-46692008 | HOXB8 | Island | Body |
| GM09256 | chr17:47209660-47209749 | B4GALNT2;B4GALNT2;B4GAI | N_Shore | TSS1500;TSS200;TSS1500 |
| GM00438 | chr17:47287492-47287526 | ABI3;GNGT2;ABI3 | | TSS200;TSS1500;TSS200 |
| GM00935 | chr17:59554672-59554806 | TBX4 | | Body |
| WG1978 | chr17:60828157-60828406 | MARCH10;MARCH10 | | Body;Body |
| GM00520 | chr17:61511683-61511829 | CYB561;CYB561;CYB561 | Island | 3'UTR;3'UTR;3'UTR |
| GM09255 | chr17:62777693-62777777 | LOC146880;LOC146880 | Island | Body;TSS200 |
| GM00290 | chr17:71772827-71772869 | C17orf54 | | Body |
| GM00290 | chr17:72439044-72439217 | GPRC5C;GPRC5C | N_Shelf | Body;Body |
| GM01096 | chr17:750241-750848 | NXN | | Body |
| GM01392 | chr17:750766-751287 | NXN | | Body |
| GM00439 | chr17:75283800-75283979 | SEPT9;SEPT9 | | TSS200;Body |
| WG2275 | chr17:75283832-75283979 | SEPT9;SEPT9 | | TSS200;Body |
| GM09256 | chr17:77037916-77037965 | C1QTNF1;C1QTNF1;C1QTNF1 | | 5'UTR;5'UTR;5'UTR |
| GM02175 | chr17:77952290-77952488 | TBC1D16 | Island | Body |
| WG1658 | chr17:78735268-78735324 | RPTOR;RPTOR | | Body;Body |
| GM00289 | chr17:78894364-78894477 | RPTOR;RPTOR | N_Shelf | Body;Body |
| GM00520 | chr17:79010458-79010831 | BAIAP2;BAIAP2;BAIAP2;BAIAI | S_Shore | Body;Body;Body;Body |
| WG1513 | chr17:79099481-79099819 | MIR657;AATK | N_Shelf | TSS1500;Body |
| WG1721 | chr17:79109729-79109808 | AATK | Island | Body |
| GM01096 | chr17:79109777-79109910 | AATK | Island | Body |
| GM00042 | chr17:79377548-79378207 | BAHCC1 | Island | Body |
| GM00439 | chr17:79881468-79881543 | MAFG;MAFG | S_Shore | 5'UTR;TSS200 |
| WG1978 | chr17:79905236-79905263 | MYADML2 | | TSS200 |
| GM00031 | chr17:80292079-80292252 | SECTM1 | Island | TSS200 |
| WG2415 | chr17:80358829-80358876 | C17orf101;C17orf101 | N_Shelf | Body;Body |
| GM00935 | chr17:80806135-80806435 | TBCD | | Body |
| GM01096 | chr17:80875998-80876074 | TBCD | | Body |
| GM00520 | chr17:80943851-80944193 | B3GNTL1 | | Body |
| GM00520 | chr17:81040663-81040906 | METRNL | N_Shore | Body |
| GM00520 | chr17:81045495-81045781 | METRNL | Island | Body |
| GM00290 | chr17:8383621-8383800 | MYH10 | S_Shelf | Body |
| WG1977 | chr17:872762-873452 | NXN | | Body |
| GM00022 | chr17:873255-873452 | NXN | | Body |
| GM02456 | chr17:933047-933155 | ABR;ABR;ABR | Island | Body;Body;Body |

| | | | | |
|---|---|---|---|---|
| GM00031 | chr18:11850139-11850153 | CHMP1B;GNAL;GNAL;GNAL | N_Shore | TSS1500;Body;Body;Body |
| GM00941 | chr18:19756582-19757016 | GATA6 | N_Shore | Body |
| GM00940 | chr18:19756582-19757022 | GATA6 | N_Shore | Body |
| GM00935 | chr18:19997912-19997969 | CTAGE1 | | TSS200 |
| GM01393 | chr18:24131115-24131604 | KCTD1 | Island | 5'UTR |
| WG1977 | chr18:30051538-30051572 | FAM59A | S_Shore | TSS1500 |
| GM02456 | chr18:46386267-46386453 | KIAA0427;KIAA0427 | Island | 3'UTR;3'UTR |
| GM00935 | chr18:56296449-56296544 | ALPK2 | | TSS1500 |
| GM00520 | chr18:56934834-56935199 | RAX | Island | 3'UTR |
| GM03073 | chr18:56936434-56936882 | RAX | Island | Body |
| WG1977 | chr18:67069959-67070120 | DOK6 | Island | Body |
| WG1513 | chr18:67070045-67070170 | DOK6 | Island | Body |
| GM02173 | chr18:7011268-7011463 | LAMA1 | Island | Body |
| GM01096 | chr18:74770402-74770589 | MBP;MBP | Island | Body;Body |
| WG1977 | chr18:74770402-74770815 | MBP;MBP | Island | Body;Body |
| GM00042 | chr18:77905298-77905947 | LOC100130522;LOC100130! | Island | TSS1500;TSS1500 |
| WG1513 | chr18:77905663-77905751 | LOC100130522;LOC100130! | S_Shore | TSS200;TSS200 |
| GM09256 | chr18:77917615-77918142 | LOC100130522;LOC100130! | N_Shore | Body;Body;3'UTR |
| WG2275 | chr19:11289349-11289376 | KANK2;KANK2 | S_Shore | Body;Body |
| GM03073 | chr19:11289356-11289376 | KANK2;KANK2 | S_Shore | Body;Body |
| WG2414 | chr19:12876846-12877000 | HOOK2;HOOK2 | N_Shore | Body;Body |
| WG2120 | chr19:12876846-12877188 | HOOK2;HOOK2 | N_Shore | Body;Body |
| GM00290 | chr19:12957979-12958340 | MAST1 | N_Shore | Body |
| GM02706 | chr19:1466647-1467032 | APC2 | Island | Body |
| GM01096 | chr19:16178091-16178099 | TPM4 | N_Shore | TSS1500 |
| WG1722 | chr19:1854549-1854724 | KLF16 | Island | Body |
| GM00935 | chr19:18771079-18771327 | KLHL26 | Island | Body |
| GM00935 | chr19:19626576-19626605 | NDUFA13;TSSK6 | Island | TSS1500;TSS200 |
| GM01096 | chr19:19639553-19639596 | YJEFN3 | Island | TSS200 |
| GM00940 | chr19:19639568-19639596 | YJEFN3 | Island | TSS200 |
| WG1657 | chr19:21264896-21265421 | ZNF714 | N_Shore | TSS200 |
| WG1977 | chr19:21265264-21265421 | ZNF714 | Island | 5'UTR |
| GM02706 | chr19:2282010-2282207 | C19orf35 | S_Shelf | 5'UTR |
| GM02555 | chr19:2282439-2282729 | C19orf35 | S_Shelf | TSS1500 |
| GM00438 | chr19:2291020-2291872 | LINGO3 | Island | Body |
| GM00439 | chr19:31640401-31640849 | DKFZp566F0947 | | TSS1500 |
| WG1513 | chr19:31640642-31640711 | DKFZp566F0947 | | TSS200 |
| GM00935 | chr19:31640642-31640849 | DKFZp566F0947 | | TSS200 |
| GM00941 | chr19:35633669-35633972 | FXYD7;FXYD1;FXYD1 | Island | TSS1500;3'UTR;3'UTR |
| GM01653 | chr19:36801408-36801503 | LOC100134317 | S_Shelf | TSS1500 |
| GM00042 | chr19:38886554-38886789 | SPRED3 | Island | Body |
| GM02175 | chr19:39695011-39695065 | SYCN | S_Shore | TSS200 |
| GM01096 | chr19:40904691-40904974 | PRX;PRX | Island | Body;Body |
| GM01392 | chr19:427022-427263 | SHC2 | Island | Body |
| GM02316 | chr19:43857861-43858174 | CD177;CD177 | | 1stExon;5'UTR |
| GM00290 | chr19:45260501-45261004 | BCL3 | Island | Body |
| WG2276 | chr19:48947560-48947578 | GRIN2D;GRWD1 | Island | 3'UTR;TSS1500 |

| | | | | |
|---|---|---|---|---|
| GM00031 | chr19:49340489-49340593 | PLEKHA4;PLEKHA4;HSD17B1 | Island | 3'UTR;3'UTR;TSS1500 |
| GM02706 | chr19:49891270-49891574 | CCDC155 | Island | TSS1500 |
| GM00031 | chr19:50931343-50931515 | SPIB | Island | Body |
| GM00042 | chr19:5147855-5148409 | KDM4B | Island | Body |
| GM09256 | chr19:52264411-52264429 | FPR2 | | TSS200 |
| GM02175 | chr19:54876728-54876752 | LAIR1;LAIR1 | | TSS200;TSS200 |
| GM00941 | chr19:55972791-55973099 | ISOC2;ISOC2;ISOC2 | Island | 5'UTR;5'UTR;5'UTR |
| GM00520 | chr19:57742421-57742444 | AURKC;AURKC;AURKC;AURK | Island | 5'UTR;TSS200;1stExon;5'UTR;1stExon |
| GM00438 | chr19:7953054-7953081 | LRRC8E | N_Shore | TSS1500 |
| GM00520 | chr19:827776-827843 | AZU1 | | TSS200 |
| WG2414 | chr2:10567628-10567843 | HPCAL1;HPCAL1 | S_Shore | 3'UTR;3'UTR |
| WG1977 | chr2:106055153-106055458 | FHL2;FHL2 | | 1stExon;5'UTR |
| GM02555 | chr2:113992843-113992930 | PAX8;PAX8;PAX8;PAX8;PAX8 | N_Shore | Body;Body;Body;Body;Body;Body;TSS |
| GM02175 | chr2:113992843-113993052 | PAX8;PAX8;PAX8;PAX8;PAX8 | N_Shore | Body;Body;Body;Body;Body;Body;TSS |
| GM09256 | chr2:114033490-114033830 | PAX8;PAX8;PAX8;PAX8;PAX8 | Island | Body;Body;Body;Body;Body;Body |
| WG2415 | chr2:121554810-121554926 | GLI2 | | TSS200 |
| GM03075 | chr2:121554817-121554926 | GLI2 | | TSS200 |
| WG1721 | chr2:142889283-142889392 | LRP1B | S_Shore | TSS200 |
| GM02149 | chr2:142889367-142889392 | LRP1B | S_Shore | TSS200 |
| GM00022 | chr2:152830189-152830521 | CACNB4;CACNB4;CACNB4;CACNB4 | | Body;Body;1stExon;Body |
| GM03073 | chr2:171676553-171676925 | GAD1;GAD1 | Island | Body;Body |
| GM02641 | chr2:176956396-176956678 | HOXD13 | N_Shore | TSS1500 |
| GM00940 | chr2:176964137-176964502 | HOXD12 | Island | TSS1500 |
| GM01096 | chr2:176964137-176964720 | HOXD12 | Island | TSS1500 |
| WG2194 | chr2:176964512-176964720 | HOXD12 | Island | TSS200 |
| GM01393 | chr2:176973275-176974060 | HOXD11 | N_Shore | Body |
| WG1721 | chr2:176973688-176973835 | HOXD11 | Island | Body |
| GM01393 | chr2:176981422-176981654 | HOXD10 | Island | TSS200 |
| GM09256 | chr2:176982108-176982342 | HOXD10 | Island | 1stExon |
| WG1310 | chr2:176993017-176993115 | HOXD8 | Island | TSS1500 |
| GM00439 | chr2:177016167-177016490 | HOXD4;HOXD4 | N_Shore | 5'UTR;1stExon |
| GM02555 | chr2:177016167-177017449 | HOXD4;HOXD4 | N_Shore | 5'UTR;1stExon |
| GM02175 | chr2:1926888-1927209 | MYT1L | | Body |
| WG1282 | chr2:19551750-19551860 | OSR1 | Island | 3'UTR |
| GM01096 | chr2:200320455-200321016 | SATB2 | N_Shore | Body |
| WG1513 | chr2:200320455-200321348 | SATB2 | N_Shore | Body |
| GM00520 | chr2:200334402-200334584 | FLJ32063 | N_Shore | Body |
| GM00935 | chr2:200334854-200335101 | FLJ32063 | N_Shore | Body |
| GM00520 | chr2:202508694-202508813 | ALS2CR4;ALS2CR4 | S_Shore | TSS1500;TSS1500 |
| GM00042 | chr2:202901288-202901470 | FZD7;FZD7 | S_Shore | 1stExon;3'UTR |
| GM02706 | chr2:215675429-215675637 | BARD1 | S_Shore | TSS1500 |
| GM02706 | chr2:220406470-220406756 | CHPF | Island | Body |
| GM09255 | chr2:223155925-223156136 | PAX3;PAX3;PAX3;PAX3;PAX3 | Island | Body;Body;Body;Body;Body;Body |
| GM03075 | chr2:223161771-223162049 | PAX3;PAX3;PAX3;CCDC140;I | Island | Body;Body;Body;TSS1500;Body;Body; |
| GM02706 | chr2:223161771-223162128 | PAX3;PAX3;PAX3;CCDC140;I | Island | Body;Body;Body;TSS1500;Body;Body; |
| GM02706 | chr2:223162666-223162800 | PAX3;PAX3;PAX3;PAX3;PAX3 | N_Shore | Body;Body;Body;Body;Body;Body;Bod |
| GM02640 | chr2:223164459-223170370 | PAX3;PAX3;PAX3;PAX3;PAX3 | S_Shore | TSS1500;TSS1500;TSS1500;TSS1500; |

| | | | | |
|---|---|---|---|---|
| GM00042 | chr2:223164747-223164869 | PAX3;PAX3;PAX3;PAX3;PAX3; | S_Shore | TSS1500;TSS1500;TSS1500;TSS1500; |
| GM01650 | chr2:223164831-223164869 | PAX3;PAX3;PAX3;PAX3;PAX3; | S_Shore | TSS1500;TSS1500;TSS1500;TSS1500; |
| GM00289 | chr2:223164854-223164869 | PAX3;PAX3;PAX3;PAX3;PAX3; | S_Shore | TSS1500;TSS1500;TSS1500;TSS1500; |
| GM01096 | chr2:235935754-235936596 | SH3BP4 | | 5'UTR |
| WG1983 | chr2:23746926-23747197 | KLHL29 | N_Shelf | 5'UTR |
| GM00935 | chr2:238394933-238395125 | MLPH;MLPH | N_Shore | TSS1500;TSS1500 |
| GM01376 | chr2:238647755-238647913 | LRRFIP1;LRRFIP1;LRRFIP1;LR | S_Shelf | Body;Body;Body;Body;Body |
| WG2414 | chr2:240020028-240020174 | HDAC4 | | Body |
| GM02555 | chr2:240233043-240233240 | HDAC4 | | Body |
| GM02706 | chr2:240291426-240291657 | HDAC4 | | 5'UTR |
| GM00438 | chr2:25391830-25391911 | POMC;POMC | Island | TSS1500;TSS1500 |
| GM00290 | chr2:27665139-27665150 | KRTCAP3;KRTCAP3 | Island | TSS200;TSS200 |
| WG2120 | chr2:30669597-30669863 | LCLAT1;LCLAT1 | N_Shore | TSS1500;TSS1500 |
| GM00438 | chr2:30669597-30669952 | LCLAT1;LCLAT1 | N_Shore | TSS1500;TSS1500 |
| WG2121 | chr2:3286338-3286479 | TSSC1 | Island | Body |
| GM01375 | chr2:3452347-3452563 | TTC15 | Island | Body |
| WG2121 | chr2:3452347-3452624 | TTC15 | Island | Body |
| GM01653 | chr2:3452543-3452624 | TTC15 | Island | Body |
| GM02640 | chr2:3642400-3642634 | COLEC11;COLEC11 | N_Shore | TSS1500;TSS1500 |
| GM00022 | chr2:3642447-3642634 | COLEC11;COLEC11 | N_Shore | TSS200;TSS200 |
| GM09256 | chr2:3714936-3715083 | ALLC | | 5'UTR |
| GM00935 | chr2:37571677-37571901 | QPCT | Island | TSS200 |
| WG1310 | chr2:54087173-54087189 | LOC100302652;GPR75 | Island | TSS200;TSS200 |
| GM02147 | chr2:54087178-54087189 | LOC100302652;GPR75 | Island | TSS200;TSS200 |
| GM01095 | chr2:54785178-54785795 | SPTBN1;SPTBN1 | Island | TSS1500;Body |
| GM09255 | chr2:56410011-56410087 | CCDC85A | N_Shore | TSS1500 |
| GM00438 | chr2:6122798-6122814 | LOC400940 | | Body |
| GM02640 | chr2:65217528-65217623 | SLC1A4;SLC1A4 | S_Shore | Body;5'UTR |
| GM02555 | chr2:66671478-66671727 | MEIS1 | N_Shore | Body |
| GM01392 | chr2:71134524-71134630 | VAX2 | N_Shore | Body |
| GM00042 | chr2:71211981-71212157 | ANKRD53;ANKRD53 | | 3'UTR;Body |
| GM01096 | chr2:72371255-72371477 | CYP26B1 | Island | Body |
| GM00520 | chr2:73964768-73964834 | TPRKB | S_Shore | TSS1500 |
| GM00520 | chr2:74669349-74669380 | RTKN | Island | TSS1500 |
| GM02456 | chr2:74780268-74780617 | LOXL3 | N_Shore | 5'UTR |
| GM01096 | chr2:74780272-74780617 | LOXL3 | N_Shore | 5'UTR |
| GM00439 | chr2:85361959-85362480 | TCF7L1 | Island | Body |
| GM00935 | chr2:85637834-85638535 | CAPG | N_Shelf | TSS200 |
| GM09256 | chr2:947167-947589 | SNTG2 | Island | Body |
| WG1721 | chr2:947515-947634 | SNTG2 | S_Shore | Body |
| GM00520 | chr2:95537314-95537475 | TEKT4;TEKT4 | Island | 1stExon;5'UTR |
| GM00439 | chr20:23067126-23067159 | CD93 | | TSS200 |
| GM02555 | chr20:25128770-25128805 | LOC284798;LOC284798;LOC | Island | TSS200;Body;Body |
| GM02706 | chr20:25128770-25128807 | LOC284798;LOC284798;LOC | Island | TSS200;Body;Body |
| GM00940 | chr20:2516256-2516592 | TMC2 | | TSS1500 |
| GM01650 | chr20:32856747-32856898 | ASIP | Island | Body |
| GM01650 | chr20:35169539-35169609 | MYL9;MYL9 | Island | TSS1500;TSS1500 |

| | | | | |
|---|---|---|---|---|
| GM00520 | chr20:35274627-35274655 | SLA2;SLA2 | | TSS200;TSS200 |
| GM00438 | chr20:36011999-36012016 | SRC;SRC | N_Shore | 5'UTR;5'UTR |
| GM00935 | chr20:36932397-36932553 | BPI | | TSS200 |
| GM01376 | chr20:43935460-43935471 | MATN4;MATN4;RBPJL;MATN4 | Island | 5'UTR;5'UTR;TSS200;5'UTR |
| GM02175 | chr20:47896448-47896888 | SNORD12B;C20orf199;MIR1: | S_Shore | TSS1500;Body;TSS1500;Body;TSS150 |
| GM00935 | chr20:49253364-49253710 | FAM65C;FAM65C | | 1stExon;5'UTR |
| GM00042 | chr20:50312386-50312632 | ATP9A | | Body |
| GM09256 | chr20:5485144-5485284 | LOC149837 | | Body |
| WG1310 | chr20:5485245-5485284 | LOC149837 | | TSS200 |
| GM02706 | chr20:5485268-5485284 | LOC149837 | | TSS200 |
| GM01392 | chr20:55925570-55925603 | RAE1;RAE1 | N_Shore | TSS1500;TSS1500 |
| GM00935 | chr20:57465439-57465448 | GNAS;GNAS;GNAS;GNAS;GN | Island | 3'UTR;TSS1500;TSS1500;Body;TSS150 |
| GM00520 | chr20:58179847-58180241 | PHACTR3;PHACTR3 | Island | 5'UTR;1stExon |
| WG1513 | chr20:58630954-58631038 | C20orf197 | | TSS200 |
| GM02175 | chr20:61447330-61447394 | COL9A3 | N_Shore | TSS1500 |
| GM00439 | chr20:61953801-61954258 | COL20A1 | N_Shore | Body |
| GM01650 | chr20:62318426-62318588 | RTEL1;RTEL1 | Island | Body;Body |
| GM00935 | chr20:62679255-62679713 | SOX18 | N_Shore | 3'UTR |
| GM02706 | chr20:62688717-62688896 | TCEA2;TCEA2 | Island | 1stExon;5'UTR |
| GM00439 | chr21:17961737-17961996 | C21orf34;MIR125B2;C21orf34 | | Body;TSS1500;Body |
| GM00940 | chr21:35831871-35832180 | KCNE1;KCNE1;KCNE1;KCNE | Island | 5'UTR;5'UTR;1stExon;1stExon;5'UTR |
| GM02147 | chr21:35832141-35832180 | KCNE1;KCNE1;KCNE1 | Island | 5'UTR;TSS1500;TSS1500 |
| GM00520 | chr21:36421467-36421503 | RUNX1;RUNX1 | | 5'UTR;1stExon |
| GM01096 | chr21:40145923-40146046 | NCRNA00114 | | TSS1500 |
| GM01096 | chr21:40759534-40760258 | WRB;WRB | N_Shore | Body;TSS200 |
| GM02555 | chr21:45148879-45149141 | PDXK | Island | Body |
| GM01095 | chr21:45705618-45705742 | AIRE | Island | TSS200 |
| WG2121 | chr21:47604052-47604654 | C21orf56;C21orf56 | S_Shore | 5'UTR;5'UTR |
| GM00031 | chr21:47604291-47604654 | C21orf56;C21orf56;C21orf56 | S_Shore | 5'UTR;5'UTR;1stExon;1stExon |
| GM00940 | chr22:19746984-19747140 | TBX1;TBX1;TBX1 | Island | 5'UTR;5'UTR;5'UTR |
| WG2276 | chr22:19754125-19754448 | TBX1;TBX1;TBX1 | Island | Body;Body;Body |
| GM00290 | chr22:20861470-20861479 | MED15;MED15 | N_Shore | TSS1500;TSS1500 |
| WG2120 | chr22:22058080-22058271 | YPEL1 | Island | Body |
| GM01393 | chr22:24181263-24181270 | DERL3;DERL3;DERL3 | Island | TSS200;TSS200;TSS200 |
| GM02456 | chr22:24348549-24348715 | GSTTP1 | | TSS1500 |
| WG1283 | chr22:24384105-24384294 | GSTT1 | N_Shore | Body |
| GM00289 | chr22:24384159-24384294 | GSTT1 | Island | 1stExon |
| WG1977 | chr22:24384400-24384944 | GSTT1 | Island | TSS200 |
| WG1977 | chr22:24979554-24979588 | GGT1 | | TSS200 |
| GM09256 | chr22:32027580-32027605 | PISD | S_Shore | TSS1500 |
| GM02175 | chr22:42828411-42828418 | NFAM1 | | TSS200 |
| GM00022 | chr22:44568337-44568423 | PARVG;PARVG | | TSS1500;TSS1500 |
| GM00439 | chr22:45125218-45126040 | PRR5;PRR5;PRR5;PRR5-ARH( | Island | Body;Body;Body;Body;Body;Body |
| WG2414 | chr22:45609402-45609421 | C22orf9;C22orf9 | | Body;TSS1500 |
| GM02149 | chr22:50629038-50629224 | TRABD | Island | 5'UTR |
| WG1283 | chr3:114790243-114790596 | ZBTB20;ZBTB20;ZBTB20 | | 1stExon;5'UTR;5'UTR |
| GM02175 | chr3:11746183-11746765 | VGLL4 | N_Shore | 5'UTR |

| | | | | |
|---|---|---|---|---|
| GM00438 | chr3:128204813-128205143 | GATA2;GATA2;GATA2 | N_Shore | Body;Body;Body |
| GM00289 | chr3:129407542-129407548 | TMCC1;TMCC1;TMCC1 | | TSS200;Body;Body |
| GM00940 | chr3:138152837-138153127 | ESYT3 | N_Shore | TSS1500 |
| GM01096 | chr3:139257775-139257907 | RBP1;RBP1;RBP1 | Island | Body;Body;Body |
| WG1977 | chr3:14443428-14443473 | SLC6A6;SLC6A6;SLC6A6 | N_Shore | TSS1500;TSS1500;TSS1500 |
| GM00941 | chr3:147112438-147112605 | ZIC4;ZIC4;ZIC4 | N_Shore | Body;Body;Body |
| WG1722 | chr3:15311021-15311269 | SH3BP5;SH3BP5 | | Body;Body |
| WG1977 | chr3:155394109-155394303 | PLCH1;PLCH1;PLCH1 | | 5'UTR;TSS200;TSS200 |
| GM00439 | chr3:155570275-155570809 | SLC33A1 | N_Shelf | Body |
| GM01653 | chr3:157821262-157822072 | SHOX2;SHOX2;SHOX2 | Island | Body;Body;Body |
| GM02706 | chr3:160475035-160475128 | PPM1L | S_Shore | Body |
| GM00439 | chr3:169377725-169377946 | MECOM | N_Shore | Body |
| GM00940 | chr3:169483122-169483295 | TERC | S_Shore | TSS1500 |
| GM00439 | chr3:169683976-169684239 | LOC100128164;SEC62;LOC1 | N_Shore | Body;TSS1500;Body |
| GM02706 | chr3:183894727-183894778 | AP2M1;AP2M1 | S_Shore | 5'UTR;5'UTR |
| WG2415 | chr3:192289245-192289293 | FGF12 | | Body |
| WG1977 | chr3:194833887-194834008 | C3orf21 | | Body |
| GM02704 | chr3:195489306-195489789 | MUC4;MUC4;MUC4 | Island | Body;Body;Body |
| WG1310 | chr3:196065357-196065506 | TM4SF19 | | TSS200 |
| GM00941 | chr3:24537277-24537309 | THRB;THRB;THRB | Island | TSS1500;TSS1500;TSS1500 |
| WG2193 | chr3:36949957-36950004 | TRANK1 | | Body |
| GM00520 | chr3:37033373-37033632 | EPM2AIP1;MLH1 | N_Shore | 1stExon;TSS1500 |
| GM00935 | chr3:42742993-42743203 | HHATL;HHATL | | Body;5'UTR |
| GM01376 | chr3:42947263-42947385 | ZNF662;ZNF662 | N_Shore | TSS1500;TSS200 |
| GM00439 | chr3:48694451-48694534 | CELSR3 | Island | Body |
| GM02175 | chr3:49203973-49204239 | CCDC71 | S_Shore | TSS200 |
| WG2121 | chr3:50488020-50488230 | CACNA2D2;CACNA2D2 | | Body;Body |
| GM00290 | chr3:52487733-52488229 | TNNC1 | N_Shore | Body |
| GM00935 | chr3:56809796-56810320 | ARHGEF3;ARHGEF3;ARHGEF3 | | TSS200;Body;Body |
| GM02456 | chr3:64547108-64547373 | ADAMTS9 | | Body |
| GM02706 | chr3:64547310-64547373 | ADAMTS9 | | Body |
| GM00042 | chr3:99357437-99357613 | MIR548G;COL8A1;COL8A1 | | Body;TSS200;TSS200 |
| GM02149 | chr4:10459219-10459352 | ZNF518B | Island | TSS200 |
| GM00520 | chr4:111397885-111398146 | ENPEP | Island | 1stExon |
| WG1722 | chr4:111398042-111398146 | ENPEP | Island | 1stExon |
| GM00520 | chr4:111544016-111544271 | PITX2;PITX2;PITX2;PITX2 | Island | 1stExon;5'UTR;Body;Body |
| GM00520 | chr4:1195845-1196179 | LOC100130872;LOC100130872-SPON2 | | Body;Body |
| GM02456 | chr4:119948360-119948542 | SYNPO2;SYNPO2;SYNPO2 | | Body;Body;Body |
| GM00940 | chr4:1243980-1244086 | CTBP1;CTBP1;C4orf42 | Island | TSS1500;TSS1500;TSS200 |
| WG1722 | chr4:154711512-154711563 | SFRP2 | N_Shore | TSS1500 |
| GM00439 | chr4:160188727-160189254 | RAPGEF2 | | TSS1500 |
| GM02149 | chr4:169799087-169799624 | PALLD;PALLD;PALLD;PALLD | Island | Body;Body;Body;Body |
| WG2193 | chr4:174447847-174448341 | HAND2 | N_Shore | 3'UTR |
| GM00520 | chr4:1804007-1804492 | FGFR3;FGFR3;FGFR3 | Island | Body;Body;Body |
| GM00520 | chr4:1844724-1844878 | LETM1 | S_Shore | Body |
| GM09255 | chr4:24818471-24818615 | CCDC149 | | Body |
| GM01096 | chr4:3372123-3372272 | RGS12;RGS12;RGS12 | N_Shelf | Body;Body;Body |

| | | | | |
|---|---|---|---|---|
| GM09256 | chr4:40858965-40859293 | APBB2;APBB2;APBB2;APBB2 | Island | Body;5'UTR;5'UTR;5'UTR;Body;5'UTR;E |
| WG1721 | chr4:40858984-40859293 | APBB2;APBB2;APBB2;APBB2 | Island | Body;5'UTR;5'UTR;5'UTR;Body;5'UTR;E |
| WG1310 | chr4:40859249-40859293 | APBB2;APBB2;APBB2;APBB2 | S_Shore | Body;TSS200;TSS200;Body;TSS200;Bo |
| GM00520 | chr4:5021084-5021113 | CYTL1 | Island | 1stExon |
| WG1721 | chr4:5021268-5021328 | CYTL1 | Island | TSS200 |
| WG1978 | chr4:52918457-52918489 | SPATA18 | S_Shore | Body |
| WG2120 | chr4:54965829-54965904 | GSX2 | N_Shore | TSS1500 |
| WG2415 | chr4:55143269-55143557 | PDGFRA | | Body |
| GM00520 | chr4:57522762-57522881 | HOPX;HOPX;HOPX;HOPX;HO | S_Shore | 5'UTR;Body;Body;TSS200;TSS200 |
| WG1282 | chr4:6052527-6052554 | JAKMIP1 | | Body |
| GM00940 | chr4:683148-683240 | MFSD7 | S_Shore | TSS200 |
| WG1513 | chr4:71587722-71587904 | RUFY3;RUFY3;RUFY3;RUFY3;RUFY3 | | 1stExon;Body;5'UTR;1stExon;5'UTR |
| GM00439 | chr4:7282819-7283184 | SORCS2 | | Body |
| GM00022 | chr4:7287492-7287737 | SORCS2 | | Body |
| GM01376 | chr4:74847646-74847829 | PF4 | Island | 1stExon |
| GM00016 | chr4:77507102-77507411 | SHROOM3 | Island | Body |
| WG1310 | chr4:94124971-94125068 | GRID2 | | Body |
| GM00022 | chr5:102898463-102898733 | NUDT12;NUDT12 | | 5'UTR;1stExon |
| GM02706 | chr5:1053545-1053774 | SLC12A7 | Island | Body |
| GM01393 | chr5:1053643-1054113 | SLC12A7 | Island | Body |
| GM03075 | chr5:1053643-1054262 | SLC12A7 | Island | Body |
| GM02706 | chr5:110062384-110062417 | TMEM232 | | TSS200 |
| GM00940 | chr5:114516368-114516422 | TRIM36;TRIM36;TRIM36 | S_Shore | TSS200;TSS200;TSS200 |
| GM00289 | chr5:126409310-126409454 | FLJ44606;FLJ44606 | Island | TSS200;TSS200 |
| GM00940 | chr5:127871927-127872329 | FBN2 | N_Shore | Body |
| GM00290 | chr5:134870506-134870517 | NEUROG1;NEUROG1 | N_Shore | 1stExon;3'UTR |
| GM01095 | chr5:135416205-135416613 | MIR886 | Island | Body |
| GM00289 | chr5:140097950-140098250 | VTRNA1-2 | | TSS1500 |
| WG1658 | chr5:140098392-140098576 | VTRNA1-2 | | TSS200 |
| GM00438 | chr5:140579296-140579303 | PCDHB11 | N_Shore | TSS200 |
| WG1658 | chr5:14406585-14406732 | TRIO | | Body |
| GM00935 | chr5:148520854-148520894 | ABLIM3 | N_Shore | TSS200 |
| GM02555 | chr5:149980486-149980682 | SYNPO | | TSS200 |
| GM01393 | chr5:149980526-149980682 | SYNPO | | TSS200 |
| WG1977 | chr5:149980671-149980682 | SYNPO;SYNPO | | 1stExon;5'UTR |
| WG1722 | chr5:150004773-150004918 | SYNPO;SYNPO | Island | Body;Body |
| GM02640 | chr5:153858816-153858936 | HAND1 | Island | TSS1500 |
| GM00439 | chr5:156886982-156887005 | NIPAL4 | Island | TSS200 |
| GM01375 | chr5:156886990-156887005 | NIPAL4 | Island | TSS200 |
| GM02555 | chr5:1594282-1594863 | SDHAP3 | Island | Body |
| GM01653 | chr5:171534678-171534792 | STK10 | | Body |
| GM00016 | chr5:175109098-175109114 | HRH2;HRH2;HRH2 | | 5'UTR;5'UTR;1stExon |
| GM00438 | chr5:176057924-176057996 | EIF4E1B;SNCB;SNCB;EIF4E1 | S_Shore | 5'UTR;TSS1500;TSS1500;1stExon |
| GM00935 | chr5:176784688-176784719 | RGS14 | | TSS200 |
| GM00289 | chr5:176829109-176829198 | PFN3 | N_Shore | TSS1500 |
| GM02555 | chr5:177548398-177548913 | N4BP3 | | Body |
| WG1722 | chr5:177548398-177549014 | N4BP3 | | Body |

| | | | | |
|---|---|---|---|---|
| WG2414 | chr5:177708304-177708528 | COL23A1 | | Body |
| GM00935 | chr5:178684416-178684658 | ADAMTS2;ADAMTS2 | Island | Body;Body |
| WG1722 | chr5:191127-191296 | LRRC14B | N_Shore | TSS1500 |
| GM00940 | chr5:23507573-23507617 | PRDM9 | | TSS200 |
| GM02173 | chr5:23507573-23507656 | PRDM9 | | TSS200 |
| WG1978 | chr5:323320-323969 | AHRR | S_Shore | Body |
| GM00439 | chr5:368394-369088 | AHRR | N_Shore | Body |
| GM03075 | chr5:368804-369088 | AHRR | N_Shore | Body |
| GM00935 | chr5:38257516-38257555 | EGFLAM | N_Shore | TSS1500 |
| GM00940 | chr5:40841315-40841590 | CARD6 | | TSS200 |
| WG1984 | chr5:40841493-40841590 | CARD6 | | 1stExon |
| GM00439 | chr5:415575-415885 | AHRR | | Body |
| GM00520 | chr5:423894-424371 | AHRR | S_Shelf | Body |
| GM00438 | chr5:42756786-42756860 | CCDC152 | | TSS200 |
| GM00438 | chr5:66462293-66462662 | MAST4;MAST4 | S_Shelf | Body;Body |
| WG1722 | chr5:68710808-68711277 | MARVELD2;MARVELD2 | Island | TSS200;TSS200 |
| GM03075 | chr5:68710818-68711277 | MARVELD2;MARVELD2 | Island | TSS200;TSS200 |
| GM00439 | chr5:68787747-68787919 | OCLN | N_Shore | TSS1500 |
| WG2120 | chr5:76010444-76010658 | F2R | N_Shore | TSS1500 |
| GM00935 | chr5:78281793-78281964 | ARSB;ARSB;ARSB | S_Shore | TSS200;1stExon;5'UTR |
| GM01095 | chr5:92908988-92909429 | FLJ42709;FLJ42709 | S_Shore | Body;Body |
| GM00935 | chr5:92908988-92909434 | FLJ42709;FLJ42709 | S_Shore | Body;Body |
| GM01393 | chr6:100911240-100911964 | SIM1 | N_Shore | 1stExon |
| WG1978 | chr6:100911240-100912072 | SIM1 | N_Shore | 1stExon |
| GM09255 | chr6:100911701-100911727 | SIM1 | N_Shore | TSS200 |
| GM09255 | chr6:100911949-100912072 | SIM1 | N_Shore | TSS1500 |
| GM00042 | chr6:101846650-101846767 | GRIK2;GRIK2;GRIK2 | N_Shore | TSS1500;TSS1500;TSS1500 |
| GM00520 | chr6:101846767-101846872 | GRIK2;GRIK2;GRIK2 | Island | TSS200;TSS200;TSS200 |
| GM02317 | chr6:101846791-101846805 | GRIK2;GRIK2;GRIK2 | Island | TSS200;TSS200;TSS200 |
| WG1282 | chr6:101846791-101846872 | GRIK2;GRIK2;GRIK2 | Island | TSS200;TSS200;TSS200 |
| WG1722 | chr6:109274429-109274496 | ARMC2 | | Body |
| GM01393 | chr6:11279656-11279727 | NEDD9 | | Body |
| GM00022 | chr6:116381609-116381957 | FRK;FRK | | 1stExon;5'UTR |
| GM02149 | chr6:134210273-134210307 | TCF21;TCF21;TCF21;TCF21 | N_Shore | 1stExon;5'UTR;1stExon;5'UTR |
| GM00438 | chr6:138200136-138200387 | TNFAIP3 | | Body |
| WG1977 | chr6:139613145-139613456 | TXLNB;TXLNB | | 5'UTR;1stExon |
| GM02555 | chr6:148869461-148869535 | SASH1 | | Body |
| GM00439 | chr6:149358743-149358802 | UST | | Body |
| WG1513 | chr6:149803087-149803444 | ZC3H12D | | 5'UTR |
| GM01393 | chr6:149805995-149806131 | ZC3H12D;ZC3H12D | | 5'UTR;1stExon |
| GM01096 | chr6:149806077-149806131 | ZC3H12D;ZC3H12D | | 5'UTR;1stExon |
| GM00042 | chr6:151186411-151186511 | MTHFD1L | N_Shore | TSS1500 |
| GM00520 | chr6:151560831-151560927 | AKAP12 | Island | TSS1500 |
| GM02706 | chr6:152085565-152085641 | ESR1 | | 5'UTR |
| WG1983 | chr6:152125861-152126080 | ESR1;ESR1 | N_Shelf | 5'UTR;TSS1500 |
| GM00290 | chr6:152125861-152126092 | ESR1;ESR1 | N_Shelf | 5'UTR;TSS1500 |
| GM01095 | chr6:152125965-152126092 | ESR1;ESR1 | N_Shelf | 5'UTR;TSS1500 |

| | | | | |
|---|---|---|---|---|
| WG1310 | chr6:152127812-152127887 | ESR1;ESR1;ESR1;ESR1 | N_Shore | TSS1500;5'UTR;TSS1500;5'UTR |
| GM02706 | chr6:152127812-152128024 | ESR1;ESR1;ESR1;ESR1 | N_Shore | TSS1500;5'UTR;TSS1500;5'UTR |
| GM00935 | chr6:158018055-158018184 | ZDHHC14;ZDHHC14 | S_Shelf | Body;Body |
| WG2276 | chr6:161796785-161796854 | PARK2;PARK2;PARK2 | | Body;Body;Body |
| WG1657 | chr6:161860203-161860359 | PARK2;PARK2;PARK2 | | Body;Body;Body |
| GM02704 | chr6:168435914-168436353 | KIF25;KIF25 | Island | Body;Body |
| GM00289 | chr6:168859941-168859997 | SMOC2;SMOC2 | | Body;Body |
| GM02555 | chr6:21667166-21667407 | FLJ22536 | S_Shore | Body |
| GM00520 | chr6:24583477-24583614 | KIAA0319;KIAA0319;KIAA0319;KIAA0319 | | Body;Body;Body;Body;Body |
| GM00438 | chr6:28493651-28493664 | GPX5;GPX5 | | TSS200;TSS200 |
| WG1513 | chr6:28555601-28555664 | SCAND3 | S_Shore | TSS1500 |
| GM00022 | chr6:291687-292823 | DUSP22 | N_Shore | TSS1500 |
| WG1513 | chr6:291882-292823 | DUSP22 | N_Shore | TSS1500 |
| GM00520 | chr6:29594830-29595016 | GABBR1;GABBR1;GABBR1 | N_Shore | Body;Body;Body |
| GM00940 | chr6:29595002-29595016 | GABBR1;GABBR1;GABBR1 | N_Shore | Body;Body;Body |
| GM02706 | chr6:29690766-29690893 | HLA-F;HLA-F;HLA-F | N_Shore | TSS1500;TSS1500;TSS1500 |
| WG1722 | chr6:29690822-29690893 | HLA-F;HLA-F;HLA-F | N_Shore | TSS1500;TSS1500;TSS1500 |
| WG2120 | chr6:29894050-29894197 | HCG4P6 | N_Shore | TSS1500 |
| GM00941 | chr6:29911091-29911104 | HLA-A | Island | Body |
| GM02555 | chr6:29943401-29943414 | HCG9 | N_Shore | Body |
| WG1978 | chr6:29943401-29943425 | HCG9 | N_Shore | Body |
| GM01096 | chr6:29943401-29943455 | HCG9 | N_Shore | Body |
| GM00940 | chr6:29943455-29943480 | HCG9 | N_Shore | Body |
| GM00439 | chr6:29944093-29944146 | HCG9 | N_Shore | Body |
| GM09256 | chr6:30038712-30038754 | RNF39;RNF39 | N_Shore | 3'UTR;3'UTR |
| WG1977 | chr6:30039432-30039442 | RNF39;RNF39 | Island | Body;Body |
| WG1309 | chr6:30079203-30079280 | TRIM31 | | Body |
| GM02555 | chr6:30079256-30079280 | TRIM31 | | Body |
| GM00290 | chr6:30113978-30114011 | TRIM40 | | Body |
| GM00438 | chr6:30131189-30131219 | TRIM15;TRIM15 | | 5'UTR;1stExon |
| GM02706 | chr6:30458158-30458237 | HLA-E | Island | Body |
| GM00520 | chr6:30510294-30510459 | GNL1 | | 3'UTR |
| GM00439 | chr6:30657882-30658014 | NRM | S_Shelf | Body |
| GM00439 | chr6:30854164-30854233 | DDR1;DDR1 | S_Shore | 5'UTR;5'UTR |
| GM02175 | chr6:30921592-30921757 | DPCR1 | N_Shore | 3'UTR |
| GM02706 | chr6:31025933-31025967 | HCG22 | | Body |
| GM00016 | chr6:31164914-31164947 | HCG27 | N_Shore | TSS1500 |
| WG2414 | chr6:31237013-31237034 | HLA-C | N_Shore | Body |
| WG2414 | chr6:31237405-31237722 | HLA-C | N_Shore | Body |
| GM00439 | chr6:31649619-31649728 | LY6G5C | N_Shore | TSS1500 |
| GM00520 | chr6:31760593-31760616 | VARS | N_Shelf | Body |
| GM01376 | chr6:31829888-31829914 | NEU1 | N_Shore | Body |
| GM02175 | chr6:31846769-31846813 | SLC44A4;SLC44A4 | | 5'UTR;1stExon |
| WG1977 | chr6:32055447-32055525 | TNXB | Island | Body |
| GM00289 | chr6:32097439-32097507 | ATF6B;ATF6B;FKBPL | N_Shore | TSS1500;TSS1500;Body |
| GM02456 | chr6:32297670-32297822 | C6orf10 | | Body |
| WG1977 | chr6:32489750-32489963 | HLA-DRB5 | Island | Body |

| | | | | |
|---|---|---|---|---|
| GM02641 | chr6:32489963-32489991 | HLA-DRB5 | Island | Body |
| GM02147 | chr6:32489991-32490043 | HLA-DRB5 | Island | Body |
| WG1658 | chr6:32526021-32526260 | HLA-DRB6 | | Body |
| GM00042 | chr6:32526027-32526342 | HLA-DRB6 | | Body |
| GM00439 | chr6:32526260-32526342 | HLA-DRB6 | | Body |
| GM03073 | chr6:32526260-32526364 | HLA-DRB6 | | Body |
| WG2193 | chr6:32551891-32551954 | HLA-DRB1 | Island | Body |
| WG2194 | chr6:32552152-32552246 | HLA-DRB1 | Island | Body |
| WG1282 | chr6:32552246-32552453 | HLA-DRB1 | Island | Body |
| GM03075 | chr6:32634245-32634316 | HLA-DQB1 | S_Shore | Body |
| GM00022 | chr6:32729545-32729590 | HLA-DQB2 | Island | Body |
| GM02175 | chr6:32729545-32729596 | HLA-DQB2 | Island | Body |
| GM00520 | chr6:32847762-32847788 | PPP1R2P1 | Island | Body |
| GM02175 | chr6:32847788-32847830 | PPP1R2P1 | Island | Body |
| GM01376 | chr6:32908567-32908642 | HLA-DMB | | 1stExon |
| GM01650 | chr6:32920880-32921142 | HLA-DMA;HLA-DMA | | 5'UTR;1stExon |
| GM01392 | chr6:33048875-33048919 | HLA-DPB1 | S_Shore | Body |
| GM09256 | chr6:33131222-33131336 | COL11A2;COL11A2;COL11A | S_Shore | 3'UTR;3'UTR;3'UTR |
| GM02316 | chr6:33288180-33288569 | DAXX;DAXX;DAXX;DAXX | N_Shore | Body;Body;Body;Body |
| GM00042 | chr6:33288366-33288569 | DAXX;DAXX;DAXX;DAXX | N_Shore | Body;Body;Body;Body |
| GM02149 | chr6:33288599-33288656 | DAXX;DAXX;DAXX;DAXX | N_Shore | Body;Body;Body;Body |
| GM00042 | chr6:33358866-33359001 | KIFC1 | N_Shore | TSS1500 |
| GM01392 | chr6:34024362-34024479 | GRM4 | Island | Body |
| GM00289 | chr6:35265534-35265555 | DEF6 | | TSS200 |
| GM00935 | chr6:35265616-35265643 | DEF6;DEF6 | | 1stExon;5'UTR |
| GM00439 | chr6:41528395-41528531 | FOXP4;FOXP4;FOXP4 | Island | 5'UTR;5'UTR;5'UTR |
| GM00289 | chr6:41528395-41528785 | FOXP4;FOXP4;FOXP4 | Island | 5'UTR;5'UTR;5'UTR |
| GM00940 | chr6:45387257-45387395 | RUNX2;RUNX2 | N_Shore | Body;Body |
| GM00042 | chr6:47624115-47624247 | GPR111 | | TSS1500 |
| WG1513 | chr6:6002865-6003303 | NRN1 | Island | Body |
| GM02173 | chr6:6614447-6614798 | LOC285780;LY86 | | Body;Body |
| WG1513 | chr6:70576168-70576321 | COL19A1 | N_Shore | TSS1500 |
| GM00520 | chr6:75912547-75912711 | COL12A1;COL12A1 | S_Shore | 5'UTR;5'UTR |
| GM00290 | chr6:91004430-91004482 | BACH2;BACH2 | N_Shore | 5'UTR;5'UTR |
| GM00520 | chr7:100880751-100881286 | CLDN15 | | 1stExon |
| GM02706 | chr7:101500014-101500514 | CUX1;CUX1;CUX1 | | Body;Body;Body |
| GM00520 | chr7:101768610-101768874 | CUX1;CUX1;CUX1 | | Body;Body;Body |
| GM00439 | chr7:105319437-105319819 | ATXN7L1;ATXN7L1 | | Body;5'UTR |
| GM00438 | chr7:1098625-1098703 | C7orf50;GPR146;C7orf50;C7 | S_Shore | Body;3'UTR;Body;Body;1stExon |
| GM03073 | chr7:114055074-114055419 | FOXP2;FOXP2;FOXP2;FOXP2;FOXP2;FOX | | 1stExon;5'UTR;5'UTR;1stExon;5'UTR;1 |
| GM09256 | chr7:114055123-114055137 | FOXP2;FOXP2;FOXP2;FOXP2;FOXP2;FOX | | 1stExon;5'UTR;5'UTR;1stExon;5'UTR;1 |
| WG2276 | chr7:114055123-114055419 | FOXP2;FOXP2;FOXP2;FOXP2;FOXP2;FOX | | 1stExon;5'UTR;5'UTR;1stExon;5'UTR;1 |
| GM09255 | chr7:115862648-115863026 | TES;TES | | TSS1500;Body |
| GM00439 | chr7:1163456-1164063 | C7orf50;C7orf50;C7orf50 | | Body;Body;Body |
| GM00520 | chr7:128550860-128550902 | KCP;KCP | Island | TSS200;TSS200 |
| GM01095 | chr7:128550862-128550902 | KCP;KCP | Island | TSS200;TSS200 |
| GM00439 | chr7:134832850-134833299 | TMEM140;TMEM140 | | 1stExon;5'UTR |

| | | | | |
|---|---|---|---|---|
| GM00290 | chr7:150434471-150434541 | GIMAP5;GIMAP5 | | 1stExon;5'UTR |
| GM00438 | chr7:151442351-151442967 | PRKAG2;PRKAG2 | | Body;Body |
| GM01096 | chr7:155299624-155299687 | CNPY1 | N_Shelf | Body |
| GM02456 | chr7:157198482-157198869 | DNAJB6 | N_Shelf | Body |
| WG1310 | chr7:158336016-158336049 | PTPRN2;PTPRN2;PTPRN2 | | Body;Body;Body |
| GM00022 | chr7:1952518-1952600 | MAD1L1;MAD1L1;MAD1L1 | S_Shelf | Body;Body;Body |
| GM02706 | chr7:1952518-1952807 | MAD1L1;MAD1L1;MAD1L1 | S_Shelf | Body;Body;Body |
| GM09256 | chr7:2109244-2109890 | MAD1L1;MAD1L1;MAD1L1 | S_Shore | Body;Body;Body |
| GM02456 | chr7:2133441-2133480 | MAD1L1;MAD1L1;MAD1L1 | | Body;Body;Body |
| WG1513 | chr7:2139216-2139259 | MAD1L1;MAD1L1;MAD1L1 | | Body;Body;Body |
| GM00439 | chr7:2563478-2563909 | LFNG;LFNG;LFNG;LFNG | N_Shore | Body;Body;Body;Body |
| GM02706 | chr7:2563611-2563909 | LFNG;LFNG;LFNG;LFNG | Island | Body;Body;Body;Body |
| GM00016 | chr7:2606716-2606824 | IQCE;IQCE | | Body;Body |
| WG1283 | chr7:2646782-2647333 | IQCE;IQCE | S_Shore | Body;Body |
| GM00940 | chr7:27143334-27143403 | HOXA2 | Island | TSS1500 |
| GM00290 | chr7:27143334-27143478 | HOXA2 | Island | TSS1500 |
| GM09255 | chr7:27150262-27150598 | HOXA3;HOXA3;HOXA3 | Island | 5'UTR;5'UTR;5'UTR |
| GM00940 | chr7:27182493-27183196 | HOXA5 | N_Shore | Body |
| GM02641 | chr7:27182493-27183436 | HOXA5 | N_Shore | Body |
| GM00439 | chr7:27183274-27183436 | HOXA5;HOXA5 | Island | 5'UTR;1stExon |
| GM02706 | chr7:27187943-27188465 | HOXA6 | S_Shore | TSS1500 |
| GM00940 | chr7:27187943-27188770 | HOXA6 | S_Shore | TSS1500 |
| GM00439 | chr7:27188364-27188770 | HOXA6 | N_Shore | TSS1500 |
| GM00031 | chr7:27225058-27225078 | HOXA11AS;HOXA11 | Island | Body;TSS1500 |
| GM00940 | chr7:27225058-27225123 | HOXA11AS;HOXA11 | Island | Body;TSS1500 |
| GM00439 | chr7:27225070-27225222 | HOXA11AS;HOXA11 | Island | Body;TSS1500 |
| GM00022 | chr7:27225127-27225222 | HOXA11AS;HOXA11 | Island | Body;TSS1500 |
| GM09256 | chr7:27225299-27225342 | HOXA11AS;HOXA11 | Island | Body;TSS1500 |
| WG1513 | chr7:27225523-27225543 | HOXA11AS;HOXA11 | Island | Body;TSS1500 |
| GM00439 | chr7:27281315-27281346 | EVX1 | N_Shore | TSS1500 |
| GM00439 | chr7:2728776-2728912 | AMZ1 | S_Shore | 5'UTR |
| GM00935 | chr7:2800701-2801248 | GNA12 | N_Shore | Body |
| WG1722 | chr7:2802374-2802560 | GNA12 | N_Shore | Body |
| GM02147 | chr7:2802374-2802976 | GNA12 | N_Shore | Body |
| GM02175 | chr7:28452066-28452289 | CREB5;CREB5 | S_Shelf | 5'UTR;TSS200 |
| WG2414 | chr7:2847517-2847575 | GNA12 | | Body |
| WG2275 | chr7:28998577-28998860 | TRIL | S_Shore | TSS1500 |
| GM00940 | chr7:29186629-29187019 | CPVL;CPVL | | TSS1500;TSS1500 |
| GM02175 | chr7:29519409-29519550 | CHN2;CHN2 | | TSS200;Body |
| GM02149 | chr7:29602763-29603253 | PRR15 | N_Shore | TSS1500 |
| GM00438 | chr7:31726692-31726912 | C7orf16;C7orf16 | | TSS200;TSS200 |
| GM01095 | chr7:35294949-35295198 | TBX20;TBX20 | N_Shore | TSS1500;TSS1500 |
| WG1513 | chr7:4065931-4066127 | SDK1 | | Body |
| GM00438 | chr7:41739678-41739801 | LOC285954;LOC285954;INHBA | | Body;Body;Body |
| GM00042 | chr7:42267547-42267747 | GLI3 | Island | 5'UTR |
| WG1722 | chr7:42267547-42268109 | GLI3 | Island | 5'UTR |
| GM03073 | chr7:4266872-4267364 | SDK1;SDK1 | | Body;Body |

| | | | | |
|---|---|---|---|---|
| WG1282 | chr7:43288635-43289504 | HECW1 | Island | Body |
| GM00439 | chr7:4729417-4729920 | FOXK1 | | Body |
| GM09255 | chr7:4762742-4763182 | FOXK1 | N_Shore | Body |
| GM00289 | chr7:4778839-4779342 | FOXK1 | | Body |
| WG1977 | chr7:4885521-4885643 | RADIL | | Body |
| GM02706 | chr7:5336543-5336811 | SLC29A4;SLC29A4 | Island | Body;Body |
| GM00935 | chr7:5518392-5518918 | FBXL18 | N_Shelf | 3'UTR |
| GM00941 | chr7:553412-553693 | PDGFA;PDGFA | N_Shelf | Body;Body |
| GM00438 | chr7:630473-630914 | PRKAR1B;PRKAR1B;PRKAR1E | N_Shore | Body;Body;Body;Body;Body;Body |
| GM00935 | chr7:644219-644417 | PRKAR1B;PRKAR1B;PRKAR1E | Island | Body;Body;Body;Body;Body;Body |
| WG1721 | chr7:70096029-70096378 | AUTS2;AUTS2 | N_Shore | Body;Body |
| GM00935 | chr7:73894573-73895278 | GTF2IRD1;GTF2IRD1 | N_Shore | 5'UTR;5'UTR |
| GM02706 | chr7:766100-766119 | PRKAR1B;PRKAR1B;PRKAR1E | Island | 5'UTR;5'UTR;5'UTR;TSS1500 |
| GM00520 | chr7:906726-906799 | UNC84A;UNC84A | S_Shore | Body;Body |
| WG2414 | chr7:944607-944776 | ADAP1 | S_Shore | Body |
| GM00042 | chr7:96650096-96650192 | DLX5 | N_Shore | Body |
| WG2121 | chr7:96650096-96650407 | DLX5 | N_Shore | Body |
| GM09256 | chr8:104032819-104032876 | ATP6V1C1 | N_Shore | TSS1500 |
| WG1722 | chr8:10586959-10587055 | SOX7 | N_Shore | Body |
| GM00438 | chr8:10907718-10907788 | XKR6 | | Body |
| GM09255 | chr8:11141365-11141424 | MTMR9 | N_Shore | TSS1500 |
| GM03075 | chr8:116660311-116661063 | TRPS1 | N_Shore | 5'UTR |
| WG1309 | chr8:11839764-11839844 | DEFB135 | | TSS200 |
| GM00940 | chr8:135490694-135490821 | ZFAT;ZFAT | N_Shore | 3'UTR;3'UTR |
| GM00042 | chr8:143324660-143324707 | TSNARE1 | | Body |
| GM02706 | chr8:143545390-143545574 | BAI1;BAI1 | N_Shore | 5'UTR;1stExon |
| GM01375 | chr8:143545478-143545789 | BAI1;BAI1 | Island | 5'UTR;1stExon |
| GM02175 | chr8:143545940-143546893 | BAI1 | Island | 1stExon |
| GM00935 | chr8:143763326-143763565 | PSCA | | Body |
| GM01650 | chr8:145048235-145048363 | PLEC1;PLEC1 | Island | Body;TSS1500 |
| GM02706 | chr8:145654565-145654635 | VPS28;NFKBIL2;VPS28 | Island | TSS1500;Body;TSS1500 |
| GM00520 | chr8:1460569-1460718 | DLGAP2 | | 5'UTR |
| GM00439 | chr8:1786207-1786322 | ARHGEF10 | | 5'UTR |
| GM03073 | chr8:1847999-1848143 | ARHGEF10 | | Body |
| GM00438 | chr8:1847999-1848407 | ARHGEF10 | | Body |
| WG1310 | chr8:1848040-1848407 | ARHGEF10 | | Body |
| GM03075 | chr8:19539991-19540272 | CSGALNACT1;CSGALNACT1;CSGALNAC | | Body;5'UTR;1stExon |
| WG1283 | chr8:2075469-2075820 | MYOM2 | Island | Body |
| GM00439 | chr8:21961713-21961788 | FAM160B2 | | 3'UTR |
| GM09256 | chr8:22132874-22132959 | PIWIL2;PIWIL2;PIWIL2 | Island | 5'UTR;1stExon;TSS1500 |
| GM00439 | chr8:23162162-23162383 | LOXL2 | | Body |
| GM09255 | chr8:28560467-28560723 | EXTL3 | S_Shore | 5'UTR |
| WG1977 | chr8:40010443-40010823 | C8orf4 | | TSS1500 |
| GM01393 | chr8:42356637-42356871 | SLC20A2 | | 5'UTR |
| WG1977 | chr8:59058585-59058660 | FAM110B | Island | 5'UTR |
| GM00016 | chr8:6420242-6420726 | ANGPT2;ANGPT2;ANGPT2;M( | N_Shelf | 1stExon;1stExon;1stExon;Body |
| GM00520 | chr8:6456182-6456465 | MCPH1 | | Body |

| | | | | |
|---|---|---|---|---|
| GM00438 | chr8:67405405-67405742 | C8orf46 | | TSS200 |
| GM01376 | chr8:72754469-72755162 | MSC | Island | 3'UTR |
| WG2276 | chr8:95962371-95962463 | TP53INP1;TP53INP1 | Island | TSS1500;TSS1500 |
| GM02641 | chr8:9952926-9953022 | MSRA;MSRA;MSRA | | Body;TSS200;Body |
| GM01376 | chr9:130955135-130955436 | CIZ1;CIZ1;CIZ1;CIZ1;CIZ1 | N_Shore | TSS1500;5'UTR;TSS1500;5'UTR;TSS15 |
| GM02456 | chr9:139237461-139237889 | GPSM1 | Island | Body |
| GM00016 | chr9:139715701-139716215 | C9orf86;C9orf86 | Island | Body;Body |
| GM02175 | chr9:140401382-140401455 | PNPLA7;PNPLA7 | Island | Body;Body |
| GM02456 | chr9:140401382-140401576 | PNPLA7;PNPLA7 | Island | Body;Body |
| GM00290 | chr9:92220388-92220771 | GADD45G | Island | Body |
| GM02704 | chr9:92220617-92221282 | GADD45G | Island | Body |
| WG1657 | chr1:108734992-108735549 | SLC25A24;SLC25A24 | N_Shore | Body;Body |
| GM00290 | chr1:109941043-109941072 | SORT1 | S_Shore | TSS1500 |
| WG1983 | chr1:110254662-110254692 | GSTM5 | | TSS1500 |
| GM00941 | chr1:110254662-110254835 | GSTM5 | | TSS1500 |
| GM00016 | chr1:110254692-110254854 | GSTM5 | | TSS200 |
| GM02175 | chr1:110610899-110611823 | ALX3 | Island | Body |
| GM00289 | chr1:110611218-110611712 | ALX3 | Island | Body |
| WG1978 | chr1:1109832-1110107 | TTLL10 | Island | 5'UTR |
| WG1977 | chr1:111218079-111218282 | KCNA3 | S_Shore | TSS1500 |
| GM00520 | chr1:112531263-112531488 | KCND3;KCND3 | N_Shore | 5'UTR;5'UTR |
| GM02317 | chr1:115238457-115238559 | AMPD1 | | TSS1500 |
| GM02175 | chr1:115881044-115881132 | NGF | Island | TSS200 |
| GM09255 | chr1:11761296-11761416 | C1orf187 | | 5'UTR |
| GM02316 | chr1:119521928-119522386 | TBX15 | N_Shore | 5'UTR |
| WG2275 | chr1:119526060-119526437 | TBX15 | N_Shore | 5'UTR |
| GM09256 | chr1:119526060-119526783 | TBX15 | N_Shore | 5'UTR |
| GM02316 | chr1:119531625-119532925 | TBX15 | N_Shore | 5'UTR |
| WG1721 | chr1:119531718-119532044 | TBX15 | N_Shore | 5'UTR |
| GM00042 | chr1:119531718-119532093 | TBX15 | N_Shore | 5'UTR |
| GM01650 | chr1:119532093-119532542 | TBX15;TBX15 | Island | 1stExon;5'UTR |
| WG2275 | chr1:119532320-119532542 | TBX15 | S_Shore | TSS200 |
| GM02456 | chr1:119532736-119532925 | TBX15 | S_Shore | TSS1500 |
| WG3465 | chr1:1266178-1266657 | TAS1R3 | N_Shore | TSS1500 |
| GM09255 | chr1:1266468-1266548 | TAS1R3 | N_Shore | TSS1500 |
| GM02175 | chr1:14027521-14027581 | PRDM2 | S_Shore | 5'UTR |
| GM00935 | chr1:150947896-150948030 | LASS2;LASS2 | Island | TSS1500;TSS1500 |
| GM01376 | chr1:150947949-150948030 | LASS2;LASS2 | Island | TSS1500;TSS1500 |
| GM00439 | chr1:15175204-15175356 | KIAA1026;KIAA1026 | | Body;Body |
| GM00935 | chr1:151811923-151812435 | LOC100132111;C2CD4D | N_Shore | Body;5'UTR |
| GM03075 | chr1:151812421-151812524 | LOC100132111;C2CD4D | Island | Body;5'UTR |
| WG1721 | chr1:15256054-15256136 | KIAA1026;KIAA1026;KIAA1026;KIAA1026 | | Body;TSS1500;Body;Body |
| WG3465 | chr1:15256054-15256222 | KIAA1026;KIAA1026;KIAA1026;KIAA1026 | | Body;TSS1500;Body;Body |
| GM00941 | chr1:15256123-15256136 | KIAA1026;KIAA1026;KIAA1026;KIAA1026 | | Body;TSS200;Body;Body |
| WG1283 | chr1:153581494-153581793 | S100A16 | | 5'UTR |
| WG1513 | chr1:154127952-154128041 | TPM3;TPM3;NUP210L;TPM3;TPM3;NUP2 | | 3'UTR;3'UTR;TSS1500;3'UTR;3'UTR;TS |
| GM00438 | chr1:155290813-155291224 | RUSC1;RUSC1;C1orf104 | Island | 5'UTR;5'UTR;Body |

| | | | | |
|---|---|---|---|---|
| GM00289 | chr1:156261200-156261351 | TMEM79;TMEM79 | Island | Body;Body |
| GM00940 | chr1:156786601-156786653 | SH2D2A;NTRK1;SH2D2A;SH2 | S_Shelf | 5'UTR;Body;1stExon;TSS1500;5'UTR;5 |
| GM02149 | chr1:158390649-158390842 | OR10K2 | | 1stExon |
| GM02456 | chr1:159796617-159796658 | SLAMF8;SLAMF8 | | 5'UTR;1stExon |
| WG2194 | chr1:161008705-161008826 | TSTD1;TSTD1;TSTD1;TSTD1;T | Island | 5'UTR;1stExon;5'UTR;1stExon;1stExon |
| GM00520 | chr1:161049443-161049579 | PVRL4 | Island | Body |
| GM00042 | chr1:165204410-165204512 | LMX1A | N_Shore | Body |
| WG1283 | chr1:165322099-165322784 | LMX1A | Island | Body |
| GM02706 | chr1:165323232-165323768 | LMX1A | N_Shore | Body |
| GM00290 | chr1:165324100-165324810 | LMX1A | N_Shore | Body |
| GM00439 | chr1:165324100-165324990 | LMX1A | N_Shore | Body |
| WG2121 | chr1:170638807-170639289 | PRRX1;PRRX1 | | Body;Body |
| GM01096 | chr1:17091946-17092279 | MST1P9 | | TSS1500 |
| GM00438 | chr1:172413492-172413542 | C1orf105;PIGC;PIGC | S_Shore | Body;TSS1500;TSS1500 |
| WG1514 | chr1:173176362-173176523 | TNFSF4;TNFSF4 | | 5'UTR;1stExon |
| WG1657 | chr1:173176366-173176523 | TNFSF4;TNFSF4 | | 5'UTR;1stExon |
| WG1978 | chr1:17944356-17944660 | ARHGEF10L;ARHGEF10L | | Body;Body |
| WG2414 | chr1:182921851-182922263 | C1orf14 | Island | Body |
| GM02175 | chr1:1957073-1957385 | GABRD | Island | Body |
| GM02175 | chr1:19600673-19600719 | AKR7L;AKR7L | Island | TSS200;TSS200 |
| WG1513 | chr1:19600673-19600730 | AKR7L;AKR7L | Island | TSS200;TSS200 |
| GM00520 | chr1:19600673-19600817 | AKR7L;AKR7L | Island | TSS200;TSS200 |
| GM01393 | chr1:19600702-19600730 | AKR7L;AKR7L | Island | TSS200;TSS200 |
| WG1513 | chr1:2005260-2005564 | PRKCZ;PRKCZ;PRKCZ | Island | 1stExon;Body;5'UTR |
| GM00042 | chr1:2005489-2005564 | PRKCZ;PRKCZ | Island | Body;5'UTR |
| GM00042 | chr1:201368946-201369031 | LAD1 | Island | TSS1500 |
| GM00439 | chr1:201709336-201709675 | NAV1;NAV1 | Island | Body;Body |
| WG1513 | chr1:202091865-202091880 | GPR37L1 | | TSS200 |
| GM00290 | chr1:2046702-2046864 | PRKCZ;PRKCZ;PRKCZ | Island | 5'UTR;Body;5'UTR |
| WG1722 | chr1:2046702-2046928 | PRKCZ;PRKCZ;PRKCZ | Island | 5'UTR;Body;5'UTR |
| GM01650 | chr1:2046783-2046928 | PRKCZ;PRKCZ;PRKCZ | Island | 5'UTR;Body;5'UTR |
| WG1977 | chr1:205602864-205603068 | ELK4;ELK4 | S_Shore | TSS1500;TSS1500 |
| WG1721 | chr1:205818956-205819463 | PM20D1 | Island | Body |
| WG1722 | chr1:205818956-205819492 | PM20D1 | Island | Body |
| GM00439 | chr1:205818956-205819609 | PM20D1 | Island | Body |
| GM02706 | chr1:20617283-20617456 | VWA5B1 | | TSS200 |
| GM09256 | chr1:206786170-206786180 | LGTN | S_Shore | TSS1500 |
| WG2120 | chr1:207992403-207992548 | LOC148696 | | Body |
| WG1722 | chr1:20988590-20988763 | DDOST | S_Shore | TSS1500 |
| GM00439 | chr1:211499620-211499648 | TRAF5;TRAF5 | N_Shore | TSS1500;TSS1500 |
| WG1658 | chr1:2120985-2121724 | C1orf86;LOC100128003 | N_Shore | Body;Body |
| WG1283 | chr1:214477063-214477280 | SMYD2 | | Body |
| WG1310 | chr1:21586831-21587174 | ECE1;ECE1;ECE1;ECE1 | | Body;Body;Body;Body |
| WG1513 | chr1:217263507-217263847 | ESRRG;ESRRG;ESRRG | S_Shore | TSS1500;TSS1500;5'UTR |
| GM00935 | chr1:217309568-217309970 | ESRRG | N_Shore | 5'UTR |
| GM00290 | chr1:217309794-217309970 | ESRRG | N_Shore | 5'UTR |
| GM02175 | chr1:217309906-217310174 | ESRRG | N_Shore | 5'UTR |

| | | | | |
|---|---|---|---|---|
| GM02456 | chr1:2174725-2174895 | SKI | | Body |
| WG1310 | chr1:221051758-221052023 | HLX | N_Shore | TSS1500 |
| WG2415 | chr1:221053841-221055097 | HLX | N_Shore | Body |
| WG1984 | chr1:221053904-221054273 | HLX | N_Shore | Body |
| GM02640 | chr1:221053904-221055097 | HLX | N_Shore | Body |
| GM01653 | chr1:221054201-221055097 | HLX | N_Shore | Body |
| WG2121 | chr1:221054615-221055097 | HLX | Island | Body |
| GM09255 | chr1:223566643-223566669 | C1orf65 | Island | TSS200 |
| WG1722 | chr1:223566643-223566709 | C1orf65 | Island | TSS200 |
| GM00438 | chr1:223566643-223567173 | C1orf65 | Island | TSS200 |
| GM00935 | chr1:223566646-223567173 | C1orf65 | Island | TSS200 |
| GM01392 | chr1:223566761-223567173 | C1orf65;C1orf65 | Island | 5'UTR;1stExon |
| GM00042 | chr1:223936343-223936799 | CAPN2;CAPN2 | Island | Body;Body |
| GM00438 | chr1:223936343-223936812 | CAPN2;CAPN2 | Island | Body;Body |
| GM02555 | chr1:223936508-223936812 | CAPN2;CAPN2 | Island | Body;Body |
| WG1722 | chr1:2257894-2258373 | MORN1 | N_Shore | Body |
| WG1722 | chr1:226825435-226825606 | ITPKB | | Body |
| GM00520 | chr1:227962723-227963171 | SNAP47 | | Body |
| GM01392 | chr1:227962873-227963171 | SNAP47 | | Body |
| WG1977 | chr1:228291591-228291682 | C1orf35 | S_Shore | TSS1500 |
| GM02704 | chr1:228291668-228291705 | C1orf35 | S_Shore | TSS1500 |
| GM00289 | chr1:228297864-228297975 | MRPL55;MRPL55;MRPL55;MI | S_Shore | TSS1500;TSS1500;TSS1500;TSS1500; |
| WG2415 | chr1:228395756-228395858 | OBSCN;OBSCN | N_Shelf | TSS200;TSS200 |
| WG2194 | chr1:23521272-23521417 | HTR1D | | TSS200 |
| GM00289 | chr1:236686564-236686618 | LGALS8;LGALS8;LGALS8;LGA | N_Shore | TSS200;TSS200;5'UTR;TSS1500 |
| WG1977 | chr1:236686598-236686671 | LGALS8;LGALS8;LGALS8;LGA | N_Shore | TSS200;TSS200;5'UTR;TSS1500 |
| GM00439 | chr1:236925877-236925965 | ACTN2 | | Body |
| WG3465 | chr1:240776162-240776211 | GREM2 | | TSS1500 |
| GM02456 | chr1:241912510-241912768 | WDR64 | | Body |
| GM09256 | chr1:2425860-2426035 | PLCH2 | | Body |
| GM02456 | chr1:243646402-243646787 | SDCCAG8 | Island | Body |
| WG1309 | chr1:247694271-247694406 | LOC148824;OR2C3 | Island | TSS200;3'UTR |
| WG1977 | chr1:25944712-25945042 | MAN1C1 | Island | 1stExon |
| WG1513 | chr1:26233565-26233623 | STMN1;STMN1;STMN1;STMN | Island | TSS1500;TSS1500;TSS1500;TSS200 |
| GM00290 | chr1:26797576-26797619 | DHDDS;HMGN2;DHDDS | N_Shore | 3'UTR;TSS1500;3'UTR |
| WG1513 | chr1:2987645-2987961 | PRDM16;PRDM16 | Island | Body;Body |
| GM00022 | chr1:3028230-3028457 | PRDM16;PRDM16 | Island | Body;Body |
| WG1977 | chr1:3028230-3028485 | PRDM16;PRDM16 | Island | Body;Body |
| GM02175 | chr1:3028338-3028485 | PRDM16;PRDM16 | Island | Body;Body |
| GM00940 | chr1:3038707-3039210 | PRDM16;PRDM16 | S_Shore | Body;Body |
| GM02175 | chr1:32714146-32714308 | FAM167B | Island | Body |
| GM00439 | chr1:32741152-32741518 | LCK;LCK | Island | Body;Body |
| GM02706 | chr1:3277525-3277623 | PRDM16;PRDM16 | S_Shelf | Body;Body |
| GM00439 | chr1:32802291-32802312 | MARCKSL1 | S_Shore | TSS1500 |
| WG2120 | chr1:3282449-3282870 | PRDM16;PRDM16 | | Body;Body |
| GM00290 | chr1:33648656-33648819 | TRIM62 | S_Shore | TSS1500 |
| WG1722 | chr1:3397061-3397125 | ARHGEF16 | S_Shore | Body |

| | | | | |
|---|---|---|---|---|
| WG2120 | chr1:3459949-3460268 | MEGF6 | Island | Body |
| WG1721 | chr1:3473665-3474202 | MEGF6 | | Body |
| GM00935 | chr1:3473923-3474376 | MEGF6 | | Body |
| GM00940 | chr1:3514921-3515194 | MEGF6 | S_Shelf | Body |
| WG2414 | chr1:36039655-36039885 | TFAP2E | Island | Body |
| GM00941 | chr1:36042986-36043014 | TFAP2E | Island | Body |
| GM00520 | chr1:36108206-36108233 | PSMB2 | S_Shore | TSS1500 |
| GM00438 | chr1:39546511-39546656 | MACF1 | N_Shore | TSS1500 |
| GM02706 | chr1:40024971-40025415 | LOC728448 | | Body |
| WG3465 | chr1:40105667-40105706 | HEYL | Island | TSS1500 |
| GM00940 | chr1:40769669-40769744 | COL9A2 | Island | Body |
| GM02706 | chr1:41250062-41250117 | KCNQ4;KCNQ4 | Island | 1stExon;1stExon |
| WG2120 | chr1:43751363-43751376 | C1orf210;C1orf210 | | TSS200;TSS200 |
| GM01096 | chr1:43751363-43751469 | C1orf210;C1orf210 | | TSS200;TSS200 |
| WG2120 | chr1:44016014-44016049 | PTPRF;PTPRF | | Body;Body |
| GM00935 | chr1:44703750-44704073 | ERI3 | | Body |
| GM00438 | chr1:4770676-4770935 | AJAP1;AJAP1 | N_Shore | Body;Body |
| GM02456 | chr1:4794912-4795114 | AJAP1;AJAP1 | | Body;Body |
| GM00438 | chr1:4841913-4841959 | AJAP1 | | 3'UTR |
| WG1513 | chr1:53600741-53600889 | SLC1A7 | | Body |
| GM00439 | chr1:54199696-54200420 | GLIS1;GLIS1 | N_Shelf | 5'UTR;1stExon |
| WG1657 | chr1:54199825-54200115 | GLIS1;GLIS1 | N_Shelf | 5'UTR;1stExon |
| WG1977 | chr1:54199825-54200420 | GLIS1;GLIS1 | N_Shelf | 5'UTR;1stExon |
| GM02555 | chr1:54200078-54200420 | GLIS1 | N_Shelf | TSS1500 |
| WG1977 | chr1:54856035-54856511 | SSBP3;SSBP3;SSBP3 | N_Shore | Body;Body;Body |
| GM02704 | chr1:54856177-54856511 | SSBP3;SSBP3;SSBP3 | N_Shore | Body;Body;Body |
| WG1513 | chr1:55267152-55267293 | TTC22;TTC22 | S_Shore | TSS1500;TSS1500 |
| GM00289 | chr1:59612408-59612710 | LOC729467 | Island | Body |
| WG1513 | chr1:6085995-6086148 | KCNAB2;KCNAB2 | N_Shore | TSS1500;TSS1500 |
| GM09256 | chr1:63249197-63249213 | ATG4C;ATG4C | | TSS1500;TSS1500 |
| GM02175 | chr1:63249199-63249213 | ATG4C;ATG4C | | TSS1500;TSS1500 |
| GM00042 | chr1:6519923-6520354 | ESPN | N_Shore | Body |
| GM02456 | chr1:6526182-6526319 | TNFRSF25;TNFRSF25;PLEKH | Island | 1stExon;1stExon;3'UTR;3'UTR;5'UTR;5 |
| WG2415 | chr1:6558085-6558113 | PLEKHG5;PLEKHG5 | S_Shore | Body;TSS1500 |
| WG1309 | chr1:7692321-7692367 | CAMTA1 | | Body |
| WG1514 | chr1:8403631-8403712 | SLC45A1 | N_Shore | Body |
| GM00290 | chr1:8403631-8403925 | SLC45A1 | N_Shore | Body |
| GM00022 | chr1:870791-870958 | SAMD11 | Island | Body |
| GM00289 | chr1:89663868-89664582 | GBP4 | | Body |
| GM00940 | chr1:89664260-89664407 | GBP4 | | Body |
| WG2121 | chr1:89664260-89664582 | GBP4 | | Body |
| GM01376 | chr1:89664338-89664407 | GBP4 | | Body |
| GM00439 | chr1:9129487-9129791 | SLC2A5;SLC2A5;SLC2A5 | N_Shelf | Body;Body;Body |
| GM02706 | chr1:9131622-9131783 | SLC2A5 | Island | Body |
| WG1513 | chr1:92352404-92352481 | TGFBR3 | S_Shore | TSS1500 |
| GM01653 | chr1:92414344-92414722 | BRDT | N_Shore | TSS1500 |
| GM02704 | chr1:92414344-92414782 | BRDT | N_Shore | TSS1500 |

| | | | | |
|---|---|---|---|---|
| GM01096 | chr1:92414520-92414737 | BRDT | N_Shore | TSS1500 |
| GM00935 | chr1:9400692-9400775 | SPSB1 | | 5'UTR |
| GM00438 | chr1:949392-949634 | ISG15 | Island | Body |
| WG1722 | chr1:95285405-95285531 | SLC44A3;SLC44A3 | N_Shore | TSS1500;TSS1500 |
| GM02173 | chr1:969825-970294 | AGRN | Island | Body |
| GM00289 | chr10:100991745-100992068 | HPSE2;HPSE2;HPSE2;HPSE2 | N_Shelf | Body;Body;Body;Body |
| GM01096 | chr10:105239164-105239205 | CALHM3 | | TSS200 |
| GM00042 | chr10:105978651-105978702 | MIR609;C10orf79 | | Body |
| GM00940 | chr10:106028373-106028519 | GSTO2 | N_Shore | TSS1500 |
| GM02173 | chr10:106088702-106088755 | ITPRIP | | 5'UTR |
| WG3465 | chr10:114877616-114877689 | TCF7L2;TCF7L2;TCF7L2;TCF7L2;TCF7L2 | | Body;Body;Body;Body;Body;Body |
| GM00935 | chr10:116164640-116164714 | AFAP1L2;AFAP1L2 | Island | TSS200;TSS200 |
| GM01376 | chr10:116418181-116418597 | ABLIM1;ABLIM1;ABLIM1 | | Body;Body;1stExon |
| GM01392 | chr10:11783823-11783974 | ECHDC3 | N_Shore | TSS1500 |
| GM02706 | chr10:119295400-119296182 | EMX2OS | Island | Body |
| GM00935 | chr10:119295619-119296182 | EMX2OS | Island | Body |
| GM02175 | chr10:119295770-119296182 | EMX2OS | Island | Body |
| GM02555 | chr10:121265587-121265893 | RGS10;RGS10 | | Body;Body |
| GM00520 | chr10:123923518-123924042 | TACC2;TACC2;TACC2;TACC2 | Island | Body;1stExon;Body;1stExon |
| GM02706 | chr10:1248770-1248818 | ADARB2 | N_Shelf | Body |
| GM00042 | chr10:125651526-125651588 | CPXM2 | S_Shore | 1stExon;5'UTR |
| GM00520 | chr10:125651526-125651621 | CPXM2 | S_Shore | 1stExon;5'UTR |
| WG1977 | chr10:126278272-126278502 | LHPP;LHPP | N_Shelf | Body;Body |
| GM00941 | chr10:126278372-126278502 | LHPP;LHPP | Island | Body;Body |
| WG1282 | chr10:12648032-12648338 | CAMK1D;CAMK1D | | Body;Body |
| GM02456 | chr10:12648032-12648526 | CAMK1D;CAMK1D | | Body;Body |
| GM02640 | chr10:12648176-12648526 | CAMK1D;CAMK1D | | Body;Body |
| GM00940 | chr10:126703991-126704259 | CTBP2;CTBP2;CTBP2 | | Body;Body;Body |
| WG1309 | chr10:131425198-131425282 | MGMT | | Body |
| WG1977 | chr10:131669406-131669630 | EBF3 | S_Shore | Body |
| WG1983 | chr10:131697080-131697345 | EBF3 | N_Shore | Body |
| WG1310 | chr10:131761206-131761386 | EBF3 | Island | Body |
| GM09256 | chr10:132891319-132891371 | TCERG1L | N_Shelf | 3'UTR |
| GM01392 | chr10:133929429-133929519 | JAKMIP3 | N_Shore | Body |
| GM00520 | chr10:133946674-133946881 | JAKMIP3 | N_Shelf | Body |
| WG1722 | chr10:133985911-133985960 | JAKMIP3 | | 3'UTR |
| GM02175 | chr10:133995286-133995387 | JAKMIP3 | | 3'UTR |
| GM00042 | chr10:134937967-134938469 | GPR123 | N_Shore | Body |
| GM00290 | chr10:135278901-135279010 | LOC619207 | Island | Body |
| GM00042 | chr10:135278976-135279147 | LOC619207 | Island | Body |
| WG2275 | chr10:135341528-135342218 | CYP2E1 | N_Shore | 1stExon |
| WG1310 | chr10:135342560-135343280 | CYP2E1 | Island | Body |
| GM01393 | chr10:135343193-135343280 | CYP2E1 | S_Shore | Body |
| WG1977 | chr10:135379213-135379242 | SYCE1;SYCE1;SYCE1 | N_Shore | Body;5'UTR;Body |
| GM00935 | chr10:13749193-13749625 | FRMD4A | | Body |
| GM02555 | chr10:13749285-13749625 | FRMD4A | | Body |
| WG1722 | chr10:13749382-13749625 | FRMD4A | | Body |

| | | | | |
|---|---|---|---|---|
| GM00941 | chr10:13932842-13933021 | FRMD4A | N_Shore | Body |
| WG2121 | chr10:13932842-13933247 | FRMD4A | N_Shore | Body |
| GM02706 | chr10:1416546-1416920 | ADARB2 | S_Shore | Body |
| GM00289 | chr10:1707523-1707576 | ADARB2 | | Body |
| WG1977 | chr10:1767482-1767511 | ADARB2 | | Body |
| GM09256 | chr10:1769480-1769577 | ADARB2 | | Body |
| WG1513 | chr10:18550033-18550223 | CACNB2;CACNB2;CACNB2;CACNB2;CA | | 1stExon;Body;Body;5'UTR;Body;Body; |
| WG1513 | chr10:22605982-22606072 | COMMD3 | S_Shore | Body |
| GM03075 | chr10:26855907-26856105 | APBB1IP | Island | Body |
| GM00439 | chr10:27541331-27541877 | LOC387646 | Island | TSS200 |
| GM01096 | chr10:28035631-28036151 | MKX | S_Shore | TSS1500 |
| WG1513 | chr10:28035731-28036151 | MKX | S_Shore | TSS1500 |
| GM02555 | chr10:29698368-29698449 | LOC387647 | Island | TSS200 |
| GM03075 | chr10:29811530-29811659 | SVIL;SVIL | | Body;Body |
| WG2415 | chr10:3138418-3138534 | PFKP | | Body |
| GM00289 | chr10:3149673-3149852 | PFKP | S_Shore | Body |
| GM00438 | chr10:31607150-31607160 | ZEB1;ZEB1;ZEB1;LOC22093( | N_Shore | TSS1500;TSS1500;TSS1500;Body;TSS |
| WG1513 | chr10:384506-384652 | DIP2C | N_Shelf | Body |
| GM00022 | chr10:38691787-38691801 | CDC10L | Island | TSS200 |
| GM00042 | chr10:42862978-42863594 | LOC441666 | N_Shore | Body |
| WG1513 | chr10:45495739-45495981 | C10orf25;ZNF22 | N_Shore | 3'UTR;TSS1500 |
| GM00935 | chr10:45495739-45496087 | C10orf25;ZNF22 | N_Shore | 3'UTR;TSS1500 |
| WG1977 | chr10:45495782-45495981 | C10orf25;ZNF22 | N_Shore | 3'UTR;TSS1500 |
| WG1977 | chr10:46993502-46993542 | GPRIN2 | Island | TSS1500 |
| GM02175 | chr10:4868328-4868333 | AKR1E2 | Island | TSS200 |
| GM09255 | chr10:50506748-50506885 | C10orf71;C10orf71 | | TSS1500;TSS1500 |
| GM09256 | chr10:50506748-50506896 | C10orf71;C10orf71 | | TSS1500;TSS1500 |
| GM00520 | chr10:50506875-50506896 | C10orf71;C10orf71 | | TSS1500;TSS1500 |
| GM00290 | chr10:50507071-50507214 | C10orf71;C10orf71 | | TSS200;TSS200 |
| GM00042 | chr10:50507071-50507298 | C10orf71;C10orf71 | | TSS200;TSS200 |
| GM02149 | chr10:530635-531569 | DIP2C | N_Shore | Body |
| GM09255 | chr10:530951-531465 | DIP2C | Island | Body |
| WG2276 | chr10:556470-556762 | DIP2C | | Body |
| GM00438 | chr10:616959-617105 | DIP2C | | Body |
| GM02456 | chr10:62671136-62671228 | RHOBTB1;RHOBTB1;RHOBTB1;RHOBTB1 | | Body;Body;Body;Body |
| GM01375 | chr10:661009-661159 | DIP2C | | Body |
| GM00935 | chr10:661009-661730 | DIP2C | | Body |
| WG2276 | chr10:670815-671628 | DIP2C | S_Shore | Body |
| WG1978 | chr10:682125-682152 | DIP2C | N_Shelf | Body |
| GM00289 | chr10:71087834-71088038 | HK1;HK1;HK1;HK1;HK1 | | Body;Body;Body;Body;Body |
| WG1513 | chr10:72201600-72201622 | NODAL | S_Shore | TSS200 |
| WG1722 | chr10:72254314-72254335 | KIAA1274 | S_Shelf | 5'UTR |
| WG2414 | chr10:72311914-72312093 | KIAA1274 | | Body |
| GM00289 | chr10:7290492-7290545 | SFMBT2 | | Body |
| GM02175 | chr10:73497514-73497629 | C10orf105;C10orf105;CDH23 | | 5'UTR;Body |
| WG1722 | chr10:73848764-73848823 | SPOCK2;SPOCK2;SPOCK2;S | S_Shore | 5'UTR;5'UTR |
| GM02706 | chr10:73976790-73976821 | ASCC1;C10orf104 | S_Shore | TSS1500;5'UTR |

| | | | | |
|---|---|---|---|---|
| GM00438 | chr10:77871618-77871958 | C10orf11 | N_Shelf | Body |
| GM00935 | chr10:77871958-77872170 | C10orf11 | N_Shore | Body |
| GM00935 | chr10:8096633-8096650 | FLJ45983;GATA3;GATA3;FLJ4 | Island | TSS1500;TSS200;TSS200;TSS1500 |
| WG2121 | chr10:82167757-82167774 | C10orf58 | N_Shelf | |
| GM00438 | chr10:88024557-88024572 | MIR346;GRID1 | S_Shore | TSS200;Body |
| GM00042 | chr10:88124936-88125351 | GRID1 | Island | Body |
| GM00042 | chr10:90712478-90712739 | ACTA2;ACTA2 | | Body;Body |
| GM00022 | chr10:90712478-90712948 | ACTA2;ACTA2 | | Body;Body |
| GM03075 | chr10:90712684-90712767 | ACTA2;ACTA2 | | 5'UTR;5'UTR |
| GM02555 | chr10:90712684-90712948 | ACTA2;ACTA2 | | 5'UTR;5'UTR |
| WG1283 | chr10:90712748-90712948 | ACTA2;ACTA2 | | TSS200;5'UTR |
| GM00289 | chr10:94451351-94451736 | HHEX | N_Shore | Body |
| WG2121 | chr10:95198200-95198374 | MYOF;MYOF | | Body;Body |
| GM00520 | chr10:95848649-95848841 | PLCE1;PLCE1 | | Body;TSS1500 |
| WG2415 | chr10:97175272-97175479 | SORBS1;SORBS1;SORBS1;SORBS1;SORI | | Body;Body;Body;Body;Body;Body;Bod |
| GM02555 | chr10:99080756-99081016 | FRAT1;FRAT1 | Island | 1stExon;3'UTR |
| GM01653 | chr10:99338056-99338188 | ANKRD2;ANKRD2 | N_Shore | Body;Body |
| WG1722 | chr10:99338056-99338240 | ANKRD2;ANKRD2 | N_Shore | Body;Body |
| GM00439 | chr10:99734081-99734154 | CRTAC1 | | Body |
| GM01096 | chr10:99735010-99735202 | CRTAC1 | | Body |
| GM01096 | chr10:99894013-99894087 | C10orf28 | N_Shelf | |
| WG2120 | chr11:1028291-1028379 | MUC6 | N_Shore | Body |
| WG1977 | chr11:10325567-10325809 | ADM | N_Shore | TSS1500 |
| WG2120 | chr11:1036727-1036766 | MUC6 | Island | TSS200 |
| WG1722 | chr11:1036754-1036865 | MUC6 | Island | TSS200 |
| WG2276 | chr11:10476976-10477452 | AMPD3;AMPD3;AMPD3;AMPD3 | | 1stExon;TSS1500;5'UTR;Body |
| GM00022 | chr11:10715175-10715595 | MRVI1;MRVI1;MRVI1;MRVI1;MRVI1 | | 1stExon;5'UTR;1stExon;5'UTR;TSS200 |
| GM00290 | chr11:107582804-107582876 | SLN | | TSS200 |
| GM02704 | chr11:109292789-109292894 | C11orf87 | N_Shore | TSS200 |
| GM00289 | chr11:111250129-111250201 | POU2AF1;POU2AF1 | | 1stExon;5'UTR |
| GM02175 | chr11:117748132-117748189 | FXYD6;FXYD6;FXYD6;FXYD6; | S_Shore | TSS1500;1stExon;5'UTR;TSS1500;TSS |
| GM09256 | chr11:118842387-118842392 | FOXR1 | Island | TSS200 |
| GM03073 | chr11:12031046-12031099 | DKK3;DKK3;DKK3 | S_Shore | TSS1500;TSS200;TSS1500 |
| WG1310 | chr11:12031093-12031099 | DKK3;DKK3;DKK3 | S_Shore | TSS1500;TSS200;TSS1500 |
| WG1513 | chr11:123066529-123066693 | ASAM | Island | TSS1500 |
| WG1283 | chr11:124767455-124767840 | ROBO4 | | Body |
| WG1978 | chr11:125037226-125037348 | PKNOX2 | S_Shore | 5'UTR |
| GM02175 | chr11:128646953-128647107 | FLI1;FLI1 | | Body;Body |
| GM00935 | chr11:128729883-128730011 | KCNJ1;KCNJ1;KCNJ1;KCNJ1 | | 5'UTR;5'UTR;5'UTR;5'UTR |
| WG1722 | chr11:128812978-128813008 | TP53AIP1 | | TSS200 |
| WG1977 | chr11:129991445-129992368 | APLP2;APLP2;APLP2;APLP2;A | N_Shelf | Body;Body;Body;Body;Body;Body |
| GM01392 | chr11:129992297-129992365 | APLP2;APLP2;APLP2;APLP2;A | N_Shore | Body;Body;Body;Body;Body;Body |
| WG2121 | chr11:129992297-129992368 | APLP2;APLP2;APLP2;APLP2;A | N_Shore | Body;Body;Body;Body;Body;Body |
| WG2414 | chr11:130319691-130320219 | ADAMTS15 | Island | 1stExon |
| GM00520 | chr11:131779604-131779859 | NTM;NTM;NTM;NTM | N_Shore | TSS1500;TSS1500;TSS1500;Body |
| WG2193 | chr11:132951838-132951950 | OPCML | Island | Body |
| WG1722 | chr11:133715076-133715511 | SPATA19 | | Body |

| | | | | |
|---|---|---|---|---|
| GM00042 | chr11:133800685-133800913 | IGSF9B | Island | Body |
| WG1977 | chr11:133994930-133995182 | JAM3 | Island | Body |
| GM00438 | chr11:134176692-134176861 | GLB1L3 | | Body |
| GM02149 | chr11:13985911-13986348 | SPON1 | Island | Body |
| WG1514 | chr11:1750302-1750381 | HCCA2 | | Body |
| GM01096 | chr11:1769289-1769390 | HCCA2;LOC402778;LOC402 | Island | Body;5'UTR;1stExon |
| WG2844 | chr11:18433500-18433564 | LDHC;LDHC | | TSS1500;TSS1500 |
| GM02175 | chr11:18477280-18477534 | LDHAL6A;LDHAL6A | N_Shore | TSS200;TSS200 |
| GM00940 | chr11:18477299-18477379 | LDHAL6A;LDHAL6A | Island | TSS200;TSS200 |
| GM02149 | chr11:18477299-18477534 | LDHAL6A;LDHAL6A | Island | TSS200;TSS200 |
| GM00042 | chr11:20692709-20692867 | NELL1;NELL1 | S_Shore | Body;Body |
| GM00520 | chr11:2162536-2162582 | IGF2AS;INS-IGF2;IGF2;IGF2A | S_Shore | Body;Body;5'UTR;Body;TSS200 |
| GM00042 | chr11:2172019-2172098 | INS-IGF2;IGF2;INS-IGF2 | | Body;TSS1500;Body |
| GM00042 | chr11:2293117-2293171 | ASCL2 | S_Shore | TSS1500 |
| GM02706 | chr11:2441045-2441141 | TRPM5 | N_Shore | Body |
| WG1722 | chr11:2464822-2464851 | KCNQ1 | N_Shore | TSS1500 |
| GM02175 | chr11:2464970-2465103 | KCNQ1 | N_Shore | TSS1500 |
| GM02706 | chr11:2554410-2554583 | KCNQ1;KCNQ1 | N_Shore | Body;Body |
| GM09256 | chr11:2907670-2907674 | CDKN1C;CDKN1C;CDKN1C | Island | TSS1500;TSS1500;TSS1500 |
| GM00031 | chr11:299564-299629 | IFITM5 | Island | TSS200 |
| WG1310 | chr11:315102-315262 | IFITM1 | N_Shore | Body |
| GM02456 | chr11:3164631-3164851 | OSBPL5;OSBPL5;OSBPL5 | | 5'UTR;5'UTR;5'UTR |
| WG1657 | chr11:31819444-31819678 | PAX6;PAX6;PAX6 | N_Shore | Body;Body;Body |
| GM02456 | chr11:31819444-31820658 | PAX6;PAX6;PAX6 | N_Shore | Body;Body;Body |
| GM02555 | chr11:31821590-31822302 | PAX6;PAX6;PAX6 | S_Shore | Body;Body;Body |
| WG2194 | chr11:31821764-31822302 | PAX6;PAX6;PAX6 | S_Shore | Body;Body;Body |
| WG1722 | chr11:31824262-31825226 | PAX6;PAX6;PAX6 | N_Shore | Body;Body;Body |
| GM01095 | chr11:31825166-31825226 | PAX6;PAX6;PAX6 | N_Shore | Body;Body;Body |
| GM01375 | chr11:31825605-31825792 | PAX6;PAX6;PAX6 | N_Shore | Body;Body;Body |
| GM02317 | chr11:31825756-31825833 | PAX6;PAX6;PAX6 | Island | Body;Body;Body |
| WG2414 | chr11:31827084-31827757 | PAX6;PAX6;PAX6 | N_Shore | Body;Body;Body |
| GM00022 | chr11:31827084-31827920 | PAX6;PAX6;PAX6 | N_Shore | Body;Body;Body |
| GM02456 | chr11:31827084-31828040 | PAX6;PAX6;PAX6 | N_Shore | Body;Body;Body |
| GM01095 | chr11:31827899-31828040 | PAX6;PAX6;PAX6 | Island | Body;Body;Body |
| GM00439 | chr11:31835929-31836334 | PAX6 | Island | 5'UTR |
| WG1977 | chr11:31837104-31837144 | PAX6 | Island | 5'UTR |
| GM01650 | chr11:32421514-32421635 | WT1;WT1;WT1;WT1 | | Body;Body;Body;Body |
| WG2120 | chr11:3253838-3254065 | MRGPRE | Island | TSS1500 |
| GM01650 | chr11:3253896-3254065 | MRGPRE | Island | TSS1500 |
| GM02555 | chr11:34460336-34460789 | CAT | Island | TSS200 |
| GM02175 | chr11:415080-415111 | SIGIRR;SIGIRR;SIGIRR | N_Shore | 5'UTR;TSS200;5'UTR |
| GM00439 | chr11:4415193-4415245 | TRIM21 | | TSS1500 |
| WG1514 | chr11:44325687-44325960 | ALX4 | Island | Body |
| GM00290 | chr11:44325687-44326516 | ALX4 | Island | Body |
| GM02175 | chr11:44325823-44326516 | ALX4 | Island | Body |
| GM02456 | chr11:44326260-44326516 | ALX4 | Island | Body |
| GM02173 | chr11:44327015-44327572 | ALX4 | N_Shore | Body |

| | | | | |
|---|---|---|---|---|
| WG1984 | chr11:44327267-44327572 | ALX4 | Island | Body |
| WG1514 | chr11:44332624-44332664 | ALX4 | Island | TSS1500 |
| GM03075 | chr11:44332624-44332685 | ALX4 | Island | TSS1500 |
| GM09255 | chr11:44332636-44332664 | ALX4 | Island | TSS1500 |
| GM01376 | chr11:44332653-44332685 | ALX4 | Island | TSS1500 |
| WG1514 | chr11:44332838-44333067 | ALX4 | S_Shore | TSS1500 |
| GM00520 | chr11:44332891-44332938 | ALX4 | S_Shore | TSS1500 |
| GM02555 | chr11:44332938-44333067 | ALX4 | S_Shore | TSS1500 |
| GM02173 | chr11:44332972-44333067 | ALX4 | S_Shore | TSS1500 |
| GM00289 | chr11:44333072-44333192 | ALX4 | S_Shore | TSS1500 |
| GM02456 | chr11:45825469-45825485 | SLC35C1;SLC35C1;SLC35C1 | N_Shore | TSS200;TSS1500;TSS200 |
| GM02149 | chr11:4902464-4903155 | OR51T1 | | TSS1500 |
| GM00438 | chr11:60048004-60048390 | MS4A4A | | TSS200 |
| GM02149 | chr11:60282420-60282742 | MS4A13;MS4A13 | N_Shore | TSS1500;TSS1500 |
| GM01393 | chr11:60719947-60720229 | SLC15A3;SLC15A3 | S_Shore | TSS1500;TSS1500 |
| WG1513 | chr11:61159602-61159687 | TMEM216 | N_Shore | TSS1500 |
| GM02456 | chr11:61248435-61248556 | C11orf66;C11orf66 | | TSS200;TSS200 |
| GM00935 | chr11:61544854-61545348 | C11orf9;C11orf9 | Island | Body;Body |
| WG1978 | chr11:628603-628677 | SCT | S_Shore | TSS1500 |
| GM00520 | chr11:63655895-63656088 | MARK2;MARK2;MARK2;MARK2;MARK2 | | Body;TSS200;Body;Body;Body |
| WG2415 | chr11:63655903-63656090 | MARK2;MARK2;MARK2;MARK2;MARK2 | | Body;TSS200;Body;Body;Body |
| WG1978 | chr11:63775226-63775436 | MACROD1 | N_Shore | Body |
| GM00289 | chr11:63784063-63784126 | MACROD1 | | Body |
| WG2844 | chr11:63974772-63975026 | FERMT3;FERMT3 | N_Shore | 5'UTR;5'UTR |
| GM00290 | chr11:64081545-64081631 | ESRRA | N_Shelf | Body |
| GM09256 | chr11:64428252-64428434 | NRXN2;NRXN2 | Island | Body;Body |
| GM00940 | chr11:66311334-66311422 | ZDHHC24 | N_Shore | Body |
| GM02175 | chr11:66650500-66650560 | PC;PC;PC | | 5'UTR;5'UTR;5'UTR |
| WG1977 | chr11:66885240-66885281 | KDM2A;KDM2A | N_Shore | TSS1500;TSS1500 |
| GM02456 | chr11:68138505-68138908 | LRP5 | | Body |
| WG1310 | chr11:68147880-68148079 | LRP5 | | Body |
| GM00940 | chr11:68579970-68580041 | CPT1A;CPT1A | | Body;Body |
| GM00520 | chr11:68610186-68610381 | CPT1A;CPT1A | N_Shore | TSS1500;TSS1500 |
| GM09255 | chr11:68610706-68611016 | CPT1A;CPT1A | N_Shore | TSS1500;TSS1500 |
| GM00439 | chr11:68610706-68611260 | CPT1A;CPT1A | N_Shore | TSS1500;TSS1500 |
| WG1283 | chr11:68782049-68782202 | MRGPRF;MRGPRF | S_Shelf | TSS1500;TSS1500 |
| GM02456 | chr11:68782049-68782211 | MRGPRF;MRGPRF | S_Shelf | TSS1500;TSS1500 |
| GM00438 | chr11:70048796-70048845 | FADD | N_Shore | TSS1500 |
| GM00520 | chr11:70048796-70048852 | FADD | N_Shore | TSS1500 |
| GM02175 | chr11:70048801-70048852 | FADD | N_Shore | TSS1500 |
| GM00520 | chr11:70559053-70559104 | SHANK2 | | Body |
| GM00438 | chr11:70644110-70644185 | SHANK2 | | Body |
| GM02706 | chr11:71709888-71709978 | IL18BP;IL18BP;IL18BP;IL18BP;IL18BP | | TSS200;TSS1500;TSS200;TSS1500;TS |
| GM00042 | chr11:71952131-71952431 | PHOX2A | Island | Body |
| GM01096 | chr11:72295425-72295716 | PDE2A;PDE2A;PDE2A;PDE2A | Island | Body;Body;Body;Body |
| WG1978 | chr11:72301235-72301724 | PDE2A;PDE2A;PDE2A;PDE2A | N_Shore | Body;Body;Body;Body |
| GM00439 | chr11:73356316-73356744 | PLEKHB1;PLEKHB1 | | TSS1500;TSS1500 |

| | | | | |
|---|---|---|---|---|
| GM02706 | chr11:74178761-74178800 | KCNE3 | Island | TSS200 |
| GM00935 | chr11:74178795-74178883 | KCNE3 | Island | TSS200 |
| GM00940 | chr11:75139390-75139736 | KLHL35 | N_Shore | Body |
| GM02456 | chr11:75898132-75898203 | WNT11 | | Body |
| WG1977 | chr11:77299805-77299962 | AQP11 | N_Shore | TSS1500 |
| WG1283 | chr11:85430629-85430657 | SYTL2;SYTL2;SYTL2;SYTL2;SYTL2;SYTL2; | | Body;TSS1500;TSS1500;Body;TSS150 |
| WG2275 | chr11:86383696-86383723 | ME3;ME3;ME3 | S_Shore | TSS1500;TSS200;TSS1500 |
| GM02175 | chr11:86383696-86383761 | ME3;ME3;ME3 | S_Shore | TSS1500;TSS200;TSS1500 |
| GM02555 | chr11:87908785-87908938 | RAB38 | S_Shore | TSS200 |
| GM00520 | chr11:87908805-87908938 | RAB38 | S_Shore | TSS1500 |
| GM02706 | chr11:87908817-87908938 | RAB38 | S_Shore | TSS1500 |
| WG1722 | chr11:93277161-93277191 | C11orf75 | S_Shore | TSS1500 |
| GM02640 | chr11:94278457-94279068 | FUT4 | Island | 1stExon |
| GM02706 | chr12:10084936-10085174 | CLEC2A;CLEC2A | | 5'UTR;1stExon |
| GM02555 | chr12:10251536-10251638 | CLEC1A;CLEC1A | | 5'UTR;1stExon |
| GM00935 | chr12:10251536-10251879 | CLEC1A;CLEC1A | | 5'UTR;1stExon |
| WG1513 | chr12:106988340-106988413 | RFX4 | | Body |
| GM02706 | chr12:107348824-107348944 | C12orf23 | N_Shore | TSS1500 |
| GM02555 | chr12:110172208-110172346 | MGC14436;MGC14436;C12orf34 | | Body;Body;5'UTR |
| GM01096 | chr12:11151478-11151622 | PRR4;TAS2R20;PRH1 | | Body;TSS1500;5'UTR |
| GM00520 | chr12:114833303-114834201 | TBX5;TBX5;TBX5;TBX5 | N_Shore | Body;Body;Body;Body |
| WG1977 | chr12:114833837-114834201 | TBX5;TBX5;TBX5;TBX5 | N_Shore | Body;Body;Body;Body |
| WG2120 | chr12:114833837-114834713 | TBX5;TBX5;TBX5;TBX5 | N_Shore | Body;Body;Body;Body |
| WG1513 | chr12:114837572-114837887 | TBX5;TBX5;TBX5;TBX5 | N_Shore | Body;Body;Body;Body |
| GM02704 | chr12:114837887-114838772 | TBX5;TBX5;TBX5;TBX5 | N_Shore | Body;Body;Body;Body |
| WG1514 | chr12:114841536-114841708 | TBX5;TBX5;TBX5;TBX5 | N_Shore | Body;Body;5'UTR;Body |
| GM01096 | chr12:114845180-114846503 | TBX5;TBX5;TBX5 | N_Shore | TSS1500;5'UTR;5'UTR |
| GM03073 | chr12:115109452-115109934 | TBX3;TBX3 | N_Shore | 3'UTR;3'UTR |
| GM00290 | chr12:116997069-116997079 | MAP1LC3B2 | | TSS200 |
| WG2276 | chr12:117155382-117155636 | C12orf49 | | 3'UTR |
| GM02456 | chr12:117407879-117408163 | FBXW8;FBXW8 | | Body;Body |
| GM01375 | chr12:122356033-122356852 | WDR66 | N_Shore | TSS1500 |
| WG1513 | chr12:122356056-122356598 | WDR66 | N_Shore | TSS1500 |
| GM00520 | chr12:122356056-122356781 | WDR66 | N_Shore | TSS1500 |
| GM03075 | chr12:122356056-122356852 | WDR66 | N_Shore | TSS1500 |
| WG2193 | chr12:122356316-122356598 | WDR66 | Island | TSS200 |
| GM02173 | chr12:122356390-122356598 | WDR66 | Island | TSS200 |
| GM00940 | chr12:123380870-123380891 | VPS37B | Island | TSS200 |
| GM00935 | chr12:124773668-124773870 | FAM101A | | TSS200 |
| WG1283 | chr12:124773668-124774078 | FAM101A | | TSS200 |
| WG1657 | chr12:124773870-124774078 | FAM101A;FAM101A | | 1stExon;5'UTR |
| WG1977 | chr12:124870112-124870323 | NCOR2;NCOR2 | N_Shelf | Body;Body |
| GM00016 | chr12:129299088-129299462 | SLC15A4;MGC16384 | | Body;Body |
| GM01392 | chr12:130316015-130316058 | TMEM132D | | Body |
| GM00289 | chr12:130823838-130824082 | PIWIL1 | Island | 5'UTR |
| GM09256 | chr12:131002386-131002918 | RIMBP2;RIMBP2 | | 5'UTR;1stExon |
| GM09256 | chr12:131303194-131303518 | STX2;STX2 | Island | Body;Body |

| | | | | |
|---|---|---|---|---|
| GM00042 | chr12:131303247-131303518 | STX2;STX2 | Island | Body;Body |
| GM00439 | chr12:131622284-131623508 | GPR133 | N_Shore | Body |
| GM00520 | chr12:131622584-131623340 | GPR133 | N_Shore | Body |
| GM00289 | chr12:132278430-132278510 | SFRS8 | S_Shore | Body |
| GM00439 | chr12:133065724-133065776 | FBRSL1 | N_Shore | TSS1500 |
| GM00042 | chr12:133159472-133159939 | FBRSL1 | Island | Body |
| GM00940 | chr12:14720938-14720974 | PLBD1 | Island | TSS200 |
| WG2276 | chr12:15114233-15114393 | ARHGDIB | | 5'UTR |
| GM00290 | chr12:15114494-15114710 | ARHGDIB;ARHGDIB | | 5'UTR;1stExon |
| GM00022 | chr12:15114494-15114732 | ARHGDIB;ARHGDIB | | 5'UTR;1stExon |
| WG1658 | chr12:15114703-15114732 | ARHGDIB | | TSS200 |
| GM00940 | chr12:15114703-15114904 | ARHGDIB | | TSS200 |
| WG1722 | chr12:1609245-1609369 | LOC100292680 | | TSS1500 |
| GM02149 | chr12:1609245-1609470 | LOC100292680 | | TSS1500 |
| GM01376 | chr12:1609346-1609470 | LOC100292680 | | TSS1500 |
| GM00290 | chr12:213776-214224 | IQSEC3;IQSEC3 | Island | Body;5'UTR |
| GM00520 | chr12:214000-214224 | IQSEC3;IQSEC3 | Island | Body;5'UTR |
| WG2193 | chr12:2339439-2339614 | CACNA1C;CACNA1C;CACNA | Island | Body;Body;Body;Body;Body;Body;Bod |
| GM00031 | chr12:25054873-25055108 | BCAT1 | N_Shore | Body |
| GM01392 | chr12:25054873-25055304 | BCAT1 | N_Shore | Body |
| GM01096 | chr12:25054873-25055421 | BCAT1 | N_Shore | Body |
| GM00520 | chr12:25055214-25055381 | BCAT1 | N_Shore | Body |
| GM01376 | chr12:25055262-25055381 | BCAT1 | N_Shore | Body |
| GM02641 | chr12:25055262-25055421 | BCAT1 | N_Shore | Body |
| WG2121 | chr12:25150530-25150626 | C12orf77 | | TSS200 |
| GM02173 | chr12:28124011-28124233 | PTHLH;PTHLH;PTHLH;PTHLH | S_Shore | TSS1500;5'UTR;5'UTR;TSS1500 |
| GM00520 | chr12:29376483-29376872 | FAR2 | | TSS200 |
| WG1513 | chr12:312592-313104 | SLC6A12;SLC6A12;SLC6A12 | Island | Body;Body;Body |
| GM02147 | chr12:45443808-45443996 | DBX2 | N_Shore | Body |
| GM02175 | chr12:46767665-46767747 | SLC38A2 | S_Shore | TSS1500 |
| GM02706 | chr12:51786038-51786547 | GALNT6 | S_Shore | TSS1500 |
| GM02706 | chr12:52281810-52282079 | ANKRD33;ANKRD33;ANKRD33;ANKRD33 | | 1stExon;5'UTR;1stExon;5'UTR |
| GM00941 | chr12:53183773-53184086 | KRT3 | Island | 3'UTR |
| GM01392 | chr12:53730072-53730277 | SP7 | | TSS1500 |
| WG2121 | chr12:54392734-54392884 | HOXC9 | N_Shore | TSS1500 |
| GM02640 | chr12:54392734-54393100 | HOXC9 | N_Shore | TSS1500 |
| GM01650 | chr12:54412269-54412565 | HOXC4;HOXC5;HOXC6 | N_Shore | 5'UTR;Body;5'UTR |
| GM01392 | chr12:54413000-54413384 | HOXC4;HOXC5;HOXC6 | Island | 5'UTR;Body;5'UTR |
| GM02147 | chr12:54423428-54423920 | HOXC4;HOXC6;HOXC5;HOX | Island | 5'UTR;Body;Body;Body |
| WG1722 | chr12:54423428-54424679 | HOXC4;HOXC6;HOXC5;HOX | Island | 5'UTR;Body;Body;Body |
| WG2120 | chr12:54423428-54425418 | HOXC4;HOXC6;HOXC5;HOX | Island | 5'UTR;Body;Body;Body |
| WG1283 | chr12:54424964-54425418 | HOXC4;HOXC5 | Island | 5'UTR;Body |
| WG2120 | chr12:54440773-54441509 | HOXC4 | Island | 5'UTR |
| WG2193 | chr12:54440806-54441213 | HOXC4 | Island | 5'UTR |
| GM00520 | chr12:54446019-54446100 | HOXC4 | Island | 5'UTR |
| WG1513 | chr12:54446019-54446253 | HOXC4 | Island | 5'UTR |
| GM00290 | chr12:54446019-54446537 | HOXC4 | Island | 5'UTR |

| | | | | |
|---|---|---|---|---|
| GM02704 | chr12:54446019-54446576 | HOXC4 | Island | 5'UTR |
| GM02173 | chr12:54446308-54446576 | HOXC4;HOXC4 | N_Shore | 5'UTR;TSS1500 |
| GM00042 | chr12:54446505-54446576 | HOXC4;HOXC4 | N_Shore | 5'UTR;TSS1500 |
| WG2414 | chr12:54448729-54448913 | HOXC4;HOXC4 | S_Shore | Body;Body |
| GM01096 | chr12:54454163-54454445 | FLJ12825 | N_Shore | Body |
| GM02173 | chr12:54785286-54785320 | ZNF385A;ZNF385A | S_Shore | TSS1500;TSS1500 |
| WG1513 | chr12:56329615-56329903 | DGKA;DGKA;DGKA;DGKA | Island | 5'UTR;5'UTR;5'UTR;5'UTR |
| WG1722 | chr12:56732615-56732723 | IL23A | | TSS200 |
| GM01096 | chr12:57915555-57915636 | MBD6;DDIT3 | N_Shore | TSS1500;TSS1500 |
| GM02704 | chr12:58013475-58013539 | SLC26A10 | Island | TSS1500 |
| GM00940 | chr12:58013487-58013651 | SLC26A10 | Island | TSS1500 |
| GM00439 | chr12:58013636-58013687 | SLC26A10 | Island | TSS200 |
| GM00520 | chr12:58021295-58021713 | B4GALNT1 | Island | Body |
| WG1310 | chr12:58021295-58021917 | B4GALNT1 | Island | Body |
| GM01096 | chr12:5937031-5937409 | ANO2 | Island | Body |
| GM09255 | chr12:6165816-6166173 | VWF | N_Shore | Body |
| GM01376 | chr12:6166874-6167002 | VWF | Island | Body |
| WG1722 | chr12:62583974-62584454 | FAM19A2 | N_Shore | 5'UTR |
| WG1978 | chr12:6442873-6443045 | TNFRSF1A | S_Shelf | Body |
| GM00289 | chr12:7070317-7070679 | PTPN6;PTPN6;PTPN6 | | 3'UTR;3'UTR;3'UTR |
| WG1977 | chr12:7281566-7281783 | CLSTN3;RBP5 | | TSS1500;TSS200 |
| GM02706 | chr12:75785230-75785295 | GLIPR1L2 | Island | Body |
| GM02555 | chr12:772861-772982 | NINJ2 | | TSS200 |
| GM00022 | chr12:81110706-81110776 | MYF5 | N_Shore | TSS200 |
| GM00042 | chr12:85306916-85307131 | SLC6A15;SLC6A15;SLC6A15 | S_Shore | TSS1500;TSS1500;TSS1500 |
| GM00520 | chr12:8815308-8815634 | MFAP5;MFAP5 | | 1stExon;5'UTR |
| GM02706 | chr12:89744471-89744621 | DUSP6;DUSP6 | N_Shore | Body;Body |
| GM01375 | chr12:89744488-89744621 | DUSP6;DUSP6 | N_Shore | Body;Body |
| GM02316 | chr12:89744621-89744701 | DUSP6;DUSP6 | N_Shore | Body;Body |
| WG2415 | chr12:97300410-97300429 | NEDD1;NEDD1;NEDD1;NEDI | N_Shore | TSS1500;TSS1500;TSS1500;TSS1500 |
| GM02149 | chr12:97885353-97886312 | RMST;MIR1251 | | Body;TSS1500 |
| WG1977 | chr13:100310162-100310405 | CLYBL | N_Shore | Body |
| WG1977 | chr13:103019936-103020405 | FGF14 | | Body |
| WG1310 | chr13:106116963-106117425 | DAOA;DAOA | | TSS1500;TSS1500 |
| WG1722 | chr13:110874267-110874326 | COL4A1 | | Body |
| GM02555 | chr13:110918331-110918683 | COL4A1 | Island | Body |
| GM01393 | chr13:110961404-110961606 | COL4A2 | S_Shore | Body |
| WG3465 | chr13:110993474-110993558 | COL4A2 | | Body |
| GM00940 | chr13:111062247-111062287 | COL4A2 | | Body |
| GM02555 | chr13:111521981-111522314 | C13orf29 | | Body |
| WG1282 | chr13:111522932-111522985 | C13orf29 | | TSS1500 |
| WG3465 | chr13:113436309-113436651 | ATP11A;ATP11A | N_Shelf | Body;Body |
| GM00042 | chr13:113436400-113436568 | ATP11A;ATP11A | N_Shelf | Body;Body |
| WG1722 | chr13:113436564-113436651 | ATP11A;ATP11A | N_Shelf | Body;Body |
| GM00042 | chr13:113540400-113540631 | ATP11A;ATP11A | Island | 3'UTR;3'UTR |
| GM09256 | chr13:113540421-113540557 | ATP11A;ATP11A | Island | 3'UTR;3'UTR |
| WG1722 | chr13:113636679-113636953 | MCF2L;MCF2L | S_Shore | Body;Body |

| | | | | |
|---|---|---|---|---|
| GM00520 | chr13:113698408-113698621 | MCF2L;MCF2L | N_Shelf | Body;Body |
| GM00935 | chr13:113698931-113699016 | MCF2L;MCF2L | N_Shelf | Body;Body |
| GM00022 | chr13:113698994-113699016 | MCF2L;MCF2L | N_Shelf | Body;Body |
| WG1978 | chr13:113784263-113784663 | F10 | Island | Body |
| GM00940 | chr13:114077033-114077172 | ADPRHL1;ADPRHL1 | | 3'UTR;3'UTR |
| WG1513 | chr13:114103646-114103796 | ADPRHL1;ADPRHL1 | | Body;TSS200 |
| GM01095 | chr13:114108052-114108652 | ADPRHL1 | | TSS1500 |
| GM00941 | chr13:114108433-114108515 | ADPRHL1 | | TSS1500 |
| WG1978 | chr13:114108433-114108652 | ADPRHL1 | | TSS1500 |
| GM00439 | chr13:114108433-114108713 | ADPRHL1 | | TSS1500 |
| WG1983 | chr13:114108471-114108652 | ADPRHL1 | | TSS1500 |
| WG1310 | chr13:114308619-114308749 | ATP4B | | Body |
| GM02555 | chr13:114828067-114828455 | RASA3 | | Body |
| WG2276 | chr13:114871422-114871575 | RASA3 | S_Shore | Body |
| WG1978 | chr13:114880761-114880889 | RASA3 | S_Shore | Body |
| GM09256 | chr13:20805196-20805385 | GJB6;GJB6;GJB6;GJB6;GJB6 | N_Shore | 5'UTR;5'UTR;5'UTR;5'UTR;1stExon |
| GM00438 | chr13:20805487-20805679 | GJB6;GJB6;GJB6;GJB6 | N_Shore | 5'UTR;5'UTR;5'UTR;TSS200 |
| GM01650 | chr13:21295941-21296465 | IL17D | Island | Body |
| GM00031 | chr13:25670170-25670228 | PABPC3 | Island | TSS200 |
| GM00520 | chr13:26586552-26586713 | ATP8A2 | Island | Body |
| GM00940 | chr13:28562819-28562850 | PRHOXNB | | TSS200 |
| GM00290 | chr13:30077315-30077489 | MTUS2;MTUS2 | Island | Body;Body |
| GM00520 | chr13:30088615-30088771 | SLC7A1 | Island | 3'UTR |
| GM02173 | chr13:30088640-30088771 | SLC7A1 | Island | Body |
| WG1977 | chr13:33590002-33590083 | KL | Island | TSS1500 |
| GM02704 | chr13:36044768-36044886 | NBEA | N_Shore | Body |
| WG1977 | chr13:36045201-36045352 | NBEA | Island | Body |
| GM02641 | chr13:36048892-36049380 | MAB21L1;MIR548F5;NBEA;M | N_Shore | 1stExon;Body;Body;3'UTR |
| WG2121 | chr13:36048892-36051073 | MAB21L1;MIR548F5;NBEA;M | N_Shore | 1stExon;Body;Body;3'UTR |
| WG2120 | chr13:36050788-36051073 | MAB21L1;MIR548F5;NBEA;M | N_Shore | 1stExon;Body;Body;5'UTR |
| GM01095 | chr13:36050881-36051073 | MIR548F5;NBEA;MAB21L1 | N_Shore | Body;Body;TSS200 |
| GM01392 | chr13:36050889-36051073 | MIR548F5;NBEA;MAB21L1 | N_Shore | Body;Body;TSS200 |
| GM02641 | chr13:36050901-36050993 | MIR548F5;NBEA;MAB21L1 | N_Shore | Body;Body;TSS200 |
| WG1722 | chr13:36050901-36051073 | MIR548F5;NBEA;MAB21L1 | N_Shore | Body;Body;TSS200 |
| GM00022 | chr13:36871646-36871878 | C13orf38;C13orf38;C13orf38 | N_Shore | Body;5'UTR;5'UTR;5'UTR;5'UTR;5'UTR |
| GM00438 | chr13:38172803-38173002 | POSTN;POSTN;POSTN;POSTN | | 1stExon;1stExon;1stExon;1stExon |
| GM00935 | chr13:41593385-41593519 | ELF1;ELF1 | | 5'UTR;1stExon |
| GM01392 | chr13:43566633-43566902 | EPSTI1;EPSTI1 | S_Shore | TSS1500;TSS1500 |
| GM01650 | chr13:43597565-43598038 | DNAJC15;DNAJC15 | Island | 5'UTR;1stExon |
| GM00940 | chr13:45149661-45149706 | TSC22D1 | N_Shore | 1stExon |
| GM00290 | chr13:45945197-45945362 | LOC100190939 | | Body |
| GM09256 | chr13:48575859-48575927 | SUCLA2 | S_Shore | TSS1500 |
| WG1283 | chr13:76334583-76334728 | LMO7;LMO7 | | TSS1500;Body |
| GM00439 | chr13:76334707-76334866 | LMO7;LMO7 | | TSS200;Body |
| GM02175 | chr13:79234435-79234847 | RNF219 | S_Shore | TSS1500 |
| GM01096 | chr13:88330200-88330614 | SLITRK5 | S_Shore | Body |
| GM01376 | chr13:95248771-95248888 | TGDS | S_Shore | TSS1500 |

| | | | | |
|---|---|---|---|---|
| GM01096 | chr13:98605770-98606196 | IPO5 | | TSS200 |
| WG1310 | chr13:98605892-98606196 | IPO5 | | TSS200 |
| GM02706 | chr13:98794353-98794593 | FARP1;FARP1 | N_Shore | TSS1500;TSS1500 |
| GM00042 | chr14:101036192-101036322 | BEGAIN | S_Shore | TSS200 |
| WG1722 | chr14:103593088-103593520 | TNFAIP2 | N_Shore | Body |
| GM02555 | chr14:103593235-103593520 | TNFAIP2 | Island | Body |
| WG2121 | chr14:103593503-103593520 | TNFAIP2 | Island | Body |
| GM02555 | chr14:104621867-104622128 | KIF26A | Island | Body |
| WG1513 | chr14:105635706-105635763 | JAG2;JAG2 | | TSS1500;TSS1500 |
| GM00940 | chr14:105714379-105714404 | BRF1;BTBD6;BRF1 | N_Shore | TSS200;TSS1500;Body |
| GM00520 | chr14:105796713-105796964 | PACS2;PACS2 | Island | Body;Body |
| WG1722 | chr14:23904678-23904930 | MYH7 | | 5'UTR |
| GM09256 | chr14:24641021-24641155 | REC8;REC8 | N_Shore | TSS1500;TSS1500 |
| GM01393 | chr14:24780167-24780557 | LTB4R2;LTB4R2;LTB4R2;LTB4 | Island | 1stExon;1stExon;Body;TSS1500;1stEx |
| GM02173 | chr14:24780540-24780557 | LTB4R2;LTB4R2;LTB4R2;CIDI | Island | Body;1stExon;Body;1stExon;5'UTR;TS |
| GM00940 | chr14:24780682-24780734 | LTB4R2;LTB4R2;LTB4R2;CIDI | Island | Body;1stExon;Body;TSS200;TSS200 |
| GM00935 | chr14:24785310-24785847 | LTB4R;LTB4R | Island | Body;Body |
| WG1283 | chr14:34269637-34270113 | NPAS3;NPAS3;NPAS3;NPAS3 | Island | Body;Body;Body;Body |
| GM01095 | chr14:37125430-37126582 | PAX9 | Island | TSS1500 |
| GM09255 | chr14:37125914-37126283 | PAX9 | N_Shore | TSS1500 |
| WG2414 | chr14:37135898-37136391 | PAX9 | Island | Body |
| WG2415 | chr14:37136192-37136391 | PAX9 | Island | Body |
| GM01392 | chr14:45432309-45432516 | KLHL28;FAM179B | S_Shore | TSS1500;1stExon |
| WG2844 | chr14:54418728-54418851 | BMP4;BMP4;BMP4 | Island | Body;Body;Body |
| GM00022 | chr14:54423807-54423845 | BMP4;BMP4 | S_Shore | TSS1500;TSS1500 |
| GM00940 | chr14:57277178-57277366 | OTX2;OTX2 | N_Shore | 1stExon;5'UTR |
| GM00289 | chr14:58862417-58862457 | TOMM20L | N_Shore | TSS1500 |
| GM02640 | chr14:73361747-73361964 | DPF3 | S_Shore | TSS1500 |
| GM02147 | chr14:75936582-75936789 | JDP2;JDP2;JDP2;JDP2 | | 3'UTR;3'UTR;3'UTR;3'UTR |
| GM09255 | chr14:80677608-80677688 | DIO2;DIO2;DIO2 | | 1stExon;1stExon;Body |
| GM01096 | chr14:81422098-81422160 | TSHR;TSHR;TSHR | S_Shore | 1stExon;1stExon;1stExon |
| GM09256 | chr14:94462309-94462337 | C14orf48;C14orf48;C14orf48 | | TSS1500;TSS1500;TSS1500 |
| GM01392 | chr14:95234822-95234965 | GSC | Island | 3'UTR |
| WG1310 | chr14:95235026-95235127 | GSC | Island | Body |
| GM02706 | chr14:96670628-96670738 | BDKRB2 | | TSS1500 |
| WG1309 | chr14:99728984-99729088 | BCL11B;BCL11B | | Body;Body |
| WG1513 | chr15:101419262-101419333 | ALDH1A3 | Island | TSS1500 |
| WG1978 | chr15:101419275-101419296 | ALDH1A3 | Island | TSS1500 |
| GM00290 | chr15:101419293-101419333 | ALDH1A3 | Island | TSS1500 |
| GM00022 | chr15:101420636-101420989 | ALDH1A3 | Island | Body |
| WG2414 | chr15:101728611-101728653 | CHSY1 | | Body |
| GM02456 | chr15:102193228-102193416 | TM2D3;TM2D3 | Island | TSS1500;TSS1500 |
| GM00438 | chr15:23810280-23810378 | MKRN3 | | TSS200 |
| WG1283 | chr15:26109215-26109261 | ATP10A | S_Shore | TSS1500 |
| GM02555 | chr15:26109252-26109394 | ATP10A | S_Shore | TSS1500 |
| GM02456 | chr15:27184766-27185073 | GABRA5;GABRA5 | | Body;Body |
| WG1977 | chr15:27215952-27215971 | GABRG3 | Island | TSS1500 |

| | | | | |
|---|---|---|---|---|
| GM00290 | chr15:28339873-28340213 | OCA2 | Island | 5'UTR |
| WG2120 | chr15:32964155-32964277 | SCG5;SCG5 | | Body;Body |
| WG1983 | chr15:32964155-32964500 | SCG5;SCG5 | | Body;Body |
| GM01650 | chr15:32964208-32964500 | SCG5;SCG5 | | Body;Body |
| GM02456 | chr15:33009409-33009531 | GREM1 | N_Shore | TSS1500 |
| GM09256 | chr15:34516488-34516777 | TMEM85 | | TSS1500 |
| GM02173 | chr15:35086890-35086985 | ACTC1 | | Body |
| GM03075 | chr15:36871588-36872085 | C15orf41 | Island | TSS1500 |
| GM02317 | chr15:37170381-37170554 | LOC145845 | N_Shelf | Body |
| WG1721 | chr15:40268610-40268777 | EIF2AK4 | Island | Body |
| WG2276 | chr15:40268610-40269214 | EIF2AK4 | Island | Body |
| WG1977 | chr15:40544174-40544427 | PAK6;C15orf56;PAK6;PAK6 | N_Shore | 5'UTR;3'UTR;5'UTR;5'UTR |
| GM00290 | chr15:41795087-41795475 | ITPKA | Island | Body |
| GM00438 | chr15:41806135-41806234 | LTK;LTK;LTK | S_Shore | TSS200;TSS200;TSS200 |
| GM00439 | chr15:42371511-42371653 | PLA2G4D | N_Shore | Body |
| WG2275 | chr15:44086617-44087089 | SERF2;MIR1282 | S_Shore | 3'UTR;TSS1500 |
| WG2415 | chr15:45314789-45314933 | SORD | N_Shore | TSS1500 |
| WG1513 | chr15:45671155-45671246 | LOC145663;GATM | S_Shore | Body;TSS200 |
| GM02555 | chr15:49716247-49716645 | C15orf33;FGF7 | | Body;5'UTR |
| WG3465 | chr15:51369862-51370019 | TNFAIP8L3 | | Body |
| WG1513 | chr15:51385891-51385970 | TNFAIP8L3 | Island | Body |
| WG1977 | chr15:55881599-55881706 | PYGO1 | S_Shore | TSS1500 |
| GM00520 | chr15:62359103-62359139 | C2CD4A | N_Shore | TSS200 |
| GM01393 | chr15:68125197-68125599 | LBXCOR1 | N_Shore | Body |
| WG2276 | chr15:70354647-70354825 | TLE3;TLE3;TLE3 | | Body;Body;Body |
| GM02704 | chr15:70994435-70994672 | UACA;UACA;UACA | | 5'UTR;Body;1stExon |
| WG1977 | chr15:70994435-70995014 | UACA;UACA;UACA | | 5'UTR;Body;1stExon |
| WG2414 | chr15:71408603-71408847 | CT62 | S_Shore | TSS1500 |
| GM00290 | chr15:74427234-74427499 | ISLR2;ISLR2;ISLR2;ISLR2 | Island | Body;Body;Body;Body |
| GM00439 | chr15:74494861-74494909 | STRA6;STRA6;STRA6;STRA6;STRA6 | | 5'UTR;5'UTR;5'UTR;5'UTR;5'UTR |
| WG1722 | chr15:74592566-74592786 | CCDC33 | | Body |
| GM02456 | chr15:76005861-76006143 | CSPG4 | | TSS1500 |
| GM00439 | chr15:81666512-81666671 | TMC3 | | TSS200 |
| WG2276 | chr15:83953690-83953744 | BNC1 | Island | TSS1500 |
| WG1310 | chr15:83953864-83953883 | BNC1 | Island | TSS1500 |
| GM00520 | chr15:84115433-84115529 | SH3GL3;SH3GL3 | N_Shore | TSS1500;TSS1500 |
| WG2844 | chr15:88576296-88576320 | NTRK3;NTRK3;NTRK3 | | Body;Body;Body |
| GM00520 | chr15:89153667-89154092 | MIR7-2 | | TSS1500 |
| GM02175 | chr15:92944664-92944756 | ST8SIA2 | | Body |
| GM01392 | chr15:93616943-93616959 | RGMA;RGMA;RGMA;RGMA;R | Island | Body;TSS200;TSS200;TSS1500;5'UTR; |
| GM09256 | chr15:93632896-93633172 | RGMA;RGMA | Island | TSS1500;TSS1500 |
| GM01096 | chr15:93632896-93633278 | RGMA;RGMA | Island | TSS1500;TSS1500 |
| GM01096 | chr15:96868935-96869132 | NR2F2 | S_Shelf | TSS1500 |
| GM02175 | chr15:96868935-96869220 | NR2F2 | S_Shelf | TSS1500 |
| GM02706 | chr15:96869027-96869220 | NR2F2 | S_Shelf | TSS200 |
| WG1513 | chr16:10912696-10912718 | FAM18A | Island | TSS200 |
| GM02149 | chr16:1128698-1128740 | SSTR5;LOC146336 | Island | TSS200;Body |

| | | | | |
|---|---|---|---|---|
| WG1513 | chr16:1429863-1430087 | UNKL | Island | TSS200 |
| GM09255 | chr16:14403004-14403040 | MIR365-1 | | TSS200 |
| GM00520 | chr16:15595987-15596215 | C16orf45;C16orf45 | | TSS200;Body |
| GM02456 | chr16:1581834-1582219 | IFT140 | N_Shore | Body |
| GM00935 | chr16:2082777-2083168 | SLC9A3R2;SLC9A3R2 | N_Shelf | Body;Body |
| GM00438 | chr16:23724835-23724943 | ERN2 | S_Shore | TSS200 |
| GM01393 | chr16:27791167-27791248 | KIAA0556 | | 3'UTR |
| GM02706 | chr16:2848577-2848830 | TESSP1 | Island | 1stExon |
| GM00941 | chr16:28634766-28635053 | SULT1A1;SULT1A1 | Island | 5'UTR;1stExon |
| WG1977 | chr16:2867434-2867683 | PRSS21;PRSS21;PRSS21 | Island | Body;Body;Body |
| WG2194 | chr16:2892766-2892782 | TMPRSS8 | Island | TSS200 |
| GM01392 | chr16:2892766-2892809 | TMPRSS8 | Island | TSS200 |
| WG3465 | chr16:2908554-2908934 | PRSS22 | S_Shore | TSS1500 |
| GM00439 | chr16:31022351-31022521 | STX1B | | TSS1500 |
| GM02555 | chr16:31147017-31147054 | PRSS8;PRSS8 | | 1stExon;5'UTR |
| GM00289 | chr16:3142911-3143041 | ZSCAN10 | S_Shelf | TSS200 |
| WG2844 | chr16:4103161-4103225 | ADCY9 | | Body |
| WG1310 | chr16:4163819-4164087 | ADCY9 | N_Shore | Body |
| GM00289 | chr16:450837-450970 | DECR2 | Island | TSS1500 |
| GM00289 | chr16:46603015-46603059 | ANKRD26P1 | | TSS200 |
| GM09256 | chr16:50347766-50347895 | ADCY7 | | Body |
| GM00940 | chr16:51183988-51184205 | SALL1;SALL1 | Island | Body;5'UTR |
| GM01393 | chr16:51184355-51184392 | SALL1;SALL1;SALL1 | Island | Body;5'UTR;1stExon |
| GM00935 | chr16:55794478-55794910 | CES4 | N_Shore | TSS200 |
| WG1282 | chr16:55794595-55794910 | CES4 | N_Shore | Body |
| GM03075 | chr16:56709853-56710256 | MT1IP | Island | Body |
| WG1721 | chr16:58019866-58019984 | TEPP;TEPP | Island | Body;Body |
| GM02706 | chr16:58533743-58534708 | NDRG4;NDRG4;NDRG4 | N_Shore | Body;Body;Body |
| WG1983 | chr16:65105516-65105714 | CDH11 | | 5'UTR |
| GM00022 | chr16:66613220-66613334 | CMTM2 | Island | TSS200 |
| GM01393 | chr16:66613266-66613278 | CMTM2 | Island | TSS200 |
| GM01095 | chr16:66613266-66613334 | CMTM2 | Island | TSS200 |
| GM01096 | chr16:67197084-67197186 | HSF4;FBXL8;HSF4 | Island | TSS1500;Body;TSS1500 |
| GM00289 | chr16:67197084-67197640 | HSF4;FBXL8;HSF4 | Island | TSS1500;Body;TSS1500 |
| GM00935 | chr16:67197129-67197640 | HSF4;HSF4;FBXL8 | Island | TSS200;TSS200;Body |
| GM02706 | chr16:67433258-67433636 | ZDHHC1 | Island | Body |
| GM01375 | chr16:67515916-67516493 | ATP6V0D1 | S_Shore | TSS1500 |
| GM00016 | chr16:68269295-68269694 | ESRP2 | Island | Body |
| WG1513 | chr16:68270252-68270282 | ESRP2 | Island | TSS200 |
| WG1977 | chr16:69958626-69958964 | WWP2;WWP2 | | TSS1500;Body |
| GM02175 | chr16:69958792-69958964 | WWP2;WWP2 | | TSS200;Body |
| GM09256 | chr16:70680320-70680380 | IL34 | | TSS200 |
| WG1722 | chr16:746048-746132 | FBXL16 | S_Shore | Body |
| WG1658 | chr16:81527427-81527504 | CMIP | | Body |
| WG2415 | chr16:83171068-83171314 | CDH13 | | Body |
| GM02706 | chr16:854168-854439 | PRR25 | N_Shelf | TSS1500 |
| GM02704 | chr16:857454-857981 | PRR25 | Island | Body |

| | | | | |
|---|---|---|---|---|
| GM00439 | chr16:8619531-8619943 | TMEM114 | N_Shore | 3'UTR |
| GM02706 | chr16:86371130-86371363 | LOC732275 | | Body |
| GM02706 | chr16:86546628-86546938 | FOXF1 | Island | Body |
| GM00290 | chr16:86547386-86547544 | FOXF1 | N_Shore | 3'UTR |
| GM00520 | chr16:87441441-87441659 | ZCCHC14 | Island | 3'UTR |
| GM00520 | chr16:87468206-87468247 | ZCCHC14 | | Body |
| GM00935 | chr16:87769432-87769974 | KLHDC4 | | Body |
| WG1310 | chr16:87769500-87769974 | KLHDC4 | | Body |
| GM00935 | chr16:87904491-87904580 | SLC7A5 | S_Shore | TSS1500 |
| WG2194 | chr16:88052956-88053104 | BANP;BANP | Island | Body;Body |
| GM02706 | chr16:88058182-88058326 | BANP;BANP | N_Shelf | Body;Body |
| GM01376 | chr16:88496788-88497293 | ZNF469 | Island | 1stExon |
| GM02555 | chr16:88590325-88590404 | ZFPM1 | Island | Body |
| WG2844 | chr16:88590325-88590409 | ZFPM1 | Island | Body |
| WG1513 | chr16:88706241-88706426 | IL17C | Island | Body |
| WG1513 | chr16:89445433-89445503 | ANKRD11 | | 5'UTR |
| GM00935 | chr16:89462292-89462418 | ANKRD11 | | 5'UTR |
| GM00520 | chr16:90016004-90016061 | DEF8;DEF8 | Island | 5'UTR;Body |
| WG2276 | chr16:968219-968265 | LMF1 | S_Shore | Body |
| WG1722 | chr17:1028394-1028676 | ABR;ABR | N_Shore | Body;Body |
| GM00520 | chr17:1028394-1028765 | ABR;ABR | N_Shore | Body;Body |
| GM09256 | chr17:1028394-1029394 | ABR;ABR | N_Shore | Body;Body |
| WG2194 | chr17:1028673-1028931 | ABR;ABR | Island | Body;5'UTR |
| GM02175 | chr17:10421635-10422077 | MYH1 | | 5'UTR |
| GM00439 | chr17:14206901-14207530 | HS3ST3B1;MGC12916 | Island | Body;TSS200 |
| GM00520 | chr17:14206981-14207530 | HS3ST3B1;MGC12916 | Island | Body;TSS200 |
| GM00520 | chr17:1466709-1466715 | PITPNA | S_Shore | TSS1500 |
| GM00439 | chr17:15244441-15244519 | TEKT3 | N_Shore | 5'UTR |
| GM00520 | chr17:15244482-15244887 | TEKT3 | N_Shore | 5'UTR |
| GM02706 | chr17:15244482-15244970 | TEKT3 | N_Shore | 5'UTR |
| WG1282 | chr17:154410-154671 | RPH3AL | S_Shore | Body |
| WG1721 | chr17:154499-155045 | RPH3AL | N_Shelf | Body |
| GM00520 | chr17:16318668-16318972 | TRPV2 | | TSS1500 |
| GM01096 | chr17:1634018-1634335 | WDR81;WDR81;WDR81;WDF | N_Shelf | Body;Body;Body;Body |
| GM00022 | chr17:16593601-16593885 | CCDC144A | Island | TSS200 |
| GM02706 | chr17:183931-184106 | RPH3AL | N_Shore | 5'UTR |
| WG2194 | chr17:1961286-1962236 | HIC1;HIC1 | Island | Body;Body |
| WG1513 | chr17:19648718-19648855 | ALDH3A1;ALDH3A1;ALDH3A | S_Shore | 5'UTR;5'UTR;5'UTR;1stExon |
| GM01095 | chr17:20799470-20799511 | CCDC144NL | Island | TSS200 |
| WG2844 | chr17:21300735-21300791 | KCNJ12 | | 5'UTR |
| WG2121 | chr17:26732191-26732272 | SLC46A1 | Island | Body |
| GM00520 | chr17:26732249-26732374 | SLC46A1 | Island | Body |
| GM00042 | chr17:29248604-29248725 | ADAP2 | N_Shore | TSS200 |
| GM03073 | chr17:29297380-29297414 | RNF135;RNF135 | N_Shore | TSS1500;TSS1500 |
| WG2414 | chr17:30822561-30823006 | MYO1D | N_Shore | Body |
| GM09256 | chr17:3117953-3118291 | OR1A1 | | TSS1500 |
| WG1722 | chr17:33416834-33417086 | RFFL | S_Shore | TSS1500 |

| | | | | |
|---|---|---|---|---|
| GM00042 | chr17:33759484-33759965 | SLFN12;SLFN12 | | 1stExon;5'UTR |
| WG1513 | chr17:33759484-33759986 | SLFN12;SLFN12 | | 1stExon;5'UTR |
| GM02555 | chr17:33759512-33759648 | SLFN12;SLFN12 | | 1stExon;5'UTR |
| WG1283 | chr17:33759929-33759965 | SLFN12 | | TSS1500 |
| GM02555 | chr17:33759957-33759986 | SLFN12 | | TSS1500 |
| GM02706 | chr17:3433428-3433970 | TRPV3 | N_Shelf | Body |
| GM00289 | chr17:34345136-34345392 | CCL23;CCL23 | | TSS200;TSS200 |
| WG1513 | chr17:36715787-36715953 | SRCIN1 | Island | Body |
| GM00439 | chr17:37028105-37028650 | LASP1 | S_Shore | Body |
| WG1978 | chr17:3705631-3705875 | ITGAE | | TSS1500 |
| GM02706 | chr17:37123767-37123843 | FBXO47 | | TSS200 |
| GM03075 | chr17:37894397-37894413 | GRB7;GRB7 | | TSS200;5'UTR |
| GM02175 | chr17:37894397-37894636 | GRB7;GRB7 | | TSS200;5'UTR |
| WG1722 | chr17:38249430-38249592 | THRA;NR1D1 | Island | Body;Body |
| GM00042 | chr17:38334161-38334439 | RAPGEFL1 | Island | TSS200 |
| GM00290 | chr17:38717206-38717275 | CCR7 | | Body |
| GM00290 | chr17:39728425-39728478 | KRT9 | | TSS200 |
| GM00935 | chr17:40259724-40260053 | DHX58 | Island | Body |
| GM02706 | chr17:40489584-40489785 | STAT3;STAT3;STAT3 | | Body;Body;Body |
| WG2415 | chr17:40687596-40687830 | NAGLU | N_Shore | TSS1500 |
| GM09256 | chr17:40835849-40836087 | CNTNAP1 | Island | Body |
| GM02706 | chr17:40935998-40936150 | WNK4 | Island | Body |
| WG1977 | chr17:40935998-40936498 | WNK4 | Island | Body |
| GM01392 | chr17:41277974-41278241 | BRCA1;NBR2;BRCA1;BRCA1; | N_Shore | TSS1500;Body;TSS1500;TSS1500;TSS |
| GM02706 | chr17:41278179-41278241 | BRCA1;NBR2;BRCA1;BRCA1; | Island | TSS1500;Body;TSS1500;TSS1500;TSS |
| GM00042 | chr17:41738893-41739195 | MEOX1;MEOX1;MEOX1 | | 1stExon;1stExon;5'UTR |
| GM02555 | chr17:41832753-41833161 | SOST | Island | Body |
| GM02706 | chr17:41909709-41909716 | MPP3;MPP3 | N_Shore | 5'UTR;Body |
| WG1722 | chr17:43319137-43319382 | FMNL1 | Island | Body |
| GM00940 | chr17:43482735-43483453 | ARHGAP27;ARHGAP27 | N_Shore | Body;Body |
| GM02706 | chr17:43483116-43483453 | ARHGAP27;ARHGAP27;ARHG | Island | 1stExon;5'UTR;5'UTR |
| GM01096 | chr17:43974869-43974975 | MAPT;MAPT;MAPT;MAPT;LOC | Island | 5'UTR;5'UTR;5'UTR;5'UTR;Body;5'UTR; |
| GM01376 | chr17:43974869-43975063 | MAPT;MAPT;MAPT;MAPT;LOC | Island | 5'UTR;5'UTR;5'UTR;5'UTR;Body;5'UTR; |
| WG1721 | chr17:43974919-43975063 | MAPT;MAPT;MAPT;MAPT;LOC | Island | 5'UTR;5'UTR;5'UTR;5'UTR;Body;5'UTR; |
| WG1722 | chr17:44847220-44847427 | WNT3 | Island | Body |
| WG1513 | chr17:45501242-45501697 | LOC100272146;C17orf57 | Island | Body;Body |
| GM00042 | chr17:46623729-46623819 | HOXB2 | S_Shelf | TSS1500 |
| WG2415 | chr17:46645972-46646359 | HOXB3 | S_Shelf | 5'UTR |
| GM02555 | chr17:46645972-46646444 | HOXB3 | S_Shelf | 5'UTR |
| WG1983 | chr17:46648014-46648533 | HOXB3 | | 5'UTR |
| GM00022 | chr17:46648014-46648582 | HOXB3 | | 5'UTR |
| GM00439 | chr17:46651722-46651920 | HOXB3;HOXB3 | N_Shelf | 5'UTR;1stExon |
| GM01375 | chr17:46651722-46652179 | HOXB3;HOXB3 | N_Shelf | 5'UTR;1stExon |
| WG1978 | chr17:46651945-46652179 | HOXB3 | N_Shelf | TSS200 |
| GM02149 | chr17:46653615-46654330 | HOXB4 | N_Shore | 3'UTR |
| WG1984 | chr17:46656543-46656690 | HOXB4 | S_Shore | TSS1500 |
| WG1722 | chr17:46656543-46657504 | HOXB4 | S_Shore | TSS1500 |

| | | | | |
|---|---|---|---|---|
| WG1513 | chr17:46657274-46657504 | MIR10A | N_Shore | Body |
| WG2844 | chr17:46669485-46669492 | LOC404266;HOXB5;LOC404 | Island | TSS200;3'UTR;Body;TSS200;Body |
| GM01650 | chr17:46669485-46669566 | LOC404266;HOXB5;LOC404 | Island | TSS200;3'UTR;Body;TSS200;Body |
| GM09256 | chr17:46669485-46669644 | LOC404266;HOXB5;LOC404 | Island | TSS200;3'UTR;Body;TSS200;Body |
| WG1722 | chr17:46669485-46670029 | LOC404266;HOXB5;LOC404 | Island | TSS200;3'UTR;Body;TSS200;Body |
| GM02704 | chr17:46671861-46672255 | LOC404266;LOC404266;LO( | N_Shore | Body;Body;Body;TSS1500;TSS1500;Bo |
| WG1978 | chr17:46673442-46674514 | LOC404266;LOC404266;LO( | N_Shore | Body;Body;Body;Body;3'UTR;Body |
| GM02175 | chr17:46673625-46674041 | LOC404266;LOC404266;LO( | Island | Body;Body;Body;Body;3'UTR;Body |
| GM00042 | chr17:46673625-46674438 | LOC404266;LOC404266;LO( | Island | Body;Body;Body;Body;3'UTR;Body |
| WG1722 | chr17:46673625-46674514 | LOC404266;LOC404266;LO( | Island | Body;Body;Body;Body;3'UTR;Body |
| WG1283 | chr17:46674041-46674438 | LOC404266;LOC404266;LO( | Island | Body;Body;Body;Body;Body;Body |
| GM03075 | chr17:46674041-46674514 | LOC404266;LOC404266;LO( | Island | Body;Body;Body;Body;Body;Body |
| GM01096 | chr17:46674395-46674514 | LOC404266;LOC404266;LO( | N_Shore | Body;Body;Body;Body;Body;Body |
| GM00520 | chr17:46676099-46676375 | LOC404266;LOC404266;LO( | S_Shore | Body;Body;Body;Body;5'UTR;Body |
| GM09255 | chr17:46681111-46681154 | LOC404266;LOC404266;HOXB6 | | Body;Body;5'UTR |
| GM00016 | chr17:46681111-46681362 | LOC404266;LOC404266;HOXB6 | | Body;Body;5'UTR |
| WG2120 | chr17:46681111-46683606 | LOC404266;LOC404266;HOXB6 | | Body;Body;5'UTR |
| GM00438 | chr17:46681135-46681362 | LOC404266;LOC404266;HOXB6 | | Body;Body;5'UTR |
| WG1513 | chr17:46681154-46681362 | LOC404266;LOC404266;HOXB6 | | Body;Body;5'UTR |
| GM02173 | chr17:46682308-46682398 | LOC404266;LOC404266;HO | N_Shelf | Body;Body;1stExon;5'UTR |
| GM01392 | chr17:46682308-46683047 | LOC404266;LOC404266;HO | N_Shelf | Body;Body;1stExon;5'UTR |
| GM00438 | chr17:46682308-46683409 | LOC404266;LOC404266;HO | N_Shelf | Body;Body;1stExon;5'UTR |
| GM00940 | chr17:46682319-46682398 | LOC404266;LOC404266;HO | N_Shelf | Body;Body;1stExon;5'UTR |
| GM09256 | chr17:46682319-46682413 | LOC404266;LOC404266;HO | N_Shelf | Body;Body;1stExon;5'UTR |
| WG1282 | chr17:46682390-46682398 | LOC404266;LOC404266;HO | N_Shelf | Body;Body;TSS200 |
| GM00031 | chr17:46682390-46682413 | LOC404266;LOC404266;HO | N_Shelf | Body;Body;TSS200 |
| GM00290 | chr17:46682390-46683606 | LOC404266;LOC404266;HO | N_Shelf | Body;Body;TSS200 |
| WG2121 | chr17:46685327-46685448 | HOXB7 | Island | Body |
| WG2415 | chr17:46685353-46685448 | HOXB7 | Island | Body |
| GM02640 | chr17:46689426-46689639 | HOXB7 | N_Shore | TSS1500 |
| GM01096 | chr17:46691543-46692008 | HOXB8 | Island | Body |
| GM02456 | chr17:47092072-47092178 | IGF2BP1;IGF2BP1 | Island | Body;Body |
| GM00520 | chr17:47113141-47113489 | IGF2BP1;IGF2BP1 | N_Shore | Body;Body |
| GM09256 | chr17:47113407-47113489 | IGF2BP1;IGF2BP1 | Island | Body;Body |
| WG2415 | chr17:47209660-47209749 | B4GALNT2;B4GALNT2;B4GAI | N_Shore | TSS1500;TSS200;TSS1500 |
| GM03075 | chr17:47287492-47287526 | ABI3;GNGT2;ABI3 | | TSS200;TSS1500;TSS200 |
| WG1983 | chr17:48263173-48263204 | COL1A1 | | Body |
| GM01376 | chr17:48912265-48912483 | WFIKKN2 | | TSS1500 |
| GM00042 | chr17:54893150-54893317 | C17orf67 | | 1stExon |
| GM01393 | chr17:56082780-56082885 | SFRS1;SFRS1 | N_Shore | 3'UTR;Body |
| GM02706 | chr17:56565286-56565644 | HSF5 | Island | 1stExon |
| GM00042 | chr17:56744332-56744490 | TEX14;TEX14 | | 5'UTR;5'UTR |
| GM01095 | chr17:59554672-59554806 | TBX4 | | Body |
| WG2276 | chr17:60828157-60828406 | MARCH10;MARCH10 | | Body;Body |
| GM02175 | chr17:61511683-61511829 | CYB561;CYB561;CYB561 | Island | 3'UTR;3'UTR;3'UTR |
| GM00438 | chr17:62097918-62097953 | ICAM2;ICAM2;ICAM2;ICAM2;ICAM2;ICAI | | 5'UTR;5'UTR;5'UTR;1stExon;1stExon;5 |
| WG2844 | chr17:62777693-62777777 | LOC146880;LOC146880 | Island | Body;TSS200 |

| | | | | |
|---|---|---|---|---|
| GM02175 | chr17:65040527-65040581 | CACNG1 | N_Shore | TSS200 |
| WG1513 | chr17:65471303-65471507 | PITPNC1;PITPNC1 | | Body;Body |
| GM02173 | chr17:6899207-6899332 | ALOX12 | Island | TSS200 |
| GM00935 | chr17:71772827-71772869 | C17orf54 | | Body |
| GM01096 | chr17:71824806-71825500 | C17orf54 | | TSS200 |
| WG1977 | chr17:72439044-72439217 | GPRC5C;GPRC5C | N_Shelf | Body;Body |
| WG1978 | chr17:74075092-74075189 | ZACN | S_Shore | TSS200 |
| WG1658 | chr17:74868561-74868670 | MGAT5B;MGAT5B | S_Shore | Body;TSS200 |
| WG1977 | chr17:74868561-74868725 | MGAT5B;MGAT5B | S_Shore | Body;TSS200 |
| GM02706 | chr17:750241-750848 | NXN | | Body |
| GM02456 | chr17:750766-751287 | NXN | | Body |
| GM02173 | chr17:75283431-75283832 | SEPT9;SEPT9 | | Body;TSS1500 |
| GM00520 | chr17:75283800-75283979 | SEPT9;SEPT9 | | TSS200;Body |
| WG2276 | chr17:75283832-75283979 | SEPT9;SEPT9 | | TSS200;Body |
| GM00438 | chr17:76128556-76128634 | TMC8;TMC6 | S_Shore | Body;TSS200 |
| GM00042 | chr17:76128634-76128683 | TMC8;TMC6 | S_Shore | Body;TSS200 |
| GM02173 | chr17:76355146-76355288 | SOCS3 | Island | Body |
| WG2275 | chr17:76976010-76976091 | LGALS3BP;LGALS3BP | | 1stExon;5'UTR |
| WG1513 | chr17:77029308-77029518 | C1QTNF1;C1QTNF1;C1QTNF1 | | 5'UTR;TSS1500;5'UTR |
| WG1658 | chr17:77037916-77037965 | C1QTNF1;C1QTNF1;C1QTNF1 | | 5'UTR;5'UTR;5'UTR |
| GM00438 | chr17:77952247-77952488 | TBC1D16 | Island | Body |
| GM09256 | chr17:77952290-77952488 | TBC1D16 | Island | Body |
| GM02317 | chr17:77952453-77952488 | TBC1D16 | S_Shore | Body |
| GM00290 | chr17:77997833-77997997 | TBC1D16 | Island | 5'UTR |
| GM00289 | chr17:78161110-78161227 | CARD14;CARD14 | N_Shelf | TSS1500;Body |
| GM00438 | chr17:7832680-7832852 | KCNAB3 | Island | 1stExon |
| WG1978 | chr17:78735268-78735324 | RPTOR;RPTOR | | Body;Body |
| GM01095 | chr17:78808060-78808570 | RPTOR;RPTOR | | Body;Body |
| WG1657 | chr17:78865263-78865373 | RPTOR;RPTOR | S_Shore | Body;Body |
| WG1984 | chr17:78894364-78894477 | RPTOR;RPTOR | N_Shelf | Body;Body |
| GM02456 | chr17:79010458-79010831 | BAIAP2;BAIAP2;BAIAP2;BAIA | S_Shore | Body;Body;Body;Body |
| GM00289 | chr17:79010458-79010968 | BAIAP2;BAIAP2;BAIAP2;BAIA | S_Shore | Body;Body;Body;Body |
| GM00289 | chr17:79011277-79011542 | BAIAP2;BAIAP2;BAIAP2;BAIA | S_Shore | Body;Body;Body;Body |
| GM02704 | chr17:79095700-79096151 | AATK | Island | Body |
| GM00520 | chr17:79099481-79099785 | MIR657;AATK | N_Shelf | TSS1500;Body |
| WG1977 | chr17:79099481-79099819 | MIR657;AATK | N_Shelf | TSS1500;Body |
| GM00042 | chr17:79099481-79099942 | MIR657;AATK | N_Shelf | TSS1500;Body |
| WG2120 | chr17:79109729-79109808 | AATK | Island | Body |
| GM09255 | chr17:79109777-79109910 | AATK | Island | Body |
| GM01096 | chr17:79259040-79259699 | SLC38A10;SLC38A10 | N_Shore | Body;Body |
| GM01096 | chr17:79377548-79378207 | BAHCC1 | Island | Body |
| GM01653 | chr17:79881468-79881543 | MAFG;MAFG | S_Shore | 5'UTR;TSS200 |
| GM03075 | chr17:79881523-79881543 | MAFG;MAFG | S_Shore | 5'UTR;TSS200 |
| GM02149 | chr17:79905219-79905263 | MYADML2 | | TSS200 |
| WG2121 | chr17:79905236-79905263 | MYADML2 | | TSS200 |
| GM02704 | chr17:800084-800749 | NXN | | Body |
| GM00439 | chr17:800100-800431 | NXN | | Body |

| | | | | |
|---|---|---|---|---|
| GM02149 | chr17:80186273-80186336 | SLC16A3;SLC16A3 | Island | TSS200;TSS1500 |
| GM02706 | chr17:80290106-80290217 | SECTM1 | N_Shore | 5'UTR |
| GM02706 | chr17:80292079-80292252 | SECTM1 | Island | TSS200 |
| WG2843 | chr17:80358829-80358876 | C17orf101;C17orf101 | N_Shelf | Body;Body |
| WG1514 | chr17:80806135-80806435 | TBCD | | Body |
| GM00290 | chr17:80875788-80876074 | TBCD | | Body |
| GM02555 | chr17:80875998-80876074 | TBCD | | Body |
| WG1977 | chr17:80943851-80944193 | B3GNTL1 | | Body |
| GM00290 | chr17:80970159-80970504 | B3GNTL1 | Island | Body |
| WG2276 | chr17:80970799-80970892 | B3GNTL1 | S_Shore | Body |
| GM00935 | chr17:80970799-80971527 | B3GNTL1 | S_Shore | Body |
| GM00935 | chr17:81040663-81040906 | METRNL | N_Shore | Body |
| GM00940 | chr17:81045495-81045781 | METRNL | Island | Body |
| GM01392 | chr17:8301166-8301192 | RNF222 | | TSS200 |
| GM00935 | chr17:8383621-8383800 | MYH10 | S_Shelf | Body |
| GM09256 | chr17:853971-854068 | NXN | S_Shelf | Body |
| WG2276 | chr17:872762-873452 | NXN | | Body |
| GM01095 | chr17:873255-873452 | NXN | | Body |
| GM00439 | chr17:8927521-8928028 | NTN1 | S_Shore | Body |
| WG1310 | chr17:933047-933155 | ABR;ABR;ABR | Island | Body;Body;Body |
| GM00940 | chr17:933047-933179 | ABR;ABR;ABR | Island | Body;Body;Body |
| GM01375 | chr17:933118-933179 | ABR;ABR;ABR | Island | Body;Body;Body |
| GM02149 | chr17:9967417-9967499 | GAS7 | | Body |
| GM02555 | chr18:11850139-11850153 | CHMP1B;GNAL;GNAL;GNAL | N_Shore | TSS1500;Body;Body;Body |
| WG1722 | chr18:19756582-19757016 | GATA6 | N_Shore | Body |
| GM09256 | chr18:19756582-19757022 | GATA6 | N_Shore | Body |
| GM01392 | chr18:19997912-19997969 | CTAGE1 | | TSS200 |
| GM00520 | chr18:20735119-20735672 | CABLES1;CABLES1;CABLES1 | | Body;Body;TSS1500 |
| GM02456 | chr18:21572622-21572748 | TTC39C | | TSS200 |
| GM02175 | chr18:24131115-24131604 | KCTD1 | Island | 5'UTR |
| WG1513 | chr18:24443729-24444196 | AQP4;AQP4 | S_Shore | TSS1500;Body |
| GM00520 | chr18:24445240-24445318 | AQP4;C18orf16 | S_Shore | Body;TSS200 |
| WG2844 | chr18:2907240-2907472 | EMILIN2 | Island | Body |
| WG2844 | chr18:30051538-30051572 | FAM59A | S_Shore | TSS1500 |
| GM02706 | chr18:37332566-37333028 | LOC647946 | | TSS1500 |
| GM00439 | chr18:45458243-45458698 | SMAD2;SMAD2;SMAD2 | S_Shore | TSS1500;TSS1500;TSS1500 |
| WG1657 | chr18:46386267-46386453 | KIAA0427;KIAA0427 | Island | 3'UTR;3'UTR |
| GM00935 | chr18:46447737-46447961 | SMAD7 | Island | 3'UTR |
| GM02149 | chr18:49866065-49866097 | DCC | N_Shelf | TSS1500 |
| GM02555 | chr18:56296449-56296544 | ALPK2 | | TSS1500 |
| GM02147 | chr18:56934834-56935199 | RAX | Island | 3'UTR |
| WG1721 | chr18:56936434-56936882 | RAX | Island | Body |
| WG1513 | chr18:61816543-61817320 | C18orf20 | | TSS1500 |
| WG2121 | chr18:67069959-67070120 | DOK6 | Island | Body |
| WG2414 | chr18:67070045-67070170 | DOK6 | Island | Body |
| GM00289 | chr18:7011217-7011463 | LAMA1 | N_Shore | Body |
| WG1513 | chr18:7011268-7011463 | LAMA1 | Island | Body |

| | | | | |
|---|---|---|---|---|
| WG2120 | chr18:74770402-74770589 | MBP;MBP | Island | Body;Body |
| WG1978 | chr18:74770402-74770815 | MBP;MBP | Island | Body;Body |
| WG1309 | chr18:74960996-74961078 | GALR1 | N_Shore | TSS1500 |
| WG1977 | chr18:77905298-77905751 | LOC100130522;LOC1001305 | Island | TSS1500;TSS1500 |
| GM00520 | chr18:77905298-77905947 | LOC100130522;LOC1001305 | Island | TSS1500;TSS1500 |
| GM02149 | chr18:77905663-77905747 | LOC100130522;LOC1001305 | S_Shore | TSS200;TSS200 |
| WG1722 | chr18:77905663-77905751 | LOC100130522;LOC1001305 | S_Shore | TSS200;TSS200 |
| WG1513 | chr18:77917615-77918142 | LOC100130522;LOC1001305 | N_Shore | Body;Body;3'UTR |
| GM00940 | chr19:1009172-1009949 | GRIN3B | Island | Body |
| WG2276 | chr19:11289349-11289376 | KANK2;KANK2 | S_Shore | Body;Body |
| WG1513 | chr19:11289356-11289376 | KANK2;KANK2 | S_Shore | Body;Body |
| GM00289 | chr19:1151960-1152189 | SBNO2 | N_Shore | Body |
| WG1722 | chr19:11689860-11689880 | ACP5;ACP5;ACP5;ACP5 | S_Shore | TSS1500;TSS1500;TSS200;TSS200 |
| WG2844 | chr19:12876846-12877000 | HOOK2;HOOK2 | N_Shore | Body;Body |
| WG2121 | chr19:12876846-12877188 | HOOK2;HOOK2 | N_Shore | Body;Body |
| GM00439 | chr19:12957979-12958340 | MAST1 | N_Shore | Body |
| WG1722 | chr19:1466647-1467032 | APC2 | Island | Body |
| WG1513 | chr19:15083795-15083842 | SLC1A6 | | TSS200 |
| GM02173 | chr19:15343350-15343395 | EPHX3;EPHX3 | N_Shore | 5'UTR;TSS200 |
| GM02175 | chr19:15582687-15582853 | PGLYRP2 | Island | Body |
| GM03075 | chr19:16178091-16178099 | TPM4 | N_Shore | TSS1500 |
| GM02175 | chr19:16999576-16999668 | F2RL3 | N_Shore | TSS1500 |
| WG1513 | chr19:16999768-16999953 | F2RL3 | N_Shore | TSS200 |
| GM00520 | chr19:17439877-17440072 | ANO8 | Island | Body |
| GM00438 | chr19:17932100-17932331 | INSL3 | Island | Body |
| GM00289 | chr19:17958892-17958961 | JAK3 | S_Shore | TSS200 |
| GM01392 | chr19:18539756-18540208 | SSBP4;SSBP4 | N_Shore | Body;Body |
| GM00941 | chr19:18539756-18540330 | SSBP4;SSBP4 | N_Shore | Body;Body |
| WG2414 | chr19:1854549-1854724 | KLF16 | Island | Body |
| GM02555 | chr19:18771079-18771327 | KLHL26 | Island | Body |
| GM00031 | chr19:18902107-18902160 | COMP;COMP | Island | 1stExon;5'UTR |
| GM00438 | chr19:18902527-18902727 | COMP | S_Shore | TSS1500 |
| GM09256 | chr19:19626576-19626605 | NDUFA13;TSSK6 | Island | TSS1500;TSS200 |
| GM01375 | chr19:19639553-19639596 | YJEFN3 | Island | TSS200 |
| GM00290 | chr19:19639553-19640075 | YJEFN3 | Island | TSS200 |
| GM09256 | chr19:19639568-19639596 | YJEFN3 | Island | TSS200 |
| GM00438 | chr19:2041985-2042592 | MKNK2;MKNK2 | Island | Body;Body |
| GM00439 | chr19:21264896-21265264 | ZNF714 | N_Shore | TSS200 |
| WG1658 | chr19:21264896-21265421 | ZNF714 | N_Shore | TSS200 |
| GM01392 | chr19:21264982-21265421 | ZNF714;ZNF714 | N_Shore | 5'UTR;1stExon |
| WG1978 | chr19:21265264-21265421 | ZNF714 | Island | 5'UTR |
| WG1722 | chr19:2253492-2253854 | JSRP1 | Island | Body |
| WG1977 | chr19:2282010-2282207 | C19orf35 | S_Shelf | 5'UTR |
| WG1309 | chr19:2282439-2282729 | C19orf35 | S_Shelf | TSS1500 |
| GM02640 | chr19:2291020-2291872 | LINGO3 | Island | Body |
| GM02704 | chr19:2302401-2302783 | LINGO3 | N_Shore | 5'UTR |
| GM02316 | chr19:23941313-23941575 | ZNF681 | Island | Body |

| | | | | |
|---|---|---|---|---|
| GM01650 | chr19:23941408-23941575 | ZNF681 | Island | Body |
| GM00940 | chr19:23942040-23942126 | ZNF681 | S_Shore | TSS1500 |
| GM00940 | chr19:23945472-23945659 | RPSAP58 | N_Shore | TSS1500 |
| GM00289 | chr19:2512999-2513436 | GNG7 | N_Shore | 3'UTR |
| WG1983 | chr19:2525384-2525634 | GNG7 | Island | 5'UTR |
| GM00520 | chr19:2588376-2588629 | GNG7 | Island | 5'UTR |
| GM00290 | chr19:2588479-2588629 | GNG7 | Island | 5'UTR |
| GM02706 | chr19:31640401-31640849 | DKFZp566F0947 | | TSS1500 |
| GM01650 | chr19:31640642-31640693 | DKFZp566F0947 | | TSS200 |
| WG2276 | chr19:31640642-31640711 | DKFZp566F0947 | | TSS200 |
| GM00940 | chr19:31640642-31640849 | DKFZp566F0947 | | TSS200 |
| GM02706 | chr19:35633553-35633972 | FXYD1;FXYD7;FXYD1 | Island | Body;TSS1500;Body |
| GM09255 | chr19:35633669-35633972 | FXYD7;FXYD1;FXYD1 | Island | TSS1500;3'UTR;3'UTR |
| GM00042 | chr19:35786821-35786961 | MAG;MAG | Island | Body;Body |
| GM00439 | chr19:36024217-36024282 | GAPDHS | Island | TSS200 |
| WG1984 | chr19:36801408-36801503 | LOC100134317 | S_Shelf | TSS1500 |
| GM00940 | chr19:38886554-38886789 | SPRED3 | Island | Body |
| GM02706 | chr19:39695011-39695065 | SYCN | S_Shore | TSS200 |
| GM03075 | chr19:39998018-39998290 | DLL3;DLL3 | Island | Body;Body |
| GM02175 | chr19:40732762-40732804 | CNTD2 | S_Shore | TSS200 |
| GM02706 | chr19:40904691-40904974 | PRX;PRX | Island | Body;Body |
| WG2121 | chr19:41119829-41120329 | LTBP4;LTBP4;LTBP4 | Island | Body;Body;Body |
| WG1513 | chr19:42636817-42637137 | POU2F2 | N_Shore | TSS200 |
| GM02456 | chr19:427022-427263 | SHC2 | Island | Body |
| WG1282 | chr19:43857861-43858174 | CD177;CD177 | | 1stExon;5'UTR |
| GM00438 | chr19:45260501-45261004 | BCL3 | Island | Body |
| WG2121 | chr19:46181221-46181330 | GIPR | Island | Body |
| GM00439 | chr19:4639844-4639992 | TNFAIP8L1;TNFAIP8L1 | S_Shore | TSS200;5'UTR |
| GM00439 | chr19:46999403-46999840 | PNMAL2 | S_Shore | TSS1500 |
| WG2844 | chr19:48947560-48947578 | GRIN2D;GRWD1 | Island | 3'UTR;TSS1500 |
| GM00289 | chr19:49055432-49055443 | SULT2B1;SULT2B1 | | 1stExon;5'UTR |
| GM02316 | chr19:49340489-49340593 | PLEKHA4;PLEKHA4;HSD17B1 | Island | 3'UTR;3'UTR;TSS1500 |
| GM09256 | chr19:49891270-49891574 | CCDC155 | Island | TSS1500 |
| WG1721 | chr19:49935823-49935948 | SLC17A7 | Island | Body |
| GM09256 | chr19:49935823-49936274 | SLC17A7 | Island | Body |
| GM00042 | chr19:50249584-50249776 | TSKS | Island | Body |
| GM00289 | chr19:50789296-50789841 | MYH14;MYH14;MYH14 | N_Shore | Body;Body;Body |
| GM09255 | chr19:50861774-50861923 | NAPSA | Island | 3'UTR |
| GM01653 | chr19:50931343-50931515 | SPIB | Island | Body |
| GM00520 | chr19:50931616-50931875 | SPIB | Island | 3'UTR |
| GM02456 | chr19:5147855-5148409 | KDM4B | Island | Body |
| WG1983 | chr19:5148023-5148409 | KDM4B | Island | Body |
| WG1977 | chr19:52264411-52264429 | FPR2 | | TSS200 |
| GM00520 | chr19:543909-544349 | GZMM | S_Shore | TSS200 |
| GM09256 | chr19:54876728-54876752 | LAIR1;LAIR1 | | TSS200;TSS200 |
| GM00935 | chr19:54876749-54876795 | LAIR1;LAIR1 | | TSS200;TSS200 |
| WG1657 | chr19:55972791-55973099 | ISOC2;ISOC2;ISOC2 | Island | 5'UTR;5'UTR;5'UTR |

| | | | | |
|---|---|---|---|---|
| WG2275 | chr19:55987366-55987415 | ZNF628 | N_Shore | TSS1500 |
| GM01096 | chr19:56686339-56686707 | GALP;GALP | | TSS1500;TSS1500 |
| GM00941 | chr19:57742421-57742444 | AURKC;AURKC;AURKC;AURK | Island | 5'UTR;TSS200;1stExon;5'UTR;1stExon |
| GM01376 | chr19:58038573-58038588 | ZNF549 | Island | TSS200 |
| GM00520 | chr19:58879022-58879059 | ZNF837;ZNF837 | Island | 3'UTR;3'UTR |
| GM09256 | chr19:710823-711001 | PALM;PALM | Island | Body;Body |
| GM02173 | chr19:7953054-7953081 | LRRC8E | N_Shore | TSS1500 |
| WG1977 | chr19:827776-827843 | AZU1 | | TSS200 |
| GM00439 | chr19:853054-853334 | ELANE | Island | Body |
| GM01392 | chr19:859499-859680 | CFD | N_Shore | TSS200 |
| GM00438 | chr19:863054-863226 | CFD | Island | Body |
| GM00439 | chr19:8642308-8642370 | MYO1F | | TSS200 |
| WG2276 | chr19:940967-941243 | ARID3A | Island | Body |
| WG2415 | chr2:10567628-10567843 | HPCAL1;HPCAL1 | S_Shore | 3'UTR;3'UTR |
| GM09256 | chr2:106015739-106015771 | FHL2;FHL2;FHL2;FHL2 | Island | TSS200;TSS200;5'UTR;TSS200 |
| WG2275 | chr2:106055153-106055458 | FHL2;FHL2 | | 1stExon;5'UTR |
| GM01392 | chr2:106055153-106055507 | FHL2;FHL2 | | 1stExon;5'UTR |
| GM02173 | chr2:112939119-112939562 | FBLN7;FBLN7 | N_Shore | Body;Body |
| WG1977 | chr2:113992843-113992930 | PAX8;PAX8;PAX8;PAX8;PAX8; | N_Shore | Body;Body;Body;Body;Body;Body;TSS |
| GM09256 | chr2:113992843-113993052 | PAX8;PAX8;PAX8;PAX8;PAX8; | N_Shore | Body;Body;Body;Body;Body;Body;TSS |
| WG1977 | chr2:114033490-114033830 | PAX8;PAX8;PAX8;PAX8;PAX8; | Island | Body;Body;Body;Body;Body;Body |
| GM09256 | chr2:11680057-11680144 | GREB1;GREB1 | | 5'UTR;TSS200 |
| WG1721 | chr2:119599748-119600002 | EN1 | Island | 3'UTR |
| GM00438 | chr2:119599748-119600948 | EN1 | Island | 3'UTR |
| GM00042 | chr2:119602480-119603489 | EN1 | N_Shore | Body |
| GM02173 | chr2:119602546-119602857 | EN1 | N_Shore | Body |
| GM01650 | chr2:119602546-119603489 | EN1 | N_Shore | Body |
| GM00289 | chr2:119603133-119603489 | EN1 | Island | Body |
| GM02456 | chr2:121554696-121554926 | GLI2 | | TSS200 |
| WG2844 | chr2:121554810-121554926 | GLI2 | | TSS200 |
| WG2275 | chr2:121554817-121554926 | GLI2 | | TSS200 |
| GM00520 | chr2:121748215-121748257 | GLI2 | S_Shore | Body |
| GM00438 | chr2:127963489-127963590 | CYP27C1 | | TSS200 |
| GM00520 | chr2:133402475-133402999 | GPR39;LYPD1;LYPD1 | N_Shore | Body;3'UTR;3'UTR |
| WG1977 | chr2:133429088-133429299 | LYPD1;LYPD1 | S_Shore | TSS1500;TSS200 |
| GM00438 | chr2:1358283-1358684 | SNTG2 | | Body |
| WG1513 | chr2:1417164-1417431 | TPO;TPO;TPO;TPO | | TSS1500;TSS200;TSS200;TSS1500 |
| WG1978 | chr2:142889283-142889392 | LRP1B | S_Shore | TSS200 |
| WG1722 | chr2:142889367-142889392 | LRP1B | S_Shore | TSS200 |
| GM00438 | chr2:1487895-1488175 | TPO;TPO;TPO;TPO | | Body;Body;Body;Body |
| GM02149 | chr2:1488252-1488516 | TPO;TPO;TPO;TPO | | Body;Body;Body;Body |
| GM00520 | chr2:150444569-150444834 | MMADHC | | TSS1500 |
| GM00290 | chr2:152830189-152830521 | CACNB4;CACNB4;CACNB4;CACNB4 | | Body;Body;1stExon;Body |
| GM00439 | chr2:157291462-157291826 | GPD2;GPD2 | N_Shore | TSS1500;TSS1500 |
| GM02456 | chr2:1647185-1647260 | PXDN | | Body |
| WG1514 | chr2:171676553-171676925 | GAD1;GAD1 | Island | Body;Body |
| GM00439 | chr2:171679906-171680337 | GAD1;GAD1 | Island | Body;Body |

| | | | | |
|---|---|---|---|---|
| GM01393 | chr2:175199259-175199349 | SP9 | N_Shore | TSS1500 |
| GM00439 | chr2:176948693-176948732 | EVX2 | N_Shore | TSS200 |
| GM00289 | chr2:176948699-176948732 | EVX2 | N_Shore | TSS200 |
| WG1722 | chr2:176948699-176948759 | EVX2 | N_Shore | TSS200 |
| GM00438 | chr2:176948728-176948759 | EVX2 | N_Shore | TSS200 |
| GM02704 | chr2:176956396-176956678 | HOXD13 | N_Shore | TSS1500 |
| WG2121 | chr2:176956396-176956841 | HOXD13 | N_Shore | TSS1500 |
| GM02641 | chr2:176964101-176964720 | HOXD12 | Island | TSS1500 |
| GM03073 | chr2:176964137-176964502 | HOXD12 | Island | TSS1500 |
| GM02640 | chr2:176964137-176964720 | HOXD12 | Island | TSS1500 |
| WG2415 | chr2:176964512-176964720 | HOXD12 | Island | TSS200 |
| GM02173 | chr2:176973275-176974060 | HOXD11 | N_Shore | Body |
| WG1977 | chr2:176973688-176973835 | HOXD11 | Island | Body |
| GM02175 | chr2:176980837-176981064 | HOXD10 | Island | TSS1500 |
| WG1978 | chr2:176981064-176981469 | HOXD10 | Island | TSS1500 |
| GM02173 | chr2:176981422-176981654 | HOXD10 | Island | TSS200 |
| WG2275 | chr2:176982108-176982342 | HOXD10 | Island | 1stExon |
| WG2276 | chr2:176988284-176988939 | HOXD9 | Island | 1stExon |
| WG1722 | chr2:176993017-176993115 | HOXD8 | Island | TSS1500 |
| GM00520 | chr2:176993017-176993320 | HOXD8 | Island | TSS1500 |
| WG1514 | chr2:177016167-177016490 | HOXD4;HOXD4 | N_Shore | 5'UTR;1stExon |
| WG3465 | chr2:177016167-177017352 | HOXD4;HOXD4 | N_Shore | 5'UTR;1stExon |
| GM00042 | chr2:177016167-177017389 | HOXD4;HOXD4 | N_Shore | 5'UTR;1stExon |
| GM02640 | chr2:177016167-177017449 | HOXD4;HOXD4 | N_Shore | 5'UTR;1stExon |
| GM00520 | chr2:177016167-177017630 | HOXD4;HOXD4 | N_Shore | 5'UTR;1stExon |
| GM00289 | chr2:177016837-177017449 | HOXD4 | N_Shore | Body |
| GM00438 | chr2:179387853-179388064 | MIR548N | | Body |
| GM02706 | chr2:1926888-1927209 | MYT1L | | Body |
| WG1658 | chr2:19551750-19551860 | OSR1 | Island | 3'UTR |
| GM00520 | chr2:19553237-19553422 | OSR1 | Island | Body |
| GM00042 | chr2:200320455-200320871 | SATB2 | N_Shore | Body |
| GM02706 | chr2:200320455-200321016 | SATB2 | N_Shore | Body |
| WG1977 | chr2:200320455-200321348 | SATB2 | N_Shore | Body |
| GM00022 | chr2:200333729-200334584 | FLJ32063 | Island | Body |
| GM02175 | chr2:200334402-200334584 | FLJ32063 | N_Shore | Body |
| WG1722 | chr2:200334854-200335101 | FLJ32063 | N_Shore | Body |
| GM00439 | chr2:200334854-200335683 | FLJ32063 | N_Shore | Body |
| GM02555 | chr2:202508694-202508813 | ALS2CR4;ALS2CR4 | S_Shore | TSS1500;TSS1500 |
| GM00289 | chr2:202901288-202901470 | FZD7;FZD7 | S_Shore | 1stExon;3'UTR |
| GM03075 | chr2:202901352-202901470 | FZD7;FZD7 | S_Shore | 1stExon;3'UTR |
| GM00042 | chr2:20870362-20871401 | GDF7 | Island | Body |
| GM09255 | chr2:211457684-211458173 | CPS1;CPS1;CPS1 | | Body;Body;TSS1500 |
| WG1513 | chr2:215675429-215675637 | BARD1 | S_Shore | TSS1500 |
| GM01392 | chr2:219735876-219736312 | WNT6 | N_Shore | Body |
| GM00290 | chr2:220253653-220253668 | DNPEP | S_Shore | TSS1500 |
| GM09256 | chr2:220253653-220253963 | DNPEP | S_Shore | TSS1500 |
| GM09256 | chr2:220406470-220406756 | CHPF | Island | Body |

| | | | | |
|---|---|---|---|---|
| WG1978 | chr2:223155925-223156136 | PAX3;PAX3;PAX3;PAX3;PAX3; | Island | Body;Body;Body;Body;Body;Body |
| WG1977 | chr2:223161771-223162049 | PAX3;PAX3;PAX3;CCDC140;P | Island | Body;Body;Body;TSS1500;Body;Body; |
| GM09256 | chr2:223161771-223162128 | PAX3;PAX3;PAX3;CCDC140;P | Island | Body;Body;Body;TSS1500;Body;Body; |
| WG1978 | chr2:223161771-223162666 | PAX3;PAX3;PAX3;CCDC140;P | Island | Body;Body;Body;TSS1500;Body;Body; |
| WG1977 | chr2:223162666-223162800 | PAX3;PAX3;PAX3;PAX3;PAX3; | N_Shore | Body;Body;Body;Body;Body;Body;Bod |
| GM00439 | chr2:223162797-223163175 | PAX3;PAX3;PAX3;PAX3;PAX3; | N_Shore | Body;Body;Body;Body;Body;Body;Bod |
| GM01095 | chr2:223164459-223166989 | PAX3;PAX3;PAX3;PAX3;PAX3; | S_Shore | TSS1500;TSS1500;TSS1500;TSS1500; |
| GM02706 | chr2:223164459-223168487 | PAX3;PAX3;PAX3;PAX3;PAX3; | S_Shore | TSS1500;TSS1500;TSS1500;TSS1500; |
| GM02641 | chr2:223164459-223170370 | PAX3;PAX3;PAX3;PAX3;PAX3; | S_Shore | TSS1500;TSS1500;TSS1500;TSS1500; |
| GM00520 | chr2:223164747-223164869 | PAX3;PAX3;PAX3;PAX3;PAX3; | S_Shore | TSS1500;TSS1500;TSS1500;TSS1500; |
| GM02555 | chr2:223164831-223164869 | PAX3;PAX3;PAX3;PAX3;PAX3; | S_Shore | TSS1500;TSS1500;TSS1500;TSS1500; |
| WG1722 | chr2:223164854-223164869 | PAX3;PAX3;PAX3;PAX3;PAX3; | S_Shore | TSS1500;TSS1500;TSS1500;TSS1500; |
| GM01392 | chr2:223166909-223166989 | CCDC140 | N_Shore | 5'UTR |
| GM01095 | chr2:223168453-223170370 | CCDC140 | N_Shore | 5'UTR |
| GM02704 | chr2:223917577-223918000 | KCNE4 | Island | Body |
| GM00940 | chr2:233323532-233323935 | ALPI | Island | Body |
| GM02706 | chr2:235935754-235936596 | SH3BP4 | | 5'UTR |
| GM00940 | chr2:235936394-235936596 | SH3BP4 | | 5'UTR |
| WG1977 | chr2:236586191-236586744 | AGAP1;AGAP1 | | Body;Body |
| GM02555 | chr2:236643527-236643760 | AGAP1;AGAP1 | | Body;Body |
| GM00290 | chr2:236674174-236674489 | AGAP1;AGAP1 | S_Shore | Body;Body |
| WG2194 | chr2:23746926-23747197 | KLHL29 | N_Shelf | 5'UTR |
| GM02175 | chr2:238394933-238395125 | MLPH;MLPH | N_Shore | TSS1500;TSS1500 |
| GM01096 | chr2:238499782-238499802 | RAB17 | | TSS200 |
| GM02456 | chr2:238647755-238647913 | LRRFIP1;LRRFIP1;LRRFIP1;LR | S_Shelf | Body;Body;Body;Body;Body |
| GM02706 | chr2:239799720-239799904 | TWIST2 | | 3'UTR |
| WG1513 | chr2:240014758-240014781 | HDAC4 | | Body |
| WG2415 | chr2:240020028-240020174 | HDAC4 | | Body |
| GM02706 | chr2:240196078-240196284 | HDAC4 | N_Shore | Body |
| WG2415 | chr2:240233043-240233240 | HDAC4 | | Body |
| WG1513 | chr2:240291426-240291657 | HDAC4 | | 5'UTR |
| WG1513 | chr2:24271827-24272044 | FKBP1B;FKBP1B | N_Shore | TSS1500;TSS1500 |
| GM00438 | chr2:25390385-25390424 | POMC;POMC | N_Shore | 5'UTR;5'UTR |
| GM00439 | chr2:25391830-25391911 | POMC;POMC | Island | TSS1500;TSS1500 |
| GM02173 | chr2:26401640-26401820 | FAM59B | N_Shore | Body |
| GM02706 | chr2:26542043-26542188 | GPR113;GPR113;GPR113 | | TSS200;TSS200;Body |
| GM00289 | chr2:27665128-27665150 | KRTCAP3;KRTCAP3 | Island | TSS200;TSS200 |
| GM01650 | chr2:27665139-27665150 | KRTCAP3;KRTCAP3 | Island | TSS200;TSS200 |
| GM01650 | chr2:287148-287395 | FAM150B | N_Shore | Body |
| WG2843 | chr2:30669597-30669863 | LCLAT1;LCLAT1 | N_Shore | TSS1500;TSS1500 |
| GM01375 | chr2:30669597-30669952 | LCLAT1;LCLAT1 | N_Shore | TSS1500;TSS1500 |
| WG3465 | chr2:3286338-3286479 | TSSC1 | Island | Body |
| GM01650 | chr2:3317860-3318039 | TSSC1 | Island | Body |
| WG1513 | chr2:3451506-3452563 | TTC15 | N_Shore | Body |
| GM01095 | chr2:3451506-3452624 | TTC15 | N_Shore | Body |
| WG1721 | chr2:3452347-3452563 | TTC15 | Island | Body |
| WG2276 | chr2:3452347-3452624 | TTC15 | Island | Body |

| | | | | |
|---|---|---|---|---|
| GM02456 | chr2:3452543-3452624 | TTC15 | Island | Body |
| GM02149 | chr2:3642211-3642634 | COLEC11;COLEC11 | N_Shore | TSS1500;TSS1500 |
| WG3465 | chr2:3642400-3642634 | COLEC11;COLEC11 | N_Shore | TSS1500;TSS1500 |
| GM02706 | chr2:3642447-3642634 | COLEC11;COLEC11 | N_Shore | TSS200;TSS200 |
| WG1282 | chr2:3714936-3715083 | ALLC | | 5'UTR |
| GM01096 | chr2:37571677-37571901 | QPCT | Island | TSS200 |
| GM00290 | chr2:43903470-43903526 | PLEKHH2;LOC728819 | | Body;TSS200 |
| GM00290 | chr2:46526785-46527098 | EPAS1 | Island | Body |
| GM00289 | chr2:48844892-48844971 | GTF2A1L;STON1-GTF2A1L;GT | Island | TSS200;Body;TSS200 |
| GM00935 | chr2:48844916-48844984 | GTF2A1L;STON1-GTF2A1L;GT | Island | TSS200;Body;TSS200 |
| GM00031 | chr2:50201413-50201505 | NRXN1;NRXN1;NRXN1 | | Body;Body;Body |
| WG1513 | chr2:54087173-54087189 | LOC100302652;GPR75 | Island | TSS200;TSS200 |
| WG1977 | chr2:54087178-54087189 | LOC100302652;GPR75 | Island | TSS200;TSS200 |
| GM01096 | chr2:54785178-54785795 | SPTBN1;SPTBN1 | Island | TSS1500;Body |
| WG2414 | chr2:56410011-56410087 | CCDC85A | N_Shore | TSS1500 |
| WG1722 | chr2:6122798-6122814 | LOC400940 | | Body |
| WG3465 | chr2:63275034-63275178 | LOC100132215 | Island | TSS200 |
| WG1983 | chr2:65217528-65217623 | SLC1A4;SLC1A4 | S_Shore | Body;5'UTR |
| WG1310 | chr2:66671478-66671727 | MEIS1 | N_Shore | Body |
| GM00439 | chr2:7057153-7057245 | RNF144A | N_Shore | TSS1500 |
| GM02706 | chr2:71134524-71134630 | VAX2 | N_Shore | Body |
| GM01392 | chr2:71211981-71212157 | ANKRD53;ANKRD53 | | 3'UTR;Body |
| GM02706 | chr2:72371255-72371477 | CYP26B1 | Island | Body |
| GM01375 | chr2:73964768-73964834 | TPRKB | S_Shore | TSS1500 |
| GM01376 | chr2:74669349-74669380 | RTKN | Island | TSS1500 |
| GM09256 | chr2:74729429-74729592 | LOC151534;LBX2 | Island | TSS1500;Body |
| GM02706 | chr2:74780268-74780617 | LOXL3 | N_Shore | 5'UTR |
| WG1283 | chr2:74780272-74780617 | LOXL3 | N_Shore | 5'UTR |
| GM00520 | chr2:8118122-8118404 | LOC339788 | | TSS1500 |
| GM00935 | chr2:85361959-85362480 | TCF7L1 | Island | Body |
| WG2276 | chr2:85637834-85638535 | CAPG | N_Shelf | TSS200 |
| GM00438 | chr2:88427581-88427637 | FABP1 | | TSS200 |
| WG1977 | chr2:947167-947589 | SNTG2 | Island | Body |
| WG2194 | chr2:947515-947634 | SNTG2 | S_Shore | Body |
| GM02175 | chr2:95537314-95537475 | TEKT4;TEKT4 | Island | 1stExon;5'UTR |
| GM00935 | chr2:98962444-98962564 | CNGA3;CNGA3 | N_Shore | TSS200;TSS200 |
| GM00520 | chr20:19915716-19915769 | RIN2 | | Body |
| GM01392 | chr20:23067126-23067159 | CD93 | | TSS200 |
| GM03075 | chr20:25128770-25128805 | LOC284798;LOC284798;LOC | Island | TSS200;Body;Body |
| GM09255 | chr20:25128770-25128807 | LOC284798;LOC284798;LOC | Island | TSS200;Body;Body |
| WG1513 | chr20:2516256-2516592 | TMC2 | | TSS1500 |
| GM00042 | chr20:30073521-30073576 | NCRNA00028 | S_Shore | TSS200 |
| GM02555 | chr20:30225666-30225685 | COX4I2 | Island | TSS200 |
| WG1977 | chr20:3051954-3052262 | OXT | N_Shore | TSS1500 |
| GM00520 | chr20:3052115-3052221 | OXT | Island | TSS200 |
| GM00289 | chr20:3052115-3052224 | OXT | Island | TSS200 |
| GM00439 | chr20:3149249-3149278 | ProSAPiP1 | S_Shore | TSS200 |

| | | | | |
|---|---|---|---|---|
| WG1513 | chr20:32856747-32856898 | ASIP | Island | Body |
| GM02706 | chr20:35169539-35169609 | MYL9;MYL9 | Island | TSS1500;TSS1500 |
| GM01096 | chr20:35274627-35274655 | SLA2;SLA2 | | TSS200;TSS200 |
| GM00439 | chr20:35274627-35274715 | SLA2;SLA2 | | TSS200;TSS200 |
| WG1722 | chr20:36011999-36012016 | SRC;SRC | N_Shore | 5'UTR;5'UTR |
| GM02456 | chr20:36011999-36012596 | SRC;SRC | N_Shore | 5'UTR;5'UTR |
| GM02456 | chr20:36932397-36932553 | BPI | | TSS200 |
| WG1513 | chr20:43729861-43729897 | KCNS1 | S_Shore | TSS200 |
| GM02147 | chr20:43935222-43935291 | MATN4;MATN4;MATN4;RBPJL | Island | 5'UTR;5'UTR;5'UTR;TSS1500 |
| WG1977 | chr20:43935460-43935471 | MATN4;MATN4;RBPJL;MATN4 | Island | 5'UTR;5'UTR;TSS200;5'UTR |
| GM01096 | chr20:44875334-44875586 | CDH22 | Island | Body |
| GM00438 | chr20:45179345-45179413 | C20orf123 | | TSS200 |
| GM09255 | chr20:47896448-47896888 | SNORD12B;C20orf199;MIR1 | S_Shore | TSS1500;Body;TSS1500;Body;TSS150 |
| GM02456 | chr20:49253364-49253710 | FAM65C;FAM65C | | 1stExon;5'UTR |
| WG1978 | chr20:50312386-50312632 | ATP9A | | Body |
| WG1309 | chr20:5485144-5485284 | LOC149837 | | Body |
| GM00940 | chr20:5485245-5485270 | LOC149837 | | TSS200 |
| WG1513 | chr20:5485245-5485284 | LOC149837 | | TSS200 |
| GM00438 | chr20:5485245-5485294 | LOC149837 | | TSS200 |
| WG1657 | chr20:5485268-5485284 | LOC149837 | | TSS200 |
| GM00439 | chr20:55203291-55203849 | TFAP2C | Island | TSS1500 |
| GM00520 | chr20:55203439-55203849 | TFAP2C | Island | TSS1500 |
| GM02175 | chr20:55925570-55925603 | RAE1;RAE1 | N_Shore | TSS1500;TSS1500 |
| GM01375 | chr20:57427146-57427173 | GNAS;GNAS;GNAS;GNASAS | N_Shore | TSS1500;TSS1500;3'UTR;TSS1500 |
| GM09255 | chr20:57465439-57465448 | GNAS;GNAS;GNAS;GNAS;GN | Island | 3'UTR;TSS1500;TSS1500;Body;TSS15 |
| GM02706 | chr20:57471654-57471672 | GNAS;GNAS;GNAS;GNAS;GN | S_Shelf | Body;Body;3'UTR;Body;Body;3'UTR;Bo |
| GM02175 | chr20:58179540-58180241 | PHACTR3 | N_Shore | TSS200 |
| WG1977 | chr20:58179847-58180241 | PHACTR3;PHACTR3 | Island | 5'UTR;1stExon |
| WG1977 | chr20:58630954-58631038 | C20orf197 | | TSS200 |
| GM02555 | chr20:6034694-6035120 | LRRN4 | S_Shore | TSS200 |
| GM00439 | chr20:61151446-61151480 | C20orf166;MIR1-1 | N_Shore | Body;TSS200 |
| GM00940 | chr20:61151467-61151480 | C20orf166;MIR1-1 | N_Shore | Body;TSS200 |
| WG1977 | chr20:61447330-61447394 | COL9A3 | N_Shore | TSS1500 |
| GM09255 | chr20:61583830-61583924 | SLC17A9 | N_Shore | TSS200 |
| GM00520 | chr20:61924405-61924802 | COL20A1 | N_Shelf | TSS200 |
| GM00042 | chr20:61953801-61954200 | COL20A1 | N_Shore | Body |
| WG1513 | chr20:61953801-61954258 | COL20A1 | N_Shore | Body |
| GM02175 | chr20:61953935-61954258 | COL20A1 | Island | Body |
| GM00438 | chr20:62153067-62153239 | PPDPF | Island | Body |
| GM02706 | chr20:62317768-62318482 | RTEL1;RTEL1 | Island | Body;Body |
| GM02555 | chr20:62318426-62318588 | RTEL1;RTEL1 | Island | Body;Body |
| GM02555 | chr20:62327968-62328427 | TNFRSF6B;TNFRSF6B | N_Shore | 5'UTR;TSS200 |
| GM02456 | chr20:62368956-62369462 | LIME1 | Island | Body |
| GM00940 | chr20:62679255-62679713 | SOX18 | N_Shore | 3'UTR |
| WG1513 | chr20:62688717-62688896 | TCEA2;TCEA2 | Island | 1stExon;5'UTR |
| GM00042 | chr20:62693943-62693971 | TCEA2;TCEA2 | Island | TSS200;5'UTR |
| GM02175 | chr20:62715942-62716485 | OPRL1;C20orf201;OPRL1 | N_Shore | 5'UTR;TSS1500;5'UTR |

| | | | | |
|---|---|---|---|---|
| GM01376 | chr21:17961737-17961996 | C21orf34;MIR125B2;C21orf34 | | Body;TSS1500;Body |
| WG1977 | chr21:17961737-17962124 | C21orf34;MIR125B2;C21orf34 | | Body;TSS1500;Body |
| GM00289 | chr21:28337594-28337869 | ADAMTS5 | N_Shore | Body |
| GM01096 | chr21:35831871-35832180 | KCNE1;KCNE1;KCNE1;KCNE | Island | 5'UTR;5'UTR;1stExon;1stExon;5'UTR |
| GM09255 | chr21:35832141-35832180 | KCNE1;KCNE1;KCNE1 | Island | 5'UTR;TSS1500;TSS1500 |
| GM01096 | chr21:36399146-36399258 | RUNX1 | Island | Body |
| GM00935 | chr21:36421467-36421503 | RUNX1;RUNX1 | | 5'UTR;1stExon |
| GM01393 | chr21:40145923-40146046 | NCRNA00114 | | TSS1500 |
| GM02704 | chr21:40759534-40760258 | WRB;WRB | N_Shore | Body;TSS200 |
| GM00438 | chr21:44104950-44105474 | PDE9A;PDE9A;PDE9A;PDE9A | N_Shore | Body;Body;5'UTR;Body;Body;Body;5'U |
| GM03075 | chr21:45148879-45149141 | PDXK | Island | Body |
| GM00439 | chr21:45148879-45149398 | PDXK | Island | Body |
| GM01376 | chr21:45705618-45705742 | AIRE | Island | TSS200 |
| GM00042 | chr21:45789122-45789379 | TRPM2 | Island | Body |
| GM00520 | chr21:45811432-45811549 | TRPM2 | Island | Body |
| GM02147 | chr21:47294739-47295215 | PCBP3;PCBP3 | N_Shore | Body;Body |
| WG2276 | chr21:47604052-47604654 | C21orf56;C21orf56 | S_Shore | 5'UTR;5'UTR |
| WG1977 | chr21:47604291-47604654 | C21orf56;C21orf56;C21orf56 | S_Shore | 5'UTR;5'UTR;1stExon;1stExon |
| GM00022 | chr21:47604361-47605174 | C21orf56;C21orf56;C21orf56 | S_Shore | 5'UTR;5'UTR;1stExon;1stExon |
| WG2415 | chr21:47813364-47813461 | PCNT | Island | Body |
| WG1514 | chr22:18632429-18632476 | USP18 | N_Shore | TSS1500 |
| WG1513 | chr22:19118165-19118202 | DGCR14;TSSK2 | N_Shore | 3'UTR;TSS200 |
| GM03073 | chr22:19746984-19747140 | TBX1;TBX1;TBX1 | Island | 5'UTR;5'UTR;5'UTR |
| WG2414 | chr22:19754125-19754448 | TBX1;TBX1;TBX1 | Island | Body;Body;Body |
| GM00439 | chr22:20861470-20861479 | MED15;MED15 | N_Shore | TSS1500;TSS1500 |
| GM00940 | chr22:22020012-22020076 | PPIL2;PPIL2;PPIL2 | N_Shore | TSS1500;TSS1500;TSS1500 |
| WG2415 | chr22:22058080-22058271 | YPEL1 | Island | Body |
| GM00022 | chr22:22901648-22901697 | PRAME;PRAME;PRAME;PRAM | Island | TSS200;5'UTR;1stExon;1stExon;TSS20 |
| GM03075 | chr22:24181263-24181270 | DERL3;DERL3;DERL3 | Island | TSS200;TSS200;TSS200 |
| WG1721 | chr22:24348549-24348715 | GSTTP1 | | TSS1500 |
| WG1721 | chr22:24384105-24384294 | GSTT1 | N_Shore | Body |
| GM00031 | chr22:24384105-24384944 | GSTT1 | N_Shore | Body |
| GM01392 | chr22:24384159-24384294 | GSTT1 | Island | 1stExon |
| GM02149 | chr22:24384397-24384525 | GSTT1 | Island | TSS200 |
| WG2414 | chr22:24384400-24384944 | GSTT1 | Island | TSS200 |
| GM00031 | chr22:24891153-24891166 | C22orf45;C22orf45;UPB1 | Island | TSS1500;TSS1500;TSS200 |
| GM02555 | chr22:24979436-24979588 | GGT1 | | TSS1500 |
| WG2415 | chr22:24979554-24979588 | GGT1 | | TSS200 |
| GM00439 | chr22:24979633-24979660 | GGT1 | | TSS200 |
| GM00935 | chr22:25159879-25160098 | TOP1P2;PIWIL3 | Island | TSS1500;5'UTR |
| GM02706 | chr22:31318147-31318181 | C22orf27 | N_Shore | TSS200 |
| GM03075 | chr22:31318181-31318373 | C22orf27 | N_Shore | TSS200 |
| WG1722 | chr22:32027580-32027605 | PISD | S_Shore | TSS1500 |
| GM00520 | chr22:32599511-32599648 | RFPL2;RFPL2;RFPL2 | | TSS200;5'UTR;TSS1500 |
| GM02706 | chr22:33196425-33196627 | SYN3;TIMP3;SYN3;SYN3 | N_Shore | Body;TSS1500;Body;Body |
| GM00520 | chr22:36681296-36681321 | MYH9 | | Body |
| GM00290 | chr22:37499649-37499859 | TMPRSS6;TMPRSS6 | | 1stExon;5'UTR |

| | | | | |
|---|---|---|---|---|
| GM00520 | chr22:37824118-37824617 | ELFN2 | Island | TSS1500 |
| GM00290 | chr22:37976063-37976192 | LGALS2 | | TSS200 |
| GM00022 | chr22:42828386-42828418 | NFAM1;NFAM1 | | 1stExon;5'UTR |
| GM02704 | chr22:42828411-42828418 | NFAM1 | | TSS200 |
| GM01096 | chr22:44568337-44568423 | PARVG;PARVG | | TSS1500;TSS1500 |
| GM00935 | chr22:44568725-44568812 | PARVG;PARVG | | TSS200;TSS200 |
| GM00289 | chr22:44708669-44708908 | KIAA1644;KIAA1644 | | 5'UTR;1stExon |
| GM00935 | chr22:45125218-45126040 | PRR5;PRR5;PRR5;PRR5-ARH( | Island | Body;Body;Body;Body;Body;Body |
| WG2415 | chr22:45609402-45609421 | C22orf9;C22orf9 | | Body;TSS1500 |
| WG1513 | chr22:45809319-45809396 | SMC1B;RIBC2 | Island | Body;TSS1500 |
| GM02317 | chr22:50584184-50585427 | MOV10L1;MOV10L1;MOV10L1;MOV10L1 | | Body;TSS1500;Body;Body |
| GM00022 | chr22:50585197-50585427 | MOV10L1;MOV10L1;MOV10L1;MOV10L1 | | Body;TSS1500;Body;Body |
| GM02175 | chr22:50629038-50629224 | TRABD | Island | 5'UTR |
| GM02173 | chr22:50985681-50986031 | KLHDC7B | Island | TSS1500 |
| WG1721 | chr3:10149882-10149963 | C3orf24;C3orf24;C3orf24;C3 | Island | 1stExon;1stExon;5'UTR;5'UTR |
| GM01392 | chr3:112358351-112358468 | CCDC80;CCDC80 | | Body;Body |
| WG1983 | chr3:11267020-11267098 | HRH1;HRH1;HRH1 | | 5'UTR;TSS1500;5'UTR |
| WG1310 | chr3:11267020-11267525 | HRH1;HRH1;HRH1 | | 5'UTR;TSS1500;5'UTR |
| WG1657 | chr3:114790243-114790596 | ZBTB20;ZBTB20;ZBTB20 | | 1stExon;5'UTR;5'UTR |
| WG3465 | chr3:11746183-11746765 | VGLL4 | N_Shore | 5'UTR |
| WG1721 | chr3:121379745-121379791 | HCLS1;HCLS1 | | 1stExon;5'UTR |
| GM00438 | chr3:121949083-121949308 | CASR | N_Shore | 5'UTR |
| GM00016 | chr3:122640778-122641029 | SEMA5B | Island | Body |
| GM00439 | chr3:128204813-128205143 | GATA2;GATA2;GATA2 | N_Shore | Body;Body;Body |
| GM09255 | chr3:129407542-129407548 | TMCC1;TMCC1;TMCC1 | | TSS200;Body;Body |
| GM00042 | chr3:137893707-137893743 | DBR1;DBR1 | Island | 1stExon;5'UTR |
| GM02706 | chr3:138152837-138153127 | ESYT3 | N_Shore | TSS1500 |
| WG2414 | chr3:138668832-138669322 | C3orf72 | Island | Body |
| WG1722 | chr3:139257775-139257907 | RBP1;RBP1;RBP1 | Island | Body;Body;Body |
| GM02173 | chr3:139257866-139258166 | RBP1;RBP1;RBP1 | Island | Body;Body;Body |
| WG2415 | chr3:14443428-14443473 | SLC6A6;SLC6A6;SLC6A6 | N_Shore | TSS1500;TSS1500;TSS1500 |
| WG1283 | chr3:147112081-147112605 | ZIC4;ZIC4;ZIC4 | N_Shore | Body;Body;Body |
| WG2120 | chr3:147112438-147112605 | ZIC4;ZIC4;ZIC4 | N_Shore | Body;Body;Body |
| WG2121 | chr3:147124363-147124523 | ZIC4;ZIC4;ZIC4 | N_Shelf | 1stExon;5'UTR;TSS1500 |
| GM01376 | chr3:147129778-147129952 | ZIC1 | N_Shore | Body |
| WG1977 | chr3:15311021-15311269 | SH3BP5;SH3BP5 | | Body;Body |
| WG1984 | chr3:155394109-155394303 | PLCH1;PLCH1;PLCH1 | | 5'UTR;TSS200;TSS200 |
| GM00940 | chr3:155570275-155570809 | SLC33A1 | N_Shelf | Body |
| GM00439 | chr3:156273297-156273458 | SSR3 | S_Shore | TSS1500 |
| GM00042 | chr3:157217556-157217569 | VEPH1;VEPH1;VEPH1;VEPH1;VEPH1 | | TSS200;5'UTR;5'UTR;5'UTR;5'UTR |
| WG1722 | chr3:157821262-157822072 | SHOX2;SHOX2;SHOX2 | Island | Body;Body;Body |
| GM02175 | chr3:157821987-157822072 | SHOX2;SHOX2;SHOX2 | N_Shore | Body;Body;Body |
| GM02149 | chr3:158390332-158390525 | LXN;LXN;GFM1 | | 1stExon;5'UTR;Body |
| GM01393 | chr3:158518888-158519176 | MFSD1;MFSD1 | N_Shore | TSS1500;TSS1500 |
| WG1977 | chr3:160475035-160475128 | PPM1L | S_Shore | Body |
| GM09255 | chr3:169377725-169377946 | MECOM | N_Shore | Body |
| WG1722 | chr3:169483122-169483295 | TERC | S_Shore | TSS1500 |

| | | | | |
|---|---|---|---|---|
| GM02706 | chr3:169683976-169684239 | LOC100128164;SEC62;LOC1 | N_Shore | Body;TSS1500;Body |
| GM00290 | chr3:169781622-169781717 | GPR160 | | 5'UTR |
| WG1984 | chr3:181437145-181437548 | SOX2OT | N_Shore | Body |
| GM09256 | chr3:183894727-183894778 | AP2M1;AP2M1 | S_Shore | 5'UTR;5'UTR |
| GM09255 | chr3:183903129-183903315 | ABCF3 | N_Shore | TSS1500 |
| WG1283 | chr3:189348936-189349021 | TP63;TP63;TP63 | | TSS1500;TSS1500;TSS1500 |
| GM00940 | chr3:189348936-189349129 | TP63;TP63;TP63 | | TSS1500;TSS1500;TSS1500 |
| WG1282 | chr3:191048377-191048439 | UTS2D;CCDC50;CCDC50 | S_Shore | TSS200;Body;Body |
| WG3465 | chr3:192289245-192289293 | FGF12 | | Body |
| GM00031 | chr3:194319644-194319931 | TMEM44;TMEM44;TMEM44;TMEM44 | | Body;Body;Body;Body |
| WG1978 | chr3:194833887-194834008 | C3orf21 | | Body |
| GM00935 | chr3:195488922-195489789 | MUC4;MUC4;MUC4 | Island | Body;Body;Body |
| GM00941 | chr3:195489306-195489782 | MUC4;MUC4;MUC4 | Island | Body;Body;Body |
| WG1310 | chr3:195489306-195489789 | MUC4;MUC4;MUC4 | Island | Body;Body;Body |
| GM00290 | chr3:195528460-195528709 | MUC4;MUC4;MUC4 | | Body;Body;Body |
| GM00439 | chr3:195530133-195530532 | MUC4;MUC4;MUC4 | | Body;Body;Body |
| GM01096 | chr3:196065289-196065318 | TM4SF19 | | TSS200 |
| WG1977 | chr3:196065357-196065506 | TM4SF19 | | TSS200 |
| GM03073 | chr3:24537277-24537309 | THRB;THRB;THRB | Island | TSS1500;TSS1500;TSS1500 |
| GM03075 | chr3:24537309-24537407 | THRB;THRB;THRB | Island | TSS1500;TSS1500;TSS1500 |
| GM00289 | chr3:24537338-24537407 | THRB;THRB;THRB | Island | TSS1500;TSS1500;TSS1500 |
| WG2120 | chr3:26665815-26666151 | LRRC3B | N_Shore | 5'UTR |
| GM00520 | chr3:32023402-32023473 | ZNF860;OSBPL10 | S_Shore | 5'UTR;TSS200 |
| GM01096 | chr3:32861150-32861407 | TRIM71 | Island | Body |
| GM01095 | chr3:350682-350892 | CHL1 | Island | 5'UTR |
| WG2275 | chr3:36949957-36950004 | TRANK1 | | Body |
| WG2414 | chr3:37033373-37033632 | EPM2AIP1;MLH1 | N_Shore | 1stExon;TSS1500 |
| GM02175 | chr3:39543515-39543776 | MOBP;MOBP | N_Shore | 5'UTR;Body |
| GM00935 | chr3:42727425-42727843 | KBTBD5 | Island | 1stExon |
| GM02555 | chr3:42742993-42743203 | HHATL;HHATL | | Body;5'UTR |
| GM02706 | chr3:42947263-42947385 | ZNF662;ZNF662 | N_Shore | TSS1500;TSS200 |
| GM00940 | chr3:44802549-44802604 | KIF15;KIAA1143 | N_Shore | TSS1500;Body |
| WG2120 | chr3:45077254-45077649 | CLEC3B | Island | Body |
| GM02173 | chr3:48694451-48694534 | CELSR3 | Island | Body |
| GM09256 | chr3:49203973-49204239 | CCDC71 | S_Shore | TSS200 |
| GM00042 | chr3:49459909-49459940 | AMT;AMT;AMT;NICN1;AMT;AMT;AMT;AMT | | Body;5'UTR;1stExon;3'UTR;1stExon;1s |
| GM00439 | chr3:50487955-50488230 | CACNA2D2;CACNA2D2 | | Body;Body |
| WG2415 | chr3:50488020-50488230 | CACNA2D2;CACNA2D2 | | Body;Body |
| WG1513 | chr3:51990575-51990929 | GPR62 | Island | 1stExon |
| GM02456 | chr3:52487733-52488229 | TNNC1 | N_Shore | Body |
| WG1513 | chr3:53290327-53290343 | TKT;TKT;TKT | S_Shore | TSS1500;TSS1500;TSS1500 |
| GM02456 | chr3:56809796-56810320 | ARHGEF3;ARHGEF3;ARHGEF3 | | TSS200;Body;Body |
| GM09256 | chr3:63087235-63087266 | LOC285401 | | TSS1500 |
| GM02555 | chr3:64547108-64547373 | ADAMTS9 | | Body |
| WG1977 | chr3:64547310-64547373 | ADAMTS9 | | Body |
| GM02706 | chr3:65342644-65342971 | MAGI1;MAGI1 | Island | Body;3'UTR |
| GM00042 | chr3:71180032-71180071 | FOXP1 | | Body |

| | | | | |
|---|---|---|---|---|
| GM00289 | chr3:99357437-99357613 | MIR548G;COL8A1;COL8A1 | | Body;TSS200;TSS200 |
| GM09255 | chr4:10459219-10459352 | ZNF518B | Island | TSS200 |
| GM00042 | chr4:10686394-10686554 | CLNK | | TSS200 |
| GM02706 | chr4:1107706-1107983 | RNF212;RNF212 | Island | TSS200;TSS200 |
| GM01393 | chr4:111397885-111398146 | ENPEP | Island | 1stExon |
| WG2193 | chr4:111398042-111398146 | ENPEP | Island | 1stExon |
| GM01096 | chr4:111544016-111544271 | PITX2;PITX2;PITX2;PITX2 | Island | 1stExon;5'UTR;Body;Body |
| GM00520 | chr4:111544291-111545062 | PITX2;PITX2;PITX2 | Island | Body;TSS200;Body |
| GM00042 | chr4:111544387-111544463 | PITX2;PITX2;PITX2 | Island | Body;TSS200;Body |
| GM01653 | chr4:113970660-113970745 | ANK2;ANK2;ANK2 | | TSS200;Body;TSS200 |
| GM00935 | chr4:1195845-1196179 | LOC100130872;LOC100130872-SPON2 | | Body;Body |
| GM02555 | chr4:119948360-119948542 | SYNPO2;SYNPO2;SYNPO2 | | Body;Body;Body |
| GM00520 | chr4:1243849-1244086 | C4orf42;CTBP1;CTBP1 | Island | TSS1500;TSS1500;TSS1500 |
| GM02173 | chr4:1243980-1244017 | CTBP1;CTBP1;C4orf42 | Island | TSS1500;TSS1500;TSS200 |
| GM01096 | chr4:1243980-1244024 | CTBP1;CTBP1;C4orf42 | Island | TSS1500;TSS1500;TSS200 |
| GM01375 | chr4:1243980-1244086 | CTBP1;CTBP1;C4orf42 | Island | TSS1500;TSS1500;TSS200 |
| GM01393 | chr4:140201537-140201574 | C4orf49 | S_Shore | TSS200 |
| WG2276 | chr4:140201537-140201652 | C4orf49 | S_Shore | TSS200 |
| GM02706 | chr4:140586881-140586988 | MGST2 | | TSS200 |
| GM00042 | chr4:151177137-151177553 | DCLK2;DCLK2 | N_Shore | Body;Body |
| WG1978 | chr4:154711512-154711563 | SFRP2 | N_Shore | TSS1500 |
| GM00289 | chr4:155702931-155703429 | RBM46 | Island | 5'UTR |
| GM00438 | chr4:156680007-156680050 | GUCY1B3 | N_Shore | TSS200 |
| GM00941 | chr4:157997064-157997086 | GLRB;GLRB;GLRB | N_Shore | TSS1500;TSS1500;TSS1500 |
| WG1310 | chr4:159092089-159092553 | FAM198B;FAM198B;FAM198B | | Body;Body;Body |
| WG1513 | chr4:160188727-160189254 | RAPGEF2 | | TSS1500 |
| GM02173 | chr4:165898707-165898835 | TRIM61;TRIM61 | | 1stExon;5'UTR |
| GM01376 | chr4:165898825-165898848 | TRIM61 | | TSS200 |
| GM00520 | chr4:169418081-169418224 | PALLD;PALLD | | TSS200;TSS200 |
| WG2414 | chr4:169799087-169799624 | PALLD;PALLD;PALLD;PALLD | Island | Body;Body;Body;Body |
| WG2415 | chr4:174447847-174448341 | HAND2 | N_Shore | 3'UTR |
| WG1978 | chr4:174448128-174448705 | HAND2 | N_Shore | 3'UTR |
| GM02173 | chr4:174452985-174453287 | NBLA00301 | S_Shore | Body |
| WG2276 | chr4:1804007-1804492 | FGFR3;FGFR3;FGFR3 | Island | Body;Body;Body |
| GM00940 | chr4:183247132-183247234 | ODZ3 | | Body |
| WG2275 | chr4:1844724-1844878 | LETM1 | S_Shore | Body |
| GM00290 | chr4:184938291-184938359 | STOX2 | | Body |
| GM00439 | chr4:184938291-184938470 | STOX2 | | Body |
| WG1977 | chr4:186732942-186733060 | SORBS2;SORBS2;SORBS2;SORBS2;SOR | | 5'UTR;5'UTR;5'UTR;1stExon;TSS1500; |
| WG1978 | chr4:186733060-186733428 | SORBS2;SORBS2;SORBS2;SORBS2;SOR | | 5'UTR;5'UTR;5'UTR;1stExon;TSS1500; |
| GM00438 | chr4:187621972-187622164 | FAT1 | | Body |
| WG2120 | chr4:24818471-24818615 | CCDC149 | | Body |
| GM00439 | chr4:2813542-2814064 | SH3BP2;SH3BP2 | | TSS1500;5'UTR |
| GM02641 | chr4:3372123-3372272 | RGS12;RGS12;RGS12 | N_Shelf | Body;Body;Body |
| WG2276 | chr4:3374767-3374909 | RGS12;RGS12;RGS12 | Island | Body;Body;Body |
| WG1513 | chr4:3486039-3486204 | DOK7;DOK7 | Island | Body;Body |
| GM01096 | chr4:40518024-40518065 | RBM47;RBM47 | | 5'UTR;TSS200 |

| | | | | |
|---|---|---|---|---|
| WG1283 | chr4:40858965-40859293 | APBB2;APBB2;APBB2;APBB2 | Island | Body;5'UTR;5'UTR;5'UTR;Body;5'UTR;E |
| WG2121 | chr4:40858984-40859293 | APBB2;APBB2;APBB2;APBB2 | Island | Body;5'UTR;5'UTR;5'UTR;Body;5'UTR;E |
| GM02175 | chr4:40858984-40859344 | APBB2;APBB2;APBB2;APBB2 | Island | Body;5'UTR;5'UTR;5'UTR;Body;5'UTR;E |
| GM00290 | chr4:40859249-40859259 | APBB2;APBB2;APBB2;APBB2 | S_Shore | Body;TSS200;TSS200;Body;TSS200;Bo |
| WG1977 | chr4:40859249-40859293 | APBB2;APBB2;APBB2;APBB2 | S_Shore | Body;TSS200;TSS200;Body;TSS200;Bo |
| GM01375 | chr4:40859259-40859293 | APBB2;APBB2;APBB2;APBB2 | S_Shore | Body;TSS200;TSS200;Body;TSS200;Bo |
| GM02640 | chr4:5021084-5021113 | CYTL1 | Island | 1stExon |
| WG1722 | chr4:5021268-5021328 | CYTL1 | Island | TSS200 |
| WG2121 | chr4:52918457-52918489 | SPATA18 | S_Shore | Body |
| GM00935 | chr4:52942997-52943197 | SPATA18 | Island | Body |
| GM00290 | chr4:54424280-54424764 | LNX1;LNX1;LNX1 | | 1stExon;Body;5'UTR |
| GM00016 | chr4:54424280-54424787 | LNX1;LNX1;LNX1 | | 1stExon;Body;5'UTR |
| WG2844 | chr4:54965829-54965904 | GSX2 | N_Shore | TSS1500 |
| WG2844 | chr4:55143269-55143557 | PDGFRA | | Body |
| GM00439 | chr4:57522632-57522881 | HOPX;HOPX;HOPX;HOPX;HO | Island | 1stExon;5'UTR;5'UTR;Body;Body;5'UTI |
| GM00935 | chr4:57522762-57522881 | HOPX;HOPX;HOPX;HOPX;HO | S_Shore | 5'UTR;Body;Body;TSS200;TSS200 |
| GM00031 | chr4:57547347-57547579 | HOPX;HOPX;HOPX | | 5'UTR;5'UTR;5'UTR |
| GM00289 | chr4:5976063-5976157 | C4orf50 | | Body |
| WG2844 | chr4:6052527-6052554 | JAKMIP1 | | Body |
| GM00940 | chr4:6694923-6695319 | S100P | | TSS1500 |
| WG1722 | chr4:683148-683240 | MFSD7 | S_Shore | TSS200 |
| WG2276 | chr4:71587722-71587904 | RUFY3;RUFY3;RUFY3;RUFY3;RUFY3 | | 1stExon;Body;5'UTR;1stExon;5'UTR |
| GM00520 | chr4:7282819-7283184 | SORCS2 | | Body |
| GM01096 | chr4:7287492-7287737 | SORCS2 | | Body |
| GM00290 | chr4:74719046-74719172 | PF4V1;PF4V1 | N_Shore | 1stExon;5'UTR |
| WG1722 | chr4:74847646-74847829 | PF4 | Island | 1stExon |
| GM00935 | chr4:74847766-74848016 | PF4 | Island | TSS200 |
| GM00439 | chr4:77507102-77507411 | SHROOM3 | Island | Body |
| GM00439 | chr4:7941894-7942031 | AFAP1;AFAP1 | S_Shore | TSS1500;TSS1500 |
| GM00042 | chr4:81106857-81107185 | PRDM8 | S_Shore | 5'UTR |
| GM00016 | chr4:81118500-81119473 | PRDM8;PRDM8 | Island | 5'UTR;TSS200 |
| WG1977 | chr4:94124971-94125068 | GRID2 | | Body |
| GM09256 | chr5:100240048-100240223 | ST8SIA4;ST8SIA4 | S_Shore | TSS1500;TSS1500 |
| GM00042 | chr5:1027884-1028394 | NKD2 | S_Shelf | Body |
| GM00438 | chr5:102898463-102898733 | NUDT12;NUDT12 | | 5'UTR;1stExon |
| WG1722 | chr5:1053545-1053774 | SLC12A7 | Island | Body |
| GM00290 | chr5:1053545-1054262 | SLC12A7 | Island | Body |
| WG1310 | chr5:1053643-1054113 | SLC12A7 | Island | Body |
| WG1977 | chr5:1053643-1054262 | SLC12A7 | Island | Body |
| GM00289 | chr5:1090619-1090936 | SLC12A7 | S_Shore | Body |
| GM01653 | chr5:110062343-110062729 | TMEM232;TMEM232 | | 1stExon;5'UTR |
| WG1514 | chr5:110062384-110062417 | TMEM232 | | TSS200 |
| GM00289 | chr5:111753803-111754296 | EPB41L4A;FLJ11235 | N_Shore | Body;TSS1500 |
| GM00438 | chr5:112824614-112824700 | MCC | S_Shore | TSS200 |
| GM02555 | chr5:114516368-114516422 | TRIM36;TRIM36;TRIM36 | S_Shore | TSS200;TSS200;TSS200 |
| GM00042 | chr5:115298704-115299475 | LVRN | Island | 1stExon |
| GM02706 | chr5:115298920-115299475 | LVRN | Island | 1stExon |

| | | | | |
|---|---|---|---|---|
| GM01653 | chr5:115299071-115299475 | LVRN | Island | Body |
| GM01393 | chr5:126409198-126409227 | FLJ44606;FLJ44606 | Island | TSS200;TSS200 |
| GM00290 | chr5:126409310-126409454 | FLJ44606;FLJ44606 | Island | TSS200;TSS200 |
| GM00289 | chr5:127871927-127872074 | FBN2 | N_Shore | Body |
| WG3465 | chr5:127871927-127872329 | FBN2 | N_Shore | Body |
| WG1310 | chr5:134870506-134870517 | NEUROG1;NEUROG1 | N_Shore | 1stExon;3'UTR |
| GM00520 | chr5:135364328-135364451 | TGFBI | N_Shore | TSS1500 |
| GM01393 | chr5:135416205-135416613 | MIR886 | Island | Body |
| GM00016 | chr5:135416412-135416613 | MIR886 | Island | TSS200 |
| GM02555 | chr5:138714136-138714337 | SLC23A1;SLC23A1 | Island | Body;Body |
| GM00438 | chr5:138714337-138714654 | SLC23A1;SLC23A1 | Island | Body;Body |
| GM00290 | chr5:140018644-140018709 | TMCO6 | N_Shore | TSS1500 |
| GM00941 | chr5:140097950-140098250 | VTRNA1-2 | | TSS1500 |
| WG1977 | chr5:140098392-140098576 | VTRNA1-2 | | TSS200 |
| GM00935 | chr5:140220686-140220822 | PCDHA6;PCDHA2;PCDHA1;P | N_Shore | Body;Body;Body;Body;Body;TSS1500; |
| WG2276 | chr5:140552992-140553150 | PCDHB7 | N_Shore | 1stExon |
| GM09255 | chr5:140579296-140579303 | PCDHB11 | N_Shore | TSS200 |
| WG2276 | chr5:140730092-140730516 | PCDHGB1;PCDHGA2;PCDHG | N_Shore | 1stExon;Body;Body;1stExon;Body |
| WG1977 | chr5:14406585-14406732 | TRIO | | Body |
| GM02175 | chr5:148520854-148520894 | ABLIM3 | N_Shore | TSS200 |
| GM02706 | chr5:149980486-149980682 | SYNPO | | TSS200 |
| GM09255 | chr5:149980526-149980682 | SYNPO | | TSS200 |
| WG2275 | chr5:149980671-149980682 | SYNPO;SYNPO | | 1stExon;5'UTR |
| WG3465 | chr5:150004773-150004918 | SYNPO;SYNPO | Island | Body;Body |
| GM00438 | chr5:150227510-150227654 | IRGM | | 5'UTR |
| GM02641 | chr5:153858816-153858936 | HAND1 | Island | TSS1500 |
| GM00520 | chr5:156886982-156887005 | NIPAL4 | Island | TSS200 |
| GM03075 | chr5:156886990-156887005 | NIPAL4 | Island | TSS200 |
| WG1983 | chr5:158522427-158522603 | EBF1 | N_Shore | Body |
| GM02175 | chr5:158524270-158524649 | EBF1 | Island | Body |
| GM02704 | chr5:1594282-1594863 | SDHAP3 | Island | Body |
| GM00042 | chr5:169063935-169064071 | DOCK2 | N_Shore | TSS1500 |
| WG1513 | chr5:171534678-171534792 | STK10 | | Body |
| WG1721 | chr5:175109098-175109114 | HRH2;HRH2;HRH2 | | 5'UTR;5'UTR;1stExon |
| GM01392 | chr5:176057924-176057996 | EIF4E1B;SNCB;SNCB;EIF4E1 | S_Shore | 5'UTR;TSS1500;TSS1500;1stExon |
| GM02706 | chr5:176559334-176559563 | NSD1;NSD1 | Island | TSS1500;TSS1500 |
| GM01392 | chr5:176784688-176784719 | RGS14 | | TSS200 |
| GM00438 | chr5:176828872-176829112 | PFN3 | N_Shore | TSS1500 |
| WG1513 | chr5:176829109-176829198 | PFN3 | N_Shore | TSS1500 |
| WG1721 | chr5:177548398-177548913 | N4BP3 | | Body |
| WG1977 | chr5:177548398-177549014 | N4BP3 | | Body |
| WG3465 | chr5:177708304-177708528 | COL23A1 | | Body |
| GM02149 | chr5:177764172-177764408 | COL23A1 | Island | Body |
| GM02640 | chr5:178004012-178004204 | COL23A1 | Island | Body |
| GM02147 | chr5:178684416-178684658 | ADAMTS2;ADAMTS2 | Island | Body;Body |
| GM00042 | chr5:179588440-179588673 | RASGEF1C | Island | 5'UTR |
| GM00935 | chr5:179588440-179588838 | RASGEF1C | Island | 5'UTR |

| | | | | |
|---|---|---|---|---|
| GM00439 | chr5:179588551-179588838 | RASGEF1C | Island | 5'UTR |
| GM01096 | chr5:179634578-179634688 | RASGEF1C | Island | 5'UTR |
| WG1722 | chr5:180230767-180230959 | MGAT1;MGAT1;MGAT1;MGAT | S_Shore | 1stExon;5'UTR;5'UTR;5'UTR;TSS1500;5 |
| WG3465 | chr5:191127-191296 | LRRC14B | N_Shore | TSS1500 |
| GM00438 | chr5:23507562-23507594 | PRDM9 | | TSS200 |
| GM09256 | chr5:23507573-23507617 | PRDM9 | | TSS200 |
| WG1513 | chr5:23507573-23507656 | PRDM9 | | TSS200 |
| GM02704 | chr5:2750163-2750263 | IRX2;IRX2 | Island | Body;Body |
| GM00016 | chr5:2753852-2754074 | C5orf38 | Island | Body |
| WG2275 | chr5:323320-323969 | AHRR | S_Shore | Body |
| GM02456 | chr5:368394-368804 | AHRR | N_Shore | Body |
| GM00520 | chr5:368394-369088 | AHRR | N_Shore | Body |
| GM09255 | chr5:368804-369088 | AHRR | N_Shore | Body |
| WG2193 | chr5:37834850-37834958 | GDNF;GDNF;GDNF | Island | 1stExon;Body;Body |
| GM01096 | chr5:38257516-38257555 | EGFLAM | N_Shore | TSS1500 |
| WG2276 | chr5:39219245-39219698 | FYB;FYB | | 5'UTR;5'UTR |
| GM01096 | chr5:40841315-40841590 | CARD6 | | TSS200 |
| WG2415 | chr5:40841493-40841590 | CARD6 | | 1stExon |
| GM02456 | chr5:415575-415885 | AHRR | | Body |
| GM00935 | chr5:423894-424371 | AHRR | S_Shelf | Body |
| GM00940 | chr5:42756786-42756860 | CCDC152 | | TSS200 |
| GM00290 | chr5:42756786-42757815 | CCDC152 | | TSS200 |
| GM00520 | chr5:502291-502571 | SLC9A3 | S_Shore | Body |
| WG1722 | chr5:54826611-54826768 | PPAP2A;PPAP2A;RNF138P1 | Island | Body;Body;Body |
| WG2415 | chr5:66462293-66462662 | MAST4;MAST4 | S_Shelf | Body;Body |
| GM00439 | chr5:68710808-68710912 | MARVELD2;MARVELD2 | Island | TSS200;TSS200 |
| WG2415 | chr5:68710808-68711277 | MARVELD2;MARVELD2 | Island | TSS200;TSS200 |
| GM09256 | chr5:68710818-68711277 | MARVELD2;MARVELD2 | Island | TSS200;TSS200 |
| GM00520 | chr5:68787747-68787919 | OCLN | N_Shore | TSS1500 |
| WG1513 | chr5:72415429-72415770 | TMEM171;TMEM171 | N_Shore | TSS1500;TSS1500 |
| WG1513 | chr5:7394927-7395395 | ADCY2 | N_Shore | TSS1500 |
| WG2415 | chr5:76010444-76010658 | F2R | N_Shore | TSS1500 |
| GM00289 | chr5:76373022-76373719 | ZBED3 | Island | Body |
| GM00290 | chr5:77805555-77806312 | LHFPL2 | N_Shore | Body |
| GM00438 | chr5:78281793-78281827 | ARSB;ARSB;ARSB | S_Shore | TSS200;1stExon;5'UTR |
| GM01376 | chr5:78281793-78281964 | ARSB;ARSB;ARSB | S_Shore | TSS200;1stExon;5'UTR |
| GM00935 | chr5:83016630-83016779 | HAPLN1;HAPLN1 | N_Shore | 1stExon;5'UTR |
| WG2275 | chr5:92907851-92908051 | FLJ42709;FLJ42709;FLJ4270 | Island | TSS1500;Body;Body |
| GM01095 | chr5:92907851-92908392 | FLJ42709;FLJ42709;FLJ4270 | Island | TSS1500;Body;Body |
| GM01096 | chr5:92907881-92908392 | FLJ42709;FLJ42709;FLJ4270 | Island | TSS1500;Body;Body |
| GM01096 | chr5:92908988-92909429 | FLJ42709;FLJ42709 | S_Shore | Body;Body |
| WG1722 | chr5:92908988-92909434 | FLJ42709;FLJ42709 | S_Shore | Body;Body |
| GM02456 | chr5:92929319-92929679 | NR2F1 | Island | Body |
| GM00289 | chr6:100911240-100911709 | SIM1 | N_Shore | 1stExon |
| GM02706 | chr6:100911240-100911964 | SIM1 | N_Shore | 1stExon |
| WG2415 | chr6:100911240-100912072 | SIM1 | N_Shore | 1stExon |
| GM00042 | chr6:100911687-100911727 | SIM1 | N_Shore | TSS200 |

| | | | | |
|---|---|---|---|---|
| WG1984 | chr6:100911701-100911727 | SIM1 | N_Shore | TSS200 |
| GM00289 | chr6:100911744-100912168 | SIM1 | N_Shore | TSS200 |
| GM00042 | chr6:100911746-100911964 | SIM1 | N_Shore | TSS200 |
| WG1722 | chr6:100911949-100912072 | SIM1 | N_Shore | TSS1500 |
| GM01375 | chr6:101846650-101846767 | GRIK2;GRIK2;GRIK2 | N_Shore | TSS1500;TSS1500;TSS1500 |
| GM00439 | chr6:101846650-101846872 | GRIK2;GRIK2;GRIK2 | N_Shore | TSS1500;TSS1500;TSS1500 |
| GM02555 | chr6:101846650-101846967 | GRIK2;GRIK2;GRIK2 | N_Shore | TSS1500;TSS1500;TSS1500 |
| WG1283 | chr6:101846767-101846872 | GRIK2;GRIK2;GRIK2 | Island | TSS200;TSS200;TSS200 |
| GM09255 | chr6:101846791-101846805 | GRIK2;GRIK2;GRIK2 | Island | TSS200;TSS200;TSS200 |
| WG1984 | chr6:101846791-101846872 | GRIK2;GRIK2;GRIK2 | Island | TSS200;TSS200;TSS200 |
| WG1513 | chr6:101846791-101846905 | GRIK2;GRIK2;GRIK2 | Island | TSS200;TSS200;TSS200 |
| GM01375 | chr6:101846791-101847388 | GRIK2;GRIK2;GRIK2 | Island | TSS200;TSS200;TSS200 |
| GM01095 | chr6:101846791-101848236 | GRIK2;GRIK2;GRIK2 | Island | TSS200;TSS200;TSS200 |
| GM00022 | chr6:101846916-101847058 | GRIK2;GRIK2;GRIK2;GRIK2;G | Island | 1stExon;1stExon;5'UTR;1stExon;5'UTR |
| GM00439 | chr6:101846916-101847388 | GRIK2;GRIK2;GRIK2;GRIK2;G | Island | 1stExon;1stExon;5'UTR;1stExon;5'UTR |
| GM00940 | chr6:101847349-101847656 | GRIK2;GRIK2;GRIK2 | S_Shore | Body;Body;Body |
| GM00438 | chr6:10416090-10416373 | TFAP2A;TFAP2A | N_Shore | Body;TSS1500 |
| WG1513 | chr6:106546747-106546824 | PRDM1;PRDM1 | | Body;TSS200 |
| WG2275 | chr6:109274429-109274496 | ARMC2 | | Body |
| WG1977 | chr6:11279656-11279727 | NEDD9 | | Body |
| GM02173 | chr6:116381609-116381957 | FRK;FRK | | 1stExon;5'UTR |
| GM02641 | chr6:117591534-117592358 | VGLL2;VGLL2 | Island | Body;Body |
| GM00438 | chr6:117869027-117869453 | DCBLD1 | N_Shore | Body |
| GM00439 | chr6:123109391-123109837 | SMPDL3A | N_Shore | TSS1500 |
| GM00042 | chr6:127796683-127797022 | C6orf174 | Island | Body |
| GM01096 | chr6:127835750-127836053 | C6orf174 | N_Shore | Body |
| GM02149 | chr6:132931714-132931736 | TAAR3 | | TSS1500 |
| WG1722 | chr6:133035150-133035266 | VNN1 | | 1stExon |
| WG1978 | chr6:134210273-134210307 | TCF21;TCF21;TCF21;TCF21 | N_Shore | 1stExon;5'UTR;1stExon;5'UTR |
| WG1722 | chr6:138200071-138200309 | TNFAIP3 | | Body |
| GM00439 | chr6:138200136-138200387 | TNFAIP3 | | Body |
| WG1977 | chr6:139351113-139351353 | C6orf115 | S_Shore | 5'UTR |
| WG1978 | chr6:139613145-139613456 | TXLNB;TXLNB | | 5'UTR;1stExon |
| GM00520 | chr6:144386416-144386422 | PLAGL1;PLAGL1 | S_Shore | TSS1500;TSS1500 |
| GM00439 | chr6:146864328-146864352 | RAB32 | N_Shore | TSS1500 |
| WG2276 | chr6:148869461-148869535 | SASH1 | | Body |
| GM02555 | chr6:149358743-149358802 | UST | | Body |
| GM00042 | chr6:149803087-149803206 | ZC3H12D | | 5'UTR |
| WG1977 | chr6:149803087-149803444 | ZC3H12D | | 5'UTR |
| WG1283 | chr6:149803111-149803444 | ZC3H12D | | 5'UTR |
| WG1978 | chr6:149805995-149806131 | ZC3H12D;ZC3H12D | | 5'UTR;1stExon |
| GM01653 | chr6:149806077-149806131 | ZC3H12D;ZC3H12D | | 5'UTR;1stExon |
| GM02706 | chr6:151186411-151186511 | MTHFD1L | N_Shore | TSS1500 |
| WG1513 | chr6:151560831-151560927 | AKAP12 | Island | TSS1500 |
| WG1977 | chr6:152085565-152085641 | ESR1 | | 5'UTR |
| WG2276 | chr6:152125861-152126080 | ESR1;ESR1 | N_Shelf | 5'UTR;TSS1500 |
| WG1310 | chr6:152125861-152126092 | ESR1;ESR1 | N_Shelf | 5'UTR;TSS1500 |

| | | | | |
|---|---|---|---|---|
| GM02555 | chr6:152125965-152126092 | ESR1;ESR1 | N_Shelf | 5'UTR;TSS1500 |
| WG1983 | chr6:152126180-152126441 | ESR1;ESR1 | N_Shelf | 5'UTR;TSS1500 |
| WG1514 | chr6:152127812-152127887 | ESR1;ESR1;ESR1;ESR1 | N_Shore | TSS1500;5'UTR;TSS1500;5'UTR |
| WG1977 | chr6:152127812-152128024 | ESR1;ESR1;ESR1;ESR1 | N_Shore | TSS1500;5'UTR;TSS1500;5'UTR |
| GM00940 | chr6:152128328-152128366 | ESR1;ESR1;ESR1;ESR1 | N_Shore | TSS1500;TSS200;5'UTR;5'UTR |
| GM00022 | chr6:152128328-152128411 | ESR1;ESR1;ESR1;ESR1 | N_Shore | TSS1500;TSS200;5'UTR;5'UTR |
| WG1657 | chr6:152130058-152130332 | ESR1;ESR1;ESR1;ESR1 | S_Shore | Body;Body;Body;Body |
| GM02456 | chr6:158018055-158018184 | ZDHHC14;ZDHHC14 | S_Shelf | Body;Body |
| GM09255 | chr6:159070959-159071076 | SYTL3 | | TSS200 |
| GM02456 | chr6:159238463-159238744 | EZR;EZR | N_Shore | Body;Body |
| GM02175 | chr6:160182438-160182624 | ACAT2 | Island | TSS1500 |
| WG2415 | chr6:161796785-161796854 | PARK2;PARK2;PARK2 | | Body;Body;Body |
| WG1658 | chr6:161860203-161860359 | PARK2;PARK2;PARK2 | | Body;Body;Body |
| GM00935 | chr6:167529765-167530253 | CCR6 | | 5'UTR |
| WG1282 | chr6:168435914-168436353 | KIF25;KIF25 | Island | Body;Body |
| GM01392 | chr6:168859941-168859997 | SMOC2;SMOC2 | | Body;Body |
| GM00289 | chr6:169026695-169027033 | SMOC2;SMOC2 | | Body;Body |
| GM02175 | chr6:18387077-18387360 | RNF144B | N_Shore | TSS1500 |
| GM02706 | chr6:21667166-21667407 | FLJ22536 | S_Shore | Body |
| GM01376 | chr6:24583477-24583614 | KIAA0319;KIAA0319;KIAA0319;KIAA0319 | | Body;Body;Body;Body;Body |
| GM00438 | chr6:2698001-2698037 | MYLK4 | | Body |
| GM00016 | chr6:27107588-27107756 | HIST1H2BK | S_Shore | 3'UTR |
| GM00022 | chr6:28058724-28058802 | ZSCAN12L1 | | TSS1500 |
| GM01393 | chr6:28493651-28493664 | GPX5;GPX5 | | TSS200;TSS200 |
| GM00438 | chr6:28497338-28497370 | GPX5;GPX5 | | Body;Body |
| WG1977 | chr6:28555601-28555664 | SCAND3 | S_Shore | TSS1500 |
| GM00438 | chr6:28875354-28875370 | TRIM27 | | Body |
| GM01392 | chr6:291687-292823 | DUSP22 | N_Shore | TSS1500 |
| WG1658 | chr6:291882-292823 | DUSP22 | N_Shore | TSS1500 |
| GM02149 | chr6:29274096-29274617 | OR14J1 | | TSS1500 |
| GM02149 | chr6:29395103-29395221 | OR11A1 | | 1stExon |
| GM02149 | chr6:29407689-29407834 | OR10C1 | | TSS200 |
| GM02149 | chr6:29408443-29408587 | OR10C1 | | 1stExon |
| GM02149 | chr6:29455302-29455331 | MAS1L | | 1stExon |
| GM02706 | chr6:29592854-29593382 | GABBR1;GABBR1;GABBR1 | N_Shelf | Body;Body;Body |
| GM02175 | chr6:29594830-29595016 | GABBR1;GABBR1;GABBR1 | N_Shore | Body;Body;Body |
| GM01096 | chr6:29595002-29595016 | GABBR1;GABBR1;GABBR1 | N_Shore | Body;Body;Body |
| GM00935 | chr6:29690766-29690889 | HLA-F;HLA-F;HLA-F | N_Shore | TSS1500;TSS1500;TSS1500 |
| WG1513 | chr6:29690766-29690893 | HLA-F;HLA-F;HLA-F | N_Shore | TSS1500;TSS1500;TSS1500 |
| WG1977 | chr6:29690822-29690893 | HLA-F;HLA-F;HLA-F | N_Shore | TSS1500;TSS1500;TSS1500 |
| GM09256 | chr6:29692082-29692092 | HLA-F;HLA-F;HLA-F | Island | Body;Body;Body |
| GM00438 | chr6:29894050-29894152 | HCG4P6 | N_Shore | TSS1500 |
| WG2121 | chr6:29894050-29894197 | HCG4P6 | N_Shore | TSS1500 |
| WG1283 | chr6:29911087-29911104 | HLA-A | Island | Body |
| GM01096 | chr6:29911091-29911104 | HLA-A | Island | Body |
| GM02641 | chr6:29943401-29943414 | HCG9 | N_Shore | Body |
| WG2415 | chr6:29943401-29943425 | HCG9 | N_Shore | Body |

| | | | | |
|---|---|---|---|---|
| GM09256 | chr6:29943401-29943455 | HCG9 | N_Shore | Body |
| WG2414 | chr6:29943455-29943480 | HCG9 | N_Shore | Body |
| GM00935 | chr6:29944093-29944146 | HCG9 | N_Shore | Body |
| WG1722 | chr6:30038712-30038754 | RNF39;RNF39 | N_Shore | 3'UTR;3'UTR |
| WG2193 | chr6:30039432-30039442 | RNF39;RNF39 | Island | Body;Body |
| WG1983 | chr6:30079203-30079280 | TRIM31 | | Body |
| WG1977 | chr6:30079256-30079280 | TRIM31 | | Body |
| WG1722 | chr6:30079256-30079307 | TRIM31 | | Body |
| GM00289 | chr6:30113774-30113825 | TRIM40 | | Body |
| WG2194 | chr6:30113978-30114011 | TRIM40 | | Body |
| GM02175 | chr6:30131189-30131219 | TRIM15;TRIM15 | | 5'UTR;1stExon |
| GM02706 | chr6:30228243-30228254 | HLA-L | Island | Body |
| GM09255 | chr6:30458158-30458237 | HLA-E | Island | Body |
| GM09255 | chr6:30510294-30510459 | GNL1 | | 3'UTR |
| GM09256 | chr6:30657882-30658014 | NRM | S_Shelf | Body |
| WG1977 | chr6:30698777-30698784 | FLOT1 | | Body |
| WG1513 | chr6:30854164-30854233 | DDR1;DDR1 | S_Shore | 5'UTR;5'UTR |
| WG1977 | chr6:30856386-30856491 | DDR1;DDR1;DDR1 | S_Shelf | TSS200;5'UTR;5'UTR |
| WG2415 | chr6:30921592-30921757 | DPCR1 | N_Shore | 3'UTR |
| GM09256 | chr6:31025933-31025967 | HCG22 | | Body |
| WG1310 | chr6:31025933-31026105 | HCG22 | | Body |
| GM00016 | chr6:31111076-31111270 | CCHCR1;CCHCR1;CCHCR1 | | Body;Body;Body |
| GM00439 | chr6:31143710-31144068 | PSORS1C3 | | Body |
| GM01650 | chr6:31164914-31164947 | HCG27 | N_Shore | TSS1500 |
| WG2415 | chr6:31237013-31237034 | HLA-C | N_Shore | Body |
| WG2415 | chr6:31237405-31237722 | HLA-C | N_Shore | Body |
| GM01096 | chr6:31238788-31238908 | HLA-C | N_Shore | Body |
| GM00290 | chr6:31323985-31324024 | HLA-B | Island | Body |
| GM01375 | chr6:31466174-31466187 | MICB | Island | Body |
| GM00042 | chr6:31507879-31508137 | BAT1;BAT1 | N_Shore | Body;Body |
| GM00439 | chr6:31549563-31549631 | LTB;LTB | S_Shore | Body;Body |
| GM00438 | chr6:31649619-31649638 | LY6G5C | N_Shore | TSS1500 |
| GM02149 | chr6:31649619-31649728 | LY6G5C | N_Shore | TSS1500 |
| WG1514 | chr6:31744628-31744735 | C6orf27 | | 5'UTR |
| GM00940 | chr6:31760593-31760616 | VARS | N_Shelf | Body |
| GM01392 | chr6:31829888-31829914 | NEU1 | N_Shore | Body |
| GM02706 | chr6:31829888-31829926 | NEU1 | N_Shore | Body |
| WG3465 | chr6:31846769-31846813 | SLC44A4;SLC44A4 | | 5'UTR;1stExon |
| GM00940 | chr6:31895464-31895493 | C2;C2;C2;C2 | | 1stExon;5'UTR;1stExon;5'UTR |
| GM00042 | chr6:31926072-31926205 | RDBP;SKIV2L;MIR1236 | N_Shore | Body;TSS1500;TSS1500 |
| GM02706 | chr6:32016214-32016247 | TNXB | | Body |
| GM02147 | chr6:32050927-32051108 | TNXB | S_Shelf | Body |
| GM00438 | chr6:32054771-32054867 | TNXB | N_Shore | Body |
| WG2194 | chr6:32055447-32055525 | TNXB | Island | Body |
| WG1513 | chr6:32097439-32097507 | ATF6B;ATF6B;FKBPL | N_Shore | TSS1500;TSS1500;Body |
| GM00042 | chr6:32120625-32120783 | PPT2;PRRT1;PPT2 | N_Shore | TSS1500;TSS1500;TSS1500 |
| WG1657 | chr6:32297670-32297822 | C6orf10 | | Body |

| | | | | |
|---|---|---|---|---|
| GM00438 | chr6:32371488-32371538 | BTNL2 | | Body |
| GM00438 | chr6:32410339-32410365 | HLA-DRA | | Body |
| WG2193 | chr6:32489750-32489963 | HLA-DRB5 | Island | Body |
| WG1984 | chr6:32489963-32489991 | HLA-DRB5 | Island | Body |
| WG2193 | chr6:32489991-32490043 | HLA-DRB5 | Island | Body |
| WG2415 | chr6:32526021-32526260 | HLA-DRB6 | | Body |
| GM00935 | chr6:32526021-32526263 | HLA-DRB6 | | Body |
| GM00520 | chr6:32526021-32526342 | HLA-DRB6 | | Body |
| GM02147 | chr6:32526027-32526342 | HLA-DRB6 | | Body |
| GM02173 | chr6:32526260-32526342 | HLA-DRB6 | | Body |
| GM09255 | chr6:32526260-32526364 | HLA-DRB6 | | Body |
| GM03075 | chr6:32526263-32526364 | HLA-DRB6 | | Body |
| WG1282 | chr6:32551888-32551954 | HLA-DRB1 | Island | Body |
| WG2194 | chr6:32551891-32551954 | HLA-DRB1 | Island | Body |
| WG2415 | chr6:32552152-32552246 | HLA-DRB1 | Island | Body |
| WG1283 | chr6:32552246-32552453 | HLA-DRB1 | Island | Body |
| WG1658 | chr6:32557970-32558459 | HLA-DRB1 | | TSS1500 |
| WG1513 | chr6:32633157-32633354 | HLA-DQB1 | S_Shore | Body |
| WG1309 | chr6:32634245-32634316 | HLA-DQB1 | S_Shore | Body |
| GM01653 | chr6:32634245-32634362 | HLA-DQB1 | S_Shore | Body |
| GM00290 | chr6:32634276-32634344 | HLA-DQB1 | S_Shore | 1stExon |
| GM02175 | chr6:32729442-32729470 | HLA-DQB2 | Island | Body |
| GM00289 | chr6:32729465-32729498 | HLA-DQB2 | Island | Body |
| GM00289 | chr6:32729545-32729590 | HLA-DQB2 | Island | Body |
| WG1513 | chr6:32729545-32729596 | HLA-DQB2 | Island | Body |
| GM00935 | chr6:32729563-32729590 | HLA-DQB2 | Island | Body |
| GM01095 | chr6:32847585-32847596 | PPP1R2P1 | Island | Body |
| GM00290 | chr6:32847585-32847605 | PPP1R2P1 | Island | Body |
| GM09256 | chr6:32847702-32847762 | PPP1R2P1 | Island | Body |
| WG1310 | chr6:32847762-32847788 | PPP1R2P1 | Island | Body |
| WG2120 | chr6:32847788-32847830 | PPP1R2P1 | Island | Body |
| WG1513 | chr6:32904061-32904317 | HLA-DMB | | Body |
| WG1513 | chr6:32908567-32908642 | HLA-DMB | | 1stExon |
| GM02640 | chr6:32920880-32921142 | HLA-DMA;HLA-DMA | | 5'UTR;1stExon |
| WG1310 | chr6:33048875-33048919 | HLA-DPB1 | S_Shore | Body |
| WG1513 | chr6:33131222-33131336 | COL11A2;COL11A2;COL11A | S_Shore | 3'UTR;3'UTR;3'UTR |
| GM00439 | chr6:33132611-33132650 | COL11A2;COL11A2;COL11A | S_Shelf | Body;Body;Body |
| GM00289 | chr6:33169731-33170101 | SLC39A7;RXRB;SLC39A7 | S_Shore | Body;TSS1500;Body |
| GM00438 | chr6:33169975-33170177 | SLC39A7;SLC39A7 | S_Shore | Body;Body |
| GM01096 | chr6:33282997-33283015 | ZBTB22;TAPBP;TAPBP;TAPBP | Island | Body;TSS1500;TSS1500;TSS1500;Bod |
| GM00940 | chr6:33282997-33283021 | ZBTB22;TAPBP;TAPBP;TAPBP | Island | Body;TSS1500;TSS1500;TSS1500;Bod |
| WG1282 | chr6:33288180-33288561 | DAXX;DAXX;DAXX;DAXX | N_Shore | Body;Body;Body;Body |
| WG1984 | chr6:33288180-33288569 | DAXX;DAXX;DAXX;DAXX | N_Shore | Body;Body;Body;Body |
| GM00289 | chr6:33288332-33288561 | DAXX;DAXX;DAXX;DAXX | N_Shore | Body;Body;Body;Body |
| GM01096 | chr6:33288366-33288569 | DAXX;DAXX;DAXX;DAXX | N_Shore | Body;Body;Body;Body |
| WG1282 | chr6:33288599-33288656 | DAXX;DAXX;DAXX;DAXX | N_Shore | Body;Body;Body;Body |
| GM02175 | chr6:33358866-33359001 | KIFC1 | N_Shore | TSS1500 |

| | | | | |
|---|---|---|---|---|
| WG1722 | chr6:34024362-34024479 | GRM4 | Island | Body |
| GM02706 | chr6:34203564-34203887 | HMGA1;HMGA1;HMGA1;HM( | Island | TSS1500;TSS1500;TSS1500;TSS1500; |
| GM00290 | chr6:35109121-35109298 | TCP11;TCP11;TCP11;TCP11 | Island | 5'UTR;1stExon;5'UTR;1stExon |
| GM00290 | chr6:35265534-35265555 | DEF6 | | TSS200 |
| GM02555 | chr6:35265616-35265643 | DEF6;DEF6 | | 1stExon;5'UTR |
| GM01653 | chr6:37461033-37461350 | C6orf129 | | Body |
| GM00042 | chr6:37617864-37618009 | MDGA1 | Island | Body |
| GM01376 | chr6:3848898-3849028 | FAM50B | N_Shore | TSS1500 |
| GM00438 | chr6:39290526-39290715 | KCNK16;KCNK16;KCNK16;KCNK16 | | TSS200;TSS200;TSS200;TSS200 |
| WG1309 | chr6:41528395-41528531 | FOXP4;FOXP4;FOXP4 | Island | 5'UTR;5'UTR;5'UTR |
| GM01095 | chr6:41528395-41528785 | FOXP4;FOXP4;FOXP4 | Island | 5'UTR;5'UTR;5'UTR |
| GM09256 | chr6:42927975-42927993 | GNMT | N_Shore | TSS1500 |
| WG2415 | chr6:43149356-43149629 | CUL9 | N_Shore | TSS1500 |
| GM02175 | chr6:45387257-45387395 | RUNX2;RUNX2 | N_Shore | Body;Body |
| GM02706 | chr6:47624115-47624247 | GPR111 | | TSS1500 |
| GM00438 | chr6:49605181-49605401 | RHAG | | TSS1500 |
| GM00289 | chr6:49754848-49755009 | PGK2 | | 1stExon |
| GM00438 | chr6:49916931-49917175 | DEFB133 | | Body |
| GM00520 | chr6:49917145-49917254 | DEFB133 | | 1stExon |
| WG2414 | chr6:6002865-6003303 | NRN1 | Island | Body |
| GM00941 | chr6:64281439-64281639 | PTP4A1 | N_Shore | TSS1500 |
| WG1310 | chr6:6614447-6614798 | LOC285780;LY86 | | Body;Body |
| WG1722 | chr6:70576168-70576321 | COL19A1 | N_Shore | TSS1500 |
| GM00042 | chr6:74008688-74009041 | C6orf147 | N_Shore | Body |
| WG1514 | chr6:74063982-74064064 | DPPA5 | Island | TSS200 |
| GM00289 | chr6:74104426-74104600 | DDX43;DDX43 | Island | 1stExon;5'UTR |
| GM01376 | chr6:75911716-75911956 | COL12A1;COL12A1 | Island | Body;Body |
| GM00438 | chr6:75911956-75912633 | COL12A1;COL12A1 | Island | Body;Body |
| GM00941 | chr6:75912410-75912633 | COL12A1;COL12A1 | S_Shore | Body;Body |
| GM00289 | chr6:75912410-75912711 | COL12A1;COL12A1 | S_Shore | Body;Body |
| WG2121 | chr6:75912547-75912711 | COL12A1;COL12A1 | S_Shore | 5'UTR;5'UTR |
| GM00520 | chr6:76059726-76059773 | FILIP1 | | Body |
| GM01393 | chr6:88757302-88757703 | SPACA1 | Island | TSS1500 |
| GM00438 | chr6:88757358-88757371 | SPACA1 | Island | TSS200 |
| GM02706 | chr6:88757371-88757703 | SPACA1 | Island | TSS200 |
| WG1513 | chr6:91004430-91004482 | BACH2;BACH2 | N_Shore | 5'UTR;5'UTR |
| GM02706 | chr7:100208533-100208950 | MOSPD3 | N_Shore | TSS1500 |
| GM00520 | chr7:100463206-100463583 | SLC12A9 | N_Shore | Body |
| GM00289 | chr7:100463416-100463583 | SLC12A9 | N_Shore | Body |
| GM01392 | chr7:100880751-100881286 | CLDN15 | | 1stExon |
| GM00016 | chr7:100881007-100881429 | CLDN15;CLDN15 | | 1stExon;5'UTR |
| GM00520 | chr7:101273741-101274037 | MYL10 | | TSS1500 |
| WG1283 | chr7:101500014-101500514 | CUX1;CUX1;CUX1 | | Body;Body;Body |
| GM01096 | chr7:1015418-1015501 | COX19 | S_Shore | TSS200 |
| GM00935 | chr7:101768610-101768874 | CUX1;CUX1;CUX1 | | Body;Body;Body |
| GM02175 | chr7:101768676-101768874 | CUX1;CUX1;CUX1 | | Body;Body;Body |
| GM00935 | chr7:105319437-105319819 | ATXN7L1;ATXN7L1 | | Body;5'UTR |

| | | | | |
|---|---|---|---|---|
| GM01376 | chr7:107205157-107205392 | COG5;COG5;COG5;DUS4L | S_Shore | TSS200;TSS200;TSS200;5'UTR |
| GM00439 | chr7:1098625-1098703 | C7orf50;GPR146;C7orf50;C7 | S_Shore | Body;3'UTR;Body;Body;1stExon |
| WG1977 | chr7:114055074-114055419 | FOXP2;FOXP2;FOXP2;FOXP2;FOXP2;FOX | | 1stExon;5'UTR;5'UTR;1stExon;5'UTR;1 |
| WG1721 | chr7:114055123-114055137 | FOXP2;FOXP2;FOXP2;FOXP2;FOXP2;FOX | | 1stExon;5'UTR;5'UTR;1stExon;5'UTR;1 |
| GM02175 | chr7:114055123-114055210 | FOXP2;FOXP2;FOXP2;FOXP2;FOXP2;FOX | | 1stExon;5'UTR;5'UTR;1stExon;5'UTR;1 |
| WG2415 | chr7:114055123-114055419 | FOXP2;FOXP2;FOXP2;FOXP2;FOXP2;FOX | | 1stExon;5'UTR;5'UTR;1stExon;5'UTR;1 |
| WG1983 | chr7:115862648-115863026 | TES;TES | | TSS1500;Body |
| GM00520 | chr7:1163456-1164063 | C7orf50;C7orf50;C7orf50 | | Body;Body;Body |
| GM00439 | chr7:12444013-12444115 | VWDE | S_Shore | TSS200 |
| GM00289 | chr7:127225891-127225949 | GCC1 | Island | TSS1500 |
| GM02706 | chr7:128550860-128550902 | KCP;KCP | Island | TSS200;TSS200 |
| GM01096 | chr7:128550860-128550946 | KCP;KCP | Island | TSS200;TSS200 |
| WG2121 | chr7:128550862-128550902 | KCP;KCP | Island | TSS200;TSS200 |
| WG1513 | chr7:134832850-134833299 | TMEM140;TMEM140 | | 1stExon;5'UTR |
| GM00042 | chr7:134833031-134833299 | TMEM140 | | 5'UTR |
| GM00042 | chr7:148480080-148480326 | CUL1 | N_Shore | Body |
| GM09255 | chr7:150147922-150147988 | GIMAP8 | | TSS200 |
| GM00520 | chr7:150434452-150434541 | GIMAP5;GIMAP5 | | 1stExon;5'UTR |
| GM00438 | chr7:150434471-150434541 | GIMAP5;GIMAP5 | | 1stExon;5'UTR |
| GM00439 | chr7:150705760-150705896 | NOS3 | Island | Body |
| GM00520 | chr7:150786044-150786082 | AGAP3;AGAP3 | S_Shore | Body;Body |
| GM00042 | chr7:151078646-151079501 | WDR86 | Island | 3'UTR |
| WG1977 | chr7:151330008-151330230 | PRKAG2;PRKAG2;PRKAG2 | | Body;TSS1500;Body |
| GM01653 | chr7:151442351-151442967 | PRKAG2;PRKAG2 | | Body;Body |
| GM00042 | chr7:1514455-1514623 | INTS1 | Island | Body |
| WG2844 | chr7:151493629-151493820 | PRKAG2;PRKAG2 | | 5'UTR;Body |
| GM02149 | chr7:154003215-154003309 | DPP6;DPP6;DPP6 | S_Shore | Body;Body;Body |
| GM00520 | chr7:154684051-154684433 | DPP6;DPP6;DPP6 | N_Shore | Body;Body;Body |
| WG2121 | chr7:155299624-155299687 | CNPY1 | N_Shelf | Body |
| GM00438 | chr7:156741514-156741538 | NOM1 | Island | TSS1500 |
| GM02456 | chr7:157185279-157185402 | DNAJB6 | | Body |
| WG1310 | chr7:157198482-157198869 | DNAJB6 | N_Shelf | Body |
| GM00439 | chr7:157333405-157333584 | PTPRN2;PTPRN2;PTPRN2 | | 3'UTR;3'UTR;3'UTR |
| GM00042 | chr7:157345746-157346224 | PTPRN2;PTPRN2;PTPRN2 | N_Shore | Body;Body;Body |
| GM01096 | chr7:157353097-157353191 | PTPRN2;PTPRN2;PTPRN2 | Island | Body;Body;Body |
| GM00438 | chr7:157402529-157402673 | PTPRN2;PTPRN2;PTPRN2 | N_Shelf | Body;Body;Body |
| GM00438 | chr7:157447178-157447249 | PTPRN2;PTPRN2;PTPRN2 | N_Shore | Body;Body;Body |
| GM02555 | chr7:157454062-157454207 | PTPRN2;PTPRN2;PTPRN2 | Island | Body;Body;Body |
| WG2844 | chr7:157475548-157476575 | PTPRN2;PTPRN2;PTPRN2 | N_Shore | Body;Body;Body |
| GM00438 | chr7:157778531-157778645 | PTPRN2;PTPRN2;PTPRN2 | | Body;Body;Body |
| GM00438 | chr7:157932921-157933354 | PTPRN2;PTPRN2;PTPRN2 | | Body;Body;Body |
| GM00289 | chr7:157985748-157985818 | PTPRN2;PTPRN2;PTPRN2 | | Body;Body;Body |
| GM02149 | chr7:158075388-158075705 | PTPRN2;PTPRN2;PTPRN2 | | Body;Body;Body |
| GM00438 | chr7:158109339-158109694 | PTPRN2;PTPRN2;PTPRN2 | Island | Body;Body;Body |
| WG1977 | chr7:158336016-158336049 | PTPRN2;PTPRN2;PTPRN2 | | Body;Body;Body |
| GM09255 | chr7:16505094-16505589 | SOSTDC1 | | 1stExon |
| WG1977 | chr7:16505094-16505602 | SOSTDC1 | | 1stExon |

| | | | | |
|---|---|---|---|---|
| GM00520 | chr7:16505409-16505592 | SOSTDC1;SOSTDC1 | | 1stExon;5'UTR |
| GM00289 | chr7:16505589-16505602 | SOSTDC1 | | TSS200 |
| GM00289 | chr7:19184555-19185139 | FERD3L | Island | 1stExon |
| GM00520 | chr7:19184555-19185165 | FERD3L | Island | 1stExon |
| GM09256 | chr7:19185260-19185407 | FERD3L | S_Shore | TSS1500 |
| GM09256 | chr7:1938450-1938791 | MAD1L1;MAD1L1;MAD1L1 | S_Shore | Body;Body;Body |
| GM00438 | chr7:1952518-1952600 | MAD1L1;MAD1L1;MAD1L1 | S_Shelf | Body;Body;Body |
| WG1722 | chr7:1952518-1952807 | MAD1L1;MAD1L1;MAD1L1 | S_Shelf | Body;Body;Body |
| WG1984 | chr7:1961785-1961968 | MAD1L1;MAD1L1;MAD1L1 | | Body;Body;Body |
| GM00290 | chr7:1980117-1980273 | MAD1L1;MAD1L1;MAD1L1 | S_Shore | Body;Body;Body |
| GM01650 | chr7:2060040-2060130 | MAD1L1;MAD1L1;MAD1L1 | Island | Body;Body;Body |
| WG1977 | chr7:2109244-2109890 | MAD1L1;MAD1L1;MAD1L1 | S_Shore | Body;Body;Body |
| GM02555 | chr7:2133441-2133480 | MAD1L1;MAD1L1;MAD1L1 | | Body;Body;Body |
| WG2415 | chr7:2139216-2139259 | MAD1L1;MAD1L1;MAD1L1 | | Body;Body;Body |
| GM00520 | chr7:226444-226563 | FAM20C | S_Shore | Body |
| WG1977 | chr7:2560976-2561116 | LFNG;LFNG;LFNG;LFNG | S_Shore | Body;Body;Body;Body |
| GM00520 | chr7:2563184-2563909 | LFNG;LFNG;LFNG;LFNG | N_Shore | Body;Body;Body;Body |
| GM01376 | chr7:2563478-2563909 | LFNG;LFNG;LFNG;LFNG | N_Shore | Body;Body;Body;Body |
| WG1977 | chr7:2563611-2563909 | LFNG;LFNG;LFNG;LFNG | Island | Body;Body;Body;Body |
| GM00940 | chr7:2563670-2563909 | LFNG;LFNG;LFNG;LFNG | Island | Body;Body;Body;Body |
| GM00042 | chr7:2606716-2606824 | IQCE;IQCE | | Body;Body |
| WG2276 | chr7:2646782-2647333 | IQCE;IQCE | S_Shore | Body;Body |
| GM01095 | chr7:26897182-26897714 | SKAP2 | | Body |
| GM02640 | chr7:26897612-26897714 | SKAP2 | | Body |
| GM02640 | chr7:27142427-27142527 | HOXA2 | N_Shore | TSS200 |
| GM02173 | chr7:27142700-27142799 | HOXA2 | N_Shore | TSS1500 |
| GM00290 | chr7:27143046-27143252 | HOXA2 | N_Shore | TSS1500 |
| GM02640 | chr7:27143046-27143287 | HOXA2 | N_Shore | TSS1500 |
| WG1977 | chr7:27143334-27143403 | HOXA2 | Island | TSS1500 |
| GM02706 | chr7:27143334-27143478 | HOXA2 | Island | TSS1500 |
| GM02641 | chr7:27146237-27146445 | HOXA3;HOXA3;HOXA3 | Island | 3'UTR;3'UTR;3'UTR |
| WG1722 | chr7:27150262-27150598 | HOXA3;HOXA3;HOXA3 | Island | 5'UTR;5'UTR;5'UTR |
| GM02149 | chr7:27153212-27153847 | HOXA3;HOXA3;HOXA3 | Island | 5'UTR;5'UTR;5'UTR |
| GM02555 | chr7:27153655-27153847 | HOXA3;HOXA3;HOXA3 | N_Shore | 5'UTR;5'UTR;TSS200 |
| GM00042 | chr7:27153655-27153944 | HOXA3;HOXA3;HOXA3 | N_Shore | 5'UTR;5'UTR;TSS200 |
| GM03075 | chr7:27182493-27183196 | HOXA5 | N_Shore | Body |
| GM09255 | chr7:27182493-27183436 | HOXA5 | N_Shore | Body |
| GM00520 | chr7:27183274-27183401 | HOXA5;HOXA5 | Island | 5'UTR;1stExon |
| GM02706 | chr7:27183274-27183436 | HOXA5;HOXA5 | Island | 5'UTR;1stExon |
| GM00940 | chr7:27183375-27183436 | HOXA5 | Island | TSS200 |
| GM02641 | chr7:27183643-27183694 | HOXA5 | Island | TSS1500 |
| GM02704 | chr7:27183794-27183861 | HOXA5 | Island | TSS1500 |
| GM02149 | chr7:27187269-27187374 | HOXA6 | Island | 1stExon |
| GM01376 | chr7:27187372-27187560 | HOXA6 | Island | TSS200 |
| WG1513 | chr7:27187943-27188465 | HOXA6 | S_Shore | TSS1500 |
| GM03075 | chr7:27187943-27188770 | HOXA6 | S_Shore | TSS1500 |
| GM01376 | chr7:27188364-27188770 | HOXA6 | N_Shore | TSS1500 |

| | | | | |
|---|---|---|---|---|
| GM02149 | chr7:27225058-27225078 | HOXA11AS;HOXA11 | Island | Body;TSS1500 |
| GM02706 | chr7:27225058-27225123 | HOXA11AS;HOXA11 | Island | Body;TSS1500 |
| GM00935 | chr7:27225070-27225222 | HOXA11AS;HOXA11 | Island | Body;TSS1500 |
| GM02175 | chr7:27225078-27225222 | HOXA11AS;HOXA11 | Island | Body;TSS1500 |
| WG1283 | chr7:27225127-27225222 | HOXA11AS;HOXA11 | Island | Body;TSS1500 |
| WG1513 | chr7:27225299-27225342 | HOXA11AS;HOXA11 | Island | Body;TSS1500 |
| GM00022 | chr7:27225448-27225485 | HOXA11AS;HOXA11 | Island | Body;TSS1500 |
| GM00940 | chr7:27225448-27225543 | HOXA11AS;HOXA11 | Island | Body;TSS1500 |
| WG1721 | chr7:27225523-27225543 | HOXA11AS;HOXA11 | Island | Body;TSS1500 |
| GM02173 | chr7:27281315-27281346 | EVX1 | N_Shore | TSS1500 |
| GM02173 | chr7:27281687-27281964 | EVX1 | N_Shore | TSS1500 |
| WG1513 | chr7:2728776-2728912 | AMZ1 | S_Shore | 5'UTR |
| GM00520 | chr7:2738875-2738961 | AMZ1 | N_Shore | 5'UTR |
| WG2120 | chr7:2800701-2801248 | GNA12 | N_Shore | Body |
| WG3465 | chr7:2802374-2802560 | GNA12 | N_Shore | Body |
| GM02641 | chr7:2802374-2802976 | GNA12 | N_Shore | Body |
| GM00022 | chr7:2842593-2842746 | GNA12 | | Body |
| GM02317 | chr7:28452066-28452289 | CREB5;CREB5 | S_Shelf | 5'UTR;TSS200 |
| WG2843 | chr7:2847517-2847575 | GNA12 | | Body |
| WG3465 | chr7:28998577-28998860 | TRIL | S_Shore | TSS1500 |
| GM01376 | chr7:29186629-29187019 | CPVL;CPVL | | TSS1500;TSS1500 |
| GM02456 | chr7:29519409-29519550 | CHN2;CHN2 | | TSS200;Body |
| WG1283 | chr7:29602763-29603253 | PRR15 | N_Shore | TSS1500 |
| GM00940 | chr7:30951381-30951425 | AQP1 | | TSS200 |
| WG1283 | chr7:30951709-30951801 | AQP1 | | 1stExon |
| GM00042 | chr7:31726654-31726912 | C7orf16;C7orf16 | | TSS200;TSS200 |
| WG2415 | chr7:31726692-31726912 | C7orf16;C7orf16 | | TSS200;TSS200 |
| WG1978 | chr7:35294949-35295198 | TBX20;TBX20 | N_Shore | TSS1500;TSS1500 |
| GM00520 | chr7:36700744-36700939 | AOAH | | Body |
| GM02149 | chr7:37959686-37959966 | EPDR1 | N_Shore | TSS1500 |
| WG2275 | chr7:4065931-4066127 | SDK1 | | Body |
| GM00520 | chr7:41739678-41739801 | LOC285954;LOC285954;INHBA | | Body;Body;Body |
| GM02706 | chr7:42267547-42267747 | GLI3 | Island | 5'UTR |
| WG2415 | chr7:42267547-42268109 | GLI3 | Island | 5'UTR |
| GM00022 | chr7:42267719-42268072 | GLI3 | Island | 5'UTR |
| GM00940 | chr7:42267719-42268109 | GLI3 | Island | 5'UTR |
| WG1722 | chr7:4266872-4267364 | SDK1;SDK1 | | Body;Body |
| WG1658 | chr7:43288635-43289504 | HECW1 | Island | Body |
| GM00290 | chr7:44801378-44801589 | ZMIZ2;ZMIZ2 | | Body;Body |
| GM00935 | chr7:4729417-4729920 | FOXK1 | | Body |
| GM09256 | chr7:4762742-4763182 | FOXK1 | N_Shore | Body |
| WG2276 | chr7:4778839-4779342 | FOXK1 | | Body |
| WG1978 | chr7:4885521-4885643 | RADIL | | Body |
| GM01392 | chr7:4901449-4901750 | PAPOLB;RADIL;PAPOLB | Island | 5'UTR;Body;1stExon |
| GM02175 | chr7:4901449-4901798 | PAPOLB;RADIL;PAPOLB | Island | 5'UTR;Body;1stExon |
| GM00439 | chr7:50518350-50518573 | FIGNL1;FIGNL1 | Island | TSS1500;TSS1500 |
| WG2194 | chr7:5336543-5336811 | SLC29A4;SLC29A4 | Island | Body;Body |

| | | | | |
|---|---|---|---|---|
| GM01095 | chr7:537740-537828 | PDGFA;PDGFA | S_Shelf | 3'UTR;3'UTR |
| GM02555 | chr7:5518392-5518918 | FBXL18 | N_Shelf | 3'UTR |
| GM01393 | chr7:553412-553693 | PDGFA;PDGFA | N_Shelf | Body;Body |
| GM09256 | chr7:56515752-56516076 | LOC650226 | Island | Body |
| WG1513 | chr7:630473-630823 | PRKAR1B;PRKAR1B;PRKAR1E | N_Shore | Body;Body;Body;Body;Body;Body |
| GM01376 | chr7:630473-630914 | PRKAR1B;PRKAR1B;PRKAR1E | N_Shore | Body;Body;Body;Body;Body;Body |
| GM01393 | chr7:644219-644417 | PRKAR1B;PRKAR1B;PRKAR1E | Island | Body;Body;Body;Body;Body;Body |
| WG3465 | chr7:70096029-70096378 | AUTS2;AUTS2 | N_Shore | Body;Body |
| GM02173 | chr7:73894573-73895278 | GTF2IRD1;GTF2IRD1 | N_Shore | 5'UTR;5'UTR |
| WG1514 | chr7:766100-766119 | PRKAR1B;PRKAR1B;PRKAR1E | Island | 5'UTR;5'UTR;5'UTR;TSS1500 |
| GM00520 | chr7:8476098-8476450 | NXPH1 | S_Shore | Body |
| GM02706 | chr7:89873797-89873891 | C7orf63;C7orf63 | N_Shore | TSS1500;TSS1500 |
| WG2121 | chr7:906726-906799 | UNC84A;UNC84A | S_Shore | Body;Body |
| GM02456 | chr7:907797-908060 | UNC84A;UNC84A | S_Shore | Body;Body |
| WG1513 | chr7:917662-917805 | C7orf20 | Island | Body |
| WG2415 | chr7:944607-944776 | ADAP1 | S_Shore | Body |
| WG1513 | chr7:94953956-94954144 | PON1 | Island | TSS200 |
| GM00439 | chr7:95025855-95025924 | PON3 | Island | TSS200 |
| WG1722 | chr7:95025955-95026073 | PON3 | Island | TSS1500 |
| WG2415 | chr7:95026121-95026193 | PON3 | Island | TSS1500 |
| GM00042 | chr7:96631580-96632460 | DLX6AS | Island | Body |
| WG1310 | chr7:96650096-96650192 | DLX5 | N_Shore | Body |
| WG2193 | chr7:96650096-96650407 | DLX5 | N_Shore | Body |
| GM02175 | chr7:96651537-96651915 | DLX5 | Island | Body |
| GM00290 | chr7:96652115-96652153 | DLX5 | Island | Body |
| GM00042 | chr7:96655044-96655889 | DLX5 | S_Shore | TSS1500 |
| GM09256 | chr7:96655501-96655798 | DLX5 | S_Shore | TSS1500 |
| GM00439 | chr7:96655501-96655981 | DLX5 | S_Shore | TSS1500 |
| WG1722 | chr8:104032819-104032876 | ATP6V1C1 | N_Shore | TSS1500 |
| WG1977 | chr8:10586959-10587055 | SOX7 | N_Shore | Body |
| GM00935 | chr8:108510286-108510343 | ANGPT1 | | TSS200 |
| GM00439 | chr8:10907718-10907788 | XKR6 | | Body |
| GM02641 | chr8:10982332-10982836 | XKR6 | | Body |
| GM01096 | chr8:110693195-110693289 | GOLSYN;GOLSYN;GOLSYN;GOLSYN;GO | | 5'UTR;5'UTR;1stExon;1stExon;5'UTR;5 |
| WG1722 | chr8:11141365-11141424 | MTMR9 | N_Shore | TSS1500 |
| GM09256 | chr8:11141415-11141680 | MTMR9 | N_Shore | TSS1500 |
| GM09255 | chr8:116660311-116661063 | TRPS1 | N_Shore | 5'UTR |
| WG1977 | chr8:11839764-11839844 | DEFB135 | | TSS200 |
| GM00935 | chr8:12237626-12237900 | FAM66A | | Body |
| GM00439 | chr8:123875036-123875223 | ZHX2 | | 5'UTR |
| GM02456 | chr8:135490694-135490821 | ZFAT;ZFAT | N_Shore | 3'UTR;3'UTR |
| GM00289 | chr8:141359539-141359786 | TRAPPC9;TRAPPC9 | Island | Body;Body |
| GM00289 | chr8:143324660-143324707 | TSNARE1 | | Body |
| GM09256 | chr8:143545390-143545574 | BAI1;BAI1 | N_Shore | 5'UTR;1stExon |
| GM02456 | chr8:143545478-143545789 | BAI1;BAI1 | Island | 5'UTR;1stExon |
| WG2415 | chr8:143545940-143546893 | BAI1 | Island | 1stExon |
| WG1977 | chr8:143763326-143763565 | PSCA | | Body |

| | | | | |
|---|---|---|---|---|
| WG1513 | chr8:143781644-143781985 | LY6K;LY6K;LY6K;LY6K;LY6K;L | Island | 1stExon;1stExon;5'UTR;1stExon;5'UTR |
| GM02456 | chr8:144344397-144344793 | ZFP41 | | 3'UTR |
| GM01392 | chr8:144513697-144513907 | MAFA | Island | TSS1500 |
| GM00042 | chr8:144513898-144513980 | MAFA | Island | TSS1500 |
| GM00941 | chr8:144659831-144660157 | NAPRT1 | Island | Body |
| GM02149 | chr8:1448750-1449108 | DLGAP2 | | TSS1500 |
| GM02456 | chr8:145019032-145019116 | PLEC1;PLEC1;PLEC1;PLEC1; | Island | Body;Body;Body;TSS1500;Body;TSS20 |
| WG1283 | chr8:145028093-145028170 | PLEC1;PLEC1;PLEC1 | Island | Body;Body;TSS200 |
| GM02706 | chr8:145048235-145048363 | PLEC1;PLEC1 | Island | Body;TSS1500 |
| GM00520 | chr8:145086243-145086427 | SPATC1;SPATC1 | | TSS1500;TSS1500 |
| GM00042 | chr8:145162529-145163136 | KIAA1875 | N_Shore | TSS200 |
| GM00439 | chr8:145163034-145163136 | KIAA1875 | N_Shore | Body |
| WG1978 | chr8:145654565-145654635 | VPS28;NFKBIL2;VPS28 | Island | TSS1500;Body;TSS1500 |
| GM01392 | chr8:145654565-145654854 | VPS28;NFKBIL2;VPS28 | Island | TSS1500;Body;TSS1500 |
| WG1978 | chr8:1460569-1460718 | DLGAP2 | | 5'UTR |
| GM02149 | chr8:1570715-1571105 | DLGAP2 | | Body |
| GM00520 | chr8:16859833-16860121 | FGF20 | S_Shore | TSS200 |
| WG1309 | chr8:17271067-17271105 | MTMR7 | Island | TSS200 |
| WG2275 | chr8:1765217-1765358 | MIR596 | S_Shore | TSS200 |
| GM00520 | chr8:1786207-1786322 | ARHGEF10 | | 5'UTR |
| WG1657 | chr8:1847999-1848143 | ARHGEF10 | | Body |
| GM02555 | chr8:1847999-1848407 | ARHGEF10 | | Body |
| WG1722 | chr8:1848040-1848407 | ARHGEF10 | | Body |
| WG3465 | chr8:18541497-18541988 | PSD3;PSD3 | | Body;Body |
| GM09256 | chr8:19539991-19540272 | CSGALNACT1;CSGALNACT1;CSGALNAC | | Body;5'UTR;1stExon |
| GM00520 | chr8:1954777-1955196 | KBTBD11 | N_Shelf | 3'UTR |
| GM00940 | chr8:2075209-2075820 | MYOM2 | N_Shore | Body |
| WG1722 | chr8:2075469-2075820 | MYOM2 | Island | Body |
| GM00520 | chr8:21961713-21961788 | FAM160B2 | | 3'UTR |
| WG1309 | chr8:22132874-22132959 | PIWIL2;PIWIL2;PIWIL2 | Island | 5'UTR;1stExon;TSS1500 |
| GM00022 | chr8:22132874-22132992 | PIWIL2;PIWIL2;PIWIL2 | Island | 5'UTR;1stExon;TSS1500 |
| GM00935 | chr8:22548400-22548828 | EGR3 | Island | Body |
| GM01393 | chr8:23162162-23162383 | LOXL2 | | Body |
| WG1310 | chr8:28560467-28560723 | EXTL3 | S_Shore | 5'UTR |
| GM02149 | chr8:37797632-37797731 | GOT1L1;GOT1L1 | | 5'UTR;1stExon |
| WG2194 | chr8:40010443-40010823 | C8orf4 | | TSS1500 |
| GM01096 | chr8:41583321-41583523 | ANK1;ANK1;ANK1;ANK1;ANK | Island | Body;Body;Body;Body;Body |
| GM02640 | chr8:41655673-41655894 | ANK1;ANK1;ANK1;ANK1;ANK | Island | TSS1500;TSS1500;TSS1500;TSS1500; |
| WG1978 | chr8:42356637-42356871 | SLC20A2 | | 5'UTR |
| GM00439 | chr8:57359956-57359992 | PENK;PENK | Island | TSS1500;TSS1500 |
| GM01392 | chr8:58192502-58192883 | C8orf71 | Island | Body |
| WG3465 | chr8:59058585-59058660 | FAM110B | Island | 5'UTR |
| GM00290 | chr8:61626185-61626646 | CHD7 | N_Shore | 5'UTR |
| GM01096 | chr8:6420242-6420726 | ANGPT2;ANGPT2;ANGPT2;M | N_Shelf | 1stExon;1stExon;1stExon;Body |
| GM00520 | chr8:6420242-6420770 | ANGPT2;ANGPT2;ANGPT2;M | N_Shelf | 1stExon;1stExon;1stExon;Body |
| GM00935 | chr8:6456182-6456465 | MCPH1 | | Body |
| GM00439 | chr8:67405405-67405742 | C8orf46 | | TSS200 |

| | | | | |
|---|---|---|---|---|
| GM09255 | chr8:72754469-72755162 | MSC | Island | 3'UTR |
| GM02704 | chr8:77585222-77585381 | LOC100192378 | | Body |
| GM00438 | chr8:77596069-77596382 | ZFHX4;LOC100192378 | S_Shore | 5'UTR;TSS1500 |
| WG2415 | chr8:95962371-95962463 | TP53INP1;TP53INP1 | Island | TSS1500;TSS1500 |
| GM02704 | chr8:9952636-9953022 | MSRA;MSRA;MSRA | | Body;TSS1500;Body |
| GM02706 | chr8:9952926-9953022 | MSRA;MSRA;MSRA | | Body;TSS200;Body |
| GM00016 | chr9:101568975-101569334 | GALNT12 | N_Shore | TSS1500 |
| GM02147 | chr9:120466493-120466837 | TLR4;TLR4;TLR4;TLR4 | | Body;Body;1stExon;5'UTR |
| GM03073 | chr9:130955135-130955436 | CIZ1;CIZ1;CIZ1;CIZ1;CIZ1 | N_Shore | TSS1500;5'UTR;TSS1500;5'UTR;TSS15 |
| GM00520 | chr9:136566991-136567339 | SARDH;SARDH | N_Shore | Body;Body |
| GM02555 | chr9:139237461-139237889 | GPSM1 | Island | Body |
| GM00042 | chr9:139715701-139716215 | C9orf86;C9orf86 | Island | Body;Body |
| GM00289 | chr9:140172089-140172631 | C9orf167 | N_Shore | TSS200 |
| GM09255 | chr9:140401382-140401455 | PNPLA7;PNPLA7 | Island | Body;Body |
| WG1722 | chr9:140401382-140401576 | PNPLA7;PNPLA7 | Island | Body;Body |
| GM02149 | chr9:92220388-92220771 | GADD45G | Island | Body |
| WG1513 | chr9:92220388-92221282 | GADD45G | Island | Body |
| GM01376 | chr9:92220617-92221133 | GADD45G | Island | Body |
| GM02706 | chr9:92220617-92221282 | GADD45G | Island | Body |
| GM02175 | chr9:975825-975997 | DMRT3 | N_Shore | TSS1500 |
| WG3465 | chr1:10056837-10056853 | RBP7 | N_Shore | TSS1500 |
| WG1722 | chr1:10056853-10057059 | RBP7 | N_Shore | TSS1500 |
| GM00289 | chr1:103574523-103574619 | COL11A1;COL11A1;COL11A1;COL11A1 | | TSS1500;TSS1500;TSS1500;TSS1500 |
| WG1977 | chr1:10490025-10490063 | APITD1;APITD1;APITD1 | N_Shore | TSS200;TSS1500;TSS200 |
| WG1513 | chr1:10732744-10732900 | CASZ1;CASZ1 | | Body;Body |
| WG2414 | chr1:10738569-10738769 | CASZ1;CASZ1 | | Body;Body |
| GM00042 | chr1:108023249-108023441 | NTNG1;NTNG1;NTNG1 | Island | Body;Body;Body |
| GM02704 | chr1:108023249-108023482 | NTNG1;NTNG1;NTNG1 | Island | Body;Body;Body |
| WG1978 | chr1:108734992-108735312 | SLC25A24;SLC25A24 | N_Shore | Body;Body |
| WG1721 | chr1:108734992-108735549 | SLC25A24;SLC25A24 | N_Shore | Body;Body |
| GM01376 | chr1:109941043-109941072 | SORT1 | S_Shore | TSS1500 |
| GM09256 | chr1:109941043-109941118 | SORT1 | S_Shore | TSS1500 |
| WG3465 | chr1:109941043-109941201 | SORT1 | S_Shore | TSS1500 |
| GM00439 | chr1:109941060-109941118 | SORT1 | S_Shore | TSS1500 |
| GM02704 | chr1:110166563-110166664 | AMPD2;AMPD2;AMPD2 | S_Shelf | Body;Body;TSS1500 |
| GM00016 | chr1:110230252-110230456 | GSTM1;GSTM1 | Island | TSS200;TSS200 |
| GM02173 | chr1:110230401-110230473 | GSTM1;GSTM1 | Island | TSS200;TSS200 |
| WG3466 | chr1:110254662-110254692 | GSTM5 | | TSS1500 |
| GM02173 | chr1:110254662-110254835 | GSTM5 | | TSS1500 |
| GM02704 | chr1:110254692-110254854 | GSTM5 | | TSS200 |
| GM00439 | chr1:110254828-110255096 | GSTM5 | | TSS200 |
| GM03075 | chr1:110254896-110255096 | GSTM5;GSTM5 | | 1stExon;5'UTR |
| GM00438 | chr1:1103225-1103312 | MIR200A;MIR429 | S_Shelf | TSS200;TSS1500 |
| GM00520 | chr1:1103225-1104366 | MIR200A;MIR429 | S_Shelf | TSS200;TSS1500 |
| GM00042 | chr1:1104257-1104399 | MIR429 | S_Shelf | TSS200 |
| WG2844 | chr1:110452372-110452616 | CSF1;CSF1;CSF1;CSF1 | N_Shore | TSS1500;TSS1500;TSS1500;TSS1500 |
| GM02555 | chr1:110610899-110611807 | ALX3 | Island | Body |

| | | | | |
|---|---|---|---|---|
| WG1977 | chr1:110610899-110611823 | ALX3 | Island | Body |
| GM01392 | chr1:110611218-110611712 | ALX3 | Island | Body |
| GM00520 | chr1:110612044-110612474 | ALX3 | Island | Body |
| WG2121 | chr1:1108820-1108924 | TTLL10 | N_Shore | TSS1500 |
| WG1977 | chr1:1109486-1109832 | TTLL10 | Island | 5'UTR |
| GM00520 | chr1:1109676-1110107 | TTLL10 | Island | 5'UTR |
| WG3465 | chr1:1109832-1110107 | TTLL10 | Island | 5'UTR |
| WG3466 | chr1:111218079-111218282 | KCNA3 | S_Shore | TSS1500 |
| WG1513 | chr1:111218079-111218287 | KCNA3 | S_Shore | TSS1500 |
| GM00042 | chr1:1114912-1114963 | TTLL10;TTLL10 | N_Shore | Body;TSS200 |
| GM02706 | chr1:112531263-112531488 | KCND3;KCND3 | N_Shore | 5'UTR;5'UTR |
| GM00290 | chr1:1141790-1141934 | TNFRSF18;TNFRSF18;TNFRS | Island | 1stExon;1stExon;1stExon |
| GM02175 | chr1:1142254-1142617 | TNFRSF18;TNFRSF18;TNFRS | S_Shore | TSS200;TSS200;TSS200 |
| GM00290 | chr1:1149091-1149211 | TNFRSF4 | Island | Body |
| WG1658 | chr1:115238457-115238559 | AMPD1 | | TSS1500 |
| GM00439 | chr1:115397374-115397449 | SYCP1 | N_Shore | TSS200 |
| GM02555 | chr1:115881044-115881132 | NGF | Island | TSS200 |
| GM01392 | chr1:115881130-115881214 | NGF | Island | TSS1500 |
| GM09255 | chr1:117213980-117214228 | MIR320B1 | S_Shelf | TSS1500 |
| WG3465 | chr1:11761296-11761416 | C1orf187 | | 5'UTR |
| GM00042 | chr1:117753591-117753616 | VTCN1 | | TSS200 |
| GM09255 | chr1:118727658-118727840 | SPAG17 | N_Shore | Body |
| WG1282 | chr1:119521928-119522386 | TBX15 | N_Shore | 5'UTR |
| WG3465 | chr1:119526060-119526437 | TBX15 | N_Shore | 5'UTR |
| WG1282 | chr1:119526060-119526783 | TBX15 | N_Shore | 5'UTR |
| GM00520 | chr1:119526060-119527156 | TBX15 | N_Shore | 5'UTR |
| WG1309 | chr1:119527008-119527156 | TBX15 | Island | 5'UTR |
| GM02316 | chr1:119527638-119528018 | TBX15 | S_Shore | 5'UTR |
| WG1309 | chr1:119531625-119532925 | TBX15 | N_Shore | 5'UTR |
| WG2120 | chr1:119531718-119532044 | TBX15 | N_Shore | 5'UTR |
| GM00520 | chr1:119531718-119532093 | TBX15 | N_Shore | 5'UTR |
| WG1722 | chr1:119531718-119532195 | TBX15 | N_Shore | 5'UTR |
| GM00289 | chr1:119531718-119532352 | TBX15 | N_Shore | 5'UTR |
| WG2415 | chr1:119531718-119532542 | TBX15 | N_Shore | 5'UTR |
| GM09255 | chr1:119531718-119532736 | TBX15 | N_Shore | 5'UTR |
| WG1721 | chr1:119532093-119532542 | TBX15;TBX15 | Island | 1stExon;5'UTR |
| GM02175 | chr1:119532093-119532736 | TBX15;TBX15 | Island | 1stExon;5'UTR |
| WG1282 | chr1:119532093-119532925 | TBX15;TBX15 | Island | 1stExon;5'UTR |
| GM00520 | chr1:119532189-119532736 | TBX15 | Island | TSS200 |
| WG3465 | chr1:119532320-119532542 | TBX15 | S_Shore | TSS200 |
| GM00290 | chr1:119532320-119532925 | TBX15 | S_Shore | TSS200 |
| GM00042 | chr1:119532352-119532925 | TBX15 | S_Shore | TSS200 |
| GM00289 | chr1:119532655-119532850 | TBX15 | S_Shore | TSS1500 |
| WG1721 | chr1:119532736-119532850 | TBX15 | S_Shore | TSS1500 |
| WG1722 | chr1:119532736-119532925 | TBX15 | S_Shore | TSS1500 |
| GM01393 | chr1:119532773-119532925 | TBX15 | S_Shore | TSS1500 |
| WG3465 | chr1:1217103-1217159 | SCNN1D;SCNN1D | | TSS1500;TSS1500 |

| | | | | |
|---|---|---|---|---|
| WG1977 | chr1:1217511-1217664 | SCNN1D;SCNN1D;SCNN1D | | 5'UTR;TSS200;1stExon |
| WG2276 | chr1:12203571-12204096 | TNFRSF8;TNFRSF8 | | 3'UTR;3'UTR |
| GM00289 | chr1:12538341-12538678 | VPS13D;VPS13D | | Body;Body |
| GM00042 | chr1:1266178-1266548 | TAS1R3 | N_Shore | TSS1500 |
| WG3466 | chr1:1266178-1266657 | TAS1R3 | N_Shore | TSS1500 |
| WG1513 | chr1:1266468-1266548 | TAS1R3 | N_Shore | TSS1500 |
| WG1722 | chr1:1266548-1266700 | TAS1R3 | N_Shore | TSS200 |
| GM02175 | chr1:1267193-1267252 | TAS1R3 | Island | Body |
| WG1513 | chr1:12703864-12704007 | AADACL4 | | TSS1500 |
| GM02555 | chr1:14027521-14027581 | PRDM2 | S_Shore | 5'UTR |
| WG1513 | chr1:145727247-145727720 | PDZK1 | | TSS1500 |
| GM00438 | chr1:146713916-146713996 | CHD1L | N_Shore | TSS1500 |
| WG3466 | chr1:147245168-147245494 | GJA5 | | 5'UTR |
| GM01392 | chr1:1490676-1490906 | SSU72 | S_Shore | Body |
| GM02706 | chr1:150947832-150947949 | LASS2;LASS2 | Island | TSS1500;TSS1500 |
| WG1514 | chr1:150947896-150948030 | LASS2;LASS2 | Island | TSS1500;TSS1500 |
| GM01392 | chr1:150947949-150948030 | LASS2;LASS2 | Island | TSS1500;TSS1500 |
| WG2120 | chr1:151344806-151345181 | SELENBP1 | | Body |
| WG1977 | chr1:151734928-151735194 | OAZ3;MRPL9 | N_Shore | TSS1500;Body |
| GM02456 | chr1:15175204-15175356 | KIAA1026;KIAA1026 | | Body;Body |
| GM02175 | chr1:151811620-151812710 | LOC100132111;C2CD4D | Island | Body;5'UTR |
| WG1978 | chr1:151811923-151812435 | LOC100132111;C2CD4D | N_Shore | Body;5'UTR |
| WG1310 | chr1:151812421-151812524 | LOC100132111;C2CD4D | Island | Body;5'UTR |
| GM09255 | chr1:151812421-151812710 | LOC100132111;C2CD4D | Island | Body;5'UTR |
| WG3466 | chr1:151825974-151826205 | THEM5 | | 1stExon |
| GM02173 | chr1:152196106-152196674 | HRNR | | 5'UTR |
| GM02173 | chr1:152298884-152299078 | FLG | | TSS1500 |
| GM00940 | chr1:152333423-152333922 | FLG2 | | TSS1500 |
| WG1513 | chr1:152538380-152538857 | LCE3E | | 3'UTR |
| WG2120 | chr1:15256054-15256127 | KIAA1026;KIAA1026;KIAA1026;KIAA1026 | | Body;TSS1500;Body;Body |
| WG1722 | chr1:15256054-15256136 | KIAA1026;KIAA1026;KIAA1026;KIAA1026 | | Body;TSS1500;Body;Body |
| WG3466 | chr1:15256054-15256222 | KIAA1026;KIAA1026;KIAA1026;KIAA1026 | | Body;TSS1500;Body;Body |
| GM01393 | chr1:15256123-15256136 | KIAA1026;KIAA1026;KIAA1026;KIAA1026 | | Body;TSS200;Body;Body |
| GM00520 | chr1:15256123-15256481 | KIAA1026;KIAA1026;KIAA1026;KIAA1026 | | Body;TSS200;Body;Body |
| GM00289 | chr1:15256127-15256222 | KIAA1026;KIAA1026;KIAA1026;KIAA1026 | | Body;TSS200;Body;Body |
| GM00042 | chr1:152595842-152595982 | LCE3A | | TSS1500 |
| GM09256 | chr1:152730027-152731117 | KPRP | N_Shelf | TSS1500 |
| GM00438 | chr1:152733118-152733223 | KPRP | Island | Body |
| WG1513 | chr1:152733140-152733337 | KPRP | Island | Body |
| WG3465 | chr1:152747472-152747562 | LCE1F | | TSS1500 |
| WG1513 | chr1:152815068-152815459 | LCE6A | | TSS1500 |
| GM02149 | chr1:152942401-152943306 | SPRR4 | | TSS1500 |
| WG1513 | chr1:153030514-153030751 | SPRR2A | | TSS1500 |
| WG3465 | chr1:153234268-153234809 | LOR | | Body |
| GM09256 | chr1:153514114-153514376 | S100A5;S100A5 | | 1stExon;5'UTR |
| WG1513 | chr1:153514114-153514442 | S100A5;S100A5 | | 1stExon;5'UTR |
| GM00289 | chr1:153514376-153514482 | S100A5 | | TSS200 |

| | | | | |
|---|---|---|---|---|
| WG1514 | chr1:153581494-153581750 | S100A16 | | 5'UTR |
| WG1977 | chr1:153581494-153581793 | S100A16 | | 5'UTR |
| WG3465 | chr1:153589781-153590243 | S100A14 | | TSS1500 |
| WG3466 | chr1:154127629-154128041 | NUP210L;NUP210L | | TSS200;TSS200 |
| WG2415 | chr1:154127952-154128041 | TPM3;TPM3;NUP210L;TPM3;TPM3;NUP2 | | 3'UTR;3'UTR;TSS1500;3'UTR;3'UTR;TS |
| GM02173 | chr1:154301468-154301964 | ATP8B2;ATP8B2 | Island | Body;Body |
| WG3465 | chr1:154929317-154929734 | PBXIP1 | | TSS1500 |
| GM00290 | chr1:154929734-154929791 | PBXIP1;PYGO2 | | TSS1500;3'UTR |
| GM00439 | chr1:1549699-1549818 | MIB2;MIB2;MIB2;MIB2;MIB2 | N_Shore | TSS1500;TSS1500;TSS1500;TSS1500; |
| WG1722 | chr1:155290813-155291224 | RUSC1;RUSC1;C1orf104 | Island | 5'UTR;5'UTR;Body |
| GM09256 | chr1:15541182-15541280 | TMEM51;TMEM51;TMEM51;T | Island | 5'UTR;5'UTR;5'UTR;5'UTR |
| GM02175 | chr1:15573050-15573525 | FHAD1 | N_Shore | TSS1500 |
| WG1977 | chr1:155931629-155931858 | ARHGEF2;ARHGEF2;ARHGEF2 | | Body;Body;Body |
| WG2120 | chr1:156215418-156215655 | PAQR6;PAQR6 | Island | Body;Body |
| GM02317 | chr1:156260918-156261207 | TMEM79;TMEM79 | N_Shore | Body;Body |
| WG1984 | chr1:156261200-156261298 | TMEM79;TMEM79 | Island | Body;Body |
| GM01392 | chr1:156261200-156261351 | TMEM79;TMEM79 | Island | Body;Body |
| WG1722 | chr1:156261200-156261841 | TMEM79;TMEM79 | Island | Body;Body |
| GM09255 | chr1:156611087-156611425 | BCAN;BCAN | N_Shore | TSS1500;TSS1500 |
| GM00439 | chr1:156786010-156786653 | SH2D2A;NTRK1;SH2D2A;SH2 | S_Shore | Body;Body;TSS200;Body;Body;Body |
| WG3466 | chr1:156786601-156786653 | SH2D2A;NTRK1;SH2D2A;SH2 | S_Shelf | 5'UTR;Body;1stExon;TSS1500;5'UTR;5 |
| GM01096 | chr1:156818749-156818961 | NTRK1;INSRR | Island | Body;Body |
| WG1513 | chr1:157670328-157670869 | FCRL3 | | 5'UTR |
| WG1513 | chr1:158390649-158390842 | OR10K2 | | 1stExon |
| GM00438 | chr1:158670124-158670617 | OR6K2 | | 1stExon |
| WG1513 | chr1:158723939-158724362 | OR6K6 | | TSS1500 |
| WG1513 | chr1:159683373-159683855 | CRP | | Body |
| WG1513 | chr1:159684337-159684486 | CRP;CRP | | 1stExon;5'UTR |
| GM00438 | chr1:159684431-159684684 | CRP | | TSS200 |
| WG1978 | chr1:159796617-159796658 | SLAMF8;SLAMF8 | | 5'UTR;1stExon |
| GM09255 | chr1:159870077-159870134 | CCDC19 | Island | TSS200 |
| GM02173 | chr1:160053931-160054219 | KCNJ9 | Island | Body |
| GM00438 | chr1:160616604-160617057 | SLAMF1 | | Body |
| GM01376 | chr1:16084445-16084758 | FBLIM1 | N_Shore | TSS1500 |
| GM00438 | chr1:16084493-16084830 | FBLIM1 | N_Shore | TSS1500 |
| GM00520 | chr1:161008297-161008644 | TSTD1;TSTD1;TSTD1 | N_Shore | Body;Body;Body |
| WG3465 | chr1:161008297-161008977 | TSTD1;TSTD1;TSTD1 | N_Shore | Body;Body;Body |
| WG2415 | chr1:161008644-161008811 | TSTD1;TSTD1;TSTD1 | Island | Body;Body;Body |
| WG2414 | chr1:161008705-161008826 | TSTD1;TSTD1;TSTD1;TSTD1;T | Island | 5'UTR;1stExon;5'UTR;1stExon;1stExon |
| GM03075 | chr1:161008705-161008977 | TSTD1;TSTD1;TSTD1;TSTD1;T | Island | 5'UTR;1stExon;5'UTR;1stExon;1stExon |
| GM02555 | chr1:161049443-161049579 | PVRL4 | Island | Body |
| GM00940 | chr1:161067927-161068039 | KLHDC9;KLHDC9;KLHDC9 | N_Shore | TSS1500;TSS1500;TSS1500 |
| GM01393 | chr1:161128567-161128720 | USP21;UFC1;USP21 | N_Shore | TSS1500;3'UTR;TSS1500 |
| GM02456 | chr1:16133715-16134184 | UQCRHL;UQCRHL | | 3'UTR;1stExon |
| GM02706 | chr1:161337603-161337703 | C1orf192 | | Body |
| WG1722 | chr1:161492666-161493011 | HSPA6 | N_Shore | TSS1500 |
| GM00016 | chr1:16163775-16164109 | FLJ37453 | Island | Body |

| | | | | |
|---|---|---|---|---|
| WG1977 | chr1:16163967-16164122 | FLJ37453 | Island | Body |
| WG2276 | chr1:162337134-162337375 | NOS1AP;NOS1AP;NOS1AP | Island | Body;Body;Body |
| WG1657 | chr1:162337248-162337375 | NOS1AP;NOS1AP;NOS1AP | Island | Body;Body;Body |
| WG2121 | chr1:162467180-162467363 | UHMK1 | N_Shore | TSS1500 |
| WG2415 | chr1:1630992-1631100 | MMP23A;CDK11B;CDK11B;C | N_Shore | TSS1500;Body;Body;Body;Body;Body; |
| GM00940 | chr1:163172914-163173262 | RGS5 | | TSS200 |
| WG1978 | chr1:16346120-16346317 | HSPB7 | | TSS1500 |
| WG1658 | chr1:16346299-16346434 | HSPB7 | | TSS1500 |
| GM00520 | chr1:16370198-16370243 | CLCNKB | | TSS200 |
| GM00520 | chr1:164545699-164546143 | PBX1 | Island | Body |
| GM00290 | chr1:165204410-165204512 | LMX1A | N_Shore | Body |
| GM01392 | chr1:165321741-165322199 | LMX1A | Island | Body |
| WG2276 | chr1:165322099-165322784 | LMX1A | Island | Body |
| WG1513 | chr1:165323232-165323768 | LMX1A | N_Shore | Body |
| GM02149 | chr1:165323232-165323896 | LMX1A | N_Shore | Body |
| GM09256 | chr1:165323692-165323896 | LMX1A | Island | Body |
| WG3465 | chr1:165324100-165324810 | LMX1A | N_Shore | Body |
| GM02706 | chr1:165324100-165324990 | LMX1A | N_Shore | Body |
| GM00520 | chr1:165324411-165324810 | LMX1A | Island | Body |
| GM00042 | chr1:165324411-165324990 | LMX1A | Island | Body |
| WG1283 | chr1:165324679-165324990 | LMX1A | Island | Body |
| GM01650 | chr1:16533891-16534157 | ARHGEF19 | Island | Body |
| WG1722 | chr1:16534283-16534579 | ARHGEF19 | Island | Body |
| GM02175 | chr1:166958401-166958439 | MAEL | Island | TSS200 |
| WG1984 | chr1:167789455-167789646 | ADCY10;ADCY10 | Island | Body;Body |
| GM01375 | chr1:170635960-170636363 | PRRX1;PRRX1 | S_Shore | Body;Body |
| WG2414 | chr1:170638807-170639289 | PRRX1;PRRX1 | | Body;Body |
| WG1977 | chr1:17091946-17092279 | MST1P9 | | TSS1500 |
| WG3465 | chr1:172413492-172413542 | C1orf105;PIGC;PIGC | S_Shore | Body;TSS1500;TSS1500 |
| GM02173 | chr1:172413492-172413612 | C1orf105;PIGC;PIGC | S_Shore | Body;TSS1500;TSS1500 |
| WG1977 | chr1:173176362-173176523 | TNFSF4;TNFSF4 | | 5'UTR;1stExon |
| WG2844 | chr1:173176366-173176523 | TNFSF4;TNFSF4 | | 5'UTR;1stExon |
| WG1310 | chr1:173835989-173836307 | GAS5;SNORD78;SNORD76;S | N_Shore | Body;TSS1500;TSS200;TSS1500;TSS1 |
| WG1977 | chr1:175162515-175162553 | KIAA0040;KIAA0040;KIAA004 | S_Shore | TSS1500;TSS1500;TSS1500;TSS1500 |
| WG1513 | chr1:177001903-177002422 | ASTN1;ASTN1 | | Body;Body |
| WG1513 | chr1:17746491-17746840 | RCC2;RCC2 | Island | Body;Body |
| WG1513 | chr1:178693972-178694013 | RALGPS2 | Island | TSS1500 |
| WG2415 | chr1:17944356-17944660 | ARHGEF10L;ARHGEF10L | | Body;Body |
| WG3465 | chr1:1795408-1795945 | GNB1 | | 5'UTR |
| GM01376 | chr1:179560700-179560754 | TDRD5 | N_Shore | TSS1500 |
| WG1513 | chr1:181767609-181767774 | CACNA1E | Island | Body |
| WG2844 | chr1:182026044-182026510 | ZNF648 | Island | Body |
| WG2415 | chr1:182921851-182922263 | C1orf14 | Island | Body |
| GM00935 | chr1:182921851-182922427 | C1orf14 | Island | Body |
| GM00520 | chr1:18806854-18807388 | KLHDC7A | N_Shelf | TSS1500 |
| GM00438 | chr1:190068053-190068180 | FAM5C | | Body |
| GM02175 | chr1:1905080-1905618 | KIAA1751 | | Body |

| | | | | |
|---|---|---|---|---|
| GM00290 | chr1:19052179-19052235 | PAX7;PAX7;PAX7 | | Body;Body;Body |
| GM09256 | chr1:1957073-1957385 | GABRD | Island | Body |
| WG1978 | chr1:19600673-19600719 | AKR7L;AKR7L | Island | TSS200;TSS200 |
| WG2121 | chr1:19600673-19600730 | AKR7L;AKR7L | Island | TSS200;TSS200 |
| GM00935 | chr1:19600673-19600817 | AKR7L;AKR7L | Island | TSS200;TSS200 |
| GM02706 | chr1:19600702-19600730 | AKR7L;AKR7L | Island | TSS200;TSS200 |
| GM01096 | chr1:19600702-19600912 | AKR7L;AKR7L | Island | TSS200;TSS200 |
| WG1722 | chr1:19614851-19615382 | AKR7A3 | N_Shore | Body |
| GM01096 | chr1:19615080-19615358 | AKR7A3 | Island | 1stExon |
| WG2120 | chr1:19615303-19615382 | AKR7A3 | Island | TSS200 |
| GM02704 | chr1:19615358-19615382 | AKR7A3 | Island | TSS200 |
| GM09256 | chr1:197744782-197744884 | DENND1B;DENND1B | Island | TSS200;TSS200 |
| WG3465 | chr1:197744884-197745025 | DENND1B;DENND1B | S_Shore | TSS1500;TSS1500 |
| GM09255 | chr1:197890479-197890812 | LHX9;LHX9 | Island | Body;Body |
| WG1722 | chr1:200005219-200005635 | NR5A2;NR5A2 | S_Shore | Body;Body |
| WG3466 | chr1:2005260-2005564 | PRKCZ;PRKCZ;PRKCZ | Island | 1stExon;Body;5'UTR |
| GM00438 | chr1:2005260-2006032 | PRKCZ;PRKCZ;PRKCZ | Island | 1stExon;Body;5'UTR |
| GM00439 | chr1:2005489-2005564 | PRKCZ;PRKCZ | Island | Body;5'UTR |
| GM00940 | chr1:201252948-201253259 | PKP1;PKP1 | Island | 1stExon;1stExon |
| GM00520 | chr1:201368843-201369031 | LAD1 | Island | TSS200 |
| GM09255 | chr1:201368914-201369031 | LAD1 | Island | TSS1500 |
| WG1721 | chr1:201368946-201369031 | LAD1 | Island | TSS1500 |
| WG1978 | chr1:20142497-20142706 | RNF186 | | TSS1500 |
| WG1513 | chr1:201476606-201476775 | CSRP1;CSRP1 | S_Shore | TSS1500;TSS1500 |
| WG2276 | chr1:201708500-201708558 | NAV1;NAV1 | N_Shore | TSS1500;Body |
| GM00520 | chr1:201708836-201709135 | NAV1;NAV1 | Island | TSS200;Body |
| WG2276 | chr1:201708836-201709336 | NAV1;NAV1 | Island | TSS200;Body |
| GM00289 | chr1:201709135-201709675 | NAV1;NAV1;NAV1 | Island | 1stExon;Body;5'UTR |
| GM09256 | chr1:201709336-201709675 | NAV1;NAV1 | Island | Body;Body |
| WG3465 | chr1:202091175-202091880 | GPR37L1 | | TSS1500 |
| WG2844 | chr1:202091865-202091880 | GPR37L1 | | TSS200 |
| GM02706 | chr1:202091944-202092005 | GPR37L1 | | TSS200 |
| WG2844 | chr1:202129854-202129894 | PTPN7;PTPN7 | | TSS1500;TSS200 |
| WG3465 | chr1:20250186-20250308 | PLA2G2E | | TSS200 |
| GM09255 | chr1:203044677-203045229 | PPFIA4 | N_Shore | Body |
| GM01392 | chr1:203829775-203830200 | SNRPE | N_Shore | TSS1500 |
| WG3466 | chr1:204183060-204183116 | GOLT1A;GOLT1A | | 5'UTR;1stExon |
| WG1977 | chr1:2046702-2046864 | PRKCZ;PRKCZ;PRKCZ | Island | 5'UTR;Body;5'UTR |
| WG3466 | chr1:2046702-2046928 | PRKCZ;PRKCZ;PRKCZ | Island | 5'UTR;Body;5'UTR |
| WG2121 | chr1:2046783-2046928 | PRKCZ;PRKCZ;PRKCZ | Island | 5'UTR;Body;5'UTR |
| GM00935 | chr1:204966197-204966235 | NFASC;NFASC;NFASC;NFASC | | Body;Body;Body;Body |
| GM00935 | chr1:205497713-205498551 | CDK18;CDK18;CDK18 | N_Shore | Body;Body;Body |
| WG2415 | chr1:205602864-205603068 | ELK4;ELK4 | S_Shore | TSS1500;TSS1500 |
| WG1513 | chr1:205818668-205819609 | PM20D1 | N_Shore | Body |
| GM00031 | chr1:205818956-205819179 | PM20D1 | Island | Body |
| GM01375 | chr1:205818956-205819406 | PM20D1 | Island | Body |
| WG1977 | chr1:205818956-205819463 | PM20D1 | Island | Body |

| | | | | |
|---|---|---|---|---|
| WG2843 | chr1:205818956-205819492 | PM20D1 | Island | Body |
| GM02149 | chr1:205818956-205819609 | PM20D1 | Island | Body |
| GM00940 | chr1:205819088-205819251 | PM20D1 | Island | 1stExon |
| GM02173 | chr1:205819251-205819463 | PM20D1;PM20D1 | S_Shore | 5'UTR;1stExon |
| WG3466 | chr1:20617283-20617456 | VWA5B1 | | TSS200 |
| GM02149 | chr1:20617329-20617456 | VWA5B1 | | TSS200 |
| GM00439 | chr1:206729034-206729099 | RASSF5;RASSF5;RASSF5 | N_Shore | Body;TSS1500;Body |
| WG2415 | chr1:206786170-206786180 | LGTN | S_Shore | TSS1500 |
| GM00289 | chr1:207224549-207225128 | YOD1 | N_Shore | TSS200 |
| WG1978 | chr1:207224567-207224982 | YOD1 | N_Shore | TSS200 |
| GM00520 | chr1:207669116-207669324 | CR1;CR1 | N_Shore | TSS1500;TSS1500 |
| WG3466 | chr1:207992403-207992548 | LOC148696 | | Body |
| WG2276 | chr1:208083913-208084071 | CD34;CD34 | N_Shore | Body;Body |
| GM00290 | chr1:20880461-20880663 | FAM43B;FAM43B | Island | 1stExon;3'UTR |
| GM01096 | chr1:209759145-209759344 | CAMK1G | | 5'UTR |
| WG2415 | chr1:209799191-209799330 | LAMB3;LAMB3;LAMB3 | Island | Body;Body;Body |
| GM00439 | chr1:209799248-209799353 | LAMB3;LAMB3;LAMB3 | Island | Body;Body;Body |
| WG2276 | chr1:209877941-209878178 | HSD11B1;HSD11B1 | | 5'UTR;TSS1500 |
| WG1977 | chr1:20988590-20988763 | DDOST | S_Shore | TSS1500 |
| WG3466 | chr1:210407556-210407677 | C1orf133;SERTAD4 | Island | TSS200;5'UTR |
| GM00438 | chr1:210407677-210407896 | C1orf133;SERTAD4 | S_Shore | TSS1500;5'UTR |
| GM02456 | chr1:210407677-210407973 | C1orf133;SERTAD4 | S_Shore | TSS1500;5'UTR |
| WG3465 | chr1:210407755-210407973 | C1orf133;SERTAD4 | S_Shore | TSS1500;5'UTR |
| WG1513 | chr1:2105154-2105392 | PRKCZ;PRKCZ;PRKCZ | | Body;Body;Body |
| WG1513 | chr1:210857111-210857246 | KCNH1;KCNH1 | Island | Body;Body |
| GM01392 | chr1:211499620-211499648 | TRAF5;TRAF5 | N_Shore | TSS1500;TSS1500 |
| WG3466 | chr1:2120985-2121724 | C1orf86;LOC100128003 | N_Shore | Body;Body |
| WG1310 | chr1:2121039-2121521 | C1orf86;LOC100128003;C1o | Island | Body;Body;3'UTR |
| GM01096 | chr1:2138442-2139657 | C1orf86 | S_Shore | 5'UTR |
| GM01095 | chr1:2138953-2139235 | C1orf86 | S_Shore | 5'UTR |
| GM09255 | chr1:214170220-214170376 | PROX1 | | Body |
| WG3465 | chr1:214477063-214477280 | SMYD2 | | Body |
| WG1977 | chr1:215255110-215255572 | KCNK2;KCNK2;KCNK2 | N_Shore | TSS1500;TSS1500;Body |
| WG2121 | chr1:215255110-215255716 | KCNK2;KCNK2;KCNK2 | N_Shore | TSS1500;TSS1500;Body |
| WG2843 | chr1:21586831-21587174 | ECE1;ECE1;ECE1;ECE1 | | Body;Body;Body;Body |
| GM00439 | chr1:217263213-217263621 | ESRRG;ESRRG;ESRRG | Island | TSS1500;TSS1500;5'UTR |
| WG1977 | chr1:217263507-217263847 | ESRRG;ESRRG;ESRRG | S_Shore | TSS1500;TSS1500;5'UTR |
| WG3466 | chr1:217306590-217306763 | ESRRG | N_Shore | 5'UTR |
| GM02555 | chr1:217309568-217309970 | ESRRG | N_Shore | 5'UTR |
| GM02173 | chr1:217309794-217309970 | ESRRG | N_Shore | 5'UTR |
| GM02704 | chr1:217309906-217310174 | ESRRG | N_Shore | 5'UTR |
| GM00022 | chr1:217309906-217310572 | ESRRG | N_Shore | 5'UTR |
| GM01095 | chr1:217309970-217310383 | ESRRG | N_Shore | 5'UTR |
| WG2415 | chr1:217311993-217312478 | ESRRG | S_Shore | TSS1500 |
| GM02706 | chr1:2174725-2174895 | SKI | | Body |
| GM00439 | chr1:2188767-2189027 | SKI | | Body |
| GM02706 | chr1:2191299-2191402 | SKI | | Body |

| | | | | |
|---|---|---|---|---|
| WG1983 | chr1:2191299-2191558 | SKI | | Body |
| WG2275 | chr1:2191354-2191558 | SKI | | Body |
| GM09256 | chr1:2191354-2191658 | SKI | | Body |
| WG2844 | chr1:221051758-221052023 | HLX | N_Shore | TSS1500 |
| GM00935 | chr1:221051758-221052143 | HLX | N_Shore | TSS1500 |
| WG2121 | chr1:221053841-221054201 | HLX | N_Shore | Body |
| GM09256 | chr1:221053841-221054273 | HLX | N_Shore | Body |
| GM02175 | chr1:221053841-221054615 | HLX | N_Shore | Body |
| WG2843 | chr1:221053841-221055097 | HLX | N_Shore | Body |
| WG3466 | chr1:221053904-221054273 | HLX | N_Shore | Body |
| GM01393 | chr1:221053904-221054825 | HLX | N_Shore | Body |
| WG1722 | chr1:221053904-221055097 | HLX | N_Shore | Body |
| GM00290 | chr1:221053904-221055401 | HLX | N_Shore | Body |
| WG1283 | chr1:221053904-221055719 | HLX | N_Shore | Body |
| WG1977 | chr1:221054201-221055097 | HLX | N_Shore | Body |
| WG2194 | chr1:221054201-221055964 | HLX | N_Shore | Body |
| GM01095 | chr1:221054273-221054615 | HLX | N_Shore | Body |
| GM01376 | chr1:221054273-221055097 | HLX | N_Shore | Body |
| GM00022 | chr1:221054365-221055097 | HLX | N_Shore | Body |
| WG3466 | chr1:221054615-221055097 | HLX | Island | Body |
| WG2275 | chr1:221055401-221055719 | HLX | N_Shore | Body |
| GM00290 | chr1:221055665-221055790 | HLX | Island | Body |
| WG2415 | chr1:223566643-223566669 | C1orf65 | Island | TSS200 |
| WG1977 | chr1:223566643-223566709 | C1orf65 | Island | TSS200 |
| GM00042 | chr1:223566643-223566794 | C1orf65 | Island | TSS200 |
| GM00290 | chr1:223566643-223567002 | C1orf65 | Island | TSS200 |
| GM00520 | chr1:223566643-223567173 | C1orf65 | Island | TSS200 |
| GM00289 | chr1:223566646-223566761 | C1orf65 | Island | TSS200 |
| WG3465 | chr1:223566646-223566794 | C1orf65 | Island | TSS200 |
| WG1514 | chr1:223566646-223567173 | C1orf65 | Island | TSS200 |
| WG3466 | chr1:223566669-223567002 | C1orf65 | Island | TSS200 |
| GM00940 | chr1:223566669-223567173 | C1orf65 | Island | TSS200 |
| WG2415 | chr1:223566761-223567173 | C1orf65;C1orf65 | Island | 5'UTR;1stExon |
| WG1513 | chr1:223566794-223567173 | C1orf65;C1orf65 | Island | 5'UTR;1stExon |
| WG1310 | chr1:223936343-223936799 | CAPN2;CAPN2 | Island | Body;Body |
| GM02706 | chr1:223936343-223936812 | CAPN2;CAPN2 | Island | Body;Body |
| WG2844 | chr1:223936508-223936812 | CAPN2;CAPN2 | Island | Body;Body |
| GM09256 | chr1:223936508-223936838 | CAPN2;CAPN2 | Island | Body;Body |
| WG2415 | chr1:2257894-2258373 | MORN1 | N_Shore | Body |
| WG3465 | chr1:2259162-2259325 | MORN1 | Island | Body |
| GM02555 | chr1:226187852-226188006 | C1orf55 | S_Shore | TSS1500 |
| WG3465 | chr1:226825435-226825606 | ITPKB | | Body |
| GM09256 | chr1:227962723-227963171 | SNAP47 | | Body |
| WG2276 | chr1:227962793-227963296 | SNAP47 | | Body |
| WG2414 | chr1:227962873-227963171 | SNAP47 | | Body |
| WG1977 | chr1:227963164-227963296 | SNAP47 | | Body |
| WG1721 | chr1:228246263-228246937 | WNT3A | N_Shore | Body |

| | | | | |
|---|---|---|---|---|
| WG2415 | chr1:228291591-228291682 | C1orf35 | S_Shore | TSS1500 |
| GM00042 | chr1:228291591-228291705 | C1orf35 | S_Shore | TSS1500 |
| GM09255 | chr1:228291668-228291705 | C1orf35 | S_Shore | TSS1500 |
| GM03075 | chr1:228297864-228297975 | MRPL55;MRPL55;MRPL55;MI | S_Shore | TSS1500;TSS1500;TSS1500;TSS1500; |
| GM02706 | chr1:228337311-228337379 | GJC2 | | TSS1500 |
| GM02555 | chr1:228337379-228337626 | GJC2 | | TSS200 |
| GM09256 | chr1:228346014-228346347 | GJC2 | Island | Body |
| WG2415 | chr1:228394559-228394900 | OBSCN;OBSCN | | TSS1500;TSS1500 |
| WG1977 | chr1:228394559-228395145 | OBSCN;OBSCN | | TSS1500;TSS1500 |
| GM00439 | chr1:228395450-228395760 | OBSCN;OBSCN | | TSS1500;TSS1500 |
| GM00940 | chr1:228395660-228395760 | OBSCN;OBSCN | N_Shelf | TSS200;TSS200 |
| WG2844 | chr1:228395756-228395858 | OBSCN;OBSCN | N_Shelf | TSS200;TSS200 |
| GM00438 | chr1:22902941-22903406 | EPHA8;EPHA8 | Island | Body;Body |
| GM02456 | chr1:22962506-22962986 | C1QA | | TSS1500 |
| GM00940 | chr1:230513337-230513566 | PGBD5 | | 1stExon |
| GM00031 | chr1:230513522-230513809 | PGBD5 | | TSS200 |
| WG2276 | chr1:231761688-231761845 | TSNAX-DISC1;DISC1;DISC1;I | N_Shore | Body;TSS1500;TSS1500;TSS1500;TSS |
| GM09256 | chr1:2324205-2324224 | MORN1;RER1 | S_Shore | TSS1500;5'UTR |
| GM00289 | chr1:233497542-233497916 | KIAA1804 | N_Shore | Body |
| GM00520 | chr1:2337334-2337711 | PEX10;PEX10 | N_Shelf | Body;Body |
| GM01376 | chr1:233765199-233765350 | KCNK1 | | Body |
| GM02173 | chr1:234350834-234350988 | SLC35F3 | S_Shore | Body |
| WG1513 | chr1:2344979-2345152 | PEX10;PEX10 | N_Shore | TSS1500;TSS1500 |
| GM09255 | chr1:2345333-2345374 | PEX10;PEX10 | N_Shore | TSS1500;TSS1500 |
| GM00520 | chr1:2345368-2345376 | PEX10;PEX10 | N_Shore | TSS1500;TSS1500 |
| WG2415 | chr1:2345398-2345410 | PEX10;PEX10 | N_Shore | TSS1500;TSS1500 |
| WG3466 | chr1:2345398-2345475 | PEX10;PEX10 | N_Shore | TSS1500;TSS1500 |
| WG1283 | chr1:23521272-23521332 | HTR1D | | TSS200 |
| WG3465 | chr1:23521272-23521417 | HTR1D | | TSS200 |
| GM02173 | chr1:235814326-235814365 | GNG4;GNG4;GNG4 | Island | TSS1500;TSS1500;TSS1500 |
| GM00520 | chr1:236557473-236557682 | EDARADD;EDARADD | N_Shore | TSS1500;TSS1500 |
| WG2844 | chr1:236557473-236557758 | EDARADD;EDARADD | N_Shore | TSS1500;TSS1500 |
| GM03075 | chr1:236686564-236686618 | LGALS8;LGALS8;LGALS8;LGA | N_Shore | TSS200;TSS200;5'UTR;TSS1500 |
| WG2121 | chr1:236686598-236686671 | LGALS8;LGALS8;LGALS8;LGA | N_Shore | TSS200;TSS200;5'UTR;TSS1500 |
| GM00438 | chr1:236902424-236902765 | ACTN2 | | Body |
| GM02704 | chr1:236925843-236925965 | ACTN2 | | Body |
| GM01096 | chr1:236925877-236925965 | ACTN2 | | Body |
| GM00438 | chr1:237628962-237629007 | RYR2 | | Body |
| GM02149 | chr1:237946534-237947435 | RYR2 | N_Shore | Body |
| WG1513 | chr1:240071723-240071966 | CHRM3 | | Body |
| WG3466 | chr1:240776162-240776211 | GREM2 | | TSS1500 |
| WG1978 | chr1:240776162-240776316 | GREM2 | | TSS1500 |
| WG1722 | chr1:2411594-2411643 | PLCH2 | S_Shore | Body |
| GM02555 | chr1:241912510-241912768 | WDR64 | | Body |
| GM00289 | chr1:242161780-242161881 | MAP1LC3C | | Body |
| WG2121 | chr1:2425860-2426035 | PLCH2 | | Body |
| GM00438 | chr1:2428864-2429005 | PLCH2 | | Body |

| | | | | |
|---|---|---|---|---|
| GM00940 | chr1:2435236-2435827 | PLCH2 | Island | Body |
| GM02706 | chr1:243646235-243646691 | SDCCAG8 | N_Shore | Body |
| WG3466 | chr1:243646235-243646787 | SDCCAG8 | N_Shore | Body |
| GM03075 | chr1:243646318-243646402 | SDCCAG8 | N_Shore | Body |
| GM02555 | chr1:243646402-243646787 | SDCCAG8 | Island | Body |
| WG2276 | chr1:244213358-244213618 | ZNF238 | N_Shore | TSS1500 |
| WG3466 | chr1:24438607-24438941 | MYOM3;MYOM3 | | 1stExon;5'UTR |
| GM02706 | chr1:24438778-24439075 | MYOM3 | | TSS200 |
| GM00290 | chr1:24438778-24439287 | MYOM3 | | TSS200 |
| GM00289 | chr1:245753302-245753539 | KIF26B | N_Shore | Body |
| GM02175 | chr1:24646191-24646392 | GRHL3 | N_Shelf | Body |
| WG1513 | chr1:247578953-247579608 | NLRP3;NLRP3 | | TSS1500;TSS1500 |
| GM00940 | chr1:247611842-247611931 | NLRP3;NLRP3;NLRP3;NLRP3 | N_Shelf | 3'UTR;3'UTR;3'UTR;3'UTR;3'UTR |
| WG3466 | chr1:247694041-247694275 | LOC148824;OR2C3 | Island | Body;3'UTR |
| WG2414 | chr1:247694041-247694531 | LOC148824;OR2C3 | Island | Body;3'UTR |
| WG2121 | chr1:247694061-247694275 | LOC148824;OR2C3 | Island | Body;3'UTR |
| WG3465 | chr1:247694061-247694406 | LOC148824;OR2C3 | Island | Body;3'UTR |
| GM01392 | chr1:247694061-247694531 | LOC148824;OR2C3 | Island | Body;3'UTR |
| WG1722 | chr1:247694271-247694406 | LOC148824;OR2C3 | Island | TSS200;3'UTR |
| GM00042 | chr1:247694271-247694531 | LOC148824;OR2C3 | Island | TSS200;3'UTR |
| WG2844 | chr1:248100228-248101009 | OR2L13 | N_Shore | TSS1500 |
| GM02706 | chr1:248100345-248100614 | OR2L13 | Island | TSS200 |
| GM00940 | chr1:248100345-248101009 | OR2L13 | Island | TSS200 |
| WG1513 | chr1:248100407-248100585 | OR2L13 | Island | TSS200 |
| GM00438 | chr1:248903100-248903325 | LOC646627 | | 1stExon |
| GM00016 | chr1:25257931-25258236 | RUNX3;RUNX3 | Island | TSS1500;Body |
| GM02175 | chr1:25944514-25944802 | MAN1C1 | Island | 1stExon |
| WG2415 | chr1:25944712-25945042 | MAN1C1 | Island | 1stExon |
| WG1977 | chr1:26233332-26233404 | STMN1;STMN1;STMN1;STMN | Island | 1stExon;TSS1500;TSS1500;5'UTR;TSS |
| GM02706 | chr1:26233376-26233459 | STMN1;STMN1;STMN1;STMN | Island | TSS1500;TSS1500;TSS1500;TSS200 |
| WG2414 | chr1:26233565-26233623 | STMN1;STMN1;STMN1;STMN | Island | TSS1500;TSS1500;TSS1500;TSS200 |
| GM00439 | chr1:26394083-26394300 | TRIM63;TRIM63 | | 1stExon;5'UTR |
| GM02706 | chr1:26394201-26394300 | TRIM63 | | TSS200 |
| GM01392 | chr1:26503776-26503847 | CNKSR1;CNKSR1 | | TSS1500;TSS1500 |
| WG3465 | chr1:26503979-26504019 | CNKSR1;CNKSR1 | | TSS200;TSS200 |
| WG3465 | chr1:26797145-26797585 | DHDDS;DHDDS | N_Shore | 3'UTR;3'UTR |
| GM02706 | chr1:26797145-26797619 | DHDDS;DHDDS | N_Shore | 3'UTR;3'UTR |
| GM02149 | chr1:26797576-26797619 | DHDDS;HMGN2;DHDDS | N_Shore | 3'UTR;TSS1500;3'UTR |
| WG3465 | chr1:27287282-27287541 | C1orf172 | S_Shore | TSS1500 |
| WG2844 | chr1:27683351-27683501 | MAP3K6 | Island | Body |
| GM00520 | chr1:27687074-27687454 | MAP3K6 | Island | Body |
| GM02149 | chr1:27718221-27718277 | GPR3 | N_Shore | TSS1500 |
| WG2844 | chr1:27961796-27962037 | FGR | Island | TSS200 |
| GM09256 | chr1:29564615-29565110 | PTPRU;PTPRU;PTPRU | Island | Body;Body;Body |
| WG1984 | chr1:2984218-2984470 | FLJ42875;FLJ42875 | Island | Body;Body |
| WG1978 | chr1:2987645-2987961 | PRDM16;PRDM16 | Island | Body;Body |
| GM01392 | chr1:3028230-3028457 | PRDM16;PRDM16 | Island | Body;Body |

| | | | | |
|---|---|---|---|---|
| WG2415 | chr1:3028230-3028485 | PRDM16;PRDM16 | Island | Body;Body |
| WG2275 | chr1:3028338-3028485 | PRDM16;PRDM16 | Island | Body;Body |
| GM00042 | chr1:3038564-3039210 | PRDM16;PRDM16 | S_Shore | Body;Body |
| GM02175 | chr1:3038707-3039210 | PRDM16;PRDM16 | S_Shore | Body;Body |
| WG3466 | chr1:3055699-3056152 | PRDM16;PRDM16 | N_Shore | Body;Body |
| WG3466 | chr1:3056742-3057366 | PRDM16;PRDM16 | Island | Body;Body |
| WG2121 | chr1:3057893-3058199 | PRDM16;PRDM16 | N_Shore | Body;Body |
| WG1984 | chr1:3058822-3059137 | PRDM16;PRDM16 | N_Shore | Body;Body |
| WG2121 | chr1:3059267-3059626 | PRDM16;PRDM16 | Island | Body;Body |
| GM01095 | chr1:3059517-3059774 | PRDM16;PRDM16 | S_Shore | Body;Body |
| GM00438 | chr1:3078839-3079212 | PRDM16;PRDM16 | N_Shelf | Body;Body |
| GM02706 | chr1:3079886-3080336 | PRDM16;PRDM16 | N_Shore | Body;Body |
| WG2120 | chr1:3100259-3100497 | PRDM16;PRDM16 | N_Shelf | Body;Body |
| GM00042 | chr1:3105189-3105326 | PRDM16;PRDM16 | Island | Body;Body |
| GM00439 | chr1:3130485-3130550 | PRDM16;PRDM16 | | Body;Body |
| WG1984 | chr1:3142925-3143965 | PRDM16;PRDM16 | | Body;Body |
| GM01096 | chr1:3159568-3159645 | PRDM16;PRDM16 | S_Shore | Body;Body |
| GM00940 | chr1:3162104-3162173 | PRDM16;PRDM16 | N_Shore | Body;Body |
| GM00520 | chr1:3162173-3162454 | PRDM16;PRDM16 | N_Shore | Body;Body |
| GM01392 | chr1:3162173-3162676 | PRDM16;PRDM16 | N_Shore | Body;Body |
| WG3465 | chr1:3162404-3162676 | PRDM16;PRDM16 | N_Shore | Body;Body |
| WG1977 | chr1:3163710-3163985 | PRDM16;PRDM16 | N_Shore | Body;Body |
| WG3465 | chr1:31654355-31654792 | NKAIN1 | N_Shore | 3'UTR |
| GM09255 | chr1:31654355-31655009 | NKAIN1 | N_Shore | 3'UTR |
| WG2415 | chr1:31654453-31655009 | NKAIN1 | Island | 3'UTR |
| WG2275 | chr1:31988357-31988579 | LOC284551 | | Body |
| WG2121 | chr1:32281460-32281665 | SPOCD1 | | 5'UTR |
| WG1722 | chr1:3229075-3229579 | PRDM16;PRDM16 | | Body;Body |
| GM02706 | chr1:3229579-3230250 | PRDM16;PRDM16 | | Body;Body |
| GM09255 | chr1:3229579-3230357 | PRDM16;PRDM16 | | Body;Body |
| GM02175 | chr1:3230250-3230609 | PRDM16;PRDM16 | | Body;Body |
| WG2415 | chr1:3230424-3230647 | PRDM16;PRDM16 | | Body;Body |
| GM00439 | chr1:32665450-32665584 | CCDC28B | N_Shore | TSS1500 |
| WG3466 | chr1:3269252-3269478 | PRDM16;PRDM16 | | Body;Body |
| GM02173 | chr1:32708234-32708276 | MTMR9L | S_Shore | TSS1500 |
| GM02706 | chr1:32714146-32714308 | FAM167B | Island | Body |
| WG2415 | chr1:32739049-32739493 | LCK;LCK | N_Shore | 5'UTR;TSS1500 |
| WG3465 | chr1:32741152-32741518 | LCK;LCK | Island | Body;Body |
| GM00520 | chr1:32741454-32741543 | LCK;LCK | Island | Body;Body |
| GM01393 | chr1:3276413-3276753 | PRDM16;PRDM16 | S_Shore | Body;Body |
| WG3466 | chr1:3276413-3276786 | PRDM16;PRDM16 | S_Shore | Body;Body |
| WG1977 | chr1:3277525-3277623 | PRDM16;PRDM16 | S_Shelf | Body;Body |
| GM00935 | chr1:32802291-32802312 | MARCKSL1 | S_Shore | TSS1500 |
| WG3465 | chr1:3282449-3282870 | PRDM16;PRDM16 | | Body;Body |
| WG2121 | chr1:32826969-32827389 | LOC100128071;TSSK3 | Island | 3'UTR;TSS1500 |
| GM01096 | chr1:3289798-3289994 | PRDM16;PRDM16 | | Body;Body |
| GM00940 | chr1:3308286-3308568 | PRDM16;PRDM16 | N_Shore | Body;Body |

| | | | | |
|---|---|---|---|---|
| WG1977 | chr1:33169558-33169850 | SYNC;SYNC | S_Shore | TSS1500;TSS1500 |
| WG2275 | chr1:3320431-3320738 | PRDM16;PRDM16 | N_Shore | Body;Body |
| WG2415 | chr1:3320446-3320738 | PRDM16;PRDM16 | N_Shore | Body;Body |
| GM00935 | chr1:3332125-3332188 | PRDM16;PRDM16 | Island | Body;Body |
| WG1513 | chr1:3332125-3332264 | PRDM16;PRDM16 | Island | Body;Body |
| GM00290 | chr1:3332160-3332264 | PRDM16;PRDM16 | Island | Body;Body |
| WG3466 | chr1:3340086-3340187 | PRDM16;PRDM16 | N_Shore | Body;Body |
| GM02149 | chr1:3352257-3352830 | PRDM16;PRDM16 | S_Shore | 3'UTR;3'UTR |
| GM02175 | chr1:33648656-33648819 | TRIM62 | S_Shore | TSS1500 |
| WG1977 | chr1:3397061-3397125 | ARHGEF16 | S_Shore | Body |
| GM00935 | chr1:34329069-34329788 | HMGB4;CSMD2;HMGB4 | | 5'UTR;Body;TSS1500 |
| WG3466 | chr1:3459949-3460268 | MEGF6 | Island | Body |
| GM00520 | chr1:34642396-34642609 | C1orf94;C1orf94 | Island | 5'UTR;TSS200 |
| WG2415 | chr1:3473665-3474202 | MEGF6 | | Body |
| GM00042 | chr1:3473665-3474376 | MEGF6 | | Body |
| WG1513 | chr1:3473923-3474376 | MEGF6 | | Body |
| GM02173 | chr1:3477168-3477302 | MEGF6 | | Body |
| WG2844 | chr1:3514921-3515194 | MEGF6 | S_Shelf | Body |
| WG1722 | chr1:35220224-35220710 | GJB5 | | TSS1500 |
| GM02175 | chr1:3569386-3569899 | TP73 | Island | 5'UTR |
| WG1513 | chr1:3574507-3575147 | TP73 | | 5'UTR |
| GM01392 | chr1:3574704-3575147 | TP73 | | 5'UTR |
| GM09255 | chr1:36038514-36038708 | TFAP2E | N_Shore | TSS1500 |
| WG3466 | chr1:36038701-36038775 | TFAP2E | N_Shore | TSS1500 |
| WG3466 | chr1:36039655-36039885 | TFAP2E | Island | Body |
| WG2276 | chr1:36042986-36043014 | TFAP2E | Island | Body |
| GM09256 | chr1:3606777-3607116 | TP73;TP73;TP73;TP73 | N_Shore | TSS1500;TSS1500;Body;TSS1500 |
| GM00439 | chr1:3607097-3607222 | TP73;TP73;TP73;TP73 | Island | Body;TSS200;TSS200;TSS200 |
| WG1722 | chr1:36108206-36108233 | PSMB2 | S_Shore | TSS1500 |
| GM01393 | chr1:3634867-3635100 | TP73;TP73;TP73;TP73 | Island | Body;Body;Body;Body |
| WG1513 | chr1:3649771-3649971 | TP73;TP73;TP73;TP73 | S_Shore | 3'UTR;3'UTR;3'UTR;3'UTR |
| GM00042 | chr1:37939320-37939724 | ZC3H12A | N_Shore | TSS1500 |
| GM02555 | chr1:38022586-38022657 | DNALI1 | Island | 1stExon |
| GM02706 | chr1:38261052-38261192 | MANEAL;MANEAL;MANEAL | S_Shore | Body;Body;TSS200 |
| WG3465 | chr1:3827065-3827428 | LOC100133612 | S_Shelf | Body |
| WG1722 | chr1:38412519-38412744 | INPP5B | | 5'UTR |
| GM02706 | chr1:39546511-39546656 | MACF1 | N_Shore | TSS1500 |
| WG1722 | chr1:40024971-40025415 | LOC728448 | | Body |
| WG3466 | chr1:40105667-40105706 | HEYL | Island | TSS1500 |
| WG3466 | chr1:40769669-40769744 | COL9A2 | Island | Body |
| GM02706 | chr1:40783060-40783234 | COL9A2 | Island | TSS200 |
| GM09256 | chr1:41250062-41250117 | KCNQ4;KCNQ4 | Island | 1stExon;1stExon |
| WG1513 | chr1:41444812-41444854 | CTPS | N_Shore | TSS200 |
| GM09255 | chr1:42384002-42384390 | HIVEP3;HIVEP3 | Island | 5'UTR;5'UTR |
| WG3465 | chr1:43751363-43751376 | C1orf210;C1orf210 | | TSS200;TSS200 |
| GM01653 | chr1:43751363-43751469 | C1orf210;C1orf210 | | TSS200;TSS200 |
| GM00935 | chr1:43751497-43751563 | C1orf210;C1orf210 | | TSS1500;TSS1500 |

| | | | | |
|---|---|---|---|---|
| GM00042 | chr1:43770523-43770851 | TIE1 | Island | Body |
| WG1514 | chr1:43770557-43770851 | TIE1 | Island | Body |
| GM02149 | chr1:43920056-43920103 | HYI | S_Shore | TSS200 |
| WG1721 | chr1:43920217-43920264 | HYI | S_Shore | TSS1500 |
| WG2844 | chr1:44016014-44016049 | PTPRF;PTPRF | | Body;Body |
| GM00290 | chr1:44401857-44402353 | ARTN;ARTN;ARTN;ARTN;ARTN | Island | 1stExon;Body;Body;Body;Body |
| WG1513 | chr1:44584417-44584446 | KLF17 | | TSS200 |
| WG2844 | chr1:44703750-44704073 | ERI3 | | Body |
| WG1513 | chr1:4471552-4471975 | LOC284661 | | TSS1500 |
| GM09255 | chr1:45187551-45187810 | C1orf228 | N_Shelf | Body |
| GM00439 | chr1:45187551-45187946 | C1orf228 | N_Shelf | Body |
| GM00016 | chr1:45187630-45187946 | C1orf228 | N_Shelf | Body |
| WG2415 | chr1:45272255-45272633 | TCTEX1D4 | Island | Body |
| GM00940 | chr1:45272261-45272637 | TCTEX1D4 | Island | Body |
| GM02175 | chr1:46088336-46089045 | CCDC17 | Island | Body |
| GM01392 | chr1:46088541-46089045 | CCDC17 | Island | Body |
| GM09255 | chr1:46859671-46859774 | FAAH | N_Shore | TSS1500 |
| GM00290 | chr1:4736099-4736187 | AJAP1;AJAP1 | | Body;Body |
| GM02149 | chr1:4770676-4770935 | AJAP1;AJAP1 | N_Shore | Body;Body |
| GM00935 | chr1:47898239-47898435 | MGC12982 | N_Shore | Body |
| WG1722 | chr1:47898911-47899230 | MGC12982 | N_Shore | Body |
| WG2415 | chr1:47902269-47902310 | FOXD2;FOXD2 | N_Shore | 1stExon;5'UTR |
| GM00290 | chr1:47902269-47902371 | FOXD2;FOXD2 | N_Shore | 1stExon;5'UTR |
| WG2415 | chr1:4794912-4795114 | AJAP1;AJAP1 | | Body;Body |
| GM00940 | chr1:4832432-4832593 | AJAP1;AJAP1 | Island | Body;Body |
| GM09256 | chr1:4832432-4832865 | AJAP1;AJAP1 | Island | Body;Body |
| GM09256 | chr1:4841913-4841959 | AJAP1 | | 3'UTR |
| GM01096 | chr1:50573970-50574383 | ELAVL4;ELAVL4;ELAVL4;ELAVL4;ELAVL4 | | Body;TSS1500;Body;Body;TSS1500 |
| GM00042 | chr1:50574383-50574517 | ELAVL4;ELAVL4;ELAVL4;ELAVL4;ELAVL4 | | Body;TSS1500;Body;Body;TSS1500 |
| GM02706 | chr1:50574507-50574591 | ELAVL4;ELAVL4;ELAVL4;ELAVL4;ELAVL4 | | Body;Body;TSS200;Body;TSS200 |
| GM09255 | chr1:50887633-50888024 | DMRTA2 | Island | 5'UTR |
| GM02555 | chr1:51888289-51888775 | EPS15;EPS15 | | TSS1500;Body |
| WG1513 | chr1:51888289-51888961 | EPS15;EPS15 | | TSS1500;Body |
| WG2415 | chr1:51888574-51888961 | EPS15;EPS15 | | TSS1500;Body |
| WG1722 | chr1:52195423-52195686 | OSBPL9;OSBPL9;OSBPL9;OS | Island | TSS200;Body;5'UTR;Body;Body;TSS20 |
| WG3465 | chr1:53600741-53600889 | SLC1A7 | | Body |
| GM00935 | chr1:53661346-53661861 | CPT2 | N_Shore | TSS1500 |
| GM00438 | chr1:53704498-53704641 | MAGOH | Island | TSS1500 |
| GM02175 | chr1:53704558-53704978 | MAGOH | Island | TSS1500 |
| GM00438 | chr1:53904065-53904319 | FLJ40434 | N_Shore | Body |
| GM00520 | chr1:54199696-54200420 | GLIS1;GLIS1 | N_Shelf | 5'UTR;1stExon |
| WG1658 | chr1:54199825-54200115 | GLIS1;GLIS1 | N_Shelf | 5'UTR;1stExon |
| WG2415 | chr1:54199825-54200420 | GLIS1;GLIS1 | N_Shelf | 5'UTR;1stExon |
| WG3466 | chr1:54199825-54200587 | GLIS1;GLIS1 | N_Shelf | 5'UTR;1stExon |
| GM03075 | chr1:54200078-54200420 | GLIS1 | N_Shelf | TSS1500 |
| WG3465 | chr1:54200115-54200587 | GLIS1 | N_Shelf | TSS1500 |
| WG3465 | chr1:54856035-54856511 | SSBP3;SSBP3;SSBP3 | N_Shore | Body;Body;Body |

| | | | | |
|---|---|---|---|---|
| GM02706 | chr1:54856177-54856511 | SSBP3;SSBP3;SSBP3 | N_Shore | Body;Body;Body |
| GM00940 | chr1:55089182-55089372 | ACOT11;FAM151A;FAM151A | | Body;1stExon;5'UTR |
| WG1977 | chr1:55267046-55267285 | TTC22;TTC22 | Island | TSS200;TSS200 |
| GM09256 | chr1:55267125-55267266 | TTC22;TTC22 | S_Shore | TSS200;TSS200 |
| GM00935 | chr1:55267152-55267285 | TTC22;TTC22 | S_Shore | TSS1500;TSS1500 |
| WG2415 | chr1:55267152-55267293 | TTC22;TTC22 | S_Shore | TSS1500;TSS1500 |
| GM09256 | chr1:55505061-55505073 | PCSK9 | Island | TSS200 |
| WG2275 | chr1:57285278-57285466 | C1orf168;C1orf168 | | 1stExon;5'UTR |
| GM09255 | chr1:59612408-59612710 | LOC729467 | Island | Body |
| WG2276 | chr1:6085872-6086018 | KCNAB2;KCNAB2 | N_Shore | TSS1500;TSS1500 |
| WG3465 | chr1:6085995-6086148 | KCNAB2;KCNAB2 | N_Shore | TSS1500;TSS1500 |
| GM00439 | chr1:6094205-6094316 | KCNAB2;KCNAB2 | | 5'UTR;5'UTR |
| WG1513 | chr1:6191821-6191879 | CHD5 | S_Shelf | Body |
| GM02149 | chr1:6191870-6192121 | CHD5 | S_Shelf | Body |
| WG1513 | chr1:6296335-6296630 | ICMT | S_Shore | TSS1500 |
| GM03073 | chr1:6309300-6309724 | GPR153 | Island | 3'UTR |
| WG1721 | chr1:63249197-63249213 | ATG4C;ATG4C | | TSS1500;TSS1500 |
| WG1513 | chr1:63249199-63249213 | ATG4C;ATG4C | | TSS1500;TSS1500 |
| GM02456 | chr1:6514605-6514781 | ESPN | Island | Body |
| GM02706 | chr1:6519923-6520354 | ESPN | N_Shore | Body |
| WG3465 | chr1:65209475-65209572 | RAVER2 | N_Shore | TSS1500 |
| GM00290 | chr1:6526049-6526319 | TNFRSF25;TNFRSF25;TNFRS | Island | Body;Body;Body;Body;Body;Body |
| WG1722 | chr1:6526049-6526469 | TNFRSF25;TNFRSF25;TNFRS | Island | Body;Body;Body;Body;Body;Body |
| WG3465 | chr1:6526051-6526398 | TNFRSF25;TNFRSF25;TNFRS | Island | Body;Body;Body;Body;Body;Body |
| GM00022 | chr1:6526129-6526309 | TNFRSF25;TNFRSF25;TNFRS | Island | 1stExon;1stExon;1stExon;1stExon;1st |
| GM02706 | chr1:6526182-6526319 | TNFRSF25;TNFRSF25;PLEKH | Island | 1stExon;1stExon;3'UTR;3'UTR;5'UTR;5 |
| GM03075 | chr1:6526309-6526398 | TNFRSF25;PLEKHG5;TNFRSF | S_Shore | TSS200;3'UTR;TSS200;3'UTR;3'UTR;TS |
| WG3465 | chr1:6546646-6546783 | PLEKHG5;PLEKHG5;PLEKHG | S_Shore | 5'UTR;TSS1500;Body;5'UTR;Body |
| WG3466 | chr1:6558085-6558113 | PLEKHG5;PLEKHG5 | S_Shore | Body;TSS1500 |
| GM02456 | chr1:6558085-6558497 | PLEKHG5;PLEKHG5 | S_Shore | Body;TSS1500 |
| WG3466 | chr1:66999753-66999929 | SGIP1 | S_Shore | TSS200 |
| GM01393 | chr1:67217673-67217826 | TCTEX1D1 | N_Shore | TSS1500 |
| GM02706 | chr1:67773020-67773128 | IL12RB2 | N_Shore | TSS200 |
| GM00439 | chr1:68517177-68517273 | DIRAS3 | N_Shore | TSS1500 |
| GM02706 | chr1:68517205-68517273 | DIRAS3 | N_Shore | TSS1500 |
| GM02706 | chr1:7120893-7120926 | CAMTA1 | N_Shore | Body |
| WG2276 | chr1:7408793-7409003 | CAMTA1 | | Body |
| GM02149 | chr1:75199075-75199117 | CRYZ;CRYZ;CRYZ;TYW3;TYW3;CRYZ;CRY | | 5'UTR;1stExon;1stExon;1stExon;Body; |
| WG1978 | chr1:75593476-75593804 | LHX8 | N_Shelf | TSS1500 |
| WG1282 | chr1:75593865-75594066 | LHX8 | N_Shore | TSS1500 |
| GM00290 | chr1:75602846-75602925 | LHX8 | Island | Body |
| GM00042 | chr1:76189707-76189801 | ACADM;ACADM | N_Shore | TSS1500;TSS1500 |
| GM00042 | chr1:76262302-76262711 | MSH4 | N_Shore | TSS1500 |
| GM02555 | chr1:76262537-76262711 | MSH4 | Island | TSS200 |
| WG3465 | chr1:76262537-76262857 | MSH4 | Island | TSS200 |
| WG3466 | chr1:7692321-7692367 | CAMTA1 | | Body |
| GM00289 | chr1:77747374-77747387 | AK5;AK5 | Island | TSS1500;TSS1500 |

| | | | | |
|---|---|---|---|---|
| WG2415 | chr1:78511140-78511433 | GIPC2 | N_Shore | TSS1500 |
| WG3465 | chr1:78511235-78511433 | GIPC2 | N_Shore | TSS1500 |
| WG1722 | chr1:78511344-78511433 | GIPC2 | N_Shore | TSS1500 |
| GM00935 | chr1:7887271-7887455 | PER3 | Island | Body |
| GM02704 | chr1:79472361-79472536 | ELTD1;ELTD1 | Island | 5'UTR;1stExon |
| WG2276 | chr1:8086721-8086959 | ERRFI1 | Island | TSS1500 |
| WG2276 | chr1:8383899-8384718 | SLC45A1 | N_Shore | TSS1500 |
| WG2194 | chr1:8403631-8403712 | SLC45A1 | N_Shore | Body |
| GM02175 | chr1:8403631-8403925 | SLC45A1 | N_Shore | Body |
| WG1283 | chr1:8403631-8404050 | SLC45A1 | N_Shore | Body |
| WG1513 | chr1:8403680-8403925 | SLC45A1 | N_Shore | Body |
| GM00438 | chr1:854766-854918 | FLJ39609 | Island | Body |
| WG1722 | chr1:854838-854966 | FLJ39609 | Island | TSS200 |
| GM02706 | chr1:854918-854966 | FLJ39609 | Island | TSS200 |
| WG1513 | chr1:85514610-85514708 | MCOLN3 | S_Shore | TSS1500 |
| GM09256 | chr1:855425-855649 | FLJ39609 | S_Shore | TSS1500 |
| GM00438 | chr1:86048479-86048585 | CYR61 | S_Shore | Body |
| GM01650 | chr1:86622624-86622682 | COL24A1 | Island | TSS1500 |
| GM01096 | chr1:86622624-86622737 | COL24A1 | Island | TSS1500 |
| WG1977 | chr1:870791-870958 | SAMD11 | Island | Body |
| GM01095 | chr1:870958-871308 | SAMD11 | Island | Body |
| WG1283 | chr1:871033-871308 | SAMD11 | Island | Body |
| WG1977 | chr1:871057-871546 | SAMD11 | Island | Body |
| GM00016 | chr1:87617710-87617916 | LOC339524 | Island | Body |
| WG2276 | chr1:87617710-87617972 | LOC339524 | Island | Body |
| GM03075 | chr1:89357908-89357988 | GTF2B | S_Shore | TSS1500 |
| GM00042 | chr1:89357908-89358043 | GTF2B | S_Shore | TSS1500 |
| GM01096 | chr1:89357911-89358043 | GTF2B | S_Shore | TSS1500 |
| GM00438 | chr1:89663868-89664582 | GBP4 | | Body |
| WG1513 | chr1:89664260-89664340 | GBP4 | | Body |
| WG2414 | chr1:89664260-89664407 | GBP4 | | Body |
| WG2415 | chr1:89664260-89664545 | GBP4 | | Body |
| WG3465 | chr1:89664260-89664582 | GBP4 | | Body |
| GM01392 | chr1:89664338-89664407 | GBP4 | | Body |
| GM02149 | chr1:89664407-89664582 | GBP4 | | Body |
| GM00520 | chr1:9129487-9129791 | SLC2A5;SLC2A5;SLC2A5 | N_Shelf | Body;Body;Body |
| WG1657 | chr1:9131622-9131783 | SLC2A5 | Island | Body |
| WG3465 | chr1:917655-917819 | C1orf170 | N_Shelf | TSS200 |
| GM01096 | chr1:92352293-92352481 | TGFBR3 | S_Shore | TSS1500 |
| WG3465 | chr1:92352404-92352481 | TGFBR3 | S_Shore | TSS1500 |
| GM00520 | chr1:92414295-92414520 | BRDT | N_Shore | TSS1500 |
| WG1978 | chr1:92414344-92414722 | BRDT | N_Shore | TSS1500 |
| WG2120 | chr1:92414344-92414782 | BRDT | N_Shore | TSS1500 |
| GM02706 | chr1:92414520-92414737 | BRDT | N_Shore | TSS1500 |
| GM01095 | chr1:92414520-92414782 | BRDT | N_Shore | TSS1500 |
| GM02706 | chr1:92545580-92545662 | BTBD8 | N_Shore | TSS1500 |
| WG1310 | chr1:9400692-9400775 | SPSB1 | | 5'UTR |

| | | | | |
|---|---|---|---|---|
| WG1283 | chr1:9409349-9409810 | SPSB1 | | 5'UTR |
| WG3465 | chr1:949392-949634 | ISG15 | Island | Body |
| GM00935 | chr1:949449-949893 | ISG15 | Island | Body |
| WG3465 | chr1:95006322-95006539 | F3 | N_Shore | Body |
| GM09255 | chr1:95006322-95006729 | F3 | N_Shore | Body |
| GM09256 | chr1:95285349-95285450 | SLC44A3;SLC44A3 | N_Shore | TSS1500;TSS1500 |
| WG2415 | chr1:95285405-95285531 | SLC44A3;SLC44A3 | N_Shore | TSS1500;TSS1500 |
| GM09255 | chr1:95698924-95699097 | RWDD3;RWDD3 | N_Shore | TSS1500;TSS1500 |
| WG3465 | chr1:95698924-95699313 | RWDD3;RWDD3 | N_Shore | TSS1500;TSS1500 |
| GM00940 | chr1:95698954-95699097 | RWDD3;RWDD3 | N_Shore | TSS1500;TSS1500 |
| WG1977 | chr1:969825-970294 | AGRN | Island | Body |
| GM02175 | chr10:100991745-100992068 | HPSE2;HPSE2;HPSE2;HPSE2 | N_Shelf | Body;Body;Body;Body |
| GM00520 | chr10:100991745-100992222 | HPSE2;HPSE2;HPSE2;HPSE2 | N_Shelf | Body;Body;Body;Body |
| WG2415 | chr10:100991959-100992222 | HPSE2;HPSE2;HPSE2;HPSE2 | N_Shore | Body;Body;Body;Body |
| WG1977 | chr10:100993025-100993556 | HPSE2;HPSE2;HPSE2;HPSE2 | Island | Body;Body;Body;Body |
| WG1977 | chr10:100993589-100993662 | HPSE2;HPSE2;HPSE2;HPSE2 | N_Shore | Body;Body;Body;Body |
| WG2844 | chr10:102896102-102896475 | TLX1 | N_Shore | Body |
| GM02317 | chr10:102896102-102896640 | TLX1 | N_Shore | Body |
| WG3465 | chr10:102997382-102997865 | FLJ41350 | S_Shore | Body |
| GM00520 | chr10:103588914-103589245 | KCNIP2;KCNIP2;KCNIP2;KCN | N_Shore | Body;Body;Body;Body;Body;Body;Bod |
| WG1722 | chr10:104613767-104613787 | C10orf32;C10orf32 | N_Shore | TSS1500;TSS1500 |
| GM00289 | chr10:105212680-105212813 | CALHM2;CALHM2 | S_Shore | TSS1500;TSS1500 |
| WG3465 | chr10:105233022-105233106 | CALHM3 | N_Shore | 3'UTR |
| WG1977 | chr10:105238873-105239085 | CALHM3;CALHM3 | | 1stExon |
| WG2121 | chr10:105238873-105239164 | CALHM3;CALHM3 | | 1stExon |
| WG2276 | chr10:105239164-105239205 | CALHM3 | | TSS200 |
| WG1309 | chr10:105978651-105978702 | MIR609;C10orf79 | | Body |
| WG1310 | chr10:106028373-106028519 | GSTO2 | N_Shore | TSS1500 |
| GM02173 | chr10:106028373-106028550 | GSTO2 | N_Shore | TSS1500 |
| GM09256 | chr10:106028543-106028939 | GSTO2 | N_Shore | TSS200 |
| WG1310 | chr10:106028703-106028928 | GSTO2;GSTO2 | Island | TSS200 |
| GM02173 | chr10:106028703-106029358 | GSTO2;GSTO2 | Island | TSS200 |
| WG2415 | chr10:106088702-106088755 | ITPRIP | | 5'UTR |
| GM00520 | chr10:111683554-111683654 | XPNPEP1;XPNPEP1;XPNPEP1 | Island | TSS200;TSS200;TSS200 |
| GM02456 | chr10:111966029-111966094 | MXI1 | N_Shelf | |
| WG1984 | chr10:114712695-114713108 | TCF7L2;TCF7L2;TCF7L2;TCF7 | Island | Body;Body;Body;Body;Body;Body |
| GM00289 | chr10:114713024-114713187 | TCF7L2;TCF7L2;TCF7L2;TCF7 | S_Shore | Body;Body;Body;Body;Body;Body |
| WG3466 | chr10:114877616-114877689 | TCF7L2;TCF7L2;TCF7L2;TCF7L2;TCF7L2 | | Body;Body;Body;Body;Body;Body |
| GM01392 | chr10:115939018-115939115 | TDRD1 | Island | TSS200 |
| GM02706 | chr10:116164640-116164714 | AFAP1L2;AFAP1L2 | Island | TSS200;TSS200 |
| GM02149 | chr10:116418181-116418597 | ABLIM1;ABLIM1;ABLIM1 | | Body;Body;1stExon |
| GM03075 | chr10:1166782-1166994 | WDR37 | | Body |
| GM00520 | chr10:1166782-1167544 | WDR37 | | Body |
| GM02175 | chr10:11783823-11783974 | ECHDC3 | N_Shore | TSS1500 |
| WG3466 | chr10:11784140-11784253 | ECHDC3 | N_Shore | TSS1500 |
| WG1513 | chr10:118084109-118084243 | C10orf96;C10orf96 | | TSS200 |
| GM02706 | chr10:118429452-118429485 | C10orf82;C10orf82 | N_Shore | 5'UTR |

| | | | | |
|---|---|---|---|---|
| WG3466 | chr10:118429481-118429648 | C10orf82 | Island | 1stExon;5'UTR |
| GM00520 | chr10:119000083-119000236 | SLC18A2 | N_Shelf | |
| WG1977 | chr10:11912126-11912597 | C10orf47 | Island | Body |
| GM02173 | chr10:119291766-119292130 | EMX2OS | | Body |
| GM00438 | chr10:119291766-119292262 | EMX2OS | | Body |
| WG2844 | chr10:119292547-119293800 | EMX2OS | N_Shore | Body |
| WG1310 | chr10:119292549-119293083 | EMX2OS | N_Shore | Body |
| WG2276 | chr10:119292549-119294332 | EMX2OS | N_Shore | Body |
| GM00438 | chr10:119294055-119294332 | EMX2OS | Island | Body |
| WG2415 | chr10:119295400-119296182 | EMX2OS | Island | Body |
| GM01393 | chr10:119295400-119296305 | EMX2OS | Island | Body |
| WG1983 | chr10:119295400-119296612 | EMX2OS | Island | Body |
| GM02704 | chr10:119295619-119295827 | EMX2OS | Island | Body |
| WG1977 | chr10:119295619-119296182 | EMX2OS | Island | Body |
| WG2844 | chr10:119295619-119296612 | EMX2OS | Island | Body |
| WG1310 | chr10:119295770-119295875 | EMX2OS | Island | Body |
| WG1513 | chr10:119295770-119296182 | EMX2OS | Island | Body |
| GM01096 | chr10:119295827-119296305 | EMX2OS | Island | Body |
| GM02173 | chr10:119296182-119296305 | EMX2OS | Island | Body |
| GM00439 | chr10:119296297-119296612 | EMX2OS | Island | Body |
| WG1310 | chr10:119305317-119305798 | EMX2;EMX2;EMX2OS | Island | Body;Body;TSS1500 |
| WG1283 | chr10:119307692-119307931 | EMX2;EMX2 | S_Shore | Body;Body |
| GM09256 | chr10:1209379-1209455 | NCRNA00200 | Island | Body |
| WG2843 | chr10:121265587-121265893 | RGS10;RGS10 | | Body;Body |
| GM00289 | chr10:121300857-121301343 | RGS10 | N_Shelf | Body |
| WG1513 | chr10:1228893-1229050 | ADARB2 | N_Shore | 3'UTR |
| GM02706 | chr10:123355268-123355576 | FGFR2;FGFR2;FGFR2;FGFR2 | N_Shelf | 5'UTR;5'UTR;TSS1500;5'UTR;TSS1500 |
| GM00016 | chr10:123355268-123356236 | FGFR2;FGFR2;FGFR2;FGFR2 | N_Shelf | 5'UTR;5'UTR;TSS1500;5'UTR;TSS1500 |
| WG2844 | chr10:123355268-123356336 | FGFR2;FGFR2;FGFR2;FGFR2 | N_Shelf | 5'UTR;5'UTR;TSS1500;5'UTR;TSS1500 |
| WG2276 | chr10:123355454-123355748 | FGFR2;FGFR2;FGFR2;FGFR2 | N_Shore | 5'UTR;5'UTR;5'UTR;5'UTR;TSS1500;5'U |
| WG3465 | chr10:123781519-123781602 | TACC2;TACC2 | | 5'UTR;5'UTR |
| GM02173 | chr10:123922863-123923099 | TACC2;TACC2;TACC2;TACC2 | N_Shore | TSS1500;Body;TSS1500;Body |
| GM02149 | chr10:123922863-123923450 | TACC2;TACC2;TACC2;TACC2 | N_Shore | TSS1500;Body;TSS1500;Body |
| WG1657 | chr10:123922970-123923273 | TACC2;TACC2;TACC2;TACC2 | Island | TSS1500;Body;TSS1500;Body |
| GM00935 | chr10:123923518-123924042 | TACC2;TACC2;TACC2;TACC2 | Island | Body;1stExon;Body;1stExon |
| WG2844 | chr10:123923518-123924085 | TACC2;TACC2;TACC2;TACC2 | Island | Body;1stExon;Body;1stExon |
| GM00439 | chr10:123923881-123924085 | TACC2;TACC2;TACC2;TACC2 | Island | Body;Body;Body;Body |
| WG2415 | chr10:124030740-124030801 | BTBD16 | | TSS1500 |
| GM01650 | chr10:124220359-124220396 | HTRA1 | N_Shore | TSS1500 |
| WG1722 | chr10:124319733-124319990 | DMBT1;DMBT1;DMBT1 | | TSS1500;TSS1500;TSS1500 |
| WG1513 | chr10:1245836-1246035 | ADARB2 | | Body |
| GM00289 | chr10:124638756-124638983 | FAM24B;LOC399815 | N_Shore | 5'UTR;TSS1500 |
| WG2276 | chr10:124638874-124638991 | FAM24B;LOC399815 | Island | 5'UTR;TSS1500 |
| GM02147 | chr10:124638976-124638991 | LOC399815;FAM24B | Island | 5'UTR;TSS1500 |
| GM02147 | chr10:124639110-124639157 | LOC399815;FAM24B;FAM24I | Island | TSS200;5'UTR;1stExon |
| GM09256 | chr10:1248770-1248818 | ADARB2 | N_Shelf | Body |
| WG1722 | chr10:124909889-124910217 | HMX2 | Island | 3'UTR |

| | | | | |
|---|---|---|---|---|
| WG3465 | chr10:1256339-1256574 | ADARB2 | S_Shelf | Body |
| WG1514 | chr10:125650957-125651085 | CPXM2 | N_Shore | Body |
| GM02149 | chr10:125651526-125651588 | CPXM2 | S_Shore | 1stExon;5'UTR |
| WG2415 | chr10:125651526-125651621 | CPXM2 | S_Shore | 1stExon;5'UTR |
| GM00439 | chr10:125651526-125651697 | CPXM2 | S_Shore | 1stExon;5'UTR |
| GM00935 | chr10:125651549-125651726 | CPXM2 | S_Shore | TSS200 |
| GM02149 | chr10:125767795-125767965 | CHST15 | | 3'UTR |
| WG1513 | chr10:125805493-125807047 | CHST15 | Island | Body |
| GM02149 | chr10:125805554-125806214 | CHST15 | S_Shore | 1stExon |
| WG2844 | chr10:126278272-126278502 | LHPP;LHPP | N_Shelf | Body;Body |
| WG1513 | chr10:126278299-126278495 | LHPP;LHPP | N_Shore | Body;Body |
| WG2415 | chr10:126278299-126278502 | LHPP;LHPP | N_Shore | Body;Body |
| GM01376 | chr10:126278372-126278502 | LHPP;LHPP | Island | Body;Body |
| WG2844 | chr10:126308034-126308552 | FAM53B | S_Shore | |
| WG1283 | chr10:12648032-12648338 | CAMK1D;CAMK1D | | Body;Body |
| WG2415 | chr10:12648032-12648526 | CAMK1D;CAMK1D | | Body;Body |
| GM02706 | chr10:12648176-12648526 | CAMK1D;CAMK1D | | Body;Body |
| WG2415 | chr10:126703991-126704259 | CTBP2;CTBP2;CTBP2 | | Body;Body;Body |
| WG1977 | chr10:126712074-126712482 | CTBP2;CTBP2;CTBP2 | | Body;Body;Body |
| GM02555 | chr10:126751950-126752037 | CTBP2;CTBP2 | | 5'UTR;5'UTR |
| GM00438 | chr10:126782538-126782592 | CTBP2;CTBP2 | | 5'UTR;5'UTR |
| GM02706 | chr10:128152075-128152124 | C10orf90 | | Body |
| GM01096 | chr10:128192896-128193332 | C10orf90 | N_Shelf | Body |
| GM00940 | chr10:128209980-128210076 | C10orf90;C10orf90 | | Body |
| GM00289 | chr10:1285332-1285535 | ADARB2 | S_Shore | Body |
| GM00438 | chr10:12999550-12999618 | CCDC3 | | Body |
| WG2414 | chr10:131265059-131265073 | MGMT | N_Shore | TSS1500 |
| GM02173 | chr10:131265575-131265796 | MGMT | Island | 1stExon;5'UTR |
| GM03075 | chr10:131412507-131412643 | MGMT | | Body |
| WG2844 | chr10:131425198-131425282 | MGMT | | Body |
| GM01096 | chr10:131640139-131640509 | EBF3 | S_Shore | Body |
| WG1978 | chr10:131669406-131669630 | EBF3 | S_Shore | Body |
| GM01096 | chr10:131682269-131682452 | EBF3 | | Body |
| WG2194 | chr10:131697080-131697345 | EBF3 | N_Shore | Body |
| GM00438 | chr10:131761184-131761386 | EBF3 | Island | Body |
| WG3465 | chr10:131761206-131761386 | EBF3 | Island | Body |
| WG2120 | chr10:131761386-131761660 | EBF3 | Island | Body |
| WG1513 | chr10:132891319-132891371 | TCERG1L | N_Shelf | 3'UTR |
| GM00438 | chr10:132891341-132892510 | TCERG1L | N_Shelf | 3'UTR |
| GM02149 | chr10:132909213-132909470 | TCERG1L | | Body |
| GM02149 | chr10:132999672-133000153 | TCERG1L | | Body |
| WG1513 | chr10:132999672-133000405 | TCERG1L | | Body |
| GM09256 | chr10:133793442-133793734 | BNIP3 | N_Shore | Body |
| GM09256 | chr10:133929429-133929519 | JAKMIP3 | N_Shore | Body |
| WG1513 | chr10:133932254-133932445 | JAKMIP3 | S_Shore | Body |
| WG1513 | chr10:133946674-133946881 | JAKMIP3 | N_Shelf | Body |
| WG2275 | chr10:133985911-133985960 | JAKMIP3 | | 3'UTR |

| | | | | |
|---|---|---|---|---|
| WG1977 | chr10:133995286-133995387 | JAKMIP3 | | 3'UTR |
| GM00940 | chr10:134043618-134043755 | STK32C | S_Shore | Body |
| GM09256 | chr10:134062171-134062614 | STK32C | | Body |
| GM00438 | chr10:134096650-134096880 | STK32C | S_Shore | Body |
| WG1722 | chr10:134119433-134120029 | STK32C | N_Shore | Body |
| WG1977 | chr10:134150489-134150760 | LRRC27;LRRC27;LRRC27;LRRC27;LRRC | | Body;TSS200;Body;Body;Body |
| GM00289 | chr10:134221503-134222218 | PWWP2B;PWWP2B | S_Shore | 3'UTR;Body |
| WG1310 | chr10:134226361-134226547 | PWWP2B;PWWP2B | | 3'UTR;Body |
| WG2276 | chr10:134377343-134377481 | INPP5A | | Body |
| WG2844 | chr10:134420036-134420623 | INPP5A | | Body |
| GM00438 | chr10:134751673-134752183 | C10orf93 | | Body |
| WG2415 | chr10:134751800-134752183 | C10orf93 | N_Shelf | Body |
| WG3465 | chr10:134910271-134910540 | GPR123 | N_Shelf | Body |
| GM09256 | chr10:134918781-134918927 | GPR123 | N_Shore | Body |
| WG2844 | chr10:134918781-134918974 | GPR123 | N_Shore | Body |
| WG1977 | chr10:134928138-134928457 | GPR123 | S_Shore | Body |
| GM09256 | chr10:134937798-134938144 | GPR123 | N_Shore | Body |
| GM02149 | chr10:134937798-134938469 | GPR123 | N_Shore | Body |
| WG2121 | chr10:134937967-134938469 | GPR123 | N_Shore | Body |
| GM00438 | chr10:134983383-134983667 | KNDC1 | S_Shore | Body |
| WG1513 | chr10:134983667-134983838 | KNDC1 | S_Shelf | Body |
| GM01096 | chr10:134994186-134994252 | KNDC1 | S_Shore | Body |
| WG1513 | chr10:135038107-135038446 | KNDC1 | N_Shore | Body |
| WG1977 | chr10:135061015-135061259 | MIR202 | | |
| WG2415 | chr10:135171713-135171746 | C10orf125;C10orf125 | Island | TSS200;TSS200 |
| GM02706 | chr10:135278901-135279010 | LOC619207 | Island | Body |
| WG3465 | chr10:135278976-135279147 | LOC619207 | Island | Body |
| GM00520 | chr10:135340721-135340785 | CYP2E1 | N_Shore | TSS1500 |
| WG2844 | chr10:135341528-135342218 | CYP2E1 | N_Shore | 1stExon |
| WG2121 | chr10:135342560-135342936 | CYP2E1 | Island | Body |
| WG2414 | chr10:135342560-135343248 | CYP2E1 | Island | Body |
| WG2275 | chr10:135342560-135343280 | CYP2E1 | Island | Body |
| WG2844 | chr10:135342936-135343248 | CYP2E1 | S_Shore | Body |
| GM02175 | chr10:135343193-135343280 | CYP2E1 | S_Shore | Body |
| WG2415 | chr10:135379213-135379242 | SYCE1;SYCE1;SYCE1 | N_Shore | Body;5'UTR;Body |
| WG1977 | chr10:13629624-13629984 | PRPF18 | S_Shore | Body |
| WG2276 | chr10:13749193-13749285 | FRMD4A | | Body |
| WG2844 | chr10:13749193-13749625 | FRMD4A | | Body |
| GM00940 | chr10:13749245-13749382 | FRMD4A | | Body |
| GM00042 | chr10:13749245-13749405 | FRMD4A | | Body |
| GM00520 | chr10:13749245-13749625 | FRMD4A | | Body |
| WG2843 | chr10:13749285-13749405 | FRMD4A | | Body |
| WG1310 | chr10:13749285-13749625 | FRMD4A | | Body |
| WG1977 | chr10:13749382-13749625 | FRMD4A | | Body |
| GM03075 | chr10:13749405-13749625 | FRMD4A | | Body |
| WG2120 | chr10:13932842-13933021 | FRMD4A | N_Shore | Body |
| WG3466 | chr10:13932842-13933247 | FRMD4A | N_Shore | Body |

| | | | | |
|---|---|---|---|---|
| GM01096 | chr10:13932842-13933452 | FRMD4A | N_Shore | Body |
| GM02704 | chr10:13933021-13933247 | FRMD4A | N_Shore | Body |
| GM00935 | chr10:14051636-14051679 | FRMD4A | | Body |
| GM09256 | chr10:1410519-1410671 | ADARB2 | | Body |
| WG1977 | chr10:1416546-1416920 | ADARB2 | S_Shore | Body |
| GM00289 | chr10:1416776-1416920 | ADARB2 | S_Shore | Body |
| GM01096 | chr10:14372879-14372913 | FRMD4A | | TSS200 |
| GM09256 | chr10:1444784-1444808 | ADARB2 | | Body |
| WG3465 | chr10:1454265-1454387 | ADARB2 | | Body |
| GM01096 | chr10:1505595-1506175 | ADARB2 | N_Shore | Body |
| GM02149 | chr10:1507708-1508061 | ADARB2 | S_Shore | Body |
| GM01650 | chr10:1509675-1509859 | ADARB2 | S_Shelf | Body |
| WG1722 | chr10:1509675-1510077 | ADARB2 | S_Shelf | Body |
| GM00438 | chr10:1567746-1567897 | ADARB2 | | Body |
| GM09256 | chr10:16478471-16478613 | PTER;PTER | N_Shore | TSS1500;TSS1500 |
| GM00438 | chr10:1707523-1707576 | ADARB2 | | Body |
| GM01095 | chr10:1712918-1713065 | ADARB2 | | Body |
| GM02706 | chr10:1712918-1713384 | ADARB2 | | Body |
| WG2120 | chr10:1712988-1713335 | ADARB2 | | Body |
| WG1978 | chr10:1767482-1767511 | ADARB2 | | Body |
| GM01096 | chr10:1767511-1767830 | ADARB2 | | Body |
| WG1513 | chr10:1769480-1769577 | ADARB2 | | Body |
| GM01650 | chr10:18549536-18550033 | CACNB2;CACNB2;CACNB2;CACNB2;CA | | Body;Body;Body;Body;Body;TSS200;B |
| WG2415 | chr10:18550033-18550223 | CACNB2;CACNB2;CACNB2;CACNB2;CA | | 1stExon;Body;Body;5'UTR;Body;Body; |
| GM09256 | chr10:18689378-18689440 | CACNB2;CACNB2;CACNB2;CACNB2;CA | | Body;Body;Body;Body;Body;TSS200;B |
| WG3466 | chr10:18689378-18689503 | CACNB2;CACNB2;CACNB2;CACNB2;CA | | Body;Body;Body;Body;Body;TSS200;B |
| WG1984 | chr10:18689430-18689471 | CACNB2;CACNB2;CACNB2;CACNB2;CA | | Body;Body;Body;Body;Body;TSS200;B |
| GM03075 | chr10:21807166-21807252 | C10orf140 | S_Shore | 5'UTR |
| WG1722 | chr10:22605982-22606072 | COMMD3 | S_Shore | Body |
| WG2276 | chr10:22634145-22634218 | SPAG6;SPAG6 | Island | TSS1500;TSS1500 |
| WG2415 | chr10:26855907-26856105 | APBB1IP | Island | Body |
| WG2415 | chr10:27541298-27541331 | LOC387646 | Island | TSS200 |
| WG1722 | chr10:27541331-27541626 | LOC387646 | Island | TSS200 |
| WG3466 | chr10:27541331-27541877 | LOC387646 | Island | TSS200 |
| WG3466 | chr10:28030398-28030839 | MKX | Island | Body |
| WG3465 | chr10:28030628-28030839 | MKX | Island | Body |
| WG3466 | chr10:28035631-28036151 | MKX | S_Shore | TSS1500 |
| GM02706 | chr10:28035719-28036151 | MKX | S_Shore | TSS1500 |
| WG1722 | chr10:28035731-28036151 | MKX | S_Shore | TSS1500 |
| WG1984 | chr10:29698368-29698449 | LOC387647 | Island | TSS200 |
| GM00439 | chr10:29698368-29698470 | LOC387647 | Island | TSS200 |
| WG1513 | chr10:29811530-29811659 | SVIL;SVIL | | Body;Body |
| GM00520 | chr10:29834201-29834266 | SVIL;MIR604;SVIL | | Body;TSS200;Body |
| WG3466 | chr10:3138418-3138534 | PFKP | | Body |
| GM00289 | chr10:3146486-3147056 | PFKP | N_Shore | Body |
| WG2415 | chr10:3149673-3149796 | PFKP | S_Shore | Body |
| WG1977 | chr10:3149673-3149852 | PFKP | S_Shore | Body |

| | | | | |
|---|---|---|---|---|
| GM02175 | chr10:31607150-31607160 | ZEB1;ZEB1;ZEB1;LOC22093( | N_Shore | TSS1500;TSS1500;TSS1500;Body;TSS |
| GM00520 | chr10:35894012-35894222 | GJD4 | N_Shelf | TSS1500 |
| GM00289 | chr10:37413782-37414261 | ANKRD30A | N_Shore | TSS1500 |
| WG1282 | chr10:3823844-3824016 | KLF6;KLF6;KLF6;KLF6 | Island | Body;Body;Body;Body |
| WG1722 | chr10:38382849-38383098 | ZNF37A;ZNF37A | N_Shore | TSS1500;TSS1500 |
| WG1722 | chr10:384506-384652 | DIP2C | N_Shelf | Body |
| GM01375 | chr10:38645376-38645450 | HSD17B7P2 | Island | Body |
| WG1722 | chr10:38691787-38691801 | CDC10L | Island | TSS200 |
| GM01392 | chr10:42862978-42863550 | LOC441666 | N_Shore | Body |
| GM02175 | chr10:42862978-42863594 | LOC441666 | N_Shore | Body |
| GM01375 | chr10:42863173-42863539 | LOC441666 | Island | Body |
| WG1722 | chr10:42863444-42863539 | LOC441666 | Island | Body |
| GM00290 | chr10:42863539-42863594 | LOC441666 | S_Shore | TSS200 |
| GM00438 | chr10:42970351-42970785 | LOC84856 | N_Shore | TSS1500 |
| GM02456 | chr10:438227-438317 | DIP2C | | Body |
| WG1722 | chr10:44101714-44101728 | ZNF485 | N_Shore | TSS1500 |
| GM00290 | chr10:45495677-45496087 | C10orf25;ZNF22 | N_Shore | 3'UTR;TSS1500 |
| WG1657 | chr10:45495739-45495981 | C10orf25;ZNF22 | N_Shore | 3'UTR;TSS1500 |
| GM02555 | chr10:45495739-45496087 | C10orf25;ZNF22 | N_Shore | 3'UTR;TSS1500 |
| WG3466 | chr10:45495782-45495981 | C10orf25;ZNF22 | N_Shore | 3'UTR;TSS1500 |
| GM00520 | chr10:45923083-45923705 | ALOX5 | Island | Body |
| GM02175 | chr10:46971222-46971495 | SYT15;SYT15 | Island | TSS1500;TSS1500 |
| GM02706 | chr10:46992448-46993586 | GPRIN2 | N_Shore | TSS1500 |
| WG3466 | chr10:46993502-46993542 | GPRIN2 | Island | TSS1500 |
| GM02317 | chr10:4868328-4868333 | AKR1E2 | Island | TSS200 |
| GM03075 | chr10:4868328-4868335 | AKR1E2 | Island | TSS200 |
| WG1513 | chr10:49482826-49483356 | FRMPD2;FRMPD2;FRMPD2 | | 5'UTR;TSS200;1stExon |
| GM00438 | chr10:50122337-50123344 | WDFY4;LRRC18 | | 1stExon;Body |
| WG1977 | chr10:50143337-50143397 | WDFY4 | | TSS1500;Body |
| GM00935 | chr10:50323825-50323832 | C10orf72;C10orf72 | Island | TSS200;TSS200 |
| WG1722 | chr10:50506748-50506885 | C10orf71;C10orf71 | | TSS1500;TSS1500 |
| WG2415 | chr10:50506748-50506896 | C10orf71;C10orf71 | | TSS1500;TSS1500 |
| WG2844 | chr10:50506748-50507092 | C10orf71;C10orf71 | | TSS1500;TSS1500 |
| WG2121 | chr10:50506875-50506896 | C10orf71;C10orf71 | | TSS1500;TSS1500 |
| GM02175 | chr10:50507071-50507214 | C10orf71;C10orf71 | | TSS200;TSS200 |
| WG2415 | chr10:50507071-50507298 | C10orf71;C10orf71 | | TSS200;TSS200 |
| WG2415 | chr10:530635-530805 | DIP2C | N_Shore | Body |
| GM00940 | chr10:530635-530836 | DIP2C | N_Shore | Body |
| WG1977 | chr10:530635-531569 | DIP2C | N_Shore | Body |
| GM09256 | chr10:530635-531584 | DIP2C | N_Shore | Body |
| GM02704 | chr10:530635-531914 | DIP2C | N_Shore | Body |
| WG1513 | chr10:530635-532791 | DIP2C | N_Shore | Body |
| GM02555 | chr10:530805-530951 | DIP2C | Island | Body |
| GM01376 | chr10:530805-531465 | DIP2C | Island | Body |
| WG3465 | chr10:530951-531320 | DIP2C | Island | Body |
| WG1722 | chr10:530951-531465 | DIP2C | Island | Body |
| GM02175 | chr10:530951-531569 | DIP2C | Island | Body |

| | | | | |
|---|---|---|---|---|
| WG1721 | chr10:531098-531320 | DIP2C | Island | Body |
| GM00940 | chr10:531098-531569 | DIP2C | Island | Body |
| WG2276 | chr10:531914-532409 | DIP2C | S_Shore | Body |
| GM09255 | chr10:54073047-54073189 | DKK1 | N_Shelf | |
| GM00439 | chr10:54073047-54073281 | DKK1 | N_Shelf | |
| WG2844 | chr10:556470-556762 | DIP2C | | Body |
| WG3465 | chr10:5734571-5735135 | C10orf18 | | 5'UTR |
| WG3465 | chr10:616959-617105 | DIP2C | | Body |
| GM02555 | chr10:62671136-62671228 | RHOBTB1;RHOBTB1;RHOBTB1;RHOBTB1 | | Body;Body;Body;Body |
| WG3466 | chr10:62760791-62761027 | RHOBTB1;RHOBTB1 | N_Shelf | Body;Body |
| GM00941 | chr10:63213339-63213391 | TMEM26 | S_Shore | 1stExon;5'UTR |
| GM00042 | chr10:63213385-63213396 | TMEM26 | S_Shore | TSS200 |
| WG2844 | chr10:639935-640085 | DIP2C | | Body |
| WG2844 | chr10:64572581-64573476 | EGR2;EGR2;EGR2;EGR2 | S_Shelf | 3'UTR;1stExon |
| GM00439 | chr10:65280473-65280795 | REEP3 | N_Shelf | |
| GM02706 | chr10:661009-661159 | DIP2C | | Body |
| GM02456 | chr10:661009-661388 | DIP2C | | Body |
| GM02555 | chr10:661009-661730 | DIP2C | | Body |
| GM00290 | chr10:661087-661319 | DIP2C | | Body |
| GM00439 | chr10:661159-661388 | DIP2C | | Body |
| WG2415 | chr10:661568-661730 | DIP2C | | Body |
| WG2844 | chr10:670815-671628 | DIP2C | S_Shore | Body |
| GM02704 | chr10:671007-671628 | DIP2C | S_Shore | Body |
| WG2120 | chr10:682125-682152 | DIP2C | N_Shelf | Body |
| GM00439 | chr10:68687951-68688167 | CTNNA3;LRRTM3;CTNNA3 | | Body;Body;5'UTR;1stExon |
| GM03075 | chr10:71087834-71087979 | HK1;HK1;HK1;HK1;HK1 | | Body;Body;Body;Body;Body |
| GM02706 | chr10:71087834-71088038 | HK1;HK1;HK1;HK1;HK1 | | Body;Body;Body;Body;Body |
| GM09256 | chr10:71087924-71088038 | HK1;HK1;HK1;HK1;HK1 | | Body;Body;Body;Body;Body |
| WG1513 | chr10:711556-712442 | DIP2C | N_Shore | Body |
| GM02173 | chr10:711676-712032 | DIP2C | N_Shore | Body |
| GM02149 | chr10:71176850-71177158 | TACR2 | | TSS200 |
| WG1977 | chr10:72201600-72201622 | NODAL | S_Shore | TSS200 |
| WG1978 | chr10:72254314-72254335 | KIAA1274 | S_Shelf | 5'UTR |
| WG2843 | chr10:72311914-72312093 | KIAA1274 | | Body |
| WG2844 | chr10:72432797-72433218 | ADAMTS14;ADAMTS14 | Island | 1stExon;1stExon |
| GM00016 | chr10:729001-729226 | DIP2C | | Body |
| WG2276 | chr10:729001-729956 | DIP2C | | Body |
| WG1513 | chr10:7290492-7290545 | SFMBT2 | | Body |
| GM02456 | chr10:73497514-73497629 | C10orf105;C10orf105;CDH23 | | 5'UTR;Body |
| GM02706 | chr10:73497514-73497772 | C10orf105;C10orf105;CDH23 | | 5'UTR;Body |
| GM02149 | chr10:73565625-73565812 | CDH23 | N_Shore | Body |
| GM00438 | chr10:73848463-73848817 | SPOCK2;SPOCK2 | S_Shore | 5'UTR;5'UTR |
| WG1977 | chr10:73848764-73848823 | SPOCK2;SPOCK2;SPOCK2;S | S_Shore | 5'UTR;5'UTR |
| GM00520 | chr10:73848764-73848825 | SPOCK2;SPOCK2;SPOCK2;S | S_Shore | 5'UTR;5'UTR |
| GM00439 | chr10:73848764-73848830 | SPOCK2;SPOCK2;SPOCK2;S | S_Shore | 5'UTR;5'UTR |
| WG1310 | chr10:73848825-73848915 | SPOCK2;SPOCK2 | S_Shore | TSS200;TSS200 |
| WG2415 | chr10:73976790-73976821 | ASCC1;C10orf104 | S_Shore | TSS1500;5'UTR |

| | | | | |
|---|---|---|---|---|
| WG1309 | chr10:77163983-77164227 | C10orf41;C10orf41 | Island | Body;TSS200 |
| WG3466 | chr10:77165167-77165750 | C10orf41;C10orf41 | Island | Body;Body |
| GM00935 | chr10:77542423-77542585 | C10orf11 | | TSS200 |
| GM03075 | chr10:77871618-77871675 | C10orf11 | N_Shelf | Body |
| GM01392 | chr10:77871618-77871958 | C10orf11 | N_Shelf | Body |
| WG3465 | chr10:77871618-77872170 | C10orf11 | N_Shelf | Body |
| GM00042 | chr10:77871644-77871958 | C10orf11 | N_Shore | Body |
| GM01096 | chr10:77871644-77872084 | C10orf11 | N_Shore | Body |
| GM00940 | chr10:77871644-77872170 | C10orf11 | N_Shore | Body |
| GM01375 | chr10:77871958-77872170 | C10orf11 | N_Shore | Body |
| GM00290 | chr10:77871958-77872288 | C10orf11 | N_Shore | Body |
| WG3465 | chr10:78870177-78870221 | KCNMA1;KCNMA1;KCNMA1;KCNMA1 | | Body;Body;Body;Body |
| WG1978 | chr10:80853937-80854035 | ZMIZ1 | | 5'UTR |
| GM02175 | chr10:8096633-8096650 | FLJ45983;GATA3;GATA3;FLJ4 | Island | TSS1500;TSS200;TSS200;TSS1500 |
| WG3466 | chr10:8100286-8100761 | GATA3;GATA3 | N_Shore | Body;Body |
| GM00042 | chr10:8101307-8101513 | GATA3;GATA3 | S_Shore | Body;Body |
| WG1513 | chr10:8106003-8106035 | GATA3;GATA3 | | Body;Body |
| WG3465 | chr10:82167757-82167774 | C10orf58 | N_Shelf | |
| GM02706 | chr10:85974018-85974623 | PCDH21 | | Body |
| GM00940 | chr10:88022641-88022859 | GRID1 | N_Shore | Body |
| GM00290 | chr10:88022641-88023185 | GRID1 | N_Shore | Body |
| WG3465 | chr10:88023052-88023185 | GRID1 | Island | Body |
| GM00042 | chr10:88024529-88024569 | MIR346;GRID1 | Island | Body |
| WG1513 | chr10:88024557-88024572 | MIR346;GRID1 | S_Shore | TSS200;Body |
| GM02704 | chr10:88123169-88123687 | GRID1 | N_Shore | Body |
| GM02173 | chr10:88124936-88125351 | GRID1 | Island | Body |
| WG1513 | chr10:88124936-88125377 | GRID1 | Island | Body |
| GM00935 | chr10:89263900-89263949 | MINPP1 | N_Shelf | |
| WG2844 | chr10:89418898-89419292 | PAPSS2;PAPSS2 | N_Shore | TSS1500;TSS1500 |
| WG1978 | chr10:90343208-90343285 | RNLS;RNLS | Island | TSS200;TSS200 |
| WG3466 | chr10:90611604-90611911 | ANKRD22;ANKRD22 | | Body |
| GM02641 | chr10:90712478-90712739 | ACTA2;ACTA2 | | Body;Body |
| WG1513 | chr10:90712478-90712748 | ACTA2;ACTA2 | | Body;Body |
| GM02149 | chr10:90712478-90712767 | ACTA2;ACTA2 | | Body;Body |
| WG1722 | chr10:90712478-90712948 | ACTA2;ACTA2 | | Body;Body |
| GM09256 | chr10:90712684-90712767 | ACTA2;ACTA2 | | 5'UTR;5'UTR |
| WG2121 | chr10:90712684-90712948 | ACTA2;ACTA2 | | 5'UTR;5'UTR |
| WG2276 | chr10:90712732-90712767 | ACTA2;ACTA2 | | TSS200;5'UTR |
| GM01095 | chr10:90712732-90712948 | ACTA2;ACTA2 | | TSS200;5'UTR |
| WG1977 | chr10:90712739-90712948 | ACTA2;ACTA2 | | TSS200;5'UTR |
| WG1310 | chr10:90712748-90712948 | ACTA2;ACTA2 | | TSS200;5'UTR |
| GM00520 | chr10:91295643-91295855 | SLC16A12 | Island | TSS200 |
| GM02149 | chr10:93392836-93393509 | PPP1R3C;PPP1R3C | Island | Body |
| WG2414 | chr10:94451351-94451736 | HHEX | N_Shore | Body |
| GM09255 | chr10:94451518-94451736 | HHEX | Island | Body |
| GM00520 | chr10:94594240-94594494 | EXOC6 | | TSS1500 |
| GM00941 | chr10:94820945-94820988 | CYP26C1 | Island | TSS200 |

| | | | | |
|---|---|---|---|---|
| WG2844 | chr10:95198200-95198374 | MYOF;MYOF | | Body;Body |
| WG1977 | chr10:95848649-95848841 | PLCE1;PLCE1 | | Body;TSS1500 |
| WG3465 | chr10:97175272-97175479 | SORBS1;SORBS1;SORBS1;SORBS1;SORI | | Body;Body;Body;Body;Body;Body;Bod |
| GM02175 | chr10:97515222-97515398 | ENTPD1;ENTPD1;ENTPD1;ENTPD1;ENTP | | TSS1500;TSS1500;TSS1500;TSS1500; |
| WG1513 | chr10:97515372-97515398 | ENTPD1;ENTPD1;ENTPD1;ENTPD1;ENTP | | TSS1500;TSS200;TSS1500;TSS1500;T |
| GM01096 | chr10:98031750-98031964 | BLNK;BLNK | | TSS200;TSS200 |
| WG1309 | chr10:99080756-99081016 | FRAT1;FRAT1 | Island | 1stExon;3'UTR |
| WG3465 | chr10:99338056-99338188 | ANKRD2;ANKRD2 | N_Shore | Body;Body |
| WG2121 | chr10:99338056-99338240 | ANKRD2;ANKRD2 | N_Shore | Body;Body |
| GM02175 | chr10:99734081-99734154 | CRTAC1 | | Body |
| WG2121 | chr10:99734120-99734416 | CRTAC1 | | Body |
| WG2121 | chr10:99734912-99735133 | CRTAC1 | | Body |
| WG1513 | chr10:99734912-99735202 | CRTAC1 | | Body |
| GM02555 | chr10:99735010-99735202 | CRTAC1 | | Body |
| GM01392 | chr10:99894013-99894087 | C10orf28 | N_Shelf | |
| GM00520 | chr11:102496104-102496159 | MMP20 | | TSS200 |
| WG2415 | chr11:1028291-1028379 | MUC6 | N_Shore | Body |
| WG1978 | chr11:10325567-10325809 | ADM | N_Shore | TSS1500 |
| WG2121 | chr11:10325778-10326100 | ADM | N_Shore | TSS1500 |
| GM00935 | chr11:1036471-1036679 | MUC6 | Island | Body |
| GM00438 | chr11:1036714-1036766 | MUC6 | Island | TSS200 |
| WG3465 | chr11:1036727-1036766 | MUC6 | Island | TSS200 |
| WG3466 | chr11:1036754-1036865 | MUC6 | Island | TSS200 |
| GM01653 | chr11:10472149-10472159 | AMPD3 | Island | TSS200 |
| WG1978 | chr11:10476494-10476976 | AMPD3;AMPD3;AMPD3 | S_Shelf | TSS200;TSS1500;Body |
| WG2844 | chr11:10476976-10477452 | AMPD3;AMPD3;AMPD3;AMPD3 | | 1stExon;TSS1500;5'UTR;Body |
| GM00940 | chr11:10476976-10477461 | AMPD3;AMPD3;AMPD3;AMPD3 | | 1stExon;TSS1500;5'UTR;Body |
| GM00438 | chr11:10477399-10477461 | AMPD3;AMPD3;AMPD3 | | TSS200;5'UTR;Body |
| GM02706 | chr11:10477399-10477847 | AMPD3;AMPD3;AMPD3 | | TSS200;5'UTR;Body |
| WG1977 | chr11:10477461-10477847 | AMPD3;AMPD3;AMPD3 | | TSS200;5'UTR;Body |
| GM02706 | chr11:10715175-10715595 | MRVI1;MRVI1;MRVI1;MRVI1;MRVI1 | | 1stExon;5'UTR;1stExon;5'UTR;TSS200 |
| WG1977 | chr11:10715360-10715595 | MRVI1;MRVI1;MRVI1;MRVI1;MRVI1 | | TSS1500;1stExon;5'UTR;1stExon;5'UT |
| GM09256 | chr11:1074749-1074869 | MUC2 | | TSS200 |
| GM00940 | chr11:107582804-107582876 | SLN | | TSS200 |
| GM02175 | chr11:107582818-107582884 | SLN | | TSS200 |
| WG2415 | chr11:108408907-108408962 | EXPH5;EXPH5;EXPH5;EXPH5 | | Body;1stExon;5'UTR;Body |
| GM01653 | chr11:109292789-109292809 | C11orf87 | N_Shore | TSS200 |
| WG1977 | chr11:109292789-109292894 | C11orf87 | N_Shore | TSS200 |
| WG2194 | chr11:111154826-111155021 | C11orf53 | | Body |
| GM00016 | chr11:111154934-111155021 | C11orf53 | | Body |
| GM02706 | chr11:111250093-111250201 | POU2AF1;POU2AF1 | | 1stExon;5'UTR |
| WG1977 | chr11:111250129-111250201 | POU2AF1;POU2AF1 | | 1stExon;5'UTR |
| GM00520 | chr11:111250129-111250338 | POU2AF1;POU2AF1 | | 1stExon;5'UTR |
| GM01096 | chr11:111789452-111789613 | C11orf52 | | TSS200 |
| WG3465 | chr11:111789452-111789693 | C11orf52 | | TSS200 |
| WG1983 | chr11:111847475-111847747 | DIXDC1;DIXDC1 | N_Shore | TSS1500;Body |
| GM02706 | chr11:111895234-111895341 | DLAT | N_Shore | TSS1500 |

| | | | | |
|---|---|---|---|---|
| GM09255 | chr11:111895279-111895455 | DLAT | N_Shore | TSS1500 |
| GM00940 | chr11:112831681-112831841 | NCAM1;NCAM1;NCAM1 | N_Shore | TSS1500;TSS1500;TSS1500 |
| GM00520 | chr11:113184990-113185089 | TTC12 | N_Shore | TSS1500 |
| WG1513 | chr11:113185561-113185879 | TTC12 | Island | 5'UTR |
| WG2415 | chr11:113947036-113947359 | ZBTB16;ZBTB16 | | Body;Body |
| WG3465 | chr11:115372874-115372975 | CADM1;CADM1 | N_Shore | Body;Body |
| WG2844 | chr11:117668140-117668288 | DSCAML1 | Island | TSS200 |
| WG1722 | chr11:117698911-117699019 | FXYD2 | | TSS200 |
| GM00940 | chr11:117747934-117748189 | FXYD6;FXYD6;FXYD6;FXYD6; | S_Shore | 5'UTR;TSS200;TSS200;TSS200;TSS20 |
| GM09255 | chr11:117747940-117748169 | FXYD6;FXYD6;FXYD6;FXYD6; | S_Shore | 5'UTR;TSS200;TSS200;TSS200;TSS20 |
| GM00042 | chr11:117748132-117748187 | FXYD6;FXYD6;FXYD6;FXYD6; | S_Shore | TSS1500;1stExon;5'UTR;TSS1500;TSS |
| WG3465 | chr11:117748132-117748189 | FXYD6;FXYD6;FXYD6;FXYD6; | S_Shore | TSS1500;1stExon;5'UTR;TSS1500;TSS |
| WG2276 | chr11:117748169-117748189 | FXYD6;FXYD6;FXYD6;FXYD6; | S_Shore | TSS1500;TSS1500;TSS1500;TSS1500; |
| WG1977 | chr11:117800255-117800467 | TMPRSS13 | | TSS200 |
| WG1722 | chr11:117800255-117800571 | TMPRSS13 | | TSS200 |
| WG1722 | chr11:118022607-118023038 | SCN4B;SCN4B | N_Shore | Body;Body |
| GM00439 | chr11:11862730-11862879 | USP47 | N_Shore | TSS1500 |
| WG1722 | chr11:118842387-118842392 | FOXR1 | Island | TSS200 |
| GM00520 | chr11:118842387-118842413 | FOXR1 | Island | TSS200 |
| GM00439 | chr11:119019361-119019412 | ABCG4;ABCG4 | N_Shore | TSS1500;TSS1500 |
| GM03075 | chr11:119294322-119294441 | THY1 | S_Shore | TSS200 |
| WG2275 | chr11:119294408-119294441 | THY1 | S_Shore | TSS200 |
| WG1977 | chr11:12031046-12031097 | DKK3;DKK3;DKK3 | S_Shore | TSS1500;TSS200;TSS1500 |
| WG2121 | chr11:12031046-12031099 | DKK3;DKK3;DKK3 | S_Shore | TSS1500;TSS200;TSS1500 |
| GM02706 | chr11:12031046-12031113 | DKK3;DKK3;DKK3 | S_Shore | TSS1500;TSS200;TSS1500 |
| WG1722 | chr11:12031093-12031099 | DKK3;DKK3;DKK3 | S_Shore | TSS1500;TSS200;TSS1500 |
| GM00520 | chr11:12031093-12031113 | DKK3;DKK3;DKK3 | S_Shore | TSS1500;TSS200;TSS1500 |
| GM02149 | chr11:121986736-121987014 | LOC399959;BLID;BLID | | Body;1stExon;5'UTR |
| WG1977 | chr11:123066529-123066693 | ASAM | Island | TSS1500 |
| GM02149 | chr11:123066529-123066956 | ASAM | Island | TSS1500 |
| WG1658 | chr11:123066560-123066956 | ASAM | Island | TSS1500 |
| GM00520 | chr11:12307450-12307924 | MICALCL | | TSS1500 |
| WG1513 | chr11:123396039-123396299 | GRAMD1B | | TSS1500 |
| GM00290 | chr11:123396039-123396370 | GRAMD1B | | TSS1500 |
| WG3466 | chr11:123396505-123396593 | GRAMD1B | | TSS200 |
| GM02149 | chr11:123814620-123814972 | OR6T1 | | TSS200 |
| WG1513 | chr11:124310939-124311399 | OR8B8 | | 1stExon |
| GM00438 | chr11:124311153-124311423 | OR8B8 | | TSS200 |
| WG2415 | chr11:124311218-124311423 | OR8B8 | | TSS1500 |
| WG1657 | chr11:124767455-124767840 | ROBO4 | | Body |
| GM00016 | chr11:124767455-124767933 | ROBO4 | | Body |
| WG2415 | chr11:1248676-1248959 | MUC5B | S_Shore | Body |
| WG2275 | chr11:125037226-125037348 | PKNOX2 | S_Shore | 5'UTR |
| GM02317 | chr11:126173516-126173568 | DCPS | N_Shore | TSS200 |
| GM02706 | chr11:126581947-126582183 | KIRREL3;KIRREL3 | | Body;Body |
| WG3465 | chr11:126581947-126582242 | KIRREL3;KIRREL3 | | Body;Body |
| WG2415 | chr11:128457400-128457682 | ETS1;ETS1 | | 1stExon;5'UTR |

| | | | | |
|---|---|---|---|---|
| WG3465 | chr11:128646953-128647107 | FLI1;FLI1 | | Body;Body |
| WG3465 | chr11:128729883-128730011 | KCNJ1;KCNJ1;KCNJ1;KCNJ1 | | 5'UTR;5'UTR;5'UTR;5'UTR |
| GM00439 | chr11:128812825-128812875 | TP53AIP1 | | TSS200 |
| WG2844 | chr11:128812846-128813008 | TP53AIP1 | | TSS200 |
| WG2415 | chr11:128812913-128813183 | TP53AIP1 | | TSS200 |
| WG2120 | chr11:128812978-128813008 | TP53AIP1 | | TSS200 |
| GM00289 | chr11:129306366-129306737 | BARX2 | N_Shore | Body |
| WG2275 | chr11:129991445-129992368 | APLP2;APLP2;APLP2;APLP2;A | N_Shelf | Body;Body;Body;Body;Body;Body |
| WG1721 | chr11:129992297-129992365 | APLP2;APLP2;APLP2;APLP2;A | N_Shore | Body;Body;Body;Body;Body;Body |
| WG3465 | chr11:129992297-129992368 | APLP2;APLP2;APLP2;APLP2;A | N_Shore | Body;Body;Body;Body;Body;Body |
| GM01392 | chr11:130029515-130029684 | ST14 | N_Shore | TSS200 |
| WG2415 | chr11:130319691-130320219 | ADAMTS15 | Island | 1stExon |
| WG1513 | chr11:131239030-131239703 | NTM | | TSS1500 |
| GM00439 | chr11:131779469-131780094 | NTM;NTM;NTM;NTM | N_Shore | TSS1500;TSS1500;TSS1500;Body |
| GM02555 | chr11:131779583-131779935 | NTM;NTM;NTM;NTM | N_Shore | TSS1500;TSS1500;TSS1500;Body |
| GM00935 | chr11:131779604-131779859 | NTM;NTM;NTM;NTM | N_Shore | TSS1500;TSS1500;TSS1500;Body |
| WG2844 | chr11:131779750-131779935 | NTM;NTM;NTM;NTM | N_Shore | TSS1500;TSS1500;TSS1500;Body |
| WG1310 | chr11:131779920-131780094 | NTM;NTM;NTM;NTM | N_Shore | TSS1500;TSS1500;TSS1500;Body |
| WG2276 | chr11:1320329-1320634 | TOLLIP | S_Shelf | Body |
| GM02704 | chr11:132526947-132527438 | OPCML;OPCML | | Body;Body |
| WG2844 | chr11:1325718-1325852 | TOLLIP | | Body |
| WG1513 | chr11:132662772-132662963 | OPCML;OPCML | | Body;Body |
| WG1977 | chr11:132912205-132912368 | OPCML | | Body |
| WG3465 | chr11:132951838-132951950 | OPCML | Island | Body |
| WG3465 | chr11:133715076-133715511 | SPATA19 | | Body |
| GM00520 | chr11:133800473-133800913 | IGSF9B | N_Shore | Body |
| WG3465 | chr11:133800685-133800913 | IGSF9B | Island | Body |
| WG1513 | chr11:133800774-133800913 | IGSF9B | Island | Body |
| GM00438 | chr11:133800800-133801225 | IGSF9B | Island | Body |
| GM02175 | chr11:133820837-133821252 | IGSF9B | | Body |
| WG3465 | chr11:133994930-133995182 | JAM3 | Island | Body |
| WG1513 | chr11:134176692-134176861 | GLB1L3 | | Body |
| GM09256 | chr11:134253609-134254128 | B3GAT1;B3GAT1 | Island | Body;Body |
| GM00520 | chr11:13983818-13983986 | SPON1 | N_Shore | TSS200 |
| GM09255 | chr11:13983986-13984067 | SPON1;SPON1 | N_Shore | 1stExon;5'UTR |
| WG1978 | chr11:13985911-13986348 | SPON1 | Island | Body |
| WG1513 | chr11:1444768-1445041 | BRSK2 | | Body |
| GM00289 | chr11:1481925-1482428 | BRSK2 | Island | Body |
| GM01376 | chr11:14994045-14994179 | CALCA;CALCA;CALCA | N_Shore | TSS1500;TSS1500;TSS1500 |
| GM00042 | chr11:14994179-14994222 | CALCA;CALCA;CALCA | N_Shore | TSS1500;TSS1500;TSS1500 |
| GM09256 | chr11:14994397-14994606 | CALCA;CALCA;CALCA | N_Shore | TSS1500;TSS1500;TSS1500 |
| GM00042 | chr11:15095017-15095068 | CALCB | Island | TSS200 |
| GM00042 | chr11:15095178-15095978 | CALCB;CALCB | Island | 1stExon;5'UTR |
| WG2844 | chr11:1542313-1542394 | HCCA2 | | Body |
| GM02175 | chr11:1673766-1673899 | HCCA2 | N_Shore | Body |
| GM00438 | chr11:1704507-1704707 | HCCA2;FAM99B | | Body;Body |
| WG1721 | chr11:1750302-1750381 | HCCA2 | | Body |

| | | | | |
|---|---|---|---|---|
| GM02706 | chr11:1769289-1769390 | HCCA2;LOC402778;LOC402 | Island | Body;5'UTR;1stExon |
| WG1283 | chr11:1769382-1769409 | HCCA2;LOC402778 | Island | Body;TSS200 |
| GM01650 | chr11:1769382-1769443 | HCCA2;LOC402778 | Island | Body;TSS200 |
| GM03075 | chr11:1769382-1769522 | HCCA2;LOC402778 | Island | Body;TSS200 |
| WG1513 | chr11:1769409-1769462 | HCCA2;LOC402778 | Island | Body;TSS200 |
| GM02456 | chr11:1769443-1769522 | HCCA2;LOC402778 | Island | Body;TSS200 |
| GM00042 | chr11:17760574-17760956 | KCNC1;KCNC1 | S_Shelf | Body;Body |
| GM02706 | chr11:1785618-1785683 | CTSD;HCCA2 | S_Shore | TSS1500;TSS200 |
| GM09256 | chr11:18127958-18128177 | SAAL1 | S_Shore | TSS1500 |
| WG3465 | chr11:18433500-18433564 | LDHC;LDHC | | TSS1500;TSS1500 |
| WG1977 | chr11:18433500-18433683 | LDHC;LDHC | | TSS1500;TSS1500 |
| WG1722 | chr11:18433564-18433745 | LDHC;LDHC | | TSS1500;TSS1500 |
| GM01375 | chr11:18433683-18433745 | LDHC;LDHC | | TSS200;TSS200 |
| WG1977 | chr11:18477254-18477299 | LDHAL6A;LDHAL6A | N_Shore | TSS200;TSS200 |
| GM02706 | chr11:18477280-18477534 | LDHAL6A;LDHAL6A | N_Shore | TSS200;TSS200 |
| GM02316 | chr11:18477299-18477379 | LDHAL6A;LDHAL6A | Island | TSS200;TSS200 |
| WG3465 | chr11:18477299-18477534 | LDHAL6A;LDHAL6A | Island | TSS200;TSS200 |
| GM00439 | chr11:1854224-1854368 | SYT8 | | TSS1500 |
| GM01653 | chr11:18727321-18727774 | IGSF22 | Island | Body |
| WG2276 | chr11:1890901-1891947 | LSP1;LSP1;LSP1;LSP1 | N_Shore | 5'UTR;TSS1500;TSS1500;Body |
| GM02555 | chr11:1891699-1891932 | LSP1;LSP1;LSP1;LSP1 | N_Shore | 5'UTR;TSS1500;TSS1500;Body |
| WG2276 | chr11:1892092-1892392 | LSP1;LSP1;LSP1;LSP1 | Island | 5'UTR;TSS200;TSS200;Body |
| WG3466 | chr11:2016513-2016747 | H19 | N_Shore | Body |
| GM00438 | chr11:2019116-2019167 | H19;MIR675 | N_Shore | TSS200;TSS1500 |
| WG1513 | chr11:2020101-2020129 | H19 | S_Shore | TSS1500 |
| GM03075 | chr11:20384886-20385172 | HTATIP2;HTATIP2;HTATIP2;HT | N_Shore | TSS1500;TSS1500;TSS1500;TSS1500; |
| GM00940 | chr11:20692709-20692867 | NELL1;NELL1 | S_Shore | Body;Body |
| GM00016 | chr11:2161892-2162183 | IGF2AS;INS-IGF2;IGF2;IGF2;I | Island | Body;Body;5'UTR;5'UTR;Body |
| WG3466 | chr11:2162406-2162445 | IGF2AS;INS-IGF2;IGF2;IGF2A | Island | Body;Body;5'UTR;Body;TSS200 |
| WG3465 | chr11:2162478-2162510 | IGF2AS;INS-IGF2;IGF2;IGF2A | Island | Body;Body;5'UTR;Body;TSS200 |
| WG1513 | chr11:2162510-2162783 | IGF2AS;INS-IGF2;IGF2;IGF2A | S_Shore | Body;Body;5'UTR;Body;TSS200 |
| GM09256 | chr11:2162536-2162582 | IGF2AS;INS-IGF2;IGF2;IGF2A | S_Shore | Body;Body;5'UTR;Body;TSS200 |
| GM00042 | chr11:2165656-2165961 | IGF2AS;INS-IGF2;IGF2;IGF2A | Island | Body;Body;5'UTR;Body |
| WG1513 | chr11:2170412-2170831 | INS-IGF2;INS-IGF2;IGF2;IGF2 | | Body;Body;1stExon;5'UTR |
| GM00290 | chr11:2170870-2171026 | IGF2;INS-IGF2;INS-IGF2 | | TSS200;Body;Body |
| GM09255 | chr11:2172019-2172098 | INS-IGF2;IGF2;INS-IGF2 | | Body;TSS1500;Body |
| WG2844 | chr11:2176877-2176921 | INS-IGF2;INS-IGF2 | Island | Body;Body |
| WG3465 | chr11:2181866-2182457 | INS-IGF2;INS-IGF2;INS | | Body;Body;Body |
| GM02706 | chr11:2292890-2292902 | ASCL2 | Island | TSS1500 |
| GM00016 | chr11:2292890-2292905 | ASCL2 | Island | TSS1500 |
| WG3466 | chr11:2292890-2292914 | ASCL2 | Island | TSS1500 |
| GM02175 | chr11:2292905-2293048 | ASCL2 | Island | TSS1500 |
| WG2415 | chr11:2293074-2293164 | ASCL2 | S_Shore | TSS1500 |
| GM01096 | chr11:2293074-2293171 | ASCL2 | S_Shore | TSS1500 |
| GM09256 | chr11:2293117-2293171 | ASCL2 | S_Shore | TSS1500 |
| WG1722 | chr11:2293164-2293173 | ASCL2 | S_Shore | TSS1500 |
| GM09255 | chr11:2293181-2293211 | ASCL2 | S_Shore | TSS1500 |

| | | | | |
|---|---|---|---|---|
| WG1513 | chr11:2293605-2293665 | ASCL2 | S_Shore | TSS1500 |
| WG1513 | chr11:2321770-2321803 | C11orf21;TSPAN32;C11orf21 | | Body;TSS1500;Body |
| WG1721 | chr11:2422730-2423045 | TSSC4 | S_Shore | TSS1500 |
| GM02175 | chr11:2423043-2423196 | TSSC4 | S_Shore | TSS1500 |
| WG1310 | chr11:2441045-2441141 | TRPM5 | N_Shore | Body |
| GM00289 | chr11:2444462-2444553 | TRPM5 | S_Shelf | TSS200 |
| WG3465 | chr11:2464822-2464851 | KCNQ1 | N_Shore | TSS1500 |
| WG2415 | chr11:2464822-2464954 | KCNQ1 | N_Shore | TSS1500 |
| GM09256 | chr11:2464845-2464970 | KCNQ1 | N_Shore | TSS1500 |
| GM09255 | chr11:2464851-2464959 | KCNQ1 | N_Shore | TSS1500 |
| GM00439 | chr11:2464954-2464970 | KCNQ1 | N_Shore | TSS1500 |
| WG1722 | chr11:2464970-2465103 | KCNQ1 | N_Shore | TSS1500 |
| GM00290 | chr11:2466788-2466909 | KCNQ1 | Island | Body |
| GM00438 | chr11:2482233-2482303 | KCNQ1;KCNQ1 | | TSS1500;Body |
| WG1513 | chr11:2482240-2482442 | KCNQ1;KCNQ1 | | TSS1500;Body |
| WG1722 | chr11:2482394-2482442 | KCNQ1;KCNQ1 | | TSS1500;Body |
| WG1513 | chr11:2554410-2554583 | KCNQ1;KCNQ1 | N_Shore | Body;Body |
| GM01096 | chr11:26743403-26743888 | SLC5A12;SLC5A12 | | 5'UTR;1stExon |
| WG2275 | chr11:27015519-27016671 | FIBIN | | TSS200 |
| WG2121 | chr11:27015656-27016671 | FIBIN;FIBIN | | 5'UTR;1stExon |
| WG1309 | chr11:27015808-27015991 | FIBIN;FIBIN | | 5'UTR;1stExon |
| WG2414 | chr11:27015813-27015991 | FIBIN;FIBIN | | 5'UTR;1stExon |
| WG1722 | chr11:2722062-2722076 | KCNQ1OT1;KCNQ1;KCNQ1 | Island | TSS1500;Body;Body |
| WG1722 | chr11:2722086-2722258 | KCNQ1OT1;KCNQ1;KCNQ1 | Island | TSS1500;Body;Body |
| GM01392 | chr11:2722440-2722625 | KCNQ1OT1;KCNQ1;KCNQ1 | S_Shore | TSS1500;Body;Body |
| GM00022 | chr11:27528430-27528587 | LIN7C;BDNFOS | Island | TSS200;Body |
| GM02704 | chr11:2812987-2813298 | KCNQ1;KCNQ1 | Island | Body;Body |
| GM09255 | chr11:2828274-2828568 | KCNQ1;KCNQ1 | N_Shore | Body;Body |
| GM00438 | chr11:2828525-2828652 | KCNQ1;KCNQ1 | Island | Body;Body |
| GM00940 | chr11:2889809-2889875 | KCNQ1DN | N_Shore | TSS1500 |
| WG3465 | chr11:2890258-2890389 | KCNQ1DN | N_Shore | TSS1500 |
| GM00520 | chr11:2890450-2890577 | KCNQ1DN | Island | TSS1500 |
| GM00290 | chr11:2890457-2890575 | KCNQ1DN | Island | TSS1500 |
| GM02149 | chr11:2890617-2890670 | KCNQ1DN | Island | TSS1500 |
| GM00290 | chr11:2890644-2890670 | KCNQ1DN | Island | TSS1500 |
| GM02706 | chr11:2904663-2904951 | CDKN1C;CDKN1C;CDKN1C | N_Shore | 3'UTR;3'UTR;3'UTR |
| WG3465 | chr11:2907670-2907674 | CDKN1C;CDKN1C;CDKN1C | Island | TSS1500;TSS1500;TSS1500 |
| WG1722 | chr11:2907670-2907819 | CDKN1C;CDKN1C;CDKN1C | Island | TSS1500;TSS1500;TSS1500 |
| GM02149 | chr11:2907754-2907884 | CDKN1C;CDKN1C;CDKN1C | S_Shore | TSS1500;TSS1500;TSS1500 |
| GM09256 | chr11:2907819-2907930 | CDKN1C;CDKN1C;CDKN1C | S_Shore | TSS1500;TSS1500;TSS1500 |
| WG3465 | chr11:2907930-2908116 | CDKN1C;CDKN1C;CDKN1C | S_Shore | TSS1500;TSS1500;TSS1500 |
| WG1978 | chr11:2920052-2920209 | SLC22A18AS;SLC22A18 | N_Shelf | Body;TSS1500 |
| GM01392 | chr11:2925470-2925570 | SLC22A18;SLC22A18;SLC22 | S_Shore | Body;Body;TSS1500 |
| GM01650 | chr11:2930485-2930995 | SLC22A18;SLC22A18 | Island | Body;Body |
| WG3465 | chr11:299503-299564 | IFITM5;IFITM5 | Island | 1stExon;5'UTR |
| GM02706 | chr11:299564-299629 | IFITM5 | Island | TSS200 |
| GM02175 | chr11:30039004-30039113 | KCNA4 | Island | TSS1500 |

| | | | | |
|---|---|---|---|---|
| GM00935 | chr11:30605411-30605797 | MPPED2 | N_Shore | 5'UTR |
| GM02173 | chr11:30605581-30605787 | MPPED2 | Island | 5'UTR |
| WG2276 | chr11:313267-313478 | IFITM1 | S_Shore | TSS1500 |
| WG2276 | chr11:3144685-3144774 | OSBPL5;OSBPL5;OSBPL5 | S_Shelf | Body;Body;Body |
| WG2415 | chr11:315102-315262 | IFITM1 | N_Shore | Body |
| GM02555 | chr11:3164631-3164851 | OSBPL5;OSBPL5;OSBPL5 | | 5'UTR;5'UTR;5'UTR |
| WG2844 | chr11:3173907-3174128 | OSBPL5;OSBPL5;OSBPL5 | | 5'UTR;5'UTR;5'UTR |
| GM00935 | chr11:3175007-3175636 | OSBPL5;OSBPL5;OSBPL5 | | 5'UTR;5'UTR;5'UTR |
| WG2276 | chr11:3177622-3178024 | OSBPL5;OSBPL5;OSBPL5 | N_Shelf | 5'UTR;5'UTR;5'UTR |
| GM00016 | chr11:31819148-31819219 | PAX6;PAX6;PAX6 | N_Shore | Body;Body;Body |
| WG3465 | chr11:31819219-31819678 | PAX6;PAX6;PAX6 | N_Shore | Body;Body;Body |
| GM03075 | chr11:31819380-31819678 | PAX6;PAX6;PAX6 | N_Shore | Body;Body;Body |
| WG3466 | chr11:31819444-31819678 | PAX6;PAX6;PAX6 | N_Shore | Body;Body;Body |
| GM02555 | chr11:31819444-31820658 | PAX6;PAX6;PAX6 | N_Shore | Body;Body;Body |
| GM09256 | chr11:31821590-31822302 | PAX6;PAX6;PAX6 | S_Shore | Body;Body;Body |
| WG3465 | chr11:31821590-31822341 | PAX6;PAX6;PAX6 | S_Shore | Body;Body;Body |
| WG2415 | chr11:31821764-31822302 | PAX6;PAX6;PAX6 | S_Shore | Body;Body;Body |
| WG2120 | chr11:31822302-31823238 | PAX6;PAX6;PAX6 | S_Shore | Body;Body;Body |
| WG1978 | chr11:31824262-31825226 | PAX6;PAX6;PAX6 | N_Shore | Body;Body;Body |
| WG2415 | chr11:31824262-31825833 | PAX6;PAX6;PAX6 | N_Shore | Body;Body;Body |
| GM02456 | chr11:31824262-31825969 | PAX6;PAX6;PAX6 | N_Shore | Body;Body;Body |
| GM02175 | chr11:31824262-31826421 | PAX6;PAX6;PAX6 | N_Shore | Body;Body;Body |
| GM01096 | chr11:31824327-31825226 | PAX6;PAX6;PAX6 | N_Shore | Body;Body;Body |
| GM00439 | chr11:31824973-31825226 | PAX6;PAX6;PAX6 | N_Shore | Body;Body;Body |
| GM09255 | chr11:31824973-31825605 | PAX6;PAX6;PAX6 | N_Shore | Body;Body;Body |
| WG2121 | chr11:31825166-31825226 | PAX6;PAX6;PAX6 | N_Shore | Body;Body;Body |
| GM00438 | chr11:31825214-31825833 | PAX6;PAX6;PAX6 | N_Shore | Body;Body;Body |
| WG1513 | chr11:31825605-31825792 | PAX6;PAX6;PAX6 | N_Shore | Body;Body;Body |
| WG2120 | chr11:31825605-31825833 | PAX6;PAX6;PAX6 | N_Shore | Body;Body;Body |
| WG1978 | chr11:31825756-31825833 | PAX6;PAX6;PAX6 | Island | Body;Body;Body |
| WG1722 | chr11:31825756-31827998 | PAX6;PAX6;PAX6 | Island | Body;Body;Body |
| GM02555 | chr11:31825756-31828040 | PAX6;PAX6;PAX6 | Island | Body;Body;Body |
| GM01375 | chr11:31827084-31827593 | PAX6;PAX6;PAX6 | N_Shore | Body;Body;Body |
| WG2415 | chr11:31827084-31827757 | PAX6;PAX6;PAX6 | N_Shore | Body;Body;Body |
| GM09256 | chr11:31827084-31827920 | PAX6;PAX6;PAX6 | N_Shore | Body;Body;Body |
| WG3465 | chr11:31827084-31827998 | PAX6;PAX6;PAX6 | N_Shore | Body;Body;Body |
| WG1978 | chr11:31827084-31828040 | PAX6;PAX6;PAX6 | N_Shore | Body;Body;Body |
| WG1977 | chr11:31827379-31828040 | PAX6;PAX6;PAX6 | N_Shore | Body;Body;Body |
| WG2193 | chr11:31827757-31827899 | PAX6;PAX6;PAX6 | Island | Body;Body;Body |
| GM00940 | chr11:31827757-31827998 | PAX6;PAX6;PAX6 | Island | Body;Body;Body |
| WG2120 | chr11:31827757-31828040 | PAX6;PAX6;PAX6 | Island | Body;Body;Body |
| WG3466 | chr11:31827899-31827998 | PAX6;PAX6;PAX6 | Island | Body;Body;Body |
| GM01096 | chr11:31827899-31828040 | PAX6;PAX6;PAX6 | Island | Body;Body;Body |
| GM02555 | chr11:31835534-31835929 | PAX6 | Island | 5'UTR |
| GM01375 | chr11:31835929-31836334 | PAX6 | Island | 5'UTR |
| GM00935 | chr11:31836673-31837124 | PAX6 | Island | 5'UTR |
| WG3466 | chr11:31837104-31837144 | PAX6 | Island | 5'UTR |

| | | | | |
|---|---|---|---|---|
| GM02555 | chr11:31839628-31839947 | PAX6 | Island | TSS200 |
| WG1977 | chr11:31840628-31840862 | PAX6 | N_Shore | TSS1500 |
| WG2415 | chr11:3239218-3239509 | MRGPRG;C11orf36 | Island | 1stExon;TSS1500 |
| WG1513 | chr11:32421514-32421635 | WT1;WT1;WT1;WT1 | | Body;Body;Body;Body |
| WG3465 | chr11:32421752-32421845 | WT1;WT1;WT1;WT1 | | Body;Body;Body;Body |
| WG2194 | chr11:32449821-32450104 | WT1;WT1;WT1;WT1 | S_Shore | Body;Body;Body;Body |
| GM00289 | chr11:32454718-32454768 | WT1;WT1;WT1;WT1 | N_Shore | Body;Body;Body;Body |
| WG2414 | chr11:3253838-3254065 | MRGPRE | Island | TSS1500 |
| WG3465 | chr11:3253896-3254065 | MRGPRE | Island | TSS1500 |
| WG2120 | chr11:34460336-34460386 | CAT | Island | TSS200 |
| WG1721 | chr11:34460336-34460789 | CAT | Island | TSS200 |
| GM03075 | chr11:34460516-34460789 | CAT;CAT | Island | 1stExon;5'UTR |
| GM02706 | chr11:35439994-35440252 | SLC1A2 | N_Shore | Body |
| GM01096 | chr11:396686-397077 | PKP3 | Island | Body |
| WG1657 | chr11:405846-405970 | SIGIRR;SIGIRR;SIGIRR | N_Shore | 3'UTR;3'UTR;3'UTR |
| WG3465 | chr11:405846-406019 | SIGIRR;SIGIRR;SIGIRR | N_Shore | 3'UTR;3'UTR;3'UTR |
| GM01392 | chr11:405846-406492 | SIGIRR;SIGIRR;SIGIRR | N_Shore | 3'UTR;3'UTR;3'UTR |
| GM00042 | chr11:405970-406492 | SIGIRR;SIGIRR;SIGIRR | N_Shore | Body;Body;Body |
| WG1977 | chr11:415080-415111 | SIGIRR;SIGIRR;SIGIRR | N_Shore | 5'UTR;TSS200;5'UTR |
| GM02149 | chr11:4389408-4389757 | OR52B4 | | 1stExon |
| GM00438 | chr11:4389638-4389856 | OR52B4 | | TSS200 |
| GM02149 | chr11:4390195-4390263 | OR52B4 | | TSS1500 |
| WG1983 | chr11:4415193-4415245 | TRIM21 | | TSS1500 |
| WG3465 | chr11:44324976-44326516 | ALX4 | Island | Body |
| WG2194 | chr11:44325687-44325960 | ALX4 | Island | Body |
| WG2193 | chr11:44325687-44326260 | ALX4 | Island | Body |
| GM09256 | chr11:44325687-44326516 | ALX4 | Island | Body |
| WG2275 | chr11:44325823-44326260 | ALX4 | Island | Body |
| GM02706 | chr11:44325823-44326516 | ALX4 | Island | Body |
| GM02173 | chr11:44325960-44326516 | ALX4 | Island | Body |
| GM02555 | chr11:44326260-44326516 | ALX4 | Island | Body |
| WG2120 | chr11:44327015-44327572 | ALX4 | N_Shore | Body |
| WG3465 | chr11:44327267-44327572 | ALX4 | Island | Body |
| GM09255 | chr11:44327267-44327869 | ALX4 | Island | Body |
| GM02555 | chr11:44327267-44328154 | ALX4 | Island | Body |
| WG1514 | chr11:44327399-44327869 | ALX4 | Island | Body |
| GM00289 | chr11:44332332-44332369 | ALX4 | Island | TSS1500 |
| WG1984 | chr11:44332340-44332685 | ALX4 | Island | TSS1500 |
| GM02706 | chr11:44332340-44332719 | ALX4 | Island | TSS1500 |
| GM00289 | chr11:44332507-44332636 | ALX4 | Island | TSS1500 |
| WG1722 | chr11:44332507-44332685 | ALX4 | Island | TSS1500 |
| GM00439 | chr11:44332507-44332891 | ALX4 | Island | TSS1500 |
| WG3465 | chr11:44332540-44332653 | ALX4 | Island | TSS1500 |
| GM00042 | chr11:44332540-44332685 | ALX4 | Island | TSS1500 |
| WG2415 | chr11:44332540-44332719 | ALX4 | Island | TSS1500 |
| WG1513 | chr11:44332540-44332754 | ALX4 | Island | TSS1500 |
| GM00520 | chr11:44332558-44332685 | ALX4 | Island | TSS1500 |

| | | | | |
|---|---|---|---|---|
| GM00290 | chr11:44332558-44332719 | ALX4 | Island | TSS1500 |
| WG2194 | chr11:44332624-44332664 | ALX4 | Island | TSS1500 |
| WG2414 | chr11:44332624-44332685 | ALX4 | Island | TSS1500 |
| WG2193 | chr11:44332636-44332664 | ALX4 | Island | TSS1500 |
| GM01653 | chr11:44332653-44332685 | ALX4 | Island | TSS1500 |
| GM02456 | chr11:44332754-44332790 | ALX4 | Island | TSS1500 |
| GM02706 | chr11:44332756-44332891 | ALX4 | Island | TSS1500 |
| WG1984 | chr11:44332790-44332891 | ALX4 | S_Shore | TSS1500 |
| GM00042 | chr11:44332790-44332913 | ALX4 | S_Shore | TSS1500 |
| WG1513 | chr11:44332790-44333070 | ALX4 | S_Shore | TSS1500 |
| WG1722 | chr11:44332838-44333067 | ALX4 | S_Shore | TSS1500 |
| GM00290 | chr11:44332838-44335005 | ALX4 | S_Shore | TSS1500 |
| WG1977 | chr11:44332891-44332938 | ALX4 | S_Shore | TSS1500 |
| GM01650 | chr11:44332891-44332940 | ALX4 | S_Shore | TSS1500 |
| GM09256 | chr11:44332913-44332958 | ALX4 | S_Shore | TSS1500 |
| WG2844 | chr11:44332938-44332958 | ALX4 | S_Shore | TSS1500 |
| WG2415 | chr11:44332938-44332972 | ALX4 | S_Shore | TSS1500 |
| GM09255 | chr11:44332938-44333067 | ALX4 | S_Shore | TSS1500 |
| GM00439 | chr11:44332938-44333192 | ALX4 | S_Shore | TSS1500 |
| GM01096 | chr11:44332958-44333067 | ALX4 | S_Shore | TSS1500 |
| GM00042 | chr11:44332958-44333131 | ALX4 | S_Shore | TSS1500 |
| GM02456 | chr11:44332972-44333067 | ALX4 | S_Shore | TSS1500 |
| WG2844 | chr11:44333030-44333192 | ALX4 | S_Shore | TSS1500 |
| WG1984 | chr11:44333067-44333192 | ALX4 | S_Shore | TSS1500 |
| WG1722 | chr11:44333072-44333192 | ALX4 | S_Shore | TSS1500 |
| GM00438 | chr11:44972679-44972744 | TP53I11;TP53I11 | Island | TSS200;TSS200 |
| WG2121 | chr11:45825469-45825485 | SLC35C1;SLC35C1;SLC35C1 | N_Shore | TSS200;TSS1500;TSS200 |
| WG3465 | chr11:45827475-45827763 | SLC35C1;SLC35C1;SLC35C1 | S_Shore | Body;Body;1stExon |
| WG3465 | chr11:47470768-47470803 | RAPSN;RAPSN | | TSS200;TSS200 |
| WG1513 | chr11:4902464-4903155 | OR51T1 | | TSS1500 |
| GM00289 | chr11:50257256-50257625 | LOC441601 | Island | Body |
| GM02175 | chr11:50257256-50257752 | LOC441601 | Island | Body |
| GM00439 | chr11:50257625-50257752 | LOC441601 | Island | Body |
| WG2415 | chr11:5291298-5291692 | HBE1;HBE1 | | 1stExon;5'UTR |
| WG2415 | chr11:536299-536817 | HRAS;HRAS;LRRC56;HRAS | Island | TSS1500;TSS1500;TSS1500;TSS1500 |
| WG1977 | chr11:537638-537737 | LRRC56 | Island | 5'UTR |
| GM02175 | chr11:5705995-5706253 | TRIM5;TRIM5;TRIM5 | | 5'UTR;5'UTR;5'UTR |
| GM02175 | chr11:5706381-5706435 | TRIM5;TRIM5;TRIM5 | | TSS200;TSS200;TSS200 |
| GM02706 | chr11:57408514-57408751 | MIR130A | | TSS200 |
| WG3466 | chr11:57508416-57508451 | TMX2;TMX2;TMX2;C11orf31 | N_Shore | 3'UTR;3'UTR;Body;TSS1500 |
| GM09255 | chr11:57508416-57508545 | TMX2;TMX2;TMX2;C11orf31 | N_Shore | 3'UTR;3'UTR;Body;TSS1500 |
| GM02456 | chr11:57508449-57508545 | C11orf31 | N_Shore | TSS1500 |
| GM00042 | chr11:57519364-57519470 | BTBD18 | N_Shore | TSS200 |
| WG1977 | chr11:57957575-57957939 | OR9Q2 | | TSS1500 |
| GM00290 | chr11:59578490-59578540 | MRPL16 | S_Shore | TSS200 |
| WG1513 | chr11:59633874-59634276 | TCN1 | | 1stExon |
| WG2120 | chr11:59634191-59634276 | TCN1 | | TSS200 |

| | | | | |
|---|---|---|---|---|
| GM00438 | chr11:59824038-59824161 | MS4A3;MS4A3;MS4A3 | | TSS200;TSS200;TSS200 |
| GM02149 | chr11:59824089-59824161 | MS4A3;MS4A3;MS4A3 | | TSS200;TSS200;TSS200 |
| GM02149 | chr11:60048004-60048390 | MS4A4A | | TSS200 |
| GM09256 | chr11:60282420-60282742 | MS4A13;MS4A13 | N_Shore | TSS1500;TSS1500 |
| WG1513 | chr11:60282420-60284021 | MS4A13;MS4A13 | N_Shore | TSS1500;TSS1500 |
| WG1722 | chr11:60414615-60414767 | C11orf64 | N_Shore | Body |
| WG2121 | chr11:60534911-60534991 | MS4A15;MS4A15 | Island | 5'UTR;Body |
| GM02175 | chr11:60719947-60720229 | SLC15A3;SLC15A3 | S_Shore | TSS1500;TSS1500 |
| GM02706 | chr11:61062902-61063062 | VWCE | Island | TSS200 |
| WG2415 | chr11:61159602-61159687 | TMEM216 | N_Shore | TSS1500 |
| WG3466 | chr11:61159602-61159736 | TMEM216 | N_Shore | TSS1500 |
| GM02555 | chr11:61248435-61248556 | C11orf66;C11orf66 | | TSS200;TSS200 |
| WG2844 | chr11:61309985-61310204 | SYT7 | | Body |
| WG2844 | chr11:61449886-61450294 | DAGLA | S_Shore | 5'UTR |
| GM02706 | chr11:61544854-61545348 | C11orf9;C11orf9 | Island | Body;Body |
| GM00290 | chr11:61717225-61717684 | BEST1;BEST1 | | TSS200;TSS200 |
| WG3465 | chr11:62272972-62273418 | AHNAK | | Body |
| GM00935 | chr11:624908-625147 | MUPCDH;MUPCDH;MUPCDH | Island | 5'UTR;5'UTR;1stExon;1stExon |
| GM01392 | chr11:625023-625147 | MUPCDH;MUPCDH | Island | TSS200;TSS200 |
| WG1722 | chr11:62783361-62783394 | SLC22A8 | | TSS200 |
| WG3465 | chr11:628603-628677 | SCT | S_Shore | TSS1500 |
| WG3465 | chr11:6292311-6292511 | CCKBR | Island | Body |
| GM09256 | chr11:62996801-62997285 | SLC22A25 | | 1stExon |
| GM02706 | chr11:63136783-63137152 | SLC22A9 | | TSS1500 |
| WG1513 | chr11:63136783-63137555 | SLC22A9 | | TSS1500 |
| GM01650 | chr11:6342214-6342510 | PRKCDBP | S_Shore | TSS1500 |
| GM02706 | chr11:63655895-63656088 | MARK2;MARK2;MARK2;MARK2;MARK2 | | Body;TSS200;Body;Body;Body |
| WG1513 | chr11:63655895-63656090 | MARK2;MARK2;MARK2;MARK2;MARK2 | | Body;TSS200;Body;Body;Body |
| WG2844 | chr11:63655903-63656090 | MARK2;MARK2;MARK2;MARK2;MARK2 | | Body;TSS200;Body;Body;Body |
| WG3465 | chr11:63753020-63753203 | OTUB1;OTUB1 | N_Shore | TSS1500;TSS1500 |
| GM00520 | chr11:63753169-63753210 | OTUB1;OTUB1 | N_Shore | TSS200;TSS200 |
| GM09255 | chr11:63753169-63753228 | OTUB1;OTUB1 | N_Shore | TSS200;TSS200 |
| WG2193 | chr11:63775226-63775436 | MACROD1 | N_Shore | Body |
| GM02175 | chr11:63775226-63775612 | MACROD1 | N_Shore | Body |
| GM02555 | chr11:63775413-63775612 | MACROD1 | Island | Body |
| GM00940 | chr11:63784063-63784126 | MACROD1 | | Body |
| WG3465 | chr11:638076-638939 | DRD4 | Island | Body |
| GM01653 | chr11:63828569-63828713 | MACROD1 | | Body |
| WG1977 | chr11:638404-638507 | DRD4 | Island | Body |
| GM02706 | chr11:638404-638939 | DRD4 | Island | Body |
| WG3465 | chr11:63974772-63975026 | FERMT3;FERMT3 | N_Shore | 5'UTR;5'UTR |
| GM00439 | chr11:63974860-63975026 | FERMT3;FERMT3 | Island | Body;Body |
| WG3466 | chr11:64058550-64058878 | KCNK4 | N_Shore | TSS1500 |
| WG2415 | chr11:64064727-64065031 | KCNK4 | N_Shelf | Body |
| GM00439 | chr11:64081430-64081631 | ESRRA | N_Shelf | Body |
| GM02175 | chr11:64081545-64081631 | ESRRA | N_Shelf | Body |
| WG2415 | chr11:64108440-64108790 | CCDC88B | N_Shore | Body |

| | | | | |
|---|---|---|---|---|
| WG3465 | chr11:64116165-64116279 | CCDC88B | S_Shelf | Body |
| WG2276 | chr11:64397734-64398015 | NRXN2;NRXN2;NRXN2 | Island | Body;Body;Body |
| GM09256 | chr11:64405452-64405919 | NRXN2;NRXN2;NRXN2 | Island | Body;Body;Body |
| WG3465 | chr11:64428252-64428434 | NRXN2;NRXN2 | Island | Body;Body |
| WG2276 | chr11:64479015-64479496 | NRXN2;NRXN2 | Island | Body;Body |
| GM09256 | chr11:64599019-64599084 | CDC42BPG | S_Shore | Body |
| GM00289 | chr11:64981081-64981227 | SLC22A20 | | TSS1500 |
| GM00439 | chr11:64981125-64981297 | SLC22A20 | | TSS1500 |
| GM03075 | chr11:6518438-6518577 | DNHD1;DNHD1 | | TSS200;TSS200 |
| GM00520 | chr11:65560635-65560763 | OVOL1 | | Body |
| WG3465 | chr11:65647270-65647288 | CTSW | | TSS200 |
| GM00016 | chr11:6592066-6592585 | DNHD1 | Island | Body |
| GM00042 | chr11:66055170-66055327 | YIF1A | N_Shore | Body |
| GM02149 | chr11:66206894-66207047 | MRPL11;MRPL11;MRPL11 | | TSS1500;TSS1500;TSS1500 |
| WG1977 | chr11:66311334-66311422 | ZDHHC24 | N_Shore | Body |
| GM02706 | chr11:66650500-66650560 | PC;PC;PC | | 5'UTR;5'UTR;5'UTR |
| WG1977 | chr11:66823587-66823684 | RHOD | N_Shore | TSS1500 |
| WG2275 | chr11:66839085-66839191 | RHOD | Island | 3'UTR |
| WG3466 | chr11:66885240-66885281 | KDM2A;KDM2A | N_Shore | TSS1500;TSS1500 |
| GM00042 | chr11:67231878-67232316 | TMEM134;TMEM134;TMEM13 | N_Shore | 3'UTR;3'UTR;3'UTR |
| WG3466 | chr11:67232269-67232403 | TMEM134;TMEM134;TMEM13 | N_Shore | Body;Body;Body |
| GM01096 | chr11:67232269-67232511 | TMEM134;TMEM134;TMEM13 | N_Shore | Body;Body;Body |
| WG3465 | chr11:67373639-67373841 | NDUFV1;NDUFV1 | N_Shore | TSS1500;TSS1500 |
| WG3465 | chr11:67418045-67418113 | ACY3 | | 5'UTR |
| GM01393 | chr11:67418291-67418358 | ACY3 | | TSS200 |
| WG1513 | chr11:67432882-67433133 | ALDH3B2;ALDH3B2 | Island | Body;Body |
| WG3465 | chr11:67442067-67442249 | ALDH3B2;ALDH3B2;ALDH3B2 | | 1stExon;5'UTR;5'UTR |
| GM00016 | chr11:67810742-67811641 | TCIRG1;TCIRG1;TCIRG1 | | Body;1stExon;5'UTR |
| WG3465 | chr11:67810905-67811265 | TCIRG1;TCIRG1;TCIRG1 | | Body;1stExon;5'UTR |
| WG1513 | chr11:67811114-67811265 | TCIRG1;TCIRG1 | | Body;Body |
| WG1657 | chr11:67811194-67811641 | TCIRG1;TCIRG1 | | Body;Body |
| GM02555 | chr11:68138505-68138908 | LRP5 | | Body |
| WG2844 | chr11:68147880-68148079 | LRP5 | | Body |
| WG2276 | chr11:68579970-68580041 | CPT1A;CPT1A | | Body;Body |
| GM01096 | chr11:68610186-68610381 | CPT1A;CPT1A | N_Shore | TSS1500;TSS1500 |
| GM02706 | chr11:68610381-68611260 | CPT1A;CPT1A | N_Shore | TSS1500;TSS1500 |
| WG1722 | chr11:68610706-68611016 | CPT1A;CPT1A | N_Shore | TSS1500;TSS1500 |
| WG3466 | chr11:68610706-68611260 | CPT1A;CPT1A | N_Shore | TSS1500;TSS1500 |
| GM00520 | chr11:68610706-68611479 | CPT1A;CPT1A | N_Shore | TSS1500;TSS1500 |
| GM02706 | chr11:68781976-68782211 | MRGPRF;MRGPRF | S_Shelf | TSS1500;TSS1500 |
| WG1722 | chr11:68782049-68782202 | MRGPRF;MRGPRF | S_Shelf | TSS1500;TSS1500 |
| WG2275 | chr11:68782049-68782211 | MRGPRF;MRGPRF | S_Shelf | TSS1500;TSS1500 |
| WG1977 | chr11:68782106-68782211 | MRGPRF;MRGPRF | S_Shelf | TSS1500;TSS1500 |
| GM01393 | chr11:69516939-69517299 | FGF19 | Island | Body |
| WG2276 | chr11:69517471-69518003 | FGF19 | N_Shore | Body |
| GM00290 | chr11:69519974-69520418 | FGF19 | S_Shore | TSS1500 |
| WG2276 | chr11:69520186-69520418 | FGF19 | S_Shore | TSS1500 |

| | | | | |
|---|---|---|---|---|
| WG3465 | chr11:69934031-69934128 | ANO1;ANO1 | Island | Body;Body |
| WG1513 | chr11:70048796-70048845 | FADD | N_Shore | TSS1500 |
| GM09255 | chr11:70048796-70048852 | FADD | N_Shore | TSS1500 |
| WG3465 | chr11:70048801-70048852 | FADD | N_Shore | TSS1500 |
| GM00439 | chr11:70261100-70261373 | CTTN;CTTN | S_Shelf | Body;Body |
| WG1514 | chr11:7041870-7041987 | NLRP14;ZNF214;NLRP14 | Island | 1stExon;TSS1500;5'UTR |
| WG2415 | chr11:70458719-70458964 | SHANK2;SHANK2 | | Body;Body |
| GM00520 | chr11:70458964-70459053 | SHANK2;SHANK2 | | Body;Body |
| GM01392 | chr11:70559053-70559104 | SHANK2 | | Body |
| GM00438 | chr11:70563264-70563792 | SHANK2 | N_Shore | Body |
| WG1513 | chr11:70563580-70563792 | SHANK2 | N_Shore | Body |
| GM02149 | chr11:70565177-70565226 | SHANK2 | S_Shore | Body |
| GM02149 | chr11:70644110-70644185 | SHANK2 | | Body |
| GM02175 | chr11:70672835-70672878 | SHANK2 | Island | Body |
| GM00438 | chr11:70829968-70830058 | SHANK2 | | Body |
| GM00289 | chr11:70935949-70936560 | SHANK2 | | TSS200 |
| WG3465 | chr11:7110074-7110561 | RBMXL2 | N_Shore | TSS200 |
| WG1722 | chr11:7110148-7110196 | RBMXL2 | Island | TSS200 |
| GM02555 | chr11:71163428-71163555 | NADSYN1 | N_Shore | TSS1500 |
| GM00438 | chr11:71259251-71260323 | KRTAP5-9 | | TSS1500 |
| WG1722 | chr11:71709837-71709978 | IL18BP;IL18BP;IL18BP;IL18BP;IL18BP | | TSS200;TSS1500;TSS200;TSS1500;TS |
| WG3466 | chr11:71709888-71709978 | IL18BP;IL18BP;IL18BP;IL18BP;IL18BP | | TSS200;TSS1500;TSS200;TSS1500;TS |
| GM00289 | chr11:71710643-71710982 | IL18BP;IL18BP;IL18BP;IL18BP;IL18BP;IL | | TSS200;5'UTR;TSS1500;5'UTR;5'UTR;1 |
| WG1513 | chr11:71724287-71724432 | NUMA1 | | Body |
| GM09256 | chr11:71724287-71724583 | NUMA1 | | Body |
| WG3465 | chr11:71724432-71724677 | NUMA1 | | Body |
| GM00439 | chr11:71724504-71725590 | NUMA1 | | Body |
| WG1978 | chr11:71899966-71900411 | FOLR1;FOLR1;FOLR1 | | TSS1500;TSS1500;TSS1500 |
| GM02456 | chr11:71952131-71952431 | PHOX2A | Island | Body |
| GM01393 | chr11:72295425-72295716 | PDE2A;PDE2A;PDE2A;PDE2A | Island | Body;Body;Body;Body |
| WG2120 | chr11:72301235-72301724 | PDE2A;PDE2A;PDE2A;PDE2A | N_Shore | Body;Body;Body;Body |
| WG2844 | chr11:72408186-72408394 | ARAP1;ARAP1;ARAP1 | Island | Body;Body;Body |
| WG1722 | chr11:73356316-73356744 | PLEKHB1;PLEKHB1 | | TSS1500;TSS1500 |
| WG2276 | chr11:73498278-73498718 | MRPL48 | N_Shore | TSS1500 |
| GM01392 | chr11:73668626-73669290 | DNAJB13 | | Body |
| GM09256 | chr11:74022875-74022930 | P4HA3 | S_Shore | TSS200 |
| WG1722 | chr11:74178761-74178800 | KCNE3 | Island | TSS200 |
| GM01376 | chr11:74178795-74178883 | KCNE3 | Island | TSS200 |
| WG1977 | chr11:75139390-75139736 | KLHL35 | N_Shore | Body |
| GM00940 | chr11:75237636-75237739 | GDPD5 | Island | TSS1500 |
| WG2843 | chr11:75898132-75898203 | WNT11 | | Body |
| GM02706 | chr11:7597349-7597814 | PPFIBP2 | | Body |
| WG2844 | chr11:77299805-77299962 | AQP11 | N_Shore | TSS1500 |
| WG2415 | chr11:77907224-77907656 | USP35 | Island | 5'UTR |
| GM09256 | chr11:78673073-78673290 | ODZ4 | Island | Body |
| GM02149 | chr11:7870238-7870558 | OR5E1P | | TSS1500 |
| WG1513 | chr11:787971-788406 | CEND1 | N_Shore | 3'UTR |

| | | | | |
|---|---|---|---|---|
| GM09256 | chr11:790293-790795 | CEND1 | N_Shore | TSS200 |
| GM00289 | chr11:7985183-7985563 | NLRP10 | | TSS200 |
| WG1722 | chr11:818834-818917 | PNPLA2 | N_Shore | TSS200 |
| WG2415 | chr11:84634409-84634615 | DLG2;DLG2;DLG2 | | 1stExon;Body;5'UTR |
| WG1513 | chr11:84634581-84634620 | DLG2;DLG2 | | TSS200;Body |
| GM00289 | chr11:84634615-84634735 | DLG2;DLG2 | | TSS200;Body |
| GM09256 | chr11:85397109-85397684 | CCDC89 | | 1stExon |
| GM00439 | chr11:85397216-85397684 | CCDC89;CCDC89 | | 1stExon;5'UTR |
| GM00289 | chr11:85430070-85430657 | SYTL2;SYTL2;SYTL2;SYTL2;SYTL2;SYTL2; | | 5'UTR;Body;5'UTR;1stExon;Body;1stE |
| WG1513 | chr11:85430629-85430657 | SYTL2;SYTL2;SYTL2;SYTL2;SYTL2;SYTL2; | | Body;TSS1500;TSS1500;Body;TSS150 |
| GM09255 | chr11:859532-859873 | TSPAN4;TSPAN4;TSPAN4;TSP | S_Shore | 5'UTR;Body;Body;Body;Body;Body;Bo |
| GM02706 | chr11:859915-860157 | TSPAN4;TSPAN4;TSPAN4;TSP | S_Shore | 5'UTR;Body;Body;Body;Body;Body;Bo |
| WG2276 | chr11:86383696-86383723 | ME3;ME3;ME3 | S_Shore | TSS1500;TSS200;TSS1500 |
| WG2415 | chr11:86383696-86383761 | ME3;ME3;ME3 | S_Shore | TSS1500;TSS200;TSS1500 |
| WG3466 | chr11:8680333-8680469 | TRIM66;TRIM66 | | 5'UTR;1stExon |
| GM00520 | chr11:8680333-8680517 | TRIM66;TRIM66 | | 5'UTR;1stExon |
| WG3465 | chr11:8680333-8680729 | TRIM66;TRIM66 | | 5'UTR;1stExon |
| WG1283 | chr11:87908783-87908817 | RAB38 | S_Shore | TSS200 |
| WG1722 | chr11:87908785-87908938 | RAB38 | S_Shore | TSS200 |
| WG1513 | chr11:87908805-87908938 | RAB38 | S_Shore | TSS1500 |
| WG3466 | chr11:87908817-87908938 | RAB38 | S_Shore | TSS1500 |
| WG3465 | chr11:88242253-88242604 | GRM5;GRM5 | Island | Body;Body |
| GM02173 | chr11:8832956-8833095 | ST5;ST5;ST5 | | TSS1500;TSS200;5'UTR |
| WG2415 | chr11:915170-915239 | CHID1 | S_Shelf | TSS200 |
| WG1513 | chr11:92615932-92616378 | FAT3 | Island | Body |
| WG2415 | chr11:93277161-93277191 | C11orf75 | S_Shore | TSS1500 |
| GM01392 | chr11:93394605-93394624 | KIAA1731 | N_Shore | TSS1500 |
| WG2844 | chr11:94278413-94279068 | FUT4 | Island | 1stExon |
| GM02149 | chr11:94278457-94278912 | FUT4 | Island | 1stExon |
| WG2276 | chr11:94278457-94279068 | FUT4 | Island | 1stExon |
| GM02704 | chr11:94278581-94278603 | FUT4 | Island | 1stExon |
| GM03073 | chr11:94278581-94278912 | FUT4 | Island | 1stExon |
| GM00941 | chr11:94278595-94278912 | FUT4 | Island | 1stExon |
| GM02706 | chr11:94965364-94965610 | SESN3 | Island | TSS1500 |
| GM02706 | chr11:9684886-9685299 | SWAP70 | N_Shore | TSS1500 |
| WG1310 | chr12:10084936-10085174 | CLEC2A;CLEC2A | | 5'UTR;1stExon |
| GM02641 | chr12:101988604-101988803 | MYBPC1;MYBPC1;MYBPC1;MYBPC1 | | TSS200;TSS200;TSS200;TSS200 |
| WG2121 | chr12:102036256-102036400 | MYBPC1;MYBPC1;MYBPC1;M | Island | Body;Body;Body;Body |
| WG3466 | chr12:102036309-102036563 | MYBPC1;MYBPC1;MYBPC1;M | Island | Body;Body;Body;Body |
| GM03073 | chr12:10251536-10251638 | CLEC1A;CLEC1A | | 5'UTR;1stExon |
| WG3465 | chr12:10251536-10251879 | CLEC1A;CLEC1A | | 5'UTR;1stExon |
| GM00438 | chr12:103980767-103981113 | STAB2 | | TSS1500 |
| GM00520 | chr12:104609476-104609797 | TXNRD1 | Island | TSS200 |
| WG3465 | chr12:106988340-106988413 | RFX4 | | Body |
| WG1722 | chr12:107348689-107348855 | C12orf23 | N_Shore | TSS1500 |
| WG2844 | chr12:107348689-107348944 | C12orf23 | N_Shore | TSS1500 |
| WG1977 | chr12:107348824-107348944 | C12orf23 | N_Shore | TSS1500 |

| | | | | |
|---|---|---|---|---|
| GM00016 | chr12:108039569-108039856 | BTBD11;BTBD11 | | Body;Body |
| GM00438 | chr12:108603943-108604021 | WSCD2 | | Body |
| GM00016 | chr12:109181748-109182032 | SSH1 | | 3'UTR |
| GM00940 | chr12:110156245-110156459 | C12orf34 | N_Shore | 5'UTR |
| WG1977 | chr12:110172208-110172346 | MGC14436;MGC14436;C12orf34 | | Body;Body;5'UTR |
| GM02175 | chr12:110211413-110211701 | MGC14436;MGC14436 | | TSS200;TSS200 |
| GM00935 | chr12:111284228-111284585 | CCDC63 | | TSS1500 |
| GM00438 | chr12:111358470-111358601 | MYL2 | | TSS200 |
| WG1977 | chr12:11151478-11151622 | PRR4;TAS2R20;PRH1 | | Body;TSS1500;5'UTR |
| GM09256 | chr12:111664582-111664723 | CUX2 | | Body |
| GM01392 | chr12:113229584-113229647 | RPH3A;RPH3A;RPH3A;RPH3A | | 5'UTR;5'UTR;1stExon;1stExon |
| GM02706 | chr12:113515332-113515969 | DTX1 | Island | Body |
| WG2415 | chr12:113587240-113587340 | CCDC42B | N_Shelf | TSS1500 |
| WG3466 | chr12:114312919-114313256 | RBM19;RBM19;RBM19 | | Body;Body;Body |
| GM09256 | chr12:114833303-114834201 | TBX5;TBX5;TBX5;TBX5 | N_Shore | Body;Body;Body;Body |
| WG1978 | chr12:114833837-114833950 | TBX5;TBX5;TBX5;TBX5 | N_Shore | Body;Body;Body;Body |
| WG3465 | chr12:114833837-114834201 | TBX5;TBX5;TBX5;TBX5 | N_Shore | Body;Body;Body;Body |
| WG2121 | chr12:114833837-114834713 | TBX5;TBX5;TBX5;TBX5 | N_Shore | Body;Body;Body;Body |
| GM01393 | chr12:114833912-114834201 | TBX5;TBX5;TBX5;TBX5 | Island | Body;Body;Body;Body |
| WG2193 | chr12:114837572-114837887 | TBX5;TBX5;TBX5;TBX5 | N_Shore | Body;Body;Body;Body |
| WG2121 | chr12:114837887-114838772 | TBX5;TBX5;TBX5;TBX5 | N_Shore | Body;Body;Body;Body |
| GM01096 | chr12:114837887-114838825 | TBX5;TBX5;TBX5;TBX5 | N_Shore | Body;Body;Body;Body |
| WG2276 | chr12:114838359-114838772 | TBX5;TBX5;TBX5;TBX5 | Island | Body;Body;Body;Body |
| GM02704 | chr12:114840854-114842031 | TBX5;TBX5;TBX5;TBX5 | Island | Body;Body;5'UTR;Body |
| WG2276 | chr12:114841202-114841708 | TBX5;TBX5;TBX5;TBX5 | N_Shore | Body;Body;5'UTR;Body |
| WG1721 | chr12:114841536-114841708 | TBX5;TBX5;TBX5;TBX5 | N_Shore | Body;Body;5'UTR;Body |
| WG2276 | chr12:114841870-114842700 | TBX5;TBX5;TBX5;TBX5 | N_Shore | 5'UTR;5'UTR;TSS200;5'UTR |
| WG2276 | chr12:114843583-114846431 | TBX5;TBX5;TBX5;TBX5 | Island | 1stExon;5'UTR;5'UTR;5'UTR |
| WG2121 | chr12:114845180-114846347 | TBX5;TBX5;TBX5 | N_Shore | TSS1500;5'UTR;5'UTR |
| GM02641 | chr12:114845180-114846399 | TBX5;TBX5;TBX5 | N_Shore | TSS1500;5'UTR;5'UTR |
| GM02640 | chr12:114845180-114846431 | TBX5;TBX5;TBX5 | N_Shore | TSS1500;5'UTR;5'UTR |
| WG3466 | chr12:114845180-114846503 | TBX5;TBX5;TBX5 | N_Shore | TSS1500;5'UTR;5'UTR |
| WG2120 | chr12:114846321-114846431 | TBX5;TBX5 | Island | TSS200;TSS200 |
| GM01096 | chr12:114846968-114847438 | TBX5;TBX5 | Island | TSS1500;TSS1500 |
| WG3466 | chr12:114847438-114847578 | TBX5;TBX5 | Island | TSS1500;TSS1500 |
| WG1513 | chr12:115109452-115109934 | TBX3;TBX3 | N_Shore | 3'UTR;3'UTR |
| GM09256 | chr12:115109504-115109934 | TBX3;TBX3 | Island | 3'UTR;3'UTR |
| WG2844 | chr12:116996773-116997185 | MAP1LC3B2 | | TSS1500 |
| GM02456 | chr12:116996931-116997185 | MAP1LC3B2 | | TSS1500 |
| WG1722 | chr12:116997022-116997072 | MAP1LC3B2 | | TSS200 |
| GM00439 | chr12:116997022-116997079 | MAP1LC3B2 | | TSS200 |
| WG3466 | chr12:116997069-116997079 | MAP1LC3B2 | | TSS200 |
| WG1977 | chr12:116997069-116997095 | MAP1LC3B2 | | TSS200 |
| WG1513 | chr12:116997079-116997185 | MAP1LC3B2 | | TSS200 |
| WG2844 | chr12:117155382-117155636 | C12orf49 | | 3'UTR |
| GM00935 | chr12:117407879-117408122 | FBXW8;FBXW8 | | Body;Body |
| GM09256 | chr12:117407879-117408163 | FBXW8;FBXW8 | | Body;Body |

| | | | | |
|---|---|---|---|---|
| WG2843 | chr12:117407970-117408163 | FBXW8;FBXW8 | | Body;Body |
| WG1722 | chr12:117536182-117536531 | TESC;TESC;TESC | N_Shore | Body;Body;Body |
| GM00289 | chr12:117797056-117797635 | NOS1 | N_Shore | 5'UTR |
| GM02175 | chr12:119591664-119592094 | SRRM4 | N_Shore | Body |
| GM00438 | chr12:122019006-122019076 | KDM2B;KDM2B | S_Shore | TSS200;TSS1500 |
| GM02706 | chr12:122019006-122019760 | KDM2B;KDM2B | S_Shore | TSS200;TSS1500 |
| GM00042 | chr12:122019076-122019117 | KDM2B;KDM2B | S_Shore | TSS200;TSS1500 |
| WG2844 | chr12:122297080-122297408 | HPD | | TSS1500 |
| GM00439 | chr12:122356033-122356390 | WDR66 | N_Shore | TSS1500 |
| WG1721 | chr12:122356033-122356852 | WDR66 | N_Shore | TSS1500 |
| GM02316 | chr12:122356056-122356390 | WDR66 | N_Shore | TSS1500 |
| GM01393 | chr12:122356056-122356400 | WDR66 | N_Shore | TSS1500 |
| WG1658 | chr12:122356056-122356598 | WDR66 | N_Shore | TSS1500 |
| GM01376 | chr12:122356056-122356781 | WDR66 | N_Shore | TSS1500 |
| WG2194 | chr12:122356056-122356852 | WDR66 | N_Shore | TSS1500 |
| WG2414 | chr12:122356316-122356400 | WDR66 | Island | TSS200 |
| WG2844 | chr12:122356316-122356598 | WDR66 | Island | TSS200 |
| GM00935 | chr12:122356316-122356852 | WDR66 | Island | TSS200 |
| GM02317 | chr12:122356390-122356598 | WDR66 | Island | TSS200 |
| GM01392 | chr12:123215010-123215288 | GPR81;GPR81 | | 1stExon;5'UTR |
| GM00940 | chr12:123215248-123215308 | GPR81 | | TSS200 |
| WG3465 | chr12:123215301-123215471 | GPR81 | | TSS200 |
| GM02175 | chr12:123215301-123215524 | GPR81 | | TSS200 |
| WG3466 | chr12:123380870-123380891 | VPS37B | Island | TSS200 |
| GM02706 | chr12:123380878-123380891 | VPS37B | Island | TSS200 |
| GM02706 | chr12:123753186-123753272 | CDK2AP1 | N_Shore | Body |
| WG1309 | chr12:123757474-123757860 | CDK2AP1 | S_Shore | TSS1500 |
| WG1722 | chr12:124773668-124773870 | FAM101A | | TSS200 |
| GM00438 | chr12:124773668-124774055 | FAM101A | | TSS200 |
| WG2276 | chr12:124773668-124774078 | FAM101A | | TSS200 |
| GM02704 | chr12:124773870-124774055 | FAM101A;FAM101A | | 1stExon;5'UTR |
| WG1658 | chr12:124773870-124774078 | FAM101A;FAM101A | | 1stExon;5'UTR |
| WG3465 | chr12:124773995-124774078 | FAM101A | | 5'UTR |
| WG2276 | chr12:124870112-124870323 | NCOR2;NCOR2 | N_Shelf | Body;Body |
| GM02456 | chr12:124876433-124876650 | NCOR2;NCOR2 | S_Shelf | Body;Body |
| WG3466 | chr12:124880022-124880260 | NCOR2;NCOR2 | | Body;Body |
| GM01096 | chr12:124942016-124942248 | NCOR2;NCOR2 | S_Shore | Body;Body |
| GM00042 | chr12:124950720-124950990 | NCOR2;NCOR2 | Island | Body;Body |
| WG3466 | chr12:125333267-125333358 | SCARB1;SCARB1 | | Body;Body |
| GM02706 | chr12:126926510-126926925 | LOC100128554 | | TSS1500 |
| WG2193 | chr12:129299088-129299462 | SLC15A4;MGC16384 | | Body;Body |
| GM01392 | chr12:129309156-129309330 | SLC15A4 | S_Shore | TSS1500 |
| WG2276 | chr12:12938082-12938425 | APOLD1;APOLD1 | N_Shore | Body;TSS1500 |
| GM02149 | chr12:129417623-129417818 | GLT1D1 | | Body |
| GM02149 | chr12:130184740-130185016 | TMEM132D | Island | Body |
| WG1513 | chr12:130195357-130195777 | TMEM132D | | Body |
| WG1513 | chr12:130316015-130316058 | TMEM132D | | Body |

| | | | | |
|---|---|---|---|---|
| GM02706 | chr12:130821530-130822674 | PIWIL1 | N_Shore | TSS1500 |
| WG1984 | chr12:130821962-130822674 | PIWIL1 | N_Shore | TSS1500 |
| WG2843 | chr12:130822603-130822674 | PIWIL1 | Island | TSS200 |
| GM02175 | chr12:130823838-130824082 | PIWIL1 | Island | 5'UTR |
| WG1513 | chr12:130958340-130958790 | RIMBP2 | | Body |
| WG1513 | chr12:131002386-131002918 | RIMBP2;RIMBP2 | | 5'UTR;1stExon |
| WG1722 | chr12:131303194-131303518 | STX2;STX2 | Island | Body;Body |
| WG3465 | chr12:131303194-131303621 | STX2;STX2 | Island | Body;Body |
| GM02706 | chr12:131303194-131303774 | STX2;STX2 | Island | Body;Body |
| GM02456 | chr12:131303247-131303518 | STX2;STX2 | Island | Body;Body |
| GM01096 | chr12:131303247-131303621 | STX2;STX2 | Island | Body;Body |
| WG2415 | chr12:131303478-131303993 | STX2;STX2 | Island | Body;Body |
| WG1513 | chr12:131355340-131355465 | RAN | N_Shore | TSS1500 |
| WG2415 | chr12:131355424-131355834 | RAN | N_Shore | TSS1500 |
| GM00439 | chr12:131561080-131561270 | GPR133 | | Body |
| WG2844 | chr12:131622284-131623508 | GPR133 | N_Shore | Body |
| WG2276 | chr12:131622584-131623340 | GPR133 | N_Shore | Body |
| GM00289 | chr12:132269643-132270218 | SFRS8 | | Body |
| GM00935 | chr12:132278430-132278510 | SFRS8 | S_Shore | Body |
| GM03073 | chr12:132689907-132690196 | GALNT9;GALNT9 | Island | Body;5'UTR |
| GM00022 | chr12:132690570-132690651 | GALNT9;GALNT9;GALNT9 | Island | Body;1stExon;5'UTR |
| GM09255 | chr12:132690643-132690692 | GALNT9;GALNT9 | S_Shore | Body;TSS200 |
| GM01096 | chr12:132696035-132696192 | GALNT9 | N_Shore | Body |
| WG1513 | chr12:132698423-132698620 | GALNT9 | S_Shore | Body |
| GM02706 | chr12:132703301-132703598 | GALNT9 | | Body |
| GM09255 | chr12:132845336-132845530 | GALNT9 | N_Shore | Body |
| GM01392 | chr12:132853954-132854189 | GALNT9;LOC100130238 | Island | Body;Body |
| GM00520 | chr12:132896199-132896405 | GALNT9 | N_Shore | Body |
| GM00289 | chr12:132896743-132896939 | GALNT9 | S_Shore | Body |
| WG3465 | chr12:132897371-132897840 | GALNT9 | S_Shore | Body |
| WG1721 | chr12:132900274-132900480 | GALNT9 | Island | Body |
| WG1977 | chr12:132900274-132900620 | GALNT9 | Island | Body |
| WG1513 | chr12:133065724-133065776 | FBRSL1 | N_Shore | TSS1500 |
| GM02706 | chr12:133065806-133065939 | FBRSL1 | N_Shore | TSS1500 |
| GM00935 | chr12:133135361-133135698 | FBRSL1 | Island | Body |
| WG1977 | chr12:133159472-133159939 | FBRSL1 | Island | Body |
| WG2844 | chr12:133380384-133380627 | GOLGA3 | S_Shore | Body |
| GM00940 | chr12:133464892-133465013 | CHFR;CHFR;CHFR;CHFR;CH | S_Shore | TSS1500;TSS1500;TSS1500;TSS1500; |
| WG1513 | chr12:14109211-14109584 | GRIN2B | | 5'UTR |
| GM01096 | chr12:14720938-14720974 | PLBD1 | Island | TSS200 |
| WG1513 | chr12:14849394-14849579 | GUCY2C;GUCY2C | | 1stExon;5'UTR |
| GM02555 | chr12:15039025-15039432 | MGP | | TSS200 |
| WG2414 | chr12:15114233-15114393 | ARHGDIB | | 5'UTR |
| GM00520 | chr12:15114233-15114732 | ARHGDIB | | 5'UTR |
| GM02456 | chr12:15114494-15114710 | ARHGDIB;ARHGDIB | | 5'UTR;1stExon |
| GM01096 | chr12:15114494-15114732 | ARHGDIB;ARHGDIB | | 5'UTR;1stExon |
| WG3465 | chr12:15114703-15114732 | ARHGDIB | | TSS200 |

| | | | | |
|---|---|---|---|---|
| GM02175 | chr12:15114703-15114904 | ARHGDIB | | TSS200 |
| WG3465 | chr12:1609245-1609369 | LOC100292680 | | TSS1500 |
| WG1978 | chr12:1609245-1609470 | LOC100292680 | | TSS1500 |
| WG1983 | chr12:1609346-1609470 | LOC100292680 | | TSS1500 |
| GM02456 | chr12:1609369-1609470 | LOC100292680 | | TSS1500 |
| WG2121 | chr12:16757954-16758047 | LMO3;LMO3 | | 5'UTR;5'UTR |
| WG1977 | chr12:16757985-16758057 | LMO3;LMO3 | | 5'UTR;5'UTR |
| GM00438 | chr12:213776-214224 | IQSEC3;IQSEC3 | Island | Body;5'UTR |
| GM02175 | chr12:214000-214224 | IQSEC3;IQSEC3 | Island | Body;5'UTR |
| GM09255 | chr12:22777409-22777465 | ETNK1;ETNK1 | N_Shore | TSS1500;TSS1500 |
| WG2844 | chr12:2339439-2339614 | CACNA1C;CACNA1C;CACNA | Island | Body;Body;Body;Body;Body;Body;Bod |
| GM01376 | chr12:23998997-23999045 | SOX5;SOX5 | | Body;Body |
| WG2276 | chr12:24102581-24102832 | SOX5;SOX5;SOX5 | | 5'UTR;1stExon;5'UTR |
| GM01095 | chr12:25054873-25055108 | BCAT1 | N_Shore | Body |
| WG1310 | chr12:25054873-25055304 | BCAT1 | N_Shore | Body |
| GM09256 | chr12:25054873-25055381 | BCAT1 | N_Shore | Body |
| WG3466 | chr12:25054873-25055421 | BCAT1 | N_Shore | Body |
| GM02706 | chr12:25054873-25055518 | BCAT1 | N_Shore | Body |
| WG2414 | chr12:25054905-25055381 | BCAT1 | N_Shore | Body |
| WG1722 | chr12:25055108-25055304 | BCAT1 | N_Shore | Body |
| GM00439 | chr12:25055108-25055381 | BCAT1 | N_Shore | Body |
| WG2194 | chr12:25055108-25055421 | BCAT1 | N_Shore | Body |
| WG2121 | chr12:25055214-25055304 | BCAT1 | N_Shore | Body |
| GM00935 | chr12:25055214-25055381 | BCAT1 | N_Shore | Body |
| WG1721 | chr12:25055262-25055381 | BCAT1 | N_Shore | Body |
| WG1657 | chr12:25055262-25055421 | BCAT1 | N_Shore | Body |
| WG1513 | chr12:25055304-25055421 | BCAT1 | N_Shore | Body |
| WG2844 | chr12:25150530-25150626 | C12orf77 | | TSS200 |
| WG1722 | chr12:25204113-25204649 | LRMP | | TSS1500 |
| GM02555 | chr12:25801455-25801621 | IFLTD1 | | 1stExon |
| WG1977 | chr12:258355-258436 | IQSEC3;IQSEC3 | | Body;Body |
| WG2844 | chr12:2800182-2800755 | CACNA1C;CACNA1C;CACNA | Island | Body;Body;Body;Body;Body;Body;Bod |
| WG3465 | chr12:2800382-2800919 | CACNA1C;CACNA1C;CACNA | Island | 3'UTR;3'UTR;3'UTR;3'UTR;3'UTR;3'UTF |
| WG1722 | chr12:2800463-2800500 | CACNA1C;CACNA1C;CACNA | Island | 3'UTR;3'UTR;3'UTR;3'UTR;3'UTR;3'UTF |
| GM00438 | chr12:2800463-2800919 | CACNA1C;CACNA1C;CACNA | Island | 3'UTR;3'UTR;3'UTR;3'UTR;3'UTR;3'UTF |
| GM01392 | chr12:2800755-2800919 | CACNA1C;CACNA1C;CACNA | Island | 3'UTR;3'UTR;3'UTR;3'UTR;3'UTR;3'UTF |
| GM00935 | chr12:2800755-2801158 | CACNA1C;CACNA1C;CACNA | Island | 3'UTR;3'UTR;3'UTR;3'UTR;3'UTR;3'UTF |
| WG2844 | chr12:2800919-2801158 | CACNA1C;CACNA1C;CACNA | Island | 3'UTR;3'UTR;3'UTR;3'UTR;3'UTR;3'UTF |
| WG2844 | chr12:28124011-28124233 | PTHLH;PTHLH;PTHLH;PTHLH | S_Shore | TSS1500;5'UTR;5'UTR;TSS1500 |
| GM00438 | chr12:29376421-29376779 | FAR2 | | TSS200 |
| GM09256 | chr12:29376483-29376872 | FAR2 | | TSS200 |
| WG3466 | chr12:29650767-29650954 | OVCH1 | | TSS200 |
| WG1978 | chr12:312592-313104 | SLC6A12;SLC6A12;SLC6A12 | Island | Body;Body;Body |
| GM02175 | chr12:322587-322659 | SLC6A12;SLC6A12;SLC6A12 | | 5'UTR;TSS200;5'UTR |
| GM00439 | chr12:322587-322831 | SLC6A12;SLC6A12;SLC6A12 | | 5'UTR;TSS200;5'UTR |
| GM00520 | chr12:322641-322659 | SLC6A12;SLC6A12;SLC6A12 | | 5'UTR;TSS200;5'UTR |
| GM02706 | chr12:322649-322831 | SLC6A12;SLC6A12;SLC6A12 | | 5'UTR;TSS200;5'UTR |

| | | | | |
|---|---|---|---|---|
| WG2844 | chr12:32654992-32655192 | FGD4 | | TSS200 |
| WG2276 | chr12:3862221-3862597 | EFCAB4B;EFCAB4B;EFCAB4E | Island | 1stExon;1stExon;5'UTR;1stExon;5'UTR |
| WG2276 | chr12:4381788-4381800 | CCND2 | Island | TSS1500 |
| WG1513 | chr12:43946493-43946880 | ADAMTS20 | S_Shore | TSS1500 |
| WG2275 | chr12:44152509-44152632 | PUS7L;PUS7L;PUS7L;IRAK4;I | Island | 1stExon;5'UTR;1stExon;TSS1500;TSS1 |
| GM02704 | chr12:44152565-44152632 | PUS7L;PUS7L;PUS7L;IRAK4;I | Island | 1stExon;5'UTR;TSS200;TSS200;TSS20 |
| WG2275 | chr12:44152807-44153217 | IRAK4;PUS7L;IRAK4;IRAK4;PU | Island | 5'UTR;TSS1500;5'UTR;1stExon;TSS15( |
| WG2415 | chr12:45443808-45443996 | DBX2 | N_Shore | Body |
| WG1722 | chr12:46767665-46767747 | SLC38A2 | S_Shore | TSS1500 |
| WG2276 | chr12:46767665-46768016 | SLC38A2 | S_Shore | TSS1500 |
| GM01096 | chr12:47219705-47219797 | SLC38A4;SLC38A4;SLC38A4;SLC38A4 | | 1stExon;5'UTR;1stExon;5'UTR |
| GM09255 | chr12:47219737-47219797 | SLC38A4;SLC38A4;SLC38A4;SLC38A4 | | 1stExon;5'UTR;1stExon;5'UTR |
| WG1978 | chr12:47219797-47219841 | SLC38A4;SLC38A4 | | TSS200;TSS200 |
| GM00438 | chr12:47219797-47219877 | SLC38A4;SLC38A4 | | TSS200;TSS200 |
| WG1978 | chr12:47219920-47219958 | SLC38A4;SLC38A4 | | TSS200;TSS200 |
| WG2415 | chr12:47219920-47220092 | SLC38A4;SLC38A4 | | TSS200;TSS200 |
| GM00940 | chr12:47219956-47220092 | SLC38A4;SLC38A4 | | TSS200;TSS200 |
| WG1513 | chr12:48722990-48723324 | H1FNT;H1FNT | N_Shore | 5'UTR;1stExon |
| GM01096 | chr12:49504967-49505053 | LMBR1L | S_Shore | TSS1500 |
| GM00289 | chr12:49740942-49740976 | DNAJC22 | S_Shore | TSS200 |
| GM02706 | chr12:50349307-50349522 | AQP2 | Island | Body |
| WG2415 | chr12:51786038-51786547 | GALNT6 | S_Shore | TSS1500 |
| GM03075 | chr12:52214320-52215216 | FIGNL2 | Island | 1stExon |
| GM09256 | chr12:52215034-52215216 | FIGNL2 | Island | 1stExon |
| GM00042 | chr12:52281674-52281997 | ANKRD33;ANKRD33 | | TSS200;TSS200 |
| WG2415 | chr12:52281810-52281997 | ANKRD33;ANKRD33;ANKRD33;ANKRD33 | | 1stExon;5'UTR;1stExon;5'UTR |
| WG1722 | chr12:52281810-52282079 | ANKRD33;ANKRD33;ANKRD33;ANKRD33 | | 1stExon;5'UTR;1stExon;5'UTR |
| GM00520 | chr12:52585844-52585892 | KRT80;KRT80 | | TSS200;TSS200 |
| GM02149 | chr12:52994933-52995295 | KRT72;KRT72;KRT72 | Island | 1stExon;1stExon;1stExon |
| WG2415 | chr12:53183773-53184086 | KRT3 | Island | 3'UTR |
| WG2276 | chr12:53448085-53448180 | TENC1;TENC1;TENC1 | Island | Body;Body;Body |
| WG1977 | chr12:53591490-53591766 | ITGB7 | Island | Body |
| WG3465 | chr12:53729629-53730072 | SP7 | | TSS200 |
| GM00941 | chr12:53729629-53730163 | SP7 | | TSS200 |
| WG1722 | chr12:53730072-53730277 | SP7 | | TSS1500 |
| GM02706 | chr12:54360262-54360511 | HOTAIR | N_Shore | Body |
| GM00042 | chr12:54360460-54360836 | HOTAIR | Island | Body |
| GM09255 | chr12:54365966-54366434 | HOXC11 | N_Shore | TSS1500 |
| GM02706 | chr12:54369987-54370458 | HOXC11 | S_Shore | 3'UTR |
| GM00439 | chr12:54383639-54383692 | HOXC10 | S_Shelf | 3'UTR |
| GM00016 | chr12:54385436-54385526 | MIR196A2 | N_Shelf | TSS200 |
| WG2415 | chr12:54392734-54392884 | HOXC9 | N_Shore | TSS1500 |
| WG1722 | chr12:54392734-54393100 | HOXC9 | N_Shore | TSS1500 |
| GM02149 | chr12:54392867-54392988 | HOXC9 | N_Shore | TSS1500 |
| GM00438 | chr12:54392867-54393100 | HOXC9 | N_Shore | TSS1500 |
| GM00438 | chr12:54409229-54409657 | HOXC4;HOXC6;HOXC5 | S_Shore | TSS1500;TSS1500;TSS1500 |
| GM00042 | chr12:54409257-54409525 | HOXC4;HOXC6;HOXC5 | S_Shore | TSS1500;TSS1500;TSS1500 |

| | | | | |
|---|---|---|---|---|
| WG1513 | chr12:54409257-54409599 | HOXC4;HOXC6;HOXC5 | S_Shore | TSS1500;TSS1500;TSS1500 |
| WG2843 | chr12:54409485-54409525 | HOXC4;HOXC6;HOXC5 | S_Shore | TSS1500;TSS1500;TSS1500 |
| GM02173 | chr12:54409491-54409599 | HOXC4;HOXC6;HOXC5 | S_Shore | TSS1500;TSS1500;TSS1500 |
| WG1283 | chr12:54412269-54412565 | HOXC4;HOXC5;HOXC6 | N_Shore | 5'UTR;Body;5'UTR |
| GM02149 | chr12:54413000-54413345 | HOXC4;HOXC5;HOXC6 | Island | 5'UTR;Body;5'UTR |
| WG2121 | chr12:54413000-54413384 | HOXC4;HOXC5;HOXC6 | Island | 5'UTR;Body;5'UTR |
| GM02173 | chr12:54420837-54421335 | HOXC4;HOXC5;HOXC6;HOX | N_Shelf | 5'UTR;Body;TSS1500;5'UTR |
| GM00439 | chr12:54423428-54423566 | HOXC4;HOXC6;HOXC5;HOX | Island | 5'UTR;Body;Body;Body |
| GM09256 | chr12:54423428-54423920 | HOXC4;HOXC6;HOXC5;HOX | Island | 5'UTR;Body;Body;Body |
| GM00289 | chr12:54423428-54424085 | HOXC4;HOXC6;HOXC5;HOX | Island | 5'UTR;Body;Body;Body |
| WG2843 | chr12:54423428-54424679 | HOXC4;HOXC6;HOXC5;HOX | Island | 5'UTR;Body;Body;Body |
| GM00031 | chr12:54423428-54424964 | HOXC4;HOXC6;HOXC5;HOX | Island | 5'UTR;Body;Body;Body |
| WG3466 | chr12:54423428-54425418 | HOXC4;HOXC6;HOXC5;HOX | Island | 5'UTR;Body;Body;Body |
| WG1721 | chr12:54423549-54423920 | HOXC4;HOXC6;HOXC5;HOX | Island | 5'UTR;Body;Body;Body |
| GM00438 | chr12:54423549-54424203 | HOXC4;HOXC6;HOXC5;HOX | Island | 5'UTR;Body;Body;Body |
| GM02555 | chr12:54423920-54424203 | HOXC4;HOXC5;HOXC6;HOX | N_Shore | 5'UTR;Body;3'UTR;3'UTR |
| GM02175 | chr12:54424679-54425156 | HOXC4;HOXC5 | Island | 5'UTR;Body |
| GM00438 | chr12:54424902-54425156 | HOXC4;HOXC5 | Island | 5'UTR;Body |
| GM00935 | chr12:54424902-54425418 | HOXC4;HOXC5 | Island | 5'UTR;Body |
| WG2843 | chr12:54424964-54425418 | HOXC4;HOXC5 | Island | 5'UTR;Body |
| WG2121 | chr12:54440773-54441509 | HOXC4 | Island | 5'UTR |
| WG3466 | chr12:54440806-54441213 | HOXC4 | Island | 5'UTR |
| GM02640 | chr12:54441213-54441509 | HOXC4 | Island | 5'UTR |
| GM01096 | chr12:54444985-54446537 | HOXC4 | N_Shore | 5'UTR |
| WG1978 | chr12:54444985-54446576 | HOXC4 | N_Shore | 5'UTR |
| WG2415 | chr12:54446019-54446100 | HOXC4 | Island | 5'UTR |
| WG3465 | chr12:54446019-54446253 | HOXC4 | Island | 5'UTR |
| WG3466 | chr12:54446019-54446308 | HOXC4 | Island | 5'UTR |
| GM09255 | chr12:54446019-54446505 | HOXC4 | Island | 5'UTR |
| GM01095 | chr12:54446019-54446537 | HOXC4 | Island | 5'UTR |
| WG1977 | chr12:54446019-54446576 | HOXC4 | Island | 5'UTR |
| GM02316 | chr12:54446308-54446576 | HOXC4;HOXC4 | N_Shore | 5'UTR;TSS1500 |
| GM00438 | chr12:54446308-54446996 | HOXC4;HOXC4 | N_Shore | 5'UTR;TSS1500 |
| WG1283 | chr12:54446505-54446576 | HOXC4;HOXC4 | N_Shore | 5'UTR;TSS1500 |
| WG3466 | chr12:54446944-54448913 | HOXC4;HOXC4 | N_Shore | 5'UTR;TSS1500 |
| GM01376 | chr12:54447136-54447283 | HOXC4;HOXC4 | N_Shore | 5'UTR;TSS1500 |
| GM01376 | chr12:54447349-54447584 | HOXC4;HOXC4 | N_Shore | 5'UTR;TSS1500 |
| GM01376 | chr12:54448090-54448769 | HOXC4;HOXC4 | Island | 1stExon;Body |
| GM02555 | chr12:54448090-54448913 | HOXC4;HOXC4 | Island | 1stExon;Body |
| WG2276 | chr12:54448265-54448913 | HOXC4;HOXC4 | S_Shore | Body;Body |
| WG2415 | chr12:54448729-54448913 | HOXC4;HOXC4 | S_Shore | Body;Body |
| WG1977 | chr12:54454163-54454445 | FLJ12825 | N_Shore | Body |
| GM02704 | chr12:54454250-54454445 | FLJ12825 | Island | Body |
| GM02640 | chr12:54454252-54454445 | FLJ12825 | Island | Body |
| GM09255 | chr12:54763211-54763433 | ZNF385A;ZNF385A;ZNF385A | Island | 3'UTR;3'UTR;3'UTR |
| WG1977 | chr12:54785286-54785320 | ZNF385A;ZNF385A | S_Shore | TSS1500;TSS1500 |
| GM00439 | chr12:54954337-54954400 | PDE1B;PDE1B | | TSS1500;Body |

| | | | | |
|---|---|---|---|---|
| GM00935 | chr12:5539832-5540426 | NTF3 | N_Shore | TSS1500 |
| GM00290 | chr12:5540116-5540426 | NTF3 | Island | TSS1500 |
| WG2844 | chr12:56105981-56106386 | ITGA7 | N_Shelf | Body |
| WG2415 | chr12:56324915-56325149 | DGKA;DGKA;DGKA;DGKA | N_Shore | TSS1500;TSS1500;TSS1500;TSS200 |
| WG1283 | chr12:56329615-56329731 | DGKA;DGKA;DGKA;DGKA | Island | 5'UTR;5'UTR;5'UTR;5'UTR |
| WG2415 | chr12:56329615-56329903 | DGKA;DGKA;DGKA;DGKA | Island | 5'UTR;5'UTR;5'UTR;5'UTR |
| GM02175 | chr12:56329641-56329903 | DGKA;DGKA;DGKA;DGKA | Island | 5'UTR;5'UTR;5'UTR;5'UTR |
| GM00438 | chr12:56365823-56365978 | CDK2;CDK2 | N_Shore | 3'UTR;3'UTR |
| WG2415 | chr12:56732615-56732723 | IL23A | | TSS200 |
| WG2415 | chr12:57410233-57410604 | TAC3;TAC3 | | 5'UTR;1stExon |
| GM02706 | chr12:57915555-57915618 | MBD6;DDIT3 | N_Shore | TSS1500;TSS1500 |
| WG1977 | chr12:57915555-57915636 | MBD6;DDIT3 | N_Shore | TSS1500;TSS1500 |
| GM02149 | chr12:57915613-57915636 | MBD6;DDIT3 | N_Shore | TSS1500;TSS1500 |
| WG1513 | chr12:58003151-58003385 | DTX3;GEFT | N_Shore | 3'UTR;TSS1500 |
| WG1977 | chr12:58013172-58013241 | SLC26A10 | Island | TSS1500 |
| WG2414 | chr12:58013174-58013267 | SLC26A10 | Island | TSS1500 |
| GM00439 | chr12:58013414-58013487 | SLC26A10 | Island | TSS1500 |
| GM00042 | chr12:58013414-58013645 | SLC26A10 | Island | TSS1500 |
| GM02706 | chr12:58013414-58013651 | SLC26A10 | Island | TSS1500 |
| WG3466 | chr12:58013414-58013687 | SLC26A10 | Island | TSS1500 |
| GM09255 | chr12:58013458-58013487 | SLC26A10 | Island | TSS1500 |
| GM01096 | chr12:58013458-58013539 | SLC26A10 | Island | TSS1500 |
| WG1722 | chr12:58013475-58013539 | SLC26A10 | Island | TSS1500 |
| GM00520 | chr12:58013475-58013651 | SLC26A10 | Island | TSS1500 |
| WG2121 | chr12:58013487-58013539 | SLC26A10 | Island | TSS1500 |
| WG1977 | chr12:58013487-58013651 | SLC26A10 | Island | TSS1500 |
| WG1309 | chr12:58013636-58013651 | SLC26A10 | Island | TSS200 |
| GM02149 | chr12:58013636-58013687 | SLC26A10 | Island | TSS200 |
| WG1722 | chr12:58021295-58021569 | B4GALNT1 | Island | Body |
| GM02173 | chr12:58021295-58021713 | B4GALNT1 | Island | Body |
| WG1977 | chr12:58021295-58021917 | B4GALNT1 | Island | Body |
| GM00438 | chr12:58135973-58136245 | AGAP2 | S_Shelf | TSS200 |
| GM09256 | chr12:5937031-5937409 | ANO2 | Island | Body |
| WG3465 | chr12:6165816-6166173 | VWF | N_Shore | Body |
| WG1722 | chr12:6166874-6167002 | VWF | Island | Body |
| WG1977 | chr12:62583974-62584454 | FAM19A2 | N_Shore | 5'UTR |
| WG2120 | chr12:6442873-6443045 | TNFRSF1A | S_Shelf | Body |
| WG1514 | chr12:6442954-6443045 | TNFRSF1A | | Body |
| GM02456 | chr12:6657744-6657949 | IFFO1;IFFO1;IFFO1 | Island | Body;Body;Body |
| WG2415 | chr12:670520-670712 | B4GALNT3 | | Body |
| GM00439 | chr12:6756377-6756751 | ACRBP | Island | Body |
| GM02149 | chr12:678853-679099 | NINJ2 | N_Shore | Body |
| GM02641 | chr12:678960-679287 | NINJ2 | Island | Body |
| WG2844 | chr12:6881601-6881629 | LAG3 | N_Shore | TSS200 |
| WG1978 | chr12:6933322-6933689 | GPR162;GPR162 | Island | Body;Body |
| WG3466 | chr12:7060257-7060368 | PTPN6;PTPN6;PTPN6 | | TSS200;Body;TSS200 |
| WG1513 | chr12:7060257-7060569 | PTPN6;PTPN6;PTPN6 | | TSS200;Body;TSS200 |

| | | | | |
|---|---|---|---|---|
| WG2415 | chr12:7070317-7070679 | PTPN6;PTPN6;PTPN6 | | 3'UTR;3'UTR;3'UTR |
| GM01376 | chr12:71551926-71552336 | TSPAN8 | | TSS200 |
| GM02175 | chr12:71552188-71552457 | TSPAN8 | | TSS1500 |
| GM09255 | chr12:72233240-72233266 | TBC1D15;TBC1D15;TBC1D15 | N_Shore | TSS1500;TSS1500;TSS1500;TSS1500 |
| WG1978 | chr12:7281566-7281783 | CLSTN3;RBP5 | | TSS1500;TSS200 |
| WG1514 | chr12:739980-740338 | NINJ2 | | Body |
| GM01096 | chr12:75728469-75728752 | GLIPR1L1;GLIPR1L1 | | 5'UTR;1stExon |
| WG1722 | chr12:75785230-75785295 | GLIPR1L2 | Island | Body |
| GM09256 | chr12:772861-772982 | NINJ2 | | TSS200 |
| WG3465 | chr12:7863853-7864047 | DPPA3 | | TSS1500 |
| WG1513 | chr12:7863853-7864076 | DPPA3 | | TSS1500 |
| GM02149 | chr12:7864013-7864076 | DPPA3 | | TSS200 |
| GM09255 | chr12:7864047-7864076 | DPPA3 | | TSS200 |
| WG1513 | chr12:7902721-7903170 | CLEC4C;CLEC4C | | TSS1500;TSS1500 |
| GM09255 | chr12:7940546-7941043 | NANOG | | TSS1500 |
| GM00935 | chr12:81102035-81102486 | MYF6 | Island | Body |
| GM00022 | chr12:81102486-81102749 | MYF6 | Island | Body |
| GM09256 | chr12:81110706-81110776 | MYF5 | N_Shore | TSS200 |
| WG1513 | chr12:85306916-85307131 | SLC6A15;SLC6A15;SLC6A15 | S_Shore | TSS1500;TSS1500;TSS1500 |
| WG3465 | chr12:85307080-85307424 | SLC6A15;SLC6A15;SLC6A15 | S_Shore | TSS1500;TSS1500;TSS1500 |
| GM00042 | chr12:85673200-85673347 | ALX1 | N_Shore | TSS1500 |
| GM01393 | chr12:8815308-8815634 | MFAP5;MFAP5 | | 1stExon;5'UTR |
| WG2844 | chr12:89742886-89744877 | DUSP6;DUSP6 | N_Shelf | 3'UTR;3'UTR |
| GM00935 | chr12:89743788-89744621 | DUSP6;DUSP6 | N_Shore | Body;Body |
| GM00520 | chr12:89743788-89744877 | DUSP6;DUSP6 | N_Shore | Body;Body |
| GM09255 | chr12:89744471-89744621 | DUSP6;DUSP6 | N_Shore | Body;Body |
| GM02456 | chr12:89744471-89744701 | DUSP6;DUSP6 | N_Shore | Body;Body |
| GM00439 | chr12:89744471-89744877 | DUSP6;DUSP6 | N_Shore | Body;Body |
| WG1978 | chr12:89744488-89744609 | DUSP6;DUSP6 | N_Shore | Body;Body |
| GM02555 | chr12:89744488-89744621 | DUSP6;DUSP6 | N_Shore | Body;Body |
| GM00289 | chr12:89744488-89744701 | DUSP6;DUSP6 | N_Shore | Body;Body |
| WG2414 | chr12:89744524-89744701 | DUSP6;DUSP6 | N_Shore | Body;Body |
| WG2276 | chr12:89744609-89744662 | DUSP6;DUSP6 | N_Shore | Body;Body |
| GM00042 | chr12:89744609-89744701 | DUSP6;DUSP6 | N_Shore | Body;Body |
| GM02175 | chr12:89744609-89744877 | DUSP6;DUSP6 | N_Shore | Body;Body |
| WG2415 | chr12:89744621-89744701 | DUSP6;DUSP6 | N_Shore | Body;Body |
| GM01393 | chr12:91573135-91573372 | DCN;DCN;DCN;DCN;DCN;DCN | | 5'UTR;TSS1500;TSS1500;TSS1500;TSS |
| WG1310 | chr12:9217763-9217859 | LOC144571 | S_Shore | TSS200 |
| WG2415 | chr12:95945277-95945384 | USP44 | S_Shelf | TSS200 |
| GM00520 | chr12:96251729-96251912 | SNRPF | N_Shore | TSS1500 |
| WG3466 | chr12:97300410-97300429 | NEDD1;NEDD1;NEDD1;NEDD | N_Shore | TSS1500;TSS1500;TSS1500;TSS1500 |
| GM00941 | chr12:97300410-97300582 | NEDD1;NEDD1;NEDD1;NEDD | N_Shore | TSS1500;TSS1500;TSS1500;TSS1500 |
| WG1513 | chr12:97885353-97886312 | RMST;MIR1251 | | Body;TSS1500 |
| GM02706 | chr12:99548329-99548504 | ANKS1B;ANKS1B;ANKS1B;ANKS1B;ANKS | | 1stExon;5'UTR;5'UTR;Body;1stExon |
| WG1513 | chr12:99548329-99548950 | ANKS1B;ANKS1B;ANKS1B;ANKS1B;ANKS | | 1stExon;5'UTR;5'UTR;Body;1stExon |
| WG3465 | chr12:99548437-99548546 | ANKS1B;ANKS1B;ANKS1B;ANKS1B;ANKS | | 1stExon;5'UTR;5'UTR;Body;1stExon |
| GM00290 | chr12:99548437-99548950 | ANKS1B;ANKS1B;ANKS1B;ANKS1B;ANKS | | 1stExon;5'UTR;5'UTR;Body;1stExon |

| | | | | |
|---|---|---|---|---|
| WG1977 | chr12:99548546-99548757 | ANKS1B;ANKS1B;ANKS1B | | TSS200;TSS200;Body |
| WG3466 | chr13:100310162-100310405 | CLYBL | N_Shore | Body |
| GM00941 | chr13:100310205-100310405 | CLYBL | Island | Body |
| WG1722 | chr13:100547292-100547728 | CLYBL | Island | 3'UTR |
| GM09255 | chr13:100624279-100624373 | ZIC5 | Island | TSS200 |
| WG1722 | chr13:100636252-100636740 | ZIC2 | N_Shore | Body |
| GM00935 | chr13:100637191-100637258 | ZIC2 | Island | Body |
| WG1513 | chr13:100637241-100637560 | ZIC2 | Island | Body |
| WG2275 | chr13:103019936-103020405 | FGF14 | | Body |
| WG2844 | chr13:106116963-106117425 | DAOA;DAOA | | TSS1500;TSS1500 |
| WG2843 | chr13:110874267-110874326 | COL4A1 | | Body |
| WG2276 | chr13:110899996-110900693 | COL4A1 | | Body |
| WG2276 | chr13:110918331-110918683 | COL4A1 | Island | Body |
| GM01393 | chr13:110918441-110918683 | COL4A1 | Island | Body |
| WG2844 | chr13:110951025-110951418 | COL4A1 | | Body |
| GM02706 | chr13:110961330-110961606 | COL4A2 | S_Shore | Body |
| WG1722 | chr13:110961404-110961606 | COL4A2 | S_Shore | Body |
| GM00438 | chr13:110992996-110993558 | COL4A2 | | Body |
| WG3466 | chr13:110993474-110993558 | COL4A2 | | Body |
| WG2276 | chr13:111061859-111062287 | COL4A2 | | Body |
| GM01393 | chr13:111062247-111062287 | COL4A2 | | Body |
| GM00935 | chr13:111301317-111301487 | CARS2 | Island | Body |
| WG3466 | chr13:111521981-111522314 | C13orf29 | | Body |
| WG1283 | chr13:111522932-111522985 | C13orf29 | | TSS1500 |
| GM00438 | chr13:112725903-112726124 | SOX1;SOX1 | N_Shore | 1stExon;3'UTR |
| GM00935 | chr13:113356724-113357112 | ATP11A;ATP11A | | Body;Body |
| WG2844 | chr13:113430153-113430397 | ATP11A;ATP11A | S_Shore | Body;Body |
| WG1977 | chr13:113435942-113437118 | ATP11A;ATP11A | | Body;Body |
| WG2415 | chr13:113436309-113436568 | ATP11A;ATP11A | N_Shelf | Body;Body |
| WG3466 | chr13:113436309-113436651 | ATP11A;ATP11A | N_Shelf | Body;Body |
| GM03075 | chr13:113436400-113436568 | ATP11A;ATP11A | N_Shelf | Body;Body |
| GM01095 | chr13:113436400-113437118 | ATP11A;ATP11A | N_Shelf | Body;Body |
| WG2276 | chr13:113436564-113436651 | ATP11A;ATP11A | N_Shelf | Body;Body |
| GM09255 | chr13:113491819-113492103 | ATP11A;ATP11A | N_Shore | Body;Body |
| GM01376 | chr13:113540189-113540557 | ATP11A;ATP11A | N_Shore | 3'UTR;3'UTR |
| GM02704 | chr13:113540400-113540631 | ATP11A;ATP11A | Island | 3'UTR;3'UTR |
| WG1283 | chr13:113540421-113540557 | ATP11A;ATP11A | Island | 3'UTR;3'UTR |
| WG1984 | chr13:113540512-113540631 | ATP11A;ATP11A | Island | 3'UTR;3'UTR |
| GM00520 | chr13:113622539-113622686 | MCF2L | N_Shore | TSS1500 |
| WG2415 | chr13:113636679-113636953 | MCF2L;MCF2L | S_Shore | Body;Body |
| GM01096 | chr13:113642457-113642703 | MCF2L;MCF2L | | Body;Body |
| WG1310 | chr13:113698408-113698621 | MCF2L;MCF2L | N_Shelf | Body;Body |
| WG1983 | chr13:113698688-113698946 | MCF2L;MCF2L | N_Shelf | Body;Body |
| GM02706 | chr13:113698931-113699016 | MCF2L;MCF2L | N_Shelf | Body;Body |
| GM02175 | chr13:113698931-113699440 | MCF2L;MCF2L | N_Shelf | Body;Body |
| GM00289 | chr13:113698946-113699440 | MCF2L;MCF2L | N_Shelf | Body;Body |
| GM01392 | chr13:113698994-113699014 | MCF2L;MCF2L | N_Shelf | Body;Body |

| | | | | |
|---|---|---|---|---|
| WG1722 | chr13:113698994-113699016 | MCF2L;MCF2L | N_Shelf | Body;Body |
| WG1310 | chr13:113699004-113699016 | MCF2L;MCF2L | N_Shelf | Body;Body |
| WG1513 | chr13:113700818-113701360 | MCF2L;MCF2L | N_Shore | Body;Body |
| WG2415 | chr13:113718667-113719172 | MCF2L;MCF2L | N_Shelf | Body;Body |
| WG3465 | chr13:113784263-113784663 | F10 | Island | Body |
| GM00520 | chr13:113832355-113832595 | PCID2;PCID2;PCID2 | | 3'UTR;3'UTR;3'UTR |
| WG2276 | chr13:113832412-113832595 | PCID2;PCID2;PCID2 | | 3'UTR;3'UTR;3'UTR |
| GM02555 | chr13:114077033-114077172 | ADPRHL1;ADPRHL1 | | 3'UTR;3'UTR |
| GM00042 | chr13:114103452-114103578 | ADPRHL1;ADPRHL1;ADPRHL1 | | 1stExon;5'UTR;Body |
| WG3465 | chr13:114103646-114103796 | ADPRHL1;ADPRHL1 | | Body;TSS200 |
| GM00289 | chr13:114107554-114107876 | ADPRHL1 | | 1stExon |
| GM00520 | chr13:114107710-114107876 | ADPRHL1 | | 1stExon |
| WG1513 | chr13:114108012-114108433 | ADPRHL1 | | TSS200 |
| GM00520 | chr13:114108012-114108652 | ADPRHL1 | | TSS200 |
| GM02175 | chr13:114108052-114108652 | ADPRHL1 | | TSS1500 |
| GM00290 | chr13:114108052-114108713 | ADPRHL1 | | TSS1500 |
| WG2275 | chr13:114108433-114108515 | ADPRHL1 | | TSS1500 |
| WG1984 | chr13:114108433-114108652 | ADPRHL1 | | TSS1500 |
| WG1310 | chr13:114108433-114108676 | ADPRHL1 | | TSS1500 |
| WG2843 | chr13:114108433-114108713 | ADPRHL1 | | TSS1500 |
| WG3466 | chr13:114108471-114108652 | ADPRHL1 | | TSS1500 |
| WG1513 | chr13:114202683-114202861 | TMCO3 | Island | Body |
| WG1977 | chr13:114308619-114308749 | ATP4B | | Body |
| WG2276 | chr13:114321390-114321473 | GRK1 | | TSS1500 |
| WG1977 | chr13:114321514-114321641 | GRK1 | | TSS200 |
| GM03075 | chr13:114321595-114321698 | GRK1 | | TSS200 |
| WG2276 | chr13:114800309-114800549 | RASA3 | N_Shelf | Body |
| WG1310 | chr13:114828067-114828455 | RASA3 | | Body |
| WG2844 | chr13:114837654-114837807 | RASA3 | N_Shore | Body |
| WG2844 | chr13:114871422-114871575 | RASA3 | S_Shore | Body |
| WG1310 | chr13:114875001-114875199 | RASA3 | N_Shore | Body |
| GM00438 | chr13:114875170-114875412 | RASA3 | N_Shore | Body |
| WG2120 | chr13:114880761-114880889 | RASA3 | S_Shore | Body |
| WG1658 | chr13:114880761-114881019 | RASA3 | S_Shore | Body |
| WG1977 | chr13:114880761-114881103 | RASA3 | S_Shore | Body |
| WG1513 | chr13:114886717-114886883 | RASA3 | | Body |
| GM02149 | chr13:19756050-19756122 | TUBA3C | | TSS200 |
| WG3465 | chr13:19919156-19919283 | LOC100101938 | Island | TSS200 |
| WG2415 | chr13:19919208-19919394 | LOC100101938 | Island | TSS200 |
| GM02555 | chr13:20716432-20716728 | GJA3 | Island | Body |
| GM00289 | chr13:20735967-20736075 | GJA3 | Island | TSS1500 |
| GM02706 | chr13:20736009-20736156 | GJA3 | Island | TSS1500 |
| GM00520 | chr13:20768523-20768560 | GJB2 | S_Shore | TSS1500 |
| WG1977 | chr13:20805196-20805385 | GJB6;GJB6;GJB6;GJB6;GJB6 | N_Shore | 5'UTR;5'UTR;5'UTR;5'UTR;1stExon |
| WG1977 | chr13:20805391-20805487 | GJB6;GJB6;GJB6;GJB6 | N_Shore | 5'UTR;5'UTR;5'UTR;TSS200 |
| GM02175 | chr13:20805391-20805656 | GJB6;GJB6;GJB6;GJB6 | N_Shore | 5'UTR;5'UTR;5'UTR;TSS200 |
| WG2121 | chr13:20805487-20805679 | GJB6;GJB6;GJB6;GJB6 | N_Shore | 5'UTR;5'UTR;5'UTR;TSS200 |

| | | | | |
|---|---|---|---|---|
| GM00042 | chr13:21295590-21296465 | IL17D | N_Shore | Body |
| WG1722 | chr13:21295941-21296465 | IL17D | Island | Body |
| GM03075 | chr13:23898364-23898864 | SGCG | | Body |
| WG3466 | chr13:24152135-24152357 | TNFRSF19 | N_Shore | 5'UTR |
| GM02149 | chr13:25338113-25338146 | RNF17 | | TSS200 |
| WG1513 | chr13:25338113-25338234 | RNF17 | | TSS200 |
| GM00042 | chr13:25469147-25469406 | CENPJ | Island | Body |
| GM00941 | chr13:25506131-25506340 | LOC646405 | Island | Body |
| WG1984 | chr13:25670170-25670187 | PABPC3 | Island | TSS200 |
| GM02704 | chr13:25670170-25670228 | PABPC3 | Island | TSS200 |
| GM09255 | chr13:25670170-25670238 | PABPC3 | Island | TSS200 |
| WG1722 | chr13:25670183-25670228 | PABPC3 | Island | TSS200 |
| GM01392 | chr13:25670228-25670319 | PABPC3 | Island | TSS200 |
| WG2415 | chr13:25746495-25746644 | FAM123A;FAM123A | S_Shore | TSS1500;TSS1500 |
| GM00290 | chr13:26586254-26586353 | ATP8A2 | N_Shore | Body |
| GM09255 | chr13:26586353-26586651 | ATP8A2 | Island | Body |
| GM01375 | chr13:26586552-26586713 | ATP8A2 | Island | Body |
| GM00438 | chr13:27335096-27335204 | GPR12 | Island | TSS200 |
| WG1513 | chr13:27335144-27335204 | GPR12 | Island | TSS1500 |
| GM02706 | chr13:28493913-28494004 | PDX1 | N_Shore | TSS1500 |
| WG1722 | chr13:28562819-28562850 | PRHOXNB | | TSS200 |
| GM02555 | chr13:28562850-28562902 | PRHOXNB | | TSS200 |
| GM09256 | chr13:28562886-28563578 | PRHOXNB | | TSS200 |
| GM01096 | chr13:30077315-30077489 | MTUS2;MTUS2 | Island | Body;Body |
| GM00940 | chr13:30088615-30088771 | SLC7A1 | Island | 3'UTR |
| WG1722 | chr13:30088640-30088771 | SLC7A1 | Island | Body |
| GM00290 | chr13:31479935-31480128 | C13orf33 | N_Shore | TSS1500 |
| GM00935 | chr13:31479935-31480225 | C13orf33 | N_Shore | TSS1500 |
| WG1310 | chr13:31480128-31480273 | C13orf33 | N_Shore | TSS200 |
| GM02175 | chr13:31506685-31506710 | C13orf26 | | TSS200 |
| WG3465 | chr13:33590002-33590083 | KL | Island | TSS1500 |
| WG2844 | chr13:33760215-33760693 | STARD13;STARD13;STARD13;STARD13 | | 1stExon;5'UTR;5'UTR;Body |
| GM02706 | chr13:33780078-33780283 | STARD13;STARD13;STARD13 | | 1stExon;5'UTR;Body |
| GM00042 | chr13:33780205-33780307 | STARD13;STARD13 | | TSS200;Body |
| WG1513 | chr13:33780205-33780362 | STARD13;STARD13 | | TSS200;Body |
| GM00935 | chr13:33780283-33780362 | STARD13;STARD13 | | TSS200;Body |
| WG2121 | chr13:36044768-36044886 | NBEA | N_Shore | Body |
| GM01096 | chr13:36044768-36045201 | NBEA | N_Shore | Body |
| GM02175 | chr13:36044768-36045352 | NBEA | N_Shore | Body |
| WG2415 | chr13:36045201-36045352 | NBEA | Island | Body |
| WG2120 | chr13:36048892-36049380 | MAB21L1;MIR548F5;NBEA;M | N_Shore | 1stExon;Body;Body;3'UTR |
| WG3466 | chr13:36048892-36051073 | MAB21L1;MIR548F5;NBEA;M | N_Shore | 1stExon;Body;Body;3'UTR |
| GM01096 | chr13:36050522-36051073 | MAB21L1;MIR548F5;NBEA;M | S_Shore | 1stExon;Body;Body;5'UTR |
| WG3465 | chr13:36050788-36051073 | MAB21L1;MIR548F5;NBEA;M | N_Shore | 1stExon;Body;Body;5'UTR |
| WG1977 | chr13:36050881-36051073 | MIR548F5;NBEA;MAB21L1 | N_Shore | Body;Body;TSS200 |
| GM00016 | chr13:36050889-36050993 | MIR548F5;NBEA;MAB21L1 | N_Shore | Body;Body;TSS200 |
| WG2193 | chr13:36050889-36051073 | MIR548F5;NBEA;MAB21L1 | N_Shore | Body;Body;TSS200 |

| | | | | |
|---|---|---|---|---|
| GM03073 | chr13:36050901-36050993 | MIR548F5;NBEA;MAB21L1 | N_Shore | Body;Body;TSS200 |
| WG2415 | chr13:36050901-36051073 | MIR548F5;NBEA;MAB21L1 | N_Shore | Body;Body;TSS200 |
| WG1721 | chr13:36050930-36051073 | MIR548F5;NBEA;MAB21L1 | N_Shore | Body;Body;TSS200 |
| WG1977 | chr13:36429936-36430134 | MIR548F5;DCLK1 | | Body;Body |
| WG1721 | chr13:36871646-36871878 | C13orf38;C13orf38;C13orf38 | N_Shore | Body;5'UTR;5'UTR;5'UTR;5'UTR;5'UTR |
| WG1283 | chr13:36871646-36871948 | C13orf38;C13orf38;C13orf38 | N_Shore | Body;5'UTR;5'UTR;5'UTR;5'UTR;5'UTR |
| WG1283 | chr13:36872048-36872148 | C13orf38;C13orf38;C13orf38 | Island | TSS200;TSS200;TSS200;TSS200;TSS2 |
| GM00940 | chr13:37004601-37004620 | CCNA1;CCNA1 | N_Shore | TSS1500;TSS1500 |
| WG2121 | chr13:37004721-37004747 | CCNA1;CCNA1 | N_Shore | TSS1500;TSS1500 |
| GM00940 | chr13:37005273-37005452 | CCNA1;CCNA1;CCNA1;CCN | N_Shore | TSS1500;TSS1500;TSS1500;TSS1500 |
| WG2275 | chr13:37005289-37005452 | CCNA1;CCNA1;CCNA1;CCN | N_Shore | TSS1500;TSS1500;TSS1500;TSS1500 |
| GM00439 | chr13:37005452-37005501 | CCNA1;CCNA1;CCNA1;CCN | N_Shore | TSS1500;TSS1500;TSS1500;TSS1500 |
| GM00520 | chr13:37249426-37249481 | C13orf36 | S_Shore | 5'UTR |
| WG1977 | chr13:38172803-38173002 | POSTN;POSTN;POSTN;POSTN | | 1stExon;1stExon;1stExon;1stExon |
| WG1283 | chr13:41593385-41593519 | ELF1;ELF1 | | 5'UTR;1stExon |
| WG1978 | chr13:41593385-41593582 | ELF1;ELF1 | | 5'UTR;1stExon |
| WG3465 | chr13:41593416-41593582 | ELF1;ELF1 | | 5'UTR;1stExon |
| GM00438 | chr13:43148278-43148436 | TNFSF11;TNFSF11 | Island | TSS200;5'UTR |
| WG2276 | chr13:43566618-43566642 | EPSTI1;EPSTI1 | S_Shore | TSS1500;TSS1500 |
| WG1722 | chr13:43566633-43566902 | EPSTI1;EPSTI1 | S_Shore | TSS1500;TSS1500 |
| WG1514 | chr13:43597565-43597736 | DNAJC15;DNAJC15 | Island | 5'UTR;1stExon |
| WG2120 | chr13:43597565-43598038 | DNAJC15;DNAJC15 | Island | 5'UTR;1stExon |
| GM01375 | chr13:43597657-43598038 | DNAJC15;DNAJC15 | Island | 5'UTR;1stExon |
| WG1977 | chr13:45149661-45149706 | TSC22D1 | N_Shore | 1stExon |
| WG1722 | chr13:45945197-45945362 | LOC100190939 | | Body |
| GM00016 | chr13:46425837-46425874 | SIAH3;SIAH3 | | 5'UTR;1stExon |
| GM09256 | chr13:46425904-46425999 | SIAH3 | | TSS200 |
| WG2415 | chr13:48575859-48575927 | SUCLA2 | S_Shore | TSS1500 |
| GM00520 | chr13:50202547-50202658 | ARL11 | | TSS200 |
| GM09255 | chr13:51417977-51418041 | DLEU7 | Island | TSS200 |
| WG1722 | chr13:52769662-52769875 | THSD1P | Island | TSS1500 |
| GM02706 | chr13:61989597-61989724 | PCDH20;PCDH20 | | 5'UTR;1stExon |
| GM02149 | chr13:70681276-70681317 | KLHL1;ATXN8OS | N_Shore | Body;TSS200 |
| WG3466 | chr13:76334583-76334728 | LMO7;LMO7 | | TSS1500;Body |
| GM09256 | chr13:76334583-76334866 | LMO7;LMO7 | | TSS1500;Body |
| WG2415 | chr13:76334707-76334866 | LMO7;LMO7 | | TSS200;Body |
| GM02706 | chr13:77461335-77461426 | KCTD12 | S_Shore | TSS1500 |
| WG3465 | chr13:78493920-78494010 | EDNRB;EDNRB;EDNRB | S_Shore | TSS1500;5'UTR;TSS1500 |
| WG1513 | chr13:78493958-78494064 | EDNRB;EDNRB;EDNRB | S_Shore | TSS1500;5'UTR;TSS1500 |
| GM03075 | chr13:78494067-78494272 | EDNRB;EDNRB;EDNRB | S_Shore | TSS1500;5'UTR;TSS1500 |
| GM00289 | chr13:78494171-78494330 | EDNRB;EDNRB;EDNRB | S_Shore | TSS1500;5'UTR;TSS1500 |
| GM02706 | chr13:79234251-79234435 | RNF219 | S_Shore | TSS1500 |
| WG1977 | chr13:79234251-79234847 | RNF219 | S_Shore | TSS1500 |
| WG2415 | chr13:79234435-79234847 | RNF219 | S_Shore | TSS1500 |
| GM01096 | chr13:80911626-80911808 | SPRY2 | N_Shore | Body |
| GM01096 | chr13:88326635-88326918 | SLITRK5 | Island | 5'UTR |
| GM09256 | chr13:88328251-88328471 | SLITRK5 | N_Shore | Body |

| | | | | |
|---|---|---|---|---|
| GM01375 | chr13:88328294-88328813 | SLITRK5 | N_Shore | Body |
| WG3466 | chr13:88330200-88330614 | SLITRK5 | S_Shore | Body |
| WG2120 | chr13:92050720-92050731 | GPC5 | N_Shore | TSS1500 |
| WG2414 | chr13:95248771-95248888 | TGDS | S_Shore | TSS1500 |
| GM00940 | chr13:96205679-96205960 | CLDN10;CLDN10;CLDN10 | S_Shore | Body;Body;Body |
| WG1722 | chr13:96297197-96297338 | DZIP1;DZIP1 | S_Shore | TSS1500;TSS1500 |
| GM02149 | chr13:97873877-97873992 | MBNL2;MBNL2 | | TSS1500;TSS1500 |
| WG1513 | chr13:97873947-97874279 | MBNL2;MBNL2 | | TSS1500;TSS1500 |
| WG2121 | chr13:98605770-98606196 | IPO5 | | TSS200 |
| WG3465 | chr13:98605886-98605951 | IPO5 | | TSS200 |
| GM00520 | chr13:98605886-98606196 | IPO5 | | TSS200 |
| WG1514 | chr13:98605892-98606196 | IPO5 | | TSS200 |
| WG2275 | chr13:98794353-98794593 | FARP1;FARP1 | N_Shore | TSS1500;TSS1500 |
| WG2415 | chr13:99100712-99100829 | FARP1 | | 3'UTR |
| WG1513 | chr13:99174312-99174494 | STK24;STK24;STK24 | | Body;5'UTR;1stExon |
| GM02175 | chr14:100069535-100069840 | CCDC85C | Island | 1stExon |
| GM00438 | chr14:100125683-100126196 | HHIPL1;HHIPL1 | Island | Body;Body |
| GM01650 | chr14:100125898-100126358 | HHIPL1;HHIPL1 | Island | Body;Body |
| GM00016 | chr14:100126358-100126741 | HHIPL1;HHIPL1 | Island | Body;Body |
| WG2844 | chr14:101035818-101036202 | BEGAIN;BEGAIN | S_Shore | TSS1500;5'UTR |
| WG1513 | chr14:101035818-101036470 | BEGAIN;BEGAIN | S_Shore | TSS1500;5'UTR |
| GM09256 | chr14:101036192-101036322 | BEGAIN | S_Shore | TSS200 |
| WG2415 | chr14:101036192-101036470 | BEGAIN | S_Shore | TSS200 |
| WG2844 | chr14:101123390-101123682 | C14orf70 | S_Shelf | TSS1500 |
| GM00439 | chr14:101192852-101192913 | DLK1 | Island | TSS1500 |
| GM00438 | chr14:101350731-101350872 | MIR136;RTL1;MIR432 | S_Shore | TSS1500;1stExon;TSS200 |
| GM02175 | chr14:102510281-102510478 | DYNC1H1 | | Body |
| GM00520 | chr14:103415458-103415882 | CDC42BPB | | Body |
| GM03075 | chr14:103592642-103593520 | TNFAIP2 | N_Shore | TSS200 |
| WG1977 | chr14:103593088-103593520 | TNFAIP2 | N_Shore | Body |
| GM00289 | chr14:103593235-103593505 | TNFAIP2 | Island | Body |
| WG2275 | chr14:103593235-103593520 | TNFAIP2 | Island | Body |
| WG3465 | chr14:103593503-103593520 | TNFAIP2 | Island | Body |
| WG1722 | chr14:104178670-104179135 | XRCC3;XRCC3;XRCC3 | N_Shelf | 5'UTR;5'UTR;5'UTR |
| WG2276 | chr14:104315145-104315223 | PPP1R13B | S_Shore | TSS1500 |
| WG1977 | chr14:104394621-104395165 | TDRD9 | Island | TSS200 |
| WG2844 | chr14:104621467-104622128 | KIF26A | N_Shore | Body |
| GM01392 | chr14:104621601-104622028 | KIF26A | N_Shore | Body |
| GM00439 | chr14:104621601-104622128 | KIF26A | N_Shore | Body |
| WG3465 | chr14:104621867-104622128 | KIF26A | Island | Body |
| GM00438 | chr14:104645687-104646121 | KIF26A | Island | Body |
| GM02706 | chr14:105154689-105154858 | INF2;INF2;INF2 | N_Shore | TSS1500;TSS1500;TSS1500 |
| WG2276 | chr14:105173681-105173801 | INF2;INF2 | | Body;Body |
| GM02706 | chr14:105287326-105287427 | MGC23270 | | TSS1500 |
| WG1722 | chr14:105635706-105635763 | JAG2;JAG2 | | TSS1500;TSS1500 |
| GM02706 | chr14:105635708-105635769 | JAG2;JAG2 | | TSS1500;TSS1500 |
| WG1513 | chr14:105714379-105714404 | BRF1;BTBD6;BRF1 | N_Shore | TSS200;TSS1500;Body |

| | | | | |
|---|---|---|---|---|
| WG1513 | chr14:105715741-105716052 | BTBD6;BRF1 | Island | Body;Body |
| WG1977 | chr14:105779826-105779952 | PACS2;PACS2 | N_Shore | TSS1500;TSS1500 |
| GM02456 | chr14:105796713-105796964 | PACS2;PACS2 | Island | Body;Body |
| GM02704 | chr14:105944604-105944942 | CRIP2 | Island | Body |
| WG3466 | chr14:105944604-105945287 | CRIP2 | Island | Body |
| GM02706 | chr14:105944604-105945699 | CRIP2 | Island | Body |
| WG1513 | chr14:106382726-106383155 | KIAA0125 | | TSS1500 |
| WG1513 | chr14:106438118-106438465 | ADAM6 | | Body |
| GM00438 | chr14:106937310-106938335 | LOC100133469 | N_Shore | TSS1500 |
| WG1513 | chr14:106937310-106938589 | LOC100133469 | N_Shore | TSS1500 |
| GM00289 | chr14:106937869-106938294 | LOC100133469 | N_Shore | TSS1500 |
| GM00289 | chr14:106938335-106938509 | LOC100133469 | Island | TSS200 |
| WG1513 | chr14:22103082-22103504 | OR10G2 | | TSS200 |
| GM02149 | chr14:22103310-22103726 | OR10G2 | | TSS1500 |
| GM00438 | chr14:23351487-23351876 | REM2 | | TSS1500 |
| GM02175 | chr14:23511078-23511306 | PSMB11 | | TSS1500 |
| WG1513 | chr14:23845819-23846456 | CMTM5;CMTM5 | | TSS200;TSS200 |
| WG2415 | chr14:23904678-23904930 | MYH7 | | 5'UTR |
| GM00042 | chr14:24046374-24047021 | JPH4;JPH4 | Island | Body;Body |
| GM02706 | chr14:24457915-24458099 | DHRS4L2 | Island | TSS200 |
| WG1977 | chr14:24457915-24458345 | DHRS4L2 | Island | TSS200 |
| WG1283 | chr14:24458000-24458099 | DHRS4L2 | Island | TSS200 |
| GM01650 | chr14:24458000-24458208 | DHRS4L2 | Island | TSS200 |
| WG1513 | chr14:24458208-24458345 | DHRS4L2 | S_Shore | 1stExon |
| WG1283 | chr14:24505537-24505554 | DHRS4L1 | N_Shore | TSS200 |
| GM01650 | chr14:24505537-24505653 | DHRS4L1 | N_Shore | TSS200 |
| GM02175 | chr14:24540174-24540415 | CPNE6 | | TSS1500 |
| GM00289 | chr14:24540235-24540452 | CPNE6 | | TSS1500 |
| WG2121 | chr14:24641021-24641155 | REC8;REC8 | N_Shore | TSS1500;TSS1500 |
| GM02706 | chr14:24641077-24641194 | REC8;REC8 | Island | TSS200;TSS200 |
| GM09256 | chr14:24641189-24641201 | REC8;REC8 | Island | TSS200;TSS200 |
| WG1977 | chr14:24779793-24780551 | LTB4R2;LTB4R2;LTB4R2;LTB4 | N_Shore | 1stExon;1stExon;5'UTR;TSS1500;5'UT |
| GM02704 | chr14:24779793-24780557 | LTB4R2;LTB4R2;LTB4R2;LTB4 | N_Shore | 1stExon;1stExon;5'UTR;TSS1500;5'UT |
| GM03075 | chr14:24779959-24780557 | LTB4R2;LTB4R2;LTB4R2;LTB4 | Island | 1stExon;1stExon;Body;TSS1500;1stEx |
| GM00016 | chr14:24780167-24780540 | LTB4R2;LTB4R2;LTB4R2;LTB4 | Island | 1stExon;1stExon;Body;TSS1500;1stEx |
| WG1983 | chr14:24780167-24780557 | LTB4R2;LTB4R2;LTB4R2;LTB4 | Island | 1stExon;1stExon;Body;TSS1500;1stEx |
| WG1721 | chr14:24780540-24780557 | LTB4R2;LTB4R2;LTB4R2;CIDI | Island | Body;1stExon;Body;1stExon;5'UTR;TS |
| WG3466 | chr14:24780682-24780734 | LTB4R2;LTB4R2;LTB4R2;CIDI | Island | Body;1stExon;Body;TSS200;TSS200 |
| GM02175 | chr14:24780734-24780890 | LTB4R;LTB4R2;LTB4R2;LTB4F | Island | 5'UTR;Body;1stExon;Body;1stExon;TS |
| GM02175 | chr14:24785310-24785847 | LTB4R;LTB4R | Island | Body;Body |
| WG1513 | chr14:25045607-25046121 | CTSG | | TSS200 |
| WG1721 | chr14:29235904-29235928 | FOXG1 | Island | TSS1500 |
| WG2276 | chr14:34269416-34269637 | NPAS3;NPAS3;NPAS3;NPAS3 | Island | Body;Body;Body;Body |
| WG2121 | chr14:34269637-34270113 | NPAS3;NPAS3;NPAS3;NPAS3 | Island | Body;Body;Body;Body |
| GM02706 | chr14:37125430-37126582 | PAX9 | Island | TSS1500 |
| WG1310 | chr14:37125768-37125997 | PAX9 | N_Shore | TSS1500 |
| GM00935 | chr14:37125872-37125997 | PAX9 | N_Shore | TSS1500 |

| | | | | |
|---|---|---|---|---|
| WG1513 | chr14:37125914-37126283 | PAX9 | N_Shore | TSS1500 |
| GM00520 | chr14:37127671-37128378 | PAX9 | Island | 5'UTR |
| GM02704 | chr14:37128378-37128790 | PAX9 | S_Shore | 5'UTR |
| WG2193 | chr14:37135586-37136391 | PAX9 | Island | Body |
| WG2843 | chr14:37135898-37136391 | PAX9 | Island | Body |
| WG3466 | chr14:37136192-37136391 | PAX9 | Island | Body |
| GM00439 | chr14:38725312-38725750 | CLEC14A;CLEC14A | Island | 1stExon;5'UTR |
| GM00935 | chr14:38725312-38726189 | CLEC14A;CLEC14A | Island | 1stExon;5'UTR |
| GM09256 | chr14:38725656-38725750 | CLEC14A | S_Shore | TSS200 |
| GM03075 | chr14:45432309-45432516 | KLHL28;FAM179B | S_Shore | TSS1500;1stExon |
| GM09256 | chr14:47121132-47121158 | RPL10L | Island | TSS200 |
| GM00289 | chr14:51296586-51296728 | NIN;NIN;NIN | N_Shore | 5'UTR;5'UTR;5'UTR |
| GM02456 | chr14:52325747-52326169 | GNG2 | | TSS1500 |
| WG3466 | chr14:54418728-54418851 | BMP4;BMP4;BMP4 | Island | Body;Body;Body |
| GM02147 | chr14:54423807-54423845 | BMP4;BMP4 | S_Shore | TSS1500;TSS1500 |
| GM00439 | chr14:54423811-54424149 | BMP4;BMP4 | S_Shore | TSS1500;TSS1500 |
| GM00289 | chr14:55907122-55907374 | TBPL2 | N_Shore | 1stExon |
| WG1722 | chr14:55907122-55907427 | TBPL2 | N_Shore | 1stExon |
| GM02706 | chr14:55907374-55907417 | TBPL2 | Island | TSS200 |
| GM00940 | chr14:55907374-55907427 | TBPL2 | Island | TSS200 |
| WG1513 | chr14:55907417-55907501 | TBPL2 | Island | TSS200 |
| GM00438 | chr14:56045827-56046001 | KTN1;C14orf33;KTN1;KTN1;K | N_Shore | TSS1500;Body;TSS1500;TSS1500;TSS |
| WG1513 | chr14:57277178-57277366 | OTX2;OTX2 | N_Shore | 1stExon;5'UTR |
| GM00290 | chr14:57277178-57277761 | OTX2;OTX2 | N_Shore | 1stExon;5'UTR |
| GM09256 | chr14:57277227-57277739 | OTX2 | N_Shore | TSS200 |
| GM00940 | chr14:58862417-58862457 | TOMM20L | N_Shore | TSS1500 |
| GM02175 | chr14:59932273-59932321 | GPR135 | S_Shore | TSS1500 |
| GM02706 | chr14:61113963-61114476 | SIX1 | N_Shore | Body |
| GM00439 | chr14:61748243-61748354 | TMEM30B;TMEM30B | S_Shore | 1stExon;5'UTR |
| GM02173 | chr14:61748243-61748440 | TMEM30B;TMEM30B | S_Shore | 1stExon;5'UTR |
| GM02149 | chr14:61748336-61748440 | TMEM30B;TMEM30B | S_Shore | 1stExon;5'UTR |
| GM01095 | chr14:61789466-61789603 | PRKCH | Island | Body |
| GM02706 | chr14:61789503-61790037 | PRKCH | S_Shore | Body |
| WG2414 | chr14:64319245-64319306 | SYNE2;SYNE2 | N_Shore | TSS1500;TSS1500 |
| WG1657 | chr14:64676792-64677007 | SYNE2;SYNE2 | | Body;Body |
| WG1513 | chr14:65005951-65006041 | HSPA2 | N_Shore | TSS1500 |
| WG1514 | chr14:65006041-65006170 | HSPA2 | N_Shore | TSS1500 |
| GM02175 | chr14:65006170-65006222 | HSPA2 | N_Shore | TSS1500 |
| GM01096 | chr14:65016517-65016591 | C14orf50 | Island | TSS200 |
| WG1977 | chr14:65016517-65016595 | C14orf50 | Island | TSS200 |
| GM00042 | chr14:65016545-65016595 | C14orf50 | Island | TSS200 |
| GM00290 | chr14:65016591-65016602 | C14orf50 | Island | TSS200 |
| GM02149 | chr14:65016595-65016802 | C14orf50 | Island | TSS200 |
| WG2415 | chr14:69341430-69341905 | ACTN1;ACTN1;ACTN1 | Island | 3'UTR;3'UTR;3'UTR |
| GM00438 | chr14:71276874-71277141 | MAP3K9 | S_Shore | TSS1500 |
| GM02706 | chr14:73361747-73361964 | DPF3 | S_Shore | TSS1500 |
| WG1977 | chr14:73712289-73712902 | PAPLN | Island | Body |

| | | | | |
|---|---|---|---|---|
| GM01096 | chr14:73712289-73712967 | PAPLN | Island | Body |
| GM00941 | chr14:73712491-73712902 | PAPLN | Island | Body |
| GM02706 | chr14:73712491-73712967 | PAPLN | Island | Body |
| GM00940 | chr14:73712574-73712967 | PAPLN | Island | Body |
| GM00438 | chr14:74083455-74083501 | ACOT6 | S_Shelf | TSS200 |
| WG2415 | chr14:75389222-75389618 | RPS6KL1 | N_Shore | TSS200 |
| GM02706 | chr14:75389480-75389716 | RPS6KL1 | N_Shore | TSS1500 |
| WG2415 | chr14:75936582-75936789 | JDP2;JDP2;JDP2;JDP2 | | 3'UTR;3'UTR;3'UTR;3'UTR |
| WG2276 | chr14:76447327-76447431 | TGFB3;TGFB3 | N_Shore | 1stExon;5'UTR |
| WG1513 | chr14:77737771-77738080 | NGB | Island | TSS200 |
| WG1513 | chr14:79111512-79111719 | NRXN3 | | 5'UTR |
| WG2844 | chr14:80677608-80677688 | DIO2;DIO2;DIO2 | | 1stExon;1stExon;Body |
| GM02706 | chr14:81421983-81422160 | TSHR;TSHR;TSHR;TSHR;TSHI | S_Shore | 5'UTR;5'UTR;1stExon;1stExon;5'UTR;1 |
| WG3466 | chr14:81422098-81422160 | TSHR;TSHR;TSHR | S_Shore | 1stExon;1stExon;1stExon |
| GM00520 | chr14:88788422-88788597 | KCNK10;KCNK10 | N_Shore | Body;Body |
| GM00439 | chr14:90042201-90042364 | PRO1768;FOXN3 | | TSS1500;5'UTR |
| WG2415 | chr14:91818497-91818668 | CCDC88C | | Body |
| GM00289 | chr14:91862813-91862907 | CCDC88C | | Body |
| GM02706 | chr14:94254131-94254586 | PRIMA1 | Island | 5'UTR |
| WG1513 | chr14:94462309-94462337 | C14orf48;C14orf48;C14orf48 | | TSS1500;TSS1500;TSS1500 |
| WG1513 | chr14:95027382-95027859 | SERPINA4 | | TSS1500 |
| WG1977 | chr14:95234658-95234965 | GSC | Island | 3'UTR |
| WG1722 | chr14:95234822-95234965 | GSC | Island | 3'UTR |
| WG2414 | chr14:95234826-95235127 | GSC | Island | 3'UTR |
| WG1721 | chr14:95235026-95235127 | GSC | Island | Body |
| GM00520 | chr14:96180643-96180653 | TCL1A;TCL1A | Island | TSS200;TSS200 |
| WG1722 | chr14:96670628-96670738 | BDKRB2 | | TSS1500 |
| GM09255 | chr14:99655593-99655747 | BCL11B;BCL11B | | Body;Body |
| WG1513 | chr14:99728984-99729088 | BCL11B;BCL11B | | Body;Body |
| GM00935 | chr15:100537865-100538145 | ADAMTS17 | | Body |
| WG1513 | chr15:100654734-100654807 | ADAMTS17 | | Body |
| GM00439 | chr15:100672282-100672379 | ADAMTS17 | | Body |
| GM02175 | chr15:100889683-100889719 | FLJ42289;FLJ42289 | N_Shore | Body;Body |
| WG2415 | chr15:101419262-101419333 | ALDH1A3 | Island | TSS1500 |
| WG1309 | chr15:101419269-101419296 | ALDH1A3 | Island | TSS1500 |
| WG2193 | chr15:101419275-101419296 | ALDH1A3 | Island | TSS1500 |
| GM02706 | chr15:101419275-101419333 | ALDH1A3 | Island | TSS1500 |
| GM02147 | chr15:101419293-101419333 | ALDH1A3 | Island | TSS1500 |
| WG1977 | chr15:101420636-101420989 | ALDH1A3 | Island | Body |
| WG3466 | chr15:101728611-101728653 | CHSY1 | | Body |
| GM02456 | chr15:101869493-101869560 | PCSK6;PCSK6;PCSK6;PCSK6 | | Body;Body;Body;Body |
| GM00289 | chr15:101991560-101991886 | PCSK6;PCSK6;PCSK6;PCSK6;PCSK6;PCS | | Body;Body;Body;Body;Body;Body;Bod |
| GM01096 | chr15:102010125-102010195 | PCSK6;PCSK6;PCSK6;PCSK6;PCSK6;PCS | | Body;Body;Body;Body;Body;Body;Bod |
| WG3466 | chr15:102193228-102193416 | TM2D3;TM2D3 | Island | TSS1500;TSS1500 |
| WG2275 | chr15:22833149-22833288 | TUBGCP5;TUBGCP5 | N_Shore | TSS1500;TSS1500 |
| WG2193 | chr15:22833288-22833335 | TUBGCP5;TUBGCP5 | Island | TSS200;TSS200 |
| GM03073 | chr15:23378429-23378571 | HERC2P2 | Island | TSS200 |

| | | | | |
|---|---|---|---|---|
| WG1513 | chr15:23810280-23810378 | MKRN3 | | TSS200 |
| WG2415 | chr15:23810652-23810861 | MKRN3;MKRN3 | | 5'UTR;1stExon |
| GM02706 | chr15:23932370-23932412 | NDN;NDN | Island | 1stExon;5'UTR |
| GM00438 | chr15:24920990-24921752 | C15orf2;C15orf2 | Island | 1stExon;5'UTR |
| GM00438 | chr15:25198793-25199164 | SNRPN;SNURF;SNRPN;SNRF | N_Shore | 5'UTR;TSS1500;TSS1500;5'UTR;5'UTR |
| GM00438 | chr15:25201224-25201732 | SNRPN;SNRPN;SNRPN;SNRF | S_Shore | 5'UTR;5'UTR;5'UTR;5'UTR;Body;Body;5 |
| WG1513 | chr15:25425326-25425615 | SNORD115-6 | | TSS1500 |
| WG1513 | chr15:25427435-25427509 | SNORD115-7 | | TSS200 |
| WG1513 | chr15:25429307-25429408 | SNORD115-5;SNORD115-12;SNORD115 | | TSS1500;TSS1500;TSS1500;TSS200;T |
| WG1513 | chr15:25432236-25432321 | SNORD115-10 | | TSS1500 |
| GM00438 | chr15:25434150-25434475 | SNORD115-29;SNORD115-36;SNORD11 | | TSS1500;TSS1500;TSS1500;TSS1500 |
| GM09256 | chr15:25434289-25434492 | SNORD115-29;SNORD115-36;SNORD11 | | TSS1500;TSS1500;TSS1500;TSS1500 |
| WG1513 | chr15:25434475-25434533 | SNORD115-11;SNORD115-36;SNORD11 | | TSS200;TSS200;TSS200;TSS200 |
| GM00438 | chr15:25438254-25438860 | SNORD115-13 | | TSS1500 |
| WG1513 | chr15:25449918-25450131 | SNORD115-21;SNORD115-20;SNORD11 | | TSS1500;TSS1500;TSS1500 |
| GM09256 | chr15:26047382-26047653 | ATP10A | | Body |
| GM00289 | chr15:26109204-26109252 | ATP10A | S_Shore | TSS1500 |
| WG2276 | chr15:26109204-26109261 | ATP10A | S_Shore | TSS1500 |
| WG3466 | chr15:26109215-26109261 | ATP10A | S_Shore | TSS1500 |
| WG1513 | chr15:26109215-26109543 | ATP10A | S_Shore | TSS1500 |
| WG1984 | chr15:26109245-26109261 | ATP10A | S_Shore | TSS1500 |
| GM03073 | chr15:26109249-26109261 | ATP10A | S_Shore | TSS1500 |
| WG2121 | chr15:26109252-26109394 | ATP10A | S_Shore | TSS1500 |
| GM00289 | chr15:26109394-26109429 | ATP10A | S_Shore | TSS1500 |
| GM00439 | chr15:26109429-26109614 | ATP10A | S_Shore | TSS1500 |
| WG1722 | chr15:26915257-26915752 | GABRB3;GABRB3 | Island | Body;Body |
| GM00016 | chr15:27111435-27111476 | GABRA5;GABRA5 | N_Shore | TSS1500;TSS1500 |
| GM00935 | chr15:27111478-27111572 | GABRA5;GABRA5 | N_Shore | TSS1500;TSS1500 |
| WG1309 | chr15:27111541-27111604 | GABRA5;GABRA5 | N_Shore | TSS1500;TSS1500 |
| WG1513 | chr15:27111597-27111749 | GABRA5;GABRA5 | N_Shore | TSS1500;TSS1500 |
| GM00520 | chr15:27111604-27111699 | GABRA5;GABRA5 | N_Shore | TSS1500;TSS1500 |
| WG2844 | chr15:27184766-27185073 | GABRA5;GABRA5 | | Body;Body |
| WG1513 | chr15:27215681-27215971 | GABRG3 | N_Shore | TSS1500 |
| WG2415 | chr15:27215952-27215971 | GABRG3 | Island | TSS1500 |
| WG1513 | chr15:28339873-28340213 | OCA2 | Island | 5'UTR |
| WG3465 | chr15:29213626-29213748 | APBA2;APBA2 | | TSS1500;TSS1500 |
| GM00290 | chr15:29213640-29213748 | APBA2;APBA2 | | TSS200;TSS200 |
| WG2844 | chr15:29409410-29409632 | APBA2;APBA2 | S_Shore | 3'UTR;3'UTR |
| WG1513 | chr15:31357535-31358057 | MIR211;TRPM1 | | TSS200;Body |
| WG3465 | chr15:32933783-32934185 | SCG5;SCG5 | | TSS200;TSS200 |
| WG1978 | chr15:32933919-32934185 | SCG5;SCG5;SCG5;SCG5 | | 1stExon;5'UTR;5'UTR;1stExon |
| WG2194 | chr15:32964155-32964277 | SCG5;SCG5 | | Body;Body |
| WG3466 | chr15:32964155-32964500 | SCG5;SCG5 | | Body;Body |
| GM03075 | chr15:32964208-32964500 | SCG5;SCG5 | | Body;Body |
| WG1722 | chr15:33009409-33009531 | GREM1 | N_Shore | TSS1500 |
| WG1722 | chr15:34516488-34516777 | TMEM85 | | TSS1500 |
| GM00290 | chr15:34610829-34611069 | SLC12A6;SLC12A6;SLC12A6;SLC12A6;S | | Body;Body;Body;1stExon;Body;Body |

| | | | | |
|---|---|---|---|---|
| GM02175 | chr15:34876295-34876471 | GOLGA8B | Island | TSS1500 |
| WG1977 | chr15:35086890-35086967 | ACTC1 | | Body |
| WG2415 | chr15:35086890-35086985 | ACTC1 | | Body |
| WG2276 | chr15:36871420-36871849 | C15orf41 | N_Shore | TSS1500 |
| GM00290 | chr15:36871588-36871852 | C15orf41 | Island | TSS1500 |
| WG2844 | chr15:36871588-36872085 | C15orf41 | Island | TSS1500 |
| GM02706 | chr15:37170381-37170554 | LOC145845 | N_Shelf | Body |
| GM02706 | chr15:37174661-37175117 | LOC145845 | S_Shore | Body |
| GM00438 | chr15:37179603-37179779 | LOC145845 | Island | TSS1500 |
| GM02706 | chr15:40268421-40268777 | EIF2AK4 | N_Shore | Body |
| WG2415 | chr15:40268610-40268777 | EIF2AK4 | Island | Body |
| WG2414 | chr15:40268610-40269214 | EIF2AK4 | Island | Body |
| GM00289 | chr15:40268687-40269214 | EIF2AK4 | Island | Body |
| WG3465 | chr15:40544174-40544427 | PAK6;C15orf56;PAK6;PAK6 | N_Shore | 5'UTR;3'UTR;5'UTR;5'UTR |
| WG2844 | chr15:41062113-41062273 | DNAJC17;C15orf62 | | Body;TSS200 |
| WG1722 | chr15:41135432-41135900 | SPINT1;SPINT1;SPINT1 | N_Shore | TSS1500;TSS1500;TSS1500 |
| WG3465 | chr15:41135847-41135900 | SPINT1;SPINT1;SPINT1 | Island | TSS1500;TSS1500;TSS1500 |
| WG1310 | chr15:41795087-41795475 | ITPKA | Island | Body |
| GM00940 | chr15:41803428-41803772 | LTK;LTK;LTK | N_Shore | Body;Body;Body |
| GM00935 | chr15:41803428-41803850 | LTK;LTK;LTK | N_Shore | Body;Body;Body |
| GM00290 | chr15:41803428-41804283 | LTK;LTK;LTK | N_Shore | Body;Body;Body |
| GM00520 | chr15:41803592-41804283 | LTK;LTK;LTK | Island | Body;Body;Body |
| WG1977 | chr15:41806135-41806234 | LTK;LTK;LTK | S_Shore | TSS200;TSS200;TSS200 |
| WG1283 | chr15:41850280-41850612 | TYRO3 | N_Shore | TSS1500 |
| WG3465 | chr15:42302379-42303032 | PLA2G4E | | 1stExon |
| WG2276 | chr15:42371511-42371653 | PLA2G4D | N_Shore | Body |
| GM00289 | chr15:42371721-42371886 | PLA2G4D | Island | Body |
| WG3465 | chr15:42371808-42371967 | PLA2G4D | Island | Body |
| GM00940 | chr15:42500697-42500737 | VPS39 | S_Shore | TSS200 |
| GM01650 | chr15:44085947-44086625 | SERF2;MIR1282 | S_Shore | Body;Body |
| WG3465 | chr15:44086617-44087089 | SERF2;MIR1282 | S_Shore | 3'UTR;TSS1500 |
| GM00289 | chr15:44092606-44092937 | SERINC4;C15orf63 | N_Shore | TSS1500;TSS200 |
| WG3465 | chr15:45314789-45314933 | SORD | N_Shore | TSS1500 |
| GM02706 | chr15:45427034-45427734 | DUOX1;DUOX1 | N_Shore | Body;Body |
| WG1977 | chr15:45671148-45671246 | LOC145663;GATM | S_Shore | Body;TSS200 |
| GM00290 | chr15:45671148-45671279 | LOC145663;GATM | S_Shore | Body;TSS200 |
| WG2121 | chr15:45671155-45671246 | LOC145663;GATM | S_Shore | Body;TSS200 |
| WG1310 | chr15:49716247-49716645 | C15orf33;FGF7 | | Body;5'UTR |
| WG1722 | chr15:50474069-50474163 | SLC27A2;SLC27A2 | N_Shore | TSS1500;TSS1500 |
| WG3466 | chr15:51369862-51370019 | TNFAIP8L3 | | Body |
| WG2844 | chr15:51385891-51385970 | TNFAIP8L3 | Island | Body |
| GM00022 | chr15:51385916-51386254 | TNFAIP8L3 | Island | Body |
| WG1513 | chr15:51387651-51387921 | TNFAIP8L3 | Island | Body |
| WG3466 | chr15:51633430-51633821 | GLDN | N_Shore | TSS1500 |
| GM09256 | chr15:51972699-51973224 | SCG3;SCG3 | N_Shore | TSS1500;TSS1500 |
| WG2415 | chr15:53075421-53075533 | ONECUT1 | N_Shore | Body |
| WG3465 | chr15:55881599-55881706 | PYGO1 | S_Shore | TSS1500 |

| | | | | |
|---|---|---|---|---|
| GM02706 | chr15:56033606-56034020 | PRTG | N_Shore | Body |
| GM00290 | chr15:56033606-56034245 | PRTG | N_Shore | Body |
| WG2414 | chr15:57026014-57026064 | ZNF280D;ZNF280D;ZNF280D | Island | TSS1500;TSS1500;TSS1500 |
| GM00520 | chr15:58358775-58358879 | ALDH1A2;ALDH1A2 | Island | TSS1500;TSS1500 |
| GM01376 | chr15:58844359-58844583 | LIPC | | Body |
| GM02175 | chr15:62358920-62359103 | C2CD4A | N_Shore | TSS1500 |
| GM01375 | chr15:62359103-62359139 | C2CD4A | N_Shore | TSS200 |
| WG3466 | chr15:62359130-62359275 | C2CD4A | N_Shore | TSS200 |
| WG1310 | chr15:64673455-64673792 | KIAA0101;KIAA0101 | | Body;Body |
| WG1978 | chr15:64673601-64673792 | KIAA0101;KIAA0101;KIAA0101;KIAA0101 | | 1stExon;1stExon;5'UTR;5'UTR |
| WG1978 | chr15:64673827-64673872 | KIAA0101;KIAA0101 | | TSS200;TSS200 |
| GM02555 | chr15:65116194-65116557 | PIF1 | Island | Body |
| WG3465 | chr15:65204370-65204428 | ANKDD1A | Island | Body |
| GM02706 | chr15:65341736-65342244 | OSTBETA | | 5'UTR |
| WG1513 | chr15:65342244-65342522 | OSTBETA | | 5'UTR |
| GM00439 | chr15:65342270-65342634 | OSTBETA | | 5'UTR |
| GM09255 | chr15:65715664-65715951 | IGDCC4 | S_Shore | TSS1500 |
| WG2276 | chr15:66858212-66858446 | LCTL | | TSS1500 |
| WG2276 | chr15:67457584-67458398 | SMAD3;SMAD3;SMAD3;SMAD3 | | Body;Body;TSS1500;Body |
| WG1283 | chr15:67813132-67813145 | C15orf61 | N_Shore | TSS1500 |
| WG2120 | chr15:68125197-68125599 | LBXCOR1 | N_Shore | Body |
| GM00042 | chr15:69222592-69223018 | MIR548H4;SPESP1;NOX5 | Island | Body;TSS1500;TSS1500 |
| WG2120 | chr15:69323995-69324156 | MIR548H4;NOX5 | Island | Body;Body |
| GM09256 | chr15:69325271-69325560 | MIR548H4;NOX5 | N_Shore | Body;Body |
| GM00520 | chr15:69744390-69744528 | RPLP1;RPLP1 | N_Shore | TSS1500;TSS1500 |
| GM02173 | chr15:69744390-69744684 | RPLP1;RPLP1 | N_Shore | TSS1500;TSS1500 |
| WG2844 | chr15:70354647-70354825 | TLE3;TLE3;TLE3 | | Body;Body;Body |
| WG1513 | chr15:70994435-70994672 | UACA;UACA;UACA | | 5'UTR;Body;1stExon |
| WG2415 | chr15:70994435-70995014 | UACA;UACA;UACA | | 5'UTR;Body;1stExon |
| WG3465 | chr15:71408603-71408847 | CT62 | S_Shore | TSS1500 |
| WG1513 | chr15:73735476-73735531 | C15orf60 | Island | TSS200 |
| GM09256 | chr15:74421523-74421584 | LOC283731;ISLR2;ISLR2;ISLI | Island | Body;TSS1500;TSS200;TSS1500;TSS1 |
| WG2121 | chr15:74421523-74421635 | LOC283731;ISLR2;ISLR2;ISLI | Island | Body;TSS1500;TSS200;TSS1500;TSS1 |
| WG3465 | chr15:74427234-74427499 | ISLR2;ISLR2;ISLR2;ISLR2 | Island | Body;Body;Body;Body |
| WG2276 | chr15:74466337-74466794 | ISLR;ISLR;ISLR | N_Shore | 1stExon;TSS1500;5'UTR |
| WG2844 | chr15:74494781-74494861 | STRA6;STRA6;STRA6;STRA6;STRA6 | | 5'UTR;5'UTR;5'UTR;5'UTR;5'UTR |
| GM00290 | chr15:74494854-74494909 | STRA6;STRA6;STRA6;STRA6;STRA6 | | 5'UTR;5'UTR;5'UTR;5'UTR;5'UTR |
| WG2415 | chr15:74494861-74494909 | STRA6;STRA6;STRA6;STRA6;STRA6 | | 5'UTR;5'UTR;5'UTR;5'UTR;5'UTR |
| WG1513 | chr15:74494909-74495354 | STRA6;STRA6;STRA6;STRA6;STRA6 | | 5'UTR;5'UTR;5'UTR;5'UTR;5'UTR |
| WG2415 | chr15:74592566-74592786 | CCDC33 | | Body |
| GM00439 | chr15:75019087-75019203 | CYP1A1 | Island | TSS1500 |
| WG2276 | chr15:75019087-75019251 | CYP1A1 | Island | TSS1500 |
| WG2844 | chr15:76005861-76006143 | CSPG4 | | TSS1500 |
| GM02149 | chr15:78631626-78631878 | CRABP1 | N_Shore | TSS1500 |
| GM09256 | chr15:78631963-78632081 | CRABP1 | N_Shore | TSS1500 |
| GM00290 | chr15:78633437-78633662 | CRABP1 | Island | Body |
| GM09256 | chr15:79502065-79502137 | MIR184 | | TSS200 |

| | | | | |
|---|---|---|---|---|
| GM02706 | chr15:81666393-81666528 | TMC3 | | 1stExon |
| GM01096 | chr15:81666393-81666671 | TMC3 | | 1stExon |
| WG2276 | chr15:81666512-81666671 | TMC3 | | TSS200 |
| GM00042 | chr15:83317306-83317526 | CPEB1 | Island | TSS1500 |
| GM02706 | chr15:83317398-83317632 | CPEB1 | Island | TSS1500 |
| WG2276 | chr15:83654517-83654607 | FAM103A1 | N_Shore | TSS1500 |
| WG2844 | chr15:83953690-83953744 | BNC1 | Island | TSS1500 |
| GM02706 | chr15:83953690-83953864 | BNC1 | Island | TSS1500 |
| GM00935 | chr15:83953690-83953929 | BNC1 | Island | TSS1500 |
| GM00520 | chr15:83953690-83954150 | BNC1 | Island | TSS1500 |
| GM01096 | chr15:83953744-83953775 | BNC1 | Island | TSS1500 |
| GM09256 | chr15:83953765-83953780 | BNC1 | Island | TSS1500 |
| GM00290 | chr15:83953775-83953929 | BNC1 | Island | TSS1500 |
| WG2844 | chr15:83953780-83953880 | BNC1 | Island | TSS1500 |
| WG2276 | chr15:83953864-83953883 | BNC1 | Island | TSS1500 |
| WG1977 | chr15:84115433-84115529 | SH3GL3;SH3GL3 | N_Shore | TSS1500;TSS1500 |
| GM01096 | chr15:85360319-85360691 | ALPK3 | Island | 1stExon |
| WG3466 | chr15:85360655-85360691 | ALPK3 | S_Shore | 1stExon |
| WG2844 | chr15:86315115-86315211 | KLHL25;MIR1276 | S_Shelf | 5'UTR;TSS1500 |
| WG3465 | chr15:88576296-88576320 | NTRK3;NTRK3;NTRK3 | | Body;Body;Body |
| WG3466 | chr15:89153667-89154092 | MIR7-2 | | TSS1500 |
| GM01392 | chr15:89154058-89154196 | MIR7-2 | | TSS1500 |
| GM02175 | chr15:89920824-89921156 | LOC254559 | Island | TSS1500 |
| GM00290 | chr15:89920887-89921158 | LOC254559 | Island | TSS1500 |
| WG1721 | chr15:89921156-89921182 | LOC254559 | Island | TSS200 |
| GM00438 | chr15:91427184-91427363 | FES;FES;FES;FES | N_Shore | TSS1500;TSS1500;TSS1500;TSS1500 |
| WG2844 | chr15:91473059-91473291 | UNC45A | N_Shore | TSS1500 |
| WG2194 | chr15:91473059-91473365 | UNC45A | N_Shore | TSS1500 |
| GM00290 | chr15:91473059-91473569 | UNC45A | N_Shore | TSS1500 |
| WG1722 | chr15:91473475-91473569 | UNC45A;UNC45A | Island | 5'UTR;1stExon |
| WG1978 | chr15:92944664-92944756 | ST8SIA2 | | Body |
| WG2415 | chr15:93616894-93616950 | RGMA;RGMA;RGMA;RGMA;R | Island | 1stExon;Body;5'UTR;Body;1stExon;5'U |
| GM00042 | chr15:93616894-93616959 | RGMA;RGMA;RGMA;RGMA;R | Island | 1stExon;Body;5'UTR;Body;1stExon;5'U |
| GM09255 | chr15:93616943-93616959 | RGMA;RGMA;RGMA;RGMA;R | Island | Body;TSS200;TSS200;TSS1500;5'UTR; |
| GM09256 | chr15:93616959-93617030 | RGMA;RGMA;RGMA;RGMA;R | Island | 1stExon;Body;TSS200;TSS200;TSS150 |
| WG2844 | chr15:93617030-93618028 | RGMA;RGMA;RGMA;RGMA;R | Island | 1stExon;Body;5'UTR;TSS200;TSS200;T |
| WG2121 | chr15:93617134-93617146 | RGMA;RGMA;RGMA;RGMA;R | Island | 1stExon;Body;5'UTR;TSS200;TSS200;T |
| WG1978 | chr15:93617141-93617168 | RGMA;RGMA;RGMA;RGMA;R | Island | 1stExon;TSS1500;Body;5'UTR;TSS150 |
| WG2844 | chr15:93632896-93633172 | RGMA;RGMA | Island | TSS1500;TSS1500 |
| GM02173 | chr15:93632896-93633249 | RGMA;RGMA | Island | TSS1500;TSS1500 |
| GM01376 | chr15:93632896-93633278 | RGMA;RGMA | Island | TSS1500;TSS1500 |
| WG2415 | chr15:93633172-93633278 | RGMA;RGMA | Island | TSS1500;TSS1500 |
| WG2275 | chr15:96868857-96869220 | NR2F2 | S_Shelf | TSS1500 |
| WG1722 | chr15:96868935-96869132 | NR2F2 | S_Shelf | TSS1500 |
| WG2415 | chr15:96868935-96869220 | NR2F2 | S_Shelf | TSS1500 |
| GM09256 | chr15:96869027-96869132 | NR2F2 | S_Shelf | TSS200 |
| GM03075 | chr15:96869027-96869220 | NR2F2 | S_Shelf | TSS200 |

| | | | | |
|---|---|---|---|---|
| GM01653 | chr15:96869043-96869220 | NR2F2 | S_Shelf | TSS200 |
| WG3465 | chr15:99408636-99409506 | IGF1R | | Body |
| GM00940 | chr15:99408804-99409506 | IGF1R | | Body |
| WG1977 | chr15:99408958-99409506 | IGF1R | | Body |
| GM01392 | chr15:99409194-99409506 | IGF1R | | Body |
| GM01653 | chr15:99409942-99409957 | IGF1R | | Body |
| GM00022 | chr15:99789622-99789855 | TTC23;TTC23;TTC23;TTC23;T | N_Shore | 1stExon;5'UTR;5'UTR;1stExon;1stExon |
| GM00289 | chr16:10274064-10274415 | GRIN2A;GRIN2A;GRIN2A | Island | Body;Body;Body |
| WG1977 | chr16:1030383-1030401 | SOX8 | Island | TSS1500 |
| GM09255 | chr16:1030388-1030444 | SOX8 | Island | TSS1500 |
| GM03075 | chr16:1030561-1030586 | SOX8 | Island | TSS1500 |
| GM03073 | chr16:1030568-1030619 | SOX8 | Island | TSS1500 |
| WG1977 | chr16:10912696-10912718 | FAM18A | Island | TSS200 |
| WG2120 | chr16:10972898-10973301 | CIITA | Island | Body |
| GM02173 | chr16:1128689-1128721 | SSTR5;LOC146336 | Island | TSS200;Body |
| WG1977 | chr16:1128698-1128740 | SSTR5;LOC146336 | Island | TSS200;Body |
| WG1513 | chr16:1138415-1138797 | C1QTNF8 | N_Shore | 3'UTR |
| WG1977 | chr16:1143566-1143887 | C1QTNF8 | Island | Body |
| GM00439 | chr16:11761803-11761964 | SNN | N_Shore | TSS1500 |
| GM09256 | chr16:11835360-11835562 | TXNDC11 | N_Shore | Body |
| GM02706 | chr16:1204876-1205018 | CACNA1H;CACNA1H | S_Shore | Body;Body |
| WG3465 | chr16:1206750-1206844 | CACNA1H;CACNA1H | N_Shore | Body;Body |
| GM02149 | chr16:1214867-1215411 | CACNA1H;CACNA1H | N_Shelf | Body;Body |
| WG2844 | chr16:12183938-12184133 | SNX29 | | Body |
| WG3465 | chr16:1393584-1393797 | BAIAP3 | Island | Body |
| WG3465 | chr16:1429863-1430087 | UNKL | Island | TSS200 |
| WG1978 | chr16:14403004-14403040 | MIR365-1 | | TSS200 |
| GM02149 | chr16:14403138-14403405 | MIR365-1 | | TSS200 |
| GM09255 | chr16:14403144-14403425 | MIR365-1 | | Body |
| WG2121 | chr16:1524174-1524521 | CLCN7;CLCN7 | N_Shore | Body;Body |
| GM00935 | chr16:15595987-15596215 | C16orf45;C16orf45 | | TSS200;Body |
| WG2415 | chr16:15596053-15596215 | C16orf45;C16orf45 | | TSS200;Body |
| WG1977 | chr16:15596082-15596423 | C16orf45;C16orf45 | | TSS200;Body |
| WG1977 | chr16:1581834-1582219 | IFT140 | N_Shore | Body |
| GM02704 | chr16:1584118-1584516 | IFT140;TMEM204 | Island | Body;TSS200 |
| GM02555 | chr16:1585644-1586043 | IFT140;TMEM204 | S_Shore | Body;Body |
| GM00520 | chr16:1593415-1593766 | IFT140;TMEM204 | S_Shore | Body;Body |
| WG1513 | chr16:1844962-1845096 | IGFALS;IGFALS;IGFALS | S_Shelf | TSS200;TSS1500;TSS1500 |
| GM00935 | chr16:2047871-2048574 | ZNF598 | N_Shore | 3'UTR |
| WG2415 | chr16:2054944-2055403 | ZNF598 | S_Shore | Body |
| GM02456 | chr16:2082777-2083168 | SLC9A3R2;SLC9A3R2 | N_Shelf | Body;Body |
| GM03075 | chr16:2082777-2083393 | SLC9A3R2;SLC9A3R2 | N_Shelf | Body;Body |
| WG2844 | chr16:2131718-2132315 | TSC2;TSC2;TSC2 | N_Shelf | Body;Body;Body |
| WG1513 | chr16:2256384-2256716 | MLST8 | S_Shore | Body |
| GM00439 | chr16:23724835-23724901 | ERN2 | S_Shore | TSS200 |
| WG2415 | chr16:23724835-23724943 | ERN2 | S_Shore | TSS200 |
| GM01096 | chr16:23724854-23724943 | ERN2 | S_Shore | TSS200 |

| | | | | |
|---|---|---|---|---|
| GM02175 | chr16:23724901-23725375 | ERN2 | S_Shore | TSS200 |
| WG2414 | chr16:2653280-2653839 | LOC652276 | Island | TSS200 |
| GM01650 | chr16:27461333-27461638 | IL21R;IL21R;IL21R | | 3'UTR;3'UTR;3'UTR |
| GM09255 | chr16:27791167-27791248 | KIAA0556 | | 3'UTR |
| GM00520 | chr16:2801793-2801952 | LOC100128788;SRRM2 | N_Shore | Body;TSS1500 |
| GM09255 | chr16:2848577-2848830 | TESSP1 | Island | 1stExon |
| GM01393 | chr16:28634766-28635053 | SULT1A1;SULT1A1 | Island | 5'UTR;1stExon |
| WG3465 | chr16:2867434-2867683 | PRSS21;PRSS21;PRSS21 | Island | Body;Body;Body |
| GM02173 | chr16:2867434-2867773 | PRSS21;PRSS21;PRSS21 | Island | Body;Body;Body |
| GM00940 | chr16:28889630-28890100 | ATP2A1;ATP2A1 | N_Shore | TSS200;TSS200 |
| GM02175 | chr16:2892518-2892766 | TMPRSS8 | N_Shore | Body |
| GM00520 | chr16:2892518-2892769 | TMPRSS8 | N_Shore | Body |
| WG3465 | chr16:2892766-2892782 | TMPRSS8 | Island | TSS200 |
| GM02555 | chr16:2892766-2892809 | TMPRSS8 | Island | TSS200 |
| GM00438 | chr16:28995994-28996053 | LAT;LAT;LAT;LAT | | TSS200;TSS1500;TSS1500;TSS1500 |
| WG2844 | chr16:28996329-28996362 | LAT;LAT;LAT;LAT | | Body;TSS200;TSS200;TSS200 |
| GM02173 | chr16:2907517-2907859 | PRSS22 | N_Shore | Body |
| GM02175 | chr16:2907819-2908245 | PRSS22 | Island | Body |
| WG3466 | chr16:2908554-2908934 | PRSS22 | S_Shore | TSS1500 |
| GM09256 | chr16:29086271-29086456 | RRN3P2 | Island | Body |
| WG1310 | chr16:2908892-2908934 | PRSS22 | S_Shore | TSS1500 |
| GM00935 | chr16:29625989-29626401 | SLC7A5P1 | S_Shore | TSS1500 |
| GM00438 | chr16:29675846-29676071 | SPN;SPN | Island | Body;Body |
| WG2415 | chr16:29875145-29875187 | LOC440356;LOC440356;CDI | S_Shore | Body;TSS200;TSS1500 |
| GM02175 | chr16:29912460-29912904 | ASPHD1 | N_Shore | 1stExon |
| WG2415 | chr16:30017582-30018105 | DOC2A | | Body |
| GM00520 | chr16:30124803-30125047 | GDPD3;GDPD3;LOC100271831 | | 1stExon;5'UTR;TSS1500 |
| GM02175 | chr16:3017902-3018104 | KREMEN2;PAQR4;KREMEN2 | Island | Body;TSS1500;Body |
| GM00439 | chr16:3018104-3018470 | KREMEN2;KREMEN2;PAQR4 | Island | 3'UTR;3'UTR;TSS1500 |
| GM00031 | chr16:30485485-30485684 | ITGAL;ITGAL | Island | Body;Body |
| GM09256 | chr16:31022351-31022521 | STX1B | | TSS1500 |
| GM00042 | chr16:31146999-31147054 | PRSS8;PRSS8 | | 1stExon;5'UTR |
| GM02706 | chr16:31147017-31147054 | PRSS8;PRSS8 | | 1stExon;5'UTR |
| WG3466 | chr16:31147017-31147177 | PRSS8;PRSS8 | | 1stExon;5'UTR |
| GM02555 | chr16:31214307-31214346 | PYCARD;PYCARD | S_Shore | TSS200;TSS200 |
| GM02555 | chr16:31235535-31235824 | TRIM72 | Island | Body |
| WG1722 | chr16:31366377-31366429 | ITGAX | | TSS200 |
| WG1513 | chr16:3142911-3143041 | ZSCAN10 | S_Shelf | TSS200 |
| WG3466 | chr16:31439106-31439393 | COX6A2 | Island | Body |
| WG1513 | chr16:31581150-31581309 | CSDAP1 | S_Shore | TSS1500 |
| GM00935 | chr16:3265228-3265467 | OR1F2P | | TSS1500 |
| WG1513 | chr16:33964783-33965322 | MIR1826 | Island | TSS1500 |
| GM00520 | chr16:3627489-3627730 | NLRC3 | | TSS200 |
| WG2415 | chr16:3724734-3724848 | TRAP1 | Island | Body |
| GM02704 | chr16:3843012-3843415 | CREBBP;CREBBP | | Body;Body |
| GM00289 | chr16:4102293-4102366 | ADCY9 | | Body |
| WG3465 | chr16:4103161-4103225 | ADCY9 | | Body |

| | | | | |
|---|---|---|---|---|
| WG3466 | chr16:4163819-4164087 | ADCY9 | N_Shore | Body |
| GM09255 | chr16:419975-420112 | MRPL28 | Island | Body |
| GM02175 | chr16:420108-420230 | MRPL28 | Island | Body |
| WG2844 | chr16:4307906-4308192 | TFAP4 | N_Shelf | 3'UTR |
| GM00290 | chr16:4311799-4311941 | TFAP4 | Island | Body |
| GM02149 | chr16:450837-450970 | DECR2 | Island | TSS1500 |
| GM02173 | chr16:450837-451495 | DECR2 | Island | TSS1500 |
| GM00438 | chr16:46603015-46603059 | ANKRD26P1 | | TSS200 |
| WG1722 | chr16:4665298-4665704 | FAM100A | Island | TSS1500 |
| GM00042 | chr16:4689950-4690020 | MGRN1;MGRN1;MGRN1;MGRN1 | | Body;Body;Body;Body |
| WG1722 | chr16:46919021-46919112 | GPT2;GPT2 | S_Shore | Body;TSS200 |
| WG2844 | chr16:4729905-4730657 | MGRN1;MGRN1;MGRN1;MGI | N_Shore | Body;Body;Body;Body |
| WG1513 | chr16:4838559-4838722 | SEPT12;LOC440335;SEPT12;LOC440335 | | TSS200;Body;TSS200;Body |
| GM02555 | chr16:49525139-49525304 | ZNF423 | N_Shore | 3'UTR |
| WG1513 | chr16:50347766-50347895 | ADCY7 | | Body |
| WG1977 | chr16:51183988-51184205 | SALL1;SALL1 | Island | Body;5'UTR |
| GM01392 | chr16:51184152-51184392 | SALL1;SALL1 | Island | Body;5'UTR |
| GM01653 | chr16:51184355-51184392 | SALL1;SALL1;SALL1 | Island | Body;5'UTR;1stExon |
| GM00438 | chr16:52582072-52582262 | TOX3;TOX3 | S_Shore | TSS1500;TSS1500 |
| WG1513 | chr16:54318044-54318221 | IRX3 | Island | Body |
| GM02706 | chr16:54318115-54318314 | IRX3 | Island | Body |
| GM00289 | chr16:54318772-54318935 | IRX3 | Island | Body |
| WG1309 | chr16:55794478-55794910 | CES4 | N_Shore | TSS200 |
| WG1978 | chr16:55794595-55794910 | CES4 | N_Shore | Body |
| GM00438 | chr16:55908870-55909393 | CES7;CES7 | | Body;Body |
| GM09256 | chr16:56554249-56554313 | BBS2 | S_Shore | TSS1500 |
| GM00520 | chr16:56677683-56677855 | MT1DP;MT1DP | S_Shore | Body;TSS200 |
| WG3466 | chr16:56703400-56703833 | MT1G;MT1H | Island | TSS1500;TSS1500 |
| WG1513 | chr16:56709853-56710256 | MT1IP | Island | Body |
| GM00290 | chr16:57317679-57318087 | PLLP | N_Shore | Body |
| WG2415 | chr16:58019866-58019984 | TEPP;TEPP | Island | Body;Body |
| GM00042 | chr16:58497230-58497239 | NDRG4;NDRG4 | Island | TSS1500;TSS1500 |
| GM00935 | chr16:58533743-58534157 | NDRG4;NDRG4;NDRG4 | N_Shore | Body;Body;Body |
| WG1977 | chr16:58533743-58534708 | NDRG4;NDRG4;NDRG4 | N_Shore | Body;Body;Body |
| GM00439 | chr16:58535416-58535556 | NDRG4;NDRG4;NDRG4 | Island | Body;Body;Body |
| WG2844 | chr16:65105516-65105714 | CDH11 | | 5'UTR |
| WG2276 | chr16:66400320-66400599 | CDH5 | | TSS1500 |
| GM01096 | chr16:66613096-66613334 | CMTM2 | Island | TSS1500 |
| GM02706 | chr16:66613220-66613334 | CMTM2 | Island | TSS200 |
| WG3466 | chr16:66613220-66613355 | CMTM2 | Island | TSS200 |
| GM02704 | chr16:66613266-66613278 | CMTM2 | Island | TSS200 |
| GM03075 | chr16:66613266-66613334 | CMTM2 | Island | TSS200 |
| GM00935 | chr16:66613273-66613334 | CMTM2 | Island | TSS200 |
| GM01650 | chr16:67197084-67197186 | HSF4;FBXL8;HSF4 | Island | TSS1500;Body;TSS1500 |
| GM02706 | chr16:67197084-67197396 | HSF4;FBXL8;HSF4 | Island | TSS1500;Body;TSS1500 |
| WG3465 | chr16:67197084-67197640 | HSF4;FBXL8;HSF4 | Island | TSS1500;Body;TSS1500 |
| WG1977 | chr16:67197084-67198029 | HSF4;FBXL8;HSF4 | Island | TSS1500;Body;TSS1500 |

| | | | | |
|---|---|---|---|---|
| WG2415 | chr16:67197129-67197640 | HSF4;HSF4;FBXL8 | Island | TSS200;TSS200;Body |
| GM02175 | chr16:67197180-67197640 | HSF4;HSF4;FBXL8 | Island | TSS200;TSS200;Body |
| WG1513 | chr16:67223733-67224199 | EXOC3L | | 5'UTR |
| GM00042 | chr16:67223955-67224341 | EXOC3L;EXOC3L | | 5'UTR;1stExon |
| GM00439 | chr16:67224199-67224629 | EXOC3L | | TSS200 |
| GM02706 | chr16:67428155-67428324 | TPPP3;TPPP3 | Island | TSS1500;TSS1500 |
| WG1722 | chr16:67428155-67428497 | TPPP3;TPPP3 | Island | TSS1500;TSS1500 |
| GM01650 | chr16:67428315-67428497 | TPPP3;TPPP3 | Island | TSS1500;TSS1500 |
| GM02555 | chr16:67433184-67433636 | ZDHHC1 | Island | Body |
| GM03075 | chr16:67433258-67433636 | ZDHHC1 | Island | Body |
| GM02555 | chr16:67515916-67516493 | ATP6V0D1 | S_Shore | TSS1500 |
| GM00935 | chr16:67516162-67516493 | ATP6V0D1 | S_Shore | TSS1500 |
| WG2121 | chr16:67516286-67516546 | ATP6V0D1 | S_Shore | TSS1500 |
| GM03075 | chr16:67695257-67695272 | PARD6A;ACD;ACD;PARD6A;A | Island | Body;TSS1500;TSS1500;Body;TSS150 |
| GM09255 | chr16:67918680-67918965 | NRN1L | Island | TSS200 |
| WG1513 | chr16:68033509-68034002 | DPEP2 | | TSS200 |
| GM00042 | chr16:68269295-68269694 | ESRP2 | Island | Body |
| WG3465 | chr16:68270252-68270282 | ESRP2 | Island | TSS200 |
| GM00438 | chr16:68481049-68481342 | SMPD3 | Island | 5'UTR |
| GM01653 | chr16:68481342-68481489 | SMPD3 | Island | 5'UTR |
| WG3465 | chr16:68677997-68678092 | CDH3 | N_Shore | TSS200 |
| WG1977 | chr16:691444-691570 | FAM195A | N_Shore | TSS1500 |
| WG2414 | chr16:69958626-69958964 | WWP2;WWP2 | | TSS1500;Body |
| WG2276 | chr16:69958792-69958964 | WWP2;WWP2 | | TSS200;Body |
| WG1721 | chr16:70680320-70680380 | IL34 | | TSS200 |
| GM01653 | chr16:70680380-70680631 | IL34 | | TSS200 |
| GM00289 | chr16:72821141-72821498 | ZFHX3;ZFHX3 | Island | Body;Body |
| WG1513 | chr16:72821353-72821665 | ZFHX3;ZFHX3 | Island | Body;Body |
| WG1513 | chr16:72981203-72981599 | ZFHX3;ZFHX3 | | Body;Body |
| WG2415 | chr16:73126608-73126909 | HTA | | Body |
| WG3466 | chr16:746048-746132 | FBXL16 | S_Shore | Body |
| GM00940 | chr16:75468057-75468297 | CFDP1 | S_Shore | TSS1500 |
| GM02706 | chr16:75468057-75468318 | CFDP1 | S_Shore | TSS1500 |
| WG1513 | chr16:75568999-75569459 | CHST5 | | 5'UTR |
| GM02149 | chr16:7615516-7615568 | A2BP1;A2BP1;A2BP1;A2BP1;A2BP1;A2BI | | Body;Body;Body;Body;Body;Body |
| GM02149 | chr16:76342377-76342678 | CNTNAP4;CNTNAP4 | | Body;TSS1500 |
| GM02706 | chr16:77469538-77469998 | ADAMTS18 | S_Shore | TSS1500 |
| WG3465 | chr16:77469583-77469710 | ADAMTS18 | S_Shore | TSS1500 |
| WG1513 | chr16:77469667-77469998 | ADAMTS18 | S_Shore | TSS1500 |
| GM00935 | chr16:78539954-78540172 | WWOX | | Body |
| WG1977 | chr16:81527360-81527504 | CMIP | | Body |
| WG1978 | chr16:81527427-81527504 | CMIP | | Body |
| GM00289 | chr16:81726721-81727056 | CMIP;CMIP | | Body;Body |
| GM02175 | chr16:82045140-82045222 | SDR42E1 | Island | TSS200 |
| GM00439 | chr16:82045183-82045222 | SDR42E1 | Island | TSS200 |
| WG3466 | chr16:82660206-82660328 | CDH13 | N_Shore | TSS1500 |
| WG3465 | chr16:83171068-83171314 | CDH13 | | Body |

| | | | | |
|---|---|---|---|---|
| GM09255 | chr16:832942-832963 | MSLNL | Island | TSS200 |
| WG1657 | chr16:84058601-84059138 | SLC38A8 | | Body |
| GM09255 | chr16:84328104-84328208 | WFDC1 | N_Shore | TSS1500 |
| WG1513 | chr16:84341323-84341364 | WFDC1 | | Body |
| WG2276 | chr16:84346722-84346930 | WFDC1 | Island | Body |
| WG1514 | chr16:84346910-84346930 | WFDC1 | Island | Body |
| WG3466 | chr16:84860766-84860918 | CRISPLD2 | | 5'UTR |
| WG3465 | chr16:85074868-85075076 | KIAA0513 | | 5'UTR |
| WG2415 | chr16:854168-854439 | PRR25 | N_Shelf | TSS1500 |
| GM02456 | chr16:85684468-85684853 | KIAA0182;KIAA0182 | Island | Body;Body |
| GM01392 | chr16:85723150-85723539 | GINS2 | S_Shore | TSS1500 |
| GM00935 | chr16:857454-857863 | PRR25 | Island | Body |
| WG1282 | chr16:857454-857981 | PRR25 | Island | Body |
| GM03075 | chr16:857576-857981 | PRR25 | Island | Body |
| WG1513 | chr16:8619531-8619943 | TMEM114 | N_Shore | 3'UTR |
| GM09255 | chr16:86370962-86371130 | LOC732275 | | Body |
| WG1977 | chr16:86371130-86371363 | LOC732275 | | Body |
| GM02149 | chr16:86371248-86371442 | LOC732275 | | Body |
| GM00022 | chr16:86546374-86546631 | FOXF1 | Island | Body |
| WG2275 | chr16:86546374-86546938 | FOXF1 | Island | Body |
| GM00520 | chr16:86546374-86546979 | FOXF1 | Island | Body |
| WG2276 | chr16:86546374-86547544 | FOXF1 | Island | Body |
| GM00438 | chr16:86546628-86546785 | FOXF1 | Island | Body |
| WG1657 | chr16:86546628-86546938 | FOXF1 | Island | Body |
| WG1977 | chr16:86546628-86547203 | FOXF1 | Island | Body |
| GM01096 | chr16:86546628-86547530 | FOXF1 | Island | Body |
| GM00935 | chr16:86546631-86546938 | FOXF1 | Island | Body |
| GM01095 | chr16:86546785-86547530 | FOXF1 | S_Shore | 3'UTR |
| GM02704 | chr16:86546979-86547530 | FOXF1 | S_Shore | 3'UTR |
| WG1657 | chr16:86547203-86547386 | FOXF1 | N_Shore | 3'UTR |
| WG2844 | chr16:86547203-86547530 | FOXF1 | N_Shore | 3'UTR |
| GM09256 | chr16:86547203-86547544 | FOXF1 | N_Shore | 3'UTR |
| GM01375 | chr16:86547386-86547544 | FOXF1 | N_Shore | 3'UTR |
| GM01392 | chr16:87441441-87441659 | ZCCHC14 | Island | 3'UTR |
| WG3466 | chr16:87467986-87469199 | ZCCHC14 | | Body |
| GM00935 | chr16:87468206-87468247 | ZCCHC14 | | Body |
| WG1977 | chr16:87637999-87638484 | JPH3 | S_Shore | Body |
| GM02555 | chr16:87769432-87769974 | KLHDC4 | | Body |
| WG2843 | chr16:87769500-87769974 | KLHDC4 | | Body |
| GM02555 | chr16:87904491-87904580 | SLC7A5 | S_Shore | TSS1500 |
| WG1977 | chr16:87996731-87996870 | BANP;BANP | | 5'UTR;5'UTR |
| WG2844 | chr16:88052956-88053104 | BANP;BANP | Island | Body;Body |
| WG1513 | chr16:88058182-88058326 | BANP;BANP | N_Shelf | Body;Body |
| GM02175 | chr16:8806966-8807018 | ABAT;ABAT;ABAT | | 5'UTR;1stExon;5'UTR |
| GM00042 | chr16:8806966-8807043 | ABAT;ABAT;ABAT | | 5'UTR;1stExon;5'UTR |
| WG2275 | chr16:8807011-8807043 | ABAT;ABAT;ABAT | | 5'UTR;1stExon;5'UTR |
| GM01096 | chr16:88496788-88497077 | ZNF469 | Island | 1stExon |

| | | | | |
|---|---|---|---|---|
| GM02173 | chr16:88496788-88497293 | ZNF469 | Island | 1stExon |
| GM03075 | chr16:88590325-88590404 | ZFPM1 | Island | Body |
| WG3466 | chr16:88590325-88590409 | ZFPM1 | Island | Body |
| WG3465 | chr16:88590340-88590409 | ZFPM1 | Island | Body |
| WG1977 | chr16:88706241-88706426 | IL17C | Island | Body |
| WG1977 | chr16:88933068-88933182 | PABPN1L | N_Shelf | TSS200 |
| WG2844 | chr16:88943319-88943533 | CBFA2T3;CBFA2T3 | N_Shore | 3'UTR;3'UTR |
| WG2844 | chr16:88990161-88990232 | CBFA2T3;CBFA2T3 | N_Shelf | 5'UTR;Body |
| WG2415 | chr16:89019403-89019632 | CBFA2T3 | | Body |
| GM00520 | chr16:89043527-89043707 | CBFA2T3 | | TSS200 |
| WG1977 | chr16:89225592-89225639 | C16orf81 | N_Shore | TSS200 |
| WG3465 | chr16:89228970-89229439 | C16orf81 | N_Shelf | Body |
| GM00439 | chr16:89381904-89382131 | ANKRD11 | | Body |
| WG2415 | chr16:89387225-89387569 | ANKRD11 | | 5'UTR |
| GM01392 | chr16:89387418-89387805 | ANKRD11 | | 5'UTR |
| WG2276 | chr16:89439771-89440324 | ANKRD11 | S_Shore | 5'UTR |
| WG2844 | chr16:89445433-89445503 | ANKRD11 | | 5'UTR |
| GM02555 | chr16:89462292-89462418 | ANKRD11 | | 5'UTR |
| GM09256 | chr16:89573369-89573562 | SPG7;SPG7 | N_Shore | TSS1500;TSS1500 |
| GM00022 | chr16:89573535-89573631 | SPG7;SPG7 | N_Shore | TSS1500;TSS1500 |
| GM09256 | chr16:89686618-89686885 | DPEP1;DPEP1 | | 5'UTR;TSS1500 |
| WG2844 | chr16:89761017-89761424 | CDK10;CDK10;CDK10;CDK1 | Island | Body;Body;Body;Body;Body;Body |
| WG2120 | chr16:90016004-90016061 | DEF8;DEF8 | Island | 5'UTR;Body |
| WG1722 | chr16:90016004-90016551 | DEF8;DEF8 | Island | 5'UTR;Body |
| WG3465 | chr16:90114523-90115013 | LOC100130015;LOC100130015 | | TSS1500;TSS1500 |
| GM01096 | chr16:90143751-90143815 | PRDM7 | Island | TSS1500 |
| WG2844 | chr16:968219-968265 | LMF1 | S_Shore | Body |
| WG1978 | chr17:1028394-1028676 | ABR;ABR | N_Shore | Body;Body |
| GM02555 | chr17:1028394-1028765 | ABR;ABR | N_Shore | Body;Body |
| WG2276 | chr17:1028394-1028931 | ABR;ABR | N_Shore | Body;Body |
| WG3466 | chr17:1028394-1029394 | ABR;ABR | N_Shore | Body;Body |
| WG2415 | chr17:1028673-1028931 | ABR;ABR | Island | Body;5'UTR |
| WG1722 | chr17:10421635-10422077 | MYH1 | | 5'UTR |
| WG1977 | chr17:10421635-10422189 | MYH1 | | 5'UTR |
| GM00016 | chr17:1062865-1063138 | ABR;ABR | N_Shore | Body;5'UTR |
| GM09256 | chr17:1202015-1202156 | TUSC5 | | 3'UTR |
| GM02175 | chr17:12567873-12568092 | MYOCD;MYOCD | N_Shore | TSS1500;TSS1500 |
| GM02706 | chr17:1390182-1390671 | MYO1C;MYO1C;MYO1C | N_Shore | 5'UTR;Body;TSS1500 |
| GM01392 | chr17:14206774-14206874 | HS3ST3B1;MGC12916 | Island | Body;TSS1500 |
| GM02555 | chr17:14206886-14207530 | HS3ST3B1;MGC12916 | Island | Body;TSS200 |
| GM00935 | chr17:14206901-14207241 | HS3ST3B1;MGC12916 | Island | Body;TSS200 |
| WG1977 | chr17:14206901-14207530 | HS3ST3B1;MGC12916 | Island | Body;TSS200 |
| GM02640 | chr17:14206981-14207241 | HS3ST3B1;MGC12916 | Island | Body;TSS200 |
| GM02456 | chr17:14206981-14207530 | HS3ST3B1;MGC12916 | Island | Body;TSS200 |
| WG1513 | chr17:14207241-14207890 | HS3ST3B1;MGC12916 | Island | Body;Body |
| WG1977 | chr17:1466709-1466715 | PITPNA | S_Shore | TSS1500 |
| WG2276 | chr17:1509883-1510080 | SLC43A2 | Island | Body |

| | | | | |
|---|---|---|---|---|
| WG2844 | chr17:15168483-15168658 | PMP22;PMP22 | S_Shore | 1stExon;5'UTR |
| GM00439 | chr17:151914-152350 | RPH3AL | N_Shore | Body |
| WG1513 | chr17:152259-152350 | RPH3AL | Island | Body |
| WG3466 | chr17:15244441-15244519 | TEKT3 | N_Shore | 5'UTR |
| WG2414 | chr17:15244441-15244970 | TEKT3 | N_Shore | 5'UTR |
| WG1722 | chr17:15244482-15244887 | TEKT3 | N_Shore | 5'UTR |
| WG1977 | chr17:15244482-15244970 | TEKT3 | N_Shore | 5'UTR |
| GM00935 | chr17:15244482-15245089 | TEKT3 | N_Shore | 5'UTR |
| GM00290 | chr17:15244519-15244970 | TEKT3 | N_Shore | 5'UTR |
| GM01393 | chr17:15244887-15245076 | TEKT3;TEKT3 | Island | 5'UTR;1stExon |
| GM02456 | chr17:15244970-15245185 | TEKT3 | Island | TSS200 |
| GM02175 | chr17:15245076-15245091 | TEKT3 | Island | TSS200 |
| GM00439 | chr17:15245089-15245196 | TEKT3 | Island | TSS200 |
| WG3465 | chr17:154410-154671 | RPH3AL | S_Shore | Body |
| WG2193 | chr17:154410-155045 | RPH3AL | S_Shore | Body |
| WG3466 | chr17:154499-155045 | RPH3AL | N_Shelf | Body |
| WG1978 | chr17:1553415-1553502 | RILP | S_Shore | TSS200 |
| GM00439 | chr17:16120399-16120890 | PIGL | S_Shore | TSS200 |
| WG1977 | chr17:1626477-1626575 | WDR81;WDR81;WDR81 | N_Shore | TSS1500;Body;5'UTR |
| WG1722 | chr17:16318668-16318930 | TRPV2 | | TSS1500 |
| GM09256 | chr17:16318668-16318972 | TRPV2 | | TSS1500 |
| GM00042 | chr17:16318860-16318972 | TRPV2 | | TSS200 |
| WG3465 | chr17:1633714-1634018 | WDR81;WDR81;WDR81;WDR | N_Shelf | Body;Body;Body;Body |
| WG3466 | chr17:1634018-1634335 | WDR81;WDR81;WDR81;WDR | N_Shelf | Body;Body;Body;Body |
| GM00042 | chr17:16593582-16593601 | CCDC144A | Island | TSS200 |
| GM01393 | chr17:16593582-16593613 | CCDC144A | Island | TSS200 |
| WG1977 | chr17:16593589-16593613 | CCDC144A | Island | TSS200 |
| WG2276 | chr17:16593601-16593885 | CCDC144A | Island | TSS200 |
| WG1513 | chr17:1664077-1664339 | SERPINF1 | | TSS1500 |
| WG3465 | chr17:1665119-1665172 | SERPINF1 | | TSS200 |
| WG1283 | chr17:1686627-1686737 | SMYD4 | | Body |
| GM00438 | chr17:17109691-17110207 | PLD6 | Island | TSS200 |
| WG2276 | chr17:17110120-17110353 | PLD6 | Island | TSS1500 |
| GM00935 | chr17:17465162-17465682 | PEMT;PEMT;PEMT | Island | Body;Body;Body |
| GM00438 | chr17:17603531-17604146 | RAI1 | N_Shore | 5'UTR |
| GM09256 | chr17:17713364-17713886 | RAI1 | Island | 3'UTR |
| GM00290 | chr17:180856-181288 | RPH3AL | | 5'UTR |
| WG2276 | chr17:183931-184106 | RPH3AL | N_Shore | 5'UTR |
| WG1977 | chr17:18528568-18528638 | CCDC144B | Island | 1stExon |
| GM00289 | chr17:1880631-1880941 | RTN4RL1 | N_Shore | Body |
| WG2844 | chr17:19247837-19247920 | MIR1180;B9D1 | Island | Body;Body |
| WG1722 | chr17:19314299-19314328 | RNF112 | | TSS200 |
| WG1977 | chr17:1957147-1957206 | HIC1 | Island | TSS1500 |
| WG1977 | chr17:1957370-1957605 | HIC1 | Island | TSS1500 |
| GM01650 | chr17:1957625-1957677 | HIC1 | Island | TSS1500 |
| WG1977 | chr17:1957625-1957794 | HIC1 | Island | TSS1500 |
| WG1977 | chr17:1957866-1957926 | HIC1 | Island | TSS1500 |

| | | | | |
|---|---|---|---|---|
| WG2276 | chr17:1961109-1962236 | HIC1;HIC1 | Island | Body;Body |
| WG3466 | chr17:1961286-1962236 | HIC1;HIC1 | Island | Body;Body |
| WG1983 | chr17:1961440-1962236 | HIC1;HIC1 | Island | Body;Body |
| GM02704 | chr17:1961778-1962236 | HIC1;HIC1 | Island | Body;Body |
| WG1722 | chr17:19648718-19648855 | ALDH3A1;ALDH3A1;ALDH3A | S_Shore | 5'UTR;5'UTR;5'UTR;1stExon |
| WG2276 | chr17:19651940-19652056 | ALDH3A1 | S_Shelf | TSS200 |
| GM02175 | chr17:20799470-20799511 | CCDC144NL | Island | TSS200 |
| WG3465 | chr17:20799511-20799676 | CCDC144NL | Island | TSS200 |
| GM00438 | chr17:20799644-20799714 | CCDC144NL | Island | TSS200 |
| GM09256 | chr17:20799644-20799769 | CCDC144NL | Island | TSS200 |
| WG3465 | chr17:21300735-21300791 | KCNJ12 | | 5'UTR |
| WG1513 | chr17:21825321-21825355 | FAM27L | | TSS200 |
| WG2415 | chr17:25958267-25958303 | LGALS9;LGALS9;LGALS9;LGALS9;LGALS | | Body;1stExon;1stExon;5'UTR;5'UTR |
| WG1513 | chr17:26127480-26127537 | NOS2;NOS2 | | 1stExon;5'UTR |
| GM00520 | chr17:26711393-26711803 | SARM1 | Island | Body |
| WG3465 | chr17:26732191-26732272 | SLC46A1 | Island | Body |
| GM00940 | chr17:26732249-26732374 | SLC46A1 | Island | Body |
| GM00520 | chr17:2699542-2699689 | RAP1GAP2;RAP1GAP2 | | TSS200;TSS200 |
| GM02175 | chr17:2699689-2699718 | RAP1GAP2;RAP1GAP2 | | TSS200;TSS200 |
| WG1722 | chr17:2699689-2699802 | RAP1GAP2;RAP1GAP2 | | TSS200;TSS200 |
| GM02175 | chr17:27049557-27050042 | RPL23A;SNORD42A;SNORD4 | N_Shelf | Body;TSS1500;TSS1500;TSS200 |
| WG2276 | chr17:28705147-28705364 | CPD | N_Shore | TSS1500 |
| GM00940 | chr17:28803808-28803927 | GOSR1;GOSR1;GOSR1 | N_Shore | TSS1500;TSS1500;TSS1500 |
| WG2844 | chr17:2886250-2886453 | RAP1GAP2;RAP1GAP2 | | Body;Body |
| WG2414 | chr17:29248604-29248725 | ADAP2 | N_Shore | TSS200 |
| GM09255 | chr17:29297380-29297414 | RNF135;RNF135 | N_Shore | TSS1500;TSS1500 |
| GM00941 | chr17:29886885-29887376 | MIR193A | Island | TSS200 |
| WG2415 | chr17:3031354-3031874 | OR1G1 | | TSS1500 |
| WG3465 | chr17:30591837-30592033 | RHBDL3 | N_Shore | TSS1500 |
| WG3466 | chr17:30822561-30823006 | MYO1D | N_Shore | Body |
| WG1310 | chr17:30822561-30823087 | MYO1D | N_Shore | Body |
| GM02555 | chr17:30822561-30823145 | MYO1D | N_Shore | Body |
| WG1977 | chr17:30822961-30823087 | MYO1D | Island | Body |
| WG1513 | chr17:3117953-3118291 | OR1A1 | | TSS1500 |
| GM00439 | chr17:32483560-32483711 | ACCN1;ACCN1 | N_Shore | 1stExon;5'UTR |
| GM02706 | chr17:32909164-32909375 | TMEM132E | Island | Body |
| WG3465 | chr17:32964265-32964615 | TMEM132E | N_Shore | Body |
| WG1977 | chr17:32964596-32964846 | TMEM132E | Island | Body |
| GM00290 | chr17:32964596-32965324 | TMEM132E | Island | Body |
| WG3465 | chr17:33416834-33417086 | RFFL | S_Shore | TSS1500 |
| WG1722 | chr17:33759484-33759573 | SLFN12;SLFN12 | | 1stExon;5'UTR |
| GM00289 | chr17:33759484-33759648 | SLFN12;SLFN12 | | 1stExon;5'UTR |
| GM00022 | chr17:33759484-33759929 | SLFN12;SLFN12 | | 1stExon;5'UTR |
| WG3465 | chr17:33759484-33759957 | SLFN12;SLFN12 | | 1stExon;5'UTR |
| WG1977 | chr17:33759484-33759965 | SLFN12;SLFN12 | | 1stExon;5'UTR |
| WG3466 | chr17:33759484-33759986 | SLFN12;SLFN12 | | 1stExon;5'UTR |
| GM09256 | chr17:33759512-33759648 | SLFN12;SLFN12 | | 1stExon;5'UTR |

| | | | | |
|---|---|---|---|---|
| GM00935 | chr17:33759648-33759986 | SLFN12 | | TSS200 |
| GM00520 | chr17:33759648-33760293 | SLFN12 | | TSS200 |
| WG1721 | chr17:33759929-33759965 | SLFN12 | | TSS1500 |
| GM02706 | chr17:33759929-33759986 | SLFN12 | | TSS1500 |
| GM03075 | chr17:33759929-33760249 | SLFN12 | | TSS1500 |
| GM09256 | chr17:33759957-33759986 | SLFN12 | | TSS1500 |
| WG2415 | chr17:33775952-33775963 | SLFN13 | N_Shore | TSS200 |
| WG1977 | chr17:33775952-33776116 | SLFN13 | N_Shore | TSS200 |
| WG1513 | chr17:33776116-33776554 | SLFN13 | N_Shore | TSS1500 |
| WG1983 | chr17:33825172-33825374 | SLFN12L | | Body |
| WG3466 | chr17:33842181-33842301 | SLFN12L | | Body |
| GM00935 | chr17:33866067-33866073 | SLFN12L | | TSS1500 |
| WG1513 | chr17:34308137-34308567 | CCL16 | | Body |
| WG3465 | chr17:3433428-3433970 | TRPV3 | N_Shelf | Body |
| WG2415 | chr17:34344995-34345172 | CCL23;CCL23;CCL23;CCL23 | | 1stExon;5'UTR;1stExon;5'UTR |
| GM09256 | chr17:34345136-34345392 | CCL23;CCL23 | | TSS200;TSS200 |
| WG1513 | chr17:34417074-34417575 | CCL3 | | Body |
| GM02555 | chr17:3461140-3461398 | TRPV3;TRPV3 | | 1stExon;5'UTR |
| WG1513 | chr17:3461140-3461450 | TRPV3;TRPV3 | | 1stExon;5'UTR |
| WG2415 | chr17:3461140-3462082 | TRPV3;TRPV3 | | 1stExon;5'UTR |
| GM03075 | chr17:3461143-3461480 | TRPV3;TRPV3 | | 1stExon;5'UTR |
| GM00016 | chr17:3461398-3461480 | TRPV3 | | TSS200 |
| GM00439 | chr17:35656860-35657046 | ACACA;ACACA;ACACA;ACACA;ACACA | | Body;TSS200;TSS200;Body;Body |
| WG1513 | chr17:36499775-36500237 | GPR179 | | TSS200 |
| GM02706 | chr17:36666462-36666953 | ARHGAP23 | Island | Body |
| WG1721 | chr17:36715787-36715953 | SRCIN1 | Island | Body |
| GM00022 | chr17:36760878-36761029 | SRCIN1 | N_Shore | Body |
| GM00520 | chr17:36997420-36997563 | C17orf98 | N_Shore | Body |
| GM09255 | chr17:37028105-37028650 | LASP1 | S_Shore | Body |
| WG3465 | chr17:3704471-3704494 | ITGAE;ITGAE | | 1stExon;5'UTR |
| WG2120 | chr17:3705631-3705875 | ITGAE | | TSS1500 |
| GM02149 | chr17:37123638-37123671 | FBXO47;FBXO47 | | 1stExon;5'UTR |
| GM09256 | chr17:37123638-37123909 | FBXO47;FBXO47 | | 1stExon;5'UTR |
| WG2275 | chr17:37123669-37123711 | FBXO47 | | TSS200 |
| WG1513 | chr17:37123767-37123843 | FBXO47 | | TSS200 |
| GM02149 | chr17:37123767-37123909 | FBXO47 | | TSS200 |
| GM09255 | chr17:37123835-37123909 | FBXO47 | | TSS200 |
| WG3465 | chr17:37821478-37821536 | TCAP | N_Shelf | TSS200 |
| WG3465 | chr17:37843591-37843818 | ERBB2;PGAP3 | N_Shore | TSS1500;Body |
| WG2194 | chr17:37894014-37894165 | GRB7;GRB7 | | TSS1500;TSS200 |
| WG1722 | chr17:37894165-37894636 | GRB7;GRB7 | | TSS1500;TSS200 |
| GM00290 | chr17:37894258-37894636 | GRB7;GRB7;GRB7 | | 5'UTR;TSS1500;1stExon |
| WG1513 | chr17:37894397-37894413 | GRB7;GRB7 | | TSS200;5'UTR |
| WG1977 | chr17:37894397-37894636 | GRB7;GRB7 | | TSS200;5'UTR |
| GM00438 | chr17:37896063-37896175 | GRB7;GRB7 | | 5'UTR;5'UTR |
| GM03075 | chr17:3791902-3792074 | CAMKK1;CAMKK1;CAMKK1 | N_Shore | 5'UTR;5'UTR;5'UTR |
| GM00290 | chr17:3794118-3794573 | CAMKK1;CAMKK1;CAMKK1 | N_Shore | 5'UTR;TSS200;5'UTR |

| | | | | |
|---|---|---|---|---|
| WG3465 | chr17:38170862-38171697 | CSF3;CSF3;CSF3 | | TSS1500;TSS1500;TSS1500 |
| GM02555 | chr17:38249266-38249592 | NR1D1;THRA | Island | 3'UTR;Body |
| GM00935 | chr17:38249352-38249592 | THRA;NR1D1 | Island | Body;Body |
| WG1977 | chr17:38249430-38249592 | THRA;NR1D1 | Island | Body;Body |
| GM00520 | chr17:38249486-38249729 | THRA;NR1D1 | Island | Body;Body |
| WG3465 | chr17:3828008-3828268 | ATP2A3;ATP2A3;ATP2A3;ATP2 | Island | 3'UTR;3'UTR;3'UTR;3'UTR;3'UTR;3'UTR |
| GM02706 | chr17:38334161-38334439 | RAPGEFL1 | Island | TSS200 |
| GM01096 | chr17:38717206-38717275 | CCR7 | | Body |
| GM02706 | chr17:38821390-38821420 | KRT222;KRT222 | | 5'UTR;1stExon |
| WG1513 | chr17:39041199-39041602 | KRT20 | | 1stExon |
| WG1658 | chr17:39204767-39204922 | KRTAP2-1 | | TSS1500 |
| WG1977 | chr17:39471738-39471940 | KRTAP17-1 | | 1stExon |
| GM00439 | chr17:39728425-39728478 | KRT9 | | TSS200 |
| WG2843 | chr17:40259724-40260053 | DHX58 | Island | Body |
| WG2121 | chr17:40274703-40274811 | HSPB9;KAT2A | Island | TSS200;TSS1500 |
| GM01393 | chr17:40274722-40274740 | HSPB9;KAT2A | Island | TSS200;TSS1500 |
| WG2276 | chr17:40438312-40438511 | STAT5A | N_Shore | TSS1500 |
| WG1977 | chr17:40440904-40441155 | STAT5A | Island | 5'UTR |
| WG2844 | chr17:40489513-40489785 | STAT3;STAT3;STAT3 | | Body;Body;Body |
| WG1657 | chr17:40489584-40489785 | STAT3;STAT3;STAT3 | | Body;Body;Body |
| WG3465 | chr17:40687596-40687830 | NAGLU | N_Shore | TSS1500 |
| GM00042 | chr17:40705273-40705708 | HSD17B1 | Island | Body |
| WG2121 | chr17:40715222-40715281 | COASY;COASY;COASY;COAS | Island | 1stExon;Body;Body;Body;5'UTR |
| GM00289 | chr17:40715236-40715281 | COASY;COASY;COASY;COAS | Island | 1stExon;Body;Body;Body;5'UTR |
| WG2414 | chr17:40835849-40836087 | CNTNAP1 | Island | Body |
| GM00289 | chr17:40838861-40839022 | CNTNAP1 | Island | Body |
| WG3466 | chr17:40935998-40936150 | WNK4 | Island | Body |
| WG1978 | chr17:40935998-40936498 | WNK4 | Island | Body |
| GM00935 | chr17:40936078-40936498 | WNK4 | Island | Body |
| GM02555 | chr17:40936150-40936658 | WNK4 | Island | Body |
| WG1977 | chr17:40936658-40936820 | WNK4 | Island | Body |
| GM02706 | chr17:41131966-41132080 | RUNDC1;AARSD1;AARSD1;A | N_Shore | TSS1500;5'UTR;1stExon;Body;Body;Bo |
| WG1977 | chr17:41131966-41132138 | RUNDC1;AARSD1;AARSD1;A | N_Shore | TSS1500;5'UTR;1stExon;Body;Body;Bo |
| GM00042 | chr17:41277974-41278141 | BRCA1;NBR2;BRCA1;BRCA1; | N_Shore | TSS1500;Body;TSS1500;TSS1500;TSS |
| WG1722 | chr17:41277974-41278241 | BRCA1;NBR2;BRCA1;BRCA1; | N_Shore | TSS1500;Body;TSS1500;TSS1500;TSS |
| GM00290 | chr17:41278135-41278197 | BRCA1;NBR2;BRCA1;BRCA1; | Island | TSS1500;Body;TSS1500;TSS1500;TSS |
| GM09255 | chr17:41278135-41278380 | BRCA1;NBR2;BRCA1;BRCA1; | Island | TSS1500;Body;TSS1500;TSS1500;TSS |
| WG1721 | chr17:41278141-41278197 | BRCA1;NBR2;BRCA1;BRCA1; | Island | TSS1500;Body;TSS1500;TSS1500;TSS |
| WG1513 | chr17:41278179-41278241 | BRCA1;NBR2;BRCA1;BRCA1; | Island | TSS1500;Body;TSS1500;TSS1500;TSS |
| GM01392 | chr17:41278281-41278357 | BRCA1;NBR2;BRCA1;BRCA1; | Island | TSS1500;Body;TSS1500;TSS1500;TSS |
| GM01392 | chr17:41278429-41278506 | BRCA1;NBR2;BRCA1;BRCA1; | Island | TSS1500;Body;TSS1500;TSS1500;TSS |
| WG2415 | chr17:41477271-41477948 | ARL4D | Island | Body |
| GM00439 | chr17:41738893-41739195 | MEOX1;MEOX1;MEOX1 | | 1stExon;1stExon;5'UTR |
| GM02173 | chr17:41739130-41739246 | MEOX1;MEOX1;MEOX1 | | TSS200;5'UTR;TSS200 |
| WG1513 | chr17:41832753-41832882 | SOST | Island | Body |
| GM02706 | chr17:41832753-41833161 | SOST | Island | Body |
| WG1722 | chr17:41832873-41832975 | SOST | Island | Body |

| | | | | |
|---|---|---|---|---|
| GM01650 | chr17:41832882-41833161 | SOST | Island | Body |
| GM00016 | chr17:41832975-41833324 | SOST | Island | Body |
| WG2415 | chr17:41909709-41909716 | MPP3;MPP3 | N_Shore | 5'UTR;Body |
| WG1978 | chr17:41924123-41924399 | CD300LG;CD300LG;CD300LG;CD300LG | | TSS1500;TSS1500;TSS1500;TSS1500 |
| GM02149 | chr17:42218908-42218924 | C17orf53;C17orf53 | N_Shore | TSS1500;TSS1500 |
| GM02704 | chr17:42275334-42275551 | ATXN7L3;ATXN7L3 | N_Shore | Body;Body |
| WG3466 | chr17:42432046-42432488 | FAM171A2 | Island | Body |
| GM01096 | chr17:42856773-42857157 | ADAM11 | Island | Body |
| WG3465 | chr17:43025585-43025655 | KIF18B | S_Shore | TSS1500 |
| GM01096 | chr17:43318760-43319382 | FMNL1 | Island | Body |
| WG2844 | chr17:43319137-43319382 | FMNL1 | Island | Body |
| GM02706 | chr17:43339328-43339512 | C17orf46;LOC100133991;LC | Island | Body;Body;TSS1500 |
| GM02706 | chr17:43339589-43339831 | LOC100133991;LOC100133 | Island | Body;Body;TSS200 |
| GM00935 | chr17:43482735-43483251 | ARHGAP27;ARHGAP27 | N_Shore | Body;Body |
| WG1977 | chr17:43482735-43483453 | ARHGAP27;ARHGAP27 | N_Shore | Body;Body |
| GM00289 | chr17:43482735-43483527 | ARHGAP27;ARHGAP27 | N_Shore | Body;Body |
| WG1513 | chr17:43483116-43483453 | ARHGAP27;ARHGAP27;ARHC | Island | 1stExon;5'UTR;5'UTR |
| GM00439 | chr17:43483116-43483527 | ARHGAP27;ARHGAP27;ARHC | Island | 1stExon;5'UTR;5'UTR |
| GM02706 | chr17:43974344-43974804 | MAPT;MAPT;LOC100128977; | Island | 5'UTR;5'UTR;TSS1500;Body;5'UTR;5'U |
| GM02175 | chr17:43974344-43975063 | MAPT;MAPT;LOC100128977; | Island | 5'UTR;5'UTR;TSS1500;Body;5'UTR;5'U |
| WG1977 | chr17:43974446-43975063 | MAPT;MAPT;MAPT;MAPT;LOC | Island | 5'UTR;5'UTR;5'UTR;5'UTR;Body;5'UTR; |
| WG2415 | chr17:43974869-43974975 | MAPT;MAPT;MAPT;MAPT;LOC | Island | 5'UTR;5'UTR;5'UTR;5'UTR;Body;5'UTR; |
| WG1513 | chr17:43974869-43975063 | MAPT;MAPT;MAPT;MAPT;LOC | Island | 5'UTR;5'UTR;5'UTR;5'UTR;Body;5'UTR; |
| WG1983 | chr17:43974919-43975063 | MAPT;MAPT;MAPT;MAPT;LOC | Island | 5'UTR;5'UTR;5'UTR;5'UTR;Body;5'UTR; |
| WG1984 | chr17:44847220-44847427 | WNT3 | Island | Body |
| WG2121 | chr17:45501030-45501697 | LOC100272146;C17orf57 | Island | Body;Body |
| WG1977 | chr17:45501242-45501697 | LOC100272146;C17orf57 | Island | Body;Body |
| WG1977 | chr17:45786010-45786512 | TBKBP1 | N_Shore | Body |
| WG3466 | chr17:45786439-45786737 | TBKBP1 | Island | Body |
| WG2120 | chr17:46018875-46019184 | PNPO | | TSS200 |
| GM00438 | chr17:46507763-46507857 | SKAP1;SKAP1 | Island | TSS200;TSS200 |
| WG1513 | chr17:46507777-46507857 | SKAP1;SKAP1 | Island | TSS200;TSS200 |
| GM00935 | chr17:46507777-46507895 | SKAP1;SKAP1 | Island | TSS200;TSS200 |
| GM00439 | chr17:46507830-46507895 | SKAP1;SKAP1 | S_Shore | TSS1500;TSS1500 |
| GM02316 | chr17:46607828-46608268 | HOXB1 | Island | 1stExon |
| GM09256 | chr17:46608277-46608293 | HOXB1 | Island | TSS200 |
| WG1721 | chr17:46620347-46620460 | HOXB2 | N_Shore | 3'UTR |
| GM01650 | chr17:46622454-46622516 | HOXB2 | S_Shore | TSS200 |
| GM02149 | chr17:46622454-46622536 | HOXB2 | S_Shore | TSS200 |
| WG1283 | chr17:46622454-46622607 | HOXB2 | S_Shore | TSS200 |
| GM00042 | chr17:46622536-46622715 | HOXB2 | S_Shore | TSS200 |
| WG2121 | chr17:46623729-46623819 | HOXB2 | S_Shelf | TSS1500 |
| GM02640 | chr17:46627384-46628717 | HOXB3 | N_Shore | 3'UTR |
| WG1977 | chr17:46627384-46629306 | HOXB3 | N_Shore | 3'UTR |
| WG2120 | chr17:46628404-46629018 | HOXB3 | Island | Body |
| WG1283 | chr17:46631910-46632154 | HOXB3 | Island | 5'UTR |
| WG3465 | chr17:46645972-46646359 | HOXB3 | S_Shelf | 5'UTR |

| | | | | |
|---|---|---|---|---|
| WG1722 | chr17:46645972-46646444 | HOXB3 | S_Shelf | 5'UTR |
| WG2844 | chr17:46648014-46648533 | HOXB3 | | 5'UTR |
| GM00520 | chr17:46648014-46648582 | HOXB3 | | 5'UTR |
| WG1722 | chr17:46651186-46651867 | HOXB3 | N_Shelf | 5'UTR |
| GM00520 | chr17:46651360-46651867 | HOXB3;HOXB3 | N_Shelf | 5'UTR;1stExon |
| GM00289 | chr17:46651722-46651822 | HOXB3;HOXB3 | N_Shelf | 5'UTR;1stExon |
| GM02149 | chr17:46651722-46651867 | HOXB3;HOXB3 | N_Shelf | 5'UTR;1stExon |
| GM02555 | chr17:46651722-46651920 | HOXB3;HOXB3 | N_Shelf | 5'UTR;1stExon |
| GM09255 | chr17:46651722-46652179 | HOXB3;HOXB3 | N_Shelf | 5'UTR;1stExon |
| WG1514 | chr17:46651745-46651867 | HOXB3;HOXB3 | N_Shelf | 5'UTR;1stExon |
| GM01376 | chr17:46651822-46652179 | HOXB3 | N_Shelf | TSS200 |
| WG1984 | chr17:46651945-46652179 | HOXB3 | N_Shelf | TSS200 |
| WG1513 | chr17:46653615-46654330 | HOXB4 | N_Shore | 3'UTR |
| WG2193 | chr17:46656543-46656690 | HOXB4 | S_Shore | TSS1500 |
| WG3465 | chr17:46656543-46657504 | HOXB4 | S_Shore | TSS1500 |
| GM03073 | chr17:46656572-46656690 | HOXB4 | S_Shore | TSS1500 |
| WG2415 | chr17:46657274-46657504 | MIR10A | N_Shore | Body |
| WG3465 | chr17:46666926-46669644 | LOC404266;LOC404266 | N_Shelf | TSS1500;TSS1500 |
| WG3466 | chr17:46667587-46667812 | LOC404266;LOC404266 | N_Shore | TSS1500;TSS1500 |
| GM02706 | chr17:46669455-46669644 | HOXB5;LOC404266;LOC404 | Island | 3'UTR;Body;TSS1500;TSS200;Body |
| WG3466 | chr17:46669485-46669492 | LOC404266;HOXB5;LOC404 | Island | TSS200;3'UTR;Body;TSS200;Body |
| WG2414 | chr17:46669485-46669566 | LOC404266;HOXB5;LOC404 | Island | TSS200;3'UTR;Body;TSS200;Body |
| WG1514 | chr17:46669485-46669644 | LOC404266;HOXB5;LOC404 | Island | TSS200;3'UTR;Body;TSS200;Body |
| WG2415 | chr17:46669485-46670029 | LOC404266;HOXB5;LOC404 | Island | TSS200;3'UTR;Body;TSS200;Body |
| WG2121 | chr17:46671861-46672255 | LOC404266;LOC404266;LO | N_Shore | Body;Body;Body;TSS1500;TSS1500;Bo |
| WG1283 | chr17:46672924-46673625 | LOC404266;LOC404266;LO | N_Shore | Body;Body;Body;TSS1500;Body |
| GM01653 | chr17:46672924-46674514 | LOC404266;LOC404266;LO | N_Shore | Body;Body;Body;TSS1500;Body |
| WG2121 | chr17:46673442-46674514 | LOC404266;LOC404266;LO | N_Shore | Body;Body;Body;Body;3'UTR;Body |
| WG2194 | chr17:46673625-46674041 | LOC404266;LOC404266;LO | Island | Body;Body;Body;Body;3'UTR;Body |
| GM00940 | chr17:46673625-46674438 | LOC404266;LOC404266;LO | Island | Body;Body;Body;Body;3'UTR;Body |
| WG1977 | chr17:46673625-46674514 | LOC404266;LOC404266;LO | Island | Body;Body;Body;Body;3'UTR;Body |
| WG1513 | chr17:46674041-46674438 | LOC404266;LOC404266;LO | Island | Body;Body;Body;Body;Body;Body |
| WG3466 | chr17:46674041-46674514 | LOC404266;LOC404266;LO | Island | Body;Body;Body;Body;Body;Body |
| WG2120 | chr17:46674395-46674514 | LOC404266;LOC404266;LO | N_Shore | Body;Body;Body;Body;Body;Body |
| WG1977 | chr17:46675892-46676215 | LOC404266;LOC404266;LO | S_Shore | Body;Body;Body;Body;5'UTR;Body |
| GM00940 | chr17:46676099-46676375 | LOC404266;LOC404266;LO | S_Shore | Body;Body;Body;Body;5'UTR;Body |
| WG1721 | chr17:46681111-46681154 | LOC404266;LOC404266;HOXB6 | | Body;Body;5'UTR |
| GM00289 | chr17:46681111-46681362 | LOC404266;LOC404266;HOXB6 | | Body;Body;5'UTR |
| GM00520 | chr17:46681111-46683409 | LOC404266;LOC404266;HOXB6 | | Body;Body;5'UTR |
| WG3465 | chr17:46681111-46683606 | LOC404266;LOC404266;HOXB6 | | Body;Body;5'UTR |
| WG1283 | chr17:46681135-46681362 | LOC404266;LOC404266;HOXB6 | | Body;Body;5'UTR |
| WG2415 | chr17:46681154-46681362 | LOC404266;LOC404266;HOXB6 | | Body;Body;5'UTR |
| GM01393 | chr17:46681362-46682413 | LOC404266;LOC404266;HO | N_Shelf | Body;Body;5'UTR |
| WG1721 | chr17:46681401-46682413 | LOC404266;LOC404266;HO | N_Shelf | Body;Body;5'UTR |
| WG2275 | chr17:46682308-46682398 | LOC404266;LOC404266;HO | N_Shelf | Body;Body;1stExon;5'UTR |
| WG1722 | chr17:46682308-46682413 | LOC404266;LOC404266;HO | N_Shelf | Body;Body;1stExon;5'UTR |
| WG1283 | chr17:46682308-46683047 | LOC404266;LOC404266;HO | N_Shelf | Body;Body;1stExon;5'UTR |

| | | | | |
|---|---|---|---|---|
| WG2415 | chr17:46682308-46683409 | LOC404266;LOC404266;HO | N_Shelf | Body;Body;1stExon;5'UTR |
| WG1657 | chr17:46682319-46682394 | LOC404266;LOC404266;HO | N_Shelf | Body;Body;1stExon;5'UTR |
| WG2843 | chr17:46682319-46682398 | LOC404266;LOC404266;HO | N_Shelf | Body;Body;1stExon;5'UTR |
| WG3466 | chr17:46682319-46682413 | LOC404266;LOC404266;HO | N_Shelf | Body;Body;1stExon;5'UTR |
| WG1513 | chr17:46682319-46683047 | LOC404266;LOC404266;HO | N_Shelf | Body;Body;1stExon;5'UTR |
| GM00935 | chr17:46682319-46683409 | LOC404266;LOC404266;HO | N_Shelf | Body;Body;1stExon;5'UTR |
| WG2193 | chr17:46682390-46682398 | LOC404266;LOC404266;HO | N_Shelf | Body;Body;TSS200 |
| GM09255 | chr17:46682390-46682413 | LOC404266;LOC404266;HO | N_Shelf | Body;Body;TSS200 |
| GM03073 | chr17:46682390-46683606 | LOC404266;LOC404266;HO | N_Shelf | Body;Body;TSS200 |
| GM00016 | chr17:46682390-46683806 | LOC404266;LOC404266;HO | N_Shelf | Body;Body;TSS200 |
| GM02175 | chr17:46685292-46685353 | HOXB7 | Island | Body |
| WG3466 | chr17:46685327-46685448 | HOXB7 | Island | Body |
| WG2843 | chr17:46685353-46685448 | HOXB7 | Island | Body |
| WG2121 | chr17:46689426-46689639 | HOXB7 | N_Shore | TSS1500 |
| WG3465 | chr17:46691543-46692008 | HOXB8 | Island | Body |
| GM02640 | chr17:46691543-46692021 | HOXB8 | Island | Body |
| WG3465 | chr17:46799479-46799745 | C17orf93;PRAC | N_Shore | TSS1500;Body |
| WG3466 | chr17:46799479-46800028 | C17orf93;PRAC | N_Shore | TSS1500;Body |
| WG1513 | chr17:46799745-46799845 | PRAC;C17orf93 | N_Shore | 1stExon;TSS1500 |
| WG2275 | chr17:46799845-46800028 | PRAC;PRAC;C17orf93 | N_Shore | 1stExon;5'UTR;TSS1500 |
| GM09255 | chr17:46800639-46800680 | C17orf93;PRAC | Island | Body;TSS1500 |
| GM00439 | chr17:47091978-47092178 | IGF2BP1;IGF2BP1 | Island | Body;Body |
| WG1282 | chr17:47092072-47092178 | IGF2BP1;IGF2BP1 | Island | Body;Body |
| GM02175 | chr17:47113141-47113489 | IGF2BP1;IGF2BP1 | N_Shore | Body;Body |
| WG3465 | chr17:47113407-47113489 | IGF2BP1;IGF2BP1 | Island | Body;Body |
| WG1513 | chr17:47113457-47113596 | IGF2BP1;IGF2BP1 | Island | Body;Body |
| WG3465 | chr17:47209660-47209749 | B4GALNT2;B4GALNT2;B4GAI | N_Shore | TSS1500;TSS200;TSS1500 |
| WG1513 | chr17:47287492-47287526 | ABI3;GNGT2;ABI3 | | TSS200;TSS1500;TSS200 |
| GM02175 | chr17:47287521-47287577 | ABI3;GNGT2;ABI3 | | TSS200;TSS1500;TSS200 |
| WG1513 | chr17:47296970-47297268 | ABI3;ABI3 | Island | Body;Body |
| GM03075 | chr17:4802847-4802906 | CHRNE;C17orf107 | Island | Body;TSS200 |
| GM00290 | chr17:48049970-48049979 | DLX4;DLX4 | Island | TSS200;Body |
| GM02706 | chr17:48206283-48206749 | SAMD14 | N_Shore | 5'UTR |
| WG1978 | chr17:48250505-48250645 | HILS1;HILS1;SGCA;SGCA | | TSS1500;TSS1500;Body;Body |
| WG1978 | chr17:48263150-48263189 | COL1A1 | | Body |
| GM09256 | chr17:48263159-48263189 | COL1A1 | | Body |
| WG2844 | chr17:48263159-48263204 | COL1A1 | | Body |
| GM01393 | chr17:48263173-48263198 | COL1A1 | | Body |
| WG2276 | chr17:48263173-48263204 | COL1A1 | | Body |
| GM00016 | chr17:4872326-4872499 | CAMTA2;CAMTA2;CAMTA2;C | S_Shore | Body;Body;Body;Body;TSS1500 |
| WG2843 | chr17:4872326-4872603 | CAMTA2;CAMTA2;CAMTA2;C | S_Shore | Body;Body;Body;Body;TSS1500 |
| WG2276 | chr17:4872488-4872603 | CAMTA2;CAMTA2;CAMTA2;C | S_Shore | Body;Body;Body;Body;TSS1500 |
| GM00016 | chr17:48912164-48912327 | WFIKKN2 | | TSS1500 |
| GM01392 | chr17:48912265-48912483 | WFIKKN2 | | TSS1500 |
| GM09255 | chr17:48912487-48912545 | WFIKKN2 | | TSS200 |
| GM00439 | chr17:49242784-49242847 | NME2;NME2;NME2;NME1-NI | N_Shore | TSS200;TSS1500;TSS1500;Body;TSS1 |
| WG1658 | chr17:5019452-5019761 | ZNF232 | Island | 5'UTR |

| | | | | |
|---|---|---|---|---|
| GM02173 | chr17:5019638-5019761 | ZNF232 | Island | 5'UTR |
| GM00042 | chr17:5026759-5026840 | ZNF232 | Island | TSS1500 |
| WG2120 | chr17:53341659-53342029 | HLF | N_Shore | TSS1500 |
| WG2276 | chr17:53810321-53810813 | TMEM100 | | TSS1500 |
| WG1977 | chr17:54230793-54230996 | ANKFN1 | | TSS200 |
| GM02175 | chr17:54893150-54893317 | C17orf67 | | 1stExon |
| WG2276 | chr17:55663225-55663284 | MSI2;MSI2 | | Body;Body |
| WG2415 | chr17:56082780-56082885 | SFRS1;SFRS1 | N_Shore | 3'UTR;Body |
| WG3465 | chr17:56296980-56297193 | MKS1;MKS1 | S_Shore | TSS200;TSS1500 |
| GM00438 | chr17:56297193-56297443 | MKS1;MKS1 | S_Shore | TSS1500;TSS1500 |
| WG1977 | chr17:56565286-56565644 | HSF5 | Island | 1stExon |
| GM00940 | chr17:56606842-56607383 | SEPT4;SEPT4;SEPT4 | N_Shelf | TSS200;Body;Body |
| WG3465 | chr17:56744332-56744490 | TEX14;TEX14 | | 5'UTR;5'UTR |
| GM00042 | chr17:56769387-56769628 | TEX14;TEX14;RAD51C;TEX14 | N_Shore | 1stExon;5'UTR;TSS1500;5'UTR;1stExo |
| GM00042 | chr17:56769696-56769767 | TEX14;RAD51C;TEX14;RAD51 | N_Shore | TSS1500;TSS1500;TSS1500;TSS1500 |
| GM02149 | chr17:57642721-57642752 | DHX40;DHX40 | Island | TSS200;TSS200 |
| WG2414 | chr17:57642749-57642763 | DHX40;DHX40 | Island | TSS200;TSS200 |
| WG2844 | chr17:58499679-58499911 | C17orf64 | S_Shore | TSS200 |
| WG1513 | chr17:59488116-59488443 | C17orf82 | Island | TSS1500 |
| WG1978 | chr17:59554672-59554806 | TBX4 | | Body |
| WG2844 | chr17:60828157-60828406 | MARCH10;MARCH10 | | Body;Body |
| WG1722 | chr17:61511683-61511829 | CYB561;CYB561;CYB561 | Island | 3'UTR;3'UTR;3'UTR |
| WG1977 | chr17:62051709-62051773 | SCN4A | | TSS1500 |
| WG2276 | chr17:62084205-62084322 | ICAM2;ICAM2;ICAM2;ICAM2;ICAM2;ICAM | | 5'UTR;1stExon;5'UTR;5'UTR;5'UTR;5'U |
| GM02175 | chr17:62084205-62084359 | ICAM2;ICAM2;ICAM2;ICAM2;ICAM2;ICAM | | 5'UTR;1stExon;5'UTR;5'UTR;5'UTR;5'U |
| GM00935 | chr17:62097918-62097953 | ICAM2;ICAM2;ICAM2;ICAM2;ICAM2;ICAM | | 5'UTR;5'UTR;5'UTR;1stExon;1stExon;5 |
| GM00042 | chr17:62097951-62098222 | ICAM2;ICAM2;ICAM2;ICAM2;ICAM2;ICAM | | 5'UTR;5'UTR;5'UTR;1stExon;1stExon;5 |
| GM00438 | chr17:62493599-62493832 | POLG2 | S_Shore | TSS1500 |
| GM01650 | chr17:62774939-62775415 | LOC146880;LOC146880 | Island | Body;Body |
| GM02706 | chr17:62777690-62777777 | LOC146880;LOC146880 | Island | Body;TSS200 |
| WG3465 | chr17:62777693-62777777 | LOC146880;LOC146880 | Island | Body;TSS200 |
| GM00290 | chr17:62777744-62777777 | LOC146880;LOC146880 | Island | Body;TSS200 |
| GM00042 | chr17:6338554-6338571 | AIPL1;AIPL1;AIPL1 | | TSS200;TSS200;TSS200 |
| GM00520 | chr17:63532630-63533152 | AXIN2 | N_Shore | Body |
| WG2843 | chr17:63532630-63533303 | AXIN2 | N_Shore | Body |
| WG2121 | chr17:6358415-6358830 | PITPNM3;PITPNM3 | Island | 3'UTR;3'UTR |
| GM00290 | chr17:6358599-6358928 | PITPNM3;PITPNM3 | Island | 3'UTR;3'UTR |
| WG1513 | chr17:65040527-65040581 | CACNG1 | N_Shore | TSS200 |
| GM09256 | chr17:65040581-65040616 | CACNG1 | N_Shore | TSS200 |
| WG2415 | chr17:65471303-65471507 | PITPNC1;PITPNC1 | | Body;Body |
| GM00520 | chr17:65990431-65990817 | C17orf58;C17orf58 | S_Shore | TSS1500;TSS1500 |
| GM02706 | chr17:66255210-66255265 | ARSG | Island | TSS200 |
| WG1722 | chr17:66286298-66286394 | ARSG;SLC16A6 | N_Shore | 5'UTR;5'UTR |
| WG1513 | chr17:6756633-6756860 | ALOX12P2 | | TSS1500 |
| GM09255 | chr17:6899095-6899332 | ALOX12 | Island | TSS1500 |
| WG2121 | chr17:6899207-6899332 | ALOX12 | Island | TSS200 |
| GM09256 | chr17:6899332-6899364 | ALOX12 | Island | TSS200 |

| | | | | |
|---|---|---|---|---|
| WG2121 | chr17:6899364-6899577 | ALOX12 | Island | TSS200 |
| GM02706 | chr17:6947323-6947337 | SLC16A11 | Island | TSS200 |
| GM00438 | chr17:70116185-70116298 | SOX9 | N_Shore | TSS1500 |
| WG1977 | chr17:7077335-7077709 | ASGR1 | Island | Body |
| WG2415 | chr17:7082562-7083015 | ASGR1;ASGR1 | | 5'UTR;1stExon |
| WG1283 | chr17:71772827-71772869 | C17orf54 | | Body |
| GM02706 | chr17:71824806-71825281 | C17orf54 | | TSS200 |
| GM09256 | chr17:71824806-71825500 | C17orf54 | | TSS200 |
| GM02706 | chr17:7227499-7227662 | NEURL4;NEURL4 | Island | Body;Body |
| GM02175 | chr17:72350354-72350710 | KIF19 | Island | Body |
| WG1721 | chr17:72350487-72350710 | KIF19 | Island | Body |
| GM00042 | chr17:72438885-72439217 | GPRC5C;GPRC5C | | Body;Body |
| WG3465 | chr17:72439044-72439217 | GPRC5C;GPRC5C | N_Shelf | Body;Body |
| WG2415 | chr17:72541932-72542430 | CD300C;CD300C | | 1stExon;5'UTR |
| GM03075 | chr17:7258070-7258501 | KCTD11;TMEM95;KCTD11 | S_Shelf | 3'UTR;TSS1500;1stExon |
| GM02175 | chr17:72580962-72581014 | C17orf77;CD300LD | | TSS200;Body |
| GM01096 | chr17:7287271-7287652 | TNK1 | Island | Body |
| WG1978 | chr17:7287271-7287740 | TNK1 | Island | Body |
| WG2121 | chr17:7287449-7287740 | TNK1 | Island | Body |
| WG1977 | chr17:7287449-7287922 | TNK1 | Island | Body |
| WG2844 | chr17:7287740-7288001 | TNK1 | Island | Body |
| GM00289 | chr17:73056104-73056187 | KCTD2 | | Body |
| GM00520 | chr17:73316496-73316721 | GRB2;GRB2 | | Body;Body |
| GM00290 | chr17:7341440-7341641 | FGF11 | N_Shore | TSS1500 |
| GM00022 | chr17:7341616-7341936 | FGF11 | N_Shore | TSS1500 |
| GM00289 | chr17:7348316-7348339 | CHRNB1 | Island | TSS200 |
| GM00290 | chr17:7348379-7348438 | CHRNB1 | Island | TSS200 |
| WG3465 | chr17:73498127-73498302 | CASKIN2;CASKIN2 | Island | Body;Body |
| GM00290 | chr17:7350001-7350282 | CHRNB1 | S_Shore | Body |
| GM02555 | chr17:73559363-73559562 | LLGL2;LLGL2;LLGL2 | Island | Body;Body;Body |
| WG3465 | chr17:73642547-73642628 | RECQL5;LOC100130933 | | Body;TSS200 |
| WG3465 | chr17:73780246-73780396 | UNK | N_Shore | TSS1500 |
| GM02706 | chr17:739341-739405 | NXN | Island | Body |
| WG2415 | chr17:74075092-74075189 | ZACN | S_Shore | TSS200 |
| WG2844 | chr17:746502-746641 | NXN | S_Shore | Body |
| WG1978 | chr17:74868561-74868670 | MGAT5B;MGAT5B | S_Shore | Body;TSS200 |
| WG2120 | chr17:74868561-74868725 | MGAT5B;MGAT5B | S_Shore | Body;TSS200 |
| WG1657 | chr17:74868604-74868725 | MGAT5B;MGAT5B | S_Shore | Body;TSS200 |
| WG3466 | chr17:750241-750848 | NXN | | Body |
| GM02704 | chr17:750306-750848 | NXN | | Body |
| WG1310 | chr17:750766-751287 | NXN | | Body |
| WG2415 | chr17:7516888-7517138 | SHBG;FXR2 | N_Shore | TSS1500;Body |
| GM02456 | chr17:75283431-75283832 | SEPT9;SEPT9 | | Body;TSS1500 |
| WG2844 | chr17:75283800-75283979 | SEPT9;SEPT9 | | TSS200;Body |
| WG3466 | chr17:75283832-75283979 | SEPT9;SEPT9 | | TSS200;Body |
| GM02706 | chr17:75319751-75320035 | SEPT9;SEPT9;SEPT9 | | 5'UTR;Body;Body |
| WG2844 | chr17:75373040-75373327 | SEPT9;SEPT9;SEPT9;SEPT9;S | S_Shelf | 5'UTR;5'UTR;Body;Body;Body |

| | | | | |
|---|---|---|---|---|
| WG2844 | chr17:75451779-75452044 | SEPT9;SEPT9;SEPT9;SEPT9;S | S_Shelf | Body;5'UTR;Body;Body;Body;Body |
| WG2276 | chr17:75451809-75452100 | SEPT9;SEPT9;SEPT9;SEPT9;S | S_Shelf | Body;5'UTR;Body;Body;Body;Body |
| WG3465 | chr17:75471308-75471491 | SEPT9;SEPT9;SEPT9;SEPT9;SEPT9;SEPT9 | | Body;TSS200;5'UTR;Body;Body;Body;I |
| WG1977 | chr17:76128481-76128556 | TMC8;TMC6;TMC6 | S_Shore | Body;5'UTR;1stExon |
| GM02706 | chr17:76128522-76128621 | TMC8;TMC6 | S_Shore | Body;TSS200 |
| GM00520 | chr17:76128556-76128634 | TMC8;TMC6 | S_Shore | Body;TSS200 |
| WG2415 | chr17:76128634-76128683 | TMC8;TMC6 | S_Shore | Body;TSS200 |
| GM00289 | chr17:76220608-76220955 | BIRC5;BIRC5;BIRC5;EPR1 | | 3'UTR;3'UTR;3'UTR;Body |
| WG3466 | chr17:76355146-76355288 | SOCS3 | Island | Body |
| WG2415 | chr17:76975944-76976057 | LGALS3BP;LGALS3BP | | 1stExon;5'UTR |
| WG3465 | chr17:76976010-76976091 | LGALS3BP;LGALS3BP | | 1stExon;5'UTR |
| WG3465 | chr17:7701546-7701636 | DNAH2 | | Body |
| WG1722 | chr17:77029308-77029518 | C1QTNF1;C1QTNF1;C1QTNF1 | | 5'UTR;TSS1500;5'UTR |
| WG3465 | chr17:77037916-77037965 | C1QTNF1;C1QTNF1;C1QTNF1 | | 5'UTR;5'UTR;5'UTR |
| GM02173 | chr17:77043179-77043544 | C1QTNF1;C1QTNF1;C1QTNF | N_Shore | Body;Body;Body |
| WG2844 | chr17:77078091-77078518 | ENGASE | | Body |
| GM02706 | chr17:77078833-77079031 | ENGASE | | Body |
| GM02704 | chr17:77086350-77086532 | HRNBP3 | | 3'UTR |
| GM00520 | chr17:77126866-77127426 | HRNBP3 | N_Shore | 5'UTR |
| GM09256 | chr17:77127220-77127487 | HRNBP3 | N_Shore | 5'UTR |
| GM00290 | chr17:77478569-77478637 | HRNBP3 | | TSS200 |
| WG2276 | chr17:77810509-77810912 | CBX4 | N_Shore | Body |
| WG1977 | chr17:7791882-7792061 | CHD3;CHD3;CHD3 | S_Shelf | Body;TSS1500;TSS1500 |
| GM02175 | chr17:7792059-7792134 | CHD3;CHD3;CHD3 | S_Shelf | Body;TSS200;TSS200 |
| GM00439 | chr17:77924665-77925136 | TBC1D16 | S_Shore | Body |
| GM02555 | chr17:77924665-77925317 | TBC1D16 | S_Shore | Body |
| WG1310 | chr17:77951408-77952488 | TBC1D16 | N_Shore | Body |
| GM00520 | chr17:77951707-77952488 | TBC1D16 | N_Shore | Body |
| WG1513 | chr17:77952109-77952290 | TBC1D16 | Island | Body |
| WG3465 | chr17:77952109-77952488 | TBC1D16 | Island | Body |
| GM02555 | chr17:77952247-77952463 | TBC1D16 | Island | Body |
| GM02640 | chr17:77952247-77952488 | TBC1D16 | Island | Body |
| WG1983 | chr17:77952290-77952488 | TBC1D16 | Island | Body |
| WG1984 | chr17:77952453-77952488 | TBC1D16 | S_Shore | Body |
| GM00439 | chr17:77997198-77997997 | TBC1D16 | N_Shore | 5'UTR |
| WG2276 | chr17:77997760-77997997 | TBC1D16 | Island | 5'UTR |
| GM02456 | chr17:77997833-77997997 | TBC1D16 | Island | 5'UTR |
| GM02706 | chr17:78079481-78079668 | GAA;GAA;GAA | N_Shore | Body;Body;Body |
| GM01096 | chr17:78161110-78161227 | CARD14;CARD14 | N_Shelf | TSS1500;Body |
| WG2276 | chr17:78161178-78161467 | CARD14;CARD14 | N_Shelf | TSS1500;Body |
| GM02173 | chr17:78166148-78166385 | CARD14;CARD14 | S_Shore | Body;Body |
| GM00438 | chr17:78237389-78237543 | RNF213;RNF213 | S_Shelf | 5'UTR;5'UTR |
| WG2415 | chr17:782910-783451 | NXN | | Body |
| GM01650 | chr17:7832680-7832769 | KCNAB3 | Island | 1stExon |
| WG1657 | chr17:7832680-7832852 | KCNAB3 | Island | 1stExon |
| GM02706 | chr17:7832764-7833237 | KCNAB3 | Island | TSS200 |
| WG1657 | chr17:7832943-7833237 | KCNAB3 | Island | TSS200 |

| | | | | |
|---|---|---|---|---|
| GM02706 | chr17:78560478-78560916 | RPTOR;RPTOR | | Body;Body |
| WG2844 | chr17:78724995-78725201 | RPTOR;RPTOR | | Body;Body |
| GM00016 | chr17:78725201-78725470 | RPTOR;RPTOR | | Body;Body |
| WG2843 | chr17:78735268-78735324 | RPTOR;RPTOR | | Body;Body |
| WG1310 | chr17:78735268-78735550 | RPTOR;RPTOR | | Body;Body |
| WG1977 | chr17:78735302-78735550 | RPTOR;RPTOR | | Body;Body |
| WG3466 | chr17:78808060-78808570 | RPTOR;RPTOR | | Body;Body |
| GM02555 | chr17:78865087-78865373 | RPTOR;RPTOR | S_Shore | Body;Body |
| WG1983 | chr17:78865263-78865373 | RPTOR;RPTOR | S_Shore | Body;Body |
| WG2276 | chr17:78892212-78892885 | RPTOR;RPTOR | | Body;Body |
| WG2276 | chr17:78894364-78894477 | RPTOR;RPTOR | N_Shelf | Body;Body |
| WG1310 | chr17:79010458-79010831 | BAIAP2;BAIAP2;BAIAP2;BAIAI | S_Shore | Body;Body;Body;Body |
| GM02555 | chr17:79010458-79010968 | BAIAP2;BAIAP2;BAIAP2;BAIAI | S_Shore | Body;Body;Body;Body |
| GM02456 | chr17:79011277-79011542 | BAIAP2;BAIAP2;BAIAP2;BAIAI | S_Shore | Body;Body;Body;Body |
| GM02706 | chr17:7905636-7905840 | GUCY2D | N_Shore | TSS1500 |
| GM00520 | chr17:790735-791217 | NXN | | Body |
| WG2844 | chr17:79095700-79096151 | AATK | Island | Body |
| GM00022 | chr17:79099481-79099777 | MIR657;AATK | N_Shelf | TSS1500;Body |
| GM09256 | chr17:79099481-79099785 | MIR657;AATK | N_Shelf | TSS1500;Body |
| WG2844 | chr17:79099481-79099819 | MIR657;AATK | N_Shelf | TSS1500;Body |
| GM01096 | chr17:79099481-79099942 | MIR657;AATK | N_Shelf | TSS1500;Body |
| WG2415 | chr17:79099711-79099819 | MIR657;AATK;MIR338 | N_Shelf | TSS1500;Body;Body |
| WG2120 | chr17:79099777-79099819 | MIR657;AATK;MIR338 | N_Shelf | TSS1500;Body;TSS200 |
| GM02555 | chr17:79100124-79100218 | MIR657;AATK;MIR338 | N_Shelf | TSS1500;Body;TSS1500 |
| WG2121 | chr17:79109729-79109808 | AATK | Island | Body |
| GM00520 | chr17:79109729-79109910 | AATK | Island | Body |
| WG2415 | chr17:79109777-79109910 | AATK | Island | Body |
| GM09255 | chr17:79140481-79140589 | AATK;LOC388428 | S_Shore | TSS1500;Body |
| WG2844 | chr17:79259040-79259699 | SLC38A10;SLC38A10 | N_Shore | Body;Body |
| GM00439 | chr17:79259283-79259699 | SLC38A10;SLC38A10 | Island | Body;Body |
| WG1977 | chr17:79259383-79259699 | SLC38A10;SLC38A10 | Island | Body;Body |
| GM09255 | chr17:79377548-79377850 | BAHCC1 | Island | Body |
| GM02149 | chr17:79377548-79378207 | BAHCC1 | Island | Body |
| GM02706 | chr17:79390726-79391071 | BAHCC1 | S_Shore | Body |
| WG1513 | chr17:79495335-79495519 | FSCN2;FSCN2 | Island | TSS200;TSS200 |
| GM01376 | chr17:79503701-79503877 | FSCN2;FSCN2 | Island | Body;Body |
| WG1977 | chr17:79609095-79609346 | TSPAN10 | | TSS1500 |
| GM01650 | chr17:79881468-79881523 | MAFG;MAFG | S_Shore | 5'UTR;TSS200 |
| GM02706 | chr17:79881468-79881543 | MAFG;MAFG | S_Shore | 5'UTR;TSS200 |
| WG2120 | chr17:79881523-79881543 | MAFG;MAFG | S_Shore | 5'UTR;TSS200 |
| WG1722 | chr17:79905219-79905263 | MYADML2 | | TSS200 |
| WG3465 | chr17:79905236-79905263 | MYADML2 | | TSS200 |
| GM02706 | chr17:79950061-79950366 | ASPSCR1 | N_Shelf | Body |
| GM02173 | chr17:800084-800431 | NXN | | Body |
| WG2276 | chr17:800084-800749 | NXN | | Body |
| WG1513 | chr17:800100-800431 | NXN | | Body |
| WG2844 | chr17:800263-800717 | NXN | | Body |

| | | | | |
|---|---|---|---|---|
| WG3465 | chr17:80186266-80186275 | SLC16A3;SLC16A3 | Island | TSS200;TSS1500 |
| WG1722 | chr17:80186273-80186336 | SLC16A3;SLC16A3 | Island | TSS200;TSS1500 |
| GM00042 | chr17:80189381-80190154 | SLC16A3;SLC16A3;SLC16A3 | Island | 5'UTR;TSS1500;5'UTR |
| GM01650 | chr17:80189973-80190054 | SLC16A3;SLC16A3;SLC16A3 | Island | TSS200;5'UTR;5'UTR |
| GM00022 | chr17:80190129-80190154 | SLC16A3;SLC16A3;SLC16A3 | S_Shore | TSS200;5'UTR;5'UTR |
| WG2844 | chr17:80196719-80197062 | SLC16A3;SLC16A3;SLC16A3 | S_Shore | Body;Body;Body |
| WG3465 | chr17:80290106-80290217 | SECTM1 | N_Shore | 5'UTR |
| WG3466 | chr17:80292079-80292252 | SECTM1 | Island | TSS200 |
| WG3465 | chr17:80358829-80358876 | C17orf101;C17orf101 | N_Shelf | Body;Body |
| WG2120 | chr17:80407590-80407779 | C17orf62;C17orf62;C17orf62 | N_Shore | 5'UTR;5'UTR;5'UTR |
| GM00935 | chr17:80708367-80708632 | FN3K | Island | Body |
| GM00439 | chr17:807121-807555 | NXN | | Body |
| WG2276 | chr17:80744833-80745235 | TBCD | S_Shelf | Body |
| WG3466 | chr17:80763035-80763117 | TBCD | N_Shore | Body |
| WG2415 | chr17:80806135-80806435 | TBCD | | Body |
| GM00520 | chr17:80875739-80876074 | TBCD | | Body |
| WG1721 | chr17:80875788-80876074 | TBCD | | Body |
| WG3465 | chr17:80875998-80876074 | TBCD | | Body |
| WG1722 | chr17:80929076-80929274 | B3GNTL1 | Island | Body |
| WG3466 | chr17:80943851-80944134 | B3GNTL1 | | Body |
| WG2844 | chr17:80943851-80944193 | B3GNTL1 | | Body |
| GM00439 | chr17:80969109-80970892 | B3GNTL1 | N_Shore | Body |
| GM02456 | chr17:80969515-80969547 | B3GNTL1 | N_Shore | Body |
| GM00520 | chr17:80969515-80969749 | B3GNTL1 | N_Shore | Body |
| GM01376 | chr17:80969525-80970434 | B3GNTL1 | N_Shore | Body |
| WG1722 | chr17:80969749-80970892 | B3GNTL1 | Island | Body |
| WG3465 | chr17:80970159-80970504 | B3GNTL1 | Island | Body |
| WG2844 | chr17:80970799-80970892 | B3GNTL1 | S_Shore | Body |
| GM02555 | chr17:80970799-80971527 | B3GNTL1 | S_Shore | Body |
| WG1658 | chr17:81006485-81006524 | B3GNTL1 | N_Shelf | Body |
| GM02555 | chr17:81040663-81040906 | METRNL | N_Shore | Body |
| WG2415 | chr17:81042782-81043176 | METRNL | Island | Body |
| GM03075 | chr17:81042947-81043176 | METRNL | Island | Body |
| WG3465 | chr17:81045495-81045781 | METRNL | Island | Body |
| GM02149 | chr17:8301166-8301192 | RNF222 | | TSS200 |
| WG1722 | chr17:8383621-8383800 | MYH10 | S_Shelf | Body |
| WG2276 | chr17:853971-854068 | NXN | S_Shelf | Body |
| WG1658 | chr17:855187-855312 | NXN | | Body |
| WG3466 | chr17:872762-873452 | NXN | | Body |
| WG3465 | chr17:873255-873452 | NXN | | Body |
| GM01095 | chr17:8927521-8928028 | NTN1 | S_Shore | Body |
| GM02706 | chr17:932700-933179 | ABR;ABR;ABR | N_Shore | Body;Body;Body |
| WG2275 | chr17:933047-933155 | ABR;ABR;ABR | Island | Body;Body;Body |
| GM03075 | chr17:933047-933179 | ABR;ABR;ABR | Island | Body;Body;Body |
| WG1983 | chr17:933118-933179 | ABR;ABR;ABR | Island | Body;Body;Body |
| GM00438 | chr17:934534-935017 | ABR;ABR;ABR | Island | Body;Body;Body |
| WG1309 | chr17:9550137-9550545 | USP43 | S_Shore | Body |

| | | | | |
|---|---|---|---|---|
| GM02149 | chr17:9559049-9559568 | USP43 | | Body |
| WG1722 | chr17:9559558-9559589 | USP43 | | Body |
| GM01096 | chr17:96503-96684 | RPH3AL | N_Shelf | Body |
| GM02175 | chr17:9694619-9694993 | DHRS7C | | TSS200 |
| GM03075 | chr17:9862752-9862873 | GAS7;GAS7;GAS7;GAS7;GAS7 | | Body;Body;Body;5'UTR;1stExon |
| GM03075 | chr17:9862924-9863081 | GAS7;GAS7;GAS7;GAS7 | | Body;Body;Body;TSS200 |
| WG2415 | chr17:9967417-9967499 | GAS7 | | Body |
| WG1657 | chr18:11850139-11850153 | CHMP1B;GNAL;GNAL;GNAL | N_Shore | TSS1500;Body;Body;Body |
| GM00439 | chr18:14748285-14748439 | ANKRD30B;ANKRD30B | S_Shore | 1stExon;5'UTR |
| WG3466 | chr18:19756582-19757016 | GATA6 | N_Shore | Body |
| WG3465 | chr18:19756582-19757022 | GATA6 | N_Shore | Body |
| GM00935 | chr18:19780587-19780878 | GATA6 | N_Shore | Body |
| WG2415 | chr18:19997803-19997924 | CTAGE1;CTAGE1 | | 1stExon;5'UTR |
| GM00520 | chr18:19997864-19997969 | CTAGE1;CTAGE1 | | 1stExon;5'UTR |
| GM02706 | chr18:19997912-19997969 | CTAGE1 | | TSS200 |
| WG2415 | chr18:20713759-20714399 | CABLES1 | N_Shore | TSS1500 |
| GM00289 | chr18:20714399-20714496 | CABLES1;CABLES1 | N_Shore | TSS1500;TSS200 |
| GM02456 | chr18:20714428-20714496 | CABLES1;CABLES1 | N_Shore | TSS1500;TSS200 |
| WG1310 | chr18:20735119-20735672 | CABLES1;CABLES1;CABLES1 | | Body;Body;TSS1500 |
| GM00290 | chr18:20735648-20735693 | CABLES1;CABLES1;CABLES1 | | Body;Body;TSS200 |
| WG1513 | chr18:21018217-21018724 | C18orf45 | S_Shore | TSS1500 |
| GM02555 | chr18:21572622-21572748 | TTC39C | | TSS200 |
| GM01650 | chr18:22006446-22006496 | IMPACT | Island | TSS200 |
| GM00042 | chr18:23713729-23713928 | PSMA8;PSMA8;PSMA8 | Island | TSS200;TSS200;TSS200 |
| GM01096 | chr18:24130839-24131115 | KCTD1;KCTD1 | Island | 5'UTR;TSS1500 |
| WG1977 | chr18:24130851-24131379 | KCTD1;KCTD1 | Island | 5'UTR;TSS1500 |
| WG2194 | chr18:24131115-24131604 | KCTD1 | Island | 5'UTR |
| WG1513 | chr18:24283669-24283696 | LOC728606 | | TSS200 |
| WG3465 | chr18:24443729-24444196 | AQP4;AQP4 | S_Shore | TSS1500;Body |
| GM02149 | chr18:24445240-24445318 | AQP4;C18orf16 | S_Shore | Body;TSS200 |
| WG3466 | chr18:24765490-24765781 | CHST9 | Island | TSS1500 |
| WG3466 | chr18:2907240-2907472 | EMILIN2 | Island | Body |
| WG2415 | chr18:29340788-29340919 | MCART2;MCART2 | | 1stExon;5'UTR |
| GM02706 | chr18:30051327-30051521 | FAM59A | Island | TSS1500 |
| GM02175 | chr18:30051521-30051538 | FAM59A | S_Shore | TSS1500 |
| WG3465 | chr18:30051538-30051572 | FAM59A | S_Shore | TSS1500 |
| WG2121 | chr18:30352782-30353125 | KLHL14;KLHL14 | S_Shore | 5'UTR;1stExon |
| WG1722 | chr18:30352975-30353000 | KLHL14 | S_Shore | TSS200 |
| GM02706 | chr18:30352980-30353283 | KLHL14 | S_Shore | TSS200 |
| WG1977 | chr18:30353000-30353283 | KLHL14 | S_Shore | TSS200 |
| GM00042 | chr18:30353073-30353240 | KLHL14 | S_Shore | TSS200 |
| WG2415 | chr18:30353073-30353283 | KLHL14 | S_Shore | TSS200 |
| GM00520 | chr18:30353073-30353699 | KLHL14 | S_Shore | TSS200 |
| GM09256 | chr18:37332566-37333028 | LOC647946 | | TSS1500 |
| WG1722 | chr18:3845348-3845703 | DLGAP1;DLGAP1 | | TSS200;Body |
| GM00438 | chr18:40695633-40696075 | RIT2;RIT2 | | 5'UTR;1stExon |
| GM00520 | chr18:45458021-45458698 | SMAD2;SMAD2;SMAD2 | Island | TSS1500;TSS1500;TSS1500 |

| | | | | |
|---|---|---|---|---|
| GM02456 | chr18:45458243-45458698 | SMAD2;SMAD2;SMAD2 | S_Shore | TSS1500;TSS1500;TSS1500 |
| WG2415 | chr18:46386267-46386453 | KIAA0427;KIAA0427 | Island | 3'UTR;3'UTR |
| GM00520 | chr18:46447634-46447961 | SMAD7 | N_Shore | 3'UTR |
| WG1310 | chr18:46447737-46447840 | SMAD7 | Island | 3'UTR |
| GM02456 | chr18:46447737-46447961 | SMAD7 | Island | 3'UTR |
| GM02706 | chr18:48346660-48346843 | MRO;MRO;MRO;MRO | | TSS1500;TSS1500;5'UTR;TSS1500 |
| GM02149 | chr18:49865270-49865971 | DCC | N_Shelf | TSS1500 |
| WG2120 | chr18:49865971-49866065 | DCC | N_Shelf | TSS1500 |
| GM02173 | chr18:49866065-49866097 | DCC | N_Shelf | TSS1500 |
| GM02706 | chr18:501109-501234 | COLEC12 | S_Shore | TSS1500 |
| WG1722 | chr18:55862577-55862676 | NEDD4L;NEDD4L;NEDD4L;N | N_Shore | 5'UTR;TSS200;Body;Body;Body;5'UTR; |
| WG1978 | chr18:55862577-55863675 | NEDD4L;NEDD4L;NEDD4L;N | N_Shore | 5'UTR;TSS200;Body;Body;Body;5'UTR; |
| WG1977 | chr18:56296243-56296449 | ALPK2 | | TSS200 |
| WG3466 | chr18:56296243-56296544 | ALPK2 | | TSS200 |
| GM02706 | chr18:56296449-56296544 | ALPK2 | | TSS1500 |
| WG2276 | chr18:56887002-56887194 | GRP;GRP;GRP | N_Shore | TSS1500;TSS1500;TSS1500 |
| WG2193 | chr18:56934834-56935199 | RAX | Island | 3'UTR |
| WG1513 | chr18:56935013-56935446 | RAX | Island | 3'UTR |
| WG3466 | chr18:56936434-56936882 | RAX | Island | Body |
| WG3466 | chr18:5890496-5891001 | TMEM200C | Island | 1stExon |
| GM00438 | chr18:60903834-60904418 | BCL2 | | Body |
| GM02706 | chr18:60904237-60904418 | BCL2 | | Body |
| GM02706 | chr18:60988017-60988058 | BCL2;BCL2 | Island | TSS1500;TSS1500 |
| WG2276 | chr18:60988056-60988085 | BCL2;BCL2 | Island | TSS1500;TSS1500 |
| GM01393 | chr18:60988056-60988096 | BCL2;BCL2 | Island | TSS1500;TSS1500 |
| GM00520 | chr18:61616451-61616526 | HMSD | | TSS200 |
| WG2121 | chr18:61616492-61616581 | HMSD | | TSS200 |
| GM02706 | chr18:61616492-61616718 | HMSD | | TSS200 |
| WG2415 | chr18:61816543-61817320 | C18orf20 | | TSS1500 |
| GM09256 | chr18:61817088-61817320 | C18orf20 | | TSS1500 |
| WG2415 | chr18:67069959-67070120 | DOK6 | Island | Body |
| WG3465 | chr18:67070045-67070170 | DOK6 | Island | Body |
| GM01392 | chr18:7011217-7011463 | LAMA1 | N_Shore | Body |
| WG3466 | chr18:7011268-7011463 | LAMA1 | Island | Body |
| GM00438 | chr18:72201590-72201716 | CNDP1 | | TSS200 |
| WG1513 | chr18:72201663-72201775 | CNDP1 | | TSS200 |
| GM00290 | chr18:72201764-72201785 | CNDP1;CNDP1 | | 1stExon;5'UTR |
| GM00042 | chr18:74535946-74535991 | ZNF236 | S_Shore | TSS200 |
| GM00520 | chr18:74729481-74729551 | MBP;MBP;MBP;MBP;MBP;MBP | | TSS1500;TSS1500;Body;Body;TSS150 |
| WG3465 | chr18:74770402-74770589 | MBP;MBP | Island | Body;Body |
| WG2844 | chr18:74770402-74770815 | MBP;MBP | Island | Body;Body |
| GM02706 | chr18:74960996-74961002 | GALR1 | N_Shore | TSS1500 |
| WG3465 | chr18:74960996-74961078 | GALR1 | N_Shore | TSS1500 |
| GM01392 | chr18:74961002-74961196 | GALR1 | N_Shore | TSS1500 |
| GM02706 | chr18:77623475-77623544 | KCNG2 | Island | TSS200 |
| GM00940 | chr18:77659572-77659949 | KCNG2 | Island | Body |
| WG2415 | chr18:77905298-77905751 | LOC100130522;LOC100130! | Island | TSS1500;TSS1500 |

| | | | | |
|---|---|---|---|---|
| GM02175 | chr18:77905298-77905947 | LOC100130522;LOC100130 | Island | TSS1500;TSS1500 |
| GM02706 | chr18:77905663-77905747 | LOC100130522;LOC100130 | S_Shore | TSS200;TSS200 |
| WG3465 | chr18:77905663-77905751 | LOC100130522;LOC100130 | S_Shore | TSS200;TSS200 |
| GM09256 | chr18:77905663-77905947 | LOC100130522;LOC100130 | S_Shore | TSS200;TSS200 |
| GM01650 | chr18:77905699-77905751 | LOC100130522;LOC100130 | S_Shore | TSS200;TSS200 |
| GM01392 | chr18:77905699-77905947 | LOC100130522;LOC100130 | S_Shore | TSS200;TSS200 |
| WG3465 | chr18:77917615-77918142 | LOC100130522;LOC100130 | N_Shore | Body;Body;3'UTR |
| GM02316 | chr19:1009172-1009764 | GRIN3B | Island | Body |
| GM01392 | chr19:1009172-1009949 | GRIN3B | Island | Body |
| GM00042 | chr19:10381063-10381393 | ICAM1 | Island | TSS1500 |
| WG1977 | chr19:10381355-10381393 | ICAM1 | Island | TSS200 |
| GM02706 | chr19:10397612-10397642 | ICAM4;ICAM4;ICAM4 | Island | TSS200;TSS200;TSS200 |
| WG3465 | chr19:10662582-10662619 | ATG4D;MIR1238 | | Body;TSS1500 |
| GM00940 | chr19:10735548-10735594 | SLC44A2;SLC44A2 | N_Shore | TSS1500;Body |
| GM00042 | chr19:11289211-11289376 | KANK2;KANK2 | Island | Body;Body |
| WG3465 | chr19:11289349-11289376 | KANK2;KANK2 | S_Shore | Body;Body |
| WG1978 | chr19:11289356-11289376 | KANK2;KANK2 | S_Shore | Body;Body |
| GM02706 | chr19:11353961-11354132 | DOCK6 | Island | Body |
| WG3465 | chr19:11406114-11406175 | TSPAN16 | S_Shore | TSS1500 |
| WG1513 | chr19:11484448-11484842 | C19orf39 | N_Shore | TSS1500 |
| GM01393 | chr19:11517079-11517260 | RGL3;RGL3 | Island | Body;Body |
| GM03075 | chr19:11517079-11517268 | RGL3;RGL3 | Island | Body;Body |
| GM02555 | chr19:1151960-1152189 | SBNO2 | N_Shore | Body |
| WG2415 | chr19:1152022-1152189 | SBNO2 | Island | Body |
| WG1513 | chr19:11565385-11565399 | ELAVL3;ELAVL3 | Island | Body;Body |
| GM01393 | chr19:11689715-11689813 | ACP5;ACP5;ACP5;ACP5;ACP | Island | TSS1500;5'UTR;1stExon;TSS200;5'UTI |
| GM01376 | chr19:11689813-11689860 | ACP5;ACP5;ACP5;ACP5 | S_Shore | TSS1500;TSS1500;TSS200;TSS200 |
| GM09255 | chr19:11689855-11689880 | ACP5;ACP5;ACP5;ACP5 | S_Shore | TSS1500;TSS1500;TSS200;TSS200 |
| WG3465 | chr19:11689860-11689880 | ACP5;ACP5;ACP5;ACP5 | S_Shore | TSS1500;TSS1500;TSS200;TSS200 |
| GM01095 | chr19:11784514-11784774 | ZNF833 | N_Shore | TSS1500 |
| GM02149 | chr19:11784647-11784761 | ZNF833 | N_Shore | TSS200 |
| WG1513 | chr19:12707946-12708001 | ZNF490 | Island | Body |
| GM00042 | chr19:1271057-1271307 | C19orf23;CIRBP;CIRBP;CIRB | S_Shore | TSS1500;Body;Body;Body |
| GM01096 | chr19:12779057-12779373 | MAN2B1;C19orf56;MORG1;N | N_Shore | TSS1500;3'UTR;TSS1500;5'UTR;Body |
| GM00935 | chr19:12831659-12832077 | TNPO2;TNPO2;TNPO2 | N_Shore | Body;Body;Body |
| GM00438 | chr19:12835001-12835134 | TNPO2 | S_Shore | TSS200 |
| WG3466 | chr19:12876846-12877000 | HOOK2;HOOK2 | N_Shore | Body;Body |
| WG3465 | chr19:12876846-12877188 | HOOK2;HOOK2 | N_Shore | Body;Body |
| GM02706 | chr19:12957979-12958340 | MAST1 | N_Shore | Body |
| WG1513 | chr19:12998135-12998435 | KLF1 | S_Shore | TSS200 |
| GM02173 | chr19:13113482-13113668 | NFIX | Island | Body |
| WG2276 | chr19:13124604-13125281 | NFIX | Island | Body |
| GM01393 | chr19:13127389-13127873 | NFIX | S_Shelf | Body |
| WG2121 | chr19:13209306-13209548 | NFIX | Island | 3'UTR |
| GM02555 | chr19:13209548-13209981 | NFIX | Island | 3'UTR |
| GM02706 | chr19:1356269-1356444 | MUM1;MUM1 | S_Shore | TSS200;5'UTR |
| WG2275 | chr19:13875014-13875137 | MRI1;MRI1 | N_Shore | TSS1500;TSS1500 |

| | | | | |
|---|---|---|---|---|
| GM02706 | chr19:1401767-1401781 | GAMT;GAMT | Island | TSS1500;TSS1500 |
| GM02173 | chr19:14260587-14260651 | LPHN1;LPHN1 | N_Shore | 3'UTR;3'UTR |
| WG3465 | chr19:1466647-1467032 | APC2 | Island | Body |
| GM09256 | chr19:1496416-1496580 | REEP6 | Island | Body |
| WG1977 | chr19:15083795-15083842 | SLC1A6 | | TSS200 |
| GM00289 | chr19:15083818-15083868 | SLC1A6 | | TSS200 |
| GM00438 | chr19:15083818-15083916 | SLC1A6 | | TSS200 |
| GM02706 | chr19:15311990-15312129 | NOTCH3 | Island | TSS200 |
| WG2276 | chr19:15343350-15343395 | EPHX3;EPHX3 | N_Shore | 5'UTR;TSS200 |
| WG2276 | chr19:15343786-15344364 | EPHX3;EPHX3;EPHX3 | N_Shore | 5'UTR;TSS1500;1stExon |
| WG1513 | chr19:15375283-15375374 | BRD4;BRD4 | | Body;Body |
| GM09256 | chr19:15580310-15580445 | PGLYRP2 | N_Shore | Body |
| WG1977 | chr19:15582687-15582853 | PGLYRP2 | Island | Body |
| GM00438 | chr19:15838375-15838756 | OR10H2 | N_Shore | TSS1500 |
| WG1513 | chr19:15838375-15838850 | OR10H2 | N_Shore | TSS1500 |
| WG1513 | chr19:15851571-15852002 | OR10H3 | | TSS1500 |
| WG2415 | chr19:16178091-16178099 | TPM4 | N_Shore | TSS1500 |
| WG1722 | chr19:16254474-16254516 | HSH2D | | 5'UTR |
| WG2415 | chr19:16284242-16284421 | CIB3 | | 1stExon |
| GM00439 | chr19:16284256-16284421 | CIB3 | | 1stExon |
| GM02706 | chr19:16999576-16999668 | F2RL3 | N_Shore | TSS1500 |
| GM00520 | chr19:16999641-16999800 | F2RL3 | N_Shore | TSS200 |
| WG3466 | chr19:16999668-16999800 | F2RL3 | N_Shore | TSS200 |
| WG1977 | chr19:16999768-16999953 | F2RL3 | N_Shore | TSS200 |
| GM02706 | chr19:17357587-17357641 | NR2F6 | S_Shore | TSS1500 |
| GM00935 | chr19:17439208-17440072 | ANO8 | Island | Body |
| GM02555 | chr19:17439877-17440072 | ANO8 | Island | Body |
| GM09256 | chr19:17858101-17858298 | FCHO1;FCHO1 | N_Shore | TSS1500;TSS1500 |
| WG2275 | chr19:17905589-17905626 | B3GNT3 | Island | TSS1500 |
| WG2415 | chr19:17918795-17919173 | B3GNT3 | Island | Body |
| GM02175 | chr19:17932100-17932331 | INSL3 | Island | Body |
| GM00520 | chr19:17932100-17932341 | INSL3 | Island | Body |
| WG3465 | chr19:17932244-17932331 | INSL3 | Island | 1stExon |
| WG2414 | chr19:17932328-17932341 | INSL3 | Island | TSS200 |
| WG1977 | chr19:17932328-17932369 | INSL3 | Island | TSS200 |
| GM00438 | chr19:17942272-17942411 | JAK3 | Island | Body |
| GM02706 | chr19:17958892-17958961 | JAK3 | S_Shore | TSS200 |
| WG3465 | chr19:18284120-18284235 | IFI30 | N_Shore | TSS1500 |
| WG1722 | chr19:18331075-18331394 | PDE4C;PDE4C;PDE4C | Island | Body;Body;Body |
| GM00438 | chr19:18344195-18344520 | PDE4C | Island | 5'UTR |
| GM02706 | chr19:18496756-18496953 | GDF15 | N_Shore | TSS1500 |
| WG3465 | chr19:18496932-18496984 | GDF15 | N_Shore | TSS200 |
| GM03073 | chr19:18539756-18540208 | SSBP4;SSBP4 | N_Shore | Body;Body |
| GM09255 | chr19:18539756-18540330 | SSBP4;SSBP4 | N_Shore | Body;Body |
| WG1309 | chr19:18539841-18540208 | SSBP4;SSBP4 | Island | Body;Body |
| WG1657 | chr19:18539841-18540330 | SSBP4;SSBP4 | Island | Body;Body |
| WG2415 | chr19:1854549-1854724 | KLF16 | Island | Body |

| | | | | |
|---|---|---|---|---|
| GM02706 | chr19:1854549-1854819 | KLF16 | Island | Body |
| GM02175 | chr19:18549131-18549274 | ISYNA1;ISYNA1;ISYNA1 | Island | TSS200;TSS1500;TSS200 |
| GM00439 | chr19:18549689-18550061 | ISYNA1;ISYNA1;ISYNA1 | S_Shore | TSS1500;TSS1500;TSS1500 |
| GM01392 | chr19:18761132-18761402 | KLHL26 | N_Shore | Body |
| GM00439 | chr19:18770726-18771327 | KLHL26 | N_Shore | Body |
| WG2276 | chr19:18771079-18771327 | KLHL26 | Island | Body |
| GM00520 | chr19:18771126-18771327 | KLHL26 | Island | Body |
| GM01376 | chr19:18902107-18902160 | COMP;COMP | Island | 1stExon;5'UTR |
| WG2415 | chr19:18902527-18902727 | COMP | S_Shore | TSS1500 |
| WG1977 | chr19:19221316-19221574 | SLC25A42 | Island | Body |
| GM02175 | chr19:19221392-19221574 | SLC25A42 | Island | Body |
| WG3465 | chr19:19626576-19626605 | NDUFA13;TSSK6 | Island | TSS1500;TSS200 |
| WG3466 | chr19:19639553-19639596 | YJEFN3 | Island | TSS200 |
| GM00935 | chr19:19639553-19639970 | YJEFN3 | Island | TSS200 |
| GM02555 | chr19:19639553-19640075 | YJEFN3 | Island | TSS200 |
| WG1977 | chr19:19639568-19639596 | YJEFN3 | Island | TSS200 |
| GM00439 | chr19:19639568-19640075 | YJEFN3 | Island | TSS200 |
| GM00935 | chr19:2041985-2042592 | MKNK2;MKNK2 | Island | Body;Body |
| GM00520 | chr19:2085286-2085483 | MOBKL2A | | 5'UTR |
| GM02555 | chr19:2095364-2095560 | MOBKL2A | N_Shore | 5'UTR |
| GM01393 | chr19:21264896-21265264 | ZNF714 | N_Shore | TSS200 |
| GM01096 | chr19:21264896-21265364 | ZNF714 | N_Shore | TSS200 |
| WG2121 | chr19:21264896-21265421 | ZNF714 | N_Shore | TSS200 |
| GM02317 | chr19:21264982-21265421 | ZNF714;ZNF714 | N_Shore | 5'UTR;1stExon |
| WG2844 | chr19:21265264-21265421 | ZNF714 | Island | 5'UTR |
| GM02706 | chr19:22193724-22193911 | ZNF208;ZNF208 | S_Shore | 5'UTR;1stExon |
| WG1977 | chr19:2250561-2251588 | AMH | Island | Body |
| WG1977 | chr19:2252069-2252432 | AMH | Island | 3'UTR |
| GM01096 | chr19:2252961-2253492 | JSRP1 | Island | Body |
| WG1977 | chr19:2252961-2253854 | JSRP1 | Island | Body |
| WG2120 | chr19:2253492-2253854 | JSRP1 | Island | Body |
| WG2844 | chr19:2276322-2276471 | C19orf35 | Island | Body |
| WG3466 | chr19:2282010-2282207 | C19orf35 | S_Shelf | 5'UTR |
| GM01650 | chr19:2282010-2282439 | C19orf35 | S_Shelf | 5'UTR |
| GM01095 | chr19:2282010-2282568 | C19orf35 | S_Shelf | 5'UTR |
| WG1978 | chr19:2282161-2282568 | C19orf35;C19orf35 | S_Shelf | 5'UTR;1stExon |
| WG2844 | chr19:2282439-2282729 | C19orf35 | S_Shelf | TSS1500 |
| WG1977 | chr19:2291020-2291872 | LINGO3 | Island | Body |
| GM01096 | chr19:2291613-2291872 | LINGO3 | Island | Body |
| WG1984 | chr19:2302401-2302783 | LINGO3 | N_Shore | 5'UTR |
| WG1722 | chr19:23941313-23941575 | ZNF681 | Island | Body |
| WG1282 | chr19:23941408-23941575 | ZNF681 | Island | Body |
| GM01096 | chr19:23942040-23942126 | ZNF681 | S_Shore | TSS1500 |
| GM01392 | chr19:23945472-23945659 | RPSAP58 | N_Shore | TSS1500 |
| GM01393 | chr19:23945659-23945996 | RPSAP58 | N_Shore | TSS200 |
| WG3466 | chr19:2428350-2429209 | LMNB2;TIMM13 | Island | 3'UTR;TSS1500 |
| WG1513 | chr19:2428514-2428677 | LMNB2;TIMM13 | Island | 3'UTR;TSS1500 |

| | | | | |
|---|---|---|---|---|
| GM00439 | chr19:2428514-2428889 | LMNB2;TIMM13 | Island | 3'UTR;TSS1500 |
| WG3465 | chr19:2428567-2428889 | LMNB2;TIMM13 | Island | 3'UTR;TSS1500 |
| WG2415 | chr19:2428677-2429209 | LMNB2;TIMM13 | Island | 3'UTR;TSS1500 |
| GM00940 | chr19:2512999-2513436 | GNG7 | N_Shore | 3'UTR |
| WG2276 | chr19:2525384-2525634 | GNG7 | Island | 5'UTR |
| GM01392 | chr19:2546598-2546877 | GNG7 | N_Shore | 5'UTR |
| WG2415 | chr19:2588241-2588553 | GNG7 | N_Shore | 5'UTR |
| WG3465 | chr19:2588376-2588629 | GNG7 | Island | 5'UTR |
| GM01653 | chr19:2588479-2588629 | GNG7 | Island | 5'UTR |
| WG2415 | chr19:2650755-2650863 | GNG7 | Island | 5'UTR |
| WG2415 | chr19:292131-292179 | PPAP2C;PPAP2C;PPAP2C | Island | TSS1500;TSS1500;TSS1500 |
| GM02706 | chr19:2977603-2977843 | TLE6;TLE6;TLE6;TLE6 | S_Shore | 1stExon;1stExon;5'UTR;5'UTR |
| GM02704 | chr19:3097565-3097728 | GNA11 | Island | Body |
| GM00042 | chr19:3097570-3097728 | GNA11 | Island | Body |
| WG2415 | chr19:31640249-31640711 | DKFZp566F0947 | | TSS1500 |
| GM00520 | chr19:31640249-31640849 | DKFZp566F0947 | | TSS1500 |
| WG1721 | chr19:31640401-31640693 | DKFZp566F0947 | | TSS1500 |
| WG3465 | chr19:31640401-31640849 | DKFZp566F0947 | | TSS1500 |
| GM03075 | chr19:31640642-31640693 | DKFZp566F0947 | | TSS200 |
| WG1983 | chr19:31640642-31640706 | DKFZp566F0947 | | TSS200 |
| WG2414 | chr19:31640642-31640711 | DKFZp566F0947 | | TSS200 |
| WG2120 | chr19:31640642-31640849 | DKFZp566F0947 | | TSS200 |
| WG1282 | chr19:31640691-31640706 | DKFZp566F0947 | | TSS200 |
| GM01393 | chr19:31640691-31640711 | DKFZp566F0947 | | TSS200 |
| WG1514 | chr19:31640691-31640849 | DKFZp566F0947 | | TSS200 |
| WG1977 | chr19:31640693-31640849 | DKFZp566F0947 | | TSS200 |
| GM00940 | chr19:34112306-34112325 | CHST8;CHST8 | Island | TSS1500;TSS1500 |
| GM09256 | chr19:34174345-34174665 | CHST8;CHST8;CHST8 | N_Shore | 5'UTR;5'UTR;TSS1500 |
| GM00520 | chr19:3480561-3480672 | C19orf77 | N_Shore | TSS200 |
| GM00042 | chr19:35531859-35532097 | HPN;HPN | Island | 5'UTR;5'UTR |
| WG3465 | chr19:35633553-35633972 | FXYD1;FXYD7;FXYD1 | Island | Body;TSS1500;Body |
| WG1309 | chr19:35633669-35633972 | FXYD7;FXYD1;FXYD1 | Island | TSS1500;3'UTR;3'UTR |
| WG3465 | chr19:35786580-35786932 | MAG;MAG | Island | Body;Body |
| WG2275 | chr19:35786821-35786961 | MAG;MAG | Island | Body;Body |
| WG1722 | chr19:35940461-35940593 | FFAR2 | | TSS200 |
| WG1513 | chr19:35981371-35981549 | KRTDAP | | TSS200 |
| WG2415 | chr19:35981517-35981549 | KRTDAP | | TSS200 |
| WG1722 | chr19:36024217-36024282 | GAPDHS | Island | TSS200 |
| WG3465 | chr19:36054621-36055013 | ATP4A | | TSS200 |
| WG3465 | chr19:36157677-36157825 | UPK1A | | TSS200 |
| WG3465 | chr19:36334925-36335320 | NPHS1 | N_Shore | Body |
| GM09256 | chr19:36484913-36485360 | SDHAF1 | Island | TSS1500 |
| GM00520 | chr19:36499666-36499839 | C19orf46;C19orf46 | N_Shore | 5'UTR;1stExon |
| GM09256 | chr19:3671903-3672228 | PIP5K1C | Island | Body |
| WG2194 | chr19:36801408-36801503 | LOC100134317 | S_Shelf | TSS1500 |
| WG2415 | chr19:37096010-37096329 | ZNF529;ZNF529;ZNF529;ZNF | Island | Body;5'UTR;5'UTR;TSS1500 |
| WG1513 | chr19:3786643-3786957 | MATK;MATK;MATK | Island | TSS1500;Body;TSS1500 |

| | | | | |
|---|---|---|---|---|
| GM00289 | chr19:37958707-37959104 | ZNF570;ZNF569 | N_Shore | TSS1500;TSS1500 |
| WG3466 | chr19:38747355-38747378 | PPP1R14A | Island | TSS200 |
| WG3465 | chr19:38794598-38794814 | C19orf33;YIF1B;YIF1B;YIF1B;YIF1B | | TSS1500;3'UTR;3'UTR;3'UTR;3'UTR;3'U |
| WG3465 | chr19:38886554-38886667 | SPRED3 | Island | Body |
| GM02706 | chr19:38886554-38886789 | SPRED3 | Island | Body |
| GM01096 | chr19:39056084-39056216 | RYR1;RYR1 | Island | Body;Body |
| WG1722 | chr19:39369067-39369277 | RINL | | TSS200 |
| WG2276 | chr19:39523309-39523395 | FBXO27 | S_Shore | TSS200 |
| GM00520 | chr19:39694993-39695034 | SYCN | Island | TSS200 |
| WG1977 | chr19:39695011-39695065 | SYCN | S_Shore | TSS200 |
| GM00438 | chr19:39695011-39695237 | SYCN | S_Shore | TSS200 |
| GM00439 | chr19:39695034-39695237 | SYCN | S_Shore | TSS200 |
| WG1513 | chr19:39785679-39785941 | IL29 | | TSS1500 |
| GM01376 | chr19:39997622-39998290 | DLL3;DLL3 | Island | Body;Body |
| WG2276 | chr19:39998018-39998290 | DLL3;DLL3 | Island | Body;Body |
| WG3465 | chr19:40229215-40229766 | CLC | | TSS1500 |
| GM01096 | chr19:406256-406560 | C2CD4C | Island | 3'UTR |
| GM00438 | chr19:40729157-40729386 | CNTD2 | Island | Body |
| GM02555 | chr19:40732762-40732804 | CNTD2 | S_Shore | TSS200 |
| WG3466 | chr19:40904691-40904974 | PRX;PRX | Island | Body;Body |
| GM02706 | chr19:41025339-41025959 | SPTBN4 | Island | Body |
| GM02704 | chr19:41025574-41025863 | SPTBN4 | Island | Body |
| GM02706 | chr19:41060190-41060475 | SPTBN4;SPTBN4 | Island | Body;Body |
| WG3466 | chr19:41119829-41120329 | LTBP4;LTBP4;LTBP4 | Island | Body;Body;Body |
| GM00042 | chr19:41510026-41510059 | CYP2B6 | | Body |
| WG3465 | chr19:4153671-4153713 | CREB3L3;CREB3L3 | | 1stExon;5'UTR |
| WG1722 | chr19:42636817-42637137 | POU2F2 | N_Shore | TSS200 |
| WG3466 | chr19:427022-427263 | SHC2 | Island | Body |
| GM00289 | chr19:4338769-4338852 | STAP2;STAP2;STAP2;STAP2 | | 1stExon;5'UTR;5'UTR;1stExon |
| WG1283 | chr19:43857861-43858174 | CD177;CD177 | | 1stExon;5'UTR |
| GM02175 | chr19:43969769-43969884 | LYPD3;LYPD3 | S_Shore | 1stExon;5'UTR |
| WG3465 | chr19:44306947-44307449 | LYPD5;LYPD5 | S_Shelf | TSS1500;5'UTR |
| GM02175 | chr19:44324481-44324856 | LYPD5 | Island | 5'UTR |
| GM09256 | chr19:44330910-44331239 | ZNF283 | | TSS1500 |
| GM02555 | chr19:4455863-4455957 | UBXN6;UBXN6 | N_Shore | Body;TSS1500 |
| GM00438 | chr19:44952725-44952766 | ZNF229 | Island | TSS200 |
| GM00438 | chr19:45033454-45033572 | CEACAM20;CEACAM20;CEACAM20;CEA | | Body;Body;Body;Body |
| GM02706 | chr19:45260501-45261004 | BCL3 | Island | Body |
| GM01096 | chr19:45579986-45580017 | ZNF296 | Island | TSS1500 |
| WG2414 | chr19:45594410-45594670 | GEMIN7;GEMIN7;GEMIN7 | Island | 3'UTR;3'UTR;3'UTR |
| WG2844 | chr19:45720771-45721076 | EXOC3L2 | Island | Body |
| WG1722 | chr19:45737208-45737485 | EXOC3L2 | N_Shore | 5'UTR |
| WG2844 | chr19:45737483-45738115 | EXOC3L2 | Island | TSS200 |
| GM02706 | chr19:45843694-45843871 | KLC3 | N_Shore | TSS1500 |
| GM09255 | chr19:45905621-45906156 | PPP1R13L;PPP1R13L | N_Shelf | 5'UTR;5'UTR |
| WG2121 | chr19:45996644-45996901 | RTN2;RTN2;RTN2 | N_Shore | Body;Body;TSS200 |
| GM02175 | chr19:46001605-46001962 | RTN2;FLJ40125;RTN2 | N_Shore | TSS1500;TSS200;TSS1500 |

| | | | | |
|---|---|---|---|---|
| GM02706 | chr19:46142212-46142357 | EML2 | N_Shelf | Body |
| WG3466 | chr19:46181221-46181330 | GIPR | Island | Body |
| GM00940 | chr19:46318439-46318633 | RSPH6A;RSPH6A | N_Shore | 1stExon;5'UTR |
| GM00935 | chr19:46318514-46318762 | RSPH6A;RSPH6A | Island | 1stExon;5'UTR |
| GM02456 | chr19:46318514-46319153 | RSPH6A;RSPH6A | Island | 1stExon;5'UTR |
| GM00520 | chr19:4639844-4639992 | TNFAIP8L1;TNFAIP8L1 | S_Shore | TSS200;5'UTR |
| GM01096 | chr19:46418849-46419054 | NANOS2 | S_Shore | TSS1500 |
| GM00042 | chr19:46443657-46444011 | NOVA2 | N_Shore | Body |
| GM00941 | chr19:46526483-46526508 | PGLYRP1 | S_Shore | TSS200 |
| WG1513 | chr19:46544662-46544802 | IGFL4 | | TSS1500 |
| GM01392 | chr19:46627896-46628137 | IGFL3 | | 1stExon |
| GM03073 | chr19:46801557-46801672 | HIF3A;HIF3A | S_Shore | Body;TSS200 |
| WG2276 | chr19:46974972-46975072 | PNMAL1;PNMAL1 | Island | TSS200;TSS200 |
| GM03075 | chr19:46999055-46999118 | PNMAL2;PNMAL2 | Island | 1stExon;5'UTR |
| GM00438 | chr19:46999055-46999307 | PNMAL2;PNMAL2 | Island | 1stExon;5'UTR |
| GM09255 | chr19:46999403-46999840 | PNMAL2 | S_Shore | TSS1500 |
| GM02706 | chr19:474973-474994 | ODF3L2;ODF3L2 | | 1stExon;5'UTR |
| GM02149 | chr19:47507399-47507461 | GRLF1 | Island | 3'UTR |
| WG1983 | chr19:47507399-47507691 | GRLF1 | Island | 3'UTR |
| GM00438 | chr19:4769206-4769240 | MIR7-3;C19orf30 | | TSS1500;Body |
| WG3466 | chr19:48947560-48947578 | GRIN2D;GRWD1 | Island | 3'UTR;TSS1500 |
| WG1514 | chr19:48983508-48984071 | CYTH2;CYTH2 | N_Shore | 3'UTR;3'UTR |
| WG1722 | chr19:48983508-48984318 | CYTH2;CYTH2 | N_Shore | 3'UTR;3'UTR |
| GM00439 | chr19:49016564-49016629 | LMTK3 | N_Shore | TSS200 |
| WG1513 | chr19:49055432-49055443 | SULT2B1;SULT2B1 | | 1stExon;5'UTR |
| GM09256 | chr19:49133421-49134105 | SPHK2 | Island | 3'UTR |
| WG3466 | chr19:49133647-49134105 | SPHK2 | Island | 3'UTR |
| GM01096 | chr19:49259265-49259452 | FGF21;FUT1 | S_Shelf | TSS200;TSS1500 |
| WG2276 | chr19:49339085-49339789 | HSD17B14 | N_Shore | Body |
| GM02706 | chr19:49339776-49340593 | HSD17B14;HSD17B14 | N_Shore | 5'UTR;1stExon |
| WG1283 | chr19:49340489-49340593 | PLEKHA4;PLEKHA4;HSD17B1 | Island | 3'UTR;3'UTR;TSS1500 |
| GM01392 | chr19:49340574-49340765 | PLEKHA4;HSD17B14;PLEKHA | Island | 3'UTR;TSS1500;Body |
| WG3465 | chr19:49891270-49891574 | CCDC155 | Island | TSS1500 |
| GM00042 | chr19:49891270-49891603 | CCDC155 | Island | TSS1500 |
| WG2415 | chr19:49935823-49935948 | SLC17A7 | Island | Body |
| WG1513 | chr19:49935823-49936274 | SLC17A7 | Island | Body |
| GM00042 | chr19:50004281-50004625 | MIR150 | | TSS200 |
| GM02706 | chr19:50192625-50193021 | C19orf76;C19orf76 | Island | 1stExon;5'UTR |
| GM09256 | chr19:50249584-50249776 | TSKS | Island | Body |
| GM00016 | chr19:5036609-5037219 | KDM4B | N_Shore | Body |
| WG2194 | chr19:5037004-5037219 | KDM4B | Island | Body |
| GM02149 | chr19:50789296-50789841 | MYH14;MYH14;MYH14 | N_Shore | Body;Body;Body |
| WG1513 | chr19:50789296-50789951 | MYH14;MYH14;MYH14 | N_Shore | Body;Body;Body |
| GM00438 | chr19:50789808-50789951 | MYH14;MYH14;MYH14 | Island | Body;Body;Body |
| WG3465 | chr19:50861774-50861923 | NAPSA | Island | 3'UTR |
| GM01650 | chr19:50931222-50931515 | SPIB | N_Shore | Body |
| WG2120 | chr19:50931343-50931515 | SPIB | Island | Body |

| | | | | |
|---|---|---|---|---|
| WG1513 | chr19:50931432-50931622 | SPIB | Island | Body |
| WG1722 | chr19:50931616-50931875 | SPIB | Island | 3'UTR |
| GM02175 | chr19:5093491-5093659 | KDM4B | Island | Body |
| GM00438 | chr19:51017123-51017311 | ASPDH;ASPDH;ASPDH | N_Shelf | 5'UTR;1stExon;Body |
| GM00520 | chr19:51017855-51018014 | ASPDH;ASPDH;ASPDH | N_Shelf | 1stExon;TSS1500;5'UTR |
| GM01376 | chr19:51107558-51107781 | SNAR-F | Island | TSS1500 |
| WG2415 | chr19:51141227-51141361 | SYT3;SYT3;SYT3;SYT3 | N_Shore | 5'UTR;1stExon;5'UTR;5'UTR |
| GM09255 | chr19:51141269-51141413 | SYT3;SYT3;SYT3;SYT3 | N_Shore | 5'UTR;1stExon;5'UTR;5'UTR |
| WG3465 | chr19:51165404-51165632 | SHANK1 | Island | Body |
| GM01376 | chr19:51165441-51165752 | SHANK1 | Island | Body |
| GM02175 | chr19:51330299-51330469 | KLK15;KLK15;KLK15 | Island | Body;Body;Body |
| GM00438 | chr19:51376305-51376711 | KLK2;KLK2 | | TSS1500;TSS1500 |
| GM02149 | chr19:51411782-51411940 | KLK4 | Island | Body |
| GM09256 | chr19:51415170-51415452 | KLK4 | N_Shore | TSS1500 |
| GM00438 | chr19:51473087-51473480 | KLK6;KLK6;KLK6 | | TSS1500;TSS200;TSS1500 |
| WG1977 | chr19:5147855-5148130 | KDM4B | Island | Body |
| WG1978 | chr19:5147855-5148409 | KDM4B | Island | Body |
| WG2276 | chr19:5148023-5148409 | KDM4B | Island | Body |
| WG2415 | chr19:51486901-51487136 | KLK7;KLK7 | | 5'UTR;5'UTR |
| GM03075 | chr19:51487229-51487305 | KLK7;KLK7 | | TSS200;TSS200 |
| WG2415 | chr19:51505779-51506273 | KLK9;KLK8;KLK8;KLK8;KLK8 | N_Shore | 3'UTR;TSS1500;TSS1500;TSS1500;TS[illegible] |
| GM02149 | chr19:51512895-51513611 | KLK9 | | TSS200 |
| WG1978 | chr19:51531318-51531484 | KLK11;KLK11;KLK11;KLK11 | N_Shelf | TSS1500;TSS200;TSS1500;TSS200 |
| GM00438 | chr19:52004992-52005108 | SIGLEC12;SIGLEC12;SIGLEC12 | | 1stExon;5'UTR;TSS1500 |
| WG1722 | chr19:52005153-52005188 | SIGLEC12 | | TSS200 |
| GM01095 | chr19:52222431-52223099 | HAS1 | Island | Body |
| WG2121 | chr19:52227309-52227357 | HAS1 | | TSS200 |
| GM09256 | chr19:52227967-52228400 | HAS1 | | TSS1500 |
| WG3465 | chr19:52264411-52264429 | FPR2 | | TSS200 |
| GM00935 | chr19:52996083-52996548 | ZNF578 | Island | 5'UTR |
| GM00289 | chr19:5335159-5335260 | PTPRS;PTPRS;PTPRS;PTPRS | Island | 5'UTR;5'UTR;5'UTR;5'UTR |
| GM00438 | chr19:54134782-54135140 | DPRX | | TSS1500 |
| WG1513 | chr19:54200863-54201247 | MIR525;MIR523 | | Body;TSS1500 |
| GM02149 | chr19:54204449-54204773 | MIR520B | | TSS200 |
| GM09256 | chr19:54219381-54219607 | MIR521-2 | | TSS1500 |
| GM00438 | chr19:54219477-54219607 | MIR521-2 | | TSS1500 |
| GM00520 | chr19:54290762-54291017 | MIR372;MIR371;MIR373 | | TSS1500;TSS200;TSS1500 |
| WG3465 | chr19:543909-544349 | GZMM | S_Shore | TSS200 |
| WG1513 | chr19:54485321-54485460 | CACNG8;MIR935 | Island | Body;TSS1500 |
| WG1977 | chr19:54515245-54515341 | CACNG6;CACNG6;CACNG6 | Island | Body;Body;Body |
| WG1513 | chr19:54850444-54850645 | LILRA4 | | TSS200 |
| GM02555 | chr19:54876664-54876749 | LAIR1;LAIR1;LAIR1;LAIR1 | | 5'UTR;1stExon;1stExon;5'UTR |
| WG1513 | chr19:54876664-54876795 | LAIR1;LAIR1;LAIR1;LAIR1 | | 5'UTR;1stExon;1stExon;5'UTR |
| WG2121 | chr19:54876728-54876752 | LAIR1;LAIR1 | | TSS200;TSS200 |
| WG2415 | chr19:54876749-54876795 | LAIR1;LAIR1 | | TSS200;TSS200 |
| GM00935 | chr19:54976211-54976506 | LENG9;CDC42EP5 | Island | TSS1500;3'UTR |
| GM00438 | chr19:55141723-55142181 | LILRB1;LILRB1;LILRB1;LILRB1 | | TSS200;5'UTR;TSS200;TSS1500 |

| | | | | |
|---|---|---|---|---|
| WG1722 | chr19:55416928-55417390 | NCR1;NCR1;NCR1;NCR1;NCR1 | | TSS1500;TSS1500;TSS1500;TSS1500; |
| GM00438 | chr19:55416928-55417500 | NCR1;NCR1;NCR1;NCR1;NCR1 | | TSS1500;TSS1500;TSS1500;TSS1500; |
| GM00438 | chr19:55458886-55459282 | NLRP7;NLRP7;NLRP7 | | TSS200;TSS200;TSS200 |
| GM00289 | chr19:55458943-55459137 | NLRP7;NLRP7;NLRP7 | | TSS200;TSS200;TSS200 |
| GM02149 | chr19:55458958-55459282 | NLRP7;NLRP7;NLRP7 | | TSS200;TSS200;TSS200 |
| WG1977 | chr19:55477653-55477759 | NLRP2 | | TSS200 |
| GM00438 | chr19:55493930-55494710 | NLRP2 | Island | Body |
| GM02149 | chr19:55549799-55549826 | GP6;GP6 | | TSS200;TSS200 |
| GM01096 | chr19:55591148-55591437 | EPS8L1;EPS8L1 | N_Shore | TSS1500;Body |
| WG1513 | chr19:55692003-55692526 | SYT5 | S_Shore | TSS1500 |
| WG2275 | chr19:55887799-55888003 | TMEM190 | N_Shore | TSS1500 |
| WG1721 | chr19:55944864-55945233 | SHISA7 | Island | Body |
| WG2414 | chr19:55972791-55973099 | ISOC2;ISOC2;ISOC2 | Island | 5'UTR;5'UTR;5'UTR |
| GM09255 | chr19:55972791-55973338 | ISOC2;ISOC2;ISOC2 | Island | 5'UTR;5'UTR;5'UTR |
| GM02704 | chr19:55972855-55973099 | ISOC2;ISOC2;ISOC2 | Island | 5'UTR;5'UTR;5'UTR |
| GM00941 | chr19:55973234-55973338 | ISOC2;ISOC2;ISOC2 | Island | TSS200;TSS200;TSS200 |
| WG2276 | chr19:55987366-55987415 | ZNF628 | N_Shore | TSS1500 |
| WG2844 | chr19:56042469-56042579 | SBK2 | Island | Body |
| GM09255 | chr19:56187475-56187915 | EPN1;EPN1;EPN1 | S_Shore | 5'UTR;5'UTR;TSS1500 |
| GM02149 | chr19:56458926-56459108 | NLRP8 | | TSS1500 |
| GM02149 | chr19:56686339-56686707 | GALP;GALP | | TSS1500;TSS1500 |
| WG1513 | chr19:57630153-57630202 | USP29 | N_Shore | TSS1500 |
| WG1722 | chr19:57630340-57630524 | USP29 | Island | TSS1500 |
| WG1513 | chr19:57630446-57630710 | USP29 | Island | TSS1500 |
| GM00438 | chr19:57630524-57630710 | USP29 | Island | TSS1500 |
| WG3465 | chr19:57630536-57630628 | USP29 | Island | TSS1500 |
| GM00438 | chr19:57631189-57631405 | USP29 | S_Shore | TSS1500 |
| GM00290 | chr19:57742217-57742444 | AURKC;AURKC;AURKC | Island | TSS200;TSS1500;TSS200 |
| WG3465 | chr19:57742421-57742444 | AURKC;AURKC;AURKC;AURK | Island | 5'UTR;TSS200;1stExon;5'UTR;1stExon |
| WG2120 | chr19:57988497-57989117 | ZNF772;ZNF772 | | Body;Body |
| GM09256 | chr19:58038573-58038588 | ZNF549 | Island | TSS200 |
| GM02175 | chr19:5822318-5822514 | NRTN | Island | TSS1500 |
| WG2415 | chr19:5838735-5838779 | FUT6;FUT6;FUT6;FUT6 | | 1stExon;5'UTR;1stExon;5'UTR |
| GM01375 | chr19:58400504-58400574 | ZNF814 | S_Shore | TSS200 |
| GM09256 | chr19:58446770-58446787 | ZNF418 | Island | TSS200 |
| GM00042 | chr19:58554401-58554527 | ZSCAN1 | Island | Body |
| WG3465 | chr19:58864846-58864929 | NCRNA00181;A1BG;A1BG | N_Shelf | Body;5'UTR;1stExon |
| WG1978 | chr19:58879022-58879059 | ZNF837;ZNF837 | Island | 3'UTR;3'UTR |
| GM02175 | chr19:59025345-59025573 | ZBTB45 | Island | 3'UTR |
| GM02706 | chr19:613111-613818 | HCN2 | Island | Body |
| WG3466 | chr19:6481819-6481857 | DENND1C | | TSS200 |
| WG1977 | chr19:695447-695568 | PRSSL1;PRSSL1 | | 5'UTR;1stExon |
| WG2276 | chr19:710823-711001 | PALM;PALM | Island | Body;Body |
| WG1513 | chr19:750564-751098 | C19orf21 | S_Shelf | TSS1500 |
| GM00042 | chr19:751098-751159 | C19orf21 | S_Shelf | TSS200 |
| GM02456 | chr19:7580209-7580275 | ZNF358 | N_Shore | TSS1500 |
| GM00438 | chr19:7701212-7701753 | STXBP2;STXBP2 | N_Shore | TSS1500;TSS1500 |

| | | | | |
|---|---|---|---|---|
| GM00438 | chr19:7741809-7741955 | C19orf59 | N_Shelf | TSS200 |
| WG1513 | chr19:7813422-7813963 | CD209;CD209;CD209;CD209;CD | | TSS1500;TSS1500;TSS1500;TSS1500; |
| WG2415 | chr19:7852051-7852238 | CLEC4GP1 | N_Shore | TSS1500 |
| GM00520 | chr19:7852204-7852214 | CLEC4GP1 | N_Shore | TSS200 |
| GM00042 | chr19:7926970-7927374 | EVI5L;EVI5L | Island | Body;Body |
| GM09255 | chr19:7953054-7953081 | LRRC8E | N_Shore | TSS1500 |
| WG1977 | chr19:7984832-7985019 | SNAPC2;SNAPC2 | N_Shore | TSS1500;TSS1500 |
| WG3465 | chr19:827715-827843 | AZU1 | | TSS200 |
| WG2414 | chr19:827776-827843 | AZU1 | | TSS200 |
| GM01096 | chr19:827776-828170 | AZU1 | | TSS200 |
| WG3465 | chr19:840873-840985 | PRTN3 | N_Shelf | TSS200 |
| GM02704 | chr19:853054-853334 | ELANE | Island | Body |
| WG1722 | chr19:859330-859680 | CFD | N_Shore | TSS1500 |
| WG2276 | chr19:859499-859680 | CFD | N_Shore | TSS200 |
| GM01096 | chr19:863054-863226 | CFD | Island | Body |
| WG2276 | chr19:863054-863244 | CFD | Island | Body |
| WG1977 | chr19:8642308-8642370 | MYO1F | | TSS200 |
| GM00289 | chr19:940724-941571 | ARID3A | Island | Body |
| WG2844 | chr19:940967-941243 | ARID3A | Island | Body |
| GM02175 | chr19:9785295-9786115 | ZNF562;ZNF562;ZNF562 | N_Shore | 5'UTR;5'UTR;5'UTR |
| GM02316 | chr19:9785511-9785779 | ZNF562;ZNF562;ZNF562 | Island | 5'UTR;5'UTR;5'UTR |
| GM09255 | chr2:100722022-100722408 | AFF3;AFF3;AFF3 | S_Shore | 5'UTR;1stExon;5'UTR |
| GM00042 | chr2:100939045-100939477 | LONRF2;LONRF2 | Island | 5'UTR;1stExon |
| GM02706 | chr2:10182878-10183115 | KLF11 | Island | TSS1500 |
| GM00016 | chr2:10302391-10302462 | C2orf48 | | Body |
| WG3465 | chr2:10567628-10567843 | HPCAL1;HPCAL1 | S_Shore | 3'UTR;3'UTR |
| WG3466 | chr2:106015739-106015771 | FHL2;FHL2;FHL2;FHL2 | Island | TSS200;TSS200;5'UTR;TSS200 |
| WG2415 | chr2:106055153-106055458 | FHL2;FHL2 | | 1stExon;5'UTR |
| GM02456 | chr2:106055153-106055507 | FHL2;FHL2 | | 1stExon;5'UTR |
| GM00940 | chr2:106055450-106055507 | FHL2 | | TSS1500 |
| GM00439 | chr2:10830698-10830764 | NOL10 | S_Shore | TSS1500 |
| WG3466 | chr2:108993844-108994311 | SULT1C4 | | TSS1500 |
| WG3466 | chr2:109788051-109788778 | SH3RF3 | | Body |
| GM00520 | chr2:110373002-110373509 | SEPT10;SEPT10;ANKRD57 | Island | TSS1500;TSS1500;1stExon |
| GM00042 | chr2:110872620-110872729 | MALL | N_Shore | Body |
| WG2194 | chr2:110969641-110970097 | NCRNA00116 | N_Shore | Body |
| GM02317 | chr2:110969641-110970204 | NCRNA00116 | N_Shore | Body |
| WG3465 | chr2:110969641-110970249 | NCRNA00116 | N_Shore | Body |
| WG1722 | chr2:112939119-112939562 | FBLN7;FBLN7 | N_Shore | Body;Body |
| GM00941 | chr2:113992762-113993304 | PAX8;PAX8;PAX8;PAX8;PAX8 | N_Shore | Body;Body;Body;Body;Body;Body;TSS |
| WG2844 | chr2:113992843-113992930 | PAX8;PAX8;PAX8;PAX8;PAX8 | N_Shore | Body;Body;Body;Body;Body;Body;TSS |
| WG2121 | chr2:113992843-113993052 | PAX8;PAX8;PAX8;PAX8;PAX8 | N_Shore | Body;Body;Body;Body;Body;Body;TSS |
| GM00042 | chr2:114033360-114033830 | PAX8;PAX8;PAX8;PAX8;PAX8 | Island | Body;Body;Body;Body;Body;Body |
| WG2193 | chr2:114033490-114033830 | PAX8;PAX8;PAX8;PAX8;PAX8 | Island | Body;Body;Body;Body;Body;Body |
| GM00042 | chr2:115419537-115420053 | DPP10 | N_Shore | Body |
| GM02149 | chr2:115419846-115420215 | DPP10 | Island | Body |
| GM01653 | chr2:11679845-11679879 | GREB1;GREB1 | | TSS1500;5'UTR |

| | | | | |
|---|---|---|---|---|
| WG1513 | chr2:11680057-11680144 | GREB1;GREB1 | | 5'UTR;TSS200 |
| WG1722 | chr2:119599748-119600002 | EN1 | Island | 3'UTR |
| WG2844 | chr2:119599748-119600408 | EN1 | Island | 3'UTR |
| GM00439 | chr2:119599748-119600627 | EN1 | Island | 3'UTR |
| GM02147 | chr2:119599748-119600948 | EN1 | Island | 3'UTR |
| WG2194 | chr2:119599819-119600948 | EN1 | Island | 3'UTR |
| WG2193 | chr2:119600002-119600627 | EN1 | Island | 3'UTR |
| WG1721 | chr2:119600408-119600948 | EN1 | Island | 3'UTR |
| WG2843 | chr2:119602212-119602857 | EN1 | N_Shore | Body |
| GM00439 | chr2:119602212-119603383 | EN1 | N_Shore | Body |
| GM00940 | chr2:119602212-119603489 | EN1 | N_Shore | Body |
| GM00290 | chr2:119602212-119603673 | EN1 | N_Shore | Body |
| GM00438 | chr2:119602480-119603383 | EN1 | N_Shore | Body |
| WG1722 | chr2:119602480-119603489 | EN1 | N_Shore | Body |
| GM03073 | chr2:119602546-119602857 | EN1 | N_Shore | Body |
| WG3465 | chr2:119602546-119603489 | EN1 | N_Shore | Body |
| GM03073 | chr2:119602901-119603383 | EN1 | Island | Body |
| WG2843 | chr2:119602901-119603489 | EN1 | Island | Body |
| GM02173 | chr2:119603133-119603489 | EN1 | Island | Body |
| WG1721 | chr2:119603383-119603969 | EN1 | Island | Body |
| WG2121 | chr2:119605959-119606151 | EN1 | N_Shore | TSS1500 |
| GM02704 | chr2:119605959-119607379 | EN1 | N_Shore | TSS1500 |
| GM02640 | chr2:119606055-119606302 | EN1 | Island | TSS1500 |
| GM01095 | chr2:119606055-119606501 | EN1 | Island | TSS1500 |
| GM00290 | chr2:119606631-119607379 | EN1 | Island | TSS1500 |
| GM01095 | chr2:119606695-119607097 | EN1 | Island | TSS1500 |
| WG2121 | chr2:119606695-119607603 | EN1 | Island | TSS1500 |
| WG3466 | chr2:119606746-119607379 | EN1 | Island | TSS1500 |
| GM00439 | chr2:120189867-120190202 | TMEM37 | N_Shore | Body |
| WG1513 | chr2:120190031-120190297 | TMEM37 | Island | Body |
| GM09256 | chr2:1204376-1204826 | SNTG2 | | Body |
| WG1283 | chr2:121105259-121105535 | INHBB | N_Shore | Body |
| GM02456 | chr2:121224327-121224679 | LOC84931 | | TSS1500 |
| WG1310 | chr2:121554696-121554817 | GLI2 | | TSS200 |
| WG1514 | chr2:121554696-121554926 | GLI2 | | TSS200 |
| WG3466 | chr2:121554810-121554926 | GLI2 | | TSS200 |
| WG2843 | chr2:121554817-121554926 | GLI2 | | TSS200 |
| GM02456 | chr2:121748215-121748257 | GLI2 | S_Shore | Body |
| WG1513 | chr2:1234178-1234687 | SNTG2 | | Body |
| GM01095 | chr2:127414802-127414883 | GYPC;GYPC | S_Shore | Body;Body |
| GM00520 | chr2:127963489-127963590 | CYP27C1 | | TSS200 |
| GM02706 | chr2:128421828-128422224 | LIMS2;LIMS2;LIMS2;LIMS2 | Island | Body;Body;Body;5'UTR |
| GM00290 | chr2:128438947-128439345 | LIMS2;LIMS2 | | 1stExon;5'UTR |
| WG2414 | chr2:128458240-128458556 | SFT2D3 | N_Shore | TSS1500 |
| WG2121 | chr2:1286216-1286397 | SNTG2 | N_Shore | Body |
| GM01096 | chr2:1287787-1287978 | SNTG2 | S_Shore | Body |
| GM00520 | chr2:131512663-131512773 | FAM123C;FAM123C;FAM123 | N_Shore | TSS1500;TSS1500;TSS1500;TSS1500 |

| | | | | |
|---|---|---|---|---|
| WG3465 | chr2:131673963-131674175 | ARHGEF4;ARHGEF4 | Island | TSS1500;TSS1500 |
| GM09256 | chr2:131674221-131674376 | ARHGEF4;ARHGEF4 | S_Shore | TSS200;TSS200 |
| WG1977 | chr2:132202427-132202537 | LOC401010 | Island | Body |
| GM00438 | chr2:133342838-133343308 | GPR39 | N_Shore | Body |
| GM00438 | chr2:133343618-133343657 | GPR39 | S_Shore | Body |
| GM00289 | chr2:133402475-133402940 | GPR39;LYPD1;LYPD1 | N_Shore | Body;3'UTR;3'UTR |
| WG1977 | chr2:133402475-133402999 | GPR39;LYPD1;LYPD1 | N_Shore | Body;3'UTR;3'UTR |
| GM00290 | chr2:133402827-133402999 | GPR39;LYPD1;LYPD1 | Island | Body;3'UTR;3'UTR |
| GM02555 | chr2:133402827-133403009 | GPR39;LYPD1;LYPD1 | Island | Body;3'UTR;3'UTR |
| WG1977 | chr2:133426298-133426653 | LYPD1;LYPD1 | N_Shore | 5'UTR;Body |
| GM00940 | chr2:133428422-133428510 | LYPD1;LYPD1;LYPD1 | Island | 1stExon;5'UTR;5'UTR |
| GM02149 | chr2:133428502-133428523 | LYPD1;LYPD1 | Island | TSS200;5'UTR |
| WG1513 | chr2:133428502-133428652 | LYPD1;LYPD1 | Island | TSS200;5'UTR |
| GM00935 | chr2:133428523-133428993 | LYPD1;LYPD1 | Island | TSS200;5'UTR |
| WG2844 | chr2:133428652-133428993 | LYPD1;LYPD1 | Island | TSS200;5'UTR |
| WG1978 | chr2:133429088-133429299 | LYPD1;LYPD1 | S_Shore | TSS1500;TSS200 |
| GM09256 | chr2:1358283-1358684 | SNTG2 | | Body |
| WG1722 | chr2:137747982-137748382 | THSD7B | | TSS1500 |
| WG1513 | chr2:137748366-137748460 | THSD7B | | TSS200 |
| GM01096 | chr2:137748382-137748460 | THSD7B | | TSS200 |
| WG1722 | chr2:1417164-1417431 | TPO;TPO;TPO;TPO | | TSS1500;TSS200;TSS200;TSS1500 |
| WG1513 | chr2:1425035-1426094 | TPO;TPO;TPO;TPO | | Body;Body;Body;Body |
| GM09256 | chr2:1426068-1426181 | TPO;TPO;TPO;TPO | | Body;Body;Body;Body |
| WG2276 | chr2:142889283-142889384 | LRP1B | S_Shore | TSS200 |
| WG2120 | chr2:142889283-142889392 | LRP1B | S_Shore | TSS200 |
| WG2415 | chr2:142889367-142889392 | LRP1B | S_Shore | TSS200 |
| GM00520 | chr2:145146928-145147291 | ZEB2;ZEB2 | N_Shore | 3'UTR;3'UTR |
| GM00935 | chr2:145146928-145147558 | ZEB2;ZEB2 | N_Shore | 3'UTR;3'UTR |
| GM09256 | chr2:1487895-1488175 | TPO;TPO;TPO;TPO | | Body;Body;Body;Body |
| WG1513 | chr2:1488252-1488516 | TPO;TPO;TPO;TPO | | Body;Body;Body;Body |
| WG1309 | chr2:150444569-150444834 | MMADHC | | TSS1500 |
| GM00520 | chr2:152830189-152830521 | CACNB4;CACNB4;CACNB4;CACNB4 | | Body;Body;1stExon;Body |
| GM02175 | chr2:152830505-152830572 | CACNB4;CACNB4;CACNB4;CACNB4 | | Body;Body;Body;TSS200 |
| GM00520 | chr2:157291462-157291826 | GPD2;GPD2 | N_Shore | TSS1500;TSS1500 |
| GM02706 | chr2:159651813-159652019 | DAPL1 | | TSS200 |
| GM00439 | chr2:160761427-160761622 | LY75 | Island | TSS200 |
| WG2414 | chr2:161265247-161265366 | RBMS1;RBMS1 | S_Shore | Body;Body |
| GM02706 | chr2:162271801-162272107 | TBR1 | N_Shore | TSS1500 |
| WG1722 | chr2:1647185-1647260 | PXDN | | Body |
| GM01653 | chr2:165811816-165812146 | SLC38A11;SLC38A11 | | 1stExon;5'UTR |
| WG1722 | chr2:165812079-165812159 | SLC38A11 | | TSS200 |
| GM00438 | chr2:165812146-165812297 | SLC38A11 | | TSS200 |
| WG3466 | chr2:169658981-169659047 | NOSTRIN;NOSTRIN;NOSTRIN;NOSTRIN | | 5'UTR;TSS200;TSS200;TSS200 |
| WG1657 | chr2:171573839-171574141 | SP5 | Island | Body |
| GM00016 | chr2:171573839-171574592 | SP5 | Island | Body |
| WG2415 | chr2:171676553-171676925 | GAD1;GAD1 | Island | Body;Body |
| WG3465 | chr2:171678751-171679402 | GAD1;GAD1 | Island | Body;Body |

| | | | | |
|---|---|---|---|---|
| WG1722 | chr2:171678751-171679591 | GAD1;GAD1 | Island | Body;Body |
| GM00935 | chr2:171679906-171680337 | GAD1;GAD1 | Island | Body;Body |
| WG1977 | chr2:172952883-172953134 | DLX1;DLX1 | Island | Body;3'UTR |
| GM01650 | chr2:172952883-172953270 | DLX1;DLX1 | Island | Body;3'UTR |
| GM00042 | chr2:172953032-172953270 | DLX1;DLX1 | Island | 3'UTR;3'UTR |
| WG3466 | chr2:172965650-172965890 | DLX2 | Island | Body |
| GM00042 | chr2:173330296-173330395 | ITGA6;ITGA6 | | Body;Body |
| WG2414 | chr2:173601103-173601360 | RAPGEF4;LOC91149 | S_Shore | Body;TSS200 |
| WG2120 | chr2:173601133-173601413 | RAPGEF4;LOC91149 | S_Shore | Body;TSS200 |
| WG2120 | chr2:1746509-1746647 | PXDN | N_Shore | Body |
| WG1978 | chr2:175198952-175199349 | SP9 | N_Shore | TSS1500 |
| WG1722 | chr2:175199259-175199349 | SP9 | N_Shore | TSS1500 |
| GM00289 | chr2:175499283-175499377 | WIPF1;WIPF1;WIPF1 | | 1stExon;5'UTR;5'UTR |
| WG2844 | chr2:175629219-175629917 | CHRNA1;CHRNA1 | | TSS200;TSS200 |
| GM09256 | chr2:176948596-176948759 | EVX2;EVX2 | N_Shore | 5'UTR;1stExon |
| WG1513 | chr2:176948596-176949017 | EVX2;EVX2 | N_Shore | 5'UTR;1stExon |
| WG1978 | chr2:176948693-176948732 | EVX2 | N_Shore | TSS200 |
| WG2415 | chr2:176948693-176948759 | EVX2 | N_Shore | TSS200 |
| GM00520 | chr2:176948699-176948732 | EVX2 | N_Shore | TSS200 |
| WG1984 | chr2:176948699-176948759 | EVX2 | N_Shore | TSS200 |
| GM02149 | chr2:176948699-176948864 | EVX2 | N_Shore | TSS200 |
| GM00940 | chr2:176948699-176949017 | EVX2 | N_Shore | TSS200 |
| GM02175 | chr2:176948728-176948759 | EVX2 | N_Shore | TSS200 |
| WG3465 | chr2:176948728-176948864 | EVX2 | N_Shore | TSS200 |
| GM03075 | chr2:176956396-176956678 | HOXD13 | N_Shore | TSS1500 |
| WG3466 | chr2:176956396-176956841 | HOXD13 | N_Shore | TSS1500 |
| GM01096 | chr2:176956573-176956761 | HOXD13 | Island | TSS1500 |
| GM09255 | chr2:176956573-176956841 | HOXD13 | Island | TSS1500 |
| WG1513 | chr2:176963315-176963583 | HOXD12 | N_Shore | TSS1500 |
| WG2121 | chr2:176963583-176964502 | HOXD12 | N_Shore | TSS1500 |
| GM01095 | chr2:176963583-176964720 | HOXD12 | N_Shore | TSS1500 |
| WG2276 | chr2:176964101-176964404 | HOXD12 | Island | TSS1500 |
| WG1977 | chr2:176964101-176964456 | HOXD12 | Island | TSS1500 |
| GM02704 | chr2:176964101-176964720 | HOXD12 | Island | TSS1500 |
| WG1978 | chr2:176964137-176964502 | HOXD12 | Island | TSS1500 |
| WG2120 | chr2:176964137-176964720 | HOXD12 | Island | TSS1500 |
| GM02706 | chr2:176964506-176964720 | HOXD12 | Island | TSS200 |
| WG3465 | chr2:176964512-176964720 | HOXD12 | Island | TSS200 |
| GM00290 | chr2:176973275-176973835 | HOXD11 | N_Shore | Body |
| WG2275 | chr2:176973275-176974060 | HOXD11 | N_Shore | Body |
| WG2414 | chr2:176973688-176973835 | HOXD11 | Island | Body |
| WG2415 | chr2:176973835-176974060 | HOXD11 | S_Shore | Body |
| WG1977 | chr2:176980837-176981064 | HOXD10 | Island | TSS1500 |
| GM00439 | chr2:176980837-176981328 | HOXD10 | Island | TSS1500 |
| GM00042 | chr2:176980837-176981469 | HOXD10 | Island | TSS1500 |
| WG2120 | chr2:176981064-176981469 | HOXD10 | Island | TSS1500 |
| WG2275 | chr2:176981064-176981654 | HOXD10 | Island | TSS1500 |

| | | | | |
|---|---|---|---|---|
| GM09256 | chr2:176981328-176981469 | HOXD10 | Island | TSS200 |
| WG3466 | chr2:176981328-176981654 | HOXD10 | Island | TSS200 |
| GM02316 | chr2:176981422-176981654 | HOXD10 | Island | TSS200 |
| GM00439 | chr2:176981919-176982342 | HOXD10 | N_Shore | 1stExon |
| WG2415 | chr2:176982108-176982342 | HOXD10 | Island | 1stExon |
| GM02640 | chr2:176983519-176983949 | HOXD10 | S_Shore | Body |
| WG1514 | chr2:176985953-176986659 | HOXD9 | N_Shore | TSS1500 |
| GM00439 | chr2:176986460-176986659 | HOXD9 | Island | TSS1500 |
| WG2844 | chr2:176988284-176988939 | HOXD9 | Island | 1stExon |
| WG1309 | chr2:176988505-176988792 | HOXD9 | N_Shore | Body |
| WG1978 | chr2:176993017-176993115 | HOXD8 | Island | TSS1500 |
| WG3466 | chr2:176993017-176993221 | HOXD8 | Island | TSS1500 |
| GM00935 | chr2:176993017-176993320 | HOXD8 | Island | TSS1500 |
| WG2120 | chr2:177016167-177016490 | HOXD4;HOXD4 | N_Shore | 5'UTR;1stExon |
| WG1513 | chr2:177016167-177016619 | HOXD4;HOXD4 | N_Shore | 5'UTR;1stExon |
| WG3466 | chr2:177016167-177017352 | HOXD4;HOXD4 | N_Shore | 5'UTR;1stExon |
| WG2414 | chr2:177016167-177017389 | HOXD4;HOXD4 | N_Shore | 5'UTR;1stExon |
| WG1977 | chr2:177016167-177017449 | HOXD4;HOXD4 | N_Shore | 5'UTR;1stExon |
| GM02706 | chr2:177016167-177017630 | HOXD4;HOXD4 | N_Shore | 5'UTR;1stExon |
| GM00439 | chr2:177016837-177017449 | HOXD4 | N_Shore | Body |
| GM00940 | chr2:179387853-179388064 | MIR548N | | Body |
| GM02555 | chr2:179672121-179672691 | TTN;TTN;TTN;TTN;TTN;TTN;TTN;TTN;TTN;T | | 1stExon;5'UTR;5'UTR;5'UTR;1stExon;1 |
| GM00289 | chr2:18059103-18059130 | KCNS3 | Island | TSS1500 |
| WG1658 | chr2:1817263-1817409 | MYT1L | | Body |
| WG2276 | chr2:1820169-1820433 | MYT1L | | Body |
| GM00520 | chr2:183943181-183943938 | DUSP19;DUSP19 | | TSS200;TSS200 |
| GM00935 | chr2:18766022-18766294 | NT5C1B;NT5C1B | Island | Body;Body |
| GM03075 | chr2:191398360-191398769 | TMEM194B | N_Shore | Body |
| WG1513 | chr2:1926366-1926671 | MYT1L | | Body |
| GM09256 | chr2:1926888-1927209 | MYT1L | | Body |
| WG1513 | chr2:1941133-1941523 | MYT1L | Island | Body |
| WG1513 | chr2:1942814-1942942 | MYT1L | S_Shore | Body |
| WG1983 | chr2:19551750-19551860 | OSR1 | Island | 3'UTR |
| WG3466 | chr2:19553237-19553422 | OSR1 | Island | Body |
| GM01096 | chr2:19553237-19553546 | OSR1 | Island | Body |
| GM02706 | chr2:19553237-19553950 | OSR1 | Island | Body |
| WG1722 | chr2:19553237-19554237 | OSR1 | Island | Body |
| GM00520 | chr2:197503878-197504046 | CCDC150 | N_Shore | TSS1500 |
| GM00022 | chr2:198029141-198029332 | ANKRD44 | Island | Body |
| WG3465 | chr2:198650752-198650987 | BOLL;BOLL;BOLL | Island | 1stExon;5'UTR;TSS1500 |
| GM02175 | chr2:200320455-200320871 | SATB2 | N_Shore | Body |
| GM09256 | chr2:200320455-200321016 | SATB2 | N_Shore | Body |
| WG2414 | chr2:200320455-200321348 | SATB2 | N_Shore | Body |
| WG3465 | chr2:200320750-200321348 | SATB2 | Island | Body |
| GM09255 | chr2:200321016-200321348 | SATB2 | Island | 5'UTR |
| GM00022 | chr2:200326721-200327336 | SATB2 | Island | TSS1500 |
| WG1657 | chr2:200331675-200332380 | FLJ32063 | S_Shore | TSS1500 |

| | | | | |
|---|---|---|---|---|
| GM00022 | chr2:200331977-200332380 | FLJ32063 | N_Shore | TSS1500 |
| WG1657 | chr2:200333729-200334439 | FLJ32063 | Island | Body |
| GM00935 | chr2:200333729-200334558 | FLJ32063 | Island | Body |
| GM02640 | chr2:200333729-200334584 | FLJ32063 | Island | Body |
| GM00289 | chr2:200333729-200335683 | FLJ32063 | Island | Body |
| GM01376 | chr2:200334001-200334402 | FLJ32063 | Island | Body |
| WG2276 | chr2:200334137-200334558 | FLJ32063 | Island | Body |
| WG3465 | chr2:200334402-200334584 | FLJ32063 | N_Shore | Body |
| GM02555 | chr2:200334402-200334805 | FLJ32063 | N_Shore | Body |
| GM00042 | chr2:200334439-200334584 | FLJ32063 | N_Shore | Body |
| WG2276 | chr2:200334805-200335874 | FLJ32063 | N_Shore | Body |
| WG3465 | chr2:200334854-200335101 | FLJ32063 | N_Shore | Body |
| WG1984 | chr2:200334854-200335558 | FLJ32063 | N_Shore | Body |
| GM02640 | chr2:200334854-200335683 | FLJ32063 | N_Shore | Body |
| GM00520 | chr2:200335148-200335683 | FLJ32063 | N_Shore | Body |
| GM02149 | chr2:202483868-202484156 | ALS2CR11;ALS2CR11;ALS2C | Island | 1stExon;5'UTR;5'UTR;1stExon;1stExon |
| WG2276 | chr2:202508694-202508813 | ALS2CR4;ALS2CR4 | S_Shore | TSS1500;TSS1500 |
| WG2121 | chr2:202671158-202671583 | CDK15 | | TSS200 |
| GM02706 | chr2:202901288-202901428 | FZD7;FZD7 | S_Shore | 1stExon;3'UTR |
| GM00520 | chr2:202901288-202901470 | FZD7;FZD7 | S_Shore | 1stExon;3'UTR |
| WG2276 | chr2:202901352-202901470 | FZD7;FZD7 | S_Shore | 1stExon;3'UTR |
| GM02706 | chr2:206628553-206628692 | NRP2;NRP2;NRP2;NRP2;NRP2 | | Body;Body;Body;Body;Body |
| GM02555 | chr2:208635549-208635637 | FZD5 | S_Shore | TSS1500 |
| GM00290 | chr2:20865651-20865847 | GDF7 | Island | TSS1500 |
| GM00940 | chr2:20870087-20870362 | GDF7 | Island | Body |
| WG1722 | chr2:20870087-20870812 | GDF7 | Island | Body |
| WG1977 | chr2:20870087-20871195 | GDF7 | Island | Body |
| GM00520 | chr2:20870362-20871401 | GDF7 | Island | Body |
| GM02175 | chr2:20871173-20871401 | GDF7 | Island | Body |
| GM00438 | chr2:20871195-20871401 | GDF7 | Island | 3'UTR |
| GM00438 | chr2:209010986-209011053 | CRYGB | | TSS200 |
| WG3466 | chr2:209054670-209055151 | C2orf80 | | 5'UTR |
| GM02555 | chr2:211457632-211457684 | CPS1;CPS1;CPS1 | | Body;Body;TSS1500 |
| GM09256 | chr2:211457661-211458004 | CPS1;CPS1;CPS1 | | Body;Body;TSS1500 |
| WG1513 | chr2:211457684-211458173 | CPS1;CPS1;CPS1 | | Body;Body;TSS1500 |
| GM02149 | chr2:21229220-21229872 | APOB | | Body |
| WG1513 | chr2:21229220-21229936 | APOB | | Body |
| WG1977 | chr2:21266947-21267113 | APOB | Island | TSS200 |
| WG1978 | chr2:215675105-215675637 | BARD1 | S_Shore | TSS1500 |
| WG2415 | chr2:215675429-215675637 | BARD1 | S_Shore | TSS1500 |
| WG1513 | chr2:2193511-2193665 | MYT1L | Island | 5'UTR |
| GM01393 | chr2:219735876-219736312 | WNT6 | N_Shore | Body |
| GM02555 | chr2:219735876-219736549 | WNT6 | N_Shore | Body |
| GM02147 | chr2:219736167-219736312 | WNT6 | Island | Body |
| WG1513 | chr2:220253653-220253668 | DNPEP | S_Shore | TSS1500 |
| WG1977 | chr2:220253653-220253963 | DNPEP | S_Shore | TSS1500 |
| WG2121 | chr2:220349208-220349733 | SPEG | Island | Body |

| | | | | |
|---|---|---|---|---|
| WG2844 | chr2:220406470-220406756 | CHPF | Island | Body |
| GM00940 | chr2:223154140-223154201 | PAX3;PAX3;PAX3;PAX3;PAX3 | N_Shore | Body;Body;Body;Body;Body;Body |
| WG3465 | chr2:223155925-223156136 | PAX3;PAX3;PAX3;PAX3;PAX3 | Island | Body;Body;Body;Body;Body;Body |
| GM02175 | chr2:223160175-223160486 | PAX3;PAX3;PAX3;PAX3;PAX3 | Island | Body;Body;Body;Body;Body;Body;Bod |
| WG2276 | chr2:223160175-223160569 | PAX3;PAX3;PAX3;PAX3;PAX3 | Island | Body;Body;Body;Body;Body;Body;Bod |
| WG2415 | chr2:223161771-223162049 | PAX3;PAX3;PAX3;CCDC140; | Island | Body;Body;Body;TSS1500;Body;Body; |
| WG2120 | chr2:223161771-223162128 | PAX3;PAX3;PAX3;CCDC140; | Island | Body;Body;Body;TSS1500;Body;Body; |
| WG3466 | chr2:223161771-223162666 | PAX3;PAX3;PAX3;CCDC140; | Island | Body;Body;Body;TSS1500;Body;Body; |
| GM02704 | chr2:223162128-223162800 | PAX3;PAX3;PAX3;CCDC140; | N_Shore | Body;Body;Body;TSS1500;Body;Body; |
| WG2121 | chr2:223162666-223162800 | PAX3;PAX3;PAX3;PAX3;PAX3 | N_Shore | Body;Body;Body;Body;Body;Body;Bod |
| WG1978 | chr2:223162797-223163175 | PAX3;PAX3;PAX3;PAX3;PAX3 | N_Shore | Body;Body;Body;Body;Body;Body;Bod |
| GM01096 | chr2:223162881-223163175 | PAX3;PAX3;PAX3;CCDC140; | N_Shore | Body;Body;Body;5'UTR;1stExon;Body; |
| GM09256 | chr2:223162881-223163573 | PAX3;PAX3;PAX3;CCDC140; | N_Shore | Body;Body;Body;5'UTR;1stExon;Body; |
| GM02175 | chr2:223164459-223164747 | PAX3;PAX3;PAX3;PAX3;PAX3 | S_Shore | TSS1500;TSS1500;TSS1500;TSS1500; |
| GM02704 | chr2:223164459-223166989 | PAX3;PAX3;PAX3;PAX3;PAX3 | S_Shore | TSS1500;TSS1500;TSS1500;TSS1500; |
| WG1977 | chr2:223164459-223168487 | PAX3;PAX3;PAX3;PAX3;PAX3 | S_Shore | TSS1500;TSS1500;TSS1500;TSS1500; |
| WG2121 | chr2:223164459-223170370 | PAX3;PAX3;PAX3;PAX3;PAX3 | S_Shore | TSS1500;TSS1500;TSS1500;TSS1500; |
| WG2120 | chr2:223164635-223167617 | PAX3;PAX3;PAX3;PAX3;PAX3 | S_Shore | TSS1500;TSS1500;TSS1500;TSS1500; |
| GM09256 | chr2:223164747-223164867 | PAX3;PAX3;PAX3;PAX3;PAX3 | S_Shore | TSS1500;TSS1500;TSS1500;TSS1500; |
| WG2275 | chr2:223164747-223164869 | PAX3;PAX3;PAX3;PAX3;PAX3 | S_Shore | TSS1500;TSS1500;TSS1500;TSS1500; |
| GM00016 | chr2:223164831-223164867 | PAX3;PAX3;PAX3;PAX3;PAX3 | S_Shore | TSS1500;TSS1500;TSS1500;TSS1500; |
| WG1978 | chr2:223164831-223164869 | PAX3;PAX3;PAX3;PAX3;PAX3 | S_Shore | TSS1500;TSS1500;TSS1500;TSS1500; |
| WG3465 | chr2:223164854-223164869 | PAX3;PAX3;PAX3;PAX3;PAX3 | S_Shore | TSS1500;TSS1500;TSS1500;TSS1500; |
| WG3466 | chr2:223164854-223164925 | PAX3;PAX3;PAX3;PAX3;PAX3 | S_Shore | TSS1500;TSS1500;TSS1500;TSS1500; |
| WG2275 | chr2:223165753-223166094 | CCDC140 | N_Shore | 5'UTR |
| GM02555 | chr2:223165753-223166989 | CCDC140 | N_Shore | 5'UTR |
| WG3466 | chr2:223165753-223167170 | CCDC140 | N_Shore | 5'UTR |
| WG3465 | chr2:223166094-223167170 | CCDC140 | N_Shore | 5'UTR |
| GM03075 | chr2:223166909-223166989 | CCDC140 | N_Shore | 5'UTR |
| GM01095 | chr2:223167280-223167617 | CCDC140 | Island | 5'UTR |
| GM02704 | chr2:223167280-223170370 | CCDC140 | Island | 5'UTR |
| WG2120 | chr2:223168453-223170370 | CCDC140 | N_Shore | 5'UTR |
| WG1513 | chr2:223168487-223168727 | CCDC140 | N_Shore | 5'UTR |
| GM00016 | chr2:223168575-223168861 | CCDC140 | N_Shore | 5'UTR |
| WG3466 | chr2:223169586-223169965 | CCDC140 | N_Shore | 3'UTR |
| WG3466 | chr2:223290168-223290515 | SGPP2 | S_Shore | Body |
| WG3465 | chr2:223917577-223918000 | KCNE4 | Island | Body |
| GM02641 | chr2:224904683-224904837 | SERPINE2;SERPINE2;SERPIN | Island | TSS1500;TSS1500;TSS1500 |
| GM00016 | chr2:227661213-227661656 | IRS1 | N_Shore | 1stExon |
| GM00016 | chr2:227662704-227663151 | IRS1 | Island | 1stExon |
| WG3465 | chr2:233274144-233274723 | ALPPL2 | Island | Body |
| GM00289 | chr2:233323381-233323718 | ALPI | Island | Body |
| GM02173 | chr2:233323532-233323935 | ALPI | Island | Body |
| WG1978 | chr2:233765872-233766118 | NGEF;NGEF | Island | Body;Body |
| GM02706 | chr2:233925030-233925181 | INPP5D;INPP5D | N_Shore | TSS200;TSS200 |
| WG1513 | chr2:234599988-234600149 | UGT1A6;UGT1A10;UGT1A9;UGT1A7;UGT | | TSS1500;Body;Body;Body;Body |
| GM02149 | chr2:234637253-234637574 | UGT1A10;UGT1A6;UGT1A8;UGT1A4;UGT | | Body;Body;Body;Body;Body;Body;Bod |

| | | | | |
|---|---|---|---|---|
| GM02175 | chr2:234847554-234847856 | TRPM8 | N_Shore | Body |
| WG1983 | chr2:235883549-235883680 | SH3BP4 | | 5'UTR |
| WG3466 | chr2:235935754-235936596 | SH3BP4 | | 5'UTR |
| GM01095 | chr2:235936137-235936596 | SH3BP4 | | 5'UTR |
| WG2194 | chr2:235936394-235936596 | SH3BP4 | | 5'UTR |
| WG3465 | chr2:236586191-236586744 | AGAP1;AGAP1 | | Body;Body |
| WG2276 | chr2:236643527-236643760 | AGAP1;AGAP1 | | Body;Body |
| GM02555 | chr2:236674174-236674489 | AGAP1;AGAP1 | S_Shore | Body;Body |
| GM00439 | chr2:236715747-236716052 | AGAP1;AGAP1 | | Body;Body |
| GM00042 | chr2:236773924-236774164 | AGAP1;AGAP1 | Island | Body;Body |
| GM09255 | chr2:236826245-236826420 | AGAP1;AGAP1 | | Body;Body |
| GM02175 | chr2:236867278-236867905 | AGAP1;AGAP1 | N_Shore | Body;Body |
| WG3466 | chr2:237122857-237123218 | ASB18 | N_Shore | Body |
| WG2415 | chr2:23746926-23747197 | KLHL29 | N_Shelf | 5'UTR |
| GM02706 | chr2:238394933-238395125 | MLPH;MLPH | N_Shore | TSS1500;TSS1500 |
| WG2276 | chr2:23840268-23840555 | KLHL29 | | Body |
| WG1977 | chr2:238446240-238446774 | MLPH;MLPH | | Body;Body |
| GM02706 | chr2:238499782-238499802 | RAB17 | | TSS200 |
| WG1722 | chr2:238499782-238500081 | RAB17 | | TSS200 |
| WG2276 | chr2:238599761-238600061 | LRRFIP1;LRRFIP1;LRRFIP1;LF | N_Shore | TSS1500;TSS1500;TSS1500;TSS1500; |
| WG2415 | chr2:238647755-238647913 | LRRFIP1;LRRFIP1;LRRFIP1;LF | S_Shelf | Body;Body;Body;Body;Body |
| GM00941 | chr2:239755455-239755703 | TWIST2 | Island | TSS1500 |
| GM01095 | chr2:239799511-239799825 | TWIST2 | | 3'UTR |
| WG2415 | chr2:239799720-239799904 | TWIST2 | | 3'UTR |
| GM02456 | chr2:239799720-239799954 | TWIST2 | | 3'UTR |
| WG1722 | chr2:240014758-240014781 | HDAC4 | | Body |
| GM02175 | chr2:240019466-240020174 | HDAC4 | | Body |
| WG3466 | chr2:240020028-240020174 | HDAC4 | | Body |
| WG2844 | chr2:240040089-240040290 | HDAC4 | | Body |
| GM00520 | chr2:240152257-240152476 | HDAC4 | N_Shore | Body |
| GM02456 | chr2:240161619-240162323 | HDAC4 | | Body |
| GM00520 | chr2:240161794-240162323 | HDAC4 | | Body |
| WG3465 | chr2:240196078-240196284 | HDAC4 | N_Shore | Body |
| GM00520 | chr2:240212763-240213173 | HDAC4 | | Body |
| WG2844 | chr2:240233043-240233240 | HDAC4 | | Body |
| WG2276 | chr2:240291068-240291426 | HDAC4 | | 5'UTR |
| WG3466 | chr2:240291426-240291657 | HDAC4 | | 5'UTR |
| GM00042 | chr2:240313541-240313668 | HDAC4 | | 5'UTR |
| GM01096 | chr2:241076189-241076208 | MYEOV2;MYEOV2 | S_Shore | TSS1500;TSS1500 |
| WG2276 | chr2:241076340-241076416 | MYEOV2;MYEOV2 | S_Shore | TSS1500;TSS1500 |
| WG1513 | chr2:241076526-241076551 | MYEOV2;MYEOV2 | S_Shore | TSS1500;TSS1500 |
| GM09256 | chr2:241076601-241076777 | MYEOV2;MYEOV2 | S_Shore | TSS1500;TSS1500 |
| GM00016 | chr2:241395288-241395678 | MIR149;PP14571;GPC1 | Island | TSS200;Body;Body |
| WG1977 | chr2:241719496-241719655 | KIF1A | Island | Body |
| GM01392 | chr2:242212884-242212901 | HDLBP;HDLBP | S_Shore | 5'UTR;TSS1500 |
| WG1722 | chr2:242448543-242448802 | STK25 | Island | TSS1500 |
| WG2844 | chr2:242715019-242715041 | GAL3ST2 | S_Shelf | TSS1500 |

| | | | | |
|---|---|---|---|---|
| WG1722 | chr2:24271827-24272044 | FKBP1B;FKBP1B | N_Shore | TSS1500;TSS1500 |
| WG1977 | chr2:242751796-242751949 | NEU4;NEU4;NEU4;NEU4;NEU | N_Shelf | TSS1500;5'UTR;5'UTR;TSS1500;TSS15 |
| GM00520 | chr2:242810748-242810927 | C2orf85 | S_Shelf | TSS1500 |
| GM01392 | chr2:24397810-24398170 | C2orf84 | Island | TSS200 |
| WG2414 | chr2:24397810-24398427 | C2orf84 | Island | TSS200 |
| GM03075 | chr2:24398170-24398519 | C2orf84 | Island | 5'UTR |
| WG3466 | chr2:24398427-24398859 | C2orf84 | Island | Body |
| WG2276 | chr2:24398668-24398859 | C2orf84 | Island | Body |
| WG2276 | chr2:25141532-25141735 | ADCY3 | N_Shore | 1stExon |
| GM02555 | chr2:25141532-25142016 | ADCY3 | N_Shore | 1stExon |
| WG1310 | chr2:25141532-25142117 | ADCY3 | N_Shore | 1stExon |
| GM00016 | chr2:25354610-25354986 | EFR3B | Island | Body |
| WG3465 | chr2:25383851-25384293 | POMC;POMC | N_Shore | 3'UTR;3'UTR |
| GM01393 | chr2:25390385-25390424 | POMC;POMC | N_Shore | 5'UTR;5'UTR |
| WG2414 | chr2:25391656-25391670 | POMC;POMC | Island | TSS200;TSS200 |
| GM00042 | chr2:25391773-25391839 | POMC;POMC | Island | TSS1500;TSS1500 |
| WG1977 | chr2:25391830-25391911 | POMC;POMC | Island | TSS1500;TSS1500 |
| WG2844 | chr2:25473781-25474327 | DNMT3A;DNMT3A;DNMT3A | N_Shore | Body;Body;Body |
| GM00290 | chr2:25499764-25500125 | DNMT3A;DNMT3A | Island | Body;Body |
| GM02706 | chr2:25500046-25500416 | DNMT3A;DNMT3A | Island | Body;Body |
| GM00520 | chr2:26045189-26045287 | ASXL2 | | Body |
| WG1282 | chr2:26401640-26401820 | FAM59B | N_Shore | Body |
| WG1722 | chr2:26541960-26542188 | GPR113;GPR113;GPR113;GPR113 | | 5'UTR;1stExon;TSS200;Body |
| WG2121 | chr2:26542043-26542188 | GPR113;GPR113;GPR113 | | TSS200;TSS200;Body |
| WG1977 | chr2:26947126-26947234 | KCNK3 | N_Shelf | Body |
| WG3465 | chr2:27371818-27371840 | TCF23 | N_Shore | TSS200 |
| GM00520 | chr2:27530884-27531163 | UCN | Island | 5'UTR |
| WG3466 | chr2:27665128-27665150 | KRTCAP3;KRTCAP3 | Island | TSS200;TSS200 |
| WG2121 | chr2:27665139-27665150 | KRTCAP3;KRTCAP3 | Island | TSS200;TSS200 |
| GM01392 | chr2:27665139-27665306 | KRTCAP3;KRTCAP3 | Island | TSS200;TSS200 |
| GM09255 | chr2:27665139-27665507 | KRTCAP3;KRTCAP3 | Island | TSS200;TSS200 |
| GM00940 | chr2:27665141-27665306 | KRTCAP3;KRTCAP3 | Island | TSS200;TSS200 |
| WG3465 | chr2:27665141-27665507 | KRTCAP3;KRTCAP3 | Island | TSS200;TSS200 |
| WG1722 | chr2:27665150-27665507 | KRTCAP3;KRTCAP3 | Island | TSS200;TSS200 |
| GM09256 | chr2:27665306-27665543 | KRTCAP3;KRTCAP3 | Island | 1stExon;1stExon |
| GM00438 | chr2:27798900-27799411 | C2orf16 | | TSS1500 |
| GM09255 | chr2:286830-287395 | FAM150B | N_Shore | Body |
| WG2121 | chr2:287148-287395 | FAM150B | N_Shore | Body |
| WG3465 | chr2:29204057-29204142 | FAM179A | | TSS200 |
| GM00520 | chr2:29204057-29204200 | FAM179A | | TSS200 |
| WG3465 | chr2:30669597-30669863 | LCLAT1;LCLAT1 | N_Shore | TSS1500;TSS1500 |
| WG1282 | chr2:30669597-30669940 | LCLAT1;LCLAT1 | N_Shore | TSS1500;TSS1500 |
| WG1977 | chr2:30669597-30669952 | LCLAT1;LCLAT1 | N_Shore | TSS1500;TSS1500 |
| WG1513 | chr2:31806183-31806551 | SRD5A2 | Island | TSS200 |
| WG1978 | chr2:31806234-31806352 | SRD5A2 | Island | TSS200 |
| WG3466 | chr2:3286338-3286479 | TSSC1 | Island | Body |
| WG1977 | chr2:3317860-3318039 | TSSC1 | Island | Body |

| | | | | |
|---|---|---|---|---|
| WG2843 | chr2:3381687-3381734 | TSSC1 | Island | TSS200 |
| WG1977 | chr2:3451506-3452563 | TTC15 | N_Shore | Body |
| GM01096 | chr2:3451506-3452624 | TTC15 | N_Shore | Body |
| GM02706 | chr2:3452347-3452437 | TTC15 | Island | Body |
| WG2414 | chr2:3452347-3452563 | TTC15 | Island | Body |
| WG2415 | chr2:3452347-3452624 | TTC15 | Island | Body |
| WG1283 | chr2:3452543-3452624 | TTC15 | Island | Body |
| GM00042 | chr2:3642211-3642447 | COLEC11;COLEC11 | N_Shore | TSS1500;TSS1500 |
| WG2415 | chr2:3642211-3642634 | COLEC11;COLEC11 | N_Shore | TSS1500;TSS1500 |
| WG3466 | chr2:3642400-3642634 | COLEC11;COLEC11 | N_Shore | TSS1500;TSS1500 |
| WG1513 | chr2:3642400-3642847 | COLEC11;COLEC11 | N_Shore | TSS1500;TSS1500 |
| WG2275 | chr2:3642447-3642634 | COLEC11;COLEC11 | N_Shore | TSS200;TSS200 |
| GM00438 | chr2:3642629-3642710 | COLEC11;COLEC11 | Island | TSS200;TSS200 |
| GM09255 | chr2:3642629-3642732 | COLEC11;COLEC11 | Island | TSS200;TSS200 |
| GM00520 | chr2:3642732-3642867 | COLEC11;COLEC11;COLEC1 | Island | 1stExon;5'UTR;1stExon;5'UTR |
| GM02149 | chr2:3648528-3648554 | COLEC11;COLEC11 | | 5'UTR;5'UTR |
| GM00042 | chr2:3654242-3654582 | COLEC11;COLEC11 | | Body;Body |
| WG3466 | chr2:3683030-3683186 | COLEC11;COLEC11 | Island | Body;Body |
| GM02640 | chr2:3683030-3683757 | COLEC11;COLEC11 | Island | Body;Body |
| GM09256 | chr2:3683119-3683757 | COLEC11;COLEC11 | Island | Body;Body |
| WG3465 | chr2:3714278-3715083 | ALLC | | 5'UTR |
| WG1977 | chr2:3714379-3715083 | ALLC | | 5'UTR |
| WG3466 | chr2:3714936-3715083 | ALLC | | 5'UTR |
| GM02175 | chr2:37545562-37545668 | PRKD3 | | TSS1500 |
| GM02706 | chr2:37571677-37571901 | QPCT | Island | TSS200 |
| WG2121 | chr2:37571696-37571732 | QPCT | Island | TSS200 |
| WG1978 | chr2:37571696-37571901 | QPCT | Island | TSS200 |
| WG1977 | chr2:38300537-38301438 | CYP1B1 | N_Shore | Body |
| GM02706 | chr2:38302487-38302892 | CYP1B1 | Island | Body |
| WG1983 | chr2:39005158-39005834 | GEMIN6 | | TSS200 |
| WG1722 | chr2:42719785-42720326 | KCNG3;KCNG3 | Island | Body;Body |
| WG1722 | chr2:43903227-43903526 | PLEKHH2;LOC728819 | | Body;1stExon |
| GM00520 | chr2:43903227-43903900 | PLEKHH2;LOC728819 | | Body;1stExon |
| GM09255 | chr2:43903470-43903526 | PLEKHH2;LOC728819 | | Body;TSS200 |
| GM01376 | chr2:46526785-46527098 | EPAS1 | Island | Body |
| GM09255 | chr2:47382427-47382462 | C2orf61;C2orf61;C2orf61;C2orf61 | | 1stExon;1stExon;5'UTR;5'UTR |
| WG1513 | chr2:47452-47716 | FAM110C | S_Shore | TSS1500 |
| GM02706 | chr2:47596912-47597188 | EPCAM | Island | Body |
| GM00290 | chr2:47597118-47597331 | EPCAM | Island | Body |
| GM00940 | chr2:48844892-48844971 | GTF2A1L;STON1-GTF2A1L;G | Island | TSS200;Body;TSS200 |
| GM01376 | chr2:48844916-48844984 | GTF2A1L;STON1-GTF2A1L;G | Island | TSS200;Body;TSS200 |
| GM09255 | chr2:48982754-48982950 | LHCGR | Island | 1stExon |
| GM02175 | chr2:50201382-50201511 | NRXN1;NRXN1;NRXN1 | | Body;Body;Body |
| GM00290 | chr2:50201413-50201505 | NRXN1;NRXN1;NRXN1 | | Body;Body;Body |
| WG3465 | chr2:54086854-54087189 | GPR75;LOC100302652 | Island | 5'UTR;Body |
| WG1722 | chr2:54087028-54087189 | GPR75;LOC100302652;GPR | Island | 5'UTR;1stExon;1stExon |
| GM02706 | chr2:54087028-54087343 | GPR75;LOC100302652;GPR | Island | 5'UTR;1stExon;1stExon |

| | | | | |
|---|---|---|---|---|
| WG2121 | chr2:54087066-54087209 | GPR75;LOC100302652;LOC1 | Island | 5'UTR;1stExon;5'UTR;1stExon |
| WG2414 | chr2:54087173-54087189 | LOC100302652;GPR75 | Island | TSS200;TSS200 |
| WG2415 | chr2:54087178-54087189 | LOC100302652;GPR75 | Island | TSS200;TSS200 |
| WG3465 | chr2:54087209-54087552 | LOC100302652;GPR75 | Island | TSS200;TSS200 |
| GM00940 | chr2:54785178-54785307 | SPTBN1;SPTBN1 | Island | TSS1500;Body |
| GM02706 | chr2:54785178-54785795 | SPTBN1;SPTBN1 | Island | TSS1500;Body |
| GM02173 | chr2:54785406-54785476 | SPTBN1;SPTBN1 | Island | TSS200;Body |
| GM00438 | chr2:54785473-54785795 | SPTBN1;SPTBN1 | Island | TSS200;Body |
| WG1283 | chr2:54785476-54785795 | SPTBN1;SPTBN1 | Island | TSS200;Body |
| GM02317 | chr2:55450577-55450874 | C2orf63;C2orf63 | N_Shore | 5'UTR;5'UTR |
| WG3466 | chr2:56410011-56410087 | CCDC85A | N_Shore | TSS1500 |
| WG1513 | chr2:56410011-56410143 | CCDC85A | N_Shore | TSS1500 |
| WG2275 | chr2:56410071-56410114 | CCDC85A | N_Shore | TSS1500 |
| WG1978 | chr2:6122798-6122814 | LOC400940 | | Body |
| WG2276 | chr2:61372165-61372256 | C2orf74 | | TSS200 |
| GM00520 | chr2:63274857-63275178 | LOC100132215 | Island | TSS200 |
| WG3466 | chr2:63275034-63275178 | LOC100132215 | Island | TSS200 |
| GM02706 | chr2:63275509-63275954 | LOC100132215 | Island | TSS1500 |
| GM00290 | chr2:63276565-63276723 | OTX1 | Island | TSS1500 |
| GM00439 | chr2:63276700-63276779 | OTX1 | Island | TSS1500 |
| GM00016 | chr2:63283939-63284132 | OTX1 | Island | 3'UTR |
| WG1984 | chr2:65217528-65217623 | SLC1A4;SLC1A4 | S_Shore | Body;5'UTR |
| GM00016 | chr2:66664798-66665428 | MEIS1 | S_Shelf | Body |
| GM00022 | chr2:66667101-66668012 | MEIS1 | | Body |
| WG1282 | chr2:66667852-66668012 | MEIS1 | | Body |
| WG1977 | chr2:66671478-66671727 | MEIS1 | N_Shore | Body |
| WG1722 | chr2:66672337-66673428 | MEIS1 | N_Shore | Body |
| GM00439 | chr2:66672553-66673428 | MEIS1 | Island | Body |
| GM02555 | chr2:66673316-66673985 | MEIS1 | Island | Body |
| GM02173 | chr2:68592345-68592394 | PLEK;PLEK | | 1stExon;5'UTR |
| GM09256 | chr2:69001825-69001870 | ARHGAP25;ARHGAP25;ARHGAP25;ARHG | | Body;TSS200;TSS200;Body |
| WG1513 | chr2:69001854-69001991 | ARHGAP25;ARHGAP25;ARHGAP25;ARHG | | Body;TSS200;TSS200;Body |
| GM02706 | chr2:69002111-69002378 | ARHGAP25;ARHGAP25;ARHGAP25;ARHG | | 5'UTR;Body;1stExon;5'UTR;1stExon;Bo |
| GM00042 | chr2:70529494-70529513 | FAM136A | S_Shore | TSS1500 |
| WG3465 | chr2:7057153-7057245 | RNF144A | N_Shore | TSS1500 |
| WG2415 | chr2:71134524-71134630 | VAX2 | N_Shore | Body |
| WG1977 | chr2:71134524-71134694 | VAX2 | N_Shore | Body |
| GM00016 | chr2:71134524-71134936 | VAX2 | N_Shore | Body |
| WG1722 | chr2:71134630-71134936 | VAX2 | Island | Body |
| WG1977 | chr2:71162693-71163123 | ATP6V1B1 | | TSS1500 |
| WG3466 | chr2:71211981-71212157 | ANKRD53;ANKRD53 | | 3'UTR;Body |
| GM00520 | chr2:71212056-71212504 | ANKRD53;ANKRD53 | | 3'UTR;Body |
| WG3465 | chr2:72371255-72371477 | CYP26B1 | Island | Body |
| GM09255 | chr2:73964768-73964834 | TPRKB | S_Shore | TSS1500 |
| GM00439 | chr2:74119781-74120092 | ACTG2 | | TSS1500 |
| GM00289 | chr2:74668072-74668672 | RTKN;RTKN;RTKN | Island | Body;TSS1500;TSS1500 |
| WG2121 | chr2:74668477-74668976 | RTKN;RTKN;RTKN | Island | Body;TSS1500;TSS1500 |

| | | | | |
|---|---|---|---|---|
| WG1658 | chr2:74669349-74669380 | RTKN | Island | TSS1500 |
| WG3465 | chr2:74669349-74669387 | RTKN | Island | TSS1500 |
| WG1977 | chr2:74669349-74669516 | RTKN | Island | TSS1500 |
| GM02173 | chr2:74700039-74700246 | CCDC142;MRPL53 | S_Shore | 3'UTR;TSS200 |
| WG3465 | chr2:74729429-74729592 | LOC151534;LBX2 | Island | TSS1500;Body |
| GM02706 | chr2:74729429-74729710 | LOC151534;LBX2 | Island | TSS1500;Body |
| GM02706 | chr2:74730630-74730948 | LOC151534;LBX2 | Island | Body;TSS200 |
| GM01392 | chr2:74780268-74780467 | LOXL3 | N_Shore | 5'UTR |
| GM09256 | chr2:74780268-74780617 | LOXL3 | N_Shore | 5'UTR |
| WG1978 | chr2:74780272-74780617 | LOXL3 | N_Shore | 5'UTR |
| GM09256 | chr2:74875263-74875387 | C2orf65 | Island | TSS200 |
| GM02706 | chr2:79347474-79347666 | REG1A | | TSS200 |
| WG1513 | chr2:79347474-79347699 | REG1A | | TSS200 |
| GM02456 | chr2:8118122-8118404 | LOC339788 | | TSS1500 |
| GM02706 | chr2:84517738-84517763 | LOC388965 | Island | TSS200 |
| WG1513 | chr2:84517738-84517877 | LOC388965 | Island | TSS200 |
| GM00290 | chr2:85106968-85107320 | C2orf89 | N_Shore | Body |
| GM02456 | chr2:85361959-85362480 | TCF7L1 | Island | Body |
| GM02173 | chr2:85637834-85638053 | CAPG | N_Shelf | TSS200 |
| WG3466 | chr2:85637834-85638535 | CAPG | N_Shelf | TSS200 |
| GM00289 | chr2:87036626-87037038 | CD8A | Island | TSS1500 |
| WG2415 | chr2:88427581-88427637 | FABP1 | | TSS200 |
| WG1513 | chr2:88427630-88427827 | FABP1 | | TSS200 |
| WG1513 | chr2:906313-906974 | C2orf90 | | TSS1500 |
| GM00941 | chr2:9144618-9144764 | MBOAT2 | Island | TSS1500 |
| WG2121 | chr2:947167-947589 | SNTG2 | Island | Body |
| GM00290 | chr2:947167-947634 | SNTG2 | Island | Body |
| WG2414 | chr2:947515-947634 | SNTG2 | S_Shore | Body |
| GM02706 | chr2:95537314-95537475 | TEKT4;TEKT4 | Island | 1stExon;5'UTR |
| GM00289 | chr2:95971840-95971994 | KCNIP3 | | Body |
| WG1513 | chr2:96990800-96990861 | ITPRIPL1;ITPRIPL1;ITPRIPL1;I | N_Shore | TSS1500;TSS1500;TSS1500;TSS1500 |
| GM00438 | chr2:97174815-97175065 | NEURL3 | S_Shore | TSS1500 |
| WG1513 | chr2:97651503-97651765 | FAM178B | N_Shore | 5'UTR |
| WG2120 | chr2:98340511-98340921 | ZAP70 | Island | Body |
| WG2415 | chr2:98962444-98962564 | CNGA3;CNGA3 | N_Shore | TSS200;TSS200 |
| GM00438 | chr2:98993614-98993730 | CNGA3;CNGA3 | | Body;Body |
| GM00940 | chr20:13975439-13975853 | MACROD2 | N_Shore | TSS1500 |
| GM09256 | chr20:1472071-1472098 | SIRPB2;SIRPB2;SIRPB2;SIRPB2 | | 1stExon;1stExon;5'UTR;5'UTR |
| GM00290 | chr20:16728895-16729319 | OTOR | | TSS200 |
| WG1310 | chr20:17295815-17296161 | PCSK2 | N_Shore | Body |
| GM00438 | chr20:17296161-17296450 | PCSK2 | Island | Body |
| WG1977 | chr20:17595355-17595472 | RRBP1;RRBP1 | Island | Body;Body |
| GM00289 | chr20:1974738-1974878 | PDYN | | TSS200 |
| GM01392 | chr20:1974924-1975372 | PDYN | | TSS1500 |
| WG2844 | chr20:19915716-19915769 | RIN2 | | Body |
| WG2844 | chr20:19915857-19916087 | RIN2 | | Body |
| GM00935 | chr20:20350367-20350813 | INSM1 | Island | 1stExon |

| | | | | |
|---|---|---|---|---|
| GM02175 | chr20:207144-207338 | DEFB129 | | TSS1500 |
| WG2276 | chr20:21685401-21685925 | PAX1 | N_Shore | TSS1500 |
| WG2276 | chr20:21694827-21695343 | PAX1 | Island | Body |
| WG3465 | chr20:23066944-23067159 | CD93;CD93 | | 5'UTR;1stExon |
| WG3466 | chr20:23067033-23067159 | CD93 | | TSS200 |
| GM02149 | chr20:23067126-23067159 | CD93 | | TSS200 |
| GM01096 | chr20:2308779-2308988 | TGM3 | Island | Body |
| WG1513 | chr20:23471737-23472171 | CST8 | | TSS200 |
| GM00438 | chr20:23498937-23498993 | CSTT | | TSS1500 |
| WG1513 | chr20:23549106-23549495 | CST9L;CST9L | | 1stExon;5'UTR |
| GM02706 | chr20:2361431-2361461 | TGM6 | | TSS200 |
| GM00940 | chr20:24452007-24452131 | TMEM90B | Island | 5'UTR |
| WG1513 | chr20:24452036-24452131 | TMEM90B | Island | 5'UTR |
| GM00941 | chr20:2490030-2490109 | ZNF343 | S_Shore | TSS1500 |
| WG2120 | chr20:24929687-24929716 | CST7 | | TSS200 |
| WG1722 | chr20:25128770-25128805 | LOC284798;LOC284798;LO( | Island | TSS200;Body;Body |
| WG1977 | chr20:25128770-25128807 | LOC284798;LOC284798;LO( | Island | TSS200;Body;Body |
| WG2844 | chr20:2516256-2516592 | TMC2 | | TSS1500 |
| WG1513 | chr20:2634697-2634874 | NOP56;SNORD110;NOP56;S | S_Shore | Body;TSS200;Body;TSS1500 |
| WG1722 | chr20:30028223-30028241 | DEFB123 | | TSS200 |
| WG1513 | chr20:30028241-30028418 | DEFB123 | | TSS200 |
| GM02706 | chr20:30060822-30060884 | DEFB124 | N_Shelf | TSS200 |
| GM09256 | chr20:30073209-30073576 | NCRNA00028 | S_Shore | TSS1500 |
| WG1722 | chr20:30073521-30073576 | NCRNA00028 | S_Shore | TSS200 |
| WG1977 | chr20:30073537-30073576 | NCRNA00028 | S_Shore | TSS200 |
| WG1514 | chr20:30134929-30135362 | HM13;HM13;HM13;PSIMCT-1 | N_Shore | Body;Body;Body;TSS1500 |
| GM00022 | chr20:30225517-30225666 | COX4I2 | N_Shore | TSS200 |
| GM02173 | chr20:30225573-30225685 | COX4I2 | Island | TSS200 |
| WG2120 | chr20:30225666-30225685 | COX4I2 | Island | TSS200 |
| WG1978 | chr20:30225666-30225706 | COX4I2 | Island | TSS200 |
| WG1977 | chr20:30225685-30225851 | COX4I2 | Island | TSS200 |
| GM02706 | chr20:30433409-30433673 | FOXS1;FOXS1 | S_Shore | 5'UTR;1stExon |
| GM02555 | chr20:3051954-3052151 | OXT | N_Shore | TSS1500 |
| WG2120 | chr20:3051954-3052262 | OXT | N_Shore | TSS1500 |
| WG2276 | chr20:3052115-3052221 | OXT | Island | TSS200 |
| GM09256 | chr20:3052115-3052224 | OXT | Island | TSS200 |
| WG1514 | chr20:3052115-3052262 | OXT | Island | TSS200 |
| GM00290 | chr20:3052151-3052224 | OXT | Island | TSS200 |
| WG2414 | chr20:3052221-3052262 | OXT | Island | TSS200 |
| WG2276 | chr20:3052253-3052274 | OXT | Island | TSS200 |
| GM00289 | chr20:3052262-3052345 | OXT | Island | TSS200 |
| GM01393 | chr20:30582838-30583001 | XKR7 | Island | Body |
| GM00439 | chr20:31331973-31332353 | COMMD7;COMMD7 | S_Shore | TSS200;TSS200 |
| WG3466 | chr20:3148785-3148934 | ProSAPiP1;ProSAPiP1 | Island | 1stExon;5'UTR |
| GM09256 | chr20:3149249-3149278 | ProSAPiP1 | S_Shore | TSS200 |
| WG3465 | chr20:3149259-3149320 | ProSAPiP1 | S_Shore | TSS200 |
| WG1722 | chr20:3149278-3149495 | ProSAPiP1 | S_Shore | TSS200 |

| | | | | |
|---|---|---|---|---|
| WG1513 | chr20:31755784-31756024 | C20orf70 | | TSS200 |
| GM02706 | chr20:31755943-31756024 | C20orf70 | | TSS200 |
| WG1513 | chr20:31781232-31781393 | BASE | | TSS200 |
| GM01653 | chr20:32031936-32032443 | SNTA1 | Island | TSS1500 |
| WG1513 | chr20:3218393-3218487 | SLC4A11 | N_Shore | TSS200 |
| GM00289 | chr20:3218476-3218500 | SLC4A11 | N_Shore | TSS200 |
| WG3465 | chr20:32856747-32856898 | ASIP | Island | Body |
| GM02147 | chr20:32856747-32857072 | ASIP | Island | Body |
| WG2121 | chr20:32856747-32857151 | ASIP | Island | Body |
| GM01393 | chr20:32856846-32857151 | ASIP | Island | Body |
| GM09255 | chr20:32857072-32857227 | ASIP | Island | 3'UTR |
| GM00289 | chr20:33585040-33585435 | MYH7B | Island | Body |
| GM02706 | chr20:33762474-33762673 | PROCR | Island | Body |
| WG3466 | chr20:33762474-33762943 | PROCR | Island | Body |
| GM00439 | chr20:33762631-33762943 | PROCR | Island | Body |
| WG1977 | chr20:35169539-35169609 | MYL9;MYL9 | Island | TSS1500;TSS1500 |
| WG1977 | chr20:35274595-35274639 | SLA2;SLA2 | | TSS200;TSS200 |
| GM00438 | chr20:35274595-35274655 | SLA2;SLA2 | | TSS200;TSS200 |
| GM02175 | chr20:35274627-35274655 | SLA2;SLA2 | | TSS200;TSS200 |
| WG2415 | chr20:35274627-35274715 | SLA2;SLA2 | | TSS200;TSS200 |
| WG2121 | chr20:35580436-35580555 | SAMHD1 | Island | TSS1500 |
| WG2415 | chr20:36011999-36012016 | SRC;SRC | N_Shore | 5'UTR;5'UTR |
| WG3466 | chr20:36011999-36012596 | SRC;SRC | N_Shore | 5'UTR;5'UTR |
| GM00940 | chr20:36153642-36153947 | BLCAP;BLCAP;BLCAP;BLCAP | N_Shelf | 5'UTR;TSS1500;5'UTR;5'UTR;5'UTR |
| WG1722 | chr20:36156919-36157016 | BLCAP;BLCAP;BLCAP;BLCAP | S_Shore | TSS1500;TSS1500;TSS1500;TSS1500 |
| WG1722 | chr20:3662512-3662948 | ADAM33;ADAM33 | Island | Body;Body |
| GM02555 | chr20:36932397-36932553 | BPI | | TSS200 |
| GM02706 | chr20:37434158-37434231 | PPP1R16B | N_Shore | TSS200 |
| GM00439 | chr20:3762422-3762653 | SPEF1 | N_Shelf | TSS1500 |
| WG1977 | chr20:39969481-39969604 | LPIN3 | S_Shore | TSS200 |
| GM02704 | chr20:39969536-39969604 | LPIN3 | S_Shore | TSS200 |
| GM00439 | chr20:39995649-39995654 | EMILIN3 | Island | TSS200 |
| GM00935 | chr20:39995649-39995747 | EMILIN3 | Island | TSS200 |
| GM00438 | chr20:42142005-42142236 | L3MBTL;L3MBTL | N_Shore | TSS1500;TSS1500 |
| GM03075 | chr20:43160249-43160301 | PKIG;PKIG;PKIG | N_Shore | TSS200;TSS200;TSS200 |
| GM00042 | chr20:43729410-43729808 | KCNS1 | N_Shore | 5'UTR |
| WG3466 | chr20:43729527-43729874 | KCNS1 | Island | 5'UTR |
| GM01392 | chr20:43729861-43729874 | KCNS1 | S_Shore | TSS200 |
| WG1722 | chr20:43729861-43729897 | KCNS1 | S_Shore | TSS200 |
| GM00520 | chr20:43729869-43729897 | KCNS1 | S_Shore | TSS200 |
| GM00042 | chr20:43729869-43729910 | KCNS1 | S_Shore | TSS200 |
| WG2415 | chr20:43935222-43935291 | MATN4;MATN4;MATN4;RBPJL | Island | 5'UTR;5'UTR;5'UTR;TSS1500 |
| WG1977 | chr20:43935283-43935361 | MATN4;MATN4;MATN4;RBPJL | Island | 5'UTR;5'UTR;5'UTR;TSS1500 |
| GM02706 | chr20:43935283-43935391 | MATN4;MATN4;MATN4;RBPJL | Island | 5'UTR;5'UTR;5'UTR;TSS1500 |
| WG2414 | chr20:43935460-43935471 | MATN4;MATN4;RBPJL;MATN4 | Island | 5'UTR;5'UTR;TSS200;5'UTR |
| GM00438 | chr20:44144582-44144711 | SPINT3 | | TSS1500 |
| WG3465 | chr20:44746499-44746825 | CD40;CD40 | N_Shore | TSS1500;TSS1500 |

| | | | | |
|---|---|---|---|---|
| WG1977 | chr20:44829602-44829821 | CDH22 | Island | Body |
| GM00520 | chr20:44838776-44839187 | CDH22 | N_Shore | Body |
| GM02706 | chr20:44875334-44875586 | CDH22 | Island | Body |
| WG3465 | chr20:44880263-44880525 | CDH22;CDH22 | Island | 1stExon;5'UTR |
| GM00520 | chr20:45179345-45179413 | C20orf123 | | TSS200 |
| GM00940 | chr20:45313200-45313266 | TP53RK;SLC13A3 | | 3'UTR;TSS200 |
| WG1722 | chr20:47896448-47896888 | SNORD12B;C20orf199;MIR1 | S_Shore | TSS1500;Body;TSS1500;Body;TSS150 |
| GM02704 | chr20:48530454-48530654 | SPATA2;SPATA2 | N_Shore | 5'UTR;TSS200 |
| GM02555 | chr20:49253364-49253710 | FAM65C;FAM65C | | 1stExon;5'UTR |
| GM00520 | chr20:49253552-49253710 | FAM65C | | TSS200 |
| WG1984 | chr20:50312386-50312632 | ATP9A | | Body |
| WG1977 | chr20:51589730-51589985 | TSHZ2;TSHZ2 | Island | 1stExon;5'UTR |
| WG1977 | chr20:52199594-52199778 | ZNF217;ZNF217 | S_Shore | 5'UTR;1stExon |
| WG1978 | chr20:52199729-52199778 | ZNF217 | S_Shore | TSS200 |
| GM02149 | chr20:5450926-5451030 | LOC643406 | | TSS1500 |
| WG1513 | chr20:5451661-5451800 | LOC643406 | | TSS200 |
| WG3466 | chr20:5485144-5485270 | LOC149837 | | Body |
| WG1722 | chr20:5485144-5485284 | LOC149837 | | Body |
| GM02149 | chr20:5485144-5485294 | LOC149837 | | Body |
| WG2194 | chr20:5485144-5485475 | LOC149837 | | Body |
| GM02175 | chr20:5485144-5485511 | LOC149837 | | Body |
| WG2415 | chr20:5485245-5485270 | LOC149837 | | TSS200 |
| WG1977 | chr20:5485245-5485284 | LOC149837 | | TSS200 |
| WG1283 | chr20:5485245-5485294 | LOC149837 | | TSS200 |
| GM00042 | chr20:5485245-5485511 | LOC149837 | | TSS200 |
| WG2121 | chr20:5485268-5485284 | LOC149837 | | TSS200 |
| GM01096 | chr20:5485268-5485294 | LOC149837 | | TSS200 |
| WG3465 | chr20:5485268-5485511 | LOC149837 | | TSS200 |
| WG2414 | chr20:55203291-55203849 | TFAP2C | Island | TSS1500 |
| WG2121 | chr20:55203439-55203849 | TFAP2C | Island | TSS1500 |
| WG2276 | chr20:55839288-55839663 | BMP7 | Island | Body |
| WG3465 | chr20:55839524-55839663 | BMP7 | Island | Body |
| WG3465 | chr20:55904677-55904856 | SPO11;SPO11 | Island | TSS200;TSS200 |
| GM02555 | chr20:55925570-55925603 | RAE1;RAE1 | N_Shore | TSS1500;TSS1500 |
| WG2844 | chr20:55964998-55965497 | RBM38;RBM38 | Island | TSS1500;TSS1500 |
| WG2276 | chr20:56286391-56286567 | PMEPA1;PMEPA1;PMEPA1;PI | S_Shore | 5'UTR;1stExon;TSS1500;TSS1500;TSS |
| GM02175 | chr20:57267493-57267680 | NPEPL1 | N_Shore | TSS1500 |
| GM00289 | chr20:57426759-57426835 | GNAS;GNAS;GNAS;GNASAS | Island | TSS1500;TSS1500;3'UTR;TSS1500 |
| GM02706 | chr20:57426950-57427017 | GNAS;GNAS;GNAS;GNASAS | Island | TSS1500;TSS1500;3'UTR;TSS1500 |
| GM02706 | chr20:57427146-57427173 | GNAS;GNAS;GNAS;GNASAS | N_Shore | TSS1500;TSS1500;3'UTR;TSS1500 |
| WG2121 | chr20:57427472-57427495 | GNAS;GNAS;GNAS | N_Shore | TSS1500;TSS1500;3'UTR |
| GM02175 | chr20:57427821-57427973 | GNAS;GNAS;GNAS | Island | TSS1500;TSS1500;3'UTR |
| WG1513 | chr20:57463325-57463357 | GNAS;GNAS;GNAS;GNAS | N_Shore | 3'UTR;3'UTR;TSS1500;Body |
| WG1722 | chr20:57465439-57465448 | GNAS;GNAS;GNAS;GNAS;GN | Island | 3'UTR;TSS1500;TSS1500;Body;TSS15 |
| WG1722 | chr20:57471654-57471672 | GNAS;GNAS;GNAS;GNAS;GN | S_Shelf | Body;Body;3'UTR;Body;Body;3'UTR;Bo |
| GM02173 | chr20:57617083-57617431 | SLMO2 | N_Shore | Body |
| WG1513 | chr20:57765489-57765943 | ZNF831 | N_Shore | TSS1500 |

| | | | | |
|---|---|---|---|---|
| GM00520 | chr20:57765815-57766071 | ZNF831 | N_Shore | TSS1500 |
| GM00940 | chr20:58179536-58179540 | PHACTR3 | N_Shore | TSS200 |
| GM09256 | chr20:58179540-58180241 | PHACTR3 | N_Shore | TSS200 |
| WG1513 | chr20:58179540-58180357 | PHACTR3 | N_Shore | TSS200 |
| WG3466 | chr20:58179847-58180241 | PHACTR3;PHACTR3 | Island | 5'UTR;1stExon |
| WG1978 | chr20:58179847-58180357 | PHACTR3;PHACTR3 | Island | 5'UTR;1stExon |
| WG3466 | chr20:58630954-58631038 | C20orf197 | | TSS200 |
| WG2414 | chr20:5891781-5891968 | CHGB | N_Shore | TSS200 |
| WG1513 | chr20:6034694-6035120 | LRRN4 | S_Shore | TSS200 |
| GM00935 | chr20:6034694-6035139 | LRRN4 | S_Shore | TSS200 |
| WG1977 | chr20:60347821-60348277 | CDH4 | N_Shore | Body |
| GM00439 | chr20:6104215-6104267 | FERMT1 | S_Shore | TSS200 |
| WG1977 | chr20:61151391-61151480 | C20orf166;MIR1-1 | N_Shore | Body;TSS200 |
| GM01096 | chr20:61151446-61151480 | C20orf166;MIR1-1 | N_Shore | Body;TSS200 |
| WG3465 | chr20:61151467-61151480 | C20orf166;MIR1-1 | N_Shore | Body;TSS200 |
| GM03073 | chr20:61151467-61151721 | C20orf166;MIR1-1 | N_Shore | Body;TSS200 |
| GM02175 | chr20:61297564-61298096 | SLCO4A1;LOC100127888 | | Body;Body |
| GM09256 | chr20:61446962-61447053 | COL9A3 | N_Shore | TSS1500 |
| WG1721 | chr20:61447110-61447272 | COL9A3 | N_Shore | TSS1500 |
| WG1977 | chr20:61447174-61447194 | COL9A3 | N_Shore | TSS1500 |
| GM01376 | chr20:61447268-61447330 | COL9A3 | N_Shore | TSS1500 |
| WG2415 | chr20:61447330-61447394 | COL9A3 | N_Shore | TSS1500 |
| WG1513 | chr20:61447614-61447661 | COL9A3 | N_Shore | TSS1500 |
| GM00289 | chr20:61583686-61583910 | SLC17A9 | N_Shore | TSS1500 |
| GM00022 | chr20:61583736-61583910 | SLC17A9 | N_Shore | TSS1500 |
| GM02706 | chr20:61583736-61584850 | SLC17A9 | N_Shore | TSS1500 |
| WG1722 | chr20:61583830-61583924 | SLC17A9 | N_Shore | TSS200 |
| GM00520 | chr20:61583910-61583987 | SLC17A9 | Island | TSS200 |
| GM00042 | chr20:61867178-61867212 | BIRC7;BIRC7 | N_Shelf | TSS200;TSS200 |
| GM00935 | chr20:61924405-61924802 | COL20A1 | N_Shelf | TSS200 |
| GM09256 | chr20:61953801-61954200 | COL20A1 | N_Shore | Body |
| WG1722 | chr20:61953801-61954258 | COL20A1 | N_Shore | Body |
| GM09255 | chr20:61953935-61954258 | COL20A1 | Island | Body |
| WG3465 | chr20:62104476-62104613 | KCNQ2;KCNQ2;KCNQ2;KCN | S_Shore | TSS1500;TSS1500;TSS1500;TSS1500; |
| GM09256 | chr20:62153067-62153239 | PPDPF | Island | Body |
| GM00940 | chr20:62183924-62183966 | C20orf195 | N_Shore | TSS1500 |
| WG1977 | chr20:62317768-62318482 | RTEL1;RTEL1 | Island | Body;Body |
| GM00022 | chr20:62317768-62318588 | RTEL1;RTEL1 | Island | Body;Body |
| WG1722 | chr20:62318426-62318588 | RTEL1;RTEL1 | Island | Body;Body |
| WG1978 | chr20:62318433-62318588 | RTEL1;RTEL1 | Island | Body;Body |
| GM00520 | chr20:62327732-62328084 | TNFRSF6B;TNFRSF6B | N_Shore | 5'UTR;TSS1500 |
| WG2415 | chr20:62327968-62328094 | TNFRSF6B;TNFRSF6B | N_Shore | 5'UTR;TSS200 |
| WG1977 | chr20:62327968-62328427 | TNFRSF6B;TNFRSF6B | N_Shore | 5'UTR;TSS200 |
| WG1977 | chr20:62368956-62369462 | LIME1 | Island | Body |
| GM00438 | chr20:62368956-62369504 | LIME1 | Island | Body |
| GM00438 | chr20:62433998-62434140 | ZBTB46 | Island | 5'UTR |
| WG2415 | chr20:62679255-62679713 | SOX18 | N_Shore | 3'UTR |

| | | | | |
|---|---|---|---|---|
| GM00439 | chr20:62688272-62688329 | TCEA2 | N_Shore | TSS200 |
| WG1977 | chr20:62688717-62688896 | TCEA2;TCEA2 | Island | 1stExon;5'UTR |
| WG2121 | chr20:62693943-62693971 | TCEA2;TCEA2 | Island | TSS200;5'UTR |
| WG1722 | chr20:62693951-62693973 | TCEA2;TCEA2 | Island | TSS200;5'UTR |
| WG3466 | chr20:62693971-62694005 | TCEA2;TCEA2 | Island | TSS200;5'UTR |
| WG2415 | chr20:62715942-62716443 | OPRL1;C20orf201;OPRL1 | N_Shore | 5'UTR;TSS1500;5'UTR |
| GM02706 | chr20:62715942-62716485 | OPRL1;C20orf201;OPRL1 | N_Shore | 5'UTR;TSS1500;5'UTR |
| GM01392 | chr20:62716332-62716606 | OPRL1;C20orf201;OPRL1 | Island | 5'UTR;TSS1500;5'UTR |
| WG1513 | chr20:62716443-62716606 | OPRL1;C20orf201;OPRL1 | Island | 5'UTR;TSS1500;5'UTR |
| WG1977 | chr20:62738880-62739073 | NPBWR2 | S_Shore | TSS1500 |
| GM00520 | chr20:749264-749360 | C20orf54 | | TSS200 |
| WG1722 | chr20:825408-825489 | FAM110A;FAM110A;FAM110A | Island | 5'UTR;5'UTR;5'UTR;1stExon |
| GM09256 | chr20:9494838-9494997 | C20orf103 | N_Shore | TSS1500 |
| GM09256 | chr21:15352213-15352641 | C21orf81 | Island | Body |
| GM02555 | chr21:17961737-17961996 | C21orf34;MIR125B2;C21orf34 | | Body;TSS1500;Body |
| WG3465 | chr21:17961737-17962124 | C21orf34;MIR125B2;C21orf34 | | Body;TSS1500;Body |
| WG1722 | chr21:17961829-17962124 | C21orf34;MIR125B2;C21orf34 | | Body;TSS1500;Body |
| WG1977 | chr21:28337594-28337869 | ADAMTS5 | N_Shore | Body |
| WG2121 | chr21:30391563-30391784 | RWDD2B | Island | 1stExon |
| GM09256 | chr21:31538714-31539057 | CLDN17 | | 1stExon |
| GM02149 | chr21:31655415-31655822 | KRTAP24-1 | | TSS200 |
| WG1513 | chr21:31655654-31655982 | KRTAP24-1 | | TSS1500 |
| GM09256 | chr21:31864089-31864374 | KRTAP19-3 | | 1stExon |
| GM02706 | chr21:32185331-32185576 | KRTAP8-1;KRTAP8-1 | | 1stExon;3'UTR |
| GM02175 | chr21:34396944-34397135 | OLIG2 | Island | TSS1500 |
| GM01653 | chr21:35831871-35832008 | KCNE1;KCNE1;KCNE1;KCNE | Island | 5'UTR;5'UTR;1stExon;1stExon;5'UTR |
| WG3465 | chr21:35831871-35832180 | KCNE1;KCNE1;KCNE1;KCNE | Island | 5'UTR;5'UTR;1stExon;1stExon;5'UTR |
| GM00935 | chr21:35831996-35832164 | KCNE1;KCNE1;KCNE1 | Island | TSS200;5'UTR;TSS200 |
| WG2844 | chr21:35831996-35832180 | KCNE1;KCNE1;KCNE1 | Island | TSS200;5'UTR;TSS200 |
| GM02555 | chr21:35832028-35832180 | KCNE1;KCNE1;KCNE1 | Island | TSS200;5'UTR;TSS200 |
| WG2275 | chr21:35832141-35832180 | KCNE1;KCNE1;KCNE1 | Island | 5'UTR;TSS1500;TSS1500 |
| GM02641 | chr21:36041334-36041579 | CLIC6 | Island | TSS1500 |
| WG2121 | chr21:36041334-36041605 | CLIC6 | Island | TSS1500 |
| GM00022 | chr21:36041527-36041579 | CLIC6 | Island | TSS200 |
| WG3466 | chr21:36263347-36263808 | RUNX1 | Island | Body |
| WG3465 | chr21:36399146-36399258 | RUNX1 | Island | Body |
| GM02456 | chr21:36421467-36421503 | RUNX1;RUNX1 | | 5'UTR;1stExon |
| GM00031 | chr21:37442104-37442476 | CBR1 | N_Shore | TSS200 |
| WG2194 | chr21:37442289-37442476 | CBR1;CBR1 | Island | 1stExon;5'UTR |
| WG3466 | chr21:38119920-38120350 | SIM2 | Island | Body |
| GM00940 | chr21:38362725-38362754 | HLCS | Island | TSS200 |
| WG2414 | chr21:38362727-38362754 | HLCS | Island | TSS200 |
| WG2276 | chr21:38580129-38580812 | DSCR9 | | TSS1500 |
| GM00940 | chr21:39644216-39644332 | KCNJ15;KCNJ15;KCNJ15 | | 5'UTR;5'UTR;TSS1500 |
| GM02706 | chr21:40145923-40146046 | NCRNA00114 | | TSS1500 |
| WG2276 | chr21:40145923-40146149 | NCRNA00114 | | TSS1500 |
| GM01650 | chr21:40176950-40177165 | ETS2 | N_Shore | TSS1500 |

| | | | | |
|---|---|---|---|---|
| WG3465 | chr21:40757691-40758208 | WRB | Island | Body |
| WG1309 | chr21:40759534-40760258 | WRB;WRB | N_Shore | Body;TSS200 |
| GM00289 | chr21:42219202-42219221 | DSCAM | Island | TSS200 |
| GM02706 | chr21:42880554-42880582 | TMPRSS2;TMPRSS2 | Island | TSS1500;TSS1500 |
| GM00520 | chr21:43221380-43221684 | PRDM15;PRDM15 | N_Shore | 3'UTR;3'UTR |
| WG2276 | chr21:43372922-43373136 | C2CD2 | N_Shore | Body |
| WG3465 | chr21:43442057-43442254 | C21orf121 | | TSS200 |
| GM01376 | chr21:43823809-43823863 | UBASH3A;UBASH3A | | TSS1500;TSS1500 |
| WG1513 | chr21:43919316-43919801 | SLC37A1 | S_Shelf | TSS1500 |
| WG2276 | chr21:44088823-44089014 | PDE9A;PDE9A;PDE9A;PDE9A | Island | Body;Body;5'UTR;Body;Body;Body;5'U |
| WG2121 | chr21:44104950-44105474 | PDE9A;PDE9A;PDE9A;PDE9A | N_Shore | Body;Body;5'UTR;Body;Body;Body;5'U |
| GM02706 | chr21:45148864-45149141 | PDXK | Island | Body |
| WG3466 | chr21:45148879-45149141 | PDXK | Island | Body |
| GM00935 | chr21:45148879-45149398 | PDXK | Island | Body |
| WG2844 | chr21:45379489-45379721 | AGPAT3;AGPAT3 | N_Shore | 5'UTR;5'UTR |
| WG1513 | chr21:45683302-45683528 | DNMT3L;DNMT3L | | TSS1500;TSS1500 |
| GM01650 | chr21:45705618-45705742 | AIRE | Island | TSS200 |
| WG1282 | chr21:45705699-45705742 | AIRE | Island | TSS200 |
| GM02175 | chr21:45773290-45773400 | TRPM2 | | TSS200 |
| GM00439 | chr21:45789122-45789379 | TRPM2 | Island | Body |
| GM00042 | chr21:45811363-45811549 | TRPM2 | Island | Body |
| WG2415 | chr21:45811432-45811549 | TRPM2 | Island | Body |
| WG1513 | chr21:45936016-45936702 | C21orf90;C21orf90;C21orf29 | | TSS1500;TSS1500;Body |
| WG1513 | chr21:45960718-45961033 | KRTAP10-1;C21orf29 | | TSS1500;Body |
| WG1722 | chr21:46077715-46077731 | KRTAP12-3;C21orf29 | | TSS200;Body |
| WG3465 | chr21:46238781-46238957 | SUMO3 | Island | TSS1500 |
| GM02456 | chr21:46367939-46368090 | C21orf70 | N_Shore | Body |
| WG1983 | chr21:47294739-47295215 | PCBP3;PCBP3 | N_Shore | Body;Body |
| WG2276 | chr21:47404165-47404268 | COL6A1 | Island | Body |
| WG1721 | chr21:47604052-47604291 | C21orf56;C21orf56 | S_Shore | 5'UTR;5'UTR |
| WG2415 | chr21:47604052-47604361 | C21orf56;C21orf56 | S_Shore | 5'UTR;5'UTR |
| WG3465 | chr21:47604052-47604654 | C21orf56;C21orf56 | S_Shore | 5'UTR;5'UTR |
| GM03075 | chr21:47604052-47605174 | C21orf56;C21orf56 | S_Shore | 5'UTR;5'UTR |
| WG2844 | chr21:47604291-47604654 | C21orf56;C21orf56;C21orf56 | S_Shore | 5'UTR;5'UTR;1stExon;1stExon |
| WG1983 | chr21:47604291-47604986 | C21orf56;C21orf56;C21orf56 | S_Shore | 5'UTR;5'UTR;1stExon;1stExon |
| GM02175 | chr21:47604291-47605072 | C21orf56;C21orf56;C21orf56 | S_Shore | 5'UTR;5'UTR;1stExon;1stExon |
| GM01650 | chr21:47604291-47605174 | C21orf56;C21orf56;C21orf56 | S_Shore | 5'UTR;5'UTR;1stExon;1stExon |
| GM09255 | chr21:47604361-47605174 | C21orf56;C21orf56;C21orf56 | S_Shore | 5'UTR;5'UTR;1stExon;1stExon |
| WG1309 | chr21:47604654-47605174 | C21orf56;C21orf56 | S_Shore | TSS1500;TSS1500 |
| GM02706 | chr21:47717575-47718080 | C21orf57 | Island | 3'UTR |
| WG3465 | chr21:47813364-47813461 | PCNT | Island | Body |
| WG2844 | chr21:47813364-47813491 | PCNT | Island | Body |
| GM02704 | chr22:17090586-17090918 | psiTPTE22 | Island | Body |
| WG1513 | chr22:17489112-17489187 | GAB4 | Island | TSS200 |
| WG3465 | chr22:18063945-18064224 | SLC25A18 | Island | Body |
| GM02706 | chr22:18064052-18064224 | SLC25A18 | Island | Body |
| WG2275 | chr22:18632429-18632476 | USP18 | N_Shore | TSS1500 |

| | | | | |
|---|---|---|---|---|
| GM00438 | chr22:19003848-19004088 | DGCR9 | | TSS1500 |
| WG2844 | chr22:19118165-19118202 | DGCR14;TSSK2 | N_Shore | 3'UTR;TSS200 |
| WG2415 | chr22:19118165-19118539 | DGCR14;TSSK2 | N_Shore | 3'UTR;TSS200 |
| GM00940 | chr22:19743892-19744078 | TBX1;TBX1;TBX1 | Island | TSS1500;TSS1500;TSS1500 |
| GM00439 | chr22:19744026-19744078 | TBX1;TBX1;TBX1 | Island | TSS200;TSS200;TSS200 |
| WG1978 | chr22:19746156-19746368 | TBX1;TBX1;TBX1 | Island | 5'UTR;5'UTR;5'UTR |
| GM00941 | chr22:19746984-19747026 | TBX1;TBX1;TBX1 | Island | 5'UTR;5'UTR;5'UTR |
| WG3465 | chr22:19746984-19747140 | TBX1;TBX1;TBX1 | Island | 5'UTR;5'UTR;5'UTR |
| WG3466 | chr22:19754125-19754448 | TBX1;TBX1;TBX1 | Island | Body;Body;Body |
| GM02706 | chr22:19754125-19754815 | TBX1;TBX1;TBX1 | Island | Body;Body;Body |
| WG2844 | chr22:19879696-19880135 | TXNRD2 | | Body |
| GM00438 | chr22:19950026-19950166 | COMT;COMT;COMT;COMT | | 5'UTR;5'UTR;5'UTR;TSS200 |
| GM00520 | chr22:20861470-20861479 | MED15;MED15 | N_Shore | TSS1500;TSS1500 |
| WG2844 | chr22:21321253-21321434 | AIFM3;AIFM3;AIFM3;AIFM3 | S_Shore | 5'UTR;5'UTR;TSS200;TSS200 |
| WG2121 | chr22:21987032-21987190 | CCDC116 | S_Shelf | TSS200 |
| GM09256 | chr22:22007154-22007528 | MIR130B;MIR301B | S_Shore | TSS1500;TSS200 |
| GM09255 | chr22:22020012-22020076 | PPIL2;PPIL2;PPIL2 | N_Shore | TSS1500;TSS1500;TSS1500 |
| WG3465 | chr22:22058080-22058271 | YPEL1 | Island | Body |
| GM09255 | chr22:22090796-22090816 | YPEL1 | Island | TSS1500 |
| GM02149 | chr22:22901648-22901697 | PRAME;PRAME;PRAME;PRAM | Island | TSS200;5'UTR;1stExon;1stExon;TSS20 |
| WG1310 | chr22:24181263-24181270 | DERL3;DERL3;DERL3 | Island | TSS200;TSS200;TSS200 |
| WG2843 | chr22:24348549-24348715 | GSTTP1 | | TSS1500 |
| GM02456 | chr22:24372913-24372926 | LOC391322 | Island | TSS1500 |
| WG2843 | chr22:24373054-24373322 | LOC391322 | Island | TSS200 |
| GM00042 | chr22:24384105-24384159 | GSTT1 | N_Shore | Body |
| WG2414 | chr22:24384105-24384294 | GSTT1 | N_Shore | Body |
| GM00439 | chr22:24384105-24384944 | GSTT1 | N_Shore | Body |
| GM02704 | chr22:24384159-24384294 | GSTT1 | Island | 1stExon |
| WG1977 | chr22:24384159-24384315 | GSTT1 | Island | 1stExon |
| WG3465 | chr22:24384245-24384315 | GSTT1;GSTT1 | Island | 5'UTR;1stExon |
| GM00042 | chr22:24384294-24384397 | GSTT1 | Island | TSS200 |
| WG2843 | chr22:24384340-24384397 | GSTT1 | Island | TSS200 |
| GM02704 | chr22:24384340-24384400 | GSTT1 | Island | TSS200 |
| GM02456 | chr22:24384340-24384525 | GSTT1 | Island | TSS200 |
| GM01392 | chr22:24384340-24384944 | GSTT1 | Island | TSS200 |
| GM00289 | chr22:24384393-24384525 | GSTT1 | Island | TSS200 |
| WG1283 | chr22:24384397-24384525 | GSTT1 | Island | TSS200 |
| WG3465 | chr22:24384400-24384944 | GSTT1 | Island | TSS200 |
| GM00042 | chr22:24384525-24384944 | GSTT1 | S_Shore | TSS1500 |
| WG2844 | chr22:24577142-24577315 | SUSD2 | | TSS1500 |
| GM02706 | chr22:24577223-24577288 | SUSD2 | | TSS1500 |
| WG1977 | chr22:24577223-24577315 | SUSD2 | | TSS1500 |
| WG3465 | chr22:24577288-24577438 | SUSD2 | | TSS200 |
| WG1721 | chr22:24891153-24891166 | C22orf45;C22orf45;UPB1 | Island | TSS1500;TSS1500;TSS200 |
| WG1722 | chr22:24891158-24891220 | C22orf45;C22orf45;UPB1 | Island | TSS1500;TSS1500;TSS200 |
| GM00940 | chr22:24979254-24979436 | GGT1 | | TSS1500 |
| WG2844 | chr22:24979254-24979554 | GGT1 | | TSS1500 |

| | | | | |
|---|---|---|---|---|
| WG1310 | chr22:24979254-24979588 | GGT1 | | TSS1500 |
| WG1983 | chr22:24979367-24979554 | GGT1 | | TSS1500 |
| WG2276 | chr22:24979436-24979588 | GGT1 | | TSS1500 |
| WG2120 | chr22:24979436-24979633 | GGT1 | | TSS1500 |
| WG3465 | chr22:24979554-24979588 | GGT1 | | TSS200 |
| GM02706 | chr22:24979554-24979633 | GGT1 | | TSS200 |
| WG1722 | chr22:24979554-24979964 | GGT1 | | TSS200 |
| GM00520 | chr22:24979633-24979660 | GGT1 | | TSS200 |
| WG3465 | chr22:24979660-24979964 | GGT1 | | TSS200 |
| WG1514 | chr22:25159879-25160098 | TOP1P2;PIWIL3 | Island | TSS1500;5'UTR |
| GM00940 | chr22:25160033-25160098 | TOP1P2;PIWIL3 | Island | TSS1500;5'UTR |
| GM00042 | chr22:25595697-25595814 | CRYBB3 | | TSS200 |
| WG1977 | chr22:29426511-29426832 | ZNRF3 | Island | Body |
| WG1722 | chr22:29426575-29426832 | ZNRF3 | Island | Body |
| WG2415 | chr22:29712045-29712472 | RASL10A;RASL10A | S_Shore | TSS1500;TSS1500 |
| GM03075 | chr22:30476089-30476285 | HORMAD2 | Island | TSS1500 |
| WG2121 | chr22:30476254-30476285 | HORMAD2 | Island | TSS200 |
| WG1722 | chr22:30476254-30476345 | HORMAD2 | Island | TSS200 |
| GM02175 | chr22:31318103-31318181 | C22orf27 | N_Shore | TSS200 |
| WG1722 | chr22:31318147-31318181 | C22orf27 | N_Shore | TSS200 |
| WG2276 | chr22:31318147-31318373 | C22orf27 | N_Shore | TSS200 |
| GM00042 | chr22:31318179-31318373 | C22orf27 | N_Shore | TSS200 |
| WG3465 | chr22:31318181-31318373 | C22orf27 | N_Shore | TSS200 |
| WG2415 | chr22:32027580-32027605 | PISD | S_Shore | TSS1500 |
| WG2276 | chr22:32438947-32439254 | SLC5A1 | N_Shore | TSS200 |
| GM00289 | chr22:32438947-32439259 | SLC5A1 | N_Shore | TSS200 |
| WG3465 | chr22:32438993-32439391 | SLC5A1 | N_Shore | TSS200 |
| GM01096 | chr22:32554996-32555275 | C22orf42 | | 1stExon |
| WG3465 | chr22:32599511-32599648 | RFPL2;RFPL2;RFPL2 | | TSS200;5'UTR;TSS1500 |
| GM01392 | chr22:32599516-32599613 | RFPL2;RFPL2;RFPL2 | | TSS200;5'UTR;TSS1500 |
| WG2121 | chr22:32599596-32599648 | RFPL2;RFPL2;RFPL2 | | TSS200;5'UTR;TSS1500 |
| GM00042 | chr22:32808489-32808786 | C22orf28 | S_Shore | TSS1500 |
| GM02456 | chr22:33196103-33196328 | SYN3;TIMP3;SYN3;SYN3 | N_Shore | Body;TSS1500;Body;Body |
| WG2276 | chr22:33196425-33196627 | SYN3;TIMP3;SYN3;SYN3 | N_Shore | Body;TSS1500;Body;Body |
| GM00289 | chr22:33196425-33197034 | SYN3;TIMP3;SYN3;SYN3 | N_Shore | Body;TSS1500;Body;Body |
| GM00520 | chr22:33454616-33454632 | SYN3 | S_Shore | TSS1500 |
| WG3465 | chr22:34046427-34046529 | LARGE;LARGE | Island | Body;Body |
| WG3465 | chr22:35795274-35795394 | MCM5 | N_Shore | TSS1500 |
| WG3465 | chr22:36013432-36013560 | MB;MB;MB;MB | | 1stExon;TSS200;5'UTR;5'UTR |
| GM02456 | chr22:36681296-36681321 | MYH9 | | Body |
| WG2415 | chr22:36960499-36960937 | CACNG2 | Island | Body |
| WG3465 | chr22:37215656-37216030 | PVALB | S_Shelf | TSS200 |
| WG2415 | chr22:37499649-37499859 | TMPRSS6;TMPRSS6 | | 1stExon;5'UTR |
| WG3465 | chr22:37545969-37546220 | IL2RB | | TSS200 |
| WG1722 | chr22:37640402-37640514 | RAC2 | | TSS200 |
| WG1977 | chr22:37824118-37824617 | ELFN2 | Island | TSS1500 |
| GM00520 | chr22:37976063-37976192 | LGALS2 | | TSS200 |

| | | | | |
|---|---|---|---|---|
| GM02706 | chr22:38035109-38035456 | SH3BP1 | N_Shore | TSS1500 |
| WG1513 | chr22:38218945-38219336 | GALR3 | N_Shore | TSS1500 |
| GM09256 | chr22:38713453-38713608 | CSNK1E;CSNK1E | S_Shore | 5'UTR;TSS200 |
| GM09256 | chr22:38714166-38714416 | CSNK1E;CSNK1E | S_Shore | TSS200;TSS1500 |
| WG2415 | chr22:38714287-38714426 | CSNK1E;CSNK1E | S_Shore | TSS200;TSS1500 |
| GM02175 | chr22:39377790-39377930 | APOBEC3B | | TSS1500 |
| WG1722 | chr22:39966585-39966711 | CACNA1I;CACNA1I | | TSS200;TSS200 |
| GM02704 | chr22:40057942-40058706 | CACNA1I;CACNA1I | Island | Body;Body |
| GM00941 | chr22:40058150-40058706 | CACNA1I;CACNA1I | Island | Body;Body |
| GM09256 | chr22:40296767-40297048 | GRAP2 | | TSS1500 |
| WG2276 | chr22:41956810-41956987 | CSDC2 | | TSS1500 |
| WG3465 | chr22:42078217-42078723 | NHP2L1;NHP2L1 | Island | Body;Body |
| GM00439 | chr22:42394590-42394659 | WBP2NL | Island | TSS200 |
| WG1978 | chr22:42394659-42395067 | WBP2NL | Island | TSS200 |
| WG1984 | chr22:42469976-42470106 | FAM109B | N_Shore | TSS1500 |
| WG1310 | chr22:42524984-42525279 | CYP2D6;CYP2D6 | Island | Body;Body |
| GM01096 | chr22:42828386-42828418 | NFAM1;NFAM1 | | 1stExon;5'UTR |
| WG3465 | chr22:42828411-42828418 | NFAM1 | | TSS200 |
| WG2844 | chr22:42828411-42828516 | NFAM1 | | TSS200 |
| GM00042 | chr22:43817258-43817384 | MPPED1 | Island | 5'UTR |
| GM02456 | chr22:44568337-44568423 | PARVG;PARVG | | TSS1500;TSS1500 |
| WG2415 | chr22:44568699-44568753 | PARVG;PARVG | | TSS200;TSS200 |
| GM02706 | chr22:44568725-44568770 | PARVG;PARVG | | TSS200;TSS200 |
| WG1722 | chr22:44568725-44568775 | PARVG;PARVG | | TSS200;TSS200 |
| GM02555 | chr22:44568725-44568812 | PARVG;PARVG | | TSS200;TSS200 |
| WG1310 | chr22:44568770-44568812 | PARVG;PARVG | | TSS200;TSS200 |
| WG2415 | chr22:44568775-44568913 | PARVG;PARVG | | TSS200;TSS200 |
| GM02456 | chr22:44708669-44708908 | KIAA1644;KIAA1644 | | 5'UTR;1stExon |
| GM00439 | chr22:44708794-44708908 | KIAA1644 | | TSS200 |
| WG2844 | chr22:45125218-45125480 | PRR5;PRR5;PRR5;PRR5-ARH | Island | Body;Body;Body;Body;Body;Body |
| WG2414 | chr22:45125218-45125666 | PRR5;PRR5;PRR5;PRR5-ARH | Island | Body;Body;Body;Body;Body;Body |
| GM02175 | chr22:45125218-45126040 | PRR5;PRR5;PRR5;PRR5-ARH | Island | Body;Body;Body;Body;Body;Body |
| WG2415 | chr22:45125395-45125666 | PRR5;PRR5;PRR5;PRR5-ARH | Island | Body;Body;Body;Body;Body;Body |
| WG2414 | chr22:45148218-45148229 | ARHGAP8;PRR5-ARHGAP8;ARHGAP8 | | TSS1500;Body;TSS1500 |
| GM00940 | chr22:45596948-45597727 | C22orf9;C22orf9;MIR1249 | N_Shore | Body;Body;TSS200 |
| WG3465 | chr22:45609402-45609421 | C22orf9;C22orf9 | | Body;TSS1500 |
| WG3466 | chr22:45680606-45680687 | UPK3A;UPK3A | N_Shore | TSS1500;TSS1500 |
| GM02706 | chr22:45809244-45809375 | SMC1B;RIBC2 | Island | Body;TSS1500 |
| WG2121 | chr22:45809244-45809396 | SMC1B;RIBC2 | Island | Body;TSS1500 |
| WG1977 | chr22:45809319-45809396 | SMC1B;RIBC2 | Island | Body;TSS1500 |
| GM09256 | chr22:45809319-45809402 | SMC1B;RIBC2 | Island | Body;TSS1500 |
| GM00940 | chr22:45809319-45809454 | SMC1B;RIBC2 | Island | Body;TSS1500 |
| WG2415 | chr22:46770063-46770440 | CELSR1 | N_Shore | Body |
| WG1513 | chr22:46805673-46805820 | CELSR1 | Island | Body |
| WG2844 | chr22:47022462-47022618 | GRAMD4 | Island | TSS200 |
| GM00520 | chr22:47069200-47069632 | GRAMD4 | N_Shore | Body |
| GM02555 | chr22:47069200-47069799 | GRAMD4 | N_Shore | Body |

| | | | | |
|---|---|---|---|---|
| GM00438 | chr22:50483123-50483545 | TTLL8 | N_Shore | Body |
| WG2415 | chr22:50483351-50483545 | TTLL8 | Island | Body |
| GM00520 | chr22:50523790-50523955 | MLC1;MLC1;MLC1 | | 1stExon;5'UTR;TSS200 |
| WG1977 | chr22:50523801-50524124 | MLC1;MLC1;MLC1 | | 1stExon;5'UTR;TSS200 |
| GM00439 | chr22:50523955-50524410 | MLC1;MLC1;MLC1 | | 1stExon;5'UTR;TSS200 |
| GM00042 | chr22:50524124-50524541 | MLC1;MLC1;MLC1 | | TSS1500;1stExon;5'UTR |
| GM00438 | chr22:50524374-50524541 | MLC1;MLC1 | N_Shelf | TSS1500;TSS200 |
| GM02317 | chr22:50528213-50528333 | MOV10L1;MOV10L1;MOV10L | Island | TSS1500;TSS1500;TSS1500 |
| WG1977 | chr22:50528282-50528333 | MOV10L1;MOV10L1;MOV10L | Island | TSS200;TSS200;TSS1500 |
| GM00520 | chr22:50528298-50528333 | MOV10L1;MOV10L1;MOV10L | Island | TSS200;TSS200;TSS1500 |
| WG2415 | chr22:50528333-50528641 | MOV10L1;MOV10L1;MOV10L | Island | TSS200;TSS200;TSS1500 |
| GM02175 | chr22:50584184-50585400 | MOV10L1;MOV10L1;MOV10L1;MOV10L1 | | Body;TSS1500;Body;Body |
| WG1977 | chr22:50584184-50585427 | MOV10L1;MOV10L1;MOV10L1;MOV10L1 | | Body;TSS1500;Body;Body |
| WG1722 | chr22:50585197-50585427 | MOV10L1;MOV10L1;MOV10L1;MOV10L1 | | Body;TSS1500;Body;Body |
| GM02173 | chr22:50585216-50585393 | MOV10L1;MOV10L1;MOV10L1;MOV10L1 | | Body;TSS1500;Body;Body |
| GM02706 | chr22:50585216-50585427 | MOV10L1;MOV10L1;MOV10L1;MOV10L1 | | Body;TSS1500;Body;Body |
| GM00290 | chr22:50617003-50617460 | PANX2;PANX2;PANX2 | Island | Body;Body;Body |
| WG3466 | chr22:50629038-50629224 | TRABD | Island | 5'UTR |
| GM02555 | chr22:50985681-50986031 | KLHDC7B | Island | TSS1500 |
| WG3466 | chr22:51016638-51016646 | CPT1B;CPT1B;CPT1B;CHKB-( | Island | 5'UTR;5'UTR;TSS200;Body;5'UTR;5'UT |
| GM00289 | chr22:51016638-51016703 | CPT1B;CPT1B;CPT1B;CHKB-( | Island | 5'UTR;5'UTR;TSS200;Body;5'UTR;5'UT |
| GM00520 | chr22:51017151-51017166 | CPT1B;CPT1B;CPT1B;CPT1B | S_Shore | TSS200;TSS200;TSS1500;TSS1500;Bo |
| GM01096 | chr22:51041213-51041732 | MAPK8IP2;MAPK8IP2 | N_Shore | TSS1500;Body |
| GM02175 | chr3:100053070-100053188 | NIT2 | N_Shore | TSS1500 |
| GM09255 | chr3:101406220-101406543 | RPL24 | S_Shore | TSS1500 |
| GM02706 | chr3:101497475-101497779 | FAM55C;FAM55C | N_Shore | TSS1500;TSS1500 |
| WG1513 | chr3:10149803-10149918 | C3orf24;C3orf24;C3orf24;C3 | Island | 1stExon;1stExon;5'UTR;5'UTR |
| WG1977 | chr3:10149882-10149963 | C3orf24;C3orf24;C3orf24;C3 | Island | 1stExon;1stExon;5'UTR;5'UTR |
| WG1978 | chr3:10149882-10150083 | C3orf24;C3orf24;C3orf24;C3 | Island | 1stExon;1stExon;5'UTR;5'UTR |
| GM00438 | chr3:10149963-10149979 | C3orf24;C3orf24 | Island | TSS200;TSS200 |
| WG1513 | chr3:10149974-10150083 | C3orf24;C3orf24 | Island | TSS200;TSS200 |
| WG1721 | chr3:10149979-10150083 | C3orf24;C3orf24 | Island | TSS200;TSS200 |
| WG1513 | chr3:10206254-10206344 | IRAK2 | N_Shore | TSS1500 |
| WG2415 | chr3:10370507-10370704 | ATP2B2;ATP2B2 | Island | Body;Body |
| WG3465 | chr3:107810507-107810687 | CD47;CD47;CD47 | Island | TSS1500;TSS1500;TSS1500 |
| GM02175 | chr3:10806022-10806175 | LOC285370 | | TSS200 |
| WG3465 | chr3:10806047-10806133 | LOC285370 | | TSS200 |
| GM02149 | chr3:108475765-108476163 | RETNLB | | Body |
| GM01096 | chr3:108837052-108837069 | MORC1 | Island | TSS200 |
| WG1513 | chr3:11058836-11058924 | SLC6A1 | Island | 5'UTR |
| GM00935 | chr3:111577634-111577743 | PHLDB2;PHLDB2;PHLDB2;PH | N_Shore | TSS1500;Body;TSS1500;TSS1500 |
| GM01096 | chr3:112013079-112013231 | SLC9A10 | | TSS200 |
| WG2843 | chr3:112358351-112358468 | CCDC80;CCDC80 | | Body;Body |
| WG2276 | chr3:11267020-11267098 | HRH1;HRH1;HRH1 | | 5'UTR;TSS1500;5'UTR |
| WG2844 | chr3:11267020-11267525 | HRH1;HRH1;HRH1 | | 5'UTR;TSS1500;5'UTR |
| GM01375 | chr3:113160183-113160623 | WDR52;WDR52 | N_Shore | 5'UTR;5'UTR |
| WG2276 | chr3:113160183-113160637 | WDR52;WDR52 | N_Shore | 5'UTR;5'UTR |

| | | | | |
|---|---|---|---|---|
| GM00289 | chr3:113160313-113160437 | WDR52;WDR52;WDR52;WDF | Island | 5'UTR;1stExon;5'UTR;1stExon |
| WG3465 | chr3:1133865-1134366 | CNTN6 | | TSS1500 |
| WG2415 | chr3:114102565-114102615 | ZBTB20;ZBTB20;ZBTB20;ZBTB20;ZBTB20 | | 5'UTR;5'UTR;5'UTR;5'UTR;5'UTR;5'UTF |
| WG2194 | chr3:114790243-114790596 | ZBTB20;ZBTB20;ZBTB20 | | 1stExon;5'UTR;5'UTR |
| WG1309 | chr3:11610239-11610613 | VGLL4;VGLL4;VGLL4;VGLL4; | Island | Body;Body;1stExon;Body;5'UTR |
| WG3466 | chr3:11746183-11746765 | VGLL4 | N_Shore | 5'UTR |
| GM01376 | chr3:11746226-11746765 | VGLL4 | N_Shore | 5'UTR |
| WG1984 | chr3:11746491-11746765 | VGLL4 | N_Shore | 5'UTR |
| GM02149 | chr3:118663470-118663642 | IGSF11;IGSF11 | | Body;Body |
| GM00289 | chr3:118864870-118864952 | C3orf30;IGSF11;IGSF11 | | TSS200;1stExon;5'UTR |
| GM00042 | chr3:118864974-118865028 | C3orf30;IGSF11 | | TSS200;TSS200 |
| GM02704 | chr3:119298195-119298236 | ADPRH | N_Shore | TSS1500 |
| WG3465 | chr3:121379745-121379791 | HCLS1;HCLS1 | | 1stExon;5'UTR |
| WG3465 | chr3:121795768-121796627 | CD86;CD86 | | TSS1500;Body |
| GM02706 | chr3:121902497-121902510 | CASR | N_Shore | TSS200 |
| WG1513 | chr3:121949083-121949308 | CASR | N_Shore | 5'UTR |
| GM00439 | chr3:12199836-12200324 | SYN2;TIMP4;SYN2 | N_Shore | Body;Body;Body |
| GM01096 | chr3:12200236-12200427 | TIMP4;SYN2;SYN2 | Island | 1stExon;Body;Body |
| WG1513 | chr3:122281881-122281975 | PARP9;PARP9;DTX3L;PARP9; | N_Shore | 5'UTR;5'UTR;TSS1500;5'UTR;5'UTR;5'U |
| WG2276 | chr3:122640778-122641029 | SEMA5B | Island | Body |
| GM01095 | chr3:123339334-123339568 | MYLK;MYLK;MYLK;MYLK;MYLK;MYLK;MYI | | 1stExon;5'UTR;Body;Body;Body;5'UTR |
| WG1722 | chr3:123699208-123699389 | ROPN1 | | Body |
| GM02456 | chr3:128204413-128204927 | GATA2;GATA2;GATA2 | N_Shore | Body;Body;Body |
| WG2120 | chr3:128204813-128205143 | GATA2;GATA2;GATA2 | N_Shore | Body;Body;Body |
| GM00520 | chr3:128204813-128205274 | GATA2;GATA2;GATA2 | N_Shore | Body;Body;Body |
| WG1513 | chr3:128292593-128292718 | C3orf27 | | 5'UTR |
| WG2276 | chr3:128722515-128723005 | CCDC48 | Island | Body |
| WG1513 | chr3:129407542-129407548 | TMCC1;TMCC1;TMCC1 | | TSS200;Body;Body |
| GM02456 | chr3:13060741-13060974 | IQSEC1 | | Body |
| WG1978 | chr3:13060915-13060974 | IQSEC1 | | Body |
| WG2276 | chr3:130745442-130745497 | NEK11;ASTE1;ASTE1;NEK11; | Island | TSS1500;1stExon;5'UTR;TSS1500;TSS |
| WG2120 | chr3:130745442-130745839 | NEK11;ASTE1;ASTE1;NEK11; | Island | TSS1500;1stExon;5'UTR;TSS1500;TSS |
| WG2276 | chr3:130745654-130745839 | NEK11;NEK11;NEK11;ASTE1 | Island | TSS200;TSS200;TSS200;TSS200 |
| GM02706 | chr3:133191270-133191407 | BFSP2 | | Body |
| WG1283 | chr3:136537169-136537752 | TMEM22;TMEM22;TMEM22 | N_Shore | TSS1500;TSS1500;TSS1500 |
| WG2415 | chr3:136537267-136537709 | TMEM22;TMEM22;TMEM22 | N_Shore | TSS1500;TSS1500;TSS1500 |
| GM02706 | chr3:136537607-136537752 | TMEM22;TMEM22;TMEM22 | Island | TSS1500;TSS1500;TSS1500 |
| GM00520 | chr3:13663398-13663538 | FBLN2;FBLN2;FBLN2 | | Body;Body;Body |
| GM01095 | chr3:137893707-137893743 | DBR1;DBR1 | Island | 1stExon;5'UTR |
| WG3465 | chr3:138048926-138049011 | TXNDC6 | S_Shore | TSS200 |
| GM09256 | chr3:138152837-138153127 | ESYT3 | N_Shore | TSS1500 |
| GM02175 | chr3:138152837-138153170 | ESYT3 | N_Shore | TSS1500 |
| WG2415 | chr3:138668832-138669322 | C3orf72 | Island | Body |
| GM09255 | chr3:138725189-138725214 | PRR23A | Island | TSS200 |
| GM01096 | chr3:138739874-138739916 | PRR23B | Island | TSS200 |
| WG3465 | chr3:138739874-138739922 | PRR23B | Island | TSS200 |
| GM00289 | chr3:138739900-138739922 | PRR23B | Island | TSS200 |

| | | | | |
|---|---|---|---|---|
| WG1977 | chr3:139257775-139257907 | RBP1;RBP1;RBP1 | Island | Body;Body;Body |
| WG1513 | chr3:139257799-139257907 | RBP1;RBP1;RBP1 | Island | Body;Body;Body |
| WG2121 | chr3:139257866-139258166 | RBP1;RBP1;RBP1 | Island | Body;Body;Body |
| WG3465 | chr3:139346734-139346818 | NMNAT3 | | 5'UTR |
| WG1513 | chr3:140401765-140401955 | TRIM42 | Island | Body |
| WG3465 | chr3:14443428-14443473 | SLC6A6;SLC6A6;SLC6A6 | N_Shore | TSS1500;TSS1500;TSS1500 |
| GM00031 | chr3:146261991-146262421 | PLSCR1 | N_Shore | 5'UTR |
| WG1309 | chr3:146261991-146262761 | PLSCR1 | N_Shore | 5'UTR |
| WG3466 | chr3:147105899-147106208 | ZIC4;ZIC4;ZIC4;ZIC4;ZIC4 | N_Shelf | 3'UTR;Body;3'UTR;Body;3'UTR |
| WG2120 | chr3:147105899-147106890 | ZIC4;ZIC4;ZIC4;ZIC4;ZIC4 | N_Shelf | 3'UTR;Body;3'UTR;Body;3'UTR |
| WG2121 | chr3:147106489-147106890 | ZIC4;ZIC4;ZIC4;ZIC4;ZIC4 | N_Shelf | 3'UTR;Body;3'UTR;Body;3'UTR |
| WG3466 | chr3:147106561-147107109 | ZIC4;ZIC4;ZIC4;ZIC4;ZIC4 | N_Shore | 3'UTR;Body;3'UTR;Body;3'UTR |
| WG1513 | chr3:147110667-147112605 | ZIC4;ZIC4;ZIC4;ZIC4;ZIC4 | Island | Body;TSS1500;Body;Body;TSS1500 |
| WG2120 | chr3:147112081-147112143 | ZIC4;ZIC4;ZIC4 | N_Shore | Body;Body;Body |
| WG2415 | chr3:147112081-147112605 | ZIC4;ZIC4;ZIC4 | N_Shore | Body;Body;Body |
| WG2121 | chr3:147112096-147112605 | ZIC4;ZIC4;ZIC4 | N_Shore | Body;Body;Body |
| GM01376 | chr3:147112143-147112605 | ZIC4;ZIC4;ZIC4 | N_Shore | Body;Body;Body |
| WG2275 | chr3:147112438-147112605 | ZIC4;ZIC4;ZIC4 | N_Shore | Body;Body;Body |
| GM00935 | chr3:147113700-147113918 | ZIC4;ZIC4;ZIC4 | Island | Body;Body;Body |
| WG3466 | chr3:147123199-147123457 | ZIC4;ZIC4;ZIC4 | N_Shelf | 5'UTR;TSS1500;1stExon |
| WG2120 | chr3:147123457-147124543 | ZIC4;ZIC4;ZIC4 | N_Shelf | 5'UTR;TSS1500;TSS200 |
| WG2415 | chr3:147124363-147124523 | ZIC4;ZIC4;ZIC4 | N_Shelf | 1stExon;5'UTR;TSS1500 |
| WG3466 | chr3:147124363-147124543 | ZIC4;ZIC4;ZIC4 | N_Shelf | 1stExon;5'UTR;TSS1500 |
| WG2120 | chr3:147124827-147125287 | ZIC4 | N_Shelf | TSS1500 |
| WG3466 | chr3:147125114-147125955 | ZIC4 | N_Shore | TSS1500 |
| WG2121 | chr3:147125712-147125924 | ZIC4;ZIC1 | N_Shore | TSS1500;TSS1500 |
| WG2120 | chr3:147125758-147125955 | ZIC4;ZIC1 | N_Shore | TSS1500;TSS1500 |
| WG2121 | chr3:147125942-147126244 | ZIC1 | N_Shore | TSS1500 |
| WG2276 | chr3:147129778-147129952 | ZIC1 | N_Shore | Body |
| WG3466 | chr3:147130536-147131332 | ZIC1 | Island | Body |
| WG2121 | chr3:147131146-147131860 | ZIC1 | Island | Body |
| WG1309 | chr3:148847273-148847295 | HPS3 | Island | TSS200 |
| GM00289 | chr3:150690357-150690456 | CLRN1;CLRN1OS | | 1stExon;TSS200 |
| GM01376 | chr3:150690403-150690585 | CLRN1;CLRN1OS | | 1stExon;TSS200 |
| GM00520 | chr3:150690456-150690810 | CLRN1;CLRN1OS | | 1stExon;TSS200 |
| WG3465 | chr3:150996078-150996563 | P2RY14;MED12L;P2RY14 | | 1stExon;Body;5'UTR |
| WG1310 | chr3:15311021-15311183 | SH3BP5;SH3BP5 | | Body;Body |
| WG3466 | chr3:15311021-15311269 | SH3BP5;SH3BP5 | | Body;Body |
| GM02175 | chr3:15311154-15311269 | SH3BP5;SH3BP5 | | Body;Body |
| WG2194 | chr3:154042278-154042375 | DHX36;DHX36;DHX36;DHX3( | Island | 1stExon;5'UTR;1stExon;5'UTR |
| WG1977 | chr3:154042611-154042778 | DHX36;DHX36 | S_Shore | TSS1500;TSS1500 |
| WG3465 | chr3:155394109-155394303 | PLCH1;PLCH1;PLCH1 | | 5'UTR;TSS200;TSS200 |
| WG2415 | chr3:155394109-155394428 | PLCH1;PLCH1;PLCH1 | | 5'UTR;TSS200;TSS200 |
| WG1721 | chr3:155394118-155394303 | PLCH1;PLCH1;PLCH1 | | 5'UTR;TSS200;TSS200 |
| WG1721 | chr3:155570275-155570809 | SLC33A1 | N_Shelf | Body |
| WG1721 | chr3:156273297-156273458 | SSR3 | S_Shore | TSS1500 |
| WG1977 | chr3:157217187-157217414 | VEPH1;VEPH1;VEPH1;VEPH1;VEPH1;VEI | | 5'UTR;5'UTR;5'UTR;1stExon;5'UTR;5'U |

| | | | | |
|---|---|---|---|---|
| GM02706 | chr3:157217187-157217435 | VEPH1;VEPH1;VEPH1;VEPH1;VEPH1;VEI | | 5'UTR;5'UTR;5'UTR;1stExon;5'UTR;5'U |
| GM01096 | chr3:157217414-157217607 | VEPH1;VEPH1;VEPH1;VEPH1;VEPH1;VEI | | 5'UTR;5'UTR;5'UTR;1stExon;5'UTR;5'U |
| WG1977 | chr3:157217556-157217569 | VEPH1;VEPH1;VEPH1;VEPH1;VEPH1 | | TSS200;5'UTR;5'UTR;5'UTR;5'UTR |
| GM02706 | chr3:157217569-157217842 | VEPH1;VEPH1;VEPH1;VEPH1;VEPH1 | | TSS200;5'UTR;5'UTR;5'UTR;5'UTR |
| GM02175 | chr3:157821262-157821536 | SHOX2;SHOX2;SHOX2 | Island | Body;Body;Body |
| WG3466 | chr3:157821262-157821994 | SHOX2;SHOX2;SHOX2 | Island | Body;Body;Body |
| WG2121 | chr3:157821262-157822072 | SHOX2;SHOX2;SHOX2 | Island | Body;Body;Body |
| GM09255 | chr3:157821407-157822072 | SHOX2;SHOX2;SHOX2 | Island | Body;Body;Body |
| WG3465 | chr3:157821987-157822072 | SHOX2;SHOX2;SHOX2 | N_Shore | Body;Body;Body |
| GM03075 | chr3:157824714-157825196 | SHOX2;SHOX2;SHOX2 | S_Shore | TSS1500;TSS1500;TSS1500 |
| WG1722 | chr3:158390074-158390336 | GFM1;LXN | | Body;Body |
| GM00042 | chr3:158390074-158390525 | GFM1;LXN | | Body;Body |
| WG2276 | chr3:158390329-158390336 | LXN;LXN;GFM1 | | 1stExon;5'UTR;Body |
| GM00016 | chr3:158390329-158390632 | LXN;LXN;GFM1 | | 1stExon;5'UTR;Body |
| WG2194 | chr3:158390332-158390525 | LXN;LXN;GFM1 | | 1stExon;5'UTR;Body |
| GM09255 | chr3:158390336-158390525 | LXN;LXN;GFM1 | | 1stExon;5'UTR;Body |
| GM09256 | chr3:158518888-158519176 | MFSD1;MFSD1 | N_Shore | TSS1500;TSS1500 |
| WG2275 | chr3:158786187-158786242 | IQCJ;IQCJ | | TSS1500;TSS1500 |
| WG1722 | chr3:160475035-160475102 | PPM1L | S_Shore | Body |
| WG2415 | chr3:160475035-160475128 | PPM1L | S_Shore | Body |
| WG1513 | chr3:160475090-160475128 | PPM1L | S_Shore | Body |
| GM02706 | chr3:168863860-168864101 | MECOM;MECOM;MECOM;MI | N_Shore | 5'UTR;Body;5'UTR;5'UTR;5'UTR;5'UTR |
| WG3465 | chr3:169377725-169377946 | MECOM | N_Shore | Body |
| GM03075 | chr3:169378675-169378764 | MECOM | N_Shore | Body |
| WG2121 | chr3:169483122-169483295 | TERC | S_Shore | TSS1500 |
| WG1721 | chr3:169683976-169684239 | LOC100128164;SEC62;LOC1 | N_Shore | Body;TSS1500;Body |
| GM03075 | chr3:169781404-169781622 | GPR160 | | 5'UTR |
| WG3465 | chr3:169781495-169781717 | GPR160 | | 5'UTR |
| GM01095 | chr3:169781622-169781717 | GPR160 | | 5'UTR |
| GM02706 | chr3:170137240-170138205 | CLDN11 | Island | Body |
| GM00290 | chr3:170137321-170138205 | CLDN11 | Island | Body |
| WG3466 | chr3:170746265-170746586 | SLC2A2 | Island | TSS1500 |
| WG1722 | chr3:171175866-171176016 | TNIK;TNIK;TNIK;TNIK;TNIK;TN | N_Shore | Body;Body;Body;Body;Body;Body;Bod |
| WG1977 | chr3:172241069-172241384 | TNFSF10 | | 1stExon |
| GM00289 | chr3:172858917-172859049 | SPATA16;SPATA16 | | 1stExon;5'UTR |
| WG1513 | chr3:172858969-172859699 | SPATA16;SPATA16 | | 1stExon;5'UTR |
| GM00940 | chr3:178275369-178275698 | KCNMB2;KCNMB2 | | TSS1500;5'UTR |
| WG1283 | chr3:178978922-178978993 | KCNMB3;KCNMB3;KCNMB3; | N_Shore | TSS1500;Body;TSS1500;Body |
| WG1514 | chr3:178978985-178979013 | KCNMB3;KCNMB3;KCNMB3; | Island | TSS1500;Body;TSS1500;Body |
| GM02149 | chr3:178984747-178984910 | KCNMB3;KCNMB3;KCNMB3;KCNMB3 | | 5'UTR;5'UTR;1stExon;1stExon |
| WG2193 | chr3:178984841-178984973 | KCNMB3;KCNMB3 | | TSS200;TSS200 |
| WG1984 | chr3:181421548-181421951 | SOX2OT | | Body |
| WG1984 | chr3:181428280-181428329 | SOX2OT;SOX2 | N_Shore | Body;TSS1500 |
| GM02147 | chr3:181428307-181428377 | SOX2OT;SOX2 | N_Shore | Body;TSS1500 |
| WG1984 | chr3:181428462-181428563 | SOX2OT;SOX2 | N_Shore | Body;TSS1500 |
| WG3466 | chr3:181437145-181437383 | SOX2OT | N_Shore | Body |
| WG2844 | chr3:181437145-181437548 | SOX2OT | N_Shore | Body |

| | | | | |
|---|---|---|---|---|
| WG3466 | chr3:181437712-181438328 | SOX2OT | S_Shore | Body |
| GM02704 | chr3:181441384-181441571 | SOX2OT | N_Shelf | Body |
| WG1984 | chr3:181441501-181441806 | SOX2OT | N_Shelf | Body |
| WG1984 | chr3:181442676-181443009 | SOX2OT | N_Shore | Body |
| WG1984 | chr3:181444757-181445385 | SOX2OT | Island | Body |
| GM02704 | chr3:181444999-181445385 | SOX2OT | Island | Body |
| GM02640 | chr3:182817338-182817389 | MCCC1;MCCC1 | Island | 1stExon;5'UTR |
| WG3465 | chr3:182970675-182970728 | MCF2L2;B3GNT5 | N_Shore | Body;TSS1500 |
| WG1721 | chr3:183273442-183273758 | KLHL6;KLHL6 | | 5'UTR;1stExon |
| GM02175 | chr3:183273557-183273807 | KLHL6 | | TSS200 |
| WG1513 | chr3:183894727-183894778 | AP2M1;AP2M1 | S_Shore | 5'UTR;5'UTR |
| WG1983 | chr3:183903129-183903315 | ABCF3 | N_Shore | TSS1500 |
| GM00290 | chr3:183958478-183959073 | VWA5B2;MIR1224 | N_Shore | Body;TSS1500 |
| WG3465 | chr3:183959062-183959088 | VWA5B2;MIR1224 | Island | Body;TSS200 |
| WG2844 | chr3:184099096-184099432 | CHRD | Island | Body |
| WG3466 | chr3:185080218-185080423 | MAP3K13 | | TSS1500 |
| GM00941 | chr3:185911702-185911885 | DGKG;DGKG;DGKG | Island | Body;Body;Body |
| GM00290 | chr3:185912227-185912486 | DGKG;DGKG;DGKG | Island | Body;Body;Body |
| WG2415 | chr3:186383386-186383874 | HRG | | TSS1500 |
| WG1977 | chr3:189348936-189349021 | TP63;TP63;TP63 | | TSS1500;TSS1500;TSS1500 |
| GM02456 | chr3:189348936-189349129 | TP63;TP63;TP63 | | TSS1500;TSS1500;TSS1500 |
| WG1657 | chr3:191048377-191048439 | UTS2D;CCDC50;CCDC50 | S_Shore | TSS200;Body;Body |
| GM00940 | chr3:191048377-191048753 | UTS2D;CCDC50;CCDC50 | S_Shore | TSS200;Body;Body |
| GM02149 | chr3:191178384-191178924 | PYDC2 | | TSS1500 |
| WG3466 | chr3:192289245-192289293 | FGF12 | | Body |
| WG1513 | chr3:193096009-193096626 | ATP13A5 | | Body |
| GM00940 | chr3:194319463-194319725 | TMEM44;TMEM44;TMEM44;TMEM44 | | Body;Body;Body;Body |
| WG2275 | chr3:194319463-194319931 | TMEM44;TMEM44;TMEM44;TMEM44 | | Body;Body;Body;Body |
| WG1983 | chr3:194319644-194319931 | TMEM44;TMEM44;TMEM44;TMEM44 | | Body;Body;Body;Body |
| GM01095 | chr3:194319644-194320017 | TMEM44;TMEM44;TMEM44;TMEM44 | | Body;Body;Body;Body |
| WG2415 | chr3:194833887-194834008 | C3orf21 | | Body |
| WG2193 | chr3:195488725-195489789 | MUC4;MUC4;MUC4 | Island | Body;Body;Body |
| GM02456 | chr3:195488922-195489789 | MUC4;MUC4;MUC4 | Island | Body;Body;Body |
| WG1283 | chr3:195489306-195489782 | MUC4;MUC4;MUC4 | Island | Body;Body;Body |
| WG2415 | chr3:195489306-195489789 | MUC4;MUC4;MUC4 | Island | Body;Body;Body |
| WG1978 | chr3:195489789-195490033 | MUC4;MUC4;MUC4 | Island | Body;Body;Body |
| GM09256 | chr3:195528460-195528709 | MUC4;MUC4;MUC4 | | Body;Body;Body |
| GM03075 | chr3:195530091-195530151 | MUC4;MUC4;MUC4 | | Body;Body;Body |
| GM00520 | chr3:195530133-195530532 | MUC4;MUC4;MUC4 | | Body;Body;Body |
| GM00438 | chr3:195929389-195929544 | ZDHHC19 | | Body |
| WG2276 | chr3:196065289-196065318 | TM4SF19 | | TSS200 |
| WG2275 | chr3:196065357-196065506 | TM4SF19 | | TSS200 |
| GM00935 | chr3:196065357-196065569 | TM4SF19 | | TSS200 |
| GM09256 | chr3:196750506-196750939 | MFI2;MFI2 | | Body;Body |
| GM00438 | chr3:24537050-24537182 | THRB;THRB;THRB | Island | TSS1500;TSS1500;TSS1500 |
| WG1283 | chr3:24537277-24537309 | THRB;THRB;THRB | Island | TSS1500;TSS1500;TSS1500 |
| GM01095 | chr3:24537277-24537338 | THRB;THRB;THRB | Island | TSS1500;TSS1500;TSS1500 |

| | | | | |
|---|---|---|---|---|
| GM01650 | chr3:24537277-24537407 | THRB;THRB;THRB | Island | TSS1500;TSS1500;TSS1500 |
| WG3465 | chr3:24537309-24537349 | THRB;THRB;THRB | Island | TSS1500;TSS1500;TSS1500 |
| WG3466 | chr3:24537309-24537407 | THRB;THRB;THRB | Island | TSS1500;TSS1500;TSS1500 |
| GM00438 | chr3:24537338-24537407 | THRB;THRB;THRB | Island | TSS1500;TSS1500;TSS1500 |
| GM00439 | chr3:25469860-25469991 | RARB;RARB;RARB;RARB | | 5'UTR;1stExon;1stExon;5'UTR |
| GM00290 | chr3:25469914-25469991 | RARB;RARB;RARB;RARB | | 5'UTR;1stExon;1stExon;5'UTR |
| WG3465 | chr3:26665815-26666151 | LRRC3B | N_Shore | 5'UTR |
| WG3465 | chr3:30647540-30647672 | TGFBR2;TGFBR2 | N_Shore | TSS1500;TSS1500 |
| WG2415 | chr3:3080327-3080621 | CNTN4;CNTN4;CNTN4 | | Body;TSS1500;Body |
| GM02149 | chr3:30936070-30936318 | GADL1 | Island | 1stExon |
| WG1722 | chr3:3151899-3152374 | IL5RA;IL5RA;IL5RA;IL5RA;IL5RA;IL5RA;IL | | 5'UTR;5'UTR;1stExon;5'UTR;1stExon;1 |
| WG3466 | chr3:32023402-32023473 | ZNF860;OSBPL10 | S_Shore | 5'UTR;TSS200 |
| WG1513 | chr3:32860909-32861318 | TRIM71 | N_Shore | Body |
| WG2193 | chr3:32861150-32861407 | TRIM71 | Island | Body |
| GM02706 | chr3:350682-350892 | CHL1 | Island | 5'UTR |
| WG2120 | chr3:350682-351003 | CHL1 | Island | 5'UTR |
| GM00438 | chr3:35720994-35721166 | ARPP-21;ARPP-21;ARPP-21;ARPP-21 | | 5'UTR;TSS200;TSS1500;TSS200 |
| GM02706 | chr3:35785735-35785897 | MIR128-2;ARPP-21 | | TSS1500;Body |
| WG1513 | chr3:35785735-35786480 | MIR128-2;ARPP-21 | | TSS1500;Body |
| WG1722 | chr3:35785890-35786480 | MIR128-2;ARPP-21 | | TSS200;Body |
| WG1977 | chr3:35785958-35786480 | MIR128-2;ARPP-21 | | TSS200;Body |
| WG2415 | chr3:36949957-36950004 | TRANK1 | | Body |
| WG2415 | chr3:37033373-37033632 | EPM2AIP1;MLH1 | N_Shore | 1stExon;TSS1500 |
| GM00438 | chr3:38080635-38080704 | DLEC1;DLEC1 | Island | TSS200;TSS200 |
| GM02706 | chr3:38080635-38080790 | DLEC1;DLEC1 | Island | TSS200;TSS200 |
| GM01096 | chr3:39424697-39424992 | SLC25A38 | Island | TSS200 |
| WG1513 | chr3:39543515-39543776 | MOBP;MOBP | N_Shore | 5'UTR;Body |
| GM03073 | chr3:39543547-39544192 | MOBP;MOBP | N_Shore | 5'UTR;Body |
| WG3466 | chr3:39543776-39544192 | MOBP;MOBP | Island | Body;Body |
| GM09256 | chr3:39543967-39544326 | MOBP;MOBP | Island | Body;Body |
| GM02706 | chr3:40428643-40428807 | ENTPD3 | N_Shore | TSS200 |
| WG1983 | chr3:42306493-42306580 | CCK | Island | TSS200 |
| WG3465 | chr3:42307761-42307866 | CCK | S_Shore | TSS1500 |
| WG3465 | chr3:42700203-42700354 | ZBTB47 | Island | Body |
| GM00520 | chr3:42727014-42727489 | KBTBD5;KBTBD5 | N_Shore | 5'UTR;1stExon |
| GM02555 | chr3:42727425-42727843 | KBTBD5 | Island | 1stExon |
| WG1513 | chr3:42742993-42743203 | HHATL;HHATL | | Body;5'UTR |
| WG3465 | chr3:42947263-42947385 | ZNF662;ZNF662 | N_Shore | TSS1500;TSS200 |
| WG1513 | chr3:44802549-44802604 | KIF15;KIAA1143 | N_Shore | TSS1500;Body |
| GM00439 | chr3:44802566-44802754 | KIF15;KIAA1143 | N_Shore | TSS1500;Body |
| WG3466 | chr3:45077254-45077649 | CLEC3B | Island | Body |
| GM02149 | chr3:45077254-45077920 | CLEC3B | Island | Body |
| GM00289 | chr3:45077369-45077765 | CLEC3B | Island | Body |
| GM02173 | chr3:45077369-45077920 | CLEC3B | Island | Body |
| GM00289 | chr3:45187717-45188186 | CDCP1;CDCP1 | Island | 1stExon;1stExon |
| GM00520 | chr3:46448579-46448671 | CCRL2;CCRL2 | | TSS1500;TSS200 |
| GM00289 | chr3:46448671-46448868 | CCRL2;CCRL2 | | TSS1500;TSS200 |

| | | | | |
|---|---|---|---|---|
| WG1977 | chr3:46448753-46448963 | CCRL2;CCRL2;CCRL2 | | TSS1500;5'UTR;1stExon |
| WG1722 | chr3:46506552-46506864 | LTF | S_Shore | TSS200 |
| WG2844 | chr3:46718369-46719086 | ALS2CL;ALS2CL | | Body;5'UTR |
| WG1721 | chr3:46759472-46759588 | PRSS50 | Island | TSS200 |
| GM02456 | chr3:46906118-46906385 | MYL3 | | TSS1500 |
| WG2844 | chr3:48310544-48310897 | ZNF589 | | 3'UTR |
| GM02149 | chr3:48632484-48632582 | COL7A1 | Island | Body |
| WG1513 | chr3:48694451-48694534 | CELSR3 | Island | Body |
| WG3465 | chr3:48694454-48694673 | CELSR3 | Island | Body |
| WG1513 | chr3:49057761-49057813 | NDUFAF3;DALRD3;MIR425;N | N_Shore | TSS200;5'UTR;TSS200;TSS1500;TSS15 |
| GM00042 | chr3:49057884-49058043 | NDUFAF3;DALRD3;NDUFAF3 | N_Shore | TSS200;5'UTR;TSS1500;TSS1500;TSS1 |
| WG1722 | chr3:49203973-49204239 | CCDC71 | S_Shore | TSS200 |
| GM00438 | chr3:49203973-49204548 | CCDC71 | S_Shore | TSS200 |
| WG1309 | chr3:49314670-49314814 | USP4;USP4;C3orf62 | Island | 3'UTR;3'UTR;TSS200 |
| WG1513 | chr3:49459909-49459940 | AMT;AMT;AMT;NICN1;AMT;AMT;AMT;AMT | | Body;5'UTR;1stExon;3'UTR;1stExon;1s |
| WG3465 | chr3:49940245-49941151 | MST1R | Island | 1stExon |
| GM01376 | chr3:49940853-49941151 | MST1R | N_Shore | 1stExon |
| GM03075 | chr3:49941378-49941470 | MST1R | Island | TSS200 |
| GM00940 | chr3:49941408-49941425 | MST1R | Island | TSS200 |
| GM00289 | chr3:50231845-50232343 | GNAT1;GNAT1 | Island | Body;Body |
| GM02147 | chr3:50304179-50304250 | SEMA3B;SEMA3B | | TSS1500;TSS1500 |
| GM00520 | chr3:50375459-50375483 | RASSF1;RASSF1;RASSF1;RAS | Island | TSS1500;Body;Body;5'UTR |
| GM09255 | chr3:50487955-50488230 | CACNA2D2;CACNA2D2 | | Body;Body |
| WG2844 | chr3:50488020-50488230 | CACNA2D2;CACNA2D2 | | Body;Body |
| WG2844 | chr3:51990575-51990929 | GPR62 | Island | 1stExon |
| GM02706 | chr3:52351355-52351680 | DNAH1 | | 5'UTR |
| WG3466 | chr3:52478965-52479105 | SEMA3G | | 1stExon |
| GM02149 | chr3:52479098-52479108 | SEMA3G | | TSS200 |
| GM00520 | chr3:52487733-52488097 | TNNC1 | N_Shore | Body |
| GM02555 | chr3:52487733-52488229 | TNNC1 | N_Shore | Body |
| GM02640 | chr3:52569026-52569097 | NT5DC2;NT5DC2;NT5DC2 | N_Shore | 5'UTR;1stExon;TSS1500 |
| WG2276 | chr3:52828680-52828818 | ITIH3 | | TSS200 |
| GM02175 | chr3:52869266-52869521 | MUSTN1 | | TSS200 |
| WG1722 | chr3:53290327-53290343 | TKT;TKT;TKT | S_Shore | TSS1500;TSS1500;TSS1500 |
| GM00520 | chr3:53857939-53857989 | CHDH | Island | Body |
| GM01392 | chr3:55517101-55517496 | WNT5A | N_Shore | Body |
| WG1722 | chr3:56809796-56810320 | ARHGEF3;ARHGEF3;ARHGEF3 | | TSS200;Body;Body |
| GM02456 | chr3:57743377-57743599 | SLMAP;SLMAP | S_Shore | 5'UTR;1stExon |
| WG2276 | chr3:57994941-57995180 | FLNB;FLNB;FLNB;FLNB | Island | Body;Body;Body;Body |
| WG1983 | chr3:58563609-58563674 | FAM107A;FAM107A | | TSS200;TSS200 |
| GM00439 | chr3:58563666-58563693 | FAM107A;FAM107A | | TSS200;TSS200 |
| GM02173 | chr3:62355443-62355525 | FEZF2 | Island | 3'UTR |
| WG1978 | chr3:63087235-63087266 | LOC285401 | | TSS1500 |
| WG2276 | chr3:64547108-64547373 | ADAMTS9 | | Body |
| WG3465 | chr3:64547310-64547373 | ADAMTS9 | | Body |
| GM00520 | chr3:64670013-64670459 | ADAMTS9 | N_Shore | Body |
| GM01096 | chr3:64670017-64670515 | ADAMTS9 | N_Shore | Body |

| | | | | |
|---|---|---|---|---|
| WG3466 | chr3:65342216-65342971 | MAGI1;MAGI1 | Island | Body;3'UTR |
| WG2121 | chr3:65342644-65342971 | MAGI1;MAGI1 | Island | Body;3'UTR |
| GM09256 | chr3:71113124-71113603 | FOXP1 | N_Shore | Body |
| WG2415 | chr3:71180032-71180064 | FOXP1 | | Body |
| WG2414 | chr3:71180032-71180071 | FOXP1 | | Body |
| GM00935 | chr3:71633826-71634044 | FOXP1;FOXP1 | S_Shore | TSS1500;TSS1500 |
| WG1310 | chr3:73045440-73045686 | PPP4R2 | N_Shore | TSS1500 |
| GM02149 | chr3:77147238-77147364 | ROBO2;ROBO2 | Island | 1stExon;Body |
| WG1978 | chr3:8809306-8810139 | OXTR | Island | Body |
| WG2121 | chr3:9258385-9258512 | SRGAP3;SRGAP3 | | Body;Body |
| WG2276 | chr3:9258385-9258573 | SRGAP3;SRGAP3 | | Body;Body |
| GM00016 | chr3:9258433-9258512 | SRGAP3;SRGAP3 | | Body;Body |
| WG1983 | chr3:9258481-9258573 | SRGAP3;SRGAP3 | | Body;Body |
| WG2276 | chr3:9289747-9289893 | SRGAP3;SRGAP3 | | Body;Body |
| GM02706 | chr3:99357437-99357613 | MIR548G;COL8A1;COL8A1 | | Body;TSS200;TSS200 |
| GM01650 | chr4:1006237-1006270 | FGFRL1;FGFRL1;FGFRL1 | Island | 5'UTR;TSS200;5'UTR |
| WG1722 | chr4:101439954-101440278 | EMCN;EMCN | | TSS1500;TSS1500 |
| WG2414 | chr4:10459219-10459352 | ZNF518B | Island | TSS200 |
| GM00520 | chr4:10459435-10460009 | ZNF518B | S_Shore | TSS1500 |
| GM00520 | chr4:104640489-104641033 | TACR3 | N_Shore | 1stExon |
| GM00520 | chr4:104641124-104641548 | TACR3 | S_Shore | TSS200 |
| GM00289 | chr4:10686394-10686554 | CLNK | | TSS200 |
| GM01096 | chr4:1107585-1107720 | RNF212;RNF212 | Island | TSS200;TSS200 |
| GM09255 | chr4:1107706-1107983 | RNF212;RNF212 | Island | TSS200;TSS200 |
| WG2415 | chr4:111397885-111398146 | ENPEP | Island | 1stExon |
| WG3465 | chr4:111398042-111398146 | ENPEP | Island | 1stExon |
| WG2276 | chr4:111539291-111539479 | PITX2;PITX2;PITX2 | Island | Body;Body;Body |
| GM00016 | chr4:111542366-111543401 | PITX2;PITX2;PITX2 | Island | Body;Body;Body |
| WG3465 | chr4:111543401-111545062 | PITX2;PITX2;PITX2 | Island | Body;Body;Body |
| GM02175 | chr4:111544016-111544271 | PITX2;PITX2;PITX2;PITX2 | Island | 1stExon;5'UTR;Body;Body |
| GM00438 | chr4:111544016-111544286 | PITX2;PITX2;PITX2;PITX2 | Island | 1stExon;5'UTR;Body;Body |
| GM00940 | chr4:111544016-111545119 | PITX2;PITX2;PITX2;PITX2 | Island | 1stExon;5'UTR;Body;Body |
| WG2415 | chr4:111544235-111544463 | PITX2;PITX2;PITX2;PITX2 | Island | 1stExon;5'UTR;Body;Body |
| GM02175 | chr4:111544291-111545062 | PITX2;PITX2;PITX2 | Island | Body;TSS200;Body |
| GM00438 | chr4:111544299-111545062 | PITX2;PITX2;PITX2 | Island | Body;TSS200;Body |
| GM00290 | chr4:111544387-111544463 | PITX2;PITX2;PITX2 | Island | Body;TSS200;Body |
| WG2276 | chr4:111555102-111555503 | PITX2;PITX2 | Island | 5'UTR;5'UTR |
| GM00438 | chr4:113065795-113066298 | C4orf32 | N_Shore | TSS1500 |
| WG2844 | chr4:113970660-113970745 | ANK2;ANK2;ANK2 | | TSS200;Body;TSS200 |
| GM09256 | chr4:113970745-113971324 | ANK2;ANK2;ANK2 | | TSS200;Body;TSS200 |
| WG1513 | chr4:11401662-11401872 | HS3ST1 | Island | 5'UTR |
| WG3466 | chr4:1160842-1161157 | SPON2;SPON2 | N_Shore | 3'UTR;3'UTR |
| WG2276 | chr4:1166755-1166824 | LOC100130872-SPON2;SPO | S_Shore | Body;TSS200;5'UTR |
| GM02706 | chr4:1167369-1167423 | LOC100130872-SPON2;SPO | S_Shore | Body;TSS1500;TSS1500 |
| GM01096 | chr4:1195845-1196179 | LOC100130872;LOC100130872-SPON2 | | Body;Body |
| WG3466 | chr4:119948360-119948505 | SYNPO2;SYNPO2;SYNPO2 | | Body;Body;Body |
| WG1978 | chr4:119948360-119948542 | SYNPO2;SYNPO2;SYNPO2 | | Body;Body;Body |

| | | | | |
|---|---|---|---|---|
| GM00042 | chr4:119948388-119948542 | SYNPO2;SYNPO2;SYNPO2 | | Body;Body;Body |
| WG2844 | chr4:1202509-1203653 | LOC100130872;LOC100130 | N_Shore | Body;Body |
| GM00438 | chr4:122685570-122685651 | LOC100192379;TMEM155 | N_Shore | TSS200;5'UTR |
| WG1722 | chr4:122685570-122685675 | LOC100192379;TMEM155 | N_Shore | TSS200;5'UTR |
| GM00438 | chr4:122853693-122853963 | TRPC3;TRPC3 | Island | 1stExon;Body |
| WG1513 | chr4:123300380-123300410 | ADAD1;ADAD1;ADAD1 | N_Shore | 5'UTR;TSS1500;TSS1500 |
| GM00438 | chr4:123300408-123300456 | ADAD1;ADAD1;ADAD1 | Island | 5'UTR;TSS200;TSS200 |
| GM09255 | chr4:124317779-124318523 | SPRY1 | N_Shore | TSS200 |
| WG2120 | chr4:1243849-1244086 | C4orf42;CTBP1;CTBP1 | Island | TSS1500;TSS1500;TSS1500 |
| GM02175 | chr4:1243980-1244017 | CTBP1;CTBP1;C4orf42 | Island | TSS1500;TSS1500;TSS200 |
| GM02149 | chr4:1243980-1244024 | CTBP1;CTBP1;C4orf42 | Island | TSS1500;TSS1500;TSS200 |
| WG1513 | chr4:1243980-1244086 | CTBP1;CTBP1;C4orf42 | Island | TSS1500;TSS1500;TSS200 |
| WG1722 | chr4:1244024-1244086 | CTBP1;CTBP1;C4orf42 | Island | TSS1500;TSS1500;TSS200 |
| GM00520 | chr4:1294996-1295078 | MAEA;MAEA | | Body;Body |
| GM01650 | chr4:1303317-1304048 | MAEA;MAEA | N_Shore | Body;Body |
| GM00520 | chr4:1352787-1353499 | KIAA1530 | | Body |
| GM01375 | chr4:1353105-1353422 | KIAA1530 | | Body |
| GM02555 | chr4:13542719-13543119 | NKX3-2 | N_Shore | 3'UTR |
| WG1722 | chr4:140201537-140201574 | C4orf49 | S_Shore | TSS200 |
| WG3465 | chr4:140201537-140201652 | C4orf49 | S_Shore | TSS200 |
| WG1983 | chr4:140586881-140586988 | MGST2 | | TSS200 |
| WG2415 | chr4:140586881-140587179 | MGST2 | | TSS200 |
| WG2121 | chr4:140586881-140587536 | MGST2 | | TSS200 |
| WG1983 | chr4:140587143-140587179 | MGST2;MGST2 | | 1stExon;5'UTR |
| GM01096 | chr4:140656749-140656926 | MAML3 | Island | Body |
| GM01096 | chr4:141264456-141264602 | SCOC;SCOC;SCOC;SCOC;SCOC | | TSS200;5'UTR;TSS200;TSS200;TSS20 |
| GM09255 | chr4:141264558-141264721 | SCOC;SCOC;SCOC;SCOC;SCOC | | TSS200;5'UTR;TSS200;TSS200;TSS20 |
| WG2844 | chr4:148401273-148401828 | EDNRA;EDNRA;EDNRA | N_Shore | TSS1500;TSS1500;TSS1500 |
| WG2415 | chr4:151177137-151177553 | DCLK2;DCLK2 | N_Shore | Body;Body |
| GM00439 | chr4:151504725-151505084 | LRBA;MAB21L2 | Island | Body;1stExon |
| GM02147 | chr4:15376072-15376119 | C1QTNF7;C1QTNF7 | | Body;TSS200 |
| GM00289 | chr4:15376072-15376135 | C1QTNF7;C1QTNF7 | | Body;TSS200 |
| GM01376 | chr4:15376072-15376180 | C1QTNF7;C1QTNF7 | | Body;TSS200 |
| GM01096 | chr4:15376072-15376215 | C1QTNF7;C1QTNF7 | | Body;TSS200 |
| GM01393 | chr4:15376116-15376135 | C1QTNF7;C1QTNF7 | | Body;TSS200 |
| WG1977 | chr4:15376116-15376180 | C1QTNF7;C1QTNF7 | | Body;TSS200 |
| WG1722 | chr4:15376119-15376215 | C1QTNF7;C1QTNF7 | | Body;TSS200 |
| WG2415 | chr4:15376129-15376180 | C1QTNF7;C1QTNF7 | | Body;TSS200 |
| GM00438 | chr4:15376135-15376180 | C1QTNF7;C1QTNF7 | | Body;TSS200 |
| WG1658 | chr4:15376135-15376215 | C1QTNF7;C1QTNF7 | | Body;TSS200 |
| GM00940 | chr4:154125208-154125453 | TRIM2;TRIM2 | | TSS1500;5'UTR |
| WG1978 | chr4:154125535-154125637 | TRIM2;TRIM2 | | TSS200;5'UTR |
| GM00289 | chr4:154125637-154125720 | TRIM2;TRIM2;TRIM2 | | 5'UTR;1stExon;5'UTR |
| WG2276 | chr4:154710961-154711183 | SFRP2 | N_Shore | TSS1500 |
| WG1983 | chr4:154711070-154711183 | SFRP2 | N_Shore | TSS1500 |
| WG2844 | chr4:154711081-154711183 | SFRP2 | N_Shore | TSS1500 |
| WG1978 | chr4:154711115-154711183 | SFRP2 | N_Shore | TSS1500 |

| | | | | |
|---|---|---|---|---|
| WG3465 | chr4:154711280-154711347 | SFRP2 | N_Shore | TSS1500 |
| GM00290 | chr4:154711470-154711563 | SFRP2 | N_Shore | TSS1500 |
| WG2276 | chr4:154711470-154711906 | SFRP2 | N_Shore | TSS1500 |
| WG3465 | chr4:154711512-154711563 | SFRP2 | N_Shore | TSS1500 |
| GM01392 | chr4:154711563-154711620 | SFRP2 | N_Shore | TSS1500 |
| GM00290 | chr4:154711620-154711675 | SFRP2 | N_Shore | TSS1500 |
| WG3465 | chr4:154711620-154711906 | SFRP2 | N_Shore | TSS1500 |
| GM00289 | chr4:154711629-154711906 | SFRP2 | N_Shore | TSS1500 |
| GM02173 | chr4:15471214-15471301 | CC2D2A;CC2D2A;CC2D2A | | TSS1500;TSS1500;TSS1500 |
| GM00016 | chr4:155253864-155254602 | DCHS2;DCHS2 | N_Shore | Body;Body |
| WG3465 | chr4:155254194-155254497 | DCHS2;DCHS2 | Island | Body;Body |
| WG2121 | chr4:155702533-155703138 | RBM46;RBM46 | Island | 1stExon;5'UTR |
| GM00290 | chr4:155702931-155703429 | RBM46 | Island | 5'UTR |
| WG2120 | chr4:156298289-156298332 | MAP9 | S_Shore | TSS200 |
| GM00289 | chr4:156298325-156298427 | MAP9 | S_Shore | TSS1500 |
| GM01650 | chr4:156680007-156680050 | GUCY1B3 | N_Shore | TSS200 |
| GM02149 | chr4:15704393-15704453 | BST1 | N_Shore | TSS200 |
| GM02706 | chr4:157996959-157997086 | GLRB;GLRB;GLRB | N_Shore | TSS1500;TSS1500;TSS1500 |
| GM01393 | chr4:157997064-157997086 | GLRB;GLRB;GLRB | N_Shore | TSS1500;TSS1500;TSS1500 |
| WG2844 | chr4:159092089-159092553 | FAM198B;FAM198B;FAM198B | | Body;Body;Body |
| GM00438 | chr4:159131324-159131393 | TMEM144 | N_Shore | TSS200 |
| WG2415 | chr4:160188727-160189254 | RAPGEF2 | | TSS1500 |
| GM01376 | chr4:165878055-165878317 | C4orf39;TRIM61 | Island | TSS200;Body |
| WG1658 | chr4:165898666-165898967 | TRIM61 | | 5'UTR |
| GM02706 | chr4:165898707-165898835 | TRIM61;TRIM61 | | 1stExon;5'UTR |
| GM00290 | chr4:165898707-165898848 | TRIM61;TRIM61 | | 1stExon;5'UTR |
| GM01653 | chr4:165898825-165898848 | TRIM61 | | TSS200 |
| GM03075 | chr4:165898835-165898912 | TRIM61 | | TSS200 |
| WG1513 | chr4:165952839-165953028 | TRIM60 | | TSS1500 |
| WG2844 | chr4:166301076-166301256 | CPE | S_Shore | Body |
| GM01096 | chr4:169418081-169418224 | PALLD;PALLD | | TSS200;TSS200 |
| WG3465 | chr4:169798931-169799624 | PALLD;PALLD;PALLD;PALLD | N_Shore | 5'UTR;Body;Body;Body |
| WG2415 | chr4:169799087-169799624 | PALLD;PALLD;PALLD;PALLD | Island | Body;Body;Body;Body |
| GM00289 | chr4:174447847-174448128 | HAND2 | N_Shore | 3'UTR |
| WG3465 | chr4:174447847-174448341 | HAND2 | N_Shore | 3'UTR |
| WG3466 | chr4:174448128-174448705 | HAND2 | N_Shore | 3'UTR |
| GM02175 | chr4:174452026-174452159 | NBLA00301;HAND2 | Island | Body;TSS1500 |
| GM00438 | chr4:174452133-174452612 | NBLA00301;HAND2 | Island | Body;TSS1500 |
| WG2193 | chr4:174452133-174453287 | NBLA00301;HAND2 | Island | Body;TSS1500 |
| WG2121 | chr4:174452159-174452612 | NBLA00301;HAND2 | Island | Body;TSS1500 |
| WG1978 | chr4:174452461-174453166 | NBLA00301;HAND2 | Island | Body;TSS1500 |
| WG1513 | chr4:174452985-174453287 | NBLA00301 | S_Shore | Body |
| GM00289 | chr4:175839468-175839610 | ADAM29;ADAM29;ADAM29;ADAM29 | | TSS200;TSS200;TSS200;TSS200 |
| WG3466 | chr4:1804007-1804492 | FGFR3;FGFR3;FGFR3 | Island | Body;Body;Body |
| WG2194 | chr4:183247132-183247234 | ODZ3 | | Body |
| WG2276 | chr4:1844724-1844878 | LETM1 | S_Shore | Body |
| GM00042 | chr4:184908510-184908573 | STOX2 | | Body |

| | | | | |
|---|---|---|---|---|
| GM02555 | chr4:184922382-184922433 | STOX2 | | Body |
| WG1722 | chr4:184930883-184930913 | STOX2 | | Body |
| GM09256 | chr4:184938291-184938359 | STOX2 | | Body |
| GM02555 | chr4:184938291-184938470 | STOX2 | | Body |
| GM00042 | chr4:184938316-184938470 | STOX2 | | Body |
| WG1513 | chr4:186131234-186131280 | SNX25 | Island | TSS200 |
| WG2121 | chr4:186559707-186560083 | SORBS2;SORBS2;SORBS2;S( | N_Shore | Body;Body;Body;Body;Body;Body;Bod |
| WG3465 | chr4:186732942-186733060 | SORBS2;SORBS2;SORBS2;SORBS2;SORI | | 5'UTR;5'UTR;5'UTR;1stExon;TSS1500; |
| WG2121 | chr4:186733060-186733428 | SORBS2;SORBS2;SORBS2;SORBS2;SORI | | 5'UTR;5'UTR;5'UTR;1stExon;TSS1500; |
| WG2276 | chr4:186877497-186877657 | SORBS2;SORBS2 | | 5'UTR;1stExon |
| GM00438 | chr4:186877550-186877657 | SORBS2;SORBS2 | | 5'UTR;1stExon |
| WG1721 | chr4:187621790-187621972 | FAT1 | | Body |
| WG2121 | chr4:187621790-187622164 | FAT1 | | Body |
| GM00935 | chr4:187621972-187622164 | FAT1 | | Body |
| GM00439 | chr4:188916865-188917251 | ZFP42 | Island | TSS200 |
| GM02149 | chr4:189060646-189061030 | TRIML1;TRIML1 | | 1stExon;5'UTR |
| WG1513 | chr4:190906077-190906123 | TUBB4Q | Island | TSS200 |
| GM00935 | chr4:20884321-20884431 | KCNIP4;KCNIP4;KCNIP4;KCNIP4;KCNIP4 | | Body;Body;1stExon;5'UTR;5'UTR |
| WG1657 | chr4:21699209-21699426 | KCNIP4;KCNIP4;KCNIP4 | | 5'UTR;1stExon;5'UTR |
| WG1983 | chr4:21699336-21699745 | KCNIP4;KCNIP4 | | 5'UTR;TSS200 |
| GM02555 | chr4:2366485-2366831 | ZFYVE28 | Island | Body |
| GM00520 | chr4:2366485-2367091 | ZFYVE28 | Island | Body |
| GM02149 | chr4:2366485-2367137 | ZFYVE28 | Island | Body |
| GM00439 | chr4:2366555-2366831 | ZFYVE28 | Island | Body |
| WG2415 | chr4:2366831-2367137 | ZFYVE28 | Island | Body |
| WG2275 | chr4:24818471-24818615 | CCDC149 | | Body |
| GM00438 | chr4:25656830-25657160 | SLC34A2 | N_Shore | TSS1500 |
| WG2844 | chr4:26491963-26492488 | CCKAR;CCKAR | | 1stExon;5'UTR |
| GM09256 | chr4:26492206-26492378 | CCKAR | | TSS200 |
| WG1310 | chr4:2813458-2814064 | SH3BP2;SH3BP2 | | TSS1500;5'UTR |
| GM02456 | chr4:2813542-2814064 | SH3BP2;SH3BP2 | | TSS1500;5'UTR |
| GM01392 | chr4:2813542-2814122 | SH3BP2;SH3BP2 | | TSS1500;5'UTR |
| GM00520 | chr4:2813724-2814122 | SH3BP2;SH3BP2 | | TSS1500;5'UTR |
| GM02175 | chr4:2964211-2964239 | GRK4;GRK4;GRK4;NOP14 | N_Shore | TSS1500;TSS1500;TSS1500;Body |
| WG3466 | chr4:30724564-30724915 | PCDH7;PCDH7;PCDH7 | Island | 1stExon;Body;1stExon |
| WG3465 | chr4:3265080-3265280 | C4orf44 | | Body |
| GM02706 | chr4:3372123-3372272 | RGS12;RGS12;RGS12 | N_Shelf | Body;Body;Body |
| WG2844 | chr4:3374767-3374909 | RGS12;RGS12;RGS12 | Island | Body;Body;Body |
| WG1983 | chr4:3486039-3486204 | DOK7;DOK7 | Island | Body;Body |
| WG2844 | chr4:3487164-3487436 | DOK7;DOK7 | S_Shore | Body;Body |
| GM02641 | chr4:36283184-36283275 | DTHD1 | | TSS200 |
| WG2415 | chr4:38807259-38807382 | TLR1 | | TSS1500 |
| GM02706 | chr4:40518024-40518065 | RBM47;RBM47 | | 5'UTR;TSS200 |
| WG1513 | chr4:40858965-40859293 | APBB2;APBB2;APBB2;APBB2 | Island | Body;5'UTR;5'UTR;5'UTR;Body;5'UTR;I |
| GM09255 | chr4:40858984-40859249 | APBB2;APBB2;APBB2;APBB2 | Island | Body;5'UTR;5'UTR;5'UTR;Body;5'UTR;I |
| GM00439 | chr4:40858984-40859259 | APBB2;APBB2;APBB2;APBB2 | Island | Body;5'UTR;5'UTR;5'UTR;Body;5'UTR;I |
| WG2276 | chr4:40858984-40859293 | APBB2;APBB2;APBB2;APBB2 | Island | Body;5'UTR;5'UTR;5'UTR;Body;5'UTR;I |

| | | | | |
|---|---|---|---|---|
| GM02555 | chr4:40858984-40859344 | APBB2;APBB2;APBB2;APBB2 | Island | Body;5'UTR;5'UTR;5'UTR;Body;5'UTR;E |
| WG1722 | chr4:40859095-40859259 | APBB2;APBB2;APBB2;APBB2 | Island | Body;5'UTR;5'UTR;5'UTR;1stExon;1stE |
| GM00935 | chr4:40859095-40859293 | APBB2;APBB2;APBB2;APBB2 | Island | Body;5'UTR;5'UTR;5'UTR;1stExon;1stE |
| WG1978 | chr4:40859249-40859259 | APBB2;APBB2;APBB2;APBB2 | S_Shore | Body;TSS200;TSS200;Body;TSS200;Bo |
| GM00941 | chr4:40859249-40859280 | APBB2;APBB2;APBB2;APBB2 | S_Shore | Body;TSS200;TSS200;Body;TSS200;Bo |
| WG3465 | chr4:40859249-40859293 | APBB2;APBB2;APBB2;APBB2 | S_Shore | Body;TSS200;TSS200;Body;TSS200;Bo |
| GM02456 | chr4:40859251-40859280 | APBB2;APBB2;APBB2;APBB2 | S_Shore | Body;TSS200;TSS200;Body;TSS200;Bo |
| WG2193 | chr4:40859251-40859293 | APBB2;APBB2;APBB2;APBB2 | S_Shore | Body;TSS200;TSS200;Body;TSS200;Bo |
| WG1658 | chr4:40859259-40859293 | APBB2;APBB2;APBB2;APBB2 | S_Shore | Body;TSS200;TSS200;Body;TSS200;Bo |
| GM00289 | chr4:42154949-42154983 | BEND4;BEND4 | Island | TSS200;TSS200 |
| GM00290 | chr4:42401056-42401242 | SHISA3 | S_Shore | Body |
| GM01376 | chr4:42401056-42401610 | SHISA3 | S_Shore | Body |
| GM09256 | chr4:4250606-4250746 | TMEM128 | S_Shore | TSS1500 |
| GM00438 | chr4:4405913-4406132 | D4S234E;D4S234E | | Body;Body |
| GM00438 | chr4:48908833-48908849 | OCIAD2;OCIAD2 | Island | TSS200;TSS200 |
| WG1513 | chr4:5021084-5021113 | CYTL1 | Island | 1stExon |
| WG2121 | chr4:5021084-5021236 | CYTL1 | Island | 1stExon |
| GM02704 | chr4:5021113-5021236 | CYTL1 | Island | 1stExon |
| WG2120 | chr4:5021203-5021328 | CYTL1 | Island | TSS200 |
| WG2844 | chr4:5021268-5021328 | CYTL1 | Island | TSS200 |
| WG2415 | chr4:52918457-52918489 | SPATA18 | S_Shore | Body |
| WG1983 | chr4:52942997-52943197 | SPATA18 | Island | Body |
| WG2276 | chr4:52942997-52943247 | SPATA18 | Island | Body |
| WG2844 | chr4:52943099-52943247 | SPATA18 | Island | Body |
| WG3465 | chr4:53728999-53729461 | RASL11B | Island | Body |
| WG2276 | chr4:54424280-54424764 | LNX1;LNX1;LNX1 | | 1stExon;Body;5'UTR |
| WG2194 | chr4:54424280-54424787 | LNX1;LNX1;LNX1 | | 1stExon;Body;5'UTR |
| GM00438 | chr4:54965109-54965999 | GSX2 | N_Shore | TSS1500 |
| WG3465 | chr4:54965829-54965904 | GSX2 | N_Shore | TSS1500 |
| WG3466 | chr4:55143269-55143557 | PDGFRA | | Body |
| GM00290 | chr4:57522493-57522762 | HOPX;HOPX;HOPX;HOPX;HC | Island | 1stExon;5'UTR;5'UTR;Body;Body;5'UTI |
| GM02555 | chr4:57522493-57522795 | HOPX;HOPX;HOPX;HOPX;HC | Island | 1stExon;5'UTR;5'UTR;Body;Body;5'UTI |
| WG1310 | chr4:57522632-57522881 | HOPX;HOPX;HOPX;HOPX;HC | Island | 1stExon;5'UTR;5'UTR;Body;Body;5'UTI |
| GM09256 | chr4:57522762-57522801 | HOPX;HOPX;HOPX;HOPX;HC | S_Shore | 5'UTR;Body;Body;TSS200;TSS200 |
| WG1513 | chr4:57522762-57522881 | HOPX;HOPX;HOPX;HOPX;HC | S_Shore | 5'UTR;Body;Body;TSS200;TSS200 |
| WG3465 | chr4:57522762-57523055 | HOPX;HOPX;HOPX;HOPX;HC | S_Shore | 5'UTR;Body;Body;TSS200;TSS200 |
| GM00438 | chr4:57547347-57547579 | HOPX;HOPX;HOPX | | 5'UTR;5'UTR;5'UTR |
| WG3465 | chr4:57547525-57547699 | HOPX;HOPX;HOPX;HOPX;HOPX;HOPX | | 5'UTR;1stExon;1stExon;5'UTR;5'UTR;1 |
| GM00042 | chr4:57547525-57547872 | HOPX;HOPX;HOPX;HOPX;HOPX;HOPX | | 5'UTR;1stExon;1stExon;5'UTR;5'UTR;1 |
| GM00290 | chr4:5976063-5976157 | C4orf50 | | Body |
| WG1722 | chr4:5990227-5990590 | C4orf50 | | TSS200 |
| WG3465 | chr4:6052527-6052554 | JAKMIP1 | | Body |
| WG1977 | chr4:6107021-6107339 | JAKMIP1;JAKMIP1 | N_Shore | Body;Body |
| WG3465 | chr4:6107131-6107320 | JAKMIP1;JAKMIP1 | Island | Body;Body |
| GM01392 | chr4:618204-618497 | PDE6B;PDE6B | N_Shore | TSS1500;TSS1500 |
| GM00042 | chr4:618753-619167 | PDE6B;PDE6B | N_Shore | TSS1500;TSS1500 |
| GM00438 | chr4:619392-619659 | PDE6B;PDE6B;PDE6B;PDE6E | N_Shore | 1stExon;1stExon;5'UTR;5'UTR |

| | | | | |
|---|---|---|---|---|
| GM09256 | chr4:6423008-6423283 | PPP2R2C | | Body |
| WG1513 | chr4:645256-645551 | PDE6B;PDE6B | Island | Body;Body |
| WG1513 | chr4:646502-646577 | PDE6B;PDE6B;PDE6B | N_Shore | TSS1500;Body;Body |
| GM02175 | chr4:66536550-66536772 | EPHA5;EPHA5 | Island | TSS1500;TSS1500 |
| GM09255 | chr4:6694923-6695319 | S100P | | TSS1500 |
| GM09255 | chr4:6695464-6695614 | S100P | | TSS200 |
| WG3465 | chr4:683148-683240 | MFSD7 | S_Shore | TSS200 |
| WG1514 | chr4:69435250-69435601 | UGT2B15;UGT2B17 | | TSS1500;TSS1500 |
| WG2844 | chr4:715865-715950 | PCGF3 | N_Shore | 5'UTR |
| WG2415 | chr4:71587722-71587904 | RUFY3;RUFY3;RUFY3;RUFY3;RUFY3 | | 1stExon;Body;5'UTR;1stExon;5'UTR |
| GM00439 | chr4:7228268-7228442 | SORCS2 | | Body |
| GM02456 | chr4:7282819-7283184 | SORCS2 | | Body |
| WG1283 | chr4:7287492-7287737 | SORCS2 | | Body |
| GM00940 | chr4:7289482-7289507 | SORCS2 | | Body |
| GM02706 | chr4:72897281-72897565 | NPFFR2 | N_Shore | TSS1500 |
| GM00016 | chr4:7373343-7373451 | SORCS2 | | Body |
| GM00290 | chr4:7436701-7437027 | PSAPL1;SORCS2 | | TSS200;Body |
| GM09256 | chr4:7436861-7437027 | PSAPL1;SORCS2 | | TSS200;Body |
| WG1514 | chr4:74718978-74719172 | PF4V1 | N_Shore | TSS200 |
| WG2275 | chr4:74718978-74719289 | PF4V1 | N_Shore | TSS200 |
| GM00940 | chr4:74719046-74719172 | PF4V1;PF4V1 | N_Shore | 1stExon;5'UTR |
| GM02555 | chr4:74847646-74847766 | PF4 | Island | 1stExon |
| WG3465 | chr4:74847646-74847824 | PF4 | Island | 1stExon |
| WG2194 | chr4:74847646-74847829 | PF4 | Island | 1stExon |
| GM01393 | chr4:74847710-74847761 | PF4;PF4 | Island | 1stExon;5'UTR |
| GM00520 | chr4:74847758-74847829 | PF4 | Island | TSS200 |
| WG2121 | chr4:74847758-74848016 | PF4 | Island | TSS200 |
| GM02149 | chr4:74847761-74847829 | PF4 | Island | TSS200 |
| GM02175 | chr4:74847766-74848016 | PF4 | Island | TSS200 |
| WG2844 | chr4:74847824-74848016 | PF4 | Island | TSS200 |
| GM09256 | chr4:74864239-74864709 | CXCL5 | Island | 1stExon |
| WG1977 | chr4:74864612-74864725 | CXCL5 | S_Shore | TSS200 |
| GM02175 | chr4:75895596-75895858 | PARM1 | | Body |
| GM09256 | chr4:7593370-7593493 | SORCS2 | Island | Body |
| WG1309 | chr4:76481053-76481269 | C4orf26 | | TSS1500 |
| WG2415 | chr4:77356008-77356180 | SHROOM3 | | TSS1500 |
| GM00940 | chr4:77507102-77507338 | SHROOM3 | Island | Body |
| WG1977 | chr4:77507102-77507411 | SHROOM3 | Island | Body |
| GM00438 | chr4:77507338-77507411 | SHROOM3 | Island | Body |
| GM00520 | chr4:7774727-7774950 | AFAP1;AFAP1;LOC84740 | Island | Body;Body;Body |
| GM02456 | chr4:7813475-7813911 | AFAP1;AFAP1 | | Body;Body |
| GM00439 | chr4:7813475-7814047 | AFAP1;AFAP1 | | Body;Body |
| WG2121 | chr4:7941894-7942031 | AFAP1;AFAP1 | S_Shore | TSS1500;TSS1500 |
| GM09256 | chr4:80246713-80247088 | NAA11 | | 1stExon |
| GM00940 | chr4:8098178-8098319 | ABLIM2;ABLIM2;ABLIM2;ABLI | N_Shore | Body;Body;Body;Body;Body;Body;Bod |
| GM02175 | chr4:81106857-81107185 | PRDM8 | S_Shore | 5'UTR |
| WG1977 | chr4:81117647-81117853 | PRDM8;PRDM8 | N_Shore | TSS1500;5'UTR |

| | | | | |
|---|---|---|---|---|
| GM02706 | chr4:81118188-81119473 | PRDM8;PRDM8 | Island | TSS1500;5'UTR |
| WG2844 | chr4:81118343-81119473 | PRDM8;PRDM8 | Island | TSS1500;5'UTR |
| WG1983 | chr4:81118500-81119473 | PRDM8;PRDM8 | Island | 5'UTR;TSS200 |
| WG1977 | chr4:81119178-81119473 | PRDM8;PRDM8 | Island | 5'UTR;5'UTR |
| WG1722 | chr4:83674615-83674996 | SCD5;MIR575;SCD5 | | Body;TSS200;Body |
| GM02555 | chr4:8386188-8386266 | ACOX3;ACOX3 | N_Shelf | Body;Body |
| GM01393 | chr4:8395941-8396163 | ACOX3;ACOX3 | | Body;Body |
| WG1310 | chr4:85417312-85417755 | NKX6-1 | N_Shore | Body |
| GM02706 | chr4:86748944-86749040 | ARHGAP24;ARHGAP24 | | Body;5'UTR |
| WG2414 | chr4:87281412-87281511 | MAPK10;MAPK10;MAPK10 | | TSS200;TSS200;5'UTR |
| WG3466 | chr4:87770586-87770782 | SLC10A6 | | TSS200 |
| WG2120 | chr4:899882-899984 | GAK | Island | Body |
| WG3465 | chr4:940614-941054 | TMEM175 | Island | 5'UTR |
| WG2844 | chr4:94124971-94125068 | GRID2 | | Body |
| WG1513 | chr4:96761229-96761796 | PDHA2 | | TSS200 |
| GM02175 | chr4:99850801-99851008 | EIF4E;EIF4E;EIF4E;EIF4E;EIF4 | S_Shore | 5'UTR;1stExon;5'UTR;TSS1500;1stExo |
| WG1658 | chr4:99850885-99851115 | EIF4E;EIF4E;EIF4E;EIF4E;EIF4 | S_Shore | 5'UTR;1stExon;5'UTR;TSS1500;1stExo |
| WG1977 | chr4:99851003-99851060 | EIF4E;EIF4E;EIF4E;EIF4E;EIF4 | S_Shore | 5'UTR;1stExon;5'UTR;TSS1500;1stExo |
| WG1977 | chr5:100240048-100240223 | ST8SIA4;ST8SIA4 | S_Shore | TSS1500;TSS1500 |
| GM02706 | chr5:101632310-101632321 | SLCO4C1 | Island | TSS200 |
| GM00520 | chr5:1027884-1028394 | NKD2 | S_Shelf | Body |
| GM01096 | chr5:102898223-102898648 | NUDT12 | | 5'UTR |
| GM01392 | chr5:102898463-102898733 | NUDT12;NUDT12 | | 5'UTR;1stExon |
| GM02555 | chr5:1052828-1054262 | SLC12A7 | N_Shore | Body |
| GM00935 | chr5:1052828-1054427 | SLC12A7 | N_Shore | Body |
| WG3465 | chr5:1053545-1053774 | SLC12A7 | Island | Body |
| GM00439 | chr5:1053545-1054262 | SLC12A7 | Island | Body |
| GM00520 | chr5:1053545-1054427 | SLC12A7 | Island | Body |
| WG2276 | chr5:1053643-1054113 | SLC12A7 | Island | Body |
| WG2843 | chr5:1053643-1054262 | SLC12A7 | Island | Body |
| GM02706 | chr5:10563092-10563501 | ANKRD33B | Island | TSS1500 |
| GM00289 | chr5:1060334-1060679 | SLC12A7 | N_Shore | Body |
| WG1978 | chr5:1068772-1068999 | SLC12A7 | Island | Body |
| GM00935 | chr5:1076587-1076758 | SLC12A7 | N_Shore | Body |
| WG2415 | chr5:1090619-1090936 | SLC12A7 | S_Shore | Body |
| GM02555 | chr5:1100209-1100544 | SLC12A7 | Island | Body |
| WG1722 | chr5:110062343-110062729 | TMEM232;TMEM232 | | 1stExon;5'UTR |
| WG2414 | chr5:110062384-110062417 | TMEM232 | | TSS200 |
| WG1513 | chr5:110062384-110062729 | TMEM232 | | TSS200 |
| GM00042 | chr5:110062417-110062729 | TMEM232 | | TSS200 |
| GM00438 | chr5:111495974-111496180 | NCRNA00219;SNORA13 | N_Shore | TSS1500;TSS1500 |
| GM01393 | chr5:111753754-111753938 | EPB41L4A | N_Shore | Body |
| GM09256 | chr5:111753803-111754296 | EPB41L4A;FLJ11235 | N_Shore | Body;TSS1500 |
| GM01096 | chr5:112824614-112824700 | MCC | S_Shore | TSS200 |
| WG1977 | chr5:112824646-112824920 | MCC | S_Shore | TSS200 |
| WG1977 | chr5:114514899-114515341 | TRIM36;TRIM36;TRIM36 | Island | Body;Body;Body |
| WG1722 | chr5:114516368-114516422 | TRIM36;TRIM36;TRIM36 | S_Shore | TSS200;TSS200;TSS200 |

| | | | | |
|---|---|---|---|---|
| WG1722 | chr5:115152785-115152938 | CDO1 | S_Shore | TSS1500 |
| GM01376 | chr5:115298704-115299071 | LVRN | Island | 1stExon |
| WG1722 | chr5:115298704-115299475 | LVRN | Island | 1stExon |
| GM00940 | chr5:115298920-115299088 | LVRN | Island | 1stExon |
| WG2121 | chr5:115298920-115299475 | LVRN | Island | 1stExon |
| GM02175 | chr5:115299071-115299475 | LVRN | Island | Body |
| GM00438 | chr5:118965093-118965202 | FAM170A;FAM170A | | TSS200;TSS200 |
| GM02175 | chr5:1201602-1201629 | SLC6A19 | Island | TSS200 |
| GM09255 | chr5:1201608-1201639 | SLC6A19 | Island | TSS200 |
| WG2415 | chr5:1201639-1201713 | SLC6A19 | Island | TSS200 |
| GM00042 | chr5:1217576-1217797 | SLC6A19 | N_Shore | Body |
| WG1513 | chr5:1218507-1218653 | SLC6A19 | Island | Body |
| GM01096 | chr5:1218561-1218653 | SLC6A19 | Island | Body |
| WG2121 | chr5:1223167-1223262 | SLC6A19 | N_Shelf | 3'UTR |
| GM00042 | chr5:122434544-122435550 | PRDM6 | Island | Body |
| WG1722 | chr5:1225328-1225355 | SLC6A18 | N_Shore | TSS200 |
| WG1513 | chr5:1225331-1225397 | SLC6A18 | N_Shore | TSS200 |
| GM00940 | chr5:1228081-1228443 | SLC6A18 | S_Shore | Body |
| GM00042 | chr5:1243714-1243809 | SLC6A18 | Island | Body |
| GM03075 | chr5:1245902-1246291 | SLC6A18 | Island | Body |
| GM09256 | chr5:1257676-1257903 | TERT;TERT | S_Shore | Body;Body |
| GM00042 | chr5:125929132-125929671 | ALDH7A1 | N_Shore | Body |
| GM09255 | chr5:126408756-126409061 | FLJ44606;FLJ44606 | N_Shore | 5'UTR;5'UTR |
| WG2276 | chr5:126408756-126409227 | FLJ44606;FLJ44606 | N_Shore | 5'UTR;5'UTR |
| WG3466 | chr5:126409061-126409198 | FLJ44606;FLJ44606 | Island | 5'UTR;5'UTR |
| GM00022 | chr5:126409061-126409227 | FLJ44606;FLJ44606 | Island | 5'UTR;5'UTR |
| GM09255 | chr5:126409198-126409227 | FLJ44606;FLJ44606 | Island | TSS200;TSS200 |
| WG3466 | chr5:126409227-126409372 | FLJ44606;FLJ44606 | Island | TSS200;TSS200 |
| GM02175 | chr5:126409308-126409372 | FLJ44606;FLJ44606 | Island | TSS200;TSS200 |
| GM02706 | chr5:126409308-126409553 | FLJ44606;FLJ44606 | Island | TSS200;TSS200 |
| GM09255 | chr5:126409310-126409454 | FLJ44606;FLJ44606 | Island | TSS200;TSS200 |
| GM02706 | chr5:126626348-126626364 | MEGF10 | Island | TSS200 |
| GM00438 | chr5:126984564-126984994 | CTXN3 | | TSS200 |
| WG2415 | chr5:127871927-127872074 | FBN2 | N_Shore | Body |
| WG3466 | chr5:127871927-127872329 | FBN2 | N_Shore | Body |
| GM00438 | chr5:127872037-127872329 | FBN2 | Island | Body |
| GM00042 | chr5:127872074-127872329 | FBN2 | Island | Body |
| WG1513 | chr5:1286162-1286354 | TERT;TERT | Island | Body;Body |
| GM00016 | chr5:132158625-132159003 | SHROOM1 | Island | Body |
| WG1310 | chr5:134363562-134364387 | PITX1 | Island | 3'UTR |
| GM02173 | chr5:134363823-134364387 | PITX1 | Island | 3'UTR |
| GM02173 | chr5:134366534-134367099 | PITX1 | N_Shore | Body |
| GM01096 | chr5:134370282-134370530 | PITX1 | Island | TSS1500 |
| WG2276 | chr5:134735544-134735613 | H2AFY;H2AFY;H2AFY;H2AFY | S_Shore | TSS1500;5'UTR;TSS1500;TSS1500;1st |
| WG3466 | chr5:134787907-134788128 | TIFAB;TIFAB | | 1stExon;5'UTR |
| WG3465 | chr5:134870506-134870517 | NEUROG1;NEUROG1 | N_Shore | 1stExon;3'UTR |
| WG3465 | chr5:134872315-134872659 | NEUROG1 | S_Shore | TSS1500 |

| | | | | |
|---|---|---|---|---|
| GM00289 | chr5:134872639-134872659 | NEUROG1 | S_Shore | TSS1500 |
| WG1722 | chr5:135364328-135364451 | TGFBI | N_Shore | TSS1500 |
| GM01650 | chr5:135416205-135416613 | MIR886 | Island | Body |
| GM02147 | chr5:135416412-135416613 | MIR886 | Island | TSS200 |
| GM02706 | chr5:135416529-135416613 | MIR886 | S_Shore | TSS1500 |
| GM02706 | chr5:136983354-136983373 | MIR874;KLHL3 | | TSS200;Body |
| GM00438 | chr5:136983354-136984054 | MIR874;KLHL3 | | TSS200;Body |
| GM01376 | chr5:137224904-137225191 | PKD2L2 | N_Shore | TSS1500 |
| GM02706 | chr5:137475115-137475288 | NME5;NME5 | | 1stExon;5'UTR |
| GM01392 | chr5:137593418-137593583 | GFRA3 | Island | Body |
| GM00439 | chr5:137610372-137610396 | GFRA3 | S_Shore | TSS200 |
| GM00941 | chr5:13810143-13810279 | DNAH5 | Island | Body |
| WG1514 | chr5:138714136-138714337 | SLC23A1;SLC23A1 | Island | Body;Body |
| GM00520 | chr5:138714322-138714339 | SLC23A1;SLC23A1 | Island | Body;Body |
| GM01392 | chr5:138714337-138714654 | SLC23A1;SLC23A1 | Island | Body;Body |
| GM01393 | chr5:138728367-138728704 | LOC389333 | Island | 1stExon |
| WG2276 | chr5:138728685-138729283 | LOC389333 | Island | 1stExon |
| GM00438 | chr5:139056667-139056804 | CXXC5 | Island | 5'UTR |
| WG2844 | chr5:1394296-1394484 | SLC6A3 | | 3'UTR |
| GM02149 | chr5:140018644-140018709 | TMCO6 | N_Shore | TSS1500 |
| WG3465 | chr5:140052017-140052475 | DND1 | N_Shore | Body |
| GM02147 | chr5:140097950-140098250 | VTRNA1-2 | | TSS1500 |
| WG1978 | chr5:140098392-140098576 | VTRNA1-2 | | TSS200 |
| WG2120 | chr5:140105405-140105629 | VTRNA1-3 | | TSS1500 |
| GM01392 | chr5:140207460-140207498 | PCDHA2;PCDHA1;PCDHA1;P | N_Shore | Body;Body;Body;TSS200;Body;TSS200 |
| GM09256 | chr5:140220686-140220822 | PCDHA6;PCDHA2;PCDHA1;P | N_Shore | Body;Body;Body;Body;Body;TSS1500; |
| WG1513 | chr5:140220803-140220822 | PCDHA6;PCDHA2;PCDHA1;P | N_Shore | Body;Body;Body;TSS200;Body;Body;B |
| GM01096 | chr5:140344290-140344620 | PCDHA7;PCDHAC2;PCDHA1 | N_Shore | Body;TSS1500;Body;Body;Body;Body; |
| GM02175 | chr5:140430569-140430713 | PCDHB1 | | TSS1500 |
| WG2276 | chr5:140480200-140480223 | PCDHB3 | N_Shore | TSS200 |
| WG1722 | chr5:140480218-140480597 | PCDHB3 | N_Shore | TSS200 |
| GM02175 | chr5:140480507-140480941 | PCDHB3 | N_Shore | 1stExon |
| GM00289 | chr5:140480597-140480872 | PCDHB3 | N_Shore | 1stExon |
| WG2276 | chr5:140480597-140480941 | PCDHB3 | N_Shore | 1stExon |
| WG2415 | chr5:140514711-140514824 | PCDHB5 | N_Shore | TSS200 |
| WG2276 | chr5:140535759-140536238 | PCDHB17 | N_Shore | Body |
| GM00940 | chr5:140552369-140552600 | PCDHB7;PCDHB7 | N_Shore | 1stExon;5'UTR |
| GM02175 | chr5:140552600-140553150 | PCDHB7 | N_Shore | 1stExon |
| WG3465 | chr5:140552992-140553150 | PCDHB7 | N_Shore | 1stExon |
| WG1513 | chr5:140557227-140557256 | PCDHB8 | N_Shore | TSS1500 |
| WG2276 | chr5:140562218-140562819 | PCDHB16 | N_Shore | 1stExon |
| WG2276 | chr5:140563213-140563400 | PCDHB16 | N_Shore | 1stExon |
| WG1722 | chr5:140579296-140579303 | PCDHB11 | N_Shore | TSS200 |
| GM00520 | chr5:140619446-140619586 | PCDHB19P | N_Shore | TSS1500 |
| WG2276 | chr5:140625750-140625848 | PCDHB15 | N_Shore | 1stExon |
| GM00042 | chr5:140683467-140683670 | SLC25A2;SLC25A2 | Island | 1stExon;5'UTR |
| WG1513 | chr5:140683467-140683681 | SLC25A2;SLC25A2 | Island | 1stExon;5'UTR |

| | | | | |
|---|---|---|---|---|
| GM09256 | chr5:140683699-140683767 | SLC25A2 | Island | TSS200 |
| GM00520 | chr5:140711852-140712459 | PCDHGA1;PCDHGA1 | Island | 1stExon;1stExon |
| GM03075 | chr5:140723509-140723568 | PCDHGA2;PCDHGA3;PCDH( | N_Shore | Body;TSS200;Body;TSS200 |
| WG2121 | chr5:140723509-140723583 | PCDHGA2;PCDHGA3;PCDH( | N_Shore | Body;TSS200;Body;TSS200 |
| GM00439 | chr5:140723549-140723583 | PCDHGA2;PCDHGA3;PCDH( | Island | Body;TSS200;Body;TSS200 |
| GM02175 | chr5:140729608-140729653 | PCDHGA2;PCDHGA1;PCDH( | N_Shore | Body;Body;Body;TSS1500;TSS1500 |
| WG3465 | chr5:140730092-140730516 | PCDHGB1;PCDHGA2;PCDH( | N_Shore | 1stExon;Body;Body;1stExon;Body |
| GM00042 | chr5:140741220-140741737 | PCDHGA4;PCDHGA2;PCDH( | Island | Body;Body;1stExon;Body;Body;Body;1 |
| GM09255 | chr5:140768213-140768251 | PCDHGB4;PCDHGA4;PCDH( | N_Shore | 1stExon;Body;Body;Body;Body;Body;E |
| GM02175 | chr5:140782367-140782755 | PCDHGA4;PCDHGA6;PCDH( | S_Shelf | Body;Body;TSS200;Body;Body;Body;B |
| GM00941 | chr5:140810106-140810123 | PCDHGA4;PCDHGA11;PCDH | N_Shore | Body;Body;Body;Body;Body;Body;Bod |
| GM02175 | chr5:140810106-140810260 | PCDHGA4;PCDHGA11;PCDH | N_Shore | Body;Body;Body;Body;Body;Body;Bod |
| WG3466 | chr5:140810109-140810137 | PCDHGA4;PCDHGA11;PCDH | N_Shore | Body;Body;Body;Body;Body;Body;Bod |
| GM01376 | chr5:140810109-140810260 | PCDHGA4;PCDHGA11;PCDH | N_Shore | Body;Body;Body;Body;Body;Body;Bod |
| WG2414 | chr5:140810805-140811102 | PCDHGA4;PCDHGA11;PCDH | Island | Body;Body;Body;Body;Body;Body;Bod |
| WG2276 | chr5:140857647-140857889 | PCDHGA4;PCDHGA12;PCDH | N_Shore | Body;Body;Body;Body;Body;Body;Bod |
| GM01392 | chr5:142065539-142065676 | FGF1;FGF1;FGF1;FGF1;FGF1;FGF1;FGF1 | | 5'UTR;Body;Body;5'UTR;1stExon;5'UTI |
| WG1978 | chr5:14406585-14406732 | TRIO | | Body |
| GM00290 | chr5:146832182-146832298 | DPYSL3 | N_Shore | Body |
| GM00935 | chr5:146832182-146832356 | DPYSL3 | N_Shore | Body |
| WG1984 | chr5:147257219-147257456 | SCGB3A2 | | TSS1500 |
| GM00031 | chr5:147443252-147443347 | SPINK5;SPINK5;SPINK5 | | TSS1500;TSS1500;TSS1500 |
| WG2414 | chr5:148442694-148442857 | SH3TC2;SH3TC2 | | 1stExon;5'UTR |
| GM00042 | chr5:148520817-148520894 | ABLIM3 | N_Shore | TSS1500 |
| GM09256 | chr5:148520854-148520894 | ABLIM3 | N_Shore | TSS200 |
| WG1722 | chr5:148785250-148785846 | LOC728264 | | TSS1500 |
| GM02555 | chr5:148810061-148810103 | MIR145;LOC728264 | | TSS200;Body |
| GM02456 | chr5:148810061-148810203 | MIR145;LOC728264 | | TSS200;Body |
| WG1310 | chr5:148810177-148810203 | MIR145;LOC728264 | | TSS200;Body |
| GM02706 | chr5:149792249-149792348 | CD74;CD74;CD74 | | 1stExon;1stExon;1stExon |
| GM00520 | chr5:149792463-149792783 | CD74;CD74;CD74 | | TSS200;TSS200;TSS200 |
| WG1722 | chr5:149980486-149980682 | SYNPO | | TSS200 |
| WG1310 | chr5:149980526-149980682 | SYNPO | | TSS200 |
| WG3466 | chr5:149980671-149980682 | SYNPO;SYNPO | | 1stExon;5'UTR |
| WG3466 | chr5:149999033-149999249 | SYNPO;SYNPO | | Body;Body |
| WG3466 | chr5:150004773-150004918 | SYNPO;SYNPO | Island | Body;Body |
| WG1513 | chr5:150227510-150227654 | IRGM | | 5'UTR |
| GM00438 | chr5:150284416-150284504 | ZNF300 | Island | TSS200 |
| WG3466 | chr5:153858585-153859854 | HAND1 | Island | TSS1500 |
| WG2121 | chr5:153858816-153858936 | HAND1 | Island | TSS1500 |
| WG3465 | chr5:156594400-156594725 | FAM71B | | TSS1500 |
| GM02555 | chr5:156886816-156887005 | NIPAL4 | N_Shore | TSS1500 |
| GM01096 | chr5:156886970-156887005 | NIPAL4 | Island | TSS200 |
| GM09255 | chr5:156886982-156887005 | NIPAL4 | Island | TSS200 |
| WG2276 | chr5:156886990-156887005 | NIPAL4 | Island | TSS200 |
| WG2844 | chr5:158522427-158522603 | EBF1 | N_Shore | Body |
| GM00438 | chr5:158523907-158524404 | EBF1 | Island | Body |

| | | | | |
|---|---|---|---|---|
| GM02555 | chr5:158524270-158524649 | EBF1 | Island | Body |
| GM03075 | chr5:158689566-158689727 | UBLCP1 | N_Shore | TSS1500 |
| GM02149 | chr5:1594282-1594676 | SDHAP3 | Island | Body |
| WG1984 | chr5:1594282-1594808 | SDHAP3 | Island | Body |
| WG1513 | chr5:1594282-1594863 | SDHAP3 | Island | Body |
| GM02149 | chr5:1594715-1594863 | SDHAP3 | Island | TSS200 |
| GM02149 | chr5:1634460-1634505 | LOC728613 | S_Shore | TSS1500 |
| WG1513 | chr5:16508920-16509123 | FAM134B;FAM134B;FAM134B | | Body;1stExon;5'UTR |
| WG1983 | chr5:16742020-16742179 | MYO10 | | Body |
| GM02706 | chr5:168728586-168729015 | SLIT3 | S_Shore | TSS1500 |
| GM01096 | chr5:168728793-168729267 | SLIT3 | S_Shore | TSS1500 |
| GM00438 | chr5:169063935-169064071 | DOCK2 | N_Shore | TSS1500 |
| WG2844 | chr5:169434-169797 | PLEKHG4B | N_Shore | Body |
| GM00520 | chr5:169434-170081 | PLEKHG4B | N_Shore | Body |
| GM09255 | chr5:169532752-169532775 | FOXI1;FOXI1 | N_Shore | TSS200;TSS200 |
| GM00520 | chr5:169757754-169758055 | LOC257358 | | TSS1500 |
| WG1657 | chr5:169815771-169816472 | KCNIP1;KCNMB1 | | Body;5'UTR |
| GM00016 | chr5:169816214-169816472 | KCNIP1;KCNMB1 | | Body;5'UTR |
| GM02149 | chr5:170210586-170210672 | GABRP | | TSS200 |
| GM02149 | chr5:170288742-170288766 | RANBP17 | N_Shore | TSS1500 |
| WG3466 | chr5:171534678-171534792 | STK10 | | Body |
| GM00438 | chr5:171708886-171709304 | UBTD2 | N_Shore | Body |
| GM01393 | chr5:172199244-172199278 | DUSP1 | Island | TSS1500 |
| GM00940 | chr5:172385677-172385817 | LOC100268168;LOC100268 | Island | Body;Body;TSS1500 |
| GM00438 | chr5:172385810-172385821 | LOC100268168;LOC100268 | Island | Body;TSS200;TSS1500 |
| GM02641 | chr5:172661803-172662315 | NKX2-5;NKX2-5;NKX2-5 | Island | 1stExon;1stExon;1stExon |
| GM02704 | chr5:172661859-172662315 | NKX2-5;NKX2-5;NKX2-5 | Island | 1stExon;1stExon;1stExon |
| GM02149 | chr5:175108269-175108315 | HRH2;HRH2 | | 5'UTR;TSS200 |
| WG2276 | chr5:175109098-175109114 | HRH2;HRH2;HRH2 | | 5'UTR;5'UTR;1stExon |
| WG2415 | chr5:176057924-176057996 | EIF4E1B;SNCB;SNCB;EIF4E1 | S_Shore | 5'UTR;TSS1500;TSS1500;1stExon |
| GM00439 | chr5:176270730-176270936 | UNC5A | | Body |
| WG2120 | chr5:176304988-176305077 | UNC5A | Island | Body |
| WG1977 | chr5:176559334-176559563 | NSD1;NSD1 | Island | TSS1500;TSS1500 |
| WG2415 | chr5:176784448-176784715 | RGS14 | | TSS1500 |
| GM00438 | chr5:176784550-176784719 | RGS14 | | TSS1500 |
| WG1978 | chr5:176784688-176784719 | RGS14 | | TSS200 |
| GM09256 | chr5:176828872-176829112 | PFN3 | N_Shore | TSS1500 |
| WG3465 | chr5:176829109-176829198 | PFN3 | N_Shore | TSS1500 |
| WG1721 | chr5:176981888-176982116 | FAM193B;FAM193B | S_Shore | TSS1500;TSS1500 |
| GM00438 | chr5:176981910-176982070 | FAM193B;FAM193B | S_Shore | TSS1500;TSS1500 |
| GM02175 | chr5:176981910-176982116 | FAM193B;FAM193B | S_Shore | TSS1500;TSS1500 |
| GM00520 | chr5:177434336-177434457 | FAM153C | S_Shore | TSS1500 |
| WG3466 | chr5:177548398-177548913 | N4BP3 | | Body |
| WG3465 | chr5:177548398-177549014 | N4BP3 | | Body |
| GM02706 | chr5:177548615-177549014 | N4BP3 | | Body |
| GM00935 | chr5:177614205-177614292 | GMCL1L | Island | Body |
| WG3466 | chr5:177708304-177708528 | COL23A1 | | Body |

| | | | | |
|---|---|---|---|---|
| WG1513 | chr5:177764172-177764408 | COL23A1 | Island | Body |
| WG3466 | chr5:177980691-177980944 | COL23A1 | | Body |
| WG3466 | chr5:178004012-178004204 | COL23A1 | Island | Body |
| GM02456 | chr5:178622572-178623035 | ADAMTS2;ADAMTS2 | | Body;Body |
| WG2415 | chr5:178684000-178684498 | ADAMTS2;ADAMTS2 | N_Shore | Body;Body |
| GM03075 | chr5:178684000-178684658 | ADAMTS2;ADAMTS2 | N_Shore | Body;Body |
| WG1722 | chr5:178684416-178684652 | ADAMTS2;ADAMTS2 | Island | Body;Body |
| WG2844 | chr5:178684416-178684658 | ADAMTS2;ADAMTS2 | Island | Body;Body |
| GM02149 | chr5:178986131-178986404 | RUFY1;RUFY1;RUFY1 | N_Shore | TSS1500;Body;TSS1500 |
| GM02706 | chr5:179588440-179588673 | RASGEF1C | Island | 5'UTR |
| WG3465 | chr5:179588440-179588838 | RASGEF1C | Island | 5'UTR |
| GM02175 | chr5:179588551-179588838 | RASGEF1C | Island | 5'UTR |
| GM01096 | chr5:179597395-179597526 | RASGEF1C | | 5'UTR |
| WG1721 | chr5:179634578-179634688 | RASGEF1C | Island | 5'UTR |
| WG2120 | chr5:179634671-179634789 | RASGEF1C | Island | 5'UTR |
| WG1977 | chr5:1800606-1800782 | NDUFS6;MRPL36 | Island | TSS1500;TSS1500 |
| GM00520 | chr5:180071709-180071816 | FLT4;FLT4 | N_Shelf | Body;Body |
| WG1983 | chr5:180230767-180230959 | MGAT1;MGAT1;MGAT1;MGAT | S_Shore | 1stExon;5'UTR;5'UTR;5'UTR;TSS1500; |
| GM02555 | chr5:180230767-180231026 | MGAT1;MGAT1;MGAT1;MGAT | S_Shore | 1stExon;5'UTR;5'UTR;5'UTR;TSS1500; |
| GM01096 | chr5:180231026-180231155 | MGAT1;MGAT1;MGAT1;MGAT | S_Shore | 5'UTR;TSS200;TSS1500;5'UTR;TSS200 |
| WG1513 | chr5:180325254-180326186 | BTNL8;BTNL8;BTNL8;BTNL8;BTNL8 | | TSS1500;TSS1500;TSS1500;TSS1500; |
| GM00438 | chr5:180325954-180326142 | BTNL8;BTNL8;BTNL8;BTNL8;BTNL8 | | TSS200;TSS200;TSS200;TSS200;TSS2 |
| GM09256 | chr5:180466679-180467044 | BTNL9 | | TSS1500 |
| GM00520 | chr5:180632948-180633326 | TRIM7;TRIM7 | S_Shore | TSS1500;TSS1500 |
| WG3466 | chr5:191127-191296 | LRRC14B | N_Shore | TSS1500 |
| GM00042 | chr5:191793-192103 | LRRC14B | Island | 1stExon |
| WG1978 | chr5:22853703-22853901 | CDH12;CDH12 | | 5'UTR;1stExon |
| WG3465 | chr5:23507450-23507469 | PRDM9 | | TSS1500 |
| WG1722 | chr5:23507562-23507594 | PRDM9 | | TSS200 |
| WG2415 | chr5:23507573-23507617 | PRDM9 | | TSS200 |
| GM00289 | chr5:23507573-23507644 | PRDM9 | | TSS200 |
| WG3465 | chr5:23507573-23507656 | PRDM9 | | TSS200 |
| GM01376 | chr5:24488007-24488076 | CDH10 | | Body |
| WG1657 | chr5:2750163-2750263 | IRX2;IRX2 | Island | Body;Body |
| WG2844 | chr5:2752882-2754148 | IRX2;IRX2;C5orf38 | Island | TSS1500;TSS1500;Body |
| WG1983 | chr5:2753852-2754074 | C5orf38 | Island | Body |
| GM00941 | chr5:2753876-2754074 | C5orf38 | Island | Body |
| GM00290 | chr5:2754148-2754753 | C5orf38 | Island | Body |
| WG2844 | chr5:2754274-2754753 | C5orf38 | Island | Body |
| WG2276 | chr5:323320-323969 | AHRR | S_Shore | Body |
| WG3465 | chr5:323794-323969 | AHRR | S_Shore | Body |
| GM01376 | chr5:33937673-33938213 | RXFP3 | Island | 1stExon |
| WG2276 | chr5:34042976-34043284 | C1QTNF3;C1QTNF3 | | Body;1stExon |
| GM00439 | chr5:34042976-34043333 | C1QTNF3;C1QTNF3 | | Body;1stExon |
| WG2844 | chr5:34042976-34043481 | C1QTNF3;C1QTNF3 | | Body;1stExon |
| GM01095 | chr5:34043140-34043333 | C1QTNF3;C1QTNF3 | | Body;1stExon |
| GM00520 | chr5:35990997-35991382 | UGT3A1 | | Body |

| | | | | |
|---|---|---|---|---|
| WG2276 | chr5:368394-368804 | AHRR | N_Shore | Body |
| GM02555 | chr5:368394-369088 | AHRR | N_Shore | Body |
| WG3466 | chr5:368804-369088 | AHRR | N_Shore | Body |
| GM09255 | chr5:37208976-37209440 | C5orf42 | N_Shore | Body |
| GM00016 | chr5:37834672-37835109 | GDNF;GDNF;GDNF | Island | Body;Body;Body |
| WG2194 | chr5:37834850-37834958 | GDNF;GDNF;GDNF | Island | 1stExon;Body;Body |
| WG2120 | chr5:37834850-37835109 | GDNF;GDNF;GDNF | Island | 1stExon;Body;Body |
| WG1977 | chr5:38257516-38257555 | EGFLAM | N_Shore | TSS1500 |
| WG2276 | chr5:38445220-38445563 | EGFLAM;EGFLAM;EGFLAM;EGFLAM | | Body;Body;TSS1500;Body |
| GM00016 | chr5:38445563-38445614 | EGFLAM;EGFLAM;EGFLAM;EGFLAM | | TSS200;Body;Body;Body |
| WG2276 | chr5:38445834-38446192 | EGFLAM;EGFLAM;EGFLAM;EGFLAM | | Body;5'UTR;Body;Body |
| WG2844 | chr5:39219245-39219698 | FYB;FYB | | 5'UTR;5'UTR |
| GM09256 | chr5:40841298-40841493 | CARD6 | | TSS200 |
| GM02175 | chr5:40841315-40841590 | CARD6 | | TSS200 |
| WG3465 | chr5:40841493-40841590 | CARD6 | | 1stExon |
| WG1309 | chr5:415575-415885 | AHRR | | Body |
| GM02456 | chr5:423894-424371 | AHRR | S_Shelf | Body |
| GM09255 | chr5:42756786-42756860 | CCDC152 | | TSS200 |
| GM00439 | chr5:42756786-42757171 | CCDC152 | | TSS200 |
| WG2415 | chr5:42756786-42757815 | CCDC152 | | TSS200 |
| WG1978 | chr5:42756851-42756876 | CCDC152 | | TSS200 |
| WG2121 | chr5:43044573-43044675 | LOC153684 | S_Shore | Body |
| GM00520 | chr5:471171-471385 | LOC25845 | N_Shore | Body |
| WG1283 | chr5:493262-493448 | SLC9A3 | N_Shore | Body |
| GM02173 | chr5:493308-493613 | SLC9A3 | N_Shore | Body |
| GM01376 | chr5:496299-496476 | SLC9A3 | N_Shore | Body |
| GM00438 | chr5:497397-497572 | SLC9A3 | N_Shore | Body |
| GM02175 | chr5:501863-502074 | SLC9A3 | Island | Body |
| WG1310 | chr5:501938-502074 | SLC9A3 | Island | Body |
| GM00439 | chr5:502074-502436 | SLC9A3 | Island | Body |
| GM02173 | chr5:502291-502571 | SLC9A3 | S_Shore | Body |
| GM00438 | chr5:5139458-5139512 | ADAMTS16 | Island | TSS1500 |
| WG3465 | chr5:54281198-54281362 | ESM1;ESM1 | | 1stExon;1stExon |
| GM02173 | chr5:54527320-54527671 | CCNO | Island | Body |
| GM01095 | chr5:54530279-54530436 | CCNO | S_Shore | TSS1500 |
| WG3466 | chr5:54826611-54826768 | PPAP2A;PPAP2A;RNF138P1 | Island | Body;Body;Body |
| GM02555 | chr5:55529213-55529375 | ANKRD55 | | TSS200 |
| GM02149 | chr5:56204884-56204925 | C5orf35 | N_Shore | TSS1500 |
| GM02555 | chr5:59188920-59189375 | PDE4D;PDE4D | N_Shore | Body;Body |
| GM00940 | chr5:6491443-6491550 | UBE2QL1 | | Body |
| WG1722 | chr5:6491472-6491754 | UBE2QL1 | | 3'UTR |
| WG3465 | chr5:66462293-66462662 | MAST4;MAST4 | S_Shelf | Body;Body |
| GM00439 | chr5:669733-669925 | TPPP | S_Shore | Body |
| GM00520 | chr5:67583598-67583627 | PIK3R1;PIK3R1 | N_Shore | Body;TSS1500 |
| GM00935 | chr5:68710808-68710912 | MARVELD2;MARVELD2 | Island | TSS200;TSS200 |
| WG3465 | chr5:68710808-68711277 | MARVELD2;MARVELD2 | Island | TSS200;TSS200 |
| GM09255 | chr5:68710813-68711277 | MARVELD2;MARVELD2 | Island | TSS200;TSS200 |

| | | | | |
|---|---|---|---|---|
| GM02456 | chr5:68710818-68710907 | MARVELD2;MARVELD2 | Island | TSS200;TSS200 |
| GM02175 | chr5:68710818-68710912 | MARVELD2;MARVELD2 | Island | TSS200;TSS200 |
| WG1977 | chr5:68710818-68711277 | MARVELD2;MARVELD2 | Island | TSS200;TSS200 |
| WG2414 | chr5:68710831-68710912 | MARVELD2;MARVELD2 | Island | TSS200;TSS200 |
| WG1722 | chr5:68787747-68787919 | OCLN | N_Shore | TSS1500 |
| WG1977 | chr5:72415429-72415770 | TMEM171;TMEM171 | N_Shore | TSS1500;TSS1500 |
| WG1977 | chr5:7394927-7395395 | ADCY2 | N_Shore | TSS1500 |
| WG2275 | chr5:74162602-74162623 | FAM169A;FAM169A | Island | 1stExon;5'UTR |
| WG3465 | chr5:76010444-76010658 | F2R | N_Shore | TSS1500 |
| WG1722 | chr5:76010472-76010658 | F2R | N_Shore | TSS1500 |
| WG1310 | chr5:76373022-76373719 | ZBED3 | Island | Body |
| WG2415 | chr5:76932016-76932062 | OTP | N_Shore | Body |
| GM00520 | chr5:77805555-77806312 | LHFPL2 | N_Shore | Body |
| GM00935 | chr5:77805555-77806538 | LHFPL2 | N_Shore | Body |
| GM02175 | chr5:77806051-77806312 | LHFPL2 | Island | 5'UTR |
| GM09256 | chr5:77806051-77806538 | LHFPL2 | Island | 5'UTR |
| GM00520 | chr5:7827486-7827964 | ADCY2 | | 3'UTR |
| WG2415 | chr5:78281793-78281827 | ARSB;ARSB;ARSB | S_Shore | TSS200;1stExon;5'UTR |
| GM01392 | chr5:78281793-78281964 | ARSB;ARSB;ARSB | S_Shore | TSS200;1stExon;5'UTR |
| GM00439 | chr5:78281793-78282099 | ARSB;ARSB;ARSB | S_Shore | TSS200;1stExon;5'UTR |
| GM09256 | chr5:78281819-78281964 | ARSB;ARSB;ARSB | S_Shore | TSS200;1stExon;5'UTR |
| WG1721 | chr5:78365647-78365710 | DMGDH;BHMT2 | Island | TSS200;Body |
| GM02704 | chr5:78365687-78365801 | BHMT2;DMGDH | Island | Body;TSS1500 |
| GM02706 | chr5:78407509-78407566 | BHMT | | TSS200 |
| WG1977 | chr5:78407562-78407566 | BHMT | | TSS200 |
| GM02706 | chr5:7851231-7851407 | C5orf49;C5orf49 | Island | 1stExon;5'UTR |
| WG2275 | chr5:83016630-83016779 | HAPLN1;HAPLN1 | N_Shore | 1stExon;5'UTR |
| WG3466 | chr5:83017009-83017184 | HAPLN1 | N_Shore | TSS200 |
| WG2415 | chr5:851225-851797 | ZDHHC11 | S_Shelf | TSS200 |
| GM00290 | chr5:87564247-87564941 | TMEM161B | N_Shore | Body |
| GM00290 | chr5:87565030-87565552 | TMEM161B | S_Shore | TSS1500 |
| GM09256 | chr5:87968118-87968528 | LOC645323;LOC645323;LO( | N_Shore | Body;Body;Body |
| WG2276 | chr5:912712-912860 | TRIP13 | Island | Body |
| GM02456 | chr5:92907151-92908051 | FLJ42709;FLJ42709;FLJ4270 | Island | TSS1500;Body;Body |
| GM00520 | chr5:92907553-92908051 | FLJ42709;FLJ42709;FLJ4270 | Island | TSS1500;Body;Body |
| WG2415 | chr5:92907851-92908051 | FLJ42709;FLJ42709;FLJ4270 | Island | TSS1500;Body;Body |
| WG2844 | chr5:92907851-92908392 | FLJ42709;FLJ42709;FLJ4270 | Island | TSS1500;Body;Body |
| GM02706 | chr5:92907851-92909069 | FLJ42709;FLJ42709;FLJ4270 | Island | TSS1500;Body;Body |
| WG2121 | chr5:92907881-92908392 | FLJ42709;FLJ42709;FLJ4270 | Island | TSS1500;Body;Body |
| GM01376 | chr5:92908988-92909429 | FLJ42709;FLJ42709 | S_Shore | Body;Body |
| WG2415 | chr5:92908988-92909434 | FLJ42709;FLJ42709 | S_Shore | Body;Body |
| WG2276 | chr5:92914202-92914319 | FLJ42709;FLJ42709 | N_Shore | Body;Body |
| GM00290 | chr5:92918912-92918943 | NR2F1 | N_Shore | TSS200 |
| GM00935 | chr5:92929319-92929573 | NR2F1 | Island | Body |
| WG1513 | chr5:92929319-92929679 | NR2F1 | Island | Body |
| GM00438 | chr5:93447570-93447700 | FAM172A;FAM172A;FAM172/ | S_Shore | TSS200;TSS200;TSS200;TSS200 |
| WG3465 | chr5:96078678-96079175 | CAST;CAST;CAST;CAST;CAST;CAST;CAS | | TSS1500;Body;Body;Body;Body;Body; |

| | | | | |
|---|---|---|---|---|
| WG1513 | chr6:100054118-100054515 | PRDM13 | N_Shore | TSS1500 |
| GM00935 | chr6:100894513-100895050 | SIM1 | N_Shore | Body |
| WG2121 | chr6:100903817-100903909 | SIM1 | N_Shore | Body |
| WG2120 | chr6:100903839-100904055 | SIM1 | N_Shore | Body |
| GM00940 | chr6:100911240-100911550 | SIM1 | N_Shore | 1stExon |
| GM00941 | chr6:100911240-100911701 | SIM1 | N_Shore | 1stExon |
| WG1977 | chr6:100911240-100911709 | SIM1 | N_Shore | 1stExon |
| WG2120 | chr6:100911240-100911727 | SIM1 | N_Shore | 1stExon |
| GM01392 | chr6:100911240-100911746 | SIM1 | N_Shore | 1stExon |
| WG2275 | chr6:100911240-100911964 | SIM1 | N_Shore | 1stExon |
| WG3465 | chr6:100911240-100912072 | SIM1 | N_Shore | 1stExon |
| WG1514 | chr6:100911240-100912168 | SIM1 | N_Shore | 1stExon |
| WG1722 | chr6:100911526-100911687 | SIM1;SIM1 | N_Shore | 1stExon;5'UTR |
| GM03075 | chr6:100911526-100911701 | SIM1;SIM1 | N_Shore | 1stExon;5'UTR |
| GM02640 | chr6:100911550-100911727 | SIM1;SIM1 | N_Shore | 1stExon;5'UTR |
| WG3466 | chr6:100911687-100911727 | SIM1 | N_Shore | TSS200 |
| WG2414 | chr6:100911701-100911727 | SIM1 | N_Shore | TSS200 |
| GM00940 | chr6:100911701-100912168 | SIM1 | N_Shore | TSS200 |
| WG1977 | chr6:100911744-100912168 | SIM1 | N_Shore | TSS200 |
| GM02640 | chr6:100911746-100911964 | SIM1 | N_Shore | TSS200 |
| WG2120 | chr6:100911746-100912072 | SIM1 | N_Shore | TSS200 |
| WG3466 | chr6:100911949-100912072 | SIM1 | N_Shore | TSS1500 |
| WG1513 | chr6:100911949-100912168 | SIM1 | N_Shore | TSS1500 |
| GM00290 | chr6:101846409-101846872 | GRIK2;GRIK2;GRIK2 | N_Shore | TSS1500;TSS1500;TSS1500 |
| GM02149 | chr6:101846409-101848236 | GRIK2;GRIK2;GRIK2 | N_Shore | TSS1500;TSS1500;TSS1500 |
| GM01392 | chr6:101846650-101846767 | GRIK2;GRIK2;GRIK2 | N_Shore | TSS1500;TSS1500;TSS1500 |
| WG3465 | chr6:101846650-101846872 | GRIK2;GRIK2;GRIK2 | N_Shore | TSS1500;TSS1500;TSS1500 |
| GM00289 | chr6:101846650-101846905 | GRIK2;GRIK2;GRIK2 | N_Shore | TSS1500;TSS1500;TSS1500 |
| WG1977 | chr6:101846650-101846967 | GRIK2;GRIK2;GRIK2 | N_Shore | TSS1500;TSS1500;TSS1500 |
| WG1310 | chr6:101846650-101847058 | GRIK2;GRIK2;GRIK2 | N_Shore | TSS1500;TSS1500;TSS1500 |
| WG1721 | chr6:101846707-101846779 | GRIK2;GRIK2;GRIK2 | N_Shore | TSS200;TSS200;TSS200 |
| GM02456 | chr6:101846707-101848236 | GRIK2;GRIK2;GRIK2 | N_Shore | TSS200;TSS200;TSS200 |
| GM00940 | chr6:101846767-101846805 | GRIK2;GRIK2;GRIK2 | Island | TSS200;TSS200;TSS200 |
| WG1657 | chr6:101846767-101846872 | GRIK2;GRIK2;GRIK2 | Island | TSS200;TSS200;TSS200 |
| GM02173 | chr6:101846767-101847058 | GRIK2;GRIK2;GRIK2 | Island | TSS200;TSS200;TSS200 |
| GM00935 | chr6:101846767-101848236 | GRIK2;GRIK2;GRIK2 | Island | TSS200;TSS200;TSS200 |
| WG3466 | chr6:101846791-101846805 | GRIK2;GRIK2;GRIK2 | Island | TSS200;TSS200;TSS200 |
| WG2843 | chr6:101846791-101846872 | GRIK2;GRIK2;GRIK2 | Island | TSS200;TSS200;TSS200 |
| WG1722 | chr6:101846791-101846905 | GRIK2;GRIK2;GRIK2 | Island | TSS200;TSS200;TSS200 |
| GM02147 | chr6:101846791-101847388 | GRIK2;GRIK2;GRIK2 | Island | TSS200;TSS200;TSS200 |
| WG2415 | chr6:101846791-101848236 | GRIK2;GRIK2;GRIK2 | Island | TSS200;TSS200;TSS200 |
| WG1978 | chr6:101846797-101847706 | GRIK2;GRIK2;GRIK2 | Island | TSS200;TSS200;TSS200 |
| GM00940 | chr6:101846916-101847058 | GRIK2;GRIK2;GRIK2;GRIK2;G | Island | 1stExon;1stExon;5'UTR;1stExon;5'UTR |
| WG2843 | chr6:101846916-101847318 | GRIK2;GRIK2;GRIK2;GRIK2;G | Island | 1stExon;1stExon;5'UTR;1stExon;5'UTR |
| WG3465 | chr6:101846916-101847388 | GRIK2;GRIK2;GRIK2;GRIK2;G | Island | 1stExon;1stExon;5'UTR;1stExon;5'UTR |
| GM00290 | chr6:101846916-101848236 | GRIK2;GRIK2;GRIK2;GRIK2;G | Island | 1stExon;1stExon;5'UTR;1stExon;5'UTR |
| GM00289 | chr6:101846967-101847318 | GRIK2;GRIK2;GRIK2;GRIK2;G | Island | 1stExon;1stExon;5'UTR;1stExon;5'UTR |

| | | | | |
|---|---|---|---|---|
| WG1282 | chr6:101846967-101847656 | GRIK2;GRIK2;GRIK2;GRIK2;G | Island | 1stExon;1stExon;5'UTR;1stExon;5'UTR |
| GM00520 | chr6:101846967-101848236 | GRIK2;GRIK2;GRIK2;GRIK2;G | Island | 1stExon;1stExon;5'UTR;1stExon;5'UTR |
| WG1722 | chr6:101847058-101847656 | GRIK2;GRIK2;GRIK2;GRIK2;G | Island | 1stExon;1stExon;5'UTR;1stExon;5'UTR |
| WG1283 | chr6:101847058-101847706 | GRIK2;GRIK2;GRIK2;GRIK2;G | Island | 1stExon;1stExon;5'UTR;1stExon;5'UTR |
| GM02555 | chr6:101847318-101847388 | GRIK2;GRIK2;GRIK2 | S_Shore | Body;Body;Body |
| WG1977 | chr6:101847318-101847656 | GRIK2;GRIK2;GRIK2 | S_Shore | Body;Body;Body |
| GM00042 | chr6:101847318-101847706 | GRIK2;GRIK2;GRIK2 | S_Shore | Body;Body;Body |
| WG1310 | chr6:101847349-101847541 | GRIK2;GRIK2;GRIK2 | S_Shore | Body;Body;Body |
| GM02173 | chr6:101847349-101847656 | GRIK2;GRIK2;GRIK2 | S_Shore | Body;Body;Body |
| GM00289 | chr6:101847388-101848236 | GRIK2;GRIK2;GRIK2 | S_Shore | Body;Body;Body |
| WG1658 | chr6:101847541-101847706 | GRIK2;GRIK2;GRIK2 | S_Shore | Body;Body;Body |
| GM09256 | chr6:10398494-10398698 | TFAP2A;TFAP2A;TFAP2A | N_Shore | 3'UTR;3'UTR;3'UTR |
| WG1978 | chr6:10398601-10398754 | TFAP2A;TFAP2A;TFAP2A | Island | 3'UTR;3'UTR;3'UTR |
| GM09256 | chr6:10410284-10410544 | TFAP2A;TFAP2A;TFAP2A | Island | Body;Body;Body |
| GM00438 | chr6:10410494-10410916 | TFAP2A;TFAP2A;TFAP2A | Island | Body;Body;Body |
| WG2414 | chr6:10415873-10416034 | TFAP2A;TFAP2A | N_Shore | Body;TSS1500 |
| GM09256 | chr6:10415873-10416090 | TFAP2A;TFAP2A | N_Shore | Body;TSS1500 |
| GM09255 | chr6:10416034-10416531 | TFAP2A;TFAP2A | N_Shore | Body;TSS1500 |
| WG1513 | chr6:10416090-10416373 | TFAP2A;TFAP2A | N_Shore | Body;TSS1500 |
| GM00439 | chr6:10416090-10416531 | TFAP2A;TFAP2A | N_Shore | Body;TSS1500 |
| GM09256 | chr6:10416373-10416531 | TFAP2A;TFAP2A | N_Shore | Body;TSS1500 |
| WG2844 | chr6:10420904-10421069 | TFAP2A | N_Shore | TSS1500 |
| GM00520 | chr6:106546074-106546824 | PRDM1;PRDM1 | | Body;TSS1500 |
| GM01393 | chr6:106546704-106546754 | PRDM1;PRDM1 | | Body;TSS200 |
| WG3465 | chr6:106546704-106546824 | PRDM1;PRDM1 | | Body;TSS200 |
| GM00935 | chr6:106546744-106546754 | PRDM1;PRDM1 | | Body;TSS200 |
| WG1978 | chr6:106546747-106546824 | PRDM1;PRDM1 | | Body;TSS200 |
| GM00438 | chr6:108145374-108145539 | SCML4;SCML4 | | 1stExon;5'UTR |
| GM01392 | chr6:108488335-108488751 | NR2E1 | Island | Body |
| GM02175 | chr6:108492653-108492769 | NR2E1 | S_Shore | Body |
| GM00290 | chr6:10881782-10881925 | GCM2 | N_Shore | Body |
| WG2276 | chr6:109274429-109274496 | ARMC2 | | Body |
| WG2415 | chr6:11044888-11044974 | ELOVL2 | Island | TSS1500 |
| GM00520 | chr6:110677913-110678179 | C6orf186 | N_Shore | Body |
| WG1977 | chr6:110720918-110721305 | DDO;DDO | N_Shore | Body;Body |
| WG1513 | chr6:110721305-110721629 | DDO;DDO | Island | Body;Body |
| GM00438 | chr6:110736865-110737053 | DDO;DDO | | TSS200;TSS200 |
| WG2415 | chr6:111888567-111888652 | TRAF3IP2;TRAF3IP2;TRAF3IP2;TRAF3IP2; | | Body;Body;Body;TSS200;Body |
| WG2276 | chr6:111888620-111888683 | TRAF3IP2;TRAF3IP2;TRAF3IP2;TRAF3IP2; | | Body;Body;Body;TSS200;Body |
| WG1983 | chr6:11279656-11279727 | NEDD9 | | Body |
| GM03075 | chr6:114181482-114181798 | MARCKS | Island | Body |
| WG1658 | chr6:114291911-114292459 | HDAC2 | Island | Body |
| WG1658 | chr6:114292723-114292751 | HDAC2 | Island | TSS1500 |
| WG1658 | chr6:116381609-116381957 | FRK;FRK | | 1stExon;5'UTR |
| WG1983 | chr6:116381904-116381976 | FRK;FRK | | 1stExon;5'UTR |
| GM02175 | chr6:117086996-117087056 | FAM162B | S_Shore | TSS200 |
| GM01650 | chr6:117591433-117592358 | VGLL2;VGLL2 | N_Shore | Body;Body |

| | | | | |
|---|---|---|---|---|
| WG3466 | chr6:117591534-117592358 | VGLL2;VGLL2 | Island | Body;Body |
| WG1722 | chr6:11771493-11771840 | C6orf105;C6orf105 | | Body;Body |
| GM02706 | chr6:117869027-117869453 | DCBLD1 | N_Shore | Body |
| GM00940 | chr6:118228493-118228514 | SLC35F1 | Island | TSS200 |
| GM00022 | chr6:118973136-118973337 | C6orf204;C6orf204 | Island | TSS200;TSS200 |
| WG2121 | chr6:122931154-122931192 | PKIB;PKIB;PKIB | Island | 5'UTR;TSS1500;TSS1500 |
| GM02706 | chr6:123109391-123109837 | SMPDL3A | N_Shore | TSS1500 |
| WG3465 | chr6:127441605-127441993 | RSPO3 | Island | Body |
| GM00438 | chr6:127796683-127797022 | C6orf174 | Island | Body |
| GM01376 | chr6:127796683-127797208 | C6orf174 | Island | Body |
| GM00935 | chr6:127796989-127797208 | C6orf174 | Island | Body |
| GM02706 | chr6:127835750-127836053 | C6orf174 | N_Shore | Body |
| WG1513 | chr6:132931714-132931736 | TAAR3 | | TSS1500 |
| WG2415 | chr6:133035150-133035266 | VNN1 | | 1stExon |
| WG2415 | chr6:133137810-133137893 | RPS12;SNORD100;SNORA33 | S_Shore | Body;TSS200;TSS1500 |
| WG2275 | chr6:133561677-133561814 | EYA4;EYA4;EYA4 | N_Shore | TSS1500;TSS1500;TSS1500 |
| GM00042 | chr6:133561814-133561886 | EYA4;EYA4;EYA4 | N_Shore | TSS1500;TSS1500;TSS1500 |
| GM00289 | chr6:133561814-133561932 | EYA4;EYA4;EYA4 | N_Shore | TSS1500;TSS1500;TSS1500 |
| WG3466 | chr6:134210273-134210307 | TCF21;TCF21;TCF21;TCF21 | N_Shore | 1stExon;5'UTR;1stExon;5'UTR |
| WG2121 | chr6:134210667-134210946 | TCF21;TCF21 | Island | 1stExon;1stExon |
| WG1722 | chr6:134213992-134214134 | TCF21 | Island | 3'UTR |
| GM02706 | chr6:134497247-134497755 | SGK1;SGK1;SGK1;SGK1 | N_Shore | Body;TSS200;Body;TSS1500 |
| GM01095 | chr6:134497379-134497627 | SGK1;SGK1;SGK1;SGK1 | N_Shore | TSS1500;Body;Body;TSS1500 |
| GM01096 | chr6:134497542-134497755 | SGK1;SGK1;SGK1 | Island | TSS1500;Body;Body |
| WG3466 | chr6:135271333-135271801 | ALDH8A1;ALDH8A1 | | TSS200;TSS200 |
| WG3465 | chr6:135271518-135271801 | ALDH8A1;ALDH8A1 | | TSS1500;TSS1500 |
| WG1513 | chr6:137494767-137495383 | IL22RA2;IL22RA2;IL22RA2;IL22RA2;IL22I | | 1stExon;5'UTR;5'UTR;1stExon;5'UTR;1 |
| WG3465 | chr6:138200071-138200309 | TNFAIP3 | | Body |
| WG2120 | chr6:138200136-138200334 | TNFAIP3 | | Body |
| GM09256 | chr6:138200136-138200387 | TNFAIP3 | | Body |
| GM01393 | chr6:138866882-138867046 | NHSL1 | | Body |
| WG1978 | chr6:1388806-1389167 | FOXF2 | N_Shore | TSS1500 |
| WG3465 | chr6:1388858-1389167 | FOXF2 | N_Shore | TSS1500 |
| GM09256 | chr6:1389146-1389367 | FOXF2 | Island | TSS1500 |
| WG2844 | chr6:139351113-139351353 | C6orf115 | S_Shore | 5'UTR |
| WG3465 | chr6:139613145-139613456 | TXLNB;TXLNB | | 5'UTR;1stExon |
| GM00042 | chr6:14118033-14118415 | CD83;CD83;CD83;CD83 | Island | 5'UTR;5'UTR;1stExon;1stExon |
| GM01392 | chr6:144386231-144386280 | PLAGL1;PLAGL1 | S_Shore | TSS1500;TSS1500 |
| WG1722 | chr6:144386275-144386416 | PLAGL1;PLAGL1 | S_Shore | TSS1500;TSS1500 |
| WG1513 | chr6:144386416-144386422 | PLAGL1;PLAGL1 | S_Shore | TSS1500;TSS1500 |
| GM01392 | chr6:146864328-146864352 | RAB32 | N_Shore | TSS1500 |
| GM02706 | chr6:147124881-147124996 | C6orf103;LOC729176 | | Body;Body |
| GM00935 | chr6:147124984-147124988 | LOC729176;C6orf103 | | TSS200;Body |
| WG2414 | chr6:148869461-148869535 | SASH1 | | Body |
| WG2194 | chr6:148869489-148869895 | SASH1 | | Body |
| WG1722 | chr6:149358743-149358802 | UST | | Body |
| WG3466 | chr6:149772707-149772854 | ZC3H12D | Island | Body |

| | | | | |
|---|---|---|---|---|
| GM02706 | chr6:149772716-149772892 | ZC3H12D | Island | Body |
| GM00290 | chr6:149803087-149803206 | ZC3H12D | | 5'UTR |
| WG2415 | chr6:149803087-149803444 | ZC3H12D | | 5'UTR |
| WG2844 | chr6:149803111-149803444 | ZC3H12D | | 5'UTR |
| WG2276 | chr6:149805995-149806131 | ZC3H12D;ZC3H12D | | 5'UTR;1stExon |
| WG3465 | chr6:149805995-149806635 | ZC3H12D;ZC3H12D | | 5'UTR;1stExon |
| WG1282 | chr6:149806077-149806131 | ZC3H12D;ZC3H12D | | 5'UTR;1stExon |
| WG1984 | chr6:149806131-149806331 | ZC3H12D;ZC3H12D | | 5'UTR;1stExon |
| WG1722 | chr6:149806131-149806635 | ZC3H12D;ZC3H12D | | 5'UTR;1stExon |
| WG1282 | chr6:149806273-149806502 | ZC3H12D | | TSS200 |
| GM01393 | chr6:149806273-149806635 | ZC3H12D | | TSS200 |
| GM02555 | chr6:149806331-149806635 | ZC3H12D | | TSS200 |
| WG2415 | chr6:150286157-150286646 | ULBP1 | Island | Body |
| WG3466 | chr6:151186411-151186511 | MTHFD1L | N_Shore | TSS1500 |
| WG3465 | chr6:151258337-151258520 | MTHFD1L | | Body |
| WG1722 | chr6:151560831-151560927 | AKAP12 | Island | TSS1500 |
| GM01096 | chr6:151560831-151560981 | AKAP12 | Island | TSS1500 |
| GM02706 | chr6:151560831-151561055 | AKAP12 | Island | TSS1500 |
| GM00042 | chr6:151560981-151561032 | AKAP12 | Island | TSS200 |
| WG2276 | chr6:152011103-152011415 | ESR1 | | TSS1500 |
| GM00940 | chr6:152011178-152011399 | ESR1 | | TSS1500 |
| GM02555 | chr6:152011399-152011656 | ESR1 | | TSS1500 |
| WG3465 | chr6:152085565-152085641 | ESR1 | | 5'UTR |
| WG2844 | chr6:152125861-152126080 | ESR1;ESR1 | N_Shelf | 5'UTR;TSS1500 |
| WG3465 | chr6:152125861-152126092 | ESR1;ESR1 | N_Shelf | 5'UTR;TSS1500 |
| WG2415 | chr6:152125861-152126180 | ESR1;ESR1 | N_Shelf | 5'UTR;TSS1500 |
| GM02706 | chr6:152125965-152126092 | ESR1;ESR1 | N_Shelf | 5'UTR;TSS1500 |
| GM00022 | chr6:152125965-152126180 | ESR1;ESR1 | N_Shelf | 5'UTR;TSS1500 |
| WG2275 | chr6:152126080-152126250 | ESR1;ESR1 | N_Shelf | 5'UTR;TSS1500 |
| WG2276 | chr6:152126180-152126441 | ESR1;ESR1 | N_Shelf | 5'UTR;TSS1500 |
| WG2844 | chr6:152127812-152127887 | ESR1;ESR1;ESR1;ESR1 | N_Shore | TSS1500;5'UTR;TSS1500;5'UTR |
| WG3465 | chr6:152127812-152128024 | ESR1;ESR1;ESR1;ESR1 | N_Shore | TSS1500;5'UTR;TSS1500;5'UTR |
| WG3466 | chr6:152127812-152128338 | ESR1;ESR1;ESR1;ESR1 | N_Shore | TSS1500;5'UTR;TSS1500;5'UTR |
| GM00042 | chr6:152128024-152128366 | ESR1;ESR1;ESR1;ESR1 | N_Shore | TSS1500;5'UTR;TSS1500;5'UTR |
| GM09256 | chr6:152128024-152128426 | ESR1;ESR1;ESR1;ESR1 | N_Shore | TSS1500;5'UTR;TSS1500;5'UTR |
| WG3465 | chr6:152128328-152128366 | ESR1;ESR1;ESR1;ESR1 | N_Shore | TSS1500;TSS200;5'UTR;5'UTR |
| GM00935 | chr6:152128328-152128411 | ESR1;ESR1;ESR1;ESR1 | N_Shore | TSS1500;TSS200;5'UTR;5'UTR |
| GM00290 | chr6:152128328-152128483 | ESR1;ESR1;ESR1;ESR1 | N_Shore | TSS1500;TSS200;5'UTR;5'UTR |
| WG1514 | chr6:152128328-152128515 | ESR1;ESR1;ESR1;ESR1 | N_Shore | TSS1500;TSS200;5'UTR;5'UTR |
| GM02173 | chr6:152128328-152128675 | ESR1;ESR1;ESR1;ESR1 | N_Shore | TSS1500;TSS200;5'UTR;5'UTR |
| GM09256 | chr6:152128515-152128584 | ESR1;ESR1;ESR1;ESR1;ESR1 | N_Shore | TSS1500;5'UTR;1stExon;5'UTR;5'UTR |
| GM09256 | chr6:152128634-152130332 | ESR1;ESR1;ESR1;ESR1;ESR1 | N_Shore | 5'UTR;TSS200;1stExon;5'UTR;5'UTR |
| GM00289 | chr6:152128675-152128865 | ESR1;ESR1;ESR1;ESR1 | N_Shore | 5'UTR;TSS200;5'UTR;5'UTR |
| GM02704 | chr6:152128865-152129400 | ESR1;ESR1;ESR1;ESR1;ESR1 | Island | 5'UTR;5'UTR;1stExon;5'UTR;5'UTR |
| GM00289 | chr6:152129400-152130207 | ESR1;ESR1;ESR1;ESR1 | Island | Body;Body;1stExon;Body |
| WG1282 | chr6:152129749-152130207 | ESR1;ESR1;ESR1;ESR1 | Island | Body;Body;Body;Body |
| GM02704 | chr6:152129791-152130332 | ESR1;ESR1;ESR1;ESR1 | S_Shore | Body;Body;Body;Body |

| | | | | |
|---|---|---|---|---|
| WG1983 | chr6:152130058-152130332 | ESR1;ESR1;ESR1;ESR1 | S_Shore | Body;Body;Body;Body |
| WG2121 | chr6:155577805-155578034 | TIAM2;TIAM2 | | Body;Body |
| GM02555 | chr6:158018055-158018184 | ZDHHC14;ZDHHC14 | S_Shelf | Body;Body |
| WG2276 | chr6:158404061-158404121 | SYNJ2 | S_Shore | Body |
| GM02456 | chr6:158464617-158464852 | SYNJ2 | | Body |
| GM00022 | chr6:159070607-159070959 | SYTL3 | | TSS1500 |
| GM00438 | chr6:159070607-159070994 | SYTL3 | | TSS1500 |
| GM09256 | chr6:159070959-159071008 | SYTL3 | | TSS200 |
| WG3466 | chr6:159070959-159071076 | SYTL3 | | TSS200 |
| GM00042 | chr6:159070959-159071106 | SYTL3 | | TSS200 |
| WG1310 | chr6:159084370-159084599 | SYTL3 | | Body |
| WG1310 | chr6:159238463-159238744 | EZR;EZR | N_Shore | Body;Body |
| WG3466 | chr6:160182184-160182805 | ACAT2 | N_Shore | TSS1500 |
| WG2194 | chr6:160182438-160182554 | ACAT2 | Island | TSS1500 |
| WG3465 | chr6:160182438-160182624 | ACAT2 | Island | TSS1500 |
| GM00940 | chr6:160783785-160784044 | SLC22A3 | Island | Body |
| WG2415 | chr6:161558075-161558105 | AGPAT4 | | Body |
| GM02706 | chr6:161561031-161561121 | AGPAT4 | | Body |
| WG3465 | chr6:161796785-161796854 | PARK2;PARK2;PARK2 | | Body;Body;Body |
| WG3466 | chr6:161860203-161860359 | PARK2;PARK2;PARK2 | | Body;Body;Body |
| WG1513 | chr6:165715586-165715757 | C6orf118 | | Body |
| GM02706 | chr6:166354015-166354232 | C6orf176;C6orf176 | Island | Body;Body |
| WG3465 | chr6:167062104-167062226 | RPS6KA2 | | Body |
| WG1513 | chr6:167529765-167530253 | CCR6 | | 5'UTR |
| WG3466 | chr6:167530185-167530343 | CCR6 | | 5'UTR |
| WG1513 | chr6:167535171-167536086 | CCR6;CCR6 | | TSS1500;5'UTR |
| GM00438 | chr6:167535909-167536063 | CCR6;CCR6 | | TSS1500;5'UTR |
| GM00042 | chr6:167704673-167704808 | UNC93A;UNC93A | | TSS200;TSS200 |
| GM00940 | chr6:168197699-168197921 | C6orf123 | Island | TSS200 |
| WG3466 | chr6:168418276-168418398 | KIF25;KIF25 | | TSS1500;TSS1500 |
| WG2193 | chr6:168435914-168436353 | KIF25;KIF25 | Island | Body;Body |
| GM00439 | chr6:168435923-168436353 | KIF25;KIF25 | Island | Body;Body |
| WG2275 | chr6:168436019-168436353 | KIF25;KIF25 | Island | Body;Body |
| GM02149 | chr6:168859941-168859997 | SMOC2;SMOC2 | | Body;Body |
| GM00438 | chr6:168897991-168898334 | SMOC2;SMOC2 | | Body;Body |
| GM00438 | chr6:168956418-168956903 | SMOC2;SMOC2 | Island | Body;Body |
| WG1513 | chr6:169013530-169013732 | SMOC2;SMOC2 | | Body;Body |
| WG1513 | chr6:169018751-169018877 | SMOC2;SMOC2 | | Body;Body |
| GM00438 | chr6:169026027-169026352 | SMOC2;SMOC2 | | Body;Body |
| GM02149 | chr6:169026695-169027033 | SMOC2;SMOC2 | | Body;Body |
| GM00438 | chr6:169027019-169027218 | SMOC2;SMOC2 | | Body;Body |
| GM02456 | chr6:170048365-170048424 | WDR27 | | Body |
| WG1977 | chr6:170564889-170565151 | LOC154449 | Island | Body |
| WG1977 | chr6:170571686-170571763 | LOC154449 | | TSS200 |
| GM09256 | chr6:18123047-18123164 | NHLRC1 | S_Shore | TSS200 |
| WG2275 | chr6:18387077-18387360 | RNF144B | N_Shore | TSS1500 |
| WG3465 | chr6:18387313-18387407 | RNF144B | N_Shore | TSS1500 |

| | | | | |
|---|---|---|---|---|
| WG2415 | chr6:21667166-21667407 | FLJ22536 | S_Shore | Body |
| GM00439 | chr6:21667166-21668049 | FLJ22536 | S_Shore | Body |
| WG2844 | chr6:21667333-21668049 | FLJ22536 | S_Shore | Body |
| GM00439 | chr6:24357720-24358236 | KAAG1;DCDC2;KAAG1 | Island | 1stExon;1stExon;5'UTR |
| WG2276 | chr6:24358306-24358681 | KAAG1;KAAG1;DCDC2 | Island | 1stExon;3'UTR;TSS200 |
| WG2415 | chr6:24583477-24583614 | KIAA0319;KIAA0319;KIAA0319;KIAA0319 | | Body;Body;Body;Body;Body |
| GM00438 | chr6:25137864-25138169 | CMAH;CMAH | N_Shelf | Body;Body |
| GM00042 | chr6:25726884-25726912 | HIST1H2AA;HIST1H2BA | S_Shore | TSS200;TSS1500 |
| WG3466 | chr6:26018084-26018185 | HIST1H1A | N_Shelf | TSS200 |
| WG3466 | chr6:26235311-26235609 | HIST1H1D | | TSS200 |
| GM01096 | chr6:26247246-26247270 | HIST1H4G | S_Shore | TSS200 |
| WG1722 | chr6:26522136-26522578 | HCG11 | Island | Body |
| GM02149 | chr6:2698001-2698037 | MYLK4 | | Body |
| GM01650 | chr6:27106988-27107019 | HIST1H2BK;HIST1H4I | N_Shore | 3'UTR;TSS200 |
| GM09256 | chr6:27107393-27107670 | HIST1H2BK;HIST1H4I | Island | 3'UTR;1stExon |
| GM02175 | chr6:27107393-27107756 | HIST1H2BK;HIST1H4I | Island | 3'UTR;1stExon |
| WG1513 | chr6:27107588-27107718 | HIST1H2BK | S_Shore | 3'UTR |
| GM00289 | chr6:27107588-27107756 | HIST1H2BK | S_Shore | 3'UTR |
| GM00439 | chr6:27215365-27215509 | PRSS16 | N_Shelf | TSS200 |
| WG3465 | chr6:27799360-27799380 | HIST1H4K | Island | TSS200 |
| GM09256 | chr6:27799360-27799385 | HIST1H4K | Island | TSS200 |
| GM00042 | chr6:27831894-27832098 | HIST1H2AL | N_Shore | TSS1500 |
| GM09256 | chr6:27841230-27841683 | HIST1H4L;HIST1H3I | Island | 1stExon;TSS1500 |
| GM03073 | chr6:28058724-28058802 | ZSCAN12L1 | | TSS1500 |
| GM00520 | chr6:28058789-28058856 | ZSCAN12L1 | | TSS200 |
| GM00042 | chr6:28227091-28227127 | NKAPL | Island | TSS200 |
| GM00438 | chr6:28478052-28478432 | GPX6 | | Body |
| GM00438 | chr6:28480628-28480984 | GPX6 | | Body |
| WG1513 | chr6:28481839-28482073 | GPX6 | | Body |
| GM00438 | chr6:28481862-28482073 | GPX6 | | Body |
| GM00438 | chr6:28493284-28493500 | GPX5;GPX5 | | TSS1500;TSS1500 |
| WG1513 | chr6:28493284-28494516 | GPX5;GPX5 | | TSS1500;TSS1500 |
| GM02175 | chr6:28493312-28493601 | GPX5;GPX5 | | TSS1500;TSS1500 |
| WG1977 | chr6:28493651-28493664 | GPX5;GPX5 | | TSS200;TSS200 |
| WG1513 | chr6:28497278-28497370 | GPX5;GPX5 | | Body;Body |
| GM09256 | chr6:28497338-28497370 | GPX5;GPX5 | | Body;Body |
| GM02706 | chr6:28501867-28501916 | GPX5;GPX5 | | Body;3'UTR |
| GM09256 | chr6:28551669-28551812 | SCAND3 | N_Shelf | Body |
| WG1977 | chr6:28554497-28554741 | SCAND3;SCAND3 | N_Shore | 1stExon;5'UTR |
| GM09255 | chr6:28554680-28554774 | SCAND3;SCAND3 | N_Shore | 1stExon;5'UTR |
| GM00439 | chr6:28555440-28555653 | SCAND3 | S_Shore | TSS1500 |
| WG3465 | chr6:28555601-28555664 | SCAND3 | S_Shore | TSS1500 |
| GM01392 | chr6:28875354-28875370 | TRIM27 | | Body |
| WG1722 | chr6:28963745-28963820 | ZNF311 | | Body |
| WG1513 | chr6:29079939-29080040 | OR2J3 | | 1stExon |
| WG2415 | chr6:291687-292823 | DUSP22 | N_Shore | TSS1500 |
| WG2844 | chr6:291882-292823 | DUSP22 | N_Shore | TSS1500 |

| | | | | |
|---|---|---|---|---|
| GM02706 | chr6:29274096-29274617 | OR14J1 | | TSS1500 |
| GM00438 | chr6:29342605-29342679 | OR12D3 | | 1stExon |
| WG1513 | chr6:29364174-29364435 | OR12D2 | | TSS1500 |
| GM00438 | chr6:29394654-29394863 | OR11A1 | | 1stExon |
| WG1513 | chr6:29395103-29395221 | OR11A1 | | 1stExon |
| GM02149 | chr6:29395421-29395657 | OR11A1;OR11A1 | | 1stExon;5'UTR |
| WG1513 | chr6:29395421-29396182 | OR11A1;OR11A1 | | 1stExon;5'UTR |
| WG1513 | chr6:29407689-29407834 | OR10C1 | | TSS200 |
| WG1513 | chr6:29408118-29408229 | OR10C1 | | 1stExon |
| WG1513 | chr6:29408443-29408587 | OR10C1 | | 1stExon |
| WG1513 | chr6:29425885-29426119 | OR2H1 | | TSS1500 |
| GM00438 | chr6:29425885-29426132 | OR2H1 | | TSS1500 |
| WG1978 | chr6:29455302-29455331 | MAS1L | | 1stExon |
| WG1513 | chr6:29523592-29523835 | UBD | S_Shore | 3'UTR |
| GM02555 | chr6:29589545-29589631 | GABBR1;GABBR1;GABBR1 | | Body;Body;Body |
| WG1722 | chr6:29592854-29593382 | GABBR1;GABBR1;GABBR1 | N_Shelf | Body;Body;Body |
| WG1977 | chr6:29594830-29595016 | GABBR1;GABBR1;GABBR1 | N_Shore | Body;Body;Body |
| WG1310 | chr6:29595002-29595016 | GABBR1;GABBR1;GABBR1 | N_Shore | Body;Body;Body |
| GM00935 | chr6:29595011-29595194 | GABBR1;GABBR1;GABBR1 | N_Shore | Body;Body;Body |
| GM02706 | chr6:29595194-29595315 | GABBR1;GABBR1;GABBR1 | N_Shore | Body;Body;Body |
| GM02175 | chr6:29595299-29595335 | GABBR1;GABBR1;GABBR1 | Island | Body;1stExon;Body |
| WG2844 | chr6:29599248-29599259 | GABBR1;GABBR1 | N_Shore | Body;Body |
| WG1513 | chr6:29627290-29627395 | MOG;MOG;MOG;MOG;MOG;MOG;MOG; | | Body;Body;Body;Body;Body;Body;Bod |
| GM09256 | chr6:29635692-29636011 | MOG;MOG;MOG;MOG;MOG;MOG;MOG; | | 3'UTR;Body;3'UTR;Body;Body;Body;Bo |
| WG2415 | chr6:29636003-29636366 | MOG;MOG;MOG;MOG;MOG;MOG;MOG; | | 3'UTR;Body;3'UTR;Body;Body;Body;Bo |
| GM02640 | chr6:29690766-29690889 | HLA-F;HLA-F;HLA-F | N_Shore | TSS1500;TSS1500;TSS1500 |
| WG2194 | chr6:29690766-29690893 | HLA-F;HLA-F;HLA-F | N_Shore | TSS1500;TSS1500;TSS1500 |
| WG3465 | chr6:29690822-29690893 | HLA-F;HLA-F;HLA-F | N_Shore | TSS1500;TSS1500;TSS1500 |
| WG3466 | chr6:29692035-29692085 | HLA-F;HLA-F;HLA-F | Island | Body;Body;Body |
| WG2415 | chr6:29692082-29692092 | HLA-F;HLA-F;HLA-F | Island | Body;Body;Body |
| GM02706 | chr6:29717368-29717475 | LOC285830;LOC285830 | S_Shore | TSS1500;TSS1500 |
| GM00290 | chr6:29759947-29759955 | HCG4 | Island | Body |
| GM01653 | chr6:29759947-29759972 | HCG4 | Island | Body |
| WG2121 | chr6:29760832-29760864 | HCG4 | Island | Body |
| WG2415 | chr6:29760909-29761075 | HCG4 | S_Shore | TSS200 |
| GM01095 | chr6:29795411-29795501 | HLA-G | N_Shore | 5'UTR |
| GM00290 | chr6:29855435-29855467 | HLA-H | Island | Body |
| GM00290 | chr6:29855636-29855970 | HLA-H | Island | Body |
| GM00290 | chr6:29856278-29856294 | HLA-H | Island | Body |
| GM02706 | chr6:29856430-29856494 | HLA-H | Island | Body |
| GM00290 | chr6:29869936-29870062 | HCG2P7 | | Body |
| GM00520 | chr6:29893168-29893605 | HCG4P6 | N_Shore | Body |
| WG2414 | chr6:29894005-29894253 | HCG4P6 | N_Shore | TSS1500 |
| WG1310 | chr6:29894050-29894152 | HCG4P6 | N_Shore | TSS1500 |
| WG3466 | chr6:29894050-29894162 | HCG4P6 | N_Shore | TSS1500 |
| WG2194 | chr6:29894050-29894197 | HCG4P6 | N_Shore | TSS1500 |
| GM01095 | chr6:29894050-29894228 | HCG4P6 | N_Shore | TSS1500 |

| | | | | |
|---|---|---|---|---|
| GM01376 | chr6:29894141-29894197 | HCG4P6 | Island | TSS1500 |
| GM00022 | chr6:29894152-29894248 | HCG4P6 | Island | TSS1500 |
| WG1722 | chr6:29894162-29894197 | HCG4P6 | Island | TSS1500 |
| GM01096 | chr6:29894162-29894253 | HCG4P6 | Island | TSS1500 |
| WG3465 | chr6:29894195-29894228 | HCG4P6 | Island | TSS1500 |
| GM00290 | chr6:29894619-29894679 | HCG4P6 | Island | TSS1500 |
| GM02316 | chr6:29911036-29911104 | HLA-A | Island | Body |
| WG3466 | chr6:29911087-29911104 | HLA-A | Island | Body |
| WG2275 | chr6:29911091-29911104 | HLA-A | Island | Body |
| GM00289 | chr6:29911261-29911295 | HLA-A | Island | Body |
| WG2276 | chr6:29911542-29911558 | HLA-A | S_Shore | Body |
| GM00520 | chr6:29943056-29943209 | HCG9 | N_Shore | Body |
| WG1513 | chr6:29943268-29943408 | HCG9 | N_Shore | Body |
| GM09255 | chr6:29943401-29943414 | HCG9 | N_Shore | Body |
| WG3465 | chr6:29943401-29943425 | HCG9 | N_Shore | Body |
| WG1984 | chr6:29943401-29943455 | HCG9 | N_Shore | Body |
| GM02175 | chr6:29943401-29943677 | HCG9 | N_Shore | Body |
| WG2120 | chr6:29943455-29943464 | HCG9 | N_Shore | Body |
| WG3466 | chr6:29943455-29943480 | HCG9 | N_Shore | Body |
| GM02555 | chr6:29943480-29943841 | HCG9 | N_Shore | Body |
| WG1722 | chr6:29944093-29944146 | HCG9 | N_Shore | Body |
| GM00520 | chr6:29945409-29945726 | HCG9 | S_Shore | Body |
| WG1977 | chr6:29975345-29975416 | HLA-J;NCRNA00171 | Island | Body;Body |
| WG3466 | chr6:29975366-29975675 | HLA-J;NCRNA00171 | Island | Body;Body |
| WG2415 | chr6:30038712-30038754 | RNF39;RNF39 | N_Shore | 3'UTR;3'UTR |
| GM02555 | chr6:30038915-30038929 | RNF39;RNF39 | Island | Body;Body |
| GM00289 | chr6:30038922-30038975 | RNF39;RNF39 | Island | Body;Body |
| WG3466 | chr6:30038922-30039151 | RNF39;RNF39 | Island | Body;Body |
| GM02147 | chr6:30039403-30039524 | RNF39;RNF39 | Island | Body;Body |
| WG3466 | chr6:30039403-30039548 | RNF39;RNF39 | Island | Body;Body |
| WG2844 | chr6:30039432-30039442 | RNF39;RNF39 | Island | Body;Body |
| GM00935 | chr6:30039432-30039466 | RNF39;RNF39 | Island | Body;Body |
| GM01376 | chr6:30039442-30039476 | RNF39;RNF39 | Island | Body;Body |
| WG3465 | chr6:30039524-30039600 | RNF39;RNF39 | S_Shore | Body;Body |
| GM02706 | chr6:30039600-30039900 | RNF39;RNF39 | S_Shore | Body;Body |
| GM01096 | chr6:30042260-30042295 | RNF39;RNF39 | N_Shore | Body;Body |
| WG1722 | chr6:30043820-30043924 | RNF39;RNF39 | S_Shore | TSS200;TSS200 |
| GM09255 | chr6:30071503-30071601 | TRIM31 | S_Shore | Body |
| WG2415 | chr6:30076659-30076980 | TRIM31 | | Body |
| WG2120 | chr6:30079102-30079155 | TRIM31 | | Body |
| GM00439 | chr6:30079102-30079192 | TRIM31 | | Body |
| GM02175 | chr6:30079155-30079307 | TRIM31 | | Body |
| WG2415 | chr6:30079203-30079280 | TRIM31 | | Body |
| WG1513 | chr6:30079203-30079307 | TRIM31 | | Body |
| WG2120 | chr6:30079256-30079280 | TRIM31 | | Body |
| WG3465 | chr6:30079256-30079307 | TRIM31 | | Body |
| GM00520 | chr6:30079265-30079307 | TRIM31 | | Body |

| | | | | |
|---|---|---|---|---|
| GM00438 | chr6:30080230-30080295 | TRIM31 | | Body |
| GM02555 | chr6:30113774-30113786 | TRIM40 | | Body |
| WG3465 | chr6:30113774-30113825 | TRIM40 | | Body |
| GM00520 | chr6:30113783-30113825 | TRIM40 | | Body |
| WG3465 | chr6:30113978-30114011 | TRIM40 | | Body |
| WG1978 | chr6:30122523-30122565 | TRIM10;TRIM10 | | Body;Body |
| WG1722 | chr6:30122523-30122593 | TRIM10;TRIM10 | | Body;Body |
| WG1721 | chr6:30122523-30122947 | TRIM10;TRIM10 | | Body;Body |
| GM02456 | chr6:30122544-30122947 | TRIM10;TRIM10 | | Body;Body |
| WG1721 | chr6:30123191-30123226 | TRIM10;TRIM10 | | Body;Body |
| GM02175 | chr6:30124240-30124299 | TRIM10;TRIM10 | | Body;Body |
| GM09256 | chr6:30130819-30131189 | TRIM15 | | TSS200 |
| WG1977 | chr6:30131189-30131219 | TRIM15;TRIM15 | | 5'UTR;1stExon |
| GM09256 | chr6:30131691-30131755 | TRIM15 | | 1stExon |
| WG1514 | chr6:30140145-30140305 | TRIM15 | Island | 3'UTR |
| WG2415 | chr6:30140305-30140409 | TRIM15 | S_Shore | 3'UTR |
| GM00439 | chr6:30140305-30140417 | TRIM15 | S_Shore | 3'UTR |
| GM00940 | chr6:30140325-30140372 | TRIM15 | S_Shore | 3'UTR |
| GM00438 | chr6:30140366-30140417 | TRIM15 | S_Shore | 3'UTR |
| GM00520 | chr6:30162942-30163098 | TRIM26 | | Body |
| GM00438 | chr6:30173974-30174704 | TRIM26 | | 5'UTR |
| WG2415 | chr6:30175299-30175792 | TRIM26 | | 5'UTR |
| GM09256 | chr6:30175305-30175792 | TRIM26 | | 5'UTR |
| WG3465 | chr6:30228243-30228254 | HLA-L | Island | Body |
| GM09256 | chr6:30231639-30231887 | HLA-L | S_Shelf | Body |
| WG1513 | chr6:30231639-30232015 | HLA-L | S_Shelf | Body |
| WG2414 | chr6:30231675-30231887 | HLA-L | S_Shelf | Body |
| WG1722 | chr6:30458147-30458161 | HLA-E | Island | Body |
| GM01096 | chr6:30458149-30458161 | HLA-E | Island | Body |
| WG3466 | chr6:30458158-30458237 | HLA-E | Island | Body |
| WG1513 | chr6:30510294-30510459 | GNL1 | | 3'UTR |
| GM02706 | chr6:30586293-30586468 | MRPS18B;PPP1R10 | S_Shore | Body;TSS1500 |
| GM09255 | chr6:30646794-30646860 | KIAA1949;KIAA1949 | | Body;Body |
| GM09256 | chr6:30647083-30647128 | KIAA1949;KIAA1949 | | Body;Body |
| WG1977 | chr6:30647561-30647649 | KIAA1949;KIAA1949 | | Body;Body |
| GM00520 | chr6:30652202-30652237 | KIAA1949;KIAA1949 | N_Shelf | Body;1stExon |
| WG1514 | chr6:30652237-30652282 | KIAA1949;KIAA1949 | N_Shelf | Body;1stExon |
| GM00520 | chr6:30657219-30657384 | NRM | S_Shore | Body |
| WG2415 | chr6:30657469-30658014 | NRM | S_Shelf | Body |
| WG3465 | chr6:30657882-30658014 | NRM | S_Shelf | Body |
| GM00438 | chr6:30684406-30684572 | MDC1 | N_Shore | 5'UTR |
| GM09256 | chr6:30695686-30696029 | FLOT1 | | 3'UTR |
| WG2844 | chr6:30695962-30696029 | FLOT1 | | Body |
| WG2844 | chr6:30698584-30698843 | FLOT1 | | Body |
| WG2415 | chr6:30698777-30698784 | FLOT1 | | Body |
| GM00289 | chr6:30698843-30698936 | FLOT1 | | Body |
| GM02706 | chr6:30711022-30711399 | IER3;FLOT1 | Island | 3'UTR;TSS1500 |

| | | | | |
|---|---|---|---|---|
| GM02706 | chr6:30711561-30711680 | IER3;FLOT1 | Island | 3'UTR;TSS1500 |
| GM00940 | chr6:30850868-30850913 | DDR1;DDR1 | N_Shore | TSS1500;TSS1500 |
| GM00290 | chr6:30850868-30850977 | DDR1;DDR1 | N_Shore | TSS1500;TSS1500 |
| GM01096 | chr6:30853255-30853333 | DDR1;DDR1 | S_Shore | 5'UTR;5'UTR |
| GM00940 | chr6:30854109-30854164 | DDR1;DDR1 | S_Shore | 5'UTR;5'UTR |
| GM00520 | chr6:30854161-30854186 | DDR1;DDR1 | S_Shore | 5'UTR;5'UTR |
| WG1977 | chr6:30854164-30854233 | DDR1;DDR1 | S_Shore | 5'UTR;5'UTR |
| WG1978 | chr6:30856386-30856491 | DDR1;DDR1;DDR1 | S_Shelf | TSS200;5'UTR;5'UTR |
| GM00289 | chr6:30882694-30882994 | VARS2;VARS2;VARS2 | S_Shore | Body;5'UTR;Body |
| WG3465 | chr6:30883762-30884124 | VARS2;VARS2;VARS2 | S_Shore | Body;Body;Body |
| WG2844 | chr6:30921592-30921757 | DPCR1 | N_Shore | 3'UTR |
| GM01096 | chr6:30955412-30955425 | MUC21 | | Body |
| GM02706 | chr6:30956440-30956603 | MUC21 | | 3'UTR |
| WG2275 | chr6:31025933-31025967 | HCG22 | | Body |
| WG1977 | chr6:31025933-31026105 | HCG22 | | Body |
| WG1309 | chr6:31025933-31026287 | HCG22 | | Body |
| GM00520 | chr6:31081323-31081617 | PSORS1C1;C6orf15 | | TSS1500;TSS1500 |
| WG2844 | chr6:31082187-31082200 | PSORS1C1 | | TSS1500 |
| GM02706 | chr6:31095908-31096102 | PSORS1C1 | | 5'UTR |
| WG1513 | chr6:31105711-31105777 | PSORS1C1;PSORS1C2 | | Body;3'UTR |
| GM01096 | chr6:31105744-31105844 | PSORS1C1;PSORS1C2 | | Body;Body |
| WG1513 | chr6:31111076-31111270 | CCHCR1;CCHCR1;CCHCR1 | | Body;Body;Body |
| WG1722 | chr6:31111088-31111270 | CCHCR1;CCHCR1;CCHCR1 | | Body;Body;Body |
| GM00520 | chr6:31139359-31139796 | POU5F1 | | TSS1500 |
| WG1513 | chr6:31143710-31144068 | PSORS1C3 | | Body |
| WG1977 | chr6:31146471-31146845 | PSORS1C3 | N_Shore | TSS1500 |
| GM00289 | chr6:31146889-31148524 | PSORS1C3 | N_Shore | TSS1500 |
| WG2275 | chr6:31164804-31164914 | HCG27 | N_Shore | TSS1500 |
| WG3466 | chr6:31164914-31164947 | HCG27 | N_Shore | TSS1500 |
| GM02316 | chr6:31164914-31164952 | HCG27 | N_Shore | TSS1500 |
| GM01376 | chr6:31164914-31164994 | HCG27 | N_Shore | TSS1500 |
| WG2275 | chr6:31164921-31164968 | HCG27 | N_Shore | TSS1500 |
| GM01096 | chr6:312026-312105 | DUSP22 | | Body |
| WG2843 | chr6:31237013-31237034 | HLA-C | N_Shore | Body |
| WG3466 | chr6:31237405-31237722 | HLA-C | N_Shore | Body |
| GM09255 | chr6:31238751-31238803 | HLA-C | N_Shore | Body |
| GM02175 | chr6:31238788-31238908 | HLA-C | N_Shore | Body |
| WG1722 | chr6:31240651-31241000 | HLA-C | S_Shore | TSS1500 |
| WG2276 | chr6:31323506-31323760 | HLA-B | N_Shore | Body |
| GM02555 | chr6:31323677-31323856 | HLA-B | N_Shore | Body |
| GM02706 | chr6:31323760-31323985 | HLA-B | N_Shore | Body |
| GM00042 | chr6:31323856-31324024 | HLA-B | N_Shore | Body |
| GM02175 | chr6:31323985-31324024 | HLA-B | Island | Body |
| GM00290 | chr6:31324500-31324551 | HLA-B | Island | Body |
| WG1513 | chr6:31379097-31379358 | MICA | | Body |
| WG3465 | chr6:31466174-31466187 | MICB | Island | Body |
| WG1977 | chr6:31468571-31468957 | MICB | S_Shelf | Body |

| | | | | |
|---|---|---|---|---|
| WG1977 | chr6:31475183-31475943 | MICB | | Body |
| GM00290 | chr6:31496733-31497233 | MCCD1 | | TSS200 |
| WG3465 | chr6:31507879-31508137 | BAT1;BAT1 | N_Shore | Body;Body |
| WG1513 | chr6:31508126-31508167 | BAT1;BAT1 | N_Shore | Body;Body |
| WG2414 | chr6:31508126-31508297 | BAT1;BAT1 | N_Shore | Body;Body |
| GM02175 | chr6:31510729-31510903 | BAT1;BAT1 | S_Shore | TSS1500;TSS1500 |
| GM01096 | chr6:31513198-31513267 | ATP6V1G2;NFKBIL1;NFKBIL1 | S_Shelf | Body;TSS1500;TSS1500;Body |
| WG1513 | chr6:31513261-31513306 | ATP6V1G2;NFKBIL1;NFKBIL1 | S_Shelf | Body;TSS1500;TSS1500;Body |
| GM09256 | chr6:31540399-31540750 | LTA;LTA | | 5'UTR;5'UTR |
| WG2415 | chr6:31543289-31543557 | TNF | | TSS200 |
| WG1722 | chr6:31543540-31543557 | TNF | | 1stExon |
| GM02706 | chr6:31549262-31549480 | LTB;LTB | Island | Body;3'UTR |
| GM02175 | chr6:31549563-31549631 | LTB;LTB | S_Shore | Body;Body |
| WG1513 | chr6:31549563-31549853 | LTB;LTB | S_Shore | Body;Body |
| GM01392 | chr6:31554548-31554829 | LST1;LST1;LST1;LST1;LST1;LST1;LST1;LS | | Body;TSS1500;TSS1500;5'UTR;TSS15( |
| WG1513 | chr6:31554848-31555016 | LST1;LST1;LST1;LST1;LST1;LST1;LST1;LS | | Body;TSS200;5'UTR;5'UTR;TSS200;TS |
| WG1513 | chr6:31599526-31599537 | BAT2 | Island | Body |
| GM00439 | chr6:31607340-31607374 | BAT3;BAT3;BAT3;BAT3 | | Body;Body;Body;Body |
| GM02175 | chr6:31607374-31607648 | BAT3;BAT3;BAT3;BAT3 | | Body;Body;Body;Body |
| WG1282 | chr6:31607634-31607674 | BAT3;BAT3;BAT3;BAT3 | | Body;Body;Body;Body |
| GM00520 | chr6:31627653-31627711 | C6orf47 | N_Shore | 1stExon |
| WG1977 | chr6:31627653-31627714 | C6orf47 | N_Shore | 1stExon |
| WG2275 | chr6:31627678-31627714 | C6orf47 | N_Shore | 1stExon |
| WG2414 | chr6:31649619-31649638 | LY6G5C | N_Shore | TSS1500 |
| WG2120 | chr6:31649619-31649728 | LY6G5C | N_Shore | TSS1500 |
| GM00439 | chr6:31673800-31673926 | LY6G6F | S_Shelf | TSS1500 |
| WG2414 | chr6:31685392-31685440 | LY6G6D | | Body |
| WG1513 | chr6:31685430-31685575 | LY6G6D | | Body |
| GM00042 | chr6:31685491-31685542 | LY6G6D | | Body |
| WG3465 | chr6:31691050-31691252 | C6orf25;C6orf25;C6orf25;C6 | N_Shore | TSS200;TSS200;TSS200;TSS200;TSS2 |
| WG2121 | chr6:31691227-31691354 | C6orf25;C6orf25;C6orf25;C6 | N_Shore | Body;Body;Body;Body;Body;Body |
| WG2276 | chr6:31696223-31696240 | DDAH2 | Island | Body |
| WG2276 | chr6:31696309-31696710 | DDAH2 | Island | Body |
| GM01393 | chr6:31696423-31696482 | DDAH2 | Island | Body |
| GM00520 | chr6:31698687-31698702 | CLIC1;DDAH2 | S_Shore | Body;TSS1500 |
| WG3465 | chr6:31701110-31701288 | CLIC1 | S_Shelf | Body |
| WG3465 | chr6:31704774-31704844 | CLIC1 | N_Shelf | TSS1500 |
| WG1978 | chr6:31707019-31707202 | MSH5;MSH5;MSH5;MSH5 | N_Shore | TSS1500;TSS1500;TSS1500;TSS1500 |
| WG3465 | chr6:31733688-31733894 | C6orf27 | | Body |
| GM01392 | chr6:31733894-31734106 | C6orf27 | | Body |
| GM02706 | chr6:31734147-31734471 | C6orf27 | | Body |
| GM01096 | chr6:31734217-31734292 | C6orf27 | | Body |
| WG1978 | chr6:31734232-31734401 | C6orf27 | | Body |
| GM02555 | chr6:31734471-31734580 | C6orf27 | | Body |
| WG3466 | chr6:31744628-31744735 | C6orf27 | | 5'UTR |
| WG2415 | chr6:31744636-31744816 | C6orf27 | | 5'UTR |
| WG2120 | chr6:31744735-31744831 | C6orf27 | | 5'UTR |

| | | | | |
|---|---|---|---|---|
| GM01392 | chr6:31760593-31760616 | VARS | N_Shelf | Body |
| WG2275 | chr6:31760593-31760796 | VARS | N_Shelf | Body |
| GM01096 | chr6:31760608-31760629 | VARS | N_Shelf | Body |
| WG3465 | chr6:31765083-31765590 | VARS | S_Shore | TSS1500 |
| WG2121 | chr6:31829644-31829800 | NEU1 | N_Shore | Body |
| GM00940 | chr6:31829644-31829840 | NEU1 | N_Shore | Body |
| WG2120 | chr6:31829888-31829914 | NEU1 | N_Shore | Body |
| WG1977 | chr6:31829888-31829926 | NEU1 | N_Shore | Body |
| GM00438 | chr6:31829895-31829926 | NEU1 | N_Shore | Body |
| GM09256 | chr6:31832962-31833028 | SLC44A4 | S_Shelf | Body |
| WG3466 | chr6:31846769-31846813 | SLC44A4;SLC44A4 | | 5'UTR;1stExon |
| WG2275 | chr6:31856367-31856431 | EHMT2;EHMT2 | N_Shore | Body;Body |
| WG1977 | chr6:31895464-31895493 | C2;C2;C2;C2 | | 1stExon;5'UTR;1stExon;5'UTR |
| GM02555 | chr6:31913759-31913814 | CFB;CFB | | 1stExon;5'UTR |
| WG1977 | chr6:31913759-31913995 | CFB;CFB | | 1stExon;5'UTR |
| GM01096 | chr6:31926014-31926205 | RDBP;SKIV2L;MIR1236 | N_Shore | Body;TSS1500;TSS1500 |
| GM09255 | chr6:31926072-31926205 | RDBP;SKIV2L;MIR1236 | N_Shore | Body;TSS1500;TSS1500 |
| WG1513 | chr6:31937470-31937731 | STK19;SKIV2L | N_Shelf | TSS1500;Body |
| WG1513 | chr6:31938207-31938372 | STK19;STK19;STK19;DOM3Z | N_Shore | TSS1500;TSS1500;TSS1500;Body |
| GM09256 | chr6:31938984-31939106 | STK19;STK19;DOM3Z;STK19 | N_Shore | TSS1500;TSS1500;Body;Body |
| GM00289 | chr6:31939025-31939085 | STK19;STK19;DOM3Z;STK19 | N_Shore | TSS1500;TSS1500;Body;Body |
| GM09255 | chr6:31973162-31973255 | CYP21A2;CYP21A2 | | TSS200;TSS200 |
| WG1983 | chr6:31980836-31980856 | TNXB;TNXB;TNXA;STK19;STK19 | | 5'UTR;1stExon;TSS200;TSS1500;TSS1 |
| WG1722 | chr6:32013699-32013845 | TNXB;TNXB;TNXB | | 1stExon;5'UTR;Body |
| WG1983 | chr6:32013699-32013974 | TNXB;TNXB;TNXB | | 1stExon;5'UTR;Body |
| WG1983 | chr6:32014100-32014300 | TNXB;TNXB | | TSS200;Body |
| GM00940 | chr6:32014381-32014484 | TNXB;TNXB | | Body;TSS1500 |
| GM00439 | chr6:32014739-32014926 | TNXB;TNXB | | Body;TSS1500 |
| WG3465 | chr6:32015175-32015226 | TNXB;TNXB | | Body;TSS1500 |
| GM00290 | chr6:32015215-32015297 | TNXB;TNXB | | Body;TSS1500 |
| WG2414 | chr6:32016214-32016247 | TNXB | | Body |
| WG2276 | chr6:32016236-32016247 | TNXB | | Body |
| GM00439 | chr6:32016336-32016368 | TNXB | | Body |
| GM00016 | chr6:32026350-32026656 | TNXB | | Body |
| WG3465 | chr6:32038177-32038230 | TNXB | | Body |
| GM02704 | chr6:32044104-32044179 | TNXB | N_Shelf | Body |
| GM00520 | chr6:32044163-32044203 | TNXB | N_Shelf | Body |
| GM00520 | chr6:32049373-32049516 | TNXB | S_Shelf | Body |
| GM01376 | chr6:32049376-32049516 | TNXB | S_Shelf | Body |
| WG2121 | chr6:32050927-32051108 | TNXB | S_Shelf | Body |
| WG1978 | chr6:32054561-32054790 | TNXB | N_Shore | Body |
| WG1977 | chr6:32054561-32054867 | TNXB | N_Shore | Body |
| GM00520 | chr6:32054771-32054867 | TNXB | N_Shore | Body |
| GM00439 | chr6:32055135-32055146 | TNXB | Island | Body |
| WG2276 | chr6:32055135-32055385 | TNXB | Island | Body |
| WG2844 | chr6:32055135-32055517 | TNXB | Island | Body |
| GM02555 | chr6:32055135-32055525 | TNXB | Island | Body |

| | | | | |
|---|---|---|---|---|
| GM00520 | chr6:32055316-32055385 | TNXB | Island | Body |
| WG2276 | chr6:32055402-32055711 | TNXB | Island | Body |
| GM03075 | chr6:32055447-32055517 | TNXB | Island | Body |
| WG3466 | chr6:32055447-32055525 | TNXB | Island | Body |
| GM00935 | chr6:32055474-32055525 | TNXB | Island | Body |
| GM00290 | chr6:32055534-32055738 | TNXB | Island | Body |
| GM02555 | chr6:32064923-32064956 | TNXB | Island | Body |
| GM00520 | chr6:32064956-32065023 | TNXB | Island | Body |
| GM00520 | chr6:32094729-32094915 | ATF6B;ATF6B | N_Shelf | Body;Body |
| WG3465 | chr6:32097439-32097507 | ATF6B;ATF6B;FKBPL | N_Shore | TSS1500;TSS1500;Body |
| WG1977 | chr6:32116207-32116225 | PRRT1 | N_Shore | 3'UTR |
| WG1977 | chr6:32116653-32116841 | PRRT1 | Island | 3'UTR |
| WG3465 | chr6:32116653-32116853 | PRRT1 | Island | 3'UTR |
| WG2276 | chr6:32116841-32116905 | PRRT1 | Island | 3'UTR |
| WG3465 | chr6:32116931-32116963 | PRRT1 | Island | 3'UTR |
| WG1513 | chr6:32118213-32118298 | PRRT1 | Island | Body |
| WG2844 | chr6:32118312-32118375 | PRRT1 | Island | Body |
| GM00520 | chr6:32118375-32118420 | PRRT1 | Island | Body |
| GM02175 | chr6:32118420-32118441 | PRRT1 | Island | Body |
| GM03075 | chr6:32118420-32118457 | PRRT1 | Island | Body |
| WG1977 | chr6:32120077-32120625 | PRRT1;PPT2 | N_Shore | TSS1500;TSS1500 |
| GM00289 | chr6:32120584-32120625 | PPT2;PRRT1;PPT2 | N_Shore | TSS1500;TSS1500;TSS1500 |
| GM00941 | chr6:32120625-32120783 | PPT2;PRRT1;PPT2 | N_Shore | TSS1500;TSS1500;TSS1500 |
| GM09256 | chr6:32122864-32122955 | PPT2;PPT2 | S_Shore | Body;Body |
| WG1513 | chr6:32122966-32123258 | PPT2;PPT2 | S_Shore | Body;Body |
| GM00520 | chr6:32123479-32123651 | PPT2;PPT2 | S_Shore | Body;Body |
| GM02175 | chr6:32160605-32160909 | GPSM3 | N_Shelf | 5'UTR |
| GM00438 | chr6:32164553-32164569 | GPSM3;NOTCH4 | S_Shore | TSS1500;Body |
| WG2276 | chr6:32164927-32165200 | NOTCH4 | S_Shore | Body |
| WG1977 | chr6:32165176-32165200 | NOTCH4 | S_Shore | Body |
| WG1722 | chr6:32165176-32165237 | NOTCH4 | S_Shore | Body |
| WG2415 | chr6:32181814-32182013 | NOTCH4 | | Body |
| GM00940 | chr6:32190281-32190325 | NOTCH4 | | Body |
| GM00042 | chr6:32191479-32191821 | NOTCH4 | | Body |
| GM00520 | chr6:32191479-32191895 | NOTCH4 | | Body |
| WG2120 | chr6:32191496-32191840 | NOTCH4 | | Body |
| WG1513 | chr6:32191821-32191895 | NOTCH4;NOTCH4 | | 1stExon;5'UTR |
| GM02149 | chr6:32294260-32294503 | C6orf10 | | Body |
| WG2120 | chr6:32297670-32297822 | C6orf10 | | Body |
| WG2276 | chr6:32305990-32306089 | C6orf10 | | Body |
| GM02149 | chr6:32305990-32306373 | C6orf10 | | Body |
| GM02706 | chr6:32371488-32371538 | BTNL2 | | Body |
| GM00042 | chr6:32407714-32407838 | HLA-DRA;HLA-DRA | | 5'UTR;1stExon |
| GM09256 | chr6:32410339-32410365 | HLA-DRA | | Body |
| WG1513 | chr6:32410347-32410459 | HLA-DRA | | Body |
| GM09256 | chr6:32411067-32411144 | HLA-DRA | | Body |
| GM00031 | chr6:32489203-32489555 | HLA-DRB5 | N_Shore | Body |

| | | | | |
|---|---|---|---|---|
| WG2194 | chr6:32489750-32489963 | HLA-DRB5 | Island | Body |
| WG3465 | chr6:32489963-32489991 | HLA-DRB5 | Island | Body |
| GM02706 | chr6:32489984-32490012 | HLA-DRB5 | Island | Body |
| WG2844 | chr6:32489991-32490043 | HLA-DRB5 | Island | Body |
| WG2844 | chr6:32526021-32526260 | HLA-DRB6 | | Body |
| GM01392 | chr6:32526021-32526263 | HLA-DRB6 | | Body |
| GM01375 | chr6:32526021-32526342 | HLA-DRB6 | | Body |
| WG1309 | chr6:32526027-32526263 | HLA-DRB6 | | Body |
| WG2193 | chr6:32526027-32526342 | HLA-DRB6 | | Body |
| WG1722 | chr6:32526260-32526342 | HLA-DRB6 | | Body |
| WG1514 | chr6:32526260-32526364 | HLA-DRB6 | | Body |
| WG2276 | chr6:32526263-32526364 | HLA-DRB6 | | Body |
| GM00031 | chr6:32526342-32526366 | HLA-DRB6 | | Body |
| WG1978 | chr6:32549361-32549631 | HLA-DRB1 | N_Shelf | Body |
| GM02175 | chr6:32551888-32551949 | HLA-DRB1 | Island | Body |
| WG2415 | chr6:32551888-32551954 | HLA-DRB1 | Island | Body |
| WG2844 | chr6:32551891-32551954 | HLA-DRB1 | Island | Body |
| WG2844 | chr6:32552152-32552246 | HLA-DRB1 | Island | Body |
| WG2120 | chr6:32552246-32552453 | HLA-DRB1 | Island | Body |
| GM00042 | chr6:32552443-32552515 | HLA-DRB1 | S_Shore | Body |
| WG2193 | chr6:32557970-32558459 | HLA-DRB1 | | TSS1500 |
| GM00439 | chr6:32608858-32609094 | HLA-DQA1 | | Body |
| WG1983 | chr6:32632565-32632639 | HLA-DQB1 | Island | Body |
| WG2276 | chr6:32633102-32633157 | HLA-DQB1 | S_Shore | Body |
| WG2415 | chr6:32633157-32633354 | HLA-DQB1 | S_Shore | Body |
| WG1514 | chr6:32634245-32634316 | HLA-DQB1 | S_Shore | Body |
| GM00042 | chr6:32634245-32634344 | HLA-DQB1 | S_Shore | Body |
| WG2193 | chr6:32634245-32634362 | HLA-DQB1 | S_Shore | Body |
| GM00941 | chr6:32634276-32634344 | HLA-DQB1 | S_Shore | 1stExon |
| WG1282 | chr6:32634276-32634362 | HLA-DQB1 | S_Shore | 1stExon |
| GM00935 | chr6:32634316-32634362 | HLA-DQB1 | S_Shore | 1stExon |
| GM00438 | chr6:32710209-32710583 | HLA-DQA2 | | Body |
| GM02149 | chr6:32710506-32710583 | HLA-DQA2 | | Body |
| GM02149 | chr6:32711889-32712093 | HLA-DQA2 | | Body |
| GM02149 | chr6:32713018-32713607 | HLA-DQA2 | | Body |
| GM02706 | chr6:32729358-32729465 | HLA-DQB2 | Island | Body |
| GM09255 | chr6:32729358-32729563 | HLA-DQB2 | Island | Body |
| WG3466 | chr6:32729442-32729470 | HLA-DQB2 | Island | Body |
| GM00438 | chr6:32729442-32729498 | HLA-DQB2 | Island | Body |
| GM00439 | chr6:32729442-32729563 | HLA-DQB2 | Island | Body |
| WG3465 | chr6:32729442-32729590 | HLA-DQB2 | Island | Body |
| GM02456 | chr6:32729442-32729643 | HLA-DQB2 | Island | Body |
| WG1309 | chr6:32729465-32729498 | HLA-DQB2 | Island | Body |
| GM00520 | chr6:32729465-32729500 | HLA-DQB2 | Island | Body |
| WG1721 | chr6:32729465-32729587 | HLA-DQB2 | Island | Body |
| GM02706 | chr6:32729498-32729596 | HLA-DQB2 | Island | Body |
| GM00940 | chr6:32729498-32729643 | HLA-DQB2 | Island | Body |

| | | | | |
|---|---|---|---|---|
| GM00042 | chr6:32729545-32729587 | HLA-DQB2 | Island | Body |
| WG1309 | chr6:32729545-32729590 | HLA-DQB2 | Island | Body |
| WG3466 | chr6:32729545-32729596 | HLA-DQB2 | Island | Body |
| WG2120 | chr6:32729563-32729590 | HLA-DQB2 | Island | Body |
| GM00438 | chr6:32729563-32729596 | HLA-DQB2 | Island | Body |
| WG1977 | chr6:32729587-32729596 | HLA-DQB2 | Island | Body |
| GM09256 | chr6:32729587-32729643 | HLA-DQB2 | Island | Body |
| GM00439 | chr6:32729590-32729643 | HLA-DQB2 | Island | Body |
| GM00289 | chr6:32729643-32729659 | HLA-DQB2 | Island | Body |
| GM00289 | chr6:32729764-32729847 | HLA-DQB2 | Island | Body |
| WG1513 | chr6:32732247-32732585 | HLA-DQB2 | S_Shelf | TSS1500 |
| WG2415 | chr6:32781171-32781307 | HLA-DOB | | Body |
| GM09256 | chr6:32805223-32805554 | TAP2;TAP2 | N_Shore | Body;Body |
| GM02175 | chr6:32805398-32805570 | TAP2;TAP2 | N_Shore | Body;Body |
| GM00439 | chr6:32805398-32805692 | TAP2;TAP2 | N_Shore | Body;Body |
| GM02706 | chr6:32805548-32805570 | TAP2;TAP2 | N_Shore | Body;Body |
| GM00520 | chr6:32805548-32805692 | TAP2;TAP2 | N_Shore | Body;Body |
| WG1513 | chr6:32805570-32805692 | TAP2;TAP2 | N_Shore | Body;Body |
| GM00042 | chr6:32813154-32813337 | PSMB8;PSMB8;TAP1 | S_Shore | TSS1500;TSS1500;3'UTR |
| GM00520 | chr6:32820201-32820355 | TAP1 | N_Shore | Body |
| GM02456 | chr6:32820214-32820355 | TAP1 | N_Shore | Body |
| GM09256 | chr6:32822565-32822911 | PSMB9;PSMB9;TAP1 | S_Shore | Body;Body;TSS1500 |
| GM00438 | chr6:32847499-32847530 | PPP1R2P1 | Island | Body |
| WG3465 | chr6:32847513-32847530 | PPP1R2P1 | Island | Body |
| GM02175 | chr6:32847585-32847596 | PPP1R2P1 | Island | Body |
| WG3465 | chr6:32847585-32847605 | PPP1R2P1 | Island | Body |
| GM00439 | chr6:32847675-32847704 | PPP1R2P1 | Island | Body |
| WG3465 | chr6:32847702-32847762 | PPP1R2P1 | Island | Body |
| WG1513 | chr6:32847702-32847830 | PPP1R2P1 | Island | Body |
| WG1722 | chr6:32847762-32847788 | PPP1R2P1 | Island | Body |
| WG1977 | chr6:32847762-32847845 | PPP1R2P1 | Island | Body |
| WG2414 | chr6:32847788-32847830 | PPP1R2P1 | Island | Body |
| GM00438 | chr6:32847830-32847845 | PPP1R2P1 | Island | Body |
| WG2415 | chr6:32904061-32904317 | HLA-DMB | | Body |
| WG1722 | chr6:32904317-32904419 | HLA-DMB | | Body |
| WG1983 | chr6:32904402-32904621 | HLA-DMB | | Body |
| GM02175 | chr6:32908466-32908642 | HLA-DMB | | Body |
| WG1977 | chr6:32908567-32908642 | HLA-DMB | | 1stExon |
| WG3465 | chr6:32908567-32908718 | HLA-DMB | | 1stExon |
| GM02704 | chr6:32920880-32920995 | HLA-DMA;HLA-DMA | | 5'UTR;1stExon |
| WG1658 | chr6:32920880-32921142 | HLA-DMA;HLA-DMA | | 5'UTR;1stExon |
| WG1983 | chr6:32920880-32921233 | HLA-DMA;HLA-DMA | | 5'UTR;1stExon |
| GM03075 | chr6:32920995-32921233 | HLA-DMA | | TSS200 |
| GM09256 | chr6:32921142-32921233 | HLA-DMA | | TSS1500 |
| WG3465 | chr6:32945336-32945631 | BRD2;BRD2 | | Body;Body |
| WG1977 | chr6:32974639-32974906 | HLA-DOA | N_Shore | Body |
| GM00438 | chr6:32974843-32974906 | HLA-DOA | N_Shore | Body |

| | | | | |
|---|---|---|---|---|
| WG1513 | chr6:32975590-32975702 | HLA-DOA | N_Shore | Body |
| WG2415 | chr6:32975875-32975925 | HLA-DOA | Island | Body |
| WG2121 | chr6:32975986-32976018 | HLA-DOA | S_Shore | Body |
| GM01392 | chr6:32976805-32976858 | HLA-DOA | S_Shore | Body |
| GM09256 | chr6:32978053-32978140 | HLA-DOA | S_Shelf | TSS1500 |
| WG2275 | chr6:32978129-32978196 | HLA-DOA | S_Shelf | TSS1500 |
| GM00022 | chr6:33032830-33033312 | HLA-DPA1 | | 3'UTR |
| GM02175 | chr6:33039481-33039804 | HLA-DPA1 | | Body |
| GM02175 | chr6:33040610-33040769 | HLA-DPA1 | | Body |
| GM00289 | chr6:33040914-33041221 | HLA-DPA1 | | Body |
| WG3466 | chr6:33048687-33048752 | HLA-DPB1 | Island | Body |
| GM02706 | chr6:33048706-33048735 | HLA-DPB1 | Island | Body |
| WG1722 | chr6:33048875-33048919 | HLA-DPB1 | S_Shore | Body |
| WG2275 | chr6:33052354-33052802 | HLA-DPB1 | S_Shelf | Body |
| GM02173 | chr6:33084549-33084608 | HLA-DPB2 | N_Shore | Body |
| GM00042 | chr6:33085590-33085755 | HLA-DPB2 | S_Shore | Body |
| GM09256 | chr6:33091357-33091567 | HLA-DPB2 | | Body |
| GM00438 | chr6:33091751-33091870 | HLA-DPB2 | | Body |
| GM01096 | chr6:33092494-33092802 | HLA-DPB2 | | Body |
| GM02175 | chr6:33096303-33096738 | HLA-DPB2 | | Body |
| WG2415 | chr6:33131222-33131319 | COL11A2;COL11A2;COL11A | S_Shore | 3'UTR;3'UTR;3'UTR |
| WG1722 | chr6:33131222-33131336 | COL11A2;COL11A2;COL11A | S_Shore | 3'UTR;3'UTR;3'UTR |
| WG3465 | chr6:33131893-33132192 | COL11A2;COL11A2;COL11A | S_Shelf | Body;Body;Body |
| GM01393 | chr6:33132611-33132637 | COL11A2;COL11A2;COL11A | S_Shelf | Body;Body;Body |
| WG1977 | chr6:33132611-33132650 | COL11A2;COL11A2;COL11A | S_Shelf | Body;Body;Body |
| WG1513 | chr6:33132611-33132693 | COL11A2;COL11A2;COL11A | S_Shelf | Body;Body;Body |
| WG2415 | chr6:33132632-33132650 | COL11A2;COL11A2;COL11A | S_Shelf | Body;Body;Body |
| WG1513 | chr6:33132727-33132996 | COL11A2;COL11A2;COL11A | S_Shelf | Body;Body;Body |
| WG2120 | chr6:33141279-33141305 | COL11A2;COL11A2;COL11A2 | | Body;Body;Body |
| GM00438 | chr6:33145955-33146120 | COL11A2;COL11A2;COL11A2 | | Body;Body;Body |
| GM00520 | chr6:33153052-33153191 | COL11A2;COL11A2;COL11A2 | | Body;Body;Body |
| GM02149 | chr6:33161336-33161394 | COL11A2;COL11A2;COL11A | Island | TSS1500;TSS1500;TSS1500;TSS1500 |
| GM02175 | chr6:33169731-33170101 | SLC39A7;RXRB;SLC39A7 | S_Shore | Body;TSS1500;Body |
| WG2415 | chr6:33169975-33170177 | SLC39A7;SLC39A7 | S_Shore | Body;Body |
| GM01376 | chr6:33170779-33170838 | SLC39A7;SLC39A7 | N_Shore | Body;Body |
| GM00042 | chr6:33175140-33175156 | RING1;MIR219-1 | N_Shore | TSS1500;TSS1500 |
| GM00438 | chr6:33219335-33219410 | VPS52 | S_Shore | Body |
| GM09256 | chr6:33240820-33241226 | VPS52;RPS18 | S_Shore | TSS1500;Body |
| GM00290 | chr6:33245488-33245516 | B3GALT4 | Island | 1stExon |
| GM00935 | chr6:33245488-33245537 | B3GALT4 | Island | 1stExon |
| GM00940 | chr6:33245490-33245537 | B3GALT4 | Island | 1stExon |
| WG1722 | chr6:33245516-33245541 | B3GALT4 | Island | 1stExon |
| GM00935 | chr6:33245895-33246008 | B3GALT4 | S_Shore | 1stExon |
| GM09256 | chr6:33256165-33256222 | PFDN6;WDR46;WDR46 | N_Shore | TSS1500;Body;Body |
| GM02555 | chr6:33265386-33265424 | RGL2;RGL2 | N_Shore | Body;Body |
| WG3465 | chr6:33273477-33273965 | TAPBP;TAPBP;TAPBP | | Body;Body;Body |
| GM02706 | chr6:33280159-33280228 | TAPBP;TAPBP;TAPBP | N_Shore | Body;Body;Body |

| | | | | |
|---|---|---|---|---|
| GM00439 | chr6:33280212-33280390 | TAPBP;TAPBP;TAPBP | N_Shore | Body;Body;Body |
| GM02175 | chr6:33280400-33280436 | TAPBP;TAPBP;TAPBP | N_Shore | Body;Body;Body |
| WG3465 | chr6:33282997-33283015 | ZBTB22;TAPBP;TAPBP;TAPBP | Island | Body;TSS1500;TSS1500;TSS1500;Bod |
| GM09255 | chr6:33282997-33283021 | ZBTB22;TAPBP;TAPBP;TAPBP | Island | Body;TSS1500;TSS1500;TSS1500;Bod |
| WG1977 | chr6:33283015-33283055 | ZBTB22;TAPBP;TAPBP;TAPBP | Island | Body;TSS1500;TSS1500;TSS1500;Bod |
| WG1722 | chr6:33283015-33283064 | ZBTB22;TAPBP;TAPBP;TAPBP | Island | Body;TSS1500;TSS1500;TSS1500;Bod |
| WG2415 | chr6:33283162-33283189 | ZBTB22;TAPBP;TAPBP;TAPBP | Island | Body;TSS1500;TSS1500;TSS1500;Bod |
| WG1513 | chr6:33283987-33284107 | ZBTB22;ZBTB22 | N_Shore | Body;Body |
| WG1977 | chr6:33288180-33288561 | DAXX;DAXX;DAXX;DAXX | N_Shore | Body;Body;Body;Body |
| WG3466 | chr6:33288180-33288569 | DAXX;DAXX;DAXX;DAXX | N_Shore | Body;Body;Body;Body |
| GM00520 | chr6:33288296-33288372 | DAXX;DAXX;DAXX;DAXX | N_Shore | Body;Body;Body;Body |
| WG1978 | chr6:33288296-33288569 | DAXX;DAXX;DAXX;DAXX | N_Shore | Body;Body;Body;Body |
| GM01375 | chr6:33288323-33288561 | DAXX;DAXX;DAXX;DAXX | N_Shore | Body;Body;Body;Body |
| WG1722 | chr6:33288329-33288366 | DAXX;DAXX;DAXX;DAXX | N_Shore | Body;Body;Body;Body |
| WG2120 | chr6:33288329-33288561 | DAXX;DAXX;DAXX;DAXX | N_Shore | Body;Body;Body;Body |
| GM09255 | chr6:33288332-33288372 | DAXX;DAXX;DAXX;DAXX | N_Shore | Body;Body;Body;Body |
| GM02175 | chr6:33288332-33288561 | DAXX;DAXX;DAXX;DAXX | N_Shore | Body;Body;Body;Body |
| GM00935 | chr6:33288366-33288561 | DAXX;DAXX;DAXX;DAXX | N_Shore | Body;Body;Body;Body |
| GM03075 | chr6:33288366-33288569 | DAXX;DAXX;DAXX;DAXX | N_Shore | Body;Body;Body;Body |
| WG1978 | chr6:33288599-33288656 | DAXX;DAXX;DAXX;DAXX | N_Shore | Body;Body;Body;Body |
| WG2276 | chr6:33288964-33288998 | DAXX;DAXX;DAXX;DAXX | Island | Body;Body;Body;Body |
| GM02175 | chr6:33291921-33291947 | DAXX;DAXX;DAXX;DAXX | S_Shore | TSS1500;TSS1500;TSS1500;TSS1500 |
| WG1513 | chr6:33291937-33292029 | DAXX;DAXX;DAXX;DAXX | S_Shore | TSS1500;TSS1500;TSS1500;TSS1500 |
| GM00438 | chr6:33358785-33358872 | KIFC1 | N_Shore | TSS1500 |
| WG1513 | chr6:33358866-33359001 | KIFC1 | N_Shore | TSS1500 |
| GM00438 | chr6:33396295-33396407 | SYNGAP1 | Island | Body |
| WG1513 | chr6:33396405-33396610 | SYNGAP1 | S_Shore | Body |
| WG2415 | chr6:33397233-33397710 | SYNGAP1 | S_Shore | Body |
| GM00438 | chr6:33401480-33401520 | SYNGAP1 | | Body |
| GM02706 | chr6:33401480-33401536 | SYNGAP1 | | Body |
| WG3465 | chr6:33418814-33419140 | SYNGAP1 | N_Shelf | Body |
| GM00940 | chr6:33422868-33422910 | ZBTB9 | S_Shore | 5'UTR |
| GM09255 | chr6:33422868-33422918 | ZBTB9 | S_Shore | 5'UTR |
| GM00935 | chr6:33560953-33561181 | C6orf227 | Island | Body |
| GM02706 | chr6:33560953-33561186 | C6orf227 | Island | Body |
| GM09256 | chr6:33561099-33561186 | C6orf227 | Island | Body |
| WG1722 | chr6:33561099-33561188 | C6orf227 | Island | Body |
| GM00289 | chr6:33561181-33561188 | C6orf227 | Island | TSS200 |
| GM01096 | chr6:33561181-33561253 | C6orf227 | Island | TSS200 |
| WG3465 | chr6:34024362-34024479 | GRM4 | Island | Body |
| GM09256 | chr6:34031208-34031694 | GRM4 | | Body |
| GM09256 | chr6:34203564-34203887 | HMGA1;HMGA1;HMGA1;HM( | Island | TSS1500;TSS1500;TSS1500;TSS1500; |
| WG1657 | chr6:34499314-34499504 | PACSIN1 | Island | Body |
| WG2414 | chr6:34524095-34524278 | SPDEF | | TSS200 |
| GM09256 | chr6:3458053-3458177 | SLC22A23 | S_Shore | TSS1500 |
| GM01096 | chr6:35027147-35027324 | ANKS1A | | Body |
| GM09256 | chr6:35109121-35109298 | TCP11;TCP11;TCP11;TCP11 | Island | 5'UTR;1stExon;5'UTR;1stExon |

| | | | | |
|---|---|---|---|---|
| WG1977 | chr6:35265308-35265411 | DEF6 | | TSS1500 |
| GM00935 | chr6:35265411-35265555 | DEF6 | | TSS200 |
| GM02555 | chr6:35265534-35265555 | DEF6 | | TSS200 |
| GM01650 | chr6:35265534-35265624 | DEF6 | | TSS200 |
| WG1977 | chr6:35265534-35265759 | DEF6 | | TSS200 |
| GM09255 | chr6:35265616-35265643 | DEF6;DEF6 | | 1stExon;5'UTR |
| GM01392 | chr6:35265616-35265759 | DEF6;DEF6 | | 1stExon;5'UTR |
| GM00935 | chr6:35754758-35754906 | C6orf127 | Island | Body |
| WG1722 | chr6:35991802-35992520 | SLC26A8;SLC26A8 | N_Shelf | 5'UTR;5'UTR |
| GM00520 | chr6:36410500-36410515 | PXT1;KCTD20;PXT1 | Island | 5'UTR;TSS200;1stExon |
| GM02706 | chr6:37400392-37400668 | FTSJD2 | N_Shore | TSS1500 |
| GM01096 | chr6:37460940-37461350 | C6orf129 | | Body |
| WG2121 | chr6:37461033-37461350 | C6orf129 | | Body |
| WG1309 | chr6:37617864-37618009 | MDGA1 | Island | Body |
| GM01393 | chr6:3848790-3848866 | FAM50B | N_Shore | TSS1500 |
| WG2121 | chr6:3848898-3849028 | FAM50B | N_Shore | TSS1500 |
| GM01393 | chr6:3849294-3849331 | FAM50B | Island | TSS1500 |
| GM00520 | chr6:3849381-3849475 | FAM50B | Island | TSS1500 |
| GM02149 | chr6:39290526-39290715 | KCNK16;KCNK16;KCNK16;KCNK16 | | TSS200;TSS200;TSS200;TSS200 |
| WG1977 | chr6:41168941-41168972 | TREML2 | | TSS200 |
| GM00290 | chr6:41528198-41528395 | FOXP4;FOXP4;FOXP4 | N_Shore | 5'UTR;5'UTR;5'UTR |
| GM09255 | chr6:41528345-41528623 | FOXP4;FOXP4;FOXP4 | Island | 5'UTR;5'UTR;5'UTR |
| WG3465 | chr6:41528395-41528531 | FOXP4;FOXP4;FOXP4 | Island | 5'UTR;5'UTR;5'UTR |
| WG1977 | chr6:41528395-41528750 | FOXP4;FOXP4;FOXP4 | Island | 5'UTR;5'UTR;5'UTR |
| GM01392 | chr6:41528395-41528785 | FOXP4;FOXP4;FOXP4 | Island | 5'UTR;5'UTR;5'UTR |
| GM00940 | chr6:41528502-41528785 | FOXP4;FOXP4;FOXP4 | Island | 5'UTR;5'UTR;5'UTR |
| WG2415 | chr6:41528531-41528785 | FOXP4;FOXP4;FOXP4 | Island | 5'UTR;5'UTR;5'UTR |
| GM01095 | chr6:41605343-41605784 | MDFI | Island | TSS1500 |
| WG2415 | chr6:41703970-41704208 | TFEB;TFEB;TFEB | S_Shore | 5'UTR;TSS1500;1stExon |
| WG2844 | chr6:42226895-42227422 | TRERF1 | N_Shore | Body |
| WG1721 | chr6:42927959-42927986 | GNMT | N_Shore | TSS1500 |
| WG2414 | chr6:42927975-42927993 | GNMT | N_Shore | TSS1500 |
| GM02706 | chr6:42928200-42928345 | GNMT | N_Shore | TSS1500 |
| WG3465 | chr6:43149356-43149629 | CUL9 | N_Shore | TSS1500 |
| GM02706 | chr6:43197544-43197675 | C6orf108;C6orf108 | S_Shore | TSS1500;TSS1500 |
| GM00438 | chr6:43214396-43214590 | TTBK1 | Island | 5'UTR |
| WG3465 | chr6:43265619-43266151 | SLC22A7;SLC22A7 | | TSS1500;TSS1500 |
| GM00935 | chr6:43456984-43457177 | TJAP1;TJAP1;TJAP1;TJAP1;TJAP1;TJAP1;TJ | | 5'UTR;5'UTR;5'UTR;5'UTR;Body;TSS15 |
| GM00935 | chr6:43478455-43478500 | C6orf154 | Island | TSS200 |
| GM00439 | chr6:43612520-43612764 | RSPH9 | N_Shore | TSS1500 |
| WG2415 | chr6:45387257-45387395 | RUNX2;RUNX2 | N_Shore | Body;Body |
| GM09256 | chr6:45387257-45387854 | RUNX2;RUNX2 | N_Shore | Body;Body |
| WG3465 | chr6:45387257-45387917 | RUNX2;RUNX2 | N_Shore | Body;Body |
| GM09255 | chr6:45387257-45387932 | RUNX2;RUNX2 | N_Shore | Body;Body |
| WG1977 | chr6:45387257-45388583 | RUNX2;RUNX2 | N_Shore | Body;Body |
| WG1978 | chr6:45387347-45388522 | RUNX2;RUNX2 | Island | Body;Body |
| GM00935 | chr6:45387854-45388522 | RUNX2;RUNX2 | Island | Body;Body |

| | | | | |
|---|---|---|---|---|
| WG1310 | chr6:45391645-45391916 | RUNX2;RUNX2;RUNX2 | Island | Body;Body;Body |
| WG2844 | chr6:45391645-45391973 | RUNX2;RUNX2;RUNX2 | Island | Body;Body;Body |
| GM00016 | chr6:45391852-45391973 | RUNX2;RUNX2;RUNX2 | S_Shore | Body;Body;Body |
| WG3465 | chr6:46703468-46703504 | PLA2G7;PLA2G7 | S_Shore | TSS1500;TSS200 |
| WG2276 | chr6:46703504-46703670 | PLA2G7;PLA2G7 | S_Shore | TSS1500;TSS200 |
| GM00438 | chr6:46761035-46761090 | MEP1A | | TSS200 |
| WG1513 | chr6:47253910-47254042 | TNFRSF21 | | Body |
| WG2121 | chr6:47624115-47624247 | GPR111 | | TSS1500 |
| WG1513 | chr6:47665926-47666329 | GPR115 | | TSS1500 |
| GM09256 | chr6:49605181-49605401 | RHAG | | TSS1500 |
| GM00439 | chr6:49681178-49681391 | CRISP2;CRISP2;CRISP2;CRISP2;CRISP2 | | 5'UTR;5'UTR;5'UTR;5'UTR;5'UTR |
| WG1513 | chr6:49754848-49755009 | PGK2 | | 1stExon |
| GM02149 | chr6:49754848-49755043 | PGK2 | | 1stExon |
| WG1513 | chr6:49916931-49917175 | DEFB133 | | Body |
| WG1310 | chr6:49917145-49917254 | DEFB133 | | 1stExon |
| GM02175 | chr6:50803820-50803903 | TFAP2B | | Body |
| WG3466 | chr6:50803820-50804174 | TFAP2B | | Body |
| WG2120 | chr6:50810432-50810993 | TFAP2B | N_Shore | Body |
| WG1513 | chr6:50814011-50814305 | TFAP2B | S_Shore | 3'UTR |
| WG2275 | chr6:51953096-51953258 | PKHD1;PKHD1 | | TSS1500;TSS1500 |
| GM09256 | chr6:52008713-52009023 | MIR206 | | TSS1500 |
| GM00438 | chr6:52009019-52009064 | MIR206 | | TSS200 |
| WG1513 | chr6:53214008-53214076 | ELOVL5 | Island | TSS200 |
| GM00940 | chr6:53530503-53530570 | KLHL31;KLHL31 | | 1stExon;5'UTR |
| GM00289 | chr6:53530538-53530600 | KLHL31 | | TSS200 |
| WG3465 | chr6:55444153-55444488 | HMGCLL1;HMGCLL1 | S_Shore | TSS200;TSS200 |
| GM00289 | chr6:6002543-6003303 | NRN1 | Island | Body |
| WG3465 | chr6:6002865-6003303 | NRN1 | Island | Body |
| WG1309 | chr6:6320324-6320895 | F13A1 | | 5'UTR |
| GM00022 | chr6:6320324-6321061 | F13A1 | | 5'UTR |
| WG1514 | chr6:6320710-6320895 | F13A1 | | 5'UTR |
| WG2415 | chr6:64281439-64281639 | PTP4A1 | N_Shore | TSS1500 |
| GM00016 | chr6:6546375-6546851 | LOC285780 | Island | Body |
| GM00940 | chr6:6547015-6547273 | LOC285780 | Island | Body |
| WG2844 | chr6:6614447-6614798 | LOC285780;LY86 | | Body;Body |
| WG3465 | chr6:70576168-70576321 | COL19A1 | N_Shore | TSS1500 |
| WG1978 | chr6:70576168-70576502 | COL19A1 | N_Shore | TSS1500 |
| GM02706 | chr6:70576168-70576541 | COL19A1 | N_Shore | TSS1500 |
| GM02641 | chr6:70576315-70576388 | COL19A1 | N_Shore | TSS200 |
| GM01096 | chr6:70576388-70576541 | COL19A1 | N_Shore | TSS200 |
| WG1513 | chr6:71012765-71013139 | COL9A1;COL9A1 | | 5'UTR;1stExon |
| WG2415 | chr6:72926121-72926485 | RIMS1;RIMS1;RIMS1;RIMS1;RIMS1 | | TSS1500;TSS200;Body;Body;Body |
| WG2415 | chr6:74008688-74009041 | C6orf147 | N_Shore | Body |
| WG3465 | chr6:74063982-74064064 | DPPA5 | Island | TSS200 |
| GM00042 | chr6:74072261-74072447 | C6orf221 | N_Shore | TSS200 |
| GM00438 | chr6:74079465-74079540 | OOEP | Island | 1stExon |
| GM00042 | chr6:74104388-74104868 | DDX43;DDX43 | N_Shore | 1stExon;5'UTR |

| | | | | |
|---|---|---|---|---|
| GM02316 | chr6:74104426-74104600 | DDX43;DDX43 | Island | 1stExon;5'UTR |
| GM02706 | chr6:74404526-74405048 | CD109;CD109;CD109 | N_Shore | TSS1500;TSS1500;TSS1500 |
| GM00042 | chr6:7540834-7541066 | DSP;DSP | N_Shore | TSS1500;TSS1500 |
| GM00940 | chr6:7541066-7541519 | DSP;DSP | N_Shore | TSS1500;TSS1500 |
| GM02555 | chr6:75911716-75911956 | COL12A1;COL12A1 | Island | Body;Body |
| GM01095 | chr6:75911716-75912132 | COL12A1;COL12A1 | Island | Body;Body |
| GM03075 | chr6:75911716-75912410 | COL12A1;COL12A1 | Island | Body;Body |
| GM01096 | chr6:75911716-75912547 | COL12A1;COL12A1 | Island | Body;Body |
| GM02149 | chr6:75911956-75912410 | COL12A1;COL12A1 | Island | Body;Body |
| WG1978 | chr6:75911956-75912633 | COL12A1;COL12A1 | Island | Body;Body |
| GM00031 | chr6:75912132-75912633 | COL12A1;COL12A1 | Island | Body;Body |
| WG1514 | chr6:75912410-75912633 | COL12A1;COL12A1 | S_Shore | Body;Body |
| WG1658 | chr6:75912410-75912711 | COL12A1;COL12A1 | S_Shore | Body;Body |
| WG3465 | chr6:75912547-75912711 | COL12A1;COL12A1 | S_Shore | 5'UTR;5'UTR |
| GM01376 | chr6:76059726-76059773 | FILIP1 | | Body |
| WG1513 | chr6:84742695-84743025 | MRAP2 | N_Shore | TSS1500 |
| GM00289 | chr6:84743033-84743154 | MRAP2 | N_Shore | TSS1500 |
| WG1978 | chr6:84743086-84743236 | MRAP2 | N_Shore | TSS1500 |
| WG3465 | chr6:88757302-88757392 | SPACA1 | Island | TSS1500 |
| WG2415 | chr6:88757302-88757703 | SPACA1 | Island | TSS1500 |
| WG1977 | chr6:88757358-88757371 | SPACA1 | Island | TSS200 |
| GM00520 | chr6:88757358-88757392 | SPACA1 | Island | TSS200 |
| WG2120 | chr6:88757371-88757703 | SPACA1 | Island | TSS200 |
| WG2415 | chr6:91004430-91004482 | BACH2;BACH2 | N_Shore | 5'UTR;5'UTR |
| WG2121 | chr6:99842351-99842368 | COQ3 | S_Shore | TSS1500 |
| GM00439 | chr7:100091181-100091364 | C7orf51 | Island | Body |
| GM02175 | chr7:100091181-100091575 | C7orf51 | Island | Body |
| WG2844 | chr7:100181701-100182085 | LRCH4 | N_Shore | Body |
| WG2415 | chr7:100208533-100208950 | MOSPD3 | N_Shore | TSS1500 |
| GM00935 | chr7:100463206-100463583 | SLC12A9 | N_Shore | Body |
| WG3466 | chr7:100463416-100463583 | SLC12A9 | N_Shore | Body |
| GM01096 | chr7:100488527-100488942 | UFSP1;ACHE;ACHE | N_Shore | TSS1500;Body;3'UTR |
| WG1283 | chr7:100880751-100881286 | CLDN15 | | 1stExon |
| GM00940 | chr7:100880751-100881429 | CLDN15 | | 1stExon |
| GM00022 | chr7:100881007-100881429 | CLDN15;CLDN15 | | 1stExon;5'UTR |
| WG1977 | chr7:101005832-101006019 | EMID2 | N_Shore | TSS1500 |
| WG1513 | chr7:101273741-101274037 | MYL10 | | TSS1500 |
| WG1657 | chr7:101500014-101500514 | CUX1;CUX1;CUX1 | | Body;Body;Body |
| WG2275 | chr7:1015418-1015501 | COX19 | S_Shore | TSS200 |
| GM01096 | chr7:101558400-101558461 | CUX1;CUX1;CUX1 | N_Shore | Body;Body;Body |
| GM02456 | chr7:101558400-101558597 | CUX1;CUX1;CUX1 | N_Shore | Body;Body;Body |
| WG2276 | chr7:101768610-101768874 | CUX1;CUX1;CUX1 | | Body;Body;Body |
| WG1977 | chr7:101768676-101768874 | CUX1;CUX1;CUX1 | | Body;Body;Body |
| WG2844 | chr7:101813842-101814109 | CUX1;CUX1;CUX1 | | Body;Body;Body |
| WG2121 | chr7:101936318-101936430 | SH2B2 | Island | Body |
| GM01393 | chr7:101936318-101936527 | SH2B2 | Island | Body |
| WG3465 | chr7:1022213-1022643 | CYP2W1 | N_Shore | TSS1500 |

| | | | | |
|---|---|---|---|---|
| WG1722 | chr7:1022471-1022675 | CYP2W1 | N_Shore | TSS1500 |
| GM02175 | chr7:1022675-1022797 | CYP2W1 | N_Shore | TSS200 |
| WG2276 | chr7:1022970-1023156 | CYP2W1 | N_Shore | 1stExon |
| GM00438 | chr7:1026168-1026363 | CYP2W1 | Island | Body |
| GM02704 | chr7:105221998-105222024 | EFCAB10 | N_Shore | TSS200 |
| GM02555 | chr7:105319437-105319615 | ATXN7L1;ATXN7L1 | | Body;5'UTR |
| WG1310 | chr7:105319437-105319819 | ATXN7L1;ATXN7L1 | | Body;5'UTR |
| GM02456 | chr7:105319486-105319615 | ATXN7L1;ATXN7L1 | | Body;5'UTR |
| GM00935 | chr7:107205105-107205253 | COG5;COG5;COG5;DUS4L | S_Shore | TSS200;TSS200;TSS200;5'UTR |
| WG2275 | chr7:107205157-107205392 | COG5;COG5;COG5;DUS4L | S_Shore | TSS200;TSS200;TSS200;5'UTR |
| WG2121 | chr7:107300957-107301010 | SLC26A4;LOC286002 | N_Shore | TSS200;Body |
| WG1513 | chr7:1080836-1081068 | C7orf50;C7orf50;C7orf50 | Island | Body;Body;Body |
| WG2276 | chr7:10979247-10979586 | NDUFA4 | N_Shore | Body |
| WG3466 | chr7:1098625-1098703 | C7orf50;GPR146;C7orf50;C7 | S_Shore | Body;3'UTR;Body;Body;1stExon |
| GM00520 | chr7:1125817-1126326 | GPER;C7orf50;C7orf50;GPEI | N_Shore | TSS1500;Body;Body;TSS1500;Body |
| GM00439 | chr7:112725899-112726396 | GPR85;GPR85;GPR85;GPR85 | | 5'UTR;5'UTR;5'UTR;5'UTR |
| GM02706 | chr7:114055074-114055137 | FOXP2;FOXP2;FOXP2;FOXP2;FOXP2;FOX | | 1stExon;5'UTR;5'UTR;1stExon;5'UTR;1 |
| WG2120 | chr7:114055074-114055419 | FOXP2;FOXP2;FOXP2;FOXP2;FOXP2;FOX | | 1stExon;5'UTR;5'UTR;1stExon;5'UTR;1 |
| WG2275 | chr7:114055123-114055137 | FOXP2;FOXP2;FOXP2;FOXP2;FOXP2;FOX | | 1stExon;5'UTR;5'UTR;1stExon;5'UTR;1 |
| GM02704 | chr7:114055123-114055204 | FOXP2;FOXP2;FOXP2;FOXP2;FOXP2;FOX | | 1stExon;5'UTR;5'UTR;1stExon;5'UTR;1 |
| WG1722 | chr7:114055123-114055210 | FOXP2;FOXP2;FOXP2;FOXP2;FOXP2;FOX | | 1stExon;5'UTR;5'UTR;1stExon;5'UTR;1 |
| WG3466 | chr7:114055123-114055419 | FOXP2;FOXP2;FOXP2;FOXP2;FOXP2;FOX | | 1stExon;5'UTR;5'UTR;1stExon;5'UTR;1 |
| WG3465 | chr7:114055204-114055419 | FOXP2;FOXP2;FOXP2;FOXP2;FOXP2;FOX | | 1stExon;5'UTR;5'UTR;1stExon;5'UTR;1 |
| GM00439 | chr7:114561804-114561988 | MDFIC;MDFIC;MDFIC | N_Shore | TSS1500;TSS1500;TSS1500 |
| GM00016 | chr7:115862648-115862891 | TES;TES | | TSS1500;Body |
| WG2844 | chr7:115862648-115863026 | TES;TES | | TSS1500;Body |
| WG1978 | chr7:1163456-1163642 | C7orf50;C7orf50;C7orf50 | | Body;Body;Body |
| GM00935 | chr7:1163456-1164063 | C7orf50;C7orf50;C7orf50 | | Body;Body;Body |
| GM02555 | chr7:1163642-1164063 | C7orf50;C7orf50;C7orf50 | | Body;Body;Body |
| WG2276 | chr7:116592956-116592968 | ST7OT4;ST7OT1;ST7;ST7 | N_Shore | TSS1500;Body;TSS1500;TSS1500 |
| GM00940 | chr7:116963492-116963502 | WNT2;WNT2 | Island | TSS200;TSS200 |
| WG1513 | chr7:117067616-117067978 | ASZ1;ASZ1 | Island | TSS200;TSS200 |
| WG1513 | chr7:117864720-117864927 | ANKRD7;ANKRD7 | | 1stExon;5'UTR |
| WG1722 | chr7:1198522-1198841 | ZFAND2A | N_Shore | 5'UTR |
| GM02555 | chr7:1198662-1198926 | ZFAND2A | N_Shore | 5'UTR |
| WG3466 | chr7:120968919-120969174 | WNT16;WNT16 | N_Shore | TSS200;Body |
| WG2844 | chr7:120969033-120969079 | WNT16;WNT16 | N_Shore | TSS200;Body |
| WG1978 | chr7:12444013-12444115 | VWDE | S_Shore | TSS200 |
| WG1513 | chr7:12609907-12609934 | SCIN | N_Shore | TSS1500 |
| GM00935 | chr7:127225891-127225949 | GCC1 | Island | TSS1500 |
| WG3465 | chr7:127255591-127255970 | PAX4;PAX4 | | 5'UTR;1stExon |
| GM00289 | chr7:127256705-127256920 | PAX4 | | TSS1500 |
| GM00439 | chr7:1273835-1274166 | UNCX | Island | Body |
| WG1310 | chr7:127881280-127881344 | LEP | Island | TSS200 |
| WG1722 | chr7:128550860-128550902 | KCP;KCP | Island | TSS200;TSS200 |
| GM01650 | chr7:128550860-128550932 | KCP;KCP | Island | TSS200;TSS200 |
| GM02555 | chr7:128550860-128550946 | KCP;KCP | Island | TSS200;TSS200 |

| | | | | |
|---|---|---|---|---|
| WG2415 | chr7:128550862-128550902 | KCP;KCP | Island | TSS200;TSS200 |
| WG1978 | chr7:128550862-128550932 | KCP;KCP | Island | TSS200;TSS200 |
| GM00940 | chr7:128550862-128550946 | KCP;KCP | Island | TSS200;TSS200 |
| WG2844 | chr7:128550898-128550932 | KCP;KCP | Island | TSS200;TSS200 |
| GM00439 | chr7:129007902-129008179 | AHCYL2;AHCYL2;AHCYL2;AHCYL2 | | Body;TSS200;Body;TSS200 |
| GM00439 | chr7:129414697-129414746 | MIR96 | N_Shelf | TSS200 |
| WG2844 | chr7:129845574-129845832 | TMEM209 | S_Shore | TSS1500 |
| WG3466 | chr7:130020011-130020130 | CPA1 | N_Shore | TSS1500 |
| GM00520 | chr7:130126024-130126182 | MEST | Island | TSS200 |
| WG1977 | chr7:130126024-130126513 | MEST | Island | TSS200 |
| GM02175 | chr7:130126182-130126513 | MEST;MEST | Island | 1stExon;5'UTR |
| WG2414 | chr7:130126368-130126637 | MEST | Island | 5'UTR |
| GM02173 | chr7:130419136-130419173 | KLF14 | Island | TSS1500 |
| WG2121 | chr7:134575187-134575456 | CALD1;CALD1;CALD1;CALD1;CALD1 | | TSS1500;Body;TSS1500;Body;Body |
| GM09255 | chr7:134575271-134575456 | CALD1;CALD1;CALD1;CALD1;CALD1 | | TSS1500;Body;TSS1500;Body;Body |
| GM01096 | chr7:134832221-134832584 | TMEM140 | | TSS1500 |
| GM00022 | chr7:134832221-134832671 | TMEM140 | | TSS1500 |
| WG3465 | chr7:134832221-134832770 | TMEM140 | | TSS1500 |
| WG1283 | chr7:134832221-134833299 | TMEM140 | | TSS1500 |
| WG3466 | chr7:134832671-134832850 | TMEM140 | | TSS200 |
| WG1722 | chr7:134832671-134832939 | TMEM140 | | TSS200 |
| GM00289 | chr7:134832770-134833031 | TMEM140;TMEM140 | | 1stExon;5'UTR |
| WG2194 | chr7:134832844-134832939 | TMEM140;TMEM140 | | 1stExon;5'UTR |
| GM00022 | chr7:134832850-134833031 | TMEM140;TMEM140 | | 1stExon;5'UTR |
| WG3465 | chr7:134832850-134833299 | TMEM140;TMEM140 | | 1stExon;5'UTR |
| GM02175 | chr7:134833031-134833299 | TMEM140 | | 5'UTR |
| GM00042 | chr7:134856562-134856681 | C7orf49;C7orf49;C7orf49 | S_Shore | TSS1500;TSS1500;TSS1500 |
| GM02706 | chr7:136553088-136553170 | CHRM2;CHRM2;CHRM2;CHF | N_Shore | TSS1500;TSS1500;TSS1500;TSS1500; |
| WG2844 | chr7:140103671-140103756 | RAB19 | | TSS200 |
| GM02149 | chr7:141673649-141673903 | TAS2R38 | | TSS200 |
| GM02706 | chr7:142636470-142636518 | C7orf34 | | TSS200 |
| GM00438 | chr7:143013161-143013444 | CLCN1 | | TSS200 |
| WG1513 | chr7:143655651-143656153 | OR2F1 | | TSS1500 |
| GM02704 | chr7:148480080-148480326 | CUL1 | N_Shore | Body |
| GM02706 | chr7:148480080-148480503 | CUL1 | N_Shore | Body |
| GM00940 | chr7:1491831-1491921 | MICALL2 | Island | Body |
| WG2415 | chr7:149471884-149472134 | SSPO | S_Shore | TSS1500 |
| GM02706 | chr7:150020025-150020136 | LRRC61;ACTR3C;LRRC61 | Island | TSS1500;5'UTR;TSS1500 |
| GM09256 | chr7:150147553-150147988 | GIMAP8 | | TSS1500 |
| WG2414 | chr7:150147922-150147988 | GIMAP8 | | TSS200 |
| GM02175 | chr7:150147922-150148022 | GIMAP8 | | TSS200 |
| GM02149 | chr7:150210785-150211061 | GIMAP7 | | TSS1500 |
| WG1513 | chr7:150210980-150211820 | GIMAP7 | | TSS1500 |
| GM01375 | chr7:150434452-150434480 | GIMAP5;GIMAP5 | | 1stExon;5'UTR |
| GM02175 | chr7:150434452-150434541 | GIMAP5;GIMAP5 | | 1stExon;5'UTR |
| GM00935 | chr7:150434471-150434541 | GIMAP5;GIMAP5 | | 1stExon;5'UTR |
| WG2415 | chr7:150498205-150498843 | TMEM176B;TMEM176B;TMEN | Island | 1stExon;5'UTR;TSS1500;TSS1500;TSS |

| | | | | |
|---|---|---|---|---|
| WG1513 | chr7:150549370-150549472 | ABP1 | | TSS1500 |
| WG1513 | chr7:150554420-150554674 | ABP1 | Island | Body |
| GM09256 | chr7:150705760-150705896 | NOS3 | Island | Body |
| GM00439 | chr7:150744274-150744599 | ACCN3;ACCN3;ABCB8;ACCN | N_Shore | TSS1500;TSS1500;3'UTR;TSS1500 |
| GM01650 | chr7:150786044-150786082 | AGAP3;AGAP3 | S_Shore | Body;Body |
| GM09255 | chr7:150822655-150822987 | AGAP3 | Island | Body |
| GM00935 | chr7:151078646-151079167 | WDR86 | Island | 3'UTR |
| WG1977 | chr7:151078646-151079501 | WDR86 | Island | 3'UTR |
| GM00520 | chr7:151078995-151079501 | WDR86 | Island | Body |
| GM00439 | chr7:151106022-151106082 | WDR86 | N_Shore | Body |
| WG3466 | chr7:151136940-151137048 | CRYGN | Island | Body |
| WG2414 | chr7:151330008-151330230 | PRKAG2;PRKAG2;PRKAG2 | | Body;TSS1500;Body |
| WG3465 | chr7:151442351-151442967 | PRKAG2;PRKAG2 | | Body;Body |
| WG3465 | chr7:1514455-1514623 | INTS1 | Island | Body |
| WG3465 | chr7:151493629-151493820 | PRKAG2;PRKAG2 | | 5'UTR;Body |
| WG3465 | chr7:151653454-151653890 | GALNTL5;GALNTL5 | | TSS200;TSS200 |
| GM09255 | chr7:153586022-153586207 | DPP6 | S_Shore | Body |
| WG1513 | chr7:154003215-154003309 | DPP6;DPP6;DPP6 | S_Shore | Body;Body;Body |
| GM00289 | chr7:154542060-154542313 | DPP6;DPP6;DPP6 | Island | Body;Body;Body |
| WG2415 | chr7:154684051-154684433 | DPP6;DPP6;DPP6 | N_Shore | Body;Body;Body |
| GM00438 | chr7:154685388-154685726 | DPP6;DPP6;DPP6 | S_Shore | 3'UTR;3'UTR;3'UTR |
| GM02175 | chr7:154862969-154863244 | HTR5A | Island | 1stExon |
| WG1977 | chr7:155255310-155255791 | EN2 | Island | Body |
| WG3466 | chr7:155299624-155299687 | CNPY1 | N_Shelf | Body |
| GM01392 | chr7:156741514-156741538 | NOM1 | Island | TSS1500 |
| GM00439 | chr7:156802325-156802502 | MNX1;MNX1 | Island | TSS200;Body |
| GM00289 | chr7:157179773-157179830 | DNAJB6 | | Body |
| WG2843 | chr7:157185279-157185402 | DNAJB6 | | Body |
| WG2843 | chr7:157198482-157198869 | DNAJB6 | N_Shelf | Body |
| WG3465 | chr7:157333405-157333584 | PTPRN2;PTPRN2;PTPRN2 | | 3'UTR;3'UTR;3'UTR |
| GM00438 | chr7:157345746-157346224 | PTPRN2;PTPRN2;PTPRN2 | N_Shore | Body;Body;Body |
| GM02149 | chr7:157353097-157353191 | PTPRN2;PTPRN2;PTPRN2 | Island | Body;Body;Body |
| GM00438 | chr7:157368901-157368976 | PTPRN2;PTPRN2;PTPRN2 | N_Shore | Body;Body;Body |
| WG1513 | chr7:157369121-157369376 | PTPRN2;PTPRN2;PTPRN2 | Island | Body;Body;Body |
| WG1977 | chr7:157370486-157370851 | PTPRN2;PTPRN2;PTPRN2 | S_Shore | Body;Body;Body |
| WG1513 | chr7:157386714-157387156 | PTPRN2;PTPRN2;PTPRN2 | N_Shore | Body;Body;Body |
| WG1513 | chr7:157402529-157402673 | PTPRN2;PTPRN2;PTPRN2 | N_Shelf | Body;Body;Body |
| WG1513 | chr7:157404440-157404705 | PTPRN2;PTPRN2;PTPRN2 | N_Shore | Body;Body;Body |
| GM00438 | chr7:157405965-157406096 | PTPRN2;PTPRN2;PTPRN2 | Island | Body;Body;Body |
| WG1513 | chr7:157412195-157413057 | PTPRN2;PTPRN2;PTPRN2 | N_Shore | Body;Body;Body |
| GM00438 | chr7:157412886-157413287 | PTPRN2;PTPRN2;PTPRN2 | N_Shore | Body;Body;Body |
| GM02149 | chr7:157414012-157414339 | PTPRN2;PTPRN2;PTPRN2 | Island | Body;Body;Body |
| GM00438 | chr7:157443760-157444239 | PTPRN2;PTPRN2;PTPRN2 | S_Shore | Body;Body;Body |
| WG1513 | chr7:157444235-157444423 | PTPRN2;PTPRN2;PTPRN2 | S_Shore | Body;Body;Body |
| WG1978 | chr7:157447178-157447249 | PTPRN2;PTPRN2;PTPRN2 | N_Shore | Body;Body;Body |
| GM02149 | chr7:157447244-157447781 | PTPRN2;PTPRN2;PTPRN2 | N_Shore | Body;Body;Body |
| GM09256 | chr7:157454062-157454207 | PTPRN2;PTPRN2;PTPRN2 | Island | Body;Body;Body |

| | | | | |
|---|---|---|---|---|
| GM00042 | chr7:157461081-157461125 | PTPRN2;PTPRN2;PTPRN2 | | Body;Body;Body |
| GM00438 | chr7:157461897-157462264 | PTPRN2;PTPRN2;PTPRN2 | | Body;Body;Body |
| WG3466 | chr7:157475548-157476575 | PTPRN2;PTPRN2;PTPRN2 | N_Shore | Body;Body;Body |
| GM00438 | chr7:157493703-157493965 | PTPRN2;PTPRN2;PTPRN2 | N_Shore | Body;Body;Body |
| WG1977 | chr7:157493965-157494424 | PTPRN2;PTPRN2;PTPRN2 | N_Shore | Body;Body;Body |
| GM02149 | chr7:157494836-157495282 | PTPRN2;PTPRN2;PTPRN2 | S_Shore | Body;Body;Body |
| GM00438 | chr7:157494836-157495532 | PTPRN2;PTPRN2;PTPRN2 | S_Shore | Body;Body;Body |
| WG1513 | chr7:157495481-157495882 | PTPRN2;PTPRN2;PTPRN2 | S_Shore | Body;Body;Body |
| GM02149 | chr7:157495532-157495969 | PTPRN2;PTPRN2;PTPRN2 | S_Shore | Body;Body;Body |
| WG1513 | chr7:157513205-157513707 | PTPRN2;PTPRN2;PTPRN2 | | Body;Body;Body |
| WG1513 | chr7:157529335-157530177 | PTPRN2;PTPRN2;PTPRN2 | S_Shelf | Body;Body;Body |
| GM02706 | chr7:157561593-157561786 | PTPRN2;PTPRN2;PTPRN2 | S_Shore | Body;Body;Body |
| WG1513 | chr7:157561684-157561848 | PTPRN2;PTPRN2;PTPRN2 | S_Shore | Body;Body;Body |
| WG1282 | chr7:157561786-157561925 | PTPRN2;PTPRN2;PTPRN2 | S_Shore | Body;Body;Body |
| GM02149 | chr7:157586219-157586345 | PTPRN2;PTPRN2;PTPRN2 | N_Shore | Body;Body;Body |
| WG1513 | chr7:157605569-157605970 | PTPRN2;PTPRN2;PTPRN2 | | Body;Body;Body |
| WG1513 | chr7:157637416-157638062 | PTPRN2;PTPRN2;PTPRN2 | | Body;Body;Body |
| GM02175 | chr7:157644273-157644555 | PTPRN2;PTPRN2;PTPRN2 | N_Shelf | Body;Body;Body |
| WG2415 | chr7:157644471-157644665 | PTPRN2;PTPRN2;PTPRN2 | N_Shelf | Body;Body;Body |
| WG1513 | chr7:157646605-157646701 | PTPRN2;PTPRN2;PTPRN2 | N_Shore | Body;Body;Body |
| WG1513 | chr7:157647847-157648006 | PTPRN2;PTPRN2;PTPRN2 | Island | Body;Body;Body |
| WG1513 | chr7:157653809-157654170 | PTPRN2;PTPRN2;PTPRN2 | | Body;Body;Body |
| GM00438 | chr7:157690246-157690657 | PTPRN2;PTPRN2;PTPRN2 | N_Shore | Body;Body;Body |
| GM09256 | chr7:157690609-157690732 | PTPRN2;PTPRN2;PTPRN2 | Island | Body;Body;Body |
| WG1513 | chr7:157690609-157691541 | PTPRN2;PTPRN2;PTPRN2 | Island | Body;Body;Body |
| GM00438 | chr7:157690769-157691376 | PTPRN2;PTPRN2;PTPRN2 | Island | Body;Body;Body |
| WG1513 | chr7:157694404-157695026 | PTPRN2;PTPRN2;PTPRN2 | Island | Body;Body;Body |
| WG1513 | chr7:157695357-157695878 | PTPRN2;PTPRN2;PTPRN2 | Island | Body;Body;Body |
| WG2415 | chr7:157695747-157695878 | PTPRN2;PTPRN2;PTPRN2 | S_Shore | Body;Body;Body |
| WG2415 | chr7:157696589-157697240 | PTPRN2;PTPRN2;PTPRN2 | S_Shore | Body;Body;Body |
| GM00438 | chr7:157730162-157730264 | PTPRN2;PTPRN2;PTPRN2 | | Body;Body;Body |
| WG1513 | chr7:157778531-157778645 | PTPRN2;PTPRN2;PTPRN2 | | Body;Body;Body |
| WG1513 | chr7:157785628-157785759 | PTPRN2;PTPRN2;PTPRN2 | | Body;Body;Body |
| GM00438 | chr7:157785628-157786056 | PTPRN2;PTPRN2;PTPRN2 | | Body;Body;Body |
| GM02149 | chr7:157785721-157786056 | PTPRN2;PTPRN2;PTPRN2 | | Body;Body;Body |
| WG1513 | chr7:157786157-157786631 | PTPRN2;PTPRN2;PTPRN2 | | Body;Body;Body |
| GM00438 | chr7:157866704-157867264 | PTPRN2;PTPRN2;PTPRN2 | | Body;Body;Body |
| GM02149 | chr7:157866922-157867264 | PTPRN2;PTPRN2;PTPRN2 | | Body;Body;Body |
| WG1513 | chr7:157867300-157867695 | PTPRN2;PTPRN2;PTPRN2 | | Body;Body;Body |
| GM01096 | chr7:157873499-157873854 | PTPRN2;PTPRN2;PTPRN2 | | Body;Body;Body |
| WG1513 | chr7:157916077-157916687 | PTPRN2;PTPRN2;PTPRN2 | N_Shelf | Body;Body;Body |
| GM01653 | chr7:157932921-157932971 | PTPRN2;PTPRN2;PTPRN2 | | Body;Body;Body |
| GM00289 | chr7:157932921-157933172 | PTPRN2;PTPRN2;PTPRN2 | | Body;Body;Body |
| GM02149 | chr7:157932921-157933354 | PTPRN2;PTPRN2;PTPRN2 | | Body;Body;Body |
| WG1513 | chr7:157932921-157933417 | PTPRN2;PTPRN2;PTPRN2 | | Body;Body;Body |
| WG1513 | chr7:157937468-157937891 | PTPRN2;PTPRN2;PTPRN2 | N_Shore | Body;Body;Body |
| WG1513 | chr7:157956042-157956161 | PTPRN2;PTPRN2;PTPRN2 | N_Shore | Body;Body;Body |

| | | | | |
|---|---|---|---|---|
| WG1513 | chr7:157957646-157958299 | PTPRN2;PTPRN2;PTPRN2 | S_Shore | Body;Body;Body |
| GM00289 | chr7:157980803-157981333 | PTPRN2;PTPRN2;PTPRN2 | Island | Body;Body;Body |
| GM09256 | chr7:157982161-157982364 | PTPRN2;PTPRN2;PTPRN2 | S_Shore | Body;Body;Body |
| WG1513 | chr7:157982224-157982427 | PTPRN2;PTPRN2;PTPRN2 | S_Shore | Body;Body;Body |
| WG1513 | chr7:157985748-157985818 | PTPRN2;PTPRN2;PTPRN2 | | Body;Body;Body |
| GM00438 | chr7:158038697-158039076 | PTPRN2;PTPRN2;PTPRN2 | S_Shore | Body;Body;Body |
| WG2415 | chr7:158045980-158046025 | PTPRN2;PTPRN2;PTPRN2 | | Body;Body;Body |
| WG1513 | chr7:158059396-158059944 | PTPRN2;PTPRN2;PTPRN2 | N_Shore | Body;Body;Body |
| GM00289 | chr7:158059790-158060003 | PTPRN2;PTPRN2;PTPRN2 | N_Shore | Body;Body;Body |
| GM01096 | chr7:158059944-158060003 | PTPRN2;PTPRN2;PTPRN2 | N_Shore | Body;Body;Body |
| GM02175 | chr7:158075388-158075705 | PTPRN2;PTPRN2;PTPRN2 | | Body;Body;Body |
| GM02149 | chr7:158092647-158093277 | PTPRN2;PTPRN2;PTPRN2 | | Body;Body;Body |
| GM00289 | chr7:158093198-158093277 | PTPRN2;PTPRN2;PTPRN2 | | Body;Body;Body |
| WG1513 | chr7:158106772-158107284 | PTPRN2;PTPRN2;PTPRN2 | N_Shelf | Body;Body;Body |
| GM01096 | chr7:158109339-158109694 | PTPRN2;PTPRN2;PTPRN2 | Island | Body;Body;Body |
| GM02149 | chr7:158109339-158110152 | PTPRN2;PTPRN2;PTPRN2 | Island | Body;Body;Body |
| GM00289 | chr7:158109492-158110152 | PTPRN2;PTPRN2;PTPRN2 | Island | Body;Body;Body |
| GM02175 | chr7:158109694-158110152 | PTPRN2;PTPRN2;PTPRN2 | N_Shore | Body;Body;Body |
| WG1722 | chr7:158110405-158110685 | PTPRN2;PTPRN2;PTPRN2 | N_Shore | Body;Body;Body |
| WG1513 | chr7:158114558-158114659 | PTPRN2;PTPRN2;PTPRN2 | S_Shelf | Body;Body;Body |
| WG1513 | chr7:158157967-158158218 | PTPRN2;PTPRN2;PTPRN2 | | Body;Body;Body |
| GM00438 | chr7:158235423-158235949 | PTPRN2;PTPRN2;PTPRN2 | | Body;Body;Body |
| WG1513 | chr7:158235423-158236160 | PTPRN2;PTPRN2;PTPRN2 | | Body;Body;Body |
| GM02149 | chr7:158235891-158236103 | PTPRN2;PTPRN2;PTPRN2 | | Body;Body;Body |
| WG1513 | chr7:158250310-158250524 | PTPRN2;PTPRN2;PTPRN2 | Island | Body;Body;Body |
| WG1513 | chr7:158263204-158263277 | PTPRN2;PTPRN2;PTPRN2 | | Body;Body;Body |
| WG1513 | chr7:158264915-158265561 | PTPRN2;PTPRN2;PTPRN2 | | Body;Body;Body |
| GM02706 | chr7:158278869-158278954 | PTPRN2;PTPRN2;PTPRN2 | S_Shore | Body;Body;Body |
| WG1513 | chr7:158279818-158279891 | PTPRN2;PTPRN2;PTPRN2 | N_Shore | Body;Body;Body |
| WG1978 | chr7:158336016-158336049 | PTPRN2;PTPRN2;PTPRN2 | | Body;Body;Body |
| WG2415 | chr7:158336016-158336562 | PTPRN2;PTPRN2;PTPRN2 | | Body;Body;Body |
| WG1513 | chr7:158349441-158349960 | PTPRN2;PTPRN2;PTPRN2 | Island | Body;Body;Body |
| GM02147 | chr7:158362729-158362907 | PTPRN2;PTPRN2;PTPRN2 | Island | Body;Body;Body |
| GM09255 | chr7:158379132-158379223 | PTPRN2;PTPRN2;PTPRN2 | N_Shore | Body;Body;Body |
| GM00438 | chr7:158656429-158656620 | WDR60 | Island | Body |
| WG1513 | chr7:158800547-158800888 | LOC154822 | S_Shore | TSS1500 |
| WG2415 | chr7:158818053-158818164 | LOC154822 | S_Shore | Body |
| GM00438 | chr7:1588319-1588579 | TMEM184A | S_Shore | Body |
| GM00289 | chr7:158854564-158854856 | VIPR2 | Island | Body |
| GM02149 | chr7:158858173-158858508 | VIPR2 | S_Shore | Body |
| WG1513 | chr7:158858498-158858862 | VIPR2 | S_Shore | Body |
| GM02555 | chr7:158905015-158905377 | VIPR2 | N_Shore | Body |
| GM02706 | chr7:158905317-158905536 | VIPR2 | Island | Body |
| GM00290 | chr7:1607247-1607601 | PSMG3;PSMG3 | N_Shelf | 3'UTR;3'UTR |
| WG1658 | chr7:1607247-1607663 | PSMG3;PSMG3 | N_Shelf | 3'UTR;3'UTR |
| WG2415 | chr7:16505094-16505589 | SOSTDC1 | | 1stExon |
| WG3465 | chr7:16505094-16505602 | SOSTDC1 | | 1stExon |

| | | | | |
|---|---|---|---|---|
| GM02706 | chr7:16505409-16505592 | SOSTDC1;SOSTDC1 | | 1stExon;5'UTR |
| GM02555 | chr7:16505409-16505602 | SOSTDC1;SOSTDC1 | | 1stExon;5'UTR |
| GM02149 | chr7:16505409-16505617 | SOSTDC1;SOSTDC1 | | 1stExon;5'UTR |
| WG1722 | chr7:16505589-16505602 | SOSTDC1 | | TSS200 |
| WG1977 | chr7:1753396-1753659 | ELFN1 | N_Shore | 5'UTR |
| GM02175 | chr7:1753573-1753659 | ELFN1 | Island | 5'UTR |
| GM00289 | chr7:1753573-1753909 | ELFN1 | Island | 5'UTR |
| WG2415 | chr7:1775904-1776076 | ELFN1 | | 5'UTR |
| WG2844 | chr7:1787310-1787831 | ELFN1 | S_Shore | 3'UTR |
| GM00520 | chr7:1882776-1883251 | MAD1L1;MAD1L1;MAD1L1 | S_Shore | Body;Body;Body |
| GM00016 | chr7:19158647-19158666 | TWIST1 | S_Shore | TSS1500 |
| GM00016 | chr7:19158721-19158954 | TWIST1 | S_Shore | TSS1500 |
| GM01393 | chr7:19184555-19185139 | FERD3L | Island | 1stExon |
| GM02555 | chr7:19184555-19185165 | FERD3L | Island | 1stExon |
| WG1513 | chr7:19184950-19185139 | FERD3L | Island | 1stExon |
| WG1283 | chr7:19185260-19185407 | FERD3L | S_Shore | TSS1500 |
| WG3465 | chr7:1938450-1938791 | MAD1L1;MAD1L1;MAD1L1 | S_Shore | Body;Body;Body |
| GM01095 | chr7:1952518-1952600 | MAD1L1;MAD1L1;MAD1L1 | S_Shelf | Body;Body;Body |
| WG1977 | chr7:1952518-1952807 | MAD1L1;MAD1L1;MAD1L1 | S_Shelf | Body;Body;Body |
| GM00520 | chr7:195475-196130 | FAM20C | Island | Body |
| WG2276 | chr7:1961785-1961968 | MAD1L1;MAD1L1;MAD1L1 | | Body;Body;Body |
| GM00520 | chr7:1980117-1980273 | MAD1L1;MAD1L1;MAD1L1 | S_Shore | Body;Body;Body |
| GM00935 | chr7:1980117-1980479 | MAD1L1;MAD1L1;MAD1L1 | S_Shore | Body;Body;Body |
| WG2276 | chr7:2058865-2059212 | MAD1L1;MAD1L1;MAD1L1 | N_Shore | Body;Body;Body |
| GM01393 | chr7:2059796-2060040 | MAD1L1;MAD1L1;MAD1L1 | N_Shore | Body;Body;Body |
| WG2120 | chr7:2060040-2060130 | MAD1L1;MAD1L1;MAD1L1 | Island | Body;Body;Body |
| WG2415 | chr7:2060040-2060179 | MAD1L1;MAD1L1;MAD1L1 | Island | Body;Body;Body |
| GM00520 | chr7:2060116-2060179 | MAD1L1;MAD1L1;MAD1L1 | Island | Body;Body;Body |
| GM02456 | chr7:2077057-2077345 | MAD1L1;MAD1L1;MAD1L1 | | Body;Body;Body |
| WG2276 | chr7:2078650-2079008 | MAD1L1;MAD1L1;MAD1L1 | | Body;Body;Body |
| GM00290 | chr7:20825824-20826128 | SP8;SP8 | S_Shore | 5'UTR;Body |
| GM00290 | chr7:2109244-2109450 | MAD1L1;MAD1L1;MAD1L1 | S_Shore | Body;Body;Body |
| WG3465 | chr7:2109244-2109890 | MAD1L1;MAD1L1;MAD1L1 | S_Shore | Body;Body;Body |
| GM09255 | chr7:2109353-2109890 | MAD1L1;MAD1L1;MAD1L1 | S_Shore | Body;Body;Body |
| WG1310 | chr7:2133441-2133480 | MAD1L1;MAD1L1;MAD1L1 | | Body;Body;Body |
| WG3465 | chr7:2139216-2139259 | MAD1L1;MAD1L1;MAD1L1 | | Body;Body;Body |
| GM00520 | chr7:214577-214779 | FAM20C | N_Shelf | Body |
| WG2276 | chr7:2150534-2151007 | MAD1L1;MAD1L1;MAD1L1 | | Body;Body;Body |
| WG2844 | chr7:2214759-2214790 | MAD1L1;MAD1L1;MAD1L1 | | Body;Body;Body |
| GM02706 | chr7:22539822-22540090 | MGC87042;MGC87042;MGC | Island | 5'UTR;1stExon;1stExon;5'UTR |
| WG2414 | chr7:226444-226563 | FAM20C | S_Shore | Body |
| WG2415 | chr7:226540-226959 | FAM20C | S_Shore | Body |
| WG2415 | chr7:22767384-22767571 | IL6 | | Body |
| GM00438 | chr7:23749667-23749740 | STK31;STK31;STK31 | | TSS200;TSS200;TSS1500 |
| GM02555 | chr7:24323128-24323559 | NPY | N_Shore | TSS1500 |
| GM02555 | chr7:24324570-24325079 | NPY | Island | 5'UTR |
| WG2844 | chr7:2557358-2557715 | LFNG;LFNG | N_Shore | Body;TSS200 |

| | | | | |
|---|---|---|---|---|
| WG3465 | chr7:2560976-2561116 | LFNG;LFNG;LFNG;LFNG | S_Shore | Body;Body;Body;Body |
| WG2276 | chr7:2563184-2563611 | LFNG;LFNG;LFNG;LFNG | N_Shore | Body;Body;Body;Body |
| GM02555 | chr7:2563184-2563909 | LFNG;LFNG;LFNG;LFNG | N_Shore | Body;Body;Body;Body |
| GM03075 | chr7:2563478-2563909 | LFNG;LFNG;LFNG;LFNG | N_Shore | Body;Body;Body;Body |
| WG2275 | chr7:2563611-2563909 | LFNG;LFNG;LFNG;LFNG | Island | Body;Body;Body;Body |
| WG1310 | chr7:2563670-2563909 | LFNG;LFNG;LFNG;LFNG | Island | Body;Body;Body;Body |
| WG1984 | chr7:2606716-2606824 | IQCE;IQCE | | Body;Body |
| GM02149 | chr7:26191458-26191599 | NFE2L3 | N_Shore | TSS1500 |
| GM00439 | chr7:26192756-26193032 | NFE2L3 | Island | Body |
| WG3466 | chr7:2646782-2647333 | IQCE;IQCE | S_Shore | Body;Body |
| GM02456 | chr7:2647039-2647414 | IQCE;IQCE | S_Shore | Body;Body |
| GM00438 | chr7:26897182-26897253 | SKAP2 | | Body |
| WG1978 | chr7:26897182-26897714 | SKAP2 | | Body |
| GM00022 | chr7:26897597-26897685 | SKAP2 | | Body |
| GM02704 | chr7:26897612-26897714 | SKAP2 | | Body |
| GM02706 | chr7:27140942-27141139 | HOXA2 | N_Shelf | Body |
| WG1978 | chr7:27140942-27141774 | HOXA2 | N_Shelf | Body |
| GM02175 | chr7:27142100-27142204 | HOXA2 | N_Shore | 1stExon |
| WG1978 | chr7:27142200-27142708 | HOXA2;HOXA2 | N_Shore | 1stExon;5'UTR |
| WG3466 | chr7:27142427-27142527 | HOXA2 | N_Shore | TSS200 |
| GM00941 | chr7:27142427-27142547 | HOXA2 | N_Shore | TSS200 |
| GM02173 | chr7:27142427-27142618 | HOXA2 | N_Shore | TSS200 |
| GM02706 | chr7:27142427-27142810 | HOXA2 | N_Shore | TSS200 |
| GM01096 | chr7:27142427-27143478 | HOXA2 | N_Shore | TSS200 |
| WG1722 | chr7:27142437-27142535 | HOXA2 | N_Shore | TSS200 |
| GM02704 | chr7:27142527-27142618 | HOXA2 | N_Shore | TSS200 |
| GM02175 | chr7:27142700-27142799 | HOXA2 | N_Shore | TSS1500 |
| GM02704 | chr7:27142700-27143287 | HOXA2 | N_Shore | TSS1500 |
| GM09255 | chr7:27142700-27143478 | HOXA2 | N_Shore | TSS1500 |
| WG1513 | chr7:27142810-27143235 | HOXA2 | N_Shore | TSS1500 |
| GM02175 | chr7:27143046-27143252 | HOXA2 | N_Shore | TSS1500 |
| GM09256 | chr7:27143046-27143287 | HOXA2 | N_Shore | TSS1500 |
| GM02641 | chr7:27143046-27143403 | HOXA2 | N_Shore | TSS1500 |
| WG1978 | chr7:27143235-27144595 | HOXA2 | Island | TSS1500 |
| WG1514 | chr7:27143252-27143403 | HOXA2 | Island | TSS1500 |
| WG2415 | chr7:27143334-27143403 | HOXA2 | Island | TSS1500 |
| WG3465 | chr7:27143334-27143478 | HOXA2 | Island | TSS1500 |
| GM02704 | chr7:27143370-27143788 | HOXA2 | Island | TSS1500 |
| GM02640 | chr7:27143403-27143585 | HOXA2 | Island | TSS1500 |
| GM02706 | chr7:27145972-27146430 | HOXA3;HOXA3;HOXA3 | N_Shore | 3'UTR;3'UTR;3'UTR |
| WG1721 | chr7:27146237-27146445 | HOXA3;HOXA3;HOXA3 | Island | 3'UTR;3'UTR;3'UTR |
| WG1978 | chr7:27147513-27148204 | HOXA3;HOXA3;HOXA3 | N_Shore | 3'UTR;3'UTR;3'UTR |
| WG3466 | chr7:27150262-27150598 | HOXA3;HOXA3;HOXA3 | Island | 5'UTR;5'UTR;5'UTR |
| WG1722 | chr7:27153212-27153605 | HOXA3;HOXA3;HOXA3 | Island | 5'UTR;5'UTR;5'UTR |
| WG1978 | chr7:27153212-27153847 | HOXA3;HOXA3;HOXA3 | Island | 5'UTR;5'UTR;5'UTR |
| WG1977 | chr7:27153605-27153663 | HOXA3;HOXA3;HOXA3;HOXA | Island | 5'UTR;1stExon;5'UTR;5'UTR |
| WG3465 | chr7:27153605-27153847 | HOXA3;HOXA3;HOXA3;HOXA | Island | 5'UTR;1stExon;5'UTR;5'UTR |

| | | | | |
|---|---|---|---|---|
| GM09255 | chr7:27153605-27153944 | HOXA3;HOXA3;HOXA3;HOXA | Island | 5'UTR;1stExon;5'UTR;5'UTR |
| GM09256 | chr7:27153605-27154262 | HOXA3;HOXA3;HOXA3;HOXA | Island | 5'UTR;1stExon;5'UTR;5'UTR |
| WG1513 | chr7:27153636-27153663 | HOXA3;HOXA3;HOXA3 | Island | 5'UTR;5'UTR;TSS200 |
| WG1721 | chr7:27153636-27153847 | HOXA3;HOXA3;HOXA3 | Island | 5'UTR;5'UTR;TSS200 |
| GM02704 | chr7:27153655-27153847 | HOXA3;HOXA3;HOXA3 | N_Shore | 5'UTR;5'UTR;TSS200 |
| GM02175 | chr7:27153655-27153944 | HOXA3;HOXA3;HOXA3 | N_Shore | 5'UTR;5'UTR;TSS200 |
| WG1978 | chr7:27154264-27154387 | HOXA3;HOXA3;HOXA3 | N_Shore | 5'UTR;TSS1500;5'UTR |
| WG1978 | chr7:27154914-27155358 | HOXA3;HOXA3;HOXA3 | N_Shore | 5'UTR;TSS1500;5'UTR |
| GM02704 | chr7:27155002-27155358 | HOXA3;HOXA3;HOXA3 | Island | 5'UTR;TSS1500;5'UTR |
| WG1513 | chr7:27160520-27160960 | HOXA3;HOXA3 | N_Shore | 5'UTR;TSS1500 |
| WG3465 | chr7:27160586-27160960 | HOXA3;HOXA3 | N_Shore | 5'UTR;TSS1500 |
| GM02706 | chr7:27170819-27170880 | HOXA4 | S_Shore | TSS1500 |
| GM02706 | chr7:27170961-27171203 | HOXA4 | S_Shore | TSS1500 |
| GM00042 | chr7:27171154-27171391 | HOXA4 | S_Shore | TSS1500 |
| WG1513 | chr7:27182493-27183196 | HOXA5 | N_Shore | Body |
| WG1977 | chr7:27182493-27183436 | HOXA5 | N_Shore | Body |
| WG1978 | chr7:27182493-27183591 | HOXA5 | N_Shore | Body |
| GM02704 | chr7:27182493-27183694 | HOXA5 | N_Shore | Body |
| GM01096 | chr7:27182493-27183816 | HOXA5 | N_Shore | Body |
| GM02175 | chr7:27182637-27183436 | HOXA5 | Island | Body |
| GM09256 | chr7:27183133-27183436 | HOXA5 | Island | 1stExon |
| WG1721 | chr7:27183133-27183694 | HOXA5 | Island | 1stExon |
| WG1513 | chr7:27183274-27183375 | HOXA5;HOXA5 | Island | 5'UTR;1stExon |
| GM02173 | chr7:27183274-27183401 | HOXA5;HOXA5 | Island | 5'UTR;1stExon |
| WG2415 | chr7:27183274-27183436 | HOXA5;HOXA5 | Island | 5'UTR;1stExon |
| GM01376 | chr7:27183375-27183436 | HOXA5 | Island | TSS200 |
| GM09255 | chr7:27183643-27183694 | HOXA5 | Island | TSS1500 |
| WG1977 | chr7:27183643-27183861 | HOXA5 | Island | TSS1500 |
| WG1978 | chr7:27183686-27183806 | HOXA5 | Island | TSS1500 |
| GM02175 | chr7:27183694-27183861 | HOXA5 | Island | TSS1500 |
| GM09255 | chr7:27183794-27183861 | HOXA5 | Island | TSS1500 |
| GM01096 | chr7:27183973-27184030 | HOXA5 | Island | TSS1500 |
| GM02704 | chr7:27183990-27184059 | HOXA5 | Island | TSS1500 |
| GM02704 | chr7:27184176-27184264 | HOXA5 | Island | TSS1500 |
| GM02704 | chr7:27184316-27184737 | HOXA5 | Island | TSS1500 |
| WG1978 | chr7:27184521-27184712 | HOXA5 | Island | TSS1500 |
| WG1310 | chr7:27187269-27187374 | HOXA6 | Island | 1stExon |
| WG1722 | chr7:27187372-27187560 | HOXA6 | Island | TSS200 |
| GM02175 | chr7:27187372-27188770 | HOXA6 | Island | TSS200 |
| GM01375 | chr7:27187553-27187560 | HOXA6 | Island | TSS200 |
| GM02706 | chr7:27187553-27187670 | HOXA6 | Island | TSS200 |
| GM02640 | chr7:27187670-27187943 | HOXA6 | Island | TSS1500 |
| WG1722 | chr7:27187691-27188364 | HOXA6 | Island | TSS1500 |
| GM01095 | chr7:27187691-27188770 | HOXA6 | Island | TSS1500 |
| WG2121 | chr7:27187943-27188465 | HOXA6 | S_Shore | TSS1500 |
| WG3465 | chr7:27187943-27188770 | HOXA6 | S_Shore | TSS1500 |
| GM02640 | chr7:27188364-27188770 | HOXA6 | N_Shore | TSS1500 |

| | | | | |
|---|---|---|---|---|
| GM02175 | chr7:27196759-27196825 | HOXA7 | N_Shore | TSS1500 |
| GM02704 | chr7:27204005-27204728 | HOXA9 | Island | Body |
| GM02704 | chr7:27204803-27205114 | HOXA9 | Island | 1stExon |
| WG1978 | chr7:27209195-27209247 | MIR196B | Island | TSS200 |
| WG1978 | chr7:27209338-27209582 | MIR196B | Island | TSS200 |
| GM00940 | chr7:27220034-27220398 | HOXA10 | S_Shore | TSS200 |
| GM00940 | chr7:27222869-27223219 | HOXA11 | N_Shore | Body |
| WG1984 | chr7:27224568-27224795 | HOXA11AS;HOXA11 | Island | TSS1500;1stExon |
| WG2121 | chr7:27224795-27225078 | HOXA11AS;HOXA11;HOXA11 | N_Shore | TSS1500;1stExon;5'UTR |
| WG2844 | chr7:27225058-27225078 | HOXA11AS;HOXA11 | Island | Body;TSS1500 |
| WG1984 | chr7:27225058-27225123 | HOXA11AS;HOXA11 | Island | Body;TSS1500 |
| GM01095 | chr7:27225058-27225222 | HOXA11AS;HOXA11 | Island | Body;TSS1500 |
| WG1722 | chr7:27225070-27225123 | HOXA11AS;HOXA11 | Island | Body;TSS1500 |
| GM02456 | chr7:27225070-27225222 | HOXA11AS;HOXA11 | Island | Body;TSS1500 |
| GM00520 | chr7:27225070-27225299 | HOXA11AS;HOXA11 | Island | Body;TSS1500 |
| WG2275 | chr7:27225078-27225222 | HOXA11AS;HOXA11 | Island | Body;TSS1500 |
| WG2414 | chr7:27225078-27225865 | HOXA11AS;HOXA11 | Island | Body;TSS1500 |
| GM03073 | chr7:27225123-27225222 | HOXA11AS;HOXA11 | Island | Body;TSS1500 |
| WG2844 | chr7:27225127-27225222 | HOXA11AS;HOXA11 | Island | Body;TSS1500 |
| WG1722 | chr7:27225143-27225299 | HOXA11AS;HOXA11 | Island | Body;TSS1500 |
| GM02706 | chr7:27225143-27225342 | HOXA11AS;HOXA11 | Island | Body;TSS1500 |
| WG1514 | chr7:27225299-27225342 | HOXA11AS;HOXA11 | Island | Body;TSS1500 |
| GM01392 | chr7:27225299-27225476 | HOXA11AS;HOXA11 | Island | Body;TSS1500 |
| WG1310 | chr7:27225299-27225543 | HOXA11AS;HOXA11 | Island | Body;TSS1500 |
| GM00438 | chr7:27225319-27226148 | HOXA11AS;HOXA11 | Island | Body;TSS1500 |
| GM00520 | chr7:27225342-27225485 | HOXA11AS;HOXA11 | Island | Body;TSS1500 |
| GM09256 | chr7:27225448-27225485 | HOXA11AS;HOXA11 | Island | Body;TSS1500 |
| GM09255 | chr7:27225448-27225543 | HOXA11AS;HOXA11 | Island | Body;TSS1500 |
| GM00042 | chr7:27225448-27225733 | HOXA11AS;HOXA11 | Island | Body;TSS1500 |
| WG3465 | chr7:27225476-27225528 | HOXA11AS;HOXA11 | Island | Body;TSS1500 |
| GM02706 | chr7:27225485-27225543 | HOXA11AS;HOXA11 | Island | Body;TSS1500 |
| WG1977 | chr7:27225523-27225543 | HOXA11AS;HOXA11 | Island | Body;TSS1500 |
| GM01096 | chr7:27225523-27225733 | HOXA11AS;HOXA11 | Island | Body;TSS1500 |
| WG2415 | chr7:27225772-27225897 | HOXA11AS;HOXA11 | N_Shore | Body;TSS1500 |
| GM00439 | chr7:27225855-27225897 | HOXA11AS;HOXA11 | N_Shore | Body;TSS1500 |
| GM01393 | chr7:27225855-27226148 | HOXA11AS;HOXA11 | N_Shore | Body;TSS1500 |
| GM02456 | chr7:27225855-27226329 | HOXA11AS;HOXA11 | N_Shore | Body;TSS1500 |
| WG1310 | chr7:27225865-27226148 | HOXA11AS;HOXA11 | N_Shore | Body;TSS1500 |
| GM01392 | chr7:27225897-27226329 | HOXA11AS;HOXA11 | N_Shore | Body;TSS1500 |
| GM00438 | chr7:27227340-27227472 | HOXA11AS | N_Shore | Body |
| WG2121 | chr7:27227340-27228112 | HOXA11AS | N_Shore | Body |
| GM02149 | chr7:27240181-27240251 | HOXA13 | Island | TSS1500 |
| WG2415 | chr7:27281315-27281346 | EVX1 | N_Shore | TSS1500 |
| GM02149 | chr7:27281315-27281404 | EVX1 | N_Shore | TSS1500 |
| WG3466 | chr7:27281344-27281404 | EVX1 | N_Shore | TSS1500 |
| GM02149 | chr7:27281465-27281569 | EVX1 | N_Shore | TSS1500 |
| WG2414 | chr7:27281687-27281964 | EVX1 | N_Shore | TSS1500 |

| | | | | |
|---|---|---|---|---|
| GM00935 | chr7:27285433-27286075 | EVX1 | Island | Body |
| WG3465 | chr7:2728776-2728912 | AMZ1 | S_Shore | 5'UTR |
| GM00935 | chr7:2738875-2738961 | AMZ1 | N_Shore | 5'UTR |
| GM02706 | chr7:2800436-2801248 | GNA12 | N_Shelf | Body |
| WG3466 | chr7:2800701-2801248 | GNA12 | N_Shore | Body |
| GM00438 | chr7:2800791-2801248 | GNA12 | N_Shore | Body |
| WG2415 | chr7:2800976-2801509 | GNA12 | N_Shore | Body |
| WG3466 | chr7:2802374-2802560 | GNA12 | N_Shore | Body |
| WG2276 | chr7:2802374-2802697 | GNA12 | N_Shore | Body |
| WG2844 | chr7:2802374-2802976 | GNA12 | N_Shore | Body |
| WG2276 | chr7:2842593-2842746 | GNA12 | | Body |
| GM02149 | chr7:28452066-28452131 | CREB5;CREB5 | S_Shelf | 5'UTR;TSS200 |
| WG3465 | chr7:28452066-28452289 | CREB5;CREB5 | S_Shelf | 5'UTR;TSS200 |
| GM00289 | chr7:28452093-28452131 | CREB5;CREB5 | S_Shelf | 5'UTR;TSS200 |
| WG1513 | chr7:28474589-28474889 | CREB5;CREB5;CREB5 | | TSS1500;5'UTR;Body |
| WG2844 | chr7:2847517-2847575 | GNA12 | | Body |
| WG3466 | chr7:28838086-28838234 | CREB5;CREB5;CREB5;CREB5 | | Body;Body;Body;Body |
| GM02706 | chr7:28998540-28998860 | TRIL | Island | TSS1500 |
| WG3466 | chr7:28998577-28998860 | TRIL | S_Shore | TSS1500 |
| GM01653 | chr7:29186629-29187019 | CPVL;CPVL | | TSS1500;TSS1500 |
| GM00520 | chr7:29519297-29519409 | CHN2;CHN2 | | TSS200;Body |
| GM00290 | chr7:29519323-29519550 | CHN2;CHN2 | | TSS200;Body |
| WG1977 | chr7:29519409-29519550 | CHN2;CHN2 | | TSS200;Body |
| GM09256 | chr7:2953836-2953930 | CARD11 | | Body |
| WG1722 | chr7:29602763-29603253 | PRR15 | N_Shore | TSS1500 |
| GM09256 | chr7:29603370-29603407 | PRR15 | N_Shore | TSS200 |
| GM00022 | chr7:29605897-29606349 | PRR15 | Island | 5'UTR |
| GM01650 | chr7:29606082-29606349 | PRR15 | Island | Body |
| WG1282 | chr7:2968406-2968627 | CARD11 | Island | Body |
| WG3466 | chr7:29923120-29924165 | WIPF3 | N_Shore | Body |
| GM01096 | chr7:30029754-30029776 | SCRN1;SCRN1;SCRN1;SCRN | Island | TSS1500;TSS200;Body;TSS1500 |
| WG1977 | chr7:30492141-30492171 | NOD1 | | Body |
| GM02706 | chr7:3082006-3082233 | CARD11 | | 5'UTR |
| GM09255 | chr7:3083721-3083773 | CARD11 | | TSS200 |
| GM01096 | chr7:30951328-30951709 | AQP1 | | TSS200 |
| WG1722 | chr7:30951381-30951425 | AQP1 | | TSS200 |
| WG1983 | chr7:30951709-30951801 | AQP1 | | 1stExon |
| GM00289 | chr7:31726654-31726912 | C7orf16;C7orf16 | | TSS200;TSS200 |
| WG3466 | chr7:31726692-31726912 | C7orf16;C7orf16 | | TSS200;TSS200 |
| GM02175 | chr7:32111062-32111068 | PDE1C | S_Shore | TSS200 |
| GM00520 | chr7:34697855-34698188 | NPSR1;NPSR1 | | TSS200;TSS200 |
| GM00438 | chr7:34873912-34874533 | AAA1;NPSR1;NPSR1 | | Body;Body;Body |
| GM02706 | chr7:35294430-35295198 | TBX20;TBX20 | Island | TSS1500;TSS1500 |
| GM02641 | chr7:35294640-35295198 | TBX20;TBX20 | Island | TSS1500;TSS1500 |
| WG2194 | chr7:35294949-35295198 | TBX20;TBX20 | N_Shore | TSS1500;TSS1500 |
| GM09256 | chr7:36700744-36700939 | AOAH | | Body |
| WG1722 | chr7:36765142-36765609 | AOAH | | TSS1500 |

| | | | | |
|---|---|---|---|---|
| GM00940 | chr7:37959686-37959782 | EPDR1 | N_Shore | TSS1500 |
| WG1722 | chr7:37959686-37959966 | EPDR1 | N_Shore | TSS1500 |
| GM00438 | chr7:37959755-37959966 | EPDR1 | N_Shore | TSS1500 |
| WG1722 | chr7:38468944-38469273 | AMPH;AMPH | | Body;Body |
| GM09255 | chr7:39170497-39170763 | POU6F2;POU6F2 | | Body;Body |
| WG3465 | chr7:39170497-39171034 | POU6F2;POU6F2 | | Body;Body |
| GM01393 | chr7:3988772-3989097 | SDK1 | | Body |
| GM02706 | chr7:3988772-3989167 | SDK1 | | Body |
| WG1513 | chr7:3990479-3991031 | SDK1 | | Body |
| WG1977 | chr7:4049531-4050114 | SDK1 | | Body |
| GM00940 | chr7:4049718-4050114 | SDK1 | | Body |
| WG2415 | chr7:4065931-4066127 | SDK1 | | Body |
| WG1513 | chr7:4118609-4119142 | SDK1 | | Body |
| WG2120 | chr7:41739678-41739801 | LOC285954;LOC285954;INHBA | | Body;Body;Body |
| GM00438 | chr7:41742630-41743001 | INHBA;LOC285954;INHBA;L( | N_Shelf | 5'UTR;Body;1stExon;Body |
| WG1513 | chr7:4174482-4174655 | SDK1;SDK1 | | Body;Body |
| GM00289 | chr7:4201510-4201594 | SDK1;SDK1 | | Body;Body |
| WG1513 | chr7:4210949-4211212 | SDK1;SDK1 | | Body;Body |
| GM00520 | chr7:4211174-4211395 | SDK1;SDK1 | | Body;Body |
| WG1513 | chr7:4218743-4218990 | SDK1;SDK1 | | Body;Body |
| GM09256 | chr7:42267547-42267747 | GLI3 | Island | 5'UTR |
| WG3465 | chr7:42267547-42268109 | GLI3 | Island | 5'UTR |
| GM00439 | chr7:42267719-42268072 | GLI3 | Island | 5'UTR |
| WG1513 | chr7:42267719-42268109 | GLI3 | Island | 5'UTR |
| GM00438 | chr7:42278077-42278096 | GLI3 | S_Shore | TSS1500 |
| WG1977 | chr7:4266872-4267364 | SDK1;SDK1 | | Body;Body |
| WG2415 | chr7:4266872-4267412 | SDK1;SDK1 | | Body;Body |
| WG2414 | chr7:4267364-4267412 | SDK1;SDK1 | | Body;Body |
| WG1513 | chr7:4307392-4307490 | SDK1;SDK1 | S_Shelf | Body;3'UTR |
| WG1513 | chr7:4308166-4308372 | SDK1;SDK1 | S_Shelf | Body;3'UTR |
| WG1977 | chr7:43288635-43288889 | HECW1 | Island | Body |
| WG1983 | chr7:43288635-43289504 | HECW1 | Island | Body |
| GM01376 | chr7:44104584-44104935 | PGAM2 | N_Shore | Body |
| GM00940 | chr7:44261120-44261293 | CAMK2B;CAMK2B;CAMK2B;( | S_Shore | Body;Body;Body;Body;Body;Body;Bod |
| WG1513 | chr7:44279680-44279938 | CAMK2B;CAMK2B;CAMK2B;( | S_Shore | Body;Body;Body;Body;Body;Body;Bod |
| GM00439 | chr7:44349704-44349955 | CAMK2B;CAMK2B;CAMK2B;( | Island | Body;Body;Body;Body;Body;Body;Bod |
| GM09255 | chr7:44614305-44614475 | DDX56 | S_Shore | TSS200 |
| WG1310 | chr7:44801378-44801589 | ZMIZ2;ZMIZ2 | | Body;Body |
| GM00439 | chr7:45066895-45067319 | CCM2;CCM2;CCM2;CCM2;CCM2 | | Body;TSS1500;Body;Body;Body |
| GM02555 | chr7:4729417-4729920 | FOXK1 | | Body |
| WG2276 | chr7:4755010-4755415 | FOXK1 | N_Shelf | Body |
| WG2415 | chr7:4762742-4763182 | FOXK1 | N_Shore | Body |
| WG1722 | chr7:47694695-47694837 | C7orf65 | | TSS200 |
| WG2844 | chr7:4778839-4779342 | FOXK1 | | Body |
| GM00940 | chr7:48019441-48019713 | HUS1 | S_Shore | TSS200 |
| GM00439 | chr7:48068321-48068758 | SUNC1;SUNC1 | | Body;Body |
| WG2194 | chr7:48129814-48129992 | UPP1;UPP1 | Island | 5'UTR;5'UTR |

| | | | | |
|---|---|---|---|---|
| GM00289 | chr7:48494579-48494734 | ABCA13 | Island | Body |
| GM00289 | chr7:4856859-4857070 | RADIL | Island | Body |
| GM00520 | chr7:4859229-4859667 | RADIL | S_Shelf | Body |
| GM02175 | chr7:4877433-4877609 | RADIL | S_Shore | Body |
| WG2194 | chr7:4885521-4885643 | RADIL | | Body |
| WG1722 | chr7:4901337-4901628 | RADIL;PAPOLB | Island | Body;1stExon |
| GM00289 | chr7:4901449-4901628 | PAPOLB;RADIL;PAPOLB | Island | 5'UTR;Body;1stExon |
| GM02149 | chr7:4901449-4901750 | PAPOLB;RADIL;PAPOLB | Island | 5'UTR;Body;1stExon |
| WG3465 | chr7:4901449-4901798 | PAPOLB;RADIL;PAPOLB | Island | 5'UTR;Body;1stExon |
| WG3465 | chr7:50132653-50132903 | ZPBP;ZPBP | Island | Body;Body |
| WG1513 | chr7:50450266-50450374 | IKZF1 | | Body |
| GM00438 | chr7:50468399-50468722 | IKZF1 | Island | 3'UTR |
| GM09255 | chr7:50518350-50518573 | FIGNL1;FIGNL1 | Island | TSS1500;TSS1500 |
| WG1513 | chr7:50634170-50634611 | DDC | | TSS1500 |
| GM02706 | chr7:50861607-50861750 | GRB10 | Island | TSS1500 |
| WG1977 | chr7:5111488-5111576 | LOC389458 | N_Shore | TSS1500 |
| GM00439 | chr7:5111621-5111641 | LOC389458 | Island | TSS200 |
| WG1977 | chr7:5111641-5111916 | LOC389458 | Island | TSS200 |
| WG1513 | chr7:5112572-5112654 | LOC389458 | S_Shore | Body |
| WG1513 | chr7:53103214-53103576 | POM121L12 | N_Shore | TSS200 |
| GM00438 | chr7:53103287-53103312 | POM121L12 | Island | TSS200 |
| WG3465 | chr7:5336543-5336811 | SLC29A4;SLC29A4 | Island | Body;Body |
| WG2120 | chr7:537740-537828 | PDGFA;PDGFA | S_Shelf | 3'UTR;3'UTR |
| WG1977 | chr7:5518392-5518918 | FBXL18 | N_Shelf | 3'UTR |
| WG2276 | chr7:553412-553693 | PDGFA;PDGFA | N_Shelf | Body;Body |
| GM02456 | chr7:5535302-5535748 | FBXL18 | S_Shore | Body |
| GM00042 | chr7:5535692-5535740 | MIR589;FBXL18 | S_Shore | TSS200;Body |
| GM09256 | chr7:55432402-55432692 | LANCL2 | N_Shore | TSS1500 |
| WG1513 | chr7:56514965-56516129 | LOC650226 | N_Shore | Body |
| WG1722 | chr7:56515510-56515666 | LOC650226 | N_Shore | Body |
| GM02175 | chr7:56515510-56515846 | LOC650226 | N_Shore | Body |
| GM00520 | chr7:56515666-56516076 | LOC650226 | Island | Body |
| WG1977 | chr7:56515752-56516076 | LOC650226 | Island | Body |
| WG3466 | chr7:56515752-56516129 | LOC650226 | Island | Body |
| GM09256 | chr7:56516129-56516248 | LOC650226 | S_Shore | TSS200 |
| WG1513 | chr7:57247707-57247912 | LOC642006 | Island | Body |
| GM09256 | chr7:57247957-57248018 | LOC642006 | S_Shore | TSS200 |
| WG1722 | chr7:5862449-5862609 | ZNF815 | N_Shore | TSS1500 |
| WG1722 | chr7:5862740-5862996 | ZNF815 | N_Shore | TSS200 |
| WG2276 | chr7:625028-625136 | PRKAR1B;PRKAR1B;PRKAR1B;PRKAR1B; | | Body;Body;Body;Body;Body;Body |
| WG3465 | chr7:630473-630581 | PRKAR1B;PRKAR1B;PRKAR1B | N_Shore | Body;Body;Body;Body;Body;Body |
| WG2121 | chr7:630473-630823 | PRKAR1B;PRKAR1B;PRKAR1B | N_Shore | Body;Body;Body;Body;Body;Body |
| WG2415 | chr7:630473-630914 | PRKAR1B;PRKAR1B;PRKAR1B | N_Shore | Body;Body;Body;Body;Body;Body |
| GM00289 | chr7:630581-630914 | PRKAR1B;PRKAR1B;PRKAR1B | N_Shore | Body;Body;Body;Body;Body;Body |
| GM00439 | chr7:63505714-63506261 | ZNF727 | N_Shore | TSS200 |
| WG1721 | chr7:63505811-63506261 | ZNF727 | N_Shore | TSS200 |
| WG1513 | chr7:63667452-63667594 | ZNF735 | | TSS200 |

| | | | | |
|---|---|---|---|---|
| GM02149 | chr7:63687600-63688007 | ZNF679 | | TSS1500 |
| GM02706 | chr7:644219-644417 | PRKAR1B;PRKAR1B;PRKAR1E | Island | Body;Body;Body;Body;Body;Body |
| WG3466 | chr7:70096029-70096378 | AUTS2;AUTS2 | N_Shore | Body;Body |
| WG2276 | chr7:70096264-70096458 | AUTS2;AUTS2 | Island | Body;Body |
| GM00520 | chr7:73245457-73245752 | CLDN4;CLDN4 | Island | 1stExon;5'UTR |
| WG2844 | chr7:736792-737039 | PRKAR1B;PRKAR1B;PRKAR1B;PRKAR1B;I | | Body;Body;Body;Body;Body;Body |
| GM02175 | chr7:73894573-73895278 | GTF2IRD1;GTF2IRD1 | N_Shore | 5'UTR;5'UTR |
| WG3465 | chr7:76027001-76027101 | ZP3;ZP3;SRCRB4D | Island | 5'UTR;1stExon;Body |
| WG1721 | chr7:766100-766119 | PRKAR1B;PRKAR1B;PRKAR1E | Island | 5'UTR;5'UTR;5'UTR;TSS1500 |
| WG2415 | chr7:808081-809109 | HEATR2 | Island | Body |
| GM00289 | chr7:809048-809109 | HEATR2 | S_Shore | Body |
| WG2844 | chr7:817139-817576 | HEATR2 | S_Shelf | Body |
| WG1513 | chr7:82580938-82581607 | PCLO;PCLO | | Body;Body |
| GM00520 | chr7:8302235-8302271 | ICA1;ICA1;ICA1 | Island | TSS200;TSS1500;TSS1500 |
| GM09256 | chr7:8476098-8476450 | NXPH1 | S_Shore | Body |
| WG1977 | chr7:855365-855632 | UNC84A;UNC84A | Island | TSS1500;TSS1500 |
| WG1310 | chr7:89873797-89873891 | C7orf63;C7orf63 | N_Shore | TSS1500;TSS1500 |
| WG2276 | chr7:906726-906799 | UNC84A;UNC84A | S_Shore | Body;Body |
| GM02555 | chr7:907797-908060 | UNC84A;UNC84A | S_Shore | Body;Body |
| GM01095 | chr7:90896329-90896701 | FZD1;FZD1 | Island | 1stExon;3'UTR |
| GM00935 | chr7:91510314-91510405 | MTERF | Island | TSS1500 |
| GM02149 | chr7:91764339-91764605 | CYP51A1;CYP51A1 | Island | TSS1500;TSS1500 |
| WG3465 | chr7:917662-917805 | C7orf20 | Island | Body |
| WG1513 | chr7:92238086-92238207 | CDK6;CDK6 | | 3'UTR;3'UTR |
| GM01096 | chr7:92238156-92238248 | CDK6;CDK6 | | 3'UTR;3'UTR |
| GM01392 | chr7:93520445-93520527 | TFPI2 | S_Shore | TSS1500 |
| GM02704 | chr7:94285286-94285343 | SGCE;SGCE;PEG10;SGCE;PE | Island | Body;Body;TSS1500;Body;TSS1500 |
| GM01393 | chr7:94285288-94285352 | SGCE;SGCE;PEG10;SGCE;PE | Island | Body;Body;TSS1500;Body;TSS1500 |
| GM02704 | chr7:94285397-94285642 | SGCE;PEG10;SGCE;PEG10;S | Island | 1stExon;TSS1500;1stExon;TSS1500;1s |
| WG3465 | chr7:944607-944776 | ADAP1 | S_Shore | Body |
| WG2276 | chr7:94538165-94538233 | PPP1R9A;PPP1R9A;PPP1R9A | S_Shore | TSS1500;5'UTR;TSS1500;TSS1500;5'U |
| WG1513 | chr7:94953653-94953810 | PON1 | N_Shore | Body |
| WG2415 | chr7:94953956-94954144 | PON1 | Island | TSS200 |
| GM00935 | chr7:94953956-94954438 | PON1 | Island | TSS200 |
| WG3466 | chr7:94954144-94954438 | PON1 | S_Shore | TSS1500 |
| WG1722 | chr7:95025611-95025740 | PON3 | Island | 1stExon |
| GM02175 | chr7:95025855-95025924 | PON3 | Island | TSS200 |
| WG1984 | chr7:95025955-95026073 | PON3 | Island | TSS1500 |
| GM01095 | chr7:95026090-95026106 | PON3 | Island | TSS1500 |
| WG1721 | chr7:95026095-95026106 | PON3 | Island | TSS1500 |
| GM09255 | chr7:95026106-95026193 | PON3 | Island | TSS1500 |
| GM00439 | chr7:95026106-95026248 | PON3 | Island | TSS1500 |
| WG3465 | chr7:95026121-95026193 | PON3 | Island | TSS1500 |
| WG1513 | chr7:95026121-95026248 | PON3 | Island | TSS1500 |
| GM00935 | chr7:95026181-95026248 | PON3 | S_Shore | TSS1500 |
| WG2844 | chr7:95115163-95115354 | ASB4;ASB4 | | TSS200;TSS200 |
| GM01096 | chr7:95402431-95402733 | DYNC1I1;DYNC1I1;DYNC1I1 | Island | 5'UTR;5'UTR;5'UTR |

| | | | | |
|---|---|---|---|---|
| GM02149 | chr7:96631384-96631680 | DLX6AS | Island | Body |
| GM02175 | chr7:96631580-96631902 | DLX6AS | Island | Body |
| WG1513 | chr7:96631580-96632460 | DLX6AS | Island | Body |
| GM02704 | chr7:96649956-96650171 | DLX5 | N_Shore | 3'UTR |
| WG2415 | chr7:96649956-96650192 | DLX5 | N_Shore | 3'UTR |
| WG1722 | chr7:96650096-96650192 | DLX5 | N_Shore | Body |
| WG3465 | chr7:96650096-96650407 | DLX5 | N_Shore | Body |
| GM00439 | chr7:96650096-96650668 | DLX5 | N_Shore | Body |
| WG3466 | chr7:96650096-96651537 | DLX5 | N_Shore | Body |
| GM00289 | chr7:96650096-96651915 | DLX5 | N_Shore | Body |
| GM00520 | chr7:96650096-96652153 | DLX5 | N_Shore | Body |
| WG1513 | chr7:96650171-96650668 | DLX5 | N_Shore | Body |
| WG2276 | chr7:96650192-96650407 | DLX5 | N_Shore | Body |
| GM00935 | chr7:96650323-96651111 | DLX5 | Island | Body |
| WG2415 | chr7:96650407-96650668 | DLX5 | Island | Body |
| GM00439 | chr7:96651281-96651915 | DLX5 | Island | Body |
| WG1722 | chr7:96651537-96651915 | DLX5 | Island | Body |
| GM00935 | chr7:96651537-96651983 | DLX5 | Island | Body |
| WG3466 | chr7:96651915-96652115 | DLX5 | N_Shore | Body |
| GM01392 | chr7:96651983-96652153 | DLX5 | Island | Body |
| WG2276 | chr7:96651983-96652894 | DLX5 | Island | Body |
| WG1513 | chr7:96652115-96652153 | DLX5 | Island | Body |
| WG1310 | chr7:96654548-96654572 | DLX5 | S_Shore | TSS1500 |
| GM03075 | chr7:96654572-96654788 | DLX5 | S_Shore | TSS1500 |
| GM01650 | chr7:96654788-96654931 | DLX5 | S_Shore | TSS1500 |
| GM00935 | chr7:96654832-96655044 | DLX5 | S_Shore | TSS1500 |
| GM09256 | chr7:96654931-96655105 | DLX5 | S_Shore | TSS1500 |
| WG2193 | chr7:96655044-96655501 | DLX5 | S_Shore | TSS1500 |
| WG3465 | chr7:96655044-96655889 | DLX5 | S_Shore | TSS1500 |
| GM00520 | chr7:96655444-96655981 | DLX5 | S_Shore | TSS1500 |
| WG1978 | chr7:96655501-96655798 | DLX5 | S_Shore | TSS1500 |
| GM00935 | chr7:96655501-96655981 | DLX5 | S_Shore | TSS1500 |
| GM00940 | chr7:96655566-96655889 | DLX5 | S_Shore | TSS1500 |
| GM00042 | chr7:974745-975358 | ADAP1 | N_Shore | Body |
| WG3465 | chr7:97841001-97841334 | BHLHA15 | N_Shore | TSS1500 |
| WG3465 | chr7:985630-985784 | ADAP1 | Island | Body |
| WG1977 | chr7:99516603-99517460 | TRIM4;TRIM4 | Island | Body;Body |
| GM01650 | chr7:99517279-99517295 | TRIM4;TRIM4 | Island | TSS200;TSS200 |
| GM00940 | chr7:99680278-99680296 | ZNF3;ZNF3 | S_Shore | TSS1500;TSS1500 |
| GM02555 | chr7:99680278-99680324 | ZNF3;ZNF3 | S_Shore | TSS1500;TSS1500 |
| GM00439 | chr8:101117949-101118083 | RGS22 | Island | Body |
| WG1310 | chr8:101117958-101118377 | RGS22 | Island | Body |
| GM00940 | chr8:101225078-101225361 | SPAG1;SPAG1 | Island | Body;Body |
| GM00022 | chr8:101225078-101225902 | SPAG1;SPAG1 | Island | Body;Body |
| WG1977 | chr8:101661478-101661788 | SNX31 | N_Shore | Body |
| GM00439 | chr8:101661767-101661918 | SNX31;SNX31 | Island | 1stExon;5'UTR |
| GM09256 | chr8:101661976-101662053 | SNX31 | Island | TSS200 |

| | | | | |
|---|---|---|---|---|
| GM02173 | chr8:102503892-102503970 | GRHL2 | N_Shore | TSS1500 |
| WG1513 | chr8:102504859-102505556 | GRHL2;GRHL2 | N_Shore | 1stExon;5'UTR |
| GM00520 | chr8:103137173-103137274 | NCALD;NCALD;NCALD;NCAL | S_Shore | TSS1500;TSS1500;TSS1500;TSS1500; |
| WG2415 | chr8:104032819-104032876 | ATP6V1C1 | N_Shore | TSS1500 |
| WG2121 | chr8:104153592-104153637 | C8orf56;BAALC;BAALC | S_Shore | TSS200;Body;Body |
| WG2414 | chr8:104153627-104153643 | C8orf56;BAALC;BAALC | S_Shore | TSS200;Body;Body |
| GM02173 | chr8:104153753-104153829 | C8orf56;BAALC;BAALC | S_Shore | TSS200;Body;Body |
| GM02173 | chr8:104310606-104310770 | FZD6;FZD6;FZD6 | N_Shore | TSS1500;TSS1500;TSS200 |
| WG1513 | chr8:10469596-10470053 | RP1L1 | N_Shore | Body |
| GM00520 | chr8:10530053-10530148 | C8orf74 | | TSS200 |
| WG1309 | chr8:10586790-10586882 | SOX7 | Island | Body |
| WG3465 | chr8:10586959-10587055 | SOX7 | N_Shore | Body |
| GM09256 | chr8:106810858-106811143 | ZFPM2 | | Body |
| GM00290 | chr8:107782513-107782632 | ABRA | | TSS200 |
| GM02456 | chr8:107782538-107782632 | ABRA | | TSS200 |
| WG1514 | chr8:108510286-108510343 | ANGPT1 | | TSS200 |
| WG1978 | chr8:10907718-10907788 | XKR6 | | Body |
| WG1513 | chr8:10929105-10929599 | XKR6 | Island | Body |
| GM02706 | chr8:10982332-10982836 | XKR6 | | Body |
| GM00290 | chr8:110098844-110098923 | TRHR | | TSS1500 |
| GM02706 | chr8:110693195-110693289 | GOLSYN;GOLSYN;GOLSYN;GOLSYN;GO | | 5'UTR;5'UTR;1stExon;1stExon;5'UTR;5 |
| GM00438 | chr8:110988043-110988248 | KCNV1 | Island | TSS1500 |
| GM09256 | chr8:110988071-110988256 | KCNV1 | Island | TSS1500 |
| WG3465 | chr8:11141365-11141424 | MTMR9 | N_Shore | TSS1500 |
| WG2414 | chr8:11141415-11141680 | MTMR9 | N_Shore | TSS1500 |
| GM09255 | chr8:11567691-11567896 | GATA4 | S_Shore | Body |
| WG3465 | chr8:11616798-11617305 | GATA4 | | 3'UTR |
| WG2275 | chr8:116660311-116661063 | TRPS1 | N_Shore | 5'UTR |
| WG1722 | chr8:11831823-11832261 | DEFB136 | | Body |
| WG1978 | chr8:11839764-11839844 | DEFB135 | | TSS200 |
| GM01096 | chr8:120220050-120220094 | MAL2 | N_Shore | TSS1500 |
| WG1977 | chr8:120220073-120220131 | MAL2 | N_Shore | TSS1500 |
| GM02149 | chr8:12237626-12237900 | FAM66A | | Body |
| GM00520 | chr8:123875036-123875223 | ZHX2 | | 5'UTR |
| GM00042 | chr8:124219425-124219699 | FAM83A;FAM83A | Island | Body;Body |
| WG2415 | chr8:125462982-125463682 | TRMT12 | | TSS200 |
| WG1721 | chr8:125463007-125463310 | TRMT12 | | TSS200 |
| WG2844 | chr8:125463024-125463310 | TRMT12 | | TSS200 |
| GM02149 | chr8:133573933-133574073 | HPYR1 | | TSS1500 |
| WG1310 | chr8:135490694-135490821 | ZFAT;ZFAT | N_Shore | 3'UTR;3'UTR |
| GM09256 | chr8:135490787-135490958 | ZFAT;ZFAT | Island | Body;Body |
| WG1977 | chr8:139613712-139614020 | COL22A1 | | Body |
| WG3465 | chr8:140714489-140714610 | KCNK9 | N_Shore | Body |
| WG2844 | chr8:140937286-140937410 | TRAPPC9;TRAPPC9 | | Body;Body |
| WG2844 | chr8:140945782-140945843 | TRAPPC9;TRAPPC9 | | Body;Body |
| WG3466 | chr8:141057453-141057827 | TRAPPC9;TRAPPC9 | Island | Body;Body |
| WG2276 | chr8:141248582-141248667 | TRAPPC9;TRAPPC9 | | Body;Body |

| | | | | |
|---|---|---|---|---|
| GM00289 | chr8:141248582-141249094 | TRAPPC9;TRAPPC9 | | Body;Body |
| WG1977 | chr8:141359539-141359674 | TRAPPC9;TRAPPC9 | Island | Body;Body |
| WG1513 | chr8:141359539-141359786 | TRAPPC9;TRAPPC9 | Island | Body;Body |
| GM01392 | chr8:141600099-141600316 | EIF2C2;EIF2C2 | | Body;Body |
| WG2844 | chr8:142232952-142233436 | SLC45A4 | S_Shore | Body |
| WG2276 | chr8:142238691-142238770 | SLC45A4 | S_Shore | TSS200 |
| GM00520 | chr8:143324660-143324707 | TSNARE1 | | Body |
| WG3465 | chr8:143545390-143545574 | BAI1;BAI1 | N_Shore | 5'UTR;1stExon |
| WG2843 | chr8:143545478-143545789 | BAI1;BAI1 | Island | 5'UTR;1stExon |
| GM00520 | chr8:143545478-143546524 | BAI1;BAI1 | Island | 5'UTR;1stExon |
| WG1721 | chr8:143545574-143545949 | BAI1 | Island | 1stExon |
| WG3465 | chr8:143545940-143546893 | BAI1 | Island | 1stExon |
| GM00042 | chr8:143560651-143561039 | BAI1 | S_Shore | Body |
| GM00042 | chr8:143580823-143581070 | BAI1 | Island | Body |
| WG1977 | chr8:143580823-143581481 | BAI1 | Island | Body |
| WG2120 | chr8:143763326-143763565 | PSCA | | Body |
| WG1977 | chr8:143781644-143781985 | LY6K;LY6K;LY6K;LY6K;LY6K;L | Island | 1stExon;1stExon;5'UTR;1stExon;5'UTR |
| WG1513 | chr8:143823734-143823898 | SLURP1 | S_Shelf | Body |
| GM01096 | chr8:143851287-143851427 | LYNX1;LYNX1 | | TSS200;Body |
| WG1513 | chr8:143868844-143868917 | LY6D | | TSS1500 |
| GM02555 | chr8:144344397-144344793 | ZFP41 | | 3'UTR |
| WG1513 | chr8:144513697-144513907 | MAFA | Island | TSS1500 |
| WG1722 | chr8:144513898-144513980 | MAFA | Island | TSS1500 |
| WG2194 | chr8:144635260-144635496 | GSDMD | N_Shore | TSS1500 |
| WG2194 | chr8:144635610-144636462 | GSDMD;GSDMD | Island | 5'UTR;1stExon |
| WG1283 | chr8:144654954-144655030 | C8orf73 | S_Shore | TSS200 |
| WG1513 | chr8:144659831-144660157 | NAPRT1 | Island | Body |
| WG2121 | chr8:144660157-144660631 | NAPRT1 | Island | Body |
| WG3466 | chr8:144660590-144660631 | NAPRT1 | Island | TSS200 |
| GM09256 | chr8:144790187-144790395 | LOC100130274 | Island | 1stExon |
| WG1513 | chr8:1448750-1449108 | DLGAP2 | | TSS1500 |
| GM00438 | chr8:1449108-1449655 | DLGAP2 | | TSS1500 |
| WG1513 | chr8:1449401-1450031 | DLGAP2 | | TSS200 |
| GM00439 | chr8:144948315-144948781 | EPPK1 | S_Shore | TSS1500 |
| GM00289 | chr8:1449634-1450031 | DLGAP2 | | 5'UTR |
| WG3466 | chr8:145018816-145019041 | PLEC1;PLEC1;PLEC1;PLEC1; | Island | Body;Body;1stExon;Body;TSS1500;Bo |
| GM01653 | chr8:145019032-145019067 | PLEC1;PLEC1;PLEC1;PLEC1; | Island | Body;Body;Body;TSS1500;Body;TSS20 |
| GM02706 | chr8:145019032-145019116 | PLEC1;PLEC1;PLEC1;PLEC1; | Island | Body;Body;Body;TSS1500;Body;TSS20 |
| GM02706 | chr8:145019475-145019639 | PLEC1;PLEC1;PLEC1;MIR661 | S_Shore | Body;Body;TSS1500;TSS200;Body;TSS |
| GM02456 | chr8:145019563-145019639 | PLEC1;PLEC1;PLEC1;MIR661 | S_Shore | Body;Body;TSS1500;TSS200;Body;TSS |
| WG2415 | chr8:145028093-145028170 | PLEC1;PLEC1;PLEC1 | Island | Body;Body;TSS200 |
| WG1722 | chr8:145028170-145028230 | PLEC1;PLEC1;PLEC1 | Island | Body;Body;TSS200 |
| WG2415 | chr8:145048235-145048363 | PLEC1;PLEC1 | Island | Body;TSS1500 |
| GM02706 | chr8:145049280-145049472 | PLEC1 | Island | Body |
| GM00439 | chr8:145085954-145086427 | SPATC1;SPATC1 | | TSS1500;TSS1500 |
| GM00935 | chr8:145086243-145086427 | SPATC1;SPATC1 | | TSS1500;TSS1500 |
| GM00935 | chr8:145162483-145162592 | MAF1;KIAA1875 | N_Shore | 3'UTR;TSS200 |

| | | | | |
|---|---|---|---|---|
| GM02175 | chr8:145162529-145163034 | KIAA1875 | N_Shore | TSS200 |
| WG2415 | chr8:145162529-145163136 | KIAA1875 | N_Shore | TSS200 |
| WG1721 | chr8:145162592-145163034 | KIAA1875 | N_Shore | TSS200 |
| GM02555 | chr8:145162592-145163102 | KIAA1875 | N_Shore | TSS200 |
| GM02317 | chr8:145162905-145163136 | KIAA1875 | N_Shore | Body |
| WG1513 | chr8:145163034-145163136 | KIAA1875 | N_Shore | Body |
| GM09256 | chr8:145577701-145578141 | C8ORFK29 | Island | Body |
| WG1984 | chr8:145638881-145639181 | SLC39A4;SLC39A4 | Island | Body;Body |
| WG2121 | chr8:145654565-145654635 | VPS28;NFKBIL2;VPS28 | Island | TSS1500;Body;TSS1500 |
| GM00520 | chr8:145654565-145654780 | VPS28;NFKBIL2;VPS28 | Island | TSS1500;Body;TSS1500 |
| WG3465 | chr8:145654565-145654854 | VPS28;NFKBIL2;VPS28 | Island | TSS1500;Body;TSS1500 |
| GM01376 | chr8:145728507-145728543 | GPT | N_Shore | TSS1500 |
| GM02706 | chr8:145728543-145728630 | GPT | N_Shore | TSS1500 |
| GM02706 | chr8:145729135-145729196 | GPT | N_Shore | TSS1500 |
| GM02706 | chr8:145747659-145748059 | LRRC14 | Island | 3'UTR |
| WG1513 | chr8:145749887-145750363 | LRRC24;LRRC14 | Island | Body;3'UTR |
| WG1309 | chr8:145755756-145755974 | MGC70857;KIAA1688 | Island | TSS1500;3'UTR |
| WG2120 | chr8:1460569-1460718 | DLGAP2 | | 5'UTR |
| GM00289 | chr8:1495025-1495185 | DLGAP2 | N_Shore | 5'UTR |
| GM00438 | chr8:1495169-1495289 | DLGAP2 | Island | 5'UTR |
| GM09256 | chr8:1496347-1496732 | DLGAP2 | N_Shore | 5'UTR |
| WG2415 | chr8:1500367-1500789 | DLGAP2 | N_Shore | Body |
| GM01096 | chr8:15093686-15094114 | SGCZ | N_Shore | Body |
| WG1513 | chr8:1570715-1571105 | DLGAP2 | | Body |
| GM02149 | chr8:1579039-1579180 | DLGAP2 | Island | Body |
| GM09256 | chr8:1616209-1616422 | DLGAP2 | N_Shore | Body |
| WG1513 | chr8:1617616-1617868 | DLGAP2 | Island | Body |
| WG1513 | chr8:1651101-1651197 | DLGAP2 | S_Shore | 3'UTR |
| GM00438 | chr8:16859636-16859833 | FGF20;FGF20 | S_Shore | 1stExon;5'UTR |
| WG1978 | chr8:16859671-16859924 | FGF20;FGF20 | S_Shore | 1stExon;5'UTR |
| GM09256 | chr8:16859754-16860088 | FGF20 | S_Shore | TSS200 |
| WG1513 | chr8:16859833-16860121 | FGF20 | S_Shore | TSS200 |
| WG2415 | chr8:16859924-16860121 | FGF20 | S_Shore | TSS1500 |
| GM02706 | chr8:17270834-17271093 | MTMR7;MTMR7 | Island | 5'UTR;1stExon |
| WG1978 | chr8:17271067-17271105 | MTMR7 | Island | TSS200 |
| GM00935 | chr8:1765066-1765477 | MIR596 | Island | TSS1500 |
| WG2844 | chr8:1765217-1765358 | MIR596 | S_Shore | TSS200 |
| WG2276 | chr8:1765217-1765421 | MIR596 | S_Shore | TSS200 |
| GM00439 | chr8:1765296-1765387 | MIR596 | S_Shore | TSS200 |
| WG2843 | chr8:1765312-1765358 | MIR596 | S_Shore | TSS200 |
| WG1721 | chr8:1780242-1780401 | ARHGEF10 | S_Shore | 5'UTR |
| WG2844 | chr8:1786207-1786322 | ARHGEF10 | | 5'UTR |
| WG1513 | chr8:18027367-18028071 | NAT1 | | TSS1500 |
| GM00935 | chr8:1809425-1809936 | ARHGEF10 | | Body |
| WG2275 | chr8:1847999-1848143 | ARHGEF10 | | Body |
| WG3466 | chr8:1847999-1848407 | ARHGEF10 | | Body |
| GM02706 | chr8:1847999-1848614 | ARHGEF10 | | Body |

| | | | | |
|---|---|---|---|---|
| WG2120 | chr8:1848040-1848407 | ARHGEF10 | | Body |
| GM02456 | chr8:1848040-1848614 | ARHGEF10 | | Body |
| WG3466 | chr8:18541497-18541988 | PSD3;PSD3 | | Body;Body |
| WG2276 | chr8:1894604-1895010 | ARHGEF10 | S_Shore | Body |
| GM02456 | chr8:1900089-1900400 | ARHGEF10 | | Body |
| WG1977 | chr8:19539991-19540272 | CSGALNACT1;CSGALNACT1;CSGALNAC | | Body;5'UTR;1stExon |
| GM00438 | chr8:19540210-19540396 | CSGALNACT1;CSGALNACT1 | | TSS200;Body |
| WG1721 | chr8:1954777-1955117 | KBTBD11 | N_Shelf | 3'UTR |
| GM02456 | chr8:1954777-1955196 | KBTBD11 | N_Shelf | 3'UTR |
| GM02555 | chr8:1955069-1955196 | KBTBD11 | N_Shore | 3'UTR |
| GM09256 | chr8:20112763-20112890 | LZTS1;LZTS1 | | 5'UTR;1stExon |
| GM00520 | chr8:20112763-20112950 | LZTS1;LZTS1 | | 5'UTR;1stExon |
| GM02706 | chr8:20112950-20113038 | LZTS1 | | TSS200 |
| WG1513 | chr8:2019935-2020566 | MYOM2 | | Body |
| WG1282 | chr8:2075209-2075820 | MYOM2 | N_Shore | Body |
| WG2121 | chr8:2075469-2075820 | MYOM2 | Island | Body |
| GM01392 | chr8:21771059-21771258 | DOK2 | S_Shore | 1stExon |
| WG1978 | chr8:21905462-21905756 | FGF17 | Island | Body |
| GM02555 | chr8:21961713-21961788 | FAM160B2 | | 3'UTR |
| WG2276 | chr8:22079107-22079330 | PHYHIP;PHYHIP | Island | Body;Body |
| WG1722 | chr8:22084861-22085227 | PHYHIP;PHYHIP | | Body;Body |
| GM00289 | chr8:22132714-22132932 | PIWIL2;PIWIL2 | N_Shore | TSS200;TSS1500 |
| GM03073 | chr8:22132874-22132932 | PIWIL2;PIWIL2;PIWIL2 | Island | 5'UTR;1stExon;TSS1500 |
| WG1513 | chr8:22132874-22132959 | PIWIL2;PIWIL2;PIWIL2 | Island | 5'UTR;1stExon;TSS1500 |
| WG3466 | chr8:22132874-22132992 | PIWIL2;PIWIL2;PIWIL2 | Island | 5'UTR;1stExon;TSS1500 |
| WG1657 | chr8:22132874-22133076 | PIWIL2;PIWIL2;PIWIL2 | Island | 5'UTR;1stExon;TSS1500 |
| GM09255 | chr8:22423014-22423026 | SORBS3;SORBS3 | Island | TSS200;Body |
| WG2844 | chr8:22548006-22548546 | EGR3 | Island | Body |
| WG1978 | chr8:22548400-22548828 | EGR3 | Island | Body |
| GM02173 | chr8:22548400-22548952 | EGR3 | Island | Body |
| GM00520 | chr8:22642322-22642638 | PEBP4 | | Body |
| GM01096 | chr8:22785712-22786374 | PEBP4 | | TSS1500 |
| WG1977 | chr8:22960257-22960427 | TNFRSF10C | N_Shore | TSS200 |
| GM00289 | chr8:23082810-23082876 | TNFRSF10A | Island | TSS200 |
| GM02456 | chr8:23162162-23162383 | LOXL2 | | Body |
| WG3466 | chr8:25902551-25902594 | EBF2 | Island | TSS200 |
| GM02706 | chr8:27115867-27116213 | STMN4;STMN4 | | 1stExon;5'UTR |
| GM00290 | chr8:27338183-27338248 | CHRNA2 | | TSS1500 |
| WG3466 | chr8:27468684-27469001 | CLU;CLU;CLU;CLU | N_Shelf | 1stExon;Body;Body;5'UTR |
| WG1977 | chr8:28174350-28174814 | PNOC | | TSS1500 |
| WG3465 | chr8:28560467-28560637 | EXTL3 | S_Shore | 5'UTR |
| WG3466 | chr8:28560467-28560658 | EXTL3 | S_Shore | 5'UTR |
| WG1513 | chr8:28560467-28560723 | EXTL3 | S_Shore | 5'UTR |
| WG1978 | chr8:28560486-28560658 | EXTL3 | S_Shore | 5'UTR |
| GM00520 | chr8:28560637-28560723 | EXTL3 | S_Shore | 5'UTR |
| WG1513 | chr8:3429382-3429480 | CSMD1 | | Body |
| GM00042 | chr8:37605359-37605783 | LOC728024;ERLIN2 | | Body;Body |

| | | | | |
|---|---|---|---|---|
| GM09255 | chr8:37605359-37605978 | LOC728024;ERLIN2 | | Body;Body |
| GM02706 | chr8:37605359-37606579 | LOC728024;ERLIN2 | | Body;Body |
| WG2414 | chr8:37605717-37605978 | LOC728024;ERLIN2 | | TSS200;Body |
| WG3465 | chr8:37797632-37797731 | GOT1L1;GOT1L1 | | 5'UTR;1stExon |
| GM09256 | chr8:39171620-39171866 | ADAM5P | | TSS1500 |
| GM00940 | chr8:39172020-39172120 | ADAM5P | | TSS200 |
| WG2415 | chr8:39172097-39172120 | ADAM5P | | TSS200 |
| GM09256 | chr8:39441968-39442089 | ADAM18 | | TSS200 |
| WG1513 | chr8:39441975-39442080 | ADAM18 | | TSS200 |
| WG1513 | chr8:39695862-39696018 | ADAM2 | | TSS200 |
| GM00289 | chr8:39695907-39696062 | ADAM2 | | TSS200 |
| GM09256 | chr8:39835944-39836064 | IDO2 | | Body |
| WG3465 | chr8:40010443-40010823 | C8orf4 | | TSS1500 |
| GM00438 | chr8:41168186-41168264 | SFRP1 | S_Shore | TSS1500 |
| GM00042 | chr8:41504890-41505042 | NKX6-3 | S_Shore | TSS200 |
| WG1513 | chr8:41522857-41522951 | ANK1;ANK1;ANK1;ANK1;ANK1;ANK1;ANI | | TSS200;TSS200;Body;Body;Body;Body |
| WG3465 | chr8:41583136-41583523 | ANK1;ANK1;ANK1;ANK1;ANK | Island | Body;Body;Body;Body;Body |
| GM09255 | chr8:41583321-41583523 | ANK1;ANK1;ANK1;ANK1;ANK | Island | Body;Body;Body;Body;Body |
| WG2415 | chr8:41655078-41655789 | ANK1;ANK1;ANK1;ANK1;ANK | Island | 1stExon;1stExon;5'UTR;5'UTR;5'UTR;1 |
| GM01095 | chr8:41655154-41655826 | ANK1;ANK1;ANK1;ANK1;ANK | Island | TSS200;TSS200;TSS200;TSS200;Body |
| GM02641 | chr8:41655154-41655894 | ANK1;ANK1;ANK1;ANK1;ANK | Island | TSS200;TSS200;TSS200;TSS200;Body |
| WG1978 | chr8:41655154-41655983 | ANK1;ANK1;ANK1;ANK1;ANK | Island | TSS200;TSS200;TSS200;TSS200;Body |
| WG1513 | chr8:41655673-41655894 | ANK1;ANK1;ANK1;ANK1;ANK | Island | TSS1500;TSS1500;TSS1500;TSS1500; |
| GM03075 | chr8:42234614-42234732 | DKK4;DKK4 | | 5'UTR;1stExon |
| WG2415 | chr8:42234669-42234732 | DKK4;DKK4 | | 5'UTR;1stExon |
| WG3465 | chr8:42356637-42356871 | SLC20A2 | | 5'UTR |
| WG1977 | chr8:42623718-42623786 | CHRNA6 | | TSS200 |
| GM01096 | chr8:42623730-42623786 | CHRNA6 | | TSS200 |
| WG1513 | chr8:43147211-43147554 | POTEA;POTEA | | TSS1500;TSS1500 |
| GM09255 | chr8:49647579-49647717 | EFCAB1;EFCAB1;EFCAB1 | N_Shore | Body;Body;Body |
| WG2415 | chr8:52322134-52322169 | PXDNL | Island | Body |
| WG1513 | chr8:53300817-53301179 | ST18 | | 5'UTR |
| GM02706 | chr8:53300830-53301169 | ST18 | | 5'UTR |
| WG1722 | chr8:53321682-53321751 | ST18 | | 5'UTR |
| WG1513 | chr8:56368017-56368202 | SBF1P1;XKR4 | | TSS200;Body |
| WG1513 | chr8:56436393-56436589 | XKR4 | | Body |
| WG2276 | chr8:56852148-56852290 | LYN;LYN | | 5'UTR;5'UTR |
| WG2844 | chr8:57233269-57233373 | SDR16C5 | S_Shore | TSS200 |
| WG2120 | chr8:57359956-57359992 | PENK;PENK | Island | TSS1500;TSS1500 |
| GM02706 | chr8:58192502-58192883 | C8orf71 | Island | Body |
| WG2121 | chr8:59058254-59058660 | FAM110B | Island | 5'UTR |
| WG3466 | chr8:59058585-59058660 | FAM110B | Island | 5'UTR |
| WG2415 | chr8:59058585-59059119 | FAM110B | Island | 5'UTR |
| GM09256 | chr8:61626185-61626646 | CHD7 | N_Shore | 5'UTR |
| WG2415 | chr8:61626185-61626832 | CHD7 | N_Shore | 5'UTR |
| WG2276 | chr8:6400815-6401045 | ANGPT2;ANGPT2;ANGPT2;MCPH1 | | Body;Body;Body;Body |
| GM02175 | chr8:6407273-6407812 | ANGPT2;ANGPT2;ANGPT2;MCPH1 | | Body;Body;Body;Body |

| | | | | |
|---|---|---|---|---|
| WG2415 | chr8:6420242-6420726 | ANGPT2;ANGPT2;ANGPT2;M( | N_Shelf | 1stExon;1stExon;1stExon;Body |
| GM02706 | chr8:6420242-6420770 | ANGPT2;ANGPT2;ANGPT2;M( | N_Shelf | 1stExon;1stExon;1stExon;Body |
| WG1513 | chr8:6420242-6420858 | ANGPT2;ANGPT2;ANGPT2;M( | N_Shelf | 1stExon;1stExon;1stExon;Body |
| GM00935 | chr8:6420770-6421063 | ANGPT2;ANGPT2;ANGPT2;AN | N_Shelf | 5'UTR;1stExon;5'UTR;1stExon;5'UTR;1 |
| GM00042 | chr8:644521-644663 | ERICH1 | | Body |
| WG3466 | chr8:6456182-6456465 | MCPH1 | | Body |
| WG2276 | chr8:6484293-6484687 | MCPH1 | | Body |
| WG3466 | chr8:65489881-65489937 | LOC401463 | N_Shore | TSS200 |
| GM00439 | chr8:65490613-65491152 | LOC401463 | N_Shore | TSS1500 |
| GM01392 | chr8:65490710-65491152 | LOC401463 | N_Shore | TSS1500 |
| GM02706 | chr8:65491906-65492280 | BHLHE22 | N_Shore | TSS1500 |
| GM09256 | chr8:6693367-6693540 | XKR5 | S_Shore | TSS1500 |
| WG3465 | chr8:67039041-67039461 | TRIM55;TRIM55;TRIM55;TRIM55 | | TSS1500;TSS1500;TSS1500;TSS1500 |
| GM02175 | chr8:6735472-6735559 | DEFB1;DEFB1 | | 5'UTR;1stExon |
| WG2415 | chr8:67405405-67405742 | C8orf46 | | TSS200 |
| GM00520 | chr8:67940776-67940913 | LRRC67;LRRC67 | Island | 1stExon;5'UTR |
| GM00520 | chr8:67940950-67941077 | LRRC67 | Island | TSS200 |
| GM02149 | chr8:6827800-6828103 | DEFA10P | | TSS1500 |
| WG1310 | chr8:72274548-72274615 | EYA1 | | TSS200 |
| WG1977 | chr8:72754469-72755162 | MSC | Island | 3'UTR |
| GM00935 | chr8:72757456-72758095 | MSC | S_Shore | TSS1500 |
| WG1513 | chr8:73848484-73849114 | KCNB2 | | Body |
| GM02706 | chr8:77585222-77585381 | LOC100192378 | | Body |
| WG3465 | chr8:77596069-77596382 | ZFHX4;LOC100192378 | S_Shore | 5'UTR;TSS1500 |
| WG3465 | chr8:80678770-80679314 | HEY1;HEY1 | N_Shore | Body;Body |
| WG1514 | chr8:81599484-81599569 | ZNF704 | | Body |
| WG2844 | chr8:8182454-8182747 | PRAGMIN | | Body |
| WG2275 | chr8:82192601-82192606 | FABP5 | Island | TSS200 |
| GM00438 | chr8:82644747-82644943 | CHMP4C;CHMP4C | Island | 1stExon;5'UTR |
| WG1977 | chr8:85097157-85097429 | RALYL;RALYL;RALYL;RALYL;R | Island | 5'UTR;5'UTR;1stExon;5'UTR;5'UTR |
| WG1513 | chr8:86290219-86290689 | CA1;CA1;CA1;CA1 | | 5'UTR;5'UTR;5'UTR;5'UTR |
| GM09256 | chr8:88884985-88885186 | DCAF4L2;DCAF4L2 | N_Shore | 3'UTR;1stExon |
| GM00941 | chr8:93107298-93107627 | RUNX1T1;RUNX1T1 | | 5'UTR;1stExon |
| WG2414 | chr8:93107298-93108262 | RUNX1T1;RUNX1T1 | | 5'UTR;1stExon |
| GM00042 | chr8:95653874-95654019 | ESRP1;ESRP1;ESRP1;ESRP1; | N_Shore | Body;Body;Body;Body;Body |
| GM02149 | chr8:95962084-95962367 | TP53INP1;TP53INP1 | Island | TSS1500;TSS1500 |
| WG1310 | chr8:95962084-95962371 | TP53INP1;TP53INP1 | Island | TSS1500;TSS1500 |
| GM01376 | chr8:95962352-95962383 | TP53INP1;TP53INP1 | Island | TSS1500;TSS1500 |
| GM02555 | chr8:95962367-95962383 | TP53INP1;TP53INP1 | Island | TSS1500;TSS1500 |
| WG3465 | chr8:95962371-95962463 | TP53INP1;TP53INP1 | Island | TSS1500;TSS1500 |
| WG1977 | chr8:96036568-96036905 | C8orf38 | N_Shore | TSS1500 |
| WG1977 | chr8:97157453-97158052 | GDF6 | Island | Body |
| GM02704 | chr8:97157856-97158015 | GDF6 | Island | Body |
| WG2415 | chr8:97165770-97166291 | GDF6 | N_Shelf | Body |
| GM01376 | chr8:97169621-97169767 | GDF6 | N_Shore | Body |
| GM00016 | chr8:97169920-97170364 | GDF6 | Island | Body |
| GM02175 | chr8:99076526-99076605 | C8orf47;C8orf47 | N_Shore | TSS1500;TSS1500 |

| | | | | |
|---|---|---|---|---|
| WG1978 | chr8:9952636-9953022 | MSRA;MSRA;MSRA | | Body;TSS1500;Body |
| WG2121 | chr8:9952926-9953022 | MSRA;MSRA;MSRA | | Body;TSS200;Body |
| GM00438 | chr8:99959430-99959602 | OSR2;OSR2 | N_Shore | 5'UTR;5'UTR |
| WG2276 | chr8:99959430-99959723 | OSR2;OSR2 | N_Shore | 5'UTR;5'UTR |
| WG2844 | chr8:99961545-99962118 | OSR2;OSR2 | S_Shore | Body;Body |
| WG2120 | chr8:99961768-99962013 | OSR2;OSR2 | S_Shore | Body;Body |
| WG1722 | chr9:100747473-100747767 | ANP32B | Island | Body |
| GM00439 | chr9:101568975-101569334 | GALNT12 | N_Shore | TSS1500 |
| WG1978 | chr9:120466493-120466837 | TLR4;TLR4;TLR4;TLR4 | | Body;Body;1stExon;5'UTR |
| WG1984 | chr9:124461998-124462182 | DAB2IP | Island | Body |
| WG2276 | chr9:124979184-124979356 | LHX6;LHX6 | N_Shore | Body;Body |
| GM01393 | chr9:124982018-124982413 | LHX6;LHX6 | Island | Body;Body |
| WG2275 | chr9:126135610-126135998 | CRB2 | Island | Body |
| GM00941 | chr9:130516695-130517095 | SH2D3C;SH2D3C;SH2D3C;S | Island | Body;5'UTR;5'UTR;Body;Body;Body |
| GM03075 | chr9:130955135-130955436 | CIZ1;CIZ1;CIZ1;CIZ1;CIZ1 | N_Shore | TSS1500;5'UTR;TSS1500;5'UTR;TSS15 |
| WG1977 | chr9:135904966-135905092 | GTF3C5;GTF3C5 | N_Shore | TSS1500;TSS1500 |
| GM02175 | chr9:136522662-136522927 | DBH | | Body |
| GM02555 | chr9:136566991-136567339 | SARDH;SARDH | N_Shore | Body;Body |
| GM02175 | chr9:137772544-137772580 | FCN2;FCN2 | | TSS200;TSS200 |
| GM00289 | chr9:139091875-139092501 | LHX3;LHX3 | Island | Body;Body |
| GM00290 | chr9:139094185-139094581 | LHX3;LHX3 | Island | Body;Body |
| GM02456 | chr9:139236663-139236777 | GPSM1 | N_Shore | Body |
| WG2843 | chr9:139237461-139237889 | GPSM1 | Island | Body |
| WG1722 | chr9:139715701-139716215 | C9orf86;C9orf86 | Island | Body;Body |
| WG1722 | chr9:140115806-140116137 | RNF208 | N_Shore | TSS200 |
| GM02704 | chr9:140171765-140172277 | C9orf167 | N_Shore | TSS1500 |
| GM00290 | chr9:140172089-140172631 | C9orf167 | N_Shore | TSS200 |
| WG1310 | chr9:140401382-140401455 | PNPLA7;PNPLA7 | Island | Body;Body |
| WG1978 | chr9:140401382-140401576 | PNPLA7;PNPLA7 | Island | Body;Body |
| WG1983 | chr9:140401437-140401576 | PNPLA7;PNPLA7 | Island | Body;Body |
| GM02706 | chr9:27109111-27109492 | TEK | | TSS200 |
| GM02706 | chr9:34370781-34370835 | KIAA1161 | Island | 3'UTR |
| GM02706 | chr9:90113998-90114156 | DAPK1 | S_Shore | Body |
| GM02175 | chr9:92220388-92220771 | GADD45G | Island | Body |
| WG1977 | chr9:92220388-92221282 | GADD45G | Island | Body |
| WG3465 | chr9:92220617-92221133 | GADD45G | Island | Body |
| WG2415 | chr9:92220617-92221282 | GADD45G | Island | Body |
| GM00042 | chr9:92220771-92221282 | GADD45G | Island | Body |
| GM00290 | chr9:96715687-96716091 | BARX1 | Island | Body |
| WG2276 | chr9:96715843-96716209 | BARX1 | Island | Body |
| WG2414 | chr9:975825-975997 | DMRT3 | N_Shore | TSS1500 |

| Supplementary Table 18: Methylation levels and expression levels for 16 genes showing significant correlation in untransformed skin fibroblasts. | | | | | | | |
|---|---|---|---|---|---|---|---|
| Gene | Coordinate | Position | CGI | $R^2$ | R | p value | Correlation type |
| AHRR | chr5:323320-323969 | Gene body | No | 0.3275 | 0.5723 | 1.00E-06 | Negative |
| C21orf56 | chr21:47604052-47604654 | Promoter | No | 0.4371 | 0.6611 | < 1.00E-06 | Negative |
| FMNL1 | chr17:43319137-43319382 | Gene body | Yes | 0.2190 | 0.4679 | 1.26E-04 | Positive |
| HLA-C | chr6:31237405-31237722 | Gene body | No | 0.3366 | 0.5802 | 1.00E-06 | Positive |
| HOXB7 | chr17:46685327-46685448 | Gene body | Yes | 0.6068 | 0.7790 | < 1.00E-06 | Positive |
| HOXC4 | chr12:54423428-54425418 | Gene body | Yes | 0.7019 | 0.8378 | < 1.00E-06 | Negative |
| HOXC5 | chr12:54423428-54425418 | Upstream | Yes | 0.595984 | 0.7720 | < 1.00E-06 | Negative |
| HOXC6 | chr12:54423428-54425418 | Promoter | Yes | 0.706776 | 0.8407 | < 1.00E-06 | Negative |
| HOXD8 | chr2:176993017-176993115 | Promoter | Yes | 0.081968 | 0.2863 | 2.41E-02 | Negative |
| HOXB5 | chr17:46669485-46669644 | Gene body | Yes | 0.753771 | 0.8682 | < 1.00E-06 | Negative |
| MAB21L1 | chr13:36050901-36051073 | Promoter | No | 0.275835 | 0.5252 | 1.20E-05 | Negative |
| PAX9 | chr14:37136192-37136391 | Gene body | Yes | 0.077173 | 0.2778 | 2.88E-02 | Positive |
| PKNOX2 | chr11:125037226-125037348 | Gene body | No | 0.097095 | 0.3116 | 1.37E-02 | Positive |
| TBX15 | chr1:119532320-119532542 | Promoter | No | 0.304594 | 0.5519 | 3.00E-06 | Negative |
| TBX5 | chr12:114833837-114834713 | Gene body | Yes | 0.376873 | 0.6139 | < 1.00E-06 | Negative |
| ZNF714 | chr19:21264896-21265421 | Downstream | Yes | 0.133152 | 0.3649 | 3.55E-03 | Negative |

**Supplementary Table 19: Correlation values between the frequency of MeVars observed in each tissue and the number of SNPs identified in dbSNP155**

| Tissue | Correlation value | No. of MeVars | p value |
|---|---|---|---|
| Cerebellum | 0.1487 | 730 | 0.000056 |
| Cortex | 0.3251 | 1320 | < 0.00001 |
| Skin | 0.0656 | 5239 | < 0.00001 |
| Liver | 0.0266 | 2607 | 0.1754 |
| Pancreas | 0.0141 | 1412 | 0.5963 |
| Thyroid | 0.1049 | 664 | 0.0068 |
| Blood | 0.5871 | 254 | < 0.00001 |
| Blood-African | 0.0985 | 193 | 0.1729 |
| Blood-Caucasian | 0.0762 | 6353 | < 0.00001 |
| Blood-Mongoloid | 0.1597 | 1315 | < 0.00001 |
| Blood-Indian | 0.1225 | 3331 | < 0.00001 |
| Blood-USA-Caucasian | 0.2900 | 1803 | < 0.00001 |
| Blood-USA-Hispanic | 0.2000 | 203 | 0.0042 |
